# A cytosine-to-uracil change within the programmed -1 ribosomal frameshift signal of SARS-CoV-2 results in structural similarities with the MERS-CoV signal

**DOI:** 10.1101/2020.06.26.174193

**Authors:** Dominique Fourmy, Satoko Yoshizawa

**Affiliations:** Université Paris-Saclay, CEA, CNRS, Institute for Integrative Biology of the Cell (I2BC), 91198, Gif-sur-Yvette, France

**Keywords:** COVID-19, coronavirus, sequence conservation, programmed ribosomal frameshifting, RNA pseudoknot, Nsp12 RNA-dependent RNA polymerase

## Abstract

The severe acute respiratory syndrome coronavirus 2 (SARS-CoV-2), the cause of the ongoing COVID-19 pandemic, like many other viruses, uses programmed ribosomal frameshifting (PRF) to enable synthesis of multiple proteins from its compact genome. In independent analyses, we evaluated the PRF regions of all SARS-CoV-2 sequences available in GenBank and from the Global Initiative on Sharing All Influenza Data for variations. Of the 5,156 and 27,153 sequences analyzed, respectively, the PRF regions were identical in 95.7% and 97.2% of isolates. The most common change from the reference sequence was from C to U at position 13,536, which lies in the three-stemmed pseudoknot known to stimulate frameshifting. With the conversion of the G_13493_-C_13536_ Watson-Crick pair to G-U, the SARS-CoV-2 PRF closely resembles its counterpart in the Middle East respiratory syndrome coronavirus. The occurrence of this change increased from 0.5 to 3% during the period of March to May 2020.

---

In many viruses, programmed ribosomal frameshifting (PRF) is necessary for the expression of essential viral proteins. For instance, a -1 frameshift results in production of the HIV polyprotein Gag-Pol, which is necessary for viral replication^1,2^. Coronaviruses also depend on -1 PRF to express a polyprotein that encodes the RNA-dependent RNA polymerase^3,4^. In SARS-CoV-2, the polymerase is an antiviral drug target^5,6^. The 30-kb positive-sense, single-stranded RNA genome of SARS-CoV-2 has an organization similar to that of other members of genus *Betacorona* virus^7^. The genomic RNA of SARS-CoV-2 encodes 14 open reading frames (ORFs) including the very long (more than 20 kb) ORF1a/b. This ORF encodes a polyprotein processed into 16 non-structural proteins (Nsp1-16). ORFs 1a and 1b are not in the same frame, and expression of proteins from ORF1b requires the ribosome to slip backwards in the 5’ direction by one nucleotide such that the translation proceeds in a new reading frame^8,9^.

In the PRF events characterized, frameshifting is dependent on three mRNA elements: a heptameric slippery sequence where the frameshift occurs, a *cis*-acting stimulatory RNA structure located downstream of the slippery sequence, and a linker of a few nucleotides separating these two elements^9–11^. This mRNA stimulating structure is either a pseudoknot^8,9^ or a hairpin^12–15^.

Among the *cis*-acting structures that induce -1 PRF, the hairpin-type (H-type) pseudoknot^16^ is observed very frequently. These pseudoknots usually fold as two double-stranded helices (stems 1 and 2) separated by two or three single-stranded connecting loops^17^. Detailed structural analyses of frameshifting pseudoknots from the beet western yellow virus, the sugarcane yellow leaf virus, and the simian retrovirus type-1 revealed how nucleotides from the connecting loop 2 have extensive contacts with the minor groove of stem 1^18–20^. In the SARS-CoV PRF region, a third internal stem-loop element contributes to the formation of a complex three-stemmed pseudoknot structure^21–23^.

Protein or RNA *trans*-acting factors can enhance or repress PRF efficiency. MicroRNAs bind and stabilize the pseudoknot of the human *CCR5* mRNA to increase -1 PRF efficiency^24^. The protein 2A from the encephalomyocarditis virus binds to the stimulatory RNA signal and as a consequence increases -1 PRF efficiency in the viral ORF^25^. Recently, the interferon-stimulated gene product Shiftless has been shown to be a host factor that inhibits the -1 PRF of HIV-1^26^. RNA elements that act in *cis* can also repress PRF. An attenuator RNA hairpin located immediately upstream of the slippery sequence represses -1 PRF in SARS-CoV^27,28^.

Changes in the sequence of the SARS-CoV pseudoknot affect frameshifting efficiency with drastic consequences on viral propagation, making the pseudoknot a potential drug target for antiviral treatment^29,30^. *In silico* screening identified a small ligand 2-{[4-(2-methylthiazol-4-ylmethyl)- [1,4]diazepane-1 carbonyl]amino}benzoic acid ethyl ester (MTDB) that binds the SARS-CoV pseudoknot. This ligand inhibits -1 PRF both in cell-free and cell-based assays^31^. A mechanical unfolding investigation of the pseudoknot in presence of the ligand suggests that rather than increasing pseudoknot stability the ligand reduces conformational plasticity of the pseudoknot leading to -1 PRF inhibition^32^.

As the structure of the three-stemmed pseudoknot is conserved in SARS-CoV and SARS-CoV-2^33^, it is of interest to evaluate how this sequence has evolved during the COVID-19 pandemic. Understanding how the -1 PRF sequence might change during adaptation to humans^34^ is important for antiviral therapy development as drugs should target conserved regions of the RNA to avoid emergence of resistance. In addition, host RNA dependent editing of SARS-CoV-2 genome may have important consequences on the fate of the virus and the patient^35^.

We independently analyzed 5,156 SARS-CoV-2 sequences available in GenBank and 27,153 sequences available in GISAID for sequence variation in the -1 PRF region. We found that the sequence has remained largely unchanged during the pandemic, with observed changes maintaining the overall architecture of the -1 PRF signals. In the small percentage of in GenBank and GISAID entries with changes in the PRF region relative to the reference sequence, we identified a recurrent C to U change at position 13,536. This position lies in the functional center of the three-stemmed pseudoknot. Interestingly, the conversion of the G_13493_-C_13536_ Watson-Crick pair into a wobble G-U pair results in a pseudoknot structure that closely resembles its counterpart in the Middle East respiratory syndrome coronavirus (MERS-CoV).

## RESULTS

### Comparative analysis of -1 PRF SARS-CoV-2 sequences obtained from GenBank and GISAID reveals a frequent change

We initially assessed the frequency of mutations in -1 frameshift signal region of SARS-CoV-2 (Figure 1) by analyzing sequences that were available in GenBank as of June 6, 2020. Only sequences longer than 29 kb were analyzed. Of the 5,156 sequences, 4,947 had -1 PRF signal identical to that of the Wuhan-Hu-1 reference sequence (NC_045512.2)^7^. The alignment of the sequences that did not contain an identical -1 PRF signal showed that within the pseudoknot region, position 13536 was most variable; it was changed from C in the reference strain to U in most cases (Supplementary Figure 1). This change was found in 12 isolates (0.23% frequency) from various origins. We refer to the C_13536_-to-U change as a type 1 mutation.

**Figure 1.**
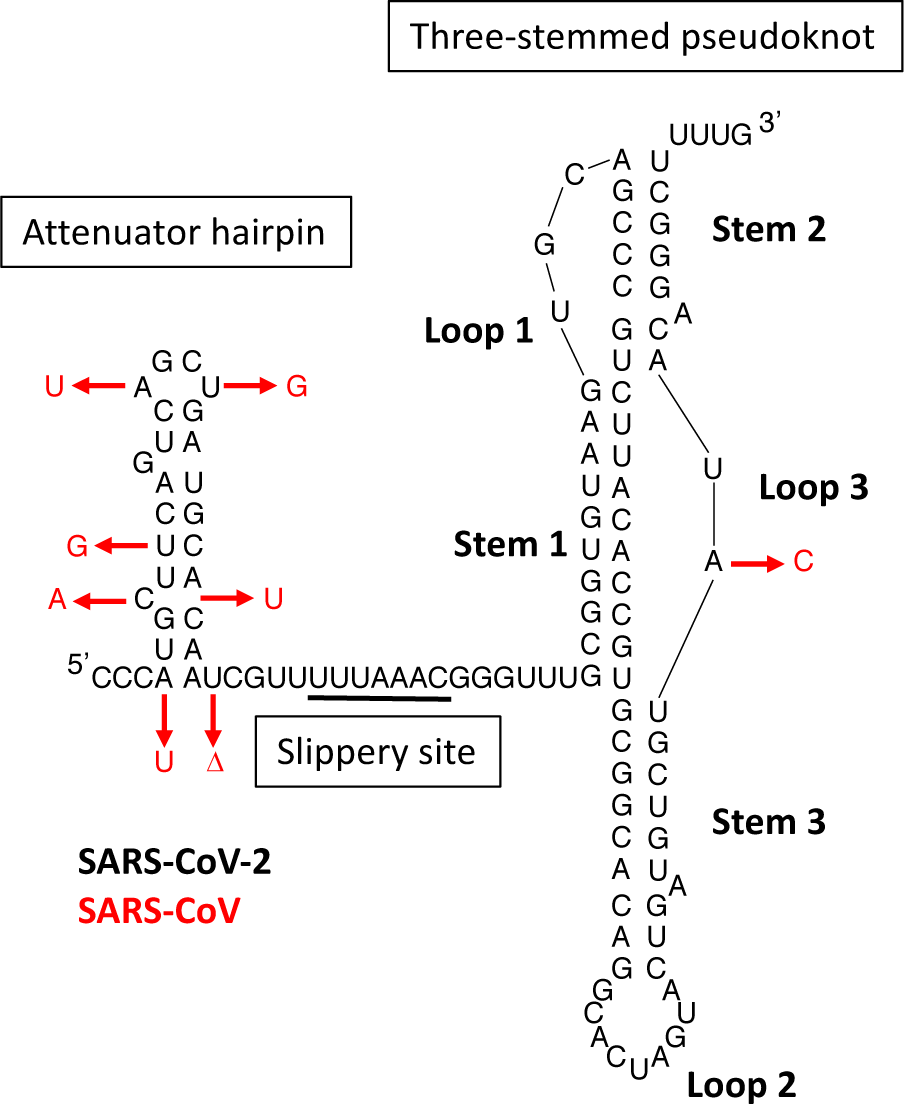
Secondary structure of the -1 PRF attenuator, slippery site (underlined), and three-stemmed pseudoknot elements of SARS-CoV-2. Differences with the SARS-CoV sequence are indicated in red (deletion: Δ).

We next extended this analysis to the larger collection of sequence data available from GISAID (https://www.epicov.org)^36,37^. Data were downloaded using “complete”, “high coverage only”, and “low coverage excluded” filtering functions. In this ensemble of 27,153 entries, conservation of the -1 PRF signal of reference strain was found for 26,405 strains, which corresponds to 97.2% (Figure 2). This agreed with the observation that any two genomes contain pairwise differences of 9.6 single nucleotide polymorphisms on average when considering the 85-nucleotide length of the -1 PRF signal^34^. The alignment of 748 sequences that did not have PRF sequence identical to that of the reference sequence revealed 418 entries with fewer than 5 ambiguous nucleotides, 93.3% of these have a single nucleotide mutation in the PRF signal. In 281 entries (1% of all sequences analyzed), the PRF region sequence was identical to that of the type 1 mutant (Figure 2). A nucleotide variation plot clearly shows that the variation is concentrated at position 13536 (Figure 3).

**Figure 2.**
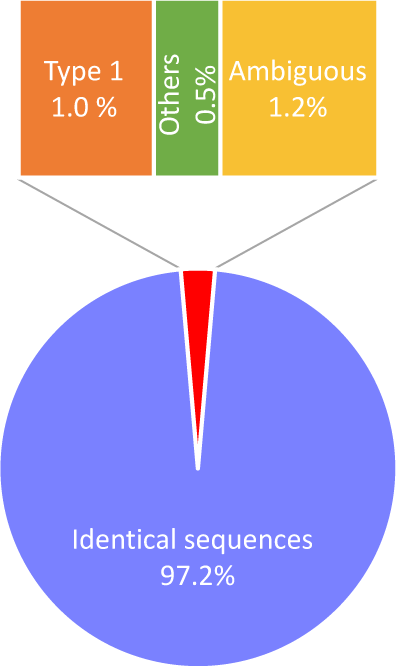
Conservation of -1 PRF region sequences among SARS-CoV-2 isolates from GISAID (n=27,153).

**Figure 3.**
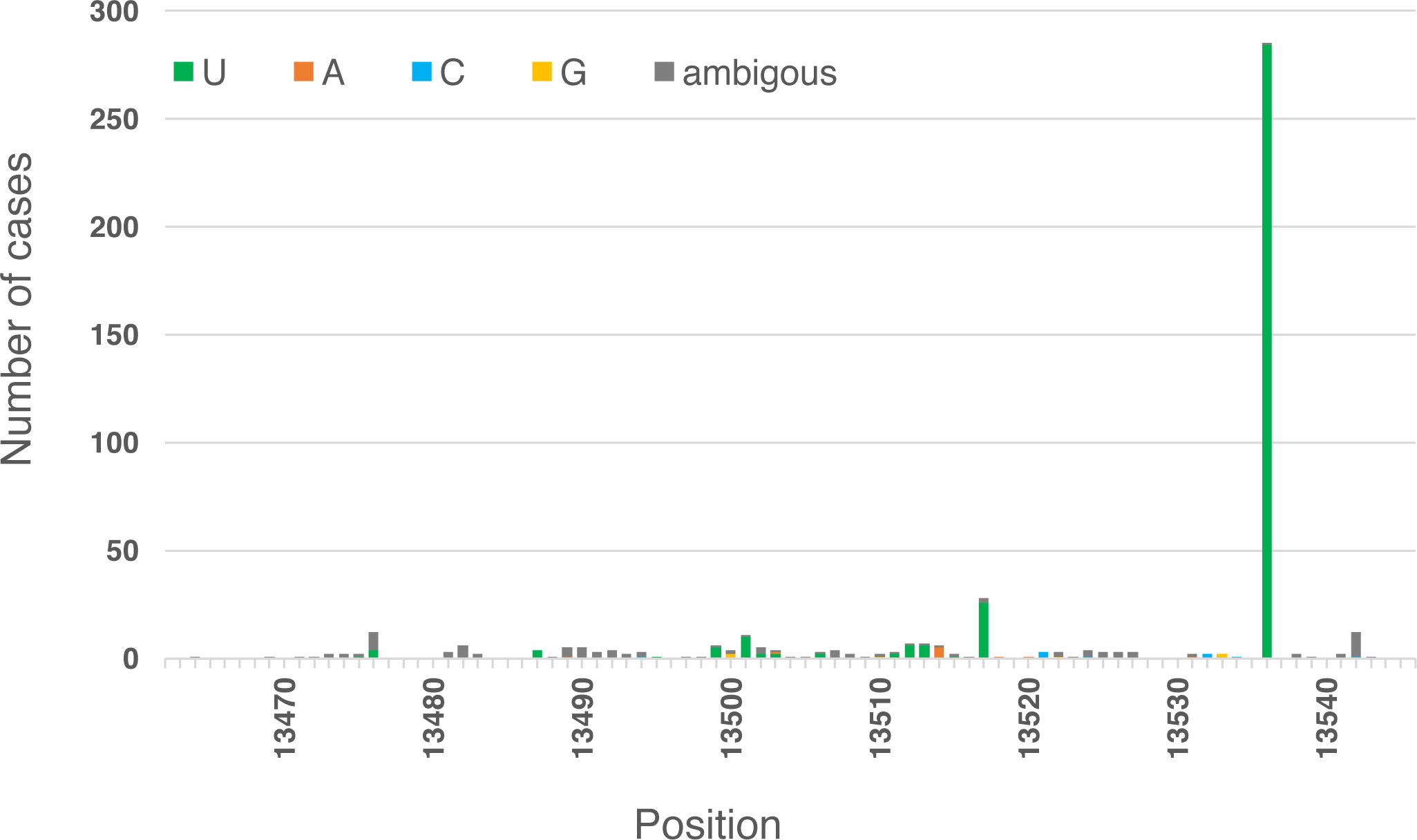
Nucleotide variations found in three-stemmed pseudoknot sequence of SARS-CoV-2 among SARS-CoV-2 isolates from GISAID (n=27,153). Number of mutations found at each position of the three-stemmed pseudoknot sequence are shown with changes indicated by bar color. Positions refer to that of the Wuhan-Hu-1 reference sequence.

There is only one nucleotide difference in the three-stemmed pseudoknot structure between SARS-CoV and SARS-CoV-233. In SARS-CoV-2, position 13533 is A, and in SARS-CoV this position is C (Figure 4). None of the SARS-CoV-2 strains analyzed had C at this position (Figure 3).

**Figure 4.**
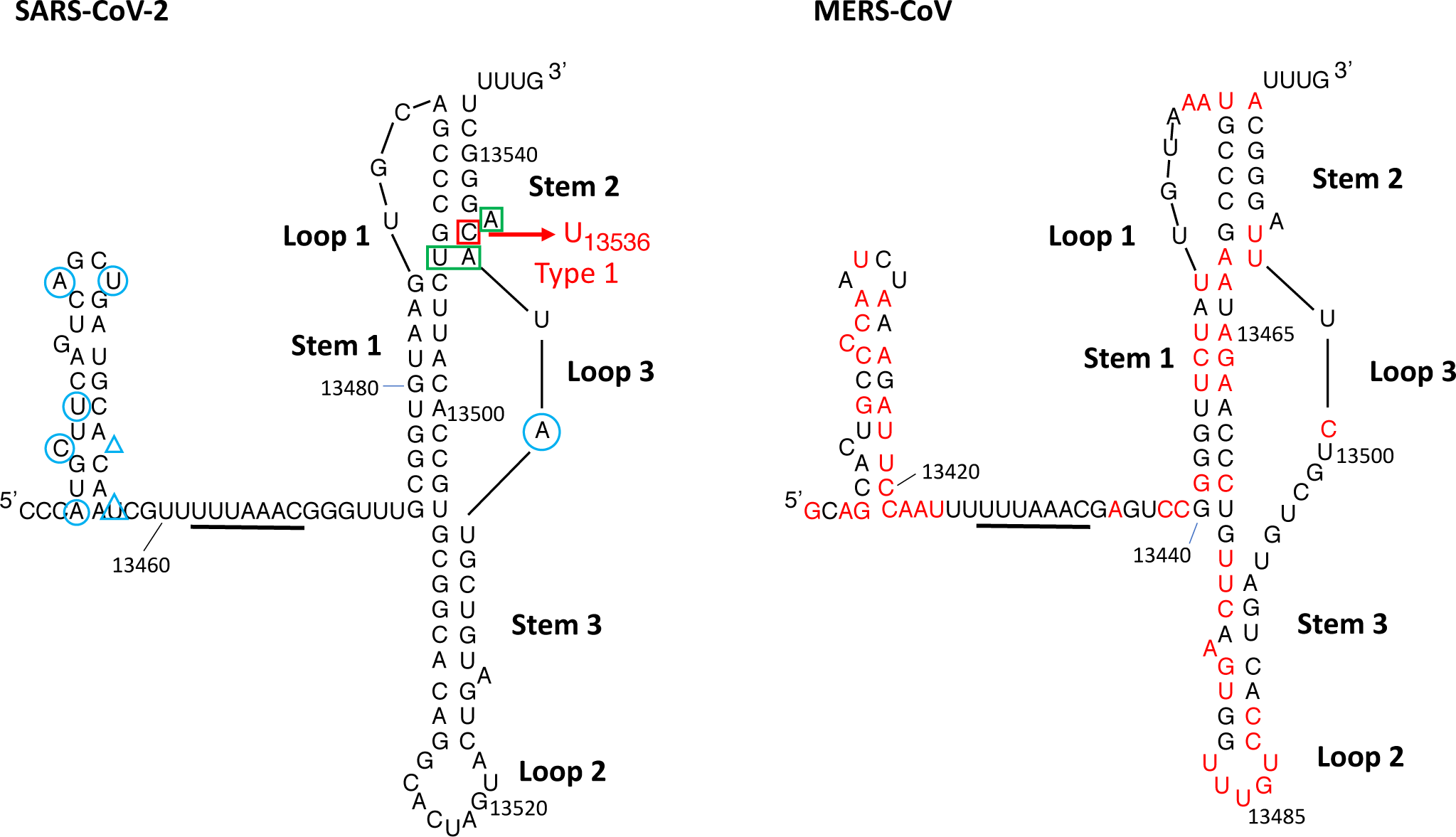
Secondary structures of the -1 PRF elements in SARS-CoV-2 and MERS-CoV. Bases that are different between the two sequences are highlighted in red in the MERS-CoV sequence (NC_019843, GenBank)^47^. Bases that differ between SARS-CoV (NC_004718, GenBank)^48^ and SARS-CoV-2 are indicated on the SARS-CoV-2 sequence with blue circles; deletions and insertions are indicated with blue triangles. Bases important for -1 PRF efficiency in SARS-CoV are marked by green boxes. The type 1 change that we have identified in SARS-CoV-2 is indicated by a red box. The type 1 mutation likely results in a non-canonical G-U pair as is observed in MERS-CoV.

### No consistent changes are observed in the 5’ attenuator stem-loop

In SARS-CoV, a short attenuator stem-loop was identified just upstream of the slippery sequence^27,28^. A comparative sequence analysis showed that in SARS-CoV-2, this stem-loop is not as highly conserved as the three-stemmed pseudoknot sequence^33^ (Figure 4). In SARS-CoV, the hairpin is capped by a UNCG tetraloop^38,39^ whereas it is an AGNN tetraloop in SARS-CoV-2^40,41^ (Figure 1). In SARS-CoVs, the stem consists of 8 to 10 base pairs. In the 35-nucleotide attenuator region of the SARS-CoV-2 isolates from the GenBank dataset, only 2.6% of the strains had attenuator hairpin sequences that differed from that of the Wuhan-Hu-1 strain (Supplementary Figure 2). Most of these strains had ambiguities in this region (more than 5 ambiguous bases in the 35-nucleotide region analyzed). Only 21 entries that had no ambiguities had mutations in the attenuator stem-loop, and none of the mutations were observed more than once. Therefore, in this hairpin, we did not identify a specific nucleotide with a strong frequency of change.

### Occurrence of type 1 changes increased over time

GISAID allows rapid data sharing, and changes in sequence over the time can be monitored. There was a considerable increase in the frequency of the type 1 change over time (Figure 5). The frequency increased 6 fold from 0.5% in March 2020 to 3% in May 2020.

**Figure 5.**
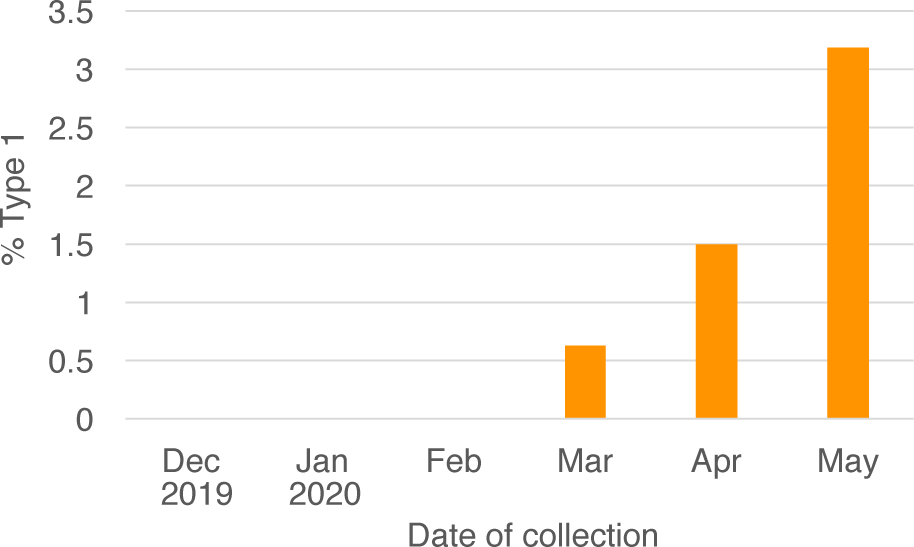
Occurrences of the SARS-CoV-2 type 1 change during the COVID-19 pandemic.

### The type 1 mutation alters the pseudoknot functional center

In the secondary structure of the Wuhan-Hu-1 reference sequence, the Watson-Crick pair G_13493_-C_13536_ lies in the vicinity of the junction between stems 1 and 2 and loop 3. The type 1 mutation likely results in the formation of a non-canonical G-U pair in the upper part of stem 2. When we compared the SARS-CoV-2 pseudoknot structure with that of related coronaviruses, we found that there is a G-U pair at the same position in the MERS-CoV pseudoknot (Figure 4). This G-U pair is flanked by two structural elements identified as important for -1 PRF efficiency in SARS-CoV: the first Watson-Crick base pair of stem 2 (U_13424_-A_13465_ in SARS-CoV and A_13462_-U_13503_ in MERS-CoV) and a bulged adenosine residue (A_13467_ and A_13505_, respectively)^22^. Consequently, the type 1 change is located in the functional center of the -1 PRF pseudoknot and the C to U substitution at position 13,536 results in interesting similarities with MERS-CoV.

### The type 1 change may have resulted from RNA editing

The deamination of cytosine into uracil is mediated by the cytidine deaminase APOBEC (Apolipoprotein B mRNA Editing Catalytic Polypeptide-like) family of proteins^42,43^. APOBECs are responsible for widespread C-to-U RNA editing of cellular transcripts. Different family members act in different tissues and cell types, and the enzyme APOBEC3A has activity in pro-inflammatory macrophages and in monocytes exposed to hypoxia and/or interferons^44^. APOBEC1 is expressed in the liver, and a shorter isoform that results from mRNA editing is detected in the small intestine. APOBECs display different sequence requirements around the target C. In SARS-CoV-2, bases flanking the C_13536_ to U mutation site match the consensus sequence of APOBEC1 (A/U-C-A/U) (Figure 4)^42,45^. A second RNA element important for editing is an 11-nucleotide mooring sequence downstream of the target C. In SARS-CoV-2, the similarity with the consensus mooring sequence of APOBEC1 is only partial. The spacer between the C_13536_ and the mooring sequence is 8 nucleotides long, within the 2- to 8-nucleotide length commonly found^46^. Thus, it is possible that the C_13536_ to U change that we identified resulted from an RNA editing event.

## DISCUSSION

As in all sequenced coronaviruses, a -1 frameshift is necessary for expression of SARS-CoV-2 early proteins such as the RNA-dependent RNA polymerase. Ribosomal frameshifting is stimulated by a three-stemmed RNA pseudoknot. As the COVID-19 pandemic continues, SARS-CoV-2 will likely evolve as it adapts to its human host^34^. Here we focused on the sequence variability of the -1 PRF signals by analysis of 5,156 SARS-CoV-2 sequences that were available in the GenBank as of June 6, 2020 and 27,153 entries from GISAID available as of June 7, 2020. We found that the 85-nucleotide -1 PRF region is highly conserved with 95.7% and 97.2% of isolates, respectively, identical in this region to the reference sequence, Wuhan-Hu-1 (NC_045512.2). Among the isolates with differences from the reference sequence in the -1 PRF region, there is a recurrent change. This change, which we refer to as a type 1 mutation, corresponds to substitution of C with U in stem 2 of the pseudoknot. SARS-CoV-2 encodes a 3’-to-5’ exoribonuclease in nonstructural protein 14 (nsp14-ExoN) which ensures high-fidelity replication. When nsp14-ExoN is inactivated, the spectrum of mutations introduced by the Nsp12 RNA-dependent RNA polymerase becomes apparent^49^. As the C-to-U transition is not a frequent change introduced by the polymerase, it is possible that it originates from RNA editing by a cytosine deaminase^35,42,43^.

Biophysical single-molecule assays and kinetics studies on ensembles revealed how the interplay between the conformational plasticity of RNA-inducing frameshifting structures and the dynamics of the ribosome play a critical role in the -1 PRF mechanism^50–54^. Interestingly, the type 1 change results in the substitution of a G-C Watson-Crick pair in the functional core of the SARS-CoV-2 pseudoknot with a wobble G-U pair. This likely alters the conformational plasticity of the pseudoknot with possible functional consequences as the dynamics of this region of the RNA are critical for frameshifting efficiency^22^.

There is a lack of clinically effective antiviral drugs for treatment of highly pathogenic coronaviruses^55^. Since frameshifts are essential, the PRF region is a target for drug development^32,33^. The small molecule MTDB binds to the SARS-CoV pseudoknot and inhibits ribosomal frameshifting^31^. Since the functional center of SARS-CoV pseudoknot was identified as the drug binding site, by analogy the same regions in SARS-CoV-2 and MERS-CoV might as well form sites that could be bound by small molecules to inhibit viral replication.

Our analysis showed that during the COVID-19 pandemic, the sequence of the stimulator pseudoknot has been highly conserved. However, it is interesting to note that the tolerated type 1 change modifies the functional center of the SARS-CoV-2 pseudoknot to closely resemble its counterpart in MERS-CoV.

## METHODS

### Sequence analysis

Genomic sequences of SARS-CoV-2 isolates were downloaded from the SARS-CoV-2 data hub of the NCBI Virus website^56^. All the sequences longer than 29,000 bases that were released by June 6, 2020 were analyzed (5,156 sequences). Sequences that did not contain a region identical to the -1 frameshift signal of Wuhan-Hu-1 strain(5’-TTTAAACGGGTTTGCGGTGTAAGTGCAGCCCGTCTTACACCGTGCGGCACAGGCACTAGTACTGATGTCGTATACAGGGCTTTTG-3’)^7,33^ were selected. From the selected sequences, a 2,500-base region (11,500 to 14,000 bases from the 5’ end) containing the 85 nucleotides of the -1 PRF region of ORF1a/b were extracted and aligned using ClustalX software^57^. For attenuator hairpin, the same analysis was performed using the 35 nucleotides including this hairpin-forming sequence:(5’-CCCATGCTTCAGTCAGCTGATGCACAATCGTTTTT-3’).

For the analysis of data from GISAID, sequences available as of June 7, 2020 were downloaded from GISAID Initiative EpiCoV platform using “complete”, “high coverage only”, and “low coverage excluded” filtering functions. Sequences derived from animals (bat and pangolin) were excluded. A total of 27,153 sequences were analyzed as described for the GenBank sequence analysis. A full acknowledgements table of the laboratories where the clinical specimens and/or virus isolates were obtained and laboratories where data have been generated was obtained from the GISAID website and is included as Supplementary Table 1.

