## Supplementary-Figures-1-2 for "A cytosine-to-uracil change within the programmed -1 ribosomal frameshift signal of SARS-CoV-2 results in structural similarities with the MERS-CoV signal"

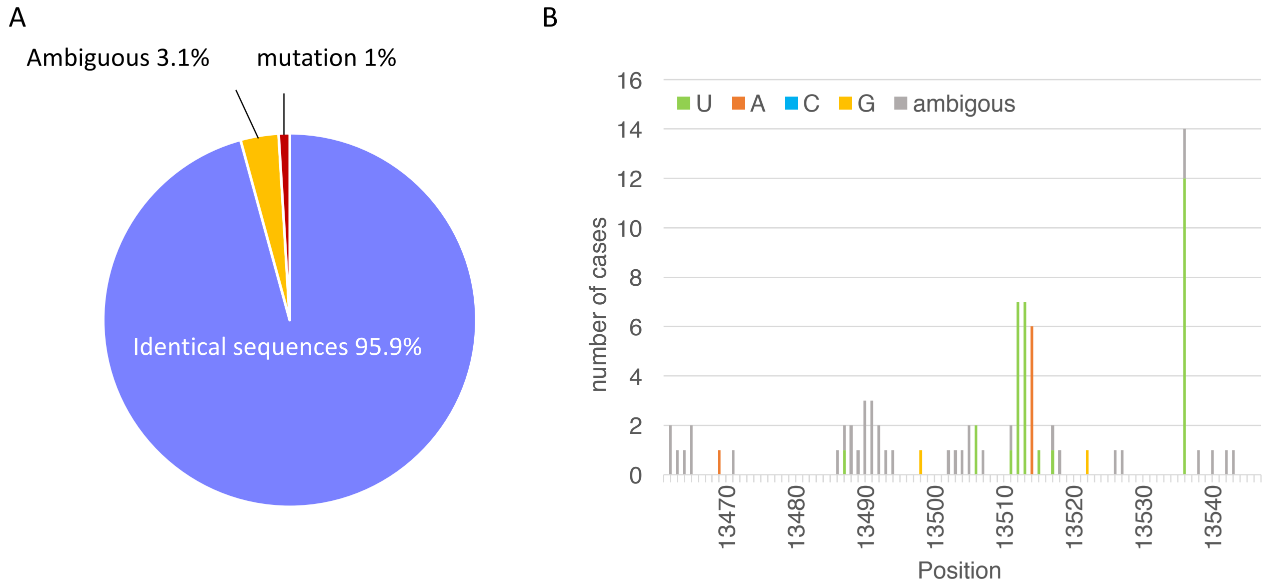


**Supplementary Figure 1.** (A) Conservation of -1 PRF region sequences among SARS-CoV-2 isolates in GenBank database (n= 5,156). (B) Nucleotide variations found in three-stemmed pseudoknot sequence of SARS-CoV-2. Frequencies and identities of the mutations found at each position of the three-stemmed pseudoknot sequence. Positions refer to that of the Wuhan-Hu-1 reference sequence.

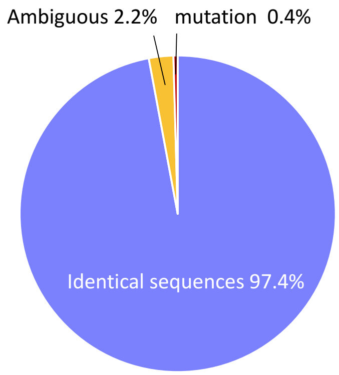


**Supplementary Figure 2.** Conservation of the attenuator sequences among SARS-CoV-2 isolates in GenBank (n= 5,156).
