## Supplementary-Table1-1 for "A cytosine-to-uracil change within the programmed -1 ribosomal frameshift signal of SARS-CoV-2 results in structural similarities with the MERS-CoV signal"

**We gratefully acknowledge the authors, originating and submitting laboratories of the sequences from GISAID's EpiFlu™ Database on which this research is based. The list is detailed below.**
**All submitters of data may be contacted directly via www.gisaid.org**

| Accession ID | Virus name | Location | Collection date | Originating lab | Submitting lab | Authors |
|---|---|---|---|---|---|---|
| EPI_ISL_404227 | hCoV-19/Zhejiang/WZ-01/2020 | Asia / China / Zhejiang | 2020-01-16 | Zhejiang Provincial Center for Disease Control and Prevention | Department of Microbiology, Zhejiang Provincial Center for Disease Control and Prevention | Yin Chen, Yanjun Zhang, Haiyan Mao, Junhang Pan, Xiuyu Lou, Yiyu Lu, Juying Yan, Hanping Zhu, Jian Gao, Yan Feng, Yi Sun, Hao Yan, Zhen Li, Yisheng Sun, Liming Gong, Qiong Ge, Wen Shi, Xinying Wang, Wenwu Yao, Zhangnv Yang, Fang Xu, Chen Chen, Enfu Chen, Zhen Wang, Zhiping Chen, Jianmin Jiang, Chonggao Hu |
| EPI_ISL_404228 | hCoV-19/Zhejiang/WZ-02/2020 | Asia / China / Zhejiang | 2020-01-17 | Zhejiang Provincial Center for Disease Control and Prevention | Department of Microbiology, Zhejiang Provincial Center for Disease Control and Prevention | Yanjun Zhang, Yin Chen, Haiyan Mao, Junhang Pan, Xiuyu Lou, Yiyu Lu, Juying Yan, Hanping Zhu, Jian Gao, Yan Feng, Yi Sun, Hao Yan, Zhen Li, Yisheng Sun, Liming Gong, Qiong Ge, Wen Shi, Xinying Wang, Wenwu Yao, Zhangnv Yang, Fang Xu, Chen Chen, Enfu Chen, Zhen Wang, Zhiping Chen, Jianmin Jiang, Chonggao Hu |
| EPI_ISL_404253 | hCoV-19/USA/IL1/2020 | North America / USA / Illinois / Chicago | 2020-01-21 | IL Department of Public Health Chicago Laboratory | Pathogen Discovery, Respiratory Viruses Branch, Division of Viral Diseases, Centers for Diseas Control and Prevention | Ying Tao, Krista Queen, Clinton R. Paden, Jing Zhang, Yan Li, Anna Uehara, Xiaoyan Lu, Brian Lynch, Senthil Kumar K. Sakthivel, Brett L. Whitaker, Shifaq Kamili, Lijuan Wang, Janna' R. Murray, Susan I. Gerber, Stephen Lindstrom, Suxiang Tong |
| EPI_ISL_406030 | hCoV-19/Shenzhen/HKU-SZ-002/2020 | Asia / China / Guangdong / Shenzhen | 2020-01-10 | The University of Hong Kong - Shenzhen Hospital | Li Ka Shing Faculty of Medicine, The University of Hong Kong | Chan,J.F.-W., Yuan,S., Kok,K.H., To,K.K.-W., Chu,H., Yang,J., Xing,F., Liu,J., Yip,C.C.-Y., Poon,R.W.-S., Tsai,H.W., Lo,S.K.-F., Chan,K.H., Poon,V.K.-M., Chan,W.M., Ip,J.D., Cai,J.P., Cheng,V.C.-C., Chen,H., Hui,C.K.-M. and Yuen,K.Y. |
| EPI_ISL_406031 | hCoV-19/Taiwan/2/2020 | Asia / Taiwan / Kaohsiung | 2020-01-23 | Centers for Disease Control, R.O.C. (Taiwan) | Centers for Disease Control, R.O.C. (Taiwan) | Ji-Rong Yang, Yu-Chi Lin, Jung-Jung Mu, Ming-Tsan Liu, Shu-Ying Li |
| EPI_ISL_406034 | hCoV-19/USA/CA1/2020 | North America / USA / California / Los An | 2020-01-23 | California Department of Public Health | Pathogen Discovery, Respiratory Viruses Branch, Division of Viral Diseases, Centers for Diseas Control and Prevention | Anna Uehara, Krista Queen, Ying Tao, Yan Li, Clinton R. Paden, Jing Zhang, Xiaoyan Lu, Brian Lynch, Senthil Kumar K. Sakthivel, Brett L. Whitaker, Shifaq Kamili, Lijuan Wang, Janna' R. Murray, Susan I. Gerber, Stephen Lindstrom, Suxiang Tong |
| EPI_ISL_406036 | hCoV-19/USA/CA2/2020 | North America / USA / California / Orange | 2020-01-22 | California Department of Public Health | Pathogen Discovery, Respiratory Viruses Branch, Division of Viral Diseases, Centers for Diseas Control and Prevention | Anna Uehara, Krista Queen, Ying Tao, Yan Li, Clinton R. Paden, Jing Zhang, Xiaoyan Lu, Brian Lynch, Senthil Kumar K. Sakthivel, Brett L. Whitaker, Shifaq Kamili, Lijuan Wang, Janna' R. Murray, Susan I. Gerber, Stephen Lindstrom, Suxiang Tong |
| EPI_ISL_406531 | hCoV-19/Guangdong/20SF174/2020 | Asia / China / Guangdong / Zhuhai | 2020-01-22 | Guangdong Provincial Center for Diseases Contorl and Prevention; Guangdong Provincial Public Health | Guangdong Provincial Center for Disease Control and Prevention | Min Kang, Jie Wu, Jing Lu, Tao Liu, Baisheng Li, Shujiang Mei, Feng Ruan, Lifeng Lin, Changwen Ke, Haojie Zhong, Yingtao Zhang, Lirong Zou, Xuguang Chen, Qi Zhu, Jianpeng Xiao, Jianxiang Geng, Zhe Liu, Jianxiong Hu, Weilin Zeng, Xing Li, Yuhuang Liao, Xiujuan Tang, Songjian Xiao, Ying Wang, Yingchao Song, Xue Zhuang, Lijun Liang, Guanhao He, Huihong Deng, Tie Song, Jianfeng He, Wenjun Ma |
| EPI_ISL_406533 | hCoV-19/Guangzhou/20SF206/2020 | Asia / China / Guangdong / Guangzhou | 2020-01-22 | Guangdong Provincial Center for Diseases Contorl and Prevention; Guangdong Provincial Public Health | Guangdong Provincial Center for Diseases Control and Prevention | Min Kang, Jie Wu, Jing Lu, Tao Liu, Baisheng Li, Shujiang Mei, Feng Ruan, Lifeng Lin, Changwen Ke, Haojie Zhong, Yingtao Zhang, Lirong Zou, Xuguang Chen, Qi Zhu, Jianpeng Xiao, Jianxiang Geng, Zhe Liu, Jianxiong Hu, Weilin Zeng, Xing Li, Yuhuang Liao, Xiujuan Tang, Songjian Xiao, Ying Wang, Yingchao Song, Xue Zhuang, Lijun Liang, Guanhao He, Huihong Deng, Tie Song, Jianfeng He, Wenjun Ma |
| EPI_ISL_406534 | hCoV-19/Foshan/20SF207/2020 | Asia / China / Guangdong / Foshan | 2020-01-22 | Guangdong Provincial Center for Diseases Control and Prevention; Guangdong Provincial Public Health | Guangdong Provincial Center for Diseases Control and Prevention | Min Kang, Jie Wu, Jing Lu, Tao Liu, Baisheng Li, Shujiang Mei, Feng Ruan, Lifeng Lin, Changwen Ke, Haojie Zhong, Yingtao Zhang, Lirong Zou, Xuguang Chen, Qi Zhu, Jianpeng Xiao, Jianxiang Geng, Zhe Liu, Jianxiong Hu, Weilin Zeng, Xing Li, Yuhuang Liao, Xiujuan Tang, Songjian Xiao, Ying Wang, Yingchao Song, Xue Zhuang, Lijun Liang, Guanhao He, Huihong Deng, Tie Song, Jianfeng He, Wenjun Ma |
| EPI_ISL_406535 | hCoV-19/Foshan/20SF210/2020 | Asia / China / Guangdong / Foshan | 2020-01-22 | Guangdong Provincial Center for Diseases Control and Prevention; Guangdong Provincial Public Health | Guangdong Provincial Center for Diseases Control and Prevention | Min Kang, Jie Wu, Jing Lu, Tao Liu, Baisheng Li, Shujiang Mei, Feng Ruan, Lifeng Lin, Changwen Ke, Haojie Zhong, Yingtao Zhang, Lirong Zou, Xuguang Chen, Qi Zhu, Jianpeng Xiao, Jianxiang Geng, Zhe Liu, Jianxiong Hu, Weilin Zeng, Xing Li, Yuhuang Liao, Xiujuan Tang, Songjian Xiao, Ying Wang, Yingchao Song, Xue Zhuang, Lijun Liang, Guanhao He, Huihong Deng, Tie Song, Jianfeng He, Wenjun Ma |
| EPI_ISL_406536 | hCoV-19/Foshan/20SF211/2020 | Asia / China / Guangdong / Foshan | 2020-01-22 | Guangdong Provincial Center for Diseases Control and Prevention; Guangdong Provincial Public Health | Guangdong Provincial Center for Diseases Control and Prevention | Min Kang, Jie Wu, Jing Lu, Tao Liu, Baisheng Li, Shujiang Mei, Feng Ruan, Lifeng Lin, Changwen Ke, Haojie Zhong, Yingtao Zhang, Lirong Zou, Xuguang Chen, Qi Zhu, Jianpeng Xiao, Jianxiang Geng, Zhe Liu, Jianxiong Hu, Weilin Zeng, Xing Li, Yuhuang Liao, Xiujuan Tang, Songjian Xiao, Ying Wang, Yingchao Song, Xue Zhuang, Lijun Liang, Guanhao He, Huihong Deng, Tie Song, Jianfeng He, Wenjun Ma |
| EPI_ISL_406538 | hCoV-19/Guangdong/20SF201/2020 | Asia / China / Guangdong | 2020-01-23 | Guangdong Provincial Center for Diseases Control and Prevention;Guangdong Provincial Institute of Public Health | Guangdong Provincial Center for Diseases Control and Prevention | Min Kang, Jie Wu, Jing Lu, Tao Liu, Baisheng Li, Shujiang Mei, Feng Ruan, Lifeng Lin, Changwen Ke, Haojie Zhong, Yingtao Zhang, Lirong Zou, Xuguang Chen, Qi Zhu, Jianpeng Xiao, Jianxiang Geng, Zhe Liu, Jianxiong Hu, Weilin Zeng, Xing Li, Yuhuang Liao, Xiujuan Tang, Songjian Xiao, Ying Wang, Yingchao Song, Xue Zhuang, Lijun Liang, Guanhao He, Huihong Deng, Tie Song, Jianfeng He, Wenjun Ma |
| EPI_ISL_406592 | hCoV-19/Shenzhen/SZTH-001/2020 | Asia / China / Guangdong / Shenzhen | 2020-01-13 | Shenzhen Third People's Hospital | Shenzhen Key Laboratory of Pathogen and Immunity, National Clinical Research Center for Infectious Disease,Shenzhen Third People's Hospital | Yang Yang, Chenguang Shen, Li Xing, Zhixiang Xu, Haixia Zheng, Yingxia Liu |
| EPI_ISL_406593 | hCoV-19/Shenzhen/SZTH-002/2020 | Asia / China / Guangdong / Shenzhen | 2020-01-13 | Shenzhen Key Laboratory of Pathogen and Immunity, National Clinical Research Center for Infectious Disease, Shenzhen Third People's Hospital | Shenzhen Key Laboratory of Pathogen and Immunity, National Clinical Research Center for Infectious Disease, Shenzhen Third People's Hospital | Yang Yang, Chenguang Shen, Li Xing, Zhixiang Xu, Haixia Zheng, Yingxia Liu |
| EPI_ISL_406594 | hCoV-19/Shenzhen/SZTH-003/2020 | Asia / China / Guangdong / Shenzhen | 2020-01-16 | Shenzhen Key Laboratory of Pathogen and Immunity, National Clinical Research Center for Infectious Disease, Shenzhen Third People's Hospital | Shenzhen Key Laboratory of Pathogen and Immunity, National Clinical Research Center for Infectious Disease, Shenzhen Third People's Hospital | Yang Yang, Chenguang Shen, Li Xing, Zhixiang Xu, Haixia Zheng, Yingxia Liu |
| EPI_ISL_406595 | hCoV-19/Shenzhen/SZTH-004/2020 | Asia / China / Guangdong / Shenzhen | 2020-01-16 | Shenzhen Key Laboratory of Pathogen and Immunity, National Clinical Research Center for Infectious Disease, Shenzhen Third People's Hospital | Shenzhen Key Laboratory of Pathogen and Immunity, National Clinical Research Center for Infectious Disease, Shenzhen Third People's Hospital | Yang Yang, Chenguang Shen, Li Xing, Zhixiang Xu, Haixia Zheng, Yingxia Liu |
| EPI_ISL_406596 | hCoV-19/France/IDF0372/2020 | Europe / France / Ile-de-France / Paris | 2020-01-23 | Department of Infectious and Tropical Diseases, Bichat Claude Bernard Hospital, Paris | National Reference Center for Viruses of Respiratory Infections, Institut Pasteur, Paris | Mélanie Albert, Marion Barbet, Sylvie Behillil, Méline Bizard, Angela Brisebarre, Flora Donati, Vincent Enouf, Maud Vanpeene, Sylvie van der Werf, Yazdan Yazdanpanah, Xavier Lescure. |
| EPI_ISL_406597 | hCoV-19/France/IDF0373/2020 | Europe / France / Ile-de-France / Paris | 2020-01-23 | Department of Infectious and Tropical Diseases, Bichat Claude Bernard Hospital, Paris | National Reference Center for Viruses of Respiratory Infections, Institut Pasteur, Paris | Mélanie Albert, Marion Barbet, Sylvie Behillil, Méline Bizard, Angela Brisebarre, Flora Donati, Vincent Enouf, Maud Vanpeene, Sylvie van der Werf, Yazdan Yazdanpanah, Xavier Lescure. |
| EPI_ISL_406800 | hCoV-19/Wuhan/WH03/2020 | Asia / China / Hubei / Wuhan | 2020-01-01 | General Hospital of Central Theater Command of People's Liberation Army of China | BGI & Institute of Microbiology, Chinese Academy of Sciences & Shandong First Medical University & Shandong Academy of Medical Sciences & General Hospital of Central Theater Command of People's Liberation Army of China | Weijun Chen, Yuhai Bi, Weifeng Shi and Zhenhong Hu |
| EPI_ISL_406801 | hCoV-19/Wuhan/WH04/2020 | Asia / China / Hubei / Wuhan | 2020-01-05 | General Hospital of Central Theater Command of People's Liberation Army of China | BGI & Institute of Microbiology, Chinese Academy of Sciences & Shandong First Medical University & Shandong Academy of Medical Sciences & General Hospital of Central Theater Command of People's Liberation Army of China | Weijun Chen, Yuhai Bi, Weifeng Shi and Zhenhong Hu |
| EPI_ISL_406862 | hCoV-19/Germany/BavPat1/2020 | Europe / Germany / Bavaria / Munich | 2020-01-28 | Charité Universitätsmedizin Berlin, Institute of Virology; Institut für Mikrobiologie der Bundeswehr, Munich | Charité Universitätsmedizin Berlin, Institute of Virology | Victor M Corman, Julia Schneider, Talitha Veith, Barbara Mühlemann, Markus Antwerpen, Christian Drosten, Roman Wölfel |
| EPI_ISL_407313 | hCoV-19/Hangzhou/HZCDC0001/2020 | Asia / China / Zhejiang / Hangzhou | 2020-01-19 | Hangzhou Center for Disease Control and Prevention | Hangzhou Center for Disease Control and Prevention | Jun Li, Haoqiu Wang, Hua Yu, Lingfeng Mao, Xinfen Yu, Zhou Sun, Qingxin Kong, Xin Qian, Shuchang Chen, Xuchu Wang |
| EPI_ISL_407893 | hCoV-19/Australia/NSW01/2020 | Oceania / Australia / New South Wales / | 2020-01-24 | Centre for Infectious Diseases and Microbiology Laboratory Services | NSW Health Pathology - Institute of Clinical Pathology and Medical Research; Westmead Hospital; University of Sydney | Eden J-S, Carter I, Rahman H, Holmes EC, Rockett R, O'Sullivan MV, Sintchenko V, Chen SC, Maddocks S, Kok J and Dwyer DE for the 2019-nCoV Study Group |
| EPI_ISL_407894 | hCoV-19/Australia/QLD01/2020 | Oceania / Australia / Queensland / Gold C | 2020-01-28 | Pathology Queensland | Public Health Virology Laboratory | Ben Huang, Alyssa Pyke, Amanda De Jong, Andrew Van Den Hurk, Carmel Taylor, David Warrilow, Doris Genge, Elisabeth Gamez, Glen Hewitson, Ian Maxwell Mackay, Inga Sultana, Jamie McMahon, Jean Barcelon, Judy Northill, Mitchell Finger, Natalie Simpson, Neelima Nair, Peter Burtonclay, Peter Moore, Sarah Wheatley, Sean Moody, Sonja Hall-Mendelin, Timothy Gardam, and Frederick Moore. |
| EPI_ISL_407896 | hCoV-19/Australia/QLD02/2020 | Oceania / Australia / Queensland / Gold C | 2020-01-30 | Pathology Queensland | Public Health Virology Laboratory | Ben Huang, Alyssa Pyke, Amanda De Jong, Andrew Van Den Hurk, Carmel Taylor, David Warrilow, Doris Genge, Elisabeth Gamez, Glen Hewitson, Ian Maxwell Mackay, Inga Sultana, Jamie McMahon, Jean Barcelon, Judy Northill, Mitchell Finger, Natalie Simpson, Neelima Nair, Peter Burtonclay, Peter Moore, Sarah Wheatley, Sean Moody, Sonja Hall-Mendelin, Timothy Gardam, and Frederick Moore. |
| EPI_ISL_408665 | hCoV-19/Japan/TY-WK-012/2020 | Asia / Japan / Tokyo | 2020-01-29 | Dept. of Virology III, National Institute of Infectious Diseases | Pathogen Genomics Center, National Institute of Infectious Diseases | Tsuyoshi Sekizuka, Shutoku Matsuyama, Naganori Nao, Kazuya Shirato, Makoto Takeda, Makoto Kuroda |
| EPI_ISL_408666 | hCoV-19/Japan/TY-WK-501/2020 | Asia / Japan / Tokyo | 2020-01-31 | Dept. of Virology III, National Institute of Infectious Diseases | Pathogen Genomics Center, National Institute of Infectious Diseases | Tsuyoshi Sekizuka, Shutoku Matsuyama, Naganori Nao, Kazuya Shirato, Makoto Takeda, Makoto Kuroda |
| EPI_ISL_408667 | hCoV-19/Japan/TY-WK-521/2020 | Asia / Japan / Tokyo | 2020-01-31 | Dept. of Virology III, National Institute of Infectious Diseases | Pathogen Genomics Center, National Institute of Infectious Diseases | Tsuyoshi Sekizuka, Shutoku Matsuyama, Naganori Nao, Kazuya Shirato, Makoto Takeda, Makoto Kuroda |
| EPI_ISL_408668 | hCoV-19/Vietnam/VR03-38142/2020 | Asia / Vietnam / Thanh Hoa | 2020-01-24 | National Influenza Center - National Institute of Hygiene and Epidemiology (NIHE) | National Influenza Center - National Institute of Hygiene and Epidemiology (NIHE) | Ung Thi Hong Trang, Hoang Vu Mai Phuong, Nguyen Le Khanh Hang, Nguyen Vu Son, Le Thi Thanh, Vuong Duc Cuong, Nguyen Phuong Anh, Pham Thi Hien, Tran Thu Huong, Le Thi Quynh Mai, |
| EPI_ISL_408669 | hCoV-19/Japan/KY-V-029/2020 | Asia / Japan / Kyoto | 2020-01-29 | Dept. of Virology III, National Institute of Infectious Diseases | Pathogen Genomics Center, National Institute of Infectious Diseases | Tsuyoshi Sekizuka, Shutoku Matsuyama, Naganori Nao, Kazuya Shirato, Makoto Takeda, Makoto Kuroda |
| EPI_ISL_408670 | hCoV-19/USA/WI1/2020 | North America / USA / Wisconsin | 2020-01-31 | Wisconsin Department of Health Services | Pathogen Discovery, Respiratory Viruses Branch, Division of Viral Diseases, Centers for Diseas Control and Prevention | Jing Zhang, Anna Uehara, Krista Queen, Yan Li, Ying Tao, Clinton R. Paden, Xiaoyan Lu, Brian Lynch, Senthil Kumar K. Sakthivel, Brett L. Whitaker, Shifaq Kamili, Lijuan Wang, Janna' R. Murray, Susan I. Gerber, Stephen Lindstrom, Suxiang Tong |
| EPI_ISL_409067 | hCoV-19/USA/MA1/2020 | North America / USA / Massachusetts | 2020-01-29 | Massachusetts Department of Public Health | Pathogen Discovery, Respiratory Viruses Branch, Division of Viral Diseases, Centers for Diseas Control and Prevention | Clinton R. Paden, Jing Zhang, Krista Queen, Yan Li, Ying Tao, Anna Uehara, Xiaoyan Lu, Brian Lynch, Senthil Kumar K. Sakthivel, Brett L. Whitaker, Shifaq Kamili, Lijuan Wang, Janna' R. Murray, Susan I. Gerber, Stephen Lindstrom, Suxiang Tong |
| EPI_ISL_410044 | hCoV-19/USA/CA6/2020 | North America / USA / California | 2020-01-27 | California Department of Public Health | Pathogen Discovery, Respiratory Viruses Branch, Division of Viral Diseases, Centers for Diseas Control and Prevention | Jing Zhang, Krista Queen, Yan Li, Ying Tao, Anna Uehara, Clinton R. Paden, Xiaoyan Lu, Brian Lynch, Senthil Kumar K. Sakthivel, Brett L. Whitaker, Shifaq Kamili, Lijuan Wang, Janna' R. Murray, Susan I. Gerber, Stephen Lindstrom, Suxiang Tong |
| EPI_ISL_410045 | hCoV-19/USA/IL2/2020 | North America / USA / Illinois | 2020-01-28 | IL Department of Public Health Chicago Laboratory | Pathogen Discovery, Respiratory Viruses Branch, Division of Viral Diseases, Centers for Diseas Control and Prevention | Yan Li, Jing Zhang, Krista Queen, Ying Tao, Anna Uehara, Clinton R. Paden, Xiaoyan Lu, Brian Lynch, Senthil Kumar K. Sakthivel, Brett L. Whitaker, Shifaq Kamili, Lijuan Wang, Janna' R. Murray, Susan I. Gerber, Stephen Lindstrom, Suxiang Tong |
| EPI_ISL_410486 | hCoV-19/France/RA739/2020 | Europe / France / Rhone-Alpes / Contami | 2020-02-08 | CNR Virus des Infections Respiratoires - France SUD | CNR Virus des Infections Respiratoires - France SUD | Bal, Antonin; Destras, Gregory; Gaymard, Alexandre; Bouscambert-Duchamp, Maude; Cheynet, Valérie; Brengel-Pesce, Karen; Morfin-Sherpa, Florence; Valette, Martine; Josset, Laurence; Lina, Bruno. |
| EPI_ISL_410531 | hCoV-19/Japan/NA-20-05-1/2020 | Asia / Japan / Nara | 2020-01-25 | Dept. of Pathology, National Institute of Infectious Diseases | Pathogen Genomics Center, National Institute of Infectious Diseases | Tsuyoshi Sekizuka, Harutaka Katano, Shutoku Matsuyama, Naganori Nao, Kazuya Shirato, Motoi Suzuki, Hideki Hasegawa, Takaji Wakita, Makoto Takeda, Tadaki Suzuki, Makoto Kuroda |
| EPI_ISL_410532 | hCoV-19/Japan/OS-20-07-1/2020 | Asia / Japan / Osaka | 2020-01-23 | Dept. of Pathology, National Institute of Infectious Diseases | Pathogen Genomics Center, National Institute of Infectious Diseases | Tsuyoshi Sekizuka, Harutaka Katano, Shutoku Matsuyama, Naganori Nao, Kazuya Shirato, Motoi Suzuki, Hideki Hasegawa, Takaji Wakita, Makoto Takeda, Tadaki Suzuki, Makoto Kuroda |
| EPI_ISL_410535 | hCoV-19/Singapore/4/2020 | Asia / Singapore | 2020-02-03 | National Centre for Infectious Diseases | Programme in Emerging Infectious Diseases, Duke-NUS Medical School | Danielle E Anderson, Martin Linster, Yan Zhuang, Jayanthi Jayakumar, David CB Lye, Yee Sin Leo, Barnaby E Young, Yvonne CF Su, Gavin JD Smith |
| EPI_ISL_410536 | hCoV-19/Singapore/5/2020 | Asia / Singapore | 2020-02-06 | Singapore General Hospital, Molecular Laboratory, Division of Pathology | Programme in Emerging Infectious Diseases, Duke-NUS Medical School | Danielle E Anderson, Martin Linster, Yan Zhuang, Jayanthi Jayakumar, Kian Sing Chan, Lynette LE Oon, Shirin Kalimuddin, Jenny GH Low, Yvonne CF Su, Gavin JD Smith |
| EPI_ISL_410537 | hCoV-19/Singapore/6/2020 | Asia / Singapore | 2020-02-09 | Singapore General Hospital, Molecular Laboratory, Division of Pathology | Programme in Emerging Infectious Diseases, Duke-NUS Medical School | Danielle E Anderson, Martin Linster, Yan Zhuang, Jayanthi Jayakumar, Kian Sing Chan, Lynette LE Oon, Shirin Kalimuddin, Jenny GH Low, Yvonne CF Su, Gavin JD Smith |
| EPI_ISL_410545 | hCoV-19/Italy/INMI1-isl/2020 | Europe / Italy / Rome | 2020-01-29 | INMI Lazzaro Spallanzani IRCCS | Laboratory of Virology, INMI Lazzaro Spallanzani IRCCS | Maria R. Capobianchi, Cesare E. M. Gruber, Martina Rueca, Barbara Bartolini, Francesco Messina, Emanuela Giombini, Francesca Colavita, Concetta Castilletti, Eleonora Lalle, Fabrizio Carletti, Emanuele Nicastri, Giuseppe Ippolito. |
| EPI_ISL_410546 | hCoV-19/Italy/INMI1-cs/2020 | Europe / Italy / Rome | 2020-01-31 | INMI Lazzaro Spallanzani IRCCS | Laboratory of Virology, INMI Lazzaro Spallanzani IRCCS | Maria R. Capobianchi, Cesare E. M. Gruber, Martina Rueca, Fabrizio Carletti, Barbara Bartolini, Francesco Messina, Emanuela Giombini, Francesca Colavita, Concetta Castilletti, Eleonora Lalle, Emanuele Nicastri, Giuseppe Ippolito. |
| EPI_ISL_410984 | hCoV-19/France/IDF0515-isl/2020 | Europe / France / Ile-de-France / Paris | 2020-01-29 | Department of Infectious and Tropical Diseases, Bichat Claude Bernard Hospital, Paris | National Reference Center for Viruses of Respiratory Infections, Institut Pasteur, Paris | Mélanie Albert, Marion Barbet, Sylvie Behillil, Méline Bizard, Angela Brisebarre, Flora Donati, Vincent Enouf, Maud Vanpeene, Sylvie van der Werf, Yazdan Yazdanpanah, Xavier Lescure |
| EPI_ISL_411902 | hCoV-19/Cambodia/0012/2020 | Asia / Cambodia / Sihanoukville | 2020-01-27 | Virology Unit, Institut Pasteur du Cambodge. | Virology Unit, Institut Pasteur du Cambodge (Sequencing done by: Jessica E Manning/Jennifer A Bohl at Malaria and Vector Research Laboratory, National Institute of Allergy and Infectious Diseases and Vida Ahyong from Chan-Zuckerberg Biohub) | Erik A Karlsson, Jennifer A Bohl, Vida Ahyong, Veasna Duong, Philippe Dussart, Jessica E Manning. |
| EPI_ISL_411915 | hCoV-19/Taiwan/CGMH-CGU-01/2020 | Asia / Taiwan / Taoyuan | 2020-01-25 | Laboratory Medicine | Department of Laboratory Medicine, Lin-Kou Chang Gung Memorial Hospital, Taoyuan, Taiwan. | Kuo-Chien Tsao, Yu-Nong Gong, Shu-Li Yang, Yi-Chun Li, Chung-Guei Huang, Yhu-Chering Huang, Shin-Ru Shih |
| EPI_ISL_411926 | hCoV-19/Taiwan/3/2020 | Asia / Taiwan / Taipei | 2020-01-24 | Taiwan Centers for Disease Control | Taiwan Centers for Disease Control | Ji-Rong Yang, Yu-Chi-Lin, Jung-Jung Mu, Ming-Tsan-Liu |
| EPI_ISL_411927 | hCoV-19/Taiwan/4/2020 | Asia / Taiwan / Taipei | 2020-01-28 | Taiwan Centers for Disease Control | Taiwan Centers for Disease Control | Ji-Rong Yang, Yu-Chi-Lin, Jung-Jung Mu, Ming-Tsan-Liu |
| EPI_ISL_411929 | hCoV-19/South Korea/SNU01/2020 | Asia / South Korea | 2020-01 | unknown | Department of Clinical Diagnostics | Park,W.B., Kwon,N.-J., Choi,S.-J., Kang,C.K., Choe,P.G., Kim,J.Y., Yun,J., Lee,G.-W., Seong,M.-W., Kim,N., Seo,J.-S. and Oh,M.-D. |
| EPI_ISL_411950 | hCoV-19/Jiangsu/JS01/2020 | Asia / China / Jiangsu | 2020-01-23 | NHC Key laboratory of Enteric Pathogenic Microbiology, Institute of Pathogenic Microbiology | Jiangsu Provincial Center for Disease Control & Prevention | Lunbiao Cui,Kangchen Zhao,Xiaojuan Zhu,Yiyue Ge,Tao Wu,Bin Wu,Yin Chen,Fengcai Zhu,Baoli Zhu,Ming Wu |
| EPI_ISL_411951 | hCoV-19/Sweden/01/2020 | Europe / Sweden | 2020-02-07 | unknown | Unit for Laboratory Development and Technology Transfer, Public Health Agency of Sweden | Bengner,M., Palmerus,M., Lindsjo,O., Lind Karlberg,M., Monteil,V., Appelberg,S., Brave,A., Muradrasoli,S. and Tegmark-Wisell,K. |
| EPI_ISL_411952 | hCoV-19/Jiangsu/JS02/2020 | Asia / China / Jiangsu | 2020-01-24 | NHC Key laboratory of Enteric Pathogenic Microbiology, Institute of Pathogenic Microbiology | Jiangsu Provincial Center for Disease Control & Prevention | Kangchen Zhao, Xiaojuan Zhu, Lunbiao Cui, Tao Wu, Yiyue Ge, Bin Wu, Yin Chen, Fengcai Zhu, Baoli Zhu, Ming Wu |
| EPI_ISL_411953 | hCoV-19/Jiangsu/JS03/2020 | Asia / China / Jiangsu | 2020-01-24 | NHC Key laboratory of Enteric Pathogenic Microbiology, Institute of Pathogenic Microbiology | Jiangsu Provincial Center for Disease Control & Prevention | Kangchen Zhao, Xiaojuan Zhu, Lunbiao Cui, Tao Wu, Yiyue Ge, Bin Wu, Yin Chen, Fengcai Zhu, Baoli Zhu, Ming Wu |
| EPI_ISL_411954 | hCoV-19/USA/CA7/2020 | North America / USA / California | 2020-02-06 | California Department of Public Health | Pathogen Discovery, Respiratory Viruses Branch, Division of Viral Diseases, Centers for Diseas Control and Prevention | Krista Queen, Anna Uehara, Jing Zhang, Yan Li, Ying Tao, Clinton R. Paden, Haibin Wang, Shifaq Kamili, Xiaoyan Lu, Brian Lynch, Senthil Kumar K. Sakthivel, Brett L. Whitaker, Lijuan Wang, Janna' R. Murray, Susan I. Gerber, Stephen Lindstrom, Suxiang Tong |
| EPI_ISL_411955 | hCoV-19/USA/CA8/2020 | North America / USA / California | 2020-02-10 | California Department of Public Health | Pathogen Discovery, Respiratory Viruses Branch, Division of Viral Diseases, Centers for Diseas Control and Prevention | Krista Queen, Anna Uehara, Jing Zhang, Yan Li, Ying Tao, Clinton R. Paden, Haibin Wang, Shifaq Kamili, Xiaoyan Lu, Brian Lynch, Senthil Kumar K. Sakthivel, Brett L. Whitaker, Lijuan Wang, Janna' R. Murray, Susan I. Gerber, Stephen Lindstrom, Suxiang Tong |
| EPI_ISL_411956 | hCoV-19/USA/TX1/2020 | North America / USA / Texas | 2020-02-11 | Texas Department of State Health Services | Pathogen Discovery, Respiratory Viruses Branch, Division of Viral Diseases, Centers for Diseas Control and Prevention | Krista Queen, Anna Uehara, Jing Zhang, Yan Li, Ying Tao, Clinton R. Paden, Haibin Wang, Shifaq Kamili, Xiaoyan Lu, Brian Lynch, Senthil Kumar K. Sakthivel, Brett L. Whitaker, Lijuan Wang, Janna' R. Murray, Susan I. Gerber, Stephen Lindstrom, Suxiang Tong |
| EPI_ISL_411957 | hCoV-19/China/WH-09/2020 | Asia / China | 2020-01-08 | unknown | Key Laboratory of Human Diseases, Comparative Medicine, Institute of Laboratory Animal Science | Linlin,B., Lili,R., Shuran,G., Jiangning,L., Feifei,Q., Qi,L., Fengdi,L., Jing,X., Wei,D., Pin,Y., Yanfeng,X., Yajin,Q., Hong,G., Qiang,W., Mingya,L., Guanpeng,W., Shunyi,W., Zhiqi,S., Li,G., Lan,C., Conghui,W., Ying,W., Xinming,W., Yan,X., Qi,J. and Chuan,Q. |
| EPI_ISL_412862 | hCoV-19/USA/CA9/2020 | North America / USA / California / Solano | 2020-02-23 | California Department of Public Health | Pathogen Discovery, Respiratory Viruses Branch, Division of Viral Diseases, Centers for Disease Control and Prevention | Krista Queen, Anna Uehara, Jing Zhang, Yan Li, Ying Tao, Clinton R. Paden, Haibin Wang, Shifaq Kamili, Xiaoyan Lu, Brian Lynch, Senthil Kumar K. Sakthivel, Brett L. Whitaker, Lijuan Wang, Janna' R. Murray, Jasmine Padilla, Justin Lee, Susan I. Gerber, Stephen Lindstrom, Suxiang Tong |
| EPI_ISL_412869 | hCoV-19/South Korea/KCDC05/2020 | Asia / South Korea / Seoul | 2020-01-30 | Division of Viral Diseases, Center for Laboratory Control of Infectious Diseases, Korea Centers for Diseases Control and Prevention | Division of Viral Diseases, Center for Laboratory Control of Infectious Diseases, Korea Centers for Diseases Control and Prevention | Jeong-Min Kim, Yoon-Seok Chung, Namjoo Lee, Mi-Seon Kim, Sang Hee Woo, Hye-Jun Jo, Sehee Park, Heui Man Kim, Myung Guk Han |
| EPI_ISL_412870 | hCoV-19/South Korea/KCDC06/2020 | Asia / South Korea / Seoul | 2020-01-30 | Division of Viral Diseases, Center for Laboratory Control of Infectious Diseases, Korea Centers for Diseases Control and Prevention | Division of Viral Diseases, Center for Laboratory Control of Infectious Diseases, Korea Centers for Diseases Control and Prevention | Jeong-Min Kim, Yoon-Seok Chung, Namjoo Lee, Mi-Seon Kim, Sang Hee Woo, Hye-Jun Jo, Sehee Park, Heui Man Kim, Myung Guk Han |
| EPI_ISL_412871 | hCoV-19/South Korea/KCDC07/2020 | Asia / South Korea / Seoul | 2020-01-31 | Division of Viral Diseases, Center for Laboratory Control of Infectious Diseases, Korea Centers for Diseases Control and Prevention | Division of Viral Diseases, Center for Laboratory Control of Infectious Diseases, Korea Centers for Diseases Control and Prevention | Jeong-Min Kim, Yoon-Seok Chung, Namjoo Lee, Mi-Seon Kim, Sang Hee Woo, Hye-Jun Jo, Sehee Park, Heui Man Kim, Myung Guk Han |
| EPI_ISL_412872 | hCoV-19/South Korea/KCDC12/2020 | Asia / South Korea / Gyeonggi-do | 2020-02-01 | Division of Viral Diseases, Center for Laboratory Control of Infectious Diseases, Korea Centers for Diseases Control and Prevention | Division of Viral Diseases, Center for Laboratory Control of Infectious Diseases, Korea Centers for Diseases Control and Prevention | Jeong-Min Kim, Yoon-Seok Chung, Namjoo Lee, Mi-Seon Kim, Sang Hee Woo, Hye-Jun Jo, Sehee Park, Heui Man Kim, Myung Guk Han |
| EPI_ISL_412873 | hCoV-19/South Korea/KCDC24/2020 | Asia / South Korea / Chungcheongnam-d | 2020-02-06 | Division of Viral Diseases, Center for Laboratory Control of Infectious Diseases, Korea Centers for Diseases Control and Prevention | Division of Viral Diseases, Center for Laboratory Control of Infectious Diseases, Korea Centers for Diseases Control and Prevention | Jeong-Min Kim, Yoon-Seok Chung, Namjoo Lee, Mi-Seon Kim, Sang Hee Woo, Hye-Jun Jo, Sehee Park, Heui Man Kim, Myung Guk Han |
| EPI_ISL_413221 | hCoV-19/Scotland/CVR01/2020 | Europe / United Kingdom / Scotland | 2020-02-28 | West of Scotland Specialist Virology Centre, NHSGGC | MRC-University of Glasgow Centre for Virus Research | Emma Thomson, Antonia Ho; James Shepherd, Shirin Ashraf; Kathy Smollett, Daniel Mair, Stephen Carmichael, Ana da Silva Filipe; Richard Orton, Josh Singer, David L Robertson; Andrew Rambaut; Alasdair MacLean, Rory Gunson. |
| EPI_ISL_413513 | hCoV-19/South Korea/KUMC03/2020 | Asia / South Korea | 2020-02-27 | Division of Infectious Diseases, Department of Internal Medicine, Korea University College of Medicine | Department of Microbiology, Institute for Viral Diseases, College of Medicine, Korea University | Changmin Kang, Joon-Yong Bae, Jungmin Lee, Jin Gu Yoon, Heedo Park, Juyoung Cho, Jeonghun Kim, Gee Eun Lee, Cui Chunguang, Kyeong-ryeol Shin, Ji Yun Noh, Joon Young Song, Hee Jin Cheong, Woo Joo Kim, Jin Il Kim, Man-Seong Park |
| EPI_ISL_413514 | hCoV-19/South Korea/KUMC04/2020 | Asia / South Korea | 2020-02-27 | Department of Microbiology, Institute for Viral Diseases, College of Medicine, Korea University | Department of Microbiology, Institute for Viral Diseases, College of Medicine, Korea University | Changmin Kang, Joon-Yong Bae, Jungmin Lee, Jin Gu Yoon, Heedo Park, Juyoung Cho, Jeonghun Kim, Gee Eun Lee, Cui Chunguang, Kyeong-ryeol Shin, Ji Yun Noh, Joon Young Song, Hee Jin Cheong, Woo Joo Kim, Jin Il Kim, Man-Seong Park |
| EPI_ISL_413515 | hCoV-19/South Korea/KUMC05/2020 | Asia / South Korea | 2020-02-27 | Division of Infectious Diseases, Department of Internal Medicine, Korea University College of Medicine | Department of Microbiology, Institute for Viral Diseases, College of Medicine, Korea University | Changmin Kang, Joon-Yong Bae, Jungmin Lee, Jin Gu Yoon, Heedo Park, Juyoung Cho, Jeonghun Kim, Gee Eun Lee, Cui Chunguang, Kyeong-ryeol Shin, Ji Yun Noh, Joon Young Song, Hee Jin Cheong, Woo Joo Kim, Jin Il Kim, Man-Seong Park |
| EPI_ISL_413516 | hCoV-19/South Korea/KUMC06/2020 | Asia / South Korea | 2020-02-27 | Department of Microbiology, Institute for Viral Diseases, College of Medicine, Korea University | Department of Microbiology, Institute for Viral Diseases, College of Medicine, Korea University | Changmin Kang, Joon-Yong Bae, Jungmin Lee, Jin Gu Yoon, Heedo Park, Juyoung Cho, Jeonghun Kim, Gee Eun Lee, Cui Chunguang, Kyeong-ryeol Shin, Ji Yun Noh, Joon Young Song, Hee Jin Cheong, Woo Joo Kim, Jin Il Kim, Man-Seong Park |
| EPI_ISL_413518 | hCoV-19/Beijing/105/2020 | Asia / China / Beijing | 2020-01-26 | unknown | Infectious Disease Control Center | Li,J., Li,L., Li,Z., Qiu,S., Song,H., Li,P. and Li,P. |
| EPI_ISL_413519 | hCoV-19/Beijing/231/2020 | Asia / China / Beijing | 2020-01-28 | unknown | Infectious Disease Control Center | Li,J., Li,L., Li,Z., Qiu,S., Song,H., Li,P. and Li,P. |
| EPI_ISL_413520 | hCoV-19/Beijing/233/2020 | Asia / China / Beijing | 2020-01-28 | unknown | Infectious Disease Control Center | Li,J., Li,L., Li,Z., Qiu,S., Song,H., Li,P. and Li,P. |
| EPI_ISL_413521 | hCoV-19/Beijing/235/2020 | Asia / China / Beijing | 2020-01-28 | unknown | Infectious Disease Control Center | Li,J., Li,L., Li,Z., Qiu,S., Song,H., Li,P. and Li,P. |
| EPI_ISL_413522 | hCoV-19/India/1-27/2020 | Asia / India / Kerala | 2020-01-27 | Indian Council of Medical Research - National Institute of Virology | National Influenza Center, Indian Council of Medical Research - National Institute of Virology | Potdar V, Yadav PD, Choudhary ML, Shete-Aich A |
| EPI_ISL_413523 | hCoV-19/India/1-31/2020 | Asia / India / Kerala | 2020-01-31 | Indian Council of Medical Research-National Institute of Virology | National Influenza Center, Indian Council of Medical Research-National Institute of Virology | Potdar V, Yadav PD, Choudhary ML, Shete-Aich A |
| EPI_ISL_413555 | hCoV-19/Wales/PHW1/2020 | Europe / United Kingdom / Wales | 2020-02-27 | Wales Specialist Virology Centre | Public Health Wales Microbiology Cardiff | Catherine Moore, Cen Sabu, Joanne Watkins, Sally Corden, Tom Connor |
| EPI_ISL_413556 | hCoV-19/Wales/PHW2/2020 | Europe / United Kingdom / Wales | 2020-03-04 | Wales Specialist Virology Centre | Public Health Wales Microbiology Cardiff | Catherine Moore, Tim Jones, Joanne Watkins, Sally Corden, Tom Connor |
| EPI_ISL_413557 | hCoV-19/USA/CA-CDPH-UC1/2020 | North America / USA / California / Sonom | 2020-02-28 | California Department of Public Health | Chiu Laboratory, University of California, San Francisco | Xianding Deng, Scot Federman, Chao-Yang Pan, Hugo Guevara,Wei Gu, Debra A. Wadford, and Charles Y. Chiu |
| EPI_ISL_413558 | hCoV-19/USA/CA-CDPH-UC2/2020 | North America / USA / California / Solano | 2020-02-27 | California Department of Public Health | Chiu Laboratory, University of California, San Francisco | Xianding Deng, Scot Federman, Chao-Yang Pan, Hugo Guevara,Wei Gu, Debra A. Wadford, and Charles Y. Chiu |
| EPI_ISL_413559 | hCoV-19/USA/CA-CDPH-UC3/2020 | North America / USA / California / Solano | 2020-02-27 | California Department of Public Health | Chiu Laboratory, University of California, San Francisco | Xianding Deng, Scot Federman, Chao-Yang Pan, Hugo Guevara,Wei Gu, Debra A. Wadford, and Charles Y. Chiu |
| EPI_ISL_413560 | hCoV-19/USA/WA-S3/2020 | North America / USA / Washington | 2020-02-28 | Seattle Flu Study | Seattle Flu Study | Chu et al |
| EPI_ISL_413561 | hCoV-19/USA/CA-CDPH-UC4/2020 | North America / USA / California / Solano | 2020-02-27 | California Department of Public Health | Chiu Laboratory, University of California, San Francisco | Xianding Deng, Scot Federman, Chao-Yang Pan, Hugo Guevara,Wei Gu, Debra A. Wadford, and Charles Y. Chiu |
| EPI_ISL_413562 | hCoV-19/USA/WA11-UW7/2020 | North America / USA / Washington | 2020-03-02 | UW Virology Lab | UW Virology Lab | Pavitra Roychoudhury, Hong Xie, Keith Jerome, Alexander Greninger |
| EPI_ISL_413563 | hCoV-19/USA/WA12-UW8/2020 | North America / USA / Washington | 2020-03-03 | UW Virology Lab | UW Virology Lab | Pavitra Roychoudhury, Hong Xie, Keith Jerome, Alexander Greninger |
| EPI_ISL_413564 | hCoV-19/Netherlands/Andel_1365066/2020 | Europe / Netherlands / Andel | 2020-03-01 | MHC West-Brabant | Erasmus Medical Center | David Nieuwenhuijse, Bas Oude Munnink, Reina Sikkema, Claudia Schapendonk, Irina Chestakova, Anne van der Linden, Mark Pronk, Pascal Lexmond, Corien Swaan, Manon Haverkate, Madelief Mollers, Mart Stein, Sandra Kengne Kamga Mobou, Jeroen van Kampen, Jolanda Voermans, Aura Timen, Corine GeurtsvanKessel, Annemiek van der Eijk, Richard Molenkamp, Marion Koopmans, on behalf of the Dutch national COVID-19 response team. |
| EPI_ISL_413566 | hCoV-19/Netherlands/Blaricum_1364780/2020 | Europe / Netherlands / Blaricum | 2020-03-02 | MHC Gooi & Vechtstreek | Erasmus Medical Center | David Nieuwenhuijse, Bas Oude Munnink, Reina Sikkema, Claudia Schapendonk, Irina Chestakova, Anne van der Linden, Mark Pronk, Pascal Lexmond, Corien Swaan, Manon Haverkate, Madelief Mollers, Mart Stein, Sandra Kengne Kamga Mobou, Jeroen van Kampen, Jolanda Voermans, Aura Timen, Corine GeurtsvanKessel, Annemiek van der Eijk, Richard Molenkamp, Marion Koopmans, on behalf of the Dutch national COVID-19 response team. |
| EPI_ISL_413568 | hCoV-19/Netherlands/Dalen_1363624/2020 | Europe / Netherlands / Dalen | 2020-03-01 | MHC Drente | Erasmus Medical Center | David Nieuwenhuijse, Bas Oude Munnink, Reina Sikkema, Claudia Schapendonk, Irina Chestakova, Anne van der Linden, Mark Pronk, Pascal Lexmond, Corien Swaan, Manon Haverkate, Madelief Mollers, Mart Stein, Sandra Kengne Kamga Mobou, Jeroen van Kampen, Jolanda Voermans, Aura Timen, Corine GeurtsvanKessel, Annemiek van der Eijk, Richard Molenkamp, Marion Koopmans, on behalf of the Dutch national COVID-19 response team. |
| EPI_ISL_413571 | hCoV-19/Netherlands/Eindhoven_1363782/2020 | Europe / Netherlands / Eindhoven | 2020-03-02 | MHC Brabant Zuidoost | Erasmus Medical Center | David Nieuwenhuijse, Bas Oude Munnink, Reina Sikkema, Claudia Schapendonk, Irina Chestakova, Anne van der Linden, Mark Pronk, Pascal Lexmond, Corien Swaan, Manon Haverkate, Madelief Mollers, Mart Stein, Sandra Kengne Kamga Mobou, Jeroen van Kampen, Jolanda Voermans, Aura Timen, Corine GeurtsvanKessel, Annemiek van der Eijk, Richard Molenkamp, Marion Koopmans, on behalf of the Dutch national COVID-19 response team. |
| EPI_ISL_413572 | hCoV-19/Netherlands/Haarlem_1363688/2020 | Europe / Netherlands / Haarlem | 2020-03-01 | MHC Kennemerland | Erasmus Medical Center | David Nieuwenhuijse, Bas Oude Munnink, Reina Sikkema, Claudia Schapendonk, Irina Chestakova, Anne van der Linden, Mark Pronk, Pascal Lexmond, Corien Swaan, Manon Haverkate, Madelief Mollers, Mart Stein, Sandra Kengne Kamga Mobou, Jeroen van Kampen, Jolanda Voermans, Aura Timen, Corine GeurtsvanKessel, Annemiek van der Eijk, Richard Molenkamp, Marion Koopmans, on behalf of the Dutch national COVID-19 response team. |
| EPI_ISL_413573 | hCoV-19/Netherlands/Hardinxveld_Giessendam_ | Europe / Netherlands / Hardinxveld Giess | 2020-03-02 | Dienst Gezondheid & Jeugd Zuid-Holland Zuid | Erasmus Medical Center | David Nieuwenhuijse, Bas Oude Munnink, Reina Sikkema, Claudia Schapendonk, Irina Chestakova, Anne van der Linden, Mark Pronk, Pascal Lexmond, Corien Swaan, Manon Haverkate, Madelief Mollers, Mart Stein, Sandra Kengne Kamga Mobou, Jeroen van Kampen, Jolanda Voermans, Aura Timen, Corine GeurtsvanKessel, Annemiek van der Eijk, Richard Molenkamp, Marion Koopmans, on behalf of the Dutch national COVID-19 response team. |
| EPI_ISL_413574 | hCoV-19/Netherlands/Helmond_1363548/2020 | Europe / Netherlands / Helmond | 2020-02-29 | MHC West-Brabant | Erasmus Medical Center | David Nieuwenhuijse, Bas Oude Munnink, Reina Sikkema, Claudia Schapendonk, Irina Chestakova, Anne van der Linden, Mark Pronk, Pascal Lexmond, Corien Swaan, Manon Haverkate, Madelief Mollers, Mart Stein, Sandra Kengne Kamga Mobou, Jeroen van Kampen, Jolanda Voermans, Aura Timen, Corine GeurtsvanKessel, Annemiek van der Eijk, Richard Molenkamp, Marion Koopmans, on behalf of the Dutch national COVID-19 response team. |
| EPI_ISL_413577 | hCoV-19/Netherlands/Naarden_1364774/2020 | Europe / Netherlands / Naarden | 2020-03-02 | MHC Gooi & Vechtstreek | Erasmus Medical Center | David Nieuwenhuijse, Bas Oude Munnink, Reina Sikkema, Claudia Schapendonk, Irina Chestakova, Anne van der Linden, Mark Pronk, Pascal Lexmond, Corien Swaan, Manon Haverkate, Madelief Mollers, Mart Stein, Sandra Kengne Kamga Mobou, Jeroen van Kampen, Jolanda Voermans, Aura Timen, Corine GeurtsvanKessel, Annemiek van der Eijk, Richard Molenkamp, Marion Koopmans, on behalf of the Dutch national COVID-19 response team. |
| EPI_ISL_413579 | hCoV-19/Netherlands/Nootdorp_1364222/2020 | Europe / Netherlands / Nootdorp | 2020-03-03 | MHC Haaglanden | Erasmus Medical Center | David Nieuwenhuijse, Bas Oude Munnink, Reina Sikkema, Claudia Schapendonk, Irina Chestakova, Anne van der Linden, Mark Pronk, Pascal Lexmond, Corien Swaan, Manon Haverkate, Madelief Mollers, Mart Stein, Sandra Kengne Kamga Mobou, Jeroen van Kampen, Jolanda Voermans, Aura Timen, Corine GeurtsvanKessel, Annemiek van der Eijk, Richard Molenkamp, Marion Koopmans, on behalf of the Dutch national COVID-19 response team. |
| EPI_ISL_413580 | hCoV-19/Netherlands/Oisterwijk_1364072/2020 | Europe / Netherlands / Oisterwijk | 2020-03-02 | MHC Hart voor Brabant | Erasmus Medical Center | David Nieuwenhuijse, Bas Oude Munnink, Reina Sikkema, Claudia Schapendonk, Irina Chestakova, Anne van der Linden, Mark Pronk, Pascal Lexmond, Corien Swaan, Manon Haverkate, Madelief Mollers, Mart Stein, Sandra Kengne Kamga Mobou, Jeroen van Kampen, Jolanda Voermans, Aura Timen, Corine GeurtsvanKessel, Annemiek van der Eijk, Richard Molenkamp, Marion Koopmans, on behalf of the Dutch national COVID-19 response team. |
| EPI_ISL_413582 | hCoV-19/Netherlands/Rotterdam_1363790/2020 | Europe / Netherlands / Rotterdam | 2020-03-01 | ErasmusMC | Erasmus Medical Center | David Nieuwenhuijse, Bas Oude Munnink, Reina Sikkema, Claudia Schapendonk, Irina Chestakova, Anne van der Linden, Mark Pronk, Pascal Lexmond, Corien Swaan, Manon Haverkate, Madelief Mollers, Mart Stein, Sandra Kengne Kamga Mobou, Jeroen van Kampen, Jolanda Voermans, Aura Timen, Corine GeurtsvanKessel, Annemiek van der Eijk, Richard Molenkamp, Marion Koopmans, on behalf of the Dutch national COVID-19 response team. |
| EPI_ISL_413583 | hCoV-19/Netherlands/Rotterdam_1364040/2020 | Europe / Netherlands / Rotterdam | 2020-03-02 | MHC Rotterdam-Rijnmond | Erasmus Medical Center | David Nieuwenhuijse, Bas Oude Munnink, Reina Sikkema, Claudia Schapendonk, Irina Chestakova, Anne van der Linden, Mark Pronk, Pascal Lexmond, Corien Swaan, Manon Haverkate, Madelief Mollers, Mart Stein, Sandra Kengne Kamga Mobou, Jeroen van Kampen, Jolanda Voermans, Aura Timen, Corine GeurtsvanKessel, Annemiek van der Eijk, Richard Molenkamp, Marion Koopmans, on behalf of the Dutch national COVID-19 response team. |
| EPI_ISL_413584 | hCoV-19/Netherlands/Rotterdam_1364740/2020 | Europe / Netherlands / Rotterdam | 2020-03-03 | unknown | Erasmus Medical Center | David Nieuwenhuijse, Bas Oude Munnink, Reina Sikkema, Claudia Schapendonk, Irina Chestakova, Anne van der Linden, Mark Pronk, Pascal Lexmond, Corien Swaan, Manon Haverkate, Madelief Mollers, Mart Stein, Sandra Kengne Kamga Mobou, Jeroen van Kampen, Jolanda Voermans, Aura Timen, Corine GeurtsvanKessel, Annemiek van der Eijk, Richard Molenkamp, Marion Koopmans, on behalf of the Dutch national COVID-19 response team. |
| EPI_ISL_413587 | hCoV-19/Netherlands/Tilburg_1364286/2020 | Europe / Netherlands / Tilburg | 2020-03-03 | Foundation Elisabeth-Tweesteden Ziekenhuis | Erasmus Medical Center | David Nieuwenhuijse, Bas Oude Munnink, Reina Sikkema, Claudia Schapendonk, Irina Chestakova, Anne van der Linden, Mark Pronk, Pascal Lexmond, Corien Swaan, Manon Haverkate, Madelief Mollers, Mart Stein, Sandra Kengne Kamga Mobou, Jeroen van Kampen, Jolanda Voermans, Aura Timen, Corine GeurtsvanKessel, Annemiek van der Eijk, Richard Molenkamp, Marion Koopmans, on behalf of the Dutch national COVID-19 response team. |
| EPI_ISL_413588 | hCoV-19/Netherlands/Utrecht_1363564/2020 | Europe / Netherlands / Utrecht | 2020-03-01 | MHC Utrecht | Erasmus Medical Center | David Nieuwenhuijse, Bas Oude Munnink, Reina Sikkema, Claudia Schapendonk, Irina Chestakova, Anne van der Linden, Mark Pronk, Pascal Lexmond, Corien Swaan, Manon Haverkate, Madelief Mollers, Mart Stein, Sandra Kengne Kamga Mobou, Jeroen van Kampen, Jolanda Voermans, Aura Timen, Corine GeurtsvanKessel, Annemiek van der Eijk, Richard Molenkamp, Marion Koopmans, on behalf of the Dutch national COVID-19 response team. |
| EPI_ISL_413589 | hCoV-19/Netherlands/Utrecht_1363628/2020 | Europe / Netherlands / Utrecht | 2020-03-01 | MHC Utrecht | Erasmus Medical Center | David Nieuwenhuijse, Bas Oude Munnink, Reina Sikkema, Claudia Schapendonk, Irina Chestakova, Anne van der Linden, Mark Pronk, Pascal Lexmond, Corien Swaan, Manon Haverkate, Madelief Mollers, Mart Stein, Sandra Kengne Kamga Mobou, Jeroen van Kampen, Jolanda Voermans, Aura Timen, Corine GeurtsvanKessel, Annemiek van der Eijk, Richard Molenkamp, Marion Koopmans, on behalf of the Dutch national COVID-19 response team. |
| EPI_ISL_413590 | hCoV-19/Netherlands/Utrecht_1364066/2020 | Europe / Netherlands / Utrecht | 2020-03-02 | MHC Utrecht | Erasmus Medical Center | David Nieuwenhuijse, Bas Oude Munnink, Reina Sikkema, Claudia Schapendonk, Irina Chestakova, Anne van der Linden, Mark Pronk, Pascal Lexmond, Corien Swaan, Manon Haverkate, Madelief Mollers, Mart Stein, Sandra Kengne Kamga Mobou, Jeroen van Kampen, Jolanda Voermans, Aura Timen, Corine GeurtsvanKessel, Annemiek van der Eijk, Richard Molenkamp, Marion Koopmans, on behalf of the Dutch national COVID-19 response team. |
| EPI_ISL_413591 | hCoV-19/Netherlands/Zeewolde_1365080/2020 | Europe / Netherlands / Zeewolde | 2020-03-02 | MHC Flevoland | Erasmus Medical Center | David Nieuwenhuijse, Bas Oude Munnink, Reina Sikkema, Claudia Schapendonk, Irina Chestakova, Anne van der Linden, Mark Pronk, Pascal Lexmond, Corien Swaan, Manon Haverkate, Madelief Mollers, Mart Stein, Sandra Kengne Kamga Mobou, Jeroen van Kampen, Jolanda Voermans, Aura Timen, Corine GeurtsvanKessel, Annemiek van der Eijk, Richard Molenkamp, Marion Koopmans, on behalf of the Dutch national COVID-19 response team. |
| EPI_ISL_413592 | hCoV-19/Taiwan/NTU03/2020 | Asia / Taiwan / Taipei | 2020-03-02 | Department of Laboratory Medicine, National Taiwan University Hospital | Microbial Genomics Core Lab, National Taiwan University Centers of Genomic and Precision Medicine | Shiou-Hwei Yeh, You-Yu Lin, Ya-Yun Lai, Chiao-Ling Li, Shan-Chwen Chang, Pei-Jer Chen, Sui-Yuan Chang |
| EPI_ISL_413593 | hCoV-19/Luxembourg/Lux1/2020 | Europe / Luxembourg | 2020-02-29 | Laboratoire National de Santé | Erasmus Medical Center | David Nieuwenhuijse, Bas Oude Munnink, Reina Sikkema, Claudia Schapendonk, Irina Chestakova, Anne van der Linden, Mark Pronk, Pascal Lexmond, T. Abdelrahman, G. Fournier, J. Mossong, T. Nguyen, Jeroen van Kampen, Jolanda Voermans, Corine GeurtsvanKessel, Annemiek van der Eijk, Richard Molenkamp, Marion Koopmans, on behalf of the Dutch national COVID-19 response team. |
| EPI_ISL_413594 | hCoV-19/Australia/NSW08/2020 | Oceania / Australia / NSW / Sydney | 2020-02-28 | Centre for Infectious Diseases and Microbiology Laboratory Services | NSW Health Pathology - Institute of Clinical Pathology and Medical Research; Westmead Hospital; University of Sydney | Rockett R, Eden J-S, Lam C, Gray K, Timms, V, Gall, M, Alicia, A, Carter I, Rahman H, Holmes EC, , O'Sullivan MV, Sintchenko V, Chen SC, Maddocks S, Kok J and Dwyer DE for the 2019-nCoV Study Group" |
| EPI_ISL_413595 | hCoV-19/Australia/NSW09/2020 | Oceania / Australia / NSW / Sydney | 2020-02-28 | Centre for Infectious Diseases and Microbiology Laboratory Services | NSW Health Pathology - Institute of Clinical Pathology and Medical Research; Westmead Hospital; University of Sydney | Rockett R, Eden J-S, Lam C, Gray K, Timms, V, Gall, M, Carter I, Rahman H, Holmes EC, O'Sullivan MV, Sintchenko V, Chen SC, Maddocks S, Kok J and Dwyer DE for the 2019-nCoV Study Group" |
| EPI_ISL_413596 | hCoV-19/Australia/NSW10/2020 | Oceania / Australia / NSW / Sydney | 2020-02-28 | Centre for Infectious Diseases and Microbiology - Public Health | NSW Health Pathology - Institute of Clinical Pathology and Medical Research; Westmead Hospital; University of Sydney | Rockett R, Eden J-S, Lam C, Gray K, Timms, V, Gall, M, Carter I, Rahman H, Holmes EC, O'Sullivan MV, Sintchenko V, Chen SC, Maddocks S, Kok J and Dwyer DE for the 2019-nCoV Study Group" |
| EPI_ISL_413597 | hCoV-19/Australia/NSW11/2020 | Oceania / Australia / NSW / Sydney | 2020-03-02 | Centre for Infectious Diseases and Microbiology- Public Health | NSW Health Pathology - Institute of Clinical Pathology and Medical Research; Westmead Hospital; University of Sydney | Lam C, Eden J-S, Rockett R, Gray K, Timms, V, Gall, M, Carter I, Rahman H, Holmes EC, O'Sullivan MV, Sintchenko V, Chen SC, Maddocks S, Kok J and Dwyer DE for the 2019-nCoV Study Group" |
| EPI_ISL_413598 | hCoV-19/Australia/NSW12/2020 | Oceania / Australia / NSW / Sydney | 2020-03-04 | Centre for Infectious Diseases and Microbiology - Public Health | NSW Health Pathology - Institute of Clinical Pathology and Medical Research; Westmead Hospital; University of Sydney | Gray K, Eden J-S, Lam C, Rockett R, Timms, V, Gall, M, Carter I, Rahman H, Holmes EC, O'Sullivan MV, Sintchenko V, Chen SC, Maddocks S, Kok J and Dwyer DE for the 2019-nCoV Study Group" |
| EPI_ISL_413599 | hCoV-19/Australia/NSW13/2020 | Oceania / Australia / NSW / Sydney | 2020-03-04 | Centre for Infectious Diseases and Microbiology - Public Health | NSW Health Pathology - Institute of Clinical Pathology and Medical Research; Westmead Hospital; University of Sydney | Timms, V, Eden J-S, Lam C, Gray K, Rockett R, Gall, M, Carter I, Rahman H, Holmes EC, O'Sullivan MV, Sintchenko V, Chen SC, Maddocks S, Kok J and Dwyer DE for the 2019-nCoV Study Group" |
| EPI_ISL_413600 | hCoV-19/Australia/NSW14/2020 | Oceania / Australia / NSW / Sydney | 2020-03-03 | Centre for Infectious Diseases and Microbiology - Public Health | NSW Health Pathology - Institute of Clinical Pathology and Medical Research; Westmead Hospital; University of Sydney | Gall, M, Eden J-S, Lam C, Gray K, Timms, V, Rockett R, Carter I, Rahman H, Holmes EC, O'Sullivan MV, Sintchenko V, Chen SC, Maddocks S, Kok J and Dwyer DE for the 2019-nCoV Study Group" |
| EPI_ISL_413601 | hCoV-19/USA/WA13-UW9/2020 | North America / USA / Washington | 2020-03-02 | UW Virology Lab | UW Virology Lab | Pavitra Roychoudhury, Hong Xie, Keith Jerome, Alexander Greninger |
| EPI_ISL_413711 | hCoV-19/China/WF0014/2020 | Asia / China | 2020-02 | Weifang Center for Disease Control and Prevention | Weifang Center for Disease Control and Prevention & BGI-Shenzhen | Qing Nie, Xingguang Li, Erik M Volz, Han Fu, Haowei Wang, Xiaoyue Xi, Wei Chen, Dehui Liu, Yingying Chen, Mengmeng Tian, Wei Tan, Junjie Zai, Wanying Sun, Jiandong Li, Junhua Li |
| EPI_ISL_413996 | hCoV-19/Switzerland/TI9486/2020 | Europe / Switzerland / Tessin | 2020-02-24 | Laboratoire de Virologie, HUG | Swiss National Reference Centre for Influenza | LAUBSCHER Florian et al. |
| EPI_ISL_413997 | hCoV-19/Switzerland/GE3895/2020 | Europe / Switzerland / Geneva | 2020-02-26 | Laboratoire de Virologie, HUG | Swiss National Reference Centre for Influenza | LAUBSCHER Florian et al. |
| EPI_ISL_413999 | hCoV-19/Switzerland/AG0361/2020 | Europe / Switzerland / Argovie | 2020-02-27 | Laboratoire de Virologie, HUG | Swiss National Reference Centre for Influenza | LAUBSCHER Florian et al. |
| EPI_ISL_414005 | hCoV-19/England/200940527/2020 | Europe / United Kingdom / England | 2020-02-25 | Respiratory Virus Unit, Microbiology Services Colindale, Public Health England | Respiratory Virus Unit, Microbiology Services Colindale, Public Health England | Monica Galiano, Shahjahan Miah, Angie Lackenby, Omolola Akinbami, Tiina Talts, Leena Bhaw, Richard Myers, Steven Platt, Kirstin Edwards, Jonathan Hubb, Joanna Ellis, Maria Zambon |
| EPI_ISL_414006 | hCoV-19/England/200990724/2020 | Europe / United Kingdom / England | 2020-02-28 | Respiratory Virus Unit, Microbiology Services Colindale, Public Health England | Respiratory Virus Unit, Microbiology Services Colindale, Public Health England | Monica Galiano, Shahjahan Miah, Angie Lackenby, Omolola Akinbami, Tiina Talts, Leena Bhaw, Richard Myers, Steven Platt, Kirstin Edwards, Jonathan Hubb, Joanna Ellis, Maria Zambon |
| EPI_ISL_414007 | hCoV-19/England/200990725/2020 | Europe / United Kingdom / England | 2020-02-28 | Respiratory Virus Unit, Microbiology Services Colindale, Public Health England | Respiratory Virus Unit, Microbiology Services Colindale, Public Health England | Monica Galiano, Shahjahan Miah, Angie Lackenby, Omolola Akinbami, Tiina Talts, Leena Bhaw, Richard Myers, Steven Platt, Kirstin Edwards, Jonathan Hubb, Joanna Ellis, Maria Zambon |
| EPI_ISL_414008 | hCoV-19/England/200960041/2020 | Europe / United Kingdom / England | 2020-02-27 | Respiratory Virus Unit, Microbiology Services Colindale, Public Health England | Respiratory Virus Unit, Microbiology Services Colindale, Public Health England | Monica Galiano, Shahjahan Miah, Angie Lackenby, Omolola Akinbami, Tiina Talts, Leena Bhaw, Richard Myers, Steven Platt, Kirstin Edwards, Jonathan Hubb, Joanna Ellis, Maria Zambon |
| EPI_ISL_414009 | hCoV-19/England/200960515/2020 | Europe / United Kingdom / England | 2020-02-25 | Respiratory Virus Unit, Microbiology Services Colindale, Public Health England | Respiratory Virus Unit, Microbiology Services Colindale, Public Health England | Monica Galiano, Shahjahan Miah, Angie Lackenby, Omolola Akinbami, Tiina Talts, Leena Bhaw, Richard Myers, Steven Platt, Kirstin Edwards, Jonathan Hubb, Joanna Ellis, Maria Zambon |
| EPI_ISL_414010 | hCoV-19/England/200981386/2020 | Europe / United Kingdom / England | 2020-02-26 | Respiratory Virus Unit, Microbiology Services Colindale, Public Health England | Respiratory Virus Unit, Microbiology Services Colindale, Public Health England | Monica Galiano, Shahjahan Miah, Angie Lackenby, Omolola Akinbami, Tiina Talts, Leena Bhaw, Richard Myers, Steven Platt, Kirstin Edwards, Jonathan Hubb, Joanna Ellis, Maria Zambon |
| EPI_ISL_414011 | hCoV-19/England/200990006/2020 | Europe / United Kingdom / England | 2020-02-26 | Respiratory Virus Unit, Microbiology Services Colindale, Public Health England | Respiratory Virus Unit, Microbiology Services Colindale, Public Health England | Monica Galiano, Shahjahan Miah, Angie Lackenby, Omolola Akinbami, Tiina Talts, Leena Bhaw, Richard Myers, Steven Platt, Kirstin Edwards, Jonathan Hubb, Joanna Ellis, Maria Zambon |
| EPI_ISL_414012 | hCoV-19/England/200990723/2020 | Europe / United Kingdom / England | 2020-02-27 | Respiratory Virus Unit, Microbiology Services Colindale, Public Health England | Respiratory Virus Unit, Microbiology Services Colindale, Public Health England | Monica Galiano, Shahjahan Miah, Angie Lackenby, Omolola Akinbami, Tiina Talts, Leena Bhaw, Richard Myers, Steven Platt, Kirstin Edwards, Jonathan Hubb, Joanna Ellis, Maria Zambon |
| EPI_ISL_414013 | hCoV-19/England/201000003/2020 | Europe / United Kingdom / England | 2020-03-01 | Respiratory Virus Unit, Microbiology Services Colindale, Public Health England | Respiratory Virus Unit, Microbiology Services Colindale, Public Health England | Monica Galiano, Shahjahan Miah, Angie Lackenby, Omolola Akinbami, Tiina Talts, Leena Bhaw, Richard Myers, Steven Platt, Kirstin Edwards, Jonathan Hubb, Joanna Ellis, Maria Zambon |
| EPI_ISL_414014 | hCoV-19/Brazil/SPBR-03/2020 | South America / Brazil / Sao Paulo | 2020-03-02 | Hospital Israelita Albert Einstein | Instituto Adolfo Lutz, Interdiciplinary Procedures Center, Strategic Laboratory | Claudio Tavares Sacchi, Claudia Regina Gonçalves, Katia Correia dos Santos, Carlos Henrique Camargo, Maria do Carmo Sampaio Tavares Timenetsky, Terezinha Maria de Paiva, Ester Cerdeira Sabino |
| EPI_ISL_414015 | hCoV-19/Brazil/SPBR-06/2020 | South America / Brazil / Sao Paulo / Sao | 2020-02-29 | Hospital São Joaquim Beneficencia Portuguesa | Instituto Adolfo Lutz, Interdiciplinary Procedures Center, Strategic Laboratory | Claudio Tavares Sacchi, Claudia Regina Gonçalves, SimoneGuadagnucci Morillo, Carlos Henrique Camargo, Maria do Carmo Sampaio Tavares Timenetsky, Fabiana Cristina Pereira dos Santos Terezinha Maria de Paiva, Ester Cerdeira Sabino |
| EPI_ISL_414016 | hCoV-19/Brazil/SPBR-05/2020 | South America / Brazil / Sao Paulo / Sao | 2020-02-29 | Hospital São Joaquim Beneficencia Portuguesa | Instituto Adolfo Lutz, Interdiciplinary Procedures Center, Strategic Laboratory | Claudio Tavares Sacchi, Claudia Regina Gonçalves, Audrey Cilli, Carlos Henrique Camargo, Maria do Carmo Sampaio Tavares Timenetsky, Daniela Bernardes Borges da Silva, Terezinha Maria de Paiva, Ester Cerdeira Sabino |
| EPI_ISL_414017 | hCoV-19/Brazil/SPBR-04/2020 | South America / Brazil / Sao Paulo | 2020-03-04 | Hospital São Joaquim Beneficencia Portuguesa | Instituto Adolfo Lutz, Interdiciplinary Procedures Center, Strategic Laboratory | Claudio Tavares Sacchi, Claudia Regina Gonçalves, Fabiana Cristina Pereira dos Santos, Carlos Henrique Camargo, Maria do Carmo Sampaio Tavares Timenetsky, Daniela Bernardes Borges da Silva, Terezinha Maria de Paiva, Ester Cerdeira Sabino |
| EPI_ISL_414019 | hCoV-19/Switzerland/GE3121/2020 | Europe / Switzerland / Geneva | 2020-02-27 | Laboratoire de Virologie, HUG | Swiss National Reference Centre for Influenza | LAUBSCHER Florian et al. |
| EPI_ISL_414020 | hCoV-19/Switzerland/GE5373/2020 | Europe / Switzerland / Geneva | 2020-02-27 | Laboratoire de Virologie, HUG | Swiss National Reference Centre for Influenza | LAUBSCHER Florian et al. |
| EPI_ISL_414021 | hCoV-19/Switzerland/BL0902/2020 | Europe / Switzerland / Basel | 2020-02-27 | Laboratoire de Virologie, HUG | Swiss National Reference Centre for Influenza | LAUBSCHER Florian et al. |
| EPI_ISL_414022 | hCoV-19/Switzerland/GE9586/2020 | Europe / Switzerland / Geneva | 2020-02-27 | Laboratoire de Virologie, HUG | Swiss National Reference Centre for Influenza | LAUBSCHER Florian et al. |
| EPI_ISL_414023 | hCoV-19/Switzerland/VD5615/2020 | Europe / Switzerland / Vaud | 2020-03-01 | Laboratoire de Virologie, HUG | Swiss National Reference Centre for Influenza | LAUBSCHER Florian et al. |
| EPI_ISL_414027 | hCoV-19/Scotland/CVR05/2020 | Europe / United Kingdom / Scotland | 2020-03-04 | West of Scotland Specialist Virology Centre, NHSGGC | MRC-University of Glasgow Centre for Virus Research | Emma Thomson, Antonia Ho; Kathy Smollett, Daniel Mair, Stephen Carmichael, Ana da Silva Filipe; Richard Orton, David L Robertson; Alasdair MacLean, Rory Gunson. |
| EPI_ISL_414040 | hCoV-19/England/200641094/2020 | Europe / United Kingdom / England | 2020-02-05 | Respiratory Virus Unit, Microbiology Services Colindale, Public Health England | Respiratory Virus Unit, Microbiology Services Colindale, Public Health England | Monica Galiano, Shahjahan Miah, Angie Lackenby, Omolola Akinbami, Tiina Talts, Leena Bhaw, Richard Myers, Steven Platt, Kirstin Edwards, Jonathan Hubb, Joanna Ellis, Maria Zambon |
| EPI_ISL_414041 | hCoV-19/England/200690245/2020 | Europe / United Kingdom / England | 2020-02-08 | Respiratory Virus Unit, Microbiology Services Colindale, Public Health England | Respiratory Virus Unit, Microbiology Services Colindale, Public Health England | Monica Galiano, Shahjahan Miah, Angie Lackenby, Omolola Akinbami, Tiina Talts, Leena Bhaw, Richard Myers, Steven Platt, Kirstin Edwards, Jonathan Hubb, Joanna Ellis, Maria Zambon |
| EPI_ISL_414042 | hCoV-19/England/200690300/2020 | Europe / United Kingdom / England | 2020-02-08 | Respiratory Virus Unit, Microbiology Services Colindale, Public Health England | Respiratory Virus Unit, Microbiology Services Colindale, Public Health England | Monica Galiano, Shahjahan Miah, Angie Lackenby, Omolola Akinbami, Tiina Talts, Leena Bhaw, Richard Myers, Steven Platt, Kirstin Edwards, Jonathan Hubb, Joanna Ellis, Maria Zambon |
| EPI_ISL_414043 | hCoV-19/England/200690306/2020 | Europe / United Kingdom / England | 2020-02-07 | Respiratory Virus Unit, Microbiology Services Colindale, Public Health England | Respiratory Virus Unit, Microbiology Services Colindale, Public Health England | Monica Galiano, Shahjahan Miah, Angie Lackenby, Omolola Akinbami, Tiina Talts, Leena Bhaw, Richard Myers, Steven Platt, Kirstin Edwards, Jonathan Hubb, Joanna Ellis, Maria Zambon |
| EPI_ISL_414044 | hCoV-19/England/200690756/2020 | Europe / United Kingdom / England | 2020-02-08 | Respiratory Virus Unit, Microbiology Services Colindale, Public Health England | Respiratory Virus Unit, Microbiology Services Colindale, Public Health England | Monica Galiano, Shahjahan Miah, Angie Lackenby, Omolola Akinbami, Tiina Talts, Leena Bhaw, Richard Myers, Steven Platt, Kirstin Edwards, Jonathan Hubb, Joanna Ellis, Maria Zambon |
| EPI_ISL_414045 | hCoV-19/Brazil/RJ-314/2020 | South America / Brazil / Rio de Janeiro / I | 2020-03-04 | LACEN RJ - Laboratório Central de Saúde Pública Noel Nutels | Instituto Oswaldo Cruz FIOCRUZ - Laboratory of Respiratory Viruses and Measles (LVRS) | Paola Resende, Aliason Fabri, Joilson Xavier, Sunando Roy, Fernando Motta, Aline Mattos, Milene Miranda, Cristiana Garcia, Braulia Caetano, Maria Ogrzewalska, Jonathan Lopes, Luciana Appolinario, Maria Nóbrega, Marilda Siqueira |
| EPI_ISL_414414 | hCoV-19/Australia/QLD09/2020 | Oceania / Australia / Queensland / Gold C | 2020-02-29 | Pathology Queensland | Public Health Virology Laboratory | Bixing Huang, Alyssa Pyke, Amanda De Jong, Andrew Van Den Hurk, Carmel Taylor, David Warrilow, Doris Genge, Elisabeth Gamez, Glen Hewitson, Ian Maxwell Mackay, Inga Sultana, Jamie McMahon, Jean Barcelon, Judy Northill, Mitchell Finger, Natalie Simpson, Neelima Nair, Peter Burtonclay, Peter Moore, Sarah Wheatley, Sean Moody, Sonja Hall-Mendelin, Timothy Gardam, and Frederick Moore |
| EPI_ISL_414423 | hCoV-19/Netherlands/Gelderland_1/2020 | Europe / Netherlands / Gelderland | 2020-03-02 | Dutch COVID-19 response team | Erasmus Medical Center | David Nieuwenhuijse, Bas Oude Munnink, Reina Sikkema, Claudia Schapendonk, Irina Chestakova, Anne van der Linden, Mark Pronk, Pascal Lexmond, Corien Swaan, Manon Haverkate, Madelief Mollers, Mart Stein, Sandra Kengne Kamga Mobou, Jeroen van Kampen, Jolanda Voermans, Aura Timen, Corine GeurtsvanKessel, Annemiek van der Eijk, Richard Molenkamp, Marion Koopmans, on behalf of the Dutch national COVID-19 response team. |
| EPI_ISL_414424 | hCoV-19/Netherlands/Limburg_2/2020 | Europe / Netherlands / Limburg | 2020-03-03 | Dutch COVID-19 response team | Erasmus Medical Center | David Nieuwenhuijse, Bas Oude Munnink, Reina Sikkema, Claudia Schapendonk, Irina Chestakova, Anne van der Linden, Mark Pronk, Pascal Lexmond, Corien Swaan, Manon Haverkate, Madelief Mollers, Mart Stein, Sandra Kengne Kamga Mobou, Jeroen van Kampen, Jolanda Voermans, Aura Timen, Corine GeurtsvanKessel, Annemiek van der Eijk, Richard Molenkamp, Marion Koopmans, on behalf of the Dutch national COVID-19 response team. |
| EPI_ISL_414425 | hCoV-19/Netherlands/Limburg_3/2020 | Europe / Netherlands / Limburg | 2020-03-03 | Dutch COVID-19 response team | Erasmus Medical Center | David Nieuwenhuijse, Bas Oude Munnink, Reina Sikkema, Claudia Schapendonk, Irina Chestakova, Anne van der Linden, Mark Pronk, Pascal Lexmond, Corien Swaan, Manon Haverkate, Madelief Mollers, Mart Stein, Sandra Kengne Kamga Mobou, Jeroen van Kampen, Jolanda Voermans, Aura Timen, Corine GeurtsvanKessel, Annemiek van der Eijk, Richard Molenkamp, Marion Koopmans, on behalf of the Dutch national COVID-19 response team. |
| EPI_ISL_414426 | hCoV-19/Netherlands/Limburg_4/2020 | Europe / Netherlands / Limburg | 2020-03-03 | Dutch COVID-19 response team | Erasmus Medical Center | David Nieuwenhuijse, Bas Oude Munnink, Reina Sikkema, Claudia Schapendonk, Irina Chestakova, Anne van der Linden, Mark Pronk, Pascal Lexmond, Corien Swaan, Manon Haverkate, Madelief Mollers, Mart Stein, Sandra Kengne Kamga Mobou, Jeroen van Kampen, Jolanda Voermans, Aura Timen, Corine GeurtsvanKessel, Annemiek van der Eijk, Richard Molenkamp, Marion Koopmans, on behalf of the Dutch national COVID-19 response team. |
| EPI_ISL_414428 | hCoV-19/Netherlands/NoordBrabant_1/2020 | Europe / Netherlands / Noord Brabant | 2020-03-02 | Dutch COVID-19 response team | Erasmus Medical Center | David Nieuwenhuijse, Bas Oude Munnink, Reina Sikkema, Claudia Schapendonk, Irina Chestakova, Anne van der Linden, Mark Pronk, Pascal Lexmond, Corien Swaan, Manon Haverkate, Madelief Mollers, Mart Stein, Sandra Kengne Kamga Mobou, Jeroen van Kampen, Jolanda Voermans, Aura Timen, Corine GeurtsvanKessel, Annemiek van der Eijk, Richard Molenkamp, Marion Koopmans, on behalf of the Dutch national COVID-19 response team. |
| EPI_ISL_414429 | hCoV-19/Netherlands/NoordBrabant_3/2020 | Europe / Netherlands / Noord Brabant | 2020-03-02 | Dutch COVID-19 response team | Erasmus Medical Center | David Nieuwenhuijse, Bas Oude Munnink, Reina Sikkema, Claudia Schapendonk, Irina Chestakova, Anne van der Linden, Mark Pronk, Pascal Lexmond, Corien Swaan, Manon Haverkate, Madelief Mollers, Mart Stein, Sandra Kengne Kamga Mobou, Jeroen van Kampen, Jolanda Voermans, Aura Timen, Corine GeurtsvanKessel, Annemiek van der Eijk, Richard Molenkamp, Marion Koopmans, on behalf of the Dutch national COVID-19 response team. |
| EPI_ISL_414430 | hCoV-19/Netherlands/NoordBrabant_5/2020 | Europe / Netherlands / Noord Brabant | 2020-03-05 | Dutch COVID-19 response team | Erasmus Medical Center | David Nieuwenhuijse, Bas Oude Munnink, Reina Sikkema, Claudia Schapendonk, Irina Chestakova, Anne van der Linden, Mark Pronk, Pascal Lexmond, Corien Swaan, Manon Haverkate, Madelief Mollers, Mart Stein, Sandra Kengne Kamga Mobou, Jeroen van Kampen, Jolanda Voermans, Aura Timen, Corine GeurtsvanKessel, Annemiek van der Eijk, Richard Molenkamp, Marion Koopmans, on behalf of the Dutch national COVID-19 response team. |
| EPI_ISL_414432 | hCoV-19/Netherlands/NoordBrabant_11/2020 | Europe / Netherlands / Noord Brabant | 2020-03-02 | Dutch COVID-19 response team | Erasmus Medical Center | David Nieuwenhuijse, Bas Oude Munnink, Reina Sikkema, Claudia Schapendonk, Irina Chestakova, Anne van der Linden, Mark Pronk, Pascal Lexmond, Corien Swaan, Manon Haverkate, Madelief Mollers, Mart Stein, Sandra Kengne Kamga Mobou, Jeroen van Kampen, Jolanda Voermans, Aura Timen, Corine GeurtsvanKessel, Annemiek van der Eijk, Richard Molenkamp, Marion Koopmans, on behalf of the Dutch national COVID-19 response team. |
| EPI_ISL_414433 | hCoV-19/Netherlands/NoordHolland_1/2020 | Europe / Netherlands / Noord Holland | 2020-03-03 | Dutch COVID-19 response team | Erasmus Medical Center | David Nieuwenhuijse, Bas Oude Munnink, Reina Sikkema, Claudia Schapendonk, Irina Chestakova, Anne van der Linden, Mark Pronk, Pascal Lexmond, Corien Swaan, Manon Haverkate, Madelief Mollers, Mart Stein, Sandra Kengne Kamga Mobou, Jeroen van Kampen, Jolanda Voermans, Aura Timen, Corine GeurtsvanKessel, Annemiek van der Eijk, Richard Molenkamp, Marion Koopmans, on behalf of the Dutch national COVID-19 response team. |
| EPI_ISL_414434 | hCoV-19/Netherlands/Overijssel_1/2020 | Europe / Netherlands / Overijssel | 2020-03-03 | Dutch COVID-19 response team | Erasmus Medical Center | David Nieuwenhuijse, Bas Oude Munnink, Reina Sikkema, Claudia Schapendonk, Irina Chestakova, Anne van der Linden, Mark Pronk, Pascal Lexmond, Corien Swaan, Manon Haverkate, Madelief Mollers, Mart Stein, Sandra Kengne Kamga Mobou, Jeroen van Kampen, Jolanda Voermans, Aura Timen, Corine GeurtsvanKessel, Annemiek van der Eijk, Richard Molenkamp, Marion Koopmans, on behalf of the Dutch national COVID-19 response team. |
| EPI_ISL_414435 | hCoV-19/Netherlands/Utrecht_1/2020 | Europe / Netherlands / Utrecht | 2020-03-03 | Dutch COVID-19 response team | Erasmus Medical Center | David Nieuwenhuijse, Bas Oude Munnink, Reina Sikkema, Claudia Schapendonk, Irina Chestakova, Anne van der Linden, Mark Pronk, Pascal Lexmond, Corien Swaan, Manon Haverkate, Madelief Mollers, Mart Stein, Sandra Kengne Kamga Mobou, Jeroen van Kampen, Jolanda Voermans, Aura Timen, Corine GeurtsvanKessel, Annemiek van der Eijk, Richard Molenkamp, Marion Koopmans, on behalf of the Dutch national COVID-19 response team. |
| EPI_ISL_414436 | hCoV-19/Netherlands/Utrecht_2/2020 | Europe / Netherlands / Utrecht | 2020-03-03 | Dutch COVID-19 response team | Erasmus Medical Center | David Nieuwenhuijse, Bas Oude Munnink, Reina Sikkema, Claudia Schapendonk, Irina Chestakova, Anne van der Linden, Mark Pronk, Pascal Lexmond, Corien Swaan, Manon Haverkate, Madelief Mollers, Mart Stein, Sandra Kengne Kamga Mobou, Jeroen van Kampen, Jolanda Voermans, Aura Timen, Corine GeurtsvanKessel, Annemiek van der Eijk, Richard Molenkamp, Marion Koopmans, on behalf of the Dutch national COVID-19 response team. |
| EPI_ISL_414437 | hCoV-19/Netherlands/Utrecht_3/2020 | Europe / Netherlands / Utrecht | 2020-03-03 | Dutch COVID-19 response team | Erasmus Medical Center | David Nieuwenhuijse, Bas Oude Munnink, Reina Sikkema, Claudia Schapendonk, Irina Chestakova, Anne van der Linden, Mark Pronk, Pascal Lexmond, Corien Swaan, Manon Haverkate, Madelief Mollers, Mart Stein, Sandra Kengne Kamga Mobou, Jeroen van Kampen, Jolanda Voermans, Aura Timen, Corine GeurtsvanKessel, Annemiek van der Eijk, Richard Molenkamp, Marion Koopmans, on behalf of the Dutch national COVID-19 response team. |
| EPI_ISL_414438 | hCoV-19/Netherlands/Utrecht_4/2020 | Europe / Netherlands / Utrecht | 2020-03-03 | Dutch COVID-19 response team | Erasmus Medical Center | David Nieuwenhuijse, Bas Oude Munnink, Reina Sikkema, Claudia Schapendonk, Irina Chestakova, Anne van der Linden, Mark Pronk, Pascal Lexmond, Corien Swaan, Manon Haverkate, Madelief Mollers, Mart Stein, Sandra Kengne Kamga Mobou, Jeroen van Kampen, Jolanda Voermans, Aura Timen, Corine GeurtsvanKessel, Annemiek van der Eijk, Richard Molenkamp, Marion Koopmans, on behalf of the Dutch national COVID-19 response team. |
| EPI_ISL_414439 | hCoV-19/Netherlands/Utrecht_5/2020 | Europe / Netherlands / Utrecht | 2020-03-02 | Dutch COVID-19 response team | Erasmus Medical Center | David Nieuwenhuijse, Bas Oude Munnink, Reina Sikkema, Claudia Schapendonk, Irina Chestakova, Anne van der Linden, Mark Pronk, Pascal Lexmond, Corien Swaan, Manon Haverkate, Madelief Mollers, Mart Stein, Sandra Kengne Kamga Mobou, Jeroen van Kampen, Jolanda Voermans, Aura Timen, Corine GeurtsvanKessel, Annemiek van der Eijk, Richard Molenkamp, Marion Koopmans, on behalf of the Dutch national COVID-19 response team. |
| EPI_ISL_414440 | hCoV-19/Netherlands/Utrecht_7/2020 | Europe / Netherlands / Utrecht | 2020-03-03 | Dutch COVID-19 response team | Erasmus Medical Center | David Nieuwenhuijse, Bas Oude Munnink, Reina Sikkema, Claudia Schapendonk, Irina Chestakova, Anne van der Linden, Mark Pronk, Pascal Lexmond, Corien Swaan, Manon Haverkate, Madelief Mollers, Mart Stein, Sandra Kengne Kamga Mobou, Jeroen van Kampen, Jolanda Voermans, Aura Timen, Corine GeurtsvanKessel, Annemiek van der Eijk, Richard Molenkamp, Marion Koopmans, on behalf of the Dutch national COVID-19 response team. |
| EPI_ISL_414441 | hCoV-19/Netherlands/Utrecht_8/2020 | Europe / Netherlands / Utrecht | 2020-03-03 | Dutch COVID-19 response team | Erasmus Medical Center | David Nieuwenhuijse, Bas Oude Munnink, Reina Sikkema, Claudia Schapendonk, Irina Chestakova, Anne van der Linden, Mark Pronk, Pascal Lexmond, Corien Swaan, Manon Haverkate, Madelief Mollers, Mart Stein, Sandra Kengne Kamga Mobou, Jeroen van Kampen, Jolanda Voermans, Aura Timen, Corine GeurtsvanKessel, Annemiek van der Eijk, Richard Molenkamp, Marion Koopmans, on behalf of the Dutch national COVID-19 response team. |
| EPI_ISL_414442 | hCoV-19/Netherlands/Utrecht_10/2020 | Europe / Netherlands / Utrecht | 2020-03-03 | Dutch COVID-19 response team | Erasmus Medical Center | David Nieuwenhuijse, Bas Oude Munnink, Reina Sikkema, Claudia Schapendonk, Irina Chestakova, Anne van der Linden, Mark Pronk, Pascal Lexmond, Corien Swaan, Manon Haverkate, Madelief Mollers, Mart Stein, Sandra Kengne Kamga Mobou, Jeroen van Kampen, Jolanda Voermans, Aura Timen, Corine GeurtsvanKessel, Annemiek van der Eijk, Richard Molenkamp, Marion Koopmans, on behalf of the Dutch national COVID-19 response team. |
| EPI_ISL_414443 | hCoV-19/Netherlands/Utrecht_11/2020 | Europe / Netherlands / Utrecht | 2020-03-03 | Dutch COVID-19 response team | Erasmus Medical Center | David Nieuwenhuijse, Bas Oude Munnink, Reina Sikkema, Claudia Schapendonk, Irina Chestakova, Anne van der Linden, Mark Pronk, Pascal Lexmond, Corien Swaan, Manon Haverkate, Madelief Mollers, Mart Stein, Sandra Kengne Kamga Mobou, Jeroen van Kampen, Jolanda Voermans, Aura Timen, Corine GeurtsvanKessel, Annemiek van der Eijk, Richard Molenkamp, Marion Koopmans, on behalf of the Dutch national COVID-19 response team. |

| | | | | | | |
|---|---|---|---|---|---|---|
| EPI_ISL_414444 | hCoV-19/Netherlands/ZuidHolland_1/2020 | Europe / Netherlands / Zuid Holland | 2020-03-02 | Dutch COVID-19 response team | Erasmus Medical Center | David Nieuwenhuijse, Bas Oude Munnink, Reina Sikkema, Claudia Schapendonk, Irina Chestakova, Anne van der Linden, Mark Pronk, Pascal Lexmond, Corien Swaan, Manon Haverkate, Madelief Mollers, Mart Stein, Sandra Kengne Kamga Mobou, Jeroen van Kampen, Jolanda Voermans, Aura Timen, Corine GeurtsvanKessel, Annemiek van der Eijk, Richard Molenkamp, Marion Koopmans, on behalf of the Dutch national COVID-19 response team. |
| EPI_ISL_414445 | hCoV-19/Netherlands/ZuidHolland_9/2020 | Europe / Netherlands / Zuid Holland | 2020-03-03 | Dutch COVID-19 response team | Erasmus Medical Center | David Nieuwenhuijse, Bas Oude Munnink, Reina Sikkema, Claudia Schapendonk, Irina Chestakova, Anne van der Linden, Mark Pronk, Pascal Lexmond, Corien Swaan, Manon Haverkate, Madelief Mollers, Mart Stein, Sandra Kengne Kamga Mobou, Jeroen van Kampen, Jolanda Voermans, Aura Timen, Corine GeurtsvanKessel, Annemiek van der Eijk, Richard Molenkamp, Marion Koopmans, on behalf of the Dutch national COVID-19 response team. |
| EPI_ISL_414446 | hCoV-19/Netherlands/ZuidHolland_10/2020 | Europe / Netherlands / Zuid Holland | 2020-03-03 | Dutch COVID-19 response team | Erasmus Medical Center | David Nieuwenhuijse, Bas Oude Munnink, Reina Sikkema, Claudia Schapendonk, Irina Chestakova, Anne van der Linden, Mark Pronk, Pascal Lexmond, Corien Swaan, Manon Haverkate, Madelief Mollers, Mart Stein, Sandra Kengne Kamga Mobou, Jeroen van Kampen, Jolanda Voermans, Aura Timen, Corine GeurtsvanKessel, Annemiek van der Eijk, Richard Molenkamp, Marion Koopmans, on behalf of the Dutch national COVID-19 response team. |
| EPI_ISL_414448 | hCoV-19/Netherlands/Limburg_5/2020 | Europe / Netherlands / Limburg | 2020-03-04 | Dutch COVID-19 response team | Erasmus Medical Center | David Nieuwenhuijse, Bas Oude Munnink, Reina Sikkema, Claudia Schapendonk, Irina Chestakova, Anne van der Linden, Mark Pronk, Pascal Lexmond, Corien Swaan, Manon Haverkate, Madelief Mollers, Mart Stein, Sandra Kengne Kamga Mobou, Jeroen van Kampen, Jolanda Voermans, Aura Timen, Corine GeurtsvanKessel, Annemiek van der Eijk, Richard Molenkamp, Marion Koopmans, on behalf of the Dutch national COVID-19 response team. |
| EPI_ISL_414449 | hCoV-19/Netherlands/NoordBrabant_2/2020 | Europe / Netherlands / Noord Brabant | 2020-03-03 | Dutch COVID-19 response team | Erasmus Medical Center | David Nieuwenhuijse, Bas Oude Munnink, Reina Sikkema, Claudia Schapendonk, Irina Chestakova, Anne van der Linden, Mark Pronk, Pascal Lexmond, Corien Swaan, Manon Haverkate, Madelief Mollers, Mart Stein, Sandra Kengne Kamga Mobou, Jeroen van Kampen, Jolanda Voermans, Aura Timen, Corine GeurtsvanKessel, Annemiek van der Eijk, Richard Molenkamp, Marion Koopmans, on behalf of the Dutch national COVID-19 response team. |
| EPI_ISL_414451 | hCoV-19/Netherlands/NoordBrabant_6/2020 | Europe / Netherlands / Noord Brabant | 2020-03-06 | Dutch COVID-19 response team | Erasmus Medical Center | David Nieuwenhuijse, Bas Oude Munnink, Reina Sikkema, Claudia Schapendonk, Irina Chestakova, Anne van der Linden, Mark Pronk, Pascal Lexmond, Corien Swaan, Manon Haverkate, Madelief Mollers, Mart Stein, Sandra Kengne Kamga Mobou, Jeroen van Kampen, Jolanda Voermans, Aura Timen, Corine GeurtsvanKessel, Annemiek van der Eijk, Richard Molenkamp, Marion Koopmans, on behalf of the Dutch national COVID-19 response team. |
| EPI_ISL_414452 | hCoV-19/Netherlands/NoordBrabant_12/2020 | Europe / Netherlands / Noord Brabant | 2020-03-06 | Dutch COVID-19 response team | Erasmus Medical Center | David Nieuwenhuijse, Bas Oude Munnink, Reina Sikkema, Claudia Schapendonk, Irina Chestakova, Anne van der Linden, Mark Pronk, Pascal Lexmond, Corien Swaan, Manon Haverkate, Madelief Mollers, Mart Stein, Sandra Kengne Kamga Mobou, Jeroen van Kampen, Jolanda Voermans, Aura Timen, Corine GeurtsvanKessel, Annemiek van der Eijk, Richard Molenkamp, Marion Koopmans, on behalf of the Dutch national COVID-19 response team. |
| EPI_ISL_414454 | hCoV-19/Netherlands/NoordBrabant_14/2020 | Europe / Netherlands / Noord Brabant | 2020-03-05 | Dutch COVID-19 response team | Erasmus Medical Center | David Nieuwenhuijse, Bas Oude Munnink, Reina Sikkema, Claudia Schapendonk, Irina Chestakova, Anne van der Linden, Mark Pronk, Pascal Lexmond, Corien Swaan, Manon Haverkate, Madelief Mollers, Mart Stein, Sandra Kengne Kamga Mobou, Jeroen van Kampen, Jolanda Voermans, Aura Timen, Corine GeurtsvanKessel, Annemiek van der Eijk, Richard Molenkamp, Marion Koopmans, on behalf of the Dutch national COVID-19 response team. |
| EPI_ISL_414456 | hCoV-19/Netherlands/NoordBrabant_16/2020 | Europe / Netherlands / Noord Brabant | 2020-03-04 | Dutch COVID-19 response team | Erasmus Medical Center | David Nieuwenhuijse, Bas Oude Munnink, Reina Sikkema, Claudia Schapendonk, Irina Chestakova, Anne van der Linden, Mark Pronk, Pascal Lexmond, Corien Swaan, Manon Haverkate, Madelief Mollers, Mart Stein, Sandra Kengne Kamga Mobou, Jeroen van Kampen, Jolanda Voermans, Aura Timen, Corine GeurtsvanKessel, Annemiek van der Eijk, Richard Molenkamp, Marion Koopmans, on behalf of the Dutch national COVID-19 response team. |
| EPI_ISL_414457 | hCoV-19/Netherlands/NoordBrabant_17/2020 | Europe / Netherlands / Noord Brabant | 2020-03-06 | Dutch COVID-19 response team | Erasmus Medical Center | David Nieuwenhuijse, Bas Oude Munnink, Reina Sikkema, Claudia Schapendonk, Irina Chestakova, Anne van der Linden, Mark Pronk, Pascal Lexmond, Corien Swaan, Manon Haverkate, Madelief Mollers, Mart Stein, Sandra Kengne Kamga Mobou, Jeroen van Kampen, Jolanda Voermans, Aura Timen, Corine GeurtsvanKessel, Annemiek van der Eijk, Richard Molenkamp, Marion Koopmans, on behalf of the Dutch national COVID-19 response team. |
| EPI_ISL_414458 | hCoV-19/Netherlands/NoordBrabant_18/2020 | Europe / Netherlands / Noord Brabant | 2020-03-05 | Dutch COVID-19 response team | Erasmus Medical Center | David Nieuwenhuijse, Bas Oude Munnink, Reina Sikkema, Claudia Schapendonk, Irina Chestakova, Anne van der Linden, Mark Pronk, Pascal Lexmond, Corien Swaan, Manon Haverkate, Madelief Mollers, Mart Stein, Sandra Kengne Kamga Mobou, Jeroen van Kampen, Jolanda Voermans, Aura Timen, Corine GeurtsvanKessel, Annemiek van der Eijk, Richard Molenkamp, Marion Koopmans, on behalf of the Dutch national COVID-19 response team. |
| EPI_ISL_414460 | hCoV-19/Netherlands/Overijssel_2/2020 | Europe / Netherlands / Overijssel | 2020-03-03 | Dutch COVID-19 response team | Erasmus Medical Center | David Nieuwenhuijse, Bas Oude Munnink, Reina Sikkema, Claudia Schapendonk, Irina Chestakova, Anne van der Linden, Mark Pronk, Pascal Lexmond, Corien Swaan, Manon Haverkate, Madelief Mollers, Mart Stein, Sandra Kengne Kamga Mobou, Jeroen van Kampen, Jolanda Voermans, Aura Timen, Corine GeurtsvanKessel, Annemiek van der Eijk, Richard Molenkamp, Marion Koopmans, on behalf of the Dutch national COVID-19 response team. |
| EPI_ISL_414461 | hCoV-19/Netherlands/Utrecht_6/2020 | Europe / Netherlands / Utrecht | 2020-03-04 | Dutch COVID-19 response team | Erasmus Medical Center | David Nieuwenhuijse, Bas Oude Munnink, Reina Sikkema, Claudia Schapendonk, Irina Chestakova, Anne van der Linden, Mark Pronk, Pascal Lexmond, Corien Swaan, Manon Haverkate, Madelief Mollers, Mart Stein, Sandra Kengne Kamga Mobou, Jeroen van Kampen, Jolanda Voermans, Aura Timen, Corine GeurtsvanKessel, Annemiek van der Eijk, Richard Molenkamp, Marion Koopmans, on behalf of the Dutch national COVID-19 response team. |
| EPI_ISL_414462 | hCoV-19/Netherlands/Utrecht_12/2020 | Europe / Netherlands / Utrecht | 2020-03-04 | Dutch COVID-19 response team | Erasmus Medical Center | David Nieuwenhuijse, Bas Oude Munnink, Reina Sikkema, Claudia Schapendonk, Irina Chestakova, Anne van der Linden, Mark Pronk, Pascal Lexmond, Corien Swaan, Manon Haverkate, Madelief Mollers, Mart Stein, Sandra Kengne Kamga Mobou, Jeroen van Kampen, Jolanda Voermans, Aura Timen, Corine GeurtsvanKessel, Annemiek van der Eijk, Richard Molenkamp, Marion Koopmans, on behalf of the Dutch national COVID-19 response team. |
| EPI_ISL_414463 | hCoV-19/Netherlands/Utrecht_13/2020 | Europe / Netherlands / Utrecht | 2020-03-04 | Dutch COVID-19 response team | Erasmus Medical Center | David Nieuwenhuijse, Bas Oude Munnink, Reina Sikkema, Claudia Schapendonk, Irina Chestakova, Anne van der Linden, Mark Pronk, Pascal Lexmond, Corien Swaan, Manon Haverkate, Madelief Mollers, Mart Stein, Sandra Kengne Kamga Mobou, Jeroen van Kampen, Jolanda Voermans, Aura Timen, Corine GeurtsvanKessel, Annemiek van der Eijk, Richard Molenkamp, Marion Koopmans, on behalf of the Dutch national COVID-19 response team. |
| EPI_ISL_414464 | hCoV-19/Netherlands/Utrecht_14/2020 | Europe / Netherlands / Utrecht | 2020-03-04 | Dutch COVID-19 response team | Erasmus Medical Center | David Nieuwenhuijse, Bas Oude Munnink, Reina Sikkema, Claudia Schapendonk, Irina Chestakova, Anne van der Linden, Mark Pronk, Pascal Lexmond, Corien Swaan, Manon Haverkate, Madelief Mollers, Mart Stein, Sandra Kengne Kamga Mobou, Jeroen van Kampen, Jolanda Voermans, Aura Timen, Corine GeurtsvanKessel, Annemiek van der Eijk, Richard Molenkamp, Marion Koopmans, on behalf of the Dutch national COVID-19 response team. |
| EPI_ISL_414467 | hCoV-19/Netherlands/ZuidHolland_7/2020 | Europe / Netherlands / Zuid Holland | 2020-03-05 | Dutch COVID-19 response team | Erasmus Medical Center | David Nieuwenhuijse, Bas Oude Munnink, Reina Sikkema, Claudia Schapendonk, Irina Chestakova, Anne van der Linden, Mark Pronk, Pascal Lexmond, Corien Swaan, Manon Haverkate, Madelief Mollers, Mart Stein, Sandra Kengne Kamga Mobou, Jeroen van Kampen, Jolanda Voermans, Aura Timen, Corine GeurtsvanKessel, Annemiek van der Eijk, Richard Molenkamp, Marion Koopmans, on behalf of the Dutch national COVID-19 response team. |
| EPI_ISL_414468 | hCoV-19/Netherlands/ZuidHolland_8/2020 | Europe / Netherlands / Zuid Holland | 2020-03-06 | Dutch COVID-19 response team | Erasmus Medical Center | David Nieuwenhuijse, Bas Oude Munnink, Reina Sikkema, Claudia Schapendonk, Irina Chestakova, Anne van der Linden, Mark Pronk, Pascal Lexmond, Corien Swaan, Manon Haverkate, Madelief Mollers, Mart Stein, Sandra Kengne Kamga Mobou, Jeroen van Kampen, Jolanda Voermans, Aura Timen, Corine GeurtsvanKessel, Annemiek van der Eijk, Richard Molenkamp, Marion Koopmans, on behalf of the Dutch national COVID-19 response team. |
| EPI_ISL_414469 | hCoV-19/Netherlands/ZuidHolland_11/2020 | Europe / Netherlands / Zuid Holland | 2020-03-04 | Dutch COVID-19 response team | Erasmus Medical Center | David Nieuwenhuijse, Bas Oude Munnink, Reina Sikkema, Claudia Schapendonk, Irina Chestakova, Anne van der Linden, Mark Pronk, Pascal Lexmond, Corien Swaan, Manon Haverkate, Madelief Mollers, Mart Stein, Sandra Kengne Kamga Mobou, Jeroen van Kampen, Jolanda Voermans, Aura Timen, Corine GeurtsvanKessel, Annemiek van der Eijk, Richard Molenkamp, Marion Koopmans, on behalf of the Dutch national COVID-19 response team. |
| EPI_ISL_414470 | hCoV-19/Netherlands/ZuidHolland_13/2020 | Europe / Netherlands / Zuid Holland | 2020-03-06 | Dutch COVID-19 response team | Erasmus Medical Center | David Nieuwenhuijse, Bas Oude Munnink, Reina Sikkema, Claudia Schapendonk, Irina Chestakova, Anne van der Linden, Mark Pronk, Pascal Lexmond, Corien Swaan, Manon Haverkate, Madelief Mollers, Mart Stein, Sandra Kengne Kamga Mobou, Jeroen van Kampen, Jolanda Voermans, Aura Timen, Corine GeurtsvanKessel, Annemiek van der Eijk, Richard Molenkamp, Marion Koopmans, on behalf of the Dutch national COVID-19 response team. |
| EPI_ISL_414471 | hCoV-19/Netherlands/ZuidHolland_14/2020 | Europe / Netherlands / Zuid Holland | 2020-03-05 | Dutch COVID-19 response team | Erasmus Medical Center | David Nieuwenhuijse, Bas Oude Munnink, Reina Sikkema, Claudia Schapendonk, Irina Chestakova, Anne van der Linden, Mark Pronk, Pascal Lexmond, Corien Swaan, Manon Haverkate, Madelief Mollers, Mart Stein, Sandra Kengne Kamga Mobou, Jeroen van Kampen, Jolanda Voermans, Aura Timen, Corine GeurtsvanKessel, Annemiek van der Eijk, Richard Molenkamp, Marion Koopmans, on behalf of the Dutch national COVID-19 response team. |
| EPI_ISL_414476 | hCoV-19/USA/NY1-PV08001/2020 | North America / USA / New York | 2020-02-29 | MSHS Clinical Microbiology Laboratories | MSHS Pathogen Surveillance Program | Gopi Patel, Emilia Sordillo, Melissa Gitman, Alberto Paniz-mondolfi, Matthew Hernandez, Shelcie Fabre, Jose Polanco, Ana Sylvia Gonzalez-Reiche, Zenab Khan, Nancy Francoeur, Melissa Smith, Robert Sebra, Lisa Miorin, Wen-chun Liu, Randy Albrecht, Judith Aberg, Florian Krammer, Adolfo Garcia-Sastre, Viviana Simon, Harm van Bakel |
| EPI_ISL_414477 | hCoV-19/Czech Republic/951/2020 | Europe / Czech Republic / Usti nad Labe | 2020-03-01 | The National Institute of Public Health Center for Epidemiology and Microbiology | State Veterinary Institute Prague | Alexander Nagy, Oldrich Bartos, Helena Jirincova, Klara Labska, Ludmila Novakova, Olga Storkanova, Dusan Trnka, Jaromira Vecerova |
| EPI_ISL_414479 | hCoV-19/USA/CruiseA-19/2020 | North America / USA | 2020-02-18 | unknown | Pathogen Discovery, Respiratory Viruses Branch, Division of Viral Diseases, Centers for Disease Control and Prevention | Ying Tao, Krista Queen, Clinton R. Paden, Anna Uehara, Jing Zhang, Yan Li, Mary S. Keckler, Alison S. Laufer Halpin, Haibin Wang, Jasmine Padilla, Justin Lee, Christopher A. Elkins, Susan I. Gerber, Suxiang Tong |
| EPI_ISL_414480 | hCoV-19/USA/CruiseA-21/2020 | North America / USA | 2020-02-21 | unknown | Pathogen Discovery, Respiratory Viruses Branch, Division of Viral Diseases, Centers for Disease Control and Prevention | Ying Tao, Krista Queen, Clinton R. Paden, Anna Uehara, Jing Zhang, Yan Li, Mary S. Keckler, Alison S. Laufer Halpin, Haibin Wang, Jasmine Padilla, Justin Lee, Christopher A. Elkins, Susan I. Gerber, Suxiang Tong |
| EPI_ISL_414481 | hCoV-19/USA/CruiseA-22/2020 | North America / USA | 2020-02-21 | unknown | Pathogen Discovery, Respiratory Viruses Branch, Division of Viral Diseases, Centers for Disease Control and Prevention | Ying Tao, Krista Queen, Clinton R. Paden, Anna Uehara, Jing Zhang, Yan Li, Mary S. Keckler, Alison S. Laufer Halpin, Haibin Wang, Jasmine Padilla, Justin Lee, Christopher A. Elkins, Susan I. Gerber, Suxiang Tong |
| EPI_ISL_414482 | hCoV-19/USA/CruiseA-23/2020 | North America / USA | 2020-02-18 | unknown | Pathogen Discovery, Respiratory Viruses Branch, Division of Viral Diseases, Centers for Disease Control and Prevention | Krista Queen, Anna Uehara, Ying Tao, Clinton R. Paden, Jing Zhang, Yan Li, Haibin Wang, Shifaq Kamili, Xiaoyan Lu, Brian Lynch, Senthil Kumar K. Sakthivel, Brett L. Whitaker, Lijuan Wang, Janna' R. Murray, Jasmine Padilla, Justin Lee, Susan I. Gerber, Stephen Lindstrom, Suxiang Tong |
| EPI_ISL_414483 | hCoV-19/USA/CruiseA-24/2020 | North America / USA | 2020-02-17 | unknown | Pathogen Discovery, Respiratory Viruses Branch, Division of Viral Diseases, Centers for Disease Control and Prevention | Krista Queen, Anna Uehara, Ying Tao, Clinton R. Paden, Jing Zhang, Yan Li, Haibin Wang, Shifaq Kamili, Xiaoyan Lu, Brian Lynch, Senthil Kumar K. Sakthivel, Brett L. Whitaker, Lijuan Wang, Janna' R. Murray, Jasmine Padilla, Justin Lee, Susan I. Gerber, Stephen Lindstrom, Suxiang Tong |
| EPI_ISL_414484 | hCoV-19/USA/CruiseA-25/2020 | North America / USA | 2020-02-17 | unknown | Pathogen Discovery, Respiratory Viruses Branch, Division of Viral Diseases, Centers for Disease Control and Prevention | Krista Queen, Anna Uehara, Ying Tao, Clinton R. Paden, Jing Zhang, Yan Li, Haibin Wang, Shifaq Kamili, Xiaoyan Lu, Brian Lynch, Senthil Kumar K. Sakthivel, Brett L. Whitaker, Lijuan Wang, Janna' R. Murray, Jasmine Padilla, Justin Lee, Susan I. Gerber, Stephen Lindstrom, Suxiang Tong |
| EPI_ISL_414485 | hCoV-19/USA/CruiseA-26/2020 | North America / USA | 2020-02-24 | unknown | Pathogen Discovery, Respiratory Viruses Branch, Division of Viral Diseases, Centers for Disease Control and Prevention | Krista Queen, Anna Uehara, Ying Tao, Clinton R. Paden, Jing Zhang, Yan Li, Haibin Wang, Shifaq Kamili, Xiaoyan Lu, Brian Lynch, Senthil Kumar K. Sakthivel, Brett L. Whitaker, Lijuan Wang, Janna' R. Murray, Jasmine Padilla, Justin Lee, Susan I. Gerber, Stephen Lindstrom, Suxiang Tong |
| EPI_ISL_414487 | hCoV-19/Ireland/COR-20134/2020 | Europe / Ireland / Cork | 2020-03-04 | UCD National Virus Reference Laboratory | UCD National Virus Reference Laboratory | Michael Carr, Gabriel Gonzalez, Jonathan Dean, Suzie Coughlan, Alison Murphy, Kevin Byrne, Ken Wolfe, Jeff Connell, Brendan Loftus, Cillian F De Gascun |
| EPI_ISL_414495 | hCoV-19/Spain/Valencia1/2020 | Europe / Spain / Comunitat Valenciana / | 2020-03-08 | Servicio Microbiología. Hospital Clínico Universitario. Valencia. | Sequencing and Bioinformatics Service. Molecular Epidemiology Laboratory. FISABIO-Public Health | David Navarro, Maria Alma Bracho, Giuseppe D'Auria, Griselda De Marco, Neris Garcia-Gonzalez, Fernando Gonzalez-Candelas |
| EPI_ISL_414496 | hCoV-19/Spain/Valencia2/2020 | Europe / Spain / Comunitat Valenciana / | 2020-03-04 | Servicio Microbiología. Hospital Clínico Universitario. Valencia. | Sequencing and Bioinformatics Service. Molecular Epidemiology Laboratory. FISABIO-Public Health | David Navarro, Maria Alma Bracho, Giuseppe D'Auria, Griselda De Marco, Neris Garcia-Gonzalez, Fernando Gonzalez-Candelas |
| EPI_ISL_414497 | hCoV-19/Germany/NRW-02-1/2020 | Europe / Germany / North Rhine Westph | 2020-02-25 | Center of Medical Microbiology, Virology, and Hospital Hygiene, University of Duesseldorf | Center of Medical Microbiology, Virology, and Hospital Hygiene, University of Duesseldorf | Ortwin Adams, Marcel Andree, Alexander Dilthey, Torsten Feldt, Sandra Hauka, Torsten Houwaart, Björn-Erik Jensen, Detlef Kindgen-Milles, Malte Kohns Vasconcelos, Klaus Pfeffer, Tina Senff, Daniel Strelow, Jörg Timm, Andreas Walker, Tobias Wienemann |
| EPI_ISL_414498 | hCoV-19/Germany/NRW-03/2020 | Europe / Germany / North Rhine Westph | 2020-02-26 | Center of Medical Microbiology, Virology, and Hospital Hygiene, University of Duesseldorf | Center of Medical Microbiology, Virology, and Hospital Hygiene, University of Duesseldorf | Ortwin Adams, Marcel Andree, Alexander Dilthey, Torsten Feldt, Sandra Hauka, Torsten Houwaart, Björn-Erik Jensen, Detlef Kindgen-Milles, Malte Kohns Vasconcelos, Klaus Pfeffer, Tina Senff, Daniel Strelow, Jörg Timm, Andreas Walker, Tobias Wienemann |
| EPI_ISL_414499 | hCoV-19/Germany/NRW-04/2020 | Europe / Germany / North Rhine Westph | 2020-02-26 | Center of Medical Microbiology, Virology, and Hospital Hygiene, University of Duesseldorf | Center of Medical Microbiology, Virology, and Hospital Hygiene, University of Duesseldorf | Ortwin Adams, Marcel Andree, Alexander Dilthey, Torsten Feldt, Sandra Hauka, Torsten Houwaart, Björn-Erik Jensen, Detlef Kindgen-Milles, Malte Kohns Vasconcelos, Klaus Pfeffer, Tina Senff, Daniel Strelow, Jörg Timm, Andreas Walker, Tobias Wienemann |
| EPI_ISL_414551 | hCoV-19/Netherlands/Utrecht_47/2020 | Europe / Netherlands / Utrecht | 2020-03-07 | Dutch COVID-19 response team | Erasmus Medical Center | David Nieuwenhuijse, Bas Oude Munnink, Reina Sikkema, Claudia Schapendonk, Irina Chestakova, Anne van der Linden, Mark Pronk, Pascal Lexmond, Corien Swaan, Manon Haverkate, Madelief Mollers, Mart Stein, Sandra Kengne Kamga Mobou, Jeroen van Kampen, Jolanda Voermans, Aura Timen, Corine GeurtsvanKessel, Annemiek van der Eijk, Richard Molenkamp, Marion Koopmans, on behalf of the Dutch national COVID-19 response team. |
| EPI_ISL_414552 | hCoV-19/Netherlands/Utrecht_46/2020 | Europe / Netherlands / Utrecht | 2020-03-07 | Dutch COVID-19 response team | Erasmus Medical Center | David Nieuwenhuijse, Bas Oude Munnink, Reina Sikkema, Claudia Schapendonk, Irina Chestakova, Anne van der Linden, Mark Pronk, Pascal Lexmond, Corien Swaan, Manon Haverkate, Madelief Mollers, Mart Stein, Sandra Kengne Kamga Mobou, Jeroen van Kampen, Jolanda Voermans, Aura Timen, Corine GeurtsvanKessel, Annemiek van der Eijk, Richard Molenkamp, Marion Koopmans, on behalf of the Dutch national COVID-19 response team. |
| EPI_ISL_414554 | hCoV-19/Netherlands/Utrecht_15/2020 | Europe / Netherlands / Utrecht | 2020-03-08 | Dutch COVID-19 response team | Erasmus Medical Center | David Nieuwenhuijse, Bas Oude Munnink, Reina Sikkema, Claudia Schapendonk, Irina Chestakova, Anne van der Linden, Mark Pronk, Pascal Lexmond, Corien Swaan, Manon Haverkate, Madelief Mollers, Mart Stein, Sandra Kengne Kamga Mobou, Jeroen van Kampen, Jolanda Voermans, Aura Timen, Corine GeurtsvanKessel, Annemiek van der Eijk, Richard Molenkamp, Marion Koopmans, on behalf of the Dutch national COVID-19 response team. |
| EPI_ISL_414556 | hCoV-19/Netherlands/ZuidHolland_2/2020 | Europe / Netherlands / Zuid Holland | 2020-03-06 | Dutch COVID-19 response team | Erasmus Medical Center | David Nieuwenhuijse, Bas Oude Munnink, Reina Sikkema, Claudia Schapendonk, Irina Chestakova, Anne van der Linden, Mark Pronk, Pascal Lexmond, Corien Swaan, Manon Haverkate, Madelief Mollers, Mart Stein, Sandra Kengne Kamga Mobou, Jeroen van Kampen, Jolanda Voermans, Aura Timen, Corine GeurtsvanKessel, Annemiek van der Eijk, Richard Molenkamp, Marion Koopmans, on behalf of the Dutch national COVID-19 response team. |
| EPI_ISL_415041 | hCoV-19/Wales/PHW04/2020 | Europe / United Kingdom / Wales | 2020-03-07 | Wales Specialist Virology Centre | Public Health Wales Microbiology Cardiff | Catherine Moore, Joanne Watkins, Sally Corden, Tom Connor |
| EPI_ISL_415470 | hCoV-19/Netherlands/NA_14/2020 | Europe / Netherlands | 2020-03-10 | Dutch COVID-19 response team | Erasmus Medical Center | David Nieuwenhuijse, Bas Oude Munnink, Reina Sikkema, Claudia Schapendonk, Irina Chestakova, Anne van der Linden, Mark Pronk, Pascal Lexmond, Corien Swaan, Manon Haverkate, Madelief Mollers, Mart Stein, Sandra Kengne Kamga Mobou, Jeroen van Kampen, Jolanda Voermans, Aura Timen, Corine GeurtsvanKessel, Annemiek van der Eijk, Richard Molenkamp, Marion Koopmans, on behalf of the Dutch national COVID-19 response team. |
| EPI_ISL_415471 | hCoV-19/Netherlands/NA_15/2020 | Europe / Netherlands | 2020-03-11 | Dutch COVID-19 response team | Erasmus Medical Center | David Nieuwenhuijse, Bas Oude Munnink, Reina Sikkema, Claudia Schapendonk, Irina Chestakova, Anne van der Linden, Mark Pronk, Pascal Lexmond, Corien Swaan, Manon Haverkate, Madelief Mollers, Mart Stein, Sandra Kengne Kamga Mobou, Jeroen van Kampen, Jolanda Voermans, Aura Timen, Corine GeurtsvanKessel, Annemiek van der Eijk, Richard Molenkamp, Marion Koopmans, on behalf of the Dutch national COVID-19 response team. |
| EPI_ISL_415472 | hCoV-19/Netherlands/NA_16/2020 | Europe / Netherlands | 2020-03-11 | Dutch COVID-19 response team | Erasmus Medical Center | David Nieuwenhuijse, Bas Oude Munnink, Reina Sikkema, Claudia Schapendonk, Irina Chestakova, Anne van der Linden, Mark Pronk, Pascal Lexmond, Corien Swaan, Manon Haverkate, Madelief Mollers, Mart Stein, Sandra Kengne Kamga Mobou, Jeroen van Kampen, Jolanda Voermans, Aura Timen, Corine GeurtsvanKessel, Annemiek van der Eijk, Richard Molenkamp, Marion Koopmans, on behalf of the Dutch national COVID-19 response team. |
| EPI_ISL_415473 | hCoV-19/Netherlands/NA_17/2020 | Europe / Netherlands | 2020-03-09 | Dutch COVID-19 response team | Erasmus Medical Center | David Nieuwenhuijse, Bas Oude Munnink, Reina Sikkema, Claudia Schapendonk, Irina Chestakova, Anne van der Linden, Mark Pronk, Pascal Lexmond, Corien Swaan, Manon Haverkate, Madelief Mollers, Mart Stein, Sandra Kengne Kamga Mobou, Jeroen van Kampen, Jolanda Voermans, Aura Timen, Corine GeurtsvanKessel, Annemiek van der Eijk, Richard Molenkamp, Marion Koopmans, on behalf of the Dutch national COVID-19 response team. |
| EPI_ISL_415474 | hCoV-19/Netherlands/NA_18/2020 | Europe / Netherlands | 2020-03-09 | Dutch COVID-19 response team | Erasmus Medical Center | David Nieuwenhuijse, Bas Oude Munnink, Reina Sikkema, Claudia Schapendonk, Irina Chestakova, Anne van der Linden, Mark Pronk, Pascal Lexmond, Corien Swaan, Manon Haverkate, Madelief Mollers, Mart Stein, Sandra Kengne Kamga Mobou, Jeroen van Kampen, Jolanda Voermans, Aura Timen, Corine GeurtsvanKessel, Annemiek van der Eijk, Richard Molenkamp, Marion Koopmans, on behalf of the Dutch national COVID-19 response team. |
| EPI_ISL_415475 | hCoV-19/Netherlands/NA_19/2020 | Europe / Netherlands | 2020-03-12 | Dutch COVID-19 response team | Erasmus Medical Center | David Nieuwenhuijse, Bas Oude Munnink, Reina Sikkema, Claudia Schapendonk, Irina Chestakova, Anne van der Linden, Mark Pronk, Pascal Lexmond, Corien Swaan, Manon Haverkate, Madelief Mollers, Mart Stein, Sandra Kengne Kamga Mobou, Jeroen van Kampen, Jolanda Voermans, Aura Timen, Corine GeurtsvanKessel, Annemiek van der Eijk, Richard Molenkamp, Marion Koopmans, on behalf of the Dutch national COVID-19 response team. |
| EPI_ISL_415476 | hCoV-19/Netherlands/NA_2/2020 | Europe / Netherlands | 2020-03-10 | Dutch COVID-19 response team | Erasmus Medical Center | David Nieuwenhuijse, Bas Oude Munnink, Reina Sikkema, Claudia Schapendonk, Irina Chestakova, Anne van der Linden, Mark Pronk, Pascal Lexmond, Corien Swaan, Manon Haverkate, Madelief Mollers, Mart Stein, Sandra Kengne Kamga Mobou, Jeroen van Kampen, Jolanda Voermans, Aura Timen, Corine GeurtsvanKessel, Annemiek van der Eijk, Richard Molenkamp, Marion Koopmans, on behalf of the Dutch national COVID-19 response team. |
| EPI_ISL_415478 | hCoV-19/Netherlands/NA_21/2020 | Europe / Netherlands | 2020-03-08 | Dutch COVID-19 response team | Erasmus Medical Center | David Nieuwenhuijse, Bas Oude Munnink, Reina Sikkema, Claudia Schapendonk, Irina Chestakova, Anne van der Linden, Mark Pronk, Pascal Lexmond, Corien Swaan, Manon Haverkate, Madelief Mollers, Mart Stein, Sandra Kengne Kamga Mobou, Jeroen van Kampen, Jolanda Voermans, Aura Timen, Corine GeurtsvanKessel, Annemiek van der Eijk, Richard Molenkamp, Marion Koopmans, on behalf of the Dutch national COVID-19 response team. |
| EPI_ISL_415479 | hCoV-19/Netherlands/NA_22/2020 | Europe / Netherlands | 2020-03-08 | Dutch COVID-19 response team | Erasmus Medical Center | David Nieuwenhuijse, Bas Oude Munnink, Reina Sikkema, Claudia Schapendonk, Irina Chestakova, Anne van der Linden, Mark Pronk, Pascal Lexmond, Corien Swaan, Manon Haverkate, Madelief Mollers, Mart Stein, Sandra Kengne Kamga Mobou, Jeroen van Kampen, Jolanda Voermans, Aura Timen, Corine GeurtsvanKessel, Annemiek van der Eijk, Richard Molenkamp, Marion Koopmans, on behalf of the Dutch national COVID-19 response team. |
| EPI_ISL_415480 | hCoV-19/Netherlands/NA_23/2020 | Europe / Netherlands | 2020-03-09 | Dutch COVID-19 response team | Erasmus Medical Center | David Nieuwenhuijse, Bas Oude Munnink, Reina Sikkema, Claudia Schapendonk, Irina Chestakova, Anne van der Linden, Mark Pronk, Pascal Lexmond, Corien Swaan, Manon Haverkate, Madelief Mollers, Mart Stein, Sandra Kengne Kamga Mobou, Jeroen van Kampen, Jolanda Voermans, Aura Timen, Corine GeurtsvanKessel, Annemiek van der Eijk, Richard Molenkamp, Marion Koopmans, on behalf of the Dutch national COVID-19 response team. |
| EPI_ISL_415481 | hCoV-19/Netherlands/NA_24/2020 | Europe / Netherlands | 2020-03-08 | Dutch COVID-19 response team | Erasmus Medical Center | David Nieuwenhuijse, Bas Oude Munnink, Reina Sikkema, Claudia Schapendonk, Irina Chestakova, Anne van der Linden, Mark Pronk, Pascal Lexmond, Corien Swaan, Manon Haverkate, Madelief Mollers, Mart Stein, Sandra Kengne Kamga Mobou, Jeroen van Kampen, Jolanda Voermans, Aura Timen, Corine GeurtsvanKessel, Annemiek van der Eijk, Richard Molenkamp, Marion Koopmans, on behalf of the Dutch national COVID-19 response team. |
| EPI_ISL_415482 | hCoV-19/Netherlands/NA_25/2020 | Europe / Netherlands | 2020-03-09 | Dutch COVID-19 response team | Erasmus Medical Center | David Nieuwenhuijse, Bas Oude Munnink, Reina Sikkema, Claudia Schapendonk, Irina Chestakova, Anne van der Linden, Mark Pronk, Pascal Lexmond, Corien Swaan, Manon Haverkate, Madelief Mollers, Mart Stein, Sandra Kengne Kamga Mobou, Jeroen van Kampen, Jolanda Voermans, Aura Timen, Corine GeurtsvanKessel, Annemiek van der Eijk, Richard Molenkamp, Marion Koopmans, on behalf of the Dutch national COVID-19 response team. |
| EPI_ISL_415483 | hCoV-19/Netherlands/NA_26/2020 | Europe / Netherlands | 2020-03-09 | Dutch COVID-19 response team | Erasmus Medical Center | David Nieuwenhuijse, Bas Oude Munnink, Reina Sikkema, Claudia Schapendonk, Irina Chestakova, Anne van der Linden, Mark Pronk, Pascal Lexmond, Corien Swaan, Manon Haverkate, Madelief Mollers, Mart Stein, Sandra Kengne Kamga Mobou, Jeroen van Kampen, Jolanda Voermans, Aura Timen, Corine GeurtsvanKessel, Annemiek van der Eijk, Richard Molenkamp, Marion Koopmans, on behalf of the Dutch national COVID-19 response team. |
| EPI_ISL_415484 | hCoV-19/Netherlands/NA_27/2020 | Europe / Netherlands | 2020-03-13 | Dutch COVID-19 response team | Erasmus Medical Center | David Nieuwenhuijse, Bas Oude Munnink, Reina Sikkema, Claudia Schapendonk, Irina Chestakova, Anne van der Linden, Mark Pronk, Pascal Lexmond, Corien Swaan, Manon Haverkate, Madelief Mollers, Mart Stein, Sandra Kengne Kamga Mobou, Jeroen van Kampen, Jolanda Voermans, Aura Timen, Corine GeurtsvanKessel, Annemiek van der Eijk, Richard Molenkamp, Marion Koopmans, on behalf of the Dutch national COVID-19 response team. |
| EPI_ISL_415485 | hCoV-19/Netherlands/NA_28/2020 | Europe / Netherlands | 2020-03-12 | Dutch COVID-19 response team | Erasmus Medical Center | David Nieuwenhuijse, Bas Oude Munnink, Reina Sikkema, Claudia Schapendonk, Irina Chestakova, Anne van der Linden, Mark Pronk, Pascal Lexmond, Corien Swaan, Manon Haverkate, Madelief Mollers, Mart Stein, Sandra Kengne Kamga Mobou, Jeroen van Kampen, Jolanda Voermans, Aura Timen, Corine GeurtsvanKessel, Annemiek van der Eijk, Richard Molenkamp, Marion Koopmans, on behalf of the Dutch national COVID-19 response team. |
| EPI_ISL_415486 | hCoV-19/Netherlands/NA_29/2020 | Europe / Netherlands | 2020-03-13 | Dutch COVID-19 response team | Erasmus Medical Center | David Nieuwenhuijse, Bas Oude Munnink, Reina Sikkema, Claudia Schapendonk, Irina Chestakova, Anne van der Linden, Mark Pronk, Pascal Lexmond, Corien Swaan, Manon Haverkate, Madelief Mollers, Mart Stein, Sandra Kengne Kamga Mobou, Jeroen van Kampen, Jolanda Voermans, Aura Timen, Corine GeurtsvanKessel, Annemiek van der Eijk, Richard Molenkamp, Marion Koopmans, on behalf of the Dutch national COVID-19 response team. |
| EPI_ISL_415487 | hCoV-19/Netherlands/NA_30/2020 | Europe / Netherlands | 2020-03-13 | Dutch COVID-19 response team | Erasmus Medical Center | David Nieuwenhuijse, Bas Oude Munnink, Reina Sikkema, Claudia Schapendonk, Irina Chestakova, Anne van der Linden, Mark Pronk, Pascal Lexmond, Corien Swaan, Manon Haverkate, Madelief Mollers, Mart Stein, Sandra Kengne Kamga Mobou, Jeroen van Kampen, Jolanda Voermans, Aura Timen, Corine GeurtsvanKessel, Annemiek van der Eijk, Richard Molenkamp, Marion Koopmans, on behalf of the Dutch national COVID-19 response team. |
| EPI_ISL_415488 | hCoV-19/Netherlands/NA_31/2020 | Europe / Netherlands | 2020-03-13 | Dutch COVID-19 response team | Erasmus Medical Center | David Nieuwenhuijse, Bas Oude Munnink, Reina Sikkema, Claudia Schapendonk, Irina Chestakova, Anne van der Linden, Mark Pronk, Pascal Lexmond, Corien Swaan, Manon Haverkate, Madelief Mollers, Mart Stein, Sandra Kengne Kamga Mobou, Jeroen van Kampen, Jolanda Voermans, Aura Timen, Corine GeurtsvanKessel, Annemiek van der Eijk, Richard Molenkamp, Marion Koopmans, on behalf of the Dutch national COVID-19 response team. |
| EPI_ISL_415489 | hCoV-19/Netherlands/NA_32/2020 | Europe / Netherlands | 2020-03-13 | Dutch COVID-19 response team | Erasmus Medical Center | David Nieuwenhuijse, Bas Oude Munnink, Reina Sikkema, Claudia Schapendonk, Irina Chestakova, Anne van der Linden, Mark Pronk, Pascal Lexmond, Corien Swaan, Manon Haverkate, Madelief Mollers, Mart Stein, Sandra Kengne Kamga Mobou, Jeroen van Kampen, Jolanda Voermans, Aura Timen, Corine GeurtsvanKessel, Annemiek van der Eijk, Richard Molenkamp, Marion Koopmans, on behalf of the Dutch national COVID-19 response team. |
| EPI_ISL_415491 | hCoV-19/Netherlands/NA_34/2020 | Europe / Netherlands | 2020-03-07 | Dutch COVID-19 response team | Erasmus Medical Center | David Nieuwenhuijse, Bas Oude Munnink, Reina Sikkema, Claudia Schapendonk, Irina Chestakova, Anne van der Linden, Mark Pronk, Pascal Lexmond, Corien Swaan, Manon Haverkate, Madelief Mollers, Mart Stein, Sandra Kengne Kamga Mobou, Jeroen van Kampen, Jolanda Voermans, Aura Timen, Corine GeurtsvanKessel, Annemiek van der Eijk, Richard Molenkamp, Marion Koopmans, on behalf of the Dutch national COVID-19 response team. |
| EPI_ISL_415492 | hCoV-19/Netherlands/NA_35/2020 | Europe / Netherlands | 2020-03-10 | Dutch COVID-19 response team | Erasmus Medical Center | David Nieuwenhuijse, Bas Oude Munnink, Reina Sikkema, Claudia Schapendonk, Irina Chestakova, Anne van der Linden, Mark Pronk, Pascal Lexmond, Corien Swaan, Manon Haverkate, Madelief Mollers, Mart Stein, Sandra Kengne Kamga Mobou, Jeroen van Kampen, Jolanda Voermans, Aura Timen, Corine GeurtsvanKessel, Annemiek van der Eijk, Richard Molenkamp, Marion Koopmans, on behalf of the Dutch national COVID-19 response team. |
| EPI_ISL_415493 | hCoV-19/Netherlands/NA_4/2020 | Europe / Netherlands | 2020 | Dutch COVID-19 response team | Erasmus Medical Center | David Nieuwenhuijse, Bas Oude Munnink, Reina Sikkema, Claudia Schapendonk, Irina Chestakova, Anne van der Linden, Mark Pronk, Pascal Lexmond, Corien Swaan, Manon Haverkate, Madelief Mollers, Mart Stein, Sandra Kengne Kamga Mobou, Jeroen van Kampen, Jolanda Voermans, Aura Timen, Corine GeurtsvanKessel, Annemiek van der Eijk, Richard Molenkamp, Marion Koopmans, on behalf of the Dutch national COVID-19 response team. |
| EPI_ISL_415494 | hCoV-19/Netherlands/NA_5/2020 | Europe / Netherlands | 2020 | Dutch COVID-19 response team | Erasmus Medical Center | David Nieuwenhuijse, Bas Oude Munnink, Reina Sikkema, Claudia Schapendonk, Irina Chestakova, Anne van der Linden, Mark Pronk, Pascal Lexmond, Corien Swaan, Manon Haverkate, Madelief Mollers, Mart Stein, Sandra Kengne Kamga Mobou, Jeroen van Kampen, Jolanda Voermans, Aura Timen, Corine GeurtsvanKessel, Annemiek van der Eijk, Richard Molenkamp, Marion Koopmans, on behalf of the Dutch national COVID-19 response team. |
| EPI_ISL_415495 | hCoV-19/Netherlands/NA_6/2020 | Europe / Netherlands | 2020-03-10 | Dutch COVID-19 response team | Erasmus Medical Center | David Nieuwenhuijse, Bas Oude Munnink, Reina Sikkema, Claudia Schapendonk, Irina Chestakova, Anne van der Linden, Mark Pronk, Pascal Lexmond, Corien Swaan, Manon Haverkate, Madelief Mollers, Mart Stein, Sandra Kengne Kamga Mobou, Jeroen van Kampen, Jolanda Voermans, Aura Timen, Corine GeurtsvanKessel, Annemiek van der Eijk, Richard Molenkamp, Marion Koopmans, on behalf of the Dutch national COVID-19 response team. |
| EPI_ISL_415496 | hCoV-19/Netherlands/NA_7/2020 | Europe / Netherlands | 2020-03-09 | Dutch COVID-19 response team | Erasmus Medical Center | David Nieuwenhuijse, Bas Oude Munnink, Reina Sikkema, Claudia Schapendonk, Irina Chestakova, Anne van der Linden, Mark Pronk, Pascal Lexmond, Corien Swaan, Manon Haverkate, Madelief Mollers, Mart Stein, Sandra Kengne Kamga Mobou, Jeroen van Kampen, Jolanda Voermans, Aura Timen, Corine GeurtsvanKessel, Annemiek van der Eijk, Richard Molenkamp, Marion Koopmans, on behalf of the Dutch national COVID-19 response team. |
| EPI_ISL_415497 | hCoV-19/Netherlands/NA_8/2020 | Europe / Netherlands | 2020-03-09 | Dutch COVID-19 response team | Erasmus Medical Center | David Nieuwenhuijse, Bas Oude Munnink, Reina Sikkema, Claudia Schapendonk, Irina Chestakova, Anne van der Linden, Mark Pronk, Pascal Lexmond, Corien Swaan, Manon Haverkate, Madelief Mollers, Mart Stein, Sandra Kengne Kamga Mobou, Jeroen van Kampen, Jolanda Voermans, Aura Timen, Corine GeurtsvanKessel, Annemiek van der Eijk, Richard Molenkamp, Marion Koopmans, on behalf of the Dutch national COVID-19 response team. |
| EPI_ISL_415498 | hCoV-19/Netherlands/NA_9/2020 | Europe / Netherlands | 2020-03-09 | Dutch COVID-19 response team | Erasmus Medical Center | David Nieuwenhuijse, Bas Oude Munnink, Reina Sikkema, Claudia Schapendonk, Irina Chestakova, Anne van der Linden, Mark Pronk, Pascal Lexmond, Corien Swaan, Manon Haverkate, Madelief Mollers, Mart Stein, Sandra Kengne Kamga Mobou, Jeroen van Kampen, Jolanda Voermans, Aura Timen, Corine GeurtsvanKessel, Annemiek van der Eijk, Richard Molenkamp, Marion Koopmans, on behalf of the Dutch national COVID-19 response team. |
| EPI_ISL_415499 | hCoV-19/Netherlands/NoordBrabant_41/2020 | Europe / Netherlands / Noord Brabant | 2020 | Dutch COVID-19 response team | Erasmus Medical Center | David Nieuwenhuijse, Bas Oude Munnink, Reina Sikkema, Claudia Schapendonk, Irina Chestakova, Anne van der Linden, Mark Pronk, Pascal Lexmond, Corien Swaan, Manon Haverkate, Madelief Mollers, Mart Stein, Sandra Kengne Kamga Mobou, Jeroen van Kampen, Jolanda Voermans, Aura Timen, Corine GeurtsvanKessel, Annemiek van der Eijk, Richard Molenkamp, Marion Koopmans, on behalf of the Dutch national COVID-19 response team. |
| EPI_ISL_415700 | hCoV-19/Switzerland/GE1402/2020 | Europe / Switzerland | 2020-02-28 | Hôpitaux universitaires de Genève Laboratoire de Virologie | Hôpitaux universitaires de Genève Laboratoire de Virologie | Laubscher F. |
| EPI_ISL_415701 | hCoV-19/Switzerland/BS0914/2020 | Europe / Switzerland | 2020-03-02 | Hôpitaux universitaires de Genève Laboratoire de Virologie | Hôpitaux universitaires de Genève Laboratoire de Virologie | Laubscher F. |
| EPI_ISL_415702 | hCoV-19/Switzerland/SZ1417/2020 | Europe / Switzerland | 2020-03-02 | Hôpitaux universitaires de Genève Laboratoire de Virologie | Hôpitaux universitaires de Genève Laboratoire de Virologie | Laubscher F. |
| EPI_ISL_415703 | hCoV-19/Switzerland/TI2045/2020 | Europe / Switzerland | 2020-03-01 | Hôpitaux universitaires de Genève Laboratoire de Virologie | Hôpitaux universitaires de Genève Laboratoire de Virologie | Laubscher F. |
| EPI_ISL_415704 | hCoV-19/Switzerland/BE2536/2020 | Europe / Switzerland | 2020-03-04 | Hôpitaux universitaires de Genève Laboratoire de Virologie | Hôpitaux universitaires de Genève Laboratoire de Virologie | Laubscher F. |
| EPI_ISL_415705 | hCoV-19/Switzerland/GE4135/2020 | Europe / Switzerland | 2020-03-06 | Hôpitaux universitaires de Genève Laboratoire de Virologie | Hôpitaux universitaires de Genève Laboratoire de Virologie | Laubscher F. |
| EPI_ISL_415706 | hCoV-19/Switzerland/GE06207/2020 | Europe / Switzerland | 2020-03-06 | Hôpitaux universitaires de Genève Laboratoire de Virologie | Hôpitaux universitaires de Genève Laboratoire de Virologie | Laubscher F. |
| EPI_ISL_415707 | hCoV-19/Switzerland/GE6679/2020 | Europe / Switzerland | 2020-03-08 | Hôpitaux universitaires de Genève Laboratoire de Virologie | Hôpitaux universitaires de Genève Laboratoire de Virologie | Laubscher F. |
| EPI_ISL_415708 | hCoV-19/Switzerland/GE4984/2020 | Europe / Switzerland | 2020-03-07 | Hôpitaux universitaires de Genève Laboratoire de Virologie | Hôpitaux universitaires de Genève Laboratoire de Virologie | Laubscher F. |
| EPI_ISL_415709 | hCoV-19/Hangzhou/ZJU-01/2020 | Asia / China / Hangzhou | 2020-01-25 | State Key Laboratory for Diagnosis and Treatment of Infectious Diseases, National Clinical Research Center for Infectious Diseases, First Affiliated Hospital, Zhejiang University School of Medicine, Hangzhou, China. 310003 | State Key Laboratory for Diagnosis and Treatment of Infectious Diseases, National Clinical Research Center for Infectious Diseases, First Affiliated Hospital, Zhejiang University School of Medicine, Hangzhou, China. 310003 | Hangping Yao, Nanping Wu, Chao Jiang, Xiangyun Lu, Linfang Cheng, Fumin Liu, Zhigang Wu, Haibo Wu, Changzhong Jin, Min Zheng, Lanjuan Li |
| EPI_ISL_415710 | hCoV-19/Russia/StPetersburg-3524/2020 | Europe / Russia / Saint Petersburg | 2020-03-15 | WHO National Influenza Centre Russian Federation | WHO National Influenza Centre Russian Federation | Andrey Komissarov, Artem Fadeev, Anna Ivanova, Daria Danilenko |
| EPI_ISL_415711 | hCoV-19/Hangzhou/ZJU-05/2020 | Asia / China / Hangzhou | 2020-01-22 | State Key Laboratory for Diagnosis and Treatment of Infectious Diseases, National Clinical Research Center for Infectious Diseases, First Affiliated Hospital, Zhejiang University School of Medicine, Hangzhou, China. 310003 | State Key Laboratory for Diagnosis and Treatment of Infectious Diseases, National Clinical Research Center for Infectious Diseases, First Affiliated Hospital, Zhejiang University School of Medicine, Hangzhou, China. 310003 | Hangping Yao, Nanping Wu, Chao Jiang, Xiangyun Lu, Linfang Cheng, Fumin Liu, Zhigang Wu, Haibo Wu, Changzhong Jin, Min Zheng, Lanjuan Li |
| EPI_ISL_415741 | hCoV-19/Taiwan/CGMH-CGU-03/2020 | Asia / Taiwan / Taoyuan | 2020-02-26 | Laboratory Medicine | Department of Laboratory Medicine, Lin-Kou Chang Gung Memorial Hospital, Taoyuan, Taiwan | Kuo-Chien Tsao, Yu-Nong Gong, Shu-Li Yang, Yi-Chun Liu, Chung-Guei Huang, Po-Wei Huang, Mei-Jen Hsiao, Cheng-Ta Yang, Cheng-Hsun Chiu, Chi-Hsien Huang, Kuang-Tao Le, Shu-Min Lin, Peng-Nien Huang, Kuo-Ming Lee, Guang-Wu Chen, Shin-Ru Shih |
| EPI_ISL_415742 | hCoV-19/Taiwan/CGMH-CGU-04/2020 | Asia / Taiwan / Taoyuan | 2020-02-27 | Laboratory Medicine | Department of Laboratory Medicine, Lin-Kou Chang Gung Memorial Hospital, Taoyuan, Taiwan | Kuo-Chien Tsao, Yu-Nong Gong, Shu-Li Yang, Yi-Chun Liu, Chung-Guei Huang, Po-Wei Huang, Mei-Jen Hsiao, Cheng-Ta Yang, Cheng-Hsun Chiu, Chi-Hsien Huang, Kuang-Tao Le, Shu-Min Lin, Peng-Nien Huang, Kuo-Ming Lee, Guang-Wu Chen, Shin-Ru Shih |
| EPI_ISL_415743 | hCoV-19/Taiwan/CGMH-CGU-05/2020 | Asia / Taiwan / Taoyuan | 2020-02-27 | Laboratory Medicine | Department of Laboratory Medicine, Lin-Kou Chang Gung Memorial Hospital, Taoyuan, Taiwan | Kuo-Chien Tsao, Yu-Nong Gong, Shu-Li Yang, Yi-Chun Liu, Chung-Guei Huang, Po-Wei Huang, Mei-Jen Hsiao, Cheng-Ta Yang, Cheng-Hsun Chiu, Chi-Hsien Huang, Kuang-Tao Le, Shu-Min Lin, Peng-Nien Huang, Kuo-Ming Lee, Guang-Wu Chen, Shin-Ru Shih |
| EPI_ISL_415787 | hCoV-19/Peru/010/2020 | South America / Peru / Lima | 2020-03-10 | Laboratorio de Referencia Nacional de Virus Respiratorio. Instituto Nacional de Salud. Peru | Laboratorio de Referencia Nacional de Biotecnologia y Biologia Molecular Instituto Nacional de Salud.Peru | Carlos Padilla Rojas, Priscila Lope Pari, Karolyn Vega Chozo, Johanna Balbuena Torres, Omar Caceres Rey, Henri Bailon Calderon, Maribel Huaringa Nuñez, Nancy Rojas Serrano |
| EPI_ISL_416319 | hCoV-19/Shanghai/SH0005/2020 | Asia / China / Shanghai | 2020-01-28 | Shanghai Public Health Clinical Center, Shanghai Medical College, Fudan University | National Research Center for Translational Medicine (Shanghai), Ruijin Hospital affiliated to Shanghai Jiao Tong University School of Medicine & Shanghai Public Health Clinical Center | Shengyue Wang, Xiaonan Zhang, Gang Lu, Yun Tan, Yun Ling, Hongzhou Lu, Saijuan Chen |
| EPI_ISL_416320 | hCoV-19/Shanghai/SH0007/2020 | Asia / China / Shanghai | 2020-01-28 | Shanghai Public Health Clinical Center, Shanghai Medical College, Fudan University | National Research Center for Translational Medicine (Shanghai), Ruijin Hospital affiliated to Shanghai Jiao Tong University School of Medicine & Shanghai Public Health Clinical Center | Shengyue Wang, Xiaonan Zhang, Gang Lu, Yun Tan, Yun Ling, Hongzhou Lu, Saijuan Chen |
| EPI_ISL_416321 | hCoV-19/Shanghai/SH0008/2020 | Asia / China / Shanghai | 2020-01-28 | Shanghai Public Health Clinical Center, Shanghai Medical College, Fudan University | National Research Center for Translational Medicine (Shanghai), Ruijin Hospital affiliated to Shanghai Jiao Tong University School of Medicine & Shanghai Public Health Clinical Center | Shengyue Wang, Xiaonan Zhang, Gang Lu, Yun Tan, Yun Ling, Hongzhou Lu, Saijuan Chen |
| EPI_ISL_416322 | hCoV-19/Shanghai/SH0009/2020 | Asia / China / Shanghai | 2020-01-29 | Shanghai Public Health Clinical Center, Shanghai Medical College, Fudan University | National Research Center for Translational Medicine (Shanghai), Ruijin Hospital affiliated to Shanghai Jiao Tong University School of Medicine & Shanghai Public Health Clinical Center | Shengyue Wang, Xiaonan Zhang, Gang Lu, Yun Tan, Yun Ling, Hongzhou Lu, Saijuan Chen |
| EPI_ISL_416323 | hCoV-19/Shanghai/SH0010/2020 | Asia / China / Shanghai | 2020-01-29 | Shanghai Public Health Clinical Center, Shanghai Medical College, Fudan University | National Research Center for Translational Medicine (Shanghai), Ruijin Hospital affiliated to Shanghai Jiao Tong University School of Medicine & Shanghai Public Health Clinical Center | Shengyue Wang, Xiaonan Zhang, Gang Lu, Yun Tan, Yun Ling, Hongzhou Lu, Saijuan Chen |
| EPI_ISL_416324 | hCoV-19/Shanghai/SH0011/2020 | Asia / China / Shanghai | 2020-01-29 | Shanghai Public Health Clinical Center, Shanghai Medical College, Fudan University | National Research Center for Translational Medicine (Shanghai), Ruijin Hospital affiliated to Shanghai Jiao Tong University School of Medicine & Shanghai Public Health Clinical Center | Shengyue Wang, Xiaonan Zhang, Gang Lu, Yun Tan, Yun Ling, Hongzhou Lu, Saijuan Chen |
| EPI_ISL_416325 | hCoV-19/Shanghai/SH0012/2020 | Asia / China / Shanghai | 2020-02-02 | Shanghai Public Health Clinical Center, Shanghai Medical College, Fudan University | National Research Center for Translational Medicine (Shanghai), Ruijin Hospital affiliated to Shanghai Jiao Tong University School of Medicine & Shanghai Public Health Clinical Center | Shengyue Wang, Xiaonan Zhang, Gang Lu, Yun Tan, Yun Ling, Hongzhou Lu, Saijuan Chen |
| EPI_ISL_416326 | hCoV-19/Shanghai/SH0013/2020 | Asia / China / Shanghai | 2020-01-30 | Shanghai Public Health Clinical Center, Shanghai Medical College, Fudan University | National Research Center for Translational Medicine (Shanghai), Ruijin Hospital affiliated to Shanghai Jiao Tong University School of Medicine & Shanghai Public Health Clinical Center | Shengyue Wang, Xiaonan Zhang, Gang Lu, Yun Tan, Yun Ling, Hongzhou Lu, Saijuan Chen |
| EPI_ISL_416329 | hCoV-19/Shanghai/SH0020/2020 | Asia / China / Shanghai | 2020-02-01 | Shanghai Public Health Clinical Center, Shanghai Medical College, Fudan University | National Research Center for Translational Medicine (Shanghai), Ruijin Hospital affiliated to Shanghai Jiao Tong University School of Medicine & Shanghai Public Health Clinical Center | Shengyue Wang, Xiaonan Zhang, Gang Lu, Yun Tan, Yun Ling, Hongzhou Lu, Saijuan Chen |
| EPI_ISL_416330 | hCoV-19/Shanghai/SH0021/2020 | Asia / China / Shanghai | 2020-02-02 | Shanghai Public Health Clinical Center, Shanghai Medical College, Fudan University | National Research Center for Translational Medicine (Shanghai), Ruijin Hospital affiliated to Shanghai Jiao Tong University School of Medicine & Shanghai Public Health Clinical Center | Shengyue Wang, Xiaonan Zhang, Gang Lu, Yun Tan, Yun Ling, Hongzhou Lu, Saijuan Chen |
| EPI_ISL_416331 | hCoV-19/Shanghai/SH0022/2020 | Asia / China / Shanghai | 2020-01-30 | Shanghai Public Health Clinical Center, Shanghai Medical College, Fudan University | National Research Center for Translational Medicine (Shanghai), Ruijin Hospital affiliated to Shanghai Jiao Tong University School of Medicine & Shanghai Public Health Clinical Center | Shengyue Wang, Xiaonan Zhang, Gang Lu, Yun Tan, Yun Ling, Hongzhou Lu, Saijuan Chen |
| EPI_ISL_416332 | hCoV-19/Shanghai/SH0023/2020 | Asia / China / Shanghai | 2020-01-30 | Shanghai Public Health Clinical Center, Shanghai Medical College, Fudan University | National Research Center for Translational Medicine (Shanghai), Ruijin Hospital affiliated to Shanghai Jiao Tong University School of Medicine & Shanghai Public Health Clinical Center | Shengyue Wang, Xiaonan Zhang, Gang Lu, Yun Tan, Yun Ling, Hongzhou Lu, Saijuan Chen |
| EPI_ISL_416333 | hCoV-19/Shanghai/SH0024/2020 | Asia / China / Shanghai | 2020-01-30 | Shanghai Public Health Clinical Center, Shanghai Medical College, Fudan University | National Research Center for Translational Medicine (Shanghai), Ruijin Hospital affiliated to Shanghai Jiao Tong University School of Medicine & Shanghai Public Health Clinical Center | Shengyue Wang, Xiaonan Zhang, Gang Lu, Yun Tan, Yun Ling, Hongzhou Lu, Saijuan Chen |
| EPI_ISL_416334 | hCoV-19/Shanghai/SH0025/2020 | Asia / China / Shanghai | 2020-02-06 | Shanghai Public Health Clinical Center, Shanghai Medical College, Fudan University | National Research Center for Translational Medicine (Shanghai), Ruijin Hospital affiliated to Shanghai Jiao Tong University School of Medicine & Shanghai Public Health Clinical Center | Shengyue Wang, Xiaonan Zhang, Gang Lu, Yun Tan, Yun Ling, Hongzhou Lu, Saijuan Chen |
| EPI_ISL_416335 | hCoV-19/Shanghai/SH0026/2020 | Asia / China / Shanghai | 2020-02-02 | Shanghai Public Health Clinical Center, Shanghai Medical College, Fudan University | National Research Center for Translational Medicine (Shanghai), Ruijin Hospital affiliated to Shanghai Jiao Tong University School of Medicine & Shanghai Public Health Clinical Center | Shengyue Wang, Xiaonan Zhang, Gang Lu, Yun Tan, Yun Ling, Hongzhou Lu, Saijuan Chen |
| EPI_ISL_416336 | hCoV-19/Shanghai/SH0027/2020 | Asia / China / Shanghai | 2020-02-01 | Shanghai Public Health Clinical Center, Shanghai Medical College, Fudan University | National Research Center for Translational Medicine (Shanghai), Ruijin Hospital affiliated to Shanghai Jiao Tong University School of Medicine & Shanghai Public Health Clinical Center | Shengyue Wang, Xiaonan Zhang, Gang Lu, Yun Tan, Yun Ling, Hongzhou Lu, Saijuan Chen |
| EPI_ISL_416337 | hCoV-19/Shanghai/SH0028/2020 | Asia / China / Shanghai | 2020-02-04 | Shanghai Public Health Clinical Center, Shanghai Medical College, Fudan University | National Research Center for Translational Medicine (Shanghai), Ruijin Hospital affiliated to Shanghai Jiao Tong University School of Medicine & Shanghai Public Health Clinical Center | Shengyue Wang, Xiaonan Zhang, Gang Lu, Yun Tan, Yun Ling, Hongzhou Lu, Saijuan Chen |
| EPI_ISL_416338 | hCoV-19/Shanghai/SH0029/2020 | Asia / China / Shanghai | 2020-02-01 | Shanghai Public Health Clinical Center, Shanghai Medical College, Fudan University | National Research Center for Translational Medicine (Shanghai), Ruijin Hospital affiliated to Shanghai Jiao Tong University School of Medicine & Shanghai Public Health Clinical Center | Shengyue Wang, Xiaonan Zhang, Gang Lu, Yun Tan, Yun Ling, Hongzhou Lu, Saijuan Chen |
| EPI_ISL_416339 | hCoV-19/Shanghai/SH0030/2020 | Asia / China / Shanghai | 2020-02-03 | Shanghai Public Health Clinical Center, Shanghai Medical College, Fudan University | National Research Center for Translational Medicine (Shanghai), Ruijin Hospital affiliated to Shanghai Jiao Tong University School of Medicine & Shanghai Public Health Clinical Center | Shengyue Wang, Xiaonan Zhang, Gang Lu, Yun Tan, Yun Ling, Hongzhou Lu, Saijuan Chen |
| EPI_ISL_416340 | hCoV-19/Shanghai/SH0031/2020 | Asia / China / Shanghai | 2020-02-02 | Shanghai Public Health Clinical Center, Shanghai Medical College, Fudan University | National Research Center for Translational Medicine (Shanghai), Ruijin Hospital affiliated to Shanghai Jiao Tong University School of Medicine & Shanghai Public Health Clinical Center | Shengyue Wang, Xiaonan Zhang, Gang Lu, Yun Tan, Yun Ling, Hongzhou Lu, Saijuan Chen |
| EPI_ISL_416341 | hCoV-19/Shanghai/SH0032/2020 | Asia / China / Shanghai | 2020-02-01 | Shanghai Public Health Clinical Center, Shanghai Medical College, Fudan University | National Research Center for Translational Medicine (Shanghai), Ruijin Hospital affiliated to Shanghai Jiao Tong University School of Medicine & Shanghai Public Health Clinical Center | Shengyue Wang, Xiaonan Zhang, Gang Lu, Yun Tan, Yun Ling, Hongzhou Lu, Saijuan Chen |
| EPI_ISL_416342 | hCoV-19/Shanghai/SH0033/2020 | Asia / China / Shanghai | 2020-02-04 | Shanghai Public Health Clinical Center, Shanghai Medical College, Fudan University | National Research Center for Translational Medicine (Shanghai), Ruijin Hospital affiliated to Shanghai Jiao Tong University School of Medicine & Shanghai Public Health Clinical Center | Shengyue Wang, Xiaonan Zhang, Gang Lu, Yun Tan, Yun Ling, Hongzhou Lu, Saijuan Chen |
| EPI_ISL_416348 | hCoV-19/Shanghai/SH0039/2020 | Asia / China / Shanghai | 2020-02-06 | Shanghai Public Health Clinical Center, Shanghai Medical College, Fudan University | National Research Center for Translational Medicine (Shanghai), Ruijin Hospital affiliated to Shanghai Jiao Tong University School of Medicine & Shanghai Public Health Clinical Center | Shengyue Wang, Xiaonan Zhang, Gang Lu, Yun Tan, Yun Ling, Hongzhou Lu, Saijuan Chen |
| EPI_ISL_416349 | hCoV-19/Shanghai/SH0040/2020 | Asia / China / Shanghai | 2020-01-30 | Shanghai Public Health Clinical Center, Shanghai Medical College, Fudan University | National Research Center for Translational Medicine (Shanghai), Ruijin Hospital affiliated to Shanghai Jiao Tong University School of Medicine & Shanghai Public Health Clinical Center | Shengyue Wang, Xiaonan Zhang, Gang Lu, Yun Tan, Yun Ling, Hongzhou Lu, Saijuan Chen |
| EPI_ISL_416350 | hCoV-19/Shanghai/SH0041/2020 | Asia / China / Shanghai | 2020-01-29 | Shanghai Public Health Clinical Center, Shanghai Medical College, Fudan University | National Research Center for Translational Medicine (Shanghai), Ruijin Hospital affiliated to Shanghai Jiao Tong University School of Medicine & Shanghai Public Health Clinical Center | Shengyue Wang, Xiaonan Zhang, Gang Lu, Yun Tan, Yun Ling, Hongzhou Lu, Saijuan Chen |
| EPI_ISL_416352 | hCoV-19/Shanghai/SH0043/2020 | Asia / China / Shanghai | 2020-01-30 | Shanghai Public Health Clinical Center, Shanghai Medical College, Fudan University | National Research Center for Translational Medicine (Shanghai), Ruijin Hospital affiliated to Shanghai Jiao Tong University School of Medicine & Shanghai Public Health Clinical Center | Shengyue Wang, Xiaonan Zhang, Gang Lu, Yun Tan, Yun Ling, Hongzhou Lu, Saijuan Chen |
| EPI_ISL_416353 | hCoV-19/Shanghai/SH0044/2020 | Asia / China / Shanghai | 2020-02-04 | Shanghai Public Health Clinical Center, Shanghai Medical College, Fudan University | National Research Center for Translational Medicine (Shanghai), Ruijin Hospital affiliated to Shanghai Jiao Tong University School of Medicine & Shanghai Public Health Clinical Center | Shengyue Wang, Xiaonan Zhang, Gang Lu, Yun Tan, Yun Ling, Hongzhou Lu, Saijuan Chen |
| EPI_ISL_416354 | hCoV-19/Shanghai/SH0045/2020 | Asia / China / Shanghai | 2020-02-04 | Shanghai Public Health Clinical Center, Shanghai Medical College, Fudan University | National Research Center for Translational Medicine (Shanghai), Ruijin Hospital affiliated to Shanghai Jiao Tong University School of Medicine & Shanghai Public Health Clinical Center | Shengyue Wang, Xiaonan Zhang, Gang Lu, Yun Tan, Yun Ling, Hongzhou Lu, Saijuan Chen |
| EPI_ISL_416355 | hCoV-19/Shanghai/SH0047/2020 | Asia / China / Shanghai | 2020-02-04 | Shanghai Public Health Clinical Center, Shanghai Medical College, Fudan University | National Research Center for Translational Medicine (Shanghai), Ruijin Hospital affiliated to Shanghai Jiao Tong University School of Medicine & Shanghai Public Health Clinical Center | Shengyue Wang, Xiaonan Zhang, Gang Lu, Yun Tan, Yun Ling, Hongzhou Lu, Saijuan Chen |
| EPI_ISL_416356 | hCoV-19/Shanghai/SH0048/2020 | Asia / China / Shanghai | 2020-02-01 | Shanghai Public Health Clinical Center, Shanghai Medical College, Fudan University | National Research Center for Translational Medicine (Shanghai), Ruijin Hospital affiliated to Shanghai Jiao Tong University School of Medicine & Shanghai Public Health Clinical Center | Shengyue Wang, Xiaonan Zhang, Gang Lu, Yun Tan, Yun Ling, Hongzhou Lu, Saijuan Chen |
| EPI_ISL_416358 | hCoV-19/Shanghai/SH0050/2020 | Asia / China / Shanghai | 2020-02-05 | Shanghai Public Health Clinical Center, Shanghai Medical College, Fudan University | National Research Center for Translational Medicine (Shanghai), Ruijin Hospital affiliated to Shanghai Jiao Tong University School of Medicine & Shanghai Public Health Clinical Center | Shengyue Wang, Xiaonan Zhang, Gang Lu, Yun Tan, Yun Ling, Hongzhou Lu, Saijuan Chen |
| EPI_ISL_416359 | hCoV-19/Shanghai/SH0051/2020 | Asia / China / Shanghai | 2020-02-06 | Shanghai Public Health Clinical Center, Shanghai Medical College, Fudan University | National Research Center for Translational Medicine (Shanghai), Ruijin Hospital affiliated to Shanghai Jiao Tong University School of Medicine & Shanghai Public Health Clinical Center | Shengyue Wang, Xiaonan Zhang, Gang Lu, Yun Tan, Yun Ling, Hongzhou Lu, Saijuan Chen |
| EPI_ISL_416361 | hCoV-19/Shanghai/SH0053/2020 | Asia / China / Shanghai | 2020-02-01 | Shanghai Public Health Clinical Center, Shanghai Medical College, Fudan University | National Research Center for Translational Medicine (Shanghai), Ruijin Hospital affiliated to Shanghai Jiao Tong University School of Medicine & Shanghai Public Health Clinical Center | Shengyue Wang, Xiaonan Zhang, Gang Lu, Yun Tan, Yun Ling, Hongzhou Lu, Saijuan Chen |
| EPI_ISL_416362 | hCoV-19/Shanghai/SH0054/2020 | Asia / China / Shanghai | 2020-02-02 | Shanghai Public Health Clinical Center, Shanghai Medical College, Fudan University | National Research Center for Translational Medicine (Shanghai), Ruijin Hospital affiliated to Shanghai Jiao Tong University School of Medicine & Shanghai Public Health Clinical Center | Shengyue Wang, Xiaonan Zhang, Gang Lu, Yun Tan, Yun Ling, Hongzhou Lu, Saijuan Chen |
| EPI_ISL_416363 | hCoV-19/Shanghai/SH0055/2020 | Asia / China / Shanghai | 2020-02-08 | Shanghai Public Health Clinical Center, Shanghai Medical College, Fudan University | National Research Center for Translational Medicine (Shanghai), Ruijin Hospital affiliated to Shanghai Jiao Tong University School of Medicine & Shanghai Public Health Clinical Center | Shengyue Wang, Xiaonan Zhang, Gang Lu, Yun Tan, Yun Ling, Hongzhou Lu, Saijuan Chen |
| EPI_ISL_416364 | hCoV-19/Shanghai/SH0056/2020 | Asia / China / Shanghai | 2020-02-08 | Shanghai Public Health Clinical Center, Shanghai Medical College, Fudan University | National Research Center for Translational Medicine (Shanghai), Ruijin Hospital affiliated to Shanghai Jiao Tong University School of Medicine & Shanghai Public Health Clinical Center | Shengyue Wang, Xiaonan Zhang, Gang Lu, Yun Tan, Yun Ling, Hongzhou Lu, Saijuan Chen |
| EPI_ISL_416365 | hCoV-19/Shanghai/SH0058/2020 | Asia / China / Shanghai | 2020-01-30 | Shanghai Public Health Clinical Center, Shanghai Medical College, Fudan University | National Research Center for Translational Medicine (Shanghai), Ruijin Hospital affiliated to Shanghai Jiao Tong University School of Medicine & Shanghai Public Health Clinical Center | Shengyue Wang, Xiaonan Zhang, Gang Lu, Yun Tan, Yun Ling, Hongzhou Lu, Saijuan Chen |
| EPI_ISL_416366 | hCoV-19/Shanghai/SH0059/2020 | Asia / China / Shanghai | 2020-01-30 | Shanghai Public Health Clinical Center, Shanghai Medical College, Fudan University | National Research Center for Translational Medicine (Shanghai), Ruijin Hospital affiliated to Shanghai Jiao Tong University School of Medicine & Shanghai Public Health Clinical Center | Shengyue Wang, Xiaonan Zhang, Gang Lu, Yun Tan, Yun Ling, Hongzhou Lu, Saijuan Chen |
| EPI_ISL_416367 | hCoV-19/Shanghai/SH0060/2020 | Asia / China / Shanghai | 2020-02-01 | Shanghai Public Health Clinical Center, Shanghai Medical College, Fudan University | National Research Center for Translational Medicine (Shanghai), Ruijin Hospital affiliated to Shanghai Jiao Tong University School of Medicine & Shanghai Public Health Clinical Center | Shengyue Wang, Xiaonan Zhang, Gang Lu, Yun Tan, Yun Ling, Hongzhou Lu, Saijuan Chen |
| EPI_ISL_416368 | hCoV-19/Shanghai/SH0065/2020 | Asia / China / Shanghai | 2020-02-05 | Shanghai Public Health Clinical Center, Shanghai Medical College, Fudan University | National Research Center for Translational Medicine (Shanghai), Ruijin Hospital affiliated to Shanghai Jiao Tong University School of Medicine & Shanghai Public Health Clinical Center | Shengyue Wang, Xiaonan Zhang, Gang Lu, Yun Tan, Yun Ling, Hongzhou Lu, Saijuan Chen |

| | | | | | | |
|---|---|---|---|---|---|---|
| EPI_ISL_416369 | hCoV-19/Shanghai/SH0066/2020 | Asia / China / Shanghai | 2020-02-05 | Shanghai Public Health Clinical Center, Shanghai Medical College, Fudan University | National Research Center for Translational Medicine (Shanghai), Ruijin Hospital affiliated to Shanghai Jiao Tong University School of Medicine & Shanghai Public Health Clinical Center | Shengyue Wang, Xiaonan Zhang, Gang Lu, Yun Tan, Yun Ling, Hongzhou Lu, Saijuan Chen |
| EPI_ISL_416370 | hCoV-19/Shanghai/SH0067/2020 | Asia / China / Shanghai | 2020-02-01 | Shanghai Public Health Clinical Center, Shanghai Medical College, Fudan University | National Research Center for Translational Medicine (Shanghai), Ruijin Hospital affiliated to Shanghai Jiao Tong University School of Medicine & Shanghai Public Health Clinical Center | Shengyue Wang, Xiaonan Zhang, Gang Lu, Yun Tan, Yun Ling, Hongzhou Lu, Saijuan Chen |
| EPI_ISL_416371 | hCoV-19/Shanghai/SH0068/2020 | Asia / China / Shanghai | 2020-02-05 | Shanghai Public Health Clinical Center, Shanghai Medical College, Fudan University | National Research Center for Translational Medicine (Shanghai), Ruijin Hospital affiliated to Shanghai Jiao Tong University School of Medicine & Shanghai Public Health Clinical Center | Shengyue Wang, Xiaonan Zhang, Gang Lu, Yun Tan, Yun Ling, Hongzhou Lu, Saijuan Chen |
| EPI_ISL_416372 | hCoV-19/Shanghai/SH0069/2020 | Asia / China / Shanghai | 2020-02-01 | Shanghai Public Health Clinical Center, Shanghai Medical College, Fudan University | National Research Center for Translational Medicine (Shanghai), Ruijin Hospital affiliated to Shanghai Jiao Tong University School of Medicine & Shanghai Public Health Clinical Center | Shengyue Wang, Xiaonan Zhang, Gang Lu, Yun Tan, Yun Ling, Hongzhou Lu, Saijuan Chen |
| EPI_ISL_416373 | hCoV-19/Shanghai/SH0070/2020 | Asia / China / Shanghai | 2020-02-07 | Shanghai Public Health Clinical Center, Shanghai Medical College, Fudan University | National Research Center for Translational Medicine (Shanghai), Ruijin Hospital affiliated to Shanghai Jiao Tong University School of Medicine & Shanghai Public Health Clinical Center | Shengyue Wang, Xiaonan Zhang, Gang Lu, Yun Tan, Yun Ling, Hongzhou Lu, Saijuan Chen |
| EPI_ISL_416374 | hCoV-19/Shanghai/SH0071/2020 | Asia / China / Shanghai | 2020-02-07 | Shanghai Public Health Clinical Center, Shanghai Medical College, Fudan University | National Research Center for Translational Medicine (Shanghai), Ruijin Hospital affiliated to Shanghai Jiao Tong University School of Medicine & Shanghai Public Health Clinical Center | Shengyue Wang, Xiaonan Zhang, Gang Lu, Yun Tan, Yun Ling, Hongzhou Lu, Saijuan Chen |
| EPI_ISL_416376 | hCoV-19/Shanghai/SH0073/2020 | Asia / China / Shanghai | 2020-02-05 | Shanghai Public Health Clinical Center, Shanghai Medical College, Fudan University | National Research Center for Translational Medicine (Shanghai), Ruijin Hospital affiliated to Shanghai Jiao Tong University School of Medicine & Shanghai Public Health Clinical Center | Shengyue Wang, Xiaonan Zhang, Gang Lu, Yun Tan, Yun Ling, Hongzhou Lu, Saijuan Chen |
| EPI_ISL_416377 | hCoV-19/Shanghai/SH0074/2020 | Asia / China / Shanghai | 2020-02-07 | Shanghai Public Health Clinical Center, Shanghai Medical College, Fudan University | National Research Center for Translational Medicine (Shanghai), Ruijin Hospital affiliated to Shanghai Jiao Tong University School of Medicine & Shanghai Public Health Clinical Center | Shengyue Wang, Xiaonan Zhang, Gang Lu, Yun Tan, Yun Ling, Hongzhou Lu, Saijuan Chen |
| EPI_ISL_416378 | hCoV-19/Shanghai/SH0075/2020 | Asia / China / Shanghai | 2020-01-30 | Shanghai Public Health Clinical Center, Shanghai Medical College, Fudan University | National Research Center for Translational Medicine (Shanghai), Ruijin Hospital affiliated to Shanghai Jiao Tong University School of Medicine & Shanghai Public Health Clinical Center | Shengyue Wang, Xiaonan Zhang, Gang Lu, Yun Tan, Yun Ling, Hongzhou Lu, Saijuan Chen |
| EPI_ISL_416379 | hCoV-19/Shanghai/SH0076/2020 | Asia / China / Shanghai | 2020-02-01 | Shanghai Public Health Clinical Center, Shanghai Medical College, Fudan University | National Research Center for Translational Medicine (Shanghai), Ruijin Hospital affiliated to Shanghai Jiao Tong University School of Medicine & Shanghai Public Health Clinical Center | Shengyue Wang, Xiaonan Zhang, Gang Lu, Yun Tan, Yun Ling, Hongzhou Lu, Saijuan Chen |
| EPI_ISL_416380 | hCoV-19/Shanghai/SH0077/2020 | Asia / China / Shanghai | 2020-02-09 | Shanghai Public Health Clinical Center, Shanghai Medical College, Fudan University | National Research Center for Translational Medicine (Shanghai), Ruijin Hospital affiliated to Shanghai Jiao Tong University School of Medicine & Shanghai Public Health Clinical Center | Shengyue Wang, Xiaonan Zhang, Gang Lu, Yun Tan, Yun Ling, Hongzhou Lu, Saijuan Chen |
| EPI_ISL_416381 | hCoV-19/Shanghai/SH0079/2020 | Asia / China / Shanghai | 2020-02-01 | Shanghai Public Health Clinical Center, Shanghai Medical College, Fudan University | National Research Center for Translational Medicine (Shanghai), Ruijin Hospital affiliated to Shanghai Jiao Tong University School of Medicine & Shanghai Public Health Clinical Center | Shengyue Wang, Xiaonan Zhang, Gang Lu, Yun Tan, Yun Ling, Hongzhou Lu, Saijuan Chen |
| EPI_ISL_416382 | hCoV-19/Shanghai/SH0080/2020 | Asia / China / Shanghai | 2020-02-08 | Shanghai Public Health Clinical Center, Shanghai Medical College, Fudan University | National Research Center for Translational Medicine (Shanghai), Ruijin Hospital affiliated to Shanghai Jiao Tong University School of Medicine & Shanghai Public Health Clinical Center | Shengyue Wang, Xiaonan Zhang, Gang Lu, Yun Tan, Yun Ling, Hongzhou Lu, Saijuan Chen |
| EPI_ISL_416384 | hCoV-19/Shanghai/SH0083/2020 | Asia / China / Shanghai | 2020-02-04 | Shanghai Public Health Clinical Center, Shanghai Medical College, Fudan University | National Research Center for Translational Medicine (Shanghai), Ruijin Hospital affiliated to Shanghai Jiao Tong University School of Medicine & Shanghai Public Health Clinical Center | Shengyue Wang, Xiaonan Zhang, Gang Lu, Yun Tan, Yun Ling, Hongzhou Lu, Saijuan Chen |
| EPI_ISL_416387 | hCoV-19/Shanghai/SH0088/2020 | Asia / China / Shanghai | 2020-02-01 | Shanghai Public Health Clinical Center, Shanghai Medical College, Fudan University | National Research Center for Translational Medicine (Shanghai), Ruijin Hospital affiliated to Shanghai Jiao Tong University School of Medicine & Shanghai Public Health Clinical Center | Shengyue Wang, Xiaonan Zhang, Gang Lu, Yun Tan, Yun Ling, Hongzhou Lu, Saijuan Chen |
| EPI_ISL_416389 | hCoV-19/Shanghai/SH0093/2020 | Asia / China / Shanghai | 2020-01-21 | Shanghai Public Health Clinical Center, Shanghai Medical College, Fudan University | National Research Center for Translational Medicine (Shanghai), Ruijin Hospital affiliated to Shanghai Jiao Tong University School of Medicine & Shanghai Public Health Clinical Center | Shengyue Wang, Xiaonan Zhang, Gang Lu, Yun Tan, Yun Ling, Hongzhou Lu, Saijuan Chen |
| EPI_ISL_416390 | hCoV-19/Shanghai/SH0094/2020 | Asia / China / Shanghai | 2020-01-31 | Shanghai Public Health Clinical Center, Shanghai Medical College, Fudan University | National Research Center for Translational Medicine (Shanghai), Ruijin Hospital affiliated to Shanghai Jiao Tong University School of Medicine & Shanghai Public Health Clinical Center | Shengyue Wang, Xiaonan Zhang, Gang Lu, Yun Tan, Yun Ling, Hongzhou Lu, Saijuan Chen |
| EPI_ISL_416393 | hCoV-19/Shanghai/SH0101/2020 | Asia / China / Shanghai | 2020-02-09 | Shanghai Public Health Clinical Center, Shanghai Medical College, Fudan University | National Research Center for Translational Medicine (Shanghai), Ruijin Hospital affiliated to Shanghai Jiao Tong University School of Medicine & Shanghai Public Health Clinical Center | Shengyue Wang, Xiaonan Zhang, Gang Lu, Yun Tan, Yun Ling, Hongzhou Lu, Saijuan Chen |
| EPI_ISL_416394 | hCoV-19/Shanghai/SH0102/2020 | Asia / China / Shanghai | 2020-02-14 | Shanghai Public Health Clinical Center, Shanghai Medical College, Fudan University | National Research Center for Translational Medicine (Shanghai), Ruijin Hospital affiliated to Shanghai Jiao Tong University School of Medicine & Shanghai Public Health Clinical Center | Shengyue Wang, Xiaonan Zhang, Gang Lu, Yun Tan, Yun Ling, Hongzhou Lu, Saijuan Chen |
| EPI_ISL_416395 | hCoV-19/Shanghai/SH0104/2020 | Asia / China / Shanghai | 2020-02-09 | Shanghai Public Health Clinical Center, Shanghai Medical College, Fudan University | National Research Center for Translational Medicine (Shanghai), Ruijin Hospital affiliated to Shanghai Jiao Tong University School of Medicine & Shanghai Public Health Clinical Center | Shengyue Wang, Xiaonan Zhang, Gang Lu, Yun Tan, Yun Ling, Hongzhou Lu, Saijuan Chen |
| EPI_ISL_416396 | hCoV-19/Shanghai/SH0106/2020 | Asia / China / Shanghai | 2020-02-10 | Shanghai Public Health Clinical Center, Shanghai Medical College, Fudan University | National Research Center for Translational Medicine (Shanghai), Ruijin Hospital affiliated to Shanghai Jiao Tong University School of Medicine & Shanghai Public Health Clinical Center | Shengyue Wang, Xiaonan Zhang, Gang Lu, Yun Tan, Yun Ling, Hongzhou Lu, Saijuan Chen |
| EPI_ISL_416397 | hCoV-19/Shanghai/SH0107/2020 | Asia / China / Shanghai | 2020-02-02 | Shanghai Public Health Clinical Center, Shanghai Medical College, Fudan University | National Research Center for Translational Medicine (Shanghai), Ruijin Hospital affiliated to Shanghai Jiao Tong University School of Medicine & Shanghai Public Health Clinical Center | Shengyue Wang, Xiaonan Zhang, Gang Lu, Yun Tan, Yun Ling, Hongzhou Lu, Saijuan Chen |
| EPI_ISL_416398 | hCoV-19/Shanghai/SH0109/2020 | Asia / China / Shanghai | 2020-02-02 | Shanghai Public Health Clinical Center, Shanghai Medical College, Fudan University | National Research Center for Translational Medicine (Shanghai), Ruijin Hospital affiliated to Shanghai Jiao Tong University School of Medicine & Shanghai Public Health Clinical Center | Shengyue Wang, Xiaonan Zhang, Gang Lu, Yun Tan, Yun Ling, Hongzhou Lu, Saijuan Chen |
| EPI_ISL_416399 | hCoV-19/Shanghai/SH0110/2020 | Asia / China / Shanghai | 2020-02-13 | Shanghai Public Health Clinical Center, Shanghai Medical College, Fudan University | National Research Center for Translational Medicine (Shanghai), Ruijin Hospital affiliated to Shanghai Jiao Tong University School of Medicine & Shanghai Public Health Clinical Center | Shengyue Wang, Xiaonan Zhang, Gang Lu, Yun Tan, Yun Ling, Hongzhou Lu, Saijuan Chen |
| EPI_ISL_416600 | hCoV-19/Japan/DP0438/2020 | Asia / Japan / unknown | 2020-02-16 | Japanese Quarantine Stations | Pathogen Genomics Center, National Institute of Infectious Diseases | Tsuyoshi Sekizuka, Kentaro Itokawa, Rina Tanaka, Masanori Hashino, Tsutomu Kageyama, Shinji Saito, Ikuyo Takayama, Hideki Hasegawa, Takuri Takahashi, Hajime Kamiya, Takuya Yamagishi, Motoi Suzuki, Takaji Wakita, Makoto Kuroda |
| EPI_ISL_416601 | hCoV-19/Japan/DP0457/2020 | Asia / Japan / unknown | 2020-02-16 | Japanese Quarantine Stations | Pathogen Genomics Center, National Institute of Infectious Diseases | Tsuyoshi Sekizuka, Kentaro Itokawa, Rina Tanaka, Masanori Hashino, Tsutomu Kageyama, Shinji Saito, Ikuyo Takayama, Hideki Hasegawa, Takuri Takahashi, Hajime Kamiya, Takuya Yamagishi, Motoi Suzuki, Takaji Wakita, Makoto Kuroda |
| EPI_ISL_416602 | hCoV-19/Japan/DP0462/2020 | Asia / Japan / unknown | 2020-02-16 | Japanese Quarantine Stations | Pathogen Genomics Center, National Institute of Infectious Diseases | Tsuyoshi Sekizuka, Kentaro Itokawa, Rina Tanaka, Masanori Hashino, Tsutomu Kageyama, Shinji Saito, Ikuyo Takayama, Hideki Hasegawa, Takuri Takahashi, Hajime Kamiya, Takuya Yamagishi, Motoi Suzuki, Takaji Wakita, Makoto Kuroda |
| EPI_ISL_416603 | hCoV-19/Japan/DP0464/2020 | Asia / Japan / unknown | 2020-02-16 | Japanese Quarantine Stations | Pathogen Genomics Center, National Institute of Infectious Diseases | Tsuyoshi Sekizuka, Kentaro Itokawa, Rina Tanaka, Masanori Hashino, Tsutomu Kageyama, Shinji Saito, Ikuyo Takayama, Hideki Hasegawa, Takuri Takahashi, Hajime Kamiya, Takuya Yamagishi, Motoi Suzuki, Takaji Wakita, Makoto Kuroda |
| EPI_ISL_416604 | hCoV-19/Japan/DP0476/2020 | Asia / Japan / unknown | 2020-02-16 | Japanese Quarantine Stations | Pathogen Genomics Center, National Institute of Infectious Diseases | Tsuyoshi Sekizuka, Kentaro Itokawa, Rina Tanaka, Masanori Hashino, Tsutomu Kageyama, Shinji Saito, Ikuyo Takayama, Hideki Hasegawa, Takuri Takahashi, Hajime Kamiya, Takuya Yamagishi, Motoi Suzuki, Takaji Wakita, Makoto Kuroda |
| EPI_ISL_416605 | hCoV-19/Japan/DP0481/2020 | Asia / Japan / unknown | 2020-02-16 | Japanese Quarantine Stations | Pathogen Genomics Center, National Institute of Infectious Diseases | Tsuyoshi Sekizuka, Kentaro Itokawa, Rina Tanaka, Masanori Hashino, Tsutomu Kageyama, Shinji Saito, Ikuyo Takayama, Hideki Hasegawa, Takuri Takahashi, Hajime Kamiya, Takuya Yamagishi, Motoi Suzuki, Takaji Wakita, Makoto Kuroda |
| EPI_ISL_416606 | hCoV-19/Japan/DP0482/2020 | Asia / Japan / unknown | 2020-02-16 | Japanese Quarantine Stations | Pathogen Genomics Center, National Institute of Infectious Diseases | Tsuyoshi Sekizuka, Kentaro Itokawa, Rina Tanaka, Masanori Hashino, Tsutomu Kageyama, Shinji Saito, Ikuyo Takayama, Hideki Hasegawa, Takuri Takahashi, Hajime Kamiya, Takuya Yamagishi, Motoi Suzuki, Takaji Wakita, Makoto Kuroda |
| EPI_ISL_416607 | hCoV-19/Japan/DP0543/2020 | Asia / Japan / unknown | 2020-02-17 | Japanese Quarantine Stations | Pathogen Genomics Center, National Institute of Infectious Diseases | Tsuyoshi Sekizuka, Kentaro Itokawa, Rina Tanaka, Masanori Hashino, Tsutomu Kageyama, Shinji Saito, Ikuyo Takayama, Hideki Hasegawa, Takuri Takahashi, Hajime Kamiya, Takuya Yamagishi, Motoi Suzuki, Takaji Wakita, Makoto Kuroda |
| EPI_ISL_416608 | hCoV-19/Japan/DP0544/2020 | Asia / Japan / unknown | 2020-02-17 | Japanese Quarantine Stations | Pathogen Genomics Center, National Institute of Infectious Diseases | Tsuyoshi Sekizuka, Kentaro Itokawa, Rina Tanaka, Masanori Hashino, Tsutomu Kageyama, Shinji Saito, Ikuyo Takayama, Hideki Hasegawa, Takuri Takahashi, Hajime Kamiya, Takuya Yamagishi, Motoi Suzuki, Takaji Wakita, Makoto Kuroda |
| EPI_ISL_416609 | hCoV-19/Japan/DP0568/2020 | Asia / Japan / unknown | 2020-02-17 | Japanese Quarantine Stations | Pathogen Genomics Center, National Institute of Infectious Diseases | Tsuyoshi Sekizuka, Kentaro Itokawa, Rina Tanaka, Masanori Hashino, Tsutomu Kageyama, Shinji Saito, Ikuyo Takayama, Hideki Hasegawa, Takuri Takahashi, Hajime Kamiya, Takuya Yamagishi, Motoi Suzuki, Takaji Wakita, Makoto Kuroda |
| EPI_ISL_416610 | hCoV-19/Japan/DP0588/2020 | Asia / Japan / unknown | 2020-02-17 | Japanese Quarantine Stations | Pathogen Genomics Center, National Institute of Infectious Diseases | Tsuyoshi Sekizuka, Kentaro Itokawa, Rina Tanaka, Masanori Hashino, Tsutomu Kageyama, Shinji Saito, Ikuyo Takayama, Hideki Hasegawa, Takuri Takahashi, Hajime Kamiya, Takuya Yamagishi, Motoi Suzuki, Takaji Wakita, Makoto Kuroda |
| EPI_ISL_416611 | hCoV-19/Japan/DP0644/2020 | Asia / Japan / unknown | 2020-02-17 | Japanese Quarantine Stations | Pathogen Genomics Center, National Institute of Infectious Diseases | Tsuyoshi Sekizuka, Kentaro Itokawa, Rina Tanaka, Masanori Hashino, Tsutomu Kageyama, Shinji Saito, Ikuyo Takayama, Hideki Hasegawa, Takuri Takahashi, Hajime Kamiya, Takuya Yamagishi, Motoi Suzuki, Takaji Wakita, Makoto Kuroda |
| EPI_ISL_416612 | hCoV-19/Japan/DP0645/2020 | Asia / Japan / unknown | 2020-02-17 | Japanese Quarantine Stations | Pathogen Genomics Center, National Institute of Infectious Diseases | Tsuyoshi Sekizuka, Kentaro Itokawa, Rina Tanaka, Masanori Hashino, Tsutomu Kageyama, Shinji Saito, Ikuyo Takayama, Hideki Hasegawa, Takuri Takahashi, Hajime Kamiya, Takuya Yamagishi, Motoi Suzuki, Takaji Wakita, Makoto Kuroda |
| EPI_ISL_416613 | hCoV-19/Japan/DP0654/2020 | Asia / Japan / unknown | 2020-02-17 | Japanese Quarantine Stations | Pathogen Genomics Center, National Institute of Infectious Diseases | Tsuyoshi Sekizuka, Kentaro Itokawa, Rina Tanaka, Masanori Hashino, Tsutomu Kageyama, Shinji Saito, Ikuyo Takayama, Hideki Hasegawa, Takuri Takahashi, Hajime Kamiya, Takuya Yamagishi, Motoi Suzuki, Takaji Wakita, Makoto Kuroda |
| EPI_ISL_416614 | hCoV-19/Japan/DP0687/2020 | Asia / Japan / unknown | 2020-02-17 | Japanese Quarantine Stations | Pathogen Genomics Center, National Institute of Infectious Diseases | Tsuyoshi Sekizuka, Kentaro Itokawa, Rina Tanaka, Masanori Hashino, Tsutomu Kageyama, Shinji Saito, Ikuyo Takayama, Hideki Hasegawa, Takuri Takahashi, Hajime Kamiya, Takuya Yamagishi, Motoi Suzuki, Takaji Wakita, Makoto Kuroda |
| EPI_ISL_416615 | hCoV-19/Japan/DP0690/2020 | Asia / Japan / unknown | 2020-02-17 | Japanese Quarantine Stations | Pathogen Genomics Center, National Institute of Infectious Diseases | Tsuyoshi Sekizuka, Kentaro Itokawa, Rina Tanaka, Masanori Hashino, Tsutomu Kageyama, Shinji Saito, Ikuyo Takayama, Hideki Hasegawa, Takuri Takahashi, Hajime Kamiya, Takuya Yamagishi, Motoi Suzuki, Takaji Wakita, Makoto Kuroda |
| EPI_ISL_416617 | hCoV-19/Japan/DP0699/2020 | Asia / Japan / unknown | 2020-02-17 | Japanese Quarantine Stations | Pathogen Genomics Center, National Institute of Infectious Diseases | Tsuyoshi Sekizuka, Kentaro Itokawa, Rina Tanaka, Masanori Hashino, Tsutomu Kageyama, Shinji Saito, Ikuyo Takayama, Hideki Hasegawa, Takuri Takahashi, Hajime Kamiya, Takuya Yamagishi, Motoi Suzuki, Takaji Wakita, Makoto Kuroda |
| EPI_ISL_416618 | hCoV-19/Japan/DP0700/2020 | Asia / Japan / unknown | 2020-02-17 | Japanese Quarantine Stations | Pathogen Genomics Center, National Institute of Infectious Diseases | Tsuyoshi Sekizuka, Kentaro Itokawa, Rina Tanaka, Masanori Hashino, Tsutomu Kageyama, Shinji Saito, Ikuyo Takayama, Hideki Hasegawa, Takuri Takahashi, Hajime Kamiya, Takuya Yamagishi, Motoi Suzuki, Takaji Wakita, Makoto Kuroda |
| EPI_ISL_416619 | hCoV-19/Japan/DP0703/2020 | Asia / Japan / unknown | 2020-02-17 | Japanese Quarantine Stations | Pathogen Genomics Center, National Institute of Infectious Diseases | Tsuyoshi Sekizuka, Kentaro Itokawa, Rina Tanaka, Masanori Hashino, Tsutomu Kageyama, Shinji Saito, Ikuyo Takayama, Hideki Hasegawa, Takuri Takahashi, Hajime Kamiya, Takuya Yamagishi, Motoi Suzuki, Takaji Wakita, Makoto Kuroda |
| EPI_ISL_416620 | hCoV-19/Japan/DP0724/2020 | Asia / Japan / unknown | 2020-02-17 | Japanese Quarantine Stations | Pathogen Genomics Center, National Institute of Infectious Diseases | Tsuyoshi Sekizuka, Kentaro Itokawa, Rina Tanaka, Masanori Hashino, Tsutomu Kageyama, Shinji Saito, Ikuyo Takayama, Hideki Hasegawa, Takuri Takahashi, Hajime Kamiya, Takuya Yamagishi, Motoi Suzuki, Takaji Wakita, Makoto Kuroda |
| EPI_ISL_416621 | hCoV-19/Japan/DP0743/2020 | Asia / Japan / unknown | 2020-02-17 | Japanese Quarantine Stations | Pathogen Genomics Center, National Institute of Infectious Diseases | Tsuyoshi Sekizuka, Kentaro Itokawa, Rina Tanaka, Masanori Hashino, Tsutomu Kageyama, Shinji Saito, Ikuyo Takayama, Hideki Hasegawa, Takuri Takahashi, Hajime Kamiya, Takuya Yamagishi, Motoi Suzuki, Takaji Wakita, Makoto Kuroda |
| EPI_ISL_416622 | hCoV-19/Japan/DP0752/2020 | Asia / Japan / unknown | 2020-02-17 | Japanese Quarantine Stations | Pathogen Genomics Center, National Institute of Infectious Diseases | Tsuyoshi Sekizuka, Kentaro Itokawa, Rina Tanaka, Masanori Hashino, Tsutomu Kageyama, Shinji Saito, Ikuyo Takayama, Hideki Hasegawa, Takuri Takahashi, Hajime Kamiya, Takuya Yamagishi, Motoi Suzuki, Takaji Wakita, Makoto Kuroda |
| EPI_ISL_416624 | hCoV-19/Japan/DP0764/2020 | Asia / Japan / unknown | 2020-02-17 | Japanese Quarantine Stations | Pathogen Genomics Center, National Institute of Infectious Diseases | Tsuyoshi Sekizuka, Kentaro Itokawa, Rina Tanaka, Masanori Hashino, Tsutomu Kageyama, Shinji Saito, Ikuyo Takayama, Hideki Hasegawa, Takuri Takahashi, Hajime Kamiya, Takuya Yamagishi, Motoi Suzuki, Takaji Wakita, Makoto Kuroda |
| EPI_ISL_416625 | hCoV-19/Japan/DP0765/2020 | Asia / Japan / unknown | 2020-02-17 | Japanese Quarantine Stations | Pathogen Genomics Center, National Institute of Infectious Diseases | Tsuyoshi Sekizuka, Kentaro Itokawa, Rina Tanaka, Masanori Hashino, Tsutomu Kageyama, Shinji Saito, Ikuyo Takayama, Hideki Hasegawa, Takuri Takahashi, Hajime Kamiya, Takuya Yamagishi, Motoi Suzuki, Takaji Wakita, Makoto Kuroda |
| EPI_ISL_416626 | hCoV-19/Japan/DP0779/2020 | Asia / Japan / unknown | 2020-02-17 | Japanese Quarantine Stations | Pathogen Genomics Center, National Institute of Infectious Diseases | Tsuyoshi Sekizuka, Kentaro Itokawa, Rina Tanaka, Masanori Hashino, Tsutomu Kageyama, Shinji Saito, Ikuyo Takayama, Hideki Hasegawa, Takuri Takahashi, Hajime Kamiya, Takuya Yamagishi, Motoi Suzuki, Takaji Wakita, Makoto Kuroda |
| EPI_ISL_416627 | hCoV-19/Japan/DP0785/2020 | Asia / Japan / unknown | 2020-02-17 | Japanese Quarantine Stations | Pathogen Genomics Center, National Institute of Infectious Diseases | Tsuyoshi Sekizuka, Kentaro Itokawa, Rina Tanaka, Masanori Hashino, Tsutomu Kageyama, Shinji Saito, Ikuyo Takayama, Hideki Hasegawa, Takuri Takahashi, Hajime Kamiya, Takuya Yamagishi, Motoi Suzuki, Takaji Wakita, Makoto Kuroda |
| EPI_ISL_416628 | hCoV-19/Japan/DP0786/2020 | Asia / Japan / unknown | 2020-02-17 | Japanese Quarantine Stations | Pathogen Genomics Center, National Institute of Infectious Diseases | Tsuyoshi Sekizuka, Kentaro Itokawa, Rina Tanaka, Masanori Hashino, Tsutomu Kageyama, Shinji Saito, Ikuyo Takayama, Hideki Hasegawa, Takuri Takahashi, Hajime Kamiya, Takuya Yamagishi, Motoi Suzuki, Takaji Wakita, Makoto Kuroda |
| EPI_ISL_416629 | hCoV-19/Japan/DP0802/2020 | Asia / Japan / unknown | 2020-02-17 | Japanese Quarantine Stations | Pathogen Genomics Center, National Institute of Infectious Diseases | Tsuyoshi Sekizuka, Kentaro Itokawa, Rina Tanaka, Masanori Hashino, Tsutomu Kageyama, Shinji Saito, Ikuyo Takayama, Hideki Hasegawa, Takuri Takahashi, Hajime Kamiya, Takuya Yamagishi, Motoi Suzuki, Takaji Wakita, Makoto Kuroda |
| EPI_ISL_416630 | hCoV-19/Japan/DP0803/2020 | Asia / Japan / unknown | 2020-02-17 | Japanese Quarantine Stations | Pathogen Genomics Center, National Institute of Infectious Diseases | Tsuyoshi Sekizuka, Kentaro Itokawa, Rina Tanaka, Masanori Hashino, Tsutomu Kageyama, Shinji Saito, Ikuyo Takayama, Hideki Hasegawa, Takuri Takahashi, Hajime Kamiya, Takuya Yamagishi, Motoi Suzuki, Takaji Wakita, Makoto Kuroda |
| EPI_ISL_416631 | hCoV-19/Japan/DP0804/2020 | Asia / Japan / unknown | 2020-02-17 | Japanese Quarantine Stations | Pathogen Genomics Center, National Institute of Infectious Diseases | Tsuyoshi Sekizuka, Kentaro Itokawa, Rina Tanaka, Masanori Hashino, Tsutomu Kageyama, Shinji Saito, Ikuyo Takayama, Hideki Hasegawa, Takuri Takahashi, Hajime Kamiya, Takuya Yamagishi, Motoi Suzuki, Takaji Wakita, Makoto Kuroda |
| EPI_ISL_416632 | hCoV-19/Japan/DP0827/2020 | Asia / Japan / unknown | 2020-02-17 | Japanese Quarantine Stations | Pathogen Genomics Center, National Institute of Infectious Diseases | Tsuyoshi Sekizuka, Kentaro Itokawa, Rina Tanaka, Masanori Hashino, Tsutomu Kageyama, Shinji Saito, Ikuyo Takayama, Hideki Hasegawa, Takuri Takahashi, Hajime Kamiya, Takuya Yamagishi, Motoi Suzuki, Takaji Wakita, Makoto Kuroda |
| EPI_ISL_416633 | hCoV-19/Japan/DP0880/2020 | Asia / Japan / unknown | 2020-02-17 | Japanese Quarantine Stations | Pathogen Genomics Center, National Institute of Infectious Diseases | Tsuyoshi Sekizuka, Kentaro Itokawa, Rina Tanaka, Masanori Hashino, Tsutomu Kageyama, Shinji Saito, Ikuyo Takayama, Hideki Hasegawa, Takuri Takahashi, Hajime Kamiya, Takuya Yamagishi, Motoi Suzuki, Takaji Wakita, Makoto Kuroda |
| EPI_ISL_416634 | hCoV-19/Japan/DP0890/2020 | Asia / Japan / unknown | 2020-02-17 | Japanese Quarantine Stations | Pathogen Genomics Center, National Institute of Infectious Diseases | Tsuyoshi Sekizuka, Kentaro Itokawa, Rina Tanaka, Masanori Hashino, Tsutomu Kageyama, Shinji Saito, Ikuyo Takayama, Hideki Hasegawa, Takuri Takahashi, Hajime Kamiya, Takuya Yamagishi, Motoi Suzuki, Takaji Wakita, Makoto Kuroda |
| EPI_ISL_416635 | hCoV-19/USA/WA-UW97/2020 | North America / USA / Washington | 2020-03-12 | UW Virology Lab | UW Virology Lab | Pavitra Roychoudhury, Hong Xie, Keith Jerome, Alexander Greninger |
| EPI_ISL_416636 | hCoV-19/USA/WA-UW98/2020 | North America / USA / Washington | 2020-03-12 | UW Virology Lab | UW Virology Lab | Pavitra Roychoudhury, Hong Xie, Keith Jerome, Alexander Greninger |
| EPI_ISL_416637 | hCoV-19/USA/WA-UW99/2020 | North America / USA / Washington | 2020-03-12 | UW Virology Lab | UW Virology Lab | Pavitra Roychoudhury, Hong Xie, Keith Jerome, Alexander Greninger |
| EPI_ISL_416638 | hCoV-19/USA/WA-UW100/2020 | North America / USA / Washington | 2020-03-12 | UW Virology Lab | UW Virology Lab | Pavitra Roychoudhury, Hong Xie, Keith Jerome, Alexander Greninger |
| EPI_ISL_416639 | hCoV-19/USA/WA-UW101/2020 | North America / USA / Washington | 2020-03-10 | UW Virology Lab | UW Virology Lab | Pavitra Roychoudhury, Hong Xie, Keith Jerome, Alexander Greninger |
| EPI_ISL_416641 | hCoV-19/USA/WA-UW103/2020 | North America / USA / Washington | 2020-03-11 | UW Virology Lab | UW Virology Lab | Pavitra Roychoudhury, Hong Xie, Keith Jerome, Alexander Greninger |
| EPI_ISL_416642 | hCoV-19/USA/WA-UW104/2020 | North America / USA / Washington | 2020-03-11 | UW Virology Lab | UW Virology Lab | Pavitra Roychoudhury, Hong Xie, Keith Jerome, Alexander Greninger |
| EPI_ISL_416643 | hCoV-19/USA/WA-UW105/2020 | North America / USA / Washington | 2020-03-11 | UW Virology Lab | UW Virology Lab | Pavitra Roychoudhury, Hong Xie, Keith Jerome, Alexander Greninger |
| EPI_ISL_416644 | hCoV-19/USA/WA-UW106/2020 | North America / USA / Washington | 2020-03-11 | UW Virology Lab | UW Virology Lab | Pavitra Roychoudhury, Hong Xie, Keith Jerome, Alexander Greninger |
| EPI_ISL_416646 | hCoV-19/USA/WA-UW108/2020 | North America / USA / Washington | 2020-03-11 | UW Virology Lab | UW Virology Lab | Pavitra Roychoudhury, Hong Xie, Keith Jerome, Alexander Greninger |
| EPI_ISL_416647 | hCoV-19/USA/WA-UW109/2020 | North America / USA / Washington | 2020-03-11 | UW Virology Lab | UW Virology Lab | Pavitra Roychoudhury, Hong Xie, Keith Jerome, Alexander Greninger |
| EPI_ISL_416648 | hCoV-19/USA/WA-UW110/2020 | North America / USA / Washington | 2020-03-11 | UW Virology Lab | UW Virology Lab | Pavitra Roychoudhury, Hong Xie, Keith Jerome, Alexander Greninger |
| EPI_ISL_416649 | hCoV-19/USA/WA-UW111/2020 | North America / USA / Washington | 2020-03-11 | UW Virology Lab | UW Virology Lab | Pavitra Roychoudhury, Hong Xie, Keith Jerome, Alexander Greninger |
| EPI_ISL_416650 | hCoV-19/USA/WA-UW112/2020 | North America / USA | 2020-03-10 | UW Virology Lab | UW Virology Lab | Pavitra Roychoudhury, Hong Xie, Keith Jerome, Alexander Greninger |
| EPI_ISL_416651 | hCoV-19/USA/WA-UW113/2020 | North America / USA | 2020-03-11 | UW Virology Lab | UW Virology Lab | Pavitra Roychoudhury, Hong Xie, Keith Jerome, Alexander Greninger |
| EPI_ISL_416652 | hCoV-19/USA/WA-UW114/2020 | North America / USA / Washington | 2020-03-11 | UW Virology Lab | UW Virology Lab | Pavitra Roychoudhury, Hong Xie, Keith Jerome, Alexander Greninger |
| EPI_ISL_416653 | hCoV-19/USA/WA-UW115/2020 | North America / USA / Washington | 2020-03-11 | UW Virology Lab | UW Virology Lab | Pavitra Roychoudhury, Hong Xie, Keith Jerome, Alexander Greninger |
| EPI_ISL_416654 | hCoV-19/USA/WA-UW116/2020 | North America / USA / Washington | 2020-03-11 | UW Virology Lab | UW Virology Lab | Pavitra Roychoudhury, Hong Xie, Keith Jerome, Alexander Greninger |
| EPI_ISL_416655 | hCoV-19/USA/WA-UW117/2020 | North America / USA / Washington | 2020-03-11 | UW Virology Lab | UW Virology Lab | Pavitra Roychoudhury, Hong Xie, Keith Jerome, Alexander Greninger |
| EPI_ISL_416656 | hCoV-19/USA/WA-UW118/2020 | North America / USA / Washington | 2020-03-11 | UW Virology Lab | UW Virology Lab | Pavitra Roychoudhury, Hong Xie, Keith Jerome, Alexander Greninger |
| EPI_ISL_416657 | hCoV-19/USA/WA-UW119/2020 | North America / USA / Washington | 2020-03-11 | UW Virology Lab | UW Virology Lab | Pavitra Roychoudhury, Hong Xie, Keith Jerome, Alexander Greninger |
| EPI_ISL_416658 | hCoV-19/USA/WA-UW120/2020 | North America / USA / Washington | 2020-03-11 | UW Virology Lab | UW Virology Lab | Pavitra Roychoudhury, Hong Xie, Keith Jerome, Alexander Greninger |
| EPI_ISL_416659 | hCoV-19/USA/WA-UW121/2020 | North America / USA / Washington | 2020-03-11 | UW Virology Lab | UW Virology Lab | Pavitra Roychoudhury, Hong Xie, Keith Jerome, Alexander Greninger |
| EPI_ISL_416660 | hCoV-19/USA/WA-UW122/2020 | North America / USA | 2020-03-10 | UW Virology Lab | UW Virology Lab | Pavitra Roychoudhury, Hong Xie, Keith Jerome, Alexander Greninger |
| EPI_ISL_416661 | hCoV-19/USA/WA-UW123/2020 | North America / USA / Washington | 2020-03-12 | UW Virology Lab | UW Virology Lab | Pavitra Roychoudhury, Hong Xie, Keith Jerome, Alexander Greninger |
| EPI_ISL_416662 | hCoV-19/USA/WA-UW124/2020 | North America / USA / Washington | 2020-03-12 | UW Virology Lab | UW Virology Lab | Pavitra Roychoudhury, Hong Xie, Keith Jerome, Alexander Greninger |
| EPI_ISL_416663 | hCoV-19/USA/WA-UW125/2020 | North America / USA / Washington | 2020-03-12 | UW Virology Lab | UW Virology Lab | Pavitra Roychoudhury, Hong Xie, Keith Jerome, Alexander Greninger |
| EPI_ISL_416664 | hCoV-19/USA/WA-UW126/2020 | North America / USA / Washington | 2020-03-12 | UW Virology Lab | UW Virology Lab | Pavitra Roychoudhury, Hong Xie, Keith Jerome, Alexander Greninger |
| EPI_ISL_416665 | hCoV-19/USA/WA-UW127/2020 | North America / USA / Washington | 2020-03-12 | UW Virology Lab | UW Virology Lab | Pavitra Roychoudhury, Hong Xie, Keith Jerome, Alexander Greninger |
| EPI_ISL_416666 | hCoV-19/USA/WA-UW128/2020 | North America / USA / Washington | 2020-03-12 | UW Virology Lab | UW Virology Lab | Pavitra Roychoudhury, Hong Xie, Keith Jerome, Alexander Greninger |
| EPI_ISL_416667 | hCoV-19/USA/WA-UW129/2020 | North America / USA / Washington | 2020-03-12 | UW Virology Lab | UW Virology Lab | Pavitra Roychoudhury, Hong Xie, Keith Jerome, Alexander Greninger |
| EPI_ISL_416668 | hCoV-19/USA/WA-UW130/2020 | North America / USA / Washington | 2020-03-11 | UW Virology Lab | UW Virology Lab | Pavitra Roychoudhury, Hong Xie, Keith Jerome, Alexander Greninger |
| EPI_ISL_416669 | hCoV-19/USA/WA-UW131/2020 | North America / USA / Washington | 2020-03-11 | UW Virology Lab | UW Virology Lab | Pavitra Roychoudhury, Hong Xie, Keith Jerome, Alexander Greninger |
| EPI_ISL_416671 | hCoV-19/USA/WA-UW133/2020 | North America / USA / Washington | 2020-03-12 | UW Virology Lab | UW Virology Lab | Pavitra Roychoudhury, Hong Xie, Keith Jerome, Alexander Greninger |
| EPI_ISL_416672 | hCoV-19/USA/WA-UW134/2020 | North America / USA | 2020-03-10 | UW Virology Lab | UW Virology Lab | Pavitra Roychoudhury, Hong Xie, Keith Jerome, Alexander Greninger |
| EPI_ISL_416673 | hCoV-19/USA/OR-UW135/2020 | North America / USA / Oregon | 2020-03-11 | UW Virology Lab | UW Virology Lab | Pavitra Roychoudhury, Hong Xie, Keith Jerome, Alexander Greninger |
| EPI_ISL_416674 | hCoV-19/USA/WA-UW136/2020 | North America / USA | 2020-03-11 | UW Virology Lab | UW Virology Lab | Pavitra Roychoudhury, Hong Xie, Keith Jerome, Alexander Greninger |
| EPI_ISL_416675 | hCoV-19/USA/WA-UW137/2020 | North America / USA | 2020-03-11 | UW Virology Lab | UW Virology Lab | Pavitra Roychoudhury, Hong Xie, Keith Jerome, Alexander Greninger |
| EPI_ISL_416676 | hCoV-19/USA/WA-UW138/2020 | North America / USA | 2020-03-11 | UW Virology Lab | UW Virology Lab | Pavitra Roychoudhury, Hong Xie, Keith Jerome, Alexander Greninger |
| EPI_ISL_416677 | hCoV-19/USA/WA-UW139/2020 | North America / USA / Washington | 2020-03-12 | UW Virology Lab | UW Virology Lab | Pavitra Roychoudhury, Hong Xie, Keith Jerome, Alexander Greninger |
| EPI_ISL_416678 | hCoV-19/USA/WA-UW140/2020 | North America / USA / Washington | 2020-03-11 | UW Virology Lab | UW Virology Lab | Pavitra Roychoudhury, Hong Xie, Keith Jerome, Alexander Greninger |
| EPI_ISL_416679 | hCoV-19/USA/WA-UW141/2020 | North America / USA / Washington | 2020-03-11 | UW Virology Lab | UW Virology Lab | Pavitra Roychoudhury, Hong Xie, Keith Jerome, Alexander Greninger |
| EPI_ISL_416680 | hCoV-19/USA/WA-UW142/2020 | North America / USA / Washington | 2020-03-11 | UW Virology Lab | UW Virology Lab | Pavitra Roychoudhury, Hong Xie, Keith Jerome, Alexander Greninger |
| EPI_ISL_416681 | hCoV-19/USA/WA-UW143/2020 | North America / USA / Washington | 2020-03-11 | UW Virology Lab | UW Virology Lab | Pavitra Roychoudhury, Hong Xie, Keith Jerome, Alexander Greninger |
| EPI_ISL_416682 | hCoV-19/USA/WA-UW144/2020 | North America / USA | 2020-03-09 | UW Virology Lab | UW Virology Lab | Pavitra Roychoudhury, Hong Xie, Keith Jerome, Alexander Greninger |
| EPI_ISL_416683 | hCoV-19/USA/WA-UW145/2020 | North America / USA / Washington | 2020-03-15 | UW Virology Lab | UW Virology Lab | Pavitra Roychoudhury, Hong Xie, Keith Jerome, Alexander Greninger |
| EPI_ISL_416684 | hCoV-19/USA/WA-UW146/2020 | North America / USA / Washington | 2020-03-14 | UW Virology Lab | UW Virology Lab | Pavitra Roychoudhury, Hong Xie, Keith Jerome, Alexander Greninger |
| EPI_ISL_416685 | hCoV-19/USA/WA-UW147/2020 | North America / USA / Washington | 2020-03-15 | UW Virology Lab | UW Virology Lab | Pavitra Roychoudhury, Hong Xie, Keith Jerome, Alexander Greninger |
| EPI_ISL_416686 | hCoV-19/USA/WA-UW148/2020 | North America / USA / Washington | 2020-03-14 | UW Virology Lab | UW Virology Lab | Pavitra Roychoudhury, Hong Xie, Keith Jerome, Alexander Greninger |
| EPI_ISL_416687 | hCoV-19/USA/WA-UW149/2020 | North America / USA / Washington | 2020-03-14 | UW Virology Lab | UW Virology Lab | Pavitra Roychoudhury, Hong Xie, Keith Jerome, Alexander Greninger |
| EPI_ISL_416688 | hCoV-19/USA/WA-UW150/2020 | North America / USA / Washington | 2020-03-14 | UW Virology Lab | UW Virology Lab | Pavitra Roychoudhury, Hong Xie, Keith Jerome, Alexander Greninger |
| EPI_ISL_416689 | hCoV-19/USA/WA-UW151/2020 | North America / USA / Washington | 2020-03-14 | UW Virology Lab | UW Virology Lab | Pavitra Roychoudhury, Hong Xie, Keith Jerome, Alexander Greninger |
| EPI_ISL_416690 | hCoV-19/USA/WA-UW152/2020 | North America / USA / Washington | 2020-03-13 | UW Virology Lab | UW Virology Lab | Pavitra Roychoudhury, Hong Xie, Keith Jerome, Alexander Greninger |
| EPI_ISL_416691 | hCoV-19/USA/WA-UW153/2020 | North America / USA / Washington | 2020-03-13 | UW Virology Lab | UW Virology Lab | Pavitra Roychoudhury, Hong Xie, Keith Jerome, Alexander Greninger |
| EPI_ISL_416692 | hCoV-19/USA/WA-UW154/2020 | North America / USA / Washington | 2020-03-14 | UW Virology Lab | UW Virology Lab | Pavitra Roychoudhury, Hong Xie, Keith Jerome, Alexander Greninger |
| EPI_ISL_416693 | hCoV-19/USA/WA-UW155/2020 | North America / USA / Washington | 2020-03-12 | UW Virology Lab | UW Virology Lab | Pavitra Roychoudhury, Hong Xie, Keith Jerome, Alexander Greninger |
| EPI_ISL_416694 | hCoV-19/USA/WA-UW156/2020 | North America / USA / Washington | 2020-03-13 | UW Virology Lab | UW Virology Lab | Pavitra Roychoudhury, Hong Xie, Keith Jerome, Alexander Greninger |
| EPI_ISL_416696 | hCoV-19/USA/CT-UW158/2020 | North America / USA / Connecticut | 2020-03-13 | UW Virology Lab | UW Virology Lab | Pavitra Roychoudhury, Hong Xie, Keith Jerome, Alexander Greninger |
| EPI_ISL_416697 | hCoV-19/USA/CT-UW159/2020 | North America / USA / Connecticut | 2020-03-13 | UW Virology Lab | UW Virology Lab | Pavitra Roychoudhury, Hong Xie, Keith Jerome, Alexander Greninger |
| EPI_ISL_416698 | hCoV-19/USA/WA-UW160/2020 | North America / USA / Washington | 2020-03-13 | UW Virology Lab | UW Virology Lab | Pavitra Roychoudhury, Hong Xie, Keith Jerome, Alexander Greninger |
| EPI_ISL_416699 | hCoV-19/USA/WA-UW161/2020 | North America / USA / Washington | 2020-03-13 | UW Virology Lab | UW Virology Lab | Pavitra Roychoudhury, Hong Xie, Keith Jerome, Alexander Greninger |
| EPI_ISL_416730 | hCoV-19/England/SHEF-BFCB1/2020 | Europe / United Kingdom / England / Sou | 2020-03-03 | Virology Department, Sheffield Teaching Hospitals NHS Foundation Trust | Department of Infection, Immunity and Cardiovascular Disease, The Florey Institute, The Medical School, University of Sheffield | Thushan de Silva, Matthew Parker, Adri Angyal, Rebecca Brown, Matthew Wyles, Mehmet Yavuz, Mohammad Raza, Cariad Evans |
| EPI_ISL_416731 | hCoV-19/England/SHEF-BFCC0/2020 | Europe / United Kingdom / England / Nor | 2020-03-03 | Virology Department, Sheffield Teaching Hospitals NHS Foundation Trust | Department of Infection, Immunity and Cardiovascular Disease, The Florey Institute, The Medical School, University of Sheffield | Thushan de Silva, Matthew Parker, Adri Angyal, Rebecca Brown, Matthew Wyles, Mehmet Yavuz, Mohammad Raza, Cariad Evans |
| EPI_ISL_416732 | hCoV-19/England/SHEF-BFCDF/2020 | Europe / United Kingdom / England / Lon | 2020-03-03 | Virology Department, Sheffield Teaching Hospitals NHS Foundation Trust | Department of Infection, Immunity and Cardiovascular Disease, The Florey Institute, The Medical School, University of Sheffield | Thushan de Silva, Matthew Parker, Adri Angyal, Rebecca Brown, Matthew Wyles, Mehmet Yavuz, Mohammad Raza, Cariad Evans |
| EPI_ISL_416735 | hCoV-19/England/SHEF-BFD09/2020 | Europe / United Kingdom / England / Sou | 2020-03-09 | Virology Department, Sheffield Teaching Hospitals NHS Foundation Trust | Department of Infection, Immunity and Cardiovascular Disease, The Florey Institute, The Medical School, University of Sheffield | Thushan de Silva, Matthew Parker, Adri Angyal, Rebecca Brown, Matthew Wyles, Mehmet Yavuz, Mohammad Raza, Cariad Evans |
| EPI_ISL_417091 | hCoV-19/USA/WA-S38/2020 | North America / USA / Washington / Sno | 2020-03-04 | Washington State Department of Health | Seattle Flu Study | Chu et al |
| EPI_ISL_417092 | hCoV-19/USA/WA-S39/2020 | North America / USA / Washington / Sno | 2020-03-04 | Washington State Department of Health | Seattle Flu Study | Chu et al |
| EPI_ISL_417093 | hCoV-19/USA/WA-S40/2020 | North America / USA / Washington / Sno | 2020-02-28 | Washington State Department of Health | Seattle Flu Study | Chu et al |
| EPI_ISL_417095 | hCoV-19/USA/WA-S42/2020 | North America / USA / Washington / King | 2020-02-28 | Washington State Department of Health | Seattle Flu Study | Chu et al |
| EPI_ISL_417096 | hCoV-19/USA/WA-S43/2020 | North America / USA / Washington | 2020-02-27 | Washington State Department of Health | Seattle Flu Study | Chu et al |
| EPI_ISL_417097 | hCoV-19/USA/WA-S44/2020 | North America / USA / Washington | 2020-02-28 | Washington State Department of Health | Seattle Flu Study | Chu et al |
| EPI_ISL_417098 | hCoV-19/USA/WA-S45/2020 | North America / USA / Washington | 2020-02-29 | Washington State Department of Health | Seattle Flu Study | Chu et al |
| EPI_ISL_417099 | hCoV-19/USA/WA-S46/2020 | North America / USA / Washington | 2020-02-29 | Washington State Department of Health | Seattle Flu Study | Chu et al |
| EPI_ISL_417100 | hCoV-19/USA/WA-S47/2020 | North America / USA / Washington / Sno | 2020-02-29 | Washington State Department of Health | Seattle Flu Study | Chu et al |
| EPI_ISL_417101 | hCoV-19/USA/WA-S48/2020 | North America / USA / Washington / Sno | 2020-02-29 | Washington State Department of Health | Seattle Flu Study | Chu et al |
| EPI_ISL_417102 | hCoV-19/USA/WA-S49/2020 | North America / USA / Washington / King | 2020-03-04 | Washington State Department of Health | Seattle Flu Study | Chu et al |
| EPI_ISL_417103 | hCoV-19/USA/WA-S50/2020 | North America / USA / Washington / King | 2020-03-05 | Washington State Department of Health | Seattle Flu Study | Chu et al |
| EPI_ISL_417104 | hCoV-19/USA/WA-S51/2020 | North America / USA / Washington | 2020-03-03 | Washington State Department of Health | Seattle Flu Study | Chu et al |
| EPI_ISL_417105 | hCoV-19/USA/WA-S52/2020 | North America / USA / Washington / Sno | 2020-03-03 | Washington State Department of Health | Seattle Flu Study | Chu et al |
| EPI_ISL_417106 | hCoV-19/USA/WA-S53/2020 | North America / USA / Washington / Sno | 2020-03-03 | Washington State Department of Health | Seattle Flu Study | Chu et al |
| EPI_ISL_417107 | hCoV-19/USA/WA-S54/2020 | North America / USA / Washington / King | 2020-03-05 | Washington State Department of Health | Seattle Flu Study | Chu et al |
| EPI_ISL_417108 | hCoV-19/USA/WA-S55/2020 | North America / USA / Washington / King | 2020-02-29 | Washington State Department of Health | Seattle Flu Study | Chu et al |
| EPI_ISL_417109 | hCoV-19/USA/WA-S56/2020 | North America / USA / Washington / King | 2020-03-05 | Washington State Department of Health | Seattle Flu Study | Chu et al |
| EPI_ISL_417110 | hCoV-19/USA/WA-S57/2020 | North America / USA / Washington | 2020-03-03 | Washington State Department of Health | Seattle Flu Study | Chu et al |
| EPI_ISL_417111 | hCoV-19/USA/WA-S58/2020 | North America / USA / Washington / Sno | 2020-03-05 | Washington State Department of Health | Seattle Flu Study | Chu et al |
| EPI_ISL_417112 | hCoV-19/USA/WA-S59/2020 | North America / USA / Washington / King | 2020-03-02 | Washington State Department of Health | Seattle Flu Study | Chu et al |
| EPI_ISL_417113 | hCoV-19/USA/WA-S60/2020 | North America / USA / Washington / King | 2020-03-02 | Washington State Department of Health | Seattle Flu Study | Chu et al |
| EPI_ISL_417114 | hCoV-19/USA/WA-S61/2020 | North America / USA / Washington / King | 2020-03-05 | Washington State Department of Health | Seattle Flu Study | Chu et al |
| EPI_ISL_417115 | hCoV-19/USA/WA-S62/2020 | North America / USA / Washington / Sno | 2020-03-03 | Washington State Department of Health | Seattle Flu Study | Chu et al |
| EPI_ISL_417116 | hCoV-19/USA/WA-S63/2020 | North America / USA / Washington / King | 2020-03-04 | Washington State Department of Health | Seattle Flu Study | Chu et al |
| EPI_ISL_417117 | hCoV-19/USA/WA-S64/2020 | North America / USA / Washington / King | 2020-03-03 | Washington State Department of Health | Seattle Flu Study | Chu et al |
| EPI_ISL_417118 | hCoV-19/USA/WA-S65/2020 | North America / USA / Washington / King | 2020-03-03 | Washington State Department of Health | Seattle Flu Study | Chu et al |
| EPI_ISL_417119 | hCoV-19/USA/WA-S66/2020 | North America / USA / Washington / Sno | 2020-03-06 | Washington State Department of Health | Seattle Flu Study | Chu et al |
| EPI_ISL_417120 | hCoV-19/USA/WA-S67/2020 | North America / USA / Washington / King | 2020-03-04 | Washington State Department of Health | Seattle Flu Study | Chu et al |
| EPI_ISL_417121 | hCoV-19/USA/WA-S68/2020 | North America / USA / Washington / King | 2020-03-04 | Washington State Department of Health | Seattle Flu Study | Chu et al |
| EPI_ISL_417122 | hCoV-19/USA/WA-S69/2020 | North America / USA / Washington / King | 2020-03-05 | Washington State Department of Health | Seattle Flu Study | Chu et al |
| EPI_ISL_417123 | hCoV-19/USA/WA-S70/2020 | North America / USA / Washington / King | 2020-03-05 | Washington State Department of Health | Seattle Flu Study | Chu et al |
| EPI_ISL_417124 | hCoV-19/USA/WA-S71/2020 | North America / USA / Washington / King | 2020-03-05 | Washington State Department of Health | Seattle Flu Study | Chu et al |
| EPI_ISL_417125 | hCoV-19/USA/WA-S72/2020 | North America / USA / Washington / King | 2020-03-06 | Washington State Department of Health | Seattle Flu Study | Chu et al |
| EPI_ISL_417126 | hCoV-19/USA/WA-S73/2020 | North America / USA / Washington | 2020-03-05 | Washington State Department of Health | Seattle Flu Study | Chu et al |
| EPI_ISL_417127 | hCoV-19/USA/WA-S74/2020 | North America / USA / Washington / King | 2020-03-05 | Washington State Department of Health | Seattle Flu Study | Chu et al |
| EPI_ISL_417128 | hCoV-19/USA/WA-S75/2020 | North America / USA / Washington / King | 2020-03-05 | Washington State Department of Health | Seattle Flu Study | Chu et al |
| EPI_ISL_417129 | hCoV-19/USA/WA-S76/2020 | North America / USA / Washington / King | 2020-03-05 | Washington State Department of Health | Seattle Flu Study | Chu et al |
| EPI_ISL_417130 | hCoV-19/USA/WA-S77/2020 | North America / USA / Washington / King | 2020-03-05 | Washington State Department of Health | Seattle Flu Study | Chu et al |
| EPI_ISL_417132 | hCoV-19/USA/WA-S79/2020 | North America / USA / Washington / King | 2020-03-05 | Washington State Department of Health | Seattle Flu Study | Chu et al |
| EPI_ISL_417133 | hCoV-19/USA/WA-S80/2020 | North America / USA / Washington / King | 2020-03-05 | Washington State Department of Health | Seattle Flu Study | Chu et al |
| EPI_ISL_417134 | hCoV-19/USA/WA-S81/2020 | North America / USA / Washington / King | 2020-02-26 | Washington State Department of Health | Seattle Flu Study | Chu et al |
| EPI_ISL_417135 | hCoV-19/USA/WA-S82/2020 | North America / USA / Washington / King | 2020-02-22 | Washington State Department of Health | Seattle Flu Study | Chu et al |
| EPI_ISL_417136 | hCoV-19/USA/WA-S83/2020 | North America / USA / Washington / King | 2020-03-05 | Washington State Department of Health | Seattle Flu Study | Chu et al |
| EPI_ISL_417137 | hCoV-19/USA/WA-S84/2020 | North America / USA / Washington / King | 2020-02-21 | Washington State Department of Health | Seattle Flu Study | Chu et al |
| EPI_ISL_417139 | hCoV-19/USA/WA-S86/2020 | North America / USA / Washington / King | 2020-03-01 | Washington State Department of Health | Seattle Flu Study | Chu et al |
| EPI_ISL_417140 | hCoV-19/USA/WA-S87/2020 | North America / USA / Washington / King | 2020-03-01 | Washington State Department of Health | Seattle Flu Study | Chu et al |
| EPI_ISL_417141 | hCoV-19/USA/WA-S88/2020 | North America / USA / Washington / King | 2020-03-01 | Washington State Department of Health | Seattle Flu Study | Chu et al |
| EPI_ISL_417142 | hCoV-19/USA/WA-S89/2020 | North America / USA / Washington / Uma | 2020-02-29 | Washington State Department of Health | Seattle Flu Study | Chu et al |
| EPI_ISL_417143 | hCoV-19/USA/WA-S90/2020 | North America / USA / Washington / Uma | 2020-02-29 | Washington State Department of Health | Seattle Flu Study | Chu et al |
| EPI_ISL_417144 | hCoV-19/USA/WA-S91/2020 | North America / USA / Washington / Sno | 2020-03-02 | Washington State Department of Health | Seattle Flu Study | Chu et al |
| EPI_ISL_417145 | hCoV-19/USA/WA-S92/2020 | North America / USA / Washington / King | 2020-02-29 | Washington State Department of Health | Seattle Flu Study | Chu et al |
| EPI_ISL_417146 | hCoV-19/USA/WA-S93/2020 | North America / USA / Washington / King | 2020-02-29 | Washington State Department of Health | Seattle Flu Study | Chu et al |
| EPI_ISL_417147 | hCoV-19/USA/WA-S94/2020 | North America / USA / Washington / Sno | 2020-02-28 | Washington State Department of Health | Seattle Flu Study | Chu et al |
| EPI_ISL_417148 | hCoV-19/USA/WA-S95/2020 | North America / USA / Washington / King | 2020-02-28 | Washington State Department of Health | Seattle Flu Study | Chu et al |
| EPI_ISL_417149 | hCoV-19/USA/WA-S96/2020 | North America / USA / Washington / King | 2020-02-28 | Washington State Department of Health | Seattle Flu Study | Chu et al |
| EPI_ISL_417150 | hCoV-19/USA/WA-S97/2020 | North America / USA / Washington / King | 2020-02-28 | Washington State Department of Health | Seattle Flu Study | Chu et al |
| EPI_ISL_417151 | hCoV-19/USA/WA-S98/2020 | North America / USA / Washington / King | 2020-02-29 | Washington State Department of Health | Seattle Flu Study | Chu et al |
| EPI_ISL_417152 | hCoV-19/USA/WA-S99/2020 | North America / USA / Washington / King | 2020-02-28 | Washington State Department of Health | Seattle Flu Study | Chu et al |
| EPI_ISL_417153 | hCoV-19/USA/WA-S100/2020 | North America / USA / Washington / King | 2020-02-29 | Washington State Department of Health | Seattle Flu Study | Chu et al |
| EPI_ISL_417154 | hCoV-19/USA/WA-S101/2020 | North America / USA / Washington / King | 2020-02-28 | Washington State Department of Health | Seattle Flu Study | Chu et al |
| EPI_ISL_417155 | hCoV-19/USA/WA-S102/2020 | North America / USA / Washington / King | 2020-02-28 | Washington State Department of Health | Seattle Flu Study | Chu et al |
| EPI_ISL_417156 | hCoV-19/USA/WA-S103/2020 | North America / USA / Washington / King | 2020-02-28 | Washington State Department of Health | Seattle Flu Study | Chu et al |
| EPI_ISL_417157 | hCoV-19/USA/WA-S104/2020 | North America / USA / Washington / Sno | 2020-03-05 | Washington State Department of Health | Seattle Flu Study | Chu et al |
| EPI_ISL_417158 | hCoV-19/USA/WA-S105/2020 | North America / USA / Washington / King | 2020-02-28 | Washington State Department of Health | Seattle Flu Study | Chu et al |
| EPI_ISL_417159 | hCoV-19/USA/WA-S106/2020 | North America / USA / Washington / King | 2020-02-29 | Washington State Department of Health | Seattle Flu Study | Chu et al |
| EPI_ISL_417160 | hCoV-19/USA/WA-S107/2020 | North America / USA / Washington / King | 2020-02-29 | Washington State Department of Health | Seattle Flu Study | Chu et al |
| EPI_ISL_417161 | hCoV-19/USA/WA-S108/2020 | North America / USA / Washington / King | 2020-02-29 | Washington State Department of Health | Seattle Flu Study | Chu et al |
| EPI_ISL_417162 | hCoV-19/USA/WA-S109/2020 | North America / USA / Washington / Sno | 2020-03-01 | Washington State Department of Health | Seattle Flu Study | Chu et al |
| EPI_ISL_417163 | hCoV-19/USA/WA-S110/2020 | North America / USA / Washington / King | 2020-03-05 | Seattle Flu Study | Seattle Flu Study | Chu et al |
| EPI_ISL_417164 | hCoV-19/USA/WA-S111/2020 | North America / USA / Washington / King | 2020-03-07 | Seattle Flu Study | Seattle Flu Study | Chu et al |
| EPI_ISL_417165 | hCoV-19/USA/WA-S112/2020 | North America / USA / Washington / King | 2020-03-08 | Seattle Flu Study | Seattle Flu Study | Chu et al |
| EPI_ISL_417167 | hCoV-19/USA/WA-S114/2020 | North America / USA / Washington / Sno | 2020-03-05 | Washington State Department of Health | Seattle Flu Study | Chu et al |
| EPI_ISL_417168 | hCoV-19/USA/WA-S115/2020 | North America / USA / Washington / King | 2020-02-29 | Washington State Department of Health | Seattle Flu Study | Chu et al |
| EPI_ISL_417169 | hCoV-19/USA/WA-S116/2020 | North America / USA / Washington | 2020-03-02 | Washington State Department of Health | Seattle Flu Study | Chu et al |
| EPI_ISL_417170 | hCoV-19/USA/WA-S117/2020 | North America / USA / Washington | 2020-03-02 | Washington State Department of Health | Seattle Flu Study | Chu et al |
| EPI_ISL_417171 | hCoV-19/USA/WA-S118/2020 | North America / USA / Washington / King | 2020-03-01 | Washington State Department of Health | Seattle Flu Study | Chu et al |
| EPI_ISL_417172 | hCoV-19/USA/WA-S119/2020 | North America / USA / Washington / King | 2020-02-29 | Washington State Department of Health | Seattle Flu Study | Chu et al |
| EPI_ISL_417173 | hCoV-19/USA/WA-S120/2020 | North America / USA / Washington / King | 2020-03-01 | Washington State Department of Health | Seattle Flu Study | Chu et al |
| EPI_ISL_417174 | hCoV-19/USA/WA-S121/2020 | North America / USA / Washington / King | 2020-03-01 | Washington State Department of Health | Seattle Flu Study | Chu et al |
| EPI_ISL_417175 | hCoV-19/USA/WA-S122/2020 | North America / USA / Washington / Gra | 2020-03-02 | Washington State Department of Health | Seattle Flu Study | Chu et al |
| EPI_ISL_417176 | hCoV-19/Hong Kong/HKPU1_2101/2020 | Asia / Hong Kong | 2020-01-21 | Department of Pathology, Princess Margaret Hospital | Department of Health Technology and Informatics, Faculty of Health and Social Science, The Hong Kong Polytechnic University | Kenneth Siu-Sing LEUNG, Timothy Ting-Leung NG, Alan Ka-Lun WU, Miranda Chong-Yee YAU, Hiu-Yin LAO, Ming-Pan CHOI, Kingsley King-Gee TAM, Lam-Kwong LEE, Barry Kin-Chung WONG, Alex Yat-Man HO, Kam-Tong Yip, Kwok-Cheung LUNG, Raymond Wai-To LIU, Eugene Yuk-Keung TSO, Wai-Shing LEUNG, Man-Chun CHAN, Yuk-Yung NG, Kit-Man SIN, Kitty Sau-Chun FUNG, Sandy Ka-Yee CHAU, Wing-Kin TO, Tak-Lun Que, David Ho-Keung SHUM, Shea Ping YIP, Wing Cheong YAM, Gilman Kit-Hang SIU |
| EPI_ISL_417178 | hCoV-19/Hong Kong/HKPU6_2101/2020 | Asia / Hong Kong | 2020-01-25 | Department of Pathology, Princess Margaret Hospital | Department of Health Technology and Informatics, Faculty of Health and Social Science, The Hong Kong Polytechnic University | Kenneth Siu-Sing LEUNG, Timothy Ting-Leung NG, Alan Ka-Lun WU, Miranda Chong-Yee YAU, Hiu-Yin LAO, Ming-Pan CHOI, Kingsley King-Gee TAM, Lam-Kwong LEE, Barry Kin-Chung WONG, Alex Yat-Man HO, Kam-Tong Yip, Kwok-Cheung LUNG, Raymond Wai-To LIU, Eugene Yuk-Keung TSO, Wai-Shing LEUNG, Man-Chun CHAN, Yuk-Yung NG, Kit-Man SIN, Kitty Sau-Chun FUNG, Sandy Ka-Yee CHAU, Wing-Kin TO, Tak-Lun Que, David Ho-Keung SHUM, Shea Ping YIP, Wing Cheong YAM, Gilman Kit-Hang SIU |
| EPI_ISL_417179 | hCoV-19/Hong Kong/HKPU12_2201/2020 | Asia / Hong Kong | 2020-01-30 | Department of Pathology, Princess Margaret Hospital | Department of Health Technology and Informatics, Faculty of Health and Social Science, The Hong Kong Polytechnic University | Kenneth Siu-Sing LEUNG, Timothy Ting-Leung NG, Alan Ka-Lun WU, Miranda Chong-Yee YAU, Hiu-Yin LAO, Ming-Pan CHOI, Kingsley King-Gee TAM, Lam-Kwong LEE, Barry Kin-Chung WONG, Alex Yat-Man HO, Kam-Tong Yip, Kwok-Cheung LUNG, Raymond Wai-To LIU, Eugene Yuk-Keung TSO, Wai-Shing LEUNG, Man-Chun CHAN, Yuk-Yung NG, Kit-Man SIN, Kitty Sau-Chun FUNG, Sandy Ka-Yee CHAU, Wing-Kin TO, Tak-Lun Que, David Ho-Keung SHUM, Shea Ping YIP, Wing Cheong YAM, Gilman Kit-Hang SIU |
| EPI_ISL_417181 | hCoV-19/Hong Kong/HKPU19_0402/2020 | Asia / Hong Kong | 2020-02-05 | Department of Pathology, United Christian Hospital | Department of Health Technology and Informatics, Faculty of Health and Social Science, The Hong Kong Polytechnic University | Kenneth Siu-Sing LEUNG, Timothy Ting-Leung NG, Alan Ka-Lun WU, Miranda Chong-Yee YAU, Hiu-Yin LAO, Ming-Pan CHOI, Kingsley King-Gee TAM, Lam-Kwong LEE, Barry Kin-Chung WONG, Alex Yat-Man HO, Kam-Tong Yip, Kwok-Cheung LUNG, Raymond Wai-To LIU, Eugene Yuk-Keung TSO, Wai-Shing LEUNG, Man-Chun CHAN, Yuk-Yung NG, Kit-Man SIN, Kitty Sau-Chun FUNG, Sandy Ka-Yee CHAU, Wing-Kin TO, Tak-Lun Que, David Ho-Keung SHUM, Shea Ping YIP, Wing Cheong YAM, Gilman Kit-Hang SIU |
| EPI_ISL_417185 | hCoV-19/Hong Kong/HKPU28_3001/2020 | Asia / Hong Kong | 2020-02-09 | Department of Pathology, United Christian Hospital | Department of Health Technology and Informatics, Faculty of Health and Social Science, The Hong Kong Polytechnic University | Kenneth Siu-Sing LEUNG, Timothy Ting-Leung NG, Alan Ka-Lun WU, Miranda Chong-Yee YAU, Hiu-Yin LAO, Ming-Pan CHOI, Kingsley King-Gee TAM, Lam-Kwong LEE, Barry Kin-Chung WONG, Alex Yat-Man HO, Kam-Tong Yip, Kwok-Cheung LUNG, Raymond Wai-To LIU, Eugene Yuk-Keung TSO, Wai-Shing LEUNG, Man-Chun CHAN, Yuk-Yung NG, Kit-Man SIN, Kitty Sau-Chun FUNG, Sandy Ka-Yee CHAU, Wing-Kin TO, Tak-Lun Que, David Ho-Keung SHUM, Shea Ping YIP, Wing Cheong YAM, Gilman Kit-Hang SIU |
| EPI_ISL_417186 | hCoV-19/South Africa/R03006/2020 | Africa / South Africa / KZN | 2020-03-07 | National Institute for Communicable Diseases of the National Health Laboratory Service | National Institute for Communicable Diseases of the National Health Laboratory Service | Allam M, Kwenda S, van Heusden P, Khumalo Z, Mohale T, Subramoney K, von Gottberg, A, Ismail A, Bhiman JN |
| EPI_ISL_417187 | hCoV-19/Hong Kong/HKPU29_0102/2020 | Asia / Hong Kong | 2020-02-08 | Department of Clinical Pathology, Pamela Youde Nethersole Eastern Hospital | Department of Health Technology and Informatics, Faculty of Health and Social Science, The Hong Kong Polytechnic University | Kenneth Siu-Sing LEUNG, Timothy Ting-Leung NG, Alan Ka-Lun WU, Miranda Chong-Yee YAU, Hiu-Yin LAO, Ming-Pan CHOI, Kingsley King-Gee TAM, Lam-Kwong LEE, Barry Kin-Chung WONG, Alex Yat-Man HO, Kam-Tong Yip, Kwok-Cheung LUNG, Raymond Wai-To LIU, Eugene Yuk-Keung TSO, Wai-Shing LEUNG, Man-Chun CHAN, Yuk-Yung NG, Kit-Man SIN, Kitty Sau-Chun FUNG, Sandy Ka-Yee CHAU, Wing-Kin TO, Tak-Lun Que, David Ho-Keung SHUM, Shea Ping YIP, Wing Cheong YAM, Gilman Kit-Hang SIU |
| EPI_ISL_417188 | hCoV-19/Hong Kong/HKPU30_2901/2020 | Asia / Hong Kong | 2020-02-08 | Department of Clinical Pathology, Pamela Youde Nethersole Eastern Hospital | Department of Health Technology and Informatics, Faculty of Health and Social Science, The Hong Kong Polytechnic University | Kenneth Siu-Sing LEUNG, Timothy Ting-Leung NG, Alan Ka-Lun WU, Miranda Chong-Yee YAU, Hiu-Yin LAO, Ming-Pan CHOI, Kingsley King-Gee TAM, Lam-Kwong LEE, Barry Kin-Chung WONG, Alex Yat-Man HO, Kam-Tong Yip, Kwok-Cheung LUNG, Raymond Wai-To LIU, Eugene Yuk-Keung TSO, Wai-Shing LEUNG, Man-Chun CHAN, Yuk-Yung NG, Kit-Man SIN, Kitty Sau-Chun FUNG, Sandy Ka-Yee CHAU, Wing-Kin TO, Tak-Lun Que, David Ho-Keung SHUM, Shea Ping YIP, Wing Cheong YAM, Gilman Kit-Hang SIU |
| EPI_ISL_417191 | hCoV-19/USA/MN5-MDH5/2020 | North America / USA / Minnesota | 2020-03-10 | Minnesota Department of Health, Public Health Laboratory | Minnesota Department of Health, Public Health Laboratory | Matt Plumb, Jake Garfin and Xiong Wang |
| EPI_ISL_417192 | hCoV-19/USA/MN25-MDH25/2020 | North America / USA / Minnesota | 2020-03-12 | Minnesota Department of Health, Public Health Laboratory | Minnesota Department of Health, Public Health Laboratory | Matt Plumb, Jake Garfin and Xiong Wang |
| EPI_ISL_417193 | hCoV-19/Hong Kong/HKPU32_0402/2020 | Asia / Hong Kong | 2020-02-09 | Department of Clinical Pathology, Pamela Youde Nethersole Eastern Hospital | Department of Health Technology and Informatics, Faculty of Health and Social Science, The Hong Kong Polytechnic University | Kenneth Siu-Sing LEUNG, Timothy Ting-Leung NG, Alan Ka-Lun WU, Miranda Chong-Yee YAU, Hiu-Yin LAO, Ming-Pan CHOI, Kingsley King-Gee TAM, Lam-Kwong LEE, Barry Kin-Chung WONG, Alex Yat-Man HO, Kam-Tong Yip, Kwok-Cheung LUNG, Raymond Wai-To LIU, Eugene Yuk-Keung TSO, Wai-Shing LEUNG, Man-Chun CHAN, Yuk-Yung NG, Kit-Man SIN, Kitty Sau-Chun FUNG, Sandy Ka-Yee CHAU, Wing-Kin TO, Tak-Lun Que, David Ho-Keung SHUM, Shea Ping YIP, Wing Cheong YAM, Gilman Kit-Hang SIU |
| EPI_ISL_417194 | hCoV-19/USA/MN26-MDH26/2020 | North America / USA / Minnesota | 2020-03-12 | Minnesota Department of Health, Public Health Laboratory | Minnesota Department of Health, Public Health Laboratory | Matt Plumb, Jake Garfin and Xiong Wang |
| EPI_ISL_417196 | hCoV-19/USA/MN29-MDH29/2020 | North America / USA / Minnesota | 2020-03-11 | Minnesota Department of Health, Public Health Laboratory | Minnesota Department of Health, Public Health Laboratory | Matt Plumb, Jake Garfin and Xiong Wang |
| EPI_ISL_417197 | hCoV-19/Hong Kong/HKPU34_3001/2020 | Asia / Hong Kong | 2020-02-09 | Department of Clinical Pathology, Pamela Youde Nethersole Eastern Hospital | Department of Health Technology and Informatics, Faculty of Health and Social Science, The Hong Kong Polytechnic University | Kenneth Siu-Sing LEUNG, Timothy Ting-Leung NG, Alan Ka-Lun WU, Miranda Chong-Yee YAU, Hiu-Yin LAO, Ming-Pan CHOI, Kingsley King-Gee TAM, Lam-Kwong LEE, Barry Kin-Chung WONG, Alex Yat-Man HO, Kam-Tong Yip, Kwok-Cheung LUNG, Raymond Wai-To LIU, Eugene Yuk-Keung TSO, Wai-Shing LEUNG, Man-Chun CHAN, Yuk-Yung NG, Kit-Man SIN, Kitty Sau-Chun FUNG, Sandy Ka-Yee CHAU, Wing-Kin TO, Tak-Lun Que, David Ho-Keung SHUM, Shea Ping YIP, Wing Cheong YAM, Gilman Kit-Hang SIU |
| EPI_ISL_417200 | hCoV-19/USA/WI-UW-06/2020 | North America / USA / Wisconsin / Cottag | 2020-03-21 | University of Wisconsin-Madison AIDS Vaccine Research Laboratories | University of Wisconsin-Madison AIDS Vaccine Research Laboratories | Gage Moreno, Katarina Braun, et al. AIDS Vaccine Research Laboratories |
| EPI_ISL_417201 | hCoV-19/USA/WI-UW-07/2020 | North America / USA / Wisconsin / Madis | 2020-03-21 | University of Wisconsin-Madison AIDS Vaccine Research Laboratories | University of Wisconsin-Madison AIDS Vaccine Research Laboratories | Gage Moreno, Katarina Braun, et al. AIDS Vaccine Research Laboratories |
| EPI_ISL_417202 | hCoV-19/USA/WI-UW-08/2020 | North America / USA / Wisconsin / Stoug | 2020-03-21 | University of Wisconsin-Madison AIDS Vaccine Research Laboratories | University of Wisconsin-Madison AIDS Vaccine Research Laboratories | Gage Moreno, Katarina Braun, et al. AIDS Vaccine Research Laboratories |
| EPI_ISL_417203 | hCoV-19/USA/WI-UW-09/2020 | North America / USA / Wisconsin / Madis | 2020-03-21 | University of Wisconsin-Madison AIDS Vaccine Research Laboratories | University of Wisconsin-Madison AIDS Vaccine Research Laboratories | Gage Moreno, Katarina Braun, et al. AIDS Vaccine Research Laboratories |
| EPI_ISL_417204 | hCoV-19/USA/WI-UW-10/2020 | North America / USA / Wisconsin / Camb | 2020-03-21 | University of Wisconsin-Madison AIDS Vaccine Research Laboratories | University of Wisconsin-Madison AIDS Vaccine Research Laboratories | Gage Moreno, Katarina Braun, et al. AIDS Vaccine Research Laboratories |
| EPI_ISL_417205 | hCoV-19/Spain/Valencia9/2020 | Europe / Spain / Comunitat Valenciana / ' | 2020-03-02 | Servicio de Microbiologia. Consorcio Hospital General Universitario de Valencia | Sequencing and Bioinformatics Service and Molecular Epidemiology Research Group. FISABIO-Public Health | Maria Alma Bracho, Maria Dolores Ocete, Concepcion Gimeno, Giuseppe D'Auria, Griselda De Marco, Neris Garcia-Gonzalez, Fernando Gonzalez-Candelas |
| EPI_ISL_417206 | hCoV-19/Spain/Valencia10/2020 | Europe / Spain / Comunitat Valenciana / ' | 2020-03-06 | Servicio de Microbiologia. Consorcio Hospital General Universitario de Valencia | Sequencing and Bioinformatics Service and Molecular Epidemiology Research Group. FISABIO-Public Health | Maria Alma Bracho, Maria Dolores Ocete, Concepcion Gimeno, Giuseppe D'Auria, Griselda De Marco, Neris Garcia-Gonzalez, Fernando Gonzalez-Candelas |
| EPI_ISL_417211 | hCoV-19/NewZealand/CoV001/2020 | Oceania / New Zealand / Otago | 2020-03-11 | Dunedin Hospital | University of Otago | M.E. Quiñones-Mateu, B. Lawley, J. Grant, R. Harfoot, J. Ussher |
| EPI_ISL_417213 | hCoV-19/England/20102068502/2020 | Europe / United Kingdom / England | 2020-03-01 | Respiratory Virus Unit, Microbiology Services Colindale, Public Health England | Respiratory Virus Unit, Microbiology Services Colindale, Public Health England | Monica Galiano, Shahjahan Miah, Angie Lackenby, Omolola Akinbami, Tiina Talts, Leena Bhaw, Richard Myers, Steven Platt, Kirstin Edwards, Jonathan Hubb, Joanna Ellis, Maria Zambon |
| EPI_ISL_417217 | hCoV-19/England/20102098802/2020 | Europe / United Kingdom / England | 2020-03-01 | Respiratory Virus Unit, Microbiology Services Colindale, Public Health England | Respiratory Virus Unit, Microbiology Services Colindale, Public Health England | Monica Galiano, Shahjahan Miah, Angie Lackenby, Omolola Akinbami, Tiina Talts, Leena Bhaw, Richard Myers, Steven Platt, Kirstin Edwards, Jonathan Hubb, Joanna Ellis, Maria Zambon |
| EPI_ISL_417220 | hCoV-19/England/20104002606/2020 | Europe / United Kingdom / England | 2020-03-03 | Respiratory Virus Unit, Microbiology Services Colindale, Public Health England | Respiratory Virus Unit, Microbiology Services Colindale, Public Health England | Monica Galiano, Shahjahan Miah, Angie Lackenby, Omolola Akinbami, Tiina Talts, Leena Bhaw, Richard Myers, Steven Platt, Kirstin Edwards, Jonathan Hubb, Joanna Ellis, Maria Zambon |
| EPI_ISL_417222 | hCoV-19/England/20104003002/2020 | Europe / United Kingdom / England | 2020-03-04 | Respiratory Virus Unit, Microbiology Services Colindale, Public Health England | Respiratory Virus Unit, Microbiology Services Colindale, Public Health England | Monica Galiano, Shahjahan Miah, Angie Lackenby, Omolola Akinbami, Tiina Talts, Leena Bhaw, Richard Myers, Steven Platt, Kirstin Edwards, Jonathan Hubb, Joanna Ellis, Maria Zambon |
| EPI_ISL_417226 | hCoV-19/England/20104004902/2020 | Europe / United Kingdom / England | 2020-03-02 | Respiratory Virus Unit, Microbiology Services Colindale, Public Health England | Respiratory Virus Unit, Microbiology Services Colindale, Public Health England | Monica Galiano, Shahjahan Miah, Angie Lackenby, Omolola Akinbami, Tiina Talts, Leena Bhaw, Richard Myers, Steven Platt, Kirstin Edwards, Jonathan Hubb, Joanna Ellis, Maria Zambon |
| EPI_ISL_417227 | hCoV-19/England/20104007503/2020 | Europe / United Kingdom / England | 2020-03-03 | Respiratory Virus Unit, Microbiology Services Colindale, Public Health England | Respiratory Virus Unit, Microbiology Services Colindale, Public Health England | Monica Galiano, Shahjahan Miah, Angie Lackenby, Omolola Akinbami, Tiina Talts, Leena Bhaw, Richard Myers, Steven Platt, Kirstin Edwards, Jonathan Hubb, Joanna Ellis, Maria Zambon |
| EPI_ISL_417228 | hCoV-19/England/20104008402/2020 | Europe / United Kingdom / England | 2020-03-03 | Respiratory Virus Unit, Microbiology Services Colindale, Public Health England | Respiratory Virus Unit, Microbiology Services Colindale, Public Health England | Monica Galiano, Shahjahan Miah, Angie Lackenby, Omolola Akinbami, Tiina Talts, Leena Bhaw, Richard Myers, Steven Platt, Kirstin Edwards, Jonathan Hubb, Joanna Ellis, Maria Zambon |
| EPI_ISL_417230 | hCoV-19/England/20104008702/2020 | Europe / United Kingdom / England | 2020-03-02 | Respiratory Virus Unit, Microbiology Services Colindale, Public Health England | Respiratory Virus Unit, Microbiology Services Colindale, Public Health England | Monica Galiano, Shahjahan Miah, Angie Lackenby, Omolola Akinbami, Tiina Talts, Leena Bhaw, Richard Myers, Steven Platt, Kirstin Edwards, Jonathan Hubb, Joanna Ellis, Maria Zambon |
| EPI_ISL_417231 | hCoV-19/England/20104008802/2020 | Europe / United Kingdom / England | 2020-03-03 | Respiratory Virus Unit, Microbiology Services Colindale, Public Health England | Respiratory Virus Unit, Microbiology Services Colindale, Public Health England | Monica Galiano, Shahjahan Miah, Angie Lackenby, Omolola Akinbami, Tiina Talts, Leena Bhaw, Richard Myers, Steven Platt, Kirstin Edwards, Jonathan Hubb, Joanna Ellis, Maria Zambon |
| EPI_ISL_417232 | hCoV-19/England/20104008902/2020 | Europe / United Kingdom / England | 2020-03-04 | Respiratory Virus Unit, Microbiology Services Colindale, Public Health England | Respiratory Virus Unit, Microbiology Services Colindale, Public Health England | Monica Galiano, Shahjahan Miah, Angie Lackenby, Omolola Akinbami, Tiina Talts, Leena Bhaw, Richard Myers, Steven Platt, Kirstin Edwards, Jonathan Hubb, Joanna Ellis, Maria Zambon |
| EPI_ISL_417233 | hCoV-19/England/20104009002/2020 | Europe / United Kingdom / England | 2020-03-02 | Respiratory Virus Unit, Microbiology Services Colindale, Public Health England | Respiratory Virus Unit, Microbiology Services Colindale, Public Health England | Monica Galiano, Shahjahan Miah, Angie Lackenby, Omolola Akinbami, Tiina Talts, Leena Bhaw, Richard Myers, Steven Platt, Kirstin Edwards, Jonathan Hubb, Joanna Ellis, Maria Zambon |
| EPI_ISL_417234 | hCoV-19/England/20104009102/2020 | Europe / United Kingdom / England | 2020-03-02 | Respiratory Virus Unit, Microbiology Services Colindale, Public Health England | Respiratory Virus Unit, Microbiology Services Colindale, Public Health England | Monica Galiano, Shahjahan Miah, Angie Lackenby, Omolola Akinbami, Tiina Talts, Leena Bhaw, Richard Myers, Steven Platt, Kirstin Edwards, Jonathan Hubb, Joanna Ellis, Maria Zambon |
| EPI_ISL_417235 | hCoV-19/England/20104013703/2020 | Europe / United Kingdom / England | 2020-03-02 | Respiratory Virus Unit, Microbiology Services Colindale, Public Health England | Respiratory Virus Unit, Microbiology Services Colindale, Public Health England | Monica Galiano, Shahjahan Miah, Angie Lackenby, Omolola Akinbami, Tiina Talts, Leena Bhaw, Richard Myers, Steven Platt, Kirstin Edwards, Jonathan Hubb, Joanna Ellis, Maria Zambon |
| EPI_ISL_417236 | hCoV-19/England/20104015302/2020 | Europe / United Kingdom / England | 2020-03-01 | Respiratory Virus Unit, Microbiology Services Colindale, Public Health England | Respiratory Virus Unit, Microbiology Services Colindale, Public Health England | Monica Galiano, Shahjahan Miah, Angie Lackenby, Omolola Akinbami, Tiina Talts, Leena Bhaw, Richard Myers, Steven Platt, Kirstin Edwards, Jonathan Hubb, Joanna Ellis, Maria Zambon |
| EPI_ISL_417238 | hCoV-19/England/20104035803/2020 | Europe / United Kingdom / England | 2020-03-03 | Respiratory Virus Unit, Microbiology Services Colindale, Public Health England | Respiratory Virus Unit, Microbiology Services Colindale, Public Health England | Monica Galiano, Shahjahan Miah, Angie Lackenby, Omolola Akinbami, Tiina Talts, Leena Bhaw, Richard Myers, Steven Platt, Kirstin Edwards, Jonathan Hubb, Joanna Ellis, Maria Zambon |
| EPI_ISL_417239 | hCoV-19/England/20106003303/2020 | Europe / United Kingdom / England | 2020-03-03 | Respiratory Virus Unit, Microbiology Services Colindale, Public Health England | Respiratory Virus Unit, Microbiology Services Colindale, Public Health England | Monica Galiano, Shahjahan Miah, Angie Lackenby, Omolola Akinbami, Tiina Talts, Leena Bhaw, Richard Myers, Steven Platt, Kirstin Edwards, Jonathan Hubb, Joanna Ellis, Maria Zambon |
| EPI_ISL_417240 | hCoV-19/England/20106004803/2020 | Europe / United Kingdom / England | 2020-03-03 | Respiratory Virus Unit, Microbiology Services Colindale, Public Health England | Respiratory Virus Unit, Microbiology Services Colindale, Public Health England | Monica Galiano, Shahjahan Miah, Angie Lackenby, Omolola Akinbami, Tiina Talts, Leena Bhaw, Richard Myers, Steven Platt, Kirstin Edwards, Jonathan Hubb, Joanna Ellis, Maria Zambon |
| EPI_ISL_417242 | hCoV-19/England/20106005103/2020 | Europe / United Kingdom / England | 2020-03-03 | Respiratory Virus Unit, Microbiology Services Colindale, Public Health England | Respiratory Virus Unit, Microbiology Services Colindale, Public Health England | Monica Galiano, Shahjahan Miah, Angie Lackenby, Omolola Akinbami, Tiina Talts, Leena Bhaw, Richard Myers, Steven Platt, Kirstin Edwards, Jonathan Hubb, Joanna Ellis, Maria Zambon |
| EPI_ISL_417244 | hCoV-19/England/20106005403/2020 | Europe / United Kingdom / England | 2020-03-03 | Respiratory Virus Unit, Microbiology Services Colindale, Public Health England | Respiratory Virus Unit, Microbiology Services Colindale, Public Health England | Monica Galiano, Shahjahan Miah, Angie Lackenby, Omolola Akinbami, Tiina Talts, Leena Bhaw, Richard Myers, Steven Platt, Kirstin Edwards, Jonathan Hubb, Joanna Ellis, Maria Zambon |
| EPI_ISL_417246 | hCoV-19/England/20106145903/2020 | Europe / United Kingdom / England | 2020-03-03 | Respiratory Virus Unit, Microbiology Services Colindale, Public Health England | Respiratory Virus Unit, Microbiology Services Colindale, Public Health England | Monica Galiano, Shahjahan Miah, Angie Lackenby, Omolola Akinbami, Tiina Talts, Leena Bhaw, Richard Myers, Steven Platt, Kirstin Edwards, Jonathan Hubb, Joanna Ellis, Maria Zambon |
| EPI_ISL_417248 | hCoV-19/England/20108003302/2020 | Europe / United Kingdom / England | 2020-03-03 | Respiratory Virus Unit, Microbiology Services Colindale, Public Health England | Respiratory Virus Unit, Microbiology Services Colindale, Public Health England | Monica Galiano, Shahjahan Miah, Angie Lackenby, Omolola Akinbami, Tiina Talts, Leena Bhaw, Richard Myers, Steven Platt, Kirstin Edwards, Jonathan Hubb, Joanna Ellis, Maria Zambon |
| EPI_ISL_417250 | hCoV-19/England/20108004702/2020 | Europe / United Kingdom / England | 2020-03-02 | Respiratory Virus Unit, Microbiology Services Colindale, Public Health England | Respiratory Virus Unit, Microbiology Services Colindale, Public Health England | Monica Galiano, Shahjahan Miah, Angie Lackenby, Omolola Akinbami, Tiina Talts, Leena Bhaw, Richard Myers, Steven Platt, Kirstin Edwards, Jonathan Hubb, Joanna Ellis, Maria Zambon |
| EPI_ISL_417252 | hCoV-19/England/20108006003/2020 | Europe / United Kingdom / England | 2020-03-05 | Respiratory Virus Unit, Microbiology Services Colindale, Public Health England | Respiratory Virus Unit, Microbiology Services Colindale, Public Health England | Monica Galiano, Shahjahan Miah, Angie Lackenby, Omolola Akinbami, Tiina Talts, Leena Bhaw, Richard Myers, Steven Platt, Kirstin Edwards, Jonathan Hubb, Joanna Ellis, Maria Zambon |
| EPI_ISL_417254 | hCoV-19/England/20108006802/2020 | Europe / United Kingdom / England | 2020-03-04 | Respiratory Virus Unit, Microbiology Services Colindale, Public Health England | Respiratory Virus Unit, Microbiology Services Colindale, Public Health England | Monica Galiano, Shahjahan Miah, Angie Lackenby, Omolola Akinbami, Tiina Talts, Leena Bhaw, Richard Myers, Steven Platt, Kirstin Edwards, Jonathan Hubb, Joanna Ellis, Maria Zambon |
| EPI_ISL_417255 | hCoV-19/England/20108007002/2020 | Europe / United Kingdom / England | 2020-03-04 | Respiratory Virus Unit, Microbiology Services Colindale, Public Health England | Respiratory Virus Unit, Microbiology Services Colindale, Public Health England | Monica Galiano, Shahjahan Miah, Angie Lackenby, Omolola Akinbami, Tiina Talts, Leena Bhaw, Richard Myers, Steven Platt, Kirstin Edwards, Jonathan Hubb, Joanna Ellis, Maria Zambon |
| EPI_ISL_417256 | hCoV-19/England/20108007302/2020 | Europe / United Kingdom / England | 2020-03-03 | Respiratory Virus Unit, Microbiology Services Colindale, Public Health England | Respiratory Virus Unit, Microbiology Services Colindale, Public Health England | Monica Galiano, Shahjahan Miah, Angie Lackenby, Omolola Akinbami, Tiina Talts, Leena Bhaw, Richard Myers, Steven Platt, Kirstin Edwards, Jonathan Hubb, Joanna Ellis, Maria Zambon |
| EPI_ISL_417258 | hCoV-19/England/20108034006/2020 | Europe / United Kingdom / England | 2020-03-06 | Respiratory Virus Unit, Microbiology Services Colindale, Public Health England | Respiratory Virus Unit, Microbiology Services Colindale, Public Health England | Monica Galiano, Shahjahan Miah, Angie Lackenby, Omolola Akinbami, Tiina Talts, Leena Bhaw, Richard Myers, Steven Platt, Kirstin Edwards, Jonathan Hubb, Joanna Ellis, Maria Zambon |
| EPI_ISL_417260 | hCoV-19/England/20109035906/2020 | Europe / United Kingdom / England | 2020-03-05 | Respiratory Virus Unit, Microbiology Services Colindale, Public Health England | Respiratory Virus Unit, Microbiology Services Colindale, Public Health England | Monica Galiano, Shahjahan Miah, Angie Lackenby, Omolola Akinbami, Tiina Talts, Leena Bhaw, Richard Myers, Steven Platt, Kirstin Edwards, Jonathan Hubb, Joanna Ellis, Maria Zambon |
| EPI_ISL_417261 | hCoV-19/England/20109038906/2020 | Europe / United Kingdom / England | 2020-03-06 | Respiratory Virus Unit, Microbiology Services Colindale, Public Health England | Respiratory Virus Unit, Microbiology Services Colindale, Public Health England | Monica Galiano, Shahjahan Miah, Angie Lackenby, Omolola Akinbami, Tiina Talts, Leena Bhaw, Richard Myers, Steven Platt, Kirstin Edwards, Jonathan Hubb, Joanna Ellis, Maria Zambon |
| EPI_ISL_417262 | hCoV-19/England/20109039306/2020 | Europe / United Kingdom / England | 2020-03-06 | Respiratory Virus Unit, Microbiology Services Colindale, Public Health England | Respiratory Virus Unit, Microbiology Services Colindale, Public Health England | Monica Galiano, Shahjahan Miah, Angie Lackenby, Omolola Akinbami, Tiina Talts, Leena Bhaw, Richard Myers, Steven Platt, Kirstin Edwards, Jonathan Hubb, Joanna Ellis, Maria Zambon |
| EPI_ISL_417263 | hCoV-19/England/20109050106/2020 | Europe / United Kingdom / England | 2020-03-06 | Respiratory Virus Unit, Microbiology Services Colindale, Public Health England | Respiratory Virus Unit, Microbiology Services Colindale, Public Health England | Monica Galiano, Shahjahan Miah, Angie Lackenby, Omolola Akinbami, Tiina Talts, Leena Bhaw, Richard Myers, Steven Platt, Kirstin Edwards, Jonathan Hubb, Joanna Ellis, Maria Zambon |
| EPI_ISL_417264 | hCoV-19/England/20109050306/2020 | Europe / United Kingdom / England | 2020-03-04 | Respiratory Virus Unit, Microbiology Services Colindale, Public Health England | Respiratory Virus Unit, Microbiology Services Colindale, Public Health England | Monica Galiano, Shahjahan Miah, Angie Lackenby, Omolola Akinbami, Tiina Talts, Leena Bhaw, Richard Myers, Steven Platt, Kirstin Edwards, Jonathan Hubb, Joanna Ellis, Maria Zambon |
| EPI_ISL_417265 | hCoV-19/England/20109050406/2020 | Europe / United Kingdom / England | 2020-03-08 | Respiratory Virus Unit, Microbiology Services Colindale, Public Health England | Respiratory Virus Unit, Microbiology Services Colindale, Public Health England | Monica Galiano, Shahjahan Miah, Angie Lackenby, Omolola Akinbami, Tiina Talts, Leena Bhaw, Richard Myers, Steven Platt, Kirstin Edwards, Jonathan Hubb, Joanna Ellis, Maria Zambon |
| EPI_ISL_417266 | hCoV-19/England/20109050506/2020 | Europe / United Kingdom / England | 2020-03-05 | Respiratory Virus Unit, Microbiology Services Colindale, Public Health England | Respiratory Virus Unit, Microbiology Services Colindale, Public Health England | Monica Galiano, Shahjahan Miah, Angie Lackenby, Omolola Akinbami, Tiina Talts, Leena Bhaw, Richard Myers, Steven Platt, Kirstin Edwards, Jonathan Hubb, Joanna Ellis, Maria Zambon |
| EPI_ISL_417267 | hCoV-19/England/20109050606/2020 | Europe / United Kingdom / England | 2020-03-05 | Respiratory Virus Unit, Microbiology Services Colindale, Public Health England | Respiratory Virus Unit, Microbiology Services Colindale, Public Health England | Monica Galiano, Shahjahan Miah, Angie Lackenby, Omolola Akinbami, Tiina Talts, Leena Bhaw, Richard Myers, Steven Platt, Kirstin Edwards, Jonathan Hubb, Joanna Ellis, Maria Zambon |
| EPI_ISL_417268 | hCoV-19/England/20109050706/2020 | Europe / United Kingdom / England | 2020-03-05 | Respiratory Virus Unit, Microbiology Services Colindale, Public Health England | Respiratory Virus Unit, Microbiology Services Colindale, Public Health England | Monica Galiano, Shahjahan Miah, Angie Lackenby, Omolola Akinbami, Tiina Talts, Leena Bhaw, Richard Myers, Steven Platt, Kirstin Edwards, Jonathan Hubb, Joanna Ellis, Maria Zambon |
| EPI_ISL_417269 | hCoV-19/England/20109050806/2020 | Europe / United Kingdom / England | 2020-03-05 | Respiratory Virus Unit, Microbiology Services Colindale, Public Health England | Respiratory Virus Unit, Microbiology Services Colindale, Public Health England | Monica Galiano, Shahjahan Miah, Angie Lackenby, Omolola Akinbami, Tiina Talts, Leena Bhaw, Richard Myers, Steven Platt, Kirstin Edwards, Jonathan Hubb, Joanna Ellis, Maria Zambon |
| EPI_ISL_417270 | hCoV-19/England/20109051806/2020 | Europe / United Kingdom / England | 2020-03-06 | Respiratory Virus Unit, Microbiology Services Colindale, Public Health England | Respiratory Virus Unit, Microbiology Services Colindale, Public Health England | Monica Galiano, Shahjahan Miah, Angie Lackenby, Omolola Akinbami, Tiina Talts, Leena Bhaw, Richard Myers, Steven Platt, Kirstin Edwards, Jonathan Hubb, Joanna Ellis, Maria Zambon |
| EPI_ISL_417272 | hCoV-19/England/20109052006/2020 | Europe / United Kingdom / England | 2020-03-06 | Respiratory Virus Unit, Microbiology Services Colindale, Public Health England | Respiratory Virus Unit, Microbiology Services Colindale, Public Health England | Monica Galiano, Shahjahan Miah, Angie Lackenby, Omolola Akinbami, Tiina Talts, Leena Bhaw, Richard Myers, Steven Platt, Kirstin Edwards, Jonathan Hubb, Joanna Ellis, Maria Zambon |
| EPI_ISL_417276 | hCoV-19/England/20109052506/2020 | Europe / United Kingdom / England | 2020-03-07 | Respiratory Virus Unit, Microbiology Services Colindale, Public Health England | Respiratory Virus Unit, Microbiology Services Colindale, Public Health England | Monica Galiano, Shahjahan Miah, Angie Lackenby, Omolola Akinbami, Tiina Talts, Leena Bhaw, Richard Myers, Steven Platt, Kirstin Edwards, Jonathan Hubb, Joanna Ellis, Maria Zambon |
| EPI_ISL_417277 | hCoV-19/England/20109053106/2020 | Europe / United Kingdom / England | 2020-03-07 | Respiratory Virus Unit, Microbiology Services Colindale, Public Health England | Respiratory Virus Unit, Microbiology Services Colindale, Public Health England | Monica Galiano, Shahjahan Miah, Angie Lackenby, Omolola Akinbami, Tiina Talts, Leena Bhaw, Richard Myers, Steven Platt, Kirstin Edwards, Jonathan Hubb, Joanna Ellis, Maria Zambon |
| EPI_ISL_417278 | hCoV-19/England/20109053406/2020 | Europe / United Kingdom / England | 2020-03-07 | Respiratory Virus Unit, Microbiology Services Colindale, Public Health England | Respiratory Virus Unit, Microbiology Services Colindale, Public Health England | Monica Galiano, Shahjahan Miah, Angie Lackenby, Omolola Akinbami, Tiina Talts, Leena Bhaw, Richard Myers, Steven Platt, Kirstin Edwards, Jonathan Hubb, Joanna Ellis, Maria Zambon |
| EPI_ISL_417279 | hCoV-19/England/20109053606/2020 | Europe / United Kingdom / England | 2020-03-07 | Respiratory Virus Unit, Microbiology Services Colindale, Public Health England | Respiratory Virus Unit, Microbiology Services Colindale, Public Health England | Monica Galiano, Shahjahan Miah, Angie Lackenby, Omolola Akinbami, Tiina Talts, Leena Bhaw, Richard Myers, Steven Platt, Kirstin Edwards, Jonathan Hubb, Joanna Ellis, Maria Zambon |

| | | | | | | |
|---|---|---|---|---|---|---|
| EPI_ISL_417280 | hCoV-19/England/20109054806/2020 | Europe / United Kingdom / England | 2020-03-07 | Respiratory Virus Unit, Microbiology Services Colindale, Public Health England | Respiratory Virus Unit, Microbiology Services Colindale, Public Health England | Monica Galiano, Shahjahan Miah, Angie Lackenby, Omolola Akinbami, Tiina Talts, Leena Bhaw, Richard Myers, Steven Platt, Kirstin Edwards, Jonathan Hubb, Joanna Ellis, Maria Zambon |
| EPI_ISL_417282 | hCoV-19/England/20109056906/2020 | Europe / United Kingdom / England | 2020-03-05 | Respiratory Virus Unit, Microbiology Services Colindale, Public Health England | Respiratory Virus Unit, Microbiology Services Colindale, Public Health England | Monica Galiano, Shahjahan Miah, Angie Lackenby, Omolola Akinbami, Tiina Talts, Leena Bhaw, Richard Myers, Steven Platt, Kirstin Edwards, Jonathan Hubb, Joanna Ellis, Maria Zambon |
| EPI_ISL_417283 | hCoV-19/England/20109058906/2020 | Europe / United Kingdom / England | 2020-03-06 | Respiratory Virus Unit, Microbiology Services Colindale, Public Health England | Respiratory Virus Unit, Microbiology Services Colindale, Public Health England | Monica Galiano, Shahjahan Miah, Angie Lackenby, Omolola Akinbami, Tiina Talts, Leena Bhaw, Richard Myers, Steven Platt, Kirstin Edwards, Jonathan Hubb, Joanna Ellis, Maria Zambon |
| EPI_ISL_417285 | hCoV-19/England/20109093606/2020 | Europe / United Kingdom / England | 2020-03-06 | Respiratory Virus Unit, Microbiology Services Colindale, Public Health England | Respiratory Virus Unit, Microbiology Services Colindale, Public Health England | Monica Galiano, Shahjahan Miah, Angie Lackenby, Omolola Akinbami, Tiina Talts, Leena Bhaw, Richard Myers, Steven Platt, Kirstin Edwards, Jonathan Hubb, Joanna Ellis, Maria Zambon |
| EPI_ISL_417286 | hCoV-19/England/20109093706/2020 | Europe / United Kingdom / England | 2020-03-05 | Respiratory Virus Unit, Microbiology Services Colindale, Public Health England | Respiratory Virus Unit, Microbiology Services Colindale, Public Health England | Monica Galiano, Shahjahan Miah, Angie Lackenby, Omolola Akinbami, Tiina Talts, Leena Bhaw, Richard Myers, Steven Platt, Kirstin Edwards, Jonathan Hubb, Joanna Ellis, Maria Zambon |
| EPI_ISL_417287 | hCoV-19/England/20109093806/2020 | Europe / United Kingdom / England | 2020-03-06 | Respiratory Virus Unit, Microbiology Services Colindale, Public Health England | Respiratory Virus Unit, Microbiology Services Colindale, Public Health England | Monica Galiano, Shahjahan Miah, Angie Lackenby, Omolola Akinbami, Tiina Talts, Leena Bhaw, Richard Myers, Steven Platt, Kirstin Edwards, Jonathan Hubb, Joanna Ellis, Maria Zambon |
| EPI_ISL_417289 | hCoV-19/England/20109094006/2020 | Europe / United Kingdom / England | 2020-03-06 | Respiratory Virus Unit, Microbiology Services Colindale, Public Health England | Respiratory Virus Unit, Microbiology Services Colindale, Public Health England | Monica Galiano, Shahjahan Miah, Angie Lackenby, Omolola Akinbami, Tiina Talts, Leena Bhaw, Richard Myers, Steven Platt, Kirstin Edwards, Jonathan Hubb, Joanna Ellis, Maria Zambon |
| EPI_ISL_417290 | hCoV-19/England/20109094106/2020 | Europe / United Kingdom / England | 2020-03-06 | Respiratory Virus Unit, Microbiology Services Colindale, Public Health England | Respiratory Virus Unit, Microbiology Services Colindale, Public Health England | Monica Galiano, Shahjahan Miah, Angie Lackenby, Omolola Akinbami, Tiina Talts, Leena Bhaw, Richard Myers, Steven Platt, Kirstin Edwards, Jonathan Hubb, Joanna Ellis, Maria Zambon |
| EPI_ISL_417291 | hCoV-19/England/20109094206/2020 | Europe / United Kingdom / England | 2020-03-08 | Respiratory Virus Unit, Microbiology Services Colindale, Public Health England | Respiratory Virus Unit, Microbiology Services Colindale, Public Health England | Monica Galiano, Shahjahan Miah, Angie Lackenby, Omolola Akinbami, Tiina Talts, Leena Bhaw, Richard Myers, Steven Platt, Kirstin Edwards, Jonathan Hubb, Joanna Ellis, Maria Zambon |
| EPI_ISL_417292 | hCoV-19/England/20109094506/2020 | Europe / United Kingdom / England | 2020-03-08 | Respiratory Virus Unit, Microbiology Services Colindale, Public Health England | Respiratory Virus Unit, Microbiology Services Colindale, Public Health England | Monica Galiano, Shahjahan Miah, Angie Lackenby, Omolola Akinbami, Tiina Talts, Leena Bhaw, Richard Myers, Steven Platt, Kirstin Edwards, Jonathan Hubb, Joanna Ellis, Maria Zambon |
| EPI_ISL_417293 | hCoV-19/England/20109097506/2020 | Europe / United Kingdom / England | 2020-03-08 | Respiratory Virus Unit, Microbiology Services Colindale, Public Health England | Respiratory Virus Unit, Microbiology Services Colindale, Public Health England | Monica Galiano, Shahjahan Miah, Angie Lackenby, Omolola Akinbami, Tiina Talts, Leena Bhaw, Richard Myers, Steven Platt, Kirstin Edwards, Jonathan Hubb, Joanna Ellis, Maria Zambon |
| EPI_ISL_417295 | hCoV-19/England/20109098806/2020 | Europe / United Kingdom / England | 2020-03-08 | Respiratory Virus Unit, Microbiology Services Colindale, Public Health England | Respiratory Virus Unit, Microbiology Services Colindale, Public Health England | Monica Galiano, Shahjahan Miah, Angie Lackenby, Omolola Akinbami, Tiina Talts, Leena Bhaw, Richard Myers, Steven Platt, Kirstin Edwards, Jonathan Hubb, Joanna Ellis, Maria Zambon |
| EPI_ISL_417297 | hCoV-19/England/20109099006/2020 | Europe / United Kingdom / England | 2020-03-06 | Respiratory Virus Unit, Microbiology Services Colindale, Public Health England | Respiratory Virus Unit, Microbiology Services Colindale, Public Health England | Monica Galiano, Shahjahan Miah, Angie Lackenby, Omolola Akinbami, Tiina Talts, Leena Bhaw, Richard Myers, Steven Platt, Kirstin Edwards, Jonathan Hubb, Joanna Ellis, Maria Zambon |
| EPI_ISL_417298 | hCoV-19/England/20109099106/2020 | Europe / United Kingdom / England | 2020-03-08 | Respiratory Virus Unit, Microbiology Services Colindale, Public Health England | Respiratory Virus Unit, Microbiology Services Colindale, Public Health England | Monica Galiano, Shahjahan Miah, Angie Lackenby, Omolola Akinbami, Tiina Talts, Leena Bhaw, Richard Myers, Steven Platt, Kirstin Edwards, Jonathan Hubb, Joanna Ellis, Maria Zambon |
| EPI_ISL_417299 | hCoV-19/England/20109099206/2020 | Europe / United Kingdom / England | 2020-03-08 | Respiratory Virus Unit, Microbiology Services Colindale, Public Health England | Respiratory Virus Unit, Microbiology Services Colindale, Public Health England | Monica Galiano, Shahjahan Miah, Angie Lackenby, Omolola Akinbami, Tiina Talts, Leena Bhaw, Richard Myers, Steven Platt, Kirstin Edwards, Jonathan Hubb, Joanna Ellis, Maria Zambon |
| EPI_ISL_417500 | hCoV-19/USA/MN56-MDH56/2020 | North America / USA / Minnesota | 2020-03-14 | Minnesota Department of Health, Public Health Laboratory | Minnesota Department of Health, Public Health Laboratory | Matt Plumb, Jake Garfin and Xiong Wang |
| EPI_ISL_417501 | hCoV-19/USA/MN57-MDH57/2020 | North America / USA / Minnesota | 2020-03-14 | Minnesota Department of Health, Public Health Laboratory | Minnesota Department of Health, Public Health Laboratory | Matt Plumb, Jake Garfin and Xiong Wang |
| EPI_ISL_417502 | hCoV-19/USA/MN58-MDH58/2020 | North America / USA / Minnesota | 2020-03-14 | Minnesota Department of Health, Public Health Laboratory | Minnesota Department of Health, Public Health Laboratory | Matt Plumb, Jake Garfin and Xiong Wang |
| EPI_ISL_417503 | hCoV-19/USA/MN59-MDH59/2020 | North America / USA / Minnesota | 2020-03-14 | Minnesota Department of Health, Public Health Laboratory | Minnesota Department of Health, Public Health Laboratory | Matt Plumb, Jake Garfin and Xiong Wang |
| EPI_ISL_417504 | hCoV-19/USA/WI-UW-15/2020 | North America / USA / Wisconsin | 2020-03-16 | University of Wisconsin-Madison AIDS Vaccine Research Laboratories | University of Wisconsin-Madison AIDS Vaccine Research Laboratories | Gage Moreno, Katarina Braun, et al. AIDS Vaccine Research Laboratories |
| EPI_ISL_417505 | hCoV-19/USA/WI-UW-11/2020 | North America / USA / Wisconsin | 2020-03-15 | University of Wisconsin-Madison AIDS Vaccine Research Laboratories | University of Wisconsin-Madison AIDS Vaccine Research Laboratories | Gage Moreno, Katarina Braun, et al. AIDS Vaccine Research Laboratories |
| EPI_ISL_417506 | hCoV-19/USA/WI-UW-12/2020 | North America / USA / Wisconsin | 2020-03-16 | University of Wisconsin-Madison AIDS Vaccine Research Laboratories | University of Wisconsin-Madison AIDS Vaccine Research Laboratories | Gage Moreno, Katarina Braun, et al. AIDS Vaccine Research Laboratories |
| EPI_ISL_417507 | hCoV-19/USA/WI-UW-23/2020 | North America / USA / Wisconsin | 2020-03-17 | University of Wisconsin-Madison AIDS Vaccine Research Laboratories | University of Wisconsin-Madison AIDS Vaccine Research Laboratories | Gage Moreno, Katarina Braun, et al. AIDS Vaccine Research Laboratories |
| EPI_ISL_417508 | hCoV-19/USA/WI-UW-21/2020 | North America / USA / Wisconsin | 2020-03-16 | University of Wisconsin-Madison AIDS Vaccine Research Laboratories | University of Wisconsin-Madison AIDS Vaccine Research Laboratories | Gage Moreno, Katarina Braun, et al. AIDS Vaccine Research Laboratories |
| EPI_ISL_417509 | hCoV-19/USA/WI-UW-16/2020 | North America / USA / Wisconsin | 2020-03-14 | University of Wisconsin-Madison AIDS Vaccine Research Laboratories | University of Wisconsin-Madison AIDS Vaccine Research Laboratories | Gage Moreno, Katarina Braun, et al. AIDS Vaccine Research Laboratories |
| EPI_ISL_417510 | hCoV-19/USA/WI-UW-19/2020 | North America / USA / Wisconsin | 2020-03-16 | University of Wisconsin-Madison AIDS Vaccine Research Laboratories | University of Wisconsin-Madison AIDS Vaccine Research Laboratories | Gage Moreno, Katarina Braun, et al. AIDS Vaccine Research Laboratories |
| EPI_ISL_417512 | hCoV-19/USA/WI-UW-24/2020 | North America / USA / Wisconsin | 2020-03-15 | University of Wisconsin-Madison AIDS Vaccine Research Laboratories | University of Wisconsin-Madison AIDS Vaccine Research Laboratories | Gage Moreno, Katarina Braun, et al. AIDS Vaccine Research Laboratories |
| EPI_ISL_417513 | hCoV-19/USA/WI-UW-14/2020 | North America / USA / Wisconsin | 2020-03-16 | University of Wisconsin-Madison AIDS Vaccine Research Laboratories | University of Wisconsin-Madison AIDS Vaccine Research Laboratories | Gage Moreno, Katarina Braun, et al. AIDS Vaccine Research Laboratories |
| EPI_ISL_417514 | hCoV-19/USA/WI-UW-22/2020 | North America / USA / Wisconsin | 2020-03-13 | University of Wisconsin-Madison AIDS Vaccine Research Laboratories | University of Wisconsin-Madison AIDS Vaccine Research Laboratories | Gage Moreno, Katarina Braun, et al. AIDS Vaccine Research Laboratories |
| EPI_ISL_417515 | hCoV-19/USA/WI-UW-18/2020 | North America / USA / Wisconsin | 2020-03-17 | University of Wisconsin-Madison AIDS Vaccine Research Laboratories | University of Wisconsin-Madison AIDS Vaccine Research Laboratories | Gage Moreno, Katarina Braun, et al. AIDS Vaccine Research Laboratories |
| EPI_ISL_417516 | hCoV-19/USA/WI-UW-13/2020 | North America / USA / Wisconsin | 2020-03-15 | University of Wisconsin-Madison AIDS Vaccine Research Laboratories | University of Wisconsin-Madison AIDS Vaccine Research Laboratories | Gage Moreno, Katarina Braun, et al. AIDS Vaccine Research Laboratories |
| EPI_ISL_417517 | hCoV-19/USA/WI-UW-17/2020 | North America / USA / Wisconsin | 2020-03-13 | University of Wisconsin-Madison AIDS Vaccine Research Laboratories | University of Wisconsin-Madison AIDS Vaccine Research Laboratories | Gage Moreno, Katarina Braun, et al. AIDS Vaccine Research Laboratories |
| EPI_ISL_417519 | hCoV-19/Taiwan/CGMH-CGU-06/2020 | Asia / Taiwan / Taoyuan | 2020-03-05 | Laboratory Medicine | Department of Laboratory Medicine, Lin-Kou Chang Gung Memorial Hospital, Taoyuan, Taiwan | Kuo-Chien Tsao, Yu-Nong Gong, Shu-Li Yang, Yi-Chun Liu, Chung-Guei Huang, Po-Wei Huang, Mei-Jen Hsiao, Cheng-Ta Yang, Cheng-Hsun Chiu, Peng-Nien Huang, Kuo-Ming Lee, Guang-Wu Chen , Shih-Ru Shih |
| EPI_ISL_417520 | hCoV-19/Taiwan/CGMH-CGU-07/2020 | Asia / Taiwan / Taoyuan | 2020-03-09 | Laboratory Medicine | Department of Laboratory Medicine, Lin-Kou Chang Gung Memorial Hospital, Taoyuan, Taiwan | Kuo-Chien Tsao, Yu-Nong Gong, Shu-Li Yang, Yi-Chun Liu, Chung-Guei Huang, Po-Wei Huang, Mei-Jen Hsiao, Cheng-Ta Yang, Cheng-Hsun Chiu, Peng-Nien Huang, Kuo-Ming Lee, Guang-Wu Chen , Shih-Ru Shih |
| EPI_ISL_417521 | hCoV-19/Taiwan/CGMH-CGU-08/2020 | Asia / Taiwan / Taoyuan | 2020-03-10 | Laboratory Medicine | Department of Laboratory Medicine, Lin-Kou Chang Gung Memorial Hospital, Taoyuan, Taiwan | Kuo-Chien Tsao, Yu-Nong Gong, Shu-Li Yang, Yi-Chun Liu, Chung-Guei Huang, Po-Wei Huang, Mei-Jen Hsiao, Cheng-Ta Yang, Cheng-Hsun Chiu, Peng-Nien Huang, Kuo-Ming Lee, Guang-Wu Chen , Shih-Ru Shih |
| EPI_ISL_417522 | hCoV-19/Taiwan/CGMH-CGU-09/2020 | Asia / Taiwan / Taoyuan | 2020-03-13 | Laboratory Medicine | Department of Laboratory Medicine, Lin-Kou Chang Gung Memorial Hospital, Taoyuan, Taiwan | Kuo-Chien Tsao, Yu-Nong Gong, Shu-Li Yang, Yi-Chun Liu, Chung-Guei Huang, Po-Wei Huang, Mei-Jen Hsiao, Cheng-Ta Yang, Cheng-Hsun Chiu, Peng-Nien Huang, Kuo-Ming Lee, Guang-Wu Chen , Shih-Ru Shih |
| EPI_ISL_417523 | hCoV-19/Taiwan/CGMH-CGU-10/2020 | Asia / Taiwan / Taoyuan | 2020-03-13 | Laboratory Medicine | Department of Laboratory Medicine, Lin-Kou Chang Gung Memorial Hospital, Taoyuan, Taiwan | Kuo-Chien Tsao, Yu-Nong Gong, Shu-Li Yang, Yi-Chun Liu, Chung-Guei Huang, Po-Wei Huang, Mei-Jen Hsiao, Cheng-Ta Yang, Cheng-Hsun Chiu, Peng-Nien Huang, Kuo-Ming Lee, Guang-Wu Chen , Shih-Ru Shih |
| EPI_ISL_417524 | hCoV-19/Taiwan/CGMH-CGU-11/2020 | Asia / Taiwan / Taoyuan | 2020-03-14 | Laboratory Medicine | Department of Laboratory Medicine, Lin-Kou Chang Gung Memorial Hospital, Taoyuan, Taiwan | Kuo-Chien Tsao, Yu-Nong Gong, Shu-Li Yang, Yi-Chun Liu, Chung-Guei Huang, Po-Wei Huang, Mei-Jen Hsiao, Cheng-Ta Yang, Cheng-Hsun Chiu, Peng-Nien Huang, Kuo-Ming Lee, Guang-Wu Chen , Shih-Ru Shih |
| EPI_ISL_417525 | hCoV-19/Taiwan/CGMH-CGU-12/2020 | Asia / Taiwan / Taoyuan | 2020-03-14 | Laboratory Medicine | Department of Laboratory Medicine, Lin-Kou Chang Gung Memorial Hospital, Taoyuan, Taiwan | Kuo-Chien Tsao, Yu-Nong Gong, Shu-Li Yang, Yi-Chun Liu, Chung-Guei Huang, Po-Wei Huang, Mei-Jen Hsiao, Cheng-Ta Yang, Cheng-Hsun Chiu, Peng-Nien Huang, Kuo-Ming Lee, Guang-Wu Chen , Shih-Ru Shih |
| EPI_ISL_417526 | hCoV-19/Luxembourg/LNS9371718/2020 | Europe / Luxembourg | 2020-03-18 | Laboratoire Nationale de Santé, Microbiology, Virology | Laboratoire Nationale de Santé, Microbiology, Epidemiology and Microbial Genomics | Anke Wienecke-Baldacchino, Ardashel Latsuzbaia, Jessica Tapp, Catherine Ragimbeau, Guillaume Fournier, Tamir Abdelrahman, Trung Nguyen Nguyen, Joel Mossong |
| EPI_ISL_417527 | hCoV-19/Luxembourg/LNS2848109/2020 | Europe / Luxembourg | 2020-03-18 | Laboratoire Nationale de Santé, Microbiology, Virology | Laboratoire Nationale de Santé, Microbiology, Epidemiology and Microbial Genomics | Anke Wienecke-Baldacchino, Ardashel Latsuzbaia, Jessica Tapp, Catherine Ragimbeau, Guillaume Fournier, Tamir Abdelrahman, Trung Nguyen Nguyen, Joel Mossong |
| EPI_ISL_417528 | hCoV-19/Luxembourg/LNS3694003/2020 | Europe / Luxembourg | 2020-03-18 | Laboratoire Nationale de Santé, Microbiology, Virology | Laboratoire Nationale de Santé, Microbiology, Epidemiology and Microbial Genomics | Anke Wienecke-Baldacchino, Ardashel Latsuzbaia, Jessica Tapp, Catherine Ragimbeau, Guillaume Fournier, Tamir Abdelrahman, Trung Nguyen Nguyen, Joel Mossong |
| EPI_ISL_417529 | hCoV-19/Luxembourg/LNS5731562/2020 | Europe / Luxembourg | 2020-03-18 | Laboratoire Nationale de Santé, Microbiology, Virology | Laboratoire Nationale de Santé, Microbiology, Epidemiology and Microbial Genomics | Anke Wienecke-Baldacchino, Ardashel Latsuzbaia, Jessica Tapp, Catherine Ragimbeau, Guillaume Fournier, Tamir Abdelrahman, Trung Nguyen Nguyen, Joel Mossong |
| EPI_ISL_417530 | hCoV-19/Luxembourg/LNS2128808/2020 | Europe / Luxembourg | 2020-03-18 | Laboratoire Nationale de Santé, Microbiology, Virology | Laboratoire Nationale de Santé, Microbiology, Epidemiology and Microbial Genomics | Anke Wienecke-Baldacchino, Ardashel Latsuzbaia, Jessica Tapp, Catherine Ragimbeau, Guillaume Fournier, Tamir Abdelrahman, Trung Nguyen Nguyen, Joel Mossong |
| EPI_ISL_417531 | hCoV-19/Luxembourg/LNS2013896/2020 | Europe / Luxembourg | 2020-03-18 | Laboratoire Nationale de Santé, Microbiology, Virology | Laboratoire Nationale de Santé, Microbiology, Epidemiology and Microbial Genomics | Anke Wienecke-Baldacchino, Ardashel Latsuzbaia, Jessica Tapp, Catherine Ragimbeau, Guillaume Fournier, Tamir Abdelrahman, Trung Nguyen Nguyen, Joel Mossong |
| EPI_ISL_417532 | hCoV-19/Luxembourg/LNS8489624/2020 | Europe / Luxembourg | 2020-03-18 | Laboratoire Nationale de Santé, Microbiology, Virology | Laboratoire Nationale de Santé, Microbiology, Epidemiology and Microbial Genomics | Anke Wienecke-Baldacchino, Ardashel Latsuzbaia, Jessica Tapp, Catherine Ragimbeau, Guillaume Fournier, Tamir Abdelrahman, Trung Nguyen Nguyen, Joel Mossong |
| EPI_ISL_417533 | hCoV-19/Luxembourg/LNS6282845/2020 | Europe / Luxembourg | 2020-03-18 | Laboratoire Nationale de Santé, Microbiology, Virology | Laboratoire Nationale de Santé, Microbiology, Epidemiology and Microbial Genomics | Anke Wienecke-Baldacchino, Ardashel Latsuzbaia, Jessica Tapp, Catherine Ragimbeau, Guillaume Fournier, Tamir Abdelrahman, Trung Nguyen Nguyen, Joel Mossong |
| EPI_ISL_417534 | hCoV-19/Luxembourg/LNS0158952/2020 | Europe / Luxembourg | 2020-03-18 | Laboratoire Nationale de Santé, Microbiology, Virology | Laboratoire Nationale de Santé, Microbiology, Epidemiology and Microbial Genomics | Anke Wienecke-Baldacchino, Ardashel Latsuzbaia, Jessica Tapp, Catherine Ragimbeau, Guillaume Fournier, Tamir Abdelrahman, Trung Nguyen Nguyen, Joel Mossong |
| EPI_ISL_417535 | hCoV-19/Iceland/1/2020 | Europe / Iceland / Reykjavik | 2020-03-13 | deCODE genetics | deCODE genetics | Daniel F Gudbjartsson; Agnar Helgason; Hakon Jonsson; Olafur T Magnusson; Pall Melsted; Gudmundur L Norddahl; Jona Saemundsdottir; Asgeir Sigurdsson; Patrick Sulem; Arna B Agustsdottir; Berglind Eiriksdottir; Run Fridriksdottir; Elisabet E Gardarsdottir; Gudmundur Georgsson; Olafia S Gretarsdottir; Kjartan R Gudmundsson; Thora R Gunnarsdottir; Arnaldur Gylfason; Hilma Holm; Brynjar O Jensson; Aslaug Jonasdottir; Kamilla S Josefsdottir; Thordur Kristjansson; Droplaug N Magnusdottir; Louise le Roux; Gudrun Sigmundsdottir; Gardar Sveinbjornsson; Kristin E Sveinsdottir; Maney Sveinsdottir; Emil A Thorarensen; Bjarni Thorbjornsson; Gisli Masson; Ingileif Jonsdottir; Alma Moller; Thorolfur Gudnason; Karl G Kristinsson; Unnur Thorsteinsdottir; Kari Stefansson |
| EPI_ISL_417536 | hCoV-19/Iceland/10/2020 | Europe / Iceland / Reykjavik | 2020-03-16 | deCODE genetics | deCODE genetics | Daniel F Gudbjartsson; Agnar Helgason; Hakon Jonsson; Olafur T Magnusson; Pall Melsted; Gudmundur L Norddahl; Jona Saemundsdottir; Asgeir Sigurdsson; Patrick Sulem; Arna B Agustsdottir; Berglind Eiriksdottir; Run Fridriksdottir; Elisabet E Gardarsdottir; Gudmundur Georgsson; Olafia S Gretarsdottir; Kjartan R Gudmundsson; Thora R Gunnarsdottir; Arnaldur Gylfason; Hilma Holm; Brynjar O Jensson; Aslaug Jonasdottir; Kamilla S Josefsdottir; Thordur Kristjansson; Droplaug N Magnusdottir; Louise le Roux; Gudrun Sigmundsdottir; Gardar Sveinbjornsson; Kristin E Sveinsdottir; Maney Sveinsdottir; Emil A Thorarensen; Bjarni Thorbjornsson; Gisli Masson; Ingileif Jonsdottir; Alma Moller; Thorolfur Gudnason; Karl G Kristinsson; Unnur Thorsteinsdottir; Kari Stefansson |
| EPI_ISL_417537 | hCoV-19/Iceland/100/2020 | Europe / Iceland / Reykjavik | 2020-03-16 | deCODE genetics | deCODE genetics | Daniel F Gudbjartsson; Agnar Helgason; Hakon Jonsson; Olafur T Magnusson; Pall Melsted; Gudmundur L Norddahl; Jona Saemundsdottir; Asgeir Sigurdsson; Patrick Sulem; Arna B Agustsdottir; Berglind Eiriksdottir; Run Fridriksdottir; Elisabet E Gardarsdottir; Gudmundur Georgsson; Olafia S Gretarsdottir; Kjartan R Gudmundsson; Thora R Gunnarsdottir; Arnaldur Gylfason; Hilma Holm; Brynjar O Jensson; Aslaug Jonasdottir; Kamilla S Josefsdottir; Thordur Kristjansson; Droplaug N Magnusdottir; Louise le Roux; Gudrun Sigmundsdottir; Gardar Sveinbjornsson; Kristin E Sveinsdottir; Maney Sveinsdottir; Emil A Thorarensen; Bjarni Thorbjornsson; Gisli Masson; Ingileif Jonsdottir; Alma Moller; Thorolfur Gudnason; Karl G Kristinsson; Unnur Thorsteinsdottir; Kari Stefansson |
| EPI_ISL_417538 | hCoV-19/Iceland/104/2020 | Europe / Iceland / Reykjavik | 2020-03-17 | deCODE genetics | deCODE genetics | Daniel F Gudbjartsson; Agnar Helgason; Hakon Jonsson; Olafur T Magnusson; Pall Melsted; Gudmundur L Norddahl; Jona Saemundsdottir; Asgeir Sigurdsson; Patrick Sulem; Arna B Agustsdottir; Berglind Eiriksdottir; Run Fridriksdottir; Elisabet E Gardarsdottir; Gudmundur Georgsson; Olafia S Gretarsdottir; Kjartan R Gudmundsson; Thora R Gunnarsdottir; Arnaldur Gylfason; Hilma Holm; Brynjar O Jensson; Aslaug Jonasdottir; Kamilla S Josefsdottir; Thordur Kristjansson; Droplaug N Magnusdottir; Louise le Roux; Gudrun Sigmundsdottir; Gardar Sveinbjornsson; Kristin E Sveinsdottir; Maney Sveinsdottir; Emil A Thorarensen; Bjarni Thorbjornsson; Gisli Masson; Ingileif Jonsdottir; Alma Moller; Thorolfur Gudnason; Karl G Kristinsson; Unnur Thorsteinsdottir; Kari Stefansson |
| EPI_ISL_417539 | hCoV-19/Iceland/11/2020 | Europe / Iceland / Reykjavik | 2020-03-16 | deCODE genetics | deCODE genetics | Daniel F Gudbjartsson; Agnar Helgason; Hakon Jonsson; Olafur T Magnusson; Pall Melsted; Gudmundur L Norddahl; Jona Saemundsdottir; Asgeir Sigurdsson; Patrick Sulem; Arna B Agustsdottir; Berglind Eiriksdottir; Run Fridriksdottir; Elisabet E Gardarsdottir; Gudmundur Georgsson; Olafia S Gretarsdottir; Kjartan R Gudmundsson; Thora R Gunnarsdottir; Arnaldur Gylfason; Hilma Holm; Brynjar O Jensson; Aslaug Jonasdottir; Kamilla S Josefsdottir; Thordur Kristjansson; Droplaug N Magnusdottir; Louise le Roux; Gudrun Sigmundsdottir; Gardar Sveinbjornsson; Kristin E Sveinsdottir; Maney Sveinsdottir; Emil A Thorarensen; Bjarni Thorbjornsson; Gisli Masson; Ingileif Jonsdottir; Alma Moller; Thorolfur Gudnason; Karl G Kristinsson; Unnur Thorsteinsdottir; Kari Stefansson |
| EPI_ISL_417540 | hCoV-19/Iceland/115/2020 | Europe / Iceland / Reykjavik | 2020-03-17 | deCODE genetics | deCODE genetics | Daniel F Gudbjartsson; Agnar Helgason; Hakon Jonsson; Olafur T Magnusson; Pall Melsted; Gudmundur L Norddahl; Jona Saemundsdottir; Asgeir Sigurdsson; Patrick Sulem; Arna B Agustsdottir; Berglind Eiriksdottir; Run Fridriksdottir; Elisabet E Gardarsdottir; Gudmundur Georgsson; Olafia S Gretarsdottir; Kjartan R Gudmundsson; Thora R Gunnarsdottir; Arnaldur Gylfason; Hilma Holm; Brynjar O Jensson; Aslaug Jonasdottir; Kamilla S Josefsdottir; Thordur Kristjansson; Droplaug N Magnusdottir; Louise le Roux; Gudrun Sigmundsdottir; Gardar Sveinbjornsson; Kristin E Sveinsdottir; Maney Sveinsdottir; Emil A Thorarensen; Bjarni Thorbjornsson; Gisli Masson; Ingileif Jonsdottir; Alma Moller; Thorolfur Gudnason; Karl G Kristinsson; Unnur Thorsteinsdottir; Kari Stefansson |
| EPI_ISL_417541 | hCoV-19/Iceland/116/2020 | Europe / Iceland / Reykjavik | 2020-03-17 | deCODE genetics | deCODE genetics | Daniel F Gudbjartsson; Agnar Helgason; Hakon Jonsson; Olafur T Magnusson; Pall Melsted; Gudmundur L Norddahl; Jona Saemundsdottir; Asgeir Sigurdsson; Patrick Sulem; Arna B Agustsdottir; Berglind Eiriksdottir; Run Fridriksdottir; Elisabet E Gardarsdottir; Gudmundur Georgsson; Olafia S Gretarsdottir; Kjartan R Gudmundsson; Thora R Gunnarsdottir; Arnaldur Gylfason; Hilma Holm; Brynjar O Jensson; Aslaug Jonasdottir; Kamilla S Josefsdottir; Thordur Kristjansson; Droplaug N Magnusdottir; Louise le Roux; Gudrun Sigmundsdottir; Gardar Sveinbjornsson; Kristin E Sveinsdottir; Maney Sveinsdottir; Emil A Thorarensen; Bjarni Thorbjornsson; Gisli Masson; Ingileif Jonsdottir; Alma Moller; Thorolfur Gudnason; Karl G Kristinsson; Unnur Thorsteinsdottir; Kari Stefansson |
| EPI_ISL_417542 | hCoV-19/Iceland/117/2020 | Europe / Iceland / Reykjavik | 2020-03-17 | deCODE genetics | deCODE genetics | Daniel F Gudbjartsson; Agnar Helgason; Hakon Jonsson; Olafur T Magnusson; Pall Melsted; Gudmundur L Norddahl; Jona Saemundsdottir; Asgeir Sigurdsson; Patrick Sulem; Arna B Agustsdottir; Berglind Eiriksdottir; Run Fridriksdottir; Elisabet E Gardarsdottir; Gudmundur Georgsson; Olafia S Gretarsdottir; Kjartan R Gudmundsson; Thora R Gunnarsdottir; Arnaldur Gylfason; Hilma Holm; Brynjar O Jensson; Aslaug Jonasdottir; Kamilla S Josefsdottir; Thordur Kristjansson; Droplaug N Magnusdottir; Louise le Roux; Gudrun Sigmundsdottir; Gardar Sveinbjornsson; Kristin E Sveinsdottir; Maney Sveinsdottir; Emil A Thorarensen; Bjarni Thorbjornsson; Gisli Masson; Ingileif Jonsdottir; Alma Moller; Thorolfur Gudnason; Karl G Kristinsson; Unnur Thorsteinsdottir; Kari Stefansson |
| EPI_ISL_417543 | hCoV-19/Iceland/118/2020 | Europe / Iceland / Reykjavik | 2020-03-17 | deCODE genetics | deCODE genetics | Daniel F Gudbjartsson; Agnar Helgason; Hakon Jonsson; Olafur T Magnusson; Pall Melsted; Gudmundur L Norddahl; Jona Saemundsdottir; Asgeir Sigurdsson; Patrick Sulem; Arna B Agustsdottir; Berglind Eiriksdottir; Run Fridriksdottir; Elisabet E Gardarsdottir; Gudmundur Georgsson; Olafia S Gretarsdottir; Kjartan R Gudmundsson; Thora R Gunnarsdottir; Arnaldur Gylfason; Hilma Holm; Brynjar O Jensson; Aslaug Jonasdottir; Kamilla S Josefsdottir; Thordur Kristjansson; Droplaug N Magnusdottir; Louise le Roux; Gudrun Sigmundsdottir; Gardar Sveinbjornsson; Kristin E Sveinsdottir; Maney Sveinsdottir; Emil A Thorarensen; Bjarni Thorbjornsson; Gisli Masson; Ingileif Jonsdottir; Alma Moller; Thorolfur Gudnason; Karl G Kristinsson; Unnur Thorsteinsdottir; Kari Stefansson |
| EPI_ISL_417544 | hCoV-19/Iceland/119/2020 | Europe / Iceland / Reykjavik | 2020-03-17 | deCODE genetics | deCODE genetics | Daniel F Gudbjartsson; Agnar Helgason; Hakon Jonsson; Olafur T Magnusson; Pall Melsted; Gudmundur L Norddahl; Jona Saemundsdottir; Asgeir Sigurdsson; Patrick Sulem; Arna B Agustsdottir; Berglind Eiriksdottir; Run Fridriksdottir; Elisabet E Gardarsdottir; Gudmundur Georgsson; Olafia S Gretarsdottir; Kjartan R Gudmundsson; Thora R Gunnarsdottir; Arnaldur Gylfason; Hilma Holm; Brynjar O Jensson; Aslaug Jonasdottir; Kamilla S Josefsdottir; Thordur Kristjansson; Droplaug N Magnusdottir; Louise le Roux; Gudrun Sigmundsdottir; Gardar Sveinbjornsson; Kristin E Sveinsdottir; Maney Sveinsdottir; Emil A Thorarensen; Bjarni Thorbjornsson; Gisli Masson; Ingileif Jonsdottir; Alma Moller; Thorolfur Gudnason; Karl G Kristinsson; Unnur Thorsteinsdottir; Kari Stefansson |
| EPI_ISL_417545 | hCoV-19/Iceland/120/2020 | Europe / Iceland / Reykjavik | 2020-03-17 | deCODE genetics | deCODE genetics | Daniel F Gudbjartsson; Agnar Helgason; Hakon Jonsson; Olafur T Magnusson; Pall Melsted; Gudmundur L Norddahl; Jona Saemundsdottir; Asgeir Sigurdsson; Patrick Sulem; Arna B Agustsdottir; Berglind Eiriksdottir; Run Fridriksdottir; Elisabet E Gardarsdottir; Gudmundur Georgsson; Olafia S Gretarsdottir; Kjartan R Gudmundsson; Thora R Gunnarsdottir; Arnaldur Gylfason; Hilma Holm; Brynjar O Jensson; Aslaug Jonasdottir; Kamilla S Josefsdottir; Thordur Kristjansson; Droplaug N Magnusdottir; Louise le Roux; Gudrun Sigmundsdottir; Gardar Sveinbjornsson; Kristin E Sveinsdottir; Maney Sveinsdottir; Emil A Thorarensen; Bjarni Thorbjornsson; Gisli Masson; Ingileif Jonsdottir; Alma Moller; Thorolfur Gudnason; Karl G Kristinsson; Unnur Thorsteinsdottir; Kari Stefansson |
| EPI_ISL_417547 | hCoV-19/Iceland/122/2020 | Europe / Iceland / Reykjavik | 2020-03-17 | deCODE genetics | deCODE genetics | Daniel F Gudbjartsson; Agnar Helgason; Hakon Jonsson; Olafur T Magnusson; Pall Melsted; Gudmundur L Norddahl; Jona Saemundsdottir; Asgeir Sigurdsson; Patrick Sulem; Arna B Agustsdottir; Berglind Eiriksdottir; Run Fridriksdottir; Elisabet E Gardarsdottir; Gudmundur Georgsson; Olafia S Gretarsdottir; Kjartan R Gudmundsson; Thora R Gunnarsdottir; Arnaldur Gylfason; Hilma Holm; Brynjar O Jensson; Aslaug Jonasdottir; Kamilla S Josefsdottir; Thordur Kristjansson; Droplaug N Magnusdottir; Louise le Roux; Gudrun Sigmundsdottir; Gardar Sveinbjornsson; Kristin E Sveinsdottir; Maney Sveinsdottir; Emil A Thorarensen; Bjarni Thorbjornsson; Gisli Masson; Ingileif Jonsdottir; Alma Moller; Thorolfur Gudnason; Karl G Kristinsson; Unnur Thorsteinsdottir; Kari Stefansson |
| EPI_ISL_417548 | hCoV-19/Iceland/170/2020 | Europe / Iceland / Reykjavik | 2020-03-17 | deCODE genetics | deCODE genetics | Daniel F Gudbjartsson; Agnar Helgason; Hakon Jonsson; Olafur T Magnusson; Pall Melsted; Gudmundur L Norddahl; Jona Saemundsdottir; Asgeir Sigurdsson; Patrick Sulem; Arna B Agustsdottir; Berglind Eiriksdottir; Run Fridriksdottir; Elisabet E Gardarsdottir; Gudmundur Georgsson; Olafia S Gretarsdottir; Kjartan R Gudmundsson; Thora R Gunnarsdottir; Arnaldur Gylfason; Hilma Holm; Brynjar O Jensson; Aslaug Jonasdottir; Kamilla S Josefsdottir; Thordur Kristjansson; Droplaug N Magnusdottir; Louise le Roux; Gudrun Sigmundsdottir; Gardar Sveinbjornsson; Kristin E Sveinsdottir; Maney Sveinsdottir; Emil A Thorarensen; Bjarni Thorbjornsson; Gisli Masson; Ingileif Jonsdottir; Alma Moller; Thorolfur Gudnason; Karl G Kristinsson; Unnur Thorsteinsdottir; Kari Stefansson |
| EPI_ISL_417549 | hCoV-19/Iceland/171/2020 | Europe / Iceland / Reykjavik | 2020-03-14 | deCODE genetics | deCODE genetics | Daniel F Gudbjartsson; Agnar Helgason; Hakon Jonsson; Olafur T Magnusson; Pall Melsted; Gudmundur L Norddahl; Jona Saemundsdottir; Asgeir Sigurdsson; Patrick Sulem; Arna B Agustsdottir; Berglind Eiriksdottir; Run Fridriksdottir; Elisabet E Gardarsdottir; Gudmundur Georgsson; Olafia S Gretarsdottir; Kjartan R Gudmundsson; Thora R Gunnarsdottir; Arnaldur Gylfason; Hilma Holm; Brynjar O Jensson; Aslaug Jonasdottir; Kamilla S Josefsdottir; Thordur Kristjansson; Droplaug N Magnusdottir; Louise le Roux; Gudrun Sigmundsdottir; Gardar Sveinbjornsson; Kristin E Sveinsdottir; Maney Sveinsdottir; Emil A Thorarensen; Bjarni Thorbjornsson; Gisli Masson; Ingileif Jonsdottir; Alma Moller; Thorolfur Gudnason; Karl G Kristinsson; Unnur Thorsteinsdottir; Kari Stefansson |
| EPI_ISL_417550 | hCoV-19/Iceland/184/2020 | Europe / Iceland / Reykjavik | 2020-03-16 | The National University Hospital of Iceland | deCODE genetics | Daniel F Gudbjartsson; Agnar Helgason; Hakon Jonsson; Olafur T Magnusson; Pall Melsted; Gudmundur L Norddahl; Jona Saemundsdottir; Asgeir Sigurdsson; Patrick Sulem; Arna B Agustsdottir; Berglind Eiriksdottir; Run Fridriksdottir; Elisabet E Gardarsdottir; Gudmundur Georgsson; Olafia S Gretarsdottir; Kjartan R Gudmundsson; Thora R Gunnarsdottir; Arnaldur Gylfason; Hilma Holm; Brynjar O Jensson; Aslaug Jonasdottir; Kamilla S Josefsdottir; Thordur Kristjansson; Droplaug N Magnusdottir; Louise le Roux; Gudrun Sigmundsdottir; Gardar Sveinbjornsson; Kristin E Sveinsdottir; Maney Sveinsdottir; Emil A Thorarensen; Bjarni Thorbjornsson; Gisli Masson; Ingileif Jonsdottir; Alma Moller; Thorolfur Gudnason; Karl G Kristinsson; Unnur Thorsteinsdottir; Kari Stefansson |
| EPI_ISL_417551 | hCoV-19/Iceland/187/2020 | Europe / Iceland / Reykjavik | 2020-03-16 | The National University Hospital of Iceland | deCODE genetics | Daniel F Gudbjartsson; Agnar Helgason; Hakon Jonsson; Olafur T Magnusson; Pall Melsted; Gudmundur L Norddahl; Jona Saemundsdottir; Asgeir Sigurdsson; Patrick Sulem; Arna B Agustsdottir; Berglind Eiriksdottir; Run Fridriksdottir; Elisabet E Gardarsdottir; Gudmundur Georgsson; Olafia S Gretarsdottir; Kjartan R Gudmundsson; Thora R Gunnarsdottir; Arnaldur Gylfason; Hilma Holm; Brynjar O Jensson; Aslaug Jonasdottir; Kamilla S Josefsdottir; Thordur Kristjansson; Droplaug N Magnusdottir; Louise le Roux; Gudrun Sigmundsdottir; Gardar Sveinbjornsson; Kristin E Sveinsdottir; Maney Sveinsdottir; Emil A Thorarensen; Bjarni Thorbjornsson; Gisli Masson; Ingileif Jonsdottir; Alma Moller; Thorolfur Gudnason; Karl G Kristinsson; Unnur Thorsteinsdottir; Kari Stefansson |
| EPI_ISL_417552 | hCoV-19/Iceland/2/2020 | Europe / Iceland / Reykjavik | 2020-03-16 | deCODE genetics | deCODE genetics | Daniel F Gudbjartsson; Agnar Helgason; Hakon Jonsson; Olafur T Magnusson; Pall Melsted; Gudmundur L Norddahl; Jona Saemundsdottir; Asgeir Sigurdsson; Patrick Sulem; Arna B Agustsdottir; Berglind Eiriksdottir; Run Fridriksdottir; Elisabet E Gardarsdottir; Gudmundur Georgsson; Olafia S Gretarsdottir; Kjartan R Gudmundsson; Thora R Gunnarsdottir; Arnaldur Gylfason; Hilma Holm; Brynjar O Jensson; Aslaug Jonasdottir; Kamilla S Josefsdottir; Thordur Kristjansson; Droplaug N Magnusdottir; Louise le Roux; Gudrun Sigmundsdottir; Gardar Sveinbjornsson; Kristin E Sveinsdottir; Maney Sveinsdottir; Emil A Thorarensen; Bjarni Thorbjornsson; Gisli Masson; Ingileif Jonsdottir; Alma Moller; Thorolfur Gudnason; Karl G Kristinsson; Unnur Thorsteinsdottir; Kari Stefansson |
| EPI_ISL_417553 | hCoV-19/Iceland/220/2020 | Europe / Iceland / Reykjavik | 2020-03-16 | The National University Hospital of Iceland | deCODE genetics | Daniel F Gudbjartsson; Agnar Helgason; Hakon Jonsson; Olafur T Magnusson; Pall Melsted; Gudmundur L Norddahl; Jona Saemundsdottir; Asgeir Sigurdsson; Patrick Sulem; Arna B Agustsdottir; Berglind Eiriksdottir; Run Fridriksdottir; Elisabet E Gardarsdottir; Gudmundur Georgsson; Olafia S Gretarsdottir; Kjartan R Gudmundsson; Thora R Gunnarsdottir; Arnaldur Gylfason; Hilma Holm; Brynjar O Jensson; Aslaug Jonasdottir; Kamilla S Josefsdottir; Thordur Kristjansson; Droplaug N Magnusdottir; Louise le Roux; Gudrun Sigmundsdottir; Gardar Sveinbjornsson; Kristin E Sveinsdottir; Maney Sveinsdottir; Emil A Thorarensen; Bjarni Thorbjornsson; Gisli Masson; Ingileif Jonsdottir; Alma Moller; Thorolfur Gudnason; Karl G Kristinsson; Unnur Thorsteinsdottir; Kari Stefansson |
| EPI_ISL_417554 | hCoV-19/Iceland/225/2020 | Europe / Iceland / Reykjavik | 2020-03-16 | The National University Hospital of Iceland | deCODE genetics | Daniel F Gudbjartsson; Agnar Helgason; Hakon Jonsson; Olafur T Magnusson; Pall Melsted; Gudmundur L Norddahl; Jona Saemundsdottir; Asgeir Sigurdsson; Patrick Sulem; Arna B Agustsdottir; Berglind Eiriksdottir; Run Fridriksdottir; Elisabet E Gardarsdottir; Gudmundur Georgsson; Olafia S Gretarsdottir; Kjartan R Gudmundsson; Thora R Gunnarsdottir; Arnaldur Gylfason; Hilma Holm; Brynjar O Jensson; Aslaug Jonasdottir; Kamilla S Josefsdottir; Thordur Kristjansson; Droplaug N Magnusdottir; Louise le Roux; Gudrun Sigmundsdottir; Gardar Sveinbjornsson; Kristin E Sveinsdottir; Maney Sveinsdottir; Emil A Thorarensen; Bjarni Thorbjornsson; Gisli Masson; Ingileif Jonsdottir; Alma Moller; Thorolfur Gudnason; Karl G Kristinsson; Unnur Thorsteinsdottir; Kari Stefansson |
| EPI_ISL_417555 | hCoV-19/Iceland/227/2020 | Europe / Iceland / Reykjavik | 2020-03-17 | The National University Hospital of Iceland | deCODE genetics | Daniel F Gudbjartsson; Agnar Helgason; Hakon Jonsson; Olafur T Magnusson; Pall Melsted; Gudmundur L Norddahl; Jona Saemundsdottir; Asgeir Sigurdsson; Patrick Sulem; Arna B Agustsdottir; Berglind Eiriksdottir; Run Fridriksdottir; Elisabet E Gardarsdottir; Gudmundur Georgsson; Olafia S Gretarsdottir; Kjartan R Gudmundsson; Thora R Gunnarsdottir; Arnaldur Gylfason; Hilma Holm; Brynjar O Jensson; Aslaug Jonasdottir; Kamilla S Josefsdottir; Thordur Kristjansson; Droplaug N Magnusdottir; Louise le Roux; Gudrun Sigmundsdottir; Gardar Sveinbjornsson; Kristin E Sveinsdottir; Maney Sveinsdottir; Emil A Thorarensen; Bjarni Thorbjornsson; Gisli Masson; Ingileif Jonsdottir; Alma Moller; Thorolfur Gudnason; Karl G Kristinsson; Unnur Thorsteinsdottir; Kari Stefansson |
| EPI_ISL_417556 | hCoV-19/Iceland/228/2020 | Europe / Iceland / Reykjavik | 2020-03-17 | The National University Hospital of Iceland | deCODE genetics | Daniel F Gudbjartsson; Agnar Helgason; Hakon Jonsson; Olafur T Magnusson; Pall Melsted; Gudmundur L Norddahl; Jona Saemundsdottir; Asgeir Sigurdsson; Patrick Sulem; Arna B Agustsdottir; Berglind Eiriksdottir; Run Fridriksdottir; Elisabet E Gardarsdottir; Gudmundur Georgsson; Olafia S Gretarsdottir; Kjartan R Gudmundsson; Thora R Gunnarsdottir; Arnaldur Gylfason; Hilma Holm; Brynjar O Jensson; Aslaug Jonasdottir; Kamilla S Josefsdottir; Thordur Kristjansson; Droplaug N Magnusdottir; Louise le Roux; Gudrun Sigmundsdottir; Gardar Sveinbjornsson; Kristin E Sveinsdottir; Maney Sveinsdottir; Emil A Thorarensen; Bjarni Thorbjornsson; Gisli Masson; Ingileif Jonsdottir; Alma Moller; Thorolfur Gudnason; Karl G Kristinsson; Unnur Thorsteinsdottir; Kari Stefansson |
| EPI_ISL_417557 | hCoV-19/Iceland/229/2020 | Europe / Iceland / Reykjavik | 2020-03-17 | The National University Hospital of Iceland | deCODE genetics | Daniel F Gudbjartsson; Agnar Helgason; Hakon Jonsson; Olafur T Magnusson; Pall Melsted; Gudmundur L Norddahl; Jona Saemundsdottir; Asgeir Sigurdsson; Patrick Sulem; Arna B Agustsdottir; Berglind Eiriksdottir; Run Fridriksdottir; Elisabet E Gardarsdottir; Gudmundur Georgsson; Olafia S Gretarsdottir; Kjartan R Gudmundsson; Thora R Gunnarsdottir; Arnaldur Gylfason; Hilma Holm; Brynjar O Jensson; Aslaug Jonasdottir; Kamilla S Josefsdottir; Thordur Kristjansson; Droplaug N Magnusdottir; Louise le Roux; Gudrun Sigmundsdottir; Gardar Sveinbjornsson; Kristin E Sveinsdottir; Maney Sveinsdottir; Emil A Thorarensen; Bjarni Thorbjornsson; Gisli Masson; Ingileif Jonsdottir; Alma Moller; Thorolfur Gudnason; Karl G Kristinsson; Unnur Thorsteinsdottir; Kari Stefansson |
| EPI_ISL_417558 | hCoV-19/Iceland/230/2020 | Europe / Iceland / Reykjavik | 2020-03-17 | The National University Hospital of Iceland | deCODE genetics | Daniel F Gudbjartsson; Agnar Helgason; Hakon Jonsson; Olafur T Magnusson; Pall Melsted; Gudmundur L Norddahl; Jona Saemundsdottir; Asgeir Sigurdsson; Patrick Sulem; Arna B Agustsdottir; Berglind Eiriksdottir; Run Fridriksdottir; Elisabet E Gardarsdottir; Gudmundur Georgsson; Olafia S Gretarsdottir; Kjartan R Gudmundsson; Thora R Gunnarsdottir; Arnaldur Gylfason; Hilma Holm; Brynjar O Jensson; Aslaug Jonasdottir; Kamilla S Josefsdottir; Thordur Kristjansson; Droplaug N Magnusdottir; Louise le Roux; Gudrun Sigmundsdottir; Gardar Sveinbjornsson; Kristin E Sveinsdottir; Maney Sveinsdottir; Emil A Thorarensen; Bjarni Thorbjornsson; Gisli Masson; Ingileif Jonsdottir; Alma Moller; Thorolfur Gudnason; Karl G Kristinsson; Unnur Thorsteinsdottir; Kari Stefansson |
| EPI_ISL_417560 | hCoV-19/Iceland/232/2020 | Europe / Iceland / Reykjavik | 2020-03-17 | The National University Hospital of Iceland | deCODE genetics | Daniel F Gudbjartsson; Agnar Helgason; Hakon Jonsson; Olafur T Magnusson; Pall Melsted; Gudmundur L Norddahl; Jona Saemundsdottir; Asgeir Sigurdsson; Patrick Sulem; Arna B Agustsdottir; Berglind Eiriksdottir; Run Fridriksdottir; Elisabet E Gardarsdottir; Gudmundur Georgsson; Olafia S Gretarsdottir; Kjartan R Gudmundsson; Thora R Gunnarsdottir; Arnaldur Gylfason; Hilma Holm; Brynjar O Jensson; Aslaug Jonasdottir; Kamilla S Josefsdottir; Thordur Kristjansson; Droplaug N Magnusdottir; Louise le Roux; Gudrun Sigmundsdottir; Gardar Sveinbjornsson; Kristin E Sveinsdottir; Maney Sveinsdottir; Emil A Thorarensen; Bjarni Thorbjornsson; Gisli Masson; Ingileif Jonsdottir; Alma Moller; Thorolfur Gudnason; Karl G Kristinsson; Unnur Thorsteinsdottir; Kari Stefansson |
| EPI_ISL_417561 | hCoV-19/Iceland/233/2020 | Europe / Iceland / Reykjavik | 2020-03-17 | The National University Hospital of Iceland | deCODE genetics | Daniel F Gudbjartsson; Agnar Helgason; Hakon Jonsson; Olafur T Magnusson; Pall Melsted; Gudmundur L Norddahl; Jona Saemundsdottir; Asgeir Sigurdsson; Patrick Sulem; Arna B Agustsdottir; Berglind Eiriksdottir; Run Fridriksdottir; Elisabet E Gardarsdottir; Gudmundur Georgsson; Olafia S Gretarsdottir; Kjartan R Gudmundsson; Thora R Gunnarsdottir; Arnaldur Gylfason; Hilma Holm; Brynjar O Jensson; Aslaug Jonasdottir; Kamilla S Josefsdottir; Thordur Kristjansson; Droplaug N Magnusdottir; Louise le Roux; Gudrun Sigmundsdottir; Gardar Sveinbjornsson; Kristin E Sveinsdottir; Maney Sveinsdottir; Emil A Thorarensen; Bjarni Thorbjornsson; Gisli Masson; Ingileif Jonsdottir; Alma Moller; Thorolfur Gudnason; Karl G Kristinsson; Unnur Thorsteinsdottir; Kari Stefansson |
| EPI_ISL_417562 | hCoV-19/Iceland/234/2020 | Europe / Iceland / Reykjavik | 2020-03-17 | The National University Hospital of Iceland | deCODE genetics | Daniel F Gudbjartsson; Agnar Helgason; Hakon Jonsson; Olafur T Magnusson; Pall Melsted; Gudmundur L Norddahl; Jona Saemundsdottir; Asgeir Sigurdsson; Patrick Sulem; Arna B Agustsdottir; Berglind Eiriksdottir; Run Fridriksdottir; Elisabet E Gardarsdottir; Gudmundur Georgsson; Olafia S Gretarsdottir; Kjartan R Gudmundsson; Thora R Gunnarsdottir; Arnaldur Gylfason; Hilma Holm; Brynjar O Jensson; Aslaug Jonasdottir; Kamilla S Josefsdottir; Thordur Kristjansson; Droplaug N Magnusdottir; Louise le Roux; Gudrun Sigmundsdottir; Gardar Sveinbjornsson; Kristin E Sveinsdottir; Maney Sveinsdottir; Emil A Thorarensen; Bjarni Thorbjornsson; Gisli Masson; Ingileif Jonsdottir; Alma Moller; Thorolfur Gudnason; Karl G Kristinsson; Unnur Thorsteinsdottir; Kari Stefansson |
| EPI_ISL_417563 | hCoV-19/Iceland/235/2020 | Europe / Iceland / Reykjavik | 2020-03-16 | The National University Hospital of Iceland | deCODE genetics | Daniel F Gudbjartsson; Agnar Helgason; Hakon Jonsson; Olafur T Magnusson; Pall Melsted; Gudmundur L Norddahl; Jona Saemundsdottir; Asgeir Sigurdsson; Patrick Sulem; Arna B Agustsdottir; Berglind Eiriksdottir; Run Fridriksdottir; Elisabet E Gardarsdottir; Gudmundur Georgsson; Olafia S Gretarsdottir; Kjartan R Gudmundsson; Thora R Gunnarsdottir; Arnaldur Gylfason; Hilma Holm; Brynjar O Jensson; Aslaug Jonasdottir; Kamilla S Josefsdottir; Thordur Kristjansson; Droplaug N Magnusdottir; Louise le Roux; Gudrun Sigmundsdottir; Gardar Sveinbjornsson; Kristin E Sveinsdottir; Maney Sveinsdottir; Emil A Thorarensen; Bjarni Thorbjornsson; Gisli Masson; Ingileif Jonsdottir; Alma Moller; Thorolfur Gudnason; Karl G Kristinsson; Unnur Thorsteinsdottir; Kari Stefansson |
| EPI_ISL_417564 | hCoV-19/Iceland/236/2020 | Europe / Iceland / Reykjavik | 2020-03-17 | The National University Hospital of Iceland | deCODE genetics | Daniel F Gudbjartsson; Agnar Helgason; Hakon Jonsson; Olafur T Magnusson; Pall Melsted; Gudmundur L Norddahl; Jona Saemundsdottir; Asgeir Sigurdsson; Patrick Sulem; Arna B Agustsdottir; Berglind Eiriksdottir; Run Fridriksdottir; Elisabet E Gardarsdottir; Gudmundur Georgsson; Olafia S Gretarsdottir; Kjartan R Gudmundsson; Thora R Gunnarsdottir; Arnaldur Gylfason; Hilma Holm; Brynjar O Jensson; Aslaug Jonasdottir; Kamilla S Josefsdottir; Thordur Kristjansson; Droplaug N Magnusdottir; Louise le Roux; Gudrun Sigmundsdottir; Gardar Sveinbjornsson; Kristin E Sveinsdottir; Maney Sveinsdottir; Emil A Thorarensen; Bjarni Thorbjornsson; Gisli Masson; Ingileif Jonsdottir; Alma Moller; Thorolfur Gudnason; Karl G Kristinsson; Unnur Thorsteinsdottir; Kari Stefansson |
| EPI_ISL_417565 | hCoV-19/Iceland/237/2020 | Europe / Iceland / Reykjavik | 2020-03-17 | The National University Hospital of Iceland | deCODE genetics | Daniel F Gudbjartsson; Agnar Helgason; Hakon Jonsson; Olafur T Magnusson; Pall Melsted; Gudmundur L Norddahl; Jona Saemundsdottir; Asgeir Sigurdsson; Patrick Sulem; Arna B Agustsdottir; Berglind Eiriksdottir; Run Fridriksdottir; Elisabet E Gardarsdottir; Gudmundur Georgsson; Olafia S Gretarsdottir; Kjartan R Gudmundsson; Thora R Gunnarsdottir; Arnaldur Gylfason; Hilma Holm; Brynjar O Jensson; Aslaug Jonasdottir; Kamilla S Josefsdottir; Thordur Kristjansson; Droplaug N Magnusdottir; Louise le Roux; Gudrun Sigmundsdottir; Gardar Sveinbjornsson; Kristin E Sveinsdottir; Maney Sveinsdottir; Emil A Thorarensen; Bjarni Thorbjornsson; Gisli Masson; Ingileif Jonsdottir; Alma Moller; Thorolfur Gudnason; Karl G Kristinsson; Unnur Thorsteinsdottir; Kari Stefansson |
| EPI_ISL_417566 | hCoV-19/Iceland/238/2020 | Europe / Iceland / Reykjavik | 2020-03-17 | The National University Hospital of Iceland | deCODE genetics | Daniel F Gudbjartsson; Agnar Helgason; Hakon Jonsson; Olafur T Magnusson; Pall Melsted; Gudmundur L Norddahl; Jona Saemundsdottir; Asgeir Sigurdsson; Patrick Sulem; Arna B Agustsdottir; Berglind Eiriksdottir; Run Fridriksdottir; Elisabet E Gardarsdottir; Gudmundur Georgsson; Olafia S Gretarsdottir; Kjartan R Gudmundsson; Thora R Gunnarsdottir; Arnaldur Gylfason; Hilma Holm; Brynjar O Jensson; Aslaug Jonasdottir; Kamilla S Josefsdottir; Thordur Kristjansson; Droplaug N Magnusdottir; Louise le Roux; Gudrun Sigmundsdottir; Gardar Sveinbjornsson; Kristin E Sveinsdottir; Maney Sveinsdottir; Emil A Thorarensen; Bjarni Thorbjornsson; Gisli Masson; Ingileif Jonsdottir; Alma Moller; Thorolfur Gudnason; Karl G Kristinsson; Unnur Thorsteinsdottir; Kari Stefansson |
| EPI_ISL_417567 | hCoV-19/Iceland/239/2020 | Europe / Iceland / Reykjavik | 2020-03-17 | The National University Hospital of Iceland | deCODE genetics | Daniel F Gudbjartsson; Agnar Helgason; Hakon Jonsson; Olafur T Magnusson; Pall Melsted; Gudmundur L Norddahl; Jona Saemundsdottir; Asgeir Sigurdsson; Patrick Sulem; Arna B Agustsdottir; Berglind Eiriksdottir; Run Fridriksdottir; Elisabet E Gardarsdottir; Gudmundur Georgsson; Olafia S Gretarsdottir; Kjartan R Gudmundsson; Thora R Gunnarsdottir; Arnaldur Gylfason; Hilma Holm; Brynjar O Jensson; Aslaug Jonasdottir; Kamilla S Josefsdottir; Thordur Kristjansson; Droplaug N Magnusdottir; Louise le Roux; Gudrun Sigmundsdottir; Gardar Sveinbjornsson; Kristin E Sveinsdottir; Maney Sveinsdottir; Emil A Thorarensen; Bjarni Thorbjornsson; Gisli Masson; Ingileif Jonsdottir; Alma Moller; Thorolfur Gudnason; Karl G Kristinsson; Unnur Thorsteinsdottir; Kari Stefansson |
| EPI_ISL_417568 | hCoV-19/Iceland/240/2020 | Europe / Iceland / Reykjavik | 2020-03-17 | The National University Hospital of Iceland | deCODE genetics | Daniel F Gudbjartsson; Agnar Helgason; Hakon Jonsson; Olafur T Magnusson; Pall Melsted; Gudmundur L Norddahl; Jona Saemundsdottir; Asgeir Sigurdsson; Patrick Sulem; Arna B Agustsdottir; Berglind Eiriksdottir; Run Fridriksdottir; Elisabet E Gardarsdottir; Gudmundur Georgsson; Olafia S Gretarsdottir; Kjartan R Gudmundsson; Thora R Gunnarsdottir; Arnaldur Gylfason; Hilma Holm; Brynjar O Jensson; Aslaug Jonasdottir; Kamilla S Josefsdottir; Thordur Kristjansson; Droplaug N Magnusdottir; Louise le Roux; Gudrun Sigmundsdottir; Gardar Sveinbjornsson; Kristin E Sveinsdottir; Maney Sveinsdottir; Emil A Thorarensen; Bjarni Thorbjornsson; Gisli Masson; Ingileif Jonsdottir; Alma Moller; Thorolfur Gudnason; Karl G Kristinsson; Unnur Thorsteinsdottir; Kari Stefansson |
| EPI_ISL_417569 | hCoV-19/Iceland/241/2020 | Europe / Iceland / Reykjavik | 2020-03-17 | The National University Hospital of Iceland | deCODE genetics | Daniel F Gudbjartsson; Agnar Helgason; Hakon Jonsson; Olafur T Magnusson; Pall Melsted; Gudmundur L Norddahl; Jona Saemundsdottir; Asgeir Sigurdsson; Patrick Sulem; Arna B Agustsdottir; Berglind Eiriksdottir; Run Fridriksdottir; Elisabet E Gardarsdottir; Gudmundur Georgsson; Olafia S Gretarsdottir; Kjartan R Gudmundsson; Thora R Gunnarsdottir; Arnaldur Gylfason; Hilma Holm; Brynjar O Jensson; Aslaug Jonasdottir; Kamilla S Josefsdottir; Thordur Kristjansson; Droplaug N Magnusdottir; Louise le Roux; Gudrun Sigmundsdottir; Gardar Sveinbjornsson; Kristin E Sveinsdottir; Maney Sveinsdottir; Emil A Thorarensen; Bjarni Thorbjornsson; Gisli Masson; Ingileif Jonsdottir; Alma Moller; Thorolfur Gudnason; Karl G Kristinsson; Unnur Thorsteinsdottir; Kari Stefansson |
| EPI_ISL_417570 | hCoV-19/Iceland/242/2020 | Europe / Iceland / Reykjavik | 2020-03-17 | The National University Hospital of Iceland | deCODE genetics | Daniel F Gudbjartsson; Agnar Helgason; Hakon Jonsson; Olafur T Magnusson; Pall Melsted; Gudmundur L Norddahl; Jona Saemundsdottir; Asgeir Sigurdsson; Patrick Sulem; Arna B Agustsdottir; Berglind Eiriksdottir; Run Fridriksdottir; Elisabet E Gardarsdottir; Gudmundur Georgsson; Olafia S Gretarsdottir; Kjartan R Gudmundsson; Thora R Gunnarsdottir; Arnaldur Gylfason; Hilma Holm; Brynjar O Jensson; Aslaug Jonasdottir; Kamilla S Josefsdottir; Thordur Kristjansson; Droplaug N Magnusdottir; Louise le Roux; Gudrun Sigmundsdottir; Gardar Sveinbjornsson; Kristin E Sveinsdottir; Maney Sveinsdottir; Emil A Thorarensen; Bjarni Thorbjornsson; Gisli Masson; Ingileif Jonsdottir; Alma Moller; Thorolfur Gudnason; Karl G Kristinsson; Unnur Thorsteinsdottir; Kari Stefansson |
| EPI_ISL_417571 | hCoV-19/Iceland/243/2020 | Europe / Iceland / Reykjavik | 2020-03-17 | The National University Hospital of Iceland | deCODE genetics | Daniel F Gudbjartsson; Agnar Helgason; Hakon Jonsson; Olafur T Magnusson; Pall Melsted; Gudmundur L Norddahl; Jona Saemundsdottir; Asgeir Sigurdsson; Patrick Sulem; Arna B Agustsdottir; Berglind Eiriksdottir; Run Fridriksdottir; Elisabet E Gardarsdottir; Gudmundur Georgsson; Olafia S Gretarsdottir; Kjartan R Gudmundsson; Thora R Gunnarsdottir; Arnaldur Gylfason; Hilma Holm; Brynjar O Jensson; Aslaug Jonasdottir; Kamilla S Josefsdottir; Thordur Kristjansson; Droplaug N Magnusdottir; Louise le Roux; Gudrun Sigmundsdottir; Gardar Sveinbjornsson; Kristin E Sveinsdottir; Maney Sveinsdottir; Emil A Thorarensen; Bjarni Thorbjornsson; Gisli Masson; Ingileif Jonsdottir; Alma Moller; Thorolfur Gudnason; Karl G Kristinsson; Unnur Thorsteinsdottir; Kari Stefansson |
| EPI_ISL_417572 | hCoV-19/Iceland/244/2020 | Europe / Iceland / Reykjavik | 2020-03-17 | The National University Hospital of Iceland | deCODE genetics | Daniel F Gudbjartsson; Agnar Helgason; Hakon Jonsson; Olafur T Magnusson; Pall Melsted; Gudmundur L Norddahl; Jona Saemundsdottir; Asgeir Sigurdsson; Patrick Sulem; Arna B Agustsdottir; Berglind Eiriksdottir; Run Fridriksdottir; Elisabet E Gardarsdottir; Gudmundur Georgsson; Olafia S Gretarsdottir; Kjartan R Gudmundsson; Thora R Gunnarsdottir; Arnaldur Gylfason; Hilma Holm; Brynjar O Jensson; Aslaug Jonasdottir; Kamilla S Josefsdottir; Thordur Kristjansson; Droplaug N Magnusdottir; Louise le Roux; Gudrun Sigmundsdottir; Gardar Sveinbjornsson; Kristin E Sveinsdottir; Maney Sveinsdottir; Emil A Thorarensen; Bjarni Thorbjornsson; Gisli Masson; Ingileif Jonsdottir; Alma Moller; Thorolfur Gudnason; Karl G Kristinsson; Unnur Thorsteinsdottir; Kari Stefansson |
| EPI_ISL_417573 | hCoV-19/Iceland/245/2020 | Europe / Iceland / Reykjavik | 2020-03-16 | The National University Hospital of Iceland | deCODE genetics | Daniel F Gudbjartsson; Agnar Helgason; Hakon Jonsson; Olafur T Magnusson; Pall Melsted; Gudmundur L Norddahl; Jona Saemundsdottir; Asgeir Sigurdsson; Patrick Sulem; Arna B Agustsdottir; Berglind Eiriksdottir; Run Fridriksdottir; Elisabet E Gardarsdottir; Gudmundur Georgsson; Olafia S Gretarsdottir; Kjartan R Gudmundsson; Thora R Gunnarsdottir; Arnaldur Gylfason; Hilma Holm; Brynjar O Jensson; Aslaug Jonasdottir; Kamilla S Josefsdottir; Thordur Kristjansson; Droplaug N Magnusdottir; Louise le Roux; Gudrun Sigmundsdottir; Gardar Sveinbjornsson; Kristin E Sveinsdottir; Maney Sveinsdottir; Emil A Thorarensen; Bjarni Thorbjornsson; Gisli Masson; Ingileif Jonsdottir; Alma Moller; Thorolfur Gudnason; Karl G Kristinsson; Unnur Thorsteinsdottir; Kari Stefansson |
| EPI_ISL_417574 | hCoV-19/Iceland/246/2020 | Europe / Iceland / Reykjavik | 2020-03-17 | The National University Hospital of Iceland | deCODE genetics | Daniel F Gudbjartsson; Agnar Helgason; Hakon Jonsson; Olafur T Magnusson; Pall Melsted; Gudmundur L Norddahl; Jona Saemundsdottir; Asgeir Sigurdsson; Patrick Sulem; Arna B Agustsdottir; Berglind Eiriksdottir; Run Fridriksdottir; Elisabet E Gardarsdottir; Gudmundur Georgsson; Olafia S Gretarsdottir; Kjartan R Gudmundsson; Thora R Gunnarsdottir; Arnaldur Gylfason; Hilma Holm; Brynjar O Jensson; Aslaug Jonasdottir; Kamilla S Josefsdottir; Thordur Kristjansson; Droplaug N Magnusdottir; Louise le Roux; Gudrun Sigmundsdottir; Gardar Sveinbjornsson; Kristin E Sveinsdottir; Maney Sveinsdottir; Emil A Thorarensen; Bjarni Thorbjornsson; Gisli Masson; Ingileif Jonsdottir; Alma Moller; Thorolfur Gudnason; Karl G Kristinsson; Unnur Thorsteinsdottir; Kari Stefansson |
| EPI_ISL_417575 | hCoV-19/Iceland/247/2020 | Europe / Iceland / Reykjavik | 2020-03-17 | The National University Hospital of Iceland | deCODE genetics | Daniel F Gudbjartsson; Agnar Helgason; Hakon Jonsson; Olafur T Magnusson; Pall Melsted; Gudmundur L Norddahl; Jona Saemundsdottir; Asgeir Sigurdsson; Patrick Sulem; Arna B Agustsdottir; Berglind Eiriksdottir; Run Fridriksdottir; Elisabet E Gardarsdottir; Gudmundur Georgsson; Olafia S Gretarsdottir; Kjartan R Gudmundsson; Thora R Gunnarsdottir; Arnaldur Gylfason; Hilma Holm; Brynjar O Jensson; Aslaug Jonasdottir; Kamilla S Josefsdottir; Thordur Kristjansson; Droplaug N Magnusdottir; Louise le Roux; Gudrun Sigmundsdottir; Gardar Sveinbjornsson; Kristin E Sveinsdottir; Maney Sveinsdottir; Emil A Thorarensen; Bjarni Thorbjornsson; Gisli Masson; Ingileif Jonsdottir; Alma Moller; Thorolfur Gudnason; Karl G Kristinsson; Unnur Thorsteinsdottir; Kari Stefansson |
| EPI_ISL_417576 | hCoV-19/Iceland/248/2020 | Europe / Iceland / Reykjavik | 2020-03-17 | The National University Hospital of Iceland | deCODE genetics | Daniel F Gudbjartsson; Agnar Helgason; Hakon Jonsson; Olafur T Magnusson; Pall Melsted; Gudmundur L Norddahl; Jona Saemundsdottir; Asgeir Sigurdsson; Patrick Sulem; Arna B Agustsdottir; Berglind Eiriksdottir; Run Fridriksdottir; Elisabet E Gardarsdottir; Gudmundur Georgsson; Olafia S Gretarsdottir; Kjartan R Gudmundsson; Thora R Gunnarsdottir; Arnaldur Gylfason; Hilma Holm; Brynjar O Jensson; Aslaug Jonasdottir; Kamilla S Josefsdottir; Thordur Kristjansson; Droplaug N Magnusdottir; Louise le Roux; Gudrun Sigmundsdottir; Gardar Sveinbjornsson; Kristin E Sveinsdottir; Maney Sveinsdottir; Emil A Thorarensen; Bjarni Thorbjornsson; Gisli Masson; Ingileif Jonsdottir; Alma Moller; Thorolfur Gudnason; Karl G Kristinsson; Unnur Thorsteinsdottir; Kari Stefansson |
| EPI_ISL_417577 | hCoV-19/Iceland/249/2020 | Europe / Iceland / Reykjavik | 2020-03-17 | The National University Hospital of Iceland | deCODE genetics | Daniel F Gudbjartsson; Agnar Helgason; Hakon Jonsson; Olafur T Magnusson; Pall Melsted; Gudmundur L Norddahl; Jona Saemundsdottir; Asgeir Sigurdsson; Patrick Sulem; Arna B Agustsdottir; Berglind Eiriksdottir; Run Fridriksdottir; Elisabet E Gardarsdottir; Gudmundur Georgsson; Olafia S Gretarsdottir; Kjartan R Gudmundsson; Thora R Gunnarsdottir; Arnaldur Gylfason; Hilma Holm; Brynjar O Jensson; Aslaug Jonasdottir; Kamilla S Josefsdottir; Thordur Kristjansson; Droplaug N Magnusdottir; Louise le Roux; Gudrun Sigmundsdottir; Gardar Sveinbjornsson; Kristin E Sveinsdottir; Maney Sveinsdottir; Emil A Thorarensen; Bjarni Thorbjornsson; Gisli Masson; Ingileif Jonsdottir; Alma Moller; Thorolfur Gudnason; Karl G Kristinsson; Unnur Thorsteinsdottir; Kari Stefansson |
| EPI_ISL_417578 | hCoV-19/Iceland/250/2020 | Europe / Iceland / Reykjavik | 2020-03-17 | The National University Hospital of Iceland | deCODE genetics | Daniel F Gudbjartsson; Agnar Helgason; Hakon Jonsson; Olafur T Magnusson; Pall Melsted; Gudmundur L Norddahl; Jona Saemundsdottir; Asgeir Sigurdsson; Patrick Sulem; Arna B Agustsdottir; Berglind Eiriksdottir; Run Fridriksdottir; Elisabet E Gardarsdottir; Gudmundur Georgsson; Olafia S Gretarsdottir; Kjartan R Gudmundsson; Thora R Gunnarsdottir; Arnaldur Gylfason; Hilma Holm; Brynjar O Jensson; Aslaug Jonasdottir; Kamilla S Josefsdottir; Thordur Kristjansson; Droplaug N Magnusdottir; Louise le Roux; Gudrun Sigmundsdottir; Gardar Sveinbjornsson; Kristin E Sveinsdottir; Maney Sveinsdottir; Emil A Thorarensen; Bjarni Thorbjornsson; Gisli Masson; Ingileif Jonsdottir; Alma Moller; Thorolfur Gudnason; Karl G Kristinsson; Unnur Thorsteinsdottir; Kari Stefansson |
| EPI_ISL_417580 | hCoV-19/Iceland/252/2020 | Europe / Iceland / Reykjavik | 2020-03-17 | The National University Hospital of Iceland | deCODE genetics | Daniel F Gudbjartsson; Agnar Helgason; Hakon Jonsson; Olafur T Magnusson; Pall Melsted; Gudmundur L Norddahl; Jona Saemundsdottir; Asgeir Sigurdsson; Patrick Sulem; Arna B Agustsdottir; Berglind Eiriksdottir; Run Fridriksdottir; Elisabet E Gardarsdottir; Gudmundur Georgsson; Olafia S Gretarsdottir; Kjartan R Gudmundsson; Thora R Gunnarsdottir; Arnaldur Gylfason; Hilma Holm; Brynjar O Jensson; Aslaug Jonasdottir; Kamilla S Josefsdottir; Thordur Kristjansson; Droplaug N Magnusdottir; Louise le Roux; Gudrun Sigmundsdottir; Gardar Sveinbjornsson; Kristin E Sveinsdottir; Maney Sveinsdottir; Emil A Thorarensen; Bjarni Thorbjornsson; Gisli Masson; Ingileif Jonsdottir; Alma Moller; Thorolfur Gudnason; Karl G Kristinsson; Unnur Thorsteinsdottir; Kari Stefansson |
| EPI_ISL_417581 | hCoV-19/Iceland/253/2020 | Europe / Iceland / Reykjavik | 2020-03-17 | The National University Hospital of Iceland | deCODE genetics | Daniel F Gudbjartsson; Agnar Helgason; Hakon Jonsson; Olafur T Magnusson; Pall Melsted; Gudmundur L Norddahl; Jona Saemundsdottir; Asgeir Sigurdsson; Patrick Sulem; Arna B Agustsdottir; Berglind Eiriksdottir; Run Fridriksdottir; Elisabet E Gardarsdottir; Gudmundur Georgsson; Olafia S Gretarsdottir; Kjartan R Gudmundsson; Thora R Gunnarsdottir; Arnaldur Gylfason; Hilma Holm; Brynjar O Jensson; Aslaug Jonasdottir; Kamilla S Josefsdottir; Thordur Kristjansson; Droplaug N Magnusdottir; Louise le Roux; Gudrun Sigmundsdottir; Gardar Sveinbjornsson; Kristin E Sveinsdottir; Maney Sveinsdottir; Emil A Thorarensen; Bjarni Thorbjornsson; Gisli Masson; Ingileif Jonsdottir; Alma Moller; Thorolfur Gudnason; Karl G Kristinsson; Unnur Thorsteinsdottir; Kari Stefansson |

| | | | | | | |
|---|---|---|---|---|---|---|
| EPI_ISL_417582 | hCoV-19/Iceland/254/2020 | Europe / Iceland / Reykjavik | 2020-03-17 | The National University Hospital of Iceland | deCODE genetics | Daniel F Gudbjartsson; Agnar Helgason; Hakon Jonsson; Olafur T Magnusson; Pall Melsted; Gudmundur L Norddahl; Jona Saemundsdottir; Asgeir Sigurdsson; Patrick Sulem; Arna B Agustsdottir; Berglind Eiriksdottir; Run Fridriksdottir; Elisabet E Gardarsdottir; Gudmundur Georgsson; Olafia S Gretarsdottir; Kjartan R Gudmundsson; Thora R Gunnarsdottir; Arnaldur Gylfason; Hilma Holm; Brynjar O Jensson; Aslaug Jonasdottir; Kamilla S Josefsdottir; Thordur Kristjansson; Droplaug N Magnusdottir; Louise le Roux; Gudrun Sigmundsdottir; Gardar Sveinbjornsson; Kristin E Sveinsdottir; Maney Sveinsdottir; Emil A Thorarensen; Bjarni Thorbjornsson; Gisli Masson; Ingileif Jonsdottir; Alma Moller; Thorolfur Gudnason; Karl G Kristinsson; Unnur Thorsteinsdottir; Kari Stefansson |
| EPI_ISL_417583 | hCoV-19/Iceland/255/2020 | Europe / Iceland / Reykjavik | 2020-03-17 | The National University Hospital of Iceland | deCODE genetics | Daniel F Gudbjartsson; Agnar Helgason; Hakon Jonsson; Olafur T Magnusson; Pall Melsted; Gudmundur L Norddahl; Jona Saemundsdottir; Asgeir Sigurdsson; Patrick Sulem; Arna B Agustsdottir; Berglind Eiriksdottir; Run Fridriksdottir; Elisabet E Gardarsdottir; Gudmundur Georgsson; Olafia S Gretarsdottir; Kjartan R Gudmundsson; Thora R Gunnarsdottir; Arnaldur Gylfason; Hilma Holm; Brynjar O Jensson; Aslaug Jonasdottir; Kamilla S Josefsdottir; Thordur Kristjansson; Droplaug N Magnusdottir; Louise le Roux; Gudrun Sigmundsdottir; Gardar Sveinbjornsson; Kristin E Sveinsdottir; Maney Sveinsdottir; Emil A Thorarensen; Bjarni Thorbjornsson; Gisli Masson; Ingileif Jonsdottir; Alma Moller; Thorolfur Gudnason; Karl G Kristinsson; Unnur Thorsteinsdottir; Kari Stefansson |
| EPI_ISL_417584 | hCoV-19/Iceland/256/2020 | Europe / Iceland / Reykjavik | 2020-03-17 | The National University Hospital of Iceland | deCODE genetics | Daniel F Gudbjartsson; Agnar Helgason; Hakon Jonsson; Olafur T Magnusson; Pall Melsted; Gudmundur L Norddahl; Jona Saemundsdottir; Asgeir Sigurdsson; Patrick Sulem; Arna B Agustsdottir; Berglind Eiriksdottir; Run Fridriksdottir; Elisabet E Gardarsdottir; Gudmundur Georgsson; Olafia S Gretarsdottir; Kjartan R Gudmundsson; Thora R Gunnarsdottir; Arnaldur Gylfason; Hilma Holm; Brynjar O Jensson; Aslaug Jonasdottir; Kamilla S Josefsdottir; Thordur Kristjansson; Droplaug N Magnusdottir; Louise le Roux; Gudrun Sigmundsdottir; Gardar Sveinbjornsson; Kristin E Sveinsdottir; Maney Sveinsdottir; Emil A Thorarensen; Bjarni Thorbjornsson; Gisli Masson; Ingileif Jonsdottir; Alma Moller; Thorolfur Gudnason; Karl G Kristinsson; Unnur Thorsteinsdottir; Kari Stefansson |
| EPI_ISL_417585 | hCoV-19/Iceland/257/2020 | Europe / Iceland / Reykjavik | 2020-03-17 | The National University Hospital of Iceland | deCODE genetics | Daniel F Gudbjartsson; Agnar Helgason; Hakon Jonsson; Olafur T Magnusson; Pall Melsted; Gudmundur L Norddahl; Jona Saemundsdottir; Asgeir Sigurdsson; Patrick Sulem; Arna B Agustsdottir; Berglind Eiriksdottir; Run Fridriksdottir; Elisabet E Gardarsdottir; Gudmundur Georgsson; Olafia S Gretarsdottir; Kjartan R Gudmundsson; Thora R Gunnarsdottir; Arnaldur Gylfason; Hilma Holm; Brynjar O Jensson; Aslaug Jonasdottir; Kamilla S Josefsdottir; Thordur Kristjansson; Droplaug N Magnusdottir; Louise le Roux; Gudrun Sigmundsdottir; Gardar Sveinbjornsson; Kristin E Sveinsdottir; Maney Sveinsdottir; Emil A Thorarensen; Bjarni Thorbjornsson; Gisli Masson; Ingileif Jonsdottir; Alma Moller; Thorolfur Gudnason; Karl G Kristinsson; Unnur Thorsteinsdottir; Kari Stefansson |
| EPI_ISL_417586 | hCoV-19/Iceland/258/2020 | Europe / Iceland / Reykjavik | 2020-03-17 | The National University Hospital of Iceland | deCODE genetics | Daniel F Gudbjartsson; Agnar Helgason; Hakon Jonsson; Olafur T Magnusson; Pall Melsted; Gudmundur L Norddahl; Jona Saemundsdottir; Asgeir Sigurdsson; Patrick Sulem; Arna B Agustsdottir; Berglind Eiriksdottir; Run Fridriksdottir; Elisabet E Gardarsdottir; Gudmundur Georgsson; Olafia S Gretarsdottir; Kjartan R Gudmundsson; Thora R Gunnarsdottir; Arnaldur Gylfason; Hilma Holm; Brynjar O Jensson; Aslaug Jonasdottir; Kamilla S Josefsdottir; Thordur Kristjansson; Droplaug N Magnusdottir; Louise le Roux; Gudrun Sigmundsdottir; Gardar Sveinbjornsson; Kristin E Sveinsdottir; Maney Sveinsdottir; Emil A Thorarensen; Bjarni Thorbjornsson; Gisli Masson; Ingileif Jonsdottir; Alma Moller; Thorolfur Gudnason; Karl G Kristinsson; Unnur Thorsteinsdottir; Kari Stefansson |
| EPI_ISL_417587 | hCoV-19/Iceland/259/2020 | Europe / Iceland / Reykjavik | 2020-03-17 | The National University Hospital of Iceland | deCODE genetics | Daniel F Gudbjartsson; Agnar Helgason; Hakon Jonsson; Olafur T Magnusson; Pall Melsted; Gudmundur L Norddahl; Jona Saemundsdottir; Asgeir Sigurdsson; Patrick Sulem; Arna B Agustsdottir; Berglind Eiriksdottir; Run Fridriksdottir; Elisabet E Gardarsdottir; Gudmundur Georgsson; Olafia S Gretarsdottir; Kjartan R Gudmundsson; Thora R Gunnarsdottir; Arnaldur Gylfason; Hilma Holm; Brynjar O Jensson; Aslaug Jonasdottir; Kamilla S Josefsdottir; Thordur Kristjansson; Droplaug N Magnusdottir; Louise le Roux; Gudrun Sigmundsdottir; Gardar Sveinbjornsson; Kristin E Sveinsdottir; Maney Sveinsdottir; Emil A Thorarensen; Bjarni Thorbjornsson; Gisli Masson; Ingileif Jonsdottir; Alma Moller; Thorolfur Gudnason; Karl G Kristinsson; Unnur Thorsteinsdottir; Kari Stefansson |
| EPI_ISL_417588 | hCoV-19/Iceland/260/2020 | Europe / Iceland / Reykjavik | 2020-03-17 | The National University Hospital of Iceland | deCODE genetics | Daniel F Gudbjartsson; Agnar Helgason; Hakon Jonsson; Olafur T Magnusson; Pall Melsted; Gudmundur L Norddahl; Jona Saemundsdottir; Asgeir Sigurdsson; Patrick Sulem; Arna B Agustsdottir; Berglind Eiriksdottir; Run Fridriksdottir; Elisabet E Gardarsdottir; Gudmundur Georgsson; Olafia S Gretarsdottir; Kjartan R Gudmundsson; Thora R Gunnarsdottir; Arnaldur Gylfason; Hilma Holm; Brynjar O Jensson; Aslaug Jonasdottir; Kamilla S Josefsdottir; Thordur Kristjansson; Droplaug N Magnusdottir; Louise le Roux; Gudrun Sigmundsdottir; Gardar Sveinbjornsson; Kristin E Sveinsdottir; Maney Sveinsdottir; Emil A Thorarensen; Bjarni Thorbjornsson; Gisli Masson; Ingileif Jonsdottir; Alma Moller; Thorolfur Gudnason; Karl G Kristinsson; Unnur Thorsteinsdottir; Kari Stefansson |
| EPI_ISL_417589 | hCoV-19/Iceland/261/2020 | Europe / Iceland / Reykjavik | 2020-03-17 | The National University Hospital of Iceland | deCODE genetics | Daniel F Gudbjartsson; Agnar Helgason; Hakon Jonsson; Olafur T Magnusson; Pall Melsted; Gudmundur L Norddahl; Jona Saemundsdottir; Asgeir Sigurdsson; Patrick Sulem; Arna B Agustsdottir; Berglind Eiriksdottir; Run Fridriksdottir; Elisabet E Gardarsdottir; Gudmundur Georgsson; Olafia S Gretarsdottir; Kjartan R Gudmundsson; Thora R Gunnarsdottir; Arnaldur Gylfason; Hilma Holm; Brynjar O Jensson; Aslaug Jonasdottir; Kamilla S Josefsdottir; Thordur Kristjansson; Droplaug N Magnusdottir; Louise le Roux; Gudrun Sigmundsdottir; Gardar Sveinbjornsson; Kristin E Sveinsdottir; Maney Sveinsdottir; Emil A Thorarensen; Bjarni Thorbjornsson; Gisli Masson; Ingileif Jonsdottir; Alma Moller; Thorolfur Gudnason; Karl G Kristinsson; Unnur Thorsteinsdottir; Kari Stefansson |
| EPI_ISL_417590 | hCoV-19/Iceland/262/2020 | Europe / Iceland / Reykjavik | 2020-03-18 | deCODE genetics | deCODE genetics | Daniel F Gudbjartsson; Agnar Helgason; Hakon Jonsson; Olafur T Magnusson; Pall Melsted; Gudmundur L Norddahl; Jona Saemundsdottir; Asgeir Sigurdsson; Patrick Sulem; Arna B Agustsdottir; Berglind Eiriksdottir; Run Fridriksdottir; Elisabet E Gardarsdottir; Gudmundur Georgsson; Olafia S Gretarsdottir; Kjartan R Gudmundsson; Thora R Gunnarsdottir; Arnaldur Gylfason; Hilma Holm; Brynjar O Jensson; Aslaug Jonasdottir; Kamilla S Josefsdottir; Thordur Kristjansson; Droplaug N Magnusdottir; Louise le Roux; Gudrun Sigmundsdottir; Gardar Sveinbjornsson; Kristin E Sveinsdottir; Maney Sveinsdottir; Emil A Thorarensen; Bjarni Thorbjornsson; Gisli Masson; Ingileif Jonsdottir; Alma Moller; Thorolfur Gudnason; Karl G Kristinsson; Unnur Thorsteinsdottir; Kari Stefansson |
| EPI_ISL_417591 | hCoV-19/Iceland/263/2020 | Europe / Iceland / Reykjavik | 2020-03-17 | The National University Hospital of Iceland | deCODE genetics | Daniel F Gudbjartsson; Agnar Helgason; Hakon Jonsson; Olafur T Magnusson; Pall Melsted; Gudmundur L Norddahl; Jona Saemundsdottir; Asgeir Sigurdsson; Patrick Sulem; Arna B Agustsdottir; Berglind Eiriksdottir; Run Fridriksdottir; Elisabet E Gardarsdottir; Gudmundur Georgsson; Olafia S Gretarsdottir; Kjartan R Gudmundsson; Thora R Gunnarsdottir; Arnaldur Gylfason; Hilma Holm; Brynjar O Jensson; Aslaug Jonasdottir; Kamilla S Josefsdottir; Thordur Kristjansson; Droplaug N Magnusdottir; Louise le Roux; Gudrun Sigmundsdottir; Gardar Sveinbjornsson; Kristin E Sveinsdottir; Maney Sveinsdottir; Emil A Thorarensen; Bjarni Thorbjornsson; Gisli Masson; Ingileif Jonsdottir; Alma Moller; Thorolfur Gudnason; Karl G Kristinsson; Unnur Thorsteinsdottir; Kari Stefansson |
| EPI_ISL_417592 | hCoV-19/Iceland/264/2020 | Europe / Iceland / Reykjavik | 2020-03-17 | The National University Hospital of Iceland | deCODE genetics | Daniel F Gudbjartsson; Agnar Helgason; Hakon Jonsson; Olafur T Magnusson; Pall Melsted; Gudmundur L Norddahl; Jona Saemundsdottir; Asgeir Sigurdsson; Patrick Sulem; Arna B Agustsdottir; Berglind Eiriksdottir; Run Fridriksdottir; Elisabet E Gardarsdottir; Gudmundur Georgsson; Olafia S Gretarsdottir; Kjartan R Gudmundsson; Thora R Gunnarsdottir; Arnaldur Gylfason; Hilma Holm; Brynjar O Jensson; Aslaug Jonasdottir; Kamilla S Josefsdottir; Thordur Kristjansson; Droplaug N Magnusdottir; Louise le Roux; Gudrun Sigmundsdottir; Gardar Sveinbjornsson; Kristin E Sveinsdottir; Maney Sveinsdottir; Emil A Thorarensen; Bjarni Thorbjornsson; Gisli Masson; Ingileif Jonsdottir; Alma Moller; Thorolfur Gudnason; Karl G Kristinsson; Unnur Thorsteinsdottir; Kari Stefansson |
| EPI_ISL_417595 | hCoV-19/Iceland/267/2020 | Europe / Iceland / Reykjavik | 2020-03-17 | The National University Hospital of Iceland | deCODE genetics | Daniel F Gudbjartsson; Agnar Helgason; Hakon Jonsson; Olafur T Magnusson; Pall Melsted; Gudmundur L Norddahl; Jona Saemundsdottir; Asgeir Sigurdsson; Patrick Sulem; Arna B Agustsdottir; Berglind Eiriksdottir; Run Fridriksdottir; Elisabet E Gardarsdottir; Gudmundur Georgsson; Olafia S Gretarsdottir; Kjartan R Gudmundsson; Thora R Gunnarsdottir; Arnaldur Gylfason; Hilma Holm; Brynjar O Jensson; Aslaug Jonasdottir; Kamilla S Josefsdottir; Thordur Kristjansson; Droplaug N Magnusdottir; Louise le Roux; Gudrun Sigmundsdottir; Gardar Sveinbjornsson; Kristin E Sveinsdottir; Maney Sveinsdottir; Emil A Thorarensen; Bjarni Thorbjornsson; Gisli Masson; Ingileif Jonsdottir; Alma Moller; Thorolfur Gudnason; Karl G Kristinsson; Unnur Thorsteinsdottir; Kari Stefansson |
| EPI_ISL_417597 | hCoV-19/Iceland/269/2020 | Europe / Iceland / Reykjavik | 2020-03-17 | The National University Hospital of Iceland | deCODE genetics | Daniel F Gudbjartsson; Agnar Helgason; Hakon Jonsson; Olafur T Magnusson; Pall Melsted; Gudmundur L Norddahl; Jona Saemundsdottir; Asgeir Sigurdsson; Patrick Sulem; Arna B Agustsdottir; Berglind Eiriksdottir; Run Fridriksdottir; Elisabet E Gardarsdottir; Gudmundur Georgsson; Olafia S Gretarsdottir; Kjartan R Gudmundsson; Thora R Gunnarsdottir; Arnaldur Gylfason; Hilma Holm; Brynjar O Jensson; Aslaug Jonasdottir; Kamilla S Josefsdottir; Thordur Kristjansson; Droplaug N Magnusdottir; Louise le Roux; Gudrun Sigmundsdottir; Gardar Sveinbjornsson; Kristin E Sveinsdottir; Maney Sveinsdottir; Emil A Thorarensen; Bjarni Thorbjornsson; Gisli Masson; Ingileif Jonsdottir; Alma Moller; Thorolfur Gudnason; Karl G Kristinsson; Unnur Thorsteinsdottir; Kari Stefansson |
| EPI_ISL_417598 | hCoV-19/Iceland/270/2020 | Europe / Iceland / Reykjavik | 2020-03-17 | The National University Hospital of Iceland | deCODE genetics | Daniel F Gudbjartsson; Agnar Helgason; Hakon Jonsson; Olafur T Magnusson; Pall Melsted; Gudmundur L Norddahl; Jona Saemundsdottir; Asgeir Sigurdsson; Patrick Sulem; Arna B Agustsdottir; Berglind Eiriksdottir; Run Fridriksdottir; Elisabet E Gardarsdottir; Gudmundur Georgsson; Olafia S Gretarsdottir; Kjartan R Gudmundsson; Thora R Gunnarsdottir; Arnaldur Gylfason; Hilma Holm; Brynjar O Jensson; Aslaug Jonasdottir; Kamilla S Josefsdottir; Thordur Kristjansson; Droplaug N Magnusdottir; Louise le Roux; Gudrun Sigmundsdottir; Gardar Sveinbjornsson; Kristin E Sveinsdottir; Maney Sveinsdottir; Emil A Thorarensen; Bjarni Thorbjornsson; Gisli Masson; Ingileif Jonsdottir; Alma Moller; Thorolfur Gudnason; Karl G Kristinsson; Unnur Thorsteinsdottir; Kari Stefansson |
| EPI_ISL_417650 | hCoV-19/Iceland/321/2020 | Europe / Iceland / Reykjavik | 2020-03-18 | The National University Hospital of Iceland | deCODE genetics | Daniel F Gudbjartsson; Agnar Helgason; Hakon Jonsson; Olafur T Magnusson; Pall Melsted; Gudmundur L Norddahl; Jona Saemundsdottir; Asgeir Sigurdsson; Patrick Sulem; Arna B Agustsdottir; Berglind Eiriksdottir; Run Fridriksdottir; Elisabet E Gardarsdottir; Gudmundur Georgsson; Olafia S Gretarsdottir; Kjartan R Gudmundsson; Thora R Gunnarsdottir; Arnaldur Gylfason; Hilma Holm; Brynjar O Jensson; Aslaug Jonasdottir; Kamilla S Josefsdottir; Thordur Kristjansson; Droplaug N Magnusdottir; Louise le Roux; Gudrun Sigmundsdottir; Gardar Sveinbjornsson; Kristin E Sveinsdottir; Maney Sveinsdottir; Emil A Thorarensen; Bjarni Thorbjornsson; Gisli Masson; Ingileif Jonsdottir; Alma Moller; Thorolfur Gudnason; Karl G Kristinsson; Unnur Thorsteinsdottir; Kari Stefansson |
| EPI_ISL_417651 | hCoV-19/Iceland/322/2020 | Europe / Iceland / Reykjavik | 2020-03-18 | The National University Hospital of Iceland | deCODE genetics | Daniel F Gudbjartsson; Agnar Helgason; Hakon Jonsson; Olafur T Magnusson; Pall Melsted; Gudmundur L Norddahl; Jona Saemundsdottir; Asgeir Sigurdsson; Patrick Sulem; Arna B Agustsdottir; Berglind Eiriksdottir; Run Fridriksdottir; Elisabet E Gardarsdottir; Gudmundur Georgsson; Olafia S Gretarsdottir; Kjartan R Gudmundsson; Thora R Gunnarsdottir; Arnaldur Gylfason; Hilma Holm; Brynjar O Jensson; Aslaug Jonasdottir; Kamilla S Josefsdottir; Thordur Kristjansson; Droplaug N Magnusdottir; Louise le Roux; Gudrun Sigmundsdottir; Gardar Sveinbjornsson; Kristin E Sveinsdottir; Maney Sveinsdottir; Emil A Thorarensen; Bjarni Thorbjornsson; Gisli Masson; Ingileif Jonsdottir; Alma Moller; Thorolfur Gudnason; Karl G Kristinsson; Unnur Thorsteinsdottir; Kari Stefansson |
| EPI_ISL_417652 | hCoV-19/Iceland/323/2020 | Europe / Iceland / Reykjavik | 2020-03-18 | The National University Hospital of Iceland | deCODE genetics | Daniel F Gudbjartsson; Agnar Helgason; Hakon Jonsson; Olafur T Magnusson; Pall Melsted; Gudmundur L Norddahl; Jona Saemundsdottir; Asgeir Sigurdsson; Patrick Sulem; Arna B Agustsdottir; Berglind Eiriksdottir; Run Fridriksdottir; Elisabet E Gardarsdottir; Gudmundur Georgsson; Olafia S Gretarsdottir; Kjartan R Gudmundsson; Thora R Gunnarsdottir; Arnaldur Gylfason; Hilma Holm; Brynjar O Jensson; Aslaug Jonasdottir; Kamilla S Josefsdottir; Thordur Kristjansson; Droplaug N Magnusdottir; Louise le Roux; Gudrun Sigmundsdottir; Gardar Sveinbjornsson; Kristin E Sveinsdottir; Maney Sveinsdottir; Emil A Thorarensen; Bjarni Thorbjornsson; Gisli Masson; Ingileif Jonsdottir; Alma Moller; Thorolfur Gudnason; Karl G Kristinsson; Unnur Thorsteinsdottir; Kari Stefansson |
| EPI_ISL_417653 | hCoV-19/Iceland/324/2020 | Europe / Iceland / Reykjavik | 2020-03-18 | The National University Hospital of Iceland | deCODE genetics | Daniel F Gudbjartsson; Agnar Helgason; Hakon Jonsson; Olafur T Magnusson; Pall Melsted; Gudmundur L Norddahl; Jona Saemundsdottir; Asgeir Sigurdsson; Patrick Sulem; Arna B Agustsdottir; Berglind Eiriksdottir; Run Fridriksdottir; Elisabet E Gardarsdottir; Gudmundur Georgsson; Olafia S Gretarsdottir; Kjartan R Gudmundsson; Thora R Gunnarsdottir; Arnaldur Gylfason; Hilma Holm; Brynjar O Jensson; Aslaug Jonasdottir; Kamilla S Josefsdottir; Thordur Kristjansson; Droplaug N Magnusdottir; Louise le Roux; Gudrun Sigmundsdottir; Gardar Sveinbjornsson; Kristin E Sveinsdottir; Maney Sveinsdottir; Emil A Thorarensen; Bjarni Thorbjornsson; Gisli Masson; Ingileif Jonsdottir; Alma Moller; Thorolfur Gudnason; Karl G Kristinsson; Unnur Thorsteinsdottir; Kari Stefansson |
| EPI_ISL_417654 | hCoV-19/Iceland/325/2020 | Europe / Iceland / Reykjavik | 2020-03-18 | The National University Hospital of Iceland | deCODE genetics | Daniel F Gudbjartsson; Agnar Helgason; Hakon Jonsson; Olafur T Magnusson; Pall Melsted; Gudmundur L Norddahl; Jona Saemundsdottir; Asgeir Sigurdsson; Patrick Sulem; Arna B Agustsdottir; Berglind Eiriksdottir; Run Fridriksdottir; Elisabet E Gardarsdottir; Gudmundur Georgsson; Olafia S Gretarsdottir; Kjartan R Gudmundsson; Thora R Gunnarsdottir; Arnaldur Gylfason; Hilma Holm; Brynjar O Jensson; Aslaug Jonasdottir; Kamilla S Josefsdottir; Thordur Kristjansson; Droplaug N Magnusdottir; Louise le Roux; Gudrun Sigmundsdottir; Gardar Sveinbjornsson; Kristin E Sveinsdottir; Maney Sveinsdottir; Emil A Thorarensen; Bjarni Thorbjornsson; Gisli Masson; Ingileif Jonsdottir; Alma Moller; Thorolfur Gudnason; Karl G Kristinsson; Unnur Thorsteinsdottir; Kari Stefansson |
| EPI_ISL_417655 | hCoV-19/Iceland/326/2020 | Europe / Iceland / Reykjavik | 2020-03-14 | deCODE genetics | deCODE genetics | Daniel F Gudbjartsson; Agnar Helgason; Hakon Jonsson; Olafur T Magnusson; Pall Melsted; Gudmundur L Norddahl; Jona Saemundsdottir; Asgeir Sigurdsson; Patrick Sulem; Arna B Agustsdottir; Berglind Eiriksdottir; Run Fridriksdottir; Elisabet E Gardarsdottir; Gudmundur Georgsson; Olafia S Gretarsdottir; Kjartan R Gudmundsson; Thora R Gunnarsdottir; Arnaldur Gylfason; Hilma Holm; Brynjar O Jensson; Aslaug Jonasdottir; Kamilla S Josefsdottir; Thordur Kristjansson; Droplaug N Magnusdottir; Louise le Roux; Gudrun Sigmundsdottir; Gardar Sveinbjornsson; Kristin E Sveinsdottir; Maney Sveinsdottir; Emil A Thorarensen; Bjarni Thorbjornsson; Gisli Masson; Ingileif Jonsdottir; Alma Moller; Thorolfur Gudnason; Karl G Kristinsson; Unnur Thorsteinsdottir; Kari Stefansson |
| EPI_ISL_417659 | hCoV-19/Iceland/330/2020 | Europe / Iceland / Reykjavik | 2020-03-14 | deCODE genetics | deCODE genetics | Daniel F Gudbjartsson; Agnar Helgason; Hakon Jonsson; Olafur T Magnusson; Pall Melsted; Gudmundur L Norddahl; Jona Saemundsdottir; Asgeir Sigurdsson; Patrick Sulem; Arna B Agustsdottir; Berglind Eiriksdottir; Run Fridriksdottir; Elisabet E Gardarsdottir; Gudmundur Georgsson; Olafia S Gretarsdottir; Kjartan R Gudmundsson; Thora R Gunnarsdottir; Arnaldur Gylfason; Hilma Holm; Brynjar O Jensson; Aslaug Jonasdottir; Kamilla S Josefsdottir; Thordur Kristjansson; Droplaug N Magnusdottir; Louise le Roux; Gudrun Sigmundsdottir; Gardar Sveinbjornsson; Kristin E Sveinsdottir; Maney Sveinsdottir; Emil A Thorarensen; Bjarni Thorbjornsson; Gisli Masson; Ingileif Jonsdottir; Alma Moller; Thorolfur Gudnason; Karl G Kristinsson; Unnur Thorsteinsdottir; Kari Stefansson |
| EPI_ISL_417660 | hCoV-19/Iceland/331/2020 | Europe / Iceland / Reykjavik | 2020-03-14 | deCODE genetics | deCODE genetics | Daniel F Gudbjartsson; Agnar Helgason; Hakon Jonsson; Olafur T Magnusson; Pall Melsted; Gudmundur L Norddahl; Jona Saemundsdottir; Asgeir Sigurdsson; Patrick Sulem; Arna B Agustsdottir; Berglind Eiriksdottir; Run Fridriksdottir; Elisabet E Gardarsdottir; Gudmundur Georgsson; Olafia S Gretarsdottir; Kjartan R Gudmundsson; Thora R Gunnarsdottir; Arnaldur Gylfason; Hilma Holm; Brynjar O Jensson; Aslaug Jonasdottir; Kamilla S Josefsdottir; Thordur Kristjansson; Droplaug N Magnusdottir; Louise le Roux; Gudrun Sigmundsdottir; Gardar Sveinbjornsson; Kristin E Sveinsdottir; Maney Sveinsdottir; Emil A Thorarensen; Bjarni Thorbjornsson; Gisli Masson; Ingileif Jonsdottir; Alma Moller; Thorolfur Gudnason; Karl G Kristinsson; Unnur Thorsteinsdottir; Kari Stefansson |
| EPI_ISL_417662 | hCoV-19/Iceland/333/2020 | Europe / Iceland / Reykjavik | 2020-03-14 | deCODE genetics | deCODE genetics | Daniel F Gudbjartsson; Agnar Helgason; Hakon Jonsson; Olafur T Magnusson; Pall Melsted; Gudmundur L Norddahl; Jona Saemundsdottir; Asgeir Sigurdsson; Patrick Sulem; Arna B Agustsdottir; Berglind Eiriksdottir; Run Fridriksdottir; Elisabet E Gardarsdottir; Gudmundur Georgsson; Olafia S Gretarsdottir; Kjartan R Gudmundsson; Thora R Gunnarsdottir; Arnaldur Gylfason; Hilma Holm; Brynjar O Jensson; Aslaug Jonasdottir; Kamilla S Josefsdottir; Thordur Kristjansson; Droplaug N Magnusdottir; Louise le Roux; Gudrun Sigmundsdottir; Gardar Sveinbjornsson; Kristin E Sveinsdottir; Maney Sveinsdottir; Emil A Thorarensen; Bjarni Thorbjornsson; Gisli Masson; Ingileif Jonsdottir; Alma Moller; Thorolfur Gudnason; Karl G Kristinsson; Unnur Thorsteinsdottir; Kari Stefansson |
| EPI_ISL_417663 | hCoV-19/Iceland/334/2020 | Europe / Iceland / Reykjavik | 2020-03-14 | deCODE genetics | deCODE genetics | Daniel F Gudbjartsson; Agnar Helgason; Hakon Jonsson; Olafur T Magnusson; Pall Melsted; Gudmundur L Norddahl; Jona Saemundsdottir; Asgeir Sigurdsson; Patrick Sulem; Arna B Agustsdottir; Berglind Eiriksdottir; Run Fridriksdottir; Elisabet E Gardarsdottir; Gudmundur Georgsson; Olafia S Gretarsdottir; Kjartan R Gudmundsson; Thora R Gunnarsdottir; Arnaldur Gylfason; Hilma Holm; Brynjar O Jensson; Aslaug Jonasdottir; Kamilla S Josefsdottir; Thordur Kristjansson; Droplaug N Magnusdottir; Louise le Roux; Gudrun Sigmundsdottir; Gardar Sveinbjornsson; Kristin E Sveinsdottir; Maney Sveinsdottir; Emil A Thorarensen; Bjarni Thorbjornsson; Gisli Masson; Ingileif Jonsdottir; Alma Moller; Thorolfur Gudnason; Karl G Kristinsson; Unnur Thorsteinsdottir; Kari Stefansson |
| EPI_ISL_417664 | hCoV-19/Iceland/335/2020 | Europe / Iceland / Reykjavik | 2020-03-13 | deCODE genetics | deCODE genetics | Daniel F Gudbjartsson; Agnar Helgason; Hakon Jonsson; Olafur T Magnusson; Pall Melsted; Gudmundur L Norddahl; Jona Saemundsdottir; Asgeir Sigurdsson; Patrick Sulem; Arna B Agustsdottir; Berglind Eiriksdottir; Run Fridriksdottir; Elisabet E Gardarsdottir; Gudmundur Georgsson; Olafia S Gretarsdottir; Kjartan R Gudmundsson; Thora R Gunnarsdottir; Arnaldur Gylfason; Hilma Holm; Brynjar O Jensson; Aslaug Jonasdottir; Kamilla S Josefsdottir; Thordur Kristjansson; Droplaug N Magnusdottir; Louise le Roux; Gudrun Sigmundsdottir; Gardar Sveinbjornsson; Kristin E Sveinsdottir; Maney Sveinsdottir; Emil A Thorarensen; Bjarni Thorbjornsson; Gisli Masson; Ingileif Jonsdottir; Alma Moller; Thorolfur Gudnason; Karl G Kristinsson; Unnur Thorsteinsdottir; Kari Stefansson |
| EPI_ISL_417665 | hCoV-19/Iceland/336/2020 | Europe / Iceland / Reykjavik | 2020-03-14 | deCODE genetics | deCODE genetics | Daniel F Gudbjartsson; Agnar Helgason; Hakon Jonsson; Olafur T Magnusson; Pall Melsted; Gudmundur L Norddahl; Jona Saemundsdottir; Asgeir Sigurdsson; Patrick Sulem; Arna B Agustsdottir; Berglind Eiriksdottir; Run Fridriksdottir; Elisabet E Gardarsdottir; Gudmundur Georgsson; Olafia S Gretarsdottir; Kjartan R Gudmundsson; Thora R Gunnarsdottir; Arnaldur Gylfason; Hilma Holm; Brynjar O Jensson; Aslaug Jonasdottir; Kamilla S Josefsdottir; Thordur Kristjansson; Droplaug N Magnusdottir; Louise le Roux; Gudrun Sigmundsdottir; Gardar Sveinbjornsson; Kristin E Sveinsdottir; Maney Sveinsdottir; Emil A Thorarensen; Bjarni Thorbjornsson; Gisli Masson; Ingileif Jonsdottir; Alma Moller; Thorolfur Gudnason; Karl G Kristinsson; Unnur Thorsteinsdottir; Kari Stefansson |
| EPI_ISL_417666 | hCoV-19/Iceland/337/2020 | Europe / Iceland / Reykjavik | 2020-03-15 | deCODE genetics | deCODE genetics | Daniel F Gudbjartsson; Agnar Helgason; Hakon Jonsson; Olafur T Magnusson; Pall Melsted; Gudmundur L Norddahl; Jona Saemundsdottir; Asgeir Sigurdsson; Patrick Sulem; Arna B Agustsdottir; Berglind Eiriksdottir; Run Fridriksdottir; Elisabet E Gardarsdottir; Gudmundur Georgsson; Olafia S Gretarsdottir; Kjartan R Gudmundsson; Thora R Gunnarsdottir; Arnaldur Gylfason; Hilma Holm; Brynjar O Jensson; Aslaug Jonasdottir; Kamilla S Josefsdottir; Thordur Kristjansson; Droplaug N Magnusdottir; Louise le Roux; Gudrun Sigmundsdottir; Gardar Sveinbjornsson; Kristin E Sveinsdottir; Maney Sveinsdottir; Emil A Thorarensen; Bjarni Thorbjornsson; Gisli Masson; Ingileif Jonsdottir; Alma Moller; Thorolfur Gudnason; Karl G Kristinsson; Unnur Thorsteinsdottir; Kari Stefansson |
| EPI_ISL_417667 | hCoV-19/Iceland/338/2020 | Europe / Iceland / Reykjavik | 2020-03-15 | deCODE genetics | deCODE genetics | Daniel F Gudbjartsson; Agnar Helgason; Hakon Jonsson; Olafur T Magnusson; Pall Melsted; Gudmundur L Norddahl; Jona Saemundsdottir; Asgeir Sigurdsson; Patrick Sulem; Arna B Agustsdottir; Berglind Eiriksdottir; Run Fridriksdottir; Elisabet E Gardarsdottir; Gudmundur Georgsson; Olafia S Gretarsdottir; Kjartan R Gudmundsson; Thora R Gunnarsdottir; Arnaldur Gylfason; Hilma Holm; Brynjar O Jensson; Aslaug Jonasdottir; Kamilla S Josefsdottir; Thordur Kristjansson; Droplaug N Magnusdottir; Louise le Roux; Gudrun Sigmundsdottir; Gardar Sveinbjornsson; Kristin E Sveinsdottir; Maney Sveinsdottir; Emil A Thorarensen; Bjarni Thorbjornsson; Gisli Masson; Ingileif Jonsdottir; Alma Moller; Thorolfur Gudnason; Karl G Kristinsson; Unnur Thorsteinsdottir; Kari Stefansson |
| EPI_ISL_417668 | hCoV-19/Iceland/339/2020 | Europe / Iceland / Reykjavik | 2020-03-15 | deCODE genetics | deCODE genetics | Daniel F Gudbjartsson; Agnar Helgason; Hakon Jonsson; Olafur T Magnusson; Pall Melsted; Gudmundur L Norddahl; Jona Saemundsdottir; Asgeir Sigurdsson; Patrick Sulem; Arna B Agustsdottir; Berglind Eiriksdottir; Run Fridriksdottir; Elisabet E Gardarsdottir; Gudmundur Georgsson; Olafia S Gretarsdottir; Kjartan R Gudmundsson; Thora R Gunnarsdottir; Arnaldur Gylfason; Hilma Holm; Brynjar O Jensson; Aslaug Jonasdottir; Kamilla S Josefsdottir; Thordur Kristjansson; Droplaug N Magnusdottir; Louise le Roux; Gudrun Sigmundsdottir; Gardar Sveinbjornsson; Kristin E Sveinsdottir; Maney Sveinsdottir; Emil A Thorarensen; Bjarni Thorbjornsson; Gisli Masson; Ingileif Jonsdottir; Alma Moller; Thorolfur Gudnason; Karl G Kristinsson; Unnur Thorsteinsdottir; Kari Stefansson |
| EPI_ISL_417670 | hCoV-19/Iceland/341/2020 | Europe / Iceland / Reykjavik | 2020-03-15 | deCODE genetics | deCODE genetics | Daniel F Gudbjartsson; Agnar Helgason; Hakon Jonsson; Olafur T Magnusson; Pall Melsted; Gudmundur L Norddahl; Jona Saemundsdottir; Asgeir Sigurdsson; Patrick Sulem; Arna B Agustsdottir; Berglind Eiriksdottir; Run Fridriksdottir; Elisabet E Gardarsdottir; Gudmundur Georgsson; Olafia S Gretarsdottir; Kjartan R Gudmundsson; Thora R Gunnarsdottir; Arnaldur Gylfason; Hilma Holm; Brynjar O Jensson; Aslaug Jonasdottir; Kamilla S Josefsdottir; Thordur Kristjansson; Droplaug N Magnusdottir; Louise le Roux; Gudrun Sigmundsdottir; Gardar Sveinbjornsson; Kristin E Sveinsdottir; Maney Sveinsdottir; Emil A Thorarensen; Bjarni Thorbjornsson; Gisli Masson; Ingileif Jonsdottir; Alma Moller; Thorolfur Gudnason; Karl G Kristinsson; Unnur Thorsteinsdottir; Kari Stefansson |
| EPI_ISL_417671 | hCoV-19/Iceland/342/2020 | Europe / Iceland / Reykjavik | 2020-03-15 | deCODE genetics | deCODE genetics | Daniel F Gudbjartsson; Agnar Helgason; Hakon Jonsson; Olafur T Magnusson; Pall Melsted; Gudmundur L Norddahl; Jona Saemundsdottir; Asgeir Sigurdsson; Patrick Sulem; Arna B Agustsdottir; Berglind Eiriksdottir; Run Fridriksdottir; Elisabet E Gardarsdottir; Gudmundur Georgsson; Olafia S Gretarsdottir; Kjartan R Gudmundsson; Thora R Gunnarsdottir; Arnaldur Gylfason; Hilma Holm; Brynjar O Jensson; Aslaug Jonasdottir; Kamilla S Josefsdottir; Thordur Kristjansson; Droplaug N Magnusdottir; Louise le Roux; Gudrun Sigmundsdottir; Gardar Sveinbjornsson; Kristin E Sveinsdottir; Maney Sveinsdottir; Emil A Thorarensen; Bjarni Thorbjornsson; Gisli Masson; Ingileif Jonsdottir; Alma Moller; Thorolfur Gudnason; Karl G Kristinsson; Unnur Thorsteinsdottir; Kari Stefansson |
| EPI_ISL_417672 | hCoV-19/Iceland/38/2020 | Europe / Iceland / Reykjavik | 2020-03-16 | deCODE genetics | deCODE genetics | Daniel F Gudbjartsson; Agnar Helgason; Hakon Jonsson; Olafur T Magnusson; Pall Melsted; Gudmundur L Norddahl; Jona Saemundsdottir; Asgeir Sigurdsson; Patrick Sulem; Arna B Agustsdottir; Berglind Eiriksdottir; Run Fridriksdottir; Elisabet E Gardarsdottir; Gudmundur Georgsson; Olafia S Gretarsdottir; Kjartan R Gudmundsson; Thora R Gunnarsdottir; Arnaldur Gylfason; Hilma Holm; Brynjar O Jensson; Aslaug Jonasdottir; Kamilla S Josefsdottir; Thordur Kristjansson; Droplaug N Magnusdottir; Louise le Roux; Gudrun Sigmundsdottir; Gardar Sveinbjornsson; Kristin E Sveinsdottir; Maney Sveinsdottir; Emil A Thorarensen; Bjarni Thorbjornsson; Gisli Masson; Ingileif Jonsdottir; Alma Moller; Thorolfur Gudnason; Karl G Kristinsson; Unnur Thorsteinsdottir; Kari Stefansson |
| EPI_ISL_417673 | hCoV-19/Iceland/42/2020 | Europe / Iceland / Reykjavik | 2020-03-04 | The National University Hospital of Iceland | deCODE genetics | Daniel F Gudbjartsson; Agnar Helgason; Hakon Jonsson; Olafur T Magnusson; Pall Melsted; Gudmundur L Norddahl; Jona Saemundsdottir; Asgeir Sigurdsson; Patrick Sulem; Arna B Agustsdottir; Berglind Eiriksdottir; Run Fridriksdottir; Elisabet E Gardarsdottir; Gudmundur Georgsson; Olafia S Gretarsdottir; Kjartan R Gudmundsson; Thora R Gunnarsdottir; Arnaldur Gylfason; Hilma Holm; Brynjar O Jensson; Aslaug Jonasdottir; Kamilla S Josefsdottir; Thordur Kristjansson; Droplaug N Magnusdottir; Louise le Roux; Gudrun Sigmundsdottir; Gardar Sveinbjornsson; Kristin E Sveinsdottir; Maney Sveinsdottir; Emil A Thorarensen; Bjarni Thorbjornsson; Gisli Masson; Ingileif Jonsdottir; Alma Moller; Thorolfur Gudnason; Karl G Kristinsson; Unnur Thorsteinsdottir; Kari Stefansson |
| EPI_ISL_417676 | hCoV-19/Iceland/98/2020 | Europe / Iceland / Reykjavik | 2020-03-16 | deCODE genetics | deCODE genetics | Daniel F Gudbjartsson; Agnar Helgason; Hakon Jonsson; Olafur T Magnusson; Pall Melsted; Gudmundur L Norddahl; Jona Saemundsdottir; Asgeir Sigurdsson; Patrick Sulem; Arna B Agustsdottir; Berglind Eiriksdottir; Run Fridriksdottir; Elisabet E Gardarsdottir; Gudmundur Georgsson; Olafia S Gretarsdottir; Kjartan R Gudmundsson; Thora R Gunnarsdottir; Arnaldur Gylfason; Hilma Holm; Brynjar O Jensson; Aslaug Jonasdottir; Kamilla S Josefsdottir; Thordur Kristjansson; Droplaug N Magnusdottir; Louise le Roux; Gudrun Sigmundsdottir; Gardar Sveinbjornsson; Kristin E Sveinsdottir; Maney Sveinsdottir; Emil A Thorarensen; Bjarni Thorbjornsson; Gisli Masson; Ingileif Jonsdottir; Alma Moller; Thorolfur Gudnason; Karl G Kristinsson; Unnur Thorsteinsdottir; Kari Stefansson |
| EPI_ISL_417677 | hCoV-19/Iceland/39/2020 | Europe / Iceland / Reykjavik | 2020-03-04 | The National University Hospital of Iceland | deCODE genetics | Daniel F Gudbjartsson; Agnar Helgason; Hakon Jonsson; Olafur T Magnusson; Pall Melsted; Gudmundur L Norddahl; Jona Saemundsdottir; Asgeir Sigurdsson; Patrick Sulem; Arna B Agustsdottir; Berglind Eiriksdottir; Run Fridriksdottir; Elisabet E Gardarsdottir; Gudmundur Georgsson; Olafia S Gretarsdottir; Kjartan R Gudmundsson; Thora R Gunnarsdottir; Arnaldur Gylfason; Hilma Holm; Brynjar O Jensson; Aslaug Jonasdottir; Kamilla S Josefsdottir; Thordur Kristjansson; Droplaug N Magnusdottir; Louise le Roux; Gudrun Sigmundsdottir; Gardar Sveinbjornsson; Kristin E Sveinsdottir; Maney Sveinsdottir; Emil A Thorarensen; Bjarni Thorbjornsson; Gisli Masson; Ingileif Jonsdottir; Alma Moller; Thorolfur Gudnason; Karl G Kristinsson; Unnur Thorsteinsdottir; Kari Stefansson |
| EPI_ISL_417678 | hCoV-19/Iceland/185/2020 | Europe / Iceland / Reykjavik | 2020-03-16 | The National University Hospital of Iceland | deCODE genetics | Daniel F Gudbjartsson; Agnar Helgason; Hakon Jonsson; Olafur T Magnusson; Pall Melsted; Gudmundur L Norddahl; Jona Saemundsdottir; Asgeir Sigurdsson; Patrick Sulem; Arna B Agustsdottir; Berglind Eiriksdottir; Run Fridriksdottir; Elisabet E Gardarsdottir; Gudmundur Georgsson; Olafia S Gretarsdottir; Kjartan R Gudmundsson; Thora R Gunnarsdottir; Arnaldur Gylfason; Hilma Holm; Brynjar O Jensson; Aslaug Jonasdottir; Kamilla S Josefsdottir; Thordur Kristjansson; Droplaug N Magnusdottir; Louise le Roux; Gudrun Sigmundsdottir; Gardar Sveinbjornsson; Kristin E Sveinsdottir; Maney Sveinsdottir; Emil A Thorarensen; Bjarni Thorbjornsson; Gisli Masson; Ingileif Jonsdottir; Alma Moller; Thorolfur Gudnason; Karl G Kristinsson; Unnur Thorsteinsdottir; Kari Stefansson |
| EPI_ISL_417679 | hCoV-19/Iceland/79/2020 | Europe / Iceland / Reykjavik | 2020-03-09 | The National University Hospital of Iceland | deCODE genetics | Daniel F Gudbjartsson; Agnar Helgason; Hakon Jonsson; Olafur T Magnusson; Pall Melsted; Gudmundur L Norddahl; Jona Saemundsdottir; Asgeir Sigurdsson; Patrick Sulem; Arna B Agustsdottir; Berglind Eiriksdottir; Run Fridriksdottir; Elisabet E Gardarsdottir; Gudmundur Georgsson; Olafia S Gretarsdottir; Kjartan R Gudmundsson; Thora R Gunnarsdottir; Arnaldur Gylfason; Hilma Holm; Brynjar O Jensson; Aslaug Jonasdottir; Kamilla S Josefsdottir; Thordur Kristjansson; Droplaug N Magnusdottir; Louise le Roux; Gudrun Sigmundsdottir; Gardar Sveinbjornsson; Kristin E Sveinsdottir; Maney Sveinsdottir; Emil A Thorarensen; Bjarni Thorbjornsson; Gisli Masson; Ingileif Jonsdottir; Alma Moller; Thorolfur Gudnason; Karl G Kristinsson; Unnur Thorsteinsdottir; Kari Stefansson |
| EPI_ISL_417680 | hCoV-19/Iceland/177/2020 | Europe / Iceland / Reykjavik | 2020-03-15 | The National University Hospital of Iceland | deCODE genetics | Daniel F Gudbjartsson; Agnar Helgason; Hakon Jonsson; Olafur T Magnusson; Pall Melsted; Gudmundur L Norddahl; Jona Saemundsdottir; Asgeir Sigurdsson; Patrick Sulem; Arna B Agustsdottir; Berglind Eiriksdottir; Run Fridriksdottir; Elisabet E Gardarsdottir; Gudmundur Georgsson; Olafia S Gretarsdottir; Kjartan R Gudmundsson; Thora R Gunnarsdottir; Arnaldur Gylfason; Hilma Holm; Brynjar O Jensson; Aslaug Jonasdottir; Kamilla S Josefsdottir; Thordur Kristjansson; Droplaug N Magnusdottir; Louise le Roux; Gudrun Sigmundsdottir; Gardar Sveinbjornsson; Kristin E Sveinsdottir; Maney Sveinsdottir; Emil A Thorarensen; Bjarni Thorbjornsson; Gisli Masson; Ingileif Jonsdottir; Alma Moller; Thorolfur Gudnason; Karl G Kristinsson; Unnur Thorsteinsdottir; Kari Stefansson |
| EPI_ISL_417681 | hCoV-19/Iceland/37/2020 | Europe / Iceland / Reykjavik | 2020-03-04 | The National University Hospital of Iceland | deCODE genetics | Daniel F Gudbjartsson; Agnar Helgason; Hakon Jonsson; Olafur T Magnusson; Pall Melsted; Gudmundur L Norddahl; Jona Saemundsdottir; Asgeir Sigurdsson; Patrick Sulem; Arna B Agustsdottir; Berglind Eiriksdottir; Run Fridriksdottir; Elisabet E Gardarsdottir; Gudmundur Georgsson; Olafia S Gretarsdottir; Kjartan R Gudmundsson; Thora R Gunnarsdottir; Arnaldur Gylfason; Hilma Holm; Brynjar O Jensson; Aslaug Jonasdottir; Kamilla S Josefsdottir; Thordur Kristjansson; Droplaug N Magnusdottir; Louise le Roux; Gudrun Sigmundsdottir; Gardar Sveinbjornsson; Kristin E Sveinsdottir; Maney Sveinsdottir; Emil A Thorarensen; Bjarni Thorbjornsson; Gisli Masson; Ingileif Jonsdottir; Alma Moller; Thorolfur Gudnason; Karl G Kristinsson; Unnur Thorsteinsdottir; Kari Stefansson |
| EPI_ISL_417682 | hCoV-19/Iceland/101/2020 | Europe / Iceland / Reykjavik | 2020-03-10 | The National University Hospital of Iceland | deCODE genetics | Daniel F Gudbjartsson; Agnar Helgason; Hakon Jonsson; Olafur T Magnusson; Pall Melsted; Gudmundur L Norddahl; Jona Saemundsdottir; Asgeir Sigurdsson; Patrick Sulem; Arna B Agustsdottir; Berglind Eiriksdottir; Run Fridriksdottir; Elisabet E Gardarsdottir; Gudmundur Georgsson; Olafia S Gretarsdottir; Kjartan R Gudmundsson; Thora R Gunnarsdottir; Arnaldur Gylfason; Hilma Holm; Brynjar O Jensson; Aslaug Jonasdottir; Kamilla S Josefsdottir; Thordur Kristjansson; Droplaug N Magnusdottir; Louise le Roux; Gudrun Sigmundsdottir; Gardar Sveinbjornsson; Kristin E Sveinsdottir; Maney Sveinsdottir; Emil A Thorarensen; Bjarni Thorbjornsson; Gisli Masson; Ingileif Jonsdottir; Alma Moller; Thorolfur Gudnason; Karl G Kristinsson; Unnur Thorsteinsdottir; Kari Stefansson |
| EPI_ISL_417683 | hCoV-19/Iceland/47/2020 | Europe / Iceland / Reykjavik | 2020-03-05 | The National University Hospital of Iceland | deCODE genetics | Daniel F Gudbjartsson; Agnar Helgason; Hakon Jonsson; Olafur T Magnusson; Pall Melsted; Gudmundur L Norddahl; Jona Saemundsdottir; Asgeir Sigurdsson; Patrick Sulem; Arna B Agustsdottir; Berglind Eiriksdottir; Run Fridriksdottir; Elisabet E Gardarsdottir; Gudmundur Georgsson; Olafia S Gretarsdottir; Kjartan R Gudmundsson; Thora R Gunnarsdottir; Arnaldur Gylfason; Hilma Holm; Brynjar O Jensson; Aslaug Jonasdottir; Kamilla S Josefsdottir; Thordur Kristjansson; Droplaug N Magnusdottir; Louise le Roux; Gudrun Sigmundsdottir; Gardar Sveinbjornsson; Kristin E Sveinsdottir; Maney Sveinsdottir; Emil A Thorarensen; Bjarni Thorbjornsson; Gisli Masson; Ingileif Jonsdottir; Alma Moller; Thorolfur Gudnason; Karl G Kristinsson; Unnur Thorsteinsdottir; Kari Stefansson |
| EPI_ISL_417684 | hCoV-19/Iceland/54/2020 | Europe / Iceland / Reykjavik | 2020-03-06 | The National University Hospital of Iceland | deCODE genetics | Daniel F Gudbjartsson; Agnar Helgason; Hakon Jonsson; Olafur T Magnusson; Pall Melsted; Gudmundur L Norddahl; Jona Saemundsdottir; Asgeir Sigurdsson; Patrick Sulem; Arna B Agustsdottir; Berglind Eiriksdottir; Run Fridriksdottir; Elisabet E Gardarsdottir; Gudmundur Georgsson; Olafia S Gretarsdottir; Kjartan R Gudmundsson; Thora R Gunnarsdottir; Arnaldur Gylfason; Hilma Holm; Brynjar O Jensson; Aslaug Jonasdottir; Kamilla S Josefsdottir; Thordur Kristjansson; Droplaug N Magnusdottir; Louise le Roux; Gudrun Sigmundsdottir; Gardar Sveinbjornsson; Kristin E Sveinsdottir; Maney Sveinsdottir; Emil A Thorarensen; Bjarni Thorbjornsson; Gisli Masson; Ingileif Jonsdottir; Alma Moller; Thorolfur Gudnason; Karl G Kristinsson; Unnur Thorsteinsdottir; Kari Stefansson |
| EPI_ISL_417685 | hCoV-19/Iceland/215/2020 | Europe / Iceland / Reykjavik | 2020-03-16 | The National University Hospital of Iceland | deCODE genetics | Daniel F Gudbjartsson; Agnar Helgason; Hakon Jonsson; Olafur T Magnusson; Pall Melsted; Gudmundur L Norddahl; Jona Saemundsdottir; Asgeir Sigurdsson; Patrick Sulem; Arna B Agustsdottir; Berglind Eiriksdottir; Run Fridriksdottir; Elisabet E Gardarsdottir; Gudmundur Georgsson; Olafia S Gretarsdottir; Kjartan R Gudmundsson; Thora R Gunnarsdottir; Arnaldur Gylfason; Hilma Holm; Brynjar O Jensson; Aslaug Jonasdottir; Kamilla S Josefsdottir; Thordur Kristjansson; Droplaug N Magnusdottir; Louise le Roux; Gudrun Sigmundsdottir; Gardar Sveinbjornsson; Kristin E Sveinsdottir; Maney Sveinsdottir; Emil A Thorarensen; Bjarni Thorbjornsson; Gisli Masson; Ingileif Jonsdottir; Alma Moller; Thorolfur Gudnason; Karl G Kristinsson; Unnur Thorsteinsdottir; Kari Stefansson |
| EPI_ISL_417686 | hCoV-19/Iceland/43/2020 | Europe / Iceland / Reykjavik | 2020-03-04 | The National University Hospital of Iceland | deCODE genetics | Daniel F Gudbjartsson; Agnar Helgason; Hakon Jonsson; Olafur T Magnusson; Pall Melsted; Gudmundur L Norddahl; Jona Saemundsdottir; Asgeir Sigurdsson; Patrick Sulem; Arna B Agustsdottir; Berglind Eiriksdottir; Run Fridriksdottir; Elisabet E Gardarsdottir; Gudmundur Georgsson; Olafia S Gretarsdottir; Kjartan R Gudmundsson; Thora R Gunnarsdottir; Arnaldur Gylfason; Hilma Holm; Brynjar O Jensson; Aslaug Jonasdottir; Kamilla S Josefsdottir; Thordur Kristjansson; Droplaug N Magnusdottir; Louise le Roux; Gudrun Sigmundsdottir; Gardar Sveinbjornsson; Kristin E Sveinsdottir; Maney Sveinsdottir; Emil A Thorarensen; Bjarni Thorbjornsson; Gisli Masson; Ingileif Jonsdottir; Alma Moller; Thorolfur Gudnason; Karl G Kristinsson; Unnur Thorsteinsdottir; Kari Stefansson |
| EPI_ISL_417688 | hCoV-19/Iceland/12/2020 | Europe / Iceland / Reykjavik | 2020-03-01 | The National University Hospital of Iceland | deCODE genetics | Daniel F Gudbjartsson; Agnar Helgason; Hakon Jonsson; Olafur T Magnusson; Pall Melsted; Gudmundur L Norddahl; Jona Saemundsdottir; Asgeir Sigurdsson; Patrick Sulem; Arna B Agustsdottir; Berglind Eiriksdottir; Run Fridriksdottir; Elisabet E Gardarsdottir; Gudmundur Georgsson; Olafia S Gretarsdottir; Kjartan R Gudmundsson; Thora R Gunnarsdottir; Arnaldur Gylfason; Hilma Holm; Brynjar O Jensson; Aslaug Jonasdottir; Kamilla S Josefsdottir; Thordur Kristjansson; Droplaug N Magnusdottir; Louise le Roux; Gudrun Sigmundsdottir; Gardar Sveinbjornsson; Kristin E Sveinsdottir; Maney Sveinsdottir; Emil A Thorarensen; Bjarni Thorbjornsson; Gisli Masson; Ingileif Jonsdottir; Alma Moller; Thorolfur Gudnason; Karl G Kristinsson; Unnur Thorsteinsdottir; Kari Stefansson |
| EPI_ISL_417690 | hCoV-19/Iceland/125/2020 | Europe / Iceland / Reykjavik | 2020-03-11 | The National University Hospital of Iceland | deCODE genetics | Daniel F Gudbjartsson; Agnar Helgason; Hakon Jonsson; Olafur T Magnusson; Pall Melsted; Gudmundur L Norddahl; Jona Saemundsdottir; Asgeir Sigurdsson; Patrick Sulem; Arna B Agustsdottir; Berglind Eiriksdottir; Run Fridriksdottir; Elisabet E Gardarsdottir; Gudmundur Georgsson; Olafia S Gretarsdottir; Kjartan R Gudmundsson; Thora R Gunnarsdottir; Arnaldur Gylfason; Hilma Holm; Brynjar O Jensson; Aslaug Jonasdottir; Kamilla S Josefsdottir; Thordur Kristjansson; Droplaug N Magnusdottir; Louise le Roux; Gudrun Sigmundsdottir; Gardar Sveinbjornsson; Kristin E Sveinsdottir; Maney Sveinsdottir; Emil A Thorarensen; Bjarni Thorbjornsson; Gisli Masson; Ingileif Jonsdottir; Alma Moller; Thorolfur Gudnason; Karl G Kristinsson; Unnur Thorsteinsdottir; Kari Stefansson |
| EPI_ISL_417691 | hCoV-19/Iceland/111/2020 | Europe / Iceland / Reykjavik | 2020-03-11 | The National University Hospital of Iceland | deCODE genetics | Daniel F Gudbjartsson; Agnar Helgason; Hakon Jonsson; Olafur T Magnusson; Pall Melsted; Gudmundur L Norddahl; Jona Saemundsdottir; Asgeir Sigurdsson; Patrick Sulem; Arna B Agustsdottir; Berglind Eiriksdottir; Run Fridriksdottir; Elisabet E Gardarsdottir; Gudmundur Georgsson; Olafia S Gretarsdottir; Kjartan R Gudmundsson; Thora R Gunnarsdottir; Arnaldur Gylfason; Hilma Holm; Brynjar O Jensson; Aslaug Jonasdottir; Kamilla S Josefsdottir; Thordur Kristjansson; Droplaug N Magnusdottir; Louise le Roux; Gudrun Sigmundsdottir; Gardar Sveinbjornsson; Kristin E Sveinsdottir; Maney Sveinsdottir; Emil A Thorarensen; Bjarni Thorbjornsson; Gisli Masson; Ingileif Jonsdottir; Alma Moller; Thorolfur Gudnason; Karl G Kristinsson; Unnur Thorsteinsdottir; Kari Stefansson |
| EPI_ISL_417692 | hCoV-19/Iceland/57/2020 | Europe / Iceland / Reykjavik | 2020-03-06 | The National University Hospital of Iceland | deCODE genetics | Daniel F Gudbjartsson; Agnar Helgason; Hakon Jonsson; Olafur T Magnusson; Pall Melsted; Gudmundur L Norddahl; Jona Saemundsdottir; Asgeir Sigurdsson; Patrick Sulem; Arna B Agustsdottir; Berglind Eiriksdottir; Run Fridriksdottir; Elisabet E Gardarsdottir; Gudmundur Georgsson; Olafia S Gretarsdottir; Kjartan R Gudmundsson; Thora R Gunnarsdottir; Arnaldur Gylfason; Hilma Holm; Brynjar O Jensson; Aslaug Jonasdottir; Kamilla S Josefsdottir; Thordur Kristjansson; Droplaug N Magnusdottir; Louise le Roux; Gudrun Sigmundsdottir; Gardar Sveinbjornsson; Kristin E Sveinsdottir; Maney Sveinsdottir; Emil A Thorarensen; Bjarni Thorbjornsson; Gisli Masson; Ingileif Jonsdottir; Alma Moller; Thorolfur Gudnason; Karl G Kristinsson; Unnur Thorsteinsdottir; Kari Stefansson |
| EPI_ISL_417693 | hCoV-19/Iceland/25/2020 | Europe / Iceland / Reykjavik | 2020-03-02 | The National University Hospital of Iceland | deCODE genetics | Daniel F Gudbjartsson; Agnar Helgason; Hakon Jonsson; Olafur T Magnusson; Pall Melsted; Gudmundur L Norddahl; Jona Saemundsdottir; Asgeir Sigurdsson; Patrick Sulem; Arna B Agustsdottir; Berglind Eiriksdottir; Run Fridriksdottir; Elisabet E Gardarsdottir; Gudmundur Georgsson; Olafia S Gretarsdottir; Kjartan R Gudmundsson; Thora R Gunnarsdottir; Arnaldur Gylfason; Hilma Holm; Brynjar O Jensson; Aslaug Jonasdottir; Kamilla S Josefsdottir; Thordur Kristjansson; Droplaug N Magnusdottir; Louise le Roux; Gudrun Sigmundsdottir; Gardar Sveinbjornsson; Kristin E Sveinsdottir; Maney Sveinsdottir; Emil A Thorarensen; Bjarni Thorbjornsson; Gisli Masson; Ingileif Jonsdottir; Alma Moller; Thorolfur Gudnason; Karl G Kristinsson; Unnur Thorsteinsdottir; Kari Stefansson |
| EPI_ISL_417694 | hCoV-19/Iceland/3/2020 | Europe / Iceland / Reykjavik | 2020-03-13 | The National University Hospital of Iceland | deCODE genetics | Daniel F Gudbjartsson; Agnar Helgason; Hakon Jonsson; Olafur T Magnusson; Pall Melsted; Gudmundur L Norddahl; Jona Saemundsdottir; Asgeir Sigurdsson; Patrick Sulem; Arna B Agustsdottir; Berglind Eiriksdottir; Run Fridriksdottir; Elisabet E Gardarsdottir; Gudmundur Georgsson; Olafia S Gretarsdottir; Kjartan R Gudmundsson; Thora R Gunnarsdottir; Arnaldur Gylfason; Hilma Holm; Brynjar O Jensson; Aslaug Jonasdottir; Kamilla S Josefsdottir; Thordur Kristjansson; Droplaug N Magnusdottir; Louise le Roux; Gudrun Sigmundsdottir; Gardar Sveinbjornsson; Kristin E Sveinsdottir; Maney Sveinsdottir; Emil A Thorarensen; Bjarni Thorbjornsson; Gisli Masson; Ingileif Jonsdottir; Alma Moller; Thorolfur Gudnason; Karl G Kristinsson; Unnur Thorsteinsdottir; Kari Stefansson |
| EPI_ISL_417695 | hCoV-19/Iceland/20/2020 | Europe / Iceland / Reykjavik | 2020-03-02 | The National University Hospital of Iceland | deCODE genetics | Daniel F Gudbjartsson; Agnar Helgason; Hakon Jonsson; Olafur T Magnusson; Pall Melsted; Gudmundur L Norddahl; Jona Saemundsdottir; Asgeir Sigurdsson; Patrick Sulem; Arna B Agustsdottir; Berglind Eiriksdottir; Run Fridriksdottir; Elisabet E Gardarsdottir; Gudmundur Georgsson; Olafia S Gretarsdottir; Kjartan R Gudmundsson; Thora R Gunnarsdottir; Arnaldur Gylfason; Hilma Holm; Brynjar O Jensson; Aslaug Jonasdottir; Kamilla S Josefsdottir; Thordur Kristjansson; Droplaug N Magnusdottir; Louise le Roux; Gudrun Sigmundsdottir; Gardar Sveinbjornsson; Kristin E Sveinsdottir; Maney Sveinsdottir; Emil A Thorarensen; Bjarni Thorbjornsson; Gisli Masson; Ingileif Jonsdottir; Alma Moller; Thorolfur Gudnason; Karl G Kristinsson; Unnur Thorsteinsdottir; Kari Stefansson |
| EPI_ISL_417696 | hCoV-19/Iceland/135/2020 | Europe / Iceland / Reykjavik | 2020-03-12 | The National University Hospital of Iceland | deCODE genetics | Daniel F Gudbjartsson; Agnar Helgason; Hakon Jonsson; Olafur T Magnusson; Pall Melsted; Gudmundur L Norddahl; Jona Saemundsdottir; Asgeir Sigurdsson; Patrick Sulem; Arna B Agustsdottir; Berglind Eiriksdottir; Run Fridriksdottir; Elisabet E Gardarsdottir; Gudmundur Georgsson; Olafia S Gretarsdottir; Kjartan R Gudmundsson; Thora R Gunnarsdottir; Arnaldur Gylfason; Hilma Holm; Brynjar O Jensson; Aslaug Jonasdottir; Kamilla S Josefsdottir; Thordur Kristjansson; Droplaug N Magnusdottir; Louise le Roux; Gudrun Sigmundsdottir; Gardar Sveinbjornsson; Kristin E Sveinsdottir; Maney Sveinsdottir; Emil A Thorarensen; Bjarni Thorbjornsson; Gisli Masson; Ingileif Jonsdottir; Alma Moller; Thorolfur Gudnason; Karl G Kristinsson; Unnur Thorsteinsdottir; Kari Stefansson |
| EPI_ISL_417698 | hCoV-19/Iceland/103/2020 | Europe / Iceland / Reykjavik | 2020-03-10 | The National University Hospital of Iceland | deCODE genetics | Daniel F Gudbjartsson; Agnar Helgason; Hakon Jonsson; Olafur T Magnusson; Pall Melsted; Gudmundur L Norddahl; Jona Saemundsdottir; Asgeir Sigurdsson; Patrick Sulem; Arna B Agustsdottir; Berglind Eiriksdottir; Run Fridriksdottir; Elisabet E Gardarsdottir; Gudmundur Georgsson; Olafia S Gretarsdottir; Kjartan R Gudmundsson; Thora R Gunnarsdottir; Arnaldur Gylfason; Hilma Holm; Brynjar O Jensson; Aslaug Jonasdottir; Kamilla S Josefsdottir; Thordur Kristjansson; Droplaug N Magnusdottir; Louise le Roux; Gudrun Sigmundsdottir; Gardar Sveinbjornsson; Kristin E Sveinsdottir; Maney Sveinsdottir; Emil A Thorarensen; Bjarni Thorbjornsson; Gisli Masson; Ingileif Jonsdottir; Alma Moller; Thorolfur Gudnason; Karl G Kristinsson; Unnur Thorsteinsdottir; Kari Stefansson |
| EPI_ISL_417699 | hCoV-19/Iceland/206/2020 | Europe / Iceland / Reykjavik | 2020-03-16 | The National University Hospital of Iceland | deCODE genetics | Daniel F Gudbjartsson; Agnar Helgason; Hakon Jonsson; Olafur T Magnusson; Pall Melsted; Gudmundur L Norddahl; Jona Saemundsdottir; Asgeir Sigurdsson; Patrick Sulem; Arna B Agustsdottir; Berglind Eiriksdottir; Run Fridriksdottir; Elisabet E Gardarsdottir; Gudmundur Georgsson; Olafia S Gretarsdottir; Kjartan R Gudmundsson; Thora R Gunnarsdottir; Arnaldur Gylfason; Hilma Holm; Brynjar O Jensson; Aslaug Jonasdottir; Kamilla S Josefsdottir; Thordur Kristjansson; Droplaug N Magnusdottir; Louise le Roux; Gudrun Sigmundsdottir; Gardar Sveinbjornsson; Kristin E Sveinsdottir; Maney Sveinsdottir; Emil A Thorarensen; Bjarni Thorbjornsson; Gisli Masson; Ingileif Jonsdottir; Alma Moller; Thorolfur Gudnason; Karl G Kristinsson; Unnur Thorsteinsdottir; Kari Stefansson |
| EPI_ISL_417917 | hCoV-19/Malaysia/189332/2020 | Asia / Malaysia / Kuala Lumpur | 2020-03-20 | Department of Medical Microbiology, University Malaya Medical Centre | Department of Medical Microbiology | Yoong Min CHONG, Sasheela PONNAMPALAVANAR, Sharifah Faridah SYED OMAR, Adeeba KAMARULZAMAN,Vijayan MUNUSAMY, Chee Kuan WONG, Cindy Shuan Ju TEH, I-Ching SAM, Yoke Fun Chan, University Malaya Medical Centre COVID Team |
| EPI_ISL_417918 | hCoV-19/Malaysia/188407/2020 | Asia / Malaysia / Kuala Lumpur | 2020-03-18 | Department of Medical Microbiology, University Malaya Medical Centre | Department of Medical Microbiology, Faculty of Medicine, University of Malaya | Yoong Min CHONG, Sasheela PONNAMPALAVANAR, Sharifah Faridah SYED OMAR, Adeeba KAMARULZAMAN,Vijayan MUNUSAMY, Chee Kuan WONG, Cindy Shuan Ju TEH, I-Ching SAM, Yoke Fun Chan, University Malaya Medical Centre COVID Team |
| EPI_ISL_417921 | hCoV-19/Italy/INMI3/2020 | Europe / Italy / Rome | 2020-03-01 | INMI Lazzaro Spallanzani IRCCS | Laboratory of Virology, INMI Lazzaro Spallanzani IRCCS | Martina Rueca, Barbara Bartolini, Francesco Messina, Cesare E. M. Gruber, Emanuela Giombini, Maria R. Capobianchi, Fabrizio Carletti, Francesca Colavita, Concetta Castilletti, Eleonora Lalle, Daniele Lapa, Giuseppe Ippolito. |
| EPI_ISL_417922 | hCoV-19/Italy/INMI4/2020 | Europe / Italy / Rome | 2020-02-28 | INMI Lazzaro Spallanzani IRCCS | Laboratory of Virology, INMI Lazzaro Spallanzani IRCCS | Cesare E. M. Gruber, Martina Rueca, Barbara Bartolini, Francesco Messina, Emanuela Giombini, Maria R. Capobianchi, Fabrizio Carletti, Francesca Colavita, Concetta Castilletti, Eleonora Lalle, Daniele Lapa, Giuseppe Ippolito. |
| EPI_ISL_417923 | hCoV-19/Italy/INMI5/2020 | Europe / Italy / Rome | 2020-03-04 | INMI Lazzaro Spallanzani IRCCS | Laboratory of Virology, INMI Lazzaro Spallanzani IRCCS | Francesco Messina, Barbara Bartolini, Martina Rueca, Cesare E. M. Gruber, Emanuela Giombini, Maria R. Capobianchi, Fabrizio Carletti, Francesca Colavita, Concetta Castilletti, Eleonora Lalle, Daniele Lapa, Giuseppe Ippolito. |
| EPI_ISL_417924 | hCoV-19/Colombia/INS-79256/2020 | South America / Colombia / Antioquia | 2020-03-11 | Secretaria de Salud Medellin | Instituto Nacional de Salud, Universidad Cooperativa de Colombia, Instituto Alexander von Humboldt, Imperial College-London, London School of Hygiene & Tropical Medicine | Marcela Mercado-Reyes, Katherine Laiton-Donato, Diego A. Álvarez-Díaz, Carlos Franco-Muñoz, Jose A. Usme-Ciro, Gloria Puerto, Nicolás D. Franco-Sierra, Mailyn A. Gonzalez, Zulma M. Cucunubá, Christian Julian Villabona-Arenas, Liz Villabona-Arenas, Sussy Echeverria-Londoño, Astrid C. Flórez, Sergio Gomez Rangel, Luz Dary Rodríguez, Juliana Barbosa, Erika Ospitia, Diana Marcela Walteros-Acero, Martha Lucía Ospina Martínez |

| | | | | | | |
|---|---|---|---|---|---|---|
| EPI_ISL_417931 | hCoV-19/USA/CA-CZB05/2020 | North America / USA / California / San Fr | 2020-03-18 | UCSF Clinical Microbiology Laboratory | Chan-Zuckerberg Biohub | Shaun Arevalo, Josh Batson, Olga Botvinnik, Gloria Castaneda, Angela Detweiler, David Dynerman, Samantha Hao, Jack Kamm, Amy Kistler, G. Renuka Kumar, Chaz Langelier, Lucy Li, Steve Miller, Lusajo Mwakibete, Norma Neff, Angela Pisco, Maira Phelps, Michelle Tan, Chunyu Zhao |
| EPI_ISL_417932 | hCoV-19/USA/CA-CZB06/2020 | North America / USA / California / San Fr | 2020-03-18 | UCSF Clinical Microbiology Laboratory | Chan-Zuckerberg Biohub | Shaun Arevalo, Josh Batson, Olga Botvinnik, Gloria Castaneda, Angela Detweiler, David Dynerman, Samantha Hao, Jack Kamm, Amy Kistler, G. Renuka Kumar, Chaz Langelier, Lucy Li, Steve Miller, Lusajo Mwakibete, Norma Neff, Angela Pisco, Maira Phelps, Michelle Tan, Chunyu Zhao |
| EPI_ISL_417933 | hCoV-19/USA/CA-CZB07/2020 | North America / USA / California / San Fr | 2020-03-18 | UCSF Clinical Microbiology Laboratory | Chan-Zuckerberg Biohub | Shaun Arevalo, Josh Batson, Olga Botvinnik, Gloria Castaneda, Angela Detweiler, David Dynerman, Samantha Hao, Jack Kamm, Amy Kistler, G. Renuka Kumar, Chaz Langelier, Lucy Li, Steve Miller, Lusajo Mwakibete, Norma Neff, Angela Pisco, Maira Phelps, Michelle Tan, Chunyu Zhao |
| EPI_ISL_417935 | hCoV-19/USA/CA-CZB011a/2020 | North America / USA / California / San Fr | 2020-03-18 | UCSF Clinical Microbiology Laboratory | Chan-Zuckerberg Biohub | Shaun Arevalo, Josh Batson, Olga Botvinnik, Gloria Castaneda, Angela Detweiler, David Dynerman, Samantha Hao, Jack Kamm, Amy Kistler, G. Renuka Kumar, Chaz Langelier, Lucy Li, Steve Miller, Lusajo Mwakibete, Norma Neff, Angela Pisco, Maira Phelps, Michelle Tan, Chunyu Zhao |
| EPI_ISL_417937 | hCoV-19/USA/CA-CZB013/2020 | North America / USA / California / San Fr | 2020-03-18 | UCSF Clinical Microbiology Laboratory | Chan-Zuckerberg Biohub | Shaun Arevalo, Josh Batson, Olga Botvinnik, Gloria Castaneda, Angela Detweiler, David Dynerman, Samantha Hao, Jack Kamm, Amy Kistler, G. Renuka Kumar, Chaz Langelier, Lucy Li, Steve Miller, Lusajo Mwakibete, Norma Neff, Angela Pisco, Maira Phelps, Michelle Tan, Chunyu Zhao |
| EPI_ISL_417938 | hCoV-19/USA/CA-CZB014/2020 | North America / USA / California / San Fr | 2020-03-18 | UCSF Clinical Microbiology Laboratory | Chan-Zuckerberg Biohub | Shaun Arevalo, Josh Batson, Olga Botvinnik, Gloria Castaneda, Angela Detweiler, David Dynerman, Samantha Hao, Jack Kamm, Amy Kistler, G. Renuka Kumar, Chaz Langelier, Lucy Li, Steve Miller, Lusajo Mwakibete, Norma Neff, Angela Pisco, Maira Phelps, Michelle Tan, Chunyu Zhao |
| EPI_ISL_417939 | hCoV-19/USA/CA-CZB015a/2020 | North America / USA / California / San Fr | 2020-03-18 | UCSF Clinical Microbiology Laboratory | Chan-Zuckerberg Biohub | Shaun Arevalo, Josh Batson, Olga Botvinnik, Gloria Castaneda, Angela Detweiler, David Dynerman, Samantha Hao, Jack Kamm, Amy Kistler, G. Renuka Kumar, Chaz Langelier, Lucy Li, Steve Miller, Lusajo Mwakibete, Norma Neff, Angela Pisco, Maira Phelps, Michelle Tan, Chunyu Zhao |
| EPI_ISL_417941 | hCoV-19/DRC/73/2020 | Africa / Democratic Republic of the Cong | 2020-03-18 | Viral Respiratory Lab, National Institute for Biomedical Research (INRB) | Pathogen Sequencing Lab, National Institute for Biomedical Research (INRB) | Placide Mbala-Kingebeni, Edith Nkwembe, Eddy Kinganda-Lusamaki, Amuri Aziza, Catherine Pratt, Matthias Pauthner, Josh Quick, Allison Black, James Hadfield, Trevor Bedford, Ian Goodfellow, Nick Loman, Kristian Andersen, Michael Wiley, Steve Ahuka-Mundeke, Jean-Jacques Muyembe Tamfum |
| EPI_ISL_417942 | hCoV-19/DRC/80/2020 | Africa / Democratic Republic of the Cong | 2020-03-18 | Viral Respiratory Lab, National Institute for Biomedical Research (INRB) | Pathogen Sequencing Lab, National Institute for Biomedical Research (INRB) | Placide Mbala-Kingebeni, Edith Nkwembe, Eddy Kinganda-Lusamaki, Amuri Aziza, Catherine Pratt, Matthias Pauthner, Josh Quick, Allison Black, James Hadfield, Trevor Bedford, Ian Goodfellow, Nick Loman, Kristian Andersen, Michael Wiley, Steve Ahuka-Mundeke, Jean-Jacques Muyembe Tamfum |
| EPI_ISL_417944 | hCoV-19/DRC/81/2020 | Africa / Democratic Republic of the Cong | 2020-03-18 | Viral Respiratory Lab, National Institute for Biomedical Research (INRB) | Pathogen Sequencing Lab, National Institute for Biomedical Research (INRB) | Placide Mbala-Kingebeni, Edith Nkwembe, Eddy Kinganda-Lusamaki, Amuri Aziza, Catherine Pratt, Matthias Pauthner, Josh Quick, Allison Black, James Hadfield, Trevor Bedford, Ian Goodfellow, Nick Loman, Kristian Andersen, Michael Wiley, Steve Ahuka-Mundeke, Jean-Jacques Muyembe Tamfum |
| EPI_ISL_417946 | hCoV-19/DRC/82/2020 | Africa / Democratic Republic of the Cong | 2020-03-18 | Viral Respiratory Lab, National Institute for Biomedical Research (INRB) | Pathogen Sequencing Lab, National Institute for Biomedical Research (INRB) | Placide Mbala-Kingebeni, Edith Nkwembe, Eddy Kinganda-Lusamaki, Amuri Aziza, Catherine Pratt, Matthias Pauthner, Josh Quick, Allison Black, James Hadfield, Trevor Bedford, Ian Goodfellow, Nick Loman, Kristian Andersen, Michael Wiley, Steve Ahuka-Mundeke, Jean-Jacques Muyembe Tamfum |
| EPI_ISL_417947 | hCoV-19/DRC/94/2020 | Africa / Democratic Republic of the Cong | 2020-03-19 | Viral Respiratory Lab, National Institute for Biomedical Research (INRB) | Pathogen Sequencing Lab, National Institute for Biomedical Research (INRB) | Placide Mbala-Kingebeni, Edith Nkwembe, Eddy Kinganda-Lusamaki, Amuri Aziza, Catherine Pratt, Matthias Pauthner, Josh Quick, Allison Black, James Hadfield, Trevor Bedford, Ian Goodfellow, Nick Loman, Kristian Andersen, Michael Wiley, Steve Ahuka-Mundeke, Jean-Jacques Muyembe Tamfum |
| EPI_ISL_417948 | hCoV-19/DRC/108/2020 | Africa / Democratic Republic of the Cong | 2020-03-19 | Viral Respiratory Lab, National Institute for Biomedical Research (INRB) | Pathogen Sequencing Lab, National Institute for Biomedical Research (INRB) | Placide Mbala-Kingebeni, Edith Nkwembe, Eddy Kinganda-Lusamaki, Amuri Aziza, Catherine Pratt, Matthias Pauthner, Josh Quick, Allison Black, James Hadfield, Trevor Bedford, Ian Goodfellow, Nick Loman, Kristian Andersen, Michael Wiley, Steve Ahuka-Mundeke, Jean-Jacques Muyembe Tamfum |
| EPI_ISL_417950 | hCoV-19/DRC/158/2020 | Africa / Democratic Republic of the Cong | 2020-03-20 | Viral Respiratory Lab, National Institute for Biomedical Research (INRB) | Pathogen Sequencing Lab, National Institute for Biomedical Research (INRB) | Placide Mbala-Kingebeni, Edith Nkwembe, Eddy Kinganda-Lusamaki, Amuri Aziza, Catherine Pratt, Matthias Pauthner, Josh Quick, Allison Black, James Hadfield, Trevor Bedford, Ian Goodfellow, Nick Loman, Kristian Andersen, Michael Wiley, Steve Ahuka-Mundeke, Jean-Jacques Muyembe Tamfum |
| EPI_ISL_417954 | hCoV-19/Spain/Madrid_H3_10/2020 | Europe / Spain / Madrid | 2020-03-12 | Hospital Universitario 12 de Octubre | Hospital Universitario La Paz | Elías Dahdouh, Sara González, Fernando Lázaro, Esther Viedma, Natalia Stella, Julio García, Juan Carlos Galán, Rafael Cantón, Mª Dolores Folgueira, Rafael Delgado, Jesús Mingorance |
| EPI_ISL_417955 | hCoV-19/DRC/191/2020 | Africa / Democratic Republic of the Cong | 2020-03-21 | Viral Respiratory Lab, National Institute for Biomedical Research (INRB) | Pathogen Sequencing Lab, National Institute for Biomedical Research (INRB) | Placide Mbala-Kingebeni, Edith Nkwembe, Eddy Kinganda-Lusamaki, Amuri Aziza, Catherine Pratt, Matthias Pauthner, Josh Quick, Allison Black, James Hadfield, Trevor Bedford, Ian Goodfellow, Nick Loman, Kristian Andersen, Michael Wiley, Steve Ahuka-Mundeke, Jean-Jacques Muyembe Tamfum |
| EPI_ISL_417956 | hCoV-19/Spain/Madrid_H5_34/2020 | Europe / Spain / Madrid | 2020-03-11 | Hospital Universitario 12 de Octubre | Hospital Universitario La Paz | Elías Dahdouh, Sara González, Fernando Lázaro, Esther Viedma, Natalia Stella, Julio García, Juan Carlos Galán, Rafael Cantón, Mª Dolores Folgueira, Rafael Delgado, Jesús Mingorance |
| EPI_ISL_417957 | hCoV-19/Spain/Madrid_H7_36/2020 | Europe / Spain / Madrid | 2020-03-12 | Hospital Universitario 12 de Octubre | Hospital Universitario La Paz | Elías Dahdouh, Sara González, Fernando Lázaro, Esther Viedma, Natalia Stella, Julio García, Juan Carlos Galán, Rafael Cantón, Mª Dolores Folgueira, Rafael Delgado, Jesús Mingorance |
| EPI_ISL_417958 | hCoV-19/USA/UT-010/2020 | North America / USA / Utah | 2020-03-10 | Utah Public Health Laboratory | Utah Public Health Laboratory | Erin Young, Kelly Oakeson |
| EPI_ISL_417959 | hCoV-19/USA/UT-011/2020 | North America / USA / Utah | 2020-03-11 | Utah Public Health Laboratory | Utah Public Health Laboratory | Erin Young, Kelly Oakeson |
| EPI_ISL_417960 | hCoV-19/USA/UT-012/2020 | North America / USA / Utah | 2020-03-12 | Utah Public Health Laboratory | Utah Public Health Laboratory | Erin Young, Kelly Oakeson |
| EPI_ISL_417961 | hCoV-19/Spain/Madrid_H9_38/2020 | Europe / Spain / Madrid | 2020-03-12 | Hospital Universitario 12 de Octubre | Hospital Universitario La Paz | Elías Dahdouh, Sara González, Fernando Lázaro, Esther Viedma, Natalia Stella, Julio García, Juan Carlos Galán, Rafael Cantón, Mª Dolores Folgueira, Rafael Delgado, Jesús Mingorance |
| EPI_ISL_417963 | hCoV-19/Spain/Madrid_H10_39/2020 | Europe / Spain / Madrid | 2020-03-12 | Hospital Universitario 12 de Octubre | Hospital Universitario La Paz | Elías Dahdouh, Sara González, Fernando Lázaro, Esther Viedma, Natalia Stella, Julio García, Juan Carlos Galán, Rafael Cantón, Mª Dolores Folgueira, Rafael Delgado, Jesús Mingorance |
| EPI_ISL_417964 | hCoV-19/USA/UT-014/2020 | North America / USA / Utah | 2020-03-13 | Utah Public Health Laboratory | Utah Public Health Laboratory | Erin Young, Kelly Oakeson |
| EPI_ISL_417966 | hCoV-19/USA/UT-016/2020 | North America / USA / Utah | 2020-03-13 | Utah Public Health Laboratory | Utah Public Health Laboratory | Erin Young, Kelly Oakeson |
| EPI_ISL_417967 | hCoV-19/Spain/Madrid_H11_40/2020 | Europe / Spain / Madrid | 2020-03-12 | Hospital Universitario 12 de Octubre | Hospital Universitario La Paz | Elías Dahdouh, Sara González, Fernando Lázaro, Esther Viedma, Natalia Stella, Julio García, Juan Carlos Galán, Rafael Cantón, Mª Dolores Folgueira, Rafael Delgado, Jesús Mingorance |
| EPI_ISL_417969 | hCoV-19/Spain/Madrid_LP10_12/2020 | Europe / Spain / Madrid | 2020-03-09 | Hospital Universitario La Paz | Hospital Universitario La Paz | Elías Dahdouh, Sara González, Fernando Lázaro, Esther Viedma, Natalia Stella, Julio García, Juan Carlos Galán, Rafael Cantón, Mª Dolores Folgueira, Rafael Delgado, Jesús Mingorance |
| EPI_ISL_417970 | hCoV-19/USA/UT-027/2020 | North America / USA / Utah | 2020-03-16 | Utah Public Health Laboratory | Utah Public Health Laboratory | Erin Young, Kelly Oakeson |
| EPI_ISL_417971 | hCoV-19/USA/UT-028/2020 | North America / USA / Utah | 2020-03-19 | Utah Public Health Laboratory | Utah Public Health Laboratory | Erin Young, Kelly Oakeson |
| EPI_ISL_417972 | hCoV-19/Spain/Madrid_LP12_21/2020 | Europe / Spain / Madrid | 2020-03-09 | Hospital Universitario La Paz | Hospital Universitario La Paz | Elías Dahdouh, Sara González, Fernando Lázaro, Esther Viedma, Natalia Stella, Julio García, Juan Carlos Galán, Rafael Cantón, Mª Dolores Folgueira, Rafael Delgado, Jesús Mingorance |
| EPI_ISL_417973 | hCoV-19/USA/UT-031/2020 | North America / USA / Utah | 2020-03-19 | Utah Public Health Laboratory | Utah Public Health Laboratory | Erin Young, Kelly Oakeson |
| EPI_ISL_417974 | hCoV-19/USA/UT-032/2020 | North America / USA / Utah | 2020-03-19 | Utah Public Health Laboratory | Utah Public Health Laboratory | Erin Young, Kelly Oakeson |
| EPI_ISL_417975 | hCoV-19/Spain/Madrid_LP14_3/2020 | Europe / Spain / Madrid | 2020-03-09 | Hospital Universitario La Paz | Hospital Universitario La Paz | Elías Dahdouh, Sara González, Fernando Lázaro, Esther Viedma, Natalia Stella, Julio García, Juan Carlos Galán, Rafael Cantón, Mª Dolores Folgueira, Rafael Delgado, Jesús Mingorance |
| EPI_ISL_417976 | hCoV-19/USA/UT-033/2020 | North America / USA / Utah | 2020-03-19 | Utah Public Health Laboratory | Utah Public Health Laboratory | Erin Young, Kelly Oakeson |
| EPI_ISL_417977 | hCoV-19/USA/UT-034/2020 | North America / USA / Utah | 2020-03-20 | Utah Public Health Laboratory | Utah Public Health Laboratory | Erin Young, Kelly Oakeson |
| EPI_ISL_417978 | hCoV-19/Spain/Madrid_LP15_4/2020 | Europe / Spain / Madrid | 2020-03-09 | Hospital Universitario La Paz | Hospital Universitario La Paz | Elías Dahdouh, Sara González, Fernando Lázaro, Esther Viedma, Natalia Stella, Julio García, Juan Carlos Galán, Rafael Cantón, Mª Dolores Folgueira, Rafael Delgado, Jesús Mingorance |
| EPI_ISL_417979 | hCoV-19/Spain/Madrid_R2_15/2020 | Europe / Spain / Madrid | 2020-03-03 | Hospital Universitario Ramón y Cajal | Hospital Universitario La Paz | Elías Dahdouh, Sara González, Fernando Lázaro, Esther Viedma, Natalia Stella, Julio García, Juan Carlos Galán, Rafael Cantón, Mª Dolores Folgueira, Rafael Delgado, Jesús Mingorance |
| EPI_ISL_417980 | hCoV-19/Spain/Madrid_R5_8/2020 | Europe / Spain / Madrid | 2020-03-03 | Hospital Universitario Ramón y Cajal | Hospital Universitario La Paz | Elías Dahdouh, Sara González, Fernando Lázaro, Esther Viedma, Natalia Stella, Julio García, Juan Carlos Galán, Rafael Cantón, Mª Dolores Folgueira, Rafael Delgado, Jesús Mingorance |
| EPI_ISL_417981 | hCoV-19/Spain/Madrid_R10_33/2020 | Europe / Spain / Madrid | 2020-03-02 | Hospital Universitario Ramón y Cajal | Hospital Universitario La Paz | Elías Dahdouh, Sara González, Fernando Lázaro, Esther Viedma, Natalia Stella, Julio García, Juan Carlos Galán, Rafael Cantón, Mª Dolores Folgueira, Rafael Delgado, Jesús Mingorance |
| EPI_ISL_417986 | hCoV-19/Portugal/PT0001b/2020 | Europe / Portugal | 2020-03-03 | Centro Hospitalar e Universitario de Sao Joao, Porto | Instituto Nacional de Saude (INSA) | Guiomar et al |
| EPI_ISL_417987 | hCoV-19/Portugal/PT0003/2020 | Europe / Portugal | 2020-03-03 | Centro Hospitalar e Universitario de Sao Joao, Porto | Instituto Nacional de Saude (INSA) | Guiomar et al |
| EPI_ISL_417988 | hCoV-19/Portugal/PT0004/2020 | Europe / Portugal | 2020-03-05 | CHULC - H Curry Cabral | Instituto Nacional de Saude (INSA) | Guiomar et al |
| EPI_ISL_417989 | hCoV-19/Portugal/PT0005/2020 | Europe / Portugal | 2020-03-04 | Centro Hospitalar e Universitario de Sao Joao, Porto | Instituto Nacional de Saude (INSA) | Guiomar et al |
| EPI_ISL_417990 | hCoV-19/Portugal/PT0006a/2020 | Europe / Portugal | 2020-03-06 | CHULC - H Curry Cabral | Instituto Nacional de Saude (INSA) | Guiomar et al |
| EPI_ISL_417991 | hCoV-19/Portugal/PT0006b/2020 | Europe / Portugal | 2020-03-08 | CHULC - H Curry Cabral | Instituto Nacional de Saude (INSA) | Guiomar et al |
| EPI_ISL_417992 | hCoV-19/Portugal/PT0007/2020 | Europe / Portugal | 2020-03-07 | CHULC - H D Estefania | Instituto Nacional de Saude (INSA) | Guiomar et al |
| EPI_ISL_417993 | hCoV-19/Portugal/PT0008/2020 | Europe / Portugal | 2020-03-08 | CHULC - H D Estefania | Instituto Nacional de Saude (INSA) | Guiomar et al |
| EPI_ISL_417994 | hCoV-19/Portugal/PT0009/2020 | Europe / Portugal | 2020-03-08 | CHULC - H Curry Cabral | Instituto Nacional de Saude (INSA) | Guiomar et al |
| EPI_ISL_417995 | hCoV-19/Portugal/PT0010/2020 | Europe / Portugal | 2020-03-08 | CHULC - H Curry Cabral | Instituto Nacional de Saude (INSA) | Guiomar et al |
| EPI_ISL_417996 | hCoV-19/Portugal/PT0011/2020 | Europe / Portugal | 2020-03-08 | CHULC - H Curry Cabral | Instituto Nacional de Saude (INSA) | Guiomar et al |
| EPI_ISL_417997 | hCoV-19/Portugal/PT0012/2020 | Europe / Portugal | 2020-03-07 | Centro Hospital do Porto, E.P.E. - H. Geral de Santo Antonio | Instituto Nacional de Saude (INSA) | Guiomar et al |
| EPI_ISL_417998 | hCoV-19/Portugal/PT0013/2020 | Europe / Portugal | 2020-03-08 | Centro Hospital do Porto, E.P.E. - H. Geral de Santo Antonio | Instituto Nacional de Saude (INSA) | Guiomar et al |
| EPI_ISL_417999 | hCoV-19/Portugal/PT0014/2020 | Europe / Portugal | 2020-03-07 | Centro Hospital do Porto, E.P.E. - H. Geral de Santo Antonio | Instituto Nacional de Saude (INSA) | Guiomar et al |
| EPI_ISL_418000 | hCoV-19/Portugal/PT0015/2020 | Europe / Portugal | 2020-03-10 | ARS Algarve - Laboratório Laura Ayres | Instituto Nacional de Saude (INSA) | Guiomar et al |
| EPI_ISL_418001 | hCoV-19/Portugal/PT0016/2020 | Europe / Portugal | 2020-03-10 | ARS Algarve - Laboratório Laura Ayres | Instituto Nacional de Saude (INSA) | Guiomar et al |
| EPI_ISL_418002 | hCoV-19/Portugal/PT0017/2020 | Europe / Portugal | 2020-03-11 | CHU Coimbra | Instituto Nacional de Saude (INSA) | Guiomar et al |
| EPI_ISL_418003 | hCoV-19/Portugal/PT0018/2020 | Europe / Portugal | 2020-03-10 | H Braga | Instituto Nacional de Saude (INSA) | Guiomar et al |
| EPI_ISL_418004 | hCoV-19/Portugal/PT0019/2020 | Europe / Portugal | 2020-03-12 | ARS Algarve - Laboratório Laura Ayres | Instituto Nacional de Saude (INSA) | Guiomar et al |
| EPI_ISL_418005 | hCoV-19/Portugal/PT0020/2020 | Europe / Portugal | 2020-03-12 | CHU Coimbra - Pediátrico | Instituto Nacional de Saude (INSA) | Guiomar et al |
| EPI_ISL_418006 | hCoV-19/Portugal/PT0021/2020 | Europe / Portugal | 2020-03-13 | CHBarreiro Montijo | Instituto Nacional de Saude (INSA) | Guiomar et al |
| EPI_ISL_418007 | hCoV-19/Portugal/PT0022/2020 | Europe / Portugal | 2020-03-13 | H Braga | Instituto Nacional de Saude (INSA) | Guiomar et al |
| EPI_ISL_418008 | hCoV-19/Portugal/PT0023/2020 | Europe / Portugal | 2020-03-13 | H Braga | Instituto Nacional de Saude (INSA) | Guiomar et al |
| EPI_ISL_418009 | hCoV-19/Portugal/PT0024/2020 | Europe / Portugal | 2020-03-15 | HSE Ilha Terceira - Angra do Heroismo | Instituto Nacional de Saude (INSA) | Guiomar et al |
| EPI_ISL_418010 | hCoV-19/Portugal/PT0025/2020 | Europe / Portugal | 2020-03-03 | CHULC - H Curry Cabral | Instituto Nacional de Saude (INSA) | Guiomar et al |
| EPI_ISL_418011 | hCoV-19/Portugal/PT0026/2020 | Europe / Portugal | 2020-03-04 | CHULC - H Curry Cabral | Instituto Nacional de Saude (INSA) | Guiomar et al |
| EPI_ISL_418012 | hCoV-19/Portugal/PT0027/2020 | Europe / Portugal | 2020-03-06 | CHULC - H Curry Cabral | Instituto Nacional de Saude (INSA) | Guiomar et al |
| EPI_ISL_418013 | hCoV-19/Portugal/PT0028/2020 | Europe / Portugal | 2020-03-07 | CHULC - H Curry Cabral | Instituto Nacional de Saude (INSA) | Guiomar et al |
| EPI_ISL_418014 | hCoV-19/Portugal/PT0029/2020 | Europe / Portugal | 2020-03-09 | CHULC - H Curry Cabral | Instituto Nacional de Saude (INSA) | Guiomar et al |
| EPI_ISL_418015 | hCoV-19/Portugal/PT0030/2020 | Europe / Portugal | 2020-03-09 | CHULC - H Curry Cabral | Instituto Nacional de Saude (INSA) | Guiomar et al |
| EPI_ISL_418016 | hCoV-19/Portugal/PT0031/2020 | Europe / Portugal | 2020-03-10 | CHULC - H Curry Cabral | Instituto Nacional de Saude (INSA) | Guiomar et al |
| EPI_ISL_418017 | hCoV-19/Portugal/PT0032/2020 | Europe / Portugal | 2020-03-16 | CHMT | Instituto Nacional de Saude (INSA) | Guiomar et al |
| EPI_ISL_418018 | hCoV-19/Portugal/PT0033/2020 | Europe / Portugal | 2020-03-16 | H Garcia de Orta | Instituto Nacional de Saude (INSA) | Guiomar et al |
| EPI_ISL_418019 | hCoV-19/Portugal/PT0034/2020 | Europe / Portugal | 2020-03-15 | H Braga | Instituto Nacional de Saude (INSA) | Guiomar et al |
| EPI_ISL_418020 | hCoV-19/Portugal/PT0035/2020 | Europe / Portugal | 2020-03-16 | H Braga | Instituto Nacional de Saude (INSA) | Guiomar et al |
| EPI_ISL_418021 | hCoV-19/Portugal/PT0036/2020 | Europe / Portugal | 2020-03-16 | H Braga | Instituto Nacional de Saude (INSA) | Guiomar et al |
| EPI_ISL_418022 | hCoV-19/Portugal/PT0037/2020 | Europe / Portugal | 2020-03-15 | H Braga | Instituto Nacional de Saude (INSA) | Guiomar et al |
| EPI_ISL_418023 | hCoV-19/Portugal/PT0038/2020 | Europe / Portugal | 2020-03-17 | H Evora | Instituto Nacional de Saude (INSA) | Guiomar et al |
| EPI_ISL_418024 | hCoV-19/Portugal/PT0039/2020 | Europe / Portugal | 2020-03-17 | CHUA - Faro | Instituto Nacional de Saude (INSA) | Guiomar et al |
| EPI_ISL_418025 | hCoV-19/Portugal/PT0040/2020 | Europe / Portugal | 2020-03-17 | H Santarem | Instituto Nacional de Saude (INSA) | Guiomar et al |
| EPI_ISL_418026 | hCoV-19/Portugal/PT0041/2020 | Europe / Portugal | 2020-03-17 | H Dr. Nelio Mendonca - Funchal | Instituto Nacional de Saude (INSA) | Guiomar et al |
| EPI_ISL_418027 | hCoV-19/Portugal/PT0042/2020 | Europe / Portugal | 2020-03-17 | CHTMAD | Instituto Nacional de Saude (INSA) | Guiomar et al |
| EPI_ISL_418028 | hCoV-19/USA/MN-UW243/2020 | North America / USA / Minnesota | 2020-03-13 | UW Virology Lab | UW Virology Lab | Pavitra Roychoudhury, Hong Xie, Keith Jerome, Alexander Greninger |
| EPI_ISL_418029 | hCoV-19/USA/WA-UW244/2020 | North America / USA / Washington | 2020-03-16 | UW Virology Lab | UW Virology Lab | Pavitra Roychoudhury, Hong Xie, Keith Jerome, Alexander Greninger |
| EPI_ISL_418030 | hCoV-19/USA/WA-UW245/2020 | North America / USA / Washington | 2020-03-16 | UW Virology Lab | UW Virology Lab | Pavitra Roychoudhury, Hong Xie, Keith Jerome, Alexander Greninger |
| EPI_ISL_418031 | hCoV-19/USA/WA-UW246/2020 | North America / USA / Washington | 2020-03-16 | UW Virology Lab | UW Virology Lab | Pavitra Roychoudhury, Hong Xie, Keith Jerome, Alexander Greninger |
| EPI_ISL_418032 | hCoV-19/USA/WA-UW247/2020 | North America / USA / Washington | 2020-03-16 | UW Virology Lab | UW Virology Lab | Pavitra Roychoudhury, Hong Xie, Keith Jerome, Alexander Greninger |
| EPI_ISL_418033 | hCoV-19/USA/WA-UW248/2020 | North America / USA / Washington | 2020-03-16 | UW Virology Lab | UW Virology Lab | Pavitra Roychoudhury, Hong Xie, Keith Jerome, Alexander Greninger |
| EPI_ISL_418034 | hCoV-19/USA/WA-UW249/2020 | North America / USA / Washington | 2020-03-16 | UW Virology Lab | UW Virology Lab | Pavitra Roychoudhury, Hong Xie, Keith Jerome, Alexander Greninger |
| EPI_ISL_418036 | hCoV-19/USA/MN-UW251/2020 | North America / USA / Minnesota | 2020-03-13 | UW Virology Lab | UW Virology Lab | Pavitra Roychoudhury, Hong Xie, Keith Jerome, Alexander Greninger |
| EPI_ISL_418037 | hCoV-19/USA/WA-UW252/2020 | North America / USA / Washington | 2020-03-16 | UW Virology Lab | UW Virology Lab | Pavitra Roychoudhury, Hong Xie, Keith Jerome, Alexander Greninger |
| EPI_ISL_418038 | hCoV-19/USA/WA-UW253/2020 | North America / USA / Washington | 2020-03-15 | UW Virology Lab | UW Virology Lab | Pavitra Roychoudhury, Hong Xie, Keith Jerome, Alexander Greninger |
| EPI_ISL_418039 | hCoV-19/USA/CT-UW254/2020 | North America / USA / Connecticut | 2020-03-13 | UW Virology Lab | UW Virology Lab | Pavitra Roychoudhury, Hong Xie, Keith Jerome, Alexander Greninger |
| EPI_ISL_418040 | hCoV-19/USA/WA-UW255/2020 | North America / USA / Washington | 2020-03-16 | UW Virology Lab | UW Virology Lab | Pavitra Roychoudhury, Hong Xie, Keith Jerome, Alexander Greninger |
| EPI_ISL_418041 | hCoV-19/USA/CT-UW256/2020 | North America / USA / Connecticut | 2020-03-14 | UW Virology Lab | UW Virology Lab | Pavitra Roychoudhury, Hong Xie, Keith Jerome, Alexander Greninger |
| EPI_ISL_418042 | hCoV-19/USA/MN-UW257/2020 | North America / USA / Minnesota | 2020-03-13 | UW Virology Lab | UW Virology Lab | Pavitra Roychoudhury, Hong Xie, Keith Jerome, Alexander Greninger |
| EPI_ISL_418043 | hCoV-19/USA/CT-UW258/2020 | North America / USA / Connecticut | 2020-03-14 | UW Virology Lab | UW Virology Lab | Pavitra Roychoudhury, Hong Xie, Keith Jerome, Alexander Greninger |
| EPI_ISL_418044 | hCoV-19/USA/CT-UW259/2020 | North America / USA / Connecticut | 2020-03-14 | UW Virology Lab | UW Virology Lab | Pavitra Roychoudhury, Hong Xie, Keith Jerome, Alexander Greninger |
| EPI_ISL_418045 | hCoV-19/USA/CT-UW260/2020 | North America / USA / Connecticut | 2020-03-14 | UW Virology Lab | UW Virology Lab | Pavitra Roychoudhury, Hong Xie, Keith Jerome, Alexander Greninger |
| EPI_ISL_418046 | hCoV-19/USA/WA-UW261/2020 | North America / USA / Washington | 2020-03-16 | UW Virology Lab | UW Virology Lab | Pavitra Roychoudhury, Hong Xie, Keith Jerome, Alexander Greninger |
| EPI_ISL_418047 | hCoV-19/USA/WA-UW262/2020 | North America / USA / Washington | 2020-03-16 | UW Virology Lab | UW Virology Lab | Pavitra Roychoudhury, Hong Xie, Keith Jerome, Alexander Greninger |
| EPI_ISL_418049 | hCoV-19/USA/MN-UW264/2020 | North America / USA / Minnesota | 2020-03-13 | UW Virology Lab | UW Virology Lab | Pavitra Roychoudhury, Hong Xie, Keith Jerome, Alexander Greninger |
| EPI_ISL_418050 | hCoV-19/USA/WA-UW265/2020 | North America / USA / Washington | 2020-03-16 | UW Virology Lab | UW Virology Lab | Pavitra Roychoudhury, Hong Xie, Keith Jerome, Alexander Greninger |
| EPI_ISL_418051 | hCoV-19/USA/MN-UW266/2020 | North America / USA / Minnesota | 2020-03-13 | UW Virology Lab | UW Virology Lab | Pavitra Roychoudhury, Hong Xie, Keith Jerome, Alexander Greninger |
| EPI_ISL_418052 | hCoV-19/USA/WA-UW267/2020 | North America / USA / Washington | 2020-03-16 | UW Virology Lab | UW Virology Lab | Pavitra Roychoudhury, Hong Xie, Keith Jerome, Alexander Greninger |
| EPI_ISL_418053 | hCoV-19/USA/WA-UW268/2020 | North America / USA / Washington | 2020-03-16 | UW Virology Lab | UW Virology Lab | Pavitra Roychoudhury, Hong Xie, Keith Jerome, Alexander Greninger |
| EPI_ISL_418054 | hCoV-19/USA/WA-UW269/2020 | North America / USA / Washington | 2020-03-16 | UW Virology Lab | UW Virology Lab | Pavitra Roychoudhury, Hong Xie, Keith Jerome, Alexander Greninger |
| EPI_ISL_418055 | hCoV-19/USA/WA-UW270/2020 | North America / USA / Washington | 2020-03-14 | UW Virology Lab | UW Virology Lab | Pavitra Roychoudhury, Hong Xie, Keith Jerome, Alexander Greninger |
| EPI_ISL_418056 | hCoV-19/USA/CT-UW271/2020 | North America / USA / Connecticut | 2020-03-13 | UW Virology Lab | UW Virology Lab | Pavitra Roychoudhury, Hong Xie, Keith Jerome, Alexander Greninger |
| EPI_ISL_418057 | hCoV-19/USA/CT-UW272/2020 | North America / USA / Connecticut | 2020-03-13 | UW Virology Lab | UW Virology Lab | Pavitra Roychoudhury, Hong Xie, Keith Jerome, Alexander Greninger |
| EPI_ISL_418058 | hCoV-19/USA/CT-UW273/2020 | North America / USA / Connecticut | 2020-03-13 | UW Virology Lab | UW Virology Lab | Pavitra Roychoudhury, Hong Xie, Keith Jerome, Alexander Greninger |
| EPI_ISL_418059 | hCoV-19/USA/WA-UW274/2020 | North America / USA | 2020-03-13 | UW Virology Lab | UW Virology Lab | Pavitra Roychoudhury, Hong Xie, Keith Jerome, Alexander Greninger |
| EPI_ISL_418105 | hCoV-19/Wales/PHWC-23CF4/2020 | Europe / United Kingdom / Wales | 2020-03-15 | Wales Specialist Virology Centre | Public Health Wales Microbiology Cardiff | Catherine Moore, Joanne Watkins, Sally Corden, Sara Rey, Matt Bull, Tom Connor |
| EPI_ISL_418106 | hCoV-19/Wales/PHWC-23D00/2020 | Europe / United Kingdom / Wales | 2020-03-13 | Wales Specialist Virology Centre | Public Health Wales Microbiology Cardiff | Catherine Moore, Joanne Watkins, Sally Corden, Sara Rey, Matt Bull, Tom Connor |
| EPI_ISL_418108 | hCoV-19/Wales/PHWC-23D2E/2020 | Europe / United Kingdom / Wales | 2020-03-13 | Wales Specialist Virology Centre | Public Health Wales Microbiology Cardiff | Catherine Moore, Joanne Watkins, Sally Corden, Sara Rey, Matt Bull, Tom Connor |
| EPI_ISL_418109 | hCoV-19/Wales/PHWC-23D4C/2020 | Europe / United Kingdom / Wales | 2020-03-12 | Wales Specialist Virology Centre | Public Health Wales Microbiology Cardiff | Catherine Moore, Joanne Watkins, Sally Corden, Sara Rey, Matt Bull, Tom Connor |
| EPI_ISL_418110 | hCoV-19/Wales/PHWC-23D6A/2020 | Europe / United Kingdom / Wales | 2020-03-13 | Wales Specialist Virology Centre | Public Health Wales Microbiology Cardiff | Catherine Moore, Joanne Watkins, Sally Corden, Sara Rey, Matt Bull, Tom Connor |
| EPI_ISL_418111 | hCoV-19/Wales/PHWC-23D79/2020 | Europe / United Kingdom / Wales | 2020-03-13 | Wales Specialist Virology Centre | Public Health Wales Microbiology Cardiff | Catherine Moore, Joanne Watkins, Sally Corden, Sara Rey, Matt Bull, Tom Connor |
| EPI_ISL_418113 | hCoV-19/Wales/PHWC-23DB5/2020 | Europe / United Kingdom / Wales | 2020-03-13 | Wales Specialist Virology Centre | Public Health Wales Microbiology Cardiff | Catherine Moore, Joanne Watkins, Sally Corden, Sara Rey, Matt Bull, Tom Connor |
| EPI_ISL_418116 | hCoV-19/Wales/PHWC-23E1C/2020 | Europe / United Kingdom / Wales | 2020-03-13 | Wales Specialist Virology Centre | Public Health Wales Microbiology Cardiff | Catherine Moore, Joanne Watkins, Sally Corden, Sara Rey, Matt Bull, Tom Connor |
| EPI_ISL_418117 | hCoV-19/Wales/PHWC-23E2B/2020 | Europe / United Kingdom / Wales | 2020-03-14 | Wales Specialist Virology Centre | Public Health Wales Microbiology Cardiff | Catherine Moore, Joanne Watkins, Sally Corden, Sara Rey, Matt Bull, Tom Connor |
| EPI_ISL_418119 | hCoV-19/Wales/PHWC-23E49/2020 | Europe / United Kingdom / Wales | 2020-03-13 | Wales Specialist Virology Centre | Public Health Wales Microbiology Cardiff | Catherine Moore, Joanne Watkins, Sally Corden, Sara Rey, Matt Bull, Tom Connor |
| EPI_ISL_418136 | hCoV-19/Wales/PHWC-23FFB/2020 | Europe / United Kingdom / Wales | 2020-03-15 | Wales Specialist Virology Centre | Public Health Wales Microbiology Cardiff | Catherine Moore, Joanne Watkins, Sally Corden, Sara Rey, Matt Bull, Tom Connor |
| EPI_ISL_418137 | hCoV-19/Wales/PHWC-24006/2020 | Europe / United Kingdom / Wales | 2020-03-15 | Wales Specialist Virology Centre | Public Health Wales Microbiology Cardiff | Catherine Moore, Joanne Watkins, Sally Corden, Sara Rey, Matt Bull, Tom Connor |
| EPI_ISL_418139 | hCoV-19/Wales/PHWC-24024/2020 | Europe / United Kingdom / Wales | 2020-03-16 | Wales Specialist Virology Centre | Public Health Wales Microbiology Cardiff | Catherine Moore, Joanne Watkins, Sally Corden, Sara Rey, Matt Bull, Tom Connor |
| EPI_ISL_418143 | hCoV-19/Wales/PHWC-240D9/2020 | Europe / United Kingdom / Wales | 2020-03-14 | Wales Specialist Virology Centre | Public Health Wales Microbiology Cardiff | Catherine Moore, Joanne Watkins, Sally Corden, Sara Rey, Matt Bull, Tom Connor |
| EPI_ISL_418144 | hCoV-19/Wales/PHWC-240E8/2020 | Europe / United Kingdom / Wales | 2020-03-14 | Wales Specialist Virology Centre | Public Health Wales Microbiology Cardiff | Catherine Moore, Joanne Watkins, Sally Corden, Sara Rey, Matt Bull, Tom Connor |
| EPI_ISL_418145 | hCoV-19/Wales/PHWC-240F7/2020 | Europe / United Kingdom / Wales | 2020-03-14 | Wales Specialist Virology Centre | Public Health Wales Microbiology Cardiff | Catherine Moore, Joanne Watkins, Sally Corden, Sara Rey, Matt Bull, Tom Connor |
| EPI_ISL_418147 | hCoV-19/Wales/PHWC-24121/2020 | Europe / United Kingdom / Wales | 2020-03-14 | Wales Specialist Virology Centre | Public Health Wales Microbiology Cardiff | Catherine Moore, Joanne Watkins, Sally Corden, Sara Rey, Matt Bull, Tom Connor |
| EPI_ISL_418148 | hCoV-19/Wales/PHWC-2414F/2020 | Europe / United Kingdom / Wales | 2020-03-16 | Wales Specialist Virology Centre | Public Health Wales Microbiology Cardiff | Catherine Moore, Joanne Watkins, Sally Corden, Sara Rey, Matt Bull, Tom Connor |
| EPI_ISL_418149 | hCoV-19/Wales/PHWC-2415E/2020 | Europe / United Kingdom / Wales | 2020-03-16 | Wales Specialist Virology Centre | Public Health Wales Microbiology Cardiff | Catherine Moore, Joanne Watkins, Sally Corden, Sara Rey, Matt Bull, Tom Connor |
| EPI_ISL_418151 | hCoV-19/Wales/PHWC-2419A/2020 | Europe / United Kingdom / Wales | 2020-03-16 | Wales Specialist Virology Centre | Public Health Wales Microbiology Cardiff | Catherine Moore, Joanne Watkins, Sally Corden, Sara Rey, Matt Bull, Tom Connor |
| EPI_ISL_418155 | hCoV-19/Wales/PHWC-2422E/2020 | Europe / United Kingdom / Wales | 2020-03-16 | Wales Specialist Virology Centre | Public Health Wales Microbiology Cardiff | Catherine Moore, Joanne Watkins, Sally Corden, Sara Rey, Matt Bull, Tom Connor |
| EPI_ISL_418156 | hCoV-19/Wales/PHWC-2423D/2020 | Europe / United Kingdom / Wales | 2020-03-16 | Wales Specialist Virology Centre | Public Health Wales Microbiology Cardiff | Catherine Moore, Joanne Watkins, Sally Corden, Sara Rey, Matt Bull, Tom Connor |
| EPI_ISL_418158 | hCoV-19/Wales/PHWC-24279/2020 | Europe / United Kingdom / Wales | 2020-03-16 | Wales Specialist Virology Centre | Public Health Wales Microbiology Cardiff | Catherine Moore, Joanne Watkins, Sally Corden, Sara Rey, Matt Bull, Tom Connor |
| EPI_ISL_418159 | hCoV-19/Wales/PHWC-24288/2020 | Europe / United Kingdom / Wales | 2020-03-16 | Wales Specialist Virology Centre | Public Health Wales Microbiology Cardiff | Catherine Moore, Joanne Watkins, Sally Corden, Sara Rey, Matt Bull, Tom Connor |
| EPI_ISL_418163 | hCoV-19/Wales/PHWC-242C4/2020 | Europe / United Kingdom / Wales | 2020-03-16 | Wales Specialist Virology Centre | Public Health Wales Microbiology Cardiff | Catherine Moore, Joanne Watkins, Sally Corden, Sara Rey, Matt Bull, Tom Connor |
| EPI_ISL_418165 | hCoV-19/Wales/PHWC-242E2/2020 | Europe / United Kingdom / Wales | 2020-03-16 | Wales Specialist Virology Centre | Public Health Wales Microbiology Cardiff | Catherine Moore, Joanne Watkins, Sally Corden, Sara Rey, Matt Bull, Tom Connor |
| EPI_ISL_418400 | hCoV-19/Finland/13M83/2020 | Europe / Finland | 2020-03-13 | Department of Virology and Immunology, University of Helsinki and Helsinki University Hospital, Huslab Finland | Department of Virology, Faculty of Medicine, University of Helsinki, Helsinki, Finland | Teemu Smura, Hannimari Kallio-Kokko, Olli Vapalahti |
| EPI_ISL_418401 | hCoV-19/Finland/14M12/2020 | Europe / Finland | 2020-03-14 | Department of Virology and Immunology, University of Helsinki and Helsinki University Hospital, Huslab Finland | Department of Virology, Faculty of Medicine, University of Helsinki, Helsinki, Finland | Teemu Smura, Hannimari Kallio-Kokko, Olli Vapalahti |
| EPI_ISL_418402 | hCoV-19/Finland/14M13/2020 | Europe / Finland | 2020-03-14 | Department of Virology and Immunology, University of Helsinki and Helsinki University Hospital, Huslab Finland | Department of Virology, Faculty of Medicine, University of Helsinki, Helsinki, Finland | Teemu Smura, Hannimari Kallio-Kokko, Olli Vapalahti |
| EPI_ISL_418403 | hCoV-19/Finland/14M14/2020 | Europe / Finland | 2020-03-14 | Department of Virology and Immunology, University of Helsinki and Helsinki University Hospital, Huslab Finland | Department of Virology, Faculty of Medicine, University of Helsinki, Helsinki, Finland | Teemu Smura, Hannimari Kallio-Kokko, Olli Vapalahti |
| EPI_ISL_418404 | hCoV-19/Finland/14M16/2020 | Europe / Finland | 2020-03-14 | Department of Virology and Immunology, University of Helsinki and Helsinki University Hospital, Huslab Finland | Department of Virology, Faculty of Medicine, University of Helsinki, Helsinki, Finland | Teemu Smura, Hannimari Kallio-Kokko, Olli Vapalahti |
| EPI_ISL_418405 | hCoV-19/Finland/14M20/2020 | Europe / Finland | 2020-03-14 | Department of Virology and Immunology, University of Helsinki and Helsinki University Hospital, Huslab Finland | Department of Virology, Faculty of Medicine, University of Helsinki, Helsinki, Finland | Teemu Smura, Hannimari Kallio-Kokko, Olli Vapalahti |
| EPI_ISL_418406 | hCoV-19/Finland/14M26/2020 | Europe / Finland | 2020-03-14 | Department of Virology and Immunology, University of Helsinki and Helsinki University Hospital, Huslab Finland | Department of Virology, Faculty of Medicine, University of Helsinki, Helsinki, Finland | Teemu Smura, Hannimari Kallio-Kokko, Olli Vapalahti |
| EPI_ISL_418408 | hCoV-19/Finland/14M32/2020 | Europe / Finland / Helsinki | 2020-03-14 | Department of Virology and Immunology, University of Helsinki and Helsinki University Hospital, Huslab Finland | Department of Virology, Faculty of Medicine, University of Helsinki, Helsinki, Finland | Teemu Smura, Hannimari Kallio-Kokko, Olli Vapalahti |
| EPI_ISL_418409 | hCoV-19/Finland/14M74/2020 | Europe / Finland / Helsinki | 2020-03-14 | Department of Virology and Immunology, University of Helsinki and Helsinki University Hospital, Huslab Finland | Department of Virology, Faculty of Medicine, University of Helsinki, Helsinki, Finland | Teemu Smura, Hannimari Kallio-Kokko, Olli Vapalahti |
| EPI_ISL_418410 | hCoV-19/Finland/14M77/2020 | Europe / Finland / Helsinki | 2020-03-14 | Department of Virology and Immunology, University of Helsinki and Helsinki University Hospital, Huslab Finland | Department of Virology, Faculty of Medicine, University of Helsinki, Helsinki, Finland | Teemu Smura, Hannimari Kallio-Kokko, Olli Vapalahti |
| EPI_ISL_418411 | hCoV-19/Finland/14M82/2020 | Europe / Finland / Helsinki | 2020-03-14 | Department of Virology and Immunology, University of Helsinki and Helsinki University Hospital, Huslab Finland | Department of Virology, Faculty of Medicine, University of Helsinki, Helsinki, Finland | Teemu Smura, Hannimari Kallio-Kokko, Olli Vapalahti |
| EPI_ISL_418412 | hCoV-19/France/ARA094100/2020 | Europe / France / ARA / Privas | 2020-03-15 | Centre Hospitalier des Vals d'Ardeche | CNR Virus des Infections Respiratoires - France SUD | Antonin Bal, Gregory Destras, Gwendolyne Burfin, Solenne Brun, Carine Moustaud, Raphaelle Lamy, Alexandre Gaymard, Maude Bouscambert-Duchamp, Florence Morfin-Sherpa, Martine Valette, Bruno Lina, Laurence Josset |
| EPI_ISL_418413 | hCoV-19/France/ARA09428/2020 | Europe / France / ARA / Macon | 2020-03-15 | Centre Hospitalier de Macon | CNR Virus des Infections Respiratoires - France SUD | Antonin Bal, Gregory Destras, Gwendolyne Burfin, Solenne Brun, Carine Moustaud, Raphaelle Lamy, Alexandre Gaymard, Maude Bouscambert-Duchamp, Florence Morfin-Sherpa, Martine Valette, Bruno Lina, Laurence Josset |
| EPI_ISL_418414 | hCoV-19/France/ARA09434/2020 | Europe / France / ARA / Valence | 2020-03-15 | Centre Hospitalier de Valence | CNR Virus des Infections Respiratoires - France SUD | Antonin Bal, Gregory Destras, Gwendolyne Burfin, Solenne Brun, Carine Moustaud, Raphaelle Lamy, Alexandre Gaymard, Maude Bouscambert-Duchamp, Florence Morfin-Sherpa, Martine Valette, Bruno Lina, Laurence Josset |
| EPI_ISL_418416 | hCoV-19/France/ARA09588/2020 | Europe / France / ARA / Venissieux | 2020-03-16 | GH Les Portes du Sud | CNR Virus des Infections Respiratoires - France SUD | Antonin Bal, Gregory Destras, Gwendolyne Burfin, Solenne Brun, Carine Moustaud, Raphaelle Lamy, Alexandre Gaymard, Maude Bouscambert-Duchamp, Florence Morfin-Sherpa, Martine Valette, Bruno Lina, Laurence Josset |
| EPI_ISL_418417 | hCoV-19/France/ARA09686/2020 | Europe / France / ARA / Valence | 2020-03-16 | Centre Hospitalier de Valence | CNR Virus des Infections Respiratoires - France SUD | Antonin Bal, Gregory Destras, Gwendolyne Burfin, Solenne Brun, Carine Moustaud, Raphaelle Lamy, Alexandre Gaymard, Maude Bouscambert-Duchamp, Florence Morfin-Sherpa, Martine Valette, Bruno Lina, Laurence Josset |
| EPI_ISL_418418 | hCoV-19/France/ARA10163/2020 | Europe / France / ARA / Lyon | 2020-03-16 | Centre Hospitalier Saint Joseph Saint Luc | CNR Virus des Infections Respiratoires - France SUD | Antonin Bal, Gregory Destras, Gwendolyne Burfin, Solenne Brun, Carine Moustaud, Raphaelle Lamy, Alexandre Gaymard, Maude Bouscambert-Duchamp, Florence Morfin-Sherpa, Martine Valette, Bruno Lina, Laurence Josset |
| EPI_ISL_418419 | hCoV-19/France/ARA10165/2020 | Europe / France / ARA / Lyon | 2020-03-16 | Centre Hospitalier Saint Joseph Saint Luc | CNR Virus des Infections Respiratoires - France SUD | Antonin Bal, Gregory Destras, Gwendolyne Burfin, Solenne Brun, Carine Moustaud, Raphaelle Lamy, Alexandre Gaymard, Maude Bouscambert-Duchamp, Florence Morfin-Sherpa, Martine Valette, Bruno Lina, Laurence Josset |
| EPI_ISL_418420 | hCoV-19/France/ARA10170/2020 | Europe / France / ARA / Lyon | 2020-03-17 | Institut des Agents Infectieux (IAI), Hospices Civils de Lyon | CNR Virus des Infections Respiratoires - France SUD | Antonin Bal, Gregory Destras, Gwendolyne Burfin, Solenne Brun, Carine Moustaud, Raphaelle Lamy, Alexandre Gaymard, Maude Bouscambert-Duchamp, Florence Morfin-Sherpa, Martine Valette, Bruno Lina, Laurence Josset |
| EPI_ISL_418422 | hCoV-19/France/ARA10184/2020 | Europe / France / ARA / Lyon | 2020-03-17 | Institut des Agents Infectieux (IAI), Hospices Civils de Lyon | CNR Virus des Infections Respiratoires - France SUD | Antonin Bal, Gregory Destras, Gwendolyne Burfin, Solenne Brun, Carine Moustaud, Raphaelle Lamy, Alexandre Gaymard, Maude Bouscambert-Duchamp, Florence Morfin-Sherpa, Martine Valette, Bruno Lina, Laurence Josset |
| EPI_ISL_418423 | hCoV-19/France/ARA10188/2020 | Europe / France / ARA / Lyon | 2020-03-17 | Institut des Agents Infectieux (IAI), Hospices Civils de Lyon | CNR Virus des Infections Respiratoires - France SUD | Antonin Bal, Gregory Destras, Gwendolyne Burfin, Solenne Brun, Carine Moustaud, Raphaelle Lamy, Alexandre Gaymard, Maude Bouscambert-Duchamp, Florence Morfin-Sherpa, Martine Valette, Bruno Lina, Laurence Josset |
| EPI_ISL_418424 | hCoV-19/France/ARA10189/2020 | Europe / France / ARA / Lyon | 2020-03-17 | Institut des Agents Infectieux (IAI), Hospices Civils de Lyon | CNR Virus des Infections Respiratoires - France SUD | Antonin Bal, Gregory Destras, Gwendolyne Burfin, Solenne Brun, Carine Moustaud, Raphaelle Lamy, Alexandre Gaymard, Maude Bouscambert-Duchamp, Florence Morfin-Sherpa, Martine Valette, Bruno Lina, Laurence Josset |
| EPI_ISL_418425 | hCoV-19/France/ARA10192/2020 | Europe / France / ARA / Lyon | 2020-03-17 | Institut des Agents Infectieux (IAI), Hospices Civils de Lyon | CNR Virus des Infections Respiratoires - France SUD | Antonin Bal, Gregory Destras, Gwendolyne Burfin, Solenne Brun, Carine Moustaud, Raphaelle Lamy, Alexandre Gaymard, Maude Bouscambert-Duchamp, Florence Morfin-Sherpa, Martine Valette, Bruno Lina, Laurence Josset |
| EPI_ISL_418426 | hCoV-19/France/ARA10251/2020 | Europe / France / ARA / Bourg-en-Bresse | 2020-03-17 | Centre Hospitalier de Bourg en Bresse | CNR Virus des Infections Respiratoires - France SUD | Antonin Bal, Gregory Destras, Gwendolyne Burfin, Solenne Brun, Carine Moustaud, Raphaelle Lamy, Alexandre Gaymard, Maude Bouscambert-Duchamp, Florence Morfin-Sherpa, Martine Valette, Bruno Lina, Laurence Josset |
| EPI_ISL_418427 | hCoV-19/France/ARA10257/2020 | Europe / France / ARA / Saint-Priest | 2020-03-17 | Hopital Privé de l'Est Lyonnais | CNR Virus des Infections Respiratoires - France SUD | Antonin Bal, Gregory Destras, Gwendolyne Burfin, Solenne Brun, Carine Moustaud, Raphaelle Lamy, Alexandre Gaymard, Maude Bouscambert-Duchamp, Florence Morfin-Sherpa, Martine Valette, Bruno Lina, Laurence Josset |
| EPI_ISL_418428 | hCoV-19/France/ARA10282/2020 | Europe / France / ARA / Vienne | 2020-03-17 | Centre Hospitalier Lucien Hussel | CNR Virus des Infections Respiratoires - France SUD | Antonin Bal, Gregory Destras, Gwendolyne Burfin, Solenne Brun, Carine Moustaud, Raphaelle Lamy, Alexandre Gaymard, Maude Bouscambert-Duchamp, Florence Morfin-Sherpa, Martine Valette, Bruno Lina, Laurence Josset |
| EPI_ISL_418429 | hCoV-19/France/ARA10876/2020 | Europe / France / ARA / Lyon | 2020-03-18 | Institut des Agents Infectieux (IAI), Hospices Civils de Lyon | CNR Virus des Infections Respiratoires - France SUD | Antonin Bal, Gregory Destras, Gwendolyne Burfin, Solenne Brun, Carine Moustaud, Raphaelle Lamy, Alexandre Gaymard, Maude Bouscambert-Duchamp, Florence Morfin-Sherpa, Martine Valette, Bruno Lina, Laurence Josset |
| EPI_ISL_418430 | hCoV-19/France/ARA10910/2020 | Europe / France / ARA / Lyon | 2020-03-18 | Institut des Agents Infectieux (IAI), Hospices Civils de Lyon | CNR Virus des Infections Respiratoires - France SUD | Antonin Bal, Gregory Destras, Gwendolyne Burfin, Solenne Brun, Carine Moustaud, Raphaelle Lamy, Alexandre Gaymard, Maude Bouscambert-Duchamp, Florence Morfin-Sherpa, Martine Valette, Bruno Lina, Laurence Josset |
| EPI_ISL_418431 | hCoV-19/France/ARA10968/2020 | Europe / France / ARA / Lyon | 2020-03-18 | Institut des Agents Infectieux (IAI), Hospices Civils de Lyon | CNR Virus des Infections Respiratoires - France SUD | Antonin Bal, Gregory Destras, Gwendolyne Burfin, Solenne Brun, Carine Moustaud, Raphaelle Lamy, Alexandre Gaymard, Maude Bouscambert-Duchamp, Florence Morfin-Sherpa, Martine Valette, Bruno Lina, Laurence Josset |
| EPI_ISL_418432 | hCoV-19/France/ARA11036/2020 | Europe / France / ARA / Lyon | 2020-03-18 | Institut des Agents Infectieux (IAI), Hospices Civils de Lyon | CNR Virus des Infections Respiratoires - France SUD | Antonin Bal, Gregory Destras, Gwendolyne Burfin, Solenne Brun, Carine Moustaud, Raphaelle Lamy, Alexandre Gaymard, Maude Bouscambert-Duchamp, Florence Morfin-Sherpa, Martine Valette, Bruno Lina, Laurence Josset |
| EPI_ISL_418441 | hCoV-19/Hangzhou/HZ48/2020 | Asia / China / Hangzhou | 2020-01-21 | Hangzhou Center for Disease Control and Prevention | Inspection Center of Hangzhou Center for Disease Control and Prevention | Yu hua, Wang haoqiu, Li jun, Yu xinfeng, Pan jingcao |
| EPI_ISL_418442 | hCoV-19/Hangzhou/HZ49/2020 | Asia / China / Hangzhou | 2020-01-21 | Hangzhou Center for Disease Control and Prevention | Inspection Center of Hangzhou Center for Disease Control and Prevention | Yu hua, Wang haoqiu, Li jun, Yu xinfeng, Pan jingcao |
| EPI_ISL_418502 | hCoV-19/Hangzhou/HZ60/2020 | Asia / China / Hangzhou | 2020-01-22 | Hangzhou Center for Disease Control and Prevention | Inspection Center of Hangzhou Center for Disease Control and Prevention | Yu hua, Wang haoqiu, Li jun, Yu xinfeng, Pan jingcao |
| EPI_ISL_418503 | hCoV-19/Hangzhou/HZ62/2020 | Asia / China / Hangzhou | 2020-01-22 | Hangzhou Center for Disease Control and Prevention | Inspection Center of Hangzhou Center for Disease Control and Prevention | Yu hua, Wang haoqiu, Li jun, Yu xinfeng, Pan jingcao |
| EPI_ISL_418504 | hCoV-19/Hangzhou/HZ79/2020 | Asia / China / Hangzhou | 2020-01-21 | Hangzhou Center for Disease Control and Prevention | Inspection Center of Hangzhou Center for Disease Control and Prevention | Yu hua, Wang haoqiu, Li jun, Yu xinfeng, Pan jingcao |
| EPI_ISL_418506 | hCoV-19/Hangzhou/HZ90/2020 | Asia / China / Hangzhou | 2020-01-21 | Hangzhou Center for Disease Control and Prevention | Inspection Center of Hangzhou Center for Disease Control and Prevention | Yu hua, Wang haoqiu, Li jun, Yu xinfeng, Pan jingcao |
| EPI_ISL_418507 | hCoV-19/Hangzhou/HZ91/2020 | Asia/China/Zhejiang/Hangzhou | 2020-01-21 | Hangzhou Center for Disease Control and Prevention | Inspection Center of Hangzhou Center for Disease Control and Prevention | Yu hua, Wang haoqiu, Li jun, Yu xinfeng, Pan jingcao |
| EPI_ISL_418508 | hCoV-19/Hangzhou/HZ162/2020 | Asia / China / Zhejiang / Hangzhou | 2020-01-23 | Hangzhou Center for Disease Control and Prevention | Inspection Center of Hangzhou Center for Disease Control and Prevention | Yu hua, Wang haoqiu, Li jun, Yu xinfeng, Pan jingcao |
| EPI_ISL_418509 | hCoV-19/Hangzhou/HZ178/2020 | Asia / China / Zhejiang / Hangzhou | 2020-01-23 | Hangzhou Center for Disease Control and Prevention | Inspection Center of Hangzhou Center for Disease Control and Prevention | Yu hua, Wang haoqiu, Li jun, Yu xinfeng, Pan jingcao |
| EPI_ISL_418510 | hCoV-19/Hangzhou/HZ185/2020 | Asia / China / Hangzhou | 2020-01-23 | Hangzhou Center for Disease Control and Prevention | Inspection Center of Hangzhou Center for Disease Control and Prevention | Yu hua, Wang haoqiu, Li jun, Yu xinfeng, Pan jingcao |
| EPI_ISL_418511 | hCoV-19/Hangzhou/HZ477/2020 | Asia / China / Zhejiang / Hangzhou | 2020-01-24 | Hangzhou Center for Disease Control and Prevention | Inspection Center of Hangzhou Center for Disease Control and Prevention | Yu hua, Wang haoqiu, Li jun, Yu xinfeng, Pan jingcao |
| EPI_ISL_418512 | hCoV-19/Hangzhou/HZ481/2020 | Asia / China / Zhejiang / Hangzhou | 2020-01-25 | Hangzhou Center for Disease Control and Prevention | Inspection Center of Hangzhou Center for Disease Control and Prevention | Yu hua, Wang haoqiu, Li jun, Yu xinfeng, Pan jingcao |
| EPI_ISL_418513 | hCoV-19/Hangzhou/HZ551/2020 | Asia / China / Hangzhou | 2020-01-25 | Hangzhou Center for Disease Control and Prevention | Inspection Center of Hangzhou Center for Disease Control and Prevention | Yu hua, Wang haoqiu, Li jun, Yu xinfeng, Pan jingcao |
| EPI_ISL_418514 | hCoV-19/Hangzhou/HZ576/2020 | Asia / China / Zhejiang / Hangzhou | 2020-01-25 | Hangzhou Center for Disease Control and Prevention | Inspection Center of Hangzhou Center for Disease Control and Prevention | Yu hua, Wang haoqiu, Li jun, Yu xinfeng, Pan jingcao |
| EPI_ISL_418515 | hCoV-19/Hangzhou/HZ638/2020 | Asia / China / Hangzhou | 2020-01-25 | Hangzhou Center for Disease Control and Prevention | Inspection Center of Hangzhou Center for Disease Control and Prevention | Yu hua, Wang haoqiu, Li jun, Yu xinfeng, Pan jingcao |
| EPI_ISL_418516 | hCoV-19/Ireland/21023/2020 | Europe / Ireland / Tipperary | 2020-03-06 | UCD National Virus Reference Laboratory | UCD National Virus Reference Laboratory | Michael Carr, Gabriel Gonzalez, Jonathan Dean, Suzie Coughlan, Alison Murphy, Kevin Byrne, Ken Wolfe, Jeff Connell, Brendan Loftus, Cillian F De Gascun |
| EPI_ISL_418548 | hCoV-19/Ireland/21145/2020 | Europe / Ireland / Dublin | 2020-03-06 | UCD National Virus Reference Laboratory | UCD National Virus Reference Laboratory | Michael Carr, Gabriel Gonzalez, Jonathan Dean, Suzie Coughlan, Alison Murphy, Kevin Byrne, Ken Wolfe, Jeff Connell, Brendan Loftus, Cillian F De Gascun |
| EPI_ISL_418580 | hCoV-19/Ireland/Dublin-22361/2020 | Europe / Ireland / Cork | 2020-03-08 | UCD National Virus Reference Laboratory | UCD National Virus Reference Laboratory | Michael Carr, Gabriel Gonzalez, Jonathan Dean, Suzie Coughlan, Alison Murphy, Kevin Byrne, Ken Wolfe, Jeff Connell, Brendan Loftus, Cillian F De Gascun |
| EPI_ISL_418581 | hCoV-19/Ireland/Dublin-22428/2020 | Europe / Ireland / Dublin | 2020-03-08 | UCD National Virus Reference Laboratory | UCD National Virus Reference Laboratory | Michael Carr, Gabriel Gonzalez, Jonathan Dean, Suzie Coughlan, Alison Murphy, Kevin Byrne, Ken Wolfe, Jeff Connell, Brendan Loftus, Cillian F De Gascun |
| EPI_ISL_418582 | hCoV-19/Ireland/22901/2020 | Europe / Ireland / Louth | 2020-03-10 | UCD National Virus Reference Laboratory | UCD National Virus Reference Laboratory | Michael Carr, Gabriel Gonzalez, Jonathan Dean, Suzie Coughlan, Alison Murphy, Kevin Byrne, Ken Wolfe, Jeff Connell, Brendan Loftus, Cillian F De Gascun |
| EPI_ISL_418583 | hCoV-19/Ireland/24042/2020 | Europe / Ireland / Dublin | 2020-03-10 | UCD National Virus Reference Laboratory | UCD National Virus Reference Laboratory | Michael Carr, Gabriel Gonzalez, Jonathan Dean, Suzie Coughlan, Alison Murphy, Kevin Byrne, Ken Wolfe, Jeff Connell, Brendan Loftus, Cillian F De Gascun |
| EPI_ISL_418584 | hCoV-19/Ireland/24052/2020 | Europe / Ireland / Wicklow | 2020-03-10 | UCD National Virus Reference Laboratory | UCD National Virus Reference Laboratory | Michael Carr, Gabriel Gonzalez, Jonathan Dean, Suzie Coughlan, Alison Murphy, Kevin Byrne, Ken Wolfe, Jeff Connell, Brendan Loftus, Cillian F De Gascun |
| EPI_ISL_418800 | hCoV-19/Belgium/CS-031052/2020 | Europe / Belgium / Gierle | 2020-03-10 | KU Leuven, Clinical and Epidemiological Virology | KU Leuven, Clinical and Epidemiological Virology | Bert Vanmechelen, Joan Martí-Carreras, Tony Wawina, Piet Maes |
| EPI_ISL_418801 | hCoV-19/Australia/QLDID923/2020 | Oceania / Australia / Queensland / Brisba | 2020-03-14 | Mater Pathology | Public Health Virology Laboratory | Bixing Huang, Alyssa Pyke, Amanda De Jong, Andrew Van Den Hurk, Carmel Taylor, David Warrilow, Doris Genge, Elisabeth Gamez, Glen Hewitson, Ian Maxwell Mackay, Inga Sultana, Jamie McMahon, Jean Barcelon, Judy Northill, Mitchell Finger, Natalie Simpson, Neelima Nair, Peter Burtonclay, Peter Moore, Sarah Wheatley, Sean Moody, Sonja Hall-Mendelin, Timothy Gardam, and Frederick Moore |
| EPI_ISL_418802 | hCoV-19/Australia/QLDID924/2020 | Oceania / Australia / Queensland / Gold ( | 2020-03-18 | Pathology Queensland | Public Health Virology Laboratory | Bixing Huang, Alyssa Pyke, Amanda De Jong, Andrew Van Den Hurk, Carmel Taylor, David Warrilow, Doris Genge, Elisabeth Gamez, Glen Hewitson, Ian Maxwell Mackay, Inga Sultana, Jamie McMahon, Jean Barcelon, Judy Northill, Mitchell Finger, Natalie Simpson, Neelima Nair, Peter Burtonclay, Peter Moore, Sarah Wheatley, Sean Moody, Sonja Hall-Mendelin, Timothy Gardam, and Frederick Moore |
| EPI_ISL_418803 | hCoV-19/Australia/QLDID925/2020 | Oceania / Australia / Queensland / Sunsh | 2020-03-18 | Pathology Queensland | Public Health Virology Laboratory | Bixing Huang, Alyssa Pyke, Amanda De Jong, Andrew Van Den Hurk, Carmel Taylor, David Warrilow, Doris Genge, Elisabeth Gamez, Glen Hewitson, Ian Maxwell Mackay, Inga Sultana, Jamie McMahon, Jean Barcelon, Judy Northill, Mitchell Finger, Natalie Simpson, Neelima Nair, Peter Burtonclay, Peter Moore, Sarah Wheatley, Sean Moody, Sonja Hall-Mendelin, Timothy Gardam, and Frederick Moore |
| EPI_ISL_418804 | hCoV-19/Australia/QLDID926/2020 | Oceania / Australia / Queensland / Sunsh | 2020-03-18 | Pathology Queensland | Public Health Virology Laboratory | Bixing Huang, Alyssa Pyke, Amanda De Jong, Andrew Van Den Hurk, Carmel Taylor, David Warrilow, Doris Genge, Elisabeth Gamez, Glen Hewitson, Ian Maxwell Mackay, Inga Sultana, Jamie McMahon, Jean Barcelon, Judy Northill, Mitchell Finger, Natalie Simpson, Neelima Nair, Peter Burtonclay, Peter Moore, Sarah Wheatley, Sean Moody, Sonja Hall-Mendelin, Timothy Gardam, and Frederick Moore |
| EPI_ISL_418805 | hCoV-19/Belgium/VPE-030650/2020 | Europe / Belgium / Deinze | 2020-03-06 | KU Leuven, Clinical and Epidemiological Virology | KU Leuven, Clinical and Epidemiological Virology | Bert Vanmechelen, Joan Martí-Carreras, Tony Wawina, Piet Maes |
| EPI_ISL_418806 | hCoV-19/Belgium/BG-030551/2020 | Europe / Belgium / Beersel | 2020-03-05 | KU Leuven, Clinical and Epidemiological Virology | KU Leuven, Clinical and Epidemiological Virology | Bert Vanmechelen, Joan Martí-Carreras, Tony Wawina, Piet Maes |
| EPI_ISL_418807 | hCoV-19/Australia/QLDID927/2020 | Oceania / Australia / Queensland / Brisba | 2020-03-19 | Pathology Queensland | Public Health Virology Laboratory | Bixing Huang, Alyssa Pyke, Amanda De Jong, Andrew Van Den Hurk, Carmel Taylor, David Warrilow, Doris Genge, Elisabeth Gamez, Glen Hewitson, Ian Maxwell Mackay, Inga Sultana, Jamie McMahon, Jean Barcelon, Judy Northill, Mitchell Finger, Natalie Simpson, Neelima Nair, Peter Burtonclay, Peter Moore, Sarah Wheatley, Sean Moody, Sonja Hall-Mendelin, Timothy Gardam, and Frederick Moore |
| EPI_ISL_418808 | hCoV-19/Australia/QLDID928/2020 | Oceania / Australia / Queensland / Brisba | 2020-03-19 | Pathology Queensland | Public Health Virology Laboratory | Bixing Huang, Alyssa Pyke, Amanda De Jong, Andrew Van Den Hurk, Carmel Taylor, David Warrilow, Doris Genge, Elisabeth Gamez, Glen Hewitson, Ian Maxwell Mackay, Inga Sultana, Jamie McMahon, Jean Barcelon, Judy Northill, Mitchell Finger, Natalie Simpson, Neelima Nair, Peter Burtonclay, Peter Moore, Sarah Wheatley, Sean Moody, Sonja Hall-Mendelin, Timothy Gardam, and Frederick Moore |
| EPI_ISL_418809 | hCoV-19/Japan/UT-NCGM02/2020 | Asia / Japan / Tokyo | 2020-02-01 | University of Wisconsin - Madison: Influenza Research Institute | University of Wisconsin Madison, AIDS Vaccine Research Laboratories | Katarina Braun, Gage Moreno, Peter Halfmann, et al. |
| EPI_ISL_418810 | hCoV-19/Canada/SK_4/2020 | North America / Canada / Saskatchewan | 2020-03-09 | Roy Romanow Provincial Laboratory | National Microbiology Laboratory | Anna Majer, Shari Tyson, Grace Seo, Philip Mabon, Natalie Knox, Morag Graham, Ryan McDonald, Amanda Lang, Rachel DePaulo, Jessica Minion, Nathalie Bastien, Yan Li, Timothy Booth, Matthew Gilmour |
| EPI_ISL_418811 | hCoV-19/Canada/NB_6/2020 | North America / Canada / New Brunswick | 2020-03-12 | Dr. Georges-L.-Dumont University Hospital Centre | National Microbiology Laboratory | Anna Majer, Shari Tyson, Grace Seo, Philip Mabon, Natalie Knox, Morag Graham, Richard Garceau, Guillaume Desnoyers, Nathalie Bastien, Yan Li, Matthew Gilmour, Timothy Booth |
| EPI_ISL_418812 | hCoV-19/Canada/MB_8/2020 | North America / Canada / Manitoba | 2020-03-12 | Cadham Provincial Laboratory | National Microbiology Laboratory | Anna Majer, Shari Tyson, Grace Seo, Philip Mabon, Natalie Knox, Morag Graham, Paul Van Caeseele, Jared Bullard, David Alexander, Kerry Dust, Nathalie Bastien, Yan Li, Matthew Gilmour, Timothy Booth |
| EPI_ISL_418813 | hCoV-19/Canada/MB_10/2020 | North America / Canada / Manitoba | 2020-03-13 | Cadham Provincial Laboratory | National Microbiology Laboratory | Anna Majer, Shari Tyson, Grace Seo, Philip Mabon, Natalie Knox, Morag Graham, Paul Van Caeseele, Jared Bullard, David Alexander, Kerry Dust, Nathalie Bastien, Yan Li, Matthew Gilmour, Timothy Booth |
| EPI_ISL_418814 | hCoV-19/Canada/NS_13/2020 | North America / Canada / Nova Scotia | 2020-03-13 | Queen Elizabeth II Health Science Centre | National Microbiology Laboratory | Anna Majer, Shari Tyson, Grace Seo, Philip Mabon, Natalie Knox, Morag Graham, Todd Hatchette, Jason LeBlanc, Nathalie Bastien, Yan Li, Matthew Gilmour, Timothy Booth |
| EPI_ISL_418815 | hCoV-19/Hong Kong/HKPU36-0702/2020 | Asia / Hong Kong | 2020-02-09 | Department of Clinical Pathology, Pamela Youde Nethersole Eastern Hospital | Department of Health Technology and Informatics, Faculty of Health and Social Science, The Hong Kong Polytechnic University | Kenneth Siu-Sing LEUNG, Timothy Ting-Leung NG, Alan Ka-Lun WU, Miranda Chong-Yee YAU, Hiu-Yin LAO, Ming-Pan CHOI, Kingsley King-Gee TAM, Lam-Kwong LEE, Barry Kin-Chung WONG, Alex Yat-Man HO, Kam-Tong YIP, Kwok-Cheung LUNG, Raymond Wai-To LIU, Eugene Yuk-Keung TSO, Wai-Shing LEUNG, Man-Chun CHAN, Yuk-Yung NG, Kit-Man SIN, Kitty Sau-Chun FUNG, Sandy Ka-Yee CHAU, Wing-Kin TO, Tak-Lun QUE, David Ho-Keung SHUM, Shea Ping YIP, Wing Cheong YAM, Gilman Kit-Hang SIU |
| EPI_ISL_418816 | hCoV-19/Canada/BC_0443574/2020 | North America / Canada / British Columbi | 2020-03-11 | BCCDC Public Health Laboratory | BCCDC Public Health Laboratory | Harrigan, Prystajecky, Krajden, Lee, Kamelian, Lapointe, Choi, Hoang, Sekirov, Levett, Tyson, Snutch, Loman, Quick, Li, Gilmour |
| EPI_ISL_418817 | hCoV-19/Canada/BC_0554880/2020 | North America / Canada / British Columbi | 2020-03-13 | BCCDC Public Health Laboratory | BCCDC Public Health Laboratory | Harrigan, Prystajecky, Krajden, Lee, Kamelian, Lapointe, Choi, Hoang, Sekirov, Levett, Tyson, Snutch, Loman, Quick, Li, Gilmour |
| EPI_ISL_418818 | hCoV-19/Canada/BC_1318414/2020 | North America / Canada / British Columbi | 2020-03-06 | BCCDC Public Health Laboratory | BCCDC Public Health Laboratory | Harrigan, Prystajecky, Krajden, Lee, Kamelian, Lapointe, Choi, Hoang, Sekirov, Levett, Tyson, Snutch, Loman, Quick, Li, Gilmour |
| EPI_ISL_418819 | hCoV-19/Canada/BC_3808524/2020 | North America / Canada / British Columbi | 2020-03-09 | BCCDC Public Health Laboratory | BCCDC Public Health Laboratory | Harrigan, Prystajecky, Krajden, Lee, Kamelian, Lapointe, Choi, Hoang, Sekirov, Levett, Tyson, Snutch, Loman, Quick, Li, Gilmour |
| EPI_ISL_418820 | hCoV-19/Canada/BC_3842755/2020 | North America / Canada / British Columbi | 2020-03-09 | BCCDC Public Health Laboratory | BCCDC Public Health Laboratory | Harrigan, Prystajecky, Krajden, Lee, Kamelian, Lapointe, Choi, Hoang, Sekirov, Levett, Tyson, Snutch, Loman, Quick, Li, Gilmour |
| EPI_ISL_418821 | hCoV-19/Canada/BC_3968175/2020 | North America / Canada / British Columbi | 2020-03-10 | BCCDC Public Health Laboratory | BCCDC Public Health Laboratory | Harrigan, Prystajecky, Krajden, Lee, Kamelian, Lapointe, Choi, Hoang, Sekirov, Levett, Tyson, Snutch, Loman, Quick, Li, Gilmour |
| EPI_ISL_418822 | hCoV-19/Canada/BC_3972884/2020 | North America / Canada / British Columbi | 2020-03-10 | BCCDC Public Health Laboratory | BCCDC Public Health Laboratory | Harrigan, Prystajecky, Krajden, Lee, Kamelian, Lapointe, Choi, Hoang, Sekirov, Levett, Tyson, Snutch, Loman, Quick, Li, Gilmour |
| EPI_ISL_418823 | hCoV-19/Canada/BC_3989992/2020 | North America / Canada / British Columbi | 2020-03-09 | BCCDC Public Health Laboratory | BCCDC Public Health Laboratory | Harrigan, Prystajecky, Krajden, Lee, Kamelian, Lapointe, Choi, Hoang, Sekirov, Levett, Tyson, Snutch, Loman, Quick, Li, Gilmour |
| EPI_ISL_418824 | hCoV-19/Canada/BC_4078583/2020 | North America / Canada / British Columbi | 2020-03-03 | BCCDC Public Health Laboratory | BCCDC Public Health Laboratory | Harrigan, Prystajecky, Krajden, Lee, Kamelian, Lapointe, Choi, Hoang, Sekirov, Levett, Tyson, Snutch, Loman, Quick, Li, Gilmour |
| EPI_ISL_418825 | hCoV-19/Canada/BC_4118226/2020 | North America / Canada / British Columbi | 2020-03-10 | BCCDC Public Health Laboratory | BCCDC Public Health Laboratory | Harrigan, Prystajecky, Krajden, Lee, Kamelian, Lapointe, Choi, Hoang, Sekirov, Levett, Tyson, Snutch, Loman, Quick, Li, Gilmour |
| EPI_ISL_418826 | hCoV-19/Canada/BC_4122951/2020 | North America / Canada / British Columbi | 2020-03-10 | BCCDC Public Health Laboratory | BCCDC Public Health Laboratory | Harrigan, Prystajecky, Krajden, Lee, Kamelian, Lapointe, Choi, Hoang, Sekirov, Levett, Tyson, Snutch, Loman, Quick, Li, Gilmour |
| EPI_ISL_418829 | hCoV-19/Canada/BC_4540462/2020 | North America / Canada / British Columbi | 2020-03-10 | BCCDC Public Health Laboratory | BCCDC Public Health Laboratory | Harrigan, Prystajecky, Krajden, Lee, Kamelian, Lapointe, Choi, Hoang, Sekirov, Levett, Tyson, Snutch, Loman, Quick, Li, Gilmour |
| EPI_ISL_418830 | hCoV-19/Canada/BC_4799711/2020 | North America / Canada / British Columbi | 2020-03-09 | BCCDC Public Health Laboratory | BCCDC Public Health Laboratory | Harrigan, Prystajecky, Krajden, Lee, Kamelian, Lapointe, Choi, Hoang, Sekirov, Levett, Tyson, Snutch, Loman, Quick, Li, Gilmour |
| EPI_ISL_418831 | hCoV-19/Canada/BC_5275718/2020 | North America / Canada / British Columbi | 2020-03-11 | BCCDC Public Health Laboratory | BCCDC Public Health Laboratory | Harrigan, Prystajecky, Krajden, Lee, Kamelian, Lapointe, Choi, Hoang, Sekirov, Levett, Tyson, Snutch, Loman, Quick, Li, Gilmour |
| EPI_ISL_418832 | hCoV-19/Canada/BC_5282984/2020 | North America / Canada / British Columbi | 2020-03-11 | BCCDC Public Health Laboratory | BCCDC Public Health Laboratory | Harrigan, Prystajecky, Krajden, Lee, Kamelian, Lapointe, Choi, Hoang, Sekirov, Levett, Tyson, Snutch, Loman, Quick, Li, Gilmour |
| EPI_ISL_418833 | hCoV-19/Canada/BC_5306970/2020 | North America / Canada / British Columbi | 2020-03-11 | BCCDC Public Health Laboratory | BCCDC Public Health Laboratory | Harrigan, Prystajecky, Krajden, Lee, Kamelian, Lapointe, Choi, Hoang, Sekirov, Levett, Tyson, Snutch, Loman, Quick, Li, Gilmour |
| EPI_ISL_418834 | hCoV-19/Canada/BC_5522039/2020 | North America / Canada / British Columbi | 2020-03-11 | BCCDC Public Health Laboratory | BCCDC Public Health Laboratory | Harrigan, Prystajecky, Krajden, Lee, Kamelian, Lapointe, Choi, Hoang, Sekirov, Levett, Tyson, Snutch, Loman, Quick, Li, Gilmour |
| EPI_ISL_418835 | hCoV-19/Canada/BC_5979789/2020 | North America / Canada / British Columbi | 2020-03-11 | BCCDC Public Health Laboratory | BCCDC Public Health Laboratory | Harrigan, Prystajecky, Krajden, Lee, Kamelian, Lapointe, Choi, Hoang, Sekirov, Levett, Tyson, Snutch, Loman, Quick, Li, Gilmour |
| EPI_ISL_418836 | hCoV-19/Canada/BC_6004567/2020 | North America / Canada / British Columbi | 2020-03-10 | BCCDC Public Health Laboratory | BCCDC Public Health Laboratory | Harrigan, Prystajecky, Krajden, Lee, Kamelian, Lapointe, Choi, Hoang, Sekirov, Levett, Tyson, Snutch, Loman, Quick, Li, Gilmour |
| EPI_ISL_418837 | hCoV-19/Canada/BC_6129127/2020 | North America / Canada / British Columbi | 2020-03-10 | BCCDC Public Health Laboratory | BCCDC Public Health Laboratory | Harrigan, Prystajecky, Krajden, Lee, Kamelian, Lapointe, Choi, Hoang, Sekirov, Levett, Tyson, Snutch, Loman, Quick, Li, Gilmour |
| EPI_ISL_418838 | hCoV-19/Canada/BC_6404889/2020 | North America / Canada / British Columbi | 2020-03-12 | BCCDC Public Health Laboratory | BCCDC Public Health Laboratory | Harrigan, Prystajecky, Krajden, Lee, Kamelian, Lapointe, Choi, Hoang, Sekirov, Levett, Tyson, Snutch, Loman, Quick, Li, Gilmour |
| EPI_ISL_418839 | hCoV-19/Canada/BC_6489864/2020 | North America / Canada / British Columbi | 2020-03-10 | BCCDC Public Health Laboratory | BCCDC Public Health Laboratory | Harrigan, Prystajecky, Krajden, Lee, Kamelian, Lapointe, Choi, Hoang, Sekirov, Levett, Tyson, Snutch, Loman, Quick, Li, Gilmour |
| EPI_ISL_418840 | hCoV-19/Canada/BC_6502001/2020 | North America / Canada / British Columbi | 2020-03-12 | BCCDC Public Health Laboratory | BCCDC Public Health Laboratory | Harrigan, Prystajecky, Krajden, Lee, Kamelian, Lapointe, Choi, Hoang, Sekirov, Levett, Tyson, Snutch, Loman, Quick, Li, Gilmour |
| EPI_ISL_418841 | hCoV-19/Canada/BC_6981299/2020 | North America / Canada / British Columbi | 2020-03-11 | BCCDC Public Health Laboratory | BCCDC Public Health Laboratory | Harrigan, Prystajecky, Krajden, Lee, Kamelian, Lapointe, Choi, Hoang, Sekirov, Levett, Tyson, Snutch, Loman, Quick, Li, Gilmour |
| EPI_ISL_418842 | hCoV-19/Canada/BC_6997898/2020 | North America / Canada / British Columbi | 2020-03-11 | BCCDC Public Health Laboratory | BCCDC Public Health Laboratory | Harrigan, Prystajecky, Krajden, Lee, Kamelian, Lapointe, Choi, Hoang, Sekirov, Levett, Tyson, Snutch, Loman, Quick, Li, Gilmour |
| EPI_ISL_418843 | hCoV-19/Canada/BC_7277381/2020 | North America / Canada / British Columbi | 2020-03-12 | BCCDC Public Health Laboratory | BCCDC Public Health Laboratory | Harrigan, Prystajecky, Krajden, Lee, Kamelian, Lapointe, Choi, Hoang, Sekirov, Levett, Tyson, Snutch, Loman, Quick, Li, Gilmour |
| EPI_ISL_418845 | hCoV-19/Canada/BC_7515663/2020 | North America / Canada / British Columbi | 2020-03-11 | BCCDC Public Health Laboratory | BCCDC Public Health Laboratory | Harrigan, Prystajecky, Krajden, Lee, Kamelian, Lapointe, Choi, Hoang, Sekirov, Levett, Tyson, Snutch, Loman, Quick, Li, Gilmour |
| EPI_ISL_418846 | hCoV-19/Canada/BC_7553799/2020 | North America / Canada / British Columbi | 2020-03-12 | BCCDC Public Health Laboratory | BCCDC Public Health Laboratory | Harrigan, Prystajecky, Krajden, Lee, Kamelian, Lapointe, Choi, Hoang, Sekirov, Levett, Tyson, Snutch, Loman, Quick, Li, Gilmour |
| EPI_ISL_418847 | hCoV-19/Canada/BC_8150537/2020 | North America / Canada / British Columbi | 2020-03-12 | BCCDC Public Health Laboratory | BCCDC Public Health Laboratory | Harrigan, Prystajecky, Krajden, Lee, Kamelian, Lapointe, Choi, Hoang, Sekirov, Levett, Tyson, Snutch, Loman, Quick, Li, Gilmour |
| EPI_ISL_418848 | hCoV-19/Canada/BC_8159203/2020 | North America / Canada / British Columbi | 2020-03-05 | BCCDC Public Health Laboratory | BCCDC Public Health Laboratory | Harrigan, Prystajecky, Krajden, Lee, Kamelian, Lapointe, Choi, Hoang, Sekirov, Levett, Tyson, Snutch, Loman, Quick, Li, Gilmour |
| EPI_ISL_418849 | hCoV-19/Canada/BC_8486790/2020 | North America / Canada / British Columbi | 2020-03-13 | BCCDC Public Health Laboratory | BCCDC Public Health Laboratory | Harrigan, Prystajecky, Krajden, Lee, Kamelian, Lapointe, Choi, Hoang, Sekirov, Levett, Tyson, Snutch, Loman, Quick, Li, Gilmour |
| EPI_ISL_418851 | hCoV-19/Canada/BC_8622445/2020 | North America / Canada / British Columbi | 2020-03-13 | BCCDC Public Health Laboratory | BCCDC Public Health Laboratory | Harrigan, Prystajecky, Krajden, Lee, Kamelian, Lapointe, Choi, Hoang, Sekirov, Levett, Tyson, Snutch, Loman, Quick, Li, Gilmour |
| EPI_ISL_418852 | hCoV-19/Canada/BC_8718874/2020 | North America / Canada / British Columbi | 2020-03-07 | BCCDC Public Health Laboratory | BCCDC Public Health Laboratory | Harrigan, Prystajecky, Krajden, Lee, Kamelian, Lapointe, Choi, Hoang, Sekirov, Levett, Tyson, Snutch, Loman, Quick, Li, Gilmour |
| EPI_ISL_418853 | hCoV-19/Canada/BC_8894200/2020 | North America / Canada / British Columbi | 2020-03-11 | BCCDC Public Health Laboratory | BCCDC Public Health Laboratory | Harrigan, Prystajecky, Krajden, Lee, Kamelian, Lapointe, Choi, Hoang, Sekirov, Levett, Tyson, Snutch, Loman, Quick, Li, Gilmour |
| EPI_ISL_418854 | hCoV-19/Canada/BC_8896915/2020 | North America / Canada / British Columbi | 2020-03-11 | BCCDC Public Health Laboratory | BCCDC Public Health Laboratory | Harrigan, Prystajecky, Krajden, Lee, Kamelian, Lapointe, Choi, Hoang, Sekirov, Levett, Tyson, Snutch, Loman, Quick, Li, Gilmour |
| EPI_ISL_418855 | hCoV-19/Canada/BC_8897642/2020 | North America / Canada / British Columbi | 2020-03-11 | BCCDC Public Health Laboratory | BCCDC Public Health Laboratory | Harrigan, Prystajecky, Krajden, Lee, Kamelian, Lapointe, Choi, Hoang, Sekirov, Levett, Tyson, Snutch, Loman, Quick, Li, Gilmour |
| EPI_ISL_418856 | hCoV-19/Canada/BC_9345042/2020 | North America / Canada / British Columbi | 2020-03-13 | BCCDC Public Health Laboratory | BCCDC Public Health Laboratory | Harrigan, Prystajecky, Krajden, Lee, Kamelian, Lapointe, Choi, Hoang, Sekirov, Levett, Tyson, Snutch, Loman, Quick, Li, Gilmour |
| EPI_ISL_418857 | hCoV-19/Canada/BC_9345715/2020 | North America / Canada / British Columbi | 2020-03-13 | BCCDC Public Health Laboratory | BCCDC Public Health Laboratory | Harrigan, Prystajecky, Krajden, Lee, Kamelian, Lapointe, Choi, Hoang, Sekirov, Levett, Tyson, Snutch, Loman, Quick, Li, Gilmour |
| EPI_ISL_418858 | hCoV-19/Canada/BC_9446031/2020 | North America / Canada / British Columbi | 2020-03-13 | BCCDC Public Health Laboratory | BCCDC Public Health Laboratory | Harrigan, Prystajecky, Krajden, Lee, Kamelian, Lapointe, Choi, Hoang, Sekirov, Levett, Tyson, Snutch, Loman, Quick, Li, Gilmour |
| EPI_ISL_418860 | hCoV-19/Spain/VH000001133/2020 | Europe / Spain / Barcelona | 2020-03-15 | Hospital Universitari Vall d'Hebron (HUVH) - Vall d'Hebron Research Institute (VHIR) | Hospital Universitari Vall d'Hebron (HUVH) - Vall d'Hebron Research Institute (VHIR) | Cristina Andrés, Dámir García-Cehic, Maria Piñana, Mercedes Guerrero-Murillo, Ariadna Rando, Tomàs Pumarola, Maria Gema Codina, Andrés Antón, Josep Quer |
| EPI_ISL_418861 | hCoV-19/Spain/VH198152683/2020 | Europe / Spain / Barcelona | 2020-03-24 | Hospital Universitari Vall d'Hebron (HUVH) - Vall d'Hebron Research Institute (VHIR) | Hospital Universitari Vall d'Hebron (HUVH) - Vall d'Hebron Research Institute (VHIR) | Cristina Andrés, Dámir García-Cehic, Maria Piñana, Mercedes Guerrero-Murillo, Ariadna Rando, Tomàs Pumarola, Maria Gema Codina, Andrés Antón, Josep Quer |
| EPI_ISL_418863 | hCoV-19/Belgium/VS-030542/2020 | Europe / Belgium / Komen | 2020-03-05 | KU Leuven, Clinical and Epidemiological Virology | KU Leuven, Clinical and Epidemiological Virology | Bert Vanmechelen, Joan Martí-Carreras, Tony Wawina, Piet Maes |
| EPI_ISL_418864 | hCoV-19/USA/VA-DCLS-0001/2020 | North America / USA / Virginia | 2020-03-17 | Virginia DCLS | Virginia DCLS | Virginia DCLS |
| EPI_ISL_418866 | hCoV-19/USA/WA-UW298/2020 | North America / USA / Washington | 2020-03-13 | UW Virology Lab | UW Virology Lab | Pavitra Roychoudhury, Hong Xie, Keith Jerome, Alexander Greninger |
| EPI_ISL_418867 | hCoV-19/USA/WA-UW299/2020 | North America / USA / Washington | 2020-03-13 | UW Virology Lab | UW Virology Lab | Pavitra Roychoudhury, Hong Xie, Keith Jerome, Alexander Greninger |
| EPI_ISL_418868 | hCoV-19/USA/WA-UW300/2020 | North America / USA / Washington | 2020-03-13 | UW Virology Lab | UW Virology Lab | Pavitra Roychoudhury, Hong Xie, Keith Jerome, Alexander Greninger |
| EPI_ISL_418869 | hCoV-19/USA/WA-UW301/2020 | North America / USA / Washington | 2020-03-16 | UW Virology Lab | UW Virology Lab | Pavitra Roychoudhury, Hong Xie, Keith Jerome, Alexander Greninger |
| EPI_ISL_418870 | hCoV-19/USA/WA-UW302/2020 | North America / USA / Washington | 2020-03-16 | UW Virology Lab | UW Virology Lab | Pavitra Roychoudhury, Hong Xie, Keith Jerome, Alexander Greninger |
| EPI_ISL_418871 | hCoV-19/USA/CT-UW303/2020 | North America / USA / Connecticut | 2020-03-14 | UW Virology Lab | UW Virology Lab | Pavitra Roychoudhury, Hong Xie, Keith Jerome, Alexander Greninger |
| EPI_ISL_418872 | hCoV-19/USA/WA-UW304/2020 | North America / USA / Washington | 2020-03-16 | UW Virology Lab | UW Virology Lab | Pavitra Roychoudhury, Hong Xie, Keith Jerome, Alexander Greninger |
| EPI_ISL_418873 | hCoV-19/USA/WA-UW305/2020 | North America / USA / Washington | 2020-03-16 | UW Virology Lab | UW Virology Lab | Pavitra Roychoudhury, Hong Xie, Keith Jerome, Alexander Greninger |
| EPI_ISL_418875 | hCoV-19/USA/WA-UW307/2020 | North America / USA / Washington | 2020-03-16 | UW Virology Lab | UW Virology Lab | Pavitra Roychoudhury, Hong Xie, Keith Jerome, Alexander Greninger |
| EPI_ISL_418876 | hCoV-19/USA/WA-UW308/2020 | North America / USA / Washington | 2020-03-16 | UW Virology Lab | UW Virology Lab | Pavitra Roychoudhury, Hong Xie, Keith Jerome, Alexander Greninger |
| EPI_ISL_418877 | hCoV-19/USA/WA-UW309/2020 | North America / USA / Washington | 2020-03-24 | UW Virology Lab | UW Virology Lab | Pavitra Roychoudhury, Hong Xie, Keith Jerome, Alexander Greninger |
| EPI_ISL_418878 | hCoV-19/USA/WA-UW310/2020 | North America / USA / Washington | 2020-03-14 | UW Virology Lab | UW Virology Lab | Pavitra Roychoudhury, Hong Xie, Keith Jerome, Alexander Greninger |
| EPI_ISL_418880 | hCoV-19/USA/WA-UW313/2020 | North America / USA / Washington | 2020-03-14 | UW Virology Lab | UW Virology Lab | Pavitra Roychoudhury, Hong Xie, Keith Jerome, Alexander Greninger |
| EPI_ISL_418881 | hCoV-19/USA/WA-UW314/2020 | North America / USA / Washington | 2020-03-14 | UW Virology Lab | UW Virology Lab | Pavitra Roychoudhury, Hong Xie, Keith Jerome, Alexander Greninger |
| EPI_ISL_418882 | hCoV-19/USA/WA-UW315/2020 | North America / USA / Washington | 2020-03-13 | UW Virology Lab | UW Virology Lab | Pavitra Roychoudhury, Hong Xie, Keith Jerome, Alexander Greninger |
| EPI_ISL_418883 | hCoV-19/USA/WA-UW316/2020 | North America / USA / Washington | 2020-03-14 | UW Virology Lab | UW Virology Lab | Pavitra Roychoudhury, Hong Xie, Keith Jerome, Alexander Greninger |
| EPI_ISL_418884 | hCoV-19/USA/WA-UW317/2020 | North America / USA / Washington | 2020-03-15 | UW Virology Lab | UW Virology Lab | Pavitra Roychoudhury, Hong Xie, Keith Jerome, Alexander Greninger |
| EPI_ISL_418886 | hCoV-19/USA/WA-UW319/2020 | North America / USA / Washington | 2020-03-15 | UW Virology Lab | UW Virology Lab | Pavitra Roychoudhury, Hong Xie, Keith Jerome, Alexander Greninger |
| EPI_ISL_418887 | hCoV-19/USA/WA-UW320/2020 | North America / USA / Washington | 2020-03-14 | UW Virology Lab | UW Virology Lab | Pavitra Roychoudhury, Hong Xie, Keith Jerome, Alexander Greninger |
| EPI_ISL_418888 | hCoV-19/USA/WA-UW321/2020 | North America / USA / Washington | 2020-03-15 | UW Virology Lab | UW Virology Lab | Pavitra Roychoudhury, Hong Xie, Keith Jerome, Alexander Greninger |
| EPI_ISL_418889 | hCoV-19/USA/WA-UW322/2020 | North America / USA / Washington | 2020-03-16 | UW Virology Lab | UW Virology Lab | Pavitra Roychoudhury, Hong Xie, Keith Jerome, Alexander Greninger |
| EPI_ISL_418890 | hCoV-19/USA/WA-UW323/2020 | North America / USA / Washington | 2020-03-16 | UW Virology Lab | UW Virology Lab | Pavitra Roychoudhury, Hong Xie, Keith Jerome, Alexander Greninger |
| EPI_ISL_418891 | hCoV-19/USA/WA-UW324/2020 | North America / USA / Washington | 2020-03-24 | UW Virology Lab | UW Virology Lab | Pavitra Roychoudhury, Hong Xie, Keith Jerome, Alexander Greninger |
| EPI_ISL_418892 | hCoV-19/USA/WA-UW325/2020 | North America / USA / Washington | 2020-03-16 | UW Virology Lab | UW Virology Lab | Pavitra Roychoudhury, Hong Xie, Keith Jerome, Alexander Greninger |
| EPI_ISL_418893 | hCoV-19/USA/MN-UW326/2020 | North America / USA / Minnesota | 2020-03-13 | UW Virology Lab | UW Virology Lab | Pavitra Roychoudhury, Hong Xie, Keith Jerome, Alexander Greninger |
| EPI_ISL_418895 | hCoV-19/USA/CT-UW328/2020 | North America / USA / Connecticut | 2020-03-14 | UW Virology Lab | UW Virology Lab | Pavitra Roychoudhury, Hong Xie, Keith Jerome, Alexander Greninger |
| EPI_ISL_418897 | hCoV-19/USA/WA-UW330/2020 | North America / USA / Washington | 2020-03-16 | UW Virology Lab | UW Virology Lab | Pavitra Roychoudhury, Hong Xie, Keith Jerome, Alexander Greninger |
| EPI_ISL_418898 | hCoV-19/USA/CT-UW331/2020 | North America / USA / Connecticut | 2020-03-14 | UW Virology Lab | UW Virology Lab | Pavitra Roychoudhury, Hong Xie, Keith Jerome, Alexander Greninger |
| EPI_ISL_418899 | hCoV-19/USA/WA-UW332/2020 | North America / USA / Washington | 2020-03-16 | UW Virology Lab | UW Virology Lab | Pavitra Roychoudhury, Hong Xie, Keith Jerome, Alexander Greninger |
| EPI_ISL_418910 | hCoV-19/USA/WA-UW343/2020 | North America / USA / Washington | 2020-03-16 | UW Virology Lab | UW Virology Lab | Pavitra Roychoudhury, Hong Xie, Keith Jerome, Alexander Greninger |
| EPI_ISL_418980 | hCoV-19/USA/NY-NYUMC34/2020 | North America / USA / New York / Manha | 2020-03-17 | NYU Langone Health | Department of Pathology and Medicine, New York University School of Medicine | Maria Aguero-Rosenfeld, Margaret Black, John Cadley, Paolo Cotzia, John Chen, Dacia Dimartino, Xiaojun Feng, Adriana Heguy, Megan Hogan, Emily Huang, George Jour, Christian Marier, Matthew T. Maurano, Mark J. Mulligan, Peter Meyn, Jared Pinnell, Sitharam Ramaswami, Amy Rapkiewicz, Marie Samanovic-Golden, Antonio Serrano, Guomiao Shen, Matija Snuderl, Nick Vulpescu, Gael Westby, Paul Zappile, Yutong Zhang |
| EPI_ISL_418981 | hCoV-19/Belgium/VRAR-030643/2020 | Europe / Belgium / Merelbeke | 2020-03-06 | KU Leuven, Clinical and Epidemiological Virology | KU Leuven, Clinical and Epidemiological Virology | Bert Vanmechelen, Joan Martí-Carreras, Tony Wawina, Piet Maes |
| EPI_ISL_418982 | hCoV-19/Belgium/HAA-030953/2020 | Europe / Belgium / Oevel | 2020-03-09 | KU Leuven, Clinical and Epidemiological Virology | KU Leuven, Clinical and Epidemiological Virology | Bert Vanmechelen, Joan Martí-Carreras, Tony Wawina, Piet Maes |
| EPI_ISL_418983 | hCoV-19/Belgium/DWAEE-030954/2020 | Europe / Belgium / Asse | 2020-03-09 | KU Leuven, Clinical and Epidemiological Virology | KU Leuven, Clinical and Epidemiological Virology | Bert Vanmechelen, Joan Martí-Carreras, Tony Wawina, Piet Maes |
| EPI_ISL_418984 | hCoV-19/Belgium/BG-030955/2020 | Europe / Belgium / Kaulille | 2020-03-09 | KU Leuven, Clinical and Epidemiological Virology | KU Leuven, Clinical and Epidemiological Virology | Bert Vanmechelen, Joan Martí-Carreras, Tony Wawina, Piet Maes |
| EPI_ISL_418985 | hCoV-19/Belgium/LT-030956/2020 | Europe / Belgium / Ellezelles | 2020-03-09 | KU Leuven, Clinical and Epidemiological Virology | KU Leuven, Clinical and Epidemiological Virology | Bert Vanmechelen, Joan Martí-Carreras, Tony Wawina, Piet Maes |
| EPI_ISL_418986 | hCoV-19/Belgium/RS-030257/2020 | Europe / Belgium / Hoegaarden | 2020-03-02 | KU Leuven, Clinical and Epidemiological Virology | KU Leuven, Clinical and Epidemiological Virology | Bert Vanmechelen, Joan Martí-Carreras, Tony Wawina, Piet Maes |
| EPI_ISL_418987 | hCoV-19/Belgium/CG-030158/2020 | Europe / Belgium / Eupen | 2020-03-01 | KU Leuven, Clinical and Epidemiological Virology | KU Leuven, Clinical and Epidemiological Virology | Bert Vanmechelen, Joan Martí-Carreras, Tony Wawina, Piet Maes |
| EPI_ISL_418988 | hCoV-19/Belgium/JRH-030459/2020 | Europe / Belgium / Zolder | 2020-03-04 | Institute information KU Leuven, Clinical and Epidemiological Virology | Institute information KU Leuven, Clinical and Epidemiological Virology | Bert Vanmechelen, Joan Martí-Carreras, Tony Wawina, Piet Maes |
| EPI_ISL_418989 | hCoV-19/Belgium/RT-030460/2020 | Europe / Belgium / Vorst | 2020-03-04 | KU Leuven, Clinical and Epidemiological Virology | KU Leuven, Clinical and Epidemiological Virology | Bert Vanmechelen, Joan Martí-Carreras, Tony Wawina, Piet Maes |
| EPI_ISL_450500 | hCoV-19/Wuhan/OS52/2020 | Asia / China / Hubei / Wuhan | 2020-01-10 | unknown | CAS Key Laboratory of Special Pathogens and Biosafety and Center for Emerging Infectious Diseases | Si,H., Zhu,Y., Lin,H., Xie,S., Shi,Z., Zhou,P. |
| EPI_ISL_450501 | hCoV-19/Wuhan/YB012611/2020 | Asia / China / Hubei / Wuhan | 2020-01-26 | unknown | CAS Key Laboratory of Special Pathogens and Biosafety and Center for Emerging Infectious Diseases | Si,H., Zhu,Y., Lin,H., Xie,S., Shi,Z., Zhou,P. |
| EPI_ISL_450502 | hCoV-19/Wuhan/YB012605/2020 | Asia / China / Hubei / Wuhan | 2020-01-26 | unknown | CAS Key Laboratory of Special Pathogens and Biosafety and Center for Emerging Infectious Diseases | Si,H., Zhu,Y., Lin,H., Xie,S., Shi,Z., Zhou,P. |
| EPI_ISL_450503 | hCoV-19/Wuhan/YB012602/2020 | Asia / China / Hubei / Wuhan | 2020-01-26 | unknown | CAS Key Laboratory of Special Pathogens and Biosafety and Center for Emerging Infectious Diseases | Si,H., Zhu,Y., Lin,H., Xie,S., Shi,Z., Zhou,P. |
| EPI_ISL_450504 | hCoV-19/Wuhan/YB012506/2020 | Asia / China / Hubei / Wuhan | 2020-01-25 | unknown | CAS Key Laboratory of Special Pathogens and Biosafety and Center for Emerging Infectious Diseases | Si,H., Zhu,Y., Lin,H., Xie,S., Shi,Z., Zhou,P. |
| EPI_ISL_402119 | hCoV-19/Wuhan/IVDC-HB-01/2019 | Asia / China / Hubei / Wuhan | 2019-12-30 | National Institute for Viral Disease Control and Prevention, China CDC | National Institute for Viral Disease Control and Prevention, China CDC | Wenjie Tan, Xiang Zhao, Wenling Wang, Xuejun Ma, Yongzhong Jiang, Roujian Lu, Ji Wang, Weimin Zhou, Peihua Niu, Peipei Liu, Faxian Zhan, Weifeng Shi, Baoying Huang, Jun Liu, Li Zhao, Yao Meng, Xiaozhou He, Fei Ye, Na Zhu, Yang Li, Jing Chen, Wenbo Xu, George F. Gao, Guizhen Wu |
| EPI_ISL_402120 | hCoV-19/Wuhan/IVDC-HB-04/2020 | Asia / China / Hubei / Wuhan | 2020-01-01 | National Institute for Viral Disease Control and Prevention, China CDC | National Institute for Viral Disease Control and Prevention, China CDC | Wenjie Tan, Xiang Zhao, Wenling Wang, Xuejun Ma, Yongzhong Jiang, Roujian Lu, Ji Wang, Weimin Zhou, Peihua Niu, Peipei Liu, Faxian Zhan, Weifeng Shi, Baoying Huang, Jun Liu, Li Zhao, Yao Meng, Xiaozhou He, Fei Ye, Na Zhu, Yang Li, Jing Chen, Wenbo Xu, George F. Gao, Guizhen Wu |
| EPI_ISL_402121 | hCoV-19/Wuhan/IVDC-HB-05/2019 | Asia / China / Hubei / Wuhan | 2019-12-30 | National Institute for Viral Disease Control and Prevention, China CDC | National Institute for Viral Disease Control and Prevention, China CDC | Wenjie Tan, Xuejun Ma, Xiang Zhao, Wenling Wang, Yongzhong Jiang, Roujian Lu, Ji Wang, Peihua Niu, Weimin Zhou, Faxian Zhan, Weifeng Shi, Baoying Huang, Jun Liu, Li Zhao, Yao Meng, Fei Ye, Na Zhu, Xiaozhou He, Peipei Liu, Yang Li, Jing Chen, Wenbo Xu, George F. Gao, Guizhen Wu |
| EPI_ISL_402123 | hCoV-19/Wuhan/IPBCAMS-WH-01/2019 | Asia / China / Hubei / Wuhan | 2019-12-24 | Institute of Pathogen Biology, Chinese Academy of Medical Sciences & Peking Union Medical College | Institute of Pathogen Biology, Chinese Academy of Medical Sciences & Peking Union Medical College | Lili Ren, Jianwei Wang, Qi Jin, Zichun Xiang, Zhiqiang Wu, Chao Wu, Yiwei Liu |
| EPI_ISL_402124 | hCoV-19/Wuhan/WIV04/2019 | Asia / China / Hubei / Wuhan | 2019-12-30 | Wuhan Jinyintan Hospital | Wuhan Institute of Virology, Chinese Academy of Sciences | Peng Zhou, Xing-Lou Yang, Ding-Yu Zhang, Lei Zhang, Yan Zhu, Hao-Rui Si, Zhengli Shi |
| EPI_ISL_402125 | hCoV-19/Wuhan/Hu-1/2019 | Asia / China | 2019-12-31 | unknown | National Institute for Communicable Disease Control and Prevention (ICDC) Chinese Center for Disease Control and Prevention (China CDC) | Zhang,Y.-Z., Wu,F., Chen,Y.-M., Pei,Y.-Y., Xu,L., Wang,W., Zhao,S., Yu,B., Hu,Y., Tao,Z.-W., Song,Z.-G., Tian,J.-H., Zhang,Y.-L., Liu,Y., Zheng,J.-J., Dai,F.-H., Wang,Q.-M., She,J.-L. and Zhu,T.-Y. |
| EPI_ISL_402127 | hCoV-19/Wuhan/WIV02/2019 | Asia / China / Hubei / Wuhan | 2019-12-30 | Wuhan Jinyintan Hospital | Wuhan Institute of Virology, Chinese Academy of Sciences | Peng Zhou, Xing-Lou Yang, Ding-Yu Zhang, Lei Zhang, Yan Zhu, Hao-Rui Si, Zhengli Shi |
| EPI_ISL_402128 | hCoV-19/Wuhan/WIV05/2019 | Asia / China / Hubei / Wuhan | 2019-12-30 | Wuhan Jinyintan Hospital | Wuhan Institute of Virology, Chinese Academy of Sciences | Peng Zhou, Xing-Lou Yang, Ding-Yu Zhang, Lei Zhang, Yan Zhu, Hao-Rui Si, Zhengli Shi |
| EPI_ISL_402129 | hCoV-19/Wuhan/WIV06/2019 | Asia / China / Hubei / Wuhan | 2019-12-30 | Wuhan Jinyintan Hospital | Wuhan Institute of Virology, Chinese Academy of Sciences | Peng Zhou, Xing-Lou Yang, Ding-Yu Zhang, Lei Zhang, Yan Zhu, Hao-Rui Si, Zhengli Shi |
| EPI_ISL_402130 | hCoV-19/Wuhan/WIV07/2019 | Asia / China / Hubei / Wuhan | 2019-12-30 | Wuhan Jinyintan Hospital | Wuhan Institute of Virology, Chinese Academy of Sciences | Peng Zhou, Xing-Lou Yang, Ding-Yu Zhang, Lei Zhang, Yan Zhu, Hao-Rui Si, Zhengli Shi |
| EPI_ISL_402132 | hCoV-19/Wuhan/HBCDC-HB-01/2019 | Asia / China / Hubei / Wuhan | 2019-12-30 | Wuhan Jinyintan Hospital | Hubei Provincial Center for Disease Control and Prevention | Bin Fang, Xiang Li, Xiao Yu, Linlin Liu, Bo Yang, Faxian Zhan, Guojun Ye, Xixiang Huo, Junqiang Xu, Bo Yu, Kun Cai, Jing Li, Yongzhong Jiang. |
| EPI_ISL_403928 | hCoV-19/Wuhan/IPBCAMS-WH-05/2020 | Asia / China / Hubei / Wuhan | 2020-01-01 | Institute of Pathogen Biology, Chinese Academy of Medical Sciences & Peking Union Medical College | Institute of Pathogen Biology, Chinese Academy of Medical Sciences & Peking Union Medical College | Lili Ren, Jianwei Wang, Qi Jin, Zichun Xiang, Zhiqiang Wu, Chao Wu, Yiwei Liu |
| EPI_ISL_403929 | hCoV-19/Wuhan/IPBCAMS-WH-04/2019 | Asia / China / Hubei / Wuhan | 2019-12-30 | Institute of Pathogen Biology, Chinese Academy of Medical Sciences & Peking Union Medical College | Institute of Pathogen Biology, Chinese Academy of Medical Sciences & Peking Union Medical College | Lili Ren, Jianwei Wang, Qi Jin, Zichun Xiang, Zhiqiang Wu, Chao Wu, Yiwei Liu |
| EPI_ISL_403930 | hCoV-19/Wuhan/IPBCAMS-WH-03/2019 | Asia / China / Hubei / Wuhan | 2019-12-30 | Institute of Pathogen Biology, Chinese Academy of Medical Sciences & Peking Union Medical College | Institute of Pathogen Biology, Chinese Academy of Medical Sciences & Peking Union Medical College | Lili Ren, Jianwei Wang, Qi Jin, Zichun Xiang, Zhiqiang Wu, Chao Wu, Yiwei Liu |

| | | | | | | |
|---|---|---|---|---|---|---|
| EPI_ISL_403931 | hCoV-19/Wuhan/IPBCAMS-WH-02/2019 | Asia / China / Hubei / Wuhan | 2019-12-30 | Institute of Pathogen Biology, Chinese Academy of Medical Sciences & Peking Union Medical College | Institute of Pathogen Biology, Chinese Academy of Medical Sciences & Peking Union Medical College | Lili Ren, Jianwei Wang, Qi Jin, Zichun Xiang, Zhiqiang Wu, Chao Wu, Yiwei Liu |
| EPI_ISL_403932 | hCoV-19/Guangdong/20SF012/2020 | Asia / China / Guangdong / Shenzhen | 2020-01-14 | Guangdong Provincial Center for Diseases Control and Prevention; Guangdong Provincial Public Health | Department of Microbiology, Guangdong Provincial Center for Diseases Control and Prevention | Min Kang, Jie Wu, Jing Lu, Tao Liu, Baisheng Li, Shujiang Mei, Feng Ruan, Lifeng Lin, Changwen Ke, Haojie Zhong, Yingtao Zhang, Lirong Zou, Xuguang Chen, Qi Zhu, Jianpeng Xiao, Jianxiang Geng, Zhe Liu, Jianxiong Hu, Weilin Zeng, Xing Li, Yuhuang Liao, Xiujuan Tang, Songjian Xiao, Ying Wang, Yingchao Song, Xue Zhuang, Lijun Liang, Guanhao He, Huihong Deng, Tie Song, Jianfeng He, Wenjun Ma |
| EPI_ISL_403933 | hCoV-19/Guangdong/20SF013/2020 | Asia / China / Guangdong / Shenzhen | 2020-01-15 | Guangdong Provincial Center for Diseases Control and Prevention; Guangdong Provincial Public Health | Department of Microbiology, Guangdong Provincial Center for Diseases Control and Prevention | Min Kang, Jie Wu, Jing Lu, Tao Liu, Baisheng Li, Shujiang Mei, Feng Ruan, Lifeng Lin, Changwen Ke, Haojie Zhong, Yingtao Zhang, Lirong Zou, Xuguang Chen, Qi Zhu, Jianpeng Xiao, Jianxiang Geng, Zhe Liu, Jianxiong Hu, Weilin Zeng, Xing Li, Yuhuang Liao, Xiujuan Tang, Songjian Xiao, Ying Wang, Yingchao Song, Xue Zhuang, Lijun Liang, Guanhao He, Huihong Deng, Tie Song, Jianfeng He, Wenjun Ma |
| EPI_ISL_403934 | hCoV-19/Guangdong/20SF014/2020 | Asia / China / Guangdong / Shenzhen | 2020-01-15 | Guangdong Provincial Center for Diseases Control and Prevention; Guangdong Provincial Public Health | Department of Microbiology, Guangdong Provincial Center for Diseases Control and Prevention | Min Kang, Jie Wu, Jing Lu, Tao Liu, Baisheng Li, Shujiang Mei, Feng Ruan, Lifeng Lin, Changwen Ke, Haojie Zhong, Yingtao Zhang, Lirong Zou, Xuguang Chen, Qi Zhu, Jianpeng Xiao, Jianxiang Geng, Zhe Liu, Jianxiong Hu, Weilin Zeng, Xing Li, Yuhuang Liao, Xiujuan Tang, Songjian Xiao, Ying Wang, Yingchao Song, Xue Zhuang, Lijun Liang, Guanhao He, Huihong Deng, Tie Song, Jianfeng He, Wenjun Ma |
| EPI_ISL_403935 | hCoV-19/Guangdong/20SF025/2020 | Asia / China / Guangdong / Shenzhen | 2020-01-15 | Guangdong Provincial Center for Diseases Control and Prevention; Guangdong Provincial Public Health | Department of Microbiology, Guangdong Provincial Center for Diseases Control and Prevention | Min Kang, Jie Wu, Jing Lu, Tao Liu, Baisheng Li, Shujiang Mei, Feng Ruan, Lifeng Lin, Changwen Ke, Haojie Zhong, Yingtao Zhang, Lirong Zou, Xuguang Chen, Qi Zhu, Jianpeng Xiao, Jianxiang Geng, Zhe Liu, Jianxiong Hu, Weilin Zeng, Xing Li, Yuhuang Liao, Xiujuan Tang, Songjian Xiao, Ying Wang, Yingchao Song, Xue Zhuang, Lijun Liang, Guanhao He, Huihong Deng, Tie Song, Jianfeng He, Wenjun Ma |
| EPI_ISL_403936 | hCoV-19/Guangdong/20SF028/2020 | Asia / China / Guangdong / Zhuhai | 2020-01-17 | Guangdong Provincial Center for Diseases Control and Prevention; Guangdong Provincial Public Health | Department of Microbiology, Guangdong Provincial Center for Diseases Control and Prevention | Min Kang, Jie Wu, Jing Lu, Tao Liu, Baisheng Li, Shujiang Mei, Feng Ruan, Lifeng Lin, Changwen Ke, Haojie Zhong, Yingtao Zhang, Lirong Zou, Xuguang Chen, Qi Zhu, Jianpeng Xiao, Jianxiang Geng, Zhe Liu, Jianxiong Hu, Weilin Zeng, Xing Li, Yuhuang Liao, Xiujuan Tang, Songjian Xiao, Ying Wang, Yingchao Song, Xue Zhuang, Lijun Liang, Guanhao He, Huihong Deng, Tie Song, Jianfeng He, Wenjun Ma |
| EPI_ISL_403937 | hCoV-19/Guangdong/20SF040/2020 | Asia / China / Guangdong / Zhuhai | 2020-01-18 | Guangdong Provincial Center for Diseases Control and Prevention; Guangdong Provincial Public Health | Department of Microbiology, Guangdong Provincial Center for Diseases Control and Prevention | Min Kang, Jie Wu, Jing Lu, Tao Liu, Baisheng Li, Shujiang Mei, Feng Ruan, Lifeng Lin, Changwen Ke, Haojie Zhong, Yingtao Zhang, Lirong Zou, Xuguang Chen, Qi Zhu, Jianpeng Xiao, Jianxiang Geng, Zhe Liu, Jianxiong Hu, Weilin Zeng, Xing Li, Yuhuang Liao, Xiujuan Tang, Songjian Xiao, Ying Wang, Yingchao Song, Xue Zhuang, Lijun Liang, Guanhao He, Huihong Deng, Tie Song, Jianfeng He, Wenjun Ma |
| EPI_ISL_403962 | hCoV-19/Thailand/61/2020 | Asia / Thailand / Nonthaburi | 2020-01-08 | Bamrasnaradura Hospital | 1. Department of Medical Sciences, Ministry of Public Health, Thailand 2. Thai Red Cross Emerging Infectious Diseases - Health Science Centre 3. Department of Disease Control, Ministry of Public Health, Thailand | Pilailuk,Okada; Siripaporn,Phuygun; Thanutsapa,Thanadachakul; Supaporn,Wacharapluesadee; Sittiporn,Parnmen; Warawan,Wongboot; Sunthareeya,Waicharoen; Rome,Buathong; Malinee,Chittaganpitch; Nanthawan,Mekha |
| EPI_ISL_403963 | hCoV-19/Thailand/74/2020 | Asia / Thailand / Nonthaburi | 2020-01-13 | Bamrasnaradura Hospital | 1. Department of Medical Sciences, Ministry of Public Health, Thailand 2. Thai Red Cross Emerging Infectious Diseases - Health Science Centre 3. Department of Disease Control, Ministry of Public Health, Thailand | Pilailuk,Okada; Siripaporn,Phuygun; Thanutsapa,Thanadachakul; Supaporn,Wacharapluesadee; Sittiporn,Parnmen; Warawan,Wongboot; Sunthareeya,Waicharoen; Rome,Buathong; Malinee,Chittaganpitch; Nanthawan,Mekha |
| EPI_ISL_404895 | hCoV-19/USA/WA1/2020 | North America / USA / Washington / Snol | 2020-01-19 | Providence Regional Medical Center | Division of Viral Diseases, Centers for Disease Control and Prevention | Queen,K., Tao,Y., Li,Y., Paden,C.R., Lu,X., Zhang,J., Gerber,S.I., Lindstrom,S., Tong,S. |
| EPI_ISL_406223 | hCoV-19/USA/AZ1/2020 | North America / USA / Arizona / Phoenix | 2020-01-22 | Arizona Department of Health Services | Pathogen Discovery, Respiratory Viruses Branch, Division of Viral Diseases, Centers for Disease Control and Prevention | Ying Tao, Clinton R. Paden, Krista Queen, Anna Uehara, Yan Li, Jing Zhang, Xiaoyan Lu, Brian Lynch, Senthil Kumar K. Sakthivel, Brett L. Whitaker, Shifaq Kamili, Lijuan Wang, Janna' R. Murray, Susan I. Gerber, Stephen Lindstrom, Suxiang Tong |
| EPI_ISL_406970 | hCoV-19/Hangzhou/HZ-1/2020 | Asia / China / Zhejiang / Hangzhou | 2020-01-20 | Hangzhou Center for Disease and Control Microbiology Lab | Hangzhou Center for Disease and Control Microbiology Lab | Yu Hua, Wang Haoqiu, Li Jun, Yu Xinfeng |
| EPI_ISL_406973 | hCoV-19/Singapore/1/2020 | Asia / Singapore | 2020-01-23 | Singapore General Hospital | National Public Health Laboratory | Mak, TM; Octavia S; Chavatte JM; Zhou, ZY; Cui, L; Lin, RTP |
| EPI_ISL_407071 | hCoV-19/England/01/2020 | Europe / United Kingdom / England | 2020-01-29 | Respiratory Virus Unit, Microbiology Services Colindale, Public Health England | Respiratory Virus Unit, Microbiology Services Colindale, Public Health England | Monica Galiano, Shahjahan Miah, Richard Myers, Angie Lackenby, Omolola Akinbami, Tiina Talts, Leena Bhaw, Kirstin Edwards, Jonathan Hubb, Joanna Ellis, Maria Zambon |
| EPI_ISL_407073 | hCoV-19/England/02/2020 | Europe / United Kingdom / England | 2020-01-29 | Respiratory Virus Unit, Microbiology Services Colindale, Public Health England | Respiratory Virus Unit, Microbiology Services Colindale, Public Health England | Monica Galiano, Shahjahan Miah, Richard Myers, Angie Lackenby, Omolola Akinbami, Tiina Talts, Leena Bhaw, Kirstin Edwards, Jonathan Hubb, Joanna Ellis, Maria Zambon. |
| EPI_ISL_407079 | hCoV-19/Finland/1/2020 | Europe / Finland / Lapland | 2020-01-29 | Lapland Central Hospital | Department of Virology, University of Helsinki and Helsinki University Hospital, Helsinki, Finland | Teemu Smura, Suvi Kuivanen, Hannimari Kallio-Kokko, Olli Vapalahti |
| EPI_ISL_407976 | hCoV-19/Belgium/GHB-03021/2020 | Europe / Belgium / Leuven | 2020-02-03 | KU Leuven, Clinical and Epidemiological Virology | KU Leuven, Clinical and Epidemiological Virology | Bert Vanmechelen, Elke Wollants, Annabel Rector, Els Keyaerts, Lies Laenen, Marc Van Ranst, and Piet Maes |
| EPI_ISL_407987 | hCoV-19/Singapore/2/2020 | Asia / Singapore | 2020-01-25 | Singapore General Hospital | Programme in Emerging Infectious Diseases, Duke-NUS Medical School | Danielle E Anderson, Martin Linster, Yan Zhuang, Jayanthi Jayakumar, Kian Sing Chan, Lynette LE Oon, Jenny GH Low, Yvonne CF Su, Linfa Wang, Gavin JD Smith |
| EPI_ISL_407988 | hCoV-19/Singapore/3/2020 | Asia / Singapore | 2020-02-01 | National Centre for Infectious Diseases | Programme in Emerging Infectious Diseases, Duke-NUS Medical School | Danielle E Anderson, Martin Linster, Yan Zhuang, Jayanthi Jayakumar, David CB Lye, Yee Sin Leo, Barnaby E Young, Yvonne CF Su, Linfa Wang, Gavin JD Smith |
| EPI_ISL_408008 | hCoV-19/USA/CA3/2020 | North America / USA / California | 2020-01-29 | California Department of Health | Pathogen Discovery, Respiratory Viruses Branch, Division of Viral Diseases, Centers for Disease Control and Prevention | Krista Queen, Jing Zhang, Yan Li, Ying Tao, Anna Uehara, Clinton Paden, Xiaoyan Lu, Brian Lynch, Senthil Kumar K. Sakthivel, Brett L. Whitaker, Shifaq Kamili, Lijuan Wang, Janna' R. Murray, Susan I. Gerber, Stephen Lindstrom, Suxiang Tong |
| EPI_ISL_408009 | hCoV-19/USA/CA4/2020 | North America / USA / California | 2020-01-29 | California Department of Health | Pathogen Discovery, Respiratory Viruses Branch, Division of Viral Diseases, Centers for Disease Control and Prevention | Krista Queen, Jing Zhang, Yan Li, Ying Tao, Anna Uehara, Clinton Paden, Xiaoyan Lu, Brian Lynch, Senthil Kumar K. Sakthivel, Brett L. Whitaker, Shifaq Kamili, Lijuan Wang, Janna' R. Murray, Susan I. Gerber, Stephen Lindstrom, Suxiang Tong |
| EPI_ISL_408010 | hCoV-19/USA/CA5/2020 | North America / USA / California | 2020-01-29 | California Department of Health | Pathogen Discovery, Respiratory Viruses Branch, Division of Viral Diseases, Centers for Disease Control and Prevention | Ying Tao, Krista Queen, Jing Zhang, Yan Li, Anna Uehara, Clinton Paden, Xiaoyan Lu, Brian Lynch, Senthil Kumar K. Sakthivel, Brett L. Whitaker, Shifaq Kamili, Lijuan Wang, Janna' R. Murray, Susan I. Gerber, Stephen Lindstrom, Suxiang Tong |
| EPI_ISL_408430 | hCoV-19/France/IDF0515/2020 | Europe / France / Ile-de-France / Paris | 2020-01-29 | Department of Infectious and Tropical Diseases, Bichat Claude Bernard Hospital, Paris | National Reference Center for Viruses of Respiratory Infections, Institut Pasteur, Paris | Mélanie Albert, Marion Barbet, Sylvie Behillil, Méline Bizard, Angela Brisebarre, Flora Donati, Vincent Enouf, Maud Vanpeene, Sylvie van der Werf, Yazdan Yazdanpanah, Xavier Lescure |
| EPI_ISL_408431 | hCoV-19/France/IDF0626/2020 | Europe / France / Ile-de-France / Paris | 2020-01-29 | Sorbonne Université, Inserm et Assistance Publique-Hôpitaux de Paris (Pitié Salpêtrière) | National Reference Center for Viruses of Respiratory Infections, Institut Pasteur, Paris | Mélanie Albert, Marion Barbet, Sylvie Behillil, Méline Bizard, Angela Brisebarre, Flora Donati, Vincent Enouf, Maud Vanpeene, Sylvie van der Werf, Sonia Burrel, Anne-Geneviève Marcelin, Vincent Calvez, David Boutolleau, Elise Klément, Valérie Pourcher, Eric Caumes. |
| EPI_ISL_408478 | hCoV-19/Chongqing/YC01/2020 | Asia / China / Chongqing / Yongchuan | 2020-01-21 | Yongchuan District Center for Disease Control and Prevention | Chongqing Municipal Center for Disease Control and Prevention | Ye Sheng, Tang Yun, Ling Hua,Yu zhen,Chen Shuang,Tan ZhangPing, Su Kun, Li Qing, Tang Wenge, Rong Rong |
| EPI_ISL_408479 | hCoV-19/Chongqing/ZX01/2020 | Asia / China / Chongqing / Zhongxian | 2020-01-23 | Zhongxian Center for Disease Control and Prevention | Chongqing Municipal Center for Disease Control and Prevention | Ye Sheng, Tang Yun, Ling Hua, Zhang Hong, Yu zhen,Chen Shuang,Tan ZhangPing, Su Kun, Li Qin, Tang Wenge, Rong Rong |
| EPI_ISL_408480 | hCoV-19/Yunnan/IVDC-YN-003/2020 | Asia / China / Yunnan / Kunming | 2020-01-17 | National Institute for Viral Disease Control and Prevention, China CDC | National Institute for Viral Disease Control & Prevention, CCDC | Wenjie Tan, Xiaoqing Fu, Xiang Zhao, Wenling Wang, Peihua Niu, Roujian Lu,Yanhong Sun, Baoying Huang, Li Zhao, Fei Ye, Wenbo Xu, George F. Gao, Guizhen Wu |
| EPI_ISL_408481 | hCoV-19/Chongqing/IVDC-CQ-001/2020 | Asia / China / Chongqing | 2020-01-18 | National Institute for Viral Disease Control and Prevention, China CDC | National Institute for Viral Disease Control & Prevention, CCDC | Wenjie Tan, Hengqin Wang, Xiang Zhao, Wenling Wang, Peihua Niu, Roujian Lu, Sheng Ye, Baoying Huang, Li Zhao, Fei Ye, Wenbo Xu, George F. Gao, Guizhen Wu |
| EPI_ISL_408482 | hCoV-19/Shandong/IVDC-SD-001/2020 | Asia / China / Shandong / Qingdao | 2020-01-19 | National Institute for Viral Disease Control and Prevention, China CDC | National Institute for Viral Disease Control & Prevention, CCDC | Wenjie Tan, Zhaoguo Wang, Xiang Zhao, Wenling Wang, Peihua Niu, Roujian Lu, Ti Liu, Baoying Huang, Li Zhao, Fei Ye, Wenbo Xu, George F. Gao, Guizhen Wu |
| EPI_ISL_408484 | hCoV-19/Sichuan/IVDC-SC-001/2020 | Asia / China / Sichuan / Chengdu | 2020-01-15 | National Institute for Viral Disease Control and Prevention, China CDC | National Institute for Viral Disease Control & Prevention, CCDC | Wenjie Tan, Jianan Xu, Wenling Wang, Peihua Niu, Roujian Lu, Huiping Yang, Xiang Zhao, Baoying Huang, Li Zhao, Fei Ye, Wenbo Xu, George F. Gao, Guizhen Wu |
| EPI_ISL_408485 | hCoV-19/Beijing/IVDC-BJ-005/2020 | Asia / China / Beijing | 2020-01-18 | National Institute for Viral Disease Control and Prevention, China CDC | National Institute for Viral Disease Control & Prevention, CCDC | Wenjie Tan,Quanyi Wang,Wenling Wang, Peihua Niu,Roujian Lu,Yang Pan,Xiang Zhao,Baoying Huang,Li Zhao,Fei Ye,Wenbo Xu,George F. Gao,Guizhen Wu |
| EPI_ISL_408486 | hCoV-19/Jiangxi/IVDC-JX-002/2020 | Asia / China / Jiangxi / Pingxiang | 2020-01-11 | National Institute for Viral Disease Control and Prevention, China CDC | National Institute for Viral Disease Control & Prevention, CCDC | Wenjie Tan, Yong Shi, Wenling Wang, Peihua Niu, Roujian Lu, Jianxiong Li, Xiang Zhao, Baoying Huang, Li Zhao, Fei Ye, Wenbo Xu, George F. Gao, Guizhen Wu |
| EPI_ISL_408488 | hCoV-19/Jiangsu/IVDC-JS-001/2020 | Asia / China / Jiangsu / Huaian | 2020-01-19 | National Institute for Viral Disease Control and Prevention, China CDC | National Institute for Viral Disease Control & Prevention, CCDC | Wenjie Tan, Shenjiao Wang, Wenling Wang, Peihua Niu, Roujian Lu, Kangchen Zhao, Xiang Zhao, Baoying Huang, Li Zhao, Fei Ye, Wenbo Xu, George F. Gao, Guizhen Wu |
| EPI_ISL_408489 | hCoV-19/Taiwan/NTU01/2020 | Asia / Taiwan / Taipei | 2020-01-31 | Department of Laboratory Medicine, National Taiwan University Hospital | Microbial Genomics Core Lab, National Taiwan University Centers of Genomic and Precision Medicine | Shiou-Hwei Yeh, You-Yu Lin, Ya-Yun Lai, Chiao-Ling Li, Shan-Chwen Chang, Pei-Jer Chen, Sui-Yuan Chang |
| EPI_ISL_410218 | hCoV-19/Taiwan/NTU02/2020 | Asia / Taiwan / Taipei | 2020-02-05 | Department of Laboratory Medicine, National Taiwan University Hospital | Microbial Genomics Core Lab, National Taiwan University Centers of Genomic and Precision Medicine | Shiou-Hwei Yeh, You-Yu Lin, Ya-Yun Lai, Chiao-Ling Li, Shan-Chwen Chang, Pei-Jer Chen, Sui-Yuan Chang |
| EPI_ISL_410301 | hCoV-19/Nepal/61/2020 | Asia / Nepal / Kathmandu | 2020-01-13 | National Influenza Centre, National Public Health Laboratory, Kathmandu, Nepal | The University of Hong Kong | Ranjit Sah , Runa Jha, Daniel Chu, Haogao Gu, Malik Peiris, Anup Bastola, Alfonso J. Rodriguez-Morales, Bibek Kumar Lal, Basu Dev Pandey, Leo Poon |
| EPI_ISL_410713 | hCoV-19/Singapore/7/2020 | Asia / Singapore | 2020-01-27 | National Public Health Laboratory, National Centre for Infectious Diseases | National Public Health Laboratory, National Centre for Infectious Diseases | Octavia S, Mak TM, Cui L, Lin RTP |
| EPI_ISL_410714 | hCoV-19/Singapore/8/2020 | Asia / Singapore | 2020-02-03 | National Public Health Laboratory, National Centre for Infectious Diseases | National Public Health Laboratory, National Centre for Infectious Diseases | Octavia S, Mak TM, Cui L, Lin RTP |
| EPI_ISL_410715 | hCoV-19/Singapore/9/2020 | Asia / Singapore | 2020-02-04 | National Public Health Laboratory, National Centre for Infectious Diseases | National Public Health Laboratory, National Centre for Infectious Diseases | Octavia S, Mak TM, Cui L, Lin RTP |
| EPI_ISL_410716 | hCoV-19/Singapore/10/2020 | Asia / Singapore | 2020-02-04 | National Public Health Laboratory, National Centre for Infectious Diseases | National Centre for Infectious Diseases, National Centre for Infectious Diseases | Octavia S, Mak TM, Cui L, Lin RTP |
| EPI_ISL_410717 | hCoV-19/Australia/QLD03/2020 | Oceania / Australia / Queensland / Gold C | 2020-02-05 | Pathology Queensland | Public Health Virology Laboratory | Ben Huang, Alyssa Pyke, Amanda De Jong, Andrew Van Den Hurk, Carmel Taylor, David Warrilow, Doris Genge, Elisabeth Gamez, Glen Hewitson, Ian Maxwell Mackay, Inga Sultana, Jamie McMahon, Jean Barcelon, Judy Northill, Mitchell Finger, Natalie Simpson, Neelima Nair, Peter Burtonclay, Peter Moore, Sarah Wheatley, Sean Moody, Sonja Hall-Mendelin, Timothy Gardam, and Frederick Moore. |
| EPI_ISL_410718 | hCoV-19/Australia/QLD04/2020 | Oceania / Australia / Queensland / Gold C | 2020-02-05 | Pathology Queensland | Public Health Virology Laboratory | Ben Huang, Alyssa Pyke, Amanda De Jong, Andrew Van Den Hurk, Carmel Taylor, David Warrilow, Doris Genge, Elisabeth Gamez, Glen Hewitson, Ian Maxwell Mackay, Inga Sultana, Jamie McMahon, Jean Barcelon, Judy Northill, Mitchell Finger, Natalie Simpson, Neelima Nair, Peter Burtonclay, Peter Moore, Sarah Wheatley, Sean Moody, Sonja Hall-Mendelin, Timothy Gardam, and Frederick Moore. |
| EPI_ISL_410719 | hCoV-19/Singapore/11/2020 | Asia / Singapore | 2020-02-02 | National Public Health Laboratory | National Public Health Laboratory | Octavia S, Mak TM, Cui L, Lin RTP |
| EPI_ISL_410720 | hCoV-19/France/IDF0372-isl/2020 | Europe / France / Ile-de-France / Paris | 2020-01-23 | Department of Infectious and Tropical Diseases, Bichat Claude Bernard Hospital, Paris | National Reference Center for Viruses of Respiratory Infections, Institut Pasteur, Paris | Mélanie Albert, Marion Barbet, Sylvie Behillil, Méline Bizard, Angela Brisebarre, Flora Donati, Vincent Enouf, Maud Vanpeene, Sylvie van der Werf, Yazdan Yazdanpanah, Xavier Lescure. |
| EPI_ISL_411060 | hCoV-19/Fujian/8/2020 | Asia / China / Fujian | 2020-01-21 | Fujian Center for Disease Control and Prevention | Fujian Center for Disease Control and Prevention | Chen Wei, Zhang Yanhua, He Wenxiang, Weng Yuwei |
| EPI_ISL_411066 | hCoV-19/Fujian/13/2020 | Asia / China / Fujian | 2020-01-22 | Fujian Center for Disease Control and Prevention | Fujian Center for Disease Control and Prevention | Chen Wei, Zhang Yanhua, He Wenxiang, Weng Yuwei |
| EPI_ISL_412026 | hCoV-19/Hefei/2/2020 | Asia / China / Anhui / Hefei | 2020-02-23 | Second Hospital of Anhui Medical University | Second Hospital of Anhui Medical University | Changtai Wang, Zhongping Liua, Zixiang Chen, Xin Huang, Mengyuan Xua, Tengfei He, Mengji Lu, Zhenhua Zhang |
| EPI_ISL_412028 | hCoV-19/Hong Kong/VM20001061-2/2020 | Asia / Hong Kong | 2020-01-22 | Hong Kong Department of Health | School of Public Health, The University of Hon g Kong | Dominic N.C. Tsang, Daniel K.W. Chu, Leo L.M. Poon, Malik Peiris |
| EPI_ISL_412029 | hCoV-19/Hong Kong/VB20024950-2/2020 | Asia / Hong Kong | 2020-01-30 | Hong Kong Department of Health | The University of Hong Kong | Dominic N.C. Tsang, Daniel K.W. Chu, Leo L.M. Poon, Malik Peiris |
| EPI_ISL_412030 | hCoV-19/Hong Kong/VB20026565-2/2020 | Asia / Hong Kong | 2020-02-01 | Hong Kong Department of Health | School of Public Health, The University of Hon g Kong | Dominic N.C. Tsang, Daniel K.W. Chu, Leo L.M. Poon, Malik Peiris |
| EPI_ISL_412459 | hCoV-19/Jingzhou/HBCDC-HB-01/2020 | Asia / China / Hubei / Jingzhou | 2020-01-08 | Jingzhou Center for Disease Control and Prevention | Hubei Provincial Center for Disease Control and Prevention | Bin Fang, Xiang Li, Xiao Yu, Linlin Liu, Bo Yang, Faxian Zhan, Guojun Ye, Xixiang Huo, Junqiang Xu, Bo Yu, Kun Cai, Jing Li, Maoyi Chen,Jie Hu, Chunlin Mao, Yongzhong Jiang. |
| EPI_ISL_412898 | hCoV-19/Wuhan/HBCDC-HB-02/2019 | Asia / China / Hubei / Wuhan | 2019-12-30 | Wuhan Jinyintan Hospital | Hubei Provincial Center for Disease Control and Prevention | Bin Fang, Xiang Li, Xiao Yu, Linlin Liu, Bo Yang, Faxian Zhan, Guojun Ye, Xixiang Huo, Junqiang Xu, Bo Yu, Kun Cai, Jing Li, Yongzhong Jiang. |
| EPI_ISL_412899 | hCoV-19/Wuhan/HBCDC-HB-03/2019 | Asia / China / Hubei / Wuhan | 2019-12-30 | Wuhan Jinyintan Hospital | Hubei Provincial Center for Disease Control and Prevention | Bin Fang, Xiang Li, Xiao Yu, Linlin Liu, Bo Yang, Faxian Zhan, Guojun Ye, Xixiang Huo, Junqiang Xu, Bo Yu, Kun Cai, Jing Li, Yongzhong Jiang. |
| EPI_ISL_412912 | hCoV-19/Germany/Baden-Wuerttemberg-1/2020 | Europe / Germany / Baden-Wuerttemberg | 2020-02-25 | State Health Office Baden-Wuerttemberg | Charité Universitätsmedizin Berlin, Institute of Virology | Victor M Corman, Julia Schneider, Barbara Mühlemann, Talitha Veith, Jörn Beheim-Schwarzbach, Terry Jones, Rainer Oehme, Silke Fischer, Christian Drosten |
| EPI_ISL_412964 | hCoV-19/Brazil/SPBR-01/2020 | South America / Brazil / Sao Paulo / Sao | 2020-02-25 | Hospital Israelita Albert Einstein | Instituto Adolfo Lutz Interdisciplinary Procedures Center Strategic Laboratory | Jaqueline Goes de Jesus, Claudio Tavares Sacchi, Daniela Bernardes Borges da Silva, Ingra Morales Claro, Flávia Cristina da Silva Sales, Claudia Regina Gonçalves, Joshua Quick, Maria do Carmo, Sampaio Tavares Timenetsky, Nicholas James Loman, Andrew Rambaut, Ester Cerdeira Sabino, Nuno Rodrigues Faria |
| EPI_ISL_412966 | hCoV-19/China/IQTC01/2020 | Asia / China / Guangdong / Guangzhou | 2020-02-05 | unknown | Technology Centre, Guangzhou Customs | Shi,Y., Sun,J., Zheng,K., Huang,J. and Zhao,J. |
| EPI_ISL_412967 | hCoV-19/China/IQTC02/2020 | Asia / China / Guangzhou | 2020-01-29 | unknown | Technology Centre, Guangzhou Customs | Shi,Y., Zheng,K., Sun,J., Huang,J., Zhu,A., Zhuang,Z., Dai,J., Chen,Z., Sun,F., Zhang,Z., Li,X. and Wang,Y. |
| EPI_ISL_412968 | hCoV-19/Japan/Hu_DP_Kng_19-020/2020 | Asia / Japan | 2020-02-10 | unknown | Takayuki Hishiki Kanagawa Prefectural Institute of Public Health, Department of Microbiology | Hishiki,T., Suzuki,R., Sakuragi,J., Usui,K., Tanaka,Y., Kawai,J., Kogo,Y., Matsuki,Y., An,T., Hayashizaki,Y. and Takasaki,T. |
| EPI_ISL_412969 | hCoV-19/Japan/Hu_DP_Kng_19-027/2020 | Asia / Japan | 2020-02-10 | unknown | Takayuki Hishiki Kanagawa Prefectural Institute of Public Health, Department of Microbiology | Hishiki,T., Suzuki,R., Sakuragi,J., Usui,K., Tanaka,Y., Kawai,J., Kogo,Y., Matsuki,Y., An,T., Hayashizaki,Y. and Takasaki,T. |
| EPI_ISL_412970 | hCoV-19/USA/WA2/2020 | North America / USA / Washington / Snol | 2020-02-24 | Washington State Department of Health | Seattle Flu Study | Helen Chu, Michael Boeckh, Janet Englund, Michael Famulare, Barry Lutz, Deborah Nickerson, Mark Rieder, Lea Starita, Matthew Thompson, Jay Shendure, and Trevor Bedford |
| EPI_ISL_412972 | hCoV-19/Mexico/CDMX-InDRE_01/2020 | North America / Mexico / Mexico City | 2020-02-27 | Instituto Nacional de Enfermedades Respiratorias | Instituto de Diagnostico y Referencia Epidemiologicos (INDRE) | Ramirez-Gonzalez Ernesto, Garces-Ayala Fabiola, Araiza-Rodriguez Adnan, Mendieta-Condado Edgar, Rodriguez-Maldonado Abril, Wong-Arambula Claudia, Vazquez-Perez Joel, Martinez Arturo, Boukadida Celia, Munoz-Medina Esteban, Sanchez Alejandro, Isa Pavel, Taboada Blanca, Lopez Susana, Arias Carlos, Barrera-Badillo Gisela, Hernandez-Rivas Lucia, Lopez-Martinez Irma |
| EPI_ISL_412973 | hCoV-19/Italy/CDG1/2020 | Europe / Italy / Lombardy | 2020-02-20 | Department of Infectious Diseases, Istituto Superiore di Sanità, Roma , Italy | Virology Laboratory, Scientific Department, Army Medical Center | Paola Stefanelli, Stefano Fiore, Antonella Marchi, Eleonora Benedetti, Concetta Fabiani, Giovanni Faggioni, Antonella Fortunato, Riccardo De Santis, Silvia Fillo, Anna Anselmo, Andrea Ciammaruconi, Stefano Palomba, Florigio Lista |
| EPI_ISL_412974 | hCoV-19/Italy/SPL1/2020 | Europe / Italy / Rome | 2020-01-29 | Department of Infectious Diseases, Istituto Superiore di Sanità, Rome, Italy | Virology Laboratory, Scientific Department, Army Medical Center | Paola Stefanelli, Stefano Fiore, Antonella Marchi, Eleonora Benedetti, Concetta Fabiani, Giovanni Faggioni, Antonella Fortunato, Silvia Fillo, Riccardo De Santis, Andrea Ciammaruconi, Giancarlo Petrallito, Filippo Molinari, Florigio Lista |
| EPI_ISL_412975 | hCoV-19/Australia/NSW05/2020 | Oceania / Australia / New South Wales / S | 2020-02-28 | Centre for Infectious Diseases and Microbiology Laboratory Services | NSW Health Pathology - Institute of Clinical Pathology and Medical Research; Westmead Hospital; University of Sydney | Eden J-S, Carter I, Rahman H, Holmes EC, Rockett R, O'Sullivan MV, Sintchenko V, Chen SC, Maddocks S, Kok J and Dwyer DE for the 2019-nCoV Study Group |
| EPI_ISL_412978 | hCoV-19/Wuhan/HBCDC-HB-02/2020 | Asia / China / Hubei / Wuhan | 2020-01-17 | The Central Hospital Of Wuhan | Hubei Provincial Center for Disease Control and Prevention | Bin Fang, Xiang Li, Xiao Yu, Linlin Liu, Bo Yang, Faxian Zhan, Guojun Ye, Xixiang Huo, Junqiang Xu, Bo Yu, Kun Cai, Jing Li, Yongzhong Jiang. |
| EPI_ISL_412979 | hCoV-19/Wuhan/HBCDC-HB-03/2020 | Asia / China / Hubei / Wuhan | 2020-01-18 | Union Hospital of Tongji Medical College, Huazhong University of Science and Technology | Hubei Provincial Center for Disease Control and Prevention | Bin Fang, Xiang Li, Xiao Yu, Linlin Liu, Bo Yang, Faxian Zhan, Guojun Ye, Xixiang Huo, Junqiang Xu, Bo Yu, Kun Cai, Jing Li, Yongzhong Jiang. |
| EPI_ISL_412980 | hCoV-19/Wuhan/HBCDC-HB-04/2020 | Asia / China / Hubei / Wuhan | 2020-01-18 | Union Hospital of Tongji Medical College, Huazhong University of Science and Technology | Hubei Provincial Center for Disease Control and Prevention | Bin Fang, Xiang Li, Xiao Yu, Linlin Liu, Bo Yang, Faxian Zhan, Guojun Ye, Xixiang Huo, Junqiang Xu, Bo Yu, Kun Cai, Jing Li, Yongzhong Jiang. |
| EPI_ISL_412981 | hCoV-19/Wuhan/HBCDC-HB-05/2020 | Asia / China / Hubei / Wuhan | 2020-01-18 | CR&WISCO GENERAL HOSPITAL | Hubei Provincial Center for Disease Control and Prevention | Bin Fang, Xiang Li, Xiao Yu, Linlin Liu, Bo Yang, Faxian Zhan, Guojun Ye, Xixiang Huo, Junqiang Xu, Bo Yu, Kun Cai, Jing Li, Yongzhong Jiang. |
| EPI_ISL_412982 | hCoV-19/Wuhan/HBCDC-HB-06/2020 | Asia / China / Hubei / Wuhan | 2020-02-07 | Wuhan Lung Hospital | Hubei Provincial Center for Disease Control and Prevention | Bin Fang, Xiang Li, Xiao Yu, Linlin Liu, Bo Yang, Faxian Zhan, Guojun Ye, Xixiang Huo, Junqiang Xu, Bo Yu, Kun Cai, Jing Li, Yongzhong Jiang. |
| EPI_ISL_412983 | hCoV-19/Tianmen/HBCDC-HB-07/2020 | Asia / China / Hubei / Tianmen | 2020-02-08 | Tianmen Center for Disease Control and Prevention | Hubei Provincial Center for Disease Control and Prevention | Bin Fang, Xiang Li, Xiao Yu, Linlin Liu, Bo Yang, Faxian Zhan, Guojun Ye, Xixiang Huo, Junqiang Xu, Bo Yu, Kun Cai, Jing Li, YiFa Zhu, Yangyang Tao,Xierong Li,Yongzhong Jiang. |
| EPI_ISL_413213 | hCoV-19/Australia/NSW06/2020 | Oceania / Australia / New South Wales / S | 2020-02-29 | Centre for Infectious Diseases and Microbiology Laboratory Services | NSW Health Pathology - Institute of Clinical Pathology and Medical Research; Westmead Hospital; University of Sydney | Eden J-S, Carter I, Rahman H, Holmes EC, Rockett R, O'Sullivan MV, Sintchenko V, Chen SC, Maddocks S, Kok J and Dwyer DE for the 2019-nCoV Study Group" |
| EPI_ISL_413214 | hCoV-19/Australia/NSW07/2020 | Oceania / Australia / New South Wales / S | 2020-02-29 | Centre for Infectious Diseases and Microbiology Laboratory Services | NSW Health Pathology - Institute of Clinical Pathology and Medical Research; Westmead Hospital; University of Sydney | Eden J-S, Carter I, Rahman H, Holmes EC, Rockett R, O'Sullivan MV, Sintchenko V, Chen SC, Maddocks S, Kok J and Dwyer DE for the 2019-nCoV Study Group" |
| EPI_ISL_413602 | hCoV-19/Finland/FIN03032020A/2020 | Europe / Finland / Helsinki | 2020-03-03 | Department of Virology and Immunology, University of Helsinki and Helsinki University Hospital, Huslab Finland | Department of Virology, Faculty of Medicine, University of Helsinki, Helsinki, Finland | Teemu Smura, Hannimari Kallio-Kokko, Olli Vapalahti |
| EPI_ISL_413603 | hCoV-19/Finland/FIN03032020B/2020 | Europe / Finland / Helsinki | 2020-03-03 | Department of Virology and Immunology, University of Helsinki and Helsinki University Hospital, Huslab Finland | Department of Virology, Faculty of Medicine, University of Helsinki, Helsinki, Finland | Teemu Smura, Hannimari Kallio-Kokko, Olli Vapalahti |
| EPI_ISL_413604 | hCoV-19/Finland/FIN03032020C/2020 | Europe / Finland / Helsinki | 2020-03-03 | Department of Virology and Immunology, University of Helsinki and Helsinki University Hospital, Huslab Finland | Department of Virology, Faculty of Medicine, University of Helsinki, Helsinki, Finland | Teemu Smura, Hannimari Kallio-Kokko, Olli Vapalahti |
| EPI_ISL_413606 | hCoV-19/USA/CruiseA-1/2020 | North America / USA | 2020-02-17 | unknown | Pathogen Discovery, Respiratory Viruses Branch, Division of Viral Diseases, Centers for Disease Control and Prevention | Anna Uehara, Ying Tao, Clinton R. Paden, Krista Queen, Jing Zhang, Yan Li, Mary S. Keckler, Alison S Laufer Halpin, Haibin Wang, Jasmine Padilla, Justin Lee, Christopher A. Elkins, Susan I. Gerber, Suxiang Tong |
| EPI_ISL_413607 | hCoV-19/USA/CruiseA-2/2020 | North America / USA | 2020-02-18 | unknown | Pathogen Discovery, Respiratory Viruses Branch, Division of Viral Diseases, Centers for Disease Control and Prevention | Anna Uehara, Ying Tao, Clinton R. Paden, Krista Queen, Jing Zhang, Yan Li, Mary S. Keckler, Alison S Laufer Halpin, Haibin Wang, Jasmine Padilla, Justin Lee, Christopher A. Elkins, Susan I. Gerber, Suxiang Tong |
| EPI_ISL_413608 | hCoV-19/USA/CruiseA-3/2020 | North America / USA | 2020-02-18 | unknown | Pathogen Discovery, Respiratory Viruses Branch, Division of Viral Diseases, Centers for Disease Control and Prevention | Anna Uehara, Ying Tao, Clinton R. Paden, Krista Queen, Jing Zhang, Yan Li, Mary S. Keckler, Alison S Laufer Halpin, Haibin Wang, Jasmine Padilla, Justin Lee, Christopher A. Elkins, Susan I. Gerber, Suxiang Tong |
| EPI_ISL_413609 | hCoV-19/USA/CruiseA-4/2020 | North America / USA | 2020-02-21 | unknown | Pathogen Discovery, Respiratory Viruses Branch, Division of Viral Diseases, Centers for Disease Control and Prevention | Anna Uehara, Ying Tao, Clinton R. Paden, Krista Queen, Jing Zhang, Yan Li, Mary S. Keckler, Alison S Laufer Halpin, Haibin Wang, Jasmine Padilla, Justin Lee, Christopher A. Elkins, Susan I. Gerber, Suxiang Tong |
| EPI_ISL_413610 | hCoV-19/USA/CruiseA-5/2020 | North America / USA | 2020-02-21 | unknown | Pathogen Discovery, Respiratory Viruses Branch, Division of Viral Diseases, Centers for Disease Control and Prevention | Anna Uehara, Ying Tao, Clinton R. Paden, Krista Queen, Jing Zhang, Yan Li, Mary S. Keckler, Alison S Laufer Halpin, Haibin Wang, Jasmine Padilla, Justin Lee, Christopher A. Elkins, Susan I. Gerber, Suxiang Tong |
| EPI_ISL_413611 | hCoV-19/USA/CruiseA-6/2020 | North America / USA | 2020-02-21 | unknown | Pathogen Discovery, Respiratory Viruses Branch, Division of Viral Diseases, Centers for Disease Control and Prevention | Anna Uehara, Ying Tao, Clinton R. Paden, Krista Queen, Jing Zhang, Yan Li, Mary S. Keckler, Alison S Laufer Halpin, Haibin Wang, Jasmine Padilla, Justin Lee, Christopher A. Elkins, Susan I. Gerber, Suxiang Tong |
| EPI_ISL_413612 | hCoV-19/USA/CruiseA-7/2020 | North America / USA | 2020-02-17 | unknown | Pathogen Discovery, Respiratory Viruses Branch, Division of Viral Diseases, Centers for Disease Control and Prevention | Ying Tao, Clinton R. Paden, Krista Queen, Anna Uehara, Jing Zhang, Yan Li, Haibin Wang, Shifaq Kamili, Xiaoyan Lu, Brian Lynch, Senthil Kumar K. Sakthivel, Brett L. Whitaker, Lijuan Wang, Janna' R. Murray, Jasmine Padilla, Justin Lee, Susan I. Gerber, Stephen Lindstrom, Suxiang Tong |
| EPI_ISL_413613 | hCoV-19/USA/CruiseA-8/2020 | North America / USA | 2020-02-17 | unknown | Pathogen Discovery, Respiratory Viruses Branch, Division of Viral Diseases, Centers for Disease Control and Prevention | Ying Tao, Clinton R. Paden, Krista Queen, Anna Uehara, Jing Zhang, Yan Li, Haibin Wang, Shifaq Kamili, Xiaoyan Lu, Brian Lynch, Senthil Kumar K. Sakthivel, Brett L. Whitaker, Lijuan Wang, Janna' R. Murray, Jasmine Padilla, Justin Lee, Susan I. Gerber, Stephen Lindstrom, Suxiang Tong |
| EPI_ISL_413614 | hCoV-19/USA/CruiseA-9/2020 | North America / USA | 2020-02-17 | unknown | Pathogen Discovery, Respiratory Viruses Branch, Division of Viral Diseases, Centers for Disease Control and Prevention | Ying Tao, Clinton R. Paden, Krista Queen, Anna Uehara, Jing Zhang, Yan Li, Haibin Wang, Shifaq Kamili, Xiaoyan Lu, Brian Lynch, Senthil Kumar K. Sakthivel, Brett L. Whitaker, Lijuan Wang, Janna' R. Murray, Jasmine Padilla, Justin Lee, Susan I. Gerber, Stephen Lindstrom, Suxiang Tong |
| EPI_ISL_413615 | hCoV-19/USA/CruiseA-10/2020 | North America / USA | 2020-02-17 | unknown | Pathogen Discovery, Respiratory Viruses Branch, Division of Viral Diseases, Centers for Disease Control and Prevention | Ying Tao, Clinton R. Paden, Krista Queen, Anna Uehara, Jing Zhang, Yan Li, Haibin Wang, Shifaq Kamili, Xiaoyan Lu, Brian Lynch, Senthil Kumar K. Sakthivel, Brett L. Whitaker, Lijuan Wang, Janna' R. Murray, Jasmine Padilla, Justin Lee, Susan I. Gerber, Stephen Lindstrom, Suxiang Tong |
| EPI_ISL_413616 | hCoV-19/USA/CruiseA-11/2020 | North America / USA | 2020-02-17 | unknown | Pathogen Discovery, Respiratory Viruses Branch, Division of Viral Diseases, Centers for Disease Control and Prevention | Ying Tao, Clinton R. Paden, Krista Queen, Anna Uehara, Jing Zhang, Yan Li, Haibin Wang, Shifaq Kamili, Xiaoyan Lu, Brian Lynch, Senthil Kumar K. Sakthivel, Brett L. Whitaker, Lijuan Wang, Janna' R. Murray, Jasmine Padilla, Justin Lee, Susan I. Gerber, Stephen Lindstrom, Suxiang Tong |
| EPI_ISL_413617 | hCoV-19/USA/CruiseA-12/2020 | North America / USA | 2020-02-20 | unknown | Pathogen Discovery, Respiratory Viruses Branch, Division of Viral Diseases, Centers for Disease Control and Prevention | Ying Tao, Clinton R. Paden, Krista Queen, Anna Uehara, Jing Zhang, Yan Li, Haibin Wang, Shifaq Kamili, Xiaoyan Lu, Brian Lynch, Senthil Kumar K. Sakthivel, Brett L. Whitaker, Lijuan Wang, Janna' R. Murray, Jasmine Padilla, Justin Lee, Susan I. Gerber, Stephen Lindstrom, Suxiang Tong |
| EPI_ISL_413618 | hCoV-19/USA/CruiseA-13/2020 | North America / USA | 2020-02-20 | unknown | Pathogen Discovery, Respiratory Viruses Branch, Division of Viral Diseases, Centers for Disease Control and Prevention | Clinton R. Paden, Ying Tao, Krista Queen, Anna Uehara, Jing Zhang, Yan Li, Haibin Wang, Shifaq Kamili, Xiaoyan Lu, Brian Lynch, Senthil Kumar K. Sakthivel, Brett L. Whitaker, Lijuan Wang, Janna' R. Murray, Jasmine Padilla, Justin Lee, Susan I. Gerber, Stephen Lindstrom, Suxiang Tong |
| EPI_ISL_413619 | hCoV-19/USA/CruiseA-14/2020 | North America / USA | 2020-02-25 | unknown | Pathogen Discovery, Respiratory Viruses Branch, Division of Viral Diseases, Centers for Disease Control and Prevention | Clinton R. Paden, Ying Tao, Krista Queen, Anna Uehara, Jing Zhang, Yan Li, Haibin Wang, Shifaq Kamili, Xiaoyan Lu, Brian Lynch, Senthil Kumar K. Sakthivel, Brett L. Whitaker, Lijuan Wang, Janna' R. Murray, Jasmine Padilla, Justin Lee, Susan I. Gerber, Stephen Lindstrom, Suxiang Tong |
| EPI_ISL_413620 | hCoV-19/USA/CruiseA-15/2020 | North America / USA | 2020-02-18 | unknown | Pathogen Discovery, Respiratory Viruses Branch, Division of Viral Diseases, Centers for Disease Control and Prevention | Clinton R. Paden, Ying Tao, Krista Queen, Anna Uehara, Jing Zhang, Yan Li, Haibin Wang, Shifaq Kamili, Xiaoyan Lu, Brian Lynch, Senthil Kumar K. Sakthivel, Brett L. Whitaker, Lijuan Wang, Janna' R. Murray, Jasmine Padilla, Justin Lee, Susan I. Gerber, Stephen Lindstrom, Suxiang Tong |
| EPI_ISL_413621 | hCoV-19/USA/CruiseA-16/2020 | North America / USA | 2020-02-18 | unknown | Pathogen Discovery, Respiratory Viruses Branch, Division of Viral Diseases, Centers for Disease Control and Prevention | Clinton R. Paden, Ying Tao, Krista Queen, Anna Uehara, Jing Zhang, Yan Li, Haibin Wang, Shifaq Kamili, Xiaoyan Lu, Brian Lynch, Senthil Kumar K. Sakthivel, Brett L. Whitaker, Lijuan Wang, Janna' R. Murray, Jasmine Padilla, Justin Lee, Susan I. Gerber, Stephen Lindstrom, Suxiang Tong |
| EPI_ISL_413622 | hCoV-19/USA/CruiseA-17/2020 | North America / USA | 2020-02-24 | unknown | Pathogen Discovery, Respiratory Viruses Branch, Division of Viral Diseases, Centers for Disease Control and Prevention | Clinton R. Paden, Ying Tao, Krista Queen, Anna Uehara, Jing Zhang, Yan Li, Haibin Wang, Shifaq Kamili, Xiaoyan Lu, Brian Lynch, Senthil Kumar K. Sakthivel, Brett L. Whitaker, Lijuan Wang, Janna' R. Murray, Jasmine Padilla, Justin Lee, Susan I. Gerber, Stephen Lindstrom, Suxiang Tong |
| EPI_ISL_413623 | hCoV-19/USA/CruiseA-18/2020 | North America / USA | 2020-02-24 | unknown | Pathogen Discovery, Respiratory Viruses Branch, Division of Viral Diseases, Centers for Disease Control and Prevention | Clinton R. Paden, Ying Tao, Krista Queen, Anna Uehara, Jing Zhang, Yan Li, Haibin Wang, Shifaq Kamili, Xiaoyan Lu, Brian Lynch, Senthil Kumar K. Sakthivel, Brett L. Whitaker, Lijuan Wang, Janna' R. Murray, Jasmine Padilla, Justin Lee, Susan I. Gerber, Stephen Lindstrom, Suxiang Tong |
| EPI_ISL_413647 | hCoV-19/Portugal/CV62/2020 | Europe / Portugal | 2020-03-01 | Centro Hospital do Porto, E.P.E. - H. Geral de Santo Antonio | Instituto Nacional de Saude (INSA) | Raquel Guiomar, Inês Costa, Pedro Pechirra, Joana Mendonça, Luís Vieira, Helena Ramos, Joana Isidro, Vítor Borges, João Paulo Gomes |
| EPI_ISL_413648 | hCoV-19/Portugal/CV63/2020 | Europe / Portugal | 2020-03-01 | Centro Hospitalar e Universitário de Sao Joao, Porto | Instituto Nacional de Saude (INSA) | Raquel Guiomar, Inês Costa, Pedro Pechirra, Joana Mendonça, Luís Vieira, João Tiago Guimarães, Joana Isidro, Vítor Borges, João Paulo Gomes |
| EPI_ISL_413649 | hCoV-19/USA/WA14-UW10/2020 | North America / USA / Washington | 2020-03-05 | UW Virology Lab | UW Virology Lab | Pavitra Roychoudhury, Hong Xie, Keith Jerome, Alexander Greninger |
| EPI_ISL_413650 | hCoV-19/USA/WA15-UW11/2020 | North America / USA / Washington | 2020-03-05 | UW Virology Lab | UW Virology Lab | Pavitra Roychoudhury, Hong Xie, Keith Jerome, Alexander Greninger |
| EPI_ISL_413651 | hCoV-19/USA/WA16-UW12/2020 | North America / USA / Washington | 2020-03-05 | UW Virology Lab | UW Virology Lab | Pavitra Roychoudhury, Hong Xie, Keith Jerome, Alexander Greninger |
| EPI_ISL_413652 | hCoV-19/USA/WA17-UW13/2020 | North America / USA / Washington | 2020-03-05 | UW Virology Lab | UW Virology Lab | Pavitra Roychoudhury, Hong Xie, Keith Jerome, Alexander Greninger |
| EPI_ISL_413653 | hCoV-19/USA/WA18-UW14/2020 | North America / USA / Washington | 2020-03-05 | UW Virology Lab | UW Virology Lab | Pavitra Roychoudhury, Hong Xie, Keith Jerome, Alexander Greninger |
| EPI_ISL_413691 | hCoV-19/China/WF0001/2020 | Asia / China | 2020-01 | Weifang Center for Disease Control and Prevention | Weifang Center for Disease Control and Prevention & BGI-Shenzhen | Qing Nie, Xingguang Li, Erik M Volz, Han Fu, Haowei Wang, Xiaoyue Xi, Wei Chen, Dehui Liu, Yingying Chen, Mengmeng Tian, Wei Tan, Junjie Zai, Wanying Sun, Jiandong Li, Junhua Li |
| EPI_ISL_413692 | hCoV-19/China/WF0002/2020 | Asia / China | 2020-01 | Weifang Center for Disease Control and Prevention | Weifang Center for Disease Control and Prevention & BGI-Shenzhen | Qing Nie, Xingguang Li, Erik M Volz, Han Fu, Haowei Wang, Xiaoyue Xi, Wei Chen, Dehui Liu, Yingying Chen, Mengmeng Tian, Wei Tan, Junjie Zai, Wanying Sun, Jiandong Li, Junhua Li |
| EPI_ISL_413693 | hCoV-19/China/WF0003/2020 | Asia / China | 2020-01 | Weifang Center for Disease Control and Prevention | Weifang Center for Disease Control and Prevention & BGI-Shenzhen | Qing Nie, Xingguang Li, Erik M Volz, Han Fu, Haowei Wang, Xiaoyue Xi, Wei Chen, Dehui Liu, Yingying Chen, Mengmeng Tian, Wei Tan, Junjie Zai, Wanying Sun, Jiandong Li, Junhua Li |
| EPI_ISL_413694 | hCoV-19/China/WF0004/2020 | Asia / China | 2020-01 | Weifang Center for Disease Control and Prevention | Weifang Center for Disease Control and Prevention & BGI-Shenzhen | Qing Nie, Xingguang Li, Erik M Volz, Han Fu, Haowei Wang, Xiaoyue Xi, Wei Chen, Dehui Liu, Yingying Chen, Mengmeng Tian, Wei Tan, Junjie Zai, Wanying Sun, Jiandong Li, Junhua Li |
| EPI_ISL_413697 | hCoV-19/China/WF0012/2020 | Asia / China | 2020-02 | Weifang Center for Disease Control and Prevention | Weifang Center for Disease Control and Prevention & BGI-Shenzhen | Qing Nie, Xingguang Li, Erik M Volz, Han Fu, Haowei Wang, Xiaoyue Xi, Wei Chen, Dehui Liu, Yingying Chen, Mengmeng Tian, Wei Tan, Junjie Zai, Wanying Sun, Jiandong Li, Junhua Li |
| EPI_ISL_413729 | hCoV-19/China/WF0015/2020 | Asia / China | 2020-02 | Weifang Center for Disease Control and Prevention | Weifang Center for Disease Control and Prevention & BGI-Shenzhen | Qing Nie, Xingguang Li, Erik M Volz, Han Fu, Haowei Wang, Xiaoyue Xi, Wei Chen, Dehui Liu, Yingying Chen, Mengmeng Tian, Wei Tan, Junjie Zai, Wanying Sun, Jiandong Li, Junhua Li |
| EPI_ISL_413746 | hCoV-19/China/WF0016/2020 | Asia / China | 2020-02 | Weifang Center for Disease Control and Prevention | Weifang Center for Disease Control and Prevention & BGI-Shenzhen | Qing Nie, Xingguang Li, Erik M Volz, Han Fu, Haowei Wang, Xiaoyue Xi, Wei Chen, Dehui Liu, Yingying Chen, Mengmeng Tian, Wei Tan, Junjie Zai, Wanying Sun, Jiandong Li, Junhua Li |
| EPI_ISL_413748 | hCoV-19/China/WF0018/2020 | Asia / China | 2020-02 | Weifang Center for Disease Control and Prevention | Weifang Center for Disease Control and Prevention & BGI-Shenzhen | Qing Nie, Xingguang Li, Erik M Volz, Han Fu, Haowei Wang, Xiaoyue Xi, Wei Chen, Dehui Liu, Yingying Chen, Mengmeng Tian, Wei Tan, Junjie Zai, Wanying Sun, Jiandong Li, Junhua Li |
| EPI_ISL_413749 | hCoV-19/China/WF0019/2020 | Asia / China | 2020-02 | Weifang Center for Disease Control and Prevention | Weifang Center for Disease Control and Prevention & BGI-Shenzhen | Qing Nie, Xingguang Li, Erik M Volz, Han Fu, Haowei Wang, Xiaoyue Xi, Wei Chen, Dehui Liu, Yingying Chen, Mengmeng Tian, Wei Tan, Junjie Zai, Wanying Sun, Jiandong Li, Junhua Li |
| EPI_ISL_413750 | hCoV-19/China/WF0020/2020 | Asia / China | 2020-02 | Weifang Center for Disease Control and Prevention | Weifang Center for Disease Control and Prevention & BGI-Shenzhen | Qing Nie, Xingguang Li, Erik M Volz, Han Fu, Haowei Wang, Xiaoyue Xi, Wei Chen, Dehui Liu, Yingying Chen, Mengmeng Tian, Wei Tan, Junjie Zai, Wanying Sun, Jiandong Li, Junhua Li |
| EPI_ISL_413751 | hCoV-19/China/WF0021/2020 | Asia / China | 2020-02 | Weifang Center for Disease Control and Prevention | Weifang Center for Disease Control and Prevention & BGI-Shenzhen | Qing Nie, Xingguang Li, Erik M Volz, Han Fu, Haowei Wang, Xiaoyue Xi, Wei Chen, Dehui Liu, Yingying Chen, Mengmeng Tian, Wei Tan, Junjie Zai, Wanying Sun, Jiandong Li, Junhua Li |
| EPI_ISL_413753 | hCoV-19/China/WF0024/2020 | Asia / China | 2020-02 | Weifang Center for Disease Control and Prevention | Weifang Center for Disease Control and Prevention & BGI-Shenzhen | Qing Nie, Xingguang Li, Erik M Volz, Han Fu, Haowei Wang, Xiaoyue Xi, Wei Chen, Dehui Liu, Yingying Chen, Mengmeng Tian, Wei Tan, Junjie Zai, Wanying Sun, Jiandong Li, Junhua Li |
| EPI_ISL_413761 | hCoV-19/China/WF0026/2020 | Asia / China | 2020-02 | Weifang Center for Disease Control and Prevention | Weifang Center for Disease Control and Prevention & BGI-Shenzhen | Qing Nie, Xingguang Li, Erik M Volz, Han Fu, Haowei Wang, Xiaoyue Xi, Wei Chen, Dehui Liu, Yingying Chen, Mengmeng Tian, Wei Tan, Junjie Zai, Wanying Sun, Jiandong Li, Junhua Li |
| EPI_ISL_413791 | hCoV-19/China/WF0028/2020 | Asia / China | 2020-02 | Weifang Center for Disease Control and Prevention | Weifang Center for Disease Control and Prevention & BGI-Shenzhen | Qing Nie, Xingguang Li, Erik M Volz, Han Fu, Haowei Wang, Xiaoyue Xi, Wei Chen, Dehui Liu, Yingying Chen, Mengmeng Tian, Wei Tan, Junjie Zai, Wanying Sun, Jiandong Li, Junhua Li |
| EPI_ISL_413809 | hCoV-19/China/WF0029/2020 | Asia / China | 2020-02 | Weifang Center for Disease Control and Prevention | Weifang Center for Disease Control and Prevention & BGI-Shenzhen | Qing Nie, Xingguang Li, Erik M Volz, Han Fu, Haowei Wang, Xiaoyue Xi, Wei Chen, Dehui Liu, Yingying Chen, Mengmeng Tian, Wei Tan, Junjie Zai, Wanying Sun, Jiandong Li, Junhua Li |
| EPI_ISL_414500 | hCoV-19/England/Sheff01/2020 | Europe / United Kingdom / England / Yor | 2020-03-04 | Virology Department, Sheffield Teaching Hospitals NHS Foundation Trust | Department of Infection, Immunity and Cardiovascular Disease, The Florey Institute, The Medical School, University of Sheffield | Thushan de Silva, Matthew Parker, Matthew Wyles, Mehmet Yavuz, Mohammad Raza, Cariad Evans |
| EPI_ISL_414501 | hCoV-19/England/Sheff02/2020 | Europe / United Kingdom / England / Yor | 2020-03-03 | Virology Department, Sheffield Teaching Hospitals NHS Foundation Trust | Department of Infection, Immunity and Cardiovascular Disease, The Florey Institute, The Medical School, University of Sheffield | Thushan de Silva, Matthew Parker, Matthew Wyles, Mehmet Yavuz, Mohammad Raza, Cariad Evans |
| EPI_ISL_414504 | hCoV-19/Germany/NRW-05/2020 | Europe / Germany / North Rhine Westph | 2020-02-26 | Center of Medical Microbiology, Virology, and Hospital Hygiene, University of Duesseldorf | Center of Medical Microbiology, Virology, and Hospital Hygiene, University of Duesseldorf | Ortwin Adams, Marcel Andree, Alexander Dilthey, Torsten Feldt, Sandra Hauka, Torsten Houwaart, Björn-Erik Jensen, Detlef Kindgen-Milles, Malte Kohns Vasconcelos, Klaus Pfeffer, Tina Senff, Daniel Strelow, Jörg Timm, Andreas Walker, Tobias Wienemann |
| EPI_ISL_414505 | hCoV-19/Germany/NRW-06/2020 | Europe / Germany / North Rhine Westph | 2020-02-27 | Center of Medical Microbiology, Virology, and Hospital Hygiene, University of Duesseldorf | Center of Medical Microbiology, Virology, and Hospital Hygiene, University of Duesseldorf | Ortwin Adams, Marcel Andree, Alexander Dilthey, Torsten Feldt, Sandra Hauka, Torsten Houwaart, Björn-Erik Jensen, Detlef Kindgen-Milles, Malte Kohns Vasconcelos, Klaus Pfeffer, Tina Senff, Daniel Strelow, Jörg Timm, Andreas Walker, Tobias Wienemann |
| EPI_ISL_414506 | hCoV-19/Germany/NRW-07/2020 | Europe / Germany / North Rhine Westph | 2020-02-27 | Center of Medical Microbiology, Virology, and Hospital Hygiene, University of Duesseldorf | Center of Medical Microbiology, Virology, and Hospital Hygiene, University of Duesseldorf | Ortwin Adams, Marcel Andree, Alexander Dilthey, Torsten Feldt, Sandra Hauka, Torsten Houwaart, Björn-Erik Jensen, Detlef Kindgen-Milles, Malte Kohns Vasconcelos, Klaus Pfeffer, Tina Senff, Daniel Strelow, Jörg Timm, Andreas Walker, Tobias Wienemann |
| EPI_ISL_414507 | hCoV-19/Germany/NRW-011/2020 | Europe / Germany / North Rhine Westph | 2020-03-03 | Center of Medical Microbiology, Virology, and Hospital Hygiene, University of Duesseldorf | Center of Medical Microbiology, Virology, and Hospital Hygiene, University of Duesseldorf | Ortwin Adams, Marcel Andree, Alexander Dilthey, Torsten Feldt, Sandra Hauka, Torsten Houwaart, Björn-Erik Jensen, Detlef Kindgen-Milles, Malte Kohns Vasconcelos, Klaus Pfeffer, Tina Senff, Daniel Strelow, Jörg Timm, Andreas Walker, Tobias Wienemann |
| EPI_ISL_414508 | hCoV-19/Germany/NRW-08/2020 | Europe / Germany / North Rhine Westph | 2020-02-27 | Center of Medical Microbiology, Virology, and Hospital Hygiene, University of Duesseldorf | Center of Medical Microbiology, Virology, and Hospital Hygiene, University of Duesseldorf | Ortwin Adams, Marcel Andree, Alexander Dilthey, Torsten Feldt, Sandra Hauka, Torsten Houwaart, Björn-Erik Jensen, Detlef Kindgen-Milles, Malte Kohns Vasconcelos, Klaus Pfeffer, Tina Senff, Daniel Strelow, Jörg Timm, Andreas Walker, Tobias Wienemann |
| EPI_ISL_414509 | hCoV-19/Germany/NRW-09/2020 | Europe / Germany / North Rhine Westph | 2020-02-28 | Center of Medical Microbiology, Virology, and Hospital Hygiene, University of Duesseldorf | Center of Medical Microbiology, Virology, and Hospital Hygiene, University of Duesseldorf | Ortwin Adams, Marcel Andree, Alexander Dilthey, Torsten Feldt, Sandra Hauka, Torsten Houwaart, Björn-Erik Jensen, Detlef Kindgen-Milles, Malte Kohns Vasconcelos, Klaus Pfeffer, Tina Senff, Daniel Strelow, Jörg Timm, Andreas Walker, Tobias Wienemann |
| EPI_ISL_414510 | hCoV-19/Shanghai/SH01/2020 | Asia / China / Shanghai | 2020-02-02 | unknown | Key Laboratory of Medical Molecular Virology (MOE/NHC/CAMS) | Zhang,R., Yi,Z., Wang,Y., Teng,Z., Xu,W., Song,W., Cai,X., Sun,Z., Gu,C., Zhou,Y., Chen,H., Ye,R., Han,W., Zhu,Y., Feng,F., Fang,F., Li,C., Zhang,X., Qu,D., Fu,C., Xie,Y. and Yuan,Z. |
| EPI_ISL_414511 | hCoV-19/Japan/TKYE6182/2020 | Asia / Japan | 2020-01 | unknown | Ryota Kumagai Tokyo Metropolitan Institute of Public Health | Kumagai,R., Yoshida,I., Nagashima,M., Chiba,T. and Sadamasu,K. |
| EPI_ISL_414517 | hCoV-19/Hong Kong/VM20002907/2020 | Asia / Hong Kong | 2020-02-25 | Hong Kong Department of Health | School of Public Health, The University of Hong Kong | Dominic N.C. Tsang, Daniel K.W. Chu, Leo L.M. Poon, Malik Peiris |
| EPI_ISL_414519 | hCoV-19/Hong Kong/VM20002868/2020 | Asia / Hong Kong | 2020-02-24 | Hong Kong Department of Health | School of Public Health, The University of Hong Kong | Dominic N.C. Tsang, Daniel K.W. Chu, Leo L.M. Poon, Malik Peiris |
| EPI_ISL_414520 | hCoV-19/Germany/BavPat2/2020 | Europe / Germany / Munich | 2020-03-02 | Bundeswehr Institute of Microbiology | Bundeswehr Institute of Microbiology | Mathias C Walter, Markus H Antwerpen and Roman Wölfel |
| EPI_ISL_414521 | hCoV-19/Germany/BavPat3/2020 | Europe / Germany / Munich | 2020-03-02 | Bundeswehr Institute of Microbiology | Bundeswehr Institute of Microbiology | Mathias C Walter, Markus H Antwerpen and Roman Wölfel |
| EPI_ISL_414522 | hCoV-19/England/200990002/2020 | Europe / United Kingdom / England | 2020-02-28 | Respiratory Virus Unit, Microbiology Services Colindale, Public Health England | Respiratory Virus Unit, Microbiology Services Colindale, Public Health England | Monica Galiano, Shahjahan Miah, Angie Lackenby, Omolola Akinbami, Tiina Talts, Leena Bhaw, Richard Myers, Steven Platt, Kirstin Edwards, Jonathan Hubb, Joanna Ellis, Maria Zambon |
| EPI_ISL_414523 | hCoV-19/England/200990660/2020 | Europe / United Kingdom / England | 2020-02-27 | Respiratory Virus Unit, Microbiology Services Colindale, Public Health England | Respiratory Virus Unit, Microbiology Services Colindale, Public Health England | Monica Galiano, Shahjahan Miah, Angie Lackenby, Omolola Akinbami, Tiina Talts, Leena Bhaw, Richard Myers, Steven Platt, Kirstin Edwards, Jonathan Hubb, Joanna Ellis, Maria Zambon |
| EPI_ISL_414524 | hCoV-19/England/200991076/2020 | Europe / United Kingdom / England | 2020-03-01 | Respiratory Virus Unit, Microbiology Services Colindale, Public Health England | Respiratory Virus Unit, Microbiology Services Colindale, Public Health England | Monica Galiano, Shahjahan Miah, Angie Lackenby, Omolola Akinbami, Tiina Talts, Leena Bhaw, Richard Myers, Steven Platt, Kirstin Edwards, Jonathan Hubb, Joanna Ellis, Maria Zambon |
| EPI_ISL_414525 | hCoV-19/England/201040081/2020 | Europe / United Kingdom / England | 2020-03-02 | Respiratory Virus Unit, Microbiology Services Colindale, Public Health England | Respiratory Virus Unit, Microbiology Services Colindale, Public Health England | Monica Galiano, Shahjahan Miah, Angie Lackenby, Omolola Akinbami, Tiina Talts, Leena Bhaw, Richard Myers, Steven Platt, Kirstin Edwards, Jonathan Hubb, Joanna Ellis, Maria Zambon |
| EPI_ISL_414526 | hCoV-19/England/201040141/2020 | Europe / United Kingdom / England | 2020-03-03 | Respiratory Virus Unit, Microbiology Services Colindale, Public Health England | Respiratory Virus Unit, Microbiology Services Colindale, Public Health England | Monica Galiano, Shahjahan Miah, Angie Lackenby, Omolola Akinbami, Tiina Talts, Leena Bhaw, Richard Myers, Steven Platt, Kirstin Edwards, Jonathan Hubb, Joanna Ellis, Maria Zambon |
| EPI_ISL_414527 | hCoV-19/Hong Kong/VM20002493-2/2020 | Asia / Hong Kong | 2020-02-09 | Hong Kong Department of Health | School of Public Health, The University of Hong Kong | Dominic N.C. Tsang, Daniel K.W. Chu, Leo L.M. Poon, Malik Peiris |
| EPI_ISL_414528 | hCoV-19/Hong Kong/VM20002507-2/2020 | Asia / Hong Kong | 2020-02-10 | Hong Kong Department of Health | School of Public Health, The University of Hong Kong | Dominic N.C. Tsang, Daniel K.W. Chu, Leo L.M. Poon, Malik Peiris |
| EPI_ISL_414529 | hCoV-19/Netherlands/NoordBrabant_20/2020 | Europe / Netherlands / Noord Brabant | 2020-03-04 | Dutch COVID-19 response team | Erasmus Medical Center | David Nieuwenhuijse, Bas Oude Munnink, Reina Sikkema, Claudia Schapendonk, Irina Chestakova, Anne van der Linden, Mark Pronk, Pascal Lexmond, Corien Swaan, Manon Haverkate, Madelief Mollers, Mart Stein, Sandra Kengne Kamga Mobou, Jeroen van Kampen, Jolanda Voermans, Aura Timen, Corine GeurtsvanKessel, Annemiek van der Eijk, Richard Molenkamp, Marion Koopmans, on behalf of the Dutch national COVID-19 response team. |
| EPI_ISL_414530 | hCoV-19/Netherlands/NoordBrabant_21/2020 | Europe / Netherlands / Noord Brabant | 2020-03-04 | Dutch COVID-19 response team | Erasmus Medical Center | David Nieuwenhuijse, Bas Oude Munnink, Reina Sikkema, Claudia Schapendonk, Irina Chestakova, Anne van der Linden, Mark Pronk, Pascal Lexmond, Corien Swaan, Manon Haverkate, Madelief Mollers, Mart Stein, Sandra Kengne Kamga Mobou, Jeroen van Kampen, Jolanda Voermans, Aura Timen, Corine GeurtsvanKessel, Annemiek van der Eijk, Richard Molenkamp, Marion Koopmans, on behalf of the Dutch national COVID-19 response team. |
| EPI_ISL_414531 | hCoV-19/Netherlands/NoordBrabant_22/2020 | Europe / Netherlands / Noord Brabant | 2020-03-04 | Dutch COVID-19 response team | Erasmus Medical Center | David Nieuwenhuijse, Bas Oude Munnink, Reina Sikkema, Claudia Schapendonk, Irina Chestakova, Anne van der Linden, Mark Pronk, Pascal Lexmond, Corien Swaan, Manon Haverkate, Madelief Mollers, Mart Stein, Sandra Kengne Kamga Mobou, Jeroen van Kampen, Jolanda Voermans, Aura Timen, Corine GeurtsvanKessel, Annemiek van der Eijk, Richard Molenkamp, Marion Koopmans, on behalf of the Dutch national COVID-19 response team. |
| EPI_ISL_414532 | hCoV-19/Netherlands/NoordBrabant_23/2020 | Europe / Netherlands / Noord Brabant | 2020-03-05 | Dutch COVID-19 response team | Erasmus Medical Center | David Nieuwenhuijse, Bas Oude Munnink, Reina Sikkema, Claudia Schapendonk, Irina Chestakova, Anne van der Linden, Mark Pronk, Pascal Lexmond, Corien Swaan, Manon Haverkate, Madelief Mollers, Mart Stein, Sandra Kengne Kamga Mobou, Jeroen van Kampen, Jolanda Voermans, Aura Timen, Corine GeurtsvanKessel, Annemiek van der Eijk, Richard Molenkamp, Marion Koopmans, on behalf of the Dutch national COVID-19 response team. |
| EPI_ISL_414534 | hCoV-19/Netherlands/NoordBrabant_25/2020 | Europe / Netherlands / Noord Brabant | 2020-03-06 | Dutch COVID-19 response team | Erasmus Medical Center | David Nieuwenhuijse, Bas Oude Munnink, Reina Sikkema, Claudia Schapendonk, Irina Chestakova, Anne van der Linden, Mark Pronk, Pascal Lexmond, Corien Swaan, Manon Haverkate, Madelief Mollers, Mart Stein, Sandra Kengne Kamga Mobou, Jeroen van Kampen, Jolanda Voermans, Aura Timen, Corine GeurtsvanKessel, Annemiek van der Eijk, Richard Molenkamp, Marion Koopmans, on behalf of the Dutch national COVID-19 response team. |
| EPI_ISL_414535 | hCoV-19/Netherlands/NoordBrabant_26/2020 | Europe / Netherlands / Noord Brabant | 2020-03-06 | Dutch COVID-19 response team | Erasmus Medical Center | David Nieuwenhuijse, Bas Oude Munnink, Reina Sikkema, Claudia Schapendonk, Irina Chestakova, Anne van der Linden, Mark Pronk, Pascal Lexmond, Corien Swaan, Manon Haverkate, Madelief Mollers, Mart Stein, Sandra Kengne Kamga Mobou, Jeroen van Kampen, Jolanda Voermans, Aura Timen, Corine GeurtsvanKessel, Annemiek van der Eijk, Richard Molenkamp, Marion Koopmans, on behalf of the Dutch national COVID-19 response team. |
| EPI_ISL_414536 | hCoV-19/Netherlands/NoordBrabant_27/2020 | Europe / Netherlands / Noord Brabant | 2020-03-08 | Dutch COVID-19 response team | Erasmus Medical Center | David Nieuwenhuijse, Bas Oude Munnink, Reina Sikkema, Claudia Schapendonk, Irina Chestakova, Anne van der Linden, Mark Pronk, Pascal Lexmond, Corien Swaan, Manon Haverkate, Madelief Mollers, Mart Stein, Sandra Kengne Kamga Mobou, Jeroen van Kampen, Jolanda Voermans, Aura Timen, Corine GeurtsvanKessel, Annemiek van der Eijk, Richard Molenkamp, Marion Koopmans, on behalf of the Dutch national COVID-19 response team. |
| EPI_ISL_414537 | hCoV-19/Netherlands/NoordBrabant_28/2020 | Europe / Netherlands / Noord Brabant | 2020-03-08 | Dutch COVID-19 response team | Erasmus Medical Center | David Nieuwenhuijse, Bas Oude Munnink, Reina Sikkema, Claudia Schapendonk, Irina Chestakova, Anne van der Linden, Mark Pronk, Pascal Lexmond, Corien Swaan, Manon Haverkate, Madelief Mollers, Mart Stein, Sandra Kengne Kamga Mobou, Jeroen van Kampen, Jolanda Voermans, Aura Timen, Corine GeurtsvanKessel, Annemiek van der Eijk, Richard Molenkamp, Marion Koopmans, on behalf of the Dutch national COVID-19 response team. |
| EPI_ISL_414539 | hCoV-19/Netherlands/NoordBrabant_30/2020 | Europe / Netherlands / Noord Brabant | 2020-03-08 | Dutch COVID-19 response team | Erasmus Medical Center | David Nieuwenhuijse, Bas Oude Munnink, Reina Sikkema, Claudia Schapendonk, Irina Chestakova, Anne van der Linden, Mark Pronk, Pascal Lexmond, Corien Swaan, Manon Haverkate, Madelief Mollers, Mart Stein, Sandra Kengne Kamga Mobou, Jeroen van Kampen, Jolanda Voermans, Aura Timen, Corine GeurtsvanKessel, Annemiek van der Eijk, Richard Molenkamp, Marion Koopmans, on behalf of the Dutch national COVID-19 response team. |
| EPI_ISL_414541 | hCoV-19/Netherlands/NoordBrabant_32/2020 | Europe / Netherlands / Noord Brabant | 2020-03-08 | Dutch COVID-19 response team | Erasmus Medical Center | David Nieuwenhuijse, Bas Oude Munnink, Reina Sikkema, Claudia Schapendonk, Irina Chestakova, Anne van der Linden, Mark Pronk, Pascal Lexmond, Corien Swaan, Manon Haverkate, Madelief Mollers, Mart Stein, Sandra Kengne Kamga Mobou, Jeroen van Kampen, Jolanda Voermans, Aura Timen, Corine GeurtsvanKessel, Annemiek van der Eijk, Richard Molenkamp, Marion Koopmans, on behalf of the Dutch national COVID-19 response team. |
| EPI_ISL_414542 | hCoV-19/Netherlands/NoordBrabant_33/2020 | Europe / Netherlands / Noord Brabant | 2020-03 | Dutch COVID-19 response team | Erasmus Medical Center | David Nieuwenhuijse, Bas Oude Munnink, Reina Sikkema, Claudia Schapendonk, Irina Chestakova, Anne van der Linden, Mark Pronk, Pascal Lexmond, Corien Swaan, Manon Haverkate, Madelief Mollers, Mart Stein, Sandra Kengne Kamga Mobou, Jeroen van Kampen, Jolanda Voermans, Aura Timen, Corine GeurtsvanKessel, Annemiek van der Eijk, Richard Molenkamp, Marion Koopmans, on behalf of the Dutch national COVID-19 response team. |
| EPI_ISL_414543 | hCoV-19/Netherlands/NoordBrabant_34/2020 | Europe / Netherlands / Noord Brabant | 2020-03 | Dutch COVID-19 response team | Erasmus Medical Center | David Nieuwenhuijse, Bas Oude Munnink, Reina Sikkema, Claudia Schapendonk, Irina Chestakova, Anne van der Linden, Mark Pronk, Pascal Lexmond, Corien Swaan, Manon Haverkate, Madelief Mollers, Mart Stein, Sandra Kengne Kamga Mobou, Jeroen van Kampen, Jolanda Voermans, Aura Timen, Corine GeurtsvanKessel, Annemiek van der Eijk, Richard Molenkamp, Marion Koopmans, on behalf of the Dutch national COVID-19 response team. |
| EPI_ISL_414544 | hCoV-19/Netherlands/NoordBrabant_35/2020 | Europe / Netherlands / Noord Brabant | 2020-03-09 | Dutch COVID-19 response team | Erasmus Medical Center | David Nieuwenhuijse, Bas Oude Munnink, Reina Sikkema, Claudia Schapendonk, Irina Chestakova, Anne van der Linden, Mark Pronk, Pascal Lexmond, Corien Swaan, Manon Haverkate, Madelief Mollers, Mart Stein, Sandra Kengne Kamga Mobou, Jeroen van Kampen, Jolanda Voermans, Aura Timen, Corine GeurtsvanKessel, Annemiek van der Eijk, Richard Molenkamp, Marion Koopmans, on behalf of the Dutch national COVID-19 response team. |
| EPI_ISL_414545 | hCoV-19/Netherlands/NoordBrabant_36/2020 | Europe / Netherlands / Noord Brabant | 2020-03-03 | Dutch COVID-19 response team | Erasmus Medical Center | David Nieuwenhuijse, Bas Oude Munnink, Reina Sikkema, Claudia Schapendonk, Irina Chestakova, Anne van der Linden, Mark Pronk, Pascal Lexmond, Corien Swaan, Manon Haverkate, Madelief Mollers, Mart Stein, Sandra Kengne Kamga Mobou, Jeroen van Kampen, Jolanda Voermans, Aura Timen, Corine GeurtsvanKessel, Annemiek van der Eijk, Richard Molenkamp, Marion Koopmans, on behalf of the Dutch national COVID-19 response team. |
| EPI_ISL_414546 | hCoV-19/Netherlands/NoordBrabant_37/2020 | Europe / Netherlands / Noord Brabant | 2020-03-06 | Dutch COVID-19 response team | Erasmus Medical Center | David Nieuwenhuijse, Bas Oude Munnink, Reina Sikkema, Claudia Schapendonk, Irina Chestakova, Anne van der Linden, Mark Pronk, Pascal Lexmond, Corien Swaan, Manon Haverkate, Madelief Mollers, Mart Stein, Sandra Kengne Kamga Mobou, Jeroen van Kampen, Jolanda Voermans, Aura Timen, Corine GeurtsvanKessel, Annemiek van der Eijk, Richard Molenkamp, Marion Koopmans, on behalf of the Dutch national COVID-19 response team. |
| EPI_ISL_414547 | hCoV-19/Netherlands/NoordBrabant_38/2020 | Europe / Netherlands / Noord Brabant | 2020-03-06 | Dutch COVID-19 response team | Erasmus Medical Center | David Nieuwenhuijse, Bas Oude Munnink, Reina Sikkema, Claudia Schapendonk, Irina Chestakova, Anne van der Linden, Mark Pronk, Pascal Lexmond, Corien Swaan, Manon Haverkate, Madelief Mollers, Mart Stein, Sandra Kengne Kamga Mobou, Jeroen van Kampen, Jolanda Voermans, Aura Timen, Corine GeurtsvanKessel, Annemiek van der Eijk, Richard Molenkamp, Marion Koopmans, on behalf of the Dutch national COVID-19 response team. |
| EPI_ISL_414548 | hCoV-19/Netherlands/NoordBrabant_39/2020 | Europe / Netherlands / Noord Brabant | 2020-03-04 | Dutch COVID-19 response team | Erasmus Medical Center | David Nieuwenhuijse, Bas Oude Munnink, Reina Sikkema, Claudia Schapendonk, Irina Chestakova, Anne van der Linden, Mark Pronk, Pascal Lexmond, Corien Swaan, Manon Haverkate, Madelief Mollers, Mart Stein, Sandra Kengne Kamga Mobou, Jeroen van Kampen, Jolanda Voermans, Aura Timen, Corine GeurtsvanKessel, Annemiek van der Eijk, Richard Molenkamp, Marion Koopmans, on behalf of the Dutch national COVID-19 response team. |
| EPI_ISL_414549 | hCoV-19/Netherlands/NoordHolland_2/2020 | Europe / Netherlands / Noord Holland | 2020-03-03 | Dutch COVID-19 response team | Erasmus Medical Center | David Nieuwenhuijse, Bas Oude Munnink, Reina Sikkema, Claudia Schapendonk, Irina Chestakova, Anne van der Linden, Mark Pronk, Pascal Lexmond, Corien Swaan, Manon Haverkate, Madelief Mollers, Mart Stein, Sandra Kengne Kamga Mobou, Jeroen van Kampen, Jolanda Voermans, Aura Timen, Corine GeurtsvanKessel, Annemiek van der Eijk, Richard Molenkamp, Marion Koopmans, on behalf of the Dutch national COVID-19 response team. |
| EPI_ISL_414555 | hCoV-19/Netherlands/Utrecht_16/2020 | Europe / Netherlands / Utrecht | 2020-03-08 | Dutch COVID-19 response team | Erasmus Medical Center | David Nieuwenhuijse, Bas Oude Munnink, Reina Sikkema, Claudia Schapendonk, Irina Chestakova, Anne van der Linden, Mark Pronk, Pascal Lexmond, Corien Swaan, Manon Haverkate, Madelief Mollers, Mart Stein, Sandra Kengne Kamga Mobou, Jeroen van Kampen, Jolanda Voermans, Aura Timen, Corine GeurtsvanKessel, Annemiek van der Eijk, Richard Molenkamp, Marion Koopmans, on behalf of the Dutch national COVID-19 response team. |
| EPI_ISL_414557 | hCoV-19/Netherlands/ZuidHolland_15/2020 | Europe / Netherlands / Zuid Holland | 2020-03-08 | Dutch COVID-19 response team | Erasmus Medical Center | David Nieuwenhuijse, Bas Oude Munnink, Reina Sikkema, Claudia Schapendonk, Irina Chestakova, Anne van der Linden, Mark Pronk, Pascal Lexmond, Corien Swaan, Manon Haverkate, Madelief Mollers, Mart Stein, Sandra Kengne Kamga Mobou, Jeroen van Kampen, Jolanda Voermans, Aura Timen, Corine GeurtsvanKessel, Annemiek van der Eijk, Richard Molenkamp, Marion Koopmans, on behalf of the Dutch national COVID-19 response team. |
| EPI_ISL_414558 | hCoV-19/Netherlands/ZuidHolland_16/2020 | Europe / Netherlands / Zuid Holland | 2020-03-06 | Dutch COVID-19 response team | Erasmus Medical Center | David Nieuwenhuijse, Bas Oude Munnink, Reina Sikkema, Claudia Schapendonk, Irina Chestakova, Anne van der Linden, Mark Pronk, Pascal Lexmond, Corien Swaan, Manon Haverkate, Madelief Mollers, Mart Stein, Sandra Kengne Kamga Mobou, Jeroen van Kampen, Jolanda Voermans, Aura Timen, Corine GeurtsvanKessel, Annemiek van der Eijk, Richard Molenkamp, Marion Koopmans, on behalf of the Dutch national COVID-19 response team. |
| EPI_ISL_414559 | hCoV-19/Netherlands/ZuidHolland_17/2020 | Europe / Netherlands / Zuid Holland | 2020-03-07 | Dutch COVID-19 response team | Erasmus Medical Center | David Nieuwenhuijse, Bas Oude Munnink, Reina Sikkema, Claudia Schapendonk, Irina Chestakova, Anne van der Linden, Mark Pronk, Pascal Lexmond, Corien Swaan, Manon Haverkate, Madelief Mollers, Mart Stein, Sandra Kengne Kamga Mobou, Jeroen van Kampen, Jolanda Voermans, Aura Timen, Corine GeurtsvanKessel, Annemiek van der Eijk, Richard Molenkamp, Marion Koopmans, on behalf of the Dutch national COVID-19 response team. |
| EPI_ISL_414560 | hCoV-19/Netherlands/ZuidHolland_18/2020 | Europe / Netherlands / Zuid Holland | 2020-03-03 | Dutch COVID-19 response team | Erasmus Medical Center | David Nieuwenhuijse, Bas Oude Munnink, Reina Sikkema, Claudia Schapendonk, Irina Chestakova, Anne van der Linden, Mark Pronk, Pascal Lexmond, Corien Swaan, Manon Haverkate, Madelief Mollers, Mart Stein, Sandra Kengne Kamga Mobou, Jeroen van Kampen, Jolanda Voermans, Aura Timen, Corine GeurtsvanKessel, Annemiek van der Eijk, Richard Molenkamp, Marion Koopmans, on behalf of the Dutch national COVID-19 response team. |
| EPI_ISL_414561 | hCoV-19/Netherlands/ZuidHolland_19/2020 | Europe / Netherlands / Zuid Holland | 2020-03-05 | Dutch COVID-19 response team | Erasmus Medical Center | David Nieuwenhuijse, Bas Oude Munnink, Reina Sikkema, Claudia Schapendonk, Irina Chestakova, Anne van der Linden, Mark Pronk, Pascal Lexmond, Corien Swaan, Manon Haverkate, Madelief Mollers, Mart Stein, Sandra Kengne Kamga Mobou, Jeroen van Kampen, Jolanda Voermans, Aura Timen, Corine GeurtsvanKessel, Annemiek van der Eijk, Richard Molenkamp, Marion Koopmans, on behalf of the Dutch national COVID-19 response team. |
| EPI_ISL_414562 | hCoV-19/Netherlands/ZuidHolland_20/2020 | Europe / Netherlands / Zuid Holland | 2020-03-03 | Dutch COVID-19 response team | Erasmus Medical Center | David Nieuwenhuijse, Bas Oude Munnink, Reina Sikkema, Claudia Schapendonk, Irina Chestakova, Anne van der Linden, Mark Pronk, Pascal Lexmond, Corien Swaan, Manon Haverkate, Madelief Mollers, Mart Stein, Sandra Kengne Kamga Mobou, Jeroen van Kampen, Jolanda Voermans, Aura Timen, Corine GeurtsvanKessel, Annemiek van der Eijk, Richard Molenkamp, Marion Koopmans, on behalf of the Dutch national COVID-19 response team. |
| EPI_ISL_414563 | hCoV-19/Netherlands/ZuidHolland_21/2020 | Europe / Netherlands / Zuid Holland | 2020-03-03 | Dutch COVID-19 response team | Erasmus Medical Center | David Nieuwenhuijse, Bas Oude Munnink, Reina Sikkema, Claudia Schapendonk, Irina Chestakova, Anne van der Linden, Mark Pronk, Pascal Lexmond, Corien Swaan, Manon Haverkate, Madelief Mollers, Mart Stein, Sandra Kengne Kamga Mobou, Jeroen van Kampen, Jolanda Voermans, Aura Timen, Corine GeurtsvanKessel, Annemiek van der Eijk, Richard Molenkamp, Marion Koopmans, on behalf of the Dutch national COVID-19 response team. |
| EPI_ISL_414564 | hCoV-19/Netherlands/ZuidHolland_22/2020 | Europe / Netherlands / Zuid Holland | 2020-03-08 | Dutch COVID-19 response team | Erasmus Medical Center | David Nieuwenhuijse, Bas Oude Munnink, Reina Sikkema, Claudia Schapendonk, Irina Chestakova, Anne van der Linden, Mark Pronk, Pascal Lexmond, Corien Swaan, Manon Haverkate, Madelief Mollers, Mart Stein, Sandra Kengne Kamga Mobou, Jeroen van Kampen, Jolanda Voermans, Aura Timen, Corine GeurtsvanKessel, Annemiek van der Eijk, Richard Molenkamp, Marion Koopmans, on behalf of the Dutch national COVID-19 response team. |
| EPI_ISL_414565 | hCoV-19/Netherlands/ZuidHolland_23/2020 | Europe / Netherlands / Zuid Holland | 2020-03-09 | Dutch COVID-19 response team | Erasmus Medical Center | David Nieuwenhuijse, Bas Oude Munnink, Reina Sikkema, Claudia Schapendonk, Irina Chestakova, Anne van der Linden, Mark Pronk, Pascal Lexmond, Corien Swaan, Manon Haverkate, Madelief Mollers, Mart Stein, Sandra Kengne Kamga Mobou, Jeroen van Kampen, Jolanda Voermans, Aura Timen, Corine GeurtsvanKessel, Annemiek van der Eijk, Richard Molenkamp, Marion Koopmans, on behalf of the Dutch national COVID-19 response team. |
| EPI_ISL_414566 | hCoV-19/Netherlands/ZuidHolland_24/2020 | Europe / Netherlands / Zuid Holland | 2020-03-06 | Dutch COVID-19 response team | Erasmus Medical Center | David Nieuwenhuijse, Bas Oude Munnink, Reina Sikkema, Claudia Schapendonk, Irina Chestakova, Anne van der Linden, Mark Pronk, Pascal Lexmond, Corien Swaan, Manon Haverkate, Madelief Mollers, Mart Stein, Sandra Kengne Kamga Mobou, Jeroen van Kampen, Jolanda Voermans, Aura Timen, Corine GeurtsvanKessel, Annemiek van der Eijk, Richard Molenkamp, Marion Koopmans, on behalf of the Dutch national COVID-19 response team. |
| EPI_ISL_414569 | hCoV-19/Hong Kong/VM20002582-2/2020 | Asia / Hong Kong | 2020-02-12 | Hong Kong Department of Health | School of Public Health, The University of Hong Kong | Dominic N.C. Tsang, Daniel K.W. Chu, Leo L.M. Poon, Malik Peiris |
| EPI_ISL_414571 | hCoV-19/Hong Kong/VM20002849/2020 | Asia / Hong Kong | 2020-02-22 | Hong Kong Department of Health | School of Public Health, The University of Hong Kong | Dominic N.C. Tsang, Daniel K.W. Chu, Leo L.M. Poon, Malik Peiris |
| EPI_ISL_414574 | hCoV-19/Germany/NRW-10/2020 | Europe / Germany / North Rhine Westph | 2020-02-28 | Center of Medical Microbiology, Virology, and Hospital Hygiene, University of Duesseldorf | Center of Medical Microbiology, Virology, and Hospital Hygiene, University of Duesseldorf | Ortwin Adams, Marcel Andree, Alexander Dilthey, Torsten Feldt, Sandra Hauka, Torsten Houwaart, Björn-Erik Jensen, Detlef Kindgen-Milles, Malte Kohns Vasconcelos, Klaus Pfeffer, Tina Senff, Daniel Strelow, Jörg Timm, Andreas Walker, Tobias Wienemann |
| EPI_ISL_414577 | hCoV-19/Chile/Talca-1/2020 | South America / Chile / Talca | 2020-03-02 | Hospital de Talca, Chile | Instituto de Salud Publica de Chile | Andrés E. Castillo, Bárbara Parra, Paz Tapia, Alejandra Acevedo, Jaime Lagos, Winston Andrade, Loredana Arata, Gabriel Leal, Gisselle Barra, Carolina Tambley, Javier Tognarelli, Patricia Bustos, Soledad Ulloa, Rodrigo Fasce, Jorge Fernández. |
| EPI_ISL_414578 | hCoV-19/Chile/Talca-2/2020 | South America / Chile / Talca | 2020-03-04 | Hospital de Talca, Chile | Instituto de Salud Publica de Chile | Andrés E. Castillo, Bárbara Parra, Paz Tapia, Alejandra Acevedo, Jaime Lagos, Winston Andrade, Loredana Arata, Gabriel Leal, Gisselle Barra, Carolina Tambley, Javier Tognarelli, Patricia Bustos, Soledad Ulloa, Rodrigo Fasce, Jorge Fernández. |
| EPI_ISL_414579 | hCoV-19/Chile/Santiago-1/2020 | South America / Chile / Santiago | 2020-03-03 | Clinica Alemana de Santiago, Chile | Instituto de Salud Publica de Chile | Andrés E. Castillo, Bárbara Parra, Paz Tapia, Alejandra Acevedo, Jaime Lagos, Winston Andrade, Loredana Arata, Gabriel Leal, Gisselle Barra, Carolina Tambley, Javier Tognarelli, Patricia Bustos, Soledad Ulloa, Rodrigo Fasce, Jorge Fernández. |
| EPI_ISL_414580 | hCoV-19/Chile/Santiago-2/2020 | South America / Chile / Santiago | 2020-03-05 | Clinica Santa Maria, Santiago, Chile | Instituto de Salud Publica de Chile | Andrés E. Castillo, Bárbara Parra, Paz Tapia, Alejandra Acevedo, Jaime Lagos, Winston Andrade, Loredana Arata, Gabriel Leal, Gisselle Barra, Carolina Tambley, Javier Tognarelli, Patricia Bustos, Soledad Ulloa, Rodrigo Fasce, Jorge Fernández. |
| EPI_ISL_414586 | hCoV-19/Ireland/Limerick-19934/2020 | Europe / Ireland / Limerick | 2020-03-03 | UCD National Virus Reference Laboratory | UCD National Virus Reference Laboratory | Michael Carr, Gabriel Gonzalez, Jonathan Dean, Suzie Coughlan, Alison Murphy, Kevin Byrne, Ken Wolfe, Jeff Connell, Brendan Loftus, Cillian F De Gascun |
| EPI_ISL_414587 | hCoV-19/Ireland/Limerick-19935/2020 | Europe / Ireland / Limerick | 2020-03-03 | UCD National Virus Reference Laboratory | UCD National Virus Reference Laboratory | Michael Carr, Gabriel Gonzalez, Jonathan Dean, Suzie Coughlan, Alison Murphy, Kevin Byrne, Ken Wolfe, Jeff Connell, Brendan Loftus, Cillian F De Gascun |
| EPI_ISL_414589 | hCoV-19/USA/MN2-MDH2/2020 | North America / USA / Minnesota | 2020-03-07 | Minnesota Department of Health, Public Health Laboratory | Minnesota Department of Health, Public Health Laboratory | Matt Plumb, Jake Garfin and Xiong Wang |
| EPI_ISL_414590 | hCoV-19/USA/MN3-MDH3/2020 | North America / USA / Minnesota | 2020-03-09 | Minnesota Department of Health, Public Health Laboratory | Minnesota Department of Health, Public Health Laboratory | Matt Plumb, Jake Garfin and Xiong Wang |
| EPI_ISL_414591 | hCoV-19/USA/WA-UW22/2020 | North America / USA / Washington / Kirkl | 2020-03-06 | UW Virology Lab | UW Virology Lab | Pavitra Roychoudhury, Hong Xie, Keith Jerome, Alexander Greninger |
| EPI_ISL_414592 | hCoV-19/USA/WA-UW23/2020 | North America / USA / Washington / Taco | 2020-03-06 | UW Virology Lab | UW Virology Lab | Pavitra Roychoudhury, Hong Xie, Keith Jerome, Alexander Greninger |
| EPI_ISL_414593 | hCoV-19/USA/WA-UW24/2020 | North America / USA / Washington / Kirkl | 2020-03-05 | UW Virology Lab | UW Virology Lab | Pavitra Roychoudhury, Hong Xie, Keith Jerome, Alexander Greninger |

| | | | | | | |
|---|---|---|---|---|---|---|
| EPI_ISL_414594 | hCoV-19/USA/WA-UW25/2020 | North America / USA / Washington / Kirkl | 2020-03-05 | UW Virology Lab | UW Virology Lab | Pavitra Roychoudhury, Hong Xie, Keith Jerome, Alexander Greninger |
| EPI_ISL_414595 | hCoV-19/USA/WA-UW26/2020 | North America / USA / Washington / Kirkl | 2020-03-05 | UW Virology Lab | UW Virology Lab | Pavitra Roychoudhury, Hong Xie, Keith Jerome, Alexander Greninger |
| EPI_ISL_414596 | hCoV-19/USA/WA-UW27/2020 | North America / USA / Washington / Kirkl | 2020-03-04 | UW Virology Lab | UW Virology Lab | Pavitra Roychoudhury, Hong Xie, Keith Jerome, Alexander Greninger |
| EPI_ISL_414597 | hCoV-19/USA/WA-UW28/2020 | North America / USA / Washington / Kirkl | 2020-03-04 | UW Virology Lab | UW Virology Lab | Pavitra Roychoudhury, Hong Xie, Keith Jerome, Alexander Greninger |
| EPI_ISL_414598 | hCoV-19/Spain/Valencia3/2020 | Europe / Spain / Comunitat Valenciana / ' | 2020-03-05 | Servicio Microbiologia, Hospital Clinico Universitario, Valencia | Sequencing and Bioinformatics Service and Molecular Epidemiology Research Group. FISABIO-Public Health. | David Navarro, Maria Alma Bracho, Giuseppe D'Auria, Griselda De Marco, Neris Garcia-Gonzalez, Fernando Gonzalez-Candelas |
| EPI_ISL_414600 | hCoV-19/France/GE1583/2020 | Europe / France / Grand-Est / Strasbourg | 2020-02-26 | Laboratoire de Virologie Institut de Virologie - INSERM U 1109 Hôpitaux Universitaires de Strasbourg | National Reference Center for Viruses of Respiratory Infections, Institut Pasteur, Paris | Mélnie Albert, Marion Barbet, Sylvie Behillil, Méline Bizard, Angela Brisebarre, Flora Donati Vincent Enouf, Maud Vanpeene, Sylvie van der Werf, Samira Fafi-Kremer |
| EPI_ISL_414616 | hCoV-19/USA/WA-UW29/2020 | North America / USA / Washington / Kirkl | 2020-03-08 | UW Virology Lab | UW Virology Lab | Pavitra Roychoudhury, Hong Xie, Keith Jerome, Alexander Greninger |
| EPI_ISL_414617 | hCoV-19/USA/WA-UW30/2020 | North America / USA | 2020-03-08 | UW Virology Lab | UW Virology Lab | Pavitra Roychoudhury, Hong Xie, Keith Jerome, Alexander Greninger |
| EPI_ISL_414618 | hCoV-19/USA/WA-UW31/2020 | North America / USA | 2020-03-08 | UW Virology Lab | UW Virology Lab | Pavitra Roychoudhury, Hong Xie, Keith Jerome, Alexander Greninger |
| EPI_ISL_414619 | hCoV-19/USA/WA-UW32/2020 | North America / USA | 2020-03-07 | UW Virology Lab | UW Virology Lab | Pavitra Roychoudhury, Hong Xie, Keith Jerome, Alexander Greninger |
| EPI_ISL_414620 | hCoV-19/USA/WA-UW33/2020 | North America / USA | 2020-03-08 | UW Virology Lab | UW Virology Lab | Pavitra Roychoudhury, Hong Xie, Keith Jerome, Alexander Greninger |
| EPI_ISL_414621 | hCoV-19/USA/WA-UW34/2020 | North America / USA / Washington | 2020-03-07 | UW Virology Lab | UW Virology Lab | Pavitra Roychoudhury, Hong Xie, Keith Jerome, Alexander Greninger |
| EPI_ISL_414622 | hCoV-19/USA/WA-UW35/2020 | North America / USA / Washington / Seat | 2020-03-08 | UW Virology Lab | UW Virology Lab | Pavitra Roychoudhury, Hong Xie, Keith Jerome, Alexander Greninger |
| EPI_ISL_414623 | hCoV-19/France/GE1583/2020 | Europe / France / Grand-Est / Strasbourg | 2020-02-25 | Laboratoire de Virologie Institut de Virologie - INSERM U 1109 Hôpitaux Universitaires de Strasbourg | National Reference Center for Viruses of Respiratory Infections, Institut Pasteur, Paris | Mélnie Albert, Marion Barbet, Sylvie Behillil, Méline Bizard, Angela Brisebarre, Flora Donati Vincent Enouf, Maud Vanpeene, Sylvie van der Werf, Samira Fafi-Kremer |
| EPI_ISL_414624 | hCoV-19/France/N1620/2020 | Europe / France / Normandie / Rouen | 2020-02-26 | Centre Hositalier Universitaire de Rouen Laboratoire de Virologie | National Reference Center for Viruses of Respiratory Infections, Institut Pasteur, Paris | Mélnie Albert, Marion Barbet, Sylvie Behillil, Méline Bizard, Angela Brisebarre, Flora Donati Vincent Enouf, Maud Vanpeene, Sylvie van der Werf, Jean-Christophe Plantier |
| EPI_ISL_414625 | hCoV-19/France/PL1643/2020 | Europe / France / Pays de la Loire / Nant | 2020-02-26 | Centre Hospitalier Régional Universitaire de Nantes Laboratoire de Virologie | National Reference Center for Viruses of Respiratory Infections, Institut Pasteur, Paris | Mélnie Albert, Marion Barbet, Sylvie Behillil, Méline Bizard, Angela Brisebarre, Flora Donati Vincent Enouf, Maud Vanpeene, Sylvie van der Werf, Marianne Coste-Burel |
| EPI_ISL_414626 | hCoV-19/France/HF1684/2020 | Europe / France / Hauts de France / Crép | 2020-02-29 | unknown | National Reference Center for Viruses of Respiratory Infections, Institut Pasteur, Paris | Mélnie Albert, Marion Barbet, Sylvie Behillil, Méline Bizard, Angela Brisebarre, Flora Donati Vincent Enouf, Maud Vanpeene, Sylvie van der Werf |
| EPI_ISL_414627 | hCoV-19/France/HF1795/2020 | Europe / France / Hauts de France / Com | 2020-03-02 | Centre Hospitalier Compiègne Laboratoire de Biologie | National Reference Center for Viruses of Respiratory Infections, Institut Pasteur, Paris | Mélnie Albert, Marion Barbet, Sylvie Behillil, Méline Bizard, Angela Brisebarre, Flora Donati Vincent Enouf, Maud Vanpeene, Sylvie van der Werf, Raulin Olivia |
| EPI_ISL_414628 | hCoV-19/France/HF1805/2020 | Europe / France / Hauts de France / Com | 2020-03-02 | Centre Hospitalier Compiègne Laboratoire de Biologie | National Reference Center for Viruses of Respiratory Infections, Institut Pasteur, Paris | Mélnie Albert, Marion Barbet, Sylvie Behillil, Méline Bizard, Angela Brisebarre, Flora Donati Vincent Enouf, Maud Vanpeene, Sylvie van der Werf, Raulin Olivia |
| EPI_ISL_414629 | hCoV-19/France/HF1870/2020 | Europe / France / Hauts de France / Com | 2020-03-03 | Centre Hospitalier Compiègne Laboratoire de Biologie | National Reference Center for Viruses of Respiratory Infections, Institut Pasteur, Paris | Mélnie Albert, Marion Barbet, Sylvie Behillil, Méline Bizard, Angela Brisebarre, Flora Donati Vincent Enouf, Maud Vanpeene, Sylvie van der Werf, Raulin Olivia |
| EPI_ISL_414630 | hCoV-19/France/HF1871/2020 | Europe / France / Hauts de France / Com | 2020-03-03 | Centre Hospitalier Compiègne Laboratoire de Biologie | National Reference Center for Viruses of Respiratory Infections, Institut Pasteur, Paris | Mélnie Albert, Marion Barbet, Sylvie Behillil, Méline Bizard, Angela Brisebarre, Flora Donati Vincent Enouf, Maud Vanpeene, Sylvie van der Werf, Raulin Olivia |
| EPI_ISL_414631 | hCoV-19/France/GE1973/2020 | Europe / France / Grand-Est / Reims | 2020-03-04 | Hôpital Robert Debré Laboratoire de Virologie | National Reference Center for Viruses of Respiratory Infections, Institut Pasteur, Paris | Mélnie Albert, Marion Barbet, Sylvie Behillil, Méline Bizard, Angela Brisebarre, Flora Donati Vincent Enouf, Maud Vanpeene, Sylvie van der Werf, Laurent Andreoletti |
| EPI_ISL_414632 | hCoV-19/France/GE1977/2020 | Europe / France / Grand-Est / Reims | 2020-03-04 | Hôpital Robert Debré Laboratoire de Virologie | National Reference Center for Viruses of Respiratory Infections, Institut Pasteur, Paris | Mélnie Albert, Marion Barbet, Sylvie Behillil, Méline Bizard, Angela Brisebarre, Flora Donati Vincent Enouf, Maud Vanpeene, Sylvie van der Werf, Laurent Andreoletti |
| EPI_ISL_414633 | hCoV-19/France/IDF1980/2020 | Europe / France / Ile-de-France / Pontois | 2020-03-04 | Centre Hospitalier René Dubois Laboratoire de Microbiologie - Bât A | National Reference Center for Viruses of Respiratory Infections, Institut Pasteur, Paris | Mélnie Albert, Marion Barbet, Sylvie Behillil, Méline Bizard, Angela Brisebarre, Flora Donati Vincent Enouf, Maud Vanpeene, Sylvie van der Werf, Pascale Martres |
| EPI_ISL_414634 | hCoV-19/France/HF1986/2020 | Europe / France / Hauts de France / Com | 2020-03-04 | Centre Hospitalier Compiègne Laboratoire de Biologie | National Reference Center for Viruses of Respiratory Infections, Institut Pasteur, Paris | Mélnie Albert, Marion Barbet, Sylvie Behillil, Méline Bizard, Angela Brisebarre, Flora Donati Vincent Enouf, Maud Vanpeene, Sylvie van der Werf, Raulin Olivia |
| EPI_ISL_414635 | hCoV-19/France/HF1988/2020 | Europe / France / Hauts de France / Com | 2020-03-04 | Centre Hospitalier Compiègne Laboratoire de Biologie | National Reference Center for Viruses of Respiratory Infections, Institut Pasteur, Paris | Mélnie Albert, Marion Barbet, Sylvie Behillil, Méline Bizard, Angela Brisebarre, Flora Donati Vincent Enouf, Maud Vanpeene, Sylvie van der Werf, Raulin Olivia |
| EPI_ISL_414636 | hCoV-19/France/HF1989/2020 | Europe / France / Hauts de France / Com | 2020-03-04 | Centre Hospitalier Compiègne Laboratoire de Biologie | National Reference Center for Viruses of Respiratory Infections, Institut Pasteur, Paris | Mélnie Albert, Marion Barbet, Sylvie Behillil, Méline Bizard, Angela Brisebarre, Flora Donati Vincent Enouf, Maud Vanpeene, Sylvie van der Werf, Raulin Olivia |
| EPI_ISL_414637 | hCoV-19/France/HF1993/2020 | Europe / France / Hauts de France / Com | 2020-03-04 | Centre Hospitalier Compiègne Laboratoire de Biologie | National Reference Center for Viruses of Respiratory Infections, Institut Pasteur, Paris | Mélnie Albert, Marion Barbet, Sylvie Behillil, Méline Bizard, Angela Brisebarre, Flora Donati Vincent Enouf, Maud Vanpeene, Sylvie van der Werf, Raulin Olivia |
| EPI_ISL_414638 | hCoV-19/France/HF1995/2020 | Europe / France / Hauts de France / Com | 2020-03-04 | Centre Hospitalier Compiègne Laboratoire de Biologie | National Reference Center for Viruses of Respiratory Infections, Institut Pasteur, Paris | Mélnie Albert, Marion Barbet, Sylvie Behillil, Méline Bizard, Angela Brisebarre, Flora Donati Vincent Enouf, Maud Vanpeene, Sylvie van der Werf, Raulin Olivia |
| EPI_ISL_414639 | hCoV-19/USA/NY-NYUMC1/2020 | North America / USA / New York / Nassa | 2020-03-04 | NYU Langone Health | Departments of Pathology and Medicine, New York University School of Medicine | John Chen, Dacia Dimartino, Xiaojun Feng, Adriana Heguy, Megan Hogan, Emily Huang, George Jour, Christian Marier, Matthew T. Maurano, Mark J. Mulligan, Peter Meyn, Marie Samanovic-Golden, Amy Rapkiewicz, Guomiao Shen, Matija Snuderl, Gael Westby, Paul Zappile |
| EPI_ISL_414641 | hCoV-19/Finland/FIN-313/2020 | Europe / Finland | 2020-03-05 | Department of Virology and Immunology, University of Helsinki and Helsinki University Hospital, Huslab Finland | Department of Virology, Faculty of Medicine, University of Helsinki, Helsinki, Finland | Teemu Smura, Hannimari Kallio-Kokko, Olli Vapalahti |
| EPI_ISL_414642 | hCoV-19/Finland/FIN-455/2020 | Europe / Finland | 2020-03-08 | Department of Virology and Immunology, University of Helsinki and Helsinki University Hospital, Huslab Finland | Department of Virology, Faculty of Medicine, University of Helsinki, Helsinki, Finland | Teemu Smura, Hannimari Kallio-Kokko, Olli Vapalahti |
| EPI_ISL_414643 | hCoV-19/Finland/FIN-508/2020 | Europe / Finland | 2020-03-07 | Department of Virology and Immunology, University of Helsinki and Helsinki University Hospital, Huslab Finland | Department of Virology, Faculty of Medicine, University of Helsinki, Helsinki, Finland | Teemu Smura, Hannimari Kallio-Kokko, Olli Vapalahti |
| EPI_ISL_414646 | hCoV-19/Finland/FIN-266/2020 | Europe / Finland | 2020-03-04 | Department of Virology and Immunology, University of Helsinki and Helsinki University Hospital, Huslab Finland | Department of Virology, Faculty of Medicine, University of Helsinki, Helsinki, Finland | Teemu Smura, Hannimari Kallio-Kokko, Olli Vapalahti |
| EPI_ISL_414647 | hCoV-19/DRC/KN-13/2020 | Africa / Democratic Republic of the Cong | 2020-03-09 | Viral Respiratory Lab, National Institute for Biomedical Research (INRB) | Pathogen Sequencing Lab, National Institute for Biomedical Research (INRB) | Placide Mbala-Kingebeni, Edith Nkwembe, Eddy Kinganda-Lusamaki, Amuri Aziza, Catherine Pratt, Matthias Pauthner, Josh Quick, Allison Black, James Hadfield, Trevor Bedford, Ian Goodfellow, Nick Loman, Kristian Andersen, Michael Wiley, Steve Ahuka-Mundeke, Jean-Jacques Muyembe Tamfum |
| EPI_ISL_414648 | hCoV-19/USA/CA-PC101P/2020 | North America / USA / California / San Di | 2020-03-11 | Andersen Lab, The Scripps Research Institute | Andersen Lab, The Scripps Research Institute | Mark Zeller, Catie Anderson, Emily Spender, Sarah Topol, Raphaelle Klitting, Refugio Robles-Sikisaka, Karthik Gangavarapu, Laura Nicholson, Kristian Andersen |
| EPI_ISL_414663 | hCoV-19/Guangzhou/GZMU0016/2020 | Asia / China / Guangdong / Guangzhou | 2020-02-25 | State Key Laboratory of Respiratory Disease, National Clinical Research Center for Respiratory Disease, Guangzhou Institute of Respiratory Health, the First Affiliated Hospital of Guangzhou Medical University | The First Affiliated Hospital of Guangzhou Medical University & BGI-Shenzhen | Zhao et al |
| EPI_ISL_414686 | hCoV-19/Guangzhou/GZMU0030/2020 | Asia / China / Guangdong / Guangzhou | 2020-02-27 | State Key Laboratory of Respiratory Disease, National Clinical Research Center for Respiratory Disease, Guangzhou Institute of Respiratory Health, the First Affiliated Hospital of Guangzhou Medical University | The First Affiliated Hospital of Guangzhou Medical University & BGI-Shenzhen | Zhao et al |
| EPI_ISL_414687 | hCoV-19/Guangzhou/GZMU0031/2020 | Asia / China / Guangdong / Guangzhou | 2020-02-25 | State Key Laboratory of Respiratory Disease, National Clinical Research Center for Respiratory Disease, Guangzhou Institute of Respiratory Health, the First Affiliated Hospital of Guangzhou Medical University | the First Affiliated Hospital of Guangzhou Medical University & BGI-Shenzhen | Zhao et al |
| EPI_ISL_414689 | hCoV-19/Guangzhou/GZMU0044/2020 | Asia / China / Guangdong / Guangzhou | 2020-02-25 | State Key Laboratory of Respiratory Disease, National Clinical Research Center for Respiratory Disease, Guangzhou Institute of Respiratory Health, the First Affiliated Hospital of Guangzhou Medical University | The First Affiliated Hospital of Guangzhou Medical University & BGI-Shenzhen | Zhao et al |
| EPI_ISL_414690 | hCoV-19/Guangzhou/GZMU0047/2020 | Asia / China / Guangdong / Guangzhou | 2020-02-25 | State Key Laboratory of Respiratory Disease, National Clinical Research Center for Respiratory Disease, Guangzhou Institute of Respiratory Health, the First Affiliated Hospital of Guangzhou Medical University | The First Affiliated Hospital of Guangzhou Medical University & BGI-Shenzhen | Zhao et al |
| EPI_ISL_414691 | hCoV-19/Guangzhou/GZMU0048/2020 | Asia / China / Guangdong / Guangzhou | 2020-02-25 | State Key Laboratory of Respiratory Disease, National Clinical Research Center for Respiratory Disease, Guangzhou Institute of Respiratory Health, the First Affiliated Hospital of Guangzhou Medical University | The First Affiliated Hospital of Guangzhou Medical University & BGI-Shenzhen | Zhao et al |
| EPI_ISL_414692 | hCoV-19/Guangzhou/GZMU0014/2020 | Asia / China / Guangdong / Guangzhou | 2020-02-25 | State Key Laboratory of Respiratory Disease, National Clinical Research Center for Respiratory Disease, Guangzhou Institute of Respiratory Health, the First Affiliated Hospital of Guangzhou Medical University | The First Affiliated Hospital of Guangzhou Medical University & BGI-Shenzhen | Zhao et al |
| EPI_ISL_414934 | hCoV-19/Shandong/LY001-2/2020 | Asia / China / Shandong | 2020-01-21 | Shandong Provincial Center for Disease Control and Prevention | Beijing Institute of Microbiology and Epidemiology | Xiao-Lin Jiang, Xiao-Li Zhang, Xiang-Na Zhao, Cun-Bao Li, Jie Lei, Zeng-Qiang Kou, Wen-Kui Sun, Yang Hang, Feng Gao, Sheng-Xiang Ji, Can-Fang Lin, Bo Pang, Ming-Xiao Yao, Guo-Lin Wang, Lin Yao, Li-Jun Duan, Xiao Wei, Dian-Ming Kang, Mai-Juan Ma |
| EPI_ISL_414936 | hCoV-19/Shandong/LY003-2/2020 | Asia / China / Shandong | 2020-01-23 | Shandong Provincial Center for Disease Control and Prevention | Beijing Institute of Microbiology and Epidemiology | Xiao-Lin Jiang, Xiao-Li Zhang, Xiang-Na Zhao, Cun-Bao Li, Jie Lei, Zeng-Qiang Kou, Wen-Kui Sun, Yang Hang, Feng Gao, Sheng-Xiang Ji, Can-Fang Lin, Bo Pang, Ming-Xiao Yao, Guo-Lin Wang, Lin Yao, Li-Jun Duan, Xiao Wei, Dian-Ming Kang, Mai-Juan Ma |
| EPI_ISL_414937 | hCoV-19/Shandong/LY004-2/2020 | Asia / China / Shandong | 2020-01-26 | Shandong Provincial Center for Disease Control and Prevention | Beijing Institute of Microbiology and Epidemiology | Xiao-Lin Jiang, Xiao-Li Zhang, Xiang-Na Zhao, Cun-Bao Li, Jie Lei, Zeng-Qiang Kou, Wen-Kui Sun, Yang Hang, Feng Gao, Sheng-Xiang Ji, Can-Fang Lin, Bo Pang, Ming-Xiao Yao, Guo-Lin Wang, Lin Yao, Li-Jun Duan, Xiao Wei, Dian-Ming Kang, Mai-Juan Ma |
| EPI_ISL_414938 | hCoV-19/Shandong/LY005-2/2020 | Asia / China / Shandong | 2020-01-24 | Shandong Provincial Center for Disease Control and Prevention | Beijing Institute of Microbiology and Epidemiology | Xiao-Lin Jiang, Xiao-Li Zhang, Xiang-Na Zhao, Cun-Bao Li, Jie Lei, Zeng-Qiang Kou, Wen-Kui Sun, Yang Hang, Feng Gao, Sheng-Xiang Ji, Can-Fang Lin, Bo Pang, Ming-Xiao Yao, Guo-Lin Wang, Lin Yao, Li-Jun Duan, Xiao Wei, Dian-Ming Kang, Mai-Juan Ma |
| EPI_ISL_414940 | hCoV-19/Shandong/LY007-2/2020 | Asia / China / Shandong | 2020-01-25 | Shandong Provincial Center for Disease Control and Prevention | Beijing Institute of Microbiology and Epidemiology | Xiao-Lin Jiang, Xiao-Li Zhang, Xiang-Na Zhao, Cun-Bao Li, Jie Lei, Zeng-Qiang Kou, Wen-Kui Sun, Yang Hang, Feng Gao, Sheng-Xiang Ji, Can-Fang Lin, Bo Pang, Ming-Xiao Yao, Guo-Lin Wang, Lin Yao, Li-Jun Duan, Xiao Wei, Dian-Ming Kang, Mai-Juan Ma |
| EPI_ISL_414941 | hCoV-19/Shandong/LY008-2/2020 | Asia / China / Shandong | 2020-01-30 | Shandong Provincial Center for Disease Control and Prevention | Beijing Institute of Microbiology and Epidemiology | Xiao-Lin Jiang, Xiao-Li Zhang, Xiang-Na Zhao, Cun-Bao Li, Jie Lei, Zeng-Qiang Kou, Wen-Kui Sun, Yang Hang, Feng Gao, Sheng-Xiang Ji, Can-Fang Lin, Bo Pang, Ming-Xiao Yao, Guo-Lin Wang, Lin Yao, Li-Jun Duan, Xiao Wei, Dian-Ming Kang, Mai-Juan Ma |
| EPI_ISL_415105 | hCoV-19/Brazil/BA-312/2020 | South America / Brazil / Bahia / Feira de | 2020-03-04 | Laboratório Central de Saúde Pública Professor Gonçalo Moniz – LACEN/BA | Instituto Oswaldo Cruz FIOCRUZ - Laboratory of Respiratory Viruses and Measles (LVRS) | Paola Resende, Allison Fabri, Joilson Xavier, Sunando Roy, Fernando Motta, Aline Mattos, Milene Miranda, Cristiana Garcia, Braulia Caetano, Maria Ogrzewalska, Jonathan Lopes, Luciana Appolinario, Maria Nóbrega, Marilda Siqueira |
| EPI_ISL_415128 | hCoV-19/Brazil/ES-225/2020 | South America / Brazil / Espirito Santo / \ | 2020-02-29 | LACEN/ES - Laboratório Central de Saúde Pública do Espírito Santo | Instituto Oswaldo Cruz FIOCRUZ - Laboratory of Respiratory Viruses and Measles (LVRS) | Paola Resende, Allison Fabri, Joilson Xavier, Sunando Roy, Fernando Motta, Aline Mattos, Milene Miranda, Cristiana Garcia, Braulia Caetano, Maria Ogrzewalska, Jonathan Lopes, Luciana Appolinario, Maria Nóbrega, Marilda Siqueira |
| EPI_ISL_415129 | hCoV-19/England/20099038206/2020 | Europe / United Kingdom / England | 2020-02-29 | Respiratory Virus Unit, Microbiology Services Colindale, Public Health England | Respiratory Virus Unit, Microbiology Services Colindale, Public Health England | Monica Galiano, Shahjahan Miah, Angie Lackenby, Omolola Akinbami, Tiina Talts, Leena Bhaw, Richard Myers, Steven Platt, Kirstin Edwards, Jonathan Hubb, Joanna Ellis, Maria Zambon |
| EPI_ISL_415134 | hCoV-19/England/20100004806/2020 | Europe / United Kingdom / England | 2020-02-29 | Respiratory Virus Unit, Microbiology Services Colindale, Public Health England | Respiratory Virus Unit, Microbiology Services Colindale, Public Health England | Monica Galiano, Shahjahan Miah, Angie Lackenby, Omolola Akinbami, Tiina Talts, Leena Bhaw, Richard Myers, Steven Platt, Kirstin Edwards, Jonathan Hubb, Joanna Ellis, Maria Zambon |
| EPI_ISL_415136 | hCoV-19/England/20100022706/2020 | Europe / United Kingdom / England | 2020-02-29 | Respiratory Virus Unit, Microbiology Services Colindale, Public Health England | Respiratory Virus Unit, Microbiology Services Colindale, Public Health England | Monica Galiano, Shahjahan Miah, Angie Lackenby, Omolola Akinbami, Tiina Talts, Leena Bhaw, Richard Myers, Steven Platt, Kirstin Edwards, Jonathan Hubb, Joanna Ellis, Maria Zambon |
| EPI_ISL_415141 | hCoV-19/England/20100121007/2020 | Europe / United Kingdom / England | 2020-02-29 | Respiratory Virus Unit, Microbiology Services Colindale, Public Health England | Respiratory Virus Unit, Microbiology Services Colindale, Public Health England | Monica Galiano, Shahjahan Miah, Angie Lackenby, Omolola Akinbami, Tiina Talts, Leena Bhaw, Richard Myers, Steven Platt, Kirstin Edwards, Jonathan Hubb, Joanna Ellis, Maria Zambon |
| EPI_ISL_415142 | hCoV-19/England/20100122106/2020 | Europe / United Kingdom / England | 2020-03-02 | Respiratory Virus Unit, Microbiology Services Colindale, Public Health England | Respiratory Virus Unit, Microbiology Services Colindale, Public Health England | Monica Galiano, Shahjahan Miah, Angie Lackenby, Omolola Akinbami, Tiina Talts, Leena Bhaw, Richard Myers, Steven Platt, Kirstin Edwards, Jonathan Hubb, Joanna Ellis, Maria Zambon |
| EPI_ISL_415147 | hCoV-19/England/20102000506/2020 | Europe / United Kingdom / England | 2020-03-01 | Respiratory Virus Unit, Microbiology Services Colindale, Public Health England | Respiratory Virus Unit, Microbiology Services Colindale, Public Health England | Monica Galiano, Shahjahan Miah, Angie Lackenby, Omolola Akinbami, Tiina Talts, Leena Bhaw, Richard Myers, Steven Platt, Kirstin Edwards, Jonathan Hubb, Joanna Ellis, Maria Zambon |
| EPI_ISL_415148 | hCoV-19/England/20102000906/2020 | Europe / United Kingdom / England | 2020-03-03 | Respiratory Virus Unit, Microbiology Services Colindale, Public Health England | Respiratory Virus Unit, Microbiology Services Colindale, Public Health England | Monica Galiano, Shahjahan Miah, Angie Lackenby, Omolola Akinbami, Tiina Talts, Leena Bhaw, Richard Myers, Steven Platt, Kirstin Edwards, Jonathan Hubb, Joanna Ellis, Maria Zambon |
| EPI_ISL_415150 | hCoV-19/England/20110003506/2020 | Europe / United Kingdom / England | 2020-03-09 | Respiratory Virus Unit, Microbiology Services Colindale, Public Health England | Respiratory Virus Unit, Microbiology Services Colindale, Public Health England | Monica Galiano, Shahjahan Miah, Angie Lackenby, Omolola Akinbami, Tiina Talts, Leena Bhaw, Richard Myers, Steven Platt, Kirstin Edwards, Jonathan Hubb, Joanna Ellis, Maria Zambon |
| EPI_ISL_415151 | hCoV-19/USA/NY2-PV08100/2020 | North America / USA / New York | 2020-03-04 | MSHS Clinical Microbiology Laboratories | MSHS Pathogen Surveillance Program | Gopi Patel, Emilia Sordillo, Melissa Gitman, Alberto Paniz-mondolfi, Matthew Hernandez, Shelcie Fabre, Jose Polanco, Ana Silvia Gonzalez-Reiche, Zenab Khan, Nancy Francoeur, Melissa Smith, Robert Sebra, Lisa Miorin, Wen-chun Liu, Randy Albrecht, Judith Aberg, Florian Krammer, Adolfo Garcia-Sarstre, Viviana Simon, Harm van Bakel |
| EPI_ISL_415152 | hCoV-19/Panama/328677/2020 | Central America / Panama / Panama City | 2020-03-06 | Gorgas Memorial Institute for Health Studies | Gorgas Memorial Institute for Health Studies | Danilo Franco, Sandra Lopez-Verges, Elimelec Valdespino, Claudia Gonzalez, Oris Chavarria, Ambar Moreno, Yamilka Diaz, Leyda Abrego, Juan M. Pascale, Alexander A. Martinez. |
| EPI_ISL_415153 | hCoV-19/Belgium/VLM-03011/2020 | Europe / Belgium / Huldenberg | 2020-03-03 | KU Leuven, Clinical and Epidemiological Virology | KU Leuven, Clinical and Epidemiological Virology | Bert Vanmechelen, Joan Marti-Carreras, Tony Wawina, Marc Van Ranst, Piet Maes |
| EPI_ISL_415154 | hCoV-19/Belgium/BM-03012/2020 | Europe / Belgium / Kraainem | 2020-03-01 | KU Leuven, Clinical and Epidemiological Virology | KU Leuven, Clinical and Epidemiological Virology | Bert Vanmechelen, Joan Marti-Carreras, Tony Wawina, Marc Van Ranst, Piet Maes. |
| EPI_ISL_415155 | hCoV-19/Belgium/VAG-03013/2020 | Europe / Belgium / Huldenberg | 2020-03-01 | KU Leuven, Clinical and Epidemiological Virology | KU Leuven, Clinical and Epidemiological Virology | Bert Vanmechelen, Joan Marti-Carreras, Tony Wawina, Marc Van Ranst, Piet Maes |
| EPI_ISL_415156 | hCoV-19/Belgium/SH-03014/2020 | Europe / Belgium / Huldenberg | 2020-03-01 | KU Leuven, Clinical and Epidemiological Virology | KU Leuven, Clinical and Epidemiological Virology | Bert Vanmechelen, Joan Marti-Carreras, Tony Wawina, Piet Maes |
| EPI_ISL_415157 | hCoV-19/Belgium/BC-03016/2020 | Europe / Belgium / Sint-Niklaas | 2020-03-01 | KU Leuven, Clinical and Epidemiological Virology | KU Leuven, Clinical and Epidemiological Virology | Bert Vanmechelen, Joan Marti-Carreras, Tony Wawina, Piet Maes |
| EPI_ISL_415158 | hCoV-19/Belgium/QKJ-03015/2020 | Europe / Belgium / Brussels | 2020-03-01 | KU Leuven, Clinical and Epidemiological Virology | KU Leuven, Clinical and Epidemiological Virology | Bert Vanmechelen, Joan Marti-Carreras, Tony Wawina, Piet Maes |
| EPI_ISL_415159 | hCoV-19/Belgium/BA-02291/2020 | Europe / Belgium / Leuven | 2020-02-29 | KU Leuven, Clinical and Epidemiological Virology | KU Leuven, Clinical and Epidemiological Virology | Bert Vanmechelen, Joan Marti-Carreras, Tony Wawina, Piet Maes |
| EPI_ISL_415435 | hCoV-19/Wales/PHW06/2020 | Europe / United Kingdom / Wales | 2020-03-06 | Wales Specialist Virology Centre | Public Health Wales Microbiology Cardiff | Catherine Moore, Joanne Watkins, Sally Corden, Tom Connor |
| EPI_ISL_415454 | hCoV-19/Switzerland/GE1422/2020 | Europe / Switzerland | 2020-02-28 | Hôpitaux universitaires de Genève Laboratoire de Virologie | Hôpitaux universitaires de Genève Laboratoire de Virologie | Laubscher F. |
| EPI_ISL_415455 | hCoV-19/Switzerland/GE0199/2020 | Europe / Switzerland | 2020-02-28 | Hôpitaux universitaires de Genève Laboratoire de Virologie | Hôpitaux universitaires de Genève Laboratoire de Virologie | Laubscher F. |
| EPI_ISL_415456 | hCoV-19/Switzerland/BE6651/2020 | Europe / Switzerland | 2020-02-29 | Hôpitaux universitaires de Genève Laboratoire de Virologie | Hôpitaux universitaires de Genève Laboratoire de Virologie | Laubscher F. |
| EPI_ISL_415457 | hCoV-19/Switzerland/AG7120/2020 | Europe / Switzerland | 2020-02-29 | Hôpitaux universitaires de Genève Laboratoire de Virologie | Hôpitaux universitaires de Genève Laboratoire de Virologie | Laubscher F. |
| EPI_ISL_415458 | hCoV-19/Switzerland/GE8102/2020 | Europe / Switzerland | 2020-03-01 | Hôpitaux universitaires de Genève Laboratoire de Virologie | Hôpitaux universitaires de Genève Laboratoire de Virologie | Laubscher F. |
| EPI_ISL_415459 | hCoV-19/Switzerland/VD0503/2020 | Europe / Switzerland / Genève | 2020-02-29 | Hôpitaux universitaires de Genève Laboratoire de Virologie | Hôpitaux universitaires de Genève Laboratoire de Virologie | Laubscher F. |
| EPI_ISL_415460 | hCoV-19/Netherlands/Flevoland_1/2020 | Europe / Netherlands / Flevoland | 2020-03-09 | Dutch COVID-19 response team | Erasmus Medical Center | David Nieuwenhuijse, Bas Oude Munnink, Reina Sikkema, Claudia Schapendonk, Irina Chestakova, Anne van der Linden, Mark Pronk, Pascal Lexmond, Corien Swaan, Manon Haverkate, Madelief Mollers, Mart Stein, Sandra Kengne Kamga Mobou, Jeroen van Kampen, Jolanda Voermans, Aura Timen, Corine GeurtsvanKessel, Annemiek van der Eijk, Richard Molenkamp, Marion Koopmans, on behalf of the Dutch national COVID-19 response team. |
| EPI_ISL_415461 | hCoV-19/Netherlands/Gelderland_1-2/2020 | Europe / Netherlands / Gelderland | 2020-03-10 | Dutch COVID-19 response team | Erasmus Medical Center | David Nieuwenhuijse, Bas Oude Munnink, Reina Sikkema, Claudia Schapendonk, Irina Chestakova, Anne van der Linden, Mark Pronk, Pascal Lexmond, Corien Swaan, Manon Haverkate, Madelief Mollers, Mart Stein, Sandra Kengne Kamga Mobou, Jeroen van Kampen, Jolanda Voermans, Aura Timen, Corine GeurtsvanKessel, Annemiek van der Eijk, Richard Molenkamp, Marion Koopmans, on behalf of the Dutch national COVID-19 response team. |
| EPI_ISL_415462 | hCoV-19/Netherlands/Gelderland_2/2020 | Europe / Netherlands / Gelderland | 2020-03-09 | Dutch COVID-19 response team | Erasmus Medical Center | David Nieuwenhuijse, Bas Oude Munnink, Reina Sikkema, Claudia Schapendonk, Irina Chestakova, Anne van der Linden, Mark Pronk, Pascal Lexmond, Corien Swaan, Manon Haverkate, Madelief Mollers, Mart Stein, Sandra Kengne Kamga Mobou, Jeroen van Kampen, Jolanda Voermans, Aura Timen, Corine GeurtsvanKessel, Annemiek van der Eijk, Richard Molenkamp, Marion Koopmans, on behalf of the Dutch national COVID-19 response team. |
| EPI_ISL_415463 | hCoV-19/Netherlands/Gelderland_3/2020 | Europe / Netherlands / Gelderland | 2020-03-09 | Dutch COVID-19 response team | Erasmus Medical Center | David Nieuwenhuijse, Bas Oude Munnink, Reina Sikkema, Claudia Schapendonk, Irina Chestakova, Anne van der Linden, Mark Pronk, Pascal Lexmond, Corien Swaan, Manon Haverkate, Madelief Mollers, Mart Stein, Sandra Kengne Kamga Mobou, Jeroen van Kampen, Jolanda Voermans, Aura Timen, Corine GeurtsvanKessel, Annemiek van der Eijk, Richard Molenkamp, Marion Koopmans, on behalf of the Dutch national COVID-19 response team. |
| EPI_ISL_415464 | hCoV-19/Netherlands/Limburg_7/2020 | Europe / Netherlands / Limburg | 2020 | Dutch COVID-19 response team | Erasmus Medical Center | David Nieuwenhuijse, Bas Oude Munnink, Reina Sikkema, Claudia Schapendonk, Irina Chestakova, Anne van der Linden, Mark Pronk, Pascal Lexmond, Corien Swaan, Manon Haverkate, Madelief Mollers, Mart Stein, Sandra Kengne Kamga Mobou, Jeroen van Kampen, Jolanda Voermans, Aura Timen, Corine GeurtsvanKessel, Annemiek van der Eijk, Richard Molenkamp, Marion Koopmans, on behalf of the Dutch national COVID-19 response team. |
| EPI_ISL_415465 | hCoV-19/Netherlands/NA_1/2020 | Europe / Netherlands | 2020-03-10 | Dutch COVID-19 response team | Erasmus Medical Center | David Nieuwenhuijse, Bas Oude Munnink, Reina Sikkema, Claudia Schapendonk, Irina Chestakova, Anne van der Linden, Mark Pronk, Pascal Lexmond, Corien Swaan, Manon Haverkate, Madelief Mollers, Mart Stein, Sandra Kengne Kamga Mobou, Jeroen van Kampen, Jolanda Voermans, Aura Timen, Corine GeurtsvanKessel, Annemiek van der Eijk, Richard Molenkamp, Marion Koopmans, on behalf of the Dutch national COVID-19 response team. |
| EPI_ISL_415466 | hCoV-19/Netherlands/NA_10/2020 | Europe / Netherlands | 2020-03-09 | Dutch COVID-19 response team | Erasmus Medical Center | David Nieuwenhuijse, Bas Oude Munnink, Reina Sikkema, Claudia Schapendonk, Irina Chestakova, Anne van der Linden, Mark Pronk, Pascal Lexmond, Corien Swaan, Manon Haverkate, Madelief Mollers, Mart Stein, Sandra Kengne Kamga Mobou, Jeroen van Kampen, Jolanda Voermans, Aura Timen, Corine GeurtsvanKessel, Annemiek van der Eijk, Richard Molenkamp, Marion Koopmans, on behalf of the Dutch national COVID-19 response team. |
| EPI_ISL_415467 | hCoV-19/Netherlands/NA_11/2020 | Europe / Netherlands | 2020-03-10 | Dutch COVID-19 response team | Erasmus Medical Center | David Nieuwenhuijse, Bas Oude Munnink, Reina Sikkema, Claudia Schapendonk, Irina Chestakova, Anne van der Linden, Mark Pronk, Pascal Lexmond, Corien Swaan, Manon Haverkate, Madelief Mollers, Mart Stein, Sandra Kengne Kamga Mobou, Jeroen van Kampen, Jolanda Voermans, Aura Timen, Corine GeurtsvanKessel, Annemiek van der Eijk, Richard Molenkamp, Marion Koopmans, on behalf of the Dutch national COVID-19 response team. |
| EPI_ISL_415468 | hCoV-19/Netherlands/NA_12/2020 | Europe / Netherlands | 2020-03-10 | Dutch COVID-19 response team | Erasmus Medical Center | David Nieuwenhuijse, Bas Oude Munnink, Reina Sikkema, Claudia Schapendonk, Irina Chestakova, Anne van der Linden, Mark Pronk, Pascal Lexmond, Corien Swaan, Manon Haverkate, Madelief Mollers, Mart Stein, Sandra Kengne Kamga Mobou, Jeroen van Kampen, Jolanda Voermans, Aura Timen, Corine GeurtsvanKessel, Annemiek van der Eijk, Richard Molenkamp, Marion Koopmans, on behalf of the Dutch national COVID-19 response team. |
| EPI_ISL_415469 | hCoV-19/Netherlands/NA_13/2020 | Europe / Netherlands | 2020-03-10 | Dutch COVID-19 response team | Erasmus Medical Center | David Nieuwenhuijse, Bas Oude Munnink, Reina Sikkema, Claudia Schapendonk, Irina Chestakova, Anne van der Linden, Mark Pronk, Pascal Lexmond, Corien Swaan, Manon Haverkate, Madelief Mollers, Mart Stein, Sandra Kengne Kamga Mobou, Jeroen van Kampen, Jolanda Voermans, Aura Timen, Corine GeurtsvanKessel, Annemiek van der Eijk, Richard Molenkamp, Marion Koopmans, on behalf of the Dutch national COVID-19 response team. |
| EPI_ISL_415500 | hCoV-19/Netherlands/NoordBrabant_42/2020 | Europe / Netherlands / Noord Brabant | 2020-03-10 | Dutch COVID-19 response team | Erasmus Medical Center | David Nieuwenhuijse, Bas Oude Munnink, Reina Sikkema, Claudia Schapendonk, Irina Chestakova, Anne van der Linden, Mark Pronk, Pascal Lexmond, Corien Swaan, Manon Haverkate, Madelief Mollers, Mart Stein, Sandra Kengne Kamga Mobou, Jeroen van Kampen, Jolanda Voermans, Aura Timen, Corine GeurtsvanKessel, Annemiek van der Eijk, Richard Molenkamp, Marion Koopmans, on behalf of the Dutch national COVID-19 response team. |
| EPI_ISL_415501 | hCoV-19/Netherlands/NoordBrabant_44/2020 | Europe / Netherlands / Noord Brabant | 2020-03-11 | Dutch COVID-19 response team | Erasmus Medical Center | David Nieuwenhuijse, Bas Oude Munnink, Reina Sikkema, Claudia Schapendonk, Irina Chestakova, Anne van der Linden, Mark Pronk, Pascal Lexmond, Corien Swaan, Manon Haverkate, Madelief Mollers, Mart Stein, Sandra Kengne Kamga Mobou, Jeroen van Kampen, Jolanda Voermans, Aura Timen, Corine GeurtsvanKessel, Annemiek van der Eijk, Richard Molenkamp, Marion Koopmans, on behalf of the Dutch national COVID-19 response team. |
| EPI_ISL_415502 | hCoV-19/Netherlands/NoordBrabant_45/2020 | Europe / Netherlands / Noord Brabant | 2020-03-09 | Dutch COVID-19 response team | Erasmus Medical Center | David Nieuwenhuijse, Bas Oude Munnink, Reina Sikkema, Claudia Schapendonk, Irina Chestakova, Anne van der Linden, Mark Pronk, Pascal Lexmond, Corien Swaan, Manon Haverkate, Madelief Mollers, Mart Stein, Sandra Kengne Kamga Mobou, Jeroen van Kampen, Jolanda Voermans, Aura Timen, Corine GeurtsvanKessel, Annemiek van der Eijk, Richard Molenkamp, Marion Koopmans, on behalf of the Dutch national COVID-19 response team. |
| EPI_ISL_415503 | hCoV-19/Netherlands/NoordBrabant_46/2020 | Europe / Netherlands / Noord Brabant | 2020-03-11 | Dutch COVID-19 response team | Erasmus Medical Center | David Nieuwenhuijse, Bas Oude Munnink, Reina Sikkema, Claudia Schapendonk, Irina Chestakova, Anne van der Linden, Mark Pronk, Pascal Lexmond, Corien Swaan, Manon Haverkate, Madelief Mollers, Mart Stein, Sandra Kengne Kamga Mobou, Jeroen van Kampen, Jolanda Voermans, Aura Timen, Corine GeurtsvanKessel, Annemiek van der Eijk, Richard Molenkamp, Marion Koopmans, on behalf of the Dutch national COVID-19 response team. |
| EPI_ISL_415504 | hCoV-19/Netherlands/NoordBrabant_47/2020 | Europe / Netherlands / Noord Brabant | 2020-03-09 | Dutch COVID-19 response team | Erasmus Medical Center | David Nieuwenhuijse, Bas Oude Munnink, Reina Sikkema, Claudia Schapendonk, Irina Chestakova, Anne van der Linden, Mark Pronk, Pascal Lexmond, Corien Swaan, Manon Haverkate, Madelief Mollers, Mart Stein, Sandra Kengne Kamga Mobou, Jeroen van Kampen, Jolanda Voermans, Aura Timen, Corine GeurtsvanKessel, Annemiek van der Eijk, Richard Molenkamp, Marion Koopmans, on behalf of the Dutch national COVID-19 response team. |
| EPI_ISL_415505 | hCoV-19/Netherlands/NoordBrabant_48/2020 | Europe / Netherlands / Noord Brabant | 2020-03-11 | Dutch COVID-19 response team | Erasmus Medical Center | David Nieuwenhuijse, Bas Oude Munnink, Reina Sikkema, Claudia Schapendonk, Irina Chestakova, Anne van der Linden, Mark Pronk, Pascal Lexmond, Corien Swaan, Manon Haverkate, Madelief Mollers, Mart Stein, Sandra Kengne Kamga Mobou, Jeroen van Kampen, Jolanda Voermans, Aura Timen, Corine GeurtsvanKessel, Annemiek van der Eijk, Richard Molenkamp, Marion Koopmans, on behalf of the Dutch national COVID-19 response team. |
| EPI_ISL_415506 | hCoV-19/Netherlands/NoordBrabant_49/2020 | Europe / Netherlands / Noord Brabant | 2020-03-11 | Dutch COVID-19 response team | Erasmus Medical Center | David Nieuwenhuijse, Bas Oude Munnink, Reina Sikkema, Claudia Schapendonk, Irina Chestakova, Anne van der Linden, Mark Pronk, Pascal Lexmond, Corien Swaan, Manon Haverkate, Madelief Mollers, Mart Stein, Sandra Kengne Kamga Mobou, Jeroen van Kampen, Jolanda Voermans, Aura Timen, Corine GeurtsvanKessel, Annemiek van der Eijk, Richard Molenkamp, Marion Koopmans, on behalf of the Dutch national COVID-19 response team. |
| EPI_ISL_415507 | hCoV-19/Netherlands/NoordBrabant_51/2020 | Europe / Netherlands / Noord Brabant | 2020 | Dutch COVID-19 response team | Erasmus Medical Center | David Nieuwenhuijse, Bas Oude Munnink, Reina Sikkema, Claudia Schapendonk, Irina Chestakova, Anne van der Linden, Mark Pronk, Pascal Lexmond, Corien Swaan, Manon Haverkate, Madelief Mollers, Mart Stein, Sandra Kengne Kamga Mobou, Jeroen van Kampen, Jolanda Voermans, Aura Timen, Corine GeurtsvanKessel, Annemiek van der Eijk, Richard Molenkamp, Marion Koopmans, on behalf of the Dutch national COVID-19 response team. |
| EPI_ISL_415508 | hCoV-19/Netherlands/NoordBrabant_52/2020 | Europe / Netherlands / Noord Brabant | 2020-03-09 | Dutch COVID-19 response team | Erasmus Medical Center | David Nieuwenhuijse, Bas Oude Munnink, Reina Sikkema, Claudia Schapendonk, Irina Chestakova, Anne van der Linden, Mark Pronk, Pascal Lexmond, Corien Swaan, Manon Haverkate, Madelief Mollers, Mart Stein, Sandra Kengne Kamga Mobou, Jeroen van Kampen, Jolanda Voermans, Aura Timen, Corine GeurtsvanKessel, Annemiek van der Eijk, Richard Molenkamp, Marion Koopmans, on behalf of the Dutch national COVID-19 response team. |
| EPI_ISL_415509 | hCoV-19/Netherlands/NoordBrabant_53/2020 | Europe / Netherlands / Noord Brabant | 2020 | Dutch COVID-19 response team | Erasmus Medical Center | David Nieuwenhuijse, Bas Oude Munnink, Reina Sikkema, Claudia Schapendonk, Irina Chestakova, Anne van der Linden, Mark Pronk, Pascal Lexmond, Corien Swaan, Manon Haverkate, Madelief Mollers, Mart Stein, Sandra Kengne Kamga Mobou, Jeroen van Kampen, Jolanda Voermans, Aura Timen, Corine GeurtsvanKessel, Annemiek van der Eijk, Richard Molenkamp, Marion Koopmans, on behalf of the Dutch national COVID-19 response team. |
| EPI_ISL_415510 | hCoV-19/Netherlands/NoordBrabant_54/2020 | Europe / Netherlands / Noord Brabant | 2020-03-09 | Dutch COVID-19 response team | Erasmus Medical Center | David Nieuwenhuijse, Bas Oude Munnink, Reina Sikkema, Claudia Schapendonk, Irina Chestakova, Anne van der Linden, Mark Pronk, Pascal Lexmond, Corien Swaan, Manon Haverkate, Madelief Mollers, Mart Stein, Sandra Kengne Kamga Mobou, Jeroen van Kampen, Jolanda Voermans, Aura Timen, Corine GeurtsvanKessel, Annemiek van der Eijk, Richard Molenkamp, Marion Koopmans, on behalf of the Dutch national COVID-19 response team. |
| EPI_ISL_415511 | hCoV-19/Netherlands/NoordBrabant_55/2020 | Europe / Netherlands / Noord Brabant | 2020-03-09 | Dutch COVID-19 response team | Erasmus Medical Center | David Nieuwenhuijse, Bas Oude Munnink, Reina Sikkema, Claudia Schapendonk, Irina Chestakova, Anne van der Linden, Mark Pronk, Pascal Lexmond, Corien Swaan, Manon Haverkate, Madelief Mollers, Mart Stein, Sandra Kengne Kamga Mobou, Jeroen van Kampen, Jolanda Voermans, Aura Timen, Corine GeurtsvanKessel, Annemiek van der Eijk, Richard Molenkamp, Marion Koopmans, on behalf of the Dutch national COVID-19 response team. |
| EPI_ISL_415512 | hCoV-19/Netherlands/NoordBrabant_56/2020 | Europe / Netherlands / Noord Brabant | 2020-03-09 | Dutch COVID-19 response team | Erasmus Medical Center | David Nieuwenhuijse, Bas Oude Munnink, Reina Sikkema, Claudia Schapendonk, Irina Chestakova, Anne van der Linden, Mark Pronk, Pascal Lexmond, Corien Swaan, Manon Haverkate, Madelief Mollers, Mart Stein, Sandra Kengne Kamga Mobou, Jeroen van Kampen, Jolanda Voermans, Aura Timen, Corine GeurtsvanKessel, Annemiek van der Eijk, Richard Molenkamp, Marion Koopmans, on behalf of the Dutch national COVID-19 response team. |
| EPI_ISL_415513 | hCoV-19/Netherlands/NoordBrabant_57/2020 | Europe / Netherlands / Noord Brabant | 2020-03-11 | Dutch COVID-19 response team | Erasmus Medical Center | David Nieuwenhuijse, Bas Oude Munnink, Reina Sikkema, Claudia Schapendonk, Irina Chestakova, Anne van der Linden, Mark Pronk, Pascal Lexmond, Corien Swaan, Manon Haverkate, Madelief Mollers, Mart Stein, Sandra Kengne Kamga Mobou, Jeroen van Kampen, Jolanda Voermans, Aura Timen, Corine GeurtsvanKessel, Annemiek van der Eijk, Richard Molenkamp, Marion Koopmans, on behalf of the Dutch national COVID-19 response team. |
| EPI_ISL_415514 | hCoV-19/Netherlands/NoordBrabant_58/2020 | Europe / Netherlands / Noord Brabant | 2020-03-11 | Dutch COVID-19 response team | Erasmus Medical Center | David Nieuwenhuijse, Bas Oude Munnink, Reina Sikkema, Claudia Schapendonk, Irina Chestakova, Anne van der Linden, Mark Pronk, Pascal Lexmond, Corien Swaan, Manon Haverkate, Madelief Mollers, Mart Stein, Sandra Kengne Kamga Mobou, Jeroen van Kampen, Jolanda Voermans, Aura Timen, Corine GeurtsvanKessel, Annemiek van der Eijk, Richard Molenkamp, Marion Koopmans, on behalf of the Dutch national COVID-19 response team. |
| EPI_ISL_415515 | hCoV-19/Netherlands/NoordBrabant_59/2020 | Europe / Netherlands / Noord Brabant | 2020-03-11 | Dutch COVID-19 response team | Erasmus Medical Center | David Nieuwenhuijse, Bas Oude Munnink, Reina Sikkema, Claudia Schapendonk, Irina Chestakova, Anne van der Linden, Mark Pronk, Pascal Lexmond, Corien Swaan, Manon Haverkate, Madelief Mollers, Mart Stein, Sandra Kengne Kamga Mobou, Jeroen van Kampen, Jolanda Voermans, Aura Timen, Corine GeurtsvanKessel, Annemiek van der Eijk, Richard Molenkamp, Marion Koopmans, on behalf of the Dutch national COVID-19 response team. |
| EPI_ISL_415517 | hCoV-19/Netherlands/NoordBrabant_61/2020 | Europe / Netherlands / Noord Brabant | 2020 | Dutch COVID-19 response team | Erasmus Medical Center | David Nieuwenhuijse, Bas Oude Munnink, Reina Sikkema, Claudia Schapendonk, Irina Chestakova, Anne van der Linden, Mark Pronk, Pascal Lexmond, Corien Swaan, Manon Haverkate, Madelief Mollers, Mart Stein, Sandra Kengne Kamga Mobou, Jeroen van Kampen, Jolanda Voermans, Aura Timen, Corine GeurtsvanKessel, Annemiek van der Eijk, Richard Molenkamp, Marion Koopmans, on behalf of the Dutch national COVID-19 response team. |
| EPI_ISL_415518 | hCoV-19/Netherlands/NoordBrabant_62/2020 | Europe / Netherlands / Noord Brabant | 2020 | Dutch COVID-19 response team | Erasmus Medical Center | David Nieuwenhuijse, Bas Oude Munnink, Reina Sikkema, Claudia Schapendonk, Irina Chestakova, Anne van der Linden, Mark Pronk, Pascal Lexmond, Corien Swaan, Manon Haverkate, Madelief Mollers, Mart Stein, Sandra Kengne Kamga Mobou, Jeroen van Kampen, Jolanda Voermans, Aura Timen, Corine GeurtsvanKessel, Annemiek van der Eijk, Richard Molenkamp, Marion Koopmans, on behalf of the Dutch national COVID-19 response team. |
| EPI_ISL_415519 | hCoV-19/Netherlands/NoordBrabant_63/2020 | Europe / Netherlands / Noord Brabant | 2020 | Dutch COVID-19 response team | Erasmus Medical Center | David Nieuwenhuijse, Bas Oude Munnink, Reina Sikkema, Claudia Schapendonk, Irina Chestakova, Anne van der Linden, Mark Pronk, Pascal Lexmond, Corien Swaan, Manon Haverkate, Madelief Mollers, Mart Stein, Sandra Kengne Kamga Mobou, Jeroen van Kampen, Jolanda Voermans, Aura Timen, Corine GeurtsvanKessel, Annemiek van der Eijk, Richard Molenkamp, Marion Koopmans, on behalf of the Dutch national COVID-19 response team. |
| EPI_ISL_415520 | hCoV-19/Netherlands/NoordBrabant_64/2020 | Europe / Netherlands / Noord Brabant | 2020 | Dutch COVID-19 response team | Erasmus Medical Center | David Nieuwenhuijse, Bas Oude Munnink, Reina Sikkema, Claudia Schapendonk, Irina Chestakova, Anne van der Linden, Mark Pronk, Pascal Lexmond, Corien Swaan, Manon Haverkate, Madelief Mollers, Mart Stein, Sandra Kengne Kamga Mobou, Jeroen van Kampen, Jolanda Voermans, Aura Timen, Corine GeurtsvanKessel, Annemiek van der Eijk, Richard Molenkamp, Marion Koopmans, on behalf of the Dutch national COVID-19 response team. |
| EPI_ISL_415521 | hCoV-19/Netherlands/NoordBrabant_65/2020 | Europe / Netherlands / Noord Brabant | 2020 | Dutch COVID-19 response team | Erasmus Medical Center | David Nieuwenhuijse, Bas Oude Munnink, Reina Sikkema, Claudia Schapendonk, Irina Chestakova, Anne van der Linden, Mark Pronk, Pascal Lexmond, Corien Swaan, Manon Haverkate, Madelief Mollers, Mart Stein, Sandra Kengne Kamga Mobou, Jeroen van Kampen, Jolanda Voermans, Aura Timen, Corine GeurtsvanKessel, Annemiek van der Eijk, Richard Molenkamp, Marion Koopmans, on behalf of the Dutch national COVID-19 response team. |
| EPI_ISL_415522 | hCoV-19/Netherlands/NoordBrabant_66/2020 | Europe / Netherlands / Noord Brabant | 2020 | Dutch COVID-19 response team | Erasmus Medical Center | David Nieuwenhuijse, Bas Oude Munnink, Reina Sikkema, Claudia Schapendonk, Irina Chestakova, Anne van der Linden, Mark Pronk, Pascal Lexmond, Corien Swaan, Manon Haverkate, Madelief Mollers, Mart Stein, Sandra Kengne Kamga Mobou, Jeroen van Kampen, Jolanda Voermans, Aura Timen, Corine GeurtsvanKessel, Annemiek van der Eijk, Richard Molenkamp, Marion Koopmans, on behalf of the Dutch national COVID-19 response team. |
| EPI_ISL_415523 | hCoV-19/Netherlands/NoordBrabant_67/2020 | Europe / Netherlands / Noord Brabant | 2020 | Dutch COVID-19 response team | Erasmus Medical Center | David Nieuwenhuijse, Bas Oude Munnink, Reina Sikkema, Claudia Schapendonk, Irina Chestakova, Anne van der Linden, Mark Pronk, Pascal Lexmond, Corien Swaan, Manon Haverkate, Madelief Mollers, Mart Stein, Sandra Kengne Kamga Mobou, Jeroen van Kampen, Jolanda Voermans, Aura Timen, Corine GeurtsvanKessel, Annemiek van der Eijk, Richard Molenkamp, Marion Koopmans, on behalf of the Dutch national COVID-19 response team. |
| EPI_ISL_415524 | hCoV-19/Netherlands/NoordBrabant_68/2020 | Europe / Netherlands / Noord Brabant | 2020-03-12 | Dutch COVID-19 response team | Erasmus Medical Center | David Nieuwenhuijse, Bas Oude Munnink, Reina Sikkema, Claudia Schapendonk, Irina Chestakova, Anne van der Linden, Mark Pronk, Pascal Lexmond, Corien Swaan, Manon Haverkate, Madelief Mollers, Mart Stein, Sandra Kengne Kamga Mobou, Jeroen van Kampen, Jolanda Voermans, Aura Timen, Corine GeurtsvanKessel, Annemiek van der Eijk, Richard Molenkamp, Marion Koopmans, on behalf of the Dutch national COVID-19 response team. |
| EPI_ISL_415525 | hCoV-19/Netherlands/NoordHolland_3/2020 | Europe / Netherlands / Noord Holland | 2020-03-12 | Dutch COVID-19 response team | Erasmus Medical Center | David Nieuwenhuijse, Bas Oude Munnink, Reina Sikkema, Claudia Schapendonk, Irina Chestakova, Anne van der Linden, Mark Pronk, Pascal Lexmond, Corien Swaan, Manon Haverkate, Madelief Mollers, Mart Stein, Sandra Kengne Kamga Mobou, Jeroen van Kampen, Jolanda Voermans, Aura Timen, Corine GeurtsvanKessel, Annemiek van der Eijk, Richard Molenkamp, Marion Koopmans, on behalf of the Dutch national COVID-19 response team. |
| EPI_ISL_415526 | hCoV-19/Netherlands/Utrecht_17/2020 | Europe / Netherlands / Utrecht | 2020-03-10 | Dutch COVID-19 response team | Erasmus Medical Center | David Nieuwenhuijse, Bas Oude Munnink, Reina Sikkema, Claudia Schapendonk, Irina Chestakova, Anne van der Linden, Mark Pronk, Pascal Lexmond, Corien Swaan, Manon Haverkate, Madelief Mollers, Mart Stein, Sandra Kengne Kamga Mobou, Jeroen van Kampen, Jolanda Voermans, Aura Timen, Corine GeurtsvanKessel, Annemiek van der Eijk, Richard Molenkamp, Marion Koopmans, on behalf of the Dutch national COVID-19 response team. |
| EPI_ISL_415527 | hCoV-19/Netherlands/Utrecht_18/2020 | Europe / Netherlands / Utrecht | 2020-03-12 | Dutch COVID-19 response team | Erasmus Medical Center | David Nieuwenhuijse, Bas Oude Munnink, Reina Sikkema, Claudia Schapendonk, Irina Chestakova, Anne van der Linden, Mark Pronk, Pascal Lexmond, Corien Swaan, Manon Haverkate, Madelief Mollers, Mart Stein, Sandra Kengne Kamga Mobou, Jeroen van Kampen, Jolanda Voermans, Aura Timen, Corine GeurtsvanKessel, Annemiek van der Eijk, Richard Molenkamp, Marion Koopmans, on behalf of the Dutch national COVID-19 response team. |
| EPI_ISL_415529 | hCoV-19/Netherlands/ZuidHolland_25/2020 | Europe / Netherlands / Zuid Holland | 2020-03-09 | Dutch COVID-19 response team | Erasmus Medical Center | David Nieuwenhuijse, Bas Oude Munnink, Reina Sikkema, Claudia Schapendonk, Irina Chestakova, Anne van der Linden, Mark Pronk, Pascal Lexmond, Corien Swaan, Manon Haverkate, Madelief Mollers, Mart Stein, Sandra Kengne Kamga Mobou, Jeroen van Kampen, Jolanda Voermans, Aura Timen, Corine GeurtsvanKessel, Annemiek van der Eijk, Richard Molenkamp, Marion Koopmans, on behalf of the Dutch national COVID-19 response team. |
| EPI_ISL_415530 | hCoV-19/Netherlands/ZuidHolland_26/2020 | Europe / Netherlands / Zuid Holland | 2020-03-09 | Dutch COVID-19 response team | Erasmus Medical Center | David Nieuwenhuijse, Bas Oude Munnink, Reina Sikkema, Claudia Schapendonk, Irina Chestakova, Anne van der Linden, Mark Pronk, Pascal Lexmond, Corien Swaan, Manon Haverkate, Madelief Mollers, Mart Stein, Sandra Kengne Kamga Mobou, Jeroen van Kampen, Jolanda Voermans, Aura Timen, Corine GeurtsvanKessel, Annemiek van der Eijk, Richard Molenkamp, Marion Koopmans, on behalf of the Dutch national COVID-19 response team. |
| EPI_ISL_415531 | hCoV-19/Netherlands/ZuidHolland_27/2020 | Europe / Netherlands / Zuid Holland | 2020-03-09 | Dutch COVID-19 response team | Erasmus Medical Center | David Nieuwenhuijse, Bas Oude Munnink, Reina Sikkema, Claudia Schapendonk, Irina Chestakova, Anne van der Linden, Mark Pronk, Pascal Lexmond, Corien Swaan, Manon Haverkate, Madelief Mollers, Mart Stein, Sandra Kengne Kamga Mobou, Jeroen van Kampen, Jolanda Voermans, Aura Timen, Corine GeurtsvanKessel, Annemiek van der Eijk, Richard Molenkamp, Marion Koopmans, on behalf of the Dutch national COVID-19 response team. |
| EPI_ISL_415532 | hCoV-19/Netherlands/ZuidHolland_28/2020 | Europe / Netherlands / Zuid Holland | 2020-03-09 | Dutch COVID-19 response team | Erasmus Medical Center | David Nieuwenhuijse, Bas Oude Munnink, Reina Sikkema, Claudia Schapendonk, Irina Chestakova, Anne van der Linden, Mark Pronk, Pascal Lexmond, Corien Swaan, Manon Haverkate, Madelief Mollers, Mart Stein, Sandra Kengne Kamga Mobou, Jeroen van Kampen, Jolanda Voermans, Aura Timen, Corine GeurtsvanKessel, Annemiek van der Eijk, Richard Molenkamp, Marion Koopmans, on behalf of the Dutch national COVID-19 response team. |
| EPI_ISL_415533 | hCoV-19/Netherlands/ZuidHolland_29/2020 | Europe / Netherlands / Zuid Holland | 2020-03-09 | Dutch COVID-19 response team | Erasmus Medical Center | David Nieuwenhuijse, Bas Oude Munnink, Reina Sikkema, Claudia Schapendonk, Irina Chestakova, Anne van der Linden, Mark Pronk, Pascal Lexmond, Corien Swaan, Manon Haverkate, Madelief Mollers, Mart Stein, Sandra Kengne Kamga Mobou, Jeroen van Kampen, Jolanda Voermans, Aura Timen, Corine GeurtsvanKessel, Annemiek van der Eijk, Richard Molenkamp, Marion Koopmans, on behalf of the Dutch national COVID-19 response team. |
| EPI_ISL_415534 | hCoV-19/Netherlands/ZuidHolland_30/2020 | Europe / Netherlands / Zuid Holland | 2020-03-11 | Dutch COVID-19 response team | Erasmus Medical Center | David Nieuwenhuijse, Bas Oude Munnink, Reina Sikkema, Claudia Schapendonk, Irina Chestakova, Anne van der Linden, Mark Pronk, Pascal Lexmond, Corien Swaan, Manon Haverkate, Madelief Mollers, Mart Stein, Sandra Kengne Kamga Mobou, Jeroen van Kampen, Jolanda Voermans, Aura Timen, Corine GeurtsvanKessel, Annemiek van der Eijk, Richard Molenkamp, Marion Koopmans, on behalf of the Dutch national COVID-19 response team. |
| EPI_ISL_415535 | hCoV-19/Netherlands/ZuidHolland_31/2020 | Europe / Netherlands / Zuid Holland | 2020-03-12 | Dutch COVID-19 response team | Erasmus Medical Center | David Nieuwenhuijse, Bas Oude Munnink, Reina Sikkema, Claudia Schapendonk, Irina Chestakova, Anne van der Linden, Mark Pronk, Pascal Lexmond, Corien Swaan, Manon Haverkate, Madelief Mollers, Mart Stein, Sandra Kengne Kamga Mobou, Jeroen van Kampen, Jolanda Voermans, Aura Timen, Corine GeurtsvanKessel, Annemiek van der Eijk, Richard Molenkamp, Marion Koopmans, on behalf of the Dutch national COVID-19 response team. |
| EPI_ISL_415537 | hCoV-19/Wales/PHW13/2020 | Europe / United Kingdom / Wales | 2020-03-08 | Wales Specialist Virology Centre | Public Health Wales Microbiology Cardiff | Catherine Moore, Joanne Watkins, Sally Corden, Tom Connor |
| EPI_ISL_415539 | hCoV-19/USA/UT-01/2020 | North America / USA / Utah | 2020-03-13 | Utah Public Health Laboratory | Utah Public Health Laboratory | Erin Young, Kelly Oakeson |
| EPI_ISL_415541 | hCoV-19/USA/UT-03/2020 | North America / USA / Utah | 2020-03-13 | Utah Public Health Laboratory | Utah Public Health Laboratory | Erin Young, Kelly Oakeson |
| EPI_ISL_415542 | hCoV-19/USA/UT-04/2020 | North America / USA / Utah | 2020-03-13 | Utah Public Health Laboratory | Utah Public Health Laboratory | Erin Young, Kelly Oakeson |
| EPI_ISL_415543 | hCoV-19/USA/UT-05/2020 | North America / USA / Utah | 2020-03-13 | Utah Public Health Laboratory | Utah Public Health Laboratory | Erin Young, Kelly Oakeson |
| EPI_ISL_415544 | hCoV-19/USA/UT-06/2020 | North America / USA / Utah | 2020-03-13 | Utah Public Health Laboratory | Utah Public Health Laboratory | Erin Young, Kelly Oakeson |
| EPI_ISL_415581 | hCoV-19/Canada/BC_02421/2020 | North America / Canada / British Columbi | 2020-03-01 | BCCDC Public Health Laboratory | BCCDC Public Health Laboratory | Harrigan, Prystajecky, Krajden, Lee, Kamelian, Lapointe, Choi, Hoang, Sekirov, Levett, Tyson, Snutch, Loman, Quick, Li, Gilmour |
| EPI_ISL_415583 | hCoV-19/Canada/BC_40860/2020 | North America / Canada / British Columbi | 2020-03-03 | BCCDC Public Health Laboratory | BCCDC Public Health Laboratory | Harrigan, Prystajecky, Krajden, Lee, Kamelian, Lapointe, Choi, Hoang, Sekirov, Levett, Tyson, Snutch, Loman, Quick, Li, Gilmour |
| EPI_ISL_415584 | hCoV-19/Canada/BC_41851/2020 | North America / Canada / British Columbi | 2020-03-02 | BCCDC Public Health Laboratory | BCCDC Public Health Laboratory | Harrigan, Prystajecky, Krajden, Lee, Kamelian, Lapointe, Choi, Hoang, Sekirov, Levett, Tyson, Snutch, Loman, Quick, Li, Gilmour |
| EPI_ISL_415588 | hCoV-19/Canada/BC_83109/2020 | North America / Canada / British Columbi | 2020-03-05 | BCCDC Public Health Laboratory | BCCDC Public Health Laboratory | Harrigan, Prystajecky, Krajden, Lee, Kamelian, Lapointe, Choi, Hoang, Sekirov, Levett, Tyson, Snutch, Loman, Quick, Li, Gilmour |
| EPI_ISL_415591 | hCoV-19/USA/WA-UW63/2020 | North America / USA / Washington | 2020-03-10 | UW Virology Lab | UW Virology Lab | Pavitra Roychoudhury, Hong Xie, Keith Jerome, Alexander Greninger |
| EPI_ISL_415592 | hCoV-19/USA/WA-UW64/2020 | North America / USA / Washington | 2020-03-09 | UW Virology Lab | UW Virology Lab | Pavitra Roychoudhury, Hong Xie, Keith Jerome, Alexander Greninger |
| EPI_ISL_415594 | hCoV-19/USA/WA-UW66/2020 | North America / USA / Washington | 2020-03-10 | UW Virology Lab | UW Virology Lab | Pavitra Roychoudhury, Hong Xie, Keith Jerome, Alexander Greninger |
| EPI_ISL_415595 | hCoV-19/USA/WA-UW67/2020 | North America / USA / Washington | 2020-03-09 | UW Virology Lab | UW Virology Lab | Pavitra Roychoudhury, Hong Xie, Keith Jerome, Alexander Greninger |
| EPI_ISL_415596 | hCoV-19/USA/WA-UW68/2020 | North America / USA / Washington | 2020-03-09 | UW Virology Lab | UW Virology Lab | Pavitra Roychoudhury, Hong Xie, Keith Jerome, Alexander Greninger |
| EPI_ISL_415597 | hCoV-19/USA/WA-UW69/2020 | North America / USA / Washington / Seat | 2020-03-10 | UW Virology Lab | UW Virology Lab | Pavitra Roychoudhury, Hong Xie, Keith Jerome, Alexander Greninger |
| EPI_ISL_415598 | hCoV-19/USA/WA-UW70/2020 | North America / USA / Washington | 2020-03-10 | UW Virology Lab | UW Virology Lab | Pavitra Roychoudhury, Hong Xie, Keith Jerome, Alexander Greninger |
| EPI_ISL_415599 | hCoV-19/USA/WA-UW71/2020 | North America / USA / Washington | 2020-03-09 | UW Virology Lab | UW Virology Lab | Pavitra Roychoudhury, Hong Xie, Keith Jerome, Alexander Greninger |
| EPI_ISL_415920 | hCoV-19/Wales/PHW32/2020 | Europe / United Kingdom / Wales | 2020-03-12 | Wales Specialist Virology Centre | Public Health Wales Microbiology Cardiff | Catherine Moore, Joanne Watkins, Sally Corden, Tom Connor |
| EPI_ISL_416024 | hCoV-19/Wales/PHW35/2020 | Europe / United Kingdom / Wales | 2020-03-11 | Wales Specialist Virology Centre | Public Health Wales Microbiology Cardiff | Catherine Moore, Joanne Watkins, Sally Corden, Tom Connor |
| EPI_ISL_416026 | hCoV-19/Wales/PHW21/2020 | Europe / United Kingdom / Wales | 2020-03-11 | Wales Specialist Virology Centre | Public Health Wales Microbiology Cardiff | Catherine Moore, Joanne Watkins, Sally Corden, Tom Connor |
| EPI_ISL_416028 | hCoV-19/Brazil/SPBR-07/2020 | South America / Brazil / Sao Paulo / Sao | 2020-03-03 | National Influenza Center - Instituto Adolfo Lutz | Instituto Adolfo Lutz, Interdiciplinary Procedures Center, Strategic Laboratory | Claudio Tavares Sacchi, Claudia Regina Gonçalves, Carlos Henrique Camargo, Fabiana Cristina Pereira dos Santos, Daniela Bernardes Borges da Silva, Simone Guadagnucci Morillo, Adriano Abbud, Adriana Bugno, Maria do Carmo Sampaio Tavares Timenetsky, Terezinha Maria de Paiva |
| EPI_ISL_416029 | hCoV-19/Brazil/SPBR-08/2020 | South America / Brazil / Sao Paulo / Sao | 2020-03-04 | Laboratório Fleury | Instituto Adolfo Lutz, Interdiciplinary Procedures Center, Strategic Laboratory | Claudio Tavares Sacchi, Claudia Regina Gonçalves, Carlos Henrique Camargo, Fabiana Cristina Pereira dos Santos, Daniela Bernardes Borges da Silva, Simone Guadagnucci Morillo, Adriano Abbud, Adriana Bugno, Maria do Carmo Sampaio Tavares Timenetsky, Terezinha Maria de Paiva |
| EPI_ISL_416031 | hCoV-19/Brazil/SPBR-09/2020 | South America / Brazil / Sao Paulo / Sao | 2020-03-04 | National Influenza Center - Instituto Adolfo Lutz | Instituto Adolfo Lutz, Interdiciplinary Procedures Center, Strategic Laboratory | Claudio Tavares Sacchi, Claudia Regina Gonçalves, Carlos Henrique Camargo, Fabiana Cristina Pereira dos Santos, Daniela Bernardes Borges da Silva, Simone Guadagnucci Morillo, Adriano Abbud, Adriana Bugno, Maria do Carmo Sampaio Tavares Timenetsky, Terezinha Maria de Paiva |
| EPI_ISL_416032 | hCoV-19/Brazil/SPBR-10/2020 | South America / Brazil / Distrito Federal / | 2020-03-04 | National Influenza Center - Instituto Adolfo Lutz | Instituto Adolfo Lutz, Interdiciplinary Procedures Center, Strategic Laboratory | Claudio Tavares Sacchi, Claudia Regina Gonçalves, Carlos Henrique Camargo, Fabiana Cristina Pereira dos Santos, Daniela Bernardes Borges da Silva, Simone Guadagnucci Morillo, Adriano Abbud, Adriana Bugno, Maria do Carmo Sampaio Tavares Timenetsky, Terezinha Maria de Paiva |
| EPI_ISL_416033 | hCoV-19/Brazil/SPBR-11/2020 | South America / Brazil / Sao Paulo / Sao | 2020-03-03 | Hospital Israelita Albert Einstein | Instituto Adolfo Lutz, Interdiciplinary Procedures Center, Strategic Laboratory | Claudio Tavares Sacchi, Claudia Regina Gonçalves, Carlos Henrique Camargo, Erica Valessa Ramos Gomes, Fabiana Cristina Pereira dos Santos, Daniela Bernardes Borges da Silva, Simone Guadagnucci Morillo, Adriano Abbud, Adriana Bugno, Maria do Carmo Sampaio Tavares Timenetsky, Terezinha Maria de Paiva |
| EPI_ISL_416034 | hCoV-19/Brazil/SPBR-12/2020 | South America / Brazil / Sao Paulo / Sao | 2020-03-04 | Hospital Israelita Albert Einstein | Instituto Adolfo Lutz, Interdiciplinary Procedures Center, Strategic Laboratory | Claudio Tavares Sacchi, Claudia Regina Gonçalves, Carlos Henrique Camargo, Erica Valessa Ramos Gomes, Fabiana Cristina Pereira dos Santos, Daniela Bernardes Borges da Silva, Simone Guadagnucci Morillo, Adriano Abbud, Adriana Bugno, Maria do Carmo Sampaio Tavares Timenetsky, Terezinha Maria de Paiva |
| EPI_ISL_416035 | hCoV-19/Brazil/SPBR-13/2020 | South America / Brazil / Sao Paulo / Sao | 2020-03-05 | National Influenza Center - Instituto Adolfo Lutz | Instituto Adolfo Lutz, Interdiciplinary Procedures Center, Strategic Laboratory | Claudio Tavares Sacchi, Claudia Regina Gonçalves, Carlos Henrique Camargo, Erica Valessa Ramos Gomes, Fabiana Cristina Pereira dos Santos, Daniela Bernardes Borges da Silva, Simone Guadagnucci Morillo, Adriano Abbud, Adriana Bugno, Maria do Carmo Sampaio Tavares Timenetsky, Terezinha Maria de Paiva |
| EPI_ISL_416036 | hCoV-19/Brazil/SPBR-14/2020 | South America / Brazil / Sao Paulo / Sao | 2020-03-05 | National Influenza Center - Instituto Adolfo Lutz | Instituto Adolfo Lutz, Interdiciplinary Procedures Center, Strategic Laboratory | Claudio Tavares Sacchi, Claudia Regina Gonçalves, Carlos Henrique Camargo, Erica Valessa Ramos Gomes, Fabiana Cristina Pereira dos Santos, Daniela Bernardes Borges da Silva, Simone Guadagnucci Morillo, Adriano Abbud, Adriana Bugno, Maria do Carmo Sampaio Tavares Timenetsky, Terezinha Maria de Paiva |
| EPI_ISL_416042 | hCoV-19/Hangzhou/ZJU-02/2020 | Asia / China / Hangzhou | 2020-01-26 | State Key Laboratory for Diagnosis and Treatment of Infectious Diseases, National Clinical Research Center for Infectious Diseases, First Affiliated Hospital, Zhejiang University School of Medicine, Hangzhou, China. 310003 | State Key Laboratory for Diagnosis and Treatment of Infectious Diseases, National Clinical Research Center for Infectious Diseases, First Affiliated Hospital, Zhejiang University School of Medicine, Hangzhou, China. 310003 | Hangping Yao, Nanping Wu, Chao Jiang, Xiangyun Lu, Linfang Cheng, Fumin Liu, Zhigang Wu, Haibo Wu, Changzhong Jin, Min Zheng, Lanjuan Li |
| EPI_ISL_416044 | hCoV-19/Hangzhou/ZJU-03/2020 | Asia / China / Hangzhou | 2020-01-25 | State Key Laboratory for Diagnosis and Treatment of Infectious Diseases, National Clinical Research Center for Infectious Diseases, First Affiliated Hospital, Zhejiang University School of Medicine, Hangzhou, China 310003 | State Key Laboratory for Diagnosis and Treatment of Infectious Diseases, National Clinical Research Center for Infectious Diseases, First Affiliated Hospital, Zhejiang University School of Medicine, Hangzhou, China 310003 | Hangping Yao, Nanping Wu, Chao Jiang, Xiangyun Lu, Linfang Cheng, Fumin Liu, Zhigang Wu, Haibo Wu, Changzhong Jin, Min Zheng, Lanjuan Li |
| EPI_ISL_416046 | hCoV-19/Hangzhou/ZJU-04/2020 | Asia / China / Hangzhou | 2020-01-24 | State Key Laboratory for Diagnosis and Treatment of Infectious Diseases, National Clinical Research Center for Infectious Diseases, First Affiliated Hospital, Zhejiang University School of Medicine, Hangzhou, China 310003 | State Key Laboratory for Diagnosis and Treatment of Infectious Diseases, National Clinical Research Center for Infectious Diseases, First Affiliated Hospital, Zhejiang University School of Medicine, Hangzhou, China 310003 | Hangping Yao, Nanping Wu, Chao Jiang, Xiangyun Lu, Linfang Cheng, Fumin Liu, Zhigang Wu, Haibo Wu, Changzhong Jin, Min Zheng, Lanjuan Li |
| EPI_ISL_416047 | hCoV-19/Hangzhou/ZJU-06/2020 | Asia / China / Hangzhou | 2020-02-02 | State Key Laboratory for Diagnosis and Treatment of Infectious Diseases, National Clinical Research Center for Infectious Diseases, First Affiliated Hospital, Zhejiang University School of Medicine, Hangzhou, China 310003 | State Key Laboratory for Diagnosis and Treatment of Infectious Diseases, National Clinical Research Center for Infectious Diseases, First Affiliated Hospital, Zhejiang University School of Medicine, Hangzhou, China 310003 | Hangping Yao, Nanping Wu, Chao Jiang, Xiangyun Lu, Linfang Cheng, Fumin Liu, Zhigang Wu, Haibo Wu, Changzhong Jin, Min Zheng, Lanjuan Li |
| EPI_ISL_416314 | hCoV-19/Hong Kong/CUHK1/2020 | Asia / Hong Kong | 2020-02-07 | Department of Microbiology, Faculty of Medicine, The Chinese University of Hong Kong, Hong Kong SAR, China | Department of Microbiology, Faculty of Medicine, Chinese University of Hong Kong, Hong Kong SAR, China | Zigui Chen, Paul KS Chan |
| EPI_ISL_416316 | hCoV-19/Shanghai/SH0002/2020 | Asia / China / Shanghai | 2020-01-25 | Shanghai Public Health Clinical Center, Shanghai Medical College, Fudan University | National Research Center for Translational Medicine (Shanghai), Ruijin Hospital affiliated to Shanghai Jiao Tong University School of Medicine & Shanghai Public Health Clinical Center | Shengyue Wang, Xiaonan Zhang, Gang Lu, Yun Tan, Yun Ling, Hongzhou Lu, Saijuan Chen |
| EPI_ISL_416317 | hCoV-19/Shanghai/SH0003/2020 | Asia / China / Shanghai | 2020-01-25 | Shanghai Public Health Clinical Center, Shanghai Medical College, Fudan University | National Research Center for Translational Medicine (Shanghai), Ruijin Hospital affiliated to Shanghai Jiao Tong University School of Medicine & Shanghai Public Health Clinical Center | Shengyue Wang, Xiaonan Zhang, Gang Lu, Yun Tan, Yun Ling, Hongzhou Lu, Saijuan Chen |
| EPI_ISL_416318 | hCoV-19/Shanghai/SH0004/2020 | Asia / China / Shanghai | 2020-01-28 | Shanghai Public Health Clinical Center, Shanghai Medical College, Fudan University | National Research Center for Translational Medicine (Shanghai), Ruijin Hospital affiliated to Shanghai Jiao Tong University School of Medicine & Shanghai Public Health Clinical Center | Shengyue Wang, Xiaonan Zhang, Gang Lu, Yun Tan, Yun Ling, Hongzhou Lu, Saijuan Chen |
| EPI_ISL_416400 | hCoV-19/Shanghai/SH0112/2020 | Asia / China / Shanghai | 2020-02-02 | Shanghai Public Health Clinical Center, Shanghai Medical College, Fudan University | National Research Center for Translational Medicine (Shanghai), Ruijin Hospital affiliated to Shanghai Jiao Tong University School of Medicine & Shanghai Public Health Clinical Center | Shengyue Wang, Xiaonan Zhang, Gang Lu, Yun Tan, Yun Ling, Hongzhou Lu, Saijuan Chen |
| EPI_ISL_416401 | hCoV-19/Shanghai/SH0114/2020 | Asia / China / Shanghai | 2020-02-02 | Shanghai Public Health Clinical Center, Shanghai Medical College, Fudan University | National Research Center for Translational Medicine (Shanghai), Ruijin Hospital affiliated to Shanghai Jiao Tong University School of Medicine & Shanghai Public Health Clinical Center | Shengyue Wang, Xiaonan Zhang, Gang Lu, Yun Tan, Yun Ling, Hongzhou Lu, Saijuan Chen |
| EPI_ISL_416402 | hCoV-19/Shanghai/SH0115/2020 | Asia / China / Shanghai | 2020-02-11 | Shanghai Public Health Clinical Center, Shanghai Medical College, Fudan University | National Research Center for Translational Medicine (Shanghai), Ruijin Hospital affiliated to Shanghai Jiao Tong University School of Medicine & Shanghai Public Health Clinical Center | Shengyue Wang, Xiaonan Zhang, Gang Lu, Yun Tan, Yun Ling, Hongzhou Lu, Saijuan Chen |
| EPI_ISL_416403 | hCoV-19/Shanghai/SH0117/2020 | Asia / China / Shanghai | 2020-02-02 | Shanghai Public Health Clinical Center, Shanghai Medical College, Fudan University | National Research Center for Translational Medicine (Shanghai), Ruijin Hospital affiliated to Shanghai Jiao Tong University School of Medicine & Shanghai Public Health Clinical Center | Shengyue Wang, Xiaonan Zhang, Gang Lu, Yun Tan, Yun Ling, Hongzhou Lu, Saijuan Chen |
| EPI_ISL_416404 | hCoV-19/Shanghai/SH0119/2020 | Asia / China / Shanghai | 2020-02-09 | Shanghai Public Health Clinical Center, Shanghai Medical College, Fudan University | National Research Center for Translational Medicine (Shanghai), Ruijin Hospital affiliated to Shanghai Jiao Tong University School of Medicine & Shanghai Public Health Clinical Center | Shengyue Wang, Xiaonan Zhang, Gang Lu, Yun Tan, Yun Ling, Hongzhou Lu, Saijuan Chen |
| EPI_ISL_416405 | hCoV-19/Shanghai/SH0121/2020 | Asia / China / Shanghai | 2020-02-02 | Shanghai Public Health Clinical Center, Shanghai Medical College, Fudan University | National Research Center for Translational Medicine (Shanghai), Ruijin Hospital affiliated to Shanghai Jiao Tong University School of Medicine & Shanghai Public Health Clinical Center | Shengyue Wang, Xiaonan Zhang, Gang Lu, Yun Tan, Yun Ling, Hongzhou Lu, Saijuan Chen |
| EPI_ISL_416406 | hCoV-19/Shanghai/SH0125/2020 | Asia / China / Shanghai | 2020-02-15 | Shanghai Public Health Clinical Center, Shanghai Medical College, Fudan University | National Research Center for Translational Medicine (Shanghai), Ruijin Hospital affiliated to Shanghai Jiao Tong University School of Medicine & Shanghai Public Health Clinical Center | Shengyue Wang, Xiaonan Zhang, Gang Lu, Yun Tan, Yun Ling, Hongzhou Lu, Saijuan Chen |
| EPI_ISL_416407 | hCoV-19/Shanghai/SH0126/2020 | Asia / China / Shanghai | 2020-02-15 | Shanghai Public Health Clinical Center, Shanghai Medical College, Fudan University | National Research Center for Translational Medicine (Shanghai), Ruijin Hospital affiliated to Shanghai Jiao Tong University School of Medicine & Shanghai Public Health Clinical Center | Shengyue Wang, Xiaonan Zhang, Gang Lu, Yun Tan, Yun Ling, Hongzhou Lu, Saijuan Chen |
| EPI_ISL_416409 | hCoV-19/Shanghai/SH0128/2020 | Asia / China / Shanghai | 2020-02-02 | Shanghai Public Health Clinical Center, Shanghai Medical College, Fudan University | National Research Center for Translational Medicine (Shanghai), Ruijin Hospital affiliated to Shanghai Jiao Tong University School of Medicine & Shanghai Public Health Clinical Center | Shengyue Wang, Xiaonan Zhang, Gang Lu, Yun Tan, Yun Ling, Hongzhou Lu, Saijuan Chen |
| EPI_ISL_416410 | hCoV-19/Australia/VIC02/2020 | Oceania / Australia / Victoria / Melbourne | 2020-01-24 | Victorian Infectious Diseases Reference Laboratory (VIDRL) | Victorian Infectious Diseases Reference Laboratory and Microbiological Diagnostic Unit Public Health Laboratory, Doherty Institute | Caly L., Seemann T., Schultz M., Druce J., Taiaroa, G. |
| EPI_ISL_416411 | hCoV-19/Australia/VIC03/2020 | Oceania / Australia / Victoria / Melbourne | 2020-01-25 | Victorian Infectious Diseases Reference Laboratory (VIDRL) | Victorian Infectious Diseases Reference Laboratory and Microbiological Diagnostic Unit Public Health Laboratory, Doherty Institute | Caly L., Seemann T., Schultz M., Druce J., Taiaroa, G. |

| | | | | | | |
|---|---|---|---|---|---|---|
| EPI_ISL_416412 | hCoV-19/Australia/VIC04/2020 | Oceania / Australia / Victoria / Melbourne | 2020-03-02 | Victorian Infectious Diseases Reference Laboratory (VIDRL) | Victorian Infectious Diseases Reference Laboratory and Microbiological Diagnostic Unit Public Health Laboratory, Doherty Institute | Caly L., Seemann T., Schultz M., Druce J., Taiaroa, G. |
| EPI_ISL_416413 | hCoV-19/Australia/VIC05/2020 | Oceania / Australia / Victoria / Melbourne | 2020-03-05 | Victorian Infectious Diseases Reference Laboratory (VIDRL) | Victorian Infectious Diseases Reference Laboratory and Microbiological Diagnostic Unit Public Health Laboratory, Doherty Institute | Caly L., Seemann T., Schultz M., Druce J., Taiaroa, G. |
| EPI_ISL_416415 | hCoV-19/Australia/VIC07/2020 | Oceania / Australia / Victoria / Melbourne | 2020-02-08 | Victorian Infectious Diseases Reference Laboratory (VIDRL) | Victorian Infectious Diseases Reference Laboratory and Microbiological Diagnostic Unit Public Health Laboratory, Doherty Institute | Caly L., Seemann T., Schultz M., Druce J., Taiaroa, G. |
| EPI_ISL_416416 | hCoV-19/USA/CT-Yale-001/2020 | North America / USA / Connecticut | 2020-03-06 | Connecticut State Department of Public Health | Grubaugh Lab - Yale School of Public Health | Joseph Fauver, Chantal Vogels, Anderson Brito, Tara Alpert, Nagarjuna Cheemarla, Ellen Foxman, Anthony Muyombwe, Jafar Razeq, Richard Martinello, Albert Ko, Marie-Louise Landry, Nathan Grubaugh |
| EPI_ISL_416417 | hCoV-19/USA/CT-Yale-002/2020 | North America / USA / Connecticut | 2020-03-10 | Connecticut State Department of Public Health | Grubaugh Lab - Yale School of Public Health | Joseph Fauver, Chantal Vogels, Anderson Brito, Tara Alpert, Nagarjuna Cheemarla, Ellen Foxman, Anthony Muyombwe, Jafar Razeq, Richard Martinello, Albert Ko, Marie-Louise Landry, Nathan Grubaugh |
| EPI_ISL_416418 | hCoV-19/USA/CT-Yale-003/2020 | North America / USA / Connecticut | 2020-03-08 | Connecticut State Department of Public Health | Grubaugh Lab - Yale School of Public Health | Joseph Fauver, Chantal Vogels, Anderson Brito, Tara Alpert, Nagarjuna Cheemarla, Ellen Foxman, Anthony Muyombwe, Jafar Razeq, Richard Martinello, Albert Ko, Marie-Louise Landry, Nathan Grubaugh |
| EPI_ISL_416419 | hCoV-19/USA/CT-Yale-005/2020 | North America / USA / Connecticut | 2020-03-09 | Connecticut State Department of Public Health | Grubaugh Lab - Yale School of Public Health | Joseph Fauver, Chantal Vogels, Anderson Brito, Tara Alpert, Nagarjuna Cheemarla, Ellen Foxman, Anthony Muyombwe, Jafar Razeq, Richard Martinello, Albert Ko, Marie-Louise Landry, Nathan Grubaugh |
| EPI_ISL_416420 | hCoV-19/USA/CT-Yale-006/2020 | North America / USA / Connecticut | 2020-03-13 | Yale Clinical Virology Laboratory | Grubaugh Lab - Yale School of Public Health | Joseph Fauver, Chantal Vogels, Anderson Brito, Tara Alpert, Nagarjuna Cheemarla, Ellen Foxman, Anthony Muyombwe, Jafar Razeq, Richard Martinello, Albert Ko, Marie-Louise Landry, Nathan Grubaugh |
| EPI_ISL_416421 | hCoV-19/USA/CT-Yale-007/2020 | North America / USA / Connecticut | 2020-03-13 | Yale Clinical Virology Laboratory | Grubaugh Lab - Yale School of Public Health | Joseph Fauver, Chantal Vogels, Anderson Brito, Tara Alpert, Nagarjuna Cheemarla, Ellen Foxman, Anthony Muyombwe, Jafar Razeq, Richard Martinello, Albert Ko, Marie-Louise Landry, Nathan Grubaugh |
| EPI_ISL_416422 | hCoV-19/USA/CT-Yale-008/2020 | North America / USA / Connecticut | 2020-03-13 | Yale Clinical Virology Laboratory | Grubaugh Lab - Yale School of Public Health | Joseph Fauver, Chantal Vogels, Anderson Brito, Tara Alpert, Nagarjuna Cheemarla, Ellen Foxman, Anthony Muyombwe, Jafar Razeq, Richard Martinello, Albert Ko, Marie-Louise Landry, Nathan Grubaugh |
| EPI_ISL_416423 | hCoV-19/USA/CT-Yale-009/2020 | North America / USA / Connecticut | 2020-03-12 | Yale Clinical Virology Laboratory | Grubaugh Lab - Yale School of Public Health | Joseph Fauver, Chantal Vogels, Anderson Brito, Tara Alpert, Nagarjuna Cheemarla, Ellen Foxman, Anthony Muyombwe, Jafar Razeq, Richard Martinello, Albert Ko, Marie-Louise Landry, Nathan Grubaugh |
| EPI_ISL_416424 | hCoV-19/USA/CT-Yale-010/2020 | North America / USA / Connecticut | 2020-03-14 | Yale Clinical Virology Laboratory | Grubaugh Lab - Yale School of Public Health | Joseph Fauver, Chantal Vogels, Anderson Brito, Tara Alpert, Nagarjuna Cheemarla, Ellen Foxman, Anthony Muyombwe, Jafar Razeq, Richard Martinello, Albert Ko, Marie-Louise Landry, Nathan Grubaugh |
| EPI_ISL_416425 | hCoV-19/Hangzhou/ZJU-07/2020 | Asia / China / Hangzhou | 2020-02-03 | State Key Laboratory for Diagnosis and Treatment of Infectious Diseases, National Clinical Research Center for Infectious Diseases, First Affiliated Hospital, Zhejiang University School of Medicine, Hangzhou, China 310003 | State Key Laboratory for Diagnosis and Treatment of Infectious Diseases, National Clinical Research Center for Infectious Diseases, First Affiliated Hospital, Zhejiang University School of Medicine, Hangzhou, China 310003 | Hangping Yao, Nanping Wu, Chao Jiang, Xiangyun Lu, Linfang Cheng, Fumin Liu, Zhigang Wu, Haibo Wu, Changzhong Jin, Min Zheng, Lanjuan Li |
| EPI_ISL_416426 | hCoV-19/Hungary/mbl1/2020 | Europe / Hungary / Budapest | 2020-03-17 | Virological Research Group, Szentágothai Research Centre, University of Pécs | Bioinformatics Research Group, Szentágothai Research Centre, University of Pécs | Péter Urbán, Endre Gábor Tóth, Gábor Kemenesi, Róbert Herczeg, Attila Gyenesei, Ferenc Jakab |
| EPI_ISL_416427 | hCoV-19/Vietnam/38142/2020 | Asia / Vietnam / Thanhhoa | 2020-01-24 | National Influenza Center, National Institute of Hygiene and Epidemiology (NIHE) | National Influenza Center, National Institute of Hygiene and Epidemiology (NIHE) | Le Quynh Mai, Taichiro Takemura, Meng Ling Moi, Takeshi Nabeshima, Nguyen Le Khanh Hang, Hoang Vu Mai Phuong, Ung Thi Hong Trang, Le Thi Thanh, Nguyen Vu Son, Vuong Duc Cuong, Pham Thi Hien, Tran Thu Huong, Nguyen Phuong Anh, Pham Hong Quynh Anh, Kouichi Morita, Futoshi Hasebe, Dang Duc Anh |
| EPI_ISL_416428 | hCoV-19/Vietnam/39607/2020 | Asia / Vietnam / Quangning | 2020-03-07 | National Influenza Center, National Institute of Hygiene and Epidemiology (NIHE) | National Influenza Center, National Institute of Hygiene and Epidemiology (NIHE) | Le Quynh Mai, Taichiro Takemura, Meng Ling Moi, Takeshi Nabeshima, Nguyen Le Khanh Hang, Hoang Vu Mai Phuong, Ung Thi Hong Trang, Le Thi Thanh, Nguyen Vu Son, Vuong Duc Cuong, Pham Thi Hien, Tran Thu Huong, Nguyen Phuong Anh, Pham Hong Quynh Anh, Kouichi Morita, Futoshi Hasebe, Dang Duc Anh |
| EPI_ISL_416429 | hCoV-19/Vietnam/CM99/2020 | Asia / Vietnam / Vinhphuc | 2020-02-11 | National Influenza Center, National Institute of Hygiene and Epidemiology (NIHE) | National Influenza Center, National Institute of Hygiene and Epidemiology (NIHE) | Le Quynh Mai, Taichiro Takemura, Meng Ling Moi, Takeshi Nabeshima, Nguyen Le Khanh Hang, Hoang Vu Mai Phuong, Ung Thi Hong Trang, Le Thi Thanh, Nguyen Vu Son, Vuong Duc Cuong, Pham Thi Hien, Tran Thu Huong, Nguyen Phuong Anh, Pham Hong Quynh Anh, Kouichi Morita, Futoshi Hasebe, Dang Duc Anh |
| EPI_ISL_416430 | hCoV-19/Vietnam/CM295/2020 | Asia / Vietnam / Hanoi | 2020-03-06 | National Influenza Center, National Institute of Hygiene and Epidemiology (NIHE) | National Influenza Center, National Institute of Hygiene and Epidemiology (NIHE) | Le Quynh Mai, Taichiro Takemura, Meng Ling Moi, Takeshi Nabeshima, Nguyen Le Khanh Hang, Hoang Vu Mai Phuong, Ung Thi Hong Trang, Le Thi Thanh, Nguyen Vu Son, Vuong Duc Cuong, Pham Thi Hien, Tran Thu Huong, Nguyen Phuong Anh, Pham Hong Quynh Anh, Kouichi Morita, Futoshi Hasebe, Dang Duc Anh |
| EPI_ISL_416431 | hCoV-19/Vietnam/CM296/2020 | Asia / Vietnam / Hanoi | 2020-03-06 | National Influenza Center, National Institute of Hygiene and Epidemiology (NIHE) | National Influenza Center, National Institute of Hygiene and Epidemiology (NIHE) | Le Quynh Mai, Taichiro Takemura, Meng Ling Moi, Takeshi Nabeshima, Nguyen Le Khanh Hang, Hoang Vu Mai Phuong, Ung Thi Hong Trang, Le Thi Thanh, Nguyen Vu Son, Vuong Duc Cuong, Pham Thi Hien, Tran Thu Huong, Nguyen Phuong Anh, Pham Hong Quynh Anh, Kouichi Morita, Futoshi Hasebe, Dang Duc Anh |
| EPI_ISL_416432 | hCoV-19/Saudi Arabia/KAIMRC-Alghoribi/2020 | Asia / Saudi Arabia / Riyadh | 2020-03-07 | Clinical Microbiology Lab | Infectious Disease Research Department, King Abdullah International Medical Research Center (KAIMRC) | Majed Alghoribi, Sadeem Alhayli, Abdulrahman Alswaji, Liliane Okdah, Sameera Al Johani, Michel Doumith |
| EPI_ISL_416433 | hCoV-19/USA/WA-UW77/2020 | North America / USA / Washington | 2020-03-10 | UW Virology Lab | UW Virology Lab | Pavitra Roychoudhury, Hong Xie, Keith Jerome, Alexander Greninger |
| EPI_ISL_416434 | hCoV-19/USA/WA-UW78/2020 | North America / USA / Washington | 2020-03-10 | UW Virology Lab | UW Virology Lab | Pavitra Roychoudhury, Hong Xie, Keith Jerome, Alexander Greninger |
| EPI_ISL_416435 | hCoV-19/USA/WA-UW79/2020 | North America / USA / Washington | 2020-03-10 | UW Virology Lab | UW Virology Lab | Pavitra Roychoudhury, Hong Xie, Keith Jerome, Alexander Greninger |
| EPI_ISL_416436 | hCoV-19/USA/WA-UW80/2020 | North America / USA / Washington | 2020-03-10 | UW Virology Lab | UW Virology Lab | Pavitra Roychoudhury, Hong Xie, Keith Jerome, Alexander Greninger |
| EPI_ISL_416437 | hCoV-19/USA/WA-UW81/2020 | North America / USA / Washington | 2020-03-10 | UW Virology Lab | UW Virology Lab | Pavitra Roychoudhury, Hong Xie, Keith Jerome, Alexander Greninger |
| EPI_ISL_416438 | hCoV-19/USA/WA-UW82/2020 | North America / USA / Washington | 2020-03-10 | UW Virology Lab | UW Virology Lab | Pavitra Roychoudhury, Hong Xie, Keith Jerome, Alexander Greninger |
| EPI_ISL_416439 | hCoV-19/USA/WA-UW83/2020 | North America / USA / Washington | 2020-03-10 | UW Virology Lab | UW Virology Lab | Pavitra Roychoudhury, Hong Xie, Keith Jerome, Alexander Greninger |
| EPI_ISL_416440 | hCoV-19/USA/WA-UW84/2020 | North America / USA / Washington | 2020-03-10 | UW Virology Lab | UW Virology Lab | Pavitra Roychoudhury, Hong Xie, Keith Jerome, Alexander Greninger |
| EPI_ISL_416441 | hCoV-19/USA/WA-UW85/2020 | North America / USA / Washington | 2020-03-10 | UW Virology Lab | UW Virology Lab | Pavitra Roychoudhury, Hong Xie, Keith Jerome, Alexander Greninger |
| EPI_ISL_416442 | hCoV-19/USA/WA-UW86/2020 | North America / USA / Washington | 2020-03-10 | UW Virology Lab | UW Virology Lab | Pavitra Roychoudhury, Hong Xie, Keith Jerome, Alexander Greninger |
| EPI_ISL_416443 | hCoV-19/USA/WA-UW87/2020 | North America / USA / Washington | 2020-03-10 | UW Virology Lab | UW Virology Lab | Pavitra Roychoudhury, Hong Xie, Keith Jerome, Alexander Greninger |
| EPI_ISL_416444 | hCoV-19/USA/WA-UW88/2020 | North America / USA / Washington | 2020-03-10 | UW Virology Lab | UW Virology Lab | Pavitra Roychoudhury, Hong Xie, Keith Jerome, Alexander Greninger |
| EPI_ISL_416445 | hCoV-19/USA/WA-UW89/2020 | North America / USA / Washington | 2020-03-10 | UW Virology Lab | UW Virology Lab | Pavitra Roychoudhury, Hong Xie, Keith Jerome, Alexander Greninger |
| EPI_ISL_416446 | hCoV-19/USA/WA-UW90/2020 | North America / USA / Washington | 2020-03-10 | UW Virology Lab | UW Virology Lab | Pavitra Roychoudhury, Hong Xie, Keith Jerome, Alexander Greninger |
| EPI_ISL_416447 | hCoV-19/USA/WA-UW91/2020 | North America / USA / Washington | 2020-03-10 | UW Virology Lab | UW Virology Lab | Pavitra Roychoudhury, Hong Xie, Keith Jerome, Alexander Greninger |
| EPI_ISL_416448 | hCoV-19/USA/WA-UW92/2020 | North America / USA / Washington | 2020-03-11 | UW Virology Lab | UW Virology Lab | Pavitra Roychoudhury, Hong Xie, Keith Jerome, Alexander Greninger |
| EPI_ISL_416449 | hCoV-19/USA/WA-UW93/2020 | North America / USA / Washington | 2020-03-11 | UW Virology Lab | UW Virology Lab | Pavitra Roychoudhury, Hong Xie, Keith Jerome, Alexander Greninger |
| EPI_ISL_416450 | hCoV-19/USA/WA-UW94/2020 | North America / USA / Washington | 2020-03-11 | UW Virology Lab | UW Virology Lab | Pavitra Roychoudhury, Hong Xie, Keith Jerome, Alexander Greninger |
| EPI_ISL_416451 | hCoV-19/USA/WA-UW95/2020 | North America / USA / Washington | 2020-03-10 | UW Virology Lab | UW Virology Lab | Pavitra Roychoudhury, Hong Xie, Keith Jerome, Alexander Greninger |
| EPI_ISL_416452 | hCoV-19/USA/WA-UW96/2020 | North America / USA / Washington | 2020-03-10 | UW Virology Lab | UW Virology Lab | Pavitra Roychoudhury, Hong Xie, Keith Jerome, Alexander Greninger |
| EPI_ISL_416453 | hCoV-19/USA/WA-UW36/2020 | North America / USA / Washington | 2020-03-07 | UW Virology Lab | UW Virology Lab | Pavitra Roychoudhury, Hong Xie, Keith Jerome, Alexander Greninger |
| EPI_ISL_416454 | hCoV-19/USA/WA-UW37/2020 | North America / USA / Washington | 2020-03-06 | UW Virology Lab | UW Virology Lab | Pavitra Roychoudhury, Hong Xie, Keith Jerome, Alexander Greninger |
| EPI_ISL_416455 | hCoV-19/USA/WA-UW38/2020 | North America / USA / Washington | 2020-03-07 | UW Virology Lab | UW Virology Lab | Pavitra Roychoudhury, Hong Xie, Keith Jerome, Alexander Greninger |
| EPI_ISL_416456 | hCoV-19/USA/WA-UW39/2020 | North America / USA / Washington | 2020-03-06 | UW Virology Lab | UW Virology Lab | Pavitra Roychoudhury, Hong Xie, Keith Jerome, Alexander Greninger |
| EPI_ISL_416457 | hCoV-19/USA/CA-MG0987/2020 | North America / USA / California / San Di | 2020-03-18 | Andersen Lab, The Scripps Research Institute | Andersen Lab, The Scripps Research Institute | Mark Zeller, Catie Anderson, Emily Spencer, Sarah Topol, Raphaelle Klitting, Refugio Robles-Sikisaka, Karthik Gangavarapu, Laura Nicholson, Kristian Andersen |
| EPI_ISL_416458 | hCoV-19/Kuwait/KU12/2020 | Asia / Kuwait / Hawali | 2020-03-02 | Virology laboratory Ministry of Health Kuwait sequenced at Dasman Diabetes Institute | Dasman Diabetes Institute | Fahd Al-Mulla, Sumi John, Sara Alqabandi, Rasheeba Iqbal, Motasem Melhem, Ebaa alOzairi, Qais Al-Duwairi |
| EPI_ISL_416459 | hCoV-19/USA/WA-S4/2020 | North America / USA / Washington / King | 2020-03-01 | Seattle Flu Study | Seattle Flu Study | Chu et al |
| EPI_ISL_416460 | hCoV-19/USA/WA-S5/2020 | North America / USA / Washington / King | 2020-02-29 | Seattle Flu Study | Seattle Flu Study | Chu et al |
| EPI_ISL_416461 | hCoV-19/USA/WA-S6/2020 | North America / USA / Washington / King | 2020-02-29 | Seattle Flu Study | Seattle Flu Study | Chu et al |
| EPI_ISL_416462 | hCoV-19/USA/WA-S7/2020 | North America / USA / Washington | 2020-02-24 | Seattle Flu Study | Seattle Flu Study | Chu et al |
| EPI_ISL_416463 | hCoV-19/USA/WA-S8/2020 | North America / USA / Washington / King | 2020-02-24 | Seattle Flu Study | Seattle Flu Study | Chu et al |
| EPI_ISL_416464 | hCoV-19/USA/WA-S9/2020 | North America / USA / Washington / Sno | 2020-03-01 | Seattle Flu Study | Seattle Flu Study | Chu et al |
| EPI_ISL_416465 | hCoV-19/USA/WA-S10/2020 | North America / USA / Washington / King | 2020-02-29 | Seattle Flu Study | Seattle Flu Study | Chu et al |
| EPI_ISL_416466 | hCoV-19/USA/WA-S11/2020 | North America / USA / Washington / King | 2020-03-03 | Seattle Flu Study | Seattle Flu Study | Chu et al |
| EPI_ISL_416467 | hCoV-19/Belgium/MTR-03021/2020 | Europe / Belgium / Holsbeek | 2020-03-02 | KU Leuven, Clinical and Epidemiological Virology | KU Leuven, Clinical and Epidemiological Virology | Bert Vanmechelen, Tony Wawina, Joan Martí-Carreras, Piet Maes |
| EPI_ISL_416468 | hCoV-19/Belgium/GMH-03022/2020 | Europe / Belgium / Holsbeek | 2020-03-02 | KU Leuven, Clinical and Epidemiological Virology | KU Leuven, Clinical and Epidemiological Virology | Bert Vanmechelen, Tony Wawina, Joan Martí-Carreras, Piet Maes |
| EPI_ISL_416469 | hCoV-19/Belgium/SN-03031/2020 | Europe / Belgium / Kessel-Lo | 2020-03-03 | KU Leuven, Clinical and Epidemiological Virology | KU Leuven, Clinical and Epidemiological Virology | Bert Vanmechelen, Tony Wawina, Joan Martí-Carreras, Piet Maes |
| EPI_ISL_416470 | hCoV-19/Belgium/DB-03023/2020 | Europe / Belgium / Couthuin | 2020-03-02 | KU Leuven, Clinical and Epidemiological Virology | KU Leuven, Clinical and Epidemiological Virology | Bert Vanmechelen, Tony Wawina, Joan Martí-Carreras, Piet Maes |
| EPI_ISL_416471 | hCoV-19/Belgium/DBD-03024/2020 | Europe / Belgium / Kessel-Lo | 2020-03-02 | KU Leuven, Clinical and Epidemiological Virology | KU Leuven, Clinical and Epidemiological Virology | Bert Vanmechelen, Tony Wawina, Joan Martí-Carreras, Piet Maes |
| EPI_ISL_416472 | hCoV-19/Belgium/UMF-03025/2020 | Europe / Belgium / Kessel-Lo | 2020-03-02 | KU Leuven, Clinical and Epidemiological Virology | KU Leuven, Clinical and Epidemiological Virology | Bert Vanmechelen, Tony Wawina, Joan Martí-Carreras, Piet Maes |
| EPI_ISL_416473 | hCoV-19/Hangzhou/ZJU-08/2020 | Asia / China / Hangzhou | 2020-01-26 | State Key Laboratory for Diagnosis and Treatment of Infectious Diseases, National Clinical Research Center for Infectious Diseases, First Affiliated Hospital, Zhejiang University School of Medicine, Hangzhou, China 310003 | State Key Laboratory for Diagnosis and Treatment of Infectious Diseases, National Clinical Research Center for Infectious Diseases, First Affiliated Hospital, Zhejiang University School of Medicine, Hangzhou, China 310003 | Hangping Yao, Nanping Wu, Chao Jiang, Xiangyun Lu, Linfang Cheng, Fumin Liu, Zhigang Wu, Haibo Wu, Changzhong Jin, Min Zheng, Lanjuan Li |
| EPI_ISL_416474 | hCoV-19/Hangzhou/ZJU-09/2020 | Asia / China / Hangzhou | 2020-01-28 | State Key Laboratory for Diagnosis and Treatment of Infectious Diseases, National Clinical Research Center for Infectious Diseases, First Affiliated Hospital, Zhejiang University School of Medicine, Hangzhou, China 310003 | State Key Laboratory for Diagnosis and Treatment of Infectious Diseases, National Clinical Research Center for Infectious Diseases, First Affiliated Hospital, Zhejiang University School of Medicine, Hangzhou, China 310003 | Hangping Yao, Nanping Wu, Chao Jiang, Xiangyun Lu, Linfang Cheng, Fumin Liu, Zhigang Wu, Haibo Wu, Changzhong Jin, Min Zheng, Lanjuan Li |
| EPI_ISL_416475 | hCoV-19/Belgium/DBA-03032/2020 | Europe / Belgium / Leuven | 2020-03-03 | KU Leuven, Clinical and Epidemiological Virology | KU Leuven, Clinical and Epidemiological Virology | Bert Vanmechelen, Tony Wawina, Joan Martí-Carreras, Piet Maes |
| EPI_ISL_416476 | hCoV-19/Belgium/MTR-03026/2020 | Europe / Belgium / Holsbeek | 2020-03-02 | KU Leuven, Clinical and Epidemiological Virology | KU Leuven, Clinical and Epidemiological Virology | Bert Vanmechelen, Tony Wawina, Joan Martí-Carreras, Piet Maes |
| EPI_ISL_416477 | hCoV-19/Georgia/Tb-390/2020 | Asia / Georgia / Tbilisi | 2020-03-08 | R. G. Lugar Center for Public Health Research, National Center for Disease Control and Public Health (NCDC) of Georgia. | R. G. Lugar Center for Public Health Research, National Center for Disease Control and Public Health (NCDC) of Georgia. | Marine Murtskhvaladze, Nato Kotaria, Ann Machablishvili, Lela Sabadze, Mari Gavashelidze, Ana Papkiauri, Meri Pantsulaia, Gvantsa Brachveli, Tata Imnadze, Tamar Jashiashvili, Tea Tevdoradze, Ketevan Sidamonidze, Ekaterine Khmaladze, Ekaterine Zhghenti, Roena Sukhiashvili, Mariam Zakalashvili, Lela Urushadze, Magda Dgebuadze, Giorgi Tomashvili, Davit Tsaguria, Ekaterine Zangaladze, Nino Berishvili, Gvantsa Chanturia, Adam Kotorashvili, Maia Alkhazashvili, Irma Burjanadze, Anna Kasradze, Khatuna Zakhashvili, Paata Imnadze, Amiran Gamkrelidze. |
| EPI_ISL_416478 | hCoV-19/Georgia/Tb-673/2020 | Asia / Georgia / Tbilisi | 2020-03-14 | R. G. Lugar Center for Public Health Research, National Center for Disease Control and Public Health (NCDC) of Georgia. | R. G. Lugar Center for Public Health Research, National Center for Disease Control and Public Health (NCDC) of Georgia. | Marine Murtskhvaladze, Nato Kotaria, Ann Machablishvili, Lela Sabadze, Mari Gavashelidze, Ana Papkiauri, Meri Pantsulaia, Gvantsa Brachveli, Tata Imnadze, Tamar Jashiashvili, Tea Tevdoradze, Ketevan Sidamonidze, Ekaterine Khmaladze, Ekaterine Zhghenti, Roena Sukhiashvili, Mariam Zakalashvili, Lela Urushadze, Magda Dgebuadze, Giorgi Tomashvili, Davit Tsaguria, Ekaterine Zangaladze, Nino Berishvili, Gvantsa Chanturia, Adam Kotorashvili, Maia Alkhazashvili, Irma Burjanadze, Anna Kasradze, Khatuna Zakhashvili, Paata Imnadze, Amiran Gamkrelidze. |
| EPI_ISL_416479 | hCoV-19/Georgia/Tb-273/2020 | Asia / Georgia / Tbilisi | 2020-03-05 | R. G. Lugar Center for Public Health Research, National Center for Disease Control and Public Health (NCDC) of Georgia. | R. G. Lugar Center for Public Health Research, National Center for Disease Control and Public Health (NCDC) of Georgia. | Marine Murtskhvaladze, Nato Kotaria, Ann Machablishvili, Lela Sabadze, Mari Gavashelidze, Ana Papkiauri, Meri Pantsulaia, Gvantsa Brachveli, Tata Imnadze, Tamar Jashiashvili, Tea Tevdoradze, Ketevan Sidamonidze, Ekaterine Khmaladze, Ekaterine Zhghenti, Roena Sukhiashvili, Mariam Zakalashvili, Lela Urushadze, Magda Dgebuadze, Giorgi Tomashvili, Davit Tsaguria, Ekaterine Zangaladze, Nino Berishvili, Gvantsa Chanturia, Adam Kotorashvili, Maia Alkhazashvili, Irma Burjanadze, Anna Kasradze, Khatuna Zakhashvili, Paata Imnadze, Amiran Gamkrelidze. |
| EPI_ISL_416480 | hCoV-19/Georgia/Tb-537/2020 | Asia / Georgia / Tbilisi | 2020-03-11 | R. G. Lugar Center for Public Health Research, National Center for Disease Control and Public Health (NCDC) of Georgia. | R. G. Lugar Center for Public Health Research, National Center for Disease Control and Public Health (NCDC) of Georgia. | Ann Machablishvili, Nato Kotaria, Marine Murtskhvaladze, Lela Sabadze, Mari Gavashelidze, Ana Papkiauri, Meri Pantsulaia, Gvantsa Brachveli, Tata Imnadze, Tamar Jashiashvili, Tea Tevdoradze, Ketevan Sidamonidze, Ekaterine Khmaladze, Ekaterine Zhghenti, Roena Sukhiashvili, Mariam Zakalashvili, Lela Urushadze, Magda Dgebuadze, Giorgi Tomashvili, Davit Tsaguria, Ekaterine Zangaladze, Nino Berishvili, Gvantsa Chanturia, Adam Kotorashvili, Maia Alkhazashvili, Irma Burjanadze, Anna Kasradze, Khatuna Zakhashvili, Paata Imnadze, Amiran Gamkrelidze. |
| EPI_ISL_416481 | hCoV-19/Georgia/Tb-712/2020 | Asia / Georgia / Tbilisi | 2020-03-16 | R. G. Lugar Center for Public Health Research, National Center for Disease Control and Public Health (NCDC) of Georgia. | R. G. Lugar Center for Public Health Research, National Center for Disease Control and Public Health (NCDC) of Georgia. | Gvantsa Chanturia, Marine Murtskhvaladze, Nato Kotaria, Ann Machablishvili, Lela Sabadze, Mari Gavashelidze, Ana Papkiauri, Meri Pantsulaia, Gvantsa Brachveli, Tata Imnadze, Tamar Jashiashvili, Tea Tevdoradze, Ketevan Sidamonidze, Ekaterine Khmaladze, Ekaterine Zhghenti, Roena Sukhiashvili, Mariam Zakalashvili, Lela Urushadze, Magda Dgebuadze, Giorgi Tomashvili, Davit Tsaguria, Ekaterine Zangaladze, Nino Berishvili, Adam Kotorashvili, Maia Alkhazashvili, Irma Burjanadze, Anna Kasradze, Khatuna Zakhashvili, Paata Imnadze, Amiran Gamkrelidze. |
| EPI_ISL_416482 | hCoV-19/Georgia/Tb/2020 | Asia / Georgia / Tbilisi | 2020-03-13 | R. G. Lugar Center for Public Health Research, National Center for Disease Control and Public Health (NCDC) of Georgia. | R. G. Lugar Center for Public Health Research, National Center for Disease Control and Public Health (NCDC) of Georgia. | Adam Kotorashvili, Marine Murtskhvaladze, Nato Kotaria, Ann Machablishvili, Lela Sabadze, Mari Gavashelidze, Ana Papkiauri, Meri Pantsulaia, Gvantsa Brachveli, Tata Imnadze, Tamar Jashiashvili, Tea Tevdoradze, Ketevan Sidamonidze, Ekaterine Khmaladze, Ekaterine Zhghenti, Roena Sukhiashvili, Mariam Zakalashvili, Lela Urushadze, Magda Dgebuadze, Giorgi Tomashvili, Davit Tsaguria, Ekaterine Zangaladze, Nino Berishvili, Gvantsa Chanturia, Maia Alkhazashvili, Irma Burjanadze, Anna Kasradze, Khatuna Zakhashvili, Paata Imnadze, Amiran Gamkrelidze. |
| EPI_ISL_416484 | hCoV-19/Spain/Valencia5/2020 | Europe / Spain / Comunitat Valenciana / | 2020-02-27 | Servicio de Microbiología. Consorcio Hospital General Universitario de Valencia | Sequencing and Bioinformatics Service and Molecular Epidemiology Research Group. FISABIO-Public Health | María Dolores Ocete, Concepcion Gimeno, Giuseppe D'Auria, Griselda De Marco, Neris García-González, María Alma Bracho, Fernando González-Candelas |
| EPI_ISL_416485 | hCoV-19/Spain/Valencia6/2020 | Europe / Spain / Comunitat Valenciana / | 2020-02-27 | Servicio de Microbiología. Consorcio Hospital General Universitario de Valencia | Sequencing and Bioinformatics Service and Molecular Epidemiology Research Group. FISABIO-Public Health | Griselda De Marco, Neris García-González, María Alma Bracho, María Dolores Ocete, Concepcion Gimeno, Giuseppe D'Auria, Fernando González-Candelas |
| EPI_ISL_416486 | hCoV-19/Spain/Valencia7/2020 | Europe / Spain / Comunitat Valenciana / | 2020-03-02 | Servicio de Microbiología. Consorcio Hospital General Universitario de Valencia | Sequencing and Bioinformatics Service and Molecular Epidemiology Research Group. FISABIO-Public Health | Neris García-González, María Alma Bracho, María Dolores Ocete, Concepcion Gimeno, Giuseppe D'Auria, Griselda De Marco, Fernando González-Candelas |
| EPI_ISL_416487 | hCoV-19/Spain/Valencia8/2020 | Europe / Spain / Comunitat Valenciana / | 2020-03-04 | Servicio de Microbiología. Consorcio Hospital General Universitario de Valencia | Sequencing and Bioinformatics Service and Molecular Epidemiology Research Group. FISABIO-Public Health | Giuseppe D'Auria, Griselda De Marco, Neris García-González, María Alma Bracho, María Dolores Ocete, Concepcion Gimeno, Fernando González-Candelas |
| EPI_ISL_416488 | hCoV-19/Poland/PL_P1/2020 | Europe / Poland / Zielonogorskie | 2020-03-03 | ViroGenetics - BSL3 Laboratory of Virology; Human Genome Variation Research Group & Genomics Centre MCB; Bioinformatics Research Group Department of Virology | ViroGenetics - BSL3 Laboratory of Virology; Human Genome Variation Research Group & Genomics Centre MCB; Bioinformatics Research Group Department of Virology | Aleksandra Milewska, Ewelina Polpiech, Agata Jarosz, Adrianna Klajmon, Kamila Marszałek, Katarzyna Pancer, Magdalena Rzeczkowska, Tomasz Wołkowicz, Katarzyna Zacharczuk, Agnieszka Kołakowska-Kulesza, Natalia Wolaniuk, Ewelina Hallman-Szelińska, Paweł P Łabaj, Wojciech Branicki, Krzysztof Pyrć |
| EPI_ISL_416489 | hCoV-19/USA/WI-UW-02/2020 | North America / USA / Wisconsin / Madis | 2020-03-15 | University of Wisconsin-Madison AIDS Vaccine Research Laboratories | University of Wisconsin-Madison AIDS Vaccine Research Laboratories | Gage Moreno, Katarina Braun, et al. AIDS Vaccine Research Laboratories |
| EPI_ISL_416491 | hCoV-19/USA/WI-UW-04/2020 | North America / USA / Wisconsin / Waun | 2020-03-15 | University of Wisconsin-Madison AIDS Vaccine Research Laboratories | University of Wisconsin-Madison AIDS Vaccine Research Laboratories | Gage Moreno, Katarina Braun, et al. AIDS Vaccine Research Laboratories |
| EPI_ISL_416492 | hCoV-19/USA/WI-UW-05/2020 | North America / USA / Wisconsin / Fitchb | 2020-03-15 | University of Wisconsin-Madison AIDS Vaccine Research Laboratories | University of Wisconsin-Madison AIDS Vaccine Research Laboratories | Gage Moreno, Katarina Braun, et al. AIDS Vaccine Research Laboratories |
| EPI_ISL_416493 | hCoV-19/France/HF2196/2020 | Europe / France / Hauts de France / Chât | 2020-03-08 | CH Jean de Navarre Laboratoire de Biologie | National Reference Center for Viruses of Respiratory Infections, Institut Pasteur, Paris | Mélnie Albert, Marion Barbet, Sylvie Behillil, Méline Bizard, Angela Brisebarre, Flora Donati, Etienne Simon-Lorière, Vincent Enouf, Maud Vanpeene, Sylvie van der Werf |
| EPI_ISL_416494 | hCoV-19/France/N2223/2020 | Europe / France / Normandie / Rouen | 2020-03-04 | Centre Hositalier Universitaire de Rouen Laboratoire de Virologie | National Reference Center for Viruses of Respiratory Infections, Institut Pasteur, Paris | Mélnie Albert, Marion Barbet, Sylvie Behillil, Méline Bizard, Angela Brisebarre, Flora Donati, Etienne Simon-Lorière, Vincent Enouf, Maud Vanpeene, Sylvie van der Werf, Jean-Christophe Plantier |
| EPI_ISL_416495 | hCoV-19/France/HF2234/2020 | Europe / France / Hauts de France / Com | 2020-03-10 | Centre Hospitalier Compiègne Laboratoire de Biologie | National Reference Center for Viruses of Respiratory Infections, Institut Pasteur, Paris | Mélnie Albert, Marion Barbet, Sylvie Behillil, Méline Bizard, Angela Brisebarre, Flora Donati, Etienne Simon-Lorière, Vincent Enouf, Maud Vanpeene, Sylvie van der Werf, Raulin Olivia |
| EPI_ISL_416496 | hCoV-19/France/HF2237/2020 | Europe / France / Hauts de France / Com | 2020-03-10 | Centre Hospitalier Compiègne Laboratoire de Biologie | National Reference Center for Viruses of Respiratory Infections, Institut Pasteur, Paris | Mélnie Albert, Marion Barbet, Sylvie Behillil, Méline Bizard, Angela Brisebarre, Flora Donati, Etienne Simon-Lorière, Vincent Enouf, Maud Vanpeene, Sylvie van der Werf, Raulin Olivia |
| EPI_ISL_416497 | hCoV-19/France/HF2239/2020 | Europe / France / Hauts de France / Com | 2020-03-10 | Centre Hospitalier Compiègne Laboratoire de Biologie | National Reference Center for Viruses of Respiratory Infections, Institut Pasteur, Paris | Mélnie Albert, Marion Barbet, Sylvie Behillil, Méline Bizard, Angela Brisebarre, Flora Donati, Etienne Simon-Lorière, Vincent Enouf, Maud Vanpeene, Sylvie van der Werf, Raulin Olivia |
| EPI_ISL_416498 | hCoV-19/France/IDF2256/2020 | Europe / France / Ile de France / Garches | 2020-03-11 | Institut Médico légal- Hop R. Poincaré | National Reference Center for Viruses of Respiratory Infections, Institut Pasteur, Paris | Mélnie Albert, Marion Barbet, Sylvie Behillil, Méline Bizard, Angela Brisebarre, Flora Donati, Etienne Simon-Lorière, Vincent Enouf, Maud Vanpeene, Sylvie van der Werf |
| EPI_ISL_416499 | hCoV-19/France/IDF2278/2020 | Europe / France / Ile de France / Longjum | 2020-03-11 | LABM GH nord Essonne | National Reference Center for Viruses of Respiratory Infections, Institut Pasteur, Paris | Mélnie Albert, Marion Barbet, Sylvie Behillil, Méline Bizard, Angela Brisebarre, Flora Donati, Etienne Simon-Lorière, Vincent Enouf, Maud Vanpeene, Sylvie van der Werf |
| EPI_ISL_416700 | hCoV-19/USA/WA-UW162/2020 | North America / USA / Washington | 2020-03-13 | UW Virology Lab | UW Virology Lab | Pavitra Roychoudhury, Hong Xie, Keith Jerome, Alexander Greninger |
| EPI_ISL_416701 | hCoV-19/USA/WA-UW163/2020 | North America / USA / Washington | 2020-03-13 | UW Virology Lab | UW Virology Lab | Pavitra Roychoudhury, Hong Xie, Keith Jerome, Alexander Greninger |
| EPI_ISL_416702 | hCoV-19/USA/WA-UW164/2020 | North America / USA / Washington | 2020-03-13 | UW Virology Lab | UW Virology Lab | Pavitra Roychoudhury, Hong Xie, Keith Jerome, Alexander Greninger |
| EPI_ISL_416703 | hCoV-19/USA/WA-UW165/2020 | North America / USA / Washington | 2020-03-13 | UW Virology Lab | UW Virology Lab | Pavitra Roychoudhury, Hong Xie, Keith Jerome, Alexander Greninger |
| EPI_ISL_416705 | hCoV-19/USA/WA-UW167/2020 | North America / USA / Washington | 2020-03-13 | UW Virology Lab | UW Virology Lab | Pavitra Roychoudhury, Hong Xie, Keith Jerome, Alexander Greninger |
| EPI_ISL_416706 | hCoV-19/USA/WA-UW168/2020 | North America / USA / Washington | 2020-03-13 | UW Virology Lab | UW Virology Lab | Pavitra Roychoudhury, Hong Xie, Keith Jerome, Alexander Greninger |
| EPI_ISL_416707 | hCoV-19/USA/WA-UW169/2020 | North America / USA / Washington | 2020-03-14 | UW Virology Lab | UW Virology Lab | Pavitra Roychoudhury, Hong Xie, Keith Jerome, Alexander Greninger |
| EPI_ISL_416708 | hCoV-19/USA/WA-UW170/2020 | North America / USA / Washington | 2020-03-14 | UW Virology Lab | UW Virology Lab | Pavitra Roychoudhury, Hong Xie, Keith Jerome, Alexander Greninger |
| EPI_ISL_416709 | hCoV-19/USA/WA-UW171/2020 | North America / USA / Washington | 2020-03-13 | UW Virology Lab | UW Virology Lab | Pavitra Roychoudhury, Hong Xie, Keith Jerome, Alexander Greninger |
| EPI_ISL_416710 | hCoV-19/USA/WA-UW172/2020 | North America / USA / Washington | 2020-03-13 | UW Virology Lab | UW Virology Lab | Pavitra Roychoudhury, Hong Xie, Keith Jerome, Alexander Greninger |
| EPI_ISL_416711 | hCoV-19/USA/WA-UW173/2020 | North America / USA / Washington | 2020-03-15 | UW Virology Lab | UW Virology Lab | Pavitra Roychoudhury, Hong Xie, Keith Jerome, Alexander Greninger |
| EPI_ISL_416712 | hCoV-19/USA/WA-UW174/2020 | North America / USA / Washington | 2020-03-14 | UW Virology Lab | UW Virology Lab | Pavitra Roychoudhury, Hong Xie, Keith Jerome, Alexander Greninger |
| EPI_ISL_416713 | hCoV-19/USA/WA-UW175/2020 | North America / USA / Washington | 2020-03-15 | UW Virology Lab | UW Virology Lab | Pavitra Roychoudhury, Hong Xie, Keith Jerome, Alexander Greninger |
| EPI_ISL_416714 | hCoV-19/USA/WA-UW176/2020 | North America / USA / Washington | 2020-03-14 | UW Virology Lab | UW Virology Lab | Pavitra Roychoudhury, Hong Xie, Keith Jerome, Alexander Greninger |
| EPI_ISL_416715 | hCoV-19/USA/WA-UW177/2020 | North America / USA / Washington | 2020-03-15 | UW Virology Lab | UW Virology Lab | Pavitra Roychoudhury, Hong Xie, Keith Jerome, Alexander Greninger |
| EPI_ISL_416716 | hCoV-19/USA/WA-UW178/2020 | North America / USA / Washington | 2020-03-13 | UW Virology Lab | UW Virology Lab | Pavitra Roychoudhury, Hong Xie, Keith Jerome, Alexander Greninger |
| EPI_ISL_416717 | hCoV-19/USA/WA-UW179/2020 | North America / USA / Washington | 2020-03-15 | UW Virology Lab | UW Virology Lab | Pavitra Roychoudhury, Hong Xie, Keith Jerome, Alexander Greninger |
| EPI_ISL_416718 | hCoV-19/USA/WA-UW180/2020 | North America / USA / Washington | 2020-03-14 | UW Virology Lab | UW Virology Lab | Pavitra Roychoudhury, Hong Xie, Keith Jerome, Alexander Greninger |
| EPI_ISL_416719 | hCoV-19/USA/WA-UW181/2020 | North America / USA / Washington | 2020-03-14 | UW Virology Lab | UW Virology Lab | Pavitra Roychoudhury, Hong Xie, Keith Jerome, Alexander Greninger |
| EPI_ISL_416720 | hCoV-19/USA/WA-UW182/2020 | North America / USA / Washington | 2020-03-13 | UW Virology Lab | UW Virology Lab | Pavitra Roychoudhury, Hong Xie, Keith Jerome, Alexander Greninger |
| EPI_ISL_416721 | hCoV-19/USA/WA-UW183/2020 | North America / USA / Washington | 2020-03-13 | UW Virology Lab | UW Virology Lab | Pavitra Roychoudhury, Hong Xie, Keith Jerome, Alexander Greninger |
| EPI_ISL_416722 | hCoV-19/USA/WA-UW184/2020 | North America / USA / Washington | 2020-03-12 | UW Virology Lab | UW Virology Lab | Pavitra Roychoudhury, Hong Xie, Keith Jerome, Alexander Greninger |
| EPI_ISL_416723 | hCoV-19/USA/WA-UW185/2020 | North America / USA / Washington | 2020-03-14 | UW Virology Lab | UW Virology Lab | Pavitra Roychoudhury, Hong Xie, Keith Jerome, Alexander Greninger |
| EPI_ISL_416724 | hCoV-19/USA/WA-UW186/2020 | North America / USA / Washington | 2020-03-13 | UW Virology Lab | UW Virology Lab | Pavitra Roychoudhury, Hong Xie, Keith Jerome, Alexander Greninger |
| EPI_ISL_416725 | hCoV-19/USA/WA-UW187/2020 | North America / USA / Washington | 2020-03-13 | UW Virology Lab | UW Virology Lab | Pavitra Roychoudhury, Hong Xie, Keith Jerome, Alexander Greninger |
| EPI_ISL_416726 | hCoV-19/USA/WA-UW188/2020 | North America / USA / Washington | 2020-03-13 | UW Virology Lab | UW Virology Lab | Pavitra Roychoudhury, Hong Xie, Keith Jerome, Alexander Greninger |
| EPI_ISL_416727 | hCoV-19/USA/WA-UW189/2020 | North America / USA / Washington | 2020-03-13 | UW Virology Lab | UW Virology Lab | Pavitra Roychoudhury, Hong Xie, Keith Jerome, Alexander Greninger |
| EPI_ISL_416728 | hCoV-19/USA/WA-UW190/2020 | North America / USA / Washington | 2020-03-13 | UW Virology Lab | UW Virology Lab | Pavitra Roychoudhury, Hong Xie, Keith Jerome, Alexander Greninger |
| EPI_ISL_416729 | hCoV-19/USA/WA-UW191/2020 | North America / USA / Washington | 2020-03-13 | UW Virology Lab | UW Virology Lab | Pavitra Roychoudhury, Hong Xie, Keith Jerome, Alexander Greninger |
| EPI_ISL_416733 | hCoV-19/England/SHEF-BFCEE/2020 | Europe / United Kingdom / England / Sou | 2020-03-07 | Virology Department, Sheffield Teaching Hospitals NHS Foundation Trust | Department of Infection, Immunity and Cardiovascular Disease, The Florey Institute, The Medical School, University of Sheffield | Thushan de Silva, Matthew Parker, Adri Angyal, Rebecca Brown, Matthew Wyles, Mehmet Yavuz, Mohammad Raza, Cariad Evans |
| EPI_ISL_416734 | hCoV-19/England/SHEF-BFCFD/2020 | Europe / United Kingdom / England / Sou | 2020-03-09 | Virology Department, Sheffield Teaching Hospitals NHS Foundation Trust | Department of Infection, Immunity and Cardiovascular Disease, The Florey Institute, The Medical School, University of Sheffield | Thushan de Silva, Matthew Parker, Adri Angyal, Rebecca Brown, Matthew Wyles, Mehmet Yavuz, Mohammad Raza, Cariad Evans |
| EPI_ISL_416736 | hCoV-19/England/SHEF-BFD18/2020 | Europe / United Kingdom / England / Sou | 2020-03-09 | Virology Department, Sheffield Teaching Hospitals NHS Foundation Trust | Department of Infection, Immunity and Cardiovascular Disease, The Florey Institute, The Medical School, University of Sheffield | Thushan de Silva, Matthew Parker, Adri Angyal, Rebecca Brown, Matthew Wyles, Mehmet Yavuz, Mohammad Raza, Cariad Evans |
| EPI_ISL_416737 | hCoV-19/England/SHEF-BFD27/2020 | Europe / United Kingdom / England / Lon | 2020-03-03 | Virology Department, Sheffield Teaching Hospitals NHS Foundation Trust | Department of Infection, Immunity and Cardiovascular Disease, The Florey Institute, The Medical School, University of Sheffield | Thushan de Silva, Matthew Parker, Adri Angyal, Rebecca Brown, Matthew Wyles, Mehmet Yavuz, Mohammad Raza, Cariad Evans |
| EPI_ISL_416738 | hCoV-19/England/SHEF-BFD36/2020 | Europe / United Kingdom / England / Nott | 2020-03-09 | Virology Department, Sheffield Teaching Hospitals NHS Foundation Trust | Department of Infection, Immunity and Cardiovascular Disease, The Florey Institute, The Medical School, University of Sheffield | Thushan de Silva, Matthew Parker, Adri Angyal, Rebecca Brown, Matthew Wyles, Mehmet Yavuz, Mohammad Raza, Cariad Evans |
| EPI_ISL_416739 | hCoV-19/England/SHEF-BFD45/2020 | Europe / United Kingdom / England / Sou | 2020-03-09 | Virology Department, Sheffield Teaching Hospitals NHS Foundation Trust | Department of Infection, Immunity and Cardiovascular Disease, The Florey Institute, The Medical School, University of Sheffield | Thushan de Silva, Matthew Parker, Adri Angyal, Rebecca Brown, Matthew Wyles, Mehmet Yavuz, Mohammad Raza, Cariad Evans |
| EPI_ISL_416740 | hCoV-19/England/SHEF-BFD54/2020 | Europe / United Kingdom / England / Sou | 2020-03-03 | Virology Department, Sheffield Teaching Hospitals NHS Foundation Trust | Department of Infection, Immunity and Cardiovascular Disease, The Florey Institute, The Medical School, University of Sheffield | Thushan de Silva, Matthew Parker, Adri Angyal, Rebecca Brown, Matthew Wyles, Mehmet Yavuz, Mohammad Raza, Cariad Evans |
| EPI_ISL_416741 | hCoV-19/Lithuania/ChVir1632/2020 | Europe / Lithuania / Vilnius | 2020-02 | National Public Health Surveillance Laboratory, Vilnius, Lithuania | Charite Universitaetsmedizin Berlin, Institute of Virology | Victor M Corman, Julia Schneider, Jörn Beheim-Schwarzbach, Talitha Veith, Barbara Muehlemann, Terry Jones, Ana Steponkiene, Christian Drosten |
| EPI_ISL_416742 | hCoV-19/Czech Republic/ChVir1630/2020 | Europe / Czech Republic / Prague | 2020-03-01 | NRL for Influenza, Centrum Epidemiology and Microbiology of National Institute of Public Health, Czech Republic | Charite Universitaetsmedizin Berlin, Institute of Virology | Victor M Corman, Julia Schneider, Jörn Beheim-Schwarzbach, Talitha Veith, Barbara Muehlemann, Terry Jones, Alexander Nagy, Jaromira Vecerova, Dusan Trnka, Ludmila Novakova, Helena Jirincova, Christian Drosten |
| EPI_ISL_416743 | hCoV-19/Czech Republic/ChVir1912/2020 | Europe / Czech Republic / Prague | 2020-03-01 | NRL for Influenza, Centrum Epidemiology and Microbiology of National Institute of Public Health, Czech Republic | Charite Universitaetsmedizin Berlin, Institute of Virology | Victor M Corman, Julia Schneider, Jörn Beheim-Schwarzbach, Talitha Veith, Barbara Muehlemann, Terry Jones, Alexander Nagy, Jaromira Vecerova, Dusan Trnka, Ludmila Novakova, Helena Jirincova, Christian Drosten |
| EPI_ISL_416744 | hCoV-19/Hungary/mbl49/2020 | Europe / Hungary / Baranya | 2020-03-20 | Virological Research Group, Szentágothai Research Centre | Bioinformatics Research Group, Szentágothai Research Centre | Péter Urbán, Endre Gábor Tóth, Gábor Kemenesi, Róbert Herczeg, Attila Gyenesei, Ferenc Jakab |
| EPI_ISL_416745 | hCoV-19/France/Pollionay_1733/2020 | Europe / France / ARA | 2020-03-10 | CNR Virus des Infections Respiratoires - France SUD | CNR Virus des Infections Respiratoires - France SUD | Bal, Antonin; Destras, Gregory; Gaymard, Alexandre; Bouscambert-Duchamp, Maude; Cheynet, Valérie; Brengel-Pesce, Karen; Morfin-Sherpa, Florence; Valette, Martine; Josset, Laurence; Lina, Bruno. |
| EPI_ISL_416746 | hCoV-19/France/Valence_425/2020 | Europe / France / ARA | 2020-03-03 | CNR Virus des Infections Respiratoires - France SUD | CNR Virus des Infections Respiratoires - France SUD | Bal, Antonin; Destras, Gregory; Gaymard, Alexandre; Bouscambert-Duchamp, Maude; Cheynet, Valérie; Brengel-Pesce, Karen; Morfin-Sherpa, Florence; Valette, Martine; Josset, Laurence; Lina, Bruno. |
| EPI_ISL_416747 | hCoV-19/France/Lyon_487/2020 | Europe / France / ARA | 2020-03-04 | Institut des Agents Infectieux (IAI) Hospices Civils de Lyon | CNR Virus des Infections Respiratoires - France SUD | Bal, Antonin; Destras, Gregory; Gaymard, Alexandre; Bouscambert-Duchamp, Maude; Cheynet, Valérie; Brengel-Pesce, Karen; Morfin-Sherpa, Florence; Valette, Martine; Josset, Laurence; Lina, Bruno. |
| EPI_ISL_416748 | hCoV-19/France/Lyon_508/2020 | Europe / France / ARA | 2020-03-04 | Institut des Agents Infectieux (IAI) Hospices Civils de Lyon | CNR Virus des Infections Respiratoires - France SUD | Bal, Antonin; Destras, Gregory; Gaymard, Alexandre; Bouscambert-Duchamp, Maude; Cheynet, Valérie; Brengel-Pesce, Karen; Morfin-Sherpa, Florence; Valette, Martine; Josset, Laurence; Lina, Bruno. |
| EPI_ISL_416749 | hCoV-19/France/Valence_532/2020 | Europe / France / ARA | 2020-03-04 | Centre Hospitalier de Valence | CNR Virus des Infections Respiratoires - France SUD | Bal, Antonin; Destras, Gregory; Gaymard, Alexandre; Bouscambert-Duchamp, Maude; Cheynet, Valérie; Brengel-Pesce, Karen; Morfin-Sherpa, Florence; Valette, Martine; Josset, Laurence; Lina, Bruno. |
| EPI_ISL_416750 | hCoV-19/France/Lyon_683/2020 | Europe / France / ARA | 2020-03-06 | Institut des Agents Infectieux (IAI) Hospices Civils de Lyon | CNR Virus des Infections Respiratoires - France SUD | Bal, Antonin; Destras, Gregory; Gaymard, Alexandre; Bouscambert-Duchamp, Maude; Cheynet, Valérie; Brengel-Pesce, Karen; Morfin-Sherpa, Florence; Valette, Martine; Josset, Laurence; Lina, Bruno. |
| EPI_ISL_416751 | hCoV-19/France/Clermont-Ferrand_651/2020 | Europe / France / ARA | 2020-03-05 | CHU Gabriel Montpied | CNR Virus des Infections Respiratoires - France SUD | Bal, Antonin; Destras, Gregory; Gaymard, Alexandre; Bouscambert-Duchamp, Maude; Cheynet, Valérie; Brengel-Pesce, Karen; Morfin-Sherpa, Florence; Valette, Martine; Josset, Laurence; Lina, Bruno. |
| EPI_ISL_416752 | hCoV-19/France/Clermont-Ferrand_650/2020 | Europe / France / ARA | 2020-03-04 | CHU Gabriel Montpied | CNR Virus des Infections Respiratoires - France SUD | Bal, Antonin; Destras, Gregory; Gaymard, Alexandre; Bouscambert-Duchamp, Maude; Cheynet, Valérie; Brengel-Pesce, Karen; Morfin-Sherpa, Florence; Valette, Martine; Josset, Laurence; Lina, Bruno. |
| EPI_ISL_416753 | hCoV-19/France/Lyon_06464 /2020 | Europe / France / ARA | 2020-03-06 | Institut des Agents Infectieux (IAI) Hospices Civils de Lyon | CNR Virus des Infections Respiratoires - France SUD | Bal, Antonin; Destras, Gregory; Gaymard, Alexandre; Bouscambert-Duchamp, Maude; Cheynet, Valérie; Brengel-Pesce, Karen; Morfin-Sherpa, Florence; Valette, Martine; Josset, Laurence; Lina, Bruno. |
| EPI_ISL_416754 | hCoV-19/France/Lyon_06487/2020 | Europe / France / ARA | 2020-03-06 | Institut des Agents Infectieux (IAI) Hospices Civils de Lyon | CNR Virus des Infections Respiratoires - France SUD | Bal, Antonin; Destras, Gregory; Gaymard, Alexandre; Bouscambert-Duchamp, Maude; Cheynet, Valérie; Brengel-Pesce, Karen; Morfin-Sherpa, Florence; Valette, Martine; Josset, Laurence; Lina, Bruno. |
| EPI_ISL_416756 | hCoV-19/France/Lyon_06531/2020 | Europe / France / ARA | 2020-03-06 | Institut des Agents Infectieux (IAI) Hospices Civils de Lyon | CNR Virus des Infections Respiratoires - France SUD | Bal, Antonin; Destras, Gregory; Gaymard, Alexandre; Bouscambert-Duchamp, Maude; Cheynet, Valérie; Brengel-Pesce, Karen; Morfin-Sherpa, Florence; Valette, Martine; Josset, Laurence; Lina, Bruno. |
| EPI_ISL_416757 | hCoV-19/France/Bourg-en-Bresse_06678/2020 | Europe / France / ARA | 2020-03-07 | Centre Hospitalier de Bourg en Bresse | CNR Virus des Infections Respiratoires - France SUD | Bal, Antonin; Destras, Gregory; Gaymard, Alexandre; Bouscambert-Duchamp, Maude; Cheynet, Valérie; Brengel-Pesce, Karen; Morfin-Sherpa, Florence; Valette, Martine; Josset, Laurence; Lina, Bruno. |
| EPI_ISL_416758 | hCoV-19/France/Lyon_0693/2020 | Europe / France / ARA | 2020-03-08 | Institut des Agents Infectieux (IAI) Hospices Civils de Lyon | CNR Virus des Infections Respiratoires - France SUD | Bal, Antonin; Destras, Gregory; Gaymard, Alexandre; Bouscambert-Duchamp, Maude; Cheynet, Valérie; Brengel-Pesce, Karen; Morfin-Sherpa, Florence; Valette, Martine; Josset, Laurence; Lina, Bruno. |
| EPI_ISL_416829 | hCoV-19/Malaysia/MKAK-CL-2020-5045/2020 | Asia / Malaysia / Selangor | 2020-01-24 | National Public Health Laboratory | Malaysia Genome Institute | Mohd Noor Mat Isa, Irni Suhayu Sapian, Yusuf Muhammad Noor, Nurhezreen Md Iqbal, Mohd Faizal Abu Bakar, Enizza Kasim, Shamsidar Sopie, Siti Noraini Othman, Azrin Ahmad, Nor Azfa Johari, Norazimah Tajudin, Noorliza Mohamad Noordin, W Afiza W Mohd Arifin, Rehan Shuhada Abu Bakar, Yu Kie Chem, Selvanesan Sengol, Hani Mat Hussin, Shahrul Hisham Zainal Ariffin |
| EPI_ISL_416830 | hCoV-19/USA/NY-NYUMC2/2020 | North America / USA / New York City | 2020-03-16 | NYU Langone Health | Department of Pathology and Medicine, New York University School of Medicine | John Chen, Dacia Dimartino, Xiaojun Feng, Adriana Heguy, Megan Hogan, Emily Huang, George Jour, Christian Marier, Matt Maurano, Mark Mulligan, Peter Meyn, Marie Samanovic-Golden, Amy Rapkiewicz, Guomiao Shen, Matija Snuderl, Gael Westby, Paul Zappile |
| EPI_ISL_416831 | hCoV-19/USA/NY-NYUMC3/2020 | North America / USA / New York City | 2020-03-16 | NYU Langone Health | Department of Pathology and Medicine, New York University School of Medicine | John Chen, Dacia Dimartino, Xiaojun Feng, Adriana Heguy, Megan Hogan, Emily Huang, George Jour, Christian Marier, Matt Maurano, Mark Mulligan, Peter Meyn, Marie Samanovic-Golden, Amy Rapkiewicz, Guomiao Shen, Matija Snuderl, Gael Westby, Paul Zappile |
| EPI_ISL_416832 | hCoV-19/USA/NY-NYUMC4/2020 | North America / USA / New York City | 2020-03-16 | NYU Langone Health | Department of Pathology and Medicine, New York University School of Medicine | John Chen, Dacia Dimartino, Xiaojun Feng, Adriana Heguy, Megan Hogan, Emily Huang, George Jour, Christian Marier, Matt Maurano, Mark Mulligan, Peter Meyn, Marie Samanovic-Golden, Amy Rapkiewicz, Guomiao Shen, Matija Snuderl, Gael Westby, Paul Zappile |
| EPI_ISL_416866 | hCoV-19/Malaysia/MKAK-CL-2020-5047/2020 | Asia / Malaysia / Selangor | 2020-01-24 | National Public Health Laboratory | Malaysia Genome Institute | Mohd Noor Mat Isa, Irni Suhayu Sapian, Yusuf Muhammad Noor, Nurhezreen Md Iqbal, Mohd Faizal Abu Bakar, Enizza Kasim, Shamsidar Sopie, Siti Noraini Othman, Azrin Ahmad, Nor Azfa Johari, Norazimah Tajudin, Noorliza Mohamad Noordin, W Afiza W Mohd Arifin, Rehan Shuhada Abu Bakar, Yu Kie Chem, Selvanesan Sengol, Hani Mat Hussin, Shahrul Hisham Zainal Ariffin |
| EPI_ISL_416885 | hCoV-19/Malaysia/MKAK-CL-2020-5096/2020 | Asia / Malaysia / Selangor | 2020-01-30 | National Public Health Laboratory | Malaysia Genome Institute | Mohd Noor Mat Isa, Irni Suhayu Sapian, Yusuf Muhammad Noor, Nurhezreen Md Iqbal, Mohd Faizal Abu Bakar, Enizza Kasim, Shamsidar Sopie, Siti Noraini Othman, Azrin Ahmad, Nor Azfa Johari, Norazimah Tajudin, Noorliza Mohamad Noordin, W Afiza W Mohd Arifin, Rehan Shuhada Abu Bakar, Yu Kie Chem, Selvanesan Sengol, Hani Mat Hussin, Shahrul Hisham Zainal Ariffin |
| EPI_ISL_416886 | hCoV-19/Malaysia/MKAK-CL-2020-6430/2020 | Asia / Malaysia / Selangor | 2020-02-04 | National Public Health Laboratory | Malaysia Genome Institute | Mohd Noor Mat Isa, Irni Suhayu Sapian, Yusuf Muhammad Noor, Nurhezreen Md Iqbal, Mohd Faizal Abu Bakar, Enizza Kasim, Shamsidar Sopie, Siti Noraini Othman, Azrin Ahmad, Nor Azfa Johari, Norazimah Tajudin, Noorliza Mohamad Noordin, W Afiza W Mohd Arifin, Rehan Shuhada Abu Bakar, Yu Kie Chem, Selvanesan Sengol, Hani Mat Hussin, Shahrul Hisham Zainal Ariffin |
| EPI_ISL_417020 | hCoV-19/Belgium/ULG-6939/2020 | Europe / Belgium / Liège | 2020-03-15 | Department of Clinical Microbiology | GIGA Medical Genomics | Durkin Keith, Artesi Maria, Bontems Sébastien, Boreux Raphaël, Meex Cécile, Melin Pierrette, Hayette Marie-Pierre, Bours Vincent. |
| EPI_ISL_417021 | hCoV-19/Belgium/ULG-6942/2020 | Europe / Belgium / Liège | 2020-03-15 | Department of Clinical Microbiology | GIGA Medical Genomics | Durkin Keith, Artesi Maria, Bontems Sébastien, Boreux Raphaël, Meex Cécile, Melin Pierrette, Hayette Marie-Pierre, Bours Vincent. |
| EPI_ISL_417022 | hCoV-19/Belgium/ULG-6948/2020 | Europe / Belgium / Liège | 2020-03-15 | Department of Clinical Microbiology | GIGA Medical Genomics | Durkin Keith, Artesi Maria, Bontems Sébastien, Boreux Raphaël, Meex Cécile, Melin Pierrette, Hayette Marie-Pierre, Bours Vincent. |
| EPI_ISL_417023 | hCoV-19/Belgium/ULG-6950/2020 | Europe / Belgium / Liège | 2020-03-15 | Department of Clinical Microbiology | GIGA Medical Genomics | Durkin Keith, Artesi Maria, Bontems Sébastien, Boreux Raphaël, Meex Cécile, Melin Pierrette, Hayette Marie-Pierre, Bours Vincent. |
| EPI_ISL_417025 | hCoV-19/Belgium/ULG-7019/2020 | Europe / Belgium / Liège | 2020-03-15 | Department of Clinical Microbiology | GIGA Medical Genomics | Durkin Keith, Artesi Maria, Bontems Sébastien, Boreux Raphaël, Meex Cécile, Melin Pierrette, Hayette Marie-Pierre, Bours Vincent. |
| EPI_ISL_417026 | hCoV-19/USA/UT-08/2020 | North America / USA / Utah | 2020-03-20 | Utah Public Health Laboratory | Utah Public Health Laboratory | Erin Young, Kelly Oakeson |
| EPI_ISL_417028 | hCoV-19/USA/UT-020/2020 | North America / USA / Utah | 2020-03-20 | Utah Public Health Laboratory | Utah Public Health Laboratory | Erin Young, Kelly Oakeson |
| EPI_ISL_417300 | hCoV-19/England/20110000106/2020 | Europe / United Kingdom / England | 2020-03-05 | Respiratory Virus Unit, Microbiology Services Colindale, Public Health England | Respiratory Virus Unit, Microbiology Services Colindale, Public Health England | Monica Galiano, Shahjahan Miah, Angie Lackenby, Omolola Akinbami, Tiina Talts, Leena Bhaw, Richard Myers, Steven Platt, Kirstin Edwards, Jonathan Hubb, Joanna Ellis, Maria Zambon |
| EPI_ISL_417301 | hCoV-19/England/20110000606/2020 | Europe / United Kingdom / England | 2020-03-06 | Respiratory Virus Unit, Microbiology Services Colindale, Public Health England | Respiratory Virus Unit, Microbiology Services Colindale, Public Health England | Monica Galiano, Shahjahan Miah, Angie Lackenby, Omolola Akinbami, Tiina Talts, Leena Bhaw, Richard Myers, Steven Platt, Kirstin Edwards, Jonathan Hubb, Joanna Ellis, Maria Zambon |
| EPI_ISL_417302 | hCoV-19/England/20110000706/2020 | Europe / United Kingdom / England | 2020-03-05 | Respiratory Virus Unit, Microbiology Services Colindale, Public Health England | Respiratory Virus Unit, Microbiology Services Colindale, Public Health England | Monica Galiano, Shahjahan Miah, Angie Lackenby, Omolola Akinbami, Tiina Talts, Leena Bhaw, Richard Myers, Steven Platt, Kirstin Edwards, Jonathan Hubb, Joanna Ellis, Maria Zambon |
| EPI_ISL_417307 | hCoV-19/England/20110023706/2020 | Europe / United Kingdom / England | 2020-03-08 | Respiratory Virus Unit, Microbiology Services Colindale, Public Health England | Respiratory Virus Unit, Microbiology Services Colindale, Public Health England | Monica Galiano, Shahjahan Miah, Angie Lackenby, Omolola Akinbami, Tiina Talts, Leena Bhaw, Richard Myers, Steven Platt, Kirstin Edwards, Jonathan Hubb, Joanna Ellis, Maria Zambon |
| EPI_ISL_417311 | hCoV-19/England/20110058706/2020 | Europe / United Kingdom / England | 2020-03-09 | Respiratory Virus Unit, Microbiology Services Colindale, Public Health England | Respiratory Virus Unit, Microbiology Services Colindale, Public Health England | Monica Galiano, Shahjahan Miah, Angie Lackenby, Omolola Akinbami, Tiina Talts, Leena Bhaw, Richard Myers, Steven Platt, Kirstin Edwards, Jonathan Hubb, Joanna Ellis, Maria Zambon |
| EPI_ISL_417312 | hCoV-19/England/20110059306/2020 | Europe / United Kingdom / England | 2020-03-08 | Respiratory Virus Unit, Microbiology Services Colindale, Public Health England | Respiratory Virus Unit, Microbiology Services Colindale, Public Health England | Monica Galiano, Shahjahan Miah, Angie Lackenby, Omolola Akinbami, Tiina Talts, Leena Bhaw, Richard Myers, Steven Platt, Kirstin Edwards, Jonathan Hubb, Joanna Ellis, Maria Zambon |
| EPI_ISL_417313 | hCoV-19/England/20110097506/2020 | Europe / United Kingdom / England | 2020-03-08 | Respiratory Virus Unit, Microbiology Services Colindale, Public Health England | Respiratory Virus Unit, Microbiology Services Colindale, Public Health England | Monica Galiano, Shahjahan Miah, Angie Lackenby, Omolola Akinbami, Tiina Talts, Leena Bhaw, Richard Myers, Steven Platt, Kirstin Edwards, Jonathan Hubb, Joanna Ellis, Maria Zambon |
| EPI_ISL_417314 | hCoV-19/England/20112112106/2020 | Europe / United Kingdom / England | 2020-03-10 | Respiratory Virus Unit, Microbiology Services Colindale, Public Health England | Respiratory Virus Unit, Microbiology Services Colindale, Public Health England | Monica Galiano, Shahjahan Miah, Angie Lackenby, Omolola Akinbami, Tiina Talts, Leena Bhaw, Richard Myers, Steven Platt, Kirstin Edwards, Jonathan Hubb, Joanna Ellis, Maria Zambon |
| EPI_ISL_417315 | hCoV-19/England/20118044606/2020 | Europe / United Kingdom / England | 2020-03-13 | Respiratory Virus Unit, Microbiology Services Colindale, Public Health England | Respiratory Virus Unit, Microbiology Services Colindale, Public Health England | Monica Galiano, Shahjahan Miah, Angie Lackenby, Omolola Akinbami, Tiina Talts, Leena Bhaw, Richard Myers, Steven Platt, Kirstin Edwards, Jonathan Hubb, Joanna Ellis, Maria Zambon |
| EPI_ISL_417316 | hCoV-19/USA/CA-CDPH-UC12/2020 | North America / USA / California / San M | 2020-03-11 | Chiu Laboratory, University of California, San Francisco | Chiu Laboratory, University of California, San Francisco | Xianding Deng, Scot Federman, Wei Gu, and Charles Y. Chiu |
| EPI_ISL_417317 | hCoV-19/USA/CA-SCCPHD-UC13/2020 | North America / USA / California / Santa | 2020-03-02 | Santa Clara County Public Health Department | Chiu Laboratory, University of California, San Francisco | Xianding Deng, Scot Federman, Wei Gu, Elsa Villarino, Brandon Bonin, Debra A. Wadford, and Charles Y. Chiu |
| EPI_ISL_417318 | hCoV-19/USA/CA-SCCPHD-UC14/2020 | North America / USA / California / Santa | 2020-02-29 | Santa Clara County Public Health Department | Chiu Laboratory, University of California, San Francisco | Xianding Deng, Scot Federman, Wei Gu, Elsa Villarino, Brandon Bonin, Debra A. Wadford, and Charles Y. Chiu |
| EPI_ISL_417319 | hCoV-19/USA/CA-SCCPHD-UC15/2020 | North America / USA / California / Santa | 2020-03-05 | Santa Clara County Public Health Department | Chiu Laboratory, University of California, San Francisco | Xianding Deng, Scot Federman, Wei Gu, Elsa Villarino, Brandon Bonin, Debra A. Wadford, and Charles Y. Chiu |
| EPI_ISL_417320 | hCoV-19/USA/CA-SCCPHD-UC16/2020 | North America / USA / California / Santa | 2020-03-04 | Santa Clara County Public Health Department | Chiu Laboratory, University of California, San Francisco | Xianding Deng, Scot Federman, Wei Gu, Elsa Villarino, Brandon Bonin, Debra A. Wadford, and Charles Y. Chiu |
| EPI_ISL_417321 | hCoV-19/USA/CA-SCCPHD-UC17/2020 | North America / USA / California / Santa | 2020-03-08 | Santa Clara County Public Health Department | Chiu Laboratory, University of California, San Francisco | Xianding Deng, Scot Federman, Wei Gu, Elsa Villarino, Brandon Bonin, Debra A. Wadford, and Charles Y. Chiu |
| EPI_ISL_417322 | hCoV-19/USA/CA-CDPH-UC18/2020 | North America / USA / California / San M | 2020-03-04 | California Department of Public Health | Chiu Laboratory, University of California, San Francisco | Xianding Deng, Scot Federman, Chao-Yang Pan, Hugo Guevara,Wei Gu, Debra A. Wadford, and Charles Y. Chiu |
| EPI_ISL_417325 | hCoV-19/USA/CA-CDPH-UC21/2020 | North America / USA / California / Solano | 2020-03-03 | California Department of Public Health | Chiu Laboratory, University of California, San Francisco | Xianding Deng, Scot Federman, Chao-Yang Pan, Hugo Guevara,Wei Gu, Debra A. Wadford, and Charles Y. Chiu |
| EPI_ISL_417326 | hCoV-19/USA/CA-CDPH-UC22/2020 | North America / USA / California / Marin | 2020-03-04 | California Department of Public Health | Chiu Laboratory, University of California, San Francisco | Xianding Deng, Scot Federman, Chao-Yang Pan, Hugo Guevara,Wei Gu, Debra A. Wadford, and Charles Y. Chiu |
| EPI_ISL_417327 | hCoV-19/USA/CA-CDPH-UC23/2020 | North America / USA / California / San Jo | 2020-03-09 | California Department of Public Health | Chiu Laboratory, University of California, San Francisco | Xianding Deng, Scot Federman, Chao-Yang Pan, Hugo Guevara,Wei Gu, Debra A. Wadford, and Charles Y. Chiu |
| EPI_ISL_417328 | hCoV-19/USA/CA-CDPH-UC24/2020 | North America / USA / California / Sacram | 2020-03-07 | California Department of Public Health | Chiu Laboratory, University of California, San Francisco | Xianding Deng, Scot Federman, Chao-Yang Pan, Hugo Guevara,Wei Gu, Debra A. Wadford, and Charles Y. Chiu |
| EPI_ISL_417329 | hCoV-19/USA/CA-CDPH-UC25/2020 | North America / USA / California / San M | 2020-03-04 | California Department of Public Health | Chiu Laboratory, University of California, San Francisco | Xianding Deng, Scot Federman, Chao-Yang Pan, Hugo Guevara,Wei Gu, Debra A. Wadford, and Charles Y. Chiu |
| EPI_ISL_417330 | hCoV-19/USA/CA-CDPH-UC26/2020 | North America / USA / California / San Fr | 2020-03-13 | Chiu Laboratory, University of California, San Francisco | Chiu Laboratory, University of California, San Francisco | Xianding Deng, Scot Federman, Wei Gu, and Charles Y. Chiu |
| EPI_ISL_417331 | hCoV-19/USA/CA-CDPH-UC27/2020 | North America / USA / California / San Fr | 2020-03-13 | Chiu Laboratory, University of California, San Francisco | Chiu Laboratory, University of California, San Francisco | Xianding Deng, Scot Federman, Wei Gu, and Charles Y. Chiu |
| EPI_ISL_417333 | hCoV-19/France/Lyon_06042/2020 | Europe / France / ARA | 2020-03-04 | Institut des Agents Infectieux (IAI), Hospices Civils de Lyon | CNR Virus des Infections Respiratoires - France SUD | Antonin Bal, Gregory Destras, Gwendolyne Burfin, Solenne Brun, Carine Moustaud, Raphaelle Lamy, Alexandre Gaymard, Maude Bouscambert-Duchamp, Florence Morfin-Sherpa, Martine Valette, Laurence Josset, Bruno Lina |
| EPI_ISL_417334 | hCoV-19/France/Lyon_06056/2020 | Europe / France / ARA | 2020-03-04 | Institut des Agents Infectieux (IAI), Hospices Civils de Lyon | CNR Virus des Infections Respiratoires - France SUD | Antonin Bal, Gregory Destras, Gwendolyne Burfin, Solenne Brun, Carine Moustaud, Raphaelle Lamy, Alexandre Gaymard, Maude Bouscambert-Duchamp, Florence Morfin-Sherpa, Martine Valette, Laurence Josset, Bruno Lina |
| EPI_ISL_417335 | hCoV-19/France/Lyon_06573/2020 | Europe / France / ARA | 2020-03-06 | Institut des Agents Infectieux (IAI), Hospices Civils de Lyon | CNR Virus des Infections Respiratoires - France SUD | Antonin Bal, Gregory Destras, Gwendolyne Burfin, Solenne Brun, Carine Moustaud, Raphaelle Lamy, Alexandre Gaymard, Maude Bouscambert-Duchamp, Florence Morfin-Sherpa, Martine Valette, Laurence Josset, Bruno Lina |
| EPI_ISL_417336 | hCoV-19/France/Lyon_0668/2020 | Europe / France / ARA | 2020-03-06 | Institut des Agents Infectieux (IAI), Hospices Civils de Lyon | CNR Virus des Infections Respiratoires - France SUD | Antonin Bal, Gregory Destras, Gwendolyne Burfin, Solenne Brun, Carine Moustaud, Raphaelle Lamy, Alexandre Gaymard, Maude Bouscambert-Duchamp, Florence Morfin-Sherpa, Martine Valette, Laurence Josset, Bruno Lina |
| EPI_ISL_417337 | hCoV-19/France/Lyon_06625/2020 | Europe / France / ARA | 2020-03-07 | Institut des Agents Infectieux (IAI), Hospices Civils de Lyon | CNR Virus des Infections Respiratoires - France SUD | Antonin Bal, Gregory Destras, Gwendolyne Burfin, Solenne Brun, Carine Moustaud, Raphaelle Lamy, Alexandre Gaymard, Maude Bouscambert-Duchamp, Florence Morfin-Sherpa, Martine Valette, Laurence Josset, Bruno Lina |
| EPI_ISL_417338 | hCoV-19/France/Macon_06756/2020 | Europe / France / ARA | 2020-03-07 | Centre Hospitalier de Macon | CNR Virus des Infections Respiratoires - France SUD | Antonin Bal, Gregory Destras, Gwendolyne Burfin, Solenne Brun, Carine Moustaud, Raphaelle Lamy, Alexandre Gaymard, Maude Bouscambert-Duchamp, Florence Morfin-Sherpa, Martine Valette, Laurence Josset, Bruno Lina |
| EPI_ISL_417339 | hCoV-19/France/Lyon_06820/2020 | Europe / France / ARA | 2020-03-08 | Institut des Agents Infectieux (IAI), Hospices Civils de Lyon | CNR Virus des Infections Respiratoires - France SUD | Antonin Bal, Gregory Destras, Gwendolyne Burfin, Solenne Brun, Carine Moustaud, Raphaelle Lamy, Alexandre Gaymard, Maude Bouscambert-Duchamp, Florence Morfin-Sherpa, Martine Valette, Laurence Josset, Bruno Lina |
| EPI_ISL_417340 | hCoV-19/France/Bourg-en-Bresse_06813/2020 | Europe / France / ARA | 2020-03-07 | Centre Hospitalier de Bourg en Bresse | CNR Virus des Infections Respiratoires - France SUD | Antonin Bal, Gregory Destras, Gwendolyne Burfin, Solenne Brun, Carine Moustaud, Raphaelle Lamy, Alexandre Gaymard, Maude Bouscambert-Duchamp, Florence Morfin-Sherpa, Martine Valette, Laurence Josset, Bruno Lina |
| EPI_ISL_417341 | hCoV-19/USA/WA-UW192/2020 | North America / USA / Washington | 2020-03-13 | UW Virology Lab | UW Virology Lab | Pavitra Roychoudhury, Hong Xie, Keith Jerome, Alexander Greninger |
| EPI_ISL_417342 | hCoV-19/USA/WA-UW193/2020 | North America / USA / Washington | 2020-03-13 | UW Virology Lab | UW Virology Lab | Pavitra Roychoudhury, Hong Xie, Keith Jerome, Alexander Greninger |
| EPI_ISL_417343 | hCoV-19/USA/WA-UW194/2020 | North America / USA / Washington | 2020-03-13 | UW Virology Lab | UW Virology Lab | Pavitra Roychoudhury, Hong Xie, Keith Jerome, Alexander Greninger |
| EPI_ISL_417344 | hCoV-19/USA/WA-UW195/2020 | North America / USA / Washington | 2020-03-13 | UW Virology Lab | UW Virology Lab | Pavitra Roychoudhury, Hong Xie, Keith Jerome, Alexander Greninger |
| EPI_ISL_417345 | hCoV-19/USA/WA-UW196/2020 | North America / USA / Washington | 2020-03-13 | UW Virology Lab | UW Virology Lab | Pavitra Roychoudhury, Hong Xie, Keith Jerome, Alexander Greninger |
| EPI_ISL_417346 | hCoV-19/USA/WA-UW197/2020 | North America / USA / Washington | 2020-03-12 | UW Virology Lab | UW Virology Lab | Pavitra Roychoudhury, Hong Xie, Keith Jerome, Alexander Greninger |
| EPI_ISL_417347 | hCoV-19/USA/WA-UW198/2020 | North America / USA / Washington | 2020-03-12 | UW Virology Lab | UW Virology Lab | Pavitra Roychoudhury, Hong Xie, Keith Jerome, Alexander Greninger |
| EPI_ISL_417348 | hCoV-19/USA/WA-UW199/2020 | North America / USA / Washington | 2020-03-13 | UW Virology Lab | UW Virology Lab | Pavitra Roychoudhury, Hong Xie, Keith Jerome, Alexander Greninger |
| EPI_ISL_417349 | hCoV-19/USA/WA-UW200/2020 | North America / USA / Washington | 2020-03-12 | UW Virology Lab | UW Virology Lab | Pavitra Roychoudhury, Hong Xie, Keith Jerome, Alexander Greninger |
| EPI_ISL_417350 | hCoV-19/USA/WA-UW201/2020 | North America / USA / Washington | 2020-03-12 | UW Virology Lab | UW Virology Lab | Pavitra Roychoudhury, Hong Xie, Keith Jerome, Alexander Greninger |
| EPI_ISL_417351 | hCoV-19/USA/WA-UW202/2020 | North America / USA / Washington | 2020-03-12 | UW Virology Lab | UW Virology Lab | Pavitra Roychoudhury, Hong Xie, Keith Jerome, Alexander Greninger |
| EPI_ISL_417352 | hCoV-19/USA/WA-UW203/2020 | North America / USA / Washington | 2020-03-12 | UW Virology Lab | UW Virology Lab | Pavitra Roychoudhury, Hong Xie, Keith Jerome, Alexander Greninger |
| EPI_ISL_417353 | hCoV-19/USA/WA-UW204/2020 | North America / USA / Washington | 2020-03-13 | UW Virology Lab | UW Virology Lab | Pavitra Roychoudhury, Hong Xie, Keith Jerome, Alexander Greninger |
| EPI_ISL_417354 | hCoV-19/USA/WA-UW205/2020 | North America / USA / Washington | 2020-03-13 | UW Virology Lab | UW Virology Lab | Pavitra Roychoudhury, Hong Xie, Keith Jerome, Alexander Greninger |
| EPI_ISL_417355 | hCoV-19/USA/WA-UW206/2020 | North America / USA / Washington | 2020-03-13 | UW Virology Lab | UW Virology Lab | Pavitra Roychoudhury, Hong Xie, Keith Jerome, Alexander Greninger |
| EPI_ISL_417356 | hCoV-19/USA/WA-UW207/2020 | North America / USA / Washington | 2020-03-13 | UW Virology Lab | UW Virology Lab | Pavitra Roychoudhury, Hong Xie, Keith Jerome, Alexander Greninger |
| EPI_ISL_417358 | hCoV-19/USA/WA-UW209/2020 | North America / USA / Washington | 2020-03-13 | UW Virology Lab | UW Virology Lab | Pavitra Roychoudhury, Hong Xie, Keith Jerome, Alexander Greninger |
| EPI_ISL_417359 | hCoV-19/USA/WA-UW210/2020 | North America / USA / Washington | 2020-03-14 | UW Virology Lab | UW Virology Lab | Pavitra Roychoudhury, Hong Xie, Keith Jerome, Alexander Greninger |
| EPI_ISL_417360 | hCoV-19/USA/WA-UW211/2020 | North America / USA / Washington | 2020-03-14 | UW Virology Lab | UW Virology Lab | Pavitra Roychoudhury, Hong Xie, Keith Jerome, Alexander Greninger |
| EPI_ISL_417361 | hCoV-19/USA/WA-UW212/2020 | North America / USA / Washington | 2020-03-14 | UW Virology Lab | UW Virology Lab | Pavitra Roychoudhury, Hong Xie, Keith Jerome, Alexander Greninger |
| EPI_ISL_417362 | hCoV-19/USA/CT-UW213/2020 | North America / USA / Connecticut | 2020-03-13 | UW Virology Lab | UW Virology Lab | Pavitra Roychoudhury, Hong Xie, Keith Jerome, Alexander Greninger |
| EPI_ISL_417363 | hCoV-19/USA/WA-UW214/2020 | North America / USA / Washington | 2020-03-14 | UW Virology Lab | UW Virology Lab | Pavitra Roychoudhury, Hong Xie, Keith Jerome, Alexander Greninger |
| EPI_ISL_417364 | hCoV-19/USA/WA-UW215/2020 | North America / USA / Washington | 2020-03-12 | UW Virology Lab | UW Virology Lab | Pavitra Roychoudhury, Hong Xie, Keith Jerome, Alexander Greninger |
| EPI_ISL_417365 | hCoV-19/USA/WA-UW216/2020 | North America / USA / Washington | 2020-03-14 | UW Virology Lab | UW Virology Lab | Pavitra Roychoudhury, Hong Xie, Keith Jerome, Alexander Greninger |
| EPI_ISL_417366 | hCoV-19/USA/WA-UW217/2020 | North America / USA / Washington | 2020-03-14 | UW Virology Lab | UW Virology Lab | Pavitra Roychoudhury, Hong Xie, Keith Jerome, Alexander Greninger |
| EPI_ISL_417367 | hCoV-19/USA/WA-UW218/2020 | North America / USA / Washington | 2020-03-13 | UW Virology Lab | UW Virology Lab | Pavitra Roychoudhury, Hong Xie, Keith Jerome, Alexander Greninger |
| EPI_ISL_417368 | hCoV-19/USA/UN-UW219/2020 | North America / USA | 2020-03-13 | UW Virology Lab | UW Virology Lab | Pavitra Roychoudhury, Hong Xie, Keith Jerome, Alexander Greninger |
| EPI_ISL_417369 | hCoV-19/USA/WA-UW220/2020 | North America / USA / Washington | 2020-03-14 | UW Virology Lab | UW Virology Lab | Pavitra Roychoudhury, Hong Xie, Keith Jerome, Alexander Greninger |
| EPI_ISL_417370 | hCoV-19/USA/WA-UW221/2020 | North America / USA / Washington | 2020-03-14 | UW Virology Lab | UW Virology Lab | Pavitra Roychoudhury, Hong Xie, Keith Jerome, Alexander Greninger |
| EPI_ISL_417371 | hCoV-19/USA/WA-UW222/2020 | North America / USA / Washington | 2020-03-13 | UW Virology Lab | UW Virology Lab | Pavitra Roychoudhury, Hong Xie, Keith Jerome, Alexander Greninger |
| EPI_ISL_417372 | hCoV-19/USA/CT-UW223/2020 | North America / USA / Connecticut | 2020-03-13 | UW Virology Lab | UW Virology Lab | Pavitra Roychoudhury, Hong Xie, Keith Jerome, Alexander Greninger |
| EPI_ISL_417373 | hCoV-19/USA/CT-UW224/2020 | North America / USA / Connecticut | 2020-03-13 | UW Virology Lab | UW Virology Lab | Pavitra Roychoudhury, Hong Xie, Keith Jerome, Alexander Greninger |
| EPI_ISL_417374 | hCoV-19/USA/WA-UW225/2020 | North America / USA / Washington | 2020-03-15 | UW Virology Lab | UW Virology Lab | Pavitra Roychoudhury, Hong Xie, Keith Jerome, Alexander Greninger |
| EPI_ISL_417375 | hCoV-19/USA/WA-UW226/2020 | North America / USA / Washington | 2020-03-15 | UW Virology Lab | UW Virology Lab | Pavitra Roychoudhury, Hong Xie, Keith Jerome, Alexander Greninger |
| EPI_ISL_417376 | hCoV-19/USA/WA-UW227/2020 | North America / USA / Washington | 2020-03-15 | UW Virology Lab | UW Virology Lab | Pavitra Roychoudhury, Hong Xie, Keith Jerome, Alexander Greninger |
| EPI_ISL_417377 | hCoV-19/USA/WA-UW228/2020 | North America / USA / Washington | 2020-03-15 | UW Virology Lab | UW Virology Lab | Pavitra Roychoudhury, Hong Xie, Keith Jerome, Alexander Greninger |
| EPI_ISL_417378 | hCoV-19/USA/WA-UW229/2020 | North America / USA / Washington | 2020-03-14 | UW Virology Lab | UW Virology Lab | Pavitra Roychoudhury, Hong Xie, Keith Jerome, Alexander Greninger |
| EPI_ISL_417379 | hCoV-19/USA/WA-UW230/2020 | North America / USA / Washington | 2020-03-15 | UW Virology Lab | UW Virology Lab | Pavitra Roychoudhury, Hong Xie, Keith Jerome, Alexander Greninger |
| EPI_ISL_417380 | hCoV-19/USA/WA-UW231/2020 | North America / USA / Washington | 2020-03-14 | UW Virology Lab | UW Virology Lab | Pavitra Roychoudhury, Hong Xie, Keith Jerome, Alexander Greninger |
| EPI_ISL_417381 | hCoV-19/USA/WA-UW232/2020 | North America / USA / Washington | 2020-03-14 | UW Virology Lab | UW Virology Lab | Pavitra Roychoudhury, Hong Xie, Keith Jerome, Alexander Greninger |
| EPI_ISL_417382 | hCoV-19/USA/WA-UW233/2020 | North America / USA / Washington | 2020-03-15 | UW Virology Lab | UW Virology Lab | Pavitra Roychoudhury, Hong Xie, Keith Jerome, Alexander Greninger |
| EPI_ISL_417383 | hCoV-19/Australia/NSW15/2020 | Oceania / Australia / New South Wales / | 2020-03-03 | Centre for Infectious Diseases and Microbiology Public Health | NSW Health Pathology - Institute of Clinical Pathology and Medical Research; Westmead Hospital; University of Sydney | Rockett R, Eden J-S, Lam C, Gray K, Timms V, Gall M, Arnott A, Sadsad R, Carter I, Rahman H, Holmes EC, O'Sullivan MV, Sintchenko V, Chen SC, Maddocks S, Kok J and Dwyer DE for the 2019-nCoV Study Group |
| EPI_ISL_417384 | hCoV-19/Australia/NSW18/2020 | Oceania / Australia / New South Wales / | 2020-03-05 | Centre for Infectious Diseases and Microbiology Public Health | NSW Health Pathology - Institute of Clinical Pathology and Medical Research; Westmead Hospital; University of Sydney | Eden J-S, Lam C, Gray K, Timms V, Gall M, Arnott A, Sadsad R, Carter I, Rahman H, Holmes EC, O'Sullivan MV, Sintchenko V, Chen SC, Maddocks S, Kok J, Dwyer DE and Rockett R for the 2019-nCoV Study Group |
| EPI_ISL_417385 | hCoV-19/Australia/NSW19/2020 | Oceania / Australia / New South Wales / | 2020-03-06 | Centre for Infectious Diseases and Microbiology Public Health | NSW Health Pathology - Institute of Clinical Pathology and Medical Research; Westmead Hospital; University of Sydney | Lam C, Gray K, Timms V, Gall M, Arnott A, Sadsad R, Carter I, Rahman H, Holmes EC, O'Sullivan MV, Sintchenko V, Chen SC, Maddocks S, Kok J, Dwyer DE, Rockett R and Eden J-S for the 2019-nCoV Study Group |
| EPI_ISL_417386 | hCoV-19/Australia/NSW21/2020 | Oceania / Australia / New South Wales / | 2020-03-04 | Centre for Infectious Diseases and Microbiology Public Health | NSW Health Pathology - Institute of Clinical Pathology and Medical Research; Westmead Hospital; University of Sydney | Gray K, Timms V, Gall M, Arnott A, Sadsad R, Carter I, Rahman H, Holmes EC, O'Sullivan MV, Sintchenko V, Chen SC, Maddocks S, Kok J, Dwyer DE, Rockett R, Eden J-S and Lam C for the 2019-nCoV Study Group |
| EPI_ISL_417387 | hCoV-19/Australia/NSW24/2020 | Oceania / Australia / New South Wales / | 2020-03-06 | Centre for Infectious Diseases and Microbiology Public Health | NSW Health Pathology - Institute of Clinical Pathology and Medical Research; Westmead Hospital; University of Sydney | Timms V, Gall M, Arnott A, Sadsad R, Carter I, Rahman H, Holmes EC, O'Sullivan MV, Sintchenko V, Chen SC, Maddocks S, Kok J, Dwyer DE, Rockett R, Eden J-S, Lam C and Gray K for the 2019-nCoV Study Group |
| EPI_ISL_417388 | hCoV-19/Australia/NSW25/2020 | Oceania / Australia / New South Wales / | 2020-03-05 | Centre for Infectious Diseases and Microbiology Public Health | NSW Health Pathology - Institute of Clinical Pathology and Medical Research; Westmead Hospital; University of Sydney | Gall M, Arnott A, Sadsad R, Carter I, Rahman H, Holmes EC, O'Sullivan MV, Sintchenko V, Chen SC, Maddocks S, Kok J, Dwyer DE, Rockett R, Eden J-S, Lam C, Gray K and Timms V for the 2019-nCoV Study Group |
| EPI_ISL_417389 | hCoV-19/Australia/NSW26/2020 | Oceania / Australia / New South Wales / | 2020-03-09 | Centre for Infectious Diseases and Microbiology Public Health | NSW Health Pathology - Institute of Clinical Pathology and Medical Research; Westmead Hospital; University of Sydney | Arnott A, Sadsad R, Carter I, Rahman H, Holmes EC, O'Sullivan MV, Sintchenko V, Chen SC, Maddocks S, Kok J, Dwyer DE, Rockett R, Eden J-S, Lam C, Gray K, Timms V and Gall M for the 2019-nCoV Study Group |
| EPI_ISL_417390 | hCoV-19/Australia/NSW27/2020 | Oceania / Australia / New South Wales / | 2020-03-08 | Centre for Infectious Diseases and Microbiology Public Health | NSW Health Pathology - Institute of Clinical Pathology and Medical Research; Westmead Hospital; University of Sydney | Sadsad R, Carter I, Rahman H, Holmes EC, O'Sullivan MV, Sintchenko V, Chen SC, Maddocks S, Kok J, Dwyer DE, Rockett R, Eden J-S, Lam C, Gray K, Timms V, Gall M and Arnott A for the 2019-nCoV Study Group |
| EPI_ISL_417391 | hCoV-19/Australia/NSW28/2020 | Oceania / Australia / New South Wales / | 2020-03-08 | Centre for Infectious Diseases and Microbiology Public Health | NSW Health Pathology - Institute of Clinical Pathology and Medical Research; Westmead Hospital; University of Sydney | Carter I, Rahman H, Holmes EC, O'Sullivan MV, Sintchenko V, Chen SC, Maddocks S, Kok J, Dwyer DE, Rockett R, Eden J-S, Lam C, Gray K, Timms V, Gall M, Arnott A and Sadsad R for the 2019-nCoV Study Group |
| EPI_ISL_417392 | hCoV-19/Australia/NSW31/2020 | Oceania / Australia / New South Wales / | 2020-03-10 | Centre for Infectious Diseases and Microbiology Public Health | NSW Health Pathology - Institute of Clinical Pathology and Medical Research; Westmead Hospital; University of Sydney | Rahman H, Holmes EC, O'Sullivan MV, Sintchenko V, Chen SC, Maddocks S, Kok J, Dwyer DE, Rockett R, Eden J-S, Lam C, Gray K, Timms V, Gall M, Arnott A, Sadsad R and Carter I for the 2019-nCoV Study Group |
| EPI_ISL_417393 | hCoV-19/Australia/NSW35/2020 | Oceania / Australia / New South Wales / | 2020-03-08 | Centre for Infectious Diseases and Microbiology Public Health | NSW Health Pathology - Institute of Clinical Pathology and Medical Research; Westmead Hospital; University of Sydney | Holmes EC, O'Sullivan MV, Sintchenko V, Chen SC, Maddocks S, Kok J, Dwyer DE, Rockett R, Eden J-S, Lam C, Gray K, Timms V, Gall M, Arnott A, Sadsad R, Carter I and Rahman H for the 2019-nCoV Study Group |
| EPI_ISL_417394 | hCoV-19/Australia/NSW36/2020 | Oceania / Australia / New South Wales / | 2020-03-10 | Centre for Infectious Diseases and Microbiology Public Health | NSW Health Pathology - Institute of Clinical Pathology and Medical Research; Westmead Hospital; University of Sydney | O'Sullivan MV, Sintchenko V, Chen SC, Maddocks S, Kok J, Dwyer DE, Rockett R, Eden J-S, Lam C, Gray K, Timms V, Gall M, Arnott A, Sadsad R, Carter I, Rahman H and Holmes EC for the 2019-nCoV Study Group |
| EPI_ISL_417395 | hCoV-19/Australia/NSW37/2020 | Oceania / Australia / New South Wales / | 2020-03-10 | Centre for Infectious Diseases and Microbiology Public Health | NSW Health Pathology - Institute of Clinical Pathology and Medical Research; Westmead Hospital; University of Sydney | Sintchenko V, Chen SC, Maddocks S, Kok J, Dwyer DE, Rockett R, Eden J-S, Lam C, Gray K, Timms V, Gall M, Arnott A, Sadsad R, Carter I, Rahman H, Holmes EC and O'Sullivan MV for the 2019-nCoV Study Group |
| EPI_ISL_417396 | hCoV-19/Australia/NSW38/2020 | Oceania / Australia / New South Wales / | 2020-03-10 | Centre for Infectious Diseases and Microbiology Public Health | NSW Health Pathology - Institute of Clinical Pathology and Medical Research; Westmead Hospital; University of Sydney | Chen SC, Maddocks S, Kok J, Dwyer DE, Rockett R, Eden J-S, Lam C, Gray K, Timms V, Gall M, Arnott A, Sadsad R, Carter I, Rahman H, Holmes EC, O'Sullivan MV and Sintchenko V for the 2019-nCoV Study Group |
| EPI_ISL_417397 | hCoV-19/Australia/NSW39/2020 | Oceania / Australia / New South Wales / | 2020-03-11 | Centre for Infectious Diseases and Microbiology Public Health | NSW Health Pathology - Institute of Clinical Pathology and Medical Research; Westmead Hospital; University of Sydney | Maddocks S, Kok J, Dwyer DE, Rockett R, Eden J-S, Lam C, Gray K, Timms V, Gall M, Arnott A, Sadsad R, Carter I, Rahman H, Holmes EC, O'Sullivan MV, Sintchenko V and Chen SC for the 2019-nCoV Study Group |
| EPI_ISL_417398 | hCoV-19/Australia/NSW40/2020 | Oceania / Australia / New South Wales / | 2020-03-11 | Centre for Infectious Diseases and Microbiology Public Health | NSW Health Pathology - Institute of Clinical Pathology and Medical Research; Westmead Hospital; University of Sydney | Kok J, Dwyer DE, Rockett R, Eden J-S, Lam C, Gray K, Timms V, Gall M, Arnott A, Sadsad R, Carter I, Rahman H, Holmes EC, O'Sullivan MV, Sintchenko V, Chen SC and Maddocks S for the 2019-nCoV Study Group |
| EPI_ISL_417399 | hCoV-19/Australia/NSW41/2020 | Oceania / Australia / New South Wales / | 2020-03-10 | Centre for Infectious Diseases and Microbiology Public Health | NSW Health Pathology - Institute of Clinical Pathology and Medical Research; Westmead Hospital; University of Sydney | Dwyer DE, Rockett R, Eden J-S, Lam C, Gray K, Timms V, Gall M, Arnott A, Sadsad R, Carter I, Rahman H, Holmes EC, O'Sullivan MV, Sintchenko V, Chen SC, Maddocks S and Kok J for the 2019-nCoV Study Group |
| EPI_ISL_417600 | hCoV-19/Iceland/272/2020 | Europe / Iceland / Reykjavik | 2020-03-17 | The National University Hospital of Iceland | deCODE genetics | Daniel F Gudbjartsson; Agnar Helgason; Hakon Jonsson; Olafur T Magnusson; Pall Melsted; Gudmundur L Norddahl; Jona Saemundsdottir; Asgeir Sigurdsson; Patrick Sulem; Arna B Agustsdottir; Berglind Eiriksdottir; Run Fridriksdottir; Elisabet E Gardarsdottir; Gudmundur Georgsson; Olafia S Gretarsdottir; Kjartan R Gudmundsson; Thora R Gunnarsdottir; Arnaldur Gylfason; Hilma Holm; Brynjar O Jensson; Aslaug Jonasdottir; Kamilla S Josefsdottir; Thordur Kristjansson; Droplaug N Magnusdottir; Louise le Roux; Gudrun Sigmundsdottir; Gardar Sveinbjornsson; Kristin E Sveinsdottir; Maney Sveinsdottir; Emil A Thorarensen; Bjarni Thorbjornsson; Gisli Masson; Ingileif Jonsdottir; Alma Moller; Thorolfur Gudnason; Karl G Kristinsson; Unnur Thorsteinsdottir; Kari Stefansson |
| EPI_ISL_417601 | hCoV-19/Iceland/273/2020 | Europe / Iceland / Reykjavik | 2020-03-17 | The National University Hospital of Iceland | deCODE genetics | Daniel F Gudbjartsson; Agnar Helgason; Hakon Jonsson; Olafur T Magnusson; Pall Melsted; Gudmundur L Norddahl; Jona Saemundsdottir; Asgeir Sigurdsson; Patrick Sulem; Arna B Agustsdottir; Berglind Eiriksdottir; Run Fridriksdottir; Elisabet E Gardarsdottir; Gudmundur Georgsson; Olafia S Gretarsdottir; Kjartan R Gudmundsson; Thora R Gunnarsdottir; Arnaldur Gylfason; Hilma Holm; Brynjar O Jensson; Aslaug Jonasdottir; Kamilla S Josefsdottir; Thordur Kristjansson; Droplaug N Magnusdottir; Louise le Roux; Gudrun Sigmundsdottir; Gardar Sveinbjornsson; Kristin E Sveinsdottir; Maney Sveinsdottir; Emil A Thorarensen; Bjarni Thorbjornsson; Gisli Masson; Ingileif Jonsdottir; Alma Moller; Thorolfur Gudnason; Karl G Kristinsson; Unnur Thorsteinsdottir; Kari Stefansson |
| EPI_ISL_417602 | hCoV-19/Iceland/274/2020 | Europe / Iceland / Reykjavik | 2020-03-17 | The National University Hospital of Iceland | deCODE genetics | Daniel F Gudbjartsson; Agnar Helgason; Hakon Jonsson; Olafur T Magnusson; Pall Melsted; Gudmundur L Norddahl; Jona Saemundsdottir; Asgeir Sigurdsson; Patrick Sulem; Arna B Agustsdottir; Berglind Eiriksdottir; Run Fridriksdottir; Elisabet E Gardarsdottir; Gudmundur Georgsson; Olafia S Gretarsdottir; Kjartan R Gudmundsson; Thora R Gunnarsdottir; Arnaldur Gylfason; Hilma Holm; Brynjar O Jensson; Aslaug Jonasdottir; Kamilla S Josefsdottir; Thordur Kristjansson; Droplaug N Magnusdottir; Louise le Roux; Gudrun Sigmundsdottir; Gardar Sveinbjornsson; Kristin E Sveinsdottir; Maney Sveinsdottir; Emil A Thorarensen; Bjarni Thorbjornsson; Gisli Masson; Ingileif Jonsdottir; Alma Moller; Thorolfur Gudnason; Karl G Kristinsson; Unnur Thorsteinsdottir; Kari Stefansson |
| EPI_ISL_417603 | hCoV-19/Iceland/275/2020 | Europe / Iceland / Reykjavik | 2020-03-17 | The National University Hospital of Iceland | deCODE genetics | Daniel F Gudbjartsson; Agnar Helgason; Hakon Jonsson; Olafur T Magnusson; Pall Melsted; Gudmundur L Norddahl; Jona Saemundsdottir; Asgeir Sigurdsson; Patrick Sulem; Arna B Agustsdottir; Berglind Eiriksdottir; Run Fridriksdottir; Elisabet E Gardarsdottir; Gudmundur Georgsson; Olafia S Gretarsdottir; Kjartan R Gudmundsson; Thora R Gunnarsdottir; Arnaldur Gylfason; Hilma Holm; Brynjar O Jensson; Aslaug Jonasdottir; Kamilla S Josefsdottir; Thordur Kristjansson; Droplaug N Magnusdottir; Louise le Roux; Gudrun Sigmundsdottir; Gardar Sveinbjornsson; Kristin E Sveinsdottir; Maney Sveinsdottir; Emil A Thorarensen; Bjarni Thorbjornsson; Gisli Masson; Ingileif Jonsdottir; Alma Moller; Thorolfur Gudnason; Karl G Kristinsson; Unnur Thorsteinsdottir; Kari Stefansson |
| EPI_ISL_417604 | hCoV-19/Iceland/276/2020 | Europe / Iceland / Reykjavik | 2020-03-17 | The National University Hospital of Iceland | deCODE genetics | Daniel F Gudbjartsson; Agnar Helgason; Hakon Jonsson; Olafur T Magnusson; Pall Melsted; Gudmundur L Norddahl; Jona Saemundsdottir; Asgeir Sigurdsson; Patrick Sulem; Arna B Agustsdottir; Berglind Eiriksdottir; Run Fridriksdottir; Elisabet E Gardarsdottir; Gudmundur Georgsson; Olafia S Gretarsdottir; Kjartan R Gudmundsson; Thora R Gunnarsdottir; Arnaldur Gylfason; Hilma Holm; Brynjar O Jensson; Aslaug Jonasdottir; Kamilla S Josefsdottir; Thordur Kristjansson; Droplaug N Magnusdottir; Louise le Roux; Gudrun Sigmundsdottir; Gardar Sveinbjornsson; Kristin E Sveinsdottir; Maney Sveinsdottir; Emil A Thorarensen; Bjarni Thorbjornsson; Gisli Masson; Ingileif Jonsdottir; Alma Moller; Thorolfur Gudnason; Karl G Kristinsson; Unnur Thorsteinsdottir; Kari Stefansson |

| | | | | | | |
|---|---|---|---|---|---|---|
| EPI_ISL_417607 | hCoV-19/Iceland/279/2020 | Europe / Iceland / Reykjavik | 2020-03-17 | The National University Hospital of Iceland | deCODE genetics | Daniel F Gudbjartsson; Agnar Helgason; Hakon Jonsson; Olafur T Magnusson; Pall Melsted; Gudmundur L Norddahl; Jona Saemundsdottir; Asgeir Sigurdsson; Patrick Sulem; Arna B Agustsdottir; Berglind Eiriksdottir; Run Fridriksdottir; Elisabet E Gardarsdottir; Gudmundur Georgsson; Olafia S Gretarsdottir; Kjartan R Gudmundsson; Thora R Gunnarsdottir; Arnaldur Gylfason; Hilma Holm; Brynjar O Jensson; Aslaug Jonasdottir; Kamilla S Josefsdottir; Thordur Kristjansson; Droplaug N Magnusdottir; Louise le Roux; Gudrun Sigmundsdottir; Gardar Sveinbjornsson; Kristin E Sveinsdottir; Maney Sveinsdottir; Emil A Thorarensen; Bjarni Thorbjornsson; Gisli Masson; Ingileif Jonsdottir; Alma Moller; Thorolfur Gudnason; Karl G Kristinsson; Unnur Thorsteinsdottir; Kari Stefansson |
| EPI_ISL_417608 | hCoV-19/Iceland/280/2020 | Europe / Iceland / Reykjavik | 2020-03-17 | The National University Hospital of Iceland | deCODE genetics | Daniel F Gudbjartsson; Agnar Helgason; Hakon Jonsson; Olafur T Magnusson; Pall Melsted; Gudmundur L Norddahl; Jona Saemundsdottir; Asgeir Sigurdsson; Patrick Sulem; Arna B Agustsdottir; Berglind Eiriksdottir; Run Fridriksdottir; Elisabet E Gardarsdottir; Gudmundur Georgsson; Olafia S Gretarsdottir; Kjartan R Gudmundsson; Thora R Gunnarsdottir; Arnaldur Gylfason; Hilma Holm; Brynjar O Jensson; Aslaug Jonasdottir; Kamilla S Josefsdottir; Thordur Kristjansson; Droplaug N Magnusdottir; Louise le Roux; Gudrun Sigmundsdottir; Gardar Sveinbjornsson; Kristin E Sveinsdottir; Maney Sveinsdottir; Emil A Thorarensen; Bjarni Thorbjornsson; Gisli Masson; Ingileif Jonsdottir; Alma Moller; Thorolfur Gudnason; Karl G Kristinsson; Unnur Thorsteinsdottir; Kari Stefansson |
| EPI_ISL_417609 | hCoV-19/Iceland/281/2020 | Europe / Iceland / Reykjavik | 2020-03-17 | The National University Hospital of Iceland | deCODE genetics | Daniel F Gudbjartsson; Agnar Helgason; Hakon Jonsson; Olafur T Magnusson; Pall Melsted; Gudmundur L Norddahl; Jona Saemundsdottir; Asgeir Sigurdsson; Patrick Sulem; Arna B Agustsdottir; Berglind Eiriksdottir; Run Fridriksdottir; Elisabet E Gardarsdottir; Gudmundur Georgsson; Olafia S Gretarsdottir; Kjartan R Gudmundsson; Thora R Gunnarsdottir; Arnaldur Gylfason; Hilma Holm; Brynjar O Jensson; Aslaug Jonasdottir; Kamilla S Josefsdottir; Thordur Kristjansson; Droplaug N Magnusdottir; Louise le Roux; Gudrun Sigmundsdottir; Gardar Sveinbjornsson; Kristin E Sveinsdottir; Maney Sveinsdottir; Emil A Thorarensen; Bjarni Thorbjornsson; Gisli Masson; Ingileif Jonsdottir; Alma Moller; Thorolfur Gudnason; Karl G Kristinsson; Unnur Thorsteinsdottir; Kari Stefansson |
| EPI_ISL_417610 | hCoV-19/Iceland/282/2020 | Europe / Iceland / Reykjavik | 2020-03-17 | The National University Hospital of Iceland | deCODE genetics | Daniel F Gudbjartsson; Agnar Helgason; Hakon Jonsson; Olafur T Magnusson; Pall Melsted; Gudmundur L Norddahl; Jona Saemundsdottir; Asgeir Sigurdsson; Patrick Sulem; Arna B Agustsdottir; Berglind Eiriksdottir; Run Fridriksdottir; Elisabet E Gardarsdottir; Gudmundur Georgsson; Olafia S Gretarsdottir; Kjartan R Gudmundsson; Thora R Gunnarsdottir; Arnaldur Gylfason; Hilma Holm; Brynjar O Jensson; Aslaug Jonasdottir; Kamilla S Josefsdottir; Thordur Kristjansson; Droplaug N Magnusdottir; Louise le Roux; Gudrun Sigmundsdottir; Gardar Sveinbjornsson; Kristin E Sveinsdottir; Maney Sveinsdottir; Emil A Thorarensen; Bjarni Thorbjornsson; Gisli Masson; Ingileif Jonsdottir; Alma Moller; Thorolfur Gudnason; Karl G Kristinsson; Unnur Thorsteinsdottir; Kari Stefansson |
| EPI_ISL_417611 | hCoV-19/Iceland/283/2020 | Europe / Iceland / Reykjavik | 2020-03-17 | The National University Hospital of Iceland | deCODE genetics | Daniel F Gudbjartsson; Agnar Helgason; Hakon Jonsson; Olafur T Magnusson; Pall Melsted; Gudmundur L Norddahl; Jona Saemundsdottir; Asgeir Sigurdsson; Patrick Sulem; Arna B Agustsdottir; Berglind Eiriksdottir; Run Fridriksdottir; Elisabet E Gardarsdottir; Gudmundur Georgsson; Olafia S Gretarsdottir; Kjartan R Gudmundsson; Thora R Gunnarsdottir; Arnaldur Gylfason; Hilma Holm; Brynjar O Jensson; Aslaug Jonasdottir; Kamilla S Josefsdottir; Thordur Kristjansson; Droplaug N Magnusdottir; Louise le Roux; Gudrun Sigmundsdottir; Gardar Sveinbjornsson; Kristin E Sveinsdottir; Maney Sveinsdottir; Emil A Thorarensen; Bjarni Thorbjornsson; Gisli Masson; Ingileif Jonsdottir; Alma Moller; Thorolfur Gudnason; Karl G Kristinsson; Unnur Thorsteinsdottir; Kari Stefansson |
| EPI_ISL_417612 | hCoV-19/Iceland/284/2020 | Europe / Iceland / Reykjavik | 2020-03-17 | The National University Hospital of Iceland | deCODE genetics | Daniel F Gudbjartsson; Agnar Helgason; Hakon Jonsson; Olafur T Magnusson; Pall Melsted; Gudmundur L Norddahl; Jona Saemundsdottir; Asgeir Sigurdsson; Patrick Sulem; Arna B Agustsdottir; Berglind Eiriksdottir; Run Fridriksdottir; Elisabet E Gardarsdottir; Gudmundur Georgsson; Olafia S Gretarsdottir; Kjartan R Gudmundsson; Thora R Gunnarsdottir; Arnaldur Gylfason; Hilma Holm; Brynjar O Jensson; Aslaug Jonasdottir; Kamilla S Josefsdottir; Thordur Kristjansson; Droplaug N Magnusdottir; Louise le Roux; Gudrun Sigmundsdottir; Gardar Sveinbjornsson; Kristin E Sveinsdottir; Maney Sveinsdottir; Emil A Thorarensen; Bjarni Thorbjornsson; Gisli Masson; Ingileif Jonsdottir; Alma Moller; Thorolfur Gudnason; Karl G Kristinsson; Unnur Thorsteinsdottir; Kari Stefansson |
| EPI_ISL_417613 | hCoV-19/Iceland/285/2020 | Europe / Iceland / Reykjavik | 2020-03-17 | The National University Hospital of Iceland | deCODE genetics | Daniel F Gudbjartsson; Agnar Helgason; Hakon Jonsson; Olafur T Magnusson; Pall Melsted; Gudmundur L Norddahl; Jona Saemundsdottir; Asgeir Sigurdsson; Patrick Sulem; Arna B Agustsdottir; Berglind Eiriksdottir; Run Fridriksdottir; Elisabet E Gardarsdottir; Gudmundur Georgsson; Olafia S Gretarsdottir; Kjartan R Gudmundsson; Thora R Gunnarsdottir; Arnaldur Gylfason; Hilma Holm; Brynjar O Jensson; Aslaug Jonasdottir; Kamilla S Josefsdottir; Thordur Kristjansson; Droplaug N Magnusdottir; Louise le Roux; Gudrun Sigmundsdottir; Gardar Sveinbjornsson; Kristin E Sveinsdottir; Maney Sveinsdottir; Emil A Thorarensen; Bjarni Thorbjornsson; Gisli Masson; Ingileif Jonsdottir; Alma Moller; Thorolfur Gudnason; Karl G Kristinsson; Unnur Thorsteinsdottir; Kari Stefansson |
| EPI_ISL_417614 | hCoV-19/Iceland/286/2020 | Europe / Iceland / Reykjavik | 2020-03-18 | The National University Hospital of Iceland | deCODE genetics | Daniel F Gudbjartsson; Agnar Helgason; Hakon Jonsson; Olafur T Magnusson; Pall Melsted; Gudmundur L Norddahl; Jona Saemundsdottir; Asgeir Sigurdsson; Patrick Sulem; Arna B Agustsdottir; Berglind Eiriksdottir; Run Fridriksdottir; Elisabet E Gardarsdottir; Gudmundur Georgsson; Olafia S Gretarsdottir; Kjartan R Gudmundsson; Thora R Gunnarsdottir; Arnaldur Gylfason; Hilma Holm; Brynjar O Jensson; Aslaug Jonasdottir; Kamilla S Josefsdottir; Thordur Kristjansson; Droplaug N Magnusdottir; Louise le Roux; Gudrun Sigmundsdottir; Gardar Sveinbjornsson; Kristin E Sveinsdottir; Maney Sveinsdottir; Emil A Thorarensen; Bjarni Thorbjornsson; Gisli Masson; Ingileif Jonsdottir; Alma Moller; Thorolfur Gudnason; Karl G Kristinsson; Unnur Thorsteinsdottir; Kari Stefansson |
| EPI_ISL_417615 | hCoV-19/Iceland/287/2020 | Europe / Iceland / Reykjavik | 2020-03-18 | The National University Hospital of Iceland | deCODE genetics | Daniel F Gudbjartsson; Agnar Helgason; Hakon Jonsson; Olafur T Magnusson; Pall Melsted; Gudmundur L Norddahl; Jona Saemundsdottir; Asgeir Sigurdsson; Patrick Sulem; Arna B Agustsdottir; Berglind Eiriksdottir; Run Fridriksdottir; Elisabet E Gardarsdottir; Gudmundur Georgsson; Olafia S Gretarsdottir; Kjartan R Gudmundsson; Thora R Gunnarsdottir; Arnaldur Gylfason; Hilma Holm; Brynjar O Jensson; Aslaug Jonasdottir; Kamilla S Josefsdottir; Thordur Kristjansson; Droplaug N Magnusdottir; Louise le Roux; Gudrun Sigmundsdottir; Gardar Sveinbjornsson; Kristin E Sveinsdottir; Maney Sveinsdottir; Emil A Thorarensen; Bjarni Thorbjornsson; Gisli Masson; Ingileif Jonsdottir; Alma Moller; Thorolfur Gudnason; Karl G Kristinsson; Unnur Thorsteinsdottir; Kari Stefansson |
| EPI_ISL_417616 | hCoV-19/Iceland/288/2020 | Europe / Iceland / Reykjavik | 2020-03-18 | The National University Hospital of Iceland | deCODE genetics | Daniel F Gudbjartsson; Agnar Helgason; Hakon Jonsson; Olafur T Magnusson; Pall Melsted; Gudmundur L Norddahl; Jona Saemundsdottir; Asgeir Sigurdsson; Patrick Sulem; Arna B Agustsdottir; Berglind Eiriksdottir; Run Fridriksdottir; Elisabet E Gardarsdottir; Gudmundur Georgsson; Olafia S Gretarsdottir; Kjartan R Gudmundsson; Thora R Gunnarsdottir; Arnaldur Gylfason; Hilma Holm; Brynjar O Jensson; Aslaug Jonasdottir; Kamilla S Josefsdottir; Thordur Kristjansson; Droplaug N Magnusdottir; Louise le Roux; Gudrun Sigmundsdottir; Gardar Sveinbjornsson; Kristin E Sveinsdottir; Maney Sveinsdottir; Emil A Thorarensen; Bjarni Thorbjornsson; Gisli Masson; Ingileif Jonsdottir; Alma Moller; Thorolfur Gudnason; Karl G Kristinsson; Unnur Thorsteinsdottir; Kari Stefansson |
| EPI_ISL_417618 | hCoV-19/Iceland/29/2020 | Europe / Iceland / Reykjavik | 2020-03-16 | deCODE genetics | deCODE genetics | Daniel F Gudbjartsson; Agnar Helgason; Hakon Jonsson; Olafur T Magnusson; Pall Melsted; Gudmundur L Norddahl; Jona Saemundsdottir; Asgeir Sigurdsson; Patrick Sulem; Arna B Agustsdottir; Berglind Eiriksdottir; Run Fridriksdottir; Elisabet E Gardarsdottir; Gudmundur Georgsson; Olafia S Gretarsdottir; Kjartan R Gudmundsson; Thora R Gunnarsdottir; Arnaldur Gylfason; Hilma Holm; Brynjar O Jensson; Aslaug Jonasdottir; Kamilla S Josefsdottir; Thordur Kristjansson; Droplaug N Magnusdottir; Louise le Roux; Gudrun Sigmundsdottir; Gardar Sveinbjornsson; Kristin E Sveinsdottir; Maney Sveinsdottir; Emil A Thorarensen; Bjarni Thorbjornsson; Gisli Masson; Ingileif Jonsdottir; Alma Moller; Thorolfur Gudnason; Karl G Kristinsson; Unnur Thorsteinsdottir; Kari Stefansson |
| EPI_ISL_417619 | hCoV-19/Iceland/290/2020 | Europe / Iceland / Reykjavik | 2020-03-18 | The National University Hospital of Iceland | deCODE genetics | Daniel F Gudbjartsson; Agnar Helgason; Hakon Jonsson; Olafur T Magnusson; Pall Melsted; Gudmundur L Norddahl; Jona Saemundsdottir; Asgeir Sigurdsson; Patrick Sulem; Arna B Agustsdottir; Berglind Eiriksdottir; Run Fridriksdottir; Elisabet E Gardarsdottir; Gudmundur Georgsson; Olafia S Gretarsdottir; Kjartan R Gudmundsson; Thora R Gunnarsdottir; Arnaldur Gylfason; Hilma Holm; Brynjar O Jensson; Aslaug Jonasdottir; Kamilla S Josefsdottir; Thordur Kristjansson; Droplaug N Magnusdottir; Louise le Roux; Gudrun Sigmundsdottir; Gardar Sveinbjornsson; Kristin E Sveinsdottir; Maney Sveinsdottir; Emil A Thorarensen; Bjarni Thorbjornsson; Gisli Masson; Ingileif Jonsdottir; Alma Moller; Thorolfur Gudnason; Karl G Kristinsson; Unnur Thorsteinsdottir; Kari Stefansson |
| EPI_ISL_417620 | hCoV-19/Iceland/291/2020 | Europe / Iceland / Reykjavik | 2020-03-18 | The National University Hospital of Iceland | deCODE genetics | Daniel F Gudbjartsson; Agnar Helgason; Hakon Jonsson; Olafur T Magnusson; Pall Melsted; Gudmundur L Norddahl; Jona Saemundsdottir; Asgeir Sigurdsson; Patrick Sulem; Arna B Agustsdottir; Berglind Eiriksdottir; Run Fridriksdottir; Elisabet E Gardarsdottir; Gudmundur Georgsson; Olafia S Gretarsdottir; Kjartan R Gudmundsson; Thora R Gunnarsdottir; Arnaldur Gylfason; Hilma Holm; Brynjar O Jensson; Aslaug Jonasdottir; Kamilla S Josefsdottir; Thordur Kristjansson; Droplaug N Magnusdottir; Louise le Roux; Gudrun Sigmundsdottir; Gardar Sveinbjornsson; Kristin E Sveinsdottir; Maney Sveinsdottir; Emil A Thorarensen; Bjarni Thorbjornsson; Gisli Masson; Ingileif Jonsdottir; Alma Moller; Thorolfur Gudnason; Karl G Kristinsson; Unnur Thorsteinsdottir; Kari Stefansson |
| EPI_ISL_417621 | hCoV-19/Iceland/292/2020 | Europe / Iceland / Reykjavik | 2020-03-18 | The National University Hospital of Iceland | deCODE genetics | Daniel F Gudbjartsson; Agnar Helgason; Hakon Jonsson; Olafur T Magnusson; Pall Melsted; Gudmundur L Norddahl; Jona Saemundsdottir; Asgeir Sigurdsson; Patrick Sulem; Arna B Agustsdottir; Berglind Eiriksdottir; Run Fridriksdottir; Elisabet E Gardarsdottir; Gudmundur Georgsson; Olafia S Gretarsdottir; Kjartan R Gudmundsson; Thora R Gunnarsdottir; Arnaldur Gylfason; Hilma Holm; Brynjar O Jensson; Aslaug Jonasdottir; Kamilla S Josefsdottir; Thordur Kristjansson; Droplaug N Magnusdottir; Louise le Roux; Gudrun Sigmundsdottir; Gardar Sveinbjornsson; Kristin E Sveinsdottir; Maney Sveinsdottir; Emil A Thorarensen; Bjarni Thorbjornsson; Gisli Masson; Ingileif Jonsdottir; Alma Moller; Thorolfur Gudnason; Karl G Kristinsson; Unnur Thorsteinsdottir; Kari Stefansson |
| EPI_ISL_417622 | hCoV-19/Iceland/293/2020 | Europe / Iceland / Reykjavik | 2020-03-18 | The National University Hospital of Iceland | deCODE genetics | Daniel F Gudbjartsson; Agnar Helgason; Hakon Jonsson; Olafur T Magnusson; Pall Melsted; Gudmundur L Norddahl; Jona Saemundsdottir; Asgeir Sigurdsson; Patrick Sulem; Arna B Agustsdottir; Berglind Eiriksdottir; Run Fridriksdottir; Elisabet E Gardarsdottir; Gudmundur Georgsson; Olafia S Gretarsdottir; Kjartan R Gudmundsson; Thora R Gunnarsdottir; Arnaldur Gylfason; Hilma Holm; Brynjar O Jensson; Aslaug Jonasdottir; Kamilla S Josefsdottir; Thordur Kristjansson; Droplaug N Magnusdottir; Louise le Roux; Gudrun Sigmundsdottir; Gardar Sveinbjornsson; Kristin E Sveinsdottir; Maney Sveinsdottir; Emil A Thorarensen; Bjarni Thorbjornsson; Gisli Masson; Ingileif Jonsdottir; Alma Moller; Thorolfur Gudnason; Karl G Kristinsson; Unnur Thorsteinsdottir; Kari Stefansson |
| EPI_ISL_417623 | hCoV-19/Iceland/294/2020 | Europe / Iceland / Reykjavik | 2020-03-18 | The National University Hospital of Iceland | deCODE genetics | Daniel F Gudbjartsson; Agnar Helgason; Hakon Jonsson; Olafur T Magnusson; Pall Melsted; Gudmundur L Norddahl; Jona Saemundsdottir; Asgeir Sigurdsson; Patrick Sulem; Arna B Agustsdottir; Berglind Eiriksdottir; Run Fridriksdottir; Elisabet E Gardarsdottir; Gudmundur Georgsson; Olafia S Gretarsdottir; Kjartan R Gudmundsson; Thora R Gunnarsdottir; Arnaldur Gylfason; Hilma Holm; Brynjar O Jensson; Aslaug Jonasdottir; Kamilla S Josefsdottir; Thordur Kristjansson; Droplaug N Magnusdottir; Louise le Roux; Gudrun Sigmundsdottir; Gardar Sveinbjornsson; Kristin E Sveinsdottir; Maney Sveinsdottir; Emil A Thorarensen; Bjarni Thorbjornsson; Gisli Masson; Ingileif Jonsdottir; Alma Moller; Thorolfur Gudnason; Karl G Kristinsson; Unnur Thorsteinsdottir; Kari Stefansson |
| EPI_ISL_417625 | hCoV-19/Iceland/296/2020 | Europe / Iceland / Reykjavik | 2020-03-18 | The National University Hospital of Iceland | deCODE genetics | Daniel F Gudbjartsson; Agnar Helgason; Hakon Jonsson; Olafur T Magnusson; Pall Melsted; Gudmundur L Norddahl; Jona Saemundsdottir; Asgeir Sigurdsson; Patrick Sulem; Arna B Agustsdottir; Berglind Eiriksdottir; Run Fridriksdottir; Elisabet E Gardarsdottir; Gudmundur Georgsson; Olafia S Gretarsdottir; Kjartan R Gudmundsson; Thora R Gunnarsdottir; Arnaldur Gylfason; Hilma Holm; Brynjar O Jensson; Aslaug Jonasdottir; Kamilla S Josefsdottir; Thordur Kristjansson; Droplaug N Magnusdottir; Louise le Roux; Gudrun Sigmundsdottir; Gardar Sveinbjornsson; Kristin E Sveinsdottir; Maney Sveinsdottir; Emil A Thorarensen; Bjarni Thorbjornsson; Gisli Masson; Ingileif Jonsdottir; Alma Moller; Thorolfur Gudnason; Karl G Kristinsson; Unnur Thorsteinsdottir; Kari Stefansson |
| EPI_ISL_417626 | hCoV-19/Iceland/297/2020 | Europe / Iceland / Reykjavik | 2020-03-18 | The National University Hospital of Iceland | deCODE genetics | Daniel F Gudbjartsson; Agnar Helgason; Hakon Jonsson; Olafur T Magnusson; Pall Melsted; Gudmundur L Norddahl; Jona Saemundsdottir; Asgeir Sigurdsson; Patrick Sulem; Arna B Agustsdottir; Berglind Eiriksdottir; Run Fridriksdottir; Elisabet E Gardarsdottir; Gudmundur Georgsson; Olafia S Gretarsdottir; Kjartan R Gudmundsson; Thora R Gunnarsdottir; Arnaldur Gylfason; Hilma Holm; Brynjar O Jensson; Aslaug Jonasdottir; Kamilla S Josefsdottir; Thordur Kristjansson; Droplaug N Magnusdottir; Louise le Roux; Gudrun Sigmundsdottir; Gardar Sveinbjornsson; Kristin E Sveinsdottir; Maney Sveinsdottir; Emil A Thorarensen; Bjarni Thorbjornsson; Gisli Masson; Ingileif Jonsdottir; Alma Moller; Thorolfur Gudnason; Karl G Kristinsson; Unnur Thorsteinsdottir; Kari Stefansson |
| EPI_ISL_417627 | hCoV-19/Iceland/298/2020 | Europe / Iceland / Reykjavik | 2020-03-18 | The National University Hospital of Iceland | deCODE genetics | Daniel F Gudbjartsson; Agnar Helgason; Hakon Jonsson; Olafur T Magnusson; Pall Melsted; Gudmundur L Norddahl; Jona Saemundsdottir; Asgeir Sigurdsson; Patrick Sulem; Arna B Agustsdottir; Berglind Eiriksdottir; Run Fridriksdottir; Elisabet E Gardarsdottir; Gudmundur Georgsson; Olafia S Gretarsdottir; Kjartan R Gudmundsson; Thora R Gunnarsdottir; Arnaldur Gylfason; Hilma Holm; Brynjar O Jensson; Aslaug Jonasdottir; Kamilla S Josefsdottir; Thordur Kristjansson; Droplaug N Magnusdottir; Louise le Roux; Gudrun Sigmundsdottir; Gardar Sveinbjornsson; Kristin E Sveinsdottir; Maney Sveinsdottir; Emil A Thorarensen; Bjarni Thorbjornsson; Gisli Masson; Ingileif Jonsdottir; Alma Moller; Thorolfur Gudnason; Karl G Kristinsson; Unnur Thorsteinsdottir; Kari Stefansson |
| EPI_ISL_417628 | hCoV-19/Iceland/299/2020 | Europe / Iceland / Reykjavik | 2020-03-18 | The National University Hospital of Iceland | deCODE genetics | Daniel F Gudbjartsson; Agnar Helgason; Hakon Jonsson; Olafur T Magnusson; Pall Melsted; Gudmundur L Norddahl; Jona Saemundsdottir; Asgeir Sigurdsson; Patrick Sulem; Arna B Agustsdottir; Berglind Eiriksdottir; Run Fridriksdottir; Elisabet E Gardarsdottir; Gudmundur Georgsson; Olafia S Gretarsdottir; Kjartan R Gudmundsson; Thora R Gunnarsdottir; Arnaldur Gylfason; Hilma Holm; Brynjar O Jensson; Aslaug Jonasdottir; Kamilla S Josefsdottir; Thordur Kristjansson; Droplaug N Magnusdottir; Louise le Roux; Gudrun Sigmundsdottir; Gardar Sveinbjornsson; Kristin E Sveinsdottir; Maney Sveinsdottir; Emil A Thorarensen; Bjarni Thorbjornsson; Gisli Masson; Ingileif Jonsdottir; Alma Moller; Thorolfur Gudnason; Karl G Kristinsson; Unnur Thorsteinsdottir; Kari Stefansson |
| EPI_ISL_417629 | hCoV-19/Iceland/300/2020 | Europe / Iceland / Reykjavik | 2020-03-18 | The National University Hospital of Iceland | deCODE genetics | Daniel F Gudbjartsson; Agnar Helgason; Hakon Jonsson; Olafur T Magnusson; Pall Melsted; Gudmundur L Norddahl; Jona Saemundsdottir; Asgeir Sigurdsson; Patrick Sulem; Arna B Agustsdottir; Berglind Eiriksdottir; Run Fridriksdottir; Elisabet E Gardarsdottir; Gudmundur Georgsson; Olafia S Gretarsdottir; Kjartan R Gudmundsson; Thora R Gunnarsdottir; Arnaldur Gylfason; Hilma Holm; Brynjar O Jensson; Aslaug Jonasdottir; Kamilla S Josefsdottir; Thordur Kristjansson; Droplaug N Magnusdottir; Louise le Roux; Gudrun Sigmundsdottir; Gardar Sveinbjornsson; Kristin E Sveinsdottir; Maney Sveinsdottir; Emil A Thorarensen; Bjarni Thorbjornsson; Gisli Masson; Ingileif Jonsdottir; Alma Moller; Thorolfur Gudnason; Karl G Kristinsson; Unnur Thorsteinsdottir; Kari Stefansson |
| EPI_ISL_417630 | hCoV-19/Iceland/301/2020 | Europe / Iceland / Reykjavik | 2020-03-18 | The National University Hospital of Iceland | deCODE genetics | Daniel F Gudbjartsson; Agnar Helgason; Hakon Jonsson; Olafur T Magnusson; Pall Melsted; Gudmundur L Norddahl; Jona Saemundsdottir; Asgeir Sigurdsson; Patrick Sulem; Arna B Agustsdottir; Berglind Eiriksdottir; Run Fridriksdottir; Elisabet E Gardarsdottir; Gudmundur Georgsson; Olafia S Gretarsdottir; Kjartan R Gudmundsson; Thora R Gunnarsdottir; Arnaldur Gylfason; Hilma Holm; Brynjar O Jensson; Aslaug Jonasdottir; Kamilla S Josefsdottir; Thordur Kristjansson; Droplaug N Magnusdottir; Louise le Roux; Gudrun Sigmundsdottir; Gardar Sveinbjornsson; Kristin E Sveinsdottir; Maney Sveinsdottir; Emil A Thorarensen; Bjarni Thorbjornsson; Gisli Masson; Ingileif Jonsdottir; Alma Moller; Thorolfur Gudnason; Karl G Kristinsson; Unnur Thorsteinsdottir; Kari Stefansson |
| EPI_ISL_417632 | hCoV-19/Iceland/303/2020 | Europe / Iceland / Reykjavik | 2020-03-18 | The National University Hospital of Iceland | deCODE genetics | Daniel F Gudbjartsson; Agnar Helgason; Hakon Jonsson; Olafur T Magnusson; Pall Melsted; Gudmundur L Norddahl; Jona Saemundsdottir; Asgeir Sigurdsson; Patrick Sulem; Arna B Agustsdottir; Berglind Eiriksdottir; Run Fridriksdottir; Elisabet E Gardarsdottir; Gudmundur Georgsson; Olafia S Gretarsdottir; Kjartan R Gudmundsson; Thora R Gunnarsdottir; Arnaldur Gylfason; Hilma Holm; Brynjar O Jensson; Aslaug Jonasdottir; Kamilla S Josefsdottir; Thordur Kristjansson; Droplaug N Magnusdottir; Louise le Roux; Gudrun Sigmundsdottir; Gardar Sveinbjornsson; Kristin E Sveinsdottir; Maney Sveinsdottir; Emil A Thorarensen; Bjarni Thorbjornsson; Gisli Masson; Ingileif Jonsdottir; Alma Moller; Thorolfur Gudnason; Karl G Kristinsson; Unnur Thorsteinsdottir; Kari Stefansson |
| EPI_ISL_417633 | hCoV-19/Iceland/304/2020 | Europe / Iceland / Reykjavik | 2020-03-17 | The National University Hospital of Iceland | deCODE genetics | Daniel F Gudbjartsson; Agnar Helgason; Hakon Jonsson; Olafur T Magnusson; Pall Melsted; Gudmundur L Norddahl; Jona Saemundsdottir; Asgeir Sigurdsson; Patrick Sulem; Arna B Agustsdottir; Berglind Eiriksdottir; Run Fridriksdottir; Elisabet E Gardarsdottir; Gudmundur Georgsson; Olafia S Gretarsdottir; Kjartan R Gudmundsson; Thora R Gunnarsdottir; Arnaldur Gylfason; Hilma Holm; Brynjar O Jensson; Aslaug Jonasdottir; Kamilla S Josefsdottir; Thordur Kristjansson; Droplaug N Magnusdottir; Louise le Roux; Gudrun Sigmundsdottir; Gardar Sveinbjornsson; Kristin E Sveinsdottir; Maney Sveinsdottir; Emil A Thorarensen; Bjarni Thorbjornsson; Gisli Masson; Ingileif Jonsdottir; Alma Moller; Thorolfur Gudnason; Karl G Kristinsson; Unnur Thorsteinsdottir; Kari Stefansson |
| EPI_ISL_417634 | hCoV-19/Iceland/305/2020 | Europe / Iceland / Reykjavik | 2020-03-18 | The National University Hospital of Iceland | deCODE genetics | Daniel F Gudbjartsson; Agnar Helgason; Hakon Jonsson; Olafur T Magnusson; Pall Melsted; Gudmundur L Norddahl; Jona Saemundsdottir; Asgeir Sigurdsson; Patrick Sulem; Arna B Agustsdottir; Berglind Eiriksdottir; Run Fridriksdottir; Elisabet E Gardarsdottir; Gudmundur Georgsson; Olafia S Gretarsdottir; Kjartan R Gudmundsson; Thora R Gunnarsdottir; Arnaldur Gylfason; Hilma Holm; Brynjar O Jensson; Aslaug Jonasdottir; Kamilla S Josefsdottir; Thordur Kristjansson; Droplaug N Magnusdottir; Louise le Roux; Gudrun Sigmundsdottir; Gardar Sveinbjornsson; Kristin E Sveinsdottir; Maney Sveinsdottir; Emil A Thorarensen; Bjarni Thorbjornsson; Gisli Masson; Ingileif Jonsdottir; Alma Moller; Thorolfur Gudnason; Karl G Kristinsson; Unnur Thorsteinsdottir; Kari Stefansson |
| EPI_ISL_417635 | hCoV-19/Iceland/306/2020 | Europe / Iceland / Reykjavik | 2020-03-18 | The National University Hospital of Iceland | deCODE genetics | Daniel F Gudbjartsson; Agnar Helgason; Hakon Jonsson; Olafur T Magnusson; Pall Melsted; Gudmundur L Norddahl; Jona Saemundsdottir; Asgeir Sigurdsson; Patrick Sulem; Arna B Agustsdottir; Berglind Eiriksdottir; Run Fridriksdottir; Elisabet E Gardarsdottir; Gudmundur Georgsson; Olafia S Gretarsdottir; Kjartan R Gudmundsson; Thora R Gunnarsdottir; Arnaldur Gylfason; Hilma Holm; Brynjar O Jensson; Aslaug Jonasdottir; Kamilla S Josefsdottir; Thordur Kristjansson; Droplaug N Magnusdottir; Louise le Roux; Gudrun Sigmundsdottir; Gardar Sveinbjornsson; Kristin E Sveinsdottir; Maney Sveinsdottir; Emil A Thorarensen; Bjarni Thorbjornsson; Gisli Masson; Ingileif Jonsdottir; Alma Moller; Thorolfur Gudnason; Karl G Kristinsson; Unnur Thorsteinsdottir; Kari Stefansson |
| EPI_ISL_417636 | hCoV-19/Iceland/307/2020 | Europe / Iceland / Reykjavik | 2020-03-18 | The National University Hospital of Iceland | deCODE genetics | Daniel F Gudbjartsson; Agnar Helgason; Hakon Jonsson; Olafur T Magnusson; Pall Melsted; Gudmundur L Norddahl; Jona Saemundsdottir; Asgeir Sigurdsson; Patrick Sulem; Arna B Agustsdottir; Berglind Eiriksdottir; Run Fridriksdottir; Elisabet E Gardarsdottir; Gudmundur Georgsson; Olafia S Gretarsdottir; Kjartan R Gudmundsson; Thora R Gunnarsdottir; Arnaldur Gylfason; Hilma Holm; Brynjar O Jensson; Aslaug Jonasdottir; Kamilla S Josefsdottir; Thordur Kristjansson; Droplaug N Magnusdottir; Louise le Roux; Gudrun Sigmundsdottir; Gardar Sveinbjornsson; Kristin E Sveinsdottir; Maney Sveinsdottir; Emil A Thorarensen; Bjarni Thorbjornsson; Gisli Masson; Ingileif Jonsdottir; Alma Moller; Thorolfur Gudnason; Karl G Kristinsson; Unnur Thorsteinsdottir; Kari Stefansson |
| EPI_ISL_417637 | hCoV-19/Iceland/308/2020 | Europe / Iceland / Reykjavik | 2020-03-18 | The National University Hospital of Iceland | deCODE genetics | Daniel F Gudbjartsson; Agnar Helgason; Hakon Jonsson; Olafur T Magnusson; Pall Melsted; Gudmundur L Norddahl; Jona Saemundsdottir; Asgeir Sigurdsson; Patrick Sulem; Arna B Agustsdottir; Berglind Eiriksdottir; Run Fridriksdottir; Elisabet E Gardarsdottir; Gudmundur Georgsson; Olafia S Gretarsdottir; Kjartan R Gudmundsson; Thora R Gunnarsdottir; Arnaldur Gylfason; Hilma Holm; Brynjar O Jensson; Aslaug Jonasdottir; Kamilla S Josefsdottir; Thordur Kristjansson; Droplaug N Magnusdottir; Louise le Roux; Gudrun Sigmundsdottir; Gardar Sveinbjornsson; Kristin E Sveinsdottir; Maney Sveinsdottir; Emil A Thorarensen; Bjarni Thorbjornsson; Gisli Masson; Ingileif Jonsdottir; Alma Moller; Thorolfur Gudnason; Karl G Kristinsson; Unnur Thorsteinsdottir; Kari Stefansson |
| EPI_ISL_417639 | hCoV-19/Iceland/310/2020 | Europe / Iceland / Reykjavik | 2020-03-18 | The National University Hospital of Iceland | deCODE genetics | Daniel F Gudbjartsson; Agnar Helgason; Hakon Jonsson; Olafur T Magnusson; Pall Melsted; Gudmundur L Norddahl; Jona Saemundsdottir; Asgeir Sigurdsson; Patrick Sulem; Arna B Agustsdottir; Berglind Eiriksdottir; Run Fridriksdottir; Elisabet E Gardarsdottir; Gudmundur Georgsson; Olafia S Gretarsdottir; Kjartan R Gudmundsson; Thora R Gunnarsdottir; Arnaldur Gylfason; Hilma Holm; Brynjar O Jensson; Aslaug Jonasdottir; Kamilla S Josefsdottir; Thordur Kristjansson; Droplaug N Magnusdottir; Louise le Roux; Gudrun Sigmundsdottir; Gardar Sveinbjornsson; Kristin E Sveinsdottir; Maney Sveinsdottir; Emil A Thorarensen; Bjarni Thorbjornsson; Gisli Masson; Ingileif Jonsdottir; Alma Moller; Thorolfur Gudnason; Karl G Kristinsson; Unnur Thorsteinsdottir; Kari Stefansson |
| EPI_ISL_417641 | hCoV-19/Iceland/312/2020 | Europe / Iceland / Reykjavik | 2020-03-18 | The National University Hospital of Iceland | deCODE genetics | Daniel F Gudbjartsson; Agnar Helgason; Hakon Jonsson; Olafur T Magnusson; Pall Melsted; Gudmundur L Norddahl; Jona Saemundsdottir; Asgeir Sigurdsson; Patrick Sulem; Arna B Agustsdottir; Berglind Eiriksdottir; Run Fridriksdottir; Elisabet E Gardarsdottir; Gudmundur Georgsson; Olafia S Gretarsdottir; Kjartan R Gudmundsson; Thora R Gunnarsdottir; Arnaldur Gylfason; Hilma Holm; Brynjar O Jensson; Aslaug Jonasdottir; Kamilla S Josefsdottir; Thordur Kristjansson; Droplaug N Magnusdottir; Louise le Roux; Gudrun Sigmundsdottir; Gardar Sveinbjornsson; Kristin E Sveinsdottir; Maney Sveinsdottir; Emil A Thorarensen; Bjarni Thorbjornsson; Gisli Masson; Ingileif Jonsdottir; Alma Moller; Thorolfur Gudnason; Karl G Kristinsson; Unnur Thorsteinsdottir; Kari Stefansson |
| EPI_ISL_417642 | hCoV-19/Iceland/313/2020 | Europe / Iceland / Reykjavik | 2020-03-18 | The National University Hospital of Iceland | deCODE genetics | Daniel F Gudbjartsson; Agnar Helgason; Hakon Jonsson; Olafur T Magnusson; Pall Melsted; Gudmundur L Norddahl; Jona Saemundsdottir; Asgeir Sigurdsson; Patrick Sulem; Arna B Agustsdottir; Berglind Eiriksdottir; Run Fridriksdottir; Elisabet E Gardarsdottir; Gudmundur Georgsson; Olafia S Gretarsdottir; Kjartan R Gudmundsson; Thora R Gunnarsdottir; Arnaldur Gylfason; Hilma Holm; Brynjar O Jensson; Aslaug Jonasdottir; Kamilla S Josefsdottir; Thordur Kristjansson; Droplaug N Magnusdottir; Louise le Roux; Gudrun Sigmundsdottir; Gardar Sveinbjornsson; Kristin E Sveinsdottir; Maney Sveinsdottir; Emil A Thorarensen; Bjarni Thorbjornsson; Gisli Masson; Ingileif Jonsdottir; Alma Moller; Thorolfur Gudnason; Karl G Kristinsson; Unnur Thorsteinsdottir; Kari Stefansson |
| EPI_ISL_417643 | hCoV-19/Iceland/314/2020 | Europe / Iceland / Reykjavik | 2020-03-18 | The National University Hospital of Iceland | deCODE genetics | Daniel F Gudbjartsson; Agnar Helgason; Hakon Jonsson; Olafur T Magnusson; Pall Melsted; Gudmundur L Norddahl; Jona Saemundsdottir; Asgeir Sigurdsson; Patrick Sulem; Arna B Agustsdottir; Berglind Eiriksdottir; Run Fridriksdottir; Elisabet E Gardarsdottir; Gudmundur Georgsson; Olafia S Gretarsdottir; Kjartan R Gudmundsson; Thora R Gunnarsdottir; Arnaldur Gylfason; Hilma Holm; Brynjar O Jensson; Aslaug Jonasdottir; Kamilla S Josefsdottir; Thordur Kristjansson; Droplaug N Magnusdottir; Louise le Roux; Gudrun Sigmundsdottir; Gardar Sveinbjornsson; Kristin E Sveinsdottir; Maney Sveinsdottir; Emil A Thorarensen; Bjarni Thorbjornsson; Gisli Masson; Ingileif Jonsdottir; Alma Moller; Thorolfur Gudnason; Karl G Kristinsson; Unnur Thorsteinsdottir; Kari Stefansson |
| EPI_ISL_417644 | hCoV-19/Iceland/315/2020 | Europe / Iceland / Reykjavik | 2020-03-18 | The National University Hospital of Iceland | deCODE genetics | Daniel F Gudbjartsson; Agnar Helgason; Hakon Jonsson; Olafur T Magnusson; Pall Melsted; Gudmundur L Norddahl; Jona Saemundsdottir; Asgeir Sigurdsson; Patrick Sulem; Arna B Agustsdottir; Berglind Eiriksdottir; Run Fridriksdottir; Elisabet E Gardarsdottir; Gudmundur Georgsson; Olafia S Gretarsdottir; Kjartan R Gudmundsson; Thora R Gunnarsdottir; Arnaldur Gylfason; Hilma Holm; Brynjar O Jensson; Aslaug Jonasdottir; Kamilla S Josefsdottir; Thordur Kristjansson; Droplaug N Magnusdottir; Louise le Roux; Gudrun Sigmundsdottir; Gardar Sveinbjornsson; Kristin E Sveinsdottir; Maney Sveinsdottir; Emil A Thorarensen; Bjarni Thorbjornsson; Gisli Masson; Ingileif Jonsdottir; Alma Moller; Thorolfur Gudnason; Karl G Kristinsson; Unnur Thorsteinsdottir; Kari Stefansson |
| EPI_ISL_417645 | hCoV-19/Iceland/316/2020 | Europe / Iceland / Reykjavik | 2020-03-18 | The National University Hospital of Iceland | deCODE genetics | Daniel F Gudbjartsson; Agnar Helgason; Hakon Jonsson; Olafur T Magnusson; Pall Melsted; Gudmundur L Norddahl; Jona Saemundsdottir; Asgeir Sigurdsson; Patrick Sulem; Arna B Agustsdottir; Berglind Eiriksdottir; Run Fridriksdottir; Elisabet E Gardarsdottir; Gudmundur Georgsson; Olafia S Gretarsdottir; Kjartan R Gudmundsson; Thora R Gunnarsdottir; Arnaldur Gylfason; Hilma Holm; Brynjar O Jensson; Aslaug Jonasdottir; Kamilla S Josefsdottir; Thordur Kristjansson; Droplaug N Magnusdottir; Louise le Roux; Gudrun Sigmundsdottir; Gardar Sveinbjornsson; Kristin E Sveinsdottir; Maney Sveinsdottir; Emil A Thorarensen; Bjarni Thorbjornsson; Gisli Masson; Ingileif Jonsdottir; Alma Moller; Thorolfur Gudnason; Karl G Kristinsson; Unnur Thorsteinsdottir; Kari Stefansson |
| EPI_ISL_417646 | hCoV-19/Iceland/317/2020 | Europe / Iceland / Reykjavik | 2020-03-18 | The National University Hospital of Iceland | deCODE genetics | Daniel F Gudbjartsson; Agnar Helgason; Hakon Jonsson; Olafur T Magnusson; Pall Melsted; Gudmundur L Norddahl; Jona Saemundsdottir; Asgeir Sigurdsson; Patrick Sulem; Arna B Agustsdottir; Berglind Eiriksdottir; Run Fridriksdottir; Elisabet E Gardarsdottir; Gudmundur Georgsson; Olafia S Gretarsdottir; Kjartan R Gudmundsson; Thora R Gunnarsdottir; Arnaldur Gylfason; Hilma Holm; Brynjar O Jensson; Aslaug Jonasdottir; Kamilla S Josefsdottir; Thordur Kristjansson; Droplaug N Magnusdottir; Louise le Roux; Gudrun Sigmundsdottir; Gardar Sveinbjornsson; Kristin E Sveinsdottir; Maney Sveinsdottir; Emil A Thorarensen; Bjarni Thorbjornsson; Gisli Masson; Ingileif Jonsdottir; Alma Moller; Thorolfur Gudnason; Karl G Kristinsson; Unnur Thorsteinsdottir; Kari Stefansson |
| EPI_ISL_417647 | hCoV-19/Iceland/318/2020 | Europe / Iceland / Reykjavik | 2020-03-18 | The National University Hospital of Iceland | deCODE genetics | Daniel F Gudbjartsson; Agnar Helgason; Hakon Jonsson; Olafur T Magnusson; Pall Melsted; Gudmundur L Norddahl; Jona Saemundsdottir; Asgeir Sigurdsson; Patrick Sulem; Arna B Agustsdottir; Berglind Eiriksdottir; Run Fridriksdottir; Elisabet E Gardarsdottir; Gudmundur Georgsson; Olafia S Gretarsdottir; Kjartan R Gudmundsson; Thora R Gunnarsdottir; Arnaldur Gylfason; Hilma Holm; Brynjar O Jensson; Aslaug Jonasdottir; Kamilla S Josefsdottir; Thordur Kristjansson; Droplaug N Magnusdottir; Louise le Roux; Gudrun Sigmundsdottir; Gardar Sveinbjornsson; Kristin E Sveinsdottir; Maney Sveinsdottir; Emil A Thorarensen; Bjarni Thorbjornsson; Gisli Masson; Ingileif Jonsdottir; Alma Moller; Thorolfur Gudnason; Karl G Kristinsson; Unnur Thorsteinsdottir; Kari Stefansson |
| EPI_ISL_417649 | hCoV-19/Iceland/320/2020 | Europe / Iceland / Reykjavik | 2020-03-18 | The National University Hospital of Iceland | deCODE genetics | Daniel F Gudbjartsson; Agnar Helgason; Hakon Jonsson; Olafur T Magnusson; Pall Melsted; Gudmundur L Norddahl; Jona Saemundsdottir; Asgeir Sigurdsson; Patrick Sulem; Arna B Agustsdottir; Berglind Eiriksdottir; Run Fridriksdottir; Elisabet E Gardarsdottir; Gudmundur Georgsson; Olafia S Gretarsdottir; Kjartan R Gudmundsson; Thora R Gunnarsdottir; Arnaldur Gylfason; Hilma Holm; Brynjar O Jensson; Aslaug Jonasdottir; Kamilla S Josefsdottir; Thordur Kristjansson; Droplaug N Magnusdottir; Louise le Roux; Gudrun Sigmundsdottir; Gardar Sveinbjornsson; Kristin E Sveinsdottir; Maney Sveinsdottir; Emil A Thorarensen; Bjarni Thorbjornsson; Gisli Masson; Ingileif Jonsdottir; Alma Moller; Thorolfur Gudnason; Karl G Kristinsson; Unnur Thorsteinsdottir; Kari Stefansson |
| EPI_ISL_417700 | hCoV-19/Iceland/173/2020 | Europe / Iceland / Reykjavik | 2020-03-15 | The National University Hospital of Iceland | deCODE genetics | Daniel F Gudbjartsson; Agnar Helgason; Hakon Jonsson; Olafur T Magnusson; Pall Melsted; Gudmundur L Norddahl; Jona Saemundsdottir; Asgeir Sigurdsson; Patrick Sulem; Arna B Agustsdottir; Berglind Eiriksdottir; Run Fridriksdottir; Elisabet E Gardarsdottir; Gudmundur Georgsson; Olafia S Gretarsdottir; Kjartan R Gudmundsson; Thora R Gunnarsdottir; Arnaldur Gylfason; Hilma Holm; Brynjar O Jensson; Aslaug Jonasdottir; Kamilla S Josefsdottir; Thordur Kristjansson; Droplaug N Magnusdottir; Louise le Roux; Gudrun Sigmundsdottir; Gardar Sveinbjornsson; Kristin E Sveinsdottir; Maney Sveinsdottir; Emil A Thorarensen; Bjarni Thorbjornsson; Gisli Masson; Ingileif Jonsdottir; Alma Moller; Thorolfur Gudnason; Karl G Kristinsson; Unnur Thorsteinsdottir; Kari Stefansson |
| EPI_ISL_417701 | hCoV-19/Iceland/74/2020 | Europe / Iceland / Reykjavik | 2020-03-09 | The National University Hospital of Iceland | deCODE genetics | Daniel F Gudbjartsson; Agnar Helgason; Hakon Jonsson; Olafur T Magnusson; Pall Melsted; Gudmundur L Norddahl; Jona Saemundsdottir; Asgeir Sigurdsson; Patrick Sulem; Arna B Agustsdottir; Berglind Eiriksdottir; Run Fridriksdottir; Elisabet E Gardarsdottir; Gudmundur Georgsson; Olafia S Gretarsdottir; Kjartan R Gudmundsson; Thora R Gunnarsdottir; Arnaldur Gylfason; Hilma Holm; Brynjar O Jensson; Aslaug Jonasdottir; Kamilla S Josefsdottir; Thordur Kristjansson; Droplaug N Magnusdottir; Louise le Roux; Gudrun Sigmundsdottir; Gardar Sveinbjornsson; Kristin E Sveinsdottir; Maney Sveinsdottir; Emil A Thorarensen; Bjarni Thorbjornsson; Gisli Masson; Ingileif Jonsdottir; Alma Moller; Thorolfur Gudnason; Karl G Kristinsson; Unnur Thorsteinsdottir; Kari Stefansson |
| EPI_ISL_417702 | hCoV-19/Iceland/33/2020 | Europe / Iceland / Reykjavik | 2020-03-03 | The National University Hospital of Iceland | deCODE genetics | Daniel F Gudbjartsson; Agnar Helgason; Hakon Jonsson; Olafur T Magnusson; Pall Melsted; Gudmundur L Norddahl; Jona Saemundsdottir; Asgeir Sigurdsson; Patrick Sulem; Arna B Agustsdottir; Berglind Eiriksdottir; Run Fridriksdottir; Elisabet E Gardarsdottir; Gudmundur Georgsson; Olafia S Gretarsdottir; Kjartan R Gudmundsson; Thora R Gunnarsdottir; Arnaldur Gylfason; Hilma Holm; Brynjar O Jensson; Aslaug Jonasdottir; Kamilla S Josefsdottir; Thordur Kristjansson; Droplaug N Magnusdottir; Louise le Roux; Gudrun Sigmundsdottir; Gardar Sveinbjornsson; Kristin E Sveinsdottir; Maney Sveinsdottir; Emil A Thorarensen; Bjarni Thorbjornsson; Gisli Masson; Ingileif Jonsdottir; Alma Moller; Thorolfur Gudnason; Karl G Kristinsson; Unnur Thorsteinsdottir; Kari Stefansson |
| EPI_ISL_417703 | hCoV-19/Iceland/179/2020 | Europe / Iceland / Reykjavik | 2020-03-16 | The National University Hospital of Iceland | deCODE genetics | Daniel F Gudbjartsson; Agnar Helgason; Hakon Jonsson; Olafur T Magnusson; Pall Melsted; Gudmundur L Norddahl; Jona Saemundsdottir; Asgeir Sigurdsson; Patrick Sulem; Arna B Agustsdottir; Berglind Eiriksdottir; Run Fridriksdottir; Elisabet E Gardarsdottir; Gudmundur Georgsson; Olafia S Gretarsdottir; Kjartan R Gudmundsson; Thora R Gunnarsdottir; Arnaldur Gylfason; Hilma Holm; Brynjar O Jensson; Aslaug Jonasdottir; Kamilla S Josefsdottir; Thordur Kristjansson; Droplaug N Magnusdottir; Louise le Roux; Gudrun Sigmundsdottir; Gardar Sveinbjornsson; Kristin E Sveinsdottir; Maney Sveinsdottir; Emil A Thorarensen; Bjarni Thorbjornsson; Gisli Masson; Ingileif Jonsdottir; Alma Moller; Thorolfur Gudnason; Karl G Kristinsson; Unnur Thorsteinsdottir; Kari Stefansson |
| EPI_ISL_417705 | hCoV-19/Iceland/65/2020 | Europe / Iceland / Reykjavik | 2020-03-07 | The National University Hospital of Iceland | deCODE genetics | Daniel F Gudbjartsson; Agnar Helgason; Hakon Jonsson; Olafur T Magnusson; Pall Melsted; Gudmundur L Norddahl; Jona Saemundsdottir; Asgeir Sigurdsson; Patrick Sulem; Arna B Agustsdottir; Berglind Eiriksdottir; Run Fridriksdottir; Elisabet E Gardarsdottir; Gudmundur Georgsson; Olafia S Gretarsdottir; Kjartan R Gudmundsson; Thora R Gunnarsdottir; Arnaldur Gylfason; Hilma Holm; Brynjar O Jensson; Aslaug Jonasdottir; Kamilla S Josefsdottir; Thordur Kristjansson; Droplaug N Magnusdottir; Louise le Roux; Gudrun Sigmundsdottir; Gardar Sveinbjornsson; Kristin E Sveinsdottir; Maney Sveinsdottir; Emil A Thorarensen; Bjarni Thorbjornsson; Gisli Masson; Ingileif Jonsdottir; Alma Moller; Thorolfur Gudnason; Karl G Kristinsson; Unnur Thorsteinsdottir; Kari Stefansson |
| EPI_ISL_417706 | hCoV-19/Iceland/199/2020 | Europe / Iceland / Reykjavik | 2020-03-16 | The National University Hospital of Iceland | deCODE genetics | Daniel F Gudbjartsson; Agnar Helgason; Hakon Jonsson; Olafur T Magnusson; Pall Melsted; Gudmundur L Norddahl; Jona Saemundsdottir; Asgeir Sigurdsson; Patrick Sulem; Arna B Agustsdottir; Berglind Eiriksdottir; Run Fridriksdottir; Elisabet E Gardarsdottir; Gudmundur Georgsson; Olafia S Gretarsdottir; Kjartan R Gudmundsson; Thora R Gunnarsdottir; Arnaldur Gylfason; Hilma Holm; Brynjar O Jensson; Aslaug Jonasdottir; Kamilla S Josefsdottir; Thordur Kristjansson; Droplaug N Magnusdottir; Louise le Roux; Gudrun Sigmundsdottir; Gardar Sveinbjornsson; Kristin E Sveinsdottir; Maney Sveinsdottir; Emil A Thorarensen; Bjarni Thorbjornsson; Gisli Masson; Ingileif Jonsdottir; Alma Moller; Thorolfur Gudnason; Karl G Kristinsson; Unnur Thorsteinsdottir; Kari Stefansson |
| EPI_ISL_417709 | hCoV-19/Iceland/189/2020 | Europe / Iceland / Reykjavik | 2020-03-16 | The National University Hospital of Iceland | deCODE genetics | Daniel F Gudbjartsson; Agnar Helgason; Hakon Jonsson; Olafur T Magnusson; Pall Melsted; Gudmundur L Norddahl; Jona Saemundsdottir; Asgeir Sigurdsson; Patrick Sulem; Arna B Agustsdottir; Berglind Eiriksdottir; Run Fridriksdottir; Elisabet E Gardarsdottir; Gudmundur Georgsson; Olafia S Gretarsdottir; Kjartan R Gudmundsson; Thora R Gunnarsdottir; Arnaldur Gylfason; Hilma Holm; Brynjar O Jensson; Aslaug Jonasdottir; Kamilla S Josefsdottir; Thordur Kristjansson; Droplaug N Magnusdottir; Louise le Roux; Gudrun Sigmundsdottir; Gardar Sveinbjornsson; Kristin E Sveinsdottir; Maney Sveinsdottir; Emil A Thorarensen; Bjarni Thorbjornsson; Gisli Masson; Ingileif Jonsdottir; Alma Moller; Thorolfur Gudnason; Karl G Kristinsson; Unnur Thorsteinsdottir; Kari Stefansson |
| EPI_ISL_417711 | hCoV-19/Iceland/150/2020 | Europe / Iceland / Reykjavik | 2020-03-13 | The National University Hospital of Iceland | deCODE genetics | Daniel F Gudbjartsson; Agnar Helgason; Hakon Jonsson; Olafur T Magnusson; Pall Melsted; Gudmundur L Norddahl; Jona Saemundsdottir; Asgeir Sigurdsson; Patrick Sulem; Arna B Agustsdottir; Berglind Eiriksdottir; Run Fridriksdottir; Elisabet E Gardarsdottir; Gudmundur Georgsson; Olafia S Gretarsdottir; Kjartan R Gudmundsson; Thora R Gunnarsdottir; Arnaldur Gylfason; Hilma Holm; Brynjar O Jensson; Aslaug Jonasdottir; Kamilla S Josefsdottir; Thordur Kristjansson; Droplaug N Magnusdottir; Louise le Roux; Gudrun Sigmundsdottir; Gardar Sveinbjornsson; Kristin E Sveinsdottir; Maney Sveinsdottir; Emil A Thorarensen; Bjarni Thorbjornsson; Gisli Masson; Ingileif Jonsdottir; Alma Moller; Thorolfur Gudnason; Karl G Kristinsson; Unnur Thorsteinsdottir; Kari Stefansson |
| EPI_ISL_417712 | hCoV-19/Iceland/172/2020 | Europe / Iceland / Reykjavik | 2020-03-15 | The National University Hospital of Iceland | deCODE genetics | Daniel F Gudbjartsson; Agnar Helgason; Hakon Jonsson; Olafur T Magnusson; Pall Melsted; Gudmundur L Norddahl; Jona Saemundsdottir; Asgeir Sigurdsson; Patrick Sulem; Arna B Agustsdottir; Berglind Eiriksdottir; Run Fridriksdottir; Elisabet E Gardarsdottir; Gudmundur Georgsson; Olafia S Gretarsdottir; Kjartan R Gudmundsson; Thora R Gunnarsdottir; Arnaldur Gylfason; Hilma Holm; Brynjar O Jensson; Aslaug Jonasdottir; Kamilla S Josefsdottir; Thordur Kristjansson; Droplaug N Magnusdottir; Louise le Roux; Gudrun Sigmundsdottir; Gardar Sveinbjornsson; Kristin E Sveinsdottir; Maney Sveinsdottir; Emil A Thorarensen; Bjarni Thorbjornsson; Gisli Masson; Ingileif Jonsdottir; Alma Moller; Thorolfur Gudnason; Karl G Kristinsson; Unnur Thorsteinsdottir; Kari Stefansson |
| EPI_ISL_417713 | hCoV-19/Iceland/130/2020 | Europe / Iceland / Reykjavik | 2020-03-12 | The National University Hospital of Iceland | deCODE genetics | Daniel F Gudbjartsson; Agnar Helgason; Hakon Jonsson; Olafur T Magnusson; Pall Melsted; Gudmundur L Norddahl; Jona Saemundsdottir; Asgeir Sigurdsson; Patrick Sulem; Arna B Agustsdottir; Berglind Eiriksdottir; Run Fridriksdottir; Elisabet E Gardarsdottir; Gudmundur Georgsson; Olafia S Gretarsdottir; Kjartan R Gudmundsson; Thora R Gunnarsdottir; Arnaldur Gylfason; Hilma Holm; Brynjar O Jensson; Aslaug Jonasdottir; Kamilla S Josefsdottir; Thordur Kristjansson; Droplaug N Magnusdottir; Louise le Roux; Gudrun Sigmundsdottir; Gardar Sveinbjornsson; Kristin E Sveinsdottir; Maney Sveinsdottir; Emil A Thorarensen; Bjarni Thorbjornsson; Gisli Masson; Ingileif Jonsdottir; Alma Moller; Thorolfur Gudnason; Karl G Kristinsson; Unnur Thorsteinsdottir; Kari Stefansson |
| EPI_ISL_417714 | hCoV-19/Iceland/129/2020 | Europe / Iceland / Reykjavik | 2020-03-12 | The National University Hospital of Iceland | deCODE genetics | Daniel F Gudbjartsson; Agnar Helgason; Hakon Jonsson; Olafur T Magnusson; Pall Melsted; Gudmundur L Norddahl; Jona Saemundsdottir; Asgeir Sigurdsson; Patrick Sulem; Arna B Agustsdottir; Berglind Eiriksdottir; Run Fridriksdottir; Elisabet E Gardarsdottir; Gudmundur Georgsson; Olafia S Gretarsdottir; Kjartan R Gudmundsson; Thora R Gunnarsdottir; Arnaldur Gylfason; Hilma Holm; Brynjar O Jensson; Aslaug Jonasdottir; Kamilla S Josefsdottir; Thordur Kristjansson; Droplaug N Magnusdottir; Louise le Roux; Gudrun Sigmundsdottir; Gardar Sveinbjornsson; Kristin E Sveinsdottir; Maney Sveinsdottir; Emil A Thorarensen; Bjarni Thorbjornsson; Gisli Masson; Ingileif Jonsdottir; Alma Moller; Thorolfur Gudnason; Karl G Kristinsson; Unnur Thorsteinsdottir; Kari Stefansson |
| EPI_ISL_417715 | hCoV-19/Iceland/45/2020 | Europe / Iceland / Reykjavik | 2020-03-04 | The National University Hospital of Iceland | deCODE genetics | Daniel F Gudbjartsson; Agnar Helgason; Hakon Jonsson; Olafur T Magnusson; Pall Melsted; Gudmundur L Norddahl; Jona Saemundsdottir; Asgeir Sigurdsson; Patrick Sulem; Arna B Agustsdottir; Berglind Eiriksdottir; Run Fridriksdottir; Elisabet E Gardarsdottir; Gudmundur Georgsson; Olafia S Gretarsdottir; Kjartan R Gudmundsson; Thora R Gunnarsdottir; Arnaldur Gylfason; Hilma Holm; Brynjar O Jensson; Aslaug Jonasdottir; Kamilla S Josefsdottir; Thordur Kristjansson; Droplaug N Magnusdottir; Louise le Roux; Gudrun Sigmundsdottir; Gardar Sveinbjornsson; Kristin E Sveinsdottir; Maney Sveinsdottir; Emil A Thorarensen; Bjarni Thorbjornsson; Gisli Masson; Ingileif Jonsdottir; Alma Moller; Thorolfur Gudnason; Karl G Kristinsson; Unnur Thorsteinsdottir; Kari Stefansson |
| EPI_ISL_417716 | hCoV-19/Iceland/174/2020 | Europe / Iceland / Reykjavik | 2020-03-16 | The National University Hospital of Iceland | deCODE genetics | Daniel F Gudbjartsson; Agnar Helgason; Hakon Jonsson; Olafur T Magnusson; Pall Melsted; Gudmundur L Norddahl; Jona Saemundsdottir; Asgeir Sigurdsson; Patrick Sulem; Arna B Agustsdottir; Berglind Eiriksdottir; Run Fridriksdottir; Elisabet E Gardarsdottir; Gudmundur Georgsson; Olafia S Gretarsdottir; Kjartan R Gudmundsson; Thora R Gunnarsdottir; Arnaldur Gylfason; Hilma Holm; Brynjar O Jensson; Aslaug Jonasdottir; Kamilla S Josefsdottir; Thordur Kristjansson; Droplaug N Magnusdottir; Louise le Roux; Gudrun Sigmundsdottir; Gardar Sveinbjornsson; Kristin E Sveinsdottir; Maney Sveinsdottir; Emil A Thorarensen; Bjarni Thorbjornsson; Gisli Masson; Ingileif Jonsdottir; Alma Moller; Thorolfur Gudnason; Karl G Kristinsson; Unnur Thorsteinsdottir; Kari Stefansson |
| EPI_ISL_417717 | hCoV-19/Iceland/195/2020 | Europe / Iceland / Reykjavik | 2020-03-16 | The National University Hospital of Iceland | deCODE genetics | Daniel F Gudbjartsson; Agnar Helgason; Hakon Jonsson; Olafur T Magnusson; Pall Melsted; Gudmundur L Norddahl; Jona Saemundsdottir; Asgeir Sigurdsson; Patrick Sulem; Arna B Agustsdottir; Berglind Eiriksdottir; Run Fridriksdottir; Elisabet E Gardarsdottir; Gudmundur Georgsson; Olafia S Gretarsdottir; Kjartan R Gudmundsson; Thora R Gunnarsdottir; Arnaldur Gylfason; Hilma Holm; Brynjar O Jensson; Aslaug Jonasdottir; Kamilla S Josefsdottir; Thordur Kristjansson; Droplaug N Magnusdottir; Louise le Roux; Gudrun Sigmundsdottir; Gardar Sveinbjornsson; Kristin E Sveinsdottir; Maney Sveinsdottir; Emil A Thorarensen; Bjarni Thorbjornsson; Gisli Masson; Ingileif Jonsdottir; Alma Moller; Thorolfur Gudnason; Karl G Kristinsson; Unnur Thorsteinsdottir; Kari Stefansson |
| EPI_ISL_417719 | hCoV-19/Iceland/154/2020 | Europe / Iceland / Reykjavik | 2020-03-13 | The National University Hospital of Iceland | deCODE genetics | Daniel F Gudbjartsson; Agnar Helgason; Hakon Jonsson; Olafur T Magnusson; Pall Melsted; Gudmundur L Norddahl; Jona Saemundsdottir; Asgeir Sigurdsson; Patrick Sulem; Arna B Agustsdottir; Berglind Eiriksdottir; Run Fridriksdottir; Elisabet E Gardarsdottir; Gudmundur Georgsson; Olafia S Gretarsdottir; Kjartan R Gudmundsson; Thora R Gunnarsdottir; Arnaldur Gylfason; Hilma Holm; Brynjar O Jensson; Aslaug Jonasdottir; Kamilla S Josefsdottir; Thordur Kristjansson; Droplaug N Magnusdottir; Louise le Roux; Gudrun Sigmundsdottir; Gardar Sveinbjornsson; Kristin E Sveinsdottir; Maney Sveinsdottir; Emil A Thorarensen; Bjarni Thorbjornsson; Gisli Masson; Ingileif Jonsdottir; Alma Moller; Thorolfur Gudnason; Karl G Kristinsson; Unnur Thorsteinsdottir; Kari Stefansson |
| EPI_ISL_417720 | hCoV-19/Iceland/138/2020 | Europe / Iceland / Reykjavik | 2020-03-12 | The National University Hospital of Iceland | deCODE genetics | Daniel F Gudbjartsson; Agnar Helgason; Hakon Jonsson; Olafur T Magnusson; Pall Melsted; Gudmundur L Norddahl; Jona Saemundsdottir; Asgeir Sigurdsson; Patrick Sulem; Arna B Agustsdottir; Berglind Eiriksdottir; Run Fridriksdottir; Elisabet E Gardarsdottir; Gudmundur Georgsson; Olafia S Gretarsdottir; Kjartan R Gudmundsson; Thora R Gunnarsdottir; Arnaldur Gylfason; Hilma Holm; Brynjar O Jensson; Aslaug Jonasdottir; Kamilla S Josefsdottir; Thordur Kristjansson; Droplaug N Magnusdottir; Louise le Roux; Gudrun Sigmundsdottir; Gardar Sveinbjornsson; Kristin E Sveinsdottir; Maney Sveinsdottir; Emil A Thorarensen; Bjarni Thorbjornsson; Gisli Masson; Ingileif Jonsdottir; Alma Moller; Thorolfur Gudnason; Karl G Kristinsson; Unnur Thorsteinsdottir; Kari Stefansson |
| EPI_ISL_417721 | hCoV-19/Iceland/27/2020 | Europe / Iceland / Reykjavik | 2020-03-03 | The National University Hospital of Iceland | deCODE genetics | Daniel F Gudbjartsson; Agnar Helgason; Hakon Jonsson; Olafur T Magnusson; Pall Melsted; Gudmundur L Norddahl; Jona Saemundsdottir; Asgeir Sigurdsson; Patrick Sulem; Arna B Agustsdottir; Berglind Eiriksdottir; Run Fridriksdottir; Elisabet E Gardarsdottir; Gudmundur Georgsson; Olafia S Gretarsdottir; Kjartan R Gudmundsson; Thora R Gunnarsdottir; Arnaldur Gylfason; Hilma Holm; Brynjar O Jensson; Aslaug Jonasdottir; Kamilla S Josefsdottir; Thordur Kristjansson; Droplaug N Magnusdottir; Louise le Roux; Gudrun Sigmundsdottir; Gardar Sveinbjornsson; Kristin E Sveinsdottir; Maney Sveinsdottir; Emil A Thorarensen; Bjarni Thorbjornsson; Gisli Masson; Ingileif Jonsdottir; Alma Moller; Thorolfur Gudnason; Karl G Kristinsson; Unnur Thorsteinsdottir; Kari Stefansson |
| EPI_ISL_417722 | hCoV-19/Iceland/110/2020 | Europe / Iceland / Reykjavik | 2020-03-11 | The National University Hospital of Iceland | deCODE genetics | Daniel F Gudbjartsson; Agnar Helgason; Hakon Jonsson; Olafur T Magnusson; Pall Melsted; Gudmundur L Norddahl; Jona Saemundsdottir; Asgeir Sigurdsson; Patrick Sulem; Arna B Agustsdottir; Berglind Eiriksdottir; Run Fridriksdottir; Elisabet E Gardarsdottir; Gudmundur Georgsson; Olafia S Gretarsdottir; Kjartan R Gudmundsson; Thora R Gunnarsdottir; Arnaldur Gylfason; Hilma Holm; Brynjar O Jensson; Aslaug Jonasdottir; Kamilla S Josefsdottir; Thordur Kristjansson; Droplaug N Magnusdottir; Louise le Roux; Gudrun Sigmundsdottir; Gardar Sveinbjornsson; Kristin E Sveinsdottir; Maney Sveinsdottir; Emil A Thorarensen; Bjarni Thorbjornsson; Gisli Masson; Ingileif Jonsdottir; Alma Moller; Thorolfur Gudnason; Karl G Kristinsson; Unnur Thorsteinsdottir; Kari Stefansson |
| EPI_ISL_417724 | hCoV-19/Iceland/182/2020 | Europe / Iceland / Reykjavik | 2020-03-16 | The National University Hospital of Iceland | deCODE genetics | Daniel F Gudbjartsson; Agnar Helgason; Hakon Jonsson; Olafur T Magnusson; Pall Melsted; Gudmundur L Norddahl; Jona Saemundsdottir; Asgeir Sigurdsson; Patrick Sulem; Arna B Agustsdottir; Berglind Eiriksdottir; Run Fridriksdottir; Elisabet E Gardarsdottir; Gudmundur Georgsson; Olafia S Gretarsdottir; Kjartan R Gudmundsson; Thora R Gunnarsdottir; Arnaldur Gylfason; Hilma Holm; Brynjar O Jensson; Aslaug Jonasdottir; Kamilla S Josefsdottir; Thordur Kristjansson; Droplaug N Magnusdottir; Louise le Roux; Gudrun Sigmundsdottir; Gardar Sveinbjornsson; Kristin E Sveinsdottir; Maney Sveinsdottir; Emil A Thorarensen; Bjarni Thorbjornsson; Gisli Masson; Ingileif Jonsdottir; Alma Moller; Thorolfur Gudnason; Karl G Kristinsson; Unnur Thorsteinsdottir; Kari Stefansson |

| | | | | | | |
|---|---|---|---|---|---|---|
| EPI_ISL_417725 | hCoV-19/Iceland/161/2020 | Europe / Iceland / Reykjavik | 2020-03-14 | The National University Hospital of Iceland | deCODE genetics | Daniel F Gudbjartsson; Agnar Helgason; Hakon Jonsson; Olafur T Magnusson; Pall Melsted; Gudmundur L Norddahl; Jona Saemundsdottir; Asgeir Sigurdsson; Patrick Sulem; Arna B Agustsdottir; Berglind Eiriksdottir; Run Fridriksdottir; Elisabet E Gardarsdottir; Gudmundur Georgsson; Olafia S Gretarsdottir; Kjartan R Gudmundsson; Thora R Gunnarsdottir; Arnaldur Gylfason; Hilma Holm; Brynjar O Jensson; Aslaug Jonasdottir; Kamilla S Josefsdottir; Thordur Kristjansson; Droplaug N Magnusdottir; Louise le Roux; Gudrun Sigmundsdottir; Gardar Sveinbjornsson; Kristin E Sveinsdottir; Maney Sveinsdottir; Emil A Thorarensen; Bjarni Thorbjornsson; Gisli Masson; Ingileif Jonsdottir; Alma Moller; Thorolfur Gudnason; Karl G Kristinsson; Unnur Thorsteinsdottir; Kari Stefansson |
| EPI_ISL_417727 | hCoV-19/Iceland/127/2020 | Europe / Iceland / Reykjavik | 2020-03-12 | The National University Hospital of Iceland | deCODE genetics | Daniel F Gudbjartsson; Agnar Helgason; Hakon Jonsson; Olafur T Magnusson; Pall Melsted; Gudmundur L Norddahl; Jona Saemundsdottir; Asgeir Sigurdsson; Patrick Sulem; Arna B Agustsdottir; Berglind Eiriksdottir; Run Fridriksdottir; Elisabet E Gardarsdottir; Gudmundur Georgsson; Olafia S Gretarsdottir; Kjartan R Gudmundsson; Thora R Gunnarsdottir; Arnaldur Gylfason; Hilma Holm; Brynjar O Jensson; Aslaug Jonasdottir; Kamilla S Josefsdottir; Thordur Kristjansson; Droplaug N Magnusdottir; Louise le Roux; Gudrun Sigmundsdottir; Gardar Sveinbjornsson; Kristin E Sveinsdottir; Maney Sveinsdottir; Emil A Thorarensen; Bjarni Thorbjornsson; Gisli Masson; Ingileif Jonsdottir; Alma Moller; Thorolfur Gudnason; Karl G Kristinsson; Unnur Thorsteinsdottir; Kari Stefansson |
| EPI_ISL_417728 | hCoV-19/Iceland/70/2020 | Europe / Iceland / Reykjavik | 2020-03-08 | The National University Hospital of Iceland | deCODE genetics | Daniel F Gudbjartsson; Agnar Helgason; Hakon Jonsson; Olafur T Magnusson; Pall Melsted; Gudmundur L Norddahl; Jona Saemundsdottir; Asgeir Sigurdsson; Patrick Sulem; Arna B Agustsdottir; Berglind Eiriksdottir; Run Fridriksdottir; Elisabet E Gardarsdottir; Gudmundur Georgsson; Olafia S Gretarsdottir; Kjartan R Gudmundsson; Thora R Gunnarsdottir; Arnaldur Gylfason; Hilma Holm; Brynjar O Jensson; Aslaug Jonasdottir; Kamilla S Josefsdottir; Thordur Kristjansson; Droplaug N Magnusdottir; Louise le Roux; Gudrun Sigmundsdottir; Gardar Sveinbjornsson; Kristin E Sveinsdottir; Maney Sveinsdottir; Emil A Thorarensen; Bjarni Thorbjornsson; Gisli Masson; Ingileif Jonsdottir; Alma Moller; Thorolfur Gudnason; Karl G Kristinsson; Unnur Thorsteinsdottir; Kari Stefansson |
| EPI_ISL_417730 | hCoV-19/Iceland/93/2020 | Europe / Iceland / Reykjavik | 2020-03-10 | The National University Hospital of Iceland | deCODE genetics | Daniel F Gudbjartsson; Agnar Helgason; Hakon Jonsson; Olafur T Magnusson; Pall Melsted; Gudmundur L Norddahl; Jona Saemundsdottir; Asgeir Sigurdsson; Patrick Sulem; Arna B Agustsdottir; Berglind Eiriksdottir; Run Fridriksdottir; Elisabet E Gardarsdottir; Gudmundur Georgsson; Olafia S Gretarsdottir; Kjartan R Gudmundsson; Thora R Gunnarsdottir; Arnaldur Gylfason; Hilma Holm; Brynjar O Jensson; Aslaug Jonasdottir; Kamilla S Josefsdottir; Thordur Kristjansson; Droplaug N Magnusdottir; Louise le Roux; Gudrun Sigmundsdottir; Gardar Sveinbjornsson; Kristin E Sveinsdottir; Maney Sveinsdottir; Emil A Thorarensen; Bjarni Thorbjornsson; Gisli Masson; Ingileif Jonsdottir; Alma Moller; Thorolfur Gudnason; Karl G Kristinsson; Unnur Thorsteinsdottir; Kari Stefansson |
| EPI_ISL_417731 | hCoV-19/Iceland/61/2020 | Europe / Iceland / Reykjavik | 2020-03-06 | The National University Hospital of Iceland | deCODE genetics | Daniel F Gudbjartsson; Agnar Helgason; Hakon Jonsson; Olafur T Magnusson; Pall Melsted; Gudmundur L Norddahl; Jona Saemundsdottir; Asgeir Sigurdsson; Patrick Sulem; Arna B Agustsdottir; Berglind Eiriksdottir; Run Fridriksdottir; Elisabet E Gardarsdottir; Gudmundur Georgsson; Olafia S Gretarsdottir; Kjartan R Gudmundsson; Thora R Gunnarsdottir; Arnaldur Gylfason; Hilma Holm; Brynjar O Jensson; Aslaug Jonasdottir; Kamilla S Josefsdottir; Thordur Kristjansson; Droplaug N Magnusdottir; Louise le Roux; Gudrun Sigmundsdottir; Gardar Sveinbjornsson; Kristin E Sveinsdottir; Maney Sveinsdottir; Emil A Thorarensen; Bjarni Thorbjornsson; Gisli Masson; Ingileif Jonsdottir; Alma Moller; Thorolfur Gudnason; Karl G Kristinsson; Unnur Thorsteinsdottir; Kari Stefansson |
| EPI_ISL_417732 | hCoV-19/Iceland/94/2020 | Europe / Iceland / Reykjavik | 2020-03-10 | The National University Hospital of Iceland | deCODE genetics | Daniel F Gudbjartsson; Agnar Helgason; Hakon Jonsson; Olafur T Magnusson; Pall Melsted; Gudmundur L Norddahl; Jona Saemundsdottir; Asgeir Sigurdsson; Patrick Sulem; Arna B Agustsdottir; Berglind Eiriksdottir; Run Fridriksdottir; Elisabet E Gardarsdottir; Gudmundur Georgsson; Olafia S Gretarsdottir; Kjartan R Gudmundsson; Thora R Gunnarsdottir; Arnaldur Gylfason; Hilma Holm; Brynjar O Jensson; Aslaug Jonasdottir; Kamilla S Josefsdottir; Thordur Kristjansson; Droplaug N Magnusdottir; Louise le Roux; Gudrun Sigmundsdottir; Gardar Sveinbjornsson; Kristin E Sveinsdottir; Maney Sveinsdottir; Emil A Thorarensen; Bjarni Thorbjornsson; Gisli Masson; Ingileif Jonsdottir; Alma Moller; Thorolfur Gudnason; Karl G Kristinsson; Unnur Thorsteinsdottir; Kari Stefansson |
| EPI_ISL_417733 | hCoV-19/Iceland/207/2020 | Europe / Iceland / Reykjavik | 2020-03-16 | The National University Hospital of Iceland | deCODE genetics | Daniel F Gudbjartsson; Agnar Helgason; Hakon Jonsson; Olafur T Magnusson; Pall Melsted; Gudmundur L Norddahl; Jona Saemundsdottir; Asgeir Sigurdsson; Patrick Sulem; Arna B Agustsdottir; Berglind Eiriksdottir; Run Fridriksdottir; Elisabet E Gardarsdottir; Gudmundur Georgsson; Olafia S Gretarsdottir; Kjartan R Gudmundsson; Thora R Gunnarsdottir; Arnaldur Gylfason; Hilma Holm; Brynjar O Jensson; Aslaug Jonasdottir; Kamilla S Josefsdottir; Thordur Kristjansson; Droplaug N Magnusdottir; Louise le Roux; Gudrun Sigmundsdottir; Gardar Sveinbjornsson; Kristin E Sveinsdottir; Maney Sveinsdottir; Emil A Thorarensen; Bjarni Thorbjornsson; Gisli Masson; Ingileif Jonsdottir; Alma Moller; Thorolfur Gudnason; Karl G Kristinsson; Unnur Thorsteinsdottir; Kari Stefansson |
| EPI_ISL_417734 | hCoV-19/Iceland/22/2020 | Europe / Iceland / Reykjavik | 2020-03-02 | The National University Hospital of Iceland | deCODE genetics | Daniel F Gudbjartsson; Agnar Helgason; Hakon Jonsson; Olafur T Magnusson; Pall Melsted; Gudmundur L Norddahl; Jona Saemundsdottir; Asgeir Sigurdsson; Patrick Sulem; Arna B Agustsdottir; Berglind Eiriksdottir; Run Fridriksdottir; Elisabet E Gardarsdottir; Gudmundur Georgsson; Olafia S Gretarsdottir; Kjartan R Gudmundsson; Thora R Gunnarsdottir; Arnaldur Gylfason; Hilma Holm; Brynjar O Jensson; Aslaug Jonasdottir; Kamilla S Josefsdottir; Thordur Kristjansson; Droplaug N Magnusdottir; Louise le Roux; Gudrun Sigmundsdottir; Gardar Sveinbjornsson; Kristin E Sveinsdottir; Maney Sveinsdottir; Emil A Thorarensen; Bjarni Thorbjornsson; Gisli Masson; Ingileif Jonsdottir; Alma Moller; Thorolfur Gudnason; Karl G Kristinsson; Unnur Thorsteinsdottir; Kari Stefansson |
| EPI_ISL_417735 | hCoV-19/Iceland/134/2020 | Europe / Iceland / Reykjavik | 2020-03-12 | The National University Hospital of Iceland | deCODE genetics | Daniel F Gudbjartsson; Agnar Helgason; Hakon Jonsson; Olafur T Magnusson; Pall Melsted; Gudmundur L Norddahl; Jona Saemundsdottir; Asgeir Sigurdsson; Patrick Sulem; Arna B Agustsdottir; Berglind Eiriksdottir; Run Fridriksdottir; Elisabet E Gardarsdottir; Gudmundur Georgsson; Olafia S Gretarsdottir; Kjartan R Gudmundsson; Thora R Gunnarsdottir; Arnaldur Gylfason; Hilma Holm; Brynjar O Jensson; Aslaug Jonasdottir; Kamilla S Josefsdottir; Thordur Kristjansson; Droplaug N Magnusdottir; Louise le Roux; Gudrun Sigmundsdottir; Gardar Sveinbjornsson; Kristin E Sveinsdottir; Maney Sveinsdottir; Emil A Thorarensen; Bjarni Thorbjornsson; Gisli Masson; Ingileif Jonsdottir; Alma Moller; Thorolfur Gudnason; Karl G Kristinsson; Unnur Thorsteinsdottir; Kari Stefansson |
| EPI_ISL_417736 | hCoV-19/Iceland/14/2020 | Europe / Iceland / Reykjavik | 2020-03-01 | The National University Hospital of Iceland | deCODE genetics | Daniel F Gudbjartsson; Agnar Helgason; Hakon Jonsson; Olafur T Magnusson; Pall Melsted; Gudmundur L Norddahl; Jona Saemundsdottir; Asgeir Sigurdsson; Patrick Sulem; Arna B Agustsdottir; Berglind Eiriksdottir; Run Fridriksdottir; Elisabet E Gardarsdottir; Gudmundur Georgsson; Olafia S Gretarsdottir; Kjartan R Gudmundsson; Thora R Gunnarsdottir; Arnaldur Gylfason; Hilma Holm; Brynjar O Jensson; Aslaug Jonasdottir; Kamilla S Josefsdottir; Thordur Kristjansson; Droplaug N Magnusdottir; Louise le Roux; Gudrun Sigmundsdottir; Gardar Sveinbjornsson; Kristin E Sveinsdottir; Maney Sveinsdottir; Emil A Thorarensen; Bjarni Thorbjornsson; Gisli Masson; Ingileif Jonsdottir; Alma Moller; Thorolfur Gudnason; Karl G Kristinsson; Unnur Thorsteinsdottir; Kari Stefansson |
| EPI_ISL_417737 | hCoV-19/Iceland/155/2020 | Europe / Iceland / Reykjavik | 2020-03-15 | The National University Hospital of Iceland | deCODE genetics | Daniel F Gudbjartsson; Agnar Helgason; Hakon Jonsson; Olafur T Magnusson; Pall Melsted; Gudmundur L Norddahl; Jona Saemundsdottir; Asgeir Sigurdsson; Patrick Sulem; Arna B Agustsdottir; Berglind Eiriksdottir; Run Fridriksdottir; Elisabet E Gardarsdottir; Gudmundur Georgsson; Olafia S Gretarsdottir; Kjartan R Gudmundsson; Thora R Gunnarsdottir; Arnaldur Gylfason; Hilma Holm; Brynjar O Jensson; Aslaug Jonasdottir; Kamilla S Josefsdottir; Thordur Kristjansson; Droplaug N Magnusdottir; Louise le Roux; Gudrun Sigmundsdottir; Gardar Sveinbjornsson; Kristin E Sveinsdottir; Maney Sveinsdottir; Emil A Thorarensen; Bjarni Thorbjornsson; Gisli Masson; Ingileif Jonsdottir; Alma Moller; Thorolfur Gudnason; Karl G Kristinsson; Unnur Thorsteinsdottir; Kari Stefansson |
| EPI_ISL_417738 | hCoV-19/Iceland/89/2020 | Europe / Iceland / Reykjavik | 2020-03-10 | The National University Hospital of Iceland | deCODE genetics | Daniel F Gudbjartsson; Agnar Helgason; Hakon Jonsson; Olafur T Magnusson; Pall Melsted; Gudmundur L Norddahl; Jona Saemundsdottir; Asgeir Sigurdsson; Patrick Sulem; Arna B Agustsdottir; Berglind Eiriksdottir; Run Fridriksdottir; Elisabet E Gardarsdottir; Gudmundur Georgsson; Olafia S Gretarsdottir; Kjartan R Gudmundsson; Thora R Gunnarsdottir; Arnaldur Gylfason; Hilma Holm; Brynjar O Jensson; Aslaug Jonasdottir; Kamilla S Josefsdottir; Thordur Kristjansson; Droplaug N Magnusdottir; Louise le Roux; Gudrun Sigmundsdottir; Gardar Sveinbjornsson; Kristin E Sveinsdottir; Maney Sveinsdottir; Emil A Thorarensen; Bjarni Thorbjornsson; Gisli Masson; Ingileif Jonsdottir; Alma Moller; Thorolfur Gudnason; Karl G Kristinsson; Unnur Thorsteinsdottir; Kari Stefansson |
| EPI_ISL_417739 | hCoV-19/Iceland/5/2020 | Europe / Iceland / Reykjavik | 2020-03-13 | The National University Hospital of Iceland | deCODE genetics | Daniel F Gudbjartsson; Agnar Helgason; Hakon Jonsson; Olafur T Magnusson; Pall Melsted; Gudmundur L Norddahl; Jona Saemundsdottir; Asgeir Sigurdsson; Patrick Sulem; Arna B Agustsdottir; Berglind Eiriksdottir; Run Fridriksdottir; Elisabet E Gardarsdottir; Gudmundur Georgsson; Olafia S Gretarsdottir; Kjartan R Gudmundsson; Thora R Gunnarsdottir; Arnaldur Gylfason; Hilma Holm; Brynjar O Jensson; Aslaug Jonasdottir; Kamilla S Josefsdottir; Thordur Kristjansson; Droplaug N Magnusdottir; Louise le Roux; Gudrun Sigmundsdottir; Gardar Sveinbjornsson; Kristin E Sveinsdottir; Maney Sveinsdottir; Emil A Thorarensen; Bjarni Thorbjornsson; Gisli Masson; Ingileif Jonsdottir; Alma Moller; Thorolfur Gudnason; Karl G Kristinsson; Unnur Thorsteinsdottir; Kari Stefansson |
| EPI_ISL_417740 | hCoV-19/Iceland/7/2020 | Europe / Iceland / Reykjavik | 2020-03-15 | The National University Hospital of Iceland | deCODE genetics | Daniel F Gudbjartsson; Agnar Helgason; Hakon Jonsson; Olafur T Magnusson; Pall Melsted; Gudmundur L Norddahl; Jona Saemundsdottir; Asgeir Sigurdsson; Patrick Sulem; Arna B Agustsdottir; Berglind Eiriksdottir; Run Fridriksdottir; Elisabet E Gardarsdottir; Gudmundur Georgsson; Olafia S Gretarsdottir; Kjartan R Gudmundsson; Thora R Gunnarsdottir; Arnaldur Gylfason; Hilma Holm; Brynjar O Jensson; Aslaug Jonasdottir; Kamilla S Josefsdottir; Thordur Kristjansson; Droplaug N Magnusdottir; Louise le Roux; Gudrun Sigmundsdottir; Gardar Sveinbjornsson; Kristin E Sveinsdottir; Maney Sveinsdottir; Emil A Thorarensen; Bjarni Thorbjornsson; Gisli Masson; Ingileif Jonsdottir; Alma Moller; Thorolfur Gudnason; Karl G Kristinsson; Unnur Thorsteinsdottir; Kari Stefansson |
| EPI_ISL_417741 | hCoV-19/Iceland/145/2020 | Europe / Iceland / Reykjavik | 2020-03-13 | The National University Hospital of Iceland | deCODE genetics | Daniel F Gudbjartsson; Agnar Helgason; Hakon Jonsson; Olafur T Magnusson; Pall Melsted; Gudmundur L Norddahl; Jona Saemundsdottir; Asgeir Sigurdsson; Patrick Sulem; Arna B Agustsdottir; Berglind Eiriksdottir; Run Fridriksdottir; Elisabet E Gardarsdottir; Gudmundur Georgsson; Olafia S Gretarsdottir; Kjartan R Gudmundsson; Thora R Gunnarsdottir; Arnaldur Gylfason; Hilma Holm; Brynjar O Jensson; Aslaug Jonasdottir; Kamilla S Josefsdottir; Thordur Kristjansson; Droplaug N Magnusdottir; Louise le Roux; Gudrun Sigmundsdottir; Gardar Sveinbjornsson; Kristin E Sveinsdottir; Maney Sveinsdottir; Emil A Thorarensen; Bjarni Thorbjornsson; Gisli Masson; Ingileif Jonsdottir; Alma Moller; Thorolfur Gudnason; Karl G Kristinsson; Unnur Thorsteinsdottir; Kari Stefansson |
| EPI_ISL_417742 | hCoV-19/Iceland/197/2020 | Europe / Iceland / Reykjavik | 2020-03-16 | The National University Hospital of Iceland | deCODE genetics | Daniel F Gudbjartsson; Agnar Helgason; Hakon Jonsson; Olafur T Magnusson; Pall Melsted; Gudmundur L Norddahl; Jona Saemundsdottir; Asgeir Sigurdsson; Patrick Sulem; Arna B Agustsdottir; Berglind Eiriksdottir; Run Fridriksdottir; Elisabet E Gardarsdottir; Gudmundur Georgsson; Olafia S Gretarsdottir; Kjartan R Gudmundsson; Thora R Gunnarsdottir; Arnaldur Gylfason; Hilma Holm; Brynjar O Jensson; Aslaug Jonasdottir; Kamilla S Josefsdottir; Thordur Kristjansson; Droplaug N Magnusdottir; Louise le Roux; Gudrun Sigmundsdottir; Gardar Sveinbjornsson; Kristin E Sveinsdottir; Maney Sveinsdottir; Emil A Thorarensen; Bjarni Thorbjornsson; Gisli Masson; Ingileif Jonsdottir; Alma Moller; Thorolfur Gudnason; Karl G Kristinsson; Unnur Thorsteinsdottir; Kari Stefansson |
| EPI_ISL_417743 | hCoV-19/Iceland/200/2020 | Europe / Iceland / Reykjavik | 2020-03-16 | The National University Hospital of Iceland | deCODE genetics | Daniel F Gudbjartsson; Agnar Helgason; Hakon Jonsson; Olafur T Magnusson; Pall Melsted; Gudmundur L Norddahl; Jona Saemundsdottir; Asgeir Sigurdsson; Patrick Sulem; Arna B Agustsdottir; Berglind Eiriksdottir; Run Fridriksdottir; Elisabet E Gardarsdottir; Gudmundur Georgsson; Olafia S Gretarsdottir; Kjartan R Gudmundsson; Thora R Gunnarsdottir; Arnaldur Gylfason; Hilma Holm; Brynjar O Jensson; Aslaug Jonasdottir; Kamilla S Josefsdottir; Thordur Kristjansson; Droplaug N Magnusdottir; Louise le Roux; Gudrun Sigmundsdottir; Gardar Sveinbjornsson; Kristin E Sveinsdottir; Maney Sveinsdottir; Emil A Thorarensen; Bjarni Thorbjornsson; Gisli Masson; Ingileif Jonsdottir; Alma Moller; Thorolfur Gudnason; Karl G Kristinsson; Unnur Thorsteinsdottir; Kari Stefansson |
| EPI_ISL_417744 | hCoV-19/Iceland/137/2020 | Europe / Iceland / Reykjavik | 2020-03-13 | The National University Hospital of Iceland | deCODE genetics | Daniel F Gudbjartsson; Agnar Helgason; Hakon Jonsson; Olafur T Magnusson; Pall Melsted; Gudmundur L Norddahl; Jona Saemundsdottir; Asgeir Sigurdsson; Patrick Sulem; Arna B Agustsdottir; Berglind Eiriksdottir; Run Fridriksdottir; Elisabet E Gardarsdottir; Gudmundur Georgsson; Olafia S Gretarsdottir; Kjartan R Gudmundsson; Thora R Gunnarsdottir; Arnaldur Gylfason; Hilma Holm; Brynjar O Jensson; Aslaug Jonasdottir; Kamilla S Josefsdottir; Thordur Kristjansson; Droplaug N Magnusdottir; Louise le Roux; Gudrun Sigmundsdottir; Gardar Sveinbjornsson; Kristin E Sveinsdottir; Maney Sveinsdottir; Emil A Thorarensen; Bjarni Thorbjornsson; Gisli Masson; Ingileif Jonsdottir; Alma Moller; Thorolfur Gudnason; Karl G Kristinsson; Unnur Thorsteinsdottir; Kari Stefansson |
| EPI_ISL_417745 | hCoV-19/Iceland/152/2020 | Europe / Iceland / Reykjavik | 2020-03-13 | The National University Hospital of Iceland | deCODE genetics | Daniel F Gudbjartsson; Agnar Helgason; Hakon Jonsson; Olafur T Magnusson; Pall Melsted; Gudmundur L Norddahl; Jona Saemundsdottir; Asgeir Sigurdsson; Patrick Sulem; Arna B Agustsdottir; Berglind Eiriksdottir; Run Fridriksdottir; Elisabet E Gardarsdottir; Gudmundur Georgsson; Olafia S Gretarsdottir; Kjartan R Gudmundsson; Thora R Gunnarsdottir; Arnaldur Gylfason; Hilma Holm; Brynjar O Jensson; Aslaug Jonasdottir; Kamilla S Josefsdottir; Thordur Kristjansson; Droplaug N Magnusdottir; Louise le Roux; Gudrun Sigmundsdottir; Gardar Sveinbjornsson; Kristin E Sveinsdottir; Maney Sveinsdottir; Emil A Thorarensen; Bjarni Thorbjornsson; Gisli Masson; Ingileif Jonsdottir; Alma Moller; Thorolfur Gudnason; Karl G Kristinsson; Unnur Thorsteinsdottir; Kari Stefansson |
| EPI_ISL_417746 | hCoV-19/Iceland/183/2020 | Europe / Iceland / Reykjavik | 2020-03-16 | The National University Hospital of Iceland | deCODE genetics | Daniel F Gudbjartsson; Agnar Helgason; Hakon Jonsson; Olafur T Magnusson; Pall Melsted; Gudmundur L Norddahl; Jona Saemundsdottir; Asgeir Sigurdsson; Patrick Sulem; Arna B Agustsdottir; Berglind Eiriksdottir; Run Fridriksdottir; Elisabet E Gardarsdottir; Gudmundur Georgsson; Olafia S Gretarsdottir; Kjartan R Gudmundsson; Thora R Gunnarsdottir; Arnaldur Gylfason; Hilma Holm; Brynjar O Jensson; Aslaug Jonasdottir; Kamilla S Josefsdottir; Thordur Kristjansson; Droplaug N Magnusdottir; Louise le Roux; Gudrun Sigmundsdottir; Gardar Sveinbjornsson; Kristin E Sveinsdottir; Maney Sveinsdottir; Emil A Thorarensen; Bjarni Thorbjornsson; Gisli Masson; Ingileif Jonsdottir; Alma Moller; Thorolfur Gudnason; Karl G Kristinsson; Unnur Thorsteinsdottir; Kari Stefansson |
| EPI_ISL_417747 | hCoV-19/Iceland/24/2020 | Europe / Iceland / Reykjavik | 2020-03-11 | The National University Hospital of Iceland | deCODE genetics | Daniel F Gudbjartsson; Agnar Helgason; Hakon Jonsson; Olafur T Magnusson; Pall Melsted; Gudmundur L Norddahl; Jona Saemundsdottir; Asgeir Sigurdsson; Patrick Sulem; Arna B Agustsdottir; Berglind Eiriksdottir; Run Fridriksdottir; Elisabet E Gardarsdottir; Gudmundur Georgsson; Olafia S Gretarsdottir; Kjartan R Gudmundsson; Thora R Gunnarsdottir; Arnaldur Gylfason; Hilma Holm; Brynjar O Jensson; Aslaug Jonasdottir; Kamilla S Josefsdottir; Thordur Kristjansson; Droplaug N Magnusdottir; Louise le Roux; Gudrun Sigmundsdottir; Gardar Sveinbjornsson; Kristin E Sveinsdottir; Maney Sveinsdottir; Emil A Thorarensen; Bjarni Thorbjornsson; Gisli Masson; Ingileif Jonsdottir; Alma Moller; Thorolfur Gudnason; Karl G Kristinsson; Unnur Thorsteinsdottir; Kari Stefansson |
| EPI_ISL_417748 | hCoV-19/Iceland/109/2020 | Europe / Iceland / Reykjavik | 2020-03-11 | The National University Hospital of Iceland | deCODE genetics | Daniel F Gudbjartsson; Agnar Helgason; Hakon Jonsson; Olafur T Magnusson; Pall Melsted; Gudmundur L Norddahl; Jona Saemundsdottir; Asgeir Sigurdsson; Patrick Sulem; Arna B Agustsdottir; Berglind Eiriksdottir; Run Fridriksdottir; Elisabet E Gardarsdottir; Gudmundur Georgsson; Olafia S Gretarsdottir; Kjartan R Gudmundsson; Thora R Gunnarsdottir; Arnaldur Gylfason; Hilma Holm; Brynjar O Jensson; Aslaug Jonasdottir; Kamilla S Josefsdottir; Thordur Kristjansson; Droplaug N Magnusdottir; Louise le Roux; Gudrun Sigmundsdottir; Gardar Sveinbjornsson; Kristin E Sveinsdottir; Maney Sveinsdottir; Emil A Thorarensen; Bjarni Thorbjornsson; Gisli Masson; Ingileif Jonsdottir; Alma Moller; Thorolfur Gudnason; Karl G Kristinsson; Unnur Thorsteinsdottir; Kari Stefansson |
| EPI_ISL_417749 | hCoV-19/Iceland/69/2020 | Europe / Iceland / Reykjavik | 2020-03-08 | The National University Hospital of Iceland | deCODE genetics | Daniel F Gudbjartsson; Agnar Helgason; Hakon Jonsson; Olafur T Magnusson; Pall Melsted; Gudmundur L Norddahl; Jona Saemundsdottir; Asgeir Sigurdsson; Patrick Sulem; Arna B Agustsdottir; Berglind Eiriksdottir; Run Fridriksdottir; Elisabet E Gardarsdottir; Gudmundur Georgsson; Olafia S Gretarsdottir; Kjartan R Gudmundsson; Thora R Gunnarsdottir; Arnaldur Gylfason; Hilma Holm; Brynjar O Jensson; Aslaug Jonasdottir; Kamilla S Josefsdottir; Thordur Kristjansson; Droplaug N Magnusdottir; Louise le Roux; Gudrun Sigmundsdottir; Gardar Sveinbjornsson; Kristin E Sveinsdottir; Maney Sveinsdottir; Emil A Thorarensen; Bjarni Thorbjornsson; Gisli Masson; Ingileif Jonsdottir; Alma Moller; Thorolfur Gudnason; Karl G Kristinsson; Unnur Thorsteinsdottir; Kari Stefansson |
| EPI_ISL_417750 | hCoV-19/Iceland/202/2020 | Europe / Iceland / Reykjavik | 2020-03-16 | The National University Hospital of Iceland | deCODE genetics | Daniel F Gudbjartsson; Agnar Helgason; Hakon Jonsson; Olafur T Magnusson; Pall Melsted; Gudmundur L Norddahl; Jona Saemundsdottir; Asgeir Sigurdsson; Patrick Sulem; Arna B Agustsdottir; Berglind Eiriksdottir; Run Fridriksdottir; Elisabet E Gardarsdottir; Gudmundur Georgsson; Olafia S Gretarsdottir; Kjartan R Gudmundsson; Thora R Gunnarsdottir; Arnaldur Gylfason; Hilma Holm; Brynjar O Jensson; Aslaug Jonasdottir; Kamilla S Josefsdottir; Thordur Kristjansson; Droplaug N Magnusdottir; Louise le Roux; Gudrun Sigmundsdottir; Gardar Sveinbjornsson; Kristin E Sveinsdottir; Maney Sveinsdottir; Emil A Thorarensen; Bjarni Thorbjornsson; Gisli Masson; Ingileif Jonsdottir; Alma Moller; Thorolfur Gudnason; Karl G Kristinsson; Unnur Thorsteinsdottir; Kari Stefansson |
| EPI_ISL_417752 | hCoV-19/Iceland/213/2020 | Europe / Iceland / Reykjavik | 2020-03-16 | The National University Hospital of Iceland | deCODE genetics | Daniel F Gudbjartsson; Agnar Helgason; Hakon Jonsson; Olafur T Magnusson; Pall Melsted; Gudmundur L Norddahl; Jona Saemundsdottir; Asgeir Sigurdsson; Patrick Sulem; Arna B Agustsdottir; Berglind Eiriksdottir; Run Fridriksdottir; Elisabet E Gardarsdottir; Gudmundur Georgsson; Olafia S Gretarsdottir; Kjartan R Gudmundsson; Thora R Gunnarsdottir; Arnaldur Gylfason; Hilma Holm; Brynjar O Jensson; Aslaug Jonasdottir; Kamilla S Josefsdottir; Thordur Kristjansson; Droplaug N Magnusdottir; Louise le Roux; Gudrun Sigmundsdottir; Gardar Sveinbjornsson; Kristin E Sveinsdottir; Maney Sveinsdottir; Emil A Thorarensen; Bjarni Thorbjornsson; Gisli Masson; Ingileif Jonsdottir; Alma Moller; Thorolfur Gudnason; Karl G Kristinsson; Unnur Thorsteinsdottir; Kari Stefansson |
| EPI_ISL_417753 | hCoV-19/Iceland/180/2020 | Europe / Iceland / Reykjavik | 2020-03-16 | The National University Hospital of Iceland | deCODE genetics | Daniel F Gudbjartsson; Agnar Helgason; Hakon Jonsson; Olafur T Magnusson; Pall Melsted; Gudmundur L Norddahl; Jona Saemundsdottir; Asgeir Sigurdsson; Patrick Sulem; Arna B Agustsdottir; Berglind Eiriksdottir; Run Fridriksdottir; Elisabet E Gardarsdottir; Gudmundur Georgsson; Olafia S Gretarsdottir; Kjartan R Gudmundsson; Thora R Gunnarsdottir; Arnaldur Gylfason; Hilma Holm; Brynjar O Jensson; Aslaug Jonasdottir; Kamilla S Josefsdottir; Thordur Kristjansson; Droplaug N Magnusdottir; Louise le Roux; Gudrun Sigmundsdottir; Gardar Sveinbjornsson; Kristin E Sveinsdottir; Maney Sveinsdottir; Emil A Thorarensen; Bjarni Thorbjornsson; Gisli Masson; Ingileif Jonsdottir; Alma Moller; Thorolfur Gudnason; Karl G Kristinsson; Unnur Thorsteinsdottir; Kari Stefansson |
| EPI_ISL_417754 | hCoV-19/Iceland/226/2020 | Europe / Iceland / Reykjavik | 2020-03-16 | The National University Hospital of Iceland | deCODE genetics | Daniel F Gudbjartsson; Agnar Helgason; Hakon Jonsson; Olafur T Magnusson; Pall Melsted; Gudmundur L Norddahl; Jona Saemundsdottir; Asgeir Sigurdsson; Patrick Sulem; Arna B Agustsdottir; Berglind Eiriksdottir; Run Fridriksdottir; Elisabet E Gardarsdottir; Gudmundur Georgsson; Olafia S Gretarsdottir; Kjartan R Gudmundsson; Thora R Gunnarsdottir; Arnaldur Gylfason; Hilma Holm; Brynjar O Jensson; Aslaug Jonasdottir; Kamilla S Josefsdottir; Thordur Kristjansson; Droplaug N Magnusdottir; Louise le Roux; Gudrun Sigmundsdottir; Gardar Sveinbjornsson; Kristin E Sveinsdottir; Maney Sveinsdottir; Emil A Thorarensen; Bjarni Thorbjornsson; Gisli Masson; Ingileif Jonsdottir; Alma Moller; Thorolfur Gudnason; Karl G Kristinsson; Unnur Thorsteinsdottir; Kari Stefansson |
| EPI_ISL_417755 | hCoV-19/Iceland/169/2020 | Europe / Iceland / Reykjavik | 2020-03-13 | The National University Hospital of Iceland | deCODE genetics | Daniel F Gudbjartsson; Agnar Helgason; Hakon Jonsson; Olafur T Magnusson; Pall Melsted; Gudmundur L Norddahl; Jona Saemundsdottir; Asgeir Sigurdsson; Patrick Sulem; Arna B Agustsdottir; Berglind Eiriksdottir; Run Fridriksdottir; Elisabet E Gardarsdottir; Gudmundur Georgsson; Olafia S Gretarsdottir; Kjartan R Gudmundsson; Thora R Gunnarsdottir; Arnaldur Gylfason; Hilma Holm; Brynjar O Jensson; Aslaug Jonasdottir; Kamilla S Josefsdottir; Thordur Kristjansson; Droplaug N Magnusdottir; Louise le Roux; Gudrun Sigmundsdottir; Gardar Sveinbjornsson; Kristin E Sveinsdottir; Maney Sveinsdottir; Emil A Thorarensen; Bjarni Thorbjornsson; Gisli Masson; Ingileif Jonsdottir; Alma Moller; Thorolfur Gudnason; Karl G Kristinsson; Unnur Thorsteinsdottir; Kari Stefansson |
| EPI_ISL_417756 | hCoV-19/Iceland/58/2020 | Europe / Iceland / Reykjavik | 2020-03-06 | The National University Hospital of Iceland | deCODE genetics | Daniel F Gudbjartsson; Agnar Helgason; Hakon Jonsson; Olafur T Magnusson; Pall Melsted; Gudmundur L Norddahl; Jona Saemundsdottir; Asgeir Sigurdsson; Patrick Sulem; Arna B Agustsdottir; Berglind Eiriksdottir; Run Fridriksdottir; Elisabet E Gardarsdottir; Gudmundur Georgsson; Olafia S Gretarsdottir; Kjartan R Gudmundsson; Thora R Gunnarsdottir; Arnaldur Gylfason; Hilma Holm; Brynjar O Jensson; Aslaug Jonasdottir; Kamilla S Josefsdottir; Thordur Kristjansson; Droplaug N Magnusdottir; Louise le Roux; Gudrun Sigmundsdottir; Gardar Sveinbjornsson; Kristin E Sveinsdottir; Maney Sveinsdottir; Emil A Thorarensen; Bjarni Thorbjornsson; Gisli Masson; Ingileif Jonsdottir; Alma Moller; Thorolfur Gudnason; Karl G Kristinsson; Unnur Thorsteinsdottir; Kari Stefansson |
| EPI_ISL_417757 | hCoV-19/Iceland/62/2020 | Europe / Iceland / Reykjavik | 2020-03-06 | The National University Hospital of Iceland | deCODE genetics | Daniel F Gudbjartsson; Agnar Helgason; Hakon Jonsson; Olafur T Magnusson; Pall Melsted; Gudmundur L Norddahl; Jona Saemundsdottir; Asgeir Sigurdsson; Patrick Sulem; Arna B Agustsdottir; Berglind Eiriksdottir; Run Fridriksdottir; Elisabet E Gardarsdottir; Gudmundur Georgsson; Olafia S Gretarsdottir; Kjartan R Gudmundsson; Thora R Gunnarsdottir; Arnaldur Gylfason; Hilma Holm; Brynjar O Jensson; Aslaug Jonasdottir; Kamilla S Josefsdottir; Thordur Kristjansson; Droplaug N Magnusdottir; Louise le Roux; Gudrun Sigmundsdottir; Gardar Sveinbjornsson; Kristin E Sveinsdottir; Maney Sveinsdottir; Emil A Thorarensen; Bjarni Thorbjornsson; Gisli Masson; Ingileif Jonsdottir; Alma Moller; Thorolfur Gudnason; Karl G Kristinsson; Unnur Thorsteinsdottir; Kari Stefansson |
| EPI_ISL_417758 | hCoV-19/Iceland/164/2020 | Europe / Iceland / Reykjavik | 2020-03-14 | The National University Hospital of Iceland | deCODE genetics | Daniel F Gudbjartsson; Agnar Helgason; Hakon Jonsson; Olafur T Magnusson; Pall Melsted; Gudmundur L Norddahl; Jona Saemundsdottir; Asgeir Sigurdsson; Patrick Sulem; Arna B Agustsdottir; Berglind Eiriksdottir; Run Fridriksdottir; Elisabet E Gardarsdottir; Gudmundur Georgsson; Olafia S Gretarsdottir; Kjartan R Gudmundsson; Thora R Gunnarsdottir; Arnaldur Gylfason; Hilma Holm; Brynjar O Jensson; Aslaug Jonasdottir; Kamilla S Josefsdottir; Thordur Kristjansson; Droplaug N Magnusdottir; Louise le Roux; Gudrun Sigmundsdottir; Gardar Sveinbjornsson; Kristin E Sveinsdottir; Maney Sveinsdottir; Emil A Thorarensen; Bjarni Thorbjornsson; Gisli Masson; Ingileif Jonsdottir; Alma Moller; Thorolfur Gudnason; Karl G Kristinsson; Unnur Thorsteinsdottir; Kari Stefansson |
| EPI_ISL_417759 | hCoV-19/Iceland/166/2020 | Europe / Iceland / Reykjavik | 2020-03-13 | The National University Hospital of Iceland | deCODE genetics | Daniel F Gudbjartsson; Agnar Helgason; Hakon Jonsson; Olafur T Magnusson; Pall Melsted; Gudmundur L Norddahl; Jona Saemundsdottir; Asgeir Sigurdsson; Patrick Sulem; Arna B Agustsdottir; Berglind Eiriksdottir; Run Fridriksdottir; Elisabet E Gardarsdottir; Gudmundur Georgsson; Olafia S Gretarsdottir; Kjartan R Gudmundsson; Thora R Gunnarsdottir; Arnaldur Gylfason; Hilma Holm; Brynjar O Jensson; Aslaug Jonasdottir; Kamilla S Josefsdottir; Thordur Kristjansson; Droplaug N Magnusdottir; Louise le Roux; Gudrun Sigmundsdottir; Gardar Sveinbjornsson; Kristin E Sveinsdottir; Maney Sveinsdottir; Emil A Thorarensen; Bjarni Thorbjornsson; Gisli Masson; Ingileif Jonsdottir; Alma Moller; Thorolfur Gudnason; Karl G Kristinsson; Unnur Thorsteinsdottir; Kari Stefansson |
| EPI_ISL_417761 | hCoV-19/Iceland/84/2020 | Europe / Iceland / Reykjavik | 2020-03-10 | The National University Hospital of Iceland | deCODE genetics | Daniel F Gudbjartsson; Agnar Helgason; Hakon Jonsson; Olafur T Magnusson; Pall Melsted; Gudmundur L Norddahl; Jona Saemundsdottir; Asgeir Sigurdsson; Patrick Sulem; Arna B Agustsdottir; Berglind Eiriksdottir; Run Fridriksdottir; Elisabet E Gardarsdottir; Gudmundur Georgsson; Olafia S Gretarsdottir; Kjartan R Gudmundsson; Thora R Gunnarsdottir; Arnaldur Gylfason; Hilma Holm; Brynjar O Jensson; Aslaug Jonasdottir; Kamilla S Josefsdottir; Thordur Kristjansson; Droplaug N Magnusdottir; Louise le Roux; Gudrun Sigmundsdottir; Gardar Sveinbjornsson; Kristin E Sveinsdottir; Maney Sveinsdottir; Emil A Thorarensen; Bjarni Thorbjornsson; Gisli Masson; Ingileif Jonsdottir; Alma Moller; Thorolfur Gudnason; Karl G Kristinsson; Unnur Thorsteinsdottir; Kari Stefansson |
| EPI_ISL_417762 | hCoV-19/Iceland/191/2020 | Europe / Iceland / Reykjavik | 2020-03-16 | The National University Hospital of Iceland | deCODE genetics | Daniel F Gudbjartsson; Agnar Helgason; Hakon Jonsson; Olafur T Magnusson; Pall Melsted; Gudmundur L Norddahl; Jona Saemundsdottir; Asgeir Sigurdsson; Patrick Sulem; Arna B Agustsdottir; Berglind Eiriksdottir; Run Fridriksdottir; Elisabet E Gardarsdottir; Gudmundur Georgsson; Olafia S Gretarsdottir; Kjartan R Gudmundsson; Thora R Gunnarsdottir; Arnaldur Gylfason; Hilma Holm; Brynjar O Jensson; Aslaug Jonasdottir; Kamilla S Josefsdottir; Thordur Kristjansson; Droplaug N Magnusdottir; Louise le Roux; Gudrun Sigmundsdottir; Gardar Sveinbjornsson; Kristin E Sveinsdottir; Maney Sveinsdottir; Emil A Thorarensen; Bjarni Thorbjornsson; Gisli Masson; Ingileif Jonsdottir; Alma Moller; Thorolfur Gudnason; Karl G Kristinsson; Unnur Thorsteinsdottir; Kari Stefansson |
| EPI_ISL_417763 | hCoV-19/Iceland/211/2020 | Europe / Iceland / Reykjavik | 2020-03-16 | The National University Hospital of Iceland | deCODE genetics | Daniel F Gudbjartsson; Agnar Helgason; Hakon Jonsson; Olafur T Magnusson; Pall Melsted; Gudmundur L Norddahl; Jona Saemundsdottir; Asgeir Sigurdsson; Patrick Sulem; Arna B Agustsdottir; Berglind Eiriksdottir; Run Fridriksdottir; Elisabet E Gardarsdottir; Gudmundur Georgsson; Olafia S Gretarsdottir; Kjartan R Gudmundsson; Thora R Gunnarsdottir; Arnaldur Gylfason; Hilma Holm; Brynjar O Jensson; Aslaug Jonasdottir; Kamilla S Josefsdottir; Thordur Kristjansson; Droplaug N Magnusdottir; Louise le Roux; Gudrun Sigmundsdottir; Gardar Sveinbjornsson; Kristin E Sveinsdottir; Maney Sveinsdottir; Emil A Thorarensen; Bjarni Thorbjornsson; Gisli Masson; Ingileif Jonsdottir; Alma Moller; Thorolfur Gudnason; Karl G Kristinsson; Unnur Thorsteinsdottir; Kari Stefansson |
| EPI_ISL_417764 | hCoV-19/Iceland/209/2020 | Europe / Iceland / Reykjavik | 2020-03-16 | The National University Hospital of Iceland | deCODE genetics | Daniel F Gudbjartsson; Agnar Helgason; Hakon Jonsson; Olafur T Magnusson; Pall Melsted; Gudmundur L Norddahl; Jona Saemundsdottir; Asgeir Sigurdsson; Patrick Sulem; Arna B Agustsdottir; Berglind Eiriksdottir; Run Fridriksdottir; Elisabet E Gardarsdottir; Gudmundur Georgsson; Olafia S Gretarsdottir; Kjartan R Gudmundsson; Thora R Gunnarsdottir; Arnaldur Gylfason; Hilma Holm; Brynjar O Jensson; Aslaug Jonasdottir; Kamilla S Josefsdottir; Thordur Kristjansson; Droplaug N Magnusdottir; Louise le Roux; Gudrun Sigmundsdottir; Gardar Sveinbjornsson; Kristin E Sveinsdottir; Maney Sveinsdottir; Emil A Thorarensen; Bjarni Thorbjornsson; Gisli Masson; Ingileif Jonsdottir; Alma Moller; Thorolfur Gudnason; Karl G Kristinsson; Unnur Thorsteinsdottir; Kari Stefansson |
| EPI_ISL_417766 | hCoV-19/Iceland/208/2020 | Europe / Iceland / Reykjavik | 2020-03-16 | The National University Hospital of Iceland | deCODE genetics | Daniel F Gudbjartsson; Agnar Helgason; Hakon Jonsson; Olafur T Magnusson; Pall Melsted; Gudmundur L Norddahl; Jona Saemundsdottir; Asgeir Sigurdsson; Patrick Sulem; Arna B Agustsdottir; Berglind Eiriksdottir; Run Fridriksdottir; Elisabet E Gardarsdottir; Gudmundur Georgsson; Olafia S Gretarsdottir; Kjartan R Gudmundsson; Thora R Gunnarsdottir; Arnaldur Gylfason; Hilma Holm; Brynjar O Jensson; Aslaug Jonasdottir; Kamilla S Josefsdottir; Thordur Kristjansson; Droplaug N Magnusdottir; Louise le Roux; Gudrun Sigmundsdottir; Gardar Sveinbjornsson; Kristin E Sveinsdottir; Maney Sveinsdottir; Emil A Thorarensen; Bjarni Thorbjornsson; Gisli Masson; Ingileif Jonsdottir; Alma Moller; Thorolfur Gudnason; Karl G Kristinsson; Unnur Thorsteinsdottir; Kari Stefansson |
| EPI_ISL_417767 | hCoV-19/Iceland/28/2020 | Europe / Iceland / Reykjavik | 2020-03-03 | The National University Hospital of Iceland | deCODE genetics | Daniel F Gudbjartsson; Agnar Helgason; Hakon Jonsson; Olafur T Magnusson; Pall Melsted; Gudmundur L Norddahl; Jona Saemundsdottir; Asgeir Sigurdsson; Patrick Sulem; Arna B Agustsdottir; Berglind Eiriksdottir; Run Fridriksdottir; Elisabet E Gardarsdottir; Gudmundur Georgsson; Olafia S Gretarsdottir; Kjartan R Gudmundsson; Thora R Gunnarsdottir; Arnaldur Gylfason; Hilma Holm; Brynjar O Jensson; Aslaug Jonasdottir; Kamilla S Josefsdottir; Thordur Kristjansson; Droplaug N Magnusdottir; Louise le Roux; Gudrun Sigmundsdottir; Gardar Sveinbjornsson; Kristin E Sveinsdottir; Maney Sveinsdottir; Emil A Thorarensen; Bjarni Thorbjornsson; Gisli Masson; Ingileif Jonsdottir; Alma Moller; Thorolfur Gudnason; Karl G Kristinsson; Unnur Thorsteinsdottir; Kari Stefansson |
| EPI_ISL_417769 | hCoV-19/Iceland/50/2020 | Europe / Iceland / Reykjavik | 2020-03-05 | The National University Hospital of Iceland | deCODE genetics | Daniel F Gudbjartsson; Agnar Helgason; Hakon Jonsson; Olafur T Magnusson; Pall Melsted; Gudmundur L Norddahl; Jona Saemundsdottir; Asgeir Sigurdsson; Patrick Sulem; Arna B Agustsdottir; Berglind Eiriksdottir; Run Fridriksdottir; Elisabet E Gardarsdottir; Gudmundur Georgsson; Olafia S Gretarsdottir; Kjartan R Gudmundsson; Thora R Gunnarsdottir; Arnaldur Gylfason; Hilma Holm; Brynjar O Jensson; Aslaug Jonasdottir; Kamilla S Josefsdottir; Thordur Kristjansson; Droplaug N Magnusdottir; Louise le Roux; Gudrun Sigmundsdottir; Gardar Sveinbjornsson; Kristin E Sveinsdottir; Maney Sveinsdottir; Emil A Thorarensen; Bjarni Thorbjornsson; Gisli Masson; Ingileif Jonsdottir; Alma Moller; Thorolfur Gudnason; Karl G Kristinsson; Unnur Thorsteinsdottir; Kari Stefansson |
| EPI_ISL_417770 | hCoV-19/Iceland/112/2020 | Europe / Iceland / Reykjavik | 2020-03-11 | The National University Hospital of Iceland | deCODE genetics | Daniel F Gudbjartsson; Agnar Helgason; Hakon Jonsson; Olafur T Magnusson; Pall Melsted; Gudmundur L Norddahl; Jona Saemundsdottir; Asgeir Sigurdsson; Patrick Sulem; Arna B Agustsdottir; Berglind Eiriksdottir; Run Fridriksdottir; Elisabet E Gardarsdottir; Gudmundur Georgsson; Olafia S Gretarsdottir; Kjartan R Gudmundsson; Thora R Gunnarsdottir; Arnaldur Gylfason; Hilma Holm; Brynjar O Jensson; Aslaug Jonasdottir; Kamilla S Josefsdottir; Thordur Kristjansson; Droplaug N Magnusdottir; Louise le Roux; Gudrun Sigmundsdottir; Gardar Sveinbjornsson; Kristin E Sveinsdottir; Maney Sveinsdottir; Emil A Thorarensen; Bjarni Thorbjornsson; Gisli Masson; Ingileif Jonsdottir; Alma Moller; Thorolfur Gudnason; Karl G Kristinsson; Unnur Thorsteinsdottir; Kari Stefansson |
| EPI_ISL_417771 | hCoV-19/Iceland/131/2020 | Europe / Iceland / Reykjavik | 2020-03-12 | The National University Hospital of Iceland | deCODE genetics | Daniel F Gudbjartsson; Agnar Helgason; Hakon Jonsson; Olafur T Magnusson; Pall Melsted; Gudmundur L Norddahl; Jona Saemundsdottir; Asgeir Sigurdsson; Patrick Sulem; Arna B Agustsdottir; Berglind Eiriksdottir; Run Fridriksdottir; Elisabet E Gardarsdottir; Gudmundur Georgsson; Olafia S Gretarsdottir; Kjartan R Gudmundsson; Thora R Gunnarsdottir; Arnaldur Gylfason; Hilma Holm; Brynjar O Jensson; Aslaug Jonasdottir; Kamilla S Josefsdottir; Thordur Kristjansson; Droplaug N Magnusdottir; Louise le Roux; Gudrun Sigmundsdottir; Gardar Sveinbjornsson; Kristin E Sveinsdottir; Maney Sveinsdottir; Emil A Thorarensen; Bjarni Thorbjornsson; Gisli Masson; Ingileif Jonsdottir; Alma Moller; Thorolfur Gudnason; Karl G Kristinsson; Unnur Thorsteinsdottir; Kari Stefansson |
| EPI_ISL_417772 | hCoV-19/Iceland/162/2020 | Europe / Iceland / Reykjavik | 2020-03-14 | The National University Hospital of Iceland | deCODE genetics | Daniel F Gudbjartsson; Agnar Helgason; Hakon Jonsson; Olafur T Magnusson; Pall Melsted; Gudmundur L Norddahl; Jona Saemundsdottir; Asgeir Sigurdsson; Patrick Sulem; Arna B Agustsdottir; Berglind Eiriksdottir; Run Fridriksdottir; Elisabet E Gardarsdottir; Gudmundur Georgsson; Olafia S Gretarsdottir; Kjartan R Gudmundsson; Thora R Gunnarsdottir; Arnaldur Gylfason; Hilma Holm; Brynjar O Jensson; Aslaug Jonasdottir; Kamilla S Josefsdottir; Thordur Kristjansson; Droplaug N Magnusdottir; Louise le Roux; Gudrun Sigmundsdottir; Gardar Sveinbjornsson; Kristin E Sveinsdottir; Maney Sveinsdottir; Emil A Thorarensen; Bjarni Thorbjornsson; Gisli Masson; Ingileif Jonsdottir; Alma Moller; Thorolfur Gudnason; Karl G Kristinsson; Unnur Thorsteinsdottir; Kari Stefansson |
| EPI_ISL_417773 | hCoV-19/Iceland/30/2020 | Europe / Iceland / Reykjavik | 2020-03-03 | The National University Hospital of Iceland | deCODE genetics | Daniel F Gudbjartsson; Agnar Helgason; Hakon Jonsson; Olafur T Magnusson; Pall Melsted; Gudmundur L Norddahl; Jona Saemundsdottir; Asgeir Sigurdsson; Patrick Sulem; Arna B Agustsdottir; Berglind Eiriksdottir; Run Fridriksdottir; Elisabet E Gardarsdottir; Gudmundur Georgsson; Olafia S Gretarsdottir; Kjartan R Gudmundsson; Thora R Gunnarsdottir; Arnaldur Gylfason; Hilma Holm; Brynjar O Jensson; Aslaug Jonasdottir; Kamilla S Josefsdottir; Thordur Kristjansson; Droplaug N Magnusdottir; Louise le Roux; Gudrun Sigmundsdottir; Gardar Sveinbjornsson; Kristin E Sveinsdottir; Maney Sveinsdottir; Emil A Thorarensen; Bjarni Thorbjornsson; Gisli Masson; Ingileif Jonsdottir; Alma Moller; Thorolfur Gudnason; Karl G Kristinsson; Unnur Thorsteinsdottir; Kari Stefansson |
| EPI_ISL_417774 | hCoV-19/Iceland/175/2020 | Europe / Iceland / Reykjavik | 2020-03-16 | The National University Hospital of Iceland | deCODE genetics | Daniel F Gudbjartsson; Agnar Helgason; Hakon Jonsson; Olafur T Magnusson; Pall Melsted; Gudmundur L Norddahl; Jona Saemundsdottir; Asgeir Sigurdsson; Patrick Sulem; Arna B Agustsdottir; Berglind Eiriksdottir; Run Fridriksdottir; Elisabet E Gardarsdottir; Gudmundur Georgsson; Olafia S Gretarsdottir; Kjartan R Gudmundsson; Thora R Gunnarsdottir; Arnaldur Gylfason; Hilma Holm; Brynjar O Jensson; Aslaug Jonasdottir; Kamilla S Josefsdottir; Thordur Kristjansson; Droplaug N Magnusdottir; Louise le Roux; Gudrun Sigmundsdottir; Gardar Sveinbjornsson; Kristin E Sveinsdottir; Maney Sveinsdottir; Emil A Thorarensen; Bjarni Thorbjornsson; Gisli Masson; Ingileif Jonsdottir; Alma Moller; Thorolfur Gudnason; Karl G Kristinsson; Unnur Thorsteinsdottir; Kari Stefansson |
| EPI_ISL_417775 | hCoV-19/Iceland/163/2020 | Europe / Iceland / Reykjavik | 2020-03-14 | The National University Hospital of Iceland | deCODE genetics | Daniel F Gudbjartsson; Agnar Helgason; Hakon Jonsson; Olafur T Magnusson; Pall Melsted; Gudmundur L Norddahl; Jona Saemundsdottir; Asgeir Sigurdsson; Patrick Sulem; Arna B Agustsdottir; Berglind Eiriksdottir; Run Fridriksdottir; Elisabet E Gardarsdottir; Gudmundur Georgsson; Olafia S Gretarsdottir; Kjartan R Gudmundsson; Thora R Gunnarsdottir; Arnaldur Gylfason; Hilma Holm; Brynjar O Jensson; Aslaug Jonasdottir; Kamilla S Josefsdottir; Thordur Kristjansson; Droplaug N Magnusdottir; Louise le Roux; Gudrun Sigmundsdottir; Gardar Sveinbjornsson; Kristin E Sveinsdottir; Maney Sveinsdottir; Emil A Thorarensen; Bjarni Thorbjornsson; Gisli Masson; Ingileif Jonsdottir; Alma Moller; Thorolfur Gudnason; Karl G Kristinsson; Unnur Thorsteinsdottir; Kari Stefansson |
| EPI_ISL_417777 | hCoV-19/Iceland/102/2020 | Europe / Iceland / Reykjavik | 2020-03-10 | The National University Hospital of Iceland | deCODE genetics | Daniel F Gudbjartsson; Agnar Helgason; Hakon Jonsson; Olafur T Magnusson; Pall Melsted; Gudmundur L Norddahl; Jona Saemundsdottir; Asgeir Sigurdsson; Patrick Sulem; Arna B Agustsdottir; Berglind Eiriksdottir; Run Fridriksdottir; Elisabet E Gardarsdottir; Gudmundur Georgsson; Olafia S Gretarsdottir; Kjartan R Gudmundsson; Thora R Gunnarsdottir; Arnaldur Gylfason; Hilma Holm; Brynjar O Jensson; Aslaug Jonasdottir; Kamilla S Josefsdottir; Thordur Kristjansson; Droplaug N Magnusdottir; Louise le Roux; Gudrun Sigmundsdottir; Gardar Sveinbjornsson; Kristin E Sveinsdottir; Maney Sveinsdottir; Emil A Thorarensen; Bjarni Thorbjornsson; Gisli Masson; Ingileif Jonsdottir; Alma Moller; Thorolfur Gudnason; Karl G Kristinsson; Unnur Thorsteinsdottir; Kari Stefansson |
| EPI_ISL_417778 | hCoV-19/Iceland/105/2020 | Europe / Iceland / Reykjavik | 2020-03-10 | The National University Hospital of Iceland | deCODE genetics | Daniel F Gudbjartsson; Agnar Helgason; Hakon Jonsson; Olafur T Magnusson; Pall Melsted; Gudmundur L Norddahl; Jona Saemundsdottir; Asgeir Sigurdsson; Patrick Sulem; Arna B Agustsdottir; Berglind Eiriksdottir; Run Fridriksdottir; Elisabet E Gardarsdottir; Gudmundur Georgsson; Olafia S Gretarsdottir; Kjartan R Gudmundsson; Thora R Gunnarsdottir; Arnaldur Gylfason; Hilma Holm; Brynjar O Jensson; Aslaug Jonasdottir; Kamilla S Josefsdottir; Thordur Kristjansson; Droplaug N Magnusdottir; Louise le Roux; Gudrun Sigmundsdottir; Gardar Sveinbjornsson; Kristin E Sveinsdottir; Maney Sveinsdottir; Emil A Thorarensen; Bjarni Thorbjornsson; Gisli Masson; Ingileif Jonsdottir; Alma Moller; Thorolfur Gudnason; Karl G Kristinsson; Unnur Thorsteinsdottir; Kari Stefansson |
| EPI_ISL_417779 | hCoV-19/Iceland/106/2020 | Europe / Iceland / Reykjavik | 2020-03-11 | The National University Hospital of Iceland | deCODE genetics | Daniel F Gudbjartsson; Agnar Helgason; Hakon Jonsson; Olafur T Magnusson; Pall Melsted; Gudmundur L Norddahl; Jona Saemundsdottir; Asgeir Sigurdsson; Patrick Sulem; Arna B Agustsdottir; Berglind Eiriksdottir; Run Fridriksdottir; Elisabet E Gardarsdottir; Gudmundur Georgsson; Olafia S Gretarsdottir; Kjartan R Gudmundsson; Thora R Gunnarsdottir; Arnaldur Gylfason; Hilma Holm; Brynjar O Jensson; Aslaug Jonasdottir; Kamilla S Josefsdottir; Thordur Kristjansson; Droplaug N Magnusdottir; Louise le Roux; Gudrun Sigmundsdottir; Gardar Sveinbjornsson; Kristin E Sveinsdottir; Maney Sveinsdottir; Emil A Thorarensen; Bjarni Thorbjornsson; Gisli Masson; Ingileif Jonsdottir; Alma Moller; Thorolfur Gudnason; Karl G Kristinsson; Unnur Thorsteinsdottir; Kari Stefansson |
| EPI_ISL_417781 | hCoV-19/Iceland/113/2020 | Europe / Iceland / Reykjavik | 2020-03-11 | The National University Hospital of Iceland | deCODE genetics | Daniel F Gudbjartsson; Agnar Helgason; Hakon Jonsson; Olafur T Magnusson; Pall Melsted; Gudmundur L Norddahl; Jona Saemundsdottir; Asgeir Sigurdsson; Patrick Sulem; Arna B Agustsdottir; Berglind Eiriksdottir; Run Fridriksdottir; Elisabet E Gardarsdottir; Gudmundur Georgsson; Olafia S Gretarsdottir; Kjartan R Gudmundsson; Thora R Gunnarsdottir; Arnaldur Gylfason; Hilma Holm; Brynjar O Jensson; Aslaug Jonasdottir; Kamilla S Josefsdottir; Thordur Kristjansson; Droplaug N Magnusdottir; Louise le Roux; Gudrun Sigmundsdottir; Gardar Sveinbjornsson; Kristin E Sveinsdottir; Maney Sveinsdottir; Emil A Thorarensen; Bjarni Thorbjornsson; Gisli Masson; Ingileif Jonsdottir; Alma Moller; Thorolfur Gudnason; Karl G Kristinsson; Unnur Thorsteinsdottir; Kari Stefansson |
| EPI_ISL_417783 | hCoV-19/Iceland/123/2020 | Europe / Iceland / Reykjavik | 2020-03-11 | The National University Hospital of Iceland | deCODE genetics | Daniel F Gudbjartsson; Agnar Helgason; Hakon Jonsson; Olafur T Magnusson; Pall Melsted; Gudmundur L Norddahl; Jona Saemundsdottir; Asgeir Sigurdsson; Patrick Sulem; Arna B Agustsdottir; Berglind Eiriksdottir; Run Fridriksdottir; Elisabet E Gardarsdottir; Gudmundur Georgsson; Olafia S Gretarsdottir; Kjartan R Gudmundsson; Thora R Gunnarsdottir; Arnaldur Gylfason; Hilma Holm; Brynjar O Jensson; Aslaug Jonasdottir; Kamilla S Josefsdottir; Thordur Kristjansson; Droplaug N Magnusdottir; Louise le Roux; Gudrun Sigmundsdottir; Gardar Sveinbjornsson; Kristin E Sveinsdottir; Maney Sveinsdottir; Emil A Thorarensen; Bjarni Thorbjornsson; Gisli Masson; Ingileif Jonsdottir; Alma Moller; Thorolfur Gudnason; Karl G Kristinsson; Unnur Thorsteinsdottir; Kari Stefansson |
| EPI_ISL_417784 | hCoV-19/Iceland/124/2020 | Europe / Iceland / Reykjavik | 2020-03-12 | The National University Hospital of Iceland | deCODE genetics | Daniel F Gudbjartsson; Agnar Helgason; Hakon Jonsson; Olafur T Magnusson; Pall Melsted; Gudmundur L Norddahl; Jona Saemundsdottir; Asgeir Sigurdsson; Patrick Sulem; Arna B Agustsdottir; Berglind Eiriksdottir; Run Fridriksdottir; Elisabet E Gardarsdottir; Gudmundur Georgsson; Olafia S Gretarsdottir; Kjartan R Gudmundsson; Thora R Gunnarsdottir; Arnaldur Gylfason; Hilma Holm; Brynjar O Jensson; Aslaug Jonasdottir; Kamilla S Josefsdottir; Thordur Kristjansson; Droplaug N Magnusdottir; Louise le Roux; Gudrun Sigmundsdottir; Gardar Sveinbjornsson; Kristin E Sveinsdottir; Maney Sveinsdottir; Emil A Thorarensen; Bjarni Thorbjornsson; Gisli Masson; Ingileif Jonsdottir; Alma Moller; Thorolfur Gudnason; Karl G Kristinsson; Unnur Thorsteinsdottir; Kari Stefansson |
| EPI_ISL_417785 | hCoV-19/Iceland/126/2020 | Europe / Iceland / Reykjavik | 2020-03-12 | The National University Hospital of Iceland | deCODE genetics | Daniel F Gudbjartsson; Agnar Helgason; Hakon Jonsson; Olafur T Magnusson; Pall Melsted; Gudmundur L Norddahl; Jona Saemundsdottir; Asgeir Sigurdsson; Patrick Sulem; Arna B Agustsdottir; Berglind Eiriksdottir; Run Fridriksdottir; Elisabet E Gardarsdottir; Gudmundur Georgsson; Olafia S Gretarsdottir; Kjartan R Gudmundsson; Thora R Gunnarsdottir; Arnaldur Gylfason; Hilma Holm; Brynjar O Jensson; Aslaug Jonasdottir; Kamilla S Josefsdottir; Thordur Kristjansson; Droplaug N Magnusdottir; Louise le Roux; Gudrun Sigmundsdottir; Gardar Sveinbjornsson; Kristin E Sveinsdottir; Maney Sveinsdottir; Emil A Thorarensen; Bjarni Thorbjornsson; Gisli Masson; Ingileif Jonsdottir; Alma Moller; Thorolfur Gudnason; Karl G Kristinsson; Unnur Thorsteinsdottir; Kari Stefansson |
| EPI_ISL_417786 | hCoV-19/Iceland/128/2020 | Europe / Iceland / Reykjavik | 2020-03-12 | The National University Hospital of Iceland | deCODE genetics | Daniel F Gudbjartsson; Agnar Helgason; Hakon Jonsson; Olafur T Magnusson; Pall Melsted; Gudmundur L Norddahl; Jona Saemundsdottir; Asgeir Sigurdsson; Patrick Sulem; Arna B Agustsdottir; Berglind Eiriksdottir; Run Fridriksdottir; Elisabet E Gardarsdottir; Gudmundur Georgsson; Olafia S Gretarsdottir; Kjartan R Gudmundsson; Thora R Gunnarsdottir; Arnaldur Gylfason; Hilma Holm; Brynjar O Jensson; Aslaug Jonasdottir; Kamilla S Josefsdottir; Thordur Kristjansson; Droplaug N Magnusdottir; Louise le Roux; Gudrun Sigmundsdottir; Gardar Sveinbjornsson; Kristin E Sveinsdottir; Maney Sveinsdottir; Emil A Thorarensen; Bjarni Thorbjornsson; Gisli Masson; Ingileif Jonsdottir; Alma Moller; Thorolfur Gudnason; Karl G Kristinsson; Unnur Thorsteinsdottir; Kari Stefansson |
| EPI_ISL_417787 | hCoV-19/Iceland/132/2020 | Europe / Iceland / Reykjavik | 2020-03-12 | The National University Hospital of Iceland | deCODE genetics | Daniel F Gudbjartsson; Agnar Helgason; Hakon Jonsson; Olafur T Magnusson; Pall Melsted; Gudmundur L Norddahl; Jona Saemundsdottir; Asgeir Sigurdsson; Patrick Sulem; Arna B Agustsdottir; Berglind Eiriksdottir; Run Fridriksdottir; Elisabet E Gardarsdottir; Gudmundur Georgsson; Olafia S Gretarsdottir; Kjartan R Gudmundsson; Thora R Gunnarsdottir; Arnaldur Gylfason; Hilma Holm; Brynjar O Jensson; Aslaug Jonasdottir; Kamilla S Josefsdottir; Thordur Kristjansson; Droplaug N Magnusdottir; Louise le Roux; Gudrun Sigmundsdottir; Gardar Sveinbjornsson; Kristin E Sveinsdottir; Maney Sveinsdottir; Emil A Thorarensen; Bjarni Thorbjornsson; Gisli Masson; Ingileif Jonsdottir; Alma Moller; Thorolfur Gudnason; Karl G Kristinsson; Unnur Thorsteinsdottir; Kari Stefansson |
| EPI_ISL_417788 | hCoV-19/Iceland/133/2020 | Europe / Iceland / Reykjavik | 2020-03-12 | The National University Hospital of Iceland | deCODE genetics | Daniel F Gudbjartsson; Agnar Helgason; Hakon Jonsson; Olafur T Magnusson; Pall Melsted; Gudmundur L Norddahl; Jona Saemundsdottir; Asgeir Sigurdsson; Patrick Sulem; Arna B Agustsdottir; Berglind Eiriksdottir; Run Fridriksdottir; Elisabet E Gardarsdottir; Gudmundur Georgsson; Olafia S Gretarsdottir; Kjartan R Gudmundsson; Thora R Gunnarsdottir; Arnaldur Gylfason; Hilma Holm; Brynjar O Jensson; Aslaug Jonasdottir; Kamilla S Josefsdottir; Thordur Kristjansson; Droplaug N Magnusdottir; Louise le Roux; Gudrun Sigmundsdottir; Gardar Sveinbjornsson; Kristin E Sveinsdottir; Maney Sveinsdottir; Emil A Thorarensen; Bjarni Thorbjornsson; Gisli Masson; Ingileif Jonsdottir; Alma Moller; Thorolfur Gudnason; Karl G Kristinsson; Unnur Thorsteinsdottir; Kari Stefansson |
| EPI_ISL_417789 | hCoV-19/Iceland/136/2020 | Europe / Iceland / Reykjavik | 2020-03-12 | The National University Hospital of Iceland | deCODE genetics | Daniel F Gudbjartsson; Agnar Helgason; Hakon Jonsson; Olafur T Magnusson; Pall Melsted; Gudmundur L Norddahl; Jona Saemundsdottir; Asgeir Sigurdsson; Patrick Sulem; Arna B Agustsdottir; Berglind Eiriksdottir; Run Fridriksdottir; Elisabet E Gardarsdottir; Gudmundur Georgsson; Olafia S Gretarsdottir; Kjartan R Gudmundsson; Thora R Gunnarsdottir; Arnaldur Gylfason; Hilma Holm; Brynjar O Jensson; Aslaug Jonasdottir; Kamilla S Josefsdottir; Thordur Kristjansson; Droplaug N Magnusdottir; Louise le Roux; Gudrun Sigmundsdottir; Gardar Sveinbjornsson; Kristin E Sveinsdottir; Maney Sveinsdottir; Emil A Thorarensen; Bjarni Thorbjornsson; Gisli Masson; Ingileif Jonsdottir; Alma Moller; Thorolfur Gudnason; Karl G Kristinsson; Unnur Thorsteinsdottir; Kari Stefansson |

| | | | | | | |
|---|---|---|---|---|---|---|
| EPI_ISL_417790 | hCoV-19/Iceland/139/2020 | Europe / Iceland / Reykjavik | 2020-03-12 | The National University Hospital of Iceland | deCODE genetics | Daniel F Gudbjartsson; Agnar Helgason; Hakon Jonsson; Olafur T Magnusson; Pall Melsted; Gudmundur L Norddahl; Jona Saemundsdottir; Asgeir Sigurdsson; Patrick Sulem; Arna B Agustsdottir; Berglind Eiriksdottir; Run Fridriksdottir; Elisabet E Gardarsdottir; Gudmundur Georgsson; Olafia S Gretarsdottir; Kjartan R Gudmundsson; Thora R Gunnarsdottir; Arnaldur Gylfason; Hilma Holm; Brynjar O Jensson; Aslaug Jonasdottir; Kamilla S Josefsdottir; Thordur Kristjansson; Droplaug N Magnusdottir; Louise le Roux; Gudrun Sigmundsdottir; Gardar Sveinbjornsson; Kristin E Sveinsdottir; Maney Sveinsdottir; Emil A Thorarensen; Bjarni Thorbjornsson; Gisli Masson; Ingileif Jonsdottir; Alma Moller; Thorolfur Gudnason; Karl G Kristinsson; Unnur Thorsteinsdottir; Kari Stefansson |
| EPI_ISL_417791 | hCoV-19/Iceland/140/2020 | Europe / Iceland / Reykjavik | 2020-03-12 | The National University Hospital of Iceland | deCODE genetics | Daniel F Gudbjartsson; Agnar Helgason; Hakon Jonsson; Olafur T Magnusson; Pall Melsted; Gudmundur L Norddahl; Jona Saemundsdottir; Asgeir Sigurdsson; Patrick Sulem; Arna B Agustsdottir; Berglind Eiriksdottir; Run Fridriksdottir; Elisabet E Gardarsdottir; Gudmundur Georgsson; Olafia S Gretarsdottir; Kjartan R Gudmundsson; Thora R Gunnarsdottir; Arnaldur Gylfason; Hilma Holm; Brynjar O Jensson; Aslaug Jonasdottir; Kamilla S Josefsdottir; Thordur Kristjansson; Droplaug N Magnusdottir; Louise le Roux; Gudrun Sigmundsdottir; Gardar Sveinbjornsson; Kristin E Sveinsdottir; Maney Sveinsdottir; Emil A Thorarensen; Bjarni Thorbjornsson; Gisli Masson; Ingileif Jonsdottir; Alma Moller; Thorolfur Gudnason; Karl G Kristinsson; Unnur Thorsteinsdottir; Kari Stefansson |
| EPI_ISL_417792 | hCoV-19/Iceland/141/2020 | Europe / Iceland / Reykjavik | 2020-03-13 | The National University Hospital of Iceland | deCODE genetics | Daniel F Gudbjartsson; Agnar Helgason; Hakon Jonsson; Olafur T Magnusson; Pall Melsted; Gudmundur L Norddahl; Jona Saemundsdottir; Asgeir Sigurdsson; Patrick Sulem; Arna B Agustsdottir; Berglind Eiriksdottir; Run Fridriksdottir; Elisabet E Gardarsdottir; Gudmundur Georgsson; Olafia S Gretarsdottir; Kjartan R Gudmundsson; Thora R Gunnarsdottir; Arnaldur Gylfason; Hilma Holm; Brynjar O Jensson; Aslaug Jonasdottir; Kamilla S Josefsdottir; Thordur Kristjansson; Droplaug N Magnusdottir; Louise le Roux; Gudrun Sigmundsdottir; Gardar Sveinbjornsson; Kristin E Sveinsdottir; Maney Sveinsdottir; Emil A Thorarensen; Bjarni Thorbjornsson; Gisli Masson; Ingileif Jonsdottir; Alma Moller; Thorolfur Gudnason; Karl G Kristinsson; Unnur Thorsteinsdottir; Kari Stefansson |
| EPI_ISL_417793 | hCoV-19/Iceland/142/2020 | Europe / Iceland / Reykjavik | 2020-03-13 | The National University Hospital of Iceland | deCODE genetics | Daniel F Gudbjartsson; Agnar Helgason; Hakon Jonsson; Olafur T Magnusson; Pall Melsted; Gudmundur L Norddahl; Jona Saemundsdottir; Asgeir Sigurdsson; Patrick Sulem; Arna B Agustsdottir; Berglind Eiriksdottir; Run Fridriksdottir; Elisabet E Gardarsdottir; Gudmundur Georgsson; Olafia S Gretarsdottir; Kjartan R Gudmundsson; Thora R Gunnarsdottir; Arnaldur Gylfason; Hilma Holm; Brynjar O Jensson; Aslaug Jonasdottir; Kamilla S Josefsdottir; Thordur Kristjansson; Droplaug N Magnusdottir; Louise le Roux; Gudrun Sigmundsdottir; Gardar Sveinbjornsson; Kristin E Sveinsdottir; Maney Sveinsdottir; Emil A Thorarensen; Bjarni Thorbjornsson; Gisli Masson; Ingileif Jonsdottir; Alma Moller; Thorolfur Gudnason; Karl G Kristinsson; Unnur Thorsteinsdottir; Kari Stefansson |
| EPI_ISL_417794 | hCoV-19/Iceland/143/2020 | Europe / Iceland / Reykjavik | 2020-03-12 | The National University Hospital of Iceland | deCODE genetics | Daniel F Gudbjartsson; Agnar Helgason; Hakon Jonsson; Olafur T Magnusson; Pall Melsted; Gudmundur L Norddahl; Jona Saemundsdottir; Asgeir Sigurdsson; Patrick Sulem; Arna B Agustsdottir; Berglind Eiriksdottir; Run Fridriksdottir; Elisabet E Gardarsdottir; Gudmundur Georgsson; Olafia S Gretarsdottir; Kjartan R Gudmundsson; Thora R Gunnarsdottir; Arnaldur Gylfason; Hilma Holm; Brynjar O Jensson; Aslaug Jonasdottir; Kamilla S Josefsdottir; Thordur Kristjansson; Droplaug N Magnusdottir; Louise le Roux; Gudrun Sigmundsdottir; Gardar Sveinbjornsson; Kristin E Sveinsdottir; Maney Sveinsdottir; Emil A Thorarensen; Bjarni Thorbjornsson; Gisli Masson; Ingileif Jonsdottir; Alma Moller; Thorolfur Gudnason; Karl G Kristinsson; Unnur Thorsteinsdottir; Kari Stefansson |
| EPI_ISL_417795 | hCoV-19/Iceland/144/2020 | Europe / Iceland / Reykjavik | 2020-03-13 | The National University Hospital of Iceland | deCODE genetics | Daniel F Gudbjartsson; Agnar Helgason; Hakon Jonsson; Olafur T Magnusson; Pall Melsted; Gudmundur L Norddahl; Jona Saemundsdottir; Asgeir Sigurdsson; Patrick Sulem; Arna B Agustsdottir; Berglind Eiriksdottir; Run Fridriksdottir; Elisabet E Gardarsdottir; Gudmundur Georgsson; Olafia S Gretarsdottir; Kjartan R Gudmundsson; Thora R Gunnarsdottir; Arnaldur Gylfason; Hilma Holm; Brynjar O Jensson; Aslaug Jonasdottir; Kamilla S Josefsdottir; Thordur Kristjansson; Droplaug N Magnusdottir; Louise le Roux; Gudrun Sigmundsdottir; Gardar Sveinbjornsson; Kristin E Sveinsdottir; Maney Sveinsdottir; Emil A Thorarensen; Bjarni Thorbjornsson; Gisli Masson; Ingileif Jonsdottir; Alma Moller; Thorolfur Gudnason; Karl G Kristinsson; Unnur Thorsteinsdottir; Kari Stefansson |
| EPI_ISL_417796 | hCoV-19/Iceland/146/2020 | Europe / Iceland / Reykjavik | 2020-03-13 | The National University Hospital of Iceland | deCODE genetics | Daniel F Gudbjartsson; Agnar Helgason; Hakon Jonsson; Olafur T Magnusson; Pall Melsted; Gudmundur L Norddahl; Jona Saemundsdottir; Asgeir Sigurdsson; Patrick Sulem; Arna B Agustsdottir; Berglind Eiriksdottir; Run Fridriksdottir; Elisabet E Gardarsdottir; Gudmundur Georgsson; Olafia S Gretarsdottir; Kjartan R Gudmundsson; Thora R Gunnarsdottir; Arnaldur Gylfason; Hilma Holm; Brynjar O Jensson; Aslaug Jonasdottir; Kamilla S Josefsdottir; Thordur Kristjansson; Droplaug N Magnusdottir; Louise le Roux; Gudrun Sigmundsdottir; Gardar Sveinbjornsson; Kristin E Sveinsdottir; Maney Sveinsdottir; Emil A Thorarensen; Bjarni Thorbjornsson; Gisli Masson; Ingileif Jonsdottir; Alma Moller; Thorolfur Gudnason; Karl G Kristinsson; Unnur Thorsteinsdottir; Kari Stefansson |
| EPI_ISL_417797 | hCoV-19/Iceland/147/2020 | Europe / Iceland / Reykjavik | 2020-03-13 | The National University Hospital of Iceland | deCODE genetics | Daniel F Gudbjartsson; Agnar Helgason; Hakon Jonsson; Olafur T Magnusson; Pall Melsted; Gudmundur L Norddahl; Jona Saemundsdottir; Asgeir Sigurdsson; Patrick Sulem; Arna B Agustsdottir; Berglind Eiriksdottir; Run Fridriksdottir; Elisabet E Gardarsdottir; Gudmundur Georgsson; Olafia S Gretarsdottir; Kjartan R Gudmundsson; Thora R Gunnarsdottir; Arnaldur Gylfason; Hilma Holm; Brynjar O Jensson; Aslaug Jonasdottir; Kamilla S Josefsdottir; Thordur Kristjansson; Droplaug N Magnusdottir; Louise le Roux; Gudrun Sigmundsdottir; Gardar Sveinbjornsson; Kristin E Sveinsdottir; Maney Sveinsdottir; Emil A Thorarensen; Bjarni Thorbjornsson; Gisli Masson; Ingileif Jonsdottir; Alma Moller; Thorolfur Gudnason; Karl G Kristinsson; Unnur Thorsteinsdottir; Kari Stefansson |
| EPI_ISL_417798 | hCoV-19/Iceland/148/2020 | Europe / Iceland / Reykjavik | 2020-03-13 | The National University Hospital of Iceland | deCODE genetics | Daniel F Gudbjartsson; Agnar Helgason; Hakon Jonsson; Olafur T Magnusson; Pall Melsted; Gudmundur L Norddahl; Jona Saemundsdottir; Asgeir Sigurdsson; Patrick Sulem; Arna B Agustsdottir; Berglind Eiriksdottir; Run Fridriksdottir; Elisabet E Gardarsdottir; Gudmundur Georgsson; Olafia S Gretarsdottir; Kjartan R Gudmundsson; Thora R Gunnarsdottir; Arnaldur Gylfason; Hilma Holm; Brynjar O Jensson; Aslaug Jonasdottir; Kamilla S Josefsdottir; Thordur Kristjansson; Droplaug N Magnusdottir; Louise le Roux; Gudrun Sigmundsdottir; Gardar Sveinbjornsson; Kristin E Sveinsdottir; Maney Sveinsdottir; Emil A Thorarensen; Bjarni Thorbjornsson; Gisli Masson; Ingileif Jonsdottir; Alma Moller; Thorolfur Gudnason; Karl G Kristinsson; Unnur Thorsteinsdottir; Kari Stefansson |
| EPI_ISL_417799 | hCoV-19/Iceland/149/2020 | Europe / Iceland / Reykjavik | 2020-03-13 | The National University Hospital of Iceland | deCODE genetics | Daniel F Gudbjartsson; Agnar Helgason; Hakon Jonsson; Olafur T Magnusson; Pall Melsted; Gudmundur L Norddahl; Jona Saemundsdottir; Asgeir Sigurdsson; Patrick Sulem; Arna B Agustsdottir; Berglind Eiriksdottir; Run Fridriksdottir; Elisabet E Gardarsdottir; Gudmundur Georgsson; Olafia S Gretarsdottir; Kjartan R Gudmundsson; Thora R Gunnarsdottir; Arnaldur Gylfason; Hilma Holm; Brynjar O Jensson; Aslaug Jonasdottir; Kamilla S Josefsdottir; Thordur Kristjansson; Droplaug N Magnusdottir; Louise le Roux; Gudrun Sigmundsdottir; Gardar Sveinbjornsson; Kristin E Sveinsdottir; Maney Sveinsdottir; Emil A Thorarensen; Bjarni Thorbjornsson; Gisli Masson; Ingileif Jonsdottir; Alma Moller; Thorolfur Gudnason; Karl G Kristinsson; Unnur Thorsteinsdottir; Kari Stefansson |
| EPI_ISL_418060 | hCoV-19/USA/WA-UW275/2020 | North America / USA | 2020-03-14 | UW Virology Lab | UW Virology Lab | Pavitra Roychoudhury, Hong Xie, Keith Jerome, Alexander Greninger |
| EPI_ISL_418061 | hCoV-19/USA/UN-UW276/2020 | North America / USA | 2020-03-13 | UW Virology Lab | UW Virology Lab | Pavitra Roychoudhury, Hong Xie, Keith Jerome, Alexander Greninger |
| EPI_ISL_418062 | hCoV-19/USA/WA-UW277/2020 | North America / USA / Washington | 2020-03-14 | UW Virology Lab | UW Virology Lab | Pavitra Roychoudhury, Hong Xie, Keith Jerome, Alexander Greninger |
| EPI_ISL_418064 | hCoV-19/USA/WA-UW279/2020 | North America / USA / Washington | 2020-03-15 | UW Virology Lab | UW Virology Lab | Pavitra Roychoudhury, Hong Xie, Keith Jerome, Alexander Greninger |
| EPI_ISL_418066 | hCoV-19/USA/WA-UW281/2020 | North America / USA / Washington | 2020-03-14 | UW Virology Lab | UW Virology Lab | Pavitra Roychoudhury, Hong Xie, Keith Jerome, Alexander Greninger |
| EPI_ISL_418067 | hCoV-19/USA/WA-UW282/2020 | North America / USA / Washington | 2020-03-15 | UW Virology Lab | UW Virology Lab | Pavitra Roychoudhury, Hong Xie, Keith Jerome, Alexander Greninger |
| EPI_ISL_418068 | hCoV-19/USA/WA-UW283/2020 | North America / USA / Washington | 2020-03-14 | UW Virology Lab | UW Virology Lab | Pavitra Roychoudhury, Hong Xie, Keith Jerome, Alexander Greninger |
| EPI_ISL_418069 | hCoV-19/USA/WA-UW284/2020 | North America / USA / Washington | 2020-03-14 | UW Virology Lab | UW Virology Lab | Pavitra Roychoudhury, Hong Xie, Keith Jerome, Alexander Greninger |
| EPI_ISL_418070 | hCoV-19/USA/WA-UW285/2020 | North America / USA / Washington | 2020-03-14 | UW Virology Lab | UW Virology Lab | Pavitra Roychoudhury, Hong Xie, Keith Jerome, Alexander Greninger |
| EPI_ISL_418071 | hCoV-19/USA/WA-UW286/2020 | North America / USA / Washington | 2020-03-15 | UW Virology Lab | UW Virology Lab | Pavitra Roychoudhury, Hong Xie, Keith Jerome, Alexander Greninger |
| EPI_ISL_418072 | hCoV-19/USA/WA-UW287/2020 | North America / USA / Washington | 2020-03-15 | UW Virology Lab | UW Virology Lab | Pavitra Roychoudhury, Hong Xie, Keith Jerome, Alexander Greninger |
| EPI_ISL_418073 | hCoV-19/USA/WA-UW288/2020 | North America / USA / Washington | 2020-03-15 | UW Virology Lab | UW Virology Lab | Pavitra Roychoudhury, Hong Xie, Keith Jerome, Alexander Greninger |
| EPI_ISL_418074 | hCoV-19/USA/WA-UW289/2020 | North America / USA / Washington | 2020-03-15 | UW Virology Lab | UW Virology Lab | Pavitra Roychoudhury, Hong Xie, Keith Jerome, Alexander Greninger |
| EPI_ISL_418075 | hCoV-19/USA/WA-UW290/2020 | North America / USA / Washington | 2020-03-15 | UW Virology Lab | UW Virology Lab | Pavitra Roychoudhury, Hong Xie, Keith Jerome, Alexander Greninger |
| EPI_ISL_418076 | hCoV-19/USA/WA-UW291/2020 | North America / USA / Washington | 2020-03-15 | UW Virology Lab | UW Virology Lab | Pavitra Roychoudhury, Hong Xie, Keith Jerome, Alexander Greninger |
| EPI_ISL_418077 | hCoV-19/USA/WA-UW292/2020 | North America / USA / Washington | 2020-03-15 | UW Virology Lab | UW Virology Lab | Pavitra Roychoudhury, Hong Xie, Keith Jerome, Alexander Greninger |
| EPI_ISL_418078 | hCoV-19/USA/WA-UW293/2020 | North America / USA / Washington | 2020-03-15 | UW Virology Lab | UW Virology Lab | Pavitra Roychoudhury, Hong Xie, Keith Jerome, Alexander Greninger |
| EPI_ISL_418079 | hCoV-19/USA/WA-UW294/2020 | North America / USA / Washington | 2020-03-15 | UW Virology Lab | UW Virology Lab | Pavitra Roychoudhury, Hong Xie, Keith Jerome, Alexander Greninger |
| EPI_ISL_418080 | hCoV-19/USA/WA-UW295/2020 | North America / USA / Washington | 2020-03-15 | UW Virology Lab | UW Virology Lab | Pavitra Roychoudhury, Hong Xie, Keith Jerome, Alexander Greninger |
| EPI_ISL_418081 | hCoV-19/USA/WA-UW296/2020 | North America / USA / Washington | 2020-03-15 | UW Virology Lab | UW Virology Lab | Pavitra Roychoudhury, Hong Xie, Keith Jerome, Alexander Greninger |
| EPI_ISL_418082 | hCoV-19/USA/WA-UW297/2020 | North America / USA / Washington | 2020-03-15 | UW Virology Lab | UW Virology Lab | Pavitra Roychoudhury, Hong Xie, Keith Jerome, Alexander Greninger |
| EPI_ISL_418084 | hCoV-19/Wales/PHWC-23A81/2020 | Europe / United Kingdom / Wales | 2020-03-12 | Wales Specialist Virology Centre | Public Health Wales Microbiology Cardiff | Catherine Moore, Joanne Watkins, Sally Corden, Sara Rey, Matt Bull, Tom Connor |
| EPI_ISL_418087 | hCoV-19/Wales/PHWC-23ADC/2020 | Europe / United Kingdom / Wales | 2020-03-12 | Wales Specialist Virology Centre | Public Health Wales Microbiology Cardiff | Catherine Moore, Joanne Watkins, Sally Corden, Sara Rey, Matt Bull, Tom Connor |
| EPI_ISL_418088 | hCoV-19/Wales/PHWC-23AEB/2020 | Europe / United Kingdom / Wales | 2020-03-12 | Wales Specialist Virology Centre | Public Health Wales Microbiology Cardiff | Catherine Moore, Joanne Watkins, Sally Corden, Sara Rey, Matt Bull, Tom Connor |
| EPI_ISL_418089 | hCoV-19/Wales/PHWC-23AFA/2020 | Europe / United Kingdom / Wales | 2020-03-12 | Wales Specialist Virology Centre | Public Health Wales Microbiology Cardiff | Catherine Moore, Joanne Watkins, Sally Corden, Sara Rey, Matt Bull, Tom Connor |
| EPI_ISL_418091 | hCoV-19/Wales/PHWC-23B24/2020 | Europe / United Kingdom / Wales | 2020-03-12 | Wales Specialist Virology Centre | Public Health Wales Microbiology Cardiff | Catherine Moore, Joanne Watkins, Sally Corden, Sara Rey, Matt Bull, Tom Connor |
| EPI_ISL_418092 | hCoV-19/Wales/PHWC-23B42/2020 | Europe / United Kingdom / Wales | 2020-03-12 | Wales Specialist Virology Centre | Public Health Wales Microbiology Cardiff | Catherine Moore, Joanne Watkins, Sally Corden, Sara Rey, Matt Bull, Tom Connor |
| EPI_ISL_418094 | hCoV-19/Wales/PHWC-23B7F/2020 | Europe / United Kingdom / Wales | 2020-03-12 | Wales Specialist Virology Centre | Public Health Wales Microbiology Cardiff | Catherine Moore, Joanne Watkins, Sally Corden, Sara Rey, Matt Bull, Tom Connor |
| EPI_ISL_418095 | hCoV-19/Wales/PHWC-23B9D/2020 | Europe / United Kingdom / Wales | 2020-03-12 | Wales Specialist Virology Centre | Public Health Wales Microbiology Cardiff | Catherine Moore, Joanne Watkins, Sally Corden, Sara Rey, Matt Bull, Tom Connor |
| EPI_ISL_418097 | hCoV-19/Wales/PHWC-23BD9/2020 | Europe / United Kingdom / Wales | 2020-03-12 | Wales Specialist Virology Centre | Public Health Wales Microbiology Cardiff | Catherine Moore, Joanne Watkins, Sally Corden, Sara Rey, Matt Bull, Tom Connor |
| EPI_ISL_418098 | hCoV-19/Wales/PHWC-23BE8/2020 | Europe / United Kingdom / Wales | 2020-03-13 | Wales Specialist Virology Centre | Public Health Wales Microbiology Cardiff | Catherine Moore, Joanne Watkins, Sally Corden, Sara Rey, Matt Bull, Tom Connor |
| EPI_ISL_418099 | hCoV-19/Wales/PHWC-23BF7/2020 | Europe / United Kingdom / Wales | 2020-03-13 | Wales Specialist Virology Centre | Public Health Wales Microbiology Cardiff | Catherine Moore, Joanne Watkins, Sally Corden, Sara Rey, Matt Bull, Tom Connor |
| EPI_ISL_418182 | hCoV-19/Spain/Madrid_H8_37/2020 | Europe / Spain / Madrid | 2020-03-12 | Hospital Universitario 12 de Octubre | Hospital Universitario La Paz | Elias Dahdouh, Sara González, Fernando Lázaro, Esther Viedma, Natalia Stella, Julio García, Juan Carlos Galán, Rafael Cantón, Mª Dolores Folgueira, Rafael Delgado, Jesús Mingorance |
| EPI_ISL_418183 | hCoV-19/Hungary/mbl2/2020 | Europe / Hungary / Baranya | 2020-03-17 | Virological Research Group, Szentágothai Research Centre | Bioinformatics Research Group, Szentágothai Research Centre | Péter Urbán, Endre Gábor Tóth, Gábor Kemenesi, Róbert Herczeg, Attila Gyenesei, Ferenc Jakab |
| EPI_ISL_418184 | hCoV-19/USA/WI-GMF-00018/2020 | North America / USA / Wisconsin / La Cr | 2020-03-18 | Gundersen Molecular Diagnostics Laboratory | Kabara Cancer Research Institute | Craig S. Richmond & Paraic A. Kenny |
| EPI_ISL_418186 | hCoV-19/USA/WI-GMF-00227/2020 | North America / USA / Wisconsin / La Cr | 2020-03-23 | Gundersen Molecular Diagnostic Laboratory | Kabara Cancer Research Institute | Craig S. Richmond & Paraic A. Kenny |
| EPI_ISL_418187 | hCoV-19/USA/WI-GMF-00228/2020 | North America / USA / Wisconsin / La Cr | 2020-03-23 | Gundersen Molecular Diagnostics Laboratory | Kabara Cancer Research Institute | Craig S. Richmond & Paraic A. Kenny |
| EPI_ISL_418188 | hCoV-19/USA/WI-GMF-00232/2020 | North America / USA / Wisconsin / Monr | 2020-03-23 | Gundersen Molecular Diagnostics Laboratory | Kabara Cancer Research Institute | Craig S. Richmond & Paraic A. Kenny |
| EPI_ISL_418191 | hCoV-19/USA/NY-NYUMC6/2020 | North America / USA / New York / Nassa | 2020-03-16 | NYU Langone Health | Department of Pathology and Medicine, New York University School of Medicine | Margaret Black, John Cadley, Paolo Cotzia, John Chen, Dacia Dimartino, Xiaojun Feng, Adriana Heguy, Megan Hogan, Emily Huang, George Jour, Christian Marier, Matthew T. Maurano, Mark J. Mulligan, Peter Meyn, Jared Pinnell, Amy Rapkiewicz, Marie Samanovic-Golden, Antonio Serrano, Guomiao Shen, Matija Snuderl, Nick Vulpescu, Gael Westby, Paul Zappile |
| EPI_ISL_418192 | hCoV-19/USA/NY-NYUMC7/2020 | North America / USA / New York / Manha | 2020-03-16 | NYU Langone Health | Department of Pathology and Medicine, New York University School of Medicine | Margaret Black, John Cadley, Paolo Cotzia, John Chen, Dacia Dimartino, Xiaojun Feng, Adriana Heguy, Megan Hogan, Emily Huang, George Jour, Christian Marier, Matthew T. Maurano, Mark J. Mulligan, Peter Meyn, Jared Pinnell, Amy Rapkiewicz, Marie Samanovic-Golden, Antonio Serrano, Guomiao Shen, Matija Snuderl, Nick Vulpescu, Gael Westby, Paul Zappile |
| EPI_ISL_418193 | hCoV-19/USA/NY-NYUMC8/2020 | North America / USA / New York / Manha | 2020-03-17 | NYU Langone Health | Department of Pathology and Medicine, New York University School of Medicine | Margaret Black, John Cadley, Paolo Cotzia, John Chen, Dacia Dimartino, Xiaojun Feng, Adriana Heguy, Megan Hogan, Emily Huang, George Jour, Christian Marier, Matthew T. Maurano, Mark J. Mulligan, Peter Meyn, Jared Pinnell, Amy Rapkiewicz, Marie Samanovic-Golden, Antonio Serrano, Guomiao Shen, Matija Snuderl, Nick Vulpescu, Gael Westby, Paul Zappile |
| EPI_ISL_418194 | hCoV-19/USA/NY-NYUMC9/2020 | North America / USA / New York / Manha | 2020-03-17 | NYU Langone Health | Department of Pathology and Medicine, New York University School of Medicine | Margaret Black, John Cadley, Paolo Cotzia, John Chen, Dacia Dimartino, Xiaojun Feng, Adriana Heguy, Megan Hogan, Emily Huang, George Jour, Christian Marier, Matthew T. Maurano, Mark J. Mulligan, Peter Meyn, Jared Pinnell, Amy Rapkiewicz, Marie Samanovic-Golden, Antonio Serrano, Guomiao Shen, Matija Snuderl, Nick Vulpescu, Gael Westby, Paul Zappile |
| EPI_ISL_418195 | hCoV-19/USA/NY-NYUMC10/2020 | North America / USA / New York / Manha | 2020-03-17 | NYU Langone Health | Department of Pathology and Medicine, New York University School of Medicine | Margaret Black, John Cadley, Paolo Cotzia, John Chen, Dacia Dimartino, Xiaojun Feng, Adriana Heguy, Megan Hogan, Emily Huang, George Jour, Christian Marier, Matthew T. Maurano, Mark J. Mulligan, Peter Meyn, Jared Pinnell, Amy Rapkiewicz, Marie Samanovic-Golden, Antonio Serrano, Guomiao Shen, Matija Snuderl, Nick Vulpescu, Gael Westby, Paul Zappile |
| EPI_ISL_418196 | hCoV-19/USA/NY-NYUMC11/2020 | North America / USA / New York / Manha | 2020-03-14 | NYU Langone Health | Department of Pathology and Medicine, New York University School of Medicine | Margaret Black, John Cadley, Paolo Cotzia, John Chen, Dacia Dimartino, Xiaojun Feng, Adriana Heguy, Megan Hogan, Emily Huang, George Jour, Christian Marier, Matthew T. Maurano, Mark J. Mulligan, Peter Meyn, Jared Pinnell, Amy Rapkiewicz, Marie Samanovic-Golden, Antonio Serrano, Guomiao Shen, Matija Snuderl, Nick Vulpescu, Gael Westby, Paul Zappile |
| EPI_ISL_418197 | hCoV-19/USA/NY-NYUMC12/2020 | North America / USA / New York / Manha | 2020-03-15 | NYU Langone Health | Department of Pathology and Medicine, New York University School of Medicine | Margaret Black, John Cadley, Paolo Cotzia, John Chen, Dacia Dimartino, Xiaojun Feng, Adriana Heguy, Megan Hogan, Emily Huang, George Jour, Christian Marier, Matthew T. Maurano, Mark J. Mulligan, Peter Meyn, Jared Pinnell, Amy Rapkiewicz, Marie Samanovic-Golden, Antonio Serrano, Guomiao Shen, Matija Snuderl, Nick Vulpescu, Gael Westby, Paul Zappile |
| EPI_ISL_418198 | hCoV-19/USA/NY-NYUMC13/2020 | North America / USA / New York / Manha | 2020-03-17 | NYU Langone Health | Department of Pathology and Medicine, New York University School of Medicine | Margaret Black, John Cadley, Paolo Cotzia, John Chen, Dacia Dimartino, Xiaojun Feng, Adriana Heguy, Megan Hogan, Emily Huang, George Jour, Christian Marier, Matthew T. Maurano, Mark J. Mulligan, Peter Meyn, Jared Pinnell, Amy Rapkiewicz, Marie Samanovic-Golden, Antonio Serrano, Guomiao Shen, Matija Snuderl, Nick Vulpescu, Gael Westby, Paul Zappile |
| EPI_ISL_418199 | hCoV-19/USA/NY-NYUMC14/2020 | North America / USA / New York / Manha | 2020-03-17 | NYU Langone Health | Department of Pathology and Medicine, New York University School of Medicine | Margaret Black, John Cadley, Paolo Cotzia, John Chen, Dacia Dimartino, Xiaojun Feng, Adriana Heguy, Megan Hogan, Emily Huang, George Jour, Christian Marier, Matthew T. Maurano, Mark J. Mulligan, Peter Meyn, Jared Pinnell, Amy Rapkiewicz, Marie Samanovic-Golden, Antonio Serrano, Guomiao Shen, Matija Snuderl, Nick Vulpescu, Gael Westby, Paul Zappile |
| EPI_ISL_418200 | hCoV-19/USA/NY-NYUMC15/2020 | North America / USA / New York / Manha | 2020-03-17 | NYU Langone Health | Department of Pathology and Medicine, New York University School of Medicine | Margaret Black, John Cadley, Paolo Cotzia, John Chen, Dacia Dimartino, Xiaojun Feng, Adriana Heguy, Megan Hogan, Emily Huang, George Jour, Christian Marier, Matthew T. Maurano, Mark J. Mulligan, Peter Meyn, Jared Pinnell, Amy Rapkiewicz, Marie Samanovic-Golden, Antonio Serrano, Guomiao Shen, Matija Snuderl, Nick Vulpescu, Gael Westby, Paul Zappile |
| EPI_ISL_418201 | hCoV-19/USA/NY-NYUMC16/2020 | North America / USA / New York / Manha | 2020-03-17 | NYU Langone Health | Department of Pathology and Medicine, New York University School of Medicine | Margaret Black, John Cadley, Paolo Cotzia, John Chen, Dacia Dimartino, Xiaojun Feng, Adriana Heguy, Megan Hogan, Emily Huang, George Jour, Christian Marier, Matthew T. Maurano, Mark J. Mulligan, Peter Meyn, Jared Pinnell, Amy Rapkiewicz, Marie Samanovic-Golden, Antonio Serrano, Guomiao Shen, Matija Snuderl, Nick Vulpescu, Gael Westby, Paul Zappile |
| EPI_ISL_418203 | hCoV-19/USA/NY-NYUMC18/2020 | North America / USA / New York / Manha | 2020-03-17 | NYU Langone Health | Department of Pathology and Medicine, New York University School of Medicine | Margaret Black, John Cadley, Paolo Cotzia, John Chen, Dacia Dimartino, Xiaojun Feng, Adriana Heguy, Megan Hogan, Emily Huang, George Jour, Christian Marier, Matthew T. Maurano, Mark J. Mulligan, Peter Meyn, Jared Pinnell, Amy Rapkiewicz, Marie Samanovic-Golden, Antonio Serrano, Guomiao Shen, Matija Snuderl, Nick Vulpescu, Gael Westby, Paul Zappile |
| EPI_ISL_418204 | hCoV-19/USA/NY-NYUMC19/2020 | North America / USA / New York / Manha | 2020-03-17 | NYU Langone Health | Department of Pathology and Medicine, New York University School of Medicine | Margaret Black, John Cadley, Paolo Cotzia, John Chen, Dacia Dimartino, Xiaojun Feng, Adriana Heguy, Megan Hogan, Emily Huang, George Jour, Christian Marier, Matthew T. Maurano, Mark J. Mulligan, Peter Meyn, Jared Pinnell, Amy Rapkiewicz, Marie Samanovic-Golden, Antonio Serrano, Guomiao Shen, Matija Snuderl, Nick Vulpescu, Gael Westby, Paul Zappile |
| EPI_ISL_418205 | hCoV-19/USA/NY-NYUMC20/2020 | North America / USA / New York / Manha | 2020-03-12 | NYU Langone Health | Department of Pathology and Medicine, New York University School of Medicine | Margaret Black, John Cadley, Paolo Cotzia, John Chen, Dacia Dimartino, Xiaojun Feng, Adriana Heguy, Megan Hogan, Emily Huang, George Jour, Christian Marier, Matthew T. Maurano, Mark J. Mulligan, Peter Meyn, Jared Pinnell, Amy Rapkiewicz, Marie Samanovic-Golden, Antonio Serrano, Guomiao Shen, Matija Snuderl, Nick Vulpescu, Gael Westby, Paul Zappile |
| EPI_ISL_418206 | hCoV-19/Senegal/003/2020 | Africa / Senegal / Dakar | 2020-02-28 | Institut Pasteur Dakar | Institut Pasteur de Dakar | Ndongo Dia, Ousmane Faye, Amadou Alpha Sall |
| EPI_ISL_418207 | hCoV-19/Senegal/016/2020 | Africa / Senegal / Dakar | 2020-03-02 | Institut Pasteur Dakar | Institut Pasteur de Dakar | Ndongo Dia, Ousmane Faye, Amadou Alpha Sall |
| EPI_ISL_418208 | hCoV-19/Senegal/020/2020 | Africa / Senegal / Dakar | 2020-03-04 | Institut Pasteur Dakar | Institut Pasteur de Dakar | Ndongo Dia, Ousmane Faye, Amadou Alpha Sall |
| EPI_ISL_418209 | hCoV-19/Senegal/026/2020 | Africa / Senegal / Dakar | 2020-03-03 | Institut Pasteur Dakar | Institut Pasteur de Dakar | Ndongo Dia, Ousmane Faye, Amadou Alpha Sall |
| EPI_ISL_418210 | hCoV-19/Senegal/073/2020 | Africa / Senegal / Touba | 2020-03-10 | Institut Pasteur Dakar | Institut Pasteur de Dakar | Ndongo Dia, Ousmane Faye, Amadou Alpha Sall |
| EPI_ISL_418211 | hCoV-19/Senegal/082/2020 | Africa / Senegal / Touba | 2020-03-11 | Institut Pasteur Dakar | Institut Pasteur de Dakar | Ndongo Dia, Ousmane Faye, Amadou Alpha Sall |
| EPI_ISL_418212 | hCoV-19/Senegal/087/2020 | Africa / Senegal / Touba | 2020-03-11 | Institut Pasteur Dakar | Institut Pasteur de Dakar | Ndongo Dia, Ousmane Faye, Amadou Alpha sall |
| EPI_ISL_418213 | hCoV-19/Senegal/094/2020 | Africa / Senegal / Touba | 2020-03-12 | Institut Pasteur Dakar | Institut Pasteur de Dakar | Ndongo Dia, Ousmane Faye, Amadou Alpha Sall |
| EPI_ISL_418215 | hCoV-19/Senegal/119/2020 | Africa / Senegal / Touba | 2020-03-12 | Institut Pasteur Dakar | Institut Pasteur de Dakar | Ndongo Dia, Ousmane Faye, Amadou Alpha Sall |
| EPI_ISL_418216 | hCoV-19/Senegal/136/2020 | Africa / Senegal / Dakar | 2020-03-13 | Institut Pasteur Dakar | Institut Pasteur de Dakar | Ndongo Dia, Ousmane Faye, Amadou Alpha Sall |
| EPI_ISL_418217 | hCoV-19/Senegal/139/2020 | Africa / Senegal / Mbour | 2020-03-13 | Institut Pasteur Dakar | Institut Pasteur de Dakar | Ndongo Dia, Ousmane Faye, Amadou Alpha Sall |
| EPI_ISL_418218 | hCoV-19/France/HF1465/2020 | Europe / France / Hauts de France / Com | 2020-02-21 | Centre Hospitalier Compiègne Laboratoire de Biologie | National Reference Center for Viruses of Respiratory Infections, Institut Pasteur, Paris | Mélanie Albert, Marion Barbet, Sylvie Behillil, Méline Bizard, Angela Brisebarre, Flora Donati, Fabiana Gambaro, Etienne Simon-Lorière, Vincent Enouf, Maud Vanpeene, Sylvie van der Werf, Raulin Olivia |
| EPI_ISL_418219 | hCoV-19/France/B1623/2020 | Europe / France / Bretagne / Brest | 2020-02-26 | CHU - Hôpital Cavale Blanche - Labo. de Virologie | National Reference Center for Viruses of Respiratory Infections, Institut Pasteur, Paris | Mélanie Albert, Marion Barbet, Sylvie Behillil, Méline Bizard, Angela Brisebarre, Flora Donati, Fabiana Gambaro, Etienne Simon-Lorière, Vincent Enouf, Maud Vanpeene, Sylvie van der Werf, Léa Pilorge |
| EPI_ISL_418220 | hCoV-19/France/HF1645/2020 | Europe / France / Hauts de France / Com | 2020-02-28 | Centre Hospitalier Compiègne Laboratoire de Biologie | National Reference Center for Viruses of Respiratory Infections, Institut Pasteur, Paris | Mélanie Albert, Marion Barbet, Sylvie Behillil, Méline Bizard, Angela Brisebarre, Flora Donati, Fabiana Gambaro, Etienne Simon-Lorière, Vincent Enouf, Maud Vanpeene, Sylvie van der Werf, Raulin Olivia |
| EPI_ISL_418222 | hCoV-19/France/CVL2000/2020 | Europe / France / Centre-Val de Loire / T | 2020-03-04 | CHRU Bretonneau - Serv. Bacterio-Virol. | National Reference Center for Viruses of Respiratory Infections, Institut Pasteur, Paris | Mélanie Albert, Marion Barbet, Sylvie Behillil, Méline Bizard, Angela Brisebarre, Flora Donati, Fabiana Gambaro, Etienne Simon-Lorière, Vincent Enouf, Maud Vanpeene, Sylvie van der Werf, Julien Marlet |
| EPI_ISL_418223 | hCoV-19/France/HF2060/2020 | Europe / France / Hauts de France / Com | 2020-03-05 | Centre Hospitalier Compiègne Laboratoire de Biologie | National Reference Center for Viruses of Respiratory Infections, Institut Pasteur, Paris | Mélanie Albert, Marion Barbet, Sylvie Behillil, Méline Bizard, Angela Brisebarre, Flora Donati, Fabiana Gambaro, Etienne Simon-Lorière, Vincent Enouf, Maud Vanpeene, Sylvie van der Werf, Raulin Olivia |
| EPI_ISL_418224 | hCoV-19/France/HF2150/2020 | Europe / France / Hauts de France / Com | 2020-03-08 | Centre Hospitalier Compiègne Laboratoire de Biologie | National Reference Center for Viruses of Respiratory Infections, Institut Pasteur, Paris | Mélanie Albert, Marion Barbet, Sylvie Behillil, Méline Bizard, Angela Brisebarre, Flora Donati, Fabiana Gambaro, Etienne Simon-Lorière, Vincent Enouf, Maud Vanpeene, Sylvie van der Werf, Raulin Olivia |
| EPI_ISL_418225 | hCoV-19/France/HF2155/2020 | Europe / France / Hauts de France / Com | 2020-03-08 | Centre Hospitalier Compiègne Laboratoire de Biologie | National Reference Center for Viruses of Respiratory Infections, Institut Pasteur, Paris | Mélanie Albert, Marion Barbet, Sylvie Behillil, Méline Bizard, Angela Brisebarre, Flora Donati, Fabiana Gambaro, Etienne Simon-Lorière, Vincent Enouf, Maud Vanpeene, Sylvie van der Werf, Raulin Olivia |
| EPI_ISL_418226 | hCoV-19/France/HF2381/2020 | Europe / France / Hauts de France / Crou | 2020-03-09 | EHPAD - Résidences les Cèdres | National Reference Center for Viruses of Respiratory Infections, Institut Pasteur, Paris | Mélanie Albert, Marion Barbet, Sylvie Behillil, Méline Bizard, Angela Brisebarre, Flora Donati, Etienne Simon-Lorière, Vincent Enouf, Maud Vanpeene, Sylvie van der Werf |
| EPI_ISL_418227 | hCoV-19/France/HF2393/2020 | Europe / France / Hauts de France / Com | 2020-03-12 | Centre Hospitalier Compiègne Laboratoire de Biologie | National Reference Center for Viruses of Respiratory Infections, Institut Pasteur, Paris | Mélanie Albert, Marion Barbet, Sylvie Behillil, Méline Bizard, Angela Brisebarre, Flora Donati, Etienne Simon-Lorière, Vincent Enouf, Maud Vanpeene, Sylvie van der Werf, Raulin Olivia |
| EPI_ISL_418228 | hCoV-19/France/HF2405/2020 | Europe / France / Hauts de France / Com | 2020-03-12 | Centre Hospitalier Compiègne Laboratoire de Biologie | National Reference Center for Viruses of Respiratory Infections, Institut Pasteur, Paris | Mélanie Albert, Marion Barbet, Sylvie Behillil, Méline Bizard, Angela Brisebarre, Flora Donati, Etienne Simon-Lorière, Vincent Enouf, Maud Vanpeene, Sylvie van der Werf, Raulin Olivia |
| EPI_ISL_418229 | hCoV-19/France/IDF2410/2020 | Europe / France / Ile de France / Levallo | 2020-03-12 | Hopital franco britannique - Laboratoire | National Reference Center for Viruses of Respiratory Infections, Institut Pasteur, Paris | Mélanie Albert, Marion Barbet, Sylvie Behillil, Méline Bizard, Angela Brisebarre, Flora Donati, Etienne Simon-Lorière, Vincent Enouf, Maud Vanpeene, Sylvie van der Werf, Marianne Asso Bonnet |
| EPI_ISL_418230 | hCoV-19/France/IDF2420/2020 | Europe / France / Ile de France / Paris | 2020-03-13 | Clinique AVERAY LA BROUSTE, Med. Polyvalente | National Reference Center for Viruses of Respiratory Infections, Institut Pasteur, Paris | Mélanie Albert, Marion Barbet, Sylvie Behillil, Méline Bizard, Angela Brisebarre, Flora Donati, Etienne Simon-Lorière, Vincent Enouf, Maud Vanpeene, Sylvie van der Werf, Elsa Ngwem |
| EPI_ISL_418231 | hCoV-19/France/HF2496/2020 | Europe / France / Hauts de France / Com | 2020-03-15 | Centre Hospitalier Compiègne Laboratoire de Biologie | National Reference Center for Viruses of Respiratory Infections, Institut Pasteur, Paris | Mélanie Albert, Marion Barbet, Sylvie Behillil, Méline Bizard, Angela Brisebarre, Flora Donati, Etienne Simon-Lorière, Vincent Enouf, Maud Vanpeene, Sylvie van der Werf, Raulin Olivia |
| EPI_ISL_418232 | hCoV-19/France/IDF2532/2020 | Europe / France / Ile de France / Meudon | 2020-03-15 | Service des Urgences | National Reference Center for Viruses of Respiratory Infections, Institut Pasteur, Paris | Mélanie Albert, Marion Barbet, Sylvie Behillil, Méline Bizard, Angela Brisebarre, Flora Donati, Etienne Simon-Lorière, Vincent Enouf, Maud Vanpeene, Sylvie van der Werf, Boubkeur |
| EPI_ISL_418233 | hCoV-19/France/IDF2533/2020 | Europe / France / Ile de France / Meudon | 2020-03-15 | Service des Urgences | National Reference Center for Viruses of Respiratory Infections, Institut Pasteur, Paris | Mélanie Albert, Marion Barbet, Sylvie Behillil, Méline Bizard, Angela Brisebarre, Flora Donati, Etienne Simon-Lorière, Vincent Enouf, Maud Vanpeene, Sylvie van der Werf, Boubkeur |
| EPI_ISL_418234 | hCoV-19/France/IDF2534/2020 | Europe / France / Ile de France / Longjun | 2020-03-14 | LABM GH nord Essonne | National Reference Center for Viruses of Respiratory Infections, Institut Pasteur, Paris | Mélanie Albert, Marion Barbet, Sylvie Behillil, Méline Bizard, Angela Brisebarre, Flora Donati, Etienne Simon-Lorière, Vincent Enouf, Maud Vanpeene, Sylvie van der Werf, Christine Lambert |
| EPI_ISL_418235 | hCoV-19/France/IDF2561/2020 | Europe / France / Ile de France / Vanves | 2020-03-16 | Cabinet médical | National Reference Center for Viruses of Respiratory Infections, Institut Pasteur, Paris | Mélanie Albert, Marion Barbet, Sylvie Behillil, Méline Bizard, Angela Brisebarre, Flora Donati, Etienne Simon-Lorière, Vincent Enouf, Maud Vanpeene, Sylvie van der Werf |
| EPI_ISL_418236 | hCoV-19/France/HF2586/2020 | Europe / France / Hauts de France / Com | 2020-03-16 | Centre Hospitalier Compiègne Laboratoire de Biologie | National Reference Center for Viruses of Respiratory Infections, Institut Pasteur, Paris | Mélanie Albert, Marion Barbet, Sylvie Behillil, Méline Bizard, Angela Brisebarre, Flora Donati, Etienne Simon-Lorière, Vincent Enouf, Maud Vanpeene, Sylvie van der Werf, Raulin Olivia |
| EPI_ISL_418237 | hCoV-19/France/HF2595/2020 | Europe / France / Hauts de France / Com | 2020-03-16 | Centre Hospitalier Compiègne Laboratoire de Biologie | National Reference Center for Viruses of Respiratory Infections, Institut Pasteur, Paris | Mélanie Albert, Marion Barbet, Sylvie Behillil, Méline Bizard, Angela Brisebarre, Flora Donati, Etienne Simon-Lorière, Vincent Enouf, Maud Vanpeene, Sylvie van der Werf, Raulin Olivia |
| EPI_ISL_418238 | hCoV-19/France/HF2597/2020 | Europe / France / Hauts de France / Com | 2020-03-16 | Centre Hospitalier Compiègne Laboratoire de Biologie | National Reference Center for Viruses of Respiratory Infections, Institut Pasteur, Paris | Mélanie Albert, Marion Barbet, Sylvie Behillil, Méline Bizard, Angela Brisebarre, Flora Donati, Etienne Simon-Lorière, Vincent Enouf, Maud Vanpeene, Sylvie van der Werf, Raulin Olivia |
| EPI_ISL_418239 | hCoV-19/France/HF2601/2020 | Europe / France / Hauts de France / Com | 2020-03-16 | Centre Hospitalier Compiègne Laboratoire de Biologie | National Reference Center for Viruses of Respiratory Infections, Institut Pasteur, Paris | Mélanie Albert, Marion Barbet, Sylvie Behillil, Méline Bizard, Angela Brisebarre, Flora Donati, Etienne Simon-Lorière, Vincent Enouf, Maud Vanpeene, Sylvie van der Werf, Raulin Olivia |
| EPI_ISL_418240 | hCoV-19/France/IDF2684/2020 | Europe / France / Ile de France / Longjun | 2020-03-16 | LABM GH nord Essonne | National Reference Center for Viruses of Respiratory Infections, Institut Pasteur, Paris | Mélanie Albert, Marion Barbet, Sylvie Behillil, Méline Bizard, Angela Brisebarre, Flora Donati, Etienne Simon-Lorière, Vincent Enouf, Maud Vanpeene, Sylvie van der Werf, Christine Lambert |
| EPI_ISL_418241 | hCoV-19/Algeria/G0638_2264/2020 | Africa / Algeria / Boufarik | 2020-03-02 | NIC Viral Respiratory Unit - Institut Pasteur of Algeria | National Reference Center for Viruses of Respiratory Infections, Institut Pasteur, Paris | Mélanie Albert, Marion Barbet, Sylvie Behillil, Méline Bizard, Angela Brisebarre, Flora Donati, Etienne Simon-Lorière, Vincent Enouf, Maud Vanpeene, Sylvie van der Werf, Fawzi Derrar |
| EPI_ISL_418242 | hCoV-19/Algeria/G0640_2265/2020 | Africa / Algeria / Blida | 2020-03-08 | NIC Viral Respiratory Unit - Institut Pasteur of Algeria | National Reference Center for Viruses of Respiratory Infections, Institut Pasteur, Paris | Mélanie Albert, Marion Barbet, Sylvie Behillil, Méline Bizard, Angela Brisebarre, Flora Donati, Etienne Simon-Lorière, Vincent Enouf, Maud Vanpeene, Sylvie van der Werf, Fawzi Derrar |
| EPI_ISL_418243 | hCoV-19/Spain/Andalucia201272/2020 | Europe / Spain / Andalusia | 2020-02-28 | HOSPITAL UNIVERSITARIO VIRGEN DE LAS NIEVES | Instituto de Salud Carlos III | Iglesias-Caballero, M. Molinero Calamita, M. González-Esguevillas, M. Camarero, S. Pozo, F. Casas, I. Jiménez, P. Jiménez, M. Zaballos, A. Monzón, S. Varona, S. Juliá, M. Cuesta, I. Sanbonmatsu S. |
| EPI_ISL_418244 | hCoV-19/Spain/Andalucia201373/2020 | Europe / Spain / Andalusia | 2020-03-02 | HOSPITAL UNIVERSITARIO VIRGEN DE LAS NIEVES | Instituto de Salud Carlos III | Iglesias-Caballero, M. Molinero Calamita, M. González-Esguevillas, M. Camarero, S. Pozo, F. Casas, I. Jiménez, P. Jiménez, M. Zaballos, A. Monzón, S. Varona, S. Juliá, M. Cuesta, I. Sanbonmatsu S. |
| EPI_ISL_418245 | hCoV-19/Spain/CastillaLaMancha201328/2020 | Europe / Spain / Castilla La Mancha | 2020-03-01 | Hospital General y Universitario de Guadalajara | Instituto de Salud Carlos III | Iglesias-Caballero, M. Molinero Calamita, M. González-Esguevillas, M. Camarero, S. Pozo, F. Casas, I. Jiménez, P. Jiménez, M. Zaballos, A. Monzón, S. Varona, S. Juliá, M. Cuesta, I. Gonzalez-Praetorius A. |
| EPI_ISL_418246 | hCoV-19/Spain/CastillaLaMancha201329/2020 | Europe / Spain / Castilla-La Mancha | 2020-03-01 | Hospital General y Universitario de Guadalajara | Instituto de Salud Carlos III | Iglesias-Caballero, M. Molinero Calamita, M. González-Esguevillas, M. Camarero, S. Pozo, F. Casas, I. Jiménez, P. Jiménez, M. Zaballos, A. Monzón, S. Varona, S. Juliá, M. Cuesta, I. Gonzalez-Praetorius A. |
| EPI_ISL_418247 | hCoV-19/Spain/CastillayLeon201061/2020 | Europe / Spain / Castilla y Leon | 2020-02-26 | HOSPITAL GENERAL DE SEGOVIA | Instituto de Salud Carlos III | Iglesias-Caballero, M. Molinero Calamita, M. González-Esguevillas, M. Camarero, S. Pozo, F. Casas, I. Jiménez, P. Jiménez, M. Zaballos, A. Monzón, S. Varona, S. Juliá, M. Cuesta, I. Hernando-Real S. |
| EPI_ISL_418248 | hCoV-19/Spain/CastillayLeon201323/2020 | Europe / Spain / Castilla y Leon | 2020-03-01 | COMPLEJO ASISTENCIAL UNIVERSITARIO DE BURGOS | Instituto de Salud Carlos III | Iglesias-Caballero, M. Molinero Calamita, M. González-Esguevillas, M. Camarero, S. Pozo, F. Casas, I. Jiménez, P. Jiménez, M. Zaballos, A. Monzón, S. Varona, S. Juliá, M. Cuesta, I. Megias-Lobon G. |
| EPI_ISL_418249 | hCoV-19/Spain/CastillayLeon201372/2020 | Europe / Spain / Castilla y Leon | 2020-03-03 | COMPLEJO ASISTENCIAL UNIVERSITARIO DE BURGOS | Instituto de Salud Carlos III | Iglesias-Caballero, M. Molinero Calamita, M. González-Esguevillas, M. Camarero, S. Pozo, F. Casas, I. Jiménez, P. Jiménez, M. Zaballos, A. Monzón, S. Varona, S. Juliá, M. Cuesta, I. Megias-Lobon G. |
| EPI_ISL_418250 | hCoV-19/Spain/Cataluna201396/2020 | Europe / Spain / Catalonia | 2020 | HOSPITAL CLINIC | Instituto de Salud Carlos III | Iglesias-Caballero, M. Molinero Calamita, M. González-Esguevillas, M. Camarero, S. Pozo, F. Casas, I. Jiménez, P. Jiménez, M. Zaballos, A. Monzón, S. Varona, S. Juliá, M. Cuesta, I. Marcos M.A |
| EPI_ISL_418251 | hCoV-19/Spain/Madrid201105/2020 | Europe / Spain / Madrid | 2020-02-25 | HOSPITAL UNIVERSITARIO LA PAZ | Instituto de Salud Carlos III | Iglesias-Caballero, M. Molinero Calamita, M. González-Esguevillas, M. Camarero, S. Pozo, F. Casas, I. Jiménez, P. Jiménez, M. Zaballos, A. Monzón, S. Varona, S. Juliá, M. Cuesta, I. Romero P. |
| EPI_ISL_418252 | hCoV-19/Spain/Madrid201449/2020 | Europe / Spain / Madrid | 2020-03-04 | FUNDACION JIMENEZ DIAZ | Instituto de Salud Carlos III | Iglesias-Caballero, M. Molinero Calamita, M. González-Esguevillas, M. Camarero, S. Pozo, F. Casas, I. Jiménez, P. Jiménez, M. Zaballos, A. Monzón, S. Varona, S. Juliá, M. Cuesta, I. Fernández Robles, R. |
| EPI_ISL_418253 | hCoV-19/Spain/PaisVasco201382/2020 | Europe / Spain / Basque Country | 2020-03-02 | HOSPITAL TXAGORRITXU | Instituto de Salud Carlos III | Iglesias-Caballero, M. Molinero Calamita, M. González-Esguevillas, M. Camarero, S. Pozo, F. Casas, I. Jiménez, P. Jiménez, M. Zaballos, A. Monzón, S. Varona, S. Juliá, M. Cuesta, I. Gomez-Gonzalez C. |
| EPI_ISL_418254 | hCoV-19/USA/NY-NYUMC21/2020 | North America / USA / New York / Nassa | 2020-03-15 | NYU Langone Health | Department of Pathology and Medicine, New York University School of Medicine | Margaret Black, John Cadley, Paolo Cotzia, John Chen, Dacia Dimartino, Xiaojun Feng, Adriana Heguy, Megan Hogan, Emily Huang, George Jour, Christian Marier, Matthew T. Maurano, Mark J. Mulligan, Peter Meyn, Jared Pinnell, Amy Rapkiewicz, Marie Samanovic-Golden, Antonio Serrano, Guomiao Shen, Matija Snuderl, Nick Vulpescu, Gael Westby, Paul Zappile |
| EPI_ISL_418255 | hCoV-19/Italy/TE4925/2020 | Europe / Italy / Abruzzo | 2020-03-14 | Presidio Ospedaliero "S. Spirito" - PESCARA | Istituto Zooprofilattico Sperimentale dell'Abruzzo e Molise "G. Caporale" | Lorusso A, Marcacci M, Cammà C, Monaco F, Puglia I, Di Pasquale A, Rinaldi A, Mangone I, Savini G |
| EPI_ISL_418256 | hCoV-19/Italy/TE4880/2020 | Europe / Italy / Abruzzo | 2020-03-14 | Ospedale "San Liberatore" di Atri | Istituto Zooprofilattico Sperimentale dell'Abruzzo e Molise "G. Caporale" | Lorusso A, Marcacci M, Di Domenico M, Puglia I, Curini V, Ancora M, Di Pasquale A, Rinaldi A, Mangone I, Cammà C, Savini G. |
| EPI_ISL_418257 | hCoV-19/Italy/TE5056/2020 | Europe / Italy / Abruzzo | 2020-03-17 | Ospedale Civile Giuseppe Mazzini, Teramo | Istituto Zooprofilattico Sperimentale dell'Abruzzo e Molise "G.Caporale" | Lorusso A, Marcacci M, Di Domenico M, Puglia I, Curini V, Ancora M, Di Pasquale A, Rinaldi A, Mangone I, Cammà C, Savini G. |
| EPI_ISL_418258 | hCoV-19/Italy/TE4953/2020 | Europe / Italy / Abruzzo | 2020-03-14 | Presidio ospedaliero "Santo Spirito" | Istituto Zooprofilattico Sperimentale dell'Abruzzo e Molise "G. Caporale" | Lorusso A, Marcacci M, Di Domenico M, Puglia I, Curini V, Ancora M, Di Pasquale A, Rinaldi A, Mangone I, Cammà C, Savini G. |
| EPI_ISL_418259 | hCoV-19/Italy/TE4959/2020 | Europe / Italy / Abruzzo | 2020-03-14 | Presidio ospedaliero "Santo Spirito" | Istituto Zooprofilattico Sperimentale dell'Abruzzo e Molise "G. Caporale" | Lorusso A, Marcacci M, Di Domenico M, Puglia I, Curini V, Ancora M, Di Pasquale A, Rinaldi A, Mangone I, Cammà C, Savini G. |
| EPI_ISL_418260 | hCoV-19/Italy/TE4836/2020 | Europe / Italy / Abruzzo | 2020-03-16 | Ospedale Civile Giuseppe Mazzini | Istituto Zooprofilattico Sperimentale dell'Abruzzo e Molise "G. Caporale" | Lorusso A, Marcacci M, Di Domenico M, Puglia I, Curini V, Ancora M, Di Pasquale A, Rinaldi A, Mangone I, Cammà C, Savini G. |
| EPI_ISL_418262 | hCoV-19/Colombia/Bogota78390/2020 | South America / Colombia / Bogota | 2020-03-06 | Instituto Nacional de Salud | Instituto Nacional de Salud Universidad Cooperativa de Colombia Instituto Alexander von Humboldt Imperial College-London London School of Hygiene & Tropical Medicine | Marcela Mercado-Reyes, Katherine Laiton-Donato, Diego A. Álvarez-Díaz, Carlos Franco-Muñoz, Jose A. Usme-Ciro, Gloria Puerto, Nicolas D. Franco-Sierra, Mailyn A.Gonzalez, Zulma M. Cucunubá, Christian Julian Villabona-Arenas, Liz Villabona-Arenas, Susey Echeverria, Astrid C. Flórez, Sergio Gomez Rangel, Luz Dary Rodriguez, Juliana Barbosa, Erika Ospitia, Diana Marcela Walteros-Acero, Nuno Rodrigues Faria, Martha Lucia Ospina Martinez |
| EPI_ISL_418263 | hCoV-19/Greece/10/2020 | Europe / Greece / Athens | 2020-03-18 | Laboratory of Microbiology, Department of Medicine, National and Kapodistrian University of Athens, Greece | Laboratory of Biology, Department of Medicine, Democritus University of Thrace, Greece | Maria Bampali, Elisavet Gatzidou, Nikolaos Dovrolis, Stavroula Veletza, Nikolaos Spanakis, Ioannis Karakasiliotis |
| EPI_ISL_418264 | hCoV-19/Greece/12/2020 | Europe / Greece / Athens | 2020-03-18 | Laboratory of Microbiology, Department of Medicine, National and Kapodistrian University of Athens, Greece | Laboratory of Biology, Department of Medicine, Democritus University of Thrace, Greece | Maria Bampali, Elisavet Gatzidou, Nikolaos Dovrolis, Stavroula Veletza, Nikolaos Spanakis, Ioannis Karakasiliotis |
| EPI_ISL_418265 | hCoV-19/Greece/16/2020 | Europe / Greece / Athens | 2020-03-18 | Laboratory of Microbiology, Department of Medicine, National and Kapodistrian University of Athens, Greece | Laboratory of Biology, Department of Medicine, Democritus University of Thrace, Greece | Maria Bampali, Elisavet Gatzidou, Nikolaos Dovrolis, Stavroula Veletza, Nikolaos Spanakis, Ioannis Karakasiliotis |
| EPI_ISL_418267 | hCoV-19/Vietnam/19-02S/2020 | Asia / Vietnam / Ho Chi Minh City | 2020-01-22 | unknown | Microbiology and Immunology department | Nguyen,H.T., Cao,T.M., Pham,H.T.T., Vu,N.P.H., Dao,M.H., Huynh,L.T.K., Nguyen,L.T., Nguyen,N.T., Nguyen,T.T.N., Nguyen,A.H., Luong,Q.C., Nguyen,T.V., Tran,K.C., Pham,Q.D., Tran,T., Hoang,C.Q., Nguyen,T.T., Le,H.Q., Phung,T.M., Vo,T.N.A., Nguyen,S.N., Pham,D.T., Nguyen,T.V. and Pham,L.T. |
| EPI_ISL_418269 | hCoV-19/Vietnam/19-01S/2020 | Asia / Vietnam / Ho Chi Minh City | 2020-01-22 | unknown | Microbiology and Immunology department | Cao,T.M., Nguyen,H.T., Pham,H.T.T., Vu,N.P.H., Dao,M.H., Huynh,L.T.K., Nguyen,L.T., Nguyen,N.T., Nguyen,T.T.N., Nguyen,A.H., Luong,Q.C., Nguyen,T.V., Tran,K.C., Pham,Q.D., Tran,T., Hoang,C.Q., Nguyen,T.T., Le,H.Q., Phung,T.M., Vo,T.N.A., Nguyen,S.N., Pham,D.T., Phan,L.T. and Nguyen,T.V. |
| EPI_ISL_418270 | hCoV-19/Belgium/JL-03044/2020 | Europe / Belgium / Antwerp | 2020-03-04 | KU Leuven, Clinical and Epidemiological Virology | KU Leuven, Clinical and Epidemiological Virology | Tony Wawina, Joan Martí-Carreras, Bert Vanmechelen, Piet Maes |
| EPI_ISL_418273 | hCoV-19/Switzerland/42169471/2020 | Europe / Switzerland / Basel-Stadt | 2020-02-28 | University Hospital Basel, Clinical Virology | University Hospital Basel, Labormedizin | Hirsch, H., Leuzinger, K., Seth-Smith, H., Mari, A., Roloff, T., Egli, A. |
| EPI_ISL_418275 | hCoV-19/Switzerland/42169310/2020 | Europe / Switzerland / Basel | 2020-02-27 | University Hospital Basel, Clinical Virology | University Hospital Basel, Clinical Bacteriology | Hirsch, H., Leuzinger, K., Seth-Smith, H., Mari, A., Roloff, T., Egli, A. |
| EPI_ISL_418278 | hCoV-19/Switzerland/42175075/2020 | Europe / Switzerland / Basel | 2020-03-03 | University Hospital Basel, Clinical Virology | University Hospital Basel, Clinical Bacteriology | Hirsch, H., Leuzinger, K., Seth-Smith, H., Mari, A., Roloff, T., Egli, A. |
| EPI_ISL_418281 | hCoV-19/Switzerland/42176216/2020 | Europe / Switzerland / Basel | 2020-03-04 | University Hospital Basel, Clinical Virology | University Hospital Basel, Clinical Bacteriology | Hirsch, H., Leuzinger, K., Seth-Smith, H., Mari, A., Roloff, T., Egli, A. |
| EPI_ISL_418286 | hCoV-19/England/SHEF-BFD63/2020 | Europe / United Kingdom / England / Sou | 2020-03-18 | Virology Department, Sheffield Teaching Hospitals NHS Foundation Trust | Department of Infection, Immunity and Cardiovascular Disease, The Florey Institute, The Medical School, University of Sheffield | Thushan de Silva, Matthew Parker, Adri Angyal, Rebecca Brown, Rachel Tucker, Paul Parsons, Danielle Groves, Alex Keeley, Dave Partridge, Matthew Wyles, Benjamin Lindsey, Mehmet Yavuz, Mohammad Raza, Cariad Evans |
| EPI_ISL_418287 | hCoV-19/England/SHEF-BFD72/2020 | Europe / United Kingdom / England / Sou | 2020-03-16 | Virology Department, Sheffield Teaching Hospitals NHS Foundation Trust | Department of Infection, Immunity and Cardiovascular Disease, The Florey Institute, The Medical School, University of Sheffield | Thushan de Silva, Matthew Parker, Adri Angyal, Rebecca Brown, Rachel Tucker, Paul Parsons, Danielle Groves, Alex Keeley, Dave Partridge, Matthew Wyles, Benjamin Lindsey, Mehmet Yavuz, Mohammad Raza, Cariad Evans |
| EPI_ISL_418288 | hCoV-19/England/SHEF-BFD81/2020 | Europe / United Kingdom / England / Sou | 2020-03-18 | Virology Department, Sheffield Teaching Hospitals NHS Foundation Trust | Department of Infection, Immunity and Cardiovascular Disease, The Florey Institute, The Medical School, University of Sheffield | Thushan de Silva, Matthew Parker, Adri Angyal, Rebecca Brown, Rachel Tucker, Paul Parsons, Danielle Groves, Alex Keeley, Dave Partridge, Matthew Wyles, Benjamin Lindsey, Mehmet Yavuz, Mohammad Raza, Cariad Evans |
| EPI_ISL_418289 | hCoV-19/England/SHEF-BFDAF/2020 | Europe / United Kingdom / England / Sou | 2020-03-18 | Virology Department, Sheffield Teaching Hospitals NHS Foundation Trust | Department of Infection, Immunity and Cardiovascular Disease, The Florey Institute, The Medical School, University of Sheffield | Thushan de Silva, Matthew Parker, Adri Angyal, Rebecca Brown, Rachel Tucker, Paul Parsons, Danielle Groves, Alex Keeley, Dave Partridge, Matthew Wyles, Benjamin Lindsey, Mehmet Yavuz, Mohammad Raza, Cariad Evans |
| EPI_ISL_418290 | hCoV-19/England/SHEF-BFDBE/2020 | Europe / United Kingdom / England / Sou | 2020-03-18 | Virology Department, Sheffield Teaching Hospitals NHS Foundation Trust | Department of Infection, Immunity and Cardiovascular Disease, The Florey Institute, The Medical School, University of Sheffield | Thushan de Silva, Matthew Parker, Adri Angyal, Rebecca Brown, Rachel Tucker, Paul Parsons, Danielle Groves, Alex Keeley, Dave Partridge, Matthew Wyles, Benjamin Lindsey, Mehmet Yavuz, Mohammad Raza, Cariad Evans |
| EPI_ISL_418291 | hCoV-19/England/SHEF-BFDCD/2020 | Europe / United Kingdom / England / Sou | 2020-03-16 | Virology Department, Sheffield Teaching Hospitals NHS Foundation Trust | Department of Infection, Immunity and Cardiovascular Disease, The Florey Institute, The Medical School, University of Sheffield | Thushan de Silva, Matthew Parker, Adri Angyal, Rebecca Brown, Rachel Tucker, Paul Parsons, Danielle Groves, Alex Keeley, Dave Partridge, Matthew Wyles, Benjamin Lindsey, Mehmet Yavuz, Mohammad Raza, Cariad Evans |
| EPI_ISL_418292 | hCoV-19/England/SHEF-BFDDC/2020 | Europe / United Kingdom / England / Sou | 2020-03-18 | Virology Department, Sheffield Teaching Hospitals NHS Foundation Trust | Department of Infection, Immunity and Cardiovascular Disease, The Florey Institute, The Medical School, University of Sheffield | Thushan de Silva, Matthew Parker, Adri Angyal, Rebecca Brown, Rachel Tucker, Paul Parsons, Danielle Groves, Alex Keeley, Dave Partridge, Matthew Wyles, Benjamin Lindsey, Mehmet Yavuz, Mohammad Raza, Cariad Evans |
| EPI_ISL_418293 | hCoV-19/England/SHEF-BFDEB/2020 | Europe / United Kingdom / England / Sou | 2020-03-16 | Virology Department, Sheffield Teaching Hospitals NHS Foundation Trust | Department of Infection, Immunity and Cardiovascular Disease, The Florey Institute, The Medical School, University of Sheffield | Thushan de Silva, Matthew Parker, Adri Angyal, Rebecca Brown, Rachel Tucker, Paul Parsons, Danielle Groves, Alex Keeley, Dave Partridge, Matthew Wyles, Benjamin Lindsey, Mehmet Yavuz, Mohammad Raza, Cariad Evans |
| EPI_ISL_418294 | hCoV-19/England/SHEF-BFDFA/2020 | Europe / United Kingdom / England / Sou | 2020-03-18 | Virology Department, Sheffield Teaching Hospitals NHS Foundation Trust | Department of Infection, Immunity and Cardiovascular Disease, The Florey Institute, The Medical School, University of Sheffield | Thushan de Silva, Matthew Parker, Adri Angyal, Rebecca Brown, Rachel Tucker, Paul Parsons, Danielle Groves, Alex Keeley, Dave Partridge, Matthew Wyles, Benjamin Lindsey, Mehmet Yavuz, Mohammad Raza, Cariad Evans |
| EPI_ISL_418295 | hCoV-19/England/SHEF-BFE06/2020 | Europe / United Kingdom / England / Sou | 2020-03-17 | Virology Department, Sheffield Teaching Hospitals NHS Foundation Trust | Department of Infection, Immunity and Cardiovascular Disease, The Florey Institute, The Medical School, University of Sheffield | Thushan de Silva, Matthew Parker, Adri Angyal, Rebecca Brown, Rachel Tucker, Paul Parsons, Danielle Groves, Alex Keeley, Dave Partridge, Matthew Wyles, Benjamin Lindsey, Mehmet Yavuz, Mohammad Raza, Cariad Evans |
| EPI_ISL_418296 | hCoV-19/England/SHEF-BFE15/2020 | Europe / United Kingdom / England / Sou | 2020-03-18 | Virology Department, Sheffield Teaching Hospitals NHS Foundation Trust | Department of Infection, Immunity and Cardiovascular Disease, The Florey Institute, The Medical School, University of Sheffield | Thushan de Silva, Matthew Parker, Adri Angyal, Rebecca Brown, Rachel Tucker, Paul Parsons, Danielle Groves, Alex Keeley, Dave Partridge, Matthew Wyles, Benjamin Lindsey, Mehmet Yavuz, Mohammad Raza, Cariad Evans |
| EPI_ISL_418297 | hCoV-19/England/SHEF-BFE24/2020 | Europe / United Kingdom / England / Sou | 2020-03-17 | Virology Department, Sheffield Teaching Hospitals NHS Foundation Trust | Department of Infection, Immunity and Cardiovascular Disease, The Florey Institute, The Medical School, University of Sheffield | Thushan de Silva, Matthew Parker, Adri Angyal, Rebecca Brown, Rachel Tucker, Paul Parsons, Danielle Groves, Alex Keeley, Dave Partridge, Matthew Wyles, Benjamin Lindsey, Mehmet Yavuz, Mohammad Raza, Cariad Evans |
| EPI_ISL_418298 | hCoV-19/England/SHEF-BFE33/2020 | Europe / United Kingdom / England / Sou | 2020-03-16 | Virology Department, Sheffield Teaching Hospitals NHS Foundation Trust | Department of Infection, Immunity and Cardiovascular Disease, The Florey Institute, The Medical School, University of Sheffield | Thushan de Silva, Matthew Parker, Adri Angyal, Rebecca Brown, Rachel Tucker, Paul Parsons, Danielle Groves, Alex Keeley, Dave Partridge, Matthew Wyles, Benjamin Lindsey, Mehmet Yavuz, Mohammad Raza, Cariad Evans |
| EPI_ISL_418299 | hCoV-19/England/SHEF-BFE42/2020 | Europe / United Kingdom / England / Sou | 2020-03-18 | Virology Department, Sheffield Teaching Hospitals NHS Foundation Trust | Department of Infection, Immunity and Cardiovascular Disease, The Florey Institute, The Medical School, University of Sheffield | Thushan de Silva, Matthew Parker, Adri Angyal, Rebecca Brown, Rachel Tucker, Paul Parsons, Danielle Groves, Alex Keeley, Dave Partridge, Matthew Wyles, Benjamin Lindsey, Mehmet Yavuz, Mohammad Raza, Cariad Evans |
| EPI_ISL_418350 | hCoV-19/Canada/ON_PHL3380/2020 | North America / Canada / Ontario | 2020-03-20 | Public Health Ontario Laboratories | Public Health Ontario Laboratories | Alireza Eshaghi, Samir N Patel, Jonathan B Gubbay, Vanessa G Allen, Christine Frantz, Aimin Li, Sandeep Nagra |
| EPI_ISL_418351 | hCoV-19/Canada/ON_PHL3575/2020 | North America / Canada / Ontario | 2020-03-12 | Public Health Ontario Laboratories | Public Health Ontario Laboratories | Alireza Eshaghi, Samir N Patel, Jonathan B Gubbay, Vanessa G Allen, Christine Frantz, Aimin Li, Sandeep Nagra |
| EPI_ISL_418352 | hCoV-19/Canada/ON_PHL5672/2020 | North America / Canada / Ontario | 2020-03-11 | Public Health Ontario Laboratories | Public Health Ontario Laboratories | Alireza Eshaghi, Samir N Patel, Jonathan B Gubbay, Vanessa G Allen, Christine Frantz, Aimin Li, Sandeep Nagra |
| EPI_ISL_418353 | hCoV-19/Canada/ON_PHL0539/2020 | North America / Canada / Ontario | 2020-03-09 | Public Health Ontario Laboratories | Public Health Ontario Laboratories | Alireza Eshaghi, Samir N Patel, Jonathan B Gubbay, Vanessa G Allen, Christine Frantz, Aimin Li, Sandeep Nagra |
| EPI_ISL_418354 | hCoV-19/Canada/ON_PHL4088/2020 | North America / Canada / Ontario | 2020-03-15 | Public Health Ontario Laboratories | Public Health Ontario Laboratories | Alireza Eshaghi, Samir N Patel, Jonathan B Gubbay, Vanessa G Allen, Christine Frantz, Aimin Li, Sandeep Nagra |
| EPI_ISL_418355 | hCoV-19/Canada/ON_PHLU8150/2020 | North America / Canada / Ontario | 2020-03-08 | Public Health Ontario Laboratories | Public Health Ontario Laboratories | Alireza Eshaghi, Samir N Patel, Jonathan B Gubbay, Vanessa G Allen, Christine Frantz, Aimin Li, Sandeep Nagra |
| EPI_ISL_418356 | hCoV-19/Canada/ON_PHL6980/2020 | North America / Canada / Ontario | 2020-03-12 | Public Health Ontario Laboratories | Public Health Ontario Laboratories | Alireza Eshaghi, Samir N Patel, Jonathan B Gubbay, Vanessa G Allen, Christine Frantz, Aimin Li, Sandeep Nagra |
| EPI_ISL_418358 | hCoV-19/Canada/ON_PHL5756/2020 | North America / Canada / Ontario | 2020-03-12 | Public Health Ontario Laboratories | Public Health Ontario Laboratories | Alireza Eshaghi, Samir N Patel, Jonathan B Gubbay, Vanessa G Allen, Christine Frantz, Aimin Li, Sandeep Nagra |
| EPI_ISL_418359 | hCoV-19/Canada/ON_PHL5757/2020 | North America / Canada / Ontario | 2020-03-12 | Public Health Ontario Laboratories | Public Health Ontario Laboratories | Alireza Eshaghi, Samir N Patel, Jonathan B Gubbay, Vanessa G Allen, Christine Frantz, Aimin Li, Sandeep Nagra |
| EPI_ISL_418624 | hCoV-19/Belgium/ULG-6972/2020 | Europe / Belgium / Liège | 2020-03-15 | Department of Clinical Microbiology | GIGA Medical Genomics | Keith Durkin, Maria Artesi, Sébastien Bontems, Raphaël Boreux, Cécile Meex, Pierrette Melin, Marie-Pierre Hayette, Vincent Bours. |
| EPI_ISL_418625 | hCoV-19/Belgium/ULG-6688/2020 | Europe / Belgium / Liège | 2020-03-14 | Department of Clinical Microbiology | GIGA Medical Genomics | Keith Durkin, Maria Artesi, Sébastien Bontems, Raphaël Boreux, Cécile Meex, Pierrette Melin, Marie-Pierre Hayette, Vincent Bours. |
| EPI_ISL_418626 | hCoV-19/Belgium/ULG-7242/2020 | Europe / Belgium / Liège | 2020-03-16 | Department of Clinical Microbiology | GIGA Medical Genomics | Keith Durkin, Maria Artesi, Sébastien Bontems, Raphaël Boreux, Cécile Meex, Pierrette Melin, Marie-Pierre Hayette, Vincent Bours. |
| EPI_ISL_418627 | hCoV-19/Belgium/ULG-6743/2020 | Europe / Belgium / Liège | 2020-03-14 | Department of Clinical Microbiology | GIGA Medical Genomics | Keith Durkin, Maria Artesi, Sébastien Bontems, Raphaël Boreux, Cécile Meex, Pierrette Melin, Marie-Pierre Hayette, Vincent Bours. |
| EPI_ISL_418628 | hCoV-19/Belgium/ULG-7134/2020 | Europe / Belgium / Liège | 2020-03-16 | Department of Clinical Microbiology | GIGA Medical Genomics | Keith Durkin, Maria Artesi, Sébastien Bontems, Raphaël Boreux, Cécile Meex, Pierrette Melin, Marie-Pierre Hayette, Vincent Bours. |
| EPI_ISL_418629 | hCoV-19/Belgium/ULG-7773/2020 | Europe / Belgium / Liège | 2020-03-17 | Department of Clinical Microbiology | GIGA Medical Genomics | Keith Durkin, Maria Artesi, Sébastien Bontems, Raphaël Boreux, Cécile Meex, Pierrette Melin, Marie-Pierre Hayette, Vincent Bours. |
| EPI_ISL_418630 | hCoV-19/Belgium/ULG-7729/2020 | Europe / Belgium / Liège | 2020-03-17 | Department of Clinical Microbiology | GIGA Medical Genomics | Keith Durkin, Maria Artesi, Sébastien Bontems, Raphaël Boreux, Cécile Meex, Pierrette Melin, Marie-Pierre Hayette, Vincent Bours. |
| EPI_ISL_418631 | hCoV-19/Belgium/ULG-6649/2020 | Europe / Belgium / Liège | 2020-03-14 | Department of Clinical Microbiology | GIGA Medical Genomics | Keith Durkin, Maria Artesi, Sébastien Bontems, Raphaël Boreux, Cécile Meex, Pierrette Melin, Marie-Pierre Hayette, Vincent Bours. |
| EPI_ISL_418632 | hCoV-19/Belgium/ULG-7500/2020 | Europe / Belgium / Liège | 2020-03-16 | Department of Clinical Microbiology | GIGA Medical Genomics | Keith Durkin, Maria Artesi, Sébastien Bontems, Raphaël Boreux, Cécile Meex, Pierrette Melin, Marie-Pierre Hayette, Vincent Bours. |
| EPI_ISL_418633 | hCoV-19/Belgium/ULG-7626/2020 | Europe / Belgium / Liège | 2020-03-17 | Department of Clinical Microbiology | GIGA Medical Genomics | Keith Durkin, Maria Artesi, Sébastien Bontems, Raphaël Boreux, Cécile Meex, Pierrette Melin, Marie-Pierre Hayette, Vincent Bours. |
| EPI_ISL_418634 | hCoV-19/Belgium/ULG-8595/2020 | Europe / Belgium / Liège | 2020-03-19 | Department of Clinical Microbiology | GIGA Medical Genomics | Keith Durkin, Maria Artesi, Sébastien Bontems, Raphaël Boreux, Cécile Meex, Pierrette Melin, Marie-Pierre Hayette, Vincent Bours. |
| EPI_ISL_418635 | hCoV-19/Belgium/ULG-8532/2020 | Europe / Belgium / Liège | 2020-03-19 | Department of Clinical Microbiology | GIGA Medical Genomics | Keith Durkin, Maria Artesi, Sébastien Bontems, Raphaël Boreux, Cécile Meex, Pierrette Melin, Marie-Pierre Hayette, Vincent Bours. |
| EPI_ISL_418636 | hCoV-19/Belgium/ULG-8541/2020 | Europe / Belgium / Liège | 2020-03-19 | Department of Clinical Microbiology | GIGA Medical Genomics | Keith Durkin, Maria Artesi, Sébastien Bontems, Raphaël Boreux, Cécile Meex, Pierrette Melin, Marie-Pierre Hayette, Vincent Bours. |
| EPI_ISL_418638 | hCoV-19/Belgium/ULG-8578/2020 | Europe / Belgium / Liège | 2020-03-19 | Department of Clinical Microbiology | GIGA Medical Genomics | Keith Durkin, Maria Artesi, Sébastien Bontems, Raphaël Boreux, Cécile Meex, Pierrette Melin, Marie-Pierre Hayette, Vincent Bours. |
| EPI_ISL_418639 | hCoV-19/Belgium/ULG-8617/2020 | Europe / Belgium / Liège | 2020-03-19 | Department of Clinical Microbiology | GIGA Medical Genomics | Keith Durkin, Maria Artesi, Sébastien Bontems, Raphaël Boreux, Cécile Meex, Pierrette Melin, Marie-Pierre Hayette, Vincent Bours. |
| EPI_ISL_418640 | hCoV-19/Belgium/ULG-8634/2020 | Europe / Belgium / Liège | 2020-03-19 | Department of Clinical Microbiology | GIGA Medical Genomics | Keith Durkin, Maria Artesi, Sébastien Bontems, Raphaël Boreux, Cécile Meex, Pierrette Melin, Marie-Pierre Hayette, Vincent Bours. |
| EPI_ISL_418645 | hCoV-19/Belgium/ULG-9558/2020 | Europe / Belgium / Liège | 2020-03-21 | Department of Clinical Microbiology | GIGA Medical Genomics | Keith Durkin, Maria Artesi, Sébastien Bontems, Raphaël Boreux, Cécile Meex, Pierrette Melin, Marie-Pierre Hayette, Vincent Bours. |
| EPI_ISL_418646 | hCoV-19/Belgium/ULG-9572/2020 | Europe / Belgium / Liège | 2020-03-21 | Department of Clinical Microbiology | GIGA Medical Genomics | Keith Durkin, Maria Artesi, Sébastien Bontems, Raphaël Boreux, Cécile Meex, Pierrette Melin, Marie-Pierre Hayette, Vincent Bours. |
| EPI_ISL_418647 | hCoV-19/Belgium/ULG-9617/2020 | Europe / Belgium / Liège | 2020-03-22 | Department of Clinical Microbiology | GIGA Medical Genomics | Keith Durkin, Maria Artesi, Sébastien Bontems, Raphaël Boreux, Cécile Meex, Pierrette Melin, Marie-Pierre Hayette, Vincent Bours. |
| EPI_ISL_418648 | hCoV-19/Belgium/ULG-9618/2020 | Europe / Belgium / Liège | 2020-03-22 | Department of Clinical Microbiology | GIGA Medical Genomics | Keith Durkin, Maria Artesi, Sébastien Bontems, Raphaël Boreux, Cécile Meex, Pierrette Melin, Marie-Pierre Hayette, Vincent Bours. |
| EPI_ISL_418649 | hCoV-19/Belgium/ULG-9619/2020 | Europe / Belgium / Liège | 2020-03-22 | Department of Clinical Microbiology | GIGA Medical Genomics | Keith Durkin, Maria Artesi, Sébastien Bontems, Raphaël Boreux, Cécile Meex, Pierrette Melin, Marie-Pierre Hayette, Vincent Bours. |
| EPI_ISL_418650 | hCoV-19/Belgium/ULG-9620/2020 | Europe / Belgium / Liège | 2020-03-22 | Department of Clinical Microbiology | GIGA Medical Genomics | Keith Durkin, Maria Artesi, Sébastien Bontems, Raphaël Boreux, Cécile Meex, Pierrette Melin, Marie-Pierre Hayette, Vincent Bours. |
| EPI_ISL_418651 | hCoV-19/Belgium/ULG-9634/2020 | Europe / Belgium / Liège | 2020-03-22 | Department of Clinical Microbiology | GIGA Medical Genomics | Keith Durkin, Maria Artesi, Sébastien Bontems, Raphaël Boreux, Cécile Meex, Pierrette Melin, Marie-Pierre Hayette, Vincent Bours. |
| EPI_ISL_418652 | hCoV-19/Belgium/ULG-9641/2020 | Europe / Belgium / Liège | 2020-03-22 | Department of Clinical Microbiology | GIGA Medical Genomics | Keith Durkin, Maria Artesi, Sébastien Bontems, Raphaël Boreux, Cécile Meex, Pierrette Melin, Marie-Pierre Hayette, Vincent Bours. |
| EPI_ISL_418653 | hCoV-19/Belgium/ULG-9647/2020 | Europe / Belgium / Liège | 2020-03-22 | Department of Clinical Microbiology | GIGA Medical Genomics | Keith Durkin, Maria Artesi, Sébastien Bontems, Raphaël Boreux, Cécile Meex, Pierrette Melin, Marie-Pierre Hayette, Vincent Bours. |
| EPI_ISL_418654 | hCoV-19/Belgium/ULG-9694/2020 | Europe / Belgium / Liège | 2020-03-22 | Department of Clinical Microbiology | GIGA Medical Genomics | Keith Durkin, Maria Artesi, Sébastien Bontems, Raphaël Boreux, Cécile Meex, Pierrette Melin, Marie-Pierre Hayette, Vincent Bours. |
| EPI_ISL_418655 | hCoV-19/Belgium/ULG-9714/2020 | Europe / Belgium / Liège | 2020-03-22 | Department of Clinical Microbiology | GIGA Medical Genomics | Keith Durkin, Maria Artesi, Sébastien Bontems, Raphaël Boreux, Cécile Meex, Pierrette Melin, Marie-Pierre Hayette, Vincent Bours. |
| EPI_ISL_418656 | hCoV-19/Belgium/ULG-9715/2020 | Europe / Belgium / Liège | 2020-03-22 | Department of Clinical Microbiology | GIGA Medical Genomics | Keith Durkin, Maria Artesi, Sébastien Bontems, Raphaël Boreux, Cécile Meex, Pierrette Melin, Marie-Pierre Hayette, Vincent Bours. |
| EPI_ISL_418657 | hCoV-19/Belgium/ULG-9716/2020 | Europe / Belgium / Liège | 2020-03-22 | Department of Clinical Microbiology | GIGA Medical Genomics | Keith Durkin, Maria Artesi, Sébastien Bontems, Raphaël Boreux, Cécile Meex, Pierrette Melin, Marie-Pierre Hayette, Vincent Bours. |
| EPI_ISL_418658 | hCoV-19/Belgium/ULG-9719/2020 | Europe / Belgium / Liège | 2020-03-22 | Department of Clinical Microbiology | GIGA Medical Genomics | Keith Durkin, Maria Artesi, Sébastien Bontems, Raphaël Boreux, Cécile Meex, Pierrette Melin, Marie-Pierre Hayette, Vincent Bours. |
| EPI_ISL_418659 | hCoV-19/Belgium/ULG-9725/2020 | Europe / Belgium / Liège | 2020-03-22 | Department of Clinical Microbiology | GIGA Medical Genomics | Keith Durkin, Maria Artesi, Sébastien Bontems, Raphaël Boreux, Cécile Meex, Pierrette Melin, Marie-Pierre Hayette, Vincent Bours. |
| EPI_ISL_418660 | hCoV-19/Belgium/ULG-9732/2020 | Europe / Belgium / Liège | 2020-03-22 | Department of Clinical Microbiology | GIGA Medical Genomics | Keith Durkin, Maria Artesi, Sébastien Bontems, Raphaël Boreux, Cécile Meex, Pierrette Melin, Marie-Pierre Hayette, Vincent Bours. |
| EPI_ISL_418661 | hCoV-19/Belgium/ULG-9735/2020 | Europe / Belgium / Liège | 2020-03-22 | Department of Clinical Microbiology | GIGA Medical Genomics | Keith Durkin, Maria Artesi, Sébastien Bontems, Raphaël Boreux, Cécile Meex, Pierrette Melin, Marie-Pierre Hayette, Vincent Bours. |
| EPI_ISL_418663 | hCoV-19/Belgium/ULG-9739/2020 | Europe / Belgium / Liège | 2020-03-22 | Department of Clinical Microbiology | GIGA Medical Genomics | Keith Durkin, Maria Artesi, Sébastien Bontems, Raphaël Boreux, Cécile Meex, Pierrette Melin, Marie-Pierre Hayette, Vincent Bours. |
| EPI_ISL_418664 | hCoV-19/Belgium/ULG-9741/2020 | Europe / Belgium / Liège | 2020-03-22 | Department of Clinical Microbiology | GIGA Medical Genomics | Keith Durkin, Maria Artesi, Sébastien Bontems, Raphaël Boreux, Cécile Meex, Pierrette Melin, Marie-Pierre Hayette, Vincent Bours. |
| EPI_ISL_418667 | hCoV-19/England/20109047803/2020 | Europe / United Kingdom / England | 2020-03-07 | Respiratory Virus Unit, Microbiology Services Colindale, Public Health England | Respiratory Virus Unit, Microbiology Services Colindale, Public Health England | Monica Galiano, Shahjahan Miah, Angie Lackenby, Omolola Akinbami, Tiina Talts, Leena Bhaw, Richard Myers, Steven Platt, Kirstin Edwards, Jonathan Hubb, Joanna Ellis, Maria Zambon |
| EPI_ISL_418668 | hCoV-19/England/20110136302/2020 | Europe / United Kingdom / England | 2020-03-09 | Respiratory Virus Unit, Microbiology Services Colindale, Public Health England | Respiratory Virus Unit, Microbiology Services Colindale, Public Health England | Monica Galiano, Shahjahan Miah, Angie Lackenby, Omolola Akinbami, Tiina Talts, Leena Bhaw, Richard Myers, Steven Platt, Kirstin Edwards, Jonathan Hubb, Joanna Ellis, Maria Zambon |
| EPI_ISL_418669 | hCoV-19/England/20116022902/2020 | Europe / United Kingdom / England | 2020-03-11 | Respiratory Virus Unit, Microbiology Services Colindale, Public Health England | Respiratory Virus Unit, Microbiology Services Colindale, Public Health England | Monica Galiano, Shahjahan Miah, Angie Lackenby, Omolola Akinbami, Tiina Talts, Leena Bhaw, Richard Myers, Steven Platt, Kirstin Edwards, Jonathan Hubb, Joanna Ellis, Maria Zambon |
| EPI_ISL_418670 | hCoV-19/England/20116025603/2020 | Europe / United Kingdom / England | 2020-03-11 | Respiratory Virus Unit, Microbiology Services Colindale, Public Health England | Respiratory Virus Unit, Microbiology Services Colindale, Public Health England | Monica Galiano, Shahjahan Miah, Angie Lackenby, Omolola Akinbami, Tiina Talts, Leena Bhaw, Richard Myers, Steven Platt, Kirstin Edwards, Jonathan Hubb, Joanna Ellis, Maria Zambon |
| EPI_ISL_418671 | hCoV-19/England/20119026102/2020 | Europe / United Kingdom / England | 2020-03-13 | Respiratory Virus Unit, Microbiology Services Colindale, Public Health England | Respiratory Virus Unit, Microbiology Services Colindale, Public Health England | Monica Galiano, Shahjahan Miah, Angie Lackenby, Omolola Akinbami, Tiina Talts, Leena Bhaw, Richard Myers, Steven Platt, Kirstin Edwards, Jonathan Hubb, Joanna Ellis, Maria Zambon |
| EPI_ISL_418672 | hCoV-19/England/20120002102/2020 | Europe / United Kingdom / England | 2020-03-15 | Respiratory Virus Unit, Microbiology Services Colindale, Public Health England | Respiratory Virus Unit, Microbiology Services Colindale, Public Health England | Monica Galiano, Shahjahan Miah, Angie Lackenby, Omolola Akinbami, Tiina Talts, Leena Bhaw, Richard Myers, Steven Platt, Kirstin Edwards, Jonathan Hubb, Joanna Ellis, Maria Zambon |
| EPI_ISL_418673 | hCoV-19/England/20122032802/2020 | Europe / United Kingdom / England | 2020-03-16 | Respiratory Virus Unit, Microbiology Services Colindale, Public Health England | Respiratory Virus Unit, Microbiology Services Colindale, Public Health England | Monica Galiano, Shahjahan Miah, Angie Lackenby, Omolola Akinbami, Tiina Talts, Leena Bhaw, Richard Myers, Steven Platt, Kirstin Edwards, Jonathan Hubb, Joanna Ellis, Maria Zambon |
| EPI_ISL_418675 | hCoV-19/England/20122073102/2020 | Europe / United Kingdom / England | 2020-03-16 | Respiratory Virus Unit, Microbiology Services Colindale, Public Health England | Respiratory Virus Unit, Microbiology Services Colindale, Public Health England | Monica Galiano, Shahjahan Miah, Angie Lackenby, Omolola Akinbami, Tiina Talts, Leena Bhaw, Richard Myers, Steven Platt, Kirstin Edwards, Jonathan Hubb, Joanna Ellis, Maria Zambon |
| EPI_ISL_418676 | hCoV-19/England/20122074602/2020 | Europe / United Kingdom / England | 2020-03-10 | Respiratory Virus Unit, Microbiology Services Colindale, Public Health England | Respiratory Virus Unit, Microbiology Services Colindale, Public Health England | Monica Galiano, Shahjahan Miah, Angie Lackenby, Omolola Akinbami, Tiina Talts, Leena Bhaw, Richard Myers, Steven Platt, Kirstin Edwards, Jonathan Hubb, Joanna Ellis, Maria Zambon |
| EPI_ISL_418677 | hCoV-19/England/20122087702/2020 | Europe / United Kingdom / England | 2020-03-15 | Respiratory Virus Unit, Microbiology Services Colindale, Public Health England | Respiratory Virus Unit, Microbiology Services Colindale, Public Health England | Monica Galiano, Shahjahan Miah, Angie Lackenby, Omolola Akinbami, Tiina Talts, Leena Bhaw, Richard Myers, Steven Platt, Kirstin Edwards, Jonathan Hubb, Joanna Ellis, Maria Zambon |
| EPI_ISL_418678 | hCoV-19/England/20122088102/2020 | Europe / United Kingdom / England | 2020-03-16 | Respiratory Virus Unit, Microbiology Services Colindale, Public Health England | Respiratory Virus Unit, Microbiology Services Colindale, Public Health England | Monica Galiano, Shahjahan Miah, Angie Lackenby, Omolola Akinbami, Tiina Talts, Leena Bhaw, Richard Myers, Steven Platt, Kirstin Edwards, Jonathan Hubb, Joanna Ellis, Maria Zambon |
| EPI_ISL_418679 | hCoV-19/England/20122119202/2020 | Europe / United Kingdom / England | 2020-03-17 | Respiratory Virus Unit, Microbiology Services Colindale, Public Health England | Respiratory Virus Unit, Microbiology Services Colindale, Public Health England | Monica Galiano, Shahjahan Miah, Angie Lackenby, Omolola Akinbami, Tiina Talts, Leena Bhaw, Richard Myers, Steven Platt, Kirstin Edwards, Jonathan Hubb, Joanna Ellis, Maria Zambon |
| EPI_ISL_418680 | hCoV-19/England/20122119302/2020 | Europe / United Kingdom / England | 2020-03-17 | Respiratory Virus Unit, Microbiology Services Colindale, Public Health England | Respiratory Virus Unit, Microbiology Services Colindale, Public Health England | Monica Galiano, Shahjahan Miah, Angie Lackenby, Omolola Akinbami, Tiina Talts, Leena Bhaw, Richard Myers, Steven Platt, Kirstin Edwards, Jonathan Hubb, Joanna Ellis, Maria Zambon |
| EPI_ISL_418681 | hCoV-19/England/20122119502/2020 | Europe / United Kingdom / England | 2020-03-17 | Respiratory Virus Unit, Microbiology Services Colindale, Public Health England | Respiratory Virus Unit, Microbiology Services Colindale, Public Health England | Monica Galiano, Shahjahan Miah, Angie Lackenby, Omolola Akinbami, Tiina Talts, Leena Bhaw, Richard Myers, Steven Platt, Kirstin Edwards, Jonathan Hubb, Joanna Ellis, Maria Zambon |
| EPI_ISL_418684 | hCoV-19/England/20124001302/2020 | Europe / United Kingdom / England | 2020-03-17 | Respiratory Virus Unit, Microbiology Services Colindale, Public Health England | Respiratory Virus Unit, Microbiology Services Colindale, Public Health England | Monica Galiano, Shahjahan Miah, Angie Lackenby, Omolola Akinbami, Tiina Talts, Leena Bhaw, Richard Myers, Steven Platt, Kirstin Edwards, Jonathan Hubb, Joanna Ellis, Maria Zambon |
| EPI_ISL_418685 | hCoV-19/England/20124001702/2020 | Europe / United Kingdom / England | 2020-03-17 | Respiratory Virus Unit, Microbiology Services Colindale, Public Health England | Respiratory Virus Unit, Microbiology Services Colindale, Public Health England | Monica Galiano, Shahjahan Miah, Angie Lackenby, Omolola Akinbami, Tiina Talts, Leena Bhaw, Richard Myers, Steven Platt, Kirstin Edwards, Jonathan Hubb, Joanna Ellis, Maria Zambon |
| EPI_ISL_418686 | hCoV-19/England/20124001902/2020 | Europe / United Kingdom / England | 2020-03-17 | Respiratory Virus Unit, Microbiology Services Colindale, Public Health England | Respiratory Virus Unit, Microbiology Services Colindale, Public Health England | Monica Galiano, Shahjahan Miah, Angie Lackenby, Omolola Akinbami, Tiina Talts, Leena Bhaw, Richard Myers, Steven Platt, Kirstin Edwards, Jonathan Hubb, Joanna Ellis, Maria Zambon |
| EPI_ISL_418687 | hCoV-19/England/20124002002/2020 | Europe / United Kingdom / England | 2020-03-17 | Respiratory Virus Unit, Microbiology Services Colindale, Public Health England | Respiratory Virus Unit, Microbiology Services Colindale, Public Health England | Monica Galiano, Shahjahan Miah, Angie Lackenby, Omolola Akinbami, Tiina Talts, Leena Bhaw, Richard Myers, Steven Platt, Kirstin Edwards, Jonathan Hubb, Joanna Ellis, Maria Zambon |
| EPI_ISL_418688 | hCoV-19/England/20124002102/2020 | Europe / United Kingdom / England | 2020-03-17 | Respiratory Virus Unit, Microbiology Services Colindale, Public Health England | Respiratory Virus Unit, Microbiology Services Colindale, Public Health England | Monica Galiano, Shahjahan Miah, Angie Lackenby, Omolola Akinbami, Tiina Talts, Leena Bhaw, Richard Myers, Steven Platt, Kirstin Edwards, Jonathan Hubb, Joanna Ellis, Maria Zambon |
| EPI_ISL_418690 | hCoV-19/England/20124003702/2020 | Europe / United Kingdom / England | 2020-03-17 | Respiratory Virus Unit, Microbiology Services Colindale, Public Health England | Respiratory Virus Unit, Microbiology Services Colindale, Public Health England | Monica Galiano, Shahjahan Miah, Angie Lackenby, Omolola Akinbami, Tiina Talts, Leena Bhaw, Richard Myers, Steven Platt, Kirstin Edwards, Jonathan Hubb, Joanna Ellis, Maria Zambon |
| EPI_ISL_418691 | hCoV-19/England/20124003902/2020 | Europe / United Kingdom / England | 2020-03-17 | Respiratory Virus Unit, Microbiology Services Colindale, Public Health England | Respiratory Virus Unit, Microbiology Services Colindale, Public Health England | Monica Galiano, Shahjahan Miah, Angie Lackenby, Omolola Akinbami, Tiina Talts, Leena Bhaw, Richard Myers, Steven Platt, Kirstin Edwards, Jonathan Hubb, Joanna Ellis, Maria Zambon |
| EPI_ISL_418692 | hCoV-19/England/20124004302/2020 | Europe / United Kingdom / England | 2020-03-17 | Respiratory Virus Unit, Microbiology Services Colindale, Public Health England | Respiratory Virus Unit, Microbiology Services Colindale, Public Health England | Monica Galiano, Shahjahan Miah, Angie Lackenby, Omolola Akinbami, Tiina Talts, Leena Bhaw, Richard Myers, Steven Platt, Kirstin Edwards, Jonathan Hubb, Joanna Ellis, Maria Zambon |

| | | | | | | |
|---|---|---|---|---|---|---|
| EPI_ISL_418894 | hCoV-19/England/20124008902/2020 | Europe / United Kingdom / England | 2020-03-16 | Respiratory Virus Unit, Microbiology Services Colindale, Public Health England | Respiratory Virus Unit, Microbiology Services Colindale, Public Health England | Monica Galiano, Shahjahan Miah, Angie Lackenby, Omolola Akinbami, Tiina Talts, Leena Bhaw, Richard Myers, Steven Platt, Kirstin Edwards, Jonathan Hubb, Joanna Ellis, Maria Zambon |
| EPI_ISL_418896 | hCoV-19/England/20124009602/2020 | Europe / United Kingdom / England | 2020-03-18 | Respiratory Virus Unit, Microbiology Services Colindale, Public Health England | Respiratory Virus Unit, Microbiology Services Colindale, Public Health England | Monica Galiano, Shahjahan Miah, Angie Lackenby, Omolola Akinbami, Tiina Talts, Leena Bhaw, Richard Myers, Steven Platt, Kirstin Edwards, Jonathan Hubb, Joanna Ellis, Maria Zambon |
| EPI_ISL_418898 | hCoV-19/England/20124012702/2020 | Europe / United Kingdom / England | 2020-03-17 | Respiratory Virus Unit, Microbiology Services Colindale, Public Health England | Respiratory Virus Unit, Microbiology Services Colindale, Public Health England | Monica Galiano, Shahjahan Miah, Angie Lackenby, Omolola Akinbami, Tiina Talts, Leena Bhaw, Richard Myers, Steven Platt, Kirstin Edwards, Jonathan Hubb, Joanna Ellis, Maria Zambon |
| EPI_ISL_418900 | hCoV-19/USA/WA-UW333/2020 | North America / USA / Washington | 2020-03-16 | UW Virology Lab | UW Virology Lab | Pavitra Roychoudhury, Hong Xie, Keith Jerome, Alexander Greninger |
| EPI_ISL_418901 | hCoV-19/USA/MN-UW334/2020 | North America / USA / Minnesota | 2020-03-13 | UW Virology Lab | UW Virology Lab | Pavitra Roychoudhury, Hong Xie, Keith Jerome, Alexander Greninger |
| EPI_ISL_418902 | hCoV-19/USA/MN-UW335/2020 | North America / USA / Minnesota | 2020-03-13 | UW Virology Lab | UW Virology Lab | Pavitra Roychoudhury, Hong Xie, Keith Jerome, Alexander Greninger |
| EPI_ISL_418903 | hCoV-19/USA/WA-UW336/2020 | North America / USA / Washington | 2020-03-16 | UW Virology Lab | UW Virology Lab | Pavitra Roychoudhury, Hong Xie, Keith Jerome, Alexander Greninger |
| EPI_ISL_418904 | hCoV-19/USA/WA-UW337/2020 | North America / USA / Washington | 2020-03-16 | UW Virology Lab | UW Virology Lab | Pavitra Roychoudhury, Hong Xie, Keith Jerome, Alexander Greninger |
| EPI_ISL_418905 | hCoV-19/USA/WA-UW338/2020 | North America / USA / Washington | 2020-03-16 | UW Virology Lab | UW Virology Lab | Pavitra Roychoudhury, Hong Xie, Keith Jerome, Alexander Greninger |
| EPI_ISL_418906 | hCoV-19/USA/WA-UW339/2020 | North America / USA / Washington | 2020-03-16 | UW Virology Lab | UW Virology Lab | Pavitra Roychoudhury, Hong Xie, Keith Jerome, Alexander Greninger |
| EPI_ISL_418907 | hCoV-19/USA/WA-UW340/2020 | North America / USA / Washington | 2020-03-16 | UW Virology Lab | UW Virology Lab | Pavitra Roychoudhury, Hong Xie, Keith Jerome, Alexander Greninger |
| EPI_ISL_418908 | hCoV-19/USA/WA-UW341/2020 | North America / USA / Washington | 2020-03-16 | UW Virology Lab | UW Virology Lab | Pavitra Roychoudhury, Hong Xie, Keith Jerome, Alexander Greninger |
| EPI_ISL_418909 | hCoV-19/USA/WA-UW342/2020 | North America / USA / Washington | 2020-03-16 | UW Virology Lab | UW Virology Lab | Pavitra Roychoudhury, Hong Xie, Keith Jerome, Alexander Greninger |
| EPI_ISL_418911 | hCoV-19/USA/WA-UW344/2020 | North America / USA / Washington | 2020-03-16 | UW Virology Lab | UW Virology Lab | Pavitra Roychoudhury, Hong Xie, Keith Jerome, Alexander Greninger |
| EPI_ISL_418912 | hCoV-19/USA/WA-UW345/2020 | North America / USA / Washington | 2020-03-16 | UW Virology Lab | UW Virology Lab | Pavitra Roychoudhury, Hong Xie, Keith Jerome, Alexander Greninger |
| EPI_ISL_418913 | hCoV-19/USA/WA-UW346/2020 | North America / USA / Washington | 2020-03-16 | UW Virology Lab | UW Virology Lab | Pavitra Roychoudhury, Hong Xie, Keith Jerome, Alexander Greninger |
| EPI_ISL_418915 | hCoV-19/USA/IL-UW348/2020 | North America / USA / Illinois | 2020-03-13 | UW Virology Lab | UW Virology Lab | Pavitra Roychoudhury, Hong Xie, Keith Jerome, Alexander Greninger |
| EPI_ISL_418916 | hCoV-19/USA/IL-UW349/2020 | North America / USA / Illinois | 2020-03-13 | UW Virology Lab | UW Virology Lab | Pavitra Roychoudhury, Hong Xie, Keith Jerome, Alexander Greninger |
| EPI_ISL_418917 | hCoV-19/USA/IL-UW350/2020 | North America / USA / Illinois | 2020-03-14 | UW Virology Lab | UW Virology Lab | Pavitra Roychoudhury, Hong Xie, Keith Jerome, Alexander Greninger |
| EPI_ISL_418918 | hCoV-19/USA/IL-UW351/2020 | North America / USA / Illinois | 2020-03-13 | UW Virology Lab | UW Virology Lab | Pavitra Roychoudhury, Hong Xie, Keith Jerome, Alexander Greninger |
| EPI_ISL_418919 | hCoV-19/USA/IL-UW352/2020 | North America / USA / Illinois | 2020-03-13 | UW Virology Lab | UW Virology Lab | Pavitra Roychoudhury, Hong Xie, Keith Jerome, Alexander Greninger |
| EPI_ISL_418920 | hCoV-19/USA/IL-UW353/2020 | North America / USA / Illinois | 2020-03-13 | UW Virology Lab | UW Virology Lab | Pavitra Roychoudhury, Hong Xie, Keith Jerome, Alexander Greninger |
| EPI_ISL_418921 | hCoV-19/USA/UN-UW354/2020 | North America / USA | 2020-03-16 | UW Virology Lab | UW Virology Lab | Pavitra Roychoudhury, Hong Xie, Keith Jerome, Alexander Greninger |
| EPI_ISL_418922 | hCoV-19/USA/WA-UW355/2020 | North America / USA / Washington | 2020-03-16 | UW Virology Lab | UW Virology Lab | Pavitra Roychoudhury, Hong Xie, Keith Jerome, Alexander Greninger |
| EPI_ISL_418923 | hCoV-19/USA/WA-UW356/2020 | North America / USA / Washington | 2020-03-16 | UW Virology Lab | UW Virology Lab | Pavitra Roychoudhury, Hong Xie, Keith Jerome, Alexander Greninger |
| EPI_ISL_418924 | hCoV-19/USA/WA-UW357/2020 | North America / USA / Washington | 2020-03-14 | UW Virology Lab | UW Virology Lab | Pavitra Roychoudhury, Hong Xie, Keith Jerome, Alexander Greninger |
| EPI_ISL_418925 | hCoV-19/USA/UN-UW358/2020 | North America / USA | 2020-03 | UW Virology Lab | UW Virology Lab | Pavitra Roychoudhury, Hong Xie, Keith Jerome, Alexander Greninger |
| EPI_ISL_418926 | hCoV-19/USA/WA-UW359/2020 | North America / USA / Washington | 2020-03-14 | UW Virology Lab | UW Virology Lab | Pavitra Roychoudhury, Hong Xie, Keith Jerome, Alexander Greninger |
| EPI_ISL_418927 | hCoV-19/USA/WA-UW360/2020 | North America / USA / Washington | 2020-03-16 | UW Virology Lab | UW Virology Lab | Pavitra Roychoudhury, Hong Xie, Keith Jerome, Alexander Greninger |
| EPI_ISL_418928 | hCoV-19/USA/WA-UW361/2020 | North America / USA / Washington | 2020-03-15 | UW Virology Lab | UW Virology Lab | Pavitra Roychoudhury, Hong Xie, Keith Jerome, Alexander Greninger |
| EPI_ISL_418929 | hCoV-19/USA/WA-UW362/2020 | North America / USA / Washington | 2020-03-17 | UW Virology Lab | UW Virology Lab | Pavitra Roychoudhury, Hong Xie, Keith Jerome, Alexander Greninger |
| EPI_ISL_418930 | hCoV-19/USA/WA-UW363/2020 | North America / USA / Washington | 2020-03-17 | UW Virology Lab | UW Virology Lab | Pavitra Roychoudhury, Hong Xie, Keith Jerome, Alexander Greninger |
| EPI_ISL_418931 | hCoV-19/USA/WA-UW364/2020 | North America / USA / Washington | 2020-03-17 | UW Virology Lab | UW Virology Lab | Pavitra Roychoudhury, Hong Xie, Keith Jerome, Alexander Greninger |
| EPI_ISL_418932 | hCoV-19/USA/WA-UW365/2020 | North America / USA / Washington | 2020-03-16 | UW Virology Lab | UW Virology Lab | Pavitra Roychoudhury, Hong Xie, Keith Jerome, Alexander Greninger |
| EPI_ISL_418933 | hCoV-19/USA/WA-UW366/2020 | North America / USA / Washington | 2020-03-17 | UW Virology Lab | UW Virology Lab | Pavitra Roychoudhury, Hong Xie, Keith Jerome, Alexander Greninger |
| EPI_ISL_418934 | hCoV-19/USA/WA-UW367/2020 | North America / USA / Washington | 2020-03-17 | UW Virology Lab | UW Virology Lab | Pavitra Roychoudhury, Hong Xie, Keith Jerome, Alexander Greninger |
| EPI_ISL_418935 | hCoV-19/USA/WA-UW368/2020 | North America / USA / Washington | 2020-03-16 | UW Virology Lab | UW Virology Lab | Pavitra Roychoudhury, Hong Xie, Keith Jerome, Alexander Greninger |
| EPI_ISL_418936 | hCoV-19/USA/WA-UW369/2020 | North America / USA / Washington | 2020-03-17 | UW Virology Lab | UW Virology Lab | Pavitra Roychoudhury, Hong Xie, Keith Jerome, Alexander Greninger |
| EPI_ISL_418937 | hCoV-19/USA/WA-UW370/2020 | North America / USA / Washington | 2020-03-17 | UW Virology Lab | UW Virology Lab | Pavitra Roychoudhury, Hong Xie, Keith Jerome, Alexander Greninger |
| EPI_ISL_418938 | hCoV-19/USA/WA-UW371/2020 | North America / USA / Washington | 2020-03-16 | UW Virology Lab | UW Virology Lab | Pavitra Roychoudhury, Hong Xie, Keith Jerome, Alexander Greninger |
| EPI_ISL_418939 | hCoV-19/USA/WA-UW372/2020 | North America / USA / Washington | 2020-03-17 | UW Virology Lab | UW Virology Lab | Pavitra Roychoudhury, Hong Xie, Keith Jerome, Alexander Greninger |
| EPI_ISL_418940 | hCoV-19/USA/WA-UW373/2020 | North America / USA / Washington | 2020-03-17 | UW Virology Lab | UW Virology Lab | Pavitra Roychoudhury, Hong Xie, Keith Jerome, Alexander Greninger |
| EPI_ISL_418941 | hCoV-19/USA/WA-UW374/2020 | North America / USA | 2020-03 | UW Virology Lab | UW Virology Lab | Pavitra Roychoudhury, Hong Xie, Keith Jerome, Alexander Greninger |
| EPI_ISL_418942 | hCoV-19/USA/WA-UW375/2020 | North America / USA / Washington | 2020-03-17 | UW Virology Lab | UW Virology Lab | Pavitra Roychoudhury, Hong Xie, Keith Jerome, Alexander Greninger |
| EPI_ISL_418943 | hCoV-19/USA/WA-UW376/2020 | North America / USA / Washington | 2020-03-17 | UW Virology Lab | UW Virology Lab | Pavitra Roychoudhury, Hong Xie, Keith Jerome, Alexander Greninger |
| EPI_ISL_418944 | hCoV-19/USA/WA-UW378/2020 | North America / USA / Washington | 2020-03-15 | UW Virology Lab | UW Virology Lab | Pavitra Roychoudhury, Hong Xie, Keith Jerome, Alexander Greninger |
| EPI_ISL_418945 | hCoV-19/USA/WA-UW379/2020 | North America / USA / Washington | 2020-03-15 | UW Virology Lab | UW Virology Lab | Pavitra Roychoudhury, Hong Xie, Keith Jerome, Alexander Greninger |
| EPI_ISL_418948 | hCoV-19/USA/WA-UW383/2020 | North America / USA / Washington | 2020-03-16 | UW Virology Lab | UW Virology Lab | Pavitra Roychoudhury, Hong Xie, Keith Jerome, Alexander Greninger |
| EPI_ISL_418949 | hCoV-19/USA/WA-UW384/2020 | North America / USA / Washington | 2020-03-16 | UW Virology Lab | UW Virology Lab | Pavitra Roychoudhury, Hong Xie, Keith Jerome, Alexander Greninger |
| EPI_ISL_418950 | hCoV-19/USA/WA-UW385/2020 | North America / USA / Washington | 2020-03-16 | UW Virology Lab | UW Virology Lab | Pavitra Roychoudhury, Hong Xie, Keith Jerome, Alexander Greninger |
| EPI_ISL_418951 | hCoV-19/USA/WA-UW386/2020 | North America / USA / Washington | 2020-03-16 | UW Virology Lab | UW Virology Lab | Pavitra Roychoudhury, Hong Xie, Keith Jerome, Alexander Greninger |
| EPI_ISL_418952 | hCoV-19/USA/WA-UW387/2020 | North America / USA / Washington | 2020-03-16 | UW Virology Lab | UW Virology Lab | Pavitra Roychoudhury, Hong Xie, Keith Jerome, Alexander Greninger |
| EPI_ISL_418953 | hCoV-19/USA/WA-UW388/2020 | North America / USA / Washington | 2020-03-16 | UW Virology Lab | UW Virology Lab | Pavitra Roychoudhury, Hong Xie, Keith Jerome, Alexander Greninger |
| EPI_ISL_418954 | hCoV-19/USA/WA-UW389/2020 | North America / USA / Washington | 2020-03-17 | UW Virology Lab | UW Virology Lab | Pavitra Roychoudhury, Hong Xie, Keith Jerome, Alexander Greninger |
| EPI_ISL_418955 | hCoV-19/USA/WA-UW390/2020 | North America / USA / Washington | 2020-03-15 | UW Virology Lab | UW Virology Lab | Pavitra Roychoudhury, Hong Xie, Keith Jerome, Alexander Greninger |
| EPI_ISL_418956 | hCoV-19/USA/VA-DCLS-0002/2020 | North America / USA / Virginia | 2020-03-17 | Virginia DCLS | Virginia DCLS | Virginia DCLS |
| EPI_ISL_418957 | hCoV-19/USA/VA-DCLS-0003/2020 | North America / USA / Virginia | 2020-03-17 | Virginia DCLS | Virginia DCLS | Virginia DCLS |
| EPI_ISL_418958 | hCoV-19/USA/VA-DCLS-0004/2020 | North America / USA / Virginia | 2020-03-17 | Virginia DCLS | Virginia DCLS | Virginia DCLS |
| EPI_ISL_418962 | hCoV-19/USA/UT-019/2020 | North America / USA / Utah | 2020-03-13 | Utah Public Health Laboratory | Utah Public Health Laboratory | Erin Young, Kelly Oakeson |
| EPI_ISL_418963 | hCoV-19/USA/UT-021/2020 | North America / USA / Utah | 2020-03-14 | Utah Public Health Laboratory | Utah Public Health Laboratory | Erin Young, Kelly Oakeson |
| EPI_ISL_418964 | hCoV-19/USA/UT-022/2020 | North America / USA / Utah | 2020-03-14 | Utah Public Health Laboratory | Utah Public Health Laboratory | Erin Young, Kelly Oakeson |
| EPI_ISL_418965 | hCoV-19/USA/UT-023/2020 | North America / USA / Utah | 2020-03-11 | Utah Public Health Laboratory | Utah Public Health Laboratory | Erin Young, Kelly Oakeson |
| EPI_ISL_418967 | hCoV-19/USA/UT-025/2020 | North America / USA / Utah | 2020-03-11 | Utah Public Health Laboratory | Utah Public Health Laboratory | Erin Young, Kelly Oakeson |
| EPI_ISL_418968 | hCoV-19/USA/NY-NYUMC22/2020 | North America / USA / New York / Manha | 2020-03-15 | NYU Langone Health | Department of Pathology and Medicine, New York University School of Medicine | Maria Aguero-Rosenfeld, Margaret Black, John Cadley, Paolo Cotzia, John Chen, Dacia Dimartino, Xiaojun Feng, Adriana Heguy, Megan Hogan, Emily Huang, George Jour, Christian Marier, Matthew T. Maurano, Mark J. Mulligan, Peter Meyn, Jared Pinnell, Sitharam Ramaswami, Amy Rapkiewicz, Marie Samanovic-Golden, Antonio Serrano, Guomiao Shen, Matija Snuderl, Nick Vulpescu, Gael Westby, Paul Zappile, Yutong Zhang |
| EPI_ISL_418969 | hCoV-19/USA/NY-NYUMC23/2020 | North America / USA / New York / Manha | 2020-03-14 | NYU Langone Health | Department of Pathology and Medicine, New York University School of Medicine | Maria Aguero-Rosenfeld, Margaret Black, John Cadley, Paolo Cotzia, John Chen, Dacia Dimartino, Xiaojun Feng, Adriana Heguy, Megan Hogan, Emily Huang, George Jour, Christian Marier, Matthew T. Maurano, Mark J. Mulligan, Peter Meyn, Jared Pinnell, Sitharam Ramaswami, Amy Rapkiewicz, Marie Samanovic-Golden, Antonio Serrano, Guomiao Shen, Matija Snuderl, Nick Vulpescu, Gael Westby, Paul Zappile, Yutong Zhang |
| EPI_ISL_418970 | hCoV-19/USA/NY-NYUMC24/2020 | North America / USA / New York / Nassa | 2020-03-16 | NYU Langone Health | Department of Pathology and Medicine, New York University School of Medicine | Maria Aguero-Rosenfeld, Margaret Black, John Cadley, Paolo Cotzia, John Chen, Dacia Dimartino, Xiaojun Feng, Adriana Heguy, Megan Hogan, Emily Huang, George Jour, Christian Marier, Matthew T. Maurano, Mark J. Mulligan, Peter Meyn, Jared Pinnell, Sitharam Ramaswami, Amy Rapkiewicz, Marie Samanovic-Golden, Antonio Serrano, Guomiao Shen, Matija Snuderl, Nick Vulpescu, Gael Westby, Paul Zappile, Yutong Zhang |
| EPI_ISL_418972 | hCoV-19/USA/NY-NYUMC26/2020 | North America / USA / New York / Manha | 2020-03-17 | NYU Langone Health | Department of Pathology and Medicine, New York University School of Medicine | Maria Aguero-Rosenfeld, Margaret Black, John Cadley, Paolo Cotzia, John Chen, Dacia Dimartino, Xiaojun Feng, Adriana Heguy, Megan Hogan, Emily Huang, George Jour, Christian Marier, Matthew T. Maurano, Mark J. Mulligan, Peter Meyn, Jared Pinnell, Sitharam Ramaswami, Amy Rapkiewicz, Marie Samanovic-Golden, Antonio Serrano, Guomiao Shen, Matija Snuderl, Nick Vulpescu, Gael Westby, Paul Zappile, Yutong Zhang |
| EPI_ISL_418973 | hCoV-19/USA/NY-NYUMC27/2020 | North America / USA / New York / Manha | 2020-03-17 | NYU Langone Health | Department of Pathology and Medicine, New York University School of Medicine | Maria Aguero-Rosenfeld, Margaret Black, John Cadley, Paolo Cotzia, John Chen, Dacia Dimartino, Xiaojun Feng, Adriana Heguy, Megan Hogan, Emily Huang, George Jour, Christian Marier, Matthew T. Maurano, Mark J. Mulligan, Peter Meyn, Jared Pinnell, Sitharam Ramaswami, Amy Rapkiewicz, Marie Samanovic-Golden, Antonio Serrano, Guomiao Shen, Matija Snuderl, Nick Vulpescu, Gael Westby, Paul Zappile, Yutong Zhang |
| EPI_ISL_418974 | hCoV-19/USA/NY-NYUMC28/2020 | North America / USA / New York / Manha | 2020-03-17 | NYU Langone Health | Department of Pathology and Medicine, New York University School of Medicine | Maria Aguero-Rosenfeld, Margaret Black, John Cadley, Paolo Cotzia, John Chen, Dacia Dimartino, Xiaojun Feng, Adriana Heguy, Megan Hogan, Emily Huang, George Jour, Christian Marier, Matthew T. Maurano, Mark J. Mulligan, Peter Meyn, Jared Pinnell, Sitharam Ramaswami, Amy Rapkiewicz, Marie Samanovic-Golden, Antonio Serrano, Guomiao Shen, Matija Snuderl, Nick Vulpescu, Gael Westby, Paul Zappile, Yutong Zhang |
| EPI_ISL_418975 | hCoV-19/USA/NY-NYUMC29/2020 | North America / USA / New York / Manha | 2020-03-17 | NYU Langone Health | Department of Pathology and Medicine, New York University School of Medicine | Maria Aguero-Rosenfeld, Margaret Black, John Cadley, Paolo Cotzia, John Chen, Dacia Dimartino, Xiaojun Feng, Adriana Heguy, Megan Hogan, Emily Huang, George Jour, Christian Marier, Matthew T. Maurano, Mark J. Mulligan, Peter Meyn, Jared Pinnell, Sitharam Ramaswami, Amy Rapkiewicz, Marie Samanovic-Golden, Antonio Serrano, Guomiao Shen, Matija Snuderl, Nick Vulpescu, Gael Westby, Paul Zappile, Yutong Zhang |
| EPI_ISL_418976 | hCoV-19/USA/NY-NYUMC30/2020 | North America / USA / New York / Manha | 2020-03-17 | NYU Langone Health | Department of Pathology and Medicine, New York University School of Medicine | Maria Aguero-Rosenfeld, Margaret Black, John Cadley, Paolo Cotzia, John Chen, Dacia Dimartino, Xiaojun Feng, Adriana Heguy, Megan Hogan, Emily Huang, George Jour, Christian Marier, Matthew T. Maurano, Mark J. Mulligan, Peter Meyn, Jared Pinnell, Sitharam Ramaswami, Amy Rapkiewicz, Marie Samanovic-Golden, Antonio Serrano, Guomiao Shen, Matija Snuderl, Nick Vulpescu, Gael Westby, Paul Zappile, Yutong Zhang |
| EPI_ISL_418977 | hCoV-19/USA/NY-NYUMC31/2020 | North America / USA / New York / Manha | 2020-03-17 | NYU Langone Health | Department of Pathology and Medicine, New York University School of Medicine | Maria Aguero-Rosenfeld, Margaret Black, John Cadley, Paolo Cotzia, John Chen, Dacia Dimartino, Xiaojun Feng, Adriana Heguy, Megan Hogan, Emily Huang, George Jour, Christian Marier, Matthew T. Maurano, Mark J. Mulligan, Peter Meyn, Jared Pinnell, Sitharam Ramaswami, Amy Rapkiewicz, Marie Samanovic-Golden, Antonio Serrano, Guomiao Shen, Matija Snuderl, Nick Vulpescu, Gael Westby, Paul Zappile, Yutong Zhang |
| EPI_ISL_418978 | hCoV-19/USA/NY-NYUMC32/2020 | North America / USA / New York / Manha | 2020-03-17 | NYU Langone Health | Department of Pathology and Medicine, New York University School of Medicine | Maria Aguero-Rosenfeld, Margaret Black, John Cadley, Paolo Cotzia, John Chen, Dacia Dimartino, Xiaojun Feng, Adriana Heguy, Megan Hogan, Emily Huang, George Jour, Christian Marier, Matthew T. Maurano, Mark J. Mulligan, Peter Meyn, Jared Pinnell, Sitharam Ramaswami, Amy Rapkiewicz, Marie Samanovic-Golden, Antonio Serrano, Guomiao Shen, Matija Snuderl, Nick Vulpescu, Gael Westby, Paul Zappile, Yutong Zhang |
| EPI_ISL_418979 | hCoV-19/USA/NY-NYUMC33/2020 | North America / USA / New York / Manha | 2020-03-17 | NYU Langone Health | Department of Pathology and Medicine, New York University School of Medicine | Maria Aguero-Rosenfeld, Margaret Black, John Cadley, Paolo Cotzia, John Chen, Dacia Dimartino, Xiaojun Feng, Adriana Heguy, Megan Hogan, Emily Huang, George Jour, Christian Marier, Matthew T. Maurano, Mark J. Mulligan, Peter Meyn, Jared Pinnell, Sitharam Ramaswami, Amy Rapkiewicz, Marie Samanovic-Golden, Antonio Serrano, Guomiao Shen, Matija Snuderl, Nick Vulpescu, Gael Westby, Paul Zappile, Yutong Zhang |
| EPI_ISL_405839 | hCoV-19/Shenzhen/HKU-SZ-005/2020 | Asia / China / Guangdong / Shenzhen | 2020-01-11 | The University of Hong Kong - Shenzhen Hospital | Li Ka Shing Faculty of Medicine, The University of Hong Kong | Chan,J.F.-W., Yuan,S., Kok,K.H., To,K.K.-W., Chu,H., Yang,J., Xing,F., Liu,J., Yip,C.C.-Y., Poon,R.W.-S., Tsai,H.W., Lo,S.K.-F., Chan,K.H., Poon,V.K.-M., Chan,W.M., Ip,J.D., Cai,J.P., Cheng,V.C.-C., Chen,H., Hui,C.K.-M. and Yuen,K.Y. |
| EPI_ISL_406716 | hCoV-19/Wuhan/WHU01/2020 | Asia / China / Hubei / Wuhan | 2020-01-02 | unknown | State Key Laboratory of Virology, Wuhan University | Chen,L., Liu,W., Zhang,Q., Xu,K., Ye,G., Wu,W., Sun,Z., Liu,F., Wu,K., Mei,Y., Zhang,W., Chen,Y., Li,Y., Shi,M., Lan,K. and Liu,Y. |
| EPI_ISL_406717 | hCoV-19/Wuhan/WHU02/2020 | Asia / China / Hubei / Wuhan | 2020-01-02 | unknown | State Key Laboratory of Virology, Wuhan University | Chen,L., Liu,W., Zhang,Q., Xu,K., Ye,G., Wu,W., Sun,Z., Liu,F., Wu,K., Mei,Y., Zhang,W., Chen,Y., Li,Y., Shi,M., Lan,K. and Liu,Y. |
| EPI_ISL_406798 | hCoV-19/Wuhan/WH01/2019 | Asia / China / Hubei / Wuhan | 2019-12-26 | General Hospital of Central Theater Command of People's Liberation Army of China | BGI & Institute of Microbiology, Chinese Academy of Sciences & Shandong First Medical University & Shandong Academy of Medical Sciences & General Hospital of Central Theater Command of People's Liberation Army of China | Weijun Chen, Yuhai Bi, Weifeng Shi and Zhenhong Hu |
| EPI_ISL_406799 | hCoV-19/Wuhan/WH02/2019 | Asia / China / Wuhan | 2019-12-31 | General Hospital of Central Theater Command of People's Liberation Army of China | BGI & Institute of Microbiology, Chinese Academy of Sciences & Shandong First Medical University & Shandong Academy of Medical Sciences & General Hospital of Central Theater Command of People's Liberation Army of China | Weijun Chen, Yuhai Bi, Weifeng Shi and Zhenhong Hu |
| EPI_ISL_407193 | hCoV-19/South Korea/KCDC03/2020 | Asia / South Korea / Gyeonggi-do | 2020-01-25 | Korea Centers for Disease Control & Prevention (KCDC) Center for Laboratory Control of Infectious Diseases Division of Viral Diseases | Korea Centers for Disease Control & Prevention (KCDC) Center for Laboratory Control of Infectious Diseases Division of Viral Diseases | Jeong-Min Kim, Yoon-Seok Chung, Namjoo Lee, Mi-Seon Kim, SangHee Woo, Hye-Joon Jo, Sehee Park, Heui Man Kim, Myung Guk Han |
| EPI_ISL_407214 | hCoV-19/USA/WA1-A12/2020 | North America / USA / Washington | 2020-01-25 | WA State Department of Health | Pathogen Discovery, Respiratory Viruses Branch, Division of Viral Diseases, Centers for Diseases Control and Prevention | Krista Queen, Azaibi Tamin, Jennifer Harcourt, Ying Tao, Clinton R. Paden, Jing Zhang, Yan Li, Anna Uehara, Xiaoyan Lu, Shifaq Kamili, Rashi Gautam, Haibin Wang, Janna' R. Murray, Susan I. Gerber, Stephen Lindstrom, Natalie Thornburg, Suxiang Tong |
| EPI_ISL_407215 | hCoV-19/USA/WA1-F6/2020 | North America / USA / Washington | 2020-01-25 | Washington State Department of Health | Pathogen Discovery, Respiratory Viruses Branch, Division of Viral Diseases, Centers for Diseases Control and Prevention | Krista Queen, Azaibi Tamin, Jennifer Harcourt, Ying Tao, Clinton R. Paden, Jing Zhang, Yan Li, Anna Uehara, Xiaoyan Lu, Shifaq Kamili, Rashi Gautam, Haibin Wang, Janna' R. Murray, Susan I. Gerber, Stephen Lindstrom, Natalie Thornburg, Suxiang Tong |
| EPI_ISL_408514 | hCoV-19/env/Wuhan/IVDC-HB-envF13-20/2020 | Asia / China / Hubei / Wuhan | 2020-01-01 | Institute of Viral Disease Control and Prevention, China CDC | Institute of Viral Disease Control and Prevention, China CDC | William J. Liu, Peipei Liu, Xiang Zhao, Peihua Niu, Yingze Zhao, Wenwen Lei, Ziqian Xu, Shumei Zou, Wei Zhen, Beiwei Ye, Mengjie Yang, Weifeng Shi, Roujian Lu, Wenjie Tan, Zhixiao Chen, Yuchao Wu, Juan Song, Weimin Zhou, Dayan Wang, Jun Han, Wenbo Xu, George F. Gao, Guizhen Wu |
| EPI_ISL_408515 | hCoV-19/env/Wuhan/IVDC-HB-envF13-21/2020 | Asia / China / Hubei / Wuhan | 2020-01-01 | Institute of Viral Disease Control and Prevention, China CDC | Institute of Viral Disease Control and Prevention, China CDC | William J. Liu, Peipei Liu, Xiang Zhao, Peihua Niu, Yingze Zhao, Wenwen Lei, Ziqian Xu, Shumei Zou, Wei Zhen, Beiwei Ye, Mengjie Yang, Weifeng Shi, Roujian Lu, Wenjie Tan, Zhixiao Chen, Yuchao Wu, Juan Song, Weimin Zhou, Dayan Wang, Jun Han, Wenbo Xu, George F. Gao, Guizhen Wu |
| EPI_ISL_408976 | hCoV-19/Australia/NSW02/2020 | Oceania / Australia / New South Wales / | 2020-01-22 | Centre for Infectious Diseases and Microbiology Laboratory Services | NSW Health Pathology - Institute of Clinical Pathology and Medical Research; Westmead Hospital; University of Sydney | Rockett R, Sadsad R, Eden J-S, Carter I, Rahman H, Holmes EC, O'Sullivan MV, Sintchenko V, Chen SC, Maddocks S, Kok J and Dwyer DE for the 2019-nCoV Study Group" |
| EPI_ISL_408977 | hCoV-19/Australia/NSW03/2020 | Oceania / Australia / New South Wales / | 2020-01-25 | Serology, Virology and OTDS Laboratories (SAViD), NSW Health Pathology Randwick | NSW Health Pathology - Institute of Clinical Pathology and Medical Research; Centre for Infectious Diseases and Microbiology Laboratory Services; Westmead Hospital; University of Sydney | Eden J-S, Carter I, Rahman H, Rawlinson W, Holmes EC, Rockett R, O'Sullivan MV, Sintchenko V, Chen SC, Maddocks S, Kok J and Dwyer DE for the 2019-nCoV Study Group" |
| EPI_ISL_408978 | hCoV-19/Wuhan/WH05/2020 | Asia / China / Hubei / Wuhan | 2020-02-07 | Wuhan Fourth Hospital | Beijing Genomics Institute (BGI) | Weijun Chen |
| EPI_ISL_411218 | hCoV-19/France/IDF0571/2020 | Europe / France / Ile-de-France / Paris | 2020-02-02 | Department of Infectious and Tropical Diseases, Bichat Claude Bernard Hospital, Paris | Laboratoire Virpath, CIRI U111, UCBL1, INSERM, CNRS, ENS Lyon | Olivier Terrier, Aurélien Traversier, Julien Fouret, Yazdan Yazdanpanah, Xavier Lescure, Catherine Legras-Lachuer, Alexandre Gaymard, Bruno Lina, Manuel Rosa-Calatrava |
| EPI_ISL_411219 | hCoV-19/France/IDF0386-islP1/2020 | Europe / France / Ile-de-France / Paris | 2020-01-28 | Department of Infectious and Tropical Diseases, Bichat Claude Bernard Hospital, Paris | Laboratoire Virpath, CIRI U111, UCBL1, INSERM, CNRS, ENS Lyon | Olivier Terrier, Aurélien Traversier, Julien Fouret, Yazdan Yazdanpanah, Xavier Lescure, Alexandre Gaymard, Bruno Lina, Manuel Rosa-Calatrava |
| EPI_ISL_411220 | hCoV-19/France/IDF0386-islP3/2020 | Europe / France / Ile-de-France / Paris | 2020-01-28 | Department of Infectious and Tropical Diseases, Bichat Claude Bernard Hospital, Paris | Laboratoire Virpath, CIRI U111, UCBL1, INSERM, CNRS, ENS Lyon | Olivier Terrier, Aurélien Traversier, Julien Fouret, Yazdan Yazdanpanah, Xavier Lescure, Alexandre Gaymard, Bruno Lina, Manuel Rosa-Calatrava |
| EPI_ISL_412116 | hCoV-19/England/09c/2020 | Europe / United Kingdom / England | 2020-02-09 | Respiratory Virus Unit, Microbiology Services Colindale, Public Health England | Respiratory Virus Unit, Microbiology Services Colindale, Public Health England | Monica Galiano, Shahjahan Miah, Angie Lackenby, Omolola Akinbami, Tiina Talts, Leena Bhaw, Richard Myers, Steven Platt, Kirstin Edwards, Jonathan Hubb, Joanna Ellis, Maria Zambon |
| EPI_ISL_413014 | hCoV-19/Canada/ON_PHL2445/2020 | North America / Canada / Ontario | 2020-01-25 | Public Health Ontario Laboratory | Ontario Agency for Health Protection and Promotion (OAHPP) | Alireza Eshaghi, Samir N Patel, Jonathan B Gubbay, Vanessa G Allen, Christine Frantz, Aimin Li, Sandeep Nagra |
| EPI_ISL_413015 | hCoV-19/Canada/ON_VIDO-01/2020 | North America / Canada / Ontario | 2020-01-23 | Public Health Ontario Laboratory | National Microbiology Laboratory | Shari Tyson, Anna Majer, Erika Landry, Morag Graham, Grace Seo, Philip Mabon, Natalie Knox, Adrian Zetner, Samira Mubareka, Rob Kozak, Jocelyne Lew, Darryl Falzarano, Gerdts Volker, Jonathan Gubbay, Stephanie Booth, Guillaume Poliquin, Tom Graefenhan, Matthew Gilmour, Nathalie Bastien, Yan Li, Timothy Booth |
| EPI_ISL_413016 | hCoV-19/Brazil/SPBR-02/2020 | South America / Brazil / Sao Paulo / Sao | 2020-02-28 | Hospital Israelita Albert Einstein | Instituto Adolfo Lutz, Interdisciplinary Procedures Center, Strategic Laboratory | Jaqueline Goes de Jesus, Claudio Tavares Sacchi, Fabiana Cristina Pereira dos Santos, Ingra Morales Claro, Flávia Cristina da Silva Sales, Claudia Regina Gonçalves, Joshua Quick, Maria do Carmo Sampaio Tavares Timenetsky, Nicholas James Loman, Andrew Rambaut, Ester Cerdeira Sabino, Nuno Rodrigues Faria |
| EPI_ISL_413017 | hCoV-19/South Korea/KUMC01/2020 | Asia / South Korea | 2020-02-06 | Department of Microbiology, Institute for Viral Diseases, College of Medicine, Korea University | Department of Microbiology, Institute for Viral Diseases, College of Medicine, Korea University | Changmin Kang, Joon-Yong Bae, Jungmin Lee, Heedo Park, Juyoung Cho, Jeonghun Kim, Gee eun Lee, Cui Chunguang, Kyeong-ryeol Shin, Dong Min Kim, Jin Il Kim, Man-Seong Park |
| EPI_ISL_413018 | hCoV-19/South Korea/KUMC02/2020 | Asia / South Korea | 2020-02-06 | Department of Microbiology, Institute for Viral Diseases, College of Medicine, Korea University | Department of Microbiology, Institute for Viral Diseases, College of Medicine, Korea University | Changmin Kang, Joon-Yong Bae, Jungmin Lee, Heedo Park, Juyoung Cho, Jeonghun Kim, Gee eun Lee, Cui Chunguang, Kyeong-ryeol Shin, Dong Min Kim, Jin Il Kim, Man-Seong Park |
| EPI_ISL_413019 | hCoV-19/Switzerland/1000477102/2020 | Europe / Switzerland / Zurich | 2020-02-26 | Department of Internal Medicine, Triemli Hospital | Institute of Medical Virology, University of Zurich | Stefan Schmutz, Maryam Zaheri, Verena Kufner, Patrick Redli, Fiona Steiner, Jon Huder, Riccarda Capaul, Andrea Zbinden, Jürg Böni, Michael Huber, Gerhard Eich, Alexandra Trkola |
| EPI_ISL_413020 | hCoV-19/Switzerland/1000477377/2020 | Europe / Switzerland / Zurich | 2020-02-27 | Department of Internal Medicine, Triemli Hospital | Institute of Medical Virology, University of Zurich | Stefan Schmutz, Maryam Zaheri, Verena Kufner, Patrick Redli, Fiona Steiner, Jon Huder, Riccarda Capaul, Andrea Zbinden, Jürg Böni, Michael Huber, Gerhard Eich, Alexandra Trkola |
| EPI_ISL_413021 | hCoV-19/Switzerland/1000477757/2020 | Europe / Switzerland / Zurich | 2020-02-29 | Klinik Hirslanden Zurich | Institute of Medical Virology, University of Zurich | Stefan Schmutz, Maryam Zaheri, Verena Kufner, Gabriela Zitlener, Patrick Redli, Fiona Steiner, Jon Huder, Riccarda Capaul, Andrea Zbinden, Jürg Böni, Michael Huber, Christian Ruef, Alexandra Trkola |
| EPI_ISL_413022 | hCoV-19/Switzerland/1000477796/2020 | Europe / Switzerland / Zurich | 2020-02-29 | Division of Infectious Diseases, University Hospital Zurich | Institute of Medical Virology, University of Zurich | Stefan Schmutz, Maryam Zaheri, Verena Kufner, Gabriela Zitlener, Patrick Redli, Fiona Steiner, Jon Huder, Riccarda Capaul, Andrea Zbinden, Jürg Böni, Michael Huber, Roberto Speck, Alexandra Trkola |
| EPI_ISL_413023 | hCoV-19/Switzerland/1000477797/2020 | Europe / Switzerland / Zurich | 2020-02-29 | Division of Infectious Diseases, University Hospital Zurich | Institute of Medical Virology, University of Zurich | Stefan Schmutz, Maryam Zaheri, Verena Kufner, Gabriela Zitlener, Patrick Redli, Fiona Steiner, Jon Huder, Riccarda Capaul, Andrea Zbinden, Jürg Böni, Michael Huber, Roberto Speck, Alexandra Trkola |
| EPI_ISL_413024 | hCoV-19/Switzerland/1000477806/2020 | Europe / Switzerland / Zurich | 2020-02-29 | Division of Infectious Diseases, University Hospital Zurich | Institute of Medical Virology, University of Zurich | Stefan Schmutz, Maryam Zaheri, Verena Kufner, Gabriela Zitlener, Patrick Redli, Fiona Steiner, Jon Huder, Riccarda Capaul, Andrea Zbinden, Jürg Böni, Michael Huber, Roberto Speck, Alexandra Trkola |
| EPI_ISL_413025 | hCoV-19/USA/WA3-UW1/2020 | North America / USA / Washington | 2020-02-27 | Harborview Medical Center | UW Virology Lab | Pavitra Roychoudhury, Arun Nalla, Hong Xie, Keith Jerome, Alexander Greninger |
| EPI_ISL_413455 | hCoV-19/USA/WA4-UW2/2020 | North America / USA / Washington | 2020-02-28 | Washington State Public Health Lab | University of Washington Virology Lab | Pavitra Roychoudhury, Arun Nalla, Hong Xie, Keith Jerome, Alexander Greninger |
| EPI_ISL_413456 | hCoV-19/USA/WA-S2/2020 | North America / USA / Washington / King | 2020-02-20 | Seattle Flu Study | Seattle Flu Study | Chu et al |
| EPI_ISL_413457 | hCoV-19/USA/WA6-UW3/2020 | North America / USA / Washington | 2020-02-29 | Washington State Public Health Lab | UW Virology Lab | Pavitra Roychoudhury, Arun Nalla, Hong Xie, Keith Jerome, Alexander Greninger |
| EPI_ISL_413458 | hCoV-19/USA/WA7-UW4/2020 | North America / USA / Washington | 2020-03-01 | Washington State Public Health Lab | UW Virology Lab | Pavitra Roychoudhury, Arun Nalla, Hong Xie, Keith Jerome, Alexander Greninger |
| EPI_ISL_413459 | hCoV-19/Japan/TK-20-31-3/2020 | Asia / Japan / Tokyo | 2020-02-20 | Department of Pathology, Toshima Hospital | Pathogen Genomics Center, National Institute of Infectious Diseases | Tsuyoshi Sekizuka, Kentaro Itokawa, Takuya Adachi, Masahiro Sano, Jun Yamazaki, Ippei Miyamoto, Haruka Nishioka, Ja-Mun Chong, Noriko Nakajima, Yuko Sato, Minoru Tobiume, Harutaka Katano, Tadaki Suzuki, Makoto Kuroda |
| EPI_ISL_413486 | hCoV-19/USA/WA8-UW5/2020 | North America / USA / Washington | 2020-03-01 | Valley Medical Center | University of Washington Virology Lab | Pavitra Roychoudhury, Arun Nalla, Hong Xie, Keith Jerome, Alexander Greninger |
| EPI_ISL_413487 | hCoV-19/USA/WA9-UW6/2020 | North America / USA / Washington | 2020-03-01 | Harborview Medical Center | University of Washington Virology Lab | Pavitra Roychoudhury, Arun Nalla, Hong Xie, Keith Jerome, Alexander Greninger |
| EPI_ISL_413488 | hCoV-19/Germany/NRW-01/2020 | Europe / Germany / North Rhine Westph | 2020-02-28 | Center of Medical Microbiology, Virology, and Hospital Hygiene, University of Duesseldorf | Center of Medical Microbiology, Virology, and Hospital Hygiene, University of Duesseldorf | Ortwin Adams, Marcel Andree, Alexander Dilthey, Torsten Feldt, Sandra Hauka, Torsten Houwaart, Björn-Erik Jensen, Detlef Kindgen-Milles, Malte Kohns Vasconcelos, Klaus Pfeffer, Tina Senff, Daniel Strelow, Jörg Timm, Andreas Walker, Tobias Wienemann |
| EPI_ISL_413489 | hCoV-19/Italy/UniSR1/2020 | Europe / Italy / Lombardy / Milan | 2020-03-03 | Laboratorio di Microbiologia e Virologia, Università Vita-Salute San Raffaele, Milano | Laboratorio di Microbiologia e Virologia, Università Vita-Salute San Raffaele, Milano | R.A Diotti, E. Criscuolo, M. Castelli, V. Caputo, R. Ferrarese, M. Sampaolo, E. Boeri, I. Negri, V. Amato, G. Lo Raso, C. Di Resta, R. Burioni, M. Clementi, N. Mancini & N. Clementi |
| EPI_ISL_413490 | hCoV-19/New Zealand/01/2020 | Oceania / New Zealand / Auckland | 2020-02-27 | Auckland Hospital | Institute of Environmental Science and Research (ESR) | Matt Storey, Xiaoyun Ren, Gary McAuliffe, Sally Roberts, Matthew Blakiston, Erasmus Smit, Lauren Jelly, Joep de Ligt |
| EPI_ISL_413850 | hCoV-19/Guangdong/GDFS2020054-P0005/2020 | Asia / China / Guangdong | 2020-02-10 | Guangdong Provincial Institution of Public Health, Guangdong Provincial Center for Disease Control and Prevention | Guangdong Provincial Institution of Public Health | Jing Lu, Louis du Plessis, Liu Zhe, Jiufeng Sun, Sarah François, Huifang Lin, Moritz Kraemer, Jingju Peng, Qianlin Xiong, Runyu Yuan, Lilian Zeng, Pingping Zhou, Chuming Liang, Tao Liu, Wei Li, Juan Su, Huanying Zheng, Kang Min, Song Tie, Bo Peng, Shisong Fang, Wenzhe Su, Kuibiao Li, Ruilin Sun, Ru bai, Xi Tang, Minfeng Liang, Nuno Faria, Josh Quick, Andrew Rambaut, Verity Hill, Wenjun Ma, Nick Loman, Oliver Pybus, Changwen Ke |
| EPI_ISL_413851 | hCoV-19/Guangdong/2020XN4373-P0039/2020 | Asia / China / Guangdong | 2020-01-30 | Guangdong Provincial Institution of Public Health, Guangdong Provincial Center for Disease Control and Prevention | Guangdong Provincial Institution of Public Health | Jing Lu, Louis du Plessis, Liu Zhe, Jiufeng Sun, Sarah François, Huifang Lin, Moritz Kraemer, Jingju Peng, Qianlin Xiong, Runyu Yuan, Lilian Zeng, Pingping Zhou, Chuming Liang, Tao Liu, Wei Li, Juan Su, Huanying Zheng, Kang Min, Song Tie, Bo Peng, Shisong Fang, Wenzhe Su, Kuibiao Li, Ruilin Sun, Ru bai, Xi Tang, Minfeng Liang, Nuno Faria, Josh Quick, Andrew Rambaut, Verity Hill, Wenjun Ma, Nick Loman, Oliver Pybus, Changwen Ke |
| EPI_ISL_413852 | hCoV-19/Guangdong/2020XN4433-P0040/2020 | Asia / China / Guangdong | 2020-01-30 | Guangdong Provincial Institution of Public Health, Guangdong Provincial Center for Disease Control and Prevention | Guangdong Provincial Institution of Public Health | Jing Lu, Louis du Plessis, Liu Zhe, Jiufeng Sun, Sarah François, Huifang Lin, Moritz Kraemer, Jingju Peng, Qianlin Xiong, Runyu Yuan, Lilian Zeng, Pingping Zhou, Chuming Liang, Tao Liu, Wei Li, Juan Su, Huanying Zheng, Kang Min, Song Tie, Bo Peng, Shisong Fang, Wenzhe Su, Kuibiao Li, Ruilin Sun, Ru bai, Xi Tang, Minfeng Liang, Nuno Faria, Josh Quick, Andrew Rambaut, Verity Hill, Wenjun Ma, Nick Loman, Oliver Pybus, Changwen Ke |
| EPI_ISL_413853 | hCoV-19/Guangdong/2020XN4243-P0035/2020 | Asia / China / Guangdong | 2020-01-30 | Guangdong Provincial Institution of Public Health, Guangdong Provincial Center for Disease Control and Prevention | Guangdong Provincial Institution of Public Health | Jing Lu, Louis du Plessis, Liu Zhe, Jiufeng Sun, Sarah François, Huifang Lin, Moritz Kraemer, Jingju Peng, Qianlin Xiong, Runyu Yuan, Lilian Zeng, Pingping Zhou, Chuming Liang, Tao Liu, Wei Li, Juan Su, Huanying Zheng, Kang Min, Song Tie, Bo Peng, Shisong Fang, Wenzhe Su, Kuibiao Li, Ruilin Sun, Ru bai, Xi Tang, Minfeng Liang, Nuno Faria, Josh Quick, Andrew Rambaut, Verity Hill, Wenjun Ma, Nick Loman, Oliver Pybus, Changwen Ke |
| EPI_ISL_413854 | hCoV-19/Guangdong/2020XN4475-P0042/2020 | Asia / China / Guangdong | 2020-01-30 | Guangdong Provincial Institution of Public Health, Guangdong Provincial Center for Disease Control and Prevention | Guangdong Provincial Institution of Public Health | Jing Lu, Louis du Plessis, Liu Zhe, Jiufeng Sun, Sarah François, Huifang Lin, Moritz Kraemer, Jingju Peng, Qianlin Xiong, Runyu Yuan, Lilian Zeng, Pingping Zhou, Chuming Liang, Tao Liu, Wei Li, Juan Su, Huanying Zheng, Kang Min, Song Tie, Bo Peng, Shisong Fang, Wenzhe Su, Kuibiao Li, Ruilin Sun, Ru bai, Xi Tang, Minfeng Liang, Nuno Faria, Josh Quick, Andrew Rambaut, Verity Hill, Wenjun Ma, Nick Loman, Oliver Pybus, Changwen Ke |
| EPI_ISL_413855 | hCoV-19/Guangdong/GD2020234-P0023/2020 | Asia / China / Guangdong | 2020-02-07 | Guangdong Provincial Institution of Public Health, Guangdong Provincial Center for Disease Control and Prevention | Guangdong Provincial Institution of Public Health | Jing Lu, Louis du Plessis, Liu Zhe, Jiufeng Sun, Sarah François, Huifang Lin, Moritz Kraemer, Jingju Peng, Qianlin Xiong, Runyu Yuan, Lilian Zeng, Pingping Zhou, Chuming Liang, Tao Liu, Wei Li, Juan Su, Huanying Zheng, Kang Min, Song Tie, Bo Peng, Shisong Fang, Wenzhe Su, Kuibiao Li, Ruilin Sun, Ru bai, Xi Tang, Minfeng Liang, Nuno Faria, Josh Quick, Andrew Rambaut, Verity Hill, Wenjun Ma, Nick Loman, Oliver Pybus, Changwen Ke |
| EPI_ISL_413856 | hCoV-19/Guangdong/GD2020233-P0027/2020 | Asia / China / Guangdong | 2020-02-07 | Guangdong Provincial Institution of Public Health, Guangdong Provincial Center for Disease Control and Prevention | Guangdong Provincial Institution of Public Health | Jing Lu, Louis du Plessis, Liu Zhe, Jiufeng Sun, Sarah François, Huifang Lin, Moritz Kraemer, Jingju Peng, Qianlin Xiong, Runyu Yuan, Lilian Zeng, Pingping Zhou, Chuming Liang, Tao Liu, Wei Li, Juan Su, Huanying Zheng, Kang Min, Song Tie, Bo Peng, Shisong Fang, Wenzhe Su, Kuibiao Li, Ruilin Sun, Ru bai, Xi Tang, Minfeng Liang, Nuno Faria, Josh Quick, Andrew Rambaut, Verity Hill, Wenjun Ma, Nick Loman, Oliver Pybus, Changwen Ke |
| EPI_ISL_413857 | hCoV-19/Guangdong/2020XN4448-P0002/2020 | Asia / China / Guangdong | 2020-01-31 | Guangdong Provincial Institution of Public Health, Guangdong Provincial Center for Disease Control and Prevention | Guangdong Provincial Institution of Public Health | Jing Lu, Louis du Plessis, Liu Zhe, Jiufeng Sun, Sarah François, Huifang Lin, Moritz Kraemer, Jingju Peng, Qianlin Xiong, Runyu Yuan, Lilian Zeng, Pingping Zhou, Chuming Liang, Tao Liu, Wei Li, Juan Su, Huanying Zheng, Kang Min, Song Tie, Bo Peng, Shisong Fang, Wenzhe Su, Kuibiao Li, Ruilin Sun, Ru bai, Xi Tang, Minfeng Liang, Nuno Faria, Josh Quick, Andrew Rambaut, Verity Hill, Wenjun Ma, Nick Loman, Oliver Pybus, Changwen Ke |
| EPI_ISL_413858 | hCoV-19/Guangdong/2020XN4459-P0041/2020 | Asia / China / Guangdong | 2020-01-30 | Guangdong Provincial Institution of Public Health, Guangdong Provincial Center for Disease Control and Prevention | Guangdong Provincial Institution of Public Health | Jing Lu, Louis du Plessis, Liu Zhe, Jiufeng Sun, Sarah François, Huifang Lin, Moritz Kraemer, Jingju Peng, Qianlin Xiong, Runyu Yuan, Lilian Zeng, Pingping Zhou, Chuming Liang, Tao Liu, Wei Li, Juan Su, Huanying Zheng, Kang Min, Song Tie, Bo Peng, Shisong Fang, Wenzhe Su, Kuibiao Li, Ruilin Sun, Ru bai, Xi Tang, Minfeng Liang, Nuno Faria, Josh Quick, Andrew Rambaut, Verity Hill, Wenjun Ma, Nick Loman, Oliver Pybus, Changwen Ke |
| EPI_ISL_413859 | hCoV-19/Guangdong/2020XN4276-P0037/2020 | Asia / China / Guangdong | 2020-01-31 | Guangdong Provincial Institution of Public Health, Guangdong Provincial Center for Disease Control and Prevention | Guangdong Provincial Institution of Public Health | Jing Lu, Louis du Plessis, Liu Zhe, Jiufeng Sun, Sarah François, Huifang Lin, Moritz Kraemer, Jingju Peng, Qianlin Xiong, Runyu Yuan, Lilian Zeng, Pingping Zhou, Chuming Liang, Tao Liu, Wei Li, Juan Su, Huanying Zheng, Kang Min, Song Tie, Bo Peng, Shisong Fang, Wenzhe Su, Kuibiao Li, Ruilin Sun, Ru bai, Xi Tang, Minfeng Liang, Nuno Faria, Josh Quick, Andrew Rambaut, Verity Hill, Wenjun Ma, Nick Loman, Oliver Pybus, Changwen Ke |
| EPI_ISL_413860 | hCoV-19/Guangdong/2020XN4273-P0036/2020 | Asia / China / Guangdong | 2020-01-30 | Guangdong Provincial Institution of Public Health, Guangdong Provincial Center for Disease Control and Prevention | Guangdong Provincial Institution of Public Health | Jing Lu, Louis du Plessis, Liu Zhe, Jiufeng Sun, Sarah François, Huifang Lin, Moritz Kraemer, Jingju Peng, Qianlin Xiong, Runyu Yuan, Lilian Zeng, Pingping Zhou, Chuming Liang, Tao Liu, Wei Li, Juan Su, Huanying Zheng, Kang Min, Song Tie, Bo Peng, Shisong Fang, Wenzhe Su, Kuibiao Li, Ruilin Sun, Ru bai, Xi Tang, Minfeng Liang, Nuno Faria, Josh Quick, Andrew Rambaut, Verity Hill, Wenjun Ma, Nick Loman, Oliver Pybus, Changwen Ke |
| EPI_ISL_413861 | hCoV-19/Guangdong/GD2020080-P0010/2020 | Asia / China / Guangdong | 2020-02-01 | Guangdong Provincial Institution of Public Health, Guangdong Provincial Center for Disease Control and Prevention | Guangdong Provincial Institution of Public Health | Jing Lu, Louis du Plessis, Liu Zhe, Jiufeng Sun, Sarah François, Huifang Lin, Moritz Kraemer, Jingju Peng, Qianlin Xiong, Runyu Yuan, Lilian Zeng, Pingping Zhou, Chuming Liang, Tao Liu, Wei Li, Juan Su, Huanying Zheng, Kang Min, Song Tie, Bo Peng, Shisong Fang, Wenzhe Su, Kuibiao Li, Ruilin Sun, Ru bai, Xi Tang, Minfeng Liang, Nuno Faria, Josh Quick, Andrew Rambaut, Verity Hill, Wenjun Ma, Nick Loman, Oliver Pybus, Changwen Ke |
| EPI_ISL_413862 | hCoV-19/Guangdong/GD2020086-P0021/2020 | Asia / China / Guangdong | 2020-02-01 | Guangdong Provincial Institution of Public Health, Guangdong Provincial Center for Disease Control and Prevention | Guangdong Provincial Institution of Public Health | Jing Lu, Louis du Plessis, Liu Zhe, Jiufeng Sun, Sarah François, Huifang Lin, Moritz Kraemer, Jingju Peng, Qianlin Xiong, Runyu Yuan, Lilian Zeng, Pingping Zhou, Chuming Liang, Tao Liu, Wei Li, Juan Su, Huanying Zheng, Kang Min, Song Tie, Bo Peng, Shisong Fang, Wenzhe Su, Kuibiao Li, Ruilin Sun, Ru bai, Xi Tang, Minfeng Liang, Nuno Faria, Josh Quick, Andrew Rambaut, Verity Hill, Wenjun Ma, Nick Loman, Oliver Pybus, Changwen Ke |
| EPI_ISL_413863 | hCoV-19/Guangdong/GD2020087-P0008/2020 | Asia / China / Guangdong | 2020-02-01 | Guangdong Provincial Institution of Public Health, Guangdong Provincial Center for Disease Control and Prevention | Guangdong Provincial Institution of Public Health | Jing Lu, Louis du Plessis, Liu Zhe, Jiufeng Sun, Sarah François, Huifang Lin, Moritz Kraemer, Jingju Peng, Qianlin Xiong, Runyu Yuan, Lilian Zeng, Pingping Zhou, Chuming Liang, Tao Liu, Wei Li, Juan Su, Huanying Zheng, Kang Min, Song Tie, Bo Peng, Shisong Fang, Wenzhe Su, Kuibiao Li, Ruilin Sun, Ru bai, Xi Tang, Minfeng Liang, Nuno Faria, Josh Quick, Andrew Rambaut, Verity Hill, Wenjun Ma, Nick Loman, Oliver Pybus, Changwen Ke |
| EPI_ISL_413864 | hCoV-19/Guangdong/GD2020246-P0028/2020 | Asia / China / Guangdong | 2020-02-09 | Guangdong Provincial Institution of Public Health, Guangdong Provincial Center for Disease Control and Prevention | Guangdong Provincial Institution of Public Health | Jing Lu, Louis du Plessis, Liu Zhe, Jiufeng Sun, Sarah François, Huifang Lin, Moritz Kraemer, Jingju Peng, Qianlin Xiong, Runyu Yuan, Lilian Zeng, Pingping Zhou, Chuming Liang, Tao Liu, Wei Li, Juan Su, Huanying Zheng, Kang Min, Song Tie, Bo Peng, Shisong Fang, Wenzhe Su, Kuibiao Li, Ruilin Sun, Ru bai, Xi Tang, Minfeng Liang, Nuno Faria, Josh Quick, Andrew Rambaut, Verity Hill, Wenjun Ma, Nick Loman, Oliver Pybus, Changwen Ke |
| EPI_ISL_413865 | hCoV-19/Guangdong/GDSZ202013-P0014/2020 | Asia / China / Guangdong | 2020-02-05 | Guangdong Provincial Institution of Public Health, Guangdong Provincial Center for Disease Control and Prevention | Guangdong Provincial Institution of Public Health | Jing Lu, Louis du Plessis, Liu Zhe, Jiufeng Sun, Sarah François, Huifang Lin, Moritz Kraemer, Jingju Peng, Qianlin Xiong, Runyu Yuan, Lilian Zeng, Pingping Zhou, Chuming Liang, Tao Liu, Wei Li, Juan Su, Huanying Zheng, Kang Min, Song Tie, Bo Peng, Shisong Fang, Wenzhe Su, Kuibiao Li, Ruilin Sun, Ru bai, Xi Tang, Minfeng Liang, Nuno Faria, Josh Quick, Andrew Rambaut, Verity Hill, Wenjun Ma, Nick Loman, Oliver Pybus, Changwen Ke |
| EPI_ISL_413866 | hCoV-19/Guangdong/GDSZ202008-P0020/2020 | Asia / China / Guangdong | 2020-02-05 | Guangdong Provincial Institution of Public Health, Guangdong Provincial Center for Disease Control and Prevention | Guangdong Provincial Institution of Public Health | Jing Lu, Louis du Plessis, Liu Zhe, Jiufeng Sun, Sarah François, Huifang Lin, Moritz Kraemer, Jingju Peng, Qianlin Xiong, Runyu Yuan, Lilian Zeng, Pingping Zhou, Chuming Liang, Tao Liu, Wei Li, Juan Su, Huanying Zheng, Kang Min, Song Tie, Bo Peng, Shisong Fang, Wenzhe Su, Kuibiao Li, Ruilin Sun, Ru bai, Xi Tang, Minfeng Liang, Nuno Faria, Josh Quick, Andrew Rambaut, Verity Hill, Wenjun Ma, Nick Loman, Oliver Pybus, Changwen Ke |
| EPI_ISL_413867 | hCoV-19/Guangdong/GDSZ202015-P0019/2020 | Asia / China / Guangdong | 2020-02-05 | Guangdong Provincial Institution of Public Health, Guangdong Provincial Center for Disease Control and Prevention | Guangdong Provincial Institution of Public Health | Jing Lu, Louis du Plessis, Liu Zhe, Jiufeng Sun, Sarah François, Huifang Lin, Moritz Kraemer, Jingju Peng, Qianlin Xiong, Runyu Yuan, Lilian Zeng, Pingping Zhou, Chuming Liang, Tao Liu, Wei Li, Juan Su, Huanying Zheng, Kang Min, Song Tie, Bo Peng, Shisong Fang, Wenzhe Su, Kuibiao Li, Ruilin Sun, Ru bai, Xi Tang, Minfeng Liang, Nuno Faria, Josh Quick, Andrew Rambaut, Verity Hill, Wenjun Ma, Nick Loman, Oliver Pybus, Changwen Ke |
| EPI_ISL_413868 | hCoV-19/Guangdong/DG-S9-P0045/2020 | Asia / China / Guangdong | 2020-02-28 | Guangdong Provincial Institution of Public Health, Guangdong Provincial Center for Disease Control and Prevention | Guangdong Provincial Institution of Public Health | Jing Lu, Louis du Plessis, Liu Zhe, Jiufeng Sun, Sarah François, Huifang Lin, Moritz Kraemer, Jingju Peng, Qianlin Xiong, Runyu Yuan, Lilian Zeng, Pingping Zhou, Chuming Liang, Tao Liu, Wei Li, Juan Su, Huanying Zheng, Kang Min, Song Tie, Bo Peng, Shisong Fang, Wenzhe Su, Kuibiao Li, Ruilin Sun, Ru bai, Xi Tang, Minfeng Liang, Nuno Faria, Josh Quick, Andrew Rambaut, Verity Hill, Wenjun Ma, Nick Loman, Oliver Pybus, Changwen Ke |
| EPI_ISL_413869 | hCoV-19/Guangdong/FS-S42-P0046/2020 | Asia / China / Guangdong | 2020-02-28 | Guangdong Provincial Institution of Public Health, Guangdong Provincial Center for Disease Control and Prevention | Guangdong Provincial Institution of Public Health | Jing Lu, Louis du Plessis, Liu Zhe, Jiufeng Sun, Sarah François, Huifang Lin, Moritz Kraemer, Jingju Peng, Qianlin Xiong, Runyu Yuan, Lilian Zeng, Pingping Zhou, Chuming Liang, Tao Liu, Wei Li, Juan Su, Huanying Zheng, Kang Min, Song Tie, Bo Peng, Shisong Fang, Wenzhe Su, Kuibiao Li, Ruilin Sun, Ru bai, Xi Tang, Minfeng Liang, Nuno Faria, Josh Quick, Andrew Rambaut, Verity Hill, Wenjun Ma, Nick Loman, Oliver Pybus, Changwen Ke |
| EPI_ISL_413870 | hCoV-19/Guangdong/FS-S48-P0047/2020 | Asia / China / Guangdong | 2020-02-28 | Guangdong Provincial Institution of Public Health, Guangdong Provincial Center for Disease Control and Prevention | Guangdong Provincial Institution of Public Health | Jing Lu, Louis du Plessis, Liu Zhe, Jiufeng Sun, Sarah François, Huifang Lin, Moritz Kraemer, Jingju Peng, Qianlin Xiong, Runyu Yuan, Lilian Zeng, Pingping Zhou, Chuming Liang, Tao Liu, Wei Li, Juan Su, Huanying Zheng, Kang Min, Song Tie, Bo Peng, Shisong Fang, Wenzhe Su, Kuibiao Li, Ruilin Sun, Ru bai, Xi Tang, Minfeng Liang, Nuno Faria, Josh Quick, Andrew Rambaut, Verity Hill, Wenjun Ma, Nick Loman, Oliver Pybus, Changwen Ke |
| EPI_ISL_413871 | hCoV-19/Guangdong/MM-S1-P0048/2020 | Asia / China / Guangdong | 2020-02-28 | Guangdong Provincial Institution of Public Health, Guangdong Provincial Center for Disease Control and Prevention | Guangdong Provincial Institution of Public Health | Jing Lu, Louis du Plessis, Liu Zhe, Jiufeng Sun, Sarah François, Huifang Lin, Moritz Kraemer, Jingju Peng, Qianlin Xiong, Runyu Yuan, Lilian Zeng, Pingping Zhou, Chuming Liang, Tao Liu, Wei Li, Juan Su, Huanying Zheng, Kang Min, Song Tie, Bo Peng, Shisong Fang, Wenzhe Su, Kuibiao Li, Ruilin Sun, Ru bai, Xi Tang, Minfeng Liang, Nuno Faria, Josh Quick, Andrew Rambaut, Verity Hill, Wenjun Ma, Nick Loman, Oliver Pybus, Changwen Ke |
| EPI_ISL_413872 | hCoV-19/Guangdong/SZ-N59-P0049/2020 | Asia / China / Guangdong | 2020-02-28 | Guangdong Provincial Institution of Public Health, Guangdong Provincial Center for Disease Control and Prevention | Guangdong Provincial Institution of Public Health | Jing Lu, Louis du Plessis, Liu Zhe, Jiufeng Sun, Sarah François, Huifang Lin, Moritz Kraemer, Jingju Peng, Qianlin Xiong, Runyu Yuan, Lilian Zeng, Pingping Zhou, Chuming Liang, Tao Liu, Wei Li, Juan Su, Huanying Zheng, Kang Min, Song Tie, Bo Peng, Shisong Fang, Wenzhe Su, Kuibiao Li, Ruilin Sun, Ru bai, Xi Tang, Minfeng Liang, Nuno Faria, Josh Quick, Andrew Rambaut, Verity Hill, Wenjun Ma, Nick Loman, Oliver Pybus, Changwen Ke |
| EPI_ISL_413873 | hCoV-19/Guangdong/GZ-S6-P0050/2020 | Asia / China / Guangdong | 2020-02-28 | Guangdong Provincial Institution of Public Health, Guangdong Provincial Center for Disease Control and Prevention | Guangdong Provincial Institution of Public Health | Jing Lu, Louis du Plessis, Liu Zhe, Jiufeng Sun, Sarah François, Huifang Lin, Moritz Kraemer, Jingju Peng, Qianlin Xiong, Runyu Yuan, Lilian Zeng, Pingping Zhou, Chuming Liang, Tao Liu, Wei Li, Juan Su, Huanying Zheng, Kang Min, Song Tie, Bo Peng, Shisong Fang, Wenzhe Su, Kuibiao Li, Ruilin Sun, Ru bai, Xi Tang, Minfeng Liang, Nuno Faria, Josh Quick, Andrew Rambaut, Verity Hill, Wenjun Ma, Nick Loman, Oliver Pybus, Changwen Ke |
| EPI_ISL_413874 | hCoV-19/Guangdong/GDFS2020127-P0026/2020 | Asia / China / Guangdong | 2020-02-12 | Guangdong Provincial Institution of Public Health, Guangdong Provincial Center for Disease Control and Prevention | Guangdong Provincial Institution of Public Health | Jing Lu, Louis du Plessis, Liu Zhe, Jiufeng Sun, Sarah François, Huifang Lin, Moritz Kraemer, Jingju Peng, Qianlin Xiong, Runyu Yuan, Lilian Zeng, Pingping Zhou, Chuming Liang, Tao Liu, Wei Li, Juan Su, Huanying Zheng, Kang Min, Song Tie, Bo Peng, Shisong Fang, Wenzhe Su, Kuibiao Li, Ruilin Sun, Ru bai, Xi Tang, Minfeng Liang, Nuno Faria, Josh Quick, Andrew Rambaut, Verity Hill, Wenjun Ma, Nick Loman, Oliver Pybus, Changwen Ke |
| EPI_ISL_413875 | hCoV-19/Guangdong/GDFS2020052-P0025/2020 | Asia / China / Guangdong | 2020-02-10 | Guangdong Provincial Institution of Public Health, Guangdong Provincial Center for Disease Control and Prevention | Guangdong Provincial Institution of Public Health | Jing Lu, Louis du Plessis, Liu Zhe, Jiufeng Sun, Sarah François, Huifang Lin, Moritz Kraemer, Jingju Peng, Qianlin Xiong, Runyu Yuan, Lilian Zeng, Pingping Zhou, Chuming Liang, Tao Liu, Wei Li, Juan Su, Huanying Zheng, Kang Min, Song Tie, Bo Peng, Shisong Fang, Wenzhe Su, Kuibiao Li, Ruilin Sun, Ru bai, Xi Tang, Minfeng Liang, Nuno Faria, Josh Quick, Andrew Rambaut, Verity Hill, Wenjun Ma, Nick Loman, Oliver Pybus, Changwen Ke |
| EPI_ISL_413876 | hCoV-19/Guangdong/GDFS2020056-P0044/2020 | Asia / China / Guangdong | 2020-02-10 | Guangdong Provincial Institution of Public Health, Guangdong Provincial Center for Disease Control and Prevention | Guangdong Provincial Institution of Public Health | Jing Lu, Louis du Plessis, Liu Zhe, Jiufeng Sun, Sarah François, Huifang Lin, Moritz Kraemer, Jingju Peng, Qianlin Xiong, Runyu Yuan, Lilian Zeng, Pingping Zhou, Chuming Liang, Tao Liu, Wei Li, Juan Su, Huanying Zheng, Kang Min, Song Tie, Bo Peng, Shisong Fang, Wenzhe Su, Kuibiao Li, Ruilin Sun, Ru bai, Xi Tang, Minfeng Liang, Nuno Faria, Josh Quick, Andrew Rambaut, Verity Hill, Wenjun Ma, Nick Loman, Oliver Pybus, Changwen Ke |
| EPI_ISL_413877 | hCoV-19/Guangdong/GD2020241-P0013/2020 | Asia / China / Guangdong | 2020-02-09 | Guangdong Provincial Institution of Public Health, Guangdong Provincial Center for Disease Control and Prevention | Guangdong Provincial Institution of Public Health | Jing Lu, Louis du Plessis, Liu Zhe, Jiufeng Sun, Sarah François, Huifang Lin, Moritz Kraemer, Jingju Peng, Qianlin Xiong, Runyu Yuan, Lilian Zeng, Pingping Zhou, Chuming Liang, Tao Liu, Wei Li, Juan Su, Huanying Zheng, Kang Min, Song Tie, Bo Peng, Shisong Fang, Wenzhe Su, Kuibiao Li, Ruilin Sun, Ru bai, Xi Tang, Minfeng Liang, Nuno Faria, Josh Quick, Andrew Rambaut, Verity Hill, Wenjun Ma, Nick Loman, Oliver Pybus, Changwen Ke |
| EPI_ISL_413878 | hCoV-19/Guangdong/GD2020258-P0018/2020 | Asia / China / Guangdong | 2020-02-09 | Guangdong Provincial Institution of Public Health, Guangdong Provincial Center for Disease Control and Prevention | Guangdong Provincial Institution of Public Health | Jing Lu, Louis du Plessis, Liu Zhe, Jiufeng Sun, Sarah François, Huifang Lin, Moritz Kraemer, Jingju Peng, Qianlin Xiong, Runyu Yuan, Lilian Zeng, Pingping Zhou, Chuming Liang, Tao Liu, Wei Li, Juan Su, Huanying Zheng, Kang Min, Song Tie, Bo Peng, Shisong Fang, Wenzhe Su, Kuibiao Li, Ruilin Sun, Ru bai, Xi Tang, Minfeng Liang, Nuno Faria, Josh Quick, Andrew Rambaut, Verity Hill, Wenjun Ma, Nick Loman, Oliver Pybus, Changwen Ke |
| EPI_ISL_413879 | hCoV-19/Guangdong/GDSZ202004-P0004/2020 | Asia / China / Guangdong | 2020-02-05 | Guangdong Provincial Institution of Public Health, Guangdong Provincial Center for Disease Control and Prevention | Guangdong Provincial Institution of Public Health | Jing Lu, Louis du Plessis, Liu Zhe, Jiufeng Sun, Sarah François, Huifang Lin, Moritz Kraemer, Jingju Peng, Qianlin Xiong, Runyu Yuan, Lilian Zeng, Pingping Zhou, Chuming Liang, Tao Liu, Wei Li, Juan Su, Huanying Zheng, Kang Min, Song Tie, Bo Peng, Shisong Fang, Wenzhe Su, Kuibiao Li, Ruilin Sun, Ru bai, Xi Tang, Minfeng Liang, Nuno Faria, Josh Quick, Andrew Rambaut, Verity Hill, Wenjun Ma, Nick Loman, Oliver Pybus, Changwen Ke |
| EPI_ISL_413880 | hCoV-19/Guangdong/GD2020227-P0029/2020 | Asia / China / Guangdong | 2020-02-05 | Guangdong Provincial Institution of Public Health, Guangdong Provincial Center for Disease Control and Prevention | Guangdong Provincial Institution of Public Health | Jing Lu, Louis du Plessis, Liu Zhe, Jiufeng Sun, Sarah François, Huifang Lin, Moritz Kraemer, Jingju Peng, Qianlin Xiong, Runyu Yuan, Lilian Zeng, Pingping Zhou, Chuming Liang, Tao Liu, Wei Li, Juan Su, Huanying Zheng, Kang Min, Song Tie, Bo Peng, Shisong Fang, Wenzhe Su, Kuibiao Li, Ruilin Sun, Ru bai, Xi Tang, Minfeng Liang, Nuno Faria, Josh Quick, Andrew Rambaut, Verity Hill, Wenjun Ma, Nick Loman, Oliver Pybus, Changwen Ke |
| EPI_ISL_413881 | hCoV-19/Guangdong/GDSZ202009-P0032/2020 | Asia / China / Guangdong | 2020-02-05 | Guangdong Provincial Institution of Public Health, Guangdong Provincial Center for Disease Control and Prevention | Guangdong Provincial Institution of Public Health | Jing Lu, Louis du Plessis, Liu Zhe, Jiufeng Sun, Sarah François, Huifang Lin, Moritz Kraemer, Jingju Peng, Qianlin Xiong, Runyu Yuan, Lilian Zeng, Pingping Zhou, Chuming Liang, Tao Liu, Wei Li, Juan Su, Huanying Zheng, Kang Min, Song Tie, Bo Peng, Shisong Fang, Wenzhe Su, Kuibiao Li, Ruilin Sun, Ru bai, Xi Tang, Minfeng Liang, Nuno Faria, Josh Quick, Andrew Rambaut, Verity Hill, Wenjun Ma, Nick Loman, Oliver Pybus, Changwen Ke |
| EPI_ISL_413882 | hCoV-19/Guangdong/GD2020139-P0007/2020 | Asia / China / Guangdong | 2020-02-02 | Guangdong Provincial Institution of Public Health, Guangdong Provincial Center for Disease Control and Prevention | Guangdong Provincial Institution of Public Health | Jing Lu, Louis du Plessis, Liu Zhe, Jiufeng Sun, Sarah François, Huifang Lin, Moritz Kraemer, Jingju Peng, Qianlin Xiong, Runyu Yuan, Lilian Zeng, Pingping Zhou, Chuming Liang, Tao Liu, Wei Li, Juan Su, Huanying Zheng, Kang Min, Song Tie, Bo Peng, Shisong Fang, Wenzhe Su, Kuibiao Li, Ruilin Sun, Ru bai, Xi Tang, Minfeng Liang, Nuno Faria, Josh Quick, Andrew Rambaut, Verity Hill, Wenjun Ma, Nick Loman, Oliver Pybus, Changwen Ke |
| EPI_ISL_413883 | hCoV-19/Guangdong/GD2020134-P0031/2020 | Asia / China / Guangdong | 2020-02-02 | Guangdong Provincial Institution of Public Health, Guangdong Provincial Center for Disease Control and Prevention | Guangdong Provincial Institution of Public Health | Jing Lu, Louis du Plessis, Liu Zhe, Jiufeng Sun, Sarah François, Huifang Lin, Moritz Kraemer, Jingju Peng, Qianlin Xiong, Runyu Yuan, Lilian Zeng, Pingping Zhou, Chuming Liang, Tao Liu, Wei Li, Juan Su, Huanying Zheng, Kang Min, Song Tie, Bo Peng, Shisong Fang, Wenzhe Su, Kuibiao Li, Ruilin Sun, Ru bai, Xi Tang, Minfeng Liang, Nuno Faria, Josh Quick, Andrew Rambaut, Verity Hill, Wenjun Ma, Nick Loman, Oliver Pybus, Changwen Ke |
| EPI_ISL_413884 | hCoV-19/Guangdong/GD2020115-P0009/2020 | Asia / China / Guangdong | 2020-02-01 | Guangdong Provincial Institution of Public Health, Guangdong Provincial Center for Disease Control and Prevention | Guangdong Provincial Institution of Public Health | Jing Lu, Louis du Plessis, Liu Zhe, Jiufeng Sun, Sarah François, Huifang Lin, Moritz Kraemer, Jingju Peng, Qianlin Xiong, Runyu Yuan, Lilian Zeng, Pingping Zhou, Chuming Liang, Tao Liu, Wei Li, Juan Su, Huanying Zheng, Kang Min, Song Tie, Bo Peng, Shisong Fang, Wenzhe Su, Kuibiao Li, Ruilin Sun, Ru bai, Xi Tang, Minfeng Liang, Nuno Faria, Josh Quick, Andrew Rambaut, Verity Hill, Wenjun Ma, Nick Loman, Oliver Pybus, Changwen Ke |
| EPI_ISL_413885 | hCoV-19/Guangdong/GD2020016-P0011/2020 | Asia / China / Guangdong | 2020-02-01 | Guangdong Provincial Institution of Public Health, Guangdong Provincial Center for Disease Control and Prevention | Guangdong Provincial Institution of Public Health | Jing Lu, Louis du Plessis, Liu Zhe, Jiufeng Sun, Sarah François, Huifang Lin, Moritz Kraemer, Jingju Peng, Qianlin Xiong, Runyu Yuan, Lilian Zeng, Pingping Zhou, Chuming Liang, Tao Liu, Wei Li, Juan Su, Huanying Zheng, Kang Min, Song Tie, Bo Peng, Shisong Fang, Wenzhe Su, Kuibiao Li, Ruilin Sun, Ru bai, Xi Tang, Minfeng Liang, Nuno Faria, Josh Quick, Andrew Rambaut, Verity Hill, Wenjun Ma, Nick Loman, Oliver Pybus, Changwen Ke |
| EPI_ISL_413886 | hCoV-19/Guangdong/GD2020012-P0022/2020 | Asia / China / Guangdong | 2020-02-01 | Guangdong Provincial Institution of Public Health, Guangdong Provincial Center for Disease Control and Prevention | Guangdong Provincial Institution of Public Health | Jing Lu, Louis du Plessis, Liu Zhe, Jiufeng Sun, Sarah François, Huifang Lin, Moritz Kraemer, Jingju Peng, Qianlin Xiong, Runyu Yuan, Lilian Zeng, Pingping Zhou, Chuming Liang, Tao Liu, Wei Li, Juan Su, Huanying Zheng, Kang Min, Song Tie, Bo Peng, Shisong Fang, Wenzhe Su, Kuibiao Li, Ruilin Sun, Ru bai, Xi Tang, Minfeng Liang, Nuno Faria, Josh Quick, Andrew Rambaut, Verity Hill, Wenjun Ma, Nick Loman, Oliver Pybus, Changwen Ke |
| EPI_ISL_413887 | hCoV-19/Guangdong/GD2020085-P0043/2020 | Asia / China / Guangdong | 2020-02-01 | Guangdong Provincial Institution of Public Health, Guangdong Provincial Center for Disease Control and Prevention | Guangdong Provincial Institution of Public Health | Jing Lu, Louis du Plessis, Liu Zhe, Jiufeng Sun, Sarah François, Huifang Lin, Moritz Kraemer, Jingju Peng, Qianlin Xiong, Runyu Yuan, Lilian Zeng, Pingping Zhou, Chuming Liang, Tao Liu, Wei Li, Juan Su, Huanying Zheng, Kang Min, Song Tie, Bo Peng, Shisong Fang, Wenzhe Su, Kuibiao Li, Ruilin Sun, Ru bai, Xi Tang, Minfeng Liang, Nuno Faria, Josh Quick, Andrew Rambaut, Verity Hill, Wenjun Ma, Nick Loman, Oliver Pybus, Changwen Ke |
| EPI_ISL_413888 | hCoV-19/Guangdong/2020XN4239-P0034/2020 | Asia / China / Guangdong | 2020-01-30 | Guangdong Provincial Institution of Public Health, Guangdong Provincial Center for Disease Control and Prevention | Guangdong Provincial Institution of Public Health | Jing Lu, Louis du Plessis, Liu Zhe, Jiufeng Sun, Sarah François, Huifang Lin, Moritz Kraemer, Jingju Peng, Qianlin Xiong, Runyu Yuan, Lilian Zeng, Pingping Zhou, Chuming Liang, Tao Liu, Wei Li, Juan Su, Huanying Zheng, Kang Min, Song Tie, Bo Peng, Shisong Fang, Wenzhe Su, Kuibiao Li, Ruilin Sun, Ru bai, Xi Tang, Minfeng Liang, Nuno Faria, Josh Quick, Andrew Rambaut, Verity Hill, Wenjun Ma, Nick Loman, Oliver Pybus, Changwen Ke |
| EPI_ISL_413889 | hCoV-19/Guangdong/2020XN4291-P0038/2020 | Asia / China / Guangdong | 2020-01-30 | Guangdong Provincial Institution of Public Health, Guangdong Provincial Center for Disease Control and Prevention | Guangdong Provincial Institution of Public Health | Jing Lu, Louis du Plessis, Liu Zhe, Jiufeng Sun, Sarah François, Huifang Lin, Moritz Kraemer, Jingju Peng, Qianlin Xiong, Runyu Yuan, Lilian Zeng, Pingping Zhou, Chuming Liang, Tao Liu, Wei Li, Juan Su, Huanying Zheng, Kang Min, Song Tie, Bo Peng, Shisong Fang, Wenzhe Su, Kuibiao Li, Ruilin Sun, Ru bai, Xi Tang, Minfeng Liang, Nuno Faria, Josh Quick, Andrew Rambaut, Verity Hill, Wenjun Ma, Nick Loman, Oliver Pybus, Changwen Ke |
| EPI_ISL_413890 | hCoV-19/Guangdong/FS-S34-P0015/2020 | Asia / China / Guangdong | 2020-02-28 | Guangdong Provincial Institution of Public Health, Guangdong Provincial Center for Disease Control and Prevention | Guangdong Provincial Institution of Public Health | Jing Lu, Louis du Plessis, Liu Zhe, Jiufeng Sun, Sarah François, Huifang Lin, Moritz Kraemer, Jingju Peng, Qianlin Xiong, Runyu Yuan, Lilian Zeng, Pingping Zhou, Chuming Liang, Tao Liu, Wei Li, Juan Su, Huanying Zheng, Kang Min, Song Tie, Bo Peng, Shisong Fang, Wenzhe Su, Kuibiao Li, Ruilin Sun, Ru bai, Xi Tang, Minfeng Liang, Nuno Faria, Josh Quick, Andrew Rambaut, Verity Hill, Wenjun Ma, Nick Loman, Oliver Pybus, Changwen Ke |
| EPI_ISL_413891 | hCoV-19/Guangdong/FS-S29-P0051/2020 | Asia / China / Guangdong | 2020-02-28 | Guangdong Provincial Institution of Public Health, Guangdong Provincial Center for Disease Control and Prevention | Guangdong Provincial Institution of Public Health | Jing Lu, Louis du Plessis, Liu Zhe, Jiufeng Sun, Sarah François, Huifang Lin, Moritz Kraemer, Jingju Peng, Qianlin Xiong, Runyu Yuan, Lilian Zeng, Pingping Zhou, Chuming Liang, Tao Liu, Wei Li, Juan Su, Huanying Zheng, Kang Min, Song Tie, Bo Peng, Shisong Fang, Wenzhe Su, Kuibiao Li, Ruilin Sun, Ru bai, Xi Tang, Minfeng Liang, Nuno Faria, Josh Quick, Andrew Rambaut, Verity Hill, Wenjun Ma, Nick Loman, Oliver Pybus, Changwen Ke |
| EPI_ISL_413892 | hCoV-19/Guangdong/FS-S30-P0052/2020 | Asia / China / Guangdong | 2020-02-28 | Guangdong Provincial Institution of Public Health, Guangdong Provincial Center for Disease Control and Prevention | Guangdong Provincial Institution of Public Health | Jing Lu, Louis du Plessis, Liu Zhe, Jiufeng Sun, Sarah François, Huifang Lin, Moritz Kraemer, Jingju Peng, Qianlin Xiong, Runyu Yuan, Lilian Zeng, Pingping Zhou, Chuming Liang, Tao Liu, Wei Li, Juan Su, Huanying Zheng, Kang Min, Song Tie, Bo Peng, Shisong Fang, Wenzhe Su, Kuibiao Li, Ruilin Sun, Ru bai, Xi Tang, Minfeng Liang, Nuno Faria, Josh Quick, Andrew Rambaut, Verity Hill, Wenjun Ma, Nick Loman, Oliver Pybus, Changwen Ke |
| EPI_ISL_413893 | hCoV-19/Guangdong/FS-S50-P0053/2020 | Asia / China / Guangdong | 2020-02-28 | Guangdong Provincial Institution of Public Health, Guangdong Provincial Center for Disease Control and Prevention | Guangdong Provincial Institution of Public Health | Jing Lu, Louis du Plessis, Liu Zhe, Jiufeng Sun, Sarah François, Huifang Lin, Moritz Kraemer, Jingju Peng, Qianlin Xiong, Runyu Yuan, Lilian Zeng, Pingping Zhou, Chuming Liang, Tao Liu, Wei Li, Juan Su, Huanying Zheng, Kang Min, Song Tie, Bo Peng, Shisong Fang, Wenzhe Su, Kuibiao Li, Ruilin Sun, Ru bai, Xi Tang, Minfeng Liang, Nuno Faria, Josh Quick, Andrew Rambaut, Verity Hill, Wenjun Ma, Nick Loman, Oliver Pybus, Changwen Ke |
| EPI_ISL_413894 | hCoV-19/Guangdong/DG-S2-P0054/2020 | Asia / China / Guangdong | 2020-02-28 | Guangdong Provincial Institution of Public Health, Guangdong Provincial Center for Disease Control and Prevention | Guangdong Provincial Institution of Public Health | Jing Lu, Louis du Plessis, Liu Zhe, Jiufeng Sun, Sarah François, Huifang Lin, Moritz Kraemer, Jingju Peng, Qianlin Xiong, Runyu Yuan, Lilian Zeng, Pingping Zhou, Chuming Liang, Tao Liu, Wei Li, Juan Su, Huanying Zheng, Kang Min, Song Tie, Bo Peng, Shisong Fang, Wenzhe Su, Kuibiao Li, Ruilin Sun, Ru bai, Xi Tang, Minfeng Liang, Nuno Faria, Josh Quick, Andrew Rambaut, Verity Hill, Wenjun Ma, Nick Loman, Oliver Pybus, Changwen Ke |
| EPI_ISL_413895 | hCoV-19/Guangdong/DG-S6-P0055/2020 | Asia / China / Guangdong | 2020-02-28 | Guangdong Provincial Institution of Public Health, Guangdong Provincial Center for Disease Control and Prevention | Guangdong Provincial Institution of Public Health | Jing Lu, Louis du Plessis, Liu Zhe, Jiufeng Sun, Sarah François, Huifang Lin, Moritz Kraemer, Jingju Peng, Qianlin Xiong, Runyu Yuan, Lilian Zeng, Pingping Zhou, Chuming Liang, Tao Liu, Wei Li, Juan Su, Huanying Zheng, Kang Min, Song Tie, Bo Peng, Shisong Fang, Wenzhe Su, Kuibiao Li, Ruilin Sun, Ru bai, Xi Tang, Minfeng Liang, Nuno Faria, Josh Quick, Andrew Rambaut, Verity Hill, Wenjun Ma, Nick Loman, Oliver Pybus, Changwen Ke |
| EPI_ISL_413896 | hCoV-19/Guangdong/DG-S41-P0056/2020 | Asia / China / Guangdong | 2020-02-28 | Guangdong Provincial Institution of Public Health, Guangdong Provincial Center for Disease Control and Prevention | Guangdong Provincial Institution of Public Health | Jing Lu, Louis du Plessis, Liu Zhe, Jiufeng Sun, Sarah François, Huifang Lin, Moritz Kraemer, Jingju Peng, Qianlin Xiong, Runyu Yuan, Lilian Zeng, Pingping Zhou, Chuming Liang, Tao Liu, Wei Li, Juan Su, Huanying Zheng, Kang Min, Song Tie, Bo Peng, Shisong Fang, Wenzhe Su, Kuibiao Li, Ruilin Sun, Ru bai, Xi Tang, Minfeng Liang, Nuno Faria, Josh Quick, Andrew Rambaut, Verity Hill, Wenjun Ma, Nick Loman, Oliver Pybus, Changwen Ke |
| EPI_ISL_413897 | hCoV-19/Guangdong/SZ-N128-P0057/2020 | Asia / China / Guangdong | 2020-02-28 | Guangdong Provincial Institution of Public Health, Guangdong Provincial Center for Disease Control and Prevention | Guangdong Provincial Institution of Public Health | Jing Lu, Louis du Plessis, Liu Zhe, Jiufeng Sun, Sarah François, Huifang Lin, Moritz Kraemer, Jingju Peng, Qianlin Xiong, Runyu Yuan, Lilian Zeng, Pingping Zhou, Chuming Liang, Tao Liu, Wei Li, Juan Su, Huanying Zheng, Kang Min, Song Tie, Bo Peng, Shisong Fang, Wenzhe Su, Kuibiao Li, Ruilin Sun, Ru bai, Xi Tang, Minfeng Liang, Nuno Faria, Josh Quick, Andrew Rambaut, Verity Hill, Wenjun Ma, Nick Loman, Oliver Pybus, Changwen Ke |
| EPI_ISL_413898 | hCoV-19/Guangdong/ZH-S33-P0058/2020 | Asia / China / Guangdong | 2020-02-28 | Guangdong Provincial Institution of Public Health, Guangdong Provincial Center for Disease Control and Prevention | Guangdong Provincial Institution of Public Health | Jing Lu, Louis du Plessis, Liu Zhe, Jiufeng Sun, Sarah François, Huifang Lin, Moritz Kraemer, Jingju Peng, Qianlin Xiong, Runyu Yuan, Lilian Zeng, Pingping Zhou, Chuming Liang, Tao Liu, Wei Li, Juan Su, Huanying Zheng, Kang Min, Song Tie, Bo Peng, Shisong Fang, Wenzhe Su, Kuibiao Li, Ruilin Sun, Ru bai, Xi Tang, Minfeng Liang, Nuno Faria, Josh Quick, Andrew Rambaut, Verity Hill, Wenjun Ma, Nick Loman, Oliver Pybus, Changwen Ke |
| EPI_ISL_413899 | hCoV-19/Guangdong/ZH-N22-P0059/2020 | Asia / China / Guangdong | 2020-02-28 | Guangdong Provincial Institution of Public Health, Guangdong Provincial Center for Disease Control and Prevention | Guangdong Provincial Institution of Public Health | Jing Lu, Louis du Plessis, Liu Zhe, Jiufeng Sun, Sarah François, Huifang Lin, Moritz Kraemer, Jingju Peng, Qianlin Xiong, Runyu Yuan, Lilian Zeng, Pingping Zhou, Chuming Liang, Tao Liu, Wei Li, Juan Su, Huanying Zheng, Kang Min, Song Tie, Bo Peng, Shisong Fang, Wenzhe Su, Kuibiao Li, Ruilin Sun, Ru bai, Xi Tang, Minfeng Liang, Nuno Faria, Josh Quick, Andrew Rambaut, Verity Hill, Wenjun Ma, Nick Loman, Oliver Pybus, Changwen Ke |
| EPI_ISL_413900 | hCoV-19/Guangdong/ZS-S6-P0060/2020 | Asia / China / Guangdong | 2020-02-28 | Guangdong Provincial Institution of Public Health, Guangdong Provincial Center for Disease Control and Prevention | Guangdong Provincial Institution of Public Health | Jing Lu, Louis du Plessis, Liu Zhe, Jiufeng Sun, Sarah François, Huifang Lin, Moritz Kraemer, Jingju Peng, Qianlin Xiong, Runyu Yuan, Lilian Zeng, Pingping Zhou, Chuming Liang, Tao Liu, Wei Li, Juan Su, Huanying Zheng, Kang Min, Song Tie, Bo Peng, Shisong Fang, Wenzhe Su, Kuibiao Li, Ruilin Sun, Ru bai, Xi Tang, Minfeng Liang, Nuno Faria, Josh Quick, Andrew Rambaut, Verity Hill, Wenjun Ma, Nick Loman, Oliver Pybus, Changwen Ke |
| EPI_ISL_413901 | hCoV-19/Guangdong/ZQ-S2-P0061/2020 | Asia / China / Guangdong | 2020-02-28 | Guangdong Provincial Institution of Public Health, Guangdong Provincial Center for Disease Control and Prevention | Guangdong Provincial Institution of Public Health | Jing Lu, Louis du Plessis, Liu Zhe, Jiufeng Sun, Sarah François, Huifang Lin, Moritz Kraemer, Jingju Peng, Qianlin Xiong, Runyu Yuan, Lilian Zeng, Pingping Zhou, Chuming Liang, Tao Liu, Wei Li, Juan Su, Huanying Zheng, Kang Min, Song Tie, Bo Peng, Shisong Fang, Wenzhe Su, Kuibiao Li, Ruilin Sun, Ru bai, Xi Tang, Minfeng Liang, Nuno Faria, Josh Quick, Andrew Rambaut, Verity Hill, Wenjun Ma, Nick Loman, Oliver Pybus, Changwen Ke |
| EPI_ISL_413902 | hCoV-19/Guangdong/JM-S1-P0062/2020 | Asia / China / Guangdong | 2020-02-28 | Guangdong Provincial Institution of Public Health, Guangdong Provincial Center for Disease Control and Prevention | Guangdong Provincial Institution of Public Health | Jing Lu, Louis du Plessis, Liu Zhe, Jiufeng Sun, Sarah François, Huifang Lin, Moritz Kraemer, Jingju Peng, Qianlin Xiong, Runyu Yuan, Lilian Zeng, Pingping Zhou, Chuming Liang, Tao Liu, Wei Li, Juan Su, Huanying Zheng, Kang Min, Song Tie, Bo Peng, Shisong Fang, Wenzhe Su, Kuibiao Li, Ruilin Sun, Ru bai, Xi Tang, Minfeng Liang, Nuno Faria, Josh Quick, Andrew Rambaut, Verity Hill, Wenjun Ma, Nick Loman, Oliver Pybus, Changwen Ke |
| EPI_ISL_413924 | hCoV-19/USA/CA-CDPH-UC6/2020 | North America / USA / California / Grand | 2020-03-05 | California Department of Public Health | Chiu Laboratory, University of California, San Francisco | Xianding Deng, Scot Federman, Chao-Yang Pan, Hugo Guevara,Wei Gu, Debra A. Wadford, and Charles Y. Chiu |
| EPI_ISL_413925 | hCoV-19/USA/CA-CDPH-UC7/2020 | North America / USA / California / Grand | 2020-03-05 | California Department of Public Health | Chiu Laboratory, University of California, San Francisco | Xianding Deng, Scot Federman, Chao-Yang Pan, Hugo Guevara,Wei Gu, Debra A. Wadford, and Charles Y. Chiu |
| EPI_ISL_413928 | hCoV-19/USA/CA-CDPH-UC9/2020 | North America / USA / California / Grand | 2020-03-05 | California Department of Public Health | Chiu Laboratory, University of California, San Francisco | Xianding Deng, Scot Federman, Chao-Yang Pan, Hugo Guevara,Wei Gu, Debra A. Wadford, and Charles Y. Chiu |

| | | | | | | |
|---|---|---|---|---|---|---|
| EPI_ISL_413931 | hCoV-19/USA/CA-CDPH-UC11/2020 | North America / USA / California / Grand | 2020-03-05 | California Department of Public Health | Chiu Laboratory, University of California, San Francisco | Xianding Deng, Scot Federman, Chao-Yang Pan, Hugo Guevara,Wei Gu, Debra A. Wadford, and Charles Y. Chiu |
| EPI_ISL_414363 | hCoV-19/USA/WA-UW15/2020 | North America / USA / Washington | 2020-03-04 | UW Virology Lab | UW Virology Lab | Pavitra Roychoudhury, Hong Xie, Keith Jerome, Alexander Greninger |
| EPI_ISL_414364 | hCoV-19/USA/WA-UW16/2020 | North America / USA / Washington | 2020-03-05 | UW Virology Lab | UW Virology Lab | Pavitra Roychoudhury, Hong Xie, Keith Jerome, Alexander Greninger |
| EPI_ISL_414365 | hCoV-19/USA/WA-UW17/2020 | North America / USA / Washington | 2020-03-06 | UW Virology Lab | UW Virology Lab | Pavitra Roychoudhury, Hong Xie, Keith Jerome, Alexander Greninger |
| EPI_ISL_414366 | hCoV-19/USA/WA-UW18/2020 | North America / USA / Washington | 2020-03-05 | UW Virology Lab | UW Virology Lab | Pavitra Roychoudhury, Hong Xie, Keith Jerome, Alexander Greninger |
| EPI_ISL_414367 | hCoV-19/USA/WA-UW19/2020 | North America / USA / Washington | 2020-03-05 | UW Virology Lab | UW Virology Lab | Pavitra Roychoudhury, Hong Xie, Keith Jerome, Alexander Greninger |
| EPI_ISL_414368 | hCoV-19/USA/WA-UW20/2020 | North America / USA / Washington | 2020-03-05 | UW Virology Lab | UW Virology Lab | Pavitra Roychoudhury, Hong Xie, Keith Jerome, Alexander Greninger |
| EPI_ISL_414369 | hCoV-19/USA/WA-UW21/2020 | North America / USA / Washington | 2020-03-05 | UW Virology Lab | UW Virology Lab | Pavitra Roychoudhury, Hong Xie, Keith Jerome, Alexander Greninger |
| EPI_ISL_414378 | hCoV-19/Singapore/12/2020 | Asia / Singapore | 2020-02-17 | National Centre for Infectious Diseases | Programme in Emerging Infectious Diseases, Duke-NUS Medical School | Danielle E Anderson, Martin Linster, Yan Zhuang, Jayanthi Jayakumar, Louisa Sun, David CB Lye, Yee Sin Leo, Barnaby E Young, Yvonne CF Su, Gavin JD Smith |
| EPI_ISL_414379 | hCoV-19/Singapore/13/2020 | Asia / Singapore | 2020-02-18 | National Centre for Infectious Diseases | Programme in Emerging Infectious Diseases, Duke-NUS Medical School | Danielle E Anderson, Martin Linster, Yan Zhuang, Jayanthi Jayakumar, David CB Lye, Yee Sin Leo, Barnaby E Young, Yvonne CF Su, Gavin JD Smith |
| EPI_ISL_414380 | hCoV-19/Singapore/14/2020 | Asia / Singapore | 2020-02-13 | National Centre for Infectious Diseases | Programme in Emerging Infectious Diseases, Duke-NUS Medical School | Danielle E Anderson, Martin Linster, Yan Zhuang, Jayanthi Jayakumar, David CB Lye, Yee Sin Leo, Barnaby E Young, Yvonne CF Su, Gavin JD Smith |
| EPI_ISL_415600 | hCoV-19/USA/WA-UW72/2020 | North America / USA | 2020-03-09 | UW Virology Lab | UW Virology Lab | Pavitra Roychoudhury, Hong Xie, Keith Jerome, Alexander Greninger |
| EPI_ISL_415601 | hCoV-19/USA/WA-UW73/2020 | North America / USA / Washington | 2020-03-10 | UW Virology Lab | UW Virology Lab | Pavitra Roychoudhury, Hong Xie, Keith Jerome, Alexander Greninger |
| EPI_ISL_415602 | hCoV-19/USA/WA-UW74/2020 | North America / USA / Washington / Seat | 2020-03-10 | UW Virology Lab | UW Virology Lab | Pavitra Roychoudhury, Hong Xie, Keith Jerome, Alexander Greninger |
| EPI_ISL_415603 | hCoV-19/USA/WA-UW75/2020 | North America / USA / Washington / Seat | 2020-03-10 | UW Virology Lab | UW Virology Lab | Pavitra Roychoudhury, Hong Xie, Keith Jerome, Alexander Greninger |
| EPI_ISL_415604 | hCoV-19/USA/WA-UW76/2020 | North America / USA / Washington | 2020-03-10 | UW Virology Lab | UW Virology Lab | Pavitra Roychoudhury, Hong Xie, Keith Jerome, Alexander Greninger |
| EPI_ISL_415605 | hCoV-19/USA/WA-UW40/2020 | North America / USA / Washington | 2020-03-05 | UW Virology Lab | UW Virology Lab | Pavitra Roychoudhury, Hong Xie, Keith Jerome, Alexander Greninger |
| EPI_ISL_415606 | hCoV-19/USA/WA-UW41/2020 | North America / USA / Washington | 2020-03-08 | UW Virology Lab | UW Virology Lab | Pavitra Roychoudhury, Hong Xie, Keith Jerome, Alexander Greninger |
| EPI_ISL_415607 | hCoV-19/USA/WA-UW42/2020 | North America / USA / Washington | 2020-03-09 | UW Virology Lab | UW Virology Lab | Pavitra Roychoudhury, Hong Xie, Keith Jerome, Alexander Greninger |
| EPI_ISL_415608 | hCoV-19/USA/WA-UW43/2020 | North America / USA / Washington | 2020-03-08 | UW Virology Lab | UW Virology Lab | Pavitra Roychoudhury, Hong Xie, Keith Jerome, Alexander Greninger |
| EPI_ISL_415609 | hCoV-19/USA/WA-UW44/2020 | North America / USA / Washington | 2020-03-08 | UW Virology Lab | UW Virology Lab | Pavitra Roychoudhury, Hong Xie, Keith Jerome, Alexander Greninger |
| EPI_ISL_415610 | hCoV-19/USA/WA-UW45/2020 | North America / USA / Washington | 2020-03-08 | UW Virology Lab | UW Virology Lab | Pavitra Roychoudhury, Hong Xie, Keith Jerome, Alexander Greninger |
| EPI_ISL_415611 | hCoV-19/USA/WA-UW46/2020 | North America / USA / Washington | 2020-03-09 | UW Virology Lab | UW Virology Lab | Pavitra Roychoudhury, Hong Xie, Keith Jerome, Alexander Greninger |
| EPI_ISL_415612 | hCoV-19/USA/WA-UW47/2020 | North America / USA / Washington / Seat | 2020-03-09 | UW Virology Lab | UW Virology Lab | Pavitra Roychoudhury, Hong Xie, Keith Jerome, Alexander Greninger |
| EPI_ISL_415613 | hCoV-19/USA/WA-UW48/2020 | North America / USA / Washington | 2020-03-09 | UW Virology Lab | UW Virology Lab | Pavitra Roychoudhury, Hong Xie, Keith Jerome, Alexander Greninger |
| EPI_ISL_415614 | hCoV-19/USA/WA-UW49/2020 | North America / USA / Washington | 2020-03-09 | UW Virology Lab | UW Virology Lab | Pavitra Roychoudhury, Hong Xie, Keith Jerome, Alexander Greninger |
| EPI_ISL_415615 | hCoV-19/USA/WA-UW50/2020 | North America / USA / Washington | 2020-03-08 | UW Virology Lab | UW Virology Lab | Pavitra Roychoudhury, Hong Xie, Keith Jerome, Alexander Greninger |
| EPI_ISL_415616 | hCoV-19/USA/WA-UW51/2020 | North America / USA / Washington | 2020-03-08 | UW Virology Lab | UW Virology Lab | Pavitra Roychoudhury, Hong Xie, Keith Jerome, Alexander Greninger |
| EPI_ISL_415617 | hCoV-19/USA/WA-UW52/2020 | North America / USA / Washington | 2020-03-09 | UW Virology Lab | UW Virology Lab | Pavitra Roychoudhury, Hong Xie, Keith Jerome, Alexander Greninger |
| EPI_ISL_415619 | hCoV-19/USA/WA-UW54/2020 | North America / USA / Washington | 2020-03-09 | UW Virology Lab | UW Virology Lab | Pavitra Roychoudhury, Hong Xie, Keith Jerome, Alexander Greninger |
| EPI_ISL_415620 | hCoV-19/USA/WA-UW55/2020 | North America / USA / Washington | 2020-03-09 | UW Virology Lab | UW Virology Lab | Pavitra Roychoudhury, Hong Xie, Keith Jerome, Alexander Greninger |
| EPI_ISL_415621 | hCoV-19/USA/WA-UW56/2020 | North America / USA / Washington / Seat | 2020-03-09 | UW Virology Lab | UW Virology Lab | Pavitra Roychoudhury, Hong Xie, Keith Jerome, Alexander Greninger |
| EPI_ISL_415622 | hCoV-19/USA/WA-UW57/2020 | North America / USA / Washington | 2020-03-09 | UW Virology Lab | UW Virology Lab | Pavitra Roychoudhury, Hong Xie, Keith Jerome, Alexander Greninger |
| EPI_ISL_415624 | hCoV-19/USA/WA-UW59/2020 | North America / USA / Washington | 2020-03-09 | UW Virology Lab | UW Virology Lab | Pavitra Roychoudhury, Hong Xie, Keith Jerome, Alexander Greninger |
| EPI_ISL_415625 | hCoV-19/USA/WA-UW60/2020 | North America / USA / Washington | 2020-03-09 | UW Virology Lab | UW Virology Lab | Pavitra Roychoudhury, Hong Xie, Keith Jerome, Alexander Greninger |
| EPI_ISL_415626 | hCoV-19/USA/WA-UW61/2020 | North America / USA / Washington | 2020-03-10 | UW Virology Lab | UW Virology Lab | Pavitra Roychoudhury, Hong Xie, Keith Jerome, Alexander Greninger |
| EPI_ISL_415627 | hCoV-19/USA/WA-UW62/2020 | North America / USA / Washington | 2020-03-09 | UW Virology Lab | UW Virology Lab | Pavitra Roychoudhury, Hong Xie, Keith Jerome, Alexander Greninger |
| EPI_ISL_415629 | hCoV-19/Scotland/EDB004/2020 | Europe / United Kingdom / Scotland | 2020-03-04 | Virology Department, Royal Infirmary of Edinburgh, NHS Lothian | Virology Department, Royal Infirmary of Edinburgh, NHS Lothian | McHugh M, Dewar R, O'Toole Á, Rambaut A, Williams TC, Templeton K |
| EPI_ISL_415630 | hCoV-19/Scotland/CVR07/2020 | Europe / United Kingdom / Scotland | 2020-03-09 | West of Scotland Specialist Virology Centre, NHSGGC | MRC-University of Glasgow Centre for Virus Research | Kathy Smollett, Daniel Mair, Stephen Carmichael, Ana da Silva Filipe; Richard Orton, David L Robertson; Alasdair MacLean, Rory Gunson; Natasha Jesudason, Kathy Li, Antonia Ho; Emma Thomson. |
| EPI_ISL_415631 | hCoV-19/Scotland/CVR10/2020 | Europe / United Kingdom / Scotland | 2020-03-10 | West of Scotland Specialist Virology Centre, NHSGGC | MRC-University of Glasgow Centre for Virus Research | Kathy Smollett, Daniel Mair, Stephen Carmichael, Ana da Silva Filipe; Richard Orton, David L Robertson; Alasdair MacLean, Rory Gunson; Natasha Jesudason, Kathy Li, Antonia Ho; Emma Thomson. |
| EPI_ISL_415640 | hCoV-19/Scotland/EDB003/2020 | Europe / United Kingdom / Scotland | 2020-03-08 | Virology Department, Royal Infirmary of Edinburgh, NHS Lothian | Virology Department, Royal Infirmary of Edinburgh, NHS Lothian | McHugh M, Dewar R, O'Toole Á, Rambaut A, Williams TC, Templeton K |
| EPI_ISL_415641 | hCoV-19/Georgia/Tb-54/2020 | Asia / Georgia / Tbilisi | 2020-02-27 | R. G. Lugar Center for Public Health Research, National Center for Disease Control and Public Health (NCDC) of Georgia. | R. G. Lugar Center for Public Health Research, National Center for Disease Control and Public Health (NCDC) of Georgia. | Nato Kotaria, Marine Murtskhvaladze, Ann Machablishvili, Lela Sabadze, Mari Gavashelidze, Ana Papkiauri, Meri Pantsulaia, Gvantsa Brachveli, Tata Imnadze, Tamar Jashiashvili, Tea Tevdoradze, Ketevan Sidamonidze, Ekaterine Khmaladze, Ekaterine Zhghenti, Roena Sukhiashvili, Mariam Zakalashvili, Lela Urushadze, Magda Dgebuadze, Giorgi Tomashvili, Davit Tsaguria, Ekaterine Zangaladze, Nino Berishvili, Gvantsa Chanturia, Adam Kotorashvili, Maia Alkhazashvili, Irma Burjanadze, Anna Kasradze, Khatuna Zakhashvili, Paata Imnadze, Amiran Gamkrelidze. |
| EPI_ISL_415642 | hCoV-19/Georgia/Tb-477/2020 | Asia / Georgia / Tbilisi | 2020-03-10 | R. G. Lugar Center for Public Health Research, National Center for Disease Control and Public Health (NCDC) of Georgia. | R. G. Lugar Center for Public Health Research, National Center for Disease Control and Public Health (NCDC) of Georgia. | Nato Kotaria, Marine Murtskhvaladze, Ann Machablishvili, Lela Sabadze, Mari Gavashelidze, Ana Papkiauri, Meri Pantsulaia, Gvantsa Brachveli, Tata Imnadze, Tamar Jashiashvili, Tea Tevdoradze, Ketevan Sidamonidze, Ekaterine Khmaladze, Ekaterine Zhghenti, Roena Sukhiashvili, Mariam Zakalashvili, Lela Urushadze, Magda Dgebuadze, Giorgi Tomashvili, Davit Tsaguria, Ekaterine Zangaladze, Nino Berishvili, Gvantsa Chanturia, Adam Kotorashvili, Maia Alkhazashvili, Irma Burjanadze, Anna Kasradze, Khatuna Zakhashvili, Paata Imnadze, Amiran Gamkrelidze. |
| EPI_ISL_415643 | hCoV-19/Georgia/Tb-468/2020 | Asia / Georgia / Tbilisi | 2020-03-10 | R. G. Lugar Center for Public Health Research, National Center for Disease Control and Public Health (NCDC) of Georgia. | R. G. Lugar Center for Public Health Research, National Center for Disease Control and Public Health (NCDC) of Georgia. | Nato Kotaria, Marine Murtskhvaladze, Ann Machablishvili, Lela Sabadze, Mari Gavashelidze, Ana Papkiauri, Meri Pantsulaia, Gvantsa Brachveli, Tata Imnadze, Tamar Jashiashvili, Tea Tevdoradze, Ketevan Sidamonidze, Ekaterine Khmaladze, Ekaterine Zhghenti, Roena Sukhiashvili, Mariam Zakalashvili, Lela Urushadze, Magda Dgebuadze, Giorgi Tomashvili, Davit Tsaguria, Ekaterine Zangaladze, Nino Berishvili, Gvantsa Chanturia, Adam Kotorashvili, Maia Alkhazashvili, Irma Burjanadze, Anna Kasradze, Khatuna Zakhashvili, Paata Imnadze, Amiran Gamkrelidze. |
| EPI_ISL_415644 | hCoV-19/Georgia/Tb-82/2020 | Asia / Georgia / Tbilisi | 2020-02-28 | R. G. Lugar Center for Public Health Research, National Center for Disease Control and Public Health (NCDC) of Georgia. | R. G. Lugar Center for Public Health Research, National Center for Disease Control and Public Health (NCDC) of Georgia. | Nato Kotaria, Marine Murtskhvaladze, Ann Machablishvili, Lela Sabadze, Mari Gavashelidze, Ana Papkiauri, Meri Pantsulaia, Gvantsa Brachveli, Tata Imnadze, Tamar Jashiashvili, Tea Tevdoradze, Ketevan Sidamonidze, Ekaterine Khmaladze, Ekaterine Zhghenti, Roena Sukhiashvili, Mariam Zakalashvili, Lela Urushadze, Magda Dgebuadze, Giorgi Tomashvili, Davit Tsaguria, Ekaterine Zangaladze, Nino Berishvili, Gvantsa Chanturia, Adam Kotorashvili, Maia Alkhazashvili, Irma Burjanadze, Anna Kasradze, Khatuna Zakhashvili, Paata Imnadze, Amiran Gamkrelidze. |
| EPI_ISL_415646 | hCoV-19/Denmark/SSI-101/2020 | Europe / Denmark | 2020-03-03 | Department of Virus and Microbiological Special diagnostics, Statens Serum Institut, Copenhagen, Denmark. | VIFU | Morten Rasmussen, Maiken Worsoe Rosenstierne , Anders Fomsgaard |
| EPI_ISL_415647 | hCoV-19/Denmark/SSI-102/2020 | Europe / Denmark / Copenhagen | 2020-03-03 | Department of Virus and Microbiological Special diagnostics, Statens Serum Institut, Copenhagen, Denmark. | Statens Serum Institute | Morten Rasmussen, Maiken Worsoe Rosenstierne , Anders Fomsgaard |
| EPI_ISL_415648 | hCoV-19/Denmark/SSI-104/2020 | Europe / Denmark | 2020-03-03 | Department of Virus and Microbiological Special diagnostics, Statens Serum Institut, Copenhagen, Denmark. | VIFU | Morten Rasmussen, Maiken Worsoe Rosenstierne , Anders Fomsgaard |
| EPI_ISL_415649 | hCoV-19/France/HF2039/2020 | Europe / France / Hauts-de-France / Crép | 2020-03-05 | unknown | National Reference Center for Viruses of Respiratory Infections, Institut Pasteur, Paris | Mélnie Albert, Marion Barbet, Sylvie Behillil, Méline Bizard, Angela Brisebarre, Flora Donati Vincent Enouf, Maud Vanpeene, Sylvie van der Werf |
| EPI_ISL_415650 | hCoV-19/France/IDF2075/2020 | Europe / France / IDF | 2020-03-02 | Hôpital Instruction des Armées - BEGIN | National Reference Center for Viruses of Respiratory Infections, Institut Pasteur, Paris | Mélnie Albert, Marion Barbet, Sylvie Behillil, Méline Bizard, Angela Brisebarre, Flora Donati Vincent Enouf, Maud Vanpeene, Sylvie van der Werf, Christine Bigaillon |
| EPI_ISL_415651 | hCoV-19/France/BFC2094/2020 | Europe / France / Bourgogne-France-Com | 2020-03-05 | Unknown | National Reference Center for Viruses of Respiratory Infections, Institut Pasteur, Paris | Mélnie Albert, Marion Barbet, Sylvie Behillil, Méline Bizard, Angela Brisebarre, Flora Donati Vincent Enouf, Maud Vanpeene, Sylvie van der Werf |
| EPI_ISL_415652 | hCoV-19/France/BFC2147/2020 | Europe / France / Bourgogne-France-Com | 2020-03-05 | unknown | National Reference Center for Viruses of Respiratory Infections, Institut Pasteur, Paris | Mélnie Albert, Marion Barbet, Sylvie Behillil, Méline Bizard, Angela Brisebarre, Flora Donati Vincent Enouf, Maud Vanpeene, Sylvie van der Werf |
| EPI_ISL_415654 | hCoV-19/France/HF2174/2020 | Europe / France / Hauts de France / Com | 2020-03-09 | Centre Hospitalier Compiègne Laboratoire de Biologie | National Reference Center for Viruses of Respiratory Infections, Institut Pasteur, Paris | Mélnie Albert, Marion Barbet, Sylvie Behillil, Méline Bizard, Angela Brisebarre, Flora Donati Vincent Enouf, Maud Vanpeene, Sylvie van der Werf, Raulin Olivia |
| EPI_ISL_415655 | hCoV-19/Wales/PHW27/2020 | Europe / United Kingdom / Wales | 2020-03-12 | Wales Specialist Virology Centre | Public Health Wales Microbiology Cardiff | Catherine Moore, Joanne Watkins, Sally Corden, Tom Connor |
| EPI_ISL_415656 | hCoV-19/Wales/PHW33/2020 | Europe / United Kingdom / Wales | 2020-03-12 | Wales Specialist Virology Centre | Public Health Wales Microbiology Cardiff | Catherine Moore, Joanne Watkins, Sally Corden, Tom Connor |
| EPI_ISL_415657 | hCoV-19/Wales/PHW37/2020 | Europe / United Kingdom / Wales | 2020-03-12 | Wales Specialist Virology Centre | Public Health Wales Microbiology Cardiff | Catherine Moore, Joanne watkins, Sally Corden, Tom Connor |
| EPI_ISL_415658 | hCoV-19/Chile/Santiago_op2d1/2020 | South America / Chile / Santiago | 2020-03-06 | Laboratory of Molecular Virology, Pontificia Universidad Católica de Chile | MSHS Pathogen Surveillance Program | Rafael A. Medina, Pablo Vial, Tamara Garcia, Eileen Serrano, Ana Silvia Gonzalez-Reiche, Zenab Khan, Mitchell Sullivan, Ajay Obla, Matthew Hernandez, Hala Alshammary, Juan Soto, Shwetha Sridhar Hara, Ying-Chih Wang, Melissa Smith, Robert Sebra, Viviana Simon, Harm van Bakel |
| EPI_ISL_415659 | hCoV-19/Wales/PHW28/2020 | Europe / United Kingdom / Wales | 2020-03-12 | Wales Specialist Virology Centre | Public Health Wales Microbiology Cardiff | Catherine Moore, Joanne Watkins, Sally Corden, Tom Connor |
| EPI_ISL_415660 | hCoV-19/Chile/Santiago_op3d1/2020 | South America / Chile / Santiago | 2020-03-07 | Laboratory of Molecular Virology, Pontificia Universidad Católica de Chile | MSHS Pathogen Surveillance Program | Rafael A. Medina, Pablo Vial, Tamara Garcia, Eileen Serrano, Ana Silvia Gonzalez-Reiche, Zenab Khan, Mitchell Sullivan, Ajay Obla, Matthew Hernandez, Hala Alshammary, Juan Soto, Shwetha Sridhar Hara, Ying-Chih Wang, Melissa Smith, Robert Sebra, Viviana Simon, Harm van Bakel |
| EPI_ISL_415661 | hCoV-19/Chile/Santiago_op4d1/2020 | South America / Chile / Santiago | 2020-03-08 | Laboratory of Molecular Virology, Pontificia Universidad Católica de Chile | MSHS Pathogen Surveillance Program | Rafael A. Medina, Pablo Vial, Tamara Garcia, Eileen Serrano, Ana Silvia Gonzalez-Reiche, Zenab Khan, Mitchell Sullivan, Ajay Obla, Matthew Hernandez, Hala Alshammary, Juan Soto, Shwetha Sridhar Hara, Ying-Chih Wang, Melissa Smith, Robert Sebra, Viviana Simon, Harm van Bakel |
| EPI_ISL_415698 | hCoV-19/Switzerland/GR2988/2020 | Europe / Switzerland | 2020-02-27 | Hôpitaux universitaires de Genève Laboratoire de Virologie | Hôpitaux universitaires de Genève Laboratoire de Virologie | Laubscher F. |
| EPI_ISL_415699 | hCoV-19/Switzerland/GR3043/2020 | Europe / Switzerland | 2020-02-27 | Hôpitaux universitaires de Genève Laboratoire de Virologie | Hôpitaux universitaires de Genève Laboratoire de Virologie | Laubscher F. |
| EPI_ISL_416140 | hCoV-19/Denmark/SSI-05/2020 | Europe / Denmark / Copenhagen | 2020-03-02 | Department of Virus and Microbiological Special diagnostics, Statens Serum Institut, Copenhagen, Denmark. | Statens Serum Institute | Morten Rasmussen, Maiken Worsoe Rosenstierne , Anders Fomsgaard |
| EPI_ISL_416141 | hCoV-19/Denmark/SSI-09/2020 | Europe / Denmark / Copenhagen | 2020-03-03 | Department of Virus and Microbiological Special diagnostics, Statens Serum Institut, Copenhagen, Denmark. | Statens Serum Institute | Morten Rasmussen, Maiken Worsoe Rosenstierne , Anders Fomsgaard |
| EPI_ISL_416142 | hCoV-19/Denmark/SSI-01/2020 | Europe / Denmark / Copenhagen | 2020-02-26 | Department of Virus and Microbiological Special diagnostics, Statens Serum Institut, Copenhagen, Denmark. | Statens Serum Institute | Morten Rasmussen, Maiken Worsoe Rosenstierne , Anders Fomsgaard |
| EPI_ISL_416143 | hCoV-19/Denmark/SSI-02/2020 | Europe / Denmark / Copenhagen | 2020-02-28 | Department of Virus and Microbiological Special diagnostics, Statens Serum Institut, Copenhagen, Denmark. | VIFU | Morten Rasmussen, Maiken Worsoe Rosenstierne , Anders Fomsgaard |
| EPI_ISL_416144 | hCoV-19/Denmark/SSI-03/2020 | Europe / Denmark / Copenhagen | 2020-03-01 | Department of Virus and Microbiological Special diagnostics, Statens Serum Institut, Copenhagen, Denmark. | VIFU | Morten Rasmussen, Maiken Worsoe Rosenstierne , Anders Fomsgaard |
| EPI_ISL_416153 | hCoV-19/Denmark/SSI-04/2020 | Europe / Denmark / Copenhagen | 2020-03-02 | Department of Virus and Microbiological Special diagnostics, Statens Serum Institut, Copenhagen, Denmark. | VIFU | Morten Rasmussen, Maiken Worsoe Rosenstierne , Anders Fomsgaard |
| EPI_ISL_416501 | hCoV-19/France/IDF2284/2020 | Europe / France / Ile de France / Levallois | 2020-03-10 | Hopital franco britannique - Service des Urgences | National Reference Center for Viruses of Respiratory Infections, Institut Pasteur, Paris | Mélnie Albert, Marion Barbet, Sylvie Behillil, Méline Bizard, Angela Brisebarre, Flora Donati, Etienne Simon-Lorière, Vincent Enouf, Maud Vanpeene, Sylvie van der Werf |
| EPI_ISL_416502 | hCoV-19/France/B2330/2020 | Europe / France / Bretagne / Rennes | 2020-02-26 | CHRU Pontchaillou - Laboratoire de Virologie | National Reference Center for Viruses of Respiratory Infections, Institut Pasteur, Paris | Mélnie Albert, Marion Barbet, Sylvie Behillil, Méline Bizard, Angela Brisebarre, Flora Donati, Etienne Simon-Lorière, Vincent Enouf, Maud Vanpeene, Sylvie van der Werf, Gisèle Lagathu |
| EPI_ISL_416503 | hCoV-19/France/B2334/2020 | Europe / France / Bretagne / Rennes | 2020-03-01 | CHRU Pontchaillou - Laboratoire de Virologie | National Reference Center for Viruses of Respiratory Infections, Institut Pasteur, Paris | Mélnie Albert, Marion Barbet, Sylvie Behillil, Méline Bizard, Angela Brisebarre, Flora Donati, Etienne Simon-Lorière, Vincent Enouf, Maud Vanpeene, Sylvie van der Werf, Gisèle Lagathu |
| EPI_ISL_416504 | hCoV-19/France/B2335/2020 | Europe / France / Bretagne / Rennes | 2020-03-02 | CHRU Pontchaillou - Laboratoire de Virologie | National Reference Center for Viruses of Respiratory Infections, Institut Pasteur, Paris | Mélnie Albert, Marion Barbet, Sylvie Behillil, Méline Bizard, Angela Brisebarre, Flora Donati, Etienne Simon-Lorière, Vincent Enouf, Maud Vanpeene, Sylvie van der Werf, Gisèle Lagathu |
| EPI_ISL_416505 | hCoV-19/France/B2336/2020 | Europe / France / Bretagne / Rennes | 2020-03-02 | CHRU Pontchaillou - Laboratoire de Virologie | National Reference Center for Viruses of Respiratory Infections, Institut Pasteur, Paris | Mélnie Albert, Marion Barbet, Sylvie Behillil, Méline Bizard, Angela Brisebarre, Flora Donati, Etienne Simon-Lorière, Vincent Enouf, Maud Vanpeene, Sylvie van der Werf, Gisèle Lagathu |
| EPI_ISL_416506 | hCoV-19/France/B2337/2020 | Europe / France / Bretagne / Rennes | 2020-03-03 | CHRU Pontchaillou - Laboratoire de Virologie | National Reference Center for Viruses of Respiratory Infections, Institut Pasteur, Paris | Mélnie Albert, Marion Barbet, Sylvie Behillil, Méline Bizard, Angela Brisebarre, Flora Donati, Etienne Simon-Lorière, Vincent Enouf, Maud Vanpeene, Sylvie van der Werf, Gisèle Lagathu |
| EPI_ISL_416507 | hCoV-19/France/B2340/2020 | Europe / France / Bretagne / Rennes | 2020-03-05 | CHRU Pontchaillou - Laboratoire de Virologie | National Reference Center for Viruses of Respiratory Infections, Institut Pasteur, Paris | Mélnie Albert, Marion Barbet, Sylvie Behillil, Méline Bizard, Angela Brisebarre, Flora Donati, Etienne Simon-Lorière, Vincent Enouf, Maud Vanpeene, Sylvie van der Werf, Gisèle Lagathu |
| EPI_ISL_416508 | hCoV-19/France/B2343/2020 | Europe / France / Bretagne / Rennes | 2020-03-06 | CHRU Pontchaillou - Laboratoire de Virologie | National Reference Center for Viruses of Respiratory Infections, Institut Pasteur, Paris | Mélnie Albert, Marion Barbet, Sylvie Behillil, Méline Bizard, Angela Brisebarre, Flora Donati, Etienne Simon-Lorière, Vincent Enouf, Maud Vanpeene, Sylvie van der Werf, Gisèle Lagathu |
| EPI_ISL_416509 | hCoV-19/France/B2344/2020 | Europe / France / Bretagne / Rennes | 2020-03-06 | CHRU Pontchaillou - Laboratoire de Virologie | National Reference Center for Viruses of Respiratory Infections, Institut Pasteur, Paris | Mélnie Albert, Marion Barbet, Sylvie Behillil, Méline Bizard, Angela Brisebarre, Flora Donati, Etienne Simon-Lorière, Vincent Enouf, Maud Vanpeene, Sylvie van der Werf, Gisèle Lagathu |
| EPI_ISL_416510 | hCoV-19/France/B2346/2020 | Europe / France / Bretagne / Rennes | 2020-03-06 | CHRU Pontchaillou - Laboratoire de Virologie | National Reference Center for Viruses of Respiratory Infections, Institut Pasteur, Paris | Mélnie Albert, Marion Barbet, Sylvie Behillil, Méline Bizard, Angela Brisebarre, Flora Donati, Etienne Simon-Lorière, Vincent Enouf, Maud Vanpeene, Sylvie van der Werf, Gisèle Lagathu |
| EPI_ISL_416511 | hCoV-19/France/B2348/2020 | Europe / France / Bretagne / Rennes | 2020-03-07 | CHRU Pontchaillou - Laboratoire de Virologie | National Reference Center for Viruses of Respiratory Infections, Institut Pasteur, Paris | Mélnie Albert, Marion Barbet, Sylvie Behillil, Méline Bizard, Angela Brisebarre, Flora Donati, Etienne Simon-Lorière, Vincent Enouf, Maud Vanpeene, Sylvie van der Werf, Gisèle Lagathu |
| EPI_ISL_416512 | hCoV-19/France/B2349/2020 | Europe / France / Bretagne / Rennes | 2020-03-07 | CHRU Pontchaillou - Laboratoire de Virologie | National Reference Center for Viruses of Respiratory Infections, Institut Pasteur, Paris | Mélnie Albert, Marion Barbet, Sylvie Behillil, Méline Bizard, Angela Brisebarre, Flora Donati, Etienne Simon-Lorière, Vincent Enouf, Maud Vanpeene, Sylvie van der Werf, Gisèle Lagathu |
| EPI_ISL_416513 | hCoV-19/France/B2351/2020 | Europe / France / Bretagne / Rennes | 2020-03-07 | CHRU Pontchaillou - Laboratoire de Virologie | National Reference Center for Viruses of Respiratory Infections, Institut Pasteur, Paris | Mélnie Albert, Marion Barbet, Sylvie Behillil, Méline Bizard, Angela Brisebarre, Flora Donati, Etienne Simon-Lorière, Vincent Enouf, Maud Vanpeene, Sylvie van der Werf, Gisèle Lagathu |
| EPI_ISL_416514 | hCoV-19/Australia/VIC08/2020 | Oceania / Australia / Victoria / Melbourne | 2020-03-15 | Victorian Infectious Diseases Reference Laboratory (VIDRL) | Victorian Infectious Diseases Reference Laboratory and Microbiological Diagnostic Unit Public Health Laboratory, Doherty Institute | Caly L., Seemann T., Schultz M., Taiaroa, G., Druce J. |
| EPI_ISL_416515 | hCoV-19/Australia/VIC09/2020 | Oceania / Australia / Victoria / Melbourne | 2020-03-15 | Victorian Infectious Diseases Reference Laboratory (VIDRL) | Victorian Infectious Diseases Reference Laboratory and Microbiological Diagnostic Unit Public Health Laboratory, Doherty Institute | Caly L., Seemann T., Schultz M., Taiaroa, G., Druce J. |
| EPI_ISL_416516 | hCoV-19/Australia/VIC10/2020 | Oceania / Australia / Victoria / Melbourne | 2020-03-16 | Victorian Infectious Diseases Reference Laboratory (VIDRL) | Victorian Infectious Diseases Reference Laboratory and Microbiological Diagnostic Unit Public Health Laboratory, Doherty Institute | Caly L., Seemann T., Schultz M., Taiaroa, G., Druce J. |
| EPI_ISL_416519 | hCoV-19/New Zealand/20VR0189/2020 | Oceania / New Zealand / Auckland | 2020-03-02 | Auckland Hospital | Institute of Environmental Science and Research (ESR) | Matt Storey, Xiaoyun Ren, Gary McAuliffe, Sally Roberts, Matthew Blakiston, Erasmus Smit, Lauren Jelly, Joep de Ligt |
| EPI_ISL_416521 | hCoV-19/Saudi Arabia/SCDC-3321/2020 | Asia / Saudi Arabia | 2020-03-10 | Public Health Laboratory | Public Health Laboratory, Saudi CDC | Albarrag, A |
| EPI_ISL_416522 | hCoV-19/Saudi Arabia/SCDC-3324/2020 | Asia / Saudi Arabia | 2020-03-10 | Public Health Laboratory, Saudi CDC | Public Health Laboratory, Saudi CDC | Albarrag,A |
| EPI_ISL_416523 | hCoV-19/USA/WI-UW-03/2020 | North America / USA / Wisconsin / Madis | 2020-03-14 | University of Wisconsin-Madison AIDS Vaccine Research Laboratories | University of Wisconsin-Madison AIDS Vaccine Research Laboratories | Gage Moreno, Katarina Braun, et al. AIDS Vaccine Research Laboratories |
| EPI_ISL_416524 | hCoV-19/Japan/SMU-0311S2/2020 | Asia / Japan / Saitama | 2020-03-11 | Saitama Medical University Hospital | Saitama Medical University | Kazuo Imai |
| EPI_ISL_416525 | hCoV-19/Japan/SMU-0311S3/2020 | Asia / Japan / Saitama | 2020-03-11 | Saitama Medical University | Saitama Medical University | Kazuo Imai |
| EPI_ISL_416538 | hCoV-19/New Zealand/20VR0275/2020 | Oceania / New Zealand / Wellington | 2020-03-15 | Wellington Hospital | Institute of Environmental Science and Research (ESR) | Wellington SCL, Wellington Hospital, Riddiford Street, Newtown, Wellington 6021, New Zealand |
| EPI_ISL_416539 | hCoV-19/New Zealand/20VR0276/2020 | Oceania / New Zealand / Wellington | 2020-03-15 | Wellington Hospital | Institute of Environmental Science and Research (ESR) | Matt Storey, Xiaoyun Ren, Craig Thornley, Maxim Bloomfield, Erasmus Smit, Lauren Jelly, Joep de Ligt |
| EPI_ISL_416541 | hCoV-19/Kuwait/KU09/2020 | Asia / Kuwait / Dasman | 2020-03-02 | Dasman Diabetes Institute and Virology Laboratory Ministry of Health | Dasman Diabetes Institute | Fahd Al-Mulla, Sumi John, Rasheeba Iqbal, Motasem Melhem, Ebaa AlOzairi, Sara Al-Qabandi, Qais Al-Duwairi |
| EPI_ISL_416542 | hCoV-19/Kuwait/KU17/2020 | Asia / Kuwait / Dasman | 2020-03-02 | Dasman Diabetes Institute | Dasman Diabetes Institute | Fahd Al-Mulla, Sumi John, Rasheeba Iqbal, Motasem Melhem, Ebaa AlOzairi, Sara Al-Qabandi, Qais Al-Duwairi |
| EPI_ISL_416543 | hCoV-19/Kuwait/KU18/2020 | Asia / Kuwait / Dasman | 2020-03-02 | Dasman Diabetes Institute | Dasman Diabetes Institute | Fahd Al-Mulla, Rasheeba Iqbal, Sumi John, Motasem Melhem, Ebaa AlOzairi, Sara Al-Qabandi, Qais Al-Duwairi |
| EPI_ISL_416565 | hCoV-19/Japan/DP0005/2020 | Asia / Japan / unknown | 2020-02-15 | Japanese Quarantine Stations | Pathogen Genomics Center, National Institute of Infectious Diseases | Tsuyoshi Sekizuka, Kentaro Itokawa, Rina Tanaka, Masanori Hashino, Tsutomu Kageyama, Shinji Saito, Ikuyo Takayama, Hideki Hasegawa, Takuri Takahashi, Hajime Kamiya, Takuya Yamagishi, Motoi Suzuki, Takaji Wakita, Makoto Kuroda |
| EPI_ISL_416566 | hCoV-19/Japan/DP0027/2020 | Asia / Japan / unknown | 2020-02-15 | Japanese Quarantine Stations | Pathogen Genomics Center, National Institute of Infectious Diseases | Tsuyoshi Sekizuka, Kentaro Itokawa, Rina Tanaka, Masanori Hashino, Tsutomu Kageyama, Shinji Saito, Ikuyo Takayama, Hideki Hasegawa, Takuri Takahashi, Hajime Kamiya, Takuya Yamagishi, Motoi Suzuki, Takaji Wakita, Makoto Kuroda |
| EPI_ISL_416567 | hCoV-19/Japan/DP0037/2020 | Asia / Japan / unknown | 2020-02-15 | Japanese Quarantine Stations | Pathogen Genomics Center, National Institute of Infectious Diseases | Tsuyoshi Sekizuka, Kentaro Itokawa, Rina Tanaka, Masanori Hashino, Tsutomu Kageyama, Shinji Saito, Ikuyo Takayama, Hideki Hasegawa, Takuri Takahashi, Hajime Kamiya, Takuya Yamagishi, Motoi Suzuki, Takaji Wakita, Makoto Kuroda |
| EPI_ISL_416569 | hCoV-19/Japan/DP0059/2020 | Asia / Japan / unknown | 2020-02-15 | Japanese Quarantine Stations | Pathogen Genomics Center, National Institute of Infectious Diseases | Tsuyoshi Sekizuka, Kentaro Itokawa, Rina Tanaka, Masanori Hashino, Tsutomu Kageyama, Shinji Saito, Ikuyo Takayama, Hideki Hasegawa, Takuri Takahashi, Hajime Kamiya, Takuya Yamagishi, Motoi Suzuki, Takaji Wakita, Makoto Kuroda |
| EPI_ISL_416570 | hCoV-19/Japan/DP0065/2020 | Asia / Japan / unknown | 2020-02-15 | Japanese Quarantine Stations | Pathogen Genomics Center, National Institute of Infectious Diseases | Tsuyoshi Sekizuka, Kentaro Itokawa, Rina Tanaka, Masanori Hashino, Tsutomu Kageyama, Shinji Saito, Ikuyo Takayama, Hideki Hasegawa, Takuri Takahashi, Hajime Kamiya, Takuya Yamagishi, Motoi Suzuki, Takaji Wakita, Makoto Kuroda |
| EPI_ISL_416571 | hCoV-19/Japan/DP0077/2020 | Asia / Japan / unknown | 2020-02-15 | Japanese Quarantine Stations | Pathogen Genomics Center, National Institute of Infectious Diseases | Tsuyoshi Sekizuka, Kentaro Itokawa, Rina Tanaka, Masanori Hashino, Tsutomu Kageyama, Shinji Saito, Ikuyo Takayama, Hideki Hasegawa, Takuri Takahashi, Hajime Kamiya, Takuya Yamagishi, Motoi Suzuki, Takaji Wakita, Makoto Kuroda |
| EPI_ISL_416572 | hCoV-19/Japan/DP0078/2020 | Asia / Japan / unknown | 2020-02-15 | Japanese Quarantine Stations | Pathogen Genomics Center, National Institute of Infectious Diseases | Tsuyoshi Sekizuka, Kentaro Itokawa, Rina Tanaka, Masanori Hashino, Tsutomu Kageyama, Shinji Saito, Ikuyo Takayama, Hideki Hasegawa, Takuri Takahashi, Hajime Kamiya, Takuya Yamagishi, Motoi Suzuki, Takaji Wakita, Makoto Kuroda |
| EPI_ISL_416573 | hCoV-19/Japan/DP0104/2020 | Asia / Japan / unknown | 2020-02-15 | Japanese Quarantine Stations | Pathogen Genomics Center, National Institute of Infectious Diseases | Tsuyoshi Sekizuka, Kentaro Itokawa, Rina Tanaka, Masanori Hashino, Tsutomu Kageyama, Shinji Saito, Ikuyo Takayama, Hideki Hasegawa, Takuri Takahashi, Hajime Kamiya, Takuya Yamagishi, Motoi Suzuki, Takaji Wakita, Makoto Kuroda |
| EPI_ISL_416574 | hCoV-19/Japan/DP0107/2020 | Asia / Japan / unknown | 2020-02-15 | Japanese Quarantine Stations | Pathogen Genomics Center, National Institute of Infectious Diseases | Tsuyoshi Sekizuka, Kentaro Itokawa, Rina Tanaka, Masanori Hashino, Tsutomu Kageyama, Shinji Saito, Ikuyo Takayama, Hideki Hasegawa, Takuri Takahashi, Hajime Kamiya, Takuya Yamagishi, Motoi Suzuki, Takaji Wakita, Makoto Kuroda |
| EPI_ISL_416575 | hCoV-19/Japan/DP0121/2020 | Asia / Japan / unknown | 2020-02-15 | Japanese Quarantine Stations | Pathogen Genomics Center, National Institute of Infectious Diseases | Tsuyoshi Sekizuka, Kentaro Itokawa, Rina Tanaka, Masanori Hashino, Tsutomu Kageyama, Shinji Saito, Ikuyo Takayama, Hideki Hasegawa, Takuri Takahashi, Hajime Kamiya, Takuya Yamagishi, Motoi Suzuki, Takaji Wakita, Makoto Kuroda |
| EPI_ISL_416576 | hCoV-19/Japan/DP0133/2020 | Asia / Japan / unknown | 2020-02-15 | Japanese Quarantine Stations | Pathogen Genomics Center, National Institute of Infectious Diseases | Tsuyoshi Sekizuka, Kentaro Itokawa, Rina Tanaka, Masanori Hashino, Tsutomu Kageyama, Shinji Saito, Ikuyo Takayama, Hideki Hasegawa, Takuri Takahashi, Hajime Kamiya, Takuya Yamagishi, Motoi Suzuki, Takaji Wakita, Makoto Kuroda |
| EPI_ISL_416577 | hCoV-19/Japan/DP0134/2020 | Asia / Japan / unknown | 2020-02-15 | Japanese Quarantine Stations | Pathogen Genomics Center, National Institute of Infectious Diseases | Tsuyoshi Sekizuka, Kentaro Itokawa, Rina Tanaka, Masanori Hashino, Tsutomu Kageyama, Shinji Saito, Ikuyo Takayama, Hideki Hasegawa, Takuri Takahashi, Hajime Kamiya, Takuya Yamagishi, Motoi Suzuki, Takaji Wakita, Makoto Kuroda |
| EPI_ISL_416578 | hCoV-19/Japan/DP0152/2020 | Asia / Japan / unknown | 2020-02-15 | Japanese Quarantine Stations | Pathogen Genomics Center, National Institute of Infectious Diseases | Tsuyoshi Sekizuka, Kentaro Itokawa, Rina Tanaka, Masanori Hashino, Tsutomu Kageyama, Shinji Saito, Ikuyo Takayama, Hideki Hasegawa, Takuri Takahashi, Hajime Kamiya, Takuya Yamagishi, Motoi Suzuki, Takaji Wakita, Makoto Kuroda |
| EPI_ISL_416579 | hCoV-19/Japan/DP0158/2020 | Asia / Japan / unknown | 2020-02-15 | Japanese Quarantine Stations | Pathogen Genomics Center, National Institute of Infectious Diseases | Tsuyoshi Sekizuka, Kentaro Itokawa, Rina Tanaka, Masanori Hashino, Tsutomu Kageyama, Shinji Saito, Ikuyo Takayama, Hideki Hasegawa, Takuri Takahashi, Hajime Kamiya, Takuya Yamagishi, Motoi Suzuki, Takaji Wakita, Makoto Kuroda |
| EPI_ISL_416580 | hCoV-19/Japan/DP0184/2020 | Asia / Japan / unknown | 2020-02-15 | Japanese Quarantine Stations | Pathogen Genomics Center, National Institute of Infectious Diseases | Tsuyoshi Sekizuka, Kentaro Itokawa, Rina Tanaka, Masanori Hashino, Tsutomu Kageyama, Shinji Saito, Ikuyo Takayama, Hideki Hasegawa, Takuri Takahashi, Hajime Kamiya, Takuya Yamagishi, Motoi Suzuki, Takaji Wakita, Makoto Kuroda |
| EPI_ISL_416581 | hCoV-19/Japan/DP0190/2020 | Asia / Japan / unknown | 2020-02-15 | Japanese Quarantine Stations | Pathogen Genomics Center, National Institute of Infectious Diseases | Tsuyoshi Sekizuka, Kentaro Itokawa, Rina Tanaka, Masanori Hashino, Tsutomu Kageyama, Shinji Saito, Ikuyo Takayama, Hideki Hasegawa, Takuri Takahashi, Hajime Kamiya, Takuya Yamagishi, Motoi Suzuki, Takaji Wakita, Makoto Kuroda |
| EPI_ISL_416582 | hCoV-19/Japan/DP0191/2020 | Asia / Japan / unknown | 2020-02-15 | Japanese Quarantine Stations | Pathogen Genomics Center, National Institute of Infectious Diseases | Tsuyoshi Sekizuka, Kentaro Itokawa, Rina Tanaka, Masanori Hashino, Tsutomu Kageyama, Shinji Saito, Ikuyo Takayama, Hideki Hasegawa, Takuri Takahashi, Hajime Kamiya, Takuya Yamagishi, Motoi Suzuki, Takaji Wakita, Makoto Kuroda |
| EPI_ISL_416583 | hCoV-19/Japan/DP0196/2020 | Asia / Japan / unknown | 2020-02-15 | Japanese Quarantine Stations | Pathogen Genomics Center, National Institute of Infectious Diseases | Tsuyoshi Sekizuka, Kentaro Itokawa, Rina Tanaka, Masanori Hashino, Tsutomu Kageyama, Shinji Saito, Ikuyo Takayama, Hideki Hasegawa, Takuri Takahashi, Hajime Kamiya, Takuya Yamagishi, Motoi Suzuki, Takaji Wakita, Makoto Kuroda |
| EPI_ISL_416584 | hCoV-19/Japan/DP0200/2020 | Asia / Japan / unknown | 2020-02-15 | Japanese Quarantine Stations | Pathogen Genomics Center, National Institute of Infectious Diseases | Tsuyoshi Sekizuka, Kentaro Itokawa, Rina Tanaka, Masanori Hashino, Tsutomu Kageyama, Shinji Saito, Ikuyo Takayama, Hideki Hasegawa, Takuri Takahashi, Hajime Kamiya, Takuya Yamagishi, Motoi Suzuki, Takaji Wakita, Makoto Kuroda |
| EPI_ISL_416585 | hCoV-19/Japan/DP0236/2020 | Asia / Japan / unknown | 2020-02-16 | Japanese Quarantine Stations | Pathogen Genomics Center, National Institute of Infectious Diseases | Tsuyoshi Sekizuka, Kentaro Itokawa, Rina Tanaka, Masanori Hashino, Tsutomu Kageyama, Shinji Saito, Ikuyo Takayama, Hideki Hasegawa, Takuri Takahashi, Hajime Kamiya, Takuya Yamagishi, Motoi Suzuki, Takaji Wakita, Makoto Kuroda |
| EPI_ISL_416586 | hCoV-19/Japan/DP0274/2020 | Asia / Japan / unknown | 2020-02-16 | Japanese Quarantine Stations | Pathogen Genomics Center, National Institute of Infectious Diseases | Tsuyoshi Sekizuka, Kentaro Itokawa, Rina Tanaka, Masanori Hashino, Tsutomu Kageyama, Shinji Saito, Ikuyo Takayama, Hideki Hasegawa, Takuri Takahashi, Hajime Kamiya, Takuya Yamagishi, Motoi Suzuki, Takaji Wakita, Makoto Kuroda |
| EPI_ISL_416587 | hCoV-19/Japan/DP0278/2020 | Asia / Japan / unknown | 2020-02-16 | Japanese Quarantine Stations | Pathogen Genomics Center, National Institute of Infectious Diseases | Tsuyoshi Sekizuka, Kentaro Itokawa, Rina Tanaka, Masanori Hashino, Tsutomu Kageyama, Shinji Saito, Ikuyo Takayama, Hideki Hasegawa, Takuri Takahashi, Hajime Kamiya, Takuya Yamagishi, Motoi Suzuki, Takaji Wakita, Makoto Kuroda |
| EPI_ISL_416589 | hCoV-19/Japan/DP0287/2020 | Asia / Japan / unknown | 2020-02-16 | Japanese Quarantine Stations | Pathogen Genomics Center, National Institute of Infectious Diseases | Tsuyoshi Sekizuka, Kentaro Itokawa, Rina Tanaka, Masanori Hashino, Tsutomu Kageyama, Shinji Saito, Ikuyo Takayama, Hideki Hasegawa, Takuri Takahashi, Hajime Kamiya, Takuya Yamagishi, Motoi Suzuki, Takaji Wakita, Makoto Kuroda |
| EPI_ISL_416590 | hCoV-19/Japan/DP0289/2020 | Asia / Japan / unknown | 2020-02-16 | Japanese Quarantine Stations | Pathogen Genomics Center, National Institute of Infectious Diseases | Tsuyoshi Sekizuka, Kentaro Itokawa, Rina Tanaka, Masanori Hashino, Tsutomu Kageyama, Shinji Saito, Ikuyo Takayama, Hideki Hasegawa, Takuri Takahashi, Hajime Kamiya, Takuya Yamagishi, Motoi Suzuki, Takaji Wakita, Makoto Kuroda |
| EPI_ISL_416591 | hCoV-19/Japan/DP0290/2020 | Asia / Japan / unknown | 2020-02-16 | Japanese Quarantine Stations | Pathogen Genomics Center, National Institute of Infectious Diseases | Tsuyoshi Sekizuka, Kentaro Itokawa, Rina Tanaka, Masanori Hashino, Tsutomu Kageyama, Shinji Saito, Ikuyo Takayama, Hideki Hasegawa, Takuri Takahashi, Hajime Kamiya, Takuya Yamagishi, Motoi Suzuki, Takaji Wakita, Makoto Kuroda |
| EPI_ISL_416592 | hCoV-19/Japan/DP0294/2020 | Asia / Japan / unknown | 2020-02-16 | Japanese Quarantine Stations | Pathogen Genomics Center, National Institute of Infectious Diseases | Tsuyoshi Sekizuka, Kentaro Itokawa, Rina Tanaka, Masanori Hashino, Tsutomu Kageyama, Shinji Saito, Ikuyo Takayama, Hideki Hasegawa, Takuri Takahashi, Hajime Kamiya, Takuya Yamagishi, Motoi Suzuki, Takaji Wakita, Makoto Kuroda |
| EPI_ISL_416593 | hCoV-19/Japan/DP0311/2020 | Asia / Japan / unknown | 2020-02-16 | Japanese Quarantine Stations | Pathogen Genomics Center, National Institute of Infectious Diseases | Tsuyoshi Sekizuka, Kentaro Itokawa, Rina Tanaka, Masanori Hashino, Tsutomu Kageyama, Shinji Saito, Ikuyo Takayama, Hideki Hasegawa, Takuri Takahashi, Hajime Kamiya, Takuya Yamagishi, Motoi Suzuki, Takaji Wakita, Makoto Kuroda |
| EPI_ISL_416594 | hCoV-19/Japan/DP0319/2020 | Asia / Japan / unknown | 2020-02-16 | Japanese Quarantine Stations | Pathogen Genomics Center, National Institute of Infectious Diseases | Tsuyoshi Sekizuka, Kentaro Itokawa, Rina Tanaka, Masanori Hashino, Tsutomu Kageyama, Shinji Saito, Ikuyo Takayama, Hideki Hasegawa, Takuri Takahashi, Hajime Kamiya, Takuya Yamagishi, Motoi Suzuki, Takaji Wakita, Makoto Kuroda |
| EPI_ISL_416595 | hCoV-19/Japan/DP0328/2020 | Asia / Japan / unknown | 2020-02-16 | Japanese Quarantine Stations | Pathogen Genomics Center, National Institute of Infectious Diseases | Tsuyoshi Sekizuka, Kentaro Itokawa, Rina Tanaka, Masanori Hashino, Tsutomu Kageyama, Shinji Saito, Ikuyo Takayama, Hideki Hasegawa, Takuri Takahashi, Hajime Kamiya, Takuya Yamagishi, Motoi Suzuki, Takaji Wakita, Makoto Kuroda |
| EPI_ISL_416596 | hCoV-19/Japan/DP0344/2020 | Asia / Japan / unknown | 2020-02-16 | Japanese Quarantine Stations | Pathogen Genomics Center, National Institute of Infectious Diseases | Tsuyoshi Sekizuka, Kentaro Itokawa, Rina Tanaka, Masanori Hashino, Tsutomu Kageyama, Shinji Saito, Ikuyo Takayama, Hideki Hasegawa, Takuri Takahashi, Hajime Kamiya, Takuya Yamagishi, Motoi Suzuki, Takaji Wakita, Makoto Kuroda |
| EPI_ISL_416597 | hCoV-19/Japan/DP0346/2020 | Asia / Japan / unknown | 2020-02-16 | Japanese Quarantine Stations | Pathogen Genomics Center, National Institute of Infectious Diseases | Tsuyoshi Sekizuka, Kentaro Itokawa, Rina Tanaka, Masanori Hashino, Tsutomu Kageyama, Shinji Saito, Ikuyo Takayama, Hideki Hasegawa, Takuri Takahashi, Hajime Kamiya, Takuya Yamagishi, Motoi Suzuki, Takaji Wakita, Makoto Kuroda |
| EPI_ISL_416598 | hCoV-19/Japan/DP0357/2020 | Asia / Japan / unknown | 2020-02-16 | Japanese Quarantine Stations | Pathogen Genomics Center, National Institute of Infectious Diseases | Tsuyoshi Sekizuka, Kentaro Itokawa, Rina Tanaka, Masanori Hashino, Tsutomu Kageyama, Shinji Saito, Ikuyo Takayama, Hideki Hasegawa, Takuri Takahashi, Hajime Kamiya, Takuya Yamagishi, Motoi Suzuki, Takaji Wakita, Makoto Kuroda |
| EPI_ISL_416599 | hCoV-19/Japan/DP0361/2020 | Asia / Japan / unknown | 2020-02-16 | Japanese Quarantine Stations | Pathogen Genomics Center, National Institute of Infectious Diseases | Tsuyoshi Sekizuka, Kentaro Itokawa, Rina Tanaka, Masanori Hashino, Tsutomu Kageyama, Shinji Saito, Ikuyo Takayama, Hideki Hasegawa, Takuri Takahashi, Hajime Kamiya, Takuya Yamagishi, Motoi Suzuki, Takaji Wakita, Makoto Kuroda |
| EPI_ISL_416907 | hCoV-19/Malaysia/MKAK-CL-2020-7554/2020 | Asia / Malaysia / Selangor | 2020-02-06 | National Public Health Laboratory | Malaysia Genome Institute | Mohd Noor Mat Isa, Irni Suhayu Sapian, Yusuf Muhammad Noor, Nurhezreen Md Iqbal, Mohd Faizal Abu Bakar, Enizza Kasim, Shamsidar Sopie, Siti Noraini Othman, Azrin Ahmad, Nor Azfa Johari, Norazimah Tajudin, Noorliza Mohamad Noordin, W Afiza W Mohd Arifin, Rehan Shuhada Abu Bakar, Yu Kie Chem, Selvanesan Sengol, Hani Mat Hussin, Shahrul Hisham Zainal Ariffin |
| EPI_ISL_416994 | hCoV-19/Spain/CastillayLeon201437/2020 | Europe / Spain / Castilla y León | 2020-03-04 | COMPLEJO ASISTENCIAL UNIVERSITARIO DE BURGOS | Instituto de Salud Carlos III | Iglesias-Caballero, M. Molinero Calamita, M. González-Esguevillas, M. Camarero S. Pozo F. Casas I. Jiménez P. Jiménez M. Zaballos A. Monzón, S. Varona, S. Juliá M. Cuesta I. Megias Lobón, G. Hospital: ------ |
| EPI_ISL_416997 | hCoV-19/Belgium/ULG-2971/2020 | Europe / Belgium / Liège | 2020-03-05 | Department of Clinical Microbiology | GIGA Medical Genomics | Durkin Keith, Artesi Maria, Bontems Sébastien, Boreux Raphaël, Meex Cécile, Melin Pierrette, Hayette Marie-Pierre, Bours Vincent. |
| EPI_ISL_417004 | hCoV-19/Belgium/ULG-3000/2020 | Europe / Belgium / Liège | 2020-03-05 | Department of Clinical Microbiology | GIGA Medical Genomics | Durkin Keith, Artesi Maria, Bontems Sébastien, Boreux Raphaël, Meex Cécile, Melin Pierrette, Hayette Marie-Pierre, Bours Vincent. |
| EPI_ISL_417005 | hCoV-19/Belgium/ULG-3162/2020 | Europe / Belgium / Liège | 2020-03-05 | Department of Clinical Microbiology | GIGA Medical Genomics | Durkin Keith, Artesi Maria, Bontems Sébastien, Boreux Raphaël, Meex Cécile, Melin Pierrette, Hayette Marie-Pierre, Bours Vincent. |
| EPI_ISL_417006 | hCoV-19/Belgium/ULG-3163/2020 | Europe / Belgium / Liège | 2020-03-05 | Department of Clinical Microbiology | GIGA Medical Genomics | Durkin Keith, Artesi Maria, Bontems Sébastien, Boreux Raphaël, Meex Cécile, Melin Pierrette, Hayette Marie-Pierre, Bours Vincent. |
| EPI_ISL_417007 | hCoV-19/Spain/Galicia201663/2020 | Europe / Spain / Galicia | 2020-03-07 | HOSPITAL SANTA MARIA NAI | Instituto de Salud Carlos III | Iglesias-Caballero, M. Molinero Calamita, M. González-Esguevillas, M. Camarero S. Pozo F. Casas I. Jiménez, P. Jiménez, M. Zaballos, A. Monzón, S. Varona, S. Juliá, M. Cuesta, I. García Costa, J. |
| EPI_ISL_417008 | hCoV-19/Belgium/ULG-3662/2020 | Europe / Belgium / Liège | 2020-03-07 | Department of Clinical Microbiology | GIGA Medical Genomics | Durkin Keith, Artesi Maria, Bontems Sébastien, Boreux Raphaël, Meex Cécile, Melin Pierrette, Hayette Marie-Pierre, Bours Vincent. |
| EPI_ISL_417009 | hCoV-19/Belgium/ULG-3665/2020 | Europe / Belgium / Liège | 2020-03-07 | Department of Clinical Microbiology | GIGA Medical Genomics | Durkin Keith, Artesi Maria, Bontems Sébastien, Boreux Raphaël, Meex Cécile, Melin Pierrette, Hayette Marie-Pierre, Bours Vincent. |
| EPI_ISL_417010 | hCoV-19/Spain/Madrid201442/2020 | Europe / Spain / Madrid | 2020-03-04 | FUNDACION JIMENEZ DIAZ | Instituto de Salud Carlos III | Iglesias-Caballero, M. Molinero Calamita, M. González-Esguevillas, M. Camarero, S. Pozo, F. Casas, I. Jiménez, P. Jiménez, M. Zaballos, A. Monzón, S. Varona, S. Juliá, M. Cuesta, I. Fernández Robles, R. |
| EPI_ISL_417012 | hCoV-19/Belgium/ULG-3843/2020 | Europe / Belgium / Liège | 2020-03-08 | Department of Clinical Microbiology | GIGA Medical Genomics | Durkin Keith, Artesi Maria, Bontems Sébastien, Boreux Raphaël, Meex Cécile, Melin Pierrette, Hayette Marie-Pierre, Bours Vincent. |
| EPI_ISL_417013 | hCoV-19/Belgium/ULG-4163/2020 | Europe / Belgium / Liège | 2020-03-09 | Department of Clinical Microbiology | GIGA Medical Genomics | Durkin Keith, Artesi Maria, Bontems Sébastien, Boreux Raphaël, Meex Cécile, Melin Pierrette, Hayette Marie-Pierre, Bours Vincent. |
| EPI_ISL_417014 | hCoV-19/Belgium/ULG-6216/2020 | Europe / Belgium / Liège | 2020-03-13 | Department of Clinical Microbiology | GIGA Medical Genomics | Durkin Keith, Artesi Maria, Bontems Sébastien, Boreux Raphaël, Meex Cécile, Melin Pierrette, Hayette Marie-Pierre, Bours Vincent. |
| EPI_ISL_417015 | hCoV-19/Belgium/ULG-6457/2020 | Europe / Belgium / Liège | 2020-03-13 | Department of Clinical Microbiology | GIGA Medical Genomics | Durkin Keith, Artesi Maria, Bontems Sébastien, Boreux Raphaël, Meex Cécile, Melin Pierrette, Hayette Marie-Pierre, Bours Vincent. |
| EPI_ISL_417016 | hCoV-19/Belgium/ULG-6503/2020 | Europe / Belgium / Liège | 2020-03-13 | Department of Clinical Microbiology | GIGA Medical Genomics | Durkin Keith, Artesi Maria, Bontems Sébastien, Boreux Raphaël, Meex Cécile, Melin Pierrette, Hayette Marie-Pierre, Bours Vincent. |
| EPI_ISL_417017 | hCoV-19/Belgium/ULG-6638/2020 | Europe / Belgium / Liège | 2020-03-14 | Department of Clinical Microbiology | GIGA Medical Genomics | Durkin Keith, Artesi Maria, Bontems Sébastien, Boreux Raphaël, Meex Cécile, Melin Pierrette, Hayette Marie-Pierre, Bours Vincent. |
| EPI_ISL_417018 | hCoV-19/Belgium/ULG-6670/2020 | Europe / Belgium / Liège | 2020-03-14 | Department of Clinical Microbiology | GIGA Medical Genomics | Durkin Keith, Artesi Maria, Bontems Sébastien, Boreux Raphaël, Meex Cécile, Melin Pierrette, Hayette Marie-Pierre, Bours Vincent. |
| EPI_ISL_417019 | hCoV-19/Belgium/ULG-6754/2020 | Europe / Belgium / Liege | 2020-03-14 | Department of Clinical Microbiology | GIGA Medical Genomics | Durkin Keith, Artesi Maria, Bontems Sébastien, Boreux Raphaël, Meex Cécile, Melin Pierrette, Hayette Marie-Pierre, Bours Vincent. |
| EPI_ISL_417027 | hCoV-19/USA/UT-09/2020 | North America / USA / Utah | 2020-03-20 | Utah Public Health Laboratory | Utah Public Health Laboratory | Erin Young, Kelly Oakeson |
| EPI_ISL_417030 | hCoV-19/Australia/NSW04/2020 | Oceania / Australia / New South Wales / | 2020-01-24 | Centre for Infectious Diseases and Microbiology Laboratory Services | NSW Health Pathology - Institute of Clinical Pathology and Medical Research; Westmead Hospital; University of Sydney | Eden J-S, Rockett R, Carter I, Rahman H, Holmes EC, O'Sullivan MV, Sintchenko V, Chen SC, Maddocks S, Kok J and Dwyer DE for the 2019-nCoV Study Group* |
| EPI_ISL_417031 | hCoV-19/Australia/QLDID919/2020 | Oceania / Australia / Queensland / Gold C | 2020-03-11 | Pathology Queensland | Public Health Virology Laboratory | Bixing Huang, Alyssa Pyke, Amanda De Jong, Andrew Van Den Hurk, Carmel Taylor, David Warrilow, Doris Genge, Elisabeth Gamez, Glen Hewitson, Ian Maxwell Mackay, Inga Sultana, Jamie McMahon, Jean Barcelon, Judy Northill, Mitchell Finger, Natalie Simpson, Neelima Nair, Peter Burtonclay, Peter Moore, Sarah Wheatley, Sean Moody, Sonja Hall-Mendelin, Timothy Gardam, and Frederick Moore |
| EPI_ISL_417032 | hCoV-19/Australia/QLDID920/2020 | Oceania / Australia / Queensland / Rockh | 2020-03-11 | Rockhampton Base Hospital | Public Health Virology Laboratory | Bixing Huang, Alyssa Pyke, Amanda De Jong, Andrew Van Den Hurk, Carmel Taylor, David Warrilow, Doris Genge, Elisabeth Gamez, Glen Hewitson, Ian Maxwell Mackay, Inga Sultana, Jamie McMahon, Jean Barcelon, Judy Northill, Mitchell Finger, Natalie Simpson, Neelima Nair, Peter Burtonclay, Peter Moore, Sarah Wheatley, Sean Moody, Sonja Hall-Mendelin, Timothy Gardam, and Frederick Moore |
| EPI_ISL_417033 | hCoV-19/Australia/QLDID921/2020 | Oceania / Australia / Queensland / Brisba | 2020-03-11 | Sullivan Nicolaides Pathology | Public Health Virology Laboratory | Bixing Huang, Alyssa Pyke, Amanda De Jong, Andrew Van Den Hurk, Carmel Taylor, David Warrilow, Doris Genge, Elisabeth Gamez, Glen Hewitson, Ian Maxwell Mackay, Inga Sultana, Jamie McMahon, Jean Barcelon, Judy Northill, Mitchell Finger, Natalie Simpson, Neelima Nair, Peter Burtonclay, Peter Moore, Sarah Wheatley, Sean Moody, Sonja Hall-Mendelin, Timothy Gardam, and Frederick Moore |
| EPI_ISL_417034 | hCoV-19/Brazil/AMBR-02/2020 | South America / Brazil / Amazonas State | 2020-03-16 | Laboratorio de Ecologia de Doencas Transmissiveis na Amazonia, Instituto Leonidas e Maria Deane - Fiocruz Amazonia | Laboratorio de Ecologia de Doencas Transmissiveis na Amazonia, Instituto Leonidas e Maria Deane - Fiocruz Amazonia | Valdinete Nascimento, André Corado, Fernanda Nascimento, Ágatha Costa, Debora Duarte, Luciana Gonçalves, Michele Jesus, Sérgio Luz, Felipe Naveca |
| EPI_ISL_417064 | hCoV-19/Hong Kong/VB20017970-2/2020 | Asia / Hong Kong | 2020-01-21 | Prince of Wales Hospital | Hong Kong Department of Health | Alan K.L. Tsang, Peter C.W. Yip, Edman T.K. Lam, Rickjason C.W. Chan, Dominic N.C. Tsang |
| EPI_ISL_417065 | hCoV-19/USA/WA-S12/2020 | North America / USA / Washington | 2020-03-03 | Washington State Department of Health | Seattle Flu Study | Chu etl al |
| EPI_ISL_417066 | hCoV-19/USA/WA-S13/2020 | North America / USA / Washington / King | 2020-03-03 | Washington State Department of Health | Seattle Flu Study | Chu etl al |
| EPI_ISL_417068 | hCoV-19/USA/WA-S15/2020 | North America / USA / Washington / King | 2020-03-02 | Washington State Department of Health | Seattle Flu Study | Chu etl al |
| EPI_ISL_417069 | hCoV-19/USA/WA-S16/2020 | North America / USA / Washington | 2020-03-03 | Washington State Department of Health | Seattle Flu Study | Chu etl al |
| EPI_ISL_417070 | hCoV-19/USA/WA-S17/2020 | North America / USA / Washington / King | 2020-03-03 | Washington State Department of Health | Seattle Flu Study | Chu etl al |
| EPI_ISL_417071 | hCoV-19/USA/WA-S18/2020 | North America / USA / Washington / King | 2020-03-03 | Washington State Department of Health | Seattle Flu Study | Chu etl al |
| EPI_ISL_417072 | hCoV-19/USA/WA-S19/2020 | North America / USA / Washington / King | 2020-03-02 | Washington State Department of Health | Seattle Flu Study | Chu etl al |
| EPI_ISL_417073 | hCoV-19/USA/WA-S20/2020 | North America / USA / Washington | 2020-03-02 | Washington State Department of Health | Seattle Flu Study | Chu etl al |
| EPI_ISL_417074 | hCoV-19/USA/WA-S21/2020 | North America / USA / Washington / King | 2020-03-02 | Washington State Department of Health | Seattle Flu Study | Chu etl al |
| EPI_ISL_417075 | hCoV-19/USA/WA-S22/2020 | North America / USA / Washington / Snoh | 2020-03-02 | Washington State Department of Health | Seattle Flu Study | Chu etl al |
| EPI_ISL_417076 | hCoV-19/USA/WA-S23/2020 | North America / USA / Washington / Snoh | 2020-03-02 | Washington State Department of Health | Seattle Flu Study | Chu etl al |
| EPI_ISL_417077 | hCoV-19/USA/WA-S24/2020 | North America / USA / Washington / Snoh | 2020-03-02 | Washington State Department of Health | Seattle Flu Study | Chu etl al |
| EPI_ISL_417079 | hCoV-19/USA/WA-S26/2020 | North America / USA / Washington / Snoh | 2020-03-02 | Washington State Department of Health | Seattle Flu Study | Chu etl al |
| EPI_ISL_417081 | hCoV-19/USA/WA-S28/2020 | North America / USA / Washington / Snoh | 2020-03-02 | Washington State Department of Health | Seattle Flu Study | Chu etl al |
| EPI_ISL_417082 | hCoV-19/USA/WA-S29/2020 | North America / USA / Washington / Gran | 2020-03-02 | Washington State Department of Health | Seattle Flu Study | Chu etl al |
| EPI_ISL_417083 | hCoV-19/USA/WA-S30/2020 | North America / USA / Washington / King | 2020-02-28 | Washington State Department of Health | Seattle Flu Study | Chu etl al |
| EPI_ISL_417085 | hCoV-19/USA/WA-S32/2020 | North America / USA / Washington / Snoh | 2020-03-02 | Washington State Department of Health | Seattle Flu Study | Chu etl al |
| EPI_ISL_417086 | hCoV-19/USA/WA-S33/2020 | North America / USA / Washington / Clark | 2020-03-01 | Washington State Department of Health | Seattle Flu Study | Chu etl al |
| EPI_ISL_417087 | hCoV-19/USA/WA-S34/2020 | North America / USA / Washington | 2020-03-02 | Washington State Department of Health | Seattle Flu Study | Chu etl al |
| EPI_ISL_417088 | hCoV-19/USA/WA-S35/2020 | North America / USA / Washington | 2020-03-02 | Washington State Department of Health | Seattle Flu Study | Chu etl al |
| EPI_ISL_417089 | hCoV-19/USA/WA-S36/2020 | North America / USA / Washington | 2020-03-02 | Washington State Department of Health | Seattle Flu Study | Chu etl al |
| EPI_ISL_417400 | hCoV-19/Australia/NSW44/2020 | Oceania / Australia / New South Wales / | 2020-03-10 | Centre for Infectious Diseases and Microbiology Public Health | NSW Health Pathology - Institute of Clinical Pathology and Medical Research; Westmead Hospital; University of Sydney | Rockett R, Eden J-S, Lam C, Gray K, Timms V, Gall M, Arnott A, Sadsad R, Carter I, Rahman H, Holmes EC, O'Sullivan MV, Sintchenko V, Chen SC, Maddocks S, Kok J and Dwyer DE for the 2019-nCoV Study Group |
| EPI_ISL_417401 | hCoV-19/Australia/NSW45/2020 | Oceania / Australia / New South Wales / | 2020-03-13 | Centre for Infectious Diseases and Microbiology Public Health | NSW Health Pathology - Institute of Clinical Pathology and Medical Research; Westmead Hospital; University of Sydney | Eden J-S, Lam C, Gray K, Timms V, Gall M, Arnott A, Sadsad R, Carter I, Rahman H, Holmes EC, O'Sullivan MV, Sintchenko V, Chen SC, Maddocks S, Kok J, Dwyer DE and Rockett R for the 2019-nCoV Study Group |
| EPI_ISL_417402 | hCoV-19/Australia/NSW47/2020 | Oceania / Australia / New South Wales / | 2020-03-13 | Centre for Infectious Diseases and Microbiology Public Health | NSW Health Pathology - Institute of Clinical Pathology and Medical Research; Westmead Hospital; University of Sydney | Lam C, Gray K, Timms V, Gall M, Arnott A, Sadsad R, Carter I, Rahman H, Holmes EC, O'Sullivan MV, Sintchenko V, Chen SC, Maddocks S, Kok J, Dwyer DE, Rockett R and Eden J-S for the 2019-nCoV Study Group |
| EPI_ISL_417403 | hCoV-19/Australia/NSW48/2020 | Oceania / Australia / New South Wales / | 2020-03-14 | Centre for Infectious Diseases and Microbiology Public Health | NSW Health Pathology - Institute of Clinical Pathology and Medical Research; Westmead Hospital; University of Sydney | Gray K, Timms V, Gall M, Arnott A, Sadsad R, Carter I, Rahman H, Holmes EC, O'Sullivan MV, Sintchenko V, Chen SC, Maddocks S, Kok J, Dwyer DE, Rockett R, Eden J-S and Lam C for the 2019-nCoV Study Group |
| EPI_ISL_417404 | hCoV-19/Australia/NSW50/2020 | Oceania / Australia / New South Wales / | 2020-03-12 | Centre for Infectious Diseases and Microbiology Public Health | NSW Health Pathology - Institute of Clinical Pathology and Medical Research; Westmead Hospital; University of Sydney | Timms V, Gall M, Arnott A, Sadsad R, Carter I, Rahman H, Holmes EC, O'Sullivan MV, Sintchenko V, Chen SC, Maddocks S, Kok J, Dwyer DE, Rockett R, Eden J-S, Lam C and Gray K for the 2019-nCoV Study Group |
| EPI_ISL_417405 | hCoV-19/Australia/NSW52/2020 | Oceania / Australia / New South Wales / | 2020-03-11 | Centre for Infectious Diseases and Microbiology Public Health | NSW Health Pathology - Institute of Clinical Pathology and Medical Research; Westmead Hospital; University of Sydney | Gall M, Arnott A, Sadsad R, Carter I, Rahman H, Holmes EC, O'Sullivan MV, Sintchenko V, Chen SC, Maddocks S, Kok J, Dwyer DE, Rockett R, Eden J-S, Lam C, Gray K and Timms V for the 2019-nCoV Study Group |
| EPI_ISL_417406 | hCoV-19/Australia/NSW53/2020 | Oceania / Australia / New South Wales / | 2020-03-11 | Centre for Infectious Diseases and Microbiology Public Health | NSW Health Pathology - Institute of Clinical Pathology and Medical Research; Westmead Hospital; University of Sydney | Arnott A, Sadsad R, Carter I, Rahman H, Holmes EC, O'Sullivan MV, Sintchenko V, Chen SC, Maddocks S, Kok J, Dwyer DE, Rockett R, Eden J-S, Lam C, Gray K, Timms V and Gall M for the 2019-nCoV Study Group |
| EPI_ISL_417407 | hCoV-19/Australia/NSW54/2020 | Oceania / Australia / New South Wales / | 2020-03-14 | Centre for Infectious Diseases and Microbiology Public Health | NSW Health Pathology - Institute of Clinical Pathology and Medical Research; Westmead Hospital; University of Sydney | Sadsad R, Carter I, Rahman H, Holmes EC, O'Sullivan MV, Sintchenko V, Chen SC, Maddocks S, Kok J, Dwyer DE, Rockett R, Eden J-S, Lam C, Gray K, Timms V, Gall M and Arnott A for the 2019-nCoV Study Group |
| EPI_ISL_417408 | hCoV-19/Australia/NSW56/2020 | Oceania / Australia / New South Wales / | 2020-03-17 | Centre for Infectious Diseases and Microbiology Public Health | NSW Health Pathology - Institute of Clinical Pathology and Medical Research; Westmead Hospital; University of Sydney | Carter I, Rahman H, Holmes EC, O'Sullivan MV, Sintchenko V, Chen SC, Maddocks S, Kok J, Dwyer DE, Rockett R, Eden J-S, Lam C, Gray K, Timms V, Gall M, Arnott A and Sadsad R for the 2019-nCoV Study Group |
| EPI_ISL_417409 | hCoV-19/Australia/NSW57/2020 | Oceania / Australia / New South Wales / | 2020-03-16 | Centre for Infectious Diseases and Microbiology Public Health | NSW Health Pathology - Institute of Clinical Pathology and Medical Research; Westmead Hospital; University of Sydney | Rahman H, Holmes EC, O'Sullivan MV, Sintchenko V, Chen SC, Maddocks S, Kok J, Dwyer DE, Rockett R, Eden J-S, Lam C, Gray K, Timms V, Gall M, Arnott A, Sadsad R and Carter I for the 2019-nCoV Study Group |
| EPI_ISL_417410 | hCoV-19/Australia/NSW58/2020 | Oceania / Australia / New South Wales / | 2020-03-16 | Centre for Infectious Diseases and Microbiology Public Health | NSW Health Pathology - Institute of Clinical Pathology and Medical Research; Westmead Hospital; University of Sydney | Holmes EC, O'Sullivan MV, Sintchenko V, Chen SC, Maddocks S, Kok J, Dwyer DE, Rockett R, Eden J-S, Lam C, Gray K, Timms V, Gall M, Arnott A, Sadsad R, Carter I and Rahman H for the 2019-nCoV Study Group |
| EPI_ISL_417411 | hCoV-19/Australia/NSW61/2020 | Oceania / Australia / New South Wales / | 2020-03-17 | Centre for Infectious Diseases and Microbiology Public Health | NSW Health Pathology - Institute of Clinical Pathology and Medical Research; Westmead Hospital; University of Sydney | O'Sullivan MV, Sintchenko V, Chen SC, Maddocks S, Kok J, Dwyer DE, Rockett R, Eden J-S, Lam C, Gray K, Timms V, Gall M, Arnott A, Sadsad R, Carter I, Rahman H and Holmes EC for the 2019-nCoV Study Group |
| EPI_ISL_417412 | hCoV-19/Australia/NSW62/2020 | Oceania / Australia / New South Wales / | 2020-03-17 | Centre for Infectious Diseases and Microbiology Public Health | NSW Health Pathology - Institute of Clinical Pathology and Medical Research; Westmead Hospital; University of Sydney | Sintchenko V, Chen SC, Maddocks S, Kok J, Dwyer DE, Rockett R, Eden J-S, Lam C, Gray K, Timms V, Gall M, Arnott A, Sadsad R, Carter I, Rahman H, Holmes EC and O'Sullivan MV for the 2019-nCoV Study Group |
| EPI_ISL_417418 | hCoV-19/Italy/FVG-ICGEB_S1/2020 | Europe / Italy / Friuli Venezia Giulia | 2020-03-01 | Laboratory of Molecular Virology International Center fro Genetic Engineering and Biotechnology (ICGEB) | ARGO Open Lab Platform for Genome sequencing | Licastro D, Rajasekharan S, Dal Monego S, Segat L, D'Agaro P, Marcello A |
| EPI_ISL_417419 | hCoV-19/Italy/FVG-ICGEB_S5/2020 | Europe / Italy / Friuli Venezia Giulia | 2020-03-01 | Laboratory of Molecular Virology International Center for Genetic Engineering and Biotechnology (ICGEB) | ARGO Open Lab Platform for Genome sequencing | Licastro D, Rajasekharan S, Dal Monego S, Segat L, D'Agaro P, Marcello A |
| EPI_ISL_417420 | hCoV-19/NanChang/JX216/2020 | Asia / China / NanChang | 2020-03-23 | Jiangxi province Center for Disease Control and Prevention | Jiangxi province Center for Disease Control and Prevention | Li jian Xiong |
| EPI_ISL_417421 | hCoV-19/Italy/FVG-ICGEB_S8/2020 | Europe / Italy / Friuli Venezia Giulia | 2020-03-01 | Laboratory of Molecular Virology International Center for Genetic Engineering and Biotechnology (ICGEB) | ARGO Open Lab Platform for Genome sequencing | Licastro D, Rajasekharan S, Dal Monego S, Segat L, D'Agaro P, Marcello A |
| EPI_ISL_417422 | hCoV-19/Belgium/DHWM-03041/2020 | Europe / Belgium / Dilbeek | 2020-03-04 | KU Leuven, Clinical and Epidemiological Virology | KU Leuven, Clinical and Epidemiological Virology | Joan Marti-Carreras, Tony Wawina, Bert Vanmechelen, Piet Maes |
| EPI_ISL_417423 | hCoV-19/Italy/FVG-ICGEB_S9/2020 | Europe / Italy / Friuli Venezia Giulia | 2020-03-01 | Laboratory of Molecular Virology International Center for Genetic Engineering and Biotechnology (ICGEB) | ARGO Open Lab Platform for Genome sequencing | Licastro D, Rajasekharan, Dal Monego S, Segat L, D'Agaro P, Marcello A |
| EPI_ISL_417424 | hCoV-19/Belgium/KA-03042/2020 | Europe / Belgium / Kasterlee | 2020-03-04 | KU Leuven, Clinical and Epidemiological Virology | KU Leuven, Clinical and Epidemiological Virology | Joan Marti-Carreras, Tony Wawina, Bert Vanmechelen, Piet Maes |
| EPI_ISL_417425 | hCoV-19/Belgium/VBK-03061/2020 | Europe / Belgium / Schoten | 2020-03-06 | KU Leuven, Clinical and Epidemiological Virology | KU Leuven, Clinical and Epidemiological Virology | Joan Marti-Carreras, Tony Wawina, Bert Vanmechelen, Piet Maes |
| EPI_ISL_417426 | hCoV-19/Belgium/SQ-03043/2020 | Europe / Belgium / Leuven | 2020-03-04 | KU Leuven, Clinical and Epidemiological Virology | KU Leuven, Clinical and Epidemiological Virology | Joan Marti-Carreras, Tony Wawina, Bert Vanmechelen, Piet Maes |
| EPI_ISL_417427 | hCoV-19/Belgium/VI-03027/2020 | Europe / Belgium / Sint-Pieters-Woluwe | 2020-03-02 | KU Leuven, Clinical and Epidemiological Virology | KU Leuven, Clinical and Epidemiological Virology | Joan Marti-Carreras, Tony Wawina, Bert Vanmechelen, Piet Maes |
| EPI_ISL_417428 | hCoV-19/Belgium/FR-03029/2020 | Europe / Belgium / Rixensart | 2020-03-02 | KU Leuven, Clinical and Epidemiological Virology | KU Leuven, Clinical and Epidemiological Virology | Joan Marti-Carreras, Tony Wawina, Bert Vanmechelen, Piet Maes |
| EPI_ISL_417429 | hCoV-19/Belgium/MMJ-03034/2020 | Europe / Belgium / Holsbeek | 2020-03-03 | KU Leuven, Clinical and Epidemiological Virology | KU Leuven, Clinical and Epidemiological Virology | Joan Marti-Carreras, Tony Wawina, Bert Vanmechelen, Piet Maes |
| EPI_ISL_417430 | hCoV-19/Belgium/RT-03035/2020 | Europe / Belgium / Vorst | 2020-03-03 | KU Leuven, Clinical and Epidemiological Virology | KU Leuven, Clinical and Epidemiological Virology | Joan Marti-Carreras, Tony Wawina, Bert Vanmechelen, Piet Maes |
| EPI_ISL_417432 | hCoV-19/Ireland/Cork2/2020 | Europe / Ireland / Cork | 2020-03-08 | UCD National Virus Reference Laboratory | Public Health Wales Microbiology Cardiff | Catherine Dempsey, Olive Murphy, Michael Prentice, Cillian F De Gascun, |
| EPI_ISL_417433 | hCoV-19/DRC/KN-0017/2020 | Africa / Democratic Republic of the Cong | 2020-03-11 | Viral Respiratory Lab, National Institute for Biomedical Research (INRB) | Pathogen Sequencing Lab, National Institute for Biomedical Research (INRB) | Placide Mbala-Kingebeni, Edith Nkwembe, Eddy Kinganda-Lusamaki, Amuri Aziza, Catherine Pratt, Matthias Pauthner, Josh Quick, Allison Black, James Hadfield, Trevor Bedford, Ian Goodfellow, Nick Loman, Kristian Andersen, Michael Wiley, Steve Ahuka-Mundeke, Jean-Jacques Muyembe Tamfum |
| EPI_ISL_417434 | hCoV-19/DRC/KN-0038/2020 | Africa / Democratic Republic of the Cong | 2020-03-14 | Viral Respiratory Lab, National Institute for Biomedical Research (INRB) | Pathogen Sequencing Lab, National Institute for Biomedical Research (INRB) | Placide Mbala-Kingebeni, Edith Nkwembe, Eddy Kinganda-Lusamaki, Amuri Aziza, Catherine Pratt, Matthias Pauthner, Josh Quick, Allison Black, James Hadfield, Trevor Bedford, Ian Goodfellow, Nick Loman, Kristian Andersen, Michael Wiley, Steve Ahuka-Mundeke, Jean-Jacques Muyembe Tamfum |
| EPI_ISL_417435 | hCoV-19/DRC/KN-0043/2020 | Africa / Democratic Republic of the Cong | 2020-03-15 | Viral Respiratory Lab, National Institute for Biomedical Research (INRB) | Pathogen Sequencing Lab, National Institute for Biomedical Research (INRB) | Placide Mbala-Kingebeni, Edith Nkwembe, Eddy Kinganda-Lusamaki, Amuri Aziza, Catherine Pratt, Matthias Pauthner, Josh Quick, Allison Black, James Hadfield, Trevor Bedford, Ian Goodfellow, Nick Loman, Kristian Andersen, Michael Wiley, Steve Ahuka-Mundeke, Jean-Jacques Muyembe Tamfum |
| EPI_ISL_417436 | hCoV-19/DRC/KN-0051/2020 | Africa / Democratic Republic of the Cong | 2020-03-17 | Viral Respiratory Lab, National Institute for Biomedical Research (INRB) | Pathogen Sequencing Lab, National Institute for Biomedical Research (INRB) | Placide Mbala-Kingebeni, Edith Nkwembe, Eddy Kinganda-Lusamaki, Amuri Aziza, Catherine Pratt, Matthias Pauthner, Josh Quick, Allison Black, James Hadfield, Trevor Bedford, Ian Goodfellow, Nick Loman, Kristian Andersen, Michael Wiley, Steve Ahuka-Mundeke, Jean-Jacques Muyembe Tamfum |
| EPI_ISL_417437 | hCoV-19/DRC/KN-0054/2020 | Africa / Democratic Republic of the Cong | 2020-03-17 | Viral Respiratory Lab, National Institute for Biomedical Research (INRB) | Pathogen Sequencing Lab, National Institute for Biomedical Research (INRB) | Placide Mbala-Kingebeni, Edith Nkwembe, Eddy Kinganda-Lusamaki, Amuri Aziza, Catherine Pratt, Matthias Pauthner, Josh Quick, Allison Black, James Hadfield, Trevor Bedford, Ian Goodfellow, Nick Loman, Kristian Andersen, Michael Wiley, Steve Ahuka-Mundeke, Jean-Jacques Muyembe Tamfum |
| EPI_ISL_417438 | hCoV-19/DRC/KN-0058/2020 | Africa / Democratic Republic of the Cong | 2020-03-17 | Viral Respiratory Lab, National Institute for Biomedical Research (INRB) | Pathogen Sequencing Lab, National Institute for Biomedical Research (INRB) | Placide Mbala-Kingebeni, Edith Nkwembe, Eddy Kinganda-Lusamaki, Amuri Aziza, Catherine Pratt, Matthias Pauthner, Josh Quick, Allison Black, James Hadfield, Trevor Bedford, Ian Goodfellow, Nick Loman, Kristian Andersen, Michael Wiley, Steve Ahuka-Mundeke, Jean-Jacques Muyembe Tamfum |
| EPI_ISL_417439 | hCoV-19/DRC/KN-0059/2020 | Africa / Democratic Republic of the Cong | 2020-03-17 | Viral Respiratory Lab, National Institute for Biomedical Research (INRB) | Pathogen Sequencing Lab, National Institute for Biomedical Research (INRB) | Placide Mbala-Kingebeni, Edith Nkwembe, Eddy Kinganda-Lusamaki, Amuri Aziza, Catherine Pratt, Matthias Pauthner, Josh Quick, Allison Black, James Hadfield, Trevor Bedford, Ian Goodfellow, Nick Loman, Kristian Andersen, Michael Wiley, Steve Ahuka-Mundeke, Jean-Jacques Muyembe Tamfum |
| EPI_ISL_417440 | hCoV-19/DRC/KN-0060/2020 | Africa / Democratic Republic of the Cong | 2020-03-17 | Viral Respiratory Lab, National Institute for Biomedical Research (INRB) | Pathogen Sequencing Lab, National Institute for Biomedical Research (INRB) | Placide Mbala-Kingebeni, Edith Nkwembe, Eddy Kinganda-Lusamaki, Amuri Aziza, Catherine Pratt, Matthias Pauthner, Josh Quick, Allison Black, James Hadfield, Trevor Bedford, Ian Goodfellow, Nick Loman, Kristian Andersen, Michael Wiley, Steve Ahuka-Mundeke, Jean-Jacques Muyembe Tamfum |
| EPI_ISL_417441 | hCoV-19/DRC/KN-0070/2020 | Africa / Democratic Republic of the Cong | 2020-03-17 | Viral Respiratory Lab, National Institute for Biomedical Research (INRB) | Pathogen Sequencing Lab, National Institute for Biomedical Research (INRB) | Placide Mbala-Kingebeni, Edith Nkwembe, Eddy Kinganda-Lusamaki, Amuri Aziza, Catherine Pratt, Matthias Pauthner, Josh Quick, Allison Black, James Hadfield, Trevor Bedford, Ian Goodfellow, Nick Loman, Kristian Andersen, Michael Wiley, Steve Ahuka-Mundeke, Jean-Jacques Muyembe Tamfum |
| EPI_ISL_417442 | hCoV-19/DRC/KN-0072/2020 | Africa / Democratic Republic of the Cong | 2020-03-18 | Viral Respiratory Lab, National Institute for Biomedical Research (INRB) | Pathogen Sequencing Lab, National Institute for Biomedical Research (INRB) | Placide Mbala-Kingebeni, Edith Nkwembe, Eddy Kinganda-Lusamaki, Amuri Aziza, Catherine Pratt, Matthias Pauthner, Josh Quick, Allison Black, James Hadfield, Trevor Bedford, Ian Goodfellow, Nick Loman, Kristian Andersen, Michael Wiley, Steve Ahuka-Mundeke, Jean-Jacques Muyembe Tamfum |
| EPI_ISL_417443 | hCoV-19/Hong Kong/XM-PII-S4/2020 | Asia / Hong Kong | 2020-01-22 | State Key Laboratory for Emerging Infectious Diseases Department of Microbiology Li Ka Shing Faculty of Medicine The University of Hong Kong | State Key Laboratory for Emerging Infectious Diseases Department of Microbiology Li Ka Shing Faculty of Medicine The University of Hong Kong | Pui Wang, Siu-Ying Lau, Shaofeng Deng, Bobo Wing-Yee Mok, Wenjun Song, Kwok-Yung Yuen, Honglin Chen |
| EPI_ISL_417444 | hCoV-19/Pakistan/Gilgit1/2020 | Asia / Pakistan / Gilgit | 2020-03-04 | unknown | Department of Healthcare Biotechnology | Javed,A., Niazi,S.K., Ghani,E., Saqib,M., Janjua,H.A., Corman,V.M. and Zohaib,A. |
| EPI_ISL_417445 | hCoV-19/Italy/UniMI01/2020 | Europe / Italy / Lombardy / Milan | 2020-02-24 | Laboratory of Infectious Diseases, Department of Biomedical and Clinical Sciences L. Sacco, University of Milan | Laboratory of Infectious Diseases, Department of Biomedical and Clinical Sciences L. Sacco, University of Milan | Gianguglielmo Zehender, Alessia Lai, Annalisa Bergna, Luca Meroni, Agostino Riva, Claudia Balotta, Maciej Tarkowski, Arianna Gabrieli, Dario Bernacchia, Stefano Rusconi, Giuliano Rizzardini, Spinello Antinori, Massimo Galli |
| EPI_ISL_417447 | hCoV-19/Italy/UniMI03/2020 | Europe / Italy / Lombardy / Milan | 2020-02-24 | Laboratory of Infectious Diseases, Department of Biomedical and Clinical Sciences L. Sacco, University of Milan | Laboratory of Infectious Diseases, Department of Biomedical and Clinical Sciences L. Sacco, University of Milan | Gianguglielmo Zehender, Alessia Lai, Annalisa Bergna, Luca Meroni, Agostino Riva, Claudia Balotta, Maciej Tarkowski, Arianna Gabrieli, Dario Bernacchia, Stefano Rusconi, Giuliano Rizzardini, Spinello Antinori, Massimo Galli |
| EPI_ISL_417448 | hCoV-19/USA/WA-UW234/2020 | North America / USA / Washington | 2020-03-14 | UW Virology Lab | UW Virology Lab | Pavitra Roychoudhury, Hong Xie, Keith Jerome, Alexander Greninger |
| EPI_ISL_417449 | hCoV-19/USA/WA-UW235/2020 | North America / USA / Washington | 2020-03-14 | UW Virology Lab | UW Virology Lab | Pavitra Roychoudhury, Hong Xie, Keith Jerome, Alexander Greninger |
| EPI_ISL_417450 | hCoV-19/USA/WA-UW236/2020 | North America / USA / Washington | 2020-03-14 | UW Virology Lab | UW Virology Lab | Pavitra Roychoudhury, Hong Xie, Keith Jerome, Alexander Greninger |
| EPI_ISL_417451 | hCoV-19/USA/WA-UW237/2020 | North America / USA / Washington | 2020-03-14 | UW Virology Lab | UW Virology Lab | Pavitra Roychoudhury, Hong Xie, Keith Jerome, Alexander Greninger |
| EPI_ISL_417452 | hCoV-19/USA/CT-UW238/2020 | North America / USA / Connecticut | 2020-03-12 | UW Virology Lab | UW Virology Lab | Pavitra Roychoudhury, Hong Xie, Keith Jerome, Alexander Greninger |
| EPI_ISL_417453 | hCoV-19/USA/WA-UW239/2020 | North America / USA / Washington | 2020-03-14 | UW Virology Lab | UW Virology Lab | Pavitra Roychoudhury, Hong Xie, Keith Jerome, Alexander Greninger |
| EPI_ISL_417454 | hCoV-19/USA/WA-UW240/2020 | North America / USA / Washington | 2020-03-14 | UW Virology Lab | UW Virology Lab | Pavitra Roychoudhury, Hong Xie, Keith Jerome, Alexander Greninger |
| EPI_ISL_417455 | hCoV-19/USA/WA-UW241/2020 | North America / USA / Washington | 2020-03-14 | UW Virology Lab | UW Virology Lab | Pavitra Roychoudhury, Hong Xie, Keith Jerome, Alexander Greninger |
| EPI_ISL_417456 | hCoV-19/USA/WA-UW242/2020 | North America / USA / Washington | 2020-03-13 | UW Virology Lab | UW Virology Lab | Pavitra Roychoudhury, Hong Xie, Keith Jerome, Alexander Greninger |
| EPI_ISL_417457 | hCoV-19/Germany/NRW-12/2020 | Europe / Germany / North Rhine Westph | 2020-03-10 | Center of Medical Microbiology, Virology, and Hospital Hygiene, University of Duesseldorf | Center of Medical Microbiology, Virology, and Hospital Hygiene, University of Duesseldorf | Ortwin Adams, Marcel Andree, Alexander Dilthey, Torsten Feldt, Sandra Hauka, Torsten Houwaart, Björn-Erik Jensen, Detlef Kindgen-Milles, Malte Kohns Vasconcelos, Klaus Pfeffer, Tina Senff, Daniel Strelow, Jörg Timm, Andreas Walker, Tobias Wienemann |
| EPI_ISL_417458 | hCoV-19/Germany/NRW-13/2020 | Europe / Germany / North Rhine Westph | 2020-03-11 | Center of Medical Microbiology, Virology, and Hospital Hygiene, University of Duesseldorf | Center of Medical Microbiology, Virology, and Hospital Hygiene, University of Duesseldorf | Ortwin Adams, Marcel Andree, Alexander Dilthey, Torsten Feldt, Sandra Hauka, Torsten Houwaart, Björn-Erik Jensen, Detlef Kindgen-Milles, Malte Kohns Vasconcelos, Klaus Pfeffer, Tina Senff, Daniel Strelow, Jörg Timm, Andreas Walker, Tobias Wienemann |
| EPI_ISL_417459 | hCoV-19/Germany/NRW-14/2020 | Europe / Germany / North Rhine Westph | 2020-03-11 | Center of Medical Microbiology, Virology, and Hospital Hygiene, University of Duesseldorf | Center of Medical Microbiology, Virology, and Hospital Hygiene, University of Duesseldorf | Ortwin Adams, Marcel Andree, Alexander Dilthey, Torsten Feldt, Sandra Hauka, Torsten Houwaart, Björn-Erik Jensen, Detlef Kindgen-Milles, Malte Kohns Vasconcelos, Klaus Pfeffer, Tina Senff, Daniel Strelow, Jörg Timm, Andreas Walker, Tobias Wienemann |
| EPI_ISL_417460 | hCoV-19/Germany/NRW-15/2020 | Europe / Germany / North Rhine Westph | 2020-03-11 | Center of Medical Microbiology, Virology, and Hospital Hygiene, University of Duesseldorf | Center of Medical Microbiology, Virology, and Hospital Hygiene, University of Duesseldorf | Ortwin Adams, Marcel Andree, Alexander Dilthey, Torsten Feldt, Sandra Hauka, Torsten Houwaart, Björn-Erik Jensen, Detlef Kindgen-Milles, Malte Kohns Vasconcelos, Klaus Pfeffer, Tina Senff, Daniel Strelow, Jörg Timm, Andreas Walker, Tobias Wienemann |
| EPI_ISL_417461 | hCoV-19/Germany/NRW-16/2020 | Europe / Germany / North Rhine Westph | 2020-03-11 | Center of Medical Microbiology, Virology, and Hospital Hygiene, University of Duesseldorf | Center of Medical Microbiology, Virology, and Hospital Hygiene, University of Duesseldorf | Ortwin Adams, Marcel Andree, Alexander Dilthey, Torsten Feldt, Sandra Hauka, Torsten Houwaart, Björn-Erik Jensen, Detlef Kindgen-Milles, Malte Kohns Vasconcelos, Klaus Pfeffer, Tina Senff, Daniel Strelow, Jörg Timm, Andreas Walker, Tobias Wienemann |
| EPI_ISL_417462 | hCoV-19/Germany/NRW-17/2020 | Europe / Germany / North Rhine Westph | 2020-03-11 | Center of Medical Microbiology, Virology, and Hospital Hygiene, University of Duesseldorf | Center of Medical Microbiology, Virology, and Hospital Hygiene, University of Duesseldorf | Ortwin Adams, Marcel Andree, Alexander Dilthey, Torsten Feldt, Sandra Hauka, Torsten Houwaart, Björn-Erik Jensen, Detlef Kindgen-Milles, Malte Kohns Vasconcelos, Klaus Pfeffer, Tina Senff, Daniel Strelow, Jörg Timm, Andreas Walker, Tobias Wienemann |
| EPI_ISL_417463 | hCoV-19/Germany/NRW-18/2020 | Europe / Germany / North Rhine Westph | 2020-03-13 | Center of Medical Microbiology, Virology, and Hospital Hygiene, University of Duesseldorf | Center of Medical Microbiology, Virology, and Hospital Hygiene, University of Duesseldorf | Ortwin Adams, Marcel Andree, Alexander Dilthey, Torsten Feldt, Sandra Hauka, Torsten Houwaart, Björn-Erik Jensen, Detlef Kindgen-Milles, Malte Kohns Vasconcelos, Klaus Pfeffer, Tina Senff, Daniel Strelow, Jörg Timm, Andreas Walker, Tobias Wienemann |
| EPI_ISL_417464 | hCoV-19/Germany/NRW-19/2020 | Europe / Germany / North Rhine Westph | 2020-03-14 | Center of Medical Microbiology, Virology, and Hospital Hygiene, University of Duesseldorf | Center of Medical Microbiology, Virology, and Hospital Hygiene, University of Duesseldorf | Ortwin Adams, Marcel Andree, Alexander Dilthey, Torsten Feldt, Sandra Hauka, Torsten Houwaart, Björn-Erik Jensen, Detlef Kindgen-Milles, Malte Kohns Vasconcelos, Klaus Pfeffer, Tina Senff, Daniel Strelow, Jörg Timm, Andreas Walker, Tobias Wienemann |
| EPI_ISL_417465 | hCoV-19/Germany/NRW-20/2020 | Europe / Germany / North Rhine Westph | 2020-03-14 | Center of Medical Microbiology, Virology, and Hospital Hygiene, University of Duesseldorf | Center of Medical Microbiology, Virology, and Hospital Hygiene, University of Duesseldorf | Ortwin Adams, Marcel Andree, Alexander Dilthey, Torsten Feldt, Sandra Hauka, Torsten Houwaart, Björn-Erik Jensen, Detlef Kindgen-Milles, Malte Kohns Vasconcelos, Klaus Pfeffer, Tina Senff, Daniel Strelow, Jörg Timm, Andreas Walker, Tobias Wienemann |
| EPI_ISL_417466 | hCoV-19/Germany/NRW-21/2020 | Europe / Germany / North Rhine Westph | 2020-03-14 | Center of Medical Microbiology, Virology, and Hospital Hygiene, University of Duesseldorf | Center of Medical Microbiology, Virology, and Hospital Hygiene, University of Duesseldorf | Ortwin Adams, Marcel Andree, Alexander Dilthey, Torsten Feldt, Sandra Hauka, Torsten Houwaart, Björn-Erik Jensen, Detlef Kindgen-Milles, Malte Kohns Vasconcelos, Klaus Pfeffer, Tina Senff, Daniel Strelow, Jörg Timm, Andreas Walker, Tobias Wienemann |
| EPI_ISL_417467 | hCoV-19/Germany/NRW-22/2020 | Europe / Germany / North Rhine Westph | 2020-03-15 | Center of Medical Microbiology, Virology, and Hospital Hygiene, University of Duesseldorf | Center of Medical Microbiology, Virology, and Hospital Hygiene, University of Duesseldorf | Ortwin Adams, Marcel Andree, Alexander Dilthey, Torsten Feldt, Sandra Hauka, Torsten Houwaart, Björn-Erik Jensen, Detlef Kindgen-Milles, Malte Kohns Vasconcelos, Klaus Pfeffer, Tina Senff, Daniel Strelow, Jörg Timm, Andreas Walker, Tobias Wienemann |
| EPI_ISL_417468 | hCoV-19/Germany/NRW-23/2020 | Europe / Germany / North Rhine Westph | 2020-03-16 | Center of Medical Microbiology, Virology, and Hospital Hygiene, University of Duesseldorf | Center of Medical Microbiology, Virology, and Hospital Hygiene, University of Duesseldorf | Ortwin Adams, Marcel Andree, Alexander Dilthey, Torsten Feldt, Sandra Hauka, Torsten Houwaart, Björn-Erik Jensen, Detlef Kindgen-Milles, Malte Kohns Vasconcelos, Klaus Pfeffer, Tina Senff, Daniel Strelow, Jörg Timm, Andreas Walker, Tobias Wienemann |
| EPI_ISL_417469 | hCoV-19/USA/MN31-MDH31/2020 | North America / USA / Minnesota | 2020-03-12 | Minnesota Department of Health, Public Health Laboratory | Minnesota Department of Health, Public Health Laboratory | Matt Plumb, Jake Garfin and Xiong Wang |
| EPI_ISL_417470 | hCoV-19/USA/MN32-MDH32/2020 | North America / USA / Minnesota | 2020-03-12 | Minnesota Department of Health, Public Health Laboratory | Minnesota Department of Health, Public Health Laboratory | Matt Plumb, Jake Garfin and Xiong Wang |
| EPI_ISL_417471 | hCoV-19/USA/MN34-MDH34/2020 | North America / USA / Minnesota | 2020-03-12 | Minnesota Department of Health, Public Health Laboratory | Minnesota Department of Health, Public Health Laboratory | Matt Plumb, Jake Garfin and Xiong Wang |
| EPI_ISL_417472 | hCoV-19/USA/MN35-MDH35/2020 | North America / USA / Minnesota | 2020-03-12 | Minnesota Department of Health, Public Health Laboratory | Minnesota Department of Health, Public Health Laboratory | Matt Plumb, Jake Garfin and Xiong Wang |
| EPI_ISL_417473 | hCoV-19/USA/MN36-MDH36/2020 | North America / USA / Minnesota | 2020-03-13 | Minnesota Department of Health, Public Health Laboratory | Minnesota Department of Health, Public Health Laboratory | Matt Plumb, Jake Garfin and Xiong Wang |
| EPI_ISL_417474 | hCoV-19/USA/MN38-MDH38/2020 | North America / USA / Minnesota | 2020-03-12 | Minnesota Department of Health, Public Health Laboratory | Minnesota Department of Health, Public Health Laboratory | Matt Plumb, Jake Garfin and Xiong Wang |
| EPI_ISL_417475 | hCoV-19/USA/MN39-MDH39/2020 | North America / USA / Minnesota | 2020-03-12 | Minnesota Department of Health, Public Health Laboratory | Minnesota Department of Health, Public Health Laboratory | Matt Plumb, Jake Garfin and Xiong Wang |
| EPI_ISL_417476 | hCoV-19/USA/MN40-MDH40/2020 | North America / USA / Minnesota | 2020-03-12 | Minnesota Department of Health, Public Health Laboratory | Minnesota Department of Health, Public Health Laboratory | Matt Plumb, Jake Garfin and Xiong Wang |
| EPI_ISL_417477 | hCoV-19/USA/MN41-MDH41/2020 | North America / USA / Minnesota | 2020-03-12 | Minnesota Department of Health, Public Health Laboratory | Minnesota Department of Health, Public Health Laboratory | Matt Plumb, Jake Garfin and Xiong Wang |
| EPI_ISL_417478 | hCoV-19/USA/MN42-MDH42/2020 | North America / USA / Minnesota | 2020-03-13 | Minnesota Department of Health, Public Health Laboratory | Minnesota Department of Health, Public Health Laboratory | Matt Plumb, Jake Garfin and Xiong Wang |
| EPI_ISL_417479 | hCoV-19/USA/MN43-MDH43/2020 | North America / USA / Minnesota | 2020-03-12 | Minnesota Department of Health, Public Health Laboratory | Minnesota Department of Health, Public Health Laboratory | Matt Plumb, Jake Garfin and Xiong Wang |
| EPI_ISL_417480 | hCoV-19/USA/MN44-MDH44/2020 | North America / USA / Minnesota | 2020-03-12 | Minnesota Department of Health, Public Health Laboratory | Minnesota Department of Health, Public Health Laboratory | Matt Plumb, Jake Garfin and Xiong Wang |
| EPI_ISL_417481 | hCoV-19/Iceland/LUH-01/2020 | Europe / Iceland / Reykjavik | 2020-03-13 | deCODE genetics | deCODE genetics | Daniel F Gudbjartsson, Agnar Helgason, Hakon Jonsson, Olafur T Magnusson, Pall Melsted, Gudmundur L Norddahl, Jona Saemundsdottir, Asgeir Sigurdsson, Patrick Sulem, Arna B Agustsdottir, Berglind Eiriksdottir, Run Fridriksdottir, Elisabet E Gardarsdottir, Gudmundur Georgsson, Olafia S Gretarsdottir, Kjartan R Gudmundsson, Thora R Gunnarsdottir, Arnaldur Gylfason, Hilma Holm, Brynjar O Jensson, Aslaug Jonasdottir, Kamilla S Josefsdottir, Thordur Kristjansson, Droplaug N Magnusdottir, Louise le Roux, Gudrun Sigmundsdottir, Gardar Sveinbjornsson, Kristin E Sveinsdottir, Maney Sveinsdottir, Emil A Thorarensen, Bjarni Thorbjornsson, Gisli Masson, Ingileif Jonsdottir, Alma Moller, Thorolfur Gudnason, Karl G Kristinsson, Unnur Thorsteinsdottir, Kari Stefansson |
| EPI_ISL_417483 | hCoV-19/Norway/1379/2020 | Europe / Norway / Oslo | 2020-02-27 | Oslo University Hospital, Department of Medical Microbiology | Norwegian Institute of Public Health | Kathrine Stene-Johansen, Kamilla Heddeland Instefjord, Hilde Elshaug, Karoline Bragstad, Olav Hungnes |

| | | | | | | |
|---|---|---|---|---|---|---|
| EPI_ISL_417484 | hCoV-19/Norway/1380/2020 | Europe / Norway / Oslo | 2020-02-26 | Oslo University Hospital, Department of Medical Microbiology | Norwegian Institute of Public Health, Department of Virology | Kathrine Stene-Johansen, Kamilla Heddeland Instefjord, Hilde Elshaug, Karoline Bragstad, Olav Hungnes |
| EPI_ISL_417485 | hCoV-19/Norway/1526/2020 | Europe / Norway / Oslo | 2020-03-02 | University Hospital of Northern Norway, Department for Microbiology and Infectious Disease Control | Norwegian Institute of Public Health, Department of Virology | Kathrine Stene-Johansen, Kamilla Heddeland Instefjord, Hilde Elshaug, Karoline Bragstad, Olav Hungnes |
| EPI_ISL_417486 | hCoV-19/Norway/1538/2020 | Europe / Norway / Oslo | 2020-03-01 | Hospital of Southern Norway - Kristiansand, Department of Medical Microbiology | Norwegian Institute of Public Health, Department of Virology | Kathrine Stene-Johansen, Kamilla Heddeland Instefjord, Hilde Elshaug, Karoline Bragstad, Olav Hungnes |
| EPI_ISL_417487 | hCoV-19/Norway/1539/2020 | Europe / Norway / Oslo | 2020-03-01 | Hospital of Southern Norway - Kristiansand, Department of Medical Microbiology | Norwegian Institute of Public Health, Department of Virology | Kathrine Stene-Johansen, Kamilla Heddeland Instefjord, Hilde Elshaug, Karoline Bragstad, Olav Hungnes |
| EPI_ISL_417488 | hCoV-19/Norway/1493/2020 | Europe / Norway / Oslo | 2020-02-29 | Oslo University Hospital, Department of Medical Microbiology | Norwegian Institute of Public Health, Department of Virology | Kathrine Stene-Johansen, Kamilla Heddeland Instefjord, Hilde Elshaug, Karoline Bragstad, Olav Hungnes |
| EPI_ISL_417491 | hCoV-19/Italy/UnivPM1/2020 | Europe / Italy / Marche | 2020-03-03 | Virology Laboratory, Department of Biomedical Sciences and Public Health, University Politecnica delle Marche | Virology and Legal Medicine Laboratories, Department of Biomedical Sciences and Public Health, University Politecnica delle Marche | Bagnarelli,P., Caucci,S., Di Sante,L., Menzo,S., Alessandrini,F., Onofri,V., Turchi,C., Tagliabracci,A. |
| EPI_ISL_417492 | hCoV-19/USA/MN45-MDH45/2020 | North America / USA / Minnesota | 2020-03-13 | Minnesota Department of Health, Public Health Laboratory | Minnesota Department of Health, Public Health Laboratory | Matt Plumb, Jake Garfin and Xiong Wang |
| EPI_ISL_417493 | hCoV-19/USA/MN46-MDH46/2020 | North America / USA / Minnesota | 2020-03-13 | Minnesota Department of Health, Public Health Laboratory | Minnesota Department of Health, Public Health Laboratory | Matt Plumb, Jake Garfin and Xiong Wang |
| EPI_ISL_417494 | hCoV-19/USA/MN47-MDH47/2020 | North America / USA / Minnesota | 2020-03-13 | Minnesota Department of Health, Public Health Laboratory | Minnesota Department of Health, Public Health Laboratory | Matt Plumb, Jake Garfin and Xiong Wang |
| EPI_ISL_417495 | hCoV-19/USA/MN48-MDH48/2020 | North America / USA / Minnesota | 2020-03-13 | Minnesota Department of Health, Public Health Laboratory | Minnesota Department of Health, Public Health Laboratory | Matt Plumb, Jake Garfin and Xiong Wang |
| EPI_ISL_417496 | hCoV-19/USA/MN51-MDH51/2020 | North America / USA / Minnesota | 2020-03-13 | Minnesota Department of Health, Public Health Laboratory | Minnesota Department of Health, Public Health Laboratory | Matt Plumb, Jake Garfin and Xiong Wang |
| EPI_ISL_417497 | hCoV-19/USA/MN52-MDH52/2020 | North America / USA / Minnesota | 2020-03-13 | Minnesota Department of Health, Public Health Laboratory | Minnesota Department of Health, Public Health Laboratory | Matt Plumb, Jake Garfin and Xiong Wang |
| EPI_ISL_417498 | hCoV-19/USA/MN53-MDH53/2020 | North America / USA / Minnesota | 2020-03-13 | Minnesota Department of Health, Public Health Laboratory | Minnesota Department of Health, Public Health Laboratory | Matt Plumb, Jake Garfin and Xiong Wang |
| EPI_ISL_417499 | hCoV-19/USA/MN55-MDH55/2020 | North America / USA / Minnesota | 2020-03-13 | Minnesota Department of Health, Public Health Laboratory | Minnesota Department of Health, Public Health Laboratory | Matt Plumb, Jake Garfin and Xiong Wang |
| EPI_ISL_417800 | hCoV-19/Iceland/15/2020 | Europe / Iceland / Reykjavik | 2020-03-02 | The National University Hospital of Iceland | deCODE genetics | Daniel F Gudbjartsson; Agnar Helgason; Hakon Jonsson; Olafur T Magnusson; Pall Melsted; Gudmundur L Norddahl; Jona Saemundsdottir; Asgeir Sigurdsson; Patrick Sulem; Arna B Agustsdottir; Berglind Eiriksdottir; Run Fridriksdottir; Elisabet E Gardarsdottir; Gudmundur Georgsson; Olafia S Gretarsdottir; Kjartan R Gudmundsson; Thora R Gunnarsdottir; Arnaldur Gylfason; Hilma Holm; Brynjar O Jensson; Aslaug Jonasdottir; Kamilla S Josefsdottir; Thordur Kristjansson; Droplaug N Magnusdottir; Louise le Roux; Gudrun Sigmundsdottir; Gardar Sveinbjornsson; Kristin E Sveinsdottir; Maney Sveinsdottir; Emil A Thorarensen; Bjarni Thorbjornsson; Gisli Masson; Ingileif Jonsdottir; Alma Moller; Thorolfur Gudnason; Karl G Kristinsson; Unnur Thorsteinsdottir; Kari Stefansson |
| EPI_ISL_417801 | hCoV-19/Iceland/151/2020 | Europe / Iceland / Reykjavik | 2020-03-13 | The National University Hospital of Iceland | deCODE genetics | Daniel F Gudbjartsson; Agnar Helgason; Hakon Jonsson; Olafur T Magnusson; Pall Melsted; Gudmundur L Norddahl; Jona Saemundsdottir; Asgeir Sigurdsson; Patrick Sulem; Arna B Agustsdottir; Berglind Eiriksdottir; Run Fridriksdottir; Elisabet E Gardarsdottir; Gudmundur Georgsson; Olafia S Gretarsdottir; Kjartan R Gudmundsson; Thora R Gunnarsdottir; Arnaldur Gylfason; Hilma Holm; Brynjar O Jensson; Aslaug Jonasdottir; Kamilla S Josefsdottir; Thordur Kristjansson; Droplaug N Magnusdottir; Louise le Roux; Gudrun Sigmundsdottir; Gardar Sveinbjornsson; Kristin E Sveinsdottir; Maney Sveinsdottir; Emil A Thorarensen; Bjarni Thorbjornsson; Gisli Masson; Ingileif Jonsdottir; Alma Moller; Thorolfur Gudnason; Karl G Kristinsson; Unnur Thorsteinsdottir; Kari Stefansson |
| EPI_ISL_417802 | hCoV-19/Iceland/153/2020 | Europe / Iceland / Reykjavik | 2020-03-13 | The National University Hospital of Iceland | deCODE genetics | Daniel F Gudbjartsson; Agnar Helgason; Hakon Jonsson; Olafur T Magnusson; Pall Melsted; Gudmundur L Norddahl; Jona Saemundsdottir; Asgeir Sigurdsson; Patrick Sulem; Arna B Agustsdottir; Berglind Eiriksdottir; Run Fridriksdottir; Elisabet E Gardarsdottir; Gudmundur Georgsson; Olafia S Gretarsdottir; Kjartan R Gudmundsson; Thora R Gunnarsdottir; Arnaldur Gylfason; Hilma Holm; Brynjar O Jensson; Aslaug Jonasdottir; Kamilla S Josefsdottir; Thordur Kristjansson; Droplaug N Magnusdottir; Louise le Roux; Gudrun Sigmundsdottir; Gardar Sveinbjornsson; Kristin E Sveinsdottir; Maney Sveinsdottir; Emil A Thorarensen; Bjarni Thorbjornsson; Gisli Masson; Ingileif Jonsdottir; Alma Moller; Thorolfur Gudnason; Karl G Kristinsson; Unnur Thorsteinsdottir; Kari Stefansson |
| EPI_ISL_417803 | hCoV-19/Iceland/156/2020 | Europe / Iceland / Reykjavik | 2020-03-13 | The National University Hospital of Iceland | deCODE genetics | Daniel F Gudbjartsson; Agnar Helgason; Hakon Jonsson; Olafur T Magnusson; Pall Melsted; Gudmundur L Norddahl; Jona Saemundsdottir; Asgeir Sigurdsson; Patrick Sulem; Arna B Agustsdottir; Berglind Eiriksdottir; Run Fridriksdottir; Elisabet E Gardarsdottir; Gudmundur Georgsson; Olafia S Gretarsdottir; Kjartan R Gudmundsson; Thora R Gunnarsdottir; Arnaldur Gylfason; Hilma Holm; Brynjar O Jensson; Aslaug Jonasdottir; Kamilla S Josefsdottir; Thordur Kristjansson; Droplaug N Magnusdottir; Louise le Roux; Gudrun Sigmundsdottir; Gardar Sveinbjornsson; Kristin E Sveinsdottir; Maney Sveinsdottir; Emil A Thorarensen; Bjarni Thorbjornsson; Gisli Masson; Ingileif Jonsdottir; Alma Moller; Thorolfur Gudnason; Karl G Kristinsson; Unnur Thorsteinsdottir; Kari Stefansson |
| EPI_ISL_417804 | hCoV-19/Iceland/157/2020 | Europe / Iceland / Reykjavik | 2020-03-13 | The National University Hospital of Iceland | deCODE genetics | Daniel F Gudbjartsson; Agnar Helgason; Hakon Jonsson; Olafur T Magnusson; Pall Melsted; Gudmundur L Norddahl; Jona Saemundsdottir; Asgeir Sigurdsson; Patrick Sulem; Arna B Agustsdottir; Berglind Eiriksdottir; Run Fridriksdottir; Elisabet E Gardarsdottir; Gudmundur Georgsson; Olafia S Gretarsdottir; Kjartan R Gudmundsson; Thora R Gunnarsdottir; Arnaldur Gylfason; Hilma Holm; Brynjar O Jensson; Aslaug Jonasdottir; Kamilla S Josefsdottir; Thordur Kristjansson; Droplaug N Magnusdottir; Louise le Roux; Gudrun Sigmundsdottir; Gardar Sveinbjornsson; Kristin E Sveinsdottir; Maney Sveinsdottir; Emil A Thorarensen; Bjarni Thorbjornsson; Gisli Masson; Ingileif Jonsdottir; Alma Moller; Thorolfur Gudnason; Karl G Kristinsson; Unnur Thorsteinsdottir; Kari Stefansson |
| EPI_ISL_417805 | hCoV-19/Iceland/158/2020 | Europe / Iceland / Reykjavik | 2020-03-13 | The National University Hospital of Iceland | deCODE genetics | Daniel F Gudbjartsson; Agnar Helgason; Hakon Jonsson; Olafur T Magnusson; Pall Melsted; Gudmundur L Norddahl; Jona Saemundsdottir; Asgeir Sigurdsson; Patrick Sulem; Arna B Agustsdottir; Berglind Eiriksdottir; Run Fridriksdottir; Elisabet E Gardarsdottir; Gudmundur Georgsson; Olafia S Gretarsdottir; Kjartan R Gudmundsson; Thora R Gunnarsdottir; Arnaldur Gylfason; Hilma Holm; Brynjar O Jensson; Aslaug Jonasdottir; Kamilla S Josefsdottir; Thordur Kristjansson; Droplaug N Magnusdottir; Louise le Roux; Gudrun Sigmundsdottir; Gardar Sveinbjornsson; Kristin E Sveinsdottir; Maney Sveinsdottir; Emil A Thorarensen; Bjarni Thorbjornsson; Gisli Masson; Ingileif Jonsdottir; Alma Moller; Thorolfur Gudnason; Karl G Kristinsson; Unnur Thorsteinsdottir; Kari Stefansson |
| EPI_ISL_417806 | hCoV-19/Iceland/159/2020 | Europe / Iceland / Reykjavik | 2020-03-13 | The National University Hospital of Iceland | deCODE genetics | Daniel F Gudbjartsson; Agnar Helgason; Hakon Jonsson; Olafur T Magnusson; Pall Melsted; Gudmundur L Norddahl; Jona Saemundsdottir; Asgeir Sigurdsson; Patrick Sulem; Arna B Agustsdottir; Berglind Eiriksdottir; Run Fridriksdottir; Elisabet E Gardarsdottir; Gudmundur Georgsson; Olafia S Gretarsdottir; Kjartan R Gudmundsson; Thora R Gunnarsdottir; Arnaldur Gylfason; Hilma Holm; Brynjar O Jensson; Aslaug Jonasdottir; Kamilla S Josefsdottir; Thordur Kristjansson; Droplaug N Magnusdottir; Louise le Roux; Gudrun Sigmundsdottir; Gardar Sveinbjornsson; Kristin E Sveinsdottir; Maney Sveinsdottir; Emil A Thorarensen; Bjarni Thorbjornsson; Gisli Masson; Ingileif Jonsdottir; Alma Moller; Thorolfur Gudnason; Karl G Kristinsson; Unnur Thorsteinsdottir; Kari Stefansson |
| EPI_ISL_417808 | hCoV-19/Iceland/165/2020 | Europe / Iceland / Reykjavik | 2020-03-15 | The National University Hospital of Iceland | deCODE genetics | Daniel F Gudbjartsson; Agnar Helgason; Hakon Jonsson; Olafur T Magnusson; Pall Melsted; Gudmundur L Norddahl; Jona Saemundsdottir; Asgeir Sigurdsson; Patrick Sulem; Arna B Agustsdottir; Berglind Eiriksdottir; Run Fridriksdottir; Elisabet E Gardarsdottir; Gudmundur Georgsson; Olafia S Gretarsdottir; Kjartan R Gudmundsson; Thora R Gunnarsdottir; Arnaldur Gylfason; Hilma Holm; Brynjar O Jensson; Aslaug Jonasdottir; Kamilla S Josefsdottir; Thordur Kristjansson; Droplaug N Magnusdottir; Louise le Roux; Gudrun Sigmundsdottir; Gardar Sveinbjornsson; Kristin E Sveinsdottir; Maney Sveinsdottir; Emil A Thorarensen; Bjarni Thorbjornsson; Gisli Masson; Ingileif Jonsdottir; Alma Moller; Thorolfur Gudnason; Karl G Kristinsson; Unnur Thorsteinsdottir; Kari Stefansson |
| EPI_ISL_417809 | hCoV-19/Iceland/167/2020 | Europe / Iceland / Reykjavik | 2020-03-15 | The National University Hospital of Iceland | deCODE genetics | Daniel F Gudbjartsson; Agnar Helgason; Hakon Jonsson; Olafur T Magnusson; Pall Melsted; Gudmundur L Norddahl; Jona Saemundsdottir; Asgeir Sigurdsson; Patrick Sulem; Arna B Agustsdottir; Berglind Eiriksdottir; Run Fridriksdottir; Elisabet E Gardarsdottir; Gudmundur Georgsson; Olafia S Gretarsdottir; Kjartan R Gudmundsson; Thora R Gunnarsdottir; Arnaldur Gylfason; Hilma Holm; Brynjar O Jensson; Aslaug Jonasdottir; Kamilla S Josefsdottir; Thordur Kristjansson; Droplaug N Magnusdottir; Louise le Roux; Gudrun Sigmundsdottir; Gardar Sveinbjornsson; Kristin E Sveinsdottir; Maney Sveinsdottir; Emil A Thorarensen; Bjarni Thorbjornsson; Gisli Masson; Ingileif Jonsdottir; Alma Moller; Thorolfur Gudnason; Karl G Kristinsson; Unnur Thorsteinsdottir; Kari Stefansson |
| EPI_ISL_417810 | hCoV-19/Iceland/168/2020 | Europe / Iceland / Reykjavik | 2020-03-13 | The National University Hospital of Iceland | deCODE genetics | Daniel F Gudbjartsson; Agnar Helgason; Hakon Jonsson; Olafur T Magnusson; Pall Melsted; Gudmundur L Norddahl; Jona Saemundsdottir; Asgeir Sigurdsson; Patrick Sulem; Arna B Agustsdottir; Berglind Eiriksdottir; Run Fridriksdottir; Elisabet E Gardarsdottir; Gudmundur Georgsson; Olafia S Gretarsdottir; Kjartan R Gudmundsson; Thora R Gunnarsdottir; Arnaldur Gylfason; Hilma Holm; Brynjar O Jensson; Aslaug Jonasdottir; Kamilla S Josefsdottir; Thordur Kristjansson; Droplaug N Magnusdottir; Louise le Roux; Gudrun Sigmundsdottir; Gardar Sveinbjornsson; Kristin E Sveinsdottir; Maney Sveinsdottir; Emil A Thorarensen; Bjarni Thorbjornsson; Gisli Masson; Ingileif Jonsdottir; Alma Moller; Thorolfur Gudnason; Karl G Kristinsson; Unnur Thorsteinsdottir; Kari Stefansson |
| EPI_ISL_417812 | hCoV-19/Iceland/176/2020 | Europe / Iceland / Reykjavik | 2020-03-14 | The National University Hospital of Iceland | deCODE genetics | Daniel F Gudbjartsson; Agnar Helgason; Hakon Jonsson; Olafur T Magnusson; Pall Melsted; Gudmundur L Norddahl; Jona Saemundsdottir; Asgeir Sigurdsson; Patrick Sulem; Arna B Agustsdottir; Berglind Eiriksdottir; Run Fridriksdottir; Elisabet E Gardarsdottir; Gudmundur Georgsson; Olafia S Gretarsdottir; Kjartan R Gudmundsson; Thora R Gunnarsdottir; Arnaldur Gylfason; Hilma Holm; Brynjar O Jensson; Aslaug Jonasdottir; Kamilla S Josefsdottir; Thordur Kristjansson; Droplaug N Magnusdottir; Louise le Roux; Gudrun Sigmundsdottir; Gardar Sveinbjornsson; Kristin E Sveinsdottir; Maney Sveinsdottir; Emil A Thorarensen; Bjarni Thorbjornsson; Gisli Masson; Ingileif Jonsdottir; Alma Moller; Thorolfur Gudnason; Karl G Kristinsson; Unnur Thorsteinsdottir; Kari Stefansson |
| EPI_ISL_417813 | hCoV-19/Iceland/178/2020 | Europe / Iceland / Reykjavik | 2020-03-15 | The National University Hospital of Iceland | deCODE genetics | Daniel F Gudbjartsson; Agnar Helgason; Hakon Jonsson; Olafur T Magnusson; Pall Melsted; Gudmundur L Norddahl; Jona Saemundsdottir; Asgeir Sigurdsson; Patrick Sulem; Arna B Agustsdottir; Berglind Eiriksdottir; Run Fridriksdottir; Elisabet E Gardarsdottir; Gudmundur Georgsson; Olafia S Gretarsdottir; Kjartan R Gudmundsson; Thora R Gunnarsdottir; Arnaldur Gylfason; Hilma Holm; Brynjar O Jensson; Aslaug Jonasdottir; Kamilla S Josefsdottir; Thordur Kristjansson; Droplaug N Magnusdottir; Louise le Roux; Gudrun Sigmundsdottir; Gardar Sveinbjornsson; Kristin E Sveinsdottir; Maney Sveinsdottir; Emil A Thorarensen; Bjarni Thorbjornsson; Gisli Masson; Ingileif Jonsdottir; Alma Moller; Thorolfur Gudnason; Karl G Kristinsson; Unnur Thorsteinsdottir; Kari Stefansson |
| EPI_ISL_417814 | hCoV-19/Iceland/181/2020 | Europe / Iceland / Reykjavik | 2020-03-16 | The National University Hospital of Iceland | deCODE genetics | Daniel F Gudbjartsson; Agnar Helgason; Hakon Jonsson; Olafur T Magnusson; Pall Melsted; Gudmundur L Norddahl; Jona Saemundsdottir; Asgeir Sigurdsson; Patrick Sulem; Arna B Agustsdottir; Berglind Eiriksdottir; Run Fridriksdottir; Elisabet E Gardarsdottir; Gudmundur Georgsson; Olafia S Gretarsdottir; Kjartan R Gudmundsson; Thora R Gunnarsdottir; Arnaldur Gylfason; Hilma Holm; Brynjar O Jensson; Aslaug Jonasdottir; Kamilla S Josefsdottir; Thordur Kristjansson; Droplaug N Magnusdottir; Louise le Roux; Gudrun Sigmundsdottir; Gardar Sveinbjornsson; Kristin E Sveinsdottir; Maney Sveinsdottir; Emil A Thorarensen; Bjarni Thorbjornsson; Gisli Masson; Ingileif Jonsdottir; Alma Moller; Thorolfur Gudnason; Karl G Kristinsson; Unnur Thorsteinsdottir; Kari Stefansson |
| EPI_ISL_417815 | hCoV-19/Iceland/186/2020 | Europe / Iceland / Reykjavik | 2020-03-16 | The National University Hospital of Iceland | deCODE genetics | Daniel F Gudbjartsson; Agnar Helgason; Hakon Jonsson; Olafur T Magnusson; Pall Melsted; Gudmundur L Norddahl; Jona Saemundsdottir; Asgeir Sigurdsson; Patrick Sulem; Arna B Agustsdottir; Berglind Eiriksdottir; Run Fridriksdottir; Elisabet E Gardarsdottir; Gudmundur Georgsson; Olafia S Gretarsdottir; Kjartan R Gudmundsson; Thora R Gunnarsdottir; Arnaldur Gylfason; Hilma Holm; Brynjar O Jensson; Aslaug Jonasdottir; Kamilla S Josefsdottir; Thordur Kristjansson; Droplaug N Magnusdottir; Louise le Roux; Gudrun Sigmundsdottir; Gardar Sveinbjornsson; Kristin E Sveinsdottir; Maney Sveinsdottir; Emil A Thorarensen; Bjarni Thorbjornsson; Gisli Masson; Ingileif Jonsdottir; Alma Moller; Thorolfur Gudnason; Karl G Kristinsson; Unnur Thorsteinsdottir; Kari Stefansson |
| EPI_ISL_417817 | hCoV-19/Iceland/19/2020 | Europe / Iceland / Reykjavik | 2020-03-02 | The National University Hospital of Iceland | deCODE genetics | Daniel F Gudbjartsson; Agnar Helgason; Hakon Jonsson; Olafur T Magnusson; Pall Melsted; Gudmundur L Norddahl; Jona Saemundsdottir; Asgeir Sigurdsson; Patrick Sulem; Arna B Agustsdottir; Berglind Eiriksdottir; Run Fridriksdottir; Elisabet E Gardarsdottir; Gudmundur Georgsson; Olafia S Gretarsdottir; Kjartan R Gudmundsson; Thora R Gunnarsdottir; Arnaldur Gylfason; Hilma Holm; Brynjar O Jensson; Aslaug Jonasdottir; Kamilla S Josefsdottir; Thordur Kristjansson; Droplaug N Magnusdottir; Louise le Roux; Gudrun Sigmundsdottir; Gardar Sveinbjornsson; Kristin E Sveinsdottir; Maney Sveinsdottir; Emil A Thorarensen; Bjarni Thorbjornsson; Gisli Masson; Ingileif Jonsdottir; Alma Moller; Thorolfur Gudnason; Karl G Kristinsson; Unnur Thorsteinsdottir; Kari Stefansson |
| EPI_ISL_417818 | hCoV-19/Iceland/190/2020 | Europe / Iceland / Reykjavik | 2020-03-16 | The National University Hospital of Iceland | deCODE genetics | Daniel F Gudbjartsson; Agnar Helgason; Hakon Jonsson; Olafur T Magnusson; Pall Melsted; Gudmundur L Norddahl; Jona Saemundsdottir; Asgeir Sigurdsson; Patrick Sulem; Arna B Agustsdottir; Berglind Eiriksdottir; Run Fridriksdottir; Elisabet E Gardarsdottir; Gudmundur Georgsson; Olafia S Gretarsdottir; Kjartan R Gudmundsson; Thora R Gunnarsdottir; Arnaldur Gylfason; Hilma Holm; Brynjar O Jensson; Aslaug Jonasdottir; Kamilla S Josefsdottir; Thordur Kristjansson; Droplaug N Magnusdottir; Louise le Roux; Gudrun Sigmundsdottir; Gardar Sveinbjornsson; Kristin E Sveinsdottir; Maney Sveinsdottir; Emil A Thorarensen; Bjarni Thorbjornsson; Gisli Masson; Ingileif Jonsdottir; Alma Moller; Thorolfur Gudnason; Karl G Kristinsson; Unnur Thorsteinsdottir; Kari Stefansson |
| EPI_ISL_417819 | hCoV-19/Iceland/192/2020 | Europe / Iceland / Reykjavik | 2020-03-15 | The National University Hospital of Iceland | deCODE genetics | Daniel F Gudbjartsson; Agnar Helgason; Hakon Jonsson; Olafur T Magnusson; Pall Melsted; Gudmundur L Norddahl; Jona Saemundsdottir; Asgeir Sigurdsson; Patrick Sulem; Arna B Agustsdottir; Berglind Eiriksdottir; Run Fridriksdottir; Elisabet E Gardarsdottir; Gudmundur Georgsson; Olafia S Gretarsdottir; Kjartan R Gudmundsson; Thora R Gunnarsdottir; Arnaldur Gylfason; Hilma Holm; Brynjar O Jensson; Aslaug Jonasdottir; Kamilla S Josefsdottir; Thordur Kristjansson; Droplaug N Magnusdottir; Louise le Roux; Gudrun Sigmundsdottir; Gardar Sveinbjornsson; Kristin E Sveinsdottir; Maney Sveinsdottir; Emil A Thorarensen; Bjarni Thorbjornsson; Gisli Masson; Ingileif Jonsdottir; Alma Moller; Thorolfur Gudnason; Karl G Kristinsson; Unnur Thorsteinsdottir; Kari Stefansson |
| EPI_ISL_417820 | hCoV-19/Iceland/193/2020 | Europe / Iceland / Reykjavik | 2020-03-16 | The National University Hospital of Iceland | deCODE genetics | Daniel F Gudbjartsson; Agnar Helgason; Hakon Jonsson; Olafur T Magnusson; Pall Melsted; Gudmundur L Norddahl; Jona Saemundsdottir; Asgeir Sigurdsson; Patrick Sulem; Arna B Agustsdottir; Berglind Eiriksdottir; Run Fridriksdottir; Elisabet E Gardarsdottir; Gudmundur Georgsson; Olafia S Gretarsdottir; Kjartan R Gudmundsson; Thora R Gunnarsdottir; Arnaldur Gylfason; Hilma Holm; Brynjar O Jensson; Aslaug Jonasdottir; Kamilla S Josefsdottir; Thordur Kristjansson; Droplaug N Magnusdottir; Louise le Roux; Gudrun Sigmundsdottir; Gardar Sveinbjornsson; Kristin E Sveinsdottir; Maney Sveinsdottir; Emil A Thorarensen; Bjarni Thorbjornsson; Gisli Masson; Ingileif Jonsdottir; Alma Moller; Thorolfur Gudnason; Karl G Kristinsson; Unnur Thorsteinsdottir; Kari Stefansson |
| EPI_ISL_417821 | hCoV-19/Iceland/194/2020 | Europe / Iceland / Reykjavik | 2020-03-16 | The National University Hospital of Iceland | deCODE genetics | Daniel F Gudbjartsson; Agnar Helgason; Hakon Jonsson; Olafur T Magnusson; Pall Melsted; Gudmundur L Norddahl; Jona Saemundsdottir; Asgeir Sigurdsson; Patrick Sulem; Arna B Agustsdottir; Berglind Eiriksdottir; Run Fridriksdottir; Elisabet E Gardarsdottir; Gudmundur Georgsson; Olafia S Gretarsdottir; Kjartan R Gudmundsson; Thora R Gunnarsdottir; Arnaldur Gylfason; Hilma Holm; Brynjar O Jensson; Aslaug Jonasdottir; Kamilla S Josefsdottir; Thordur Kristjansson; Droplaug N Magnusdottir; Louise le Roux; Gudrun Sigmundsdottir; Gardar Sveinbjornsson; Kristin E Sveinsdottir; Maney Sveinsdottir; Emil A Thorarensen; Bjarni Thorbjornsson; Gisli Masson; Ingileif Jonsdottir; Alma Moller; Thorolfur Gudnason; Karl G Kristinsson; Unnur Thorsteinsdottir; Kari Stefansson |
| EPI_ISL_417822 | hCoV-19/Iceland/196/2020 | Europe / Iceland / Reykjavik | 2020-03-16 | The National University Hospital of Iceland | deCODE genetics | Daniel F Gudbjartsson; Agnar Helgason; Hakon Jonsson; Olafur T Magnusson; Pall Melsted; Gudmundur L Norddahl; Jona Saemundsdottir; Asgeir Sigurdsson; Patrick Sulem; Arna B Agustsdottir; Berglind Eiriksdottir; Run Fridriksdottir; Elisabet E Gardarsdottir; Gudmundur Georgsson; Olafia S Gretarsdottir; Kjartan R Gudmundsson; Thora R Gunnarsdottir; Arnaldur Gylfason; Hilma Holm; Brynjar O Jensson; Aslaug Jonasdottir; Kamilla S Josefsdottir; Thordur Kristjansson; Droplaug N Magnusdottir; Louise le Roux; Gudrun Sigmundsdottir; Gardar Sveinbjornsson; Kristin E Sveinsdottir; Maney Sveinsdottir; Emil A Thorarensen; Bjarni Thorbjornsson; Gisli Masson; Ingileif Jonsdottir; Alma Moller; Thorolfur Gudnason; Karl G Kristinsson; Unnur Thorsteinsdottir; Kari Stefansson |
| EPI_ISL_417823 | hCoV-19/Iceland/198/2020 | Europe / Iceland / Reykjavik | 2020-03-16 | The National University Hospital of Iceland | deCODE genetics | Daniel F Gudbjartsson; Agnar Helgason; Hakon Jonsson; Olafur T Magnusson; Pall Melsted; Gudmundur L Norddahl; Jona Saemundsdottir; Asgeir Sigurdsson; Patrick Sulem; Arna B Agustsdottir; Berglind Eiriksdottir; Run Fridriksdottir; Elisabet E Gardarsdottir; Gudmundur Georgsson; Olafia S Gretarsdottir; Kjartan R Gudmundsson; Thora R Gunnarsdottir; Arnaldur Gylfason; Hilma Holm; Brynjar O Jensson; Aslaug Jonasdottir; Kamilla S Josefsdottir; Thordur Kristjansson; Droplaug N Magnusdottir; Louise le Roux; Gudrun Sigmundsdottir; Gardar Sveinbjornsson; Kristin E Sveinsdottir; Maney Sveinsdottir; Emil A Thorarensen; Bjarni Thorbjornsson; Gisli Masson; Ingileif Jonsdottir; Alma Moller; Thorolfur Gudnason; Karl G Kristinsson; Unnur Thorsteinsdottir; Kari Stefansson |
| EPI_ISL_417824 | hCoV-19/Iceland/201/2020 | Europe / Iceland / Reykjavik | 2020-03-16 | The National University Hospital of Iceland | deCODE genetics | Daniel F Gudbjartsson; Agnar Helgason; Hakon Jonsson; Olafur T Magnusson; Pall Melsted; Gudmundur L Norddahl; Jona Saemundsdottir; Asgeir Sigurdsson; Patrick Sulem; Arna B Agustsdottir; Berglind Eiriksdottir; Run Fridriksdottir; Elisabet E Gardarsdottir; Gudmundur Georgsson; Olafia S Gretarsdottir; Kjartan R Gudmundsson; Thora R Gunnarsdottir; Arnaldur Gylfason; Hilma Holm; Brynjar O Jensson; Aslaug Jonasdottir; Kamilla S Josefsdottir; Thordur Kristjansson; Droplaug N Magnusdottir; Louise le Roux; Gudrun Sigmundsdottir; Gardar Sveinbjornsson; Kristin E Sveinsdottir; Maney Sveinsdottir; Emil A Thorarensen; Bjarni Thorbjornsson; Gisli Masson; Ingileif Jonsdottir; Alma Moller; Thorolfur Gudnason; Karl G Kristinsson; Unnur Thorsteinsdottir; Kari Stefansson |
| EPI_ISL_417825 | hCoV-19/Iceland/203/2020 | Europe / Iceland / Reykjavik | 2020-03-16 | The National University Hospital of Iceland | deCODE genetics | Daniel F Gudbjartsson; Agnar Helgason; Hakon Jonsson; Olafur T Magnusson; Pall Melsted; Gudmundur L Norddahl; Jona Saemundsdottir; Asgeir Sigurdsson; Patrick Sulem; Arna B Agustsdottir; Berglind Eiriksdottir; Run Fridriksdottir; Elisabet E Gardarsdottir; Gudmundur Georgsson; Olafia S Gretarsdottir; Kjartan R Gudmundsson; Thora R Gunnarsdottir; Arnaldur Gylfason; Hilma Holm; Brynjar O Jensson; Aslaug Jonasdottir; Kamilla S Josefsdottir; Thordur Kristjansson; Droplaug N Magnusdottir; Louise le Roux; Gudrun Sigmundsdottir; Gardar Sveinbjornsson; Kristin E Sveinsdottir; Maney Sveinsdottir; Emil A Thorarensen; Bjarni Thorbjornsson; Gisli Masson; Ingileif Jonsdottir; Alma Moller; Thorolfur Gudnason; Karl G Kristinsson; Unnur Thorsteinsdottir; Kari Stefansson |
| EPI_ISL_417826 | hCoV-19/Iceland/204/2020 | Europe / Iceland / Reykjavik | 2020-03-16 | The National University Hospital of Iceland | deCODE genetics | Daniel F Gudbjartsson; Agnar Helgason; Hakon Jonsson; Olafur T Magnusson; Pall Melsted; Gudmundur L Norddahl; Jona Saemundsdottir; Asgeir Sigurdsson; Patrick Sulem; Arna B Agustsdottir; Berglind Eiriksdottir; Run Fridriksdottir; Elisabet E Gardarsdottir; Gudmundur Georgsson; Olafia S Gretarsdottir; Kjartan R Gudmundsson; Thora R Gunnarsdottir; Arnaldur Gylfason; Hilma Holm; Brynjar O Jensson; Aslaug Jonasdottir; Kamilla S Josefsdottir; Thordur Kristjansson; Droplaug N Magnusdottir; Louise le Roux; Gudrun Sigmundsdottir; Gardar Sveinbjornsson; Kristin E Sveinsdottir; Maney Sveinsdottir; Emil A Thorarensen; Bjarni Thorbjornsson; Gisli Masson; Ingileif Jonsdottir; Alma Moller; Thorolfur Gudnason; Karl G Kristinsson; Unnur Thorsteinsdottir; Kari Stefansson |
| EPI_ISL_417827 | hCoV-19/Iceland/205/2020 | Europe / Iceland / Reykjavik | 2020-03-16 | The National University Hospital of Iceland | deCODE genetics | Daniel F Gudbjartsson; Agnar Helgason; Hakon Jonsson; Olafur T Magnusson; Pall Melsted; Gudmundur L Norddahl; Jona Saemundsdottir; Asgeir Sigurdsson; Patrick Sulem; Arna B Agustsdottir; Berglind Eiriksdottir; Run Fridriksdottir; Elisabet E Gardarsdottir; Gudmundur Georgsson; Olafia S Gretarsdottir; Kjartan R Gudmundsson; Thora R Gunnarsdottir; Arnaldur Gylfason; Hilma Holm; Brynjar O Jensson; Aslaug Jonasdottir; Kamilla S Josefsdottir; Thordur Kristjansson; Droplaug N Magnusdottir; Louise le Roux; Gudrun Sigmundsdottir; Gardar Sveinbjornsson; Kristin E Sveinsdottir; Maney Sveinsdottir; Emil A Thorarensen; Bjarni Thorbjornsson; Gisli Masson; Ingileif Jonsdottir; Alma Moller; Thorolfur Gudnason; Karl G Kristinsson; Unnur Thorsteinsdottir; Kari Stefansson |
| EPI_ISL_417828 | hCoV-19/Iceland/21/2020 | Europe / Iceland / Reykjavik | 2020-03-02 | The National University Hospital of Iceland | deCODE genetics | Daniel F Gudbjartsson; Agnar Helgason; Hakon Jonsson; Olafur T Magnusson; Pall Melsted; Gudmundur L Norddahl; Jona Saemundsdottir; Asgeir Sigurdsson; Patrick Sulem; Arna B Agustsdottir; Berglind Eiriksdottir; Run Fridriksdottir; Elisabet E Gardarsdottir; Gudmundur Georgsson; Olafia S Gretarsdottir; Kjartan R Gudmundsson; Thora R Gunnarsdottir; Arnaldur Gylfason; Hilma Holm; Brynjar O Jensson; Aslaug Jonasdottir; Kamilla S Josefsdottir; Thordur Kristjansson; Droplaug N Magnusdottir; Louise le Roux; Gudrun Sigmundsdottir; Gardar Sveinbjornsson; Kristin E Sveinsdottir; Maney Sveinsdottir; Emil A Thorarensen; Bjarni Thorbjornsson; Gisli Masson; Ingileif Jonsdottir; Alma Moller; Thorolfur Gudnason; Karl G Kristinsson; Unnur Thorsteinsdottir; Kari Stefansson |
| EPI_ISL_417829 | hCoV-19/Iceland/210/2020 | Europe / Iceland / Reykjavik | 2020-03-16 | The National University Hospital of Iceland | deCODE genetics | Daniel F Gudbjartsson; Agnar Helgason; Hakon Jonsson; Olafur T Magnusson; Pall Melsted; Gudmundur L Norddahl; Jona Saemundsdottir; Asgeir Sigurdsson; Patrick Sulem; Arna B Agustsdottir; Berglind Eiriksdottir; Run Fridriksdottir; Elisabet E Gardarsdottir; Gudmundur Georgsson; Olafia S Gretarsdottir; Kjartan R Gudmundsson; Thora R Gunnarsdottir; Arnaldur Gylfason; Hilma Holm; Brynjar O Jensson; Aslaug Jonasdottir; Kamilla S Josefsdottir; Thordur Kristjansson; Droplaug N Magnusdottir; Louise le Roux; Gudrun Sigmundsdottir; Gardar Sveinbjornsson; Kristin E Sveinsdottir; Maney Sveinsdottir; Emil A Thorarensen; Bjarni Thorbjornsson; Gisli Masson; Ingileif Jonsdottir; Alma Moller; Thorolfur Gudnason; Karl G Kristinsson; Unnur Thorsteinsdottir; Kari Stefansson |
| EPI_ISL_417830 | hCoV-19/Iceland/212/2020 | Europe / Iceland / Reykjavik | 2020-03-16 | The National University Hospital of Iceland | deCODE genetics | Daniel F Gudbjartsson; Agnar Helgason; Hakon Jonsson; Olafur T Magnusson; Pall Melsted; Gudmundur L Norddahl; Jona Saemundsdottir; Asgeir Sigurdsson; Patrick Sulem; Arna B Agustsdottir; Berglind Eiriksdottir; Run Fridriksdottir; Elisabet E Gardarsdottir; Gudmundur Georgsson; Olafia S Gretarsdottir; Kjartan R Gudmundsson; Thora R Gunnarsdottir; Arnaldur Gylfason; Hilma Holm; Brynjar O Jensson; Aslaug Jonasdottir; Kamilla S Josefsdottir; Thordur Kristjansson; Droplaug N Magnusdottir; Louise le Roux; Gudrun Sigmundsdottir; Gardar Sveinbjornsson; Kristin E Sveinsdottir; Maney Sveinsdottir; Emil A Thorarensen; Bjarni Thorbjornsson; Gisli Masson; Ingileif Jonsdottir; Alma Moller; Thorolfur Gudnason; Karl G Kristinsson; Unnur Thorsteinsdottir; Kari Stefansson |
| EPI_ISL_417831 | hCoV-19/Iceland/214/2020 | Europe / Iceland / Reykjavik | 2020-03-16 | The National University Hospital of Iceland | deCODE genetics | Daniel F Gudbjartsson; Agnar Helgason; Hakon Jonsson; Olafur T Magnusson; Pall Melsted; Gudmundur L Norddahl; Jona Saemundsdottir; Asgeir Sigurdsson; Patrick Sulem; Arna B Agustsdottir; Berglind Eiriksdottir; Run Fridriksdottir; Elisabet E Gardarsdottir; Gudmundur Georgsson; Olafia S Gretarsdottir; Kjartan R Gudmundsson; Thora R Gunnarsdottir; Arnaldur Gylfason; Hilma Holm; Brynjar O Jensson; Aslaug Jonasdottir; Kamilla S Josefsdottir; Thordur Kristjansson; Droplaug N Magnusdottir; Louise le Roux; Gudrun Sigmundsdottir; Gardar Sveinbjornsson; Kristin E Sveinsdottir; Maney Sveinsdottir; Emil A Thorarensen; Bjarni Thorbjornsson; Gisli Masson; Ingileif Jonsdottir; Alma Moller; Thorolfur Gudnason; Karl G Kristinsson; Unnur Thorsteinsdottir; Kari Stefansson |
| EPI_ISL_417832 | hCoV-19/Iceland/216/2020 | Europe / Iceland / Reykjavik | 2020-03-16 | The National University Hospital of Iceland | deCODE genetics | Daniel F Gudbjartsson; Agnar Helgason; Hakon Jonsson; Olafur T Magnusson; Pall Melsted; Gudmundur L Norddahl; Jona Saemundsdottir; Asgeir Sigurdsson; Patrick Sulem; Arna B Agustsdottir; Berglind Eiriksdottir; Run Fridriksdottir; Elisabet E Gardarsdottir; Gudmundur Georgsson; Olafia S Gretarsdottir; Kjartan R Gudmundsson; Thora R Gunnarsdottir; Arnaldur Gylfason; Hilma Holm; Brynjar O Jensson; Aslaug Jonasdottir; Kamilla S Josefsdottir; Thordur Kristjansson; Droplaug N Magnusdottir; Louise le Roux; Gudrun Sigmundsdottir; Gardar Sveinbjornsson; Kristin E Sveinsdottir; Maney Sveinsdottir; Emil A Thorarensen; Bjarni Thorbjornsson; Gisli Masson; Ingileif Jonsdottir; Alma Moller; Thorolfur Gudnason; Karl G Kristinsson; Unnur Thorsteinsdottir; Kari Stefansson |
| EPI_ISL_417833 | hCoV-19/Iceland/217/2020 | Europe / Iceland / Reykjavik | 2020-03-16 | The National University Hospital of Iceland | deCODE genetics | Daniel F Gudbjartsson; Agnar Helgason; Hakon Jonsson; Olafur T Magnusson; Pall Melsted; Gudmundur L Norddahl; Jona Saemundsdottir; Asgeir Sigurdsson; Patrick Sulem; Arna B Agustsdottir; Berglind Eiriksdottir; Run Fridriksdottir; Elisabet E Gardarsdottir; Gudmundur Georgsson; Olafia S Gretarsdottir; Kjartan R Gudmundsson; Thora R Gunnarsdottir; Arnaldur Gylfason; Hilma Holm; Brynjar O Jensson; Aslaug Jonasdottir; Kamilla S Josefsdottir; Thordur Kristjansson; Droplaug N Magnusdottir; Louise le Roux; Gudrun Sigmundsdottir; Gardar Sveinbjornsson; Kristin E Sveinsdottir; Maney Sveinsdottir; Emil A Thorarensen; Bjarni Thorbjornsson; Gisli Masson; Ingileif Jonsdottir; Alma Moller; Thorolfur Gudnason; Karl G Kristinsson; Unnur Thorsteinsdottir; Kari Stefansson |
| EPI_ISL_417834 | hCoV-19/Iceland/218/2020 | Europe / Iceland / Reykjavik | 2020-03-16 | The National University Hospital of Iceland | deCODE genetics | Daniel F Gudbjartsson; Agnar Helgason; Hakon Jonsson; Olafur T Magnusson; Pall Melsted; Gudmundur L Norddahl; Jona Saemundsdottir; Asgeir Sigurdsson; Patrick Sulem; Arna B Agustsdottir; Berglind Eiriksdottir; Run Fridriksdottir; Elisabet E Gardarsdottir; Gudmundur Georgsson; Olafia S Gretarsdottir; Kjartan R Gudmundsson; Thora R Gunnarsdottir; Arnaldur Gylfason; Hilma Holm; Brynjar O Jensson; Aslaug Jonasdottir; Kamilla S Josefsdottir; Thordur Kristjansson; Droplaug N Magnusdottir; Louise le Roux; Gudrun Sigmundsdottir; Gardar Sveinbjornsson; Kristin E Sveinsdottir; Maney Sveinsdottir; Emil A Thorarensen; Bjarni Thorbjornsson; Gisli Masson; Ingileif Jonsdottir; Alma Moller; Thorolfur Gudnason; Karl G Kristinsson; Unnur Thorsteinsdottir; Kari Stefansson |
| EPI_ISL_417835 | hCoV-19/Iceland/219/2020 | Europe / Iceland / Reykjavik | 2020-03-16 | The National University Hospital of Iceland | deCODE genetics | Daniel F Gudbjartsson; Agnar Helgason; Hakon Jonsson; Olafur T Magnusson; Pall Melsted; Gudmundur L Norddahl; Jona Saemundsdottir; Asgeir Sigurdsson; Patrick Sulem; Arna B Agustsdottir; Berglind Eiriksdottir; Run Fridriksdottir; Elisabet E Gardarsdottir; Gudmundur Georgsson; Olafia S Gretarsdottir; Kjartan R Gudmundsson; Thora R Gunnarsdottir; Arnaldur Gylfason; Hilma Holm; Brynjar O Jensson; Aslaug Jonasdottir; Kamilla S Josefsdottir; Thordur Kristjansson; Droplaug N Magnusdottir; Louise le Roux; Gudrun Sigmundsdottir; Gardar Sveinbjornsson; Kristin E Sveinsdottir; Maney Sveinsdottir; Emil A Thorarensen; Bjarni Thorbjornsson; Gisli Masson; Ingileif Jonsdottir; Alma Moller; Thorolfur Gudnason; Karl G Kristinsson; Unnur Thorsteinsdottir; Kari Stefansson |
| EPI_ISL_417836 | hCoV-19/Iceland/221/2020 | Europe / Iceland / Reykjavik | 2020-03-16 | The National University Hospital of Iceland | deCODE genetics | Daniel F Gudbjartsson; Agnar Helgason; Hakon Jonsson; Olafur T Magnusson; Pall Melsted; Gudmundur L Norddahl; Jona Saemundsdottir; Asgeir Sigurdsson; Patrick Sulem; Arna B Agustsdottir; Berglind Eiriksdottir; Run Fridriksdottir; Elisabet E Gardarsdottir; Gudmundur Georgsson; Olafia S Gretarsdottir; Kjartan R Gudmundsson; Thora R Gunnarsdottir; Arnaldur Gylfason; Hilma Holm; Brynjar O Jensson; Aslaug Jonasdottir; Kamilla S Josefsdottir; Thordur Kristjansson; Droplaug N Magnusdottir; Louise le Roux; Gudrun Sigmundsdottir; Gardar Sveinbjornsson; Kristin E Sveinsdottir; Maney Sveinsdottir; Emil A Thorarensen; Bjarni Thorbjornsson; Gisli Masson; Ingileif Jonsdottir; Alma Moller; Thorolfur Gudnason; Karl G Kristinsson; Unnur Thorsteinsdottir; Kari Stefansson |
| EPI_ISL_417837 | hCoV-19/Iceland/222/2020 | Europe / Iceland / Reykjavik | 2020-03-16 | The National University Hospital of Iceland | deCODE genetics | Daniel F Gudbjartsson; Agnar Helgason; Hakon Jonsson; Olafur T Magnusson; Pall Melsted; Gudmundur L Norddahl; Jona Saemundsdottir; Asgeir Sigurdsson; Patrick Sulem; Arna B Agustsdottir; Berglind Eiriksdottir; Run Fridriksdottir; Elisabet E Gardarsdottir; Gudmundur Georgsson; Olafia S Gretarsdottir; Kjartan R Gudmundsson; Thora R Gunnarsdottir; Arnaldur Gylfason; Hilma Holm; Brynjar O Jensson; Aslaug Jonasdottir; Kamilla S Josefsdottir; Thordur Kristjansson; Droplaug N Magnusdottir; Louise le Roux; Gudrun Sigmundsdottir; Gardar Sveinbjornsson; Kristin E Sveinsdottir; Maney Sveinsdottir; Emil A Thorarensen; Bjarni Thorbjornsson; Gisli Masson; Ingileif Jonsdottir; Alma Moller; Thorolfur Gudnason; Karl G Kristinsson; Unnur Thorsteinsdottir; Kari Stefansson |
| EPI_ISL_417838 | hCoV-19/Iceland/223/2020 | Europe / Iceland / Reykjavik | 2020-03-16 | The National University Hospital of Iceland | deCODE genetics | Daniel F Gudbjartsson; Agnar Helgason; Hakon Jonsson; Olafur T Magnusson; Pall Melsted; Gudmundur L Norddahl; Jona Saemundsdottir; Asgeir Sigurdsson; Patrick Sulem; Arna B Agustsdottir; Berglind Eiriksdottir; Run Fridriksdottir; Elisabet E Gardarsdottir; Gudmundur Georgsson; Olafia S Gretarsdottir; Kjartan R Gudmundsson; Thora R Gunnarsdottir; Arnaldur Gylfason; Hilma Holm; Brynjar O Jensson; Aslaug Jonasdottir; Kamilla S Josefsdottir; Thordur Kristjansson; Droplaug N Magnusdottir; Louise le Roux; Gudrun Sigmundsdottir; Gardar Sveinbjornsson; Kristin E Sveinsdottir; Maney Sveinsdottir; Emil A Thorarensen; Bjarni Thorbjornsson; Gisli Masson; Ingileif Jonsdottir; Alma Moller; Thorolfur Gudnason; Karl G Kristinsson; Unnur Thorsteinsdottir; Kari Stefansson |
| EPI_ISL_417839 | hCoV-19/Iceland/224/2020 | Europe / Iceland / Reykjavik | 2020-03-16 | The National University Hospital of Iceland | deCODE genetics | Daniel F Gudbjartsson; Agnar Helgason; Hakon Jonsson; Olafur T Magnusson; Pall Melsted; Gudmundur L Norddahl; Jona Saemundsdottir; Asgeir Sigurdsson; Patrick Sulem; Arna B Agustsdottir; Berglind Eiriksdottir; Run Fridriksdottir; Elisabet E Gardarsdottir; Gudmundur Georgsson; Olafia S Gretarsdottir; Kjartan R Gudmundsson; Thora R Gunnarsdottir; Arnaldur Gylfason; Hilma Holm; Brynjar O Jensson; Aslaug Jonasdottir; Kamilla S Josefsdottir; Thordur Kristjansson; Droplaug N Magnusdottir; Louise le Roux; Gudrun Sigmundsdottir; Gardar Sveinbjornsson; Kristin E Sveinsdottir; Maney Sveinsdottir; Emil A Thorarensen; Bjarni Thorbjornsson; Gisli Masson; Ingileif Jonsdottir; Alma Moller; Thorolfur Gudnason; Karl G Kristinsson; Unnur Thorsteinsdottir; Kari Stefansson |
| EPI_ISL_417840 | hCoV-19/Iceland/23/2020 | Europe / Iceland / Reykjavik | 2020-03-02 | The National University Hospital of Iceland | deCODE genetics | Daniel F Gudbjartsson; Agnar Helgason; Hakon Jonsson; Olafur T Magnusson; Pall Melsted; Gudmundur L Norddahl; Jona Saemundsdottir; Asgeir Sigurdsson; Patrick Sulem; Arna B Agustsdottir; Berglind Eiriksdottir; Run Fridriksdottir; Elisabet E Gardarsdottir; Gudmundur Georgsson; Olafia S Gretarsdottir; Kjartan R Gudmundsson; Thora R Gunnarsdottir; Arnaldur Gylfason; Hilma Holm; Brynjar O Jensson; Aslaug Jonasdottir; Kamilla S Josefsdottir; Thordur Kristjansson; Droplaug N Magnusdottir; Louise le Roux; Gudrun Sigmundsdottir; Gardar Sveinbjornsson; Kristin E Sveinsdottir; Maney Sveinsdottir; Emil A Thorarensen; Bjarni Thorbjornsson; Gisli Masson; Ingileif Jonsdottir; Alma Moller; Thorolfur Gudnason; Karl G Kristinsson; Unnur Thorsteinsdottir; Kari Stefansson |
| EPI_ISL_417842 | hCoV-19/Iceland/31/2020 | Europe / Iceland / Reykjavik | 2020-03-03 | The National University Hospital of Iceland | deCODE genetics | Daniel F Gudbjartsson; Agnar Helgason; Hakon Jonsson; Olafur T Magnusson; Pall Melsted; Gudmundur L Norddahl; Jona Saemundsdottir; Asgeir Sigurdsson; Patrick Sulem; Arna B Agustsdottir; Berglind Eiriksdottir; Run Fridriksdottir; Elisabet E Gardarsdottir; Gudmundur Georgsson; Olafia S Gretarsdottir; Kjartan R Gudmundsson; Thora R Gunnarsdottir; Arnaldur Gylfason; Hilma Holm; Brynjar O Jensson; Aslaug Jonasdottir; Kamilla S Josefsdottir; Thordur Kristjansson; Droplaug N Magnusdottir; Louise le Roux; Gudrun Sigmundsdottir; Gardar Sveinbjornsson; Kristin E Sveinsdottir; Maney Sveinsdottir; Emil A Thorarensen; Bjarni Thorbjornsson; Gisli Masson; Ingileif Jonsdottir; Alma Moller; Thorolfur Gudnason; Karl G Kristinsson; Unnur Thorsteinsdottir; Kari Stefansson |
| EPI_ISL_417844 | hCoV-19/Iceland/36/2020 | Europe / Iceland / Reykjavik | 2020-03-04 | The National University Hospital of Iceland | deCODE genetics | Daniel F Gudbjartsson; Agnar Helgason; Hakon Jonsson; Olafur T Magnusson; Pall Melsted; Gudmundur L Norddahl; Jona Saemundsdottir; Asgeir Sigurdsson; Patrick Sulem; Arna B Agustsdottir; Berglind Eiriksdottir; Run Fridriksdottir; Elisabet E Gardarsdottir; Gudmundur Georgsson; Olafia S Gretarsdottir; Kjartan R Gudmundsson; Thora R Gunnarsdottir; Arnaldur Gylfason; Hilma Holm; Brynjar O Jensson; Aslaug Jonasdottir; Kamilla S Josefsdottir; Thordur Kristjansson; Droplaug N Magnusdottir; Louise le Roux; Gudrun Sigmundsdottir; Gardar Sveinbjornsson; Kristin E Sveinsdottir; Maney Sveinsdottir; Emil A Thorarensen; Bjarni Thorbjornsson; Gisli Masson; Ingileif Jonsdottir; Alma Moller; Thorolfur Gudnason; Karl G Kristinsson; Unnur Thorsteinsdottir; Kari Stefansson |
| EPI_ISL_417846 | hCoV-19/Iceland/40/2020 | Europe / Iceland / Reykjavik | 2020-03-04 | The National University Hospital of Iceland | deCODE genetics | Daniel F Gudbjartsson; Agnar Helgason; Hakon Jonsson; Olafur T Magnusson; Pall Melsted; Gudmundur L Norddahl; Jona Saemundsdottir; Asgeir Sigurdsson; Patrick Sulem; Arna B Agustsdottir; Berglind Eiriksdottir; Run Fridriksdottir; Elisabet E Gardarsdottir; Gudmundur Georgsson; Olafia S Gretarsdottir; Kjartan R Gudmundsson; Thora R Gunnarsdottir; Arnaldur Gylfason; Hilma Holm; Brynjar O Jensson; Aslaug Jonasdottir; Kamilla S Josefsdottir; Thordur Kristjansson; Droplaug N Magnusdottir; Louise le Roux; Gudrun Sigmundsdottir; Gardar Sveinbjornsson; Kristin E Sveinsdottir; Maney Sveinsdottir; Emil A Thorarensen; Bjarni Thorbjornsson; Gisli Masson; Ingileif Jonsdottir; Alma Moller; Thorolfur Gudnason; Karl G Kristinsson; Unnur Thorsteinsdottir; Kari Stefansson |
| EPI_ISL_417847 | hCoV-19/Iceland/41/2020 | Europe / Iceland / Reykjavik | 2020-03-04 | The National University Hospital of Iceland | deCODE genetics | Daniel F Gudbjartsson; Agnar Helgason; Hakon Jonsson; Olafur T Magnusson; Pall Melsted; Gudmundur L Norddahl; Jona Saemundsdottir; Asgeir Sigurdsson; Patrick Sulem; Arna B Agustsdottir; Berglind Eiriksdottir; Run Fridriksdottir; Elisabet E Gardarsdottir; Gudmundur Georgsson; Olafia S Gretarsdottir; Kjartan R Gudmundsson; Thora R Gunnarsdottir; Arnaldur Gylfason; Hilma Holm; Brynjar O Jensson; Aslaug Jonasdottir; Kamilla S Josefsdottir; Thordur Kristjansson; Droplaug N Magnusdottir; Louise le Roux; Gudrun Sigmundsdottir; Gardar Sveinbjornsson; Kristin E Sveinsdottir; Maney Sveinsdottir; Emil A Thorarensen; Bjarni Thorbjornsson; Gisli Masson; Ingileif Jonsdottir; Alma Moller; Thorolfur Gudnason; Karl G Kristinsson; Unnur Thorsteinsdottir; Kari Stefansson |
| EPI_ISL_417850 | hCoV-19/Iceland/52/2020 | Europe / Iceland / Reykjavik | 2020-03-05 | The National University Hospital of Iceland | deCODE genetics | Daniel F Gudbjartsson; Agnar Helgason; Hakon Jonsson; Olafur T Magnusson; Pall Melsted; Gudmundur L Norddahl; Jona Saemundsdottir; Asgeir Sigurdsson; Patrick Sulem; Arna B Agustsdottir; Berglind Eiriksdottir; Run Fridriksdottir; Elisabet E Gardarsdottir; Gudmundur Georgsson; Olafia S Gretarsdottir; Kjartan R Gudmundsson; Thora R Gunnarsdottir; Arnaldur Gylfason; Hilma Holm; Brynjar O Jensson; Aslaug Jonasdottir; Kamilla S Josefsdottir; Thordur Kristjansson; Droplaug N Magnusdottir; Louise le Roux; Gudrun Sigmundsdottir; Gardar Sveinbjornsson; Kristin E Sveinsdottir; Maney Sveinsdottir; Emil A Thorarensen; Bjarni Thorbjornsson; Gisli Masson; Ingileif Jonsdottir; Alma Moller; Thorolfur Gudnason; Karl G Kristinsson; Unnur Thorsteinsdottir; Kari Stefansson |
| EPI_ISL_417851 | hCoV-19/Iceland/53/2020 | Europe / Iceland / Reykjavik | 2020-03-05 | The National University Hospital of Iceland | deCODE genetics | Daniel F Gudbjartsson; Agnar Helgason; Hakon Jonsson; Olafur T Magnusson; Pall Melsted; Gudmundur L Norddahl; Jona Saemundsdottir; Asgeir Sigurdsson; Patrick Sulem; Arna B Agustsdottir; Berglind Eiriksdottir; Run Fridriksdottir; Elisabet E Gardarsdottir; Gudmundur Georgsson; Olafia S Gretarsdottir; Kjartan R Gudmundsson; Thora R Gunnarsdottir; Arnaldur Gylfason; Hilma Holm; Brynjar O Jensson; Aslaug Jonasdottir; Kamilla S Josefsdottir; Thordur Kristjansson; Droplaug N Magnusdottir; Louise le Roux; Gudrun Sigmundsdottir; Gardar Sveinbjornsson; Kristin E Sveinsdottir; Maney Sveinsdottir; Emil A Thorarensen; Bjarni Thorbjornsson; Gisli Masson; Ingileif Jonsdottir; Alma Moller; Thorolfur Gudnason; Karl G Kristinsson; Unnur Thorsteinsdottir; Kari Stefansson |
| EPI_ISL_417852 | hCoV-19/Iceland/56/2020 | Europe / Iceland / Reykjavik | 2020-03-06 | The National University Hospital of Iceland | deCODE genetics | Daniel F Gudbjartsson; Agnar Helgason; Hakon Jonsson; Olafur T Magnusson; Pall Melsted; Gudmundur L Norddahl; Jona Saemundsdottir; Asgeir Sigurdsson; Patrick Sulem; Arna B Agustsdottir; Berglind Eiriksdottir; Run Fridriksdottir; Elisabet E Gardarsdottir; Gudmundur Georgsson; Olafia S Gretarsdottir; Kjartan R Gudmundsson; Thora R Gunnarsdottir; Arnaldur Gylfason; Hilma Holm; Brynjar O Jensson; Aslaug Jonasdottir; Kamilla S Josefsdottir; Thordur Kristjansson; Droplaug N Magnusdottir; Louise le Roux; Gudrun Sigmundsdottir; Gardar Sveinbjornsson; Kristin E Sveinsdottir; Maney Sveinsdottir; Emil A Thorarensen; Bjarni Thorbjornsson; Gisli Masson; Ingileif Jonsdottir; Alma Moller; Thorolfur Gudnason; Karl G Kristinsson; Unnur Thorsteinsdottir; Kari Stefansson |
| EPI_ISL_417853 | hCoV-19/Iceland/59/2020 | Europe / Iceland / Reykjavik | 2020-03-06 | The National University Hospital of Iceland | deCODE genetics | Daniel F Gudbjartsson; Agnar Helgason; Hakon Jonsson; Olafur T Magnusson; Pall Melsted; Gudmundur L Norddahl; Jona Saemundsdottir; Asgeir Sigurdsson; Patrick Sulem; Arna B Agustsdottir; Berglind Eiriksdottir; Run Fridriksdottir; Elisabet E Gardarsdottir; Gudmundur Georgsson; Olafia S Gretarsdottir; Kjartan R Gudmundsson; Thora R Gunnarsdottir; Arnaldur Gylfason; Hilma Holm; Brynjar O Jensson; Aslaug Jonasdottir; Kamilla S Josefsdottir; Thordur Kristjansson; Droplaug N Magnusdottir; Louise le Roux; Gudrun Sigmundsdottir; Gardar Sveinbjornsson; Kristin E Sveinsdottir; Maney Sveinsdottir; Emil A Thorarensen; Bjarni Thorbjornsson; Gisli Masson; Ingileif Jonsdottir; Alma Moller; Thorolfur Gudnason; Karl G Kristinsson; Unnur Thorsteinsdottir; Kari Stefansson |
| EPI_ISL_417854 | hCoV-19/Iceland/6/2020 | Europe / Iceland / Reykjavik | 2020-03-13 | The National University Hospital of Iceland | deCODE genetics | Daniel F Gudbjartsson; Agnar Helgason; Hakon Jonsson; Olafur T Magnusson; Pall Melsted; Gudmundur L Norddahl; Jona Saemundsdottir; Asgeir Sigurdsson; Patrick Sulem; Arna B Agustsdottir; Berglind Eiriksdottir; Run Fridriksdottir; Elisabet E Gardarsdottir; Gudmundur Georgsson; Olafia S Gretarsdottir; Kjartan R Gudmundsson; Thora R Gunnarsdottir; Arnaldur Gylfason; Hilma Holm; Brynjar O Jensson; Aslaug Jonasdottir; Kamilla S Josefsdottir; Thordur Kristjansson; Droplaug N Magnusdottir; Louise le Roux; Gudrun Sigmundsdottir; Gardar Sveinbjornsson; Kristin E Sveinsdottir; Maney Sveinsdottir; Emil A Thorarensen; Bjarni Thorbjornsson; Gisli Masson; Ingileif Jonsdottir; Alma Moller; Thorolfur Gudnason; Karl G Kristinsson; Unnur Thorsteinsdottir; Kari Stefansson |
| EPI_ISL_417855 | hCoV-19/Iceland/63/2020 | Europe / Iceland / Reykjavik | 2020-03-07 | The National University Hospital of Iceland | deCODE genetics | Daniel F Gudbjartsson; Agnar Helgason; Hakon Jonsson; Olafur T Magnusson; Pall Melsted; Gudmundur L Norddahl; Jona Saemundsdottir; Asgeir Sigurdsson; Patrick Sulem; Arna B Agustsdottir; Berglind Eiriksdottir; Run Fridriksdottir; Elisabet E Gardarsdottir; Gudmundur Georgsson; Olafia S Gretarsdottir; Kjartan R Gudmundsson; Thora R Gunnarsdottir; Arnaldur Gylfason; Hilma Holm; Brynjar O Jensson; Aslaug Jonasdottir; Kamilla S Josefsdottir; Thordur Kristjansson; Droplaug N Magnusdottir; Louise le Roux; Gudrun Sigmundsdottir; Gardar Sveinbjornsson; Kristin E Sveinsdottir; Maney Sveinsdottir; Emil A Thorarensen; Bjarni Thorbjornsson; Gisli Masson; Ingileif Jonsdottir; Alma Moller; Thorolfur Gudnason; Karl G Kristinsson; Unnur Thorsteinsdottir; Kari Stefansson |
| EPI_ISL_417856 | hCoV-19/Iceland/64/2020 | Europe / Iceland / Reykjavik | 2020-03-07 | The National University Hospital of Iceland | deCODE genetics | Daniel F Gudbjartsson; Agnar Helgason; Hakon Jonsson; Olafur T Magnusson; Pall Melsted; Gudmundur L Norddahl; Jona Saemundsdottir; Asgeir Sigurdsson; Patrick Sulem; Arna B Agustsdottir; Berglind Eiriksdottir; Run Fridriksdottir; Elisabet E Gardarsdottir; Gudmundur Georgsson; Olafia S Gretarsdottir; Kjartan R Gudmundsson; Thora R Gunnarsdottir; Arnaldur Gylfason; Hilma Holm; Brynjar O Jensson; Aslaug Jonasdottir; Kamilla S Josefsdottir; Thordur Kristjansson; Droplaug N Magnusdottir; Louise le Roux; Gudrun Sigmundsdottir; Gardar Sveinbjornsson; Kristin E Sveinsdottir; Maney Sveinsdottir; Emil A Thorarensen; Bjarni Thorbjornsson; Gisli Masson; Ingileif Jonsdottir; Alma Moller; Thorolfur Gudnason; Karl G Kristinsson; Unnur Thorsteinsdottir; Kari Stefansson |
| EPI_ISL_417857 | hCoV-19/Iceland/66/2020 | Europe / Iceland / Reykjavik | 2020-03-07 | The National University Hospital of Iceland | deCODE genetics | Daniel F Gudbjartsson; Agnar Helgason; Hakon Jonsson; Olafur T Magnusson; Pall Melsted; Gudmundur L Norddahl; Jona Saemundsdottir; Asgeir Sigurdsson; Patrick Sulem; Arna B Agustsdottir; Berglind Eiriksdottir; Run Fridriksdottir; Elisabet E Gardarsdottir; Gudmundur Georgsson; Olafia S Gretarsdottir; Kjartan R Gudmundsson; Thora R Gunnarsdottir; Arnaldur Gylfason; Hilma Holm; Brynjar O Jensson; Aslaug Jonasdottir; Kamilla S Josefsdottir; Thordur Kristjansson; Droplaug N Magnusdottir; Louise le Roux; Gudrun Sigmundsdottir; Gardar Sveinbjornsson; Kristin E Sveinsdottir; Maney Sveinsdottir; Emil A Thorarensen; Bjarni Thorbjornsson; Gisli Masson; Ingileif Jonsdottir; Alma Moller; Thorolfur Gudnason; Karl G Kristinsson; Unnur Thorsteinsdottir; Kari Stefansson |
| EPI_ISL_417858 | hCoV-19/Iceland/68/2020 | Europe / Iceland / Reykjavik | 2020-03-08 | The National University Hospital of Iceland | deCODE genetics | Daniel F Gudbjartsson; Agnar Helgason; Hakon Jonsson; Olafur T Magnusson; Pall Melsted; Gudmundur L Norddahl; Jona Saemundsdottir; Asgeir Sigurdsson; Patrick Sulem; Arna B Agustsdottir; Berglind Eiriksdottir; Run Fridriksdottir; Elisabet E Gardarsdottir; Gudmundur Georgsson; Olafia S Gretarsdottir; Kjartan R Gudmundsson; Thora R Gunnarsdottir; Arnaldur Gylfason; Hilma Holm; Brynjar O Jensson; Aslaug Jonasdottir; Kamilla S Josefsdottir; Thordur Kristjansson; Droplaug N Magnusdottir; Louise le Roux; Gudrun Sigmundsdottir; Gardar Sveinbjornsson; Kristin E Sveinsdottir; Maney Sveinsdottir; Emil A Thorarensen; Bjarni Thorbjornsson; Gisli Masson; Ingileif Jonsdottir; Alma Moller; Thorolfur Gudnason; Karl G Kristinsson; Unnur Thorsteinsdottir; Kari Stefansson |
| EPI_ISL_417859 | hCoV-19/Iceland/71/2020 | Europe / Iceland / Reykjavik | 2020-03-08 | The National University Hospital of Iceland | deCODE genetics | Daniel F Gudbjartsson; Agnar Helgason; Hakon Jonsson; Olafur T Magnusson; Pall Melsted; Gudmundur L Norddahl; Jona Saemundsdottir; Asgeir Sigurdsson; Patrick Sulem; Arna B Agustsdottir; Berglind Eiriksdottir; Run Fridriksdottir; Elisabet E Gardarsdottir; Gudmundur Georgsson; Olafia S Gretarsdottir; Kjartan R Gudmundsson; Thora R Gunnarsdottir; Arnaldur Gylfason; Hilma Holm; Brynjar O Jensson; Aslaug Jonasdottir; Kamilla S Josefsdottir; Thordur Kristjansson; Droplaug N Magnusdottir; Louise le Roux; Gudrun Sigmundsdottir; Gardar Sveinbjornsson; Kristin E Sveinsdottir; Maney Sveinsdottir; Emil A Thorarensen; Bjarni Thorbjornsson; Gisli Masson; Ingileif Jonsdottir; Alma Moller; Thorolfur Gudnason; Karl G Kristinsson; Unnur Thorsteinsdottir; Kari Stefansson |
| EPI_ISL_417860 | hCoV-19/Iceland/72/2020 | Europe / Iceland / Reykjavik | 2020-03-08 | The National University Hospital of Iceland | deCODE genetics | Daniel F Gudbjartsson; Agnar Helgason; Hakon Jonsson; Olafur T Magnusson; Pall Melsted; Gudmundur L Norddahl; Jona Saemundsdottir; Asgeir Sigurdsson; Patrick Sulem; Arna B Agustsdottir; Berglind Eiriksdottir; Run Fridriksdottir; Elisabet E Gardarsdottir; Gudmundur Georgsson; Olafia S Gretarsdottir; Kjartan R Gudmundsson; Thora R Gunnarsdottir; Arnaldur Gylfason; Hilma Holm; Brynjar O Jensson; Aslaug Jonasdottir; Kamilla S Josefsdottir; Thordur Kristjansson; Droplaug N Magnusdottir; Louise le Roux; Gudrun Sigmundsdottir; Gardar Sveinbjornsson; Kristin E Sveinsdottir; Maney Sveinsdottir; Emil A Thorarensen; Bjarni Thorbjornsson; Gisli Masson; Ingileif Jonsdottir; Alma Moller; Thorolfur Gudnason; Karl G Kristinsson; Unnur Thorsteinsdottir; Kari Stefansson |
| EPI_ISL_417861 | hCoV-19/Iceland/73/2020 | Europe / Iceland / Reykjavik | 2020-03-09 | The National University Hospital of Iceland | deCODE genetics | Daniel F Gudbjartsson; Agnar Helgason; Hakon Jonsson; Olafur T Magnusson; Pall Melsted; Gudmundur L Norddahl; Jona Saemundsdottir; Asgeir Sigurdsson; Patrick Sulem; Arna B Agustsdottir; Berglind Eiriksdottir; Run Fridriksdottir; Elisabet E Gardarsdottir; Gudmundur Georgsson; Olafia S Gretarsdottir; Kjartan R Gudmundsson; Thora R Gunnarsdottir; Arnaldur Gylfason; Hilma Holm; Brynjar O Jensson; Aslaug Jonasdottir; Kamilla S Josefsdottir; Thordur Kristjansson; Droplaug N Magnusdottir; Louise le Roux; Gudrun Sigmundsdottir; Gardar Sveinbjornsson; Kristin E Sveinsdottir; Maney Sveinsdottir; Emil A Thorarensen; Bjarni Thorbjornsson; Gisli Masson; Ingileif Jonsdottir; Alma Moller; Thorolfur Gudnason; Karl G Kristinsson; Unnur Thorsteinsdottir; Kari Stefansson |

| | | | | | | |
|---|---|---|---|---|---|---|
| EPI_ISL_417862 | hCoV-19/Iceland/77/2020 | Europe / Iceland / Reykjavik | 2020-03-09 | The National University Hospital of Iceland | deCODE genetics | Daniel F Gudbjartsson; Agnar Helgason; Hakon Jonsson; Olafur T Magnusson; Pall Melsted; Gudmundur L Norddahl; Jona Saemundsdottir; Asgeir Sigurdsson; Patrick Sulem; Arna B Agustsdottir; Berglind Eiriksdottir; Run Fridriksdottir; Elisabet E Gardarsdottir; Gudmundur Georgsson; Olafia S Gretarsdottir; Kjartan R Gudmundsson; Thora R Gunnarsdottir; Arnaldur Gylfason; Hilma Holm; Brynjar O Jensson; Aslaug Jonasdottir; Kamilla S Josefsdottir; Thordur Kristjansson; Droplaug N Magnusdottir; Louise le Roux; Gudrun Sigmundsdottir; Gardar Sveinbjornsson; Kristin E Sveinsdottir; Maney Sveinsdottir; Emil A Thorarensen; Bjarni Thorbjornsson; Gisli Masson; Ingileif Jonsdottir; Alma Moller; Thorolfur Gudnason; Karl G Kristinsson; Unnur Thorsteinsdottir; Kari Stefansson |
| EPI_ISL_417863 | hCoV-19/Iceland/78/2020 | Europe / Iceland / Reykjavik | 2020-03-09 | The National University Hospital of Iceland | deCODE genetics | Daniel F Gudbjartsson; Agnar Helgason; Hakon Jonsson; Olafur T Magnusson; Pall Melsted; Gudmundur L Norddahl; Jona Saemundsdottir; Asgeir Sigurdsson; Patrick Sulem; Arna B Agustsdottir; Berglind Eiriksdottir; Run Fridriksdottir; Elisabet E Gardarsdottir; Gudmundur Georgsson; Olafia S Gretarsdottir; Kjartan R Gudmundsson; Thora R Gunnarsdottir; Arnaldur Gylfason; Hilma Holm; Brynjar O Jensson; Aslaug Jonasdottir; Kamilla S Josefsdottir; Thordur Kristjansson; Droplaug N Magnusdottir; Louise le Roux; Gudrun Sigmundsdottir; Gardar Sveinbjornsson; Kristin E Sveinsdottir; Maney Sveinsdottir; Emil A Thorarensen; Bjarni Thorbjornsson; Gisli Masson; Ingileif Jonsdottir; Alma Moller; Thorolfur Gudnason; Karl G Kristinsson; Unnur Thorsteinsdottir; Kari Stefansson |
| EPI_ISL_417864 | hCoV-19/Iceland/8/2020 | Europe / Iceland / Reykjavik | 2020-03-15 | The National University Hospital of Iceland | deCODE genetics | Daniel F Gudbjartsson; Agnar Helgason; Hakon Jonsson; Olafur T Magnusson; Pall Melsted; Gudmundur L Norddahl; Jona Saemundsdottir; Asgeir Sigurdsson; Patrick Sulem; Arna B Agustsdottir; Berglind Eiriksdottir; Run Fridriksdottir; Elisabet E Gardarsdottir; Gudmundur Georgsson; Olafia S Gretarsdottir; Kjartan R Gudmundsson; Thora R Gunnarsdottir; Arnaldur Gylfason; Hilma Holm; Brynjar O Jensson; Aslaug Jonasdottir; Kamilla S Josefsdottir; Thordur Kristjansson; Droplaug N Magnusdottir; Louise le Roux; Gudrun Sigmundsdottir; Gardar Sveinbjornsson; Kristin E Sveinsdottir; Maney Sveinsdottir; Emil A Thorarensen; Bjarni Thorbjornsson; Gisli Masson; Ingileif Jonsdottir; Alma Moller; Thorolfur Gudnason; Karl G Kristinsson; Unnur Thorsteinsdottir; Kari Stefansson |
| EPI_ISL_417865 | hCoV-19/Iceland/80/2020 | Europe / Iceland / Reykjavik | 2020-03-09 | The National University Hospital of Iceland | deCODE genetics | Daniel F Gudbjartsson; Agnar Helgason; Hakon Jonsson; Olafur T Magnusson; Pall Melsted; Gudmundur L Norddahl; Jona Saemundsdottir; Asgeir Sigurdsson; Patrick Sulem; Arna B Agustsdottir; Berglind Eiriksdottir; Run Fridriksdottir; Elisabet E Gardarsdottir; Gudmundur Georgsson; Olafia S Gretarsdottir; Kjartan R Gudmundsson; Thora R Gunnarsdottir; Arnaldur Gylfason; Hilma Holm; Brynjar O Jensson; Aslaug Jonasdottir; Kamilla S Josefsdottir; Thordur Kristjansson; Droplaug N Magnusdottir; Louise le Roux; Gudrun Sigmundsdottir; Gardar Sveinbjornsson; Kristin E Sveinsdottir; Maney Sveinsdottir; Emil A Thorarensen; Bjarni Thorbjornsson; Gisli Masson; Ingileif Jonsdottir; Alma Moller; Thorolfur Gudnason; Karl G Kristinsson; Unnur Thorsteinsdottir; Kari Stefansson |
| EPI_ISL_417866 | hCoV-19/Iceland/81/2020 | Europe / Iceland / Reykjavik | 2020-03-09 | The National University Hospital of Iceland | deCODE genetics | Daniel F Gudbjartsson; Agnar Helgason; Hakon Jonsson; Olafur T Magnusson; Pall Melsted; Gudmundur L Norddahl; Jona Saemundsdottir; Asgeir Sigurdsson; Patrick Sulem; Arna B Agustsdottir; Berglind Eiriksdottir; Run Fridriksdottir; Elisabet E Gardarsdottir; Gudmundur Georgsson; Olafia S Gretarsdottir; Kjartan R Gudmundsson; Thora R Gunnarsdottir; Arnaldur Gylfason; Hilma Holm; Brynjar O Jensson; Aslaug Jonasdottir; Kamilla S Josefsdottir; Thordur Kristjansson; Droplaug N Magnusdottir; Louise le Roux; Gudrun Sigmundsdottir; Gardar Sveinbjornsson; Kristin E Sveinsdottir; Maney Sveinsdottir; Emil A Thorarensen; Bjarni Thorbjornsson; Gisli Masson; Ingileif Jonsdottir; Alma Moller; Thorolfur Gudnason; Karl G Kristinsson; Unnur Thorsteinsdottir; Kari Stefansson |
| EPI_ISL_417867 | hCoV-19/Iceland/82/2020 | Europe / Iceland / Reykjavik | 2020-03-09 | The National University Hospital of Iceland | deCODE genetics | Daniel F Gudbjartsson; Agnar Helgason; Hakon Jonsson; Olafur T Magnusson; Pall Melsted; Gudmundur L Norddahl; Jona Saemundsdottir; Asgeir Sigurdsson; Patrick Sulem; Arna B Agustsdottir; Berglind Eiriksdottir; Run Fridriksdottir; Elisabet E Gardarsdottir; Gudmundur Georgsson; Olafia S Gretarsdottir; Kjartan R Gudmundsson; Thora R Gunnarsdottir; Arnaldur Gylfason; Hilma Holm; Brynjar O Jensson; Aslaug Jonasdottir; Kamilla S Josefsdottir; Thordur Kristjansson; Droplaug N Magnusdottir; Louise le Roux; Gudrun Sigmundsdottir; Gardar Sveinbjornsson; Kristin E Sveinsdottir; Maney Sveinsdottir; Emil A Thorarensen; Bjarni Thorbjornsson; Gisli Masson; Ingileif Jonsdottir; Alma Moller; Thorolfur Gudnason; Karl G Kristinsson; Unnur Thorsteinsdottir; Kari Stefansson |
| EPI_ISL_417868 | hCoV-19/Iceland/83/2020 | Europe / Iceland / Reykjavik | 2020-03-09 | The National University Hospital of Iceland | deCODE genetics | Daniel F Gudbjartsson; Agnar Helgason; Hakon Jonsson; Olafur T Magnusson; Pall Melsted; Gudmundur L Norddahl; Jona Saemundsdottir; Asgeir Sigurdsson; Patrick Sulem; Arna B Agustsdottir; Berglind Eiriksdottir; Run Fridriksdottir; Elisabet E Gardarsdottir; Gudmundur Georgsson; Olafia S Gretarsdottir; Kjartan R Gudmundsson; Thora R Gunnarsdottir; Arnaldur Gylfason; Hilma Holm; Brynjar O Jensson; Aslaug Jonasdottir; Kamilla S Josefsdottir; Thordur Kristjansson; Droplaug N Magnusdottir; Louise le Roux; Gudrun Sigmundsdottir; Gardar Sveinbjornsson; Kristin E Sveinsdottir; Maney Sveinsdottir; Emil A Thorarensen; Bjarni Thorbjornsson; Gisli Masson; Ingileif Jonsdottir; Alma Moller; Thorolfur Gudnason; Karl G Kristinsson; Unnur Thorsteinsdottir; Kari Stefansson |
| EPI_ISL_417869 | hCoV-19/Iceland/85/2020 | Europe / Iceland / Reykjavik | 2020-03-10 | The National University Hospital of Iceland | deCODE genetics | Daniel F Gudbjartsson; Agnar Helgason; Hakon Jonsson; Olafur T Magnusson; Pall Melsted; Gudmundur L Norddahl; Jona Saemundsdottir; Asgeir Sigurdsson; Patrick Sulem; Arna B Agustsdottir; Berglind Eiriksdottir; Run Fridriksdottir; Elisabet E Gardarsdottir; Gudmundur Georgsson; Olafia S Gretarsdottir; Kjartan R Gudmundsson; Thora R Gunnarsdottir; Arnaldur Gylfason; Hilma Holm; Brynjar O Jensson; Aslaug Jonasdottir; Kamilla S Josefsdottir; Thordur Kristjansson; Droplaug N Magnusdottir; Louise le Roux; Gudrun Sigmundsdottir; Gardar Sveinbjornsson; Kristin E Sveinsdottir; Maney Sveinsdottir; Emil A Thorarensen; Bjarni Thorbjornsson; Gisli Masson; Ingileif Jonsdottir; Alma Moller; Thorolfur Gudnason; Karl G Kristinsson; Unnur Thorsteinsdottir; Kari Stefansson |
| EPI_ISL_417870 | hCoV-19/Iceland/86/2020 | Europe / Iceland / Reykjavik | 2020-03-10 | The National University Hospital of Iceland | deCODE genetics | Daniel F Gudbjartsson; Agnar Helgason; Hakon Jonsson; Olafur T Magnusson; Pall Melsted; Gudmundur L Norddahl; Jona Saemundsdottir; Asgeir Sigurdsson; Patrick Sulem; Arna B Agustsdottir; Berglind Eiriksdottir; Run Fridriksdottir; Elisabet E Gardarsdottir; Gudmundur Georgsson; Olafia S Gretarsdottir; Kjartan R Gudmundsson; Thora R Gunnarsdottir; Arnaldur Gylfason; Hilma Holm; Brynjar O Jensson; Aslaug Jonasdottir; Kamilla S Josefsdottir; Thordur Kristjansson; Droplaug N Magnusdottir; Louise le Roux; Gudrun Sigmundsdottir; Gardar Sveinbjornsson; Kristin E Sveinsdottir; Maney Sveinsdottir; Emil A Thorarensen; Bjarni Thorbjornsson; Gisli Masson; Ingileif Jonsdottir; Alma Moller; Thorolfur Gudnason; Karl G Kristinsson; Unnur Thorsteinsdottir; Kari Stefansson |
| EPI_ISL_417871 | hCoV-19/Iceland/9/2020 | Europe / Iceland / Reykjavik | 2020-03-11 | The National University Hospital of Iceland | deCODE genetics | Daniel F Gudbjartsson; Agnar Helgason; Hakon Jonsson; Olafur T Magnusson; Pall Melsted; Gudmundur L Norddahl; Jona Saemundsdottir; Asgeir Sigurdsson; Patrick Sulem; Arna B Agustsdottir; Berglind Eiriksdottir; Run Fridriksdottir; Elisabet E Gardarsdottir; Gudmundur Georgsson; Olafia S Gretarsdottir; Kjartan R Gudmundsson; Thora R Gunnarsdottir; Arnaldur Gylfason; Hilma Holm; Brynjar O Jensson; Aslaug Jonasdottir; Kamilla S Josefsdottir; Thordur Kristjansson; Droplaug N Magnusdottir; Louise le Roux; Gudrun Sigmundsdottir; Gardar Sveinbjornsson; Kristin E Sveinsdottir; Maney Sveinsdottir; Emil A Thorarensen; Bjarni Thorbjornsson; Gisli Masson; Ingileif Jonsdottir; Alma Moller; Thorolfur Gudnason; Karl G Kristinsson; Unnur Thorsteinsdottir; Kari Stefansson |
| EPI_ISL_417872 | hCoV-19/Iceland/90/2020 | Europe / Iceland / Reykjavik | 2020-03-10 | The National University Hospital of Iceland | deCODE genetics | Daniel F Gudbjartsson; Agnar Helgason; Hakon Jonsson; Olafur T Magnusson; Pall Melsted; Gudmundur L Norddahl; Jona Saemundsdottir; Asgeir Sigurdsson; Patrick Sulem; Arna B Agustsdottir; Berglind Eiriksdottir; Run Fridriksdottir; Elisabet E Gardarsdottir; Gudmundur Georgsson; Olafia S Gretarsdottir; Kjartan R Gudmundsson; Thora R Gunnarsdottir; Arnaldur Gylfason; Hilma Holm; Brynjar O Jensson; Aslaug Jonasdottir; Kamilla S Josefsdottir; Thordur Kristjansson; Droplaug N Magnusdottir; Louise le Roux; Gudrun Sigmundsdottir; Gardar Sveinbjornsson; Kristin E Sveinsdottir; Maney Sveinsdottir; Emil A Thorarensen; Bjarni Thorbjornsson; Gisli Masson; Ingileif Jonsdottir; Alma Moller; Thorolfur Gudnason; Karl G Kristinsson; Unnur Thorsteinsdottir; Kari Stefansson |
| EPI_ISL_417873 | hCoV-19/Iceland/91/2020 | Europe / Iceland / Reykjavik | 2020-03-10 | The National University Hospital of Iceland | deCODE genetics | Daniel F Gudbjartsson; Agnar Helgason; Hakon Jonsson; Olafur T Magnusson; Pall Melsted; Gudmundur L Norddahl; Jona Saemundsdottir; Asgeir Sigurdsson; Patrick Sulem; Arna B Agustsdottir; Berglind Eiriksdottir; Run Fridriksdottir; Elisabet E Gardarsdottir; Gudmundur Georgsson; Olafia S Gretarsdottir; Kjartan R Gudmundsson; Thora R Gunnarsdottir; Arnaldur Gylfason; Hilma Holm; Brynjar O Jensson; Aslaug Jonasdottir; Kamilla S Josefsdottir; Thordur Kristjansson; Droplaug N Magnusdottir; Louise le Roux; Gudrun Sigmundsdottir; Gardar Sveinbjornsson; Kristin E Sveinsdottir; Maney Sveinsdottir; Emil A Thorarensen; Bjarni Thorbjornsson; Gisli Masson; Ingileif Jonsdottir; Alma Moller; Thorolfur Gudnason; Karl G Kristinsson; Unnur Thorsteinsdottir; Kari Stefansson |
| EPI_ISL_417874 | hCoV-19/Iceland/95/2020 | Europe / Iceland / Reykjavik | 2020-03-10 | The National University Hospital of Iceland | deCODE genetics | Daniel F Gudbjartsson; Agnar Helgason; Hakon Jonsson; Olafur T Magnusson; Pall Melsted; Gudmundur L Norddahl; Jona Saemundsdottir; Asgeir Sigurdsson; Patrick Sulem; Arna B Agustsdottir; Berglind Eiriksdottir; Run Fridriksdottir; Elisabet E Gardarsdottir; Gudmundur Georgsson; Olafia S Gretarsdottir; Kjartan R Gudmundsson; Thora R Gunnarsdottir; Arnaldur Gylfason; Hilma Holm; Brynjar O Jensson; Aslaug Jonasdottir; Kamilla S Josefsdottir; Thordur Kristjansson; Droplaug N Magnusdottir; Louise le Roux; Gudrun Sigmundsdottir; Gardar Sveinbjornsson; Kristin E Sveinsdottir; Maney Sveinsdottir; Emil A Thorarensen; Bjarni Thorbjornsson; Gisli Masson; Ingileif Jonsdottir; Alma Moller; Thorolfur Gudnason; Karl G Kristinsson; Unnur Thorsteinsdottir; Kari Stefansson |
| EPI_ISL_417875 | hCoV-19/Iceland/96/2020 | Europe / Iceland / Reykjavik | 2020-03-10 | The National University Hospital of Iceland | deCODE genetics | Daniel F Gudbjartsson; Agnar Helgason; Hakon Jonsson; Olafur T Magnusson; Pall Melsted; Gudmundur L Norddahl; Jona Saemundsdottir; Asgeir Sigurdsson; Patrick Sulem; Arna B Agustsdottir; Berglind Eiriksdottir; Run Fridriksdottir; Elisabet E Gardarsdottir; Gudmundur Georgsson; Olafia S Gretarsdottir; Kjartan R Gudmundsson; Thora R Gunnarsdottir; Arnaldur Gylfason; Hilma Holm; Brynjar O Jensson; Aslaug Jonasdottir; Kamilla S Josefsdottir; Thordur Kristjansson; Droplaug N Magnusdottir; Louise le Roux; Gudrun Sigmundsdottir; Gardar Sveinbjornsson; Kristin E Sveinsdottir; Maney Sveinsdottir; Emil A Thorarensen; Bjarni Thorbjornsson; Gisli Masson; Ingileif Jonsdottir; Alma Moller; Thorolfur Gudnason; Karl G Kristinsson; Unnur Thorsteinsdottir; Kari Stefansson |
| EPI_ISL_417876 | hCoV-19/Iceland/97/2020 | Europe / Iceland / Reykjavik | 2020-03-10 | The National University Hospital of Iceland | deCODE genetics | Daniel F Gudbjartsson; Agnar Helgason; Hakon Jonsson; Olafur T Magnusson; Pall Melsted; Gudmundur L Norddahl; Jona Saemundsdottir; Asgeir Sigurdsson; Patrick Sulem; Arna B Agustsdottir; Berglind Eiriksdottir; Run Fridriksdottir; Elisabet E Gardarsdottir; Gudmundur Georgsson; Olafia S Gretarsdottir; Kjartan R Gudmundsson; Thora R Gunnarsdottir; Arnaldur Gylfason; Hilma Holm; Brynjar O Jensson; Aslaug Jonasdottir; Kamilla S Josefsdottir; Thordur Kristjansson; Droplaug N Magnusdottir; Louise le Roux; Gudrun Sigmundsdottir; Gardar Sveinbjornsson; Kristin E Sveinsdottir; Maney Sveinsdottir; Emil A Thorarensen; Bjarni Thorbjornsson; Gisli Masson; Ingileif Jonsdottir; Alma Moller; Thorolfur Gudnason; Karl G Kristinsson; Unnur Thorsteinsdottir; Kari Stefansson |
| EPI_ISL_417877 | hCoV-19/Slovakia/SK-BMC1/2020 | Europe / Slovakia / Kostoliste | 2020-03-06 | Institute of Virology, Biomedical Research Center of the Slovak Academy of Sciences, Bratislava; Public Health Authority of the Slovak Republic, Bratislava | Institute of Virology, Biomedical Research Center of the Slovak Academy of Sciences, Bratislava; Comenius University Science Park, Bratislava | Monika Slávikova, Martina Ličková, Sabina Fumačová Havlíková, Juraj Koči, Juraj Kopáček, Elena Tichá, Edita Staroňová, Jaroslav Budiš, Werner Krampl, Miroslav Böhmer, Diana Rusňáková, Tomáš Szemeš, Boris Klempa |
| EPI_ISL_417878 | hCoV-19/Slovakia/SK-BMC2/2020 | Europe / Slovakia / Bratislava | 2020-03-07 | Institute of Virology, Biomedical Research Center of the Slovak Academy of Sciences, Bratislava; Public Health Authority of the Slovak Republic, Bratislava | Institute of Virology, Biomedical Research Center of the Slovak Academy of Sciences, Bratislava; Comenius University Science Park, Bratislava | Monika Slávikova, Martina Ličková, Sabina Fumačová Havlíková, Juraj Koči, Juraj Kopáček, Elena Tichá, Edita Staroňová, Jaroslav Budiš, Werner Krampl, Miroslav Böhmer, Diana Rusňáková, Tomáš Szemeš, Boris Klempa |
| EPI_ISL_417879 | hCoV-19/Slovakia/SK-BMC5/2020 | Europe / Slovakia / Kosice | 2020-03-06 | Institute of Virology, Biomedical Research Center of the Slovak Academy of Sciences, Bratislava; Public Health Authority of the Slovak Republic, Bratislava | Institute of Virology, Biomedical Research Center of the Slovak Academy of Sciences, Bratislava; Comenius University Science Park, Bratislava | Monika Slávikova, Martina Ličková, Sabina Fumačová Havlíková, Juraj Koči, Juraj Kopáček, Elena Tichá, Edita Staroňová, Jaroslav Budiš, Werner Krampl, Miroslav Böhmer, Diana Rusňáková, Tomáš Szemeš, Boris Klempa |
| EPI_ISL_417880 | hCoV-19/Slovakia/SK-BMC6/2020 | Europe / Slovakia / Martin | 2020-03-08 | Institute of Virology, Biomedical Research Center of the Slovak Academy of Sciences, Bratislava; Public Health Authority of the Slovak Republic, Bratislava | Institute of Virology, Biomedical Research Center of the Slovak Academy of Sciences, Bratislava; Comenius University Science Park, Bratislava | Monika Slávikova, Martina Ličková, Sabina Fumačová Havlíková, Juraj Koči, Juraj Kopáček, Elena Tichá, Edita Staroňová, Jaroslav Budiš, Werner Krampl, Miroslav Böhmer, Diana Rusňáková, Tomáš Szemeš, Boris Klempa |
| EPI_ISL_418300 | hCoV-19/England/SHEF-8FE51/2020 | Europe / United Kingdom / England / Sou | 2020-03-15 | Virology Department, Sheffield Teaching Hospitals NHS Foundation Trust | Department of Infection, Immunity and Cardiovascular Disease, The Florey Institute, The Medical School, University of Sheffield | Thushan de Silva, Matthew Parker, Adri Angyal, Rebecca Brown, Rachel Tucker, Paul Parsons, Danielle Groves, Alex Keeley, Dave Partridge, Matthew Wyles, Benjamin Lindsey, Mehmet Yavuz, Mohammad Raza, Cariad Evans |
| EPI_ISL_418301 | hCoV-19/England/SHEF-8FE60/2020 | Europe / United Kingdom / England / Sou | 2020-03-05 | Virology Department, Sheffield Teaching Hospitals NHS Foundation Trust | Department of Infection, Immunity and Cardiovascular Disease, The Florey Institute, The Medical School, University of Sheffield | Thushan de Silva, Matthew Parker, Adri Angyal, Rebecca Brown, Rachel Tucker, Paul Parsons, Danielle Groves, Alex Keeley, Dave Partridge, Matthew Wyles, Benjamin Lindsey, Mehmet Yavuz, Mohammad Raza, Cariad Evans |
| EPI_ISL_418302 | hCoV-19/England/SHEF-8FE9D/2020 | Europe / United Kingdom / England / Sou | 2020-03-24 | Virology Department, Sheffield Teaching Hospitals NHS Foundation Trust | Department of Infection, Immunity and Cardiovascular Disease, The Florey Institute, The Medical School, University of Sheffield | Thushan de Silva, Matthew Parker, Adri Angyal, Rebecca Brown, Rachel Tucker, Paul Parsons, Danielle Groves, Alex Keeley, Dave Partridge, Matthew Wyles, Benjamin Lindsey, Mehmet Yavuz, Mohammad Raza, Cariad Evans |
| EPI_ISL_418303 | hCoV-19/England/SHEF-8FEAC/2020 | Europe / United Kingdom / England / Der | 2020-03-25 | Virology Department, Sheffield Teaching Hospitals NHS Foundation Trust | Department of Infection, Immunity and Cardiovascular Disease, The Florey Institute, The Medical School, University of Sheffield | Thushan de Silva, Matthew Parker, Adri Angyal, Rebecca Brown, Rachel Tucker, Paul Parsons, Danielle Groves, Alex Keeley, Dave Partridge, Matthew Wyles, Benjamin Lindsey, Mehmet Yavuz, Mohammad Raza, Cariad Evans |
| EPI_ISL_418304 | hCoV-19/England/SHEF-8FEB8/2020 | Europe / United Kingdom / England / Der | 2020-03-24 | Virology Department, Sheffield Teaching Hospitals NHS Foundation Trust | Department of Infection, Immunity and Cardiovascular Disease, The Florey Institute, The Medical School, University of Sheffield | Thushan de Silva, Matthew Parker, Adri Angyal, Rebecca Brown, Rachel Tucker, Paul Parsons, Danielle Groves, Alex Keeley, Dave Partridge, Matthew Wyles, Benjamin Lindsey, Mehmet Yavuz, Mohammad Raza, Cariad Evans |
| EPI_ISL_418305 | hCoV-19/England/SHEF-8FED9/2020 | Europe / United Kingdom / England / Sou | 2020-03-25 | Virology Department, Sheffield Teaching Hospitals NHS Foundation Trust | Department of Infection, Immunity and Cardiovascular Disease, The Florey Institute, The Medical School, University of Sheffield | Thushan de Silva, Matthew Parker, Adri Angyal, Rebecca Brown, Rachel Tucker, Paul Parsons, Danielle Groves, Alex Keeley, Dave Partridge, Matthew Wyles, Benjamin Lindsey, Mehmet Yavuz, Mohammad Raza, Cariad Evans |
| EPI_ISL_418306 | hCoV-19/England/SHEF-8FEE8/2020 | Europe / United Kingdom / England / Sou | 2020-03-25 | Virology Department, Sheffield Teaching Hospitals NHS Foundation Trust | Department of Infection, Immunity and Cardiovascular Disease, The Florey Institute, The Medical School, University of Sheffield | Thushan de Silva, Matthew Parker, Adri Angyal, Rebecca Brown, Rachel Tucker, Paul Parsons, Danielle Groves, Alex Keeley, Dave Partridge, Matthew Wyles, Benjamin Lindsey, Mehmet Yavuz, Mohammad Raza, Cariad Evans |
| EPI_ISL_418307 | hCoV-19/England/SHEF-8FF03/2020 | Europe / United Kingdom / England / Der | 2020-03-24 | Virology Department, Sheffield Teaching Hospitals NHS Foundation Trust | Department of Infection, Immunity and Cardiovascular Disease, The Florey Institute, The Medical School, University of Sheffield | Thushan de Silva, Matthew Parker, Adri Angyal, Rebecca Brown, Rachel Tucker, Paul Parsons, Danielle Groves, Alex Keeley, Dave Partridge, Matthew Wyles, Benjamin Lindsey, Mehmet Yavuz, Mohammad Raza, Cariad Evans |
| EPI_ISL_418308 | hCoV-19/England/SHEF-8FF12/2020 | Europe / United Kingdom / England / Sou | 2020-03-25 | Virology Department, Sheffield Teaching Hospitals NHS Foundation Trust | Department of Infection, Immunity and Cardiovascular Disease, The Florey Institute, The Medical School, University of Sheffield | Thushan de Silva, Matthew Parker, Adri Angyal, Rebecca Brown, Rachel Tucker, Paul Parsons, Danielle Groves, Alex Keeley, Dave Partridge, Matthew Wyles, Benjamin Lindsey, Mehmet Yavuz, Mohammad Raza, Cariad Evans |
| EPI_ISL_418309 | hCoV-19/England/SHEF-8FF21/2020 | Europe / United Kingdom / England / Der | 2020-03-25 | Virology Department, Sheffield Teaching Hospitals NHS Foundation Trust | Department of Infection, Immunity and Cardiovascular Disease, The Florey Institute, The Medical School, University of Sheffield | Thushan de Silva, Matthew Parker, Adri Angyal, Rebecca Brown, Rachel Tucker, Paul Parsons, Danielle Groves, Alex Keeley, Dave Partridge, Matthew Wyles, Benjamin Lindsey, Mehmet Yavuz, Mohammad Raza, Cariad Evans |
| EPI_ISL_418310 | hCoV-19/England/SHEF-8FF30/2020 | Europe / United Kingdom / England / Sou | 2020-03-25 | Virology Department, Sheffield Teaching Hospitals NHS Foundation Trust | Department of Infection, Immunity and Cardiovascular Disease, The Florey Institute, The Medical School, University of Sheffield | Thushan de Silva, Matthew Parker, Adri Angyal, Rebecca Brown, Rachel Tucker, Paul Parsons, Danielle Groves, Alex Keeley, Dave Partridge, Matthew Wyles, Benjamin Lindsey, Mehmet Yavuz, Mohammad Raza, Cariad Evans |
| EPI_ISL_418311 | hCoV-19/England/SHEF-8FF4F/2020 | Europe / United Kingdom / England / Der | 2020-03-25 | Virology Department, Sheffield Teaching Hospitals NHS Foundation Trust | Department of Infection, Immunity and Cardiovascular Disease, The Florey Institute, The Medical School, University of Sheffield | Thushan de Silva, Matthew Parker, Adri Angyal, Rebecca Brown, Rachel Tucker, Paul Parsons, Danielle Groves, Alex Keeley, Dave Partridge, Matthew Wyles, Benjamin Lindsey, Mehmet Yavuz, Mohammad Raza, Cariad Evans |
| EPI_ISL_418312 | hCoV-19/England/SHEF-8FF5E/2020 | Europe / United Kingdom / England / Sou | 2020-03-25 | Virology Department, Sheffield Teaching Hospitals NHS Foundation Trust | Department of Infection, Immunity and Cardiovascular Disease, The Florey Institute, The Medical School, University of Sheffield | Thushan de Silva, Matthew Parker, Adri Angyal, Rebecca Brown, Rachel Tucker, Paul Parsons, Danielle Groves, Alex Keeley, Dave Partridge, Matthew Wyles, Benjamin Lindsey, Mehmet Yavuz, Mohammad Raza, Cariad Evans |
| EPI_ISL_418313 | hCoV-19/England/SHEF-8FF6D/2020 | Europe / United Kingdom / England / Der | 2020-03-24 | Virology Department, Sheffield Teaching Hospitals NHS Foundation Trust | Department of Infection, Immunity and Cardiovascular Disease, The Florey Institute, The Medical School, University of Sheffield | Thushan de Silva, Matthew Parker, Adri Angyal, Rebecca Brown, Rachel Tucker, Paul Parsons, Danielle Groves, Alex Keeley, Dave Partridge, Matthew Wyles, Benjamin Lindsey, Mehmet Yavuz, Mohammad Raza, Cariad Evans |
| EPI_ISL_418314 | hCoV-19/England/SHEF-8FF7C/2020 | Europe / United Kingdom / England / Sou | 2020-03-25 | Virology Department, Sheffield Teaching Hospitals NHS Foundation Trust | Department of Infection, Immunity and Cardiovascular Disease, The Florey Institute, The Medical School, University of Sheffield | Thushan de Silva, Matthew Parker, Adri Angyal, Rebecca Brown, Rachel Tucker, Paul Parsons, Danielle Groves, Alex Keeley, Dave Partridge, Matthew Wyles, Benjamin Lindsey, Mehmet Yavuz, Mohammad Raza, Cariad Evans |
| EPI_ISL_418315 | hCoV-19/England/SHEF-8FF8B/2020 | Europe / United Kingdom / England / Der | 2020-03-25 | Virology Department, Sheffield Teaching Hospitals NHS Foundation Trust | Department of Infection, Immunity and Cardiovascular Disease, The Florey Institute, The Medical School, University of Sheffield | Thushan de Silva, Matthew Parker, Adri Angyal, Rebecca Brown, Rachel Tucker, Paul Parsons, Danielle Groves, Alex Keeley, Dave Partridge, Matthew Wyles, Benjamin Lindsey, Mehmet Yavuz, Mohammad Raza, Cariad Evans |
| EPI_ISL_418316 | hCoV-19/England/SHEF-8FF9A/2020 | Europe / United Kingdom / England / Sou | 2020-03-25 | Virology Department, Sheffield Teaching Hospitals NHS Foundation Trust | Department of Infection, Immunity and Cardiovascular Disease, The Florey Institute, The Medical School, University of Sheffield | Thushan de Silva, Matthew Parker, Adri Angyal, Rebecca Brown, Rachel Tucker, Paul Parsons, Danielle Groves, Alex Keeley, Dave Partridge, Matthew Wyles, Benjamin Lindsey, Mehmet Yavuz, Mohammad Raza, Cariad Evans |
| EPI_ISL_418317 | hCoV-19/England/SHEF-8FFA9/2020 | Europe / United Kingdom / England / Sou | 2020-03-23 | Virology Department, Sheffield Teaching Hospitals NHS Foundation Trust | Department of Infection, Immunity and Cardiovascular Disease, The Florey Institute, The Medical School, University of Sheffield | Thushan de Silva, Matthew Parker, Adri Angyal, Rebecca Brown, Rachel Tucker, Paul Parsons, Danielle Groves, Alex Keeley, Dave Partridge, Matthew Wyles, Benjamin Lindsey, Mehmet Yavuz, Mohammad Raza, Cariad Evans |
| EPI_ISL_418318 | hCoV-19/England/SHEF-8FFB8/2020 | Europe / United Kingdom / England / Sou | 2020-03-25 | Virology Department, Sheffield Teaching Hospitals NHS Foundation Trust | Department of Infection, Immunity and Cardiovascular Disease, The Florey Institute, The Medical School, University of Sheffield | Thushan de Silva, Matthew Parker, Adri Angyal, Rebecca Brown, Rachel Tucker, Paul Parsons, Danielle Groves, Alex Keeley, Dave Partridge, Matthew Wyles, Benjamin Lindsey, Mehmet Yavuz, Mohammad Raza, Cariad Evans |
| EPI_ISL_418319 | hCoV-19/England/SHEF-8FFC7/2020 | Europe / United Kingdom / England / Der | 2020-03-25 | Virology Department, Sheffield Teaching Hospitals NHS Foundation Trust | Department of Infection, Immunity and Cardiovascular Disease, The Florey Institute, The Medical School, University of Sheffield | Thushan de Silva, Matthew Parker, Adri Angyal, Rebecca Brown, Rachel Tucker, Paul Parsons, Danielle Groves, Alex Keeley, Dave Partridge, Matthew Wyles, Benjamin Lindsey, Mehmet Yavuz, Mohammad Raza, Cariad Evans |
| EPI_ISL_418320 | hCoV-19/England/SHEF-8FFD6/2020 | Europe / United Kingdom / England / Sou | 2020-03-25 | Virology Department, Sheffield Teaching Hospitals NHS Foundation Trust | Department of Infection, Immunity and Cardiovascular Disease, The Florey Institute, The Medical School, University of Sheffield | Thushan de Silva, Matthew Parker, Adri Angyal, Rebecca Brown, Rachel Tucker, Paul Parsons, Danielle Groves, Alex Keeley, Dave Partridge, Matthew Wyles, Benjamin Lindsey, Mehmet Yavuz, Mohammad Raza, Cariad Evans |
| EPI_ISL_418321 | hCoV-19/England/SHEF-8FFE5/2020 | Europe / United Kingdom / England / Sou | 2020-03-25 | Virology Department, Sheffield Teaching Hospitals NHS Foundation Trust | Department of Infection, Immunity and Cardiovascular Disease, The Florey Institute, The Medical School, University of Sheffield | Thushan de Silva, Matthew Parker, Adri Angyal, Rebecca Brown, Rachel Tucker, Paul Parsons, Danielle Groves, Alex Keeley, Dave Partridge, Matthew Wyles, Benjamin Lindsey, Mehmet Yavuz, Mohammad Raza, Cariad Evans |
| EPI_ISL_418322 | hCoV-19/Canada/ON_PHL3917/2020 | North America / Canada / Ontario | 2020-03-08 | Public Health Ontario Laboratories | Public Health Ontario Laboratories | Alireza Eshaghi, Samir N Patel, Jonathan B Gubbay, Vanessa G Allen, Christine Frantz, Aimin Li, Sandeep Nagra |
| EPI_ISL_418323 | hCoV-19/Canada/ON_PHL3919/2020 | North America / Canada / Ontario | 2020-03-14 | Public Health Ontario Laboratories | Public Health Ontario Laboratories | Alireza Eshaghi, Samir N Patel, Jonathan B Gubbay, Vanessa G Allen, Christine Frantz, Aimin Li, Sandeep Nagra |
| EPI_ISL_418324 | hCoV-19/Canada/ON_PHL3318/2020 | North America / Canada / Ontario | 2020-03-11 | Public Health Ontario Laboratories | Public Health Ontario Laboratories | Alireza Eshaghi, Samir N Patel, Jonathan B Gubbay, Vanessa G Allen, Christine Frantz, Aimin Li, Sandeep Nagra |
| EPI_ISL_418325 | hCoV-19/Canada/ON_PHL5472/2020 | North America / Canada / Ontario | 2020-02-20 | Public Health Ontario Laboratories | Public Health Ontario Laboratories | Alireza Eshaghi, Samir N Patel, Jonathan B Gubbay, Vanessa G Allen, Christine Frantz, Aimin Li, Sandeep Nagra |
| EPI_ISL_418326 | hCoV-19/Canada/ON_PHL7972/2020 | North America / Canada / Ontario | 2020-03-12 | Public Health Ontario Laboratories | Public Health Ontario Laboratories | Alireza Eshaghi, Samir N Patel, Jonathan B Gubbay, Vanessa G Allen, Christine Frantz, Aimin Li, Sandeep Nagra |
| EPI_ISL_418327 | hCoV-19/Canada/ON_PHL4181/2020 | North America / Canada / Ontario | 2020-01-25 | Public Health Ontario Laboratories | Public Health Ontario Laboratories | Alireza Eshaghi, Samir N Patel, Jonathan B Gubbay, Vanessa G Allen, Christine Frantz, Aimin Li, Sandeep Nagra |
| EPI_ISL_418328 | hCoV-19/Canada/ON_PHL3670/2020 | North America / Canada / Ontario | 2020-03-20 | Public Health Ontario Laboratories | Public Health Ontario Laboratories | Alireza Eshaghi, Samir N Patel, Jonathan B Gubbay, Vanessa G Allen, Christine Frantz, Aimin Li, Sandeep Nagra |
| EPI_ISL_418329 | hCoV-19/Canada/ON_PHL5694/2020 | North America / Canada / Ontario | 2020-03-12 | Public Health Ontario Laboratories | Public Health Ontario Laboratories | Alireza Eshaghi, Samir N Patel, Jonathan B Gubbay, Vanessa G Allen, Christine Frantz, Aimin Li, Sandeep Nagra |
| EPI_ISL_418330 | hCoV-19/Canada/ON_PHL8580/2020 | North America / Canada / Ontario | 2020-03-05 | Public Health Ontario Laboratories | Public Health Ontario Laboratories | Alireza Eshaghi, Samir N Patel, Jonathan B Gubbay, Vanessa G Allen, Christine Frantz, Aimin Li, Sandeep Nagra |
| EPI_ISL_418331 | hCoV-19/Canada/ON_PHL6883/2020 | North America / Canada / Ontario | 2020-03-08 | Public Health Ontario Laboratories | Public Health Ontario Laboratories | Alireza Eshaghi, Samir N Patel, Jonathan B Gubbay, Vanessa G Allen, Christine Frantz, Aimin Li, Sandeep Nagra |
| EPI_ISL_418332 | hCoV-19/Canada/ON_PHL0052/2020 | North America / Canada / Ontario | 2020-03-11 | Public Health Ontario Laboratories | Public Health Ontario Laboratories | Alireza Eshaghi, Samir N Patel, Jonathan B Gubbay, Vanessa G Allen, Christine Frantz, Aimin Li, Sandeep Nagra |
| EPI_ISL_418333 | hCoV-19/Canada/ON_PHL3501/2020 | North America / Canada / Ontario | 2020-03-10 | Public Health Ontario Laboratories | Public Health Ontario Laboratories | Alireza Eshaghi, Samir N Patel, Jonathan B Gubbay, Vanessa G Allen, Christine Frantz, Aimin Li, Sandeep Nagra |
| EPI_ISL_418334 | hCoV-19/Canada/ON_PHL4069/2020 | North America / Canada / Ontario | 2020-03-07 | Public Health Ontario Laboratories | Public Health Ontario Laboratories | Alireza Eshaghi, Samir N Patel, Jonathan B Gubbay, Vanessa G Allen, Christine Frantz, Aimin Li, Sandeep Nagra |
| EPI_ISL_418335 | hCoV-19/Canada/ON_PHL4464/2020 | North America / Canada / Ontario | 2020-03-14 | Public Health Ontario Laboratories | Public Health Ontario Laboratories | Alireza Eshaghi, Samir N Patel, Jonathan B Gubbay, Vanessa G Allen, Christine Frantz, Aimin Li, Sandeep Nagra |
| EPI_ISL_418336 | hCoV-19/Canada/ON_PHL5705/2020 | North America / Canada / Ontario | 2020-03-12 | Public Health Ontario Laboratories | Public Health Ontario Laboratories | Alireza Eshaghi, Samir N Patel, Jonathan B Gubbay, Vanessa G Allen, Christine Frantz, Aimin Li, Sandeep Nagra |
| EPI_ISL_418337 | hCoV-19/Canada/ON_PHL3802/2020 | North America / Canada / Ontario | 2020-03-12 | Public Health Ontario Laboratories | Public Health Ontario Laboratories | Alireza Eshaghi, Samir N Patel, Jonathan B Gubbay, Vanessa G Allen, Christine Frantz, Aimin Li, Sandeep Nagra |
| EPI_ISL_418338 | hCoV-19/Canada/ON_PHL5710/2020 | North America / Canada / Ontario | 2020-03-12 | Public Health Ontario Laboratories | Public Health Ontario Laboratories | Alireza Eshaghi, Samir N Patel, Jonathan B Gubbay, Vanessa G Allen, Christine Frantz, Aimin Li, Sandeep Nagra |
| EPI_ISL_418339 | hCoV-19/Canada/ON_PHL3877/2020 | North America / Canada / Ontario | 2020-03-09 | Public Health Ontario Laboratories | Public Health Ontario Laboratories | Alireza Eshaghi, Samir N Patel, Jonathan B Gubbay, Vanessa G Allen, Christine Frantz, Aimin Li, Sandeep Nagra |
| EPI_ISL_418340 | hCoV-19/Canada/ON_PHL1083/2020 | North America / Canada / Ontario | 2020-03-09 | Public Health Ontario Laboratories | Public Health Ontario Laboratories | Alireza Eshaghi, Samir N Patel, Jonathan B Gubbay, Vanessa G Allen, Christine Frantz, Aimin Li, Sandeep Nagra |
| EPI_ISL_418341 | hCoV-19/Canada/ON_PHL0142/2020 | North America / Canada / Ontario | 2020-03-12 | Public Health Ontario Laboratories | Public Health Ontario Laboratories | Alireza Eshaghi, Samir N Patel, Jonathan B Gubbay, Vanessa G Allen, Christine Frantz, Aimin Li, Sandeep Nagra |
| EPI_ISL_418342 | hCoV-19/Canada/ON_PHL0178/2020 | North America / Canada / Ontario | 2020-03-11 | Public Health Ontario Laboratories | Public Health Ontario Laboratories | Alireza Eshaghi, Samir N Patel, Jonathan B Gubbay, Vanessa G Allen, Christine Frantz, Aimin Li, Sandeep Nagra |
| EPI_ISL_418343 | hCoV-19/Canada/ON_PHL6884/2020 | North America / Canada / Ontario | 2020-03-10 | Public Health Ontario Laboratories | Public Health Ontario Laboratories | Alireza Eshaghi, Samir N Patel, Jonathan B Gubbay, Vanessa G Allen, Christine Frantz, Aimin Li, Sandeep Nagra |
| EPI_ISL_418344 | hCoV-19/Canada/ON_PHL2259/2020 | North America / Canada / Ontario | 2020-02-29 | Public Health Ontario Laboratories | Public Health Ontario Laboratories | Alireza Eshaghi, Samir N Patel, Jonathan B Gubbay, Vanessa G Allen, Christine Frantz, Aimin Li, Sandeep Nagra |
| EPI_ISL_418345 | hCoV-19/Canada/ON_PHL8751/2020 | North America / Canada / Ontario | 2020-02-29 | Public Health Ontario Laboratories | Public Health Ontario Laboratories | Alireza Eshaghi, Samir N Patel, Jonathan B Gubbay, Vanessa G Allen, Christine Frantz, Aimin Li, Sandeep Nagra |
| EPI_ISL_418346 | hCoV-19/Canada/ON_PHL0743/2020 | North America / Canada / Ontario | 2020-03-07 | Public Health Ontario Laboratories | Public Health Ontario Laboratories | Alireza Eshaghi, Samir N Patel, Jonathan B Gubbay, Vanessa G Allen, Christine Frantz, Aimin Li, Sandeep Nagra |
| EPI_ISL_418347 | hCoV-19/Canada/ON_PHL3741/2020 | North America / Canada / Ontario | 2020-03-11 | Public Health Ontario Laboratories | Public Health Ontario Laboratories | Alireza Eshaghi, Samir N Patel, Jonathan B Gubbay, Vanessa G Allen, Christine Frantz, Aimin Li, Sandeep Nagra |
| EPI_ISL_418348 | hCoV-19/Canada/ON_PHL3680/2020 | North America / Canada / Ontario | 2020-03-08 | Public Health Ontario Laboratories | Public Health Ontario Laboratories | Alireza Eshaghi, Samir N Patel, Jonathan B Gubbay, Vanessa G Allen, Christine Frantz, Aimin Li, Sandeep Nagra |
| EPI_ISL_418349 | hCoV-19/Canada/ON_PHL3650/2020 | North America / Canada / Ontario | 2020-03-07 | Public Health Ontario Laboratories | Public Health Ontario Laboratories | Alireza Eshaghi, Samir N Patel, Jonathan B Gubbay, Vanessa G Allen, Christine Frantz, Aimin Li, Sandeep Nagra |
| EPI_ISL_418357 | hCoV-19/Canada/ON_PHLH6415/2020 | North America / Canada / Ontario | 2020-03-12 | Public Health Ontario Laboratories | Public Health Ontario Laboratories | Alireza Eshaghi, Samir N Patel, Jonathan B Gubbay, Vanessa G Allen, Christine Frantz, Aimin Li, Sandeep Nagra |
| EPI_ISL_418360 | hCoV-19/Canada/ON_PHL3692/2020 | North America / Canada / Ontario | 2020-03-11 | Public Health Ontario Laboratories | Public Health Ontario Laboratories | Alireza Eshaghi, Samir N Patel, Jonathan B Gubbay, Vanessa G Allen, Christine Frantz, Aimin Li, Sandeep Nagra |
| EPI_ISL_418361 | hCoV-19/Canada/ON_PHL1095/2020 | North America / Canada / Ontario | 2020-03-12 | Public Health Ontario Laboratories | Public Health Ontario Laboratories | Alireza Eshaghi, Samir N Patel, Jonathan B Gubbay, Vanessa G Allen, Christine Frantz, Aimin Li, Sandeep Nagra |
| EPI_ISL_418362 | hCoV-19/Canada/ON_PHL3536/2020 | North America / Canada / Ontario | 2020-03-10 | Public Health Ontario Laboratories | Public Health Ontario Laboratories | Alireza Eshaghi, Samir N Patel, Jonathan B Gubbay, Vanessa G Allen, Christine Frantz, Aimin Li, Sandeep Nagra |
| EPI_ISL_418363 | hCoV-19/Canada/ON_PHL3695/2020 | North America / Canada / Ontario | 2020-03-11 | Public Health Ontario Laboratories | Public Health Ontario Laboratories | Alireza Eshaghi, Samir N Patel, Jonathan B Gubbay, Vanessa G Allen, Christine Frantz, Aimin Li, Sandeep Nagra |
| EPI_ISL_418364 | hCoV-19/Canada/ON_PHL6922/2020 | North America / Canada / Ontario | 2020-03-13 | Public Health Ontario Laboratories | Public Health Ontario Laboratories | Alireza Eshaghi, Samir N Patel, Jonathan B Gubbay, Vanessa G Allen, Christine Frantz, Aimin Li, Sandeep Nagra |
| EPI_ISL_418365 | hCoV-19/Canada/ON_PHL0977/2020 | North America / Canada / Ontario | 2020-03-10 | Public Health Ontario Laboratories | Public Health Ontario Laboratories | Alireza Eshaghi, Samir N Patel, Jonathan B Gubbay, Vanessa G Allen, Christine Frantz, Aimin Li, Sandeep Nagra |
| EPI_ISL_418366 | hCoV-19/Canada/ON_PHL3350/2020 | North America / Canada / Ontario | 2020-03-05 | Public Health Ontario Laboratories | Public Health Ontario Laboratories | Alireza Eshaghi, Samir N Patel, Jonathan B Gubbay, Vanessa G Allen, Christine Frantz, Aimin Li, Sandeep Nagra |
| EPI_ISL_418367 | hCoV-19/Canada/ON_PHL7513/2020 | North America / Canada / Ontario | 2020-03-12 | Public Health Ontario Laboratories | Public Health Ontario Laboratories | Alireza Eshaghi, Samir N Patel, Jonathan B Gubbay, Vanessa G Allen, Christine Frantz, Aimin Li, Sandeep Nagra |
| EPI_ISL_418368 | hCoV-19/Canada/ON_PHL3458/2020 | North America / Canada / Ontario | 2020-03-12 | Public Health Ontario Laboratories | Public Health Ontario Laboratories | Alireza Eshaghi, Samir N Patel, Jonathan B Gubbay, Vanessa G Allen, Christine Frantz, Aimin Li, Sandeep Nagra |
| EPI_ISL_418369 | hCoV-19/Canada/ON_PHL8539/2020 | North America / Canada / Ontario | 2020-03-10 | Public Health Ontario Laboratories | Public Health Ontario Laboratories | Alireza Eshaghi, Samir N Patel, Jonathan B Gubbay, Vanessa G Allen, Christine Frantz, Aimin Li, Sandeep Nagra |
| EPI_ISL_418370 | hCoV-19/Canada/ON_PHL0654/2020 | North America / Canada / Ontario | 2020-03-11 | Public Health Ontario Laboratories | Public Health Ontario Laboratories | Alireza Eshaghi, Samir N Patel, Jonathan B Gubbay, Vanessa G Allen, Christine Frantz, Aimin Li, Sandeep Nagra |
| EPI_ISL_418371 | hCoV-19/Canada/ON_PHL2653/2020 | North America / Canada / Ontario | 2020-03-07 | Public Health Ontario Laboratories | Public Health Ontario Laboratories | Alireza Eshaghi, Samir N Patel, Jonathan B Gubbay, Vanessa G Allen, Christine Frantz, Aimin Li, Sandeep Nagra |
| EPI_ISL_418372 | hCoV-19/Canada/ON_PHL8458/2020 | North America / Canada / Ontario | 2020-03-12 | Public Health Ontario Laboratories | Public Health Ontario Laboratories | Alireza Eshaghi, Samir N Patel, Jonathan B Gubbay, Vanessa G Allen, Christine Frantz, Aimin Li, Sandeep Nagra |
| EPI_ISL_418373 | hCoV-19/Canada/ON_PHL7590/2020 | North America / Canada / Ontario | 2020-03-14 | Public Health Ontario Laboratories | Public Health Ontario Laboratories | Alireza Eshaghi, Samir N Patel, Jonathan B Gubbay, Vanessa G Allen, Christine Frantz, Aimin Li, Sandeep Nagra |
| EPI_ISL_418374 | hCoV-19/Canada/ON_PHL4232/2020 | North America / Canada / Ontario | 2020-03-11 | Public Health Ontario Laboratories | Public Health Ontario Laboratories | Alireza Eshaghi, Samir N Patel, Jonathan B Gubbay, Vanessa G Allen, Christine Frantz, Aimin Li, Sandeep Nagra |
| EPI_ISL_418375 | hCoV-19/Canada/ON_PHL0141/2020 | North America / Canada / Ontario | 2020-03-12 | Public Health Ontario Laboratories | Public Health Ontario Laboratories | Alireza Eshaghi, Samir N Patel, Jonathan B Gubbay, Vanessa G Allen, Christine Frantz, Aimin Li, Sandeep Nagra |
| EPI_ISL_418376 | hCoV-19/Canada/ON_PHL3459/2020 | North America / Canada / Ontario | 2020-03-12 | Public Health Ontario Laboratories | Public Health Ontario Laboratories | Alireza Eshaghi, Samir N Patel, Jonathan B Gubbay, Vanessa G Allen, Christine Frantz, Aimin Li, Sandeep Nagra |
| EPI_ISL_418377 | hCoV-19/Canada/ON_PHL1898/2020 | North America / Canada / Ontario | 2020-03-13 | Public Health Ontario Laboratories | Public Health Ontario Laboratories | Alireza Eshaghi, Samir N Patel, Jonathan B Gubbay, Vanessa G Allen, Christine Frantz, Aimin Li, Sandeep Nagra |
| EPI_ISL_418378 | hCoV-19/Canada/ON_PHL0976/2020 | North America / Canada / Ontario | 2020-03-13 | Public Health Ontario Laboratories | Public Health Ontario Laboratories | Alireza Eshaghi, Samir N Patel, Jonathan B Gubbay, Vanessa G Allen, Christine Frantz, Aimin Li, Sandeep Nagra |
| EPI_ISL_418379 | hCoV-19/Canada/ON_PHL7512/2020 | North America / Canada / Ontario | 2020-03-13 | Public Health Ontario Laboratories | Public Health Ontario Laboratories | Alireza Eshaghi, Samir N Patel, Jonathan B Gubbay, Vanessa G Allen, Christine Frantz, Aimin Li, Sandeep Nagra |
| EPI_ISL_418380 | hCoV-19/Canada/ON_PHL3476/2020 | North America / Canada / Ontario | 2020-03-13 | Public Health Ontario Laboratories | Public Health Ontario Laboratories | Alireza Eshaghi, Samir N Patel, Jonathan B Gubbay, Vanessa G Allen, Christine Frantz, Aimin Li, Sandeep Nagra |
| EPI_ISL_418381 | hCoV-19/Canada/ON_PHL2223/2020 | North America / Canada / Ontario | 2020-03-19 | Public Health Ontario Laboratories | Public Health Ontario Laboratories | Alireza Eshaghi, Samir N Patel, Jonathan B Gubbay, Vanessa G Allen, Christine Frantz, Aimin Li, Sandeep Nagra |
| EPI_ISL_418382 | hCoV-19/Canada/ON_PHL5930/2020 | North America / Canada / Ontario | 2020-02-28 | Public Health Ontario Laboratories | Public Health Ontario Laboratories | Alireza Eshaghi, Samir N Patel, Jonathan B Gubbay, Vanessa G Allen, Christine Frantz, Aimin Li, Sandeep Nagra |
| EPI_ISL_418383 | hCoV-19/Canada/ON_PHL2273/2020 | North America / Canada / Ontario | 2020-02-29 | Public Health Ontario Laboratories | Public Health Ontario Laboratories | Alireza Eshaghi, Samir N Patel, Jonathan B Gubbay, Vanessa G Allen, Christine Frantz, Aimin Li, Sandeep Nagra |
| EPI_ISL_418385 | hCoV-19/Finland/13M19/2020 | Europe / Finland | 2020-03-13 | Department of Virology and Immunology, University of Helsinki and Helsinki University Hospital, Huslab Finland | Department of Virology, Faculty of Medicine, University of Helsinki, Helsinki, Finland | Teemu Smura, Hannimari Kallio-Kokko, Olli Vapalahti |
| EPI_ISL_418386 | hCoV-19/Finland/13M26/2020 | Europe / Finland | 2020-03-13 | Department of Virology and Immunology, University of Helsinki and Helsinki University Hospital, Huslab Finland | Department of Virology, Faculty of Medicine, University of Helsinki, Helsinki, Finland | Teemu Smura, Hannimari Kallio-Kokko, Olli Vapalahti |
| EPI_ISL_418387 | hCoV-19/Finland/13M27/2020 | Europe / Finland | 2020-03-13 | Department of Virology and Immunology, University of Helsinki and Helsinki University Hospital, Huslab Finland | Department of Virology, Faculty of Medicine, University of Helsinki, Helsinki, Finland | Teemu Smura, Hannimari Kallio-Kokko, Olli Vapalahti |
| EPI_ISL_418388 | hCoV-19/Finland/13M29/2020 | Europe / Finland | 2020-03-13 | Department of Virology and Immunology, University of Helsinki and Helsinki University Hospital, Huslab Finland | Department of Virology, Faculty of Medicine, University of Helsinki, Helsinki, Finland | Teemu Smura, Hannimari Kallio-Kokko, Olli Vapalahti |
| EPI_ISL_418389 | hCoV-19/Finland/13M3/2020 | Europe / Finland | 2020-03-13 | Department of Virology and Immunology, University of Helsinki and Helsinki University Hospital, Huslab Finland | Department of Virology, Faculty of Medicine, University of Helsinki, Helsinki, Finland | Teemu Smura, Hannimari Kallio-Kokko, Olli Vapalahti |
| EPI_ISL_418390 | hCoV-19/Finland/13M33/2020 | Europe / Finland | 2020-03-13 | Department of Virology and Immunology, University of Helsinki and Helsinki University Hospital, Huslab Finland | Department of Virology, Faculty of Medicine, University of Helsinki, Helsinki, Finland | Teemu Smura, Hannimari Kallio-Kokko, Olli Vapalahti |
| EPI_ISL_418391 | hCoV-19/Finland/13M57/2020 | Europe / Finland | 2020-03-13 | Department of Virology and Immunology, University of Helsinki and Helsinki University Hospital, Huslab Finland | Department of Virology, Faculty of Medicine, University of Helsinki, Helsinki, Finland | Teemu Smura, Hannimari Kallio-Kokko, Olli Vapalahti |
| EPI_ISL_418392 | hCoV-19/Finland/13M58/2020 | Europe / Finland | 2020-03-13 | Department of Virology and Immunology, University of Helsinki and Helsinki University Hospital, Huslab Finland | Department of Virology, Faculty of Medicine, University of Helsinki, Helsinki, Finland | Teemu Smura, Hannimari Kallio-Kokko, Olli Vapalahti |
| EPI_ISL_418393 | hCoV-19/Finland/13M60/2020 | Europe / Finland | 2020-03-13 | Department of Virology and Immunology, University of Helsinki and Helsinki University Hospital, Huslab Finland | Department of Virology, Faculty of Medicine, University of Helsinki, Helsinki, Finland | Teemu Smura, Hannimari Kallio-Kokko, Olli Vapalahti |
| EPI_ISL_418394 | hCoV-19/Finland/13M64/2020 | Europe / Finland | 2020-03-13 | Department of Virology and Immunology, University of Helsinki and Helsinki University Hospital, Huslab Finland | Department of Virology, Faculty of Medicine, University of Helsinki, Helsinki, Finland | Teemu Smura, Hannimari Kallio-Kokko, Olli Vapalahti |
| EPI_ISL_418395 | hCoV-19/Finland/13M65/2020 | Europe / Finland | 2020-03-13 | Department of Virology and Immunology, University of Helsinki and Helsinki University Hospital, Huslab Finland | Department of Virology, Faculty of Medicine, University of Helsinki, Helsinki, Finland | Teemu Smura, Hannimari Kallio-Kokko, Olli Vapalahti |
| EPI_ISL_418396 | hCoV-19/Finland/13M69/2020 | Europe / Finland | 2020-03-13 | Department of Virology and Immunology, University of Helsinki and Helsinki University Hospital, Huslab Finland | Department of Virology, Faculty of Medicine, University of Helsinki, Helsinki, Finland | Teemu Smura, Hannimari Kallio-Kokko, Olli Vapalahti |
| EPI_ISL_418397 | hCoV-19/Finland/13M77/2020 | Europe / Finland | 2020-03-13 | Department of Virology and Immunology, University of Helsinki and Helsinki University Hospital, Huslab Finland | Department of Virology, Faculty of Medicine, University of Helsinki, Helsinki, Finland | Teemu Smura, Hannimari Kallio-Kokko, Olli Vapalahti |
| EPI_ISL_418399 | hCoV-19/Finland/13M82/2020 | Europe / Finland | 2020-03-13 | Department of Virology and Immunology, University of Helsinki and Helsinki University Hospital, Huslab Finland | Department of Virology, Faculty of Medicine, University of Helsinki, Helsinki, Finland | Teemu Smura, Hannimari Kallio-Kokko, Olli Vapalahti |
| EPI_ISL_418700 | hCoV-19/England/20124013202/2020 | Europe / United Kingdom / England | 2020-03-17 | Respiratory Virus Unit, Microbiology Services Colindale, Public Health England | Respiratory Virus Unit, Microbiology Services Colindale, Public Health England | Monica Galiano, Shahjahan Miah, Angie Lackenby, Omolola Akinbami, Tiina Talts, Leena Bhaw, Richard Myers, Steven Platt, Kirstin Edwards, Jonathan Hubb, Joanna Ellis, Maria Zambon |
| EPI_ISL_418701 | hCoV-19/England/20124020402/2020 | Europe / United Kingdom / England | 2020-03-17 | Respiratory Virus Unit, Microbiology Services Colindale, Public Health England | Respiratory Virus Unit, Microbiology Services Colindale, Public Health England | Monica Galiano, Shahjahan Miah, Angie Lackenby, Omolola Akinbami, Tiina Talts, Leena Bhaw, Richard Myers, Steven Platt, Kirstin Edwards, Jonathan Hubb, Joanna Ellis, Maria Zambon |
| EPI_ISL_418702 | hCoV-19/England/20124020802/2020 | Europe / United Kingdom / England | 2020-03-16 | Respiratory Virus Unit, Microbiology Services Colindale, Public Health England | Respiratory Virus Unit, Microbiology Services Colindale, Public Health England | Monica Galiano, Shahjahan Miah, Angie Lackenby, Omolola Akinbami, Tiina Talts, Leena Bhaw, Richard Myers, Steven Platt, Kirstin Edwards, Jonathan Hubb, Joanna Ellis, Maria Zambon |
| EPI_ISL_418703 | hCoV-19/England/20124021002/2020 | Europe / United Kingdom / England | 2020-03-17 | Respiratory Virus Unit, Microbiology Services Colindale, Public Health England | Respiratory Virus Unit, Microbiology Services Colindale, Public Health England | Monica Galiano, Shahjahan Miah, Angie Lackenby, Omolola Akinbami, Tiina Talts, Leena Bhaw, Richard Myers, Steven Platt, Kirstin Edwards, Jonathan Hubb, Joanna Ellis, Maria Zambon |
| EPI_ISL_418704 | hCoV-19/England/20124021402/2020 | Europe / United Kingdom / England | 2020-03-17 | Respiratory Virus Unit, Microbiology Services Colindale, Public Health England | Respiratory Virus Unit, Microbiology Services Colindale, Public Health England | Monica Galiano, Shahjahan Miah, Angie Lackenby, Omolola Akinbami, Tiina Talts, Leena Bhaw, Richard Myers, Steven Platt, Kirstin Edwards, Jonathan Hubb, Joanna Ellis, Maria Zambon |
| EPI_ISL_418706 | hCoV-19/England/20124029802/2020 | Europe / United Kingdom / England | 2020-03-17 | Respiratory Virus Unit, Microbiology Services Colindale, Public Health England | Respiratory Virus Unit, Microbiology Services Colindale, Public Health England | Monica Galiano, Shahjahan Miah, Angie Lackenby, Omolola Akinbami, Tiina Talts, Leena Bhaw, Richard Myers, Steven Platt, Kirstin Edwards, Jonathan Hubb, Joanna Ellis, Maria Zambon |
| EPI_ISL_418707 | hCoV-19/England/20124030003/2020 | Europe / United Kingdom / England | 2020-03-17 | Respiratory Virus Unit, Microbiology Services Colindale, Public Health England | Respiratory Virus Unit, Microbiology Services Colindale, Public Health England | Monica Galiano, Shahjahan Miah, Angie Lackenby, Omolola Akinbami, Tiina Talts, Leena Bhaw, Richard Myers, Steven Platt, Kirstin Edwards, Jonathan Hubb, Joanna Ellis, Maria Zambon |
| EPI_ISL_418708 | hCoV-19/England/20124030802/2020 | Europe / United Kingdom / England | 2020-03-17 | Respiratory Virus Unit, Microbiology Services Colindale, Public Health England | Respiratory Virus Unit, Microbiology Services Colindale, Public Health England | Monica Galiano, Shahjahan Miah, Angie Lackenby, Omolola Akinbami, Tiina Talts, Leena Bhaw, Richard Myers, Steven Platt, Kirstin Edwards, Jonathan Hubb, Joanna Ellis, Maria Zambon |
| EPI_ISL_418709 | hCoV-19/England/20124034002/2020 | Europe / United Kingdom / England | 2020-03-16 | Respiratory Virus Unit, Microbiology Services Colindale, Public Health England | Respiratory Virus Unit, Microbiology Services Colindale, Public Health England | Monica Galiano, Shahjahan Miah, Angie Lackenby, Omolola Akinbami, Tiina Talts, Leena Bhaw, Richard Myers, Steven Platt, Kirstin Edwards, Jonathan Hubb, Joanna Ellis, Maria Zambon |
| EPI_ISL_418711 | hCoV-19/England/20124034702/2020 | Europe / United Kingdom / England | 2020-03-16 | Respiratory Virus Unit, Microbiology Services Colindale, Public Health England | Respiratory Virus Unit, Microbiology Services Colindale, Public Health England | Monica Galiano, Shahjahan Miah, Angie Lackenby, Omolola Akinbami, Tiina Talts, Leena Bhaw, Richard Myers, Steven Platt, Kirstin Edwards, Jonathan Hubb, Joanna Ellis, Maria Zambon |
| EPI_ISL_418713 | hCoV-19/England/20124035502/2020 | Europe / United Kingdom / England | 2020-03-17 | Respiratory Virus Unit, Microbiology Services Colindale, Public Health England | Respiratory Virus Unit, Microbiology Services Colindale, Public Health England | Monica Galiano, Shahjahan Miah, Angie Lackenby, Omolola Akinbami, Tiina Talts, Leena Bhaw, Richard Myers, Steven Platt, Kirstin Edwards, Jonathan Hubb, Joanna Ellis, Maria Zambon |
| EPI_ISL_418715 | hCoV-19/England/20124035802/2020 | Europe / United Kingdom / England | 2020-03-17 | Respiratory Virus Unit, Microbiology Services Colindale, Public Health England | Respiratory Virus Unit, Microbiology Services Colindale, Public Health England | Monica Galiano, Shahjahan Miah, Angie Lackenby, Omolola Akinbami, Tiina Talts, Leena Bhaw, Richard Myers, Steven Platt, Kirstin Edwards, Jonathan Hubb, Joanna Ellis, Maria Zambon |
| EPI_ISL_418716 | hCoV-19/England/20124036002/2020 | Europe / United Kingdom / England | 2020-03-17 | Respiratory Virus Unit, Microbiology Services Colindale, Public Health England | Respiratory Virus Unit, Microbiology Services Colindale, Public Health England | Monica Galiano, Shahjahan Miah, Angie Lackenby, Omolola Akinbami, Tiina Talts, Leena Bhaw, Richard Myers, Steven Platt, Kirstin Edwards, Jonathan Hubb, Joanna Ellis, Maria Zambon |
| EPI_ISL_418718 | hCoV-19/England/20124036902/2020 | Europe / United Kingdom / England | 2020-03-18 | Respiratory Virus Unit, Microbiology Services Colindale, Public Health England | Respiratory Virus Unit, Microbiology Services Colindale, Public Health England | Monica Galiano, Shahjahan Miah, Angie Lackenby, Omolola Akinbami, Tiina Talts, Leena Bhaw, Richard Myers, Steven Platt, Kirstin Edwards, Jonathan Hubb, Joanna Ellis, Maria Zambon |
| EPI_ISL_418720 | hCoV-19/England/20124037302/2020 | Europe / United Kingdom / England | 2020-03-17 | Respiratory Virus Unit, Microbiology Services Colindale, Public Health England | Respiratory Virus Unit, Microbiology Services Colindale, Public Health England | Monica Galiano, Shahjahan Miah, Angie Lackenby, Omolola Akinbami, Tiina Talts, Leena Bhaw, Richard Myers, Steven Platt, Kirstin Edwards, Jonathan Hubb, Joanna Ellis, Maria Zambon |
| EPI_ISL_418722 | hCoV-19/England/20124048202/2020 | Europe / United Kingdom / England | 2020-03-17 | Respiratory Virus Unit, Microbiology Services Colindale, Public Health England | Respiratory Virus Unit, Microbiology Services Colindale, Public Health England | Monica Galiano, Shahjahan Miah, Angie Lackenby, Omolola Akinbami, Tiina Talts, Leena Bhaw, Richard Myers, Steven Platt, Kirstin Edwards, Jonathan Hubb, Joanna Ellis, Maria Zambon |
| EPI_ISL_418723 | hCoV-19/England/20124049402/2020 | Europe / United Kingdom / England | 2020-03-16 | Respiratory Virus Unit, Microbiology Services Colindale, Public Health England | Respiratory Virus Unit, Microbiology Services Colindale, Public Health England | Monica Galiano, Shahjahan Miah, Angie Lackenby, Omolola Akinbami, Tiina Talts, Leena Bhaw, Richard Myers, Steven Platt, Kirstin Edwards, Jonathan Hubb, Joanna Ellis, Maria Zambon |
| EPI_ISL_418729 | hCoV-19/England/20124095502/2020 | Europe / United Kingdom / England | 2020-03-18 | Respiratory Virus Unit, Microbiology Services Colindale, Public Health England | Respiratory Virus Unit, Microbiology Services Colindale, Public Health England | Monica Galiano, Shahjahan Miah, Angie Lackenby, Omolola Akinbami, Tiina Talts, Leena Bhaw, Richard Myers, Steven Platt, Kirstin Edwards, Jonathan Hubb, Joanna Ellis, Maria Zambon |
| EPI_ISL_418733 | hCoV-19/England/20124096802/2020 | Europe / United Kingdom / England | 2020-03-18 | Respiratory Virus Unit, Microbiology Services Colindale, Public Health England | Respiratory Virus Unit, Microbiology Services Colindale, Public Health England | Monica Galiano, Shahjahan Miah, Angie Lackenby, Omolola Akinbami, Tiina Talts, Leena Bhaw, Richard Myers, Steven Platt, Kirstin Edwards, Jonathan Hubb, Joanna Ellis, Maria Zambon |
| EPI_ISL_418734 | hCoV-19/England/20124097102/2020 | Europe / United Kingdom / England | 2020-03-13 | Respiratory Virus Unit, Microbiology Services Colindale, Public Health England | Respiratory Virus Unit, Microbiology Services Colindale, Public Health England | Monica Galiano, Shahjahan Miah, Angie Lackenby, Omolola Akinbami, Tiina Talts, Leena Bhaw, Richard Myers, Steven Platt, Kirstin Edwards, Jonathan Hubb, Joanna Ellis, Maria Zambon |
| EPI_ISL_418737 | hCoV-19/England/20124100002/2020 | Europe / United Kingdom / England | 2020-03-17 | Respiratory Virus Unit, Microbiology Services Colindale, Public Health England | Respiratory Virus Unit, Microbiology Services Colindale, Public Health England | Monica Galiano, Shahjahan Miah, Angie Lackenby, Omolola Akinbami, Tiina Talts, Leena Bhaw, Richard Myers, Steven Platt, Kirstin Edwards, Jonathan Hubb, Joanna Ellis, Maria Zambon |
| EPI_ISL_418738 | hCoV-19/England/20124100602/2020 | Europe / United Kingdom / England | 2020-03-17 | Respiratory Virus Unit, Microbiology Services Colindale, Public Health England | Respiratory Virus Unit, Microbiology Services Colindale, Public Health England | Monica Galiano, Shahjahan Miah, Angie Lackenby, Omolola Akinbami, Tiina Talts, Leena Bhaw, Richard Myers, Steven Platt, Kirstin Edwards, Jonathan Hubb, Joanna Ellis, Maria Zambon |
| EPI_ISL_418739 | hCoV-19/England/20124101102/2020 | Europe / United Kingdom / England | 2020-03-18 | Respiratory Virus Unit, Microbiology Services Colindale, Public Health England | Respiratory Virus Unit, Microbiology Services Colindale, Public Health England | Monica Galiano, Shahjahan Miah, Angie Lackenby, Omolola Akinbami, Tiina Talts, Leena Bhaw, Richard Myers, Steven Platt, Kirstin Edwards, Jonathan Hubb, Joanna Ellis, Maria Zambon |
| EPI_ISL_418742 | hCoV-19/England/20126003302/2020 | Europe / United Kingdom / England | 2020-03-19 | Respiratory Virus Unit, Microbiology Services Colindale, Public Health England | Respiratory Virus Unit, Microbiology Services Colindale, Public Health England | Monica Galiano, Shahjahan Miah, Angie Lackenby, Omolola Akinbami, Tiina Talts, Leena Bhaw, Richard Myers, Steven Platt, Kirstin Edwards, Jonathan Hubb, Joanna Ellis, Maria Zambon |
| EPI_ISL_418749 | hCoV-19/England/20126006802/2020 | Europe / United Kingdom / England | 2020-03-18 | Respiratory Virus Unit, Microbiology Services Colindale, Public Health England | Respiratory Virus Unit, Microbiology Services Colindale, Public Health England | Monica Galiano, Shahjahan Miah, Angie Lackenby, Omolola Akinbami, Tiina Talts, Leena Bhaw, Richard Myers, Steven Platt, Kirstin Edwards, Jonathan Hubb, Joanna Ellis, Maria Zambon |
| EPI_ISL_418751 | hCoV-19/England/20126007102/2020 | Europe / United Kingdom / England | 2020-03-18 | Respiratory Virus Unit, Microbiology Services Colindale, Public Health England | Respiratory Virus Unit, Microbiology Services Colindale, Public Health England | Monica Galiano, Shahjahan Miah, Angie Lackenby, Omolola Akinbami, Tiina Talts, Leena Bhaw, Richard Myers, Steven Platt, Kirstin Edwards, Jonathan Hubb, Joanna Ellis, Maria Zambon |
| EPI_ISL_418755 | hCoV-19/England/20126012302/2020 | Europe / United Kingdom / England | 2020-03-17 | Respiratory Virus Unit, Microbiology Services Colindale, Public Health England | Respiratory Virus Unit, Microbiology Services Colindale, Public Health England | Monica Galiano, Shahjahan Miah, Angie Lackenby, Omolola Akinbami, Tiina Talts, Leena Bhaw, Richard Myers, Steven Platt, Kirstin Edwards, Jonathan Hubb, Joanna Ellis, Maria Zambon |
| EPI_ISL_418756 | hCoV-19/England/20126012702/2020 | Europe / United Kingdom / England | 2020-03-17 | Respiratory Virus Unit, Microbiology Services Colindale, Public Health England | Respiratory Virus Unit, Microbiology Services Colindale, Public Health England | Monica Galiano, Shahjahan Miah, Angie Lackenby, Omolola Akinbami, Tiina Talts, Leena Bhaw, Richard Myers, Steven Platt, Kirstin Edwards, Jonathan Hubb, Joanna Ellis, Maria Zambon |
| EPI_ISL_418764 | hCoV-19/England/20126035502/2020 | Europe / United Kingdom / England | 2020-03-18 | Respiratory Virus Unit, Microbiology Services Colindale, Public Health England | Respiratory Virus Unit, Microbiology Services Colindale, Public Health England | Monica Galiano, Shahjahan Miah, Angie Lackenby, Omolola Akinbami, Tiina Talts, Leena Bhaw, Richard Myers, Steven Platt, Kirstin Edwards, Jonathan Hubb, Joanna Ellis, Maria Zambon |
| EPI_ISL_418770 | hCoV-19/England/20126096602/2020 | Europe / United Kingdom / England | 2020-03-18 | Respiratory Virus Unit, Microbiology Services Colindale, Public Health England | Respiratory Virus Unit, Microbiology Services Colindale, Public Health England | Monica Galiano, Shahjahan Miah, Angie Lackenby, Omolola Akinbami, Tiina Talts, Leena Bhaw, Richard Myers, Steven Platt, Kirstin Edwards, Jonathan Hubb, Joanna Ellis, Maria Zambon |
| EPI_ISL_418771 | hCoV-19/USA/WA-NH2/2020 | North America / USA / Washington | 2020-03-13 | WA State Department of Health | Pathogen Discovery, Respiratory Viruses Branch, Division of Viral Diseases, Centers for Disease Control and Prevention | Jing Zhang, Ying Tao, Clinton R. Paden, Krista Queen, Anna Uehara, Yan Li, Haibin Wang, Jessica Jacobs, Denny Russell, Brian Hiatt, Jessica Gant, Suxiang Tong |
| EPI_ISL_418772 | hCoV-19/USA/WA-NH3/2020 | North America / USA / Washington | 2020-03-13 | WA State Department of Health | Pathogen Discovery, Respiratory Viruses Branch, Division of Viral Diseases, Centers for Disease Control and Prevention | Jing Zhang, Ying Tao, Clinton R. Paden, Krista Queen, Anna Uehara, Yan Li, Haibin Wang, Jessica Jacobs, Denny Russell, Brian Hiatt, Jessica Gant, Suxiang Tong |
| EPI_ISL_418773 | hCoV-19/USA/WA-NH4/2020 | North America / USA / Washington | 2020-03-13 | WA State Department of Health | Pathogen Discovery, Respiratory Viruses Branch, Division of Viral Diseases, Centers for Disease Control and Prevention | Jing Zhang, Ying Tao, Clinton R. Paden, Krista Queen, Anna Uehara, Yan Li, Haibin Wang, Jessica Jacobs, Denny Russell, Brian Hiatt, Jessica Gant, Suxiang Tong |
| EPI_ISL_418774 | hCoV-19/USA/WA-NH5/2020 | North America / USA / Washington | 2020-03-13 | WA State Department of Health | Pathogen Discovery, Respiratory Viruses Branch, Division of Viral Diseases, Centers for Disease Control and Prevention | Jing Zhang, Ying Tao, Clinton R. Paden, Krista Queen, Anna Uehara, Yan Li, Haibin Wang, Jessica Jacobs, Denny Russell, Brian Hiatt, Jessica Gant, Suxiang Tong |
| EPI_ISL_418775 | hCoV-19/USA/WA-NH6/2020 | North America / USA / Washington | 2020-03-13 | WA State Department of Health | Pathogen Discovery, Respiratory Viruses Branch, Division of Viral Diseases, Centers for Disease Control and Prevention | Jing Zhang, Ying Tao, Clinton R. Paden, Krista Queen, Anna Uehara, Yan Li, Haibin Wang, Jessica Jacobs, Denny Russell, Brian Hiatt, Jessica Gant, Suxiang Tong |
| EPI_ISL_418776 | hCoV-19/USA/WA-NH7/2020 | North America / USA / Washington | 2020-03-13 | WA State Department of Health | Pathogen Discovery, Respiratory Viruses Branch, Division of Viral Diseases, Centers for Disease Control and Prevention | Jing Zhang, Ying Tao, Clinton R. Paden, Krista Queen, Anna Uehara, Yan Li, Haibin Wang, Jessica Jacobs, Denny Russell, Brian Hiatt, Jessica Gant, Suxiang Tong |
| EPI_ISL_418777 | hCoV-19/USA/WA-NH8/2020 | North America / USA / Washington | 2020-03-13 | WA State Department of Health | Pathogen Discovery, Respiratory Viruses Branch, Division of Viral Diseases, Centers for Disease Control and Prevention | Jing Zhang, Ying Tao, Clinton R. Paden, Krista Queen, Anna Uehara, Yan Li, Haibin Wang, Jessica Jacobs, Denny Russell, Brian Hiatt, Jessica Gant, Suxiang Tong |
| EPI_ISL_418778 | hCoV-19/USA/WA-NH9/2020 | North America / USA / Washington | 2020-03-13 | WA State Department of Health | Pathogen Discovery, Respiratory Viruses Branch, Division of Viral Diseases, Centers for Disease Control and Prevention | Ying Tao, Jing Zhang, Clinton R. Paden, Krista Queen, Anna Uehara, Yan Li, Haibin Wang, Jessica Jacobs, Denny Russell, Brian Hiatt, Jessica Gant, Suxiang Tong |
| EPI_ISL_418779 | hCoV-19/USA/WA-NH10/2020 | North America / USA / Washington | 2020-03-13 | WA State Department of Health | Pathogen Discovery, Respiratory Viruses Branch, Division of Viral Diseases, Centers for Disease Control and Prevention | Ying Tao, Jing Zhang, Clinton R. Paden, Krista Queen, Anna Uehara, Yan Li, Haibin Wang, Jessica Jacobs, Denny Russell, Brian Hiatt, Jessica Gant, Suxiang Tong |
| EPI_ISL_418780 | hCoV-19/USA/WA-NH11/2020 | North America / USA / Washington | 2020-03-13 | WA State Department of Health | Pathogen Discovery, Respiratory Viruses Branch, Division of Viral Diseases, Centers for Disease Control and Prevention | Ying Tao, Jing Zhang, Clinton R. Paden, Krista Queen, Anna Uehara, Yan Li, Haibin Wang, Jessica Jacobs, Denny Russell, Brian Hiatt, Jessica Gant, Suxiang Tong |
| EPI_ISL_418781 | hCoV-19/USA/WA-NH12/2020 | North America / USA / Washington | 2020-03-13 | WA State Department of Health | Pathogen Discovery, Respiratory Viruses Branch, Division of Viral Diseases, Centers for Disease Control and Prevention | Ying Tao, Jing Zhang, Clinton R. Paden, Krista Queen, Anna Uehara, Yan Li, Haibin Wang, Jessica Jacobs, Denny Russell, Brian Hiatt, Jessica Gant, Suxiang Tong |
| EPI_ISL_418782 | hCoV-19/USA/WA-NH13/2020 | North America / USA / Washington | 2020-03-13 | WA State Department of Health | Pathogen Discovery, Respiratory Viruses Branch, Division of Viral Diseases, Centers for Disease Control and Prevention | Ying Tao, Jing Zhang, Clinton R. Paden, Krista Queen, Anna Uehara, Yan Li, Haibin Wang, Jessica Jacobs, Denny Russell, Brian Hiatt, Jessica Gant, Suxiang Tong |
| EPI_ISL_418783 | hCoV-19/USA/WA-NH14/2020 | North America / USA / Washington | 2020-03-13 | WA State Department of Health | Pathogen Discovery, Respiratory Viruses Branch, Division of Viral Diseases, Centers for Disease Control and Prevention | Ying Tao, Jing Zhang, Clinton R. Paden, Krista Queen, Anna Uehara, Yan Li, Haibin Wang, Jessica Jacobs, Denny Russell, Brian Hiatt, Jessica Gant, Suxiang Tong |
| EPI_ISL_418784 | hCoV-19/USA/WA-NH17/2020 | North America / USA / Washington | 2020-03-13 | WA State Department of Health | Pathogen Discovery, Respiratory Viruses Branch, Division of Viral Diseases, Centers for Disease Control and Prevention | Ying Tao, Jing Zhang, Clinton R. Paden, Krista Queen, Anna Uehara, Yan Li, Haibin Wang, Jessica Jacobs, Denny Russell, Brian Hiatt, Jessica Gant, Suxiang Tong |
| EPI_ISL_418785 | hCoV-19/USA/WA-NH18/2020 | North America / USA / Washington | 2020-03-13 | WA State Department of Health | Pathogen Discovery, Respiratory Viruses Branch, Division of Viral Diseases, Centers for Disease Control and Prevention | Ying Tao, Jing Zhang, Clinton R. Paden, Krista Queen, Anna Uehara, Yan Li, Haibin Wang, Jessica Jacobs, Denny Russell, Brian Hiatt, Jessica Gant, Suxiang Tong |
| EPI_ISL_418786 | hCoV-19/USA/WA-NH19/2020 | North America / USA / Washington | 2020-03-13 | WA State Department of Health | Pathogen Discovery, Respiratory Viruses Branch, Division of Viral Diseases, Centers for Disease Control and Prevention | Ying Tao, Jing Zhang, Clinton R. Paden, Krista Queen, Anna Uehara, Yan Li, Haibin Wang, Jessica Jacobs, Denny Russell, Brian Hiatt, Jessica Gant, Suxiang Tong |
| EPI_ISL_418787 | hCoV-19/USA/WA-NH20/2020 | North America / USA / Washington | 2020-03-13 | WA State Department of Health | Pathogen Discovery, Respiratory Viruses Branch, Division of Viral Diseases, Centers for Disease Control and Prevention | Ying Tao, Jing Zhang, Clinton R. Paden, Krista Queen, Anna Uehara, Yan Li, Haibin Wang, Jessica Jacobs, Denny Russell, Brian Hiatt, Jessica Gant, Suxiang Tong |
| EPI_ISL_418788 | hCoV-19/USA/WA-NH21/2020 | North America / USA / Washington | 2020-03-13 | WA State Department of Health | Pathogen Discovery, Respiratory Viruses Branch, Division of Viral Diseases, Centers for Disease Control and Prevention | Ying Tao, Jing Zhang, Clinton R. Paden, Krista Queen, Anna Uehara, Yan Li, Haibin Wang, Jessica Jacobs, Denny Russell, Brian Hiatt, Jessica Gant, Suxiang Tong |
| EPI_ISL_418789 | hCoV-19/USA/WA-NH22/2020 | North America / USA / Washington | 2020-03-13 | WA State Department of Health | Pathogen Discovery, Respiratory Viruses Branch, Division of Viral Diseases, Centers for Disease Control and Prevention | Ying Tao, Jing Zhang, Clinton R. Paden, Krista Queen, Anna Uehara, Yan Li, Haibin Wang, Jessica Jacobs, Denny Russell, Brian Hiatt, Jessica Gant, Suxiang Tong |
| EPI_ISL_418790 | hCoV-19/USA/WA-NH23/2020 | North America / USA / Washington | 2020-03-13 | WA State Department of Health | Pathogen Discovery, Respiratory Viruses Branch, Division of Viral Diseases, Centers for Disease Control and Prevention | Ying Tao, Jing Zhang, Clinton R. Paden, Krista Queen, Anna Uehara, Yan Li, Haibin Wang, Jessica Jacobs, Denny Russell, Brian Hiatt, Jessica Gant, Suxiang Tong |
| EPI_ISL_418791 | hCoV-19/USA/WA-NH24/2020 | North America / USA / Washington | 2020-03-13 | WA State Department of Health | Pathogen Discovery, Respiratory Viruses Branch, Division of Viral Diseases, Centers for Disease Control and Prevention | Ying Tao, Jing Zhang, Clinton R. Paden, Krista Queen, Anna Uehara, Yan Li, Haibin Wang, Jessica Jacobs, Denny Russell, Brian Hiatt, Jessica Gant, Suxiang Tong |
| EPI_ISL_418792 | hCoV-19/Belgium/GL-030546/2020 | Europe / Belgium / Houthalen-Helchteren | 2020-03-05 | KU Leuven, Clinical and Epidemiological Virology | KU Leuven, Clinical and Epidemiological Virology | Bert Vanmechelen, Tony Wawina, Joan Martí-Carreras, Piet Maes |
| EPI_ISL_418793 | hCoV-19/Belgium/SS-031047/2020 | Europe / Belgium / Zonhoven | 2020-03-10 | KU Leuven, Clinical and Epidemiological Virology | KU Leuven, Clinical and Epidemiological Virology | Bert Vanmechelen, Tony Wawina, Joan Martí-Carreras, Piet Maes |
| EPI_ISL_418794 | hCoV-19/Belgium/VHRA-030644/2020 | Europe / Belgium / Nevele | 2020-03-06 | KU Leuven, Clinical and Epidemiological Virology | KU Leuven, Clinical and Epidemiological Virology | Bert Vanmechelen, Joan Martí-Carreras, Tony Wawina, Piet Maes |
| EPI_ISL_418795 | hCoV-19/Belgium/NKR-030645/2020 | Europe / Belgium / Ganshoren | 2020-03-06 | KU Leuven, Clinical and Epidemiological Virology | KU Leuven, Clinical and Epidemiological Virology | Bert Vanmechelen, Joan Martí-Carreras, Tony Wawina, Piet Maes |
| EPI_ISL_418796 | hCoV-19/Belgium/VDLH-030548/2020 | Europe / Belgium / Kalmthout | 2020-03-05 | KU Leuven, Clinical and Epidemiological Virology | KU Leuven, Clinical and Epidemiological Virology | Bert Vanmechelen, Joan Martí-Carreras, Tony Wawina, Piet Maes |
| EPI_ISL_418797 | hCoV-19/Belgium/VDLH-030548/2020 | Europe / Belgium / Kalmthout | 2020-03-05 | KU Leuven, Clinical and Epidemiological Virology | KU Leuven, Clinical and Epidemiological Virology | Bert Vanmechelen, Joan Martí-Carreras, Tony Wawina, Piet Maes |
| EPI_ISL_418798 | hCoV-19/Belgium/GS-030549/2020 | Europe / Belgium / Chaineux | 2020-03-05 | KU Leuven, Clinical and Epidemiological Virology | KU Leuven, Clinical and Epidemiological Virology | Bert Vanmechelen, Joan Martí-Carreras, Tony Wawina, Piet Maes |
| EPI_ISL_418799 | hCoV-19/Australia/QLDID922/2020 | Oceania / Australia / Queensland / Brisba | 2020-02-28 | Mater Pathology | Public Health Virology Laboratory | Bixing Huang, Alyssa Pyke, Amanda De Jong, Andrew Van Den Hurk, Carmel Taylor, David Warrilow, Doris Genge, Elisabeth Gamez, Glen Hewitson, Ian Maxwell Mackay, Inga Sultana, Jamie McMahon, Jean Barcelon, Judy Northill, Mitchell Finger, Natalie Simpson, Neelima Nair, Peter Burtonclay, Peter Moore, Sarah Wheatley, Sean Moody, Sonja Hall-Mendelin, Timothy Gardam, and Frederick Moore |
| EPI_ISL_450404 | hCoV-19/Hong Kong/VM20009579/2020 | Asia / Hong Kong | 2020-03-16 | unknown | School of Public Health | Sit,T.H.S., Brackman,C.J., Sims,L.D., Tsang,D.N.C., Chu,D.K.W., Perera,R.A.P.M., Poon,L.L.M. and Peiris,M. |
| EPI_ISL_450489 | hCoV-19/Wuhan/YB012504/2020 | Asia / China / Wuhan / Hubei | 2020-01-25 | unknown | CAS Key Laboratory of Special Pathogens and Biosafety and Center for Emerging Infectious Diseases | Si,H., Zhu,Y., Lin,H., Xie,S., Shi,Z. and Zhou,P. |
