## Supplementary-Table1-3 for "A cytosine-to-uracil change within the programmed -1 ribosomal frameshift signal of SARS-CoV-2 results in structural similarities with the MERS-CoV signal"

**We gratefully acknowledge the authors, originating and submitting laboratories of the sequences from GISAID's EpiFlu™ Database on which this research is based. The list is detailed below.**
**All submitters of data may be contacted directly via www.gisaid.org**

| Accession ID | Virus name | Location | Collection date | Originating lab | Submitting lab | Authors |
|---|---|---|---|---|---|---|
| EPI_ISL_426159 | hCoV-19/USA/WI-GMF-00986/2020 | North America / USA / Minnesota / Winor | 2020-04-06 | Gundersen Molecular Diagnostics Laboratory | Kabara Cancer Research Institute | Craig S. Richmond, Paraic A. Kenny |
| EPI_ISL_426160 | hCoV-19/USA/WI-GMF-00921/2020 | North America / USA / Iowa / Allamakee ( | 2020-04-03 | Gundersen Molecular Diagnostics Laboratory | Kabara Cancer Research Institute | Craig S. Richmond, Paraic A. Kenny |
| EPI_ISL_426161 | hCoV-19/USA/WI-GMF-00928/2020 | North America / USA / Iowa | 2020-04-06 | Gundersen Molecular Diagnostics Laboratory | Kabara Cancer Research Institute | Craig S. Richmond, Paraic A. Kenny |
| EPI_ISL_426163 | hCoV-19/Korea/KCDC2003/2020 | Asia / South Korea | 2020-02-23 | Division of Viral Diseases, Center for Laboratory Control of Infectious Diseases, Korea Centers for Diseases Control and Prevention | Division of Viral Diseases, Center for Laboratory Control of Infectious Diseases, Korea Centers for Diseases Control and Prevention | Jeong-Min Kim, Yoon-Seok Chung, Namjoo Lee, Mi-Seon Kim, Sang Hee Woo, Hye-Jun Jo, Sehee Park, Heui Man Kim, Jun-Sub Kim, Junhyeong Jang, Myung Guk Han |
| EPI_ISL_426164 | hCoV-19/Korea/KCDC2004/2020 | Asia / South Korea | 2020-02-02 | Division of Viral Diseases, Center for Laboratory Control of Infectious Diseases, Korea Centers for Diseases Control and Prevention | Division of Viral Diseases, Center for Laboratory Control of Infectious Diseases, Korea Centers for Diseases Control and Prevention | Jeong-Min Kim, Yoon-Seok Chung, Namjoo Lee, Mi-Seon Kim, Sang Hee Woo, Hye-Jun Jo, Sehee Park, Heui Man Kim, Jun-Sub Kim, Junhyeong Jang, Dong Hyun Song, Daesang Lee, Seong Tae Jeong, Myung Guk Han |
| EPI_ISL_426166 | hCoV-19/Korea/KCDC2005/2020 | Asia / South Korea | 2020-02-04 | Division of Viral Diseases, Center for Laboratory Control of Infectious Diseases, Korea Centers for Diseases Control and Prevention | Division of Viral Diseases, Center for Laboratory Control of Infectious Diseases, Korea Centers for Diseases Control and Prevention | Jeong-Min Kim, Yoon-Seok Chung, Namjoo Lee, Mi-Seon Kim, Sang Hee Woo, Hye-Jun Jo, Sehee Park, Heui Man Kim, Jun-Sub Kim, Junhyeong Jang, Myung Guk Han |
| EPI_ISL_426167 | hCoV-19/Korea/KCDC2003/2020 | Asia / South Korea | 2020-02-23 | Division of Viral Diseases, Center for Laboratory Control of Infectious Diseases, Korea Centers for Diseases Control and Prevention | Division of Viral Diseases, Center for Laboratory Control of Infectious Diseases, Korea Centers for Diseases Control and Prevention | Jeong-Min Kim, Yoon-Seok Chung, Namjoo Lee, Mi-Seon Kim, Sang Hee Woo, Hye-Jun Jo, Sehee Park, Heui Man Kim, Jun-Sub Kim, Junhyeong Jang, Myung Guk Han |
| EPI_ISL_426168 | hCoV-19/Korea/KCDC2006/2020 | Asia / South Korea | 2020-02-02 | Division of Viral Diseases, Center for Laboratory Control of Infectious Diseases, Korea Centers for Diseases Control and Prevention | Division of Viral Diseases, Center for Laboratory Control of Infectious Diseases, Korea Centers for Diseases Control and Prevention | Jeong-Min Kim, Yoon-Seok Chung, Namjoo Lee, Mi-Seon Kim, Sang Hee Woo, Hye-Jun Jo, Sehee Park, Heui Man Kim, Jun-Sub Kim, Junhyeong Jang, Myung Guk Han |
| EPI_ISL_426169 | hCoV-19/Korea/KCDC2007/2020 | Asia / South Korea | 2020-02-05 | Division of Viral Diseases, Center for Laboratory Control of Infectious Diseases, Korea Centers for Diseases Control and Prevention | Division of Viral Diseases, Center for Laboratory Control of Infectious Diseases, Korea Centers for Diseases Control and Prevention | Jeong-Min Kim, Yoon-Seok Chung, Namjoo Lee, Mi-Seon Kim, Sang Hee Woo, Hye-Jun Jo, Sehee Park, Heui Man Kim, Jun-Sub Kim, Junhyeong Jang, Dong Hyun Song, Daesang Lee, Seong Tae Jeong, Myung Guk Han |
| EPI_ISL_426171 | hCoV-19/Korea/KCDC2008/2020 | Asia / South Korea | 2020-02-05 | Division of Viral Diseases, Center for Laboratory Control of Infectious Diseases, Korea Centers for Diseases Control and Prevention | Division of Viral Diseases, Center for Laboratory Control of Infectious Diseases, Korea Centers for Diseases Control and Prevention | Jeong-Min Kim, Yoon-Seok Chung, Namjoo Lee, Mi-Seon Kim, Sang Hee Woo, Hye-Jun Jo, Sehee Park, Heui Man Kim, Jun-Sub Kim, Junhyeong Jang, Dong Hyun Song, Daesang Lee, Seong Tae Jeong, Myung Guk Han |
| EPI_ISL_426173 | hCoV-19/Korea/KCDC2009/2020 | Asia / South Korea | 2020-02-07 | Division of Viral Diseases, Center for Laboratory Control of Infectious Diseases, Korea Centers for Diseases Control and Prevention | Division of Viral Diseases, Center for Laboratory Control of Infectious Diseases, Korea Centers for Diseases Control and Prevention | Jeong-Min Kim, Yoon-Seok Chung, Namjoo Lee, Mi-Seon Kim, Sang Hee Woo, Hye-Jun Jo, Sehee Park, Heui Man Kim, Jun-Sub Kim, Junhyeong Jang, Myung Guk Han |
| EPI_ISL_426179 | hCoV-19/India/c31/2020 | Asia / India | 2020-03-02 | National Influenza Center, Indian Council of Medical Research - National Institute of Virology | Indian Council of Medical Research-National Institute of Virology, Microbial Containment Complex | Pragya D. Yadav, Varsha Potdar, Savita Patil, Dimpal A. Nyayanit, Triparna Majumdar, Manohar. L. Chaudhary, Gururaj Deshpande, Padinjaremattathil Thankappan Ullas, Anita Shete-Aich, Hitesh Dighe, Sreelekshmy Mohandas, Gajanan Sapkal, Atanu Basu, Amita Jain, Bharti Malhotra, Deepika Chaudhary, Sarah Cherian, Priya Abraham |
| EPI_ISL_426180 | hCoV-19/Korea/KCDC2010/2020 | Asia / South Korea | 2020-02-05 | Division of Viral Diseases, Center for Laboratory Control of Infectious Diseases, Korea Centers for Diseases Control and Prevention | Division of Viral Diseases, Center for Laboratory Control of Infectious Diseases, Korea Centers for Diseases Control and Prevention | Jeong-Min Kim, Yoon-Seok Chung, Namjoo Lee, Mi-Seon Kim, Sang Hee Woo, Hye-Jun Jo, Sehee Park, Heui Man Kim, Jun-Sub Kim, Junhyeong Jang, Myung Guk Han |
| EPI_ISL_426181 | hCoV-19/Korea/KCDC2011/2020 | Asia / South Korea | 2020-02-23 | Division of Viral Diseases, Center for Laboratory Control of Infectious Diseases, Korea Centers for Diseases Control and Prevention | Division of Viral Diseases, Center for Laboratory Control of Infectious Diseases, Korea Centers for Diseases Control and Prevention | Jeong-Min Kim, Yoon-Seok Chung, Namjoo Lee, Mi-Seon Kim, Sang Hee Woo, Hye-Jun Jo, Sehee Park, Heui Man Kim, Jun-Sub Kim, Junhyeong Jang, Myung Guk Han |
| EPI_ISL_426182 | hCoV-19/Korea/KCDC2012/2020 | Asia / South Korea | 2020-02-09 | Division of Viral Diseases, Center for Laboratory Control of Infectious Diseases, Korea Centers for Diseases Control and Prevention | Division of Viral Diseases, Center for Laboratory Control of Infectious Diseases, Korea Centers for Diseases Control and Prevention | Jeong-Min Kim, Yoon-Seok Chung, Namjoo Lee, Mi-Seon Kim, Sang Hee Woo, Hye-Jun Jo, Sehee Park, Heui Man Kim, Jun-Sub Kim, Junhyeong Jang, Myung Guk Han |
| EPI_ISL_426183 | hCoV-19/Korea/KCDC2013/2020 | Asia / South Korea | 2020-02-23 | Division of Viral Diseases, Center for Laboratory Control of Infectious Diseases, Korea Centers for Diseases Control and Prevention | Division of Viral Diseases, Center for Laboratory Control of Infectious Diseases, Korea Centers for Diseases Control and Prevention | Jeong-Min Kim, Yoon-Seok Chung, Namjoo Lee, Mi-Seon Kim, Sang Hee Woo, Hye-Jun Jo, Sehee Park, Heui Man Kim, Jun-Sub Kim, Junhyeong Jang, Myung Guk Han |
| EPI_ISL_426187 | hCoV-19/Korea/KCDC2014/2020 | Asia / South Korea | 2020-02-12 | Division of Viral Diseases, Center for Laboratory Control of Infectious Diseases, Korea Centers for Diseases Control and Prevention | Division of Viral Diseases, Center for Laboratory Control of Infectious Diseases, Korea Centers for Diseases Control and Prevention | Jeong-Min Kim, Yoon-Seok Chung, Namjoo Lee, Mi-Seon Kim, Sang Hee Woo, Hye-Jun Jo, Sehee Park, Heui Man Kim, Jun-Sub Kim, Junhyeong Jang, Myung Guk Han |
| EPI_ISL_426800 | hCoV-19/Australia/VIC534/2020 | Oceania / Australia / Victoria | 2020-03-25 | Victorian Infectious Diseases Reference Laboratory (VIDRL) | Microbiological Diagnostic Unit Public Health Laboratory and Victorian Infectious Diseases Reference Laboratory, Doherty Institute | Caly L., Seemann T., Sait, M., Schultz M., Druce J., Sherry, N. |
| EPI_ISL_426801 | hCoV-19/Australia/VIC535/2020 | Oceania / Australia / Victoria | 2020-03-25 | Victorian Infectious Diseases Reference Laboratory (VIDRL) | Microbiological Diagnostic Unit Public Health Laboratory and Victorian Infectious Diseases Reference Laboratory, Doherty Institute | Caly L., Seemann T., Sait, M., Schultz M., Druce J., Sherry, N. |
| EPI_ISL_426802 | hCoV-19/Australia/VIC536/2020 | Oceania / Australia / Victoria | 2020-03-24 | Victorian Infectious Diseases Reference Laboratory (VIDRL) | Microbiological Diagnostic Unit Public Health Laboratory and Victorian Infectious Diseases Reference Laboratory, Doherty Institute | Caly L., Seemann T., Sait, M., Schultz M., Druce J., Sherry, N. |
| EPI_ISL_426803 | hCoV-19/Australia/VIC537/2020 | Oceania / Australia / Victoria | 2020-03-24 | Victorian Infectious Diseases Reference Laboratory (VIDRL) | Microbiological Diagnostic Unit Public Health Laboratory and Victorian Infectious Diseases Reference Laboratory, Doherty Institute | Caly L., Seemann T., Sait, M., Schultz M., Druce J., Sherry, N. |
| EPI_ISL_426804 | hCoV-19/Australia/VIC538/2020 | Oceania / Australia / Victoria | 2020-03-23 | Victorian Infectious Diseases Reference Laboratory (VIDRL) | Microbiological Diagnostic Unit Public Health Laboratory and Victorian Infectious Diseases Reference Laboratory, Doherty Institute | Caly L., Seemann T., Sait, M., Schultz M., Druce J., Sherry, N. |
| EPI_ISL_426805 | hCoV-19/Australia/VIC540/2020 | Oceania / Australia / Victoria | 2020-03-24 | Victorian Infectious Diseases Reference Laboratory (VIDRL) | Microbiological Diagnostic Unit Public Health Laboratory and Victorian Infectious Diseases Reference Laboratory, Doherty Institute | Caly L., Seemann T., Sait, M., Schultz M., Druce J., Sherry, N. |
| EPI_ISL_426806 | hCoV-19/Australia/VIC541/2020 | Oceania / Australia / Victoria | 2020-03-24 | Victorian Infectious Diseases Reference Laboratory (VIDRL) | Microbiological Diagnostic Unit Public Health Laboratory and Victorian Infectious Diseases Reference Laboratory, Doherty Institute | Caly L., Seemann T., Sait, M., Schultz M., Druce J., Sherry, N. |
| EPI_ISL_426807 | hCoV-19/Australia/VIC544/2020 | Oceania / Australia / Victoria | 2020-03-23 | Victorian Infectious Diseases Reference Laboratory (VIDRL) | Microbiological Diagnostic Unit Public Health Laboratory and Victorian Infectious Diseases Reference Laboratory, Doherty Institute | Caly L., Seemann T., Sait, M., Schultz M., Druce J., Sherry, N. |
| EPI_ISL_426809 | hCoV-19/Australia/VIC546/2020 | Oceania / Australia / Victoria | 2020-03-24 | Victorian Infectious Diseases Reference Laboratory (VIDRL) | Microbiological Diagnostic Unit Public Health Laboratory and Victorian Infectious Diseases Reference Laboratory, Doherty Institute | Caly L., Seemann T., Sait, M., Schultz M., Druce J., Sherry, N. |
| EPI_ISL_426810 | hCoV-19/Australia/VIC547/2020 | Oceania / Australia / Victoria | 2020-03-25 | Victorian Infectious Diseases Reference Laboratory (VIDRL) | Microbiological Diagnostic Unit Public Health Laboratory and Victorian Infectious Diseases Reference Laboratory, Doherty Institute | Caly L., Seemann T., Sait, M., Schultz M., Druce J., Sherry, N. |
| EPI_ISL_426811 | hCoV-19/Australia/VIC548/2020 | Oceania / Australia / Victoria | 2020-03-25 | Victorian Infectious Diseases Reference Laboratory (VIDRL) | Microbiological Diagnostic Unit Public Health Laboratory and Victorian Infectious Diseases Reference Laboratory, Doherty Institute | Caly L., Seemann T., Sait, M., Schultz M., Druce J., Sherry, N. |
| EPI_ISL_426812 | hCoV-19/Australia/VIC549/2020 | Oceania / Australia / Victoria | 2020-03-23 | Victorian Infectious Diseases Reference Laboratory (VIDRL) | Microbiological Diagnostic Unit Public Health Laboratory and Victorian Infectious Diseases Reference Laboratory, Doherty Institute | Caly L., Seemann T., Sait, M., Schultz M., Druce J., Sherry, N. |
| EPI_ISL_426813 | hCoV-19/Australia/VIC551/2020 | Oceania / Australia / Victoria | 2020-03-25 | Victorian Infectious Diseases Reference Laboratory (VIDRL) | Microbiological Diagnostic Unit Public Health Laboratory and Victorian Infectious Diseases Reference Laboratory, Doherty Institute | Caly L., Seemann T., Sait, M., Schultz M., Druce J., Sherry, N. |
| EPI_ISL_426816 | hCoV-19/Australia/VIC555/2020 | Oceania / Australia / Victoria | 2020-03-25 | Victorian Infectious Diseases Reference Laboratory (VIDRL) | Microbiological Diagnostic Unit Public Health Laboratory and Victorian Infectious Diseases Reference Laboratory, Doherty Institute | Caly L., Seemann T., Sait, M., Schultz M., Druce J., Sherry, N. |
| EPI_ISL_426817 | hCoV-19/Australia/VIC556/2020 | Oceania / Australia / Victoria | 2020-03-25 | Victorian Infectious Diseases Reference Laboratory (VIDRL) | Microbiological Diagnostic Unit Public Health Laboratory and Victorian Infectious Diseases Reference Laboratory, Doherty Institute | Caly L., Seemann T., Sait, M., Schultz M., Druce J., Sherry, N. |
| EPI_ISL_426819 | hCoV-19/Australia/VIC558/2020 | Oceania / Australia / Victoria | 2020-03-25 | Victorian Infectious Diseases Reference Laboratory (VIDRL) | Microbiological Diagnostic Unit Public Health Laboratory and Victorian Infectious Diseases Reference Laboratory, Doherty Institute | Caly L., Seemann T., Sait, M., Schultz M., Druce J., Sherry, N. |
| EPI_ISL_426821 | hCoV-19/Australia/VIC560/2020 | Oceania / Australia / Victoria | 2020-03-25 | Victorian Infectious Diseases Reference Laboratory (VIDRL) | Microbiological Diagnostic Unit Public Health Laboratory and Victorian Infectious Diseases Reference Laboratory, Doherty Institute | Caly L., Seemann T., Sait, M., Schultz M., Druce J., Sherry, N. |
| EPI_ISL_426822 | hCoV-19/Australia/VIC561/2020 | Oceania / Australia / Victoria | 2020-03-25 | Victorian Infectious Diseases Reference Laboratory (VIDRL) | Microbiological Diagnostic Unit Public Health Laboratory and Victorian Infectious Diseases Reference Laboratory, Doherty Institute | Caly L., Seemann T., Sait, M., Schultz M., Druce J., Sherry, N. |
| EPI_ISL_426824 | hCoV-19/Australia/VIC563/2020 | Oceania / Australia / Victoria | 2020-03-24 | Victorian Infectious Diseases Reference Laboratory (VIDRL) | Microbiological Diagnostic Unit Public Health Laboratory and Victorian Infectious Diseases Reference Laboratory, Doherty Institute | Caly L., Seemann T., Sait, M., Schultz M., Druce J., Sherry, N. |
| EPI_ISL_426825 | hCoV-19/Australia/VIC565/2020 | Oceania / Australia / Victoria | 2020-03-25 | Victorian Infectious Diseases Reference Laboratory (VIDRL) | Microbiological Diagnostic Unit Public Health Laboratory and Victorian Infectious Diseases Reference Laboratory, Doherty Institute | Caly L., Seemann T., Sait, M., Schultz M., Druce J., Sherry, N. |
| EPI_ISL_426826 | hCoV-19/Australia/VIC566/2020 | Oceania / Australia / Victoria | 2020-03-25 | Victorian Infectious Diseases Reference Laboratory (VIDRL) | Microbiological Diagnostic Unit Public Health Laboratory and Victorian Infectious Diseases Reference Laboratory, Doherty Institute | Caly L., Seemann T., Sait, M., Schultz M., Druce J., Sherry, N. |
| EPI_ISL_426827 | hCoV-19/Australia/VIC567/2020 | Oceania / Australia / Victoria | 2020-03-25 | Victorian Infectious Diseases Reference Laboratory (VIDRL) | Microbiological Diagnostic Unit Public Health Laboratory and Victorian Infectious Diseases Reference Laboratory, Doherty Institute | Caly L., Seemann T., Sait, M., Schultz M., Druce J., Sherry, N. |
| EPI_ISL_426828 | hCoV-19/Australia/VIC568/2020 | Oceania / Australia / Victoria | 2020-03-25 | Victorian Infectious Diseases Reference Laboratory (VIDRL) | Microbiological Diagnostic Unit Public Health Laboratory and Victorian Infectious Diseases Reference Laboratory, Doherty Institute | Caly L., Seemann T., Sait, M., Schultz M., Druce J., Sherry, N. |
| EPI_ISL_426831 | hCoV-19/Australia/VIC571/2020 | Oceania / Australia / Victoria | 2020-03-25 | Victorian Infectious Diseases Reference Laboratory (VIDRL) | Microbiological Diagnostic Unit Public Health Laboratory and Victorian Infectious Diseases Reference Laboratory, Doherty Institute | Caly L., Seemann T., Sait, M., Schultz M., Druce J., Sherry, N. |
| EPI_ISL_426832 | hCoV-19/Australia/VIC572/2020 | Oceania / Australia / Victoria | 2020-03-25 | Victorian Infectious Diseases Reference Laboratory (VIDRL) | Microbiological Diagnostic Unit Public Health Laboratory and Victorian Infectious Diseases Reference Laboratory, Doherty Institute | Caly L., Seemann T., Sait, M., Schultz M., Druce J., Sherry, N. |
| EPI_ISL_426833 | hCoV-19/Australia/VIC574/2020 | Oceania / Australia / Victoria | 2020-03-26 | Victorian Infectious Diseases Reference Laboratory (VIDRL) | Microbiological Diagnostic Unit Public Health Laboratory and Victorian Infectious Diseases Reference Laboratory, Doherty Institute | Caly L., Seemann T., Sait, M., Schultz M., Druce J., Sherry, N. |
| EPI_ISL_426835 | hCoV-19/Australia/VIC576/2020 | Oceania / Australia / Victoria | 2020-03-26 | Victorian Infectious Diseases Reference Laboratory (VIDRL) | Microbiological Diagnostic Unit Public Health Laboratory and Victorian Infectious Diseases Reference Laboratory, Doherty Institute | Caly L., Seemann T., Sait, M., Schultz M., Druce J., Sherry, N. |
| EPI_ISL_426836 | hCoV-19/Australia/VIC578/2020 | Oceania / Australia / Victoria | 2020-03-26 | Victorian Infectious Diseases Reference Laboratory (VIDRL) | Microbiological Diagnostic Unit Public Health Laboratory and Victorian Infectious Diseases Reference Laboratory, Doherty Institute | Caly L., Seemann T., Sait, M., Schultz M., Druce J., Sherry, N. |
| EPI_ISL_426839 | hCoV-19/Australia/VIC582/2020 | Oceania / Australia / Victoria | 2020-03-25 | Victorian Infectious Diseases Reference Laboratory (VIDRL) | Microbiological Diagnostic Unit Public Health Laboratory and Victorian Infectious Diseases Reference Laboratory, Doherty Institute | Caly L., Seemann T., Sait, M., Schultz M., Druce J., Sherry, N. |
| EPI_ISL_426840 | hCoV-19/Australia/VIC583/2020 | Oceania / Australia / Victoria | 2020-03-25 | Victorian Infectious Diseases Reference Laboratory (VIDRL) | Microbiological Diagnostic Unit Public Health Laboratory and Victorian Infectious Diseases Reference Laboratory, Doherty Institute | Caly L., Seemann T., Sait, M., Schultz M., Druce J., Sherry, N. |
| EPI_ISL_426841 | hCoV-19/Australia/VIC584/2020 | Oceania / Australia / Victoria | 2020-03-25 | Victorian Infectious Diseases Reference Laboratory (VIDRL) | Microbiological Diagnostic Unit Public Health Laboratory and Victorian Infectious Diseases Reference Laboratory, Doherty Institute | Caly L., Seemann T., Sait, M., Schultz M., Druce J., Sherry, N. |
| EPI_ISL_426842 | hCoV-19/Australia/VIC585/2020 | Oceania / Australia / Victoria | 2020-03-25 | Victorian Infectious Diseases Reference Laboratory (VIDRL) | Microbiological Diagnostic Unit Public Health Laboratory and Victorian Infectious Diseases Reference Laboratory, Doherty Institute | Caly L., Seemann T., Sait, M., Schultz M., Druce J., Sherry, N. |
| EPI_ISL_426844 | hCoV-19/Australia/VIC587/2020 | Oceania / Australia / Victoria | 2020-03-25 | Victorian Infectious Diseases Reference Laboratory (VIDRL) | Microbiological Diagnostic Unit Public Health Laboratory and Victorian Infectious Diseases Reference Laboratory, Doherty Institute | Caly L., Seemann T., Sait, M., Schultz M., Druce J., Sherry, N. |
| EPI_ISL_426846 | hCoV-19/Australia/VIC589/2020 | Oceania / Australia / Victoria | 2020-03-22 | Victorian Infectious Diseases Reference Laboratory (VIDRL) | Microbiological Diagnostic Unit Public Health Laboratory and Victorian Infectious Diseases Reference Laboratory, Doherty Institute | Caly L., Seemann T., Sait, M., Schultz M., Druce J., Sherry, N. |
| EPI_ISL_426847 | hCoV-19/Australia/VIC590/2020 | Oceania / Australia / Victoria | 2020-03-22 | Victorian Infectious Diseases Reference Laboratory (VIDRL) | Microbiological Diagnostic Unit Public Health Laboratory and Victorian Infectious Diseases Reference Laboratory, Doherty Institute | Caly L., Seemann T., Sait, M., Schultz M., Druce J., Sherry, N. |
| EPI_ISL_426848 | hCoV-19/Australia/VIC591/2020 | Oceania / Australia / Victoria | 2020-03-22 | Victorian Infectious Diseases Reference Laboratory (VIDRL) | Microbiological Diagnostic Unit Public Health Laboratory and Victorian Infectious Diseases Reference Laboratory, Doherty Institute | Caly L., Seemann T., Sait, M., Schultz M., Druce J., Sherry, N. |
| EPI_ISL_426850 | hCoV-19/Australia/VIC593/2020 | Oceania / Australia / Victoria | 2020-03-23 | Victorian Infectious Diseases Reference Laboratory (VIDRL) | Microbiological Diagnostic Unit Public Health Laboratory and Victorian Infectious Diseases Reference Laboratory, Doherty Institute | Caly L., Seemann T., Sait, M., Schultz M., Druce J., Sherry, N. |
| EPI_ISL_426851 | hCoV-19/Australia/VIC594/2020 | Oceania / Australia / Victoria | 2020-03-25 | Victorian Infectious Diseases Reference Laboratory (VIDRL) | Microbiological Diagnostic Unit Public Health Laboratory and Victorian Infectious Diseases Reference Laboratory, Doherty Institute | Caly L., Seemann T., Sait, M., Schultz M., Druce J., Sherry, N. |
| EPI_ISL_426852 | hCoV-19/Australia/VIC595/2020 | Oceania / Australia / Victoria | 2020-03-25 | Victorian Infectious Diseases Reference Laboratory (VIDRL) | Microbiological Diagnostic Unit Public Health Laboratory and Victorian Infectious Diseases Reference Laboratory, Doherty Institute | Caly L., Seemann T., Sait, M., Schultz M., Druce J., Sherry, N. |
| EPI_ISL_426853 | hCoV-19/Australia/VIC596/2020 | Oceania / Australia / Victoria | 2020-03-25 | Victorian Infectious Diseases Reference Laboratory (VIDRL) | Microbiological Diagnostic Unit Public Health Laboratory and Victorian Infectious Diseases Reference Laboratory, Doherty Institute | Caly L., Seemann T., Sait, M., Schultz M., Druce J., Sherry, N. |
| EPI_ISL_426854 | hCoV-19/Australia/VIC597/2020 | Oceania / Australia / Victoria | 2020-03-25 | Victorian Infectious Diseases Reference Laboratory (VIDRL) | Microbiological Diagnostic Unit Public Health Laboratory and Victorian Infectious Diseases Reference Laboratory, Doherty Institute | Caly L., Seemann T., Sait, M., Schultz M., Druce J., Sherry, N. |
| EPI_ISL_426855 | hCoV-19/Australia/VIC598/2020 | Oceania / Australia / Victoria | 2020-03-25 | Victorian Infectious Diseases Reference Laboratory (VIDRL) | Microbiological Diagnostic Unit Public Health Laboratory and Victorian Infectious Diseases Reference Laboratory, Doherty Institute | Caly L., Seemann T., Sait, M., Schultz M., Druce J., Sherry, N. |
| EPI_ISL_426856 | hCoV-19/Australia/VIC599/2020 | Oceania / Australia / Victoria | 2020-03-25 | Victorian Infectious Diseases Reference Laboratory (VIDRL) | Microbiological Diagnostic Unit Public Health Laboratory and Victorian Infectious Diseases Reference Laboratory, Doherty Institute | Caly L., Seemann T., Sait, M., Schultz M., Druce J., Sherry, N. |
| EPI_ISL_426858 | hCoV-19/Australia/VIC603/2020 | Oceania / Australia / Victoria | 2020-03-26 | Victorian Infectious Diseases Reference Laboratory (VIDRL) | Microbiological Diagnostic Unit Public Health Laboratory and Victorian Infectious Diseases Reference Laboratory, Doherty Institute | Caly L., Seemann T., Sait, M., Schultz M., Druce J., Sherry, N. |
| EPI_ISL_426860 | hCoV-19/Australia/VIC605/2020 | Oceania / Australia / Victoria | 2020-03-27 | Victorian Infectious Diseases Reference Laboratory (VIDRL) | Microbiological Diagnostic Unit Public Health Laboratory and Victorian Infectious Diseases Reference Laboratory, Doherty Institute | Caly L., Seemann T., Sait, M., Schultz M., Druce J., Sherry, N. |
| EPI_ISL_426861 | hCoV-19/Australia/VIC606/2020 | Oceania / Australia / Victoria | 2020-03-27 | Victorian Infectious Diseases Reference Laboratory (VIDRL) | Microbiological Diagnostic Unit Public Health Laboratory and Victorian Infectious Diseases Reference Laboratory, Doherty Institute | Caly L., Seemann T., Sait, M., Schultz M., Druce J., Sherry, N. |
| EPI_ISL_426862 | hCoV-19/Australia/VIC607/2020 | Oceania / Australia / Victoria | 2020-03-27 | Victorian Infectious Diseases Reference Laboratory (VIDRL) | Microbiological Diagnostic Unit Public Health Laboratory and Victorian Infectious Diseases Reference Laboratory, Doherty Institute | Caly L., Seemann T., Sait, M., Schultz M., Druce J., Sherry, N. |
| EPI_ISL_426864 | hCoV-19/Australia/VIC610/2020 | Oceania / Australia / Victoria | 2020-03-26 | Victorian Infectious Diseases Reference Laboratory (VIDRL) | Microbiological Diagnostic Unit Public Health Laboratory and Victorian Infectious Diseases Reference Laboratory, Doherty Institute | Caly L., Seemann T., Sait, M., Schultz M., Druce J., Sherry, N. |
| EPI_ISL_426865 | hCoV-19/Australia/VIC611/2020 | Oceania / Australia / Victoria | 2020-03-27 | Victorian Infectious Diseases Reference Laboratory (VIDRL) | Microbiological Diagnostic Unit Public Health Laboratory and Victorian Infectious Diseases Reference Laboratory, Doherty Institute | Caly L., Seemann T., Sait, M., Schultz M., Druce J., Sherry, N. |
| EPI_ISL_426866 | hCoV-19/Australia/VIC612/2020 | Oceania / Australia / Victoria | 2020-03-26 | Victorian Infectious Diseases Reference Laboratory (VIDRL) | Microbiological Diagnostic Unit Public Health Laboratory and Victorian Infectious Diseases Reference Laboratory, Doherty Institute | Caly L., Seemann T., Sait, M., Schultz M., Druce J., Sherry, N. |
| EPI_ISL_426870 | hCoV-19/Australia/VIC615/2020 | Oceania / Australia / Victoria | 2020-03-28 | Victorian Infectious Diseases Reference Laboratory (VIDRL) | Microbiological Diagnostic Unit Public Health Laboratory and Victorian Infectious Diseases Reference Laboratory, Doherty Institute | Caly L., Seemann T., Sait, M., Schultz M., Druce J., Sherry, N. |
| EPI_ISL_426872 | hCoV-19/Australia/VIC617/2020 | Oceania / Australia / Victoria | 2020-03-26 | Victorian Infectious Diseases Reference Laboratory (VIDRL) | Microbiological Diagnostic Unit Public Health Laboratory and Victorian Infectious Diseases Reference Laboratory, Doherty Institute | Caly L., Seemann T., Sait, M., Schultz M., Druce J., Sherry, N. |
| EPI_ISL_426873 | hCoV-19/Australia/VIC629/2020 | Oceania / Australia / Victoria | 2020-03-25 | Victorian Infectious Diseases Reference Laboratory (VIDRL) | Microbiological Diagnostic Unit Public Health Laboratory and Victorian Infectious Diseases Reference Laboratory, Doherty Institute | Caly L., Seemann T., Sait, M., Schultz M., Druce J., Sherry, N. |
| EPI_ISL_426874 | hCoV-19/Australia/VIC630/2020 | Oceania / Australia / Victoria | 2020-03-26 | Victorian Infectious Diseases Reference Laboratory (VIDRL) | Microbiological Diagnostic Unit Public Health Laboratory and Victorian Infectious Diseases Reference Laboratory, Doherty Institute | Caly L., Seemann T., Sait, M., Schultz M., Druce J., Sherry, N. |
| EPI_ISL_426875 | hCoV-19/Australia/VIC633/2020 | Oceania / Australia / Victoria | 2020-03-27 | Victorian Infectious Diseases Reference Laboratory (VIDRL) | Microbiological Diagnostic Unit Public Health Laboratory and Victorian Infectious Diseases Reference Laboratory, Doherty Institute | Caly L., Seemann T., Sait, M., Schultz M., Druce J., Sherry, N. |
| EPI_ISL_426876 | hCoV-19/Australia/VIC635/2020 | Oceania / Australia / Victoria | 2020-03-28 | Victorian Infectious Diseases Reference Laboratory (VIDRL) | Microbiological Diagnostic Unit Public Health Laboratory and Victorian Infectious Diseases Reference Laboratory, Doherty Institute | Caly L., Seemann T., Sait, M., Schultz M., Druce J., Sherry, N. |
| EPI_ISL_426877 | hCoV-19/Australia/VIC636/2020 | Oceania / Australia / Victoria | 2020-03-28 | Victorian Infectious Diseases Reference Laboratory (VIDRL) | Microbiological Diagnostic Unit Public Health Laboratory and Victorian Infectious Diseases Reference Laboratory, Doherty Institute | Caly L., Seemann T., Sait, M., Schultz M., Druce J., Sherry, N. |
| EPI_ISL_426878 | hCoV-19/Australia/VIC701/2020 | Oceania / Australia / Victoria | 2020-03-21 | Victorian Infectious Diseases Reference Laboratory (VIDRL) | Microbiological Diagnostic Unit Public Health Laboratory and Victorian Infectious Diseases Reference Laboratory, Doherty Institute | Caly L., Seemann T., Sait, M., Schultz M., Druce J., Sherry, N. |
| EPI_ISL_426879 | hCoV-19/Australia/VIC908/2020 | Oceania / Australia / Victoria | 2020-03-23 | Victorian Infectious Diseases Reference Laboratory (VIDRL) | Microbiological Diagnostic Unit Public Health Laboratory and Victorian Infectious Diseases Reference Laboratory, Doherty Institute | Caly L., Seemann T., Sait, M., Schultz M., Druce J., Sherry, N. |
| EPI_ISL_426880 | hCoV-19/Australia/VIC909/2020 | Oceania / Australia / Victoria | 2020-03-23 | Victorian Infectious Diseases Reference Laboratory (VIDRL) | Microbiological Diagnostic Unit Public Health Laboratory and Victorian Infectious Diseases Reference Laboratory, Doherty Institute | Caly L., Seemann T., Sait, M., Schultz M., Druce J., Sherry, N. |
| EPI_ISL_426881 | hCoV-19/Australia/VIC912/2020 | Oceania / Australia / Victoria | 2020-03-24 | Victorian Infectious Diseases Reference Laboratory (VIDRL) | Microbiological Diagnostic Unit Public Health Laboratory and Victorian Infectious Diseases Reference Laboratory, Doherty Institute | Caly L., Seemann T., Sait, M., Schultz M., Druce J., Sherry, N. |
| EPI_ISL_426882 | hCoV-19/Australia/VIC913/2020 | Oceania / Australia / Victoria | 2020-03-24 | Victorian Infectious Diseases Reference Laboratory (VIDRL) | Microbiological Diagnostic Unit Public Health Laboratory and Victorian Infectious Diseases Reference Laboratory, Doherty Institute | Caly L., Seemann T., Sait, M., Schultz M., Druce J., Sherry, N. |
| EPI_ISL_426883 | hCoV-19/Czech Republic/IAB_1/2020 | Europe / Czech Republic | 2020-03-27 | Motol University Hospital | Institute of Applied Biotechnologies a.s. | Petr Brož, Jan Geryk, Petr Klempt, Martin Kalný, Adam Novotný, Kateřina Kvapilová, Pavel Dřevínek, Petr Kvapil, Milan Macek |
| EPI_ISL_426884 | hCoV-19/Czech Republic/IAB_4/2020 | Europe / Czech Republic | 2020-03-27 | Motol University Hospital | Institute of Applied Biotechnologies a.s. | Petr Brož, Jan Geryk, Petr Klempt, Martin Kalný, Adam Novotný, Kateřina Kvapilová, Pavel Dřevínek, Petr Kvapil, Milan Macek |
| EPI_ISL_426885 | hCoV-19/Czech Republic/IAB_8/2020 | Europe / Czech Republic | 2020-03-27 | Motol University Hospital | Institute of Applied Biotechnologies a.s. | Petr Brož, Jan Geryk, Petr Klempt, Martin Kalný, Adam Novotný, Kateřina Kvapilová, Pavel Dřevínek, Petr Kvapil, Milan Macek |
| EPI_ISL_426886 | hCoV-19/Czech Republic/IAB_9/2020 | Europe / Czech Republic | 2020-03-28 | Motol University Hospital | Institute of Applied Biotechnologies a.s. | Petr Brož, Jan Geryk, Petr Klempt, Martin Kalný, Adam Novotný, Kateřina Kvapilová, Pavel Dřevínek, Petr Kvapil, Milan Macek |
| EPI_ISL_426887 | hCoV-19/Czech Republic/IAB_10/2020 | Europe / Czech Republic | 2020-03-31 | Motol University Hospital | Institute of Applied Biotechnologies a.s. | Petr Brož, Jan Geryk, Petr Klempt, Martin Kalný, Adam Novotný, Kateřina Kvapilová, Pavel Dřevínek, Petr Kvapil, Milan Macek |
| EPI_ISL_426888 | hCoV-19/Czech Republic/IAB_12/2020 | Europe / Czech Republic | 2020-03-31 | Motol University Hospital | Institute of Applied Biotechnologies a.s. | Petr Brož, Jan Geryk, Petr Klempt, Martin Kalný, Adam Novotný, Kateřina Kvapilová, Pavel Dřevínek, Petr Kvapil, Milan Macek |
| EPI_ISL_426889 | hCoV-19/Czech Republic/IAB_14/2020 | Europe / Czech Republic | 2020-04-02 | Motol University Hospital | Institute of Applied Biotechnologies a.s. | Petr Brož, Jan Geryk, Petr Klempt, Martin Kalný, Adam Novotný, Kateřina Kvapilová, Pavel Dřevínek, Petr Kvapil, Milan Macek |
| EPI_ISL_426890 | hCoV-19/Czech Republic/IAB_15/2020 | Europe / Czech Republic | 2020-03-26 | Motol University Hospital | Institute of Applied Biotechnologies a.s. | Petr Brož, Jan Geryk, Petr Klempt, Martin Kalný, Adam Novotný, Kateřina Kvapilová, Pavel Dřevínek, Petr Kvapil, Milan Macek |
| EPI_ISL_426891 | hCoV-19/Czech Republic/IAB_16/2020 | Europe / Czech Republic | 2020-03-26 | Motol University Hospital | Institute of Applied Biotechnologies a.s. | Petr Brož, Jan Geryk, Petr Klempt, Martin Kalný, Adam Novotný, Kateřina Kvapilová, Pavel Dřevínek, Petr Kvapil, Milan Macek |
| EPI_ISL_426892 | hCoV-19/Czech Republic/IAB_17/2020 | Europe / Czech Republic | 2020-03-27 | Motol University Hospital | Institute of Applied Biotechnologies a.s. | Petr Brož, Jan Geryk, Petr Klempt, Martin Kalný, Adam Novotný, Kateřina Kvapilová, Pavel Dřevínek, Petr Kvapil, Milan Macek |
| EPI_ISL_426893 | hCoV-19/Czech Republic/IAB_18/2020 | Europe / Czech Republic | 2020-03-27 | Motol University Hospital | Institute of Applied Biotechnologies a.s. | Petr Brož, Jan Geryk, Petr Klempt, Martin Kalný, Adam Novotný, Kateřina Kvapilová, Pavel Dřevínek, Petr Kvapil, Milan Macek |
| EPI_ISL_426894 | hCoV-19/Czech Republic/IAB_20/2020 | Europe / Czech Republic | 2020-03-27 | Motol University Hospital | Institute of Applied Biotechnologies a.s. | Petr Brož, Jan Geryk, Petr Klempt, Martin Kalný, Adam Novotný, Kateřina Kvapilová, Pavel Dřevínek, Petr Kvapil, Milan Macek |
| EPI_ISL_426895 | hCoV-19/Czech Republic/IAB_21/2020 | Europe / Czech Republic | 2020-03-27 | Motol University Hospital | Institute of Applied Biotechnologies a.s. | Petr Brož, Jan Geryk, Petr Klempt, Martin Kalný, Adam Novotný, Kateřina Kvapilová, Pavel Dřevínek, Petr Kvapil, Milan Macek |
| EPI_ISL_426896 | hCoV-19/Czech Republic/IAB_22/2020 | Europe / Czech Republic | 2020-03-28 | Motol University Hospital | Institute of Applied Biotechnologies a.s. | Petr Brož, Jan Geryk, Petr Klempt, Martin Kalný, Adam Novotný, Kateřina Kvapilová, Pavel Dřevínek, Petr Kvapil, Milan Macek |
| EPI_ISL_426897 | hCoV-19/Czech Republic/IAB_23/2020 | Europe / Czech Republic | 2020-03-29 | Motol University Hospital | Institute of Applied Biotechnologies a.s. | Petr Brož, Jan Geryk, Petr Klempt, Martin Kalný, Adam Novotný, Kateřina Kvapilová, Pavel Dřevínek, Petr Kvapil, Milan Macek |
| EPI_ISL_426898 | hCoV-19/Australia/NT10/2020 | Oceania / Australia / Northern territory | 2020-03-26 | Royal Darwin Hospital Pathology | Microbiological Diagnostic Unit Public Health Laboratory and Victorian Infectious Diseases Reference Laboratory, Doherty Institute | Meumann, E., Caly L., Seemann T., Sait, M., Schultz M., Druce J., Sherry, N. |
| EPI_ISL_426899 | hCoV-19/Australia/NT11/2020 | Oceania / Australia / Northern territory | 2020-03-27 | Royal Darwin Hospital Pathology | Microbiological Diagnostic Unit Public Health Laboratory and Victorian Infectious Diseases Reference Laboratory, Doherty Institute | Meumann, E., Caly L., Seemann T., Sait, M., Schultz M., Druce J., Sherry, N. |
| EPI_ISL_426900 | hCoV-19/Australia/NT12/2020 | Oceania / Australia / Northern territory | 2020-03-25 | Royal Darwin Hospital Pathology | Microbiological Diagnostic Unit Public Health Laboratory and Victorian Infectious Diseases Reference Laboratory, Doherty Institute | Meumann, E., Caly L., Seemann T., Sait, M., Schultz M., Druce J., Sherry, N. |
| EPI_ISL_426901 | hCoV-19/Australia/NT13/2020 | Oceania / Australia / Northern territory | 2020-03-25 | Royal Darwin Hospital Pathology | Microbiological Diagnostic Unit Public Health Laboratory and Victorian Infectious Diseases Reference Laboratory, Doherty Institute | Meumann, E., Caly L., Seemann T., Sait, M., Schultz M., Druce J., Sherry, N. |
| EPI_ISL_426904 | hCoV-19/Australia/NT17/2020 | Oceania / Australia / Northern territory | 2020-03-24 | Royal Darwin Hospital Pathology | Microbiological Diagnostic Unit Public Health Laboratory and Victorian Infectious Diseases Reference Laboratory, Doherty Institute | Meumann, E., Caly L., Seemann T., Sait, M., Schultz M., Druce J., Sherry, N. |
| EPI_ISL_426906 | hCoV-19/Australia/VIC473/2020 | Oceania / Australia / Victoria | 2020-03-28 | Microbiological Diagnostic Unit Public Health Laboratory | Microbiological Diagnostic Unit Public Health Laboratory | Seemann T., Schultz M., Sait, M., Sherry, N. |
| EPI_ISL_426907 | hCoV-19/Australia/VIC475/2020 | Oceania / Australia / Victoria | 2020-03-28 | Microbiological Diagnostic Unit Public Health Laboratory | Microbiological Diagnostic Unit Public Health Laboratory | Seemann T., Schultz M., Sait, M., Sherry, N. |
| EPI_ISL_426910 | hCoV-19/Australia/VIC478/2020 | Oceania / Australia / Victoria | 2020-03-28 | Microbiological Diagnostic Unit Public Health Laboratory | Microbiological Diagnostic Unit Public Health Laboratory | Seemann T., Schultz M., Sait, M., Sherry, N. |
| EPI_ISL_427300 | hCoV-19/Brazil/RJ-477/2020 | South America / Brazil / Rio de Janeiro / I | 2020-03-11 | Instituto Oswaldo Cruz FIOCRUZ - Laboratory of Respiratory Viruses and Measles (LVRS) | Instituto Oswaldo Cruz FIOCRUZ - Laboratory of Respiratory Viruses and Measles (LVRS) | Paola Resende, Fernando Motta, Luciana Appolinario, Sunando Roy, Aline Mattos, Milene Miranda, Cristiana Garcia, Braulia Caetano, Maria Ogrzewalska, Priscila Born, Jonathan Lopes, Marilda Siqueira |
| EPI_ISL_427301 | hCoV-19/Brazil/RJ-477V/2020 | South America / Brazil / Rio de Janeiro / I | 2020-03-11 | Instituto Oswaldo Cruz FIOCRUZ - Laboratory of Respiratory Viruses and Measles (LVRS) | Instituto Oswaldo Cruz FIOCRUZ - Laboratory of Respiratory Viruses and Measles (LVRS) | Paola Resende, Fernando Motta, Luciana Appolinario, Sunando Roy, Aline Mattos, Milene Miranda, Cristiana Garcia, Braulia Caetano, Maria Ogrzewalska, Priscila Born, Jonathan Lopes, Marilda Siqueira |
| EPI_ISL_427302 | hCoV-19/Brazil/RJ-763/2020 | South America / Brazil / Rio de Janeiro / I | 2020-03-20 | Instituto Oswaldo Cruz FIOCRUZ - Laboratory of Respiratory Viruses and Measles (LVRS) | Instituto Oswaldo Cruz FIOCRUZ - Laboratory of Respiratory Viruses and Measles (LVRS) | Paola Resende, Fernando Motta, Luciana Appolinario, Sunando Roy, Aline Mattos, Milene Miranda, Cristiana Garcia, Braulia Caetano, Maria Ogrzewalska, Priscila Born, Jonathan Lopes, Marilda Siqueira |
| EPI_ISL_427303 | hCoV-19/Brazil/RJ-818/2020 | South America / Brazil / Rio de Janeiro / I | 2020-03-25 | Instituto Oswaldo Cruz FIOCRUZ - Laboratory of Respiratory Viruses and Measles (LVRS) | Instituto Oswaldo Cruz FIOCRUZ - Laboratory of Respiratory Viruses and Measles (LVRS) | Paola Resende, Fernando Motta, Luciana Appolinario, Sunando Roy, Aline Mattos, Milene Miranda, Cristiana Garcia, Braulia Caetano, Maria Ogrzewalska, Priscila Born, Jonathan Lopes, Marilda Siqueira |
| EPI_ISL_427304 | hCoV-19/Brazil/RJ-872/2020 | South America / Brazil / Rio de Janeiro / I | 2020-03-26 | Instituto Oswaldo Cruz FIOCRUZ - Laboratory of Respiratory Viruses and Measles (LVRS) | Instituto Oswaldo Cruz FIOCRUZ - Laboratory of Respiratory Viruses and Measles (LVRS) | Paola Resende, Fernando Motta, Luciana Appolinario, Sunando Roy, Aline Mattos, Milene Miranda, Cristiana Garcia, Braulia Caetano, Maria Ogrzewalska, Priscila Born, Jonathan Lopes, Marilda Siqueira |
| EPI_ISL_427305 | hCoV-19/Brazil/SC-766/2020 | South America / Brazil / Santa Catarina / | 2020-03-10 | LACEN-SC - Laboratorio Central de Santa Catarina | Instituto Oswaldo Cruz FIOCRUZ - Laboratory of Respiratory Viruses and Measles (LVRS) | Paola Resende, Fernando Motta, Luciana Appolinario, Sunando Roy, Aline Mattos, Milene Miranda, Cristiana Garcia, Braulia Caetano, Maria Ogrzewalska, Priscila Born, Jonathan Lopes, Marilda Siqueira |
| EPI_ISL_427306 | hCoV-19/Brazil/SC-769/2020 | South America / Brazil / Santa Catarina / | 2020-03-10 | LACEN-SC - Laboratorio Central de Santa Catarina | Instituto Oswaldo Cruz FIOCRUZ - Laboratory of Respiratory Viruses and Measles (LVRS) | Paola Resende, Fernando Motta, Luciana Appolinario, Sunando Roy, Aline Mattos, Milene Miranda, Cristiana Garcia, Braulia Caetano, Maria Ogrzewalska, Priscila Born, Jonathan Lopes, Marilda Siqueira |
| EPI_ISL_427307 | hCoV-19/Russia/StPetersburg-RII4144S/2020 | Europe / Russia / Saint-Petersburg | 2020-03-20 | WHO National Influenza Centre Russian Federation | WHO National Influenza Centre Russian Federation | Andrey Komissarov, Artem Fadeev, Mariia Sergeeva, Anna Ivanova, Daria Danilenko |
| EPI_ISL_427308 | hCoV-19/Russia/StPetersburg-RII4307V/2020 | Europe / Russia / Saint-Petersburg | 2020-03-24 | WHO National Influenza Centre Russian Federation | WHO National Influenza Centre Russian Federation | Andrey Komissarov, Artem Fadeev, Mariia Sergeeva, Anna Ivanova, Daria Danilenko |
| EPI_ISL_427309 | hCoV-19/Russia/StPetersburg-RII4308V/2020 | Europe / Russia / Saint-Petersburg | 2020-03-24 | WHO National Influenza Centre Russian Federation | WHO National Influenza Centre Russian Federation | Andrey Komissarov, Artem Fadeev, Mariia Sergeeva, Anna Ivanova, Daria Danilenko |
| EPI_ISL_427310 | hCoV-19/Russia/StPetersburg-RII4382V/2020 | Europe / Russia / Saint-Petersburg | 2020-03-29 | WHO National Influenza Centre Russian Federation | WHO National Influenza Centre Russian Federation | Andrey Komissarov, Artem Fadeev, Mariia Sergeeva, Anna Ivanova, Daria Danilenko |
| EPI_ISL_427311 | hCoV-19/Russia/StPetersburg-RII4384V/2020 | Europe / Russia / Saint-Petersburg | 2020-03-29 | WHO National Influenza Centre Russian Federation | WHO National Influenza Centre Russian Federation | Andrey Komissarov, Artem Fadeev, Mariia Sergeeva, Anna Ivanova, Daria Danilenko |
| EPI_ISL_427312 | hCoV-19/Russia/StPetersburg-RII4386V/2020 | Europe / Russia / Saint-Petersburg | 2020-03-29 | WHO National Influenza Centre Russian Federation | WHO National Influenza Centre Russian Federation | Andrey Komissarov, Artem Fadeev, Mariia Sergeeva, Anna Ivanova, Daria Danilenko |
| EPI_ISL_427313 | hCoV-19/Russia/StPetersburg-RII4532V/2020 | Europe / Russia / Saint-Petersburg | 2020-04-03 | WHO National Influenza Centre Russian Federation | WHO National Influenza Centre Russian Federation | Andrey Komissarov, Artem Fadeev, Mariia Sergeeva, Anna Ivanova, Daria Danilenko |
| EPI_ISL_427314 | hCoV-19/Russia/StPetersburg-RII4546V/2020 | Europe / Russia / Saint-Petersburg | 2020-04-02 | WHO National Influenza Centre Russian Federation | WHO National Influenza Centre Russian Federation | Andrey Komissarov, Artem Fadeev, Mariia Sergeeva, Anna Ivanova, Daria Danilenko |
| EPI_ISL_427316 | hCoV-19/Russia/StPetersburg-RII4584S/2020 | Europe / Russia / Saint-Petersburg | 2020-04-03 | WHO National Influenza Centre Russian Federation | WHO National Influenza Centre Russian Federation | Andrey Komissarov, Artem Fadeev, Mariia Sergeeva, Anna Ivanova, Daria Danilenko |
| EPI_ISL_427317 | hCoV-19/Russia/StPetersburg-RII4585S/2020 | Europe / Russia / Saint-Petersburg | 2020-04-03 | WHO National Influenza Centre Russian Federation | WHO National Influenza Centre Russian Federation | Andrey Komissarov, Artem Fadeev, Mariia Sergeeva, Anna Ivanova, Daria Danilenko |
| EPI_ISL_427318 | hCoV-19/Russia/StPetersburg-RII4591S/2020 | Europe / Russia / Saint-Petersburg | 2020-04-03 | WHO National Influenza Centre Russian Federation | WHO National Influenza Centre Russian Federation | Andrey Komissarov, Artem Fadeev, Mariia Sergeeva, Anna Ivanova, Daria Danilenko |
| EPI_ISL_427319 | hCoV-19/Russia/StPetersburg-RII4597S/2020 | Europe / Russia / Saint-Petersburg | 2020-04-04 | WHO National Influenza Centre Russian Federation | WHO National Influenza Centre Russian Federation | Andrey Komissarov, Artem Fadeev, Mariia Sergeeva, Anna Ivanova, Daria Danilenko |
| EPI_ISL_427320 | hCoV-19/Russia/StPetersburg-RII4637S/2020 | Europe / Russia / Saint-Petersburg | 2020-04-05 | WHO National Influenza Centre Russian Federation | WHO National Influenza Centre Russian Federation | Andrey Komissarov, Artem Fadeev, Mariia Sergeeva, Anna Ivanova, Daria Danilenko |
| EPI_ISL_427321 | hCoV-19/Russia/StPetersburg-RII4649S/2020 | Europe / Russia / Saint-Petersburg | 2020-04-05 | WHO National Influenza Centre Russian Federation | WHO National Influenza Centre Russian Federation | Andrey Komissarov, Artem Fadeev, Mariia Sergeeva, Anna Ivanova, Daria Danilenko |
| EPI_ISL_427322 | hCoV-19/Russia/StPetersburg-RII4655S/2020 | Europe / Russia / Saint-Petersburg | 2020-04-05 | WHO National Influenza Centre Russian Federation | WHO National Influenza Centre Russian Federation | Andrey Komissarov, Artem Fadeev, Mariia Sergeeva, Anna Ivanova, Daria Danilenko |
| EPI_ISL_427323 | hCoV-19/Russia/StPetersburg-RII4678S/2020 | Europe / Russia / Saint-Petersburg | 2020-04-06 | WHO National Influenza Centre Russian Federation | WHO National Influenza Centre Russian Federation | Andrey Komissarov, Artem Fadeev, Mariia Sergeeva, Anna Ivanova, Daria Danilenko |
| EPI_ISL_427324 | hCoV-19/Russia/StPetersburg-RII4686S/2020 | Europe / Russia / Saint-Petersburg | 2020-04-07 | WHO National Influenza Centre Russian Federation | WHO National Influenza Centre Russian Federation | Andrey Komissarov, Artem Fadeev, Mariia Sergeeva, Anna Ivanova, Daria Danilenko |
| EPI_ISL_427325 | hCoV-19/Russia/StPetersburg-RII4693S/2020 | Europe / Russia / Saint-Petersburg | 2020-04-07 | WHO National Influenza Centre Russian Federation | WHO National Influenza Centre Russian Federation | Andrey Komissarov, Artem Fadeev, Mariia Sergeeva, Anna Ivanova, Daria Danilenko |
| EPI_ISL_427326 | hCoV-19/Russia/StPetersburg-RII4694S/2020 | Europe / Russia / Saint-Petersburg | 2020-04-07 | WHO National Influenza Centre Russian Federation | WHO National Influenza Centre Russian Federation | Andrey Komissarov, Artem Fadeev, Mariia Sergeeva, Anna Ivanova, Daria Danilenko |
| EPI_ISL_427327 | hCoV-19/Russia/StPetersburg-RII4696S/2020 | Europe / Russia / Saint-Petersburg | 2020-04-07 | WHO National Influenza Centre Russian Federation | WHO National Influenza Centre Russian Federation | Andrey Komissarov, Artem Fadeev, Mariia Sergeeva, Anna Ivanova, Daria Danilenko |
| EPI_ISL_427328 | hCoV-19/Russia/StPetersburg-RII4697S/2020 | Europe / Russia / Saint-Petersburg | 2020-04-07 | WHO National Influenza Centre Russian Federation | WHO National Influenza Centre Russian Federation | Andrey Komissarov, Artem Fadeev, Mariia Sergeeva, Anna Ivanova, Daria Danilenko |
| EPI_ISL_427329 | hCoV-19/Russia/StPetersburg-RII4707S/2020 | Europe / Russia / Saint-Petersburg | 2020-04-07 | WHO National Influenza Centre Russian Federation | WHO National Influenza Centre Russian Federation | Andrey Komissarov, Artem Fadeev, Mariia Sergeeva, Anna Ivanova, Daria Danilenko |
| EPI_ISL_427330 | hCoV-19/Russia/StPetersburg-RII4711S/2020 | Europe / Russia / Saint-Petersburg | 2020-04-07 | WHO National Influenza Centre Russian Federation | WHO National Influenza Centre Russian Federation | Andrey Komissarov, Artem Fadeev, Mariia Sergeeva, Anna Ivanova, Daria Danilenko |
| EPI_ISL_427331 | hCoV-19/Russia/StPetersburg-RII4713S/2020 | Europe / Russia / Saint-Petersburg | 2020-04-07 | WHO National Influenza Centre Russian Federation | WHO National Influenza Centre Russian Federation | Andrey Komissarov, Artem Fadeev, Mariia Sergeeva, Anna Ivanova, Daria Danilenko |
| EPI_ISL_427332 | hCoV-19/Russia/StPetersburg-RII4723S/2020 | Europe / Russia / Saint-Petersburg | 2020-04-07 | WHO National Influenza Centre Russian Federation | WHO National Influenza Centre Russian Federation | Andrey Komissarov, Artem Fadeev, Mariia Sergeeva, Anna Ivanova, Daria Danilenko |
| EPI_ISL_427333 | hCoV-19/Russia/StPetersburg-RII4724S/2020 | Europe / Russia / Saint-Petersburg | 2020-04-07 | WHO National Influenza Centre Russian Federation | WHO National Influenza Centre Russian Federation | Andrey Komissarov, Artem Fadeev, Mariia Sergeeva, Anna Ivanova, Daria Danilenko |
| EPI_ISL_427334 | hCoV-19/Russia/StPetersburg-RII4726S/2020 | Europe / Russia / Saint-Petersburg | 2020-04-07 | WHO National Influenza Centre Russian Federation | WHO National Influenza Centre Russian Federation | Andrey Komissarov, Artem Fadeev, Mariia Sergeeva, Anna Ivanova, Daria Danilenko |
| EPI_ISL_427335 | hCoV-19/Russia/StPetersburg-RII4728S/2020 | Europe / Russia / Saint-Petersburg | 2020-04-07 | WHO National Influenza Centre Russian Federation | WHO National Influenza Centre Russian Federation | Andrey Komissarov, Artem Fadeev, Mariia Sergeeva, Anna Ivanova, Daria Danilenko |
| EPI_ISL_427336 | hCoV-19/Russia/StPetersburg-RII4781S/2020 | Europe / Russia / Saint-Petersburg | 2020-04-07 | WHO National Influenza Centre Russian Federation | WHO National Influenza Centre Russian Federation | Andrey Komissarov, Artem Fadeev, Mariia Sergeeva, Anna Ivanova, Daria Danilenko |
| EPI_ISL_427337 | hCoV-19/Russia/StPetersburg-RII4326S/2020 | Europe / Russia / Saint-Petersburg | 2020-03-25 | WHO National Influenza Centre Russian Federation | WHO National Influenza Centre Russian Federation | Andrey Komissarov, Artem Fadeev, Mariia Sergeeva, Anna Ivanova, Daria Danilenko |
| EPI_ISL_427338 | hCoV-19/Russia/StPetersburg-RII4328S/2020 | Europe / Russia / Saint-Petersburg | 2020-03-25 | WHO National Influenza Centre Russian Federation | WHO National Influenza Centre Russian Federation | Andrey Komissarov, Artem Fadeev, Mariia Sergeeva, Anna Ivanova, Daria Danilenko |
| EPI_ISL_427339 | hCoV-19/Russia/StPetersburg-RII4332S/2020 | Europe / Russia / Saint-Petersburg | 2020-03-25 | WHO National Influenza Centre Russian Federation | WHO National Influenza Centre Russian Federation | Andrey Komissarov, Artem Fadeev, Mariia Sergeeva, Anna Ivanova, Daria Danilenko |
| EPI_ISL_427342 | hCoV-19/Belgium/ULG-10034/2020 | Europe / Belgium / Liege | 2020-03-31 | Department of Clinical Microbiology | GIGA Medical Genomics | Keith Durkin, Maria Artesi, Sébastien Bontems, Raphaël Boreux, Cécile Meex, Pierrette Melin, Marie-Pierre Hayette, Vincent Bours. |
| EPI_ISL_427349 | hCoV-19/Belgium/ULG-10098/2020 | Europe / Belgium / Liege | 2020-04-06 | Department of Clinical Microbiology | GIGA Medical Genomics | Keith Durkin, Maria Artesi, Sébastien Bontems, Raphaël Boreux, Cécile Meex, Pierrette Melin, Marie-Pierre Hayette, Vincent Bours. |
| EPI_ISL_427350 | hCoV-19/Belgium/ULG-10099/2020 | Europe / Belgium / Liege | 2020-04-06 | Department of Clinical Microbiology | GIGA Medical Genomics | Keith Durkin, Maria Artesi, Sébastien Bontems, Raphaël Boreux, Cécile Meex, Pierrette Melin, Marie-Pierre Hayette, Vincent Bours. |
| EPI_ISL_427351 | hCoV-19/Belgium/ULG-10100/2020 | Europe / Belgium / Liege | 2020-04-07 | Department of Clinical Microbiology | GIGA Medical Genomics | Keith Durkin, Maria Artesi, Sébastien Bontems, Raphaël Boreux, Cécile Meex, Pierrette Melin, Marie-Pierre Hayette, Vincent Bours. |
| EPI_ISL_427352 | hCoV-19/Belgium/ULG-10101/2020 | Europe / Belgium / Liege | 2020-04-06 | Department of Clinical Microbiology | GIGA Medical Genomics | Keith Durkin, Maria Artesi, Sébastien Bontems, Raphaël Boreux, Cécile Meex, Pierrette Melin, Marie-Pierre Hayette, Vincent Bours. |
| EPI_ISL_427354 | hCoV-19/Belgium/ULG-10103/2020 | Europe / Belgium / Liege | 2020-04-06 | Department of Clinical Microbiology | GIGA Medical Genomics | Keith Durkin, Maria Artesi, Sébastien Bontems, Raphaël Boreux, Cécile Meex, Pierrette Melin, Marie-Pierre Hayette, Vincent Bours. |
| EPI_ISL_427357 | hCoV-19/Belgium/ULG-10106/2020 | Europe / Belgium / Liege | 2020-04-07 | Department of Clinical Microbiology | GIGA Medical Genomics | Keith Durkin, Maria Artesi, Sébastien Bontems, Raphaël Boreux, Cécile Meex, Pierrette Melin, Marie-Pierre Hayette, Vincent Bours. |
| EPI_ISL_427358 | hCoV-19/Belgium/ULG-10107/2020 | Europe / Belgium / Liege | 2020-04-06 | Department of Clinical Microbiology | GIGA Medical Genomics | Keith Durkin, Maria Artesi, Sébastien Bontems, Raphaël Boreux, Cécile Meex, Pierrette Melin, Marie-Pierre Hayette, Vincent Bours. |
| EPI_ISL_427359 | hCoV-19/Belgium/ULG-10108/2020 | Europe / Belgium / Liege | 2020-04-06 | Department of Clinical Microbiology | GIGA Medical Genomics | Keith Durkin, Maria Artesi, Sébastien Bontems, Raphaël Boreux, Cécile Meex, Pierrette Melin, Marie-Pierre Hayette, Vincent Bours. |
| EPI_ISL_427364 | hCoV-19/Belgium/ULG-10115/2020 | Europe / Belgium / Liege | 2020-04-07 | Department of Clinical Microbiology | GIGA Medical Genomics | Keith Durkin, Maria Artesi, Sébastien Bontems, Raphaël Boreux, Cécile Meex, Pierrette Melin, Marie-Pierre Hayette, Vincent Bours. |
| EPI_ISL_427366 | hCoV-19/Belgium/ULG-10118/2020 | Europe / Belgium / Liege | 2020-04-07 | Department of Clinical Microbiology | GIGA Medical Genomics | Keith Durkin, Maria Artesi, Sébastien Bontems, Raphaël Boreux, Cécile Meex, Pierrette Melin, Marie-Pierre Hayette, Vincent Bours. |
| EPI_ISL_427370 | hCoV-19/Belgium/ULG-10122/2020 | Europe / Belgium / Liege | 2020-04-07 | Department of Clinical Microbiology | GIGA Medical Genomics | Keith Durkin, Maria Artesi, Sébastien Bontems, Raphaël Boreux, Cécile Meex, Pierrette Melin, Marie-Pierre Hayette, Vincent Bours. |
| EPI_ISL_427371 | hCoV-19/Belgium/ULG-10124/2020 | Europe / Belgium / Liege | 2020-04-07 | Department of Clinical Microbiology | GIGA Medical Genomics | Keith Durkin, Maria Artesi, Sébastien Bontems, Raphaël Boreux, Cécile Meex, Pierrette Melin, Marie-Pierre Hayette, Vincent Bours. |
| EPI_ISL_427372 | hCoV-19/Belgium/ULG-10126/2020 | Europe / Belgium / Liege | 2020-04-07 | Department of Clinical Microbiology | GIGA Medical Genomics | Keith Durkin, Maria Artesi, Sébastien Bontems, Raphaël Boreux, Cécile Meex, Pierrette Melin, Marie-Pierre Hayette, Vincent Bours. |
| EPI_ISL_427379 | hCoV-19/Belgium/ULG-10134/2020 | Europe / Belgium / Liege | 2020-04-08 | Department of Clinical Microbiology | GIGA Medical Genomics | Keith Durkin, Maria Artesi, Sébastien Bontems, Raphaël Boreux, Cécile Meex, Pierrette Melin, Marie-Pierre Hayette, Vincent Bours. |
| EPI_ISL_427383 | hCoV-19/Belgium/ULG-10138/2020 | Europe / Belgium / Liege | 2020-04-08 | Department of Clinical Microbiology | GIGA Medical Genomics | Keith Durkin, Maria Artesi, Sébastien Bontems, Raphaël Boreux, Cécile Meex, Pierrette Melin, Marie-Pierre Hayette, Vincent Bours. |
| EPI_ISL_427385 | hCoV-19/Belgium/ULG-10142/2020 | Europe / Belgium / Liege | 2020-04-08 | Department of Clinical Microbiology | GIGA Medical Genomics | Keith Durkin, Maria Artesi, Sébastien Bontems, Raphaël Boreux, Cécile Meex, Pierrette Melin, Marie-Pierre Hayette, Vincent Bours. |
| EPI_ISL_427388 | hCoV-19/Belgium/ULG-10144/2020 | Europe / Belgium / Liege | 2020-04-07 | Department of Clinical Microbiology | GIGA Medical Genomics | Keith Durkin, Maria Artesi, Sébastien Bontems, Raphaël Boreux, Cécile Meex, Pierrette Melin, Marie-Pierre Hayette, Vincent Bours. |
| EPI_ISL_427389 | hCoV-19/Belgium/ULG-10096/2020 | Europe / Belgium / Liege | 2020-04-05 | Department of Clinical Microbiology | GIGA Medical Genomics | Keith Durkin, Maria Artesi, Sébastien Bontems, Raphaël Boreux, Cécile Meex, Pierrette Melin, Marie-Pierre Hayette, Vincent Bours. |
| EPI_ISL_427390 | hCoV-19/Belgium/ULG-10125/2020 | Europe / Belgium / Liege | 2020-04-07 | Department of Clinical Microbiology | GIGA Medical Genomics | Keith Durkin, Maria Artesi, Sébastien Bontems, Raphaël Boreux, Cécile Meex, Pierrette Melin, Marie-Pierre Hayette, Vincent Bours. |
| EPI_ISL_427391 | hCoV-19/Turkey/GLAB-CoV008/2020 | Europe / Turkey / Istanbul | 2020-04-13 | Genomic Laboratory (GLAB) (Conjoint lab of Health Directorate of Istanbul and Istanbul Technical University) | Genomic Laboratory (GLAB), Istanbul Technical University | Ilker Karacan, Tugba Kizilboga Akgun, Bugra Agaoglu, Gizem Alkurt, Jale Yildiz, Betsi Köse, Elifnaz Çelik, Mehtap Aydin, Levent Doganay, Gizem Dinler Doganay |
| EPI_ISL_427392 | hCoV-19/Taiwan/TSGH-05/2020 | Asia / Taiwan / New Taipei City | 2020-03-14 | TSGH-CP molecular lab | TSGH-CP molecular lab | Cherng-Lih Perng, Ming-Jr Jian, Chih-Kai Chang, Jung-Chung Lin, Kuo-Ming Yeh, Chien-Wen Chen, Sheng-Kang Chiu, Hsing-Yi Chung, Shih-Hung Tsai, Kuo-Sheng Hung, Tien-Yao Chang, Feng-Yee Chang, Hung-Sheng Shang |
| EPI_ISL_427393 | hCoV-19/Taiwan/TSGH-06/2020 | Asia / Taiwan / Taipei | 2020-03-15 | TSGH-CP molecular lab | TSGH-CP molecular lab | Cherng-Lih Perng, Ming-Jr Jian, Chih-Kai Chang, Jung-Chung Lin, Kuo-Ming Yeh, Chien-Wen Chen, Sheng-Kang Chiu, Hsing-Yi Chung, Shih-Hung Tsai, Kuo-Sheng Hung, Tien-Yao Chang, Feng-Yee Chang, Hung-Sheng Shang |
| EPI_ISL_427394 | hCoV-19/Taiwan/TSGH-07/2020 | Asia / Taiwan / New Taipei City | 2020-03-22 | TSGH-CP molecular lab | TSGH-CP molecular lab | Cherng-Lih Perng, Ming-Jr Jian, Chih-Kai Chang, Jung-Chung Lin, Kuo-Ming Yeh, Chien-Wen Chen, Sheng-Kang Chiu, Hsing-Yi Chung, Shih-Hung Tsai, Kuo-Sheng Hung, Tien-Yao Chang, Feng-Yee Chang, Hung-Sheng Shang |
| EPI_ISL_427395 | hCoV-19/Taiwan/TSGH-08/2020 | Asia / Taiwan / Keelung | 2020-03-23 | TSGH-CP molecular lab | TSGH-CP molecular lab | Cherng-Lih Perng, Ming-Jr Jian, Chih-Kai Chang, Jung-Chung Lin, Kuo-Ming Yeh, Chien-Wen Chen, Sheng-Kang Chiu, Hsing-Yi Chung, Shih-Hung Tsai, Kuo-Sheng Hung, Tien-Yao Chang, Feng-Yee Chang, Hung-Sheng Shang |
| EPI_ISL_427396 | hCoV-19/Taiwan/TSGH-09/2020 | Asia / Taiwan / New Taipei City | 2020-03-23 | TSGH-CP molecular lab | TSGH-CP molecular lab | Cherng-Lih Perng, Ming-Jr Jian, Chih-Kai Chang, Jung-Chung Lin, Kuo-Ming Yeh, Chien-Wen Chen, Sheng-Kang Chiu, Hsing-Yi Chung, Shih-Hung Tsai, Kuo-Sheng Hung, Tien-Yao Chang, Feng-Yee Chang, Hung-Sheng Shang |
| EPI_ISL_427397 | hCoV-19/Taiwan/TSGH-10/2020 | Asia / Taiwan / Taipei | 2020-03-24 | TSGH-CP molecular lab | TSGH-CP molecular lab | Cherng-Lih Perng, Ming-Jr Jian, Chih-Kai Chang, Jung-Chung Lin, Kuo-Ming Yeh, Chien-Wen Chen, Sheng-Kang Chiu, Hsing-Yi Chung, Shih-Hung Tsai, Kuo-Sheng Hung, Tien-Yao Chang, Feng-Yee Chang, Hung-Sheng Shang |
| EPI_ISL_427398 | hCoV-19/Taiwan/TSGH-11/2020 | Asia / Taiwan / New Taipei City | 2020-03-25 | TSGH-CP molecular lab | TSGH-CP molecular lab | Cherng-Lih Perng, Ming-Jr Jian, Chih-Kai Chang, Jung-Chung Lin, Kuo-Ming Yeh, Chien-Wen Chen, Sheng-Kang Chiu, Hsing-Yi Chung, Shih-Hung Tsai, Kuo-Sheng Hung, Tien-Yao Chang, Feng-Yee Chang, Hung-Sheng Shang |
| EPI_ISL_427450 | hCoV-19/USA/WI-UW-143/2020 | North America / USA / Illinois | 2020-04-13 | University of Wisconsin-Madison AIDS Vaccine Research Laboratories | University of Wisconsin-Madison AIDS Vaccine Research Laboratories | Gage Moreno, Katarina Braun, et al. AIDS Vaccine Research Laboratories |
| EPI_ISL_427451 | hCoV-19/USA/WI-UW-144/2020 | North America / USA / Wisconsin / Madis | 2020-04-06 | University of Wisconsin-Madison AIDS Vaccine Research Laboratories | University of Wisconsin-Madison AIDS Vaccine Research Laboratories | Gage Moreno, Katarina Braun, et al. AIDS Vaccine Research Laboratories |
| EPI_ISL_427452 | hCoV-19/USA/WI-UW-145/2020 | North America / USA / Wisconsin / DeFor | 2020-04-10 | University of Wisconsin-Madison AIDS Vaccine Research Laboratories | University of Wisconsin-Madison AIDS Vaccine Research Laboratories | Gage Moreno, Katarina Braun, et al. AIDS Vaccine Research Laboratories |
| EPI_ISL_427453 | hCoV-19/USA/WI-UW-146/2020 | North America / USA / Wisconsin / Madis | 2020-04-06 | University of Wisconsin-Madison AIDS Vaccine Research Laboratories | University of Wisconsin-Madison AIDS Vaccine Research Laboratories | Gage Moreno, Katarina Braun, et al. AIDS Vaccine Research Laboratories |
| EPI_ISL_427454 | hCoV-19/USA/WI-UW-147/2020 | North America / USA / Wisconsin / Madis | 2020-04-10 | University of Wisconsin-Madison AIDS Vaccine Research Laboratories | University of Wisconsin-Madison AIDS Vaccine Research Laboratories | Gage Moreno, Katarina Braun, et al. AIDS Vaccine Research Laboratories |
| EPI_ISL_427455 | hCoV-19/USA/WI-UW-148/2020 | North America / USA / Wisconsin / Waun | 2020-04-07 | University of Wisconsin-Madison AIDS Vaccine Research Laboratories | University of Wisconsin-Madison AIDS Vaccine Research Laboratories | Gage Moreno, Katarina Braun, et al. AIDS Vaccine Research Laboratories |
| EPI_ISL_427456 | hCoV-19/USA/WI-UW-149/2020 | North America / USA / Illinois | 2020-04-07 | University of Wisconsin-Madison AIDS Vaccine Research Laboratories | University of Wisconsin-Madison AIDS Vaccine Research Laboratories | Gage Moreno, Katarina Braun, et al. AIDS Vaccine Research Laboratories |
| EPI_ISL_427457 | hCoV-19/USA/WI-UW-150/2020 | North America / USA / Wisconsin / Madis | 2020-04-13 | University of Wisconsin-Madison AIDS Vaccine Research Laboratories | University of Wisconsin-Madison AIDS Vaccine Research Laboratories | Gage Moreno, Katarina Braun, et al. AIDS Vaccine Research Laboratories |
| EPI_ISL_427458 | hCoV-19/USA/WI-UW-151/2020 | North America / USA / Wisconsin / Sun P | 2020-04-08 | University of Wisconsin-Madison AIDS Vaccine Research Laboratories | University of Wisconsin-Madison AIDS Vaccine Research Laboratories | Gage Moreno, Katarina Braun, et al. AIDS Vaccine Research Laboratories |
| EPI_ISL_427459 | hCoV-19/USA/WI-UW-152/2020 | North America / USA / Wisconsin / Madis | 2020-04-06 | University of Wisconsin-Madison AIDS Vaccine Research Laboratories | University of Wisconsin-Madison AIDS Vaccine Research Laboratories | Gage Moreno, Katarina Braun, et al. AIDS Vaccine Research Laboratories |
| EPI_ISL_427460 | hCoV-19/USA/WI-UW-153/2020 | North America / USA / Wisconsin / Madis | 2020-04-09 | University of Wisconsin-Madison AIDS Vaccine Research Laboratories | University of Wisconsin-Madison AIDS Vaccine Research Laboratories | Gage Moreno, Katarina Braun, et al. AIDS Vaccine Research Laboratories |
| EPI_ISL_427461 | hCoV-19/USA/WI-UW-154/2020 | North America / USA / Wisconsin / Tomal | 2020-04-12 | University of Wisconsin-Madison AIDS Vaccine Research Laboratories | University of Wisconsin-Madison AIDS Vaccine Research Laboratories | Gage Moreno, Katarina Braun, et al. AIDS Vaccine Research Laboratories |
| EPI_ISL_427462 | hCoV-19/USA/WI-UW-155/2020 | North America / USA / Wisconsin / Madis | 2020-04-10 | University of Wisconsin-Madison AIDS Vaccine Research Laboratories | University of Wisconsin-Madison AIDS Vaccine Research Laboratories | Gage Moreno, Katarina Braun, et al. AIDS Vaccine Research Laboratories |
| EPI_ISL_427469 | hCoV-19/USA/NY-NYUMC172/2020 | North America / USA / New York / Manha | 2020-04-01 | NYU Langone Health | Departments of Pathology and Medicine, New York University School of Medicine | Maria Aguero-Rosenfeld, Brendan Belovarac, Margaret Black, Ludovic Boytard, John Cadley, Paolo Cotzia, John Chen, Dacia Dimartino, Xiaojun Feng, Tatyana Gindin, Emily Guzman, Adriana Heguy, Megan Hogan, Emily Huang, George Jour, Andrew Lytle, Christian Marier, Matthew T. Maurano, Mark J. Mulligan, Peter Meyn, Iman Osman, Jared Pinnell, Vanessa Raabe, Sitharam Ramaswami, Amy Rapkiewicz, Marie Samanovic-Golden, Antonio Serrano, Guomiao Shen, Matija Snuderl, Theodore Vougiouklakis, Nick Vulpescu, Gael Westby, Paul Zappile, Yutong Zhang |
| EPI_ISL_427470 | hCoV-19/USA/NY-NYUMC173/2020 | North America / USA / New York / Staten | 2020-04-01 | NYU Langone Health | Departments of Pathology and Medicine, New York University School of Medicine | Maria Aguero-Rosenfeld, Brendan Belovarac, Margaret Black, Ludovic Boytard, John Cadley, Paolo Cotzia, John Chen, Dacia Dimartino, Xiaojun Feng, Tatyana Gindin, Emily Guzman, Adriana Heguy, Megan Hogan, Emily Huang, George Jour, Andrew Lytle, Christian Marier, Matthew T. Maurano, Mark J. Mulligan, Peter Meyn, Iman Osman, Jared Pinnell, Vanessa Raabe, Sitharam Ramaswami, Amy Rapkiewicz, Marie Samanovic-Golden, Antonio Serrano, Guomiao Shen, Matija Snuderl, Theodore Vougiouklakis, Nick Vulpescu, Gael Westby, Paul Zappile, Yutong Zhang |
| EPI_ISL_427471 | hCoV-19/USA/NY-NYUMC174/2020 | North America / USA / New York / Nassa | 2020-04-01 | NYU Langone Health | Departments of Pathology and Medicine, New York University School of Medicine | Maria Aguero-Rosenfeld, Brendan Belovarac, Margaret Black, Ludovic Boytard, John Cadley, Paolo Cotzia, John Chen, Dacia Dimartino, Xiaojun Feng, Tatyana Gindin, Emily Guzman, Adriana Heguy, Megan Hogan, Emily Huang, George Jour, Andrew Lytle, Christian Marier, Matthew T. Maurano, Mark J. Mulligan, Peter Meyn, Iman Osman, Jared Pinnell, Vanessa Raabe, Sitharam Ramaswami, Amy Rapkiewicz, Marie Samanovic-Golden, Antonio Serrano, Guomiao Shen, Matija Snuderl, Theodore Vougiouklakis, Nick Vulpescu, Gael Westby, Paul Zappile, Yutong Zhang |
| EPI_ISL_427472 | hCoV-19/USA/NY-NYUMC175/2020 | North America / USA / New York / Nassa | 2020-04-01 | NYU Langone Health | Departments of Pathology and Medicine, New York University School of Medicine | Maria Aguero-Rosenfeld, Brendan Belovarac, Margaret Black, Ludovic Boytard, John Cadley, Paolo Cotzia, John Chen, Dacia Dimartino, Xiaojun Feng, Tatyana Gindin, Emily Guzman, Adriana Heguy, Megan Hogan, Emily Huang, George Jour, Andrew Lytle, Christian Marier, Matthew T. Maurano, Mark J. Mulligan, Peter Meyn, Iman Osman, Jared Pinnell, Vanessa Raabe, Sitharam Ramaswami, Amy Rapkiewicz, Marie Samanovic-Golden, Antonio Serrano, Guomiao Shen, Matija Snuderl, Theodore Vougiouklakis, Nick Vulpescu, Gael Westby, Paul Zappile, Yutong Zhang |
| EPI_ISL_427473 | hCoV-19/USA/NY-NYUMC177/2020 | North America / USA / New York / Brookl | 2020-04-01 | NYU Langone Health | Departments of Pathology and Medicine, New York University School of Medicine | Maria Aguero-Rosenfeld, Brendan Belovarac, Margaret Black, Ludovic Boytard, John Cadley, Paolo Cotzia, John Chen, Dacia Dimartino, Xiaojun Feng, Tatyana Gindin, Emily Guzman, Adriana Heguy, Megan Hogan, Emily Huang, George Jour, Andrew Lytle, Christian Marier, Matthew T. Maurano, Mark J. Mulligan, Peter Meyn, Iman Osman, Jared Pinnell, Vanessa Raabe, Sitharam Ramaswami, Amy Rapkiewicz, Marie Samanovic-Golden, Antonio Serrano, Guomiao Shen, Matija Snuderl, Theodore Vougiouklakis, Nick Vulpescu, Gael Westby, Paul Zappile, Yutong Zhang |
| EPI_ISL_427474 | hCoV-19/USA/NY-NYUMC178/2020 | North America / USA / New York / Manha | 2020-04-01 | NYU Langone Health | Departments of Pathology and Medicine, New York University School of Medicine | Maria Aguero-Rosenfeld, Brendan Belovarac, Margaret Black, Ludovic Boytard, John Cadley, Paolo Cotzia, John Chen, Dacia Dimartino, Xiaojun Feng, Tatyana Gindin, Emily Guzman, Adriana Heguy, Megan Hogan, Emily Huang, George Jour, Andrew Lytle, Christian Marier, Matthew T. Maurano, Mark J. Mulligan, Peter Meyn, Iman Osman, Jared Pinnell, Vanessa Raabe, Sitharam Ramaswami, Amy Rapkiewicz, Marie Samanovic-Golden, Antonio Serrano, Guomiao Shen, Matija Snuderl, Theodore Vougiouklakis, Nick Vulpescu, Gael Westby, Paul Zappile, Yutong Zhang |
| EPI_ISL_427475 | hCoV-19/USA/NY-NYUMC179/2020 | North America / USA / New York / Staten | 2020-04-01 | NYU Langone Health | Departments of Pathology and Medicine, New York University School of Medicine | Maria Aguero-Rosenfeld, Brendan Belovarac, Margaret Black, Ludovic Boytard, John Cadley, Paolo Cotzia, John Chen, Dacia Dimartino, Xiaojun Feng, Tatyana Gindin, Emily Guzman, Adriana Heguy, Megan Hogan, Emily Huang, George Jour, Andrew Lytle, Christian Marier, Matthew T. Maurano, Mark J. Mulligan, Peter Meyn, Iman Osman, Jared Pinnell, Vanessa Raabe, Sitharam Ramaswami, Amy Rapkiewicz, Marie Samanovic-Golden, Antonio Serrano, Guomiao Shen, Matija Snuderl, Theodore Vougiouklakis, Nick Vulpescu, Gael Westby, Paul Zappile, Yutong Zhang |
| EPI_ISL_427476 | hCoV-19/USA/NY-NYUMC180/2020 | North America / USA / New York / Brookl | 2020-04-01 | NYU Langone Health | Departments of Pathology and Medicine, New York University School of Medicine | Maria Aguero-Rosenfeld, Brendan Belovarac, Margaret Black, Ludovic Boytard, John Cadley, Paolo Cotzia, John Chen, Dacia Dimartino, Xiaojun Feng, Tatyana Gindin, Emily Guzman, Adriana Heguy, Megan Hogan, Emily Huang, George Jour, Andrew Lytle, Christian Marier, Matthew T. Maurano, Mark J. Mulligan, Peter Meyn, Iman Osman, Jared Pinnell, Vanessa Raabe, Sitharam Ramaswami, Amy Rapkiewicz, Marie Samanovic-Golden, Antonio Serrano, Guomiao Shen, Matija Snuderl, Theodore Vougiouklakis, Nick Vulpescu, Gael Westby, Paul Zappile, Yutong Zhang |
| EPI_ISL_427477 | hCoV-19/USA/NY-NYUMC181/2020 | North America / USA / New York / Queen | 2020-04-01 | NYU Langone Health | Departments of Pathology and Medicine, New York University School of Medicine | Maria Aguero-Rosenfeld, Brendan Belovarac, Margaret Black, Ludovic Boytard, John Cadley, Paolo Cotzia, John Chen, Dacia Dimartino, Xiaojun Feng, Tatyana Gindin, Emily Guzman, Adriana Heguy, Megan Hogan, Emily Huang, George Jour, Andrew Lytle, Christian Marier, Matthew T. Maurano, Mark J. Mulligan, Peter Meyn, Iman Osman, Jared Pinnell, Vanessa Raabe, Sitharam Ramaswami, Amy Rapkiewicz, Marie Samanovic-Golden, Antonio Serrano, Guomiao Shen, Matija Snuderl, Theodore Vougiouklakis, Nick Vulpescu, Gael Westby, Paul Zappile, Yutong Zhang |
| EPI_ISL_427478 | hCoV-19/USA/NY-NYUMC182/2020 | North America / USA / New York / Manha | 2020-04-01 | NYU Langone Health | Departments of Pathology and Medicine, New York University School of Medicine | Maria Aguero-Rosenfeld, Brendan Belovarac, Margaret Black, Ludovic Boytard, John Cadley, Paolo Cotzia, John Chen, Dacia Dimartino, Xiaojun Feng, Tatyana Gindin, Emily Guzman, Adriana Heguy, Megan Hogan, Emily Huang, George Jour, Andrew Lytle, Christian Marier, Matthew T. Maurano, Mark J. Mulligan, Peter Meyn, Iman Osman, Jared Pinnell, Vanessa Raabe, Sitharam Ramaswami, Amy Rapkiewicz, Marie Samanovic-Golden, Antonio Serrano, Guomiao Shen, Matija Snuderl, Theodore Vougiouklakis, Nick Vulpescu, Gael Westby, Paul Zappile, Yutong Zhang |
| EPI_ISL_427479 | hCoV-19/USA/NY-NYUMC183/2020 | North America / USA / New York / Manha | 2020-04-01 | NYU Langone Health | Departments of Pathology and Medicine, New York University School of Medicine | Maria Aguero-Rosenfeld, Brendan Belovarac, Margaret Black, Ludovic Boytard, John Cadley, Paolo Cotzia, John Chen, Dacia Dimartino, Xiaojun Feng, Tatyana Gindin, Emily Guzman, Adriana Heguy, Megan Hogan, Emily Huang, George Jour, Andrew Lytle, Christian Marier, Matthew T. Maurano, Mark J. Mulligan, Peter Meyn, Iman Osman, Jared Pinnell, Vanessa Raabe, Sitharam Ramaswami, Amy Rapkiewicz, Marie Samanovic-Golden, Antonio Serrano, Guomiao Shen, Matija Snuderl, Theodore Vougiouklakis, Nick Vulpescu, Gael Westby, Paul Zappile, Yutong Zhang |
| EPI_ISL_427480 | hCoV-19/USA/NY-NYUMC184/2020 | North America / USA / New York / Queen | 2020-04-01 | NYU Langone Health | Departments of Pathology and Medicine, New York University School of Medicine | Maria Aguero-Rosenfeld, Brendan Belovarac, Margaret Black, Ludovic Boytard, John Cadley, Paolo Cotzia, John Chen, Dacia Dimartino, Xiaojun Feng, Tatyana Gindin, Emily Guzman, Adriana Heguy, Megan Hogan, Emily Huang, George Jour, Andrew Lytle, Christian Marier, Matthew T. Maurano, Mark J. Mulligan, Peter Meyn, Iman Osman, Jared Pinnell, Vanessa Raabe, Sitharam Ramaswami, Amy Rapkiewicz, Marie Samanovic-Golden, Antonio Serrano, Guomiao Shen, Matija Snuderl, Theodore Vougiouklakis, Nick Vulpescu, Gael Westby, Paul Zappile, Yutong Zhang |
| EPI_ISL_427481 | hCoV-19/USA/NY-NYUMC185/2020 | North America / USA / New York / Manha | 2020-04-01 | NYU Langone Health | Departments of Pathology and Medicine, New York University School of Medicine | Maria Aguero-Rosenfeld, Brendan Belovarac, Margaret Black, Ludovic Boytard, John Cadley, Paolo Cotzia, John Chen, Dacia Dimartino, Xiaojun Feng, Tatyana Gindin, Emily Guzman, Adriana Heguy, Megan Hogan, Emily Huang, George Jour, Andrew Lytle, Christian Marier, Matthew T. Maurano, Mark J. Mulligan, Peter Meyn, Iman Osman, Jared Pinnell, Vanessa Raabe, Sitharam Ramaswami, Amy Rapkiewicz, Marie Samanovic-Golden, Antonio Serrano, Guomiao Shen, Matija Snuderl, Theodore Vougiouklakis, Nick Vulpescu, Gael Westby, Paul Zappile, Yutong Zhang |
| EPI_ISL_427482 | hCoV-19/USA/NY-NYUMC186/2020 | North America / USA / New York / Brookl | 2020-04-01 | NYU Langone Health | Departments of Pathology and Medicine, New York University School of Medicine | Maria Aguero-Rosenfeld, Brendan Belovarac, Margaret Black, Ludovic Boytard, John Cadley, Paolo Cotzia, John Chen, Dacia Dimartino, Xiaojun Feng, Tatyana Gindin, Emily Guzman, Adriana Heguy, Megan Hogan, Emily Huang, George Jour, Andrew Lytle, Christian Marier, Matthew T. Maurano, Mark J. Mulligan, Peter Meyn, Iman Osman, Jared Pinnell, Vanessa Raabe, Sitharam Ramaswami, Amy Rapkiewicz, Marie Samanovic-Golden, Antonio Serrano, Guomiao Shen, Matija Snuderl, Theodore Vougiouklakis, Nick Vulpescu, Gael Westby, Paul Zappile, Yutong Zhang |
| EPI_ISL_427484 | hCoV-19/USA/NY-NYUMC188/2020 | North America / USA / New York / Brookl | 2020-04-01 | NYU Langone Health | Departments of Pathology and Medicine, New York University School of Medicine | Maria Aguero-Rosenfeld, Brendan Belovarac, Margaret Black, Ludovic Boytard, John Cadley, Paolo Cotzia, John Chen, Dacia Dimartino, Xiaojun Feng, Tatyana Gindin, Emily Guzman, Adriana Heguy, Megan Hogan, Emily Huang, George Jour, Andrew Lytle, Christian Marier, Matthew T. Maurano, Mark J. Mulligan, Peter Meyn, Iman Osman, Jared Pinnell, Vanessa Raabe, Sitharam Ramaswami, Amy Rapkiewicz, Marie Samanovic-Golden, Antonio Serrano, Guomiao Shen, Matija Snuderl, Theodore Vougiouklakis, Nick Vulpescu, Gael Westby, Paul Zappile, Yutong Zhang |
| EPI_ISL_427485 | hCoV-19/USA/NY-NYUMC189/2020 | North America / USA / New York / Brookl | 2020-04-01 | NYU Langone Health | Departments of Pathology and Medicine, New York University School of Medicine | Maria Aguero-Rosenfeld, Brendan Belovarac, Margaret Black, Ludovic Boytard, John Cadley, Paolo Cotzia, John Chen, Dacia Dimartino, Xiaojun Feng, Tatyana Gindin, Emily Guzman, Adriana Heguy, Megan Hogan, Emily Huang, George Jour, Andrew Lytle, Christian Marier, Matthew T. Maurano, Mark J. Mulligan, Peter Meyn, Iman Osman, Jared Pinnell, Vanessa Raabe, Sitharam Ramaswami, Amy Rapkiewicz, Marie Samanovic-Golden, Antonio Serrano, Guomiao Shen, Matija Snuderl, Theodore Vougiouklakis, Nick Vulpescu, Gael Westby, Paul Zappile, Yutong Zhang |
| EPI_ISL_427487 | hCoV-19/USA/NY-NYUMC192/2020 | North America / USA / New York / Brookl | 2020-04-01 | NYU Langone Health | Departments of Pathology and Medicine, New York University School of Medicine | Maria Aguero-Rosenfeld, Brendan Belovarac, Margaret Black, Ludovic Boytard, John Cadley, Paolo Cotzia, John Chen, Dacia Dimartino, Xiaojun Feng, Tatyana Gindin, Emily Guzman, Adriana Heguy, Megan Hogan, Emily Huang, George Jour, Andrew Lytle, Christian Marier, Matthew T. Maurano, Mark J. Mulligan, Peter Meyn, Iman Osman, Jared Pinnell, Vanessa Raabe, Sitharam Ramaswami, Amy Rapkiewicz, Marie Samanovic-Golden, Antonio Serrano, Guomiao Shen, Matija Snuderl, Theodore Vougiouklakis, Nick Vulpescu, Gael Westby, Paul Zappile, Yutong Zhang |
| EPI_ISL_427488 | hCoV-19/USA/NY-NYUMC193/2020 | North America / USA / New York / Queen | 2020-04-01 | NYU Langone Health | Departments of Pathology and Medicine, New York University School of Medicine | Maria Aguero-Rosenfeld, Brendan Belovarac, Margaret Black, Ludovic Boytard, John Cadley, Paolo Cotzia, John Chen, Dacia Dimartino, Xiaojun Feng, Tatyana Gindin, Emily Guzman, Adriana Heguy, Megan Hogan, Emily Huang, George Jour, Andrew Lytle, Christian Marier, Matthew T. Maurano, Mark J. Mulligan, Peter Meyn, Iman Osman, Jared Pinnell, Vanessa Raabe, Sitharam Ramaswami, Amy Rapkiewicz, Marie Samanovic-Golden, Antonio Serrano, Guomiao Shen, Matija Snuderl, Theodore Vougiouklakis, Nick Vulpescu, Gael Westby, Paul Zappile, Yutong Zhang |
| EPI_ISL_427489 | hCoV-19/USA/NY-NYUMC194/2020 | North America / USA / New York / Brookl | 2020-04-01 | NYU Langone Health | Departments of Pathology and Medicine, New York University School of Medicine | Maria Aguero-Rosenfeld, Brendan Belovarac, Margaret Black, Ludovic Boytard, John Cadley, Paolo Cotzia, John Chen, Dacia Dimartino, Xiaojun Feng, Tatyana Gindin, Emily Guzman, Adriana Heguy, Megan Hogan, Emily Huang, George Jour, Andrew Lytle, Christian Marier, Matthew T. Maurano, Mark J. Mulligan, Peter Meyn, Iman Osman, Jared Pinnell, Vanessa Raabe, Sitharam Ramaswami, Amy Rapkiewicz, Marie Samanovic-Golden, Antonio Serrano, Guomiao Shen, Matija Snuderl, Theodore Vougiouklakis, Nick Vulpescu, Gael Westby, Paul Zappile, Yutong Zhang |

| | | | | | | |
|---|---|---|---|---|---|---|
| EPI_ISL_427490 | hCoV-19/USA/NY-NYUMC195/2020 | North America / USA / New York / Manha | 2020-04-01 | NYU Langone Health | Departments of Pathology and Medicine, New York University School of Medicine | Maria Aguero-Rosenfeld, Brendan Belovarac, Margaret Black, Ludovic Boytard, John Cadley, Paolo Cotzia, John Chen, Dacia Dimartino, Xiaojun Feng, Tatyana Gindin, Emily Guzman, Adriana Heguy, Megan Hogan, Emily Huang, George Jour, Andrew Lytle, Christian Marier, Matthew T. Maurano, Mark J. Mulligan, Peter Meyn, Iman Osman, Jared Pinnell, Vanessa Raabe, Sitharam Ramaswami, Amy Rapkiewicz, Marie Samanovic-Golden, Antonio Serrano, Guomiao Shen, Matija Snuderl, Theodore Vougiouklakis, Nick Vulpescu, Gael Westby, Paul Zappile, Yutong Zhang |
| EPI_ISL_427491 | hCoV-19/USA/NY-NYUMC196/2020 | North America / USA / New York / Brookl | 2020-04-01 | NYU Langone Health | Departments of Pathology and Medicine, New York University School of Medicine | Maria Aguero-Rosenfeld, Brendan Belovarac, Margaret Black, Ludovic Boytard, John Cadley, Paolo Cotzia, John Chen, Dacia Dimartino, Xiaojun Feng, Tatyana Gindin, Emily Guzman, Adriana Heguy, Megan Hogan, Emily Huang, George Jour, Andrew Lytle, Christian Marier, Matthew T. Maurano, Mark J. Mulligan, Peter Meyn, Iman Osman, Jared Pinnell, Vanessa Raabe, Sitharam Ramaswami, Amy Rapkiewicz, Marie Samanovic-Golden, Antonio Serrano, Guomiao Shen, Matija Snuderl, Theodore Vougiouklakis, Nick Vulpescu, Gael Westby, Paul Zappile, Yutong Zhang |
| EPI_ISL_427492 | hCoV-19/USA/NY-NYUMC197/2020 | North America / USA / New York / Queen | 2020-04-01 | NYU Langone Health | Departments of Pathology and Medicine, New York University School of Medicine | Maria Aguero-Rosenfeld, Brendan Belovarac, Margaret Black, Ludovic Boytard, John Cadley, Paolo Cotzia, John Chen, Dacia Dimartino, Xiaojun Feng, Tatyana Gindin, Emily Guzman, Adriana Heguy, Megan Hogan, Emily Huang, George Jour, Andrew Lytle, Christian Marier, Matthew T. Maurano, Mark J. Mulligan, Peter Meyn, Iman Osman, Jared Pinnell, Vanessa Raabe, Sitharam Ramaswami, Amy Rapkiewicz, Marie Samanovic-Golden, Antonio Serrano, Guomiao Shen, Matija Snuderl, Theodore Vougiouklakis, Nick Vulpescu, Gael Westby, Paul Zappile, Yutong Zhang |
| EPI_ISL_427493 | hCoV-19/USA/NY-NYUMC198/2020 | North America / USA / New York / Brookl | 2020-04-01 | NYU Langone Health | Departments of Pathology and Medicine, New York University School of Medicine | Maria Aguero-Rosenfeld, Brendan Belovarac, Margaret Black, Ludovic Boytard, John Cadley, Paolo Cotzia, John Chen, Dacia Dimartino, Xiaojun Feng, Tatyana Gindin, Emily Guzman, Adriana Heguy, Megan Hogan, Emily Huang, George Jour, Andrew Lytle, Christian Marier, Matthew T. Maurano, Mark J. Mulligan, Peter Meyn, Iman Osman, Jared Pinnell, Vanessa Raabe, Sitharam Ramaswami, Amy Rapkiewicz, Marie Samanovic-Golden, Antonio Serrano, Guomiao Shen, Matija Snuderl, Theodore Vougiouklakis, Nick Vulpescu, Gael Westby, Paul Zappile, Yutong Zhang |
| EPI_ISL_427494 | hCoV-19/USA/NY-NYUMC199/2020 | North America / USA / New York / Bronx | 2020-04-01 | NYU Langone Health | Departments of Pathology and Medicine, New York University School of Medicine | Maria Aguero-Rosenfeld, Brendan Belovarac, Margaret Black, Ludovic Boytard, John Cadley, Paolo Cotzia, John Chen, Dacia Dimartino, Xiaojun Feng, Tatyana Gindin, Emily Guzman, Adriana Heguy, Megan Hogan, Emily Huang, George Jour, Andrew Lytle, Christian Marier, Matthew T. Maurano, Mark J. Mulligan, Peter Meyn, Iman Osman, Jared Pinnell, Vanessa Raabe, Sitharam Ramaswami, Amy Rapkiewicz, Marie Samanovic-Golden, Antonio Serrano, Guomiao Shen, Matija Snuderl, Theodore Vougiouklakis, Nick Vulpescu, Gael Westby, Paul Zappile, Yutong Zhang |
| EPI_ISL_427495 | hCoV-19/USA/NY-NYUMC200/2020 | North America / USA / New York / Brookl | 2020-04-01 | NYU Langone Health | Departments of Pathology and Medicine, New York University School of Medicine | Maria Aguero-Rosenfeld, Brendan Belovarac, Margaret Black, Ludovic Boytard, John Cadley, Paolo Cotzia, John Chen, Dacia Dimartino, Xiaojun Feng, Tatyana Gindin, Emily Guzman, Adriana Heguy, Megan Hogan, Emily Huang, George Jour, Andrew Lytle, Christian Marier, Matthew T. Maurano, Mark J. Mulligan, Peter Meyn, Iman Osman, Jared Pinnell, Vanessa Raabe, Sitharam Ramaswami, Amy Rapkiewicz, Marie Samanovic-Golden, Antonio Serrano, Guomiao Shen, Matija Snuderl, Theodore Vougiouklakis, Nick Vulpescu, Gael Westby, Paul Zappile, Yutong Zhang |
| EPI_ISL_427496 | hCoV-19/USA/NY-NYUMC201/2020 | North America / USA / New York / Brookl | 2020-04-01 | NYU Langone Health | Departments of Pathology and Medicine, New York University School of Medicine | Maria Aguero-Rosenfeld, Brendan Belovarac, Margaret Black, Ludovic Boytard, John Cadley, Paolo Cotzia, John Chen, Dacia Dimartino, Xiaojun Feng, Tatyana Gindin, Emily Guzman, Adriana Heguy, Megan Hogan, Emily Huang, George Jour, Andrew Lytle, Christian Marier, Matthew T. Maurano, Mark J. Mulligan, Peter Meyn, Iman Osman, Jared Pinnell, Vanessa Raabe, Sitharam Ramaswami, Amy Rapkiewicz, Marie Samanovic-Golden, Antonio Serrano, Guomiao Shen, Matija Snuderl, Theodore Vougiouklakis, Nick Vulpescu, Gael Westby, Paul Zappile, Yutong Zhang |
| EPI_ISL_427498 | hCoV-19/USA/NY-NYUMC203/2020 | North America / USA / New York / Brookl | 2020-04-01 | NYU Langone Health | Departments of Pathology and Medicine, New York University School of Medicine | Maria Aguero-Rosenfeld, Brendan Belovarac, Margaret Black, Ludovic Boytard, John Cadley, Paolo Cotzia, John Chen, Dacia Dimartino, Xiaojun Feng, Tatyana Gindin, Emily Guzman, Adriana Heguy, Megan Hogan, Emily Huang, George Jour, Andrew Lytle, Christian Marier, Matthew T. Maurano, Mark J. Mulligan, Peter Meyn, Iman Osman, Jared Pinnell, Vanessa Raabe, Sitharam Ramaswami, Amy Rapkiewicz, Marie Samanovic-Golden, Antonio Serrano, Guomiao Shen, Matija Snuderl, Theodore Vougiouklakis, Nick Vulpescu, Gael Westby, Paul Zappile, Yutong Zhang |
| EPI_ISL_427499 | hCoV-19/USA/NY-NYUMC204/2020 | North America / USA / New York / Manha | 2020-04-01 | NYU Langone Health | Departments of Pathology and Medicine, New York University School of Medicine | Maria Aguero-Rosenfeld, Brendan Belovarac, Margaret Black, Ludovic Boytard, John Cadley, Paolo Cotzia, John Chen, Dacia Dimartino, Xiaojun Feng, Tatyana Gindin, Emily Guzman, Adriana Heguy, Megan Hogan, Emily Huang, George Jour, Andrew Lytle, Christian Marier, Matthew T. Maurano, Mark J. Mulligan, Peter Meyn, Iman Osman, Jared Pinnell, Vanessa Raabe, Sitharam Ramaswami, Amy Rapkiewicz, Marie Samanovic-Golden, Antonio Serrano, Guomiao Shen, Matija Snuderl, Theodore Vougiouklakis, Nick Vulpescu, Gael Westby, Paul Zappile, Yutong Zhang |
| EPI_ISL_428670 | hCoV-19/Sri Lanka/COV38/2020 | Asia / Sri Lanka | 2020-03-16 | Centre for Dengue Research | Centre for Dengue Research | Chandima Jeewandara, Dinuka Ariyaratne, Laksiri Gomes, Deshni Jayathilaka, Ananda Wijewickrama, Eranga Narangoda, Damayanthi Idampitiya, Neelika Malaige |
| EPI_ISL_428671 | hCoV-19/Sri Lanka/COV53/2020 | Asia / Sri Lanka | 2020-03-10 | Centre for Dengue Research | Centre for Dengue Research | Chandima Jeewandara, Dinuka Ariyatane, Laksiri Gomes, Deshni Jayathilaka, Diyanath Ranasinghe, Ananda Wijewickrama, Eranga Narangoda, Damayanthi Tdampitiya, Neelika Malavige |
| EPI_ISL_428672 | hCoV-19/Sri Lanka/COV91/2020 | Asia / Sri Lanka | 2020-03-19 | Centre for Dengue Research | Centre for Dengue Research | Chandima Jeewandara, Dinuka Ariyaratne, Laksiri Gomes, Deshni Jayathilaka, Diyanath Ranasinghe, Ananda Wijewickrama, Eranga Narangoda, Damayanthi Idampitiya, Neelika Malavige |
| EPI_ISL_428673 | hCoV-19/Sri Lanka/COV486/2020 | Asia / Sri Lanka | 2020-03-31 | Centre for Dengue Research | Centre for Dengue Research | Chandima Jeewandara, Dinuka Ariyaratne, Laksiri Gomes, Deshni Jayathilaka, Diyanath Ranasinghe, Ananda Wijewickrama, Eranga Narangoda, Damayanthi Idampitiya, Neelika Malavige |
| EPI_ISL_428674 | hCoV-19/Spain/Madrid_LP11_2271/2020 | Europe / Spain / Madrid | 2020-03-09 | Hospital Universitario La Paz | Hospital Universitario 12 de Octubre | Elias Dahdouh, Sara González, Raúl Recio, Fernando Lázaro, Esther Viedma, Natalia Stella, Julio García, Juan Carlos Galán, Rafael Cantón, Mª Dolores Folgueira, Rafael Delgado, Jesús Mingorance |
| EPI_ISL_428675 | hCoV-19/Spain/Madrid_LP16_6193/2020 | Europe / Spain / Madrid | 2020-03-09 | Hospital Universitario La Paz | Hospital Universitario 12 de Octubre | Elias Dahdouh, Sara González, Raúl Recio, Fernando Lázaro, Esther Viedma, Natalia Stella, Julio García, Juan Carlos Galán, Rafael Cantón, Mª Dolores Folgueira, Rafael Delgado, Jesús Mingorance |
| EPI_ISL_428676 | hCoV-19/Spain/Madrid_LP19_4952/2020 | Europe / Spain / Madrid | 2020-03-07 | Hospital Universitario La Paz | Hospital Universitario 12 de Octubre | Elias Dahdouh, Sara González, Raúl Recio, Fernando Lázaro, Esther Viedma, Natalia Stella, Julio García, Juan Carlos Galán, Rafael Cantón, Mª Dolores Folgueira, Rafael Delgado, Jesús Mingorance |
| EPI_ISL_428677 | hCoV-19/Spain/Madrid_LP20_2327/2020 | Europe / Spain / Madrid | 2020-03-11 | Hospital Universitario La Paz | Hospital Universitario 12 de Octubre | Elias Dahdouh, Sara González, Raúl Recio, Fernando Lázaro, Esther Viedma, Natalia Stella, Julio García, Juan Carlos Galán, Rafael Cantón, Mª Dolores Folgueira, Rafael Delgado, Jesús Mingorance |
| EPI_ISL_428678 | hCoV-19/Spain/Madrid_LP22_5885/2020 | Europe / Spain / Madrid | 2020-03-08 | Hospital Universitario La Paz | Hospital Universitario 12 de Octubre | Elias Dahdouh, Sara González, Raúl Recio, Fernando Lázaro, Esther Viedma, Natalia Stella, Julio García, Juan Carlos Galán, Rafael Cantón, Mª Dolores Folgueira, Rafael Delgado, Jesús Mingorance |
| EPI_ISL_428679 | hCoV-19/Spain/Madrid_LP23_5852/2020 | Europe / Spain / Madrid | 2020-03-08 | Hospital Universitario La Paz | Hospital Universitario 12 de Octubre | Elias Dahdouh, Sara González, Raúl Recio, Fernando Lázaro, Esther Viedma, Natalia Stella, Julio García, Juan Carlos Galán, Rafael Cantón, Mª Dolores Folgueira, Rafael Delgado, Jesús Mingorance |
| EPI_ISL_428680 | hCoV-19/Spain/Madrid_LP24_5999/2020 | Europe / Spain / Madrid | 2020-03-09 | Hospital Universitario La Paz | Hospital Universitario 12 de Octubre | Elias Dahdouh, Sara González, Raúl Recio, Fernando Lázaro, Esther Viedma, Natalia Stella, Julio García, Juan Carlos Galán, Rafael Cantón, Mª Dolores Folgueira, Rafael Delgado, Jesús Mingorance |
| EPI_ISL_428681 | hCoV-19/Spain/Madrid_LP27_2548/2020 | Europe / Spain / Madrid | 2020-03-11 | Hospital Universitario La Paz | Hospital Universitario 12 de Octubre | Elias Dahdouh, Sara González, Raúl Recio, Fernando Lázaro, Esther Viedma, Natalia Stella, Julio García, Juan Carlos Galán, Rafael Cantón, Mª Dolores Folgueira, Rafael Delgado, Jesús Mingorance |
| EPI_ISL_428682 | hCoV-19/Spain/Madrid_LP30_2226/2020 | Europe / Spain / Madrid | 2020-03-11 | Hospital Universitario La Paz | Hospital Universitario 12 de Octubre | Elias Dahdouh, Sara González, Raúl Recio, Fernando Lázaro, Esther Viedma, Natalia Stella, Julio García, Juan Carlos Galán, Rafael Cantón, Mª Dolores Folgueira, Rafael Delgado, Jesús Mingorance |
| EPI_ISL_428683 | hCoV-19/Spain/Madrid_H12_13/2020 | Europe / Spain / Madrid | 2020-03-03 | Hospital Universitario 12 de Octubre | Hospital Universitario 12 de Octubre | Sara González, Raúl Recio,Elias Dahdouh, Fernando Lázaro, Esther Viedma, Natalia Stella, Julio García, Juan Carlos Galán, Rafael Cantón, Mª Dolores Folgueira, Rafael Delgado, Jesús Mingorance |
| EPI_ISL_428684 | hCoV-19/Spain/Madrid_H12_20/2020 | Europe / Spain / Madrid | 2020-03-27 | Hospital Universitario 12 de Octubre | Hospital Universitario 12 de Octubre | Sara González, Raúl Recio,Elias Dahdouh, Fernando Lázaro, Esther Viedma, Natalia Stella, Julio García, Juan Carlos Galán, Rafael Cantón, Mª Dolores Folgueira, Rafael Delgado, Jesús Mingorance |
| EPI_ISL_428686 | hCoV-19/Spain/Madrid_H12_23/2020 | Europe / Spain / Madrid | 2020-03-22 | Hospital Universitario 12 de Octubre | Hospital Universitario 12 de Octubre | Sara González, Raúl Recio,Elias Dahdouh, Fernando Lázaro, Esther Viedma, Natalia Stella, Julio García, Juan Carlos Galán, Rafael Cantón, Mª Dolores Folgueira, Rafael Delgado, Jesús Mingorance |
| EPI_ISL_428687 | hCoV-19/Spain/Madrid_H12_24/2020 | Europe / Spain / Madrid | 2020-03-22 | Hospital Universitario 12 de Octubre | Hospital Universitario 12 de Octubre | Sara González, Raúl Recio,Elias Dahdouh, Fernando Lázaro, Esther Viedma, Natalia Stella, Julio García, Juan Carlos Galán, Rafael Cantón, Mª Dolores Folgueira, Rafael Delgado, Jesús Mingorance |
| EPI_ISL_428688 | hCoV-19/Spain/Madrid_H12_25/2020 | Europe / Spain / Madrid | 2020-03-22 | Hospital Universitario 12 de Octubre | Hospital Universitario 12 de Octubre | Sara González, Raúl Recio,Elias Dahdouh, Fernando Lázaro, Esther Viedma, Natalia Stella, Julio García, Juan Carlos Galán, Rafael Cantón, Mª Dolores Folgueira, Rafael Delgado, Jesús Mingorance |
| EPI_ISL_428689 | hCoV-19/Spain/Madrid_H12_26/2020 | Europe / Spain / Madrid | 2020-03-26 | Hospital Universitario 12 de Octubre | Hospital Universitario 12 de Octubre | Sara González, Raúl Recio,Elias Dahdouh, Fernando Lázaro, Esther Viedma, Natalia Stella, Julio García, Juan Carlos Galán, Rafael Cantón, Mª Dolores Folgueira, Rafael Delgado, Jesús Mingorance |
| EPI_ISL_428690 | hCoV-19/Spain/Madrid_H12_27/2020 | Europe / Spain / Madrid | 2020-03-21 | Hospital Universitario 12 de Octubre | Hospital Universitario 12 de Octubre | Sara González, Raúl Recio,Elias Dahdouh, Fernando Lázaro, Esther Viedma, Natalia Stella, Julio García, Juan Carlos Galán, Rafael Cantón, Mª Dolores Folgueira, Rafael Delgado, Jesús Mingorance |
| EPI_ISL_428691 | hCoV-19/Spain/Madrid_H12_28/2020 | Europe / Spain / Madrid | 2020-03-21 | Hospital Universitario 12 de Octubre | Hospital Universitario 12 de Octubre | Sara González, Raúl Recio,Elias Dahdouh, Fernando Lázaro, Esther Viedma, Natalia Stella, Julio García, Juan Carlos Galán, Rafael Cantón, Mª Dolores Folgueira, Rafael Delgado, Jesús Mingorance |
| EPI_ISL_428692 | hCoV-19/Spain/Madrid_H12_29/2020 | Europe / Spain / Madrid | 2020-03-12 | Hospital Universitario 12 de Octubre | Hospital Universitario 12 de Octubre | Sara González, Raúl Recio,Elias Dahdouh, Fernando Lázaro, Esther Viedma, Natalia Stella, Julio García, Juan Carlos Galán, Rafael Cantón, Mª Dolores Folgueira, Rafael Delgado, Jesús Mingorance |
| EPI_ISL_428693 | hCoV-19/Spain/Madrid_H12_30/2020 | Europe / Spain / Madrid | 2020-03-12 | Hospital Universitario 12 de Octubre | Hospital Universitario 12 de Octubre | Sara González, Raúl Recio,Elias Dahdouh, Fernando Lázaro, Esther Viedma, Natalia Stella, Julio García, Juan Carlos Galán, Rafael Cantón, Mª Dolores Folgueira, Rafael Delgado, Jesús Mingorance |
| EPI_ISL_428694 | hCoV-19/Spain/Madrid_H12_31/2020 | Europe / Spain / Madrid | 2020-03-12 | Hospital Universitario 12 de Octubre | Hospital Universitario 12 de Octubre | Sara González, Raúl Recio,Elias Dahdouh, Fernando Lázaro, Esther Viedma, Natalia Stella, Julio García, Juan Carlos Galán, Rafael Cantón, Mª Dolores Folgueira, Rafael Delgado, Jesús Mingorance |
| EPI_ISL_428695 | hCoV-19/Spain/Madrid_H12_32/2020 | Europe / Spain / Madrid | 2020-03-12 | Hospital Universitario 12 de Octubre | Hospital Universitario 12 de Octubre | Raúl Recio, Sara González, Elias Dahdouh, Fernando Lázaro, Esther Viedma, Natalia Stella, Julio García, Juan Carlos Galán, Rafael Cantón, Mª Dolores Folgueira, Rafael Delgado, Jesús Mingorance |
| EPI_ISL_428696 | hCoV-19/Spain/Madrid_H12_33/2020 | Europe / Spain / Madrid | 2020-03-12 | Hospital Universitario 12 de Octubre | Hospital Universitario 12 de Octubre | Raúl Recio, Sara González, Elias Dahdouh, Fernando Lázaro, Esther Viedma, Natalia Stella, Julio García, Juan Carlos Galán, Rafael Cantón, Mª Dolores Folgueira, Rafael Delgado, Jesús Mingorance |
| EPI_ISL_428697 | hCoV-19/Spain/Madrid_H12_34/2020 | Europe / Spain / Madrid | 2020-03-12 | Hospital Universitario 12 de Octubre | Hospital Universitario 12 de Octubre | Raúl Recio, Sara González, Elias Dahdouh, Fernando Lázaro, Esther Viedma, Natalia Stella, Julio García, Juan Carlos Galán, Rafael Cantón, Mª Dolores Folgueira, Rafael Delgado, Jesús Mingorance |
| EPI_ISL_428698 | hCoV-19/Spain/Madrid_H12_35/2020 | Europe / Spain / Madrid | 2020-03-12 | Hospital Universitario 12 de Octubre | Hospital Universitario 12 de Octubre | Raúl Recio, Sara González, Elias Dahdouh, Fernando Lázaro, Esther Viedma, Natalia Stella, Julio García, Juan Carlos Galán, Rafael Cantón, Mª Dolores Folgueira, Rafael Delgado, Jesús Mingorance |
| EPI_ISL_428699 | hCoV-19/Spain/Madrid_H12_36/2020 | Europe / Spain / Madrid | 2020-03-09 | Hospital Universitario 12 de Octubre | Hospital Universitario 12 de Octubre | Raúl Recio, Sara González, Elias Dahdouh, Fernando Lázaro, Esther Viedma, Natalia Stella, Julio García, Juan Carlos Galán, Rafael Cantón, Mª Dolores Folgueira, Rafael Delgado, Jesús Mingorance |
| EPI_ISL_428716 | hCoV-19/Turkey/HSGM-5711/2020 | Europe / Turkey / Ankara | 2020-03-18 | Ministry of Health Turkey | Ministry of Health Turkey | Fatma Bayrakdar,Ayşe Başak Altaş,Yasemin Coşgun,Gülay Korukluoğlu,Selçuk Kılıç |
| EPI_ISL_428717 | hCoV-19/Turkey/HSGM-5769/2020 | Europe / Turkey / Kocaeli | 2020-03-19 | Ministry of Health Turkey | Ministry of Health Turkey | Fatma Bayrakdar,Ayşe Başak Altaş,Yasemin Coşgun,Gülay Korukluoğlu,Selçuk Kılıç |
| EPI_ISL_428718 | hCoV-19/Turkey/HSGM-5770/2020 | Europe / Turkey / Kocaeli | 2020-03-19 | Ministry of Health Turkey | Ministry of Health Turkey | Fatma Bayrakdar,Ayşe Başak Altaş,Yasemin Coşgun,Gülay Korukluoğlu,Selçuk Kılıç |
| EPI_ISL_428719 | hCoV-19/Turkey/HSGM-7718/2020 | Europe / Turkey / Siirt | 2020-03-21 | Ministry of Health Turkey | Ministry of Health Turkey | Fatma Bayrakdar,Ayşe Başak Altaş,Yasemin Coşgun,Gülay Korukluoğlu,Selçuk Kılıç |
| EPI_ISL_428720 | hCoV-19/Turkey/HSGM-8001/2020 | Europe / Turkey / Ankara | 2020-03-21 | Ministry of Health Turkey | Ministry of Health Turkey | Fatma Bayrakdar,Ayşe Başak Altaş,Yasemin Coşgun,Gülay Korukluoğlu,Selçuk Kılıç |
| EPI_ISL_428721 | hCoV-19/Turkey/HSGM-8003/2020 | Europe / Turkey / Ankara | 2020-03-21 | Ministry of Health Turkey | Ministry of Health Turkey | Fatma Bayrakdar,Ayşe Başak Altaş,Yasemin Coşgun,Gülay Korukluoğlu,Selçuk Kılıç |
| EPI_ISL_428722 | hCoV-19/Turkey/HSGM-8010/2020 | Europe / Turkey / Balikesir | 2020-03-22 | Ministry of Health Turkey | Ministry of Health Turkey | Fatma Bayrakdar,Ayşe Başak Altaş,Yasemin Coşgun,Gülay Korukluoğlu,Selçuk Kılıç |
| EPI_ISL_428724 | hCoV-19/USA/WI-UW-250/2020 | North America / USA / Wisconsin | 2020-03-13 | University of Wisconsin-Madison AIDS Vaccine Research Laboratories | University of Wisconsin-Madison AIDS Vaccine Research Laboratories | Gage Moreno, Katarina Braun, et al. AIDS Vaccine Research Laboratories |
| EPI_ISL_428780 | hCoV-19/USA/NY-NYUMC266/2020 | North America / USA / New York / Brookl | 2020-04-06 | NYU Langone Health | Departments of Pathology and Medicine, New York University School of Medicine | Maria Aguero-Rosenfeld, Brendan Belovarac, Margaret Black, Ludovic Boytard, John Cadley, Paolo Cotzia, John Chen, Dacia Dimartino, Xiaojun Feng, Tatyana Gindin, Emily Guzman, Adriana Heguy, Megan Hogan, Emily Huang, George Jour, Andrew Lytle, Christian Marier, Matthew T. Maurano, Mark J. Mulligan, Peter Meyn, Iman Osman, Jared Pinnell, Vanessa Raabe, Sitharam Ramaswami, Amy Rapkiewicz, Marie Samanovic-Golden, Antonio Serrano, Guomiao Shen, Matija Snuderl, Theodore Vougiouklakis, Nick Vulpescu, Gael Westby, Paul Zappile, Yutong Zhang |
| EPI_ISL_428781 | hCoV-19/USA/NY-NYUMC267/2020 | North America / USA / New York / Brookl | 2020-04-06 | NYU Langone Health | Departments of Pathology and Medicine, New York University School of Medicine | Maria Aguero-Rosenfeld, Brendan Belovarac, Margaret Black, Ludovic Boytard, John Cadley, Paolo Cotzia, John Chen, Dacia Dimartino, Xiaojun Feng, Tatyana Gindin, Emily Guzman, Adriana Heguy, Megan Hogan, Emily Huang, George Jour, Andrew Lytle, Christian Marier, Matthew T. Maurano, Mark J. Mulligan, Peter Meyn, Iman Osman, Jared Pinnell, Vanessa Raabe, Sitharam Ramaswami, Amy Rapkiewicz, Marie Samanovic-Golden, Antonio Serrano, Guomiao Shen, Matija Snuderl, Theodore Vougiouklakis, Nick Vulpescu, Gael Westby, Paul Zappile, Yutong Zhang |
| EPI_ISL_428783 | hCoV-19/USA/NY-NYUMC269/2020 | North America / USA / New York / Brookl | 2020-04-06 | NYU Langone Health | Departments of Pathology and Medicine, New York University School of Medicine | Maria Aguero-Rosenfeld, Brendan Belovarac, Margaret Black, Ludovic Boytard, John Cadley, Paolo Cotzia, John Chen, Dacia Dimartino, Xiaojun Feng, Tatyana Gindin, Emily Guzman, Adriana Heguy, Megan Hogan, Emily Huang, George Jour, Andrew Lytle, Christian Marier, Matthew T. Maurano, Mark J. Mulligan, Peter Meyn, Iman Osman, Jared Pinnell, Vanessa Raabe, Sitharam Ramaswami, Amy Rapkiewicz, Marie Samanovic-Golden, Antonio Serrano, Guomiao Shen, Matija Snuderl, Theodore Vougiouklakis, Nick Vulpescu, Gael Westby, Paul Zappile, Yutong Zhang |
| EPI_ISL_428784 | hCoV-19/USA/NY-NYUMC270/2020 | North America / USA / New York / Brookl | 2020-04-06 | NYU Langone Health | Departments of Pathology and Medicine, New York University School of Medicine | Maria Aguero-Rosenfeld, Brendan Belovarac, Margaret Black, Ludovic Boytard, John Cadley, Paolo Cotzia, John Chen, Dacia Dimartino, Xiaojun Feng, Tatyana Gindin, Emily Guzman, Adriana Heguy, Megan Hogan, Emily Huang, George Jour, Andrew Lytle, Christian Marier, Matthew T. Maurano, Mark J. Mulligan, Peter Meyn, Iman Osman, Jared Pinnell, Vanessa Raabe, Sitharam Ramaswami, Amy Rapkiewicz, Marie Samanovic-Golden, Antonio Serrano, Guomiao Shen, Matija Snuderl, Theodore Vougiouklakis, Nick Vulpescu, Gael Westby, Paul Zappile, Yutong Zhang |
| EPI_ISL_428785 | hCoV-19/USA/NY-NYUMC271/2020 | North America / USA / New York / Brookl | 2020-04-06 | NYU Langone Health | Departments of Pathology and Medicine, New York University School of Medicine | Maria Aguero-Rosenfeld, Brendan Belovarac, Margaret Black, Ludovic Boytard, John Cadley, Paolo Cotzia, John Chen, Dacia Dimartino, Xiaojun Feng, Tatyana Gindin, Emily Guzman, Adriana Heguy, Megan Hogan, Emily Huang, George Jour, Andrew Lytle, Christian Marier, Matthew T. Maurano, Mark J. Mulligan, Peter Meyn, Iman Osman, Jared Pinnell, Vanessa Raabe, Sitharam Ramaswami, Amy Rapkiewicz, Marie Samanovic-Golden, Antonio Serrano, Guomiao Shen, Matija Snuderl, Theodore Vougiouklakis, Nick Vulpescu, Gael Westby, Paul Zappile, Yutong Zhang |
| EPI_ISL_428786 | hCoV-19/USA/NY-NYUMC272/2020 | North America / USA / New York / Brookl | 2020-04-06 | NYU Langone Health | Departments of Pathology and Medicine, New York University School of Medicine | Maria Aguero-Rosenfeld, Brendan Belovarac, Margaret Black, Ludovic Boytard, John Cadley, Paolo Cotzia, John Chen, Dacia Dimartino, Xiaojun Feng, Tatyana Gindin, Emily Guzman, Adriana Heguy, Megan Hogan, Emily Huang, George Jour, Andrew Lytle, Christian Marier, Matthew T. Maurano, Mark J. Mulligan, Peter Meyn, Iman Osman, Jared Pinnell, Vanessa Raabe, Sitharam Ramaswami, Amy Rapkiewicz, Marie Samanovic-Golden, Antonio Serrano, Guomiao Shen, Matija Snuderl, Theodore Vougiouklakis, Nick Vulpescu, Gael Westby, Paul Zappile, Yutong Zhang |
| EPI_ISL_428787 | hCoV-19/USA/NY-NYUMC273/2020 | North America / USA / New York / Brookl | 2020-04-06 | NYU Langone Health | Departments of Pathology and Medicine, New York University School of Medicine | Maria Aguero-Rosenfeld, Brendan Belovarac, Margaret Black, Ludovic Boytard, John Cadley, Paolo Cotzia, John Chen, Dacia Dimartino, Xiaojun Feng, Tatyana Gindin, Emily Guzman, Adriana Heguy, Megan Hogan, Emily Huang, George Jour, Andrew Lytle, Christian Marier, Matthew T. Maurano, Mark J. Mulligan, Peter Meyn, Iman Osman, Jared Pinnell, Vanessa Raabe, Sitharam Ramaswami, Amy Rapkiewicz, Marie Samanovic-Golden, Antonio Serrano, Guomiao Shen, Matija Snuderl, Theodore Vougiouklakis, Nick Vulpescu, Gael Westby, Paul Zappile, Yutong Zhang |
| EPI_ISL_428788 | hCoV-19/USA/NY-NYUMC274/2020 | North America / USA / New York / Brookl | 2020-04-06 | NYU Langone Health | Departments of Pathology and Medicine, New York University School of Medicine | Maria Aguero-Rosenfeld, Brendan Belovarac, Margaret Black, Ludovic Boytard, John Cadley, Paolo Cotzia, John Chen, Dacia Dimartino, Xiaojun Feng, Tatyana Gindin, Emily Guzman, Adriana Heguy, Megan Hogan, Emily Huang, George Jour, Andrew Lytle, Christian Marier, Matthew T. Maurano, Mark J. Mulligan, Peter Meyn, Iman Osman, Jared Pinnell, Vanessa Raabe, Sitharam Ramaswami, Amy Rapkiewicz, Marie Samanovic-Golden, Antonio Serrano, Guomiao Shen, Matija Snuderl, Theodore Vougiouklakis, Nick Vulpescu, Gael Westby, Paul Zappile, Yutong Zhang |
| EPI_ISL_428789 | hCoV-19/USA/NY-NYUMC275/2020 | North America / USA / New York / Brookl | 2020-04-06 | NYU Langone Health | Departments of Pathology and Medicine, New York University School of Medicine | Maria Aguero-Rosenfeld, Brendan Belovarac, Margaret Black, Ludovic Boytard, John Cadley, Paolo Cotzia, John Chen, Dacia Dimartino, Xiaojun Feng, Tatyana Gindin, Emily Guzman, Adriana Heguy, Megan Hogan, Emily Huang, George Jour, Andrew Lytle, Christian Marier, Matthew T. Maurano, Mark J. Mulligan, Peter Meyn, Iman Osman, Jared Pinnell, Vanessa Raabe, Sitharam Ramaswami, Amy Rapkiewicz, Marie Samanovic-Golden, Antonio Serrano, Guomiao Shen, Matija Snuderl, Theodore Vougiouklakis, Nick Vulpescu, Gael Westby, Paul Zappile, Yutong Zhang |
| EPI_ISL_428790 | hCoV-19/USA/NY-NYUMC276/2020 | North America / USA / New York / Brookl | 2020-04-06 | NYU Langone Health | Departments of Pathology and Medicine, New York University School of Medicine | Maria Aguero-Rosenfeld, Brendan Belovarac, Margaret Black, Ludovic Boytard, John Cadley, Paolo Cotzia, John Chen, Dacia Dimartino, Xiaojun Feng, Tatyana Gindin, Emily Guzman, Adriana Heguy, Megan Hogan, Emily Huang, George Jour, Andrew Lytle, Christian Marier, Matthew T. Maurano, Mark J. Mulligan, Peter Meyn, Iman Osman, Jared Pinnell, Vanessa Raabe, Sitharam Ramaswami, Amy Rapkiewicz, Marie Samanovic-Golden, Antonio Serrano, Guomiao Shen, Matija Snuderl, Theodore Vougiouklakis, Nick Vulpescu, Gael Westby, Paul Zappile, Yutong Zhang |
| EPI_ISL_428791 | hCoV-19/USA/NY-NYUMC277/2020 | North America / USA / New York / Brookl | 2020-04-04 | NYU Langone Health | Departments of Pathology and Medicine, New York University School of Medicine | Maria Aguero-Rosenfeld, Brendan Belovarac, Margaret Black, Ludovic Boytard, John Cadley, Paolo Cotzia, John Chen, Dacia Dimartino, Xiaojun Feng, Tatyana Gindin, Emily Guzman, Adriana Heguy, Megan Hogan, Emily Huang, George Jour, Andrew Lytle, Christian Marier, Matthew T. Maurano, Mark J. Mulligan, Peter Meyn, Iman Osman, Jared Pinnell, Vanessa Raabe, Sitharam Ramaswami, Amy Rapkiewicz, Marie Samanovic-Golden, Antonio Serrano, Guomiao Shen, Matija Snuderl, Theodore Vougiouklakis, Nick Vulpescu, Gael Westby, Paul Zappile, Yutong Zhang |
| EPI_ISL_428792 | hCoV-19/USA/NY-NYUMC278/2020 | North America / USA / New York / Brookl | 2020-04-04 | NYU Langone Health | Departments of Pathology and Medicine, New York University School of Medicine | Maria Aguero-Rosenfeld, Brendan Belovarac, Margaret Black, Ludovic Boytard, John Cadley, Paolo Cotzia, John Chen, Dacia Dimartino, Xiaojun Feng, Tatyana Gindin, Emily Guzman, Adriana Heguy, Megan Hogan, Emily Huang, George Jour, Andrew Lytle, Christian Marier, Matthew T. Maurano, Mark J. Mulligan, Peter Meyn, Iman Osman, Jared Pinnell, Vanessa Raabe, Sitharam Ramaswami, Amy Rapkiewicz, Marie Samanovic-Golden, Antonio Serrano, Guomiao Shen, Matija Snuderl, Theodore Vougiouklakis, Nick Vulpescu, Gael Westby, Paul Zappile, Yutong Zhang |
| EPI_ISL_428793 | hCoV-19/USA/NY-NYUMC279/2020 | North America / USA / New York / Brookl | 2020-04-05 | NYU Langone Health | Departments of Pathology and Medicine, New York University School of Medicine | Maria Aguero-Rosenfeld, Brendan Belovarac, Margaret Black, Ludovic Boytard, John Cadley, Paolo Cotzia, John Chen, Dacia Dimartino, Xiaojun Feng, Tatyana Gindin, Emily Guzman, Adriana Heguy, Megan Hogan, Emily Huang, George Jour, Andrew Lytle, Christian Marier, Matthew T. Maurano, Mark J. Mulligan, Peter Meyn, Iman Osman, Jared Pinnell, Vanessa Raabe, Sitharam Ramaswami, Amy Rapkiewicz, Marie Samanovic-Golden, Antonio Serrano, Guomiao Shen, Matija Snuderl, Theodore Vougiouklakis, Nick Vulpescu, Gael Westby, Paul Zappile, Yutong Zhang |
| EPI_ISL_428794 | hCoV-19/USA/NY-NYUMC280/2020 | North America / USA / New York / Brookl | 2020-04-05 | NYU Langone Health | Departments of Pathology and Medicine, New York University School of Medicine | Maria Aguero-Rosenfeld, Brendan Belovarac, Margaret Black, Ludovic Boytard, John Cadley, Paolo Cotzia, John Chen, Dacia Dimartino, Xiaojun Feng, Tatyana Gindin, Emily Guzman, Adriana Heguy, Megan Hogan, Emily Huang, George Jour, Andrew Lytle, Christian Marier, Matthew T. Maurano, Mark J. Mulligan, Peter Meyn, Iman Osman, Jared Pinnell, Vanessa Raabe, Sitharam Ramaswami, Amy Rapkiewicz, Marie Samanovic-Golden, Antonio Serrano, Guomiao Shen, Matija Snuderl, Theodore Vougiouklakis, Nick Vulpescu, Gael Westby, Paul Zappile, Yutong Zhang |
| EPI_ISL_428795 | hCoV-19/USA/NY-NYUMC281/2020 | North America / USA / New York / Brookl | 2020-04-05 | NYU Langone Health | Departments of Pathology and Medicine, New York University School of Medicine | Maria Aguero-Rosenfeld, Brendan Belovarac, Margaret Black, Ludovic Boytard, John Cadley, Paolo Cotzia, John Chen, Dacia Dimartino, Xiaojun Feng, Tatyana Gindin, Emily Guzman, Adriana Heguy, Megan Hogan, Emily Huang, George Jour, Andrew Lytle, Christian Marier, Matthew T. Maurano, Mark J. Mulligan, Peter Meyn, Iman Osman, Jared Pinnell, Vanessa Raabe, Sitharam Ramaswami, Amy Rapkiewicz, Marie Samanovic-Golden, Antonio Serrano, Guomiao Shen, Matija Snuderl, Theodore Vougiouklakis, Nick Vulpescu, Gael Westby, Paul Zappile, Yutong Zhang |
| EPI_ISL_428797 | hCoV-19/USA/NY-NYUMC283/2020 | North America / USA / New York / Brookl | 2020-04-05 | NYU Langone Health | Departments of Pathology and Medicine, New York University School of Medicine | Maria Aguero-Rosenfeld, Brendan Belovarac, Margaret Black, Ludovic Boytard, John Cadley, Paolo Cotzia, John Chen, Dacia Dimartino, Xiaojun Feng, Tatyana Gindin, Emily Guzman, Adriana Heguy, Megan Hogan, Emily Huang, George Jour, Andrew Lytle, Christian Marier, Matthew T. Maurano, Mark J. Mulligan, Peter Meyn, Iman Osman, Jared Pinnell, Vanessa Raabe, Sitharam Ramaswami, Amy Rapkiewicz, Marie Samanovic-Golden, Antonio Serrano, Guomiao Shen, Matija Snuderl, Theodore Vougiouklakis, Nick Vulpescu, Gael Westby, Paul Zappile, Yutong Zhang |
| EPI_ISL_428798 | hCoV-19/USA/NY-NYUMC284/2020 | North America / USA / New York / Brookl | 2020-04-05 | NYU Langone Health | Departments of Pathology and Medicine, New York University School of Medicine | Maria Aguero-Rosenfeld, Brendan Belovarac, Margaret Black, Ludovic Boytard, John Cadley, Paolo Cotzia, John Chen, Dacia Dimartino, Xiaojun Feng, Tatyana Gindin, Emily Guzman, Adriana Heguy, Megan Hogan, Emily Huang, George Jour, Andrew Lytle, Christian Marier, Matthew T. Maurano, Mark J. Mulligan, Peter Meyn, Iman Osman, Jared Pinnell, Vanessa Raabe, Sitharam Ramaswami, Amy Rapkiewicz, Marie Samanovic-Golden, Antonio Serrano, Guomiao Shen, Matija Snuderl, Theodore Vougiouklakis, Nick Vulpescu, Gael Westby, Paul Zappile, Yutong Zhang |
| EPI_ISL_428799 | hCoV-19/USA/NY-NYUMC285/2020 | North America / USA / New York / Brookl | 2020-04-05 | NYU Langone Health | Departments of Pathology and Medicine, New York University School of Medicine | Maria Aguero-Rosenfeld, Brendan Belovarac, Margaret Black, Ludovic Boytard, John Cadley, Paolo Cotzia, John Chen, Dacia Dimartino, Xiaojun Feng, Tatyana Gindin, Emily Guzman, Adriana Heguy, Megan Hogan, Emily Huang, George Jour, Andrew Lytle, Christian Marier, Matthew T. Maurano, Mark J. Mulligan, Peter Meyn, Iman Osman, Jared Pinnell, Vanessa Raabe, Sitharam Ramaswami, Amy Rapkiewicz, Marie Samanovic-Golden, Antonio Serrano, Guomiao Shen, Matija Snuderl, Theodore Vougiouklakis, Nick Vulpescu, Gael Westby, Paul Zappile, Yutong Zhang |
| EPI_ISL_429070 | hCoV-19/USA/CA-CZB0700/2020 | North America / USA / California / San Fr | 2020-03-23 | UCSF Clinical Microbiology Laboratory | Chan-Zuckerberg Biohub | CZB Cliahub Consortium |
| EPI_ISL_429071 | hCoV-19/USA/CA-CZB099/2020 | North America / USA / California / San Fr | 2020-03-28 | UCSF Clinical Microbiology Laboratory | Chan-Zuckerberg Biohub | CZB Cliahub Consortium |
| EPI_ISL_429072 | hCoV-19/USA/CA-CZB0719/2020 | North America / USA / California / San Fr | 2020-03-31 | UCSF Clinical Microbiology Laboratory | Chan-Zuckerberg Biohub | CZB Cliahub Consortium |
| EPI_ISL_429073 | hCoV-19/USA/CA-CZB017/2020 | North America / USA / California / San Fr | 2020-03-25 | UCSF Clinical Microbiology Laboratory | Chan-Zuckerberg Biohub | CZB Cliahub Consortium |
| EPI_ISL_429074 | hCoV-19/Guangzhou/GZMU0003/2020 | Asia / China / Guangzhou | 2020-02-07 | The First Affiliated Hospital of Guangzhou Medical University | BGI-shenzhen & The First Affiliated Hospital of Guangzhou Medical University | Yanqun Wang, Daxi Wang, Lu Zhang, Wanying Sun, Zhaoyong Zhang et al. |
| EPI_ISL_429075 | hCoV-19/Guangzhou/GZMU0008/2020 | Asia / China / Guangzhou | 2020-02-07 | The First Affiliated Hospital of Guangzhou Medical University | BGI-shenzhen & The First Affiliated Hospital of Guangzhou Medical University | Yanqun Wang, Daxi Wang, Lu Zhang, Wanying Sun, Zhaoyong Zhang et al. |
| EPI_ISL_429076 | hCoV-19/Guangzhou/GZMU0009/2020 | Asia / China / Guangzhou | 2020-02-07 | The First Affiliated Hospital of Guangzhou Medical University | BGI-shenzhen & The First Affiliated Hospital of Guangzhou Medical University | |
| EPI_ISL_429077 | hCoV-19/Guangzhou/GZMU0010/2020 | Asia / China / Guangzhou | 2020-02-07 | The First Affiliated Hospital of Guangzhou Medical University | BGI-shenzhen & The First Affiliated Hospital of Guangzhou Medical University | Yanqun Wang, Daxi Wang, Lu Zhang, Wanying Sun, Zhaoyong Zhang et al. |
| EPI_ISL_429078 | hCoV-19/Guangzhou/GZMU0013/2020 | Asia / China / Guangzhou | 2020-02-07 | The First Affiliated Hospital of Guangzhou Medical University | BGI-shenzhen & The First Affiliated Hospital of Guangzhou Medical University | |
| EPI_ISL_429079 | hCoV-19/Guangzhou/GZMU0017/2020 | Asia / China / Guangzhou | 2020-02-05 | The First Affiliated Hospital of Guangzhou Medical University | BGI-shenzhen & The First Affiliated Hospital of Guangzhou Medical University | |
| EPI_ISL_429080 | hCoV-19/Guangzhou/GZMU0019/2020 | Asia / China / Guangzhou | 2020-02-05 | The First Affiliated Hospital of Guangzhou Medical University | BGI-shenzhen & The First Affiliated Hospital of Guangzhou Medical University | |
| EPI_ISL_429081 | hCoV-19/Guangzhou/GZMU0020/2020 | Asia / China / Guangzhou | 2020-02-05 | The First Affiliated Hospital of Guangzhou Medical University | BGI-shenzhen & The First Affiliated Hospital of Guangzhou Medical University | |
| EPI_ISL_429082 | hCoV-19/Guangzhou/GZMU0023/2020 | Asia / China / Guangzhou | 2020-02-05 | The First Affiliated Hospital of Guangzhou Medical University | BGI-shenzhen & The First Affiliated Hospital of Guangzhou Medical University | Yanqun Wang, Daxi Wang, Lu Zhang, Wanying Sun, Zhaoyong Zhang et al. |
| EPI_ISL_429083 | hCoV-19/Guangzhou/GZMU0024/2020 | Asia / China / Guangzhou | 2020-02-05 | The First Affiliated Hospital of Guangzhou Medical University | BGI-shenzhen & The First Affiliated Hospital of Guangzhou Medical University | Yanqun Wang, Daxi Wang, Lu Zhang, Wanying Sun, Zhaoyong Zhang et al. |
| EPI_ISL_429084 | hCoV-19/Guangzhou/GZMU0025/2020 | Asia / China / Guangzhou | 2020-02-05 | The First Affiliated Hospital of Guangzhou Medical University | BGI-shenzhen & The First Affiliated Hospital of Guangzhou Medical University | |
| EPI_ISL_429085 | hCoV-19/Guangzhou/GZMU0033/2020 | Asia / China / Guangzhou | 2020-02-01 | The First Affiliated Hospital of Guangzhou Medical University | BGI-shenzhen & The First Affiliated Hospital of Guangzhou Medical University | Yanqun Wang, Daxi Wang, Lu Zhang, Wanying Sun, Zhaoyong Zhang et al. |
| EPI_ISL_429086 | hCoV-19/Guangzhou/GZMU0034/2020 | Asia / China / Guangzhou | 2020-02-01 | The First Affiliated Hospital of Guangzhou Medical University | BGI-shenzhen & The First Affiliated Hospital of Guangzhou Medical University | |
| EPI_ISL_429088 | hCoV-19/Guangzhou/GZMU0035/2020 | Asia / China / Guangzhou | 2020-02-01 | The First Affiliated Hospital of Guangzhou Medical University | BGI-shenzhen & The First Affiliated Hospital of Guangzhou Medical University | Yanqun Wang, Daxi Wang, Lu Zhang, Wanying Sun, Zhaoyong Zhang et al. |
| EPI_ISL_429089 | hCoV-19/Guangzhou/GZMU0037/2020 | Asia / China / Guangzhou | 2020-02-08 | The First Affiliated Hospital of Guangzhou Medical University | BGI-shenzhen & The First Affiliated Hospital of Guangzhou Medical University | Yanqun Wang, Daxi Wang, Lu Zhang, Wanying Sun, Zhaoyong Zhang et al. |
| EPI_ISL_429090 | hCoV-19/Guangzhou/GZMU0039/2020 | Asia / China / Guangzhou | 2020-02-08 | The First Affiliated Hospital of Guangzhou Medical University | BGI-shenzhen & The First Affiliated Hospital of Guangzhou Medical University | Yanqun Wang, Daxi Wang, Lu Zhang, Wanying Sun, Zhaoyong Zhang et al. |
| EPI_ISL_429091 | hCoV-19/Guangzhou/GZMU0041/2020 | Asia / China / Guangzhou | 2020-02-08 | The First Affiliated Hospital of Guangzhou Medical University | BGI-shenzhen & The First Affiliated Hospital of Guangzhou Medical University | Yanqun Wang, Daxi Wang, Lu Zhang, Wanying Sun, Zhaoyong Zhang et al. |
| EPI_ISL_429092 | hCoV-19/Guangzhou/GZMU0045/2020 | Asia / China / Guangzhou | 2020-02-08 | The First Affiliated Hospital of Guangzhou Medical University | BGI-shenzhen & The First Affiliated Hospital of Guangzhou Medical University | Yanqun Wang, Daxi Wang, Lu Zhang, Wanying Sun, Zhaoyong Zhang et al. |
| EPI_ISL_429093 | hCoV-19/Guangzhou/GZMU0046/2020 | Asia / China / Guangzhou | 2020-02-08 | The First Affiliated Hospital of Guangzhou Medical University | BGI-shenzhen & The First Affiliated Hospital of Guangzhou Medical University | Yanqun Wang, Daxi Wang, Lu Zhang, Wanying Sun, Zhaoyong Zhang et al. |
| EPI_ISL_429094 | hCoV-19/Guangzhou/GZMU0049/2020 | Asia / China / Guangzhou | 2020-02-08 | The First Affiliated Hospital of Guangzhou Medical University | BGI-shenzhen & The First Affiliated Hospital of Guangzhou Medical University | |
| EPI_ISL_429095 | hCoV-19/Guangzhou/GZMU0050/2020 | Asia / China / Guangzhou | 2020-02-08 | The First Affiliated Hospital of Guangzhou Medical University | BGI-shenzhen & The First Affiliated Hospital of Guangzhou Medical University | |
| EPI_ISL_429096 | hCoV-19/Guangzhou/GZMU0051/2020 | Asia / China / Guangzhou | 2020-01-29 | The First Affiliated Hospital of Guangzhou Medical University | BGI-shenzhen & The First Affiliated Hospital of Guangzhou Medical University | Yanqun Wang, Daxi Wang, Lu Zhang, Wanying Sun, Zhaoyong Zhang et al. |
| EPI_ISL_429097 | hCoV-19/Guangzhou/GZMU0052/2020 | Asia / China / Guangzhou | 2020-01-29 | The First Affiliated Hospital of Guangzhou Medical University | BGI-shenzhen & The First Affiliated Hospital of Guangzhou Medical University | Yanqun Wang, Daxi Wang, Lu Zhang, Wanying Sun, Zhaoyong Zhang et al. |
| EPI_ISL_429098 | hCoV-19/Guangzhou/GZMU0054/2020 | Asia / China / Guangzhou | 2020-01-29 | The First Affiliated Hospital of Guangzhou Medical University | BGI-shenzhen & The First Affiliated Hospital of Guangzhou Medical University | Yanqun Wang, Daxi Wang, Lu Zhang, Wanying Sun, Zhaoyong Zhang et al. |
| EPI_ISL_429100 | hCoV-19/Guangzhou/GZMU0057/2020 | Asia / China / Guangzhou | 2020-01-27 | The First Affiliated Hospital of Guangzhou Medical University | BGI-shenzhen & The First Affiliated Hospital of Guangzhou Medical University | |
| EPI_ISL_429101 | hCoV-19/Guangzhou/GZMU0058/2020 | Asia / China / Guangzhou | 2020-01-27 | The First Affiliated Hospital of Guangzhou Medical University | BGI-shenzhen & The First Affiliated Hospital of Guangzhou Medical University | |
| EPI_ISL_429102 | hCoV-19/Guangzhou/GZMU0059/2020 | Asia / China / Guangzhou | 2020-01-27 | The First Affiliated Hospital of Guangzhou Medical University | BGI-shenzhen & The First Affiliated Hospital of Guangzhou Medical University | |
| EPI_ISL_429103 | hCoV-19/Guangzhou/GZMU0060/2020 | Asia / China / Guangzhou | 2020-02-09 | The First Affiliated Hospital of Guangzhou Medical University | BGI-shenzhen & The First Affiliated Hospital of Guangzhou Medical University | |
| EPI_ISL_429104 | hCoV-19/Guangzhou/GZMU0036/2020 | Asia / China / Guangzhou | 2020-02-01 | The First Affiliated Hospital of Guangzhou Medical University | BGI-shenzhen & The First Affiliated Hospital of Guangzhou Medical University | |
| EPI_ISL_429105 | hCoV-19/Guangzhou/GZMU0055/2020 | Asia / China / Guangzhou | 2020-01-29 | The First Affiliated Hospital of Guangzhou Medical University | BGI-shenzhen & The First Affiliated Hospital of Guangzhou Medical University | |
| EPI_ISL_429115 | hCoV-19/Sweden/20-08142/2020 | Europe / Sweden / Halland | 2020-03-11 | Klinisk mikrobiologi och vardhygien Halmstad | The Public Health Agency of Sweden | Arne Kotz, Olov Svartstrom, Maria Lind Karlberg, Anna-Malin Linde, Oskar Karlsson Lindsjo, Anna Risberg, Shaman Muradrasoli, Karin Tegmark-Wisell |
| EPI_ISL_429116 | hCoV-19/Sweden/20-08143/2020 | Europe / Sweden / Halland | 2020-03-15 | Klinisk mikrobiologi och vardhygien Halmstad | The Public Health Agency of Sweden | Arne Kotz, Olov Svartstrom, Maria Lind Karlberg, Anna-Malin Linde, Oskar Karlsson Lindsjo, Anna Risberg, Shaman Muradrasoli, Karin Tegmark-Wisell |
| EPI_ISL_429117 | hCoV-19/Sweden/20-08145/2020 | Europe / Sweden / Halland | 2020-03-11 | Klinisk mikrobiologi och vardhygien Halmstad | The Public Health Agency of Sweden | Arne Kotz, Olov Svartstrom, Maria Lind Karlberg, Anna-Malin Linde, Oskar Karlsson Lindsjo, Anna Risberg, Shaman Muradrasoli, Karin Tegmark-Wisell |
| EPI_ISL_429118 | hCoV-19/Sweden/20-08146/2020 | Europe / Sweden / Halland | 2020-03-11 | Klinisk mikrobiologi och vardhygien Halmstad | The Public Health Agency of Sweden | Arne Kotz, Olov Svartstrom, Maria Lind Karlberg, Anna-Malin Linde, Oskar Karlsson Lindsjo, Anna Risberg, Shaman Muradrasoli, Karin Tegmark-Wisell |
| EPI_ISL_429119 | hCoV-19/Sweden/20-08148/2020 | Europe / Sweden / Halland | 2020-03-11 | Klinisk mikrobiologi och vardhygien Halmstad | The Public Health Agency of Sweden | Arne Kotz, Olov Svartstrom, Maria Lind Karlberg, Anna-Malin Linde, Oskar Karlsson Lindsjo, Anna Risberg, Shaman Muradrasoli, Karin Tegmark-Wisell |
| EPI_ISL_429120 | hCoV-19/Sweden/20-08149/2020 | Europe / Sweden / Halland | 2020-03-11 | Klinisk mikrobiologi och vardhygien Halmstad | The Public Health Agency of Sweden | Arne Kotz, Olov Svartstrom, Maria Lind Karlberg, Anna-Malin Linde, Oskar Karlsson Lindsjo, Anna Risberg, Shaman Muradrasoli, Karin Tegmark-Wisell |
| EPI_ISL_429121 | hCoV-19/Sweden/20-08151/2020 | Europe / Sweden / Halland | 2020-03-11 | Klinisk mikrobiologi och vardhygien Halmstad | The Public Health Agency of Sweden | Arne Kotz, Olov Svartstrom, Maria Lind Karlberg, Anna-Malin Linde, Oskar Karlsson Lindsjo, Anna Risberg, Shaman Muradrasoli, Karin Tegmark-Wisell |
| EPI_ISL_429122 | hCoV-19/Sweden/20-08155/2020 | Europe / Sweden / Halland | 2020-03-11 | Klinisk mikrobiologi och vardhygien Halmstad | The Public Health Agency of Sweden | Arne Kotz, Olov Svartstrom, Maria Lind Karlberg, Anna-Malin Linde, Oskar Karlsson Lindsjo, Anna Risberg, Shaman Muradrasoli, Karin Tegmark-Wisell |
| EPI_ISL_429123 | hCoV-19/Sweden/20-08169/2020 | Europe / Sweden / Halland | 2020-03-12 | Klinisk mikrobiologi och vardhygien Halmstad | The Public Health Agency of Sweden | Arne Kotz, Olov Svartstrom, Maria Lind Karlberg, Anna-Malin Linde, Oskar Karlsson Lindsjo, Anna Risberg, Shaman Muradrasoli, Karin Tegmark-Wisell |
| EPI_ISL_429124 | hCoV-19/Sweden/20-08170/2020 | Europe / Sweden / Halland | 2020-03-12 | Klinisk mikrobiologi och vardhygien Halmstad | The Public Health Agency of Sweden | Arne Kotz, Olov Svartstrom, Maria Lind Karlberg, Anna-Malin Linde, Oskar Karlsson Lindsjo, Anna Risberg, Shaman Muradrasoli, Karin Tegmark-Wisell |
| EPI_ISL_429125 | hCoV-19/Sweden/20-08173/2020 | Europe / Sweden / Halland | 2020-03-12 | Klinisk mikrobiologi och vardhygien Halmstad | The Public Health Agency of Sweden | Arne Kotz, Olov Svartstrom, Maria Lind Karlberg, Anna-Malin Linde, Oskar Karlsson Lindsjo, Anna Risberg, Shaman Muradrasoli, Karin Tegmark-Wisell |
| EPI_ISL_429126 | hCoV-19/Sweden/20-08320/2020 | Europe / Sweden / Sormland | 2020-03-21 | Unilabs Skovde | The Public Health Agency of Sweden | Tobias Kollberg, Helena Enroth, Olov Svartstrom, Maria Lind Karlberg, Anna-Malin Linde, Oskar Karlsson Lindsjo, Anna Risberg, Shaman Muradrasoli, Karin Tegmark-Wisell |
| EPI_ISL_429127 | hCoV-19/Sweden/20-08332/2020 | Europe / Sweden / Sormland | 2020-03-20 | Unilabs Skovde | The Public Health Agency of Sweden | Tobias Kollberg, Helena Enroth, Olov Svartstrom, Maria Lind Karlberg, Anna-Malin Linde, Oskar Karlsson Lindsjo, Anna Risberg, Shaman Muradrasoli, Karin Tegmark-Wisell |
| EPI_ISL_429129 | hCoV-19/Sweden/20-08681/2020 | Europe / Sweden / Norrbotten | 2020-03-27 | Laboratoriemedicin | The Public Health Agency of Sweden | Olov Svartstrom, Maria Lind Karlberg, Anna-Malin Linde, Oskar Karlsson Lindsjo, Anna Risberg, Shaman Muradrasoli, Karin Tegmark-Wisell |
| EPI_ISL_429130 | hCoV-19/Sweden/20-08683/2020 | Europe / Sweden / Norrbotten | 2020-03-23 | Laboratoriemedicin | The Public Health Agency of Sweden | Olov Svartstrom, Maria Lind Karlberg, Anna-Malin Linde, Oskar Karlsson Lindsjo, Anna Risberg, Shaman Muradrasoli, Karin Tegmark-Wisell |
| EPI_ISL_429131 | hCoV-19/Sweden/20-08684/2020 | Europe / Sweden / Norrbotten | 2020-03-27 | Laboratoriemedicin | The Public Health Agency of Sweden | Olov Svartstrom, Maria Lind Karlberg, Anna-Malin Linde, Oskar Karlsson Lindsjo, Anna Risberg, Shaman Muradrasoli, Karin Tegmark-Wisell |
| EPI_ISL_429132 | hCoV-19/Sweden/20-08685/2020 | Europe / Sweden / Norrbotten | 2020-03-27 | Laboratoriemedicin | The Public Health Agency of Sweden | Olov Svartstrom, Maria Lind Karlberg, Anna-Malin Linde, Oskar Karlsson Lindsjo, Anna Risberg, Shaman Muradrasoli, Karin Tegmark-Wisell |
| EPI_ISL_429133 | hCoV-19/Sweden/20-08690/2020 | Europe / Sweden / Norrbotten | 2020-03-26 | Laboratoriemedicin | The Public Health Agency of Sweden | Olov Svartstrom, Maria Lind Karlberg, Anna-Malin Linde, Oskar Karlsson Lindsjo, Anna Risberg, Shaman Muradrasoli, Karin Tegmark-Wisell |
| EPI_ISL_429134 | hCoV-19/Sweden/20-08693/2020 | Europe / Sweden / Norrbotten | 2020-03-24 | Laboratoriemedicin | The Public Health Agency of Sweden | Olov Svartstrom, Maria Lind Karlberg, Anna-Malin Linde, Oskar Karlsson Lindsjo, Anna Risberg, Shaman Muradrasoli, Karin Tegmark-Wisell |
| EPI_ISL_429135 | hCoV-19/Sweden/20-08719/2020 | Europe / Sweden / Norrbotten | 2020-04-02 | Laboratoriemedicin | The Public Health Agency of Sweden | Olov Svartstrom, Maria Lind Karlberg, Anna-Malin Linde, Oskar Karlsson Lindsjo, Anna Risberg, Shaman Muradrasoli, Karin Tegmark-Wisell |
| EPI_ISL_429136 | hCoV-19/Sweden/20-50048/2020 | Europe / Sweden / Varmland | 2020-03-04 | Klinisk mikrobiologi Orebro | The Public Health Agency of Sweden | Martin Sundqvist, Olov Svartstrom, Maria Lind Karlberg, Anna-Malin Linde, Oskar Karlsson Lindsjo, Anna Risberg, Shaman Muradrasoli, Karin Tegmark-Wisell |
| EPI_ISL_429137 | hCoV-19/Sweden/20-50076/2020 | Europe / Sweden / Orebro lan | 2020-03-04 | Klinisk mikrobiologi Orebro | The Public Health Agency of Sweden | Martin Sundqvist, Olov Svartstrom, Maria Lind Karlberg, Anna-Malin Linde, Oskar Karlsson Lindsjo, Anna Risberg, Shaman Muradrasoli, Karin Tegmark-Wisell |
| EPI_ISL_429138 | hCoV-19/Sweden/20-50087/2020 | Europe / Sweden / Varmland | 2020-03-05 | Klinisk mikrobiologi Orebro | The Public Health Agency of Sweden | Martin Sundqvist, Olov Svartstrom, Maria Lind Karlberg, Anna-Malin Linde, Oskar Karlsson Lindsjo, Anna Risberg, Shaman Muradrasoli, Karin Tegmark-Wisell |
| EPI_ISL_429139 | hCoV-19/Sweden/20-50123/2020 | Europe / Sweden / Varmland | 2020-03-06 | Klinisk mikrobiologi Orebro | The Public Health Agency of Sweden | Martin Sundqvist, Olov Svartstrom, Maria Lind Karlberg, Anna-Malin Linde, Oskar Karlsson Lindsjo, Anna Risberg, Shaman Muradrasoli, Karin Tegmark-Wisell |
| EPI_ISL_429140 | hCoV-19/Sweden/20-50125/2020 | Europe / Sweden / Varmland | 2020-03-06 | Klinisk mikrobiologi Orebro | The Public Health Agency of Sweden | Martin Sundqvist, Olov Svartstrom, Maria Lind Karlberg, Anna-Malin Linde, Oskar Karlsson Lindsjo, Anna Risberg, Shaman Muradrasoli, Karin Tegmark-Wisell |
| EPI_ISL_429141 | hCoV-19/Sweden/20-50130/2020 | Europe / Sweden / Varmland | 2020-03-06 | Klinisk mikrobiologi Orebro | The Public Health Agency of Sweden | Martin Sundqvist, Olov Svartstrom, Maria Lind Karlberg, Anna-Malin Linde, Oskar Karlsson Lindsjo, Anna Risberg, Shaman Muradrasoli, Karin Tegmark-Wisell |
| EPI_ISL_429142 | hCoV-19/Sweden/20-50131/2020 | Europe / Sweden / Varmland | 2020-03-06 | Klinisk mikrobiologi Orebro | The Public Health Agency of Sweden | Martin Sundqvist, Olov Svartstrom, Maria Lind Karlberg, Anna-Malin Linde, Oskar Karlsson Lindsjo, Anna Risberg, Shaman Muradrasoli, Karin Tegmark-Wisell |
| EPI_ISL_429143 | hCoV-19/Sweden/20-50132/2020 | Europe / Sweden / Varmland | 2020-03-06 | Klinisk mikrobiologi Orebro | The Public Health Agency of Sweden | Martin Sundqvist, Olov Svartstrom, Maria Lind Karlberg, Anna-Malin Linde, Oskar Karlsson Lindsjo, Anna Risberg, Shaman Muradrasoli, Karin Tegmark-Wisell |
| EPI_ISL_429144 | hCoV-19/Sweden/20-50133/2020 | Europe / Sweden / Varmland | 2020-03-06 | Klinisk mikrobiologi Orebro | The Public Health Agency of Sweden | Martin Sundqvist, Olov Svartstrom, Maria Lind Karlberg, Anna-Malin Linde, Oskar Karlsson Lindsjo, Anna Risberg, Shaman Muradrasoli, Karin Tegmark-Wisell |
| EPI_ISL_429145 | hCoV-19/Sweden/20-50170/2020 | Europe / Sweden / Varmland | 2020-03-08 | Klinisk mikrobiologi Orebro | The Public Health Agency of Sweden | Martin Sundqvist, Olov Svartstrom, Maria Lind Karlberg, Anna-Malin Linde, Oskar Karlsson Lindsjo, Anna Risberg, Shaman Muradrasoli, Karin Tegmark-Wisell |
| EPI_ISL_429146 | hCoV-19/Sweden/20-50171/2020 | Europe / Sweden / Varmland | 2020-03-08 | Klinisk mikrobiologi Orebro | The Public Health Agency of Sweden | Martin Sundqvist, Olov Svartstrom, Maria Lind Karlberg, Anna-Malin Linde, Oskar Karlsson Lindsjo, Anna Risberg, Shaman Muradrasoli, Karin Tegmark-Wisell |
| EPI_ISL_429147 | hCoV-19/Sweden/20-50172/2020 | Europe / Sweden / Varmland | 2020-03-08 | Klinisk mikrobiologi Orebro | The Public Health Agency of Sweden | Martin Sundqvist, Olov Svartstrom, Maria Lind Karlberg, Anna-Malin Linde, Oskar Karlsson Lindsjo, Anna Risberg, Shaman Muradrasoli, Karin Tegmark-Wisell |
| EPI_ISL_429148 | hCoV-19/Sweden/20-50173/2020 | Europe / Sweden / Varmland | 2020-03-08 | Klinisk mikrobiologi Orebro | The Public Health Agency of Sweden | Martin Sundqvist, Olov Svartstrom, Maria Lind Karlberg, Anna-Malin Linde, Oskar Karlsson Lindsjo, Anna Risberg, Shaman Muradrasoli, Karin Tegmark-Wisell |
| EPI_ISL_429150 | hCoV-19/Sweden/20-50175/2020 | Europe / Sweden / Varmland | 2020-03-08 | Klinisk mikrobiologi Orebro | The Public Health Agency of Sweden | Martin Sundqvist, Olov Svartstrom, Maria Lind Karlberg, Anna-Malin Linde, Oskar Karlsson Lindsjo, Anna Risberg, Shaman Muradrasoli, Karin Tegmark-Wisell |
| EPI_ISL_429151 | hCoV-19/Sweden/20-50176/2020 | Europe / Sweden / Varmland | 2020-03-08 | Klinisk mikrobiologi Orebro | The Public Health Agency of Sweden | Martin Sundqvist, Olov Svartstrom, Maria Lind Karlberg, Anna-Malin Linde, Oskar Karlsson Lindsjo, Anna Risberg, Shaman Muradrasoli, Karin Tegmark-Wisell |
| EPI_ISL_429152 | hCoV-19/Sweden/20-50179/2020 | Europe / Sweden / Orebro lan | 2020-03-08 | Klinisk mikrobiologi Orebro | The Public Health Agency of Sweden | Martin Sundqvist, Olov Svartstrom, Maria Lind Karlberg, Anna-Malin Linde, Oskar Karlsson Lindsjo, Anna Risberg, Shaman Muradrasoli, Karin Tegmark-Wisell |
| EPI_ISL_429153 | hCoV-19/Sweden/20-50180/2020 | Europe / Sweden / Varmland | 2020-03-08 | Klinisk mikrobiologi Orebro | The Public Health Agency of Sweden | Martin Sundqvist, Olov Svartstrom, Maria Lind Karlberg, Anna-Malin Linde, Oskar Karlsson Lindsjo, Anna Risberg, Shaman Muradrasoli, Karin Tegmark-Wisell |
| EPI_ISL_429154 | hCoV-19/Sweden/20-50181/2020 | Europe / Sweden / Varmland | 2020-03-08 | Klinisk mikrobiologi Orebro | The Public Health Agency of Sweden | Martin Sundqvist, Olov Svartstrom, Maria Lind Karlberg, Anna-Malin Linde, Oskar Karlsson Lindsjo, Anna Risberg, Shaman Muradrasoli, Karin Tegmark-Wisell |
| EPI_ISL_429155 | hCoV-19/Sweden/20-50230/2020 | Europe / Sweden / Orebro lan | 2020-03-09 | Klinisk mikrobiologi Orebro | The Public Health Agency of Sweden | Martin Sundqvist, Olov Svartstrom, Maria Lind Karlberg, Anna-Malin Linde, Oskar Karlsson Lindsjo, Anna Risberg, Shaman Muradrasoli, Karin Tegmark-Wisell |
| EPI_ISL_429156 | hCoV-19/Sweden/20-50232/2020 | Europe / Sweden / Varmland | 2020-03-09 | Klinisk mikrobiologi Orebro | The Public Health Agency of Sweden | Martin Sundqvist, Olov Svartstrom, Maria Lind Karlberg, Anna-Malin Linde, Oskar Karlsson Lindsjo, Anna Risberg, Shaman Muradrasoli, Karin Tegmark-Wisell |
| EPI_ISL_429157 | hCoV-19/Sweden/20-50234/2020 | Europe / Sweden / Orebro lan | 2020-03-10 | Klinisk mikrobiologi Orebro | The Public Health Agency of Sweden | Martin Sundqvist, Olov Svartstrom, Maria Lind Karlberg, Anna-Malin Linde, Oskar Karlsson Lindsjo, Anna Risberg, Shaman Muradrasoli, Karin Tegmark-Wisell |
| EPI_ISL_429158 | hCoV-19/Sweden/20-50235/2020 | Europe / Sweden / Varmland | 2020-03-10 | Klinisk mikrobiologi Orebro | The Public Health Agency of Sweden | Martin Sundqvist, Olov Svartstrom, Maria Lind Karlberg, Anna-Malin Linde, Oskar Karlsson Lindsjo, Anna Risberg, Shaman Muradrasoli, Karin Tegmark-Wisell |
| EPI_ISL_429160 | hCoV-19/Sweden/20-50312/2020 | Europe / Sweden / Ostergotland | 2020-03-12 | Klinisk mikrobiologi Orebro | The Public Health Agency of Sweden | Martin Sundqvist, Olov Svartstrom, Maria Lind Karlberg, Anna-Malin Linde, Oskar Karlsson Lindsjo, Anna Risberg, Shaman Muradrasoli, Karin Tegmark-Wisell |
| EPI_ISL_429161 | hCoV-19/Sweden/20-50315/2020 | Europe / Sweden / Stockholm | 2020-03-12 | The Public Health Agency of Sweden | The Public Health Agency of Sweden | Olov Svartstrom, Maria Lind Karlberg, Anna-Malin Linde, Oskar Karlsson Lindsjo, Anna Risberg, Shaman Muradrasoli, Karin Tegmark-Wisell |
| EPI_ISL_429162 | hCoV-19/Sweden/20-50357/2020 | Europe / Sweden / Stockholm | 2020-03-15 | The Public Health Agency of Sweden | The Public Health Agency of Sweden | Olov Svartstrom, Maria Lind Karlberg, Anna-Malin Linde, Oskar Karlsson Lindsjo, Anna Risberg, Shaman Muradrasoli, Karin Tegmark-Wisell |
| EPI_ISL_429163 | hCoV-19/Sweden/20-50361/2020 | Europe / Sweden / Gavleborgs lan | 2020-03-16 | The Public Health Agency of Sweden | The Public Health Agency of Sweden | Olov Svartstrom, Maria Lind Karlberg, Anna-Malin Linde, Oskar Karlsson Lindsjo, Anna Risberg, Shaman Muradrasoli, Karin Tegmark-Wisell |
| EPI_ISL_429164 | hCoV-19/Thailand/Bangkok-0025/2020 | Asia / Thailand / Bangkok | 2020-03-13 | Ramathibodi Hospital | COVID-19 Network Investigations (CONI) Alliance | Elizabeth Batty, Wasun Chantratita, Thanat Chookajorn, Stefan Fernandez, Angkana Huang, Poramate Jiaranai, Anthony R. Jones, Khajohn Joonsalak, Chonticha Klungtong, Theerarat Kochakarn, Namfon Kotanan, Kritikorn Kumpornsin, Wudtichai Manasatienkij, Bhakbhoom Panthan, Ekawat Pasomsub, Kingkan Rakmanee, Insee Sensorn, Janjira Thaipadungpanit, Arporn Wangwiwatsin,Treewat Watthanachockchai |
| EPI_ISL_429165 | hCoV-19/Thailand/Bangkok-0026/2020 | Asia / Thailand / Bangkok | 2020-03-13 | Ramathibodi Hospital | COVID-19 Network Investigations (CONI) Alliance | Elizabeth Batty, Wasun Chantratita, Thanat Chookajorn, Stefan Fernandez, Angkana Huang, Poramate Jiaranai, Anthony R. Jones, Khajohn Joonsalak, Chonticha Klungtong, Theerarat Kochakarn, Namfon Kotanan, Kritikorn Kumpornsin, Wudtichai Manasatienkij, Bhakbhoom Panthan, Ekawat Pasomsub, Kingkan Rakmanee, Insee Sensorn, Janjira Thaipadungpanit, Arporn Wangwiwatsin,Treewat Watthanachockchai |
| EPI_ISL_429166 | hCoV-19/Thailand/Bangkok-0028/2020 | Asia / Thailand / Bangkok | 2020-03-13 | Ramathibodi Hospital | COVID-19 Network Investigations (CONI) Alliance | Elizabeth Batty, Wasun Chantratita, Thanat Chookajorn, Stefan Fernandez, Angkana Huang, Poramate Jiaranai, Anthony R. Jones, Khajohn Joonsalak, Chonticha Klungtong, Theerarat Kochakarn, Namfon Kotanan, Kritikorn Kumpornsin, Wudtichai Manasatienkij, Bhakbhoom Panthan, Ekawat Pasomsub, Kingkan Rakmanee, Insee Sensorn, Janjira Thaipadungpanit, Arporn Wangwiwatsin,Treewat Watthanachockchai |
| EPI_ISL_429167 | hCoV-19/Thailand/Bangkok-0029/2020 | Asia / Thailand / Bangkok | 2020-03-13 | Ramathibodi Hospital | COVID-19 Network Investigations (CONI) Alliance | Elizabeth Batty, Wasun Chantratita, Thanat Chookajorn, Stefan Fernandez, Angkana Huang, Poramate Jiaranai, Anthony R. Jones, Khajohn Joonsalak, Chonticha Klungtong, Theerarat Kochakarn, Namfon Kotanan, Kritikorn Kumpornsin, Wudtichai Manasatienkij, Bhakbhoom Panthan, Ekawat Pasomsub, Kingkan Rakmanee, Insee Sensorn, Janjira Thaipadungpanit, Arporn Wangwiwatsin,Treewat Watthanachockchai |
| EPI_ISL_429168 | hCoV-19/Thailand/Bangkok-0030/2020 | Asia / Thailand / Bangkok | 2020-03-17 | Ramathibodi Hospital | COVID-19 Network Investigations (CONI) Alliance | Elizabeth Batty, Wasun Chantratita, Thanat Chookajorn, Stefan Fernandez, Angkana Huang, Poramate Jiaranai, Anthony R. Jones, Khajohn Joonsalak, Chonticha Klungtong, Theerarat Kochakarn, Namfon Kotanan, Kritikorn Kumpornsin, Wudtichai Manasatienkij, Bhakbhoom Panthan, Ekawat Pasomsub, Kingkan Rakmanee, Insee Sensorn, Janjira Thaipadungpanit, Arporn Wangwiwatsin,Treewat Watthanachockchai |
| EPI_ISL_429169 | hCoV-19/Thailand/Bangkok-0033/2020 | Asia / Thailand / Bangkok | 2020-03-17 | Ramathibodi Hospital | COVID-19 Network Investigations (CONI) Alliance | Elizabeth Batty, Wasun Chantratita, Thanat Chookajorn, Stefan Fernandez, Angkana Huang, Poramate Jiaranai, Anthony R. Jones, Khajohn Joonsalak, Chonticha Klungtong, Theerarat Kochakarn, Namfon Kotanan, Kritikorn Kumpornsin, Wudtichai Manasatienkij, Bhakbhoom Panthan, Ekawat Pasomsub, Kingkan Rakmanee, Insee Sensorn, Janjira Thaipadungpanit, Arporn Wangwiwatsin,Treewat Watthanachockchai |
| EPI_ISL_429170 | hCoV-19/Thailand/Bangkok-0034/2020 | Asia / Thailand / Bangkok | 2020-03-17 | Ramathibodi Hospital | COVID-19 Network Investigations (CONI) Alliance | Elizabeth Batty, Wasun Chantratita, Thanat Chookajorn, Stefan Fernandez, Angkana Huang, Poramate Jiaranai, Anthony R. Jones, Khajohn Joonsalak, Chonticha Klungtong, Theerarat Kochakarn, Namfon Kotanan, Kritikorn Kumpornsin, Wudtichai Manasatienkij, Bhakbhoom Panthan, Ekawat Pasomsub, Kingkan Rakmanee, Insee Sensorn, Janjira Thaipadungpanit, Arporn Wangwiwatsin,Treewat Watthanachockchai |
| EPI_ISL_429171 | hCoV-19/Thailand/Bangkok-0035/2020 | Asia / Thailand / Bangkok | 2020-03-18 | Ramathibodi Hospital | COVID-19 Network Investigations (CONI) Alliance | Elizabeth Batty, Wasun Chantratita, Thanat Chookajorn, Stefan Fernandez, Angkana Huang, Poramate Jiaranai, Anthony R. Jones, Khajohn Joonsalak, Chonticha Klungtong, Theerarat Kochakarn, Namfon Kotanan, Kritikorn Kumpornsin, Wudtichai Manasatienkij, Bhakbhoom Panthan, Ekawat Pasomsub, Kingkan Rakmanee, Insee Sensorn, Janjira Thaipadungpanit, Arporn Wangwiwatsin,Treewat Watthanachockchai |
| EPI_ISL_429172 | hCoV-19/Thailand/Bangkok-0036/2020 | Asia / Thailand / Bangkok | 2020-03-21 | Ramathibodi Hospital | COVID-19 Network Investigations (CONI) Alliance | Elizabeth Batty, Wasun Chantratita, Thanat Chookajorn, Stefan Fernandez, Angkana Huang, Poramate Jiaranai, Anthony R. Jones, Khajohn Joonsalak, Chonticha Klungtong, Theerarat Kochakarn, Namfon Kotanan, Kritikorn Kumpornsin, Wudtichai Manasatienkij, Bhakbhoom Panthan, Ekawat Pasomsub, Kingkan Rakmanee, Insee Sensorn, Janjira Thaipadungpanit, Arporn Wangwiwatsin,Treewat Watthanachockchai |
| EPI_ISL_429173 | hCoV-19/Thailand/Bangkok-0037/2020 | Asia / Thailand / Bangkok | 2020-03-21 | Ramathibodi Hospital | COVID-19 Network Investigations (CONI) Alliance | Elizabeth Batty, Wasun Chantratita, Thanat Chookajorn, Stefan Fernandez, Angkana Huang, Poramate Jiaranai, Anthony R. Jones, Khajohn Joonsalak, Chonticha Klungtong, Theerarat Kochakarn, Namfon Kotanan, Kritikorn Kumpornsin, Wudtichai Manasatienkij, Bhakbhoom Panthan, Ekawat Pasomsub, Kingkan Rakmanee, Insee Sensorn, Janjira Thaipadungpanit, Arporn Wangwiwatsin,Treewat Watthanachockchai |
| EPI_ISL_429174 | hCoV-19/Thailand/Bangkok-0038/2020 | Asia / Thailand / Bangkok | 2020-03-25 | Ramathibodi Hospital | COVID-19 Network Investigations (CONI) Alliance | Elizabeth Batty, Wasun Chantratita, Thanat Chookajorn, Stefan Fernandez, Angkana Huang, Poramate Jiaranai, Anthony R. Jones, Khajohn Joonsalak, Chonticha Klungtong, Theerarat Kochakarn, Namfon Kotanan, Kritikorn Kumpornsin, Wudtichai Manasatienkij, Bhakbhoom Panthan, Ekawat Pasomsub, Kingkan Rakmanee, Insee Sensorn, Janjira Thaipadungpanit, Arporn Wangwiwatsin,Treewat Watthanachockchai |
| EPI_ISL_429175 | hCoV-19/Thailand/Bangkok-0039/2020 | Asia / Thailand / Bangkok | 2020-03-28 | Ramathibodi Hospital | COVID-19 Network Investigations (CONI) Alliance | Elizabeth Batty, Wasun Chantratita, Thanat Chookajorn, Stefan Fernandez, Angkana Huang, Poramate Jiaranai, Anthony R. Jones, Khajohn Joonsalak, Chonticha Klungtong, Theerarat Kochakarn, Namfon Kotanan, Kritikorn Kumpornsin, Wudtichai Manasatienkij, Bhakbhoom Panthan, Ekawat Pasomsub, Kingkan Rakmanee, Insee Sensorn, Janjira Thaipadungpanit, Arporn Wangwiwatsin,Treewat Watthanachockchai |
| EPI_ISL_429178 | hCoV-19/Thailand/Bangkok-0042/2020 | Asia / Thailand / Bangkok | 2020-03-28 | Ramathibodi Hospital | COVID-19 Network Investigations (CONI) Alliance | Elizabeth Batty, Wasun Chantratita, Thanat Chookajorn, Stefan Fernandez, Angkana Huang, Poramate Jiaranai, Anthony R. Jones, Khajohn Joonsalak, Chonticha Klungtong, Theerarat Kochakarn, Namfon Kotanan, Kritikorn Kumpornsin, Wudtichai Manasatienkij, Bhakbhoom Panthan, Ekawat Pasomsub, Kingkan Rakmanee, Insee Sensorn, Janjira Thaipadungpanit, Arporn Wangwiwatsin,Treewat Watthanachockchai |
| EPI_ISL_429179 | hCoV-19/Thailand/Bangkok-0043/2020 | Asia / Thailand / Bangkok | 2020-03-28 | Ramathibodi Hospital | COVID-19 Network Investigations (CONI) Alliance | Elizabeth Batty, Wasun Chantratita, Thanat Chookajorn, Stefan Fernandez, Angkana Huang, Poramate Jiaranai, Anthony R. Jones, Khajohn Joonsalak, Chonticha Klungtong, Theerarat Kochakarn, Namfon Kotanan, Kritikorn Kumpornsin, Wudtichai Manasatienkij, Bhakbhoom Panthan, Ekawat Pasomsub, Kingkan Rakmanee, Insee Sensorn, Janjira Thaipadungpanit, Arporn Wangwiwatsin,Treewat Watthanachockchai |
| EPI_ISL_429181 | hCoV-19/Thailand/Bangkok-0045/2020 | Asia / Thailand / Bangkok | 2020-03-28 | Ramathibodi Hospital | COVID-19 Network Investigations (CONI) Alliance | Elizabeth Batty, Wasun Chantratita, Thanat Chookajorn, Stefan Fernandez, Angkana Huang, Poramate Jiaranai, Anthony R. Jones, Khajohn Joonsalak, Chonticha Klungtong, Theerarat Kochakarn, Namfon Kotanan, Kritikorn Kumpornsin, Wudtichai Manasatienkij, Bhakbhoom Panthan, Ekawat Pasomsub, Kingkan Rakmanee, Insee Sensorn, Janjira Thaipadungpanit, Arporn Wangwiwatsin,Treewat Watthanachockchai |
| EPI_ISL_429182 | hCoV-19/Thailand/Bangkok-0046/2020 | Asia / Thailand / Bangkok | 2020-03-28 | Ramathibodi Hospital | COVID-19 Network Investigations (CONI) Alliance | Elizabeth Batty, Wasun Chantratita, Thanat Chookajorn, Stefan Fernandez, Angkana Huang, Poramate Jiaranai, Anthony R. Jones, Khajohn Joonsalak, Chonticha Klungtong, Theerarat Kochakarn, Namfon Kotanan, Kritikorn Kumpornsin, Wudtichai Manasatienkij, Bhakbhoom Panthan, Ekawat Pasomsub, Kingkan Rakmanee, Insee Sensorn, Janjira Thaipadungpanit, Arporn Wangwiwatsin,Treewat Watthanachockchai |
| EPI_ISL_429183 | hCoV-19/Thailand/Bangkok-0047/2020 | Asia / Thailand / Bangkok | 2020-03-28 | Ramathibodi Hospital | COVID-19 Network Investigations (CONI) Alliance | Elizabeth Batty, Wasun Chantratita, Thanat Chookajorn, Stefan Fernandez, Angkana Huang, Poramate Jiaranai, Anthony R. Jones, Khajohn Joonsalak, Chonticha Klungtong, Theerarat Kochakarn, Namfon Kotanan, Kritikorn Kumpornsin, Wudtichai Manasatienkij, Bhakbhoom Panthan, Ekawat Pasomsub, Kingkan Rakmanee, Insee Sensorn, Janjira Thaipadungpanit, Arporn Wangwiwatsin,Treewat Watthanachockchai |
| EPI_ISL_429184 | hCoV-19/Thailand/Bangkok-0048/2020 | Asia / Thailand / Bangkok | 2020-03-28 | Ramathibodi Hospital | COVID-19 Network Investigations (CONI) Alliance | Elizabeth Batty, Wasun Chantratita, Thanat Chookajorn, Stefan Fernandez, Angkana Huang, Poramate Jiaranai, Anthony R. Jones, Khajohn Joonsalak, Chonticha Klungtong, Theerarat Kochakarn, Namfon Kotanan, Kritikorn Kumpornsin, Wudtichai Manasatienkij, Bhakbhoom Panthan, Ekawat Pasomsub, Kingkan Rakmanee, Insee Sensorn, Janjira Thaipadungpanit, Arporn Wangwiwatsin,Treewat Watthanachockchai |
| EPI_ISL_429200 | hCoV-19/Switzerland/GE5737/2020 | Europe / Switzerland | 2020-03-19 | University Hospitals of Geneva Laboratory of Virology | University Hospitals of Geneva Laboratory of Virology | Laubscher F. |
| EPI_ISL_429201 | hCoV-19/Switzerland/GE2297/2020 | Europe / Switzerland | 2020-03-29 | University Hospitals of Geneva Laboratory of Virology | University Hospitals of Geneva Laboratory of Virology | Laubscher F. |
| EPI_ISL_429203 | hCoV-19/Switzerland/GE1736/2020 | Europe / Switzerland | 2020-03-21 | University Hospitals of Geneva Laboratory of Virology | University Hospitals of Geneva Laboratory of Virology | Laubscher F. |
| EPI_ISL_429204 | hCoV-19/Switzerland/GE7617/2020 | Europe / Switzerland | 2020-04-03 | University Hospitals of Geneva Laboratory of Virology | University Hospitals of Geneva Laboratory of Virology | Laubscher F. |
| EPI_ISL_429206 | hCoV-19/Switzerland/GE5007/2020 | Europe / Switzerland | 2020-04-07 | University Hospitals of Geneva Laboratory of Virology | University Hospitals of Geneva Laboratory of Virology | Laubscher F. |
| EPI_ISL_429207 | hCoV-19/Switzerland/GE1184/2020 | Europe / Switzerland | 2020-04-03 | University Hospitals of Geneva Laboratory of Virology | University Hospitals of Geneva Laboratory of Virology | Laubscher F. |
| EPI_ISL_429208 | hCoV-19/Switzerland/GE2164/2020 | Europe / Switzerland | 2020-03-17 | University Hospitals of Geneva Laboratory of Virology | University Hospitals of Geneva Laboratory of Virology | Laubscher F. |
| EPI_ISL_429209 | hCoV-19/Switzerland/GE3650/2020 | Europe / Switzerland | 2020-03-16 | University Hospitals of Geneva Laboratory of Virology | University Hospitals of Geneva Laboratory of Virology | Laubscher F. |
| EPI_ISL_429210 | hCoV-19/Switzerland/GE8147/2020 | Europe / Switzerland | 2020-03-16 | University Hospitals of Geneva Laboratory of Virology | University Hospitals of Geneva Laboratory of Virology | Laubscher F. |
| EPI_ISL_429211 | hCoV-19/Switzerland/GE5708/2020 | Europe / Switzerland | 2020-03-24 | University Hospitals of Geneva Laboratory of Virology | University Hospitals of Geneva Laboratory of Virology | Laubscher F. |
| EPI_ISL_429212 | hCoV-19/Switzerland/GE0056/2020 | Europe / Switzerland | 2020-03-24 | University Hospitals of Geneva Laboratory of Virology | University Hospitals of Geneva Laboratory of Virology | Laubscher F. |
| EPI_ISL_429213 | hCoV-19/Switzerland/GE8086/2020 | Europe / Switzerland | 2020-03-16 | University Hospitals of Geneva Laboratory of Virology | University Hospitals of Geneva Laboratory of Virology | Laubscher F. |
| EPI_ISL_429214 | hCoV-19/Switzerland/GE2759/2020 | Europe / Switzerland | 2020-03-23 | University Hospitals of Geneva Laboratory of Virology | University Hospitals of Geneva Laboratory of Virology | Laubscher F. |
| EPI_ISL_429215 | hCoV-19/Switzerland/GE4807/2020 | Europe / Switzerland | 2020-03-23 | University Hospitals of Geneva Laboratory of Virology | University Hospitals of Geneva Laboratory of Virology | Laubscher F. |
| EPI_ISL_429217 | hCoV-19/Switzerland/GE6414/2020 | Europe / Switzerland | 2020-04-06 | University Hospitals of Geneva Laboratory of Virology | University Hospitals of Geneva Laboratory of Virology | Laubscher F. |
| EPI_ISL_429218 | hCoV-19/Switzerland/GE0304/2020 | Europe / Switzerland | 2020-04-02 | University Hospitals of Geneva Laboratory of Virology | University Hospitals of Geneva Laboratory of Virology | Laubscher F. |
| EPI_ISL_429219 | hCoV-19/Switzerland/GE2453/2020 | Europe / Switzerland | 2020-04-03 | University Hospitals of Geneva Laboratory of Virology | University Hospitals of Geneva Laboratory of Virology | Laubscher F. |
| EPI_ISL_429220 | hCoV-19/Switzerland/GE6065/2020 | Europe / Switzerland | 2020-04-06 | University Hospitals of Geneva Laboratory of Virology | University Hospitals of Geneva Laboratory of Virology | Laubscher F. |
| EPI_ISL_429221 | hCoV-19/Switzerland/GE6099/2020 | Europe / Switzerland | 2020-04-06 | University Hospitals of Geneva Laboratory of Virology | University Hospitals of Geneva Laboratory of Virology | Laubscher F. |
| EPI_ISL_429222 | hCoV-19/Switzerland/GE3655/2020 | Europe / Switzerland | 2020-04-07 | University Hospitals of Geneva Laboratory of Virology | University Hospitals of Geneva Laboratory of Virology | Laubscher F. |
| EPI_ISL_429223 | hCoV-19/Switzerland/GE4644/2020 | Europe / Switzerland | 2020-04-07 | University Hospitals of Geneva Laboratory of Virology | University Hospitals of Geneva Laboratory of Virology | Laubscher F. |
| EPI_ISL_429226 | hCoV-19/Italy/TE5551/2020 | Europe / Italy / Abruzzo | 2020-03-17 | Presidio Ospedaliero Santo Spirito | Istituto Zooprofilattico Sperimentale dell'Abruzzo e Molise "G. Caporale" | Lorusso A, Marcacci M, Di Domenico M, Ancora M, Curini V, Mangone I, Rinaldi A, Di Pasquale A, Camma C, Puglia I, Savini G |
| EPI_ISL_429227 | hCoV-19/Italy/TE5553/2020 | Europe / Italy / Abruzzo | 2020-03-17 | Presidio Ospedaliero Santo Spirito | Istituto Zooprofilattico Sperimentale dell'Abruzzo e Molise "G. Caporale" | Lorusso A, Marcacci M, Di Domenico M, Ancora M, Curini V, Mangone I, Rinaldi A, Di Pasquale A, Camma C, Puglia I, Savini G |
| EPI_ISL_429228 | hCoV-19/Italy/TE5674/2020 | Europe / Italy / Abruzzo | 2020-03-20 | Ospedale Civile Giuseppe Mazzini | Istituto Zooprofilattico Sperimentale dell'Abruzzo e Molise "G. Caporale" | Lorusso A, Marcacci M, Di Domenico M, Ancora M, Curini V, Mangone I, Rinaldi A, Di Pasquale A, Camma C, Puglia I, Savini G |
| EPI_ISL_429230 | hCoV-19/Italy/TE6570/2020 | Europe / Italy / Abruzzo | 2020-03-24 | Ospedale Civile Giuseppe Mazzini | Istituto Zooprofilattico Sperimentale dell'Abruzzo e Molise "G. Caporale" | Lorusso A, Marcacci M, Di Domenico M, Ancora M, Curini V, Mangone I, Rinaldi A, Di Pasquale A, Camma C, Puglia I, Savini G |
| EPI_ISL_429231 | hCoV-19/Italy/TE6571/2020 | Europe / Italy / Abruzzo | 2020-03-24 | Ospedale Civile Giuseppe Mazzini | Istituto Zooprofilattico Sperimentale dell'Abruzzo e Molise "G. Caporale" | Lorusso A, Marcacci M, Di Domenico M, Ancora M, Curini V, Mangone I, Rinaldi A, Di Pasquale A, Camma C, Puglia I, Savini G |

| | | | | | | |
|---|---|---|---|---|---|---|
| EPI_ISL_429232 | hCoV-19/Italy/TE6572/2020 | Europe / Italy / Abruzzo | 2020-03-24 | Ospedale Civile Giuseppe Mazzini | Istituto Zooprofilattico Sperimentale dell'Abruzzo e Molise "G. Caporale" | Lorusso A, Marcacci M, Di Domenico M, Ancora M, Curini V, Mangone I, Rinaldi A, Di Pasquale A, Camma C, Puglia I, Savini G |
| EPI_ISL_429233 | hCoV-19/Italy/TE6575/2020 | Europe / Italy / Abruzzo | 2020-03-24 | Ospedale Civile Giuseppe Mazzini | Istituto Zooprofilattico Sperimentale dell'Abruzzo e Molise "G. Caporale" | Lorusso A, Marcacci M, Di Domenico M, Ancora M, Curini V, Mangone I, Rinaldi A, Di Pasquale A, Camma C, Puglia I, Savini G |
| EPI_ISL_429235 | hCoV-19/Italy/TE6580/2020 | Europe / Italy / Abruzzo | 2020-03-24 | Ospedale Civile Giuseppe Mazzini | Istituto Zooprofilattico Sperimentale dell'Abruzzo e Molise "G. Caporale" | Lorusso A, Marcacci M, Di Domenico M, Ancora M, Curini V, Mangone I, Rinaldi A, Di Pasquale A, Camma C, Puglia I, Savini G |
| EPI_ISL_429236 | hCoV-19/Italy/TE7859/2020 | Europe / Italy / Abruzzo | 2020-03-28 | Ospedale Civile S. Liberatore di Atri | Istituto Zooprofilattico Sperimentale dell'Abruzzo e Molise "G. Caporale" | Lorusso A, Marcacci M, Di Domenico M, Ancora M, Curini V, Mangone I, Rinaldi A, Di Pasquale A, Camma C, Puglia I, Savini G |
| EPI_ISL_429239 | hCoV-19/Yunnan/0306-466/2020 | Asia / China / Yunnan | 2020-03-06 | Department of Clinical Laboratory, the First People's Hospital of Yunnan Province | Department of Clinical Laboratory, the First People's Hospital of Yunnan Province | Yi Sun,Ziqin Dian,Ya Xu,Guiqian Zhang,Xin Fan,Yu Zhang |
| EPI_ISL_429254 | hCoV-19/DRC/998/2020 | Africa / Democratic Republic of the Cong | 2020-04-03 | Viral Respiratory Lab, National Institute for Biomedical Research (INRB) | Pathogen Sequencing Lab, National Institute for Biomedical Research (INRB) | Placide Mbala-Kingebeni, Edith Nkwembe, Eddy Kinganda-Lusamaki, Amuri Aziza, Catherine Pratt, Matthias Pauthner, Josh Quick, Allison Black, James Hadfield, Trevor Bedford, Ian Goodfellow, Nick Loman, Kristian Andersen, Michael Wiley, Steve Ahuka-Mundeke, Jean-Jacques Muyembe Tamfum |
| EPI_ISL_429255 | hCoV-19/DRC/1131/2020 | Africa / Democratic Republic of the Cong | 2020-04-05 | Viral Respiratory Lab, National Institute for Biomedical Research (INRB) | Pathogen Sequencing Lab, National Institute for Biomedical Research (INRB) | Placide Mbala-Kingebeni, Edith Nkwembe, Eddy Kinganda-Lusamaki, Amuri Aziza, Catherine Pratt, Matthias Pauthner, Josh Quick, Allison Black, James Hadfield, Trevor Bedford, Ian Goodfellow, Nick Loman, Kristian Andersen, Michael Wiley, Steve Ahuka-Mundeke, Jean-Jacques Muyembe Tamfum |
| EPI_ISL_429256 | hCoV-19/Spain/Vircell001/2020 | Europe / Spain / Madrid | 2020-03-12 | Health Sciences Technology Park, Avicena, 8, 18016 Granada. Spain | Sequencing and Bioinformatics Service FISABIO-Public Health | Joaquin Mendoza, Almudena Rojas, Pablo Mendoza |
| EPI_ISL_429258 | hCoV-19/DRC/1234/2020 | Africa / Democratic Republic of the Cong | 2020-04-06 | Viral Respiratory Lab, National Institute for Biomedical Research (INRB) | Pathogen Sequencing Lab, National Institute for Biomedical Research (INRB) | Placide Mbala-Kingebeni, Edith Nkwembe, Eddy Kinganda-Lusamaki, Amuri Aziza, Catherine Pratt, Matthias Pauthner, Josh Quick, Allison Black, James Hadfield, Trevor Bedford, Ian Goodfellow, Nick Loman, Kristian Andersen, Michael Wiley, Steve Ahuka-Mundeke, Jean-Jacques Muyembe Tamfum |
| EPI_ISL_429259 | hCoV-19/DRC/1249/2020 | Africa / Democratic Republic of the Cong | 2020-04-06 | Viral Respiratory Lab, National Institute for Biomedical Research (INRB) | Pathogen Sequencing Lab, National Institute for Biomedical Research (INRB) | Placide Mbala-Kingebeni, Edith Nkwembe, Eddy Kinganda-Lusamaki, Amuri Aziza, Catherine Pratt, Matthias Pauthner, Josh Quick, Allison Black, James Hadfield, Trevor Bedford, Ian Goodfellow, Nick Loman, Kristian Andersen, Michael Wiley, Steve Ahuka-Mundeke, Jean-Jacques Muyembe Tamfum |
| EPI_ISL_429262 | hCoV-19/Denmark/ALAB-HH01/2020 | Europe / Denmark / Unknown | 2020-03-10 | Department of Clinical Microbiology, Copenhagen University Hospital, Hvidovre, Kettegaard Alle 30, 2650 Hvidovre. | Albertsen lab, Department of Chemistry and Bioscience, Aalborg University, Denmark | Rasmus Kirkegaard |
| EPI_ISL_429263 | hCoV-19/Denmark/ALAB-HH02/2020 | Europe / Denmark / Unknown | 2020-03-10 | Department of Clinical Microbiology, Copenhagen University Hospital, Hvidovre, Kettegaard Alle 30, 2650 Hvidovre. | Albertsen lab, Department of Chemistry and Bioscience, Aalborg University, Denmark | Rasmus Kirkegaard |
| EPI_ISL_429264 | hCoV-19/Denmark/ALAB-HH03/2020 | Europe / Denmark / Unknown | 2020-03-10 | Department of Clinical Microbiology, Copenhagen University Hospital, Hvidovre, Kettegaard Alle 30, 2650 Hvidovre. | Albertsen lab, Department of Chemistry and Bioscience, Aalborg University, Denmark | Rasmus Kirkegaard |
| EPI_ISL_429265 | hCoV-19/Denmark/ALAB-HH04/2020 | Europe / Denmark / Unknown | 2020-03-10 | Department of Clinical Microbiology, Copenhagen University Hospital, Hvidovre, Kettegaard Alle 30, 2650 Hvidovre. | Albertsen lab, Department of Chemistry and Bioscience, Aalborg University, Denmark | Rasmus Kirkegaard |
| EPI_ISL_429266 | hCoV-19/Denmark/ALAB-HH05/2020 | Europe / Denmark / Unknown | 2020-03-10 | Department of Clinical Microbiology, Copenhagen University Hospital, Hvidovre, Kettegaard Alle 30, 2650 Hvidovre. | Albertsen lab, Department of Chemistry and Bioscience, Aalborg University, Denmark | Rasmus Kirkegaard |
| EPI_ISL_429267 | hCoV-19/Denmark/ALAB-HH06/2020 | Europe / Denmark / Unknown | 2020-03-10 | Department of Clinical Microbiology, Copenhagen University Hospital, Hvidovre, Kettegaard Alle 30, 2650 Hvidovre. | Albertsen lab, Department of Chemistry and Bioscience, Aalborg University, Denmark | Rasmus Kirkegaard |
| EPI_ISL_429268 | hCoV-19/Denmark/ALAB-HH07/2020 | Europe / Denmark / Unknown | 2020-03-10 | Department of Clinical Microbiology, Copenhagen University Hospital, Hvidovre, Kettegaard Alle 30, 2650 Hvidovre. | Albertsen lab, Department of Chemistry and Bioscience, Aalborg University, Denmark | Rasmus Kirkegaard |
| EPI_ISL_429269 | hCoV-19/Denmark/ALAB-HH08/2020 | Europe / Denmark / Unknown | 2020-03-10 | Department of Clinical Microbiology, Copenhagen University Hospital, Hvidovre, Kettegaard Alle 30, 2650 Hvidovre. | Albertsen lab, Department of Chemistry and Bioscience, Aalborg University, Denmark | Rasmus Kirkegaard |
| EPI_ISL_429270 | hCoV-19/Denmark/ALAB-HH11/2020 | Europe / Denmark / Unknown | 2020-03-10 | Department of Clinical Microbiology, Copenhagen University Hospital, Hvidovre, Kettegaard Alle 30, 2650 Hvidovre. | Albertsen lab, Department of Chemistry and Bioscience, Aalborg University, Denmark | Rasmus Kirkegaard |
| EPI_ISL_429271 | hCoV-19/Denmark/ALAB-HH13/2020 | Europe / Denmark / Unknown | 2020-03-10 | Department of Clinical Microbiology, Copenhagen University Hospital, Hvidovre, Kettegaard Alle 30, 2650 Hvidovre. | Albertsen lab, Department of Chemistry and Bioscience, Aalborg University, Denmark | Rasmus Kirkegaard |
| EPI_ISL_429272 | hCoV-19/Denmark/ALAB-HH14/2020 | Europe / Denmark / Unknown | 2020-03-10 | Department of Clinical Microbiology, Copenhagen University Hospital, Hvidovre, Kettegaard Alle 30, 2650 Hvidovre. | Albertsen lab, Department of Chemistry and Bioscience, Aalborg University, Denmark | Rasmus Kirkegaard |
| EPI_ISL_429273 | hCoV-19/Denmark/ALAB-HH15/2020 | Europe / Denmark / Unknown | 2020-03-10 | Department of Clinical Microbiology, Copenhagen University Hospital, Hvidovre, Kettegaard Alle 30, 2650 Hvidovre. | Albertsen lab, Department of Chemistry and Bioscience, Aalborg University, Denmark | Rasmus Kirkegaard |
| EPI_ISL_429274 | hCoV-19/Denmark/ALAB-HH16/2020 | Europe / Denmark / Unknown | 2020-03-10 | Department of Clinical Microbiology, Copenhagen University Hospital, Hvidovre, Kettegaard Alle 30, 2650 Hvidovre. | Albertsen lab, Department of Chemistry and Bioscience, Aalborg University, Denmark | Rasmus Kirkegaard |
| EPI_ISL_429275 | hCoV-19/Denmark/ALAB-HH17/2020 | Europe / Denmark / Unknown | 2020-03-10 | Department of Clinical Microbiology, Copenhagen University Hospital, Hvidovre, Kettegaard Alle 30, 2650 Hvidovre. | Albertsen lab, Department of Chemistry and Bioscience, Aalborg University, Denmark | Rasmus Kirkegaard |
| EPI_ISL_429276 | hCoV-19/Denmark/ALAB-HH18/2020 | Europe / Denmark / Unknown | 2020-03-10 | Department of Clinical Microbiology, Copenhagen University Hospital, Hvidovre, Kettegaard Alle 30, 2650 Hvidovre. | Albertsen lab, Department of Chemistry and Bioscience, Aalborg University, Denmark | Rasmus Kirkegaard |
| EPI_ISL_429277 | hCoV-19/Denmark/ALAB-HH19/2020 | Europe / Denmark / Unknown | 2020-03-10 | Department of Clinical Microbiology, Copenhagen University Hospital, Hvidovre, Kettegaard Alle 30, 2650 Hvidovre. | Albertsen lab, Department of Chemistry and Bioscience, Aalborg University, Denmark | Rasmus Kirkegaard |
| EPI_ISL_429278 | hCoV-19/Denmark/ALAB-HH20/2020 | Europe / Denmark / Unknown | 2020-03-10 | Department of Clinical Microbiology, Copenhagen University Hospital, Hvidovre, Kettegaard Alle 30, 2650 Hvidovre. | Albertsen lab, Department of Chemistry and Bioscience, Aalborg University, Denmark | Rasmus Kirkegaard |
| EPI_ISL_429279 | hCoV-19/Denmark/ALAB-HH21/2020 | Europe / Denmark / Unknown | 2020-03-10 | Department of Clinical Microbiology, Copenhagen University Hospital, Hvidovre, Kettegaard Alle 30, 2650 Hvidovre. | Albertsen lab, Department of Chemistry and Bioscience, Aalborg University, Denmark | Rasmus Kirkegaard |
| EPI_ISL_429280 | hCoV-19/Denmark/ALAB-HH22/2020 | Europe / Denmark / Unknown | 2020-03-10 | Department of Clinical Microbiology, Copenhagen University Hospital, Hvidovre, Kettegaard Alle 30, 2650 Hvidovre. | Albertsen lab, Department of Chemistry and Bioscience, Aalborg University, Denmark | Rasmus Kirkegaard |
| EPI_ISL_429281 | hCoV-19/Denmark/ALAB-HH23/2020 | Europe / Denmark / Unknown | 2020-03-10 | Department of Clinical Microbiology, Copenhagen University Hospital, Hvidovre, Kettegaard Alle 30, 2650 Hvidovre. | Albertsen lab, Department of Chemistry and Bioscience, Aalborg University, Denmark | Rasmus Kirkegaard |
| EPI_ISL_429282 | hCoV-19/Denmark/ALAB-HH24/2020 | Europe / Denmark / Unknown | 2020-03-11 | Department of Clinical Microbiology, Copenhagen University Hospital, Hvidovre, Kettegaard Alle 30, 2650 Hvidovre. | Albertsen lab, Department of Chemistry and Bioscience, Aalborg University, Denmark | Rasmus Kirkegaard |
| EPI_ISL_429283 | hCoV-19/Denmark/ALAB-HH25/2020 | Europe / Denmark / Unknown | 2020-03-11 | Department of Clinical Microbiology, Copenhagen University Hospital, Hvidovre, Kettegaard Alle 30, 2650 Hvidovre. | Albertsen lab, Department of Chemistry and Bioscience, Aalborg University, Denmark | Rasmus Kirkegaard |
| EPI_ISL_429284 | hCoV-19/Denmark/ALAB-HH26/2020 | Europe / Denmark / Unknown | 2020-03-11 | Department of Clinical Microbiology, Copenhagen University Hospital, Hvidovre, Kettegaard Alle 30, 2650 Hvidovre. | Albertsen lab, Department of Chemistry and Bioscience, Aalborg University, Denmark | Rasmus Kirkegaard |
| EPI_ISL_429285 | hCoV-19/Denmark/ALAB-HH27/2020 | Europe / Denmark / Unknown | 2020-03-11 | Department of Clinical Microbiology, Copenhagen University Hospital, Hvidovre, Kettegaard Alle 30, 2650 Hvidovre. | Albertsen lab, Department of Chemistry and Bioscience, Aalborg University, Denmark | Rasmus Kirkegaard |
| EPI_ISL_429287 | hCoV-19/Denmark/ALAB-HH29/2020 | Europe / Denmark / Unknown | 2020-03-11 | Department of Clinical Microbiology, Copenhagen University Hospital, Hvidovre, Kettegaard Alle 30, 2650 Hvidovre. | Albertsen lab, Department of Chemistry and Bioscience, Aalborg University, Denmark | Rasmus Kirkegaard |
| EPI_ISL_429289 | hCoV-19/Denmark/ALAB-HH31/2020 | Europe / Denmark / Unknown | 2020-03-12 | Department of Clinical Microbiology, Copenhagen University Hospital, Hvidovre, Kettegaard Alle 30, 2650 Hvidovre. | Albertsen lab, Department of Chemistry and Bioscience, Aalborg University, Denmark | Rasmus Kirkegaard |
| EPI_ISL_429290 | hCoV-19/Denmark/ALAB-HH32/2020 | Europe / Denmark / Unknown | 2020-03-11 | Department of Clinical Microbiology, Copenhagen University Hospital, Hvidovre, Kettegaard Alle 30, 2650 Hvidovre. | Albertsen lab, Department of Chemistry and Bioscience, Aalborg University, Denmark | Rasmus Kirkegaard |
| EPI_ISL_429292 | hCoV-19/Denmark/ALAB-HH36/2020 | Europe / Denmark / Unknown | 2020-03-12 | Department of Clinical Microbiology, Copenhagen University Hospital, Hvidovre, Kettegaard Alle 30, 2650 Hvidovre. | Albertsen lab, Department of Chemistry and Bioscience, Aalborg University, Denmark | Rasmus Kirkegaard |
| EPI_ISL_429293 | hCoV-19/Denmark/ALAB-HH37/2020 | Europe / Denmark / Unknown | 2020-03-13 | Department of Clinical Microbiology, Copenhagen University Hospital, Hvidovre, Kettegaard Alle 30, 2650 Hvidovre. | Albertsen lab, Department of Chemistry and Bioscience, Aalborg University, Denmark | Rasmus Kirkegaard |
| EPI_ISL_429295 | hCoV-19/Denmark/ALAB-HH39/2020 | Europe / Denmark / Unknown | 2020-03-13 | Department of Clinical Microbiology, Copenhagen University Hospital, Hvidovre, Kettegaard Alle 30, 2650 Hvidovre. | Albertsen lab, Department of Chemistry and Bioscience, Aalborg University, Denmark | Rasmus Kirkegaard |
| EPI_ISL_429296 | hCoV-19/Denmark/ALAB-HH40/2020 | Europe / Denmark / Unknown | 2020-03-14 | Department of Clinical Microbiology, Copenhagen University Hospital, Hvidovre, Kettegaard Alle 30, 2650 Hvidovre. | Albertsen lab, Department of Chemistry and Bioscience, Aalborg University, Denmark | Rasmus Kirkegaard |
| EPI_ISL_429299 | hCoV-19/Denmark/ALAB-HH43/2020 | Europe / Denmark / Unknown | 2020-03-14 | Department of Clinical Microbiology, Copenhagen University Hospital, Hvidovre, Kettegaard Alle 30, 2650 Hvidovre. | Albertsen lab, Department of Chemistry and Bioscience, Aalborg University, Denmark | Rasmus Kirkegaard |
| EPI_ISL_429968 | hCoV-19/France/HF1463/2020 | Europe / France / Hauts de France / Com | 2020-02-21 | Centre Hospitalier Compiègne Laboratoire de Biologie | National Reference Center for Viruses of Respiratory Infections, Institut Pasteur, Paris | Mélanie Albert, Marion Barbet, Sylvie Behillil, Méline Bizard, Angela Brisebarre, Flora Donati, Fabiana Gambaro, Etienne Simon-Lorière, Vincent Enouf, Maud Vanpeene, Sylvie van der Werf, Raulin Olivia |
| EPI_ISL_429969 | hCoV-19/USA/VA-DCLS-0069/2020 | North America / USA / Virginia | 2020-03-26 | Virginia DCLS | Virginia DCLS | Virginia DCLS |
| EPI_ISL_429970 | hCoV-19/USA/VA-DCLS-0070/2020 | North America / USA / Virginia | 2020-03-28 | Virginia DCLS | Virginia DCLS | Virginia DCLS |
| EPI_ISL_429971 | hCoV-19/USA/VA-DCLS-0071/2020 | North America / USA / Virginia | 2020-03-30 | Virginia DCLS | Virginia DCLS | Virginia DCLS |
| EPI_ISL_429972 | hCoV-19/USA/VA-DCLS-0072/2020 | North America / USA / Virginia | 2020-03-28 | Virginia DCLS | Virginia DCLS | Virginia DCLS |
| EPI_ISL_429973 | hCoV-19/USA/VA-DCLS-0073/2020 | North America / USA / Virginia | 2020-03-28 | Virginia DCLS | Virginia DCLS | Virginia DCLS |
| EPI_ISL_429974 | hCoV-19/USA/VA-DCLS-0075/2020 | North America / USA / Virginia | 2020-03-29 | Virginia DCLS | Virginia DCLS | Virginia DCLS |
| EPI_ISL_429975 | hCoV-19/USA/VA-DCLS-0076/2020 | North America / USA / Virginia | 2020-03-27 | Virginia DCLS | Virginia DCLS | Virginia DCLS |
| EPI_ISL_429976 | hCoV-19/USA/VA-DCLS-0077/2020 | North America / USA / Virginia | 2020-04-04 | Virginia DCLS | Virginia DCLS | Virginia DCLS |
| EPI_ISL_429977 | hCoV-19/USA/VA-DCLS-0078/2020 | North America / USA / Virginia | 2020-03-27 | Virginia DCLS | Virginia DCLS | Virginia DCLS |
| EPI_ISL_429978 | hCoV-19/USA/VA-DCLS-0079/2020 | North America / USA / Virginia | 2020-03-30 | Virginia DCLS | Virginia DCLS | Virginia DCLS |
| EPI_ISL_429979 | hCoV-19/USA/VA-DCLS-0080/2020 | North America / USA / Virginia | 2020-03-26 | Virginia DCLS | Virginia DCLS | Virginia DCLS |
| EPI_ISL_429980 | hCoV-19/USA/VA-DCLS-0081/2020 | North America / USA / Virginia | 2020-04-03 | Virginia DCLS | Virginia DCLS | Virginia DCLS |
| EPI_ISL_429981 | hCoV-19/USA/VA-DCLS-0083/2020 | North America / USA / Virginia | 2020-04-03 | Virginia DCLS | Virginia DCLS | Virginia DCLS |
| EPI_ISL_429982 | hCoV-19/USA/VA-DCLS-0084/2020 | North America / USA / Virginia | 2020-04-06 | Virginia DCLS | Virginia DCLS | Virginia DCLS |
| EPI_ISL_429983 | hCoV-19/USA/VA-DCLS-0086/2020 | North America / USA / Virginia | 2020-03-30 | Virginia DCLS | Virginia DCLS | Virginia DCLS |
| EPI_ISL_429984 | hCoV-19/USA/VA-DCLS-0087/2020 | North America / USA / Virginia | 2020-04-02 | Virginia DCLS | Virginia DCLS | Virginia DCLS |
| EPI_ISL_429985 | hCoV-19/USA/VA-DCLS-0088/2020 | North America / USA / Virginia | 2020-04-02 | Virginia DCLS | Virginia DCLS | Virginia DCLS |
| EPI_ISL_429986 | hCoV-19/USA/VA-DCLS-0089/2020 | North America / USA / Virginia | 2020-04-02 | Virginia DCLS | Virginia DCLS | Virginia DCLS |
| EPI_ISL_429987 | hCoV-19/USA/VA-DCLS-0090/2020 | North America / USA / Virginia | 2020-04-02 | Virginia DCLS | Virginia DCLS | Virginia DCLS |
| EPI_ISL_429988 | hCoV-19/USA/VA-DCLS-0091/2020 | North America / USA / Virginia | 2020-04-02 | Virginia DCLS | Virginia DCLS | Virginia DCLS |
| EPI_ISL_429989 | hCoV-19/USA/VA-DCLS-0092/2020 | North America / USA / Virginia | 2020-04-01 | Virginia DCLS | Virginia DCLS | Virginia DCLS |
| EPI_ISL_429990 | hCoV-19/USA/CA-SR007/2020 | North America / USA / California / San Di | 2020-03-21 | Rady's Childrens Hospital | Andersen lab at Scripps Research | SEARCH Alliance San Diego with Christina Clarke, Michelle Vanderpool, Teresa Mueller, Denise Malicki |
| EPI_ISL_429991 | hCoV-19/USA/CA-SR017/2020 | North America / USA / California / San Di | 2020-03-24 | Andersen lab at Scripps Research | Andersen lab at Scripps Research | SEARCH Alliance San Diego |
| EPI_ISL_429992 | hCoV-19/Jordan/SR-032/2020 | Asia / Jordan / Amman | 2020-03-22 | Biolab Diagnostic Laboratories | Andersen lab at Scripps Research | Issa Abu-Dayyeh, Ahmad Tibi, Lama Hussein, Lina Mohammad, Zein Naber, Amid Abdelnour with SEARCH Alliance San Diego |
| EPI_ISL_429993 | hCoV-19/Jordan/SR-033/2020 | Asia / Jordan / Amman | 2020-03-16 | Biolab Diagnostic Laboratories | Andersen lab at Scripps Research | Issa Abu-Dayyeh, Ahmad Tibi, Lama Hussein, Lina Mohammad, Zein Naber, Amid Abdelnour with SEARCH Alliance San Diego |
| EPI_ISL_429994 | hCoV-19/Jordan/SR-034/2020 | Asia / Jordan / Amman | 2020-03-17 | Biolab Diagnostic Laboratories | Andersen lab at Scripps Research | Issa Abu-Dayyeh, Ahmad Tibi, Lama Hussein, Lina Mohammad, Zein Naber, Amid Abdelnour with SEARCH Alliance San Diego |
| EPI_ISL_429995 | hCoV-19/Jordan/SR-035/2020 | Asia / Jordan / Amman | 2020-03-30 | Biolab Diagnostic Laboratories | Andersen lab at Scripps Research | Issa Abu-Dayyeh, Ahmad Tibi, Lama Hussein, Lina Mohammad, Zein Naber, Amid Abdelnour with SEARCH Alliance San Diego |
| EPI_ISL_429996 | hCoV-19/Jordan/SR-036/2020 | Asia / Jordan / Amman | 2020-03-23 | Biolab Diagnostic Laboratories | Andersen lab at Scripps Research | Issa Abu-Dayyeh, Ahmad Tibi, Lama Hussein, Lina Mohammad, Zein Naber, Amid Abdelnour with SEARCH Alliance San Diego |
| EPI_ISL_429997 | hCoV-19/Jordan/SR-037/2020 | Asia / Jordan / Amman | 2020-03-16 | Biolab Diagnostic Laboratories | Andersen lab at Scripps Research | Issa Abu-Dayyeh, Ahmad Tibi, Lama Hussein, Lina Mohammad, Zein Naber, Amid Abdelnour with SEARCH Alliance San Diego |
| EPI_ISL_429998 | hCoV-19/Jordan/SR-039/2020 | Asia / Jordan / Amman | 2020-03-28 | Biolab Diagnostic Laboratories | Andersen lab at Scripps Research | Issa Abu-Dayyeh, Ahmad Tibi, Lama Hussein, Lina Mohammad, Zein Naber, Amid Abdelnour with SEARCH Alliance San Diego |
| EPI_ISL_429999 | hCoV-19/Jordan/SR-041/2020 | Asia / Jordan / Amman | 2020-03-17 | Biolab Diagnostic Laboratories | Andersen lab at Scripps Research | Issa Abu-Dayyeh, Ahmad Tibi, Lama Hussein, Lina Mohammad, Zein Naber, Amid Abdelnour with SEARCH Alliance San Diego |
| EPI_ISL_430058 | hCoV-19/USA/UT-0511/2020 | North America / USA / Utah | 2020-04-03 | Utah Public Health Laboratory | Utah Public Health Laboratory | Erin Young, Kelly Oakeson |
| EPI_ISL_430059 | hCoV-19/USA/UT-0513/2020 | North America / USA / Utah | 2020-04-03 | Utah Public Health Laboratory | Utah Public Health Laboratory | Erin Young, Kelly Oakeson |
| EPI_ISL_430060 | hCoV-19/USA/UT-0514/2020 | North America / USA / Utah | 2020-04-03 | Utah Public Health Laboratory | Utah Public Health Laboratory | Erin Young, Kelly Oakeson |
| EPI_ISL_430061 | hCoV-19/USA/UT-0519/2020 | North America / USA / Utah | 2020-04-03 | Utah Public Health Laboratory | Utah Public Health Laboratory | Erin Young, Kelly Oakeson |
| EPI_ISL_430062 | hCoV-19/USA/UT-0520/2020 | North America / USA / Utah | 2020-04-03 | Utah Public Health Laboratory | Utah Public Health Laboratory | Erin Young, Kelly Oakeson |
| EPI_ISL_430064 | hCoV-19/Australia/VIC-CBA4/2020 | Oceania / Australia / Victoria | 2020-03-07 | Geelong Centre for Emerging Infectious Diseases | Geelong Centre for Emerging Infectious Diseases | Chamings A., Bhatta T.R., Alexandersen S. |
| EPI_ISL_430065 | hCoV-19/Australia/VIC-CBA5/2020 | Oceania / Australia / Victoria | 2020-04-08 | Geelong Centre for Emerging Infectious Diseases | Geelong Centre for Emerging Infectious Diseases | Chamings A., Bhatta T.R., Alexandersen S. |
| EPI_ISL_430067 | hCoV-19/Russia/StPetersburg-RII4705S/2020 | Europe / Russia / St.Petersburg | 2020-04-07 | WHO National Influenza Centre Russian Federation | WHO National Influenza Centre Russian Federation | Andrey Komissarov, Artem Fadeev, Mariia Sergeeva, Anna Ivanova, Daria Danilenko |
| EPI_ISL_430068 | hCoV-19/Russia/StPetersburg-RII4712V/2020 | Europe / Russia / St.Petersburg | 2020-04-07 | WHO National Influenza Centre Russian Federation | WHO National Influenza Centre Russian Federation | Andrey Komissarov, Artem Fadeev, Mariia Sergeeva, Anna Ivanova, Daria Danilenko |
| EPI_ISL_430069 | hCoV-19/Russia/StPetersburg-RII4780S/2020 | Europe / Russia / St.Petersburg | 2020-04-07 | WHO National Influenza Centre Russian Federation | WHO National Influenza Centre Russian Federation | Andrey Komissarov, Artem Fadeev, Mariia Sergeeva, Anna Ivanova, Daria Danilenko |
| EPI_ISL_430070 | hCoV-19/Russia/StPetersburg-RII4917S/2020 | Europe / Russia / St.Petersburg | 2020-04-07 | WHO National Influenza Centre Russian Federation | WHO National Influenza Centre Russian Federation | Andrey Komissarov, Artem Fadeev, Mariia Sergeeva, Anna Ivanova, Daria Danilenko |
| EPI_ISL_430071 | hCoV-19/Russia/StPetersburg-RII4936S/2020 | Europe / Russia / St.Petersburg | 2020-04-08 | WHO National Influenza Centre Russian Federation | WHO National Influenza Centre Russian Federation | Andrey Komissarov, Artem Fadeev, Mariia Sergeeva, Anna Ivanova, Daria Danilenko |
| EPI_ISL_430072 | hCoV-19/Russia/StPetersburg-RII4937S/2020 | Europe / Russia / St.Petersburg | 2020-04-08 | WHO National Influenza Centre Russian Federation | WHO National Influenza Centre Russian Federation | Andrey Komissarov, Artem Fadeev, Mariia Sergeeva, Anna Ivanova, Daria Danilenko |
| EPI_ISL_430073 | hCoV-19/Russia/StPetersburg-RII4938S/2020 | Europe / Russia / St.Petersburg | 2020-04-08 | WHO National Influenza Centre Russian Federation | WHO National Influenza Centre Russian Federation | Andrey Komissarov, Artem Fadeev, Mariia Sergeeva, Anna Ivanova, Daria Danilenko |
| EPI_ISL_430074 | hCoV-19/Russia/StPetersburg-RII4944S/2020 | Europe / Russia / St.Petersburg | 2020-04-08 | WHO National Influenza Centre Russian Federation | WHO National Influenza Centre Russian Federation | Andrey Komissarov, Artem Fadeev, Mariia Sergeeva, Anna Ivanova, Daria Danilenko |
| EPI_ISL_430075 | hCoV-19/Russia/StPetersburg-RII4949S/2020 | Europe / Russia / St.Petersburg | 2020-04-09 | WHO National Influenza Centre Russian Federation | WHO National Influenza Centre Russian Federation | Andrey Komissarov, Artem Fadeev, Mariia Sergeeva, Anna Ivanova, Daria Danilenko |
| EPI_ISL_430076 | hCoV-19/Russia/StPetersburg-RII4969S/2020 | Europe / Russia / St.Petersburg | 2020-04-10 | WHO National Influenza Centre Russian Federation | WHO National Influenza Centre Russian Federation | Andrey Komissarov, Artem Fadeev, Mariia Sergeeva, Anna Ivanova, Daria Danilenko |
| EPI_ISL_430077 | hCoV-19/Russia/StPetersburg-RII4970S/2020 | Europe / Russia / St.Petersburg | 2020-04-10 | WHO National Influenza Centre Russian Federation | WHO National Influenza Centre Russian Federation | Andrey Komissarov, Artem Fadeev, Mariia Sergeeva, Anna Ivanova, Daria Danilenko |
| EPI_ISL_430078 | hCoV-19/Russia/StPetersburg-RII4972S/2020 | Europe / Russia / St.Petersburg | 2020-04-10 | WHO National Influenza Centre Russian Federation | WHO National Influenza Centre Russian Federation | Andrey Komissarov, Artem Fadeev, Mariia Sergeeva, Anna Ivanova, Daria Danilenko |
| EPI_ISL_430079 | hCoV-19/Russia/StPetersburg-RII4980S/2020 | Europe / Russia / St.Petersburg | 2020-04-10 | WHO National Influenza Centre Russian Federation | WHO National Influenza Centre Russian Federation | Andrey Komissarov, Artem Fadeev, Mariia Sergeeva, Anna Ivanova, Daria Danilenko |
| EPI_ISL_430080 | hCoV-19/Russia/StPetersburg-RII4983S/2020 | Europe / Russia / St.Petersburg | 2020-04-10 | WHO National Influenza Centre Russian Federation | WHO National Influenza Centre Russian Federation | Andrey Komissarov, Artem Fadeev, Mariia Sergeeva, Anna Ivanova, Daria Danilenko |
| EPI_ISL_430081 | hCoV-19/Russia/StPetersburg-RII4984S/2020 | Europe / Russia / St.Petersburg | 2020-04-10 | WHO National Influenza Centre Russian Federation | WHO National Influenza Centre Russian Federation | Andrey Komissarov, Artem Fadeev, Mariia Sergeeva, Anna Ivanova, Daria Danilenko |
| EPI_ISL_430082 | hCoV-19/Russia/StPetersburg-RII4985S/2020 | Europe / Russia / St.Petersburg | 2020-04-10 | WHO National Influenza Centre Russian Federation | WHO National Influenza Centre Russian Federation | Andrey Komissarov, Artem Fadeev, Mariia Sergeeva, Anna Ivanova, Daria Danilenko |
| EPI_ISL_430083 | hCoV-19/Russia/StPetersburg-RII4988S/2020 | Europe / Russia / St.Petersburg | 2020-04-10 | WHO National Influenza Centre Russian Federation | WHO National Influenza Centre Russian Federation | Andrey Komissarov, Artem Fadeev, Mariia Sergeeva, Anna Ivanova, Daria Danilenko |
| EPI_ISL_430084 | hCoV-19/Russia/StPetersburg-RII5017V/2020 | Europe / Russia / St.Petersburg | 2020-04-09 | WHO National Influenza Centre Russian Federation | WHO National Influenza Centre Russian Federation | Andrey Komissarov, Artem Fadeev, Mariia Sergeeva, Anna Ivanova, Daria Danilenko |
| EPI_ISL_430085 | hCoV-19/Russia/StPetersburg-RII5029S/2020 | Europe / Russia / St.Petersburg | 2020-04-10 | WHO National Influenza Centre Russian Federation | WHO National Influenza Centre Russian Federation | Andrey Komissarov, Artem Fadeev, Mariia Sergeeva, Anna Ivanova, Daria Danilenko |
| EPI_ISL_430086 | hCoV-19/Russia/StPetersburg-RII5033S/2020 | Europe / Russia / St.Petersburg | 2020-04-10 | WHO National Influenza Centre Russian Federation | WHO National Influenza Centre Russian Federation | Andrey Komissarov, Artem Fadeev, Mariia Sergeeva, Anna Ivanova, Daria Danilenko |
| EPI_ISL_430087 | hCoV-19/Russia/StPetersburg-RII5047S/2020 | Europe / Russia / St.Petersburg | 2020-04-10 | WHO National Influenza Centre Russian Federation | WHO National Influenza Centre Russian Federation | Andrey Komissarov, Artem Fadeev, Mariia Sergeeva, Anna Ivanova, Daria Danilenko |
| EPI_ISL_430088 | hCoV-19/Russia/StPetersburg-RII5169S/2020 | Europe / Russia / St.Petersburg | 2020-04-09 | WHO National Influenza Centre Russian Federation | WHO National Influenza Centre Russian Federation | Andrey Komissarov, Artem Fadeev, Mariia Sergeeva, Anna Ivanova, Daria Danilenko |
| EPI_ISL_430089 | hCoV-19/Russia/StPetersburg-RII5170S/2020 | Europe / Russia / St.Petersburg | 2020-04-09 | WHO National Influenza Centre Russian Federation | WHO National Influenza Centre Russian Federation | Andrey Komissarov, Artem Fadeev, Mariia Sergeeva, Anna Ivanova, Daria Danilenko |
| EPI_ISL_430090 | hCoV-19/Russia/StPetersburg-RII5244S/2020 | Europe / Russia / St.Petersburg | 2020-04-10 | WHO National Influenza Centre Russian Federation | WHO National Influenza Centre Russian Federation | Andrey Komissarov, Artem Fadeev, Mariia Sergeeva, Anna Ivanova, Daria Danilenko |
| EPI_ISL_430091 | hCoV-19/Russia/StPetersburg-RII5247S/2020 | Europe / Russia / St.Petersburg | 2020-04-10 | WHO National Influenza Centre Russian Federation | WHO National Influenza Centre Russian Federation | Andrey Komissarov, Artem Fadeev, Mariia Sergeeva, Anna Ivanova, Daria Danilenko |
| EPI_ISL_430092 | hCoV-19/Russia/StPetersburg-RII5633S/2020 | Europe / Russia / St.Petersburg | 2020-04-14 | WHO National Influenza Centre Russian Federation | WHO National Influenza Centre Russian Federation | Andrey Komissarov, Artem Fadeev, Mariia Sergeeva, Anna Ivanova, Daria Danilenko |
| EPI_ISL_430093 | hCoV-19/Russia/StPetersburg-RII5634S/2020 | Europe / Russia / St.Petersburg | 2020-04-14 | WHO National Influenza Centre Russian Federation | WHO National Influenza Centre Russian Federation | Andrey Komissarov, Artem Fadeev, Mariia Sergeeva, Anna Ivanova, Daria Danilenko |
| EPI_ISL_430094 | hCoV-19/Russia/StPetersburg-RII5635S/2020 | Europe / Russia / St.Petersburg | 2020-04-14 | WHO National Influenza Centre Russian Federation | WHO National Influenza Centre Russian Federation | Andrey Komissarov, Artem Fadeev, Mariia Sergeeva, Anna Ivanova, Daria Danilenko |
| EPI_ISL_430095 | hCoV-19/Russia/StPetersburg-RII5642S/2020 | Europe / Russia / St.Petersburg | 2020-04-14 | WHO National Influenza Centre Russian Federation | WHO National Influenza Centre Russian Federation | Andrey Komissarov, Artem Fadeev, Mariia Sergeeva, Anna Ivanova, Daria Danilenko |
| EPI_ISL_430096 | hCoV-19/Russia/StPetersburg-RII5643S/2020 | Europe / Russia / St.Petersburg | 2020-04-14 | WHO National Influenza Centre Russian Federation | WHO National Influenza Centre Russian Federation | Andrey Komissarov, Artem Fadeev, Mariia Sergeeva, Anna Ivanova, Daria Danilenko |
| EPI_ISL_430097 | hCoV-19/Russia/StPetersburg-RII5644S/2020 | Europe / Russia / St.Petersburg | 2020-04-14 | WHO National Influenza Centre Russian Federation | WHO National Influenza Centre Russian Federation | Andrey Komissarov, Artem Fadeev, Mariia Sergeeva, Anna Ivanova, Daria Danilenko |
| EPI_ISL_430098 | hCoV-19/Russia/StPetersburg-RII5654S/2020 | Europe / Russia / St.Petersburg | 2020-04-14 | WHO National Influenza Centre Russian Federation | WHO National Influenza Centre Russian Federation | Andrey Komissarov, Artem Fadeev, Mariia Sergeeva, Anna Ivanova, Daria Danilenko |
| EPI_ISL_430099 | hCoV-19/Russia/StPetersburg-RII5942S/2020 | Europe / Russia / St.Petersburg | 2020-04-13 | WHO National Influenza Centre Russian Federation | WHO National Influenza Centre Russian Federation | Andrey Komissarov, Artem Fadeev, Mariia Sergeeva, Anna Ivanova, Daria Danilenko |
| EPI_ISL_430100 | hCoV-19/Russia/StPetersburg-RII6027S/2020 | Europe / Russia / St.Petersburg | 2020-04-14 | WHO National Influenza Centre Russian Federation | WHO National Influenza Centre Russian Federation | Andrey Komissarov, Artem Fadeev, Mariia Sergeeva, Anna Ivanova, Daria Danilenko |
| EPI_ISL_430101 | hCoV-19/Russia/StPetersburg-RII6031S/2020 | Europe / Russia / St.Petersburg | 2020-04-14 | WHO National Influenza Centre Russian Federation | WHO National Influenza Centre Russian Federation | Andrey Komissarov, Artem Fadeev, Mariia Sergeeva, Anna Ivanova, Daria Danilenko |
| EPI_ISL_430102 | hCoV-19/Russia/StPetersburg-RII6039S/2020 | Europe / Russia / St.Petersburg | 2020-04-14 | WHO National Influenza Centre Russian Federation | WHO National Influenza Centre Russian Federation | Andrey Komissarov, Artem Fadeev, Mariia Sergeeva, Anna Ivanova, Daria Danilenko |
| EPI_ISL_430103 | hCoV-19/Russia/StPetersburg-RII6043S/2020 | Europe / Russia / St.Petersburg | 2020-04-14 | WHO National Influenza Centre Russian Federation | WHO National Influenza Centre Russian Federation | Andrey Komissarov, Artem Fadeev, Mariia Sergeeva, Anna Ivanova, Daria Danilenko |
| EPI_ISL_430104 | hCoV-19/Russia/StPetersburg-RII6053S/2020 | Europe / Russia / St.Petersburg | 2020-04-15 | WHO National Influenza Centre Russian Federation | WHO National Influenza Centre Russian Federation | Andrey Komissarov, Artem Fadeev, Mariia Sergeeva, Anna Ivanova, Daria Danilenko |
| EPI_ISL_430105 | hCoV-19/Russia/StPetersburg-RII6055S/2020 | Europe / Russia / St.Petersburg | 2020-04-15 | WHO National Influenza Centre Russian Federation | WHO National Influenza Centre Russian Federation | Andrey Komissarov, Artem Fadeev, Mariia Sergeeva, Anna Ivanova, Daria Danilenko |
| EPI_ISL_430106 | hCoV-19/Russia/StPetersburg-RII6056S/2020 | Europe / Russia / St.Petersburg | 2020-04-15 | WHO National Influenza Centre Russian Federation | WHO National Influenza Centre Russian Federation | Andrey Komissarov, Artem Fadeev, Mariia Sergeeva, Anna Ivanova, Daria Danilenko |
| EPI_ISL_430107 | hCoV-19/Russia/StPetersburg-RII6060S/2020 | Europe / Russia / St.Petersburg | 2020-04-15 | WHO National Influenza Centre Russian Federation | WHO National Influenza Centre Russian Federation | Andrey Komissarov, Artem Fadeev, Mariia Sergeeva, Anna Ivanova, Daria Danilenko |
| EPI_ISL_430108 | hCoV-19/Russia/StPetersburg-RII6061S/2020 | Europe / Russia / St.Petersburg | 2020-04-15 | WHO National Influenza Centre Russian Federation | WHO National Influenza Centre Russian Federation | Andrey Komissarov, Artem Fadeev, Mariia Sergeeva, Anna Ivanova, Daria Danilenko |
| EPI_ISL_430109 | hCoV-19/Russia/StPetersburg-RII6063S/2020 | Europe / Russia / St.Petersburg | 2020-04-15 | WHO National Influenza Centre Russian Federation | WHO National Influenza Centre Russian Federation | Andrey Komissarov, Artem Fadeev, Mariia Sergeeva, Anna Ivanova, Daria Danilenko |
| EPI_ISL_430110 | hCoV-19/Russia/StPetersburg-RII6065S/2020 | Europe / Russia / St.Petersburg | 2020-04-15 | WHO National Influenza Centre Russian Federation | WHO National Influenza Centre Russian Federation | Andrey Komissarov, Artem Fadeev, Mariia Sergeeva, Anna Ivanova, Daria Danilenko |
| EPI_ISL_430111 | hCoV-19/Russia/StPetersburg-RII6086S/2020 | Europe / Russia / St.Petersburg | 2020-04-15 | WHO National Influenza Centre Russian Federation | WHO National Influenza Centre Russian Federation | Andrey Komissarov, Artem Fadeev, Mariia Sergeeva, Anna Ivanova, Daria Danilenko |
| EPI_ISL_430112 | hCoV-19/Russia/Ulan-Ude-RII4562V/2020 | Europe / Russia / Buryat Republic / Ulan- | 2020-03-25 | WHO National Influenza Centre Russian Federation | WHO National Influenza Centre Russian Federation | Andrey Komissarov, Artem Fadeev, Mariia Sergeeva, Anna Ivanova, Daria Danilenko |
| EPI_ISL_430113 | hCoV-19/USA/WA-S259/2020 | North America / USA / Washington / King | 2020-03-04 | Seattle Flu Study | Seattle Flu Study | Chu et al |
| EPI_ISL_430114 | hCoV-19/USA/WA-S260/2020 | North America / USA / Washington / King | 2020-03-12 | Seattle Flu Study | Seattle Flu Study | Chu et al |
| EPI_ISL_430115 | hCoV-19/USA/WA-S261/2020 | North America / USA / Washington | 2020-03-30 | Seattle Flu Study | Seattle Flu Study | Chu et al |
| EPI_ISL_430116 | hCoV-19/USA/WA-S262/2020 | North America / USA / Washington | 2020-03-16 | Seattle Flu Study | Seattle Flu Study | Chu et al |
| EPI_ISL_430117 | hCoV-19/USA/WA-S263/2020 | North America / USA / Washington / King | 2020-03-05 | Seattle Flu Study | Seattle Flu Study | Chu et al |
| EPI_ISL_430118 | hCoV-19/USA/WA-S264/2020 | North America / USA / Washington / King | 2020-03-03 | Seattle Flu Study | Seattle Flu Study | Chu et al |
| EPI_ISL_430119 | hCoV-19/USA/WA-S265/2020 | North America / USA / Washington / Snol | 2020-03-05 | Seattle Flu Study | Seattle Flu Study | Chu et al |
| EPI_ISL_430121 | hCoV-19/USA/WA-S267/2020 | North America / USA / Washington / King | 2020-03-31 | Seattle Flu Study | Seattle Flu Study | Chu et al |
| EPI_ISL_430122 | hCoV-19/USA/WA-S268/2020 | North America / USA / Washington / King | 2020-03-05 | Seattle Flu Study | Seattle Flu Study | Chu et al |
| EPI_ISL_430123 | hCoV-19/USA/WA-S269/2020 | North America / USA / Washington / King | 2020-03-10 | Seattle Flu Study | Seattle Flu Study | Chu et al |
| EPI_ISL_430124 | hCoV-19/USA/WA-S270/2020 | North America / USA / Washington / King | 2020-03-06 | Seattle Flu Study | Seattle Flu Study | Chu et al |
| EPI_ISL_430125 | hCoV-19/USA/WA-S271/2020 | North America / USA / Washington / King | 2020-03-31 | Seattle Flu Study | Seattle Flu Study | Chu et al |
| EPI_ISL_430126 | hCoV-19/USA/WA-S272/2020 | North America / USA / Washington / King | 2020-02-26 | Seattle Flu Study | Seattle Flu Study | Chu et al |
| EPI_ISL_430127 | hCoV-19/USA/WA-S273/2020 | North America / USA / Washington / King | 2020-03-12 | Seattle Flu Study | Seattle Flu Study | Chu et al |
| EPI_ISL_430128 | hCoV-19/USA/WA-S274/2020 | North America / USA / Washington / King | 2020-03-08 | Seattle Flu Study | Seattle Flu Study | Chu et al |
| EPI_ISL_430129 | hCoV-19/USA/WA-S275/2020 | North America / USA / Washington / King | 2020-03-06 | Seattle Flu Study | Seattle Flu Study | Chu et al |
| EPI_ISL_430130 | hCoV-19/USA/WA-S276/2020 | North America / USA / Washington | 2020-04-01 | Seattle Flu Study | Seattle Flu Study | Chu et al |
| EPI_ISL_430131 | hCoV-19/USA/WA-S277/2020 | North America / USA / Washington / King | 2020-02-22 | Seattle Flu Study | Seattle Flu Study | Chu et al |
| EPI_ISL_430132 | hCoV-19/USA/WA-S278/2020 | North America / USA / Washington | 2020-03-17 | Seattle Flu Study | Seattle Flu Study | Chu et al |
| EPI_ISL_430133 | hCoV-19/USA/WA-S279/2020 | North America / USA / Washington / King | 2020-03-04 | Seattle Flu Study | Seattle Flu Study | Chu et al |
| EPI_ISL_430134 | hCoV-19/USA/WA-S280/2020 | North America / USA / Washington / King | 2020-03-09 | Seattle Flu Study | Seattle Flu Study | Chu et al |
| EPI_ISL_430135 | hCoV-19/USA/WA-S281/2020 | North America / USA / Washington / King | 2020-02-29 | Seattle Flu Study | Seattle Flu Study | Chu et al |
| EPI_ISL_430136 | hCoV-19/USA/WA-S282/2020 | North America / USA / Washington / King | 2020-03-10 | Seattle Flu Study | Seattle Flu Study | Chu et al |
| EPI_ISL_430137 | hCoV-19/USA/WA-S283/2020 | North America / USA / Washington / King | 2020-03-21 | Seattle Flu Study | Seattle Flu Study | Chu et al |
| EPI_ISL_430140 | hCoV-19/USA/WA-S286/2020 | North America / USA / Washington / Pier | 2020-03-15 | Seattle Flu Study | Seattle Flu Study | Chu et al |
| EPI_ISL_430141 | hCoV-19/USA/WA-S287/2020 | North America / USA / Washington / King | 2020-03-06 | Seattle Flu Study | Seattle Flu Study | Chu et al |
| EPI_ISL_430142 | hCoV-19/USA/WA-S288/2020 | North America / USA / Washington / King | 2020-03-06 | Seattle Flu Study | Seattle Flu Study | Chu et al |
| EPI_ISL_430143 | hCoV-19/USA/WA-S289/2020 | North America / USA / Washington / King | 2020-03-23 | Seattle Flu Study | Seattle Flu Study | Chu et al |
| EPI_ISL_430146 | hCoV-19/USA/WA-S292/2020 | North America / USA / Washington / King | 2020-03-05 | Seattle Flu Study | Seattle Flu Study | Chu et al |
| EPI_ISL_430147 | hCoV-19/USA/WA-S293/2020 | North America / USA / Washington | 2020-04-01 | Seattle Flu Study | Seattle Flu Study | Chu et al |
| EPI_ISL_430148 | hCoV-19/USA/WA-S294/2020 | North America / USA / Washington / King | 2020-03-23 | Seattle Flu Study | Seattle Flu Study | Chu et al |
| EPI_ISL_430149 | hCoV-19/USA/WA-S295/2020 | North America / USA / Washington / King | 2020-03-31 | Seattle Flu Study | Seattle Flu Study | Chu et al |
| EPI_ISL_430150 | hCoV-19/USA/WA-S296/2020 | North America / USA / Washington / King | 2020-03-13 | Seattle Flu Study | Seattle Flu Study | Chu et al |
| EPI_ISL_430151 | hCoV-19/USA/WA-S297/2020 | North America / USA / Washington / King | 2020-03-13 | Seattle Flu Study | Seattle Flu Study | Chu et al |
| EPI_ISL_430152 | hCoV-19/USA/WA-S298/2020 | North America / USA / Washington / King | 2020-03-05 | Seattle Flu Study | Seattle Flu Study | Chu et al |
| EPI_ISL_430153 | hCoV-19/USA/WA-S299/2020 | North America / USA / Washington / King | 2020-03-31 | Seattle Flu Study | Seattle Flu Study | Chu et al |
| EPI_ISL_430154 | hCoV-19/USA/WA-S300/2020 | North America / USA / Washington / King | 2020-03-25 | Seattle Flu Study | Seattle Flu Study | Chu et al |
| EPI_ISL_430155 | hCoV-19/USA/WA-S301/2020 | North America / USA / Washington / Pier | 2020-03-14 | Seattle Flu Study | Seattle Flu Study | Chu et al |
| EPI_ISL_430160 | hCoV-19/USA/WA-S123/2020 | North America / USA / Washington | 2020-02-29 | Washington State Department of Health | Seattle Flu Study | Chu et al |
| EPI_ISL_430161 | hCoV-19/USA/WA-S124/2020 | North America / USA / Washington / King | 2020-03-13 | Washington State Department of Health | Seattle Flu Study | Chu et al |
| EPI_ISL_430162 | hCoV-19/USA/WA-S125/2020 | North America / USA / Washington / King | 2020-03-13 | Washington State Department of Health | Seattle Flu Study | Chu et al |
| EPI_ISL_430163 | hCoV-19/USA/WA-S126/2020 | North America / USA / Washington / King | 2020-03-13 | Washington State Department of Health | Seattle Flu Study | Chu et al |
| EPI_ISL_430164 | hCoV-19/USA/WA-S127/2020 | North America / USA / Washington / King | 2020-03-13 | Washington State Department of Health | Seattle Flu Study | Chu et al |
| EPI_ISL_430165 | hCoV-19/USA/WA-S128/2020 | North America / USA / Washington / King | 2020-03-13 | Washington State Department of Health | Seattle Flu Study | Chu et al |
| EPI_ISL_430166 | hCoV-19/USA/WA-S129/2020 | North America / USA / Washington / King | 2020-03-13 | Washington State Department of Health | Seattle Flu Study | Chu et al |
| EPI_ISL_430167 | hCoV-19/USA/WA-S130/2020 | North America / USA / Washington / King | 2020-03-13 | Washington State Department of Health | Seattle Flu Study | Chu et al |
| EPI_ISL_430168 | hCoV-19/USA/WA-S131/2020 | North America / USA / Washington / King | 2020-03-13 | Washington State Department of Health | Seattle Flu Study | Chu et al |
| EPI_ISL_430169 | hCoV-19/USA/WA-S132/2020 | North America / USA / Washington / King | 2020-03-13 | Washington State Department of Health | Seattle Flu Study | Chu et al |
| EPI_ISL_430170 | hCoV-19/USA/WA-S133/2020 | North America / USA / Washington / King | 2020-03-13 | Washington State Department of Health | Seattle Flu Study | Chu et al |
| EPI_ISL_430171 | hCoV-19/USA/WA-S134/2020 | North America / USA / Washington / King | 2020-03-13 | Washington State Department of Health | Seattle Flu Study | Chu et al |
| EPI_ISL_430172 | hCoV-19/USA/WA-S135/2020 | North America / USA / Washington / King | 2020-03-13 | Washington State Department of Health | Seattle Flu Study | Chu et al |
| EPI_ISL_430173 | hCoV-19/USA/WA-S136/2020 | North America / USA / Washington / King | 2020-03-13 | Washington State Department of Health | Seattle Flu Study | Chu et al |
| EPI_ISL_430174 | hCoV-19/USA/WA-S137/2020 | North America / USA / Washington / King | 2020-03-13 | Washington State Department of Health | Seattle Flu Study | Chu et al |
| EPI_ISL_430175 | hCoV-19/USA/WA-S138/2020 | North America / USA / Washington / Snol | 2020-03-12 | Washington State Department of Health | Seattle Flu Study | Chu et al |
| EPI_ISL_430176 | hCoV-19/USA/WA-S139/2020 | North America / USA / Washington / King | 2020-03-12 | Washington State Department of Health | Seattle Flu Study | Chu et al |
| EPI_ISL_430177 | hCoV-19/USA/WA-S140/2020 | North America / USA / Washington / King | 2020-03-12 | Washington State Department of Health | Seattle Flu Study | Chu et al |
| EPI_ISL_430178 | hCoV-19/USA/WA-S141/2020 | North America / USA / Washington / King | 2020-03-13 | Washington State Department of Health | Seattle Flu Study | Chu et al |
| EPI_ISL_430200 | hCoV-19/USA/WA-S162/2020 | North America / USA / Washington | 2020-03-04 | Washington State Department of Health | Seattle Flu Study | Chu et al |
| EPI_ISL_430201 | hCoV-19/USA/WA-S163/2020 | North America / USA / Washington | 2020-03-04 | Washington State Department of Health | Seattle Flu Study | Chu et al |
| EPI_ISL_430202 | hCoV-19/USA/WA-S164/2020 | North America / USA / Washington | 2020-03-02 | Washington State Department of Health | Seattle Flu Study | Chu et al |
| EPI_ISL_430204 | hCoV-19/USA/WA-S166/2020 | North America / USA / Washington / King | 2020-03-18 | Washington State Department of Health | Seattle Flu Study | Chu et al |
| EPI_ISL_430205 | hCoV-19/USA/WA-S167/2020 | North America / USA / Washington / Wha | 2020-03-18 | Washington State Department of Health | Seattle Flu Study | Chu et al |
| EPI_ISL_430206 | hCoV-19/USA/WA-S168/2020 | North America / USA / Washington / Wha | 2020-03-18 | Washington State Department of Health | Seattle Flu Study | Chu et al |
| EPI_ISL_430207 | hCoV-19/USA/WA-S169/2020 | North America / USA / Washington / Wha | 2020-03-18 | Washington State Department of Health | Seattle Flu Study | Chu et al |
| EPI_ISL_430208 | hCoV-19/USA/WA-S170/2020 | North America / USA / Washington / Yaki | 2020-03-17 | Washington State Department of Health | Seattle Flu Study | Chu et al |
| EPI_ISL_430209 | hCoV-19/USA/WA-S171/2020 | North America / USA / Washington | 2020-03-19 | Washington State Department of Health | Seattle Flu Study | Chu et al |
| EPI_ISL_430210 | hCoV-19/USA/WA-S172/2020 | North America / USA / Washington / King | 2020-03-19 | Washington State Department of Health | Seattle Flu Study | Chu et al |
| EPI_ISL_430211 | hCoV-19/USA/WA-S173/2020 | North America / USA / Washington | 2020-03-19 | Washington State Department of Health | Seattle Flu Study | Chu et al |
| EPI_ISL_430212 | hCoV-19/USA/WA-S174/2020 | North America / USA / Washington / King | 2020-03-19 | Washington State Department of Health | Seattle Flu Study | Chu et al |
| EPI_ISL_430213 | hCoV-19/USA/WA-S175/2020 | North America / USA / Washington / King | 2020-03-19 | Washington State Department of Health | Seattle Flu Study | Chu et al |
| EPI_ISL_430214 | hCoV-19/USA/WA-S176/2020 | North America / USA / Washington / King | 2020-03-19 | Washington State Department of Health | Seattle Flu Study | Chu et al |
| EPI_ISL_430215 | hCoV-19/USA/WA-S177/2020 | North America / USA / Washington / King | 2020-03-19 | Washington State Department of Health | Seattle Flu Study | Chu et al |
| EPI_ISL_430216 | hCoV-19/USA/WA-S178/2020 | North America / USA / Washington / King | 2020-03-20 | Washington State Department of Health | Seattle Flu Study | Chu et al |
| EPI_ISL_430217 | hCoV-19/USA/WA-S179/2020 | North America / USA / Washington / Wha | 2020-03-20 | Washington State Department of Health | Seattle Flu Study | Chu et al |
| EPI_ISL_430219 | hCoV-19/USA/WA-S181/2020 | North America / USA / Washington | 2020-03-19 | Washington State Department of Health | Seattle Flu Study | Chu et al |
| EPI_ISL_430220 | hCoV-19/USA/WA-S182/2020 | North America / USA / Washington / Wha | 2020-03-20 | Washington State Department of Health | Seattle Flu Study | Chu et al |
| EPI_ISL_430221 | hCoV-19/USA/WA-S183/2020 | North America / USA / Washington / King | 2020-03-19 | Washington State Department of Health | Seattle Flu Study | Chu et al |
| EPI_ISL_430222 | hCoV-19/USA/WA-S184/2020 | North America / USA / Washington / King | 2020-03-19 | Washington State Department of Health | Seattle Flu Study | Chu et al |
| EPI_ISL_430223 | hCoV-19/USA/WA-S185/2020 | North America / USA / Washington | 2020-03-19 | Washington State Department of Health | Seattle Flu Study | Chu et al |
| EPI_ISL_430224 | hCoV-19/USA/WA-S186/2020 | North America / USA / Washington / Wha | 2020-03-20 | Washington State Department of Health | Seattle Flu Study | Chu et al |
| EPI_ISL_430225 | hCoV-19/USA/WA-S187/2020 | North America / USA / Washington / King | 2020-03-20 | Washington State Department of Health | Seattle Flu Study | Chu et al |
| EPI_ISL_430226 | hCoV-19/USA/WA-S188/2020 | North America / USA / Washington / Wha | 2020-03-20 | Washington State Department of Health | Seattle Flu Study | Chu et al |
| EPI_ISL_430227 | hCoV-19/USA/WA-S189/2020 | North America / USA / Washington / Wha | 2020-03-20 | Washington State Department of Health | Seattle Flu Study | Chu et al |
| EPI_ISL_430228 | hCoV-19/USA/WA-S190/2020 | North America / USA / Washington / King | 2020-03-23 | Washington State Department of Health | Seattle Flu Study | Chu et al |
| EPI_ISL_430229 | hCoV-19/USA/WA-S191/2020 | North America / USA / Washington / Wha | 2020-03-20 | Washington State Department of Health | Seattle Flu Study | Chu et al |
| EPI_ISL_430230 | hCoV-19/USA/WA-S192/2020 | North America / USA / Washington / Wha | 2020-03-20 | Washington State Department of Health | Seattle Flu Study | Chu et al |
| EPI_ISL_430231 | hCoV-19/USA/WA-S193/2020 | North America / USA / Washington / Grar | 2020-03-19 | Washington State Department of Health | Seattle Flu Study | Chu et al |
| EPI_ISL_430232 | hCoV-19/USA/WA-S194/2020 | North America / USA / Washington / King | 2020-03-20 | Washington State Department of Health | Seattle Flu Study | Chu et al |
| EPI_ISL_430233 | hCoV-19/USA/WA-S195/2020 | North America / USA / Washington / Wha | 2020-03-20 | Washington State Department of Health | Seattle Flu Study | Chu et al |
| EPI_ISL_430234 | hCoV-19/USA/WA-S196/2020 | North America / USA / Washington / King | 2020-03-19 | Washington State Department of Health | Seattle Flu Study | Chu et al |
| EPI_ISL_430235 | hCoV-19/USA/WA-S197/2020 | North America / USA / Washington / Wha | 2020-03-20 | Washington State Department of Health | Seattle Flu Study | Chu et al |
| EPI_ISL_430236 | hCoV-19/USA/WA-S198/2020 | North America / USA / Washington / Wha | 2020-03-20 | Washington State Department of Health | Seattle Flu Study | Chu et al |
| EPI_ISL_430237 | hCoV-19/USA/WA-S199/2020 | North America / USA / Washington / Wha | 2020-03-20 | Washington State Department of Health | Seattle Flu Study | Chu et al |
| EPI_ISL_430238 | hCoV-19/USA/WA-S200/2020 | North America / USA / Washington / Yaki | 2020-03-20 | Washington State Department of Health | Seattle Flu Study | Chu et al |
| EPI_ISL_430239 | hCoV-19/USA/WA-S201/2020 | North America / USA / Washington / King | 2020-03-20 | Washington State Department of Health | Seattle Flu Study | Chu et al |
| EPI_ISL_430240 | hCoV-19/USA/WA-S202/2020 | North America / USA / Washington / Wha | 2020-03-20 | Washington State Department of Health | Seattle Flu Study | Chu et al |
| EPI_ISL_430241 | hCoV-19/USA/WA-S203/2020 | North America / USA / Washington / Wha | 2020-03-20 | Washington State Department of Health | Seattle Flu Study | Chu et al |
| EPI_ISL_430242 | hCoV-19/USA/WA-S204/2020 | North America / USA / Washington / Wha | 2020-03-22 | Washington State Department of Health | Seattle Flu Study | Chu et al |
| EPI_ISL_430243 | hCoV-19/USA/WA-S205/2020 | North America / USA / Washington / Wha | 2020-03-20 | Washington State Department of Health | Seattle Flu Study | Chu et al |
| EPI_ISL_430244 | hCoV-19/USA/WA-S206/2020 | North America / USA / Washington / Wha | 2020-03-20 | Washington State Department of Health | Seattle Flu Study | Chu et al |
| EPI_ISL_430245 | hCoV-19/USA/WA-S207/2020 | North America / USA / Washington | 2020-03-20 | Washington State Department of Health | Seattle Flu Study | Chu et al |
| EPI_ISL_430246 | hCoV-19/USA/WA-S208/2020 | North America / USA / Washington / Wha | 2020-03-23 | Washington State Department of Health | Seattle Flu Study | Chu et al |
| EPI_ISL_430247 | hCoV-19/USA/WA-S209/2020 | North America / USA / Washington / Wha | 2020-03-23 | Washington State Department of Health | Seattle Flu Study | Chu et al |
| EPI_ISL_430248 | hCoV-19/USA/WA-S210/2020 | North America / USA / Washington / King | 2020-03-19 | Washington State Department of Health | Seattle Flu Study | Chu et al |
| EPI_ISL_430249 | hCoV-19/USA/WA-S211/2020 | North America / USA / Washington / King | 2020-03-20 | Washington State Department of Health | Seattle Flu Study | Chu et al |
| EPI_ISL_430250 | hCoV-19/USA/WA-S212/2020 | North America / USA / Washington | 2020-03-19 | Washington State Department of Health | Seattle Flu Study | Chu et al |
| EPI_ISL_430252 | hCoV-19/USA/WA-S214/2020 | North America / USA / Washington / Unkı | 2020-03-20 | Washington State Department of Health | Seattle Flu Study | Chu et al |
| EPI_ISL_430253 | hCoV-19/USA/WA-S215/2020 | North America / USA / Washington | 2020-03-20 | Washington State Department of Health | Seattle Flu Study | Chu et al |
| EPI_ISL_430254 | hCoV-19/USA/WA-S216/2020 | North America / USA / Washington / Wha | 2020-03-20 | Washington State Department of Health | Seattle Flu Study | Chu et al |
| EPI_ISL_430255 | hCoV-19/USA/WA-S217/2020 | North America / USA / Washington / Wha | 2020-03-20 | Washington State Department of Health | Seattle Flu Study | Chu et al |
| EPI_ISL_430256 | hCoV-19/USA/WA-S218/2020 | North America / USA / Washington / Wha | 2020-03-23 | Washington State Department of Health | Seattle Flu Study | Chu et al |
| EPI_ISL_430257 | hCoV-19/USA/WA-S219/2020 | North America / USA / Washington / Pier | 2020-03-20 | Washington State Department of Health | Seattle Flu Study | Chu et al |
| EPI_ISL_430258 | hCoV-19/USA/WA-S220/2020 | North America / USA / Washington / Wha | 2020-03-20 | Washington State Department of Health | Seattle Flu Study | Chu et al |
| EPI_ISL_430259 | hCoV-19/USA/WA-S221/2020 | North America / USA / Washington / King | 2020-03-19 | Washington State Department of Health | Seattle Flu Study | Chu et al |
| EPI_ISL_430261 | hCoV-19/USA/WA-S223/2020 | North America / USA / Washington / Wha | 2020-03-20 | Washington State Department of Health | Seattle Flu Study | Chu et al |
| EPI_ISL_430262 | hCoV-19/USA/WA-S224/2020 | North America / USA / Washington / King | 2020-03-19 | Washington State Department of Health | Seattle Flu Study | Chu et al |
| EPI_ISL_430263 | hCoV-19/USA/WA-S225/2020 | North America / USA / Washington / Wha | 2020-03-20 | Washington State Department of Health | Seattle Flu Study | Chu et al |
| EPI_ISL_430264 | hCoV-19/USA/WA-S226/2020 | North America / USA / Washington / Wha | 2020-03-21 | Washington State Department of Health | Seattle Flu Study | Chu et al |
| EPI_ISL_430265 | hCoV-19/USA/WA-S227/2020 | North America / USA / Washington | 2020-03-20 | Washington State Department of Health | Seattle Flu Study | Chu et al |
| EPI_ISL_430266 | hCoV-19/USA/WA-S228/2020 | North America / USA / Washington / Wha | 2020-03-20 | Washington State Department of Health | Seattle Flu Study | Chu et al |
| EPI_ISL_430267 | hCoV-19/USA/WA-S229/2020 | North America / USA / Washington / King | 2020-03-19 | Washington State Department of Health | Seattle Flu Study | Chu et al |
| EPI_ISL_430268 | hCoV-19/USA/WA-S230/2020 | North America / USA / Washington / King | 2020-03-19 | Washington State Department of Health | Seattle Flu Study | Chu et al |
| EPI_ISL_430269 | hCoV-19/USA/WA-S231/2020 | North America / USA / Washington / Wha | 2020-03-20 | Washington State Department of Health | Seattle Flu Study | Chu et al |
| EPI_ISL_430270 | hCoV-19/USA/WA-S232/2020 | North America / USA / Washington / Clar | 2020-03-08 | Washington State Department of Health | Seattle Flu Study | Chu et al |
| EPI_ISL_430271 | hCoV-19/USA/WA-S233/2020 | North America / USA / Washington / Snol | 2020-03-03 | Washington State Department of Health | Seattle Flu Study | Chu et al |
| EPI_ISL_430272 | hCoV-19/USA/WA-S234/2020 | North America / USA / Washington / King | 2020-02-28 | Washington State Department of Health | Seattle Flu Study | Chu et al |
| EPI_ISL_430273 | hCoV-19/USA/WA-S235/2020 | North America / USA / Washington / King | 2020-02-29 | Washington State Department of Health | Seattle Flu Study | Chu et al |
| EPI_ISL_430274 | hCoV-19/USA/WA-S236/2020 | North America / USA / Washington / King | 2020-03-01 | Washington State Department of Health | Seattle Flu Study | Chu et al |
| EPI_ISL_430275 | hCoV-19/USA/WA-S237/2020 | North America / USA / Washington | 2020-03-06 | Washington State Department of Health | Seattle Flu Study | Chu et al |
| EPI_ISL_430276 | hCoV-19/USA/WA-S238/2020 | North America / USA / Washington / King | 2020-03-02 | Washington State Department of Health | Seattle Flu Study | Chu et al |
| EPI_ISL_430277 | hCoV-19/USA/WA-S239/2020 | North America / USA / Washington / King | 2020-03-07 | Washington State Department of Health | Seattle Flu Study | Chu et al |
| EPI_ISL_430278 | hCoV-19/USA/WA-S240/2020 | North America / USA / Washington / King | 2020-03-06 | Washington State Department of Health | Seattle Flu Study | Chu et al |
| EPI_ISL_430279 | hCoV-19/USA/WA-S241/2020 | North America / USA / Washington / King | 2020-03-04 | Washington State Department of Health | Seattle Flu Study | Chu et al |
| EPI_ISL_430280 | hCoV-19/USA/WA-S242/2020 | North America / USA / Washington / King | 2020-03-01 | Washington State Department of Health | Seattle Flu Study | Chu et al |
| EPI_ISL_430281 | hCoV-19/USA/WA-S243/2020 | North America / USA / Washington / King | 2020-02-29 | Washington State Department of Health | Seattle Flu Study | Chu et al |
| EPI_ISL_430282 | hCoV-19/USA/WA-S244/2020 | North America / USA / Washington / Thur | 2020-03-11 | Washington State Department of Health | Seattle Flu Study | Chu et al |
| EPI_ISL_430283 | hCoV-19/USA/WA-S245/2020 | North America / USA / Washington / Snol | 2020-03-02 | Washington State Department of Health | Seattle Flu Study | Chu et al |
| EPI_ISL_430285 | hCoV-19/USA/WA-S247/2020 | North America / USA / Washington / King | 2020-03-01 | Washington State Department of Health | Seattle Flu Study | Chu et al |
| EPI_ISL_430286 | hCoV-19/USA/WA-S248/2020 | North America / USA / Washington / King | 2020-03-10 | Washington State Department of Health | Seattle Flu Study | Chu et al |
| EPI_ISL_430287 | hCoV-19/USA/WA-S249/2020 | North America / USA / Washington / King | 2020-03-01 | Washington State Department of Health | Seattle Flu Study | Chu et al |
| EPI_ISL_430288 | hCoV-19/USA/WA-S250/2020 | North America / USA / Washington / Snol | 2020-03-03 | Washington State Department of Health | Seattle Flu Study | Chu et al |
| EPI_ISL_430289 | hCoV-19/USA/WA-S251/2020 | North America / USA / Washington / Skag | 2020-03-11 | Washington State Department of Health | Seattle Flu Study | Chu et al |
| EPI_ISL_430290 | hCoV-19/USA/WA-S252/2020 | North America / USA / Washington / King | 2020-03-02 | Washington State Department of Health | Seattle Flu Study | Chu et al |
| EPI_ISL_430291 | hCoV-19/USA/WA-S253/2020 | North America / USA / Washington / Snol | 2020-03-07 | Washington State Department of Health | Seattle Flu Study | Chu et al |
| EPI_ISL_430292 | hCoV-19/USA/WA-S254/2020 | North America / USA / Washington / Snol | 2020-02-24 | Washington State Department of Health | Seattle Flu Study | Chu et al |
| EPI_ISL_430293 | hCoV-19/USA/WA-S255/2020 | North America / USA / Washington / Clar | 2020-03-08 | Washington State Department of Health | Seattle Flu Study | Chu et al |
| EPI_ISL_430294 | hCoV-19/USA/WA-S256/2020 | North America / USA / Washington / King | 2020-03-02 | Washington State Department of Health | Seattle Flu Study | Chu et al |
| EPI_ISL_430295 | hCoV-19/USA/WA-S257/2020 | North America / USA / Washington / King | 2020-02-25 | Washington State Department of Health | Seattle Flu Study | Chu et al |
| EPI_ISL_430297 | hCoV-19/South Africa/R02827/2020 | Africa / South Africa / GP | 2020-03-06 | National Institute for Communicable Diseases of the National Health Laboratory Service | National Institute for Communicable Diseases of the National Health Laboratory Service | Allam M, Kwenda S, van Heusden P, Khumalo Z, Mohale T, Subramoney K, von Gottberg, A, Ismail A, Bhiman JN |
| EPI_ISL_430900 | hCoV-19/USA/WA-UW-5205/2020 | North America / USA / Washington | 2020-04-02 | UW Virology Lab | UW Virology Lab | Pavitra Roychoudhury, Hong Xie, Keith Jerome, Alexander Greninger |
| EPI_ISL_430901 | hCoV-19/USA/WA-UW-5359/2020 | North America / USA / Washington | 2020-04-05 | UW Virology Lab | UW Virology Lab | Pavitra Roychoudhury, Hong Xie, Keith Jerome, Alexander Greninger |
| EPI_ISL_430902 | hCoV-19/USA/WA-UW-5364/2020 | North America / USA / Washington | 2020-04-05 | UW Virology Lab | UW Virology Lab | Pavitra Roychoudhury, Hong Xie, Keith Jerome, Alexander Greninger |
| EPI_ISL_430904 | hCoV-19/USA/WA-UW-5371/2020 | North America / USA / Washington | 2020-04-05 | UW Virology Lab | UW Virology Lab | Pavitra Roychoudhury, Hong Xie, Keith Jerome, Alexander Greninger |
| EPI_ISL_430905 | hCoV-19/USA/WA-UW-5397/2020 | North America / USA / Washington | 2020-04-05 | UW Virology Lab | UW Virology Lab | Pavitra Roychoudhury, Hong Xie, Keith Jerome, Alexander Greninger |
| EPI_ISL_430906 | hCoV-19/USA/WA-UW-5400/2020 | North America / USA / Washington | 2020-04-05 | UW Virology Lab | UW Virology Lab | Pavitra Roychoudhury, Hong Xie, Keith Jerome, Alexander Greninger |
| EPI_ISL_430907 | hCoV-19/USA/CT-UW-5595/2020 | North America / USA / Connecticut | 2020-04-07 | UW Virology Lab | UW Virology Lab | Pavitra Roychoudhury, Hong Xie, Keith Jerome, Alexander Greninger |
| EPI_ISL_430908 | hCoV-19/USA/CT-UW-5599/2020 | North America / USA / Connecticut | 2020-04-07 | UW Virology Lab | UW Virology Lab | Pavitra Roychoudhury, Hong Xie, Keith Jerome, Alexander Greninger |
| EPI_ISL_430909 | hCoV-19/USA/UN-UW-5605/2020 | North America / USA | 2020-04-08 | UW Virology Lab | UW Virology Lab | Pavitra Roychoudhury, Hong Xie, Keith Jerome, Alexander Greninger |
| EPI_ISL_430910 | hCoV-19/USA/UN-UW-5614/2020 | North America / USA | 2020-04-08 | UW Virology Lab | UW Virology Lab | Pavitra Roychoudhury, Hong Xie, Keith Jerome, Alexander Greninger |
| EPI_ISL_430911 | hCoV-19/USA/WA-UW-5617/2020 | North America / USA / Washington | 2020-04-08 | UW Virology Lab | UW Virology Lab | Pavitra Roychoudhury, Hong Xie, Keith Jerome, Alexander Greninger |
| EPI_ISL_430912 | hCoV-19/USA/WA-UW-5629/2020 | North America / USA / Washington | 2020-04-08 | UW Virology Lab | UW Virology Lab | Pavitra Roychoudhury, Hong Xie, Keith Jerome, Alexander Greninger |
| EPI_ISL_430914 | hCoV-19/USA/WA-UW-5732/2020 | North America / USA / Washington | 2020-04-06 | UW Virology Lab | UW Virology Lab | Pavitra Roychoudhury, Hong Xie, Keith Jerome, Alexander Greninger |
| EPI_ISL_430915 | hCoV-19/USA/WA-UW-5737/2020 | North America / USA / Washington | 2020-04-06 | UW Virology Lab | UW Virology Lab | Pavitra Roychoudhury, Hong Xie, Keith Jerome, Alexander Greninger |
| EPI_ISL_430916 | hCoV-19/USA/UN-UW-5742/2020 | North America / USA | 2020-04-07 | UW Virology Lab | UW Virology Lab | Pavitra Roychoudhury, Hong Xie, Keith Jerome, Alexander Greninger |
| EPI_ISL_430917 | hCoV-19/USA/CT-UW-5749/2020 | North America / USA / Connecticut | 2020-04-04 | UW Virology Lab | UW Virology Lab | Pavitra Roychoudhury, Hong Xie, Keith Jerome, Alexander Greninger |
| EPI_ISL_430918 | hCoV-19/USA/WA-UW-5750/2020 | North America / USA / Washington | 2020-04-06 | UW Virology Lab | UW Virology Lab | Pavitra Roychoudhury, Hong Xie, Keith Jerome, Alexander Greninger |
| EPI_ISL_430919 | hCoV-19/USA/CT-UW-5754/2020 | North America / USA / Connecticut | 2020-04-05 | UW Virology Lab | UW Virology Lab | Pavitra Roychoudhury, Hong Xie, Keith Jerome, Alexander Greninger |
| EPI_ISL_430920 | hCoV-19/USA/WA-UW-5756/2020 | North America / USA / Washington | 2020-04-07 | UW Virology Lab | UW Virology Lab | Pavitra Roychoudhury, Hong Xie, Keith Jerome, Alexander Greninger |
| EPI_ISL_430921 | hCoV-19/USA/CT-UW-5757/2020 | North America / USA / Connecticut | 2020-04-06 | UW Virology Lab | UW Virology Lab | Pavitra Roychoudhury, Hong Xie, Keith Jerome, Alexander Greninger |
| EPI_ISL_430922 | hCoV-19/USA/CT-UW-5763/2020 | North America / USA / Connecticut | 2020-04-05 | UW Virology Lab | UW Virology Lab | Pavitra Roychoudhury, Hong Xie, Keith Jerome, Alexander Greninger |
| EPI_ISL_430923 | hCoV-19/USA/CT-UW-5764/2020 | North America / USA / Connecticut | 2020-04-05 | UW Virology Lab | UW Virology Lab | Pavitra Roychoudhury, Hong Xie, Keith Jerome, Alexander Greninger |
| EPI_ISL_430925 | hCoV-19/USA/CT-UW-5770/2020 | North America / USA / Connecticut | 2020-04-04 | UW Virology Lab | UW Virology Lab | Pavitra Roychoudhury, Hong Xie, Keith Jerome, Alexander Greninger |
| EPI_ISL_430926 | hCoV-19/USA/CT-UW-5772/2020 | North America / USA / Connecticut | 2020-04-04 | UW Virology Lab | UW Virology Lab | Pavitra Roychoudhury, Hong Xie, Keith Jerome, Alexander Greninger |
| EPI_ISL_430927 | hCoV-19/USA/CT-UW-5773/2020 | North America / USA / Connecticut | 2020-04-04 | UW Virology Lab | UW Virology Lab | Pavitra Roychoudhury, Hong Xie, Keith Jerome, Alexander Greninger |
| EPI_ISL_430928 | hCoV-19/USA/WA-UW-5879/2020 | North America / USA / Washington | 2020-04-07 | UW Virology Lab | UW Virology Lab | Pavitra Roychoudhury, Hong Xie, Keith Jerome, Alexander Greninger |
| EPI_ISL_430929 | hCoV-19/USA/WA-UW-5884/2020 | North America / USA / Washington | 2020-04-07 | UW Virology Lab | UW Virology Lab | Pavitra Roychoudhury, Hong Xie, Keith Jerome, Alexander Greninger |
| EPI_ISL_430930 | hCoV-19/USA/WA-UW-5911/2020 | North America / USA / Washington | 2020-04-07 | UW Virology Lab | UW Virology Lab | Pavitra Roychoudhury, Hong Xie, Keith Jerome, Alexander Greninger |
| EPI_ISL_430931 | hCoV-19/USA/WA-UW-5932/2020 | North America / USA / Washington | 2020-04-07 | UW Virology Lab | UW Virology Lab | Pavitra Roychoudhury, Hong Xie, Keith Jerome, Alexander Greninger |
| EPI_ISL_430932 | hCoV-19/USA/WA-UW-5947/2020 | North America / USA / Washington | 2020-04-07 | UW Virology Lab | UW Virology Lab | Pavitra Roychoudhury, Hong Xie, Keith Jerome, Alexander Greninger |
| EPI_ISL_430933 | hCoV-19/USA/WA-UW-6162/2020 | North America / USA / Washington | 2020-04-01 | UW Virology Lab | UW Virology Lab | Pavitra Roychoudhury, Hong Xie, Keith Jerome, Alexander Greninger |
| EPI_ISL_430934 | hCoV-19/USA/WA-UW-6164/2020 | North America / USA / Washington | 2020-04-01 | UW Virology Lab | UW Virology Lab | Pavitra Roychoudhury, Hong Xie, Keith Jerome, Alexander Greninger |
| EPI_ISL_430936 | hCoV-19/USA/WA-UW-6173/2020 | North America / USA / Washington | 2020-04-01 | UW Virology Lab | UW Virology Lab | Pavitra Roychoudhury, Hong Xie, Keith Jerome, Alexander Greninger |
| EPI_ISL_430937 | hCoV-19/USA/WA-UW-6178/2020 | North America / USA / Washington | 2020-04-01 | UW Virology Lab | UW Virology Lab | Pavitra Roychoudhury, Hong Xie, Keith Jerome, Alexander Greninger |
| EPI_ISL_430938 | hCoV-19/USA/WA-UW-4226/2020 | North America / USA / Washington | 2020-03-31 | UW Virology Lab | UW Virology Lab | Pavitra Roychoudhury, Hong Xie, Keith Jerome, Alexander Greninger |
| EPI_ISL_430939 | hCoV-19/USA/WA-UW-4243/2020 | North America / USA / Washington | 2020-04-01 | UW Virology Lab | UW Virology Lab | Pavitra Roychoudhury, Hong Xie, Keith Jerome, Alexander Greninger |
| EPI_ISL_430940 | hCoV-19/USA/ID-UW-4255/2020 | North America / USA / Idaho | 2020-03-30 | UW Virology Lab | UW Virology Lab | Pavitra Roychoudhury, Hong Xie, Keith Jerome, Alexander Greninger |
| EPI_ISL_430941 | hCoV-19/USA/WA-UW-4279/2020 | North America / USA / Washington | 2020-03-31 | UW Virology Lab | UW Virology Lab | Pavitra Roychoudhury, Hong Xie, Keith Jerome, Alexander Greninger |
| EPI_ISL_430942 | hCoV-19/USA/ID-UW-4307/2020 | North America / USA / Idaho | 2020-03-31 | UW Virology Lab | UW Virology Lab | Pavitra Roychoudhury, Hong Xie, Keith Jerome, Alexander Greninger |
| EPI_ISL_430943 | hCoV-19/USA/WA-UW-4338/2020 | North America / USA / Washington | 2020-03-30 | UW Virology Lab | UW Virology Lab | Pavitra Roychoudhury, Hong Xie, Keith Jerome, Alexander Greninger |
| EPI_ISL_430945 | hCoV-19/USA/CT-UW-4345/2020 | North America / USA / Connecticut | 2020-03-30 | UW Virology Lab | UW Virology Lab | Pavitra Roychoudhury, Hong Xie, Keith Jerome, Alexander Greninger |
| EPI_ISL_430946 | hCoV-19/USA/CT-UW-4348/2020 | North America / USA / Connecticut | 2020-03-30 | UW Virology Lab | UW Virology Lab | Pavitra Roychoudhury, Hong Xie, Keith Jerome, Alexander Greninger |
| EPI_ISL_430947 | hCoV-19/USA/ID-UW-4354/2020 | North America / USA / Idaho | 2020-03-27 | UW Virology Lab | UW Virology Lab | Pavitra Roychoudhury, Hong Xie, Keith Jerome, Alexander Greninger |
| EPI_ISL_430948 | hCoV-19/USA/ID-UW-4383/2020 | North America / USA / Idaho | 2020-03-29 | UW Virology Lab | UW Virology Lab | Pavitra Roychoudhury, Hong Xie, Keith Jerome, Alexander Greninger |
| EPI_ISL_430949 | hCoV-19/USA/ID-UW-4402/2020 | North America / USA / Idaho | 2020-03-29 | UW Virology Lab | UW Virology Lab | Pavitra Roychoudhury, Hong Xie, Keith Jerome, Alexander Greninger |
| EPI_ISL_430950 | hCoV-19/USA/OR-UW-4410/2020 | North America / USA / Oregon | 2020-03-29 | UW Virology Lab | UW Virology Lab | Pavitra Roychoudhury, Hong Xie, Keith Jerome, Alexander Greninger |
| EPI_ISL_430951 | hCoV-19/USA/WA-UW-4413/2020 | North America / USA / Washington | 2020-03-31 | UW Virology Lab | UW Virology Lab | Pavitra Roychoudhury, Hong Xie, Keith Jerome, Alexander Greninger |
| EPI_ISL_430952 | hCoV-19/USA/ID-UW-4418/2020 | North America / USA / Idaho | 2020-03-28 | UW Virology Lab | UW Virology Lab | Pavitra Roychoudhury, Hong Xie, Keith Jerome, Alexander Greninger |
| EPI_ISL_430953 | hCoV-19/USA/WA-UW-6185/2020 | North America / USA / Washington | 2020-04-01 | UW Virology Lab | UW Virology Lab | Pavitra Roychoudhury, Hong Xie, Keith Jerome, Alexander Greninger |
| EPI_ISL_430954 | hCoV-19/USA/WA-UW-6186/2020 | North America / USA / Washington | 2020-04-01 | UW Virology Lab | UW Virology Lab | Pavitra Roychoudhury, Hong Xie, Keith Jerome, Alexander Greninger |
| EPI_ISL_430955 | hCoV-19/USA/WA-UW-6211/2020 | North America / USA / Washington | 2020-04-01 | UW Virology Lab | UW Virology Lab | Pavitra Roychoudhury, Hong Xie, Keith Jerome, Alexander Greninger |
| EPI_ISL_430957 | hCoV-19/USA/WA-UW-6240/2020 | North America / USA / Washington | 2020-04-01 | UW Virology Lab | UW Virology Lab | Pavitra Roychoudhury, Hong Xie, Keith Jerome, Alexander Greninger |
| EPI_ISL_430958 | hCoV-19/USA/WA-UW-6241/2020 | North America / USA / Washington | 2020-04-01 | UW Virology Lab | UW Virology Lab | Pavitra Roychoudhury, Hong Xie, Keith Jerome, Alexander Greninger |
| EPI_ISL_430959 | hCoV-19/USA/WA-UW-6243/2020 | North America / USA / Washington | 2020-04-02 | UW Virology Lab | UW Virology Lab | Pavitra Roychoudhury, Hong Xie, Keith Jerome, Alexander Greninger |
| EPI_ISL_430960 | hCoV-19/USA/CT-UW-6259/2020 | North America / USA / Connecticut | 2020-04-01 | UW Virology Lab | UW Virology Lab | Pavitra Roychoudhury, Hong Xie, Keith Jerome, Alexander Greninger |
| EPI_ISL_430961 | hCoV-19/USA/CT-UW-6270/2020 | North America / USA / Connecticut | 2020-04-01 | UW Virology Lab | UW Virology Lab | Pavitra Roychoudhury, Hong Xie, Keith Jerome, Alexander Greninger |
| EPI_ISL_430962 | hCoV-19/USA/CT-UW-6271/2020 | North America / USA / Connecticut | 2020-04-01 | UW Virology Lab | UW Virology Lab | Pavitra Roychoudhury, Hong Xie, Keith Jerome, Alexander Greninger |
| EPI_ISL_430964 | hCoV-19/USA/WA-UW-6302/2020 | North America / USA / Washington | 2020-04-02 | UW Virology Lab | UW Virology Lab | Pavitra Roychoudhury, Hong Xie, Keith Jerome, Alexander Greninger |
| EPI_ISL_430965 | hCoV-19/USA/CT-UW-6314/2020 | North America / USA / Connecticut | 2020-04-01 | UW Virology Lab | UW Virology Lab | Pavitra Roychoudhury, Hong Xie, Keith Jerome, Alexander Greninger |
| EPI_ISL_430967 | hCoV-19/USA/WA-UW-6327/2020 | North America / USA / Washington | 2020-04-02 | UW Virology Lab | UW Virology Lab | Pavitra Roychoudhury, Hong Xie, Keith Jerome, Alexander Greninger |
| EPI_ISL_430968 | hCoV-19/USA/WA-UW-6331/2020 | North America / USA / Washington | 2020-04-01 | UW Virology Lab | UW Virology Lab | Pavitra Roychoudhury, Hong Xie, Keith Jerome, Alexander Greninger |
| EPI_ISL_430969 | hCoV-19/USA/WA-UW-6335/2020 | North America / USA / Washington | 2020-04-02 | UW Virology Lab | UW Virology Lab | Pavitra Roychoudhury, Hong Xie, Keith Jerome, Alexander Greninger |
| EPI_ISL_430970 | hCoV-19/USA/ID-UW-6349/2020 | North America / USA / Idaho | 2020-04-01 | UW Virology Lab | UW Virology Lab | Pavitra Roychoudhury, Hong Xie, Keith Jerome, Alexander Greninger |
| EPI_ISL_430971 | hCoV-19/USA/CT-UW-6362/2020 | North America / USA / Connecticut | 2020-04-02 | UW Virology Lab | UW Virology Lab | Pavitra Roychoudhury, Hong Xie, Keith Jerome, Alexander Greninger |
| EPI_ISL_430972 | hCoV-19/USA/CT-UW-6380/2020 | North America / USA / Connecticut | 2020-04-01 | UW Virology Lab | UW Virology Lab | Pavitra Roychoudhury, Hong Xie, Keith Jerome, Alexander Greninger |
| EPI_ISL_430973 | hCoV-19/USA/CT-UW-6385/2020 | North America / USA / Connecticut | 2020-04-01 | UW Virology Lab | UW Virology Lab | Pavitra Roychoudhury, Hong Xie, Keith Jerome, Alexander Greninger |
| EPI_ISL_430974 | hCoV-19/USA/WA-UW-6399/2020 | North America / USA / Washington | 2020-04-03 | UW Virology Lab | UW Virology Lab | Pavitra Roychoudhury, Hong Xie, Keith Jerome, Alexander Greninger |
| EPI_ISL_430975 | hCoV-19/USA/WA-UW-6400/2020 | North America / USA / Washington | 2020-04-03 | UW Virology Lab | UW Virology Lab | Pavitra Roychoudhury, Hong Xie, Keith Jerome, Alexander Greninger |
| EPI_ISL_430976 | hCoV-19/USA/WA-UW-6403/2020 | North America / USA / Washington | 2020-04-03 | UW Virology Lab | UW Virology Lab | Pavitra Roychoudhury, Hong Xie, Keith Jerome, Alexander Greninger |
| EPI_ISL_430977 | hCoV-19/USA/WA-UW-6404/2020 | North America / USA / Washington | 2020-04-03 | UW Virology Lab | UW Virology Lab | Pavitra Roychoudhury, Hong Xie, Keith Jerome, Alexander Greninger |
| EPI_ISL_430979 | hCoV-19/USA/WA-UW-6410/2020 | North America / USA / Washington | 2020-04-03 | UW Virology Lab | UW Virology Lab | Pavitra Roychoudhury, Hong Xie, Keith Jerome, Alexander Greninger |
| EPI_ISL_430980 | hCoV-19/USA/WA-UW-6417/2020 | North America / USA / Washington | 2020-04-03 | UW Virology Lab | UW Virology Lab | Pavitra Roychoudhury, Hong Xie, Keith Jerome, Alexander Greninger |
| EPI_ISL_432706 | hCoV-19/England/NOTT-10FC84/2020 | Europe / United Kingdom / England | 2020-04-08 | Queens Medical Centre, Clinical Microbiology Department / DeepSeq Nottingham | COVID-19 Genomics UK (COG-UK) Consortium | Gemma Clark, Wendy Smith, Manjinder Khakh, Hannah Howson-Wells, Jonathan Ball, Patrick McClure, Joseph Chappell, Theocharis Tsoleridis, Nadine Holmes, Matthew Carlisle, Christopher Moore, Fei Sang, Johnny Debebe, Victoria Wright, Matthew Loose |
| EPI_ISL_432712 | hCoV-19/England/SHEF-BFE8E/2020 | Europe / United Kingdom / England | 2020-03-25 | Virology Department, Sheffield Teaching Hospitals NHS Foundation Trust / Virology Department, Sheffield Teaching Hospitals NHS Foundation Trust | COVID-19 Genomics UK (COG-UK) Consortium | Thushan de Silva, Matthew Parker,Adri Angyal, Rebecca Brown, Luke Green, Rachel Tucker, Paul Parsons, Danielle Groves, Alex Keeley, Dave Partridge, Matthew Wyles, Benjamin Lindsey, Mehmet Yavuz, Mohammad Raza, Cariad Evans |
| EPI_ISL_432713 | hCoV-19/England/SHEF-BFEF7/2020 | Europe / United Kingdom / England | 2020-03-25 | Virology Department, Sheffield Teaching Hospitals NHS Foundation Trust / Virology Department, Sheffield Teaching Hospitals NHS Foundation Trust | COVID-19 Genomics UK (COG-UK) Consortium | Thushan de Silva, Matthew Parker,Adri Angyal, Rebecca Brown, Luke Green, Rachel Tucker, Paul Parsons, Danielle Groves, Alex Keeley, Dave Partridge, Matthew Wyles, Benjamin Lindsey, Mehmet Yavuz, Mohammad Raza, Cariad Evans |
| EPI_ISL_432716 | hCoV-19/England/SHEF-C0C35/2020 | Europe / United Kingdom / England | 2020-03-24 | Virology Department, Sheffield Teaching Hospitals NHS Foundation Trust / Virology Department, Sheffield Teaching Hospitals NHS Foundation Trust | COVID-19 Genomics UK (COG-UK) Consortium | Thushan de Silva, Matthew Parker,Adri Angyal, Rebecca Brown, Luke Green, Rachel Tucker, Paul Parsons, Danielle Groves, Alex Keeley, Dave Partridge, Matthew Wyles, Benjamin Lindsey, Mehmet Yavuz, Mohammad Raza, Cariad Evans |
| EPI_ISL_432720 | hCoV-19/England/SHEF-C0FB4/2020 | Europe / United Kingdom / England | 2020-03-30 | Virology Department, Sheffield Teaching Hospitals NHS Foundation Trust / Virology Department, Sheffield Teaching Hospitals NHS Foundation Trust | COVID-19 Genomics UK (COG-UK) Consortium | Thushan de Silva, Matthew Parker,Adri Angyal, Rebecca Brown, Luke Green, Rachel Tucker, Paul Parsons, Danielle Groves, Alex Keeley, Dave Partridge, Matthew Wyles, Benjamin Lindsey, Mehmet Yavuz, Mohammad Raza, Cariad Evans |
| EPI_ISL_432722 | hCoV-19/England/SHEF-C11BD/2020 | Europe / United Kingdom / England | 2020-04-07 | Virology Department, Sheffield Teaching Hospitals NHS Foundation Trust / Virology Department, Sheffield Teaching Hospitals NHS Foundation Trust | COVID-19 Genomics UK (COG-UK) Consortium | Thushan de Silva, Matthew Parker,Adri Angyal, Rebecca Brown, Luke Green, Rachel Tucker, Paul Parsons, Danielle Groves, Alex Keeley, Dave Partridge, Matthew Wyles, Benjamin Lindsey, Mehmet Yavuz, Mohammad Raza, Cariad Evans |
| EPI_ISL_432724 | hCoV-19/England/SHEF-C151B/2020 | Europe / United Kingdom / England | 2020-03-29 | Virology Department, Sheffield Teaching Hospitals NHS Foundation Trust / Virology Department, Sheffield Teaching Hospitals NHS Foundation Trust | COVID-19 Genomics UK (COG-UK) Consortium | Thushan de Silva, Matthew Parker,Adri Angyal, Rebecca Brown, Luke Green, Rachel Tucker, Paul Parsons, Danielle Groves, Alex Keeley, Dave Partridge, Matthew Wyles, Benjamin Lindsey, Mehmet Yavuz, Mohammad Raza, Cariad Evans |
| EPI_ISL_432727 | hCoV-19/England/SHEF-C1645/2020 | Europe / United Kingdom / England | 2020-04-01 | Virology Department, Sheffield Teaching Hospitals NHS Foundation Trust / Virology Department, Sheffield Teaching Hospitals NHS Foundation Trust | COVID-19 Genomics UK (COG-UK) Consortium | Thushan de Silva, Matthew Parker,Adri Angyal, Rebecca Brown, Luke Green, Rachel Tucker, Paul Parsons, Danielle Groves, Alex Keeley, Dave Partridge, Matthew Wyles, Benjamin Lindsey, Mehmet Yavuz, Mohammad Raza, Cariad Evans |
| EPI_ISL_432732 | hCoV-19/England/SHEF-C18E5/2020 | Europe / United Kingdom / England | 2020-04-01 | Virology Department, Sheffield Teaching Hospitals NHS Foundation Trust / Virology Department, Sheffield Teaching Hospitals NHS Foundation Trust | COVID-19 Genomics UK (COG-UK) Consortium | Thushan de Silva, Matthew Parker,Adri Angyal, Rebecca Brown, Luke Green, Rachel Tucker, Paul Parsons, Danielle Groves, Alex Keeley, Dave Partridge, Matthew Wyles, Benjamin Lindsey, Mehmet Yavuz, Mohammad Raza, Cariad Evans |
| EPI_ISL_432734 | hCoV-19/England/SHEF-C1B64/2020 | Europe / United Kingdom / England | 2020-03-30 | Virology Department, Sheffield Teaching Hospitals NHS Foundation Trust / Virology Department, Sheffield Teaching Hospitals NHS Foundation Trust | COVID-19 Genomics UK (COG-UK) Consortium | Thushan de Silva, Matthew Parker,Adri Angyal, Rebecca Brown, Luke Green, Rachel Tucker, Paul Parsons, Danielle Groves, Alex Keeley, Dave Partridge, Matthew Wyles, Benjamin Lindsey, Mehmet Yavuz, Mohammad Raza, Cariad Evans |
| EPI_ISL_432742 | hCoV-19/England/SHEF-C2222/2020 | Europe / United Kingdom / England | 2020-04-02 | Virology Department, Sheffield Teaching Hospitals NHS Foundation Trust / Virology Department, Sheffield Teaching Hospitals NHS Foundation Trust | COVID-19 Genomics UK (COG-UK) Consortium | Thushan de Silva, Matthew Parker,Adri Angyal, Rebecca Brown, Luke Green, Rachel Tucker, Paul Parsons, Danielle Groves, Alex Keeley, Dave Partridge, Matthew Wyles, Benjamin Lindsey, Mehmet Yavuz, Mohammad Raza, Cariad Evans |
| EPI_ISL_432744 | hCoV-19/England/SHEF-C2310/2020 | Europe / United Kingdom / England | 2020-04-01 | Virology Department, Sheffield Teaching Hospitals NHS Foundation Trust / Virology Department, Sheffield Teaching Hospitals NHS Foundation Trust | COVID-19 Genomics UK (COG-UK) Consortium | Thushan de Silva, Matthew Parker,Adri Angyal, Rebecca Brown, Luke Green, Rachel Tucker, Paul Parsons, Danielle Groves, Alex Keeley, Dave Partridge, Matthew Wyles, Benjamin Lindsey, Mehmet Yavuz, Mohammad Raza, Cariad Evans |
| EPI_ISL_432746 | hCoV-19/England/SHEF-C2626/2020 | Europe / United Kingdom / England | 2020-03-30 | Virology Department, Sheffield Teaching Hospitals NHS Foundation Trust / Virology Department, Sheffield Teaching Hospitals NHS Foundation Trust | COVID-19 Genomics UK (COG-UK) Consortium | Thushan de Silva, Matthew Parker,Adri Angyal, Rebecca Brown, Luke Green, Rachel Tucker, Paul Parsons, Danielle Groves, Alex Keeley, Dave Partridge, Matthew Wyles, Benjamin Lindsey, Mehmet Yavuz, Mohammad Raza, Cariad Evans |
| EPI_ISL_432753 | hCoV-19/England/SHEF-C2CF7/2020 | Europe / United Kingdom / England | 2020-03-29 | Virology Department, Sheffield Teaching Hospitals NHS Foundation Trust / Virology Department, Sheffield Teaching Hospitals NHS Foundation Trust | COVID-19 Genomics UK (COG-UK) Consortium | Thushan de Silva, Matthew Parker,Adri Angyal, Rebecca Brown, Luke Green, Rachel Tucker, Paul Parsons, Danielle Groves, Alex Keeley, Dave Partridge, Matthew Wyles, Benjamin Lindsey, Mehmet Yavuz, Mohammad Raza, Cariad Evans |
| EPI_ISL_432762 | hCoV-19/England/SHEF-C3115/2020 | Europe / United Kingdom / England | 2020-04-15 | Virology Department, Sheffield Teaching Hospitals NHS Foundation Trust / Virology Department, Sheffield Teaching Hospitals NHS Foundation Trust | COVID-19 Genomics UK (COG-UK) Consortium | Thushan de Silva, Matthew Parker,Adri Angyal, Rebecca Brown, Luke Green, Rachel Tucker, Paul Parsons, Danielle Groves, Alex Keeley, Dave Partridge, Matthew Wyles, Benjamin Lindsey, Mehmet Yavuz, Mohammad Raza, Cariad Evans |
| EPI_ISL_432766 | hCoV-19/England/SHEF-C3212/2020 | Europe / United Kingdom / England | 2020-04-03 | Virology Department, Sheffield Teaching Hospitals NHS Foundation Trust / Virology Department, Sheffield Teaching Hospitals NHS Foundation Trust | COVID-19 Genomics UK (COG-UK) Consortium | Thushan de Silva, Matthew Parker,Adri Angyal, Rebecca Brown, Luke Green, Rachel Tucker, Paul Parsons, Danielle Groves, Alex Keeley, Dave Partridge, Matthew Wyles, Benjamin Lindsey, Mehmet Yavuz, Mohammad Raza, Cariad Evans |
| EPI_ISL_432768 | hCoV-19/England/SHEF-C32C7/2020 | Europe / United Kingdom / England | 2020-03-24 | Virology Department, Sheffield Teaching Hospitals NHS Foundation Trust / Virology Department, Sheffield Teaching Hospitals NHS Foundation Trust | COVID-19 Genomics UK (COG-UK) Consortium | Thushan de Silva, Matthew Parker,Adri Angyal, Rebecca Brown, Luke Green, Rachel Tucker, Paul Parsons, Danielle Groves, Alex Keeley, Dave Partridge, Matthew Wyles, Benjamin Lindsey, Mehmet Yavuz, Mohammad Raza, Cariad Evans |
| EPI_ISL_432777 | hCoV-19/England/SHEF-C384D/2020 | Europe / United Kingdom / England | 2020-03-30 | Virology Department, Sheffield Teaching Hospitals NHS Foundation Trust / Virology Department, Sheffield Teaching Hospitals NHS Foundation Trust | COVID-19 Genomics UK (COG-UK) Consortium | Thushan de Silva, Matthew Parker,Adri Angyal, Rebecca Brown, Luke Green, Rachel Tucker, Paul Parsons, Danielle Groves, Alex Keeley, Dave Partridge, Matthew Wyles, Benjamin Lindsey, Mehmet Yavuz, Mohammad Raza, Cariad Evans |
| EPI_ISL_432780 | hCoV-19/England/SHEF-C3CC9/2020 | Europe / United Kingdom / England | 2020-04-15 | Virology Department, Sheffield Teaching Hospitals NHS Foundation Trust / Virology Department, Sheffield Teaching Hospitals NHS Foundation Trust | COVID-19 Genomics UK (COG-UK) Consortium | Thushan de Silva, Matthew Parker,Adri Angyal, Rebecca Brown, Luke Green, Rachel Tucker, Paul Parsons, Danielle Groves, Alex Keeley, Dave Partridge, Matthew Wyles, Benjamin Lindsey, Mehmet Yavuz, Mohammad Raza, Cariad Evans |
| EPI_ISL_432786 | hCoV-19/England/SHEF-C426C/2020 | Europe / United Kingdom / England | 2020-03-29 | Virology Department, Sheffield Teaching Hospitals NHS Foundation Trust / Virology Department, Sheffield Teaching Hospitals NHS Foundation Trust | COVID-19 Genomics UK (COG-UK) Consortium | Thushan de Silva, Matthew Parker,Adri Angyal, Rebecca Brown, Luke Green, Rachel Tucker, Paul Parsons, Danielle Groves, Alex Keeley, Dave Partridge, Matthew Wyles, Benjamin Lindsey, Mehmet Yavuz, Mohammad Raza, Cariad Evans |
| EPI_ISL_432788 | hCoV-19/England/SHEF-C4493/2020 | Europe / United Kingdom / England | 2020-03-30 | Virology Department, Sheffield Teaching Hospitals NHS Foundation Trust / Virology Department, Sheffield Teaching Hospitals NHS Foundation Trust | COVID-19 Genomics UK (COG-UK) Consortium | Thushan de Silva, Matthew Parker,Adri Angyal, Rebecca Brown, Luke Green, Rachel Tucker, Paul Parsons, Danielle Groves, Alex Keeley, Dave Partridge, Matthew Wyles, Benjamin Lindsey, Mehmet Yavuz, Mohammad Raza, Cariad Evans |
| EPI_ISL_432790 | hCoV-19/England/SHEF-C45BE/2020 | Europe / United Kingdom / England | 2020-04-01 | Virology Department, Sheffield Teaching Hospitals NHS Foundation Trust / Virology Department, Sheffield Teaching Hospitals NHS Foundation Trust | COVID-19 Genomics UK (COG-UK) Consortium | Thushan de Silva, Matthew Parker,Adri Angyal, Rebecca Brown, Luke Green, Rachel Tucker, Paul Parsons, Danielle Groves, Alex Keeley, Dave Partridge, Matthew Wyles, Benjamin Lindsey, Mehmet Yavuz, Mohammad Raza, Cariad Evans |
| EPI_ISL_432799 | hCoV-19/England/SHEF-C51E6/2020 | Europe / United Kingdom / England | 2020-03-30 | Virology Department, Sheffield Teaching Hospitals NHS Foundation Trust / Virology Department, Sheffield Teaching Hospitals NHS Foundation Trust | COVID-19 Genomics UK (COG-UK) Consortium | Thushan de Silva, Matthew Parker,Adri Angyal, Rebecca Brown, Luke Green, Rachel Tucker, Paul Parsons, Danielle Groves, Alex Keeley, Dave Partridge, Matthew Wyles, Benjamin Lindsey, Mehmet Yavuz, Mohammad Raza, Cariad Evans |
| EPI_ISL_432802 | hCoV-19/England/SHEF-C5359/2020 | Europe / United Kingdom / England | 2020-03-31 | Virology Department, Sheffield Teaching Hospitals NHS Foundation Trust / Virology Department, Sheffield Teaching Hospitals NHS Foundation Trust | COVID-19 Genomics UK (COG-UK) Consortium | Thushan de Silva, Matthew Parker,Adri Angyal, Rebecca Brown, Luke Green, Rachel Tucker, Paul Parsons, Danielle Groves, Alex Keeley, Dave Partridge, Matthew Wyles, Benjamin Lindsey, Mehmet Yavuz, Mohammad Raza, Cariad Evans |
| EPI_ISL_432804 | hCoV-19/England/SHEF-C56AB/2020 | Europe / United Kingdom / England | 2020-03-27 | Virology Department, Sheffield Teaching Hospitals NHS Foundation Trust / Virology Department, Sheffield Teaching Hospitals NHS Foundation Trust | COVID-19 Genomics UK (COG-UK) Consortium | Thushan de Silva, Matthew Parker,Adri Angyal, Rebecca Brown, Luke Green, Rachel Tucker, Paul Parsons, Danielle Groves, Alex Keeley, Dave Partridge, Matthew Wyles, Benjamin Lindsey, Mehmet Yavuz, Mohammad Raza, Cariad Evans |
| EPI_ISL_432812 | hCoV-19/England/SHEF-C5D79/2020 | Europe / United Kingdom / England | 2020-04-01 | Virology Department, Sheffield Teaching Hospitals NHS Foundation Trust / Virology Department, Sheffield Teaching Hospitals NHS Foundation Trust | COVID-19 Genomics UK (COG-UK) Consortium | Thushan de Silva, Matthew Parker,Adri Angyal, Rebecca Brown, Luke Green, Rachel Tucker, Paul Parsons, Danielle Groves, Alex Keeley, Dave Partridge, Matthew Wyles, Benjamin Lindsey, Mehmet Yavuz, Mohammad Raza, Cariad Evans |
| EPI_ISL_432813 | hCoV-19/England/SHEF-C5DD3/2020 | Europe / United Kingdom / England | 2020-03-30 | Virology Department, Sheffield Teaching Hospitals NHS Foundation Trust / Virology Department, Sheffield Teaching Hospitals NHS Foundation Trust | COVID-19 Genomics UK (COG-UK) Consortium | Thushan de Silva, Matthew Parker,Adri Angyal, Rebecca Brown, Luke Green, Rachel Tucker, Paul Parsons, Danielle Groves, Alex Keeley, Dave Partridge, Matthew Wyles, Benjamin Lindsey, Mehmet Yavuz, Mohammad Raza, Cariad Evans |
| EPI_ISL_432825 | hCoV-19/England/SHEF-C6789/2020 | Europe / United Kingdom / England | 2020-04-01 | Virology Department, Sheffield Teaching Hospitals NHS Foundation Trust / Virology Department, Sheffield Teaching Hospitals NHS Foundation Trust | COVID-19 Genomics UK (COG-UK) Consortium | Thushan de Silva, Matthew Parker,Adri Angyal, Rebecca Brown, Luke Green, Rachel Tucker, Paul Parsons, Danielle Groves, Alex Keeley, Dave Partridge, Matthew Wyles, Benjamin Lindsey, Mehmet Yavuz, Mohammad Raza, Cariad Evans |
| EPI_ISL_432828 | hCoV-19/England/SHEF-C6C11/2020 | Europe / United Kingdom / England | 2020-03-30 | Virology Department, Sheffield Teaching Hospitals NHS Foundation Trust / Virology Department, Sheffield Teaching Hospitals NHS Foundation Trust | COVID-19 Genomics UK (COG-UK) Consortium | Thushan de Silva, Matthew Parker,Adri Angyal, Rebecca Brown, Luke Green, Rachel Tucker, Paul Parsons, Danielle Groves, Alex Keeley, Dave Partridge, Matthew Wyles, Benjamin Lindsey, Mehmet Yavuz, Mohammad Raza, Cariad Evans |
| EPI_ISL_432831 | hCoV-19/England/SHEF-C6EA2/2020 | Europe / United Kingdom / England | 2020-03-30 | Virology Department, Sheffield Teaching Hospitals NHS Foundation Trust / Virology Department, Sheffield Teaching Hospitals NHS Foundation Trust | COVID-19 Genomics UK (COG-UK) Consortium | Thushan de Silva, Matthew Parker,Adri Angyal, Rebecca Brown, Luke Green, Rachel Tucker, Paul Parsons, Danielle Groves, Alex Keeley, Dave Partridge, Matthew Wyles, Benjamin Lindsey, Mehmet Yavuz, Mohammad Raza, Cariad Evans |
| EPI_ISL_432836 | hCoV-19/England/SHEF-C715D/2020 | Europe / United Kingdom / England | 2020-04-03 | Virology Department, Sheffield Teaching Hospitals NHS Foundation Trust / Virology Department, Sheffield Teaching Hospitals NHS Foundation Trust | COVID-19 Genomics UK (COG-UK) Consortium | Thushan de Silva, Matthew Parker,Adri Angyal, Rebecca Brown, Luke Green, Rachel Tucker, Paul Parsons, Danielle Groves, Alex Keeley, Dave Partridge, Matthew Wyles, Benjamin Lindsey, Mehmet Yavuz, Mohammad Raza, Cariad Evans |
| EPI_ISL_432837 | hCoV-19/England/SHEF-C7296/2020 | Europe / United Kingdom / England | 2020-03-29 | Virology Department, Sheffield Teaching Hospitals NHS Foundation Trust / Virology Department, Sheffield Teaching Hospitals NHS Foundation Trust | COVID-19 Genomics UK (COG-UK) Consortium | Thushan de Silva, Matthew Parker,Adri Angyal, Rebecca Brown, Luke Green, Rachel Tucker, Paul Parsons, Danielle Groves, Alex Keeley, Dave Partridge, Matthew Wyles, Benjamin Lindsey, Mehmet Yavuz, Mohammad Raza, Cariad Evans |
| EPI_ISL_432850 | hCoV-19/England/SHEF-C7937/2020 | Europe / United Kingdom / England | 2020-04-03 | Virology Department, Sheffield Teaching Hospitals NHS Foundation Trust / Virology Department, Sheffield Teaching Hospitals NHS Foundation Trust | COVID-19 Genomics UK (COG-UK) Consortium | Thushan de Silva, Matthew Parker,Adri Angyal, Rebecca Brown, Luke Green, Rachel Tucker, Paul Parsons, Danielle Groves, Alex Keeley, Dave Partridge, Matthew Wyles, Benjamin Lindsey, Mehmet Yavuz, Mohammad Raza, Cariad Evans |
| EPI_ISL_432859 | hCoV-19/England/SHEF-C81D4/2020 | Europe / United Kingdom / England | 2020-04-01 | Virology Department, Sheffield Teaching Hospitals NHS Foundation Trust / Virology Department, Sheffield Teaching Hospitals NHS Foundation Trust | COVID-19 Genomics UK (COG-UK) Consortium | Thushan de Silva, Matthew Parker,Adri Angyal, Rebecca Brown, Luke Green, Rachel Tucker, Paul Parsons, Danielle Groves, Alex Keeley, Dave Partridge, Matthew Wyles, Benjamin Lindsey, Mehmet Yavuz, Mohammad Raza, Cariad Evans |
| EPI_ISL_432861 | hCoV-19/England/SHEF-C8338/2020 | Europe / United Kingdom / England | 2020-04-01 | Virology Department, Sheffield Teaching Hospitals NHS Foundation Trust / Virology Department, Sheffield Teaching Hospitals NHS Foundation Trust | COVID-19 Genomics UK (COG-UK) Consortium | Thushan de Silva, Matthew Parker,Adri Angyal, Rebecca Brown, Luke Green, Rachel Tucker, Paul Parsons, Danielle Groves, Alex Keeley, Dave Partridge, Matthew Wyles, Benjamin Lindsey, Mehmet Yavuz, Mohammad Raza, Cariad Evans |
| EPI_ISL_432864 | hCoV-19/England/SHEF-C83ED/2020 | Europe / United Kingdom / England | 2020-03-23 | Virology Department, Sheffield Teaching Hospitals NHS Foundation Trust / Virology Department, Sheffield Teaching Hospitals NHS Foundation Trust | COVID-19 Genomics UK (COG-UK) Consortium | Thushan de Silva, Matthew Parker,Adri Angyal, Rebecca Brown, Luke Green, Rachel Tucker, Paul Parsons, Danielle Groves, Alex Keeley, Dave Partridge, Matthew Wyles, Benjamin Lindsey, Mehmet Yavuz, Mohammad Raza, Cariad Evans |
| EPI_ISL_432867 | hCoV-19/England/SHEF-C8532/2020 | Europe / United Kingdom / England | 2020-03-30 | Virology Department, Sheffield Teaching Hospitals NHS Foundation Trust / Virology Department, Sheffield Teaching Hospitals NHS Foundation Trust | COVID-19 Genomics UK (COG-UK) Consortium | Thushan de Silva, Matthew Parker,Adri Angyal, Rebecca Brown, Luke Green, Rachel Tucker, Paul Parsons, Danielle Groves, Alex Keeley, Dave Partridge, Matthew Wyles, Benjamin Lindsey, Mehmet Yavuz, Mohammad Raza, Cariad Evans |
| EPI_ISL_432868 | hCoV-19/Scotland/EDB894/2020 | Europe / United Kingdom / Scotland | 2020-04-05 | Virology Department, Royal Infirmary of Edinburgh, NHS Lothian / School of Biological Sciences, University of Edinburgh / Institute of Genetics and Molecular Medicine, University of Edinburgh | COVID-19 Genomics UK (COG-UK) Consortium | McHugh M, Dewar R, Rooke S, Gallagher M, Balcaza C, O'Toole A, Hill V, McCrone JT, Colquhoun R, Yu X, Jackson B, Rambaut A, Williams TC, Templeton K |
| EPI_ISL_432869 | hCoV-19/Scotland/EDB895/2020 | Europe / United Kingdom / Scotland | 2020-04-05 | Virology Department, Royal Infirmary of Edinburgh, NHS Lothian / School of Biological Sciences, University of Edinburgh / Institute of Genetics and Molecular Medicine, University of Edinburgh | COVID-19 Genomics UK (COG-UK) Consortium | McHugh M, Dewar R, Rooke S, Gallagher M, Balcaza C, O'Toole A, Hill V, McCrone JT, Colquhoun R, Yu X, Jackson B, Rambaut A, Williams TC, Templeton K |
| EPI_ISL_432870 | hCoV-19/Scotland/EDB896/2020 | Europe / United Kingdom / Scotland | 2020-04-05 | Virology Department, Royal Infirmary of Edinburgh, NHS Lothian / School of Biological Sciences, University of Edinburgh / Institute of Genetics and Molecular Medicine, University of Edinburgh | COVID-19 Genomics UK (COG-UK) Consortium | McHugh M, Dewar R, Rooke S, Gallagher M, Balcaza C, O'Toole A, Hill V, McCrone JT, Colquhoun R, Yu X, Jackson B, Rambaut A, Williams TC, Templeton K |
| EPI_ISL_432871 | hCoV-19/Scotland/EDB897/2020 | Europe / United Kingdom / Scotland | 2020-04-05 | Virology Department, Royal Infirmary of Edinburgh, NHS Lothian / School of Biological Sciences, University of Edinburgh / Institute of Genetics and Molecular Medicine, University of Edinburgh | COVID-19 Genomics UK (COG-UK) Consortium | McHugh M, Dewar R, Rooke S, Gallagher M, Balcaza C, O'Toole A, Hill V, McCrone JT, Colquhoun R, Yu X, Jackson B, Rambaut A, Williams TC, Templeton K |
| EPI_ISL_432872 | hCoV-19/Scotland/EDB899/2020 | Europe / United Kingdom / Scotland | 2020-04-05 | Virology Department, Royal Infirmary of Edinburgh, NHS Lothian / School of Biological Sciences, University of Edinburgh / Institute of Genetics and Molecular Medicine, University of Edinburgh | COVID-19 Genomics UK (COG-UK) Consortium | McHugh M, Dewar R, Rooke S, Gallagher M, Balcaza C, O'Toole A, Hill V, McCrone JT, Colquhoun R, Yu X, Jackson B, Rambaut A, Williams TC, Templeton K |
| EPI_ISL_432873 | hCoV-19/Scotland/EDB901/2020 | Europe / United Kingdom / Scotland | 2020-04-05 | Virology Department, Royal Infirmary of Edinburgh, NHS Lothian / School of Biological Sciences, University of Edinburgh / Institute of Genetics and Molecular Medicine, University of Edinburgh | COVID-19 Genomics UK (COG-UK) Consortium | McHugh M, Dewar R, Rooke S, Gallagher M, Balcaza C, O'Toole A, Hill V, McCrone JT, Colquhoun R, Yu X, Jackson B, Rambaut A, Williams TC, Templeton K |
| EPI_ISL_432874 | hCoV-19/Scotland/EDB909/2020 | Europe / United Kingdom / Scotland | 2020-04-05 | Virology Department, Royal Infirmary of Edinburgh, NHS Lothian / School of Biological Sciences, University of Edinburgh / Institute of Genetics and Molecular Medicine, University of Edinburgh | COVID-19 Genomics UK (COG-UK) Consortium | McHugh M, Dewar R, Rooke S, Gallagher M, Balcaza C, O'Toole A, Hill V, McCrone JT, Colquhoun R, Yu X, Jackson B, Rambaut A, Williams TC, Templeton K |
| EPI_ISL_432875 | hCoV-19/Scotland/EDB910/2020 | Europe / United Kingdom / Scotland | 2020-04-06 | Virology Department, Royal Infirmary of Edinburgh, NHS Lothian / School of Biological Sciences, University of Edinburgh / Institute of Genetics and Molecular Medicine, University of Edinburgh | COVID-19 Genomics UK (COG-UK) Consortium | McHugh M, Dewar R, Rooke S, Gallagher M, Balcaza C, O'Toole A, Hill V, McCrone JT, Colquhoun R, Yu X, Jackson B, Rambaut A, Williams TC, Templeton K |
| EPI_ISL_432877 | hCoV-19/Scotland/EDB914/2020 | Europe / United Kingdom / Scotland | 2020-04-05 | Virology Department, Royal Infirmary of Edinburgh, NHS Lothian / School of Biological Sciences, University of Edinburgh / Institute of Genetics and Molecular Medicine, University of Edinburgh | COVID-19 Genomics UK (COG-UK) Consortium | McHugh M, Dewar R, Rooke S, Gallagher M, Balcaza C, O'Toole A, Hill V, McCrone JT, Colquhoun R, Yu X, Jackson B, Rambaut A, Williams TC, Templeton K |
| EPI_ISL_432880 | hCoV-19/Scotland/EDB921/2020 | Europe / United Kingdom / Scotland | 2020-04-04 | Virology Department, Royal Infirmary of Edinburgh, NHS Lothian / School of Biological Sciences, University of Edinburgh / Institute of Genetics and Molecular Medicine, University of Edinburgh | COVID-19 Genomics UK (COG-UK) Consortium | McHugh M, Dewar R, Rooke S, Gallagher M, Balcaza C, O'Toole A, Hill V, McCrone JT, Colquhoun R, Yu X, Jackson B, Rambaut A, Williams TC, Templeton K |
| EPI_ISL_432881 | hCoV-19/Scotland/EDB922/2020 | Europe / United Kingdom / Scotland | 2020-04-05 | Virology Department, Royal Infirmary of Edinburgh, NHS Lothian / School of Biological Sciences, University of Edinburgh / Institute of Genetics and Molecular Medicine, University of Edinburgh | COVID-19 Genomics UK (COG-UK) Consortium | McHugh M, Dewar R, Rooke S, Gallagher M, Balcaza C, O'Toole A, Hill V, McCrone JT, Colquhoun R, Yu X, Jackson B, Rambaut A, Williams TC, Templeton K |
| EPI_ISL_432883 | hCoV-19/Scotland/EDB924/2020 | Europe / United Kingdom / Scotland | 2020-04-05 | Virology Department, Royal Infirmary of Edinburgh, NHS Lothian / School of Biological Sciences, University of Edinburgh / Institute of Genetics and Molecular Medicine, University of Edinburgh | COVID-19 Genomics UK (COG-UK) Consortium | McHugh M, Dewar R, Rooke S, Gallagher M, Balcaza C, O'Toole A, Hill V, McCrone JT, Colquhoun R, Yu X, Jackson B, Rambaut A, Williams TC, Templeton K |
| EPI_ISL_432885 | hCoV-19/Scotland/EDB926/2020 | Europe / United Kingdom / Scotland | 2020-04-04 | Virology Department, Royal Infirmary of Edinburgh, NHS Lothian / School of Biological Sciences, University of Edinburgh / Institute of Genetics and Molecular Medicine, University of Edinburgh | COVID-19 Genomics UK (COG-UK) Consortium | McHugh M, Dewar R, Rooke S, Gallagher M, Balcaza C, O'Toole A, Hill V, McCrone JT, Colquhoun R, Yu X, Jackson B, Rambaut A, Williams TC, Templeton K |
| EPI_ISL_432886 | hCoV-19/Scotland/EDB927/2020 | Europe / United Kingdom / Scotland | 2020-04-05 | Virology Department, Royal Infirmary of Edinburgh, NHS Lothian / School of Biological Sciences, University of Edinburgh / Institute of Genetics and Molecular Medicine, University of Edinburgh | COVID-19 Genomics UK (COG-UK) Consortium | McHugh M, Dewar R, Rooke S, Gallagher M, Balcaza C, O'Toole A, Hill V, McCrone JT, Colquhoun R, Yu X, Jackson B, Rambaut A, Williams TC, Templeton K |

| | | | | | | |
|---|---|---|---|---|---|---|
| EPI_ISL_432890 | hCoV-19/Scotland/EDB945/2020 | Europe / United Kingdom / Scotland | 2020-04-05 | Virology Department, Royal Infirmary of Edinburgh, NHS Lothian / School of Biological Sciences, University of Edinburgh / Institute of Genetics and Molecular Medicine, University of Edinburgh | COVID-19 Genomics UK (COG-UK) Consortium | McHugh M, Dewar R, Rooke S, Gallagher M, Balcaza C, O'Toole A, Hill V, McCrone JT, Colquhoun R, Yu X, Jackson B, Rambaut A, Williams TC, Templeton K |
| EPI_ISL_432891 | hCoV-19/Scotland/EDB964/2020 | Europe / United Kingdom / Scotland | 2020-04-06 | Virology Department, Royal Infirmary of Edinburgh, NHS Lothian / School of Biological Sciences, University of Edinburgh / Institute of Genetics and Molecular Medicine, University of Edinburgh | COVID-19 Genomics UK (COG-UK) Consortium | McHugh M, Dewar R, Rooke S, Gallagher M, Balcaza C, O'Toole A, Hill V, McCrone JT, Colquhoun R, Yu X, Jackson B, Rambaut A, Williams TC, Templeton K |
| EPI_ISL_432892 | hCoV-19/Scotland/EDB971/2020 | Europe / United Kingdom / Scotland | 2020-04-06 | Virology Department, Royal Infirmary of Edinburgh, NHS Lothian / School of Biological Sciences, University of Edinburgh / Institute of Genetics and Molecular Medicine, University of Edinburgh | COVID-19 Genomics UK (COG-UK) Consortium | McHugh M, Dewar R, Rooke S, Gallagher M, Balcaza C, O'Toole A, Hill V, McCrone JT, Colquhoun R, Yu X, Jackson B, Rambaut A, Williams TC, Templeton K |
| EPI_ISL_432893 | hCoV-19/Scotland/EDB972/2020 | Europe / United Kingdom / Scotland | 2020-04-05 | Virology Department, Royal Infirmary of Edinburgh, NHS Lothian / School of Biological Sciences, University of Edinburgh / Institute of Genetics and Molecular Medicine, University of Edinburgh | COVID-19 Genomics UK (COG-UK) Consortium | McHugh M, Dewar R, Rooke S, Gallagher M, Balcaza C, O'Toole A, Hill V, McCrone JT, Colquhoun R, Yu X, Jackson B, Rambaut A, Williams TC, Templeton K |
| EPI_ISL_432894 | hCoV-19/Scotland/EDB973/2020 | Europe / United Kingdom / Scotland | 2020-04-05 | Virology Department, Royal Infirmary of Edinburgh, NHS Lothian / School of Biological Sciences, University of Edinburgh / Institute of Genetics and Molecular Medicine, University of Edinburgh | COVID-19 Genomics UK (COG-UK) Consortium | McHugh M, Dewar R, Rooke S, Gallagher M, Balcaza C, O'Toole A, Hill V, McCrone JT, Colquhoun R, Yu X, Jackson B, Rambaut A, Williams TC, Templeton K |
| EPI_ISL_432895 | hCoV-19/Scotland/EDB986/2020 | Europe / United Kingdom / Scotland | 2020-04-06 | Virology Department, Royal Infirmary of Edinburgh, NHS Lothian / School of Biological Sciences, University of Edinburgh / Institute of Genetics and Molecular Medicine, University of Edinburgh | COVID-19 Genomics UK (COG-UK) Consortium | McHugh M, Dewar R, Rooke S, Gallagher M, Balcaza C, O'Toole A, Hill V, McCrone JT, Colquhoun R, Yu X, Jackson B, Rambaut A, Williams TC, Templeton K |
| EPI_ISL_432898 | hCoV-19/Scotland/EDB997/2020 | Europe / United Kingdom / Scotland | 2020-04-06 | Virology Department, Royal Infirmary of Edinburgh, NHS Lothian / School of Biological Sciences, University of Edinburgh / Institute of Genetics and Molecular Medicine, University of Edinburgh | COVID-19 Genomics UK (COG-UK) Consortium | McHugh M, Dewar R, Rooke S, Gallagher M, Balcaza C, O'Toole A, Hill V, McCrone JT, Colquhoun R, Yu X, Jackson B, Rambaut A, Williams TC, Templeton K |
| EPI_ISL_432899 | hCoV-19/Scotland/EDB999/2020 | Europe / United Kingdom / Scotland | 2020-04-06 | Virology Department, Royal Infirmary of Edinburgh, NHS Lothian / School of Biological Sciences, University of Edinburgh / Institute of Genetics and Molecular Medicine, University of Edinburgh | COVID-19 Genomics UK (COG-UK) Consortium | McHugh M, Dewar R, Rooke S, Gallagher M, Balcaza C, O'Toole A, Hill V, McCrone JT, Colquhoun R, Yu X, Jackson B, Rambaut A, Williams TC, Templeton K |
| EPI_ISL_433101 | hCoV-19/Scotland/EDB1642/2020 | Europe / United Kingdom / Scotland | 2020-04-14 | Virology Department, Royal Infirmary of Edinburgh, NHS Lothian / School of Biological Sciences, University of Edinburgh / Institute of Genetics and Molecular Medicine, University of Edinburgh | COVID-19 Genomics UK (COG-UK) Consortium | McHugh M, Dewar R, Rooke S, Gallagher M, Balcaza C, O'Toole A, Hill V, McCrone JT, Colquhoun R, Yu X, Jackson B, Rambaut A, Williams TC, Templeton K |
| EPI_ISL_433104 | hCoV-19/Scotland/EDB1647/2020 | Europe / United Kingdom / Scotland | 2020-04-14 | Virology Department, Royal Infirmary of Edinburgh, NHS Lothian / School of Biological Sciences, University of Edinburgh / Institute of Genetics and Molecular Medicine, University of Edinburgh | COVID-19 Genomics UK (COG-UK) Consortium | McHugh M, Dewar R, Rooke S, Gallagher M, Balcaza C, O'Toole A, Hill V, McCrone JT, Colquhoun R, Yu X, Jackson B, Rambaut A, Williams TC, Templeton K |
| EPI_ISL_433107 | hCoV-19/Scotland/EDB1655/2020 | Europe / United Kingdom / Scotland | 2020-04-14 | Virology Department, Royal Infirmary of Edinburgh, NHS Lothian / School of Biological Sciences, University of Edinburgh / Institute of Genetics and Molecular Medicine, University of Edinburgh | COVID-19 Genomics UK (COG-UK) Consortium | McHugh M, Dewar R, Rooke S, Gallagher M, Balcaza C, O'Toole A, Hill V, McCrone JT, Colquhoun R, Yu X, Jackson B, Rambaut A, Williams TC, Templeton K |
| EPI_ISL_433108 | hCoV-19/Scotland/EDB1656/2020 | Europe / United Kingdom / Scotland | 2020-04-14 | Virology Department, Royal Infirmary of Edinburgh, NHS Lothian / School of Biological Sciences, University of Edinburgh / Institute of Genetics and Molecular Medicine, University of Edinburgh | COVID-19 Genomics UK (COG-UK) Consortium | McHugh M, Dewar R, Rooke S, Gallagher M, Balcaza C, O'Toole A, Hill V, McCrone JT, Colquhoun R, Yu X, Jackson B, Rambaut A, Williams TC, Templeton K |
| EPI_ISL_433110 | hCoV-19/Scotland/EDB1663/2020 | Europe / United Kingdom / Scotland | 2020-04-15 | Virology Department, Royal Infirmary of Edinburgh, NHS Lothian / School of Biological Sciences, University of Edinburgh / Institute of Genetics and Molecular Medicine, University of Edinburgh | COVID-19 Genomics UK (COG-UK) Consortium | McHugh M, Dewar R, Rooke S, Gallagher M, Balcaza C, O'Toole A, Hill V, McCrone JT, Colquhoun R, Yu X, Jackson B, Rambaut A, Williams TC, Templeton K |
| EPI_ISL_433112 | hCoV-19/Scotland/EDB1671/2020 | Europe / United Kingdom / Scotland | 2020-04-15 | Virology Department, Royal Infirmary of Edinburgh, NHS Lothian / School of Biological Sciences, University of Edinburgh / Institute of Genetics and Molecular Medicine, University of Edinburgh | COVID-19 Genomics UK (COG-UK) Consortium | McHugh M, Dewar R, Rooke S, Gallagher M, Balcaza C, O'Toole A, Hill V, McCrone JT, Colquhoun R, Yu X, Jackson B, Rambaut A, Williams TC, Templeton K |
| EPI_ISL_433113 | hCoV-19/Scotland/EDB1674/2020 | Europe / United Kingdom / Scotland | 2020-04-15 | Virology Department, Royal Infirmary of Edinburgh, NHS Lothian / School of Biological Sciences, University of Edinburgh / Institute of Genetics and Molecular Medicine, University of Edinburgh | COVID-19 Genomics UK (COG-UK) Consortium | McHugh M, Dewar R, Rooke S, Gallagher M, Balcaza C, O'Toole A, Hill V, McCrone JT, Colquhoun R, Yu X, Jackson B, Rambaut A, Williams TC, Templeton K |
| EPI_ISL_433114 | hCoV-19/Scotland/EDB1677/2020 | Europe / United Kingdom / Scotland | 2020-04-15 | Virology Department, Royal Infirmary of Edinburgh, NHS Lothian / School of Biological Sciences, University of Edinburgh / Institute of Genetics and Molecular Medicine, University of Edinburgh | COVID-19 Genomics UK (COG-UK) Consortium | McHugh M, Dewar R, Rooke S, Gallagher M, Balcaza C, O'Toole A, Hill V, McCrone JT, Colquhoun R, Yu X, Jackson B, Rambaut A, Williams TC, Templeton K |
| EPI_ISL_433115 | hCoV-19/Scotland/EDB1679/2020 | Europe / United Kingdom / Scotland | 2020-04-15 | Virology Department, Royal Infirmary of Edinburgh, NHS Lothian / School of Biological Sciences, University of Edinburgh / Institute of Genetics and Molecular Medicine, University of Edinburgh | COVID-19 Genomics UK (COG-UK) Consortium | McHugh M, Dewar R, Rooke S, Gallagher M, Balcaza C, O'Toole A, Hill V, McCrone JT, Colquhoun R, Yu X, Jackson B, Rambaut A, Williams TC, Templeton K |
| EPI_ISL_433116 | hCoV-19/Scotland/EDB1681/2020 | Europe / United Kingdom / Scotland | 2020-04-15 | Virology Department, Royal Infirmary of Edinburgh, NHS Lothian / School of Biological Sciences, University of Edinburgh / Institute of Genetics and Molecular Medicine, University of Edinburgh | COVID-19 Genomics UK (COG-UK) Consortium | McHugh M, Dewar R, Rooke S, Gallagher M, Balcaza C, O'Toole A, Hill V, McCrone JT, Colquhoun R, Yu X, Jackson B, Rambaut A, Williams TC, Templeton K |
| EPI_ISL_433117 | hCoV-19/Scotland/EDB1683/2020 | Europe / United Kingdom / Scotland | 2020-04-16 | Virology Department, Royal Infirmary of Edinburgh, NHS Lothian / School of Biological Sciences, University of Edinburgh / Institute of Genetics and Molecular Medicine, University of Edinburgh | COVID-19 Genomics UK (COG-UK) Consortium | McHugh M, Dewar R, Rooke S, Gallagher M, Balcaza C, O'Toole A, Hill V, McCrone JT, Colquhoun R, Yu X, Jackson B, Rambaut A, Williams TC, Templeton K |
| EPI_ISL_433118 | hCoV-19/Scotland/EDB1693/2020 | Europe / United Kingdom / Scotland | 2020-04-15 | Virology Department, Royal Infirmary of Edinburgh, NHS Lothian / School of Biological Sciences, University of Edinburgh / Institute of Genetics and Molecular Medicine, University of Edinburgh | COVID-19 Genomics UK (COG-UK) Consortium | McHugh M, Dewar R, Rooke S, Gallagher M, Balcaza C, O'Toole A, Hill V, McCrone JT, Colquhoun R, Yu X, Jackson B, Rambaut A, Williams TC, Templeton K |
| EPI_ISL_433119 | hCoV-19/Scotland/EDB1694/2020 | Europe / United Kingdom / Scotland | 2020-04-15 | Virology Department, Royal Infirmary of Edinburgh, NHS Lothian / School of Biological Sciences, University of Edinburgh / Institute of Genetics and Molecular Medicine, University of Edinburgh | COVID-19 Genomics UK (COG-UK) Consortium | McHugh M, Dewar R, Rooke S, Gallagher M, Balcaza C, O'Toole A, Hill V, McCrone JT, Colquhoun R, Yu X, Jackson B, Rambaut A, Williams TC, Templeton K |
| EPI_ISL_433120 | hCoV-19/Scotland/EDB1697/2020 | Europe / United Kingdom / Scotland | 2020-04-15 | Virology Department, Royal Infirmary of Edinburgh, NHS Lothian / School of Biological Sciences, University of Edinburgh / Institute of Genetics and Molecular Medicine, University of Edinburgh | COVID-19 Genomics UK (COG-UK) Consortium | McHugh M, Dewar R, Rooke S, Gallagher M, Balcaza C, O'Toole A, Hill V, McCrone JT, Colquhoun R, Yu X, Jackson B, Rambaut A, Williams TC, Templeton K |
| EPI_ISL_433121 | hCoV-19/Scotland/EDB1700/2020 | Europe / United Kingdom / Scotland | 2020-04-16 | Virology Department, Royal Infirmary of Edinburgh, NHS Lothian / School of Biological Sciences, University of Edinburgh / Institute of Genetics and Molecular Medicine, University of Edinburgh | COVID-19 Genomics UK (COG-UK) Consortium | McHugh M, Dewar R, Rooke S, Gallagher M, Balcaza C, O'Toole A, Hill V, McCrone JT, Colquhoun R, Yu X, Jackson B, Rambaut A, Williams TC, Templeton K |
| EPI_ISL_433122 | hCoV-19/Scotland/EDB1701/2020 | Europe / United Kingdom / Scotland | 2020-04-15 | Virology Department, Royal Infirmary of Edinburgh, NHS Lothian / School of Biological Sciences, University of Edinburgh / Institute of Genetics and Molecular Medicine, University of Edinburgh | COVID-19 Genomics UK (COG-UK) Consortium | McHugh M, Dewar R, Rooke S, Gallagher M, Balcaza C, O'Toole A, Hill V, McCrone JT, Colquhoun R, Yu X, Jackson B, Rambaut A, Williams TC, Templeton K |
| EPI_ISL_433123 | hCoV-19/Scotland/EDB1704/2020 | Europe / United Kingdom / Scotland | 2020-04-15 | Virology Department, Royal Infirmary of Edinburgh, NHS Lothian / School of Biological Sciences, University of Edinburgh / Institute of Genetics and Molecular Medicine, University of Edinburgh | COVID-19 Genomics UK (COG-UK) Consortium | McHugh M, Dewar R, Rooke S, Gallagher M, Balcaza C, O'Toole A, Hill V, McCrone JT, Colquhoun R, Yu X, Jackson B, Rambaut A, Williams TC, Templeton K |
| EPI_ISL_433125 | hCoV-19/Scotland/EDB1707/2020 | Europe / United Kingdom / Scotland | 2020-04-16 | Virology Department, Royal Infirmary of Edinburgh, NHS Lothian / School of Biological Sciences, University of Edinburgh / Institute of Genetics and Molecular Medicine, University of Edinburgh | COVID-19 Genomics UK (COG-UK) Consortium | McHugh M, Dewar R, Rooke S, Gallagher M, Balcaza C, O'Toole A, Hill V, McCrone JT, Colquhoun R, Yu X, Jackson B, Rambaut A, Williams TC, Templeton K |
| EPI_ISL_433126 | hCoV-19/Scotland/EDB1711/2020 | Europe / United Kingdom / Scotland | 2020-04-15 | Virology Department, Royal Infirmary of Edinburgh, NHS Lothian / School of Biological Sciences, University of Edinburgh / Institute of Genetics and Molecular Medicine, University of Edinburgh | COVID-19 Genomics UK (COG-UK) Consortium | McHugh M, Dewar R, Rooke S, Gallagher M, Balcaza C, O'Toole A, Hill V, McCrone JT, Colquhoun R, Yu X, Jackson B, Rambaut A, Williams TC, Templeton K |
| EPI_ISL_433127 | hCoV-19/Scotland/EDB1717/2020 | Europe / United Kingdom / Scotland | 2020-04-15 | Virology Department, Royal Infirmary of Edinburgh, NHS Lothian / School of Biological Sciences, University of Edinburgh / Institute of Genetics and Molecular Medicine, University of Edinburgh | COVID-19 Genomics UK (COG-UK) Consortium | McHugh M, Dewar R, Rooke S, Gallagher M, Balcaza C, O'Toole A, Hill V, McCrone JT, Colquhoun R, Yu X, Jackson B, Rambaut A, Williams TC, Templeton K |
| EPI_ISL_433129 | hCoV-19/Scotland/EDB1725/2020 | Europe / United Kingdom / Scotland | 2020-04-15 | Virology Department, Royal Infirmary of Edinburgh, NHS Lothian / School of Biological Sciences, University of Edinburgh / Institute of Genetics and Molecular Medicine, University of Edinburgh | COVID-19 Genomics UK (COG-UK) Consortium | McHugh M, Dewar R, Rooke S, Gallagher M, Balcaza C, O'Toole A, Hill V, McCrone JT, Colquhoun R, Yu X, Jackson B, Rambaut A, Williams TC, Templeton K |
| EPI_ISL_433130 | hCoV-19/Scotland/EDB1730/2020 | Europe / United Kingdom / Scotland | 2020-04-15 | Virology Department, Royal Infirmary of Edinburgh, NHS Lothian / School of Biological Sciences, University of Edinburgh / Institute of Genetics and Molecular Medicine, University of Edinburgh | COVID-19 Genomics UK (COG-UK) Consortium | McHugh M, Dewar R, Rooke S, Gallagher M, Balcaza C, O'Toole A, Hill V, McCrone JT, Colquhoun R, Yu X, Jackson B, Rambaut A, Williams TC, Templeton K |
| EPI_ISL_433132 | hCoV-19/Scotland/EDB1738/2020 | Europe / United Kingdom / Scotland | 2020-04-16 | Virology Department, Royal Infirmary of Edinburgh, NHS Lothian / School of Biological Sciences, University of Edinburgh / Institute of Genetics and Molecular Medicine, University of Edinburgh | COVID-19 Genomics UK (COG-UK) Consortium | McHugh M, Dewar R, Rooke S, Gallagher M, Balcaza C, O'Toole A, Hill V, McCrone JT, Colquhoun R, Yu X, Jackson B, Rambaut A, Williams TC, Templeton K |
| EPI_ISL_433133 | hCoV-19/Scotland/EDB1740/2020 | Europe / United Kingdom / Scotland | 2020-04-15 | Virology Department, Royal Infirmary of Edinburgh, NHS Lothian / School of Biological Sciences, University of Edinburgh / Institute of Genetics and Molecular Medicine, University of Edinburgh | COVID-19 Genomics UK (COG-UK) Consortium | McHugh M, Dewar R, Rooke S, Gallagher M, Balcaza C, O'Toole A, Hill V, McCrone JT, Colquhoun R, Yu X, Jackson B, Rambaut A, Williams TC, Templeton K |
| EPI_ISL_433135 | hCoV-19/Scotland/EDB1811/2020 | Europe / United Kingdom / Scotland | 2020-04-16 | Virology Department, Royal Infirmary of Edinburgh, NHS Lothian / School of Biological Sciences, University of Edinburgh / Institute of Genetics and Molecular Medicine, University of Edinburgh | COVID-19 Genomics UK (COG-UK) Consortium | McHugh M, Dewar R, Rooke S, Gallagher M, Balcaza C, O'Toole A, Hill V, McCrone JT, Colquhoun R, Yu X, Jackson B, Rambaut A, Williams TC, Templeton K |
| EPI_ISL_433136 | hCoV-19/Scotland/EDB1812/2020 | Europe / United Kingdom / Scotland | 2020-04-16 | Virology Department, Royal Infirmary of Edinburgh, NHS Lothian / School of Biological Sciences, University of Edinburgh / Institute of Genetics and Molecular Medicine, University of Edinburgh | COVID-19 Genomics UK (COG-UK) Consortium | McHugh M, Dewar R, Rooke S, Gallagher M, Balcaza C, O'Toole A, Hill V, McCrone JT, Colquhoun R, Yu X, Jackson B, Rambaut A, Williams TC, Templeton K |
| EPI_ISL_433137 | hCoV-19/Scotland/EDB1813/2020 | Europe / United Kingdom / Scotland | 2020-04-16 | Virology Department, Royal Infirmary of Edinburgh, NHS Lothian / School of Biological Sciences, University of Edinburgh / Institute of Genetics and Molecular Medicine, University of Edinburgh | COVID-19 Genomics UK (COG-UK) Consortium | McHugh M, Dewar R, Rooke S, Gallagher M, Balcaza C, O'Toole A, Hill V, McCrone JT, Colquhoun R, Yu X, Jackson B, Rambaut A, Williams TC, Templeton K |
| EPI_ISL_433139 | hCoV-19/Scotland/EDB1816/2020 | Europe / United Kingdom / Scotland | 2020-04-16 | Virology Department, Royal Infirmary of Edinburgh, NHS Lothian / School of Biological Sciences, University of Edinburgh / Institute of Genetics and Molecular Medicine, University of Edinburgh | COVID-19 Genomics UK (COG-UK) Consortium | McHugh M, Dewar R, Rooke S, Gallagher M, Balcaza C, O'Toole A, Hill V, McCrone JT, Colquhoun R, Yu X, Jackson B, Rambaut A, Williams TC, Templeton K |
| EPI_ISL_433140 | hCoV-19/Scotland/EDB1817/2020 | Europe / United Kingdom / Scotland | 2020-04-16 | Virology Department, Royal Infirmary of Edinburgh, NHS Lothian / School of Biological Sciences, University of Edinburgh / Institute of Genetics and Molecular Medicine, University of Edinburgh | COVID-19 Genomics UK (COG-UK) Consortium | McHugh M, Dewar R, Rooke S, Gallagher M, Balcaza C, O'Toole A, Hill V, McCrone JT, Colquhoun R, Yu X, Jackson B, Rambaut A, Williams TC, Templeton K |
| EPI_ISL_433141 | hCoV-19/Scotland/EDB1819/2020 | Europe / United Kingdom / Scotland | 2020-04-16 | Virology Department, Royal Infirmary of Edinburgh, NHS Lothian / School of Biological Sciences, University of Edinburgh / Institute of Genetics and Molecular Medicine, University of Edinburgh | COVID-19 Genomics UK (COG-UK) Consortium | McHugh M, Dewar R, Rooke S, Gallagher M, Balcaza C, O'Toole A, Hill V, McCrone JT, Colquhoun R, Yu X, Jackson B, Rambaut A, Williams TC, Templeton K |
| EPI_ISL_433142 | hCoV-19/Scotland/EDB1820/2020 | Europe / United Kingdom / Scotland | 2020-04-16 | Virology Department, Royal Infirmary of Edinburgh, NHS Lothian / School of Biological Sciences, University of Edinburgh / Institute of Genetics and Molecular Medicine, University of Edinburgh | COVID-19 Genomics UK (COG-UK) Consortium | McHugh M, Dewar R, Rooke S, Gallagher M, Balcaza C, O'Toole A, Hill V, McCrone JT, Colquhoun R, Yu X, Jackson B, Rambaut A, Williams TC, Templeton K |
| EPI_ISL_433143 | hCoV-19/Scotland/EDB1821/2020 | Europe / United Kingdom / Scotland | 2020-04-16 | Virology Department, Royal Infirmary of Edinburgh, NHS Lothian / School of Biological Sciences, University of Edinburgh / Institute of Genetics and Molecular Medicine, University of Edinburgh | COVID-19 Genomics UK (COG-UK) Consortium | McHugh M, Dewar R, Rooke S, Gallagher M, Balcaza C, O'Toole A, Hill V, McCrone JT, Colquhoun R, Yu X, Jackson B, Rambaut A, Williams TC, Templeton K |
| EPI_ISL_433145 | hCoV-19/Scotland/EDB1825/2020 | Europe / United Kingdom / Scotland | 2020-04-16 | Virology Department, Royal Infirmary of Edinburgh, NHS Lothian / School of Biological Sciences, University of Edinburgh / Institute of Genetics and Molecular Medicine, University of Edinburgh | COVID-19 Genomics UK (COG-UK) Consortium | McHugh M, Dewar R, Rooke S, Gallagher M, Balcaza C, O'Toole A, Hill V, McCrone JT, Colquhoun R, Yu X, Jackson B, Rambaut A, Williams TC, Templeton K |
| EPI_ISL_433147 | hCoV-19/Scotland/EDB1827/2020 | Europe / United Kingdom / Scotland | 2020-04-16 | Virology Department, Royal Infirmary of Edinburgh, NHS Lothian / School of Biological Sciences, University of Edinburgh / Institute of Genetics and Molecular Medicine, University of Edinburgh | COVID-19 Genomics UK (COG-UK) Consortium | McHugh M, Dewar R, Rooke S, Gallagher M, Balcaza C, O'Toole A, Hill V, McCrone JT, Colquhoun R, Yu X, Jackson B, Rambaut A, Williams TC, Templeton K |
| EPI_ISL_433148 | hCoV-19/Scotland/EDB1828/2020 | Europe / United Kingdom / Scotland | 2020-04-16 | Virology Department, Royal Infirmary of Edinburgh, NHS Lothian / School of Biological Sciences, University of Edinburgh / Institute of Genetics and Molecular Medicine, University of Edinburgh | COVID-19 Genomics UK (COG-UK) Consortium | McHugh M, Dewar R, Rooke S, Gallagher M, Balcaza C, O'Toole A, Hill V, McCrone JT, Colquhoun R, Yu X, Jackson B, Rambaut A, Williams TC, Templeton K |
| EPI_ISL_433149 | hCoV-19/Scotland/EDB1829/2020 | Europe / United Kingdom / Scotland | 2020-04-16 | Virology Department, Royal Infirmary of Edinburgh, NHS Lothian / School of Biological Sciences, University of Edinburgh / Institute of Genetics and Molecular Medicine, University of Edinburgh | COVID-19 Genomics UK (COG-UK) Consortium | McHugh M, Dewar R, Rooke S, Gallagher M, Balcaza C, O'Toole A, Hill V, McCrone JT, Colquhoun R, Yu X, Jackson B, Rambaut A, Williams TC, Templeton K |
| EPI_ISL_433150 | hCoV-19/Scotland/EDB1830/2020 | Europe / United Kingdom / Scotland | 2020-04-16 | Virology Department, Royal Infirmary of Edinburgh, NHS Lothian / School of Biological Sciences, University of Edinburgh / Institute of Genetics and Molecular Medicine, University of Edinburgh | COVID-19 Genomics UK (COG-UK) Consortium | McHugh M, Dewar R, Rooke S, Gallagher M, Balcaza C, O'Toole A, Hill V, McCrone JT, Colquhoun R, Yu X, Jackson B, Rambaut A, Williams TC, Templeton K |
| EPI_ISL_433153 | hCoV-19/Scotland/EDB1841/2020 | Europe / United Kingdom / Scotland | 2020-04-17 | Virology Department, Royal Infirmary of Edinburgh, NHS Lothian / School of Biological Sciences, University of Edinburgh / Institute of Genetics and Molecular Medicine, University of Edinburgh | COVID-19 Genomics UK (COG-UK) Consortium | McHugh M, Dewar R, Rooke S, Gallagher M, Balcaza C, O'Toole A, Hill V, McCrone JT, Colquhoun R, Yu X, Jackson B, Rambaut A, Williams TC, Templeton K |
| EPI_ISL_433154 | hCoV-19/Scotland/EDB1842/2020 | Europe / United Kingdom / Scotland | 2020-04-17 | Virology Department, Royal Infirmary of Edinburgh, NHS Lothian / School of Biological Sciences, University of Edinburgh / Institute of Genetics and Molecular Medicine, University of Edinburgh | COVID-19 Genomics UK (COG-UK) Consortium | McHugh M, Dewar R, Rooke S, Gallagher M, Balcaza C, O'Toole A, Hill V, McCrone JT, Colquhoun R, Yu X, Jackson B, Rambaut A, Williams TC, Templeton K |
| EPI_ISL_433155 | hCoV-19/Scotland/EDB1848/2020 | Europe / United Kingdom / Scotland | 2020-04-17 | Virology Department, Royal Infirmary of Edinburgh, NHS Lothian / School of Biological Sciences, University of Edinburgh / Institute of Genetics and Molecular Medicine, University of Edinburgh | COVID-19 Genomics UK (COG-UK) Consortium | McHugh M, Dewar R, Rooke S, Gallagher M, Balcaza C, O'Toole A, Hill V, McCrone JT, Colquhoun R, Yu X, Jackson B, Rambaut A, Williams TC, Templeton K |
| EPI_ISL_433156 | hCoV-19/Scotland/EDB1897/2020 | Europe / United Kingdom / Scotland | 2020-04-17 | Virology Department, Royal Infirmary of Edinburgh, NHS Lothian / School of Biological Sciences, University of Edinburgh / Institute of Genetics and Molecular Medicine, University of Edinburgh | COVID-19 Genomics UK (COG-UK) Consortium | McHugh M, Dewar R, Rooke S, Gallagher M, Balcaza C, O'Toole A, Hill V, McCrone JT, Colquhoun R, Yu X, Jackson B, Rambaut A, Williams TC, Templeton K |
| EPI_ISL_433162 | hCoV-19/Scotland/EDB2011/2020 | Europe / United Kingdom / Scotland | 2020-04-18 | Virology Department, Royal Infirmary of Edinburgh, NHS Lothian / School of Biological Sciences, University of Edinburgh / Institute of Genetics and Molecular Medicine, University of Edinburgh | COVID-19 Genomics UK (COG-UK) Consortium | McHugh M, Dewar R, Rooke S, Gallagher M, Balcaza C, O'Toole A, Hill V, McCrone JT, Colquhoun R, Yu X, Jackson B, Rambaut A, Williams TC, Templeton K |
| EPI_ISL_433164 | hCoV-19/Scotland/EDB2048/2020 | Europe / United Kingdom / Scotland | 2020-04-18 | Virology Department, Royal Infirmary of Edinburgh, NHS Lothian / School of Biological Sciences, University of Edinburgh / Institute of Genetics and Molecular Medicine, University of Edinburgh | COVID-19 Genomics UK (COG-UK) Consortium | McHugh M, Dewar R, Rooke S, Gallagher M, Balcaza C, O'Toole A, Hill V, McCrone JT, Colquhoun R, Yu X, Jackson B, Rambaut A, Williams TC, Templeton K |
| EPI_ISL_433165 | hCoV-19/Scotland/EDB2049/2020 | Europe / United Kingdom / Scotland | 2020-04-18 | Virology Department, Royal Infirmary of Edinburgh, NHS Lothian / School of Biological Sciences, University of Edinburgh / Institute of Genetics and Molecular Medicine, University of Edinburgh | COVID-19 Genomics UK (COG-UK) Consortium | McHugh M, Dewar R, Rooke S, Gallagher M, Balcaza C, O'Toole A, Hill V, McCrone JT, Colquhoun R, Yu X, Jackson B, Rambaut A, Williams TC, Templeton K |
| EPI_ISL_433166 | hCoV-19/Scotland/EDB2050/2020 | Europe / United Kingdom / Scotland | 2020-04-18 | Virology Department, Royal Infirmary of Edinburgh, NHS Lothian / School of Biological Sciences, University of Edinburgh / Institute of Genetics and Molecular Medicine, University of Edinburgh | COVID-19 Genomics UK (COG-UK) Consortium | McHugh M, Dewar R, Rooke S, Gallagher M, Balcaza C, O'Toole A, Hill V, McCrone JT, Colquhoun R, Yu X, Jackson B, Rambaut A, Williams TC, Templeton K |
| EPI_ISL_433167 | hCoV-19/Scotland/EDB2051/2020 | Europe / United Kingdom / Scotland | 2020-04-18 | Virology Department, Royal Infirmary of Edinburgh, NHS Lothian / School of Biological Sciences, University of Edinburgh / Institute of Genetics and Molecular Medicine, University of Edinburgh | COVID-19 Genomics UK (COG-UK) Consortium | McHugh M, Dewar R, Rooke S, Gallagher M, Balcaza C, O'Toole A, Hill V, McCrone JT, Colquhoun R, Yu X, Jackson B, Rambaut A, Williams TC, Templeton K |
| EPI_ISL_433168 | hCoV-19/Scotland/EDB2052/2020 | Europe / United Kingdom / Scotland | 2020-04-18 | Virology Department, Royal Infirmary of Edinburgh, NHS Lothian / School of Biological Sciences, University of Edinburgh / Institute of Genetics and Molecular Medicine, University of Edinburgh | COVID-19 Genomics UK (COG-UK) Consortium | McHugh M, Dewar R, Rooke S, Gallagher M, Balcaza C, O'Toole A, Hill V, McCrone JT, Colquhoun R, Yu X, Jackson B, Rambaut A, Williams TC, Templeton K |
| EPI_ISL_433169 | hCoV-19/Scotland/EDB2057/2020 | Europe / United Kingdom / Scotland | 2020-04-18 | Virology Department, Royal Infirmary of Edinburgh, NHS Lothian / School of Biological Sciences, University of Edinburgh / Institute of Genetics and Molecular Medicine, University of Edinburgh | COVID-19 Genomics UK (COG-UK) Consortium | McHugh M, Dewar R, Rooke S, Gallagher M, Balcaza C, O'Toole A, Hill V, McCrone JT, Colquhoun R, Yu X, Jackson B, Rambaut A, Williams TC, Templeton K |
| EPI_ISL_433170 | hCoV-19/Scotland/EDB2062/2020 | Europe / United Kingdom / Scotland | 2020-04-18 | Virology Department, Royal Infirmary of Edinburgh, NHS Lothian / School of Biological Sciences, University of Edinburgh / Institute of Genetics and Molecular Medicine, University of Edinburgh | COVID-19 Genomics UK (COG-UK) Consortium | McHugh M, Dewar R, Rooke S, Gallagher M, Balcaza C, O'Toole A, Hill V, McCrone JT, Colquhoun R, Yu X, Jackson B, Rambaut A, Williams TC, Templeton K |
| EPI_ISL_433171 | hCoV-19/Scotland/EDB2064/2020 | Europe / United Kingdom / Scotland | 2020-04-18 | Virology Department, Royal Infirmary of Edinburgh, NHS Lothian / School of Biological Sciences, University of Edinburgh / Institute of Genetics and Molecular Medicine, University of Edinburgh | COVID-19 Genomics UK (COG-UK) Consortium | McHugh M, Dewar R, Rooke S, Gallagher M, Balcaza C, O'Toole A, Hill V, McCrone JT, Colquhoun R, Yu X, Jackson B, Rambaut A, Williams TC, Templeton K |
| EPI_ISL_433172 | hCoV-19/Scotland/EDB2065/2020 | Europe / United Kingdom / Scotland | 2020-04-18 | Virology Department, Royal Infirmary of Edinburgh, NHS Lothian / School of Biological Sciences, University of Edinburgh / Institute of Genetics and Molecular Medicine, University of Edinburgh | COVID-19 Genomics UK (COG-UK) Consortium | McHugh M, Dewar R, Rooke S, Gallagher M, Balcaza C, O'Toole A, Hill V, McCrone JT, Colquhoun R, Yu X, Jackson B, Rambaut A, Williams TC, Templeton K |
| EPI_ISL_433173 | hCoV-19/Scotland/EDB2066/2020 | Europe / United Kingdom / Scotland | 2020-04-18 | Virology Department, Royal Infirmary of Edinburgh, NHS Lothian / School of Biological Sciences, University of Edinburgh / Institute of Genetics and Molecular Medicine, University of Edinburgh | COVID-19 Genomics UK (COG-UK) Consortium | McHugh M, Dewar R, Rooke S, Gallagher M, Balcaza C, O'Toole A, Hill V, McCrone JT, Colquhoun R, Yu X, Jackson B, Rambaut A, Williams TC, Templeton K |
| EPI_ISL_433174 | hCoV-19/Scotland/EDB2071/2020 | Europe / United Kingdom / Scotland | 2020-04-19 | Virology Department, Royal Infirmary of Edinburgh, NHS Lothian / School of Biological Sciences, University of Edinburgh / Institute of Genetics and Molecular Medicine, University of Edinburgh | COVID-19 Genomics UK (COG-UK) Consortium | McHugh M, Dewar R, Rooke S, Gallagher M, Balcaza C, O'Toole A, Hill V, McCrone JT, Colquhoun R, Yu X, Jackson B, Rambaut A, Williams TC, Templeton K |
| EPI_ISL_433176 | hCoV-19/Scotland/EDB2077/2020 | Europe / United Kingdom / Scotland | 2020-04-19 | Virology Department, Royal Infirmary of Edinburgh, NHS Lothian / School of Biological Sciences, University of Edinburgh / Institute of Genetics and Molecular Medicine, University of Edinburgh | COVID-19 Genomics UK (COG-UK) Consortium | McHugh M, Dewar R, Rooke S, Gallagher M, Balcaza C, O'Toole A, Hill V, McCrone JT, Colquhoun R, Yu X, Jackson B, Rambaut A, Williams TC, Templeton K |
| EPI_ISL_433177 | hCoV-19/Scotland/EDB2078/2020 | Europe / United Kingdom / Scotland | 2020-04-19 | Virology Department, Royal Infirmary of Edinburgh, NHS Lothian / School of Biological Sciences, University of Edinburgh / Institute of Genetics and Molecular Medicine, University of Edinburgh | COVID-19 Genomics UK (COG-UK) Consortium | McHugh M, Dewar R, Rooke S, Gallagher M, Balcaza C, O'Toole A, Hill V, McCrone JT, Colquhoun R, Yu X, Jackson B, Rambaut A, Williams TC, Templeton K |
| EPI_ISL_433178 | hCoV-19/Scotland/EDB2079/2020 | Europe / United Kingdom / Scotland | 2020-04-19 | Virology Department, Royal Infirmary of Edinburgh, NHS Lothian / School of Biological Sciences, University of Edinburgh / Institute of Genetics and Molecular Medicine, University of Edinburgh | COVID-19 Genomics UK (COG-UK) Consortium | McHugh M, Dewar R, Rooke S, Gallagher M, Balcaza C, O'Toole A, Hill V, McCrone JT, Colquhoun R, Yu X, Jackson B, Rambaut A, Williams TC, Templeton K |
| EPI_ISL_433180 | hCoV-19/Scotland/EDB2081/2020 | Europe / United Kingdom / Scotland | 2020-04-19 | Virology Department, Royal Infirmary of Edinburgh, NHS Lothian / School of Biological Sciences, University of Edinburgh / Institute of Genetics and Molecular Medicine, University of Edinburgh | COVID-19 Genomics UK (COG-UK) Consortium | McHugh M, Dewar R, Rooke S, Gallagher M, Balcaza C, O'Toole A, Hill V, McCrone JT, Colquhoun R, Yu X, Jackson B, Rambaut A, Williams TC, Templeton K |
| EPI_ISL_433181 | hCoV-19/Scotland/EDB2082/2020 | Europe / United Kingdom / Scotland | 2020-04-19 | Virology Department, Royal Infirmary of Edinburgh, NHS Lothian / School of Biological Sciences, University of Edinburgh / Institute of Genetics and Molecular Medicine, University of Edinburgh | COVID-19 Genomics UK (COG-UK) Consortium | McHugh M, Dewar R, Rooke S, Gallagher M, Balcaza C, O'Toole A, Hill V, McCrone JT, Colquhoun R, Yu X, Jackson B, Rambaut A, Williams TC, Templeton K |
| EPI_ISL_433182 | hCoV-19/Scotland/EDB2083/2020 | Europe / United Kingdom / Scotland | 2020-04-19 | Virology Department, Royal Infirmary of Edinburgh, NHS Lothian / School of Biological Sciences, University of Edinburgh / Institute of Genetics and Molecular Medicine, University of Edinburgh | COVID-19 Genomics UK (COG-UK) Consortium | McHugh M, Dewar R, Rooke S, Gallagher M, Balcaza C, O'Toole A, Hill V, McCrone JT, Colquhoun R, Yu X, Jackson B, Rambaut A, Williams TC, Templeton K |
| EPI_ISL_433183 | hCoV-19/Scotland/EDB2084/2020 | Europe / United Kingdom / Scotland | 2020-04-19 | Virology Department, Royal Infirmary of Edinburgh, NHS Lothian / School of Biological Sciences, University of Edinburgh / Institute of Genetics and Molecular Medicine, University of Edinburgh | COVID-19 Genomics UK (COG-UK) Consortium | McHugh M, Dewar R, Rooke S, Gallagher M, Balcaza C, O'Toole A, Hill V, McCrone JT, Colquhoun R, Yu X, Jackson B, Rambaut A, Williams TC, Templeton K |
| EPI_ISL_433184 | hCoV-19/Scotland/EDB2085/2020 | Europe / United Kingdom / Scotland | 2020-04-19 | Virology Department, Royal Infirmary of Edinburgh, NHS Lothian / School of Biological Sciences, University of Edinburgh / Institute of Genetics and Molecular Medicine, University of Edinburgh | COVID-19 Genomics UK (COG-UK) Consortium | McHugh M, Dewar R, Rooke S, Gallagher M, Balcaza C, O'Toole A, Hill V, McCrone JT, Colquhoun R, Yu X, Jackson B, Rambaut A, Williams TC, Templeton K |
| EPI_ISL_433185 | hCoV-19/Scotland/EDB2086/2020 | Europe / United Kingdom / Scotland | 2020-04-19 | Virology Department, Royal Infirmary of Edinburgh, NHS Lothian / School of Biological Sciences, University of Edinburgh / Institute of Genetics and Molecular Medicine, University of Edinburgh | COVID-19 Genomics UK (COG-UK) Consortium | McHugh M, Dewar R, Rooke S, Gallagher M, Balcaza C, O'Toole A, Hill V, McCrone JT, Colquhoun R, Yu X, Jackson B, Rambaut A, Williams TC, Templeton K |
| EPI_ISL_433186 | hCoV-19/Scotland/EDB2087/2020 | Europe / United Kingdom / Scotland | 2020-04-19 | Virology Department, Royal Infirmary of Edinburgh, NHS Lothian / School of Biological Sciences, University of Edinburgh / Institute of Genetics and Molecular Medicine, University of Edinburgh | COVID-19 Genomics UK (COG-UK) Consortium | McHugh M, Dewar R, Rooke S, Gallagher M, Balcaza C, O'Toole A, Hill V, McCrone JT, Colquhoun R, Yu X, Jackson B, Rambaut A, Williams TC, Templeton K |
| EPI_ISL_433187 | hCoV-19/Scotland/EDB2088/2020 | Europe / United Kingdom / Scotland | 2020-04-19 | Virology Department, Royal Infirmary of Edinburgh, NHS Lothian / School of Biological Sciences, University of Edinburgh / Institute of Genetics and Molecular Medicine, University of Edinburgh | COVID-19 Genomics UK (COG-UK) Consortium | McHugh M, Dewar R, Rooke S, Gallagher M, Balcaza C, O'Toole A, Hill V, McCrone JT, Colquhoun R, Yu X, Jackson B, Rambaut A, Williams TC, Templeton K |
| EPI_ISL_433188 | hCoV-19/Scotland/EDB2089/2020 | Europe / United Kingdom / Scotland | 2020-04-19 | Virology Department, Royal Infirmary of Edinburgh, NHS Lothian / School of Biological Sciences, University of Edinburgh / Institute of Genetics and Molecular Medicine, University of Edinburgh | COVID-19 Genomics UK (COG-UK) Consortium | McHugh M, Dewar R, Rooke S, Gallagher M, Balcaza C, O'Toole A, Hill V, McCrone JT, Colquhoun R, Yu X, Jackson B, Rambaut A, Williams TC, Templeton K |
| EPI_ISL_433190 | hCoV-19/Scotland/EDB2092/2020 | Europe / United Kingdom / Scotland | 2020-04-19 | Virology Department, Royal Infirmary of Edinburgh, NHS Lothian / School of Biological Sciences, University of Edinburgh / Institute of Genetics and Molecular Medicine, University of Edinburgh | COVID-19 Genomics UK (COG-UK) Consortium | McHugh M, Dewar R, Rooke S, Gallagher M, Balcaza C, O'Toole A, Hill V, McCrone JT, Colquhoun R, Yu X, Jackson B, Rambaut A, Williams TC, Templeton K |
| EPI_ISL_433192 | hCoV-19/Scotland/EDB2094/2020 | Europe / United Kingdom / Scotland | 2020-04-18 | Virology Department, Royal Infirmary of Edinburgh, NHS Lothian / School of Biological Sciences, University of Edinburgh / Institute of Genetics and Molecular Medicine, University of Edinburgh | COVID-19 Genomics UK (COG-UK) Consortium | McHugh M, Dewar R, Rooke S, Gallagher M, Balcaza C, O'Toole A, Hill V, McCrone JT, Colquhoun R, Yu X, Jackson B, Rambaut A, Williams TC, Templeton K |
| EPI_ISL_433193 | hCoV-19/Scotland/EDB2095/2020 | Europe / United Kingdom / Scotland | 2020-04-18 | Virology Department, Royal Infirmary of Edinburgh, NHS Lothian / School of Biological Sciences, University of Edinburgh / Institute of Genetics and Molecular Medicine, University of Edinburgh | COVID-19 Genomics UK (COG-UK) Consortium | McHugh M, Dewar R, Rooke S, Gallagher M, Balcaza C, O'Toole A, Hill V, McCrone JT, Colquhoun R, Yu X, Jackson B, Rambaut A, Williams TC, Templeton K |
| EPI_ISL_433194 | hCoV-19/Scotland/EDB2096/2020 | Europe / United Kingdom / Scotland | 2020-04-19 | Virology Department, Royal Infirmary of Edinburgh, NHS Lothian / School of Biological Sciences, University of Edinburgh / Institute of Genetics and Molecular Medicine, University of Edinburgh | COVID-19 Genomics UK (COG-UK) Consortium | McHugh M, Dewar R, Rooke S, Gallagher M, Balcaza C, O'Toole A, Hill V, McCrone JT, Colquhoun R, Yu X, Jackson B, Rambaut A, Williams TC, Templeton K |
| EPI_ISL_433195 | hCoV-19/Scotland/EDB2097/2020 | Europe / United Kingdom / Scotland | 2020-04-19 | Virology Department, Royal Infirmary of Edinburgh, NHS Lothian / School of Biological Sciences, University of Edinburgh / Institute of Genetics and Molecular Medicine, University of Edinburgh | COVID-19 Genomics UK (COG-UK) Consortium | McHugh M, Dewar R, Rooke S, Gallagher M, Balcaza C, O'Toole A, Hill V, McCrone JT, Colquhoun R, Yu X, Jackson B, Rambaut A, Williams TC, Templeton K |
| EPI_ISL_433196 | hCoV-19/Scotland/EDB2098/2020 | Europe / United Kingdom / Scotland | 2020-04-19 | Virology Department, Royal Infirmary of Edinburgh, NHS Lothian / School of Biological Sciences, University of Edinburgh / Institute of Genetics and Molecular Medicine, University of Edinburgh | COVID-19 Genomics UK (COG-UK) Consortium | McHugh M, Dewar R, Rooke S, Gallagher M, Balcaza C, O'Toole A, Hill V, McCrone JT, Colquhoun R, Yu X, Jackson B, Rambaut A, Williams TC, Templeton K |
| EPI_ISL_433197 | hCoV-19/Scotland/EDB2099/2020 | Europe / United Kingdom / Scotland | 2020-04-18 | Virology Department, Royal Infirmary of Edinburgh, NHS Lothian / School of Biological Sciences, University of Edinburgh / Institute of Genetics and Molecular Medicine, University of Edinburgh | COVID-19 Genomics UK (COG-UK) Consortium | McHugh M, Dewar R, Rooke S, Gallagher M, Balcaza C, O'Toole A, Hill V, McCrone JT, Colquhoun R, Yu X, Jackson B, Rambaut A, Williams TC, Templeton K |
| EPI_ISL_433198 | hCoV-19/Scotland/EDB328/2020 | Europe / United Kingdom / Scotland | 2020-03-28 | Virology Department, Royal Infirmary of Edinburgh, NHS Lothian / School of Biological Sciences, University of Edinburgh / Institute of Genetics and Molecular Medicine, University of Edinburgh | COVID-19 Genomics UK (COG-UK) Consortium | McHugh M, Dewar R, Rooke S, Gallagher M, Balcaza C, O'Toole A, Hill V, McCrone JT, Colquhoun R, Yu X, Jackson B, Rambaut A, Williams TC, Templeton K |
| EPI_ISL_433200 | hCoV-19/Scotland/EDB573/2020 | Europe / United Kingdom / Scotland | 2020-04-01 | Virology Department, Royal Infirmary of Edinburgh, NHS Lothian / School of Biological Sciences, University of Edinburgh / Institute of Genetics and Molecular Medicine, University of Edinburgh | COVID-19 Genomics UK (COG-UK) Consortium | McHugh M, Dewar R, Rooke S, Gallagher M, Balcaza C, O'Toole A, Hill V, McCrone JT, Colquhoun R, Yu X, Jackson B, Rambaut A, Williams TC, Templeton K |
| EPI_ISL_433302 | hCoV-19/Scotland/CVR473/2020 | Europe / United Kingdom / Scotland | 2020-03-23 | West of Scotland Specialist Virology Centre, NHSGGC / MRC-University of Glasgow Centre for Virus Research | COVID-19 Genomics UK (COG-UK) Consortium | Ana da Silva Filipe, Natasha Johnson, Kathy Smollett, Daniel Mair, Stephen Carmichael, Lily Tong, Jenna Nichols, Elihu Aranday-Cortes, Kirstyn Brunker, Yasmin Parr, Kyriaki Nomikou; Sarah McDonald, Marc Niebel, Patawee Asamaphan; Richard Orton, Joseph Hughes, Sreenu Vattipally, David L Robertson; Alasdair MacLean, Rory Gunson; Kathy Li, Natasha Jesudason, Rajiv Shah, James Shepherd, Antonia Ho, Emma Thomson |
| EPI_ISL_433303 | hCoV-19/Scotland/CVR487/2020 | Europe / United Kingdom / Scotland | 2020-03-23 | West of Scotland Specialist Virology Centre, NHSGGC / MRC-University of Glasgow Centre for Virus Research | COVID-19 Genomics UK (COG-UK) Consortium | Ana da Silva Filipe, Natasha Johnson, Kathy Smollett, Daniel Mair, Stephen Carmichael, Lily Tong, Jenna Nichols, Elihu Aranday-Cortes, Kirstyn Brunker, Yasmin Parr, Kyriaki Nomikou; Sarah McDonald, Marc Niebel, Patawee Asamaphan; Richard Orton, Joseph Hughes, Sreenu Vattipally, David L Robertson; Alasdair MacLean, Rory Gunson; Kathy Li, Natasha Jesudason, Rajiv Shah, James Shepherd, Antonia Ho, Emma Thomson |
| EPI_ISL_433304 | hCoV-19/Scotland/CVR491/2020 | Europe / United Kingdom / Scotland | 2020-03-23 | West of Scotland Specialist Virology Centre, NHSGGC / MRC-University of Glasgow Centre for Virus Research | COVID-19 Genomics UK (COG-UK) Consortium | Ana da Silva Filipe, Natasha Johnson, Kathy Smollett, Daniel Mair, Stephen Carmichael, Lily Tong, Jenna Nichols, Elihu Aranday-Cortes, Kirstyn Brunker, Yasmin Parr, Kyriaki Nomikou; Sarah McDonald, Marc Niebel, Patawee Asamaphan; Richard Orton, Joseph Hughes, Sreenu Vattipally, David L Robertson; Alasdair MacLean, Rory Gunson; Kathy Li, Natasha Jesudason, Rajiv Shah, James Shepherd, Antonia Ho, Emma Thomson |
| EPI_ISL_433305 | hCoV-19/Scotland/CVR495/2020 | Europe / United Kingdom / Scotland | 2020-03-24 | West of Scotland Specialist Virology Centre, NHSGGC / MRC-University of Glasgow Centre for Virus Research | COVID-19 Genomics UK (COG-UK) Consortium | Ana da Silva Filipe, Natasha Johnson, Kathy Smollett, Daniel Mair, Stephen Carmichael, Lily Tong, Jenna Nichols, Elihu Aranday-Cortes, Kirstyn Brunker, Yasmin Parr, Kyriaki Nomikou; Sarah McDonald, Marc Niebel, Patawee Asamaphan; Richard Orton, Joseph Hughes, Sreenu Vattipally, David L Robertson; Alasdair MacLean, Rory Gunson; Kathy Li, Natasha Jesudason, Rajiv Shah, James Shepherd, Antonia Ho, Emma Thomson |
| EPI_ISL_433307 | hCoV-19/Scotland/CVR507/2020 | Europe / United Kingdom / Scotland | 2020-03-24 | West of Scotland Specialist Virology Centre, NHSGGC / MRC-University of Glasgow Centre for Virus Research | COVID-19 Genomics UK (COG-UK) Consortium | Ana da Silva Filipe, Natasha Johnson, Kathy Smollett, Daniel Mair, Stephen Carmichael, Lily Tong, Jenna Nichols, Elihu Aranday-Cortes, Kirstyn Brunker, Yasmin Parr, Kyriaki Nomikou; Sarah McDonald, Marc Niebel, Patawee Asamaphan; Richard Orton, Joseph Hughes, Sreenu Vattipally, David L Robertson; Alasdair MacLean, Rory Gunson; Kathy Li, Natasha Jesudason, Rajiv Shah, James Shepherd, Antonia Ho, Emma Thomson |
| EPI_ISL_433308 | hCoV-19/Scotland/CVR512/2020 | Europe / United Kingdom / Scotland | 2020-03-24 | West of Scotland Specialist Virology Centre, NHSGGC / MRC-University of Glasgow Centre for Virus Research | COVID-19 Genomics UK (COG-UK) Consortium | Ana da Silva Filipe, Natasha Johnson, Kathy Smollett, Daniel Mair, Stephen Carmichael, Lily Tong, Jenna Nichols, Elihu Aranday-Cortes, Kirstyn Brunker, Yasmin Parr, Kyriaki Nomikou; Sarah McDonald, Marc Niebel, Patawee Asamaphan; Richard Orton, Joseph Hughes, Sreenu Vattipally, David L Robertson; Alasdair MacLean, Rory Gunson; Kathy Li, Natasha Jesudason, Rajiv Shah, James Shepherd, Antonia Ho, Emma Thomson |
| EPI_ISL_433313 | hCoV-19/Scotland/CVR583/2020 | Europe / United Kingdom / Scotland | 2020-03-25 | West of Scotland Specialist Virology Centre, NHSGGC / MRC-University of Glasgow Centre for Virus Research | COVID-19 Genomics UK (COG-UK) Consortium | Ana da Silva Filipe, Natasha Johnson, Kathy Smollett, Daniel Mair, Stephen Carmichael, Lily Tong, Jenna Nichols, Elihu Aranday-Cortes, Kirstyn Brunker, Yasmin Parr, Kyriaki Nomikou; Sarah McDonald, Marc Niebel, Patawee Asamaphan; Richard Orton, Joseph Hughes, Sreenu Vattipally, David L Robertson; Alasdair MacLean, Rory Gunson; Kathy Li, Natasha Jesudason, Rajiv Shah, James Shepherd, Antonia Ho, Emma Thomson |
| EPI_ISL_433314 | hCoV-19/Scotland/CVR599/2020 | Europe / United Kingdom / Scotland | 2020-03-24 | West of Scotland Specialist Virology Centre, NHSGGC / MRC-University of Glasgow Centre for Virus Research | COVID-19 Genomics UK (COG-UK) Consortium | Ana da Silva Filipe, Natasha Johnson, Kathy Smollett, Daniel Mair, Stephen Carmichael, Lily Tong, Jenna Nichols, Elihu Aranday-Cortes, Kirstyn Brunker, Yasmin Parr, Kyriaki Nomikou; Sarah McDonald, Marc Niebel, Patawee Asamaphan; Richard Orton, Joseph Hughes, Sreenu Vattipally, David L Robertson; Alasdair MacLean, Rory Gunson; Kathy Li, Natasha Jesudason, Rajiv Shah, James Shepherd, Antonia Ho, Emma Thomson |
| EPI_ISL_433316 | hCoV-19/Scotland/CVR607/2020 | Europe / United Kingdom / Scotland | 2020-03-25 | West of Scotland Specialist Virology Centre, NHSGGC / MRC-University of Glasgow Centre for Virus Research | COVID-19 Genomics UK (COG-UK) Consortium | Ana da Silva Filipe, Natasha Johnson, Kathy Smollett, Daniel Mair, Stephen Carmichael, Lily Tong, Jenna Nichols, Elihu Aranday-Cortes, Kirstyn Brunker, Yasmin Parr, Kyriaki Nomikou; Sarah McDonald, Marc Niebel, Patawee Asamaphan; Richard Orton, Joseph Hughes, Sreenu Vattipally, David L Robertson; Alasdair MacLean, Rory Gunson; Kathy Li, Natasha Jesudason, Rajiv Shah, James Shepherd, Antonia Ho, Emma Thomson |
| EPI_ISL_433321 | hCoV-19/Scotland/CVR663/2020 | Europe / United Kingdom / Scotland | 2020-03-25 | West of Scotland Specialist Virology Centre, NHSGGC / MRC-University of Glasgow Centre for Virus Research | COVID-19 Genomics UK (COG-UK) Consortium | Ana da Silva Filipe, Natasha Johnson, Kathy Smollett, Daniel Mair, Stephen Carmichael, Lily Tong, Jenna Nichols, Elihu Aranday-Cortes, Kirstyn Brunker, Yasmin Parr, Kyriaki Nomikou; Sarah McDonald, Marc Niebel, Patawee Asamaphan; Richard Orton, Joseph Hughes, Sreenu Vattipally, David L Robertson; Alasdair MacLean, Rory Gunson; Kathy Li, Natasha Jesudason, Rajiv Shah, James Shepherd, Antonia Ho, Emma Thomson |
| EPI_ISL_433323 | hCoV-19/Scotland/CVR672/2020 | Europe / United Kingdom / Scotland | 2020-03-26 | West of Scotland Specialist Virology Centre, NHSGGC / MRC-University of Glasgow Centre for Virus Research | COVID-19 Genomics UK (COG-UK) Consortium | Ana da Silva Filipe, Natasha Johnson, Kathy Smollett, Daniel Mair, Stephen Carmichael, Lily Tong, Jenna Nichols, Elihu Aranday-Cortes, Kirstyn Brunker, Yasmin Parr, Kyriaki Nomikou; Sarah McDonald, Marc Niebel, Patawee Asamaphan; Richard Orton, Joseph Hughes, Sreenu Vattipally, David L Robertson; Alasdair MacLean, Rory Gunson; Kathy Li, Natasha Jesudason, Rajiv Shah, James Shepherd, Antonia Ho, Emma Thomson |
| EPI_ISL_433324 | hCoV-19/Scotland/CVR681/2020 | Europe / United Kingdom / Scotland | 2020-03-25 | West of Scotland Specialist Virology Centre, NHSGGC / MRC-University of Glasgow Centre for Virus Research | COVID-19 Genomics UK (COG-UK) Consortium | Ana da Silva Filipe, Natasha Johnson, Kathy Smollett, Daniel Mair, Stephen Carmichael, Lily Tong, Jenna Nichols, Elihu Aranday-Cortes, Kirstyn Brunker, Yasmin Parr, Kyriaki Nomikou; Sarah McDonald, Marc Niebel, Patawee Asamaphan; Richard Orton, Joseph Hughes, Sreenu Vattipally, David L Robertson; Alasdair MacLean, Rory Gunson; Kathy Li, Natasha Jesudason, Rajiv Shah, James Shepherd, Antonia Ho, Emma Thomson |
| EPI_ISL_434455 | hCoV-19/Greece/133/2020 | Europe / Greece / Athens | 2020-03-09 | Laboratory of Microbiology, Medical School, National and Kapodistrian University of Athens | Laboratory of Biology, Department of Medicine, Democritus University of Thrace | Kassela K., Bampali,M., Dovrolis,N., Gatzidou,E., Froukala,E., Stavropoulou,A., Veletza,S., Tsakris,A., Spanakis,N. and Karakasiliotis,I. |
| EPI_ISL_434456 | hCoV-19/Greece/48/2020 | Europe / Greece / Athens | 2020-03-22 | Laboratory of Microbiology, Medical School, National and Kapodistrian University of Athens | Laboratory of Biology, Department of Medicine, Democritus University of Thrace | Kassela K., Bampali,M., Dovrolis,N., Gatzidou,E., Froukala,E., Stavropoulou,A., Veletza,S., Tsakris,A., Spanakis,N. and Karakasiliotis,I. |
| EPI_ISL_434457 | hCoV-19/Greece/36/2020 | Europe / Greece / Athens | 2020-03-28 | Laboratory of Microbiology, Medical School, National and Kapodistrian University of Athens | Laboratory of Biology, Department of Medicine, Democritus University of Thrace | Kassela K., Bampali,M., Dovrolis,N., Gatzidou,E., Froukala,E., Stavropoulou,A., Veletza,S., Tsakris,A., Spanakis,N. and Karakasiliotis,I. |
| EPI_ISL_434458 | hCoV-19/Greece/37/2020 | Europe / Greece / Athens | 2020-03-20 | Laboratory of Microbiology, Medical School, National and Kapodistrian University of Athens | Laboratory of Biology, Department of Medicine, Democritus University of Thrace | Kassela K., Bampali,M., Dovrolis,N., Gatzidou,E., Froukala,E., Stavropoulou,A., Veletza,S., Tsakris,A., Spanakis,N. and Karakasiliotis,I. |
| EPI_ISL_434459 | hCoV-19/Greece/35/2020 | Europe / Greece / Athens | 2020-03-29 | Laboratory of Microbiology, Medical School, National and Kapodistrian University of Athens | Laboratory of Biology, Department of Medicine, Democritus University of Thrace | Kassela K., Bampali,M., Dovrolis,N., Gatzidou,E., Froukala,E., Stavropoulou,A., Veletza,S., Tsakris,A., Spanakis,N. and Karakasiliotis,I. |
| EPI_ISL_434460 | hCoV-19/Greece/38/2020 | Europe / Greece / Athens | 2020-03-23 | Laboratory of Microbiology, Medical School, National and Kapodistrian University of Athens | Laboratory of Biology, Department of Medicine, Democritus University of Thrace | Kassela K., Bampali,M., Dovrolis,N., Gatzidou,E., Froukala,E., Stavropoulou,A., Veletza,S., Tsakris,A., Spanakis,N. and Karakasiliotis,I. |
| EPI_ISL_434461 | hCoV-19/Greece/39/2020 | Europe / Greece / Athens | 2020-03-29 | Laboratory of Microbiology, Medical School, National and Kapodistrian University of Athens | Laboratory of Biology, Department of Medicine, Democritus University of Thrace | Kassela K., Bampali,M., Dovrolis,N., Gatzidou,E., Froukala,E., Stavropoulou,A., Veletza,S., Tsakris,A., Spanakis,N. and Karakasiliotis,I. |
| EPI_ISL_434462 | hCoV-19/Greece/47/2020 | Europe / Greece / Athens | 2020-03-23 | Laboratory of Microbiology, Medical School, National and Kapodistrian University of Athens | Laboratory of Biology, Department of Medicine, Democritus University of Thrace | Kassela K., Bampali,M., Dovrolis,N., Gatzidou,E., Froukala,E., Stavropoulou,A., Veletza,S., Tsakris,A., Spanakis,N. and Karakasiliotis,I. |
| EPI_ISL_434463 | hCoV-19/Greece/46/2020 | Europe / Greece / Athens | 2020-03-23 | Laboratory of Microbiology, Medical School, National and Kapodistrian University of Athens | Laboratory of Biology, Department of Medicine, Democritus University of Thrace | Kassela K., Bampali,M., Dovrolis,N., Gatzidou,E., Froukala,E., Stavropoulou,A., Veletza,S., Tsakris,A., Spanakis,N. and Karakasiliotis,I. |
| EPI_ISL_434464 | hCoV-19/Greece/145/2020 | Europe / Greece / Athens | 2020-03-16 | Laboratory of Microbiology, Medical School, National and Kapodistrian University of Athens | Laboratory of Biology, Department of Medicine, Democritus University of Thrace | Kassela K., Bampali,M., Dovrolis,N., Gatzidou,E., Froukala,E., Stavropoulou,A., Veletza,S., Tsakris,A., Spanakis,N. and Karakasiliotis,I. |
| EPI_ISL_434465 | hCoV-19/Greece/44/2020 | Europe / Greece / Athens | 2020-03-23 | Laboratory of Microbiology, Medical School, National and Kapodistrian University of Athens | Laboratory of Biology, Department of Medicine, Democritus University of Thrace | Kassela K., Bampali,M., Dovrolis,N., Gatzidou,E., Froukala,E., Stavropoulou,A., Veletza,S., Tsakris,A., Spanakis,N. and Karakasiliotis,I. |
| EPI_ISL_434466 | hCoV-19/Greece/53/2020 | Europe / Greece / Athens | 2020-04-04 | Laboratory of Microbiology, Medical School, National and Kapodistrian University of Athens | Laboratory of Biology, Department of Medicine, Democritus University of Thrace | Kassela K., Bampali,M., Dovrolis,N., Gatzidou,E., Froukala,E., Stavropoulou,A., Veletza,S., Tsakris,A., Spanakis,N. and Karakasiliotis,I. |
| EPI_ISL_434467 | hCoV-19/Greece/141/2020 | Europe / Greece / Athens | 2020-03-12 | Laboratory of Microbiology, Medical School, National and Kapodistrian University of Athens | Laboratory of Biology, Department of Medicine, Democritus University of Thrace | Kassela K., Bampali,M., Dovrolis,N., Gatzidou,E., Froukala,E., Stavropoulou,A., Veletza,S., Tsakris,A., Spanakis,N. and Karakasiliotis,I. |
| EPI_ISL_434468 | hCoV-19/Greece/51/2020 | Europe / Greece / Athens | 2020-03-26 | Laboratory of Microbiology, Medical School, National and Kapodistrian University of Athens | Laboratory of Biology, Department of Medicine, Democritus University of Thrace | Kassela K., Bampali,M., Dovrolis,N., Gatzidou,E., Froukala,E., Stavropoulou,A., Veletza,S., Tsakris,A., Spanakis,N. and Karakasiliotis,I. |
| EPI_ISL_434469 | hCoV-19/Greece/154/2020 | Europe / Greece / Athens | 2020-03-20 | Laboratory of Microbiology, Medical School, National and Kapodistrian University of Athens | Laboratory of Biology, Department of Medicine, Democritus University of Thrace | Kassela K., Bampali,M., Dovrolis,N., Gatzidou,E., Froukala,E., Stavropoulou,A., Veletza,S., Tsakris,A., Spanakis,N. and Karakasiliotis,I. |
| EPI_ISL_434470 | hCoV-19/Greece/155/2020 | Europe / Greece / Athens | 2020-03-19 | Laboratory of Microbiology, Medical School, National and Kapodistrian University of Athens | Laboratory of Biology, Department of Medicine, Democritus University of Thrace | Kassela K., Bampali,M., Dovrolis,N., Gatzidou,E., Froukala,E., Stavropoulou,A., Veletza,S., Tsakris,A., Spanakis,N. and Karakasiliotis,I. |
| EPI_ISL_434471 | hCoV-19/Greece/149/2020 | Europe / Greece / Athens | 2020-03-11 | Laboratory of Microbiology, Medical School, National and Kapodistrian University of Athens | Laboratory of Biology, Department of Medicine, Democritus University of Thrace | Kassela K., Bampali,M., Dovrolis,N., Gatzidou,E., Froukala,E., Stavropoulou,A., Veletza,S., Tsakris,A., Spanakis,N. and Karakasiliotis,I. |
| EPI_ISL_434472 | hCoV-19/Greece/139/2020 | Europe / Greece / Athens | 2020-03-12 | Laboratory of Microbiology, Medical School, National and Kapodistrian University of Athens | Laboratory of Biology, Department of Medicine, Democritus University of Thrace | Kassela K., Bampali,M., Dovrolis,N., Gatzidou,E., Froukala,E., Stavropoulou,A., Veletza,S., Tsakris,A., Spanakis,N. and Karakasiliotis,I. |
| EPI_ISL_434473 | hCoV-19/Greece/136/2020 | Europe / Greece / Athens | 2020-03-11 | Laboratory of Microbiology, Medical School, National and Kapodistrian University of Athens | Laboratory of Biology, Department of Medicine, Democritus University of Thrace | Kassela K., Bampali,M., Dovrolis,N., Gatzidou,E., Froukala,E., Stavropoulou,A., Veletza,S., Tsakris,A., Spanakis,N. and Karakasiliotis,I. |
| EPI_ISL_434474 | hCoV-19/Greece/137/2020 | Europe / Greece / Athens | 2020-03-11 | Laboratory of Microbiology, Medical School, National and Kapodistrian University of Athens | Laboratory of Biology, Department of Medicine, Democritus University of Thrace | Kassela K., Bampali,M., Dovrolis,N., Gatzidou,E., Froukala,E., Stavropoulou,A., Veletza,S., Tsakris,A., Spanakis,N. and Karakasiliotis,I. |
| EPI_ISL_434475 | hCoV-19/Greece/134/2020 | Europe / Greece / Athens | 2020-03-10 | Laboratory of Microbiology, Medical School, National and Kapodistrian University of Athens | Laboratory of Biology, Department of Medicine, Democritus University of Thrace | Kassela K., Bampali,M., Dovrolis,N., Gatzidou,E., Froukala,E., Stavropoulou,A., Veletza,S., Tsakris,A., Spanakis,N. and Karakasiliotis,I. |
| EPI_ISL_434476 | hCoV-19/Greece/135/2020 | Europe / Greece / Athens | 2020-03-10 | Laboratory of Microbiology, Medical School, National and Kapodistrian University of Athens | Laboratory of Biology, Department of Medicine, Democritus University of Thrace | Kassela K., Bampali,M., Dovrolis,N., Gatzidou,E., Froukala,E., Stavropoulou,A., Veletza,S., Tsakris,A., Spanakis,N. and Karakasiliotis,I. |
| EPI_ISL_434477 | hCoV-19/Greece/143/2020 | Europe / Greece / Athens | 2020-03-16 | Laboratory of Microbiology, Medical School, National and Kapodistrian University of Athens | Laboratory of Biology, Department of Medicine, Democritus University of Thrace | Kassela K., Bampali,M., Dovrolis,N., Gatzidou,E., Froukala,E., Stavropoulou,A., Veletza,S., Tsakris,A., Spanakis,N. and Karakasiliotis,I. |
| EPI_ISL_434478 | hCoV-19/Greece/142/2020 | Europe / Greece / Athens | 2020-03-14 | Laboratory of Microbiology, Medical School, National and Kapodistrian University of Athens | Laboratory of Biology, Department of Medicine, Democritus University of Thrace | Kassela K., Bampali,M., Dovrolis,N., Gatzidou,E., Froukala,E., Stavropoulou,A., Veletza,S., Tsakris,A., Spanakis,N. and Karakasiliotis,I. |
| EPI_ISL_434479 | hCoV-19/Greece/138/2020 | Europe / Greece / Athens | 2020-03-11 | Laboratory of Microbiology, Medical School, National and Kapodistrian University of Athens | Laboratory of Biology, Department of Medicine, Democritus University of Thrace | Kassela K., Bampali,M., Dovrolis,N., Gatzidou,E., Froukala,E., Stavropoulou,A., Veletza,S., Tsakris,A., Spanakis,N. and Karakasiliotis,I. |
| EPI_ISL_434480 | hCoV-19/Greece/140/2020 | Europe / Greece / Athens | 2020-03-12 | Laboratory of Microbiology, Medical School, National and Kapodistrian University of Athens | Laboratory of Biology, Department of Medicine, Democritus University of Thrace | Kassela K., Bampali,M., Dovrolis,N., Gatzidou,E., Froukala,E., Stavropoulou,A., Veletza,S., Tsakris,A., Spanakis,N. and Karakasiliotis,I. |
| EPI_ISL_434481 | hCoV-19/Greece/146/2020 | Europe / Greece / Athens | 2020-03-19 | Laboratory of Microbiology, Medical School, National and Kapodistrian University of Athens | Laboratory of Biology, Department of Medicine, Democritus University of Thrace | Kassela K., Bampali,M., Dovrolis,N., Gatzidou,E., Froukala,E., Stavropoulou,A., Veletza,S., Tsakris,A., Spanakis,N. and Karakasiliotis,I. |
| EPI_ISL_434482 | hCoV-19/Greece/156/2020 | Europe / Greece / Athens | 2020-03-16 | Laboratory of Microbiology, Medical School, National and Kapodistrian University of Athens | Laboratory of Biology, Department of Medicine, Democritus University of Thrace | Kassela K., Bampali,M., Dovrolis,N., Gatzidou,E., Froukala,E., Stavropoulou,A., Veletza,S., Tsakris,A., Spanakis,N. and Karakasiliotis,I. |
| EPI_ISL_434483 | hCoV-19/Greece/144/2020 | Europe / Greece / Athens | 2020-03-16 | Laboratory of Microbiology, Medical School, National and Kapodistrian University of Athens | Laboratory of Biology, Department of Medicine, Democritus University of Thrace | Kassela K., Bampali,M., Dovrolis,N., Gatzidou,E., Froukala,E., Stavropoulou,A., Veletza,S., Tsakris,A., Spanakis,N. and Karakasiliotis,I. |
| EPI_ISL_434484 | hCoV-19/Greece/148/2020 | Europe / Greece / Athens | 2020-03-09 | Laboratory of Microbiology, Medical School, National and Kapodistrian University of Athens | Laboratory of Biology, Department of Medicine, Democritus University of Thrace | Kassela K., Bampali,M., Dovrolis,N., Gatzidou,E., Froukala,E., Stavropoulou,A., Veletza,S., Tsakris,A., Spanakis,N. and Karakasiliotis,I. |
| EPI_ISL_434485 | hCoV-19/Greece/150/2020 | Europe / Greece / Athens | 2020-03-12 | Laboratory of Microbiology, Medical School, National and Kapodistrian University of Athens | Laboratory of Biology, Department of Medicine, Democritus University of Thrace | Kassela K., Bampali,M., Dovrolis,N., Gatzidou,E., Froukala,E., Stavropoulou,A., Veletza,S., Tsakris,A., Spanakis,N. and Karakasiliotis,I. |
| EPI_ISL_434486 | hCoV-19/Greece/151/2020 | Europe / Greece / Athens | 2020-03-13 | Laboratory of Microbiology, Medical School, National and Kapodistrian University of Athens | Laboratory of Biology, Department of Medicine, Democritus University of Thrace | Kassela K., Bampali,M., Dovrolis,N., Gatzidou,E., Froukala,E., Stavropoulou,A., Veletza,S., Tsakris,A., Spanakis,N. and Karakasiliotis,I. |
| EPI_ISL_434487 | hCoV-19/Luxembourg/LNS0972189/2020 | Europe / Luxembourg | 2020-04-13 | Laboratoire National de Sante, Microbiology, Virology | Laboratoire National de Sante, Microbiology, Epidemiology and Microbial Genomics | Anke Wienecke-Baldacchino, Ardashel Latsuzbaia, Jessica Tapp, Catherine Ragimbeau, Guillaume Fournier, Tamir Abdelrahman, Trung Nguyen Nguyen, Joel Mossong |
| EPI_ISL_434488 | hCoV-19/Luxembourg/LNS1673015/2020 | Europe / Luxembourg | 2020-04-21 | Laboratoire National de Sante, Microbiology, Virology | Laboratoire National de Sante, Microbiology, Epidemiology and Microbial Genomics | Anke Wienecke-Baldacchino, Ardashel Latsuzbaia, Jessica Tapp, Catherine Ragimbeau, Guillaume Fournier, Tamir Abdelrahman, Trung Nguyen Nguyen, Joel Mossong |
| EPI_ISL_434489 | hCoV-19/Luxembourg/LNS8742871/2020 | Europe / Luxembourg | 2020-04-13 | Laboratoire National de Sante, Microbiology, Virology | Laboratoire National de Sante, Microbiology, Epidemiology and Microbial Genomics | Anke Wienecke-Baldacchino, Ardashel Latsuzbaia, Jessica Tapp, Catherine Ragimbeau, Guillaume Fournier, Tamir Abdelrahman, Trung Nguyen Nguyen, Joel Mossong |
| EPI_ISL_434491 | hCoV-19/Luxembourg/LNS8258882/2020 | Europe / Luxembourg | 2020-04-14 | Laboratoire National de Sante, Microbiology, Virology | Laboratoire National de Sante, Microbiology, Epidemiology and Microbial Genomics | Anke Wienecke-Baldacchino, Ardashel Latsuzbaia, Jessica Tapp, Catherine Ragimbeau, Guillaume Fournier, Tamir Abdelrahman, Trung Nguyen Nguyen, Joel Mossong |
| EPI_ISL_434492 | hCoV-19/Luxembourg/LNS8655659/2020 | Europe / Luxembourg | 2020-04-12 | Laboratoire National de Sante, Microbiology, Virology | Laboratoire National de Sante, Microbiology, Epidemiology and Microbial Genomics | Anke Wienecke-Baldacchino, Ardashel Latsuzbaia, Jessica Tapp, Catherine Ragimbeau, Guillaume Fournier, Tamir Abdelrahman, Trung Nguyen Nguyen, Joel Mossong |
| EPI_ISL_434493 | hCoV-19/Luxembourg/LNS1368952/2020 | Europe / Luxembourg | 2020-04-11 | Laboratoire National de Sante, Microbiology, Virology | Laboratoire National de Sante, Microbiology, Epidemiology and Microbial Genomics | Anke Wienecke-Baldacchino, Ardashel Latsuzbaia, Jessica Tapp, Catherine Ragimbeau, Guillaume Fournier, Tamir Abdelrahman, Trung Nguyen Nguyen, Joel Mossong |
| EPI_ISL_434495 | hCoV-19/Luxembourg/LNS0188413/2020 | Europe / Luxembourg | 2020-04-12 | Laboratoire National de Sante, Microbiology, Virology | Laboratoire National de Sante, Microbiology, Epidemiology and Microbial Genomics | Anke Wienecke-Baldacchino, Ardashel Latsuzbaia, Jessica Tapp, Catherine Ragimbeau, Guillaume Fournier, Tamir Abdelrahman, Trung Nguyen Nguyen, Joel Mossong |
| EPI_ISL_434496 | hCoV-19/Luxembourg/LNS0361493/2020 | Europe / Luxembourg | 2020-04-10 | Laboratoire National de Sante, Microbiology, Virology | Laboratoire National de Sante, Microbiology, Epidemiology and Microbial Genomics | Anke Wienecke-Baldacchino, Ardashel Latsuzbaia, Jessica Tapp, Catherine Ragimbeau, Guillaume Fournier, Tamir Abdelrahman, Trung Nguyen Nguyen, Joel Mossong |
| EPI_ISL_434497 | hCoV-19/Luxembourg/LNS2209136/2020 | Europe / Luxembourg | 2020-04-12 | Laboratoire National de Sante, Microbiology, Virology | Laboratoire National de Sante, Microbiology, Epidemiology and Microbial Genomics | Anke Wienecke-Baldacchino, Ardashel Latsuzbaia, Jessica Tapp, Catherine Ragimbeau, Guillaume Fournier, Tamir Abdelrahman, Trung Nguyen Nguyen, Joel Mossong |
| EPI_ISL_434498 | hCoV-19/Luxembourg/LNS4757989/2020 | Europe / Luxembourg | 2020-04-15 | Laboratoire National de Sante, Microbiology, Virology | Laboratoire National de Sante, Microbiology, Epidemiology and Microbial Genomics | Anke Wienecke-Baldacchino, Ardashel Latsuzbaia, Jessica Tapp, Catherine Ragimbeau, Guillaume Fournier, Tamir Abdelrahman, Trung Nguyen Nguyen, Joel Mossong |
| EPI_ISL_434500 | hCoV-19/Luxembourg/LNS2818424/2020 | Europe / Luxembourg | 2020-04-15 | Laboratoire National de Sante, Microbiology, Virology | Laboratoire National de Sante, Microbiology, Epidemiology and Microbial Genomics | Anke Wienecke-Baldacchino, Ardashel Latsuzbaia, Jessica Tapp, Catherine Ragimbeau, Guillaume Fournier, Tamir Abdelrahman, Trung Nguyen Nguyen, Joel Mossong |
| EPI_ISL_434502 | hCoV-19/Luxembourg/LNS5807524/2020 | Europe / Luxembourg | 2020-04-13 | Laboratoire National de Sante, Microbiology, Virology | Laboratoire National de Sante, Microbiology, Epidemiology and Microbial Genomics | Anke Wienecke-Baldacchino, Ardashel Latsuzbaia, Jessica Tapp, Catherine Ragimbeau, Guillaume Fournier, Tamir Abdelrahman, Trung Nguyen Nguyen, Joel Mossong |
| EPI_ISL_434503 | hCoV-19/Luxembourg/LNS0364174/2020 | Europe / Luxembourg | 2020-04-12 | Laboratoire National de Sante, Microbiology, Virology | Laboratoire National de Sante, Microbiology, Epidemiology and Microbial Genomics | Anke Wienecke-Baldacchino, Ardashel Latsuzbaia, Jessica Tapp, Catherine Ragimbeau, Guillaume Fournier, Tamir Abdelrahman, Trung Nguyen Nguyen, Joel Mossong |
| EPI_ISL_434504 | hCoV-19/Luxembourg/LNS2458393/2020 | Europe / Luxembourg | 2020-04-14 | Laboratoire National de Sante, Microbiology, Virology | Laboratoire National de Sante, Microbiology, Epidemiology and Microbial Genomics | Anke Wienecke-Baldacchino, Ardashel Latsuzbaia, Jessica Tapp, Catherine Ragimbeau, Guillaume Fournier, Tamir Abdelrahman, Trung Nguyen Nguyen, Joel Mossong |
| EPI_ISL_434505 | hCoV-19/Luxembourg/LNS9470500/2020 | Europe / Luxembourg | 2020-04-08 | Laboratoire National de Sante, Microbiology, Virology | Laboratoire National de Sante, Microbiology, Epidemiology and Microbial Genomics | Anke Wienecke-Baldacchino, Ardashel Latsuzbaia, Jessica Tapp, Catherine Ragimbeau, Guillaume Fournier, Tamir Abdelrahman, Trung Nguyen Nguyen, Joel Mossong |
| EPI_ISL_434506 | hCoV-19/Luxembourg/LNS6690193/2020 | Europe / Luxembourg | 2020-04-15 | Laboratoire National de Sante, Microbiology, Virology | Laboratoire National de Sante, Microbiology, Epidemiology and Microbial Genomics | Anke Wienecke-Baldacchino, Ardashel Latsuzbaia, Jessica Tapp, Catherine Ragimbeau, Guillaume Fournier, Tamir Abdelrahman, Trung Nguyen Nguyen, Joel Mossong |
| EPI_ISL_434507 | hCoV-19/Luxembourg/LNS9175349/2020 | Europe / Luxembourg | 2020-04-12 | Laboratoire National de Sante, Microbiology, Virology | Laboratoire National de Sante, Microbiology, Epidemiology and Microbial Genomics | Anke Wienecke-Baldacchino, Ardashel Latsuzbaia, Jessica Tapp, Catherine Ragimbeau, Guillaume Fournier, Tamir Abdelrahman, Trung Nguyen Nguyen, Joel Mossong |
| EPI_ISL_434511 | hCoV-19/Luxembourg/LNS2435702/2020 | Europe / Luxembourg | 2020-04-21 | Laboratoire National de Sante, Microbiology, Virology | Laboratoire National de Sante, Microbiology, Epidemiology and Microbial Genomics | Anke Wienecke-Baldacchino, Ardashel Latsuzbaia, Jessica Tapp, Catherine Ragimbeau, Guillaume Fournier, Tamir Abdelrahman, Trung Nguyen Nguyen, Joel Mossong |
| EPI_ISL_434512 | hCoV-19/Luxembourg/LNS9272998/2020 | Europe / Luxembourg | 2020-04-17 | Laboratoire National de Sante, Microbiology, Virology | Laboratoire National de Sante, Microbiology, Epidemiology and Microbial Genomics | Anke Wienecke-Baldacchino, Ardashel Latsuzbaia, Jessica Tapp, Catherine Ragimbeau, Guillaume Fournier, Tamir Abdelrahman, Trung Nguyen Nguyen, Joel Mossong |
| EPI_ISL_434514 | hCoV-19/Luxembourg/LNS2629386/2020 | Europe / Luxembourg | 2020-04-16 | Laboratoire National de Sante, Microbiology, Virology | Laboratoire National de Sante, Microbiology, Epidemiology and Microbial Genomics | Anke Wienecke-Baldacchino, Ardashel Latsuzbaia, Jessica Tapp, Catherine Ragimbeau, Guillaume Fournier, Tamir Abdelrahman, Trung Nguyen Nguyen, Joel Mossong |
| EPI_ISL_434515 | hCoV-19/Luxembourg/LNS5953507/2020 | Europe / Luxembourg | 2020-04-15 | Laboratoire National de Sante, Microbiology, Virology | Laboratoire National de Sante, Microbiology, Epidemiology and Microbial Genomics | Anke Wienecke-Baldacchino, Ardashel Latsuzbaia, Jessica Tapp, Catherine Ragimbeau, Guillaume Fournier, Tamir Abdelrahman, Trung Nguyen Nguyen, Joel Mossong |
| EPI_ISL_434516 | hCoV-19/Jordan/SR-084/2020 | Asia / Jordan / Amman | 2020-03-16 | Biolab Diagnostic Laboratories | Andersen lab at Scripps Research | Issa Abu-Dayyeh, Ahmad Tibi, Lama Hussein, Lina Mohammad, Zein Naber, Amid Abdelnour with SEARCH Alliance San Diego |
| EPI_ISL_434517 | hCoV-19/USA/SEARCH-0058-NBG/2020 | North America / USA / Louisiana / New O | 2020-03-23 | Robert Garry lab | Andersen lab at Scripps Research | Allison Smither, Gilberto Sabino-Santos, Patricia Snarski, Lilia Melnik, Antoinette Bell, Kaylynn Genemaras, Arnaud Drouin, Dahlene Fusco, Robert Garry with SEARCH Alliance San Diego |
| EPI_ISL_434518 | hCoV-19/USA/SEARCH-0059-NBG/2020 | North America / USA / Louisiana / New O | 2020-03-23 | Robert Garry lab | Andersen lab at Scripps Research | Allison Smither, Gilberto Sabino-Santos, Patricia Snarski, Lilia Melnik, Antoinette Bell, Kaylynn Genemaras, Arnaud Drouin, Dahlene Fusco, Robert Garry with SEARCH Alliance San Diego |
| EPI_ISL_434519 | hCoV-19/USA/SEARCH-0060-NBG/2020 | North America / USA / Louisiana / New O | 2020-03-25 | Robert Garry lab | Andersen lab at Scripps Research | Allison Smither, Gilberto Sabino-Santos, Patricia Snarski, Lilia Melnik, Antoinette Bell, Kaylynn Genemaras, Arnaud Drouin, Dahlene Fusco, Robert Garry with SEARCH Alliance San Diego |
| EPI_ISL_434526 | hCoV-19/USA/SEARCH-0070-NBG/2020 | North America / USA / Louisiana / New O | 2020-03-26 | Robert Garry lab | Andersen lab at Scripps Research | Allison Smither, Gilberto Sabino-Santos, Patricia Snarski, Lilia Melnik, Antoinette Bell, Kaylynn Genemaras, Arnaud Drouin, Dahlene Fusco, Robert Garry with SEARCH Alliance San Diego |
| EPI_ISL_434527 | hCoV-19/USA/SEARCH-0071-NBG/2020 | North America / USA / Louisiana / New O | 2020-03-26 | Robert Garry lab | Andersen lab at Scripps Research | Allison Smither, Gilberto Sabino-Santos, Patricia Snarski, Lilia Melnik, Antoinette Bell, Kaylynn Genemaras, Arnaud Drouin, Dahlene Fusco, Robert Garry with SEARCH Alliance San Diego |
| EPI_ISL_434530 | hCoV-19/USA/SEARCH-0076-NBG/2020 | North America / USA / Louisiana / New O | 2020-03-31 | Robert Garry lab | Andersen lab at Scripps Research | Allison Smither, Gilberto Sabino-Santos, Patricia Snarski, Lilia Melnik, Antoinette Bell, Kaylynn Genemaras, Arnaud Drouin, Dahlene Fusco, Robert Garry with SEARCH Alliance San Diego |
| EPI_ISL_434533 | hCoV-19/Costa Rica/01/2020 | Central America / Costa Rica | 2020-03-16 | Area de Salud Alajuela Sur | Inciensa, Instituto Costarricense de Investigación y Enseñanza en Nutrición y Salud | Francisco Duarte, Hebleen Porras, Claudio Soto-Garita, Estela Cordero, Adriana Godinez & Melany Calderon |
| EPI_ISL_434534 | hCoV-19/Wuhan/IVDC-HB-GX02/2019 | Asia / China / Hubei / Wuhan | 2019-12-30 | National Institute for Viral Disease Control and Prevention, China CDC | National Institute for Viral Disease Control and Prevention, China CDC, Yunnan Provincial CDC | Wenjie Tan, Roujian Lu, Wenling Wang, Peihua Niu, Huijuan Wang, Baoying Huang, Li Zhao, Fei Ye, Guizhen Wu |
| EPI_ISL_434600 | hCoV-19/USA/VA-DCLS-0108/2020 | North America / USA / Virginia | 2020-04-14 | Virginia DCLS | Virginia DCLS | Virginia DCLS |
| EPI_ISL_434601 | hCoV-19/USA/VA-DCLS-0109/2020 | North America / USA / Virginia | 2020-04-08 | Virginia DCLS | Virginia DCLS | Virginia DCLS |
| EPI_ISL_434602 | hCoV-19/USA/VA-DCLS-0110/2020 | North America / USA / Virginia | 2020-04-08 | Virginia DCLS | Virginia DCLS | Virginia DCLS |
| EPI_ISL_434603 | hCoV-19/USA/VA-DCLS-0111/2020 | North America / USA / Virginia | 2020-04-08 | Virginia DCLS | Virginia DCLS | Virginia DCLS |
| EPI_ISL_434604 | hCoV-19/USA/VA-DCLS-0112/2020 | North America / USA / Virginia | 2020-04-08 | Virginia DCLS | Virginia DCLS | Virginia DCLS |
| EPI_ISL_434605 | hCoV-19/USA/VA-DCLS-0114/2020 | North America / USA / Virginia | 2020-04-09 | Virginia DCLS | Virginia DCLS | Virginia DCLS |
| EPI_ISL_434607 | hCoV-19/USA/WI-UW-261/2020 | North America / USA / Wisconsin | 2020-03-26 | University of Wisconsin-Madison AIDS Vaccine Research Laboratories | University of Wisconsin-Madison AIDS Vaccine Research Laboratories | Gage Moreno, Katarina Braun, et al. AIDS Vaccine Research Laboratories |
| EPI_ISL_434608 | hCoV-19/USA/WI-UW-262/2020 | North America / USA / Wisconsin | 2020-04-10 | University of Wisconsin-Madison AIDS Vaccine Research Laboratories | University of Wisconsin-Madison AIDS Vaccine Research Laboratories | Gage Moreno, Katarina Braun, et al. AIDS Vaccine Research Laboratories |
| EPI_ISL_434609 | hCoV-19/USA/WI-UW-263/2020 | North America / USA / Wisconsin | 2020-04-08 | University of Wisconsin-Madison AIDS Vaccine Research Laboratories | University of Wisconsin-Madison AIDS Vaccine Research Laboratories | Gage Moreno, Katarina Braun, et al. AIDS Vaccine Research Laboratories |
| EPI_ISL_434610 | hCoV-19/USA/WI-UW-264/2020 | North America / USA / Wisconsin | 2020-04-12 | University of Wisconsin-Madison AIDS Vaccine Research Laboratories | University of Wisconsin-Madison AIDS Vaccine Research Laboratories | Gage Moreno, Katarina Braun, et al. AIDS Vaccine Research Laboratories |
| EPI_ISL_434611 | hCoV-19/USA/WI-UW-265/2020 | North America / USA / Wisconsin | 2020-04-08 | University of Wisconsin-Madison AIDS Vaccine Research Laboratories | University of Wisconsin-Madison AIDS Vaccine Research Laboratories | Gage Moreno, Katarina Braun, et al. AIDS Vaccine Research Laboratories |
| EPI_ISL_434612 | hCoV-19/USA/WI-UW-266/2020 | North America / USA / Wisconsin | 2020-04-07 | University of Wisconsin-Madison AIDS Vaccine Research Laboratories | University of Wisconsin-Madison AIDS Vaccine Research Laboratories | Gage Moreno, Katarina Braun, et al. AIDS Vaccine Research Laboratories |
| EPI_ISL_434613 | hCoV-19/USA/WI-UW-267/2020 | North America / USA / Wisconsin | 2020-04-08 | University of Wisconsin-Madison AIDS Vaccine Research Laboratories | University of Wisconsin-Madison AIDS Vaccine Research Laboratories | Gage Moreno, Katarina Braun, et al. AIDS Vaccine Research Laboratories |
| EPI_ISL_434614 | hCoV-19/USA/WI-UW-268/2020 | North America / USA / Wisconsin | 2020-04-06 | University of Wisconsin-Madison AIDS Vaccine Research Laboratories | University of Wisconsin-Madison AIDS Vaccine Research Laboratories | Gage Moreno, Katarina Braun, et al. AIDS Vaccine Research Laboratories |
| EPI_ISL_434615 | hCoV-19/USA/WI-UW-269/2020 | North America / USA / Wisconsin | 2020-04-06 | University of Wisconsin-Madison AIDS Vaccine Research Laboratories | University of Wisconsin-Madison AIDS Vaccine Research Laboratories | Gage Moreno, Katarina Braun, et al. AIDS Vaccine Research Laboratories |
| EPI_ISL_434616 | hCoV-19/France/OCC-1/2020 | Europe / France / Occitanie | 2020-03 | CHU Purpan - Laboratoire de Virologie - Institut Fédératif de Biologie | Laboratoire de virologie - École Nationale Vétérinaire de Toulouse | Guillaume Croville, Jean-Luc Guérin, Jacques Izopet |
| EPI_ISL_434617 | hCoV-19/France/OCC-2/2020 | Europe / France / Occitanie | 2020-03 | CHU Purpan - Laboratoire de Virologie - Institut Fédératif de Biologie | Laboratoire de virologie - École Nationale Vétérinaire de Toulouse | Guillaume Croville, Jean-Luc Guérin, Jacques Izopet |
| EPI_ISL_434618 | hCoV-19/France/OCC-3/2020 | Europe / France / Occitanie | 2020-03 | CHU Purpan - Laboratoire de Virologie - Institut Fédératif de Biologie | Laboratoire de virologie - École Nationale Vétérinaire de Toulouse | Guillaume Croville, Jean-Luc Guérin, Jacques Izopet |
| EPI_ISL_434619 | hCoV-19/France/OCC-4/2020 | Europe / France / Occitanie | 2020-03 | CHU Purpan - Laboratoire de Virologie - Institut Fédératif de Biologie | Laboratoire de virologie - École Nationale Vétérinaire de Toulouse | Guillaume Croville, Jean-Luc Guérin, Jacques Izopet |
| EPI_ISL_434620 | hCoV-19/France/OCC-5/2020 | Europe / France / Occitanie | 2020-03 | CHU Purpan - Laboratoire de Virologie - Institut Fédératif de Biologie | Laboratoire de virologie - École Nationale Vétérinaire de Toulouse | Guillaume Croville, Jean-Luc Guérin, Jacques Izopet |
| EPI_ISL_434621 | hCoV-19/France/OCC-6/2020 | Europe / France / Occitanie | 2020-03 | CHU Purpan - Laboratoire de Virologie - Institut Fédératif de Biologie | Laboratoire de virologie - École Nationale Vétérinaire de Toulouse | Guillaume Croville, Jean-Luc Guérin, Jacques Izopet |
| EPI_ISL_434622 | hCoV-19/France/OCC-7/2020 | Europe / France / Occitanie | 2020-03 | CHU Purpan - Laboratoire de Virologie - Institut Fédératif de Biologie | Laboratoire de virologie - École Nationale Vétérinaire de Toulouse | Guillaume Croville, Jean-Luc Guérin, Jacques Izopet |
| EPI_ISL_434623 | hCoV-19/France/OCC-8/2020 | Europe / France / Occitanie | 2020-03 | CHU Purpan - Laboratoire de Virologie - Institut Fédératif de Biologie | Laboratoire de virologie - École Nationale Vétérinaire de Toulouse | Guillaume Croville, Jean-Luc Guérin, Jacques Izopet |
| EPI_ISL_434624 | hCoV-19/France/OCC-9/2020 | Europe / France / Occitanie | 2020-03 | CHU Purpan - Laboratoire de Virologie - Institut Fédératif de Biologie | Laboratoire de virologie - École Nationale Vétérinaire de Toulouse | Guillaume Croville, Jean-Luc Guérin, Jacques Izopet |
| EPI_ISL_434625 | hCoV-19/France/OCC-10/2020 | Europe / France / Occitanie | 2020-03 | CHU Purpan - Laboratoire de Virologie - Institut Fédératif de Biologie | Laboratoire de virologie - École Nationale Vétérinaire de Toulouse | Guillaume Croville, Jean-Luc Guérin, Jacques Izopet |
| EPI_ISL_434626 | hCoV-19/France/OCC-11/2020 | Europe / France / Occitanie | 2020-03-26 | CHU Purpan - Laboratoire de Virologie - Institut Fédératif de Biologie | Laboratoire de virologie - École Nationale Vétérinaire de Toulouse | Guillaume Croville, Jean-Luc Guérin, Jacques Izopet |
| EPI_ISL_434627 | hCoV-19/France/OCC-12/2020 | Europe / France / Occitanie | 2020-03-26 | CHU Purpan - Laboratoire de Virologie - Institut Fédératif de Biologie | Laboratoire de virologie - École Nationale Vétérinaire de Toulouse | Guillaume Croville, Jean-Luc Guérin, Jacques Izopet |
| EPI_ISL_434628 | hCoV-19/France/OCC-13/2020 | Europe / France / Occitanie | 2020-04-07 | CHU Purpan - Laboratoire de Virologie - Institut Fédératif de Biologie | Laboratoire de virologie - École Nationale Vétérinaire de Toulouse | Guillaume Croville, Jean-Luc Guérin, Jacques Izopet |
| EPI_ISL_434629 | hCoV-19/France/OCC-14/2020 | Europe / France / Occitanie | 2020-04-07 | CHU Purpan - Laboratoire de Virologie - Institut Fédératif de Biologie | Laboratoire de virologie - École Nationale Vétérinaire de Toulouse | Guillaume Croville, Jean-Luc Guérin, Jacques Izopet |
| EPI_ISL_434630 | hCoV-19/France/OCC-15/2020 | Europe / France / Occitanie | 2020-04-08 | CHU Purpan - Laboratoire de Virologie - Institut Fédératif de Biologie | Laboratoire de virologie - École Nationale Vétérinaire de Toulouse | Guillaume Croville, Jean-Luc Guérin, Jacques Izopet |
| EPI_ISL_434631 | hCoV-19/France/OCC-16/2020 | Europe / France / Occitanie | 2020-04-08 | CHU Purpan - Laboratoire de Virologie - Institut Fédératif de Biologie | Laboratoire de virologie - École Nationale Vétérinaire de Toulouse | Guillaume Croville, Jean-Luc Guérin, Jacques Izopet |
| EPI_ISL_434632 | hCoV-19/France/OCC-17/2020 | Europe / France / Occitanie | 2020-04-09 | CHU Purpan - Laboratoire de Virologie - Institut Fédératif de Biologie | Laboratoire de virologie - École Nationale Vétérinaire de Toulouse | Guillaume Croville, Jean-Luc Guérin, Jacques Izopet |
| EPI_ISL_434633 | hCoV-19/France/OCC-18/2020 | Europe / France / Occitanie | 2020-04-09 | CHU Purpan - Laboratoire de Virologie - Institut Fédératif de Biologie | Laboratoire de virologie - École Nationale Vétérinaire de Toulouse | Guillaume Croville, Jean-Luc Guérin, Jacques Izopet |
| EPI_ISL_434634 | hCoV-19/France/OCC-19/2020 | Europe / France / Occitanie | 2020-04-11 | CHU Purpan - Laboratoire de Virologie - Institut Fédératif de Biologie | Laboratoire de virologie - École Nationale Vétérinaire de Toulouse | Guillaume Croville, Jean-Luc Guérin, Jacques Izopet |
| EPI_ISL_434635 | hCoV-19/France/OCC-20/2020 | Europe / France / Occitanie | 2020-04-11 | CHU Purpan - Laboratoire de Virologie - Institut Fédératif de Biologie | Laboratoire de virologie - École Nationale Vétérinaire de Toulouse | Guillaume Croville, Jean-Luc Guérin, Jacques Izopet |
| EPI_ISL_434636 | hCoV-19/USA/FL-UF1/2020 | North America / USA / Florida | 2020-03-10 | Lednicky Laboratory, Emerging Pathogens Institute, University of Florida | Lednicky Laboratory at Emerging Pathogens Institute, University of Florida | Elbadry,M.A.; Subramaniam,K.; Waltzek,T.B.; Stephenson,C.J.;Gibson,J.C.; Alam,M.; Morris,J.G. Jr. and Lednicky,J.A. |
| EPI_ISL_434637 | hCoV-19/USA/FL-UF2/2020 | North America / USA / Florida | 2020-03-27 | Lednicky Laboratory, Emerging Pathogens Institute, University of Florida. | Lednicky Laboratory, Emerging Pathogens Institute, University of Florida. | Elbadry,M.A.; Subramaniam,K.; Waltzek,T.B.; Gibson,J.C.; Stephenson,C.J.; Morris,J.G. Jr. and Lednicky,J.A. |
| EPI_ISL_434638 | hCoV-19/USA/MD-HP00041/2020 | North America / USA / Maryland | 2020-03-19 | Johns Hopkins Hospital Department of Pathology | Johns Hopkins Hospital Department of Pathology | Peter M. Thielen, Thomas Mehoke, Shirlee Wohl, Srividya Ramakrishnan, Oluwaseun Nwulia-Falade, Amanda Emlund, Melanie Kirsche, Paul Morris, Norah Sadowski, Nídia Trovao, Victoria Gniazdowski, Michael Schatz, Stuart C. Ray, Winston Timp, Heba Mostafa |
| EPI_ISL_434639 | hCoV-19/USA/DC-HP00043/2020 | North America / USA / District of Columbi | 2020-03-22 | Johns Hopkins Hospital Department of Pathology | Johns Hopkins Hospital Department of Pathology | Peter M. Thielen, Thomas Mehoke, Shirlee Wohl, Srividya Ramakrishnan, Oluwaseun Nwulia-Falade, Amanda Emlund, Melanie Kirsche, Paul Morris, Norah Sadowski, Nídia Trovao, Victoria Gniazdowski, Michael Schatz, Stuart C. Ray, Winston Timp, Heba Mostafa |
| EPI_ISL_434640 | hCoV-19/USA/MD-HP00044/2020 | North America / USA / Maryland | 2020-03-24 | Johns Hopkins Hospital Department of Pathology | Johns Hopkins Hospital Department of Pathology | Peter M. Thielen, Thomas Mehoke, Shirlee Wohl, Srividya Ramakrishnan, Oluwaseun Nwulia-Falade, Amanda Emlund, Melanie Kirsche, Paul Morris, Norah Sadowski, Nídia Trovao, Victoria Gniazdowski, Michael Schatz, Stuart C. Ray, Winston Timp, Heba Mostafa |
| EPI_ISL_434641 | hCoV-19/Sweden/20-08682/2020 | Europe / Sweden / Norrbotten | 2020-03-25 | Laboratoriemedicin | The Public Health Agency of Sweden | Oskar Karlsson Lindsjo, Maria Lind Karlberg, Anna-Malin Linde, Olov Svartstrom, Anna Risberg, Shaman Muradrasoli, Karin Tegmark-Wisell |
| EPI_ISL_434642 | hCoV-19/Sweden/20-08686/2020 | Europe / Sweden / Norrbotten | 2020-03-27 | Laboratoriemedicin | The Public Health Agency of Sweden | Oskar Karlsson Lindsjo, Maria Lind Karlberg, Anna-Malin Linde, Olov Svartstrom, Anna Risberg, Shaman Muradrasoli, Karin Tegmark-Wisell |
| EPI_ISL_434643 | hCoV-19/Sweden/20-05448/2020 | Europe / Sweden / Uppsala | 2020-03-10 | Uppsala Narakut Aleris | The Public Health Agency of Sweden | Annika Nilsson, Oskar Karlsson Lindsjo, Maria Lind Karlberg, Anna-Malin Linde, Olov Svartstrom, Anna Risberg, Theresa Enkirch, Mia Brytting, Karin Tegmark-Wisell |
| EPI_ISL_434644 | hCoV-19/Sweden/20-05469/2020 | Europe / Sweden / Stockholm | 2020-03-10 | Kungsholmsdoktorn | The Public Health Agency of Sweden | Linus Hammar, Oskar Karlsson Lindsjo, Maria Lind Karlberg, Anna-Malin Linde, Olov Svartstrom, Anna Risberg, Theresa Enkirch, Mia Brytting, Karin Tegmark-Wisell |
| EPI_ISL_434645 | hCoV-19/Sweden/20-05627/2020 | Europe / Sweden / Dalarna | 2020-03-13 | Svardsjo VC | The Public Health Agency of Sweden | Tommy Janers, Oskar Karlsson Lindsjo, Maria Lind Karlberg, Anna-Malin Linde, Olov Svartstrom, Anna Risberg, Theresa Enkirch, Mia Brytting, Karin Tegmark-Wisell |
| EPI_ISL_434646 | hCoV-19/Sweden/20-05653/2020 | Europe / Sweden / Uppsala | 2020-03-12 | Ulltuna Vardcentral | The Public Health Agency of Sweden | Heidi Lindback, Oskar Karlsson Lindsjo, Maria Lind Karlberg, Anna-Malin Linde, Olov Svartstrom, Anna Risberg, Theresa Enkirch, Mia Brytting, Karin Tegmark-Wisell |
| EPI_ISL_434647 | hCoV-19/Sweden/20-05721/2020 | Europe / Sweden / Skane | 2020-03-16 | Victoria Vard och Halsa | The Public Health Agency of Sweden | Sarah Henriksson, Oskar Karlsson Lindsjo, Maria Lind Karlberg, Anna-Malin Linde, Olov Svartstrom, Anna Risberg, Theresa Enkirch, Mia Brytting, Karin Tegmark-Wisell |

| | | | | | | |
|---|---|---|---|---|---|---|
| EPI_ISL_434648 | hCoV-19/Sweden/20-05847/2020 | Europe / Sweden / Skane | 2020-03-16 | Victoria Vard och Halsa | The Public Health Agency of Sweden | Sarah Henriksson, Oskar Karlsson Lindsjo, Maria Lind Karlberg, Anna-Malin Linde, Olov Svartstrom, Anna Risberg, Theresa Enkirch, Mia Brytting, Karin Tegmark-Wisell |
| EPI_ISL_434649 | hCoV-19/Sweden/20-05856/2020 | Europe / Sweden / Dalarna | 2020-03-17 | Svardsjo VC | The Public Health Agency of Sweden | Tommy Janers, Oskar Karlsson Lindsjo, Maria Lind Karlberg, Anna-Malin Linde, Olov Svartstrom, Anna Risberg, Theresa Enkirch, Mia Brytting, Karin Tegmark-Wisell |
| EPI_ISL_434650 | hCoV-19/Sweden/20-06071/2020 | Europe / Sweden / Jamtland | 2020-03-20 | Follinge Halsocentral | The Public Health Agency of Sweden | Kerstin Persson Moberg, Oskar Karlsson Lindsjo, Maria Lind Karlberg, Anna-Malin Linde, Olov Svartstrom, Anna Risberg, Theresa Enkirch, Mia Brytting, Karin Tegmark-Wisell |
| EPI_ISL_434651 | hCoV-19/Sweden/20-06306/2020 | Europe / Sweden / Jamtland | 2020-03-23 | Krokoms Halsocentral | The Public Health Agency of Sweden | Martin Ensson, Oskar Karlsson Lindsjo, Maria Lind Karlberg, Anna-Malin Linde, Olov Svartstrom, Anna Risberg, Theresa Enkirch, Mia Brytting, Karin Tegmark-Wisell |
| EPI_ISL_434652 | hCoV-19/Sweden/20-06308/2020 | Europe / Sweden / Stockholm | 2020-03-20 | Ektorps Vardcentral | The Public Health Agency of Sweden | Eva Espmark, Oskar Karlsson Lindsjo, Maria Lind Karlberg, Anna-Malin Linde, Olov Svartstrom, Anna Risberg, Theresa Enkirch, Mia Brytting, Karin Tegmark-Wisell |
| EPI_ISL_434653 | hCoV-19/Sweden/20-06376/2020 | Europe / Sweden / Stockholm | 2020-03-24 | Trollbackens VC | The Public Health Agency of Sweden | Amelie Holmqvist, Oskar Karlsson Lindsjo, Maria Lind Karlberg, Anna-Malin Linde, Olov Svartstrom, Anna Risberg, Theresa Enkirch, Mia Brytting, Karin Tegmark-Wisell |
| EPI_ISL_434654 | hCoV-19/Sweden/20-06383/2020 | Europe / Sweden / Halland | 2020-03-25 | Saroledens Familjelakare | The Public Health Agency of Sweden | Katarina Jarbur, Oskar Karlsson Lindsjo, Maria Lind Karlberg, Anna-Malin Linde, Olov Svartstrom, Anna Risberg, Theresa Enkirch, Mia Brytting, Karin Tegmark-Wisell |
| EPI_ISL_434655 | hCoV-19/Sweden/20-06468/2020 | Europe / Sweden / Uppsala | 2020-03-25 | Uppsala Narakut Aleris | The Public Health Agency of Sweden | Annika Nilsson, Oskar Karlsson Lindsjo, Maria Lind Karlberg, Anna-Malin Linde, Olov Svartstrom, Anna Risberg, Theresa Enkirch, Mia Brytting, Karin Tegmark-Wisell |
| EPI_ISL_450506 | hCoV-19/Brazil/SP02cc/2020 | South America / Brazil | 2020-03-18 | unknown | Clinical Laboratory | Malta,F., Amgarten,D., de Oliveira,D.B.L., Araujo,D.B., Machado,R.R.G., Santana,R.A.F., Mangueira,C.L.P., Durigon,E.L. and Pinho,J.R.R. |
| EPI_ISL_426617 | hCoV-19/USA/NY-NYUMC162/2020 | North America / USA / New York / Suffolk | 2020-04-01 | NYU Langone Health | Departments of Pathology and Medicine, New York University School of Medicine | Maria Aguero-Rosenfeld, Brendan Belovarac, Margaret Black, Ludovic Boytard, John Cadley, Paolo Cotzia, John Chen, Dacia Dimartino, Xiaojun Feng, Tatyana Gindin, Emily Guzman, Adriana Heguy, Megan Hogan, Emily Huang, George Jour, Andrew Lytle, Christian Marier, Matthew T. Maurano, Mark J. Mulligan, Peter Meyn, Iman Osman, Jared Pinnell, Vanessa Raabe, Sitharam Ramaswami, Amy Rapkiewicz, Marie Samanovic-Golden, Antonio Serrano, Guomiao Shen, Matija Snuderl, Theodore Vougiouklakis, Nick Vulpescu, Gael Westby, Paul Zappile, Yutong Zhang |
| EPI_ISL_426618 | hCoV-19/USA/NY-NYUMC163/2020 | North America / USA / New York / Nassa | 2020-04-01 | NYU Langone Health | Departments of Pathology and Medicine, New York University School of Medicine | Maria Aguero-Rosenfeld, Brendan Belovarac, Margaret Black, Ludovic Boytard, John Cadley, Paolo Cotzia, John Chen, Dacia Dimartino, Xiaojun Feng, Tatyana Gindin, Emily Guzman, Adriana Heguy, Megan Hogan, Emily Huang, George Jour, Andrew Lytle, Christian Marier, Matthew T. Maurano, Mark J. Mulligan, Peter Meyn, Iman Osman, Jared Pinnell, Vanessa Raabe, Sitharam Ramaswami, Amy Rapkiewicz, Marie Samanovic-Golden, Antonio Serrano, Guomiao Shen, Matija Snuderl, Theodore Vougiouklakis, Nick Vulpescu, Gael Westby, Paul Zappile, Yutong Zhang |
| EPI_ISL_426619 | hCoV-19/USA/NY-NYUMC164/2020 | North America / USA / New York / Nassa | 2020-04-01 | NYU Langone Health | Departments of Pathology and Medicine, New York University School of Medicine | Maria Aguero-Rosenfeld, Brendan Belovarac, Margaret Black, Ludovic Boytard, John Cadley, Paolo Cotzia, John Chen, Dacia Dimartino, Xiaojun Feng, Tatyana Gindin, Emily Guzman, Adriana Heguy, Megan Hogan, Emily Huang, George Jour, Andrew Lytle, Christian Marier, Matthew T. Maurano, Mark J. Mulligan, Peter Meyn, Iman Osman, Jared Pinnell, Vanessa Raabe, Sitharam Ramaswami, Amy Rapkiewicz, Marie Samanovic-Golden, Antonio Serrano, Guomiao Shen, Matija Snuderl, Theodore Vougiouklakis, Nick Vulpescu, Gael Westby, Paul Zappile, Yutong Zhang |
| EPI_ISL_426620 | hCoV-19/USA/NY-NYUMC165/2020 | North America / USA / New York / Brookl | 2020-04-01 | NYU Langone Health | Departments of Pathology and Medicine, New York University School of Medicine | Maria Aguero-Rosenfeld, Brendan Belovarac, Margaret Black, Ludovic Boytard, John Cadley, Paolo Cotzia, John Chen, Dacia Dimartino, Xiaojun Feng, Tatyana Gindin, Emily Guzman, Adriana Heguy, Megan Hogan, Emily Huang, George Jour, Andrew Lytle, Christian Marier, Matthew T. Maurano, Mark J. Mulligan, Peter Meyn, Iman Osman, Jared Pinnell, Vanessa Raabe, Sitharam Ramaswami, Amy Rapkiewicz, Marie Samanovic-Golden, Antonio Serrano, Guomiao Shen, Matija Snuderl, Theodore Vougiouklakis, Nick Vulpescu, Gael Westby, Paul Zappile, Yutong Zhang |
| EPI_ISL_426621 | hCoV-19/USA/NY-NYUMC166/2020 | North America / USA / New York / Brookl | 2020-04-01 | NYU Langone Health | Departments of Pathology and Medicine, New York University School of Medicine | Maria Aguero-Rosenfeld, Brendan Belovarac, Margaret Black, Ludovic Boytard, John Cadley, Paolo Cotzia, John Chen, Dacia Dimartino, Xiaojun Feng, Tatyana Gindin, Emily Guzman, Adriana Heguy, Megan Hogan, Emily Huang, George Jour, Andrew Lytle, Christian Marier, Matthew T. Maurano, Mark J. Mulligan, Peter Meyn, Iman Osman, Jared Pinnell, Vanessa Raabe, Sitharam Ramaswami, Amy Rapkiewicz, Marie Samanovic-Golden, Antonio Serrano, Guomiao Shen, Matija Snuderl, Theodore Vougiouklakis, Nick Vulpescu, Gael Westby, Paul Zappile, Yutong Zhang |
| EPI_ISL_426622 | hCoV-19/USA/NY-NYUMC167/2020 | North America / USA / New York / Manha | 2020-04-01 | NYU Langone Health | Departments of Pathology and Medicine, New York University School of Medicine | Maria Aguero-Rosenfeld, Brendan Belovarac, Margaret Black, Ludovic Boytard, John Cadley, Paolo Cotzia, John Chen, Dacia Dimartino, Xiaojun Feng, Tatyana Gindin, Emily Guzman, Adriana Heguy, Megan Hogan, Emily Huang, George Jour, Andrew Lytle, Christian Marier, Matthew T. Maurano, Mark J. Mulligan, Peter Meyn, Iman Osman, Jared Pinnell, Vanessa Raabe, Sitharam Ramaswami, Amy Rapkiewicz, Marie Samanovic-Golden, Antonio Serrano, Guomiao Shen, Matija Snuderl, Theodore Vougiouklakis, Nick Vulpescu, Gael Westby, Paul Zappile, Yutong Zhang |
| EPI_ISL_426623 | hCoV-19/USA/NY-NYUMC168/2020 | North America / USA / New York / Manha | 2020-04-01 | NYU Langone Health | Departments of Pathology and Medicine, New York University School of Medicine | Maria Aguero-Rosenfeld, Brendan Belovarac, Margaret Black, Ludovic Boytard, John Cadley, Paolo Cotzia, John Chen, Dacia Dimartino, Xiaojun Feng, Tatyana Gindin, Emily Guzman, Adriana Heguy, Megan Hogan, Emily Huang, George Jour, Andrew Lytle, Christian Marier, Matthew T. Maurano, Mark J. Mulligan, Peter Meyn, Iman Osman, Jared Pinnell, Vanessa Raabe, Sitharam Ramaswami, Amy Rapkiewicz, Marie Samanovic-Golden, Antonio Serrano, Guomiao Shen, Matija Snuderl, Theodore Vougiouklakis, Nick Vulpescu, Gael Westby, Paul Zappile, Yutong Zhang |
| EPI_ISL_426624 | hCoV-19/USA/NY-NYUMC169/2020 | North America / USA / New York / Brookl | 2020-04-01 | NYU Langone Health | Departments of Pathology and Medicine, New York University School of Medicine | Maria Aguero-Rosenfeld, Brendan Belovarac, Margaret Black, Ludovic Boytard, John Cadley, Paolo Cotzia, John Chen, Dacia Dimartino, Xiaojun Feng, Tatyana Gindin, Emily Guzman, Adriana Heguy, Megan Hogan, Emily Huang, George Jour, Andrew Lytle, Christian Marier, Matthew T. Maurano, Mark J. Mulligan, Peter Meyn, Iman Osman, Jared Pinnell, Vanessa Raabe, Sitharam Ramaswami, Amy Rapkiewicz, Marie Samanovic-Golden, Antonio Serrano, Guomiao Shen, Matija Snuderl, Theodore Vougiouklakis, Nick Vulpescu, Gael Westby, Paul Zappile, Yutong Zhang |
| EPI_ISL_426625 | hCoV-19/USA/NY-NYUMC170/2020 | North America / USA / New York / Brookl | 2020-04-01 | NYU Langone Health | Departments of Pathology and Medicine, New York University School of Medicine | Maria Aguero-Rosenfeld, Brendan Belovarac, Margaret Black, Ludovic Boytard, John Cadley, Paolo Cotzia, John Chen, Dacia Dimartino, Xiaojun Feng, Tatyana Gindin, Emily Guzman, Adriana Heguy, Megan Hogan, Emily Huang, George Jour, Andrew Lytle, Christian Marier, Matthew T. Maurano, Mark J. Mulligan, Peter Meyn, Iman Osman, Jared Pinnell, Vanessa Raabe, Sitharam Ramaswami, Amy Rapkiewicz, Marie Samanovic-Golden, Antonio Serrano, Guomiao Shen, Matija Snuderl, Theodore Vougiouklakis, Nick Vulpescu, Gael Westby, Paul Zappile, Yutong Zhang |
| EPI_ISL_426626 | hCoV-19/USA/NY-NYUMC171/2020 | North America / USA / New York / Brookl | 2020-04-01 | NYU Langone Health | Departments of Pathology and Medicine, New York University School of Medicine | Maria Aguero-Rosenfeld, Brendan Belovarac, Margaret Black, Ludovic Boytard, John Cadley, Paolo Cotzia, John Chen, Dacia Dimartino, Xiaojun Feng, Tatyana Gindin, Emily Guzman, Adriana Heguy, Megan Hogan, Emily Huang, George Jour, Andrew Lytle, Christian Marier, Matthew T. Maurano, Mark J. Mulligan, Peter Meyn, Iman Osman, Jared Pinnell, Vanessa Raabe, Sitharam Ramaswami, Amy Rapkiewicz, Marie Samanovic-Golden, Antonio Serrano, Guomiao Shen, Matija Snuderl, Theodore Vougiouklakis, Nick Vulpescu, Gael Westby, Paul Zappile, Yutong Zhang |
| EPI_ISL_426627 | hCoV-19/USA/LA-BIE-096/2020 | North America / USA / Louisiana | 2020-04-06 | Ochsner Health | BioInfoExperts, LLC | Amy Feehan, David Nolan, Rebecca Rose, Susanna Lamers, Sissy Cross, Julia Garcia-Diaz, Tong Yang, Luke Caruso, David Moraga Amador, Wayra Navia, Lydia Von Bondel, Xiao Hui Zhou |
| EPI_ISL_426628 | hCoV-19/USA/LA-BIE-097/2020 | North America / USA / Louisiana | 2020-04-06 | Ochsner Health | BioInfoExperts, LLC | Amy Feehan, David Nolan, Rebecca Rose, Susanna Lamers, Sissy Cross, Julia Garcia-Diaz, Tong Yang, Luke Caruso, David Moraga Amador, Wayra Navia, Lydia Von Bondel, Xiao Hui Zhou |
| EPI_ISL_426629 | hCoV-19/Taiwan/TSGH-01/2020 | Asia / Taiwan / Taipei | 2020-02-07 | TSGH-CP molecular lab | TSGH-CP molecular lab | Cherng-Lih Perng, Ming-Jr Jian, Chih-Kai Chang, Jung-Chung Lin, Kuo-Ming Yeh, Chien-Wen Chen, Sheng-Kang Chiu, Hsing-Yi Chung, Shih-Hung Tsai, Kuo-Sheng Hung, Feng-Yee Chang, Hung-Sheng Shang |
| EPI_ISL_426630 | hCoV-19/Taiwan/TSGH-02/2020 | Asia / Taiwan / New Taipei city | 2020-03-17 | TSGH-CP molecular lab | TSGH-CP molecular lab | Cherng-Lih Perng, Ming-Jr Jian, Chih-Kai Chang, Jung-Chung Lin, Kuo-Ming Yeh, Chien-Wen Chen, Sheng-Kang Chiu, Hsing-Yi Chung, Shih-Hung Tsai, Kuo-Sheng Hung, Feng-Yee Chang, Hung-Sheng Shang |
| EPI_ISL_426631 | hCoV-19/Taiwan/TSGH-03/2020 | Asia / Taiwan / New Taipei City | 2020-03-17 | TSGH-CP molecular lab | TSGH-CP molecular lab | Cherng-Lih Perng, Ming-Jr Jian, Chih-Kai Chang, Jung-Chung Lin, Kuo-Ming Yeh, Chien-Wen Chen, Sheng-Kang Chiu, Hsing-Yi Chung, Shih-Hung Tsai, Kuo-Sheng Hung, Tien-Yao Chang, Feng-Yee Chang, Hung-Sheng Shang |
| EPI_ISL_426632 | hCoV-19/Taiwan/TSGH-04/2020 | Asia / Taiwan / New Taipei City | 2020-03-17 | TSGH-CP molecular lab | TSGH-CP molecular lab | Cherng-Lih Perng, Ming-Jr Jian, Chih-Kai Chang, Jung-Chung Lin, Kuo-Ming Yeh, Chien-Wen Chen, Sheng-Kang Chiu, Hsing-Yi Chung, Shih-Hung Tsai, Kuo-Sheng Hung, Tien-Yao Chang, Feng-Yee Chang, Hung-Sheng Shang |
| EPI_ISL_426634 | hCoV-19/Australia/NT08/2020 | Oceania / Australia / Northern territory | 2020-03-20 | Royal Darwin Hospital Pathology | Microbiological Diagnostic Unit Public Health Laboratory and Victorian Infectious Diseases Reference Laboratory, Doherty Institute | Meumann, E., Caly L., Seemann T., Sait, M., Schultz M., Druce J., Sherry, N. |
| EPI_ISL_426635 | hCoV-19/Australia/NT18/2020 | Oceania / Australia / Northern territory | 2020-03-23 | Royal Darwin Hospital Pathology | Microbiological Diagnostic Unit Public Health Laboratory and Victorian Infectious Diseases Reference Laboratory, Doherty Institute | Meumann, E., Caly L., Seemann T., Sait, M., Schultz M., Druce J., Sherry, N. |
| EPI_ISL_426636 | hCoV-19/Australia/NT19/2020 | Oceania / Australia / Northern territory | 2020-03-25 | Royal Darwin Hospital Pathology | Microbiological Diagnostic Unit Public Health Laboratory and Victorian Infectious Diseases Reference Laboratory, Doherty Institute | Meumann, E., Caly L., Seemann T., Sait, M., Schultz M., Druce J., Sherry, N. |
| EPI_ISL_426637 | hCoV-19/Australia/VIC327/2020 | Oceania / Australia / Victoria | 2020-03-17 | Victorian Infectious Diseases Reference Laboratory (VIDRL) | Microbiological Diagnostic Unit Public Health Laboratory and Victorian Infectious Diseases Reference Laboratory, Doherty Institute | Caly L., Seemann T., Sait, M., Schultz M., Druce J., Sherry, N. |
| EPI_ISL_426638 | hCoV-19/Australia/VIC328/2020 | Oceania / Australia / Victoria | 2020-03-17 | Victorian Infectious Diseases Reference Laboratory (VIDRL) | Microbiological Diagnostic Unit Public Health Laboratory and Victorian Infectious Diseases Reference Laboratory, Doherty Institute | Caly L., Seemann T., Sait, M., Schultz M., Druce J., Sherry, N. |
| EPI_ISL_426639 | hCoV-19/Australia/VIC329/2020 | Oceania / Australia / Victoria | 2020-03-17 | Victorian Infectious Diseases Reference Laboratory (VIDRL) | Microbiological Diagnostic Unit Public Health Laboratory and Victorian Infectious Diseases Reference Laboratory, Doherty Institute | Caly L., Seemann T., Sait, M., Schultz M., Druce J., Sherry, N. |
| EPI_ISL_426640 | hCoV-19/Australia/VIC330/2020 | Oceania / Australia / Victoria | 2020-03-18 | Victorian Infectious Diseases Reference Laboratory (VIDRL) | Microbiological Diagnostic Unit Public Health Laboratory and Victorian Infectious Diseases Reference Laboratory, Doherty Institute | Caly L., Seemann T., Sait, M., Schultz M., Druce J., Sherry, N. |
| EPI_ISL_426641 | hCoV-19/Australia/VIC332/2020 | Oceania / Australia / Victoria | 2020-03-18 | Victorian Infectious Diseases Reference Laboratory (VIDRL) | Microbiological Diagnostic Unit Public Health Laboratory and Victorian Infectious Diseases Reference Laboratory, Doherty Institute | Caly L., Seemann T., Sait, M., Schultz M., Druce J., Sherry, N. |
| EPI_ISL_426642 | hCoV-19/Australia/VIC333/2020 | Oceania / Australia / Victoria | 2020-03-18 | Victorian Infectious Diseases Reference Laboratory (VIDRL) | Microbiological Diagnostic Unit Public Health Laboratory and Victorian Infectious Diseases Reference Laboratory, Doherty Institute | Caly L., Seemann T., Sait, M., Schultz M., Druce J., Sherry, N. |
| EPI_ISL_426643 | hCoV-19/Australia/VIC334/2020 | Oceania / Australia / Victoria | 2020-03-19 | Victorian Infectious Diseases Reference Laboratory (VIDRL) | Microbiological Diagnostic Unit Public Health Laboratory and Victorian Infectious Diseases Reference Laboratory, Doherty Institute | Caly L., Seemann T., Sait, M., Schultz M., Druce J., Sherry, N. |
| EPI_ISL_426644 | hCoV-19/Australia/VIC335/2020 | Oceania / Australia / Victoria | 2020-03-20 | Victorian Infectious Diseases Reference Laboratory (VIDRL) | Microbiological Diagnostic Unit Public Health Laboratory and Victorian Infectious Diseases Reference Laboratory, Doherty Institute | Caly L., Seemann T., Sait, M., Schultz M., Druce J., Sherry, N. |
| EPI_ISL_426645 | hCoV-19/Australia/VIC336/2020 | Oceania / Australia / Victoria | 2020-03-20 | Victorian Infectious Diseases Reference Laboratory (VIDRL) | Microbiological Diagnostic Unit Public Health Laboratory and Victorian Infectious Diseases Reference Laboratory, Doherty Institute | Caly L., Seemann T., Sait, M., Schultz M., Druce J., Sherry, N. |
| EPI_ISL_426646 | hCoV-19/Australia/VIC337/2020 | Oceania / Australia / Victoria | 2020-03-20 | Victorian Infectious Diseases Reference Laboratory (VIDRL) | Microbiological Diagnostic Unit Public Health Laboratory and Victorian Infectious Diseases Reference Laboratory, Doherty Institute | Caly L., Seemann T., Sait, M., Schultz M., Druce J., Sherry, N. |
| EPI_ISL_426647 | hCoV-19/Australia/VIC338/2020 | Oceania / Australia / Victoria | 2020-03-20 | Victorian Infectious Diseases Reference Laboratory (VIDRL) | Microbiological Diagnostic Unit Public Health Laboratory and Victorian Infectious Diseases Reference Laboratory, Doherty Institute | Caly L., Seemann T., Sait, M., Schultz M., Druce J., Sherry, N. |
| EPI_ISL_426648 | hCoV-19/Australia/VIC339/2020 | Oceania / Australia / Victoria | 2020-03-20 | Victorian Infectious Diseases Reference Laboratory (VIDRL) | Microbiological Diagnostic Unit Public Health Laboratory and Victorian Infectious Diseases Reference Laboratory, Doherty Institute | Caly L., Seemann T., Sait, M., Schultz M., Druce J., Sherry, N. |
| EPI_ISL_426649 | hCoV-19/Australia/VIC340/2020 | Oceania / Australia / Victoria | 2020-03-20 | Victorian Infectious Diseases Reference Laboratory (VIDRL) | Microbiological Diagnostic Unit Public Health Laboratory and Victorian Infectious Diseases Reference Laboratory, Doherty Institute | Caly L., Seemann T., Sait, M., Schultz M., Druce J., Sherry, N. |
| EPI_ISL_426650 | hCoV-19/Australia/VIC341/2020 | Oceania / Australia / Victoria | 2020-03-20 | Victorian Infectious Diseases Reference Laboratory (VIDRL) | Microbiological Diagnostic Unit Public Health Laboratory and Victorian Infectious Diseases Reference Laboratory, Doherty Institute | Caly L., Seemann T., Sait, M., Schultz M., Druce J., Sherry, N. |
| EPI_ISL_426651 | hCoV-19/Australia/VIC342/2020 | Oceania / Australia / Victoria | 2020-03-20 | Victorian Infectious Diseases Reference Laboratory (VIDRL) | Microbiological Diagnostic Unit Public Health Laboratory and Victorian Infectious Diseases Reference Laboratory, Doherty Institute | Caly L., Seemann T., Sait, M., Schultz M., Druce J., Sherry, N. |
| EPI_ISL_426652 | hCoV-19/Australia/VIC343/2020 | Oceania / Australia / Victoria | 2020-03-20 | Victorian Infectious Diseases Reference Laboratory (VIDRL) | Microbiological Diagnostic Unit Public Health Laboratory and Victorian Infectious Diseases Reference Laboratory, Doherty Institute | Caly L., Seemann T., Sait, M., Schultz M., Druce J., Sherry, N. |
| EPI_ISL_426653 | hCoV-19/Australia/VIC344/2020 | Oceania / Australia / Victoria | 2020-03-20 | Victorian Infectious Diseases Reference Laboratory (VIDRL) | Microbiological Diagnostic Unit Public Health Laboratory and Victorian Infectious Diseases Reference Laboratory, Doherty Institute | Caly L., Seemann T., Sait, M., Schultz M., Druce J., Sherry, N. |
| EPI_ISL_426654 | hCoV-19/Australia/VIC345/2020 | Oceania / Australia / Victoria | 2020-03-20 | Victorian Infectious Diseases Reference Laboratory (VIDRL) | Microbiological Diagnostic Unit Public Health Laboratory and Victorian Infectious Diseases Reference Laboratory, Doherty Institute | Caly L., Seemann T., Sait, M., Schultz M., Druce J., Sherry, N. |
| EPI_ISL_426655 | hCoV-19/Australia/VIC346/2020 | Oceania / Australia / Victoria | 2020-03-20 | Victorian Infectious Diseases Reference Laboratory (VIDRL) | Microbiological Diagnostic Unit Public Health Laboratory and Victorian Infectious Diseases Reference Laboratory, Doherty Institute | Caly L., Seemann T., Sait, M., Schultz M., Druce J., Sherry, N. |
| EPI_ISL_426656 | hCoV-19/Australia/VIC347/2020 | Oceania / Australia / Victoria | 2020-03-20 | Victorian Infectious Diseases Reference Laboratory (VIDRL) | Microbiological Diagnostic Unit Public Health Laboratory and Victorian Infectious Diseases Reference Laboratory, Doherty Institute | Caly L., Seemann T., Sait, M., Schultz M., Druce J., Sherry, N. |
| EPI_ISL_426657 | hCoV-19/Australia/VIC348/2020 | Oceania / Australia / Victoria | 2020-03-20 | Victorian Infectious Diseases Reference Laboratory (VIDRL) | Microbiological Diagnostic Unit Public Health Laboratory and Victorian Infectious Diseases Reference Laboratory, Doherty Institute | Caly L., Seemann T., Sait, M., Schultz M., Druce J., Sherry, N. |
| EPI_ISL_426658 | hCoV-19/Australia/VIC350/2020 | Oceania / Australia / Victoria | 2020-03-20 | Victorian Infectious Diseases Reference Laboratory (VIDRL) | Microbiological Diagnostic Unit Public Health Laboratory and Victorian Infectious Diseases Reference Laboratory, Doherty Institute | Caly L., Seemann T., Sait, M., Schultz M., Druce J., Sherry, N. |
| EPI_ISL_426659 | hCoV-19/Australia/VIC351/2020 | Oceania / Australia / Victoria | 2020-03-20 | Victorian Infectious Diseases Reference Laboratory (VIDRL) | Microbiological Diagnostic Unit Public Health Laboratory and Victorian Infectious Diseases Reference Laboratory, Doherty Institute | Caly L., Seemann T., Sait, M., Schultz M., Druce J., Sherry, N. |
| EPI_ISL_426661 | hCoV-19/Australia/VIC353/2020 | Oceania / Australia / Victoria | 2020-03-21 | Victorian Infectious Diseases Reference Laboratory (VIDRL) | Microbiological Diagnostic Unit Public Health Laboratory and Victorian Infectious Diseases Reference Laboratory, Doherty Institute | Caly L., Seemann T., Sait, M., Schultz M., Druce J., Sherry, N. |
| EPI_ISL_426662 | hCoV-19/Australia/VIC354/2020 | Oceania / Australia / Victoria | 2020-03-21 | Victorian Infectious Diseases Reference Laboratory (VIDRL) | Microbiological Diagnostic Unit Public Health Laboratory and Victorian Infectious Diseases Reference Laboratory, Doherty Institute | Caly L., Seemann T., Sait, M., Schultz M., Druce J., Sherry, N. |
| EPI_ISL_426663 | hCoV-19/Australia/VIC355/2020 | Oceania / Australia / Victoria | 2020-03-21 | Victorian Infectious Diseases Reference Laboratory (VIDRL) | Microbiological Diagnostic Unit Public Health Laboratory and Victorian Infectious Diseases Reference Laboratory, Doherty Institute | Caly L., Seemann T., Sait, M., Schultz M., Druce J., Sherry, N. |
| EPI_ISL_426664 | hCoV-19/Australia/VIC356/2020 | Oceania / Australia / Victoria | 2020-03-21 | Victorian Infectious Diseases Reference Laboratory (VIDRL) | Microbiological Diagnostic Unit Public Health Laboratory and Victorian Infectious Diseases Reference Laboratory, Doherty Institute | Caly L., Seemann T., Sait, M., Schultz M., Druce J., Sherry, N. |
| EPI_ISL_426665 | hCoV-19/Australia/VIC357/2020 | Oceania / Australia / Victoria | 2020-03-21 | Victorian Infectious Diseases Reference Laboratory (VIDRL) | Microbiological Diagnostic Unit Public Health Laboratory and Victorian Infectious Diseases Reference Laboratory, Doherty Institute | Caly L., Seemann T., Sait, M., Schultz M., Druce J., Sherry, N. |
| EPI_ISL_426666 | hCoV-19/Australia/VIC358/2020 | Oceania / Australia / Victoria | 2020-03-21 | Victorian Infectious Diseases Reference Laboratory (VIDRL) | Microbiological Diagnostic Unit Public Health Laboratory and Victorian Infectious Diseases Reference Laboratory, Doherty Institute | Caly L., Seemann T., Sait, M., Schultz M., Druce J., Sherry, N. |
| EPI_ISL_426667 | hCoV-19/Australia/VIC359/2020 | Oceania / Australia / Victoria | 2020-03-22 | Victorian Infectious Diseases Reference Laboratory (VIDRL) | Microbiological Diagnostic Unit Public Health Laboratory and Victorian Infectious Diseases Reference Laboratory, Doherty Institute | Caly L., Seemann T., Sait, M., Schultz M., Druce J., Sherry, N. |
| EPI_ISL_426669 | hCoV-19/Australia/VIC361/2020 | Oceania / Australia / Victoria | 2020-03-22 | Victorian Infectious Diseases Reference Laboratory (VIDRL) | Microbiological Diagnostic Unit Public Health Laboratory and Victorian Infectious Diseases Reference Laboratory, Doherty Institute | Caly L., Seemann T., Sait, M., Schultz M., Druce J., Sherry, N. |
| EPI_ISL_426670 | hCoV-19/Australia/VIC362/2020 | Oceania / Australia / Victoria | 2020-03-22 | Victorian Infectious Diseases Reference Laboratory (VIDRL) | Microbiological Diagnostic Unit Public Health Laboratory and Victorian Infectious Diseases Reference Laboratory, Doherty Institute | Caly L., Seemann T., Sait, M., Schultz M., Druce J., Sherry, N. |
| EPI_ISL_426671 | hCoV-19/Australia/VIC363/2020 | Oceania / Australia / Victoria | 2020-03-22 | Victorian Infectious Diseases Reference Laboratory (VIDRL) | Microbiological Diagnostic Unit Public Health Laboratory and Victorian Infectious Diseases Reference Laboratory, Doherty Institute | Caly L., Seemann T., Sait, M., Schultz M., Druce J., Sherry, N. |
| EPI_ISL_426672 | hCoV-19/Australia/VIC364/2020 | Oceania / Australia / Victoria | 2020-03-22 | Victorian Infectious Diseases Reference Laboratory (VIDRL) | Microbiological Diagnostic Unit Public Health Laboratory and Victorian Infectious Diseases Reference Laboratory, Doherty Institute | Caly L., Seemann T., Sait, M., Schultz M., Druce J., Sherry, N. |
| EPI_ISL_426673 | hCoV-19/Australia/VIC365/2020 | Oceania / Australia / Victoria | 2020-03-22 | Victorian Infectious Diseases Reference Laboratory (VIDRL) | Microbiological Diagnostic Unit Public Health Laboratory and Victorian Infectious Diseases Reference Laboratory, Doherty Institute | Caly L., Seemann T., Sait, M., Schultz M., Druce J., Sherry, N. |
| EPI_ISL_426674 | hCoV-19/Australia/VIC366/2020 | Oceania / Australia / Victoria | 2020-03-22 | Victorian Infectious Diseases Reference Laboratory (VIDRL) | Microbiological Diagnostic Unit Public Health Laboratory and Victorian Infectious Diseases Reference Laboratory, Doherty Institute | Caly L., Seemann T., Sait, M., Schultz M., Druce J., Sherry, N. |
| EPI_ISL_426675 | hCoV-19/Australia/VIC367/2020 | Oceania / Australia / Victoria | 2020-03-22 | Victorian Infectious Diseases Reference Laboratory (VIDRL) | Microbiological Diagnostic Unit Public Health Laboratory and Victorian Infectious Diseases Reference Laboratory, Doherty Institute | Caly L., Seemann T., Sait, M., Schultz M., Druce J., Sherry, N. |
| EPI_ISL_426676 | hCoV-19/Australia/VIC368/2020 | Oceania / Australia / Victoria | 2020-03-22 | Victorian Infectious Diseases Reference Laboratory (VIDRL) | Microbiological Diagnostic Unit Public Health Laboratory and Victorian Infectious Diseases Reference Laboratory, Doherty Institute | Caly L., Seemann T., Sait, M., Schultz M., Druce J., Sherry, N. |
| EPI_ISL_426677 | hCoV-19/Australia/VIC369/2020 | Oceania / Australia / Victoria | 2020-03-22 | Victorian Infectious Diseases Reference Laboratory (VIDRL) | Microbiological Diagnostic Unit Public Health Laboratory and Victorian Infectious Diseases Reference Laboratory, Doherty Institute | Caly L., Seemann T., Sait, M., Schultz M., Druce J., Sherry, N. |
| EPI_ISL_426679 | hCoV-19/Australia/VIC372/2020 | Oceania / Australia / Victoria | 2020-03-22 | Victorian Infectious Diseases Reference Laboratory (VIDRL) | Microbiological Diagnostic Unit Public Health Laboratory and Victorian Infectious Diseases Reference Laboratory, Doherty Institute | Caly L., Seemann T., Sait, M., Schultz M., Druce J., Sherry, N. |
| EPI_ISL_426680 | hCoV-19/Australia/VIC373/2020 | Oceania / Australia / Victoria | 2020-03-22 | Victorian Infectious Diseases Reference Laboratory (VIDRL) | Microbiological Diagnostic Unit Public Health Laboratory and Victorian Infectious Diseases Reference Laboratory, Doherty Institute | Caly L., Seemann T., Sait, M., Schultz M., Druce J., Sherry, N. |
| EPI_ISL_426682 | hCoV-19/Australia/VIC375/2020 | Oceania / Australia / Victoria | 2020-03-22 | Victorian Infectious Diseases Reference Laboratory (VIDRL) | Microbiological Diagnostic Unit Public Health Laboratory and Victorian Infectious Diseases Reference Laboratory, Doherty Institute | Caly L., Seemann T., Sait, M., Schultz M., Druce J., Sherry, N. |
| EPI_ISL_426683 | hCoV-19/Australia/VIC376/2020 | Oceania / Australia / Victoria | 2020-03-22 | Victorian Infectious Diseases Reference Laboratory (VIDRL) | Microbiological Diagnostic Unit Public Health Laboratory and Victorian Infectious Diseases Reference Laboratory, Doherty Institute | Caly L., Seemann T., Sait, M., Schultz M., Druce J., Sherry, N. |
| EPI_ISL_426684 | hCoV-19/Australia/VIC377/2020 | Oceania / Australia / Victoria | 2020-03-22 | Victorian Infectious Diseases Reference Laboratory (VIDRL) | Microbiological Diagnostic Unit Public Health Laboratory and Victorian Infectious Diseases Reference Laboratory, Doherty Institute | Caly L., Seemann T., Sait, M., Schultz M., Druce J., Sherry, N. |
| EPI_ISL_426685 | hCoV-19/Australia/VIC378/2020 | Oceania / Australia / Victoria | 2020-03-22 | Victorian Infectious Diseases Reference Laboratory (VIDRL) | Microbiological Diagnostic Unit Public Health Laboratory and Victorian Infectious Diseases Reference Laboratory, Doherty Institute | Caly L., Seemann T., Sait, M., Schultz M., Druce J., Sherry, N. |
| EPI_ISL_426686 | hCoV-19/Australia/VIC379/2020 | Oceania / Australia / Victoria | 2020-03-22 | Victorian Infectious Diseases Reference Laboratory (VIDRL) | Microbiological Diagnostic Unit Public Health Laboratory and Victorian Infectious Diseases Reference Laboratory, Doherty Institute | Caly L., Seemann T., Sait, M., Schultz M., Druce J., Sherry, N. |
| EPI_ISL_426687 | hCoV-19/Australia/VIC380/2020 | Oceania / Australia / Victoria | 2020-03-23 | Victorian Infectious Diseases Reference Laboratory (VIDRL) | Microbiological Diagnostic Unit Public Health Laboratory and Victorian Infectious Diseases Reference Laboratory, Doherty Institute | Caly L., Seemann T., Sait, M., Schultz M., Druce J., Sherry, N. |
| EPI_ISL_426688 | hCoV-19/Australia/VIC381/2020 | Oceania / Australia / Victoria | 2020-03-23 | Victorian Infectious Diseases Reference Laboratory (VIDRL) | Microbiological Diagnostic Unit Public Health Laboratory and Victorian Infectious Diseases Reference Laboratory, Doherty Institute | Caly L., Seemann T., Sait, M., Schultz M., Druce J., Sherry, N. |
| EPI_ISL_426689 | hCoV-19/Australia/VIC382/2020 | Oceania / Australia / Victoria | 2020-03-23 | Victorian Infectious Diseases Reference Laboratory (VIDRL) | Microbiological Diagnostic Unit Public Health Laboratory and Victorian Infectious Diseases Reference Laboratory, Doherty Institute | Caly L., Seemann T., Sait, M., Schultz M., Druce J., Sherry, N. |
| EPI_ISL_426690 | hCoV-19/Australia/VIC383/2020 | Oceania / Australia / Victoria | 2020-03-23 | Victorian Infectious Diseases Reference Laboratory (VIDRL) | Microbiological Diagnostic Unit Public Health Laboratory and Victorian Infectious Diseases Reference Laboratory, Doherty Institute | Caly L., Seemann T., Sait, M., Schultz M., Druce J., Sherry, N. |
| EPI_ISL_426691 | hCoV-19/Australia/VIC384/2020 | Oceania / Australia / Victoria | 2020-03-23 | Victorian Infectious Diseases Reference Laboratory (VIDRL) | Microbiological Diagnostic Unit Public Health Laboratory and Victorian Infectious Diseases Reference Laboratory, Doherty Institute | Caly L., Seemann T., Sait, M., Schultz M., Druce J., Sherry, N. |
| EPI_ISL_426692 | hCoV-19/Australia/VIC385/2020 | Oceania / Australia / Victoria | 2020-03-23 | Victorian Infectious Diseases Reference Laboratory (VIDRL) | Microbiological Diagnostic Unit Public Health Laboratory and Victorian Infectious Diseases Reference Laboratory, Doherty Institute | Caly L., Seemann T., Sait, M., Schultz M., Druce J., Sherry, N. |
| EPI_ISL_426693 | hCoV-19/Australia/VIC386/2020 | Oceania / Australia / Victoria | 2020-03-23 | Victorian Infectious Diseases Reference Laboratory (VIDRL) | Microbiological Diagnostic Unit Public Health Laboratory and Victorian Infectious Diseases Reference Laboratory, Doherty Institute | Caly L., Seemann T., Sait, M., Schultz M., Druce J., Sherry, N. |
| EPI_ISL_426694 | hCoV-19/Australia/VIC387/2020 | Oceania / Australia / Victoria | 2020-03-23 | Victorian Infectious Diseases Reference Laboratory (VIDRL) | Microbiological Diagnostic Unit Public Health Laboratory and Victorian Infectious Diseases Reference Laboratory, Doherty Institute | Caly L., Seemann T., Sait, M., Schultz M., Druce J., Sherry, N. |
| EPI_ISL_426696 | hCoV-19/Australia/VIC389/2020 | Oceania / Australia / Victoria | 2020-03-23 | Victorian Infectious Diseases Reference Laboratory (VIDRL) | Microbiological Diagnostic Unit Public Health Laboratory and Victorian Infectious Diseases Reference Laboratory, Doherty Institute | Caly L., Seemann T., Sait, M., Schultz M., Druce J., Sherry, N. |
| EPI_ISL_426697 | hCoV-19/Australia/VIC394/2020 | Oceania / Australia / Victoria | 2020-03-24 | Victorian Infectious Diseases Reference Laboratory (VIDRL) | Microbiological Diagnostic Unit Public Health Laboratory and Victorian Infectious Diseases Reference Laboratory, Doherty Institute | Caly L., Seemann T., Sait, M., Schultz M., Druce J., Sherry, N. |
| EPI_ISL_426698 | hCoV-19/Australia/VIC395/2020 | Oceania / Australia / Victoria | 2020-03-24 | Victorian Infectious Diseases Reference Laboratory (VIDRL) | Microbiological Diagnostic Unit Public Health Laboratory and Victorian Infectious Diseases Reference Laboratory, Doherty Institute | Caly L., Seemann T., Sait, M., Schultz M., Druce J., Sherry, N. |
| EPI_ISL_426914 | hCoV-19/Australia/VIC483/2020 | Oceania / Australia / Victoria | 2020-03-29 | Microbiological Diagnostic Unit Public Health Laboratory | Microbiological Diagnostic Unit Public Health Laboratory | Seemann T., Schultz M., Sait, M., Sherry, N. |
| EPI_ISL_426917 | hCoV-19/Australia/VIC486/2020 | Oceania / Australia / Victoria | 2020-03-29 | Microbiological Diagnostic Unit Public Health Laboratory | Microbiological Diagnostic Unit Public Health Laboratory | Seemann T., Schultz M., Sait, M., Sherry, N. |
| EPI_ISL_426918 | hCoV-19/Australia/VIC487/2020 | Oceania / Australia / Victoria | 2020-03-29 | Microbiological Diagnostic Unit Public Health Laboratory | Microbiological Diagnostic Unit Public Health Laboratory | Seemann T., Schultz M., Sait, M., Sherry, N. |
| EPI_ISL_426919 | hCoV-19/Australia/VIC488/2020 | Oceania / Australia / Victoria | 2020-03-29 | Microbiological Diagnostic Unit Public Health Laboratory | Microbiological Diagnostic Unit Public Health Laboratory | Seemann T., Schultz M., Sait, M., Sherry, N. |
| EPI_ISL_426920 | hCoV-19/Australia/VIC489/2020 | Oceania / Australia / Victoria | 2020-03-29 | Microbiological Diagnostic Unit Public Health Laboratory | Microbiological Diagnostic Unit Public Health Laboratory | Seemann T., Schultz M., Sait, M., Sherry, N. |
| EPI_ISL_426921 | hCoV-19/Australia/VIC490/2020 | Oceania / Australia / Victoria | 2020-03-30 | Microbiological Diagnostic Unit Public Health Laboratory | Microbiological Diagnostic Unit Public Health Laboratory | Seemann T., Schultz M., Sait, M., Sherry, N. |
| EPI_ISL_426922 | hCoV-19/Australia/VIC491/2020 | Oceania / Australia / Victoria | 2020-03-30 | Microbiological Diagnostic Unit Public Health Laboratory | Microbiological Diagnostic Unit Public Health Laboratory | Seemann T., Schultz M., Sait, M., Sherry, N. |
| EPI_ISL_426923 | hCoV-19/Australia/VIC619/2020 | Oceania / Australia / Victoria | 2020-03-28 | Victorian Infectious Diseases Reference Laboratory (VIDRL) | Microbiological Diagnostic Unit Public Health Laboratory and Victorian Infectious Diseases Reference Laboratory, Doherty Institute | Caly L., Seemann T., Sait, M., Schultz M., Druce J., Sherry, N. |
| EPI_ISL_426924 | hCoV-19/Australia/VIC620/2020 | Oceania / Australia / Victoria | 2020-03-28 | Victorian Infectious Diseases Reference Laboratory (VIDRL) | Microbiological Diagnostic Unit Public Health Laboratory and Victorian Infectious Diseases Reference Laboratory, Doherty Institute | Caly L., Seemann T., Sait, M., Schultz M., Druce J., Sherry, N. |
| EPI_ISL_426926 | hCoV-19/Australia/VIC622/2020 | Oceania / Australia / Victoria | 2020-03-29 | Victorian Infectious Diseases Reference Laboratory (VIDRL) | Microbiological Diagnostic Unit Public Health Laboratory and Victorian Infectious Diseases Reference Laboratory, Doherty Institute | Caly L., Seemann T., Sait, M., Schultz M., Druce J., Sherry, N. |
| EPI_ISL_426927 | hCoV-19/Australia/VIC623/2020 | Oceania / Australia / Victoria | 2020-03-28 | Victorian Infectious Diseases Reference Laboratory (VIDRL) | Microbiological Diagnostic Unit Public Health Laboratory and Victorian Infectious Diseases Reference Laboratory, Doherty Institute | Caly L., Seemann T., Sait, M., Schultz M., Druce J., Sherry, N. |
| EPI_ISL_426929 | hCoV-19/Australia/VIC625/2020 | Oceania / Australia / Victoria | 2020-03-28 | Victorian Infectious Diseases Reference Laboratory (VIDRL) | Microbiological Diagnostic Unit Public Health Laboratory and Victorian Infectious Diseases Reference Laboratory, Doherty Institute | Caly L., Seemann T., Sait, M., Schultz M., Druce J., Sherry, N. |
| EPI_ISL_426930 | hCoV-19/Australia/VIC626/2020 | Oceania / Australia / Victoria | 2020-03-28 | Victorian Infectious Diseases Reference Laboratory (VIDRL) | Microbiological Diagnostic Unit Public Health Laboratory and Victorian Infectious Diseases Reference Laboratory, Doherty Institute | Caly L., Seemann T., Sait, M., Schultz M., Druce J., Sherry, N. |
| EPI_ISL_426931 | hCoV-19/Australia/VIC627/2020 | Oceania / Australia / Victoria | 2020-03-29 | Victorian Infectious Diseases Reference Laboratory (VIDRL) | Microbiological Diagnostic Unit Public Health Laboratory and Victorian Infectious Diseases Reference Laboratory, Doherty Institute | Caly L., Seemann T., Sait, M., Schultz M., Druce J., Sherry, N. |
| EPI_ISL_426932 | hCoV-19/Australia/VIC628/2020 | Oceania / Australia / Victoria | 2020-03-29 | Victorian Infectious Diseases Reference Laboratory (VIDRL) | Microbiological Diagnostic Unit Public Health Laboratory and Victorian Infectious Diseases Reference Laboratory, Doherty Institute | Caly L., Seemann T., Sait, M., Schultz M., Druce J., Sherry, N. |
| EPI_ISL_426933 | hCoV-19/Australia/VIC637/2020 | Oceania / Australia / Victoria | 2020-03-28 | Victorian Infectious Diseases Reference Laboratory (VIDRL) | Microbiological Diagnostic Unit Public Health Laboratory and Victorian Infectious Diseases Reference Laboratory, Doherty Institute | Caly L., Seemann T., Sait, M., Schultz M., Druce J., Sherry, N. |
| EPI_ISL_426934 | hCoV-19/Australia/VIC638/2020 | Oceania / Australia / Victoria | 2020-03-29 | Victorian Infectious Diseases Reference Laboratory (VIDRL) | Microbiological Diagnostic Unit Public Health Laboratory and Victorian Infectious Diseases Reference Laboratory, Doherty Institute | Caly L., Seemann T., Sait, M., Schultz M., Druce J., Sherry, N. |
| EPI_ISL_426935 | hCoV-19/Australia/VIC639/2020 | Oceania / Australia / Victoria | 2020-03-29 | Victorian Infectious Diseases Reference Laboratory (VIDRL) | Microbiological Diagnostic Unit Public Health Laboratory and Victorian Infectious Diseases Reference Laboratory, Doherty Institute | Caly L., Seemann T., Sait, M., Schultz M., Druce J., Sherry, N. |
| EPI_ISL_426940 | hCoV-19/Australia/VIC644/2020 | Oceania / Australia / Victoria | 2020-03-28 | Victorian Infectious Diseases Reference Laboratory (VIDRL) | Microbiological Diagnostic Unit Public Health Laboratory and Victorian Infectious Diseases Reference Laboratory, Doherty Institute | Caly L., Seemann T., Sait, M., Schultz M., Druce J., Sherry, N. |
| EPI_ISL_426941 | hCoV-19/Australia/VIC645/2020 | Oceania / Australia / Victoria | 2020-03-28 | Victorian Infectious Diseases Reference Laboratory (VIDRL) | Microbiological Diagnostic Unit Public Health Laboratory and Victorian Infectious Diseases Reference Laboratory, Doherty Institute | Caly L., Seemann T., Sait, M., Schultz M., Druce J., Sherry, N. |
| EPI_ISL_426942 | hCoV-19/Australia/VIC646/2020 | Oceania / Australia / Victoria | 2020-03-24 | Victorian Infectious Diseases Reference Laboratory (VIDRL) | Microbiological Diagnostic Unit Public Health Laboratory and Victorian Infectious Diseases Reference Laboratory, Doherty Institute | Caly L., Seemann T., Sait, M., Schultz M., Druce J., Sherry, N. |
| EPI_ISL_426943 | hCoV-19/Australia/VIC647/2020 | Oceania / Australia / Victoria | 2020-03-28 | Victorian Infectious Diseases Reference Laboratory (VIDRL) | Microbiological Diagnostic Unit Public Health Laboratory and Victorian Infectious Diseases Reference Laboratory, Doherty Institute | Caly L., Seemann T., Sait, M., Schultz M., Druce J., Sherry, N. |
| EPI_ISL_426944 | hCoV-19/Australia/VIC648/2020 | Oceania / Australia / Victoria | 2020-03-28 | Victorian Infectious Diseases Reference Laboratory (VIDRL) | Microbiological Diagnostic Unit Public Health Laboratory and Victorian Infectious Diseases Reference Laboratory, Doherty Institute | Caly L., Seemann T., Sait, M., Schultz M., Druce J., Sherry, N. |
| EPI_ISL_426946 | hCoV-19/Australia/VIC650/2020 | Oceania / Australia / Victoria | 2020-03-28 | Victorian Infectious Diseases Reference Laboratory (VIDRL) | Microbiological Diagnostic Unit Public Health Laboratory and Victorian Infectious Diseases Reference Laboratory, Doherty Institute | Caly L., Seemann T., Sait, M., Schultz M., Druce J., Sherry, N. |
| EPI_ISL_426947 | hCoV-19/Australia/VIC651/2020 | Oceania / Australia / Victoria | 2020-03-28 | Victorian Infectious Diseases Reference Laboratory (VIDRL) | Microbiological Diagnostic Unit Public Health Laboratory and Victorian Infectious Diseases Reference Laboratory, Doherty Institute | Caly L., Seemann T., Sait, M., Schultz M., Druce J., Sherry, N. |
| EPI_ISL_426948 | hCoV-19/Australia/VIC652/2020 | Oceania / Australia / Victoria | 2020-03-28 | Victorian Infectious Diseases Reference Laboratory (VIDRL) | Microbiological Diagnostic Unit Public Health Laboratory and Victorian Infectious Diseases Reference Laboratory, Doherty Institute | Caly L., Seemann T., Sait, M., Schultz M., Druce J., Sherry, N. |
| EPI_ISL_426951 | hCoV-19/Australia/VIC655/2020 | Oceania / Australia / Victoria | 2020-03-28 | Victorian Infectious Diseases Reference Laboratory (VIDRL) | Microbiological Diagnostic Unit Public Health Laboratory and Victorian Infectious Diseases Reference Laboratory, Doherty Institute | Caly L., Seemann T., Sait, M., Schultz M., Druce J., Sherry, N. |
| EPI_ISL_426952 | hCoV-19/Australia/VIC656/2020 | Oceania / Australia / Victoria | 2020-03-28 | Victorian Infectious Diseases Reference Laboratory (VIDRL) | Microbiological Diagnostic Unit Public Health Laboratory and Victorian Infectious Diseases Reference Laboratory, Doherty Institute | Caly L., Seemann T., Sait, M., Schultz M., Druce J., Sherry, N. |
| EPI_ISL_426953 | hCoV-19/Australia/VIC657/2020 | Oceania / Australia / Victoria | 2020-03-28 | Victorian Infectious Diseases Reference Laboratory (VIDRL) | Microbiological Diagnostic Unit Public Health Laboratory and Victorian Infectious Diseases Reference Laboratory, Doherty Institute | Caly L., Seemann T., Sait, M., Schultz M., Druce J., Sherry, N. |
| EPI_ISL_426954 | hCoV-19/Australia/VIC658/2020 | Oceania / Australia / Victoria | 2020-03-28 | Victorian Infectious Diseases Reference Laboratory (VIDRL) | Microbiological Diagnostic Unit Public Health Laboratory and Victorian Infectious Diseases Reference Laboratory, Doherty Institute | Caly L., Seemann T., Sait, M., Schultz M., Druce J., Sherry, N. |
| EPI_ISL_426955 | hCoV-19/Australia/VIC659/2020 | Oceania / Australia / Victoria | 2020-03-27 | Victorian Infectious Diseases Reference Laboratory (VIDRL) | Microbiological Diagnostic Unit Public Health Laboratory and Victorian Infectious Diseases Reference Laboratory, Doherty Institute | Caly L., Seemann T., Sait, M., Schultz M., Druce J., Sherry, N. |
| EPI_ISL_426956 | hCoV-19/Australia/VIC660/2020 | Oceania / Australia / Victoria | 2020-03-28 | Victorian Infectious Diseases Reference Laboratory (VIDRL) | Microbiological Diagnostic Unit Public Health Laboratory and Victorian Infectious Diseases Reference Laboratory, Doherty Institute | Caly L., Seemann T., Sait, M., Schultz M., Druce J., Sherry, N. |
| EPI_ISL_426957 | hCoV-19/Australia/VIC661/2020 | Oceania / Australia / Victoria | 2020-03-27 | Victorian Infectious Diseases Reference Laboratory (VIDRL) | Microbiological Diagnostic Unit Public Health Laboratory and Victorian Infectious Diseases Reference Laboratory, Doherty Institute | Caly L., Seemann T., Sait, M., Schultz M., Druce J., Sherry, N. |
| EPI_ISL_426958 | hCoV-19/Australia/VIC662/2020 | Oceania / Australia / Victoria | 2020-03-27 | Victorian Infectious Diseases Reference Laboratory (VIDRL) | Microbiological Diagnostic Unit Public Health Laboratory and Victorian Infectious Diseases Reference Laboratory, Doherty Institute | Caly L., Seemann T., Sait, M., Schultz M., Druce J., Sherry, N. |
| EPI_ISL_426959 | hCoV-19/Australia/VIC663/2020 | Oceania / Australia / Victoria | 2020-03-28 | Victorian Infectious Diseases Reference Laboratory (VIDRL) | Microbiological Diagnostic Unit Public Health Laboratory and Victorian Infectious Diseases Reference Laboratory, Doherty Institute | Caly L., Seemann T., Sait, M., Schultz M., Druce J., Sherry, N. |
| EPI_ISL_426960 | hCoV-19/Australia/VIC664/2020 | Oceania / Australia / Victoria | 2020-03-27 | Victorian Infectious Diseases Reference Laboratory (VIDRL) | Microbiological Diagnostic Unit Public Health Laboratory and Victorian Infectious Diseases Reference Laboratory, Doherty Institute | Caly L., Seemann T., Sait, M., Schultz M., Druce J., Sherry, N. |
| EPI_ISL_426961 | hCoV-19/Australia/VIC665/2020 | Oceania / Australia / Victoria | 2020-03-27 | Victorian Infectious Diseases Reference Laboratory (VIDRL) | Microbiological Diagnostic Unit Public Health Laboratory and Victorian Infectious Diseases Reference Laboratory, Doherty Institute | Caly L., Seemann T., Sait, M., Schultz M., Druce J., Sherry, N. |
| EPI_ISL_426962 | hCoV-19/Australia/VIC666/2020 | Oceania / Australia / Victoria | 2020-03-27 | Victorian Infectious Diseases Reference Laboratory (VIDRL) | Microbiological Diagnostic Unit Public Health Laboratory and Victorian Infectious Diseases Reference Laboratory, Doherty Institute | Caly L., Seemann T., Sait, M., Schultz M., Druce J., Sherry, N. |
| EPI_ISL_426963 | hCoV-19/Australia/VIC667/2020 | Oceania / Australia / Victoria | 2020-03-27 | Victorian Infectious Diseases Reference Laboratory (VIDRL) | Microbiological Diagnostic Unit Public Health Laboratory and Victorian Infectious Diseases Reference Laboratory, Doherty Institute | Caly L., Seemann T., Sait, M., Schultz M., Druce J., Sherry, N. |
| EPI_ISL_426964 | hCoV-19/Australia/VIC668/2020 | Oceania / Australia / Victoria | 2020-03-27 | Victorian Infectious Diseases Reference Laboratory (VIDRL) | Microbiological Diagnostic Unit Public Health Laboratory and Victorian Infectious Diseases Reference Laboratory, Doherty Institute | Caly L., Seemann T., Sait, M., Schultz M., Druce J., Sherry, N. |
| EPI_ISL_426968 | hCoV-19/Australia/VIC672/2020 | Oceania / Australia / Victoria | 2020-03-27 | Victorian Infectious Diseases Reference Laboratory (VIDRL) | Microbiological Diagnostic Unit Public Health Laboratory and Victorian Infectious Diseases Reference Laboratory, Doherty Institute | Caly L., Seemann T., Sait, M., Schultz M., Druce J., Sherry, N. |
| EPI_ISL_426970 | hCoV-19/Australia/VIC674/2020 | Oceania / Australia / Victoria | 2020-03-27 | Victorian Infectious Diseases Reference Laboratory (VIDRL) | Microbiological Diagnostic Unit Public Health Laboratory and Victorian Infectious Diseases Reference Laboratory, Doherty Institute | Caly L., Seemann T., Sait, M., Schultz M., Druce J., Sherry, N. |
| EPI_ISL_426971 | hCoV-19/Australia/VIC676/2020 | Oceania / Australia / Victoria | 2020-03-27 | Victorian Infectious Diseases Reference Laboratory (VIDRL) | Microbiological Diagnostic Unit Public Health Laboratory and Victorian Infectious Diseases Reference Laboratory, Doherty Institute | Caly L., Seemann T., Sait, M., Schultz M., Druce J., Sherry, N. |
| EPI_ISL_426972 | hCoV-19/Australia/VIC677/2020 | Oceania / Australia / Victoria | 2020-03-27 | Victorian Infectious Diseases Reference Laboratory (VIDRL) | Microbiological Diagnostic Unit Public Health Laboratory and Victorian Infectious Diseases Reference Laboratory, Doherty Institute | Caly L., Seemann T., Sait, M., Schultz M., Druce J., Sherry, N. |
| EPI_ISL_426973 | hCoV-19/Australia/VIC678/2020 | Oceania / Australia / Victoria | 2020-03-27 | Victorian Infectious Diseases Reference Laboratory (VIDRL) | Microbiological Diagnostic Unit Public Health Laboratory and Victorian Infectious Diseases Reference Laboratory, Doherty Institute | Caly L., Seemann T., Sait, M., Schultz M., Druce J., Sherry, N. |
| EPI_ISL_426974 | hCoV-19/Australia/VIC679/2020 | Oceania / Australia / Victoria | 2020-03-27 | Victorian Infectious Diseases Reference Laboratory (VIDRL) | Microbiological Diagnostic Unit Public Health Laboratory and Victorian Infectious Diseases Reference Laboratory, Doherty Institute | Caly L., Seemann T., Sait, M., Schultz M., Druce J., Sherry, N. |
| EPI_ISL_426975 | hCoV-19/Australia/VIC680/2020 | Oceania / Australia / Victoria | 2020-03-27 | Victorian Infectious Diseases Reference Laboratory (VIDRL) | Microbiological Diagnostic Unit Public Health Laboratory and Victorian Infectious Diseases Reference Laboratory, Doherty Institute | Caly L., Seemann T., Sait, M., Schultz M., Druce J., Sherry, N. |
| EPI_ISL_426976 | hCoV-19/Australia/VIC681/2020 | Oceania / Australia / Victoria | 2020-03-27 | Victorian Infectious Diseases Reference Laboratory (VIDRL) | Microbiological Diagnostic Unit Public Health Laboratory and Victorian Infectious Diseases Reference Laboratory, Doherty Institute | Caly L., Seemann T., Sait, M., Schultz M., Druce J., Sherry, N. |
| EPI_ISL_426977 | hCoV-19/Australia/VIC682/2020 | Oceania / Australia / Victoria | 2020-03-29 | Victorian Infectious Diseases Reference Laboratory (VIDRL) | Microbiological Diagnostic Unit Public Health Laboratory and Victorian Infectious Diseases Reference Laboratory, Doherty Institute | Caly L., Seemann T., Sait, M., Schultz M., Druce J., Sherry, N. |
| EPI_ISL_426978 | hCoV-19/Australia/VIC683/2020 | Oceania / Australia / Victoria | 2020-03-29 | Victorian Infectious Diseases Reference Laboratory (VIDRL) | Microbiological Diagnostic Unit Public Health Laboratory and Victorian Infectious Diseases Reference Laboratory, Doherty Institute | Caly L., Seemann T., Sait, M., Schultz M., Druce J., Sherry, N. |
| EPI_ISL_426980 | hCoV-19/Australia/VIC685/2020 | Oceania / Australia / Victoria | 2020-03-28 | Victorian Infectious Diseases Reference Laboratory (VIDRL) | Microbiological Diagnostic Unit Public Health Laboratory and Victorian Infectious Diseases Reference Laboratory, Doherty Institute | Caly L., Seemann T., Sait, M., Schultz M., Druce J., Sherry, N. |
| EPI_ISL_426981 | hCoV-19/Australia/VIC687/2020 | Oceania / Australia / Victoria | 2020-03-29 | Victorian Infectious Diseases Reference Laboratory (VIDRL) | Microbiological Diagnostic Unit Public Health Laboratory and Victorian Infectious Diseases Reference Laboratory, Doherty Institute | Caly L., Seemann T., Sait, M., Schultz M., Druce J., Sherry, N. |
| EPI_ISL_426986 | hCoV-19/Australia/VIC694/2020 | Oceania / Australia / Victoria | 2020-03-28 | Victorian Infectious Diseases Reference Laboratory (VIDRL) | Microbiological Diagnostic Unit Public Health Laboratory and Victorian Infectious Diseases Reference Laboratory, Doherty Institute | Caly L., Seemann T., Sait, M., Schultz M., Druce J., Sherry, N. |
| EPI_ISL_426988 | hCoV-19/Australia/VIC696/2020 | Oceania / Australia / Victoria | 2020-03-28 | Victorian Infectious Diseases Reference Laboratory (VIDRL) | Microbiological Diagnostic Unit Public Health Laboratory and Victorian Infectious Diseases Reference Laboratory, Doherty Institute | Caly L., Seemann T., Sait, M., Schultz M., Druce J., Sherry, N. |
| EPI_ISL_426989 | hCoV-19/Australia/VIC697/2020 | Oceania / Australia / Victoria | 2020-03-28 | Victorian Infectious Diseases Reference Laboratory (VIDRL) | Microbiological Diagnostic Unit Public Health Laboratory and Victorian Infectious Diseases Reference Laboratory, Doherty Institute | Caly L., Seemann T., Sait, M., Schultz M., Druce J., Sherry, N. |
| EPI_ISL_426993 | hCoV-19/Australia/VIC706/2020 | Oceania / Australia / Victoria | 2020-03-30 | Victorian Infectious Diseases Reference Laboratory (VIDRL) | Microbiological Diagnostic Unit Public Health Laboratory and Victorian Infectious Diseases Reference Laboratory, Doherty Institute | Caly L., Seemann T., Sait, M., Schultz M., Druce J., Sherry, N. |
| EPI_ISL_426994 | hCoV-19/Australia/VIC707/2020 | Oceania / Australia / Victoria | 2020-03-29 | Victorian Infectious Diseases Reference Laboratory (VIDRL) | Microbiological Diagnostic Unit Public Health Laboratory and Victorian Infectious Diseases Reference Laboratory, Doherty Institute | Caly L., Seemann T., Sait, M., Schultz M., Druce J., Sherry, N. |
| EPI_ISL_426995 | hCoV-19/Australia/VIC708/2020 | Oceania / Australia / Victoria | 2020-03-29 | Victorian Infectious Diseases Reference Laboratory (VIDRL) | Microbiological Diagnostic Unit Public Health Laboratory and Victorian Infectious Diseases Reference Laboratory, Doherty Institute | Caly L., Seemann T., Sait, M., Schultz M., Druce J., Sherry, N. |
| EPI_ISL_426997 | hCoV-19/Australia/VIC710/2020 | Oceania / Australia / Victoria | 2020-03-29 | Victorian Infectious Diseases Reference Laboratory (VIDRL) | Microbiological Diagnostic Unit Public Health Laboratory and Victorian Infectious Diseases Reference Laboratory, Doherty Institute | Caly L., Seemann T., Sait, M., Schultz M., Druce J., Sherry, N. |
| EPI_ISL_426999 | hCoV-19/Australia/VIC712/2020 | Oceania / Australia / Victoria | 2020-03-29 | Victorian Infectious Diseases Reference Laboratory (VIDRL) | Microbiological Diagnostic Unit Public Health Laboratory and Victorian Infectious Diseases Reference Laboratory, Doherty Institute | Caly L., Seemann T., Sait, M., Schultz M., Druce J., Sherry, N. |
| EPI_ISL_427100 | hCoV-19/Australia/VIC836/2020 | Oceania / Australia / Victoria | 2020-04-04 | Victorian Infectious Diseases Reference Laboratory (VIDRL) | Microbiological Diagnostic Unit Public Health Laboratory and Victorian Infectious Diseases Reference Laboratory, Doherty Institute | Caly L., Seemann T., Sait, M., Schultz M., Druce J., Sherry, N. |
| EPI_ISL_427101 | hCoV-19/Australia/VIC837/2020 | Oceania / Australia / Victoria | 2020-04-04 | Victorian Infectious Diseases Reference Laboratory (VIDRL) | Microbiological Diagnostic Unit Public Health Laboratory and Victorian Infectious Diseases Reference Laboratory, Doherty Institute | Caly L., Seemann T., Sait, M., Schultz M., Druce J., Sherry, N. |
| EPI_ISL_427102 | hCoV-19/Australia/VIC842/2020 | Oceania / Australia / Victoria | 2020-04-04 | Victorian Infectious Diseases Reference Laboratory (VIDRL) | Microbiological Diagnostic Unit Public Health Laboratory and Victorian Infectious Diseases Reference Laboratory, Doherty Institute | Caly L., Seemann T., Sait, M., Schultz M., Druce J., Sherry, N. |
| EPI_ISL_427103 | hCoV-19/Australia/VIC843/2020 | Oceania / Australia / Victoria | 2020-04-04 | Victorian Infectious Diseases Reference Laboratory (VIDRL) | Microbiological Diagnostic Unit Public Health Laboratory and Victorian Infectious Diseases Reference Laboratory, Doherty Institute | Caly L., Seemann T., Sait, M., Schultz M., Druce J., Sherry, N. |
| EPI_ISL_427105 | hCoV-19/Australia/VIC845/2020 | Oceania / Australia / Victoria | 2020-04-04 | Victorian Infectious Diseases Reference Laboratory (VIDRL) | Microbiological Diagnostic Unit Public Health Laboratory and Victorian Infectious Diseases Reference Laboratory, Doherty Institute | Caly L., Seemann T., Sait, M., Schultz M., Druce J., Sherry, N. |
| EPI_ISL_427106 | hCoV-19/Australia/VIC846/2020 | Oceania / Australia / Victoria | 2020-04-04 | Victorian Infectious Diseases Reference Laboratory (VIDRL) | Microbiological Diagnostic Unit Public Health Laboratory and Victorian Infectious Diseases Reference Laboratory, Doherty Institute | Caly L., Seemann T., Sait, M., Schultz M., Druce J., Sherry, N. |
| EPI_ISL_427107 | hCoV-19/Australia/VIC847/2020 | Oceania / Australia / Victoria | 2020-04-04 | Victorian Infectious Diseases Reference Laboratory (VIDRL) | Microbiological Diagnostic Unit Public Health Laboratory and Victorian Infectious Diseases Reference Laboratory, Doherty Institute | Caly L., Seemann T., Sait, M., Schultz M., Druce J., Sherry, N. |
| EPI_ISL_427108 | hCoV-19/Australia/VIC848/2020 | Oceania / Australia / Victoria | 2020-04-04 | Victorian Infectious Diseases Reference Laboratory (VIDRL) | Microbiological Diagnostic Unit Public Health Laboratory and Victorian Infectious Diseases Reference Laboratory, Doherty Institute | Caly L., Seemann T., Sait, M., Schultz M., Druce J., Sherry, N. |
| EPI_ISL_427109 | hCoV-19/Australia/VIC849/2020 | Oceania / Australia / Victoria | 2020-04-04 | Victorian Infectious Diseases Reference Laboratory (VIDRL) | Microbiological Diagnostic Unit Public Health Laboratory and Victorian Infectious Diseases Reference Laboratory, Doherty Institute | Caly L., Seemann T., Sait, M., Schultz M., Druce J., Sherry, N. |
| EPI_ISL_427110 | hCoV-19/Australia/VIC850/2020 | Oceania / Australia / Victoria | 2020-04-04 | Victorian Infectious Diseases Reference Laboratory (VIDRL) | Microbiological Diagnostic Unit Public Health Laboratory and Victorian Infectious Diseases Reference Laboratory, Doherty Institute | Caly L., Seemann T., Sait, M., Schultz M., Druce J., Sherry, N. |
| EPI_ISL_427111 | hCoV-19/Australia/VIC851/2020 | Oceania / Australia / Victoria | 2020-04-04 | Victorian Infectious Diseases Reference Laboratory (VIDRL) | Microbiological Diagnostic Unit Public Health Laboratory and Victorian Infectious Diseases Reference Laboratory, Doherty Institute | Caly L., Seemann T., Sait, M., Schultz M., Druce J., Sherry, N. |
| EPI_ISL_427113 | hCoV-19/Australia/VIC853/2020 | Oceania / Australia / Victoria | 2020-04-04 | Victorian Infectious Diseases Reference Laboratory (VIDRL) | Microbiological Diagnostic Unit Public Health Laboratory and Victorian Infectious Diseases Reference Laboratory, Doherty Institute | Caly L., Seemann T., Sait, M., Schultz M., Druce J., Sherry, N. |
| EPI_ISL_427115 | hCoV-19/Australia/VIC856/2020 | Oceania / Australia / Victoria | 2020-04-04 | Victorian Infectious Diseases Reference Laboratory (VIDRL) | Microbiological Diagnostic Unit Public Health Laboratory and Victorian Infectious Diseases Reference Laboratory, Doherty Institute | Caly L., Seemann T., Sait, M., Schultz M., Druce J., Sherry, N. |
| EPI_ISL_427116 | hCoV-19/Australia/VIC857/2020 | Oceania / Australia / Victoria | 2020-04-05 | Victorian Infectious Diseases Reference Laboratory (VIDRL) | Microbiological Diagnostic Unit Public Health Laboratory and Victorian Infectious Diseases Reference Laboratory, Doherty Institute | Caly L., Seemann T., Sait, M., Schultz M., Druce J., Sherry, N. |
| EPI_ISL_427117 | hCoV-19/Australia/VIC858/2020 | Oceania / Australia / Victoria | 2020-04-05 | Victorian Infectious Diseases Reference Laboratory (VIDRL) | Microbiological Diagnostic Unit Public Health Laboratory and Victorian Infectious Diseases Reference Laboratory, Doherty Institute | Caly L., Seemann T., Sait, M., Schultz M., Druce J., Sherry, N. |
| EPI_ISL_427118 | hCoV-19/Australia/VIC860/2020 | Oceania / Australia / Victoria | 2020-04-05 | Victorian Infectious Diseases Reference Laboratory (VIDRL) | Microbiological Diagnostic Unit Public Health Laboratory and Victorian Infectious Diseases Reference Laboratory, Doherty Institute | Caly L., Seemann T., Sait, M., Schultz M., Druce J., Sherry, N. |
| EPI_ISL_427119 | hCoV-19/Australia/VIC862/2020 | Oceania / Australia / Victoria | 2020-04-05 | Victorian Infectious Diseases Reference Laboratory (VIDRL) | Microbiological Diagnostic Unit Public Health Laboratory and Victorian Infectious Diseases Reference Laboratory, Doherty Institute | Caly L., Seemann T., Sait, M., Schultz M., Druce J., Sherry, N. |
| EPI_ISL_427120 | hCoV-19/Australia/VIC863/2020 | Oceania / Australia / Victoria | 2020-04-05 | Victorian Infectious Diseases Reference Laboratory (VIDRL) | Microbiological Diagnostic Unit Public Health Laboratory and Victorian Infectious Diseases Reference Laboratory, Doherty Institute | Caly L., Seemann T., Sait, M., Schultz M., Druce J., Sherry, N. |
| EPI_ISL_427122 | hCoV-19/Australia/VIC867/2020 | Oceania / Australia / Victoria | 2020-04-05 | Victorian Infectious Diseases Reference Laboratory (VIDRL) | Microbiological Diagnostic Unit Public Health Laboratory and Victorian Infectious Diseases Reference Laboratory, Doherty Institute | Caly L., Seemann T., Sait, M., Schultz M., Druce J., Sherry, N. |
| EPI_ISL_427123 | hCoV-19/Australia/VIC868/2020 | Oceania / Australia / Victoria | 2020-04-05 | Victorian Infectious Diseases Reference Laboratory (VIDRL) | Microbiological Diagnostic Unit Public Health Laboratory and Victorian Infectious Diseases Reference Laboratory, Doherty Institute | Caly L., Seemann T., Sait, M., Schultz M., Druce J., Sherry, N. |
| EPI_ISL_427124 | hCoV-19/Australia/VIC869/2020 | Oceania / Australia / Victoria | 2020-04-05 | Victorian Infectious Diseases Reference Laboratory (VIDRL) | Microbiological Diagnostic Unit Public Health Laboratory and Victorian Infectious Diseases Reference Laboratory, Doherty Institute | Caly L., Seemann T., Sait, M., Schultz M., Druce J., Sherry, N. |
| EPI_ISL_427125 | hCoV-19/Australia/VIC870/2020 | Oceania / Australia / Victoria | 2020-04-05 | Victorian Infectious Diseases Reference Laboratory (VIDRL) | Microbiological Diagnostic Unit Public Health Laboratory and Victorian Infectious Diseases Reference Laboratory, Doherty Institute | Caly L., Seemann T., Sait, M., Schultz M., Druce J., Sherry, N. |
| EPI_ISL_427126 | hCoV-19/Australia/VIC871/2020 | Oceania / Australia / Victoria | 2020-04-05 | Victorian Infectious Diseases Reference Laboratory (VIDRL) | Microbiological Diagnostic Unit Public Health Laboratory and Victorian Infectious Diseases Reference Laboratory, Doherty Institute | Caly L., Seemann T., Sait, M., Schultz M., Druce J., Sherry, N. |
| EPI_ISL_427128 | hCoV-19/Australia/VIC873/2020 | Oceania / Australia / Victoria | 2020-04-05 | Victorian Infectious Diseases Reference Laboratory (VIDRL) | Microbiological Diagnostic Unit Public Health Laboratory and Victorian Infectious Diseases Reference Laboratory, Doherty Institute | Caly L., Seemann T., Sait, M., Schultz M., Druce J., Sherry, N. |
| EPI_ISL_427129 | hCoV-19/Australia/VIC874/2020 | Oceania / Australia / Victoria | 2020-04-05 | Victorian Infectious Diseases Reference Laboratory (VIDRL) | Microbiological Diagnostic Unit Public Health Laboratory and Victorian Infectious Diseases Reference Laboratory, Doherty Institute | Caly L., Seemann T., Sait, M., Schultz M., Druce J., Sherry, N. |
| EPI_ISL_427130 | hCoV-19/Australia/VIC875/2020 | Oceania / Australia / Victoria | 2020-04-05 | Victorian Infectious Diseases Reference Laboratory (VIDRL) | Microbiological Diagnostic Unit Public Health Laboratory and Victorian Infectious Diseases Reference Laboratory, Doherty Institute | Caly L., Seemann T., Sait, M., Schultz M., Druce J., Sherry, N. |
| EPI_ISL_427131 | hCoV-19/Australia/VIC876/2020 | Oceania / Australia / Victoria | 2020-04-05 | Victorian Infectious Diseases Reference Laboratory (VIDRL) | Microbiological Diagnostic Unit Public Health Laboratory and Victorian Infectious Diseases Reference Laboratory, Doherty Institute | Caly L., Seemann T., Sait, M., Schultz M., Druce J., Sherry, N. |
| EPI_ISL_427132 | hCoV-19/Australia/VIC877/2020 | Oceania / Australia / Victoria | 2020-04-05 | Victorian Infectious Diseases Reference Laboratory (VIDRL) | Microbiological Diagnostic Unit Public Health Laboratory and Victorian Infectious Diseases Reference Laboratory, Doherty Institute | Caly L., Seemann T., Sait, M., Schultz M., Druce J., Sherry, N. |
| EPI_ISL_427133 | hCoV-19/Australia/VIC878/2020 | Oceania / Australia / Victoria | 2020-04-06 | Microbiological Diagnostic Unit Public Health Laboratory | Microbiological Diagnostic Unit Public Health Laboratory | Seemann T., Schultz M., Sait, M., Sherry, N. |
| EPI_ISL_427134 | hCoV-19/Australia/VIC879/2020 | Oceania / Australia / Victoria | 2020-04-06 | Victorian Infectious Diseases Reference Laboratory (VIDRL) | Microbiological Diagnostic Unit Public Health Laboratory and Victorian Infectious Diseases Reference Laboratory, Doherty Institute | Caly L., Seemann T., Sait, M., Schultz M., Druce J., Sherry, N. |
| EPI_ISL_427135 | hCoV-19/Australia/VIC881/2020 | Oceania / Australia / Victoria | 2020-04-06 | Victorian Infectious Diseases Reference Laboratory (VIDRL) | Microbiological Diagnostic Unit Public Health Laboratory and Victorian Infectious Diseases Reference Laboratory, Doherty Institute | Caly L., Seemann T., Sait, M., Schultz M., Druce J., Sherry, N. |
| EPI_ISL_427136 | hCoV-19/Australia/VIC883/2020 | Oceania / Australia / Victoria | 2020-04-06 | Victorian Infectious Diseases Reference Laboratory (VIDRL) | Microbiological Diagnostic Unit Public Health Laboratory and Victorian Infectious Diseases Reference Laboratory, Doherty Institute | Caly L., Seemann T., Sait, M., Schultz M., Druce J., Sherry, N. |
| EPI_ISL_427139 | hCoV-19/Australia/VIC886/2020 | Oceania / Australia / Victoria | 2020-04-06 | Victorian Infectious Diseases Reference Laboratory (VIDRL) | Microbiological Diagnostic Unit Public Health Laboratory and Victorian Infectious Diseases Reference Laboratory, Doherty Institute | Caly L., Seemann T., Sait, M., Schultz M., Druce J., Sherry, N. |
| EPI_ISL_427140 | hCoV-19/Australia/VIC887/2020 | Oceania / Australia / Victoria | 2020-04-06 | Victorian Infectious Diseases Reference Laboratory (VIDRL) | Microbiological Diagnostic Unit Public Health Laboratory and Victorian Infectious Diseases Reference Laboratory, Doherty Institute | Caly L., Seemann T., Sait, M., Schultz M., Druce J., Sherry, N. |
| EPI_ISL_427143 | hCoV-19/Australia/VIC891/2020 | Oceania / Australia / Victoria | 2020-04-06 | Victorian Infectious Diseases Reference Laboratory (VIDRL) | Microbiological Diagnostic Unit Public Health Laboratory and Victorian Infectious Diseases Reference Laboratory, Doherty Institute | Caly L., Seemann T., Sait, M., Schultz M., Druce J., Sherry, N. |
| EPI_ISL_427144 | hCoV-19/Australia/VIC892/2020 | Oceania / Australia / Victoria | 2020-04-06 | Victorian Infectious Diseases Reference Laboratory (VIDRL) | Microbiological Diagnostic Unit Public Health Laboratory and Victorian Infectious Diseases Reference Laboratory, Doherty Institute | Caly L., Seemann T., Sait, M., Schultz M., Druce J., Sherry, N. |
| EPI_ISL_427145 | hCoV-19/Australia/VIC893/2020 | Oceania / Australia / Victoria | 2020-04-06 | Victorian Infectious Diseases Reference Laboratory (VIDRL) | Microbiological Diagnostic Unit Public Health Laboratory and Victorian Infectious Diseases Reference Laboratory, Doherty Institute | Caly L., Seemann T., Sait, M., Schultz M., Druce J., Sherry, N. |
| EPI_ISL_427148 | hCoV-19/Australia/VIC898/2020 | Oceania / Australia / Victoria | 2020-04-08 | Microbiological Diagnostic Unit Public Health Laboratory | Microbiological Diagnostic Unit Public Health Laboratory | Seemann T., Schultz M., Sait, M., Sherry, N. |
| EPI_ISL_427149 | hCoV-19/Australia/VIC899/2020 | Oceania / Australia / Victoria | 2020-04-08 | Microbiological Diagnostic Unit Public Health Laboratory | Microbiological Diagnostic Unit Public Health Laboratory | Seemann T., Schultz M., Sait, M., Sherry, N. |

| | | | | | | |
|---|---|---|---|---|---|---|
| EPI_ISL_427150 | hCoV-19/Australia/VIC915/2020 | Oceania / Australia / Victoria | 2020-03-26 | Victorian Infectious Diseases Reference Laboratory (VIDRL) | Microbiological Diagnostic Unit Public Health Laboratory and Victorian Infectious Diseases Reference Laboratory, Doherty Institute | Caly L., Seemann T., Sait, M., Schultz M., Druce J., Sherry, N. |
| EPI_ISL_427151 | hCoV-19/Australia/VIC916/2020 | Oceania / Australia / Victoria | 2020-03-26 | Victorian Infectious Diseases Reference Laboratory (VIDRL) | Microbiological Diagnostic Unit Public Health Laboratory and Victorian Infectious Diseases Reference Laboratory, Doherty Institute | Caly L., Seemann T., Sait, M., Schultz M., Druce J., Sherry, N. |
| EPI_ISL_427153 | hCoV-19/Australia/VIC920/2020 | Oceania / Australia / Victoria | 2020-03-27 | Victorian Infectious Diseases Reference Laboratory (VIDRL) | Microbiological Diagnostic Unit Public Health Laboratory and Victorian Infectious Diseases Reference Laboratory, Doherty Institute | Caly L., Seemann T., Sait, M., Schultz M., Druce J., Sherry, N. |
| EPI_ISL_427154 | hCoV-19/Australia/VIC922/2020 | Oceania / Australia / Victoria | 2020-03-28 | Victorian Infectious Diseases Reference Laboratory (VIDRL) | Microbiological Diagnostic Unit Public Health Laboratory and Victorian Infectious Diseases Reference Laboratory, Doherty Institute | Caly L., Seemann T., Sait, M., Schultz M., Druce J., Sherry, N. |
| EPI_ISL_427155 | hCoV-19/Australia/VIC927/2020 | Oceania / Australia / Victoria | 2020-03-30 | Victorian Infectious Diseases Reference Laboratory (VIDRL) | Microbiological Diagnostic Unit Public Health Laboratory and Victorian Infectious Diseases Reference Laboratory, Doherty Institute | Caly L., Seemann T., Sait, M., Schultz M., Druce J., Sherry, N. |
| EPI_ISL_427157 | hCoV-19/Australia/VIC929/2020 | Oceania / Australia / Victoria | 2020-03-30 | Victorian Infectious Diseases Reference Laboratory (VIDRL) | Microbiological Diagnostic Unit Public Health Laboratory and Victorian Infectious Diseases Reference Laboratory, Doherty Institute | Caly L., Seemann T., Sait, M., Schultz M., Druce J., Sherry, N. |
| EPI_ISL_427159 | hCoV-19/Australia/VIC931/2020 | Oceania / Australia / Victoria | 2020-03-30 | Victorian Infectious Diseases Reference Laboratory (VIDRL) | Microbiological Diagnostic Unit Public Health Laboratory and Victorian Infectious Diseases Reference Laboratory, Doherty Institute | Caly L., Seemann T., Sait, M., Schultz M., Druce J., Sherry, N. |
| EPI_ISL_427160 | hCoV-19/Australia/VIC932/2020 | Oceania / Australia / Victoria | 2020-03-31 | Victorian Infectious Diseases Reference Laboratory (VIDRL) | Microbiological Diagnostic Unit Public Health Laboratory and Victorian Infectious Diseases Reference Laboratory, Doherty Institute | Caly L., Seemann T., Sait, M., Schultz M., Druce J., Sherry, N. |
| EPI_ISL_427161 | hCoV-19/USA/CT-UW-3845/2020 | North America / USA / Connecticut | 2020-03-25 | UW Virology Lab | UW Virology Lab | Pavitra Roychoudhury, Hong Xie, Keith Jerome, Alexander Greninger |
| EPI_ISL_427162 | hCoV-19/USA/CT-UW-4238/2020 | North America / USA / Connecticut | 2020-03-30 | UW Virology Lab | UW Virology Lab | Pavitra Roychoudhury, Hong Xie, Keith Jerome, Alexander Greninger |
| EPI_ISL_427163 | hCoV-19/USA/CT-UW-4244/2020 | North America / USA / Connecticut | 2020-03-31 | UW Virology Lab | UW Virology Lab | Pavitra Roychoudhury, Hong Xie, Keith Jerome, Alexander Greninger |
| EPI_ISL_427164 | hCoV-19/USA/CT-UW-4251/2020 | North America / USA / Connecticut | 2020-03-31 | UW Virology Lab | UW Virology Lab | Pavitra Roychoudhury, Hong Xie, Keith Jerome, Alexander Greninger |
| EPI_ISL_427165 | hCoV-19/USA/CT-UW-4341/2020 | North America / USA / Connecticut | 2020-03-30 | UW Virology Lab | UW Virology Lab | Pavitra Roychoudhury, Hong Xie, Keith Jerome, Alexander Greninger |
| EPI_ISL_427166 | hCoV-19/USA/ID-UW-3888/2020 | North America / USA / Idaho | 2020-03-25 | UW Virology Lab | UW Virology Lab | Pavitra Roychoudhury, Hong Xie, Keith Jerome, Alexander Greninger |
| EPI_ISL_427167 | hCoV-19/USA/ID-UW-3892/2020 | North America / USA / Idaho | 2020-03-26 | UW Virology Lab | UW Virology Lab | Pavitra Roychoudhury, Hong Xie, Keith Jerome, Alexander Greninger |
| EPI_ISL_427169 | hCoV-19/USA/ID-UW-4239/2020 | North America / USA / Idaho | 2020-03-30 | UW Virology Lab | UW Virology Lab | Pavitra Roychoudhury, Hong Xie, Keith Jerome, Alexander Greninger |
| EPI_ISL_427170 | hCoV-19/USA/ID-UW-4266/2020 | North America / USA / Idaho | 2020-03-30 | UW Virology Lab | UW Virology Lab | Pavitra Roychoudhury, Hong Xie, Keith Jerome, Alexander Greninger |
| EPI_ISL_427171 | hCoV-19/USA/ID-UW-4462/2020 | North America / USA / Idaho | 2020-03-29 | UW Virology Lab | UW Virology Lab | Pavitra Roychoudhury, Hong Xie, Keith Jerome, Alexander Greninger |
| EPI_ISL_427172 | hCoV-19/USA/ID-UW-4486/2020 | North America / USA / Idaho | 2020-03-29 | UW Virology Lab | UW Virology Lab | Pavitra Roychoudhury, Hong Xie, Keith Jerome, Alexander Greninger |
| EPI_ISL_427173 | hCoV-19/USA/OR-UW-4300/2020 | North America / USA / Oregon | 2020-03-31 | UW Virology Lab | UW Virology Lab | Pavitra Roychoudhury, Hong Xie, Keith Jerome, Alexander Greninger |
| EPI_ISL_427174 | hCoV-19/USA/WA-UW-3809/2020 | North America / USA / Washington | 2020-03-26 | UW Virology Lab | UW Virology Lab | Pavitra Roychoudhury, Hong Xie, Keith Jerome, Alexander Greninger |
| EPI_ISL_427175 | hCoV-19/USA/WA-UW-3810/2020 | North America / USA / Washington | 2020-03-26 | UW Virology Lab | UW Virology Lab | Pavitra Roychoudhury, Hong Xie, Keith Jerome, Alexander Greninger |
| EPI_ISL_427177 | hCoV-19/USA/WA-UW-3814/2020 | North America / USA / Washington | 2020-03-26 | UW Virology Lab | UW Virology Lab | Pavitra Roychoudhury, Hong Xie, Keith Jerome, Alexander Greninger |
| EPI_ISL_427178 | hCoV-19/USA/WA-UW-3815/2020 | North America / USA / Washington | 2020-03-27 | UW Virology Lab | UW Virology Lab | Pavitra Roychoudhury, Hong Xie, Keith Jerome, Alexander Greninger |
| EPI_ISL_427179 | hCoV-19/USA/WA-UW-3816/2020 | North America / USA / Washington | 2020-03-26 | UW Virology Lab | UW Virology Lab | Pavitra Roychoudhury, Hong Xie, Keith Jerome, Alexander Greninger |
| EPI_ISL_427180 | hCoV-19/USA/WA-UW-3819/2020 | North America / USA / Washington | 2020-03-26 | UW Virology Lab | UW Virology Lab | Pavitra Roychoudhury, Hong Xie, Keith Jerome, Alexander Greninger |
| EPI_ISL_427181 | hCoV-19/USA/WA-UW-3826/2020 | North America / USA / Washington | 2020-03-26 | UW Virology Lab | UW Virology Lab | Pavitra Roychoudhury, Hong Xie, Keith Jerome, Alexander Greninger |
| EPI_ISL_427182 | hCoV-19/USA/WA-UW-3827/2020 | North America / USA / Washington | 2020-03-26 | UW Virology Lab | UW Virology Lab | Pavitra Roychoudhury, Hong Xie, Keith Jerome, Alexander Greninger |
| EPI_ISL_427183 | hCoV-19/USA/WA-UW-3828/2020 | North America / USA / Washington | 2020-03-25 | UW Virology Lab | UW Virology Lab | Pavitra Roychoudhury, Hong Xie, Keith Jerome, Alexander Greninger |
| EPI_ISL_427184 | hCoV-19/USA/WA-UW-3829/2020 | North America / USA / Washington | 2020-03-25 | UW Virology Lab | UW Virology Lab | Pavitra Roychoudhury, Hong Xie, Keith Jerome, Alexander Greninger |
| EPI_ISL_427185 | hCoV-19/USA/WA-UW-3832/2020 | North America / USA / Washington | 2020-03-26 | UW Virology Lab | UW Virology Lab | Pavitra Roychoudhury, Hong Xie, Keith Jerome, Alexander Greninger |
| EPI_ISL_427187 | hCoV-19/USA/WA-UW-3834/2020 | North America / USA / Washington | 2020-03-26 | UW Virology Lab | UW Virology Lab | Pavitra Roychoudhury, Hong Xie, Keith Jerome, Alexander Greninger |
| EPI_ISL_427188 | hCoV-19/USA/WA-UW-3838/2020 | North America / USA / Washington | 2020-03-26 | UW Virology Lab | UW Virology Lab | Pavitra Roychoudhury, Hong Xie, Keith Jerome, Alexander Greninger |
| EPI_ISL_427189 | hCoV-19/USA/WA-UW-3839/2020 | North America / USA / Washington | 2020-03-26 | UW Virology Lab | UW Virology Lab | Pavitra Roychoudhury, Hong Xie, Keith Jerome, Alexander Greninger |
| EPI_ISL_427190 | hCoV-19/USA/WA-UW-3841/2020 | North America / USA / Washington | 2020-03-26 | UW Virology Lab | UW Virology Lab | Pavitra Roychoudhury, Hong Xie, Keith Jerome, Alexander Greninger |
| EPI_ISL_427191 | hCoV-19/USA/WA-UW-3843/2020 | North America / USA / Washington | 2020-03-27 | UW Virology Lab | UW Virology Lab | Pavitra Roychoudhury, Hong Xie, Keith Jerome, Alexander Greninger |
| EPI_ISL_427192 | hCoV-19/USA/WA-UW-3844/2020 | North America / USA / Washington | 2020-03-25 | UW Virology Lab | UW Virology Lab | Pavitra Roychoudhury, Hong Xie, Keith Jerome, Alexander Greninger |
| EPI_ISL_427193 | hCoV-19/USA/WA-UW-3850/2020 | North America / USA / Washington | 2020-03-26 | UW Virology Lab | UW Virology Lab | Pavitra Roychoudhury, Hong Xie, Keith Jerome, Alexander Greninger |
| EPI_ISL_427194 | hCoV-19/USA/WA-UW-3855/2020 | North America / USA / Washington | 2020-03-26 | UW Virology Lab | UW Virology Lab | Pavitra Roychoudhury, Hong Xie, Keith Jerome, Alexander Greninger |
| EPI_ISL_427195 | hCoV-19/USA/WA-UW-3858/2020 | North America / USA / Washington | 2020-03-26 | UW Virology Lab | UW Virology Lab | Pavitra Roychoudhury, Hong Xie, Keith Jerome, Alexander Greninger |
| EPI_ISL_427196 | hCoV-19/USA/WA-UW-3862/2020 | North America / USA / Washington | 2020-03-26 | UW Virology Lab | UW Virology Lab | Pavitra Roychoudhury, Hong Xie, Keith Jerome, Alexander Greninger |
| EPI_ISL_427197 | hCoV-19/USA/WA-UW-3868/2020 | North America / USA / Washington | 2020-03-25 | UW Virology Lab | UW Virology Lab | Pavitra Roychoudhury, Hong Xie, Keith Jerome, Alexander Greninger |
| EPI_ISL_427198 | hCoV-19/USA/WA-UW-3871/2020 | North America / USA / Washington | 2020-03-26 | UW Virology Lab | UW Virology Lab | Pavitra Roychoudhury, Hong Xie, Keith Jerome, Alexander Greninger |
| EPI_ISL_427199 | hCoV-19/USA/WA-UW-3880/2020 | North America / USA / Washington | 2020-03-26 | UW Virology Lab | UW Virology Lab | Pavitra Roychoudhury, Hong Xie, Keith Jerome, Alexander Greninger |
| EPI_ISL_427209 | hCoV-19/USA/WA-UW-3918/2020 | North America / USA / Washington | 2020-03-27 | UW Virology Lab | UW Virology Lab | Pavitra Roychoudhury, Hong Xie, Keith Jerome, Alexander Greninger |
| EPI_ISL_427210 | hCoV-19/USA/WA-UW-3923/2020 | North America / USA / Washington | 2020-03-27 | UW Virology Lab | UW Virology Lab | Pavitra Roychoudhury, Hong Xie, Keith Jerome, Alexander Greninger |
| EPI_ISL_427211 | hCoV-19/USA/WA-UW-3932/2020 | North America / USA / Washington | 2020-03-27 | UW Virology Lab | UW Virology Lab | Pavitra Roychoudhury, Hong Xie, Keith Jerome, Alexander Greninger |
| EPI_ISL_427230 | hCoV-19/USA/WA-UW-3990/2020 | North America / USA / Washington | 2020-03-29 | UW Virology Lab | UW Virology Lab | Pavitra Roychoudhury, Hong Xie, Keith Jerome, Alexander Greninger |
| EPI_ISL_427232 | hCoV-19/USA/WA-UW-3994/2020 | North America / USA / Washington | 2020-03-28 | UW Virology Lab | UW Virology Lab | Pavitra Roychoudhury, Hong Xie, Keith Jerome, Alexander Greninger |
| EPI_ISL_427233 | hCoV-19/USA/WA-UW-3996/2020 | North America / USA / Washington | 2020-03-28 | UW Virology Lab | UW Virology Lab | Pavitra Roychoudhury, Hong Xie, Keith Jerome, Alexander Greninger |
| EPI_ISL_427234 | hCoV-19/USA/WA-UW-3997/2020 | North America / USA / Washington | 2020-03-29 | UW Virology Lab | UW Virology Lab | Pavitra Roychoudhury, Hong Xie, Keith Jerome, Alexander Greninger |
| EPI_ISL_427235 | hCoV-19/USA/WA-UW-4035/2020 | North America / USA / Washington | 2020-03-28 | UW Virology Lab | UW Virology Lab | Pavitra Roychoudhury, Hong Xie, Keith Jerome, Alexander Greninger |
| EPI_ISL_427236 | hCoV-19/USA/WA-UW-4038/2020 | North America / USA / Washington | 2020-03-29 | UW Virology Lab | UW Virology Lab | Pavitra Roychoudhury, Hong Xie, Keith Jerome, Alexander Greninger |
| EPI_ISL_427237 | hCoV-19/USA/WA-UW-4039/2020 | North America / USA / Washington | 2020-03-29 | UW Virology Lab | UW Virology Lab | Pavitra Roychoudhury, Hong Xie, Keith Jerome, Alexander Greninger |
| EPI_ISL_427238 | hCoV-19/USA/WA-UW-4068/2020 | North America / USA / Washington | 2020-04-01 | UW Virology Lab | UW Virology Lab | Pavitra Roychoudhury, Hong Xie, Keith Jerome, Alexander Greninger |
| EPI_ISL_427239 | hCoV-19/USA/WA-UW-4082/2020 | North America / USA / Washington | 2020-04-01 | UW Virology Lab | UW Virology Lab | Pavitra Roychoudhury, Hong Xie, Keith Jerome, Alexander Greninger |
| EPI_ISL_427240 | hCoV-19/USA/WA-UW-4105/2020 | North America / USA / Washington | 2020-03-31 | UW Virology Lab | UW Virology Lab | Pavitra Roychoudhury, Hong Xie, Keith Jerome, Alexander Greninger |
| EPI_ISL_427241 | hCoV-19/USA/WA-UW-4109/2020 | North America / USA / Washington | 2020-03-31 | UW Virology Lab | UW Virology Lab | Pavitra Roychoudhury, Hong Xie, Keith Jerome, Alexander Greninger |
| EPI_ISL_427242 | hCoV-19/USA/WA-UW-4111/2020 | North America / USA / Washington | 2020-03-31 | UW Virology Lab | UW Virology Lab | Pavitra Roychoudhury, Hong Xie, Keith Jerome, Alexander Greninger |
| EPI_ISL_427245 | hCoV-19/USA/WA-UW-4180/2020 | North America / USA / Washington | 2020-04-01 | UW Virology Lab | UW Virology Lab | Pavitra Roychoudhury, Hong Xie, Keith Jerome, Alexander Greninger |
| EPI_ISL_427246 | hCoV-19/USA/WA-UW-4188/2020 | North America / USA / Washington | 2020-03-31 | UW Virology Lab | UW Virology Lab | Pavitra Roychoudhury, Hong Xie, Keith Jerome, Alexander Greninger |
| EPI_ISL_427247 | hCoV-19/USA/WA-UW-4197/2020 | North America / USA / Washington | 2020-03-31 | UW Virology Lab | UW Virology Lab | Pavitra Roychoudhury, Hong Xie, Keith Jerome, Alexander Greninger |
| EPI_ISL_427248 | hCoV-19/USA/WA-UW-4213/2020 | North America / USA / Washington | 2020-03-31 | UW Virology Lab | UW Virology Lab | Pavitra Roychoudhury, Hong Xie, Keith Jerome, Alexander Greninger |
| EPI_ISL_427249 | hCoV-19/USA/WA-UW-4217/2020 | North America / USA / Washington | 2020-03-31 | UW Virology Lab | UW Virology Lab | Pavitra Roychoudhury, Hong Xie, Keith Jerome, Alexander Greninger |
| EPI_ISL_427290 | hCoV-19/USA/OH-OSUP0017/2020 | North America / USA / Ohio / Columbus | 2020-03-31 | The Ohio State University | The Ohio State University-James Molecular Lab at Polaris | Huolin Tu, Jason Garee, Matthew Hunt, Joan-Miquel Balada-Llasat, Preeti Pancholi, Erica Vincent, Rongqin Ren, Dan Jones |
| EPI_ISL_427292 | hCoV-19/Brazil/AL-837/2020 | South America / Brazil / Alagoas / Maceic | 2020-03-18 | LACEN-AL - Laboratorio Central de Alagoas | Instituto Oswaldo Cruz FIOCRUZ - Laboratory of Respiratory Viruses and Measles (LVRS) | Paola Resende, Fernando Motta, Luciana Appolinario, Sunando Roy, Aline Mattos, Milene Miranda, Cristiana Garcia, Braulia Caetano, Maria Ogrzewalska, Priscila Born, Jonathan Lopes, Marilda Siqueira |
| EPI_ISL_427293 | hCoV-19/Brazil/BA-510/2020 | South America / Brazil / Bahia / Feira de | 2020-03-06 | LACEN-BA - Laboratório Central de Saúde Pública Professor Gonçalo Moniz | Instituto Oswaldo Cruz FIOCRUZ - Laboratory of Respiratory Viruses and Measles (LVRS) | Paola Resende, Fernando Motta, Luciana Appolinario, Sunando Roy, Aline Mattos, Milene Miranda, Cristiana Garcia, Braulia Caetano, Maria Ogrzewalska, Priscila Born, Jonathan Lopes, Marilda Siqueira |
| EPI_ISL_427294 | hCoV-19/Brazil/DF-615/2020 | South America / Brazil / Distrito Federal / | 2020-03-13 | Instituto Oswaldo Cruz FIOCRUZ - Laboratory of Respiratory Viruses and Measles (LVRS) | Instituto Oswaldo Cruz FIOCRUZ - Laboratory of Respiratory Viruses and Measles (LVRS) | Paola Resende, Fernando Motta, Luciana Appolinario, Sunando Roy, Aline Mattos, Milene Miranda, Cristiana Garcia, Braulia Caetano, Maria Ogrzewalska, Priscila Born, Jonathan Lopes, Marilda Siqueira |
| EPI_ISL_427295 | hCoV-19/Brazil/DF-619/2020 | South America / Brazil / Distrito Federal / | 2020-03-13 | Instituto Oswaldo Cruz FIOCRUZ - Laboratory of Respiratory Viruses and Measles (LVRS) | Instituto Oswaldo Cruz FIOCRUZ - Laboratory of Respiratory Viruses and Measles (LVRS) | Paola Resende, Fernando Motta, Luciana Appolinario, Sunando Roy, Aline Mattos, Milene Miranda, Cristiana Garcia, Braulia Caetano, Maria Ogrzewalska, Priscila Born, Jonathan Lopes, Marilda Siqueira |
| EPI_ISL_427296 | hCoV-19/Brazil/DF-861/2020 | South America / Brazil / Distrito Federal / | 2020-03-23 | Instituto Oswaldo Cruz FIOCRUZ - Laboratory of Respiratory Viruses and Measles (LVRS) | Instituto Oswaldo Cruz FIOCRUZ - Laboratory of Respiratory Viruses and Measles (LVRS) | Paola Resende, Fernando Motta, Luciana Appolinario, Sunando Roy, Aline Mattos, Milene Miranda, Cristiana Garcia, Braulia Caetano, Maria Ogrzewalska, Priscila Born, Jonathan Lopes, Marilda Siqueira |
| EPI_ISL_427297 | hCoV-19/Brazil/DF-862/2020 | South America / Brazil / Distrito Federal / | 2020-03-23 | Instituto Oswaldo Cruz FIOCRUZ - Laboratory of Respiratory Viruses and Measles (LVRS) | Instituto Oswaldo Cruz FIOCRUZ - Laboratory of Respiratory Viruses and Measles (LVRS) | Paola Resende, Fernando Motta, Luciana Appolinario, Sunando Roy, Aline Mattos, Milene Miranda, Cristiana Garcia, Braulia Caetano, Maria Ogrzewalska, Priscila Born, Jonathan Lopes, Marilda Siqueira |
| EPI_ISL_427298 | hCoV-19/Brazil/DF-891/2020 | South America / Brazil / Distrito Federal / | 2020-03-22 | Instituto Oswaldo Cruz FIOCRUZ - Laboratory of Respiratory Viruses and Measles (LVRS) | Instituto Oswaldo Cruz FIOCRUZ - Laboratory of Respiratory Viruses and Measles (LVRS) | Paola Resende, Fernando Motta, Luciana Appolinario, Sunando Roy, Aline Mattos, Milene Miranda, Cristiana Garcia, Braulia Caetano, Maria Ogrzewalska, Priscila Born, Jonathan Lopes, Marilda Siqueira |
| EPI_ISL_427299 | hCoV-19/Brazil/RJ-352/2020 | South America / Brazil / Rio de Janeiro / I | 2020-03-05 | Instituto Oswaldo Cruz FIOCRUZ - Laboratory of Respiratory Viruses and Measles (LVRS) | Instituto Oswaldo Cruz FIOCRUZ - Laboratory of Respiratory Viruses and Measles (LVRS) | Paola Resende, Fernando Motta, Luciana Appolinario, Sunando Roy, Aline Mattos, Milene Miranda, Cristiana Garcia, Braulia Caetano, Maria Ogrzewalska, Priscila Born, Jonathan Lopes, Marilda Siqueira |
| EPI_ISL_427408 | hCoV-19/Qatar/QA13/2020 | Asia / Qatar / Doha | 2020-03-23 | Ministry of Public Health (MoPH) | Biomedical Research Center (BRC) | Abdullatif Al-Khal, Muna A. S. Al-Maslamani, Ajaeb D. M. H. Al-Nabet, Peter V. Coyle, Einas A. E. Al-Kuwari, Nourah B. M. Younes, Hamad E. Al-Romaihi, Salih Al-Marri, Mohammed Al-Thani, Fatiha M. Benslimane, Heba A. Al-Khatib, Sonia Boughattas, Hadi M. Yassine, Asmaa A. Al-Thani. |
| EPI_ISL_427409 | hCoV-19/Qatar/QA14/2020 | Asia / Qatar / Doha | 2020-03-23 | Ministry of Public Health (MoPH) | Biomedical Research Center (BRC) | Abdullatif Al-Khal, Muna A. S. Al-Maslamani, Ajaeb D. M. H. Al-Nabet, Peter V. Coyle, Einas A. E. Al-Kuwari, Nourah B. M. Younes, Hamad E. Al-Romaihi, Salih Al-Marri, Mohammed Al-Thani, Fatiha M. Benslimane, Heba A. Al-Khatib, Sonia Boughattas, Hadi M. Yassine, Asmaa A. Al-Thani. |
| EPI_ISL_427410 | hCoV-19/Qatar/QA15/2020 | Asia / Qatar / Doha | 2020-03-23 | Ministry of Public Health (MoPH) | Biomedical Research Center (BRC) | Abdullatif Al-Khal, Muna A. S. Al-Maslamani, Ajaeb D. M. H. Al-Nabet, Peter V. Coyle, Einas A. E. Al-Kuwari, Nourah B. M. Younes, Hamad E. Al-Romaihi, Salih Al-Marri, Mohammed Al-Thani, Fatiha M. Benslimane, Heba A. Al-Khatib, Sonia Boughattas, Hadi M. Yassine, Asmaa A. Al-Thani. |
| EPI_ISL_427412 | hCoV-19/Qatar/QA17/2020 | Asia / Qatar / Doha | 2020-03-23 | Ministry of Public Health (MoPH) | Biomedical Research Center (BRC) | Abdullatif Al-Khal, Muna A. S. Al-Maslamani, Ajaeb D. M. H. Al-Nabet, Peter V. Coyle, Einas A. E. Al-Kuwari, Nourah B. M. Younes, Hamad E. Al-Romaihi, Salih Al-Marri, Mohammed Al-Thani, Fatiha M. Benslimane, Heba A. Al-Khatib, Sonia Boughattas, Hadi M. Yassine, Asmaa A. Al-Thani. |
| EPI_ISL_427414 | hCoV-19/Qatar/QA18/2020 | Asia / Qatar / Doha | 2020-03-23 | Ministry of Public Health (MoPH) | Biomedical Research Center (BRC) | Abdullatif Al-Khal, Muna A. S. Al-Maslamani, Ajaeb D. M. H. Al-Nabet, Peter V. Coyle, Einas A. E. Al-Kuwari, Nourah B. M. Younes, Hamad E. Al-Romaihi, Salih Al-Marri, Mohammed Al-Thani, Fatiha M. Benslimane, Heba A. Al-Khatib, Sonia Boughattas, Hadi M. Yassine, Asmaa A. Al-Thani. |
| EPI_ISL_427415 | hCoV-19/Qatar/QA19/2020 | Asia / Qatar / Doha | 2020-03-23 | Ministry of Public Health (MoPH) | Biomedical Research Center (BRC) | Abdullatif Al-Khal, Muna A. S. Al-Maslamani, Ajaeb D. M. H. Al-Nabet, Peter V. Coyle, Einas A. E. Al-Kuwari, Nourah B. M. Younes, Hamad E. Al-Romaihi, Salih Al-Marri, Mohammed Al-Thani, Fatiha M. Benslimane, Heba A. Al-Khatib, Sonia Boughattas, Hadi M. Yassine, Asmaa A. Al-Thani. |
| EPI_ISL_427416 | hCoV-19/Qatar/QA20/2020 | Asia / Qatar / Doha | 2020-03-23 | Ministry of Public Health (MoPH) | Biomedical Research Center (BRC) | Abdullatif Al-Khal, Muna A. S. Al-Maslamani, Ajaeb D. M. H. Al-Nabet, Peter V. Coyle, Einas A. E. Al-Kuwari, Nourah B. M. Younes, Hamad E. Al-Romaihi, Salih Al-Marri, Mohammed Al-Thani, Fatiha M. Benslimane, Heba A. Al-Khatib, Sonia Boughattas, Hadi M. Yassine, Asmaa A. Al-Thani. |
| EPI_ISL_427417 | hCoV-19/Qatar/QA21/2020 | Asia / Qatar / Doha | 2020-03-23 | Ministry of Public Health (MoPH) | Biomedical Research Center (BRC) | Abdullatif Al-Khal, Muna A. S. Al-Maslamani, Ajaeb D. M. H. Al-Nabet, Peter V. Coyle, Einas A. E. Al-Kuwari, Nourah B. M. Younes, Hamad E. Al-Romaihi, Salih Al-Marri, Mohammed Al-Thani, Fatiha M. Benslimane, Heba A. Al-Khatib, Sonia Boughattas, Hadi M. Yassine, Asmaa A. Al-Thani. |
| EPI_ISL_427418 | hCoV-19/Qatar/QA22/2020 | Asia / Qatar / Doha | 2020-03-23 | Ministry of Public Health (MoPH) | Biomedical Research Center (BRC) | Abdullatif Al-Khal, Muna A. S. Al-Maslamani, Ajaeb D. M. H. Al-Nabet, Peter V. Coyle, Einas A. E. Al-Kuwari, Nourah B. M. Younes, Hamad E. Al-Romaihi, Salih Al-Marri, Mohammed Al-Thani, Fatiha M. Benslimane, Heba A. Al-Khatib, Sonia Boughattas, Hadi M. Yassine, Asmaa A. Al-Thani. |
| EPI_ISL_427420 | hCoV-19/Qatar/QA24/2020 | Asia / Qatar / Doha | 2020-03-23 | Ministry of Public Health (MoPH) | Biomedical Research Center (BRC) | Abdullatif Al-Khal, Muna A. S. Al-Maslamani, Ajaeb D. M. H. Al-Nabet, Peter V. Coyle, Einas A. E. Al-Kuwari, Nourah B. M. Younes, Hamad E. Al-Romaihi, Salih Al-Marri, Mohammed Al-Thani, Fatiha M. Benslimane, Heba A. Al-Khatib, Sonia Boughattas, Hadi M. Yassine, Asmaa A. Al-Thani. |
| EPI_ISL_427427 | hCoV-19/USA/WI-UW-120/2020 | North America / USA / Wisconsin / Veron | 2020-04-13 | University of Wisconsin-Madison AIDS Vaccine Research Laboratories | University of Wisconsin-Madison AIDS Vaccine Research Laboratories | Gage Moreno, Katarina Braun, et al. AIDS Vaccine Research Laboratories |
| EPI_ISL_427428 | hCoV-19/USA/WI-UW-121/2020 | North America / USA / Wisconsin / Holy C | 2020-04-11 | University of Wisconsin-Madison AIDS Vaccine Research Laboratories | University of Wisconsin-Madison AIDS Vaccine Research Laboratories | Gage Moreno, Katarina Braun, et al. AIDS Vaccine Research Laboratories |
| EPI_ISL_427429 | hCoV-19/USA/WI-UW-122/2020 | North America / USA / Wisconsin / Madis | 2020-04-07 | University of Wisconsin-Madison AIDS Vaccine Research Laboratories | University of Wisconsin-Madison AIDS Vaccine Research Laboratories | Gage Moreno, Katarina Braun, et al. AIDS Vaccine Research Laboratories |
| EPI_ISL_427430 | hCoV-19/USA/WI-UW-123/2020 | North America / USA / Illinois | 2020-04-11 | University of Wisconsin-Madison AIDS Vaccine Research Laboratories | University of Wisconsin-Madison AIDS Vaccine Research Laboratories | Gage Moreno, Katarina Braun, et al. AIDS Vaccine Research Laboratories |
| EPI_ISL_427431 | hCoV-19/USA/WI-UW-124/2020 | North America / USA / Wisconsin / Madis | 2020-04-07 | University of Wisconsin-Madison AIDS Vaccine Research Laboratories | University of Wisconsin-Madison AIDS Vaccine Research Laboratories | Gage Moreno, Katarina Braun, et al. AIDS Vaccine Research Laboratories |
| EPI_ISL_427432 | hCoV-19/USA/WI-UW-125/2020 | North America / USA / Illinois | 2020-04-07 | University of Wisconsin-Madison AIDS Vaccine Research Laboratories | University of Wisconsin-Madison AIDS Vaccine Research Laboratories | Gage Moreno, Katarina Braun, et al. AIDS Vaccine Research Laboratories |
| EPI_ISL_427433 | hCoV-19/USA/WI-UW-126/2020 | North America / USA / Illinois | 2020-04-11 | University of Wisconsin-Madison AIDS Vaccine Research Laboratories | University of Wisconsin-Madison AIDS Vaccine Research Laboratories | Gage Moreno, Katarina Braun, et al. AIDS Vaccine Research Laboratories |
| EPI_ISL_427434 | hCoV-19/USA/WI-UW-127/2020 | North America / USA / Wisconsin / Madis | 2020-04-07 | University of Wisconsin-Madison AIDS Vaccine Research Laboratories | University of Wisconsin-Madison AIDS Vaccine Research Laboratories | Gage Moreno, Katarina Braun, et al. AIDS Vaccine Research Laboratories |
| EPI_ISL_427435 | hCoV-19/USA/WI-UW-128/2020 | North America / USA / Wisconsin / Sun P | 2020-04-07 | University of Wisconsin-Madison AIDS Vaccine Research Laboratories | University of Wisconsin-Madison AIDS Vaccine Research Laboratories | Gage Moreno, Katarina Braun, et al. AIDS Vaccine Research Laboratories |
| EPI_ISL_427436 | hCoV-19/USA/WI-UW-129/2020 | North America / USA / Wisconsin / Middle | 2020-04-06 | University of Wisconsin-Madison AIDS Vaccine Research Laboratories | University of Wisconsin-Madison AIDS Vaccine Research Laboratories | Gage Moreno, Katarina Braun, et al. AIDS Vaccine Research Laboratories |
| EPI_ISL_427437 | hCoV-19/USA/WI-UW-130/2020 | North America / USA / Illinois | 2020-04-13 | University of Wisconsin-Madison AIDS Vaccine Research Laboratories | University of Wisconsin-Madison AIDS Vaccine Research Laboratories | Gage Moreno, Katarina Braun, et al. AIDS Vaccine Research Laboratories |
| EPI_ISL_427438 | hCoV-19/USA/WI-UW-131/2020 | North America / USA / Wisconsin / Richla | 2020-04-08 | University of Wisconsin-Madison AIDS Vaccine Research Laboratories | University of Wisconsin-Madison AIDS Vaccine Research Laboratories | Gage Moreno, Katarina Braun, et al. AIDS Vaccine Research Laboratories |
| EPI_ISL_427439 | hCoV-19/USA/WI-UW-132/2020 | North America / USA / Wisconsin / Beloit | 2020-04-10 | University of Wisconsin-Madison AIDS Vaccine Research Laboratories | University of Wisconsin-Madison AIDS Vaccine Research Laboratories | Gage Moreno, Katarina Braun, et al. AIDS Vaccine Research Laboratories |
| EPI_ISL_427440 | hCoV-19/USA/WI-UW-133/2020 | North America / USA / Illinois | 2020-04-12 | University of Wisconsin-Madison AIDS Vaccine Research Laboratories | University of Wisconsin-Madison AIDS Vaccine Research Laboratories | Gage Moreno, Katarina Braun, et al. AIDS Vaccine Research Laboratories |
| EPI_ISL_427441 | hCoV-19/USA/WI-UW-134/2020 | North America / USA / Wisconsin | 2020-04-13 | University of Wisconsin-Madison AIDS Vaccine Research Laboratories | University of Wisconsin-Madison AIDS Vaccine Research Laboratories | Gage Moreno, Katarina Braun, et al. AIDS Vaccine Research Laboratories |
| EPI_ISL_427442 | hCoV-19/USA/WI-UW-135/2020 | North America / USA / Wisconsin / Madis | 2020-04-09 | University of Wisconsin-Madison AIDS Vaccine Research Laboratories | University of Wisconsin-Madison AIDS Vaccine Research Laboratories | Gage Moreno, Katarina Braun, et al. AIDS Vaccine Research Laboratories |
| EPI_ISL_427443 | hCoV-19/USA/WI-UW-136/2020 | North America / USA / Wisconsin / Musco | 2020-04-09 | University of Wisconsin-Madison AIDS Vaccine Research Laboratories | University of Wisconsin-Madison AIDS Vaccine Research Laboratories | Gage Moreno, Katarina Braun, et al. AIDS Vaccine Research Laboratories |
| EPI_ISL_427444 | hCoV-19/USA/WI-UW-137/2020 | North America / USA / Illinois | 2020-04-09 | University of Wisconsin-Madison AIDS Vaccine Research Laboratories | University of Wisconsin-Madison AIDS Vaccine Research Laboratories | Gage Moreno, Katarina Braun, et al. AIDS Vaccine Research Laboratories |
| EPI_ISL_427445 | hCoV-19/USA/WI-UW-138/2020 | North America / USA / Illinois | 2020-04-12 | University of Wisconsin-Madison AIDS Vaccine Research Laboratories | University of Wisconsin-Madison AIDS Vaccine Research Laboratories | Gage Moreno, Katarina Braun, et al. AIDS Vaccine Research Laboratories |
| EPI_ISL_427446 | hCoV-19/USA/WI-UW-139/2020 | North America / USA / Illinois | 2020-04-12 | University of Wisconsin-Madison AIDS Vaccine Research Laboratories | University of Wisconsin-Madison AIDS Vaccine Research Laboratories | Gage Moreno, Katarina Braun, et al. AIDS Vaccine Research Laboratories |
| EPI_ISL_427447 | hCoV-19/USA/WI-UW-140/2020 | North America / USA / Wisconsin / Waun | 2020-04-09 | University of Wisconsin-Madison AIDS Vaccine Research Laboratories | University of Wisconsin-Madison AIDS Vaccine Research Laboratories | Gage Moreno, Katarina Braun, et al. AIDS Vaccine Research Laboratories |
| EPI_ISL_427448 | hCoV-19/USA/WI-UW-141/2020 | North America / USA / Illinois | 2020-04-07 | University of Wisconsin-Madison AIDS Vaccine Research Laboratories | University of Wisconsin-Madison AIDS Vaccine Research Laboratories | Gage Moreno, Katarina Braun, et al. AIDS Vaccine Research Laboratories |
| EPI_ISL_427449 | hCoV-19/USA/WI-UW-142/2020 | North America / USA / Illinois | 2020-04-10 | University of Wisconsin-Madison AIDS Vaccine Research Laboratories | University of Wisconsin-Madison AIDS Vaccine Research Laboratories | Gage Moreno, Katarina Braun, et al. AIDS Vaccine Research Laboratories |
| EPI_ISL_428400 | hCoV-19/USA/CT-Yale-072/2020 | North America/ USA / Connecticut | 2020-03-30 | Yale COVID-19 Biorepository | Grubaugh Lab - Yale School of Public Health | Joseph Fauver, Tara Alpert, Anderson Brito, Anne Wyllie, Chantal Vogels, Mary Petrone, Cole Jensen, Chaney Kalinich, Isabel Ott, Arnau Casanovas, Catherine Muenker, Adam Moore, Alice Lu, Maria Tokuyama, Patrick Wong, Peiwen Lu, Saad Omer, Richard Martinello, Allison Nelson, Shelli Farhadian, Akiko Iwasaki, Charlese Dela Cruz, Albert Ko, Nathan Grubaugh |
| EPI_ISL_428401 | hCoV-19/USA/CT-Yale-073/2020 | North America/ USA / Connecticut | 2020-04-02 | Yale COVID-19 Biorepository | Grubaugh Lab - Yale School of Public Health | Joseph Fauver, Tara Alpert, Anderson Brito, Anne Wyllie, Chantal Vogels, Mary Petrone, Cole Jensen, Chaney Kalinich, Isabel Ott, Arnau Casanovas, Catherine Muenker, Adam Moore, Alice Lu, Maria Tokuyama, Patrick Wong, Peiwen Lu, Saad Omer, Richard Martinello, Allison Nelson, Shelli Farhadian, Akiko Iwasaki, Charlese Dela Cruz, Albert Ko, Nathan Grubaugh |
| EPI_ISL_428402 | hCoV-19/USA/CT-Yale-080/2020 | North America/ USA / Connecticut | 2020-04-06 | Yale COVID-19 Biorepository | Grubaugh Lab - Yale School of Public Health | Joseph Fauver, Tara Alpert, Anderson Brito, Anne Wyllie, Chantal Vogels, Mary Petrone, Cole Jensen, Chaney Kalinich, Isabel Ott, Arnau Casanovas, Catherine Muenker, Adam Moore, Alice Lu, Maria Tokuyama, Patrick Wong, Peiwen Lu, Saad Omer, Richard Martinello, Allison Nelson, Shelli Farhadian, Akiko Iwasaki, Charlese Dela Cruz, Albert Ko, Nathan Grubaugh |
| EPI_ISL_428403 | hCoV-19/USA/CT-Yale-107/2020 | North America/ USA / Connecticut | 2020-04-07 | Yale COVID-19 Biorepository | Grubaugh Lab - Yale School of Public Health | Joseph Fauver, Tara Alpert, Anderson Brito, Anne Wyllie, Chantal Vogels, Mary Petrone, Cole Jensen, Chaney Kalinich, Isabel Ott, Arnau Casanovas, Catherine Muenker, Adam Moore, Alice Lu, Maria Tokuyama, Patrick Wong, Peiwen Lu, Saad Omer, Richard Martinello, Allison Nelson, Shelli Farhadian, Akiko Iwasaki, Charlese Dela Cruz, Albert Ko, Nathan Grubaugh |
| EPI_ISL_428404 | hCoV-19/USA/CT-Yale-115/2020 | North America/ USA / Connecticut | 2020-04-11 | Yale COVID-19 Biorepository | Grubaugh Lab - Yale School of Public Health | Joseph Fauver, Tara Alpert, Anderson Brito, Anne Wyllie, Chantal Vogels, Mary Petrone, Cole Jensen, Chaney Kalinich, Isabel Ott, Arnau Casanovas, Catherine Muenker, Adam Moore, Alice Lu, Maria Tokuyama, Patrick Wong, Peiwen Lu, Saad Omer, Richard Martinello, Allison Nelson, Shelli Farhadian, Akiko Iwasaki, Charlese Dela Cruz, Albert Ko, Nathan Grubaugh |
| EPI_ISL_428405 | hCoV-19/USA/CT-Yale-117/2020 | North America/ USA / Connecticut | 2020-04-11 | Yale COVID-19 Biorepository | Grubaugh Lab - Yale School of Public Health | Joseph Fauver, Tara Alpert, Anderson Brito, Anne Wyllie, Chantal Vogels, Mary Petrone, Cole Jensen, Chaney Kalinich, Isabel Ott, Arnau Casanovas, Catherine Muenker, Adam Moore, Alice Lu, Maria Tokuyama, Patrick Wong, Peiwen Lu, Saad Omer, Richard Martinello, Allison Nelson, Shelli Farhadian, Akiko Iwasaki, Charlese Dela Cruz, Albert Ko, Nathan Grubaugh |
| EPI_ISL_428441 | hCoV-19/Guangdong/20SF1107/2020 | Asia / China / Guangdong / Huizhou | 2020-01-29 | Guangdong Provincial Center for Diseases Control and Prevention;Guangdong Provincial Institute of Public Health | School of Public Health, The University of Hong Kong | Bosheng Li, Haogao Gu, Lijun Liang, Zhencui Li, Hui-Ling Yen, Yao Hu, Yingchao Song , Hanri Zeng, Tie Song, Jie Wu, Leo L.M. Poon |
| EPI_ISL_428442 | hCoV-19/Guangdong/20SF1136/2020 | Asia / China / Guangdong / Shanwei | 2020-01-30 | Guangdong Provincial Center for Diseases Control and Prevention;Guangdong Provincial Institute of Public Health | School of Public Health, The University of Hong Kong | Bosheng Li, Haogao Gu, Lijun Liang, Zhencui Li, Hui-Ling Yen, Yao Hu, Yingchao Song , Hanri Zeng, Tie Song, Jie Wu, Leo L.M. Poon |
| EPI_ISL_428443 | hCoV-19/Guangdong/20SF115/2020 | Asia / China / Guangdong / Guangzhou | 2020-01-21 | Guangdong Provincial Center for Diseases Control and Prevention;Guangdong Provincial Institute of Public Health | School of Public Health, The University of Hong Kong | Bosheng Li, Haogao Gu, Lijun Liang, Zhencui Li, Hui-Ling Yen, Yao Hu, Yingchao Song , Hanri Zeng, Tie Song, Jie Wu, Leo L.M. Poon |
| EPI_ISL_428444 | hCoV-19/Guangdong/20SF1152/2020 | Asia / China / Guangdong / Huizhou | 2020-01-30 | Guangdong Provincial Center for Diseases Control and Prevention;Guangdong Provincial Institute of Public Health | School of Public Health, The University of Hong Kong | Bosheng Li, Haogao Gu, Lijun Liang, Zhencui Li, Hui-Ling Yen, Yao Hu, Yingchao Song , Hanri Zeng, Tie Song, Jie Wu, Leo L.M. Poon |
| EPI_ISL_428445 | hCoV-19/Guangdong/20SF1153/2020 | Asia / China / Guangdong / Huizhou | 2020-01-30 | Guangdong Provincial Center for Diseases Control and Prevention;Guangdong Provincial Institute of Public Health | School of Public Health, The University of Hong Kong | Bosheng Li, Haogao Gu, Lijun Liang, Zhencui Li, Hui-Ling Yen, Yao Hu, Yingchao Song , Hanri Zeng, Tie Song, Jie Wu, Leo L.M. Poon |
| EPI_ISL_428446 | hCoV-19/Guangdong/20SF1159/2020 | Asia / China / Guangdong / Zhuhai | 2020-01-31 | Guangdong Provincial Center for Diseases Control and Prevention;Guangdong Provincial Institute of Public Health | School of Public Health, The University of Hong Kong | Bosheng Li, Haogao Gu, Lijun Liang, Zhencui Li, Hui-Ling Yen, Yao Hu, Yingchao Song , Hanri Zeng, Tie Song, Jie Wu, Leo L.M. Poon |
| EPI_ISL_428447 | hCoV-19/Guangdong/20SF117/2020 | Asia / China / Guangdong / Shenzhen | 2020-01-21 | Guangdong Provincial Center for Diseases Control and Prevention;Guangdong Provincial Institute of Public Health | School of Public Health, The University of Hong Kong | Bosheng Li, Haogao Gu, Lijun Liang, Zhencui Li, Hui-Ling Yen, Yao Hu, Yingchao Song , Hanri Zeng, Tie Song, Jie Wu, Leo L.M. Poon |
| EPI_ISL_428448 | hCoV-19/Guangdong/20SF118/2020 | Asia / China / Guangdong / Shenzhen | 2020-01-21 | Guangdong Provincial Center for Diseases Control and Prevention;Guangdong Provincial Institute of Public Health | School of Public Health, The University of Hong Kong | Bosheng Li, Haogao Gu, Lijun Liang, Zhencui Li, Hui-Ling Yen, Yao Hu, Yingchao Song , Hanri Zeng, Tie Song, Jie Wu, Leo L.M. Poon |
| EPI_ISL_428449 | hCoV-19/Guangdong/20SF123/2020 | Asia / China / Guangdong / Zhanjiang | 2020-01-20 | Guangdong Provincial Center for Diseases Control and Prevention;Guangdong Provincial Institute of Public Health | School of Public Health, The University of Hong Kong | Bosheng Li, Haogao Gu, Lijun Liang, Zhencui Li, Hui-Ling Yen, Yao Hu, Yingchao Song , Hanri Zeng, Tie Song, Jie Wu, Leo L.M. Poon |
| EPI_ISL_428450 | hCoV-19/Guangdong/20SF156/2020 | Asia / China / Guangdong / Guangzhou | 2020-01-22 | Guangdong Provincial Center for Diseases Control and Prevention;Guangdong Provincial Institute of Public Health | School of Public Health, The University of Hong Kong | Bosheng Li, Haogao Gu, Lijun Liang, Zhencui Li, Hui-Ling Yen, Yao Hu, Yingchao Song , Hanri Zeng, Tie Song, Jie Wu, Leo L.M. Poon |
| EPI_ISL_428451 | hCoV-19/Guangdong/20SF179/2020 | Asia / China / Guangdong / Zhuhai | 2020-01-22 | Guangdong Provincial Center for Diseases Control and Prevention;Guangdong Provincial Institute of Public Health | School of Public Health, The University of Hong Kong | Bosheng Li, Haogao Gu, Lijun Liang, Zhencui Li, Hui-Ling Yen, Yao Hu, Yingchao Song , Hanri Zeng, Tie Song, Jie Wu, Leo L.M. Poon |
| EPI_ISL_428452 | hCoV-19/Guangdong/20SF190/2020 | Asia / China / Guangdong / Shaoguan | 2020-01-23 | Guangdong Provincial Center for Diseases Control and Prevention;Guangdong Provincial Institute of Public Health | School of Public Health, The University of Hong Kong | Bosheng Li, Haogao Gu, Lijun Liang, Zhencui Li, Hui-Ling Yen, Yao Hu, Yingchao Song , Hanri Zeng, Tie Song, Jie Wu, Leo L.M. Poon |
| EPI_ISL_428454 | hCoV-19/Guangdong/20SF198/2020 | Asia / China / Guangdong / Huizhou | 2020-01-22 | Guangdong Provincial Center for Diseases Control and Prevention;Guangdong Provincial Institute of Public Health | School of Public Health, The University of Hong Kong | Bosheng Li, Haogao Gu, Lijun Liang, Zhencui Li, Hui-Ling Yen, Yao Hu, Yingchao Song , Hanri Zeng, Tie Song, Jie Wu, Leo L.M. Poon |
| EPI_ISL_428455 | hCoV-19/Guangdong/20SF200/2020 | Asia / China / Guangdong / Yangjiang | 2020-01-23 | Guangdong Provincial Center for Diseases Control and Prevention;Guangdong Provincial Institute of Public Health | School of Public Health, The University of Hong Kong | Bosheng Li, Haogao Gu, Lijun Liang, Zhencui Li, Hui-Ling Yen, Yao Hu, Yingchao Song , Hanri Zeng, Tie Song, Jie Wu, Leo L.M. Poon |
| EPI_ISL_428458 | hCoV-19/Guangdong/20SF2546/2020 | Asia / China / Guangdong / Guangzhou | 2020-02-06 | Guangdong Provincial Center for Diseases Control and Prevention;Guangdong Provincial Institute of Public Health | School of Public Health, The University of Hong Kong | Bosheng Li, Haogao Gu, Lijun Liang, Zhencui Li, Hui-Ling Yen, Yao Hu, Yingchao Song , Hanri Zeng, Tie Song, Jie Wu, Leo L.M. Poon |
| EPI_ISL_428459 | hCoV-19/Guangdong/20SF262/2020 | Asia / China / Guangdong / Shenzhen | 2020-01-23 | Guangdong Provincial Center for Diseases Control and Prevention;Guangdong Provincial Institute of Public Health | School of Public Health, The University of Hong Kong | Bosheng Li, Haogao Gu, Lijun Liang, Zhencui Li, Hui-Ling Yen, Yao Hu, Yingchao Song , Hanri Zeng, Tie Song, Jie Wu, Leo L.M. Poon |
| EPI_ISL_428460 | hCoV-19/Guangdong/20SF273/2020 | Asia / China / Guangdong / Guangzhou | 2020-01-23 | Guangdong Provincial Center for Diseases Control and Prevention;Guangdong Provincial Institute of Public Health | School of Public Health, The University of Hong Kong | Bosheng Li, Haogao Gu, Lijun Liang, Zhencui Li, Hui-Ling Yen, Yao Hu, Yingchao Song , Hanri Zeng, Tie Song, Jie Wu, Leo L.M. Poon |
| EPI_ISL_428461 | hCoV-19/Guangdong/20SF316/2020 | Asia / China / Guangdong / Huizhou | 2020-01-24 | Guangdong Provincial Center for Diseases Control and Prevention;Guangdong Provincial Institute of Public Health | School of Public Health, The University of Hong Kong | Bosheng Li, Haogao Gu, Lijun Liang, Zhencui Li, Hui-Ling Yen, Yao Hu, Yingchao Song , Hanri Zeng, Tie Song, Jie Wu, Leo L.M. Poon |
| EPI_ISL_428462 | hCoV-19/Guangdong/20SF4047/2020 | Asia / China / Guangdong / Guangzhou | 2020-02-17 | Guangdong Provincial Center for Diseases Control and Prevention;Guangdong Provincial Institute of Public Health | School of Public Health, The University of Hong Kong | Bosheng Li, Haogao Gu, Lijun Liang, Zhencui Li, Hui-Ling Yen, Yao Hu, Yingchao Song , Hanri Zeng, Tie Song, Jie Wu, Leo L.M. Poon |
| EPI_ISL_428463 | hCoV-19/Guangdong/20SF4051/2020 | Asia / China / Guangdong / Guangzhou | 2020-02-17 | Guangdong Provincial Center for Diseases Control and Prevention;Guangdong Provincial Institute of Public Health | School of Public Health, The University of Hong Kong | Bosheng Li, Haogao Gu, Lijun Liang, Zhencui Li, Hui-Ling Yen, Yao Hu, Yingchao Song , Hanri Zeng, Tie Song, Jie Wu, Leo L.M. Poon |
| EPI_ISL_428464 | hCoV-19/Guangdong/20SF602/2020 | Asia / China / Guangdong / Zhanjiang | 2020-01-25 | Guangdong Provincial Center for Diseases Control and Prevention;Guangdong Provincial Institute of Public Health | School of Public Health, The University of Hong Kong | Bosheng Li, Haogao Gu, Lijun Liang, Zhencui Li, Hui-Ling Yen, Yao Hu, Yingchao Song , Hanri Zeng, Tie Song, Jie Wu, Leo L.M. Poon |
| EPI_ISL_428465 | hCoV-19/Guangdong/20SF616/2020 | Asia / China / Guangdong / Shenzhen | 2020-01-24 | Guangdong Provincial Center for Diseases Control and Prevention;Guangdong Provincial Institute of Public Health | School of Public Health, The University of Hong Kong | Bosheng Li, Haogao Gu, Lijun Liang, Zhencui Li, Hui-Ling Yen, Yao Hu, Yingchao Song , Hanri Zeng, Tie Song, Jie Wu, Leo L.M. Poon |
| EPI_ISL_428466 | hCoV-19/Guangdong/20SF629/2020 | Asia / China / Guangdong / Dongguan | 2020-01-26 | Guangdong Provincial Center for Diseases Control and Prevention;Guangdong Provincial Institute of Public Health | School of Public Health, The University of Hong Kong | Bosheng Li, Haogao Gu, Lijun Liang, Zhencui Li, Hui-Ling Yen, Yao Hu, Yingchao Song , Hanri Zeng, Tie Song, Jie Wu, Leo L.M. Poon |
| EPI_ISL_428467 | hCoV-19/Guangdong/20SF630/2020 | Asia / China / Guangdong / Dongguan | 2020-01-26 | Guangdong Provincial Center for Diseases Control and Prevention;Guangdong Provincial Institute of Public Health | School of Public Health, The University of Hong Kong | Bosheng Li, Haogao Gu, Lijun Liang, Zhencui Li, Hui-Ling Yen, Yao Hu, Yingchao Song , Hanri Zeng, Tie Song, Jie Wu, Leo L.M. Poon |
| EPI_ISL_428468 | hCoV-19/Guangdong/20SF632/2020 | Asia / China / Guangdong / Dongguan | 2020-01-26 | Guangdong Provincial Center for Diseases Control and Prevention;Guangdong Provincial Institute of Public Health | School of Public Health, The University of Hong Kong | Bosheng Li, Haogao Gu, Lijun Liang, Zhencui Li, Hui-Ling Yen, Yao Hu, Yingchao Song , Hanri Zeng, Tie Song, Jie Wu, Leo L.M. Poon |
| EPI_ISL_428469 | hCoV-19/Guangdong/20SF665/2020 | Asia / China / Guangdong / Zhongshan | 2020-01-27 | Guangdong Provincial Center for Diseases Control and Prevention;Guangdong Provincial Institute of Public Health | School of Public Health, The University of Hong Kong | Bosheng Li, Haogao Gu, Lijun Liang, Zhencui Li, Hui-Ling Yen, Yao Hu, Yingchao Song , Hanri Zeng, Tie Song, Jie Wu, Leo L.M. Poon |
| EPI_ISL_428470 | hCoV-19/Guangdong/20SF684/2020 | Asia / China / Guangdong / Shantou | 2020-01-27 | Guangdong Provincial Center for Diseases Control and Prevention;Guangdong Provincial Institute of Public Health | School of Public Health, The University of Hong Kong | Bosheng Li, Haogao Gu, Lijun Liang, Zhencui Li, Hui-Ling Yen, Yao Hu, Yingchao Song , Hanri Zeng, Tie Song, Jie Wu, Leo L.M. Poon |
| EPI_ISL_428471 | hCoV-19/Guangdong/20SF685/2020 | Asia / China / Guangdong / Shantou | 2020-01-27 | Guangdong Provincial Center for Diseases Control and Prevention;Guangdong Provincial Institute of Public Health | School of Public Health, The University of Hong Kong | Bosheng Li, Haogao Gu, Lijun Liang, Zhencui Li, Hui-Ling Yen, Yao Hu, Yingchao Song , Hanri Zeng, Tie Song, Jie Wu, Leo L.M. Poon |
| EPI_ISL_428472 | hCoV-19/Guangdong/20SF753/2020 | Asia / China / Guangdong / Zhuhai | 2020-01-28 | Guangdong Provincial Center for Diseases Control and Prevention;Guangdong Provincial Institute of Public Health | School of Public Health, The University of Hong Kong | Bosheng Li, Haogao Gu, Lijun Liang, Zhencui Li, Hui-Ling Yen, Yao Hu, Yingchao Song , Hanri Zeng, Tie Song, Jie Wu, Leo L.M. Poon |
| EPI_ISL_428473 | hCoV-19/Guangdong/20SF758/2020 | Asia / China / Guangdong / Zhuhai | 2020-01-27 | Guangdong Provincial Center for Diseases Control and Prevention;Guangdong Provincial Institute of Public Health | School of Public Health, The University of Hong Kong | Bosheng Li, Haogao Gu, Lijun Liang, Zhencui Li, Hui-Ling Yen, Yao Hu, Yingchao Song , Hanri Zeng, Tie Song, Jie Wu, Leo L.M. Poon |
| EPI_ISL_428474 | hCoV-19/Guangdong/20SF808/2020 | Asia / China / Guangdong / Huizhou | 2020-01-28 | Guangdong Provincial Center for Diseases Control and Prevention;Guangdong Provincial Institute of Public Health | School of Public Health, The University of Hong Kong | Bosheng Li, Haogao Gu, Lijun Liang, Zhencui Li, Hui-Ling Yen, Yao Hu, Yingchao Song , Hanri Zeng, Tie Song, Jie Wu, Leo L.M. Poon |
| EPI_ISL_428475 | hCoV-19/Guangdong/20SF812/2020 | Asia / China / Guangdong / Huizhou | 2020-01-28 | Guangdong Provincial Center for Diseases Control and Prevention;Guangdong Provincial Institute of Public Health | School of Public Health, The University of Hong Kong | Bosheng Li, Haogao Gu, Lijun Liang, Zhencui Li, Hui-Ling Yen, Yao Hu, Yingchao Song , Hanri Zeng, Tie Song, Jie Wu, Leo L.M. Poon |
| EPI_ISL_428476 | hCoV-19/Guangdong/20SF813/2020 | Asia / China / Guangdong / Huizhou | 2020-01-28 | Guangdong Provincial Center for Diseases Control and Prevention;Guangdong Provincial Institute of Public Health | School of Public Health, The University of Hong Kong | Bosheng Li, Haogao Gu, Lijun Liang, Zhencui Li, Hui-Ling Yen, Yao Hu, Yingchao Song , Hanri Zeng, Tie Song, Jie Wu, Leo L.M. Poon |
| EPI_ISL_428477 | hCoV-19/Guangdong/20SF822/2020 | Asia / China / Guangdong / Foshan | 2020-01-28 | Guangdong Provincial Center for Diseases Control and Prevention;Guangdong Provincial Institute of Public Health | School of Public Health, The University of Hong Kong | Bosheng Li, Haogao Gu, Lijun Liang, Zhencui Li, Hui-Ling Yen, Yao Hu, Yingchao Song , Hanri Zeng, Tie Song, Jie Wu, Leo L.M. Poon |
| EPI_ISL_428478 | hCoV-19/Guangdong/20SF840/2020 | Asia / China / Guangdong / Dongguan | 2020-01-29 | Guangdong Provincial Center for Diseases Control and Prevention;Guangdong Provincial Institute of Public Health | School of Public Health, The University of Hong Kong | Bosheng Li, Haogao Gu, Lijun Liang, Zhencui Li, Hui-Ling Yen, Yao Hu, Yingchao Song , Hanri Zeng, Tie Song, Jie Wu, Leo L.M. Poon |
| EPI_ISL_428479 | hCoV-19/India/nimh-0113/2020 | Asia / India / Karnataka | 2020-04-06 | District Surveillance Unit | Department of Neurovirology, National Institute of Mental Health and Neuroscience (NIMHANS) | Chitra Pattabiraman, Vijayalakshmi Reddy, Harsha PK, Risha Rasheed, Shafeeq S Hameed, Manjunatha Venkataswamy, Anita Desai, Ravi Vasanthapuram |
| EPI_ISL_428480 | hCoV-19/India/nimh-0116/2020 | Asia / India / Karnataka | 2020-04-06 | District Surveillance Unit | Department of Neurovirology, National Institute of Mental Health and Neuroscience (NIMHANS) | Chitra Pattabiraman, Vijayalakshmi Reddy, Harsha PK, Risha Rasheed, Shafeeq S Hameed, Manjunatha Venkataswamy, Anita Desai, Ravi Vasanthapuram |
| EPI_ISL_428481 | hCoV-19/India/nimh-0130/2020 | Asia / India / Karnataka | 2020-04-06 | District Surveillance Unit | Department of Neurovirology, National Institute of Mental Health and Neuroscience (NIMHANS) | Chitra Pattabiraman, Vijayalakshmi Reddy, Harsha PK, Risha Rasheed, Shafeeq S Hameed, Manjunatha Venkataswamy, Anita Desai, Ravi Vasanthapuram |
| EPI_ISL_428482 | hCoV-19/India/nimh-0182/2020 | Asia / India / Karnataka | 2020-04-08 | District Surveillance Unit | Department of Neurovirology, National Institute of Mental Health and Neuroscience (NIMHANS) | Chitra Pattabiraman, Vijayalakshmi Reddy, Harsha PK, Risha Rasheed, Shafeeq S Hameed, Manjunatha Venkataswamy, Anita Desai, Ravi Vasanthapuram |
| EPI_ISL_428483 | hCoV-19/India/nimh-0318/2020 | Asia / India / Karnataka | 2020-04-10 | District Surveillance Unit | Department of Neurovirology, National Institute of Mental Health and Neuroscience (NIMHANS) | Chitra Pattabiraman, Vijayalakshmi Reddy, Harsha PK, Risha Rasheed, Shafeeq S Hameed, Manjunatha Venkataswamy, Anita Desai, Ravi Vasanthapuram |
| EPI_ISL_428484 | hCoV-19/India/nimh-0351/2020 | Asia / India / Karnataka | 2020-04-10 | District Surveillance Unit | Department of Neurovirology, National Institute of Mental Health and Neuroscience (NIMHANS) | Chitra Pattabiraman, Vijayalakshmi Reddy, Harsha PK, Risha Rasheed, Shafeeq S Hameed, Manjunatha Venkataswamy, Anita Desai, Ravi Vasanthapuram |
| EPI_ISL_428485 | hCoV-19/India/nimh-0834/2020 | Asia / India / Karnataka | 2020-04-12 | District Surveillance Unit | Department of Neurovirology, National Institute of Mental Health and Neuroscience (NIMHANS) | Chitra Pattabiraman, Vijayalakshmi Reddy, Harsha PK, Risha Rasheed, Shafeeq S Hameed, Manjunatha Venkataswamy, Anita Desai, Ravi Vasanthapuram |
| EPI_ISL_428486 | hCoV-19/India/nimh-0996/2020 | Asia / India / Karnataka | 2020-04-14 | District Surveillance Unit | Department of Neurovirology, National Institute of Mental Health and Neuroscience (NIMHANS) | Chitra Pattabiraman, Vijayalakshmi Reddy, Harsha PK, Risha Rasheed, Shafeeq S Hameed, Manjunatha Venkataswamy, Anita Desai, Ravi Vasanthapuram |
| EPI_ISL_428487 | hCoV-19/India/nimh-1071/2020 | Asia / India / Karnataka | 2020-04-14 | District Surveillance Unit | Department of Neurovirology, National Institute of Mental Health and Neuroscience (NIMHANS) | Chitra Pattabiraman, Vijayalakshmi Reddy, Harsha PK, Risha Rasheed, Shafeeq S Hameed, Manjunatha Venkataswamy, Anita Desai, Ravi Vasanthapuram |
| EPI_ISL_428488 | hCoV-19/Taiwan/8/2020 | Asia / Taiwan | 2020-01-31 | Centers for Disease Control, R.O.C. (Taiwan) | Centers for Disease Control, R.O.C. (Taiwan) | Ji-Rong Yang, Yu-Chi Lin, Jung-Jung Mu, Ming-Tsan Liu |
| EPI_ISL_428489 | hCoV-19/Taiwan/5/2020 | Asia / Taiwan | 2020-01-31 | Centers for Disease Control, R.O.C. (Taiwan) | Centers for Disease Control, R.O.C. (Taiwan) | Ji-Rong Yang, Yu-Chi Lin, Jung-Jung Mu, Ming-Tsan Liu |
| EPI_ISL_428490 | hCoV-19/Taiwan/78/2020 | Asia / Taiwan | 2020-03-16 | Centers for Disease Control, R.O.C. (Taiwan) | Centers for Disease Control, R.O.C. (Taiwan) | Ji-Rong Yang, Yu-Chi Lin, Jung-Jung Mu, Ming-Tsan Liu |
| EPI_ISL_428491 | hCoV-19/Taiwan/58/2020 | Asia / Taiwan | 2020-03-15 | Centers for Disease Control, R.O.C. (Taiwan) | Centers for Disease Control, R.O.C. (Taiwan) | Ji-Rong Yang, Yu-Chi Lin, Jung-Jung Mu, Ming-Tsan Liu |
| EPI_ISL_428700 | hCoV-19/Spain/Madrid_H12_2501/2020 | Europe / Spain / Madrid | 2020-03-27 | Hospital Universitario 12 de Octubre | Hospital Universitario 12 de Octubre | Esther Viedma, Sara González, Raúl Recio, Elías Dahdouh, Fernando Lázaro, Julio García, Mª Dolores Folgueira, Jesús Mingorance, Rafael Delgado |
| EPI_ISL_428701 | hCoV-19/Spain/Madrid_H12_2902/2020 | Europe / Spain / Madrid | 2020-03-27 | Hospital Universitario 12 de Octubre | Hospital Universitario 12 de Octubre | Esther Viedma, Sara González, Raúl Recio, Elías Dahdouh, Fernando Lázaro, Julio García, Mª Dolores Folgueira, Jesús Mingorance, Rafael Delgado |
| EPI_ISL_428703 | hCoV-19/Spain/Madrid_H12_3304/2020 | Europe / Spain / Madrid | 2020-03-11 | Hospital Universitario 12 de Octubre | Hospital Universitario 12 de Octubre | Esther Viedma, Sara González, Raúl Recio, Elías Dahdouh, Fernando Lázaro, Julio García, Mª Dolores Folgueira, Jesús Mingorance, Rafael Delgado |
| EPI_ISL_428704 | hCoV-19/Spain/Madrid_H12_3405/2020 | Europe / Spain / Madrid | 2020-03-11 | Hospital Universitario 12 de Octubre | Hospital Universitario 12 de Octubre | Esther Viedma, Sara González, Raúl Recio, Elías Dahdouh, Fernando Lázaro, Julio García, Mª Dolores Folgueira, Jesús Mingorance, Rafael Delgado |
| EPI_ISL_428705 | hCoV-19/Spain/Madrid_H12_3506/2020 | Europe / Spain / Madrid | 2020-03-21 | Hospital Universitario 12 de Octubre | Hospital Universitario 12 de Octubre | Esther Viedma, Sara González, Raúl Recio, Elías Dahdouh, Fernando Lázaro, Julio García, Mª Dolores Folgueira, Jesús Mingorance, Rafael Delgado |
| EPI_ISL_428706 | hCoV-19/Spain/Madrid_H12_3607/2020 | Europe / Spain / Madrid | 2020-03-21 | Hospital Universitario 12 de Octubre | Hospital Universitario 12 de Octubre | Esther Viedma, Sara González, Raúl Recio, Elías Dahdouh, Fernando Lázaro, Julio García, Mª Dolores Folgueira, Jesús Mingorance, Rafael Delgado |
| EPI_ISL_428707 | hCoV-19/Spain/Madrid_H12_3708/2020 | Europe / Spain / Madrid | 2020-03-26 | Hospital Universitario 12 de Octubre | Hospital Universitario 12 de Octubre | Esther Viedma, Sara González, Raúl Recio, Elías Dahdouh, Fernando Lázaro, Julio García, Mª Dolores Folgueira, Jesús Mingorance, Rafael Delgado |
| EPI_ISL_428708 | hCoV-19/Spain/Madrid_H12_3809/2020 | Europe / Spain / Madrid | 2020-03-21 | Hospital Universitario 12 de Octubre | Hospital Universitario 12 de Octubre | Esther Viedma, Sara González, Raúl Recio, Elías Dahdouh, Fernando Lázaro, Julio García, Mª Dolores Folgueira, Jesús Mingorance, Rafael Delgado |
| EPI_ISL_428709 | hCoV-19/Spain/Madrid_H12_3910/2020 | Europe / Spain / Madrid | 2020-03-21 | Hospital Universitario 12 de Octubre | Hospital Universitario 12 de Octubre | Esther Viedma, Sara González, Raúl Recio, Elías Dahdouh, Fernando Lázaro, Julio García, Mª Dolores Folgueira, Jesús Mingorance, Rafael Delgado |
| EPI_ISL_428710 | hCoV-19/Spain/Madrid_H12_4011/2020 | Europe / Spain / Madrid | 2020-03-22 | Hospital Universitario 12 de Octubre | Hospital Universitario 12 de Octubre | Esther Viedma, Sara González, Raúl Recio, Elías Dahdouh, Fernando Lázaro, Julio García, Mª Dolores Folgueira, Jesús Mingorance, Rafael Delgado |
| EPI_ISL_428711 | hCoV-19/Spain/Madrid_H12_4112/2020 | Europe / Spain / Madrid | 2020-03-22 | Hospital Universitario 12 de Octubre | Hospital Universitario 12 de Octubre | Esther Viedma, Sara González, Raúl Recio, Elías Dahdouh, Fernando Lázaro, Julio García, Mª Dolores Folgueira, Jesús Mingorance, Rafael Delgado |
| EPI_ISL_428712 | hCoV-19/Turkey/HSGM-4706/2020 | Europe / Turkey / Karaman | 2020-03-17 | Ministry of Health Turkey | Ministry of Health Turkey | Fatma Bayrakdar,Ayşe Başak Altaş,Yasemin Coşgun,Gülay Korukluoğlu,Selçuk Kılıç |
| EPI_ISL_428713 | hCoV-19/Turkey/HSGM-5516/2020 | Europe / Turkey / Ankara | 2020-03-18 | Ministry of Health Turkey | Ministry of Health Turkey | Fatma Bayrakdar,Ayşe Başak Altaş,Yasemin Coşgun,Gülay Korukluoğlu,Selçuk Kılıç |
| EPI_ISL_428714 | hCoV-19/Turkey/HSGM-5602/2020 | Europe / Turkey / Kastamonu | 2020-03-18 | Ministry of Health Turkey | Ministry of Health Turkey | Fatma Bayrakdar,Ayşe Başak Altaş,Yasemin Coşgun,Gülay Korukluoğlu,Selçuk Kılıç |
| EPI_ISL_428715 | hCoV-19/Turkey/HSGM-5624/2020 | Europe / Turkey / Nevsehir | 2020-03-18 | Ministry of Health Turkey | Ministry of Health Turkey | Fatma Bayrakdar,Ayşe Başak Altaş,Yasemin Coşgun,Gülay Korukluoğlu,Selçuk Kılıç |
| EPI_ISL_428723 | hCoV-19/Turkey/HSGM-8964/2020 | Europe / Turkey / Aksaray | 2020-03-22 | Ministry of Health Turkey | Ministry of Health Turkey | Fatma Bayrakdar,Ayşe Başak Altaş,Yasemin Coşgun,Gülay Korukluoğlu,Selçuk Kılıç |
| EPI_ISL_428725 | hCoV-19/USA/WI-UW-251/2020 | North America / USA / Wisconsin | 2020-03-14 | University of Wisconsin-Madison AIDS Vaccine Research Laboratories | University of Wisconsin-Madison AIDS Vaccine Research Laboratories | Gage Moreno, Katarina Braun, et al. AIDS Vaccine Research Laboratories |
| EPI_ISL_428726 | hCoV-19/USA/WI-UW-252/2020 | North America / USA / Wisconsin | 2020-03-24 | University of Wisconsin-Madison AIDS Vaccine Research Laboratories | University of Wisconsin-Madison AIDS Vaccine Research Laboratories | Gage Moreno, Katarina Braun, et al. AIDS Vaccine Research Laboratories |
| EPI_ISL_428727 | hCoV-19/USA/WI-UW-253/2020 | North America / USA / Wisconsin | 2020-03-26 | University of Wisconsin-Madison AIDS Vaccine Research Laboratories | University of Wisconsin-Madison AIDS Vaccine Research Laboratories | Gage Moreno, Katarina Braun, et al. AIDS Vaccine Research Laboratories |
| EPI_ISL_428728 | hCoV-19/USA/WI-UW-254/2020 | North America / USA / Wisconsin | 2020-04-01 | University of Wisconsin-Madison AIDS Vaccine Research Laboratories | University of Wisconsin-Madison AIDS Vaccine Research Laboratories | Gage Moreno, Katarina Braun, et al. AIDS Vaccine Research Laboratories |
| EPI_ISL_428729 | hCoV-19/USA/WI-UW-255/2020 | North America / USA / Wisconsin | 2020-04-02 | University of Wisconsin-Madison AIDS Vaccine Research Laboratories | University of Wisconsin-Madison AIDS Vaccine Research Laboratories | Gage Moreno, Katarina Braun, et al. AIDS Vaccine Research Laboratories |
| EPI_ISL_428730 | hCoV-19/USA/WI-UW-256/2020 | North America / USA / Wisconsin | 2020-04-06 | University of Wisconsin-Madison AIDS Vaccine Research Laboratories | University of Wisconsin-Madison AIDS Vaccine Research Laboratories | Gage Moreno, Katarina Braun, et al. AIDS Vaccine Research Laboratories |
| EPI_ISL_428731 | hCoV-19/USA/WI-UW-257/2020 | North America / USA / Wisconsin | 2020-04-06 | University of Wisconsin-Madison AIDS Vaccine Research Laboratories | University of Wisconsin-Madison AIDS Vaccine Research Laboratories | Gage Moreno, Katarina Braun, et al. AIDS Vaccine Research Laboratories |
| EPI_ISL_428732 | hCoV-19/USA/WI-UW-258/2020 | North America / USA / Wisconsin | 2020-04-06 | University of Wisconsin-Madison AIDS Vaccine Research Laboratories | University of Wisconsin-Madison AIDS Vaccine Research Laboratories | Gage Moreno, Katarina Braun, et al. AIDS Vaccine Research Laboratories |
| EPI_ISL_428740 | hCoV-19/USA/CT-Yale-019/2020 | North America / USA / Connecticut | 2020-03-19 | Yale Clinical Virology Laboratory | Grubaugh Lab - Yale School of Public Health | Joseph Fauver, Anderson Brito, Tara Alpert, Chantal Vogels, Ellen Foxman, Albert Ko, Marie Landry, Nathan Grubaugh |
| EPI_ISL_428745 | hCoV-19/USA/CT-Yale-070/2020 | North America / USA / Connecticut | 2020-03-26 | Yale COVID-19 Biorepository | Grubaugh Lab - Yale School of Public Health | Joseph Fauver, Tara Alpert, Anderson Brito, Anne Wyllie, Chantal Vogels, Mary Petrone, Cole Jensen, Chaney Kalinich, Isabel Ott, Arnau Casanovas, Catherine Muenker, Adam Moore, Alice Lu, Maria Tokuyama, Patrick Wong, Peiwen Lu, Saad Omer, Richard Martinello, Allison Nelson, Shelli Farhadian, Akiko Iwasaki, Charlese Dela Cruz, Albert Ko, Nathan Grubaugh |
| EPI_ISL_428746 | hCoV-19/USA/CT-Yale-082/2020 | North America / USA / Connecticut | 2020-04-07 | Yale COVID-19 Biorepository | Grubaugh Lab - Yale School of Public Health | Joseph Fauver, Tara Alpert, Anderson Brito, Anne Wyllie, Chantal Vogels, Mary Petrone, Cole Jensen, Chaney Kalinich, Isabel Ott, Arnau Casanovas, Catherine Muenker, Adam Moore, Alice Lu, Maria Tokuyama, Patrick Wong, Peiwen Lu, Saad Omer, Richard Martinello, Allison Nelson, Shelli Farhadian, Akiko Iwasaki, Charlese Dela Cruz, Albert Ko, Nathan Grubaugh |
| EPI_ISL_428747 | hCoV-19/USA/CT-Yale-088/2020 | North America / USA / Connecticut | 2020-04-06 | Yale COVID-19 Biorepository | Grubaugh Lab - Yale School of Public Health | Joseph Fauver, Tara Alpert, Anderson Brito, Anne Wyllie, Chantal Vogels, Mary Petrone, Cole Jensen, Chaney Kalinich, Isabel Ott, Arnau Casanovas, Catherine Muenker, Adam Moore, Alice Lu, Maria Tokuyama, Patrick Wong, Peiwen Lu, Saad Omer, Richard Martinello, Allison Nelson, Shelli Farhadian, Akiko Iwasaki, Charlese Dela Cruz, Albert Ko, Nathan Grubaugh |
| EPI_ISL_428748 | hCoV-19/USA/CT-Yale-089/2020 | North America / USA / Connecticut | 2020-04-07 | Yale COVID-19 Biorepository | Grubaugh Lab - Yale School of Public Health | Joseph Fauver, Tara Alpert, Anderson Brito, Anne Wyllie, Chantal Vogels, Mary Petrone, Cole Jensen, Chaney Kalinich, Isabel Ott, Arnau Casanovas, Catherine Muenker, Adam Moore, Alice Lu, Maria Tokuyama, Patrick Wong, Peiwen Lu, Saad Omer, Richard Martinello, Allison Nelson, Shelli Farhadian, Akiko Iwasaki, Charlese Dela Cruz, Albert Ko, Nathan Grubaugh |
| EPI_ISL_428749 | hCoV-19/USA/CT-Yale-098/2020 | North America / USA / Connecticut | 2020-04-02 | Yale COVID-19 Biorepository | Grubaugh Lab - Yale School of Public Health | Joseph Fauver, Tara Alpert, Anderson Brito, Anne Wyllie, Chantal Vogels, Mary Petrone, Cole Jensen, Chaney Kalinich, Isabel Ott, Arnau Casanovas, Catherine Muenker, Adam Moore, Alice Lu, Maria Tokuyama, Patrick Wong, Peiwen Lu, Saad Omer, Richard Martinello, Allison Nelson, Shelli Farhadian, Akiko Iwasaki, Charlese Dela Cruz, Albert Ko, Nathan Grubaugh |
| EPI_ISL_428750 | hCoV-19/USA/CT-Yale-099/2020 | North America / USA / Connecticut | 2020-04-07 | Yale COVID-19 Biorepository | Grubaugh Lab - Yale School of Public Health | Joseph Fauver, Tara Alpert, Anderson Brito, Anne Wyllie, Chantal Vogels, Mary Petrone, Cole Jensen, Chaney Kalinich, Isabel Ott, Arnau Casanovas, Catherine Muenker, Adam Moore, Alice Lu, Maria Tokuyama, Patrick Wong, Peiwen Lu, Saad Omer, Richard Martinello, Allison Nelson, Shelli Farhadian, Akiko Iwasaki, Charlese Dela Cruz, Albert Ko, Nathan Grubaugh |
| EPI_ISL_428751 | hCoV-19/USA/CT-Yale-110/2020 | North America / USA / Connecticut | 2020-04-09 | Yale COVID-19 Biorepository | Grubaugh Lab - Yale School of Public Health | Joseph Fauver, Tara Alpert, Anderson Brito, Anne Wyllie, Chantal Vogels, Mary Petrone, Cole Jensen, Chaney Kalinich, Isabel Ott, Arnau Casanovas, Catherine Muenker, Adam Moore, Alice Lu, Maria Tokuyama, Patrick Wong, Peiwen Lu, Saad Omer, Richard Martinello, Allison Nelson, Shelli Farhadian, Akiko Iwasaki, Charlese Dela Cruz, Albert Ko, Nathan Grubaugh |
| EPI_ISL_428752 | hCoV-19/USA/CT-Yale-113/2020 | North America / USA / Connecticut | 2020-04-09 | Yale COVID-19 Biorepository | Grubaugh Lab - Yale School of Public Health | Joseph Fauver, Tara Alpert, Anderson Brito, Anne Wyllie, Chantal Vogels, Mary Petrone, Cole Jensen, Chaney Kalinich, Isabel Ott, Arnau Casanovas, Catherine Muenker, Adam Moore, Alice Lu, Maria Tokuyama, Patrick Wong, Peiwen Lu, Saad Omer, Richard Martinello, Allison Nelson, Shelli Farhadian, Akiko Iwasaki, Charlese Dela Cruz, Albert Ko, Nathan Grubaugh |
| EPI_ISL_428757 | hCoV-19/USA/NY-NYUMC243/2020 | North America / USA / New York / Brookl | 2020-03-30 | NYU Langone Health | Departments of Pathology and Medicine, New York University School of Medicine | Maria Aguero-Rosenfeld, Brendan Belovarac, Margaret Black, Ludovic Boytard, John Cadley, Paolo Cotzia, John Chen, Dacia Dimartino, Xiaojun Feng, Tatyana Gindin, Emily Guzman, Adriana Heguy, Megan Hogan, Emily Huang, George Jour, Andrew Lytle, Christian Marier, Matthew T. Maurano, Mark J. Mulligan, Peter Meyn, Iman Osman, Jared Pinnell, Vanessa Raabe, Sitharam Ramaswami, Amy Rapkiewicz, Marie Samanovic-Golden, Antonio Serrano, Guomiao Shen, Matija Snuderl, Theodore Vougiouklakis, Nick Vulpescu, Gael Westby, Paul Zappile, Yutong Zhang |
| EPI_ISL_428758 | hCoV-19/USA/NY-NYUMC244/2020 | North America / USA / New York / Manha | 2020-04-03 | NYU Langone Health | Departments of Pathology and Medicine, New York University School of Medicine | Maria Aguero-Rosenfeld, Brendan Belovarac, Margaret Black, Ludovic Boytard, John Cadley, Paolo Cotzia, John Chen, Dacia Dimartino, Xiaojun Feng, Tatyana Gindin, Emily Guzman, Adriana Heguy, Megan Hogan, Emily Huang, George Jour, Andrew Lytle, Christian Marier, Matthew T. Maurano, Mark J. Mulligan, Peter Meyn, Iman Osman, Jared Pinnell, Vanessa Raabe, Sitharam Ramaswami, Amy Rapkiewicz, Marie Samanovic-Golden, Antonio Serrano, Guomiao Shen, Matija Snuderl, Theodore Vougiouklakis, Nick Vulpescu, Gael Westby, Paul Zappile, Yutong Zhang |
| EPI_ISL_428759 | hCoV-19/USA/NY-NYUMC245/2020 | North America / USA / New York / Brookl | 2020-04-03 | NYU Langone Health | Departments of Pathology and Medicine, New York University School of Medicine | Maria Aguero-Rosenfeld, Brendan Belovarac, Margaret Black, Ludovic Boytard, John Cadley, Paolo Cotzia, John Chen, Dacia Dimartino, Xiaojun Feng, Tatyana Gindin, Emily Guzman, Adriana Heguy, Megan Hogan, Emily Huang, George Jour, Andrew Lytle, Christian Marier, Matthew T. Maurano, Mark J. Mulligan, Peter Meyn, Iman Osman, Jared Pinnell, Vanessa Raabe, Sitharam Ramaswami, Amy Rapkiewicz, Marie Samanovic-Golden, Antonio Serrano, Guomiao Shen, Matija Snuderl, Theodore Vougiouklakis, Nick Vulpescu, Gael Westby, Paul Zappile, Yutong Zhang |
| EPI_ISL_428760 | hCoV-19/USA/NY-NYUMC246/2020 | North America / USA / New York / Brookl | 2020-04-03 | NYU Langone Health | Departments of Pathology and Medicine, New York University School of Medicine | Maria Aguero-Rosenfeld, Brendan Belovarac, Margaret Black, Ludovic Boytard, John Cadley, Paolo Cotzia, John Chen, Dacia Dimartino, Xiaojun Feng, Tatyana Gindin, Emily Guzman, Adriana Heguy, Megan Hogan, Emily Huang, George Jour, Andrew Lytle, Christian Marier, Matthew T. Maurano, Mark J. Mulligan, Peter Meyn, Iman Osman, Jared Pinnell, Vanessa Raabe, Sitharam Ramaswami, Amy Rapkiewicz, Marie Samanovic-Golden, Antonio Serrano, Guomiao Shen, Matija Snuderl, Theodore Vougiouklakis, Nick Vulpescu, Gael Westby, Paul Zappile, Yutong Zhang |
| EPI_ISL_428761 | hCoV-19/USA/NY-NYUMC247/2020 | North America / USA / New York / Brookl | 2020-04-03 | NYU Langone Health | Departments of Pathology and Medicine, New York University School of Medicine | Maria Aguero-Rosenfeld, Brendan Belovarac, Margaret Black, Ludovic Boytard, John Cadley, Paolo Cotzia, John Chen, Dacia Dimartino, Xiaojun Feng, Tatyana Gindin, Emily Guzman, Adriana Heguy, Megan Hogan, Emily Huang, George Jour, Andrew Lytle, Christian Marier, Matthew T. Maurano, Mark J. Mulligan, Peter Meyn, Iman Osman, Jared Pinnell, Vanessa Raabe, Sitharam Ramaswami, Amy Rapkiewicz, Marie Samanovic-Golden, Antonio Serrano, Guomiao Shen, Matija Snuderl, Theodore Vougiouklakis, Nick Vulpescu, Gael Westby, Paul Zappile, Yutong Zhang |
| EPI_ISL_428762 | hCoV-19/USA/NY-NYUMC248/2020 | North America / USA / New York / Brookl | 2020-04-03 | NYU Langone Health | Departments of Pathology and Medicine, New York University School of Medicine | Maria Aguero-Rosenfeld, Brendan Belovarac, Margaret Black, Ludovic Boytard, John Cadley, Paolo Cotzia, John Chen, Dacia Dimartino, Xiaojun Feng, Tatyana Gindin, Emily Guzman, Adriana Heguy, Megan Hogan, Emily Huang, George Jour, Andrew Lytle, Christian Marier, Matthew T. Maurano, Mark J. Mulligan, Peter Meyn, Iman Osman, Jared Pinnell, Vanessa Raabe, Sitharam Ramaswami, Amy Rapkiewicz, Marie Samanovic-Golden, Antonio Serrano, Guomiao Shen, Matija Snuderl, Theodore Vougiouklakis, Nick Vulpescu, Gael Westby, Paul Zappile, Yutong Zhang |
| EPI_ISL_428763 | hCoV-19/USA/NY-NYUMC249/2020 | North America / USA / New York / Brookl | 2020-04-03 | NYU Langone Health | Departments of Pathology and Medicine, New York University School of Medicine | Maria Aguero-Rosenfeld, Brendan Belovarac, Margaret Black, Ludovic Boytard, John Cadley, Paolo Cotzia, John Chen, Dacia Dimartino, Xiaojun Feng, Tatyana Gindin, Emily Guzman, Adriana Heguy, Megan Hogan, Emily Huang, George Jour, Andrew Lytle, Christian Marier, Matthew T. Maurano, Mark J. Mulligan, Peter Meyn, Iman Osman, Jared Pinnell, Vanessa Raabe, Sitharam Ramaswami, Amy Rapkiewicz, Marie Samanovic-Golden, Antonio Serrano, Guomiao Shen, Matija Snuderl, Theodore Vougiouklakis, Nick Vulpescu, Gael Westby, Paul Zappile, Yutong Zhang |
| EPI_ISL_428765 | hCoV-19/USA/NY-NYUMC251/2020 | North America / USA / New York / Queen | 2020-04-03 | NYU Langone Health | Departments of Pathology and Medicine, New York University School of Medicine | Maria Aguero-Rosenfeld, Brendan Belovarac, Margaret Black, Ludovic Boytard, John Cadley, Paolo Cotzia, John Chen, Dacia Dimartino, Xiaojun Feng, Tatyana Gindin, Emily Guzman, Adriana Heguy, Megan Hogan, Emily Huang, George Jour, Andrew Lytle, Christian Marier, Matthew T. Maurano, Mark J. Mulligan, Peter Meyn, Iman Osman, Jared Pinnell, Vanessa Raabe, Sitharam Ramaswami, Amy Rapkiewicz, Marie Samanovic-Golden, Antonio Serrano, Guomiao Shen, Matija Snuderl, Theodore Vougiouklakis, Nick Vulpescu, Gael Westby, Paul Zappile, Yutong Zhang |
| EPI_ISL_428766 | hCoV-19/USA/NY-NYUMC252/2020 | North America / USA / New York / Manha | 2020-04-03 | NYU Langone Health | Departments of Pathology and Medicine, New York University School of Medicine | Maria Aguero-Rosenfeld, Brendan Belovarac, Margaret Black, Ludovic Boytard, John Cadley, Paolo Cotzia, John Chen, Dacia Dimartino, Xiaojun Feng, Tatyana Gindin, Emily Guzman, Adriana Heguy, Megan Hogan, Emily Huang, George Jour, Andrew Lytle, Christian Marier, Matthew T. Maurano, Mark J. Mulligan, Peter Meyn, Iman Osman, Jared Pinnell, Vanessa Raabe, Sitharam Ramaswami, Amy Rapkiewicz, Marie Samanovic-Golden, Antonio Serrano, Guomiao Shen, Matija Snuderl, Theodore Vougiouklakis, Nick Vulpescu, Gael Westby, Paul Zappile, Yutong Zhang |
| EPI_ISL_428767 | hCoV-19/USA/NY-NYUMC253/2020 | North America / USA / New York / Brookl | 2020-04-05 | NYU Langone Health | Departments of Pathology and Medicine, New York University School of Medicine | Maria Aguero-Rosenfeld, Brendan Belovarac, Margaret Black, Ludovic Boytard, John Cadley, Paolo Cotzia, John Chen, Dacia Dimartino, Xiaojun Feng, Tatyana Gindin, Emily Guzman, Adriana Heguy, Megan Hogan, Emily Huang, George Jour, Andrew Lytle, Christian Marier, Matthew T. Maurano, Mark J. Mulligan, Peter Meyn, Iman Osman, Jared Pinnell, Vanessa Raabe, Sitharam Ramaswami, Amy Rapkiewicz, Marie Samanovic-Golden, Antonio Serrano, Guomiao Shen, Matija Snuderl, Theodore Vougiouklakis, Nick Vulpescu, Gael Westby, Paul Zappile, Yutong Zhang |
| EPI_ISL_428768 | hCoV-19/USA/NY-NYUMC254/2020 | North America / USA / New York / Brookl | 2020-04-05 | NYU Langone Health | Departments of Pathology and Medicine, New York University School of Medicine | Maria Aguero-Rosenfeld, Brendan Belovarac, Margaret Black, Ludovic Boytard, John Cadley, Paolo Cotzia, John Chen, Dacia Dimartino, Xiaojun Feng, Tatyana Gindin, Emily Guzman, Adriana Heguy, Megan Hogan, Emily Huang, George Jour, Andrew Lytle, Christian Marier, Matthew T. Maurano, Mark J. Mulligan, Peter Meyn, Iman Osman, Jared Pinnell, Vanessa Raabe, Sitharam Ramaswami, Amy Rapkiewicz, Marie Samanovic-Golden, Antonio Serrano, Guomiao Shen, Matija Snuderl, Theodore Vougiouklakis, Nick Vulpescu, Gael Westby, Paul Zappile, Yutong Zhang |
| EPI_ISL_428769 | hCoV-19/USA/NY-NYUMC255/2020 | North America / USA / New York / Brookl | 2020-04-05 | NYU Langone Health | Departments of Pathology and Medicine, New York University School of Medicine | Maria Aguero-Rosenfeld, Brendan Belovarac, Margaret Black, Ludovic Boytard, John Cadley, Paolo Cotzia, John Chen, Dacia Dimartino, Xiaojun Feng, Tatyana Gindin, Emily Guzman, Adriana Heguy, Megan Hogan, Emily Huang, George Jour, Andrew Lytle, Christian Marier, Matthew T. Maurano, Mark J. Mulligan, Peter Meyn, Iman Osman, Jared Pinnell, Vanessa Raabe, Sitharam Ramaswami, Amy Rapkiewicz, Marie Samanovic-Golden, Antonio Serrano, Guomiao Shen, Matija Snuderl, Theodore Vougiouklakis, Nick Vulpescu, Gael Westby, Paul Zappile, Yutong Zhang |
| EPI_ISL_428770 | hCoV-19/USA/NY-NYUMC256/2020 | North America / USA / New York / Brookl | 2020-04-05 | NYU Langone Health | Departments of Pathology and Medicine, New York University School of Medicine | Maria Aguero-Rosenfeld, Brendan Belovarac, Margaret Black, Ludovic Boytard, John Cadley, Paolo Cotzia, John Chen, Dacia Dimartino, Xiaojun Feng, Tatyana Gindin, Emily Guzman, Adriana Heguy, Megan Hogan, Emily Huang, George Jour, Andrew Lytle, Christian Marier, Matthew T. Maurano, Mark J. Mulligan, Peter Meyn, Iman Osman, Jared Pinnell, Vanessa Raabe, Sitharam Ramaswami, Amy Rapkiewicz, Marie Samanovic-Golden, Antonio Serrano, Guomiao Shen, Matija Snuderl, Theodore Vougiouklakis, Nick Vulpescu, Gael Westby, Paul Zappile, Yutong Zhang |
| EPI_ISL_428772 | hCoV-19/USA/NY-NYUMC258/2020 | North America / USA / New York / Brookl | 2020-04-05 | NYU Langone Health | Departments of Pathology and Medicine, New York University School of Medicine | Maria Aguero-Rosenfeld, Brendan Belovarac, Margaret Black, Ludovic Boytard, John Cadley, Paolo Cotzia, John Chen, Dacia Dimartino, Xiaojun Feng, Tatyana Gindin, Emily Guzman, Adriana Heguy, Megan Hogan, Emily Huang, George Jour, Andrew Lytle, Christian Marier, Matthew T. Maurano, Mark J. Mulligan, Peter Meyn, Iman Osman, Jared Pinnell, Vanessa Raabe, Sitharam Ramaswami, Amy Rapkiewicz, Marie Samanovic-Golden, Antonio Serrano, Guomiao Shen, Matija Snuderl, Theodore Vougiouklakis, Nick Vulpescu, Gael Westby, Paul Zappile, Yutong Zhang |
| EPI_ISL_428773 | hCoV-19/USA/NY-NYUMC259/2020 | North America / USA / New York / Brookl | 2020-04-05 | NYU Langone Health | Departments of Pathology and Medicine, New York University School of Medicine | Maria Aguero-Rosenfeld, Brendan Belovarac, Margaret Black, Ludovic Boytard, John Cadley, Paolo Cotzia, John Chen, Dacia Dimartino, Xiaojun Feng, Tatyana Gindin, Emily Guzman, Adriana Heguy, Megan Hogan, Emily Huang, George Jour, Andrew Lytle, Christian Marier, Matthew T. Maurano, Mark J. Mulligan, Peter Meyn, Iman Osman, Jared Pinnell, Vanessa Raabe, Sitharam Ramaswami, Amy Rapkiewicz, Marie Samanovic-Golden, Antonio Serrano, Guomiao Shen, Matija Snuderl, Theodore Vougiouklakis, Nick Vulpescu, Gael Westby, Paul Zappile, Yutong Zhang |
| EPI_ISL_428774 | hCoV-19/USA/NY-NYUMC260/2020 | North America / USA / New York / Brookl | 2020-04-05 | NYU Langone Health | Departments of Pathology and Medicine, New York University School of Medicine | Maria Aguero-Rosenfeld, Brendan Belovarac, Margaret Black, Ludovic Boytard, John Cadley, Paolo Cotzia, John Chen, Dacia Dimartino, Xiaojun Feng, Tatyana Gindin, Emily Guzman, Adriana Heguy, Megan Hogan, Emily Huang, George Jour, Andrew Lytle, Christian Marier, Matthew T. Maurano, Mark J. Mulligan, Peter Meyn, Iman Osman, Jared Pinnell, Vanessa Raabe, Sitharam Ramaswami, Amy Rapkiewicz, Marie Samanovic-Golden, Antonio Serrano, Guomiao Shen, Matija Snuderl, Theodore Vougiouklakis, Nick Vulpescu, Gael Westby, Paul Zappile, Yutong Zhang |
| EPI_ISL_428775 | hCoV-19/USA/NY-NYUMC261/2020 | North America / USA / New York / Brookl | 2020-04-05 | NYU Langone Health | Departments of Pathology and Medicine, New York University School of Medicine | Maria Aguero-Rosenfeld, Brendan Belovarac, Margaret Black, Ludovic Boytard, John Cadley, Paolo Cotzia, John Chen, Dacia Dimartino, Xiaojun Feng, Tatyana Gindin, Emily Guzman, Adriana Heguy, Megan Hogan, Emily Huang, George Jour, Andrew Lytle, Christian Marier, Matthew T. Maurano, Mark J. Mulligan, Peter Meyn, Iman Osman, Jared Pinnell, Vanessa Raabe, Sitharam Ramaswami, Amy Rapkiewicz, Marie Samanovic-Golden, Antonio Serrano, Guomiao Shen, Matija Snuderl, Theodore Vougiouklakis, Nick Vulpescu, Gael Westby, Paul Zappile, Yutong Zhang |
| EPI_ISL_428776 | hCoV-19/USA/NY-NYUMC262/2020 | North America / USA / New York / Brookl | 2020-04-05 | NYU Langone Health | Departments of Pathology and Medicine, New York University School of Medicine | Maria Aguero-Rosenfeld, Brendan Belovarac, Margaret Black, Ludovic Boytard, John Cadley, Paolo Cotzia, John Chen, Dacia Dimartino, Xiaojun Feng, Tatyana Gindin, Emily Guzman, Adriana Heguy, Megan Hogan, Emily Huang, George Jour, Andrew Lytle, Christian Marier, Matthew T. Maurano, Mark J. Mulligan, Peter Meyn, Iman Osman, Jared Pinnell, Vanessa Raabe, Sitharam Ramaswami, Amy Rapkiewicz, Marie Samanovic-Golden, Antonio Serrano, Guomiao Shen, Matija Snuderl, Theodore Vougiouklakis, Nick Vulpescu, Gael Westby, Paul Zappile, Yutong Zhang |
| EPI_ISL_428777 | hCoV-19/USA/NY-NYUMC263/2020 | North America / USA / New York / Brookl | 2020-04-06 | NYU Langone Health | Departments of Pathology and Medicine, New York University School of Medicine | Maria Aguero-Rosenfeld, Brendan Belovarac, Margaret Black, Ludovic Boytard, John Cadley, Paolo Cotzia, John Chen, Dacia Dimartino, Xiaojun Feng, Tatyana Gindin, Emily Guzman, Adriana Heguy, Megan Hogan, Emily Huang, George Jour, Andrew Lytle, Christian Marier, Matthew T. Maurano, Mark J. Mulligan, Peter Meyn, Iman Osman, Jared Pinnell, Vanessa Raabe, Sitharam Ramaswami, Amy Rapkiewicz, Marie Samanovic-Golden, Antonio Serrano, Guomiao Shen, Matija Snuderl, Theodore Vougiouklakis, Nick Vulpescu, Gael Westby, Paul Zappile, Yutong Zhang |
| EPI_ISL_428778 | hCoV-19/USA/NY-NYUMC264/2020 | North America / USA / New York / Brookl | 2020-04-06 | NYU Langone Health | Departments of Pathology and Medicine, New York University School of Medicine | Maria Aguero-Rosenfeld, Brendan Belovarac, Margaret Black, Ludovic Boytard, John Cadley, Paolo Cotzia, John Chen, Dacia Dimartino, Xiaojun Feng, Tatyana Gindin, Emily Guzman, Adriana Heguy, Megan Hogan, Emily Huang, George Jour, Andrew Lytle, Christian Marier, Matthew T. Maurano, Mark J. Mulligan, Peter Meyn, Iman Osman, Jared Pinnell, Vanessa Raabe, Sitharam Ramaswami, Amy Rapkiewicz, Marie Samanovic-Golden, Antonio Serrano, Guomiao Shen, Matija Snuderl, Theodore Vougiouklakis, Nick Vulpescu, Gael Westby, Paul Zappile, Yutong Zhang |
| EPI_ISL_428779 | hCoV-19/USA/NY-NYUMC265/2020 | North America / USA / New York / Queen | 2020-04-06 | NYU Langone Health | Departments of Pathology and Medicine, New York University School of Medicine | Maria Aguero-Rosenfeld, Brendan Belovarac, Margaret Black, Ludovic Boytard, John Cadley, Paolo Cotzia, John Chen, Dacia Dimartino, Xiaojun Feng, Tatyana Gindin, Emily Guzman, Adriana Heguy, Megan Hogan, Emily Huang, George Jour, Andrew Lytle, Christian Marier, Matthew T. Maurano, Mark J. Mulligan, Peter Meyn, Iman Osman, Jared Pinnell, Vanessa Raabe, Sitharam Ramaswami, Amy Rapkiewicz, Marie Samanovic-Golden, Antonio Serrano, Guomiao Shen, Matija Snuderl, Theodore Vougiouklakis, Nick Vulpescu, Gael Westby, Paul Zappile, Yutong Zhang |
| EPI_ISL_429000 | hCoV-19/USA/CA-CZB0103/2020 | North America / USA / California / San Fr | 2020-03-28 | UCSF Clinical Microbiology Laboratory | Chan-Zuckerberg Biohub | CZB Cliahub Consortium |
| EPI_ISL_429001 | hCoV-19/USA/CA-CZB046/2020 | North America / USA / California / San Fr | 2020-03-25 | UCSF Clinical Microbiology Laboratory | Chan-Zuckerberg Biohub | CZB Cliahub Consortium |
| EPI_ISL_429002 | hCoV-19/USA/CA-CZB068/2020 | North America / USA / California / San Fr | 2020-03-28 | UCSF Clinical Microbiology Laboratory | Chan-Zuckerberg Biohub | CZB Cliahub Consortium |
| EPI_ISL_429003 | hCoV-19/USA/CA-CZB0136/2020 | North America / USA / California / San Fr | 2020-04-08 | UCSF Clinical Microbiology Laboratory | Chan-Zuckerberg Biohub | CZB Cliahub Consortium |
| EPI_ISL_429004 | hCoV-19/USA/CA-CZB085/2020 | North America / USA / California / San Fr | 2020-03-28 | UCSF Clinical Microbiology Laboratory | Chan-Zuckerberg Biohub | CZB Cliahub Consortium |
| EPI_ISL_429005 | hCoV-19/USA/CA-CZB079/2020 | North America / USA / California / San Fr | 2020-03-28 | UCSF Clinical Microbiology Laboratory | Chan-Zuckerberg Biohub | CZB Cliahub Consortium |
| EPI_ISL_429006 | hCoV-19/USA/CA-CZB095/2020 | North America / USA / California / San Fr | 2020-03-28 | UCSF Clinical Microbiology Laboratory | Chan-Zuckerberg Biohub | CZB Cliahub Consortium |
| EPI_ISL_429007 | hCoV-19/USA/CA-CZB0458/2020 | North America / USA / California / San Fr | 2020-03-18 | UCSF Clinical Microbiology Laboratory | Chan-Zuckerberg Biohub | CZB Cliahub Consortium |
| EPI_ISL_429008 | hCoV-19/USA/CA-CZB091/2020 | North America / USA / California / San Fr | 2020-03-28 | UCSF Clinical Microbiology Laboratory | Chan-Zuckerberg Biohub | CZB Cliahub Consortium |
| EPI_ISL_429009 | hCoV-19/USA/CA-CZB0122/2020 | North America / USA / California / San Fr | 2020-04-08 | UCSF Clinical Microbiology Laboratory | Chan-Zuckerberg Biohub | CZB Cliahub Consortium |
| EPI_ISL_429010 | hCoV-19/USA/CA-CZB0108/2020 | North America / USA / California / San Fr | 2020-03-28 | UCSF Clinical Microbiology Laboratory | Chan-Zuckerberg Biohub | CZB Cliahub Consortium |
| EPI_ISL_429011 | hCoV-19/USA/CA-CZB0147/2020 | North America / USA / California / San Fr | 2020-04-08 | UCSF Clinical Microbiology Laboratory | Chan-Zuckerberg Biohub | CZB Cliahub Consortium |
| EPI_ISL_429012 | hCoV-19/USA/CA-CZB016/2020 | North America / USA / California / San Fr | 2020-03-25 | UCSF Clinical Microbiology Laboratory | Chan-Zuckerberg Biohub | CZB Cliahub Consortium |
| EPI_ISL_429013 | hCoV-19/USA/CA-CZB080/2020 | North America / USA / California / San Fr | 2020-03-28 | UCSF Clinical Microbiology Laboratory | Chan-Zuckerberg Biohub | CZB Cliahub Consortium |
| EPI_ISL_429014 | hCoV-19/USA/CA-CZB0142/2020 | North America / USA / California / San Fr | 2020-04-08 | UCSF Clinical Microbiology Laboratory | Chan-Zuckerberg Biohub | CZB Cliahub Consortium |
| EPI_ISL_429015 | hCoV-19/USA/CA-CZB0100/2020 | North America / USA / California / San Fr | 2020-03-28 | UCSF Clinical Microbiology Laboratory | Chan-Zuckerberg Biohub | CZB Cliahub Consortium |
| EPI_ISL_429016 | hCoV-19/USA/CA-CZB0709/2020 | North America / USA / California / San Fr | 2020-03-23 | UCSF Clinical Microbiology Laboratory | Chan-Zuckerberg Biohub | CZB Cliahub Consortium |
| EPI_ISL_429017 | hCoV-19/USA/CA-CZB030/2020 | North America / USA / California / San Fr | 2020-03-25 | UCSF Clinical Microbiology Laboratory | Chan-Zuckerberg Biohub | CZB Cliahub Consortium |

| | | | | | | |
|---|---|---|---|---|---|---|
| EPI_ISL_429018 | hCoV-19/USA/CA-CZB0750/2020 | North America / USA / California / San Fr | 2020-04-03 | UCSF Clinical Microbiology Laboratory | Chan-Zuckerberg Biohub | CZB Cliahub Consortium |
| EPI_ISL_429019 | hCoV-19/USA/CA-CZB078/2020 | North America / USA / California / San Fr | 2020-03-28 | UCSF Clinical Microbiology Laboratory | Chan-Zuckerberg Biohub | CZB Cliahub Consortium |
| EPI_ISL_429020 | hCoV-19/USA/CA-CZB0721/2020 | North America / USA / California / San Fr | 2020-03-31 | UCSF Clinical Microbiology Laboratory | Chan-Zuckerberg Biohub | CZB Cliahub Consortium |
| EPI_ISL_429021 | hCoV-19/USA/CA-CZB045/2020 | North America / USA / California / San Fr | 2020-03-25 | UCSF Clinical Microbiology Laboratory | Chan-Zuckerberg Biohub | CZB Cliahub Consortium |
| EPI_ISL_429022 | hCoV-19/USA/CA-CZB0713/2020 | North America / USA / California / San Fr | 2020-03-23 | UCSF Clinical Microbiology Laboratory | Chan-Zuckerberg Biohub | CZB Cliahub Consortium |
| EPI_ISL_429023 | hCoV-19/USA/CA-CZB0740/2020 | North America / USA / California / San Fr | 2020-03-31 | UCSF Clinical Microbiology Laboratory | Chan-Zuckerberg Biohub | CZB Cliahub Consortium |
| EPI_ISL_429024 | hCoV-19/USA/CA-CZB0705/2020 | North America / USA / California / San Fr | 2020-03-23 | UCSF Clinical Microbiology Laboratory | Chan-Zuckerberg Biohub | CZB Cliahub Consortium |
| EPI_ISL_429025 | hCoV-19/USA/CA-CZB0737/2020 | North America / USA / California / San Fr | 2020-03-31 | UCSF Clinical Microbiology Laboratory | Chan-Zuckerberg Biohub | CZB Cliahub Consortium |
| EPI_ISL_429026 | hCoV-19/USA/CA-CZB0731/2020 | North America / USA / California / San Fr | 2020-03-31 | UCSF Clinical Microbiology Laboratory | Chan-Zuckerberg Biohub | CZB Cliahub Consortium |
| EPI_ISL_429027 | hCoV-19/USA/CA-CZB0137/2020 | North America / USA / California / San Fr | 2020-04-08 | UCSF Clinical Microbiology Laboratory | Chan-Zuckerberg Biohub | CZB Cliahub Consortium |
| EPI_ISL_429028 | hCoV-19/USA/CA-CZB0746/2020 | North America / USA / California / San Fr | 2020-04-03 | UCSF Clinical Microbiology Laboratory | Chan-Zuckerberg Biohub | CZB Cliahub Consortium |
| EPI_ISL_429029 | hCoV-19/USA/CA-CZB067/2020 | North America / USA / California / San Fr | 2020-03-28 | UCSF Clinical Microbiology Laboratory | Chan-Zuckerberg Biohub | CZB Cliahub Consortium |
| EPI_ISL_429030 | hCoV-19/USA/CA-CZB072/2020 | North America / USA / California / San Fr | 2020-03-28 | UCSF Clinical Microbiology Laboratory | Chan-Zuckerberg Biohub | CZB Cliahub Consortium |
| EPI_ISL_429031 | hCoV-19/USA/CA-CZB0139/2020 | North America / USA / California / San Fr | 2020-04-08 | UCSF Clinical Microbiology Laboratory | Chan-Zuckerberg Biohub | CZB Cliahub Consortium |
| EPI_ISL_429032 | hCoV-19/USA/CA-CZB082/2020 | North America / USA / California / San Fr | 2020-03-28 | UCSF Clinical Microbiology Laboratory | Chan-Zuckerberg Biohub | CZB Cliahub Consortium |
| EPI_ISL_429033 | hCoV-19/USA/CA-CZB086/2020 | North America / USA / California / San Fr | 2020-03-28 | UCSF Clinical Microbiology Laboratory | Chan-Zuckerberg Biohub | CZB Cliahub Consortium |
| EPI_ISL_429034 | hCoV-19/USA/CA-CZB0706/2020 | North America / USA / California / San Fr | 2020-03-23 | UCSF Clinical Microbiology Laboratory | Chan-Zuckerberg Biohub | CZB Cliahub Consortium |
| EPI_ISL_429035 | hCoV-19/USA/CA-CZB042/2020 | North America / USA / California / San Fr | 2020-03-25 | UCSF Clinical Microbiology Laboratory | Chan-Zuckerberg Biohub | CZB Cliahub Consortium |
| EPI_ISL_429036 | hCoV-19/USA/CA-CZB0107/2020 | North America / USA / California / San Fr | 2020-03-28 | UCSF Clinical Microbiology Laboratory | Chan-Zuckerberg Biohub | CZB Cliahub Consortium |
| EPI_ISL_429037 | hCoV-19/USA/CA-CZB09/2020 | North America / USA / California / San Fr | 2020-03-25 | UCSF Clinical Microbiology Laboratory | Chan-Zuckerberg Biohub | CZB Cliahub Consortium |
| EPI_ISL_429038 | hCoV-19/USA/CA-CZB0707/2020 | North America / USA / California / San Fr | 2020-03-23 | UCSF Clinical Microbiology Laboratory | Chan-Zuckerberg Biohub | CZB Cliahub Consortium |
| EPI_ISL_429039 | hCoV-19/USA/CA-CZB036/2020 | North America / USA / California / San Fr | 2020-03-25 | UCSF Clinical Microbiology Laboratory | Chan-Zuckerberg Biohub | CZB Cliahub Consortium |
| EPI_ISL_429040 | hCoV-19/USA/CA-CZB0701/2020 | North America / USA / California / San Fr | 2020-03-23 | UCSF Clinical Microbiology Laboratory | Chan-Zuckerberg Biohub | CZB Cliahub Consortium |
| EPI_ISL_429041 | hCoV-19/USA/CA-CZB096/2020 | North America / USA / California / San Fr | 2020-03-28 | UCSF Clinical Microbiology Laboratory | Chan-Zuckerberg Biohub | CZB Cliahub Consortium |
| EPI_ISL_429042 | hCoV-19/USA/CA-CZB0710/2020 | North America / USA / California / San Fr | 2020-03-23 | UCSF Clinical Microbiology Laboratory | Chan-Zuckerberg Biohub | CZB Cliahub Consortium |
| EPI_ISL_429043 | hCoV-19/USA/CA-CZB0156/2020 | North America / USA / California / San Fr | 2020-04-08 | UCSF Clinical Microbiology Laboratory | Chan-Zuckerberg Biohub | CZB Cliahub Consortium |
| EPI_ISL_429044 | hCoV-19/USA/CA-CZB0158/2020 | North America / USA / California / San Fr | 2020-04-08 | UCSF Clinical Microbiology Laboratory | Chan-Zuckerberg Biohub | CZB Cliahub Consortium |
| EPI_ISL_429045 | hCoV-19/USA/CA-CZB039/2020 | North America / USA / California / San Fr | 2020-03-25 | UCSF Clinical Microbiology Laboratory | Chan-Zuckerberg Biohub | CZB Cliahub Consortium |
| EPI_ISL_429046 | hCoV-19/USA/CA-CZB0125/2020 | North America / USA / California / San Fr | 2020-04-08 | UCSF Clinical Microbiology Laboratory | Chan-Zuckerberg Biohub | CZB Cliahub Consortium |
| EPI_ISL_429047 | hCoV-19/USA/CA-CZB0154/2020 | North America / USA / California / San Fr | 2020-04-08 | UCSF Clinical Microbiology Laboratory | Chan-Zuckerberg Biohub | CZB Cliahub Consortium |
| EPI_ISL_429048 | hCoV-19/USA/CA-CZB043/2020 | North America / USA / California / San Fr | 2020-03-25 | UCSF Clinical Microbiology Laboratory | Chan-Zuckerberg Biohub | CZB Cliahub Consortium |
| EPI_ISL_429049 | hCoV-19/USA/CA-CZB090/2020 | North America / USA / California / San Fr | 2020-03-28 | UCSF Clinical Microbiology Laboratory | Chan-Zuckerberg Biohub | CZB Cliahub Consortium |
| EPI_ISL_429050 | hCoV-19/USA/CA-CZB089/2020 | North America / USA / California / San Fr | 2020-03-28 | UCSF Clinical Microbiology Laboratory | Chan-Zuckerberg Biohub | CZB Cliahub Consortium |
| EPI_ISL_429051 | hCoV-19/USA/CA-CZB0741/2020 | North America / USA / California / San Fr | 2020-04-03 | UCSF Clinical Microbiology Laboratory | Chan-Zuckerberg Biohub | CZB Cliahub Consortium |
| EPI_ISL_429052 | hCoV-19/USA/CA-CZB0714/2020 | North America / USA / California / San Fr | 2020-03-23 | UCSF Clinical Microbiology Laboratory | Chan-Zuckerberg Biohub | CZB Cliahub Consortium |
| EPI_ISL_429053 | hCoV-19/USA/CA-CZB0167/2020 | North America / USA / California / San Fr | 2020-04-08 | UCSF Clinical Microbiology Laboratory | Chan-Zuckerberg Biohub | CZB Cliahub Consortium |
| EPI_ISL_429054 | hCoV-19/USA/CA-CZB0141/2020 | North America / USA / California / San Fr | 2020-04-08 | UCSF Clinical Microbiology Laboratory | Chan-Zuckerberg Biohub | CZB Cliahub Consortium |
| EPI_ISL_429055 | hCoV-19/USA/CA-CZB0726/2020 | North America / USA / California / San Fr | 2020-03-31 | UCSF Clinical Microbiology Laboratory | Chan-Zuckerberg Biohub | CZB Cliahub Consortium |
| EPI_ISL_429056 | hCoV-19/USA/CA-CZB0101/2020 | North America / USA / California / San Fr | 2020-03-28 | UCSF Clinical Microbiology Laboratory | Chan-Zuckerberg Biohub | CZB Cliahub Consortium |
| EPI_ISL_429057 | hCoV-19/USA/CA-CZB076/2020 | North America / USA / California / San Fr | 2020-03-28 | UCSF Clinical Microbiology Laboratory | Chan-Zuckerberg Biohub | CZB Cliahub Consortium |
| EPI_ISL_429058 | hCoV-19/USA/CA-CZB0114/2020 | North America / USA / California / San Fr | 2020-03-28 | UCSF Clinical Microbiology Laboratory | Chan-Zuckerberg Biohub | CZB Cliahub Consortium |
| EPI_ISL_429059 | hCoV-19/USA/CA-CZB0110/2020 | North America / USA / California / San Fr | 2020-03-28 | UCSF Clinical Microbiology Laboratory | Chan-Zuckerberg Biohub | CZB Cliahub Consortium |
| EPI_ISL_429060 | hCoV-19/USA/CA-CZB0725/2020 | North America / USA / California / San Fr | 2020-03-31 | UCSF Clinical Microbiology Laboratory | Chan-Zuckerberg Biohub | CZB Cliahub Consortium |
| EPI_ISL_429061 | hCoV-19/USA/CA-CZB0151/2020 | North America / USA / California / San Fr | 2020-04-08 | UCSF Clinical Microbiology Laboratory | Chan-Zuckerberg Biohub | CZB Cliahub Consortium |
| EPI_ISL_429062 | hCoV-19/USA/CA-CZB0145/2020 | North America / USA / California / San Fr | 2020-04-08 | UCSF Clinical Microbiology Laboratory | Chan-Zuckerberg Biohub | CZB Cliahub Consortium |
| EPI_ISL_429063 | hCoV-19/USA/CA-CZB011b/2020 | North America / USA / California / San Fr | 2020-03-25 | UCSF Clinical Microbiology Laboratory | Chan-Zuckerberg Biohub | CZB Cliahub Consortium |
| EPI_ISL_429064 | hCoV-19/USA/CA-CZB040/2020 | North America / USA / California / San Fr | 2020-03-25 | UCSF Clinical Microbiology Laboratory | Chan-Zuckerberg Biohub | CZB Cliahub Consortium |
| EPI_ISL_429065 | hCoV-19/USA/CA-CZB0112/2020 | North America / USA / California / San Fr | 2020-03-28 | UCSF Clinical Microbiology Laboratory | Chan-Zuckerberg Biohub | CZB Cliahub Consortium |
| EPI_ISL_429066 | hCoV-19/USA/CA-CZB0109/2020 | North America / USA / California / San Fr | 2020-03-28 | UCSF Clinical Microbiology Laboratory | Chan-Zuckerberg Biohub | CZB Cliahub Consortium |
| EPI_ISL_429067 | hCoV-19/USA/CA-CZB0735/2020 | North America / USA / California / San Fr | 2020-03-31 | UCSF Clinical Microbiology Laboratory | Chan-Zuckerberg Biohub | CZB Cliahub Consortium |
| EPI_ISL_429068 | hCoV-19/USA/CA-CZB027/2020 | North America / USA / California / San Fr | 2020-03-25 | UCSF Clinical Microbiology Laboratory | Chan-Zuckerberg Biohub | CZB Cliahub Consortium |
| EPI_ISL_429069 | hCoV-19/USA/CA-CZB04/2020 | North America / USA / California / San Fr | 2020-03-25 | UCSF Clinical Microbiology Laboratory | Chan-Zuckerberg Biohub | CZB Cliahub Consortium |
| EPI_ISL_429196 | hCoV-19/Switzerland/GE3144/2020 | Europe / Switzerland | 2020-03-18 | University Hospitals of Geneva Laboratory of Virology | University Hospitals of Geneva Laboratory of Virology | Laubscher F. |
| EPI_ISL_429197 | hCoV-19/Switzerland/GE0199-2/2020 | Europe / Switzerland | 2020-03-21 | University Hospitals of Geneva Laboratory of Virology | University Hospitals of Geneva Laboratory of Virology | Laubscher F. |
| EPI_ISL_429199 | hCoV-19/Switzerland/GE0636/2020 | Europe / Switzerland | 2020-03-22 | University Hospitals of Geneva Laboratory of Virology | University Hospitals of Geneva Laboratory of Virology | Laubscher F. |
| EPI_ISL_429702 | hCoV-19/Brazil/CV49/2020 | South America / Brazil / Minas gerais | 2020-03-22 | Central Public Health Laboratory/Octávio Magalhães Institute (IOM) from the Ezequiel Dias Foundation (FUNED) | Instituto Octávio Magalhães / Fundação Ezequiel Dias (IOM/Funed) | Talita Adelino, Joilson Xavier, Marta Giovanetti, Vagner Fonseca, Marcos Vinícius Silva, Luiz Carlos Junior Alcantara, Marluce Aparecida Assunção Oliveira |
| EPI_ISL_429705 | hCoV-19/Croatia/AU-S10/2020 | Europe / Croatia / Istria | 2020-03 | Institute for Public Health | Laboratory for advanced genomics | Filip Rokić, Lovro Trgovec-Greif, Neven Sučić, Tomislav Rukavina, Igor Jurak, Oliver Vugrek |
| EPI_ISL_429706 | hCoV-19/Luxembourg/LNS3652733/2020 | Europe / Luxembourg | 2020-04-09 | Laboratoire National de Sante, Microbiology, Virology | Laboratoire National de Sante, Microbiology, Epidemiology and Microbial Genomics | Anke Wienecke-Baldacchino, Ardashel Latsuzbaia, Jessica Tapp, Catherine Ragimbeau, Guillaume Fournier, Tamir Abdelrahman, Trung Nguyen Nguyen, Joel Mossong |
| EPI_ISL_429707 | hCoV-19/Luxembourg/LNS4641661/2020 | Europe / Luxembourg | 2020-03-18 | Laboratoire National de Sante, Microbiology, Virology | Laboratoire National de Sante, Microbiology, Epidemiology and Microbial Genomics | Anke Wienecke-Baldacchino, Ardashel Latsuzbaia, Jessica Tapp, Catherine Ragimbeau, Guillaume Fournier, Tamir Abdelrahman, Trung Nguyen Nguyen, Joel Mossong |
| EPI_ISL_429710 | hCoV-19/Luxembourg/LNS5041225/2020 | Europe / Luxembourg | 2020-03-28 | Laboratoire National de Sante, Microbiology, Virology | Laboratoire National de Sante, Microbiology, Epidemiology and Microbial Genomics | Anke Wienecke-Baldacchino, Ardashel Latsuzbaia, Jessica Tapp, Catherine Ragimbeau, Guillaume Fournier, Tamir Abdelrahman, Trung Nguyen Nguyen, Joel Mossong |
| EPI_ISL_429711 | hCoV-19/Luxembourg/LNS5519604/2020 | Europe / Luxembourg | 2020-03-20 | Laboratoire National de Sante, Microbiology, Virology | Laboratoire National de Sante, Microbiology, Epidemiology and Microbial Genomics | Anke Wienecke-Baldacchino, Ardashel Latsuzbaia, Jessica Tapp, Catherine Ragimbeau, Guillaume Fournier, Tamir Abdelrahman, Trung Nguyen Nguyen, Joel Mossong |
| EPI_ISL_429713 | hCoV-19/Luxembourg/LNS9831617/2020 | Europe / Luxembourg | 2020-04-08 | Laboratoire National de Sante, Microbiology, Virology | Laboratoire National de Sante, Microbiology, Epidemiology and Microbial Genomics | Anke Wienecke-Baldacchino, Ardashel Latsuzbaia, Jessica Tapp, Catherine Ragimbeau, Guillaume Fournier, Tamir Abdelrahman, Trung Nguyen Nguyen, Joel Mossong |
| EPI_ISL_429715 | hCoV-19/Luxembourg/LNS7469301/2020 | Europe / Luxembourg | 2020-03-23 | Laboratoire National de Sante, Microbiology, Virology | Laboratoire National de Sante, Microbiology, Epidemiology and Microbial Genomics | Anke Wienecke-Baldacchino, Ardashel Latsuzbaia, Jessica Tapp, Catherine Ragimbeau, Guillaume Fournier, Tamir Abdelrahman, Trung Nguyen Nguyen, Joel Mossong |
| EPI_ISL_429716 | hCoV-19/Luxembourg/LNS7032690/2020 | Europe / Luxembourg | 2020-03-29 | Laboratoire National de Sante, Microbiology, Virology | Laboratoire National de Sante, Microbiology, Epidemiology and Microbial Genomics | Anke Wienecke-Baldacchino, Ardashel Latsuzbaia, Jessica Tapp, Catherine Ragimbeau, Guillaume Fournier, Tamir Abdelrahman, Trung Nguyen Nguyen, Joel Mossong |
| EPI_ISL_429718 | hCoV-19/Luxembourg/LNS1995247/2020 | Europe / Luxembourg | 2020-03-19 | Laboratoire National de Sante, Microbiology, Virology | Laboratoire National de Sante, Microbiology, Epidemiology and Microbial Genomics | Anke Wienecke-Baldacchino, Ardashel Latsuzbaia, Jessica Tapp, Catherine Ragimbeau, Guillaume Fournier, Tamir Abdelrahman, Trung Nguyen Nguyen, Joel Mossong |
| EPI_ISL_429722 | hCoV-19/Luxembourg/LNS6137379/2020 | Europe / Luxembourg | 2020-04-01 | Laboratoire National de Sante, Microbiology, Virology | Laboratoire National de Sante, Microbiology, Epidemiology and Microbial Genomics | Anke Wienecke-Baldacchino, Ardashel Latsuzbaia, Jessica Tapp, Catherine Ragimbeau, Guillaume Fournier, Tamir Abdelrahman, Trung Nguyen Nguyen, Joel Mossong |
| EPI_ISL_429723 | hCoV-19/Luxembourg/LNS5277361/2020 | Europe / Luxembourg | 2020-03-22 | Laboratoire National de Sante, Microbiology, Virology | Laboratoire National de Sante, Microbiology, Epidemiology and Microbial Genomics | Anke Wienecke-Baldacchino, Ardashel Latsuzbaia, Jessica Tapp, Catherine Ragimbeau, Guillaume Fournier, Tamir Abdelrahman, Trung Nguyen Nguyen, Joel Mossong |
| EPI_ISL_429724 | hCoV-19/Luxembourg/LNS9984061/2020 | Europe / Luxembourg | 2020-03-19 | Laboratoire National de Sante, Microbiology, Virology | Laboratoire National de Sante, Microbiology, Epidemiology and Microbial Genomics | Anke Wienecke-Baldacchino, Ardashel Latsuzbaia, Jessica Tapp, Catherine Ragimbeau, Guillaume Fournier, Tamir Abdelrahman, Trung Nguyen Nguyen, Joel Mossong |
| EPI_ISL_429725 | hCoV-19/Luxembourg/LNS0756265/2020 | Europe / Luxembourg | 2020-03-30 | Laboratoire National de Sante, Microbiology, Virology | Laboratoire National de Sante, Microbiology, Epidemiology and Microbial Genomics | Anke Wienecke-Baldacchino, Ardashel Latsuzbaia, Jessica Tapp, Catherine Ragimbeau, Guillaume Fournier, Tamir Abdelrahman, Trung Nguyen Nguyen, Joel Mossong |
| EPI_ISL_429727 | hCoV-19/Luxembourg/LNS9102295/2020 | Europe / Luxembourg | 2020-03-19 | Laboratoire National de Sante, Microbiology, Virology | Laboratoire National de Sante, Microbiology, Epidemiology and Microbial Genomics | Anke Wienecke-Baldacchino, Ardashel Latsuzbaia, Jessica Tapp, Catherine Ragimbeau, Guillaume Fournier, Tamir Abdelrahman, Trung Nguyen Nguyen, Joel Mossong |
| EPI_ISL_429728 | hCoV-19/Luxembourg/LNS3212178/2020 | Europe / Luxembourg | 2020-03-26 | Laboratoire National de Sante, Microbiology, Virology | Laboratoire National de Sante, Microbiology, Epidemiology and Microbial Genomics | Anke Wienecke-Baldacchino, Ardashel Latsuzbaia, Jessica Tapp, Catherine Ragimbeau, Guillaume Fournier, Tamir Abdelrahman, Trung Nguyen Nguyen, Joel Mossong |
| EPI_ISL_429729 | hCoV-19/Luxembourg/LNS0481305/2020 | Europe / Luxembourg | 2020-03-22 | Laboratoire National de Sante, Microbiology, Virology | Laboratoire National de Sante, Microbiology, Epidemiology and Microbial Genomics | Anke Wienecke-Baldacchino, Ardashel Latsuzbaia, Jessica Tapp, Catherine Ragimbeau, Guillaume Fournier, Tamir Abdelrahman, Trung Nguyen Nguyen, Joel Mossong |
| EPI_ISL_429730 | hCoV-19/Luxembourg/LNS7438855/2020 | Europe / Luxembourg | 2020-04-09 | Laboratoire National de Sante, Microbiology, Virology | Laboratoire National de Sante, Microbiology, Epidemiology and Microbial Genomics | Anke Wienecke-Baldacchino, Ardashel Latsuzbaia, Jessica Tapp, Catherine Ragimbeau, Guillaume Fournier, Tamir Abdelrahman, Trung Nguyen Nguyen, Joel Mossong |
| EPI_ISL_429732 | hCoV-19/Luxembourg/LNS7580831/2020 | Europe / Luxembourg | 2020-03-23 | Laboratoire National de Sante, Microbiology, Virology | Laboratoire National de Sante, Microbiology, Epidemiology and Microbial Genomics | Anke Wienecke-Baldacchino, Ardashel Latsuzbaia, Jessica Tapp, Catherine Ragimbeau, Guillaume Fournier, Tamir Abdelrahman, Trung Nguyen Nguyen, Joel Mossong |
| EPI_ISL_429733 | hCoV-19/Luxembourg/LNS4423730/2020 | Europe / Luxembourg | 2020-03-22 | Laboratoire National de Sante, Microbiology, Virology | Laboratoire National de Sante, Microbiology, Epidemiology and Microbial Genomics | Anke Wienecke-Baldacchino, Ardashel Latsuzbaia, Jessica Tapp, Catherine Ragimbeau, Guillaume Fournier, Tamir Abdelrahman, Trung Nguyen Nguyen, Joel Mossong |
| EPI_ISL_429734 | hCoV-19/Luxembourg/LNS6783622/2020 | Europe / Luxembourg | 2020-03-20 | Laboratoire National de Sante, Microbiology, Virology | Laboratoire National de Sante, Microbiology, Epidemiology and Microbial Genomics | Anke Wienecke-Baldacchino, Ardashel Latsuzbaia, Jessica Tapp, Catherine Ragimbeau, Guillaume Fournier, Tamir Abdelrahman, Trung Nguyen Nguyen, Joel Mossong |
| EPI_ISL_429735 | hCoV-19/Luxembourg/LNS5358505/2020 | Europe / Luxembourg | 2020-03-20 | Laboratoire National de Sante, Microbiology, Virology | Laboratoire National de Sante, Microbiology, Epidemiology and Microbial Genomics | Anke Wienecke-Baldacchino, Ardashel Latsuzbaia, Jessica Tapp, Catherine Ragimbeau, Guillaume Fournier, Tamir Abdelrahman, Trung Nguyen Nguyen, Joel Mossong |
| EPI_ISL_429736 | hCoV-19/Luxembourg/LNS7640155/2020 | Europe / Luxembourg | 2020-03-29 | Laboratoire National de Sante, Microbiology, Virology | Laboratoire National de Sante, Microbiology, Epidemiology and Microbial Genomics | Anke Wienecke-Baldacchino, Ardashel Latsuzbaia, Jessica Tapp, Catherine Ragimbeau, Guillaume Fournier, Tamir Abdelrahman, Trung Nguyen Nguyen, Joel Mossong |
| EPI_ISL_429737 | hCoV-19/Luxembourg/LNS4254328/2020 | Europe / Luxembourg | 2020-03-24 | Laboratoire National de Sante, Microbiology, Virology | Laboratoire National de Sante, Microbiology, Epidemiology and Microbial Genomics | Anke Wienecke-Baldacchino, Ardashel Latsuzbaia, Jessica Tapp, Catherine Ragimbeau, Guillaume Fournier, Tamir Abdelrahman, Trung Nguyen Nguyen, Joel Mossong |
| EPI_ISL_429738 | hCoV-19/Luxembourg/LNS5488745/2020 | Europe / Luxembourg | 2020-03-18 | Laboratoire National de Sante, Microbiology, Virology | Laboratoire National de Sante, Microbiology, Epidemiology and Microbial Genomics | Anke Wienecke-Baldacchino, Ardashel Latsuzbaia, Jessica Tapp, Catherine Ragimbeau, Guillaume Fournier, Tamir Abdelrahman, Trung Nguyen Nguyen, Joel Mossong |
| EPI_ISL_429739 | hCoV-19/Luxembourg/LNS2696129/2020 | Europe / Luxembourg | 2020-04-09 | Laboratoire National de Sante, Microbiology, Virology | Laboratoire National de Sante, Microbiology, Epidemiology and Microbial Genomics | Anke Wienecke-Baldacchino, Ardashel Latsuzbaia, Jessica Tapp, Catherine Ragimbeau, Guillaume Fournier, Tamir Abdelrahman, Trung Nguyen Nguyen, Joel Mossong |
| EPI_ISL_429740 | hCoV-19/Luxembourg/LNS5549490/2020 | Europe / Luxembourg | 2020-03-25 | Laboratoire National de Sante, Microbiology, Virology | Laboratoire National de Sante, Microbiology, Epidemiology and Microbial Genomics | Anke Wienecke-Baldacchino, Ardashel Latsuzbaia, Jessica Tapp, Catherine Ragimbeau, Guillaume Fournier, Tamir Abdelrahman, Trung Nguyen Nguyen, Joel Mossong |
| EPI_ISL_429741 | hCoV-19/Luxembourg/LNS6533667/2020 | Europe / Luxembourg | 2020-03-28 | Laboratoire National de Sante, Microbiology, Virology | Laboratoire National de Sante, Microbiology, Epidemiology and Microbial Genomics | Anke Wienecke-Baldacchino, Ardashel Latsuzbaia, Jessica Tapp, Catherine Ragimbeau, Guillaume Fournier, Tamir Abdelrahman, Trung Nguyen Nguyen, Joel Mossong |
| EPI_ISL_429742 | hCoV-19/Luxembourg/LNS1658150/2020 | Europe / Luxembourg | 2020-04-10 | Laboratoire National de Sante, Microbiology, Virology | Laboratoire National de Sante, Microbiology, Epidemiology and Microbial Genomics | Anke Wienecke-Baldacchino, Ardashel Latsuzbaia, Jessica Tapp, Catherine Ragimbeau, Guillaume Fournier, Tamir Abdelrahman, Trung Nguyen Nguyen, Joel Mossong |
| EPI_ISL_429743 | hCoV-19/Luxembourg/LNS6325758/2020 | Europe / Luxembourg | 2020-03-29 | Laboratoire National de Sante, Microbiology, Virology | Laboratoire National de Sante, Microbiology, Epidemiology and Microbial Genomics | Anke Wienecke-Baldacchino, Ardashel Latsuzbaia, Jessica Tapp, Catherine Ragimbeau, Guillaume Fournier, Tamir Abdelrahman, Trung Nguyen Nguyen, Joel Mossong |
| EPI_ISL_429745 | hCoV-19/Luxembourg/LNS6344924/2020 | Europe / Luxembourg | 2020-03-30 | Laboratoire National de Sante, Microbiology, Virology | Laboratoire National de Sante, Microbiology, Epidemiology and Microbial Genomics | Anke Wienecke-Baldacchino, Ardashel Latsuzbaia, Jessica Tapp, Catherine Ragimbeau, Guillaume Fournier, Tamir Abdelrahman, Trung Nguyen Nguyen, Joel Mossong |
| EPI_ISL_429746 | hCoV-19/Luxembourg/LNS2913521/2020 | Europe / Luxembourg | 2020-03-26 | Laboratoire National de Sante, Microbiology, Virology | Laboratoire National de Sante, Microbiology, Epidemiology and Microbial Genomics | Anke Wienecke-Baldacchino, Ardashel Latsuzbaia, Jessica Tapp, Catherine Ragimbeau, Guillaume Fournier, Tamir Abdelrahman, Trung Nguyen Nguyen, Joel Mossong |
| EPI_ISL_429748 | hCoV-19/Luxembourg/LNS4830703/2020 | Europe / Luxembourg | 2020-03-18 | Laboratoire National de Sante, Microbiology, Virology | Laboratoire National de Sante, Microbiology, Epidemiology and Microbial Genomics | Anke Wienecke-Baldacchino, Ardashel Latsuzbaia, Jessica Tapp, Catherine Ragimbeau, Guillaume Fournier, Tamir Abdelrahman, Trung Nguyen Nguyen, Joel Mossong |
| EPI_ISL_429749 | hCoV-19/Luxembourg/LNS3989293/2020 | Europe / Luxembourg | 2020-03-24 | Laboratoire National de Sante, Microbiology, Virology | Laboratoire National de Sante, Microbiology, Epidemiology and Microbial Genomics | Anke Wienecke-Baldacchino, Ardashel Latsuzbaia, Jessica Tapp, Catherine Ragimbeau, Guillaume Fournier, Tamir Abdelrahman, Trung Nguyen Nguyen, Joel Mossong |
| EPI_ISL_429750 | hCoV-19/Luxembourg/LNS9598665/2020 | Europe / Luxembourg | 2020-03-18 | Laboratoire National de Sante, Microbiology, Virology | Laboratoire National de Sante, Microbiology, Epidemiology and Microbial Genomics | Anke Wienecke-Baldacchino, Ardashel Latsuzbaia, Jessica Tapp, Catherine Ragimbeau, Guillaume Fournier, Tamir Abdelrahman, Trung Nguyen Nguyen, Joel Mossong |
| EPI_ISL_429751 | hCoV-19/Luxembourg/LNS4576534/2020 | Europe / Luxembourg | 2020-03-28 | Laboratoire National de Sante, Microbiology, Virology | Laboratoire National de Sante, Microbiology, Epidemiology and Microbial Genomics | Anke Wienecke-Baldacchino, Ardashel Latsuzbaia, Jessica Tapp, Catherine Ragimbeau, Guillaume Fournier, Tamir Abdelrahman, Trung Nguyen Nguyen, Joel Mossong |
| EPI_ISL_429753 | hCoV-19/Luxembourg/LNS4245799/2020 | Europe / Luxembourg | 2020-03-24 | Laboratoire National de Sante, Microbiology, Virology | Laboratoire National de Sante, Microbiology, Epidemiology and Microbial Genomics | Anke Wienecke-Baldacchino, Ardashel Latsuzbaia, Jessica Tapp, Catherine Ragimbeau, Guillaume Fournier, Tamir Abdelrahman, Trung Nguyen Nguyen, Joel Mossong |
| EPI_ISL_429755 | hCoV-19/Luxembourg/LNS8772889/2020 | Europe / Luxembourg | 2020-03-29 | Laboratoire National de Sante, Microbiology, Virology | Laboratoire National de Sante, Microbiology, Epidemiology and Microbial Genomics | Anke Wienecke-Baldacchino, Ardashel Latsuzbaia, Jessica Tapp, Catherine Ragimbeau, Guillaume Fournier, Tamir Abdelrahman, Trung Nguyen Nguyen, Joel Mossong |
| EPI_ISL_429756 | hCoV-19/Luxembourg/LNS4946091/2020 | Europe / Luxembourg | 2020-03-27 | Laboratoire National de Sante, Microbiology, Virology | Laboratoire National de Sante, Microbiology, Epidemiology and Microbial Genomics | Anke Wienecke-Baldacchino, Ardashel Latsuzbaia, Jessica Tapp, Catherine Ragimbeau, Guillaume Fournier, Tamir Abdelrahman, Trung Nguyen Nguyen, Joel Mossong |
| EPI_ISL_429757 | hCoV-19/Luxembourg/LNS7006156/2020 | Europe / Luxembourg | 2020-03-24 | Laboratoire National de Sante, Microbiology, Virology | Laboratoire National de Sante, Microbiology, Epidemiology and Microbial Genomics | Anke Wienecke-Baldacchino, Ardashel Latsuzbaia, Jessica Tapp, Catherine Ragimbeau, Guillaume Fournier, Tamir Abdelrahman, Trung Nguyen Nguyen, Joel Mossong |
| EPI_ISL_429758 | hCoV-19/Luxembourg/LNS8167314/2020 | Europe / Luxembourg | 2020-03-24 | Laboratoire National de Sante, Microbiology, Virology | Laboratoire National de Sante, Microbiology, Epidemiology and Microbial Genomics | Anke Wienecke-Baldacchino, Ardashel Latsuzbaia, Jessica Tapp, Catherine Ragimbeau, Guillaume Fournier, Tamir Abdelrahman, Trung Nguyen Nguyen, Joel Mossong |
| EPI_ISL_429759 | hCoV-19/Luxembourg/LNS9455951/2020 | Europe / Luxembourg | 2020-03-24 | Laboratoire National de Sante, Microbiology, Virology | Laboratoire National de Sante, Microbiology, Epidemiology and Microbial Genomics | Anke Wienecke-Baldacchino, Ardashel Latsuzbaia, Jessica Tapp, Catherine Ragimbeau, Guillaume Fournier, Tamir Abdelrahman, Trung Nguyen Nguyen, Joel Mossong |
| EPI_ISL_429760 | hCoV-19/Luxembourg/LNS0211710/2020 | Europe / Luxembourg | 2020-03-22 | Laboratoire National de Sante, Microbiology, Virology | Laboratoire National de Sante, Microbiology, Epidemiology and Microbial Genomics | Anke Wienecke-Baldacchino, Ardashel Latsuzbaia, Jessica Tapp, Catherine Ragimbeau, Guillaume Fournier, Tamir Abdelrahman, Trung Nguyen Nguyen, Joel Mossong |
| EPI_ISL_429762 | hCoV-19/Luxembourg/LNS6277439/2020 | Europe / Luxembourg | 2020-03-22 | Laboratoire National de Sante, Microbiology, Virology | Laboratoire National de Sante, Microbiology, Epidemiology and Microbial Genomics | Anke Wienecke-Baldacchino, Ardashel Latsuzbaia, Jessica Tapp, Catherine Ragimbeau, Guillaume Fournier, Tamir Abdelrahman, Trung Nguyen Nguyen, Joel Mossong |
| EPI_ISL_429764 | hCoV-19/Luxembourg/LNS1703112/2020 | Europe / Luxembourg | 2020-04-09 | Laboratoire National de Sante, Microbiology, Virology | Laboratoire National de Sante, Microbiology, Epidemiology and Microbial Genomics | Anke Wienecke-Baldacchino, Ardashel Latsuzbaia, Jessica Tapp, Catherine Ragimbeau, Guillaume Fournier, Tamir Abdelrahman, Trung Nguyen Nguyen, Joel Mossong |
| EPI_ISL_429766 | hCoV-19/Luxembourg/LNS0814393/2020 | Europe / Luxembourg | 2020-03-23 | Laboratoire National de Sante, Microbiology, Virology | Laboratoire National de Sante, Microbiology, Epidemiology and Microbial Genomics | Anke Wienecke-Baldacchino, Ardashel Latsuzbaia, Jessica Tapp, Catherine Ragimbeau, Guillaume Fournier, Tamir Abdelrahman, Trung Nguyen Nguyen, Joel Mossong |
| EPI_ISL_429767 | hCoV-19/Luxembourg/LNS3437586/2020 | Europe / Luxembourg | 2020-03-25 | Laboratoire National de Sante, Microbiology, Virology | Laboratoire National de Sante, Microbiology, Epidemiology and Microbial Genomics | Anke Wienecke-Baldacchino, Ardashel Latsuzbaia, Jessica Tapp, Catherine Ragimbeau, Guillaume Fournier, Tamir Abdelrahman, Trung Nguyen Nguyen, Joel Mossong |
| EPI_ISL_429769 | hCoV-19/Luxembourg/LNS7721101/2020 | Europe / Luxembourg | 2020-03-28 | Laboratoire National de Sante, Microbiology, Virology | Laboratoire National de Sante, Microbiology, Epidemiology and Microbial Genomics | Anke Wienecke-Baldacchino, Ardashel Latsuzbaia, Jessica Tapp, Catherine Ragimbeau, Guillaume Fournier, Tamir Abdelrahman, Trung Nguyen Nguyen, Joel Mossong |
| EPI_ISL_429770 | hCoV-19/Luxembourg/LNS4530810/2020 | Europe / Luxembourg | 2020-03-28 | Laboratoire National de Sante, Microbiology, Virology | Laboratoire National de Sante, Microbiology, Epidemiology and Microbial Genomics | Anke Wienecke-Baldacchino, Ardashel Latsuzbaia, Jessica Tapp, Catherine Ragimbeau, Guillaume Fournier, Tamir Abdelrahman, Trung Nguyen Nguyen, Joel Mossong |
| EPI_ISL_429772 | hCoV-19/Luxembourg/LNS4107430/2020 | Europe / Luxembourg | 2020-03-28 | Laboratoire National de Sante, Microbiology, Virology | Laboratoire National de Sante, Microbiology, Epidemiology and Microbial Genomics | Anke Wienecke-Baldacchino, Ardashel Latsuzbaia, Jessica Tapp, Catherine Ragimbeau, Guillaume Fournier, Tamir Abdelrahman, Trung Nguyen Nguyen, Joel Mossong |
| EPI_ISL_429773 | hCoV-19/Luxembourg/LNS3257811/2020 | Europe / Luxembourg | 2020-04-09 | Laboratoire National de Sante, Microbiology, Virology | Laboratoire National de Sante, Microbiology, Epidemiology and Microbial Genomics | Anke Wienecke-Baldacchino, Ardashel Latsuzbaia, Jessica Tapp, Catherine Ragimbeau, Guillaume Fournier, Tamir Abdelrahman, Trung Nguyen Nguyen, Joel Mossong |
| EPI_ISL_429774 | hCoV-19/Luxembourg/LNS5611124/2020 | Europe / Luxembourg | 2020-03-23 | Laboratoire National de Sante, Microbiology, Virology | Laboratoire National de Sante, Microbiology, Epidemiology and Microbial Genomics | Anke Wienecke-Baldacchino, Ardashel Latsuzbaia, Jessica Tapp, Catherine Ragimbeau, Guillaume Fournier, Tamir Abdelrahman, Trung Nguyen Nguyen, Joel Mossong |
| EPI_ISL_429775 | hCoV-19/Luxembourg/LNS7299024/2020 | Europe / Luxembourg | 2020-03-23 | Laboratoire National de Sante, Microbiology, Virology | Laboratoire National de Sante, Microbiology, Epidemiology and Microbial Genomics | Anke Wienecke-Baldacchino, Ardashel Latsuzbaia, Jessica Tapp, Catherine Ragimbeau, Guillaume Fournier, Tamir Abdelrahman, Trung Nguyen Nguyen, Joel Mossong |
| EPI_ISL_429776 | hCoV-19/Luxembourg/LNS8726389/2020 | Europe / Luxembourg | 2020-03-23 | Laboratoire National de Sante, Microbiology, Virology | Laboratoire National de Sante, Microbiology, Epidemiology and Microbial Genomics | Anke Wienecke-Baldacchino, Ardashel Latsuzbaia, Jessica Tapp, Catherine Ragimbeau, Guillaume Fournier, Tamir Abdelrahman, Trung Nguyen Nguyen, Joel Mossong |
| EPI_ISL_429777 | hCoV-19/Luxembourg/LNS0994857/2020 | Europe / Luxembourg | 2020-03-24 | Laboratoire National de Sante, Microbiology, Virology | Laboratoire National de Sante, Microbiology, Epidemiology and Microbial Genomics | Anke Wienecke-Baldacchino, Ardashel Latsuzbaia, Jessica Tapp, Catherine Ragimbeau, Guillaume Fournier, Tamir Abdelrahman, Trung Nguyen Nguyen, Joel Mossong |
| EPI_ISL_429778 | hCoV-19/Luxembourg/LNS2580723/2020 | Europe / Luxembourg | 2020-03-22 | Laboratoire National de Sante, Microbiology, Virology | Laboratoire National de Sante, Microbiology, Epidemiology and Microbial Genomics | Anke Wienecke-Baldacchino, Ardashel Latsuzbaia, Jessica Tapp, Catherine Ragimbeau, Guillaume Fournier, Tamir Abdelrahman, Trung Nguyen Nguyen, Joel Mossong |
| EPI_ISL_429779 | hCoV-19/Luxembourg/LNS7357097/2020 | Europe / Luxembourg | 2020-03-26 | Laboratoire National de Sante, Microbiology, Virology | Laboratoire National de Sante, Microbiology, Epidemiology and Microbial Genomics | Anke Wienecke-Baldacchino, Ardashel Latsuzbaia, Jessica Tapp, Catherine Ragimbeau, Guillaume Fournier, Tamir Abdelrahman, Trung Nguyen Nguyen, Joel Mossong |
| EPI_ISL_429780 | hCoV-19/Luxembourg/LNS5680075/2020 | Europe / Luxembourg | 2020-03-26 | Laboratoire National de Sante, Microbiology, Virology | Laboratoire National de Sante, Microbiology, Epidemiology and Microbial Genomics | Anke Wienecke-Baldacchino, Ardashel Latsuzbaia, Jessica Tapp, Catherine Ragimbeau, Guillaume Fournier, Tamir Abdelrahman, Trung Nguyen Nguyen, Joel Mossong |
| EPI_ISL_429781 | hCoV-19/Luxembourg/LNS0709551/2020 | Europe / Luxembourg | 2020-03-23 | Laboratoire National de Sante, Microbiology, Virology | Laboratoire National de Sante, Microbiology, Epidemiology and Microbial Genomics | Anke Wienecke-Baldacchino, Ardashel Latsuzbaia, Jessica Tapp, Catherine Ragimbeau, Guillaume Fournier, Tamir Abdelrahman, Trung Nguyen Nguyen, Joel Mossong |
| EPI_ISL_429782 | hCoV-19/Luxembourg/LNS3169999/2020 | Europe / Luxembourg | 2020-03-29 | Laboratoire National de Sante, Microbiology, Virology | Laboratoire National de Sante, Microbiology, Epidemiology and Microbial Genomics | Anke Wienecke-Baldacchino, Ardashel Latsuzbaia, Jessica Tapp, Catherine Ragimbeau, Guillaume Fournier, Tamir Abdelrahman, Trung Nguyen Nguyen, Joel Mossong |
| EPI_ISL_429784 | hCoV-19/Luxembourg/LNS5677421/2020 | Europe / Luxembourg | 2020-03-27 | Laboratoire National de Sante, Microbiology, Virology | Laboratoire National de Sante, Microbiology, Epidemiology and Microbial Genomics | Anke Wienecke-Baldacchino, Ardashel Latsuzbaia, Jessica Tapp, Catherine Ragimbeau, Guillaume Fournier, Tamir Abdelrahman, Trung Nguyen Nguyen, Joel Mossong |
| EPI_ISL_429786 | hCoV-19/Luxembourg/LNS4818581/2020 | Europe / Luxembourg | 2020-03-31 | Laboratoire National de Sante, Microbiology, Virology | Laboratoire National de Sante, Microbiology, Epidemiology and Microbial Genomics | Anke Wienecke-Baldacchino, Ardashel Latsuzbaia, Jessica Tapp, Catherine Ragimbeau, Guillaume Fournier, Tamir Abdelrahman, Trung Nguyen Nguyen, Joel Mossong |
| EPI_ISL_429787 | hCoV-19/Luxembourg/LNS3283768/2020 | Europe / Luxembourg | 2020-03-22 | Laboratoire National de Sante, Microbiology, Virology | Laboratoire National de Sante, Microbiology, Epidemiology and Microbial Genomics | Anke Wienecke-Baldacchino, Ardashel Latsuzbaia, Jessica Tapp, Catherine Ragimbeau, Guillaume Fournier, Tamir Abdelrahman, Trung Nguyen Nguyen, Joel Mossong |
| EPI_ISL_429791 | hCoV-19/Luxembourg/LNS7007586/2020 | Europe / Luxembourg | 2020-03-28 | Laboratoire National de Sante, Microbiology, Virology | Laboratoire National de Sante, Microbiology, Epidemiology and Microbial Genomics | Anke Wienecke-Baldacchino, Ardashel Latsuzbaia, Jessica Tapp, Catherine Ragimbeau, Guillaume Fournier, Tamir Abdelrahman, Trung Nguyen Nguyen, Joel Mossong |
| EPI_ISL_429792 | hCoV-19/Croatia/LG-S2/2020 | Europe / Croatia / Istria | 2020-03 | Institute for Public Health | Laboratory for advanced genomics | Filip Rokić, Lovro Trgovec-Greif, Neven Sučić, Tomislav Rukavina, Igor Jurak, Oliver Vugrek |
| EPI_ISL_429793 | hCoV-19/Luxembourg/LNS7812353/2020 | Europe / Luxembourg | 2020-03-27 | Laboratoire National de Sante, Microbiology, Virology | Laboratoire National de Sante, Microbiology, Epidemiology and Microbial Genomics | Anke Wienecke-Baldacchino, Ardashel Latsuzbaia, Jessica Tapp, Catherine Ragimbeau, Guillaume Fournier, Tamir Abdelrahman, Trung Nguyen Nguyen, Joel Mossong |
| EPI_ISL_429794 | hCoV-19/Luxembourg/LNS6479831/2020 | Europe / Luxembourg | 2020-03-31 | Laboratoire National de Sante, Microbiology, Virology | Laboratoire National de Sante, Microbiology, Epidemiology and Microbial Genomics | Anke Wienecke-Baldacchino, Ardashel Latsuzbaia, Jessica Tapp, Catherine Ragimbeau, Guillaume Fournier, Tamir Abdelrahman, Trung Nguyen Nguyen, Joel Mossong |
| EPI_ISL_429796 | hCoV-19/Luxembourg/LNS8367209/2020 | Europe / Luxembourg | 2020-04-08 | Laboratoire National de Sante, Microbiology, Virology | Laboratoire National de Sante, Microbiology, Epidemiology and Microbial Genomics | Anke Wienecke-Baldacchino, Ardashel Latsuzbaia, Jessica Tapp, Catherine Ragimbeau, Guillaume Fournier, Tamir Abdelrahman, Trung Nguyen Nguyen, Joel Mossong |
| EPI_ISL_429797 | hCoV-19/Luxembourg/LNS0687874/2020 | Europe / Luxembourg | 2020-03-30 | Laboratoire National de Sante, Microbiology, Virology | Laboratoire National de Sante, Microbiology, Epidemiology and Microbial Genomics | Anke Wienecke-Baldacchino, Ardashel Latsuzbaia, Jessica Tapp, Catherine Ragimbeau, Guillaume Fournier, Tamir Abdelrahman, Trung Nguyen Nguyen, Joel Mossong |
| EPI_ISL_429870 | hCoV-19/Turkey/HSGM-8990/2020 | Europe / Turkey / Sakarya | 2020-03-22 | Ministry of Health Turkey | Ministry of Health Turkey | Fatma Bayrakdar,Ayşe Başak Altaş,Yasemin Coşgun,Gülay Korukluoğlu,Selçuk Kılıç |
| EPI_ISL_429871 | hCoV-19/Turkey/HSGM-10241/2020 | Europe / Turkey / Ankara | 2020-03-23 | Ministry of Health Turkey | Ministry of Health Turkey | Fatma Bayrakdar,Ayşe Başak Altaş,Yasemin Coşgun,Gülay Korukluoğlu,Selçuk Kılıç |
| EPI_ISL_429872 | hCoV-19/Turkey/HSGM-11518/2020 | Europe / Turkey / Kocaeli | 2020-03-25 | Ministry of Health Turkey | Ministry of Health Turkey | Fatma Bayrakdar,Ayşe Başak Altaş,Yasemin Coşgun,Gülay Korukluoğlu,Selçuk Kılıç |
| EPI_ISL_429873 | hCoV-19/Turkey/HSGM-11520/2020 | Europe / Turkey / Kocaeli | 2020-03-23 | Ministry of Health Turkey | Ministry of Health Turkey | Fatma Bayrakdar,Ayşe Başak Altaş,Yasemin Coşgun,Gülay Korukluoğlu,Selçuk Kılıç |
| EPI_ISL_430000 | hCoV-19/Jordan/SR-042/2020 | Asia / Jordan / Amman | 2020-03-30 | Biolab Diagnostic Laboratories | Andersen lab at Scripps Research | Issa Abu-Dayyeh, Ahmad Tibi, Lama Hussein, Lina Mohammad, Zein Naber, Amid Abdelnour with SEARCH Alliance San Diego |
| EPI_ISL_430001 | hCoV-19/Jordan/SR-043/2020 | Asia / Jordan / Amman | 2020-03-17 | Biolab Diagnostic Laboratories | Andersen lab at Scripps Research | Issa Abu-Dayyeh, Ahmad Tibi, Lama Hussein, Lina Mohammad, Zein Naber, Amid Abdelnour with SEARCH Alliance San Diego |
| EPI_ISL_430002 | hCoV-19/Jordan/SR-044/2020 | Asia / Jordan / Amman | 2020-03-16 | Biolab Diagnostic Laboratories | Andersen lab at Scripps Research | Issa Abu-Dayyeh, Ahmad Tibi, Lama Hussein, Lina Mohammad, Zein Naber, Amid Abdelnour with SEARCH Alliance San Diego |
| EPI_ISL_430003 | hCoV-19/Jordan/SR-045/2020 | Asia / Jordan / Amman | 2020-03-16 | Biolab Diagnostic Laboratories | Andersen lab at Scripps Research | Issa Abu-Dayyeh, Ahmad Tibi, Lama Hussein, Lina Mohammad, Zein Naber, Amid Abdelnour with SEARCH Alliance San Diego |
| EPI_ISL_430004 | hCoV-19/Jordan/SR-046/2020 | Asia / Jordan / Amman | 2020-03-17 | Biolab Diagnostic Laboratories | Andersen lab at Scripps Research | Issa Abu-Dayyeh, Ahmad Tibi, Lama Hussein, Lina Mohammad, Zein Naber, Amid Abdelnour with SEARCH Alliance San Diego |
| EPI_ISL_430006 | hCoV-19/Jordan/SR-048/2020 | Asia / Jordan / Amman | 2020-03-23 | Biolab Diagnostic Laboratories | Andersen lab at Scripps Research | Issa Abu-Dayyeh, Ahmad Tibi, Lama Hussein, Lina Mohammad, Zein Naber, Amid Abdelnour with SEARCH Alliance San Diego |
| EPI_ISL_430007 | hCoV-19/Jordan/SR-049/2020 | Asia / Jordan / Amman | 2020-03-19 | Biolab Diagnostic Laboratories | Andersen lab at Scripps Research | Issa Abu-Dayyeh, Ahmad Tibi, Lama Hussein, Lina Mohammad, Zein Naber, Amid Abdelnour with SEARCH Alliance San Diego |
| EPI_ISL_430009 | hCoV-19/Jordan/SR-052/2020 | Asia / Jordan / Irbid | 2020-03-30 | Biolab Diagnostic Laboratories | Andersen lab at Scripps Research | Issa Abu-Dayyeh, Ahmad Tibi, Lama Hussein, Lina Mohammad, Zein Naber, Amid Abdelnour with SEARCH Alliance San Diego |
| EPI_ISL_430011 | hCoV-19/Jordan/SR-053/2020 | Asia / Jordan / Amman | 2020-03-19 | Biolab Diagnostic Laboratories | Andersen lab at Scripps Research | Issa Abu-Dayyeh, Ahmad Tibi, Lama Hussein, Lina Mohammad, Zein Naber, Amid Abdelnour with SEARCH Alliance San Diego |
| EPI_ISL_430012 | hCoV-19/Jordan/SR-054/2020 | Asia / Jordan / Irbid | 2020-03-30 | Biolab Diagnostic Laboratories | Andersen lab at Scripps Research | Issa Abu-Dayyeh, Ahmad Tibi, Lama Hussein, Lina Mohammad, Zein Naber, Amid Abdelnour with SEARCH Alliance San Diego |
| EPI_ISL_430014 | hCoV-19/Jordan/SR-056/2020 | Asia / Jordan / Amman | 2020-03-17 | Biolab Diagnostic Laboratories | Andersen lab at Scripps Research | Issa Abu-Dayyeh, Ahmad Tibi, Lama Hussein, Lina Mohammad, Zein Naber, Amid Abdelnour with SEARCH Alliance San Diego |
| EPI_ISL_430015 | hCoV-19/Jordan/SR-057/2020 | Asia / Jordan / Amman | 2020-03-24 | Biolab Diagnostic Laboratories | Andersen lab at Scripps Research | Issa Abu-Dayyeh, Ahmad Tibi, Lama Hussein, Lina Mohammad, Zein Naber, Amid Abdelnour with SEARCH Alliance San Diego |
| EPI_ISL_430016 | hCoV-19/USA/CA-SR016/2020 | North America / USA / California / San Di | 2020-03-24 | Andersen lab at Scripps Research | Andersen lab at Scripps Research | SEARCH Alliance San Diego |
| EPI_ISL_430019 | hCoV-19/USA/UT-068/2020 | North America / USA / Utah | 2020-03-16 | Utah Public Health Laboratory | Utah Public Health Laboratory | Erin Young, Kelly Oakeson |
| EPI_ISL_430021 | hCoV-19/USA/UT-069/2020 | North America / USA / Utah | 2020-03-16 | Utah Public Health Laboratory | Utah Public Health Laboratory | Erin Young, Kelly Oakeson |
| EPI_ISL_430022 | hCoV-19/USA/UT-070/2020 | North America / USA / Utah | 2020-03-16 | Utah Public Health Laboratory | Utah Public Health Laboratory | Erin Young, Kelly Oakeson |
| EPI_ISL_430023 | hCoV-19/USA/UT-071/2020 | North America / USA / Utah | 2020-03-16 | Utah Public Health Laboratory | Utah Public Health Laboratory | Erin Young, Kelly Oakeson |
| EPI_ISL_430024 | hCoV-19/USA/UT-072/2020 | North America / USA / Utah | 2020-03-16 | Utah Public Health Laboratory | Utah Public Health Laboratory | Erin Young, Kelly Oakeson |
| EPI_ISL_430025 | hCoV-19/USA/UT-074/2020 | North America / USA / Utah | 2020-03-17 | Utah Public Health Laboratory | Utah Public Health Laboratory | Erin Young, Kelly Oakeson |
| EPI_ISL_430026 | hCoV-19/USA/UT-075/2020 | North America / USA / Utah | 2020-03-16 | Utah Public Health Laboratory | Utah Public Health Laboratory | Erin Young, Kelly Oakeson |
| EPI_ISL_430027 | hCoV-19/USA/UT-076/2020 | North America / USA / Utah | 2020-03-17 | Utah Public Health Laboratory | Utah Public Health Laboratory | Erin Young, Kelly Oakeson |
| EPI_ISL_430028 | hCoV-19/USA/UT-077/2020 | North America / USA / Utah | 2020-03-17 | Utah Public Health Laboratory | Utah Public Health Laboratory | Erin Young, Kelly Oakeson |
| EPI_ISL_430029 | hCoV-19/USA/UT-078/2020 | North America / USA / Utah | 2020-03-15 | Utah Public Health Laboratory | Utah Public Health Laboratory | Erin Young, Kelly Oakeson |
| EPI_ISL_430031 | hCoV-19/USA/UT-080/2020 | North America / USA / Utah | 2020-03-15 | Utah Public Health Laboratory | Utah Public Health Laboratory | Erin Young, Kelly Oakeson |
| EPI_ISL_430032 | hCoV-19/USA/UT-083/2020 | North America / USA / Utah | 2020-03-17 | Utah Public Health Laboratory | Utah Public Health Laboratory | Erin Young, Kelly Oakeson |
| EPI_ISL_430033 | hCoV-19/USA/UT-084/2020 | North America / USA / Utah | 2020-03-17 | Utah Public Health Laboratory | Utah Public Health Laboratory | Erin Young, Kelly Oakeson |
| EPI_ISL_430034 | hCoV-19/USA/UT-085/2020 | North America / USA / Utah | 2020-03-17 | Utah Public Health Laboratory | Utah Public Health Laboratory | Erin Young, Kelly Oakeson |
| EPI_ISL_430035 | hCoV-19/USA/UT-097/2020 | North America / USA / Utah | 2020-03-15 | Utah Public Health Laboratory | Utah Public Health Laboratory | Erin Young, Kelly Oakeson |
| EPI_ISL_430036 | hCoV-19/USA/UT-098/2020 | North America / USA / Utah | 2020-03-14 | Utah Public Health Laboratory | Utah Public Health Laboratory | Erin Young, Kelly Oakeson |
| EPI_ISL_430037 | hCoV-19/USA/UT-099/2020 | North America / USA / Utah | 2020-03-14 | Utah Public Health Laboratory | Utah Public Health Laboratory | Erin Young, Kelly Oakeson |
| EPI_ISL_430038 | hCoV-19/USA/UT-0100/2020 | North America / USA / Utah | 2020-03-15 | Utah Public Health Laboratory | Utah Public Health Laboratory | Erin Young, Kelly Oakeson |
| EPI_ISL_430039 | hCoV-19/USA/UT-0370/2020 | North America / USA / Utah | 2020-03-30 | Utah Public Health Laboratory | Utah Public Health Laboratory | Erin Young, Kelly Oakeson |
| EPI_ISL_430041 | hCoV-19/USA/UT-0443/2020 | North America / USA / Utah | 2020-03-30 | Utah Public Health Laboratory | Utah Public Health Laboratory | Erin Young, Kelly Oakeson |
| EPI_ISL_430042 | hCoV-19/USA/UT-0444/2020 | North America / USA / Utah | 2020-03-30 | Utah Public Health Laboratory | Utah Public Health Laboratory | Erin Young, Kelly Oakeson |
| EPI_ISL_430043 | hCoV-19/USA/UT-0445/2020 | North America / USA / Utah | 2020-03-30 | Utah Public Health Laboratory | Utah Public Health Laboratory | Erin Young, Kelly Oakeson |
| EPI_ISL_430044 | hCoV-19/USA/UT-0446/2020 | North America / USA / Utah | 2020-03-30 | Utah Public Health Laboratory | Utah Public Health Laboratory | Erin Young, Kelly Oakeson |
| EPI_ISL_430045 | hCoV-19/USA/UT-0447/2020 | North America / USA / Utah | 2020-03-31 | Utah Public Health Laboratory | Utah Public Health Laboratory | Erin Young, Kelly Oakeson |
| EPI_ISL_430046 | hCoV-19/USA/UT-0491/2020 | North America / USA / Utah | 2020-04-02 | Utah Public Health Laboratory | Utah Public Health Laboratory | Erin Young, Kelly Oakeson |
| EPI_ISL_430047 | hCoV-19/USA/UT-0492/2020 | North America / USA / Utah | 2020-04-02 | Utah Public Health Laboratory | Utah Public Health Laboratory | Erin Young, Kelly Oakeson |
| EPI_ISL_430048 | hCoV-19/USA/UT-0495/2020 | North America / USA / Utah | 2020-04-02 | Utah Public Health Laboratory | Utah Public Health Laboratory | Erin Young, Kelly Oakeson |
| EPI_ISL_430050 | hCoV-19/USA/UT-0500/2020 | North America / USA / Utah | 2020-03-31 | Utah Public Health Laboratory | Utah Public Health Laboratory | Erin Young, Kelly Oakeson |
| EPI_ISL_430051 | hCoV-19/USA/UT-0501/2020 | North America / USA / Utah | 2020-04-02 | Utah Public Health Laboratory | Utah Public Health Laboratory | Erin Young, Kelly Oakeson |
| EPI_ISL_430052 | hCoV-19/USA/UT-0502/2020 | North America / USA / Utah | 2020-04-03 | Utah Public Health Laboratory | Utah Public Health Laboratory | Erin Young, Kelly Oakeson |
| EPI_ISL_430053 | hCoV-19/USA/UT-0503/2020 | North America / USA / Utah | 2020-04-02 | Utah Public Health Laboratory | Utah Public Health Laboratory | Erin Young, Kelly Oakeson |
| EPI_ISL_430054 | hCoV-19/USA/UT-0506/2020 | North America / USA / Utah | 2020-04-02 | Utah Public Health Laboratory | Utah Public Health Laboratory | Erin Young, Kelly Oakeson |
| EPI_ISL_430055 | hCoV-19/USA/UT-0507/2020 | North America / USA / Utah | 2020-04-02 | Utah Public Health Laboratory | Utah Public Health Laboratory | Erin Young, Kelly Oakeson |
| EPI_ISL_430056 | hCoV-19/USA/UT-0508/2020 | North America / USA / Utah | 2020-04-02 | Utah Public Health Laboratory | Utah Public Health Laboratory | Erin Young, Kelly Oakeson |
| EPI_ISL_430057 | hCoV-19/USA/UT-0510/2020 | North America / USA / Utah | 2020-04-03 | Utah Public Health Laboratory | Utah Public Health Laboratory | Erin Young, Kelly Oakeson |
| EPI_ISL_430179 | hCoV-19/USA/WA-S142/2020 | North America / USA / Washington / King | 2020-03-13 | Washington State Department of Health | Seattle Flu Study | Chu et al |
| EPI_ISL_430180 | hCoV-19/USA/WA-S143/2020 | North America / USA / Washington / King | 2020-03-15 | Washington State Department of Health | Seattle Flu Study | Chu et al |
| EPI_ISL_430181 | hCoV-19/USA/WA-S144/2020 | North America / USA / Washington / King | 2020-03-15 | Washington State Department of Health | Seattle Flu Study | Chu et al |
| EPI_ISL_430182 | hCoV-19/USA/WA-S145/2020 | North America / USA / Washington / King | 2020-03-15 | Washington State Department of Health | Seattle Flu Study | Chu et al |
| EPI_ISL_430183 | hCoV-19/USA/WA-S146/2020 | North America / USA / Washington / King | 2020-03-14 | Washington State Department of Health | Seattle Flu Study | Chu et al |
| EPI_ISL_430184 | hCoV-19/USA/WA-S147/2020 | North America / USA / Washington / King | 2020-03-14 | Washington State Department of Health | Seattle Flu Study | Chu et al |
| EPI_ISL_430185 | hCoV-19/USA/WA-S148/2020 | North America / USA / Washington / King | 2020-03-13 | Washington State Department of Health | Seattle Flu Study | Chu et al |
| EPI_ISL_430186 | hCoV-19/USA/WA-S149/2020 | North America / USA / Washington / King | 2020-03-14 | Washington State Department of Health | Seattle Flu Study | Chu et al |
| EPI_ISL_430187 | hCoV-19/USA/WA-S150/2020 | North America / USA / Washington / King | 2020-03-14 | Washington State Department of Health | Seattle Flu Study | Chu et al |
| EPI_ISL_430188 | hCoV-19/USA/WA-S151/2020 | North America / USA / Washington / Snoh | 2020-03-16 | Washington State Department of Health | Seattle Flu Study | Chu et al |
| EPI_ISL_430189 | hCoV-19/USA/WA-S152/2020 | North America / USA / Washington / King | 2020-03-16 | Washington State Department of Health | Seattle Flu Study | Chu et al |
| EPI_ISL_430190 | hCoV-19/USA/WA-S153/2020 | North America / USA / Washington / King | 2020-03-16 | Washington State Department of Health | Seattle Flu Study | Chu et al |
| EPI_ISL_430191 | hCoV-19/USA/WA-S154/2020 | North America / USA / Washington / King | 2020-03-16 | Washington State Department of Health | Seattle Flu Study | Chu et al |
| EPI_ISL_430193 | hCoV-19/USA/WA-S156/2020 | North America / USA / Washington / King | 2020-03-16 | Washington State Department of Health | Seattle Flu Study | Chu et al |
| EPI_ISL_430196 | hCoV-19/USA/WA-S158/2020 | North America / USA / Washington / Snoh | 2020-03-16 | Washington State Department of Health | Seattle Flu Study | Chu et al |
| EPI_ISL_430197 | hCoV-19/USA/WA-S159/2020 | North America / USA / Washington / Snoh | 2020-03-16 | Washington State Department of Health | Seattle Flu Study | Chu et al |
| EPI_ISL_430198 | hCoV-19/USA/WA-S160/2020 | North America / USA / Washington / King | 2020-03-17 | Washington State Department of Health | Seattle Flu Study | Chu et al |
| EPI_ISL_430199 | hCoV-19/USA/WA-S161/2020 | North America / USA / Washington / King | 2020-03-17 | Washington State Department of Health | Seattle Flu Study | Chu et al |
| EPI_ISL_430701 | hCoV-19/Australia/VIC1284/2020 | Oceania / Australia / Victoria | 2020-04-09 | Victorian Infectious Diseases Reference Laboratory (VIDRL) | Microbiological Diagnostic Unit Public Health Laboratory and Victorian Infectious Diseases Reference Laboratory, The Peter Doherty Institute for Infection and Immunity | Caly L., Seemann T., Sait, M., Schultz M., Druce J., Sherry, N. |
| EPI_ISL_430705 | hCoV-19/Australia/VIC1305/2020 | Oceania / Australia / Victoria | 2020-04-10 | Victorian Infectious Diseases Reference Laboratory (VIDRL) | Microbiological Diagnostic Unit Public Health Laboratory and Victorian Infectious Diseases Reference Laboratory, The Peter Doherty Institute for Infection and Immunity | Caly L., Seemann T., Sait, M., Schultz M., Druce J., Sherry, N. |
| EPI_ISL_430708 | hCoV-19/Australia/VIC1345/2020 | Oceania / Australia / Victoria | 2020-04-14 | Victorian Infectious Diseases Reference Laboratory (VIDRL) | Microbiological Diagnostic Unit Public Health Laboratory and Victorian Infectious Diseases Reference Laboratory, The Peter Doherty Institute for Infection and Immunity | Caly L., Seemann T., Sait, M., Schultz M., Druce J., Sherry, N. |
| EPI_ISL_430711 | hCoV-19/Australia/VIC1292/2020 | Oceania / Australia / Victoria | 2020-04-09 | Victorian Infectious Diseases Reference Laboratory (VIDRL) | Microbiological Diagnostic Unit Public Health Laboratory and Victorian Infectious Diseases Reference Laboratory, The Peter Doherty Institute for Infection and Immunity | Caly L., Seemann T., Sait, M., Schultz M., Druce J., Sherry, N. |
| EPI_ISL_430714 | hCoV-19/Australia/VIC1313/2020 | Oceania / Australia / Victoria | 2020-04-10 | Victorian Infectious Diseases Reference Laboratory (VIDRL) | Microbiological Diagnostic Unit Public Health Laboratory and Victorian Infectious Diseases Reference Laboratory, The Peter Doherty Institute for Infection and Immunity | Caly L., Seemann T., Sait, M., Schultz M., Druce J., Sherry, N. |
| EPI_ISL_430718 | hCoV-19/Spain/Madrid_H12_16/2020 | Europe / Spain / Madrid | 2020-03-10 | Hospital Universitario 12 de Octubre | Hospital Universitario 12 de Octubre | Sara González, Raúl Recio,Elias Dahdouh, Fernando Lázaro, Esther Viedma, Natalia Stella, Julio García, Juan Carlos Galán, Rafael Cantón, Mª Dolores Folgueira, Rafael Delgado, Jesús Mingorance |
| EPI_ISL_430719 | hCoV-19/Spain/Madrid-LP29/2020 | Europe / Spain / Madrid | 2020-03-11 | Hospital Universitario La Paz | Hospital Universitario 12 de Octubre | Elias Dahdouh, Sara González, Raúl Recio, Fernando Lázaro, Esther Viedma, Natalia Stella, Julio García, Juan Carlos Galán, Rafael Cantón, Mª Dolores Folgueira, Rafael Delgado, Jesús Mingorance |
| EPI_ISL_430720 | hCoV-19/Spain/Madrid-LP32/2020 | Europe / Spain / Madrid | 2020-03-09 | Hospital Universitario La Paz | Hospital Universitario 12 de Octubre | Elias Dahdouh, Sara González, Raúl Recio, Fernando Lázaro, Esther Viedma, Natalia Stella, Julio García, Juan Carlos Galán, Rafael Cantón, Mª Dolores Folgueira, Rafael Delgado, Jesús Mingorance |
| EPI_ISL_430721 | hCoV-19/Spain/Madrid-LP33/2020 | Europe / Spain / Madrid | 2020-03-11 | Hospital Universitario La Paz | Hospital Universitario 12 de Octubre | Elias Dahdouh, Sara González, Raúl Recio, Fernando Lázaro, Esther Viedma, Natalia Stella, Julio García, Juan Carlos Galán, Rafael Cantón, Mª Dolores Folgueira, Rafael Delgado, Jesús Mingorance |
| EPI_ISL_430722 | hCoV-19/Beijing/BJ287/2020 | Asia / China / Beijing | 2020-01-29 | Chinese PLA Institute for Disease Control and Prevention | Chinese PLA Institute for Disease Control and Prevention | Peng Li, Jinhui Li, Lizhong Li |
| EPI_ISL_430724 | hCoV-19/Beijing/BJ752/2020 | Asia / China / Beijing | 2020-02-08 | Chinese PLA Institute for Disease Control and Prevention | Chinese PLA Institute for Disease Control and Prevention | Peng Li, Jinhui Li, Lizhong Li |
| EPI_ISL_430725 | hCoV-19/Beijing/BJ288/2020 | Asia / China / Beijing | 2020-01-29 | Chinese PLA Institute for Disease Control and Prevention | Chinese PLA Institute for Disease Control and Prevention | Peng Li, Jinhui Li, Lizhong Li |
| EPI_ISL_430728 | hCoV-19/Beijing/BJ492/2020 | Asia / China / Beijing | 2020-02-02 | Chinese PLA Institute for Disease Control and Prevention | Chinese PLA Institute for Disease Control and Prevention | Peng Li, Jinhui Li, Lizhong Li |
| EPI_ISL_430729 | hCoV-19/Beijing/BJ709/2020 | Asia / China / Beijing | 2020-02-07 | Chinese PLA Institute for Disease Control and Prevention | Chinese PLA Institute for Disease Control and Prevention | Peng Li, Jinhui Li, Lizhong Li |
| EPI_ISL_430730 | hCoV-19/Beijing/BJ390/2020 | Asia / China / Beijing | 2020-01-31 | Chinese PLA Institute for Disease Control and Prevention | Chinese PLA Institute for Disease Control and Prevention | Peng Li, Jinhui Li, Lizhong Li |
| EPI_ISL_430731 | hCoV-19/Beijing/BJ561/2020 | Asia / China / Beijing | 2020-02-04 | Chinese PLA Institute for Disease Control and Prevention | Chinese PLA Institute for Disease Control and Prevention | Peng Li, Jinhui Li, Lizhong Li |
| EPI_ISL_430733 | hCoV-19/Beijing/BJ556/2020 | Asia / China / Beijing | 2020-02-04 | Chinese PLA Institute for Disease Control and Prevention | Chinese PLA Institute for Disease Control and Prevention | Peng Li, Jinhui Li, Lizhong Li |
| EPI_ISL_430734 | hCoV-19/Beijing/BJ53/2020 | Asia / China / Beijing | 2020-01-24 | Chinese PLA Institute for Disease Control and Prevention | Chinese PLA Institute for Disease Control and Prevention | Peng Li, Jinhui Li, Lizhong Li |
| EPI_ISL_430735 | hCoV-19/Beijing/BJ54/2020 | Asia / China / Beijing | 2020-01-24 | Chinese PLA Institute for Disease Control and Prevention | Chinese PLA Institute for Disease Control and Prevention | Peng Li, Jinhui Li, Lizhong Li |
| EPI_ISL_430736 | hCoV-19/Beijing/BJ252/2020 | Asia / China / Beijing | 2020-01-29 | Chinese PLA Institute for Disease Control and Prevention | Chinese PLA Institute for Disease Control and Prevention | Peng Li, Jinhui Li, Lizhong Li |
| EPI_ISL_430737 | hCoV-19/Beijing/BJ1112/2020 | Asia / China / Beijing | 2020-02-25 | Chinese PLA Institute for Disease Control and Prevention | Chinese PLA Institute for Disease Control and Prevention | Peng Li, Jinhui Li, Lizhong Li |
| EPI_ISL_430738 | hCoV-19/Beijing/BJ589/2020 | Asia / China / Beijing | 2020-02-05 | Chinese PLA Institute for Disease Control and Prevention | Chinese PLA Institute for Disease Control and Prevention | Peng Li, Jinhui Li, Lizhong Li |
| EPI_ISL_430740 | hCoV-19/Beijing/BJ710/2020 | Asia / China / Beijing | 2020-02-07 | Chinese PLA Institute for Disease Control and Prevention | Chinese PLA Institute for Disease Control and Prevention | Peng Li, Jinhui Li, Lizhong Li |
| EPI_ISL_430741 | hCoV-19/Beijing/BJ782/2020 | Asia / China / Beijing | 2020-02-10 | Chinese PLA Institute for Disease Control and Prevention | Chinese PLA Institute for Disease Control and Prevention | Peng Li, Jinhui Li, Lizhong Li |
| EPI_ISL_430742 | hCoV-19/Beijing/BJ251/2020 | Asia / China / Beijing | 2020-01-29 | Chinese PLA Institute for Disease Control and Prevention | Chinese PLA Institute for Disease Control and Prevention | Peng Li, Jinhui Li, Lizhong Li |
| EPI_ISL_430743 | hCoV-19/env/Beijing/BJ2452/2020 | Asia / China / Beijing | 2020-03-14 | Chinese PLA Institute for Disease Control and Prevention | Chinese PLA Institute for Disease Control and Prevention | Peng Li, Jinhui Li, Lizhong Li |
| EPI_ISL_430744 | hCoV-19/env/Beijing/BJ2454/2020 | Asia / China / Beijing | 2020-03-14 | Chinese PLA Institute for Disease Control and Prevention | Chinese PLA Institute for Disease Control and Prevention | Peng Li, Jinhui Li, Lizhong Li |
| EPI_ISL_430745 | hCoV-19/env/Beijing/BJ2456/2020 | Asia / China / Beijing | 2020-03-14 | Chinese PLA Institute for Disease Control and Prevention | Chinese PLA Institute for Disease Control and Prevention | Peng Li, Jinhui Li, Lizhong Li |
| EPI_ISL_430746 | hCoV-19/env/Beijing/BJ2460/2020 | Asia / China / Beijing | 2020-03-14 | Chinese PLA Institute for Disease Control and Prevention | Chinese PLA Institute for Disease Control and Prevention | Peng Li, Jinhui Li, Lizhong Li |
| EPI_ISL_430791 | hCoV-19/USA/CA-CZB088/2020 | North America / USA / California / San Fr | 2020-03-28 | UCSF Clinical Microbiology Laboratory | Chan-Zuckerberg Biohub | CZB Cliahub Consortium |
| EPI_ISL_430792 | hCoV-19/USA/CA-CZB0716/2020 | North America / USA / California / San Fr | 2020-03-23 | UCSF Clinical Microbiology Laboratory | Chan-Zuckerberg Biohub | CZB Cliahub Consortium |
| EPI_ISL_430793 | hCoV-19/Argentina/PAIS_A001/2020 | South America / Argentina / Ciudad Autor | 2020-03-01 | Laboratorio Análisis Clínicos, Unidad de Servicios Diagnósticos, Swiss Medical Group | Área de Secuenciación del Laboratorio de Virología del Hospital de Niños Dr. Ricardo Gutierrez | Nabaes Jodar, MS; Goya, S; Natale, MI; Lusso, S; Sanchez, O; Guevara, D; Vicario, SM; Mistchenko, AS; Valinotto, LE; Viegas, M. |
| EPI_ISL_430796 | hCoV-19/Argentina/PAIS_A004/2020 | South America / Argentina / Ciudad Autor | 2020-03-27 | Departamento de Biología y genética molecular, IACA Laboratorios. | Área de Secuenciación del Laboratorio de Virología del Hospital de Niños Dr. Ricardo Gutierrez | Nabaes Jodar, MS; Goya, S; Natale, MI; Lusso, S; Tittarelli, E; Suárez, A; Masciovecchio MV; Streitenberger ER; Mistchenko, AS; Valinotto, LE; Viegas, M. |
| EPI_ISL_430797 | hCoV-19/Argentina/PAIS_A005/2020 | South America / Argentina / Buenos Aires | 2020-03-27 | Departamento de Biología y genética molecular, IACA Laboratorios. | Área de Secuenciación del Laboratorio de Virología del Hospital de Niños Dr. Ricardo Gutierrez | Nabaes Jodar, MS; Goya, S; Natale, MI; Lusso, S; Tittarelli, E; Suárez, A; Masciovecchio MV; Streitenberger ER; Mistchenko, AS; Valinotto, LE; Viegas, M. |
| EPI_ISL_430798 | hCoV-19/Argentina/PAIS_A006/2020 | South America / Argentina / Buenos Aires | 2020-03-28 | Departamento de Biología y genética molecular, IACA Laboratorios. | Área de Secuenciación del Laboratorio de Virología del Hospital de Niños Dr. Ricardo Gutierrez | Nabaes Jodar, MS; Goya, S; Natale, MI; Lusso, S; Tittarelli, E; Suárez, A; Masciovecchio MV; Streitenberger ER; Mistchenko, AS; Valinotto, LE; Viegas, M. |
| EPI_ISL_430799 | hCoV-19/Argentina/PAIS_A007/2020 | South America / Argentina / Ciudad Autor | 2020-03-30 | Laboratorio de Virología del Hospital de Niños Dr. Ricardo Gutierrez | Área de Secuenciación del Laboratorio de Virología del Hospital de Niños Dr. Ricardo Gutierrez | Nabaes Jodar, MS; Goya, S; Natale, MI; Lusso, S; Gravis, E; Mistchenko, AS; Valinotto, LE; Viegas, M. |
| EPI_ISL_430850 | hCoV-19/Sweden/20-08150/2020 | Europe / Sweden / Halland | 2020-03-11 | Klinisk mikrobiologi och vardhygien Halmstad | The Public Health Agency of Sweden | Arne Kotz, Oskar Karlsson Lindsjo, Maria Lind Karlberg, Anna-Malin Linde, Olov Svartstrom, Anna Riaberg, Shaman Muradrasoli, Karin Tegmark-Wisell |
| EPI_ISL_430851 | hCoV-19/Sweden/20-08152/2020 | Europe / Sweden / Halland | 2020-03-11 | Klinisk mikrobiologi och vardhygien Halmstad | The Public Health Agency of Sweden | Arne Kotz, Oskar Karlsson Lindsjo, Maria Lind Karlberg, Anna-Malin Linde, Olov Svartstrom, Anna Riaberg, Shaman Muradrasoli, Karin Tegmark-Wisell |
| EPI_ISL_430852 | hCoV-19/Sweden/20-08154/2020 | Europe / Sweden / Halland | 2020-03-11 | Klinisk mikrobiologi och vardhygien Halmstad | The Public Health Agency of Sweden | Arne Kotz, Oskar Karlsson Lindsjo, Maria Lind Karlberg, Anna-Malin Linde, Olov Svartstrom, Anna Riaberg, Shaman Muradrasoli, Karin Tegmark-Wisell |
| EPI_ISL_430853 | hCoV-19/Sweden/20-08167/2020 | Europe / Sweden / Halland | 2020-03-11 | Klinisk mikrobiologi och vardhygien Halmstad | The Public Health Agency of Sweden | Arne Kotz, Oskar Karlsson Lindsjo, Maria Lind Karlberg, Anna-Malin Linde, Olov Svartstrom, Anna Riaberg, Shaman Muradrasoli, Karin Tegmark-Wisell |
| EPI_ISL_430854 | hCoV-19/Sweden/20-08171/2020 | Europe / Sweden / Halland | 2020-03-12 | Klinisk mikrobiologi och vardhygien Halmstad | The Public Health Agency of Sweden | Arne Kotz, Oskar Karlsson Lindsjo, Maria Lind Karlberg, Anna-Malin Linde, Olov Svartstrom, Anna Riaberg, Shaman Muradrasoli, Karin Tegmark-Wisell |
| EPI_ISL_430855 | hCoV-19/Sweden/20-08172/2020 | Europe / Sweden / Halland | 2020-03-12 | Klinisk mikrobiologi och vardhygien Halmstad | The Public Health Agency of Sweden | Arne Kotz, Oskar Karlsson Lindsjo, Maria Lind Karlberg, Anna-Malin Linde, Olov Svartstrom, Anna Riaberg, Shaman Muradrasoli, Karin Tegmark-Wisell |
| EPI_ISL_430856 | hCoV-19/Sweden/20-08687/2020 | Europe / Sweden / Norrbotten | 2020-03-22 | Laboratoriemedicin | The Public Health Agency of Sweden | Oskar Karlsson Lindsjo, Maria Lind Karlberg, Anna-Malin Linde, Olov Svartstrom, Anna Riaberg, Shaman Muradrasoli, Karin Tegmark-Wisell |
| EPI_ISL_430857 | hCoV-19/Sweden/20-08691/2020 | Europe / Sweden / Norrbotten | 2020-03-22 | Laboratoriemedicin | The Public Health Agency of Sweden | Oskar Karlsson Lindsjo, Maria Lind Karlberg, Anna-Malin Linde, Olov Svartstrom, Anna Riaberg, Shaman Muradrasoli, Karin Tegmark-Wisell |
| EPI_ISL_430858 | hCoV-19/Sweden/20-08714/2020 | Europe / Sweden / Norrbotten | 2020-04-02 | Laboratoriemedicin | The Public Health Agency of Sweden | Oskar Karlsson Lindsjo, Maria Lind Karlberg, Anna-Malin Linde, Olov Svartstrom, Anna Riaberg, Shaman Muradrasoli, Karin Tegmark-Wisell |
| EPI_ISL_430859 | hCoV-19/Sweden/20-08717/2020 | Europe / Sweden / Norrbotten | 2020-04-02 | Laboratoriemedicin | The Public Health Agency of Sweden | Oskar Karlsson Lindsjo, Maria Lind Karlberg, Anna-Malin Linde, Olov Svartstrom, Anna Riaberg, Shaman Muradrasoli, Karin Tegmark-Wisell |
| EPI_ISL_430860 | hCoV-19/Sweden/20-50043/2020 | Europe / Sweden / Varmland | 2020-03-04 | Klinisk mikrobiologi Orebro | The Public Health Agency of Sweden | Martin Sundqvist, Oskar Karlsson Lindsjo, Maria Lind Karlberg, Anna-Malin Linde, Olov Svartstrom, Anna Riaberg, Shaman Muradrasoli, Karin Tegmark-Wisell |
| EPI_ISL_430872 | hCoV-19/USA/WA-UW-5182/2020 | North America / USA / Washington | 2020-04-02 | UW Virology Lab | UW Virology Lab | Pavitra Roychoudhury, Hong Xie, Keith Jerome, Alexander Greninger |
| EPI_ISL_430873 | hCoV-19/USA/WA-UW-5189/2020 | North America / USA / Washington | 2020-04-01 | UW Virology Lab | UW Virology Lab | Pavitra Roychoudhury, Hong Xie, Keith Jerome, Alexander Greninger |
| EPI_ISL_430874 | hCoV-19/USA/WA-UW-5190/2020 | North America / USA / Washington | 2020-04-01 | UW Virology Lab | UW Virology Lab | Pavitra Roychoudhury, Hong Xie, Keith Jerome, Alexander Greninger |
| EPI_ISL_430877 | hCoV-19/USA/WA-UW-524/2020 | North America / USA / Washington | 2020-03-12 | UW Virology Lab | UW Virology Lab | Pavitra Roychoudhury, Hong Xie, Keith Jerome, Alexander Greninger |
| EPI_ISL_430878 | hCoV-19/USA/WA-UW-525/2020 | North America / USA / Washington | 2020-03-11 | UW Virology Lab | UW Virology Lab | Pavitra Roychoudhury, Hong Xie, Keith Jerome, Alexander Greninger |
| EPI_ISL_430879 | hCoV-19/USA/UN-UW-526/2020 | North America / USA | 2020-03-11 | UW Virology Lab | UW Virology Lab | Pavitra Roychoudhury, Hong Xie, Keith Jerome, Alexander Greninger |
| EPI_ISL_430880 | hCoV-19/USA/WA-UW-530/2020 | North America / USA / Washington | 2020-03-12 | UW Virology Lab | UW Virology Lab | Pavitra Roychoudhury, Hong Xie, Keith Jerome, Alexander Greninger |
| EPI_ISL_430881 | hCoV-19/USA/UN-UW-531/2020 | North America / USA | 2020-03-13 | UW Virology Lab | UW Virology Lab | Pavitra Roychoudhury, Hong Xie, Keith Jerome, Alexander Greninger |
| EPI_ISL_430882 | hCoV-19/USA/UN-UW-536/2020 | North America / USA | 2020-03-11 | UW Virology Lab | UW Virology Lab | Pavitra Roychoudhury, Hong Xie, Keith Jerome, Alexander Greninger |
| EPI_ISL_430883 | hCoV-19/USA/WA-UW-542/2020 | North America / USA / Washington | 2020-03-12 | UW Virology Lab | UW Virology Lab | Pavitra Roychoudhury, Hong Xie, Keith Jerome, Alexander Greninger |
| EPI_ISL_430884 | hCoV-19/USA/WA-UW-543/2020 | North America / USA / Washington | 2020-03-12 | UW Virology Lab | UW Virology Lab | Pavitra Roychoudhury, Hong Xie, Keith Jerome, Alexander Greninger |
| EPI_ISL_430885 | hCoV-19/USA/WA-UW-546/2020 | North America / USA / Washington | 2020-03-12 | UW Virology Lab | UW Virology Lab | Pavitra Roychoudhury, Hong Xie, Keith Jerome, Alexander Greninger |
| EPI_ISL_430886 | hCoV-19/USA/UN-UW-547/2020 | North America / USA | 2020-03-10 | UW Virology Lab | UW Virology Lab | Pavitra Roychoudhury, Hong Xie, Keith Jerome, Alexander Greninger |
| EPI_ISL_430887 | hCoV-19/USA/WA-UW-555/2020 | North America / USA / Washington | 2020-03-12 | UW Virology Lab | UW Virology Lab | Pavitra Roychoudhury, Hong Xie, Keith Jerome, Alexander Greninger |
| EPI_ISL_430888 | hCoV-19/USA/WA-UW-558/2020 | North America / USA / Washington | 2020-03-13 | UW Virology Lab | UW Virology Lab | Pavitra Roychoudhury, Hong Xie, Keith Jerome, Alexander Greninger |
| EPI_ISL_430889 | hCoV-19/USA/WA-UW-572/2020 | North America / USA / Washington | 2020-03-12 | UW Virology Lab | UW Virology Lab | Pavitra Roychoudhury, Hong Xie, Keith Jerome, Alexander Greninger |
| EPI_ISL_430890 | hCoV-19/USA/UN-UW-581/2020 | North America / USA | 2020-03-12 | UW Virology Lab | UW Virology Lab | Pavitra Roychoudhury, Hong Xie, Keith Jerome, Alexander Greninger |
| EPI_ISL_430891 | hCoV-19/USA/WA-UW-586/2020 | North America / USA / Washington | 2020-03-12 | UW Virology Lab | UW Virology Lab | Pavitra Roychoudhury, Hong Xie, Keith Jerome, Alexander Greninger |
| EPI_ISL_430892 | hCoV-19/USA/WA-UW-596/2020 | North America / USA / Washington | 2020-03-12 | UW Virology Lab | UW Virology Lab | Pavitra Roychoudhury, Hong Xie, Keith Jerome, Alexander Greninger |
| EPI_ISL_430893 | hCoV-19/USA/WA-UW-600/2020 | North America / USA / Washington | 2020-03-12 | UW Virology Lab | UW Virology Lab | Pavitra Roychoudhury, Hong Xie, Keith Jerome, Alexander Greninger |
| EPI_ISL_431180 | hCoV-19/Fujian/IM3520002T/2020 | Asia / China / Fujian | 2020-03-15 | Fujian Center for Disease Control and Prevention | Fujian Center for Disease Control and Prevention | Lin Qi, Huang Zhimiao, Zhang Yanhua, Weng Yuwei |
| EPI_ISL_431240 | hCoV-19/Fujian/IM3520003T/2020 | Asia / China / Fujian | 2020-03-16 | Fujian Center for Disease Control and Prevention | Fujian Center for Disease Control and Prevention | Lin Qi, Huang Zhimiao, Zhang Yanhua, Weng Yuwei |
| EPI_ISL_432478 | hCoV-19/England/SHEF-C9E36/2020 | Europe / United Kingdom / England | 2020-04-07 | Virology Department, Sheffield Teaching Hospitals NHS Foundation Trust / Virology Department, Sheffield Teaching Hospitals NHS Foundation Trust | COVID-19 Genomics UK (COG-UK) Consortium | Thushan de Silva, Matthew Parker,Adri Angyal, Rebecca Brown, Luke Green, Rachel Tucker, Paul Parsons, Danielle Groves, Alex Keeley, Dave Partridge, Matthew Wyles, Benjamin Lindsey, Mehmet Yavuz, Mohammad Raza, Cariad Evans |
| EPI_ISL_432480 | hCoV-19/England/SHEF-CA020/2020 | Europe / United Kingdom / England | 2020-03-29 | Virology Department, Sheffield Teaching Hospitals NHS Foundation Trust / Virology Department, Sheffield Teaching Hospitals NHS Foundation Trust | COVID-19 Genomics UK (COG-UK) Consortium | Thushan de Silva, Matthew Parker,Adri Angyal, Rebecca Brown, Luke Green, Rachel Tucker, Paul Parsons, Danielle Groves, Alex Keeley, Dave Partridge, Matthew Wyles, Benjamin Lindsey, Mehmet Yavuz, Mohammad Raza, Cariad Evans |
| EPI_ISL_432481 | hCoV-19/England/SHEF-CA0B7/2020 | Europe / United Kingdom / England | 2020-03-30 | Virology Department, Sheffield Teaching Hospitals NHS Foundation Trust / Virology Department, Sheffield Teaching Hospitals NHS Foundation Trust | COVID-19 Genomics UK (COG-UK) Consortium | Thushan de Silva, Matthew Parker,Adri Angyal, Rebecca Brown, Luke Green, Rachel Tucker, Paul Parsons, Danielle Groves, Alex Keeley, Dave Partridge, Matthew Wyles, Benjamin Lindsey, Mehmet Yavuz, Mohammad Raza, Cariad Evans |
| EPI_ISL_432487 | hCoV-19/England/SHEF-CA1F0/2020 | Europe / United Kingdom / England | 2020-03-30 | Virology Department, Sheffield Teaching Hospitals NHS Foundation Trust / Virology Department, Sheffield Teaching Hospitals NHS Foundation Trust | COVID-19 Genomics UK (COG-UK) Consortium | Thushan de Silva, Matthew Parker,Adri Angyal, Rebecca Brown, Luke Green, Rachel Tucker, Paul Parsons, Danielle Groves, Alex Keeley, Dave Partridge, Matthew Wyles, Benjamin Lindsey, Mehmet Yavuz, Mohammad Raza, Cariad Evans |
| EPI_ISL_432502 | hCoV-19/England/SHEF-CB0F2/2020 | Europe / United Kingdom / England | 2020-03-25 | Virology Department, Sheffield Teaching Hospitals NHS Foundation Trust / Virology Department, Sheffield Teaching Hospitals NHS Foundation Trust | COVID-19 Genomics UK (COG-UK) Consortium | Thushan de Silva, Matthew Parker,Adri Angyal, Rebecca Brown, Luke Green, Rachel Tucker, Paul Parsons, Danielle Groves, Alex Keeley, Dave Partridge, Matthew Wyles, Benjamin Lindsey, Mehmet Yavuz, Mohammad Raza, Cariad Evans |
| EPI_ISL_432505 | hCoV-19/England/SHEF-CB317/2020 | Europe / United Kingdom / England | 2020-03-31 | Virology Department, Sheffield Teaching Hospitals NHS Foundation Trust / Virology Department, Sheffield Teaching Hospitals NHS Foundation Trust | COVID-19 Genomics UK (COG-UK) Consortium | Thushan de Silva, Matthew Parker,Adri Angyal, Rebecca Brown, Luke Green, Rachel Tucker, Paul Parsons, Danielle Groves, Alex Keeley, Dave Partridge, Matthew Wyles, Benjamin Lindsey, Mehmet Yavuz, Mohammad Raza, Cariad Evans |
| EPI_ISL_432510 | hCoV-19/England/SHEF-CB5E4/2020 | Europe / United Kingdom / England | 2020-03-27 | Virology Department, Sheffield Teaching Hospitals NHS Foundation Trust / Virology Department, Sheffield Teaching Hospitals NHS Foundation Trust | COVID-19 Genomics UK (COG-UK) Consortium | Thushan de Silva, Matthew Parker,Adri Angyal, Rebecca Brown, Luke Green, Rachel Tucker, Paul Parsons, Danielle Groves, Alex Keeley, Dave Partridge, Matthew Wyles, Benjamin Lindsey, Mehmet Yavuz, Mohammad Raza, Cariad Evans |
| EPI_ISL_432517 | hCoV-19/England/SHEF-CBECB/2020 | Europe / United Kingdom / England | 2020-03-30 | Virology Department, Sheffield Teaching Hospitals NHS Foundation Trust / Virology Department, Sheffield Teaching Hospitals NHS Foundation Trust | COVID-19 Genomics UK (COG-UK) Consortium | Thushan de Silva, Matthew Parker,Adri Angyal, Rebecca Brown, Luke Green, Rachel Tucker, Paul Parsons, Danielle Groves, Alex Keeley, Dave Partridge, Matthew Wyles, Benjamin Lindsey, Mehmet Yavuz, Mohammad Raza, Cariad Evans |
| EPI_ISL_432519 | hCoV-19/England/SHEF-CBF31/2020 | Europe / United Kingdom / England | 2020-04-02 | Virology Department, Sheffield Teaching Hospitals NHS Foundation Trust / Virology Department, Sheffield Teaching Hospitals NHS Foundation Trust | COVID-19 Genomics UK (COG-UK) Consortium | Thushan de Silva, Matthew Parker,Adri Angyal, Rebecca Brown, Luke Green, Rachel Tucker, Paul Parsons, Danielle Groves, Alex Keeley, Dave Partridge, Matthew Wyles, Benjamin Lindsey, Mehmet Yavuz, Mohammad Raza, Cariad Evans |
| EPI_ISL_432529 | hCoV-19/England/SHEF-CC905/2020 | Europe / United Kingdom / England | 2020-03-31 | Virology Department, Sheffield Teaching Hospitals NHS Foundation Trust / Virology Department, Sheffield Teaching Hospitals NHS Foundation Trust | COVID-19 Genomics UK (COG-UK) Consortium | Thushan de Silva, Matthew Parker,Adri Angyal, Rebecca Brown, Luke Green, Rachel Tucker, Paul Parsons, Danielle Groves, Alex Keeley, Dave Partridge, Matthew Wyles, Benjamin Lindsey, Mehmet Yavuz, Mohammad Raza, Cariad Evans |
| EPI_ISL_432533 | hCoV-19/England/SHEF-CCC93/2020 | Europe / United Kingdom / England | 2020-03-31 | Virology Department, Sheffield Teaching Hospitals NHS Foundation Trust / Virology Department, Sheffield Teaching Hospitals NHS Foundation Trust | COVID-19 Genomics UK (COG-UK) Consortium | Thushan de Silva, Matthew Parker,Adri Angyal, Rebecca Brown, Luke Green, Rachel Tucker, Paul Parsons, Danielle Groves, Alex Keeley, Dave Partridge, Matthew Wyles, Benjamin Lindsey, Mehmet Yavuz, Mohammad Raza, Cariad Evans |
| EPI_ISL_432539 | hCoV-19/England/SHEF-CD05A/2020 | Europe / United Kingdom / England | 2020-03-27 | Virology Department, Sheffield Teaching Hospitals NHS Foundation Trust / Virology Department, Sheffield Teaching Hospitals NHS Foundation Trust | COVID-19 Genomics UK (COG-UK) Consortium | Thushan de Silva, Matthew Parker,Adri Angyal, Rebecca Brown, Luke Green, Rachel Tucker, Paul Parsons, Danielle Groves, Alex Keeley, Dave Partridge, Matthew Wyles, Benjamin Lindsey, Mehmet Yavuz, Mohammad Raza, Cariad Evans |
| EPI_ISL_432541 | hCoV-19/England/SHEF-CD2AF/2020 | Europe / United Kingdom / England | 2020-04-03 | Virology Department, Sheffield Teaching Hospitals NHS Foundation Trust / Virology Department, Sheffield Teaching Hospitals NHS Foundation Trust | COVID-19 Genomics UK (COG-UK) Consortium | Thushan de Silva, Matthew Parker,Adri Angyal, Rebecca Brown, Luke Green, Rachel Tucker, Paul Parsons, Danielle Groves, Alex Keeley, Dave Partridge, Matthew Wyles, Benjamin Lindsey, Mehmet Yavuz, Mohammad Raza, Cariad Evans |
| EPI_ISL_432542 | hCoV-19/England/SHEF-CD315/2020 | Europe / United Kingdom / England | 2020-04-01 | Virology Department, Sheffield Teaching Hospitals NHS Foundation Trust / Virology Department, Sheffield Teaching Hospitals NHS Foundation Trust | COVID-19 Genomics UK (COG-UK) Consortium | Thushan de Silva, Matthew Parker,Adri Angyal, Rebecca Brown, Luke Green, Rachel Tucker, Paul Parsons, Danielle Groves, Alex Keeley, Dave Partridge, Matthew Wyles, Benjamin Lindsey, Mehmet Yavuz, Mohammad Raza, Cariad Evans |
| EPI_ISL_432547 | hCoV-19/England/SHEF-CD52E/2020 | Europe / United Kingdom / England | 2020-03-31 | Virology Department, Sheffield Teaching Hospitals NHS Foundation Trust / Virology Department, Sheffield Teaching Hospitals NHS Foundation Trust | COVID-19 Genomics UK (COG-UK) Consortium | Thushan de Silva, Matthew Parker,Adri Angyal, Rebecca Brown, Luke Green, Rachel Tucker, Paul Parsons, Danielle Groves, Alex Keeley, Dave Partridge, Matthew Wyles, Benjamin Lindsey, Mehmet Yavuz, Mohammad Raza, Cariad Evans |
| EPI_ISL_432548 | hCoV-19/England/SHEF-CD737/2020 | Europe / United Kingdom / England | 2020-03-29 | Virology Department, Sheffield Teaching Hospitals NHS Foundation Trust / Virology Department, Sheffield Teaching Hospitals NHS Foundation Trust | COVID-19 Genomics UK (COG-UK) Consortium | Thushan de Silva, Matthew Parker,Adri Angyal, Rebecca Brown, Luke Green, Rachel Tucker, Paul Parsons, Danielle Groves, Alex Keeley, Dave Partridge, Matthew Wyles, Benjamin Lindsey, Mehmet Yavuz, Mohammad Raza, Cariad Evans |
| EPI_ISL_432549 | hCoV-19/England/SHEF-CD89E/2020 | Europe / United Kingdom / England | 2020-03-28 | Virology Department, Sheffield Teaching Hospitals NHS Foundation Trust / Virology Department, Sheffield Teaching Hospitals NHS Foundation Trust | COVID-19 Genomics UK (COG-UK) Consortium | Thushan de Silva, Matthew Parker,Adri Angyal, Rebecca Brown, Luke Green, Rachel Tucker, Paul Parsons, Danielle Groves, Alex Keeley, Dave Partridge, Matthew Wyles, Benjamin Lindsey, Mehmet Yavuz, Mohammad Raza, Cariad Evans |
| EPI_ISL_432551 | hCoV-19/England/SHEF-CD97D/2020 | Europe / United Kingdom / England | 2020-03-30 | Virology Department, Sheffield Teaching Hospitals NHS Foundation Trust / Virology Department, Sheffield Teaching Hospitals NHS Foundation Trust | COVID-19 Genomics UK (COG-UK) Consortium | Thushan de Silva, Matthew Parker,Adri Angyal, Rebecca Brown, Luke Green, Rachel Tucker, Paul Parsons, Danielle Groves, Alex Keeley, Dave Partridge, Matthew Wyles, Benjamin Lindsey, Mehmet Yavuz, Mohammad Raza, Cariad Evans |
| EPI_ISL_432570 | hCoV-19/England/SHEF-CE89D/2020 | Europe / United Kingdom / England | 2020-04-06 | Virology Department, Sheffield Teaching Hospitals NHS Foundation Trust / Virology Department, Sheffield Teaching Hospitals NHS Foundation Trust | COVID-19 Genomics UK (COG-UK) Consortium | Thushan de Silva, Matthew Parker,Adri Angyal, Rebecca Brown, Luke Green, Rachel Tucker, Paul Parsons, Danielle Groves, Alex Keeley, Dave Partridge, Matthew Wyles, Benjamin Lindsey, Mehmet Yavuz, Mohammad Raza, Cariad Evans |
| EPI_ISL_432578 | hCoV-19/England/SHEF-CECEC/2020 | Europe / United Kingdom / England | 2020-04-04 | Virology Department, Sheffield Teaching Hospitals NHS Foundation Trust / Virology Department, Sheffield Teaching Hospitals NHS Foundation Trust | COVID-19 Genomics UK (COG-UK) Consortium | Thushan de Silva, Matthew Parker,Adri Angyal, Rebecca Brown, Luke Green, Rachel Tucker, Paul Parsons, Danielle Groves, Alex Keeley, Dave Partridge, Matthew Wyles, Benjamin Lindsey, Mehmet Yavuz, Mohammad Raza, Cariad Evans |
| EPI_ISL_432579 | hCoV-19/England/SHEF-CED9E/2020 | Europe / United Kingdom / England | 2020-04-01 | Virology Department, Sheffield Teaching Hospitals NHS Foundation Trust / Virology Department, Sheffield Teaching Hospitals NHS Foundation Trust | COVID-19 Genomics UK (COG-UK) Consortium | Thushan de Silva, Matthew Parker,Adri Angyal, Rebecca Brown, Luke Green, Rachel Tucker, Paul Parsons, Danielle Groves, Alex Keeley, Dave Partridge, Matthew Wyles, Benjamin Lindsey, Mehmet Yavuz, Mohammad Raza, Cariad Evans |
| EPI_ISL_432588 | hCoV-19/England/SHEF-CF340/2020 | Europe / United Kingdom / England | 2020-03-28 | Virology Department, Sheffield Teaching Hospitals NHS Foundation Trust / Virology Department, Sheffield Teaching Hospitals NHS Foundation Trust | COVID-19 Genomics UK (COG-UK) Consortium | Thushan de Silva, Matthew Parker,Adri Angyal, Rebecca Brown, Luke Green, Rachel Tucker, Paul Parsons, Danielle Groves, Alex Keeley, Dave Partridge, Matthew Wyles, Benjamin Lindsey, Mehmet Yavuz, Mohammad Raza, Cariad Evans |
| EPI_ISL_432642 | hCoV-19/England/SHEF-D1E2C/2020 | Europe / United Kingdom / England | 2020-04-02 | Virology Department, Sheffield Teaching Hospitals NHS Foundation Trust / Virology Department, Sheffield Teaching Hospitals NHS Foundation Trust | COVID-19 Genomics UK (COG-UK) Consortium | Thushan de Silva, Matthew Parker,Adri Angyal, Rebecca Brown, Luke Green, Rachel Tucker, Paul Parsons, Danielle Groves, Alex Keeley, Dave Partridge, Matthew Wyles, Benjamin Lindsey, Mehmet Yavuz, Mohammad Raza, Cariad Evans |
| EPI_ISL_432644 | hCoV-19/England/SHEF-D1F56/2020 | Europe / United Kingdom / England | 2020-03-30 | Virology Department, Sheffield Teaching Hospitals NHS Foundation Trust / Virology Department, Sheffield Teaching Hospitals NHS Foundation Trust | COVID-19 Genomics UK (COG-UK) Consortium | Thushan de Silva, Matthew Parker,Adri Angyal, Rebecca Brown, Luke Green, Rachel Tucker, Paul Parsons, Danielle Groves, Alex Keeley, Dave Partridge, Matthew Wyles, Benjamin Lindsey, Mehmet Yavuz, Mohammad Raza, Cariad Evans |
| EPI_ISL_432675 | hCoV-19/England/NOTT-10FA3F/2020 | Europe / United Kingdom / England | 2020-04-05 | Queens Medical Centre, Clinical Microbiology Department / DeepSeq Nottingham | COVID-19 Genomics UK (COG-UK) Consortium | Gemma Clark, Wendy Smith, Manjinder Khakh, Hannah Howson-Wells, Jonathan Ball, Patrick McClure, Joseph Chappell, Theocharis Tsoleridis, Nadine Holmes, Matthew Carlisle, Christopher Moore, Fei Sang, Johnny Debebe, Victoria Wright, Matthew Loose |
| EPI_ISL_432691 | hCoV-19/England/NOTT-10FB69/2020 | Europe / United Kingdom / England | 2020-04-07 | Queens Medical Centre, Clinical Microbiology Department / DeepSeq Nottingham | COVID-19 Genomics UK (COG-UK) Consortium | Gemma Clark, Wendy Smith, Manjinder Khakh, Hannah Howson-Wells, Jonathan Ball, Patrick McClure, Joseph Chappell, Theocharis Tsoleridis, Nadine Holmes, Matthew Carlisle, Christopher Moore, Fei Sang, Johnny Debebe, Victoria Wright, Matthew Loose |
| EPI_ISL_433204 | hCoV-19/Scotland/EDB582/2020 | Europe / United Kingdom / Scotland | 2020-04-02 | Virology Department, Royal Infirmary of Edinburgh, NHS Lothian / School of Biological Sciences, University of Edinburgh / Institute of Genetics and Molecular Medicine, University of Edinburgh | COVID-19 Genomics UK (COG-UK) Consortium | McHugh M, Dewar R, Rooke S, Gallagher M, Balcaza C, O'Toole A, Hill V, McCrone JT, Colquhoun R, Yu X, Jackson B, Rambaut A, Williams TC, Templeton K |
| EPI_ISL_433205 | hCoV-19/Scotland/EDB587/2020 | Europe / United Kingdom / Scotland | 2020-04-02 | Virology Department, Royal Infirmary of Edinburgh, NHS Lothian / School of Biological Sciences, University of Edinburgh / Institute of Genetics and Molecular Medicine, University of Edinburgh | COVID-19 Genomics UK (COG-UK) Consortium | McHugh M, Dewar R, Rooke S, Gallagher M, Balcaza C, O'Toole A, Hill V, McCrone JT, Colquhoun R, Yu X, Jackson B, Rambaut A, Williams TC, Templeton K |
| EPI_ISL_433206 | hCoV-19/Scotland/EDB588/2020 | Europe / United Kingdom / Scotland | 2020-04-02 | Virology Department, Royal Infirmary of Edinburgh, NHS Lothian / School of Biological Sciences, University of Edinburgh / Institute of Genetics and Molecular Medicine, University of Edinburgh | COVID-19 Genomics UK (COG-UK) Consortium | McHugh M, Dewar R, Rooke S, Gallagher M, Balcaza C, O'Toole A, Hill V, McCrone JT, Colquhoun R, Yu X, Jackson B, Rambaut A, Williams TC, Templeton K |
| EPI_ISL_433207 | hCoV-19/Scotland/EDB589/2020 | Europe / United Kingdom / Scotland | 2020-04-01 | Virology Department, Royal Infirmary of Edinburgh, NHS Lothian / School of Biological Sciences, University of Edinburgh / Institute of Genetics and Molecular Medicine, University of Edinburgh | COVID-19 Genomics UK (COG-UK) Consortium | McHugh M, Dewar R, Rooke S, Gallagher M, Balcaza C, O'Toole A, Hill V, McCrone JT, Colquhoun R, Yu X, Jackson B, Rambaut A, Williams TC, Templeton K |
| EPI_ISL_433208 | hCoV-19/Scotland/EDB590/2020 | Europe / United Kingdom / Scotland | 2020-04-01 | Virology Department, Royal Infirmary of Edinburgh, NHS Lothian / School of Biological Sciences, University of Edinburgh / Institute of Genetics and Molecular Medicine, University of Edinburgh | COVID-19 Genomics UK (COG-UK) Consortium | McHugh M, Dewar R, Rooke S, Gallagher M, Balcaza C, O'Toole A, Hill V, McCrone JT, Colquhoun R, Yu X, Jackson B, Rambaut A, Williams TC, Templeton K |
| EPI_ISL_433211 | hCoV-19/Scotland/EDB598/2020 | Europe / United Kingdom / Scotland | 2020-04-01 | Virology Department, Royal Infirmary of Edinburgh, NHS Lothian / School of Biological Sciences, University of Edinburgh / Institute of Genetics and Molecular Medicine, University of Edinburgh | COVID-19 Genomics UK (COG-UK) Consortium | McHugh M, Dewar R, Rooke S, Gallagher M, Balcaza C, O'Toole A, Hill V, McCrone JT, Colquhoun R, Yu X, Jackson B, Rambaut A, Williams TC, Templeton K |
| EPI_ISL_433214 | hCoV-19/Scotland/EDB607/2020 | Europe / United Kingdom / Scotland | 2020-04-01 | Virology Department, Royal Infirmary of Edinburgh, NHS Lothian / School of Biological Sciences, University of Edinburgh / Institute of Genetics and Molecular Medicine, University of Edinburgh | COVID-19 Genomics UK (COG-UK) Consortium | McHugh M, Dewar R, Rooke S, Gallagher M, Balcaza C, O'Toole A, Hill V, McCrone JT, Colquhoun R, Yu X, Jackson B, Rambaut A, Williams TC, Templeton K |
| EPI_ISL_433215 | hCoV-19/Scotland/EDB612/2020 | Europe / United Kingdom / Scotland | 2020-04-02 | Virology Department, Royal Infirmary of Edinburgh, NHS Lothian / School of Biological Sciences, University of Edinburgh / Institute of Genetics and Molecular Medicine, University of Edinburgh | COVID-19 Genomics UK (COG-UK) Consortium | McHugh M, Dewar R, Rooke S, Gallagher M, Balcaza C, O'Toole A, Hill V, McCrone JT, Colquhoun R, Yu X, Jackson B, Rambaut A, Williams TC, Templeton K |
| EPI_ISL_433216 | hCoV-19/Scotland/EDB613/2020 | Europe / United Kingdom / Scotland | 2020-03-31 | Virology Department, Royal Infirmary of Edinburgh, NHS Lothian / School of Biological Sciences, University of Edinburgh / Institute of Genetics and Molecular Medicine, University of Edinburgh | COVID-19 Genomics UK (COG-UK) Consortium | McHugh M, Dewar R, Rooke S, Gallagher M, Balcaza C, O'Toole A, Hill V, McCrone JT, Colquhoun R, Yu X, Jackson B, Rambaut A, Williams TC, Templeton K |
| EPI_ISL_433217 | hCoV-19/Scotland/EDB620/2020 | Europe / United Kingdom / Scotland | 2020-03-31 | Virology Department, Royal Infirmary of Edinburgh, NHS Lothian / School of Biological Sciences, University of Edinburgh / Institute of Genetics and Molecular Medicine, University of Edinburgh | COVID-19 Genomics UK (COG-UK) Consortium | McHugh M, Dewar R, Rooke S, Gallagher M, Balcaza C, O'Toole A, Hill V, McCrone JT, Colquhoun R, Yu X, Jackson B, Rambaut A, Williams TC, Templeton K |
| EPI_ISL_433218 | hCoV-19/Scotland/EDB625/2020 | Europe / United Kingdom / Scotland | 2020-04-01 | Virology Department, Royal Infirmary of Edinburgh, NHS Lothian / School of Biological Sciences, University of Edinburgh / Institute of Genetics and Molecular Medicine, University of Edinburgh | COVID-19 Genomics UK (COG-UK) Consortium | McHugh M, Dewar R, Rooke S, Gallagher M, Balcaza C, O'Toole A, Hill V, McCrone JT, Colquhoun R, Yu X, Jackson B, Rambaut A, Williams TC, Templeton K |
| EPI_ISL_433222 | hCoV-19/Scotland/EDB682/2020 | Europe / United Kingdom / Scotland | 2020-04-03 | Virology Department, Royal Infirmary of Edinburgh, NHS Lothian / School of Biological Sciences, University of Edinburgh / Institute of Genetics and Molecular Medicine, University of Edinburgh | COVID-19 Genomics UK (COG-UK) Consortium | McHugh M, Dewar R, Rooke S, Gallagher M, Balcaza C, O'Toole A, Hill V, McCrone JT, Colquhoun R, Yu X, Jackson B, Rambaut A, Williams TC, Templeton K |
| EPI_ISL_433223 | hCoV-19/Scotland/EDB689/2020 | Europe / United Kingdom / Scotland | 2020-04-03 | Virology Department, Royal Infirmary of Edinburgh, NHS Lothian / School of Biological Sciences, University of Edinburgh / Institute of Genetics and Molecular Medicine, University of Edinburgh | COVID-19 Genomics UK (COG-UK) Consortium | McHugh M, Dewar R, Rooke S, Gallagher M, Balcaza C, O'Toole A, Hill V, McCrone JT, Colquhoun R, Yu X, Jackson B, Rambaut A, Williams TC, Templeton K |
| EPI_ISL_433224 | hCoV-19/Scotland/EDB694/2020 | Europe / United Kingdom / Scotland | 2020-04-01 | Virology Department, Royal Infirmary of Edinburgh, NHS Lothian / School of Biological Sciences, University of Edinburgh / Institute of Genetics and Molecular Medicine, University of Edinburgh | COVID-19 Genomics UK (COG-UK) Consortium | McHugh M, Dewar R, Rooke S, Gallagher M, Balcaza C, O'Toole A, Hill V, McCrone JT, Colquhoun R, Yu X, Jackson B, Rambaut A, Williams TC, Templeton K |

| | | | | | | |
|---|---|---|---|---|---|---|
| EPI_ISL_433225 | hCoV-19/Scotland/EDB695/2020 | Europe / United Kingdom / Scotland | 2020-04-01 | Virology Department, Royal Infirmary of Edinburgh, NHS Lothian / School of Biological Sciences, University of Edinburgh / Institute of Genetics and Molecular Medicine, University of Edinburgh | COVID-19 Genomics UK (COG-UK) Consortium | McHugh M, Dewar R, Rooke S, Gallagher M, Balcaza C, O'Toole A, Hill V, McCrone JT, Colquhoun R, Yu X, Jackson B, Rambaut A, Williams TC, Templeton K |
| EPI_ISL_433226 | hCoV-19/Scotland/EDB703/2020 | Europe / United Kingdom / Scotland | 2020-04-01 | Virology Department, Royal Infirmary of Edinburgh, NHS Lothian / School of Biological Sciences, University of Edinburgh / Institute of Genetics and Molecular Medicine, University of Edinburgh | COVID-19 Genomics UK (COG-UK) Consortium | McHugh M, Dewar R, Rooke S, Gallagher M, Balcaza C, O'Toole A, Hill V, McCrone JT, Colquhoun R, Yu X, Jackson B, Rambaut A, Williams TC, Templeton K |
| EPI_ISL_433227 | hCoV-19/Scotland/EDB707/2020 | Europe / United Kingdom / Scotland | 2020-04-01 | Virology Department, Royal Infirmary of Edinburgh, NHS Lothian / School of Biological Sciences, University of Edinburgh / Institute of Genetics and Molecular Medicine, University of Edinburgh | COVID-19 Genomics UK (COG-UK) Consortium | McHugh M, Dewar R, Rooke S, Gallagher M, Balcaza C, O'Toole A, Hill V, McCrone JT, Colquhoun R, Yu X, Jackson B, Rambaut A, Williams TC, Templeton K |
| EPI_ISL_433228 | hCoV-19/Scotland/EDB721/2020 | Europe / United Kingdom / Scotland | 2020-04-02 | Virology Department, Royal Infirmary of Edinburgh, NHS Lothian / School of Biological Sciences, University of Edinburgh / Institute of Genetics and Molecular Medicine, University of Edinburgh | COVID-19 Genomics UK (COG-UK) Consortium | McHugh M, Dewar R, Rooke S, Gallagher M, Balcaza C, O'Toole A, Hill V, McCrone JT, Colquhoun R, Yu X, Jackson B, Rambaut A, Williams TC, Templeton K |
| EPI_ISL_433229 | hCoV-19/Scotland/EDB726/2020 | Europe / United Kingdom / Scotland | 2020-04-02 | Virology Department, Royal Infirmary of Edinburgh, NHS Lothian / School of Biological Sciences, University of Edinburgh / Institute of Genetics and Molecular Medicine, University of Edinburgh | COVID-19 Genomics UK (COG-UK) Consortium | McHugh M, Dewar R, Rooke S, Gallagher M, Balcaza C, O'Toole A, Hill V, McCrone JT, Colquhoun R, Yu X, Jackson B, Rambaut A, Williams TC, Templeton K |
| EPI_ISL_433233 | hCoV-19/Scotland/EDB734/2020 | Europe / United Kingdom / Scotland | 2020-04-02 | Virology Department, Royal Infirmary of Edinburgh, NHS Lothian / School of Biological Sciences, University of Edinburgh / Institute of Genetics and Molecular Medicine, University of Edinburgh | COVID-19 Genomics UK (COG-UK) Consortium | McHugh M, Dewar R, Rooke S, Gallagher M, Balcaza C, O'Toole A, Hill V, McCrone JT, Colquhoun R, Yu X, Jackson B, Rambaut A, Williams TC, Templeton K |
| EPI_ISL_433234 | hCoV-19/Scotland/EDB735/2020 | Europe / United Kingdom / Scotland | 2020-04-02 | Virology Department, Royal Infirmary of Edinburgh, NHS Lothian / School of Biological Sciences, University of Edinburgh / Institute of Genetics and Molecular Medicine, University of Edinburgh | COVID-19 Genomics UK (COG-UK) Consortium | McHugh M, Dewar R, Rooke S, Gallagher M, Balcaza C, O'Toole A, Hill V, McCrone JT, Colquhoun R, Yu X, Jackson B, Rambaut A, Williams TC, Templeton K |
| EPI_ISL_433236 | hCoV-19/Scotland/EDB753/2020 | Europe / United Kingdom / Scotland | 2020-04-03 | Virology Department, Royal Infirmary of Edinburgh, NHS Lothian / School of Biological Sciences, University of Edinburgh / Institute of Genetics and Molecular Medicine, University of Edinburgh | COVID-19 Genomics UK (COG-UK) Consortium | McHugh M, Dewar R, Rooke S, Gallagher M, Balcaza C, O'Toole A, Hill V, McCrone JT, Colquhoun R, Yu X, Jackson B, Rambaut A, Williams TC, Templeton K |
| EPI_ISL_433237 | hCoV-19/Scotland/EDB764/2020 | Europe / United Kingdom / Scotland | 2020-04-02 | Virology Department, Royal Infirmary of Edinburgh, NHS Lothian / School of Biological Sciences, University of Edinburgh / Institute of Genetics and Molecular Medicine, University of Edinburgh | COVID-19 Genomics UK (COG-UK) Consortium | McHugh M, Dewar R, Rooke S, Gallagher M, Balcaza C, O'Toole A, Hill V, McCrone JT, Colquhoun R, Yu X, Jackson B, Rambaut A, Williams TC, Templeton K |
| EPI_ISL_433238 | hCoV-19/Scotland/EDB766/2020 | Europe / United Kingdom / Scotland | 2020-04-03 | Virology Department, Royal Infirmary of Edinburgh, NHS Lothian / School of Biological Sciences, University of Edinburgh / Institute of Genetics and Molecular Medicine, University of Edinburgh | COVID-19 Genomics UK (COG-UK) Consortium | McHugh M, Dewar R, Rooke S, Gallagher M, Balcaza C, O'Toole A, Hill V, McCrone JT, Colquhoun R, Yu X, Jackson B, Rambaut A, Williams TC, Templeton K |
| EPI_ISL_433240 | hCoV-19/Scotland/EDB773/2020 | Europe / United Kingdom / Scotland | 2020-04-03 | Virology Department, Royal Infirmary of Edinburgh, NHS Lothian / School of Biological Sciences, University of Edinburgh / Institute of Genetics and Molecular Medicine, University of Edinburgh | COVID-19 Genomics UK (COG-UK) Consortium | McHugh M, Dewar R, Rooke S, Gallagher M, Balcaza C, O'Toole A, Hill V, McCrone JT, Colquhoun R, Yu X, Jackson B, Rambaut A, Williams TC, Templeton K |
| EPI_ISL_433241 | hCoV-19/Scotland/EDB774/2020 | Europe / United Kingdom / Scotland | 2020-04-03 | Virology Department, Royal Infirmary of Edinburgh, NHS Lothian / School of Biological Sciences, University of Edinburgh / Institute of Genetics and Molecular Medicine, University of Edinburgh | COVID-19 Genomics UK (COG-UK) Consortium | McHugh M, Dewar R, Rooke S, Gallagher M, Balcaza C, O'Toole A, Hill V, McCrone JT, Colquhoun R, Yu X, Jackson B, Rambaut A, Williams TC, Templeton K |
| EPI_ISL_433242 | hCoV-19/Scotland/EDB781/2020 | Europe / United Kingdom / Scotland | 2020-04-03 | Virology Department, Royal Infirmary of Edinburgh, NHS Lothian / School of Biological Sciences, University of Edinburgh / Institute of Genetics and Molecular Medicine, University of Edinburgh | COVID-19 Genomics UK (COG-UK) Consortium | McHugh M, Dewar R, Rooke S, Gallagher M, Balcaza C, O'Toole A, Hill V, McCrone JT, Colquhoun R, Yu X, Jackson B, Rambaut A, Williams TC, Templeton K |
| EPI_ISL_433243 | hCoV-19/Scotland/EDB787/2020 | Europe / United Kingdom / Scotland | 2020-04-03 | Virology Department, Royal Infirmary of Edinburgh, NHS Lothian / School of Biological Sciences, University of Edinburgh / Institute of Genetics and Molecular Medicine, University of Edinburgh | COVID-19 Genomics UK (COG-UK) Consortium | McHugh M, Dewar R, Rooke S, Gallagher M, Balcaza C, O'Toole A, Hill V, McCrone JT, Colquhoun R, Yu X, Jackson B, Rambaut A, Williams TC, Templeton K |
| EPI_ISL_433244 | hCoV-19/Scotland/EDB788/2020 | Europe / United Kingdom / Scotland | 2020-04-04 | Virology Department, Royal Infirmary of Edinburgh, NHS Lothian / School of Biological Sciences, University of Edinburgh / Institute of Genetics and Molecular Medicine, University of Edinburgh | COVID-19 Genomics UK (COG-UK) Consortium | McHugh M, Dewar R, Rooke S, Gallagher M, Balcaza C, O'Toole A, Hill V, McCrone JT, Colquhoun R, Yu X, Jackson B, Rambaut A, Williams TC, Templeton K |
| EPI_ISL_433245 | hCoV-19/Scotland/EDB789/2020 | Europe / United Kingdom / Scotland | 2020-04-03 | Virology Department, Royal Infirmary of Edinburgh, NHS Lothian / School of Biological Sciences, University of Edinburgh / Institute of Genetics and Molecular Medicine, University of Edinburgh | COVID-19 Genomics UK (COG-UK) Consortium | McHugh M, Dewar R, Rooke S, Gallagher M, Balcaza C, O'Toole A, Hill V, McCrone JT, Colquhoun R, Yu X, Jackson B, Rambaut A, Williams TC, Templeton K |
| EPI_ISL_433246 | hCoV-19/Scotland/EDB795/2020 | Europe / United Kingdom / Scotland | 2020-04-03 | Virology Department, Royal Infirmary of Edinburgh, NHS Lothian / School of Biological Sciences, University of Edinburgh / Institute of Genetics and Molecular Medicine, University of Edinburgh | COVID-19 Genomics UK (COG-UK) Consortium | McHugh M, Dewar R, Rooke S, Gallagher M, Balcaza C, O'Toole A, Hill V, McCrone JT, Colquhoun R, Yu X, Jackson B, Rambaut A, Williams TC, Templeton K |
| EPI_ISL_433247 | hCoV-19/Scotland/EDB799/2020 | Europe / United Kingdom / Scotland | 2020-04-04 | Virology Department, Royal Infirmary of Edinburgh, NHS Lothian / School of Biological Sciences, University of Edinburgh / Institute of Genetics and Molecular Medicine, University of Edinburgh | COVID-19 Genomics UK (COG-UK) Consortium | McHugh M, Dewar R, Rooke S, Gallagher M, Balcaza C, O'Toole A, Hill V, McCrone JT, Colquhoun R, Yu X, Jackson B, Rambaut A, Williams TC, Templeton K |
| EPI_ISL_433248 | hCoV-19/Scotland/EDB801/2020 | Europe / United Kingdom / Scotland | 2020-04-04 | Virology Department, Royal Infirmary of Edinburgh, NHS Lothian / School of Biological Sciences, University of Edinburgh / Institute of Genetics and Molecular Medicine, University of Edinburgh | COVID-19 Genomics UK (COG-UK) Consortium | McHugh M, Dewar R, Rooke S, Gallagher M, Balcaza C, O'Toole A, Hill V, McCrone JT, Colquhoun R, Yu X, Jackson B, Rambaut A, Williams TC, Templeton K |
| EPI_ISL_433249 | hCoV-19/Scotland/EDB807/2020 | Europe / United Kingdom / Scotland | 2020-04-03 | Virology Department, Royal Infirmary of Edinburgh, NHS Lothian / School of Biological Sciences, University of Edinburgh / Institute of Genetics and Molecular Medicine, University of Edinburgh | COVID-19 Genomics UK (COG-UK) Consortium | McHugh M, Dewar R, Rooke S, Gallagher M, Balcaza C, O'Toole A, Hill V, McCrone JT, Colquhoun R, Yu X, Jackson B, Rambaut A, Williams TC, Templeton K |
| EPI_ISL_433250 | hCoV-19/Scotland/EDB821/2020 | Europe / United Kingdom / Scotland | 2020-04-04 | Virology Department, Royal Infirmary of Edinburgh, NHS Lothian / School of Biological Sciences, University of Edinburgh / Institute of Genetics and Molecular Medicine, University of Edinburgh | COVID-19 Genomics UK (COG-UK) Consortium | McHugh M, Dewar R, Rooke S, Gallagher M, Balcaza C, O'Toole A, Hill V, McCrone JT, Colquhoun R, Yu X, Jackson B, Rambaut A, Williams TC, Templeton K |
| EPI_ISL_433253 | hCoV-19/Scotland/EDB838/2020 | Europe / United Kingdom / Scotland | 2020-04-04 | Virology Department, Royal Infirmary of Edinburgh, NHS Lothian / School of Biological Sciences, University of Edinburgh / Institute of Genetics and Molecular Medicine, University of Edinburgh | COVID-19 Genomics UK (COG-UK) Consortium | McHugh M, Dewar R, Rooke S, Gallagher M, Balcaza C, O'Toole A, Hill V, McCrone JT, Colquhoun R, Yu X, Jackson B, Rambaut A, Williams TC, Templeton K |
| EPI_ISL_433254 | hCoV-19/Scotland/EDB842/2020 | Europe / United Kingdom / Scotland | 2020-04-04 | Virology Department, Royal Infirmary of Edinburgh, NHS Lothian / School of Biological Sciences, University of Edinburgh / Institute of Genetics and Molecular Medicine, University of Edinburgh | COVID-19 Genomics UK (COG-UK) Consortium | McHugh M, Dewar R, Rooke S, Gallagher M, Balcaza C, O'Toole A, Hill V, McCrone JT, Colquhoun R, Yu X, Jackson B, Rambaut A, Williams TC, Templeton K |
| EPI_ISL_433255 | hCoV-19/Scotland/EDB848/2020 | Europe / United Kingdom / Scotland | 2020-04-04 | Virology Department, Royal Infirmary of Edinburgh, NHS Lothian / School of Biological Sciences, University of Edinburgh / Institute of Genetics and Molecular Medicine, University of Edinburgh | COVID-19 Genomics UK (COG-UK) Consortium | McHugh M, Dewar R, Rooke S, Gallagher M, Balcaza C, O'Toole A, Hill V, McCrone JT, Colquhoun R, Yu X, Jackson B, Rambaut A, Williams TC, Templeton K |
| EPI_ISL_433256 | hCoV-19/Scotland/EDB852/2020 | Europe / United Kingdom / Scotland | 2020-04-04 | Virology Department, Royal Infirmary of Edinburgh, NHS Lothian / School of Biological Sciences, University of Edinburgh / Institute of Genetics and Molecular Medicine, University of Edinburgh | COVID-19 Genomics UK (COG-UK) Consortium | McHugh M, Dewar R, Rooke S, Gallagher M, Balcaza C, O'Toole A, Hill V, McCrone JT, Colquhoun R, Yu X, Jackson B, Rambaut A, Williams TC, Templeton K |
| EPI_ISL_433257 | hCoV-19/Scotland/EDB860/2020 | Europe / United Kingdom / Scotland | 2020-04-04 | Virology Department, Royal Infirmary of Edinburgh, NHS Lothian / School of Biological Sciences, University of Edinburgh / Institute of Genetics and Molecular Medicine, University of Edinburgh | COVID-19 Genomics UK (COG-UK) Consortium | McHugh M, Dewar R, Rooke S, Gallagher M, Balcaza C, O'Toole A, Hill V, McCrone JT, Colquhoun R, Yu X, Jackson B, Rambaut A, Williams TC, Templeton K |
| EPI_ISL_433260 | hCoV-19/Scotland/EDB868/2020 | Europe / United Kingdom / Scotland | 2020-04-04 | Virology Department, Royal Infirmary of Edinburgh, NHS Lothian / School of Biological Sciences, University of Edinburgh / Institute of Genetics and Molecular Medicine, University of Edinburgh | COVID-19 Genomics UK (COG-UK) Consortium | McHugh M, Dewar R, Rooke S, Gallagher M, Balcaza C, O'Toole A, Hill V, McCrone JT, Colquhoun R, Yu X, Jackson B, Rambaut A, Williams TC, Templeton K |
| EPI_ISL_433261 | hCoV-19/Scotland/EDB877/2020 | Europe / United Kingdom / Scotland | 2020-04-04 | Virology Department, Royal Infirmary of Edinburgh, NHS Lothian / School of Biological Sciences, University of Edinburgh / Institute of Genetics and Molecular Medicine, University of Edinburgh | COVID-19 Genomics UK (COG-UK) Consortium | McHugh M, Dewar R, Rooke S, Gallagher M, Balcaza C, O'Toole A, Hill V, McCrone JT, Colquhoun R, Yu X, Jackson B, Rambaut A, Williams TC, Templeton K |
| EPI_ISL_433262 | hCoV-19/Scotland/EDB880/2020 | Europe / United Kingdom / Scotland | 2020-04-04 | Virology Department, Royal Infirmary of Edinburgh, NHS Lothian / School of Biological Sciences, University of Edinburgh / Institute of Genetics and Molecular Medicine, University of Edinburgh | COVID-19 Genomics UK (COG-UK) Consortium | McHugh M, Dewar R, Rooke S, Gallagher M, Balcaza C, O'Toole A, Hill V, McCrone JT, Colquhoun R, Yu X, Jackson B, Rambaut A, Williams TC, Templeton K |
| EPI_ISL_433263 | hCoV-19/Scotland/EDB884/2020 | Europe / United Kingdom / Scotland | 2020-04-04 | Virology Department, Royal Infirmary of Edinburgh, NHS Lothian / School of Biological Sciences, University of Edinburgh / Institute of Genetics and Molecular Medicine, University of Edinburgh | COVID-19 Genomics UK (COG-UK) Consortium | McHugh M, Dewar R, Rooke S, Gallagher M, Balcaza C, O'Toole A, Hill V, McCrone JT, Colquhoun R, Yu X, Jackson B, Rambaut A, Williams TC, Templeton K |
| EPI_ISL_433265 | hCoV-19/Scotland/EDB886/2020 | Europe / United Kingdom / Scotland | 2020-04-04 | Virology Department, Royal Infirmary of Edinburgh, NHS Lothian / School of Biological Sciences, University of Edinburgh / Institute of Genetics and Molecular Medicine, University of Edinburgh | COVID-19 Genomics UK (COG-UK) Consortium | McHugh M, Dewar R, Rooke S, Gallagher M, Balcaza C, O'Toole A, Hill V, McCrone JT, Colquhoun R, Yu X, Jackson B, Rambaut A, Williams TC, Templeton K |
| EPI_ISL_433266 | hCoV-19/Scotland/EDB888/2020 | Europe / United Kingdom / Scotland | 2020-04-04 | Virology Department, Royal Infirmary of Edinburgh, NHS Lothian / School of Biological Sciences, University of Edinburgh / Institute of Genetics and Molecular Medicine, University of Edinburgh | COVID-19 Genomics UK (COG-UK) Consortium | McHugh M, Dewar R, Rooke S, Gallagher M, Balcaza C, O'Toole A, Hill V, McCrone JT, Colquhoun R, Yu X, Jackson B, Rambaut A, Williams TC, Templeton K |
| EPI_ISL_433267 | hCoV-19/Scotland/EDB889/2020 | Europe / United Kingdom / Scotland | 2020-04-04 | Virology Department, Royal Infirmary of Edinburgh, NHS Lothian / School of Biological Sciences, University of Edinburgh / Institute of Genetics and Molecular Medicine, University of Edinburgh | COVID-19 Genomics UK (COG-UK) Consortium | McHugh M, Dewar R, Rooke S, Gallagher M, Balcaza C, O'Toole A, Hill V, McCrone JT, Colquhoun R, Yu X, Jackson B, Rambaut A, Williams TC, Templeton K |
| EPI_ISL_433270 | hCoV-19/Scotland/CVR1978/2020 | Europe / United Kingdom / Scotland | 2020-04-07 | West of Scotland Specialist Virology Centre, NHSGGC / MRC-University of Glasgow Centre for Virus Research | COVID-19 Genomics UK (COG-UK) Consortium | Ana da Silva Filipe, Natasha Johnson, Kathy Smollett, Daniel Mair, Stephen Carmichael, Lily Tong, Jenna Nichols, Elihu Aranday-Cortes, Kirstyn Brunker, Yasmin Parr, Kyriaki Nomikou; Sarah McDonald, Marc Niebel, Patawee Asamaphan; Richard Orton, Joseph Hughes, Sreenu Vattipally, David L Robertson; Alasdair MacLean, Rory Gunson; Kathy Li, Natasha Jesudason, Rajiv Shah, James Shepherd, Antonia Ho, Emma Thomson |
| EPI_ISL_433277 | hCoV-19/Scotland/CVR261/2020 | Europe / United Kingdom / Scotland | 2020-03-20 | West of Scotland Specialist Virology Centre, NHSGGC / MRC-University of Glasgow Centre for Virus Research | COVID-19 Genomics UK (COG-UK) Consortium | Ana da Silva Filipe, Natasha Johnson, Kathy Smollett, Daniel Mair, Stephen Carmichael, Lily Tong, Jenna Nichols, Elihu Aranday-Cortes, Kirstyn Brunker, Yasmin Parr, Kyriaki Nomikou; Sarah McDonald, Marc Niebel, Patawee Asamaphan; Richard Orton, Joseph Hughes, Sreenu Vattipally, David L Robertson; Alasdair MacLean, Rory Gunson; Kathy Li, Natasha Jesudason, Rajiv Shah, James Shepherd, Antonia Ho, Emma Thomson |
| EPI_ISL_433278 | hCoV-19/Scotland/CVR273/2020 | Europe / United Kingdom / Scotland | 2020-03-20 | West of Scotland Specialist Virology Centre, NHSGGC / MRC-University of Glasgow Centre for Virus Research | COVID-19 Genomics UK (COG-UK) Consortium | Ana da Silva Filipe, Natasha Johnson, Kathy Smollett, Daniel Mair, Stephen Carmichael, Lily Tong, Jenna Nichols, Elihu Aranday-Cortes, Kirstyn Brunker, Yasmin Parr, Kyriaki Nomikou; Sarah McDonald, Marc Niebel, Patawee Asamaphan; Richard Orton, Joseph Hughes, Sreenu Vattipally, David L Robertson; Alasdair MacLean, Rory Gunson; Kathy Li, Natasha Jesudason, Rajiv Shah, James Shepherd, Antonia Ho, Emma Thomson |
| EPI_ISL_433279 | hCoV-19/Scotland/CVR278/2020 | Europe / United Kingdom / Scotland | 2020-03-20 | West of Scotland Specialist Virology Centre, NHSGGC / MRC-University of Glasgow Centre for Virus Research | COVID-19 Genomics UK (COG-UK) Consortium | Ana da Silva Filipe, Natasha Johnson, Kathy Smollett, Daniel Mair, Stephen Carmichael, Lily Tong, Jenna Nichols, Elihu Aranday-Cortes, Kirstyn Brunker, Yasmin Parr, Kyriaki Nomikou; Sarah McDonald, Marc Niebel, Patawee Asamaphan; Richard Orton, Joseph Hughes, Sreenu Vattipally, David L Robertson; Alasdair MacLean, Rory Gunson; Kathy Li, Natasha Jesudason, Rajiv Shah, James Shepherd, Antonia Ho, Emma Thomson |
| EPI_ISL_433280 | hCoV-19/Scotland/CVR282/2020 | Europe / United Kingdom / Scotland | 2020-03-20 | West of Scotland Specialist Virology Centre, NHSGGC / MRC-University of Glasgow Centre for Virus Research | COVID-19 Genomics UK (COG-UK) Consortium | Ana da Silva Filipe, Natasha Johnson, Kathy Smollett, Daniel Mair, Stephen Carmichael, Lily Tong, Jenna Nichols, Elihu Aranday-Cortes, Kirstyn Brunker, Yasmin Parr, Kyriaki Nomikou; Sarah McDonald, Marc Niebel, Patawee Asamaphan; Richard Orton, Joseph Hughes, Sreenu Vattipally, David L Robertson; Alasdair MacLean, Rory Gunson; Kathy Li, Natasha Jesudason, Rajiv Shah, James Shepherd, Antonia Ho, Emma Thomson |
| EPI_ISL_433281 | hCoV-19/Scotland/CVR284/2020 | Europe / United Kingdom / Scotland | 2020-03-21 | West of Scotland Specialist Virology Centre, NHSGGC / MRC-University of Glasgow Centre for Virus Research | COVID-19 Genomics UK (COG-UK) Consortium | Ana da Silva Filipe, Natasha Johnson, Kathy Smollett, Daniel Mair, Stephen Carmichael, Lily Tong, Jenna Nichols, Elihu Aranday-Cortes, Kirstyn Brunker, Yasmin Parr, Kyriaki Nomikou; Sarah McDonald, Marc Niebel, Patawee Asamaphan; Richard Orton, Joseph Hughes, Sreenu Vattipally, David L Robertson; Alasdair MacLean, Rory Gunson; Kathy Li, Natasha Jesudason, Rajiv Shah, James Shepherd, Antonia Ho, Emma Thomson |
| EPI_ISL_433282 | hCoV-19/Scotland/CVR304/2020 | Europe / United Kingdom / Scotland | 2020-03-21 | West of Scotland Specialist Virology Centre, NHSGGC / MRC-University of Glasgow Centre for Virus Research | COVID-19 Genomics UK (COG-UK) Consortium | Ana da Silva Filipe, Natasha Johnson, Kathy Smollett, Daniel Mair, Stephen Carmichael, Lily Tong, Jenna Nichols, Elihu Aranday-Cortes, Kirstyn Brunker, Yasmin Parr, Kyriaki Nomikou; Sarah McDonald, Marc Niebel, Patawee Asamaphan; Richard Orton, Joseph Hughes, Sreenu Vattipally, David L Robertson; Alasdair MacLean, Rory Gunson; Kathy Li, Natasha Jesudason, Rajiv Shah, James Shepherd, Antonia Ho, Emma Thomson |
| EPI_ISL_433283 | hCoV-19/Scotland/CVR309/2020 | Europe / United Kingdom / Scotland | 2020-03-20 | West of Scotland Specialist Virology Centre, NHSGGC / MRC-University of Glasgow Centre for Virus Research | COVID-19 Genomics UK (COG-UK) Consortium | Ana da Silva Filipe, Natasha Johnson, Kathy Smollett, Daniel Mair, Stephen Carmichael, Lily Tong, Jenna Nichols, Elihu Aranday-Cortes, Kirstyn Brunker, Yasmin Parr, Kyriaki Nomikou; Sarah McDonald, Marc Niebel, Patawee Asamaphan; Richard Orton, Joseph Hughes, Sreenu Vattipally, David L Robertson; Alasdair MacLean, Rory Gunson; Kathy Li, Natasha Jesudason, Rajiv Shah, James Shepherd, Antonia Ho, Emma Thomson |
| EPI_ISL_433284 | hCoV-19/Scotland/CVR338/2020 | Europe / United Kingdom / Scotland | 2020-03-21 | West of Scotland Specialist Virology Centre, NHSGGC / MRC-University of Glasgow Centre for Virus Research | COVID-19 Genomics UK (COG-UK) Consortium | Ana da Silva Filipe, Natasha Johnson, Kathy Smollett, Daniel Mair, Stephen Carmichael, Lily Tong, Jenna Nichols, Elihu Aranday-Cortes, Kirstyn Brunker, Yasmin Parr, Kyriaki Nomikou; Sarah McDonald, Marc Niebel, Patawee Asamaphan; Richard Orton, Joseph Hughes, Sreenu Vattipally, David L Robertson; Alasdair MacLean, Rory Gunson; Kathy Li, Natasha Jesudason, Rajiv Shah, James Shepherd, Antonia Ho, Emma Thomson |
| EPI_ISL_433286 | hCoV-19/Scotland/CVR367/2020 | Europe / United Kingdom / Scotland | 2020-03-22 | West of Scotland Specialist Virology Centre, NHSGGC / MRC-University of Glasgow Centre for Virus Research | COVID-19 Genomics UK (COG-UK) Consortium | Ana da Silva Filipe, Natasha Johnson, Kathy Smollett, Daniel Mair, Stephen Carmichael, Lily Tong, Jenna Nichols, Elihu Aranday-Cortes, Kirstyn Brunker, Yasmin Parr, Kyriaki Nomikou; Sarah McDonald, Marc Niebel, Patawee Asamaphan; Richard Orton, Joseph Hughes, Sreenu Vattipally, David L Robertson; Alasdair MacLean, Rory Gunson; Kathy Li, Natasha Jesudason, Rajiv Shah, James Shepherd, Antonia Ho, Emma Thomson |
| EPI_ISL_433287 | hCoV-19/Scotland/CVR375/2020 | Europe / United Kingdom / Scotland | 2020-03-21 | West of Scotland Specialist Virology Centre, NHSGGC / MRC-University of Glasgow Centre for Virus Research | COVID-19 Genomics UK (COG-UK) Consortium | Ana da Silva Filipe, Natasha Johnson, Kathy Smollett, Daniel Mair, Stephen Carmichael, Lily Tong, Jenna Nichols, Elihu Aranday-Cortes, Kirstyn Brunker, Yasmin Parr, Kyriaki Nomikou; Sarah McDonald, Marc Niebel, Patawee Asamaphan; Richard Orton, Joseph Hughes, Sreenu Vattipally, David L Robertson; Alasdair MacLean, Rory Gunson; Kathy Li, Natasha Jesudason, Rajiv Shah, James Shepherd, Antonia Ho, Emma Thomson |
| EPI_ISL_433288 | hCoV-19/Scotland/CVR392/2020 | Europe / United Kingdom / Scotland | 2020-03-23 | West of Scotland Specialist Virology Centre, NHSGGC / MRC-University of Glasgow Centre for Virus Research | COVID-19 Genomics UK (COG-UK) Consortium | Ana da Silva Filipe, Natasha Johnson, Kathy Smollett, Daniel Mair, Stephen Carmichael, Lily Tong, Jenna Nichols, Elihu Aranday-Cortes, Kirstyn Brunker, Yasmin Parr, Kyriaki Nomikou; Sarah McDonald, Marc Niebel, Patawee Asamaphan; Richard Orton, Joseph Hughes, Sreenu Vattipally, David L Robertson; Alasdair MacLean, Rory Gunson; Kathy Li, Natasha Jesudason, Rajiv Shah, James Shepherd, Antonia Ho, Emma Thomson |
| EPI_ISL_433289 | hCoV-19/Scotland/CVR397/2020 | Europe / United Kingdom / Scotland | 2020-03-23 | West of Scotland Specialist Virology Centre, NHSGGC / MRC-University of Glasgow Centre for Virus Research | COVID-19 Genomics UK (COG-UK) Consortium | Ana da Silva Filipe, Natasha Johnson, Kathy Smollett, Daniel Mair, Stephen Carmichael, Lily Tong, Jenna Nichols, Elihu Aranday-Cortes, Kirstyn Brunker, Yasmin Parr, Kyriaki Nomikou; Sarah McDonald, Marc Niebel, Patawee Asamaphan; Richard Orton, Joseph Hughes, Sreenu Vattipally, David L Robertson; Alasdair MacLean, Rory Gunson; Kathy Li, Natasha Jesudason, Rajiv Shah, James Shepherd, Antonia Ho, Emma Thomson |
| EPI_ISL_433291 | hCoV-19/Scotland/CVR425/2020 | Europe / United Kingdom / Scotland | 2020-03-22 | West of Scotland Specialist Virology Centre, NHSGGC / MRC-University of Glasgow Centre for Virus Research | COVID-19 Genomics UK (COG-UK) Consortium | Ana da Silva Filipe, Natasha Johnson, Kathy Smollett, Daniel Mair, Stephen Carmichael, Lily Tong, Jenna Nichols, Elihu Aranday-Cortes, Kirstyn Brunker, Yasmin Parr, Kyriaki Nomikou; Sarah McDonald, Marc Niebel, Patawee Asamaphan; Richard Orton, Joseph Hughes, Sreenu Vattipally, David L Robertson; Alasdair MacLean, Rory Gunson; Kathy Li, Natasha Jesudason, Rajiv Shah, James Shepherd, Antonia Ho, Emma Thomson |
| EPI_ISL_433295 | hCoV-19/Scotland/CVR450/2020 | Europe / United Kingdom / Scotland | 2020-03-23 | West of Scotland Specialist Virology Centre, NHSGGC / MRC-University of Glasgow Centre for Virus Research | COVID-19 Genomics UK (COG-UK) Consortium | Ana da Silva Filipe, Natasha Johnson, Kathy Smollett, Daniel Mair, Stephen Carmichael, Lily Tong, Jenna Nichols, Elihu Aranday-Cortes, Kirstyn Brunker, Yasmin Parr, Kyriaki Nomikou; Sarah McDonald, Marc Niebel, Patawee Asamaphan; Richard Orton, Joseph Hughes, Sreenu Vattipally, David L Robertson; Alasdair MacLean, Rory Gunson; Kathy Li, Natasha Jesudason, Rajiv Shah, James Shepherd, Antonia Ho, Emma Thomson |
| EPI_ISL_433298 | hCoV-19/Scotland/CVR458/2020 | Europe / United Kingdom / Scotland | 2020-03-23 | West of Scotland Specialist Virology Centre, NHSGGC / MRC-University of Glasgow Centre for Virus Research | COVID-19 Genomics UK (COG-UK) Consortium | Ana da Silva Filipe, Natasha Johnson, Kathy Smollett, Daniel Mair, Stephen Carmichael, Lily Tong, Jenna Nichols, Elihu Aranday-Cortes, Kirstyn Brunker, Yasmin Parr, Kyriaki Nomikou; Sarah McDonald, Marc Niebel, Patawee Asamaphan; Richard Orton, Joseph Hughes, Sreenu Vattipally, David L Robertson; Alasdair MacLean, Rory Gunson; Kathy Li, Natasha Jesudason, Rajiv Shah, James Shepherd, Antonia Ho, Emma Thomson |
| EPI_ISL_433800 | hCoV-19/England/CAMB-789AE/2020 | Europe / United Kingdom / England | 2020-04-06 | Department of Pathology, University of Cambridge | COVID-19 Genomics UK (COG-UK) Consortium | Luke W Meredith, M. Estee Torok , Myra Hosmillo, William L. Hamilton, Martin D. Curran, Theresa Feltwell, Grant Hall, Anna Yakovleva, Fahad A Khokhar, Charlotte J. Houldcroft, Laura G Caller, Aminu S. Jahun, Sarah L. Caddy, Ian Goodfellow |
| EPI_ISL_433801 | hCoV-19/England/CAMB-789BD/2020 | Europe / United Kingdom / England | 2020-04-06 | Department of Pathology, University of Cambridge | COVID-19 Genomics UK (COG-UK) Consortium | Luke W Meredith, M. Estee Torok , Myra Hosmillo, William L. Hamilton, Martin D. Curran, Theresa Feltwell, Grant Hall, Anna Yakovleva, Fahad A Khokhar, Charlotte J. Houldcroft, Laura G Caller, Aminu S. Jahun, Sarah L. Caddy, Ian Goodfellow |
| EPI_ISL_433803 | hCoV-19/England/CAMB-789DB/2020 | Europe / United Kingdom / England | 2020-04-07 | Department of Pathology, University of Cambridge | COVID-19 Genomics UK (COG-UK) Consortium | Luke W Meredith, M. Estee Torok , Myra Hosmillo, William L. Hamilton, Martin D. Curran, Theresa Feltwell, Grant Hall, Anna Yakovleva, Fahad A Khokhar, Charlotte J. Houldcroft, Laura G Caller, Aminu S. Jahun, Sarah L. Caddy, Ian Goodfellow |
| EPI_ISL_433805 | hCoV-19/England/CAMB-789F9/2020 | Europe / United Kingdom / England | 2020-04-07 | Department of Pathology, University of Cambridge | COVID-19 Genomics UK (COG-UK) Consortium | Luke W Meredith, M. Estee Torok , Myra Hosmillo, William L. Hamilton, Martin D. Curran, Theresa Feltwell, Grant Hall, Anna Yakovleva, Fahad A Khokhar, Charlotte J. Houldcroft, Laura G Caller, Aminu S. Jahun, Sarah L. Caddy, Ian Goodfellow |
| EPI_ISL_433806 | hCoV-19/England/CAMB-78A05/2020 | Europe / United Kingdom / England | 2020-04-06 | Department of Pathology, University of Cambridge | COVID-19 Genomics UK (COG-UK) Consortium | Luke W Meredith, M. Estee Torok , Myra Hosmillo, William L. Hamilton, Martin D. Curran, Theresa Feltwell, Grant Hall, Anna Yakovleva, Fahad A Khokhar, Charlotte J. Houldcroft, Laura G Caller, Aminu S. Jahun, Sarah L. Caddy, Ian Goodfellow |
| EPI_ISL_433808 | hCoV-19/England/CAMB-78A32/2020 | Europe / United Kingdom / England | 2020-04-06 | Department of Pathology, University of Cambridge | COVID-19 Genomics UK (COG-UK) Consortium | Luke W Meredith, M. Estee Torok , Myra Hosmillo, William L. Hamilton, Martin D. Curran, Theresa Feltwell, Grant Hall, Anna Yakovleva, Fahad A Khokhar, Charlotte J. Houldcroft, Laura G Caller, Aminu S. Jahun, Sarah L. Caddy, Ian Goodfellow |
| EPI_ISL_433809 | hCoV-19/England/CAMB-78A50/2020 | Europe / United Kingdom / England | 2020-04-07 | Department of Pathology, University of Cambridge | COVID-19 Genomics UK (COG-UK) Consortium | Luke W Meredith, M. Estee Torok , Myra Hosmillo, William L. Hamilton, Martin D. Curran, Theresa Feltwell, Grant Hall, Anna Yakovleva, Fahad A Khokhar, Charlotte J. Houldcroft, Laura G Caller, Aminu S. Jahun, Sarah L. Caddy, Ian Goodfellow |
| EPI_ISL_433810 | hCoV-19/England/CAMB-78A7E/2020 | Europe / United Kingdom / England | 2020-04-06 | Department of Pathology, University of Cambridge | COVID-19 Genomics UK (COG-UK) Consortium | Luke W Meredith, M. Estee Torok , Myra Hosmillo, William L. Hamilton, Martin D. Curran, Theresa Feltwell, Grant Hall, Anna Yakovleva, Fahad A Khokhar, Charlotte J. Houldcroft, Laura G Caller, Aminu S. Jahun, Sarah L. Caddy, Ian Goodfellow |
| EPI_ISL_433811 | hCoV-19/England/CAMB-78A9C/2020 | Europe / United Kingdom / England | 2020-04-06 | Department of Pathology, University of Cambridge | COVID-19 Genomics UK (COG-UK) Consortium | Luke W Meredith, M. Estee Torok , Myra Hosmillo, William L. Hamilton, Martin D. Curran, Theresa Feltwell, Grant Hall, Anna Yakovleva, Fahad A Khokhar, Charlotte J. Houldcroft, Laura G Caller, Aminu S. Jahun, Sarah L. Caddy, Ian Goodfellow |
| EPI_ISL_433812 | hCoV-19/England/CAMB-78ABA/2020 | Europe / United Kingdom / England | 2020-04-07 | Department of Pathology, University of Cambridge | COVID-19 Genomics UK (COG-UK) Consortium | Luke W Meredith, M. Estee Torok , Myra Hosmillo, William L. Hamilton, Martin D. Curran, Theresa Feltwell, Grant Hall, Anna Yakovleva, Fahad A Khokhar, Charlotte J. Houldcroft, Laura G Caller, Aminu S. Jahun, Sarah L. Caddy, Ian Goodfellow |
| EPI_ISL_433813 | hCoV-19/England/CAMB-79A04/2020 | Europe / United Kingdom / England | 2020-03-26 | Department of Pathology, University of Cambridge | COVID-19 Genomics UK (COG-UK) Consortium | Luke W Meredith, M. Estee Torok , Myra Hosmillo, William L. Hamilton, Martin D. Curran, Theresa Feltwell, Grant Hall, Anna Yakovleva, Fahad A Khokhar, Charlotte J. Houldcroft, Laura G Caller, Aminu S. Jahun, Sarah L. Caddy, Ian Goodfellow |
| EPI_ISL_433814 | hCoV-19/England/CAMB-79A22/2020 | Europe / United Kingdom / England | 2020-03-26 | Department of Pathology, University of Cambridge | COVID-19 Genomics UK (COG-UK) Consortium | Luke W Meredith, M. Estee Torok , Myra Hosmillo, William L. Hamilton, Martin D. Curran, Theresa Feltwell, Grant Hall, Anna Yakovleva, Fahad A Khokhar, Charlotte J. Houldcroft, Laura G Caller, Aminu S. Jahun, Sarah L. Caddy, Ian Goodfellow |
| EPI_ISL_433815 | hCoV-19/England/CAMB-79A40/2020 | Europe / United Kingdom / England | 2020-03-26 | Department of Pathology, University of Cambridge | COVID-19 Genomics UK (COG-UK) Consortium | Luke W Meredith, M. Estee Torok , Myra Hosmillo, William L. Hamilton, Martin D. Curran, Theresa Feltwell, Grant Hall, Anna Yakovleva, Fahad A Khokhar, Charlotte J. Houldcroft, Laura G Caller, Aminu S. Jahun, Sarah L. Caddy, Ian Goodfellow |
| EPI_ISL_433816 | hCoV-19/England/CAMB-79A5F/2020 | Europe / United Kingdom / England | 2020-03-27 | Department of Pathology, University of Cambridge | COVID-19 Genomics UK (COG-UK) Consortium | Luke W Meredith, M. Estee Torok , Myra Hosmillo, William L. Hamilton, Martin D. Curran, Theresa Feltwell, Grant Hall, Anna Yakovleva, Fahad A Khokhar, Charlotte J. Houldcroft, Laura G Caller, Aminu S. Jahun, Sarah L. Caddy, Ian Goodfellow |
| EPI_ISL_433817 | hCoV-19/England/CAMB-79A8C/2020 | Europe / United Kingdom / England | 2020-03-26 | Department of Pathology, University of Cambridge | COVID-19 Genomics UK (COG-UK) Consortium | Luke W Meredith, M. Estee Torok , Myra Hosmillo, William L. Hamilton, Martin D. Curran, Theresa Feltwell, Grant Hall, Anna Yakovleva, Fahad A Khokhar, Charlotte J. Houldcroft, Laura G Caller, Aminu S. Jahun, Sarah L. Caddy, Ian Goodfellow |
| EPI_ISL_433818 | hCoV-19/England/CAMB-79A9B/2020 | Europe / United Kingdom / England | 2020-03-27 | Department of Pathology, University of Cambridge | COVID-19 Genomics UK (COG-UK) Consortium | Luke W Meredith, M. Estee Torok , Myra Hosmillo, William L. Hamilton, Martin D. Curran, Theresa Feltwell, Grant Hall, Anna Yakovleva, Fahad A Khokhar, Charlotte J. Houldcroft, Laura G Caller, Aminu S. Jahun, Sarah L. Caddy, Ian Goodfellow |
| EPI_ISL_433819 | hCoV-19/England/CAMB-79AB9/2020 | Europe / United Kingdom / England | 2020-03-27 | Department of Pathology, University of Cambridge | COVID-19 Genomics UK (COG-UK) Consortium | Luke W Meredith, M. Estee Torok , Myra Hosmillo, William L. Hamilton, Martin D. Curran, Theresa Feltwell, Grant Hall, Anna Yakovleva, Fahad A Khokhar, Charlotte J. Houldcroft, Laura G Caller, Aminu S. Jahun, Sarah L. Caddy, Ian Goodfellow |
| EPI_ISL_433820 | hCoV-19/England/CAMB-79AC8/2020 | Europe / United Kingdom / England | 2020-03-26 | Department of Pathology, University of Cambridge | COVID-19 Genomics UK (COG-UK) Consortium | Luke W Meredith, M. Estee Torok , Myra Hosmillo, William L. Hamilton, Martin D. Curran, Theresa Feltwell, Grant Hall, Anna Yakovleva, Fahad A Khokhar, Charlotte J. Houldcroft, Laura G Caller, Aminu S. Jahun, Sarah L. Caddy, Ian Goodfellow |
| EPI_ISL_433821 | hCoV-19/England/CAMB-79AD7/2020 | Europe / United Kingdom / England | 2020-03-27 | Department of Pathology, University of Cambridge | COVID-19 Genomics UK (COG-UK) Consortium | Luke W Meredith, M. Estee Torok , Myra Hosmillo, William L. Hamilton, Martin D. Curran, Theresa Feltwell, Grant Hall, Anna Yakovleva, Fahad A Khokhar, Charlotte J. Houldcroft, Laura G Caller, Aminu S. Jahun, Sarah L. Caddy, Ian Goodfellow |
| EPI_ISL_433823 | hCoV-19/England/CAMB-79AF5/2020 | Europe / United Kingdom / England | 2020-03-27 | Department of Pathology, University of Cambridge | COVID-19 Genomics UK (COG-UK) Consortium | Luke W Meredith, M. Estee Torok , Myra Hosmillo, William L. Hamilton, Martin D. Curran, Theresa Feltwell, Grant Hall, Anna Yakovleva, Fahad A Khokhar, Charlotte J. Houldcroft, Laura G Caller, Aminu S. Jahun, Sarah L. Caddy, Ian Goodfellow |
| EPI_ISL_433824 | hCoV-19/England/CAMB-79B01/2020 | Europe / United Kingdom / England | 2020-03-26 | Department of Pathology, University of Cambridge | COVID-19 Genomics UK (COG-UK) Consortium | Luke W Meredith, M. Estee Torok , Myra Hosmillo, William L. Hamilton, Martin D. Curran, Theresa Feltwell, Grant Hall, Anna Yakovleva, Fahad A Khokhar, Charlotte J. Houldcroft, Laura G Caller, Aminu S. Jahun, Sarah L. Caddy, Ian Goodfellow |
| EPI_ISL_433826 | hCoV-19/England/CAMB-7A8BE/2020 | Europe / United Kingdom / England | 2020-04-08 | Department of Pathology, University of Cambridge | COVID-19 Genomics UK (COG-UK) Consortium | Luke W Meredith, M. Estee Torok , Myra Hosmillo, William L. Hamilton, Martin D. Curran, Theresa Feltwell, Grant Hall, Anna Yakovleva, Fahad A Khokhar, Charlotte J. Houldcroft, Laura G Caller, Aminu S. Jahun, Sarah L. Caddy, Ian Goodfellow |
| EPI_ISL_433827 | hCoV-19/England/CAMB-7A8DC/2020 | Europe / United Kingdom / England | 2020-04-08 | Department of Pathology, University of Cambridge | COVID-19 Genomics UK (COG-UK) Consortium | Luke W Meredith, M. Estee Torok , Myra Hosmillo, William L. Hamilton, Martin D. Curran, Theresa Feltwell, Grant Hall, Anna Yakovleva, Fahad A Khokhar, Charlotte J. Houldcroft, Laura G Caller, Aminu S. Jahun, Sarah L. Caddy, Ian Goodfellow |
| EPI_ISL_433828 | hCoV-19/England/CAMB-7A8FA/2020 | Europe / United Kingdom / England | 2020-04-07 | Department of Pathology, University of Cambridge | COVID-19 Genomics UK (COG-UK) Consortium | Luke W Meredith, M. Estee Torok , Myra Hosmillo, William L. Hamilton, Martin D. Curran, Theresa Feltwell, Grant Hall, Anna Yakovleva, Fahad A Khokhar, Charlotte J. Houldcroft, Laura G Caller, Aminu S. Jahun, Sarah L. Caddy, Ian Goodfellow |
| EPI_ISL_433829 | hCoV-19/England/CAMB-7AA30/2020 | Europe / United Kingdom / England | 2020-04-08 | Department of Pathology, University of Cambridge | COVID-19 Genomics UK (COG-UK) Consortium | Luke W Meredith, M. Estee Torok , Myra Hosmillo, William L. Hamilton, Martin D. Curran, Theresa Feltwell, Grant Hall, Anna Yakovleva, Fahad A Khokhar, Charlotte J. Houldcroft, Laura G Caller, Aminu S. Jahun, Sarah L. Caddy, Ian Goodfellow |
| EPI_ISL_433833 | hCoV-19/England/CAMB-7AA8B/2020 | Europe / United Kingdom / England | 2020-04-08 | Department of Pathology, University of Cambridge | COVID-19 Genomics UK (COG-UK) Consortium | Luke W Meredith, M. Estee Torok , Myra Hosmillo, William L. Hamilton, Martin D. Curran, Theresa Feltwell, Grant Hall, Anna Yakovleva, Fahad A Khokhar, Charlotte J. Houldcroft, Laura G Caller, Aminu S. Jahun, Sarah L. Caddy, Ian Goodfellow |
| EPI_ISL_433834 | hCoV-19/England/CAMB-7AAA9/2020 | Europe / United Kingdom / England | 2020-04-07 | Department of Pathology, University of Cambridge | COVID-19 Genomics UK (COG-UK) Consortium | Luke W Meredith, M. Estee Torok , Myra Hosmillo, William L. Hamilton, Martin D. Curran, Theresa Feltwell, Grant Hall, Anna Yakovleva, Fahad A Khokhar, Charlotte J. Houldcroft, Laura G Caller, Aminu S. Jahun, Sarah L. Caddy, Ian Goodfellow |
| EPI_ISL_433835 | hCoV-19/England/CAMB-7AAB8/2020 | Europe / United Kingdom / England | 2020-04-07 | Department of Pathology, University of Cambridge | COVID-19 Genomics UK (COG-UK) Consortium | Luke W Meredith, M. Estee Torok , Myra Hosmillo, William L. Hamilton, Martin D. Curran, Theresa Feltwell, Grant Hall, Anna Yakovleva, Fahad A Khokhar, Charlotte J. Houldcroft, Laura G Caller, Aminu S. Jahun, Sarah L. Caddy, Ian Goodfellow |
| EPI_ISL_433837 | hCoV-19/England/CAMB-7AAD6/2020 | Europe / United Kingdom / England | 2020-04-07 | Department of Pathology, University of Cambridge | COVID-19 Genomics UK (COG-UK) Consortium | Luke W Meredith, M. Estee Torok , Myra Hosmillo, William L. Hamilton, Martin D. Curran, Theresa Feltwell, Grant Hall, Anna Yakovleva, Fahad A Khokhar, Charlotte J. Houldcroft, Laura G Caller, Aminu S. Jahun, Sarah L. Caddy, Ian Goodfellow |
| EPI_ISL_433838 | hCoV-19/England/CAMB-7AAE5/2020 | Europe / United Kingdom / England | 2020 | Department of Pathology, University of Cambridge | COVID-19 Genomics UK (COG-UK) Consortium | Luke W Meredith, M. Estee Torok , Myra Hosmillo, William L. Hamilton, Martin D. Curran, Theresa Feltwell, Grant Hall, Anna Yakovleva, Fahad A Khokhar, Charlotte J. Houldcroft, Laura G Caller, Aminu S. Jahun, Sarah L. Caddy, Ian Goodfellow |
| EPI_ISL_433839 | hCoV-19/England/CAMB-7AAF4/2020 | Europe / United Kingdom / England | 2020-04-07 | Department of Pathology, University of Cambridge | COVID-19 Genomics UK (COG-UK) Consortium | Luke W Meredith, M. Estee Torok , Myra Hosmillo, William L. Hamilton, Martin D. Curran, Theresa Feltwell, Grant Hall, Anna Yakovleva, Fahad A Khokhar, Charlotte J. Houldcroft, Laura G Caller, Aminu S. Jahun, Sarah L. Caddy, Ian Goodfellow |
| EPI_ISL_433840 | hCoV-19/England/CAMB-7AB00/2020 | Europe / United Kingdom / England | 2020-04-07 | Department of Pathology, University of Cambridge | COVID-19 Genomics UK (COG-UK) Consortium | Luke W Meredith, M. Estee Torok , Myra Hosmillo, William L. Hamilton, Martin D. Curran, Theresa Feltwell, Grant Hall, Anna Yakovleva, Fahad A Khokhar, Charlotte J. Houldcroft, Laura G Caller, Aminu S. Jahun, Sarah L. Caddy, Ian Goodfellow |
| EPI_ISL_433842 | hCoV-19/England/CAMB-7AB2E/2020 | Europe / United Kingdom / England | 2020-04-05 | Department of Pathology, University of Cambridge | COVID-19 Genomics UK (COG-UK) Consortium | Luke W Meredith, M. Estee Torok , Myra Hosmillo, William L. Hamilton, Martin D. Curran, Theresa Feltwell, Grant Hall, Anna Yakovleva, Fahad A Khokhar, Charlotte J. Houldcroft, Laura G Caller, Aminu S. Jahun, Sarah L. Caddy, Ian Goodfellow |
| EPI_ISL_433843 | hCoV-19/England/CAMB-7AB3D/2020 | Europe / United Kingdom / England | 2020-04-08 | Department of Pathology, University of Cambridge | COVID-19 Genomics UK (COG-UK) Consortium | Luke W Meredith, M. Estee Torok , Myra Hosmillo, William L. Hamilton, Martin D. Curran, Theresa Feltwell, Grant Hall, Anna Yakovleva, Fahad A Khokhar, Charlotte J. Houldcroft, Laura G Caller, Aminu S. Jahun, Sarah L. Caddy, Ian Goodfellow |
| EPI_ISL_433844 | hCoV-19/England/CAMB-7AB79/2020 | Europe / United Kingdom / England | 2020-04-07 | Department of Pathology, University of Cambridge | COVID-19 Genomics UK (COG-UK) Consortium | Luke W Meredith, M. Estee Torok , Myra Hosmillo, William L. Hamilton, Martin D. Curran, Theresa Feltwell, Grant Hall, Anna Yakovleva, Fahad A Khokhar, Charlotte J. Houldcroft, Laura G Caller, Aminu S. Jahun, Sarah L. Caddy, Ian Goodfellow |
| EPI_ISL_433846 | hCoV-19/England/CAMB-7AB97/2020 | Europe / United Kingdom / England | 2020-04-07 | Department of Pathology, University of Cambridge | COVID-19 Genomics UK (COG-UK) Consortium | Luke W Meredith, M. Estee Torok , Myra Hosmillo, William L. Hamilton, Martin D. Curran, Theresa Feltwell, Grant Hall, Anna Yakovleva, Fahad A Khokhar, Charlotte J. Houldcroft, Laura G Caller, Aminu S. Jahun, Sarah L. Caddy, Ian Goodfellow |
| EPI_ISL_433847 | hCoV-19/England/CAMB-7ABA6/2020 | Europe / United Kingdom / England | 2020-04-07 | Department of Pathology, University of Cambridge | COVID-19 Genomics UK (COG-UK) Consortium | Luke W Meredith, M. Estee Torok , Myra Hosmillo, William L. Hamilton, Martin D. Curran, Theresa Feltwell, Grant Hall, Anna Yakovleva, Fahad A Khokhar, Charlotte J. Houldcroft, Laura G Caller, Aminu S. Jahun, Sarah L. Caddy, Ian Goodfellow |
| EPI_ISL_433848 | hCoV-19/England/CAMB-7ABB5/2020 | Europe / United Kingdom / England | 2020-04-06 | Department of Pathology, University of Cambridge | COVID-19 Genomics UK (COG-UK) Consortium | Luke W Meredith, M. Estee Torok , Myra Hosmillo, William L. Hamilton, Martin D. Curran, Theresa Feltwell, Grant Hall, Anna Yakovleva, Fahad A Khokhar, Charlotte J. Houldcroft, Laura G Caller, Aminu S. Jahun, Sarah L. Caddy, Ian Goodfellow |
| EPI_ISL_433849 | hCoV-19/England/CAMB-7ABC4/2020 | Europe / United Kingdom / England | 2020-04-07 | Department of Pathology, University of Cambridge | COVID-19 Genomics UK (COG-UK) Consortium | Luke W Meredith, M. Estee Torok , Myra Hosmillo, William L. Hamilton, Martin D. Curran, Theresa Feltwell, Grant Hall, Anna Yakovleva, Fahad A Khokhar, Charlotte J. Houldcroft, Laura G Caller, Aminu S. Jahun, Sarah L. Caddy, Ian Goodfellow |
| EPI_ISL_433851 | hCoV-19/England/CAMB-7ABF1/2020 | Europe / United Kingdom / England | 2020-04-09 | Department of Pathology, University of Cambridge | COVID-19 Genomics UK (COG-UK) Consortium | Luke W Meredith, M. Estee Torok , Myra Hosmillo, William L. Hamilton, Martin D. Curran, Theresa Feltwell, Grant Hall, Anna Yakovleva, Fahad A Khokhar, Charlotte J. Houldcroft, Laura G Caller, Aminu S. Jahun, Sarah L. Caddy, Ian Goodfellow |
| EPI_ISL_433852 | hCoV-19/England/CAMB-7AC0D/2020 | Europe / United Kingdom / England | 2020-04-08 | Department of Pathology, University of Cambridge | COVID-19 Genomics UK (COG-UK) Consortium | Luke W Meredith, M. Estee Torok , Myra Hosmillo, William L. Hamilton, Martin D. Curran, Theresa Feltwell, Grant Hall, Anna Yakovleva, Fahad A Khokhar, Charlotte J. Houldcroft, Laura G Caller, Aminu S. Jahun, Sarah L. Caddy, Ian Goodfellow |
| EPI_ISL_433854 | hCoV-19/England/CAMB-7AC2B/2020 | Europe / United Kingdom / England | 2020-04-08 | Department of Pathology, University of Cambridge | COVID-19 Genomics UK (COG-UK) Consortium | Luke W Meredith, M. Estee Torok , Myra Hosmillo, William L. Hamilton, Martin D. Curran, Theresa Feltwell, Grant Hall, Anna Yakovleva, Fahad A Khokhar, Charlotte J. Houldcroft, Laura G Caller, Aminu S. Jahun, Sarah L. Caddy, Ian Goodfellow |
| EPI_ISL_433855 | hCoV-19/England/CAMB-7AC3A/2020 | Europe / United Kingdom / England | 2020-04-08 | Department of Pathology, University of Cambridge | COVID-19 Genomics UK (COG-UK) Consortium | Luke W Meredith, M. Estee Torok , Myra Hosmillo, William L. Hamilton, Martin D. Curran, Theresa Feltwell, Grant Hall, Anna Yakovleva, Fahad A Khokhar, Charlotte J. Houldcroft, Laura G Caller, Aminu S. Jahun, Sarah L. Caddy, Ian Goodfellow |
| EPI_ISL_433856 | hCoV-19/England/CAMB-7AC49/2020 | Europe / United Kingdom / England | 2020-04-08 | Department of Pathology, University of Cambridge | COVID-19 Genomics UK (COG-UK) Consortium | Luke W Meredith, M. Estee Torok , Myra Hosmillo, William L. Hamilton, Martin D. Curran, Theresa Feltwell, Grant Hall, Anna Yakovleva, Fahad A Khokhar, Charlotte J. Houldcroft, Laura G Caller, Aminu S. Jahun, Sarah L. Caddy, Ian Goodfellow |
| EPI_ISL_433857 | hCoV-19/England/CAMB-7AC58/2020 | Europe / United Kingdom / England | 2020-04-08 | Department of Pathology, University of Cambridge | COVID-19 Genomics UK (COG-UK) Consortium | Luke W Meredith, M. Estee Torok , Myra Hosmillo, William L. Hamilton, Martin D. Curran, Theresa Feltwell, Grant Hall, Anna Yakovleva, Fahad A Khokhar, Charlotte J. Houldcroft, Laura G Caller, Aminu S. Jahun, Sarah L. Caddy, Ian Goodfellow |
| EPI_ISL_433858 | hCoV-19/England/CAMB-7AC67/2020 | Europe / United Kingdom / England | 2020-04-08 | Department of Pathology, University of Cambridge | COVID-19 Genomics UK (COG-UK) Consortium | Luke W Meredith, M. Estee Torok , Myra Hosmillo, William L. Hamilton, Martin D. Curran, Theresa Feltwell, Grant Hall, Anna Yakovleva, Fahad A Khokhar, Charlotte J. Houldcroft, Laura G Caller, Aminu S. Jahun, Sarah L. Caddy, Ian Goodfellow |
| EPI_ISL_433859 | hCoV-19/England/CAMB-7AC76/2020 | Europe / United Kingdom / England | 2020-04-08 | Department of Pathology, University of Cambridge | COVID-19 Genomics UK (COG-UK) Consortium | Luke W Meredith, M. Estee Torok , Myra Hosmillo, William L. Hamilton, Martin D. Curran, Theresa Feltwell, Grant Hall, Anna Yakovleva, Fahad A Khokhar, Charlotte J. Houldcroft, Laura G Caller, Aminu S. Jahun, Sarah L. Caddy, Ian Goodfellow |
| EPI_ISL_433860 | hCoV-19/England/CAMB-7AC85/2020 | Europe / United Kingdom / England | 2020-04-08 | Department of Pathology, University of Cambridge | COVID-19 Genomics UK (COG-UK) Consortium | Luke W Meredith, M. Estee Torok , Myra Hosmillo, William L. Hamilton, Martin D. Curran, Theresa Feltwell, Grant Hall, Anna Yakovleva, Fahad A Khokhar, Charlotte J. Houldcroft, Laura G Caller, Aminu S. Jahun, Sarah L. Caddy, Ian Goodfellow |
| EPI_ISL_433861 | hCoV-19/England/CAMB-7AC94/2020 | Europe / United Kingdom / England | 2020-04-07 | Department of Pathology, University of Cambridge | COVID-19 Genomics UK (COG-UK) Consortium | Luke W Meredith, M. Estee Torok , Myra Hosmillo, William L. Hamilton, Martin D. Curran, Theresa Feltwell, Grant Hall, Anna Yakovleva, Fahad A Khokhar, Charlotte J. Houldcroft, Laura G Caller, Aminu S. Jahun, Sarah L. Caddy, Ian Goodfellow |
| EPI_ISL_433862 | hCoV-19/England/CAMB-7ACA3/2020 | Europe / United Kingdom / England | 2020-04-06 | Department of Pathology, University of Cambridge | COVID-19 Genomics UK (COG-UK) Consortium | Luke W Meredith, M. Estee Torok , Myra Hosmillo, William L. Hamilton, Martin D. Curran, Theresa Feltwell, Grant Hall, Anna Yakovleva, Fahad A Khokhar, Charlotte J. Houldcroft, Laura G Caller, Aminu S. Jahun, Sarah L. Caddy, Ian Goodfellow |
| EPI_ISL_433863 | hCoV-19/England/CAMB-7ACB2/2020 | Europe / United Kingdom / England | 2020-04-09 | Department of Pathology, University of Cambridge | COVID-19 Genomics UK (COG-UK) Consortium | Luke W Meredith, M. Estee Torok , Myra Hosmillo, William L. Hamilton, Martin D. Curran, Theresa Feltwell, Grant Hall, Anna Yakovleva, Fahad A Khokhar, Charlotte J. Houldcroft, Laura G Caller, Aminu S. Jahun, Sarah L. Caddy, Ian Goodfellow |
| EPI_ISL_433864 | hCoV-19/England/CAMB-7ACC1/2020 | Europe / United Kingdom / England | 2020-04-09 | Department of Pathology, University of Cambridge | COVID-19 Genomics UK (COG-UK) Consortium | Luke W Meredith, M. Estee Torok , Myra Hosmillo, William L. Hamilton, Martin D. Curran, Theresa Feltwell, Grant Hall, Anna Yakovleva, Fahad A Khokhar, Charlotte J. Houldcroft, Laura G Caller, Aminu S. Jahun, Sarah L. Caddy, Ian Goodfellow |
| EPI_ISL_433865 | hCoV-19/England/CAMB-7ACD0/2020 | Europe / United Kingdom / England | 2020-04-08 | Department of Pathology, University of Cambridge | COVID-19 Genomics UK (COG-UK) Consortium | Luke W Meredith, M. Estee Torok , Myra Hosmillo, William L. Hamilton, Martin D. Curran, Theresa Feltwell, Grant Hall, Anna Yakovleva, Fahad A Khokhar, Charlotte J. Houldcroft, Laura G Caller, Aminu S. Jahun, Sarah L. Caddy, Ian Goodfellow |
| EPI_ISL_433866 | hCoV-19/England/CAMB-7ACFE/2020 | Europe / United Kingdom / England | 2020-04-08 | Department of Pathology, University of Cambridge | COVID-19 Genomics UK (COG-UK) Consortium | Luke W Meredith, M. Estee Torok , Myra Hosmillo, William L. Hamilton, Martin D. Curran, Theresa Feltwell, Grant Hall, Anna Yakovleva, Fahad A Khokhar, Charlotte J. Houldcroft, Laura G Caller, Aminu S. Jahun, Sarah L. Caddy, Ian Goodfellow |
| EPI_ISL_433868 | hCoV-19/England/CAMB-7AD28/2020 | Europe / United Kingdom / England | 2020-04-08 | Department of Pathology, University of Cambridge | COVID-19 Genomics UK (COG-UK) Consortium | Luke W Meredith, M. Estee Torok , Myra Hosmillo, William L. Hamilton, Martin D. Curran, Theresa Feltwell, Grant Hall, Anna Yakovleva, Fahad A Khokhar, Charlotte J. Houldcroft, Laura G Caller, Aminu S. Jahun, Sarah L. Caddy, Ian Goodfellow |
| EPI_ISL_433870 | hCoV-19/England/CAMB-7AD46/2020 | Europe / United Kingdom / England | 2020-04-07 | Department of Pathology, University of Cambridge | COVID-19 Genomics UK (COG-UK) Consortium | Luke W Meredith, M. Estee Torok , Myra Hosmillo, William L. Hamilton, Martin D. Curran, Theresa Feltwell, Grant Hall, Anna Yakovleva, Fahad A Khokhar, Charlotte J. Houldcroft, Laura G Caller, Aminu S. Jahun, Sarah L. Caddy, Ian Goodfellow |
| EPI_ISL_433872 | hCoV-19/England/CAMB-7AD64/2020 | Europe / United Kingdom / England | 2020-04-08 | Department of Pathology, University of Cambridge | COVID-19 Genomics UK (COG-UK) Consortium | Luke W Meredith, M. Estee Torok , Myra Hosmillo, William L. Hamilton, Martin D. Curran, Theresa Feltwell, Grant Hall, Anna Yakovleva, Fahad A Khokhar, Charlotte J. Houldcroft, Laura G Caller, Aminu S. Jahun, Sarah L. Caddy, Ian Goodfellow |
| EPI_ISL_433875 | hCoV-19/England/CAMB-7ADA0/2020 | Europe / United Kingdom / England | 2020-04-08 | Department of Pathology, University of Cambridge | COVID-19 Genomics UK (COG-UK) Consortium | Luke W Meredith, M. Estee Torok , Myra Hosmillo, William L. Hamilton, Martin D. Curran, Theresa Feltwell, Grant Hall, Anna Yakovleva, Fahad A Khokhar, Charlotte J. Houldcroft, Laura G Caller, Aminu S. Jahun, Sarah L. Caddy, Ian Goodfellow |
| EPI_ISL_433876 | hCoV-19/England/CAMB-7ADBF/2020 | Europe / United Kingdom / England | 2020-04-08 | Department of Pathology, University of Cambridge | COVID-19 Genomics UK (COG-UK) Consortium | Luke W Meredith, M. Estee Torok , Myra Hosmillo, William L. Hamilton, Martin D. Curran, Theresa Feltwell, Grant Hall, Anna Yakovleva, Fahad A Khokhar, Charlotte J. Houldcroft, Laura G Caller, Aminu S. Jahun, Sarah L. Caddy, Ian Goodfellow |
| EPI_ISL_433877 | hCoV-19/England/CAMB-7ADDD/2020 | Europe / United Kingdom / England | 2020-04-09 | Department of Pathology, University of Cambridge | COVID-19 Genomics UK (COG-UK) Consortium | Luke W Meredith, M. Estee Torok , Myra Hosmillo, William L. Hamilton, Martin D. Curran, Theresa Feltwell, Grant Hall, Anna Yakovleva, Fahad A Khokhar, Charlotte J. Houldcroft, Laura G Caller, Aminu S. Jahun, Sarah L. Caddy, Ian Goodfellow |
| EPI_ISL_433878 | hCoV-19/England/CAMB-7ADEC/2020 | Europe / United Kingdom / England | 2020-04-09 | Department of Pathology, University of Cambridge | COVID-19 Genomics UK (COG-UK) Consortium | Luke W Meredith, M. Estee Torok , Myra Hosmillo, William L. Hamilton, Martin D. Curran, Theresa Feltwell, Grant Hall, Anna Yakovleva, Fahad A Khokhar, Charlotte J. Houldcroft, Laura G Caller, Aminu S. Jahun, Sarah L. Caddy, Ian Goodfellow |
| EPI_ISL_433880 | hCoV-19/England/CAMB-7AE07/2020 | Europe / United Kingdom / England | 2020-04-09 | Department of Pathology, University of Cambridge | COVID-19 Genomics UK (COG-UK) Consortium | Luke W Meredith, M. Estee Torok , Myra Hosmillo, William L. Hamilton, Martin D. Curran, Theresa Feltwell, Grant Hall, Anna Yakovleva, Fahad A Khokhar, Charlotte J. Houldcroft, Laura G Caller, Aminu S. Jahun, Sarah L. Caddy, Ian Goodfellow |
| EPI_ISL_433882 | hCoV-19/England/CAMB-7AE34/2020 | Europe / United Kingdom / England | 2020-04-08 | Department of Pathology, University of Cambridge | COVID-19 Genomics UK (COG-UK) Consortium | Luke W Meredith, M. Estee Torok , Myra Hosmillo, William L. Hamilton, Martin D. Curran, Theresa Feltwell, Grant Hall, Anna Yakovleva, Fahad A Khokhar, Charlotte J. Houldcroft, Laura G Caller, Aminu S. Jahun, Sarah L. Caddy, Ian Goodfellow |
| EPI_ISL_433883 | hCoV-19/England/CAMB-7AE43/2020 | Europe / United Kingdom / England | 2020-04-09 | Department of Pathology, University of Cambridge | COVID-19 Genomics UK (COG-UK) Consortium | Luke W Meredith, M. Estee Torok , Myra Hosmillo, William L. Hamilton, Martin D. Curran, Theresa Feltwell, Grant Hall, Anna Yakovleva, Fahad A Khokhar, Charlotte J. Houldcroft, Laura G Caller, Aminu S. Jahun, Sarah L. Caddy, Ian Goodfellow |
| EPI_ISL_433884 | hCoV-19/England/CAMB-7AE52/2020 | Europe / United Kingdom / England | 2020-04-09 | Department of Pathology, University of Cambridge | COVID-19 Genomics UK (COG-UK) Consortium | Luke W Meredith, M. Estee Torok , Myra Hosmillo, William L. Hamilton, Martin D. Curran, Theresa Feltwell, Grant Hall, Anna Yakovleva, Fahad A Khokhar, Charlotte J. Houldcroft, Laura G Caller, Aminu S. Jahun, Sarah L. Caddy, Ian Goodfellow |
| EPI_ISL_433885 | hCoV-19/England/CAMB-7AE61/2020 | Europe / United Kingdom / England | 2020-04-09 | Department of Pathology, University of Cambridge | COVID-19 Genomics UK (COG-UK) Consortium | Luke W Meredith, M. Estee Torok , Myra Hosmillo, William L. Hamilton, Martin D. Curran, Theresa Feltwell, Grant Hall, Anna Yakovleva, Fahad A Khokhar, Charlotte J. Houldcroft, Laura G Caller, Aminu S. Jahun, Sarah L. Caddy, Ian Goodfellow |
| EPI_ISL_433886 | hCoV-19/England/CAMB-7AE70/2020 | Europe / United Kingdom / England | 2020-04-09 | Department of Pathology, University of Cambridge | COVID-19 Genomics UK (COG-UK) Consortium | Luke W Meredith, M. Estee Torok , Myra Hosmillo, William L. Hamilton, Martin D. Curran, Theresa Feltwell, Grant Hall, Anna Yakovleva, Fahad A Khokhar, Charlotte J. Houldcroft, Laura G Caller, Aminu S. Jahun, Sarah L. Caddy, Ian Goodfellow |
| EPI_ISL_433888 | hCoV-19/England/CAMB-7AEAD/2020 | Europe / United Kingdom / England | 2020-04-09 | Department of Pathology, University of Cambridge | COVID-19 Genomics UK (COG-UK) Consortium | Luke W Meredith, M. Estee Torok , Myra Hosmillo, William L. Hamilton, Martin D. Curran, Theresa Feltwell, Grant Hall, Anna Yakovleva, Fahad A Khokhar, Charlotte J. Houldcroft, Laura G Caller, Aminu S. Jahun, Sarah L. Caddy, Ian Goodfellow |
| EPI_ISL_433890 | hCoV-19/England/CAMB-7AEDA/2020 | Europe / United Kingdom / England | 2020-04-09 | Department of Pathology, University of Cambridge | COVID-19 Genomics UK (COG-UK) Consortium | Luke W Meredith, M. Estee Torok , Myra Hosmillo, William L. Hamilton, Martin D. Curran, Theresa Feltwell, Grant Hall, Anna Yakovleva, Fahad A Khokhar, Charlotte J. Houldcroft, Laura G Caller, Aminu S. Jahun, Sarah L. Caddy, Ian Goodfellow |
| EPI_ISL_433891 | hCoV-19/England/CAMB-7AEE9/2020 | Europe / United Kingdom / England | 2020-04-09 | Department of Pathology, University of Cambridge | COVID-19 Genomics UK (COG-UK) Consortium | Luke W Meredith, M. Estee Torok , Myra Hosmillo, William L. Hamilton, Martin D. Curran, Theresa Feltwell, Grant Hall, Anna Yakovleva, Fahad A Khokhar, Charlotte J. Houldcroft, Laura G Caller, Aminu S. Jahun, Sarah L. Caddy, Ian Goodfellow |
| EPI_ISL_433892 | hCoV-19/England/CAMB-7AEF8/2020 | Europe / United Kingdom / England | 2020-04-09 | Department of Pathology, University of Cambridge | COVID-19 Genomics UK (COG-UK) Consortium | Luke W Meredith, M. Estee Torok , Myra Hosmillo, William L. Hamilton, Martin D. Curran, Theresa Feltwell, Grant Hall, Anna Yakovleva, Fahad A Khokhar, Charlotte J. Houldcroft, Laura G Caller, Aminu S. Jahun, Sarah L. Caddy, Ian Goodfellow |
| EPI_ISL_433893 | hCoV-19/England/CAMB-7AF04/2020 | Europe / United Kingdom / England | 2020-04-10 | Department of Pathology, University of Cambridge | COVID-19 Genomics UK (COG-UK) Consortium | Luke W Meredith, M. Estee Torok , Myra Hosmillo, William L. Hamilton, Martin D. Curran, Theresa Feltwell, Grant Hall, Anna Yakovleva, Fahad A Khokhar, Charlotte J. Houldcroft, Laura G Caller, Aminu S. Jahun, Sarah L. Caddy, Ian Goodfellow |
| EPI_ISL_433894 | hCoV-19/England/CAMB-7AF13/2020 | Europe / United Kingdom / England | 2020-04-10 | Department of Pathology, University of Cambridge | COVID-19 Genomics UK (COG-UK) Consortium | Luke W Meredith, M. Estee Torok , Myra Hosmillo, William L. Hamilton, Martin D. Curran, Theresa Feltwell, Grant Hall, Anna Yakovleva, Fahad A Khokhar, Charlotte J. Houldcroft, Laura G Caller, Aminu S. Jahun, Sarah L. Caddy, Ian Goodfellow |
| EPI_ISL_433895 | hCoV-19/England/CAMB-7AF22/2020 | Europe / United Kingdom / England | 2020-04-10 | Department of Pathology, University of Cambridge | COVID-19 Genomics UK (COG-UK) Consortium | Luke W Meredith, M. Estee Torok , Myra Hosmillo, William L. Hamilton, Martin D. Curran, Theresa Feltwell, Grant Hall, Anna Yakovleva, Fahad A Khokhar, Charlotte J. Houldcroft, Laura G Caller, Aminu S. Jahun, Sarah L. Caddy, Ian Goodfellow |
| EPI_ISL_433896 | hCoV-19/England/CAMB-7AF31/2020 | Europe / United Kingdom / England | 2020-04-10 | Department of Pathology, University of Cambridge | COVID-19 Genomics UK (COG-UK) Consortium | Luke W Meredith, M. Estee Torok , Myra Hosmillo, William L. Hamilton, Martin D. Curran, Theresa Feltwell, Grant Hall, Anna Yakovleva, Fahad A Khokhar, Charlotte J. Houldcroft, Laura G Caller, Aminu S. Jahun, Sarah L. Caddy, Ian Goodfellow |
| EPI_ISL_433897 | hCoV-19/England/CAMB-7AF40/2020 | Europe / United Kingdom / England | 2020-04-10 | Department of Pathology, University of Cambridge | COVID-19 Genomics UK (COG-UK) Consortium | Luke W Meredith, M. Estee Torok , Myra Hosmillo, William L. Hamilton, Martin D. Curran, Theresa Feltwell, Grant Hall, Anna Yakovleva, Fahad A Khokhar, Charlotte J. Houldcroft, Laura G Caller, Aminu S. Jahun, Sarah L. Caddy, Ian Goodfellow |
| EPI_ISL_433898 | hCoV-19/England/CAMB-7AF5F/2020 | Europe / United Kingdom / England | 2020-04-10 | Department of Pathology, University of Cambridge | COVID-19 Genomics UK (COG-UK) Consortium | Luke W Meredith, M. Estee Torok , Myra Hosmillo, William L. Hamilton, Martin D. Curran, Theresa Feltwell, Grant Hall, Anna Yakovleva, Fahad A Khokhar, Charlotte J. Houldcroft, Laura G Caller, Aminu S. Jahun, Sarah L. Caddy, Ian Goodfellow |
| EPI_ISL_433899 | hCoV-19/England/CAMB-7AF6E/2020 | Europe / United Kingdom / England | 2020-04-10 | Department of Pathology, University of Cambridge | COVID-19 Genomics UK (COG-UK) Consortium | Luke W Meredith, M. Estee Torok , Myra Hosmillo, William L. Hamilton, Martin D. Curran, Theresa Feltwell, Grant Hall, Anna Yakovleva, Fahad A Khokhar, Charlotte J. Houldcroft, Laura G Caller, Aminu S. Jahun, Sarah L. Caddy, Ian Goodfellow |
| EPI_ISL_433900 | hCoV-19/England/CAMB-7AF7D/2020 | Europe / United Kingdom / England | 2020-04-10 | Department of Pathology, University of Cambridge | COVID-19 Genomics UK (COG-UK) Consortium | Luke W Meredith, M. Estee Torok , Myra Hosmillo, William L. Hamilton, Martin D. Curran, Theresa Feltwell, Grant Hall, Anna Yakovleva, Fahad A Khokhar, Charlotte J. Houldcroft, Laura G Caller, Aminu S. Jahun, Sarah L. Caddy, Ian Goodfellow |
| EPI_ISL_433902 | hCoV-19/England/CAMB-7AFAA/2020 | Europe / United Kingdom / England | 2020-04-09 | Department of Pathology, University of Cambridge | COVID-19 Genomics UK (COG-UK) Consortium | Luke W Meredith, M. Estee Torok , Myra Hosmillo, William L. Hamilton, Martin D. Curran, Theresa Feltwell, Grant Hall, Anna Yakovleva, Fahad A Khokhar, Charlotte J. Houldcroft, Laura G Caller, Aminu S. Jahun, Sarah L. Caddy, Ian Goodfellow |
| EPI_ISL_433904 | hCoV-19/England/CAMB-7AFF5/2020 | Europe / United Kingdom / England | 2020-04-10 | Department of Pathology, University of Cambridge | COVID-19 Genomics UK (COG-UK) Consortium | Luke W Meredith, M. Estee Torok , Myra Hosmillo, William L. Hamilton, Martin D. Curran, Theresa Feltwell, Grant Hall, Anna Yakovleva, Fahad A Khokhar, Charlotte J. Houldcroft, Laura G Caller, Aminu S. Jahun, Sarah L. Caddy, Ian Goodfellow |
| EPI_ISL_433905 | hCoV-19/England/CAMB-7B000/2020 | Europe / United Kingdom / England | 2020-04-10 | Department of Pathology, University of Cambridge | COVID-19 Genomics UK (COG-UK) Consortium | Luke W Meredith, M. Estee Torok , Myra Hosmillo, William L. Hamilton, Martin D. Curran, Theresa Feltwell, Grant Hall, Anna Yakovleva, Fahad A Khokhar, Charlotte J. Houldcroft, Laura G Caller, Aminu S. Jahun, Sarah L. Caddy, Ian Goodfellow |
| EPI_ISL_433906 | hCoV-19/England/CAMB-7B01F/2020 | Europe / United Kingdom / England | 2020-04-11 | Department of Pathology, University of Cambridge | COVID-19 Genomics UK (COG-UK) Consortium | Luke W Meredith, M. Estee Torok , Myra Hosmillo, William L. Hamilton, Martin D. Curran, Theresa Feltwell, Grant Hall, Anna Yakovleva, Fahad A Khokhar, Charlotte J. Houldcroft, Laura G Caller, Aminu S. Jahun, Sarah L. Caddy, Ian Goodfellow |
| EPI_ISL_433907 | hCoV-19/England/CAMB-7B02E/2020 | Europe / United Kingdom / England | 2020-04-11 | Department of Pathology, University of Cambridge | COVID-19 Genomics UK (COG-UK) Consortium | Luke W Meredith, M. Estee Torok , Myra Hosmillo, William L. Hamilton, Martin D. Curran, Theresa Feltwell, Grant Hall, Anna Yakovleva, Fahad A Khokhar, Charlotte J. Houldcroft, Laura G Caller, Aminu S. Jahun, Sarah L. Caddy, Ian Goodfellow |
| EPI_ISL_433909 | hCoV-19/England/CAMB-7B06A/2020 | Europe / United Kingdom / England | 2020-04-11 | Department of Pathology, University of Cambridge | COVID-19 Genomics UK (COG-UK) Consortium | Luke W Meredith, M. Estee Torok , Myra Hosmillo, William L. Hamilton, Martin D. Curran, Theresa Feltwell, Grant Hall, Anna Yakovleva, Fahad A Khokhar, Charlotte J. Houldcroft, Laura G Caller, Aminu S. Jahun, Sarah L. Caddy, Ian Goodfellow |
| EPI_ISL_433910 | hCoV-19/England/CAMB-7B079/2020 | Europe / United Kingdom / England | 2020-04-11 | Department of Pathology, University of Cambridge | COVID-19 Genomics UK (COG-UK) Consortium | Luke W Meredith, M. Estee Torok , Myra Hosmillo, William L. Hamilton, Martin D. Curran, Theresa Feltwell, Grant Hall, Anna Yakovleva, Fahad A Khokhar, Charlotte J. Houldcroft, Laura G Caller, Aminu S. Jahun, Sarah L. Caddy, Ian Goodfellow |
| EPI_ISL_433911 | hCoV-19/England/CAMB-7B088/2020 | Europe / United Kingdom / England | 2020-04-11 | Department of Pathology, University of Cambridge | COVID-19 Genomics UK (COG-UK) Consortium | Luke W Meredith, M. Estee Torok , Myra Hosmillo, William L. Hamilton, Martin D. Curran, Theresa Feltwell, Grant Hall, Anna Yakovleva, Fahad A Khokhar, Charlotte J. Houldcroft, Laura G Caller, Aminu S. Jahun, Sarah L. Caddy, Ian Goodfellow |
| EPI_ISL_433913 | hCoV-19/England/CAMB-7B0A6/2020 | Europe / United Kingdom / England | 2020-04-11 | Department of Pathology, University of Cambridge | COVID-19 Genomics UK (COG-UK) Consortium | Luke W Meredith, M. Estee Torok , Myra Hosmillo, William L. Hamilton, Martin D. Curran, Theresa Feltwell, Grant Hall, Anna Yakovleva, Fahad A Khokhar, Charlotte J. Houldcroft, Laura G Caller, Aminu S. Jahun, Sarah L. Caddy, Ian Goodfellow |
| EPI_ISL_433914 | hCoV-19/England/CAMB-7B0B5/2020 | Europe / United Kingdom / England | 2020-04-11 | Department of Pathology, University of Cambridge | COVID-19 Genomics UK (COG-UK) Consortium | Luke W Meredith, M. Estee Torok , Myra Hosmillo, William L. Hamilton, Martin D. Curran, Theresa Feltwell, Grant Hall, Anna Yakovleva, Fahad A Khokhar, Charlotte J. Houldcroft, Laura G Caller, Aminu S. Jahun, Sarah L. Caddy, Ian Goodfellow |
| EPI_ISL_433915 | hCoV-19/England/CAMB-7B0C4/2020 | Europe / United Kingdom / England | 2020-04-11 | Department of Pathology, University of Cambridge | COVID-19 Genomics UK (COG-UK) Consortium | Luke W Meredith, M. Estee Torok , Myra Hosmillo, William L. Hamilton, Martin D. Curran, Theresa Feltwell, Grant Hall, Anna Yakovleva, Fahad A Khokhar, Charlotte J. Houldcroft, Laura G Caller, Aminu S. Jahun, Sarah L. Caddy, Ian Goodfellow |
| EPI_ISL_433918 | hCoV-19/England/CAMB-7B0F1/2020 | Europe / United Kingdom / England | 2020-04-10 | Department of Pathology, University of Cambridge | COVID-19 Genomics UK (COG-UK) Consortium | Luke W Meredith, M. Estee Torok , Myra Hosmillo, William L. Hamilton, Martin D. Curran, Theresa Feltwell, Grant Hall, Anna Yakovleva, Fahad A Khokhar, Charlotte J. Houldcroft, Laura G Caller, Aminu S. Jahun, Sarah L. Caddy, Ian Goodfellow |
| EPI_ISL_433919 | hCoV-19/England/CAMB-7B10D/2020 | Europe / United Kingdom / England | 2020-04-11 | Department of Pathology, University of Cambridge | COVID-19 Genomics UK (COG-UK) Consortium | Luke W Meredith, M. Estee Torok , Myra Hosmillo, William L. Hamilton, Martin D. Curran, Theresa Feltwell, Grant Hall, Anna Yakovleva, Fahad A Khokhar, Charlotte J. Houldcroft, Laura G Caller, Aminu S. Jahun, Sarah L. Caddy, Ian Goodfellow |
| EPI_ISL_433963 | hCoV-19/England/CAMB-7B4AA/2020 | Europe / United Kingdom / England | 2020-04-12 | Department of Pathology, University of Cambridge | COVID-19 Genomics UK (COG-UK) Consortium | Luke W Meredith, M. Estee Torok , Myra Hosmillo, William L. Hamilton, Martin D. Curran, Theresa Feltwell, Grant Hall, Anna Yakovleva, Fahad A Khokhar, Charlotte J. Houldcroft, Laura G Caller, Aminu S. Jahun, Sarah L. Caddy, Ian Goodfellow |
| EPI_ISL_433965 | hCoV-19/England/CAMB-7B4C8/2020 | Europe / United Kingdom / England | 2020-04-12 | Department of Pathology, University of Cambridge | COVID-19 Genomics UK (COG-UK) Consortium | Luke W Meredith, M. Estee Torok , Myra Hosmillo, William L. Hamilton, Martin D. Curran, Theresa Feltwell, Grant Hall, Anna Yakovleva, Fahad A Khokhar, Charlotte J. Houldcroft, Laura G Caller, Aminu S. Jahun, Sarah L. Caddy, Ian Goodfellow |
| EPI_ISL_433966 | hCoV-19/England/CAMB-7B4D7/2020 | Europe / United Kingdom / England | 2020-04-12 | Department of Pathology, University of Cambridge | COVID-19 Genomics UK (COG-UK) Consortium | Luke W Meredith, M. Estee Torok , Myra Hosmillo, William L. Hamilton, Martin D. Curran, Theresa Feltwell, Grant Hall, Anna Yakovleva, Fahad A Khokhar, Charlotte J. Houldcroft, Laura G Caller, Aminu S. Jahun, Sarah L. Caddy, Ian Goodfellow |
| EPI_ISL_433967 | hCoV-19/England/CAMB-7B4E6/2020 | Europe / United Kingdom / England | 2020-04-12 | Department of Pathology, University of Cambridge | COVID-19 Genomics UK (COG-UK) Consortium | Luke W Meredith, M. Estee Torok , Myra Hosmillo, William L. Hamilton, Martin D. Curran, Theresa Feltwell, Grant Hall, Anna Yakovleva, Fahad A Khokhar, Charlotte J. Houldcroft, Laura G Caller, Aminu S. Jahun, Sarah L. Caddy, Ian Goodfellow |
| EPI_ISL_433968 | hCoV-19/England/CAMB-7B4F5/2020 | Europe / United Kingdom / England | 2020-04-12 | Department of Pathology, University of Cambridge | COVID-19 Genomics UK (COG-UK) Consortium | Luke W Meredith, M. Estee Torok , Myra Hosmillo, William L. Hamilton, Martin D. Curran, Theresa Feltwell, Grant Hall, Anna Yakovleva, Fahad A Khokhar, Charlotte J. Houldcroft, Laura G Caller, Aminu S. Jahun, Sarah L. Caddy, Ian Goodfellow |
| EPI_ISL_433969 | hCoV-19/England/CAMB-7B501/2020 | Europe / United Kingdom / England | 2020-04-12 | Department of Pathology, University of Cambridge | COVID-19 Genomics UK (COG-UK) Consortium | Luke W Meredith, M. Estee Torok , Myra Hosmillo, William L. Hamilton, Martin D. Curran, Theresa Feltwell, Grant Hall, Anna Yakovleva, Fahad A Khokhar, Charlotte J. Houldcroft, Laura G Caller, Aminu S. Jahun, Sarah L. Caddy, Ian Goodfellow |
| EPI_ISL_433970 | hCoV-19/England/CAMB-7B510/2020 | Europe / United Kingdom / England | 2020-04-13 | Department of Pathology, University of Cambridge | COVID-19 Genomics UK (COG-UK) Consortium | Luke W Meredith, M. Estee Torok , Myra Hosmillo, William L. Hamilton, Martin D. Curran, Theresa Feltwell, Grant Hall, Anna Yakovleva, Fahad A Khokhar, Charlotte J. Houldcroft, Laura G Caller, Aminu S. Jahun, Sarah L. Caddy, Ian Goodfellow |
| EPI_ISL_433971 | hCoV-19/England/CAMB-7B52F/2020 | Europe / United Kingdom / England | 2020-04-13 | Department of Pathology, University of Cambridge | COVID-19 Genomics UK (COG-UK) Consortium | Luke W Meredith, M. Estee Torok , Myra Hosmillo, William L. Hamilton, Martin D. Curran, Theresa Feltwell, Grant Hall, Anna Yakovleva, Fahad A Khokhar, Charlotte J. Houldcroft, Laura G Caller, Aminu S. Jahun, Sarah L. Caddy, Ian Goodfellow |
| EPI_ISL_433972 | hCoV-19/England/CAMB-7B53E/2020 | Europe / United Kingdom / England | 2020 | Department of Pathology, University of Cambridge | COVID-19 Genomics UK (COG-UK) Consortium | Luke W Meredith, M. Estee Torok , Myra Hosmillo, William L. Hamilton, Martin D. Curran, Theresa Feltwell, Grant Hall, Anna Yakovleva, Fahad A Khokhar, Charlotte J. Houldcroft, Laura G Caller, Aminu S. Jahun, Sarah L. Caddy, Ian Goodfellow |
| EPI_ISL_433973 | hCoV-19/England/CAMB-7B54D/2020 | Europe / United Kingdom / England | 2020 | Department of Pathology, University of Cambridge | COVID-19 Genomics UK (COG-UK) Consortium | Luke W Meredith, M. Estee Torok , Myra Hosmillo, William L. Hamilton, Martin D. Curran, Theresa Feltwell, Grant Hall, Anna Yakovleva, Fahad A Khokhar, Charlotte J. Houldcroft, Laura G Caller, Aminu S. Jahun, Sarah L. Caddy, Ian Goodfellow |
| EPI_ISL_433975 | hCoV-19/England/CAMB-7B56B/2020 | Europe / United Kingdom / England | 2020-04-13 | Department of Pathology, University of Cambridge | COVID-19 Genomics UK (COG-UK) Consortium | Luke W Meredith, M. Estee Torok , Myra Hosmillo, William L. Hamilton, Martin D. Curran, Theresa Feltwell, Grant Hall, Anna Yakovleva, Fahad A Khokhar, Charlotte J. Houldcroft, Laura G Caller, Aminu S. Jahun, Sarah L. Caddy, Ian Goodfellow |
| EPI_ISL_433976 | hCoV-19/England/CAMB-7BFD6/2020 | Europe / United Kingdom / England | 2020-04-13 | Department of Pathology, University of Cambridge | COVID-19 Genomics UK (COG-UK) Consortium | Luke W Meredith, M. Estee Torok , Myra Hosmillo, William L. Hamilton, Martin D. Curran, Theresa Feltwell, Grant Hall, Anna Yakovleva, Fahad A Khokhar, Charlotte J. Houldcroft, Laura G Caller, Aminu S. Jahun, Sarah L. Caddy, Ian Goodfellow |
| EPI_ISL_433982 | hCoV-19/England/CAMB-7C03C/2020 | Europe / United Kingdom / England | 2020-04-13 | Department of Pathology, University of Cambridge | COVID-19 Genomics UK (COG-UK) Consortium | Luke W Meredith, M. Estee Torok , Myra Hosmillo, William L. Hamilton, Martin D. Curran, Theresa Feltwell, Grant Hall, Anna Yakovleva, Fahad A Khokhar, Charlotte J. Houldcroft, Laura G Caller, Aminu S. Jahun, Sarah L. Caddy, Ian Goodfellow |
| EPI_ISL_433999 | hCoV-19/England/CAMB-7C148/2020 | Europe / United Kingdom / England | 2020-04-13 | Department of Pathology, University of Cambridge | COVID-19 Genomics UK (COG-UK) Consortium | Luke W Meredith, M. Estee Torok , Myra Hosmillo, William L. Hamilton, Martin D. Curran, Theresa Feltwell, Grant Hall, Anna Yakovleva, Fahad A Khokhar, Charlotte J. Houldcroft, Laura G Caller, Aminu S. Jahun, Sarah L. Caddy, Ian Goodfellow |
| EPI_ISL_434200 | hCoV-19/USA/WA-S440/2020 | North America / USA / Washington / Yaki | 2020-04-01 | Washington State Department of Health | Seattle Flu Study | Chu et al |
| EPI_ISL_434201 | hCoV-19/USA/WA-S441/2020 | North America / USA / Washington / Pier | 2020-03-31 | Washington State Department of Health | Seattle Flu Study | Chu et al |
| EPI_ISL_434202 | hCoV-19/USA/WA-S442/2020 | North America / USA / Washington / Yaki | 2020-03-26 | Washington State Department of Health | Seattle Flu Study | Chu et al |
| EPI_ISL_434204 | hCoV-19/USA/WA-S444/2020 | North America / USA / Washington / Pier | 2020-04-01 | Washington State Department of Health | Seattle Flu Study | Chu et al |
| EPI_ISL_434205 | hCoV-19/USA/WA-S445/2020 | North America / USA / Washington / Yaki | 2020-04-01 | Washington State Department of Health | Seattle Flu Study | Chu et al |
| EPI_ISL_434206 | hCoV-19/USA/WA-S446/2020 | North America / USA / Washington / Yaki | 2020-04-01 | Washington State Department of Health | Seattle Flu Study | Chu et al |
| EPI_ISL_434207 | hCoV-19/USA/WA-S447/2020 | North America / USA / Washington / Fran | 2020-04-01 | Washington State Department of Health | Seattle Flu Study | Chu et al |
| EPI_ISL_434208 | hCoV-19/USA/WA-S448/2020 | North America / USA / Washington / Yaki | 2020-04-01 | Washington State Department of Health | Seattle Flu Study | Chu et al |
| EPI_ISL_434209 | hCoV-19/USA/WA-S449/2020 | North America / USA / Washington / Skag | 2020-03-31 | Washington State Department of Health | Seattle Flu Study | Chu et al |
| EPI_ISL_434211 | hCoV-19/USA/WA-S451/2020 | North America / USA / Washington / Pier | 2020-03-31 | Washington State Department of Health | Seattle Flu Study | Chu et al |
| EPI_ISL_434212 | hCoV-19/USA/WA-S452/2020 | North America / USA / Washington / Yaki | 2020-03-31 | Washington State Department of Health | Seattle Flu Study | Chu et al |
| EPI_ISL_434213 | hCoV-19/USA/WA-S453/2020 | North America / USA / Washington / Mas | 2020-03-30 | Washington State Department of Health | Seattle Flu Study | Chu et al |
| EPI_ISL_434214 | hCoV-19/USA/WA-S454/2020 | North America / USA / Washington / Pier | 2020-03-31 | Washington State Department of Health | Seattle Flu Study | Chu et al |
| EPI_ISL_434215 | hCoV-19/USA/WA-S455/2020 | North America / USA / Washington / Pier | 2020-03-31 | Washington State Department of Health | Seattle Flu Study | Chu et al |
| EPI_ISL_434216 | hCoV-19/USA/WA-S456/2020 | North America / USA / Washington / Snoh | 2020-04-06 | Washington State Department of Health | Seattle Flu Study | Chu et al |
| EPI_ISL_434217 | hCoV-19/USA/WA-S457/2020 | North America / USA / Washington / Yaki | 2020-04-02 | Washington State Department of Health | Seattle Flu Study | Chu et al |
| EPI_ISL_434218 | hCoV-19/USA/WA-S458/2020 | North America / USA / Washington / Snoh | 2020-04-06 | Washington State Department of Health | Seattle Flu Study | Chu et al |
| EPI_ISL_434219 | hCoV-19/USA/WA-S459/2020 | North America / USA / Washington / Snoh | 2020-04-06 | Washington State Department of Health | Seattle Flu Study | Chu et al |
| EPI_ISL_434220 | hCoV-19/USA/WA-S460/2020 | North America / USA / Washington / King | 2020-04-05 | Washington State Department of Health | Seattle Flu Study | Chu et al |
| EPI_ISL_434221 | hCoV-19/USA/WA-S461/2020 | North America / USA / Washington / Snoh | 2020-04-06 | Washington State Department of Health | Seattle Flu Study | Chu et al |
| EPI_ISL_434222 | hCoV-19/USA/WA-S462/2020 | North America / USA / Washington / Yaki | 2020-04-02 | Washington State Department of Health | Seattle Flu Study | Chu et al |
| EPI_ISL_434223 | hCoV-19/USA/WA-S463/2020 | North America / USA / Washington / Yaki | 2020-04-04 | Washington State Department of Health | Seattle Flu Study | Chu et al |
| EPI_ISL_434224 | hCoV-19/USA/WA-S464/2020 | North America / USA / Washington / King | 2020-04-04 | Washington State Department of Health | Seattle Flu Study | Chu et al |
| EPI_ISL_434225 | hCoV-19/USA/WA-S465/2020 | North America / USA / Washington / Pier | 2020-04-01 | Washington State Department of Health | Seattle Flu Study | Chu et al |
| EPI_ISL_434226 | hCoV-19/USA/WA-S466/2020 | North America / USA / Washington / Yaki | 2020-04-01 | Washington State Department of Health | Seattle Flu Study | Chu et al |
| EPI_ISL_434227 | hCoV-19/USA/WA-S467/2020 | North America / USA / Washington / Yaki | 2020-04-02 | Washington State Department of Health | Seattle Flu Study | Chu et al |
| EPI_ISL_434228 | hCoV-19/USA/WA-S468/2020 | North America / USA / Washington / King | 2020-04-05 | Washington State Department of Health | Seattle Flu Study | Chu et al |
| EPI_ISL_434229 | hCoV-19/USA/WA-S469/2020 | North America / USA / Washington / King | 2020-04-05 | Washington State Department of Health | Seattle Flu Study | Chu et al |
| EPI_ISL_434230 | hCoV-19/USA/WA-S470/2020 | North America / USA / Washington / King | 2020-04-05 | Washington State Department of Health | Seattle Flu Study | Chu et al |
| EPI_ISL_434231 | hCoV-19/USA/WA-S471/2020 | North America / USA / Washington / Snoh | 2020-04-06 | Washington State Department of Health | Seattle Flu Study | Chu et al |
| EPI_ISL_434232 | hCoV-19/USA/WA-S472/2020 | North America / USA / Washington / Yaki | 2020-04-05 | Washington State Department of Health | Seattle Flu Study | Chu et al |
| EPI_ISL_434233 | hCoV-19/USA/WA-S473/2020 | North America / USA / Washington / Yaki | 2020-04-03 | Washington State Department of Health | Seattle Flu Study | Chu et al |
| EPI_ISL_434234 | hCoV-19/USA/WA-S474/2020 | North America / USA / Washington / King | 2020-04-01 | Washington State Department of Health | Seattle Flu Study | Chu et al |
| EPI_ISL_434235 | hCoV-19/USA/WA-S475/2020 | North America / USA / Washington / King | 2020-04-01 | Washington State Department of Health | Seattle Flu Study | Chu et al |
| EPI_ISL_434236 | hCoV-19/USA/WA-S476/2020 | North America / USA / Washington / Yaki | 2020-04-03 | Washington State Department of Health | Seattle Flu Study | Chu et al |
| EPI_ISL_434237 | hCoV-19/USA/WA-S477/2020 | North America / USA / Washington / King | 2020-04-05 | Washington State Department of Health | Seattle Flu Study | Chu et al |
| EPI_ISL_434238 | hCoV-19/USA/WA-S478/2020 | North America / USA / Washington / Yaki | 2020-04-01 | Washington State Department of Health | Seattle Flu Study | Chu et al |
| EPI_ISL_434239 | hCoV-19/USA/WA-S479/2020 | North America / USA / Washington / Yaki | 2020-04-04 | Washington State Department of Health | Seattle Flu Study | Chu et al |
| EPI_ISL_434240 | hCoV-19/USA/WA-S480/2020 | North America / USA / Washington / King | 2020-04-05 | Washington State Department of Health | Seattle Flu Study | Chu et al |
| EPI_ISL_434241 | hCoV-19/USA/WA-S481/2020 | North America / USA / Washington / Yaki | 2020-04-02 | Washington State Department of Health | Seattle Flu Study | Chu et al |
| EPI_ISL_434242 | hCoV-19/USA/WA-S482/2020 | North America / USA / Washington / Snoh | 2020-04-06 | Washington State Department of Health | Seattle Flu Study | Chu et al |
| EPI_ISL_434243 | hCoV-19/USA/WA-S483/2020 | North America / USA / Washington / King | 2020-04-05 | Washington State Department of Health | Seattle Flu Study | Chu et al |
| EPI_ISL_434244 | hCoV-19/USA/WA-S484/2020 | North America / USA / Washington / Pier | 2020-04-02 | Washington State Department of Health | Seattle Flu Study | Chu et al |
| EPI_ISL_434245 | hCoV-19/USA/WA-S485/2020 | North America / USA / Washington / Yaki | 2020-04-01 | Washington State Department of Health | Seattle Flu Study | Chu et al |
| EPI_ISL_434246 | hCoV-19/USA/WA-S486/2020 | North America / USA / Washington / Yaki | 2020-04-01 | Washington State Department of Health | Seattle Flu Study | Chu et al |
| EPI_ISL_434247 | hCoV-19/USA/WA-S487/2020 | North America / USA / Washington / Yaki | 2020-03-31 | Washington State Department of Health | Seattle Flu Study | Chu et al |
| EPI_ISL_434248 | hCoV-19/USA/WA-S488/2020 | North America / USA / Washington / King | 2020-04-01 | Washington State Department of Health | Seattle Flu Study | Chu et al |
| EPI_ISL_434249 | hCoV-19/USA/WA-S489/2020 | North America / USA / Washington / Snoh | 2020-04-02 | Washington State Department of Health | Seattle Flu Study | Chu et al |
| EPI_ISL_434250 | hCoV-19/USA/WA-S490/2020 | North America / USA / Washington / Pier | 2020-03-31 | Washington State Department of Health | Seattle Flu Study | Chu et al |
| EPI_ISL_434251 | hCoV-19/USA/WA-S491/2020 | North America / USA / Washington / Pier | 2020-04-02 | Washington State Department of Health | Seattle Flu Study | Chu et al |
| EPI_ISL_434252 | hCoV-19/USA/WA-S492/2020 | North America / USA / Washington / Yaki | 2020-03-26 | Washington State Department of Health | Seattle Flu Study | Chu et al |

| | | | | | | |
|---|---|---|---|---|---|---|
| EPI_ISL_434253 | hCoV-19/USA/WA-S493/2020 | North America / USA / Washington / Yaki | 2020-04-01 | Washington State Department of Health | Seattle Flu Study | Chu et al |
| EPI_ISL_434254 | hCoV-19/USA/WA-S494/2020 | North America / USA / Washington / Yaki | 2020-04-01 | Washington State Department of Health | Seattle Flu Study | Chu et al |
| EPI_ISL_434255 | hCoV-19/USA/WA-S495/2020 | North America / USA / Washington / King | 2020-04-01 | Washington State Department of Health | Seattle Flu Study | Chu et al |
| EPI_ISL_434256 | hCoV-19/USA/WA-S496/2020 | North America / USA / Washington / Pier | 2020-03-31 | Washington State Department of Health | Seattle Flu Study | Chu et al |
| EPI_ISL_434257 | hCoV-19/USA/WA-S497/2020 | North America / USA / Washington / Yaki | 2020-03-31 | Washington State Department of Health | Seattle Flu Study | Chu et al |
| EPI_ISL_434258 | hCoV-19/USA/WA-S498/2020 | North America / USA / Washington / Yaki | 2020-04-01 | Washington State Department of Health | Seattle Flu Study | Chu et al |
| EPI_ISL_434259 | hCoV-19/USA/WA-S499/2020 | North America / USA / Washington / Pier | 2020-03-31 | Washington State Department of Health | Seattle Flu Study | Chu et al |
| EPI_ISL_434260 | hCoV-19/USA/WA-S500/2020 | North America / USA / Washington / Yaki | 2020-03-25 | Washington State Department of Health | Seattle Flu Study | Chu et al |
| EPI_ISL_434261 | hCoV-19/USA/WA-S501/2020 | North America / USA / Washington | 2020-03-29 | Washington State Department of Health | Seattle Flu Study | Chu et al |
| EPI_ISL_434262 | hCoV-19/USA/WA-S502/2020 | North America / USA / Washington / Yaki | 2020-03-24 | Washington State Department of Health | Seattle Flu Study | Chu et al |
| EPI_ISL_434263 | hCoV-19/USA/WA-S503/2020 | North America / USA / Washington / King | 2020-03-25 | Washington State Department of Health | Seattle Flu Study | Chu et al |
| EPI_ISL_434264 | hCoV-19/USA/WA-S504/2020 | North America / USA / Washington / Yaki | 2020-04-01 | Washington State Department of Health | Seattle Flu Study | Chu et al |
| EPI_ISL_434265 | hCoV-19/USA/WA-S505/2020 | North America / USA / Washington / Pier | 2020-03-29 | Washington State Department of Health | Seattle Flu Study | Chu et al |
| EPI_ISL_434266 | hCoV-19/USA/WA-S506/2020 | North America / USA / Washington / Yaki | 2020-04-01 | Washington State Department of Health | Seattle Flu Study | Chu et al |
| EPI_ISL_434267 | hCoV-19/USA/WA-S507/2020 | North America / USA / Washington | 2020-03-23 | Washington State Department of Health | Seattle Flu Study | Chu et al |
| EPI_ISL_434268 | hCoV-19/USA/WA-S508/2020 | North America / USA / Washington / Yaki | 2020-03-30 | Washington State Department of Health | Seattle Flu Study | Chu et al |
| EPI_ISL_434269 | hCoV-19/USA/WA-S509/2020 | North America / USA / Washington / Yaki | 2020-03-30 | Washington State Department of Health | Seattle Flu Study | Chu et al |
| EPI_ISL_434270 | hCoV-19/USA/WA-S510/2020 | North America / USA / Washington / Yaki | 2020-03-25 | Washington State Department of Health | Seattle Flu Study | Chu et al |
| EPI_ISL_434271 | hCoV-19/USA/WA-S511/2020 | North America / USA / Washington / Wha | 2020-03-24 | Washington State Department of Health | Seattle Flu Study | Chu et al |
| EPI_ISL_434272 | hCoV-19/USA/WA-S512/2020 | North America / USA / Washington / Yaki | 2020-04-01 | Washington State Department of Health | Seattle Flu Study | Chu et al |
| EPI_ISL_434273 | hCoV-19/USA/WA-S513/2020 | North America / USA / Washington / Yaki | 2020-04-01 | Washington State Department of Health | Seattle Flu Study | Chu et al |
| EPI_ISL_434274 | hCoV-19/USA/WA-S514/2020 | North America / USA / Washington / Snol | 2020-03-29 | Washington State Department of Health | Seattle Flu Study | Chu et al |
| EPI_ISL_434276 | hCoV-19/USA/WA-S516/2020 | North America / USA / Washington / Yaki | 2020-04-01 | Washington State Department of Health | Seattle Flu Study | Chu et al |
| EPI_ISL_434277 | hCoV-19/USA/WA-S517/2020 | North America / USA / Washington / Yaki | 2020-03-24 | Washington State Department of Health | Seattle Flu Study | Chu et al |
| EPI_ISL_434278 | hCoV-19/USA/WA-S518/2020 | North America / USA / Washington | 2020-03-29 | Washington State Department of Health | Seattle Flu Study | Chu et al |
| EPI_ISL_434279 | hCoV-19/USA/WA-S519/2020 | North America / USA / Washington / Adar | 2020-03-27 | Washington State Department of Health | Seattle Flu Study | Chu et al |
| EPI_ISL_434280 | hCoV-19/USA/WA-S520/2020 | North America / USA / Washington / Yaki | 2020-03-31 | Washington State Department of Health | Seattle Flu Study | Chu et al |
| EPI_ISL_434281 | hCoV-19/USA/WA-S521/2020 | North America / USA / Washington / Wha | 2020-03-25 | Washington State Department of Health | Seattle Flu Study | Chu et al |
| EPI_ISL_434282 | hCoV-19/USA/WA-S522/2020 | North America / USA / Washington / Yaki | 2020-04-01 | Washington State Department of Health | Seattle Flu Study | Chu et al |
| EPI_ISL_434283 | hCoV-19/USA/WA-S523/2020 | North America / USA / Washington / King | 2020-03-26 | Washington State Department of Health | Seattle Flu Study | Chu et al |
| EPI_ISL_434284 | hCoV-19/USA/WA-S524/2020 | North America / USA / Washington / Yaki | 2020-04-01 | Washington State Department of Health | Seattle Flu Study | Chu et al |
| EPI_ISL_434285 | hCoV-19/USA/WA-S525/2020 | North America / USA / Washington / Yaki | 2020-04-01 | Washington State Department of Health | Seattle Flu Study | Chu et al |
| EPI_ISL_434286 | hCoV-19/USA/WA-S526/2020 | North America / USA / Washington / Pier | 2020-03-31 | Washington State Department of Health | Seattle Flu Study | Chu et al |
| EPI_ISL_434287 | hCoV-19/USA/WA-S527/2020 | North America / USA / Washington / Pier | 2020-03-31 | Washington State Department of Health | Seattle Flu Study | Chu et al |
| EPI_ISL_434288 | hCoV-19/USA/WA-S528/2020 | North America / USA / Washington / King | 2020-03-31 | Washington State Department of Health | Seattle Flu Study | Chu et al |
| EPI_ISL_434289 | hCoV-19/USA/WA-S529/2020 | North America / USA / Washington / King | 2020-03-24 | Washington State Department of Health | Seattle Flu Study | Chu et al |
| EPI_ISL_434290 | hCoV-19/USA/WA-S530/2020 | North America / USA / Washington / Yaki | 2020-03-24 | Washington State Department of Health | Seattle Flu Study | Chu et al |
| EPI_ISL_434291 | hCoV-19/USA/WA-S531/2020 | North America / USA / Washington | 2020-03-24 | Washington State Department of Health | Seattle Flu Study | Chu et al |
| EPI_ISL_434292 | hCoV-19/USA/WA-S532/2020 | North America / USA / Washington | 2020-03-25 | Washington State Department of Health | Seattle Flu Study | Chu et al |
| EPI_ISL_434293 | hCoV-19/USA/WA-S533/2020 | North America / USA / Washington / Yaki | 2020-03-25 | Washington State Department of Health | Seattle Flu Study | Chu et al |
| EPI_ISL_434294 | hCoV-19/USA/WA-S534/2020 | North America / USA / Washington | 2020-03-25 | Washington State Department of Health | Seattle Flu Study | Chu et al |
| EPI_ISL_434295 | hCoV-19/USA/WA-S535/2020 | North America / USA / Washington | 2020-03-25 | Washington State Department of Health | Seattle Flu Study | Chu et al |
| EPI_ISL_434296 | hCoV-19/USA/WA-S536/2020 | North America / USA / Washington / Wha | 2020-03-24 | Washington State Department of Health | Seattle Flu Study | Chu et al |
| EPI_ISL_434297 | hCoV-19/USA/WA-S537/2020 | North America / USA / Washington | 2020-03-23 | Washington State Department of Health | Seattle Flu Study | Chu et al |
| EPI_ISL_434298 | hCoV-19/USA/WA-S538/2020 | North America / USA / Washington / Pier | 2020-03-31 | Washington State Department of Health | Seattle Flu Study | Chu et al |
| EPI_ISL_434299 | hCoV-19/USA/WA-S539/2020 | North America / USA / Washington / Pier | 2020-03-23 | Washington State Department of Health | Seattle Flu Study | Chu et al |
| EPI_ISL_434535 | hCoV-19/Costa Rica/02/2020 | Central America / Costa Rica | 2020-03-16 | Area de Salud Alajuela Sur | Inciensa, Instituto Costarricense de Investigación y Enseñanza en Nutrición y Salud | Francisco Duarte, Hebleen Porras, Claudio Soto-Garita, Estela Cordero, Adriana Godinez & Melany Calderon |
| EPI_ISL_434538 | hCoV-19/Costa Rica/05/2020 | Central America / Costa Rica | 2020-03-20 | COOPESAIN | Inciensa, Instituto Costarricense de Investigación y Enseñanza en Nutrición y Salud | Francisco Duarte, Hebleen Porras, Claudio Soto-Garita, Estela Cordero, Adriana Godinez & Melany Calderon |
| EPI_ISL_434539 | hCoV-19/Costa Rica/06/2020 | Central America / Costa Rica | 2020-03-25 | Area de Salud Orotina | Inciensa, Instituto Costarricense de Investigación y Enseñanza en Nutrición y Salud | Francisco Duarte, Hebleen Porras, Claudio Soto-Garita, Estela Cordero, Adriana Godinez & Melany Calderon |
| EPI_ISL_434540 | hCoV-19/Costa Rica/04/2020 | Central America / Costa Rica | 2020-03-18 | EBAIS Concepción Norte | Inciensa, Instituto Costarricense de Investigación y Enseñanza en Nutrición y Salud | Francisco Duarte, Hebleen Porras, Claudio Soto-Garita, Estela Cordero, Adriana Godinez & Melany Calderon |
| EPI_ISL_434541 | hCoV-19/Puerto Rico/CDC-S1/2020 | North America / USA / Puerto Rico | 2020-03-23 | Puerto Rico Department of Health | Centers for Disease Control and Prevention, Dengue Branch | Gilberto A. Santiago, Glenda Gonzalez, Betzabel Flores, Keyla Charriez, Fabiola Cruz, Chaney Kalinich, Joseph Fauver, Jessica I. Falcon, Nathan Grubaugh, Jorge L. Munoz-Jordan |
| EPI_ISL_434543 | hCoV-19/Puerto Rico/CDC-S3/2020 | North America / USA / Puerto Rico | 2020-03-23 | Puerto Rico Department of Health | Centers for Disease Control and Prevention, Dengue Branch | Gilberto A. Santiago, Glenda Gonzalez, Betzabel Flores, Keyla Charriez, Fabiola Cruz, Chaney Kalinich, Joseph Fauver, Jessica I. Falcon, Nathan Grubaugh, Jorge L. Munoz-Jordan |
| EPI_ISL_434545 | hCoV-19/Puerto Rico/CDC-S5/2020 | North America / USA / Puerto Rico | 2020-03-23 | Puerto Rico Department of Health | Centers for Disease Control and Prevention, Dengue Branch | Gilberto A. Santiago, Glenda Gonzalez, Betzabel Flores, Keyla Charriez, Fabiola Cruz, Chaney Kalinich, Joseph Fauver, Jessica I. Falcon, Nathan Grubaugh, Jorge L. Munoz-Jordan |
| EPI_ISL_434546 | hCoV-19/Puerto Rico/CDC-S6/2020 | North America / USA / Puerto Rico | 2020-03-24 | Puerto Rico Department of Health | Centers for Disease Control and Prevention, Dengue Branch | Gilberto A. Santiago, Glenda Gonzalez, Betzabel Flores, Keyla Charriez, Fabiola Cruz, Chaney Kalinich, Joseph Fauver, Jessica I. Falcon, Nathan Grubaugh, Jorge L. Munoz-Jordan |
| EPI_ISL_434547 | hCoV-19/Puerto Rico/CDC-S7/2020 | North America / USA / Puerto Rico | 2020-03-24 | Puerto Rico Department of Health | Centers for Disease Control and Prevention, Dengue Branch | Gilberto A. Santiago, Glenda Gonzalez, Betzabel Flores, Keyla Charriez, Fabiola Cruz, Chaney Kalinich, Joseph Fauver, Jessica I. Falcon, Nathan Grubaugh, Jorge L. Munoz-Jordan |
| EPI_ISL_434549 | hCoV-19/Puerto Rico/CDC-S9/2020 | North America / USA / Puerto Rico | 2020-03-23 | Puerto Rico Department of Health | Centers for Disease Control and Prevention, Dengue Branch | Gilberto A. Santiago, Glenda Gonzalez, Betzabel Flores, Keyla Charriez, Fabiola Cruz, Chaney Kalinich, Joseph Fauver, Jessica I. Falcon, Nathan Grubaugh, Jorge L. Munoz-Jordan |
| EPI_ISL_434551 | hCoV-19/Puerto Rico/CDC-S11/2020 | North America / USA / Puerto Rico | 2020-03-24 | Puerto Rico Department of Health | Centers for Disease Control and Prevention, Dengue Branch | Gilberto A. Santiago, Glenda Gonzalez, Betzabel Flores, Keyla Charriez, Fabiola Cruz, Chaney Kalinich, Joseph Fauver, Jessica I. Falcon, Nathan Grubaugh, Jorge L. Munoz-Jordan |
| EPI_ISL_434555 | hCoV-19/Philippines/PGC03/2020 | Asia / Philippines / Manila | 2020-03-26 | National Institutes of Health, University of the Philippines Manila | Philippine Genome Center | Carlo M. Lapid, Francis A. Tablizo, Benedict A. Maralit, Jan Michael C. Yap, Raul V. Destura, Marissa M. Alejandria, El King D. Morado, Joshua Gregor A. Dizon, Jo-Hannah S. Llames, Shiela Mae M. Araiza, Kris P. Punayan, Kristianne Arielle D. Gabriel, Shebna Rose D. Fabilloren, Shana F. Genavia, Jarvin E. Nipales, Alessandra C. Sanchez, Haifa L.Gaza, Joy Ann Petronio-Santos, Julius Aaron Mejia, Maribell Dollete, Sonia Salamat, Christina Tan, Bernard Demot, John Mark Velasco, Eva Maria Cutiongco-de la Paz, and Cynthia P. Saloma |
| EPI_ISL_434558 | hCoV-19/Philippines/PGC06/2020 | Asia / Philippines / Manila | 2020-03-28 | National Institutes of Health, University of the Philippines Manila | Philippine Genome Center | Carlo M. Lapid, Francis A. Tablizo, Benedict A. Maralit, Jan Michael C. Yap, Raul V. Destura, Marissa M. Alejandria, El King D. Morado, Joshua Gregor A. Dizon, Jo-Hannah S. Llames, Shiela Mae M. Araiza, Kris P. Punayan, Kristianne Arielle D. Gabriel, Shebna Rose D. Fabilloren, Shana F. Genavia, Jarvin E. Nipales, Alessandra C. Sanchez, Haifa L.Gaza, Joy Ann Petronio-Santos, Julius Aaron Mejia, Maribell Dollete, Sonia Salamat, Christina Tan, Bernard Demot, John Mark Velasco, Eva Maria Cutiongco-de la Paz, and Cynthia P. Saloma |
| EPI_ISL_434560 | hCoV-19/Hong Kong/HK20/2020 | Asia / Hong Kong | 2020-01 | unknown | Department of Microbiology | Lau,S.K.P., Luk,H.K.H., Wong,A.C.P., Li,K.S.M., Zhu,L., He,Z., Fung,J., Chan,T.T.Y., Fung,K.S.C. and Woo,P.C.Y. |
| EPI_ISL_434561 | hCoV-19/Japan/TKYE6968/2020 | Asia / Japan | 2020-02 | unknown | Ryota Kumagai Tokyo Metropolitan Institute of Public Health | Kumagai,R., Yoshida,I., Asakura,H., Nagashima,M., Chiba,T. and Sadamasu,K. |
| EPI_ISL_434562 | hCoV-19/Japan/TKYE6947/2020 | Asia / Japan | 2020-03 | unknown | Ryota Kumagai Tokyo Metropolitan Institute of Public Health | Kumagai,R., Yoshida,I., Asakura,H., Nagashima,M., Chiba,T. and Sadamasu,K. |
| EPI_ISL_434563 | hCoV-19/Hong Kong/HKU-902a/2020 | Asia / Hong Kong | 2020-01-23 | unknown | Microbiology | To,K.K.W. and Yuen,K.-Y. |
| EPI_ISL_434564 | hCoV-19/Hong Kong/HKU-902b/2020 | Asia / Hong Kong | 2020-01-24 | unknown | unknown | To,K.K.W. and Yuen,K.-Y. |
| EPI_ISL_434565 | hCoV-19/Hong Kong/HKU-903a/2020 | Asia / Hong Kong | 2020-01-24 | unknown | Microbiology | To,K.K.W. and Yuen,K.-Y. |
| EPI_ISL_434566 | hCoV-19/Hong Kong/HKU-903b/2020 | Asia / Hong Kong | 2020-01-27 | unknown | Microbiology | To,K.K.W. and Yuen,K.-Y. |
| EPI_ISL_434567 | hCoV-19/Hong Kong/HKU-907a/2020 | Asia / Hong Kong | 2020-01-27 | unknown | Microbiology | To,K.K.W. and Yuen,K.-Y. |
| EPI_ISL_434568 | hCoV-19/Hong Kong/HKU-907b/2020 | Asia / Hong Kong | 2020-01-29 | unknown | Microbiology | To,K.K.W. and Yuen,K.-Y. |
| EPI_ISL_434569 | hCoV-19/Hong Kong/HKU-908a/2020 | Asia / Hong Kong | 2020-01-27 | unknown | Microbiology | To,K.K.W. and Yuen,K.-Y. |
| EPI_ISL_434570 | hCoV-19/Hong Kong/HKU-908b/2020 | Asia / Hong Kong | 2020-01-29 | unknown | Microbiology | To,K.K.W. and Yuen,K.-Y. |
| EPI_ISL_434571 | hCoV-19/Hong Kong/HKU-001a/2020 | Asia / Hong Kong | 2020-01-22 | unknown | Microbiology | Chan,J.F.W. and Yuen,K.-Y. |
| EPI_ISL_434572 | hCoV-19/Czech Republic/2308/2020 | Europe / Czech Republic / Vysocina Regi | 2020-04-14 | The National Institute of Public Health Center for Epidemiology and Microbiology | The National Institute of Public Health Center for Epidemiology and Microbiology | Alexander Nagy, Helena Jirincova, Ludmila Novakova, Dusan Trnka, Jaromira Vecerova |
| EPI_ISL_434586 | hCoV-19/USA/MD-HP00036/2020 | North America / USA / Maryland | 2020-03-19 | Johns Hopkins Hospital Department of Pathology | Johns Hopkins Hospital Department of Pathology | Peter M. Thielen, Thomas Mehoke, Shirlee Wohl, Srividya Ramakrishnan, Oluwaseun Nwulia-Falade, Amanda Ernlund, Melanie Kirsche, Paul Morris, Norah Sadowski, Nidiá Trovao, Victoria Gniazdowski, Michael Schatz, Stuart C. Ray, Winston Timp, Heba Mostafa |
| EPI_ISL_434587 | hCoV-19/USA/MD-HP00037/2020 | North America / USA / Maryland | 2020-03-24 | Johns Hopkins Hospital Department of Pathology | Johns Hopkins Hospital Department of Pathology | Peter M. Thielen, Thomas Mehoke, Shirlee Wohl, Srividya Ramakrishnan, Oluwaseun Nwulia-Falade, Amanda Ernlund, Melanie Kirsche, Paul Morris, Norah Sadowski, Nidiá Trovao, Victoria Gniazdowski, Michael Schatz, Stuart C. Ray, Winston Timp, Heba Mostafa |
| EPI_ISL_434588 | hCoV-19/USA/MD-HP00038/2020 | North America / USA / Maryland | 2020-03-14 | Johns Hopkins Hospital Department of Pathology | Johns Hopkins Hospital Department of Pathology | Peter M. Thielen, Thomas Mehoke, Shirlee Wohl, Srividya Ramakrishnan, Oluwaseun Nwulia-Falade, Amanda Ernlund, Melanie Kirsche, Paul Morris, Norah Sadowski, Nidiá Trovao, Victoria Gniazdowski, Michael Schatz, Stuart C. Ray, Winston Timp, Heba Mostafa |
| EPI_ISL_434590 | hCoV-19/USA/VA-DCLS-0095/2020 | North America / USA / Virginia | 2020-04-09 | Virginia DCLS | Virginia DCLS | Virginia DCLS |
| EPI_ISL_434592 | hCoV-19/USA/VA-DCLS-0098/2020 | North America / USA / Virginia | 2020-04-10 | Virginia DCLS | Virginia DCLS | Virginia DCLS |
| EPI_ISL_434593 | hCoV-19/USA/VA-DCLS-0099/2020 | North America / USA / Virginia | 2020-04-11 | Virginia DCLS | Virginia DCLS | Virginia DCLS |
| EPI_ISL_434594 | hCoV-19/USA/VA-DCLS-0100/2020 | North America / USA / Virginia | 2020-04-10 | Virginia DCLS | Virginia DCLS | Virginia DCLS |
| EPI_ISL_434595 | hCoV-19/USA/VA-DCLS-0101/2020 | North America / USA / Virginia | 2020-04-10 | Virginia DCLS | Virginia DCLS | Virginia DCLS |
| EPI_ISL_434596 | hCoV-19/USA/VA-DCLS-0104/2020 | North America / USA / Virginia | 2020-04-08 | Virginia DCLS | Virginia DCLS | Virginia DCLS |
| EPI_ISL_434597 | hCoV-19/USA/VA-DCLS-0105/2020 | North America / USA / Virginia | 2020-04-14 | Virginia DCLS | Virginia DCLS | Virginia DCLS |
| EPI_ISL_434598 | hCoV-19/USA/VA-DCLS-0106/2020 | North America / USA / Virginia | 2020-04-14 | Virginia DCLS | Virginia DCLS | Virginia DCLS |
| EPI_ISL_434599 | hCoV-19/USA/VA-DCLS-0107/2020 | North America / USA / Virginia | 2020-04-14 | Virginia DCLS | Virginia DCLS | Virginia DCLS |
| EPI_ISL_426414 | hCoV-19/India/GBRC1/2020 | Asia / India / Gujarat | 2020-04-05 | Sir M P Shah Government Medical College | Gujarat Biotechnology Research Centre | Ramesh Pandit, Tejas Shah, Ankit Hinsu, Pritesh Sabara, Apurvasinh Puvar, Janvi Raval, Monika Gandhi, Pinal Trivedi, Maharshi Pandya, Amit Kanani, Akanksha Verma, Nitin Savaliya, Raghawendra Kumar, Dinesh Kumar, Zubair Saiyed, Dipa Kinariwala, Disha Patel, Binita Aring, Geeta Vaghela, Sonia Barve, Bhavesh Modi, Kairavi Joshi, Nidhi Sood, Pranay Shah, R D Dixit, Snehal Bagatharia, Madhvi Joshi, Chaitanya Joshi |
| EPI_ISL_426415 | hCoV-19/India/GBRC1a/2020 | Asia / India / Gujarat | 2020-04-05 | Sir M P Shah Government Medical College, Jamnagar | Gujarat Biotechnology Research Centre, Gandhinagar | Ramesh Pandit, Tejas Shah, Ankit Hinsu, Pritesh Sabara, Apurvasinh Puvar, Janvi Raval, Monika Gandhi, Pinal Trivedi, Maharshi Pandya, Amit Kanani, Akanksha Verma, Nitin Savaliya, Raghawendra Kumar, Dinesh Kumar, Zubair Saiyed, Dipa Kinariwala, Disha Patel, Binita Aring, Geeta Vaghela, Sonia Barve, Bhavesh Modi, Kairavi Joshi, Nidhi Sood, Pranay Shah, R D Dixit, Snehal Bagatharia, Madhvi Joshi, Chaitanya Joshi |
| EPI_ISL_426416 | hCoV-19/USA/CT_9849/2020 | North America / USA / Connecticut | 2020-03-08 | CT-Dr. Katherine A. Kelley State Public Health Lab | Pathogen Discovery, Respiratory Viruses Branch, Division of Viral Diseases, Centers for Disease Control and Prevention | Anna Uehara, Yan Li, Krista Queen, Clinton R. Paden, Rachel Marine, Ying Tao, Jing Zhang, Haibin Wang, Mary S. Keckler, Alison S. Laufer Halpin, Christopher A. Elkins, Suxiang Tong |
| EPI_ISL_426417 | hCoV-19/USA/GA_2070/2020 | North America / USA / Georgia | 2020-03-07 | GA Department of Public Health Laboratory | Pathogen Discovery, Respiratory Viruses Branch, Division of Viral Diseases, Centers for Disease Control and Prevention | Anna Uehara, Yan Li, Krista Queen, Clinton R. Paden, Rachel Marine, Ying Tao, Jing Zhang, Haibin Wang, Mary S. Keckler, Alison S. Laufer Halpin, Christopher A. Elkins, Suxiang Tong |
| EPI_ISL_426418 | hCoV-19/USA/GA_2147/2020 | North America / USA / Georgia | 2020-03 | GA Department of Public Health Laboratory | Pathogen Discovery, Respiratory Viruses Branch, Division of Viral Diseases, Centers for Disease Control and Prevention | Anna Uehara, Yan Li, Krista Queen, Clinton R. Paden, Rachel Marine, Ying Tao, Jing Zhang, Haibin Wang, Mary S. Keckler, Alison S. Laufer Halpin, Christopher A. Elkins, Suxiang Tong |
| EPI_ISL_426419 | hCoV-19/USA/GA_2275/2020 | North America / USA / Georgia | 2020-03 | GA Department of Public Health Laboratory | Pathogen Discovery, Respiratory Viruses Branch, Division of Viral Diseases, Centers for Disease Control and Prevention | Anna Uehara, Yan Li, Krista Queen, Clinton R. Paden, Rachel Marine, Ying Tao, Jing Zhang, Haibin Wang, Mary S. Keckler, Alison S. Laufer Halpin, Christopher A. Elkins, Suxiang Tong |
| EPI_ISL_426420 | hCoV-19/USA/HI_4970/2020 | North America / USA / Hawaii | 2020-03-05 | HI Dept. of Health, State Laboratories Division | Pathogen Discovery, Respiratory Viruses Branch, Division of Viral Diseases, Centers for Disease Control and Prevention | Anna Uehara, Yan Li, Krista Queen, Clinton R. Paden, Rachel Marine, Ying Tao, Jing Zhang, Haibin Wang, Mary S. Keckler, Alison S. Laufer Halpin, Christopher A. Elkins, Suxiang Tong |
| EPI_ISL_426421 | hCoV-19/USA/HI_7881/2020 | North America / USA / Hawaii | 2020-03-07 | HI Dept. of Health, State Laboratories Division | Pathogen Discovery, Respiratory Viruses Branch, Division of Viral Diseases, Centers for Disease Control and Prevention | Anna Uehara, Yan Li, Krista Queen, Clinton R. Paden, Rachel Marine, Ying Tao, Jing Zhang, Haibin Wang, Mary S. Keckler, Alison S. Laufer Halpin, Christopher A. Elkins, Suxiang Tong |
| EPI_ISL_426425 | hCoV-19/USA/MD_0027/2020 | North America / USA / Maryland | 2020-03-04 | MD DOH Laboratories Administration | Pathogen Discovery, Respiratory Viruses Branch, Division of Viral Diseases, Centers for Disease Control and Prevention | Krista Queen, Yan Li, Anna Uehara, Clinton R. Paden, Rachel Marine, Ying Tao, Jing Zhang, Haibin Wang, Mary S. Keckler, Alison S. Laufer Halpin, Christopher A. Elkins, Suxiang Tong |
| EPI_ISL_426426 | hCoV-19/USA/MN_0100/2020 | North America / USA / Minnesota | 2020-03-05 | MN PHL Division, Minnesota Department of Health | Pathogen Discovery, Respiratory Viruses Branch, Division of Viral Diseases, Centers for Disease Control and Prevention | Krista Queen, Yan Li, Anna Uehara, Clinton R. Paden, Rachel Marine, Ying Tao, Jing Zhang, Haibin Wang, Mary S. Keckler, Alison S. Laufer Halpin, Christopher A. Elkins, Suxiang Tong |
| EPI_ISL_426427 | hCoV-19/USA/MN_0101/2020 | North America / USA / Minnesota | 2020-03-07 | MN PHL Division, Minnesota Department of Health | Pathogen Discovery, Respiratory Viruses Branch, Division of Viral Diseases, Centers for Disease Control and Prevention | Krista Queen, Yan Li, Anna Uehara, Clinton R. Paden, Rachel Marine, Ying Tao, Jing Zhang, Haibin Wang, Mary S. Keckler, Alison S. Laufer Halpin, Christopher A. Elkins, Suxiang Tong |
| EPI_ISL_426428 | hCoV-19/USA/NC_0025/2020 | North America / USA / North Carolina | 2020-03-08 | NC State Laboratory of Public Health | Pathogen Discovery, Respiratory Viruses Branch, Division of Viral Diseases, Centers for Disease Control and Prevention | Krista Queen, Yan Li, Anna Uehara, Clinton R. Paden, Rachel Marine, Ying Tao, Jing Zhang, Haibin Wang, Mary S. Keckler, Alison S. Laufer Halpin, Christopher A. Elkins, Suxiang Tong |
| EPI_ISL_426429 | hCoV-19/USA/NV_0016/2020 | North America / USA / Nevada | 2020-03-07 | NV State Public Health Laboratory | Pathogen Discovery, Respiratory Viruses Branch, Division of Viral Diseases, Centers for Disease Control and Prevention | Krista Queen, Yan Li, Anna Uehara, Clinton R. Paden, Rachel Marine, Ying Tao, Jing Zhang, Haibin Wang, Mary S. Keckler, Alison S. Laufer Halpin, Christopher A. Elkins, Suxiang Tong |
| EPI_ISL_426430 | hCoV-19/USA/OH_0019/2020 | North America / USA / Ohio | 2020-03-08 | OH Department of Health Laboratory | Pathogen Discovery, Respiratory Viruses Branch, Division of Viral Diseases, Centers for Disease Control and Prevention | Krista Queen, Yan Li, Anna Uehara, Clinton R. Paden, Rachel Marine, Ying Tao, Jing Zhang, Haibin Wang, Mary S. Keckler, Alison S. Laufer Halpin, Christopher A. Elkins, Suxiang Tong |
| EPI_ISL_426431 | hCoV-19/USA/OH_0020/2020 | North America / USA / Ohio | 2020-03-07 | OH Department of Health Laboratory | Pathogen Discovery, Respiratory Viruses Branch, Division of Viral Diseases, Centers for Disease Control and Prevention | Krista Queen, Yan Li, Anna Uehara, Clinton R. Paden, Rachel Marine, Ying Tao, Jing Zhang, Haibin Wang, Mary S. Keckler, Alison S. Laufer Halpin, Christopher A. Elkins, Suxiang Tong |
| EPI_ISL_426432 | hCoV-19/USA/PA_2734/2020 | North America / USA / Pennsylvania | 2020-03-07 | PA Department of Health, Bureau of Laboratories | Pathogen Discovery, Respiratory Viruses Branch, Division of Viral Diseases, Centers for Disease Control and Prevention | Krista Queen, Yan Li, Anna Uehara, Clinton R. Paden, Rachel Marine, Ying Tao, Jing Zhang, Haibin Wang, Mary S. Keckler, Alison S. Laufer Halpin, Christopher A. Elkins, Suxiang Tong |
| EPI_ISL_426433 | hCoV-19/USA/PA_2735/2020 | North America / USA / Pennsylvania | 2020-03-07 | PA Department of Health, Bureau of Laboratories | Pathogen Discovery, Respiratory Viruses Branch, Division of Viral Diseases, Centers for Disease Control and Prevention | Krista Queen, Yan Li, Anna Uehara, Clinton R. Paden, Rachel Marine, Ying Tao, Jing Zhang, Haibin Wang, Mary S. Keckler, Alison S. Laufer Halpin, Christopher A. Elkins, Suxiang Tong |
| EPI_ISL_426434 | hCoV-19/USA/PA_2887/2020 | North America / USA / Pennsylvania | 2020-03-08 | PA Department of Health, Bureau of Laboratories | Pathogen Discovery, Respiratory Viruses Branch, Division of Viral Diseases, Centers for Disease Control and Prevention | Krista Queen, Yan Li, Anna Uehara, Clinton R. Paden, Rachel Marine, Ying Tao, Jing Zhang, Haibin Wang, Mary S. Keckler, Alison S. Laufer Halpin, Christopher A. Elkins, Suxiang Tong |
| EPI_ISL_426435 | hCoV-19/USA/RI_0882/2020 | North America / USA / Rhode Island | 2020-03-09 | RI State Health Laboratories | Pathogen Discovery, Respiratory Viruses Branch, Division of Viral Diseases, Centers for Disease Control and Prevention | Krista Queen, Yan Li, Anna Uehara, Clinton R. Paden, Rachel Marine, Ying Tao, Jing Zhang, Haibin Wang, Mary S. Keckler, Alison S. Laufer Halpin, Christopher A. Elkins, Suxiang Tong |
| EPI_ISL_426436 | hCoV-19/USA/WA_0418/2020 | North America / USA / Washington | 2020-03-19 | WA State Department of Health | Pathogen Discovery, Respiratory Viruses Branch, Division of Viral Diseases, Centers for Disease Control and Prevention | Jing Zhang, Ying Tao, Clinton R. Paden, Krista Queen, Anna Uehara, Yan Li, Haibin Wang, Jessica Jacobs, Denny Russell, Brian Hiatt, Jessica Gant, Suxiang Tong |
| EPI_ISL_426437 | hCoV-19/USA/WA_0422/2020 | North America / USA / Washington | 2020-03-19 | WA State Department of Health | Pathogen Discovery, Respiratory Viruses Branch, Division of Viral Diseases, Centers for Disease Control and Prevention | Ying Tao, Jing Zhang, Clinton R. Paden, Krista Queen, Anna Uehara, Yan Li, Haibin Wang, Jessica Jacobs, Denny Russell, Brian Hiatt, Jessica Gant, Suxiang Tong |
| EPI_ISL_426438 | hCoV-19/USA/WA_0427/2020 | North America / USA / Washington | 2020-03-19 | WA State Department of Health | Pathogen Discovery, Respiratory Viruses Branch, Division of Viral Diseases, Centers for Disease Control and Prevention | Jing Zhang, Ying Tao, Clinton R. Paden, Krista Queen, Anna Uehara, Yan Li, Haibin Wang, Jessica Jacobs, Denny Russell, Brian Hiatt, Jessica Gant, Suxiang Tong |
| EPI_ISL_426439 | hCoV-19/USA/WA_0428/2020 | North America / USA / Washington | 2020-03-19 | WA State Department of Health | Pathogen Discovery, Respiratory Viruses Branch, Division of Viral Diseases, Centers for Disease Control and Prevention | Jing Zhang, Ying Tao, Clinton R. Paden, Krista Queen, Anna Uehara, Yan Li, Haibin Wang, Jessica Jacobs, Denny Russell, Brian Hiatt, Jessica Gant, Suxiang Tong |
| EPI_ISL_426440 | hCoV-19/USA/WA_0429/2020 | North America / USA / Washington | 2020-03-19 | WA State Department of Health | Pathogen Discovery, Respiratory Viruses Branch, Division of Viral Diseases, Centers for Disease Control and Prevention | Ying Tao, Jing Zhang, Clinton R. Paden, Krista Queen, Anna Uehara, Yan Li, Haibin Wang, Jessica Jacobs, Denny Russell, Brian Hiatt, Jessica Gant, Suxiang Tong |
| EPI_ISL_426441 | hCoV-19/USA/WA_0432/2020 | North America / USA / Washington | 2020-03-19 | WA State Department of Health | Pathogen Discovery, Respiratory Viruses Branch, Division of Viral Diseases, Centers for Disease Control and Prevention | Ying Tao, Jing Zhang, Clinton R. Paden, Krista Queen, Anna Uehara, Yan Li, Haibin Wang, Jessica Jacobs, Denny Russell, Brian Hiatt, Jessica Gant, Suxiang Tong |
| EPI_ISL_426442 | hCoV-19/USA/WA_0434/2020 | North America / USA / Washington | 2020-03-19 | WA State Department of Health | Pathogen Discovery, Respiratory Viruses Branch, Division of Viral Diseases, Centers for Disease Control and Prevention | Jing Zhang, Ying Tao, Clinton R. Paden, Krista Queen, Anna Uehara, Yan Li, Haibin Wang, Jessica Jacobs, Denny Russell, Brian Hiatt, Jessica Gant, Suxiang Tong |
| EPI_ISL_426443 | hCoV-19/USA/WA_0440/2020 | North America / USA / Washington | 2020-03-19 | WA State Department of Health | Pathogen Discovery, Respiratory Viruses Branch, Division of Viral Diseases, Centers for Disease Control and Prevention | Ying Tao, Jing Zhang, Clinton R. Paden, Krista Queen, Anna Uehara, Yan Li, Haibin Wang, Jessica Jacobs, Denny Russell, Brian Hiatt, Jessica Gant, Suxiang Tong |
| EPI_ISL_426444 | hCoV-19/USA/WA_0441/2020 | North America / USA / Washington | 2020-03-19 | WA State Department of Health | Pathogen Discovery, Respiratory Viruses Branch, Division of Viral Diseases, Centers for Disease Control and Prevention | Ying Tao, Jing Zhang, Clinton R. Paden, Krista Queen, Anna Uehara, Yan Li, Haibin Wang, Jessica Jacobs, Denny Russell, Brian Hiatt, Jessica Gant, Suxiang Tong |
| EPI_ISL_426445 | hCoV-19/USA/WA_0442/2020 | North America / USA / Washington | 2020-03-19 | WA State Department of Health | Pathogen Discovery, Respiratory Viruses Branch, Division of Viral Diseases, Centers for Disease Control and Prevention | Jing Zhang, Ying Tao, Clinton R. Paden, Krista Queen, Anna Uehara, Yan Li, Haibin Wang, Jessica Jacobs, Denny Russell, Brian Hiatt, Jessica Gant, Suxiang Tong |
| EPI_ISL_426446 | hCoV-19/USA/WA_0447/2020 | North America / USA / Washington | 2020-03-19 | WA State Department of Health | Pathogen Discovery, Respiratory Viruses Branch, Division of Viral Diseases, Centers for Disease Control and Prevention | Ying Tao, Jing Zhang, Clinton R. Paden, Krista Queen, Anna Uehara, Yan Li, Haibin Wang, Jessica Jacobs, Denny Russell, Brian Hiatt, Jessica Gant, Suxiang Tong |
| EPI_ISL_426447 | hCoV-19/USA/WA_0448/2020 | North America / USA / Washington | 2020-03-19 | WA State Department of Health | Pathogen Discovery, Respiratory Viruses Branch, Division of Viral Diseases, Centers for Disease Control and Prevention | Ying Tao, Jing Zhang, Clinton R. Paden, Krista Queen, Anna Uehara, Yan Li, Haibin Wang, Jessica Jacobs, Denny Russell, Brian Hiatt, Jessica Gant, Suxiang Tong |
| EPI_ISL_426448 | hCoV-19/USA/WA_0457/2020 | North America / USA / Washington | 2020-03-20 | WA State Department of Health | Pathogen Discovery, Respiratory Viruses Branch, Division of Viral Diseases, Centers for Disease Control and Prevention | Ying Tao, Jing Zhang, Clinton R. Paden, Krista Queen, Anna Uehara, Yan Li, Haibin Wang, Jessica Jacobs, Denny Russell, Brian Hiatt, Jessica Gant, Suxiang Tong |
| EPI_ISL_426449 | hCoV-19/USA/WA_0478/2020 | North America / USA / Washington | 2020-03-19 | WA State Department of Health | Pathogen Discovery, Respiratory Viruses Branch, Division of Viral Diseases, Centers for Disease Control and Prevention | Ying Tao, Jing Zhang, Clinton R. Paden, Krista Queen, Anna Uehara, Yan Li, Haibin Wang, Jessica Jacobs, Denny Russell, Brian Hiatt, Jessica Gant, Suxiang Tong |
| EPI_ISL_426450 | hCoV-19/USA/WA_0480/2020 | North America / USA / Washington | 2020-03-19 | WA State Department of Health | Pathogen Discovery, Respiratory Viruses Branch, Division of Viral Diseases, Centers for Disease Control and Prevention | Jing Zhang, Ying Tao, Clinton R. Paden, Krista Queen, Anna Uehara, Yan Li, Haibin Wang, Jessica Jacobs, Denny Russell, Brian Hiatt, Jessica Gant, Suxiang Tong |
| EPI_ISL_426451 | hCoV-19/USA/WA_0484/2020 | North America / USA / Washington | 2020-03-19 | WA State Department of Health | Pathogen Discovery, Respiratory Viruses Branch, Division of Viral Diseases, Centers for Disease Control and Prevention | Ying Tao, Jing Zhang, Clinton R. Paden, Krista Queen, Anna Uehara, Yan Li, Haibin Wang, Jessica Jacobs, Denny Russell, Brian Hiatt, Jessica Gant, Suxiang Tong |
| EPI_ISL_426452 | hCoV-19/USA/WA_0711/2020 | North America / USA / Washington | 2020-03-19 | WA State Department of Health | Pathogen Discovery, Respiratory Viruses Branch, Division of Viral Diseases, Centers for Disease Control and Prevention | Ying Tao, Jing Zhang, Clinton R. Paden, Krista Queen, Anna Uehara, Yan Li, Haibin Wang, Jessica Jacobs, Denny Russell, Brian Hiatt, Jessica Gant, Suxiang Tong |
| EPI_ISL_426453 | hCoV-19/USA/WA_0797/2020 | North America / USA / Washington | 2020-03-20 | WA State Department of Health | Pathogen Discovery, Respiratory Viruses Branch, Division of Viral Diseases, Centers for Disease Control and Prevention | Ying Tao, Jing Zhang, Clinton R. Paden, Krista Queen, Anna Uehara, Yan Li, Haibin Wang, Jessica Jacobs, Denny Russell, Brian Hiatt, Jessica Gant, Suxiang Tong |
| EPI_ISL_426454 | hCoV-19/USA/VA-DCLS-0041/2020 | North America / USA / Virginia | 2020-03-23 | Virginia DCLS | Virginia DCLS | Virginia DCLS |
| EPI_ISL_426455 | hCoV-19/USA/VA-DCLS-0047/2020 | North America / USA / Virginia | 2020-03-27 | Virginia DCLS | Virginia DCLS | Virginia DCLS |
| EPI_ISL_426456 | hCoV-19/USA/VA-DCLS-0048/2020 | North America / USA / Virginia | 2020-03-30 | Virginia DCLS | Virginia DCLS | Virginia DCLS |
| EPI_ISL_426457 | hCoV-19/USA/VA-DCLS-0049/2020 | North America / USA / Virginia | 2020-03-30 | Virginia DCLS | Virginia DCLS | Virginia DCLS |
| EPI_ISL_426458 | hCoV-19/USA/VA-DCLS-0050/2020 | North America / USA / Virginia | 2020-03-30 | Virginia DCLS | Virginia DCLS | Virginia DCLS |
| EPI_ISL_426459 | hCoV-19/USA/VA-DCLS-0051/2020 | North America / USA / Virginia | 2020-03-28 | Virginia DCLS | Virginia DCLS | Virginia DCLS |
| EPI_ISL_426460 | hCoV-19/USA/VA-DCLS-0052/2020 | North America / USA / Virginia | 2020-03-28 | Virginia DCLS | Virginia DCLS | Virginia DCLS |
| EPI_ISL_426461 | hCoV-19/USA/VA-DCLS-0053/2020 | North America / USA / Virginia | 2020-03-27 | Virginia DCLS | Virginia DCLS | Virginia DCLS |
| EPI_ISL_426462 | hCoV-19/USA/VA-DCLS-0054/2020 | North America / USA / Virginia | 2020-03-29 | Virginia DCLS | Virginia DCLS | Virginia DCLS |
| EPI_ISL_426463 | hCoV-19/USA/VA-DCLS-0055/2020 | North America / USA / Virginia | 2020-03-29 | Virginia DCLS | Virginia DCLS | Virginia DCLS |
| EPI_ISL_426464 | hCoV-19/USA/VA-DCLS-0056/2020 | North America / USA / Virginia | 2020-04-01 | Virginia DCLS | Virginia DCLS | Virginia DCLS |
| EPI_ISL_426465 | hCoV-19/USA/VA-DCLS-0057/2020 | North America / USA / Virginia | 2020-03-31 | Virginia DCLS | Virginia DCLS | Virginia DCLS |
| EPI_ISL_426466 | hCoV-19/USA/VA-DCLS-0058/2020 | North America / USA / Virginia | 2020-03-31 | Virginia DCLS | Virginia DCLS | Virginia DCLS |
| EPI_ISL_426467 | hCoV-19/USA/VA-DCLS-0059/2020 | North America / USA / Virginia | 2020-03-31 | Virginia DCLS | Virginia DCLS | Virginia DCLS |
| EPI_ISL_426468 | hCoV-19/USA/VA-DCLS-0060/2020 | North America / USA / Virginia | 2020-03-31 | Virginia DCLS | Virginia DCLS | Virginia DCLS |
| EPI_ISL_426469 | hCoV-19/USA/VA-DCLS-0061/2020 | North America / USA / Virginia | 2020-03-30 | Virginia DCLS | Virginia DCLS | Virginia DCLS |
| EPI_ISL_426470 | hCoV-19/USA/VA-DCLS-0063/2020 | North America / USA / Virginia | 2020-03-31 | Virginia DCLS | Virginia DCLS | Virginia DCLS |
| EPI_ISL_426471 | hCoV-19/USA/VA-DCLS-0064/2020 | North America / USA / Virginia | 2020-04-02 | Virginia DCLS | Virginia DCLS | Virginia DCLS |
| EPI_ISL_426472 | hCoV-19/USA/VA-DCLS-0065/2020 | North America / USA / Virginia | 2020-04-03 | Virginia DCLS | Virginia DCLS | Virginia DCLS |
| EPI_ISL_426473 | hCoV-19/USA/VA-DCLS-0066/2020 | North America / USA / Virginia | 2020-04-03 | Virginia DCLS | Virginia DCLS | Virginia DCLS |
| EPI_ISL_426474 | hCoV-19/USA/VA-DCLS-0067/2020 | North America / USA / Virginia | 2020-04-03 | Virginia DCLS | Virginia DCLS | Virginia DCLS |
| EPI_ISL_426475 | hCoV-19/USA/VA-DCLS-0068/2020 | North America / USA / Virginia | 2020-04-04 | Virginia DCLS | Virginia DCLS | Virginia DCLS |
| EPI_ISL_426482 | hCoV-19/Uruguay/UY-7/2020 | South America / Uruguay / Montevideo | 2020-03-17 | Microbial Genomics Laboratory, Institut Pasteur Montevideo, Uruguay | Microbial Genomics Laboratory, Institut Pasteur Montevideo | Cecilia Salazar, Florencia Díaz-Viraqué, Marianoel Pereira, Pilar Moreno, Gonzalo Moratorio, Gregorio Iraola |
| EPI_ISL_426485 | hCoV-19/USA/AZ-TG271872/2020 | North America / USA / Arizona | 2020-03-24 | AZ SPHL, Arizona Department of Health Services | TGen North | Jolene Bowers, Megan Folkerts, Darrin Lemmer, Dave Engelthaler |
| EPI_ISL_426700 | hCoV-19/Australia/VIC397/2020 | Oceania / Australia / Victoria | 2020-03-24 | Victorian Infectious Diseases Reference Laboratory (VIDRL) | Microbiological Diagnostic Unit Public Health Laboratory and Victorian Infectious Diseases Reference Laboratory, Doherty Institute | Caly L., Seemann T., Sait, M., Schultz M., Druce J., Sherry, N. |
| EPI_ISL_426701 | hCoV-19/Australia/VIC398/2020 | Oceania / Australia / Victoria | 2020-03-24 | Victorian Infectious Diseases Reference Laboratory (VIDRL) | Microbiological Diagnostic Unit Public Health Laboratory and Victorian Infectious Diseases Reference Laboratory, Doherty Institute | Caly L., Seemann T., Sait, M., Schultz M., Druce J., Sherry, N. |
| EPI_ISL_426702 | hCoV-19/Australia/VIC399/2020 | Oceania / Australia / Victoria | 2020-03-24 | Victorian Infectious Diseases Reference Laboratory (VIDRL) | Microbiological Diagnostic Unit Public Health Laboratory and Victorian Infectious Diseases Reference Laboratory, Doherty Institute | Caly L., Seemann T., Sait, M., Schultz M., Druce J., Sherry, N. |
| EPI_ISL_426703 | hCoV-19/Australia/VIC400/2020 | Oceania / Australia / Victoria | 2020-03-24 | Victorian Infectious Diseases Reference Laboratory (VIDRL) | Microbiological Diagnostic Unit Public Health Laboratory and Victorian Infectious Diseases Reference Laboratory, Doherty Institute | Caly L., Seemann T., Sait, M., Schultz M., Druce J., Sherry, N. |
| EPI_ISL_426704 | hCoV-19/Australia/VIC401/2020 | Oceania / Australia / Victoria | 2020-03-24 | Victorian Infectious Diseases Reference Laboratory (VIDRL) | Microbiological Diagnostic Unit Public Health Laboratory and Victorian Infectious Diseases Reference Laboratory, Doherty Institute | Caly L., Seemann T., Sait, M., Schultz M., Druce J., Sherry, N. |
| EPI_ISL_426705 | hCoV-19/Australia/VIC402/2020 | Oceania / Australia / Victoria | 2020-03-24 | Victorian Infectious Diseases Reference Laboratory (VIDRL) | Microbiological Diagnostic Unit Public Health Laboratory and Victorian Infectious Diseases Reference Laboratory, Doherty Institute | Caly L., Seemann T., Sait, M., Schultz M., Druce J., Sherry, N. |
| EPI_ISL_426706 | hCoV-19/Australia/VIC403/2020 | Oceania / Australia / Victoria | 2020-03-22 | Victorian Infectious Diseases Reference Laboratory (VIDRL) | Microbiological Diagnostic Unit Public Health Laboratory and Victorian Infectious Diseases Reference Laboratory, Doherty Institute | Caly L., Seemann T., Sait, M., Schultz M., Druce J., Sherry, N. |
| EPI_ISL_426707 | hCoV-19/Australia/VIC404/2020 | Oceania / Australia / Victoria | 2020-03-24 | Victorian Infectious Diseases Reference Laboratory (VIDRL) | Microbiological Diagnostic Unit Public Health Laboratory and Victorian Infectious Diseases Reference Laboratory, Doherty Institute | Caly L., Seemann T., Sait, M., Schultz M., Druce J., Sherry, N. |
| EPI_ISL_426709 | hCoV-19/Australia/VIC407/2020 | Oceania / Australia / Victoria | 2020-03-25 | Victorian Infectious Diseases Reference Laboratory (VIDRL) | Microbiological Diagnostic Unit Public Health Laboratory and Victorian Infectious Diseases Reference Laboratory, Doherty Institute | Caly L., Seemann T., Sait, M., Schultz M., Druce J., Sherry, N. |
| EPI_ISL_426711 | hCoV-19/Australia/VIC409/2020 | Oceania / Australia / Victoria | 2020-03-26 | Victorian Infectious Diseases Reference Laboratory (VIDRL) | Microbiological Diagnostic Unit Public Health Laboratory and Victorian Infectious Diseases Reference Laboratory, Doherty Institute | Caly L., Seemann T., Sait, M., Schultz M., Druce J., Sherry, N. |
| EPI_ISL_426714 | hCoV-19/Australia/VIC414/2020 | Oceania / Australia / Victoria | 2020-03-26 | Victorian Infectious Diseases Reference Laboratory (VIDRL) | Microbiological Diagnostic Unit Public Health Laboratory and Victorian Infectious Diseases Reference Laboratory, Doherty Institute | Caly L., Seemann T., Sait, M., Schultz M., Druce J., Sherry, N. |
| EPI_ISL_426715 | hCoV-19/Australia/VIC415/2020 | Oceania / Australia / Victoria | 2020-03-25 | Victorian Infectious Diseases Reference Laboratory (VIDRL) | Microbiological Diagnostic Unit Public Health Laboratory and Victorian Infectious Diseases Reference Laboratory, Doherty Institute | Caly L., Seemann T., Sait, M., Schultz M., Druce J., Sherry, N. |
| EPI_ISL_426717 | hCoV-19/Australia/VIC417/2020 | Oceania / Australia / Victoria | 2020-03-25 | Victorian Infectious Diseases Reference Laboratory (VIDRL) | Microbiological Diagnostic Unit Public Health Laboratory and Victorian Infectious Diseases Reference Laboratory, Doherty Institute | Caly L., Seemann T., Sait, M., Schultz M., Druce J., Sherry, N. |
| EPI_ISL_426718 | hCoV-19/Australia/VIC418/2020 | Oceania / Australia / Victoria | 2020-03-25 | Victorian Infectious Diseases Reference Laboratory (VIDRL) | Microbiological Diagnostic Unit Public Health Laboratory and Victorian Infectious Diseases Reference Laboratory, Doherty Institute | Caly L., Seemann T., Sait, M., Schultz M., Druce J., Sherry, N. |
| EPI_ISL_426720 | hCoV-19/Australia/VIC420/2020 | Oceania / Australia / Victoria | 2020-03-25 | Victorian Infectious Diseases Reference Laboratory (VIDRL) | Microbiological Diagnostic Unit Public Health Laboratory and Victorian Infectious Diseases Reference Laboratory, Doherty Institute | Caly L., Seemann T., Sait, M., Schultz M., Druce J., Sherry, N. |
| EPI_ISL_426721 | hCoV-19/Australia/VIC421/2020 | Oceania / Australia / Victoria | 2020-03-25 | Victorian Infectious Diseases Reference Laboratory (VIDRL) | Microbiological Diagnostic Unit Public Health Laboratory and Victorian Infectious Diseases Reference Laboratory, Doherty Institute | Caly L., Seemann T., Sait, M., Schultz M., Druce J., Sherry, N. |
| EPI_ISL_426725 | hCoV-19/Australia/VIC425/2020 | Oceania / Australia / Victoria | 2020-03-25 | Victorian Infectious Diseases Reference Laboratory (VIDRL) | Microbiological Diagnostic Unit Public Health Laboratory and Victorian Infectious Diseases Reference Laboratory, Doherty Institute | Caly L., Seemann T., Sait, M., Schultz M., Druce J., Sherry, N. |
| EPI_ISL_426727 | hCoV-19/Australia/VIC427/2020 | Oceania / Australia / Victoria | 2020-03-25 | Victorian Infectious Diseases Reference Laboratory (VIDRL) | Microbiological Diagnostic Unit Public Health Laboratory and Victorian Infectious Diseases Reference Laboratory, Doherty Institute | Caly L., Seemann T., Sait, M., Schultz M., Druce J., Sherry, N. |
| EPI_ISL_426729 | hCoV-19/Australia/VIC429/2020 | Oceania / Australia / Victoria | 2020-03-26 | Victorian Infectious Diseases Reference Laboratory (VIDRL) | Microbiological Diagnostic Unit Public Health Laboratory and Victorian Infectious Diseases Reference Laboratory, Doherty Institute | Caly L., Seemann T., Sait, M., Schultz M., Druce J., Sherry, N. |
| EPI_ISL_426730 | hCoV-19/Australia/VIC430/2020 | Oceania / Australia / Victoria | 2020-03-26 | Victorian Infectious Diseases Reference Laboratory (VIDRL) | Microbiological Diagnostic Unit Public Health Laboratory and Victorian Infectious Diseases Reference Laboratory, Doherty Institute | Caly L., Seemann T., Sait, M., Schultz M., Druce J., Sherry, N. |
| EPI_ISL_426732 | hCoV-19/Australia/VIC433/2020 | Oceania / Australia / Victoria | 2020-03-25 | Victorian Infectious Diseases Reference Laboratory (VIDRL) | Microbiological Diagnostic Unit Public Health Laboratory and Victorian Infectious Diseases Reference Laboratory, Doherty Institute | Caly L., Seemann T., Sait, M., Schultz M., Druce J., Sherry, N. |
| EPI_ISL_426733 | hCoV-19/Australia/VIC434/2020 | Oceania / Australia / Victoria | 2020-03-25 | Victorian Infectious Diseases Reference Laboratory (VIDRL) | Microbiological Diagnostic Unit Public Health Laboratory and Victorian Infectious Diseases Reference Laboratory, Doherty Institute | Caly L., Seemann T., Sait, M., Schultz M., Druce J., Sherry, N. |
| EPI_ISL_426734 | hCoV-19/Australia/VIC437/2020 | Oceania / Australia / Victoria | 2020-03-26 | Victorian Infectious Diseases Reference Laboratory (VIDRL) | Microbiological Diagnostic Unit Public Health Laboratory and Victorian Infectious Diseases Reference Laboratory, Doherty Institute | Caly L., Seemann T., Sait, M., Schultz M., Druce J., Sherry, N. |
| EPI_ISL_426735 | hCoV-19/Australia/VIC438/2020 | Oceania / Australia / Victoria | 2020-03-26 | Victorian Infectious Diseases Reference Laboratory (VIDRL) | Microbiological Diagnostic Unit Public Health Laboratory and Victorian Infectious Diseases Reference Laboratory, Doherty Institute | Caly L., Seemann T., Sait, M., Schultz M., Druce J., Sherry, N. |
| EPI_ISL_426736 | hCoV-19/Australia/VIC439/2020 | Oceania / Australia / Victoria | 2020-03-26 | Victorian Infectious Diseases Reference Laboratory (VIDRL) | Microbiological Diagnostic Unit Public Health Laboratory and Victorian Infectious Diseases Reference Laboratory, Doherty Institute | Caly L., Seemann T., Sait, M., Schultz M., Druce J., Sherry, N. |
| EPI_ISL_426737 | hCoV-19/Australia/VIC440/2020 | Oceania / Australia / Victoria | 2020-03-26 | Victorian Infectious Diseases Reference Laboratory (VIDRL) | Microbiological Diagnostic Unit Public Health Laboratory and Victorian Infectious Diseases Reference Laboratory, Doherty Institute | Caly L., Seemann T., Sait, M., Schultz M., Druce J., Sherry, N. |
| EPI_ISL_426738 | hCoV-19/Australia/VIC441/2020 | Oceania / Australia / Victoria | 2020-03-27 | Victorian Infectious Diseases Reference Laboratory (VIDRL) | Microbiological Diagnostic Unit Public Health Laboratory and Victorian Infectious Diseases Reference Laboratory, Doherty Institute | Caly L., Seemann T., Sait, M., Schultz M., Druce J., Sherry, N. |
| EPI_ISL_426740 | hCoV-19/Australia/VIC443/2020 | Oceania / Australia / Victoria | 2020-03-27 | Victorian Infectious Diseases Reference Laboratory (VIDRL) | Microbiological Diagnostic Unit Public Health Laboratory and Victorian Infectious Diseases Reference Laboratory, Doherty Institute | Caly L., Seemann T., Sait, M., Schultz M., Druce J., Sherry, N. |
| EPI_ISL_426741 | hCoV-19/Australia/VIC444/2020 | Oceania / Australia / Victoria | 2020-03-27 | Victorian Infectious Diseases Reference Laboratory (VIDRL) | Microbiological Diagnostic Unit Public Health Laboratory and Victorian Infectious Diseases Reference Laboratory, Doherty Institute | Caly L., Seemann T., Sait, M., Schultz M., Druce J., Sherry, N. |
| EPI_ISL_426742 | hCoV-19/Australia/VIC446/2020 | Oceania / Australia / Victoria | 2020-03-27 | Victorian Infectious Diseases Reference Laboratory (VIDRL) | Microbiological Diagnostic Unit Public Health Laboratory and Victorian Infectious Diseases Reference Laboratory, Doherty Institute | Caly L., Seemann T., Sait, M., Schultz M., Druce J., Sherry, N. |
| EPI_ISL_426743 | hCoV-19/Australia/VIC447/2020 | Oceania / Australia / Victoria | 2020-03-27 | Victorian Infectious Diseases Reference Laboratory (VIDRL) | Microbiological Diagnostic Unit Public Health Laboratory and Victorian Infectious Diseases Reference Laboratory, Doherty Institute | Caly L., Seemann T., Sait, M., Schultz M., Druce J., Sherry, N. |
| EPI_ISL_426745 | hCoV-19/Australia/VIC449/2020 | Oceania / Australia / Victoria | 2020-03-24 | Victorian Infectious Diseases Reference Laboratory (VIDRL) | Microbiological Diagnostic Unit Public Health Laboratory and Victorian Infectious Diseases Reference Laboratory, Doherty Institute | Caly L., Seemann T., Sait, M., Schultz M., Druce J., Sherry, N. |
| EPI_ISL_426746 | hCoV-19/Australia/VIC450/2020 | Oceania / Australia / Victoria | 2020-03-25 | Victorian Infectious Diseases Reference Laboratory (VIDRL) | Microbiological Diagnostic Unit Public Health Laboratory and Victorian Infectious Diseases Reference Laboratory, Doherty Institute | Caly L., Seemann T., Sait, M., Schultz M., Druce J., Sherry, N. |
| EPI_ISL_426747 | hCoV-19/Australia/VIC451/2020 | Oceania / Australia / Victoria | 2020-03-25 | Victorian Infectious Diseases Reference Laboratory (VIDRL) | Microbiological Diagnostic Unit Public Health Laboratory and Victorian Infectious Diseases Reference Laboratory, Doherty Institute | Caly L., Seemann T., Sait, M., Schultz M., Druce J., Sherry, N. |
| EPI_ISL_426751 | hCoV-19/Australia/VIC456/2020 | Oceania / Australia / Victoria | 2020-03-25 | Victorian Infectious Diseases Reference Laboratory (VIDRL) | Microbiological Diagnostic Unit Public Health Laboratory and Victorian Infectious Diseases Reference Laboratory, Doherty Institute | Caly L., Seemann T., Sait, M., Schultz M., Druce J., Sherry, N. |
| EPI_ISL_426752 | hCoV-19/Australia/VIC458/2020 | Oceania / Australia / Victoria | 2020-03-26 | Victorian Infectious Diseases Reference Laboratory (VIDRL) | Microbiological Diagnostic Unit Public Health Laboratory and Victorian Infectious Diseases Reference Laboratory, Doherty Institute | Caly L., Seemann T., Sait, M., Schultz M., Druce J., Sherry, N. |
| EPI_ISL_426754 | hCoV-19/Australia/VIC461/2020 | Oceania / Australia / Victoria | 2020-03-27 | Victorian Infectious Diseases Reference Laboratory (VIDRL) | Microbiological Diagnostic Unit Public Health Laboratory and Victorian Infectious Diseases Reference Laboratory, Doherty Institute | Caly L., Seemann T., Sait, M., Schultz M., Druce J., Sherry, N. |
| EPI_ISL_426755 | hCoV-19/Australia/VIC462/2020 | Oceania / Australia / Victoria | 2020-03-27 | Victorian Infectious Diseases Reference Laboratory (VIDRL) | Microbiological Diagnostic Unit Public Health Laboratory and Victorian Infectious Diseases Reference Laboratory, Doherty Institute | Caly L., Seemann T., Sait, M., Schultz M., Druce J., Sherry, N. |
| EPI_ISL_426756 | hCoV-19/Australia/VIC463/2020 | Oceania / Australia / Victoria | 2020-03-26 | Victorian Infectious Diseases Reference Laboratory (VIDRL) | Microbiological Diagnostic Unit Public Health Laboratory and Victorian Infectious Diseases Reference Laboratory, Doherty Institute | Caly L., Seemann T., Sait, M., Schultz M., Druce J., Sherry, N. |
| EPI_ISL_426761 | hCoV-19/Australia/VIC468/2020 | Oceania / Australia / Victoria | 2020-03-27 | Victorian Infectious Diseases Reference Laboratory (VIDRL) | Microbiological Diagnostic Unit Public Health Laboratory and Victorian Infectious Diseases Reference Laboratory, Doherty Institute | Caly L., Seemann T., Sait, M., Schultz M., Druce J., Sherry, N. |
| EPI_ISL_426763 | hCoV-19/Australia/VIC471/2020 | Oceania / Australia / Victoria | 2020-03-26 | Victorian Infectious Diseases Reference Laboratory (VIDRL) | Microbiological Diagnostic Unit Public Health Laboratory and Victorian Infectious Diseases Reference Laboratory, Doherty Institute | Caly L., Seemann T., Sait, M., Schultz M., Druce J., Sherry, N. |
| EPI_ISL_426764 | hCoV-19/Australia/VIC472/2020 | Oceania / Australia / Victoria | 2020-03-26 | Victorian Infectious Diseases Reference Laboratory (VIDRL) | Microbiological Diagnostic Unit Public Health Laboratory and Victorian Infectious Diseases Reference Laboratory, Doherty Institute | Caly L., Seemann T., Sait, M., Schultz M., Druce J., Sherry, N. |
| EPI_ISL_426765 | hCoV-19/Australia/VIC492/2020 | Oceania / Australia / Victoria | 2020-03-23 | Victorian Infectious Diseases Reference Laboratory (VIDRL) | Microbiological Diagnostic Unit Public Health Laboratory and Victorian Infectious Diseases Reference Laboratory, Doherty Institute | Caly L., Seemann T., Sait, M., Schultz M., Druce J., Sherry, N. |
| EPI_ISL_426766 | hCoV-19/Australia/VIC493/2020 | Oceania / Australia / Victoria | 2020-03-22 | Victorian Infectious Diseases Reference Laboratory (VIDRL) | Microbiological Diagnostic Unit Public Health Laboratory and Victorian Infectious Diseases Reference Laboratory, Doherty Institute | Caly L., Seemann T., Sait, M., Schultz M., Druce J., Sherry, N. |
| EPI_ISL_426767 | hCoV-19/Australia/VIC494/2020 | Oceania / Australia / Victoria | 2020-03-23 | Victorian Infectious Diseases Reference Laboratory (VIDRL) | Microbiological Diagnostic Unit Public Health Laboratory and Victorian Infectious Diseases Reference Laboratory, Doherty Institute | Caly L., Seemann T., Sait, M., Schultz M., Druce J., Sherry, N. |
| EPI_ISL_426768 | hCoV-19/Australia/VIC495/2020 | Oceania / Australia / Victoria | 2020-03-23 | Victorian Infectious Diseases Reference Laboratory (VIDRL) | Microbiological Diagnostic Unit Public Health Laboratory and Victorian Infectious Diseases Reference Laboratory, Doherty Institute | Caly L., Seemann T., Sait, M., Schultz M., Druce J., Sherry, N. |
| EPI_ISL_426769 | hCoV-19/Australia/VIC496/2020 | Oceania / Australia / Victoria | 2020-03-23 | Victorian Infectious Diseases Reference Laboratory (VIDRL) | Microbiological Diagnostic Unit Public Health Laboratory and Victorian Infectious Diseases Reference Laboratory, Doherty Institute | Caly L., Seemann T., Sait, M., Schultz M., Druce J., Sherry, N. |
| EPI_ISL_426770 | hCoV-19/Australia/VIC497/2020 | Oceania / Australia / Victoria | 2020-03-23 | Victorian Infectious Diseases Reference Laboratory (VIDRL) | Microbiological Diagnostic Unit Public Health Laboratory and Victorian Infectious Diseases Reference Laboratory, Doherty Institute | Caly L., Seemann T., Sait, M., Schultz M., Druce J., Sherry, N. |
| EPI_ISL_426771 | hCoV-19/Australia/VIC498/2020 | Oceania / Australia / Victoria | 2020-03-23 | Victorian Infectious Diseases Reference Laboratory (VIDRL) | Microbiological Diagnostic Unit Public Health Laboratory and Victorian Infectious Diseases Reference Laboratory, Doherty Institute | Caly L., Seemann T., Sait, M., Schultz M., Druce J., Sherry, N. |
| EPI_ISL_426772 | hCoV-19/Australia/VIC500/2020 | Oceania / Australia / Victoria | 2020-03-24 | Victorian Infectious Diseases Reference Laboratory (VIDRL) | Microbiological Diagnostic Unit Public Health Laboratory and Victorian Infectious Diseases Reference Laboratory, Doherty Institute | Caly L., Seemann T., Sait, M., Schultz M., Druce J., Sherry, N. |
| EPI_ISL_426774 | hCoV-19/Australia/VIC503/2020 | Oceania / Australia / Victoria | 2020-03-23 | Victorian Infectious Diseases Reference Laboratory (VIDRL) | Microbiological Diagnostic Unit Public Health Laboratory and Victorian Infectious Diseases Reference Laboratory, Doherty Institute | Caly L., Seemann T., Sait, M., Schultz M., Druce J., Sherry, N. |
| EPI_ISL_426775 | hCoV-19/Australia/VIC504/2020 | Oceania / Australia / Victoria | 2020-03-23 | Victorian Infectious Diseases Reference Laboratory (VIDRL) | Microbiological Diagnostic Unit Public Health Laboratory and Victorian Infectious Diseases Reference Laboratory, Doherty Institute | Caly L., Seemann T., Sait, M., Schultz M., Druce J., Sherry, N. |
| EPI_ISL_426776 | hCoV-19/Australia/VIC505/2020 | Oceania / Australia / Victoria | 2020-03-22 | Victorian Infectious Diseases Reference Laboratory (VIDRL) | Microbiological Diagnostic Unit Public Health Laboratory and Victorian Infectious Diseases Reference Laboratory, Doherty Institute | Caly L., Seemann T., Sait, M., Schultz M., Druce J., Sherry, N. |
| EPI_ISL_426777 | hCoV-19/Australia/VIC506/2020 | Oceania / Australia / Victoria | 2020-03-23 | Victorian Infectious Diseases Reference Laboratory (VIDRL) | Microbiological Diagnostic Unit Public Health Laboratory and Victorian Infectious Diseases Reference Laboratory, Doherty Institute | Caly L., Seemann T., Sait, M., Schultz M., Druce J., Sherry, N. |
| EPI_ISL_426779 | hCoV-19/Australia/VIC509/2020 | Oceania / Australia / Victoria | 2020-03-22 | Victorian Infectious Diseases Reference Laboratory (VIDRL) | Microbiological Diagnostic Unit Public Health Laboratory and Victorian Infectious Diseases Reference Laboratory, Doherty Institute | Caly L., Seemann T., Sait, M., Schultz M., Druce J., Sherry, N. |
| EPI_ISL_426781 | hCoV-19/Australia/VIC511/2020 | Oceania / Australia / Victoria | 2020-03-23 | Victorian Infectious Diseases Reference Laboratory (VIDRL) | Microbiological Diagnostic Unit Public Health Laboratory and Victorian Infectious Diseases Reference Laboratory, Doherty Institute | Caly L., Seemann T., Sait, M., Schultz M., Druce J., Sherry, N. |
| EPI_ISL_426786 | hCoV-19/Australia/VIC516/2020 | Oceania / Australia / Victoria | 2020-03-23 | Victorian Infectious Diseases Reference Laboratory (VIDRL) | Microbiological Diagnostic Unit Public Health Laboratory and Victorian Infectious Diseases Reference Laboratory, Doherty Institute | Caly L., Seemann T., Sait, M., Schultz M., Druce J., Sherry, N. |
| EPI_ISL_426788 | hCoV-19/Australia/VIC518/2020 | Oceania / Australia / Victoria | 2020-03-23 | Victorian Infectious Diseases Reference Laboratory (VIDRL) | Microbiological Diagnostic Unit Public Health Laboratory and Victorian Infectious Diseases Reference Laboratory, Doherty Institute | Caly L., Seemann T., Sait, M., Schultz M., Druce J., Sherry, N. |
| EPI_ISL_426789 | hCoV-19/Australia/VIC520/2020 | Oceania / Australia / Victoria | 2020-03-23 | Victorian Infectious Diseases Reference Laboratory (VIDRL) | Microbiological Diagnostic Unit Public Health Laboratory and Victorian Infectious Diseases Reference Laboratory, Doherty Institute | Caly L., Seemann T., Sait, M., Schultz M., Druce J., Sherry, N. |
| EPI_ISL_426790 | hCoV-19/Australia/VIC522/2020 | Oceania / Australia / Victoria | 2020-03-23 | Victorian Infectious Diseases Reference Laboratory (VIDRL) | Microbiological Diagnostic Unit Public Health Laboratory and Victorian Infectious Diseases Reference Laboratory, Doherty Institute | Caly L., Seemann T., Sait, M., Schultz M., Druce J., Sherry, N. |
| EPI_ISL_426792 | hCoV-19/Australia/VIC525/2020 | Oceania / Australia / Victoria | 2020-03-24 | Victorian Infectious Diseases Reference Laboratory (VIDRL) | Microbiological Diagnostic Unit Public Health Laboratory and Victorian Infectious Diseases Reference Laboratory, Doherty Institute | Caly L., Seemann T., Sait, M., Schultz M., Druce J., Sherry, N. |
| EPI_ISL_426793 | hCoV-19/Australia/VIC527/2020 | Oceania / Australia / Victoria | 2020-03-24 | Victorian Infectious Diseases Reference Laboratory (VIDRL) | Microbiological Diagnostic Unit Public Health Laboratory and Victorian Infectious Diseases Reference Laboratory, Doherty Institute | Caly L., Seemann T., Sait, M., Schultz M., Druce J., Sherry, N. |
| EPI_ISL_426796 | hCoV-19/Australia/VIC530/2020 | Oceania / Australia / Victoria | 2020-03-23 | Victorian Infectious Diseases Reference Laboratory (VIDRL) | Microbiological Diagnostic Unit Public Health Laboratory and Victorian Infectious Diseases Reference Laboratory, Doherty Institute | Caly L., Seemann T., Sait, M., Schultz M., Druce J., Sherry, N. |
| EPI_ISL_426797 | hCoV-19/Australia/VIC531/2020 | Oceania / Australia / Victoria | 2020-03-24 | Victorian Infectious Diseases Reference Laboratory (VIDRL) | Microbiological Diagnostic Unit Public Health Laboratory and Victorian Infectious Diseases Reference Laboratory, Doherty Institute | Caly L., Seemann T., Sait, M., Schultz M., Druce J., Sherry, N. |
| EPI_ISL_426798 | hCoV-19/Australia/VIC532/2020 | Oceania / Australia / Victoria | 2020-03-23 | Victorian Infectious Diseases Reference Laboratory (VIDRL) | Microbiological Diagnostic Unit Public Health Laboratory and Victorian Infectious Diseases Reference Laboratory, Doherty Institute | Caly L., Seemann T., Sait, M., Schultz M., Druce J., Sherry, N. |
| EPI_ISL_426799 | hCoV-19/Australia/VIC533/2020 | Oceania / Australia / Victoria | 2020-03-24 | Victorian Infectious Diseases Reference Laboratory (VIDRL) | Microbiological Diagnostic Unit Public Health Laboratory and Victorian Infectious Diseases Reference Laboratory, Doherty Institute | Caly L., Seemann T., Sait, M., Schultz M., Druce J., Sherry, N. |
| EPI_ISL_427054 | hCoV-19/Australia/VIC779/2020 | Oceania / Australia / Victoria | 2020-03-30 | Microbiological Diagnostic Unit Public Health Laboratory | Microbiological Diagnostic Unit Public Health Laboratory | Seemann T., Schultz M., Sait, M., Sherry, N. |
| EPI_ISL_427055 | hCoV-19/Australia/VIC780/2020 | Oceania / Australia / Victoria | 2020-03-30 | Microbiological Diagnostic Unit Public Health Laboratory | Microbiological Diagnostic Unit Public Health Laboratory | Seemann T., Schultz M., Sait, M., Sherry, N. |
| EPI_ISL_427057 | hCoV-19/Australia/VIC782/2020 | Oceania / Australia / Victoria | 2020-03-30 | Microbiological Diagnostic Unit Public Health Laboratory | Microbiological Diagnostic Unit Public Health Laboratory | Seemann T., Schultz M., Sait, M., Sherry, N. |
| EPI_ISL_427070 | hCoV-19/Australia/VIC796/2020 | Oceania / Australia / Victoria | 2020-03-31 | Microbiological Diagnostic Unit Public Health Laboratory | Microbiological Diagnostic Unit Public Health Laboratory | Seemann T., Schultz M., Sait, M., Sherry, N. |
| EPI_ISL_427071 | hCoV-19/Australia/VIC797/2020 | Oceania / Australia / Victoria | 2020-04-01 | Microbiological Diagnostic Unit Public Health Laboratory | Microbiological Diagnostic Unit Public Health Laboratory | Seemann T., Schultz M., Sait, M., Sherry, N. |
| EPI_ISL_427072 | hCoV-19/Australia/VIC798/2020 | Oceania / Australia / Victoria | 2020-04-01 | Microbiological Diagnostic Unit Public Health Laboratory | Microbiological Diagnostic Unit Public Health Laboratory | Seemann T., Schultz M., Sait, M., Sherry, N. |
| EPI_ISL_427073 | hCoV-19/Australia/VIC799/2020 | Oceania / Australia / Victoria | 2020-04-01 | Microbiological Diagnostic Unit Public Health Laboratory | Microbiological Diagnostic Unit Public Health Laboratory | Seemann T., Schultz M., Sait, M., Sherry, N. |
| EPI_ISL_427074 | hCoV-19/Australia/VIC800/2020 | Oceania / Australia / Victoria | 2020-04-01 | Microbiological Diagnostic Unit Public Health Laboratory | Microbiological Diagnostic Unit Public Health Laboratory | Seemann T., Schultz M., Sait, M., Sherry, N. |
| EPI_ISL_427076 | hCoV-19/Australia/VIC802/2020 | Oceania / Australia / Victoria | 2020-04-01 | Microbiological Diagnostic Unit Public Health Laboratory | Microbiological Diagnostic Unit Public Health Laboratory | Seemann T., Schultz M., Sait, M., Sherry, N. |
| EPI_ISL_427079 | hCoV-19/Australia/VIC805/2020 | Oceania / Australia / Victoria | 2020-03-16 | Victorian Infectious Diseases Reference Laboratory (VIDRL) | Microbiological Diagnostic Unit Public Health Laboratory and Victorian Infectious Diseases Reference Laboratory, Doherty Institute | Caly L., Seemann T., Sait, M., Schultz M., Druce J., Sherry, N. |
| EPI_ISL_427080 | hCoV-19/Australia/VIC806/2020 | Oceania / Australia / Victoria | 2020-03-23 | Victorian Infectious Diseases Reference Laboratory (VIDRL) | Microbiological Diagnostic Unit Public Health Laboratory and Victorian Infectious Diseases Reference Laboratory, Doherty Institute | Caly L., Seemann T., Sait, M., Schultz M., Druce J., Sherry, N. |
| EPI_ISL_427081 | hCoV-19/Australia/VIC807/2020 | Oceania / Australia / Victoria | 2020-03-24 | Victorian Infectious Diseases Reference Laboratory (VIDRL) | Microbiological Diagnostic Unit Public Health Laboratory and Victorian Infectious Diseases Reference Laboratory, Doherty Institute | Caly L., Seemann T., Sait, M., Schultz M., Druce J., Sherry, N. |
| EPI_ISL_427082 | hCoV-19/Australia/VIC808/2020 | Oceania / Australia / Victoria | 2020-03-28 | Victorian Infectious Diseases Reference Laboratory (VIDRL) | Microbiological Diagnostic Unit Public Health Laboratory and Victorian Infectious Diseases Reference Laboratory, Doherty Institute | Caly L., Seemann T., Sait, M., Schultz M., Druce J., Sherry, N. |
| EPI_ISL_427085 | hCoV-19/Australia/VIC812/2020 | Oceania / Australia / Victoria | 2020-04-02 | Victorian Infectious Diseases Reference Laboratory (VIDRL) | Microbiological Diagnostic Unit Public Health Laboratory and Victorian Infectious Diseases Reference Laboratory, Doherty Institute | Caly L., Seemann T., Sait, M., Schultz M., Druce J., Sherry, N. |
| EPI_ISL_427200 | hCoV-19/USA/WA-UW-3881/2020 | North America / USA / Washington | 2020-03-25 | UW Virology Lab | UW Virology Lab | Pavitra Roychoudhury, Hong Xie, Keith Jerome, Alexander Greninger |
| EPI_ISL_427201 | hCoV-19/USA/WA-UW-3883/2020 | North America / USA / Washington | 2020-03-27 | UW Virology Lab | UW Virology Lab | Pavitra Roychoudhury, Hong Xie, Keith Jerome, Alexander Greninger |
| EPI_ISL_427202 | hCoV-19/USA/WA-UW-3889/2020 | North America / USA / Washington | 2020-03-27 | UW Virology Lab | UW Virology Lab | Pavitra Roychoudhury, Hong Xie, Keith Jerome, Alexander Greninger |
| EPI_ISL_427203 | hCoV-19/USA/WA-UW-3895/2020 | North America / USA / Washington | 2020-03-27 | UW Virology Lab | UW Virology Lab | Pavitra Roychoudhury, Hong Xie, Keith Jerome, Alexander Greninger |
| EPI_ISL_427204 | hCoV-19/USA/WA-UW-3900/2020 | North America / USA / Washington | 2020-03-27 | UW Virology Lab | UW Virology Lab | Pavitra Roychoudhury, Hong Xie, Keith Jerome, Alexander Greninger |
| EPI_ISL_427205 | hCoV-19/USA/WA-UW-3903/2020 | North America / USA / Washington | 2020-03-27 | UW Virology Lab | UW Virology Lab | Pavitra Roychoudhury, Hong Xie, Keith Jerome, Alexander Greninger |
| EPI_ISL_427206 | hCoV-19/USA/WA-UW-3908/2020 | North America / USA / Washington | 2020-03-27 | UW Virology Lab | UW Virology Lab | Pavitra Roychoudhury, Hong Xie, Keith Jerome, Alexander Greninger |
| EPI_ISL_427207 | hCoV-19/USA/WA-UW-3911/2020 | North America / USA / Washington | 2020-03-27 | UW Virology Lab | UW Virology Lab | Pavitra Roychoudhury, Hong Xie, Keith Jerome, Alexander Greninger |
| EPI_ISL_427208 | hCoV-19/USA/WA-UW-3917/2020 | North America / USA / Washington | 2020-03-27 | UW Virology Lab | UW Virology Lab | Pavitra Roychoudhury, Hong Xie, Keith Jerome, Alexander Greninger |
| EPI_ISL_427212 | hCoV-19/USA/WA-UW-3935/2020 | North America / USA / Washington | 2020-03-27 | UW Virology Lab | UW Virology Lab | Pavitra Roychoudhury, Hong Xie, Keith Jerome, Alexander Greninger |
| EPI_ISL_427213 | hCoV-19/USA/WA-UW-3939/2020 | North America / USA / Washington | 2020-03-27 | UW Virology Lab | UW Virology Lab | Pavitra Roychoudhury, Hong Xie, Keith Jerome, Alexander Greninger |
| EPI_ISL_427214 | hCoV-19/USA/WA-UW-3940/2020 | North America / USA / Washington | 2020-03-27 | UW Virology Lab | UW Virology Lab | Pavitra Roychoudhury, Hong Xie, Keith Jerome, Alexander Greninger |
| EPI_ISL_427215 | hCoV-19/USA/WA-UW-3941/2020 | North America / USA / Washington | 2020-03-27 | UW Virology Lab | UW Virology Lab | Pavitra Roychoudhury, Hong Xie, Keith Jerome, Alexander Greninger |
| EPI_ISL_427216 | hCoV-19/USA/WA-UW-3942/2020 | North America / USA / Washington | 2020-03-27 | UW Virology Lab | UW Virology Lab | Pavitra Roychoudhury, Hong Xie, Keith Jerome, Alexander Greninger |
| EPI_ISL_427217 | hCoV-19/USA/WA-UW-3946/2020 | North America / USA / Washington | 2020-03-27 | UW Virology Lab | UW Virology Lab | Pavitra Roychoudhury, Hong Xie, Keith Jerome, Alexander Greninger |
| EPI_ISL_427218 | hCoV-19/USA/WA-UW-3950/2020 | North America / USA / Washington | 2020-03-27 | UW Virology Lab | UW Virology Lab | Pavitra Roychoudhury, Hong Xie, Keith Jerome, Alexander Greninger |
| EPI_ISL_427219 | hCoV-19/USA/WA-UW-3954/2020 | North America / USA / Washington | 2020-03-27 | UW Virology Lab | UW Virology Lab | Pavitra Roychoudhury, Hong Xie, Keith Jerome, Alexander Greninger |
| EPI_ISL_427220 | hCoV-19/USA/WA-UW-3959/2020 | North America / USA / Washington | 2020-03-27 | UW Virology Lab | UW Virology Lab | Pavitra Roychoudhury, Hong Xie, Keith Jerome, Alexander Greninger |
| EPI_ISL_427222 | hCoV-19/USA/WA-UW-3964/2020 | North America / USA / Washington | 2020-03-31 | UW Virology Lab | UW Virology Lab | Pavitra Roychoudhury, Hong Xie, Keith Jerome, Alexander Greninger |
| EPI_ISL_427223 | hCoV-19/USA/WA-UW-3969/2020 | North America / USA / Washington | 2020-03-31 | UW Virology Lab | UW Virology Lab | Pavitra Roychoudhury, Hong Xie, Keith Jerome, Alexander Greninger |
| EPI_ISL_427224 | hCoV-19/USA/WA-UW-3983/2020 | North America / USA / Washington | 2020-03-29 | UW Virology Lab | UW Virology Lab | Pavitra Roychoudhury, Hong Xie, Keith Jerome, Alexander Greninger |
| EPI_ISL_427225 | hCoV-19/USA/WA-UW-3984/2020 | North America / USA / Washington | 2020-03-29 | UW Virology Lab | UW Virology Lab | Pavitra Roychoudhury, Hong Xie, Keith Jerome, Alexander Greninger |
| EPI_ISL_427226 | hCoV-19/USA/WA-UW-3985/2020 | North America / USA / Washington | 2020-03-29 | UW Virology Lab | UW Virology Lab | Pavitra Roychoudhury, Hong Xie, Keith Jerome, Alexander Greninger |
| EPI_ISL_427227 | hCoV-19/USA/WA-UW-3986/2020 | North America / USA / Washington | 2020-03-29 | UW Virology Lab | UW Virology Lab | Pavitra Roychoudhury, Hong Xie, Keith Jerome, Alexander Greninger |
| EPI_ISL_427228 | hCoV-19/USA/WA-UW-3988/2020 | North America / USA / Washington | 2020-03-29 | UW Virology Lab | UW Virology Lab | Pavitra Roychoudhury, Hong Xie, Keith Jerome, Alexander Greninger |
| EPI_ISL_427229 | hCoV-19/USA/WA-UW-3989/2020 | North America / USA / Washington | 2020-03-28 | UW Virology Lab | UW Virology Lab | Pavitra Roychoudhury, Hong Xie, Keith Jerome, Alexander Greninger |
| EPI_ISL_427250 | hCoV-19/USA/WA-UW-4227/2020 | North America / USA / Washington | 2020-03-31 | UW Virology Lab | UW Virology Lab | Pavitra Roychoudhury, Hong Xie, Keith Jerome, Alexander Greninger |
| EPI_ISL_427251 | hCoV-19/USA/WA-UW-4228/2020 | North America / USA / Washington | 2020-03-31 | UW Virology Lab | UW Virology Lab | Pavitra Roychoudhury, Hong Xie, Keith Jerome, Alexander Greninger |
| EPI_ISL_427252 | hCoV-19/USA/WA-UW-4281/2020 | North America / USA / Washington | 2020-03-31 | UW Virology Lab | UW Virology Lab | Pavitra Roychoudhury, Hong Xie, Keith Jerome, Alexander Greninger |
| EPI_ISL_427253 | hCoV-19/USA/WA-UW-4289/2020 | North America / USA / Washington | 2020-03-31 | UW Virology Lab | UW Virology Lab | Pavitra Roychoudhury, Hong Xie, Keith Jerome, Alexander Greninger |
| EPI_ISL_427254 | hCoV-19/USA/WA-UW-4290/2020 | North America / USA / Washington | 2020-03-31 | UW Virology Lab | UW Virology Lab | Pavitra Roychoudhury, Hong Xie, Keith Jerome, Alexander Greninger |
| EPI_ISL_427255 | hCoV-19/USA/WA-UW-4291/2020 | North America / USA / Washington | 2020-03-31 | UW Virology Lab | UW Virology Lab | Pavitra Roychoudhury, Hong Xie, Keith Jerome, Alexander Greninger |
| EPI_ISL_427256 | hCoV-19/USA/WA-UW-4292/2020 | North America / USA / Washington | 2020-03-31 | UW Virology Lab | UW Virology Lab | Pavitra Roychoudhury, Hong Xie, Keith Jerome, Alexander Greninger |
| EPI_ISL_427257 | hCoV-19/USA/WA-UW-4315/2020 | North America / USA / Washington | 2020-03-31 | UW Virology Lab | UW Virology Lab | Pavitra Roychoudhury, Hong Xie, Keith Jerome, Alexander Greninger |
| EPI_ISL_427258 | hCoV-19/USA/WA-UW-4316/2020 | North America / USA / Washington | 2020-03-31 | UW Virology Lab | UW Virology Lab | Pavitra Roychoudhury, Hong Xie, Keith Jerome, Alexander Greninger |
| EPI_ISL_427259 | hCoV-19/USA/WA-UW-4318/2020 | North America / USA / Washington | 2020-03-31 | UW Virology Lab | UW Virology Lab | Pavitra Roychoudhury, Hong Xie, Keith Jerome, Alexander Greninger |
| EPI_ISL_427260 | hCoV-19/USA/WA-UW-4321/2020 | North America / USA / Washington | 2020-03-31 | UW Virology Lab | UW Virology Lab | Pavitra Roychoudhury, Hong Xie, Keith Jerome, Alexander Greninger |
| EPI_ISL_427261 | hCoV-19/USA/WA-UW-4322/2020 | North America / USA / Washington | 2020-03-31 | UW Virology Lab | UW Virology Lab | Pavitra Roychoudhury, Hong Xie, Keith Jerome, Alexander Greninger |
| EPI_ISL_427262 | hCoV-19/USA/WA-UW-4324/2020 | North America / USA / Washington | 2020-03-30 | UW Virology Lab | UW Virology Lab | Pavitra Roychoudhury, Hong Xie, Keith Jerome, Alexander Greninger |
| EPI_ISL_427263 | hCoV-19/USA/WA-UW-4326/2020 | North America / USA / Washington | 2020-03-31 | UW Virology Lab | UW Virology Lab | Pavitra Roychoudhury, Hong Xie, Keith Jerome, Alexander Greninger |
| EPI_ISL_427264 | hCoV-19/USA/WA-UW-4329/2020 | North America / USA / Washington | 2020-03-29 | UW Virology Lab | UW Virology Lab | Pavitra Roychoudhury, Hong Xie, Keith Jerome, Alexander Greninger |
| EPI_ISL_427265 | hCoV-19/USA/WA-UW-4330/2020 | North America / USA / Washington | 2020-03-31 | UW Virology Lab | UW Virology Lab | Pavitra Roychoudhury, Hong Xie, Keith Jerome, Alexander Greninger |
| EPI_ISL_427266 | hCoV-19/USA/WA-UW-4335/2020 | North America / USA / Washington | 2020-03-30 | UW Virology Lab | UW Virology Lab | Pavitra Roychoudhury, Hong Xie, Keith Jerome, Alexander Greninger |
| EPI_ISL_427267 | hCoV-19/USA/WA-UW-4337/2020 | North America / USA / Washington | 2020-03-30 | UW Virology Lab | UW Virology Lab | Pavitra Roychoudhury, Hong Xie, Keith Jerome, Alexander Greninger |
| EPI_ISL_427268 | hCoV-19/USA/WA-UW-4340/2020 | North America / USA / Washington | 2020-03-30 | UW Virology Lab | UW Virology Lab | Pavitra Roychoudhury, Hong Xie, Keith Jerome, Alexander Greninger |
| EPI_ISL_427269 | hCoV-19/USA/WA-UW-4499/2020 | North America / USA / Washington | 2020-03-31 | UW Virology Lab | UW Virology Lab | Pavitra Roychoudhury, Hong Xie, Keith Jerome, Alexander Greninger |
| EPI_ISL_427270 | hCoV-19/USA/WA-UW-4500/2020 | North America / USA / Washington | 2020-03-31 | UW Virology Lab | UW Virology Lab | Pavitra Roychoudhury, Hong Xie, Keith Jerome, Alexander Greninger |
| EPI_ISL_427271 | hCoV-19/USA/AZ-TG271866/2020 | North America / USA / Arizona | 2020-03-23 | AZ SPHL, Arizona Department of Health Services | TGen North | Jolene Bowers, Megan Folkerts, Darrin Lemmer, Dave Engelthaler |
| EPI_ISL_427272 | hCoV-19/USA/AZ-TG271868/2020 | North America / USA / Arizona | 2020-03-23 | AZ SPHL, Arizona Department of Health Services | TGen North | Jolene Bowers, Megan Folkerts, Darrin Lemmer, Dave Engelthaler |
| EPI_ISL_427273 | hCoV-19/USA/MN-MDH-117/2020 | North America / USA / Minnesota | 2020-03-20 | Minnesota Department of Health, Public Health Laboratory | Minnesota Department of Health, Public Health Laboratory | Matt Plumb, Jacob Garfin and Xiong Wang |
| EPI_ISL_427274 | hCoV-19/USA/MN-MDH-118/2020 | North America / USA / Minnesota | 2020-03-20 | Minnesota Department of Health, Public Health Laboratory | Minnesota Department of Health, Public Health Laboratory | Matt Plumb, Jacob Garfin and Xiong Wang |
| EPI_ISL_427275 | hCoV-19/USA/MN-MDH-119/2020 | North America / USA / Minnesota | 2020-03-20 | Minnesota Department of Health, Public Health Laboratory | Minnesota Department of Health, Public Health Laboratory | Matt Plumb, Jacob Garfin and Xiong Wang |
| EPI_ISL_427276 | hCoV-19/USA/MN-MDH-120/2020 | North America / USA / Minnesota | 2020-03-20 | Minnesota Department of Health, Public Health Laboratory | Minnesota Department of Health, Public Health Laboratory | Matt Plumb, Jacob Garfin and Xiong Wang |
| EPI_ISL_427277 | hCoV-19/USA/MN-MDH-121/2020 | North America / USA / Minnesota | 2020-03-20 | Minnesota Department of Health, Public Health Laboratory | Minnesota Department of Health, Public Health Laboratory | Matt Plumb, Jacob Garfin and Xiong Wang |
| EPI_ISL_427278 | hCoV-19/USA/MN-MDH-138/2020 | North America / USA / Minnesota | 2020-03-22 | Minnesota Department of Health, Public Health Laboratory | Minnesota Department of Health, Public Health Laboratory | Matt Plumb, Jacob Garfin and Xiong Wang |
| EPI_ISL_427279 | hCoV-19/USA/MN-MDH-140/2020 | North America / USA / Minnesota | 2020-03-21 | Minnesota Department of Health, Public Health Laboratory | Minnesota Department of Health, Public Health Laboratory | Matt Plumb, Jacob Garfin and Xiong Wang |
| EPI_ISL_427280 | hCoV-19/USA/MN-MDH-141/2020 | North America / USA / Minnesota | 2020-03-21 | Minnesota Department of Health, Public Health Laboratory | Minnesota Department of Health, Public Health Laboratory | Matt Plumb, Jacob Garfin and Xiong Wang |
| EPI_ISL_427281 | hCoV-19/USA/MN-MDH-142/2020 | North America / USA / Minnesota | 2020-03-18 | Minnesota Department of Health, Public Health Laboratory | Minnesota Department of Health, Public Health Laboratory | Matt Plumb, Jacob Garfin and Xiong Wang |
| EPI_ISL_427282 | hCoV-19/USA/MN-MDH-143/2020 | North America / USA / Minnesota | 2020-03-22 | Minnesota Department of Health, Public Health Laboratory | Minnesota Department of Health, Public Health Laboratory | Matt Plumb, Jacob Garfin and Xiong Wang |
| EPI_ISL_427283 | hCoV-19/USA/MN-MDH-144/2020 | North America / USA / Minnesota | 2020-03-22 | Minnesota Department of Health, Public Health Laboratory | Minnesota Department of Health, Public Health Laboratory | Matt Plumb, Jacob Garfin and Xiong Wang |
| EPI_ISL_427284 | hCoV-19/USA/MN-MDH-145/2020 | North America / USA / Minnesota | 2020-03-22 | Minnesota Department of Health, Public Health Laboratory | Minnesota Department of Health, Public Health Laboratory | Matt Plumb, Jacob Garfin and Xiong Wang |
| EPI_ISL_427285 | hCoV-19/USA/MN-MDH-146/2020 | North America / USA / Minnesota | 2020-03-22 | Minnesota Department of Health, Public Health Laboratory | Minnesota Department of Health, Public Health Laboratory | Matt Plumb, Jacob Garfin and Xiong Wang |
| EPI_ISL_427286 | hCoV-19/USA/MN-MDH-147/2020 | North America / USA / Minnesota | 2020-03-23 | Minnesota Department of Health, Public Health Laboratory | Minnesota Department of Health, Public Health Laboratory | Matt Plumb, Jacob Garfin and Xiong Wang |
| EPI_ISL_427287 | hCoV-19/USA/MN-MDH-149/2020 | North America / USA / Minnesota | 2020-03-23 | Minnesota Department of Health, Public Health Laboratory | Minnesota Department of Health, Public Health Laboratory | Matt Plumb, Jacob Garfin and Xiong Wang |
| EPI_ISL_427288 | hCoV-19/USA/OH-OSUP0016/2020 | North America / USA / Ohio / Columbus | 2020-03-31 | The Ohio State University | The Ohio State University-James Molecular Lab at Polaris | Huolin Tu, Preeti Panchioli, Jason Garee, Matthew Hunt, Joan-Miquel Balada-Llasat, Erica Vincent, Weiqiang Zhao, Dan Jones |
| EPI_ISL_427700 | hCoV-19/Australia/NSW137/2020 | Oceania / Australia / New South Wales / | 2020-03-23 | Centre for Infectious Diseases and Microbiology Public Health | NSW Health Pathology - Institute of Clinical Pathology and Medical Research; Westmead Hospital; University of Sydney | Gall M, Arnott A, Sadsad R, Draper J, Sim E, Bachmann N, Rockett R, Lam C, Gray K, Timms V, Carter I, Holmes EC, O'Sullivan MV, Byun R, Sintchenko V, Chen SC, Eden JS, Maddocks S, Kok J, Propenko M, Sorrell T, Chang S, Basile K, Dwyer DE for the 2019-nCoV Study Group |
| EPI_ISL_427701 | hCoV-19/Australia/NSW180/2020 | Oceania / Australia / New South Wales / | 2020-03-22 | Centre for Infectious Diseases and Microbiology Public Health | NSW Health Pathology - Institute of Clinical Pathology and Medical Research; Westmead Hospital; University of Sydney | Draper J, Sim E, Bachmann N, Rockett R, Lam C, Gray K, Timms V, Gall M, Arnott A, Sadsad R, Carter I, Holmes EC, O'Sullivan MV, Byun R, Sintchenko V, Chen SC, Eden JS, Maddocks S, Kok J, Propenko M, Sorrell T, Chang S, Basile K, Dwyer DE for the 2019-nCoV Study Group |
| EPI_ISL_427702 | hCoV-19/Australia/NSW88/2020 | Oceania / Australia / New South Wales / | 2020-03-16 | Centre for Infectious Diseases and Microbiology Public Health | NSW Health Pathology - Institute of Clinical Pathology and Medical Research; Westmead Hospital; University of Sydney | Arnott A, Sadsad R, Draper J, Sim E, Bachmann N, Rockett R, Lam C, Gray K, Timms V, Gall M, Carter I, Holmes EC, O'Sullivan MV, Byun R, Sintchenko V, Chen SC, Eden JS, Maddocks S, Kok J, Propenko M, Sorrell T, Chang S, Basile K, Dwyer DE for the 2019-nCoV Study Group |
| EPI_ISL_427703 | hCoV-19/Australia/NSW75/2020 | Oceania / Australia / New South Wales / | 2020-03-16 | Centre for Infectious Diseases and Microbiology Public Health | NSW Health Pathology - Institute of Clinical Pathology and Medical Research; Westmead Hospital; University of Sydney | Gray K, Timms V, Gall M, Arnott A, Sadsad R, Draper J, Sim E, Bachmann N, Rockett R, Lam C, Carter I, Holmes EC, O'Sullivan MV, Byun R, Sintchenko V, Chen SC, Eden JS, Maddocks S, Kok J, Propenko M, Sorrell T, Chang S, Basile K, Dwyer DE for the 2019-nCoV Study Group |
| EPI_ISL_427704 | hCoV-19/Australia/NSW76/2020 | Oceania / Australia / New South Wales / | 2020-03-19 | Centre for Infectious Diseases and Microbiology Public Health | NSW Health Pathology - Institute of Clinical Pathology and Medical Research; Westmead Hospital; University of Sydney | Timms V, Gall M, Arnott A, Sadsad R, Draper J, Sim E, Bachmann N, Rockett R, Lam C, Gray K, Carter I, Holmes EC, O'Sullivan MV, Byun R, Sintchenko V, Chen SC, Eden JS, Maddocks S, Kok J, Propenko M, Sorrell T, Chang S, Basile K, Dwyer DE for the 2019-nCoV Study Group |
| EPI_ISL_427705 | hCoV-19/Australia/NSW69/2020 | Oceania / Australia / New South Wales / | 2020-03-20 | Centre for Infectious Diseases and Microbiology Public Health | NSW Health Pathology - Institute of Clinical Pathology and Medical Research; Westmead Hospital; University of Sydney | Sadsad R, Draper J, Sim E, Bachmann N, Rockett R, Lam C, Gray K, Timms V, Gall M, Arnott A, Carter I, Holmes EC, O'Sullivan MV, Byun R, Sintchenko V, Chen SC, Eden JS, Maddocks S, Kok J, Propenko M, Sorrell T, Chang S, Basile K, Dwyer DE for the 2019-nCoV Study Group |

| | | | | | | |
|---|---|---|---|---|---|---|
| EPI_ISL_427706 | hCoV-19/Australia/NSW74/2020 | Oceania / Australia / New South Wales / | 2020-03-19 | Centre for Infectious Diseases and Microbiology Public Health | NSW Health Pathology - Institute of Clinical Pathology and Medical Research; Westmead Hospital; University of Sydney | Lam C, Gray K, Timms, V, Gall M, Arnott A, Sadsad R, Draper J, Sim E, Bachmann N, Rockett R, Carter I, Holmes EC, O'Sullivan MV, Byun R, Sintchenko V, Chen SC, Eden JS, Maddocks S, Kok J, Propenko M, Sorrell T, Chang S, Basile K, Dwyer DE for the 2019-nCoV Study Group |
| EPI_ISL_427707 | hCoV-19/Australia/NSW198/2020 | Oceania / Australia / New South Wales / | 2020-03-23 | Centre for Infectious Diseases and Microbiology Public Health | NSW Health Pathology - Institute of Clinical Pathology and Medical Research; Westmead Hospital; University of Sydney | Arnott A, Sadsad R, Draper J, Sim E, Bachmann N, Rockett R, Lam C, Gray K, Timms V, Gall M, Carter I, Holmes EC, O'Sullivan MV, Byun R, Sintchenko V, Chen SC, Eden JS, Maddocks S, Kok J, Propenko M, Sorrell T, Chang S, Basile K, Dwyer DE for the 2019-nCoV Study Group |
| EPI_ISL_427708 | hCoV-19/Australia/NSW145/2020 | Oceania / Australia / New South Wales / | 2020-03-21 | Centre for Infectious Diseases and Microbiology Public Health | NSW Health Pathology - Institute of Clinical Pathology and Medical Research; Westmead Hospital; University of Sydney | Gray K, Timms V, Gall M, Arnott A, Sadsad R, Draper J, Sim E, Bachmann N, Rockett R, Lam C, Carter I, Holmes EC, O'Sullivan MV, Byun R, Sintchenko V, Chen SC, Eden JS, Maddocks S, Kok J, Propenko M, Sorrell T, Chang S, Basile K, Dwyer DE for the 2019-nCoV Study Group |
| EPI_ISL_427709 | hCoV-19/Australia/NSW181/2020 | Oceania / Australia / New South Wales / | 2020-03-21 | Centre for Infectious Diseases and Microbiology Public Health | NSW Health Pathology - Institute of Clinical Pathology and Medical Research; Westmead Hospital; University of Sydney | Sim E, Bachmann N, Rockett R, Lam C, Gray K, Timms V, Gall M, Arnott A, Sadsad R, Draper J, Carter I, Holmes EC, O'Sullivan MV, Byun R, Sintchenko V, Chen SC, Eden JS, Maddocks S, Kok J, Propenko M, Sorrell T, Chang S, Basile K, Dwyer DE for the 2019-nCoV Study Group |
| EPI_ISL_427710 | hCoV-19/Australia/NSW201/2020 | Oceania / Australia / New South Wales / | 2020-03-21 | Centre for Infectious Diseases and Microbiology Public Health | NSW Health Pathology - Institute of Clinical Pathology and Medical Research; Westmead Hospital; University of Sydney | Sim E, Bachmann N, Rockett R, Lam C, Gray K, Timms V, Gall M, Arnott A, Sadsad R, Draper J, Carter I, Holmes EC, O'Sullivan MV, Byun R, Sintchenko V, Chen SC, Eden JS, Maddocks S, Kok J, Propenko M, Sorrell T, Chang S, Basile K, Dwyer DE for the 2019-nCoV Study Group |
| EPI_ISL_427711 | hCoV-19/Australia/NSW83/2020 | Oceania / Australia / New South Wales / | 2020-03-21 | ACT Pathology, The Canberra Hospital | NSW Health Pathology - Institute of Clinical Pathology and Medical Research; Westmead Hospital; University of Sydney | Rockett R, Lam C, Gray K, Timms V, Gall M, Arnott A, Sadsad R, Draper J, Sim E, Bachmann N, Carter I, Holmes EC, O'Sullivan MV, Byun R, Sintchenko V, Chen SC, Eden JS, Maddocks S, Kok J, Propenko M, Sorrell T, Chang S, Basile K, Dwyer DE for the 2019-nCoV Study Group |
| EPI_ISL_427712 | hCoV-19/Australia/NSW91/2020 | Oceania / Australia / New South Wales / | 2020-03-20 | Centre for Infectious Diseases and Microbiology Public Health | NSW Health Pathology - Institute of Clinical Pathology and Medical Research; Westmead Hospital; University of Sydney | Sim E, Bachmann N, Rockett R, Lam C, Gray K, Timms V, Gall M, Arnott A, Sadsad R, Draper J, Carter I, Holmes EC, O'Sullivan MV, Byun R, Sintchenko V, Chen SC, Eden JS, Maddocks S, Kok J, Propenko M, Sorrell T, Chang S, Basile K, Dwyer DE for the 2019-nCoV Study Group |
| EPI_ISL_427713 | hCoV-19/Australia/NSW182/2020 | Oceania / Australia / New South Wales / | 2020-03-21 | Centre for Infectious Diseases and Microbiology Public Health | NSW Health Pathology - Institute of Clinical Pathology and Medical Research; Westmead Hospital; University of Sydney | Bachmann N, Rockett R, Lam C, Gray K, Timms V, Gall M, Arnott A, Sadsad R, Draper J, Sim E, Carter I, Holmes EC, O'Sullivan MV, Byun R, Sintchenko V, Chen SC, Eden JS, Maddocks S, Kok J, Propenko M, Sorrell T, Chang S, Basile K, Dwyer DE for the 2019-nCoV Study Group |
| EPI_ISL_427714 | hCoV-19/Australia/NSW183/2020 | Oceania / Australia / New South Wales / | 2020-03-21 | Centre for Infectious Diseases and Microbiology Public Health | NSW Health Pathology - Institute of Clinical Pathology and Medical Research; Westmead Hospital; University of Sydney | Rockett R, Lam C, Gray K, Timms V, Gall M, Arnott A, Sadsad R, Draper J, Sim E, Bachmann N, Carter I, Holmes EC, O'Sullivan MV, Byun R, Sintchenko V, Chen SC, Eden JS, Maddocks S, Kok J, Propenko M, Sorrell T, Chang S, Basile K, Dwyer DE for the 2019-nCoV Study Group |
| EPI_ISL_427715 | hCoV-19/Australia/NSW164/2020 | Oceania / Australia / New South Wales / | 2020-03-22 | Centre for Infectious Diseases and Microbiology Public Health | NSW Health Pathology - Institute of Clinical Pathology and Medical Research; Westmead Hospital; University of Sydney | Lam C, Gray K, Timms, V, Gall M, Arnott A, Sadsad R, Draper J, Sim E, Bachmann N, Rockett R, Carter I, Holmes EC, O'Sullivan MV, Byun R, Sintchenko V, Chen SC, Eden JS, Maddocks S, Kok J, Propenko M, Sorrell T, Chang S, Basile K, Dwyer DE for the 2019-nCoV Study Group |
| EPI_ISL_427716 | hCoV-19/Australia/NSW94/2020 | Oceania / Australia / New South Wales / | 2020-03-22 | Centre for Infectious Diseases and Microbiology Public Health | NSW Health Pathology - Institute of Clinical Pathology and Medical Research; Westmead Hospital; University of Sydney | Lam C, Gray K, Timms, V, Gall M, Arnott A, Sadsad R, Draper J, Sim E, Bachmann N, Rockett R, Carter I, Holmes EC, O'Sullivan MV, Byun R, Sintchenko V, Chen SC, Eden JS, Maddocks S, Kok J, Propenko M, Sorrell T, Chang S, Basile K, Dwyer DE for the 2019-nCoV Study Group |
| EPI_ISL_427717 | hCoV-19/Australia/NSW101/2020 | Oceania / Australia / New South Wales / | 2020-03-17 | ACT Pathology, The Canberra Hospital | NSW Health Pathology - Institute of Clinical Pathology and Medical Research; Westmead Hospital; University of Sydney | Sim E, Bachmann N, Rockett R, Lam C, Gray K, Timms V, Gall M, Arnott A, Sadsad R, Draper J, Carter I, Holmes EC, O'Sullivan MV, Byun R, Sintchenko V, Chen SC, Eden JS, Maddocks S, Kok J, Propenko M, Sorrell T, Chang S, Basile K, Dwyer DE for the 2019-nCoV Study Group |
| EPI_ISL_427718 | hCoV-19/Australia/NSW85/2020 | Oceania / Australia / New South Wales / | 2020-03-16 | ACT Pathology, The Canberra Hospital | NSW Health Pathology - Institute of Clinical Pathology and Medical Research; Westmead Hospital; University of Sydney | Gray K, Timms V, Gall M, Arnott A, Sadsad R, Draper J, Sim E, Bachmann N, Rockett R, Lam C, Carter I, Holmes EC, O'Sullivan MV, Byun R, Sintchenko V, Chen SC, Eden JS, Maddocks S, Kok J, Propenko M, Sorrell T, Chang S, Basile K, Dwyer DE for the 2019-nCoV Study Group |
| EPI_ISL_427719 | hCoV-19/Australia/NSW156/2020 | Oceania / Australia / New South Wales / | 2020-03-21 | ACT Pathology, The Canberra Hospital | NSW Health Pathology - Institute of Clinical Pathology and Medical Research; Westmead Hospital; University of Sydney | Timms V, Gall M, Arnott A, Sadsad R, Draper J, Sim E, Bachmann N, Rockett R, Lam C, Gray K, Carter I, Holmes EC, O'Sullivan MV, Byun R, Sintchenko V, Chen SC, Eden JS, Maddocks S, Kok J, Propenko M, Sorrell T, Chang S, Basile K, Dwyer DE for the 2019-nCoV Study Group |
| EPI_ISL_427720 | hCoV-19/Australia/NSW81/2020 | Oceania / Australia / New South Wales / | 2020-03-18 | Centre for Infectious Diseases and Microbiology Public Health | NSW Health Pathology - Institute of Clinical Pathology and Medical Research; Westmead Hospital; University of Sydney | Sim E, Bachmann N, Rockett R, Lam C, Gray K, Timms V, Gall M, Arnott A, Sadsad R, Draper J, Carter I, Holmes EC, O'Sullivan MV, Byun R, Sintchenko V, Chen SC, Eden JS, Maddocks S, Kok J, Propenko M, Sorrell T, Chang S, Basile K, Dwyer DE for the 2019-nCoV Study Group |
| EPI_ISL_427721 | hCoV-19/Australia/NSW151/2020 | Oceania / Australia / New South Wales / | 2020-03-19 | ACT Pathology, The Canberra Hospital | NSW Health Pathology - Institute of Clinical Pathology and Medical Research; Westmead Hospital; University of Sydney | Sim E, Bachmann N, Rockett R, Lam C, Gray K, Timms V, Gall M, Arnott A, Sadsad R, Draper J, Carter I, Holmes EC, O'Sullivan MV, Byun R, Sintchenko V, Chen SC, Eden JS, Maddocks S, Kok J, Propenko M, Sorrell T, Chang S, Basile K, Dwyer DE for the 2019-nCoV Study Group |
| EPI_ISL_427722 | hCoV-19/Australia/NSW100/2020 | Oceania / Australia / New South Wales / | 2020-03-18 | Centre for Infectious Diseases and Microbiology Public Health | NSW Health Pathology - Institute of Clinical Pathology and Medical Research; Westmead Hospital; University of Sydney | Draper J, Sim E, Bachmann N, Rockett R, Lam C, Gray K, Timms V, Gall M, Arnott A, Sadsad R, Carter I, Holmes EC, O'Sullivan MV, Byun R, Sintchenko V, Chen SC, Eden JS, Maddocks S, Kok J, Propenko M, Sorrell T, Chang S, Basile K, Dwyer DE for the 2019-nCoV Study Group |
| EPI_ISL_427723 | hCoV-19/Australia/NSW84/2020 | Oceania / Australia / New South Wales / | 2020-03-19 | Centre for Infectious Diseases and Microbiology Public Health | NSW Health Pathology - Institute of Clinical Pathology and Medical Research; Westmead Hospital; University of Sydney | Lam C, Gray K, Timms, V, Gall M, Arnott A, Sadsad R, Draper J, Sim E, Bachmann N, Rockett R, Carter I, Holmes EC, O'Sullivan MV, Byun R, Sintchenko V, Chen SC, Eden JS, Maddocks S, Kok J, Propenko M, Sorrell T, Chang S, Basile K, Dwyer DE for the 2019-nCoV Study Group |
| EPI_ISL_427724 | hCoV-19/Australia/NSW114/2020 | Oceania / Australia / New South Wales / | 2020-03-19 | Centre for Infectious Diseases and Microbiology Public Health | NSW Health Pathology - Institute of Clinical Pathology and Medical Research; Westmead Hospital; University of Sydney | Lam C, Gray K, Timms, V, Gall M, Arnott A, Sadsad R, Draper J, Sim E, Bachmann N, Rockett R, Carter I, Holmes EC, O'Sullivan MV, Byun R, Sintchenko V, Chen SC, Eden JS, Maddocks S, Kok J, Propenko M, Sorrell T, Chang S, Basile K, Dwyer DE for the 2019-nCoV Study Group |
| EPI_ISL_427725 | hCoV-19/Australia/NSW86/2020 | Oceania / Australia / New South Wales / | 2020-03-19 | Centre for Infectious Diseases and Microbiology Public Health | NSW Health Pathology - Institute of Clinical Pathology and Medical Research; Westmead Hospital; University of Sydney | Timms V, Gall M, Arnott A, Sadsad R, Draper J, Sim E, Bachmann N, Rockett R, Lam C, Gray K, Carter I, Holmes EC, O'Sullivan MV, Byun R, Sintchenko V, Chen SC, Eden JS, Maddocks S, Kok J, Propenko M, Sorrell T, Chang S, Basile K, Dwyer DE for the 2019-nCoV Study Group |
| EPI_ISL_427726 | hCoV-19/Australia/NSW213/2020 | Oceania / Australia / New South Wales / | 2020-03-19 | Centre for Infectious Diseases and Microbiology Public Health | NSW Health Pathology - Institute of Clinical Pathology and Medical Research; Westmead Hospital; University of Sydney | Rockett R, Lam C, Gray K, Timms V, Gall M, Arnott A, Sadsad R, Draper J, Sim E, Bachmann N, Carter I, Holmes EC, O'Sullivan MV, Byun R, Sintchenko V, Chen SC, Eden JS, Maddocks S, Kok J, Propenko M, Sorrell T, Chang S, Basile K, Dwyer DE for the 2019-nCoV Study Group |
| EPI_ISL_427727 | hCoV-19/Australia/NSW214/2020 | Oceania / Australia / New South Wales / | 2020-03-17 | Centre for Infectious Diseases and Microbiology Public Health | NSW Health Pathology - Institute of Clinical Pathology and Medical Research; Westmead Hospital; University of Sydney | Lam C, Gray K, Timms, V, Gall M, Arnott A, Sadsad R, Draper J, Sim E, Bachmann N, Rockett R, Carter I, Holmes EC, O'Sullivan MV, Byun R, Sintchenko V, Chen SC, Eden JS, Maddocks S, Kok J, Propenko M, Sorrell T, Chang S, Basile K, Dwyer DE for the 2019-nCoV Study Group |
| EPI_ISL_427728 | hCoV-19/Australia/NSW113/2020 | Oceania / Australia / New South Wales / | 2020-03-17 | Centre for Infectious Diseases and Microbiology Public Health | NSW Health Pathology - Institute of Clinical Pathology and Medical Research; Westmead Hospital; University of Sydney | Rockett R, Lam C, Gray K, Timms V, Gall M, Arnott A, Sadsad R, Draper J, Sim E, Bachmann N, Carter I, Holmes EC, O'Sullivan MV, Byun R, Sintchenko V, Chen SC, Eden JS, Maddocks S, Kok J, Propenko M, Sorrell T, Chang S, Basile K, Dwyer DE for the 2019-nCoV Study Group |
| EPI_ISL_427729 | hCoV-19/Australia/NSW77/2020 | Oceania / Australia / New South Wales / | 2020-03-17 | Centre for Infectious Diseases and Microbiology Public Health | NSW Health Pathology - Institute of Clinical Pathology and Medical Research; Westmead Hospital; University of Sydney | Gall M, Arnott A, Sadsad R, Draper J, Sim E, Bachmann N, Rockett R, Lam C, Gray K, Timms V, Carter I, Holmes EC, O'Sullivan MV, Byun R, Sintchenko V, Chen SC, Eden JS, Maddocks S, Kok J, Propenko M, Sorrell T, Chang S, Basile K, Dwyer DE for the 2019-nCoV Study Group |
| EPI_ISL_427730 | hCoV-19/Australia/NSW215/2020 | Oceania / Australia / New South Wales / | 2020-03-17 | Centre for Infectious Diseases and Microbiology Public Health | NSW Health Pathology - Institute of Clinical Pathology and Medical Research; Westmead Hospital; University of Sydney | Gray K, Timms V, Gall M, Arnott A, Sadsad R, Draper J, Sim E, Bachmann N, Rockett R, Lam C, Carter I, Holmes EC, O'Sullivan MV, Byun R, Sintchenko V, Chen SC, Eden JS, Maddocks S, Kok J, Propenko M, Sorrell T, Chang S, Basile K, Dwyer DE for the 2019-nCoV Study Group |
| EPI_ISL_427731 | hCoV-19/Australia/NSW165/2020 | Oceania / Australia / New South Wales / | 2020-03-17 | Centre for Infectious Diseases and Microbiology Public Health | NSW Health Pathology - Institute of Clinical Pathology and Medical Research; Westmead Hospital; University of Sydney | Gray K, Timms V, Gall M, Arnott A, Sadsad R, Draper J, Sim E, Bachmann N, Rockett R, Lam C, Carter I, Holmes EC, O'Sullivan MV, Byun R, Sintchenko V, Chen SC, Eden JS, Maddocks S, Kok J, Propenko M, Sorrell T, Chang S, Basile K, Dwyer DE for the 2019-nCoV Study Group |
| EPI_ISL_427732 | hCoV-19/Australia/NSW216/2020 | Oceania / Australia / New South Wales / | 2020-03-17 | Centre for Infectious Diseases and Microbiology Public Health | NSW Health Pathology - Institute of Clinical Pathology and Medical Research; Westmead Hospital; University of Sydney | Gray K, Timms V, Gall M, Arnott A, Sadsad R, Draper J, Sim E, Bachmann N, Rockett R, Lam C, Carter I, Holmes EC, O'Sullivan MV, Byun R, Sintchenko V, Chen SC, Eden JS, Maddocks S, Kok J, Propenko M, Sorrell T, Chang S, Basile K, Dwyer DE for the 2019-nCoV Study Group |
| EPI_ISL_427733 | hCoV-19/Australia/NSW67/2020 | Oceania / Australia / New South Wales / | 2020-03-17 | Centre for Infectious Diseases and Microbiology Public Health | NSW Health Pathology - Institute of Clinical Pathology and Medical Research; Westmead Hospital; University of Sydney | Gall M, Arnott A, Sadsad R, Draper J, Sim E, Bachmann N, Rockett R, Lam C, Gray K, Timms V, Carter I, Holmes EC, O'Sullivan MV, Byun R, Sintchenko V, Chen SC, Eden JS, Maddocks S, Kok J, Propenko M, Sorrell T, Chang S, Basile K, Dwyer DE for the 2019-nCoV Study Group |
| EPI_ISL_427734 | hCoV-19/Australia/NSW176/2020 | Oceania / Australia / New South Wales / | 2020-03-18 | Centre for Infectious Diseases and Microbiology Public Health | NSW Health Pathology - Institute of Clinical Pathology and Medical Research; Westmead Hospital; University of Sydney | Timms V, Gall M, Arnott A, Sadsad R, Draper J, Sim E, Bachmann N, Rockett R, Lam C, Gray K, Carter I, Holmes EC, O'Sullivan MV, Byun R, Sintchenko V, Chen SC, Eden JS, Maddocks S, Kok J, Propenko M, Sorrell T, Chang S, Basile K, Dwyer DE for the 2019-nCoV Study Group |
| EPI_ISL_427735 | hCoV-19/Australia/NSW158/2020 | Oceania / Australia / New South Wales / | 2020-03-18 | Centre for Infectious Diseases and Microbiology Public Health | NSW Health Pathology - Institute of Clinical Pathology and Medical Research; Westmead Hospital; University of Sydney | Arnott A, Sadsad R, Draper J, Sim E, Bachmann N, Rockett R, Lam C, Gray K, Timms V, Gall M, Carter I, Holmes EC, O'Sullivan MV, Byun R, Sintchenko V, Chen SC, Eden JS, Maddocks S, Kok J, Propenko M, Sorrell T, Chang S, Basile K, Dwyer DE for the 2019-nCoV Study Group |
| EPI_ISL_427736 | hCoV-19/Australia/NSW190/2020 | Oceania / Australia / New South Wales / | 2020-03-16 | Centre for Infectious Diseases and Microbiology Public Health | NSW Health Pathology - Institute of Clinical Pathology and Medical Research; Westmead Hospital; University of Sydney | Draper J, Sim E, Bachmann N, Rockett R, Lam C, Gray K, Timms V, Gall M, Arnott A, Sadsad R, Carter I, Holmes EC, O'Sullivan MV, Byun R, Sintchenko V, Chen SC, Eden JS, Maddocks S, Kok J, Propenko M, Sorrell T, Chang S, Basile K, Dwyer DE for the 2019-nCoV Study Group |
| EPI_ISL_427737 | hCoV-19/Australia/NSW143/2020 | Oceania / Australia / New South Wales / | 2020-03-20 | Centre for Infectious Diseases and Microbiology Public Health | NSW Health Pathology - Institute of Clinical Pathology and Medical Research; Westmead Hospital; University of Sydney | Rockett R, Lam C, Gray K, Timms V, Gall M, Arnott A, Sadsad R, Draper J, Sim E, Bachmann N, Carter I, Holmes EC, O'Sullivan MV, Byun R, Sintchenko V, Chen SC, Eden JS, Maddocks S, Kok J, Propenko M, Sorrell T, Chang S, Basile K, Dwyer DE for the 2019-nCoV Study Group |
| EPI_ISL_427738 | hCoV-19/Australia/NSW144/2020 | Oceania / Australia / New South Wales / | 2020-03-21 | Centre for Infectious Diseases and Microbiology Public Health | NSW Health Pathology - Institute of Clinical Pathology and Medical Research; Westmead Hospital; University of Sydney | Lam C, Gray K, Timms, V, Gall M, Arnott A, Sadsad R, Draper J, Sim E, Bachmann N, Rockett R, Carter I, Holmes EC, O'Sullivan MV, Byun R, Sintchenko V, Chen SC, Eden JS, Maddocks S, Kok J, Propenko M, Sorrell T, Chang S, Basile K, Dwyer DE for the 2019-nCoV Study Group |
| EPI_ISL_427739 | hCoV-19/Australia/NSW168/2020 | Oceania / Australia / New South Wales / | 2020-03-21 | Centre for Infectious Diseases and Microbiology Public Health | NSW Health Pathology - Institute of Clinical Pathology and Medical Research; Westmead Hospital; University of Sydney | Arnott A, Sadsad R, Draper J, Sim E, Bachmann N, Rockett R, Lam C, Gray K, Timms V, Gall M, Carter I, Holmes EC, O'Sullivan MV, Byun R, Sintchenko V, Chen SC, Eden JS, Maddocks S, Kok J, Propenko M, Sorrell T, Chang S, Basile K, Dwyer DE for the 2019-nCoV Study Group |
| EPI_ISL_427740 | hCoV-19/Australia/NSW189/2020 | Oceania / Australia / New South Wales / | 2020-03-21 | Centre for Infectious Diseases and Microbiology Public Health | NSW Health Pathology - Institute of Clinical Pathology and Medical Research; Westmead Hospital; University of Sydney | Sadsad R, Draper J, Sim E, Bachmann N, Rockett R, Lam C, Gray K, Timms V, Gall M, Arnott A, Carter I, Holmes EC, O'Sullivan MV, Byun R, Sintchenko V, Chen SC, Eden JS, Maddocks S, Kok J, Propenko M, Sorrell T, Chang S, Basile K, Dwyer DE for the 2019-nCoV Study Group |
| EPI_ISL_427741 | hCoV-19/Australia/NSW110/2020 | Oceania / Australia / New South Wales / | 2020-03-21 | Centre for Infectious Diseases and Microbiology Public Health | NSW Health Pathology - Institute of Clinical Pathology and Medical Research; Westmead Hospital; University of Sydney | Draper J, Sim E, Bachmann N, Rockett R, Lam C, Gray K, Timms V, Gall M, Arnott A, Sadsad R, Carter I, Holmes EC, O'Sullivan MV, Byun R, Sintchenko V, Chen SC, Eden JS, Maddocks S, Kok J, Propenko M, Sorrell T, Chang S, Basile K, Dwyer DE for the 2019-nCoV Study Group |
| EPI_ISL_427742 | hCoV-19/Australia/NSW134/2020 | Oceania / Australia / New South Wales / | 2020-03-21 | Centre for Infectious Diseases and Microbiology Public Health | NSW Health Pathology - Institute of Clinical Pathology and Medical Research; Westmead Hospital; University of Sydney | Lam C, Gray K, Timms, V, Gall M, Arnott A, Sadsad R, Draper J, Sim E, Bachmann N, Rockett R, Carter I, Holmes EC, O'Sullivan MV, Byun R, Sintchenko V, Chen SC, Eden JS, Maddocks S, Kok J, Propenko M, Sorrell T, Chang S, Basile K, Dwyer DE for the 2019-nCoV Study Group |
| EPI_ISL_427743 | hCoV-19/Australia/NSW171/2020 | Oceania / Australia / New South Wales / | 2020-03-21 | Centre for Infectious Diseases and Microbiology Public Health | NSW Health Pathology - Institute of Clinical Pathology and Medical Research; Westmead Hospital; University of Sydney | Sim E, Bachmann N, Rockett R, Lam C, Gray K, Timms V, Gall M, Arnott A, Sadsad R, Draper J, Carter I, Holmes EC, O'Sullivan MV, Byun R, Sintchenko V, Chen SC, Eden JS, Maddocks S, Kok J, Propenko M, Sorrell T, Chang S, Basile K, Dwyer DE for the 2019-nCoV Study Group |
| EPI_ISL_427744 | hCoV-19/Australia/NSW72/2020 | Oceania / Australia / New South Wales / | 2020-03-21 | Centre for Infectious Diseases and Microbiology Public Health | NSW Health Pathology - Institute of Clinical Pathology and Medical Research; Westmead Hospital; University of Sydney | Bachmann N, Rockett R, Lam C, Gray K, Timms V, Gall M, Arnott A, Sadsad R, Draper J, Sim E, Carter I, Holmes EC, O'Sullivan MV, Byun R, Sintchenko V, Chen SC, Eden JS, Maddocks S, Kok J, Propenko M, Sorrell T, Chang S, Basile K, Dwyer DE for the 2019-nCoV Study Group |
| EPI_ISL_427745 | hCoV-19/Australia/NSW205/2020 | Oceania / Australia / New South Wales / | 2020-03-21 | Centre for Infectious Diseases and Microbiology Public Health | NSW Health Pathology - Institute of Clinical Pathology and Medical Research; Westmead Hospital; University of Sydney | Gray K, Timms V, Gall M, Arnott A, Sadsad R, Draper J, Sim E, Bachmann N, Rockett R, Lam C, Carter I, Holmes EC, O'Sullivan MV, Byun R, Sintchenko V, Chen SC, Eden JS, Maddocks S, Kok J, Propenko M, Sorrell T, Chang S, Basile K, Dwyer DE for the 2019-nCoV Study Group |
| EPI_ISL_427746 | hCoV-19/Australia/NSW197/2020 | Oceania / Australia / New South Wales / | 2020-03-21 | Centre for Infectious Diseases and Microbiology Public Health | NSW Health Pathology - Institute of Clinical Pathology and Medical Research; Westmead Hospital; University of Sydney | Gall M, Arnott A, Sadsad R, Draper J, Sim E, Bachmann N, Rockett R, Lam C, Gray K, Timms V, Carter I, Holmes EC, O'Sullivan MV, Byun R, Sintchenko V, Chen SC, Eden JS, Maddocks S, Kok J, Propenko M, Sorrell T, Chang S, Basile K, Dwyer DE for the 2019-nCoV Study Group |
| EPI_ISL_427747 | hCoV-19/Australia/NSW199/2020 | Oceania / Australia / New South Wales / | 2020-03-22 | Centre for Infectious Diseases and Microbiology Public Health | NSW Health Pathology - Institute of Clinical Pathology and Medical Research; Westmead Hospital; University of Sydney | Sadsad R, Draper J, Sim E, Bachmann N, Rockett R, Lam C, Gray K, Timms V, Gall M, Arnott A, Carter I, Holmes EC, O'Sullivan MV, Byun R, Sintchenko V, Chen SC, Eden JS, Maddocks S, Kok J, Propenko M, Sorrell T, Chang S, Basile K, Dwyer DE for the 2019-nCoV Study Group |
| EPI_ISL_427748 | hCoV-19/Australia/NSW106/2020 | Oceania / Australia / New South Wales / | 2020-03-22 | Centre for Infectious Diseases and Microbiology Public Health | NSW Health Pathology - Institute of Clinical Pathology and Medical Research; Westmead Hospital; University of Sydney | Timms V, Gall M, Arnott A, Sadsad R, Draper J, Sim E, Bachmann N, Rockett R, Lam C, Gray K, Carter I, Holmes EC, O'Sullivan MV, Byun R, Sintchenko V, Chen SC, Eden JS, Maddocks S, Kok J, Propenko M, Sorrell T, Chang S, Basile K, Dwyer DE for the 2019-nCoV Study Group |
| EPI_ISL_427749 | hCoV-19/Australia/NSW142/2020 | Oceania / Australia / New South Wales / | 2020-03-23 | Centre for Infectious Diseases and Microbiology Public Health | NSW Health Pathology - Institute of Clinical Pathology and Medical Research; Westmead Hospital; University of Sydney | Bachmann N, Rockett R, Lam C, Gray K, Timms V, Gall M, Arnott A, Sadsad R, Draper J, Sim E, Carter I, Holmes EC, O'Sullivan MV, Byun R, Sintchenko V, Chen SC, Eden JS, Maddocks S, Kok J, Propenko M, Sorrell T, Chang S, Basile K, Dwyer DE for the 2019-nCoV Study Group |
| EPI_ISL_427750 | hCoV-19/Australia/NSW73/2020 | Oceania / Australia / New South Wales / | 2020-03-23 | Centre for Infectious Diseases and Microbiology Public Health | NSW Health Pathology - Institute of Clinical Pathology and Medical Research; Westmead Hospital; University of Sydney | Rockett R, Lam C, Gray K, Timms V, Gall M, Arnott A, Sadsad R, Draper J, Sim E, Bachmann N, Carter I, Holmes EC, O'Sullivan MV, Byun R, Sintchenko V, Chen SC, Eden JS, Maddocks S, Kok J, Propenko M, Sorrell T, Chang S, Basile K, Dwyer DE for the 2019-nCoV Study Group |
| EPI_ISL_427751 | hCoV-19/Australia/NSW68/2020 | Oceania / Australia / New South Wales / | 2020-03-25 | Centre for Infectious Diseases and Microbiology Public Health | NSW Health Pathology - Institute of Clinical Pathology and Medical Research; Westmead Hospital; University of Sydney | Arnott A, Sadsad R, Draper J, Sim E, Bachmann N, Rockett R, Lam C, Gray K, Timms V, Gall M, Carter I, Holmes EC, O'Sullivan MV, Byun R, Sintchenko V, Chen SC, Eden JS, Maddocks S, Kok J, Propenko M, Sorrell T, Chang S, Basile K, Dwyer DE for the 2019-nCoV Study Group |
| EPI_ISL_427752 | hCoV-19/Australia/NSW184/2020 | Oceania / Australia / New South Wales / | 2020-03-24 | Centre for Infectious Diseases and Microbiology Public Health | NSW Health Pathology - Institute of Clinical Pathology and Medical Research; Westmead Hospital; University of Sydney | Lam C, Gray K, Timms, V, Gall M, Arnott A, Sadsad R, Draper J, Sim E, Bachmann N, Rockett R, Carter I, Holmes EC, O'Sullivan MV, Byun R, Sintchenko V, Chen SC, Eden JS, Maddocks S, Kok J, Propenko M, Sorrell T, Chang S, Basile K, Dwyer DE for the 2019-nCoV Study Group |
| EPI_ISL_427753 | hCoV-19/Australia/NSW208/2020 | Oceania / Australia / New South Wales / | 2020-03-24 | Centre for Infectious Diseases and Microbiology Public Health | NSW Health Pathology - Institute of Clinical Pathology and Medical Research; Westmead Hospital; University of Sydney | Arnott A, Sadsad R, Draper J, Sim E, Bachmann N, Rockett R, Lam C, Gray K, Timms V, Gall M, Carter I, Holmes EC, O'Sullivan MV, Byun R, Sintchenko V, Chen SC, Eden JS, Maddocks S, Kok J, Propenko M, Sorrell T, Chang S, Basile K, Dwyer DE for the 2019-nCoV Study Group |
| EPI_ISL_427754 | hCoV-19/Australia/NSW109/2020 | Oceania / Australia / New South Wales / | 2020-03-24 | Centre for Infectious Diseases and Microbiology Public Health | NSW Health Pathology - Institute of Clinical Pathology and Medical Research; Westmead Hospital; University of Sydney | Sadsad R, Draper J, Sim E, Bachmann N, Rockett R, Lam C, Gray K, Timms V, Gall M, Arnott A, Carter I, Holmes EC, O'Sullivan MV, Byun R, Sintchenko V, Chen SC, Eden JS, Maddocks S, Kok J, Propenko M, Sorrell T, Chang S, Basile K, Dwyer DE for the 2019-nCoV Study Group |
| EPI_ISL_427755 | hCoV-19/Australia/NSW209/2020 | Oceania / Australia / New South Wales / | 2020-03-23 | Centre for Infectious Diseases and Microbiology Public Health | NSW Health Pathology - Institute of Clinical Pathology and Medical Research; Westmead Hospital; University of Sydney | Sadsad R, Draper J, Sim E, Bachmann N, Rockett R, Lam C, Gray K, Timms V, Gall M, Arnott A, Carter I, Holmes EC, O'Sullivan MV, Byun R, Sintchenko V, Chen SC, Eden JS, Maddocks S, Kok J, Propenko M, Sorrell T, Chang S, Basile K, Dwyer DE for the 2019-nCoV Study Group |
| EPI_ISL_427756 | hCoV-19/Australia/NSW117/2020 | Oceania / Australia / New South Wales / | 2020-03-25 | Centre for Infectious Diseases and Microbiology Public Health | NSW Health Pathology - Institute of Clinical Pathology and Medical Research; Westmead Hospital; University of Sydney | Gall M, Arnott A, Sadsad R, Draper J, Sim E, Bachmann N, Rockett R, Lam C, Gray K, Timms V, Carter I, Holmes EC, O'Sullivan MV, Byun R, Sintchenko V, Chen SC, Eden JS, Maddocks S, Kok J, Propenko M, Sorrell T, Chang S, Basile K, Dwyer DE for the 2019-nCoV Study Group |
| EPI_ISL_427757 | hCoV-19/Australia/NSW70/2020 | Oceania / Australia / New South Wales / | 2020-03-20 | Centre for Infectious Diseases and Microbiology Public Health | NSW Health Pathology - Institute of Clinical Pathology and Medical Research; Westmead Hospital; University of Sydney | Draper J, Sim E, Bachmann N, Rockett R, Lam C, Gray K, Timms V, Gall M, Arnott A, Sadsad R, Carter I, Holmes EC, O'Sullivan MV, Byun R, Sintchenko V, Chen SC, Eden JS, Maddocks S, Kok J, Propenko M, Sorrell T, Chang S, Basile K, Dwyer DE for the 2019-nCoV Study Group |
| EPI_ISL_427758 | hCoV-19/Australia/NSW162/2020 | Oceania / Australia / New South Wales / | 2020-03-20 | Centre for Infectious Diseases and Microbiology Public Health | NSW Health Pathology - Institute of Clinical Pathology and Medical Research; Westmead Hospital; University of Sydney | Bachmann N, Rockett R, Lam C, Gray K, Timms V, Gall M, Arnott A, Sadsad R, Draper J, Sim E, Carter I, Holmes EC, O'Sullivan MV, Byun R, Sintchenko V, Chen SC, Eden JS, Maddocks S, Kok J, Propenko M, Sorrell T, Chang S, Basile K, Dwyer DE for the 2019-nCoV Study Group |
| EPI_ISL_427759 | hCoV-19/Australia/NSW71/2020 | Oceania / Australia / New South Wales / | 2020-03-20 | Centre for Infectious Diseases and Microbiology Public Health | NSW Health Pathology - Institute of Clinical Pathology and Medical Research; Westmead Hospital; University of Sydney | Sim E, Bachmann N, Rockett R, Lam C, Gray K, Timms V, Gall M, Arnott A, Sadsad R, Draper J, Carter I, Holmes EC, O'Sullivan MV, Byun R, Sintchenko V, Chen SC, Eden JS, Maddocks S, Kok J, Propenko M, Sorrell T, Chang S, Basile K, Dwyer DE for the 2019-nCoV Study Group |
| EPI_ISL_427760 | hCoV-19/Australia/NSW129/2020 | Oceania / Australia / New South Wales / | 2020-03-24 | Centre for Infectious Diseases and Microbiology Public Health | NSW Health Pathology - Institute of Clinical Pathology and Medical Research; Westmead Hospital; University of Sydney | Sadsad R, Draper J, Sim E, Bachmann N, Rockett R, Lam C, Gray K, Timms V, Gall M, Arnott A, Carter I, Holmes EC, O'Sullivan MV, Byun R, Sintchenko V, Chen SC, Eden JS, Maddocks S, Kok J, Propenko M, Sorrell T, Chang S, Basile K, Dwyer DE for the 2019-nCoV Study Group |
| EPI_ISL_427761 | hCoV-19/Australia/NSW138/2020 | Oceania / Australia / New South Wales / | 2020-03-24 | Centre for Infectious Diseases and Microbiology Public Health | NSW Health Pathology - Institute of Clinical Pathology and Medical Research; Westmead Hospital; University of Sydney | Arnott A, Sadsad R, Draper J, Sim E, Bachmann N, Rockett R, Lam C, Gray K, Timms V, Gall M, Carter I, Holmes EC, O'Sullivan MV, Byun R, Sintchenko V, Chen SC, Eden JS, Maddocks S, Kok J, Propenko M, Sorrell T, Chang S, Basile K, Dwyer DE for the 2019-nCoV Study Group |
| EPI_ISL_427762 | hCoV-19/Australia/NSW202/2020 | Oceania / Australia / New South Wales / | 2020-03-22 | Centre for Infectious Diseases and Microbiology Public Health | NSW Health Pathology - Institute of Clinical Pathology and Medical Research; Westmead Hospital; University of Sydney | Bachmann N, Rockett R, Lam C, Gray K, Timms V, Gall M, Arnott A, Sadsad R, Draper J, Sim E, Carter I, Holmes EC, O'Sullivan MV, Byun R, Sintchenko V, Chen SC, Eden JS, Maddocks S, Kok J, Propenko M, Sorrell T, Chang S, Basile K, Dwyer DE for the 2019-nCoV Study Group |
| EPI_ISL_427763 | hCoV-19/Australia/NSW194/2020 | Oceania / Australia / New South Wales / | 2020-03-23 | Centre for Infectious Diseases and Microbiology Public Health | NSW Health Pathology - Institute of Clinical Pathology and Medical Research; Westmead Hospital; University of Sydney | Lam C, Gray K, Timms, V, Gall M, Arnott A, Sadsad R, Draper J, Sim E, Bachmann N, Rockett R, Carter I, Holmes EC, O'Sullivan MV, Byun R, Sintchenko V, Chen SC, Eden JS, Maddocks S, Kok J, Propenko M, Sorrell T, Chang S, Basile K, Dwyer DE for the 2019-nCoV Study Group |
| EPI_ISL_427764 | hCoV-19/Australia/NSW147/2020 | Oceania / Australia / New South Wales / | 2020-03-23 | Centre for Infectious Diseases and Microbiology Public Health | NSW Health Pathology - Institute of Clinical Pathology and Medical Research; Westmead Hospital; University of Sydney | Gall M, Arnott A, Sadsad R, Draper J, Sim E, Bachmann N, Rockett R, Lam C, Gray K, Timms V, Carter I, Holmes EC, O'Sullivan MV, Byun R, Sintchenko V, Chen SC, Eden JS, Maddocks S, Kok J, Propenko M, Sorrell T, Chang S, Basile K, Dwyer DE for the 2019-nCoV Study Group |
| EPI_ISL_427765 | hCoV-19/Australia/NSW203/2020 | Oceania / Australia / New South Wales / | 2020-03-22 | Centre for Infectious Diseases and Microbiology Public Health | NSW Health Pathology - Institute of Clinical Pathology and Medical Research; Westmead Hospital; University of Sydney | Rockett R, Lam C, Gray K, Timms V, Gall M, Arnott A, Sadsad R, Draper J, Sim E, Bachmann N, Carter I, Holmes EC, O'Sullivan MV, Byun R, Sintchenko V, Chen SC, Eden JS, Maddocks S, Kok J, Propenko M, Sorrell T, Chang S, Basile K, Dwyer DE for the 2019-nCoV Study Group |
| EPI_ISL_427766 | hCoV-19/Australia/NSW210/2020 | Oceania / Australia / New South Wales / | 2020-03-25 | Centre for Infectious Diseases and Microbiology Public Health | NSW Health Pathology - Institute of Clinical Pathology and Medical Research; Westmead Hospital; University of Sydney | Draper J, Sim E, Bachmann N, Rockett R, Lam C, Gray K, Timms V, Gall M, Arnott A, Sadsad R, Carter I, Holmes EC, O'Sullivan MV, Byun R, Sintchenko V, Chen SC, Eden JS, Maddocks S, Kok J, Propenko M, Sorrell T, Chang S, Basile K, Dwyer DE for the 2019-nCoV Study Group |
| EPI_ISL_427767 | hCoV-19/Australia/NSW195/2020 | Oceania / Australia / New South Wales / | 2020-03-22 | Centre for Infectious Diseases and Microbiology Public Health | NSW Health Pathology - Institute of Clinical Pathology and Medical Research; Westmead Hospital; University of Sydney | Gray K, Timms V, Gall M, Arnott A, Sadsad R, Draper J, Sim E, Bachmann N, Rockett R, Lam C, Carter I, Holmes EC, O'Sullivan MV, Byun R, Sintchenko V, Chen SC, Eden JS, Maddocks S, Kok J, Propenko M, Sorrell T, Chang S, Basile K, Dwyer DE for the 2019-nCoV Study Group |
| EPI_ISL_427768 | hCoV-19/Australia/NSW130/2020 | Oceania / Australia / New South Wales / | 2020-03-23 | Centre for Infectious Diseases and Microbiology Public Health | NSW Health Pathology - Institute of Clinical Pathology and Medical Research; Westmead Hospital; University of Sydney | Draper J, Sim E, Bachmann N, Rockett R, Lam C, Gray K, Timms V, Gall M, Arnott A, Sadsad R, Carter I, Holmes EC, O'Sullivan MV, Byun R, Sintchenko V, Chen SC, Eden JS, Maddocks S, Kok J, Propenko M, Sorrell T, Chang S, Basile K, Dwyer DE for the 2019-nCoV Study Group |
| EPI_ISL_427769 | hCoV-19/Australia/NSW97/2020 | Oceania / Australia / New South Wales / | 2020-03-23 | Centre for Infectious Diseases and Microbiology Public Health | NSW Health Pathology - Institute of Clinical Pathology and Medical Research; Westmead Hospital; University of Sydney | Gall M, Arnott A, Sadsad R, Draper J, Sim E, Bachmann N, Rockett R, Lam C, Gray K, Timms V, Carter I, Holmes EC, O'Sullivan MV, Byun R, Sintchenko V, Chen SC, Eden JS, Maddocks S, Kok J, Propenko M, Sorrell T, Chang S, Basile K, Dwyer DE for the 2019-nCoV Study Group |
| EPI_ISL_427770 | hCoV-19/Australia/NSW78/2020 | Oceania / Australia / New South Wales / | 2020-03-23 | Centre for Infectious Diseases and Microbiology Public Health | NSW Health Pathology - Institute of Clinical Pathology and Medical Research; Westmead Hospital; University of Sydney | Arnott A, Sadsad R, Draper J, Sim E, Bachmann N, Rockett R, Lam C, Gray K, Timms V, Gall M, Carter I, Holmes EC, O'Sullivan MV, Byun R, Sintchenko V, Chen SC, Eden JS, Maddocks S, Kok J, Propenko M, Sorrell T, Chang S, Basile K, Dwyer DE for the 2019-nCoV Study Group |
| EPI_ISL_427771 | hCoV-19/Australia/NSW166/2020 | Oceania / Australia / New South Wales / | 2020-03-23 | Centre for Infectious Diseases and Microbiology Public Health | NSW Health Pathology - Institute of Clinical Pathology and Medical Research; Westmead Hospital; University of Sydney | Timms V, Gall M, Arnott A, Sadsad R, Draper J, Sim E, Bachmann N, Rockett R, Lam C, Gray K, Carter I, Holmes EC, O'Sullivan MV, Byun R, Sintchenko V, Chen SC, Eden JS, Maddocks S, Kok J, Propenko M, Sorrell T, Chang S, Basile K, Dwyer DE for the 2019-nCoV Study Group |
| EPI_ISL_427772 | hCoV-19/Australia/NSW107/2020 | Oceania / Australia / New South Wales / | 2020-03-22 | Centre for Infectious Diseases and Microbiology Public Health | NSW Health Pathology - Institute of Clinical Pathology and Medical Research; Westmead Hospital; University of Sydney | Gall M, Arnott A, Sadsad R, Draper J, Sim E, Bachmann N, Rockett R, Lam C, Gray K, Timms V, Carter I, Holmes EC, O'Sullivan MV, Byun R, Sintchenko V, Chen SC, Eden JS, Maddocks S, Kok J, Propenko M, Sorrell T, Chang S, Basile K, Dwyer DE for the 2019-nCoV Study Group |
| EPI_ISL_427773 | hCoV-19/Australia/NSW64/2020 | Oceania / Australia / New South Wales / | 2020-03-25 | Centre for Infectious Diseases and Microbiology Public Health | NSW Health Pathology - Institute of Clinical Pathology and Medical Research; Westmead Hospital; University of Sydney | Lam C, Gray K, Timms, V, Gall M, Arnott A, Sadsad R, Draper J, Sim E, Bachmann N, Rockett R, Carter I, Holmes EC, O'Sullivan MV, Byun R, Sintchenko V, Chen SC, Eden JS, Maddocks S, Kok J, Propenko M, Sorrell T, Chang S, Basile K, Dwyer DE for the 2019-nCoV Study Group |
| EPI_ISL_427774 | hCoV-19/Australia/NSW98/2020 | Oceania / Australia / New South Wales / | 2020-03-22 | Centre for Infectious Diseases and Microbiology Public Health | NSW Health Pathology - Institute of Clinical Pathology and Medical Research; Westmead Hospital; University of Sydney | Arnott A, Sadsad R, Draper J, Sim E, Bachmann N, Rockett R, Lam C, Gray K, Timms V, Gall M, Carter I, Holmes EC, O'Sullivan MV, Byun R, Sintchenko V, Chen SC, Eden JS, Maddocks S, Kok J, Propenko M, Sorrell T, Chang S, Basile K, Dwyer DE for the 2019-nCoV Study Group |
| EPI_ISL_427775 | hCoV-19/Australia/NSW149/2020 | Oceania / Australia / New South Wales / | 2020-03-25 | Centre for Infectious Diseases and Microbiology Public Health | NSW Health Pathology - Institute of Clinical Pathology and Medical Research; Westmead Hospital; University of Sydney | Sadsad R, Draper J, Sim E, Bachmann N, Rockett R, Lam C, Gray K, Timms V, Gall M, Arnott A, Carter I, Holmes EC, O'Sullivan MV, Byun R, Sintchenko V, Chen SC, Eden JS, Maddocks S, Kok J, Propenko M, Sorrell T, Chang S, Basile K, Dwyer DE for the 2019-nCoV Study Group |
| EPI_ISL_427776 | hCoV-19/Australia/NSW89/2020 | Oceania / Australia / New South Wales / | 2020-03-23 | Centre for Infectious Diseases and Microbiology Public Health | NSW Health Pathology - Institute of Clinical Pathology and Medical Research; Westmead Hospital; University of Sydney | Sadsad R, Draper J, Sim E, Bachmann N, Rockett R, Lam C, Gray K, Timms V, Gall M, Arnott A, Carter I, Holmes EC, O'Sullivan MV, Byun R, Sintchenko V, Chen SC, Eden JS, Maddocks S, Kok J, Propenko M, Sorrell T, Chang S, Basile K, Dwyer DE for the 2019-nCoV Study Group |
| EPI_ISL_427777 | hCoV-19/Australia/NSW99/2020 | Oceania / Australia / New South Wales / | 2020-03-21 | Centre for Infectious Diseases and Microbiology Public Health | NSW Health Pathology - Institute of Clinical Pathology and Medical Research; Westmead Hospital; University of Sydney | Sadsad R, Draper J, Sim E, Bachmann N, Rockett R, Lam C, Gray K, Timms V, Gall M, Arnott A, Carter I, Holmes EC, O'Sullivan MV, Byun R, Sintchenko V, Chen SC, Eden JS, Maddocks S, Kok J, Propenko M, Sorrell T, Chang S, Basile K, Dwyer DE for the 2019-nCoV Study Group |
| EPI_ISL_427778 | hCoV-19/Australia/NSW122/2020 | Oceania / Australia / New South Wales / | 2020-03-25 | Centre for Infectious Diseases and Microbiology Public Health | NSW Health Pathology - Institute of Clinical Pathology and Medical Research; Westmead Hospital; University of Sydney | Bachmann N, Rockett R, Lam C, Gray K, Timms V, Gall M, Arnott A, Sadsad R, Draper J, Sim E, Carter I, Holmes EC, O'Sullivan MV, Byun R, Sintchenko V, Chen SC, Eden JS, Maddocks S, Kok J, Propenko M, Sorrell T, Chang S, Basile K, Dwyer DE for the 2019-nCoV Study Group |
| EPI_ISL_427779 | hCoV-19/Australia/NSW169/2020 | Oceania / Australia / New South Wales / | 2020-03-24 | Centre for Infectious Diseases and Microbiology Public Health | NSW Health Pathology - Institute of Clinical Pathology and Medical Research; Westmead Hospital; University of Sydney | Sadsad R, Draper J, Sim E, Bachmann N, Rockett R, Lam C, Gray K, Timms V, Gall M, Arnott A, Carter I, Holmes EC, O'Sullivan MV, Byun R, Sintchenko V, Chen SC, Eden JS, Maddocks S, Kok J, Propenko M, Sorrell T, Chang S, Basile K, Dwyer DE for the 2019-nCoV Study Group |
| EPI_ISL_427780 | hCoV-19/Australia/NSW123/2020 | Oceania / Australia / New South Wales / | 2020-03-23 | Centre for Infectious Diseases and Microbiology Public Health | NSW Health Pathology - Institute of Clinical Pathology and Medical Research; Westmead Hospital; University of Sydney | Rockett R, Lam C, Gray K, Timms V, Gall M, Arnott A, Sadsad R, Draper J, Sim E, Bachmann N, Carter I, Holmes EC, O'Sullivan MV, Byun R, Sintchenko V, Chen SC, Eden JS, Maddocks S, Kok J, Propenko M, Sorrell T, Chang S, Basile K, Dwyer DE for the 2019-nCoV Study Group |
| EPI_ISL_427781 | hCoV-19/Australia/NSW159/2020 | Oceania / Australia / New South Wales / | 2020-03-22 | Centre for Infectious Diseases and Microbiology Public Health | NSW Health Pathology - Institute of Clinical Pathology and Medical Research; Westmead Hospital; University of Sydney | Sadsad R, Draper J, Sim E, Bachmann N, Rockett R, Lam C, Gray K, Timms V, Gall M, Arnott A, Carter I, Holmes EC, O'Sullivan MV, Byun R, Sintchenko V, Chen SC, Eden JS, Maddocks S, Kok J, Propenko M, Sorrell T, Chang S, Basile K, Dwyer DE for the 2019-nCoV Study Group |
| EPI_ISL_427782 | hCoV-19/Australia/NSW124/2020 | Oceania / Australia / New South Wales / | 2020-03-25 | Centre for Infectious Diseases and Microbiology Public Health | NSW Health Pathology - Institute of Clinical Pathology and Medical Research; Westmead Hospital; University of Sydney | Lam C, Gray K, Timms, V, Gall M, Arnott A, Sadsad R, Draper J, Sim E, Bachmann N, Rockett R, Carter I, Holmes EC, O'Sullivan MV, Byun R, Sintchenko V, Chen SC, Eden JS, Maddocks S, Kok J, Propenko M, Sorrell T, Chang S, Basile K, Dwyer DE for the 2019-nCoV Study Group |
| EPI_ISL_427783 | hCoV-19/Australia/NSW131/2020 | Oceania / Australia / New South Wales / | 2020-03-25 | Centre for Infectious Diseases and Microbiology Public Health | NSW Health Pathology - Institute of Clinical Pathology and Medical Research; Westmead Hospital; University of Sydney | Sim E, Bachmann N, Rockett R, Lam C, Gray K, Timms V, Gall M, Arnott A, Sadsad R, Draper J, Carter I, Holmes EC, O'Sullivan MV, Byun R, Sintchenko V, Chen SC, Eden JS, Maddocks S, Kok J, Propenko M, Sorrell T, Chang S, Basile K, Dwyer DE for the 2019-nCoV Study Group |
| EPI_ISL_427784 | hCoV-19/Australia/NSW160/2020 | Oceania / Australia / New South Wales / | 2020-03-20 | Centre for Infectious Diseases and Microbiology Public Health | NSW Health Pathology - Institute of Clinical Pathology and Medical Research; Westmead Hospital; University of Sydney | Draper J, Sim E, Bachmann N, Rockett R, Lam C, Gray K, Timms V, Gall M, Arnott A, Sadsad R, Carter I, Holmes EC, O'Sullivan MV, Byun R, Sintchenko V, Chen SC, Eden JS, Maddocks S, Kok J, Propenko M, Sorrell T, Chang S, Basile K, Dwyer DE for the 2019-nCoV Study Group |
| EPI_ISL_427785 | hCoV-19/Australia/NSW125/2020 | Oceania / Australia / New South Wales / | 2020-03-23 | Centre for Infectious Diseases and Microbiology Public Health | NSW Health Pathology - Institute of Clinical Pathology and Medical Research; Westmead Hospital; University of Sydney | Gray K, Timms V, Gall M, Arnott A, Sadsad R, Draper J, Sim E, Bachmann N, Rockett R, Lam C, Carter I, Holmes EC, O'Sullivan MV, Byun R, Sintchenko V, Chen SC, Eden JS, Maddocks S, Kok J, Propenko M, Sorrell T, Chang S, Basile K, Dwyer DE for the 2019-nCoV Study Group |
| EPI_ISL_427786 | hCoV-19/Australia/NSW191/2020 | Oceania / Australia / New South Wales / | 2020-03-25 | Centre for Infectious Diseases and Microbiology Public Health | NSW Health Pathology - Institute of Clinical Pathology and Medical Research; Westmead Hospital; University of Sydney | Sim E, Bachmann N, Rockett R, Lam C, Gray K, Timms V, Gall M, Arnott A, Sadsad R, Draper J, Carter I, Holmes EC, O'Sullivan MV, Byun R, Sintchenko V, Chen SC, Eden JS, Maddocks S, Kok J, Propenko M, Sorrell T, Chang S, Basile K, Dwyer DE for the 2019-nCoV Study Group |
| EPI_ISL_427787 | hCoV-19/Australia/NSW132/2020 | Oceania / Australia / New South Wales / | 2020-03-26 | Centre for Infectious Diseases and Microbiology Public Health | NSW Health Pathology - Institute of Clinical Pathology and Medical Research; Westmead Hospital; University of Sydney | Bachmann N, Rockett R, Lam C, Gray K, Timms V, Gall M, Arnott A, Sadsad R, Draper J, Sim E, Carter I, Holmes EC, O'Sullivan MV, Byun R, Sintchenko V, Chen SC, Eden JS, Maddocks S, Kok J, Propenko M, Sorrell T, Chang S, Basile K, Dwyer DE for the 2019-nCoV Study Group |
| EPI_ISL_427788 | hCoV-19/Australia/NSW161/2020 | Oceania / Australia / New South Wales / | 2020-03-20 | Centre for Infectious Diseases and Microbiology Public Health | NSW Health Pathology - Institute of Clinical Pathology and Medical Research; Westmead Hospital; University of Sydney | Sim E, Bachmann N, Rockett R, Lam C, Gray K, Timms V, Gall M, Arnott A, Sadsad R, Draper J, Carter I, Holmes EC, O'Sullivan MV, Byun R, Sintchenko V, Chen SC, Eden JS, Maddocks S, Kok J, Propenko M, Sorrell T, Chang S, Basile K, Dwyer DE for the 2019-nCoV Study Group |
| EPI_ISL_427789 | hCoV-19/Australia/NSW163/2020 | Oceania / Australia / New South Wales / | 2020-03-20 | Centre for Infectious Diseases and Microbiology Public Health | NSW Health Pathology - Institute of Clinical Pathology and Medical Research; Westmead Hospital; University of Sydney | Rockett R, Lam C, Gray K, Timms V, Gall M, Arnott A, Sadsad R, Draper J, Sim E, Bachmann N, Carter I, Holmes EC, O'Sullivan MV, Byun R, Sintchenko V, Chen SC, Eden JS, Maddocks S, Kok J, Propenko M, Sorrell T, Chang S, Basile K, Dwyer DE for the 2019-nCoV Study Group |
| EPI_ISL_427790 | hCoV-19/Australia/NSW79/2020 | Oceania / Australia / New South Wales / | 2020-03-23 | Centre for Infectious Diseases and Microbiology Public Health | NSW Health Pathology - Institute of Clinical Pathology and Medical Research; Westmead Hospital; University of Sydney | Sadsad R, Draper J, Sim E, Bachmann N, Rockett R, Lam C, Gray K, Timms V, Gall M, Arnott A, Carter I, Holmes EC, O'Sullivan MV, Byun R, Sintchenko V, Chen SC, Eden JS, Maddocks S, Kok J, Propenko M, Sorrell T, Chang S, Basile K, Dwyer DE for the 2019-nCoV Study Group |
| EPI_ISL_427791 | hCoV-19/Australia/NSW133/2020 | Oceania / Australia / New South Wales / | 2020-03-24 | Centre for Infectious Diseases and Microbiology Public Health | NSW Health Pathology - Institute of Clinical Pathology and Medical Research; Westmead Hospital; University of Sydney | Rockett R, Lam C, Gray K, Timms V, Gall M, Arnott A, Sadsad R, Draper J, Sim E, Bachmann N, Carter I, Holmes EC, O'Sullivan MV, Byun R, Sintchenko V, Chen SC, Eden JS, Maddocks S, Kok J, Propenko M, Sorrell T, Chang S, Basile K, Dwyer DE for the 2019-nCoV Study Group |
| EPI_ISL_427792 | hCoV-19/Australia/NSW204/2020 | Oceania / Australia / New South Wales / | 2020-03-25 | Centre for Infectious Diseases and Microbiology Public Health | NSW Health Pathology - Institute of Clinical Pathology and Medical Research; Westmead Hospital; University of Sydney | Lam C, Gray K, Timms, V, Gall M, Arnott A, Sadsad R, Draper J, Sim E, Bachmann N, Rockett R, Carter I, Holmes EC, O'Sullivan MV, Byun R, Sintchenko V, Chen SC, Eden JS, Maddocks S, Kok J, Propenko M, Sorrell T, Chang S, Basile K, Dwyer DE for the 2019-nCoV Study Group |
| EPI_ISL_427793 | hCoV-19/Australia/NSW185/2020 | Oceania / Australia / New South Wales / | 2020-03-26 | Centre for Infectious Diseases and Microbiology Public Health | NSW Health Pathology - Institute of Clinical Pathology and Medical Research; Westmead Hospital; University of Sydney | Gray K, Timms V, Gall M, Arnott A, Sadsad R, Draper J, Sim E, Bachmann N, Rockett R, Lam C, Carter I, Holmes EC, O'Sullivan MV, Byun R, Sintchenko V, Chen SC, Eden JS, Maddocks S, Kok J, Propenko M, Sorrell T, Chang S, Basile K, Dwyer DE for the 2019-nCoV Study Group |
| EPI_ISL_427794 | hCoV-19/Australia/NSW126/2020 | Oceania / Australia / New South Wales / | 2020-03-23 | Centre for Infectious Diseases and Microbiology Public Health | NSW Health Pathology - Institute of Clinical Pathology and Medical Research; Westmead Hospital; University of Sydney | Timms V, Gall M, Arnott A, Sadsad R, Draper J, Sim E, Bachmann N, Rockett R, Lam C, Gray K, Carter I, Holmes EC, O'Sullivan MV, Byun R, Sintchenko V, Chen SC, Eden JS, Maddocks S, Kok J, Propenko M, Sorrell T, Chang S, Basile K, Dwyer DE for the 2019-nCoV Study Group |
| EPI_ISL_427795 | hCoV-19/Australia/NSW139/2020 | Oceania / Australia / New South Wales / | 2020-03-24 | Centre for Infectious Diseases and Microbiology Public Health | NSW Health Pathology - Institute of Clinical Pathology and Medical Research; Westmead Hospital; University of Sydney | Sadsad R, Draper J, Sim E, Bachmann N, Rockett R, Lam C, Gray K, Timms V, Gall M, Arnott A, Carter I, Holmes EC, O'Sullivan MV, Byun R, Sintchenko V, Chen SC, Eden JS, Maddocks S, Kok J, Propenko M, Sorrell T, Chang S, Basile K, Dwyer DE for the 2019-nCoV Study Group |
| EPI_ISL_427796 | hCoV-19/Australia/NSW150/2020 | Oceania / Australia / New South Wales / | 2020-03-25 | Centre for Infectious Diseases and Microbiology Public Health | NSW Health Pathology - Institute of Clinical Pathology and Medical Research; Westmead Hospital; University of Sydney | Draper J, Sim E, Bachmann N, Rockett R, Lam C, Gray K, Timms V, Gall M, Arnott A, Sadsad R, Carter I, Holmes EC, O'Sullivan MV, Byun R, Sintchenko V, Chen SC, Eden JS, Maddocks S, Kok J, Propenko M, Sorrell T, Chang S, Basile K, Dwyer DE for the 2019-nCoV Study Group |
| EPI_ISL_427797 | hCoV-19/Australia/NSW192/2020 | Oceania / Australia / New South Wales / | 2020-03-26 | Centre for Infectious Diseases and Microbiology Public Health | NSW Health Pathology - Institute of Clinical Pathology and Medical Research; Westmead Hospital; University of Sydney | Bachmann N, Rockett R, Lam C, Gray K, Timms V, Gall M, Arnott A, Sadsad R, Draper J, Sim E, Carter I, Holmes EC, O'Sullivan MV, Byun R, Sintchenko V, Chen SC, Eden JS, Maddocks S, Kok J, Propenko M, Sorrell T, Chang S, Basile K, Dwyer DE for the 2019-nCoV Study Group |
| EPI_ISL_427798 | hCoV-19/Australia/NSW193/2020 | Oceania / Australia / New South Wales / | 2020-03-20 | Centre for Infectious Diseases and Microbiology Public Health | NSW Health Pathology - Institute of Clinical Pathology and Medical Research; Westmead Hospital; University of Sydney | Rockett R, Lam C, Gray K, Timms V, Gall M, Arnott A, Sadsad R, Draper J, Sim E, Bachmann N, Carter I, Holmes EC, O'Sullivan MV, Byun R, Sintchenko V, Chen SC, Eden JS, Maddocks S, Kok J, Propenko M, Sorrell T, Chang S, Basile K, Dwyer DE for the 2019-nCoV Study Group |
| EPI_ISL_427799 | hCoV-19/Australia/NSW140/2020 | Oceania / Australia / New South Wales / | 2020-03-17 | Centre for Infectious Diseases and Microbiology Public Health | NSW Health Pathology - Institute of Clinical Pathology and Medical Research; Westmead Hospital; University of Sydney | Draper J, Sim E, Bachmann N, Rockett R, Lam C, Gray K, Timms V, Gall M, Arnott A, Sadsad R, Carter I, Holmes EC, O'Sullivan MV, Byun R, Sintchenko V, Chen SC, Eden JS, Maddocks S, Kok J, Propenko M, Sorrell T, Chang S, Basile K, Dwyer DE for the 2019-nCoV Study Group |
| EPI_ISL_428201 | hCoV-19/Sweden/RV-FOI-6/2020 | Europe / Sweden / Västerbotten | 2020-03-20 | Klinisk mikrobiologi, Region Västerbotten | Unit for Biological Agents, Department for CBRN Defence and Security, Swedish Defence Research Agency | FOI Bioinformatics team |
| EPI_ISL_428229 | hCoV-19/Taiwan/TSGH-12/2020 | Asia / Taiwan / Taipei | 2020-03-18 | TSGH-CP molecular lab | TSGH-CP molecular lab | Cherng-Lih Perng, Ming-Jr Jian, Chih-Kai Chang, Jung-Chung Lin, Kuo-Ming Yeh, Chien-Wen Chen, Sheng-Kang Chiu, Hsing-Yi Chung, Shih-Hung Tsai, Kuo-Sheng Hung, Tien-Yao Chang, Feng-Yee Chang, Hung-Sheng Shang |
| EPI_ISL_428230 | hCoV-19/Taiwan/TSGH-13/2020 | Asia / Taiwan / New Taipei City | 2020-03-17 | TSGH-CP molecular lab | TSGH-CP molecular lab | Cherng-Lih Perng, Ming-Jr Jian, Chih-Kai Chang, Jung-Chung Lin, Kuo-Ming Yeh, Chien-Wen Chen, Sheng-Kang Chiu, Hsing-Yi Chung, Shih-Hung Tsai, Kuo-Sheng Hung, Tien-Yao Chang, Feng-Yee Chang, Hung-Sheng Shang |
| EPI_ISL_428231 | hCoV-19/Taiwan/TSGH-14/2020 | Asia / Taiwan / New Taipei City | 2020-03-24 | TSGH-CP molecular lab | TSGH-CP molecular lab | Cherng-Lih Perng, Ming-Jr JIAN, Chih-Kai Chang, Jung-Chung Lin, Kuo-Ming Yeh, Chien-Wen Chen, Sheng-Kang Chiu, Hsing-Yi Chung, Shih-Hung Tsai, Kuo-Sheng Hung, Tien-Yao Chang, Feng-Yee Chang, Hung-Sheng Shang |
| EPI_ISL_428250 | hCoV-19/USA/CT-Yale-011/2020 | North America / USA / Connecticut | 2020-03-11 | Yale Clinical Virology Laboratory | Grubaugh Lab - Yale School of Public Health | Joseph Fauver, Anderson Brito, Tara Alpert, Chantal Vogels, Ellen Foxman, Albert Ko, Marie Landry, Nathan Grubaugh |
| EPI_ISL_428252 | hCoV-19/USA/WI-UW-156/2020 | North America / USA / Wisconsin / Wauw | 2020-03-25 | University of Wisconsin-Madison AIDS Vaccine Research Laboratories | University of Wisconsin-Madison AIDS Vaccine Research Laboratories | Gage Moreno, Katarina Braun, et al. AIDS Vaccine Research Laboratories |
| EPI_ISL_428253 | hCoV-19/USA/WI-UW-157/2020 | North America / USA / Wisconsin / Milwa | 2020-03-15 | University of Wisconsin-Madison AIDS Vaccine Research Laboratories | University of Wisconsin-Madison AIDS Vaccine Research Laboratories | Gage Moreno, Katarina Braun, et al. AIDS Vaccine Research Laboratories |
| EPI_ISL_428254 | hCoV-19/USA/WI-UW-158/2020 | North America / USA / Wisconsin / Oak C | 2020-03-15 | University of Wisconsin-Madison AIDS Vaccine Research Laboratories | University of Wisconsin-Madison AIDS Vaccine Research Laboratories | Gage Moreno, Katarina Braun, et al. AIDS Vaccine Research Laboratories |
| EPI_ISL_428255 | hCoV-19/USA/WI-UW-159/2020 | North America / USA / Wisconsin / Oak C | 2020-03-15 | University of Wisconsin-Madison AIDS Vaccine Research Laboratories | University of Wisconsin-Madison AIDS Vaccine Research Laboratories | Gage Moreno, Katarina Braun, et al. AIDS Vaccine Research Laboratories |
| EPI_ISL_428256 | hCoV-19/USA/WI-UW-160/2020 | North America / USA / Wisconsin / Oak C | 2020-03-15 | University of Wisconsin-Madison AIDS Vaccine Research Laboratories | University of Wisconsin-Madison AIDS Vaccine Research Laboratories | Gage Moreno, Katarina Braun, et al. AIDS Vaccine Research Laboratories |
| EPI_ISL_428257 | hCoV-19/USA/WI-UW-161/2020 | North America / USA / Wisconsin / Milwa | 2020-03-29 | University of Wisconsin-Madison AIDS Vaccine Research Laboratories | University of Wisconsin-Madison AIDS Vaccine Research Laboratories | Gage Moreno, Katarina Braun, et al. AIDS Vaccine Research Laboratories |
| EPI_ISL_428258 | hCoV-19/USA/WI-UW-162/2020 | North America / USA / Wisconsin / Milwa | 2020-03-29 | University of Wisconsin-Madison AIDS Vaccine Research Laboratories | University of Wisconsin-Madison AIDS Vaccine Research Laboratories | Gage Moreno, Katarina Braun, et al. AIDS Vaccine Research Laboratories |
| EPI_ISL_428259 | hCoV-19/USA/WI-UW-163/2020 | North America / USA / Wisconsin / Elm G | 2020-03-16 | University of Wisconsin-Madison AIDS Vaccine Research Laboratories | University of Wisconsin-Madison AIDS Vaccine Research Laboratories | Gage Moreno, Katarina Braun, et al. AIDS Vaccine Research Laboratories |
| EPI_ISL_428260 | hCoV-19/USA/WI-UW-164/2020 | North America / USA / Wisconsin / Pewa | 2020-03-16 | University of Wisconsin-Madison AIDS Vaccine Research Laboratories | University of Wisconsin-Madison AIDS Vaccine Research Laboratories | Gage Moreno, Katarina Braun, et al. AIDS Vaccine Research Laboratories |
| EPI_ISL_428261 | hCoV-19/USA/WI-UW-165/2020 | North America / USA / Wisconsin / Milwa | 2020-03-17 | University of Wisconsin-Madison AIDS Vaccine Research Laboratories | University of Wisconsin-Madison AIDS Vaccine Research Laboratories | Gage Moreno, Katarina Braun, et al. AIDS Vaccine Research Laboratories |
| EPI_ISL_428262 | hCoV-19/USA/WI-UW-166/2020 | North America / USA / Wisconsin / Green | 2020-03-17 | University of Wisconsin-Madison AIDS Vaccine Research Laboratories | University of Wisconsin-Madison AIDS Vaccine Research Laboratories | Gage Moreno, Katarina Braun, et al. AIDS Vaccine Research Laboratories |
| EPI_ISL_428263 | hCoV-19/USA/WI-UW-167/2020 | North America / USA / Wisconsin / New E | 2020-03-18 | University of Wisconsin-Madison AIDS Vaccine Research Laboratories | University of Wisconsin-Madison AIDS Vaccine Research Laboratories | Gage Moreno, Katarina Braun, et al. AIDS Vaccine Research Laboratories |
| EPI_ISL_428264 | hCoV-19/USA/WI-UW-168/2020 | North America / USA / Wisconsin / Milwa | 2020-03-19 | University of Wisconsin-Madison AIDS Vaccine Research Laboratories | University of Wisconsin-Madison AIDS Vaccine Research Laboratories | Gage Moreno, Katarina Braun, et al. AIDS Vaccine Research Laboratories |
| EPI_ISL_428265 | hCoV-19/USA/WI-UW-169/2020 | North America / USA / Wisconsin / Grafto | 2020-03-19 | University of Wisconsin-Madison AIDS Vaccine Research Laboratories | University of Wisconsin-Madison AIDS Vaccine Research Laboratories | Gage Moreno, Katarina Braun, et al. AIDS Vaccine Research Laboratories |
| EPI_ISL_428266 | hCoV-19/USA/WI-UW-170/2020 | North America / USA / Wisconsin / Milwa | 2020-03-19 | University of Wisconsin-Madison AIDS Vaccine Research Laboratories | University of Wisconsin-Madison AIDS Vaccine Research Laboratories | Gage Moreno, Katarina Braun, et al. AIDS Vaccine Research Laboratories |
| EPI_ISL_428267 | hCoV-19/USA/WI-UW-171/2020 | North America / USA / Wisconsin / Milwa | 2020-03-19 | University of Wisconsin-Madison AIDS Vaccine Research Laboratories | University of Wisconsin-Madison AIDS Vaccine Research Laboratories | Gage Moreno, Katarina Braun, et al. AIDS Vaccine Research Laboratories |
| EPI_ISL_428268 | hCoV-19/USA/WI-UW-172/2020 | North America / USA / Wisconsin / Milwa | 2020-03-19 | University of Wisconsin-Madison AIDS Vaccine Research Laboratories | University of Wisconsin-Madison AIDS Vaccine Research Laboratories | Gage Moreno, Katarina Braun, et al. AIDS Vaccine Research Laboratories |
| EPI_ISL_428269 | hCoV-19/USA/WI-UW-173/2020 | North America / USA / Wisconsin / Milwa | 2020-03-20 | University of Wisconsin-Madison AIDS Vaccine Research Laboratories | University of Wisconsin-Madison AIDS Vaccine Research Laboratories | Gage Moreno, Katarina Braun, et al. AIDS Vaccine Research Laboratories |
| EPI_ISL_428270 | hCoV-19/USA/WI-UW-174/2020 | North America / USA / Wisconsin / Grafto | 2020-03-20 | University of Wisconsin-Madison AIDS Vaccine Research Laboratories | University of Wisconsin-Madison AIDS Vaccine Research Laboratories | Gage Moreno, Katarina Braun, et al. AIDS Vaccine Research Laboratories |
| EPI_ISL_428271 | hCoV-19/USA/WI-UW-175/2020 | North America / USA / Wisconsin / Grafto | 2020-03-20 | University of Wisconsin-Madison AIDS Vaccine Research Laboratories | University of Wisconsin-Madison AIDS Vaccine Research Laboratories | Gage Moreno, Katarina Braun, et al. AIDS Vaccine Research Laboratories |
| EPI_ISL_428272 | hCoV-19/USA/WI-UW-176/2020 | North America / USA / Wisconsin / Milwa | 2020-03-20 | University of Wisconsin-Madison AIDS Vaccine Research Laboratories | University of Wisconsin-Madison AIDS Vaccine Research Laboratories | Gage Moreno, Katarina Braun, et al. AIDS Vaccine Research Laboratories |
| EPI_ISL_428273 | hCoV-19/USA/WI-UW-177/2020 | North America / USA / Wisconsin / Milwa | 2020-03-20 | University of Wisconsin-Madison AIDS Vaccine Research Laboratories | University of Wisconsin-Madison AIDS Vaccine Research Laboratories | Gage Moreno, Katarina Braun, et al. AIDS Vaccine Research Laboratories |
| EPI_ISL_428274 | hCoV-19/USA/WI-UW-178/2020 | North America / USA / Wisconsin / Milwa | 2020-03-20 | University of Wisconsin-Madison AIDS Vaccine Research Laboratories | University of Wisconsin-Madison AIDS Vaccine Research Laboratories | Gage Moreno, Katarina Braun, et al. AIDS Vaccine Research Laboratories |
| EPI_ISL_428275 | hCoV-19/USA/WI-UW-179/2020 | North America / USA / Wisconsin / Milwa | 2020-03-21 | University of Wisconsin-Madison AIDS Vaccine Research Laboratories | University of Wisconsin-Madison AIDS Vaccine Research Laboratories | Gage Moreno, Katarina Braun, et al. AIDS Vaccine Research Laboratories |
| EPI_ISL_428276 | hCoV-19/USA/WI-UW-180/2020 | North America / USA / Wisconsin / Grafto | 2020-03-21 | University of Wisconsin-Madison AIDS Vaccine Research Laboratories | University of Wisconsin-Madison AIDS Vaccine Research Laboratories | Gage Moreno, Katarina Braun, et al. AIDS Vaccine Research Laboratories |
| EPI_ISL_428277 | hCoV-19/USA/WI-UW-181/2020 | North America / USA / Wisconsin / Milwa | 2020-03-22 | University of Wisconsin-Madison AIDS Vaccine Research Laboratories | University of Wisconsin-Madison AIDS Vaccine Research Laboratories | Gage Moreno, Katarina Braun, et al. AIDS Vaccine Research Laboratories |
| EPI_ISL_428278 | hCoV-19/USA/WI-UW-182/2020 | North America / USA / Wisconsin / Milwa | 2020-03-22 | University of Wisconsin-Madison AIDS Vaccine Research Laboratories | University of Wisconsin-Madison AIDS Vaccine Research Laboratories | Gage Moreno, Katarina Braun, et al. AIDS Vaccine Research Laboratories |
| EPI_ISL_428279 | hCoV-19/USA/WI-UW-183/2020 | North America / USA / Wisconsin / Milwa | 2020-03-22 | University of Wisconsin-Madison AIDS Vaccine Research Laboratories | University of Wisconsin-Madison AIDS Vaccine Research Laboratories | Gage Moreno, Katarina Braun, et al. AIDS Vaccine Research Laboratories |
| EPI_ISL_428280 | hCoV-19/USA/WI-UW-184/2020 | North America / USA / Wisconsin / Milwa | 2020-03-22 | University of Wisconsin-Madison AIDS Vaccine Research Laboratories | University of Wisconsin-Madison AIDS Vaccine Research Laboratories | Gage Moreno, Katarina Braun, et al. AIDS Vaccine Research Laboratories |
| EPI_ISL_428281 | hCoV-19/USA/WI-UW-185/2020 | North America / USA / Wisconsin / Milwa | 2020-03-22 | University of Wisconsin-Madison AIDS Vaccine Research Laboratories | University of Wisconsin-Madison AIDS Vaccine Research Laboratories | Gage Moreno, Katarina Braun, et al. AIDS Vaccine Research Laboratories |
| EPI_ISL_428282 | hCoV-19/USA/WI-UW-186/2020 | North America / USA / Wisconsin / Milwa | 2020-03-23 | University of Wisconsin-Madison AIDS Vaccine Research Laboratories | University of Wisconsin-Madison AIDS Vaccine Research Laboratories | Gage Moreno, Katarina Braun, et al. AIDS Vaccine Research Laboratories |
| EPI_ISL_428283 | hCoV-19/USA/WI-UW-187/2020 | North America / USA / Wisconsin / Milwa | 2020-03-23 | University of Wisconsin-Madison AIDS Vaccine Research Laboratories | University of Wisconsin-Madison AIDS Vaccine Research Laboratories | Gage Moreno, Katarina Braun, et al. AIDS Vaccine Research Laboratories |
| EPI_ISL_428284 | hCoV-19/USA/WI-UW-188/2020 | North America / USA / Wisconsin / Milwa | 2020-03-23 | University of Wisconsin-Madison AIDS Vaccine Research Laboratories | University of Wisconsin-Madison AIDS Vaccine Research Laboratories | Gage Moreno, Katarina Braun, et al. AIDS Vaccine Research Laboratories |
| EPI_ISL_428285 | hCoV-19/USA/WI-UW-189/2020 | North America / USA / Wisconsin / Milwa | 2020-03-23 | University of Wisconsin-Madison AIDS Vaccine Research Laboratories | University of Wisconsin-Madison AIDS Vaccine Research Laboratories | Gage Moreno, Katarina Braun, et al. AIDS Vaccine Research Laboratories |
| EPI_ISL_428286 | hCoV-19/USA/WI-UW-190/2020 | North America / USA / Wisconsin / Wauw | 2020-03-23 | University of Wisconsin-Madison AIDS Vaccine Research Laboratories | University of Wisconsin-Madison AIDS Vaccine Research Laboratories | Gage Moreno, Katarina Braun, et al. AIDS Vaccine Research Laboratories |
| EPI_ISL_428287 | hCoV-19/USA/WI-UW-191/2020 | North America / USA / Wisconsin / Milwa | 2020-03-23 | University of Wisconsin-Madison AIDS Vaccine Research Laboratories | University of Wisconsin-Madison AIDS Vaccine Research Laboratories | Gage Moreno, Katarina Braun, et al. AIDS Vaccine Research Laboratories |
| EPI_ISL_428288 | hCoV-19/USA/WI-UW-192/2020 | North America / USA / Wisconsin / Milwa | 2020-03-23 | University of Wisconsin-Madison AIDS Vaccine Research Laboratories | University of Wisconsin-Madison AIDS Vaccine Research Laboratories | Gage Moreno, Katarina Braun, et al. AIDS Vaccine Research Laboratories |
| EPI_ISL_428289 | hCoV-19/USA/WI-UW-193/2020 | North America / USA / Wisconsin / Jacks | 2020-03-23 | University of Wisconsin-Madison AIDS Vaccine Research Laboratories | University of Wisconsin-Madison AIDS Vaccine Research Laboratories | Gage Moreno, Katarina Braun, et al. AIDS Vaccine Research Laboratories |
| EPI_ISL_428290 | hCoV-19/USA/WI-UW-194/2020 | North America / USA / Wisconsin / Grafto | 2020-03-24 | University of Wisconsin-Madison AIDS Vaccine Research Laboratories | University of Wisconsin-Madison AIDS Vaccine Research Laboratories | Gage Moreno, Katarina Braun, et al. AIDS Vaccine Research Laboratories |
| EPI_ISL_428291 | hCoV-19/USA/WI-UW-195/2020 | North America / USA / Wisconsin / Milwa | 2020-03-24 | University of Wisconsin-Madison AIDS Vaccine Research Laboratories | University of Wisconsin-Madison AIDS Vaccine Research Laboratories | Gage Moreno, Katarina Braun, et al. AIDS Vaccine Research Laboratories |
| EPI_ISL_428292 | hCoV-19/USA/WI-UW-196/2020 | North America / USA / Wisconsin / Milwa | 2020-03-21 | University of Wisconsin-Madison AIDS Vaccine Research Laboratories | University of Wisconsin-Madison AIDS Vaccine Research Laboratories | Gage Moreno, Katarina Braun, et al. AIDS Vaccine Research Laboratories |
| EPI_ISL_428293 | hCoV-19/USA/WI-UW-197/2020 | North America / USA / Wisconsin / Green | 2020-03-24 | University of Wisconsin-Madison AIDS Vaccine Research Laboratories | University of Wisconsin-Madison AIDS Vaccine Research Laboratories | Gage Moreno, Katarina Braun, et al. AIDS Vaccine Research Laboratories |
| EPI_ISL_428294 | hCoV-19/USA/WI-UW-198/2020 | North America / USA / Wisconsin / Brook | 2020-03-24 | University of Wisconsin-Madison AIDS Vaccine Research Laboratories | University of Wisconsin-Madison AIDS Vaccine Research Laboratories | Gage Moreno, Katarina Braun, et al. AIDS Vaccine Research Laboratories |
| EPI_ISL_428295 | hCoV-19/USA/WI-UW-199/2020 | North America / USA / Wisconsin / Brook | 2020-03-24 | University of Wisconsin-Madison AIDS Vaccine Research Laboratories | University of Wisconsin-Madison AIDS Vaccine Research Laboratories | Gage Moreno, Katarina Braun, et al. AIDS Vaccine Research Laboratories |
| EPI_ISL_428296 | hCoV-19/USA/WI-UW-200/2020 | North America / USA / Wisconsin / Milwa | 2020-03-24 | University of Wisconsin-Madison AIDS Vaccine Research Laboratories | University of Wisconsin-Madison AIDS Vaccine Research Laboratories | Gage Moreno, Katarina Braun, et al. AIDS Vaccine Research Laboratories |
| EPI_ISL_428297 | hCoV-19/USA/WI-UW-201/2020 | North America / USA / Wisconsin / Grafto | 2020-03-25 | University of Wisconsin-Madison AIDS Vaccine Research Laboratories | University of Wisconsin-Madison AIDS Vaccine Research Laboratories | Gage Moreno, Katarina Braun, et al. AIDS Vaccine Research Laboratories |
| EPI_ISL_428298 | hCoV-19/USA/WI-UW-202/2020 | North America / USA / Wisconsin / Grafto | 2020-03-25 | University of Wisconsin-Madison AIDS Vaccine Research Laboratories | University of Wisconsin-Madison AIDS Vaccine Research Laboratories | Gage Moreno, Katarina Braun, et al. AIDS Vaccine Research Laboratories |
| EPI_ISL_428299 | hCoV-19/USA/WI-UW-203/2020 | North America / USA / Wisconsin / Grafto | 2020-03-25 | University of Wisconsin-Madison AIDS Vaccine Research Laboratories | University of Wisconsin-Madison AIDS Vaccine Research Laboratories | Gage Moreno, Katarina Braun, et al. AIDS Vaccine Research Laboratories |
| EPI_ISL_428380 | hCoV-19/USA/CT-Yale-061/2020 | North America / USA / Connecticut | 2020-03-31 | Yale COVID-19 Biorepository | Grubaugh Lab - Yale School of Public Health | Joseph Fauver, Tara Alpert, Anderson Brito, Anne Wyllie, Chantal Vogels, Mary Petrone, Chaney Kalinich, Isabel Ott, Arnau Casanovas, Catherine Muenker, Adam Moore, Alice Lu, Maria Tokuyama, Patrick Wong, Peiwen Lu, Saad Omer, Richard Martinello, Allison Nelson, Shelli Farhadian, Akiko Iwasaki, Charlese Dela Cruz, Albert Ko, Nathan Grubaugh |
| EPI_ISL_428381 | hCoV-19/USA/CT-Yale-038/2020 | North America / USA / Connecticut | 2020-03-17 | Yale Clinical Virology Laboratory | Grubaugh Lab - Yale School of Public Health | Joseph Fauver, Anderson Brito, Tara Alpert, Chantal Vogels, Ellen Foxman, Albert Ko, Marie Landry, Nathan Grubaugh |
| EPI_ISL_428382 | hCoV-19/USA/CT-Yale-042/2020 | North America / USA / Connecticut | 2020-03-19 | Yale Clinical Virology Laboratory | Grubaugh Lab - Yale School of Public Health | Joseph Fauver, Anderson Brito, Tara Alpert, Chantal Vogels, Ellen Foxman, Albert Ko, Marie Landry, Nathan Grubaugh |
| EPI_ISL_428384 | hCoV-19/USA/CT-Yale-076/2020 | North America / USA / Connecticut | 2020-04-04 | Yale COVID-19 Biorepository | Grubaugh Lab - Yale School of Public Health | Joseph Fauver, Tara Alpert, Anderson Brito, Anne Wyllie, Chantal Vogels, Mary Petrone, Chaney Kalinich, Isabel Ott, Arnau Casanovas, Catherine Muenker, Adam Moore, Alice Lu, Maria Tokuyama, Patrick Wong, Peiwen Lu, Saad Omer, Richard Martinello, Allison Nelson, Shelli Farhadian, Akiko Iwasaki, Charlese Dela Cruz, Albert Ko, Nathan Grubaugh |
| EPI_ISL_428385 | hCoV-19/USA/CT-Yale-066/2020 | North America / USA / Connecticut | 2020-04-04 | Yale COVID-19 Biorepository | Grubaugh Lab - Yale School of Public Health | Joseph Fauver, Tara Alpert, Anderson Brito, Anne Wyllie, Chantal Vogels, Mary Petrone, Chaney Kalinich, Isabel Ott, Arnau Casanovas, Catherine Muenker, Adam Moore, Alice Lu, Maria Tokuyama, Patrick Wong, Peiwen Lu, Saad Omer, Richard Martinello, Allison Nelson, Shelli Farhadian, Akiko Iwasaki, Charlese Dela Cruz, Albert Ko, Nathan Grubaugh |
| EPI_ISL_428386 | hCoV-19/USA/CT-Yale-068/2020 | North America / USA / Connecticut | 2020-04-03 | Yale COVID-19 Biorepository | Grubaugh Lab - Yale School of Public Health | Joseph Fauver, Tara Alpert, Anderson Brito, Anne Wyllie, Chantal Vogels, Mary Petrone, Chaney Kalinich, Isabel Ott, Arnau Casanovas, Catherine Muenker, Adam Moore, Alice Lu, Maria Tokuyama, Patrick Wong, Peiwen Lu, Saad Omer, Richard Martinello, Allison Nelson, Shelli Farhadian, Akiko Iwasaki, Charlese Dela Cruz, Albert Ko, Nathan Grubaugh |
| EPI_ISL_428388 | hCoV-19/USA/CT-Yale-058/2020 | North America / USA / Connecticut | 2020-03-28 | Yale COVID-19 Biorepository | Grubaugh Lab - Yale School of Public Health | Joseph Fauver, Tara Alpert, Anderson Brito, Anne Wyllie, Chantal Vogels, Mary Petrone, Chaney Kalinich, Isabel Ott, Arnau Casanovas, Catherine Muenker, Adam Moore, Alice Lu, Maria Tokuyama, Patrick Wong, Peiwen Lu, Saad Omer, Richard Martinello, Allison Nelson, Shelli Farhadian, Akiko Iwasaki, Charlese Dela Cruz, Albert Ko, Nathan Grubaugh |
| EPI_ISL_429501 | hCoV-19/Denmark/ALAB-SSI407/2020 | Europe / Denmark / Unknown | 2020-03-10 | Department of Virus and Microbiological Special Diagnostics, Statens Serum Institut, Copenhagen, Denmark, Artillerivej 5, 2300 Copenahgen S | Albertsen lab, Department of Chemistry and Bioscience, Aalborg University, Denmark | Rasmus Kirkegaard |
| EPI_ISL_429502 | hCoV-19/Denmark/ALAB-SSI408/2020 | Europe / Denmark / Unknown | 2020-03-10 | Department of Virus and Microbiological Special Diagnostics, Statens Serum Institut, Copenhagen, Denmark, Artillerivej 5, 2300 Copenahgen S | Albertsen lab, Department of Chemistry and Bioscience, Aalborg University, Denmark | Rasmus Kirkegaard |
| EPI_ISL_429503 | hCoV-19/Denmark/ALAB-SSI409/2020 | Europe / Denmark / Unknown | 2020-03-10 | Department of Virus and Microbiological Special Diagnostics, Statens Serum Institut, Copenhagen, Denmark, Artillerivej 5, 2300 Copenahgen S | Albertsen lab, Department of Chemistry and Bioscience, Aalborg University, Denmark | Rasmus Kirkegaard |
| EPI_ISL_429505 | hCoV-19/Denmark/ALAB-SSI411/2020 | Europe / Denmark / Unknown | 2020-03-10 | Department of Virus and Microbiological Special Diagnostics, Statens Serum Institut, Copenhagen, Denmark, Artillerivej 5, 2300 Copenahgen S | Albertsen lab, Department of Chemistry and Bioscience, Aalborg University, Denmark | Rasmus Kirkegaard |
| EPI_ISL_429506 | hCoV-19/Denmark/ALAB-SSI411A/2020 | Europe / Denmark / Unknown | 2020-03-23 | Department of Virus and Microbiological Special Diagnostics, Statens Serum Institut, Copenhagen, Denmark, Artillerivej 5, 2300 Copenahgen S | Albertsen lab, Department of Chemistry and Bioscience, Aalborg University, Denmark | Rasmus Kirkegaard |
| EPI_ISL_429507 | hCoV-19/Denmark/ALAB-SSI412/2020 | Europe / Denmark / Unknown | 2020-03-10 | Department of Virus and Microbiological Special Diagnostics, Statens Serum Institut, Copenhagen, Denmark, Artillerivej 5, 2300 Copenahgen S | Albertsen lab, Department of Chemistry and Bioscience, Aalborg University, Denmark | Rasmus Kirkegaard |
| EPI_ISL_429508 | hCoV-19/Denmark/ALAB-SSI412A/2020 | Europe / Denmark / Unknown | 2020-03-23 | Department of Virus and Microbiological Special Diagnostics, Statens Serum Institut, Copenhagen, Denmark, Artillerivej 5, 2300 Copenahgen S | Albertsen lab, Department of Chemistry and Bioscience, Aalborg University, Denmark | Rasmus Kirkegaard |
| EPI_ISL_429509 | hCoV-19/Denmark/ALAB-SSI413/2020 | Europe / Denmark / Unknown | 2020-03-10 | Department of Virus and Microbiological Special Diagnostics, Statens Serum Institut, Copenhagen, Denmark, Artillerivej 5, 2300 Copenahgen S | Albertsen lab, Department of Chemistry and Bioscience, Aalborg University, Denmark | Rasmus Kirkegaard |
| EPI_ISL_429510 | hCoV-19/Denmark/ALAB-SSI413A/2020 | Europe / Denmark / Unknown | 2020-03-23 | Department of Virus and Microbiological Special Diagnostics, Statens Serum Institut, Copenhagen, Denmark, Artillerivej 5, 2300 Copenahgen S | Albertsen lab, Department of Chemistry and Bioscience, Aalborg University, Denmark | Rasmus Kirkegaard |
| EPI_ISL_429511 | hCoV-19/Denmark/ALAB-SSI414/2020 | Europe / Denmark / Unknown | 2020-03-10 | Department of Virus and Microbiological Special Diagnostics, Statens Serum Institut, Copenhagen, Denmark, Artillerivej 5, 2300 Copenahgen S | Albertsen lab, Department of Chemistry and Bioscience, Aalborg University, Denmark | Rasmus Kirkegaard |
| EPI_ISL_429512 | hCoV-19/Denmark/ALAB-SSI414A/2020 | Europe / Denmark / Unknown | 2020-03-23 | Department of Virus and Microbiological Special Diagnostics, Statens Serum Institut, Copenhagen, Denmark, Artillerivej 5, 2300 Copenahgen S | Albertsen lab, Department of Chemistry and Bioscience, Aalborg University, Denmark | Rasmus Kirkegaard |
| EPI_ISL_429513 | hCoV-19/Denmark/ALAB-SSI416/2020 | Europe / Denmark / Unknown | 2020-03-10 | Department of Virus and Microbiological Special Diagnostics, Statens Serum Institut, Copenhagen, Denmark, Artillerivej 5, 2300 Copenahgen S | Albertsen lab, Department of Chemistry and Bioscience, Aalborg University, Denmark | Rasmus Kirkegaard |
| EPI_ISL_429514 | hCoV-19/Denmark/ALAB-SSI417/2020 | Europe / Denmark / Unknown | 2020-03-10 | Department of Virus and Microbiological Special Diagnostics, Statens Serum Institut, Copenhagen, Denmark, Artillerivej 5, 2300 Copenahgen S | Albertsen lab, Department of Chemistry and Bioscience, Aalborg University, Denmark | Rasmus Kirkegaard |
| EPI_ISL_429516 | hCoV-19/Denmark/ALAB-SSI418/2020 | Europe / Denmark / Unknown | 2020-03-10 | Department of Virus and Microbiological Special Diagnostics, Statens Serum Institut, Copenhagen, Denmark, Artillerivej 5, 2300 Copenahgen S | Albertsen lab, Department of Chemistry and Bioscience, Aalborg University, Denmark | Rasmus Kirkegaard |
| EPI_ISL_429517 | hCoV-19/Denmark/ALAB-SSI419/2020 | Europe / Denmark / Unknown | 2020-03-11 | Department of Virus and Microbiological Special Diagnostics, Statens Serum Institut, Copenhagen, Denmark, Artillerivej 5, 2300 Copenahgen S | Albertsen lab, Department of Chemistry and Bioscience, Aalborg University, Denmark | Rasmus Kirkegaard |
| EPI_ISL_429518 | hCoV-19/Denmark/ALAB-SSI420/2020 | Europe / Denmark / Unknown | 2020-03-11 | Department of Virus and Microbiological Special Diagnostics, Statens Serum Institut, Copenhagen, Denmark, Artillerivej 5, 2300 Copenahgen S | Albertsen lab, Department of Chemistry and Bioscience, Aalborg University, Denmark | Rasmus Kirkegaard |

| | | | | | | |
|---|---|---|---|---|---|---|
| EPI_ISL_429519 | hCoV-19/Denmark/ALAB-SSI420A/2020 | Europe / Denmark / Unknown | 2020-03-22 | Department of Virus and Microbiological Special Diagnostics, Statens Serum Institut, Copenhagen, Denmark, Artillerivej 5, 2300 Copenahgen S | Albertsen lab, Department of Chemistry and Bioscience, Aalborg University, Denmark | Rasmus Kirkegaard |
| EPI_ISL_429520 | hCoV-19/Denmark/ALAB-SSI423/2020 | Europe / Denmark / Unknown | 2020-03-11 | Department of Virus and Microbiological Special Diagnostics, Statens Serum Institut, Copenhagen, Denmark, Artillerivej 5, 2300 Copenahgen S | Albertsen lab, Department of Chemistry and Bioscience, Aalborg University, Denmark | Rasmus Kirkegaard |
| EPI_ISL_429521 | hCoV-19/Denmark/ALAB-SSI424/2020 | Europe / Denmark / Unknown | 2020-03-11 | Department of Virus and Microbiological Special Diagnostics, Statens Serum Institut, Copenhagen, Denmark, Artillerivej 5, 2300 Copenahgen S | Albertsen lab, Department of Chemistry and Bioscience, Aalborg University, Denmark | Rasmus Kirkegaard |
| EPI_ISL_429523 | hCoV-19/Denmark/ALAB-SSI425/2020 | Europe / Denmark / Unknown | 2020-03-02 | Department of Virus and Microbiological Special Diagnostics, Statens Serum Institut, Copenhagen, Denmark, Artillerivej 5, 2300 Copenahgen S | Albertsen lab, Department of Chemistry and Bioscience, Aalborg University, Denmark | Rasmus Kirkegaard |
| EPI_ISL_429524 | hCoV-19/Denmark/ALAB-SSI425A/2020 | Europe / Denmark / Unknown | 2020-03-24 | Department of Virus and Microbiological Special Diagnostics, Statens Serum Institut, Copenhagen, Denmark, Artillerivej 5, 2300 Copenahgen S | Albertsen lab, Department of Chemistry and Bioscience, Aalborg University, Denmark | Rasmus Kirkegaard |
| EPI_ISL_429525 | hCoV-19/Denmark/ALAB-SSI426/2020 | Europe / Denmark / Unknown | 2020-03-12 | Department of Virus and Microbiological Special Diagnostics, Statens Serum Institut, Copenhagen, Denmark, Artillerivej 5, 2300 Copenahgen S | Albertsen lab, Department of Chemistry and Bioscience, Aalborg University, Denmark | Rasmus Kirkegaard |
| EPI_ISL_429526 | hCoV-19/Denmark/ALAB-SSI430/2020 | Europe / Denmark / Unknown | 2020-03-24 | Department of Virus and Microbiological Special Diagnostics, Statens Serum Institut, Copenhagen, Denmark, Artillerivej 5, 2300 Copenahgen S | Albertsen lab, Department of Chemistry and Bioscience, Aalborg University, Denmark | Rasmus Kirkegaard |
| EPI_ISL_429527 | hCoV-19/Denmark/ALAB-SSI431/2020 | Europe / Denmark / Unknown | 2020-03-24 | Department of Virus and Microbiological Special Diagnostics, Statens Serum Institut, Copenhagen, Denmark, Artillerivej 5, 2300 Copenahgen S | Albertsen lab, Department of Chemistry and Bioscience, Aalborg University, Denmark | Rasmus Kirkegaard |
| EPI_ISL_429528 | hCoV-19/Denmark/ALAB-SSI432/2020 | Europe / Denmark / Unknown | 2020-03-24 | Department of Virus and Microbiological Special Diagnostics, Statens Serum Institut, Copenhagen, Denmark, Artillerivej 5, 2300 Copenahgen S | Albertsen lab, Department of Chemistry and Bioscience, Aalborg University, Denmark | Rasmus Kirkegaard |
| EPI_ISL_429529 | hCoV-19/Denmark/ALAB-SSI434/2020 | Europe / Denmark / Unknown | 2020-03-24 | Department of Virus and Microbiological Special Diagnostics, Statens Serum Institut, Copenhagen, Denmark, Artillerivej 5, 2300 Copenahgen S | Albertsen lab, Department of Chemistry and Bioscience, Aalborg University, Denmark | Rasmus Kirkegaard |
| EPI_ISL_429530 | hCoV-19/Denmark/ALAB-SSI437/2020 | Europe / Denmark / Unknown | 2020-03-25 | Department of Virus and Microbiological Special Diagnostics, Statens Serum Institut, Copenhagen, Denmark, Artillerivej 5, 2300 Copenahgen S | Albertsen lab, Department of Chemistry and Bioscience, Aalborg University, Denmark | Rasmus Kirkegaard |
| EPI_ISL_429533 | hCoV-19/Denmark/ALAB-SSI444/2020 | Europe / Denmark / Unknown | 2020-03-25 | Department of Virus and Microbiological Special Diagnostics, Statens Serum Institut, Copenhagen, Denmark, Artillerivej 5, 2300 Copenahgen S | Albertsen lab, Department of Chemistry and Bioscience, Aalborg University, Denmark | Rasmus Kirkegaard |
| EPI_ISL_429534 | hCoV-19/Denmark/ALAB-SSI446/2020 | Europe / Denmark / Unknown | 2020-03-25 | Department of Virus and Microbiological Special Diagnostics, Statens Serum Institut, Copenhagen, Denmark, Artillerivej 5, 2300 Copenahgen S | Albertsen lab, Department of Chemistry and Bioscience, Aalborg University, Denmark | Rasmus Kirkegaard |
| EPI_ISL_429535 | hCoV-19/Denmark/ALAB-SSI448/2020 | Europe / Denmark / Unknown | 2020-03-25 | Department of Virus and Microbiological Special Diagnostics, Statens Serum Institut, Copenhagen, Denmark, Artillerivej 5, 2300 Copenahgen S | Albertsen lab, Department of Chemistry and Bioscience, Aalborg University, Denmark | Rasmus Kirkegaard |
| EPI_ISL_429536 | hCoV-19/Denmark/ALAB-SSI449/2020 | Europe / Denmark / Unknown | 2020-03-25 | Department of Virus and Microbiological Special Diagnostics, Statens Serum Institut, Copenhagen, Denmark, Artillerivej 5, 2300 Copenahgen S | Albertsen lab, Department of Chemistry and Bioscience, Aalborg University, Denmark | Rasmus Kirkegaard |
| EPI_ISL_429537 | hCoV-19/Denmark/ALAB-SSI450/2020 | Europe / Denmark / Unknown | 2020-03-24 | Department of Virus and Microbiological Special Diagnostics, Statens Serum Institut, Copenhagen, Denmark, Artillerivej 5, 2300 Copenahgen S | Albertsen lab, Department of Chemistry and Bioscience, Aalborg University, Denmark | Rasmus Kirkegaard |
| EPI_ISL_429538 | hCoV-19/Denmark/ALAB-SSI451/2020 | Europe / Denmark / Unknown | 2020-03-25 | Department of Virus and Microbiological Special Diagnostics, Statens Serum Institut, Copenhagen, Denmark, Artillerivej 5, 2300 Copenahgen S | Albertsen lab, Department of Chemistry and Bioscience, Aalborg University, Denmark | Rasmus Kirkegaard |
| EPI_ISL_429539 | hCoV-19/Denmark/ALAB-SSI454/2020 | Europe / Denmark / Unknown | 2020-03-25 | Department of Virus and Microbiological Special Diagnostics, Statens Serum Institut, Copenhagen, Denmark, Artillerivej 5, 2300 Copenahgen S | Albertsen lab, Department of Chemistry and Bioscience, Aalborg University, Denmark | Rasmus Kirkegaard |
| EPI_ISL_429540 | hCoV-19/Denmark/ALAB-SSI456/2020 | Europe / Denmark / Unknown | 2020-03-25 | Department of Virus and Microbiological Special Diagnostics, Statens Serum Institut, Copenhagen, Denmark, Artillerivej 5, 2300 Copenahgen S | Albertsen lab, Department of Chemistry and Bioscience, Aalborg University, Denmark | Rasmus Kirkegaard |
| EPI_ISL_429542 | hCoV-19/Denmark/ALAB-SSI461/2020 | Europe / Denmark / Unknown | 2020-03-25 | Department of Virus and Microbiological Special Diagnostics, Statens Serum Institut, Copenhagen, Denmark, Artillerivej 5, 2300 Copenahgen S | Albertsen lab, Department of Chemistry and Bioscience, Aalborg University, Denmark | Rasmus Kirkegaard |
| EPI_ISL_429543 | hCoV-19/Denmark/ALAB-SSI462/2020 | Europe / Denmark / Unknown | 2020-03-25 | Department of Virus and Microbiological Special Diagnostics, Statens Serum Institut, Copenhagen, Denmark, Artillerivej 5, 2300 Copenahgen S | Albertsen lab, Department of Chemistry and Bioscience, Aalborg University, Denmark | Rasmus Kirkegaard |
| EPI_ISL_429544 | hCoV-19/Denmark/ALAB-SSI463/2020 | Europe / Denmark / Unknown | 2020-03-25 | Department of Virus and Microbiological Special Diagnostics, Statens Serum Institut, Copenhagen, Denmark, Artillerivej 5, 2300 Copenahgen S | Albertsen lab, Department of Chemistry and Bioscience, Aalborg University, Denmark | Rasmus Kirkegaard |
| EPI_ISL_429545 | hCoV-19/Denmark/ALAB-SSI464/2020 | Europe / Denmark / Unknown | 2020-03-25 | Department of Virus and Microbiological Special Diagnostics, Statens Serum Institut, Copenhagen, Denmark, Artillerivej 5, 2300 Copenahgen S | Albertsen lab, Department of Chemistry and Bioscience, Aalborg University, Denmark | Rasmus Kirkegaard |
| EPI_ISL_429546 | hCoV-19/Denmark/ALAB-SSI466/2020 | Europe / Denmark / Unknown | 2020-03-25 | Department of Virus and Microbiological Special Diagnostics, Statens Serum Institut, Copenhagen, Denmark, Artillerivej 5, 2300 Copenahgen S | Albertsen lab, Department of Chemistry and Bioscience, Aalborg University, Denmark | Rasmus Kirkegaard |
| EPI_ISL_429547 | hCoV-19/Denmark/ALAB-SSI467/2020 | Europe / Denmark / Unknown | 2020-03-25 | Department of Virus and Microbiological Special Diagnostics, Statens Serum Institut, Copenhagen, Denmark, Artillerivej 5, 2300 Copenahgen S | Albertsen lab, Department of Chemistry and Bioscience, Aalborg University, Denmark | Rasmus Kirkegaard |
| EPI_ISL_429548 | hCoV-19/Denmark/ALAB-SSI468/2020 | Europe / Denmark / Unknown | 2020-03-25 | Department of Virus and Microbiological Special Diagnostics, Statens Serum Institut, Copenhagen, Denmark, Artillerivej 5, 2300 Copenahgen S | Albertsen lab, Department of Chemistry and Bioscience, Aalborg University, Denmark | Rasmus Kirkegaard |
| EPI_ISL_429549 | hCoV-19/Denmark/ALAB-SSI469/2020 | Europe / Denmark / Unknown | 2020-03-25 | Department of Virus and Microbiological Special Diagnostics, Statens Serum Institut, Copenhagen, Denmark, Artillerivej 5, 2300 Copenahgen S | Albertsen lab, Department of Chemistry and Bioscience, Aalborg University, Denmark | Rasmus Kirkegaard |
| EPI_ISL_429550 | hCoV-19/Denmark/ALAB-SSI470/2020 | Europe / Denmark / Unknown | 2020-03-25 | Department of Virus and Microbiological Special Diagnostics, Statens Serum Institut, Copenhagen, Denmark, Artillerivej 5, 2300 Copenahgen S | Albertsen lab, Department of Chemistry and Bioscience, Aalborg University, Denmark | Rasmus Kirkegaard |
| EPI_ISL_429551 | hCoV-19/Denmark/ALAB-SSI471/2020 | Europe / Denmark / Unknown | 2020-03-25 | Department of Virus and Microbiological Special Diagnostics, Statens Serum Institut, Copenhagen, Denmark, Artillerivej 5, 2300 Copenahgen S | Albertsen lab, Department of Chemistry and Bioscience, Aalborg University, Denmark | Rasmus Kirkegaard |
| EPI_ISL_429552 | hCoV-19/Denmark/ALAB-SSI472/2020 | Europe / Denmark / Unknown | 2020-03-25 | Department of Virus and Microbiological Special Diagnostics, Statens Serum Institut, Copenhagen, Denmark, Artillerivej 5, 2300 Copenahgen S | Albertsen lab, Department of Chemistry and Bioscience, Aalborg University, Denmark | Rasmus Kirkegaard |
| EPI_ISL_429553 | hCoV-19/Denmark/ALAB-SSI473/2020 | Europe / Denmark / Unknown | 2020-03-25 | Department of Virus and Microbiological Special Diagnostics, Statens Serum Institut, Copenhagen, Denmark, Artillerivej 5, 2300 Copenahgen S | Albertsen lab, Department of Chemistry and Bioscience, Aalborg University, Denmark | Rasmus Kirkegaard |
| EPI_ISL_429554 | hCoV-19/Denmark/ALAB-SSI474/2020 | Europe / Denmark / Unknown | 2020-03-25 | Department of Virus and Microbiological Special Diagnostics, Statens Serum Institut, Copenhagen, Denmark, Artillerivej 5, 2300 Copenahgen S | Albertsen lab, Department of Chemistry and Bioscience, Aalborg University, Denmark | Rasmus Kirkegaard |
| EPI_ISL_429555 | hCoV-19/Denmark/ALAB-SSI475/2020 | Europe / Denmark / Unknown | 2020-03-25 | Department of Virus and Microbiological Special Diagnostics, Statens Serum Institut, Copenhagen, Denmark, Artillerivej 5, 2300 Copenahgen S | Albertsen lab, Department of Chemistry and Bioscience, Aalborg University, Denmark | Rasmus Kirkegaard |
| EPI_ISL_429556 | hCoV-19/Denmark/ALAB-SSI477/2020 | Europe / Denmark / Unknown | 2020-03-26 | Department of Virus and Microbiological Special Diagnostics, Statens Serum Institut, Copenhagen, Denmark, Artillerivej 5, 2300 Copenahgen S | Albertsen lab, Department of Chemistry and Bioscience, Aalborg University, Denmark | Rasmus Kirkegaard |
| EPI_ISL_429557 | hCoV-19/Denmark/ALAB-SSI478/2020 | Europe / Denmark / Unknown | 2020-03-26 | Department of Virus and Microbiological Special Diagnostics, Statens Serum Institut, Copenhagen, Denmark, Artillerivej 5, 2300 Copenahgen S | Albertsen lab, Department of Chemistry and Bioscience, Aalborg University, Denmark | Rasmus Kirkegaard |
| EPI_ISL_429558 | hCoV-19/Denmark/ALAB-SSI479/2020 | Europe / Denmark / Unknown | 2020-03-25 | Department of Virus and Microbiological Special Diagnostics, Statens Serum Institut, Copenhagen, Denmark, Artillerivej 5, 2300 Copenahgen S | Albertsen lab, Department of Chemistry and Bioscience, Aalborg University, Denmark | Rasmus Kirkegaard |
| EPI_ISL_429559 | hCoV-19/Denmark/ALAB-SSI480/2020 | Europe / Denmark / Unknown | 2020-03-25 | Department of Virus and Microbiological Special Diagnostics, Statens Serum Institut, Copenhagen, Denmark, Artillerivej 5, 2300 Copenahgen S | Albertsen lab, Department of Chemistry and Bioscience, Aalborg University, Denmark | Rasmus Kirkegaard |
| EPI_ISL_429560 | hCoV-19/Denmark/ALAB-SSI481/2020 | Europe / Denmark / Unknown | 2020-03-25 | Department of Virus and Microbiological Special Diagnostics, Statens Serum Institut, Copenhagen, Denmark, Artillerivej 5, 2300 Copenahgen S | Albertsen lab, Department of Chemistry and Bioscience, Aalborg University, Denmark | Rasmus Kirkegaard |
| EPI_ISL_429561 | hCoV-19/Denmark/ALAB-SSI482/2020 | Europe / Denmark / Unknown | 2020-03-25 | Department of Virus and Microbiological Special Diagnostics, Statens Serum Institut, Copenhagen, Denmark, Artillerivej 5, 2300 Copenahgen S | Albertsen lab, Department of Chemistry and Bioscience, Aalborg University, Denmark | Rasmus Kirkegaard |
| EPI_ISL_429562 | hCoV-19/Denmark/ALAB-SSI483/2020 | Europe / Denmark / Unknown | 2020-03-26 | Department of Virus and Microbiological Special Diagnostics, Statens Serum Institut, Copenhagen, Denmark, Artillerivej 5, 2300 Copenahgen S | Albertsen lab, Department of Chemistry and Bioscience, Aalborg University, Denmark | Rasmus Kirkegaard |
| EPI_ISL_429563 | hCoV-19/Denmark/ALAB-SSI484/2020 | Europe / Denmark / Unknown | 2020-03-23 | Department of Virus and Microbiological Special Diagnostics, Statens Serum Institut, Copenhagen, Denmark, Artillerivej 5, 2300 Copenahgen S | Albertsen lab, Department of Chemistry and Bioscience, Aalborg University, Denmark | Rasmus Kirkegaard |
| EPI_ISL_429564 | hCoV-19/Denmark/ALAB-SSI486/2020 | Europe / Denmark / Unknown | 2020-03-26 | Department of Virus and Microbiological Special Diagnostics, Statens Serum Institut, Copenhagen, Denmark, Artillerivej 5, 2300 Copenahgen S | Albertsen lab, Department of Chemistry and Bioscience, Aalborg University, Denmark | Rasmus Kirkegaard |
| EPI_ISL_429565 | hCoV-19/Denmark/ALAB-SSI488/2020 | Europe / Denmark / Unknown | 2020-03-24 | Department of Virus and Microbiological Special Diagnostics, Statens Serum Institut, Copenhagen, Denmark, Artillerivej 5, 2300 Copenahgen S | Albertsen lab, Department of Chemistry and Bioscience, Aalborg University, Denmark | Rasmus Kirkegaard |
| EPI_ISL_429566 | hCoV-19/Denmark/ALAB-SSI489/2020 | Europe / Denmark / Unknown | 2020-03-23 | Department of Virus and Microbiological Special Diagnostics, Statens Serum Institut, Copenhagen, Denmark, Artillerivej 5, 2300 Copenahgen S | Albertsen lab, Department of Chemistry and Bioscience, Aalborg University, Denmark | Rasmus Kirkegaard |
| EPI_ISL_429567 | hCoV-19/Denmark/ALAB-SSI490/2020 | Europe / Denmark / Unknown | 2020-03-21 | Department of Virus and Microbiological Special Diagnostics, Statens Serum Institut, Copenhagen, Denmark, Artillerivej 5, 2300 Copenahgen S | Albertsen lab, Department of Chemistry and Bioscience, Aalborg University, Denmark | Rasmus Kirkegaard |
| EPI_ISL_429568 | hCoV-19/Denmark/ALAB-SSI491/2020 | Europe / Denmark / Unknown | 2020-03-23 | Department of Virus and Microbiological Special Diagnostics, Statens Serum Institut, Copenhagen, Denmark, Artillerivej 5, 2300 Copenahgen S | Albertsen lab, Department of Chemistry and Bioscience, Aalborg University, Denmark | Rasmus Kirkegaard |
| EPI_ISL_429569 | hCoV-19/Denmark/ALAB-SSI494/2020 | Europe / Denmark / Unknown | 2020-03-24 | Department of Virus and Microbiological Special Diagnostics, Statens Serum Institut, Copenhagen, Denmark, Artillerivej 5, 2300 Copenahgen S | Albertsen lab, Department of Chemistry and Bioscience, Aalborg University, Denmark | Rasmus Kirkegaard |
| EPI_ISL_429570 | hCoV-19/Denmark/ALAB-SSI495/2020 | Europe / Denmark / Unknown | 2020-03-24 | Department of Virus and Microbiological Special Diagnostics, Statens Serum Institut, Copenhagen, Denmark, Artillerivej 5, 2300 Copenahgen S | Albertsen lab, Department of Chemistry and Bioscience, Aalborg University, Denmark | Rasmus Kirkegaard |
| EPI_ISL_429571 | hCoV-19/Denmark/ALAB-SSI497/2020 | Europe / Denmark / Unknown | 2020-03-26 | Department of Virus and Microbiological Special Diagnostics, Statens Serum Institut, Copenhagen, Denmark, Artillerivej 5, 2300 Copenahgen S | Albertsen lab, Department of Chemistry and Bioscience, Aalborg University, Denmark | Rasmus Kirkegaard |
| EPI_ISL_429572 | hCoV-19/Denmark/ALAB-SSI501/2020 | Europe / Denmark / Unknown | 2020-03-24 | Department of Virus and Microbiological Special Diagnostics, Statens Serum Institut, Copenhagen, Denmark, Artillerivej 5, 2300 Copenahgen S | Albertsen lab, Department of Chemistry and Bioscience, Aalborg University, Denmark | Rasmus Kirkegaard |
| EPI_ISL_429573 | hCoV-19/Denmark/ALAB-SSI502/2020 | Europe / Denmark / Unknown | 2020-03-23 | Department of Virus and Microbiological Special Diagnostics, Statens Serum Institut, Copenhagen, Denmark, Artillerivej 5, 2300 Copenahgen S | Albertsen lab, Department of Chemistry and Bioscience, Aalborg University, Denmark | Rasmus Kirkegaard |
| EPI_ISL_429574 | hCoV-19/Denmark/ALAB-SSI506/2020 | Europe / Denmark / Unknown | 2020-03-26 | Department of Virus and Microbiological Special Diagnostics, Statens Serum Institut, Copenhagen, Denmark, Artillerivej 5, 2300 Copenahgen S | Albertsen lab, Department of Chemistry and Bioscience, Aalborg University, Denmark | Rasmus Kirkegaard |
| EPI_ISL_429575 | hCoV-19/Denmark/ALAB-SSI507/2020 | Europe / Denmark / Unknown | 2020-03-25 | Department of Virus and Microbiological Special Diagnostics, Statens Serum Institut, Copenhagen, Denmark, Artillerivej 5, 2300 Copenahgen S | Albertsen lab, Department of Chemistry and Bioscience, Aalborg University, Denmark | Rasmus Kirkegaard |
| EPI_ISL_429577 | hCoV-19/Denmark/ALAB-SSI509/2020 | Europe / Denmark / Unknown | 2020-03-25 | Department of Virus and Microbiological Special Diagnostics, Statens Serum Institut, Copenhagen, Denmark, Artillerivej 5, 2300 Copenahgen S | Albertsen lab, Department of Chemistry and Bioscience, Aalborg University, Denmark | Rasmus Kirkegaard |
| EPI_ISL_429578 | hCoV-19/Denmark/ALAB-SSI510/2020 | Europe / Denmark / Unknown | 2020-03-25 | Department of Virus and Microbiological Special Diagnostics, Statens Serum Institut, Copenhagen, Denmark, Artillerivej 5, 2300 Copenahgen S | Albertsen lab, Department of Chemistry and Bioscience, Aalborg University, Denmark | Rasmus Kirkegaard |
| EPI_ISL_429579 | hCoV-19/Denmark/ALAB-SSI511/2020 | Europe / Denmark / Unknown | 2020-03-26 | Department of Virus and Microbiological Special Diagnostics, Statens Serum Institut, Copenhagen, Denmark, Artillerivej 5, 2300 Copenahgen S | Albertsen lab, Department of Chemistry and Bioscience, Aalborg University, Denmark | Rasmus Kirkegaard |
| EPI_ISL_429580 | hCoV-19/Denmark/ALAB-SSI512/2020 | Europe / Denmark / Unknown | 2020-03-25 | Department of Virus and Microbiological Special Diagnostics, Statens Serum Institut, Copenhagen, Denmark, Artillerivej 5, 2300 Copenahgen S | Albertsen lab, Department of Chemistry and Bioscience, Aalborg University, Denmark | Rasmus Kirkegaard |
| EPI_ISL_429581 | hCoV-19/Denmark/ALAB-SSI513/2020 | Europe / Denmark / Unknown | 2020-03-24 | Department of Virus and Microbiological Special Diagnostics, Statens Serum Institut, Copenhagen, Denmark, Artillerivej 5, 2300 Copenahgen S | Albertsen lab, Department of Chemistry and Bioscience, Aalborg University, Denmark | Rasmus Kirkegaard |
| EPI_ISL_429582 | hCoV-19/Denmark/ALAB-SSI514/2020 | Europe / Denmark / Unknown | 2020-03-16 | Department of Virus and Microbiological Special Diagnostics, Statens Serum Institut, Copenhagen, Denmark, Artillerivej 5, 2300 Copenahgen S | Albertsen lab, Department of Chemistry and Bioscience, Aalborg University, Denmark | Rasmus Kirkegaard |
| EPI_ISL_429583 | hCoV-19/Denmark/ALAB-SSI515/2020 | Europe / Denmark / Unknown | 2020-03-23 | Department of Virus and Microbiological Special Diagnostics, Statens Serum Institut, Copenhagen, Denmark, Artillerivej 5, 2300 Copenahgen S | Albertsen lab, Department of Chemistry and Bioscience, Aalborg University, Denmark | Rasmus Kirkegaard |
| EPI_ISL_429584 | hCoV-19/Denmark/ALAB-SSI518/2020 | Europe / Denmark / Unknown | 2020-03-26 | Department of Virus and Microbiological Special Diagnostics, Statens Serum Institut, Copenhagen, Denmark, Artillerivej 5, 2300 Copenahgen S | Albertsen lab, Department of Chemistry and Bioscience, Aalborg University, Denmark | Rasmus Kirkegaard |
| EPI_ISL_429585 | hCoV-19/Denmark/ALAB-SSI520/2020 | Europe / Denmark / Unknown | 2020-03-26 | Department of Virus and Microbiological Special Diagnostics, Statens Serum Institut, Copenhagen, Denmark, Artillerivej 5, 2300 Copenahgen S | Albertsen lab, Department of Chemistry and Bioscience, Aalborg University, Denmark | Rasmus Kirkegaard |
| EPI_ISL_429586 | hCoV-19/Denmark/ALAB-SSI521/2020 | Europe / Denmark / Unknown | 2020-03-26 | Department of Virus and Microbiological Special Diagnostics, Statens Serum Institut, Copenhagen, Denmark, Artillerivej 5, 2300 Copenahgen S | Albertsen lab, Department of Chemistry and Bioscience, Aalborg University, Denmark | Rasmus Kirkegaard |
| EPI_ISL_429587 | hCoV-19/Denmark/ALAB-SSI522/2020 | Europe / Denmark / Unknown | 2020-03-26 | Department of Virus and Microbiological Special Diagnostics, Statens Serum Institut, Copenhagen, Denmark, Artillerivej 5, 2300 Copenahgen S | Albertsen lab, Department of Chemistry and Bioscience, Aalborg University, Denmark | Rasmus Kirkegaard |
| EPI_ISL_429588 | hCoV-19/Denmark/ALAB-SSI523/2020 | Europe / Denmark / Unknown | 2020-03-26 | Department of Virus and Microbiological Special Diagnostics, Statens Serum Institut, Copenhagen, Denmark, Artillerivej 5, 2300 Copenahgen S | Albertsen lab, Department of Chemistry and Bioscience, Aalborg University, Denmark | Rasmus Kirkegaard |
| EPI_ISL_429589 | hCoV-19/Denmark/ALAB-SSI592/2020 | Europe / Denmark / Unknown | 2020-03-28 | Department of Virus and Microbiological Special Diagnostics, Statens Serum Institut, Copenhagen, Denmark, Artillerivej 5, 2300 Copenahgen S | Albertsen lab, Department of Chemistry and Bioscience, Aalborg University, Denmark | Rasmus Kirkegaard |
| EPI_ISL_429590 | hCoV-19/Denmark/ALAB-SSI595/2020 | Europe / Denmark / Unknown | 2020-03-28 | Department of Virus and Microbiological Special Diagnostics, Statens Serum Institut, Copenhagen, Denmark, Artillerivej 5, 2300 Copenahgen S | Albertsen lab, Department of Chemistry and Bioscience, Aalborg University, Denmark | Rasmus Kirkegaard |
| EPI_ISL_429598 | hCoV-19/USA/WA-UW-4546/2020 | North America / USA / Washington | 2020-03-31 | UW Virology Lab | UW Virology Lab | Pavitra Roychoudhury, Hong Xie, Keith Jerome, Alexander Greninger |
| EPI_ISL_429802 | hCoV-19/Croatia/17-S7/2020 | Europe / Croatia / Istria | 2020-03 | Institute for Public Health | Laboratory for advanced genomics | Filip Rokić, Lovro Trgovec-Greif, Neven Sučić, Tomislav Rukavina, Igor Jurak, Oliver Vugrek |
| EPI_ISL_429803 | hCoV-19/Croatia/I7-S21/2020 | Europe / Croatia / Istria | 2020-03 | Institute for Public Health | Laboratory for advanced genomics | Filip Rokić, Lovro Trgovec-Greif, Neven Sučić, Tomislav Rukavina, Igor Jurak, Oliver Vugrek |
| EPI_ISL_429805 | hCoV-19/Croatia/7R-S8/2020 | Europe / Croatia / Istria | 2020-03 | Institute for Public Health | Laboratory for advanced genomics | Filip Rokić, Lovro Trgovec-Greif, Neven Sučić, Tomislav Rukavina, Igor Jurak, Oliver Vugrek |
| EPI_ISL_429806 | hCoV-19/Canada/NB_1/2020 | North America / Canada / New Brunswick | 2020-03-09 | Dr. Georges-L.-Dumont University Hospital Centre | National Microbiology Laboratory | Anna Majer, Shari Tyson, Grace Seo, Kristyn Burak, Philip Mabon, Elsie Grudeski, Rhiannon Huzarewich, Russell Mandes, Jennifer Tanner, Natalie Knox, Morag Graham, Gary Van Domselaar, Richard Garceau, Guillaume Desnoyers, Nathalie Bastien, Yan Li, Timothy Booth, Matthew Gilmour |
| EPI_ISL_429807 | hCoV-19/Canada/MB_2/2020 | North America / Canada / Manitoba | 2020-03-10 | Cadham Provincial Laboratory | National Microbiology Laboratory | Anna Majer, Shari Tyson, Grace Seo, Kristyn Burak, Philip Mabon, Elsie Grudeski, Rhiannon Huzarewich, Russell Mandes, Jennifer Tanner, Natalie Knox, Morag Graham, Gary Van Domselaar, Paul Van Caeseele, Jared Bullard, David Alexander, Kerry Dust, Nathalie Bastien, Yan Li, Timothy Booth, Matthew Gilmour |
| EPI_ISL_429809 | hCoV-19/Canada/SK_16/2020 | North America / Canada / Saskatchewan | 2020-03-13 | Roy Romanow Provincial Laboratory | National Microbiology Laboratory | Anna Majer, Shari Tyson, Grace Seo, Kristyn Burak, Philip Mabon, Elsie Grudeski, Rhiannon Huzarewich, Russell Mandes, Jennifer Tanner, Natalie Knox, Morag Graham, Gary Van Domselaar, Ryan McDonald, Amanda Lang, Rachel DePaulo, Jessica Minion, Nathalie Bastien, Yan Li, Timothy Booth, Matthew Gilmour |
| EPI_ISL_429810 | hCoV-19/Canada/SK_20/2020 | North America / Canada / Saskatchewan | 2020-03-15 | Roy Romanow Provincial Laboratory | National Microbiology Laboratory | Anna Majer, Shari Tyson, Grace Seo, Kristyn Burak, Philip Mabon, Elsie Grudeski, Rhiannon Huzarewich, Russell Mandes, Jennifer Tanner, Natalie Knox, Morag Graham, Gary Van Domselaar, Ryan McDonald, Amanda Lang, Rachel DePaulo, Jessica Minion, Nathalie Bastien, Yan Li, Timothy Booth, Matthew Gilmour |
| EPI_ISL_429811 | hCoV-19/Canada/NB_21/2020 | North America / Canada / New Brunswick | 2020-03-12 | Dr. Georges-L.-Dumont University Hospital Centre | National Microbiology Laboratory | Anna Majer, Shari Tyson, Grace Seo, Kristyn Burak, Philip Mabon, Elsie Grudeski, Rhiannon Huzarewich, Russell Mandes, Jennifer Tanner, Natalie Knox, Morag Graham, Gary Van Domselaar, Richard Garceau, Guillaume Desnoyers, Nathalie Bastien, Yan Li, Timothy Booth, Matthew Gilmour |
| EPI_ISL_429812 | hCoV-19/Canada/NS_30/2020 | North America / Canada / Nova Scotia | 2020-03-14 | Queen Elizabeth II Health Science Centre | National Microbiology Laboratory | Anna Majer, Shari Tyson, Grace Seo, Kristyn Burak, Philip Mabon, Elsie Grudeski, Rhiannon Huzarewich, Russell Mandes, Jennifer Tanner, Natalie Knox, Morag Graham, Gary Van Domselaar, Todd Hatchette, Jason LeBlanc, Nathalie Bastien, Yan Li, Timothy Booth, Matthew Gilmour |
| EPI_ISL_429813 | hCoV-19/Canada/NS_31/2020 | North America / Canada / Nova Scotia | 2020-03-15 | Queen Elizabeth II Health Science Centre | National Microbiology Laboratory | Anna Majer, Shari Tyson, Grace Seo, Kristyn Burak, Philip Mabon, Elsie Grudeski, Rhiannon Huzarewich, Russell Mandes, Jennifer Tanner, Natalie Knox, Morag Graham, Gary Van Domselaar, Todd Hatchette, Jason LeBlanc, Nathalie Bastien, Yan Li, Timothy Booth, Matthew Gilmour |
| EPI_ISL_429814 | hCoV-19/Canada/NS_32/2020 | North America / Canada / Nova Scotia | 2020-03-15 | Queen Elizabeth II Health Science Centre | National Microbiology Laboratory | Anna Majer, Shari Tyson, Grace Seo, Kristyn Burak, Philip Mabon, Elsie Grudeski, Rhiannon Huzarewich, Russell Mandes, Jennifer Tanner, Natalie Knox, Morag Graham, Gary Van Domselaar, Todd Hatchette, Jason LeBlanc, Nathalie Bastien, Yan Li, Timothy Booth, Matthew Gilmour |
| EPI_ISL_429815 | hCoV-19/Canada/NL_34/2020 | North America / Canada / Newfoundland | 2020-03-15 | Public Health Laboratory | National Microbiology Laboratory | Anna Majer, Shari Tyson, Grace Seo, Kristyn Burak, Philip Mabon, Elsie Grudeski, Rhiannon Huzarewich, Russell Mandes, Jennifer Tanner, Natalie Knox, Morag Graham, Gary Van Domselaar, Robert Needle, Yang Yu, Adel Malek, Laura Gilbert, George Zahariadis, Nathalie Bastien, Yan Li, Timothy Booth, Matthew Gilmour |
| EPI_ISL_429816 | hCoV-19/Canada/NB_35/2020 | North America / Canada / New Brunswick | 2020-03-14 | Dr. Georges-L.-Dumont University Hospital Centre | National Microbiology Laboratory | Anna Majer, Shari Tyson, Grace Seo, Kristyn Burak, Philip Mabon, Elsie Grudeski, Rhiannon Huzarewich, Russell Mandes, Jennifer Tanner, Natalie Knox, Morag Graham, Gary Van Domselaar, Richard Garceau, Guillaume Desnoyers, Nathalie Bastien, Yan Li, Timothy Booth, Matthew Gilmour |
| EPI_ISL_429817 | hCoV-19/Canada/MB_36/2020 | North America / Canada / Manitoba | 2020-03-14 | Cadham Provincial Laboratory | National Microbiology Laboratory | Anna Majer, Shari Tyson, Grace Seo, Kristyn Burak, Philip Mabon, Elsie Grudeski, Rhiannon Huzarewich, Russell Mandes, Jennifer Tanner, Natalie Knox, Morag Graham, Gary Van Domselaar, Paul Van Caeseele, Jared Bullard, David Alexander, Kerry Dust, Nathalie Bastien, Yan Li, Timothy Booth, Matthew Gilmour |
| EPI_ISL_429818 | hCoV-19/Canada/MB_37/2020 | North America / Canada / Manitoba | 2020-03-18 | Cadham Provincial Laboratory | National Microbiology Laboratory | Anna Majer, Shari Tyson, Grace Seo, Kristyn Burak, Philip Mabon, Elsie Grudeski, Rhiannon Huzarewich, Russell Mandes, Jennifer Tanner, Natalie Knox, Morag Graham, Gary Van Domselaar, Paul Van Caeseele, Jared Bullard, David Alexander, Kerry Dust, Nathalie Bastien, Yan Li, Timothy Booth, Matthew Gilmour |
| EPI_ISL_429819 | hCoV-19/Canada/MB_38/2020 | North America / Canada / Manitoba | 2020-03-18 | Cadham Provincial Laboratory | National Microbiology Laboratory | Anna Majer, Shari Tyson, Grace Seo, Kristyn Burak, Philip Mabon, Elsie Grudeski, Rhiannon Huzarewich, Russell Mandes, Jennifer Tanner, Natalie Knox, Morag Graham, Gary Van Domselaar, Paul Van Caeseele, Jared Bullard, David Alexander, Kerry Dust, Nathalie Bastien, Yan Li, Timothy Booth, Matthew Gilmour |
| EPI_ISL_429820 | hCoV-19/Canada/MB_39/2020 | North America / Canada / Manitoba | 2020-03-14 | Cadham Provincial Laboratory | National Microbiology Laboratory | Anna Majer, Shari Tyson, Grace Seo, Kristyn Burak, Philip Mabon, Elsie Grudeski, Rhiannon Huzarewich, Russell Mandez, Jennifer Tanner, Natalie Knox, Morag Graham, Gary Van Domselaar, Paul Van Caeseele, Jared Bullard, David Alexander, Kerry Dust, Nathalie Bastien, Yan Li, Timothy Booth, Matthew Gilmour |
| EPI_ISL_429821 | hCoV-19/Canada/SK_52/2020 | North America / Canada / Saskatchewan | 2020-03-16 | Roy Romanow Provincial Laboratory | National Microbiology Laboratory | Anna Majer, Shari Tyson, Grace Seo, Kristyn Burak, Philip Mabon, Elsie Grudeski, Rhiannon Huzarewich, Russell Mandes, Jennifer Tanner, Natalie Knox, Morag Graham, Gary Van Domselaar, Ryan McDonald, Amanda Lang, Rachel DePaulo, Jessica Minion, Nathalie Bastien, Yan Li, Timothy Booth, Matthew Gilmour |
| EPI_ISL_429822 | hCoV-19/Canada/SK_63/2020 | North America / Canada / Saskatchewan | 2020-03-16 | Roy Romanow Provincial Laboratory | National Microbiology Laboratory | Anna Majer, Shari Tyson, Grace Seo, Kristyn Burak, Philip Mabon, Elsie Grudeski, Rhiannon Huzarewich, Russell Mandes, Jennifer Tanner, Natalie Knox, Morag Graham, Gary Van Domselaar, Ryan McDonald, Amanda Lang, Rachel DePaulo, Jessica Minion, Nathalie Bastien, Yan Li, Timothy Booth, Matthew Gilmour |
| EPI_ISL_429843 | hCoV-19/USA/WI-GMF-00822/2020 | North America / USA / Wisconsin / La Cr | 2020-04-08 | Gundersen Molecular Diagnostics Laboratory | Kabara Cancer Research Institute | Craig S. Richmond, Paraic A. Kenny |
| EPI_ISL_429844 | hCoV-19/USA/WI-GMF-00857/2020 | North America / USA / Iowa | 2020-04-09 | Gundersen Molecular Diagnostics Laboratory | Kabara Cancer Research Institute | Craig S. Richmond, Paraic A. Kenny |
| EPI_ISL_429845 | hCoV-19/USA/WI-GMF-01047/2020 | North America / USA / Iowa / Allamakee ( | 2020-04-14 | Gundersen Molecular Diagnostics Laboratory | Kabara Cancer Research Institute | Craig S. Richmond, Paraic A. Kenny |
| EPI_ISL_429846 | hCoV-19/USA/WI-GMF-01158/2020 | North America / USA / Iowa / Allamakee ( | 2020-04-16 | Gundersen Molecular Diagnostics Laboratory | Kabara Cancer Research Institute | Craig S. Richmond, Paraic A. Kenny |
| EPI_ISL_429847 | hCoV-19/USA/WI-GMF-01159/2020 | North America / USA / Iowa / Allamakee ( | 2020-04-16 | Gundersen Molecular Diagnostics Laboratory | Kabara Cancer Research Institute | Craig S. Richmond, Paraic A. Kenny |
| EPI_ISL_429848 | hCoV-19/USA/WI-GMF-01316/2020 | North America / USA / Iowa / Winneshiek | 2020-04-06 | Gundersen Molecular Diagnostics Laboratory | Kabara Cancer Research Institute | Craig S. Richmond, Paraic A. Kenny |
| EPI_ISL_429852 | hCoV-19/Lishui/LS111/2020 | Asia / China / Zhejiang / Lishui | 2020-01-25 | Centers for Disease Control and Prevention of Lishui | Department of Inspection, Centers for Disease Control and Prevention of Lishui | Wang Xiaoguang,Ji Qiaoying,Ji Jiansong,Ye Bifeng,Ye Ling |
| EPI_ISL_429853 | hCoV-19/Lishui/LS556/2020 | Asia / China / Zhejiang / Lishui | 2020-01-28 | Centers for Disease Control and Prevention of Lishui | Department of Inspection, Centers for Disease Control and Prevention of Lishui | Wang Xiaoguang,Ji Qiaoying,Ji Jiansong,Ye Bifeng,Ye Ling |
| EPI_ISL_429854 | hCoV-19/Lishui/LS557/2020 | Asia / China / Zhejiang / Lishui | 2020-02-01 | Centers for Disease Control and Prevention of Lishui | Department of Inspection, Centers for Disease Control and Prevention of Lishui | Wang Xiaoguang,Ji Qiaoying,Ji Jiansong,Ye Bifeng,Ye Ling |
| EPI_ISL_429855 | hCoV-19/Lishui/LS559/2020 | Asia / China / Zhejiang / Lishui | 2020-01-31 | Centers for Disease Control and Prevention of Lishui | Department of Inspection, Centers for Disease Control and Prevention of Lishui | Wang Xiaoguang,Ji Qiaoying,Ji Jiansong,Ye Bifeng,Ye Ling |
| EPI_ISL_429861 | hCoV-19/Turkey/HSGM-8968/2020 | Europe / Turkey / Ankara | 2020-03-22 | Ministry of Health Turkey | Ministry of Health Turkey | Fatma Bayrakdar,Ayşe Başak Altaş,Yasemin Coşgun,Gülay Korukluoğlu,Selçuk Kılıç |
| EPI_ISL_429862 | hCoV-19/Turkey/HSGM-8970/2020 | Europe / Turkey / Ankara | 2020-03-22 | Ministry of Health Turkey | Ministry of Health Turkey | Fatma Bayrakdar,Ayşe Başak Altaş,Yasemin Coşgun,Gülay Korukluoğlu,Selçuk Kılıç |
| EPI_ISL_429863 | hCoV-19/Turkey/HSGM-8975/2020 | Europe / Turkey / Sakarya | 2020-03-22 | Ministry of Health Turkey | Ministry of Health Turkey | Fatma Bayrakdar,Ayşe Başak Altaş,Yasemin Coşgun,Gülay Korukluoğlu,Selçuk Kılıç |
| EPI_ISL_429864 | hCoV-19/Turkey/HSGM-8979/2020 | Europe / Turkey / Sakarya | 2020-03-22 | Ministry of Health Turkey | Ministry of Health Turkey | Fatma Bayrakdar,Ayşe Başak Altaş,Yasemin Coşgun,Gülay Korukluoğlu,Selçuk Kılıç |
| EPI_ISL_429865 | hCoV-19/Turkey/HSGM-8992/2020 | Europe / Turkey / Canakkale | 2020-03-18 | Ministry of Health Turkey | Ministry of Health Turkey | Fatma Bayrakdar,Ayşe Başak Altaş,Yasemin Coşgun,Gülay Korukluoğlu,Selçuk Kılıç |
| EPI_ISL_429866 | hCoV-19/Turkey/HSGM-4236/2020 | Europe / Turkey / Afyon | 2020-03-16 | Ministry of Health Turkey | Ministry of Health Turkey | Fatma Bayrakdar,Ayşe Başak Altaş,Yasemin Coşgun,Gülay Korukluoğlu,Selçuk Kılıç |
| EPI_ISL_429867 | hCoV-19/Turkey/HSGM-4505/2020 | Europe / Turkey / Balikesir | 2020-03-17 | Ministry of Health Turkey | Ministry of Health Turkey | Fatma Bayrakdar,Ayşe Başak Altaş,Yasemin Coşgun,Gülay Korukluoğlu,Selçuk Kılıç |
| EPI_ISL_429868 | hCoV-19/Turkey/HSGM-4598/2020 | Europe / Turkey / Eskisehir | 2020-03-17 | Ministry of Health Turkey | Ministry of Health Turkey | Fatma Bayrakdar,Ayşe Başak Altaş,Yasemin Coşgun,Gülay Korukluoğlu,Selçuk Kılıç |
| EPI_ISL_429869 | hCoV-19/Turkey/HSGM-4701/2020 | Europe / Turkey / Konya | 2020-03-17 | Ministry of Health Turkey | Ministry of Health Turkey | Fatma Bayrakdar,Ayşe Başak Altaş,Yasemin Coşgun,Gülay Korukluoğlu,Selçuk Kılıç |
| EPI_ISL_429874 | hCoV-19/env/Italy/HSacco-1/2020 | Europe / Italy / Lombardy | 2020-04-14 | Microbiology, Virology and Biemergency Laboratory-ASST FBF Sacco | Microbiology, Virology and Biemergency Laboratory-ASST FBF Sacco | Rimoldi SG, Stefani F |
| EPI_ISL_429875 | hCoV-19/USA/CA-CDPH-UC30/2020 | North America / USA / California / Grand | 2020-03-02 | California Department of Public Health | Chiu Laboratory, University of California, San Francisco | Xianding Deng, Scot Federman, Chao-Yang Pan, Hugo Guevara,Wei Gu, Debra A. Wadford, and Charles Y. Chiu |
| EPI_ISL_429876 | hCoV-19/USA/CA-CDPH-UC31/2020 | North America / USA / California / Grand | 2020-03-05 | California Department of Public Health | Chiu Laboratory, University of California, San Francisco | Xianding Deng, Scot Federman, Chao-Yang Pan, Hugo Guevara,Wei Gu, Debra A. Wadford, and Charles Y. Chiu |
| EPI_ISL_429881 | hCoV-19/USA/CA-UCSF-UC36/2020 | North America / USA / California / San Fr | 2020-03-20 | California Department of Public Health | Chiu Laboratory, University of California, San Francisco | Xianding Deng, Scot Federman, Wei Gu, and Charles Y. Chiu |
| EPI_ISL_429882 | hCoV-19/Taiwan/299/2020 | Asia / Taiwan | 2020-03-29 | Centers for Disease Control, R.O.C. (Taiwan) | Centers for Disease Control, R.O.C. (Taiwan) | Ji-Rong Yang, Yu-Chi Lin, Jung-Jung Mu, Ming-Tsan Liu |
| EPI_ISL_429883 | hCoV-19/Taiwan/108/2020 | Asia / Taiwan | 2020-03-23 | Centers for Disease Control, R.O.C. (Taiwan) | Centers for Disease Control, R.O.C. (Taiwan) | Ji-Rong Yang, Yu-Chi Lin, Jung-Jung Mu, Ming-Tsan Liu |
| EPI_ISL_429884 | hCoV-19/Taiwan/284/2020 | Asia / Taiwan | 2020-03-30 | Centers for Disease Control, R.O.C. (Taiwan) | Centers for Disease Control, R.O.C. (Taiwan) | Ji-Rong Yang, Yu-Chi Lin, Jung-Jung Mu, Ming-Tsan Liu |
| EPI_ISL_430500 | hCoV-19/Australia/VIC1210/2020 | Oceania / Australia / Victoria | 2020-04-01 | Victorian Infectious Diseases Reference Laboratory (VIDRL) | Microbiological Diagnostic Unit Public Health Laboratory and Victorian Infectious Diseases Reference Laboratory, The Peter Doherty Institute for Infection and Immunity | Caly L., Seemann T., Sait, M., Schultz M., Druce J., Sherry, N. |
| EPI_ISL_430512 | hCoV-19/Australia/VIC1235/2020 | Oceania / Australia / Victoria | 2020-04-02 | Victorian Infectious Diseases Reference Laboratory (VIDRL) | Microbiological Diagnostic Unit Public Health Laboratory and Victorian Infectious Diseases Reference Laboratory, The Peter Doherty Institute for Infection and Immunity | Caly L., Seemann T., Sait, M., Schultz M., Druce J., Sherry, N. |
| EPI_ISL_430519 | hCoV-19/Australia/VIC1222/2020 | Oceania / Australia / Victoria | 2020-04-01 | Victorian Infectious Diseases Reference Laboratory (VIDRL) | Microbiological Diagnostic Unit Public Health Laboratory and Victorian Infectious Diseases Reference Laboratory, The Peter Doherty Institute for Infection and Immunity | Caly L., Seemann T., Sait, M., Schultz M., Druce J., Sherry, N. |
| EPI_ISL_430522 | hCoV-19/Australia/VIC1224/2020 | Oceania / Australia / Victoria | 2020-04-01 | Victorian Infectious Diseases Reference Laboratory (VIDRL) | Microbiological Diagnostic Unit Public Health Laboratory and Victorian Infectious Diseases Reference Laboratory, The Peter Doherty Institute for Infection and Immunity | Caly L., Seemann T., Sait, M., Schultz M., Druce J., Sherry, N. |
| EPI_ISL_430528 | hCoV-19/Australia/VIC1192/2020 | Oceania / Australia / Victoria | 2020-03-30 | Victorian Infectious Diseases Reference Laboratory (VIDRL) | Microbiological Diagnostic Unit Public Health Laboratory and Victorian Infectious Diseases Reference Laboratory, The Peter Doherty Institute for Infection and Immunity | Caly L., Seemann T., Sait, M., Schultz M., Druce J., Sherry, N. |
| EPI_ISL_430532 | hCoV-19/Australia/VIC1025/2020 | Oceania / Australia / Victoria | 2020-04-02 | Victorian Infectious Diseases Reference Laboratory (VIDRL) | Microbiological Diagnostic Unit Public Health Laboratory and Victorian Infectious Diseases Reference Laboratory, The Peter Doherty Institute for Infection and Immunity | Caly L., Seemann T., Sait, M., Schultz M., Druce J., Sherry, N. |
| EPI_ISL_430533 | hCoV-19/Australia/VIC1037/2020 | Oceania / Australia / Victoria | 2020-04-03 | Victorian Infectious Diseases Reference Laboratory (VIDRL) | Microbiological Diagnostic Unit Public Health Laboratory and Victorian Infectious Diseases Reference Laboratory, The Peter Doherty Institute for Infection and Immunity | Caly L., Seemann T., Sait, M., Schultz M., Druce J., Sherry, N. |
| EPI_ISL_430535 | hCoV-19/Australia/VIC1039/2020 | Oceania / Australia / Victoria | 2020-04-03 | Victorian Infectious Diseases Reference Laboratory (VIDRL) | Microbiological Diagnostic Unit Public Health Laboratory and Victorian Infectious Diseases Reference Laboratory, The Peter Doherty Institute for Infection and Immunity | Caly L., Seemann T., Sait, M., Schultz M., Druce J., Sherry, N. |
| EPI_ISL_430536 | hCoV-19/Australia/VIC1041/2020 | Oceania / Australia / Victoria | 2020-04-03 | Victorian Infectious Diseases Reference Laboratory (VIDRL) | Microbiological Diagnostic Unit Public Health Laboratory and Victorian Infectious Diseases Reference Laboratory, The Peter Doherty Institute for Infection and Immunity | Caly L., Seemann T., Sait, M., Schultz M., Druce J., Sherry, N. |
| EPI_ISL_430538 | hCoV-19/Australia/VIC1042/2020 | Oceania / Australia / Victoria | 2020-04-03 | Victorian Infectious Diseases Reference Laboratory (VIDRL) | Microbiological Diagnostic Unit Public Health Laboratory and Victorian Infectious Diseases Reference Laboratory, The Peter Doherty Institute for Infection and Immunity | Caly L., Seemann T., Sait, M., Schultz M., Druce J., Sherry, N. |
| EPI_ISL_430539 | hCoV-19/Australia/VIC1043/2020 | Oceania / Australia / Victoria | 2020-04-03 | Victorian Infectious Diseases Reference Laboratory (VIDRL) | Microbiological Diagnostic Unit Public Health Laboratory and Victorian Infectious Diseases Reference Laboratory, The Peter Doherty Institute for Infection and Immunity | Caly L., Seemann T., Sait, M., Schultz M., Druce J., Sherry, N. |
| EPI_ISL_430542 | hCoV-19/Australia/VIC978/2020 | Oceania / Australia / Victoria | 2020-03-29 | Victorian Infectious Diseases Reference Laboratory (VIDRL) | Microbiological Diagnostic Unit Public Health Laboratory and Victorian Infectious Diseases Reference Laboratory, The Peter Doherty Institute for Infection and Immunity | Caly L., Seemann T., Sait, M., Schultz M., Druce J., Sherry, N. |
| EPI_ISL_430543 | hCoV-19/Australia/VIC982/2020 | Oceania / Australia / Victoria | 2020-03-30 | Victorian Infectious Diseases Reference Laboratory (VIDRL) | Microbiological Diagnostic Unit Public Health Laboratory and Victorian Infectious Diseases Reference Laboratory, The Peter Doherty Institute for Infection and Immunity | Caly L., Seemann T., Sait, M., Schultz M., Druce J., Sherry, N. |
| EPI_ISL_430545 | hCoV-19/Australia/VIC983/2020 | Oceania / Australia / Victoria | 2020-03-30 | Victorian Infectious Diseases Reference Laboratory (VIDRL) | Microbiological Diagnostic Unit Public Health Laboratory and Victorian Infectious Diseases Reference Laboratory, The Peter Doherty Institute for Infection and Immunity | Caly L., Seemann T., Sait, M., Schultz M., Druce J., Sherry, N. |
| EPI_ISL_430547 | hCoV-19/Australia/VIC1011/2020 | Oceania / Australia / Victoria | 2020-04-01 | Victorian Infectious Diseases Reference Laboratory (VIDRL) | Microbiological Diagnostic Unit Public Health Laboratory and Victorian Infectious Diseases Reference Laboratory, The Peter Doherty Institute for Infection and Immunity | Caly L., Seemann T., Sait, M., Schultz M., Druce J., Sherry, N. |
| EPI_ISL_430550 | hCoV-19/Australia/VIC1046/2020 | Oceania / Australia / Victoria | 2020-04-03 | Victorian Infectious Diseases Reference Laboratory (VIDRL) | Microbiological Diagnostic Unit Public Health Laboratory and Victorian Infectious Diseases Reference Laboratory, The Peter Doherty Institute for Infection and Immunity | Caly L., Seemann T., Sait, M., Schultz M., Druce J., Sherry, N. |
| EPI_ISL_430551 | hCoV-19/Australia/VIC989/2020 | Oceania / Australia / Victoria | 2020-03-31 | Victorian Infectious Diseases Reference Laboratory (VIDRL) | Microbiological Diagnostic Unit Public Health Laboratory and Victorian Infectious Diseases Reference Laboratory, The Peter Doherty Institute for Infection and Immunity | Caly L., Seemann T., Sait, M., Schultz M., Druce J., Sherry, N. |
| EPI_ISL_430552 | hCoV-19/Australia/VIC984/2020 | Oceania / Australia / Victoria | 2020-03-30 | Victorian Infectious Diseases Reference Laboratory (VIDRL) | Microbiological Diagnostic Unit Public Health Laboratory and Victorian Infectious Diseases Reference Laboratory, The Peter Doherty Institute for Infection and Immunity | Caly L., Seemann T., Sait, M., Schultz M., Druce J., Sherry, N. |
| EPI_ISL_430553 | hCoV-19/Australia/VIC1013/2020 | Oceania / Australia / Victoria | 2020-04-01 | Victorian Infectious Diseases Reference Laboratory (VIDRL) | Microbiological Diagnostic Unit Public Health Laboratory and Victorian Infectious Diseases Reference Laboratory, The Peter Doherty Institute for Infection and Immunity | Caly L., Seemann T., Sait, M., Schultz M., Druce J., Sherry, N. |
| EPI_ISL_430556 | hCoV-19/Australia/VIC990/2020 | Oceania / Australia / Victoria | 2020-03-31 | Victorian Infectious Diseases Reference Laboratory (VIDRL) | Microbiological Diagnostic Unit Public Health Laboratory and Victorian Infectious Diseases Reference Laboratory, The Peter Doherty Institute for Infection and Immunity | Caly L., Seemann T., Sait, M., Schultz M., Druce J., Sherry, N. |
| EPI_ISL_430559 | hCoV-19/Australia/VIC1015/2020 | Oceania / Australia / Victoria | 2020-04-01 | Victorian Infectious Diseases Reference Laboratory (VIDRL) | Microbiological Diagnostic Unit Public Health Laboratory and Victorian Infectious Diseases Reference Laboratory, The Peter Doherty Institute for Infection and Immunity | Caly L., Seemann T., Sait, M., Schultz M., Druce J., Sherry, N. |
| EPI_ISL_430567 | hCoV-19/Australia/VIC1119/2020 | Oceania / Australia / Victoria | 2020-04-08 | Victorian Infectious Diseases Reference Laboratory (VIDRL) | Microbiological Diagnostic Unit Public Health Laboratory and Victorian Infectious Diseases Reference Laboratory, The Peter Doherty Institute for Infection and Immunity | Caly L., Seemann T., Sait, M., Schultz M., Druce J., Sherry, N. |
| EPI_ISL_430569 | hCoV-19/Australia/VIC1110/2020 | Oceania / Australia / Victoria | 2020-04-07 | Victorian Infectious Diseases Reference Laboratory (VIDRL) | Microbiological Diagnostic Unit Public Health Laboratory and Victorian Infectious Diseases Reference Laboratory, The Peter Doherty Institute for Infection and Immunity | Caly L., Seemann T., Sait, M., Schultz M., Druce J., Sherry, N. |
| EPI_ISL_430573 | hCoV-19/Australia/VIC1112/2020 | Oceania / Australia / Victoria | 2020-04-07 | Victorian Infectious Diseases Reference Laboratory (VIDRL) | Microbiological Diagnostic Unit Public Health Laboratory and Victorian Infectious Diseases Reference Laboratory, The Peter Doherty Institute for Infection and Immunity | Caly L., Seemann T., Sait, M., Schultz M., Druce J., Sherry, N. |
| EPI_ISL_430574 | hCoV-19/Australia/VIC1087/2020 | Oceania / Australia / Victoria | 2020-04-06 | Victorian Infectious Diseases Reference Laboratory (VIDRL) | Microbiological Diagnostic Unit Public Health Laboratory and Victorian Infectious Diseases Reference Laboratory, The Peter Doherty Institute for Infection and Immunity | Caly L., Seemann T., Sait, M., Schultz M., Druce J., Sherry, N. |
| EPI_ISL_430576 | hCoV-19/Australia/VIC1114/2020 | Oceania / Australia / Victoria | 2020-04-07 | Victorian Infectious Diseases Reference Laboratory (VIDRL) | Microbiological Diagnostic Unit Public Health Laboratory and Victorian Infectious Diseases Reference Laboratory, The Peter Doherty Institute for Infection and Immunity | Caly L., Seemann T., Sait, M., Schultz M., Druce J., Sherry, N. |
| EPI_ISL_430577 | hCoV-19/Australia/VIC1115/2020 | Oceania / Australia / Victoria | 2020-04-07 | Victorian Infectious Diseases Reference Laboratory (VIDRL) | Microbiological Diagnostic Unit Public Health Laboratory and Victorian Infectious Diseases Reference Laboratory, The Peter Doherty Institute for Infection and Immunity | Caly L., Seemann T., Sait, M., Schultz M., Druce J., Sherry, N. |
| EPI_ISL_430578 | hCoV-19/Australia/VIC1062/2020 | Oceania / Australia / Victoria | 2020-04-04 | Victorian Infectious Diseases Reference Laboratory (VIDRL) | Microbiological Diagnostic Unit Public Health Laboratory and Victorian Infectious Diseases Reference Laboratory, The Peter Doherty Institute for Infection and Immunity | Caly L., Seemann T., Sait, M., Schultz M., Druce J., Sherry, N. |
| EPI_ISL_430579 | hCoV-19/Australia/VIC1080/2020 | Oceania / Australia / Victoria | 2020-04-06 | Victorian Infectious Diseases Reference Laboratory (VIDRL) | Microbiological Diagnostic Unit Public Health Laboratory and Victorian Infectious Diseases Reference Laboratory, The Peter Doherty Institute for Infection and Immunity | Caly L., Seemann T., Sait, M., Schultz M., Druce J., Sherry, N. |
| EPI_ISL_430580 | hCoV-19/Australia/VIC1081/2020 | Oceania / Australia / Victoria | 2020-04-06 | Victorian Infectious Diseases Reference Laboratory (VIDRL) | Microbiological Diagnostic Unit Public Health Laboratory and Victorian Infectious Diseases Reference Laboratory, The Peter Doherty Institute for Infection and Immunity | Caly L., Seemann T., Sait, M., Schultz M., Druce J., Sherry, N. |
| EPI_ISL_430581 | hCoV-19/Australia/VIC1071/2020 | Oceania / Australia / Victoria | 2020-04-05 | Victorian Infectious Diseases Reference Laboratory (VIDRL) | Microbiological Diagnostic Unit Public Health Laboratory and Victorian Infectious Diseases Reference Laboratory, The Peter Doherty Institute for Infection and Immunity | Caly L., Seemann T., Sait, M., Schultz M., Druce J., Sherry, N. |
| EPI_ISL_430583 | hCoV-19/Australia/VIC1094/2020 | Oceania / Australia / Victoria | 2020-04-07 | Victorian Infectious Diseases Reference Laboratory (VIDRL) | Microbiological Diagnostic Unit Public Health Laboratory and Victorian Infectious Diseases Reference Laboratory, The Peter Doherty Institute for Infection and Immunity | Caly L., Seemann T., Sait, M., Schultz M., Druce J., Sherry, N. |
| EPI_ISL_430584 | hCoV-19/Australia/VIC1103/2020 | Oceania / Australia / Victoria | 2020-04-07 | Victorian Infectious Diseases Reference Laboratory (VIDRL) | Microbiological Diagnostic Unit Public Health Laboratory and Victorian Infectious Diseases Reference Laboratory, The Peter Doherty Institute for Infection and Immunity | Caly L., Seemann T., Sait, M., Schultz M., Druce J., Sherry, N. |
| EPI_ISL_430590 | hCoV-19/Australia/VIC1107/2020 | Oceania / Australia / Victoria | 2020-04-07 | Victorian Infectious Diseases Reference Laboratory (VIDRL) | Microbiological Diagnostic Unit Public Health Laboratory and Victorian Infectious Diseases Reference Laboratory, The Peter Doherty Institute for Infection and Immunity | Caly L., Seemann T., Sait, M., Schultz M., Druce J., Sherry, N. |
| EPI_ISL_430596 | hCoV-19/Australia/VIC1133/2020 | Oceania / Australia / Victoria | 2020-04-08 | Victorian Infectious Diseases Reference Laboratory (VIDRL) | Microbiological Diagnostic Unit Public Health Laboratory and Victorian Infectious Diseases Reference Laboratory, The Peter Doherty Institute for Infection and Immunity | Caly L., Seemann T., Sait, M., Schultz M., Druce J., Sherry, N. |
| EPI_ISL_430597 | hCoV-19/Australia/VIC1086/2020 | Oceania / Australia / Victoria | 2020-04-06 | Victorian Infectious Diseases Reference Laboratory (VIDRL) | Microbiological Diagnostic Unit Public Health Laboratory and Victorian Infectious Diseases Reference Laboratory, The Peter Doherty Institute for Infection and Immunity | Caly L., Seemann T., Sait, M., Schultz M., Druce J., Sherry, N. |
| EPI_ISL_430598 | hCoV-19/Australia/VIC1157/2020 | Oceania / Australia / Victoria | 2020-04-09 | Victorian Infectious Diseases Reference Laboratory (VIDRL) | Microbiological Diagnostic Unit Public Health Laboratory and Victorian Infectious Diseases Reference Laboratory, The Peter Doherty Institute for Infection and Immunity | Caly L., Seemann T., Sait, M., Schultz M., Druce J., Sherry, N. |
| EPI_ISL_430696 | hCoV-19/Australia/VIC1329/2020 | Oceania / Australia / Victoria | 2020-04-12 | Victorian Infectious Diseases Reference Laboratory (VIDRL) | Microbiological Diagnostic Unit Public Health Laboratory and Victorian Infectious Diseases Reference Laboratory, The Peter Doherty Institute for Infection and Immunity | Caly L., Seemann T., Sait, M., Schultz M., Druce J., Sherry, N. |
| EPI_ISL_430697 | hCoV-19/Australia/VIC1344/2020 | Oceania / Australia / Victoria | 2020-04-14 | Victorian Infectious Diseases Reference Laboratory (VIDRL) | Microbiological Diagnostic Unit Public Health Laboratory and Victorian Infectious Diseases Reference Laboratory, The Peter Doherty Institute for Infection and Immunity | Caly L., Seemann T., Sait, M., Schultz M., Druce J., Sherry, N. |
| EPI_ISL_430698 | hCoV-19/Australia/VIC1336/2020 | Oceania / Australia / Victoria | 2020-04-13 | Victorian Infectious Diseases Reference Laboratory (VIDRL) | Microbiological Diagnostic Unit Public Health Laboratory and Victorian Infectious Diseases Reference Laboratory, The Peter Doherty Institute for Infection and Immunity | Caly L., Seemann T., Sait, M., Schultz M., Druce J., Sherry, N. |
| EPI_ISL_430800 | hCoV-19/Argentina/PAIS_A008/2020 | South America / Argentina / Ciudad Autor | 2020-03-30 | Laboratorio de Virología del Hospital de Niños Dr. Ricardo Gutierrez | Área de Secuenciación del Laboratorio de Virología del Hospital de Niños Dr. Ricardo Gutierrez | Nabaes Jodar, MS; Goya, S; Natale, MI; Lusso, S; Gravis, E; Mistchenko, AS; Valinotto, LE; Viegas, M. |
| EPI_ISL_430801 | hCoV-19/Argentina/PAIS_A009/2020 | South America / Argentina / Ciudad Autor | 2020-03-31 | Laboratorio de Virología del Hospital de Niños Dr. Ricardo Gutierrez | Área de Secuenciación del Laboratorio de Virología del Hospital de Niños Dr. Ricardo Gutierrez | Nabaes Jodar, MS; Goya, S; Natale, MI; Lusso, S; Gravis, E; Mistchenko, AS; Valinotto, LE; Viegas, M. |
| EPI_ISL_430802 | hCoV-19/Argentina/PAIS_A011/2020 | South America / Argentina / Buenos Aire | 2020-03-31 | Departamento de Biología y genética molecular, IACA Laboratorios. | Área de Secuenciación del Laboratorio de Virología del Hospital de Niños Dr. Ricardo Gutierrez | Nabaes Jodar, MS; Goya, S; Natale, MI; Lusso, S; Tittarelli, E; Suárez, A; Masciovecchio MV; Streitenberger ER; Mistchenko, AS; Valinotto, LE; Viegas, M. |
| EPI_ISL_430803 | hCoV-19/Argentina/PAIS_A012/2020 | South America / Argentina / Ciudad Autor | 2020-04-01 | Laboratorio de Virología del Hospital de Niños Dr. Ricardo Gutierrez | Área de Secuenciación del Laboratorio de Virología del Hospital de Niños Dr. Ricardo Gutierrez | Nabaes Jodar, MS; Goya, S; Natale, MI; Lusso, S; Gravis, E; Mistchenko, AS; Valinotto, LE; Viegas, M. |
| EPI_ISL_430804 | hCoV-19/Argentina/PAIS_A013/2020 | South America / Argentina / Ciudad Autor | 2020-04-01 | Laboratorio de Virología del Hospital de Niños Dr. Ricardo Gutierrez | Área de Secuenciación del Laboratorio de Virología del Hospital de Niños Dr. Ricardo Gutierrez | Nabaes Jodar, MS; Goya, S; Natale, MI; Lusso, S; Gravis, E; Mistchenko, AS; Valinotto, LE; Viegas, M. |
| EPI_ISL_430805 | hCoV-19/Argentina/PAIS_A014/2020 | South America / Argentina / Buenos Aire | 2020-04-01 | Departamento de Biología y genética molecular, IACA Laboratorios. | Área de Secuenciación del Laboratorio de Virología del Hospital de Niños Dr. Ricardo Gutierrez | Nabaes Jodar, MS; Goya, S; Natale, MI; Lusso, S; Tittarelli, E; Suárez, A; Masciovecchio MV; Streitenberger ER; Mistchenko, AS; Valinotto, LE; Viegas, M. |
| EPI_ISL_430806 | hCoV-19/Argentina/PAIS_A015/2020 | South America / Argentina / Buenos Aire | 2020-04-01 | Departamento de Biología y genética molecular, IACA Laboratorios. | Área de Secuenciación del Laboratorio de Virología del Hospital de Niños Dr. Ricardo Gutierrez | Nabaes Jodar, MS; Goya, S; Natale, MI; Lusso, S; Tittarelli, E; Suárez, A; Masciovecchio MV; Streitenberger ER; Mistchenko, AS; Valinotto, LE; Viegas, M. |
| EPI_ISL_430807 | hCoV-19/Argentina/PAIS_A016/2020 | South America / Argentina / Ciudad Autor | 2020-04-02 | Laboratorio de Virología del Hospital de Niños Dr. Ricardo Gutierrez | Área de Secuenciación del Laboratorio de Virología del Hospital de Niños Dr. Ricardo Gutierrez | Nabaes Jodar, MS; Goya, S; Natale, MI; Lusso, S; Gravis, E; Mistchenko, AS; Valinotto, LE; Viegas, M. |
| EPI_ISL_430808 | hCoV-19/Argentina/PAIS_A017/2020 | South America / Argentina / Rio Negro | 2020-04-03 | Departamento de Biología y genética molecular, IACA Laboratorios. | Área de Secuenciación del Laboratorio de Virología del Hospital de Niños Dr. Ricardo Gutierrez | Nabaes Jodar, MS; Goya, S; Natale, MI; Lusso, S; Tittarelli, E; Suárez, A; Masciovecchio MV; Streitenberger ER; Mistchenko, AS; Valinotto, LE; Viegas, M. |
| EPI_ISL_430809 | hCoV-19/Argentina/PAIS_A018/2020 | South America / Argentina / Ciudad Autor | 2020-04-04 | Laboratorio de Virología del Hospital de Niños Dr. Ricardo Gutierrez | Área de Secuenciación del Laboratorio de Virología del Hospital de Niños Dr. Ricardo Gutierrez | Nabaes Jodar, MS; Goya, S; Natale, MI; Lusso, S; Gravis, E; Mistchenko, AS; Valinotto, LE; Viegas, M. |
| EPI_ISL_430810 | hCoV-19/Argentina/PAIS_A019/2020 | South America / Argentina / Ciudad Autor | 2020-04-07 | Laboratorio de Virología del Hospital de Niños Dr. Ricardo Gutierrez | Área de Secuenciación del Laboratorio de Virología del Hospital de Niños Dr. Ricardo Gutierrez | Nabaes Jodar, MS; Goya, S; Natale, MI; Lusso, S; Gravis, E; Mistchenko, AS; Valinotto, LE; Viegas, M. |
| EPI_ISL_430811 | hCoV-19/Argentina/PAIS_A020/2020 | South America / Argentina / Ciudad Autor | 2020-04-11 | Laboratorio de Virología del Hospital de Niños Dr. Ricardo Gutierrez | Área de Secuenciación del Laboratorio de Virología del Hospital de Niños Dr. Ricardo Gutierrez | Nabaes Jodar, MS; Goya, S; Natale, MI; Lusso, S; Gravis, E; Mistchenko, AS; Valinotto, LE; Viegas, M. |
| EPI_ISL_430812 | hCoV-19/Argentina/PAIS_A021/2020 | South America / Argentina / Ciudad Autor | 2020-04-11 | Laboratorio de Virología del Hospital de Niños Dr. Ricardo Gutierrez | Área de Secuenciación del Laboratorio de Virología del Hospital de Niños Dr. Ricardo Gutierrez | Nabaes Jodar, MS; Goya, S; Natale, MI; Lusso, S; Gravis, E; Mistchenko, AS; Valinotto, LE; Viegas, M. |
| EPI_ISL_430814 | hCoV-19/Argentina/PAIS_A023/2020 | South America / Argentina / Ciudad Autor | 2020-04-17 | Laboratorio de Virología del Hospital de Niños Dr. Ricardo Gutierrez | Área de Secuenciación del Laboratorio de Virología del Hospital de Niños Dr. Ricardo Gutierrez | Nabaes Jodar, MS; Goya, S; Natale, MI; Lusso, S; Gravis, E; Mistchenko, AS; Valinotto, LE; Viegas, M. |
| EPI_ISL_430815 | hCoV-19/Argentina/PAIS_A024/2020 | South America / Argentina / Ciudad Autor | 2020-04-18 | Laboratorio de Virología del Hospital de Niños Dr. Ricardo Gutierrez | Área de Secuenciación del Laboratorio de Virología del Hospital de Niños Dr. Ricardo Gutierrez | Nabaes Jodar, MS; Goya, S; Natale, MI; Lusso, S; Gravis, E; Mistchenko, AS; Valinotto, LE; Viegas, M. |
| EPI_ISL_430816 | hCoV-19/Argentina/PAIS_A025/2020 | South America / Argentina / Ciudad Autor | 2020-04-18 | Laboratorio de Virología del Hospital de Niños Dr. Ricardo Gutierrez | Área de Secuenciación del Laboratorio de Virología del Hospital de Niños Dr. Ricardo Gutierrez | Nabaes Jodar, MS; Goya, S; Natale, MI; Lusso, S; Gravis, E; Mistchenko, AS; Valinotto, LE; Viegas, M. |
| EPI_ISL_430817 | hCoV-19/Argentina/PAIS_A026/2020 | South America / Argentina / Ciudad Autor | 2020-04-18 | Laboratorio de Virología del Hospital de Niños Dr. Ricardo Gutierrez | Área de Secuenciación del Laboratorio de Virología del Hospital de Niños Dr. Ricardo Gutierrez | Nabaes Jodar, MS; Goya, S; Natale, MI; Lusso, S; Gravis, E; Mistchenko, AS; Valinotto, LE; Viegas, M. |
| EPI_ISL_430818 | hCoV-19/Argentina/PAIS_A027/2020 | South America / Argentina / Ciudad Autor | 2020-04-19 | Laboratorio de Virología del Hospital de Niños Dr. Ricardo Gutierrez | Área de Secuenciación del Laboratorio de Virología del Hospital de Niños Dr. Ricardo Gutierrez | Nabaes Jodar, MS; Goya, S; Natale, MI; Lusso, S; Gravis, E; Mistchenko, AS; Valinotto, LE; Viegas, M. |
| EPI_ISL_430819 | hCoV-19/Egypt/NRC-03/2020 | Africa / Egypt | 2020-03-18 | Center of Scientific Excellence for Influenza Viruses,National Research Centre (NRC), Egypt. | Center of Scientific Excellence for Influenza Viruses,National Research Centre (NRC), Egypt. | Mohamed Ahmed Ali, Ahmed Kandeil, Ahmed Mostafa, Rabeh El-Shesheny, Mahmoud Shehata, Wael Roshdy, Shymaa Showky Ahmed , Amal Naguib, Nancy M. El Guindy, Mokhtar Gomaa, Ahmed El-Taweel, Ahmed E Kayed, Yassmin Moatasim, Omnia Kutkat, Sara Mahmoud, Mina Kamel, Abo Shama, M Noura, Mohamed El Sayes |
| EPI_ISL_430820 | hCoV-19/Egypt/NRC-01/2020 | Africa / Egypt | 2020-03-18 | Center of Scientific Excellence for Influenza Viruses, National Research Centre (NRC), Egypt. | Center of Scientific Excellence for Influenza Viruses, National Research Centre (NRC), Egypt. | Mohamed Ahmed Ali, Ahmed Kandeil, Ahmed Mostafa, Rabeh El-Shesheny, Mahmoud Shehata, Wael Roshdy, Shymaa Showky Ahmed , Amal Naguib, Mokhtar Gomaa, Ahmed El-Taweel, Ahmed E Kayed, Yassmin Moatasim, Omnia Kutkat, Sara Mahmoud, Mina Kamel, Abo Shama, M Noura, Mohamed El Sayes, Nancy M. El Guindy |
| EPI_ISL_430837 | hCoV-19/Thailand/TNIC-1889/2020 | Asia / Thailand / Nonthaburi | 2020-03-03 | n/a | Thai National Influenza Center, Department of medical Science, Ministry of Public Health, Thailand | Pilailuk,Okada; Siripaporn,Phuygun; Thanutsapa,Thanadachakul;Sittiporn,Parnmen;Warawan,Wongboot;Sunthareeya,Waicharoen; Malinee,Chittaganpitch |
| EPI_ISL_430841 | hCoV-19/Thailand/NIH-2492/2020 | Asia / Thailand / Bangkok | 2020-03-17 | Praram 9 Hospital | National Institute of Health. Department of medical Sciences, Ministry of Public Health, Thailand | Pilailuk,Okada; Siripaporn,Phuygun; Thanutsapa,Thanadachakul; Sittiporn,Parnmen;Warawan,Wongboot; Sunthareeya,Waicharoen; Malinee,Chittaganpitch |
| EPI_ISL_430842 | hCoV-19/Thailand/NIH-2982/2020 | Asia / Thailand / Nonthaburi | 2020-03-18 | Central chest Institute of Thailand | National Institute of Health. Department of medical Sciences, Ministry of Public Health, Thailand | Pilailuk,Okada; Siripaporn,Phuygun; Thanutsapa,Thanadachakul; Sittiporn,Parnmen;Warawan,Wongboot; Sunthareeya,Waicharoen; Malinee,Chittaganpitch |
| EPI_ISL_430847 | hCoV-19/Sweden/20-04631/2020 | Europe / Sweden / Stockholm | 2020-02-27 | HS mikrobiologi virus | The Public Health Agency of Sweden | Zhibing Yun, Oskar Karlsson Lindsjo, Maria Lind Karlberg, Anna-Malin Linde, Olov Svartstrom, Anna Risberg, Shaman Muradrasoli, Karin Tegmark-Wisell |
| EPI_ISL_430848 | hCoV-19/Sweden/20-08144/2020 | Europe / Sweden / Halland | 2020-03-11 | Klinisk mikrobiologi och vardhygien Halmstad | The Public Health Agency of Sweden | Arne Kotz, Oskar Karlsson Lindsjo, Maria Lind Karlberg, Anna-Malin Linde, Olov Svartstrom, Anna Risberg, Shaman Muradrasoli, Karin Tegmark-Wisell |
| EPI_ISL_430849 | hCoV-19/Sweden/20-08147/2020 | Europe / Sweden / Halland | 2020-03-11 | Klinisk mikrobiologi och vardhygien Halmstad | The Public Health Agency of Sweden | Arne Kotz, Oskar Karlsson Lindsjo, Maria Lind Karlberg, Anna-Malin Linde, Olov Svartstrom, Anna Risberg, Shaman Muradrasoli, Karin Tegmark-Wisell |
| EPI_ISL_430861 | hCoV-19/Sweden/20-50129/2020 | Europe / Sweden / Varmland | 2020-03-06 | Klinisk mikrobiologi Orebro | The Public Health Agency of Sweden | Martin Sundqvist, Oskar Karlsson Lindsjo, Maria Lind Karlberg, Anna-Malin Linde, Olov Svartstrom, Anna Risberg, Shaman Muradrasoli, Karin Tegmark-Wisell |
| EPI_ISL_430862 | hCoV-19/Sweden/20-50185/2020 | Europe / Sweden / Varmland | 2020-03-08 | The Public Health Agency of Sweden | The Public Health Agency of Sweden | Oskar Karlsson Lindsjo, Maria Lind Karlberg, Anna-Malin Linde, Olov Svartstrom, Anna Risberg, Shaman Muradrasoli, Karin Tegmark-Wisell |
| EPI_ISL_430863 | hCoV-19/Sweden/20-50231/2020 | Europe / Sweden / Orebro lan | 2020-03-09 | Klinisk mikrobiologi Orebro | The Public Health Agency of Sweden | Martin Sundqvist, Oskar Karlsson Lindsjo, Maria Lind Karlberg, Anna-Malin Linde, Olov Svartstrom, Anna Risberg, Shaman Muradrasoli, Karin Tegmark-Wisell |
| EPI_ISL_430864 | hCoV-19/Sweden/20-50360/2020 | Europe / Sweden / Gavleborgs lan | 2020-03-16 | The Public Health Agency of Sweden | The Public Health Agency of Sweden | Oskar Karlsson Lindsjo, Maria Lind Karlberg, Anna-Malin Linde, Olov Svartstrom, Anna Risberg, Shaman Muradrasoli, Karin Tegmark-Wisell |
| EPI_ISL_430867 | hCoV-19/USA/WA-UW-517/2020 | North America / USA / Washington | 2020-03-11 | UW Virology Lab | UW Virology Lab | Pavitra Roychoudhury, Hong Xie, Keith Jerome, Alexander Greninger |
| EPI_ISL_430868 | hCoV-19/USA/WA-UW-5177/2020 | North America / USA / Washington | 2020-04-01 | UW Virology Lab | UW Virology Lab | Pavitra Roychoudhury, Hong Xie, Keith Jerome, Alexander Greninger |
| EPI_ISL_430869 | hCoV-19/USA/WA-UW-5178/2020 | North America / USA / Washington | 2020-04-01 | UW Virology Lab | UW Virology Lab | Pavitra Roychoudhury, Hong Xie, Keith Jerome, Alexander Greninger |
| EPI_ISL_430870 | hCoV-19/USA/WA-UW-518/2020 | North America / USA / Washington | 2020-03-11 | UW Virology Lab | UW Virology Lab | Pavitra Roychoudhury, Hong Xie, Keith Jerome, Alexander Greninger |
| EPI_ISL_430871 | hCoV-19/USA/WA-UW-5181/2020 | North America / USA / Washington | 2020-04-02 | UW Virology Lab | UW Virology Lab | Pavitra Roychoudhury, Hong Xie, Keith Jerome, Alexander Greninger |
| EPI_ISL_430894 | hCoV-19/USA/WA-UW-602/2020 | North America / USA / Washington | 2020-03-12 | UW Virology Lab | UW Virology Lab | Pavitra Roychoudhury, Hong Xie, Keith Jerome, Alexander Greninger |
| EPI_ISL_430895 | hCoV-19/USA/WA-UW-604/2020 | North America / USA / Washington | 2020-03-13 | UW Virology Lab | UW Virology Lab | Pavitra Roychoudhury, Hong Xie, Keith Jerome, Alexander Greninger |
| EPI_ISL_430897 | hCoV-19/USA/UN-UW-5199/2020 | North America / USA | 2020-04-02 | UW Virology Lab | UW Virology Lab | Pavitra Roychoudhury, Hong Xie, Keith Jerome, Alexander Greninger |
| EPI_ISL_430898 | hCoV-19/USA/WA-UW-5200/2020 | North America / USA / Washington | 2020-04-02 | UW Virology Lab | UW Virology Lab | Pavitra Roychoudhury, Hong Xie, Keith Jerome, Alexander Greninger |
| EPI_ISL_430899 | hCoV-19/USA/WA-UW-5203/2020 | North America / USA / Washington | 2020-04-01 | UW Virology Lab | UW Virology Lab | Pavitra Roychoudhury, Hong Xie, Keith Jerome, Alexander Greninger |
| EPI_ISL_431011 | hCoV-19/DRC/1319/2020 | Africa / Democratic Republic of the Cong | 2020-04-07 | Viral Respiratory Lab, National Institute for Biomedical Research (INRB) | Pathogen Sequencing Lab, National Institute for Biomedical Research (INRB) | Placide Mbala-Kingebeni, Edith Nkwembe, Eddy Kinganda-Lusamaki, Amuri Aziza, Francisca Muyembe Mawete, Catherine Pratt, Matthias Pauthner, Josh Quick, Allison Black, James Hadfield, Trevor Bedford, Ian Goodfellow, Andrew Rambaut, Nick Loman, Kristian Andersen, Michael Wiley, Steve Ahuka-Mundeke, Jean-Jacques Muyembe Tamfum |
| EPI_ISL_431012 | hCoV-19/DRC/1376/2020 | Africa / Democratic Republic of the Cong | 2020-04-07 | Viral Respiratory Lab, National Institute for Biomedical Research (INRB) | Pathogen Sequencing Lab, National Institute for Biomedical Research (INRB) | Placide Mbala-Kingebeni, Edith Nkwembe, Eddy Kinganda-Lusamaki, Amuri Aziza, Francisca Muyembe Mawete, Catherine Pratt, Matthias Pauthner, Josh Quick, Allison Black, James Hadfield, Trevor Bedford, Ian Goodfellow, Andrew Rambaut, Nick Loman, Kristian Andersen, Michael Wiley, Steve Ahuka-Mundeke, Jean-Jacques Muyembe Tamfum |
| EPI_ISL_431013 | hCoV-19/USA/AK-PHL010/2020 | North America / USA / Alaska / Interior | 2020-03-17 | Alaska State Virology Laboratory | Alaska State Virology Laboratory | Jack Chen |
| EPI_ISL_431014 | hCoV-19/USA/AK-PHL011/2020 | North America / USA / Alaska / Southeas | 2020-03-17 | Alaska State Virology Laboratory | Alaska State Virology Laboratory | Jack Chen, Ph.D. |
| EPI_ISL_431015 | hCoV-19/USA/AK-PHL013/2020 | North America / USA / Alaska / Interior | 2020-03-19 | Alaska State Virology Laboratory | Alaska State Virology Laboratory | Jack Chen, Ph.D. |
| EPI_ISL_431016 | hCoV-19/USA/AK-PHL016/2020 | North America / USA / Alaska / Southeas | 2020-03-18 | Alaska State Virology Laboratory | Alaska State Virology Laboratory | Jack Chen, Ph.D. |
| EPI_ISL_431017 | hCoV-19/USA/AK-PHL040/2020 | North America / USA / Alaska / Interior | 2020-03-24 | Alaska State Virology Laboratory | Alaska State Virology Laboratory | Jack Chen, Ph.D. |
| EPI_ISL_431018 | hCoV-19/USA/AK-PHL052/2020 | North America / USA / Alaska / Interior | 2020-03-26 | Alaska State Virology Laboratory | Alaska State Virology Laboratory | Jack Chen, Ph.D. |
| EPI_ISL_431019 | hCoV-19/USA/AK-PHL074/2020 | North America / USA / Alaska / Southeas | 2020-03-27 | Alaska State Virology Laboratory | Alaska State Virology Laboratory | Jack Chen, Ph.D. |
| EPI_ISL_431101 | hCoV-19/India/GMC-RK100/2020 | Asia / India / Telangana / Hyderabad | 2020-03-01 | Department of Microbiology,Gandhi Medical College and Hospital | Virus Research Laboratory, Department of Zoology, Osmania University,Hyderabad,India | Muttineni Radhakrishna, Nagamani K, Thrilok Chander B, Raja Rao M, Kalyani Putty, Ravikumar P, Sunitha P, Pankaj Singh D, Anand Kumar K, Amit A. Upadhyay Steven E. Bosinger, Rama Amara |
| EPI_ISL_431102 | hCoV-19/India/GMC-KN318/2020 | Asia / India / Telangana / Hyderabad | 2020-03-11 | Department of MicroBiology,Gandhi Medical College and Hospital,Secendrabad,Hyderabad,India | Department of Microbiology, Gandhi Medical College and Hospital, Secendrabad, Hyderabad | Nagamani K, Muttineni Radhakrishna, Thrilok Chander B, Raja Rao M, Kalyani Putty, Ravikumar P, Sunitha P, Pankaj Singh D, Anand Kumar K, Amit A. Upadhyay, Steven E. Bosinger, Rama Amara |
| EPI_ISL_432236 | hCoV-19/Wales/PHWC-27D2A/2020 | Europe / United Kingdom / Wales | 2020-04-01 | Wales Specialist Virology Centre | Public Health Wales Microbiology Cardiff | Catherine Moore, Johnathan Evans, Malorie Perry, Simon Cottrell, Alec Birchley, Alexander Adams, Amy Gaskin, Bree Gatica-Wilcox, Jason Coombes, Lauren Gilbert, Lee Graham, Nicole Pacchiarini, Sara Kumziene-Summerhayes, Sarah Taylor, Sophie Jones, Sara Rey, Matthew Bull, Joanne Watkins, Sally Corden, Tom Connor |
| EPI_ISL_432237 | hCoV-19/Wales/PHWC-2A03D/2020 | Europe / United Kingdom / Wales | 2020-04-03 | Wales Specialist Virology Centre | Public Health Wales Microbiology Cardiff | Catherine Moore, Johnathan Evans, Malorie Perry, Simon Cottrell, Alec Birchley, Alexander Adams, Amy Gaskin, Bree Gatica-Wilcox, Jason Coombes, Lauren Gilbert, Lee Graham, Nicole Pacchiarini, Sara Kumziene-Summerhayes, Sarah Taylor, Sophie Jones, Sara Rey, Matthew Bull, Joanne Watkins, Sally Corden, Tom Connor |
| EPI_ISL_432238 | hCoV-19/Wales/PHWC-27C1E/2020 | Europe / United Kingdom / Wales | 2020-03-31 | Wales Specialist Virology Centre | Public Health Wales Microbiology Cardiff | Catherine Moore, Johnathan Evans, Malorie Perry, Simon Cottrell, Alec Birchley, Alexander Adams, Amy Gaskin, Bree Gatica-Wilcox, Jason Coombes, Lauren Gilbert, Lee Graham, Nicole Pacchiarini, Sara Kumziene-Summerhayes, Sarah Taylor, Sophie Jones, Sara Rey, Matthew Bull, Joanne Watkins, Sally Corden, Tom Connor |
| EPI_ISL_432239 | hCoV-19/Wales/PHWC-27D39/2020 | Europe / United Kingdom / Wales | 2020-04-01 | Wales Specialist Virology Centre | Public Health Wales Microbiology Cardiff | Catherine Moore, Johnathan Evans, Malorie Perry, Simon Cottrell, Alec Birchley, Alexander Adams, Amy Gaskin, Bree Gatica-Wilcox, Jason Coombes, Lauren Gilbert, Lee Graham, Nicole Pacchiarini, Sara Kumziene-Summerhayes, Sarah Taylor, Sophie Jones, Sara Rey, Matthew Bull, Joanne Watkins, Sally Corden, Tom Connor |
| EPI_ISL_432240 | hCoV-19/Wales/PHWC-27BA8/2020 | Europe / United Kingdom / Wales | 2020-04-01 | Wales Specialist Virology Centre | Public Health Wales Microbiology Cardiff | Catherine Moore, Johnathan Evans, Malorie Perry, Simon Cottrell, Alec Birchley, Alexander Adams, Amy Gaskin, Bree Gatica-Wilcox, Jason Coombes, Lauren Gilbert, Lee Graham, Nicole Pacchiarini, Sara Kumziene-Summerhayes, Sarah Taylor, Sophie Jones, Sara Rey, Matthew Bull, Joanne Watkins, Sally Corden, Tom Connor |
| EPI_ISL_432241 | hCoV-19/Wales/PHWC-2B306/2020 | Europe / United Kingdom / Wales | 2020-04-08 | Wales Specialist Virology Centre | Public Health Wales Microbiology Cardiff | Catherine Moore, Johnathan Evans, Malorie Perry, Simon Cottrell, Alec Birchley, Alexander Adams, Amy Gaskin, Bree Gatica-Wilcox, Jason Coombes, Lauren Gilbert, Lee Graham, Nicole Pacchiarini, Sara Kumziene-Summerhayes, Sarah Taylor, Sophie Jones, Sara Rey, Matthew Bull, Joanne Watkins, Sally Corden, Tom Connor |
| EPI_ISL_432242 | hCoV-19/Wales/PHWC-27328/2020 | Europe / United Kingdom / Wales | 2020-03-31 | Wales Specialist Virology Centre | Public Health Wales Microbiology Cardiff | Catherine Moore, Johnathan Evans, Malorie Perry, Simon Cottrell, Alec Birchley, Alexander Adams, Amy Gaskin, Bree Gatica-Wilcox, Jason Coombes, Lauren Gilbert, Lee Graham, Nicole Pacchiarini, Sara Kumziene-Summerhayes, Sarah Taylor, Sophie Jones, Sara Rey, Matthew Bull, Joanne Watkins, Sally Corden, Tom Connor |
| EPI_ISL_432243 | hCoV-19/Wales/PHWC-29BF1/2020 | Europe / United Kingdom / Wales | 2020-04-04 | Wales Specialist Virology Centre | Public Health Wales Microbiology Cardiff | Catherine Moore, Johnathan Evans, Malorie Perry, Simon Cottrell, Alec Birchley, Alexander Adams, Amy Gaskin, Bree Gatica-Wilcox, Jason Coombes, Lauren Gilbert, Lee Graham, Nicole Pacchiarini, Sara Kumziene-Summerhayes, Sarah Taylor, Sophie Jones, Sara Rey, Matthew Bull, Joanne Watkins, Sally Corden, Tom Connor |
| EPI_ISL_432244 | hCoV-19/Wales/PHWC-29D55/2020 | Europe / United Kingdom / Wales | 2020-04-04 | Wales Specialist Virology Centre | Public Health Wales Microbiology Cardiff | Catherine Moore, Johnathan Evans, Malorie Perry, Simon Cottrell, Alec Birchley, Alexander Adams, Amy Gaskin, Bree Gatica-Wilcox, Jason Coombes, Lauren Gilbert, Lee Graham, Nicole Pacchiarini, Sara Kumziene-Summerhayes, Sarah Taylor, Sophie Jones, Sara Rey, Matthew Bull, Joanne Watkins, Sally Corden, Tom Connor |

| | | | | | | |
|---|---|---|---|---|---|---|
| EPI_ISL_432245 | hCoV-19/Wales/PHWC-27A23/2020 | Europe / United Kingdom / Wales | 2020-04-01 | Wales Specialist Virology Centre | Public Health Wales Microbiology Cardiff | Catherine Moore, Johnathan Evans, Malorie Perry, Simon Cottrell, Alec Birchley, Alexander Adams, Amy Gaskin, Bree Gatica-Wilcox, Jason Coombes, Lauren Gilbert, Lee Graham, Nicole Pacchiarini, Sara Kumziene-Summerhayes, Sarah Taylor, Sophie Jones, Sara Rey, Matthew Bull, Joanne Watkins, Sally Corden, Tom Connor |
| EPI_ISL_432246 | hCoV-19/Wales/PHWC-2B1FD/2020 | Europe / United Kingdom / Wales | 2020-04-08 | Wales Specialist Virology Centre | Public Health Wales Microbiology Cardiff | Catherine Moore, Johnathan Evans, Malorie Perry, Simon Cottrell, Alec Birchley, Alexander Adams, Amy Gaskin, Bree Gatica-Wilcox, Jason Coombes, Lauren Gilbert, Lee Graham, Nicole Pacchiarini, Sara Kumziene-Summerhayes, Sarah Taylor, Sophie Jones, Sara Rey, Matthew Bull, Joanne Watkins, Sally Corden, Tom Connor |
| EPI_ISL_432247 | hCoV-19/Wales/PHWC-29C85/2020 | Europe / United Kingdom / Wales | 2020-04-05 | Wales Specialist Virology Centre | Public Health Wales Microbiology Cardiff | Catherine Moore, Johnathan Evans, Malorie Perry, Simon Cottrell, Alec Birchley, Alexander Adams, Amy Gaskin, Bree Gatica-Wilcox, Jason Coombes, Lauren Gilbert, Lee Graham, Nicole Pacchiarini, Sara Kumziene-Summerhayes, Sarah Taylor, Sophie Jones, Sara Rey, Matthew Bull, Joanne Watkins, Sally Corden, Tom Connor |
| EPI_ISL_432248 | hCoV-19/Wales/PHWC-29D73/2020 | Europe / United Kingdom / Wales | 2020-04-05 | Wales Specialist Virology Centre | Public Health Wales Microbiology Cardiff | Catherine Moore, Johnathan Evans, Malorie Perry, Simon Cottrell, Alec Birchley, Alexander Adams, Amy Gaskin, Bree Gatica-Wilcox, Jason Coombes, Lauren Gilbert, Lee Graham, Nicole Pacchiarini, Sara Kumziene-Summerhayes, Sarah Taylor, Sophie Jones, Sara Rey, Matthew Bull, Joanne Watkins, Sally Corden, Tom Connor |
| EPI_ISL_432249 | hCoV-19/Wales/PHWC-29DBF/2020 | Europe / United Kingdom / Wales | 2020-04-05 | Wales Specialist Virology Centre | Public Health Wales Microbiology Cardiff | Catherine Moore, Johnathan Evans, Malorie Perry, Simon Cottrell, Alec Birchley, Alexander Adams, Amy Gaskin, Bree Gatica-Wilcox, Jason Coombes, Lauren Gilbert, Lee Graham, Nicole Pacchiarini, Sara Kumziene-Summerhayes, Sarah Taylor, Sophie Jones, Sara Rey, Matthew Bull, Joanne Watkins, Sally Corden, Tom Connor |
| EPI_ISL_432250 | hCoV-19/Wales/PHWC-2722B/2020 | Europe / United Kingdom / Wales | 2020-03-30 | Wales Specialist Virology Centre | Public Health Wales Microbiology Cardiff | Catherine Moore, Johnathan Evans, Malorie Perry, Simon Cottrell, Alec Birchley, Alexander Adams, Amy Gaskin, Bree Gatica-Wilcox, Jason Coombes, Lauren Gilbert, Lee Graham, Nicole Pacchiarini, Sara Kumziene-Summerhayes, Sarah Taylor, Sophie Jones, Sara Rey, Matthew Bull, Joanne Watkins, Sally Corden, Tom Connor |
| EPI_ISL_432251 | hCoV-19/Wales/PHWC-2A088/2020 | Europe / United Kingdom / Wales | 2020-04-04 | Wales Specialist Virology Centre | Public Health Wales Microbiology Cardiff | Catherine Moore, Johnathan Evans, Malorie Perry, Simon Cottrell, Alec Birchley, Alexander Adams, Amy Gaskin, Bree Gatica-Wilcox, Jason Coombes, Lauren Gilbert, Lee Graham, Nicole Pacchiarini, Sara Kumziene-Summerhayes, Sarah Taylor, Sophie Jones, Sara Rey, Matthew Bull, Joanne Watkins, Sally Corden, Tom Connor |
| EPI_ISL_432252 | hCoV-19/Wales/PHWC-2A0F1/2020 | Europe / United Kingdom / Wales | 2020-04-04 | Wales Specialist Virology Centre | Public Health Wales Microbiology Cardiff | Catherine Moore, Johnathan Evans, Malorie Perry, Simon Cottrell, Alec Birchley, Alexander Adams, Amy Gaskin, Bree Gatica-Wilcox, Jason Coombes, Lauren Gilbert, Lee Graham, Nicole Pacchiarini, Sara Kumziene-Summerhayes, Sarah Taylor, Sophie Jones, Sara Rey, Matthew Bull, Joanne Watkins, Sally Corden, Tom Connor |
| EPI_ISL_432253 | hCoV-19/Wales/PHWC-29B79/2020 | Europe / United Kingdom / Wales | 2020-04-04 | Wales Specialist Virology Centre | Public Health Wales Microbiology Cardiff | Catherine Moore, Johnathan Evans, Malorie Perry, Simon Cottrell, Alec Birchley, Alexander Adams, Amy Gaskin, Bree Gatica-Wilcox, Jason Coombes, Lauren Gilbert, Lee Graham, Nicole Pacchiarini, Sara Kumziene-Summerhayes, Sarah Taylor, Sophie Jones, Sara Rey, Matthew Bull, Joanne Watkins, Sally Corden, Tom Connor |
| EPI_ISL_432254 | hCoV-19/Wales/PHWC-2748F/2020 | Europe / United Kingdom / Wales | 2020-03-31 | Wales Specialist Virology Centre | Public Health Wales Microbiology Cardiff | Catherine Moore, Johnathan Evans, Malorie Perry, Simon Cottrell, Alec Birchley, Alexander Adams, Amy Gaskin, Bree Gatica-Wilcox, Jason Coombes, Lauren Gilbert, Lee Graham, Nicole Pacchiarini, Sara Kumziene-Summerhayes, Sarah Taylor, Sophie Jones, Sara Rey, Matthew Bull, Joanne Watkins, Sally Corden, Tom Connor |
| EPI_ISL_432255 | hCoV-19/Wales/PHWC-27531/2020 | Europe / United Kingdom / Wales | 2020-03-30 | Wales Specialist Virology Centre | Public Health Wales Microbiology Cardiff | Catherine Moore, Johnathan Evans, Malorie Perry, Simon Cottrell, Alec Birchley, Alexander Adams, Amy Gaskin, Bree Gatica-Wilcox, Jason Coombes, Lauren Gilbert, Lee Graham, Nicole Pacchiarini, Sara Kumziene-Summerhayes, Sarah Taylor, Sophie Jones, Sara Rey, Matthew Bull, Joanne Watkins, Sally Corden, Tom Connor |
| EPI_ISL_432256 | hCoV-19/Wales/PHWC-275F5/2020 | Europe / United Kingdom / Wales | 2020-04-01 | Wales Specialist Virology Centre | Public Health Wales Microbiology Cardiff | Catherine Moore, Johnathan Evans, Malorie Perry, Simon Cottrell, Alec Birchley, Alexander Adams, Amy Gaskin, Bree Gatica-Wilcox, Jason Coombes, Lauren Gilbert, Lee Graham, Nicole Pacchiarini, Sara Kumziene-Summerhayes, Sarah Taylor, Sophie Jones, Sara Rey, Matthew Bull, Joanne Watkins, Sally Corden, Tom Connor |
| EPI_ISL_432257 | hCoV-19/Wales/PHWC-2762F/2020 | Europe / United Kingdom / Wales | 2020-03-30 | Wales Specialist Virology Centre | Public Health Wales Microbiology Cardiff | Catherine Moore, Johnathan Evans, Malorie Perry, Simon Cottrell, Alec Birchley, Alexander Adams, Amy Gaskin, Bree Gatica-Wilcox, Jason Coombes, Lauren Gilbert, Lee Graham, Nicole Pacchiarini, Sara Kumziene-Summerhayes, Sarah Taylor, Sophie Jones, Sara Rey, Matthew Bull, Joanne Watkins, Sally Corden, Tom Connor |
| EPI_ISL_432258 | hCoV-19/Wales/PHWC-2757D/2020 | Europe / United Kingdom / Wales | 2020-04-01 | Wales Specialist Virology Centre | Public Health Wales Microbiology Cardiff | Catherine Moore, Johnathan Evans, Malorie Perry, Simon Cottrell, Alec Birchley, Alexander Adams, Amy Gaskin, Bree Gatica-Wilcox, Jason Coombes, Lauren Gilbert, Lee Graham, Nicole Pacchiarini, Sara Kumziene-Summerhayes, Sarah Taylor, Sophie Jones, Sara Rey, Matthew Bull, Joanne Watkins, Sally Corden, Tom Connor |
| EPI_ISL_432259 | hCoV-19/Wales/PHWC-29DFB/2020 | Europe / United Kingdom / Wales | 2020-04-04 | Wales Specialist Virology Centre | Public Health Wales Microbiology Cardiff | Catherine Moore, Johnathan Evans, Malorie Perry, Simon Cottrell, Alec Birchley, Alexander Adams, Amy Gaskin, Bree Gatica-Wilcox, Jason Coombes, Lauren Gilbert, Lee Graham, Nicole Pacchiarini, Sara Kumziene-Summerhayes, Sarah Taylor, Sophie Jones, Sara Rey, Matthew Bull, Joanne Watkins, Sally Corden, Tom Connor |
| EPI_ISL_432260 | hCoV-19/Wales/PHWC-27197/2020 | Europe / United Kingdom / Wales | 2020-03-31 | Wales Specialist Virology Centre | Public Health Wales Microbiology Cardiff | Catherine Moore, Johnathan Evans, Malorie Perry, Simon Cottrell, Alec Birchley, Alexander Adams, Amy Gaskin, Bree Gatica-Wilcox, Jason Coombes, Lauren Gilbert, Lee Graham, Nicole Pacchiarini, Sara Kumziene-Summerhayes, Sarah Taylor, Sophie Jones, Sara Rey, Matthew Bull, Joanne Watkins, Sally Corden, Tom Connor |
| EPI_ISL_432261 | hCoV-19/Wales/PHWC-2B1A2/2020 | Europe / United Kingdom / Wales | 2020-04-07 | Wales Specialist Virology Centre | Public Health Wales Microbiology Cardiff | Catherine Moore, Johnathan Evans, Malorie Perry, Simon Cottrell, Alec Birchley, Alexander Adams, Amy Gaskin, Bree Gatica-Wilcox, Jason Coombes, Lauren Gilbert, Lee Graham, Nicole Pacchiarini, Sara Kumziene-Summerhayes, Sarah Taylor, Sophie Jones, Sara Rey, Matthew Bull, Joanne Watkins, Sally Corden, Tom Connor |
| EPI_ISL_432262 | hCoV-19/Wales/PHWC-2720D/2020 | Europe / United Kingdom / Wales | 2020-03-30 | Wales Specialist Virology Centre | Public Health Wales Microbiology Cardiff | Catherine Moore, Johnathan Evans, Malorie Perry, Simon Cottrell, Alec Birchley, Alexander Adams, Amy Gaskin, Bree Gatica-Wilcox, Jason Coombes, Lauren Gilbert, Lee Graham, Nicole Pacchiarini, Sara Kumziene-Summerhayes, Sarah Taylor, Sophie Jones, Sara Rey, Matthew Bull, Joanne Watkins, Sally Corden, Tom Connor |
| EPI_ISL_432263 | hCoV-19/Wales/PHWC-2B00F/2020 | Europe / United Kingdom / Wales | 2020-04-07 | Wales Specialist Virology Centre | Public Health Wales Microbiology Cardiff | Catherine Moore, Johnathan Evans, Malorie Perry, Simon Cottrell, Alec Birchley, Alexander Adams, Amy Gaskin, Bree Gatica-Wilcox, Jason Coombes, Lauren Gilbert, Lee Graham, Nicole Pacchiarini, Sara Kumziene-Summerhayes, Sarah Taylor, Sophie Jones, Sara Rey, Matthew Bull, Joanne Watkins, Sally Corden, Tom Connor |
| EPI_ISL_432264 | hCoV-19/Wales/PHWC-2B148/2020 | Europe / United Kingdom / Wales | 2020-04-07 | Wales Specialist Virology Centre | Public Health Wales Microbiology Cardiff | Catherine Moore, Johnathan Evans, Malorie Perry, Simon Cottrell, Alec Birchley, Alexander Adams, Amy Gaskin, Bree Gatica-Wilcox, Jason Coombes, Lauren Gilbert, Lee Graham, Nicole Pacchiarini, Sara Kumziene-Summerhayes, Sarah Taylor, Sophie Jones, Sara Rey, Matthew Bull, Joanne Watkins, Sally Corden, Tom Connor |
| EPI_ISL_432265 | hCoV-19/Wales/PHWC-2B01E/2020 | Europe / United Kingdom / Wales | 2020-04-08 | Wales Specialist Virology Centre | Public Health Wales Microbiology Cardiff | Catherine Moore, Johnathan Evans, Malorie Perry, Simon Cottrell, Alec Birchley, Alexander Adams, Amy Gaskin, Bree Gatica-Wilcox, Jason Coombes, Lauren Gilbert, Lee Graham, Nicole Pacchiarini, Sara Kumziene-Summerhayes, Sarah Taylor, Sophie Jones, Sara Rey, Matthew Bull, Joanne Watkins, Sally Corden, Tom Connor |
| EPI_ISL_432266 | hCoV-19/Wales/PHWC-27513/2020 | Europe / United Kingdom / Wales | 2020-03-30 | Wales Specialist Virology Centre | Public Health Wales Microbiology Cardiff | Catherine Moore, Johnathan Evans, Malorie Perry, Simon Cottrell, Alec Birchley, Alexander Adams, Amy Gaskin, Bree Gatica-Wilcox, Jason Coombes, Lauren Gilbert, Lee Graham, Nicole Pacchiarini, Sara Kumziene-Summerhayes, Sarah Taylor, Sophie Jones, Sara Rey, Matthew Bull, Joanne Watkins, Sally Corden, Tom Connor |
| EPI_ISL_432267 | hCoV-19/Wales/PHWC-29E07/2020 | Europe / United Kingdom / Wales | 2020-04-04 | Wales Specialist Virology Centre | Public Health Wales Microbiology Cardiff | Catherine Moore, Johnathan Evans, Malorie Perry, Simon Cottrell, Alec Birchley, Alexander Adams, Amy Gaskin, Bree Gatica-Wilcox, Jason Coombes, Lauren Gilbert, Lee Graham, Nicole Pacchiarini, Sara Kumziene-Summerhayes, Sarah Taylor, Sophie Jones, Sara Rey, Matthew Bull, Joanne Watkins, Sally Corden, Tom Connor |
| EPI_ISL_432268 | hCoV-19/Wales/PHWC-2B0D2/2020 | Europe / United Kingdom / Wales | 2020-04-08 | Wales Specialist Virology Centre | Public Health Wales Microbiology Cardiff | Catherine Moore, Johnathan Evans, Malorie Perry, Simon Cottrell, Alec Birchley, Alexander Adams, Amy Gaskin, Bree Gatica-Wilcox, Jason Coombes, Lauren Gilbert, Lee Graham, Nicole Pacchiarini, Sara Kumziene-Summerhayes, Sarah Taylor, Sophie Jones, Sara Rey, Matthew Bull, Joanne Watkins, Sally Corden, Tom Connor |
| EPI_ISL_432269 | hCoV-19/Wales/PHWC-2AE42/2020 | Europe / United Kingdom / Wales | 2020-04-07 | Wales Specialist Virology Centre | Public Health Wales Microbiology Cardiff | Catherine Moore, Johnathan Evans, Malorie Perry, Simon Cottrell, Alec Birchley, Alexander Adams, Amy Gaskin, Bree Gatica-Wilcox, Jason Coombes, Lauren Gilbert, Lee Graham, Nicole Pacchiarini, Sara Kumziene-Summerhayes, Sarah Taylor, Sophie Jones, Sara Rey, Matthew Bull, Joanne Watkins, Sally Corden, Tom Connor |
| EPI_ISL_432270 | hCoV-19/Wales/PHWC-27935/2020 | Europe / United Kingdom / Wales | 2020-04-01 | Wales Specialist Virology Centre | Public Health Wales Microbiology Cardiff | Catherine Moore, Johnathan Evans, Malorie Perry, Simon Cottrell, Alec Birchley, Alexander Adams, Amy Gaskin, Bree Gatica-Wilcox, Jason Coombes, Lauren Gilbert, Lee Graham, Nicole Pacchiarini, Sara Kumziene-Summerhayes, Sarah Taylor, Sophie Jones, Sara Rey, Matthew Bull, Joanne Watkins, Sally Corden, Tom Connor |
| EPI_ISL_432271 | hCoV-19/Wales/PHWC-2B04B/2020 | Europe / United Kingdom / Wales | 2020-04-08 | Wales Specialist Virology Centre | Public Health Wales Microbiology Cardiff | Catherine Moore, Johnathan Evans, Malorie Perry, Simon Cottrell, Alec Birchley, Alexander Adams, Amy Gaskin, Bree Gatica-Wilcox, Jason Coombes, Lauren Gilbert, Lee Graham, Nicole Pacchiarini, Sara Kumziene-Summerhayes, Sarah Taylor, Sophie Jones, Sara Rey, Matthew Bull, Joanne Watkins, Sally Corden, Tom Connor |
| EPI_ISL_432272 | hCoV-19/Wales/PHWC-2B1EE/2020 | Europe / United Kingdom / Wales | 2020-04-07 | Wales Specialist Virology Centre | Public Health Wales Microbiology Cardiff | Catherine Moore, Johnathan Evans, Malorie Perry, Simon Cottrell, Alec Birchley, Alexander Adams, Amy Gaskin, Bree Gatica-Wilcox, Jason Coombes, Lauren Gilbert, Lee Graham, Nicole Pacchiarini, Sara Kumziene-Summerhayes, Sarah Taylor, Sophie Jones, Sara Rey, Matthew Bull, Joanne Watkins, Sally Corden, Tom Connor |
| EPI_ISL_432273 | hCoV-19/Wales/PHWC-2B218/2020 | Europe / United Kingdom / Wales | 2020-04-08 | Wales Specialist Virology Centre | Public Health Wales Microbiology Cardiff | Catherine Moore, Johnathan Evans, Malorie Perry, Simon Cottrell, Alec Birchley, Alexander Adams, Amy Gaskin, Bree Gatica-Wilcox, Jason Coombes, Lauren Gilbert, Lee Graham, Nicole Pacchiarini, Sara Kumziene-Summerhayes, Sarah Taylor, Sophie Jones, Sara Rey, Matthew Bull, Joanne Watkins, Sally Corden, Tom Connor |
| EPI_ISL_432274 | hCoV-19/Wales/PHWC-2A11C/2020 | Europe / United Kingdom / Wales | 2020-04-04 | Wales Specialist Virology Centre | Public Health Wales Microbiology Cardiff | Catherine Moore, Johnathan Evans, Malorie Perry, Simon Cottrell, Alec Birchley, Alexander Adams, Amy Gaskin, Bree Gatica-Wilcox, Jason Coombes, Lauren Gilbert, Lee Graham, Nicole Pacchiarini, Sara Kumziene-Summerhayes, Sarah Taylor, Sophie Jones, Sara Rey, Matthew Bull, Joanne Watkins, Sally Corden, Tom Connor |
| EPI_ISL_432275 | hCoV-19/Wales/PHWC-2AE9D/2020 | Europe / United Kingdom / Wales | 2020-04-08 | Wales Specialist Virology Centre | Public Health Wales Microbiology Cardiff | Catherine Moore, Johnathan Evans, Malorie Perry, Simon Cottrell, Alec Birchley, Alexander Adams, Amy Gaskin, Bree Gatica-Wilcox, Jason Coombes, Lauren Gilbert, Lee Graham, Nicole Pacchiarini, Sara Kumziene-Summerhayes, Sarah Taylor, Sophie Jones, Sara Rey, Matthew Bull, Joanne Watkins, Sally Corden, Tom Connor |
| EPI_ISL_432276 | hCoV-19/Wales/PHWC-2AEB8/2020 | Europe / United Kingdom / Wales | 2020-04-07 | Wales Specialist Virology Centre | Public Health Wales Microbiology Cardiff | Catherine Moore, Johnathan Evans, Malorie Perry, Simon Cottrell, Alec Birchley, Alexander Adams, Amy Gaskin, Bree Gatica-Wilcox, Jason Coombes, Lauren Gilbert, Lee Graham, Nicole Pacchiarini, Sara Kumziene-Summerhayes, Sarah Taylor, Sophie Jones, Sara Rey, Matthew Bull, Joanne Watkins, Sally Corden, Tom Connor |
| EPI_ISL_432277 | hCoV-19/Wales/PHWC-27285/2020 | Europe / United Kingdom / Wales | 2020-03-31 | Wales Specialist Virology Centre | Public Health Wales Microbiology Cardiff | Catherine Moore, Johnathan Evans, Malorie Perry, Simon Cottrell, Alec Birchley, Alexander Adams, Amy Gaskin, Bree Gatica-Wilcox, Jason Coombes, Lauren Gilbert, Lee Graham, Nicole Pacchiarini, Sara Kumziene-Summerhayes, Sarah Taylor, Sophie Jones, Sara Rey, Matthew Bull, Joanne Watkins, Sally Corden, Tom Connor |
| EPI_ISL_432278 | hCoV-19/Wales/PHWC-275C8/2020 | Europe / United Kingdom / Wales | 2020-03-31 | Wales Specialist Virology Centre | Public Health Wales Microbiology Cardiff | Catherine Moore, Johnathan Evans, Malorie Perry, Simon Cottrell, Alec Birchley, Alexander Adams, Amy Gaskin, Bree Gatica-Wilcox, Jason Coombes, Lauren Gilbert, Lee Graham, Nicole Pacchiarini, Sara Kumziene-Summerhayes, Sarah Taylor, Sophie Jones, Sara Rey, Matthew Bull, Joanne Watkins, Sally Corden, Tom Connor |
| EPI_ISL_432279 | hCoV-19/Wales/PHWC-29C94/2020 | Europe / United Kingdom / Wales | 2020-04-05 | Wales Specialist Virology Centre | Public Health Wales Microbiology Cardiff | Catherine Moore, Johnathan Evans, Malorie Perry, Simon Cottrell, Alec Birchley, Alexander Adams, Amy Gaskin, Bree Gatica-Wilcox, Jason Coombes, Lauren Gilbert, Lee Graham, Nicole Pacchiarini, Sara Kumziene-Summerhayes, Sarah Taylor, Sophie Jones, Sara Rey, Matthew Bull, Joanne Watkins, Sally Corden, Tom Connor |
| EPI_ISL_432280 | hCoV-19/Wales/PHWC-2B184/2020 | Europe / United Kingdom / Wales | 2020-04-08 | Wales Specialist Virology Centre | Public Health Wales Microbiology Cardiff | Catherine Moore, Johnathan Evans, Malorie Perry, Simon Cottrell, Alec Birchley, Alexander Adams, Amy Gaskin, Bree Gatica-Wilcox, Jason Coombes, Lauren Gilbert, Lee Graham, Nicole Pacchiarini, Sara Kumziene-Summerhayes, Sarah Taylor, Sophie Jones, Sara Rey, Matthew Bull, Joanne Watkins, Sally Corden, Tom Connor |
| EPI_ISL_432281 | hCoV-19/Wales/PHWC-2B227/2020 | Europe / United Kingdom / Wales | 2020-04-08 | Wales Specialist Virology Centre | Public Health Wales Microbiology Cardiff | Catherine Moore, Johnathan Evans, Malorie Perry, Simon Cottrell, Alec Birchley, Alexander Adams, Amy Gaskin, Bree Gatica-Wilcox, Jason Coombes, Lauren Gilbert, Lee Graham, Nicole Pacchiarini, Sara Kumziene-Summerhayes, Sarah Taylor, Sophie Jones, Sara Rey, Matthew Bull, Joanne Watkins, Sally Corden, Tom Connor |
| EPI_ISL_432282 | hCoV-19/Wales/PHWC-274AD/2020 | Europe / United Kingdom / Wales | 2020-03-31 | Wales Specialist Virology Centre | Public Health Wales Microbiology Cardiff | Catherine Moore, Johnathan Evans, Malorie Perry, Simon Cottrell, Alec Birchley, Alexander Adams, Amy Gaskin, Bree Gatica-Wilcox, Jason Coombes, Lauren Gilbert, Lee Graham, Nicole Pacchiarini, Sara Kumziene-Summerhayes, Sarah Taylor, Sophie Jones, Sara Rey, Matthew Bull, Joanne Watkins, Sally Corden, Tom Connor |
| EPI_ISL_432283 | hCoV-19/Wales/PHWC-2B05A/2020 | Europe / United Kingdom / Wales | 2020-04-08 | Wales Specialist Virology Centre | Public Health Wales Microbiology Cardiff | Catherine Moore, Johnathan Evans, Malorie Perry, Simon Cottrell, Alec Birchley, Alexander Adams, Amy Gaskin, Bree Gatica-Wilcox, Jason Coombes, Lauren Gilbert, Lee Graham, Nicole Pacchiarini, Sara Kumziene-Summerhayes, Sarah Taylor, Sophie Jones, Sara Rey, Matthew Bull, Joanne Watkins, Sally Corden, Tom Connor |
| EPI_ISL_432284 | hCoV-19/Wales/PHWC-2B0B4/2020 | Europe / United Kingdom / Wales | 2020-04-07 | Wales Specialist Virology Centre | Public Health Wales Microbiology Cardiff | Catherine Moore, Johnathan Evans, Malorie Perry, Simon Cottrell, Alec Birchley, Alexander Adams, Amy Gaskin, Bree Gatica-Wilcox, Jason Coombes, Lauren Gilbert, Lee Graham, Nicole Pacchiarini, Sara Kumziene-Summerhayes, Sarah Taylor, Sophie Jones, Sara Rey, Matthew Bull, Joanne Watkins, Sally Corden, Tom Connor |
| EPI_ISL_432285 | hCoV-19/Wales/PHWC-271E2/2020 | Europe / United Kingdom / Wales | 2020-03-31 | Wales Specialist Virology Centre | Public Health Wales Microbiology Cardiff | Catherine Moore, Johnathan Evans, Malorie Perry, Simon Cottrell, Alec Birchley, Alexander Adams, Amy Gaskin, Bree Gatica-Wilcox, Jason Coombes, Lauren Gilbert, Lee Graham, Nicole Pacchiarini, Sara Kumziene-Summerhayes, Sarah Taylor, Sophie Jones, Sara Rey, Matthew Bull, Joanne Watkins, Sally Corden, Tom Connor |
| EPI_ISL_432286 | hCoV-19/Wales/PHWC-2B236/2020 | Europe / United Kingdom / Wales | 2020-04-08 | Wales Specialist Virology Centre | Public Health Wales Microbiology Cardiff | Catherine Moore, Johnathan Evans, Malorie Perry, Simon Cottrell, Alec Birchley, Alexander Adams, Amy Gaskin, Bree Gatica-Wilcox, Jason Coombes, Lauren Gilbert, Lee Graham, Nicole Pacchiarini, Sara Kumziene-Summerhayes, Sarah Taylor, Sophie Jones, Sara Rey, Matthew Bull, Joanne Watkins, Sally Corden, Tom Connor |
| EPI_ISL_432287 | hCoV-19/Wales/PHWC-2772C/2020 | Europe / United Kingdom / Wales | 2020-04-01 | Wales Specialist Virology Centre | Public Health Wales Microbiology Cardiff | Catherine Moore, Johnathan Evans, Malorie Perry, Simon Cottrell, Alec Birchley, Alexander Adams, Amy Gaskin, Bree Gatica-Wilcox, Jason Coombes, Lauren Gilbert, Lee Graham, Nicole Pacchiarini, Sara Kumziene-Summerhayes, Sarah Taylor, Sophie Jones, Sara Rey, Matthew Bull, Joanne Watkins, Sally Corden, Tom Connor |
| EPI_ISL_432288 | hCoV-19/Wales/PHWC-277A4/2020 | Europe / United Kingdom / Wales | 2020-03-31 | Wales Specialist Virology Centre | Public Health Wales Microbiology Cardiff | Catherine Moore, Johnathan Evans, Malorie Perry, Simon Cottrell, Alec Birchley, Alexander Adams, Amy Gaskin, Bree Gatica-Wilcox, Jason Coombes, Lauren Gilbert, Lee Graham, Nicole Pacchiarini, Sara Kumziene-Summerhayes, Sarah Taylor, Sophie Jones, Sara Rey, Matthew Bull, Joanne Watkins, Sally Corden, Tom Connor |
| EPI_ISL_432289 | hCoV-19/Wales/PHWC-27971/2020 | Europe / United Kingdom / Wales | 2020-04-01 | Wales Specialist Virology Centre | Public Health Wales Microbiology Cardiff | Catherine Moore, Johnathan Evans, Malorie Perry, Simon Cottrell, Alec Birchley, Alexander Adams, Amy Gaskin, Bree Gatica-Wilcox, Jason Coombes, Lauren Gilbert, Lee Graham, Nicole Pacchiarini, Sara Kumziene-Summerhayes, Sarah Taylor, Sophie Jones, Sara Rey, Matthew Bull, Joanne Watkins, Sally Corden, Tom Connor |
| EPI_ISL_432290 | hCoV-19/Wales/PHWC-2730A/2020 | Europe / United Kingdom / Wales | 2020-03-30 | Wales Specialist Virology Centre | Public Health Wales Microbiology Cardiff | Catherine Moore, Johnathan Evans, Malorie Perry, Simon Cottrell, Alec Birchley, Alexander Adams, Amy Gaskin, Bree Gatica-Wilcox, Jason Coombes, Lauren Gilbert, Lee Graham, Nicole Pacchiarini, Sara Kumziene-Summerhayes, Sarah Taylor, Sophie Jones, Sara Rey, Matthew Bull, Joanne Watkins, Sally Corden, Tom Connor |
| EPI_ISL_432291 | hCoV-19/Wales/PHWC-2B2AF/2020 | Europe / United Kingdom / Wales | 2020-04-08 | Wales Specialist Virology Centre | Public Health Wales Microbiology Cardiff | Catherine Moore, Johnathan Evans, Malorie Perry, Simon Cottrell, Alec Birchley, Alexander Adams, Amy Gaskin, Bree Gatica-Wilcox, Jason Coombes, Lauren Gilbert, Lee Graham, Nicole Pacchiarini, Sara Kumziene-Summerhayes, Sarah Taylor, Sophie Jones, Sara Rey, Matthew Bull, Joanne Watkins, Sally Corden, Tom Connor |
| EPI_ISL_432292 | hCoV-19/Wales/PHWC-29C58/2020 | Europe / United Kingdom / Wales | 2020-04-05 | Wales Specialist Virology Centre | Public Health Wales Microbiology Cardiff | Catherine Moore, Johnathan Evans, Malorie Perry, Simon Cottrell, Alec Birchley, Alexander Adams, Amy Gaskin, Bree Gatica-Wilcox, Jason Coombes, Lauren Gilbert, Lee Graham, Nicole Pacchiarini, Sara Kumziene-Summerhayes, Sarah Taylor, Sophie Jones, Sara Rey, Matthew Bull, Joanne Watkins, Sally Corden, Tom Connor |
| EPI_ISL_432293 | hCoV-19/Wales/PHWC-27504/2020 | Europe / United Kingdom / Wales | 2020-03-30 | Wales Specialist Virology Centre | Public Health Wales Microbiology Cardiff | Catherine Moore, Johnathan Evans, Malorie Perry, Simon Cottrell, Alec Birchley, Alexander Adams, Amy Gaskin, Bree Gatica-Wilcox, Jason Coombes, Lauren Gilbert, Lee Graham, Nicole Pacchiarini, Sara Kumziene-Summerhayes, Sarah Taylor, Sophie Jones, Sara Rey, Matthew Bull, Joanne Watkins, Sally Corden, Tom Connor |
| EPI_ISL_432294 | hCoV-19/Wales/PHWC-29FD7/2020 | Europe / United Kingdom / Wales | 2020-04-03 | Wales Specialist Virology Centre | Public Health Wales Microbiology Cardiff | Catherine Moore, Johnathan Evans, Malorie Perry, Simon Cottrell, Alec Birchley, Alexander Adams, Amy Gaskin, Bree Gatica-Wilcox, Jason Coombes, Lauren Gilbert, Lee Graham, Nicole Pacchiarini, Sara Kumziene-Summerhayes, Sarah Taylor, Sophie Jones, Sara Rey, Matthew Bull, Joanne Watkins, Sally Corden, Tom Connor |
| EPI_ISL_432295 | hCoV-19/Wales/PHWC-2AF9A/2020 | Europe / United Kingdom / Wales | 2020-04-08 | Wales Specialist Virology Centre | Public Health Wales Microbiology Cardiff | Catherine Moore, Johnathan Evans, Malorie Perry, Simon Cottrell, Alec Birchley, Alexander Adams, Amy Gaskin, Bree Gatica-Wilcox, Jason Coombes, Lauren Gilbert, Lee Graham, Nicole Pacchiarini, Sara Kumziene-Summerhayes, Sarah Taylor, Sophie Jones, Sara Rey, Matthew Bull, Joanne Watkins, Sally Corden, Tom Connor |
| EPI_ISL_432296 | hCoV-19/Wales/PHWC-2B078/2020 | Europe / United Kingdom / Wales | 2020-04-08 | Wales Specialist Virology Centre | Public Health Wales Microbiology Cardiff | Catherine Moore, Johnathan Evans, Malorie Perry, Simon Cottrell, Alec Birchley, Alexander Adams, Amy Gaskin, Bree Gatica-Wilcox, Jason Coombes, Lauren Gilbert, Lee Graham, Nicole Pacchiarini, Sara Kumziene-Summerhayes, Sarah Taylor, Sophie Jones, Sara Rey, Matthew Bull, Joanne Watkins, Sally Corden, Tom Connor |
| EPI_ISL_432297 | hCoV-19/Wales/PHWC-29D82/2020 | Europe / United Kingdom / Wales | 2020-04-04 | Wales Specialist Virology Centre | Public Health Wales Microbiology Cardiff | Catherine Moore, Johnathan Evans, Malorie Perry, Simon Cottrell, Alec Birchley, Alexander Adams, Amy Gaskin, Bree Gatica-Wilcox, Jason Coombes, Lauren Gilbert, Lee Graham, Nicole Pacchiarini, Sara Kumziene-Summerhayes, Sarah Taylor, Sophie Jones, Sara Rey, Matthew Bull, Joanne Watkins, Sally Corden, Tom Connor |
| EPI_ISL_432298 | hCoV-19/Wales/PHWC-27C0F/2020 | Europe / United Kingdom / Wales | 2020-03-31 | Wales Specialist Virology Centre | Public Health Wales Microbiology Cardiff | Catherine Moore, Johnathan Evans, Malorie Perry, Simon Cottrell, Alec Birchley, Alexander Adams, Amy Gaskin, Bree Gatica-Wilcox, Jason Coombes, Lauren Gilbert, Lee Graham, Nicole Pacchiarini, Sara Kumziene-Summerhayes, Sarah Taylor, Sophie Jones, Sara Rey, Matthew Bull, Joanne Watkins, Sally Corden, Tom Connor |
| EPI_ISL_432299 | hCoV-19/Wales/PHWC-27470/2020 | Europe / United Kingdom / Wales | 2020-03-31 | Wales Specialist Virology Centre | Public Health Wales Microbiology Cardiff | Catherine Moore, Johnathan Evans, Malorie Perry, Simon Cottrell, Alec Birchley, Alexander Adams, Amy Gaskin, Bree Gatica-Wilcox, Jason Coombes, Lauren Gilbert, Lee Graham, Nicole Pacchiarini, Sara Kumziene-Summerhayes, Sarah Taylor, Sophie Jones, Sara Rey, Matthew Bull, Joanne Watkins, Sally Corden, Tom Connor |
| EPI_ISL_432300 | hCoV-19/Wales/PHWC-2B096/2020 | Europe / United Kingdom / Wales | 2020-04-07 | Wales Specialist Virology Centre | Public Health Wales Microbiology Cardiff | Catherine Moore, Johnathan Evans, Malorie Perry, Simon Cottrell, Alec Birchley, Alexander Adams, Amy Gaskin, Bree Gatica-Wilcox, Jason Coombes, Lauren Gilbert, Lee Graham, Nicole Pacchiarini, Sara Kumziene-Summerhayes, Sarah Taylor, Sophie Jones, Sara Rey, Matthew Bull, Joanne Watkins, Sally Corden, Tom Connor |
| EPI_ISL_432301 | hCoV-19/Wales/PHWC-27407/2020 | Europe / United Kingdom / Wales | 2020-03-30 | Wales Specialist Virology Centre | Public Health Wales Microbiology Cardiff | Catherine Moore, Johnathan Evans, Malorie Perry, Simon Cottrell, Alec Birchley, Alexander Adams, Amy Gaskin, Bree Gatica-Wilcox, Jason Coombes, Lauren Gilbert, Lee Graham, Nicole Pacchiarini, Sara Kumziene-Summerhayes, Sarah Taylor, Sophie Jones, Sara Rey, Matthew Bull, Joanne Watkins, Sally Corden, Tom Connor |
| EPI_ISL_432302 | hCoV-19/Wales/PHWC-273A0/2020 | Europe / United Kingdom / Wales | 2020-03-31 | Wales Specialist Virology Centre | Public Health Wales Microbiology Cardiff | Catherine Moore, Johnathan Evans, Malorie Perry, Simon Cottrell, Alec Birchley, Alexander Adams, Amy Gaskin, Bree Gatica-Wilcox, Jason Coombes, Lauren Gilbert, Lee Graham, Nicole Pacchiarini, Sara Kumziene-Summerhayes, Sarah Taylor, Sophie Jones, Sara Rey, Matthew Bull, Joanne Watkins, Sally Corden, Tom Connor |
| EPI_ISL_432303 | hCoV-19/Wales/PHWC-27759/2020 | Europe / United Kingdom / Wales | 2020-03-31 | Wales Specialist Virology Centre | Public Health Wales Microbiology Cardiff | Catherine Moore, Johnathan Evans, Malorie Perry, Simon Cottrell, Alec Birchley, Alexander Adams, Amy Gaskin, Bree Gatica-Wilcox, Jason Coombes, Lauren Gilbert, Lee Graham, Nicole Pacchiarini, Sara Kumziene-Summerhayes, Sarah Taylor, Sophie Jones, Sara Rey, Matthew Bull, Joanne Watkins, Sally Corden, Tom Connor |
| EPI_ISL_432304 | hCoV-19/Wales/PHWC-2B157/2020 | Europe / United Kingdom / Wales | 2020-04-08 | Wales Specialist Virology Centre | Public Health Wales Microbiology Cardiff | Catherine Moore, Johnathan Evans, Malorie Perry, Simon Cottrell, Alec Birchley, Alexander Adams, Amy Gaskin, Bree Gatica-Wilcox, Jason Coombes, Lauren Gilbert, Lee Graham, Nicole Pacchiarini, Sara Kumziene-Summerhayes, Sarah Taylor, Sophie Jones, Sara Rey, Matthew Bull, Joanne Watkins, Sally Corden, Tom Connor |
| EPI_ISL_432305 | hCoV-19/Wales/PHWC-2B087/2020 | Europe / United Kingdom / Wales | 2020-04-07 | Wales Specialist Virology Centre | Public Health Wales Microbiology Cardiff | Catherine Moore, Johnathan Evans, Malorie Perry, Simon Cottrell, Alec Birchley, Alexander Adams, Amy Gaskin, Bree Gatica-Wilcox, Jason Coombes, Lauren Gilbert, Lee Graham, Nicole Pacchiarini, Sara Kumziene-Summerhayes, Sarah Taylor, Sophie Jones, Sara Rey, Matthew Bull, Joanne Watkins, Sally Corden, Tom Connor |
| EPI_ISL_432306 | hCoV-19/Wales/PHWC-27689/2020 | Europe / United Kingdom / Wales | 2020-03-31 | Wales Specialist Virology Centre | Public Health Wales Microbiology Cardiff | Catherine Moore, Johnathan Evans, Malorie Perry, Simon Cottrell, Alec Birchley, Alexander Adams, Amy Gaskin, Bree Gatica-Wilcox, Jason Coombes, Lauren Gilbert, Lee Graham, Nicole Pacchiarini, Sara Kumziene-Summerhayes, Sarah Taylor, Sophie Jones, Sara Rey, Matthew Bull, Joanne Watkins, Sally Corden, Tom Connor |
| EPI_ISL_432307 | hCoV-19/Wales/PHWC-27267/2020 | Europe / United Kingdom / Wales | 2020-03-31 | Wales Specialist Virology Centre | Public Health Wales Microbiology Cardiff | Catherine Moore, Johnathan Evans, Malorie Perry, Simon Cottrell, Alec Birchley, Alexander Adams, Amy Gaskin, Bree Gatica-Wilcox, Jason Coombes, Lauren Gilbert, Lee Graham, Nicole Pacchiarini, Sara Kumziene-Summerhayes, Sarah Taylor, Sophie Jones, Sara Rey, Matthew Bull, Joanne Watkins, Sally Corden, Tom Connor |
| EPI_ISL_432308 | hCoV-19/Wales/PHWC-24E39/2020 | Europe / United Kingdom / Wales | 2020-03 | Wales Specialist Virology Centre | Public Health Wales Microbiology Cardiff | Catherine Moore, Johnathan Evans, Malorie Perry, Simon Cottrell, Alec Birchley, Alexander Adams, Amy Gaskin, Bree Gatica-Wilcox, Jason Coombes, Lauren Gilbert, Lee Graham, Nicole Pacchiarini, Sara Kumziene-Summerhayes, Sarah Taylor, Sophie Jones, Sara Rey, Matthew Bull, Joanne Watkins, Sally Corden, Tom Connor |
| EPI_ISL_432309 | hCoV-19/Wales/PHWC-2B0A5/2020 | Europe / United Kingdom / Wales | 2020-04-07 | Wales Specialist Virology Centre | Public Health Wales Microbiology Cardiff | Catherine Moore, Johnathan Evans, Malorie Perry, Simon Cottrell, Alec Birchley, Alexander Adams, Amy Gaskin, Bree Gatica-Wilcox, Jason Coombes, Lauren Gilbert, Lee Graham, Nicole Pacchiarini, Sara Kumziene-Summerhayes, Sarah Taylor, Sophie Jones, Sara Rey, Matthew Bull, Joanne Watkins, Sally Corden, Tom Connor |
| EPI_ISL_432310 | hCoV-19/Wales/PHWC-27BB7/2020 | Europe / United Kingdom / Wales | 2020-04-01 | Wales Specialist Virology Centre | Public Health Wales Microbiology Cardiff | Catherine Moore, Johnathan Evans, Malorie Perry, Simon Cottrell, Alec Birchley, Alexander Adams, Amy Gaskin, Bree Gatica-Wilcox, Jason Coombes, Lauren Gilbert, Lee Graham, Nicole Pacchiarini, Sara Kumziene-Summerhayes, Sarah Taylor, Sophie Jones, Sara Rey, Matthew Bull, Joanne Watkins, Sally Corden, Tom Connor |
| EPI_ISL_432311 | hCoV-19/Wales/PHWC-26B6D/2020 | Europe / United Kingdom / Wales | 2020-03 | Wales Specialist Virology Centre | Public Health Wales Microbiology Cardiff | Catherine Moore, Johnathan Evans, Malorie Perry, Simon Cottrell, Alec Birchley, Alexander Adams, Amy Gaskin, Bree Gatica-Wilcox, Jason Coombes, Lauren Gilbert, Lee Graham, Nicole Pacchiarini, Sara Kumziene-Summerhayes, Sarah Taylor, Sophie Jones, Sara Rey, Matthew Bull, Joanne Watkins, Sally Corden, Tom Connor |
| EPI_ISL_432312 | hCoV-19/Wales/PHWC-271F1/2020 | Europe / United Kingdom / Wales | 2020-03-31 | Wales Specialist Virology Centre | Public Health Wales Microbiology Cardiff | Catherine Moore, Johnathan Evans, Malorie Perry, Simon Cottrell, Alec Birchley, Alexander Adams, Amy Gaskin, Bree Gatica-Wilcox, Jason Coombes, Lauren Gilbert, Lee Graham, Nicole Pacchiarini, Sara Kumziene-Summerhayes, Sarah Taylor, Sophie Jones, Sara Rey, Matthew Bull, Joanne Watkins, Sally Corden, Tom Connor |
| EPI_ISL_432313 | hCoV-19/Wales/PHWC-29FC8/2020 | Europe / United Kingdom / Wales | 2020-04-03 | Wales Specialist Virology Centre | Public Health Wales Microbiology Cardiff | Catherine Moore, Johnathan Evans, Malorie Perry, Simon Cottrell, Alec Birchley, Alexander Adams, Amy Gaskin, Bree Gatica-Wilcox, Jason Coombes, Lauren Gilbert, Lee Graham, Nicole Pacchiarini, Sara Kumziene-Summerhayes, Sarah Taylor, Sophie Jones, Sara Rey, Matthew Bull, Joanne Watkins, Sally Corden, Tom Connor |
| EPI_ISL_432314 | hCoV-19/Wales/PHWC-276B6/2020 | Europe / United Kingdom / Wales | 2020-03-31 | Wales Specialist Virology Centre | Public Health Wales Microbiology Cardiff | Catherine Moore, Johnathan Evans, Malorie Perry, Simon Cottrell, Alec Birchley, Alexander Adams, Amy Gaskin, Bree Gatica-Wilcox, Jason Coombes, Lauren Gilbert, Lee Graham, Nicole Pacchiarini, Sara Kumziene-Summerhayes, Sarah Taylor, Sophie Jones, Sara Rey, Matthew Bull, Joanne Watkins, Sally Corden, Tom Connor |
| EPI_ISL_432315 | hCoV-19/Wales/PHWC-2B2EB/2020 | Europe / United Kingdom / Wales | 2020-04-07 | Wales Specialist Virology Centre | Public Health Wales Microbiology Cardiff | Catherine Moore, Johnathan Evans, Malorie Perry, Simon Cottrell, Alec Birchley, Alexander Adams, Amy Gaskin, Bree Gatica-Wilcox, Jason Coombes, Lauren Gilbert, Lee Graham, Nicole Pacchiarini, Sara Kumziene-Summerhayes, Sarah Taylor, Sophie Jones, Sara Rey, Matthew Bull, Joanne Watkins, Sally Corden, Tom Connor |
| EPI_ISL_432316 | hCoV-19/Wales/PHWC-274CB/2020 | Europe / United Kingdom / Wales | 2020-04-01 | Wales Specialist Virology Centre | Public Health Wales Microbiology Cardiff | Catherine Moore, Johnathan Evans, Malorie Perry, Simon Cottrell, Alec Birchley, Alexander Adams, Amy Gaskin, Bree Gatica-Wilcox, Jason Coombes, Lauren Gilbert, Lee Graham, Nicole Pacchiarini, Sara Kumziene-Summerhayes, Sarah Taylor, Sophie Jones, Sara Rey, Matthew Bull, Joanne Watkins, Sally Corden, Tom Connor |
| EPI_ISL_432317 | hCoV-19/Wales/PHWC-2723A/2020 | Europe / United Kingdom / Wales | 2020-03-30 | Wales Specialist Virology Centre | Public Health Wales Microbiology Cardiff | Catherine Moore, Johnathan Evans, Malorie Perry, Simon Cottrell, Alec Birchley, Alexander Adams, Amy Gaskin, Bree Gatica-Wilcox, Jason Coombes, Lauren Gilbert, Lee Graham, Nicole Pacchiarini, Sara Kumziene-Summerhayes, Sarah Taylor, Sophie Jones, Sara Rey, Matthew Bull, Joanne Watkins, Sally Corden, Tom Connor |
| EPI_ISL_432318 | hCoV-19/Wales/PHWC-2B290/2020 | Europe / United Kingdom / Wales | 2020-04-08 | Wales Specialist Virology Centre | Public Health Wales Microbiology Cardiff | Catherine Moore, Johnathan Evans, Malorie Perry, Simon Cottrell, Alec Birchley, Alexander Adams, Amy Gaskin, Bree Gatica-Wilcox, Jason Coombes, Lauren Gilbert, Lee Graham, Nicole Pacchiarini, Sara Kumziene-Summerhayes, Sarah Taylor, Sophie Jones, Sara Rey, Matthew Bull, Joanne Watkins, Sally Corden, Tom Connor |
| EPI_ISL_432319 | hCoV-19/Wales/PHWC-29EDA/2020 | Europe / United Kingdom / Wales | 2020-03-30 | Wales Specialist Virology Centre | Public Health Wales Microbiology Cardiff | Catherine Moore, Johnathan Evans, Malorie Perry, Simon Cottrell, Alec Birchley, Alexander Adams, Amy Gaskin, Bree Gatica-Wilcox, Jason Coombes, Lauren Gilbert, Lee Graham, Nicole Pacchiarini, Sara Kumziene-Summerhayes, Sarah Taylor, Sophie Jones, Sara Rey, Matthew Bull, Joanne Watkins, Sally Corden, Tom Connor |
| EPI_ISL_432320 | hCoV-19/Wales/PHWC-2ADCD/2020 | Europe / United Kingdom / Wales | 2020-04-06 | Wales Specialist Virology Centre | Public Health Wales Microbiology Cardiff | Catherine Moore, Johnathan Evans, Malorie Perry, Simon Cottrell, Alec Birchley, Alexander Adams, Amy Gaskin, Bree Gatica-Wilcox, Jason Coombes, Lauren Gilbert, Lee Graham, Nicole Pacchiarini, Sara Kumziene-Summerhayes, Sarah Taylor, Sophie Jones, Sara Rey, Matthew Bull, Joanne Watkins, Sally Corden, Tom Connor |
| EPI_ISL_432321 | hCoV-19/Wales/PHWC-27B02/2020 | Europe / United Kingdom / Wales | 2020-03-30 | Wales Specialist Virology Centre | Public Health Wales Microbiology Cardiff | Catherine Moore, Johnathan Evans, Malorie Perry, Simon Cottrell, Alec Birchley, Alexander Adams, Amy Gaskin, Bree Gatica-Wilcox, Jason Coombes, Lauren Gilbert, Lee Graham, Nicole Pacchiarini, Sara Kumziene-Summerhayes, Sarah Taylor, Sophie Jones, Sara Rey, Matthew Bull, Joanne Watkins, Sally Corden, Tom Connor |
| EPI_ISL_432322 | hCoV-19/Wales/PHWC-2780B/2020 | Europe / United Kingdom / Wales | 2020-04-01 | Wales Specialist Virology Centre | Public Health Wales Microbiology Cardiff | Catherine Moore, Johnathan Evans, Malorie Perry, Simon Cottrell, Alec Birchley, Alexander Adams, Amy Gaskin, Bree Gatica-Wilcox, Jason Coombes, Lauren Gilbert, Lee Graham, Nicole Pacchiarini, Sara Kumziene-Summerhayes, Sarah Taylor, Sophie Jones, Sara Rey, Matthew Bull, Joanne Watkins, Sally Corden, Tom Connor |
| EPI_ISL_432323 | hCoV-19/Wales/PHWC-25348/2020 | Europe / United Kingdom / Wales | 2020-03-23 | Wales Specialist Virology Centre | Public Health Wales Microbiology Cardiff | Catherine Moore, Johnathan Evans, Malorie Perry, Simon Cottrell, Alec Birchley, Alexander Adams, Amy Gaskin, Bree Gatica-Wilcox, Jason Coombes, Lauren Gilbert, Lee Graham, Nicole Pacchiarini, Sara Kumziene-Summerhayes, Sarah Taylor, Sophie Jones, Sara Rey, Matthew Bull, Joanne Watkins, Sally Corden, Tom Connor |
| EPI_ISL_432324 | hCoV-19/Wales/PHWC-2AE7F/2020 | Europe / United Kingdom / Wales | 2020-04-07 | Wales Specialist Virology Centre | Public Health Wales Microbiology Cardiff | Catherine Moore, Johnathan Evans, Malorie Perry, Simon Cottrell, Alec Birchley, Alexander Adams, Amy Gaskin, Bree Gatica-Wilcox, Jason Coombes, Lauren Gilbert, Lee Graham, Nicole Pacchiarini, Sara Kumziene-Summerhayes, Sarah Taylor, Sophie Jones, Sara Rey, Matthew Bull, Joanne Watkins, Sally Corden, Tom Connor |
| EPI_ISL_432325 | hCoV-19/Wales/PHWC-27B6C/2020 | Europe / United Kingdom / Wales | 2020-04-01 | Wales Specialist Virology Centre | Public Health Wales Microbiology Cardiff | Catherine Moore, Johnathan Evans, Malorie Perry, Simon Cottrell, Alec Birchley, Alexander Adams, Amy Gaskin, Bree Gatica-Wilcox, Jason Coombes, Lauren Gilbert, Lee Graham, Nicole Pacchiarini, Sara Kumziene-Summerhayes, Sarah Taylor, Sophie Jones, Sara Rey, Matthew Bull, Joanne Watkins, Sally Corden, Tom Connor |
| EPI_ISL_432326 | hCoV-19/Wales/PHWC-247E3/2020 | Europe / United Kingdom / Wales | 2020-03 | Wales Specialist Virology Centre | Public Health Wales Microbiology Cardiff | Catherine Moore, Johnathan Evans, Malorie Perry, Simon Cottrell, Alec Birchley, Alexander Adams, Amy Gaskin, Bree Gatica-Wilcox, Jason Coombes, Lauren Gilbert, Lee Graham, Nicole Pacchiarini, Sara Kumziene-Summerhayes, Sarah Taylor, Sophie Jones, Sara Rey, Matthew Bull, Joanne Watkins, Sally Corden, Tom Connor |
| EPI_ISL_432327 | hCoV-19/Wales/PHWC-27B5D/2020 | Europe / United Kingdom / Wales | 2020-04-01 | Wales Specialist Virology Centre | Public Health Wales Microbiology Cardiff | Catherine Moore, Johnathan Evans, Malorie Perry, Simon Cottrell, Alec Birchley, Alexander Adams, Amy Gaskin, Bree Gatica-Wilcox, Jason Coombes, Lauren Gilbert, Lee Graham, Nicole Pacchiarini, Sara Kumziene-Summerhayes, Sarah Taylor, Sophie Jones, Sara Rey, Matthew Bull, Joanne Watkins, Sally Corden, Tom Connor |
| EPI_ISL_432328 | hCoV-19/Wales/PHWC-2756E/2020 | Europe / United Kingdom / Wales | 2020-03-31 | Wales Specialist Virology Centre | Public Health Wales Microbiology Cardiff | Catherine Moore, Johnathan Evans, Malorie Perry, Simon Cottrell, Alec Birchley, Alexander Adams, Amy Gaskin, Bree Gatica-Wilcox, Jason Coombes, Lauren Gilbert, Lee Graham, Nicole Pacchiarini, Sara Kumziene-Summerhayes, Sarah Taylor, Sophie Jones, Sara Rey, Matthew Bull, Joanne Watkins, Sally Corden, Tom Connor |
| EPI_ISL_432329 | hCoV-19/Wales/PHWC-27D48/2020 | Europe / United Kingdom / Wales | 2020-04-01 | Wales Specialist Virology Centre | Public Health Wales Microbiology Cardiff | Catherine Moore, Johnathan Evans, Malorie Perry, Simon Cottrell, Alec Birchley, Alexander Adams, Amy Gaskin, Bree Gatica-Wilcox, Jason Coombes, Lauren Gilbert, Lee Graham, Nicole Pacchiarini, Sara Kumziene-Summerhayes, Sarah Taylor, Sophie Jones, Sara Rey, Matthew Bull, Joanne Watkins, Sally Corden, Tom Connor |
| EPI_ISL_432330 | hCoV-19/Wales/PHWC-29CC1/2020 | Europe / United Kingdom / Wales | 2020-04-05 | Wales Specialist Virology Centre | Public Health Wales Microbiology Cardiff | Catherine Moore, Johnathan Evans, Malorie Perry, Simon Cottrell, Alec Birchley, Alexander Adams, Amy Gaskin, Bree Gatica-Wilcox, Jason Coombes, Lauren Gilbert, Lee Graham, Nicole Pacchiarini, Sara Kumziene-Summerhayes, Sarah Taylor, Sophie Jones, Sara Rey, Matthew Bull, Joanne Watkins, Sally Corden, Tom Connor |
| EPI_ISL_432331 | hCoV-19/Wales/PHWC-2861F/2020 | Europe / United Kingdom / Wales | 2020-03-30 | Wales Specialist Virology Centre | Public Health Wales Microbiology Cardiff | Catherine Moore, Johnathan Evans, Malorie Perry, Simon Cottrell, Alec Birchley, Alexander Adams, Amy Gaskin, Bree Gatica-Wilcox, Jason Coombes, Lauren Gilbert, Lee Graham, Nicole Pacchiarini, Sara Kumziene-Summerhayes, Sarah Taylor, Sophie Jones, Sara Rey, Matthew Bull, Joanne Watkins, Sally Corden, Tom Connor |
| EPI_ISL_432332 | hCoV-19/Wales/PHWC-26BE5/2020 | Europe / United Kingdom / Wales | 2020-04-08 | Wales Specialist Virology Centre | Public Health Wales Microbiology Cardiff | Catherine Moore, Johnathan Evans, Malorie Perry, Simon Cottrell, Alec Birchley, Alexander Adams, Amy Gaskin, Bree Gatica-Wilcox, Jason Coombes, Lauren Gilbert, Lee Graham, Nicole Pacchiarini, Sara Kumziene-Summerhayes, Sarah Taylor, Sophie Jones, Sara Rey, Matthew Bull, Joanne Watkins, Sally Corden, Tom Connor |
| EPI_ISL_432333 | hCoV-19/Wales/PHWC-29CB2/2020 | Europe / United Kingdom / Wales | 2020-04-05 | Wales Specialist Virology Centre | Public Health Wales Microbiology Cardiff | Catherine Moore, Johnathan Evans, Malorie Perry, Simon Cottrell, Alec Birchley, Alexander Adams, Amy Gaskin, Bree Gatica-Wilcox, Jason Coombes, Lauren Gilbert, Lee Graham, Nicole Pacchiarini, Sara Kumziene-Summerhayes, Sarah Taylor, Sophie Jones, Sara Rey, Matthew Bull, Joanne Watkins, Sally Corden, Tom Connor |
| EPI_ISL_432334 | hCoV-19/Wales/PHWC-2A079/2020 | Europe / United Kingdom / Wales | 2020-04-04 | Wales Specialist Virology Centre | Public Health Wales Microbiology Cardiff | Catherine Moore, Johnathan Evans, Malorie Perry, Simon Cottrell, Alec Birchley, Alexander Adams, Amy Gaskin, Bree Gatica-Wilcox, Jason Coombes, Lauren Gilbert, Lee Graham, Nicole Pacchiarini, Sara Kumziene-Summerhayes, Sarah Taylor, Sophie Jones, Sara Rey, Matthew Bull, Joanne Watkins, Sally Corden, Tom Connor |
| EPI_ISL_432335 | hCoV-19/Wales/PHWC-27D1B/2020 | Europe / United Kingdom / Wales | 2020-04-01 | Wales Specialist Virology Centre | Public Health Wales Microbiology Cardiff | Catherine Moore, Johnathan Evans, Malorie Perry, Simon Cottrell, Alec Birchley, Alexander Adams, Amy Gaskin, Bree Gatica-Wilcox, Jason Coombes, Lauren Gilbert, Lee Graham, Nicole Pacchiarini, Sara Kumziene-Summerhayes, Sarah Taylor, Sophie Jones, Sara Rey, Matthew Bull, Joanne Watkins, Sally Corden, Tom Connor |
| EPI_ISL_432336 | hCoV-19/Wales/PHWC-24E1B/2020 | Europe / United Kingdom / Wales | 2020-03 | Wales Specialist Virology Centre | Public Health Wales Microbiology Cardiff | Catherine Moore, Johnathan Evans, Malorie Perry, Simon Cottrell, Alec Birchley, Alexander Adams, Amy Gaskin, Bree Gatica-Wilcox, Jason Coombes, Lauren Gilbert, Lee Graham, Nicole Pacchiarini, Sara Kumziene-Summerhayes, Sarah Taylor, Sophie Jones, Sara Rey, Matthew Bull, Joanne Watkins, Sally Corden, Tom Connor |
| EPI_ISL_432337 | hCoV-19/Wales/PHWC-27B4E/2020 | Europe / United Kingdom / Wales | 2020-03-30 | Wales Specialist Virology Centre | Public Health Wales Microbiology Cardiff | Catherine Moore, Johnathan Evans, Malorie Perry, Simon Cottrell, Alec Birchley, Alexander Adams, Amy Gaskin, Bree Gatica-Wilcox, Jason Coombes, Lauren Gilbert, Lee Graham, Nicole Pacchiarini, Sara Kumziene-Summerhayes, Sarah Taylor, Sophie Jones, Sara Rey, Matthew Bull, Joanne Watkins, Sally Corden, Tom Connor |
| EPI_ISL_432338 | hCoV-19/Wales/PHWC-27425/2020 | Europe / United Kingdom / Wales | 2020-03-30 | Wales Specialist Virology Centre | Public Health Wales Microbiology Cardiff | Catherine Moore, Johnathan Evans, Malorie Perry, Simon Cottrell, Alec Birchley, Alexander Adams, Amy Gaskin, Bree Gatica-Wilcox, Jason Coombes, Lauren Gilbert, Lee Graham, Nicole Pacchiarini, Sara Kumziene-Summerhayes, Sarah Taylor, Sophie Jones, Sara Rey, Matthew Bull, Joanne Watkins, Sally Corden, Tom Connor |
| EPI_ISL_432339 | hCoV-19/Wales/PHWC-277D1/2020 | Europe / United Kingdom / Wales | 2020-03-27 | Wales Specialist Virology Centre | Public Health Wales Microbiology Cardiff | Catherine Moore, Johnathan Evans, Malorie Perry, Simon Cottrell, Alec Birchley, Alexander Adams, Amy Gaskin, Bree Gatica-Wilcox, Jason Coombes, Lauren Gilbert, Lee Graham, Nicole Pacchiarini, Sara Kumziene-Summerhayes, Sarah Taylor, Sophie Jones, Sara Rey, Matthew Bull, Joanne Watkins, Sally Corden, Tom Connor |
| EPI_ISL_432340 | hCoV-19/Wales/PHWC-27461/2020 | Europe / United Kingdom / Wales | 2020-03-31 | Wales Specialist Virology Centre | Public Health Wales Microbiology Cardiff | Catherine Moore, Johnathan Evans, Malorie Perry, Simon Cottrell, Alec Birchley, Alexander Adams, Amy Gaskin, Bree Gatica-Wilcox, Jason Coombes, Lauren Gilbert, Lee Graham, Nicole Pacchiarini, Sara Kumziene-Summerhayes, Sarah Taylor, Sophie Jones, Sara Rey, Matthew Bull, Joanne Watkins, Sally Corden, Tom Connor |
| EPI_ISL_432341 | hCoV-19/Wales/PHWC-27522/2020 | Europe / United Kingdom / Wales | 2020-03-30 | Wales Specialist Virology Centre | Public Health Wales Microbiology Cardiff | Catherine Moore, Johnathan Evans, Malorie Perry, Simon Cottrell, Alec Birchley, Alexander Adams, Amy Gaskin, Bree Gatica-Wilcox, Jason Coombes, Lauren Gilbert, Lee Graham, Nicole Pacchiarini, Sara Kumziene-Summerhayes, Sarah Taylor, Sophie Jones, Sara Rey, Matthew Bull, Joanne Watkins, Sally Corden, Tom Connor |
| EPI_ISL_432342 | hCoV-19/Wales/PHWC-28451/2020 | Europe / United Kingdom / Wales | 2020-03-28 | Wales Specialist Virology Centre | Public Health Wales Microbiology Cardiff | Catherine Moore, Johnathan Evans, Malorie Perry, Simon Cottrell, Alec Birchley, Alexander Adams, Amy Gaskin, Bree Gatica-Wilcox, Jason Coombes, Lauren Gilbert, Lee Graham, Nicole Pacchiarini, Sara Kumziene-Summerhayes, Sarah Taylor, Sophie Jones, Sara Rey, Matthew Bull, Joanne Watkins, Sally Corden, Tom Connor |
| EPI_ISL_432343 | hCoV-19/Wales/PHWC-29FAA/2020 | Europe / United Kingdom / Wales | 2020-04-03 | Wales Specialist Virology Centre | Public Health Wales Microbiology Cardiff | Catherine Moore, Johnathan Evans, Malorie Perry, Simon Cottrell, Alec Birchley, Alexander Adams, Amy Gaskin, Bree Gatica-Wilcox, Jason Coombes, Lauren Gilbert, Lee Graham, Nicole Pacchiarini, Sara Kumziene-Summerhayes, Sarah Taylor, Sophie Jones, Sara Rey, Matthew Bull, Joanne Watkins, Sally Corden, Tom Connor |
| EPI_ISL_432344 | hCoV-19/Wales/PHWC-29FF5/2020 | Europe / United Kingdom / Wales | 2020-04-03 | Wales Specialist Virology Centre | Public Health Wales Microbiology Cardiff | Catherine Moore, Johnathan Evans, Malorie Perry, Simon Cottrell, Alec Birchley, Alexander Adams, Amy Gaskin, Bree Gatica-Wilcox, Jason Coombes, Lauren Gilbert, Lee Graham, Nicole Pacchiarini, Sara Kumziene-Summerhayes, Sarah Taylor, Sophie Jones, Sara Rey, Matthew Bull, Joanne Watkins, Sally Corden, Tom Connor |
| EPI_ISL_432345 | hCoV-19/Wales/PHWC-2ADFA/2020 | Europe / United Kingdom / Wales | 2020-04-06 | Wales Specialist Virology Centre | Public Health Wales Microbiology Cardiff | Catherine Moore, Johnathan Evans, Malorie Perry, Simon Cottrell, Alec Birchley, Alexander Adams, Amy Gaskin, Bree Gatica-Wilcox, Jason Coombes, Lauren Gilbert, Lee Graham, Nicole Pacchiarini, Sara Kumziene-Summerhayes, Sarah Taylor, Sophie Jones, Sara Rey, Matthew Bull, Joanne Watkins, Sally Corden, Tom Connor |
| EPI_ISL_432346 | hCoV-19/Wales/PHWC-27BC6/2020 | Europe / United Kingdom / Wales | 2020-04-01 | Wales Specialist Virology Centre | Public Health Wales Microbiology Cardiff | Catherine Moore, Johnathan Evans, Malorie Perry, Simon Cottrell, Alec Birchley, Alexander Adams, Amy Gaskin, Bree Gatica-Wilcox, Jason Coombes, Lauren Gilbert, Lee Graham, Nicole Pacchiarini, Sara Kumziene-Summerhayes, Sarah Taylor, Sophie Jones, Sara Rey, Matthew Bull, Joanne Watkins, Sally Corden, Tom Connor |
| EPI_ISL_432347 | hCoV-19/Wales/PHWC-28460/2020 | Europe / United Kingdom / Wales | 2020-03-29 | Wales Specialist Virology Centre | Public Health Wales Microbiology Cardiff | Catherine Moore, Johnathan Evans, Malorie Perry, Simon Cottrell, Alec Birchley, Alexander Adams, Amy Gaskin, Bree Gatica-Wilcox, Jason Coombes, Lauren Gilbert, Lee Graham, Nicole Pacchiarini, Sara Kumziene-Summerhayes, Sarah Taylor, Sophie Jones, Sara Rey, Matthew Bull, Joanne Watkins, Sally Corden, Tom Connor |
| EPI_ISL_432348 | hCoV-19/Wales/PHWC-25339/2020 | Europe / United Kingdom / Wales | 2020-03-23 | Wales Specialist Virology Centre | Public Health Wales Microbiology Cardiff | Catherine Moore, Johnathan Evans, Malorie Perry, Simon Cottrell, Alec Birchley, Alexander Adams, Amy Gaskin, Bree Gatica-Wilcox, Jason Coombes, Lauren Gilbert, Lee Graham, Nicole Pacchiarini, Sara Kumziene-Summerhayes, Sarah Taylor, Sophie Jones, Sara Rey, Matthew Bull, Joanne Watkins, Sally Corden, Tom Connor |
| EPI_ISL_432349 | hCoV-19/Wales/PHWC-28891/2020 | Europe / United Kingdom / Wales | 2020-03-31 | Wales Specialist Virology Centre | Public Health Wales Microbiology Cardiff | Catherine Moore, Johnathan Evans, Malorie Perry, Simon Cottrell, Alec Birchley, Alexander Adams, Amy Gaskin, Bree Gatica-Wilcox, Jason Coombes, Lauren Gilbert, Lee Graham, Nicole Pacchiarini, Sara Kumziene-Summerhayes, Sarah Taylor, Sophie Jones, Sara Rey, Matthew Bull, Joanne Watkins, Sally Corden, Tom Connor |
| EPI_ISL_432350 | hCoV-19/Wales/PHWC-2799F/2020 | Europe / United Kingdom / Wales | 2020-04-01 | Wales Specialist Virology Centre | Public Health Wales Microbiology Cardiff | Catherine Moore, Johnathan Evans, Malorie Perry, Simon Cottrell, Alec Birchley, Alexander Adams, Amy Gaskin, Bree Gatica-Wilcox, Jason Coombes, Lauren Gilbert, Lee Graham, Nicole Pacchiarini, Sara Kumziene-Summerhayes, Sarah Taylor, Sophie Jones, Sara Rey, Matthew Bull, Joanne Watkins, Sally Corden, Tom Connor |
| EPI_ISL_432351 | hCoV-19/Wales/PHWC-2866A/2020 | Europe / United Kingdom / Wales | 2020-03-31 | Wales Specialist Virology Centre | Public Health Wales Microbiology Cardiff | Catherine Moore, Johnathan Evans, Malorie Perry, Simon Cottrell, Alec Birchley, Alexander Adams, Amy Gaskin, Bree Gatica-Wilcox, Jason Coombes, Lauren Gilbert, Lee Graham, Nicole Pacchiarini, Sara Kumziene-Summerhayes, Sarah Taylor, Sophie Jones, Sara Rey, Matthew Bull, Joanne Watkins, Sally Corden, Tom Connor |
| EPI_ISL_432352 | hCoV-19/Wales/PHWC-283CD/2020 | Europe / United Kingdom / Wales | 2020-03-27 | Wales Specialist Virology Centre | Public Health Wales Microbiology Cardiff | Catherine Moore, Johnathan Evans, Malorie Perry, Simon Cottrell, Alec Birchley, Alexander Adams, Amy Gaskin, Bree Gatica-Wilcox, Jason Coombes, Lauren Gilbert, Lee Graham, Nicole Pacchiarini, Sara Kumziene-Summerhayes, Sarah Taylor, Sophie Jones, Sara Rey, Matthew Bull, Joanne Watkins, Sally Corden, Tom Connor |
| EPI_ISL_432353 | hCoV-19/Wales/PHWC-29E25/2020 | Europe / United Kingdom / Wales | 2020-04-04 | Wales Specialist Virology Centre | Public Health Wales Microbiology Cardiff | Catherine Moore, Johnathan Evans, Malorie Perry, Simon Cottrell, Alec Birchley, Alexander Adams, Amy Gaskin, Bree Gatica-Wilcox, Jason Coombes, Lauren Gilbert, Lee Graham, Nicole Pacchiarini, Sara Kumziene-Summerhayes, Sarah Taylor, Sophie Jones, Sara Rey, Matthew Bull, Joanne Watkins, Sally Corden, Tom Connor |
| EPI_ISL_432354 | hCoV-19/Wales/PHWC-2AF6D/2020 | Europe / United Kingdom / Wales | 2020-04-07 | Wales Specialist Virology Centre | Public Health Wales Microbiology Cardiff | Catherine Moore, Johnathan Evans, Malorie Perry, Simon Cottrell, Alec Birchley, Alexander Adams, Amy Gaskin, Bree Gatica-Wilcox, Jason Coombes, Lauren Gilbert, Lee Graham, Nicole Pacchiarini, Sara Kumziene-Summerhayes, Sarah Taylor, Sophie Jones, Sara Rey, Matthew Bull, Joanne Watkins, Sally Corden, Tom Connor |
| EPI_ISL_432355 | hCoV-19/Wales/PHWC-286D3/2020 | Europe / United Kingdom / Wales | 2020-04-01 | Wales Specialist Virology Centre | Public Health Wales Microbiology Cardiff | Catherine Moore, Johnathan Evans, Malorie Perry, Simon Cottrell, Alec Birchley, Alexander Adams, Amy Gaskin, Bree Gatica-Wilcox, Jason Coombes, Lauren Gilbert, Lee Graham, Nicole Pacchiarini, Sara Kumziene-Summerhayes, Sarah Taylor, Sophie Jones, Sara Rey, Matthew Bull, Joanne Watkins, Sally Corden, Tom Connor |
| EPI_ISL_432356 | hCoV-19/Wales/PHWC-286E2/2020 | Europe / United Kingdom / Wales | 2020-04-01 | Wales Specialist Virology Centre | Public Health Wales Microbiology Cardiff | Catherine Moore, Johnathan Evans, Malorie Perry, Simon Cottrell, Alec Birchley, Alexander Adams, Amy Gaskin, Bree Gatica-Wilcox, Jason Coombes, Lauren Gilbert, Lee Graham, Nicole Pacchiarini, Sara Kumziene-Summerhayes, Sarah Taylor, Sophie Jones, Sara Rey, Matthew Bull, Joanne Watkins, Sally Corden, Tom Connor |
| EPI_ISL_432601 | hCoV-19/England/SHEF-CFE03/2020 | Europe / United Kingdom / England | 2020-03-30 | Virology Department, Sheffield Teaching Hospitals NHS Foundation Trust / Virology Department, Sheffield Teaching Hospitals NHS Foundation Trust | COVID-19 Genomics UK (COG-UK) Consortium | Thushan de Silva, Matthew Parker,Adri Angyal, Rebecca Brown, Luke Green, Rachel Tucker, Paul Parsons, Danielle Groves, Alex Keeley, Dave Partridge, Matthew Wyles, Benjamin Lindsey, Mehmet Yavuz, Mohammad Raza, Cariad Evans |
| EPI_ISL_432602 | hCoV-19/England/SHEF-CFE21/2020 | Europe / United Kingdom / England | 2020-03-30 | Virology Department, Sheffield Teaching Hospitals NHS Foundation Trust / Virology Department, Sheffield Teaching Hospitals NHS Foundation Trust | COVID-19 Genomics UK (COG-UK) Consortium | Thushan de Silva, Matthew Parker,Adri Angyal, Rebecca Brown, Luke Green, Rachel Tucker, Paul Parsons, Danielle Groves, Alex Keeley, Dave Partridge, Matthew Wyles, Benjamin Lindsey, Mehmet Yavuz, Mohammad Raza, Cariad Evans |
| EPI_ISL_432614 | hCoV-19/England/SHEF-D0731/2020 | Europe / United Kingdom / England | 2020-04-01 | Virology Department, Sheffield Teaching Hospitals NHS Foundation Trust / Virology Department, Sheffield Teaching Hospitals NHS Foundation Trust | COVID-19 Genomics UK (COG-UK) Consortium | Thushan de Silva, Matthew Parker,Adri Angyal, Rebecca Brown, Luke Green, Rachel Tucker, Paul Parsons, Danielle Groves, Alex Keeley, Dave Partridge, Matthew Wyles, Benjamin Lindsey, Mehmet Yavuz, Mohammad Raza, Cariad Evans |
| EPI_ISL_432620 | hCoV-19/England/SHEF-D0D99/2020 | Europe / United Kingdom / England | 2020-03-28 | Virology Department, Sheffield Teaching Hospitals NHS Foundation Trust / Virology Department, Sheffield Teaching Hospitals NHS Foundation Trust | COVID-19 Genomics UK (COG-UK) Consortium | Thushan de Silva, Matthew Parker,Adri Angyal, Rebecca Brown, Luke Green, Rachel Tucker, Paul Parsons, Danielle Groves, Alex Keeley, Dave Partridge, Matthew Wyles, Benjamin Lindsey, Mehmet Yavuz, Mohammad Raza, Cariad Evans |
| EPI_ISL_432622 | hCoV-19/England/SHEF-D1053/2020 | Europe / United Kingdom / England | 2020-04-02 | Virology Department, Sheffield Teaching Hospitals NHS Foundation Trust / Virology Department, Sheffield Teaching Hospitals NHS Foundation Trust | COVID-19 Genomics UK (COG-UK) Consortium | Thushan de Silva, Matthew Parker,Adri Angyal, Rebecca Brown, Luke Green, Rachel Tucker, Paul Parsons, Danielle Groves, Alex Keeley, Dave Partridge, Matthew Wyles, Benjamin Lindsey, Mehmet Yavuz, Mohammad Raza, Cariad Evans |
| EPI_ISL_432630 | hCoV-19/England/SHEF-D1536/2020 | Europe / United Kingdom / England | 2020-04-02 | Virology Department, Sheffield Teaching Hospitals NHS Foundation Trust / Virology Department, Sheffield Teaching Hospitals NHS Foundation Trust | COVID-19 Genomics UK (COG-UK) Consortium | Thushan de Silva, Matthew Parker,Adri Angyal, Rebecca Brown, Luke Green, Rachel Tucker, Paul Parsons, Danielle Groves, Alex Keeley, Dave Partridge, Matthew Wyles, Benjamin Lindsey, Mehmet Yavuz, Mohammad Raza, Cariad Evans |
| EPI_ISL_433000 | hCoV-19/England/NOTT-10F3E9/2020 | Europe / United Kingdom / England | 2020-04-02 | Queens Medical Centre, Clinical Microbiology Department / DeepSeq Nottingham | COVID-19 Genomics UK (COG-UK) Consortium | Gemma Clark, Wendy Smith, Manjinder Khakh, Hannah Howson-Wells, Jonathan Ball, Patrick McClure, Joseph Chappell, Theocharis Tsoleridis, Nadine Holmes, Matthew Carlisle, Christopher Moore, Fei Sang, Johnny Debebe, Victoria Wright, Matthew Loose |
| EPI_ISL_433002 | hCoV-19/England/NOTT-10F413/2020 | Europe / United Kingdom / England | 2020-04-02 | Queens Medical Centre, Clinical Microbiology Department / DeepSeq Nottingham | COVID-19 Genomics UK (COG-UK) Consortium | Gemma Clark, Wendy Smith, Manjinder Khakh, Hannah Howson-Wells, Jonathan Ball, Patrick McClure, Joseph Chappell, Theocharis Tsoleridis, Nadine Holmes, Matthew Carlisle, Christopher Moore, Fei Sang, Johnny Debebe, Victoria Wright, Matthew Loose |
| EPI_ISL_433012 | hCoV-19/England/NOTT-10F4F5/2020 | Europe / United Kingdom / England | 2020-04-02 | Queens Medical Centre, Clinical Microbiology Department / DeepSeq Nottingham | COVID-19 Genomics UK (COG-UK) Consortium | Gemma Clark, Wendy Smith, Manjinder Khakh, Hannah Howson-Wells, Jonathan Ball, Patrick McClure, Joseph Chappell, Theocharis Tsoleridis, Nadine Holmes, Matthew Carlisle, Christopher Moore, Fei Sang, Johnny Debebe, Victoria Wright, Matthew Loose |
| EPI_ISL_433016 | hCoV-19/England/NOTT-10F53E/2020 | Europe / United Kingdom / England | 2020-04-02 | Queens Medical Centre, Clinical Microbiology Department / DeepSeq Nottingham | COVID-19 Genomics UK (COG-UK) Consortium | Gemma Clark, Wendy Smith, Manjinder Khakh, Hannah Howson-Wells, Jonathan Ball, Patrick McClure, Joseph Chappell, Theocharis Tsoleridis, Nadine Holmes, Matthew Carlisle, Christopher Moore, Fei Sang, Johnny Debebe, Victoria Wright, Matthew Loose |
| EPI_ISL_433034 | hCoV-19/England/NOTT-10F695/2020 | Europe / United Kingdom / England | 2020-04-03 | Queens Medical Centre, Clinical Microbiology Department / DeepSeq Nottingham | COVID-19 Genomics UK (COG-UK) Consortium | Gemma Clark, Wendy Smith, Manjinder Khakh, Hannah Howson-Wells, Jonathan Ball, Patrick McClure, Joseph Chappell, Theocharis Tsoleridis, Nadine Holmes, Matthew Carlisle, Christopher Moore, Fei Sang, Johnny Debebe, Victoria Wright, Matthew Loose |
| EPI_ISL_433049 | hCoV-19/England/NOTT-10F792/2020 | Europe / United Kingdom / England | 2020-04-04 | Queens Medical Centre, Clinical Microbiology Department / DeepSeq Nottingham | COVID-19 Genomics UK (COG-UK) Consortium | Gemma Clark, Wendy Smith, Manjinder Khakh, Hannah Howson-Wells, Jonathan Ball, Patrick McClure, Joseph Chappell, Theocharis Tsoleridis, Nadine Holmes, Matthew Carlisle, Christopher Moore, Fei Sang, Johnny Debebe, Victoria Wright, Matthew Loose |
| EPI_ISL_433066 | hCoV-19/England/NOTT-10F98D/2020 | Europe / United Kingdom / England | 2020-04-05 | Queens Medical Centre, Clinical Microbiology Department / DeepSeq Nottingham | COVID-19 Genomics UK (COG-UK) Consortium | Gemma Clark, Wendy Smith, Manjinder Khakh, Hannah Howson-Wells, Jonathan Ball, Patrick McClure, Joseph Chappell, Theocharis Tsoleridis, Nadine Holmes, Matthew Carlisle, Christopher Moore, Fei Sang, Johnny Debebe, Victoria Wright, Matthew Loose |
| EPI_ISL_433068 | hCoV-19/Scotland/EDB1518/2020 | Europe / United Kingdom / Scotland | 2020-04-12 | Virology Department, Royal Infirmary of Edinburgh, NHS Lothian / School of Biological Sciences, University of Edinburgh / Institute of Genetics and Molecular Medicine, University of Edinburgh | COVID-19 Genomics UK (COG-UK) Consortium | McHugh M, Dewar R, Rooke S, Gallagher M, Balcaza C, O'Toole A, Hill V, McCrone JT, Colquhoun R, Yu X, Jackson B, Rambaut A, Williams TC, Templeton K |
| EPI_ISL_433069 | hCoV-19/Scotland/EDB1519/2020 | Europe / United Kingdom / Scotland | 2020-04-13 | Virology Department, Royal Infirmary of Edinburgh, NHS Lothian / School of Biological Sciences, University of Edinburgh / Institute of Genetics and Molecular Medicine, University of Edinburgh | COVID-19 Genomics UK (COG-UK) Consortium | McHugh M, Dewar R, Rooke S, Gallagher M, Balcaza C, O'Toole A, Hill V, McCrone JT, Colquhoun R, Yu X, Jackson B, Rambaut A, Williams TC, Templeton K |
| EPI_ISL_433070 | hCoV-19/Scotland/EDB1523/2020 | Europe / United Kingdom / Scotland | 2020-04-14 | Virology Department, Royal Infirmary of Edinburgh, NHS Lothian / School of Biological Sciences, University of Edinburgh / Institute of Genetics and Molecular Medicine, University of Edinburgh | COVID-19 Genomics UK (COG-UK) Consortium | McHugh M, Dewar R, Rooke S, Gallagher M, Balcaza C, O'Toole A, Hill V, McCrone JT, Colquhoun R, Yu X, Jackson B, Rambaut A, Williams TC, Templeton K |
| EPI_ISL_433071 | hCoV-19/Scotland/EDB1525/2020 | Europe / United Kingdom / Scotland | 2020-04-14 | Virology Department, Royal Infirmary of Edinburgh, NHS Lothian / School of Biological Sciences, University of Edinburgh / Institute of Genetics and Molecular Medicine, University of Edinburgh | COVID-19 Genomics UK (COG-UK) Consortium | McHugh M, Dewar R, Rooke S, Gallagher M, Balcaza C, O'Toole A, Hill V, McCrone JT, Colquhoun R, Yu X, Jackson B, Rambaut A, Williams TC, Templeton K |
| EPI_ISL_433072 | hCoV-19/Scotland/EDB1526/2020 | Europe / United Kingdom / Scotland | 2020-04-12 | Virology Department, Royal Infirmary of Edinburgh, NHS Lothian / School of Biological Sciences, University of Edinburgh / Institute of Genetics and Molecular Medicine, University of Edinburgh | COVID-19 Genomics UK (COG-UK) Consortium | McHugh M, Dewar R, Rooke S, Gallagher M, Balcaza C, O'Toole A, Hill V, McCrone JT, Colquhoun R, Yu X, Jackson B, Rambaut A, Williams TC, Templeton K |
| EPI_ISL_433073 | hCoV-19/Scotland/EDB1529/2020 | Europe / United Kingdom / Scotland | 2020-04-13 | Virology Department, Royal Infirmary of Edinburgh, NHS Lothian / School of Biological Sciences, University of Edinburgh / Institute of Genetics and Molecular Medicine, University of Edinburgh | COVID-19 Genomics UK (COG-UK) Consortium | McHugh M, Dewar R, Rooke S, Gallagher M, Balcaza C, O'Toole A, Hill V, McCrone JT, Colquhoun R, Yu X, Jackson B, Rambaut A, Williams TC, Templeton K |
| EPI_ISL_433074 | hCoV-19/Scotland/EDB153/2020 | Europe / United Kingdom / Scotland | 2020-03-24 | Virology Department, Royal Infirmary of Edinburgh, NHS Lothian / School of Biological Sciences, University of Edinburgh / Institute of Genetics and Molecular Medicine, University of Edinburgh | COVID-19 Genomics UK (COG-UK) Consortium | McHugh M, Dewar R, Rooke S, Gallagher M, Balcaza C, O'Toole A, Hill V, McCrone JT, Colquhoun R, Yu X, Jackson B, Rambaut A, Williams TC, Templeton K |
| EPI_ISL_433075 | hCoV-19/Scotland/EDB1536/2020 | Europe / United Kingdom / Scotland | 2020-04-12 | Virology Department, Royal Infirmary of Edinburgh, NHS Lothian / School of Biological Sciences, University of Edinburgh / Institute of Genetics and Molecular Medicine, University of Edinburgh | COVID-19 Genomics UK (COG-UK) Consortium | McHugh M, Dewar R, Rooke S, Gallagher M, Balcaza C, O'Toole A, Hill V, McCrone JT, Colquhoun R, Yu X, Jackson B, Rambaut A, Williams TC, Templeton K |
| EPI_ISL_433077 | hCoV-19/Scotland/EDB1541/2020 | Europe / United Kingdom / Scotland | 2020-04-12 | Virology Department, Royal Infirmary of Edinburgh, NHS Lothian / School of Biological Sciences, University of Edinburgh / Institute of Genetics and Molecular Medicine, University of Edinburgh | COVID-19 Genomics UK (COG-UK) Consortium | McHugh M, Dewar R, Rooke S, Gallagher M, Balcaza C, O'Toole A, Hill V, McCrone JT, Colquhoun R, Yu X, Jackson B, Rambaut A, Williams TC, Templeton K |
| EPI_ISL_433078 | hCoV-19/Scotland/EDB1544/2020 | Europe / United Kingdom / Scotland | 2020-04-12 | Virology Department, Royal Infirmary of Edinburgh, NHS Lothian / School of Biological Sciences, University of Edinburgh / Institute of Genetics and Molecular Medicine, University of Edinburgh | COVID-19 Genomics UK (COG-UK) Consortium | McHugh M, Dewar R, Rooke S, Gallagher M, Balcaza C, O'Toole A, Hill V, McCrone JT, Colquhoun R, Yu X, Jackson B, Rambaut A, Williams TC, Templeton K |
| EPI_ISL_433079 | hCoV-19/Scotland/EDB1550/2020 | Europe / United Kingdom / Scotland | 2020-04-12 | Virology Department, Royal Infirmary of Edinburgh, NHS Lothian / School of Biological Sciences, University of Edinburgh / Institute of Genetics and Molecular Medicine, University of Edinburgh | COVID-19 Genomics UK (COG-UK) Consortium | McHugh M, Dewar R, Rooke S, Gallagher M, Balcaza C, O'Toole A, Hill V, McCrone JT, Colquhoun R, Yu X, Jackson B, Rambaut A, Williams TC, Templeton K |
| EPI_ISL_433080 | hCoV-19/Scotland/EDB1554/2020 | Europe / United Kingdom / Scotland | 2020-04-13 | Virology Department, Royal Infirmary of Edinburgh, NHS Lothian / School of Biological Sciences, University of Edinburgh / Institute of Genetics and Molecular Medicine, University of Edinburgh | COVID-19 Genomics UK (COG-UK) Consortium | McHugh M, Dewar R, Rooke S, Gallagher M, Balcaza C, O'Toole A, Hill V, McCrone JT, Colquhoun R, Yu X, Jackson B, Rambaut A, Williams TC, Templeton K |
| EPI_ISL_433081 | hCoV-19/Scotland/EDB1559/2020 | Europe / United Kingdom / Scotland | 2020-04-12 | Virology Department, Royal Infirmary of Edinburgh, NHS Lothian / School of Biological Sciences, University of Edinburgh / Institute of Genetics and Molecular Medicine, University of Edinburgh | COVID-19 Genomics UK (COG-UK) Consortium | McHugh M, Dewar R, Rooke S, Gallagher M, Balcaza C, O'Toole A, Hill V, McCrone JT, Colquhoun R, Yu X, Jackson B, Rambaut A, Williams TC, Templeton K |
| EPI_ISL_433082 | hCoV-19/Scotland/EDB1565/2020 | Europe / United Kingdom / Scotland | 2020-04-12 | Virology Department, Royal Infirmary of Edinburgh, NHS Lothian / School of Biological Sciences, University of Edinburgh / Institute of Genetics and Molecular Medicine, University of Edinburgh | COVID-19 Genomics UK (COG-UK) Consortium | McHugh M, Dewar R, Rooke S, Gallagher M, Balcaza C, O'Toole A, Hill V, McCrone JT, Colquhoun R, Yu X, Jackson B, Rambaut A, Williams TC, Templeton K |
| EPI_ISL_433083 | hCoV-19/Scotland/EDB1566/2020 | Europe / United Kingdom / Scotland | 2020-04-13 | Virology Department, Royal Infirmary of Edinburgh, NHS Lothian / School of Biological Sciences, University of Edinburgh / Institute of Genetics and Molecular Medicine, University of Edinburgh | COVID-19 Genomics UK (COG-UK) Consortium | McHugh M, Dewar R, Rooke S, Gallagher M, Balcaza C, O'Toole A, Hill V, McCrone JT, Colquhoun R, Yu X, Jackson B, Rambaut A, Williams TC, Templeton K |
| EPI_ISL_433084 | hCoV-19/Scotland/EDB1568/2020 | Europe / United Kingdom / Scotland | 2020-04-12 | Virology Department, Royal Infirmary of Edinburgh, NHS Lothian / School of Biological Sciences, University of Edinburgh / Institute of Genetics and Molecular Medicine, University of Edinburgh | COVID-19 Genomics UK (COG-UK) Consortium | McHugh M, Dewar R, Rooke S, Gallagher M, Balcaza C, O'Toole A, Hill V, McCrone JT, Colquhoun R, Yu X, Jackson B, Rambaut A, Williams TC, Templeton K |
| EPI_ISL_433085 | hCoV-19/Scotland/EDB1580/2020 | Europe / United Kingdom / Scotland | 2020-04-13 | Virology Department, Royal Infirmary of Edinburgh, NHS Lothian / School of Biological Sciences, University of Edinburgh / Institute of Genetics and Molecular Medicine, University of Edinburgh | COVID-19 Genomics UK (COG-UK) Consortium | McHugh M, Dewar R, Rooke S, Gallagher M, Balcaza C, O'Toole A, Hill V, McCrone JT, Colquhoun R, Yu X, Jackson B, Rambaut A, Williams TC, Templeton K |
| EPI_ISL_433086 | hCoV-19/Scotland/EDB1581/2020 | Europe / United Kingdom / Scotland | 2020-04-14 | Virology Department, Royal Infirmary of Edinburgh, NHS Lothian / School of Biological Sciences, University of Edinburgh / Institute of Genetics and Molecular Medicine, University of Edinburgh | COVID-19 Genomics UK (COG-UK) Consortium | McHugh M, Dewar R, Rooke S, Gallagher M, Balcaza C, O'Toole A, Hill V, McCrone JT, Colquhoun R, Yu X, Jackson B, Rambaut A, Williams TC, Templeton K |
| EPI_ISL_433087 | hCoV-19/Scotland/EDB1582/2020 | Europe / United Kingdom / Scotland | 2020-04-13 | Virology Department, Royal Infirmary of Edinburgh, NHS Lothian / School of Biological Sciences, University of Edinburgh / Institute of Genetics and Molecular Medicine, University of Edinburgh | COVID-19 Genomics UK (COG-UK) Consortium | McHugh M, Dewar R, Rooke S, Gallagher M, Balcaza C, O'Toole A, Hill V, McCrone JT, Colquhoun R, Yu X, Jackson B, Rambaut A, Williams TC, Templeton K |
| EPI_ISL_433088 | hCoV-19/Scotland/EDB1583/2020 | Europe / United Kingdom / Scotland | 2020-04-13 | Virology Department, Royal Infirmary of Edinburgh, NHS Lothian / School of Biological Sciences, University of Edinburgh / Institute of Genetics and Molecular Medicine, University of Edinburgh | COVID-19 Genomics UK (COG-UK) Consortium | McHugh M, Dewar R, Rooke S, Gallagher M, Balcaza C, O'Toole A, Hill V, McCrone JT, Colquhoun R, Yu X, Jackson B, Rambaut A, Williams TC, Templeton K |
| EPI_ISL_433091 | hCoV-19/Scotland/EDB1617/2020 | Europe / United Kingdom / Scotland | 2020-04-14 | Virology Department, Royal Infirmary of Edinburgh, NHS Lothian / School of Biological Sciences, University of Edinburgh / Institute of Genetics and Molecular Medicine, University of Edinburgh | COVID-19 Genomics UK (COG-UK) Consortium | McHugh M, Dewar R, Rooke S, Gallagher M, Balcaza C, O'Toole A, Hill V, McCrone JT, Colquhoun R, Yu X, Jackson B, Rambaut A, Williams TC, Templeton K |
| EPI_ISL_433093 | hCoV-19/Scotland/EDB1629/2020 | Europe / United Kingdom / Scotland | 2020-04-14 | Virology Department, Royal Infirmary of Edinburgh, NHS Lothian / School of Biological Sciences, University of Edinburgh / Institute of Genetics and Molecular Medicine, University of Edinburgh | COVID-19 Genomics UK (COG-UK) Consortium | McHugh M, Dewar R, Rooke S, Gallagher M, Balcaza C, O'Toole A, Hill V, McCrone JT, Colquhoun R, Yu X, Jackson B, Rambaut A, Williams TC, Templeton K |
| EPI_ISL_433095 | hCoV-19/Scotland/EDB1631/2020 | Europe / United Kingdom / Scotland | 2020-04-14 | Virology Department, Royal Infirmary of Edinburgh, NHS Lothian / School of Biological Sciences, University of Edinburgh / Institute of Genetics and Molecular Medicine, University of Edinburgh | COVID-19 Genomics UK (COG-UK) Consortium | McHugh M, Dewar R, Rooke S, Gallagher M, Balcaza C, O'Toole A, Hill V, McCrone JT, Colquhoun R, Yu X, Jackson B, Rambaut A, Williams TC, Templeton K |
| EPI_ISL_433096 | hCoV-19/Scotland/EDB1632/2020 | Europe / United Kingdom / Scotland | 2020-04-14 | Virology Department, Royal Infirmary of Edinburgh, NHS Lothian / School of Biological Sciences, University of Edinburgh / Institute of Genetics and Molecular Medicine, University of Edinburgh | COVID-19 Genomics UK (COG-UK) Consortium | McHugh M, Dewar R, Rooke S, Gallagher M, Balcaza C, O'Toole A, Hill V, McCrone JT, Colquhoun R, Yu X, Jackson B, Rambaut A, Williams TC, Templeton K |
| EPI_ISL_433098 | hCoV-19/Scotland/EDB1638/2020 | Europe / United Kingdom / Scotland | 2020-04-15 | Virology Department, Royal Infirmary of Edinburgh, NHS Lothian / School of Biological Sciences, University of Edinburgh / Institute of Genetics and Molecular Medicine, University of Edinburgh | COVID-19 Genomics UK (COG-UK) Consortium | McHugh M, Dewar R, Rooke S, Gallagher M, Balcaza C, O'Toole A, Hill V, McCrone JT, Colquhoun R, Yu X, Jackson B, Rambaut A, Williams TC, Templeton K |

| | | | | | | |
|---|---|---|---|---|---|---|
| EPI_ISL_433952 | hCoV-19/England/CAMB-7B3E9/2020 | Europe / United Kingdom / England | 2020-04-12 | Department of Pathology, University of Cambridge | COVID-19 Genomics UK (COG-UK) Consortium | Luke W Meredith, M. Estee Torok , Myra Hosmillo, William L. Hamilton, Martin D. Curran, Theresa Feltwell, Grant Hall, Anna Yakovleva, Fahad A Khokhar, Charlotte J. Houldcroft, Laura G Caller, Aminu S. Jahun, Sarah L. Caddy, Ian Goodfellow |
| EPI_ISL_433953 | hCoV-19/England/CAMB-7B3F8/2020 | Europe / United Kingdom / England | 2020-04-13 | Department of Pathology, University of Cambridge | COVID-19 Genomics UK (COG-UK) Consortium | Luke W Meredith, M. Estee Torok , Myra Hosmillo, William L. Hamilton, Martin D. Curran, Theresa Feltwell, Grant Hall, Anna Yakovleva, Fahad A Khokhar, Charlotte J. Houldcroft, Laura G Caller, Aminu S. Jahun, Sarah L. Caddy, Ian Goodfellow |
| EPI_ISL_433954 | hCoV-19/England/CAMB-7B404/2020 | Europe / United Kingdom / England | 2020-04-12 | Department of Pathology, University of Cambridge | COVID-19 Genomics UK (COG-UK) Consortium | Luke W Meredith, M. Estee Torok , Myra Hosmillo, William L. Hamilton, Martin D. Curran, Theresa Feltwell, Grant Hall, Anna Yakovleva, Fahad A Khokhar, Charlotte J. Houldcroft, Laura G Caller, Aminu S. Jahun, Sarah L. Caddy, Ian Goodfellow |
| EPI_ISL_433956 | hCoV-19/England/CAMB-7B422/2020 | Europe / United Kingdom / England | 2020-04-13 | Department of Pathology, University of Cambridge | COVID-19 Genomics UK (COG-UK) Consortium | Luke W Meredith, M. Estee Torok , Myra Hosmillo, William L. Hamilton, Martin D. Curran, Theresa Feltwell, Grant Hall, Anna Yakovleva, Fahad A Khokhar, Charlotte J. Houldcroft, Laura G Caller, Aminu S. Jahun, Sarah L. Caddy, Ian Goodfellow |
| EPI_ISL_433957 | hCoV-19/England/CAMB-7B431/2020 | Europe / United Kingdom / England | 2020-04-12 | Department of Pathology, University of Cambridge | COVID-19 Genomics UK (COG-UK) Consortium | Luke W Meredith, M. Estee Torok , Myra Hosmillo, William L. Hamilton, Martin D. Curran, Theresa Feltwell, Grant Hall, Anna Yakovleva, Fahad A Khokhar, Charlotte J. Houldcroft, Laura G Caller, Aminu S. Jahun, Sarah L. Caddy, Ian Goodfellow |
| EPI_ISL_433958 | hCoV-19/England/CAMB-7B440/2020 | Europe / United Kingdom / England | 2020-04-13 | Department of Pathology, University of Cambridge | COVID-19 Genomics UK (COG-UK) Consortium | Luke W Meredith, M. Estee Torok , Myra Hosmillo, William L. Hamilton, Martin D. Curran, Theresa Feltwell, Grant Hall, Anna Yakovleva, Fahad A Khokhar, Charlotte J. Houldcroft, Laura G Caller, Aminu S. Jahun, Sarah L. Caddy, Ian Goodfellow |
| EPI_ISL_433959 | hCoV-19/England/CAMB-7B45F/2020 | Europe / United Kingdom / England | 2020-04-11 | Department of Pathology, University of Cambridge | COVID-19 Genomics UK (COG-UK) Consortium | Luke W Meredith, M. Estee Torok , Myra Hosmillo, William L. Hamilton, Martin D. Curran, Theresa Feltwell, Grant Hall, Anna Yakovleva, Fahad A Khokhar, Charlotte J. Houldcroft, Laura G Caller, Aminu S. Jahun, Sarah L. Caddy, Ian Goodfellow |
| EPI_ISL_433961 | hCoV-19/England/CAMB-7B48C/2020 | Europe / United Kingdom / England | 2020-04-12 | Department of Pathology, University of Cambridge | COVID-19 Genomics UK (COG-UK) Consortium | Luke W Meredith, M. Estee Torok , Myra Hosmillo, William L. Hamilton, Martin D. Curran, Theresa Feltwell, Grant Hall, Anna Yakovleva, Fahad A Khokhar, Charlotte J. Houldcroft, Laura G Caller, Aminu S. Jahun, Sarah L. Caddy, Ian Goodfellow |
| EPI_ISL_433962 | hCoV-19/England/CAMB-7B49B/2020 | Europe / United Kingdom / England | 2020-04-12 | Department of Pathology, University of Cambridge | COVID-19 Genomics UK (COG-UK) Consortium | Luke W Meredith, M. Estee Torok , Myra Hosmillo, William L. Hamilton, Martin D. Curran, Theresa Feltwell, Grant Hall, Anna Yakovleva, Fahad A Khokhar, Charlotte J. Houldcroft, Laura G Caller, Aminu S. Jahun, Sarah L. Caddy, Ian Goodfellow |
| EPI_ISL_434001 | hCoV-19/England/CAMB-7C166/2020 | Europe / United Kingdom / England | 2020-04-13 | Department of Pathology, University of Cambridge | COVID-19 Genomics UK (COG-UK) Consortium | Luke W Meredith, M. Estee Torok , Myra Hosmillo, William L. Hamilton, Martin D. Curran, Theresa Feltwell, Grant Hall, Anna Yakovleva, Fahad A Khokhar, Charlotte J. Houldcroft, Laura G Caller, Aminu S. Jahun, Sarah L. Caddy, Ian Goodfellow |
| EPI_ISL_434002 | hCoV-19/England/CAMB-7C175/2020 | Europe / United Kingdom / England | 2020-04-12 | Department of Pathology, University of Cambridge | COVID-19 Genomics UK (COG-UK) Consortium | Luke W Meredith, M. Estee Torok , Myra Hosmillo, William L. Hamilton, Martin D. Curran, Theresa Feltwell, Grant Hall, Anna Yakovleva, Fahad A Khokhar, Charlotte J. Houldcroft, Laura G Caller, Aminu S. Jahun, Sarah L. Caddy, Ian Goodfellow |
| EPI_ISL_434007 | hCoV-19/England/CAMB-7C1C0/2020 | Europe / United Kingdom / England | 2020 | Department of Pathology, University of Cambridge | COVID-19 Genomics UK (COG-UK) Consortium | Luke W Meredith, M. Estee Torok , Myra Hosmillo, William L. Hamilton, Martin D. Curran, Theresa Feltwell, Grant Hall, Anna Yakovleva, Fahad A Khokhar, Charlotte J. Houldcroft, Laura G Caller, Aminu S. Jahun, Sarah L. Caddy, Ian Goodfellow |
| EPI_ISL_434013 | hCoV-19/England/CAMB-7C227/2020 | Europe / United Kingdom / England | 2020-04-14 | Department of Pathology, University of Cambridge | COVID-19 Genomics UK (COG-UK) Consortium | Luke W Meredith, M. Estee Torok , Myra Hosmillo, William L. Hamilton, Martin D. Curran, Theresa Feltwell, Grant Hall, Anna Yakovleva, Fahad A Khokhar, Charlotte J. Houldcroft, Laura G Caller, Aminu S. Jahun, Sarah L. Caddy, Ian Goodfellow |
| EPI_ISL_434014 | hCoV-19/England/CAMB-7C236/2020 | Europe / United Kingdom / England | 2020-04-12 | Department of Pathology, University of Cambridge | COVID-19 Genomics UK (COG-UK) Consortium | Luke W Meredith, M. Estee Torok , Myra Hosmillo, William L. Hamilton, Martin D. Curran, Theresa Feltwell, Grant Hall, Anna Yakovleva, Fahad A Khokhar, Charlotte J. Houldcroft, Laura G Caller, Aminu S. Jahun, Sarah L. Caddy, Ian Goodfellow |
| EPI_ISL_434020 | hCoV-19/England/CAMB-7C290/2020 | Europe / United Kingdom / England | 2020-04-15 | Department of Pathology, University of Cambridge | COVID-19 Genomics UK (COG-UK) Consortium | Luke W Meredith, M. Estee Torok , Myra Hosmillo, William L. Hamilton, Martin D. Curran, Theresa Feltwell, Grant Hall, Anna Yakovleva, Fahad A Khokhar, Charlotte J. Houldcroft, Laura G Caller, Aminu S. Jahun, Sarah L. Caddy, Ian Goodfellow |
| EPI_ISL_434021 | hCoV-19/England/CAMB-7C2AF/2020 | Europe / United Kingdom / England | 2020-04-14 | Department of Pathology, University of Cambridge | COVID-19 Genomics UK (COG-UK) Consortium | Luke W Meredith, M. Estee Torok , Myra Hosmillo, William L. Hamilton, Martin D. Curran, Theresa Feltwell, Grant Hall, Anna Yakovleva, Fahad A Khokhar, Charlotte J. Houldcroft, Laura G Caller, Aminu S. Jahun, Sarah L. Caddy, Ian Goodfellow |
| EPI_ISL_434022 | hCoV-19/England/CAMB-7C2BE/2020 | Europe / United Kingdom / England | 2020-04-14 | Department of Pathology, University of Cambridge | COVID-19 Genomics UK (COG-UK) Consortium | Luke W Meredith, M. Estee Torok , Myra Hosmillo, William L. Hamilton, Martin D. Curran, Theresa Feltwell, Grant Hall, Anna Yakovleva, Fahad A Khokhar, Charlotte J. Houldcroft, Laura G Caller, Aminu S. Jahun, Sarah L. Caddy, Ian Goodfellow |
| EPI_ISL_434024 | hCoV-19/England/CAMB-7C2DC/2020 | Europe / United Kingdom / England | 2020-04-15 | Department of Pathology, University of Cambridge | COVID-19 Genomics UK (COG-UK) Consortium | Luke W Meredith, M. Estee Torok , Myra Hosmillo, William L. Hamilton, Martin D. Curran, Theresa Feltwell, Grant Hall, Anna Yakovleva, Fahad A Khokhar, Charlotte J. Houldcroft, Laura G Caller, Aminu S. Jahun, Sarah L. Caddy, Ian Goodfellow |
| EPI_ISL_434027 | hCoV-19/England/CAMB-7C315/2020 | Europe / United Kingdom / England | 2020-04-14 | Department of Pathology, University of Cambridge | COVID-19 Genomics UK (COG-UK) Consortium | Luke W Meredith, M. Estee Torok , Myra Hosmillo, William L. Hamilton, Martin D. Curran, Theresa Feltwell, Grant Hall, Anna Yakovleva, Fahad A Khokhar, Charlotte J. Houldcroft, Laura G Caller, Aminu S. Jahun, Sarah L. Caddy, Ian Goodfellow |
| EPI_ISL_434029 | hCoV-19/England/CAMB-7C333/2020 | Europe / United Kingdom / England | 2020-04-14 | Department of Pathology, University of Cambridge | COVID-19 Genomics UK (COG-UK) Consortium | Luke W Meredith, M. Estee Torok , Myra Hosmillo, William L. Hamilton, Martin D. Curran, Theresa Feltwell, Grant Hall, Anna Yakovleva, Fahad A Khokhar, Charlotte J. Houldcroft, Laura G Caller, Aminu S. Jahun, Sarah L. Caddy, Ian Goodfellow |
| EPI_ISL_434032 | hCoV-19/England/CAMB-7C360/2020 | Europe / United Kingdom / England | 2020-04-14 | Department of Pathology, University of Cambridge | COVID-19 Genomics UK (COG-UK) Consortium | Luke W Meredith, M. Estee Torok , Myra Hosmillo, William L. Hamilton, Martin D. Curran, Theresa Feltwell, Grant Hall, Anna Yakovleva, Fahad A Khokhar, Charlotte J. Houldcroft, Laura G Caller, Aminu S. Jahun, Sarah L. Caddy, Ian Goodfellow |
| EPI_ISL_434034 | hCoV-19/England/CAMB-7C38E/2020 | Europe / United Kingdom / England | 2020-04-14 | Department of Pathology, University of Cambridge | COVID-19 Genomics UK (COG-UK) Consortium | Luke W Meredith, M. Estee Torok , Myra Hosmillo, William L. Hamilton, Martin D. Curran, Theresa Feltwell, Grant Hall, Anna Yakovleva, Fahad A Khokhar, Charlotte J. Houldcroft, Laura G Caller, Aminu S. Jahun, Sarah L. Caddy, Ian Goodfellow |
| EPI_ISL_434035 | hCoV-19/England/CAMB-7C39D/2020 | Europe / United Kingdom / England | 2020-04-13 | Department of Pathology, University of Cambridge | COVID-19 Genomics UK (COG-UK) Consortium | Luke W Meredith, M. Estee Torok , Myra Hosmillo, William L. Hamilton, Martin D. Curran, Theresa Feltwell, Grant Hall, Anna Yakovleva, Fahad A Khokhar, Charlotte J. Houldcroft, Laura G Caller, Aminu S. Jahun, Sarah L. Caddy, Ian Goodfellow |
| EPI_ISL_434036 | hCoV-19/England/CAMB-7C3AC/2020 | Europe / United Kingdom / England | 2020-04-14 | Department of Pathology, University of Cambridge | COVID-19 Genomics UK (COG-UK) Consortium | Luke W Meredith, M. Estee Torok , Myra Hosmillo, William L. Hamilton, Martin D. Curran, Theresa Feltwell, Grant Hall, Anna Yakovleva, Fahad A Khokhar, Charlotte J. Houldcroft, Laura G Caller, Aminu S. Jahun, Sarah L. Caddy, Ian Goodfellow |
| EPI_ISL_434038 | hCoV-19/England/CAMB-7C3CA/2020 | Europe / United Kingdom / England | 2020-04-14 | Department of Pathology, University of Cambridge | COVID-19 Genomics UK (COG-UK) Consortium | Luke W Meredith, M. Estee Torok , Myra Hosmillo, William L. Hamilton, Martin D. Curran, Theresa Feltwell, Grant Hall, Anna Yakovleva, Fahad A Khokhar, Charlotte J. Houldcroft, Laura G Caller, Aminu S. Jahun, Sarah L. Caddy, Ian Goodfellow |
| EPI_ISL_434041 | hCoV-19/England/CAMB-7E119/2020 | Europe / United Kingdom / England | 2020-04-16 | Department of Pathology, University of Cambridge | COVID-19 Genomics UK (COG-UK) Consortium | Luke W Meredith, M. Estee Torok , Myra Hosmillo, William L. Hamilton, Martin D. Curran, Theresa Feltwell, Grant Hall, Anna Yakovleva, Fahad A Khokhar, Charlotte J. Houldcroft, Laura G Caller, Aminu S. Jahun, Sarah L. Caddy, Ian Goodfellow |
| EPI_ISL_434045 | hCoV-19/England/CAMB-7E155/2020 | Europe / United Kingdom / England | 2020-04-16 | Department of Pathology, University of Cambridge | COVID-19 Genomics UK (COG-UK) Consortium | Luke W Meredith, M. Estee Torok , Myra Hosmillo, William L. Hamilton, Martin D. Curran, Theresa Feltwell, Grant Hall, Anna Yakovleva, Fahad A Khokhar, Charlotte J. Houldcroft, Laura G Caller, Aminu S. Jahun, Sarah L. Caddy, Ian Goodfellow |
| EPI_ISL_434046 | hCoV-19/England/CAMB-7E164/2020 | Europe / United Kingdom / England | 2020-04-15 | Department of Pathology, University of Cambridge | COVID-19 Genomics UK (COG-UK) Consortium | Luke W Meredith, M. Estee Torok , Myra Hosmillo, William L. Hamilton, Martin D. Curran, Theresa Feltwell, Grant Hall, Anna Yakovleva, Fahad A Khokhar, Charlotte J. Houldcroft, Laura G Caller, Aminu S. Jahun, Sarah L. Caddy, Ian Goodfellow |
| EPI_ISL_434047 | hCoV-19/England/CAMB-7E1BF/2020 | Europe / United Kingdom / England | 2020-04-16 | Department of Pathology, University of Cambridge | COVID-19 Genomics UK (COG-UK) Consortium | Luke W Meredith, M. Estee Torok , Myra Hosmillo, William L. Hamilton, Martin D. Curran, Theresa Feltwell, Grant Hall, Anna Yakovleva, Fahad A Khokhar, Charlotte J. Houldcroft, Laura G Caller, Aminu S. Jahun, Sarah L. Caddy, Ian Goodfellow |
| EPI_ISL_434049 | hCoV-19/England/CAMB-7E1DD/2020 | Europe / United Kingdom / England | 2020-04-16 | Department of Pathology, University of Cambridge | COVID-19 Genomics UK (COG-UK) Consortium | Luke W Meredith, M. Estee Torok , Myra Hosmillo, William L. Hamilton, Martin D. Curran, Theresa Feltwell, Grant Hall, Anna Yakovleva, Fahad A Khokhar, Charlotte J. Houldcroft, Laura G Caller, Aminu S. Jahun, Sarah L. Caddy, Ian Goodfellow |
| EPI_ISL_434051 | hCoV-19/England/CAMB-7E1FB/2020 | Europe / United Kingdom / England | 2020-04-16 | Department of Pathology, University of Cambridge | COVID-19 Genomics UK (COG-UK) Consortium | Luke W Meredith, M. Estee Torok , Myra Hosmillo, William L. Hamilton, Martin D. Curran, Theresa Feltwell, Grant Hall, Anna Yakovleva, Fahad A Khokhar, Charlotte J. Houldcroft, Laura G Caller, Aminu S. Jahun, Sarah L. Caddy, Ian Goodfellow |
| EPI_ISL_434054 | hCoV-19/England/CAMB-7E234/2020 | Europe / United Kingdom / England | 2020-04-15 | Department of Pathology, University of Cambridge | COVID-19 Genomics UK (COG-UK) Consortium | Luke W Meredith, M. Estee Torok , Myra Hosmillo, William L. Hamilton, Martin D. Curran, Theresa Feltwell, Grant Hall, Anna Yakovleva, Fahad A Khokhar, Charlotte J. Houldcroft, Laura G Caller, Aminu S. Jahun, Sarah L. Caddy, Ian Goodfellow |
| EPI_ISL_434063 | hCoV-19/USA/WA-S303/2020 | North America / USA / Washington / Yaki | 2020-04-03 | Washington State Department of Health | Seattle Flu Study | Chu et al |
| EPI_ISL_434064 | hCoV-19/USA/WA-S304/2020 | North America / USA / Washington / Yaki | 2020-04-05 | Washington State Department of Health | Seattle Flu Study | Chu et al |
| EPI_ISL_434065 | hCoV-19/USA/WA-S305/2020 | North America / USA / Washington / Pier | 2020-04-01 | Washington State Department of Health | Seattle Flu Study | Chu et al |
| EPI_ISL_434066 | hCoV-19/USA/WA-S306/2020 | North America / USA / Washington / King | 2020-04-02 | Washington State Department of Health | Seattle Flu Study | Chu et al |
| EPI_ISL_434067 | hCoV-19/USA/WA-S307/2020 | North America / USA / Washington | 2020-03-28 | Washington State Department of Health | Seattle Flu Study | Chu et al |
| EPI_ISL_434068 | hCoV-19/USA/WA-S308/2020 | North America / USA / Washington / Yaki | 2020-04-02 | Washington State Department of Health | Seattle Flu Study | Chu et al |
| EPI_ISL_434069 | hCoV-19/USA/WA-S309/2020 | North America / USA / Washington / Asot | 2020-04-03 | Washington State Department of Health | Seattle Flu Study | Chu et al |
| EPI_ISL_434070 | hCoV-19/USA/WA-S310/2020 | North America / USA / Washington / Bent | 2020-04-01 | Washington State Department of Health | Seattle Flu Study | Chu et al |
| EPI_ISL_434071 | hCoV-19/USA/WA-S311/2020 | North America / USA / Washington / Yaki | 2020-04-04 | Washington State Department of Health | Seattle Flu Study | Chu et al |
| EPI_ISL_434072 | hCoV-19/USA/WA-S312/2020 | North America / USA / Washington / Yaki | 2020-04-01 | Washington State Department of Health | Seattle Flu Study | Chu et al |
| EPI_ISL_434073 | hCoV-19/USA/WA-S313/2020 | North America / USA / Washington / Yaki | 2020-04-02 | Washington State Department of Health | Seattle Flu Study | Chu et al |
| EPI_ISL_434074 | hCoV-19/USA/WA-S314/2020 | North America / USA / Washington / King | 2020-04-03 | Washington State Department of Health | Seattle Flu Study | Chu et al |
| EPI_ISL_434075 | hCoV-19/USA/WA-S315/2020 | North America / USA / Washington / Yaki | 2020-04-02 | Washington State Department of Health | Seattle Flu Study | Chu et al |
| EPI_ISL_434076 | hCoV-19/USA/WA-S316/2020 | North America / USA / Washington / Fran | 2020-04-01 | Washington State Department of Health | Seattle Flu Study | Chu et al |
| EPI_ISL_434077 | hCoV-19/USA/WA-S317/2020 | North America / USA / Washington / Yaki | 2020-04-04 | Washington State Department of Health | Seattle Flu Study | Chu et al |
| EPI_ISL_434079 | hCoV-19/USA/WA-S319/2020 | North America / USA / Washington / Yaki | 2020-04-04 | Washington State Department of Health | Seattle Flu Study | Chu et al |
| EPI_ISL_434080 | hCoV-19/USA/WA-S320/2020 | North America / USA / Washington / Fran | 2020-04-01 | Washington State Department of Health | Seattle Flu Study | Chu et al |
| EPI_ISL_434081 | hCoV-19/USA/WA-S321/2020 | North America / USA / Washington / Yaki | 2020-03-30 | Washington State Department of Health | Seattle Flu Study | Chu et al |
| EPI_ISL_434082 | hCoV-19/USA/WA-S322/2020 | North America / USA / Washington / Yaki | 2020-03-31 | Washington State Department of Health | Seattle Flu Study | Chu et al |
| EPI_ISL_434083 | hCoV-19/USA/WA-S323/2020 | North America / USA / Washington / Yaki | 2020-04-02 | Washington State Department of Health | Seattle Flu Study | Chu et al |
| EPI_ISL_434084 | hCoV-19/USA/WA-S324/2020 | North America / USA / Washington / Yaki | 2020-04-01 | Washington State Department of Health | Seattle Flu Study | Chu et al |
| EPI_ISL_434085 | hCoV-19/USA/WA-S325/2020 | North America / USA / Washington / Yaki | 2020-04-03 | Washington State Department of Health | Seattle Flu Study | Chu et al |
| EPI_ISL_434086 | hCoV-19/USA/WA-S326/2020 | North America / USA / Washington / Adar | 2020-04-01 | Washington State Department of Health | Seattle Flu Study | Chu et al |
| EPI_ISL_434087 | hCoV-19/USA/WA-S327/2020 | North America / USA / Washington / Yaki | 2020-04-05 | Washington State Department of Health | Seattle Flu Study | Chu et al |
| EPI_ISL_434088 | hCoV-19/USA/WA-S328/2020 | North America / USA / Washington / Yaki | 2020-04-03 | Washington State Department of Health | Seattle Flu Study | Chu et al |
| EPI_ISL_434089 | hCoV-19/USA/WA-S329/2020 | North America / USA / Washington / Yaki | 2020-04-01 | Washington State Department of Health | Seattle Flu Study | Chu et al |
| EPI_ISL_434090 | hCoV-19/USA/WA-S330/2020 | North America / USA / Washington / Yaki | 2020-04-02 | Washington State Department of Health | Seattle Flu Study | Chu et al |
| EPI_ISL_434091 | hCoV-19/USA/WA-S331/2020 | North America / USA / Washington / Pier | 2020-04-02 | Washington State Department of Health | Seattle Flu Study | Chu et al |
| EPI_ISL_434092 | hCoV-19/USA/WA-S332/2020 | North America / USA / Washington / Wha | 2020-04-02 | Washington State Department of Health | Seattle Flu Study | Chu et al |
| EPI_ISL_434093 | hCoV-19/USA/WA-S333/2020 | North America / USA / Washington / Yaki | 2020-04-04 | Washington State Department of Health | Seattle Flu Study | Chu et al |
| EPI_ISL_434094 | hCoV-19/USA/WA-S334/2020 | North America / USA / Washington / Yaki | 2020-04-04 | Washington State Department of Health | Seattle Flu Study | Chu et al |
| EPI_ISL_434095 | hCoV-19/USA/WA-S335/2020 | North America / USA / Washington / Yaki | 2020-04-02 | Washington State Department of Health | Seattle Flu Study | Chu et al |
| EPI_ISL_434096 | hCoV-19/USA/WA-S336/2020 | North America / USA / Washington / Pier | 2020-04-03 | Washington State Department of Health | Seattle Flu Study | Chu et al |
| EPI_ISL_434097 | hCoV-19/USA/WA-S337/2020 | North America / USA / Washington / Pier | 2020-04-01 | Washington State Department of Health | Seattle Flu Study | Chu et al |
| EPI_ISL_434098 | hCoV-19/USA/WA-S338/2020 | North America / USA / Washington / Yaki | 2020-04-02 | Washington State Department of Health | Seattle Flu Study | Chu et al |
| EPI_ISL_434099 | hCoV-19/USA/WA-S339/2020 | North America / USA / Washington / Yaki | 2020-04-02 | Washington State Department of Health | Seattle Flu Study | Chu et al |
| EPI_ISL_434300 | hCoV-19/USA/WA-S540/2020 | North America / USA / Washington / Yaki | 2020-03-24 | Washington State Department of Health | Seattle Flu Study | Chu et al |
| EPI_ISL_434301 | hCoV-19/USA/WA-S541/2020 | North America / USA / Washington / Wha | 2020-03-24 | Washington State Department of Health | Seattle Flu Study | Chu et al |
| EPI_ISL_434302 | hCoV-19/USA/WA-S542/2020 | North America / USA / Washington / Adar | 2020-03-23 | Washington State Department of Health | Seattle Flu Study | Chu et al |
| EPI_ISL_434303 | hCoV-19/USA/WA-S543/2020 | North America / USA / Washington / Yaki | 2020-03-24 | Washington State Department of Health | Seattle Flu Study | Chu et al |
| EPI_ISL_434304 | hCoV-19/USA/WA-S544/2020 | North America / USA / Washington | 2020-03-25 | Washington State Department of Health | Seattle Flu Study | Chu et al |
| EPI_ISL_434305 | hCoV-19/USA/WA-S545/2020 | North America / USA / Washington / King | 2020-03-31 | Washington State Department of Health | Seattle Flu Study | Chu et al |
| EPI_ISL_434306 | hCoV-19/USA/WA-S546/2020 | North America / USA / Washington / Adar | 2020-03-23 | Washington State Department of Health | Seattle Flu Study | Chu et al |
| EPI_ISL_434307 | hCoV-19/USA/WA-S547/2020 | North America / USA / Washington / Yaki | 2020-03-24 | Washington State Department of Health | Seattle Flu Study | Chu et al |
| EPI_ISL_434308 | hCoV-19/USA/WA-S548/2020 | North America / USA / Washington / Pier | 2020-03-31 | Washington State Department of Health | Seattle Flu Study | Chu et al |
| EPI_ISL_434309 | hCoV-19/USA/WA-S549/2020 | North America / USA / Washington / Yaki | 2020-03-30 | Washington State Department of Health | Seattle Flu Study | Chu et al |
| EPI_ISL_434310 | hCoV-19/USA/WA-S550/2020 | North America / USA / Washington / Yaki | 2020-03-24 | Washington State Department of Health | Seattle Flu Study | Chu et al |
| EPI_ISL_434311 | hCoV-19/USA/WA-S551/2020 | North America / USA / Washington / Yaki | 2020-03-24 | Washington State Department of Health | Seattle Flu Study | Chu et al |
| EPI_ISL_434312 | hCoV-19/USA/WA-S552/2020 | North America / USA / Washington | 2020-03-23 | Washington State Department of Health | Seattle Flu Study | Chu et al |
| EPI_ISL_434313 | hCoV-19/USA/WA-S553/2020 | North America / USA / Washington / Pier | 2020-03-31 | Washington State Department of Health | Seattle Flu Study | Chu et al |
| EPI_ISL_434314 | hCoV-19/USA/WA-S554/2020 | North America / USA / Washington / King | 2020-03-31 | Washington State Department of Health | Seattle Flu Study | Chu et al |
| EPI_ISL_434315 | hCoV-19/USA/WA-S555/2020 | North America / USA / Washington / Adar | 2020-03-30 | Washington State Department of Health | Seattle Flu Study | Chu et al |
| EPI_ISL_434316 | hCoV-19/USA/WA-S556/2020 | North America / USA / Washington / Yaki | 2020-03-29 | Washington State Department of Health | Seattle Flu Study | Chu et al |
| EPI_ISL_434317 | hCoV-19/USA/WA-S557/2020 | North America / USA / Washington / Yaki | 2020-04-01 | Washington State Department of Health | Seattle Flu Study | Chu et al |
| EPI_ISL_434318 | hCoV-19/USA/WA-S558/2020 | North America / USA / Washington | 2020-03-30 | Washington State Department of Health | Seattle Flu Study | Chu et al |
| EPI_ISL_434319 | hCoV-19/USA/WA-S559/2020 | North America / USA / Washington / Snol | 2020-03-27 | Washington State Department of Health | Seattle Flu Study | Chu et al |
| EPI_ISL_434320 | hCoV-19/USA/WA-S560/2020 | North America / USA / Washington / Yaki | 2020-03-24 | Washington State Department of Health | Seattle Flu Study | Chu et al |
| EPI_ISL_434321 | hCoV-19/USA/WA-S561/2020 | North America / USA / Washington / King | 2020-03-30 | Washington State Department of Health | Seattle Flu Study | Chu et al |
| EPI_ISL_434322 | hCoV-19/USA/WA-S562/2020 | North America / USA / Washington / Wha | 2020-03-24 | Washington State Department of Health | Seattle Flu Study | Chu et al |
| EPI_ISL_434323 | hCoV-19/USA/WA-S563/2020 | North America / USA / Washington / Pier | 2020-03-31 | Washington State Department of Health | Seattle Flu Study | Chu et al |
| EPI_ISL_434324 | hCoV-19/USA/WA-S564/2020 | North America / USA / Washington / Yaki | 2020-03-24 | Washington State Department of Health | Seattle Flu Study | Chu et al |
| EPI_ISL_434326 | hCoV-19/USA/WA-S566/2020 | North America / USA / Washington | 2020-03-29 | Washington State Department of Health | Seattle Flu Study | Chu et al |
| EPI_ISL_434327 | hCoV-19/USA/WA-S567/2020 | North America / USA / Washington / King | 2020-03-25 | Washington State Department of Health | Seattle Flu Study | Chu et al |
| EPI_ISL_434328 | hCoV-19/USA/WA-S568/2020 | North America / USA / Washington / Pier | 2020-03-31 | Washington State Department of Health | Seattle Flu Study | Chu et al |
| EPI_ISL_434329 | hCoV-19/USA/WA-S569/2020 | North America / USA / Washington / Grar | 2020-03-25 | Washington State Department of Health | Seattle Flu Study | Chu et al |
| EPI_ISL_434330 | hCoV-19/USA/WA-S570/2020 | North America / USA / Washington / Pier | 2020-03-24 | Washington State Department of Health | Seattle Flu Study | Chu et al |
| EPI_ISL_434331 | hCoV-19/USA/WA-S571/2020 | North America / USA / Washington / King | 2020-03-30 | Washington State Department of Health | Seattle Flu Study | Chu et al |
| EPI_ISL_434332 | hCoV-19/USA/WA-S572/2020 | North America / USA / Washington / King | 2020-03-26 | Washington State Department of Health | Seattle Flu Study | Chu et al |
| EPI_ISL_434334 | hCoV-19/USA/WA-S574/2020 | North America / USA / Washington / Pier | 2020-03-31 | Washington State Department of Health | Seattle Flu Study | Chu et al |
| EPI_ISL_434335 | hCoV-19/USA/WA-S575/2020 | North America / USA / Washington / Pier | 2020-03-31 | Washington State Department of Health | Seattle Flu Study | Chu et al |
| EPI_ISL_434336 | hCoV-19/USA/WA-S576/2020 | North America / USA / Washington / Pier | 2020-03-29 | Washington State Department of Health | Seattle Flu Study | Chu et al |
| EPI_ISL_434337 | hCoV-19/USA/WA-S577/2020 | North America / USA / Washington / Yaki | 2020-03-28 | Washington State Department of Health | Seattle Flu Study | Chu et al |
| EPI_ISL_434338 | hCoV-19/USA/WA-S578/2020 | North America / USA / Washington / Yaki | 2020-03-30 | Washington State Department of Health | Seattle Flu Study | Chu et al |
| EPI_ISL_434339 | hCoV-19/USA/WA-S579/2020 | North America / USA / Washington / Kittit | 2020-03-24 | Washington State Department of Health | Seattle Flu Study | Chu et al |
| EPI_ISL_434340 | hCoV-19/USA/WA-S580/2020 | North America / USA / Washington | 2020-03-25 | Washington State Department of Health | Seattle Flu Study | Chu et al |
| EPI_ISL_434341 | hCoV-19/USA/WA-S581/2020 | North America / USA / Washington / Yaki | 2020-03-30 | Washington State Department of Health | Seattle Flu Study | Chu et al |
| EPI_ISL_434342 | hCoV-19/USA/WA-S582/2020 | North America / USA / Washington / Yaki | 2020-03-31 | Washington State Department of Health | Seattle Flu Study | Chu et al |
| EPI_ISL_434343 | hCoV-19/USA/WA-S583/2020 | North America / USA / Washington / King | 2020-03-24 | Washington State Department of Health | Seattle Flu Study | Chu et al |
| EPI_ISL_434345 | hCoV-19/USA/WA-S585/2020 | North America / USA / Washington / Yaki | 2020-03-26 | Washington State Department of Health | Seattle Flu Study | Chu et al |
| EPI_ISL_434346 | hCoV-19/USA/WA-S586/2020 | North America / USA / Washington / Pier | 2020-03-31 | Washington State Department of Health | Seattle Flu Study | Chu et al |
| EPI_ISL_434347 | hCoV-19/Belgium/UGent-28/2020 | Europe / Belgium / Ghent | 2020-03-26 | Lab voor klinische biologie | Onderzoeksgroep Virologie | Laurens Lambrechts, Nick Vereecke, Marthe Pauwels, Jozefien De Clercq, Bruno Verhasselt, Linos Vandekerckhove, Hans Nauwynck, Sebastiaan Theuns |
| EPI_ISL_434348 | hCoV-19/Belgium/UGent-30/2020 | Europe / Belgium / Ghent | 2020-03-26 | Lab voor klinische biologie | Onderzoeksgroep Virologie | Laurens Lambrechts, Nick Vereecke, Marthe Pauwels, Jozefien De Clercq, Bruno Verhasselt, Linos Vandekerckhove, Hans Nauwynck, Sebastiaan Theuns |
| EPI_ISL_434349 | hCoV-19/Belgium/UGent-31/2020 | Europe / Belgium / Ghent | 2020-03-26 | Lab voor klinische biologie | Onderzoeksgroep Virologie | Laurens Lambrechts, Nick Vereecke, Marthe Pauwels, Jozefien De Clercq, Bruno Verhasselt, Linos Vandekerckhove, Hans Nauwynck, Sebastiaan Theuns |
| EPI_ISL_434351 | hCoV-19/Belgium/UGent-33/2020 | Europe / Belgium / Ghent | 2020-03-28 | Lab voor klinische biologie | Onderzoeksgroep Virologie | Laurens Lambrechts, Nick Vereecke, Marthe Pauwels, Jozefien De Clercq, Bruno Verhasselt, Linos Vandekerckhove, Hans Nauwynck, Sebastiaan Theuns |
| EPI_ISL_434352 | hCoV-19/Belgium/UGent-34/2020 | Europe / Belgium / Ghent | 2020-03-26 | Lab voor klinische biologie | Onderzoeksgroep Virologie | Laurens Lambrechts, Nick Vereecke, Marthe Pauwels, Jozefien De Clercq, Bruno Verhasselt, Linos Vandekerckhove, Hans Nauwynck, Sebastiaan Theuns |
| EPI_ISL_434355 | hCoV-19/Belgium/UGent-37/2020 | Europe / Belgium / Ghent | 2020-03-27 | Lab voor klinische biologie | Onderzoeksgroep Virologie | Laurens Lambrechts, Nick Vereecke, Marthe Pauwels, Jozefien De Clercq, Bruno Verhasselt, Linos Vandekerckhove, Hans Nauwynck, Sebastiaan Theuns |
| EPI_ISL_434357 | hCoV-19/Belgium/UGent-40/2020 | Europe / Belgium / Ghent | 2020-03-27 | Lab voor klinische biologie | Onderzoeksgroep Virologie | Nick Vereecke, Laurens Lambrechts, Marthe Pauwels, Jozefien De Clercq, Bruno Verhasselt, Linos Vandekerckhove, Hans Nauwynck, Sebastiaan Theuns |
| EPI_ISL_434360 | hCoV-19/Belgium/UGent-44/2020 | Europe / Belgium / Ghent | 2020-03-29 | Lab voor klinische biologie | Onderzoeksgroep Virologie | Nick Vereecke, Laurens Lambrechts, Marthe Pauwels, Jozefien De Clercq, Bruno Verhasselt, Linos Vandekerckhove, Hans Nauwynck, Sebastiaan Theuns |
| EPI_ISL_434361 | hCoV-19/Belgium/UGent-45/2020 | Europe / Belgium / Ghent | 2020-03-29 | Lab voor klinische biologie | Onderzoeksgroep Virologie | Nick Vereecke, Laurens Lambrechts, Marthe Pauwels, Jozefien De Clercq, Bruno Verhasselt, Linos Vandekerckhove, Hans Nauwynck, Sebastiaan Theuns |
| EPI_ISL_434362 | hCoV-19/Belgium/UGent-46/2020 | Europe / Belgium / Ghent | 2020-03-29 | Lab voor klinische biologie | Onderzoeksgroep Virologie | Nick Vereecke, Laurens Lambrechts, Marthe Pauwels, Jozefien De Clercq, Bruno Verhasselt, Linos Vandekerckhove, Hans Nauwynck, Sebastiaan Theuns |
| EPI_ISL_434368 | hCoV-19/Belgium/ITM_C211/2020 | Europe / Belgium / Antwerp | 2020-03-31 | Hospital AZ Rivierenland | Institute of Tropical Medicine | Philippe Selhorst, Colin Anthony |
| EPI_ISL_434369 | hCoV-19/Belgium/ITM_C262/2020 | Europe / Belgium / Antwerp | 2020-04-02 | Hospital AZ Rivierenland | Institute of Tropical Medicine | Philippe Selhorst, Colin Anthony |
| EPI_ISL_434372 | hCoV-19/Belgium/ITM_C053/2020 | Europe / Belgium / Antwerp | 2020-03-26 | Hospital AZ Rivierenland | Institute of Tropical Medicine | Philippe Selhorst, Colin Anthony, |
| EPI_ISL_434373 | hCoV-19/Belgium/ITM_C147/2020 | Europe / Belgium / Antwerp | 2020-03-28 | Hospital AZ Rivierenland | Institute of Tropical Medicine | Philippe Selhorst, Colin Anthony, |
| EPI_ISL_450442 | hCoV-19/Henan/HN03/2020 | Asia / China / Henan | 2020-02-24 | unknown | The Department of Infectious Disease Prevention and Control | Li,X., Lu,S., Wu,B., Hu,X., Li,D., Huang,X. and Guo,W. |
| EPI_ISL_426285 | hCoV-19/Latvia/05/2020 | Europe / Latvia | 2020-03-30 | E. Gulbja Laboratorija | Latvian Biomedical Research and Study Centre | Ivars Silamiķelis, Kaspars Megnis, Monta Ustinova, Ņikita Zrelovs, Vita Rovīte, Mikus Gavars, Dmitrijs Perminovs, Uga Dumpis, Jānis Kloviņš |
| EPI_ISL_426286 | hCoV-19/Latvia/06/2020 | Europe / Latvia | 2020-03-30 | E. Gulbja Laboratorija | Latvian Biomedical Research and Study Centre | Ivars Silamiķelis, Kaspars Megnis, Monta Ustinova, Ņikita Zrelovs, Vita Rovīte, Mikus Gavars, Dmitrijs Perminovs, Uga Dumpis, Jānis Kloviņš |
| EPI_ISL_426287 | hCoV-19/Latvia/07/2020 | Europe / Latvia | 2020-03-30 | E. Gulbja Laboratorija | Latvian Biomedical Research and Study Centre | Ivars Silamiķelis, Kaspars Megnis, Monta Ustinova, Ņikita Zrelovs, Vita Rovīte, Mikus Gavars, Dmitrijs Perminovs, Uga Dumpis, Jānis Kloviņš |
| EPI_ISL_426288 | hCoV-19/Latvia/08/2020 | Europe / Latvia | 2020-03-30 | E. Gulbja Laboratorija | Latvian Biomedical Research and Study Centre | Ivars Silamiķelis, Kaspars Megnis, Monta Ustinova, Ņikita Zrelovs, Vita Rovīte, Mikus Gavars, Dmitrijs Perminovs, Uga Dumpis, Jānis Kloviņš |
| EPI_ISL_426289 | hCoV-19/Latvia/09/2020 | Europe / Latvia | 2020-03-30 | E. Gulbja Laboratorija | Latvian Biomedical Research and Study Centre | Ivars Silamiķelis, Kaspars Megnis, Monta Ustinova, Ņikita Zrelovs, Vita Rovīte, Mikus Gavars, Dmitrijs Perminovs, Uga Dumpis, Jānis Kloviņš |
| EPI_ISL_426290 | hCoV-19/USA/NY-Wadsworth-11735-01/2020 | North America / USA / New York / Westc | 2020-03-08 | Wadsworth Center, New York State Department of Health | Wadsworth Center, New York State Department.of Health | Kirsten St. George, Daryl M. Lamson, Sara Griesemer, Jonathan Plitnick, Navjot Singh, Matthew D. Shudt, Erica Lasek-Nesselquist |
| EPI_ISL_426291 | hCoV-19/USA/NY-Wadsworth-11717-01/2020 | North America / USA / New York / Westc | 2020-03-08 | Wadsworth Center, New York State Department of Health | Wadsworth Center, New York State Department.of Health | Kirsten St. George, Daryl M. Lamson, Sara Griesemer, Jonathan Plitnick, Navjot Singh, Matthew D. Shudt, Erica Lasek-Nesselquist |
| EPI_ISL_426292 | hCoV-19/USA/NY-Wadsworth-11207-01/2020 | North America / USA / New York / Sarato | 2020-03-06 | Wadsworth Center, New York State Department of Health | Wadsworth Center, New York State Department.of Health | Kirsten St. George, Daryl M. Lamson, Sara Griesemer, Jonathan Plitnick, Navjot Singh, Matthew D. Shudt, Erica Lasek-Nesselquist |
| EPI_ISL_426293 | hCoV-19/USA/NY-Wadsworth-12883-02/2020 | North America / USA / New York / Nassa | 2020-03-10 | Wadsworth Center, New York State Department of Health | Wadsworth Center, New York State Department.of Health | Kirsten St. George, Daryl M. Lamson, Sara Griesemer, Jonathan Plitnick, Navjot Singh, Matthew D. Shudt, Erica Lasek-Nesselquist |
| EPI_ISL_426294 | hCoV-19/USA/NY-Wadsworth-13468-02/2020 | North America / USA / New York / Suffolk | 2020-03-11 | Wadsworth Center, New York State Department of Health | Wadsworth Center, New York State Department.of Health | Kirsten St. George, Daryl M. Lamson, Sara Griesemer, Jonathan Plitnick, Navjot Singh, Matthew D. Shudt, Erica Lasek-Nesselquist |
| EPI_ISL_426295 | hCoV-19/USA/NY-Wadsworth-27073-01/2020 | North America / USA / New York / Queen | 2020-03-22 | Wadsworth Center, New York State Department of Health | Wadsworth Center, New York State Department.of Health | Kirsten St. George, Daryl M. Lamson, Sara Griesemer, Jonathan Plitnick, Navjot Singh, Matthew D. Shudt, Erica Lasek-Nesselquist |
| EPI_ISL_426296 | hCoV-19/USA/NY-Wadsworth-16080-01/2020 | North America / USA / New York / Orang | 2020-03-14 | Wadsworth Center, New York State Department of Health | Wadsworth Center, New York State Department.of Health | Kirsten St. George, Daryl M. Lamson, Sara Griesemer, Jonathan Plitnick, Navjot Singh, Matthew D. Shudt, Erica Lasek-Nesselquist |
| EPI_ISL_426297 | hCoV-19/USA/NY-Wadsworth-27804-01/2020 | North America / USA / New York / Schen | 2020-03-16 | Wadsworth Center, New York State Department of Health | Wadsworth Center, New York State Department.of Health | Kirsten St. George, Daryl M. Lamson, Sara Griesemer, Jonathan Plitnick, Navjot Singh, Matthew D. Shudt, Erica Lasek-Nesselquist |
| EPI_ISL_426298 | hCoV-19/USA/NY-Wadsworth-31488-01/2020 | North America / USA / New York / Frankl | 2020-03-27 | Wadsworth Center, New York State Department of Health | Wadsworth Center, New York State Department.of Health | Kirsten St. George, Daryl M. Lamson, Sara Griesemer, Jonathan Plitnick, Navjot Singh, Matthew D. Shudt, Erica Lasek-Nesselquist |
| EPI_ISL_426299 | hCoV-19/USA/NY-Wadsworth-32851-01/2020 | North America / USA / New York / Onond | 2020-03-28 | Wadsworth Center, New York State Department of Health | Wadsworth Center, New York State Department.of Health | Kirsten St. George, Daryl M. Lamson, Sara Griesemer, Jonathan Plitnick, Navjot Singh, Matthew D. Shudt, Erica Lasek-Nesselquist |
| EPI_ISL_426500 | hCoV-19/USA/AZ-TG268099/2020 | North America / USA / Arizona | 2020-03-17 | TGen North | TGen North | Jolene Bowers, Megan Folkerts, Darrin Lemmer, Dave Engelthaler |
| EPI_ISL_426501 | hCoV-19/USA/AZ-TG268183/2020 | North America / USA / Arizona | 2020-03-17 | TGen North | TGen North | Jolene Bowers, Megan Folkerts, Darrin Lemmer, Dave Engelthaler |
| EPI_ISL_426502 | hCoV-19/USA/AZ-TG268188/2020 | North America / USA / Arizona | 2020-03-17 | TGen North | TGen North | Jolene Bowers, Megan Folkerts, Darrin Lemmer, Dave Engelthaler |
| EPI_ISL_426503 | hCoV-19/USA/AZ-TG268281/2020 | North America / USA / Arizona | 2020-03-17 | TGen North | TGen North | Jolene Bowers, Megan Folkerts, Darrin Lemmer, Dave Engelthaler |
| EPI_ISL_426504 | hCoV-19/USA/AZ-TG268282/2020 | North America / USA / Arizona | 2020-03-17 | TGen North | TGen North | Jolene Bowers, Megan Folkerts, Darrin Lemmer, Dave Engelthaler |
| EPI_ISL_426505 | hCoV-19/USA/AZ-TG268289/2020 | North America / USA / Arizona | 2020-03-17 | TGen North | TGen North | Jolene Bowers, Megan Folkerts, Darrin Lemmer, Dave Engelthaler |
| EPI_ISL_426506 | hCoV-19/USA/AZ-TG268350/2020 | North America / USA / Arizona | 2020-03-17 | TGen North | TGen North | Jolene Bowers, Megan Folkerts, Darrin Lemmer, Dave Engelthaler |
| EPI_ISL_426507 | hCoV-19/USA/AZ-TG268360/2020 | North America / USA / Arizona | 2020-03-18 | TGen North | TGen North | Jolene Bowers, Megan Folkerts, Darrin Lemmer, Dave Engelthaler |
| EPI_ISL_426508 | hCoV-19/USA/AZ-TG268417/2020 | North America / USA / Arizona | 2020-03 | TGen North | TGen North | Jolene Bowers, Megan Folkerts, Darrin Lemmer, Dave Engelthaler |
| EPI_ISL_426509 | hCoV-19/USA/AZ-TG268629/2020 | North America / USA / Arizona | 2020-03-18 | TGen North | TGen North | Jolene Bowers, Megan Folkerts, Darrin Lemmer, Dave Engelthaler |
| EPI_ISL_426510 | hCoV-19/USA/AZ-TG268657/2020 | North America / USA / Arizona | 2020-03-17 | TGen North | TGen North | Jolene Bowers, Megan Folkerts, Darrin Lemmer, Dave Engelthaler |
| EPI_ISL_426511 | hCoV-19/USA/AZ-TG268770/2020 | North America / USA / Arizona | 2020-03-19 | TGen North | TGen North | Jolene Bowers, Megan Folkerts, Darrin Lemmer, Dave Engelthaler |
| EPI_ISL_426512 | hCoV-19/USA/AZ-TG268893/2020 | North America / USA / Arizona | 2020-03-05 | AZ SPHL, Arizona Department of Health Services | TGen North | Jolene Bowers, Megan Folkerts, Darrin Lemmer, Dave Engelthaler |
| EPI_ISL_426513 | hCoV-19/USA/AZ-TG268903/2020 | North America / USA / Arizona | 2020-03-11 | AZ SPHL, Arizona Department of Health Services | TGen North | Jolene Bowers, Megan Folkerts, Darrin Lemmer, Dave Engelthaler |
| EPI_ISL_426514 | hCoV-19/USA/AZ-TG268905/2020 | North America / USA / Arizona | 2020-03-11 | AZ SPHL, Arizona Department of Health Services | TGen North | Jolene Bowers, Megan Folkerts, Darrin Lemmer, Dave Engelthaler |
| EPI_ISL_426515 | hCoV-19/USA/AZ-TG268909/2020 | North America / USA / Arizona | 2020-03-13 | AZ SPHL, Arizona Department of Health Services | TGen North | Jolene Bowers, Megan Folkerts, Darrin Lemmer, Dave Engelthaler |
| EPI_ISL_426516 | hCoV-19/USA/AZ-TG268911/2020 | North America / USA / Arizona | 2020-03-13 | AZ SPHL, Arizona Department of Health Services | TGen North | Jolene Bowers, Megan Folkerts, Darrin Lemmer, Dave Engelthaler |
| EPI_ISL_426517 | hCoV-19/USA/AZ-TG268913/2020 | North America / USA / Arizona | 2020-03-14 | AZ SPHL, Arizona Department of Health Services | TGen North | Jolene Bowers, Megan Folkerts, Darrin Lemmer, Dave Engelthaler |
| EPI_ISL_426518 | hCoV-19/USA/AZ-TG268915/2020 | North America / USA / Arizona | 2020-03-12 | AZ SPHL, Arizona Department of Health Services | TGen North | Jolene Bowers, Megan Folkerts, Darrin Lemmer, Dave Engelthaler |
| EPI_ISL_426519 | hCoV-19/USA/AZ-TG268917/2020 | North America / USA / Arizona | 2020-03-14 | AZ SPHL, Arizona Department of Health Services | TGen North | Jolene Bowers, Megan Folkerts, Darrin Lemmer, Dave Engelthaler |
| EPI_ISL_426520 | hCoV-19/USA/AZ-TG269027/2020 | North America / USA / Arizona | 2020-03-20 | TGen North | TGen North | Jolene Bowers, Megan Folkerts, Darrin Lemmer, Dave Engelthaler |
| EPI_ISL_426521 | hCoV-19/USA/AZ-TG269238/2020 | North America / USA / Arizona | 2020-03-20 | TGen North | TGen North | Jolene Bowers, Megan Folkerts, Darrin Lemmer, Dave Engelthaler |
| EPI_ISL_426522 | hCoV-19/USA/AZ-TG269290/2020 | North America / USA / Arizona | 2020-03-20 | TGen North | TGen North | Jolene Bowers, Megan Folkerts, Darrin Lemmer, Dave Engelthaler |
| EPI_ISL_426523 | hCoV-19/USA/AZ-TG269358/2020 | North America / USA / Arizona | 2020-03-20 | TGen North | TGen North | Jolene Bowers, Megan Folkerts, Darrin Lemmer, Dave Engelthaler |
| EPI_ISL_426524 | hCoV-19/USA/AZ-TG269439/2020 | North America / USA / Arizona | 2020-03-20 | TGen North | TGen North | Jolene Bowers, Megan Folkerts, Darrin Lemmer, Dave Engelthaler |
| EPI_ISL_426525 | hCoV-19/USA/AZ-TG269440/2020 | North America / USA / Arizona | 2020-03-19 | TGen North | TGen North | Jolene Bowers, Megan Folkerts, Darrin Lemmer, Dave Engelthaler |
| EPI_ISL_426526 | hCoV-19/USA/AZ-TG269824/2020 | North America / USA / Arizona | 2020-03-23 | TGen North | TGen North | Jolene Bowers, Megan Folkerts, Darrin Lemmer, Dave Engelthaler |
| EPI_ISL_426527 | hCoV-19/USA/AZ-TG269848/2020 | North America / USA / Arizona | 2020-03-17 | AZ SPHL, Arizona Department of Health Services | TGen North | Jolene Bowers, Megan Folkerts, Darrin Lemmer, Dave Engelthaler |
| EPI_ISL_426528 | hCoV-19/USA/AZ-TG269856/2020 | North America / USA / Arizona | 2020-03-14 | AZ SPHL, Arizona Department of Health Services | TGen North | Jolene Bowers, Megan Folkerts, Darrin Lemmer, Dave Engelthaler |
| EPI_ISL_426529 | hCoV-19/USA/AZ-TG269863/2020 | North America / USA / Arizona | 2020-03-16 | AZ SPHL, Arizona Department of Health Services | TGen North | Jolene Bowers, Megan Folkerts, Darrin Lemmer, Dave Engelthaler |
| EPI_ISL_426530 | hCoV-19/USA/AZ-TG269875/2020 | North America / USA / Arizona | 2020-03-17 | AZ SPHL, Arizona Department of Health Services | TGen North | Jolene Bowers, Megan Folkerts, Darrin Lemmer, Dave Engelthaler |
| EPI_ISL_426531 | hCoV-19/USA/AZ-TG269883/2020 | North America / USA / Arizona | 2020-03-12 | AZ SPHL, Arizona Department of Health Services | TGen North | Jolene Bowers, Megan Folkerts, Darrin Lemmer, Dave Engelthaler |
| EPI_ISL_426532 | hCoV-19/USA/AZ-TG270152/2020 | North America / USA / Arizona | 2020-03-25 | TGen North | TGen North | Jolene Bowers, Megan Folkerts, Darrin Lemmer, Dave Engelthaler |
| EPI_ISL_426533 | hCoV-19/USA/AZ-TG270200/2020 | North America / USA / Arizona | 2020-03-26 | TGen North | TGen North | Jolene Bowers, Megan Folkerts, Darrin Lemmer, Dave Engelthaler |
| EPI_ISL_426534 | hCoV-19/USA/AZ-TG271435/2020 | North America / USA / Arizona | 2020-03-27 | TGen North | TGen North | Jolene Bowers, Megan Folkerts, Darrin Lemmer, Dave Engelthaler |
| EPI_ISL_426535 | hCoV-19/USA/AZ-TG271578/2020 | North America / USA / Arizona | 2020-03-17 | TGen North | TGen North | Jolene Bowers, Megan Folkerts, Darrin Lemmer, Dave Engelthaler |
| EPI_ISL_426536 | hCoV-19/USA/AZ-TG271591/2020 | North America / USA / Arizona | 2020-03-25 | TGen North | TGen North | Jolene Bowers, Megan Folkerts, Darrin Lemmer, Dave Engelthaler |
| EPI_ISL_426537 | hCoV-19/USA/AZ-TG271854/2020 | North America / USA / Arizona | 2020-03-17 | AZ SPHL, Arizona Department of Health Services | TGen North | Jolene Bowers, Megan Folkerts, Darrin Lemmer, Dave Engelthaler |
| EPI_ISL_426538 | hCoV-19/USA/AZ-TG271856/2020 | North America / USA / Arizona | 2020-03-19 | AZ SPHL, Arizona Department of Health Services | TGen North | Jolene Bowers, Megan Folkerts, Darrin Lemmer, Dave Engelthaler |
| EPI_ISL_426540 | hCoV-19/USA/AZ-TG271864/2020 | North America / USA / Arizona | 2020-03-23 | AZ SPHL, Arizona Department of Health Services | TGen North | Jolene Bowers, Megan Folkerts, Darrin Lemmer, Dave Engelthaler |
| EPI_ISL_426541 | hCoV-19/USA/AZ-TG271870/2020 | North America / USA / Arizona | 2020-03-24 | AZ SPHL, Arizona Department of Health Services | TGen North | Jolene Bowers, Megan Folkerts, Darrin Lemmer, Dave Engelthaler |
| EPI_ISL_426542 | hCoV-19/USA/AZ-TG271878/2020 | North America / USA / Arizona | 2020-03-20 | AZ SPHL, Arizona Department of Health Services | TGen North | Jolene Bowers, Megan Folkerts, Darrin Lemmer, Dave Engelthaler |
| EPI_ISL_426543 | hCoV-19/USA/AZ-TG271880/2020 | North America / USA / Arizona | 2020-03-24 | AZ SPHL, Arizona Department of Health Services | TGen North | Jolene Bowers, Megan Folkerts, Darrin Lemmer, Dave Engelthaler |
| EPI_ISL_426544 | hCoV-19/USA/AZ-TG271882/2020 | North America / USA / Arizona | 2020-03-20 | AZ SPHL, Arizona Department of Health Services | TGen North | Jolene Bowers, Megan Folkerts, Darrin Lemmer, Dave Engelthaler |
| EPI_ISL_426545 | hCoV-19/USA/AZ-TG271884/2020 | North America / USA / Arizona | 2020-03-25 | AZ SPHL, Arizona Department of Health Services | TGen North | Jolene Bowers, Megan Folkerts, Darrin Lemmer, Dave Engelthaler |
| EPI_ISL_426546 | hCoV-19/USA/AZ-TG271886/2020 | North America / USA / Arizona | 2020-03-26 | AZ SPHL, Arizona Department of Health Services | TGen North | Jolene Bowers, Megan Folkerts, Darrin Lemmer, Dave Engelthaler |
| EPI_ISL_426547 | hCoV-19/USA/AZ-TG271888/2020 | North America / USA / Arizona | 2020-03-26 | AZ SPHL, Arizona Department of Health Services | TGen North | Jolene Bowers, Megan Folkerts, Darrin Lemmer, Dave Engelthaler |
| EPI_ISL_426548 | hCoV-19/USA/AZ-TG271890/2020 | North America / USA / Arizona | 2020-03-26 | AZ SPHL, Arizona Department of Health Services | TGen North | Jolene Bowers, Megan Folkerts, Darrin Lemmer, Dave Engelthaler |
| EPI_ISL_426549 | hCoV-19/USA/AZ-TG271892/2020 | North America / USA / Arizona | 2020-03-26 | AZ SPHL, Arizona Department of Health Services | TGen North | Jolene Bowers, Megan Folkerts, Darrin Lemmer, Dave Engelthaler |
| EPI_ISL_426550 | hCoV-19/USA/AZ-TG271894/2020 | North America / USA / Arizona | 2020-03-26 | AZ SPHL, Arizona Department of Health Services | TGen North | Jolene Bowers, Megan Folkerts, Darrin Lemmer, Dave Engelthaler |
| EPI_ISL_426551 | hCoV-19/USA/AZ-TG271896/2020 | North America / USA / Arizona | 2020-03-27 | AZ SPHL, Arizona Department of Health Services | TGen North | Jolene Bowers, Megan Folkerts, Darrin Lemmer, Dave Engelthaler |
| EPI_ISL_426552 | hCoV-19/USA/AZ-TG271898/2020 | North America / USA / Arizona | 2020-03-27 | AZ SPHL, Arizona Department of Health Services | TGen North | Jolene Bowers, Megan Folkerts, Darrin Lemmer, Dave Engelthaler |
| EPI_ISL_426553 | hCoV-19/USA/AZ-TG271900/2020 | North America / USA / Arizona | 2020-03-27 | AZ SPHL, Arizona Department of Health Services | TGen North | Jolene Bowers, Megan Folkerts, Darrin Lemmer, Dave Engelthaler |
| EPI_ISL_426554 | hCoV-19/USA/AZ-TG271902/2020 | North America / USA / Arizona | 2020-03-26 | AZ SPHL, Arizona Department of Health Services | TGen North | Jolene Bowers, Megan Folkerts, Darrin Lemmer, Dave Engelthaler |
| EPI_ISL_426555 | hCoV-19/USA/AZ-TG271922/2020 | North America / USA / Arizona | 2020-03-26 | AZ SPHL, Arizona Department of Health Services | TGen North | Jolene Bowers, Megan Folkerts, Darrin Lemmer, Dave Engelthaler |
| EPI_ISL_426556 | hCoV-19/USA/AZ-TG272213/2020 | North America / USA / Arizona | 2020-04-01 | TGen North | TGen North | Jolene Bowers, Megan Folkerts, Darrin Lemmer, Dave Engelthaler |
| EPI_ISL_426557 | hCoV-19/USA/AZ-TG272215/2020 | North America / USA / Arizona | 2020-04-01 | TGen North | TGen North | Jolene Bowers, Megan Folkerts, Darrin Lemmer, Dave Engelthaler |
| EPI_ISL_426558 | hCoV-19/USA/AZ-TG273280/2020 | North America / USA / Arizona | 2020-03-25 | AZ SPHL, Arizona Department of Health Services | TGen North | Jolene Bowers, Megan Folkerts, Darrin Lemmer, Dave Engelthaler |
| EPI_ISL_426559 | hCoV-19/USA/AZ-TG273282/2020 | North America / USA / Arizona | 2020-03-25 | AZ SPHL, Arizona Department of Health Services | TGen North | Jolene Bowers, Megan Folkerts, Darrin Lemmer, Dave Engelthaler |
| EPI_ISL_426560 | hCoV-19/USA/AZ-TG273286/2020 | North America / USA / Arizona | 2020-03-30 | AZ SPHL, Arizona Department of Health Services | TGen North | Jolene Bowers, Megan Folkerts, Darrin Lemmer, Dave Engelthaler |
| EPI_ISL_426561 | hCoV-19/USA/AZ-TG273294/2020 | North America / USA / Arizona | 2020-03-31 | AZ SPHL, Arizona Department of Health Services | TGen North | Jolene Bowers, Megan Folkerts, Darrin Lemmer, Dave Engelthaler |
| EPI_ISL_426562 | hCoV-19/USA/AZ-TG273300/2020 | North America / USA / Arizona | 2020-03-31 | AZ SPHL, Arizona Department of Health Services | TGen North | Jolene Bowers, Megan Folkerts, Darrin Lemmer, Dave Engelthaler |
| EPI_ISL_426563 | hCoV-19/USA/AZ-TG273302/2020 | North America / USA / Arizona | 2020-03-31 | AZ SPHL, Arizona Department of Health Services | TGen North | Jolene Bowers, Megan Folkerts, Darrin Lemmer, Dave Engelthaler |
| EPI_ISL_426564 | hCoV-19/USA/AZ-TG273304/2020 | North America / USA / Arizona | 2020-04-01 | AZ SPHL, Arizona Department of Health Services | TGen North | Jolene Bowers, Megan Folkerts, Darrin Lemmer, Dave Engelthaler |
| EPI_ISL_426565 | hCoV-19/USA/AZ-TG273308/2020 | North America / USA / Arizona | 2020-04-01 | AZ SPHL, Arizona Department of Health Services | TGen North | Jolene Bowers, Megan Folkerts, Darrin Lemmer, Dave Engelthaler |
| EPI_ISL_426566 | hCoV-19/USA/AZ-TG273310/2020 | North America / USA / Arizona | 2020-04-01 | AZ SPHL, Arizona Department of Health Services | TGen North | Jolene Bowers, Megan Folkerts, Darrin Lemmer, Dave Engelthaler |
| EPI_ISL_426567 | hCoV-19/USA/AZ-TG273314/2020 | North America / USA / Arizona | 2020-04-01 | AZ SPHL, Arizona Department of Health Services | TGen North | Jolene Bowers, Megan Folkerts, Darrin Lemmer, Dave Engelthaler |
| EPI_ISL_426568 | hCoV-19/USA/AZ-TG273316/2020 | North America / USA / Arizona | 2020-04-02 | AZ SPHL, Arizona Department of Health Services | TGen North | Jolene Bowers, Megan Folkerts, Darrin Lemmer, Dave Engelthaler |
| EPI_ISL_426569 | hCoV-19/USA/AZ-TG273318/2020 | North America / USA / Arizona | 2020-04-02 | AZ SPHL, Arizona Department of Health Services | TGen North | Jolene Bowers, Megan Folkerts, Darrin Lemmer, Dave Engelthaler |
| EPI_ISL_426580 | hCoV-19/Brazil/DFBR-0001/2020 | South America / Brazil / Distrito Federal / | 2020-03-13 | Instituto Sabin | Laboratory of Virology | Fernando L Melo,Gustavo Barra, Ticiane H Santa-Rita, Pedro G Mesquita, Ikaro A Andrade, Tatsuya Nagata, Bergmann M Ribeiro |
| EPI_ISL_426581 | hCoV-19/Czech Republic/IAB20-006-16/2020 | Europe / Czech Republic | 2020 | Motol University Hospital | Institute of Applied Biotechnologies a.s. | Petr Brož, Jan Geryk, Petr Klempt, Martin Kalný, Adam Novotný, Kateřina Kvapilová, Pavel Dřevínek, Petr Kvapil, Milan Macek |
| EPI_ISL_426584 | hCoV-19/Uruguay/UY-9/2020 | South America / Uruguay / Montevideo | 2020-03-19 | Microbial Genomics Laboratory, Institut Pasteur Montevideo, Uruguay | Microbial Genomics Laboratory, Institut Pasteur Montevideo, Uruguay | Cecilia Salazar, Florencia Díaz-Viraqué, Mariancel Pereira, Pilar Moreno, Gonzalo Moratorio, Gregorio Iraola |
| EPI_ISL_427001 | hCoV-19/Australia/VIC716/2020 | Oceania / Australia / Victoria | 2020-03-27 | Victorian Infectious Diseases Reference Laboratory (VIDRL) | Microbiological Diagnostic Unit Public Health Laboratory and Victorian Infectious Diseases Reference Laboratory, Doherty Institute | Caly L., Seemann T., Sait, M., Schultz M., Druce J., Sherry, N. |
| EPI_ISL_427002 | hCoV-19/Australia/VIC718/2020 | Oceania / Australia / Victoria | 2020-03-27 | Victorian Infectious Diseases Reference Laboratory (VIDRL) | Microbiological Diagnostic Unit Public Health Laboratory and Victorian Infectious Diseases Reference Laboratory, Doherty Institute | Caly L., Seemann T., Sait, M., Schultz M., Druce J., Sherry, N. |
| EPI_ISL_427003 | hCoV-19/Australia/VIC720/2020 | Oceania / Australia / Victoria | 2020-03-28 | Victorian Infectious Diseases Reference Laboratory (VIDRL) | Microbiological Diagnostic Unit Public Health Laboratory and Victorian Infectious Diseases Reference Laboratory, Doherty Institute | Caly L., Seemann T., Sait, M., Schultz M., Druce J., Sherry, N. |
| EPI_ISL_427005 | hCoV-19/Australia/VIC722/2020 | Oceania / Australia / Victoria | 2020-03-29 | Victorian Infectious Diseases Reference Laboratory (VIDRL) | Microbiological Diagnostic Unit Public Health Laboratory and Victorian Infectious Diseases Reference Laboratory, Doherty Institute | Caly L., Seemann T., Sait, M., Schultz M., Druce J., Sherry, N. |
| EPI_ISL_427006 | hCoV-19/Australia/VIC725/2020 | Oceania / Australia / Victoria | 2020-03-28 | Victorian Infectious Diseases Reference Laboratory (VIDRL) | Microbiological Diagnostic Unit Public Health Laboratory and Victorian Infectious Diseases Reference Laboratory, Doherty Institute | Caly L., Seemann T., Sait, M., Schultz M., Druce J., Sherry, N. |
| EPI_ISL_427010 | hCoV-19/Australia/VIC731/2020 | Oceania / Australia / Victoria | 2020-03-30 | Victorian Infectious Diseases Reference Laboratory (VIDRL) | Microbiological Diagnostic Unit Public Health Laboratory and Victorian Infectious Diseases Reference Laboratory, Doherty Institute | Caly L., Seemann T., Sait, M., Schultz M., Druce J., Sherry, N. |
| EPI_ISL_427011 | hCoV-19/Australia/VIC732/2020 | Oceania / Australia / Victoria | 2020-03-30 | Victorian Infectious Diseases Reference Laboratory (VIDRL) | Microbiological Diagnostic Unit Public Health Laboratory and Victorian Infectious Diseases Reference Laboratory, Doherty Institute | Caly L., Seemann T., Sait, M., Schultz M., Druce J., Sherry, N. |
| EPI_ISL_427012 | hCoV-19/Australia/VIC733/2020 | Oceania / Australia / Victoria | 2020-03-30 | Victorian Infectious Diseases Reference Laboratory (VIDRL) | Microbiological Diagnostic Unit Public Health Laboratory and Victorian Infectious Diseases Reference Laboratory, Doherty Institute | Caly L., Seemann T., Sait, M., Schultz M., Druce J., Sherry, N. |
| EPI_ISL_427013 | hCoV-19/Australia/VIC734/2020 | Oceania / Australia / Victoria | 2020-03-30 | Victorian Infectious Diseases Reference Laboratory (VIDRL) | Microbiological Diagnostic Unit Public Health Laboratory and Victorian Infectious Diseases Reference Laboratory, Doherty Institute | Caly L., Seemann T., Sait, M., Schultz M., Druce J., Sherry, N. |
| EPI_ISL_427014 | hCoV-19/Australia/VIC735/2020 | Oceania / Australia / Victoria | 2020-03-30 | Victorian Infectious Diseases Reference Laboratory (VIDRL) | Microbiological Diagnostic Unit Public Health Laboratory and Victorian Infectious Diseases Reference Laboratory, Doherty Institute | Caly L., Seemann T., Sait, M., Schultz M., Druce J., Sherry, N. |
| EPI_ISL_427015 | hCoV-19/Australia/VIC736/2020 | Oceania / Australia / Victoria | 2020-03-30 | Victorian Infectious Diseases Reference Laboratory (VIDRL) | Microbiological Diagnostic Unit Public Health Laboratory and Victorian Infectious Diseases Reference Laboratory, Doherty Institute | Caly L., Seemann T., Sait, M., Schultz M., Druce J., Sherry, N. |
| EPI_ISL_427016 | hCoV-19/Australia/VIC737/2020 | Oceania / Australia / Victoria | 2020-03-30 | Victorian Infectious Diseases Reference Laboratory (VIDRL) | Microbiological Diagnostic Unit Public Health Laboratory and Victorian Infectious Diseases Reference Laboratory, Doherty Institute | Caly L., Seemann T., Sait, M., Schultz M., Druce J., Sherry, N. |
| EPI_ISL_427017 | hCoV-19/Australia/VIC738/2020 | Oceania / Australia / Victoria | 2020-03-30 | Victorian Infectious Diseases Reference Laboratory (VIDRL) | Microbiological Diagnostic Unit Public Health Laboratory and Victorian Infectious Diseases Reference Laboratory, Doherty Institute | Caly L., Seemann T., Sait, M., Schultz M., Druce J., Sherry, N. |
| EPI_ISL_427018 | hCoV-19/Australia/VIC739/2020 | Oceania / Australia / Victoria | 2020-03-30 | Victorian Infectious Diseases Reference Laboratory (VIDRL) | Microbiological Diagnostic Unit Public Health Laboratory and Victorian Infectious Diseases Reference Laboratory, Doherty Institute | Caly L., Seemann T., Sait, M., Schultz M., Druce J., Sherry, N. |
| EPI_ISL_427019 | hCoV-19/Australia/VIC741/2020 | Oceania / Australia / Victoria | 2020-03-30 | Victorian Infectious Diseases Reference Laboratory (VIDRL) | Microbiological Diagnostic Unit Public Health Laboratory and Victorian Infectious Diseases Reference Laboratory, Doherty Institute | Caly L., Seemann T., Sait, M., Schultz M., Druce J., Sherry, N. |
| EPI_ISL_427020 | hCoV-19/Australia/VIC742/2020 | Oceania / Australia / Victoria | 2020-03-29 | Victorian Infectious Diseases Reference Laboratory (VIDRL) | Microbiological Diagnostic Unit Public Health Laboratory and Victorian Infectious Diseases Reference Laboratory, Doherty Institute | Caly L., Seemann T., Sait, M., Schultz M., Druce J., Sherry, N. |
| EPI_ISL_427021 | hCoV-19/Australia/VIC743/2020 | Oceania / Australia / Victoria | 2020-03-29 | Victorian Infectious Diseases Reference Laboratory (VIDRL) | Microbiological Diagnostic Unit Public Health Laboratory and Victorian Infectious Diseases Reference Laboratory, Doherty Institute | Caly L., Seemann T., Sait, M., Schultz M., Druce J., Sherry, N. |
| EPI_ISL_427022 | hCoV-19/Australia/VIC744/2020 | Oceania / Australia / Victoria | 2020-03-29 | Victorian Infectious Diseases Reference Laboratory (VIDRL) | Microbiological Diagnostic Unit Public Health Laboratory and Victorian Infectious Diseases Reference Laboratory, Doherty Institute | Caly L., Seemann T., Sait, M., Schultz M., Druce J., Sherry, N. |
| EPI_ISL_427023 | hCoV-19/Australia/VIC745/2020 | Oceania / Australia / Victoria | 2020-03-29 | Victorian Infectious Diseases Reference Laboratory (VIDRL) | Microbiological Diagnostic Unit Public Health Laboratory and Victorian Infectious Diseases Reference Laboratory, Doherty Institute | Caly L., Seemann T., Sait, M., Schultz M., Druce J., Sherry, N. |
| EPI_ISL_427024 | hCoV-19/Australia/VIC746/2020 | Oceania / Australia / Victoria | 2020-03-29 | Victorian Infectious Diseases Reference Laboratory (VIDRL) | Microbiological Diagnostic Unit Public Health Laboratory and Victorian Infectious Diseases Reference Laboratory, Doherty Institute | Caly L., Seemann T., Sait, M., Schultz M., Druce J., Sherry, N. |
| EPI_ISL_427025 | hCoV-19/Australia/VIC747/2020 | Oceania / Australia / Victoria | 2020-03-31 | Victorian Infectious Diseases Reference Laboratory (VIDRL) | Microbiological Diagnostic Unit Public Health Laboratory and Victorian Infectious Diseases Reference Laboratory, Doherty Institute | Caly L., Seemann T., Sait, M., Schultz M., Druce J., Sherry, N. |
| EPI_ISL_427026 | hCoV-19/Australia/VIC748/2020 | Oceania / Australia / Victoria | 2020-03-30 | Victorian Infectious Diseases Reference Laboratory (VIDRL) | Microbiological Diagnostic Unit Public Health Laboratory and Victorian Infectious Diseases Reference Laboratory, Doherty Institute | Caly L., Seemann T., Sait, M., Schultz M., Druce J., Sherry, N. |
| EPI_ISL_427027 | hCoV-19/Australia/VIC749/2020 | Oceania / Australia / Victoria | 2020-03-31 | Victorian Infectious Diseases Reference Laboratory (VIDRL) | Microbiological Diagnostic Unit Public Health Laboratory and Victorian Infectious Diseases Reference Laboratory, Doherty Institute | Caly L., Seemann T., Sait, M., Schultz M., Druce J., Sherry, N. |
| EPI_ISL_427028 | hCoV-19/Australia/VIC751/2020 | Oceania / Australia / Victoria | 2020-03-30 | Victorian Infectious Diseases Reference Laboratory (VIDRL) | Microbiological Diagnostic Unit Public Health Laboratory and Victorian Infectious Diseases Reference Laboratory, Doherty Institute | Caly L., Seemann T., Sait, M., Schultz M., Druce J., Sherry, N. |
| EPI_ISL_427029 | hCoV-19/Australia/VIC752/2020 | Oceania / Australia / Victoria | 2020-03-30 | Victorian Infectious Diseases Reference Laboratory (VIDRL) | Microbiological Diagnostic Unit Public Health Laboratory and Victorian Infectious Diseases Reference Laboratory, Doherty Institute | Caly L., Seemann T., Sait, M., Schultz M., Druce J., Sherry, N. |
| EPI_ISL_427031 | hCoV-19/Australia/VIC754/2020 | Oceania / Australia / Victoria | 2020-03-30 | Victorian Infectious Diseases Reference Laboratory (VIDRL) | Microbiological Diagnostic Unit Public Health Laboratory and Victorian Infectious Diseases Reference Laboratory, Doherty Institute | Caly L., Seemann T., Sait, M., Schultz M., Druce J., Sherry, N. |
| EPI_ISL_427033 | hCoV-19/Australia/VIC756/2020 | Oceania / Australia / Victoria | 2020-03-30 | Victorian Infectious Diseases Reference Laboratory (VIDRL) | Microbiological Diagnostic Unit Public Health Laboratory and Victorian Infectious Diseases Reference Laboratory, Doherty Institute | Caly L., Seemann T., Sait, M., Schultz M., Druce J., Sherry, N. |
| EPI_ISL_427034 | hCoV-19/Australia/VIC757/2020 | Oceania / Australia / Victoria | 2020-03-31 | Victorian Infectious Diseases Reference Laboratory (VIDRL) | Microbiological Diagnostic Unit Public Health Laboratory and Victorian Infectious Diseases Reference Laboratory, Doherty Institute | Caly L., Seemann T., Sait, M., Schultz M., Druce J., Sherry, N. |
| EPI_ISL_427037 | hCoV-19/Australia/VIC761/2020 | Oceania / Australia / Victoria | 2020-03-30 | Victorian Infectious Diseases Reference Laboratory (VIDRL) | Microbiological Diagnostic Unit Public Health Laboratory and Victorian Infectious Diseases Reference Laboratory, Doherty Institute | Caly L., Seemann T., Sait, M., Schultz M., Druce J., Sherry, N. |
| EPI_ISL_427038 | hCoV-19/Australia/VIC762/2020 | Oceania / Australia / Victoria | 2020-03-29 | Victorian Infectious Diseases Reference Laboratory (VIDRL) | Microbiological Diagnostic Unit Public Health Laboratory and Victorian Infectious Diseases Reference Laboratory, Doherty Institute | Caly L., Seemann T., Sait, M., Schultz M., Druce J., Sherry, N. |
| EPI_ISL_427040 | hCoV-19/Australia/VIC764/2020 | Oceania / Australia / Victoria | 2020-03-29 | Victorian Infectious Diseases Reference Laboratory (VIDRL) | Microbiological Diagnostic Unit Public Health Laboratory and Victorian Infectious Diseases Reference Laboratory, Doherty Institute | Caly L., Seemann T., Sait, M., Schultz M., Druce J., Sherry, N. |
| EPI_ISL_427041 | hCoV-19/Australia/VIC765/2020 | Oceania / Australia / Victoria | 2020-03-29 | Victorian Infectious Diseases Reference Laboratory (VIDRL) | Microbiological Diagnostic Unit Public Health Laboratory and Victorian Infectious Diseases Reference Laboratory, Doherty Institute | Caly L., Seemann T., Sait, M., Schultz M., Druce J., Sherry, N. |
| EPI_ISL_427043 | hCoV-19/Greece/13/2020 | Europe / Greece / Athens | 2020-03-18 | Laboratory of Microbiology, Medical School, National and Kapodistrian University of Athens | Laboratory of Biology, Department of Medicine, Democritus University of Thrace | Bampali,M., Dovrolis,N., Gatzidou,E., Froukala,E., Stavropoulou,A., Veletza,S., Tsakris,A., Spanakis,N. and Karakasiliotis,I. |
| EPI_ISL_427045 | hCoV-19/Australia/VIC768/2020 | Oceania / Australia / Victoria | 2020-03-20 | Victorian Infectious Diseases Reference Laboratory (VIDRL) | Microbiological Diagnostic Unit Public Health Laboratory and Victorian Infectious Diseases Reference Laboratory, Doherty Institute | Caly L., Seemann T., Sait, M., Schultz M., Druce J., Sherry, N. |
| EPI_ISL_427046 | hCoV-19/Australia/VIC769/2020 | Oceania / Australia / Victoria | 2020-03-20 | Victorian Infectious Diseases Reference Laboratory (VIDRL) | Microbiological Diagnostic Unit Public Health Laboratory and Victorian Infectious Diseases Reference Laboratory, Doherty Institute | Caly L., Seemann T., Sait, M., Schultz M., Druce J., Sherry, N. |
| EPI_ISL_427048 | hCoV-19/Australia/VIC772/2020 | Oceania / Australia / Victoria | 2020-03-20 | Victorian Infectious Diseases Reference Laboratory (VIDRL) | Microbiological Diagnostic Unit Public Health Laboratory and Victorian Infectious Diseases Reference Laboratory, Doherty Institute | Caly L., Seemann T., Sait, M., Schultz M., Druce J., Sherry, N. |
| EPI_ISL_427049 | hCoV-19/Australia/VIC773/2020 | Oceania / Australia / Victoria | 2020-03-20 | Victorian Infectious Diseases Reference Laboratory (VIDRL) | Microbiological Diagnostic Unit Public Health Laboratory and Victorian Infectious Diseases Reference Laboratory, Doherty Institute | Caly L., Seemann T., Sait, M., Schultz M., Druce J., Sherry, N. |
| EPI_ISL_427058 | hCoV-19/Australia/VIC783/2020 | Oceania / Australia / Victoria | 2020-03-30 | Microbiological Diagnostic Unit Public Health Laboratory | Microbiological Diagnostic Unit Public Health Laboratory | Seemann T., Schultz M., Sait, M., Sherry, N. |
| EPI_ISL_427060 | hCoV-19/Australia/VIC785/2020 | Oceania / Australia / Victoria | 2020-03-30 | Microbiological Diagnostic Unit Public Health Laboratory | Microbiological Diagnostic Unit Public Health Laboratory | Seemann T., Schultz M., Sait, M., Sherry, N. |
| EPI_ISL_427061 | hCoV-19/Australia/VIC786/2020 | Oceania / Australia / Victoria | 2020-03-30 | Microbiological Diagnostic Unit Public Health Laboratory | Microbiological Diagnostic Unit Public Health Laboratory | Seemann T., Schultz M., Sait, M., Sherry, N. |
| EPI_ISL_427062 | hCoV-19/Australia/VIC787/2020 | Oceania / Australia / Victoria | 2020-03-30 | Microbiological Diagnostic Unit Public Health Laboratory | Microbiological Diagnostic Unit Public Health Laboratory | Seemann T., Schultz M., Sait, M., Sherry, N. |
| EPI_ISL_427063 | hCoV-19/Australia/VIC788/2020 | Oceania / Australia / Victoria | 2020-03-31 | Microbiological Diagnostic Unit Public Health Laboratory | Microbiological Diagnostic Unit Public Health Laboratory | Seemann T., Schultz M., Sait, M., Sherry, N. |
| EPI_ISL_427065 | hCoV-19/Australia/VIC790/2020 | Oceania / Australia / Victoria | 2020-03-31 | Microbiological Diagnostic Unit Public Health Laboratory | Microbiological Diagnostic Unit Public Health Laboratory | Seemann T., Schultz M., Sait, M., Sherry, N. |
| EPI_ISL_427068 | hCoV-19/Australia/VIC793/2020 | Oceania / Australia / Victoria | 2020-03-31 | Microbiological Diagnostic Unit Public Health Laboratory | Microbiological Diagnostic Unit Public Health Laboratory | Seemann T., Schultz M., Sait, M., Sherry, N. |
| EPI_ISL_427069 | hCoV-19/Australia/VIC794/2020 | Oceania / Australia / Victoria | 2020-03-31 | Microbiological Diagnostic Unit Public Health Laboratory | Microbiological Diagnostic Unit Public Health Laboratory | Seemann T., Schultz M., Sait, M., Sherry, N. |
| EPI_ISL_427092 | hCoV-19/Australia/VIC820/2020 | Oceania / Australia / Victoria | 2020-04-03 | Victorian Infectious Diseases Reference Laboratory (VIDRL) | Microbiological Diagnostic Unit Public Health Laboratory and Victorian Infectious Diseases Reference Laboratory, Doherty Institute | Caly L., Seemann T., Sait, M., Schultz M., Druce J., Sherry, N. |
| EPI_ISL_427093 | hCoV-19/Australia/VIC821/2020 | Oceania / Australia / Victoria | 2020-04-03 | Victorian Infectious Diseases Reference Laboratory (VIDRL) | Microbiological Diagnostic Unit Public Health Laboratory and Victorian Infectious Diseases Reference Laboratory, Doherty Institute | Caly L., Seemann T., Sait, M., Schultz M., Druce J., Sherry, N. |
| EPI_ISL_427094 | hCoV-19/Australia/VIC823/2020 | Oceania / Australia / Victoria | 2020-04-03 | Victorian Infectious Diseases Reference Laboratory (VIDRL) | Microbiological Diagnostic Unit Public Health Laboratory and Victorian Infectious Diseases Reference Laboratory, Doherty Institute | Caly L., Seemann T., Sait, M., Schultz M., Druce J., Sherry, N. |
| EPI_ISL_427095 | hCoV-19/Australia/VIC824/2020 | Oceania / Australia / Victoria | 2020-04-03 | Victorian Infectious Diseases Reference Laboratory (VIDRL) | Microbiological Diagnostic Unit Public Health Laboratory and Victorian Infectious Diseases Reference Laboratory, Doherty Institute | Caly L., Seemann T., Sait, M., Schultz M., Druce J., Sherry, N. |
| EPI_ISL_427096 | hCoV-19/Australia/VIC828/2020 | Oceania / Australia / Victoria | 2020-04-03 | Victorian Infectious Diseases Reference Laboratory (VIDRL) | Microbiological Diagnostic Unit Public Health Laboratory and Victorian Infectious Diseases Reference Laboratory, Doherty Institute | Caly L., Seemann T., Sait, M., Schultz M., Druce J., Sherry, N. |
| EPI_ISL_427098 | hCoV-19/Australia/VIC831/2020 | Oceania / Australia / Victoria | 2020-04-04 | Victorian Infectious Diseases Reference Laboratory (VIDRL) | Microbiological Diagnostic Unit Public Health Laboratory and Victorian Infectious Diseases Reference Laboratory, Doherty Institute | Caly L., Seemann T., Sait, M., Schultz M., Druce J., Sherry, N. |
| EPI_ISL_427500 | hCoV-19/USA/NY-NYUMC205/2020 | North America / USA / New York / Brookl | 2020-04-01 | NYU Langone Health | Departments of Pathology and Medicine, New York University School of Medicine | Maria Aguero-Rosenfeld, Brendan Belovarac, Margaret Black, Ludovic Boytard, John Cadley, Paolo Cotzia, John Chen, Dacia Dimartino, Xiaojun Feng, Tatyana Gindin, Emily Guzman, Adriana Heguy, Megan Hogan, Emily Huang, George Jour, Andrew Lytle, Christian Marier, Matthew T. Maurano, Mark J. Mulligan, Peter Meyn, Iman Osman, Jared Pinnell, Vanessa Raabe, Sitharam Ramaswami, Amy Rapkiewicz, Marie Samanovic-Golden, Antonio Serrano, Guomiao Shen, Matija Snuderl, Theodore Vougiouklakis, Nick Vulpescu, Gael Westby, Paul Zappile, Yutong Zhang |
| EPI_ISL_427502 | hCoV-19/USA/NY-NYUMC207/2020 | North America / USA / New York / Brookl | 2020-04-03 | NYU Langone Health | Departments of Pathology and Medicine, New York University School of Medicine | Maria Aguero-Rosenfeld, Brendan Belovarac, Margaret Black, Ludovic Boytard, John Cadley, Paolo Cotzia, John Chen, Dacia Dimartino, Xiaojun Feng, Tatyana Gindin, Emily Guzman, Adriana Heguy, Megan Hogan, Emily Huang, George Jour, Andrew Lytle, Christian Marier, Matthew T. Maurano, Mark J. Mulligan, Peter Meyn, Iman Osman, Jared Pinnell, Vanessa Raabe, Sitharam Ramaswami, Amy Rapkiewicz, Marie Samanovic-Golden, Antonio Serrano, Guomiao Shen, Matija Snuderl, Theodore Vougiouklakis, Nick Vulpescu, Gael Westby, Paul Zappile, Yutong Zhang |
| EPI_ISL_427503 | hCoV-19/USA/NY-NYUMC208/2020 | North America / USA / New York / Brookl | 2020-04-03 | NYU Langone Health | Departments of Pathology and Medicine, New York University School of Medicine | Maria Aguero-Rosenfeld, Brendan Belovarac, Margaret Black, Ludovic Boytard, John Cadley, Paolo Cotzia, John Chen, Dacia Dimartino, Xiaojun Feng, Tatyana Gindin, Emily Guzman, Adriana Heguy, Megan Hogan, Emily Huang, George Jour, Andrew Lytle, Christian Marier, Matthew T. Maurano, Mark J. Mulligan, Peter Meyn, Iman Osman, Jared Pinnell, Vanessa Raabe, Sitharam Ramaswami, Amy Rapkiewicz, Marie Samanovic-Golden, Antonio Serrano, Guomiao Shen, Matija Snuderl, Theodore Vougiouklakis, Nick Vulpescu, Gael Westby, Paul Zappile, Yutong Zhang |
| EPI_ISL_427504 | hCoV-19/USA/NY-NYUMC209/2020 | North America / USA / New York / Brookl | 2020-04-03 | NYU Langone Health | Departments of Pathology and Medicine, New York University School of Medicine | Maria Aguero-Rosenfeld, Brendan Belovarac, Margaret Black, Ludovic Boytard, John Cadley, Paolo Cotzia, John Chen, Dacia Dimartino, Xiaojun Feng, Tatyana Gindin, Emily Guzman, Adriana Heguy, Megan Hogan, Emily Huang, George Jour, Andrew Lytle, Christian Marier, Matthew T. Maurano, Mark J. Mulligan, Peter Meyn, Iman Osman, Jared Pinnell, Vanessa Raabe, Sitharam Ramaswami, Amy Rapkiewicz, Marie Samanovic-Golden, Antonio Serrano, Guomiao Shen, Matija Snuderl, Theodore Vougiouklakis, Nick Vulpescu, Gael Westby, Paul Zappile, Yutong Zhang |
| EPI_ISL_427505 | hCoV-19/USA/NY-NYUMC210/2020 | North America / USA / New York / Brookl | 2020-04-03 | NYU Langone Health | Departments of Pathology and Medicine, New York University School of Medicine | Maria Aguero-Rosenfeld, Brendan Belovarac, Margaret Black, Ludovic Boytard, John Cadley, Paolo Cotzia, John Chen, Dacia Dimartino, Xiaojun Feng, Tatyana Gindin, Emily Guzman, Adriana Heguy, Megan Hogan, Emily Huang, George Jour, Andrew Lytle, Christian Marier, Matthew T. Maurano, Mark J. Mulligan, Peter Meyn, Iman Osman, Jared Pinnell, Vanessa Raabe, Sitharam Ramaswami, Amy Rapkiewicz, Marie Samanovic-Golden, Antonio Serrano, Guomiao Shen, Matija Snuderl, Theodore Vougiouklakis, Nick Vulpescu, Gael Westby, Paul Zappile, Yutong Zhang |
| EPI_ISL_427506 | hCoV-19/USA/NY-NYUMC211/2020 | North America / USA / New York / Manha | 2020-04-03 | NYU Langone Health | Departments of Pathology and Medicine, New York University School of Medicine | Maria Aguero-Rosenfeld, Brendan Belovarac, Margaret Black, Ludovic Boytard, John Cadley, Paolo Cotzia, John Chen, Dacia Dimartino, Xiaojun Feng, Tatyana Gindin, Emily Guzman, Adriana Heguy, Megan Hogan, Emily Huang, George Jour, Andrew Lytle, Christian Marier, Matthew T. Maurano, Mark J. Mulligan, Peter Meyn, Iman Osman, Jared Pinnell, Vanessa Raabe, Sitharam Ramaswami, Amy Rapkiewicz, Marie Samanovic-Golden, Antonio Serrano, Guomiao Shen, Matija Snuderl, Theodore Vougiouklakis, Nick Vulpescu, Gael Westby, Paul Zappile, Yutong Zhang |
| EPI_ISL_427507 | hCoV-19/USA/NY-NYUMC212/2020 | North America / USA / New York / Brookl | 2020-04-03 | NYU Langone Health | Departments of Pathology and Medicine, New York University School of Medicine | Maria Aguero-Rosenfeld, Brendan Belovarac, Margaret Black, Ludovic Boytard, John Cadley, Paolo Cotzia, John Chen, Dacia Dimartino, Xiaojun Feng, Tatyana Gindin, Emily Guzman, Adriana Heguy, Megan Hogan, Emily Huang, George Jour, Andrew Lytle, Christian Marier, Matthew T. Maurano, Mark J. Mulligan, Peter Meyn, Iman Osman, Jared Pinnell, Vanessa Raabe, Sitharam Ramaswami, Amy Rapkiewicz, Marie Samanovic-Golden, Antonio Serrano, Guomiao Shen, Matija Snuderl, Theodore Vougiouklakis, Nick Vulpescu, Gael Westby, Paul Zappile, Yutong Zhang |
| EPI_ISL_427508 | hCoV-19/USA/NY-NYUMC213/2020 | North America / USA / New York / Brookl | 2020-04-03 | NYU Langone Health | Departments of Pathology and Medicine, New York University School of Medicine | Maria Aguero-Rosenfeld, Brendan Belovarac, Margaret Black, Ludovic Boytard, John Cadley, Paolo Cotzia, John Chen, Dacia Dimartino, Xiaojun Feng, Tatyana Gindin, Emily Guzman, Adriana Heguy, Megan Hogan, Emily Huang, George Jour, Andrew Lytle, Christian Marier, Matthew T. Maurano, Mark J. Mulligan, Peter Meyn, Iman Osman, Jared Pinnell, Vanessa Raabe, Sitharam Ramaswami, Amy Rapkiewicz, Marie Samanovic-Golden, Antonio Serrano, Guomiao Shen, Matija Snuderl, Theodore Vougiouklakis, Nick Vulpescu, Gael Westby, Paul Zappile, Yutong Zhang |
| EPI_ISL_427509 | hCoV-19/USA/NY-NYUMC214/2020 | North America / USA / New York / Manha | 2020-04-03 | NYU Langone Health | Departments of Pathology and Medicine, New York University School of Medicine | Maria Aguero-Rosenfeld, Brendan Belovarac, Margaret Black, Ludovic Boytard, John Cadley, Paolo Cotzia, John Chen, Dacia Dimartino, Xiaojun Feng, Tatyana Gindin, Emily Guzman, Adriana Heguy, Megan Hogan, Emily Huang, George Jour, Andrew Lytle, Christian Marier, Matthew T. Maurano, Mark J. Mulligan, Peter Meyn, Iman Osman, Jared Pinnell, Vanessa Raabe, Sitharam Ramaswami, Amy Rapkiewicz, Marie Samanovic-Golden, Antonio Serrano, Guomiao Shen, Matija Snuderl, Theodore Vougiouklakis, Nick Vulpescu, Gael Westby, Paul Zappile, Yutong Zhang |
| EPI_ISL_427510 | hCoV-19/USA/NY-NYUMC215/2020 | North America / USA / New York / Brookl | 2020-04-03 | NYU Langone Health | Departments of Pathology and Medicine, New York University School of Medicine | Maria Aguero-Rosenfeld, Brendan Belovarac, Margaret Black, Ludovic Boytard, John Cadley, Paolo Cotzia, John Chen, Dacia Dimartino, Xiaojun Feng, Tatyana Gindin, Emily Guzman, Adriana Heguy, Megan Hogan, Emily Huang, George Jour, Andrew Lytle, Christian Marier, Matthew T. Maurano, Mark J. Mulligan, Peter Meyn, Iman Osman, Jared Pinnell, Vanessa Raabe, Sitharam Ramaswami, Amy Rapkiewicz, Marie Samanovic-Golden, Antonio Serrano, Guomiao Shen, Matija Snuderl, Theodore Vougiouklakis, Nick Vulpescu, Gael Westby, Paul Zappile, Yutong Zhang |
| EPI_ISL_427511 | hCoV-19/USA/NY-NYUMC216/2020 | North America / USA / New York | 2020-04-03 | NYU Langone Health | Departments of Pathology and Medicine, New York University School of Medicine | Maria Aguero-Rosenfeld, Brendan Belovarac, Margaret Black, Ludovic Boytard, John Cadley, Paolo Cotzia, John Chen, Dacia Dimartino, Xiaojun Feng, Tatyana Gindin, Emily Guzman, Adriana Heguy, Megan Hogan, Emily Huang, George Jour, Andrew Lytle, Christian Marier, Matthew T. Maurano, Mark J. Mulligan, Peter Meyn, Iman Osman, Jared Pinnell, Vanessa Raabe, Sitharam Ramaswami, Amy Rapkiewicz, Marie Samanovic-Golden, Antonio Serrano, Guomiao Shen, Matija Snuderl, Theodore Vougiouklakis, Nick Vulpescu, Gael Westby, Paul Zappile, Yutong Zhang |
| EPI_ISL_427512 | hCoV-19/USA/NY-NYUMC217/2020 | North America / USA / New York / Brookl | 2020-04-03 | NYU Langone Health | Departments of Pathology and Medicine, New York University School of Medicine | Maria Aguero-Rosenfeld, Brendan Belovarac, Margaret Black, Ludovic Boytard, John Cadley, Paolo Cotzia, John Chen, Dacia Dimartino, Xiaojun Feng, Tatyana Gindin, Emily Guzman, Adriana Heguy, Megan Hogan, Emily Huang, George Jour, Andrew Lytle, Christian Marier, Matthew T. Maurano, Mark J. Mulligan, Peter Meyn, Iman Osman, Jared Pinnell, Vanessa Raabe, Sitharam Ramaswami, Amy Rapkiewicz, Marie Samanovic-Golden, Antonio Serrano, Guomiao Shen, Matija Snuderl, Theodore Vougiouklakis, Nick Vulpescu, Gael Westby, Paul Zappile, Yutong Zhang |
| EPI_ISL_427513 | hCoV-19/USA/NY-NYUMC218/2020 | North America / USA / New York / Brookl | 2020-04-03 | NYU Langone Health | Departments of Pathology and Medicine, New York University School of Medicine | Maria Aguero-Rosenfeld, Brendan Belovarac, Margaret Black, Ludovic Boytard, John Cadley, Paolo Cotzia, John Chen, Dacia Dimartino, Xiaojun Feng, Tatyana Gindin, Emily Guzman, Adriana Heguy, Megan Hogan, Emily Huang, George Jour, Andrew Lytle, Christian Marier, Matthew T. Maurano, Mark J. Mulligan, Peter Meyn, Iman Osman, Jared Pinnell, Vanessa Raabe, Sitharam Ramaswami, Amy Rapkiewicz, Marie Samanovic-Golden, Antonio Serrano, Guomiao Shen, Matija Snuderl, Theodore Vougiouklakis, Nick Vulpescu, Gael Westby, Paul Zappile, Yutong Zhang |
| EPI_ISL_427514 | hCoV-19/USA/NY-NYUMC219/2020 | North America / USA / New York / Brookl | 2020-04-03 | NYU Langone Health | Departments of Pathology and Medicine, New York University School of Medicine | Maria Aguero-Rosenfeld, Brendan Belovarac, Margaret Black, Ludovic Boytard, John Cadley, Paolo Cotzia, John Chen, Dacia Dimartino, Xiaojun Feng, Tatyana Gindin, Emily Guzman, Adriana Heguy, Megan Hogan, Emily Huang, George Jour, Andrew Lytle, Christian Marier, Matthew T. Maurano, Mark J. Mulligan, Peter Meyn, Iman Osman, Jared Pinnell, Vanessa Raabe, Sitharam Ramaswami, Amy Rapkiewicz, Marie Samanovic-Golden, Antonio Serrano, Guomiao Shen, Matija Snuderl, Theodore Vougiouklakis, Nick Vulpescu, Gael Westby, Paul Zappile, Yutong Zhang |
| EPI_ISL_427515 | hCoV-19/USA/NY-NYUMC220/2020 | North America / USA / New York / Manha | 2020-03-18 | NYU Langone Health | Departments of Pathology and Medicine, New York University School of Medicine | Maria Aguero-Rosenfeld, Brendan Belovarac, Margaret Black, Ludovic Boytard, John Cadley, Paolo Cotzia, John Chen, Dacia Dimartino, Xiaojun Feng, Tatyana Gindin, Emily Guzman, Adriana Heguy, Megan Hogan, Emily Huang, George Jour, Andrew Lytle, Christian Marier, Matthew T. Maurano, Mark J. Mulligan, Peter Meyn, Iman Osman, Jared Pinnell, Vanessa Raabe, Sitharam Ramaswami, Amy Rapkiewicz, Marie Samanovic-Golden, Antonio Serrano, Guomiao Shen, Matija Snuderl, Theodore Vougiouklakis, Nick Vulpescu, Gael Westby, Paul Zappile, Yutong Zhang |
| EPI_ISL_427516 | hCoV-19/USA/NY-NYUMC221/2020 | North America / USA / New York / Manha | 2020-03-18 | NYU Langone Health | Departments of Pathology and Medicine, New York University School of Medicine | Maria Aguero-Rosenfeld, Brendan Belovarac, Margaret Black, Ludovic Boytard, John Cadley, Paolo Cotzia, John Chen, Dacia Dimartino, Xiaojun Feng, Tatyana Gindin, Emily Guzman, Adriana Heguy, Megan Hogan, Emily Huang, George Jour, Andrew Lytle, Christian Marier, Matthew T. Maurano, Mark J. Mulligan, Peter Meyn, Iman Osman, Jared Pinnell, Vanessa Raabe, Sitharam Ramaswami, Amy Rapkiewicz, Marie Samanovic-Golden, Antonio Serrano, Guomiao Shen, Matija Snuderl, Theodore Vougiouklakis, Nick Vulpescu, Gael Westby, Paul Zappile, Yutong Zhang |
| EPI_ISL_427518 | hCoV-19/USA/NY-NYUMC223/2020 | North America / USA / New York / Nassa | 2020-03-18 | NYU Langone Health | Departments of Pathology and Medicine, New York University School of Medicine | Maria Aguero-Rosenfeld, Brendan Belovarac, Margaret Black, Ludovic Boytard, John Cadley, Paolo Cotzia, John Chen, Dacia Dimartino, Xiaojun Feng, Tatyana Gindin, Emily Guzman, Adriana Heguy, Megan Hogan, Emily Huang, George Jour, Andrew Lytle, Christian Marier, Matthew T. Maurano, Mark J. Mulligan, Peter Meyn, Iman Osman, Jared Pinnell, Vanessa Raabe, Sitharam Ramaswami, Amy Rapkiewicz, Marie Samanovic-Golden, Antonio Serrano, Guomiao Shen, Matija Snuderl, Theodore Vougiouklakis, Nick Vulpescu, Gael Westby, Paul Zappile, Yutong Zhang |

| | | | | | | |
|---|---|---|---|---|---|---|
| EPI_ISL_427519 | hCoV-19/USA/NY-NYUMC224/2020 | North America / USA / New York / Passai | 2020-03-18 | NYU Langone Health | Departments of Pathology and Medicine, New York University School of Medicine | Maria Aguero-Rosenfeld, Brendan Belovarac, Margaret Black, Ludovic Boytard, John Cadley, Paolo Cotzia, John Chen, Dacia Dimartino, Xiaojun Feng, Tatyana Gindin, Emily Guzman, Adriana Heguy, Megan Hogan, Emily Huang, George Jour, Andrew Lytle, Christian Marier, Matthew T. Maurano, Mark J. Mulligan, Peter Meyn, Iman Osman, Jared Pinnell, Vanessa Raabe, Sitharam Ramaswami, Amy Rapkiewicz, Marie Samanovic-Golden, Antonio Serrano, Guomiao Shen, Matija Snuderl, Theodore Vougiouklakis, Nick Vulpescu, Gael Westby, Paul Zappile, Yutong Zhang |
| EPI_ISL_427520 | hCoV-19/USA/NY-NYUMC225/2020 | North America / USA / New York / Brookl | 2020-03-18 | NYU Langone Health | Departments of Pathology and Medicine, New York University School of Medicine | Maria Aguero-Rosenfeld, Brendan Belovarac, Margaret Black, Ludovic Boytard, John Cadley, Paolo Cotzia, John Chen, Dacia Dimartino, Xiaojun Feng, Tatyana Gindin, Emily Guzman, Adriana Heguy, Megan Hogan, Emily Huang, George Jour, Andrew Lytle, Christian Marier, Matthew T. Maurano, Mark J. Mulligan, Peter Meyn, Iman Osman, Jared Pinnell, Vanessa Raabe, Sitharam Ramaswami, Amy Rapkiewicz, Marie Samanovic-Golden, Antonio Serrano, Guomiao Shen, Matija Snuderl, Theodore Vougiouklakis, Nick Vulpescu, Gael Westby, Paul Zappile, Yutong Zhang |
| EPI_ISL_427521 | hCoV-19/USA/NY-NYUMC226/2020 | North America / USA / New York / Nassa | 2020-03-18 | NYU Langone Health | Departments of Pathology and Medicine, New York University School of Medicine | Maria Aguero-Rosenfeld, Brendan Belovarac, Margaret Black, Ludovic Boytard, John Cadley, Paolo Cotzia, John Chen, Dacia Dimartino, Xiaojun Feng, Tatyana Gindin, Emily Guzman, Adriana Heguy, Megan Hogan, Emily Huang, George Jour, Andrew Lytle, Christian Marier, Matthew T. Maurano, Mark J. Mulligan, Peter Meyn, Iman Osman, Jared Pinnell, Vanessa Raabe, Sitharam Ramaswami, Amy Rapkiewicz, Marie Samanovic-Golden, Antonio Serrano, Guomiao Shen, Matija Snuderl, Theodore Vougiouklakis, Nick Vulpescu, Gael Westby, Paul Zappile, Yutong Zhang |
| EPI_ISL_427525 | hCoV-19/USA/NY-NYUMC190/2020 | North America / USA / New York / Queen | 2020-04-01 | NYU Langone Health | Departments of Pathology and Medicine, New York University School of Medicine | Maria Aguero-Rosenfeld, Brendan Belovarac, Margaret Black, Ludovic Boytard, John Cadley, Paolo Cotzia, John Chen, Dacia Dimartino, Xiaojun Feng, Tatyana Gindin, Emily Guzman, Adriana Heguy, Megan Hogan, Emily Huang, George Jour, Andrew Lytle, Christian Marier, Matthew T. Maurano, Mark J. Mulligan, Peter Meyn, Iman Osman, Jared Pinnell, Vanessa Raabe, Sitharam Ramaswami, Amy Rapkiewicz, Marie Samanovic-Golden, Antonio Serrano, Guomiao Shen, Matija Snuderl, Theodore Vougiouklakis, Nick Vulpescu, Gael Westby, Paul Zappile, Yutong Zhang |
| EPI_ISL_427526 | hCoV-19/USA/NY-WCMP2C03P/2020 | North America / USA / New York / Manha | 2020-03-12 | NewYork-Presbyterian & Mason Lab | Mason Lab | Daniel J. Butler, Christopher Mozsary, Cem Meydan, David Danko, Jonathan Foox, Joel Rosiene, Alon Shaiber, Matthew MacKay, Ebrahim Afshinnekoo, Fritz J. Sedlazeck, Nikolay A. Ivanov, Maria Sierra, Craig D. Westover, Krista Ryon, Benjamin Young, Chandrima Bhattacharya, Phyllis Ruggiero, Justyna Gawrys, Iman Hajirasouliha, Dmitry Meleshko, Mirella Salvatore, Dong Xu, Jenny Xiang, John Sipley, Lin Cong, Arryn Craney, Priya Velu, Lars F. Westblade, Massimo Loda, Shawn Levy, Melissa Cushing, Marcin Imielinski, Hanna Rennert, Christopher E. Mason |
| EPI_ISL_427527 | hCoV-19/USA/NY-WCMP2D02P/2020 | North America / USA / New York / Manha | 2020-03-12 | NewYork-Presbyterian & Mason Lab | Mason Lab | Daniel J. Butler, Christopher Mozsary, Cem Meydan, David Danko, Jonathan Foox, Joel Rosiene, Alon Shaiber, Matthew MacKay, Ebrahim Afshinnekoo, Fritz J. Sedlazeck, Nikolay A. Ivanov, Maria Sierra, Craig D. Westover, Krista Ryon, Benjamin Young, Chandrima Bhattacharya, Phyllis Ruggiero, Justyna Gawrys, Iman Hajirasouliha, Dmitry Meleshko, Mirella Salvatore, Dong Xu, Jenny Xiang, John Sipley, Lin Cong, Arryn Craney, Priya Velu, Lars F. Westblade, Massimo Loda, Shawn Levy, Melissa Cushing, Marcin Imielinski, Hanna Rennert, Christopher E. Mason |
| EPI_ISL_427528 | hCoV-19/USA/NY-WCMP2D04P/2020 | North America / USA / New York / Manha | 2020-03-12 | NewYork-Presbyterian & Mason Lab | Mason Lab | Daniel J. Butler, Christopher Mozsary, Cem Meydan, David Danko, Jonathan Foox, Joel Rosiene, Alon Shaiber, Matthew MacKay, Ebrahim Afshinnekoo, Fritz J. Sedlazeck, Nikolay A. Ivanov, Maria Sierra, Craig D. Westover, Krista Ryon, Benjamin Young, Chandrima Bhattacharya, Phyllis Ruggiero, Justyna Gawrys, Iman Hajirasouliha, Dmitry Meleshko, Mirella Salvatore, Dong Xu, Jenny Xiang, John Sipley, Lin Cong, Arryn Craney, Priya Velu, Lars F. Westblade, Massimo Loda, Shawn Levy, Melissa Cushing, Marcin Imielinski, Hanna Rennert, Christopher E. Mason |
| EPI_ISL_427529 | hCoV-19/USA/NY-WCMP2H03P/2020 | North America / USA / New York / Manha | 2020-03-12 | NewYork-Presbyterian & Mason Lab | Mason Lab | Daniel J. Butler, Christopher Mozsary, Cem Meydan, David Danko, Jonathan Foox, Joel Rosiene, Alon Shaiber, Matthew MacKay, Ebrahim Afshinnekoo, Fritz J. Sedlazeck, Nikolay A. Ivanov, Maria Sierra, Craig D. Westover, Krista Ryon, Benjamin Young, Chandrima Bhattacharya, Phyllis Ruggiero, Justyna Gawrys, Iman Hajirasouliha, Dmitry Meleshko, Mirella Salvatore, Dong Xu, Jenny Xiang, John Sipley, Lin Cong, Arryn Craney, Priya Velu, Lars F. Westblade, Massimo Loda, Shawn Levy, Melissa Cushing, Marcin Imielinski, Hanna Rennert, Christopher E. Mason |
| EPI_ISL_427530 | hCoV-19/USA/NY-WCMP3E02P/2020 | North America / USA / New York / Manha | 2020-03-12 | NewYork-Presbyterian & Mason Lab | Mason Lab | Daniel J. Butler, Christopher Mozsary, Cem Meydan, David Danko, Jonathan Foox, Joel Rosiene, Alon Shaiber, Matthew MacKay, Ebrahim Afshinnekoo, Fritz J. Sedlazeck, Nikolay A. Ivanov, Maria Sierra, Craig D. Westover, Krista Ryon, Benjamin Young, Chandrima Bhattacharya, Phyllis Ruggiero, Justyna Gawrys, Iman Hajirasouliha, Dmitry Meleshko, Mirella Salvatore, Dong Xu, Jenny Xiang, John Sipley, Lin Cong, Arryn Craney, Priya Velu, Lars F. Westblade, Massimo Loda, Shawn Levy, Melissa Cushing, Marcin Imielinski, Hanna Rennert, Christopher E. Mason |
| EPI_ISL_427531 | hCoV-19/USA/NY-WCMP4A02N/2020 | North America / USA / New York / Manha | 2020-03-13 | NewYork-Presbyterian & Mason Lab | Mason Lab | Daniel J. Butler, Christopher Mozsary, Cem Meydan, David Danko, Jonathan Foox, Joel Rosiene, Alon Shaiber, Matthew MacKay, Ebrahim Afshinnekoo, Fritz J. Sedlazeck, Nikolay A. Ivanov, Maria Sierra, Craig D. Westover, Krista Ryon, Benjamin Young, Chandrima Bhattacharya, Phyllis Ruggiero, Justyna Gawrys, Iman Hajirasouliha, Dmitry Meleshko, Mirella Salvatore, Dong Xu, Jenny Xiang, John Sipley, Lin Cong, Arryn Craney, Priya Velu, Lars F. Westblade, Massimo Loda, Shawn Levy, Melissa Cushing, Marcin Imielinski, Hanna Rennert, Christopher E. Mason |
| EPI_ISL_427532 | hCoV-19/USA/NY-WCMP4B06P/2020 | North America / USA / New York / Manha | 2020-03-13 | NewYork-Presbyterian & Mason Lab | Mason Lab | Daniel J. Butler, Christopher Mozsary, Cem Meydan, David Danko, Jonathan Foox, Joel Rosiene, Alon Shaiber, Matthew MacKay, Ebrahim Afshinnekoo, Fritz J. Sedlazeck, Nikolay A. Ivanov, Maria Sierra, Craig D. Westover, Krista Ryon, Benjamin Young, Chandrima Bhattacharya, Phyllis Ruggiero, Justyna Gawrys, Iman Hajirasouliha, Dmitry Meleshko, Mirella Salvatore, Dong Xu, Jenny Xiang, John Sipley, Lin Cong, Arryn Craney, Priya Velu, Lars F. Westblade, Massimo Loda, Shawn Levy, Melissa Cushing, Marcin Imielinski, Hanna Rennert, Christopher E. Mason |
| EPI_ISL_427533 | hCoV-19/USA/NY-WCMP4C01P/2020 | North America / USA / New York / Manha | 2020-03-13 | NewYork-Presbyterian & Mason Lab | Mason Lab | Daniel J. Butler, Christopher Mozsary, Cem Meydan, David Danko, Jonathan Foox, Joel Rosiene, Alon Shaiber, Matthew MacKay, Ebrahim Afshinnekoo, Fritz J. Sedlazeck, Nikolay A. Ivanov, Maria Sierra, Craig D. Westover, Krista Ryon, Benjamin Young, Chandrima Bhattacharya, Phyllis Ruggiero, Justyna Gawrys, Iman Hajirasouliha, Dmitry Meleshko, Mirella Salvatore, Dong Xu, Jenny Xiang, John Sipley, Lin Cong, Arryn Craney, Priya Velu, Lars F. Westblade, Massimo Loda, Shawn Levy, Melissa Cushing, Marcin Imielinski, Hanna Rennert, Christopher E. Mason |
| EPI_ISL_427534 | hCoV-19/USA/NY-WCMP4C02N/2020 | North America / USA / New York / Manha | 2020-03-13 | NewYork-Presbyterian & Mason Lab | Mason Lab | Daniel J. Butler, Christopher Mozsary, Cem Meydan, David Danko, Jonathan Foox, Joel Rosiene, Alon Shaiber, Matthew MacKay, Ebrahim Afshinnekoo, Fritz J. Sedlazeck, Nikolay A. Ivanov, Maria Sierra, Craig D. Westover, Krista Ryon, Benjamin Young, Chandrima Bhattacharya, Phyllis Ruggiero, Justyna Gawrys, Iman Hajirasouliha, Dmitry Meleshko, Mirella Salvatore, Dong Xu, Jenny Xiang, John Sipley, Lin Cong, Arryn Craney, Priya Velu, Lars F. Westblade, Massimo Loda, Shawn Levy, Melissa Cushing, Marcin Imielinski, Hanna Rennert, Christopher E. Mason |
| EPI_ISL_427535 | hCoV-19/USA/NY-WCMP4D01P/2020 | North America / USA / New York / Manha | 2020-03-13 | NewYork-Presbyterian & Mason Lab | Mason Lab | Daniel J. Butler, Christopher Mozsary, Cem Meydan, David Danko, Jonathan Foox, Joel Rosiene, Alon Shaiber, Matthew MacKay, Ebrahim Afshinnekoo, Fritz J. Sedlazeck, Nikolay A. Ivanov, Maria Sierra, Craig D. Westover, Krista Ryon, Benjamin Young, Chandrima Bhattacharya, Phyllis Ruggiero, Justyna Gawrys, Iman Hajirasouliha, Dmitry Meleshko, Mirella Salvatore, Dong Xu, Jenny Xiang, John Sipley, Lin Cong, Arryn Craney, Priya Velu, Lars F. Westblade, Massimo Loda, Shawn Levy, Melissa Cushing, Marcin Imielinski, Hanna Rennert, Christopher E. Mason |
| EPI_ISL_427536 | hCoV-19/USA/NY-WCMP4D04P/2020 | North America / USA / New York / Manha | 2020-03-13 | NewYork-Presbyterian & Mason Lab | Mason Lab | Daniel J. Butler, Christopher Mozsary, Cem Meydan, David Danko, Jonathan Foox, Joel Rosiene, Alon Shaiber, Matthew MacKay, Ebrahim Afshinnekoo, Fritz J. Sedlazeck, Nikolay A. Ivanov, Maria Sierra, Craig D. Westover, Krista Ryon, Benjamin Young, Chandrima Bhattacharya, Phyllis Ruggiero, Justyna Gawrys, Iman Hajirasouliha, Dmitry Meleshko, Mirella Salvatore, Dong Xu, Jenny Xiang, John Sipley, Lin Cong, Arryn Craney, Priya Velu, Lars F. Westblade, Massimo Loda, Shawn Levy, Melissa Cushing, Marcin Imielinski, Hanna Rennert, Christopher E. Mason |
| EPI_ISL_427537 | hCoV-19/USA/NY-WCMP4E05P/2020 | North America / USA / New York / Manha | 2020-03-13 | NewYork-Presbyterian & Mason Lab | Mason Lab | Daniel J. Butler, Christopher Mozsary, Cem Meydan, David Danko, Jonathan Foox, Joel Rosiene, Alon Shaiber, Matthew MacKay, Ebrahim Afshinnekoo, Fritz J. Sedlazeck, Nikolay A. Ivanov, Maria Sierra, Craig D. Westover, Krista Ryon, Benjamin Young, Chandrima Bhattacharya, Phyllis Ruggiero, Justyna Gawrys, Iman Hajirasouliha, Dmitry Meleshko, Mirella Salvatore, Dong Xu, Jenny Xiang, John Sipley, Lin Cong, Arryn Craney, Priya Velu, Lars F. Westblade, Massimo Loda, Shawn Levy, Melissa Cushing, Marcin Imielinski, Hanna Rennert, Christopher E. Mason |
| EPI_ISL_427538 | hCoV-19/USA/NY-WCMP4G04P/2020 | North America / USA / New York / Manha | 2020-03-13 | NewYork-Presbyterian & Mason Lab | Mason Lab | Daniel J. Butler, Christopher Mozsary, Cem Meydan, David Danko, Jonathan Foox, Joel Rosiene, Alon Shaiber, Matthew MacKay, Ebrahim Afshinnekoo, Fritz J. Sedlazeck, Nikolay A. Ivanov, Maria Sierra, Craig D. Westover, Krista Ryon, Benjamin Young, Chandrima Bhattacharya, Phyllis Ruggiero, Justyna Gawrys, Iman Hajirasouliha, Dmitry Meleshko, Mirella Salvatore, Dong Xu, Jenny Xiang, John Sipley, Lin Cong, Arryn Craney, Priya Velu, Lars F. Westblade, Massimo Loda, Shawn Levy, Melissa Cushing, Marcin Imielinski, Hanna Rennert, Christopher E. Mason |
| EPI_ISL_427539 | hCoV-19/USA/NY-WCMP4H03P/2020 | North America / USA / New York / Manha | 2020-03-13 | NewYork-Presbyterian & Mason Lab | Mason Lab | Daniel J. Butler, Christopher Mozsary, Cem Meydan, David Danko, Jonathan Foox, Joel Rosiene, Alon Shaiber, Matthew MacKay, Ebrahim Afshinnekoo, Fritz J. Sedlazeck, Nikolay A. Ivanov, Maria Sierra, Craig D. Westover, Krista Ryon, Benjamin Young, Chandrima Bhattacharya, Phyllis Ruggiero, Justyna Gawrys, Iman Hajirasouliha, Dmitry Meleshko, Mirella Salvatore, Dong Xu, Jenny Xiang, John Sipley, Lin Cong, Arryn Craney, Priya Velu, Lars F. Westblade, Massimo Loda, Shawn Levy, Melissa Cushing, Marcin Imielinski, Hanna Rennert, Christopher E. Mason |
| EPI_ISL_427540 | hCoV-19/USA/NY-WCMP5F01P/2020 | North America / USA / New York / Manha | 2020-03-13 | NewYork-Presbyterian & Mason Lab | Mason Lab | Daniel J. Butler, Christopher Mozsary, Cem Meydan, David Danko, Jonathan Foox, Joel Rosiene, Alon Shaiber, Matthew MacKay, Ebrahim Afshinnekoo, Fritz J. Sedlazeck, Nikolay A. Ivanov, Maria Sierra, Craig D. Westover, Krista Ryon, Benjamin Young, Chandrima Bhattacharya, Phyllis Ruggiero, Justyna Gawrys, Iman Hajirasouliha, Dmitry Meleshko, Mirella Salvatore, Dong Xu, Jenny Xiang, John Sipley, Lin Cong, Arryn Craney, Priya Velu, Lars F. Westblade, Massimo Loda, Shawn Levy, Melissa Cushing, Marcin Imielinski, Hanna Rennert, Christopher E. Mason |
| EPI_ISL_427541 | hCoV-19/USA/NY-WCMP5G03P/2020 | North America / USA / New York / Manha | 2020-03-13 | NewYork-Presbyterian & Mason Lab | Mason Lab | Daniel J. Butler, Christopher Mozsary, Cem Meydan, David Danko, Jonathan Foox, Joel Rosiene, Alon Shaiber, Matthew MacKay, Ebrahim Afshinnekoo, Fritz J. Sedlazeck, Nikolay A. Ivanov, Maria Sierra, Craig D. Westover, Krista Ryon, Benjamin Young, Chandrima Bhattacharya, Phyllis Ruggiero, Justyna Gawrys, Iman Hajirasouliha, Dmitry Meleshko, Mirella Salvatore, Dong Xu, Jenny Xiang, John Sipley, Lin Cong, Arryn Craney, Priya Velu, Lars F. Westblade, Massimo Loda, Shawn Levy, Melissa Cushing, Marcin Imielinski, Hanna Rennert, Christopher E. Mason |
| EPI_ISL_427542 | hCoV-19/USA/NY-WCMP5H02P/2020 | North America / USA / New York / Manha | 2020-03-13 | NewYork-Presbyterian & Mason Lab | Mason Lab | Daniel J. Butler, Christopher Mozsary, Cem Meydan, David Danko, Jonathan Foox, Joel Rosiene, Alon Shaiber, Matthew MacKay, Ebrahim Afshinnekoo, Fritz J. Sedlazeck, Nikolay A. Ivanov, Maria Sierra, Craig D. Westover, Krista Ryon, Benjamin Young, Chandrima Bhattacharya, Phyllis Ruggiero, Justyna Gawrys, Iman Hajirasouliha, Dmitry Meleshko, Mirella Salvatore, Dong Xu, Jenny Xiang, John Sipley, Lin Cong, Arryn Craney, Priya Velu, Lars F. Westblade, Massimo Loda, Shawn Levy, Melissa Cushing, Marcin Imielinski, Hanna Rennert, Christopher E. Mason |
| EPI_ISL_427543 | hCoV-19/USA/NY-WCMP5H03P/2020 | North America / USA / New York / Manha | 2020-03-13 | NewYork-Presbyterian & Mason Lab | Mason Lab | Daniel J. Butler, Christopher Mozsary, Cem Meydan, David Danko, Jonathan Foox, Joel Rosiene, Alon Shaiber, Matthew MacKay, Ebrahim Afshinnekoo, Fritz J. Sedlazeck, Nikolay A. Ivanov, Maria Sierra, Craig D. Westover, Krista Ryon, Benjamin Young, Chandrima Bhattacharya, Phyllis Ruggiero, Justyna Gawrys, Iman Hajirasouliha, Dmitry Meleshko, Mirella Salvatore, Dong Xu, Jenny Xiang, John Sipley, Lin Cong, Arryn Craney, Priya Velu, Lars F. Westblade, Massimo Loda, Shawn Levy, Melissa Cushing, Marcin Imielinski, Hanna Rennert, Christopher E. Mason |
| EPI_ISL_427544 | hCoV-19/USA/NY-WCMP9A05P/2020 | North America / USA / New York / Manha | 2020-03-14 | NewYork-Presbyterian & Mason Lab | Mason Lab | Daniel J. Butler, Christopher Mozsary, Cem Meydan, David Danko, Jonathan Foox, Joel Rosiene, Alon Shaiber, Matthew MacKay, Ebrahim Afshinnekoo, Fritz J. Sedlazeck, Nikolay A. Ivanov, Maria Sierra, Craig D. Westover, Krista Ryon, Benjamin Young, Chandrima Bhattacharya, Phyllis Ruggiero, Justyna Gawrys, Iman Hajirasouliha, Dmitry Meleshko, Mirella Salvatore, Dong Xu, Jenny Xiang, John Sipley, Lin Cong, Arryn Craney, Priya Velu, Lars F. Westblade, Massimo Loda, Shawn Levy, Melissa Cushing, Marcin Imielinski, Hanna Rennert, Christopher E. Mason |
| EPI_ISL_427545 | hCoV-19/USA/NY-WCMP9A08P/2020 | North America / USA / New York / Manha | 2020-03-14 | NewYork-Presbyterian & Mason Lab | Mason Lab | Daniel J. Butler, Christopher Mozsary, Cem Meydan, David Danko, Jonathan Foox, Joel Rosiene, Alon Shaiber, Matthew MacKay, Ebrahim Afshinnekoo, Fritz J. Sedlazeck, Nikolay A. Ivanov, Maria Sierra, Craig D. Westover, Krista Ryon, Benjamin Young, Chandrima Bhattacharya, Phyllis Ruggiero, Justyna Gawrys, Iman Hajirasouliha, Dmitry Meleshko, Mirella Salvatore, Dong Xu, Jenny Xiang, John Sipley, Lin Cong, Arryn Craney, Priya Velu, Lars F. Westblade, Massimo Loda, Shawn Levy, Melissa Cushing, Marcin Imielinski, Hanna Rennert, Christopher E. Mason |
| EPI_ISL_427546 | hCoV-19/USA/NY-WCMP9B05P/2020 | North America / USA / New York / Manha | 2020-03-14 | NewYork-Presbyterian & Mason Lab | Mason Lab | Daniel J. Butler, Christopher Mozsary, Cem Meydan, David Danko, Jonathan Foox, Joel Rosiene, Alon Shaiber, Matthew MacKay, Ebrahim Afshinnekoo, Fritz J. Sedlazeck, Nikolay A. Ivanov, Maria Sierra, Craig D. Westover, Krista Ryon, Benjamin Young, Chandrima Bhattacharya, Phyllis Ruggiero, Justyna Gawrys, Iman Hajirasouliha, Dmitry Meleshko, Mirella Salvatore, Dong Xu, Jenny Xiang, John Sipley, Lin Cong, Arryn Craney, Priya Velu, Lars F. Westblade, Massimo Loda, Shawn Levy, Melissa Cushing, Marcin Imielinski, Hanna Rennert, Christopher E. Mason |
| EPI_ISL_427547 | hCoV-19/USA/NY-WCMP9B06P/2020 | North America / USA / New York / Manha | 2020-03-14 | NewYork-Presbyterian & Mason Lab | Mason Lab | Daniel J. Butler, Christopher Mozsary, Cem Meydan, David Danko, Jonathan Foox, Joel Rosiene, Alon Shaiber, Matthew MacKay, Ebrahim Afshinnekoo, Fritz J. Sedlazeck, Nikolay A. Ivanov, Maria Sierra, Craig D. Westover, Krista Ryon, Benjamin Young, Chandrima Bhattacharya, Phyllis Ruggiero, Justyna Gawrys, Iman Hajirasouliha, Dmitry Meleshko, Mirella Salvatore, Dong Xu, Jenny Xiang, John Sipley, Lin Cong, Arryn Craney, Priya Velu, Lars F. Westblade, Massimo Loda, Shawn Levy, Melissa Cushing, Marcin Imielinski, Hanna Rennert, Christopher E. Mason |
| EPI_ISL_427548 | hCoV-19/USA/NY-WCMP9B08P/2020 | North America / USA / New York / Manha | 2020-03-14 | NewYork-Presbyterian & Mason Lab | Mason Lab | Daniel J. Butler, Christopher Mozsary, Cem Meydan, David Danko, Jonathan Foox, Joel Rosiene, Alon Shaiber, Matthew MacKay, Ebrahim Afshinnekoo, Fritz J. Sedlazeck, Nikolay A. Ivanov, Maria Sierra, Craig D. Westover, Krista Ryon, Benjamin Young, Chandrima Bhattacharya, Phyllis Ruggiero, Justyna Gawrys, Iman Hajirasouliha, Dmitry Meleshko, Mirella Salvatore, Dong Xu, Jenny Xiang, John Sipley, Lin Cong, Arryn Craney, Priya Velu, Lars F. Westblade, Massimo Loda, Shawn Levy, Melissa Cushing, Marcin Imielinski, Hanna Rennert, Christopher E. Mason |
| EPI_ISL_427549 | hCoV-19/USA/NY-WCMP9B09P/2020 | North America / USA / New York / Manha | 2020-03-14 | NewYork-Presbyterian & Mason Lab | Mason Lab | Daniel J. Butler, Christopher Mozsary, Cem Meydan, David Danko, Jonathan Foox, Joel Rosiene, Alon Shaiber, Matthew MacKay, Ebrahim Afshinnekoo, Fritz J. Sedlazeck, Nikolay A. Ivanov, Maria Sierra, Craig D. Westover, Krista Ryon, Benjamin Young, Chandrima Bhattacharya, Phyllis Ruggiero, Justyna Gawrys, Iman Hajirasouliha, Dmitry Meleshko, Mirella Salvatore, Dong Xu, Jenny Xiang, John Sipley, Lin Cong, Arryn Craney, Priya Velu, Lars F. Westblade, Massimo Loda, Shawn Levy, Melissa Cushing, Marcin Imielinski, Hanna Rennert, Christopher E. Mason |
| EPI_ISL_427550 | hCoV-19/USA/NY-WCMP9D03P/2020 | North America / USA / New York / Manha | 2020-03-14 | NewYork-Presbyterian & Mason Lab | Mason Lab | Daniel J. Butler, Christopher Mozsary, Cem Meydan, David Danko, Jonathan Foox, Joel Rosiene, Alon Shaiber, Matthew MacKay, Ebrahim Afshinnekoo, Fritz J. Sedlazeck, Nikolay A. Ivanov, Maria Sierra, Craig D. Westover, Krista Ryon, Benjamin Young, Chandrima Bhattacharya, Phyllis Ruggiero, Justyna Gawrys, Iman Hajirasouliha, Dmitry Meleshko, Mirella Salvatore, Dong Xu, Jenny Xiang, John Sipley, Lin Cong, Arryn Craney, Priya Velu, Lars F. Westblade, Massimo Loda, Shawn Levy, Melissa Cushing, Marcin Imielinski, Hanna Rennert, Christopher E. Mason |
| EPI_ISL_427551 | hCoV-19/USA/NY-WCMP9E04P/2020 | North America / USA / New York / Manha | 2020-03-14 | NewYork-Presbyterian & Mason Lab | Mason Lab | Daniel J. Butler, Christopher Mozsary, Cem Meydan, David Danko, Jonathan Foox, Joel Rosiene, Alon Shaiber, Matthew MacKay, Ebrahim Afshinnekoo, Fritz J. Sedlazeck, Nikolay A. Ivanov, Maria Sierra, Craig D. Westover, Krista Ryon, Benjamin Young, Chandrima Bhattacharya, Phyllis Ruggiero, Justyna Gawrys, Iman Hajirasouliha, Dmitry Meleshko, Mirella Salvatore, Dong Xu, Jenny Xiang, John Sipley, Lin Cong, Arryn Craney, Priya Velu, Lars F. Westblade, Massimo Loda, Shawn Levy, Melissa Cushing, Marcin Imielinski, Hanna Rennert, Christopher E. Mason |
| EPI_ISL_427552 | hCoV-19/USA/NY-WCMP9G04P/2020 | North America / USA / New York / Manha | 2020-03-14 | NewYork-Presbyterian & Mason Lab | Mason Lab | Daniel J. Butler, Christopher Mozsary, Cem Meydan, David Danko, Jonathan Foox, Joel Rosiene, Alon Shaiber, Matthew MacKay, Ebrahim Afshinnekoo, Fritz J. Sedlazeck, Nikolay A. Ivanov, Maria Sierra, Craig D. Westover, Krista Ryon, Benjamin Young, Chandrima Bhattacharya, Phyllis Ruggiero, Justyna Gawrys, Iman Hajirasouliha, Dmitry Meleshko, Mirella Salvatore, Dong Xu, Jenny Xiang, John Sipley, Lin Cong, Arryn Craney, Priya Velu, Lars F. Westblade, Massimo Loda, Shawn Levy, Melissa Cushing, Marcin Imielinski, Hanna Rennert, Christopher E. Mason |
| EPI_ISL_427553 | hCoV-19/USA/NY-WCMP10A02P/2020 | North America / USA / New York / Manha | 2020-03-15 | NewYork-Presbyterian & Mason Lab | Mason Lab | Daniel J. Butler, Christopher Mozsary, Cem Meydan, David Danko, Jonathan Foox, Joel Rosiene, Alon Shaiber, Matthew MacKay, Ebrahim Afshinnekoo, Fritz J. Sedlazeck, Nikolay A. Ivanov, Maria Sierra, Craig D. Westover, Krista Ryon, Benjamin Young, Chandrima Bhattacharya, Phyllis Ruggiero, Justyna Gawrys, Iman Hajirasouliha, Dmitry Meleshko, Mirella Salvatore, Dong Xu, Jenny Xiang, John Sipley, Lin Cong, Arryn Craney, Priya Velu, Lars F. Westblade, Massimo Loda, Shawn Levy, Melissa Cushing, Marcin Imielinski, Hanna Rennert, Christopher E. Mason |
| EPI_ISL_427554 | hCoV-19/USA/NY-WCMP10A05P/2020 | North America / USA / New York / Manha | 2020-03-15 | NewYork-Presbyterian & Mason Lab | Mason Lab | Daniel J. Butler, Christopher Mozsary, Cem Meydan, David Danko, Jonathan Foox, Joel Rosiene, Alon Shaiber, Matthew MacKay, Ebrahim Afshinnekoo, Fritz J. Sedlazeck, Nikolay A. Ivanov, Maria Sierra, Craig D. Westover, Krista Ryon, Benjamin Young, Chandrima Bhattacharya, Phyllis Ruggiero, Justyna Gawrys, Iman Hajirasouliha, Dmitry Meleshko, Mirella Salvatore, Dong Xu, Jenny Xiang, John Sipley, Lin Cong, Arryn Craney, Priya Velu, Lars F. Westblade, Massimo Loda, Shawn Levy, Melissa Cushing, Marcin Imielinski, Hanna Rennert, Christopher E. Mason |
| EPI_ISL_427555 | hCoV-19/USA/NY-WCMP10A06P/2020 | North America / USA / New York / Manha | 2020-03-15 | NewYork-Presbyterian & Mason Lab | Mason Lab | Daniel J. Butler, Christopher Mozsary, Cem Meydan, David Danko, Jonathan Foox, Joel Rosiene, Alon Shaiber, Matthew MacKay, Ebrahim Afshinnekoo, Fritz J. Sedlazeck, Nikolay A. Ivanov, Maria Sierra, Craig D. Westover, Krista Ryon, Benjamin Young, Chandrima Bhattacharya, Phyllis Ruggiero, Justyna Gawrys, Iman Hajirasouliha, Dmitry Meleshko, Mirella Salvatore, Dong Xu, Jenny Xiang, John Sipley, Lin Cong, Arryn Craney, Priya Velu, Lars F. Westblade, Massimo Loda, Shawn Levy, Melissa Cushing, Marcin Imielinski, Hanna Rennert, Christopher E. Mason |
| EPI_ISL_427556 | hCoV-19/USA/NY-WCMP10A12P/2020 | North America / USA / New York / Manha | 2020-03-15 | NewYork-Presbyterian & Mason Lab | Mason Lab | Daniel J. Butler, Christopher Mozsary, Cem Meydan, David Danko, Jonathan Foox, Joel Rosiene, Alon Shaiber, Matthew MacKay, Ebrahim Afshinnekoo, Fritz J. Sedlazeck, Nikolay A. Ivanov, Maria Sierra, Craig D. Westover, Krista Ryon, Benjamin Young, Chandrima Bhattacharya, Phyllis Ruggiero, Justyna Gawrys, Iman Hajirasouliha, Dmitry Meleshko, Mirella Salvatore, Dong Xu, Jenny Xiang, John Sipley, Lin Cong, Arryn Craney, Priya Velu, Lars F. Westblade, Massimo Loda, Shawn Levy, Melissa Cushing, Marcin Imielinski, Hanna Rennert, Christopher E. Mason |
| EPI_ISL_427557 | hCoV-19/USA/NY-WCMP10B03P/2020 | North America / USA / New York / Manha | 2020-03-15 | NewYork-Presbyterian & Mason Lab | Mason Lab | Daniel J. Butler, Christopher Mozsary, Cem Meydan, David Danko, Jonathan Foox, Joel Rosiene, Alon Shaiber, Matthew MacKay, Ebrahim Afshinnekoo, Fritz J. Sedlazeck, Nikolay A. Ivanov, Maria Sierra, Craig D. Westover, Krista Ryon, Benjamin Young, Chandrima Bhattacharya, Phyllis Ruggiero, Justyna Gawrys, Iman Hajirasouliha, Dmitry Meleshko, Mirella Salvatore, Dong Xu, Jenny Xiang, John Sipley, Lin Cong, Arryn Craney, Priya Velu, Lars F. Westblade, Massimo Loda, Shawn Levy, Melissa Cushing, Marcin Imielinski, Hanna Rennert, Christopher E. Mason |
| EPI_ISL_427558 | hCoV-19/USA/NY-WCMP10B07N/2020 | North America / USA / New York / Manha | 2020-03-15 | NewYork-Presbyterian & Mason Lab | Mason Lab | Daniel J. Butler, Christopher Mozsary, Cem Meydan, David Danko, Jonathan Foox, Joel Rosiene, Alon Shaiber, Matthew MacKay, Ebrahim Afshinnekoo, Fritz J. Sedlazeck, Nikolay A. Ivanov, Maria Sierra, Craig D. Westover, Krista Ryon, Benjamin Young, Chandrima Bhattacharya, Phyllis Ruggiero, Justyna Gawrys, Iman Hajirasouliha, Dmitry Meleshko, Mirella Salvatore, Dong Xu, Jenny Xiang, John Sipley, Lin Cong, Arryn Craney, Priya Velu, Lars F. Westblade, Massimo Loda, Shawn Levy, Melissa Cushing, Marcin Imielinski, Hanna Rennert, Christopher E. Mason |
| EPI_ISL_427559 | hCoV-19/USA/NY-WCMP10B08P/2020 | North America / USA / New York / Manha | 2020-03-15 | NewYork-Presbyterian & Mason Lab | Mason Lab | Daniel J. Butler, Christopher Mozsary, Cem Meydan, David Danko, Jonathan Foox, Joel Rosiene, Alon Shaiber, Matthew MacKay, Ebrahim Afshinnekoo, Fritz J. Sedlazeck, Nikolay A. Ivanov, Maria Sierra, Craig D. Westover, Krista Ryon, Benjamin Young, Chandrima Bhattacharya, Phyllis Ruggiero, Justyna Gawrys, Iman Hajirasouliha, Dmitry Meleshko, Mirella Salvatore, Dong Xu, Jenny Xiang, John Sipley, Lin Cong, Arryn Craney, Priya Velu, Lars F. Westblade, Massimo Loda, Shawn Levy, Melissa Cushing, Marcin Imielinski, Hanna Rennert, Christopher E. Mason |
| EPI_ISL_427560 | hCoV-19/USA/NY-WCMP10B11P/2020 | North America / USA / New York / Manha | 2020-03-15 | NewYork-Presbyterian & Mason Lab | Mason Lab | Daniel J. Butler, Christopher Mozsary, Cem Meydan, David Danko, Jonathan Foox, Joel Rosiene, Alon Shaiber, Matthew MacKay, Ebrahim Afshinnekoo, Fritz J. Sedlazeck, Nikolay A. Ivanov, Maria Sierra, Craig D. Westover, Krista Ryon, Benjamin Young, Chandrima Bhattacharya, Phyllis Ruggiero, Justyna Gawrys, Iman Hajirasouliha, Dmitry Meleshko, Mirella Salvatore, Dong Xu, Jenny Xiang, John Sipley, Lin Cong, Arryn Craney, Priya Velu, Lars F. Westblade, Massimo Loda, Shawn Levy, Melissa Cushing, Marcin Imielinski, Hanna Rennert, Christopher E. Mason |
| EPI_ISL_427561 | hCoV-19/USA/NY-WCMP10B12P/2020 | North America / USA / New York / Manha | 2020-03-15 | NewYork-Presbyterian & Mason Lab | Mason Lab | Daniel J. Butler, Christopher Mozsary, Cem Meydan, David Danko, Jonathan Foox, Joel Rosiene, Alon Shaiber, Matthew MacKay, Ebrahim Afshinnekoo, Fritz J. Sedlazeck, Nikolay A. Ivanov, Maria Sierra, Craig D. Westover, Krista Ryon, Benjamin Young, Chandrima Bhattacharya, Phyllis Ruggiero, Justyna Gawrys, Iman Hajirasouliha, Dmitry Meleshko, Mirella Salvatore, Dong Xu, Jenny Xiang, John Sipley, Lin Cong, Arryn Craney, Priya Velu, Lars F. Westblade, Massimo Loda, Shawn Levy, Melissa Cushing, Marcin Imielinski, Hanna Rennert, Christopher E. Mason |
| EPI_ISL_427562 | hCoV-19/USA/NY-WCMP10C03P/2020 | North America / USA / New York / Manha | 2020-03-15 | NewYork-Presbyterian & Mason Lab | Mason Lab | Daniel J. Butler, Christopher Mozsary, Cem Meydan, David Danko, Jonathan Foox, Joel Rosiene, Alon Shaiber, Matthew MacKay, Ebrahim Afshinnekoo, Fritz J. Sedlazeck, Nikolay A. Ivanov, Maria Sierra, Craig D. Westover, Krista Ryon, Benjamin Young, Chandrima Bhattacharya, Phyllis Ruggiero, Justyna Gawrys, Iman Hajirasouliha, Dmitry Meleshko, Mirella Salvatore, Dong Xu, Jenny Xiang, John Sipley, Lin Cong, Arryn Craney, Priya Velu, Lars F. Westblade, Massimo Loda, Shawn Levy, Melissa Cushing, Marcin Imielinski, Hanna Rennert, Christopher E. Mason |
| EPI_ISL_427563 | hCoV-19/USA/NY-WCMP10C04P/2020 | North America / USA / New York / Manha | 2020-03-15 | NewYork-Presbyterian & Mason Lab | Mason Lab | Daniel J. Butler, Christopher Mozsary, Cem Meydan, David Danko, Jonathan Foox, Joel Rosiene, Alon Shaiber, Matthew MacKay, Ebrahim Afshinnekoo, Fritz J. Sedlazeck, Nikolay A. Ivanov, Maria Sierra, Craig D. Westover, Krista Ryon, Benjamin Young, Chandrima Bhattacharya, Phyllis Ruggiero, Justyna Gawrys, Iman Hajirasouliha, Dmitry Meleshko, Mirella Salvatore, Dong Xu, Jenny Xiang, John Sipley, Lin Cong, Arryn Craney, Priya Velu, Lars F. Westblade, Massimo Loda, Shawn Levy, Melissa Cushing, Marcin Imielinski, Hanna Rennert, Christopher E. Mason |
| EPI_ISL_427564 | hCoV-19/USA/NY-WCMP10C09P/2020 | North America / USA / New York / Manha | 2020-03-15 | NewYork-Presbyterian & Mason Lab | Mason Lab | Daniel J. Butler, Christopher Mozsary, Cem Meydan, David Danko, Jonathan Foox, Joel Rosiene, Alon Shaiber, Matthew MacKay, Ebrahim Afshinnekoo, Fritz J. Sedlazeck, Nikolay A. Ivanov, Maria Sierra, Craig D. Westover, Krista Ryon, Benjamin Young, Chandrima Bhattacharya, Phyllis Ruggiero, Justyna Gawrys, Iman Hajirasouliha, Dmitry Meleshko, Mirella Salvatore, Dong Xu, Jenny Xiang, John Sipley, Lin Cong, Arryn Craney, Priya Velu, Lars F. Westblade, Massimo Loda, Shawn Levy, Melissa Cushing, Marcin Imielinski, Hanna Rennert, Christopher E. Mason |
| EPI_ISL_427565 | hCoV-19/USA/NY-WCMP10C11P/2020 | North America / USA / New York / Manha | 2020-03-15 | NewYork-Presbyterian & Mason Lab | Mason Lab | Daniel J. Butler, Christopher Mozsary, Cem Meydan, David Danko, Jonathan Foox, Joel Rosiene, Alon Shaiber, Matthew MacKay, Ebrahim Afshinnekoo, Fritz J. Sedlazeck, Nikolay A. Ivanov, Maria Sierra, Craig D. Westover, Krista Ryon, Benjamin Young, Chandrima Bhattacharya, Phyllis Ruggiero, Justyna Gawrys, Iman Hajirasouliha, Dmitry Meleshko, Mirella Salvatore, Dong Xu, Jenny Xiang, John Sipley, Lin Cong, Arryn Craney, Priya Velu, Lars F. Westblade, Massimo Loda, Shawn Levy, Melissa Cushing, Marcin Imielinski, Hanna Rennert, Christopher E. Mason |
| EPI_ISL_427566 | hCoV-19/USA/NY-WCMP10D06P/2020 | North America / USA / New York / Manha | 2020-03-15 | NewYork-Presbyterian & Mason Lab | Mason Lab | Daniel J. Butler, Christopher Mozsary, Cem Meydan, David Danko, Jonathan Foox, Joel Rosiene, Alon Shaiber, Matthew MacKay, Ebrahim Afshinnekoo, Fritz J. Sedlazeck, Nikolay A. Ivanov, Maria Sierra, Craig D. Westover, Krista Ryon, Benjamin Young, Chandrima Bhattacharya, Phyllis Ruggiero, Justyna Gawrys, Iman Hajirasouliha, Dmitry Meleshko, Mirella Salvatore, Dong Xu, Jenny Xiang, John Sipley, Lin Cong, Arryn Craney, Priya Velu, Lars F. Westblade, Massimo Loda, Shawn Levy, Melissa Cushing, Marcin Imielinski, Hanna Rennert, Christopher E. Mason |
| EPI_ISL_427567 | hCoV-19/USA/NY-WCMP10D09P/2020 | North America / USA / New York / Manha | 2020-03-15 | NewYork-Presbyterian & Mason Lab | Mason Lab | Daniel J. Butler, Christopher Mozsary, Cem Meydan, David Danko, Jonathan Foox, Joel Rosiene, Alon Shaiber, Matthew MacKay, Ebrahim Afshinnekoo, Fritz J. Sedlazeck, Nikolay A. Ivanov, Maria Sierra, Craig D. Westover, Krista Ryon, Benjamin Young, Chandrima Bhattacharya, Phyllis Ruggiero, Justyna Gawrys, Iman Hajirasouliha, Dmitry Meleshko, Mirella Salvatore, Dong Xu, Jenny Xiang, John Sipley, Lin Cong, Arryn Craney, Priya Velu, Lars F. Westblade, Massimo Loda, Shawn Levy, Melissa Cushing, Marcin Imielinski, Hanna Rennert, Christopher E. Mason |
| EPI_ISL_427568 | hCoV-19/USA/NY-WCMP10D11P/2020 | North America / USA / New York / Manha | 2020-03-15 | NewYork-Presbyterian & Mason Lab | Mason Lab | Daniel J. Butler, Christopher Mozsary, Cem Meydan, David Danko, Jonathan Foox, Joel Rosiene, Alon Shaiber, Matthew MacKay, Ebrahim Afshinnekoo, Fritz J. Sedlazeck, Nikolay A. Ivanov, Maria Sierra, Craig D. Westover, Krista Ryon, Benjamin Young, Chandrima Bhattacharya, Phyllis Ruggiero, Justyna Gawrys, Iman Hajirasouliha, Dmitry Meleshko, Mirella Salvatore, Dong Xu, Jenny Xiang, John Sipley, Lin Cong, Arryn Craney, Priya Velu, Lars F. Westblade, Massimo Loda, Shawn Levy, Melissa Cushing, Marcin Imielinski, Hanna Rennert, Christopher E. Mason |
| EPI_ISL_427569 | hCoV-19/USA/NY-WCMP10E02P/2020 | North America / USA / New York / Manha | 2020-03-15 | NewYork-Presbyterian & Mason Lab | Mason Lab | Daniel J. Butler, Christopher Mozsary, Cem Meydan, David Danko, Jonathan Foox, Joel Rosiene, Alon Shaiber, Matthew MacKay, Ebrahim Afshinnekoo, Fritz J. Sedlazeck, Nikolay A. Ivanov, Maria Sierra, Craig D. Westover, Krista Ryon, Benjamin Young, Chandrima Bhattacharya, Phyllis Ruggiero, Justyna Gawrys, Iman Hajirasouliha, Dmitry Meleshko, Mirella Salvatore, Dong Xu, Jenny Xiang, John Sipley, Lin Cong, Arryn Craney, Priya Velu, Lars F. Westblade, Massimo Loda, Shawn Levy, Melissa Cushing, Marcin Imielinski, Hanna Rennert, Christopher E. Mason |
| EPI_ISL_427570 | hCoV-19/USA/NY-WCMP10E06P/2020 | North America / USA / New York / Manha | 2020-03-15 | NewYork-Presbyterian & Mason Lab | Mason Lab | Daniel J. Butler, Christopher Mozsary, Cem Meydan, David Danko, Jonathan Foox, Joel Rosiene, Alon Shaiber, Matthew MacKay, Ebrahim Afshinnekoo, Fritz J. Sedlazeck, Nikolay A. Ivanov, Maria Sierra, Craig D. Westover, Krista Ryon, Benjamin Young, Chandrima Bhattacharya, Phyllis Ruggiero, Justyna Gawrys, Iman Hajirasouliha, Dmitry Meleshko, Mirella Salvatore, Dong Xu, Jenny Xiang, John Sipley, Lin Cong, Arryn Craney, Priya Velu, Lars F. Westblade, Massimo Loda, Shawn Levy, Melissa Cushing, Marcin Imielinski, Hanna Rennert, Christopher E. Mason |
| EPI_ISL_427571 | hCoV-19/USA/NY-WCMP10E07P/2020 | North America / USA / New York / Manha | 2020-03-15 | NewYork-Presbyterian & Mason Lab | Mason Lab | Daniel J. Butler, Christopher Mozsary, Cem Meydan, David Danko, Jonathan Foox, Joel Rosiene, Alon Shaiber, Matthew MacKay, Ebrahim Afshinnekoo, Fritz J. Sedlazeck, Nikolay A. Ivanov, Maria Sierra, Craig D. Westover, Krista Ryon, Benjamin Young, Chandrima Bhattacharya, Phyllis Ruggiero, Justyna Gawrys, Iman Hajirasouliha, Dmitry Meleshko, Mirella Salvatore, Dong Xu, Jenny Xiang, John Sipley, Lin Cong, Arryn Craney, Priya Velu, Lars F. Westblade, Massimo Loda, Shawn Levy, Melissa Cushing, Marcin Imielinski, Hanna Rennert, Christopher E. Mason |
| EPI_ISL_427572 | hCoV-19/USA/NY-WCMP10F04P/2020 | North America / USA / New York / Manha | 2020-03-15 | NewYork-Presbyterian & Mason Lab | Mason Lab | Daniel J. Butler, Christopher Mozsary, Cem Meydan, David Danko, Jonathan Foox, Joel Rosiene, Alon Shaiber, Matthew MacKay, Ebrahim Afshinnekoo, Fritz J. Sedlazeck, Nikolay A. Ivanov, Maria Sierra, Craig D. Westover, Krista Ryon, Benjamin Young, Chandrima Bhattacharya, Phyllis Ruggiero, Justyna Gawrys, Iman Hajirasouliha, Dmitry Meleshko, Mirella Salvatore, Dong Xu, Jenny Xiang, John Sipley, Lin Cong, Arryn Craney, Priya Velu, Lars F. Westblade, Massimo Loda, Shawn Levy, Melissa Cushing, Marcin Imielinski, Hanna Rennert, Christopher E. Mason |
| EPI_ISL_427573 | hCoV-19/USA/NY-WCMP10F06P/2020 | North America / USA / New York / Manha | 2020-03-15 | NewYork-Presbyterian & Mason Lab | Mason Lab | Daniel J. Butler, Christopher Mozsary, Cem Meydan, David Danko, Jonathan Foox, Joel Rosiene, Alon Shaiber, Matthew MacKay, Ebrahim Afshinnekoo, Fritz J. Sedlazeck, Nikolay A. Ivanov, Maria Sierra, Craig D. Westover, Krista Ryon, Benjamin Young, Chandrima Bhattacharya, Phyllis Ruggiero, Justyna Gawrys, Iman Hajirasouliha, Dmitry Meleshko, Mirella Salvatore, Dong Xu, Jenny Xiang, John Sipley, Lin Cong, Arryn Craney, Priya Velu, Lars F. Westblade, Massimo Loda, Shawn Levy, Melissa Cushing, Marcin Imielinski, Hanna Rennert, Christopher E. Mason |
| EPI_ISL_427574 | hCoV-19/USA/NY-WCMP10F09P/2020 | North America / USA / New York / Manha | 2020-03-15 | NewYork-Presbyterian & Mason Lab | Mason Lab | Daniel J. Butler, Christopher Mozsary, Cem Meydan, David Danko, Jonathan Foox, Joel Rosiene, Alon Shaiber, Matthew MacKay, Ebrahim Afshinnekoo, Fritz J. Sedlazeck, Nikolay A. Ivanov, Maria Sierra, Craig D. Westover, Krista Ryon, Benjamin Young, Chandrima Bhattacharya, Phyllis Ruggiero, Justyna Gawrys, Iman Hajirasouliha, Dmitry Meleshko, Mirella Salvatore, Dong Xu, Jenny Xiang, John Sipley, Lin Cong, Arryn Craney, Priya Velu, Lars F. Westblade, Massimo Loda, Shawn Levy, Melissa Cushing, Marcin Imielinski, Hanna Rennert, Christopher E. Mason |
| EPI_ISL_427575 | hCoV-19/USA/NY-WCMP10F10P/2020 | North America / USA / New York / Manha | 2020-03-15 | NewYork-Presbyterian & Mason Lab | Mason Lab | Daniel J. Butler, Christopher Mozsary, Cem Meydan, David Danko, Jonathan Foox, Joel Rosiene, Alon Shaiber, Matthew MacKay, Ebrahim Afshinnekoo, Fritz J. Sedlazeck, Nikolay A. Ivanov, Maria Sierra, Craig D. Westover, Krista Ryon, Benjamin Young, Chandrima Bhattacharya, Phyllis Ruggiero, Justyna Gawrys, Iman Hajirasouliha, Dmitry Meleshko, Mirella Salvatore, Dong Xu, Jenny Xiang, John Sipley, Lin Cong, Arryn Craney, Priya Velu, Lars F. Westblade, Massimo Loda, Shawn Levy, Melissa Cushing, Marcin Imielinski, Hanna Rennert, Christopher E. Mason |
| EPI_ISL_427576 | hCoV-19/USA/NY-WCMP10G05P/2020 | North America / USA / New York / Manha | 2020-03-15 | NewYork-Presbyterian & Mason Lab | Mason Lab | Daniel J. Butler, Christopher Mozsary, Cem Meydan, David Danko, Jonathan Foox, Joel Rosiene, Alon Shaiber, Matthew MacKay, Ebrahim Afshinnekoo, Fritz J. Sedlazeck, Nikolay A. Ivanov, Maria Sierra, Craig D. Westover, Krista Ryon, Benjamin Young, Chandrima Bhattacharya, Phyllis Ruggiero, Justyna Gawrys, Iman Hajirasouliha, Dmitry Meleshko, Mirella Salvatore, Dong Xu, Jenny Xiang, John Sipley, Lin Cong, Arryn Craney, Priya Velu, Lars F. Westblade, Massimo Loda, Shawn Levy, Melissa Cushing, Marcin Imielinski, Hanna Rennert, Christopher E. Mason |
| EPI_ISL_427577 | hCoV-19/USA/NY-WCMP10G06P/2020 | North America / USA / New York / Manha | 2020-03-15 | NewYork-Presbyterian & Mason Lab | Mason Lab | Daniel J. Butler, Christopher Mozsary, Cem Meydan, David Danko, Jonathan Foox, Joel Rosiene, Alon Shaiber, Matthew MacKay, Ebrahim Afshinnekoo, Fritz J. Sedlazeck, Nikolay A. Ivanov, Maria Sierra, Craig D. Westover, Krista Ryon, Benjamin Young, Chandrima Bhattacharya, Phyllis Ruggiero, Justyna Gawrys, Iman Hajirasouliha, Dmitry Meleshko, Mirella Salvatore, Dong Xu, Jenny Xiang, John Sipley, Lin Cong, Arryn Craney, Priya Velu, Lars F. Westblade, Massimo Loda, Shawn Levy, Melissa Cushing, Marcin Imielinski, Hanna Rennert, Christopher E. Mason |
| EPI_ISL_427578 | hCoV-19/USA/NY-WCMP10G10P/2020 | North America / USA / New York / Manha | 2020-03-15 | NewYork-Presbyterian & Mason Lab | Mason Lab | Daniel J. Butler, Christopher Mozsary, Cem Meydan, David Danko, Jonathan Foox, Joel Rosiene, Alon Shaiber, Matthew MacKay, Ebrahim Afshinnekoo, Fritz J. Sedlazeck, Nikolay A. Ivanov, Maria Sierra, Craig D. Westover, Krista Ryon, Benjamin Young, Chandrima Bhattacharya, Phyllis Ruggiero, Justyna Gawrys, Iman Hajirasouliha, Dmitry Meleshko, Mirella Salvatore, Dong Xu, Jenny Xiang, John Sipley, Lin Cong, Arryn Craney, Priya Velu, Lars F. Westblade, Massimo Loda, Shawn Levy, Melissa Cushing, Marcin Imielinski, Hanna Rennert, Christopher E. Mason |
| EPI_ISL_427579 | hCoV-19/USA/NY-WCMP10G11P/2020 | North America / USA / New York / Manha | 2020-03-15 | NewYork-Presbyterian & Mason Lab | Mason Lab | Daniel J. Butler, Christopher Mozsary, Cem Meydan, David Danko, Jonathan Foox, Joel Rosiene, Alon Shaiber, Matthew MacKay, Ebrahim Afshinnekoo, Fritz J. Sedlazeck, Nikolay A. Ivanov, Maria Sierra, Craig D. Westover, Krista Ryon, Benjamin Young, Chandrima Bhattacharya, Phyllis Ruggiero, Justyna Gawrys, Iman Hajirasouliha, Dmitry Meleshko, Mirella Salvatore, Dong Xu, Jenny Xiang, John Sipley, Lin Cong, Arryn Craney, Priya Velu, Lars F. Westblade, Massimo Loda, Shawn Levy, Melissa Cushing, Marcin Imielinski, Hanna Rennert, Christopher E. Mason |
| EPI_ISL_427580 | hCoV-19/USA/NY-WCMP10H01P/2020 | North America / USA / New York / Manha | 2020-03-15 | NewYork-Presbyterian & Mason Lab | Mason Lab | Daniel J. Butler, Christopher Mozsary, Cem Meydan, David Danko, Jonathan Foox, Joel Rosiene, Alon Shaiber, Matthew MacKay, Ebrahim Afshinnekoo, Fritz J. Sedlazeck, Nikolay A. Ivanov, Maria Sierra, Craig D. Westover, Krista Ryon, Benjamin Young, Chandrima Bhattacharya, Phyllis Ruggiero, Justyna Gawrys, Iman Hajirasouliha, Dmitry Meleshko, Mirella Salvatore, Dong Xu, Jenny Xiang, John Sipley, Lin Cong, Arryn Craney, Priya Velu, Lars F. Westblade, Massimo Loda, Shawn Levy, Melissa Cushing, Marcin Imielinski, Hanna Rennert, Christopher E. Mason |
| EPI_ISL_427581 | hCoV-19/USA/NY-WCMP10H04P/2020 | North America / USA / New York / Manha | 2020-03-15 | NewYork-Presbyterian & Mason Lab | Mason Lab | Daniel J. Butler, Christopher Mozsary, Cem Meydan, David Danko, Jonathan Foox, Joel Rosiene, Alon Shaiber, Matthew MacKay, Ebrahim Afshinnekoo, Fritz J. Sedlazeck, Nikolay A. Ivanov, Maria Sierra, Craig D. Westover, Krista Ryon, Benjamin Young, Chandrima Bhattacharya, Phyllis Ruggiero, Justyna Gawrys, Iman Hajirasouliha, Dmitry Meleshko, Mirella Salvatore, Dong Xu, Jenny Xiang, John Sipley, Lin Cong, Arryn Craney, Priya Velu, Lars F. Westblade, Massimo Loda, Shawn Levy, Melissa Cushing, Marcin Imielinski, Hanna Rennert, Christopher E. Mason |
| EPI_ISL_427582 | hCoV-19/USA/NY-WCMP10H08P/2020 | North America / USA / New York / Manha | 2020-03-15 | NewYork-Presbyterian & Mason Lab | Mason Lab | Daniel J. Butler, Christopher Mozsary, Cem Meydan, David Danko, Jonathan Foox, Joel Rosiene, Alon Shaiber, Matthew MacKay, Ebrahim Afshinnekoo, Fritz J. Sedlazeck, Nikolay A. Ivanov, Maria Sierra, Craig D. Westover, Krista Ryon, Benjamin Young, Chandrima Bhattacharya, Phyllis Ruggiero, Justyna Gawrys, Iman Hajirasouliha, Dmitry Meleshko, Mirella Salvatore, Dong Xu, Jenny Xiang, John Sipley, Lin Cong, Arryn Craney, Priya Velu, Lars F. Westblade, Massimo Loda, Shawn Levy, Melissa Cushing, Marcin Imielinski, Hanna Rennert, Christopher E. Mason |
| EPI_ISL_427583 | hCoV-19/USA/NY-WCMP11A03P/2020 | North America / USA / New York / Manha | 2020-03-16 | NewYork-Presbyterian & Mason Lab | Mason Lab | Daniel J. Butler, Christopher Mozsary, Cem Meydan, David Danko, Jonathan Foox, Joel Rosiene, Alon Shaiber, Matthew MacKay, Ebrahim Afshinnekoo, Fritz J. Sedlazeck, Nikolay A. Ivanov, Maria Sierra, Craig D. Westover, Krista Ryon, Benjamin Young, Chandrima Bhattacharya, Phyllis Ruggiero, Justyna Gawrys, Iman Hajirasouliha, Dmitry Meleshko, Mirella Salvatore, Dong Xu, Jenny Xiang, John Sipley, Lin Cong, Arryn Craney, Priya Velu, Lars F. Westblade, Massimo Loda, Shawn Levy, Melissa Cushing, Marcin Imielinski, Hanna Rennert, Christopher E. Mason |
| EPI_ISL_427584 | hCoV-19/USA/NY-WCMP11A06P/2020 | North America / USA / New York / Manha | 2020-03-16 | NewYork-Presbyterian & Mason Lab | Mason Lab | Daniel J. Butler, Christopher Mozsary, Cem Meydan, David Danko, Jonathan Foox, Joel Rosiene, Alon Shaiber, Matthew MacKay, Ebrahim Afshinnekoo, Fritz J. Sedlazeck, Nikolay A. Ivanov, Maria Sierra, Craig D. Westover, Krista Ryon, Benjamin Young, Chandrima Bhattacharya, Phyllis Ruggiero, Justyna Gawrys, Iman Hajirasouliha, Dmitry Meleshko, Mirella Salvatore, Dong Xu, Jenny Xiang, John Sipley, Lin Cong, Arryn Craney, Priya Velu, Lars F. Westblade, Massimo Loda, Shawn Levy, Melissa Cushing, Marcin Imielinski, Hanna Rennert, Christopher E. Mason |
| EPI_ISL_427585 | hCoV-19/USA/NY-WCMP11B03P/2020 | North America / USA / New York / Manha | 2020-03-16 | NewYork-Presbyterian & Mason Lab | Mason Lab | Daniel J. Butler, Christopher Mozsary, Cem Meydan, David Danko, Jonathan Foox, Joel Rosiene, Alon Shaiber, Matthew MacKay, Ebrahim Afshinnekoo, Fritz J. Sedlazeck, Nikolay A. Ivanov, Maria Sierra, Craig D. Westover, Krista Ryon, Benjamin Young, Chandrima Bhattacharya, Phyllis Ruggiero, Justyna Gawrys, Iman Hajirasouliha, Dmitry Meleshko, Mirella Salvatore, Dong Xu, Jenny Xiang, John Sipley, Lin Cong, Arryn Craney, Priya Velu, Lars F. Westblade, Massimo Loda, Shawn Levy, Melissa Cushing, Marcin Imielinski, Hanna Rennert, Christopher E. Mason |
| EPI_ISL_427586 | hCoV-19/USA/NY-WCMP11B04P/2020 | North America / USA / New York / Manha | 2020-03-16 | NewYork-Presbyterian & Mason Lab | Mason Lab | Daniel J. Butler, Christopher Mozsary, Cem Meydan, David Danko, Jonathan Foox, Joel Rosiene, Alon Shaiber, Matthew MacKay, Ebrahim Afshinnekoo, Fritz J. Sedlazeck, Nikolay A. Ivanov, Maria Sierra, Craig D. Westover, Krista Ryon, Benjamin Young, Chandrima Bhattacharya, Phyllis Ruggiero, Justyna Gawrys, Iman Hajirasouliha, Dmitry Meleshko, Mirella Salvatore, Dong Xu, Jenny Xiang, John Sipley, Lin Cong, Arryn Craney, Priya Velu, Lars F. Westblade, Massimo Loda, Shawn Levy, Melissa Cushing, Marcin Imielinski, Hanna Rennert, Christopher E. Mason |
| EPI_ISL_427587 | hCoV-19/USA/NY-WCMP11B07P/2020 | North America / USA / New York / Manha | 2020-03-16 | NewYork-Presbyterian & Mason Lab | Mason Lab | Daniel J. Butler, Christopher Mozsary, Cem Meydan, David Danko, Jonathan Foox, Joel Rosiene, Alon Shaiber, Matthew MacKay, Ebrahim Afshinnekoo, Fritz J. Sedlazeck, Nikolay A. Ivanov, Maria Sierra, Craig D. Westover, Krista Ryon, Benjamin Young, Chandrima Bhattacharya, Phyllis Ruggiero, Justyna Gawrys, Iman Hajirasouliha, Dmitry Meleshko, Mirella Salvatore, Dong Xu, Jenny Xiang, John Sipley, Lin Cong, Arryn Craney, Priya Velu, Lars F. Westblade, Massimo Loda, Shawn Levy, Melissa Cushing, Marcin Imielinski, Hanna Rennert, Christopher E. Mason |
| EPI_ISL_427588 | hCoV-19/USA/NY-WCMP11C03P/2020 | North America / USA / New York / Manha | 2020-03-16 | NewYork-Presbyterian & Mason Lab | Mason Lab | Daniel J. Butler, Christopher Mozsary, Cem Meydan, David Danko, Jonathan Foox, Joel Rosiene, Alon Shaiber, Matthew MacKay, Ebrahim Afshinnekoo, Fritz J. Sedlazeck, Nikolay A. Ivanov, Maria Sierra, Craig D. Westover, Krista Ryon, Benjamin Young, Chandrima Bhattacharya, Phyllis Ruggiero, Justyna Gawrys, Iman Hajirasouliha, Dmitry Meleshko, Mirella Salvatore, Dong Xu, Jenny Xiang, John Sipley, Lin Cong, Arryn Craney, Priya Velu, Lars F. Westblade, Massimo Loda, Shawn Levy, Melissa Cushing, Marcin Imielinski, Hanna Rennert, Christopher E. Mason |
| EPI_ISL_427589 | hCoV-19/USA/NY-WCMP11C04P/2020 | North America / USA / New York / Manha | 2020-03-16 | NewYork-Presbyterian & Mason Lab | Mason Lab | Daniel J. Butler, Christopher Mozsary, Cem Meydan, David Danko, Jonathan Foox, Joel Rosiene, Alon Shaiber, Matthew MacKay, Ebrahim Afshinnekoo, Fritz J. Sedlazeck, Nikolay A. Ivanov, Maria Sierra, Craig D. Westover, Krista Ryon, Benjamin Young, Chandrima Bhattacharya, Phyllis Ruggiero, Justyna Gawrys, Iman Hajirasouliha, Dmitry Meleshko, Mirella Salvatore, Dong Xu, Jenny Xiang, John Sipley, Lin Cong, Arryn Craney, Priya Velu, Lars F. Westblade, Massimo Loda, Shawn Levy, Melissa Cushing, Marcin Imielinski, Hanna Rennert, Christopher E. Mason |
| EPI_ISL_427590 | hCoV-19/USA/NY-WCMP11C05P/2020 | North America / USA / New York / Manha | 2020-03-16 | NewYork-Presbyterian & Mason Lab | Mason Lab | Daniel J. Butler, Christopher Mozsary, Cem Meydan, David Danko, Jonathan Foox, Joel Rosiene, Alon Shaiber, Matthew MacKay, Ebrahim Afshinnekoo, Fritz J. Sedlazeck, Nikolay A. Ivanov, Maria Sierra, Craig D. Westover, Krista Ryon, Benjamin Young, Chandrima Bhattacharya, Phyllis Ruggiero, Justyna Gawrys, Iman Hajirasouliha, Dmitry Meleshko, Mirella Salvatore, Dong Xu, Jenny Xiang, John Sipley, Lin Cong, Arryn Craney, Priya Velu, Lars F. Westblade, Massimo Loda, Shawn Levy, Melissa Cushing, Marcin Imielinski, Hanna Rennert, Christopher E. Mason |
| EPI_ISL_427591 | hCoV-19/USA/NY-WCMP11D01N/2020 | North America / USA / New York / Manha | 2020-03-16 | NewYork-Presbyterian & Mason Lab | Mason Lab | Daniel J. Butler, Christopher Mozsary, Cem Meydan, David Danko, Jonathan Foox, Joel Rosiene, Alon Shaiber, Matthew MacKay, Ebrahim Afshinnekoo, Fritz J. Sedlazeck, Nikolay A. Ivanov, Maria Sierra, Craig D. Westover, Krista Ryon, Benjamin Young, Chandrima Bhattacharya, Phyllis Ruggiero, Justyna Gawrys, Iman Hajirasouliha, Dmitry Meleshko, Mirella Salvatore, Dong Xu, Jenny Xiang, John Sipley, Lin Cong, Arryn Craney, Priya Velu, Lars F. Westblade, Massimo Loda, Shawn Levy, Melissa Cushing, Marcin Imielinski, Hanna Rennert, Christopher E. Mason |
| EPI_ISL_427592 | hCoV-19/USA/NY-WCMP11E04P/2020 | North America / USA / New York / Manha | 2020-03-16 | NewYork-Presbyterian & Mason Lab | Mason Lab | Daniel J. Butler, Christopher Mozsary, Cem Meydan, David Danko, Jonathan Foox, Joel Rosiene, Alon Shaiber, Matthew MacKay, Ebrahim Afshinnekoo, Fritz J. Sedlazeck, Nikolay A. Ivanov, Maria Sierra, Craig D. Westover, Krista Ryon, Benjamin Young, Chandrima Bhattacharya, Phyllis Ruggiero, Justyna Gawrys, Iman Hajirasouliha, Dmitry Meleshko, Mirella Salvatore, Dong Xu, Jenny Xiang, John Sipley, Lin Cong, Arryn Craney, Priya Velu, Lars F. Westblade, Massimo Loda, Shawn Levy, Melissa Cushing, Marcin Imielinski, Hanna Rennert, Christopher E. Mason |
| EPI_ISL_427593 | hCoV-19/USA/NY-WCMP11E05P/2020 | North America / USA / New York / Manha | 2020-03-16 | NewYork-Presbyterian & Mason Lab | Mason Lab | Daniel J. Butler, Christopher Mozsary, Cem Meydan, David Danko, Jonathan Foox, Joel Rosiene, Alon Shaiber, Matthew MacKay, Ebrahim Afshinnekoo, Fritz J. Sedlazeck, Nikolay A. Ivanov, Maria Sierra, Craig D. Westover, Krista Ryon, Benjamin Young, Chandrima Bhattacharya, Phyllis Ruggiero, Justyna Gawrys, Iman Hajirasouliha, Dmitry Meleshko, Mirella Salvatore, Dong Xu, Jenny Xiang, John Sipley, Lin Cong, Arryn Craney, Priya Velu, Lars F. Westblade, Massimo Loda, Shawn Levy, Melissa Cushing, Marcin Imielinski, Hanna Rennert, Christopher E. Mason |
| EPI_ISL_427594 | hCoV-19/USA/NY-WCMP11E09P/2020 | North America / USA / New York / Manha | 2020-03-16 | NewYork-Presbyterian & Mason Lab | Mason Lab | Daniel J. Butler, Christopher Mozsary, Cem Meydan, David Danko, Jonathan Foox, Joel Rosiene, Alon Shaiber, Matthew MacKay, Ebrahim Afshinnekoo, Fritz J. Sedlazeck, Nikolay A. Ivanov, Maria Sierra, Craig D. Westover, Krista Ryon, Benjamin Young, Chandrima Bhattacharya, Phyllis Ruggiero, Justyna Gawrys, Iman Hajirasouliha, Dmitry Meleshko, Mirella Salvatore, Dong Xu, Jenny Xiang, John Sipley, Lin Cong, Arryn Craney, Priya Velu, Lars F. Westblade, Massimo Loda, Shawn Levy, Melissa Cushing, Marcin Imielinski, Hanna Rennert, Christopher E. Mason |
| EPI_ISL_427595 | hCoV-19/USA/NY-WCMP11F04P/2020 | North America / USA / New York / Manha | 2020-03-16 | NewYork-Presbyterian & Mason Lab | Mason Lab | Daniel J. Butler, Christopher Mozsary, Cem Meydan, David Danko, Jonathan Foox, Joel Rosiene, Alon Shaiber, Matthew MacKay, Ebrahim Afshinnekoo, Fritz J. Sedlazeck, Nikolay A. Ivanov, Maria Sierra, Craig D. Westover, Krista Ryon, Benjamin Young, Chandrima Bhattacharya, Phyllis Ruggiero, Justyna Gawrys, Iman Hajirasouliha, Dmitry Meleshko, Mirella Salvatore, Dong Xu, Jenny Xiang, John Sipley, Lin Cong, Arryn Craney, Priya Velu, Lars F. Westblade, Massimo Loda, Shawn Levy, Melissa Cushing, Marcin Imielinski, Hanna Rennert, Christopher E. Mason |
| EPI_ISL_427596 | hCoV-19/USA/NY-WCMP11F06P/2020 | North America / USA / New York / Manha | 2020-03-16 | NewYork-Presbyterian & Mason Lab | Mason Lab | Daniel J. Butler, Christopher Mozsary, Cem Meydan, David Danko, Jonathan Foox, Joel Rosiene, Alon Shaiber, Matthew MacKay, Ebrahim Afshinnekoo, Fritz J. Sedlazeck, Nikolay A. Ivanov, Maria Sierra, Craig D. Westover, Krista Ryon, Benjamin Young, Chandrima Bhattacharya, Phyllis Ruggiero, Justyna Gawrys, Iman Hajirasouliha, Dmitry Meleshko, Mirella Salvatore, Dong Xu, Jenny Xiang, John Sipley, Lin Cong, Arryn Craney, Priya Velu, Lars F. Westblade, Massimo Loda, Shawn Levy, Melissa Cushing, Marcin Imielinski, Hanna Rennert, Christopher E. Mason |
| EPI_ISL_427597 | hCoV-19/USA/NY-WCMP11F07P/2020 | North America / USA / New York / Manha | 2020-03-16 | NewYork-Presbyterian & Mason Lab | Mason Lab | Daniel J. Butler, Christopher Mozsary, Cem Meydan, David Danko, Jonathan Foox, Joel Rosiene, Alon Shaiber, Matthew MacKay, Ebrahim Afshinnekoo, Fritz J. Sedlazeck, Nikolay A. Ivanov, Maria Sierra, Craig D. Westover, Krista Ryon, Benjamin Young, Chandrima Bhattacharya, Phyllis Ruggiero, Justyna Gawrys, Iman Hajirasouliha, Dmitry Meleshko, Mirella Salvatore, Dong Xu, Jenny Xiang, John Sipley, Lin Cong, Arryn Craney, Priya Velu, Lars F. Westblade, Massimo Loda, Shawn Levy, Melissa Cushing, Marcin Imielinski, Hanna Rennert, Christopher E. Mason |
| EPI_ISL_427598 | hCoV-19/USA/NY-WCMP11H06P/2020 | North America / USA / New York / Manha | 2020-03-16 | NewYork-Presbyterian & Mason Lab | Mason Lab | Daniel J. Butler, Christopher Mozsary, Cem Meydan, David Danko, Jonathan Foox, Joel Rosiene, Alon Shaiber, Matthew MacKay, Ebrahim Afshinnekoo, Fritz J. Sedlazeck, Nikolay A. Ivanov, Maria Sierra, Craig D. Westover, Krista Ryon, Benjamin Young, Chandrima Bhattacharya, Phyllis Ruggiero, Justyna Gawrys, Iman Hajirasouliha, Dmitry Meleshko, Mirella Salvatore, Dong Xu, Jenny Xiang, John Sipley, Lin Cong, Arryn Craney, Priya Velu, Lars F. Westblade, Massimo Loda, Shawn Levy, Melissa Cushing, Marcin Imielinski, Hanna Rennert, Christopher E. Mason |
| EPI_ISL_427599 | hCoV-19/USA/NY-WCMP12B04P/2020 | North America / USA / New York / Manha | 2020-03-16 | NewYork-Presbyterian & Mason Lab | Mason Lab | Daniel J. Butler, Christopher Mozsary, Cem Meydan, David Danko, Jonathan Foox, Joel Rosiene, Alon Shaiber, Matthew MacKay, Ebrahim Afshinnekoo, Fritz J. Sedlazeck, Nikolay A. Ivanov, Maria Sierra, Craig D. Westover, Krista Ryon, Benjamin Young, Chandrima Bhattacharya, Phyllis Ruggiero, Justyna Gawrys, Iman Hajirasouliha, Dmitry Meleshko, Mirella Salvatore, Dong Xu, Jenny Xiang, John Sipley, Lin Cong, Arryn Craney, Priya Velu, Lars F. Westblade, Massimo Loda, Shawn Levy, Melissa Cushing, Marcin Imielinski, Hanna Rennert, Christopher E. Mason |
| EPI_ISL_427800 | hCoV-19/Australia/NSW87/2020 | Oceania / Australia / New South Wales / | 2020-03-24 | Centre for Infectious Diseases and Microbiology Public Health | NSW Health Pathology - Institute of Clinical Pathology and Medical Research; Westmead Hospital; University of Sydney | Gall M, Arnott A, Sadsad R, Draper J, Sim E, Bachmann N, Rockett R, Lam C, Gray K, Timms V, Carter I, Holmes EC, O'Sullivan MV, Byun R, Sintchenko V, Chen SC, Eden JS, Maddocks S, Kok J, Propenko M, Sorrell T, Chang S, Basile K, Dwyer DE for the 2019-nCoV Study Group |
| EPI_ISL_427801 | hCoV-19/Australia/NSW173/2020 | Oceania / Australia / New South Wales / | 2020-03-26 | Centre for Infectious Diseases and Microbiology Public Health | NSW Health Pathology - Institute of Clinical Pathology and Medical Research; Westmead Hospital; University of Sydney | Rockett R, Lam C, Gray K, Timms V, Gall M, Arnott A, Sadsad R, Draper J, Sim E, Bachmann N, Carter I, Holmes EC, O'Sullivan MV, Byun R, Sintchenko V, Chen SC, Eden JS, Maddocks S, Kok J, Propenko M, Sorrell T, Chang S, Basile K, Dwyer DE for the 2019-nCoV Study Group |
| EPI_ISL_427802 | hCoV-19/Australia/NSW141/2020 | Oceania / Australia / New South Wales / | 2020-03-18 | Centre for Infectious Diseases and Microbiology Public Health | NSW Health Pathology - Institute of Clinical Pathology and Medical Research; Westmead Hospital; University of Sydney | Sim E, Bachmann N, Rockett R, Lam C, Gray K, Timms V, Gall M, Arnott A, Sadsad R, Draper J, Carter I, Holmes EC, O'Sullivan MV, Byun R, Sintchenko V, Chen SC, Eden JS, Maddocks S, Kok J, Propenko M, Sorrell T, Chang S, Basile K, Dwyer DE for the 2019-nCoV Study Group |
| EPI_ISL_427803 | hCoV-19/Australia/NSW186/2020 | Oceania / Australia / New South Wales / | 2020-03-24 | Centre for Infectious Diseases and Microbiology Public Health | NSW Health Pathology - Institute of Clinical Pathology and Medical Research; Westmead Hospital; University of Sydney | Timms V, Gall M, Arnott A, Sadsad R, Draper J, Sim E, Bachmann N, Rockett R, Lam C, Gray K, Carter I, Holmes EC, O'Sullivan MV, Byun R, Sintchenko V, Chen SC, Eden JS, Maddocks S, Kok J, Propenko M, Sorrell T, Chang S, Basile K, Dwyer DE for the 2019-nCoV Study Group |

| | | | | | | |
|---|---|---|---|---|---|---|
| EPI_ISL_427804 | hCoV-19/Australia/NSW108/2020 | Oceania / Australia / New South Wales / | 2020-03-23 | Centre for Infectious Diseases and Microbiology Public Health | NSW Health Pathology - Institute of Clinical Pathology and Medical Research; Westmead Hospital; University of Sydney | Arnott A, Sadsad R, Draper J, Sim E, Bachmann N, Rockett R, Lam C, Gray K, Timms V, Gall M, Carter I, Holmes EC, O'Sullivan MV, Byun R, Sintchenko V, Chen SC, Eden JS, Maddocks S, Kok J, Propenko M, Sorrell T, Chang S, Basile K, Dwyer DE for the 2019-nCoV Study Group |
| EPI_ISL_427805 | hCoV-19/Australia/NSW82/2020 | Oceania / Australia / New South Wales / | 2020-03-20 | Centre for Infectious Diseases and Microbiology Public Health | NSW Health Pathology - Institute of Clinical Pathology and Medical Research; Westmead Hospital; University of Sydney | Bachmann N, Rockett R, Lam C, Gray K, Timms V, Gall M, Arnott A, Sadsad R, Draper J, Sim E, Carter I, Holmes EC, O'Sullivan MV, Byun R, Sintchenko V, Chen SC, Eden JS, Maddocks S, Kok J, Propenko M, Sorrell T, Chang S, Basile K, Dwyer DE for the 2019-nCoV Study Group |
| EPI_ISL_427806 | hCoV-19/Australia/NSW196/2020 | Oceania / Australia / New South Wales / | 2020-03-26 | Centre for Infectious Diseases and Microbiology Public Health | NSW Health Pathology - Institute of Clinical Pathology and Medical Research; Westmead Hospital; University of Sydney | Timms V, Gall M, Arnott A, Sadsad R, Draper J, Sim E, Bachmann N, Rockett R, Lam C, Gray K, Carter I, Holmes EC, O'Sullivan MV, Byun R, Sintchenko V, Chen SC, Eden JS, Maddocks S, Kok J, Propenko M, Sorrell T, Chang S, Basile K, Dwyer DE for the 2019-nCoV Study Group |
| EPI_ISL_427807 | hCoV-19/Australia/NSW207/2020 | Oceania / Australia / New South Wales / | 2020-03-23 | Centre for Infectious Diseases and Microbiology Public Health | NSW Health Pathology - Institute of Clinical Pathology and Medical Research; Westmead Hospital; University of Sydney | Gall M, Arnott A, Sadsad R, Draper J, Sim E, Bachmann N, Rockett R, Lam C, Gray K, Timms V, Carter I, Holmes EC, O'Sullivan MV, Byun R, Sintchenko V, Chen SC, Eden JS, Maddocks S, Kok J, Propenko M, Sorrell T, Chang S, Basile K, Dwyer DE for the 2019-nCoV Study Group |
| EPI_ISL_427808 | hCoV-19/Australia/NSW127/2020 | Oceania / Australia / New South Wales / | 2020-03-24 | Centre for Infectious Diseases and Microbiology Public Health | NSW Health Pathology - Institute of Clinical Pathology and Medical Research; Westmead Hospital; University of Sydney | Gall M, Arnott A, Sadsad R, Draper J, Sim E, Bachmann N, Rockett R, Lam C, Gray K, Timms V, Carter I, Holmes EC, O'Sullivan MV, Byun R, Sintchenko V, Chen SC, Eden JS, Maddocks S, Kok J, Propenko M, Sorrell T, Chang S, Basile K, Dwyer DE for the 2019-nCoV Study Group |
| EPI_ISL_427809 | hCoV-19/Korea/KCDC2015/2020 | Asia / South Korea | 2020-02-26 | Division of Viral Diseases, Center for Laboratory Control of Infectious Diseases, Korea Centers for Diseases Control and Prevention | Division of Viral Diseases, Center for Laboratory Control of Infectious Diseases, Korea Centers for Diseases Control and Prevention | Jeong-Min Kim, Yoon-Seok Chung, Namjoo Lee, Mi-Seon Kim, Sang Hee Woo, Hye-Jun Jo, Sehee Park, Heui Man Kim, Jun-Sub Kim, Junhyeong Jang, Myung Guk Han |
| EPI_ISL_427810 | hCoV-19/Korea/KCDC2016/2020 | Asia / South Korea | 2020-02-18 | Division of Viral Diseases, Center for Laboratory Control of Infectious Diseases, Korea Centers for Diseases Control and Prevention | Division of Viral Diseases, Center for Laboratory Control of Infectious Diseases, Korea Centers for Diseases Control and Prevention | Jeong-Min Kim, Yoon-Seok Chung, Namjoo Lee, Mi-Seon Kim, Sang Hee Woo, Hye-Jun Jo, Sehee Park, Heui Man Kim, Jun-Sub Kim, Junhyeong Jang, Dong Hyun Song, Daesang Lee, Seong Tae Jeong, Myung Guk Han |
| EPI_ISL_427811 | hCoV-19/Korea/KCDC2017/2020 | Asia / South Korea | 2020-02-18 | Division of Viral Diseases, Center for Laboratory Control of Infectious Diseases, Korea Centers for Diseases Control and Prevention | Division of Viral Diseases, Center for Laboratory Control of Infectious Diseases, Korea Centers for Diseases Control and Prevention | Jeong-Min Kim, Yoon-Seok Chung, Namjoo Lee, Mi-Seon Kim, Sang Hee Woo, Hye-Jun Jo, Sehee Park, Heui Man Kim, Jun-Sub Kim, Junhyeong Jang, Dong Hyun Song, Daesang Lee, Seong Tae Jeong, Myung Guk Han |
| EPI_ISL_427812 | hCoV-19/Korea/KCDC2018/2020 | Asia / South Korea | 2020-02-19 | Division of Viral Diseases, Center for Laboratory Control of Infectious Diseases, Korea Centers for Diseases Control and Prevention | Division of Viral Diseases, Center for Laboratory Control of Infectious Diseases, Korea Centers for Diseases Control and Prevention | Jeong-Min Kim, Yoon-Seok Chung, Namjoo Lee, Mi-Seon Kim, Sang Hee Woo, Hye-Jun Jo, Sehee Park, Heui Man Kim, Jun-Sub Kim, Junhyeong Jang, Dong Hyun Song, Daesang Lee, Seong Tae Jeong, Myung Guk Han |
| EPI_ISL_427813 | hCoV-19/Korea/KCDC2019/2020 | Asia / South Korea | 2020-02-18 | Division of Viral Diseases, Center for Laboratory Control of Infectious Diseases, Korea Centers for Diseases Control and Prevention | Division of Viral Diseases, Center for Laboratory Control of Infectious Diseases, Korea Centers for Diseases Control and Prevention | Jeong-Min Kim, Yoon-Seok Chung, Namjoo Lee, Mi-Seon Kim, Sang Hee Woo, Hye-Jun Jo, Sehee Park, Heui Man Kim, Jun-Sub Kim, Junhyeong Jang, Dong Hyun Song, Daesang Lee, Seong Tae Jeong, Myung Guk Han |
| EPI_ISL_427815 | hCoV-19/Russia/StPetersburg-RII4706S/2020 | Europe / Russia / Saint Petersburg | 2020-04-07 | WHO National Influenza Centre Russian Federation | WHO National Influenza Centre Russian Federation | Andrey Komissarov, Artem Fadeev, Anna Ivanova, Daria Danilenko |
| EPI_ISL_428300 | hCoV-19/USA/WI-UW-204/2020 | North America / USA / Wisconsin / Camp | 2020-03-25 | University of Wisconsin-Madison AIDS Vaccine Research Laboratories | University of Wisconsin-Madison AIDS Vaccine Research Laboratories | Gage Moreno, Katarina Braun, et al. AIDS Vaccine Research Laboratories |
| EPI_ISL_428301 | hCoV-19/USA/WI-UW-205/2020 | North America / USA / Wisconsin / Milwa | 2020-03-25 | University of Wisconsin-Madison AIDS Vaccine Research Laboratories | University of Wisconsin-Madison AIDS Vaccine Research Laboratories | Gage Moreno, Katarina Braun, et al. AIDS Vaccine Research Laboratories |
| EPI_ISL_428302 | hCoV-19/USA/WI-UW-206/2020 | North America / USA / Wisconsin / Grafto | 2020-03-25 | University of Wisconsin-Madison AIDS Vaccine Research Laboratories | University of Wisconsin-Madison AIDS Vaccine Research Laboratories | Gage Moreno, Katarina Braun, et al. AIDS Vaccine Research Laboratories |
| EPI_ISL_428303 | hCoV-19/USA/WI-UW-207/2020 | North America / USA / Wisconsin / Milwa | 2020-03-23 | University of Wisconsin-Madison AIDS Vaccine Research Laboratories | University of Wisconsin-Madison AIDS Vaccine Research Laboratories | Gage Moreno, Katarina Braun, et al. AIDS Vaccine Research Laboratories |
| EPI_ISL_428304 | hCoV-19/USA/WI-UW-208/2020 | North America / USA / Wisconsin / Milwa | 2020-03-25 | University of Wisconsin-Madison AIDS Vaccine Research Laboratories | University of Wisconsin-Madison AIDS Vaccine Research Laboratories | Gage Moreno, Katarina Braun, et al. AIDS Vaccine Research Laboratories |
| EPI_ISL_428305 | hCoV-19/USA/WI-UW-209/2020 | North America / USA / Wisconsin / Milwa | 2020-03-25 | University of Wisconsin-Madison AIDS Vaccine Research Laboratories | University of Wisconsin-Madison AIDS Vaccine Research Laboratories | Gage Moreno, Katarina Braun, et al. AIDS Vaccine Research Laboratories |
| EPI_ISL_428306 | hCoV-19/USA/WI-UW-210/2020 | North America / USA / Wisconsin / Milwa | 2020-03-24 | University of Wisconsin-Madison AIDS Vaccine Research Laboratories | University of Wisconsin-Madison AIDS Vaccine Research Laboratories | Gage Moreno, Katarina Braun, et al. AIDS Vaccine Research Laboratories |
| EPI_ISL_428307 | hCoV-19/USA/WI-UW-211/2020 | North America / USA / Wisconsin / Milwa | 2020-03-25 | University of Wisconsin-Madison AIDS Vaccine Research Laboratories | University of Wisconsin-Madison AIDS Vaccine Research Laboratories | Gage Moreno, Katarina Braun, et al. AIDS Vaccine Research Laboratories |
| EPI_ISL_428308 | hCoV-19/USA/WI-UW-212/2020 | North America / USA / Wisconsin / Milwa | 2020-03-25 | University of Wisconsin-Madison AIDS Vaccine Research Laboratories | University of Wisconsin-Madison AIDS Vaccine Research Laboratories | Gage Moreno, Katarina Braun, et al. AIDS Vaccine Research Laboratories |
| EPI_ISL_428309 | hCoV-19/USA/WI-UW-213/2020 | North America / USA / Wisconsin / Milwa | 2020-03-25 | University of Wisconsin-Madison AIDS Vaccine Research Laboratories | University of Wisconsin-Madison AIDS Vaccine Research Laboratories | Gage Moreno, Katarina Braun, et al. AIDS Vaccine Research Laboratories |
| EPI_ISL_428310 | hCoV-19/USA/WI-UW-214/2020 | North America / USA / Wisconsin / Grafto | 2020-03-25 | University of Wisconsin-Madison AIDS Vaccine Research Laboratories | University of Wisconsin-Madison AIDS Vaccine Research Laboratories | Gage Moreno, Katarina Braun, et al. AIDS Vaccine Research Laboratories |
| EPI_ISL_428311 | hCoV-19/USA/WI-UW-215/2020 | North America / USA / Wisconsin / Milwa | 2020-03-25 | University of Wisconsin-Madison AIDS Vaccine Research Laboratories | University of Wisconsin-Madison AIDS Vaccine Research Laboratories | Gage Moreno, Katarina Braun, et al. AIDS Vaccine Research Laboratories |
| EPI_ISL_428312 | hCoV-19/USA/WI-UW-216/2020 | North America / USA / Wisconsin / Milwa | 2020-03-25 | University of Wisconsin-Madison AIDS Vaccine Research Laboratories | University of Wisconsin-Madison AIDS Vaccine Research Laboratories | Gage Moreno, Katarina Braun, et al. AIDS Vaccine Research Laboratories |
| EPI_ISL_428313 | hCoV-19/USA/WI-UW-217/2020 | North America / USA / Wisconsin / Milwa | 2020-03-25 | University of Wisconsin-Madison AIDS Vaccine Research Laboratories | University of Wisconsin-Madison AIDS Vaccine Research Laboratories | Gage Moreno, Katarina Braun, et al. AIDS Vaccine Research Laboratories |
| EPI_ISL_428314 | hCoV-19/USA/WI-UW-218/2020 | North America / USA / Wisconsin / Milwa | 2020-03-25 | University of Wisconsin-Madison AIDS Vaccine Research Laboratories | University of Wisconsin-Madison AIDS Vaccine Research Laboratories | Gage Moreno, Katarina Braun, et al. AIDS Vaccine Research Laboratories |
| EPI_ISL_428315 | hCoV-19/USA/WI-UW-219/2020 | North America / USA / Wisconsin / Milwa | 2020-03-25 | University of Wisconsin-Madison AIDS Vaccine Research Laboratories | University of Wisconsin-Madison AIDS Vaccine Research Laboratories | Gage Moreno, Katarina Braun, et al. AIDS Vaccine Research Laboratories |
| EPI_ISL_428316 | hCoV-19/USA/WI-UW-220/2020 | North America / USA / Wisconsin / Mequ | 2020-03-25 | University of Wisconsin-Madison AIDS Vaccine Research Laboratories | University of Wisconsin-Madison AIDS Vaccine Research Laboratories | Gage Moreno, Katarina Braun, et al. AIDS Vaccine Research Laboratories |
| EPI_ISL_428317 | hCoV-19/USA/WI-UW-221/2020 | North America / USA / Wisconsin / Milwa | 2020-03-25 | University of Wisconsin-Madison AIDS Vaccine Research Laboratories | University of Wisconsin-Madison AIDS Vaccine Research Laboratories | Gage Moreno, Katarina Braun, et al. AIDS Vaccine Research Laboratories |
| EPI_ISL_428318 | hCoV-19/USA/WI-UW-222/2020 | North America / USA / Wisconsin / Milwa | 2020-03-25 | University of Wisconsin-Madison AIDS Vaccine Research Laboratories | University of Wisconsin-Madison AIDS Vaccine Research Laboratories | Gage Moreno, Katarina Braun, et al. AIDS Vaccine Research Laboratories |
| EPI_ISL_428319 | hCoV-19/USA/WI-UW-223/2020 | North America / USA / Wisconsin / Milwa | 2020-03-26 | University of Wisconsin-Madison AIDS Vaccine Research Laboratories | University of Wisconsin-Madison AIDS Vaccine Research Laboratories | Gage Moreno, Katarina Braun, et al. AIDS Vaccine Research Laboratories |
| EPI_ISL_428320 | hCoV-19/USA/WI-UW-224/2020 | North America / USA / Wisconsin / Brook | 2020-03-26 | University of Wisconsin-Madison AIDS Vaccine Research Laboratories | University of Wisconsin-Madison AIDS Vaccine Research Laboratories | Gage Moreno, Katarina Braun, et al. AIDS Vaccine Research Laboratories |
| EPI_ISL_428321 | hCoV-19/USA/WI-UW-225/2020 | North America / USA / Wisconsin / Frankl | 2020-03-26 | University of Wisconsin-Madison AIDS Vaccine Research Laboratories | University of Wisconsin-Madison AIDS Vaccine Research Laboratories | Gage Moreno, Katarina Braun, et al. AIDS Vaccine Research Laboratories |
| EPI_ISL_428322 | hCoV-19/USA/WI-UW-226/2020 | North America / USA / Wisconsin / South | 2020-03-26 | University of Wisconsin-Madison AIDS Vaccine Research Laboratories | University of Wisconsin-Madison AIDS Vaccine Research Laboratories | Gage Moreno, Katarina Braun, et al. AIDS Vaccine Research Laboratories |
| EPI_ISL_428323 | hCoV-19/USA/WI-UW-227/2020 | North America / USA / Wisconsin / Milwa | 2020-03-26 | University of Wisconsin-Madison AIDS Vaccine Research Laboratories | University of Wisconsin-Madison AIDS Vaccine Research Laboratories | Gage Moreno, Katarina Braun, et al. AIDS Vaccine Research Laboratories |
| EPI_ISL_428324 | hCoV-19/USA/WI-UW-228/2020 | North America / USA / Wisconsin / Green | 2020-03-26 | University of Wisconsin-Madison AIDS Vaccine Research Laboratories | University of Wisconsin-Madison AIDS Vaccine Research Laboratories | Gage Moreno, Katarina Braun, et al. AIDS Vaccine Research Laboratories |
| EPI_ISL_428325 | hCoV-19/USA/WI-UW-229/2020 | North America / USA / Wisconsin / Milwa | 2020-03-26 | University of Wisconsin-Madison AIDS Vaccine Research Laboratories | University of Wisconsin-Madison AIDS Vaccine Research Laboratories | Gage Moreno, Katarina Braun, et al. AIDS Vaccine Research Laboratories |
| EPI_ISL_428326 | hCoV-19/USA/WI-UW-230/2020 | North America / USA / Wisconsin / Milwa | 2020-03-26 | University of Wisconsin-Madison AIDS Vaccine Research Laboratories | University of Wisconsin-Madison AIDS Vaccine Research Laboratories | Gage Moreno, Katarina Braun, et al. AIDS Vaccine Research Laboratories |
| EPI_ISL_428327 | hCoV-19/USA/WI-UW-231/2020 | North America / USA / Wisconsin / Milwa | 2020-03-26 | University of Wisconsin-Madison AIDS Vaccine Research Laboratories | University of Wisconsin-Madison AIDS Vaccine Research Laboratories | Gage Moreno, Katarina Braun, et al. AIDS Vaccine Research Laboratories |
| EPI_ISL_428328 | hCoV-19/USA/WI-UW-232/2020 | North America / USA / Wisconsin / Glend | 2020-03-27 | University of Wisconsin-Madison AIDS Vaccine Research Laboratories | University of Wisconsin-Madison AIDS Vaccine Research Laboratories | Gage Moreno, Katarina Braun, et al. AIDS Vaccine Research Laboratories |
| EPI_ISL_428329 | hCoV-19/USA/WI-UW-233/2020 | North America / USA / Wisconsin / Milwa | 2020-03-27 | University of Wisconsin-Madison AIDS Vaccine Research Laboratories | University of Wisconsin-Madison AIDS Vaccine Research Laboratories | Gage Moreno, Katarina Braun, et al. AIDS Vaccine Research Laboratories |
| EPI_ISL_428330 | hCoV-19/USA/WI-UW-234/2020 | North America / USA / Wisconsin / Milwa | 2020-03-26 | University of Wisconsin-Madison AIDS Vaccine Research Laboratories | University of Wisconsin-Madison AIDS Vaccine Research Laboratories | Gage Moreno, Katarina Braun, et al. AIDS Vaccine Research Laboratories |
| EPI_ISL_428331 | hCoV-19/USA/WI-UW-235/2020 | North America / USA / Wisconsin / Cudah | 2020-03-28 | University of Wisconsin-Madison AIDS Vaccine Research Laboratories | University of Wisconsin-Madison AIDS Vaccine Research Laboratories | Gage Moreno, Katarina Braun, et al. AIDS Vaccine Research Laboratories |
| EPI_ISL_428332 | hCoV-19/USA/WI-UW-236/2020 | North America / USA / Wisconsin / Milwa | 2020-03-27 | University of Wisconsin-Madison AIDS Vaccine Research Laboratories | University of Wisconsin-Madison AIDS Vaccine Research Laboratories | Gage Moreno, Katarina Braun, et al. AIDS Vaccine Research Laboratories |
| EPI_ISL_428333 | hCoV-19/USA/WI-UW-237/2020 | North America / USA / Wisconsin / Whitef | 2020-03-27 | University of Wisconsin-Madison AIDS Vaccine Research Laboratories | University of Wisconsin-Madison AIDS Vaccine Research Laboratories | Gage Moreno, Katarina Braun, et al. AIDS Vaccine Research Laboratories |
| EPI_ISL_428334 | hCoV-19/USA/WI-UW-238/2020 | North America / USA / Wisconsin / Milwa | 2020-03-27 | University of Wisconsin-Madison AIDS Vaccine Research Laboratories | University of Wisconsin-Madison AIDS Vaccine Research Laboratories | Gage Moreno, Katarina Braun, et al. AIDS Vaccine Research Laboratories |
| EPI_ISL_428335 | hCoV-19/USA/WI-UW-239/2020 | North America / USA / Wisconsin / Mequ | 2020-03-27 | University of Wisconsin-Madison AIDS Vaccine Research Laboratories | University of Wisconsin-Madison AIDS Vaccine Research Laboratories | Gage Moreno, Katarina Braun, et al. AIDS Vaccine Research Laboratories |
| EPI_ISL_428336 | hCoV-19/USA/WI-UW-240/2020 | North America / USA / Wisconsin / Milwa | 2020-03-27 | University of Wisconsin-Madison AIDS Vaccine Research Laboratories | University of Wisconsin-Madison AIDS Vaccine Research Laboratories | Gage Moreno, Katarina Braun, et al. AIDS Vaccine Research Laboratories |
| EPI_ISL_428337 | hCoV-19/USA/WI-UW-241/2020 | North America / USA / Wisconsin / Milwa | 2020-03-27 | University of Wisconsin-Madison AIDS Vaccine Research Laboratories | University of Wisconsin-Madison AIDS Vaccine Research Laboratories | Gage Moreno, Katarina Braun, et al. AIDS Vaccine Research Laboratories |
| EPI_ISL_428338 | hCoV-19/USA/WI-UW-242/2020 | North America / USA / Wisconsin / Milwa | 2020-03-28 | University of Wisconsin-Madison AIDS Vaccine Research Laboratories | University of Wisconsin-Madison AIDS Vaccine Research Laboratories | Gage Moreno, Katarina Braun, et al. AIDS Vaccine Research Laboratories |
| EPI_ISL_428339 | hCoV-19/USA/WI-UW-243/2020 | North America / USA / Wisconsin / Grafto | 2020-03-28 | University of Wisconsin-Madison AIDS Vaccine Research Laboratories | University of Wisconsin-Madison AIDS Vaccine Research Laboratories | Gage Moreno, Katarina Braun, et al. AIDS Vaccine Research Laboratories |
| EPI_ISL_428340 | hCoV-19/USA/WI-UW-244/2020 | North America / USA / Wisconsin / Thiens | 2020-03-28 | University of Wisconsin-Madison AIDS Vaccine Research Laboratories | University of Wisconsin-Madison AIDS Vaccine Research Laboratories | Gage Moreno, Katarina Braun, et al. AIDS Vaccine Research Laboratories |
| EPI_ISL_428341 | hCoV-19/USA/WI-UW-245/2020 | North America / USA / Wisconsin / Milwa | 2020-03-28 | University of Wisconsin-Madison AIDS Vaccine Research Laboratories | University of Wisconsin-Madison AIDS Vaccine Research Laboratories | Gage Moreno, Katarina Braun, et al. AIDS Vaccine Research Laboratories |
| EPI_ISL_428342 | hCoV-19/USA/WI-UW-246/2020 | North America / USA / Wisconsin / Milwa | 2020-03-28 | University of Wisconsin-Madison AIDS Vaccine Research Laboratories | University of Wisconsin-Madison AIDS Vaccine Research Laboratories | Gage Moreno, Katarina Braun, et al. AIDS Vaccine Research Laboratories |
| EPI_ISL_428343 | hCoV-19/USA/WI-UW-247/2020 | North America / USA / Wisconsin / Port W | 2020-03-28 | University of Wisconsin-Madison AIDS Vaccine Research Laboratories | University of Wisconsin-Madison AIDS Vaccine Research Laboratories | Gage Moreno, Katarina Braun, et al. AIDS Vaccine Research Laboratories |
| EPI_ISL_428344 | hCoV-19/USA/WI-UW-248/2020 | North America / USA / Wisconsin / Milwa | 2020-03-28 | University of Wisconsin-Madison AIDS Vaccine Research Laboratories | University of Wisconsin-Madison AIDS Vaccine Research Laboratories | Gage Moreno, Katarina Braun, et al. AIDS Vaccine Research Laboratories |
| EPI_ISL_428345 | hCoV-19/USA/WI-UW-249/2020 | North America / USA / Wisconsin / Racin | 2020-03-16 | University of Wisconsin-Madison AIDS Vaccine Research Laboratories | University of Wisconsin-Madison AIDS Vaccine Research Laboratories | Gage Moreno, Katarina Braun, et al. AIDS Vaccine Research Laboratories |
| EPI_ISL_428346 | hCoV-19/Turkey/GLAB-CoV012/2020 | Europe / Turkey / Istanbul | 2020-04-17 | Genomic Laboratory (GLAB) (Conjoint lab of Health Directorate of Istanbul and Istanbul Technical University) | Genomic Laboratory (GLAB), Istanbul Technical University | Ilker Karacan, Tugba Kizilboga Akgun, Bugra Agaoglu, Gizem Alkurt, Jale Yildiz, Betsi Köse, Elifnaz Çelik, Azzu Irvem, Yasemin Kendir Demirkol, Ozlem Akgun Dogan, Mehtap Aydin, Levent Doganay, Gizem Dinler Doganay |
| EPI_ISL_428347 | hCoV-19/France/IDF3384/2020 | Europe / France / Ile De France / Longjur | 2020-03-23 | Service de Biologie Médicale - BP 125 | National Reference Center for Viruses of Respiratory Infections, Institut Pasteur, Paris | Mélanie Albert, Marion Barbet, Sylvie Behillil, Méline Bizard, Angela Brisebarre, Flora Donati, Etienne Simon-Lorière, Vincent Enouf, Maud Vanpeene, Sylvie van der Werf |
| EPI_ISL_428348 | hCoV-19/France/GE3372/2020 | Europe / France / Grand-est / Gondrecou | 2020-03-20 | Maison de Santé du Val d'Ormois | National Reference Center for Viruses of Respiratory Infections, Institut Pasteur, Paris | Mélanie Albert, Marion Barbet, Sylvie Behillil, Méline Bizard, Angela Brisebarre, Flora Donati, Etienne Simon-Lorière, Vincent Enouf, Maud Vanpeene, Sylvie van der Werf |
| EPI_ISL_428349 | hCoV-19/France/IDF3386/2020 | Europe / France / Ile De France / Longjur | 2020-03-23 | Service de Biologie Médicale - BP 125 | National Reference Center for Viruses of Respiratory Infections, Institut Pasteur, Paris | Mélanie Albert, Marion Barbet, Sylvie Behillil, Méline Bizard, Angela Brisebarre, Flora Donati, Etienne Simon-Lorière, Vincent Enouf, Maud Vanpeene, Sylvie van der Werf |
| EPI_ISL_428350 | hCoV-19/France/HF3419/2020 | Europe / France / Hauts De France / Cha | 2020-03-24 | CH Jean de Navarre Laboratoire de Biologie | National Reference Center for Viruses of Respiratory Infections, Institut Pasteur, Paris | Mélanie Albert, Marion Barbet, Sylvie Behillil, Méline Bizard, Angela Brisebarre, Flora Donati, Etienne Simon-Lorière, Vincent Enouf, Maud Vanpeene, Sylvie van der Werf |
| EPI_ISL_428351 | hCoV-19/France/IDF3509/2020 | Europe / France / Ile De France / Orsay | 2020-03-24 | GH Nord Essonne Service de Biologie clinique | National Reference Center for Viruses of Respiratory Infections, Institut Pasteur, Paris | Mélanie Albert, Marion Barbet, Sylvie Behillil, Méline Bizard, Angela Brisebarre, Flora Donati, Etienne Simon-Lorière, Vincent Enouf, Maud Vanpeene, Sylvie van der Werf |
| EPI_ISL_428352 | hCoV-19/France/IDF3518/2020 | Europe / France / Ile De France / Orsay | 2020-03-23 | GH Nord Essonne Service de Biologie clinique | National Reference Center for Viruses of Respiratory Infections, Institut Pasteur, Paris | Mélanie Albert, Marion Barbet, Sylvie Behillil, Méline Bizard, Angela Brisebarre, Flora Donati, Etienne Simon-Lorière, Vincent Enouf, Maud Vanpeene, Sylvie van der Werf |
| EPI_ISL_428353 | hCoV-19/France/HF3534/2020 | Europe / France / Hauts De France / Con | 2020-03-24 | CH Compiègne Laboratoire de Biologie | National Reference Center for Viruses of Respiratory Infections, Institut Pasteur, Paris | Mélanie Albert, Marion Barbet, Sylvie Behillil, Méline Bizard, Angela Brisebarre, Flora Donati, Etienne Simon-Lorière, Vincent Enouf, Maud Vanpeene, Sylvie van der Werf |
| EPI_ISL_428354 | hCoV-19/France/IDF3577/2020 | Europe / France / Ile De France / Longjur | 2020-03-25 | LABM GH nord Essonne de Longjumeau - BP 125 | National Reference Center for Viruses of Respiratory Infections, Institut Pasteur, Paris | Mélanie Albert, Marion Barbet, Sylvie Behillil, Méline Bizard, Angela Brisebarre, Flora Donati, Etienne Simon-Lorière, Vincent Enouf, Maud Vanpeene, Sylvie van der Werf |
| EPI_ISL_428355 | hCoV-19/France/IDF2251/2020 | Europe / France / Ile De France / Garche | 2020-03-11 | Institut Médico légal- Hop R. Poincaré | National Reference Center for Viruses of Respiratory Infections, Institut Pasteur, Paris | Mélanie Albert, Marion Barbet, Sylvie Behillil, Méline Bizard, Angela Brisebarre, Flora Donati, Etienne Simon-Lorière, Vincent Enouf, Maud Vanpeene, Sylvie van der Werf |
| EPI_ISL_428356 | hCoV-19/France/IDF2252/2020 | Europe / France / Ile De France / Garche | 2020-03-11 | Institut Médico légal- Hop R. Poincaré | National Reference Center for Viruses of Respiratory Infections, Institut Pasteur, Paris | Mélanie Albert, Marion Barbet, Sylvie Behillil, Méline Bizard, Angela Brisebarre, Flora Donati, Etienne Simon-Lorière, Vincent Enouf, Maud Vanpeene, Sylvie van der Werf |
| EPI_ISL_428357 | hCoV-19/France/IDF2253/2020 | Europe / France / Ile De France / Garche | 2020-03-11 | Institut Médico légal- Hop R. Poincaré | National Reference Center for Viruses of Respiratory Infections, Institut Pasteur, Paris | Mélanie Albert, Marion Barbet, Sylvie Behillil, Méline Bizard, Angela Brisebarre, Flora Donati, Etienne Simon-Lorière, Vincent Enouf, Maud Vanpeene, Sylvie van der Werf |
| EPI_ISL_428358 | hCoV-19/France/HF3598/2020 | Europe / France / Hauts De France / Cha | 2020-03-24 | CH Jeanne de Navarre Laboratoire de Biologie | National Reference Center for Viruses of Respiratory Infections, Institut Pasteur, Paris | Mélanie Albert, Marion Barbet, Sylvie Behillil, Méline Bizard, Angela Brisebarre, Flora Donati, Etienne Simon-Lorière, Vincent Enouf, Maud Vanpeene, Sylvie van der Werf |
| EPI_ISL_428359 | hCoV-19/France/HF3677/2020 | Europe / France / Hauts De France / Con | 2020-03-23 | CH Compiègne Laboratoire de Biologie | National Reference Center for Viruses of Respiratory Infections, Institut Pasteur, Paris | Mélanie Albert, Marion Barbet, Sylvie Behillil, Méline Bizard, Angela Brisebarre, Flora Donati, Etienne Simon-Lorière, Vincent Enouf, Maud Vanpeene, Sylvie van der Werf |
| EPI_ISL_428360 | hCoV-19/France/HF3678/2020 | Europe / France / Hauts De France / Con | 2020-03-25 | CH Compiègne Laboratoire de Biologie | National Reference Center for Viruses of Respiratory Infections, Institut Pasteur, Paris | Mélanie Albert, Marion Barbet, Sylvie Behillil, Méline Bizard, Angela Brisebarre, Flora Donati, Etienne Simon-Lorière, Vincent Enouf, Maud Vanpeene, Sylvie van der Werf |
| EPI_ISL_428361 | hCoV-19/France/IDF3703/2020 | Europe / France / Ile De France / Longjur | 2020-03-25 | LABM GH nord Essonne de Longjumeau - BP 125 | National Reference Center for Viruses of Respiratory Infections, Institut Pasteur, Paris | Mélanie Albert, Marion Barbet, Sylvie Behillil, Méline Bizard, Angela Brisebarre, Flora Donati, Etienne Simon-Lorière, Vincent Enouf, Maud Vanpeene, Sylvie van der Werf |
| EPI_ISL_428363 | hCoV-19/France/IDF3745/2020 | Europe / France / Ile De France / Orsay | 2020-03-26 | GH Nord Essonne Service de Biologie clinique | National Reference Center for Viruses of Respiratory Infections, Institut Pasteur, Paris | Mélanie Albert, Marion Barbet, Sylvie Behillil, Méline Bizard, Angela Brisebarre, Flora Donati, Etienne Simon-Lorière, Vincent Enouf, Maud Vanpeene, Sylvie van der Werf |
| EPI_ISL_428365 | hCoV-19/France/IDF3930/2020 | Europe / France / Ile De France / Longjur | 2020-03-26 | LABM GH nord Essonne de Longjumeau - BP 125 | National Reference Center for Viruses of Respiratory Infections, Institut Pasteur, Paris | Mélanie Albert, Marion Barbet, Sylvie Behillil, Méline Bizard, Angela Brisebarre, Flora Donati, Etienne Simon-Lorière, Vincent Enouf, Maud Vanpeene, Sylvie van der Werf |
| EPI_ISL_428366 | hCoV-19/France/HF4220/2020 | Europe / France / Hauts De France / Cha | 2020-03-30 | CH Jeanne de Navarre Laboratoire de Biologie | National Reference Center for Viruses of Respiratory Infections, Institut Pasteur, Paris | Mélanie Albert, Marion Barbet, Sylvie Behillil, Méline Bizard, Angela Brisebarre, Flora Donati, Etienne Simon-Lorière, Vincent Enouf, Maud Vanpeene, Sylvie van der Werf |
| EPI_ISL_428367 | hCoV-19/France/N4427/2020 | Europe / France / Normandie / Carentan | 2020-03-23 | Cabinet Médical | National Reference Center for Viruses of Respiratory Infections, Institut Pasteur, Paris | Mélanie Albert, Marion Barbet, Sylvie Behillil, Méline Bizard, Angela Brisebarre, Flora Donati, Etienne Simon-Lorière, Vincent Enouf, Maud Vanpeene, Sylvie van der Werf |
| EPI_ISL_428368 | hCoV-19/Turkey/GLAB-CoV013/2020 | Europe / Turkey / Istanbul | 2020-04-16 | Genomic Laboratory (GLAB) (Conjoint lab of Health Directorate of Istanbul and Istanbul Technical University) | Genomic Laboratory (GLAB), Istanbul Technical University | Ilker Karacan, Tugba Kizilboga Akgun, Bugra Agaoglu, Gizem Alkurt, Jale Yildiz, Betsi Köse, Elifnaz Çelik, Azzu Irvem, Yasemin Kendir Demirkol, Ozlem Akgun Dogan, Mehtap Aydin, Levent Doganay, Gizem Dinler Doganay |
| EPI_ISL_428369 | hCoV-19/USA/CT-Yale-011/2020 | North America / USA / Connecticut | 2020-03-11 | Yale Clinical Virology Laboratory | Grubaugh Lab - Yale School of Public Health | Joseph Fauver, Anderson Brito, Tara Alpert, Chantal Vogels, Ellen Foxman, Albert Ko, Marie Landry, Nathan Grubaugh |
| EPI_ISL_428370 | hCoV-19/USA/CT-Yale-032/2020 | North America / USA / Connecticut | 2020-03-18 | Yale Clinical Virology Laboratory | Grubaugh Lab - Yale School of Public Health | Joseph Fauver, Anderson Brito, Tara Alpert, Chantal Vogels, Ellen Foxman, Albert Ko, Marie Landry, Nathan Grubaugh |
| EPI_ISL_428371 | hCoV-19/USA/CT-Yale-033/2020 | North America / USA / Connecticut | 2020-03-18 | Yale Clinical Virology Laboratory | Grubaugh Lab - Yale School of Public Health | Joseph Fauver, Anderson Brito, Tara Alpert, Chantal Vogels, Ellen Foxman, Albert Ko, Marie Landry, Nathan Grubaugh |
| EPI_ISL_428372 | hCoV-19/USA/CT-Yale-041/2020 | North America / USA / Connecticut | 2020-03-16 | Yale Clinical Virology Laboratory | Grubaugh Lab - Yale School of Public Health | Joseph Fauver, Anderson Brito, Tara Alpert, Chantal Vogels, Ellen Foxman, Albert Ko, Marie Landry, Nathan Grubaugh |
| EPI_ISL_428373 | hCoV-19/USA/CT-Yale-052/2020 | North America / USA / Connecticut | 2020-03-26 | Yale COVID-19 Biorepository | Grubaugh Lab - Yale School of Public Health | Joseph Fauver, Tara Alpert, Anderson Brito, Anne Wyllie, Chantal Vogels, Mary Petrone, Chaney Kalinich, Isabel Ott, Arnau Casanovas, Catherine Muenker, Adam Moore, Alice Lu, Maria Tokuyama, Patrick Wong, Peiwen Lu, Saad Omer, Richard Martinello, Allison Nelson, Shelli Farhadian, Akiko Iwasaki, Charles Dela Cruz, Albert Ko, Nathan Grubaugh |
| EPI_ISL_428374 | hCoV-19/USA/CT-Yale-053/2020 | North America / USA / Connecticut | 2020-03-30 | Yale COVID-19 Biorepository | Grubaugh Lab - Yale School of Public Health | Joseph Fauver, Tara Alpert, Anderson Brito, Anne Wyllie, Chantal Vogels, Mary Petrone, Chaney Kalinich, Isabel Ott, Arnau Casanovas, Catherine Muenker, Adam Moore, Alice Lu, Maria Tokuyama, Patrick Wong, Peiwen Lu, Saad Omer, Richard Martinello, Allison Nelson, Shelli Farhadian, Akiko Iwasaki, Charles Dela Cruz, Albert Ko, Nathan Grubaugh |
| EPI_ISL_428375 | hCoV-19/USA/CT-Yale-054/2020 | North America / USA / Connecticut | 2020-03-27 | Yale COVID-19 Biorepository | Grubaugh Lab - Yale School of Public Health | Joseph Fauver, Tara Alpert, Anderson Brito, Anne Wyllie, Chantal Vogels, Mary Petrone, Chaney Kalinich, Isabel Ott, Arnau Casanovas, Catherine Muenker, Adam Moore, Alice Lu, Maria Tokuyama, Patrick Wong, Peiwen Lu, Saad Omer, Richard Martinello, Allison Nelson, Shelli Farhadian, Akiko Iwasaki, Charles Dela Cruz, Albert Ko, Nathan Grubaugh |
| EPI_ISL_428376 | hCoV-19/USA/CT-Yale-055/2020 | North America / USA / Connecticut | 2020-03-27 | Yale COVID-19 Biorepository | Grubaugh Lab - Yale School of Public Health | Joseph Fauver, Tara Alpert, Anderson Brito, Anne Wyllie, Chantal Vogels, Mary Petrone, Chaney Kalinich, Isabel Ott, Arnau Casanovas, Catherine Muenker, Adam Moore, Alice Lu, Maria Tokuyama, Patrick Wong, Peiwen Lu, Saad Omer, Richard Martinello, Allison Nelson, Shelli Farhadian, Akiko Iwasaki, Charles Dela Cruz, Albert Ko, Nathan Grubaugh |
| EPI_ISL_428377 | hCoV-19/USA/CT-Yale-056/2020 | North America / USA / Connecticut | 2020-03-27 | Yale COVID-19 Biorepository | Grubaugh Lab - Yale School of Public Health | Joseph Fauver, Tara Alpert, Anderson Brito, Anne Wyllie, Chantal Vogels, Mary Petrone, Chaney Kalinich, Isabel Ott, Arnau Casanovas, Catherine Muenker, Adam Moore, Alice Lu, Maria Tokuyama, Patrick Wong, Peiwen Lu, Saad Omer, Richard Martinello, Allison Nelson, Shelli Farhadian, Akiko Iwasaki, Charles Dela Cruz, Albert Ko, Nathan Grubaugh |
| EPI_ISL_428378 | hCoV-19/USA/CT-Yale-057/2020 | North America / USA / Connecticut | 2020-03-28 | Yale COVID-19 Biorepository | Grubaugh Lab - Yale School of Public Health | Joseph Fauver, Tara Alpert, Anderson Brito, Anne Wyllie, Chantal Vogels, Mary Petrone, Chaney Kalinich, Isabel Ott, Arnau Casanovas, Catherine Muenker, Adam Moore, Alice Lu, Maria Tokuyama, Patrick Wong, Peiwen Lu, Saad Omer, Richard Martinello, Allison Nelson, Shelli Farhadian, Akiko Iwasaki, Charles Dela Cruz, Albert Ko, Nathan Grubaugh |
| EPI_ISL_428379 | hCoV-19/USA/CT-Yale-059/2020 | North America / USA / Connecticut | 2020-03-28 | Yale COVID-19 Biorepository | Grubaugh Lab - Yale School of Public Health | Joseph Fauver, Tara Alpert, Anderson Brito, Anne Wyllie, Chantal Vogels, Mary Petrone, Chaney Kalinich, Isabel Ott, Arnau Casanovas, Catherine Muenker, Adam Moore, Alice Lu, Maria Tokuyama, Patrick Wong, Peiwen Lu, Saad Omer, Richard Martinello, Allison Nelson, Shelli Farhadian, Akiko Iwasaki, Charles Dela Cruz, Albert Ko, Nathan Grubaugh |
| EPI_ISL_428387 | hCoV-19/USA/CT-Yale-062/2020 | North America / USA / Connecticut | 2020-03-31 | Yale COVID-19 Biorepository | Grubaugh Lab - Yale School of Public Health | Joseph Fauver, Tara Alpert, Anderson Brito, Anne Wyllie, Chantal Vogels, Mary Petrone, Chaney Kalinich, Isabel Ott, Arnau Casanovas, Catherine Muenker, Adam Moore, Alice Lu, Maria Tokuyama, Patrick Wong, Peiwen Lu, Saad Omer, Richard Martinello, Allison Nelson, Shelli Farhadian, Akiko Iwasaki, Charles Dela Cruz, Albert Ko, Nathan Grubaugh |
| EPI_ISL_428389 | hCoV-19/USA/CT-Yale-075/2020 | North America / USA / Connecticut | 2020-04-02 | Yale COVID-19 Biorepository | Grubaugh Lab - Yale School of Public Health | Joseph Fauver, Tara Alpert, Anderson Brito, Anne Wyllie, Chantal Vogels, Mary Petrone, Chaney Kalinich, Isabel Ott, Arnau Casanovas, Catherine Muenker, Adam Moore, Alice Lu, Maria Tokuyama, Patrick Wong, Peiwen Lu, Saad Omer, Richard Martinello, Allison Nelson, Shelli Farhadian, Akiko Iwasaki, Charles Dela Cruz, Albert Ko, Nathan Grubaugh |
| EPI_ISL_428390 | hCoV-19/USA/CT-Yale-051/2020 | North America / USA / Connecticut | 2020-03-26 | Yale COVID-19 Biorepository | Grubaugh Lab - Yale School of Public Health | Joseph Fauver, Tara Alpert, Anderson Brito, Anne Wyllie, Chantal Vogels, Mary Petrone, Chaney Kalinich, Isabel Ott, Arnau Casanovas, Catherine Muenker, Adam Moore, Alice Lu, Maria Tokuyama, Patrick Wong, Peiwen Lu, Saad Omer, Richard Martinello, Allison Nelson, Shelli Farhadian, Akiko Iwasaki, Charles Dela Cruz, Albert Ko, Nathan Grubaugh |
| EPI_ISL_428391 | hCoV-19/USA/CT-Yale-060/2020 | North America / USA / Connecticut | 2020-04-01 | Yale COVID-19 Biorepository | Grubaugh Lab - Yale School of Public Health | Joseph Fauver, Tara Alpert, Anderson Brito, Anne Wyllie, Chantal Vogels, Mary Petrone, Chaney Kalinich, Isabel Ott, Arnau Casanovas, Catherine Muenker, Adam Moore, Alice Lu, Maria Tokuyama, Patrick Wong, Peiwen Lu, Saad Omer, Richard Martinello, Allison Nelson, Shelli Farhadian, Akiko Iwasaki, Charles Dela Cruz, Albert Ko, Nathan Grubaugh |
| EPI_ISL_428392 | hCoV-19/USA/CT-Yale-065/2020 | North America / USA / Connecticut | 2020-04-02 | Yale COVID-19 Biorepository | Grubaugh Lab - Yale School of Public Health | Joseph Fauver, Tara Alpert, Anderson Brito, Anne Wyllie, Chantal Vogels, Mary Petrone, Chaney Kalinich, Isabel Ott, Arnau Casanovas, Catherine Muenker, Adam Moore, Alice Lu, Maria Tokuyama, Patrick Wong, Peiwen Lu, Saad Omer, Richard Martinello, Allison Nelson, Shelli Farhadian, Akiko Iwasaki, Charles Dela Cruz, Albert Ko, Nathan Grubaugh |
| EPI_ISL_428393 | hCoV-19/USA/CT-Yale-084/2020 | North America / USA / Connecticut | 2020-04-06 | Yale COVID-19 Biorepository | Grubaugh Lab - Yale School of Public Health | Joseph Fauver, Tara Alpert, Anderson Brito, Anne Wyllie, Chantal Vogels, Mary Petrone, Chaney Kalinich, Isabel Ott, Arnau Casanovas, Catherine Muenker, Adam Moore, Alice Lu, Maria Tokuyama, Patrick Wong, Peiwen Lu, Saad Omer, Richard Martinello, Allison Nelson, Shelli Farhadian, Akiko Iwasaki, Charles Dela Cruz, Albert Ko, Nathan Grubaugh |
| EPI_ISL_428394 | hCoV-19/USA/CT-Yale-086/2020 | North America / USA / Connecticut | 2020-04-07 | Yale COVID-19 Biorepository | Grubaugh Lab - Yale School of Public Health | Joseph Fauver, Tara Alpert, Anderson Brito, Anne Wyllie, Chantal Vogels, Mary Petrone, Chaney Kalinich, Isabel Ott, Arnau Casanovas, Catherine Muenker, Adam Moore, Alice Lu, Maria Tokuyama, Patrick Wong, Peiwen Lu, Saad Omer, Richard Martinello, Allison Nelson, Shelli Farhadian, Akiko Iwasaki, Charles Dela Cruz, Albert Ko, Nathan Grubaugh |
| EPI_ISL_428395 | hCoV-19/USA/CT-Yale-091/2020 | North America / USA / Connecticut | 2020-04-07 | Yale COVID-19 Biorepository | Grubaugh Lab - Yale School of Public Health | Joseph Fauver, Tara Alpert, Anderson Brito, Anne Wyllie, Chantal Vogels, Mary Petrone, Chaney Kalinich, Isabel Ott, Arnau Casanovas, Catherine Muenker, Adam Moore, Alice Lu, Maria Tokuyama, Patrick Wong, Peiwen Lu, Saad Omer, Richard Martinello, Allison Nelson, Shelli Farhadian, Akiko Iwasaki, Charles Dela Cruz, Albert Ko, Nathan Grubaugh |
| EPI_ISL_428396 | hCoV-19/USA/CT-Yale-092/2020 | North America / USA / Connecticut | 2020-04-07 | Yale COVID-19 Biorepository | Grubaugh Lab - Yale School of Public Health | Joseph Fauver, Tara Alpert, Anderson Brito, Anne Wyllie, Chantal Vogels, Mary Petrone, Chaney Kalinich, Isabel Ott, Arnau Casanovas, Catherine Muenker, Adam Moore, Alice Lu, Maria Tokuyama, Patrick Wong, Peiwen Lu, Saad Omer, Richard Martinello, Allison Nelson, Shelli Farhadian, Akiko Iwasaki, Charles Dela Cruz, Albert Ko, Nathan Grubaugh |
| EPI_ISL_428397 | hCoV-19/USA/CT-Yale-094/2020 | North America / USA / Connecticut | 2020-04-07 | Yale COVID-19 Biorepository | Grubaugh Lab - Yale School of Public Health | Joseph Fauver, Tara Alpert, Anderson Brito, Anne Wyllie, Chantal Vogels, Mary Petrone, Chaney Kalinich, Isabel Ott, Arnau Casanovas, Catherine Muenker, Adam Moore, Alice Lu, Maria Tokuyama, Patrick Wong, Peiwen Lu, Saad Omer, Richard Martinello, Allison Nelson, Shelli Farhadian, Akiko Iwasaki, Charles Dela Cruz, Albert Ko, Nathan Grubaugh |
| EPI_ISL_428398 | hCoV-19/USA/CT-Yale-095/2020 | North America / USA / Connecticut | 2020-04-08 | Yale COVID-19 Biorepository | Grubaugh Lab - Yale School of Public Health | Joseph Fauver, Tara Alpert, Anderson Brito, Anne Wyllie, Chantal Vogels, Mary Petrone, Chaney Kalinich, Isabel Ott, Arnau Casanovas, Catherine Muenker, Adam Moore, Alice Lu, Maria Tokuyama, Patrick Wong, Peiwen Lu, Saad Omer, Richard Martinello, Allison Nelson, Shelli Farhadian, Akiko Iwasaki, Charles Dela Cruz, Albert Ko, Nathan Grubaugh |
| EPI_ISL_428399 | hCoV-19/USA/CT-Yale-067/2020 | North America/ USA / Connecticut | 2020-04-04 | Yale COVID-19 Biorepository | Grubaugh Lab - Yale School of Public Health | Joseph Fauver, Tara Alpert, Anderson Brito, Anne Wyllie, Chantal Vogels, Mary Petrone, Cole Jensen, Chaney Kalinich, Isabel Ott, Arnau Casanovas, Catherine Muenker, Adam Moore, Alice Lu, Maria Tokuyama, Patrick Wong, Peiwen Lu, Saad Omer, Richard Martinello, Allison Nelson, Shelli Farhadian, Akiko Iwasaki, Charles Dela Cruz, Albert Ko, Nathan Grubaugh |
| EPI_ISL_428822 | hCoV-19/Singapore/43/2020 | Asia / Singapore | 2020-02-16 | National Public Health Laboratory, National Centre for Infectious Diseases | National Public Health Laboratory, National Centre for Infectious Diseases | Mak TM, Octavia S, Chavatte JM, Cui L, Lin RTP |
| EPI_ISL_428823 | hCoV-19/Singapore/44/2020 | Asia / Singapore | 2020-03-08 | National Public Health Laboratory, National Centre for Infectious Diseases | National Public Health Laboratory, National Centre for Infectious Diseases | Mak TM, Octavia S, Chavatte JM, Cui L, Lin RTP |
| EPI_ISL_428824 | hCoV-19/Singapore/45/2020 | Asia / Singapore | 2020-03-16 | National Public Health Laboratory, National Centre for Infectious Diseases | National Public Health Laboratory, National Centre for Infectious Diseases | Mak TM, Octavia S, Chavatte JM, Cui L, Lin RTP |
| EPI_ISL_428825 | hCoV-19/Singapore/46/2020 | Asia / Singapore | 2020-03-16 | National Public Health Laboratory, National Centre for Infectious Diseases | National Public Health Laboratory, National Centre for Infectious Diseases | Mak TM, Octavia S, Chavatte JM, Cui L, Lin RTP |
| EPI_ISL_428826 | hCoV-19/Singapore/47/2020 | Asia / Singapore | 2020-03-17 | National Public Health Laboratory, National Centre for Infectious Diseases | National Public Health Laboratory, National Centre for Infectious Diseases | Mak TM, Octavia S, Chavatte JM, Cui L, Lin RTP |
| EPI_ISL_428827 | hCoV-19/Singapore/48/2020 | Asia / Singapore | 2020-03-17 | National Public Health Laboratory, National Centre for Infectious Diseases | National Public Health Laboratory, National Centre for Infectious Diseases | Mak TM, Octavia S, Chavatte JM, Cui L, Lin RTP |
| EPI_ISL_428828 | hCoV-19/Singapore/49/2020 | Asia / Singapore | 2020-03-08 | National Public Health Laboratory, National Centre for Infectious Diseases | National Public Health Laboratory, National Centre for Infectious Diseases | Mak TM, Octavia S, Chavatte JM, Cui L, Lin RTP |
| EPI_ISL_428829 | hCoV-19/Singapore/50/2020 | Asia / Singapore | 2020-03-08 | National Public Health Laboratory, National Centre for Infectious Diseases | National Public Health Laboratory, National Centre for Infectious Diseases | Mak TM, Octavia S, Chavatte JM, Cui L, Lin RTP |
| EPI_ISL_428830 | hCoV-19/Singapore/51/2020 | Asia / Singapore | 2020-02-27 | National Public Health Laboratory, National Centre for Infectious Diseases | National Public Health Laboratory, National Centre for Infectious Diseases | Mak TM, Octavia S, Chavatte JM, Cui L, Lin RTP |
| EPI_ISL_428831 | hCoV-19/Singapore/52/2020 | Asia / Singapore | 2020-03-13 | National Public Health Laboratory, National Centre for Infectious Diseases | National Public Health Laboratory, National Centre for Infectious Diseases | Mak TM, Octavia S, Chavatte JM, Cui L, Lin RTP |
| EPI_ISL_428832 | hCoV-19/Singapore/53/2020 | Asia / Singapore | 2020-04-08 | National Public Health Laboratory, National Centre for Infectious Diseases | National Public Health Laboratory, National Centre for Infectious Diseases | Mak TM, Octavia S, Chavatte JM, Cui L, Lin RTP |
| EPI_ISL_428833 | hCoV-19/Singapore/54/2020 | Asia / Singapore | 2020-04-08 | National Public Health Laboratory, National Centre for Infectious Diseases | National Public Health Laboratory, National Centre for Infectious Diseases | Mak TM, Octavia S, Chavatte JM, Cui L, Lin RTP |
| EPI_ISL_428834 | hCoV-19/Singapore/55/2020 | Asia / Singapore | 2020-04-08 | National Public Health Laboratory, National Centre for Infectious Diseases | National Public Health Laboratory, National Centre for Infectious Diseases | Mak TM, Octavia S, Chavatte JM, Cui L, Lin RTP |
| EPI_ISL_428835 | hCoV-19/Singapore/56/2020 | Asia / Singapore | 2020-04-08 | National Public Health Laboratory, National Centre for Infectious Diseases | National Public Health Laboratory, National Centre for Infectious Diseases | Mak TM, Octavia S, Chavatte JM, Cui L, Lin RTP |
| EPI_ISL_428836 | hCoV-19/Singapore/57/2020 | Asia / Singapore | 2020-04-08 | National Public Health Laboratory, National Centre for Infectious Diseases | National Public Health Laboratory, National Centre for Infectious Diseases | Mak TM, Octavia S, Chavatte JM, Cui L, Lin RTP |
| EPI_ISL_428837 | hCoV-19/Singapore/58/2020 | Asia / Singapore | 2020-04-08 | National Public Health Laboratory, National Centre for Infectious Diseases | National Public Health Laboratory, National Centre for Infectious Diseases | Mak TM, Octavia S, Chavatte JM, Cui L, Lin RTP |
| EPI_ISL_428838 | hCoV-19/Singapore/59/2020 | Asia / Singapore | 2020-04-08 | National Public Health Laboratory, National Centre for Infectious Diseases | National Public Health Laboratory, National Centre for Infectious Diseases | Mak TM, Octavia S, Chavatte JM, Cui L, Lin RTP |
| EPI_ISL_428839 | hCoV-19/Singapore/60/2020 | Asia / Singapore | 2020-04-08 | National Public Health Laboratory, National Centre for Infectious Diseases | National Public Health Laboratory, National Centre for Infectious Diseases | Mak TM, Octavia S, Chavatte JM, Cui L, Lin RTP |
| EPI_ISL_428840 | hCoV-19/Singapore/61/2020 | Asia / Singapore | 2020-04-07 | National Public Health Laboratory, National Centre for Infectious Diseases | National Public Health Laboratory, National Centre for Infectious Diseases | Mak TM, Octavia S, Chavatte JM, Cui L, Lin RTP |
| EPI_ISL_428841 | hCoV-19/Singapore/62/2020 | Asia / Singapore | 2020-04-08 | National Public Health Laboratory, National Centre for Infectious Diseases | National Public Health Laboratory, National Centre for Infectious Diseases | Mak TM, Octavia S, Chavatte JM, Cui L, Lin RTP |
| EPI_ISL_428842 | hCoV-19/Singapore/63/2020 | Asia / Singapore | 2020-04-08 | National Public Health Laboratory, National Centre for Infectious Diseases | National Public Health Laboratory, National Centre for Infectious Diseases | Mak TM, Octavia S, Chavatte JM, Cui L, Lin RTP |
| EPI_ISL_428843 | hCoV-19/Singapore/64/2020 | Asia / Singapore | 2020-04-08 | National Public Health Laboratory, National Centre for Infectious Diseases | National Public Health Laboratory, National Centre for Infectious Diseases | Mak TM, Octavia S, Chavatte JM, Cui L, Lin RTP |
| EPI_ISL_428844 | hCoV-19/Singapore/65/2020 | Asia / Singapore | 2020-04-08 | National Public Health Laboratory, National Centre for Infectious Diseases | National Public Health Laboratory, National Centre for Infectious Diseases | Mak TM, Octavia S, Chavatte JM, Cui L, Lin RTP |
| EPI_ISL_428845 | hCoV-19/Singapore/66/2020 | Asia / Singapore | 2020-04-08 | National Public Health Laboratory, National Centre for Infectious Diseases | National Public Health Laboratory, National Centre for Infectious Diseases | Mak TM, Octavia S, Chavatte JM, Cui L, Lin RTP |
| EPI_ISL_428846 | hCoV-19/Singapore/67/2020 | Asia / Singapore | 2020-04-15 | National Public Health Laboratory, National Centre for Infectious Diseases | National Public Health Laboratory, National Centre for Infectious Diseases | Mak TM, Octavia S, Chavatte JM, Cui L, Lin RTP |
| EPI_ISL_428847 | hCoV-19/Singapore/68/2020 | Asia / Singapore | 2020-04-15 | National Public Health Laboratory, National Centre for Infectious Diseases | National Public Health Laboratory, National Centre for Infectious Diseases | Mak TM, Octavia S, Chavatte JM, Cui L, Lin RTP |
| EPI_ISL_428848 | hCoV-19/Singapore/69/2020 | Asia / Singapore | 2020-04-15 | National Public Health Laboratory, National Centre for Infectious Diseases | National Public Health Laboratory, National Centre for Infectious Diseases | Mak TM, Octavia S, Chavatte JM, Cui L, Lin RTP |
| EPI_ISL_428849 | hCoV-19/Singapore/70/2020 | Asia / Singapore | 2020-04-15 | National Public Health Laboratory, National Centre for Infectious Diseases | National Public Health Laboratory, National Centre for Infectious Diseases | Mak TM, Octavia S, Chavatte JM, Cui L, Lin RTP |
| EPI_ISL_428850 | hCoV-19/Singapore/71/2020 | Asia / Singapore | 2020-04-15 | National Public Health Laboratory, National Centre for Infectious Diseases | National Public Health Laboratory, National Centre for Infectious Diseases | Mak TM, Octavia S, Chavatte JM, Cui L, Lin RTP |
| EPI_ISL_428851 | hCoV-19/Russia/Moscow-25l/2020 | Europe / Russia / Moscow | 2020-04-02 | FSBSI "Chumakov Federal Scientific Center for Research and Development of Immune-and-Biological Products of Russian Academy of Sciences" | FSBSI "Chumakov Federal Scientific Center for Research and Development of Immune-and-Biological Products of Russian Academy of Sciences" & NRC "Kurchatov institute" | Liubov Kozlovskaya, Anastasia Piniaeva, Georgy Ignatyev, Anna Shishova, Aydar Ishmukhametov, Mikhail Rychev, Egor Prokhorchuk, Denis Protsenko, Anastasia Berestovskaya |
| EPI_ISL_428852 | hCoV-19/Russia/Moscow-35l/2020 | Europe / Russia / Moscow | 2020-04-02 | FSBSI "Chumakov Federal Scientific Center for Research and Development of Immune-and-Biological Products of Russian Academy of Sciences" | FSBSI "Chumakov Federal Scientific Center for Research and Development of Immune-and-Biological Products of Russian Academy of Sciences" & NRC "Kurchatov institute" | Liubov Kozlovskaya, Anastasia Piniaeva, Georgy Ignatyev, Anna Shishova, Aydar Ishmukhametov, Mikhail Rychev, Egor Prokhorchuk, Denis Protsenko, Anastasia Berestovskaya |
| EPI_ISL_428853 | hCoV-19/Italy/FVG-ICGEB_S14/2020 | Europe / Italy / Friuli Venezia Giulia | 2020-03-30 | Laboratory of Molecular Virology International Center for Genetic Engineering and Biotechnology (ICGEB) | ARGO Open Lab Platform for Genome Sequencing | Licastro D, Rajasekharan S, Dal Monego S, Segat L, D'Agaro P, Marcello A |
| EPI_ISL_428854 | hCoV-19/Italy/FVG-ICGEB_S18/2020 | Europe / Italy / Friuli Venezia Giulia | 2020-03-30 | Laboratory of Molecular Virology International Center for Genetic Engineering and Biotechnology (ICGEB) | ARGO Open Lab Platform for Genome sequencing | Licastro D, Rajasekharan S, Dal Monego S, Segat L, D'Agaro P, Marcello A |
| EPI_ISL_428855 | hCoV-19/Gambia/GC19-015/2020 | Africa / Gambia / West Coast Region | 2020-03-17 | MRCG at LSHTM Geomics lab | MRCG at LSHTM Genomics lab | Sesay et al |
| EPI_ISL_428856 | hCoV-19/Gambia/GC19-026/2020 | Africa / Gambia / West Coast Region | 2020-03-21 | MRCG at LSHTM Genomics Lab | MRCG at LSHTM Genomics lab | Sesay et al |
| EPI_ISL_428857 | hCoV-19/Gambia/GC19-029/2020 | Africa / Gambia / West Coast Region | 2020-04-20 | MRCG at LSHTM Genomics lab | MRCG at LSHTM Genomics lab | Sesay et al |
| EPI_ISL_428860 | hCoV-19/Russia/Moscow-62505/2020 | Europe / Russia / Moscow | 2020-03-11 | State Research Center of Virology and Biotechnology VECTOR, Department of Collection of Microorganisms | State Research Center of Virology and Biotechnology VECTOR, Department of Collection of Microorganisms | Oleg V. Pyankov, Sergey A. Bodnev, Tatyana V. Tregubchak, Alexander N. Shvalov, Elena V. Gavrilova, Rinat A. Maksyutov |
| EPI_ISL_428861 | hCoV-19/Russia/Moscow-62501/2020 | Europe / Russia / Moscow | 2020-03-11 | State Research Center of Virology and Biotechnology VECTOR, Department of Collection of Microorganisms | State Research Center of Virology and Biotechnology VECTOR, Department of Collection of Microorganisms | Oleg V. Pyankov, Sergey A. Bodnev, Tatyana V. Tregubchak, Alexander N. Shvalov, Elena V. Gavrilova, Rinat A. Maksyutov |
| EPI_ISL_428862 | hCoV-19/Russia/Moscow-62506/2020 | Europe / Russia / Moscow | 2020-03-11 | State Research Center of Virology and Biotechnology VECTOR, Department of Collection of Microorganisms | State Research Center of Virology and Biotechnology VECTOR, Department of Collection of Microorganisms | Oleg V. Pyankov, Sergey A. Bodnev, Anastasiya M. Smirnova, Anastasiya A. Nazarenko, Tatyana V. Tregubchak, Alexander N. Shvalov, Elena V. Gavrilova, Rinat A. Maksyutov |
| EPI_ISL_428863 | hCoV-19/Russia/Moscow-62508/2020 | Europe / Russia / Moscow | 2020-03-11 | State Research Center of Virology and Biotechnology VECTOR, Department of Collection of Microorganisms | State Research Center of Virology and Biotechnology VECTOR, Department of Collection of Microorganisms | Oleg V. Pyankov, Sergey A. Bodnev, Anastasiya M. Smirnova, Anastasiya A. Nazarenko, Tatyana V. Tregubchak, Alexander N. Shvalov, Elena V. Gavrilova, Rinat A. Maksyutov |
| EPI_ISL_428864 | hCoV-19/Russia/Moscow-62502/2020 | Europe / Russia / Moscow | 2020-03-11 | State Research Center of Virology and Biotechnology VECTOR, Department of Collection of Microorganisms | State Research Center of Virology and Biotechnology VECTOR, Department of Collection of Microorganisms | Oleg V. Pyankov, Sergey A. Bodnev, Tatyana V. Tregubchak, Alexander N. Shvalov, Elena V. Gavrilova, Rinat A. Maksyutov |
| EPI_ISL_428865 | hCoV-19/Russia/Lipetsk-62704/2020 | Europe / Russia / Lipetsk Region | 2020-03-11 | State Research Center of Virology and Biotechnology VECTOR, Department of Collection of Microorganisms | State Research Center of Virology and Biotechnology VECTOR, Department of Collection of Microorganisms | Oleg V. Pyankov, Sergey A. Bodnev, Tatyana V. Tregubchak, Alexander N. Shvalov, Elena V. Gavrilova, Rinat A. Maksyutov |
| EPI_ISL_428866 | hCoV-19/Russia/Lipetsk-62707/2020 | Europe / Russia / Lipetsk Region | 2020-03-11 | State Research Center of Virology and Biotechnology VECTOR, Department of Collection of Microorganisms | State Research Center of Virology and Biotechnology VECTOR, Department of Collection of Microorganisms | Oleg V. Pyankov, Sergey A. Bodnev, Tatyana V. Tregubchak, Alexander N. Shvalov, Elena V. Gavrilova, Rinat A. Maksyutov |
| EPI_ISL_428867 | hCoV-19/Russia/Krasnodar-63401/2020 | Europe / Russia / Krasnodar Region | 2020-03-11 | State Research Center of Virology and Biotechnology VECTOR, Department of Collection of Microorganisms | State Research Center of Virology and Biotechnology VECTOR, Department of Collection of Microorganisms | Oleg V. Pyankov, Sergey A. Bodnev, Tatyana V. Tregubchak, Alexander N. Shvalov, Elena V. Gavrilova, Rinat A. Maksyutov |
| EPI_ISL_428868 | hCoV-19/Russia/StPetersburg-64304/2020 | Europe / Russia / St. Petersburg | 2020-03-13 | State Research Center of Virology and Biotechnology VECTOR, Department of Collection of Microorganisms | State Research Center of Virology and Biotechnology VECTOR, Department of Collection of Microorganisms | Oleg V. Pyankov, Sergey A. Bodnev, Tatyana V. Tregubchak, Alexander N. Shvalov, Elena V. Gavrilova, Rinat A. Maksyutov |
| EPI_ISL_428869 | hCoV-19/Russia/Moscow_64405/2020 | Europe / Russia / Moscow | 2020-03-13 | State Research Center of Virology and Biotechnology VECTOR, Department of Collection of Microorganisms | State Research Center of Virology and Biotechnology VECTOR, Department of Collection of Microorganisms | Oleg V. Pyankov, Sergey A. Bodnev, Tatyana V. Tregubchak, Alexander N. Shvalov, Elena V. Gavrilova, Rinat A. Maksyutov |
| EPI_ISL_428870 | hCoV-19/Russia/Moscow-64601/2020 | Europe / Russia / Moscow Region | 2020-03-13 | State Research Center of Virology and Biotechnology VECTOR, Department of Collection of Microorganisms | State Research Center of Virology and Biotechnology VECTOR, Department of Collection of Microorganisms | Oleg V. Pyankov, Sergey A. Bodnev, Tatyana V. Tregubchak, Alexander N. Shvalov, Elena V. Gavrilova, Rinat A. Maksyutov |
| EPI_ISL_428871 | hCoV-19/Russia/Krasnoyarsk-66501/2020 | Europe / Russia / Krasnoyarsk Region | 2020-03-13 | State Research Center of Virology and Biotechnology VECTOR, Department of Collection of Microorganisms | State Research Center of Virology and Biotechnology VECTOR, Department of Collection of Microorganisms | Oleg V. Pyankov, Sergey A. Bodnev, Tatyana V. Tregubchak, Alexander N. Shvalov, Elena V. Gavrilova, Rinat A. Maksyutov |
| EPI_ISL_428872 | hCoV-19/Russia/Moscow-71605/2020 | Europe / Russia / Moscow Region | 2020-03-19 | State Research Center of Virology and Biotechnology VECTOR, Department of Collection of Microorganisms | State Research Center of Virology and Biotechnology VECTOR, Department of Collection of Microorganisms | Sergey A. Bodnev, Oleg V. Pyankov, Tatyana V. Tregubchak, Alexander N. Shvalov, Elena V. Gavrilova, Rinat A. Maksyutov |
| EPI_ISL_428873 | hCoV-19/Russia/Sverdlovsk-71904/2020 | Europe / Russia / Sverdlovsk Region | 2020-03-19 | State Research Center of Virology and Biotechnology VECTOR, Department of Collection of Microorganisms | State Research Center of Virology and Biotechnology VECTOR, Department of Collection of Microorganisms | Sergey A. Bodnev, Oleg V. Pyankov, Tatyana V. Tregubchak, Alexander N. Shvalov, Elena V. Gavrilova, Rinat A. Maksyutov |
| EPI_ISL_428874 | hCoV-19/Russia/Yakutia-73709/2020 | Europe / Russia / Republic of Sakha (Yak | 2020-03-20 | State Research Center of Virology and Biotechnology VECTOR, Department of Collection of Microorganisms | State Research Center of Virology and Biotechnology VECTOR, Department of Collection of Microorganisms | Sergey A. Bodnev, Oleg V. Pyankov, Tatyana V. Tregubchak, Alexander N. Shvalov, Elena V. Gavrilova, Rinat A. Maksyutov |
| EPI_ISL_428875 | hCoV-19/Russia/Moscow-67117/2020 | Europe / Russia / Moscow | 2020-03-16 | State Research Center of Virology and Biotechnology VECTOR, Department of Collection of Microorganisms | State Research Center of Virology and Biotechnology VECTOR, Department of Collection of Microorganisms | Sergey A. Bodnev, Oleg V. Pyankov, Tatyana V. Tregubchak, Alexander N. Shvalov, Elena V. Gavrilova, Rinat A. Maksyutov |
| EPI_ISL_428876 | hCoV-19/Russia/Moscow-71604/2020 | Europe / Russia / Moscow Region | 2020-03-19 | State Research Center of Virology and Biotechnology VECTOR, Department of Collection of Microorganisms | State Research Center of Virology and Biotechnology VECTOR, Department of Collection of Microorganisms | Sergey A. Bodnev, Oleg V. Pyankov, Anastasiya M. Smirnova, Anastasiya A. Nazarenko, Tatyana V. Tregubchak, Alexander N. Shvalov, Elena V. Gavrilova, Rinat A. Maksyutov |
| EPI_ISL_428877 | hCoV-19/Russia/Moscow-71602/2020 | Europe / Russia / Moscow Region | 2020-03-19 | State Research Center of Virology and Biotechnology VECTOR, Department of Collection of Microorganisms | State Research Center of Virology and Biotechnology VECTOR, Department of Collection of Microorganisms | Sergey A. Bodnev, Oleg V. Pyankov, Anastasiya M. Smirnova, Anastasiya A. Nazarenko, Tatyana V. Tregubchak, Alexander N. Shvalov, Elena V. Gavrilova, Rinat A. Maksyutov |
| EPI_ISL_428878 | hCoV-19/Russia/Omsk-89001/2020 | Europe / Russia / Omsk Region | 2020-04-01 | State Research Center of Virology and Biotechnology VECTOR, Department of Collection of Microorganisms | State Research Center of Virology and Biotechnology VECTOR, Department of Collection of Microorganisms | Sergey A. Bodnev, Oleg V. Pyankov, Tatyana V. Tregubchak, Alexander N. Shvalov, Elena V. Gavrilova, Rinat A. Maksyutov |
| EPI_ISL_428879 | hCoV-19/Russia/Sverdlovsk-71905/2020 | Europe / Russia / Sverdlovsk Region | 2020-03-19 | State Research Center of Virology and Biotechnology VECTOR, Department of Collection of Microorganisms | State Research Center of Virology and Biotechnology VECTOR, Department of Collection of Microorganisms | Sergey A. Bodnev, Oleg V. Pyankov, Tatyana V. Tregubchak, Alexander N. Shvalov, Elena V. Gavrilova, Rinat A. Maksyutov |
| EPI_ISL_428880 | hCoV-19/Russia/Khabarovsk-71403/2020 | Europe / Russia / Khabarovsk Region | 2020-03-19 | State Research Center of Virology and Biotechnology VECTOR, Department of Collection of Microorganisms | State Research Center of Virology and Biotechnology VECTOR, Department of Collection of Microorganisms | Sergey A. Bodnev, Oleg V. Pyankov, Tatyana V. Tregubchak, Alexander N. Shvalov, Elena V. Gavrilova, Rinat A. Maksyutov |
| EPI_ISL_428881 | hCoV-19/Russia/Moscow-67142/2020 | Europe / Russia / Moscow | 2020-03-16 | State Research Center of Virology and Biotechnology VECTOR, Department of Collection of Microorganisms | State Research Center of Virology and Biotechnology VECTOR, Department of Collection of Microorganisms | Sergey A. Bodnev, Oleg V. Pyankov, Tatyana V. Tregubchak, Alexander N. Shvalov, Elena V. Gavrilova, Rinat A. Maksyutov |
| EPI_ISL_428882 | hCoV-19/Russia/Moscow-67609/2020 | Europe / Russia / Moscow | 2020-03-16 | State Research Center of Virology and Biotechnology VECTOR, Department of Collection of Microorganisms | State Research Center of Virology and Biotechnology VECTOR, Department of Collection of Microorganisms | Oleg V. Pyankov, Sergey A. Bodnev, Tatyana V. Tregubchak, Alexander N. Shvalov, Elena V. Gavrilova, Rinat A. Maksyutov |
| EPI_ISL_428883 | hCoV-19/Russia/StPetersburg-73603/2020 | Europe / Russia / St. Petersburg | 2020-03-20 | State Research Center of Virology and Biotechnology VECTOR, Department of Collection of Microorganisms | State Research Center of Virology and Biotechnology VECTOR, Department of Collection of Microorganisms | Oleg V. Pyankov, Sergey A. Bodnev, Tatyana V. Tregubchak, Alexander N. Shvalov, Elena V. Gavrilova, Rinat A. Maksyutov |
| EPI_ISL_428884 | hCoV-19/Russia/Moscow-77627/2020 | Europe / Russia / Moscow | 2020-03-22 | State Research Center of Virology and Biotechnology VECTOR, Department of Collection of Microorganisms | State Research Center of Virology and Biotechnology VECTOR, Department of Collection of Microorganisms | Oleg V. Pyankov, Sergey A. Bodnev, Tatyana V. Tregubchak, Alexander N. Shvalov, Elena V. Gavrilova, Rinat A. Maksyutov |
| EPI_ISL_428885 | hCoV-19/Russia/Samara-73406/2020 | Europe / Russia / Samara Region | 2020-03-20 | State Research Center of Virology and Biotechnology VECTOR, Department of Collection of Microorganisms | State Research Center of Virology and Biotechnology VECTOR, Department of Collection of Microorganisms | Oleg V. Pyankov, Sergey A. Bodnev, Tatyana V. Tregubchak, Alexander N. Shvalov, Elena V. Gavrilova, Rinat A. Maksyutov |
| EPI_ISL_428886 | hCoV-19/Russia/Samara-73403/2020 | Europe / Russia / Samara Region | 2020-03-20 | State Research Center of Virology and Biotechnology VECTOR, Department of Collection of Microorganisms | State Research Center of Virology and Biotechnology VECTOR, Department of Collection of Microorganisms | Oleg V. Pyankov, Sergey A. Bodnev, Tatyana V. Tregubchak, Alexander N. Shvalov, Elena V. Gavrilova, Rinat A. Maksyutov |
| EPI_ISL_428887 | hCoV-19/Russia/Moscow-77610/2020 | Europe / Russia / Moscow | 2020-03-22 | State Research Center of Virology and Biotechnology VECTOR, Department of Collection of Microorganisms | State Research Center of Virology and Biotechnology VECTOR, Department of Collection of Microorganisms | Oleg V. Pyankov, Sergey A. Bodnev, Tatyana V. Tregubchak, Alexander N. Shvalov, Elena V. Gavrilova, Rinat A. Maksyutov |
| EPI_ISL_428888 | hCoV-19/Russia/Yakutia-73703/2020 | Europe / Russia / Republic of Sakha (Yak | 2020-03-20 | State Research Center of Virology and Biotechnology VECTOR, Department of Collection of Microorganisms | State Research Center of Virology and Biotechnology VECTOR, Department of Collection of Microorganisms | Oleg V. Pyankov, Sergey A. Bodnev, Tatyana V. Tregubchak, Alexander N. Shvalov, Elena V. Gavrilova, Rinat A. Maksyutov |
| EPI_ISL_428889 | hCoV-19/Russia/Moscow-77620/2020 | Europe / Russia / Moscow | 2020-03-22 | State Research Center of Virology and Biotechnology VECTOR, Department of Collection of Microorganisms | State Research Center of Virology and Biotechnology VECTOR, Department of Collection of Microorganisms | Oleg V. Pyankov, Sergey A. Bodnev, Tatyana V. Tregubchak, Alexander N. Shvalov, Elena V. Gavrilova, Rinat A. Maksyutov |
| EPI_ISL_428890 | hCoV-19/Russia/Sverdlovsk-71903/2020 | Europe / Russia / Sverdlovsk Region | 2020-03-19 | State Research Center of Virology and Biotechnology VECTOR, Department of Collection of Microorganisms | State Research Center of Virology and Biotechnology VECTOR, Department of Collection of Microorganisms | Oleg V. Pyankov, Sergey A. Bodnev, Tatyana V. Tregubchak, Alexander N. Shvalov, Elena V. Gavrilova, Rinat A. Maksyutov |
| EPI_ISL_428891 | hCoV-19/Russia/Yakutia-83301/2020 | Europe / Russia / Republic of Sakha (Yak | 2020-03-25 | State Research Center of Virology and Biotechnology VECTOR, Department of Collection of Microorganisms | State Research Center of Virology and Biotechnology VECTOR, Department of Collection of Microorganisms | Oleg V. Pyankov, Sergey A. Bodnev, Tatyana V. Tregubchak, Alexander N. Shvalov, Elena V. Gavrilova, Rinat A. Maksyutov |
| EPI_ISL_428892 | hCoV-19/Russia/Krasnodar-80902/2020 | Europe / Russia / Krasnodar Region | 2020-03-23 | State Research Center of Virology and Biotechnology VECTOR, Department of Collection of Microorganisms | State Research Center of Virology and Biotechnology VECTOR, Department of Collection of Microorganisms | Oleg V. Pyankov, Sergey A. Bodnev, Tatyana V. Tregubchak, Alexander N. Shvalov, Elena V. Gavrilova, Rinat A. Maksyutov |
| EPI_ISL_428893 | hCoV-19/Russia/Kabardino-Balkaria-80501/2020 | Europe / Russia / Republic of Kabardino- | 2020-03-23 | State Research Center of Virology and Biotechnology VECTOR, Department of Collection of Microorganisms | State Research Center of Virology and Biotechnology VECTOR, Department of Collection of Microorganisms | Oleg V. Pyankov, Sergey A. Bodnev, Tatyana V. Tregubchak, Alexander N. Shvalov, Elena V. Gavrilova, Rinat A. Maksyutov |
| EPI_ISL_428894 | hCoV-19/Russia/Buryatia-84506/2020 | Europe / Russia / Republic of Buryatia | 2020-03-27 | State Research Center of Virology and Biotechnology VECTOR, Department of Collection of Microorganisms | State Research Center of Virology and Biotechnology VECTOR, Department of Collection of Microorganisms | Oleg V. Pyankov, Sergey A. Bodnev, Tatyana V. Tregubchak, Alexander N. Shvalov, Elena V. Gavrilova, Rinat A. Maksyutov |
| EPI_ISL_428895 | hCoV-19/Russia/Perm-82707/2020 | Europe / Russia / Perm Region | 2020-03-24 | State Research Center of Virology and Biotechnology VECTOR, Department of Collection of Microorganisms | State Research Center of Virology and Biotechnology VECTOR, Department of Collection of Microorganisms | Sergey A. Bodnev, Oleg V. Pyankov, Tatyana V. Tregubchak, Alexander N. Shvalov, Elena V. Gavrilova, Rinat A. Maksyutov |
| EPI_ISL_428896 | hCoV-19/Russia/Vector_84801/2020 | Europe / Russia / Republic of Sakha (Yak | 2020-03-27 | State Research Center of Virology and Biotechnology VECTOR, Department of Collection of Microorganisms | State Research Center of Virology and Biotechnology VECTOR, Department of Collection of Microorganisms | Sergey A. Bodnev, Oleg V. Pyankov, Tatyana V. Tregubchak, Alexander N. Shvalov, Elena V. Gavrilova, Rinat A. Maksyutov |
| EPI_ISL_428897 | hCoV-19/Russia/StPetersburg-82005/2020 | Europe / Russia / St. Petersburg | 2020-03-24 | State Research Center of Virology and Biotechnology VECTOR, Department of Collection of Microorganisms | State Research Center of Virology and Biotechnology VECTOR, Department of Collection of Microorganisms | Sergey A. Bodnev, Oleg V. Pyankov, Tatyana V. Tregubchak, Alexander N. Shvalov, Elena V. Gavrilova, Rinat A. Maksyutov |
| EPI_ISL_428898 | hCoV-19/Russia/Kabardino-Balkaria-80503/2020 | Europe / Russia / Republic of Kabardino- | 2020-03-23 | State Research Center of Virology and Biotechnology VECTOR, Department of Collection of Microorganisms | State Research Center of Virology and Biotechnology VECTOR, Department of Collection of Microorganisms | Sergey A. Bodnev, Oleg V. Pyankov, Tatyana V. Tregubchak, Alexander N. Shvalov, Elena V. Gavrilova, Rinat A. Maksyutov |
| EPI_ISL_428899 | hCoV-19/Russia/Buryatia-86204/2020 | Europe / Russia / Republic of Buryatia | 2020-03-29 | State Research Center of Virology and Biotechnology VECTOR, Department of Collection of Microorganisms | State Research Center of Virology and Biotechnology VECTOR, Department of Collection of Microorganisms | Sergey A. Bodnev, Oleg V. Pyankov, Tatyana V. Tregubchak, Alexander N. Shvalov, Elena V. Gavrilova, Rinat A. Maksyutov |
| EPI_ISL_429300 | hCoV-19/Denmark/ALAB-HH46/2020 | Europe / Denmark / Unknown | 2020-03-13 | Department of Clinical Microbiology, Copenhagen University Hospital, Hvidovre, Kettegaard Alle 30, 2650 Hvidovre. | Albertsen lab, Department of Chemistry and Bioscience, Aalborg University, Denmark | Rasmus Kirkegaard |
| EPI_ISL_429301 | hCoV-19/Denmark/ALAB-HH49/2020 | Europe / Denmark / Unknown | 2020-03-16 | Department of Clinical Microbiology, Copenhagen University Hospital, Hvidovre, Kettegaard Alle 30, 2650 Hvidovre. | Albertsen lab, Department of Chemistry and Bioscience, Aalborg University, Denmark | Rasmus Kirkegaard |
| EPI_ISL_429302 | hCoV-19/Denmark/ALAB-HH50/2020 | Europe / Denmark / Unknown | 2020-03-16 | Department of Clinical Microbiology, Copenhagen University Hospital, Hvidovre, Kettegaard Alle 30, 2650 Hvidovre. | Albertsen lab, Department of Chemistry and Bioscience, Aalborg University, Denmark | Rasmus Kirkegaard |
| EPI_ISL_429303 | hCoV-19/Denmark/ALAB-HH51/2020 | Europe / Denmark / Unknown | 2020-03-16 | Department of Clinical Microbiology, Copenhagen University Hospital, Hvidovre, Kettegaard Alle 30, 2650 Hvidovre. | Albertsen lab, Department of Chemistry and Bioscience, Aalborg University, Denmark | Rasmus Kirkegaard |
| EPI_ISL_429304 | hCoV-19/Denmark/ALAB-HH56/2020 | Europe / Denmark / Unknown | 2020-03-17 | Department of Clinical Microbiology, Copenhagen University Hospital, Hvidovre, Kettegaard Alle 30, 2650 Hvidovre. | Albertsen lab, Department of Chemistry and Bioscience, Aalborg University, Denmark | Rasmus Kirkegaard |
| EPI_ISL_429305 | hCoV-19/Denmark/ALAB-HH57/2020 | Europe / Denmark / Unknown | 2020-03-17 | Department of Clinical Microbiology, Copenhagen University Hospital, Hvidovre, Kettegaard Alle 30, 2650 Hvidovre. | Albertsen lab, Department of Chemistry and Bioscience, Aalborg University, Denmark | Rasmus Kirkegaard |
| EPI_ISL_429306 | hCoV-19/Denmark/ALAB-HH60/2020 | Europe / Denmark / Unknown | 2020-03-17 | Department of Clinical Microbiology, Copenhagen University Hospital, Hvidovre, Kettegaard Alle 30, 2650 Hvidovre. | Albertsen lab, Department of Chemistry and Bioscience, Aalborg University, Denmark | Rasmus Kirkegaard |
| EPI_ISL_429307 | hCoV-19/Denmark/ALAB-HH61/2020 | Europe / Denmark / Unknown | 2020-03-17 | Department of Clinical Microbiology, Copenhagen University Hospital, Hvidovre, Kettegaard Alle 30, 2650 Hvidovre. | Albertsen lab, Department of Chemistry and Bioscience, Aalborg University, Denmark | Rasmus Kirkegaard |
| EPI_ISL_429308 | hCoV-19/Denmark/ALAB-HH62/2020 | Europe / Denmark / Unknown | 2020-03-17 | Department of Clinical Microbiology, Copenhagen University Hospital, Hvidovre, Kettegaard Alle 30, 2650 Hvidovre. | Albertsen lab, Department of Chemistry and Bioscience, Aalborg University, Denmark | Rasmus Kirkegaard |
| EPI_ISL_429309 | hCoV-19/Denmark/ALAB-HH63/2020 | Europe / Denmark / Unknown | 2020-03-18 | Department of Clinical Microbiology, Copenhagen University Hospital, Hvidovre, Kettegaard Alle 30, 2650 Hvidovre. | Albertsen lab, Department of Chemistry and Bioscience, Aalborg University, Denmark | Rasmus Kirkegaard |
| EPI_ISL_429310 | hCoV-19/Denmark/ALAB-HH64/2020 | Europe / Denmark / Unknown | 2020-03-17 | Department of Clinical Microbiology, Copenhagen University Hospital, Hvidovre, Kettegaard Alle 30, 2650 Hvidovre. | Albertsen lab, Department of Chemistry and Bioscience, Aalborg University, Denmark | Rasmus Kirkegaard |
| EPI_ISL_429311 | hCoV-19/Denmark/ALAB-HH65/2020 | Europe / Denmark / Unknown | 2020-03-17 | Department of Clinical Microbiology, Copenhagen University Hospital, Hvidovre, Kettegaard Alle 30, 2650 Hvidovre. | Albertsen lab, Department of Chemistry and Bioscience, Aalborg University, Denmark | Rasmus Kirkegaard |
| EPI_ISL_429312 | hCoV-19/Denmark/ALAB-HH66/2020 | Europe / Denmark / Unknown | 2020-03-18 | Department of Clinical Microbiology, Copenhagen University Hospital, Hvidovre, Kettegaard Alle 30, 2650 Hvidovre. | Albertsen lab, Department of Chemistry and Bioscience, Aalborg University, Denmark | Rasmus Kirkegaard |
| EPI_ISL_429313 | hCoV-19/Denmark/ALAB-HH67/2020 | Europe / Denmark / Unknown | 2020-03-18 | Department of Clinical Microbiology, Copenhagen University Hospital, Hvidovre, Kettegaard Alle 30, 2650 Hvidovre. | Albertsen lab, Department of Chemistry and Bioscience, Aalborg University, Denmark | Rasmus Kirkegaard |
| EPI_ISL_429314 | hCoV-19/Denmark/ALAB-HH73/2020 | Europe / Denmark / Unknown | 2020-03-18 | Department of Clinical Microbiology, Copenhagen University Hospital, Hvidovre, Kettegaard Alle 30, 2650 Hvidovre. | Albertsen lab, Department of Chemistry and Bioscience, Aalborg University, Denmark | Rasmus Kirkegaard |
| EPI_ISL_429315 | hCoV-19/Denmark/ALAB-HH74/2020 | Europe / Denmark / Unknown | 2020-03-18 | Department of Clinical Microbiology, Copenhagen University Hospital, Hvidovre, Kettegaard Alle 30, 2650 Hvidovre. | Albertsen lab, Department of Chemistry and Bioscience, Aalborg University, Denmark | Rasmus Kirkegaard |
| EPI_ISL_429316 | hCoV-19/Denmark/ALAB-HH75/2020 | Europe / Denmark / Unknown | 2020-03-18 | Department of Clinical Microbiology, Copenhagen University Hospital, Hvidovre, Kettegaard Alle 30, 2650 Hvidovre. | Albertsen lab, Department of Chemistry and Bioscience, Aalborg University, Denmark | Rasmus Kirkegaard |
| EPI_ISL_429317 | hCoV-19/Denmark/ALAB-HH76/2020 | Europe / Denmark / Unknown | 2020-03-18 | Department of Clinical Microbiology, Copenhagen University Hospital, Hvidovre, Kettegaard Alle 30, 2650 Hvidovre. | Albertsen lab, Department of Chemistry and Bioscience, Aalborg University, Denmark | Rasmus Kirkegaard |
| EPI_ISL_429318 | hCoV-19/Denmark/ALAB-HH77/2020 | Europe / Denmark / Unknown | 2020-03-16 | Department of Clinical Microbiology, Copenhagen University Hospital, Hvidovre, Kettegaard Alle 30, 2650 Hvidovre. | Albertsen lab, Department of Chemistry and Bioscience, Aalborg University, Denmark | Rasmus Kirkegaard |
| EPI_ISL_429320 | hCoV-19/Denmark/ALAB-HH78/2020 | Europe / Denmark / Unknown | 2020-03-16 | Department of Clinical Microbiology, Copenhagen University Hospital, Hvidovre, Kettegaard Alle 30, 2650 Hvidovre. | Albertsen lab, Department of Chemistry and Bioscience, Aalborg University, Denmark | Rasmus Kirkegaard |
| EPI_ISL_429321 | hCoV-19/Denmark/ALAB-HH80/2020 | Europe / Denmark / Unknown | 2020-03-16 | Department of Clinical Microbiology, Copenhagen University Hospital, Hvidovre, Kettegaard Alle 30, 2650 Hvidovre. | Albertsen lab, Department of Chemistry and Bioscience, Aalborg University, Denmark | Rasmus Kirkegaard |
| EPI_ISL_429322 | hCoV-19/Denmark/ALAB-HH81/2020 | Europe / Denmark / Unknown | 2020-03-16 | Department of Clinical Microbiology, Copenhagen University Hospital, Hvidovre, Kettegaard Alle 30, 2650 Hvidovre. | Albertsen lab, Department of Chemistry and Bioscience, Aalborg University, Denmark | Rasmus Kirkegaard |
| EPI_ISL_429323 | hCoV-19/Denmark/ALAB-HH82/2020 | Europe / Denmark / Unknown | 2020-03-15 | Department of Clinical Microbiology, Copenhagen University Hospital, Hvidovre, Kettegaard Alle 30, 2650 Hvidovre. | Albertsen lab, Department of Chemistry and Bioscience, Aalborg University, Denmark | Rasmus Kirkegaard |
| EPI_ISL_429324 | hCoV-19/Denmark/ALAB-HH83/2020 | Europe / Denmark / Unknown | 2020-03-16 | Department of Clinical Microbiology, Copenhagen University Hospital, Hvidovre, Kettegaard Alle 30, 2650 Hvidovre. | Albertsen lab, Department of Chemistry and Bioscience, Aalborg University, Denmark | Rasmus Kirkegaard |
| EPI_ISL_429325 | hCoV-19/Denmark/ALAB-HH84/2020 | Europe / Denmark / Unknown | 2020-03-16 | Department of Clinical Microbiology, Copenhagen University Hospital, Hvidovre, Kettegaard Alle 30, 2650 Hvidovre. | Albertsen lab, Department of Chemistry and Bioscience, Aalborg University, Denmark | Rasmus Kirkegaard |
| EPI_ISL_429326 | hCoV-19/Denmark/ALAB-HH86/2020 | Europe / Denmark / Unknown | 2020-03-15 | Department of Clinical Microbiology, Copenhagen University Hospital, Hvidovre, Kettegaard Alle 30, 2650 Hvidovre. | Albertsen lab, Department of Chemistry and Bioscience, Aalborg University, Denmark | Rasmus Kirkegaard |
| EPI_ISL_429327 | hCoV-19/Denmark/ALAB-HH87/2020 | Europe / Denmark / Unknown | 2020-03-15 | Department of Clinical Microbiology, Copenhagen University Hospital, Hvidovre, Kettegaard Alle 30, 2650 Hvidovre. | Albertsen lab, Department of Chemistry and Bioscience, Aalborg University, Denmark | Rasmus Kirkegaard |
| EPI_ISL_429328 | hCoV-19/Denmark/ALAB-HH88/2020 | Europe / Denmark / Unknown | 2020-03-15 | Department of Clinical Microbiology, Copenhagen University Hospital, Hvidovre, Kettegaard Alle 30, 2650 Hvidovre. | Albertsen lab, Department of Chemistry and Bioscience, Aalborg University, Denmark | Rasmus Kirkegaard |
| EPI_ISL_429329 | hCoV-19/Denmark/ALAB-HH89/2020 | Europe / Denmark / Unknown | 2020-03-15 | Department of Clinical Microbiology, Copenhagen University Hospital, Hvidovre, Kettegaard Alle 30, 2650 Hvidovre. | Albertsen lab, Department of Chemistry and Bioscience, Aalborg University, Denmark | Rasmus Kirkegaard |
| EPI_ISL_429331 | hCoV-19/Denmark/ALAB-HH92/2020 | Europe / Denmark / Unknown | 2020-03-15 | Department of Clinical Microbiology, Copenhagen University Hospital, Hvidovre, Kettegaard Alle 30, 2650 Hvidovre. | Albertsen lab, Department of Chemistry and Bioscience, Aalborg University, Denmark | Rasmus Kirkegaard |
| EPI_ISL_429335 | hCoV-19/Denmark/ALAB-SSI103/2020 | Europe / Denmark / Unknown | 2020-03-02 | Department of Virus and Microbiological Special Diagnostics, Statens Serum Institut, Copenhagen, Denmark, Artillerivej 5, 2300 Copenahgen S | Albertsen lab, Department of Chemistry and Bioscience, Aalborg University, Denmark | Rasmus Kirkegaard |
| EPI_ISL_429337 | hCoV-19/Denmark/ALAB-SSI105/2020 | Europe / Denmark / Unknown | 2020-03-08 | Department of Virus and Microbiological Special Diagnostics, Statens Serum Institut, Copenhagen, Denmark, Artillerivej 5, 2300 Copenahgen S | Albertsen lab, Department of Chemistry and Bioscience, Aalborg University, Denmark | Rasmus Kirkegaard |
| EPI_ISL_429339 | hCoV-19/Denmark/ALAB-SSI107/2020 | Europe / Denmark / Unknown | 2020-03-07 | Department of Virus and Microbiological Special Diagnostics, Statens Serum Institut, Copenhagen, Denmark, Artillerivej 5, 2300 Copenahgen S | Albertsen lab, Department of Chemistry and Bioscience, Aalborg University, Denmark | Rasmus Kirkegaard |
| EPI_ISL_429340 | hCoV-19/Denmark/ALAB-SSI108/2020 | Europe / Denmark / Unknown | 2020-03-07 | Department of Virus and Microbiological Special Diagnostics, Statens Serum Institut, Copenhagen, Denmark, Artillerivej 5, 2300 Copenahgen S | Albertsen lab, Department of Chemistry and Bioscience, Aalborg University, Denmark | Rasmus Kirkegaard |
| EPI_ISL_429341 | hCoV-19/Denmark/ALAB-SSI109/2020 | Europe / Denmark / Unknown | 2020-03-04 | Department of Virus and Microbiological Special Diagnostics, Statens Serum Institut, Copenhagen, Denmark, Artillerivej 5, 2300 Copenahgen S | Albertsen lab, Department of Chemistry and Bioscience, Aalborg University, Denmark | Rasmus Kirkegaard |
| EPI_ISL_429342 | hCoV-19/Denmark/ALAB-SSI110/2020 | Europe / Denmark / Unknown | 2020-03-08 | Department of Virus and Microbiological Special Diagnostics, Statens Serum Institut, Copenhagen, Denmark, Artillerivej 5, 2300 Copenahgen S | Albertsen lab, Department of Chemistry and Bioscience, Aalborg University, Denmark | Rasmus Kirkegaard |
| EPI_ISL_429343 | hCoV-19/Denmark/ALAB-SSI111/2020 | Europe / Denmark / Unknown | 2020-03-09 | Department of Virus and Microbiological Special Diagnostics, Statens Serum Institut, Copenhagen, Denmark, Artillerivej 5, 2300 Copenahgen S | Albertsen lab, Department of Chemistry and Bioscience, Aalborg University, Denmark | Rasmus Kirkegaard |
| EPI_ISL_429344 | hCoV-19/Denmark/ALAB-SSI113/2020 | Europe / Denmark / Unknown | 2020-03-08 | Department of Virus and Microbiological Special Diagnostics, Statens Serum Institut, Copenhagen, Denmark, Artillerivej 5, 2300 Copenahgen S | Albertsen lab, Department of Chemistry and Bioscience, Aalborg University, Denmark | Rasmus Kirkegaard |
| EPI_ISL_429345 | hCoV-19/Denmark/ALAB-SSI114/2020 | Europe / Denmark / Unknown | 2020-03-09 | Department of Virus and Microbiological Special Diagnostics, Statens Serum Institut, Copenhagen, Denmark, Artillerivej 5, 2300 Copenahgen S | Albertsen lab, Department of Chemistry and Bioscience, Aalborg University, Denmark | Rasmus Kirkegaard |
| EPI_ISL_429346 | hCoV-19/Denmark/ALAB-SSI117/2020 | Europe / Denmark / Unknown | 2020-03-06 | Department of Virus and Microbiological Special Diagnostics, Statens Serum Institut, Copenhagen, Denmark, Artillerivej 5, 2300 Copenahgen S | Albertsen lab, Department of Chemistry and Bioscience, Aalborg University, Denmark | Rasmus Kirkegaard |
| EPI_ISL_429348 | hCoV-19/Denmark/ALAB-SSI119/2020 | Europe / Denmark / Unknown | 2020-03-08 | Department of Virus and Microbiological Special Diagnostics, Statens Serum Institut, Copenhagen, Denmark, Artillerivej 5, 2300 Copenahgen S | Albertsen lab, Department of Chemistry and Bioscience, Aalborg University, Denmark | Rasmus Kirkegaard |
| EPI_ISL_429349 | hCoV-19/Denmark/ALAB-SSI120/2020 | Europe / Denmark / Unknown | 2020-03-08 | Department of Virus and Microbiological Special Diagnostics, Statens Serum Institut, Copenhagen, Denmark, Artillerivej 5, 2300 Copenahgen S | Albertsen lab, Department of Chemistry and Bioscience, Aalborg University, Denmark | Rasmus Kirkegaard |
| EPI_ISL_429350 | hCoV-19/Denmark/ALAB-SSI121/2020 | Europe / Denmark / Unknown | 2020-03-08 | Department of Virus and Microbiological Special Diagnostics, Statens Serum Institut, Copenhagen, Denmark, Artillerivej 5, 2300 Copenahgen S | Albertsen lab, Department of Chemistry and Bioscience, Aalborg University, Denmark | Rasmus Kirkegaard |
| EPI_ISL_429351 | hCoV-19/Denmark/ALAB-SSI122/2020 | Europe / Denmark / Unknown | 2020-03-08 | Department of Virus and Microbiological Special Diagnostics, Statens Serum Institut, Copenhagen, Denmark, Artillerivej 5, 2300 Copenahgen S | Albertsen lab, Department of Chemistry and Bioscience, Aalborg University, Denmark | Rasmus Kirkegaard |
| EPI_ISL_429352 | hCoV-19/Denmark/ALAB-SSI123/2020 | Europe / Denmark / Unknown | 2020-03-08 | Department of Virus and Microbiological Special Diagnostics, Statens Serum Institut, Copenhagen, Denmark, Artillerivej 5, 2300 Copenahgen S | Albertsen lab, Department of Chemistry and Bioscience, Aalborg University, Denmark | Rasmus Kirkegaard |
| EPI_ISL_429353 | hCoV-19/Denmark/ALAB-SSI125/2020 | Europe / Denmark / Unknown | 2020-03-08 | Department of Virus and Microbiological Special Diagnostics, Statens Serum Institut, Copenhagen, Denmark, Artillerivej 5, 2300 Copenahgen S | Albertsen lab, Department of Chemistry and Bioscience, Aalborg University, Denmark | Rasmus Kirkegaard |
| EPI_ISL_429354 | hCoV-19/Denmark/ALAB-SSI126/2020 | Europe / Denmark / Unknown | 2020-03-08 | Department of Virus and Microbiological Special Diagnostics, Statens Serum Institut, Copenhagen, Denmark, Artillerivej 5, 2300 Copenahgen S | Albertsen lab, Department of Chemistry and Bioscience, Aalborg University, Denmark | Rasmus Kirkegaard |
| EPI_ISL_429355 | hCoV-19/Denmark/ALAB-SSI128/2020 | Europe / Denmark / Unknown | 2020-03-08 | Department of Virus and Microbiological Special Diagnostics, Statens Serum Institut, Copenhagen, Denmark, Artillerivej 5, 2300 Copenahgen S | Albertsen lab, Department of Chemistry and Bioscience, Aalborg University, Denmark | Rasmus Kirkegaard |
| EPI_ISL_429356 | hCoV-19/Denmark/ALAB-SSI129/2020 | Europe / Denmark / Unknown | 2020-03-08 | Department of Virus and Microbiological Special Diagnostics, Statens Serum Institut, Copenhagen, Denmark, Artillerivej 5, 2300 Copenahgen S | Albertsen lab, Department of Chemistry and Bioscience, Aalborg University, Denmark | Rasmus Kirkegaard |
| EPI_ISL_429357 | hCoV-19/Denmark/ALAB-SSI132/2020 | Europe / Denmark / Unknown | 2020-03-08 | Department of Virus and Microbiological Special Diagnostics, Statens Serum Institut, Copenhagen, Denmark, Artillerivej 5, 2300 Copenahgen S | Albertsen lab, Department of Chemistry and Bioscience, Aalborg University, Denmark | Rasmus Kirkegaard |
| EPI_ISL_429358 | hCoV-19/Denmark/ALAB-SSI139/2020 | Europe / Denmark / Unknown | 2020-03-09 | Department of Virus and Microbiological Special Diagnostics, Statens Serum Institut, Copenhagen, Denmark, Artillerivej 5, 2300 Copenahgen S | Albertsen lab, Department of Chemistry and Bioscience, Aalborg University, Denmark | Rasmus Kirkegaard |
| EPI_ISL_429360 | hCoV-19/Denmark/ALAB-SSI142/2020 | Europe / Denmark / Unknown | 2020-03-09 | Department of Virus and Microbiological Special Diagnostics, Statens Serum Institut, Copenhagen, Denmark, Artillerivej 5, 2300 Copenahgen S | Albertsen lab, Department of Chemistry and Bioscience, Aalborg University, Denmark | Rasmus Kirkegaard |
| EPI_ISL_429361 | hCoV-19/Denmark/ALAB-SSI143/2020 | Europe / Denmark / Unknown | 2020-03-09 | Department of Virus and Microbiological Special Diagnostics, Statens Serum Institut, Copenhagen, Denmark, Artillerivej 5, 2300 Copenahgen S | Albertsen lab, Department of Chemistry and Bioscience, Aalborg University, Denmark | Rasmus Kirkegaard |
| EPI_ISL_429364 | hCoV-19/Denmark/ALAB-SSI146/2020 | Europe / Denmark / Unknown | 2020-03-09 | Department of Virus and Microbiological Special Diagnostics, Statens Serum Institut, Copenhagen, Denmark, Artillerivej 5, 2300 Copenahgen S | Albertsen lab, Department of Chemistry and Bioscience, Aalborg University, Denmark | Rasmus Kirkegaard |

| | | | | | | |
|---|---|---|---|---|---|---|
| EPI_ISL_429368 | hCoV-19/Denmark/ALAB-SSI150/2020 | Europe / Denmark / Unknown | 2020-03-09 | Department of Virus and Microbiological Special Diagnostics, Statens Serum Institut, Copenhagen, Denmark, Artillerivej 5, 2300 Copenahgen S | Albertsen lab, Department of Chemistry and Bioscience, Aalborg University, Denmark | Rasmus Kirkegaard |
| EPI_ISL_429369 | hCoV-19/Denmark/ALAB-SSI151/2020 | Europe / Denmark / Unknown | 2020-03-09 | Department of Virus and Microbiological Special Diagnostics, Statens Serum Institut, Copenhagen, Denmark, Artillerivej 5, 2300 Copenahgen S | Albertsen lab, Department of Chemistry and Bioscience, Aalborg University, Denmark | Rasmus Kirkegaard |
| EPI_ISL_429370 | hCoV-19/Denmark/ALAB-SSI154/2020 | Europe / Denmark / Unknown | 2020-03-09 | Department of Virus and Microbiological Special Diagnostics, Statens Serum Institut, Copenhagen, Denmark, Artillerivej 5, 2300 Copenahgen S | Albertsen lab, Department of Chemistry and Bioscience, Aalborg University, Denmark | Rasmus Kirkegaard |
| EPI_ISL_429371 | hCoV-19/Denmark/ALAB-SSI155/2020 | Europe / Denmark / Unknown | 2020-03-10 | Department of Virus and Microbiological Special Diagnostics, Statens Serum Institut, Copenhagen, Denmark, Artillerivej 5, 2300 Copenahgen S | Albertsen lab, Department of Chemistry and Bioscience, Aalborg University, Denmark | Rasmus Kirkegaard |
| EPI_ISL_429372 | hCoV-19/Denmark/ALAB-SSI156/2020 | Europe / Denmark / Unknown | 2020-03-10 | Department of Virus and Microbiological Special Diagnostics, Statens Serum Institut, Copenhagen, Denmark, Artillerivej 5, 2300 Copenahgen S | Albertsen lab, Department of Chemistry and Bioscience, Aalborg University, Denmark | Rasmus Kirkegaard |
| EPI_ISL_429373 | hCoV-19/Denmark/ALAB-SSI157/2020 | Europe / Denmark / Unknown | 2020-03-09 | Department of Virus and Microbiological Special Diagnostics, Statens Serum Institut, Copenhagen, Denmark, Artillerivej 5, 2300 Copenahgen S | Albertsen lab, Department of Chemistry and Bioscience, Aalborg University, Denmark | Rasmus Kirkegaard |
| EPI_ISL_429374 | hCoV-19/Denmark/ALAB-SSI158/2020 | Europe / Denmark / Unknown | 2020-03-09 | Department of Virus and Microbiological Special Diagnostics, Statens Serum Institut, Copenhagen, Denmark, Artillerivej 5, 2300 Copenahgen S | Albertsen lab, Department of Chemistry and Bioscience, Aalborg University, Denmark | Rasmus Kirkegaard |
| EPI_ISL_429375 | hCoV-19/Denmark/ALAB-SSI159/2020 | Europe / Denmark / Unknown | 2020-03-09 | Department of Virus and Microbiological Special Diagnostics, Statens Serum Institut, Copenhagen, Denmark, Artillerivej 5, 2300 Copenahgen S | Albertsen lab, Department of Chemistry and Bioscience, Aalborg University, Denmark | Rasmus Kirkegaard |
| EPI_ISL_429376 | hCoV-19/Denmark/ALAB-SSI160/2020 | Europe / Denmark / Unknown | 2020-03-09 | Department of Virus and Microbiological Special Diagnostics, Statens Serum Institut, Copenhagen, Denmark, Artillerivej 5, 2300 Copenahgen S | Albertsen lab, Department of Chemistry and Bioscience, Aalborg University, Denmark | Rasmus Kirkegaard |
| EPI_ISL_429377 | hCoV-19/Denmark/ALAB-SSI162/2020 | Europe / Denmark / Unknown | 2020-03-09 | Department of Virus and Microbiological Special Diagnostics, Statens Serum Institut, Copenhagen, Denmark, Artillerivej 5, 2300 Copenahgen S | Albertsen lab, Department of Chemistry and Bioscience, Aalborg University, Denmark | Rasmus Kirkegaard |
| EPI_ISL_429378 | hCoV-19/Denmark/ALAB-SSI163/2020 | Europe / Denmark / Unknown | 2020-03-09 | Department of Virus and Microbiological Special Diagnostics, Statens Serum Institut, Copenhagen, Denmark, Artillerivej 5, 2300 Copenahgen S | Albertsen lab, Department of Chemistry and Bioscience, Aalborg University, Denmark | Rasmus Kirkegaard |
| EPI_ISL_429379 | hCoV-19/Denmark/ALAB-SSI164/2020 | Europe / Denmark / Unknown | 2020-03-09 | Department of Virus and Microbiological Special Diagnostics, Statens Serum Institut, Copenhagen, Denmark, Artillerivej 5, 2300 Copenahgen S | Albertsen lab, Department of Chemistry and Bioscience, Aalborg University, Denmark | Rasmus Kirkegaard |
| EPI_ISL_429380 | hCoV-19/Denmark/ALAB-SSI166/2020 | Europe / Denmark / Unknown | 2020-03-09 | Department of Virus and Microbiological Special Diagnostics, Statens Serum Institut, Copenhagen, Denmark, Artillerivej 5, 2300 Copenahgen S | Albertsen lab, Department of Chemistry and Bioscience, Aalborg University, Denmark | Rasmus Kirkegaard |
| EPI_ISL_429381 | hCoV-19/Denmark/ALAB-SSI167/2020 | Europe / Denmark / Unknown | 2020-03-09 | Department of Virus and Microbiological Special Diagnostics, Statens Serum Institut, Copenhagen, Denmark, Artillerivej 5, 2300 Copenahgen S | Albertsen lab, Department of Chemistry and Bioscience, Aalborg University, Denmark | Rasmus Kirkegaard |
| EPI_ISL_429382 | hCoV-19/Denmark/ALAB-SSI168/2020 | Europe / Denmark / Unknown | 2020-03-09 | Department of Virus and Microbiological Special Diagnostics, Statens Serum Institut, Copenhagen, Denmark, Artillerivej 5, 2300 Copenahgen S | Albertsen lab, Department of Chemistry and Bioscience, Aalborg University, Denmark | Rasmus Kirkegaard |
| EPI_ISL_429383 | hCoV-19/Denmark/ALAB-SSI169/2020 | Europe / Denmark / Unknown | 2020-03-09 | Department of Virus and Microbiological Special Diagnostics, Statens Serum Institut, Copenhagen, Denmark, Artillerivej 5, 2300 Copenahgen S | Albertsen lab, Department of Chemistry and Bioscience, Aalborg University, Denmark | Rasmus Kirkegaard |
| EPI_ISL_429384 | hCoV-19/Denmark/ALAB-SSI171/2020 | Europe / Denmark / Unknown | 2020-03-09 | Department of Virus and Microbiological Special Diagnostics, Statens Serum Institut, Copenhagen, Denmark, Artillerivej 5, 2300 Copenahgen S | Albertsen lab, Department of Chemistry and Bioscience, Aalborg University, Denmark | Rasmus Kirkegaard |
| EPI_ISL_429385 | hCoV-19/Denmark/ALAB-SSI172/2020 | Europe / Denmark / Unknown | 2020-03-09 | Department of Virus and Microbiological Special Diagnostics, Statens Serum Institut, Copenhagen, Denmark, Artillerivej 5, 2300 Copenahgen S | Albertsen lab, Department of Chemistry and Bioscience, Aalborg University, Denmark | Rasmus Kirkegaard |
| EPI_ISL_429386 | hCoV-19/Denmark/ALAB-SSI173/2020 | Europe / Denmark / Unknown | 2020-03-09 | Department of Virus and Microbiological Special Diagnostics, Statens Serum Institut, Copenhagen, Denmark, Artillerivej 5, 2300 Copenahgen S | Albertsen lab, Department of Chemistry and Bioscience, Aalborg University, Denmark | Rasmus Kirkegaard |
| EPI_ISL_429387 | hCoV-19/Denmark/ALAB-SSI174/2020 | Europe / Denmark / Unknown | 2020-03-09 | Department of Virus and Microbiological Special Diagnostics, Statens Serum Institut, Copenhagen, Denmark, Artillerivej 5, 2300 Copenahgen S | Albertsen lab, Department of Chemistry and Bioscience, Aalborg University, Denmark | Rasmus Kirkegaard |
| EPI_ISL_429391 | hCoV-19/Denmark/ALAB-SSI187/2020 | Europe / Denmark / Unknown | 2020-03-09 | Department of Virus and Microbiological Special Diagnostics, Statens Serum Institut, Copenhagen, Denmark, Artillerivej 5, 2300 Copenahgen S | Albertsen lab, Department of Chemistry and Bioscience, Aalborg University, Denmark | Rasmus Kirkegaard |
| EPI_ISL_429395 | hCoV-19/Denmark/ALAB-SSI194/2020 | Europe / Denmark / Unknown | 2020-03-09 | Department of Virus and Microbiological Special Diagnostics, Statens Serum Institut, Copenhagen, Denmark, Artillerivej 5, 2300 Copenahgen S | Albertsen lab, Department of Chemistry and Bioscience, Aalborg University, Denmark | Rasmus Kirkegaard |
| EPI_ISL_429396 | hCoV-19/Denmark/ALAB-SSI195/2020 | Europe / Denmark / Unknown | 2020-03-09 | Department of Virus and Microbiological Special Diagnostics, Statens Serum Institut, Copenhagen, Denmark, Artillerivej 5, 2300 Copenahgen S | Albertsen lab, Department of Chemistry and Bioscience, Aalborg University, Denmark | Rasmus Kirkegaard |
| EPI_ISL_429399 | hCoV-19/Denmark/ALAB-SSI201/2020 | Europe / Denmark / Unknown | 2020-03-09 | Department of Virus and Microbiological Special Diagnostics, Statens Serum Institut, Copenhagen, Denmark, Artillerivej 5, 2300 Copenahgen S | Albertsen lab, Department of Chemistry and Bioscience, Aalborg University, Denmark | Rasmus Kirkegaard |
| EPI_ISL_429600 | hCoV-19/USA/WA-UW-4549/2020 | North America / USA / Washington | 2020-03-31 | UW Virology Lab | UW Virology Lab | Pavitra Roychoudhury, Hong Xie, Keith Jerome, Alexander Greninger |
| EPI_ISL_429601 | hCoV-19/USA/ID-UW-4552/2020 | North America / USA / Idaho | 2020-03-29 | UW Virology Lab | UW Virology Lab | Pavitra Roychoudhury, Hong Xie, Keith Jerome, Alexander Greninger |
| EPI_ISL_429602 | hCoV-19/USA/WA-UW-4564/2020 | North America / USA / Washington | 2020-03-30 | UW Virology Lab | UW Virology Lab | Pavitra Roychoudhury, Hong Xie, Keith Jerome, Alexander Greninger |
| EPI_ISL_429603 | hCoV-19/USA/WA-UW-4565/2020 | North America / USA / Washington | 2020-03-30 | UW Virology Lab | UW Virology Lab | Pavitra Roychoudhury, Hong Xie, Keith Jerome, Alexander Greninger |
| EPI_ISL_429604 | hCoV-19/USA/WA-UW-4568/2020 | North America / USA / Washington | 2020-03-29 | UW Virology Lab | UW Virology Lab | Pavitra Roychoudhury, Hong Xie, Keith Jerome, Alexander Greninger |
| EPI_ISL_429605 | hCoV-19/USA/WA-UW-4578/2020 | North America / USA / Washington | 2020-03-31 | UW Virology Lab | UW Virology Lab | Pavitra Roychoudhury, Hong Xie, Keith Jerome, Alexander Greninger |
| EPI_ISL_429606 | hCoV-19/USA/WA-UW-4579/2020 | North America / USA / Washington | 2020-03-31 | UW Virology Lab | UW Virology Lab | Pavitra Roychoudhury, Hong Xie, Keith Jerome, Alexander Greninger |
| EPI_ISL_429608 | hCoV-19/USA/WA-UW-4593/2020 | North America / USA / Washington | 2020-03-31 | UW Virology Lab | UW Virology Lab | Pavitra Roychoudhury, Hong Xie, Keith Jerome, Alexander Greninger |
| EPI_ISL_429609 | hCoV-19/USA/WA-UW-4594/2020 | North America / USA / Washington | 2020-03-31 | UW Virology Lab | UW Virology Lab | Pavitra Roychoudhury, Hong Xie, Keith Jerome, Alexander Greninger |
| EPI_ISL_429610 | hCoV-19/USA/ID-UW-4598/2020 | North America / USA / Idaho | 2020-03-30 | UW Virology Lab | UW Virology Lab | Pavitra Roychoudhury, Hong Xie, Keith Jerome, Alexander Greninger |
| EPI_ISL_429611 | hCoV-19/USA/WA-UW-4602/2020 | North America / USA / Washington | 2020-03-31 | UW Virology Lab | UW Virology Lab | Pavitra Roychoudhury, Hong Xie, Keith Jerome, Alexander Greninger |
| EPI_ISL_429612 | hCoV-19/USA/WA-UW-4609/2020 | North America / USA / Washington | 2020-03-26 | UW Virology Lab | UW Virology Lab | Pavitra Roychoudhury, Hong Xie, Keith Jerome, Alexander Greninger |
| EPI_ISL_429613 | hCoV-19/USA/WA-UW-4611/2020 | North America / USA / Washington | 2020-03-26 | UW Virology Lab | UW Virology Lab | Pavitra Roychoudhury, Hong Xie, Keith Jerome, Alexander Greninger |
| EPI_ISL_429614 | hCoV-19/USA/WA-UW-4612/2020 | North America / USA / Washington | 2020-03-26 | UW Virology Lab | UW Virology Lab | Pavitra Roychoudhury, Hong Xie, Keith Jerome, Alexander Greninger |
| EPI_ISL_429615 | hCoV-19/USA/WA-UW-4703/2020 | North America / USA / Washington | 2020-03-29 | UW Virology Lab | UW Virology Lab | Pavitra Roychoudhury, Hong Xie, Keith Jerome, Alexander Greninger |
| EPI_ISL_429616 | hCoV-19/USA/WA-UW-4732/2020 | North America / USA / Washington | 2020-03-29 | UW Virology Lab | UW Virology Lab | Pavitra Roychoudhury, Hong Xie, Keith Jerome, Alexander Greninger |
| EPI_ISL_429617 | hCoV-19/USA/WA-UW-4734/2020 | North America / USA / Washington | 2020-03-28 | UW Virology Lab | UW Virology Lab | Pavitra Roychoudhury, Hong Xie, Keith Jerome, Alexander Greninger |
| EPI_ISL_429618 | hCoV-19/USA/WA-UW-4741/2020 | North America / USA / Washington | 2020-03-25 | UW Virology Lab | UW Virology Lab | Pavitra Roychoudhury, Hong Xie, Keith Jerome, Alexander Greninger |
| EPI_ISL_429621 | hCoV-19/USA/WA-UW-4760/2020 | North America / USA / Washington | 2020-03-28 | UW Virology Lab | UW Virology Lab | Pavitra Roychoudhury, Hong Xie, Keith Jerome, Alexander Greninger |
| EPI_ISL_429622 | hCoV-19/USA/WA-UW-4763/2020 | North America / USA / Washington | 2020-03-27 | UW Virology Lab | UW Virology Lab | Pavitra Roychoudhury, Hong Xie, Keith Jerome, Alexander Greninger |
| EPI_ISL_429623 | hCoV-19/USA/WA-UW-4766/2020 | North America / USA / Washington | 2020-03-28 | UW Virology Lab | UW Virology Lab | Pavitra Roychoudhury, Hong Xie, Keith Jerome, Alexander Greninger |
| EPI_ISL_429624 | hCoV-19/USA/WA-UW-4769/2020 | North America / USA / Washington | 2020-03-27 | UW Virology Lab | UW Virology Lab | Pavitra Roychoudhury, Hong Xie, Keith Jerome, Alexander Greninger |
| EPI_ISL_429625 | hCoV-19/USA/WA-UW-4770/2020 | North America / USA / Washington | 2020-03-27 | UW Virology Lab | UW Virology Lab | Pavitra Roychoudhury, Hong Xie, Keith Jerome, Alexander Greninger |
| EPI_ISL_429626 | hCoV-19/USA/WA-UW-4775/2020 | North America / USA / Washington | 2020-03-28 | UW Virology Lab | UW Virology Lab | Pavitra Roychoudhury, Hong Xie, Keith Jerome, Alexander Greninger |
| EPI_ISL_429627 | hCoV-19/USA/WA-UW-4777/2020 | North America / USA / Washington | 2020-03-27 | UW Virology Lab | UW Virology Lab | Pavitra Roychoudhury, Hong Xie, Keith Jerome, Alexander Greninger |
| EPI_ISL_429628 | hCoV-19/USA/UN-UW-4645/2020 | North America / USA | 2020-04-06 | UW Virology Lab | UW Virology Lab | Pavitra Roychoudhury, Hong Xie, Keith Jerome, Alexander Greninger |
| EPI_ISL_429629 | hCoV-19/USA/WA-UW-4649/2020 | North America / USA / Washington | 2020-04-06 | UW Virology Lab | UW Virology Lab | Pavitra Roychoudhury, Hong Xie, Keith Jerome, Alexander Greninger |
| EPI_ISL_429630 | hCoV-19/USA/WA-UW-4677/2020 | North America / USA / Washington | 2020-04-05 | UW Virology Lab | UW Virology Lab | Pavitra Roychoudhury, Hong Xie, Keith Jerome, Alexander Greninger |
| EPI_ISL_429631 | hCoV-19/USA/WA-UW-4695/2020 | North America / USA / Washington | 2020-04-06 | UW Virology Lab | UW Virology Lab | Pavitra Roychoudhury, Hong Xie, Keith Jerome, Alexander Greninger |
| EPI_ISL_429632 | hCoV-19/USA/WA-UW-4705/2020 | North America / USA / Washington | 2020-04-06 | UW Virology Lab | UW Virology Lab | Pavitra Roychoudhury, Hong Xie, Keith Jerome, Alexander Greninger |
| EPI_ISL_429634 | hCoV-19/USA/WA-UW-4892/2020 | North America / USA / Washington | 2020-04-07 | UW Virology Lab | UW Virology Lab | Pavitra Roychoudhury, Hong Xie, Keith Jerome, Alexander Greninger |
| EPI_ISL_429635 | hCoV-19/USA/WA-UW-4897/2020 | North America / USA / Washington | 2020-04-08 | UW Virology Lab | UW Virology Lab | Pavitra Roychoudhury, Hong Xie, Keith Jerome, Alexander Greninger |
| EPI_ISL_429636 | hCoV-19/USA/WA-UW-4921/2020 | North America / USA / Washington | 2020-04-07 | UW Virology Lab | UW Virology Lab | Pavitra Roychoudhury, Hong Xie, Keith Jerome, Alexander Greninger |
| EPI_ISL_429637 | hCoV-19/USA/WA-UW-4925/2020 | North America / USA / Washington | 2020-04-07 | UW Virology Lab | UW Virology Lab | Pavitra Roychoudhury, Hong Xie, Keith Jerome, Alexander Greninger |
| EPI_ISL_429638 | hCoV-19/USA/WA-UW-4999/2020 | North America / USA / Washington | 2020-04-07 | UW Virology Lab | UW Virology Lab | Pavitra Roychoudhury, Hong Xie, Keith Jerome, Alexander Greninger |
| EPI_ISL_429639 | hCoV-19/USA/WA-UW-5025/2020 | North America / USA / Washington | 2020-04-07 | UW Virology Lab | UW Virology Lab | Pavitra Roychoudhury, Hong Xie, Keith Jerome, Alexander Greninger |
| EPI_ISL_429641 | hCoV-19/USA/CT-UW-5038/2020 | North America / USA / Connecticut | 2020-04-06 | UW Virology Lab | UW Virology Lab | Pavitra Roychoudhury, Hong Xie, Keith Jerome, Alexander Greninger |
| EPI_ISL_429642 | hCoV-19/USA/CT-UW-5043/2020 | North America / USA / Connecticut | 2020-04-06 | UW Virology Lab | UW Virology Lab | Pavitra Roychoudhury, Hong Xie, Keith Jerome, Alexander Greninger |
| EPI_ISL_429643 | hCoV-19/USA/UN-UW-5044/2020 | North America / USA | 2020-04-08 | UW Virology Lab | UW Virology Lab | Pavitra Roychoudhury, Hong Xie, Keith Jerome, Alexander Greninger |
| EPI_ISL_429644 | hCoV-19/USA/WA-UW-5065/2020 | North America / USA / Washington | 2020-04-08 | UW Virology Lab | UW Virology Lab | Pavitra Roychoudhury, Hong Xie, Keith Jerome, Alexander Greninger |
| EPI_ISL_429645 | hCoV-19/USA/CT-UW-5073/2020 | North America / USA / Connecticut | 2020-04-06 | UW Virology Lab | UW Virology Lab | Pavitra Roychoudhury, Hong Xie, Keith Jerome, Alexander Greninger |
| EPI_ISL_429646 | hCoV-19/USA/CT-UW-5086/2020 | North America / USA / Connecticut | 2020-04-06 | UW Virology Lab | UW Virology Lab | Pavitra Roychoudhury, Hong Xie, Keith Jerome, Alexander Greninger |
| EPI_ISL_429647 | hCoV-19/USA/WA-UW-5095/2020 | North America / USA / Washington | 2020-04-08 | UW Virology Lab | UW Virology Lab | Pavitra Roychoudhury, Hong Xie, Keith Jerome, Alexander Greninger |
| EPI_ISL_429649 | hCoV-19/USA/WA-UW-5144/2020 | North America / USA / Washington | 2020-04-02 | UW Virology Lab | UW Virology Lab | Pavitra Roychoudhury, Hong Xie, Keith Jerome, Alexander Greninger |
| EPI_ISL_429650 | hCoV-19/USA/WA-UW-5152/2020 | North America / USA / Washington | 2020-04-02 | UW Virology Lab | UW Virology Lab | Pavitra Roychoudhury, Hong Xie, Keith Jerome, Alexander Greninger |
| EPI_ISL_429651 | hCoV-19/USA/WA-UW-5158/2020 | North America / USA / Washington | 2020-04-02 | UW Virology Lab | UW Virology Lab | Pavitra Roychoudhury, Hong Xie, Keith Jerome, Alexander Greninger |
| EPI_ISL_429652 | hCoV-19/USA/WA-UW-5164/2020 | North America / USA / Washington | 2020-04-02 | UW Virology Lab | UW Virology Lab | Pavitra Roychoudhury, Hong Xie, Keith Jerome, Alexander Greninger |
| EPI_ISL_429653 | hCoV-19/USA/WA-UW-5166/2020 | North America / USA / Washington | 2020-04-02 | UW Virology Lab | UW Virology Lab | Pavitra Roychoudhury, Hong Xie, Keith Jerome, Alexander Greninger |
| EPI_ISL_429654 | hCoV-19/USA/WA-UW-5169/2020 | North America / USA / Washington | 2020-04-02 | UW Virology Lab | UW Virology Lab | Pavitra Roychoudhury, Hong Xie, Keith Jerome, Alexander Greninger |
| EPI_ISL_429655 | hCoV-19/USA/WA-UW-5170/2020 | North America / USA / Washington | 2020-04-02 | UW Virology Lab | UW Virology Lab | Pavitra Roychoudhury, Hong Xie, Keith Jerome, Alexander Greninger |
| EPI_ISL_429656 | hCoV-19/USA/UN-UW-5172/2020 | North America / USA | 2020-04-02 | UW Virology Lab | UW Virology Lab | Pavitra Roychoudhury, Hong Xie, Keith Jerome, Alexander Greninger |
| EPI_ISL_429659 | hCoV-19/Croatia/OY-S1/2020 | Europe / Croatia / Istria | 2020-03 | Institute for Public Health | Laboratory for advanced genomics | Filip Rokić, Lovro Trgovec-Greif, Neven Sučić, Tomislav Rukavina, Igor Jurak, Oliver Vugrek |
| EPI_ISL_429663 | hCoV-19/Croatia/8U-S17/2020 | Europe / Croatia / Istria | 2020-03 | Institute for Public Health | Laboratory for advanced genomics | Filip Rokić, Lovro Trgovec-Greif, Neven Sučić, Tomislav Rukavina, Igor Jurak, Oliver Vugrek |
| EPI_ISL_429667 | hCoV-19/Brazil/CV4/2020 | South America / Brazil / Minas gerais | 2020-03-09 | Central Public Health Laboratory/Octávio Magalhães Institute (IOM) from the Ezequiel Dias Foundation (FUNED) | Instituto Octávio Magalhães / Fundação Ezequiel Dias (IOM/Funed) | Talita Adelino, Joilson Xavier, Marta Giovanetti, Vagner Fonseca, Marcos Vinícius Silva, Luiz Carlos Junior Alcantara, Mariuce Aparecida Assunção Oliveira |
| EPI_ISL_429669 | hCoV-19/Brazil/CV6/2020 | South America / Brazil / Minas gerais | 2020-03-13 | Central Public Health Laboratory/Octávio Magalhães Institute (IOM) from the Ezequiel Dias Foundation (FUNED) | Instituto Octávio Magalhães / Fundação Ezequiel Dias (IOM/Funed) | Talita Adelino, Joilson Xavier, Marta Giovanetti, Vagner Fonseca, Marcos Vinícius Silva, Luiz Carlos Junior Alcantara, Mariuce Aparecida Assunção Oliveira |
| EPI_ISL_429671 | hCoV-19/Brazil/CV8/2020 | South America / Brazil / Minas gerais | 2020-03-13 | Central Public Health Laboratory/Octávio Magalhães Institute (IOM) from the Ezequiel Dias Foundation (FUNED) | Instituto Octávio Magalhães / Fundação Ezequiel Dias (IOM/Funed) | Talita Adelino, Joilson Xavier, Marta Giovanetti, Vagner Fonseca, Marcos Vinícius Silva, Luiz Carlos Junior Alcantara, Mariuce Aparecida Assunção Oliveira |
| EPI_ISL_429674 | hCoV-19/Brazil/CV12/2020 | South America / Brazil / Minas gerais | 2020-03-11 | Central Public Health Laboratory/Octávio Magalhães Institute (IOM) from the Ezequiel Dias Foundation (FUNED) | Instituto Octávio Magalhães / Fundação Ezequiel Dias (IOM/Funed) | Talita Adelino, Joilson Xavier, Marta Giovanetti, Vagner Fonseca, Marcos Vinícius Silva, Luiz Carlos Junior Alcantara, Mariuce Aparecida Assunção Oliveira |
| EPI_ISL_429676 | hCoV-19/Brazil/CV16/2020 | South America / Brazil / Minas gerais | 2020-03-16 | Central Public Health Laboratory/Octávio Magalhães Institute (IOM) from the Ezequiel Dias Foundation (FUNED) | Instituto Octávio Magalhães / Fundação Ezequiel Dias (IOM/Funed) | Talita Adelino, Joilson Xavier, Marta Giovanetti, Vagner Fonseca, Marcos Vinícius Silva, Luiz Carlos Junior Alcantara, Mariuce Aparecida Assunção Oliveira |
| EPI_ISL_429679 | hCoV-19/Brazil/CV19/2020 | South America / Brazil / Minas gerais | 2020-03-16 | Central Public Health Laboratory/Octávio Magalhães Institute (IOM) from the Ezequiel Dias Foundation (FUNED) | Instituto Octávio Magalhães / Fundação Ezequiel Dias (IOM/Funed) | Talita Adelino, Joilson Xavier, Marta Giovanetti, Vagner Fonseca, Marcos Vinícius Silva, Luiz Carlos Junior Alcantara, Mariuce Aparecida Assunção Oliveira |
| EPI_ISL_429681 | hCoV-19/Brazil/CV21/2020 | South America / Brazil / Minas gerais | 2020-03-16 | Central Public Health Laboratory/Octávio Magalhães Institute (IOM) from the Ezequiel Dias Foundation (FUNED) | Instituto Octávio Magalhães / Fundação Ezequiel Dias (IOM/Funed) | Talita Adelino, Joilson Xavier, Marta Giovanetti, Vagner Fonseca, Marcos Vinícius Silva, Luiz Carlos Junior Alcantara, Mariuce Aparecida Assunção Oliveira |
| EPI_ISL_429684 | hCoV-19/Brazil/CV26/2020 | South America / Brazil / Minas gerais | 2020-03-17 | Central Public Health Laboratory/Octávio Magalhães Institute (IOM) from the Ezequiel Dias Foundation (FUNED) | Instituto Octávio Magalhães / Fundação Ezequiel Dias (IOM/Funed) | Talita Adelino, Joilson Xavier, Marta Giovanetti, Vagner Fonseca, Marcos Vinícius Silva, Luiz Carlos Junior Alcantara, Mariuce Aparecida Assunção Oliveira |
| EPI_ISL_429687 | hCoV-19/Brazil/CV31/2020 | South America / Brazil / Minas gerais | 2020-03-17 | Central Public Health Laboratory/Octávio Magalhães Institute (IOM) from the Ezequiel Dias Foundation (FUNED) | Instituto Octávio Magalhães / Fundação Ezequiel Dias (IOM/Funed) | Talita Adelino, Joilson Xavier, Marta Giovanetti, Vagner Fonseca, Marcos Vinícius Silva, Luiz Carlos Junior Alcantara, Mariuce Aparecida Assunção Oliveira |
| EPI_ISL_429688 | hCoV-19/Brazil/CV32/2020 | South America / Brazil / Minas gerais | 2020-03-17 | Central Public Health Laboratory/Octávio Magalhães Institute (IOM) from the Ezequiel Dias Foundation (FUNED) | Instituto Octávio Magalhães / Fundação Ezequiel Dias (IOM/Funed) | Talita Adelino, Joilson Xavier, Marta Giovanetti, Vagner Fonseca, Marcos Vinícius Silva, Luiz Carlos Junior Alcantara, Mariuce Aparecida Assunção Oliveira |
| EPI_ISL_429689 | hCoV-19/Brazil/CV33/2020 | South America / Brazil / Minas gerais | 2020-03-17 | Central Public Health Laboratory/Octávio Magalhães Institute (IOM) from the Ezequiel Dias Foundation (FUNED) | Instituto Octávio Magalhães / Fundação Ezequiel Dias (IOM/Funed) | Talita Adelino, Joilson Xavier, Marta Giovanetti, Vagner Fonseca, Marcos Vinícius Silva, Luiz Carlos Junior Alcantara, Mariuce Aparecida Assunção Oliveira |
| EPI_ISL_429695 | hCoV-19/Brazil/CV42/2020 | South America / Brazil / Minas gerais | 2020-03-20 | Central Public Health Laboratory/Octávio Magalhães Institute (IOM) from the Ezequiel Dias Foundation (FUNED) | Instituto Octávio Magalhães / Fundação Ezequiel Dias (IOM/Funed) | Talita Adelino, Joilson Xavier, Marta Giovanetti, Vagner Fonseca, Marcos Vinícius Silva, Luiz Carlos Junior Alcantara, Mariuce Aparecida Assunção Oliveira |
| EPI_ISL_430323 | hCoV-19/USA/NY-NYUMC296/2020 | North America / USA / New York / Queen | 2020-04-10 | NYU Langone Health | Departments of Pathology and Medicine, New York University School of Medicine | Maria Aguero-Rosenfeld, Brendan Belovarac, Margaret Black, Ludovic Boytard, John Cadley, Paolo Cotzia, John Chen, Dacia Dimartino, Xiaojun Feng, Tatyana Gindin, Emily Guzman, Adriana Heguy, Megan Hogan, Emily Huang, George Jour, Lawrence H. Lin, Raven Luther, Andrew Lytle, Christian Marier, Matthew T. Maurano, Mark J. Mulligan, Peter Meyn, Raquel Ordonez Ciriza, Iman Osman, Jared Pinnell, Vanessa Raabe, Sitharam Ramaswami, Amy Rapkiewicz, Andre M. Ribeiro-dos-Santos, Marie Samanovic-Golden, Antonio Serrano, Guomiao Shen, Matija Snuderl, Theodore Vougioukalis, Nick Vulpescu, Gael Westby, Paul Zappile, Yutong Zhang |
| EPI_ISL_430324 | hCoV-19/USA/NY-NYUMC297/2020 | North America / USA / New York / Westc | 2020-04-10 | NYU Langone Health | Departments of Pathology and Medicine, New York University School of Medicine | Maria Aguero-Rosenfeld, Brendan Belovarac, Margaret Black, Ludovic Boytard, John Cadley, Paolo Cotzia, John Chen, Dacia Dimartino, Xiaojun Feng, Tatyana Gindin, Emily Guzman, Adriana Heguy, Megan Hogan, Emily Huang, George Jour, Lawrence H. Lin, Raven Luther, Andrew Lytle, Christian Marier, Matthew T. Maurano, Mark J. Mulligan, Peter Meyn, Raquel Ordonez Ciriza, Iman Osman, Jared Pinnell, Vanessa Raabe, Sitharam Ramaswami, Amy Rapkiewicz, Andre M. Ribeiro-dos-Santos, Marie Samanovic-Golden, Antonio Serrano, Guomiao Shen, Matija Snuderl, Theodore Vougioukalis, Nick Vulpescu, Gael Westby, Paul Zappile, Yutong Zhang |
| EPI_ISL_430325 | hCoV-19/USA/NY-NYUMC298/2020 | North America / USA / New York / Brookl | 2020-04-08 | NYU Langone Health | Departments of Pathology and Medicine, New York University School of Medicine | Maria Aguero-Rosenfeld, Brendan Belovarac, Margaret Black, Ludovic Boytard, John Cadley, Paolo Cotzia, John Chen, Dacia Dimartino, Xiaojun Feng, Tatyana Gindin, Emily Guzman, Adriana Heguy, Megan Hogan, Emily Huang, George Jour, Lawrence H. Lin, Raven Luther, Andrew Lytle, Christian Marier, Matthew T. Maurano, Mark J. Mulligan, Peter Meyn, Raquel Ordonez Ciriza, Iman Osman, Jared Pinnell, Vanessa Raabe, Sitharam Ramaswami, Amy Rapkiewicz, Andre M. Ribeiro-dos-Santos, Marie Samanovic-Golden, Antonio Serrano, Guomiao Shen, Matija Snuderl, Theodore Vougioukalis, Nick Vulpescu, Gael Westby, Paul Zappile, Yutong Zhang |
| EPI_ISL_430326 | hCoV-19/USA/NY-NYUMC299/2020 | North America / USA / New York / Brookl | 2020-03-30 | NYU Langone Health | Departments of Pathology and Medicine, New York University School of Medicine | Maria Aguero-Rosenfeld, Brendan Belovarac, Margaret Black, Ludovic Boytard, John Cadley, Paolo Cotzia, John Chen, Dacia Dimartino, Xiaojun Feng, Tatyana Gindin, Emily Guzman, Adriana Heguy, Megan Hogan, Emily Huang, George Jour, Lawrence H. Lin, Raven Luther, Andrew Lytle, Christian Marier, Matthew T. Maurano, Mark J. Mulligan, Peter Meyn, Raquel Ordonez Ciriza, Iman Osman, Jared Pinnell, Vanessa Raabe, Sitharam Ramaswami, Amy Rapkiewicz, Andre M. Ribeiro-dos-Santos, Marie Samanovic-Golden, Antonio Serrano, Guomiao Shen, Matija Snuderl, Theodore Vougioukalis, Nick Vulpescu, Gael Westby, Paul Zappile, Yutong Zhang |
| EPI_ISL_430327 | hCoV-19/USA/NY-NYUMC300/2020 | North America / USA / New York / Manha | 2020-03-30 | NYU Langone Health | Departments of Pathology and Medicine, New York University School of Medicine | Maria Aguero-Rosenfeld, Brendan Belovarac, Margaret Black, Ludovic Boytard, John Cadley, Paolo Cotzia, John Chen, Dacia Dimartino, Xiaojun Feng, Tatyana Gindin, Emily Guzman, Adriana Heguy, Megan Hogan, Emily Huang, George Jour, Lawrence H. Lin, Raven Luther, Andrew Lytle, Christian Marier, Matthew T. Maurano, Mark J. Mulligan, Peter Meyn, Raquel Ordonez Ciriza, Iman Osman, Jared Pinnell, Vanessa Raabe, Sitharam Ramaswami, Amy Rapkiewicz, Andre M. Ribeiro-dos-Santos, Marie Samanovic-Golden, Antonio Serrano, Guomiao Shen, Matija Snuderl, Theodore Vougioukalis, Nick Vulpescu, Gael Westby, Paul Zappile, Yutong Zhang |
| EPI_ISL_430328 | hCoV-19/USA/NY-NYUMC301/2020 | North America / USA / New York / Brookl | 2020-03-30 | NYU Langone Health | Departments of Pathology and Medicine, New York University School of Medicine | Maria Aguero-Rosenfeld, Brendan Belovarac, Margaret Black, Ludovic Boytard, John Cadley, Paolo Cotzia, John Chen, Dacia Dimartino, Xiaojun Feng, Tatyana Gindin, Emily Guzman, Adriana Heguy, Megan Hogan, Emily Huang, George Jour, Lawrence H. Lin, Raven Luther, Andrew Lytle, Christian Marier, Matthew T. Maurano, Mark J. Mulligan, Peter Meyn, Raquel Ordonez Ciriza, Iman Osman, Jared Pinnell, Vanessa Raabe, Sitharam Ramaswami, Amy Rapkiewicz, Andre M. Ribeiro-dos-Santos, Marie Samanovic-Golden, Antonio Serrano, Guomiao Shen, Matija Snuderl, Theodore Vougioukalis, Nick Vulpescu, Gael Westby, Paul Zappile, Yutong Zhang |
| EPI_ISL_430329 | hCoV-19/USA/NY-NYUMC302/2020 | North America / USA / New York / Staten | 2020-03-30 | NYU Langone Health | Departments of Pathology and Medicine, New York University School of Medicine | Maria Aguero-Rosenfeld, Brendan Belovarac, Margaret Black, Ludovic Boytard, John Cadley, Paolo Cotzia, John Chen, Dacia Dimartino, Xiaojun Feng, Tatyana Gindin, Emily Guzman, Adriana Heguy, Megan Hogan, Emily Huang, George Jour, Lawrence H. Lin, Raven Luther, Andrew Lytle, Christian Marier, Matthew T. Maurano, Mark J. Mulligan, Peter Meyn, Raquel Ordonez Ciriza, Iman Osman, Jared Pinnell, Vanessa Raabe, Sitharam Ramaswami, Amy Rapkiewicz, Andre M. Ribeiro-dos-Santos, Marie Samanovic-Golden, Antonio Serrano, Guomiao Shen, Matija Snuderl, Theodore Vougioukalis, Nick Vulpescu, Gael Westby, Paul Zappile, Yutong Zhang |
| EPI_ISL_430330 | hCoV-19/USA/NY-NYUMC303/2020 | North America / USA / New York / Nassa | 2020-03-30 | NYU Langone Health | Departments of Pathology and Medicine, New York University School of Medicine | Maria Aguero-Rosenfeld, Brendan Belovarac, Margaret Black, Ludovic Boytard, John Cadley, Paolo Cotzia, John Chen, Dacia Dimartino, Xiaojun Feng, Tatyana Gindin, Emily Guzman, Adriana Heguy, Megan Hogan, Emily Huang, George Jour, Lawrence H. Lin, Raven Luther, Andrew Lytle, Christian Marier, Matthew T. Maurano, Mark J. Mulligan, Peter Meyn, Raquel Ordonez Ciriza, Iman Osman, Jared Pinnell, Vanessa Raabe, Sitharam Ramaswami, Amy Rapkiewicz, Andre M. Ribeiro-dos-Santos, Marie Samanovic-Golden, Antonio Serrano, Guomiao Shen, Matija Snuderl, Theodore Vougioukalis, Nick Vulpescu, Gael Westby, Paul Zappile, Yutong Zhang |
| EPI_ISL_430331 | hCoV-19/USA/NY-NYUMC304/2020 | North America / USA / New York / Manha | 2020-03-30 | NYU Langone Health | Departments of Pathology and Medicine, New York University School of Medicine | Maria Aguero-Rosenfeld, Brendan Belovarac, Margaret Black, Ludovic Boytard, John Cadley, Paolo Cotzia, John Chen, Dacia Dimartino, Xiaojun Feng, Tatyana Gindin, Emily Guzman, Adriana Heguy, Megan Hogan, Emily Huang, George Jour, Lawrence H. Lin, Raven Luther, Andrew Lytle, Christian Marier, Matthew T. Maurano, Mark J. Mulligan, Peter Meyn, Raquel Ordonez Ciriza, Iman Osman, Jared Pinnell, Vanessa Raabe, Sitharam Ramaswami, Amy Rapkiewicz, Andre M. Ribeiro-dos-Santos, Marie Samanovic-Golden, Antonio Serrano, Guomiao Shen, Matija Snuderl, Theodore Vougioukalis, Nick Vulpescu, Gael Westby, Paul Zappile, Yutong Zhang |
| EPI_ISL_430332 | hCoV-19/USA/NY-NYUMC305/2020 | North America / USA / New York / Brookl | 2020-03-30 | NYU Langone Health | Departments of Pathology and Medicine, New York University School of Medicine | Maria Aguero-Rosenfeld, Brendan Belovarac, Margaret Black, Ludovic Boytard, John Cadley, Paolo Cotzia, John Chen, Dacia Dimartino, Xiaojun Feng, Tatyana Gindin, Emily Guzman, Adriana Heguy, Megan Hogan, Emily Huang, George Jour, Lawrence H. Lin, Raven Luther, Andrew Lytle, Christian Marier, Matthew T. Maurano, Mark J. Mulligan, Peter Meyn, Raquel Ordonez Ciriza, Iman Osman, Jared Pinnell, Vanessa Raabe, Sitharam Ramaswami, Amy Rapkiewicz, Andre M. Ribeiro-dos-Santos, Marie Samanovic-Golden, Antonio Serrano, Guomiao Shen, Matija Snuderl, Theodore Vougioukalis, Nick Vulpescu, Gael Westby, Paul Zappile, Yutong Zhang |
| EPI_ISL_430333 | hCoV-19/USA/NY-NYUMC306/2020 | North America / USA / New York / Brookl | 2020-03-30 | NYU Langone Health | Departments of Pathology and Medicine, New York University School of Medicine | Maria Aguero-Rosenfeld, Brendan Belovarac, Margaret Black, Ludovic Boytard, John Cadley, Paolo Cotzia, John Chen, Dacia Dimartino, Xiaojun Feng, Tatyana Gindin, Emily Guzman, Adriana Heguy, Megan Hogan, Emily Huang, George Jour, Lawrence H. Lin, Raven Luther, Andrew Lytle, Christian Marier, Matthew T. Maurano, Mark J. Mulligan, Peter Meyn, Raquel Ordonez Ciriza, Iman Osman, Jared Pinnell, Vanessa Raabe, Sitharam Ramaswami, Amy Rapkiewicz, Andre M. Ribeiro-dos-Santos, Marie Samanovic-Golden, Antonio Serrano, Guomiao Shen, Matija Snuderl, Theodore Vougioukalis, Nick Vulpescu, Gael Westby, Paul Zappile, Yutong Zhang |
| EPI_ISL_430334 | hCoV-19/USA/NY-NYUMC307/2020 | North America / USA / New York / Brookl | 2020-03-30 | NYU Langone Health | Departments of Pathology and Medicine, New York University School of Medicine | Maria Aguero-Rosenfeld, Brendan Belovarac, Margaret Black, Ludovic Boytard, John Cadley, Paolo Cotzia, John Chen, Dacia Dimartino, Xiaojun Feng, Tatyana Gindin, Emily Guzman, Adriana Heguy, Megan Hogan, Emily Huang, George Jour, Lawrence H. Lin, Raven Luther, Andrew Lytle, Christian Marier, Matthew T. Maurano, Mark J. Mulligan, Peter Meyn, Raquel Ordonez Ciriza, Iman Osman, Jared Pinnell, Vanessa Raabe, Sitharam Ramaswami, Amy Rapkiewicz, Andre M. Ribeiro-dos-Santos, Marie Samanovic-Golden, Antonio Serrano, Guomiao Shen, Matija Snuderl, Theodore Vougioukalis, Nick Vulpescu, Gael Westby, Paul Zappile, Yutong Zhang |
| EPI_ISL_430335 | hCoV-19/USA/NY-NYUMC308/2020 | North America / USA / New York / Brookl | 2020-03-30 | NYU Langone Health | Departments of Pathology and Medicine, New York University School of Medicine | Maria Aguero-Rosenfeld, Brendan Belovarac, Margaret Black, Ludovic Boytard, John Cadley, Paolo Cotzia, John Chen, Dacia Dimartino, Xiaojun Feng, Tatyana Gindin, Emily Guzman, Adriana Heguy, Megan Hogan, Emily Huang, George Jour, Lawrence H. Lin, Raven Luther, Andrew Lytle, Christian Marier, Matthew T. Maurano, Mark J. Mulligan, Peter Meyn, Raquel Ordonez Ciriza, Iman Osman, Jared Pinnell, Vanessa Raabe, Sitharam Ramaswami, Amy Rapkiewicz, Andre M. Ribeiro-dos-Santos, Marie Samanovic-Golden, Antonio Serrano, Guomiao Shen, Matija Snuderl, Theodore Vougioukalis, Nick Vulpescu, Gael Westby, Paul Zappile, Yutong Zhang |
| EPI_ISL_430336 | hCoV-19/USA/NY-NYUMC309/2020 | North America / USA / New York / Manha | 2020-03-30 | NYU Langone Health | Departments of Pathology and Medicine, New York University School of Medicine | Maria Aguero-Rosenfeld, Brendan Belovarac, Margaret Black, Ludovic Boytard, John Cadley, Paolo Cotzia, John Chen, Dacia Dimartino, Xiaojun Feng, Tatyana Gindin, Emily Guzman, Adriana Heguy, Megan Hogan, Emily Huang, George Jour, Lawrence H. Lin, Raven Luther, Andrew Lytle, Christian Marier, Matthew T. Maurano, Mark J. Mulligan, Peter Meyn, Raquel Ordonez Ciriza, Iman Osman, Jared Pinnell, Vanessa Raabe, Sitharam Ramaswami, Amy Rapkiewicz, Andre M. Ribeiro-dos-Santos, Marie Samanovic-Golden, Antonio Serrano, Guomiao Shen, Matija Snuderl, Theodore Vougioukalis, Nick Vulpescu, Gael Westby, Paul Zappile, Yutong Zhang |
| EPI_ISL_430337 | hCoV-19/USA/NY-NYUMC310/2020 | North America / USA / New York / Manha | 2020-03-30 | NYU Langone Health | Departments of Pathology and Medicine, New York University School of Medicine | Maria Aguero-Rosenfeld, Brendan Belovarac, Margaret Black, Ludovic Boytard, John Cadley, Paolo Cotzia, John Chen, Dacia Dimartino, Xiaojun Feng, Tatyana Gindin, Emily Guzman, Adriana Heguy, Megan Hogan, Emily Huang, George Jour, Lawrence H. Lin, Raven Luther, Andrew Lytle, Christian Marier, Matthew T. Maurano, Mark J. Mulligan, Peter Meyn, Raquel Ordonez Ciriza, Iman Osman, Jared Pinnell, Vanessa Raabe, Sitharam Ramaswami, Amy Rapkiewicz, Andre M. Ribeiro-dos-Santos, Marie Samanovic-Golden, Antonio Serrano, Guomiao Shen, Matija Snuderl, Theodore Vougioukalis, Nick Vulpescu, Gael Westby, Paul Zappile, Yutong Zhang |
| EPI_ISL_430338 | hCoV-19/USA/NY-NYUMC311/2020 | North America / USA / New York / Manha | 2020-03-30 | NYU Langone Health | Departments of Pathology and Medicine, New York University School of Medicine | Maria Aguero-Rosenfeld, Brendan Belovarac, Margaret Black, Ludovic Boytard, John Cadley, Paolo Cotzia, John Chen, Dacia Dimartino, Xiaojun Feng, Tatyana Gindin, Emily Guzman, Adriana Heguy, Megan Hogan, Emily Huang, George Jour, Lawrence H. Lin, Raven Luther, Andrew Lytle, Christian Marier, Matthew T. Maurano, Mark J. Mulligan, Peter Meyn, Raquel Ordonez Ciriza, Iman Osman, Jared Pinnell, Vanessa Raabe, Sitharam Ramaswami, Amy Rapkiewicz, Andre M. Ribeiro-dos-Santos, Marie Samanovic-Golden, Antonio Serrano, Guomiao Shen, Matija Snuderl, Theodore Vougioukalis, Nick Vulpescu, Gael Westby, Paul Zappile, Yutong Zhang |
| EPI_ISL_430339 | hCoV-19/USA/NY-NYUMC312/2020 | North America / USA / New York / Manha | 2020-03-30 | NYU Langone Health | Departments of Pathology and Medicine, New York University School of Medicine | Maria Aguero-Rosenfeld, Brendan Belovarac, Margaret Black, Ludovic Boytard, John Cadley, Paolo Cotzia, John Chen, Dacia Dimartino, Xiaojun Feng, Tatyana Gindin, Emily Guzman, Adriana Heguy, Megan Hogan, Emily Huang, George Jour, Lawrence H. Lin, Raven Luther, Andrew Lytle, Christian Marier, Matthew T. Maurano, Mark J. Mulligan, Peter Meyn, Raquel Ordonez Ciriza, Iman Osman, Jared Pinnell, Vanessa Raabe, Sitharam Ramaswami, Amy Rapkiewicz, Andre M. Ribeiro-dos-Santos, Marie Samanovic-Golden, Antonio Serrano, Guomiao Shen, Matija Snuderl, Theodore Vougioukalis, Nick Vulpescu, Gael Westby, Paul Zappile, Yutong Zhang |
| EPI_ISL_430340 | hCoV-19/USA/NY-NYUMC313/2020 | North America / USA / New York / Westc | 2020-03-30 | NYU Langone Health | Departments of Pathology and Medicine, New York University School of Medicine | Maria Aguero-Rosenfeld, Brendan Belovarac, Margaret Black, Ludovic Boytard, John Cadley, Paolo Cotzia, John Chen, Dacia Dimartino, Xiaojun Feng, Tatyana Gindin, Emily Guzman, Adriana Heguy, Megan Hogan, Emily Huang, George Jour, Lawrence H. Lin, Raven Luther, Andrew Lytle, Christian Marier, Matthew T. Maurano, Mark J. Mulligan, Peter Meyn, Raquel Ordonez Ciriza, Iman Osman, Jared Pinnell, Vanessa Raabe, Sitharam Ramaswami, Amy Rapkiewicz, Andre M. Ribeiro-dos-Santos, Marie Samanovic-Golden, Antonio Serrano, Guomiao Shen, Matija Snuderl, Theodore Vougioukalis, Nick Vulpescu, Gael Westby, Paul Zappile, Yutong Zhang |
| EPI_ISL_430341 | hCoV-19/USA/NY-NYUMC314/2020 | North America / USA / New York / Brookl | 2020-03-30 | NYU Langone Health | Departments of Pathology and Medicine, New York University School of Medicine | Maria Aguero-Rosenfeld, Brendan Belovarac, Margaret Black, Ludovic Boytard, John Cadley, Paolo Cotzia, John Chen, Dacia Dimartino, Xiaojun Feng, Tatyana Gindin, Emily Guzman, Adriana Heguy, Megan Hogan, Emily Huang, George Jour, Lawrence H. Lin, Raven Luther, Andrew Lytle, Christian Marier, Matthew T. Maurano, Mark J. Mulligan, Peter Meyn, Raquel Ordonez Ciriza, Iman Osman, Jared Pinnell, Vanessa Raabe, Sitharam Ramaswami, Amy Rapkiewicz, Andre M. Ribeiro-dos-Santos, Marie Samanovic-Golden, Antonio Serrano, Guomiao Shen, Matija Snuderl, Theodore Vougioukalis, Nick Vulpescu, Gael Westby, Paul Zappile, Yutong Zhang |
| EPI_ISL_430342 | hCoV-19/USA/NY-NYUMC315/2020 | North America / USA / New York / Queen | 2020-03-30 | NYU Langone Health | Departments of Pathology and Medicine, New York University School of Medicine | Maria Aguero-Rosenfeld, Brendan Belovarac, Margaret Black, Ludovic Boytard, John Cadley, Paolo Cotzia, John Chen, Dacia Dimartino, Xiaojun Feng, Tatyana Gindin, Emily Guzman, Adriana Heguy, Megan Hogan, Emily Huang, George Jour, Lawrence H. Lin, Raven Luther, Andrew Lytle, Christian Marier, Matthew T. Maurano, Mark J. Mulligan, Peter Meyn, Raquel Ordonez Ciriza, Iman Osman, Jared Pinnell, Vanessa Raabe, Sitharam Ramaswami, Amy Rapkiewicz, Andre M. Ribeiro-dos-Santos, Marie Samanovic-Golden, Antonio Serrano, Guomiao Shen, Matija Snuderl, Theodore Vougioukalis, Nick Vulpescu, Gael Westby, Paul Zappile, Yutong Zhang |
| EPI_ISL_430343 | hCoV-19/USA/NY-NYUMC316/2020 | North America / USA / New York / Manha | 2020-03-30 | NYU Langone Health | Departments of Pathology and Medicine, New York University School of Medicine | Maria Aguero-Rosenfeld, Brendan Belovarac, Margaret Black, Ludovic Boytard, John Cadley, Paolo Cotzia, John Chen, Dacia Dimartino, Xiaojun Feng, Tatyana Gindin, Emily Guzman, Adriana Heguy, Megan Hogan, Emily Huang, George Jour, Lawrence H. Lin, Raven Luther, Andrew Lytle, Christian Marier, Matthew T. Maurano, Mark J. Mulligan, Peter Meyn, Raquel Ordonez Ciriza, Iman Osman, Jared Pinnell, Vanessa Raabe, Sitharam Ramaswami, Amy Rapkiewicz, Andre M. Ribeiro-dos-Santos, Marie Samanovic-Golden, Antonio Serrano, Guomiao Shen, Matija Snuderl, Theodore Vougioukalis, Nick Vulpescu, Gael Westby, Paul Zappile, Yutong Zhang |
| EPI_ISL_430344 | hCoV-19/USA/NY-NYUMC317/2020 | North America / USA / New York / Brookl | 2020-03-30 | NYU Langone Health | Departments of Pathology and Medicine, New York University School of Medicine | Maria Aguero-Rosenfeld, Brendan Belovarac, Margaret Black, Ludovic Boytard, John Cadley, Paolo Cotzia, John Chen, Dacia Dimartino, Xiaojun Feng, Tatyana Gindin, Emily Guzman, Adriana Heguy, Megan Hogan, Emily Huang, George Jour, Lawrence H. Lin, Raven Luther, Andrew Lytle, Christian Marier, Matthew T. Maurano, Mark J. Mulligan, Peter Meyn, Raquel Ordonez Ciriza, Iman Osman, Jared Pinnell, Vanessa Raabe, Sitharam Ramaswami, Amy Rapkiewicz, Andre M. Ribeiro-dos-Santos, Marie Samanovic-Golden, Antonio Serrano, Guomiao Shen, Matija Snuderl, Theodore Vougioukalis, Nick Vulpescu, Gael Westby, Paul Zappile, Yutong Zhang |
| EPI_ISL_430345 | hCoV-19/USA/NY-NYUMC318/2020 | North America / USA / New York / Manha | 2020-03-30 | NYU Langone Health | Departments of Pathology and Medicine, New York University School of Medicine | Maria Aguero-Rosenfeld, Brendan Belovarac, Margaret Black, Ludovic Boytard, John Cadley, Paolo Cotzia, John Chen, Dacia Dimartino, Xiaojun Feng, Tatyana Gindin, Emily Guzman, Adriana Heguy, Megan Hogan, Emily Huang, George Jour, Lawrence H. Lin, Raven Luther, Andrew Lytle, Christian Marier, Matthew T. Maurano, Mark J. Mulligan, Peter Meyn, Raquel Ordonez Ciriza, Iman Osman, Jared Pinnell, Vanessa Raabe, Sitharam Ramaswami, Amy Rapkiewicz, Andre M. Ribeiro-dos-Santos, Marie Samanovic-Golden, Antonio Serrano, Guomiao Shen, Matija Snuderl, Theodore Vougioukalis, Nick Vulpescu, Gael Westby, Paul Zappile, Yutong Zhang |
| EPI_ISL_430346 | hCoV-19/USA/NY-NYUMC319/2020 | North America / USA / New York / Brookl | 2020-03-30 | NYU Langone Health | Departments of Pathology and Medicine, New York University School of Medicine | Maria Aguero-Rosenfeld, Brendan Belovarac, Margaret Black, Ludovic Boytard, John Cadley, Paolo Cotzia, John Chen, Dacia Dimartino, Xiaojun Feng, Tatyana Gindin, Emily Guzman, Adriana Heguy, Megan Hogan, Emily Huang, George Jour, Lawrence H. Lin, Raven Luther, Andrew Lytle, Christian Marier, Matthew T. Maurano, Mark J. Mulligan, Peter Meyn, Raquel Ordonez Ciriza, Iman Osman, Jared Pinnell, Vanessa Raabe, Sitharam Ramaswami, Amy Rapkiewicz, Andre M. Ribeiro-dos-Santos, Marie Samanovic-Golden, Antonio Serrano, Guomiao Shen, Matija Snuderl, Theodore Vougioukalis, Nick Vulpescu, Gael Westby, Paul Zappile, Yutong Zhang |
| EPI_ISL_430347 | hCoV-19/USA/NY-NYUMC320/2020 | North America / USA / New York / Manha | 2020-03-30 | NYU Langone Health | Departments of Pathology and Medicine, New York University School of Medicine | Maria Aguero-Rosenfeld, Brendan Belovarac, Margaret Black, Ludovic Boytard, John Cadley, Paolo Cotzia, John Chen, Dacia Dimartino, Xiaojun Feng, Tatyana Gindin, Emily Guzman, Adriana Heguy, Megan Hogan, Emily Huang, George Jour, Lawrence H. Lin, Raven Luther, Andrew Lytle, Christian Marier, Matthew T. Maurano, Mark J. Mulligan, Peter Meyn, Raquel Ordonez Ciriza, Iman Osman, Jared Pinnell, Vanessa Raabe, Sitharam Ramaswami, Amy Rapkiewicz, Andre M. Ribeiro-dos-Santos, Marie Samanovic-Golden, Antonio Serrano, Guomiao Shen, Matija Snuderl, Theodore Vougioukalis, Nick Vulpescu, Gael Westby, Paul Zappile, Yutong Zhang |
| EPI_ISL_430348 | hCoV-19/USA/NY-NYUMC321/2020 | North America / USA / New York / Manha | 2020-03-30 | NYU Langone Health | Departments of Pathology and Medicine, New York University School of Medicine | Maria Aguero-Rosenfeld, Brendan Belovarac, Margaret Black, Ludovic Boytard, John Cadley, Paolo Cotzia, John Chen, Dacia Dimartino, Xiaojun Feng, Tatyana Gindin, Emily Guzman, Adriana Heguy, Megan Hogan, Emily Huang, George Jour, Lawrence H. Lin, Raven Luther, Andrew Lytle, Christian Marier, Matthew T. Maurano, Mark J. Mulligan, Peter Meyn, Raquel Ordonez Ciriza, Iman Osman, Jared Pinnell, Vanessa Raabe, Sitharam Ramaswami, Amy Rapkiewicz, Andre M. Ribeiro-dos-Santos, Marie Samanovic-Golden, Antonio Serrano, Guomiao Shen, Matija Snuderl, Theodore Vougioukalis, Nick Vulpescu, Gael Westby, Paul Zappile, Yutong Zhang |
| EPI_ISL_430349 | hCoV-19/USA/NY-NYUMC322/2020 | North America / USA / New York / Staten | 2020-03-30 | NYU Langone Health | Departments of Pathology and Medicine, New York University School of Medicine | Maria Aguero-Rosenfeld, Brendan Belovarac, Margaret Black, Ludovic Boytard, John Cadley, Paolo Cotzia, John Chen, Dacia Dimartino, Xiaojun Feng, Tatyana Gindin, Emily Guzman, Adriana Heguy, Megan Hogan, Emily Huang, George Jour, Lawrence H. Lin, Raven Luther, Andrew Lytle, Christian Marier, Matthew T. Maurano, Mark J. Mulligan, Peter Meyn, Raquel Ordonez Ciriza, Iman Osman, Jared Pinnell, Vanessa Raabe, Sitharam Ramaswami, Amy Rapkiewicz, Andre M. Ribeiro-dos-Santos, Marie Samanovic-Golden, Antonio Serrano, Guomiao Shen, Matija Snuderl, Theodore Vougioukalis, Nick Vulpescu, Gael Westby, Paul Zappile, Yutong Zhang |
| EPI_ISL_430350 | hCoV-19/USA/NY-NYUMC323/2020 | North America / USA / New York / Queen | 2020-04-02 | NYU Langone Health | Departments of Pathology and Medicine, New York University School of Medicine | Maria Aguero-Rosenfeld, Brendan Belovarac, Margaret Black, Ludovic Boytard, John Cadley, Paolo Cotzia, John Chen, Dacia Dimartino, Xiaojun Feng, Tatyana Gindin, Emily Guzman, Adriana Heguy, Megan Hogan, Emily Huang, George Jour, Lawrence H. Lin, Raven Luther, Andrew Lytle, Christian Marier, Matthew T. Maurano, Mark J. Mulligan, Peter Meyn, Raquel Ordonez Ciriza, Iman Osman, Jared Pinnell, Vanessa Raabe, Sitharam Ramaswami, Amy Rapkiewicz, Andre M. Ribeiro-dos-Santos, Marie Samanovic-Golden, Antonio Serrano, Guomiao Shen, Matija Snuderl, Theodore Vougioukalis, Nick Vulpescu, Gael Westby, Paul Zappile, Yutong Zhang |
| EPI_ISL_430351 | hCoV-19/USA/NY-NYUMC324/2020 | North America / USA / New York / Brookl | 2020-03-31 | NYU Langone Health | Departments of Pathology and Medicine, New York University School of Medicine | Maria Aguero-Rosenfeld, Brendan Belovarac, Margaret Black, Ludovic Boytard, John Cadley, Paolo Cotzia, John Chen, Dacia Dimartino, Xiaojun Feng, Tatyana Gindin, Emily Guzman, Adriana Heguy, Megan Hogan, Emily Huang, George Jour, Lawrence H. Lin, Raven Luther, Andrew Lytle, Christian Marier, Matthew T. Maurano, Mark J. Mulligan, Peter Meyn, Raquel Ordonez Ciriza, Iman Osman, Jared Pinnell, Vanessa Raabe, Sitharam Ramaswami, Amy Rapkiewicz, Andre M. Ribeiro-dos-Santos, Marie Samanovic-Golden, Antonio Serrano, Guomiao Shen, Matija Snuderl, Theodore Vougioukalis, Nick Vulpescu, Gael Westby, Paul Zappile, Yutong Zhang |
| EPI_ISL_430352 | hCoV-19/USA/NY-NYUMC325/2020 | North America / USA / New York / Suffolk | 2020-04-01 | NYU Langone Health | Departments of Pathology and Medicine, New York University School of Medicine | Maria Aguero-Rosenfeld, Brendan Belovarac, Margaret Black, Ludovic Boytard, John Cadley, Paolo Cotzia, John Chen, Dacia Dimartino, Xiaojun Feng, Tatyana Gindin, Emily Guzman, Adriana Heguy, Megan Hogan, Emily Huang, George Jour, Lawrence H. Lin, Raven Luther, Andrew Lytle, Christian Marier, Matthew T. Maurano, Mark J. Mulligan, Peter Meyn, Raquel Ordonez Ciriza, Iman Osman, Jared Pinnell, Vanessa Raabe, Sitharam Ramaswami, Amy Rapkiewicz, Andre M. Ribeiro-dos-Santos, Marie Samanovic-Golden, Antonio Serrano, Guomiao Shen, Matija Snuderl, Theodore Vougioukalis, Nick Vulpescu, Gael Westby, Paul Zappile, Yutong Zhang |
| EPI_ISL_430353 | hCoV-19/USA/NY-NYUMC326/2020 | North America / USA / New York / Brookl | 2020-04-01 | NYU Langone Health | Departments of Pathology and Medicine, New York University School of Medicine | Maria Aguero-Rosenfeld, Brendan Belovarac, Margaret Black, Ludovic Boytard, John Cadley, Paolo Cotzia, John Chen, Dacia Dimartino, Xiaojun Feng, Tatyana Gindin, Emily Guzman, Adriana Heguy, Megan Hogan, Emily Huang, George Jour, Lawrence H. Lin, Raven Luther, Andrew Lytle, Christian Marier, Matthew T. Maurano, Mark J. Mulligan, Peter Meyn, Raquel Ordonez Ciriza, Iman Osman, Jared Pinnell, Vanessa Raabe, Sitharam Ramaswami, Amy Rapkiewicz, Andre M. Ribeiro-dos-Santos, Marie Samanovic-Golden, Antonio Serrano, Guomiao Shen, Matija Snuderl, Theodore Vougioukalis, Nick Vulpescu, Gael Westby, Paul Zappile, Yutong Zhang |
| EPI_ISL_430354 | hCoV-19/USA/NY-NYUMC327/2020 | North America / USA / New York / Nassa | 2020-04-03 | NYU Langone Health | Departments of Pathology and Medicine, New York University School of Medicine | Maria Aguero-Rosenfeld, Brendan Belovarac, Margaret Black, Ludovic Boytard, John Cadley, Paolo Cotzia, John Chen, Dacia Dimartino, Xiaojun Feng, Tatyana Gindin, Emily Guzman, Adriana Heguy, Megan Hogan, Emily Huang, George Jour, Lawrence H. Lin, Raven Luther, Andrew Lytle, Christian Marier, Matthew T. Maurano, Mark J. Mulligan, Peter Meyn, Raquel Ordonez Ciriza, Iman Osman, Jared Pinnell, Vanessa Raabe, Sitharam Ramaswami, Amy Rapkiewicz, Andre M. Ribeiro-dos-Santos, Marie Samanovic-Golden, Antonio Serrano, Guomiao Shen, Matija Snuderl, Theodore Vougioukalis, Nick Vulpescu, Gael Westby, Paul Zappile, Yutong Zhang |
| EPI_ISL_430355 | hCoV-19/USA/NY-NYUMC328/2020 | North America / USA / New York / Nassa | 2020-04-02 | NYU Langone Health | Departments of Pathology and Medicine, New York University School of Medicine | Maria Aguero-Rosenfeld, Brendan Belovarac, Margaret Black, Ludovic Boytard, John Cadley, Paolo Cotzia, John Chen, Dacia Dimartino, Xiaojun Feng, Tatyana Gindin, Emily Guzman, Adriana Heguy, Megan Hogan, Emily Huang, George Jour, Lawrence H. Lin, Raven Luther, Andrew Lytle, Christian Marier, Matthew T. Maurano, Mark J. Mulligan, Peter Meyn, Raquel Ordonez Ciriza, Iman Osman, Jared Pinnell, Vanessa Raabe, Sitharam Ramaswami, Amy Rapkiewicz, Andre M. Ribeiro-dos-Santos, Marie Samanovic-Golden, Antonio Serrano, Guomiao Shen, Matija Snuderl, Theodore Vougioukalis, Nick Vulpescu, Gael Westby, Paul Zappile, Yutong Zhang |
| EPI_ISL_430356 | hCoV-19/USA/NJ-NYUMC329/2020 | North America / USA / New Jersey / Berg | 2020-04-02 | NYU Langone Health | Departments of Pathology and Medicine, New York University School of Medicine | Maria Aguero-Rosenfeld, Brendan Belovarac, Margaret Black, Ludovic Boytard, John Cadley, Paolo Cotzia, John Chen, Dacia Dimartino, Xiaojun Feng, Tatyana Gindin, Emily Guzman, Adriana Heguy, Megan Hogan, Emily Huang, George Jour, Lawrence H. Lin, Raven Luther, Andrew Lytle, Christian Marier, Matthew T. Maurano, Mark J. Mulligan, Peter Meyn, Raquel Ordonez Ciriza, Iman Osman, Jared Pinnell, Vanessa Raabe, Sitharam Ramaswami, Amy Rapkiewicz, Andre M. Ribeiro-dos-Santos, Marie Samanovic-Golden, Antonio Serrano, Guomiao Shen, Matija Snuderl, Theodore Vougioukalis, Nick Vulpescu, Gael Westby, Paul Zappile, Yutong Zhang |
| EPI_ISL_430357 | hCoV-19/USA/NY-NYUMC330/2020 | North America / USA / New York / Brookl | 2020-04-03 | NYU Langone Health | Departments of Pathology and Medicine, New York University School of Medicine | Maria Aguero-Rosenfeld, Brendan Belovarac, Margaret Black, Ludovic Boytard, John Cadley, Paolo Cotzia, John Chen, Dacia Dimartino, Xiaojun Feng, Tatyana Gindin, Emily Guzman, Adriana Heguy, Megan Hogan, Emily Huang, George Jour, Lawrence H. Lin, Raven Luther, Andrew Lytle, Christian Marier, Matthew T. Maurano, Mark J. Mulligan, Peter Meyn, Raquel Ordonez Ciriza, Iman Osman, Jared Pinnell, Vanessa Raabe, Sitharam Ramaswami, Amy Rapkiewicz, Andre M. Ribeiro-dos-Santos, Marie Samanovic-Golden, Antonio Serrano, Guomiao Shen, Matija Snuderl, Theodore Vougioukalis, Nick Vulpescu, Gael Westby, Paul Zappile, Yutong Zhang |
| EPI_ISL_430358 | hCoV-19/USA/NY-NYUMC331/2020 | North America / USA / New York / Manha | 2020-04-13 | NYU Langone Health | Departments of Pathology and Medicine, New York University School of Medicine | Maria Aguero-Rosenfeld, Brendan Belovarac, Margaret Black, Ludovic Boytard, John Cadley, Paolo Cotzia, John Chen, Dacia Dimartino, Xiaojun Feng, Tatyana Gindin, Emily Guzman, Adriana Heguy, Megan Hogan, Emily Huang, George Jour, Lawrence H. Lin, Raven Luther, Andrew Lytle, Christian Marier, Matthew T. Maurano, Mark J. Mulligan, Peter Meyn, Raquel Ordonez Ciriza, Iman Osman, Jared Pinnell, Vanessa Raabe, Sitharam Ramaswami, Amy Rapkiewicz, Andre M. Ribeiro-dos-Santos, Marie Samanovic-Golden, Antonio Serrano, Guomiao Shen, Matija Snuderl, Theodore Vougioukalis, Nick Vulpescu, Gael Westby, Paul Zappile, Yutong Zhang |
| EPI_ISL_430359 | hCoV-19/USA/NY-NYUMC332/2020 | North America / USA / New York / Brookl | 2020-04-08 | NYU Langone Health | Departments of Pathology and Medicine, New York University School of Medicine | Maria Aguero-Rosenfeld, Brendan Belovarac, Margaret Black, Ludovic Boytard, John Cadley, Paolo Cotzia, John Chen, Dacia Dimartino, Xiaojun Feng, Tatyana Gindin, Emily Guzman, Adriana Heguy, Megan Hogan, Emily Huang, George Jour, Lawrence H. Lin, Raven Luther, Andrew Lytle, Christian Marier, Matthew T. Maurano, Mark J. Mulligan, Peter Meyn, Raquel Ordonez Ciriza, Iman Osman, Jared Pinnell, Vanessa Raabe, Sitharam Ramaswami, Amy Rapkiewicz, Andre M. Ribeiro-dos-Santos, Marie Samanovic-Golden, Antonio Serrano, Guomiao Shen, Matija Snuderl, Theodore Vougioukalis, Nick Vulpescu, Gael Westby, Paul Zappile, Yutong Zhang |
| EPI_ISL_430360 | hCoV-19/USA/NY-NYUMC333/2020 | North America / USA / New York / Queen | 2020-04-07 | NYU Langone Health | Departments of Pathology and Medicine, New York University School of Medicine | Maria Aguero-Rosenfeld, Brendan Belovarac, Margaret Black, Ludovic Boytard, John Cadley, Paolo Cotzia, John Chen, Dacia Dimartino, Xiaojun Feng, Tatyana Gindin, Emily Guzman, Adriana Heguy, Megan Hogan, Emily Huang, George Jour, Lawrence H. Lin, Raven Luther, Andrew Lytle, Christian Marier, Matthew T. Maurano, Mark J. Mulligan, Peter Meyn, Raquel Ordonez Ciriza, Iman Osman, Jared Pinnell, Vanessa Raabe, Sitharam Ramaswami, Amy Rapkiewicz, Andre M. Ribeiro-dos-Santos, Marie Samanovic-Golden, Antonio Serrano, Guomiao Shen, Matija Snuderl, Theodore Vougioukalis, Nick Vulpescu, Gael Westby, Paul Zappile, Yutong Zhang |
| EPI_ISL_430361 | hCoV-19/USA/NY-NYUMC334/2020 | North America / USA / New York / Brookl | 2020-04-07 | NYU Langone Health | Departments of Pathology and Medicine, New York University School of Medicine | Maria Aguero-Rosenfeld, Brendan Belovarac, Margaret Black, Ludovic Boytard, John Cadley, Paolo Cotzia, John Chen, Dacia Dimartino, Xiaojun Feng, Tatyana Gindin, Emily Guzman, Adriana Heguy, Megan Hogan, Emily Huang, George Jour, Lawrence H. Lin, Raven Luther, Andrew Lytle, Christian Marier, Matthew T. Maurano, Mark J. Mulligan, Peter Meyn, Raquel Ordonez Ciriza, Iman Osman, Jared Pinnell, Vanessa Raabe, Sitharam Ramaswami, Amy Rapkiewicz, Andre M. Ribeiro-dos-Santos, Marie Samanovic-Golden, Antonio Serrano, Guomiao Shen, Matija Snuderl, Theodore Vougioukalis, Nick Vulpescu, Gael Westby, Paul Zappile, Yutong Zhang |
| EPI_ISL_430362 | hCoV-19/USA/NY-NYUMC335/2020 | North America / USA / New York / Brookl | 2020-04-08 | NYU Langone Health | Departments of Pathology and Medicine, New York University School of Medicine | Maria Aguero-Rosenfeld, Brendan Belovarac, Margaret Black, Ludovic Boytard, John Cadley, Paolo Cotzia, John Chen, Dacia Dimartino, Xiaojun Feng, Tatyana Gindin, Emily Guzman, Adriana Heguy, Megan Hogan, Emily Huang, George Jour, Lawrence H. Lin, Raven Luther, Andrew Lytle, Christian Marier, Matthew T. Maurano, Mark J. Mulligan, Peter Meyn, Raquel Ordonez Ciriza, Iman Osman, Jared Pinnell, Vanessa Raabe, Sitharam Ramaswami, Amy Rapkiewicz, Andre M. Ribeiro-dos-Santos, Marie Samanovic-Golden, Antonio Serrano, Guomiao Shen, Matija Snuderl, Theodore Vougioukalis, Nick Vulpescu, Gael Westby, Paul Zappile, Yutong Zhang |
| EPI_ISL_430363 | hCoV-19/USA/NY-NYUMC336/2020 | North America / USA / New York / Brookl | 2020-04-08 | NYU Langone Health | Departments of Pathology and Medicine, New York University School of Medicine | Maria Aguero-Rosenfeld, Brendan Belovarac, Margaret Black, Ludovic Boytard, John Cadley, Paolo Cotzia, John Chen, Dacia Dimartino, Xiaojun Feng, Tatyana Gindin, Emily Guzman, Adriana Heguy, Megan Hogan, Emily Huang, George Jour, Lawrence H. Lin, Raven Luther, Andrew Lytle, Christian Marier, Matthew T. Maurano, Mark J. Mulligan, Peter Meyn, Raquel Ordonez Ciriza, Iman Osman, Jared Pinnell, Vanessa Raabe, Sitharam Ramaswami, Amy Rapkiewicz, Andre M. Ribeiro-dos-Santos, Marie Samanovic-Golden, Antonio Serrano, Guomiao Shen, Matija Snuderl, Theodore Vougioukalis, Nick Vulpescu, Gael Westby, Paul Zappile, Yutong Zhang |
| EPI_ISL_430364 | hCoV-19/USA/NY-NYUMC337/2020 | North America / USA / New York / Brookl | 2020-04-08 | NYU Langone Health | Departments of Pathology and Medicine, New York University School of Medicine | Maria Aguero-Rosenfeld, Brendan Belovarac, Margaret Black, Ludovic Boytard, John Cadley, Paolo Cotzia, John Chen, Dacia Dimartino, Xiaojun Feng, Tatyana Gindin, Emily Guzman, Adriana Heguy, Megan Hogan, Emily Huang, George Jour, Lawrence H. Lin, Raven Luther, Andrew Lytle, Christian Marier, Matthew T. Maurano, Mark J. Mulligan, Peter Meyn, Raquel Ordonez Ciriza, Iman Osman, Jared Pinnell, Vanessa Raabe, Sitharam Ramaswami, Amy Rapkiewicz, Andre M. Ribeiro-dos-Santos, Marie Samanovic-Golden, Antonio Serrano, Guomiao Shen, Matija Snuderl, Theodore Vougioukalis, Nick Vulpescu, Gael Westby, Paul Zappile, Yutong Zhang |
| EPI_ISL_430365 | hCoV-19/USA/NY-NYUMC338/2020 | North America / USA / New York / Brookl | 2020-04-08 | NYU Langone Health | Departments of Pathology and Medicine, New York University School of Medicine | Maria Aguero-Rosenfeld, Brendan Belovarac, Margaret Black, Ludovic Boytard, John Cadley, Paolo Cotzia, John Chen, Dacia Dimartino, Xiaojun Feng, Tatyana Gindin, Emily Guzman, Adriana Heguy, Megan Hogan, Emily Huang, George Jour, Lawrence H. Lin, Raven Luther, Andrew Lytle, Christian Marier, Matthew T. Maurano, Mark J. Mulligan, Peter Meyn, Raquel Ordonez Ciriza, Iman Osman, Jared Pinnell, Vanessa Raabe, Sitharam Ramaswami, Amy Rapkiewicz, Andre M. Ribeiro-dos-Santos, Marie Samanovic-Golden, Antonio Serrano, Guomiao Shen, Matija Snuderl, Theodore Vougioukalis, Nick Vulpescu, Gael Westby, Paul Zappile, Yutong Zhang |
| EPI_ISL_430366 | hCoV-19/USA/NY-NYUMC339/2020 | North America / USA / New York / Brookl | 2020-04-08 | NYU Langone Health | Departments of Pathology and Medicine, New York University School of Medicine | Maria Aguero-Rosenfeld, Brendan Belovarac, Margaret Black, Ludovic Boytard, John Cadley, Paolo Cotzia, John Chen, Dacia Dimartino, Xiaojun Feng, Tatyana Gindin, Emily Guzman, Adriana Heguy, Megan Hogan, Emily Huang, George Jour, Lawrence H. Lin, Raven Luther, Andrew Lytle, Christian Marier, Matthew T. Maurano, Mark J. Mulligan, Peter Meyn, Raquel Ordonez Ciriza, Iman Osman, Jared Pinnell, Vanessa Raabe, Sitharam Ramaswami, Amy Rapkiewicz, Andre M. Ribeiro-dos-Santos, Marie Samanovic-Golden, Antonio Serrano, Guomiao Shen, Matija Snuderl, Theodore Vougioukalis, Nick Vulpescu, Gael Westby, Paul Zappile, Yutong Zhang |
| EPI_ISL_430367 | hCoV-19/USA/NY-NYUMC340/2020 | North America / USA / New York / Brookl | 2020-04-07 | NYU Langone Health | Departments of Pathology and Medicine, New York University School of Medicine | Maria Aguero-Rosenfeld, Brendan Belovarac, Margaret Black, Ludovic Boytard, John Cadley, Paolo Cotzia, John Chen, Dacia Dimartino, Xiaojun Feng, Tatyana Gindin, Emily Guzman, Adriana Heguy, Megan Hogan, Emily Huang, George Jour, Lawrence H. Lin, Raven Luther, Andrew Lytle, Christian Marier, Matthew T. Maurano, Mark J. Mulligan, Peter Meyn, Raquel Ordonez Ciriza, Iman Osman, Jared Pinnell, Vanessa Raabe, Sitharam Ramaswami, Amy Rapkiewicz, Andre M. Ribeiro-dos-Santos, Marie Samanovic-Golden, Antonio Serrano, Guomiao Shen, Matija Snuderl, Theodore Vougioukalis, Nick Vulpescu, Gael Westby, Paul Zappile, Yutong Zhang |
| EPI_ISL_430368 | hCoV-19/USA/NY-NYUMC341/2020 | North America / USA / New York / Brookl | 2020-04-08 | NYU Langone Health | Departments of Pathology and Medicine, New York University School of Medicine | Maria Aguero-Rosenfeld, Brendan Belovarac, Margaret Black, Ludovic Boytard, John Cadley, Paolo Cotzia, John Chen, Dacia Dimartino, Xiaojun Feng, Tatyana Gindin, Emily Guzman, Adriana Heguy, Megan Hogan, Emily Huang, George Jour, Lawrence H. Lin, Raven Luther, Andrew Lytle, Christian Marier, Matthew T. Maurano, Mark J. Mulligan, Peter Meyn, Raquel Ordonez Ciriza, Iman Osman, Jared Pinnell, Vanessa Raabe, Sitharam Ramaswami, Amy Rapkiewicz, Andre M. Ribeiro-dos-Santos, Marie Samanovic-Golden, Antonio Serrano, Guomiao Shen, Matija Snuderl, Theodore Vougioukalis, Nick Vulpescu, Gael Westby, Paul Zappile, Yutong Zhang |
| EPI_ISL_430369 | hCoV-19/USA/NY-NYUMC342/2020 | North America / USA / New York / Manha | 2020-04-08 | NYU Langone Health | Departments of Pathology and Medicine, New York University School of Medicine | Maria Aguero-Rosenfeld, Brendan Belovarac, Margaret Black, Ludovic Boytard, John Cadley, Paolo Cotzia, John Chen, Dacia Dimartino, Xiaojun Feng, Tatyana Gindin, Emily Guzman, Adriana Heguy, Megan Hogan, Emily Huang, George Jour, Lawrence H. Lin, Raven Luther, Andrew Lytle, Christian Marier, Matthew T. Maurano, Mark J. Mulligan, Peter Meyn, Raquel Ordonez Ciriza, Iman Osman, Jared Pinnell, Vanessa Raabe, Sitharam Ramaswami, Amy Rapkiewicz, Andre M. Ribeiro-dos-Santos, Marie Samanovic-Golden, Antonio Serrano, Guomiao Shen, Matija Snuderl, Theodore Vougioukalis, Nick Vulpescu, Gael Westby, Paul Zappile, Yutong Zhang |
| EPI_ISL_430370 | hCoV-19/USA/NY-NYUMC343/2020 | North America / USA / New York / Queen | 2020-04-08 | NYU Langone Health | Departments of Pathology and Medicine, New York University School of Medicine | Maria Aguero-Rosenfeld, Brendan Belovarac, Margaret Black, Ludovic Boytard, John Cadley, Paolo Cotzia, John Chen, Dacia Dimartino, Xiaojun Feng, Tatyana Gindin, Emily Guzman, Adriana Heguy, Megan Hogan, Emily Huang, George Jour, Lawrence H. Lin, Raven Luther, Andrew Lytle, Christian Marier, Matthew T. Maurano, Mark J. Mulligan, Peter Meyn, Raquel Ordonez Ciriza, Iman Osman, Jared Pinnell, Vanessa Raabe, Sitharam Ramaswami, Amy Rapkiewicz, Andre M. Ribeiro-dos-Santos, Marie Samanovic-Golden, Antonio Serrano, Guomiao Shen, Matija Snuderl, Theodore Vougioukalis, Nick Vulpescu, Gael Westby, Paul Zappile, Yutong Zhang |
| EPI_ISL_430371 | hCoV-19/USA/NJ-NYUMC344/2020 | North America / USA / New Jersey / Huds | 2020-04-08 | NYU Langone Health | Departments of Pathology and Medicine, New York University School of Medicine | Maria Aguero-Rosenfeld, Brendan Belovarac, Margaret Black, Ludovic Boytard, John Cadley, Paolo Cotzia, John Chen, Dacia Dimartino, Xiaojun Feng, Tatyana Gindin, Emily Guzman, Adriana Heguy, Megan Hogan, Emily Huang, George Jour, Lawrence H. Lin, Raven Luther, Andrew Lytle, Christian Marier, Matthew T. Maurano, Mark J. Mulligan, Peter Meyn, Raquel Ordonez Ciriza, Iman Osman, Jared Pinnell, Vanessa Raabe, Sitharam Ramaswami, Amy Rapkiewicz, Andre M. Ribeiro-dos-Santos, Marie Samanovic-Golden, Antonio Serrano, Guomiao Shen, Matija Snuderl, Theodore Vougioukalis, Nick Vulpescu, Gael Westby, Paul Zappile, Yutong Zhang |
| EPI_ISL_430372 | hCoV-19/USA/NY-NYUMC345/2020 | North America / USA / New York / Brookl | 2020-04-07 | NYU Langone Health | Departments of Pathology and Medicine, New York University School of Medicine | Maria Aguero-Rosenfeld, Brendan Belovarac, Margaret Black, Ludovic Boytard, John Cadley, Paolo Cotzia, John Chen, Dacia Dimartino, Xiaojun Feng, Tatyana Gindin, Emily Guzman, Adriana Heguy, Megan Hogan, Emily Huang, George Jour, Lawrence H. Lin, Raven Luther, Andrew Lytle, Christian Marier, Matthew T. Maurano, Mark J. Mulligan, Peter Meyn, Raquel Ordonez Ciriza, Iman Osman, Jared Pinnell, Vanessa Raabe, Sitharam Ramaswami, Amy Rapkiewicz, Andre M. Ribeiro-dos-Santos, Marie Samanovic-Golden, Antonio Serrano, Guomiao Shen, Matija Snuderl, Theodore Vougioukalis, Nick Vulpescu, Gael Westby, Paul Zappile, Yutong Zhang |
| EPI_ISL_430373 | hCoV-19/USA/NY-NYUMC346/2020 | North America / USA / New York / Brookl | 2020-04-08 | NYU Langone Health | Departments of Pathology and Medicine, New York University School of Medicine | Maria Aguero-Rosenfeld, Brendan Belovarac, Margaret Black, Ludovic Boytard, John Cadley, Paolo Cotzia, John Chen, Dacia Dimartino, Xiaojun Feng, Tatyana Gindin, Emily Guzman, Adriana Heguy, Megan Hogan, Emily Huang, George Jour, Lawrence H. Lin, Raven Luther, Andrew Lytle, Christian Marier, Matthew T. Maurano, Mark J. Mulligan, Peter Meyn, Raquel Ordonez Ciriza, Iman Osman, Jared Pinnell, Vanessa Raabe, Sitharam Ramaswami, Amy Rapkiewicz, Andre M. Ribeiro-dos-Santos, Marie Samanovic-Golden, Antonio Serrano, Guomiao Shen, Matija Snuderl, Theodore Vougioukalis, Nick Vulpescu, Gael Westby, Paul Zappile, Yutong Zhang |

| | | | | | | |
|---|---|---|---|---|---|---|
| EPI_ISL_430374 | hCoV-19/USA/NY-NYUMC347/2020 | North America / USA / New York / Brookl | 2020-04-08 | NYU Langone Health | Departments of Pathology and Medicine, New York University School of Medicine | Maria Aguero-Rosenfeld, Brendan Belovarac, Margaret Black, Ludovic Boytard, John Cadley, Paolo Cotzia, John Chen, Dacia Dimartino, Xiaojun Feng, Tatyana Gindin, Emily Guzman, Adriana Heguy, Megan Hogan, Emily Huang, George Jour, Lawrence H. Lin, Raven Luther, Andrew Lytle, Christian Marier, Matthew T. Maurano, Mark J. Mulligan, Peter Meyn, Raquel Ordonez Ciriza, Iman Osman, Jared Pinnell, Vanessa Raabe, Sitharam Ramaswami, Amy Rapkiewicz, Andre M. Ribeiro-dos-Santos, Marie Samanovic-Golden, Antonio Serrano, Guomiao Shen, Matija Snuderl, Theodore Vougiouklakis, Nick Vulpescu, Gael Westby, Paul Zappile, Yutong Zhang |
| EPI_ISL_430375 | hCoV-19/USA/NY-NYUMC348/2020 | North America / USA / New York / Brookl | 2020-04-08 | NYU Langone Health | Departments of Pathology and Medicine, New York University School of Medicine | Maria Aguero-Rosenfeld, Brendan Belovarac, Margaret Black, Ludovic Boytard, John Cadley, Paolo Cotzia, John Chen, Dacia Dimartino, Xiaojun Feng, Tatyana Gindin, Emily Guzman, Adriana Heguy, Megan Hogan, Emily Huang, George Jour, Lawrence H. Lin, Raven Luther, Andrew Lytle, Christian Marier, Matthew T. Maurano, Mark J. Mulligan, Peter Meyn, Raquel Ordonez Ciriza, Iman Osman, Jared Pinnell, Vanessa Raabe, Sitharam Ramaswami, Amy Rapkiewicz, Andre M. Ribeiro-dos-Santos, Marie Samanovic-Golden, Antonio Serrano, Guomiao Shen, Matija Snuderl, Theodore Vougiouklakis, Nick Vulpescu, Gael Westby, Paul Zappile, Yutong Zhang |
| EPI_ISL_430376 | hCoV-19/USA/NY-NYUMC349/2020 | North America / USA / New York / Brookl | 2020-04-08 | NYU Langone Health | Departments of Pathology and Medicine, New York University School of Medicine | Maria Aguero-Rosenfeld, Brendan Belovarac, Margaret Black, Ludovic Boytard, John Cadley, Paolo Cotzia, John Chen, Dacia Dimartino, Xiaojun Feng, Tatyana Gindin, Emily Guzman, Adriana Heguy, Megan Hogan, Emily Huang, George Jour, Lawrence H. Lin, Raven Luther, Andrew Lytle, Christian Marier, Matthew T. Maurano, Mark J. Mulligan, Peter Meyn, Raquel Ordonez Ciriza, Iman Osman, Jared Pinnell, Vanessa Raabe, Sitharam Ramaswami, Amy Rapkiewicz, Andre M. Ribeiro-dos-Santos, Marie Samanovic-Golden, Antonio Serrano, Guomiao Shen, Matija Snuderl, Theodore Vougiouklakis, Nick Vulpescu, Gael Westby, Paul Zappile, Yutong Zhang |
| EPI_ISL_430377 | hCoV-19/USA/NY-NYUMC350/2020 | North America / USA / New York / Brookl | 2020-04-08 | NYU Langone Health | Departments of Pathology and Medicine, New York University School of Medicine | Maria Aguero-Rosenfeld, Brendan Belovarac, Margaret Black, Ludovic Boytard, John Cadley, Paolo Cotzia, John Chen, Dacia Dimartino, Xiaojun Feng, Tatyana Gindin, Emily Guzman, Adriana Heguy, Megan Hogan, Emily Huang, George Jour, Lawrence H. Lin, Raven Luther, Andrew Lytle, Christian Marier, Matthew T. Maurano, Mark J. Mulligan, Peter Meyn, Raquel Ordonez Ciriza, Iman Osman, Jared Pinnell, Vanessa Raabe, Sitharam Ramaswami, Amy Rapkiewicz, Andre M. Ribeiro-dos-Santos, Marie Samanovic-Golden, Antonio Serrano, Guomiao Shen, Matija Snuderl, Theodore Vougiouklakis, Nick Vulpescu, Gael Westby, Paul Zappile, Yutong Zhang |
| EPI_ISL_430378 | hCoV-19/USA/NY-NYUMC351/2020 | North America / USA / New York / Brookl | 2020-04-07 | NYU Langone Health | Departments of Pathology and Medicine, New York University School of Medicine | Maria Aguero-Rosenfeld, Brendan Belovarac, Margaret Black, Ludovic Boytard, John Cadley, Paolo Cotzia, John Chen, Dacia Dimartino, Xiaojun Feng, Tatyana Gindin, Emily Guzman, Adriana Heguy, Megan Hogan, Emily Huang, George Jour, Lawrence H. Lin, Raven Luther, Andrew Lytle, Christian Marier, Matthew T. Maurano, Mark J. Mulligan, Peter Meyn, Raquel Ordonez Ciriza, Iman Osman, Jared Pinnell, Vanessa Raabe, Sitharam Ramaswami, Amy Rapkiewicz, Andre M. Ribeiro-dos-Santos, Marie Samanovic-Golden, Antonio Serrano, Guomiao Shen, Matija Snuderl, Theodore Vougiouklakis, Nick Vulpescu, Gael Westby, Paul Zappile, Yutong Zhang |
| EPI_ISL_430379 | hCoV-19/USA/NY-NYUMC352/2020 | North America / USA / New York / Brookl | 2020-04-07 | NYU Langone Health | Departments of Pathology and Medicine, New York University School of Medicine | Maria Aguero-Rosenfeld, Brendan Belovarac, Margaret Black, Ludovic Boytard, John Cadley, Paolo Cotzia, John Chen, Dacia Dimartino, Xiaojun Feng, Tatyana Gindin, Emily Guzman, Adriana Heguy, Megan Hogan, Emily Huang, George Jour, Lawrence H. Lin, Raven Luther, Andrew Lytle, Christian Marier, Matthew T. Maurano, Mark J. Mulligan, Peter Meyn, Raquel Ordonez Ciriza, Iman Osman, Jared Pinnell, Vanessa Raabe, Sitharam Ramaswami, Amy Rapkiewicz, Andre M. Ribeiro-dos-Santos, Marie Samanovic-Golden, Antonio Serrano, Guomiao Shen, Matija Snuderl, Theodore Vougiouklakis, Nick Vulpescu, Gael Westby, Paul Zappile, Yutong Zhang |
| EPI_ISL_430380 | hCoV-19/USA/NY-NYUMC353/2020 | North America / USA / New York / Queen | 2020-04-08 | NYU Langone Health | Departments of Pathology and Medicine, New York University School of Medicine | Maria Aguero-Rosenfeld, Brendan Belovarac, Margaret Black, Ludovic Boytard, John Cadley, Paolo Cotzia, John Chen, Dacia Dimartino, Xiaojun Feng, Tatyana Gindin, Emily Guzman, Adriana Heguy, Megan Hogan, Emily Huang, George Jour, Lawrence H. Lin, Raven Luther, Andrew Lytle, Christian Marier, Matthew T. Maurano, Mark J. Mulligan, Peter Meyn, Raquel Ordonez Ciriza, Iman Osman, Jared Pinnell, Vanessa Raabe, Sitharam Ramaswami, Amy Rapkiewicz, Andre M. Ribeiro-dos-Santos, Marie Samanovic-Golden, Antonio Serrano, Guomiao Shen, Matija Snuderl, Theodore Vougiouklakis, Nick Vulpescu, Gael Westby, Paul Zappile, Yutong Zhang |
| EPI_ISL_430381 | hCoV-19/USA/NY-NYUMC354/2020 | North America / USA / New York / Brookl | 2020-04-07 | NYU Langone Health | Departments of Pathology and Medicine, New York University School of Medicine | Maria Aguero-Rosenfeld, Brendan Belovarac, Margaret Black, Ludovic Boytard, John Cadley, Paolo Cotzia, John Chen, Dacia Dimartino, Xiaojun Feng, Tatyana Gindin, Emily Guzman, Adriana Heguy, Megan Hogan, Emily Huang, George Jour, Lawrence H. Lin, Raven Luther, Andrew Lytle, Christian Marier, Matthew T. Maurano, Mark J. Mulligan, Peter Meyn, Raquel Ordonez Ciriza, Iman Osman, Jared Pinnell, Vanessa Raabe, Sitharam Ramaswami, Amy Rapkiewicz, Andre M. Ribeiro-dos-Santos, Marie Samanovic-Golden, Antonio Serrano, Guomiao Shen, Matija Snuderl, Theodore Vougiouklakis, Nick Vulpescu, Gael Westby, Paul Zappile, Yutong Zhang |
| EPI_ISL_430382 | hCoV-19/USA/NY-NYUMC355/2020 | North America / USA / New York / Brookl | 2020-04-09 | NYU Langone Health | Departments of Pathology and Medicine, New York University School of Medicine | Maria Aguero-Rosenfeld, Brendan Belovarac, Margaret Black, Ludovic Boytard, John Cadley, Paolo Cotzia, John Chen, Dacia Dimartino, Xiaojun Feng, Tatyana Gindin, Emily Guzman, Adriana Heguy, Megan Hogan, Emily Huang, George Jour, Lawrence H. Lin, Raven Luther, Andrew Lytle, Christian Marier, Matthew T. Maurano, Mark J. Mulligan, Peter Meyn, Raquel Ordonez Ciriza, Iman Osman, Jared Pinnell, Vanessa Raabe, Sitharam Ramaswami, Amy Rapkiewicz, Andre M. Ribeiro-dos-Santos, Marie Samanovic-Golden, Antonio Serrano, Guomiao Shen, Matija Snuderl, Theodore Vougiouklakis, Nick Vulpescu, Gael Westby, Paul Zappile, Yutong Zhang |
| EPI_ISL_430383 | hCoV-19/USA/NY-NYUMC356/2020 | North America / USA / New York / Brookl | 2020-04-08 | NYU Langone Health | Departments of Pathology and Medicine, New York University School of Medicine | Maria Aguero-Rosenfeld, Brendan Belovarac, Margaret Black, Ludovic Boytard, John Cadley, Paolo Cotzia, John Chen, Dacia Dimartino, Xiaojun Feng, Tatyana Gindin, Emily Guzman, Adriana Heguy, Megan Hogan, Emily Huang, George Jour, Lawrence H. Lin, Raven Luther, Andrew Lytle, Christian Marier, Matthew T. Maurano, Mark J. Mulligan, Peter Meyn, Raquel Ordonez Ciriza, Iman Osman, Jared Pinnell, Vanessa Raabe, Sitharam Ramaswami, Amy Rapkiewicz, Andre M. Ribeiro-dos-Santos, Marie Samanovic-Golden, Antonio Serrano, Guomiao Shen, Matija Snuderl, Theodore Vougiouklakis, Nick Vulpescu, Gael Westby, Paul Zappile, Yutong Zhang |
| EPI_ISL_430384 | hCoV-19/USA/NY-NYUMC357/2020 | North America / USA / New York / Brookl | 2020-04-09 | NYU Langone Health | Departments of Pathology and Medicine, New York University School of Medicine | Maria Aguero-Rosenfeld, Brendan Belovarac, Margaret Black, Ludovic Boytard, John Cadley, Paolo Cotzia, John Chen, Dacia Dimartino, Xiaojun Feng, Tatyana Gindin, Emily Guzman, Adriana Heguy, Megan Hogan, Emily Huang, George Jour, Lawrence H. Lin, Raven Luther, Andrew Lytle, Christian Marier, Matthew T. Maurano, Mark J. Mulligan, Peter Meyn, Raquel Ordonez Ciriza, Iman Osman, Jared Pinnell, Vanessa Raabe, Sitharam Ramaswami, Amy Rapkiewicz, Andre M. Ribeiro-dos-Santos, Marie Samanovic-Golden, Antonio Serrano, Guomiao Shen, Matija Snuderl, Theodore Vougiouklakis, Nick Vulpescu, Gael Westby, Paul Zappile, Yutong Zhang |
| EPI_ISL_430385 | hCoV-19/USA/NY-NYUMC358/2020 | North America / USA / New York / Brookl | 2020-04-08 | NYU Langone Health | Departments of Pathology and Medicine, New York University School of Medicine | Maria Aguero-Rosenfeld, Brendan Belovarac, Margaret Black, Ludovic Boytard, John Cadley, Paolo Cotzia, John Chen, Dacia Dimartino, Xiaojun Feng, Tatyana Gindin, Emily Guzman, Adriana Heguy, Megan Hogan, Emily Huang, George Jour, Lawrence H. Lin, Raven Luther, Andrew Lytle, Christian Marier, Matthew T. Maurano, Mark J. Mulligan, Peter Meyn, Raquel Ordonez Ciriza, Iman Osman, Jared Pinnell, Vanessa Raabe, Sitharam Ramaswami, Amy Rapkiewicz, Andre M. Ribeiro-dos-Santos, Marie Samanovic-Golden, Antonio Serrano, Guomiao Shen, Matija Snuderl, Theodore Vougiouklakis, Nick Vulpescu, Gael Westby, Paul Zappile, Yutong Zhang |
| EPI_ISL_430386 | hCoV-19/USA/NY-NYUMC359/2020 | North America / USA / New York / Brookl | 2020-04-08 | NYU Langone Health | Departments of Pathology and Medicine, New York University School of Medicine | Maria Aguero-Rosenfeld, Brendan Belovarac, Margaret Black, Ludovic Boytard, John Cadley, Paolo Cotzia, John Chen, Dacia Dimartino, Xiaojun Feng, Tatyana Gindin, Emily Guzman, Adriana Heguy, Megan Hogan, Emily Huang, George Jour, Lawrence H. Lin, Raven Luther, Andrew Lytle, Christian Marier, Matthew T. Maurano, Mark J. Mulligan, Peter Meyn, Raquel Ordonez Ciriza, Iman Osman, Jared Pinnell, Vanessa Raabe, Sitharam Ramaswami, Amy Rapkiewicz, Andre M. Ribeiro-dos-Santos, Marie Samanovic-Golden, Antonio Serrano, Guomiao Shen, Matija Snuderl, Theodore Vougiouklakis, Nick Vulpescu, Gael Westby, Paul Zappile, Yutong Zhang |
| EPI_ISL_430387 | hCoV-19/USA/NY-NYUMC360/2020 | North America / USA / New York / Brookl | 2020-04-08 | NYU Langone Health | Departments of Pathology and Medicine, New York University School of Medicine | Maria Aguero-Rosenfeld, Brendan Belovarac, Margaret Black, Ludovic Boytard, John Cadley, Paolo Cotzia, John Chen, Dacia Dimartino, Xiaojun Feng, Tatyana Gindin, Emily Guzman, Adriana Heguy, Megan Hogan, Emily Huang, George Jour, Lawrence H. Lin, Raven Luther, Andrew Lytle, Christian Marier, Matthew T. Maurano, Mark J. Mulligan, Peter Meyn, Raquel Ordonez Ciriza, Iman Osman, Jared Pinnell, Vanessa Raabe, Sitharam Ramaswami, Amy Rapkiewicz, Andre M. Ribeiro-dos-Santos, Marie Samanovic-Golden, Antonio Serrano, Guomiao Shen, Matija Snuderl, Theodore Vougiouklakis, Nick Vulpescu, Gael Westby, Paul Zappile, Yutong Zhang |
| EPI_ISL_430388 | hCoV-19/USA/NY-NYUMC361/2020 | North America / USA / New York / Queen | 2020-04-09 | NYU Langone Health | Departments of Pathology and Medicine, New York University School of Medicine | Maria Aguero-Rosenfeld, Brendan Belovarac, Margaret Black, Ludovic Boytard, John Cadley, Paolo Cotzia, John Chen, Dacia Dimartino, Xiaojun Feng, Tatyana Gindin, Emily Guzman, Adriana Heguy, Megan Hogan, Emily Huang, George Jour, Lawrence H. Lin, Raven Luther, Andrew Lytle, Christian Marier, Matthew T. Maurano, Mark J. Mulligan, Peter Meyn, Raquel Ordonez Ciriza, Iman Osman, Jared Pinnell, Vanessa Raabe, Sitharam Ramaswami, Amy Rapkiewicz, Andre M. Ribeiro-dos-Santos, Marie Samanovic-Golden, Antonio Serrano, Guomiao Shen, Matija Snuderl, Theodore Vougiouklakis, Nick Vulpescu, Gael Westby, Paul Zappile, Yutong Zhang |
| EPI_ISL_430389 | hCoV-19/USA/NY-NYUMC362/2020 | North America / USA / New York / Brookl | 2020-04-08 | NYU Langone Health | Departments of Pathology and Medicine, New York University School of Medicine | Maria Aguero-Rosenfeld, Brendan Belovarac, Margaret Black, Ludovic Boytard, John Cadley, Paolo Cotzia, John Chen, Dacia Dimartino, Xiaojun Feng, Tatyana Gindin, Emily Guzman, Adriana Heguy, Megan Hogan, Emily Huang, George Jour, Lawrence H. Lin, Raven Luther, Andrew Lytle, Christian Marier, Matthew T. Maurano, Mark J. Mulligan, Peter Meyn, Raquel Ordonez Ciriza, Iman Osman, Jared Pinnell, Vanessa Raabe, Sitharam Ramaswami, Amy Rapkiewicz, Andre M. Ribeiro-dos-Santos, Marie Samanovic-Golden, Antonio Serrano, Guomiao Shen, Matija Snuderl, Theodore Vougiouklakis, Nick Vulpescu, Gael Westby, Paul Zappile, Yutong Zhang |
| EPI_ISL_430390 | hCoV-19/USA/NY-NYUMC363/2020 | North America / USA / New York / Brookl | 2020-04-08 | NYU Langone Health | Departments of Pathology and Medicine, New York University School of Medicine | Maria Aguero-Rosenfeld, Brendan Belovarac, Margaret Black, Ludovic Boytard, John Cadley, Paolo Cotzia, John Chen, Dacia Dimartino, Xiaojun Feng, Tatyana Gindin, Emily Guzman, Adriana Heguy, Megan Hogan, Emily Huang, George Jour, Lawrence H. Lin, Raven Luther, Andrew Lytle, Christian Marier, Matthew T. Maurano, Mark J. Mulligan, Peter Meyn, Raquel Ordonez Ciriza, Iman Osman, Jared Pinnell, Vanessa Raabe, Sitharam Ramaswami, Amy Rapkiewicz, Andre M. Ribeiro-dos-Santos, Marie Samanovic-Golden, Antonio Serrano, Guomiao Shen, Matija Snuderl, Theodore Vougiouklakis, Nick Vulpescu, Gael Westby, Paul Zappile, Yutong Zhang |
| EPI_ISL_430391 | hCoV-19/USA/NY-NYUMC364/2020 | North America / USA / New York / Queen | 2020-04-09 | NYU Langone Health | Departments of Pathology and Medicine, New York University School of Medicine | Maria Aguero-Rosenfeld, Brendan Belovarac, Margaret Black, Ludovic Boytard, John Cadley, Paolo Cotzia, John Chen, Dacia Dimartino, Xiaojun Feng, Tatyana Gindin, Emily Guzman, Adriana Heguy, Megan Hogan, Emily Huang, George Jour, Lawrence H. Lin, Raven Luther, Andrew Lytle, Christian Marier, Matthew T. Maurano, Mark J. Mulligan, Peter Meyn, Raquel Ordonez Ciriza, Iman Osman, Jared Pinnell, Vanessa Raabe, Sitharam Ramaswami, Amy Rapkiewicz, Andre M. Ribeiro-dos-Santos, Marie Samanovic-Golden, Antonio Serrano, Guomiao Shen, Matija Snuderl, Theodore Vougiouklakis, Nick Vulpescu, Gael Westby, Paul Zappile, Yutong Zhang |
| EPI_ISL_430392 | hCoV-19/USA/NY-NYUMC365/2020 | North America / USA / New York / Brookl | 2020-04-08 | NYU Langone Health | Departments of Pathology and Medicine, New York University School of Medicine | Maria Aguero-Rosenfeld, Brendan Belovarac, Margaret Black, Ludovic Boytard, John Cadley, Paolo Cotzia, John Chen, Dacia Dimartino, Xiaojun Feng, Tatyana Gindin, Emily Guzman, Adriana Heguy, Megan Hogan, Emily Huang, George Jour, Lawrence H. Lin, Raven Luther, Andrew Lytle, Christian Marier, Matthew T. Maurano, Mark J. Mulligan, Peter Meyn, Raquel Ordonez Ciriza, Iman Osman, Jared Pinnell, Vanessa Raabe, Sitharam Ramaswami, Amy Rapkiewicz, Andre M. Ribeiro-dos-Santos, Marie Samanovic-Golden, Antonio Serrano, Guomiao Shen, Matija Snuderl, Theodore Vougiouklakis, Nick Vulpescu, Gael Westby, Paul Zappile, Yutong Zhang |
| EPI_ISL_430393 | hCoV-19/USA/NY-NYUMC366/2020 | North America / USA / New York / Manha | 2020-04-09 | NYU Langone Health | Departments of Pathology and Medicine, New York University School of Medicine | Maria Aguero-Rosenfeld, Brendan Belovarac, Margaret Black, Ludovic Boytard, John Cadley, Paolo Cotzia, John Chen, Dacia Dimartino, Xiaojun Feng, Tatyana Gindin, Emily Guzman, Adriana Heguy, Megan Hogan, Emily Huang, George Jour, Lawrence H. Lin, Raven Luther, Andrew Lytle, Christian Marier, Matthew T. Maurano, Mark J. Mulligan, Peter Meyn, Raquel Ordonez Ciriza, Iman Osman, Jared Pinnell, Vanessa Raabe, Sitharam Ramaswami, Amy Rapkiewicz, Andre M. Ribeiro-dos-Santos, Marie Samanovic-Golden, Antonio Serrano, Guomiao Shen, Matija Snuderl, Theodore Vougiouklakis, Nick Vulpescu, Gael Westby, Paul Zappile, Yutong Zhang |
| EPI_ISL_430394 | hCoV-19/USA/NJ-NYUMC367/2020 | North America / USA / New Jersey / Hud | 2020-04-09 | NYU Langone Health | Departments of Pathology and Medicine, New York University School of Medicine | Maria Aguero-Rosenfeld, Brendan Belovarac, Margaret Black, Ludovic Boytard, John Cadley, Paolo Cotzia, John Chen, Dacia Dimartino, Xiaojun Feng, Tatyana Gindin, Emily Guzman, Adriana Heguy, Megan Hogan, Emily Huang, George Jour, Lawrence H. Lin, Raven Luther, Andrew Lytle, Christian Marier, Matthew T. Maurano, Mark J. Mulligan, Peter Meyn, Raquel Ordonez Ciriza, Iman Osman, Jared Pinnell, Vanessa Raabe, Sitharam Ramaswami, Amy Rapkiewicz, Andre M. Ribeiro-dos-Santos, Marie Samanovic-Golden, Antonio Serrano, Guomiao Shen, Matija Snuderl, Theodore Vougiouklakis, Nick Vulpescu, Gael Westby, Paul Zappile, Yutong Zhang |
| EPI_ISL_430395 | hCoV-19/USA/NY-NYUMC368/2020 | North America / USA / New York / Brookl | 2020-04-09 | NYU Langone Health | Departments of Pathology and Medicine, New York University School of Medicine | Maria Aguero-Rosenfeld, Brendan Belovarac, Margaret Black, Ludovic Boytard, John Cadley, Paolo Cotzia, John Chen, Dacia Dimartino, Xiaojun Feng, Tatyana Gindin, Emily Guzman, Adriana Heguy, Megan Hogan, Emily Huang, George Jour, Lawrence H. Lin, Raven Luther, Andrew Lytle, Christian Marier, Matthew T. Maurano, Mark J. Mulligan, Peter Meyn, Raquel Ordonez Ciriza, Iman Osman, Jared Pinnell, Vanessa Raabe, Sitharam Ramaswami, Amy Rapkiewicz, Andre M. Ribeiro-dos-Santos, Marie Samanovic-Golden, Antonio Serrano, Guomiao Shen, Matija Snuderl, Theodore Vougiouklakis, Nick Vulpescu, Gael Westby, Paul Zappile, Yutong Zhang |
| EPI_ISL_430396 | hCoV-19/USA/NY-NYUMC369/2020 | North America / USA / New York / Nassa | 2020-04-10 | NYU Langone Health | Departments of Pathology and Medicine, New York University School of Medicine | Maria Aguero-Rosenfeld, Brendan Belovarac, Margaret Black, Ludovic Boytard, John Cadley, Paolo Cotzia, John Chen, Dacia Dimartino, Xiaojun Feng, Tatyana Gindin, Emily Guzman, Adriana Heguy, Megan Hogan, Emily Huang, George Jour, Lawrence H. Lin, Raven Luther, Andrew Lytle, Christian Marier, Matthew T. Maurano, Mark J. Mulligan, Peter Meyn, Raquel Ordonez Ciriza, Iman Osman, Jared Pinnell, Vanessa Raabe, Sitharam Ramaswami, Amy Rapkiewicz, Andre M. Ribeiro-dos-Santos, Marie Samanovic-Golden, Antonio Serrano, Guomiao Shen, Matija Snuderl, Theodore Vougiouklakis, Nick Vulpescu, Gael Westby, Paul Zappile, Yutong Zhang |
| EPI_ISL_430397 | hCoV-19/USA/NY-NYUMC370/2020 | North America / USA / New York / Nassa | 2020-04-09 | NYU Langone Health | Departments of Pathology and Medicine, New York University School of Medicine | Maria Aguero-Rosenfeld, Brendan Belovarac, Margaret Black, Ludovic Boytard, John Cadley, Paolo Cotzia, John Chen, Dacia Dimartino, Xiaojun Feng, Tatyana Gindin, Emily Guzman, Adriana Heguy, Megan Hogan, Emily Huang, George Jour, Lawrence H. Lin, Raven Luther, Andrew Lytle, Christian Marier, Matthew T. Maurano, Mark J. Mulligan, Peter Meyn, Raquel Ordonez Ciriza, Iman Osman, Jared Pinnell, Vanessa Raabe, Sitharam Ramaswami, Amy Rapkiewicz, Andre M. Ribeiro-dos-Santos, Marie Samanovic-Golden, Antonio Serrano, Guomiao Shen, Matija Snuderl, Theodore Vougiouklakis, Nick Vulpescu, Gael Westby, Paul Zappile, Yutong Zhang |
| EPI_ISL_430398 | hCoV-19/USA/NY-NYUMC371/2020 | North America / USA / New York / Nassa | 2020-04-10 | NYU Langone Health | Departments of Pathology and Medicine, New York University School of Medicine | Maria Aguero-Rosenfeld, Brendan Belovarac, Margaret Black, Ludovic Boytard, John Cadley, Paolo Cotzia, John Chen, Dacia Dimartino, Xiaojun Feng, Tatyana Gindin, Emily Guzman, Adriana Heguy, Megan Hogan, Emily Huang, George Jour, Lawrence H. Lin, Raven Luther, Andrew Lytle, Christian Marier, Matthew T. Maurano, Mark J. Mulligan, Peter Meyn, Raquel Ordonez Ciriza, Iman Osman, Jared Pinnell, Vanessa Raabe, Sitharam Ramaswami, Amy Rapkiewicz, Andre M. Ribeiro-dos-Santos, Marie Samanovic-Golden, Antonio Serrano, Guomiao Shen, Matija Snuderl, Theodore Vougiouklakis, Nick Vulpescu, Gael Westby, Paul Zappile, Yutong Zhang |
| EPI_ISL_430399 | hCoV-19/USA/NY-NYUMC372/2020 | North America / USA / New York / Nassa | 2020-04-10 | NYU Langone Health | Departments of Pathology and Medicine, New York University School of Medicine | Maria Aguero-Rosenfeld, Brendan Belovarac, Margaret Black, Ludovic Boytard, John Cadley, Paolo Cotzia, John Chen, Dacia Dimartino, Xiaojun Feng, Tatyana Gindin, Emily Guzman, Adriana Heguy, Megan Hogan, Emily Huang, George Jour, Lawrence H. Lin, Raven Luther, Andrew Lytle, Christian Marier, Matthew T. Maurano, Mark J. Mulligan, Peter Meyn, Raquel Ordonez Ciriza, Iman Osman, Jared Pinnell, Vanessa Raabe, Sitharam Ramaswami, Amy Rapkiewicz, Andre M. Ribeiro-dos-Santos, Marie Samanovic-Golden, Antonio Serrano, Guomiao Shen, Matija Snuderl, Theodore Vougiouklakis, Nick Vulpescu, Gael Westby, Paul Zappile, Yutong Zhang |
| EPI_ISL_430600 | hCoV-19/Australia/VIC1134/2020 | Oceania / Australia / Victoria | 2020-04-08 | Victorian Infectious Diseases Reference Laboratory (VIDRL) | Microbiological Diagnostic Unit Public Health Laboratory and Victorian Infectious Diseases Reference Laboratory, The Peter Doherty Institute for Infection and Immunity | Caly L., Seemann T., Sait, M., Schultz M., Druce J., Sherry, N. |
| EPI_ISL_430601 | hCoV-19/Australia/VIC1135/2020 | Oceania / Australia / Victoria | 2020-04-08 | Victorian Infectious Diseases Reference Laboratory (VIDRL) | Microbiological Diagnostic Unit Public Health Laboratory and Victorian Infectious Diseases Reference Laboratory, The Peter Doherty Institute for Infection and Immunity | Caly L., Seemann T., Sait, M., Schultz M., Druce J., Sherry, N. |
| EPI_ISL_430602 | hCoV-19/Australia/VIC1158/2020 | Oceania / Australia / Victoria | 2020-04-09 | Victorian Infectious Diseases Reference Laboratory (VIDRL) | Microbiological Diagnostic Unit Public Health Laboratory and Victorian Infectious Diseases Reference Laboratory, The Peter Doherty Institute for Infection and Immunity | Caly L., Seemann T., Sait, M., Schultz M., Druce J., Sherry, N. |
| EPI_ISL_430604 | hCoV-19/Australia/VIC1159/2020 | Oceania / Australia / Victoria | 2020-04-09 | Victorian Infectious Diseases Reference Laboratory (VIDRL) | Microbiological Diagnostic Unit Public Health Laboratory and Victorian Infectious Diseases Reference Laboratory, The Peter Doherty Institute for Infection and Immunity | Caly L., Seemann T., Sait, M., Schultz M., Druce J., Sherry, N. |
| EPI_ISL_430605 | hCoV-19/Australia/VIC1160/2020 | Oceania / Australia / Victoria | 2020-04-09 | Victorian Infectious Diseases Reference Laboratory (VIDRL) | Microbiological Diagnostic Unit Public Health Laboratory and Victorian Infectious Diseases Reference Laboratory, The Peter Doherty Institute for Infection and Immunity | Caly L., Seemann T., Sait, M., Schultz M., Druce J., Sherry, N. |
| EPI_ISL_430607 | hCoV-19/Australia/VIC1137/2020 | Oceania / Australia / Victoria | 2020-04-08 | Victorian Infectious Diseases Reference Laboratory (VIDRL) | Microbiological Diagnostic Unit Public Health Laboratory and Victorian Infectious Diseases Reference Laboratory, The Peter Doherty Institute for Infection and Immunity | Caly L., Seemann T., Sait, M., Schultz M., Druce J., Sherry, N. |
| EPI_ISL_430609 | hCoV-19/Australia/VIC1138/2020 | Oceania / Australia / Victoria | 2020-04-08 | Victorian Infectious Diseases Reference Laboratory (VIDRL) | Microbiological Diagnostic Unit Public Health Laboratory and Victorian Infectious Diseases Reference Laboratory, The Peter Doherty Institute for Infection and Immunity | Caly L., Seemann T., Sait, M., Schultz M., Druce J., Sherry, N. |
| EPI_ISL_430610 | hCoV-19/Australia/VIC1139/2020 | Oceania / Australia / Victoria | 2020-04-08 | Victorian Infectious Diseases Reference Laboratory (VIDRL) | Microbiological Diagnostic Unit Public Health Laboratory and Victorian Infectious Diseases Reference Laboratory, The Peter Doherty Institute for Infection and Immunity | Caly L., Seemann T., Sait, M., Schultz M., Druce J., Sherry, N. |
| EPI_ISL_430614 | hCoV-19/Australia/VIC1143/2020 | Oceania / Australia / Victoria | 2020-04-08 | Victorian Infectious Diseases Reference Laboratory (VIDRL) | Microbiological Diagnostic Unit Public Health Laboratory and Victorian Infectious Diseases Reference Laboratory, The Peter Doherty Institute for Infection and Immunity | Caly L., Seemann T., Sait, M., Schultz M., Druce J., Sherry, N. |
| EPI_ISL_430615 | hCoV-19/Australia/VIC1161/2020 | Oceania / Australia / Victoria | 2020-04-09 | Victorian Infectious Diseases Reference Laboratory (VIDRL) | Microbiological Diagnostic Unit Public Health Laboratory and Victorian Infectious Diseases Reference Laboratory, The Peter Doherty Institute for Infection and Immunity | Caly L., Seemann T., Sait, M., Schultz M., Druce J., Sherry, N. |
| EPI_ISL_430616 | hCoV-19/Australia/VIC1162/2020 | Oceania / Australia / Victoria | 2020-04-09 | Victorian Infectious Diseases Reference Laboratory (VIDRL) | Microbiological Diagnostic Unit Public Health Laboratory and Victorian Infectious Diseases Reference Laboratory, The Peter Doherty Institute for Infection and Immunity | Caly L., Seemann T., Sait, M., Schultz M., Druce J., Sherry, N. |
| EPI_ISL_430618 | hCoV-19/Australia/VIC1109/2020 | Oceania / Australia / Victoria | 2020-04-07 | Victorian Infectious Diseases Reference Laboratory (VIDRL) | Microbiological Diagnostic Unit Public Health Laboratory and Victorian Infectious Diseases Reference Laboratory, The Peter Doherty Institute for Infection and Immunity | Caly L., Seemann T., Sait, M., Schultz M., Druce J., Sherry, N. |
| EPI_ISL_430619 | hCoV-19/Australia/VIC1144/2020 | Oceania / Australia / Victoria | 2020-04-08 | Victorian Infectious Diseases Reference Laboratory (VIDRL) | Microbiological Diagnostic Unit Public Health Laboratory and Victorian Infectious Diseases Reference Laboratory, The Peter Doherty Institute for Infection and Immunity | Caly L., Seemann T., Sait, M., Schultz M., Druce J., Sherry, N. |
| EPI_ISL_430623 | hCoV-19/Australia/VIC1174/2020 | Oceania / Australia / Victoria | 2020-04-10 | Victorian Infectious Diseases Reference Laboratory (VIDRL) | Microbiological Diagnostic Unit Public Health Laboratory and Victorian Infectious Diseases Reference Laboratory, The Peter Doherty Institute for Infection and Immunity | Caly L., Seemann T., Sait, M., Schultz M., Druce J., Sherry, N. |
| EPI_ISL_430632 | hCoV-19/Australia/NT29/2020 | Oceania / Australia / Northern Territory | 2020 | Royal Darwin Hospital Pathology | Microbiological Diagnostic Unit Public Health Laboratory and Victorian Infectious Diseases Reference Laboratory, The Peter Doherty Institute for Infection and Immunity | Meumann, E., Caly L., Seemann T., Sait, M., Schultz M., Druce J., Sherry, N. |
| EPI_ISL_430633 | hCoV-19/Australia/NT30/2020 | Oceania / Australia / Northern Territory | 2020 | Royal Darwin Hospital Pathology | Microbiological Diagnostic Unit Public Health Laboratory and Victorian Infectious Diseases Reference Laboratory, The Peter Doherty Institute for Infection and Immunity | Meumann, E., Caly L., Seemann T., Sait, M., Schultz M., Druce J., Sherry, N. |
| EPI_ISL_430635 | hCoV-19/Australia/NT31/2020 | Oceania / Australia / Northern Territory | 2020 | Royal Darwin Hospital Pathology | Microbiological Diagnostic Unit Public Health Laboratory and Victorian Infectious Diseases Reference Laboratory, The Peter Doherty Institute for Infection and Immunity | Meumann, E., Caly L., Seemann T., Sait, M., Schultz M., Druce J., Sherry, N. |
| EPI_ISL_430638 | hCoV-19/Australia/VIC1355/2020 | Oceania / Australia / Victoria | 2020-04-09 | Victorian Infectious Diseases Reference Laboratory (VIDRL) | Microbiological Diagnostic Unit Public Health Laboratory and Victorian Infectious Diseases Reference Laboratory, The Peter Doherty Institute for Infection and Immunity | Caly L., Seemann T., Sait, M., Druce J., Sherry, N. |
| EPI_ISL_430639 | hCoV-19/Australia/VIC942/2020 | Oceania / Australia / Victoria | 2020-03-26 | Microbiological Diagnostic Unit Public Health Laboratory | Microbiological Diagnostic Unit Public Health Laboratory | Seemann T., Schultz M., Sait, M., Sherry, N. |
| EPI_ISL_430640 | hCoV-19/Australia/VIC946/2020 | Oceania / Australia / Victoria | 2020-03-26 | Microbiological Diagnostic Unit Public Health Laboratory | Microbiological Diagnostic Unit Public Health Laboratory | Seemann T., Schultz M., Sait, M., Sherry, N. |
| EPI_ISL_430645 | hCoV-19/Australia/VIC953/2020 | Oceania / Australia / Victoria | 2020-03-26 | Microbiological Diagnostic Unit Public Health Laboratory | Microbiological Diagnostic Unit Public Health Laboratory | Seemann T., Schultz M., Sait, M., Sherry, N. |
| EPI_ISL_430646 | hCoV-19/Australia/VIC951/2020 | Oceania / Australia / Victoria | 2020-03-26 | Microbiological Diagnostic Unit Public Health Laboratory | Microbiological Diagnostic Unit Public Health Laboratory | Seemann T., Schultz M., Sait, M., Sherry, N. |
| EPI_ISL_430647 | hCoV-19/Australia/VIC948/2020 | Oceania / Australia / Victoria | 2020-03-26 | Microbiological Diagnostic Unit Public Health Laboratory | Microbiological Diagnostic Unit Public Health Laboratory | Seemann T., Schultz M., Sait, M., Sherry, N. |
| EPI_ISL_430649 | hCoV-19/Australia/VIC957/2020 | Oceania / Australia / Victoria | 2020-03-27 | Microbiological Diagnostic Unit Public Health Laboratory | Microbiological Diagnostic Unit Public Health Laboratory | Seemann T., Schultz M., Sait, M., Sherry, N. |
| EPI_ISL_430653 | hCoV-19/Australia/VIC966/2020 | Oceania / Australia / Victoria | 2020-03-27 | Microbiological Diagnostic Unit Public Health Laboratory | Microbiological Diagnostic Unit Public Health Laboratory | Seemann T., Schultz M., Sait, M., Sherry, N. |
| EPI_ISL_430654 | hCoV-19/Australia/VIC961/2020 | Oceania / Australia / Victoria | 2020-03-27 | Microbiological Diagnostic Unit Public Health Laboratory | Microbiological Diagnostic Unit Public Health Laboratory | Seemann T., Schultz M., Sait, M., Sherry, N. |
| EPI_ISL_430655 | hCoV-19/Australia/VIC964/2020 | Oceania / Australia / Victoria | 2020-03-27 | Microbiological Diagnostic Unit Public Health Laboratory | Microbiological Diagnostic Unit Public Health Laboratory | Seemann T., Schultz M., Sait, M., Sherry, N. |
| EPI_ISL_430658 | hCoV-19/Australia/VIC975/2020 | Oceania / Australia / Victoria | 2020-03-28 | Microbiological Diagnostic Unit Public Health Laboratory | Microbiological Diagnostic Unit Public Health Laboratory | Seemann T., Schultz M., Sait, M., Sherry, N. |
| EPI_ISL_430659 | hCoV-19/Australia/VIC974/2020 | Oceania / Australia / Victoria | 2020-03-28 | Microbiological Diagnostic Unit Public Health Laboratory | Microbiological Diagnostic Unit Public Health Laboratory | Seemann T., Schultz M., Sait, M., Sherry, N. |
| EPI_ISL_430661 | hCoV-19/Australia/VIC971/2020 | Oceania / Australia / Victoria | 2020-03-27 | Microbiological Diagnostic Unit Public Health Laboratory | Microbiological Diagnostic Unit Public Health Laboratory | Seemann T., Schultz M., Sait, M., Sherry, N. |
| EPI_ISL_430663 | hCoV-19/Australia/VIC959/2020 | Oceania / Australia / Victoria | 2020-03-27 | Microbiological Diagnostic Unit Public Health Laboratory | Microbiological Diagnostic Unit Public Health Laboratory | Seemann T., Schultz M., Sait, M., Sherry, N. |
| EPI_ISL_430669 | hCoV-19/Australia/VIC1056/2020 | Oceania / Australia / Victoria | 2020-04-03 | Microbiological Diagnostic Unit Public Health Laboratory | Microbiological Diagnostic Unit Public Health Laboratory | Seemann T., Schultz M., Sait, M., Sherry, N. |
| EPI_ISL_430670 | hCoV-19/Australia/VIC1067/2020 | Oceania / Australia / Victoria | 2020-04-04 | Microbiological Diagnostic Unit Public Health Laboratory | Microbiological Diagnostic Unit Public Health Laboratory | Seemann T., Schultz M., Sait, M., Sherry, N. |
| EPI_ISL_430674 | hCoV-19/Australia/VIC1178/2020 | Oceania / Australia / Victoria | 2020-04-10 | Microbiological Diagnostic Unit Public Health Laboratory | Microbiological Diagnostic Unit Public Health Laboratory | Seemann T., Schultz M., Sait, M., Sherry, N. |
| EPI_ISL_430676 | hCoV-19/Australia/VIC1281/2020 | Oceania / Australia / Victoria | 2020-04-08 | Microbiological Diagnostic Unit Public Health Laboratory | Microbiological Diagnostic Unit Public Health Laboratory | Seemann T., Schultz M., Sait, M., Sherry, N. |
| EPI_ISL_430677 | hCoV-19/Australia/VIC1300/2020 | Oceania / Australia / Victoria | 2020-04-10 | Microbiological Diagnostic Unit Public Health Laboratory | Microbiological Diagnostic Unit Public Health Laboratory | Seemann T., Schultz M., Sait, M., Sherry, N. |
| EPI_ISL_430678 | hCoV-19/Australia/VIC1279/2020 | Oceania / Australia / Victoria | 2020-04-07 | Microbiological Diagnostic Unit Public Health Laboratory | Microbiological Diagnostic Unit Public Health Laboratory | Seemann T., Schultz M., Sait, M., Sherry, N. |
| EPI_ISL_430679 | hCoV-19/Australia/VIC1301/2020 | Oceania / Australia / Victoria | 2020-04-10 | Microbiological Diagnostic Unit Public Health Laboratory | Microbiological Diagnostic Unit Public Health Laboratory | Seemann T., Schultz M., Sait, M., Sherry, N. |
| EPI_ISL_430680 | hCoV-19/Australia/VIC1302/2020 | Oceania / Australia / Victoria | 2020-04-10 | Microbiological Diagnostic Unit Public Health Laboratory | Microbiological Diagnostic Unit Public Health Laboratory | Seemann T., Schultz M., Sait, M., Sherry, N. |
| EPI_ISL_430681 | hCoV-19/Australia/VIC1314/2020 | Oceania / Australia / Victoria | 2020-04-11 | Microbiological Diagnostic Unit Public Health Laboratory | Microbiological Diagnostic Unit Public Health Laboratory | Seemann T., Schultz M., Sait, M., Sherry, N. |
| EPI_ISL_430682 | hCoV-19/Australia/VIC1334/2020 | Oceania / Australia / Victoria | 2020-04-13 | Microbiological Diagnostic Unit Public Health Laboratory | Microbiological Diagnostic Unit Public Health Laboratory | Seemann T., Schultz M., Sait, M., Sherry, N. |
| EPI_ISL_430683 | hCoV-19/Australia/VIC1342/2020 | Oceania / Australia / Victoria | 2020-04-14 | Microbiological Diagnostic Unit Public Health Laboratory | Microbiological Diagnostic Unit Public Health Laboratory | Seemann T., Schultz M., Sait, M., Sherry, N. |
| EPI_ISL_430684 | hCoV-19/Australia/VIC1343/2020 | Oceania / Australia / Victoria | 2020-04-14 | Microbiological Diagnostic Unit Public Health Laboratory | Microbiological Diagnostic Unit Public Health Laboratory | Seemann T., Schultz M., Sait, M., Sherry, N. |
| EPI_ISL_430686 | hCoV-19/Australia/VIC1340/2020 | Oceania / Australia / Victoria | 2020-04-13 | Microbiological Diagnostic Unit Public Health Laboratory | Microbiological Diagnostic Unit Public Health Laboratory | Seemann T., Schultz M., Sait, M., Sherry, N. |
| EPI_ISL_430689 | hCoV-19/Australia/VIC1315/2020 | Oceania / Australia / Victoria | 2020-04-11 | Victorian Infectious Diseases Reference Laboratory (VIDRL) | Microbiological Diagnostic Unit Public Health Laboratory and Victorian Infectious Diseases Reference Laboratory, The Peter Doherty Institute for Infection and Immunity | Caly L., Seemann T., Sait, M., Schultz M., Druce J., Sherry, N. |
| EPI_ISL_430690 | hCoV-19/Australia/VIC1316/2020 | Oceania / Australia / Victoria | 2020-04-11 | Victorian Infectious Diseases Reference Laboratory (VIDRL) | Microbiological Diagnostic Unit Public Health Laboratory and Victorian Infectious Diseases Reference Laboratory, The Peter Doherty Institute for Infection and Immunity | Caly L., Seemann T., Sait, M., Schultz M., Druce J., Sherry, N. |
| EPI_ISL_430691 | hCoV-19/Australia/VIC1317/2020 | Oceania / Australia / Victoria | 2020-04-11 | Victorian Infectious Diseases Reference Laboratory (VIDRL) | Microbiological Diagnostic Unit Public Health Laboratory and Victorian Infectious Diseases Reference Laboratory, The Peter Doherty Institute for Infection and Immunity | Caly L., Seemann T., Sait, M., Schultz M., Druce J., Sherry, N. |
| EPI_ISL_430693 | hCoV-19/Australia/VIC1318/2020 | Oceania / Australia / Victoria | 2020-04-11 | Victorian Infectious Diseases Reference Laboratory (VIDRL) | Microbiological Diagnostic Unit Public Health Laboratory and Victorian Infectious Diseases Reference Laboratory, The Peter Doherty Institute for Infection and Immunity | Caly L., Seemann T., Sait, M., Schultz M., Druce J., Sherry, N. |
| EPI_ISL_430694 | hCoV-19/Australia/VIC1303/2020 | Oceania / Australia / Victoria | 2020-04-10 | Victorian Infectious Diseases Reference Laboratory (VIDRL) | Microbiological Diagnostic Unit Public Health Laboratory and Victorian Infectious Diseases Reference Laboratory, The Peter Doherty Institute for Infection and Immunity | Caly L., Seemann T., Sait, M., Schultz M., Druce J., Sherry, N. |
| EPI_ISL_430695 | hCoV-19/Australia/VIC1328/2020 | Oceania / Australia / Victoria | 2020-04-12 | Victorian Infectious Diseases Reference Laboratory (VIDRL) | Microbiological Diagnostic Unit Public Health Laboratory and Victorian Infectious Diseases Reference Laboratory, The Peter Doherty Institute for Infection and Immunity | Caly L., Seemann T., Sait, M., Druce J., Sherry, N. |
| EPI_ISL_431080 | hCoV-19/USA/CT-Yale-079/2020 | North America / USA / Connecticut | 2020-03-02 | Yale COVID-19 Biorepository | Grubaugh Lab - Yale School of Public Health | Joseph Fauver, Tara Alpert, Anderson Brito, Anne Wyllie, Chantal Vogels, Mary Petrone, Cole Jensen, Chaney Kalinich, Isabel Ott, Arnau Casanovas, Catherine Muenker, Adam Moore, Alice Lu, Maria Tokuyama, Patrick Wong, Peiwen Lu, Saad Omer, Richard Martinello, Allison Nelson, Shelli Farhadian, Akiko Iwasaki, Charles Dela Cruz, Albert Ko, Nathan Grubaugh |
| EPI_ISL_431081 | hCoV-19/USA/CT-Yale-102/2020 | North America / USA / Connecticut | 2020-04-06 | Yale COVID-19 Biorepository | Grubaugh Lab - Yale School of Public Health | Joseph Fauver, Tara Alpert, Anderson Brito, Anne Wyllie, Chantal Vogels, Mary Petrone, Cole Jensen, Chaney Kalinich, Isabel Ott, Arnau Casanovas, Catherine Muenker, Adam Moore, Alice Lu, Maria Tokuyama, Patrick Wong, Peiwen Lu, Saad Omer, Richard Martinello, Allison Nelson, Shelli Farhadian, Akiko Iwasaki, Charles Dela Cruz, Albert Ko, Nathan Grubaugh |
| EPI_ISL_431082 | hCoV-19/USA/CT-Yale-114/2020 | North America / USA / Connecticut | 2020-04-10 | Yale COVID-19 Biorepository | Grubaugh Lab - Yale School of Public Health | Joseph Fauver, Tara Alpert, Anderson Brito, Anne Wyllie, Chantal Vogels, Mary Petrone, Cole Jensen, Chaney Kalinich, Isabel Ott, Arnau Casanovas, Catherine Muenker, Adam Moore, Alice Lu, Maria Tokuyama, Patrick Wong, Peiwen Lu, Saad Omer, Richard Martinello, Allison Nelson, Shelli Farhadian, Akiko Iwasaki, Charles Dela Cruz, Albert Ko, Nathan Grubaugh |
| EPI_ISL_431083 | hCoV-19/USA/CT-Yale-116/2020 | North America / USA / Connecticut | 2020-04-10 | Yale COVID-19 Biorepository | Grubaugh Lab - Yale School of Public Health | Joseph Fauver, Tara Alpert, Anderson Brito, Anne Wyllie, Chantal Vogels, Mary Petrone, Cole Jensen, Chaney Kalinich, Isabel Ott, Arnau Casanovas, Catherine Muenker, Adam Moore, Alice Lu, Maria Tokuyama, Patrick Wong, Peiwen Lu, Saad Omer, Richard Martinello, Allison Nelson, Shelli Farhadian, Akiko Iwasaki, Charles Dela Cruz, Albert Ko, Nathan Grubaugh |
| EPI_ISL_431084 | hCoV-19/USA/CT-Yale-120/2020 | North America / USA / Connecticut | 2020-04-11 | Yale COVID-19 Biorepository | Grubaugh Lab - Yale School of Public Health | Joseph Fauver, Tara Alpert, Anderson Brito, Anne Wyllie, Chantal Vogels, Mary Petrone, Cole Jensen, Chaney Kalinich, Isabel Ott, Arnau Casanovas, Catherine Muenker, Adam Moore, Alice Lu, Maria Tokuyama, Patrick Wong, Peiwen Lu, Saad Omer, Richard Martinello, Allison Nelson, Shelli Farhadian, Akiko Iwasaki, Charles Dela Cruz, Albert Ko, Nathan Grubaugh |
| EPI_ISL_431085 | hCoV-19/USA/CT-Yale-121/2020 | North America / USA / Connecticut | 2020-04-11 | Yale COVID-19 Biorepository | Grubaugh Lab - Yale School of Public Health | Joseph Fauver, Tara Alpert, Anderson Brito, Anne Wyllie, Chantal Vogels, Mary Petrone, Cole Jensen, Chaney Kalinich, Isabel Ott, Arnau Casanovas, Catherine Muenker, Adam Moore, Alice Lu, Maria Tokuyama, Patrick Wong, Peiwen Lu, Saad Omer, Richard Martinello, Allison Nelson, Shelli Farhadian, Akiko Iwasaki, Charles Dela Cruz, Albert Ko, Nathan Grubaugh |
| EPI_ISL_431086 | hCoV-19/USA/CT-Yale-123/2020 | North America / USA / Connecticut | 2020-04-11 | Yale COVID-19 Biorepository | Grubaugh Lab - Yale School of Public Health | Joseph Fauver, Tara Alpert, Anderson Brito, Anne Wyllie, Chantal Vogels, Mary Petrone, Cole Jensen, Chaney Kalinich, Isabel Ott, Arnau Casanovas, Catherine Muenker, Adam Moore, Alice Lu, Maria Tokuyama, Patrick Wong, Peiwen Lu, Saad Omer, Richard Martinello, Allison Nelson, Shelli Farhadian, Akiko Iwasaki, Charles Dela Cruz, Albert Ko, Nathan Grubaugh |
| EPI_ISL_431089 | hCoV-19/USA/CT-Yale-127/2020 | North America / USA / Connecticut | 2020-04-13 | Yale COVID-19 Biorepository | Grubaugh Lab - Yale School of Public Health | Joseph Fauver, Tara Alpert, Anderson Brito, Anne Wyllie, Chantal Vogels, Mary Petrone, Cole Jensen, Chaney Kalinich, Isabel Ott, Arnau Casanovas, Catherine Muenker, Adam Moore, Alice Lu, Maria Tokuyama, Patrick Wong, Peiwen Lu, Saad Omer, Richard Martinello, Allison Nelson, Shelli Farhadian, Akiko Iwasaki, Charles Dela Cruz, Albert Ko, Nathan Grubaugh |
| EPI_ISL_431092 | hCoV-19/USA/CT-Yale-129/2020 | North America / USA / Connecticut | 2020-04-13 | Yale COVID-19 Biorepository | Grubaugh Lab - Yale School of Public Health | Joseph Fauver, Tara Alpert, Anderson Brito, Anne Wyllie, Chantal Vogels, Mary Petrone, Cole Jensen, Chaney Kalinich, Isabel Ott, Arnau Casanovas, Catherine Muenker, Adam Moore, Alice Lu, Maria Tokuyama, Patrick Wong, Peiwen Lu, Saad Omer, Richard Martinello, Allison Nelson, Shelli Farhadian, Akiko Iwasaki, Charles Dela Cruz, Albert Ko, Nathan Grubaugh |
| EPI_ISL_431096 | hCoV-19/USA/CT-Yale-134/2020 | North America / USA / Connecticut | 2020-04-16 | Yale COVID-19 Biorepository | Grubaugh Lab - Yale School of Public Health | Joseph Fauver, Tara Alpert, Anderson Brito, Anne Wyllie, Chantal Vogels, Mary Petrone, Cole Jensen, Chaney Kalinich, Isabel Ott, Arnau Casanovas, Catherine Muenker, Adam Moore, Alice Lu, Maria Tokuyama, Patrick Wong, Peiwen Lu, Saad Omer, Richard Martinello, Allison Nelson, Shelli Farhadian, Akiko Iwasaki, Charles Dela Cruz, Albert Ko, Nathan Grubaugh |
| EPI_ISL_431103 | hCoV-19/India/GMC-KN443/2020 | Asia / India / Telangana / Hyderabad | 2020-03-16 | Department of Microbiology, Gandhi Medical College and Hospital, Secendrabad, Hyderabad, India | Department of Microbiology, Gandhi Medical College and Hospital, Secendrabad, Hyderabad, India | Nagamani K, Muttineni Radhakrishna, Thrilok Chander B, Raja Rao M, Kalyani Putty, Ravikumar P, Sunitha P, Pankaj Singh D, Anand Kumar K, Amit A. Upadhyay, Steven E. Bosinger, Rama Amara |
| EPI_ISL_431117 | hCoV-19/India/GMC-TC469/2020 | Asia / India / Telangana / Hyderabad | 2020-03-20 | Department of Microbiology, Gandhi Medical College and Hospital, Secendrabad, Hyderabad, India | Department of Microbiology, Gandhi Medical College and Hospital, Secendrabad, Hyderabad, India | Thrilok Chander B, Muttineni Radhakrishna, Nagamani K, Raja Rao M, Kalyani Putty, Ravikumar P, Sunitha P, Pankaj Singh D, Anand Kumar K, Amit A. Upadhyay, Steven E.Bosinger, Rama Amara |
| EPI_ISL_431118 | hCoV-19/Fujian/IM3520001T/2020 | Asia / China / Fujian | 2020-03-13 | Fujian Center for Disease Control and Prevention | Fujian Center for Disease Control and Prevention | Lin Qi, Huang Zhimiao, Zhang Yanhua, Weng Yuwei |
| EPI_ISL_432357 | hCoV-19/Wales/PHWC-285A9/2020 | Europe / United Kingdom / Wales | 2020-03-30 | Wales Specialist Virology Centre | Public Health Wales Microbiology Cardiff | Catherine Moore, Johnathan Evans, Malorie Perry, Simon Cottrell, Alec Birchley, Alexander Adams, Amy Gaskin, Bree Gatica-Wilcox, Jason Coombes, Lauren Gilbert, Lee Graham, Nicole Pacchiarini, Sara Kumziene-Summerhayes, Sarah Taylor, Sophie Jones, Sara Rey, Matthew Bull, Joanne Watkins, Sally Corden, Tom Connor |
| EPI_ISL_432358 | hCoV-19/Wales/PHWC-2B263/2020 | Europe / United Kingdom / Wales | 2020-04-08 | Wales Specialist Virology Centre | Public Health Wales Microbiology Cardiff | Catherine Moore, Johnathan Evans, Malorie Perry, Simon Cottrell, Alec Birchley, Alexander Adams, Amy Gaskin, Bree Gatica-Wilcox, Jason Coombes, Lauren Gilbert, Lee Graham, Nicole Pacchiarini, Sara Kumziene-Summerhayes, Sarah Taylor, Sophie Jones, Sara Rey, Matthew Bull, Joanne Watkins, Sally Corden, Tom Connor |
| EPI_ISL_432359 | hCoV-19/Wales/PHWC-2863D/2020 | Europe / United Kingdom / Wales | 2020-03-30 | Wales Specialist Virology Centre | Public Health Wales Microbiology Cardiff | Catherine Moore, Johnathan Evans, Malorie Perry, Simon Cottrell, Alec Birchley, Alexander Adams, Amy Gaskin, Bree Gatica-Wilcox, Jason Coombes, Lauren Gilbert, Lee Graham, Nicole Pacchiarini, Sara Kumziene-Summerhayes, Sarah Taylor, Sophie Jones, Sara Rey, Matthew Bull, Joanne Watkins, Sally Corden, Tom Connor |
| EPI_ISL_432360 | hCoV-19/Wales/PHWC-2849D/2020 | Europe / United Kingdom / Wales | 2020-03-29 | Wales Specialist Virology Centre | Public Health Wales Microbiology Cardiff | Catherine Moore, Johnathan Evans, Malorie Perry, Simon Cottrell, Alec Birchley, Alexander Adams, Amy Gaskin, Bree Gatica-Wilcox, Jason Coombes, Lauren Gilbert, Lee Graham, Nicole Pacchiarini, Sara Kumziene-Summerhayes, Sarah Taylor, Sophie Jones, Sara Rey, Matthew Bull, Joanne Watkins, Sally Corden, Tom Connor |
| EPI_ISL_432361 | hCoV-19/Wales/PHWC-274F8/2020 | Europe / United Kingdom / Wales | 2020-03-30 | Wales Specialist Virology Centre | Public Health Wales Microbiology Cardiff | Catherine Moore, Johnathan Evans, Malorie Perry, Simon Cottrell, Alec Birchley, Alexander Adams, Amy Gaskin, Bree Gatica-Wilcox, Jason Coombes, Lauren Gilbert, Lee Graham, Nicole Pacchiarini, Sara Kumziene-Summerhayes, Sarah Taylor, Sophie Jones, Sara Rey, Matthew Bull, Joanne Watkins, Sally Corden, Tom Connor |
| EPI_ISL_432362 | hCoV-19/Wales/PHWC-27C3C/2020 | Europe / United Kingdom / Wales | 2020-03-31 | Wales Specialist Virology Centre | Public Health Wales Microbiology Cardiff | Catherine Moore, Johnathan Evans, Malorie Perry, Simon Cottrell, Alec Birchley, Alexander Adams, Amy Gaskin, Bree Gatica-Wilcox, Jason Coombes, Lauren Gilbert, Lee Graham, Nicole Pacchiarini, Sara Kumziene-Summerhayes, Sarah Taylor, Sophie Jones, Sara Rey, Matthew Bull, Joanne Watkins, Sally Corden, Tom Connor |
| EPI_ISL_432363 | hCoV-19/Wales/PHWC-29D46/2020 | Europe / United Kingdom / Wales | 2020-04-03 | Wales Specialist Virology Centre | Public Health Wales Microbiology Cardiff | Catherine Moore, Johnathan Evans, Malorie Perry, Simon Cottrell, Alec Birchley, Alexander Adams, Amy Gaskin, Bree Gatica-Wilcox, Jason Coombes, Lauren Gilbert, Lee Graham, Nicole Pacchiarini, Sara Kumziene-Summerhayes, Sarah Taylor, Sophie Jones, Sara Rey, Matthew Bull, Joanne Watkins, Sally Corden, Tom Connor |
| EPI_ISL_432364 | hCoV-19/Wales/PHWC-2AED9/2020 | Europe / United Kingdom / Wales | 2020-04-07 | Wales Specialist Virology Centre | Public Health Wales Microbiology Cardiff | Catherine Moore, Johnathan Evans, Malorie Perry, Simon Cottrell, Alec Birchley, Alexander Adams, Amy Gaskin, Bree Gatica-Wilcox, Jason Coombes, Lauren Gilbert, Lee Graham, Nicole Pacchiarini, Sara Kumziene-Summerhayes, Sarah Taylor, Sophie Jones, Sara Rey, Matthew Bull, Joanne Watkins, Sally Corden, Tom Connor |
| EPI_ISL_432365 | hCoV-19/Wales/PHWC-28819/2020 | Europe / United Kingdom / Wales | 2020-04-01 | Wales Specialist Virology Centre | Public Health Wales Microbiology Cardiff | Catherine Moore, Johnathan Evans, Malorie Perry, Simon Cottrell, Alec Birchley, Alexander Adams, Amy Gaskin, Bree Gatica-Wilcox, Jason Coombes, Lauren Gilbert, Lee Graham, Nicole Pacchiarini, Sara Kumziene-Summerhayes, Sarah Taylor, Sophie Jones, Sara Rey, Matthew Bull, Joanne Watkins, Sally Corden, Tom Connor |
| EPI_ISL_432366 | hCoV-19/Wales/PHWC-28688/2020 | Europe / United Kingdom / Wales | 2020-03-31 | Wales Specialist Virology Centre | Public Health Wales Microbiology Cardiff | Catherine Moore, Johnathan Evans, Malorie Perry, Simon Cottrell, Alec Birchley, Alexander Adams, Amy Gaskin, Bree Gatica-Wilcox, Jason Coombes, Lauren Gilbert, Lee Graham, Nicole Pacchiarini, Sara Kumziene-Summerhayes, Sarah Taylor, Sophie Jones, Sara Rey, Matthew Bull, Joanne Watkins, Sally Corden, Tom Connor |
| EPI_ISL_432367 | hCoV-19/Wales/PHWC-27B20/2020 | Europe / United Kingdom / Wales | 2020-03-30 | Wales Specialist Virology Centre | Public Health Wales Microbiology Cardiff | Catherine Moore, Johnathan Evans, Malorie Perry, Simon Cottrell, Alec Birchley, Alexander Adams, Amy Gaskin, Bree Gatica-Wilcox, Jason Coombes, Lauren Gilbert, Lee Graham, Nicole Pacchiarini, Sara Kumziene-Summerhayes, Sarah Taylor, Sophie Jones, Sara Rey, Matthew Bull, Joanne Watkins, Sally Corden, Tom Connor |
| EPI_ISL_432368 | hCoV-19/Wales/PHWC-29ECB/2020 | Europe / United Kingdom / Wales | 2020-04-04 | Wales Specialist Virology Centre | Public Health Wales Microbiology Cardiff | Catherine Moore, Johnathan Evans, Malorie Perry, Simon Cottrell, Alec Birchley, Alexander Adams, Amy Gaskin, Bree Gatica-Wilcox, Jason Coombes, Lauren Gilbert, Lee Graham, Nicole Pacchiarini, Sara Kumziene-Summerhayes, Sarah Taylor, Sophie Jones, Sara Rey, Matthew Bull, Joanne Watkins, Sally Corden, Tom Connor |
| EPI_ISL_432369 | hCoV-19/Wales/PHWC-27AD8/2020 | Europe / United Kingdom / Wales | 2020-03-30 | Wales Specialist Virology Centre | Public Health Wales Microbiology Cardiff | Catherine Moore, Johnathan Evans, Malorie Perry, Simon Cottrell, Alec Birchley, Alexander Adams, Amy Gaskin, Bree Gatica-Wilcox, Jason Coombes, Lauren Gilbert, Lee Graham, Nicole Pacchiarini, Sara Kumziene-Summerhayes, Sarah Taylor, Sophie Jones, Sara Rey, Matthew Bull, Joanne Watkins, Sally Corden, Tom Connor |
| EPI_ISL_432370 | hCoV-19/Wales/PHWC-28776/2020 | Europe / United Kingdom / Wales | 2020-03-31 | Wales Specialist Virology Centre | Public Health Wales Microbiology Cardiff | Catherine Moore, Johnathan Evans, Malorie Perry, Simon Cottrell, Alec Birchley, Alexander Adams, Amy Gaskin, Bree Gatica-Wilcox, Jason Coombes, Lauren Gilbert, Lee Graham, Nicole Pacchiarini, Sara Kumziene-Summerhayes, Sarah Taylor, Sophie Jones, Sara Rey, Matthew Bull, Joanne Watkins, Sally Corden, Tom Connor |
| EPI_ISL_432371 | hCoV-19/Wales/PHWC-288FB/2020 | Europe / United Kingdom / Wales | 2020-03-31 | Wales Specialist Virology Centre | Public Health Wales Microbiology Cardiff | Catherine Moore, Johnathan Evans, Malorie Perry, Simon Cottrell, Alec Birchley, Alexander Adams, Amy Gaskin, Bree Gatica-Wilcox, Jason Coombes, Lauren Gilbert, Lee Graham, Nicole Pacchiarini, Sara Kumziene-Summerhayes, Sarah Taylor, Sophie Jones, Sara Rey, Matthew Bull, Joanne Watkins, Sally Corden, Tom Connor |
| EPI_ISL_432372 | hCoV-19/Wales/PHWC-29D28/2020 | Europe / United Kingdom / Wales | 2020-04-02 | Wales Specialist Virology Centre | Public Health Wales Microbiology Cardiff | Catherine Moore, Johnathan Evans, Malorie Perry, Simon Cottrell, Alec Birchley, Alexander Adams, Amy Gaskin, Bree Gatica-Wilcox, Jason Coombes, Lauren Gilbert, Lee Graham, Nicole Pacchiarini, Sara Kumziene-Summerhayes, Sarah Taylor, Sophie Jones, Sara Rey, Matthew Bull, Joanne Watkins, Sally Corden, Tom Connor |
| EPI_ISL_432373 | hCoV-19/Wales/PHWC-2A02E/2020 | Europe / United Kingdom / Wales | 2020-04-03 | Wales Specialist Virology Centre | Public Health Wales Microbiology Cardiff | Catherine Moore, Johnathan Evans, Malorie Perry, Simon Cottrell, Alec Birchley, Alexander Adams, Amy Gaskin, Bree Gatica-Wilcox, Jason Coombes, Lauren Gilbert, Lee Graham, Nicole Pacchiarini, Sara Kumziene-Summerhayes, Sarah Taylor, Sophie Jones, Sara Rey, Matthew Bull, Joanne Watkins, Sally Corden, Tom Connor |
| EPI_ISL_432374 | hCoV-19/Wales/PHWC-27CC3/2020 | Europe / United Kingdom / Wales | 2020-04-01 | Wales Specialist Virology Centre | Public Health Wales Microbiology Cardiff | Catherine Moore, Johnathan Evans, Malorie Perry, Simon Cottrell, Alec Birchley, Alexander Adams, Amy Gaskin, Bree Gatica-Wilcox, Jason Coombes, Lauren Gilbert, Lee Graham, Nicole Pacchiarini, Sara Kumziene-Summerhayes, Sarah Taylor, Sophie Jones, Sara Rey, Matthew Bull, Joanne Watkins, Sally Corden, Tom Connor |
| EPI_ISL_432375 | hCoV-19/Wales/PHWC-27E45/2020 | Europe / United Kingdom / Wales | 2020 | Wales Specialist Virology Centre | Public Health Wales Microbiology Cardiff | Catherine Moore, Johnathan Evans, Malorie Perry, Simon Cottrell, Alec Birchley, Alexander Adams, Amy Gaskin, Bree Gatica-Wilcox, Jason Coombes, Lauren Gilbert, Lee Graham, Nicole Pacchiarini, Sara Kumziene-Summerhayes, Sarah Taylor, Sophie Jones, Sara Rey, Matthew Bull, Joanne Watkins, Sally Corden, Tom Connor |
| EPI_ISL_432376 | hCoV-19/Wales/PHWC-24877/2020 | Europe / United Kingdom / Wales | 2020-03 | Wales Specialist Virology Centre | Public Health Wales Microbiology Cardiff | Catherine Moore, Johnathan Evans, Malorie Perry, Simon Cottrell, Alec Birchley, Alexander Adams, Amy Gaskin, Bree Gatica-Wilcox, Jason Coombes, Lauren Gilbert, Lee Graham, Nicole Pacchiarini, Sara Kumziene-Summerhayes, Sarah Taylor, Sophie Jones, Sara Rey, Matthew Bull, Joanne Watkins, Sally Corden, Tom Connor |
| EPI_ISL_432377 | hCoV-19/Wales/PHWC-27F7F/2020 | Europe / United Kingdom / Wales | 2020-03-24 | Wales Specialist Virology Centre | Public Health Wales Microbiology Cardiff | Catherine Moore, Johnathan Evans, Malorie Perry, Simon Cottrell, Alec Birchley, Alexander Adams, Amy Gaskin, Bree Gatica-Wilcox, Jason Coombes, Lauren Gilbert, Lee Graham, Nicole Pacchiarini, Sara Kumziene-Summerhayes, Sarah Taylor, Sophie Jones, Sara Rey, Matthew Bull, Joanne Watkins, Sally Corden, Tom Connor |
| EPI_ISL_432378 | hCoV-19/Wales/PHWC-2B1C0/2020 | Europe / United Kingdom / Wales | 2020-04-08 | Wales Specialist Virology Centre | Public Health Wales Microbiology Cardiff | Catherine Moore, Johnathan Evans, Malorie Perry, Simon Cottrell, Alec Birchley, Alexander Adams, Amy Gaskin, Bree Gatica-Wilcox, Jason Coombes, Lauren Gilbert, Lee Graham, Nicole Pacchiarini, Sara Kumziene-Summerhayes, Sarah Taylor, Sophie Jones, Sara Rey, Matthew Bull, Joanne Watkins, Sally Corden, Tom Connor |
| EPI_ISL_432379 | hCoV-19/Wales/PHWC-28424/2020 | Europe / United Kingdom / Wales | 2020-03-27 | Wales Specialist Virology Centre | Public Health Wales Microbiology Cardiff | Catherine Moore, Johnathan Evans, Malorie Perry, Simon Cottrell, Alec Birchley, Alexander Adams, Amy Gaskin, Bree Gatica-Wilcox, Jason Coombes, Lauren Gilbert, Lee Graham, Nicole Pacchiarini, Sara Kumziene-Summerhayes, Sarah Taylor, Sophie Jones, Sara Rey, Matthew Bull, Joanne Watkins, Sally Corden, Tom Connor |
| EPI_ISL_432380 | hCoV-19/Wales/PHWC-2781A/2020 | Europe / United Kingdom / Wales | 2020-04-01 | Wales Specialist Virology Centre | Public Health Wales Microbiology Cardiff | Catherine Moore, Johnathan Evans, Malorie Perry, Simon Cottrell, Alec Birchley, Alexander Adams, Amy Gaskin, Bree Gatica-Wilcox, Jason Coombes, Lauren Gilbert, Lee Graham, Nicole Pacchiarini, Sara Kumziene-Summerhayes, Sarah Taylor, Sophie Jones, Sara Rey, Matthew Bull, Joanne Watkins, Sally Corden, Tom Connor |
| EPI_ISL_432381 | hCoV-19/Wales/PHWC-276D4/2020 | Europe / United Kingdom / Wales | 2020-03-31 | Wales Specialist Virology Centre | Public Health Wales Microbiology Cardiff | Catherine Moore, Johnathan Evans, Malorie Perry, Simon Cottrell, Alec Birchley, Alexander Adams, Amy Gaskin, Bree Gatica-Wilcox, Jason Coombes, Lauren Gilbert, Lee Graham, Nicole Pacchiarini, Sara Kumziene-Summerhayes, Sarah Taylor, Sophie Jones, Sara Rey, Matthew Bull, Joanne Watkins, Sally Corden, Tom Connor |
| EPI_ISL_432382 | hCoV-19/Wales/PHWC-2B245/2020 | Europe / United Kingdom / Wales | 2020-04-08 | Wales Specialist Virology Centre | Public Health Wales Microbiology Cardiff | Catherine Moore, Johnathan Evans, Malorie Perry, Simon Cottrell, Alec Birchley, Alexander Adams, Amy Gaskin, Bree Gatica-Wilcox, Jason Coombes, Lauren Gilbert, Lee Graham, Nicole Pacchiarini, Sara Kumziene-Summerhayes, Sarah Taylor, Sophie Jones, Sara Rey, Matthew Bull, Joanne Watkins, Sally Corden, Tom Connor |
| EPI_ISL_432383 | hCoV-19/Wales/PHWC-29C67/2020 | Europe / United Kingdom / Wales | 2020-04-05 | Wales Specialist Virology Centre | Public Health Wales Microbiology Cardiff | Catherine Moore, Johnathan Evans, Malorie Perry, Simon Cottrell, Alec Birchley, Alexander Adams, Amy Gaskin, Bree Gatica-Wilcox, Jason Coombes, Lauren Gilbert, Lee Graham, Nicole Pacchiarini, Sara Kumziene-Summerhayes, Sarah Taylor, Sophie Jones, Sara Rey, Matthew Bull, Joanne Watkins, Sally Corden, Tom Connor |
| EPI_ISL_432384 | hCoV-19/Wales/PHWC-27249/2020 | Europe / United Kingdom / Wales | 2020-03-30 | Wales Specialist Virology Centre | Public Health Wales Microbiology Cardiff | Catherine Moore, Johnathan Evans, Malorie Perry, Simon Cottrell, Alec Birchley, Alexander Adams, Amy Gaskin, Bree Gatica-Wilcox, Jason Coombes, Lauren Gilbert, Lee Graham, Nicole Pacchiarini, Sara Kumziene-Summerhayes, Sarah Taylor, Sophie Jones, Sara Rey, Matthew Bull, Joanne Watkins, Sally Corden, Tom Connor |
| EPI_ISL_432385 | hCoV-19/Wales/PHWC-285F4/2020 | Europe / United Kingdom / Wales | 2020-04-01 | Wales Specialist Virology Centre | Public Health Wales Microbiology Cardiff | Catherine Moore, Johnathan Evans, Malorie Perry, Simon Cottrell, Alec Birchley, Alexander Adams, Amy Gaskin, Bree Gatica-Wilcox, Jason Coombes, Lauren Gilbert, Lee Graham, Nicole Pacchiarini, Sara Kumziene-Summerhayes, Sarah Taylor, Sophie Jones, Sara Rey, Matthew Bull, Joanne Watkins, Sally Corden, Tom Connor |
| EPI_ISL_432386 | hCoV-19/Wales/PHWC-252E1/2020 | Europe / United Kingdom / Wales | 2020-03-22 | Wales Specialist Virology Centre | Public Health Wales Microbiology Cardiff | Catherine Moore, Johnathan Evans, Malorie Perry, Simon Cottrell, Alec Birchley, Alexander Adams, Amy Gaskin, Bree Gatica-Wilcox, Jason Coombes, Lauren Gilbert, Lee Graham, Nicole Pacchiarini, Sara Kumziene-Summerhayes, Sarah Taylor, Sophie Jones, Sara Rey, Matthew Bull, Joanne Watkins, Sally Corden, Tom Connor |
| EPI_ISL_432387 | hCoV-19/Wales/PHWC-28433/2020 | Europe / United Kingdom / Wales | 2020-03-27 | Wales Specialist Virology Centre | Public Health Wales Microbiology Cardiff | Catherine Moore, Johnathan Evans, Malorie Perry, Simon Cottrell, Alec Birchley, Alexander Adams, Amy Gaskin, Bree Gatica-Wilcox, Jason Coombes, Lauren Gilbert, Lee Graham, Nicole Pacchiarini, Sara Kumziene-Summerhayes, Sarah Taylor, Sophie Jones, Sara Rey, Matthew Bull, Joanne Watkins, Sally Corden, Tom Connor |
| EPI_ISL_432388 | hCoV-19/Wales/PHWC-29CA3/2020 | Europe / United Kingdom / Wales | 2020-04-05 | Wales Specialist Virology Centre | Public Health Wales Microbiology Cardiff | Catherine Moore, Johnathan Evans, Malorie Perry, Simon Cottrell, Alec Birchley, Alexander Adams, Amy Gaskin, Bree Gatica-Wilcox, Jason Coombes, Lauren Gilbert, Lee Graham, Nicole Pacchiarini, Sara Kumziene-Summerhayes, Sarah Taylor, Sophie Jones, Sara Rey, Matthew Bull, Joanne Watkins, Sally Corden, Tom Connor |
| EPI_ISL_432389 | hCoV-19/Wales/PHWC-278B0/2020 | Europe / United Kingdom / Wales | 2020-04-01 | Wales Specialist Virology Centre | Public Health Wales Microbiology Cardiff | Catherine Moore, Johnathan Evans, Malorie Perry, Simon Cottrell, Alec Birchley, Alexander Adams, Amy Gaskin, Bree Gatica-Wilcox, Jason Coombes, Lauren Gilbert, Lee Graham, Nicole Pacchiarini, Sara Kumziene-Summerhayes, Sarah Taylor, Sophie Jones, Sara Rey, Matthew Bull, Joanne Watkins, Sally Corden, Tom Connor |
| EPI_ISL_432390 | hCoV-19/Wales/PHWC-271A6/2020 | Europe / United Kingdom / Wales | 2020-03-30 | Wales Specialist Virology Centre | Public Health Wales Microbiology Cardiff | Catherine Moore, Johnathan Evans, Malorie Perry, Simon Cottrell, Alec Birchley, Alexander Adams, Amy Gaskin, Bree Gatica-Wilcox, Jason Coombes, Lauren Gilbert, Lee Graham, Nicole Pacchiarini, Sara Kumziene-Summerhayes, Sarah Taylor, Sophie Jones, Sara Rey, Matthew Bull, Joanne Watkins, Sally Corden, Tom Connor |
| EPI_ISL_432458 | hCoV-19/England/SHEF-C8C2E/2020 | Europe / United Kingdom / England | 2020-03-28 | Virology Department, Sheffield Teaching Hospitals NHS Foundation Trust / Virology Department, Sheffield Teaching Hospitals NHS Foundation Trust | COVID-19 Genomics UK (COG-UK) Consortium | Thushan de Silva, Matthew Parker,Adri Angyal, Rebecca Brown, Luke Green, Rachel Tucker, Paul Parsons, Danielle Groves, Alex Keeley, Dave Partridge, Matthew Wyles, Benjamin Lindsey, Mehmet Yavuz, Mohammad Raza, Cariad Evans |
| EPI_ISL_432466 | hCoV-19/England/SHEF-C914C/2020 | Europe / United Kingdom / England | 2020-04-01 | Virology Department, Sheffield Teaching Hospitals NHS Foundation Trust / Virology Department, Sheffield Teaching Hospitals NHS Foundation Trust | COVID-19 Genomics UK (COG-UK) Consortium | Thushan de Silva, Matthew Parker,Adri Angyal, Rebecca Brown, Luke Green, Rachel Tucker, Paul Parsons, Danielle Groves, Alex Keeley, Dave Partridge, Matthew Wyles, Benjamin Lindsey, Mehmet Yavuz, Mohammad Raza, Cariad Evans |
| EPI_ISL_432900 | hCoV-19/England/NOTT-10DF8D/2020 | Europe / United Kingdom / England | 2020-03-02 | Queens Medical Centre, Clinical Microbiology Department / DeepSeq Nottingham | COVID-19 Genomics UK (COG-UK) Consortium | Gemma Clark, Wendy Smith, Manjinder Khakh, Hannah Howson-Wells, Jonathan Ball, Patrick McClure, Joseph Chappell, Theocharis Tsoleridis, Nadine Holmes, Matthew Carlisle, Christopher Moore, Fei Sang, Johnny Debebe, Victoria Wright, Matthew Loose |
| EPI_ISL_432916 | hCoV-19/England/NOTT-10ED28/2020 | Europe / United Kingdom / England | 2020-03-28 | Queens Medical Centre, Clinical Microbiology Department / DeepSeq Nottingham | COVID-19 Genomics UK (COG-UK) Consortium | Gemma Clark, Wendy Smith, Manjinder Khakh, Hannah Howson-Wells, Jonathan Ball, Patrick McClure, Joseph Chappell, Theocharis Tsoleridis, Nadine Holmes, Matthew Carlisle, Christopher Moore, Fei Sang, Johnny Debebe, Victoria Wright, Matthew Loose |
| EPI_ISL_432921 | hCoV-19/England/NOTT-10ED73/2020 | Europe / United Kingdom / England | 2020-03-28 | Queens Medical Centre, Clinical Microbiology Department / DeepSeq Nottingham | COVID-19 Genomics UK (COG-UK) Consortium | Gemma Clark, Wendy Smith, Manjinder Khakh, Hannah Howson-Wells, Jonathan Ball, Patrick McClure, Joseph Chappell, Theocharis Tsoleridis, Nadine Holmes, Matthew Carlisle, Christopher Moore, Fei Sang, Johnny Debebe, Victoria Wright, Matthew Loose |
| EPI_ISL_432925 | hCoV-19/England/NOTT-10EDCE/2020 | Europe / United Kingdom / England | 2020-03-28 | Queens Medical Centre, Clinical Microbiology Department / DeepSeq Nottingham | COVID-19 Genomics UK (COG-UK) Consortium | Gemma Clark, Wendy Smith, Manjinder Khakh, Hannah Howson-Wells, Jonathan Ball, Patrick McClure, Joseph Chappell, Theocharis Tsoleridis, Nadine Holmes, Matthew Carlisle, Christopher Moore, Fei Sang, Johnny Debebe, Victoria Wright, Matthew Loose |
| EPI_ISL_432930 | hCoV-19/England/NOTT-10EE16/2020 | Europe / United Kingdom / England | 2020-03-28 | Queens Medical Centre, Clinical Microbiology Department / DeepSeq Nottingham | COVID-19 Genomics UK (COG-UK) Consortium | Gemma Clark, Wendy Smith, Manjinder Khakh, Hannah Howson-Wells, Jonathan Ball, Patrick McClure, Joseph Chappell, Theocharis Tsoleridis, Nadine Holmes, Matthew Carlisle, Christopher Moore, Fei Sang, Johnny Debebe, Victoria Wright, Matthew Loose |
| EPI_ISL_432935 | hCoV-19/England/NOTT-10EE8F/2020 | Europe / United Kingdom / England | 2020-03-28 | Queens Medical Centre, Clinical Microbiology Department / DeepSeq Nottingham | COVID-19 Genomics UK (COG-UK) Consortium | Gemma Clark, Wendy Smith, Manjinder Khakh, Hannah Howson-Wells, Jonathan Ball, Patrick McClure, Joseph Chappell, Theocharis Tsoleridis, Nadine Holmes, Matthew Carlisle, Christopher Moore, Fei Sang, Johnny Debebe, Victoria Wright, Matthew Loose |
| EPI_ISL_432938 | hCoV-19/England/NOTT-10EEBC/2020 | Europe / United Kingdom / England | 2020-03-28 | Queens Medical Centre, Clinical Microbiology Department / DeepSeq Nottingham | COVID-19 Genomics UK (COG-UK) Consortium | Gemma Clark, Wendy Smith, Manjinder Khakh, Hannah Howson-Wells, Jonathan Ball, Patrick McClure, Joseph Chappell, Theocharis Tsoleridis, Nadine Holmes, Matthew Carlisle, Christopher Moore, Fei Sang, Johnny Debebe, Victoria Wright, Matthew Loose |
| EPI_ISL_432942 | hCoV-19/England/NOTT-10EF13/2020 | Europe / United Kingdom / England | 2020-03-30 | Queens Medical Centre, Clinical Microbiology Department / DeepSeq Nottingham | COVID-19 Genomics UK (COG-UK) Consortium | Gemma Clark, Wendy Smith, Manjinder Khakh, Hannah Howson-Wells, Jonathan Ball, Patrick McClure, Joseph Chappell, Theocharis Tsoleridis, Nadine Holmes, Matthew Carlisle, Christopher Moore, Fei Sang, Johnny Debebe, Victoria Wright, Matthew Loose |
| EPI_ISL_432951 | hCoV-19/England/NOTT-10EFB9/2020 | Europe / United Kingdom / England | 2020-03-30 | Queens Medical Centre, Clinical Microbiology Department / DeepSeq Nottingham | COVID-19 Genomics UK (COG-UK) Consortium | Gemma Clark, Wendy Smith, Manjinder Khakh, Hannah Howson-Wells, Jonathan Ball, Patrick McClure, Joseph Chappell, Theocharis Tsoleridis, Nadine Holmes, Matthew Carlisle, Christopher Moore, Fei Sang, Johnny Debebe, Victoria Wright, Matthew Loose |
| EPI_ISL_432954 | hCoV-19/England/NOTT-10EFF5/2020 | Europe / United Kingdom / England | 2020-03-30 | Queens Medical Centre, Clinical Microbiology Department / DeepSeq Nottingham | COVID-19 Genomics UK (COG-UK) Consortium | Gemma Clark, Wendy Smith, Manjinder Khakh, Hannah Howson-Wells, Jonathan Ball, Patrick McClure, Joseph Chappell, Theocharis Tsoleridis, Nadine Holmes, Matthew Carlisle, Christopher Moore, Fei Sang, Johnny Debebe, Victoria Wright, Matthew Loose |
| EPI_ISL_432955 | hCoV-19/England/NOTT-10F000/2020 | Europe / United Kingdom / England | 2020-03-30 | Queens Medical Centre, Clinical Microbiology Department / DeepSeq Nottingham | COVID-19 Genomics UK (COG-UK) Consortium | Gemma Clark, Wendy Smith, Manjinder Khakh, Hannah Howson-Wells, Jonathan Ball, Patrick McClure, Joseph Chappell, Theocharis Tsoleridis, Nadine Holmes, Matthew Carlisle, Christopher Moore, Fei Sang, Johnny Debebe, Victoria Wright, Matthew Loose |
| EPI_ISL_432956 | hCoV-19/England/NOTT-10F01F/2020 | Europe / United Kingdom / England | 2020-03-30 | Queens Medical Centre, Clinical Microbiology Department / DeepSeq Nottingham | COVID-19 Genomics UK (COG-UK) Consortium | Gemma Clark, Wendy Smith, Manjinder Khakh, Hannah Howson-Wells, Jonathan Ball, Patrick McClure, Joseph Chappell, Theocharis Tsoleridis, Nadine Holmes, Matthew Carlisle, Christopher Moore, Fei Sang, Johnny Debebe, Victoria Wright, Matthew Loose |
| EPI_ISL_432961 | hCoV-19/England/NOTT-10F0B5/2020 | Europe / United Kingdom / England | 2020-03-30 | Queens Medical Centre, Clinical Microbiology Department / DeepSeq Nottingham | COVID-19 Genomics UK (COG-UK) Consortium | Gemma Clark, Wendy Smith, Manjinder Khakh, Hannah Howson-Wells, Jonathan Ball, Patrick McClure, Joseph Chappell, Theocharis Tsoleridis, Nadine Holmes, Matthew Carlisle, Christopher Moore, Fei Sang, Johnny Debebe, Victoria Wright, Matthew Loose |
| EPI_ISL_432963 | hCoV-19/England/NOTT-10F0E2/2020 | Europe / United Kingdom / England | 2020-03-31 | Queens Medical Centre, Clinical Microbiology Department / DeepSeq Nottingham | COVID-19 Genomics UK (COG-UK) Consortium | Gemma Clark, Wendy Smith, Manjinder Khakh, Hannah Howson-Wells, Jonathan Ball, Patrick McClure, Joseph Chappell, Theocharis Tsoleridis, Nadine Holmes, Matthew Carlisle, Christopher Moore, Fei Sang, Johnny Debebe, Victoria Wright, Matthew Loose |
| EPI_ISL_432965 | hCoV-19/England/NOTT-10F10D/2020 | Europe / United Kingdom / England | 2020-03-31 | Queens Medical Centre, Clinical Microbiology Department / DeepSeq Nottingham | COVID-19 Genomics UK (COG-UK) Consortium | Gemma Clark, Wendy Smith, Manjinder Khakh, Hannah Howson-Wells, Jonathan Ball, Patrick McClure, Joseph Chappell, Theocharis Tsoleridis, Nadine Holmes, Matthew Carlisle, Christopher Moore, Fei Sang, Johnny Debebe, Victoria Wright, Matthew Loose |
| EPI_ISL_432967 | hCoV-19/England/NOTT-10F12B/2020 | Europe / United Kingdom / England | 2020-03-31 | Queens Medical Centre, Clinical Microbiology Department / DeepSeq Nottingham | COVID-19 Genomics UK (COG-UK) Consortium | Gemma Clark, Wendy Smith, Manjinder Khakh, Hannah Howson-Wells, Jonathan Ball, Patrick McClure, Joseph Chappell, Theocharis Tsoleridis, Nadine Holmes, Matthew Carlisle, Christopher Moore, Fei Sang, Johnny Debebe, Victoria Wright, Matthew Loose |
| EPI_ISL_432970 | hCoV-19/England/NOTT-10F176/2020 | Europe / United Kingdom / England | 2020-03-31 | Queens Medical Centre, Clinical Microbiology Department / DeepSeq Nottingham | COVID-19 Genomics UK (COG-UK) Consortium | Gemma Clark, Wendy Smith, Manjinder Khakh, Hannah Howson-Wells, Jonathan Ball, Patrick McClure, Joseph Chappell, Theocharis Tsoleridis, Nadine Holmes, Matthew Carlisle, Christopher Moore, Fei Sang, Johnny Debebe, Victoria Wright, Matthew Loose |
| EPI_ISL_432979 | hCoV-19/England/NOTT-10F237/2020 | Europe / United Kingdom / England | 2020-04-01 | Queens Medical Centre, Clinical Microbiology Department / DeepSeq Nottingham | COVID-19 Genomics UK (COG-UK) Consortium | Gemma Clark, Wendy Smith, Manjinder Khakh, Hannah Howson-Wells, Jonathan Ball, Patrick McClure, Joseph Chappell, Theocharis Tsoleridis, Nadine Holmes, Matthew Carlisle, Christopher Moore, Fei Sang, Johnny Debebe, Victoria Wright, Matthew Loose |
| EPI_ISL_432984 | hCoV-19/England/NOTT-10F282/2020 | Europe / United Kingdom / England | 2020-04-01 | Queens Medical Centre, Clinical Microbiology Department / DeepSeq Nottingham | COVID-19 Genomics UK (COG-UK) Consortium | Gemma Clark, Wendy Smith, Manjinder Khakh, Hannah Howson-Wells, Jonathan Ball, Patrick McClure, Joseph Chappell, Theocharis Tsoleridis, Nadine Holmes, Matthew Carlisle, Christopher Moore, Fei Sang, Johnny Debebe, Victoria Wright, Matthew Loose |

| | | | | | | |
|---|---|---|---|---|---|---|
| EPI_ISL_432988 | hCoV-19/England/NOTT-10F2DD/2020 | Europe / United Kingdom / England | 2020-04-01 | Queens Medical Centre, Clinical Microbiology Department / DeepSeq Nottingham | COVID-19 Genomics UK (COG-UK) Consortium | Gemma Clark, Wendy Smith, Manjinder Khakh, Hannah Howson-Wells, Jonathan Ball, Patrick McClure, Joseph Chappell, Theocharis Tsoleridis, Nadine Holmes, Matthew Carlisle, Christopher Moore, Fei Sang, Johnny Debebe, Victoria Wright, Matthew Loose |
| EPI_ISL_432990 | hCoV-19/England/NOTT-10F2FB/2020 | Europe / United Kingdom / England | 2020-04-01 | Queens Medical Centre, Clinical Microbiology Department / DeepSeq Nottingham | COVID-19 Genomics UK (COG-UK) Consortium | Gemma Clark, Wendy Smith, Manjinder Khakh, Hannah Howson-Wells, Jonathan Ball, Patrick McClure, Joseph Chappell, Theocharis Tsoleridis, Nadine Holmes, Matthew Carlisle, Christopher Moore, Fei Sang, Johnny Debebe, Victoria Wright, Matthew Loose |
| EPI_ISL_432992 | hCoV-19/England/NOTT-10F334/2020 | Europe / United Kingdom / England | 2020-04-01 | Queens Medical Centre, Clinical Microbiology Department / DeepSeq Nottingham | COVID-19 Genomics UK (COG-UK) Consortium | Gemma Clark, Wendy Smith, Manjinder Khakh, Hannah Howson-Wells, Jonathan Ball, Patrick McClure, Joseph Chappell, Theocharis Tsoleridis, Nadine Holmes, Matthew Carlisle, Christopher Moore, Fei Sang, Johnny Debebe, Victoria Wright, Matthew Loose |
| EPI_ISL_432993 | hCoV-19/England/NOTT-10F343/2020 | Europe / United Kingdom / England | 2020-04-02 | Queens Medical Centre, Clinical Microbiology Department / DeepSeq Nottingham | COVID-19 Genomics UK (COG-UK) Consortium | Gemma Clark, Wendy Smith, Manjinder Khakh, Hannah Howson-Wells, Jonathan Ball, Patrick McClure, Joseph Chappell, Theocharis Tsoleridis, Nadine Holmes, Matthew Carlisle, Christopher Moore, Fei Sang, Johnny Debebe, Victoria Wright, Matthew Loose |
| EPI_ISL_432998 | hCoV-19/England/NOTT-10F3CB/2020 | Europe / United Kingdom / England | 2020-04-02 | Queens Medical Centre, Clinical Microbiology Department / DeepSeq Nottingham | COVID-19 Genomics UK (COG-UK) Consortium | Gemma Clark, Wendy Smith, Manjinder Khakh, Hannah Howson-Wells, Jonathan Ball, Patrick McClure, Joseph Chappell, Theocharis Tsoleridis, Nadine Holmes, Matthew Carlisle, Christopher Moore, Fei Sang, Johnny Debebe, Victoria Wright, Matthew Loose |
| EPI_ISL_433326 | hCoV-19/Scotland/CVR694/2020 | Europe / United Kingdom / Scotland | 2020-03-25 | West of Scotland Specialist Virology Centre, NHSGGC / MRC-University of Glasgow Centre for Virus Research | COVID-19 Genomics UK (COG-UK) Consortium | Ana da Silva Filipe, Natasha Johnson, Kathy Smollett, Daniel Mair, Stephen Carmichael, Lily Tong, Jenna Nichols, Elihu Aranday-Cortes, Kirstyn Brunker, Yasmin Parr, Kyriaki Nomikou; Sarah McDonald, Marc Niebel, Patawee Asamaphan; Richard Orton, Joseph Hughes, Sreenu Vattipally, David L Robertson; Alasdair MacLean, Rory Gunson; Kathy Li, Natasha Jesudason, Rajiv Shah, James Shepherd, Antonia Ho, Emma Thomson |
| EPI_ISL_433327 | hCoV-19/Scotland/CVR702/2020 | Europe / United Kingdom / Scotland | 2020-03-25 | West of Scotland Specialist Virology Centre, NHSGGC / MRC-University of Glasgow Centre for Virus Research | COVID-19 Genomics UK (COG-UK) Consortium | Ana da Silva Filipe, Natasha Johnson, Kathy Smollett, Daniel Mair, Stephen Carmichael, Lily Tong, Jenna Nichols, Elihu Aranday-Cortes, Kirstyn Brunker, Yasmin Parr, Kyriaki Nomikou; Sarah McDonald, Marc Niebel, Patawee Asamaphan; Richard Orton, Joseph Hughes, Sreenu Vattipally, David L Robertson; Alasdair MacLean, Rory Gunson; Kathy Li, Natasha Jesudason, Rajiv Shah, James Shepherd, Antonia Ho, Emma Thomson |
| EPI_ISL_433329 | hCoV-19/Scotland/CVR705/2020 | Europe / United Kingdom / Scotland | 2020-03-26 | West of Scotland Specialist Virology Centre, NHSGGC / MRC-University of Glasgow Centre for Virus Research | COVID-19 Genomics UK (COG-UK) Consortium | Ana da Silva Filipe, Natasha Johnson, Kathy Smollett, Daniel Mair, Stephen Carmichael, Lily Tong, Jenna Nichols, Elihu Aranday-Cortes, Kirstyn Brunker, Yasmin Parr, Kyriaki Nomikou; Sarah McDonald, Marc Niebel, Patawee Asamaphan; Richard Orton, Joseph Hughes, Sreenu Vattipally, David L Robertson; Alasdair MacLean, Rory Gunson; Kathy Li, Natasha Jesudason, Rajiv Shah, James Shepherd, Antonia Ho, Emma Thomson |
| EPI_ISL_433330 | hCoV-19/Scotland/CVR712/2020 | Europe / United Kingdom / Scotland | 2020-03-26 | West of Scotland Specialist Virology Centre, NHSGGC / MRC-University of Glasgow Centre for Virus Research | COVID-19 Genomics UK (COG-UK) Consortium | Ana da Silva Filipe, Natasha Johnson, Kathy Smollett, Daniel Mair, Stephen Carmichael, Lily Tong, Jenna Nichols, Elihu Aranday-Cortes, Kirstyn Brunker, Yasmin Parr, Kyriaki Nomikou; Sarah McDonald, Marc Niebel, Patawee Asamaphan; Richard Orton, Joseph Hughes, Sreenu Vattipally, David L Robertson; Alasdair MacLean, Rory Gunson; Kathy Li, Natasha Jesudason, Rajiv Shah, James Shepherd, Antonia Ho, Emma Thomson |
| EPI_ISL_433333 | hCoV-19/Scotland/CVR749/2020 | Europe / United Kingdom / Scotland | 2020-03-26 | West of Scotland Specialist Virology Centre, NHSGGC / MRC-University of Glasgow Centre for Virus Research | COVID-19 Genomics UK (COG-UK) Consortium | Ana da Silva Filipe, Natasha Johnson, Kathy Smollett, Daniel Mair, Stephen Carmichael, Lily Tong, Jenna Nichols, Elihu Aranday-Cortes, Kirstyn Brunker, Yasmin Parr, Kyriaki Nomikou; Sarah McDonald, Marc Niebel, Patawee Asamaphan; Richard Orton, Joseph Hughes, Sreenu Vattipally, David L Robertson; Alasdair MacLean, Rory Gunson; Kathy Li, Natasha Jesudason, Rajiv Shah, James Shepherd, Antonia Ho, Emma Thomson |
| EPI_ISL_433334 | hCoV-19/Scotland/CVR752/2020 | Europe / United Kingdom / Scotland | 2020-03-26 | West of Scotland Specialist Virology Centre, NHSGGC / MRC-University of Glasgow Centre for Virus Research | COVID-19 Genomics UK (COG-UK) Consortium | Ana da Silva Filipe, Natasha Johnson, Kathy Smollett, Daniel Mair, Stephen Carmichael, Lily Tong, Jenna Nichols, Elihu Aranday-Cortes, Kirstyn Brunker, Yasmin Parr, Kyriaki Nomikou; Sarah McDonald, Marc Niebel, Patawee Asamaphan; Richard Orton, Joseph Hughes, Sreenu Vattipally, David L Robertson; Alasdair MacLean, Rory Gunson; Kathy Li, Natasha Jesudason, Rajiv Shah, James Shepherd, Antonia Ho, Emma Thomson |
| EPI_ISL_433335 | hCoV-19/Scotland/CVR754/2020 | Europe / United Kingdom / Scotland | 2020-03-26 | West of Scotland Specialist Virology Centre, NHSGGC / MRC-University of Glasgow Centre for Virus Research | COVID-19 Genomics UK (COG-UK) Consortium | Ana da Silva Filipe, Natasha Johnson, Kathy Smollett, Daniel Mair, Stephen Carmichael, Lily Tong, Jenna Nichols, Elihu Aranday-Cortes, Kirstyn Brunker, Yasmin Parr, Kyriaki Nomikou; Sarah McDonald, Marc Niebel, Patawee Asamaphan; Richard Orton, Joseph Hughes, Sreenu Vattipally, David L Robertson; Alasdair MacLean, Rory Gunson; Kathy Li, Natasha Jesudason, Rajiv Shah, James Shepherd, Antonia Ho, Emma Thomson |
| EPI_ISL_433338 | hCoV-19/Scotland/CVR780/2020 | Europe / United Kingdom / Scotland | 2020-03-27 | West of Scotland Specialist Virology Centre, NHSGGC / MRC-University of Glasgow Centre for Virus Research | COVID-19 Genomics UK (COG-UK) Consortium | Ana da Silva Filipe, Natasha Johnson, Kathy Smollett, Daniel Mair, Stephen Carmichael, Lily Tong, Jenna Nichols, Elihu Aranday-Cortes, Kirstyn Brunker, Yasmin Parr, Kyriaki Nomikou; Sarah McDonald, Marc Niebel, Patawee Asamaphan; Richard Orton, Joseph Hughes, Sreenu Vattipally, David L Robertson; Alasdair MacLean, Rory Gunson; Kathy Li, Natasha Jesudason, Rajiv Shah, James Shepherd, Antonia Ho, Emma Thomson |
| EPI_ISL_433339 | hCoV-19/Scotland/CVR785/2020 | Europe / United Kingdom / Scotland | 2020-03-26 | West of Scotland Specialist Virology Centre, NHSGGC / MRC-University of Glasgow Centre for Virus Research | COVID-19 Genomics UK (COG-UK) Consortium | Ana da Silva Filipe, Natasha Johnson, Kathy Smollett, Daniel Mair, Stephen Carmichael, Lily Tong, Jenna Nichols, Elihu Aranday-Cortes, Kirstyn Brunker, Yasmin Parr, Kyriaki Nomikou; Sarah McDonald, Marc Niebel, Patawee Asamaphan; Richard Orton, Joseph Hughes, Sreenu Vattipally, David L Robertson; Alasdair MacLean, Rory Gunson; Kathy Li, Natasha Jesudason, Rajiv Shah, James Shepherd, Antonia Ho, Emma Thomson |
| EPI_ISL_433340 | hCoV-19/Scotland/CVR788/2020 | Europe / United Kingdom / Scotland | 2020-03-26 | West of Scotland Specialist Virology Centre, NHSGGC / MRC-University of Glasgow Centre for Virus Research | COVID-19 Genomics UK (COG-UK) Consortium | Ana da Silva Filipe, Natasha Johnson, Kathy Smollett, Daniel Mair, Stephen Carmichael, Lily Tong, Jenna Nichols, Elihu Aranday-Cortes, Kirstyn Brunker, Yasmin Parr, Kyriaki Nomikou; Sarah McDonald, Marc Niebel, Patawee Asamaphan; Richard Orton, Joseph Hughes, Sreenu Vattipally, David L Robertson; Alasdair MacLean, Rory Gunson; Kathy Li, Natasha Jesudason, Rajiv Shah, James Shepherd, Antonia Ho, Emma Thomson |
| EPI_ISL_433341 | hCoV-19/Scotland/CVR796/2020 | Europe / United Kingdom / Scotland | 2020-03-26 | West of Scotland Specialist Virology Centre, NHSGGC / MRC-University of Glasgow Centre for Virus Research | COVID-19 Genomics UK (COG-UK) Consortium | Ana da Silva Filipe, Natasha Johnson, Kathy Smollett, Daniel Mair, Stephen Carmichael, Lily Tong, Jenna Nichols, Elihu Aranday-Cortes, Kirstyn Brunker, Yasmin Parr, Kyriaki Nomikou; Sarah McDonald, Marc Niebel, Patawee Asamaphan; Richard Orton, Joseph Hughes, Sreenu Vattipally, David L Robertson; Alasdair MacLean, Rory Gunson; Kathy Li, Natasha Jesudason, Rajiv Shah, James Shepherd, Antonia Ho, Emma Thomson |
| EPI_ISL_433342 | hCoV-19/Scotland/CVR800/2020 | Europe / United Kingdom / Scotland | 2020-03-27 | West of Scotland Specialist Virology Centre, NHSGGC / MRC-University of Glasgow Centre for Virus Research | COVID-19 Genomics UK (COG-UK) Consortium | Ana da Silva Filipe, Natasha Johnson, Kathy Smollett, Daniel Mair, Stephen Carmichael, Lily Tong, Jenna Nichols, Elihu Aranday-Cortes, Kirstyn Brunker, Yasmin Parr, Kyriaki Nomikou; Sarah McDonald, Marc Niebel, Patawee Asamaphan; Richard Orton, Joseph Hughes, Sreenu Vattipally, David L Robertson; Alasdair MacLean, Rory Gunson; Kathy Li, Natasha Jesudason, Rajiv Shah, James Shepherd, Antonia Ho, Emma Thomson |
| EPI_ISL_433344 | hCoV-19/Scotland/CVR803/2020 | Europe / United Kingdom / Scotland | 2020-03-27 | West of Scotland Specialist Virology Centre, NHSGGC / MRC-University of Glasgow Centre for Virus Research | COVID-19 Genomics UK (COG-UK) Consortium | Ana da Silva Filipe, Natasha Johnson, Kathy Smollett, Daniel Mair, Stephen Carmichael, Lily Tong, Jenna Nichols, Elihu Aranday-Cortes, Kirstyn Brunker, Yasmin Parr, Kyriaki Nomikou; Sarah McDonald, Marc Niebel, Patawee Asamaphan; Richard Orton, Joseph Hughes, Sreenu Vattipally, David L Robertson; Alasdair MacLean, Rory Gunson; Kathy Li, Natasha Jesudason, Rajiv Shah, James Shepherd, Antonia Ho, Emma Thomson |
| EPI_ISL_433346 | hCoV-19/Scotland/CVR811/2020 | Europe / United Kingdom / Scotland | 2020-03-27 | West of Scotland Specialist Virology Centre, NHSGGC / MRC-University of Glasgow Centre for Virus Research | COVID-19 Genomics UK (COG-UK) Consortium | Ana da Silva Filipe, Natasha Johnson, Kathy Smollett, Daniel Mair, Stephen Carmichael, Lily Tong, Jenna Nichols, Elihu Aranday-Cortes, Kirstyn Brunker, Yasmin Parr, Kyriaki Nomikou; Sarah McDonald, Marc Niebel, Patawee Asamaphan; Richard Orton, Joseph Hughes, Sreenu Vattipally, David L Robertson; Alasdair MacLean, Rory Gunson; Kathy Li, Natasha Jesudason, Rajiv Shah, James Shepherd, Antonia Ho, Emma Thomson |
| EPI_ISL_433347 | hCoV-19/Scotland/CVR826/2020 | Europe / United Kingdom / Scotland | 2020-03-27 | West of Scotland Specialist Virology Centre, NHSGGC / MRC-University of Glasgow Centre for Virus Research | COVID-19 Genomics UK (COG-UK) Consortium | Ana da Silva Filipe, Natasha Johnson, Kathy Smollett, Daniel Mair, Stephen Carmichael, Lily Tong, Jenna Nichols, Elihu Aranday-Cortes, Kirstyn Brunker, Yasmin Parr, Kyriaki Nomikou; Sarah McDonald, Marc Niebel, Patawee Asamaphan; Richard Orton, Joseph Hughes, Sreenu Vattipally, David L Robertson; Alasdair MacLean, Rory Gunson; Kathy Li, Natasha Jesudason, Rajiv Shah, James Shepherd, Antonia Ho, Emma Thomson |
| EPI_ISL_433348 | hCoV-19/Scotland/CVR833/2020 | Europe / United Kingdom / Scotland | 2020-03-27 | West of Scotland Specialist Virology Centre, NHSGGC / MRC-University of Glasgow Centre for Virus Research | COVID-19 Genomics UK (COG-UK) Consortium | Ana da Silva Filipe, Natasha Johnson, Kathy Smollett, Daniel Mair, Stephen Carmichael, Lily Tong, Jenna Nichols, Elihu Aranday-Cortes, Kirstyn Brunker, Yasmin Parr, Kyriaki Nomikou; Sarah McDonald, Marc Niebel, Patawee Asamaphan; Richard Orton, Joseph Hughes, Sreenu Vattipally, David L Robertson; Alasdair MacLean, Rory Gunson; Kathy Li, Natasha Jesudason, Rajiv Shah, James Shepherd, Antonia Ho, Emma Thomson |
| EPI_ISL_433350 | hCoV-19/Scotland/CVR849/2020 | Europe / United Kingdom / Scotland | 2020-03-25 | West of Scotland Specialist Virology Centre, NHSGGC / MRC-University of Glasgow Centre for Virus Research | COVID-19 Genomics UK (COG-UK) Consortium | Ana da Silva Filipe, Natasha Johnson, Kathy Smollett, Daniel Mair, Stephen Carmichael, Lily Tong, Jenna Nichols, Elihu Aranday-Cortes, Kirstyn Brunker, Yasmin Parr, Kyriaki Nomikou; Sarah McDonald, Marc Niebel, Patawee Asamaphan; Richard Orton, Joseph Hughes, Sreenu Vattipally, David L Robertson; Alasdair MacLean, Rory Gunson; Kathy Li, Natasha Jesudason, Rajiv Shah, James Shepherd, Antonia Ho, Emma Thomson |
| EPI_ISL_433351 | hCoV-19/Scotland/CVR851/2020 | Europe / United Kingdom / Scotland | 2020-03-27 | West of Scotland Specialist Virology Centre, NHSGGC / MRC-University of Glasgow Centre for Virus Research | COVID-19 Genomics UK (COG-UK) Consortium | Ana da Silva Filipe, Natasha Johnson, Kathy Smollett, Daniel Mair, Stephen Carmichael, Lily Tong, Jenna Nichols, Elihu Aranday-Cortes, Kirstyn Brunker, Yasmin Parr, Kyriaki Nomikou; Sarah McDonald, Marc Niebel, Patawee Asamaphan; Richard Orton, Joseph Hughes, Sreenu Vattipally, David L Robertson; Alasdair MacLean, Rory Gunson; Kathy Li, Natasha Jesudason, Rajiv Shah, James Shepherd, Antonia Ho, Emma Thomson |
| EPI_ISL_433353 | hCoV-19/Scotland/CVR855/2020 | Europe / United Kingdom / Scotland | 2020-03-26 | West of Scotland Specialist Virology Centre, NHSGGC / MRC-University of Glasgow Centre for Virus Research | COVID-19 Genomics UK (COG-UK) Consortium | Ana da Silva Filipe, Natasha Johnson, Kathy Smollett, Daniel Mair, Stephen Carmichael, Lily Tong, Jenna Nichols, Elihu Aranday-Cortes, Kirstyn Brunker, Yasmin Parr, Kyriaki Nomikou; Sarah McDonald, Marc Niebel, Patawee Asamaphan; Richard Orton, Joseph Hughes, Sreenu Vattipally, David L Robertson; Alasdair MacLean, Rory Gunson; Kathy Li, Natasha Jesudason, Rajiv Shah, James Shepherd, Antonia Ho, Emma Thomson |
| EPI_ISL_433355 | hCoV-19/Scotland/CVR867/2020 | Europe / United Kingdom / Scotland | 2020-03-27 | West of Scotland Specialist Virology Centre, NHSGGC / MRC-University of Glasgow Centre for Virus Research | COVID-19 Genomics UK (COG-UK) Consortium | Ana da Silva Filipe, Natasha Johnson, Kathy Smollett, Daniel Mair, Stephen Carmichael, Lily Tong, Jenna Nichols, Elihu Aranday-Cortes, Kirstyn Brunker, Yasmin Parr, Kyriaki Nomikou; Sarah McDonald, Marc Niebel, Patawee Asamaphan; Richard Orton, Joseph Hughes, Sreenu Vattipally, David L Robertson; Alasdair MacLean, Rory Gunson; Kathy Li, Natasha Jesudason, Rajiv Shah, James Shepherd, Antonia Ho, Emma Thomson |
| EPI_ISL_433356 | hCoV-19/Scotland/CVR874/2020 | Europe / United Kingdom / Scotland | 2020-03-27 | West of Scotland Specialist Virology Centre, NHSGGC / MRC-University of Glasgow Centre for Virus Research | COVID-19 Genomics UK (COG-UK) Consortium | Ana da Silva Filipe, Natasha Johnson, Kathy Smollett, Daniel Mair, Stephen Carmichael, Lily Tong, Jenna Nichols, Elihu Aranday-Cortes, Kirstyn Brunker, Yasmin Parr, Kyriaki Nomikou; Sarah McDonald, Marc Niebel, Patawee Asamaphan; Richard Orton, Joseph Hughes, Sreenu Vattipally, David L Robertson; Alasdair MacLean, Rory Gunson; Kathy Li, Natasha Jesudason, Rajiv Shah, James Shepherd, Antonia Ho, Emma Thomson |
| EPI_ISL_433357 | hCoV-19/Scotland/CVR901/2020 | Europe / United Kingdom / Scotland | 2020-03-27 | West of Scotland Specialist Virology Centre, NHSGGC / MRC-University of Glasgow Centre for Virus Research | COVID-19 Genomics UK (COG-UK) Consortium | Ana da Silva Filipe, Natasha Johnson, Kathy Smollett, Daniel Mair, Stephen Carmichael, Lily Tong, Jenna Nichols, Elihu Aranday-Cortes, Kirstyn Brunker, Yasmin Parr, Kyriaki Nomikou; Sarah McDonald, Marc Niebel, Patawee Asamaphan; Richard Orton, Joseph Hughes, Sreenu Vattipally, David L Robertson; Alasdair MacLean, Rory Gunson; Kathy Li, Natasha Jesudason, Rajiv Shah, James Shepherd, Antonia Ho, Emma Thomson |
| EPI_ISL_433358 | hCoV-19/Scotland/CVR902/2020 | Europe / United Kingdom / Scotland | 2020-03-27 | West of Scotland Specialist Virology Centre, NHSGGC / MRC-University of Glasgow Centre for Virus Research | COVID-19 Genomics UK (COG-UK) Consortium | Ana da Silva Filipe, Natasha Johnson, Kathy Smollett, Daniel Mair, Stephen Carmichael, Lily Tong, Jenna Nichols, Elihu Aranday-Cortes, Kirstyn Brunker, Yasmin Parr, Kyriaki Nomikou; Sarah McDonald, Marc Niebel, Patawee Asamaphan; Richard Orton, Joseph Hughes, Sreenu Vattipally, David L Robertson; Alasdair MacLean, Rory Gunson; Kathy Li, Natasha Jesudason, Rajiv Shah, James Shepherd, Antonia Ho, Emma Thomson |
| EPI_ISL_433359 | hCoV-19/Scotland/CVR903/2020 | Europe / United Kingdom / Scotland | 2020-03-27 | West of Scotland Specialist Virology Centre, NHSGGC / MRC-University of Glasgow Centre for Virus Research | COVID-19 Genomics UK (COG-UK) Consortium | Ana da Silva Filipe, Natasha Johnson, Kathy Smollett, Daniel Mair, Stephen Carmichael, Lily Tong, Jenna Nichols, Elihu Aranday-Cortes, Kirstyn Brunker, Yasmin Parr, Kyriaki Nomikou; Sarah McDonald, Marc Niebel, Patawee Asamaphan; Richard Orton, Joseph Hughes, Sreenu Vattipally, David L Robertson; Alasdair MacLean, Rory Gunson; Kathy Li, Natasha Jesudason, Rajiv Shah, James Shepherd, Antonia Ho, Emma Thomson |
| EPI_ISL_433360 | hCoV-19/Scotland/CVR919/2020 | Europe / United Kingdom / Scotland | 2020-03-28 | West of Scotland Specialist Virology Centre, NHSGGC / MRC-University of Glasgow Centre for Virus Research | COVID-19 Genomics UK (COG-UK) Consortium | Ana da Silva Filipe, Natasha Johnson, Kathy Smollett, Daniel Mair, Stephen Carmichael, Lily Tong, Jenna Nichols, Elihu Aranday-Cortes, Kirstyn Brunker, Yasmin Parr, Kyriaki Nomikou; Sarah McDonald, Marc Niebel, Patawee Asamaphan; Richard Orton, Joseph Hughes, Sreenu Vattipally, David L Robertson; Alasdair MacLean, Rory Gunson; Kathy Li, Natasha Jesudason, Rajiv Shah, James Shepherd, Antonia Ho, Emma Thomson |
| EPI_ISL_433361 | hCoV-19/Scotland/CVR922/2020 | Europe / United Kingdom / Scotland | 2020-03-28 | West of Scotland Specialist Virology Centre, NHSGGC / MRC-University of Glasgow Centre for Virus Research | COVID-19 Genomics UK (COG-UK) Consortium | Ana da Silva Filipe, Natasha Johnson, Kathy Smollett, Daniel Mair, Stephen Carmichael, Lily Tong, Jenna Nichols, Elihu Aranday-Cortes, Kirstyn Brunker, Yasmin Parr, Kyriaki Nomikou; Sarah McDonald, Marc Niebel, Patawee Asamaphan; Richard Orton, Joseph Hughes, Sreenu Vattipally, David L Robertson; Alasdair MacLean, Rory Gunson; Kathy Li, Natasha Jesudason, Rajiv Shah, James Shepherd, Antonia Ho, Emma Thomson |
| EPI_ISL_433362 | hCoV-19/Scotland/CVR929/2020 | Europe / United Kingdom / Scotland | 2020-03-28 | West of Scotland Specialist Virology Centre, NHSGGC / MRC-University of Glasgow Centre for Virus Research | COVID-19 Genomics UK (COG-UK) Consortium | Ana da Silva Filipe, Natasha Johnson, Kathy Smollett, Daniel Mair, Stephen Carmichael, Lily Tong, Jenna Nichols, Elihu Aranday-Cortes, Kirstyn Brunker, Yasmin Parr, Kyriaki Nomikou; Sarah McDonald, Marc Niebel, Patawee Asamaphan; Richard Orton, Joseph Hughes, Sreenu Vattipally, David L Robertson; Alasdair MacLean, Rory Gunson; Kathy Li, Natasha Jesudason, Rajiv Shah, James Shepherd, Antonia Ho, Emma Thomson |
| EPI_ISL_433364 | hCoV-19/Scotland/CVR942/2020 | Europe / United Kingdom / Scotland | 2020-03-29 | West of Scotland Specialist Virology Centre, NHSGGC / MRC-University of Glasgow Centre for Virus Research | COVID-19 Genomics UK (COG-UK) Consortium | Ana da Silva Filipe, Natasha Johnson, Kathy Smollett, Daniel Mair, Stephen Carmichael, Lily Tong, Jenna Nichols, Elihu Aranday-Cortes, Kirstyn Brunker, Yasmin Parr, Kyriaki Nomikou; Sarah McDonald, Marc Niebel, Patawee Asamaphan; Richard Orton, Joseph Hughes, Sreenu Vattipally, David L Robertson; Alasdair MacLean, Rory Gunson; Kathy Li, Natasha Jesudason, Rajiv Shah, James Shepherd, Antonia Ho, Emma Thomson |
| EPI_ISL_433366 | hCoV-19/Scotland/CVR965/2020 | Europe / United Kingdom / Scotland | 2020-03-28 | West of Scotland Specialist Virology Centre, NHSGGC / MRC-University of Glasgow Centre for Virus Research | COVID-19 Genomics UK (COG-UK) Consortium | Ana da Silva Filipe, Natasha Johnson, Kathy Smollett, Daniel Mair, Stephen Carmichael, Lily Tong, Jenna Nichols, Elihu Aranday-Cortes, Kirstyn Brunker, Yasmin Parr, Kyriaki Nomikou; Sarah McDonald, Marc Niebel, Patawee Asamaphan; Richard Orton, Joseph Hughes, Sreenu Vattipally, David L Robertson; Alasdair MacLean, Rory Gunson; Kathy Li, Natasha Jesudason, Rajiv Shah, James Shepherd, Antonia Ho, Emma Thomson |
| EPI_ISL_433367 | hCoV-19/Scotland/CVR968/2020 | Europe / United Kingdom / Scotland | 2020-03-28 | West of Scotland Specialist Virology Centre, NHSGGC / MRC-University of Glasgow Centre for Virus Research | COVID-19 Genomics UK (COG-UK) Consortium | Ana da Silva Filipe, Natasha Johnson, Kathy Smollett, Daniel Mair, Stephen Carmichael, Lily Tong, Jenna Nichols, Elihu Aranday-Cortes, Kirstyn Brunker, Yasmin Parr, Kyriaki Nomikou; Sarah McDonald, Marc Niebel, Patawee Asamaphan; Richard Orton, Joseph Hughes, Sreenu Vattipally, David L Robertson; Alasdair MacLean, Rory Gunson; Kathy Li, Natasha Jesudason, Rajiv Shah, James Shepherd, Antonia Ho, Emma Thomson |
| EPI_ISL_433368 | hCoV-19/Scotland/CVR969/2020 | Europe / United Kingdom / Scotland | 2020-03-28 | West of Scotland Specialist Virology Centre, NHSGGC / MRC-University of Glasgow Centre for Virus Research | COVID-19 Genomics UK (COG-UK) Consortium | Ana da Silva Filipe, Natasha Johnson, Kathy Smollett, Daniel Mair, Stephen Carmichael, Lily Tong, Jenna Nichols, Elihu Aranday-Cortes, Kirstyn Brunker, Yasmin Parr, Kyriaki Nomikou; Sarah McDonald, Marc Niebel, Patawee Asamaphan; Richard Orton, Joseph Hughes, Sreenu Vattipally, David L Robertson; Alasdair MacLean, Rory Gunson; Kathy Li, Natasha Jesudason, Rajiv Shah, James Shepherd, Antonia Ho, Emma Thomson |
| EPI_ISL_433399 | hCoV-19/Scotland/EDB1109/2020 | Europe / United Kingdom / Scotland | 2020-04-06 | Virology Department, Royal Infirmary of Edinburgh, NHS Lothian / School of Biological Sciences, University of Edinburgh / Institute of Genetics and Molecular Medicine, University of Edinburgh | COVID-19 Genomics UK (COG-UK) Consortium | McHugh M, Dewar R, Rooke S, Gallagher M, Balcaza C, O'Toole A, Hill V, McCrone JT, Colquhoun R, Yu X, Jackson B, Rambaut A, Williams TC, Templeton K |
| EPI_ISL_433400 | hCoV-19/Scotland/EDB1110/2020 | Europe / United Kingdom / Scotland | 2020-04-06 | Virology Department, Royal Infirmary of Edinburgh, NHS Lothian / School of Biological Sciences, University of Edinburgh / Institute of Genetics and Molecular Medicine, University of Edinburgh | COVID-19 Genomics UK (COG-UK) Consortium | McHugh M, Dewar R, Rooke S, Gallagher M, Balcaza C, O'Toole A, Hill V, McCrone JT, Colquhoun R, Yu X, Jackson B, Rambaut A, Williams TC, Templeton K |
| EPI_ISL_433401 | hCoV-19/Scotland/EDB1111/2020 | Europe / United Kingdom / Scotland | 2020-04-06 | Virology Department, Royal Infirmary of Edinburgh, NHS Lothian / School of Biological Sciences, University of Edinburgh / Institute of Genetics and Molecular Medicine, University of Edinburgh | COVID-19 Genomics UK (COG-UK) Consortium | McHugh M, Dewar R, Rooke S, Gallagher M, Balcaza C, O'Toole A, Hill V, McCrone JT, Colquhoun R, Yu X, Jackson B, Rambaut A, Williams TC, Templeton K |
| EPI_ISL_433402 | hCoV-19/Scotland/EDB1112/2020 | Europe / United Kingdom / Scotland | 2020-04-06 | Virology Department, Royal Infirmary of Edinburgh, NHS Lothian / School of Biological Sciences, University of Edinburgh / Institute of Genetics and Molecular Medicine, University of Edinburgh | COVID-19 Genomics UK (COG-UK) Consortium | McHugh M, Dewar R, Rooke S, Gallagher M, Balcaza C, O'Toole A, Hill V, McCrone JT, Colquhoun R, Yu X, Jackson B, Rambaut A, Williams TC, Templeton K |
| EPI_ISL_433403 | hCoV-19/Scotland/EDB1113/2020 | Europe / United Kingdom / Scotland | 2020-04-06 | Virology Department, Royal Infirmary of Edinburgh, NHS Lothian / School of Biological Sciences, University of Edinburgh / Institute of Genetics and Molecular Medicine, University of Edinburgh | COVID-19 Genomics UK (COG-UK) Consortium | McHugh M, Dewar R, Rooke S, Gallagher M, Balcaza C, O'Toole A, Hill V, McCrone JT, Colquhoun R, Yu X, Jackson B, Rambaut A, Williams TC, Templeton K |
| EPI_ISL_433405 | hCoV-19/Scotland/EDB1117/2020 | Europe / United Kingdom / Scotland | 2020-04-06 | Virology Department, Royal Infirmary of Edinburgh, NHS Lothian / School of Biological Sciences, University of Edinburgh / Institute of Genetics and Molecular Medicine, University of Edinburgh | COVID-19 Genomics UK (COG-UK) Consortium | McHugh M, Dewar R, Rooke S, Gallagher M, Balcaza C, O'Toole A, Hill V, McCrone JT, Colquhoun R, Yu X, Jackson B, Rambaut A, Williams TC, Templeton K |
| EPI_ISL_433406 | hCoV-19/Scotland/EDB1118/2020 | Europe / United Kingdom / Scotland | 2020-04-06 | Virology Department, Royal Infirmary of Edinburgh, NHS Lothian / School of Biological Sciences, University of Edinburgh / Institute of Genetics and Molecular Medicine, University of Edinburgh | COVID-19 Genomics UK (COG-UK) Consortium | McHugh M, Dewar R, Rooke S, Gallagher M, Balcaza C, O'Toole A, Hill V, McCrone JT, Colquhoun R, Yu X, Jackson B, Rambaut A, Williams TC, Templeton K |
| EPI_ISL_433407 | hCoV-19/Scotland/EDB112/2020 | Europe / United Kingdom / Scotland | 2020-03-22 | Virology Department, Royal Infirmary of Edinburgh, NHS Lothian / School of Biological Sciences, University of Edinburgh / Institute of Genetics and Molecular Medicine, University of Edinburgh | COVID-19 Genomics UK (COG-UK) Consortium | McHugh M, Dewar R, Rooke S, Gallagher M, Balcaza C, O'Toole A, Hill V, McCrone JT, Colquhoun R, Yu X, Jackson B, Rambaut A, Williams TC, Templeton K |
| EPI_ISL_433408 | hCoV-19/Scotland/EDB1148/2020 | Europe / United Kingdom / Scotland | 2020-04-07 | Virology Department, Royal Infirmary of Edinburgh, NHS Lothian / School of Biological Sciences, University of Edinburgh / Institute of Genetics and Molecular Medicine, University of Edinburgh | COVID-19 Genomics UK (COG-UK) Consortium | McHugh M, Dewar R, Rooke S, Gallagher M, Balcaza C, O'Toole A, Hill V, McCrone JT, Colquhoun R, Yu X, Jackson B, Rambaut A, Williams TC, Templeton K |
| EPI_ISL_433410 | hCoV-19/Scotland/EDB1151/2020 | Europe / United Kingdom / Scotland | 2020-04-07 | Virology Department, Royal Infirmary of Edinburgh, NHS Lothian / School of Biological Sciences, University of Edinburgh / Institute of Genetics and Molecular Medicine, University of Edinburgh | COVID-19 Genomics UK (COG-UK) Consortium | McHugh M, Dewar R, Rooke S, Gallagher M, Balcaza C, O'Toole A, Hill V, McCrone JT, Colquhoun R, Yu X, Jackson B, Rambaut A, Williams TC, Templeton K |
| EPI_ISL_433411 | hCoV-19/Scotland/EDB1152/2020 | Europe / United Kingdom / Scotland | 2020-04-07 | Virology Department, Royal Infirmary of Edinburgh, NHS Lothian / School of Biological Sciences, University of Edinburgh / Institute of Genetics and Molecular Medicine, University of Edinburgh | COVID-19 Genomics UK (COG-UK) Consortium | McHugh M, Dewar R, Rooke S, Gallagher M, Balcaza C, O'Toole A, Hill V, McCrone JT, Colquhoun R, Yu X, Jackson B, Rambaut A, Williams TC, Templeton K |
| EPI_ISL_433412 | hCoV-19/Scotland/EDB1158/2020 | Europe / United Kingdom / Scotland | 2020-04-07 | Virology Department, Royal Infirmary of Edinburgh, NHS Lothian / School of Biological Sciences, University of Edinburgh / Institute of Genetics and Molecular Medicine, University of Edinburgh | COVID-19 Genomics UK (COG-UK) Consortium | McHugh M, Dewar R, Rooke S, Gallagher M, Balcaza C, O'Toole A, Hill V, McCrone JT, Colquhoun R, Yu X, Jackson B, Rambaut A, Williams TC, Templeton K |
| EPI_ISL_433413 | hCoV-19/Scotland/EDB1161/2020 | Europe / United Kingdom / Scotland | 2020-04-08 | Virology Department, Royal Infirmary of Edinburgh, NHS Lothian / School of Biological Sciences, University of Edinburgh / Institute of Genetics and Molecular Medicine, University of Edinburgh | COVID-19 Genomics UK (COG-UK) Consortium | McHugh M, Dewar R, Rooke S, Gallagher M, Balcaza C, O'Toole A, Hill V, McCrone JT, Colquhoun R, Yu X, Jackson B, Rambaut A, Williams TC, Templeton K |
| EPI_ISL_433415 | hCoV-19/Scotland/EDB1164/2020 | Europe / United Kingdom / Scotland | 2020-04-08 | Virology Department, Royal Infirmary of Edinburgh, NHS Lothian / School of Biological Sciences, University of Edinburgh / Institute of Genetics and Molecular Medicine, University of Edinburgh | COVID-19 Genomics UK (COG-UK) Consortium | McHugh M, Dewar R, Rooke S, Gallagher M, Balcaza C, O'Toole A, Hill V, McCrone JT, Colquhoun R, Yu X, Jackson B, Rambaut A, Williams TC, Templeton K |
| EPI_ISL_433417 | hCoV-19/Scotland/EDB1167/2020 | Europe / United Kingdom / Scotland | 2020-04-08 | Virology Department, Royal Infirmary of Edinburgh, NHS Lothian / School of Biological Sciences, University of Edinburgh / Institute of Genetics and Molecular Medicine, University of Edinburgh | COVID-19 Genomics UK (COG-UK) Consortium | McHugh M, Dewar R, Rooke S, Gallagher M, Balcaza C, O'Toole A, Hill V, McCrone JT, Colquhoun R, Yu X, Jackson B, Rambaut A, Williams TC, Templeton K |
| EPI_ISL_433418 | hCoV-19/Scotland/EDB1168/2020 | Europe / United Kingdom / Scotland | 2020-04-08 | Virology Department, Royal Infirmary of Edinburgh, NHS Lothian / School of Biological Sciences, University of Edinburgh / Institute of Genetics and Molecular Medicine, University of Edinburgh | COVID-19 Genomics UK (COG-UK) Consortium | McHugh M, Dewar R, Rooke S, Gallagher M, Balcaza C, O'Toole A, Hill V, McCrone JT, Colquhoun R, Yu X, Jackson B, Rambaut A, Williams TC, Templeton K |
| EPI_ISL_433419 | hCoV-19/Scotland/EDB1169/2020 | Europe / United Kingdom / Scotland | 2020-04-08 | Virology Department, Royal Infirmary of Edinburgh, NHS Lothian / School of Biological Sciences, University of Edinburgh / Institute of Genetics and Molecular Medicine, University of Edinburgh | COVID-19 Genomics UK (COG-UK) Consortium | McHugh M, Dewar R, Rooke S, Gallagher M, Balcaza C, O'Toole A, Hill V, McCrone JT, Colquhoun R, Yu X, Jackson B, Rambaut A, Williams TC, Templeton K |
| EPI_ISL_433420 | hCoV-19/Scotland/EDB1170/2020 | Europe / United Kingdom / Scotland | 2020-04-09 | Virology Department, Royal Infirmary of Edinburgh, NHS Lothian / School of Biological Sciences, University of Edinburgh / Institute of Genetics and Molecular Medicine, University of Edinburgh | COVID-19 Genomics UK (COG-UK) Consortium | McHugh M, Dewar R, Rooke S, Gallagher M, Balcaza C, O'Toole A, Hill V, McCrone JT, Colquhoun R, Yu X, Jackson B, Rambaut A, Williams TC, Templeton K |
| EPI_ISL_433421 | hCoV-19/Scotland/EDB1171/2020 | Europe / United Kingdom / Scotland | 2020-04-07 | Virology Department, Royal Infirmary of Edinburgh, NHS Lothian / School of Biological Sciences, University of Edinburgh / Institute of Genetics and Molecular Medicine, University of Edinburgh | COVID-19 Genomics UK (COG-UK) Consortium | McHugh M, Dewar R, Rooke S, Gallagher M, Balcaza C, O'Toole A, Hill V, McCrone JT, Colquhoun R, Yu X, Jackson B, Rambaut A, Williams TC, Templeton K |
| EPI_ISL_433423 | hCoV-19/Scotland/EDB1173/2020 | Europe / United Kingdom / Scotland | 2020-04-08 | Virology Department, Royal Infirmary of Edinburgh, NHS Lothian / School of Biological Sciences, University of Edinburgh / Institute of Genetics and Molecular Medicine, University of Edinburgh | COVID-19 Genomics UK (COG-UK) Consortium | McHugh M, Dewar R, Rooke S, Gallagher M, Balcaza C, O'Toole A, Hill V, McCrone JT, Colquhoun R, Yu X, Jackson B, Rambaut A, Williams TC, Templeton K |
| EPI_ISL_433425 | hCoV-19/Scotland/EDB1180/2020 | Europe / United Kingdom / Scotland | 2020-04-07 | Virology Department, Royal Infirmary of Edinburgh, NHS Lothian / School of Biological Sciences, University of Edinburgh / Institute of Genetics and Molecular Medicine, University of Edinburgh | COVID-19 Genomics UK (COG-UK) Consortium | McHugh M, Dewar R, Rooke S, Gallagher M, Balcaza C, O'Toole A, Hill V, McCrone JT, Colquhoun R, Yu X, Jackson B, Rambaut A, Williams TC, Templeton K |
| EPI_ISL_433426 | hCoV-19/Scotland/EDB1182/2020 | Europe / United Kingdom / Scotland | 2020-04-08 | Virology Department, Royal Infirmary of Edinburgh, NHS Lothian / School of Biological Sciences, University of Edinburgh / Institute of Genetics and Molecular Medicine, University of Edinburgh | COVID-19 Genomics UK (COG-UK) Consortium | McHugh M, Dewar R, Rooke S, Gallagher M, Balcaza C, O'Toole A, Hill V, McCrone JT, Colquhoun R, Yu X, Jackson B, Rambaut A, Williams TC, Templeton K |
| EPI_ISL_433428 | hCoV-19/Scotland/EDB1197/2020 | Europe / United Kingdom / Scotland | 2020-04-08 | Virology Department, Royal Infirmary of Edinburgh, NHS Lothian / School of Biological Sciences, University of Edinburgh / Institute of Genetics and Molecular Medicine, University of Edinburgh | COVID-19 Genomics UK (COG-UK) Consortium | McHugh M, Dewar R, Rooke S, Gallagher M, Balcaza C, O'Toole A, Hill V, McCrone JT, Colquhoun R, Yu X, Jackson B, Rambaut A, Williams TC, Templeton K |
| EPI_ISL_433429 | hCoV-19/Scotland/EDB1198/2020 | Europe / United Kingdom / Scotland | 2020-04-08 | Virology Department, Royal Infirmary of Edinburgh, NHS Lothian / School of Biological Sciences, University of Edinburgh / Institute of Genetics and Molecular Medicine, University of Edinburgh | COVID-19 Genomics UK (COG-UK) Consortium | McHugh M, Dewar R, Rooke S, Gallagher M, Balcaza C, O'Toole A, Hill V, McCrone JT, Colquhoun R, Yu X, Jackson B, Rambaut A, Williams TC, Templeton K |
| EPI_ISL_433430 | hCoV-19/Scotland/EDB1207/2020 | Europe / United Kingdom / Scotland | 2020-04-08 | Virology Department, Royal Infirmary of Edinburgh, NHS Lothian / School of Biological Sciences, University of Edinburgh / Institute of Genetics and Molecular Medicine, University of Edinburgh | COVID-19 Genomics UK (COG-UK) Consortium | McHugh M, Dewar R, Rooke S, Gallagher M, Balcaza C, O'Toole A, Hill V, McCrone JT, Colquhoun R, Yu X, Jackson B, Rambaut A, Williams TC, Templeton K |
| EPI_ISL_433431 | hCoV-19/Scotland/EDB1216/2020 | Europe / United Kingdom / Scotland | 2020-04-08 | Virology Department, Royal Infirmary of Edinburgh, NHS Lothian / School of Biological Sciences, University of Edinburgh / Institute of Genetics and Molecular Medicine, University of Edinburgh | COVID-19 Genomics UK (COG-UK) Consortium | McHugh M, Dewar R, Rooke S, Gallagher M, Balcaza C, O'Toole A, Hill V, McCrone JT, Colquhoun R, Yu X, Jackson B, Rambaut A, Williams TC, Templeton K |
| EPI_ISL_433432 | hCoV-19/Scotland/EDB1220/2020 | Europe / United Kingdom / Scotland | 2020-04-08 | Virology Department, Royal Infirmary of Edinburgh, NHS Lothian / School of Biological Sciences, University of Edinburgh / Institute of Genetics and Molecular Medicine, University of Edinburgh | COVID-19 Genomics UK (COG-UK) Consortium | McHugh M, Dewar R, Rooke S, Gallagher M, Balcaza C, O'Toole A, Hill V, McCrone JT, Colquhoun R, Yu X, Jackson B, Rambaut A, Williams TC, Templeton K |
| EPI_ISL_433433 | hCoV-19/Scotland/EDB129/2020 | Europe / United Kingdom / Scotland | 2020-03-23 | Virology Department, Royal Infirmary of Edinburgh, NHS Lothian / School of Biological Sciences, University of Edinburgh / Institute of Genetics and Molecular Medicine, University of Edinburgh | COVID-19 Genomics UK (COG-UK) Consortium | McHugh M, Dewar R, Rooke S, Gallagher M, Balcaza C, O'Toole A, Hill V, McCrone JT, Colquhoun R, Yu X, Jackson B, Rambaut A, Williams TC, Templeton K |
| EPI_ISL_433434 | hCoV-19/Scotland/EDB1317/2020 | Europe / United Kingdom / Scotland | 2020-04-09 | Virology Department, Royal Infirmary of Edinburgh, NHS Lothian / School of Biological Sciences, University of Edinburgh / Institute of Genetics and Molecular Medicine, University of Edinburgh | COVID-19 Genomics UK (COG-UK) Consortium | McHugh M, Dewar R, Rooke S, Gallagher M, Balcaza C, O'Toole A, Hill V, McCrone JT, Colquhoun R, Yu X, Jackson B, Rambaut A, Williams TC, Templeton K |
| EPI_ISL_433435 | hCoV-19/Scotland/EDB1324/2020 | Europe / United Kingdom / Scotland | 2020-04-10 | Virology Department, Royal Infirmary of Edinburgh, NHS Lothian / School of Biological Sciences, University of Edinburgh / Institute of Genetics and Molecular Medicine, University of Edinburgh | COVID-19 Genomics UK (COG-UK) Consortium | McHugh M, Dewar R, Rooke S, Gallagher M, Balcaza C, O'Toole A, Hill V, McCrone JT, Colquhoun R, Yu X, Jackson B, Rambaut A, Williams TC, Templeton K |
| EPI_ISL_433436 | hCoV-19/Scotland/EDB1339/2020 | Europe / United Kingdom / Scotland | 2020-04-10 | Virology Department, Royal Infirmary of Edinburgh, NHS Lothian / School of Biological Sciences, University of Edinburgh / Institute of Genetics and Molecular Medicine, University of Edinburgh | COVID-19 Genomics UK (COG-UK) Consortium | McHugh M, Dewar R, Rooke S, Gallagher M, Balcaza C, O'Toole A, Hill V, McCrone JT, Colquhoun R, Yu X, Jackson B, Rambaut A, Williams TC, Templeton K |
| EPI_ISL_433437 | hCoV-19/Scotland/EDB1340/2020 | Europe / United Kingdom / Scotland | 2020-04-10 | Virology Department, Royal Infirmary of Edinburgh, NHS Lothian / School of Biological Sciences, University of Edinburgh / Institute of Genetics and Molecular Medicine, University of Edinburgh | COVID-19 Genomics UK (COG-UK) Consortium | McHugh M, Dewar R, Rooke S, Gallagher M, Balcaza C, O'Toole A, Hill V, McCrone JT, Colquhoun R, Yu X, Jackson B, Rambaut A, Williams TC, Templeton K |
| EPI_ISL_433439 | hCoV-19/Scotland/EDB1404/2020 | Europe / United Kingdom / Scotland | 2020-04-11 | Virology Department, Royal Infirmary of Edinburgh, NHS Lothian / School of Biological Sciences, University of Edinburgh / Institute of Genetics and Molecular Medicine, University of Edinburgh | COVID-19 Genomics UK (COG-UK) Consortium | McHugh M, Dewar R, Rooke S, Gallagher M, Balcaza C, O'Toole A, Hill V, McCrone JT, Colquhoun R, Yu X, Jackson B, Rambaut A, Williams TC, Templeton K |
| EPI_ISL_433440 | hCoV-19/Scotland/EDB1411/2020 | Europe / United Kingdom / Scotland | 2020-04-10 | Virology Department, Royal Infirmary of Edinburgh, NHS Lothian / School of Biological Sciences, University of Edinburgh / Institute of Genetics and Molecular Medicine, University of Edinburgh | COVID-19 Genomics UK (COG-UK) Consortium | McHugh M, Dewar R, Rooke S, Gallagher M, Balcaza C, O'Toole A, Hill V, McCrone JT, Colquhoun R, Yu X, Jackson B, Rambaut A, Williams TC, Templeton K |
| EPI_ISL_433442 | hCoV-19/Scotland/EDB1424/2020 | Europe / United Kingdom / Scotland | 2020-04-12 | Virology Department, Royal Infirmary of Edinburgh, NHS Lothian / School of Biological Sciences, University of Edinburgh / Institute of Genetics and Molecular Medicine, University of Edinburgh | COVID-19 Genomics UK (COG-UK) Consortium | McHugh M, Dewar R, Rooke S, Gallagher M, Balcaza C, O'Toole A, Hill V, McCrone JT, Colquhoun R, Yu X, Jackson B, Rambaut A, Williams TC, Templeton K |
| EPI_ISL_433443 | hCoV-19/Scotland/EDB1428/2020 | Europe / United Kingdom / Scotland | 2020-04-12 | Virology Department, Royal Infirmary of Edinburgh, NHS Lothian / School of Biological Sciences, University of Edinburgh / Institute of Genetics and Molecular Medicine, University of Edinburgh | COVID-19 Genomics UK (COG-UK) Consortium | McHugh M, Dewar R, Rooke S, Gallagher M, Balcaza C, O'Toole A, Hill V, McCrone JT, Colquhoun R, Yu X, Jackson B, Rambaut A, Williams TC, Templeton K |
| EPI_ISL_433444 | hCoV-19/Scotland/EDB1431/2020 | Europe / United Kingdom / Scotland | 2020-04-12 | Virology Department, Royal Infirmary of Edinburgh, NHS Lothian / School of Biological Sciences, University of Edinburgh / Institute of Genetics and Molecular Medicine, University of Edinburgh | COVID-19 Genomics UK (COG-UK) Consortium | McHugh M, Dewar R, Rooke S, Gallagher M, Balcaza C, O'Toole A, Hill V, McCrone JT, Colquhoun R, Yu X, Jackson B, Rambaut A, Williams TC, Templeton K |
| EPI_ISL_433688 | hCoV-19/England/CAMB-73D2F/2020 | Europe / United Kingdom / England | 2020-04-01 | Department of Pathology, University of Cambridge | COVID-19 Genomics UK (COG-UK) Consortium | Luke W Meredith, M. Estee Torok , Myra Hosmillo, William L. Hamilton, Martin D. Curran, Theresa Feltwell, Grant Hall, Anna Yakovleva, Fahad A Khokhar, Charlotte J. Houldcroft, Laura G Caller, Aminu S. Jahun, Sarah L. Caddy, Ian Goodfellow |
| EPI_ISL_433689 | hCoV-19/England/CAMB-73D3E/2020 | Europe / United Kingdom / England | 2020-04-01 | Department of Pathology, University of Cambridge | COVID-19 Genomics UK (COG-UK) Consortium | Luke W Meredith, M. Estee Torok , Myra Hosmillo, William L. Hamilton, Martin D. Curran, Theresa Feltwell, Grant Hall, Anna Yakovleva, Fahad A Khokhar, Charlotte J. Houldcroft, Laura G Caller, Aminu S. Jahun, Sarah L. Caddy, Ian Goodfellow |
| EPI_ISL_433691 | hCoV-19/England/CAMB-73D5C/2020 | Europe / United Kingdom / England | 2020-04-01 | Department of Pathology, University of Cambridge | COVID-19 Genomics UK (COG-UK) Consortium | Luke W Meredith, M. Estee Torok , Myra Hosmillo, William L. Hamilton, Martin D. Curran, Theresa Feltwell, Grant Hall, Anna Yakovleva, Fahad A Khokhar, Charlotte J. Houldcroft, Laura G Caller, Aminu S. Jahun, Sarah L. Caddy, Ian Goodfellow |
| EPI_ISL_433693 | hCoV-19/England/CAMB-73D7A/2020 | Europe / United Kingdom / England | 2020-04-01 | Department of Pathology, University of Cambridge | COVID-19 Genomics UK (COG-UK) Consortium | Luke W Meredith, M. Estee Torok , Myra Hosmillo, William L. Hamilton, Martin D. Curran, Theresa Feltwell, Grant Hall, Anna Yakovleva, Fahad A Khokhar, Charlotte J. Houldcroft, Laura G Caller, Aminu S. Jahun, Sarah L. Caddy, Ian Goodfellow |
| EPI_ISL_433695 | hCoV-19/England/CAMB-73D98/2020 | Europe / United Kingdom / England | 2020-04-02 | Department of Pathology, University of Cambridge | COVID-19 Genomics UK (COG-UK) Consortium | Luke W Meredith, M. Estee Torok , Myra Hosmillo, William L. Hamilton, Martin D. Curran, Theresa Feltwell, Grant Hall, Anna Yakovleva, Fahad A Khokhar, Charlotte J. Houldcroft, Laura G Caller, Aminu S. Jahun, Sarah L. Caddy, Ian Goodfellow |
| EPI_ISL_433697 | hCoV-19/England/CAMB-73DB6/2020 | Europe / United Kingdom / England | 2020-04-01 | Department of Pathology, University of Cambridge | COVID-19 Genomics UK (COG-UK) Consortium | Luke W Meredith, M. Estee Torok , Myra Hosmillo, William L. Hamilton, Martin D. Curran, Theresa Feltwell, Grant Hall, Anna Yakovleva, Fahad A Khokhar, Charlotte J. Houldcroft, Laura G Caller, Aminu S. Jahun, Sarah L. Caddy, Ian Goodfellow |
| EPI_ISL_433698 | hCoV-19/England/CAMB-73DC5/2020 | Europe / United Kingdom / England | 2020-04-01 | Department of Pathology, University of Cambridge | COVID-19 Genomics UK (COG-UK) Consortium | Luke W Meredith, M. Estee Torok , Myra Hosmillo, William L. Hamilton, Martin D. Curran, Theresa Feltwell, Grant Hall, Anna Yakovleva, Fahad A Khokhar, Charlotte J. Houldcroft, Laura G Caller, Aminu S. Jahun, Sarah L. Caddy, Ian Goodfellow |
| EPI_ISL_433701 | hCoV-19/England/CAMB-779FA/2020 | Europe / United Kingdom / England | 2020-04-04 | Department of Pathology, University of Cambridge | COVID-19 Genomics UK (COG-UK) Consortium | Luke W Meredith, M. Estee Torok , Myra Hosmillo, William L. Hamilton, Martin D. Curran, Theresa Feltwell, Grant Hall, Anna Yakovleva, Fahad A Khokhar, Charlotte J. Houldcroft, Laura G Caller, Aminu S. Jahun, Sarah L. Caddy, Ian Goodfellow |
| EPI_ISL_433706 | hCoV-19/England/CAMB-7820D/2020 | Europe / United Kingdom / England | 2020-04-05 | Department of Pathology, University of Cambridge | COVID-19 Genomics UK (COG-UK) Consortium | Luke W Meredith, M. Estee Torok , Myra Hosmillo, William L. Hamilton, Martin D. Curran, Theresa Feltwell, Grant Hall, Anna Yakovleva, Fahad A Khokhar, Charlotte J. Houldcroft, Laura G Caller, Aminu S. Jahun, Sarah L. Caddy, Ian Goodfellow |
| EPI_ISL_433707 | hCoV-19/England/CAMB-7822B/2020 | Europe / United Kingdom / England | 2020-04-05 | Department of Pathology, University of Cambridge | COVID-19 Genomics UK (COG-UK) Consortium | Luke W Meredith, M. Estee Torok , Myra Hosmillo, William L. Hamilton, Martin D. Curran, Theresa Feltwell, Grant Hall, Anna Yakovleva, Fahad A Khokhar, Charlotte J. Houldcroft, Laura G Caller, Aminu S. Jahun, Sarah L. Caddy, Ian Goodfellow |
| EPI_ISL_433709 | hCoV-19/England/CAMB-78285/2020 | Europe / United Kingdom / England | 2020-04-04 | Department of Pathology, University of Cambridge | COVID-19 Genomics UK (COG-UK) Consortium | Luke W Meredith, M. Estee Torok , Myra Hosmillo, William L. Hamilton, Martin D. Curran, Theresa Feltwell, Grant Hall, Anna Yakovleva, Fahad A Khokhar, Charlotte J. Houldcroft, Laura G Caller, Aminu S. Jahun, Sarah L. Caddy, Ian Goodfellow |
| EPI_ISL_433710 | hCoV-19/England/CAMB-78294/2020 | Europe / United Kingdom / England | 2020-04-04 | Department of Pathology, University of Cambridge | COVID-19 Genomics UK (COG-UK) Consortium | Luke W Meredith, M. Estee Torok , Myra Hosmillo, William L. Hamilton, Martin D. Curran, Theresa Feltwell, Grant Hall, Anna Yakovleva, Fahad A Khokhar, Charlotte J. Houldcroft, Laura G Caller, Aminu S. Jahun, Sarah L. Caddy, Ian Goodfellow |
| EPI_ISL_433711 | hCoV-19/England/CAMB-782A3/2020 | Europe / United Kingdom / England | 2020-04-04 | Department of Pathology, University of Cambridge | COVID-19 Genomics UK (COG-UK) Consortium | Luke W Meredith, M. Estee Torok , Myra Hosmillo, William L. Hamilton, Martin D. Curran, Theresa Feltwell, Grant Hall, Anna Yakovleva, Fahad A Khokhar, Charlotte J. Houldcroft, Laura G Caller, Aminu S. Jahun, Sarah L. Caddy, Ian Goodfellow |
| EPI_ISL_433712 | hCoV-19/England/CAMB-782B2/2020 | Europe / United Kingdom / England | 2020-04-05 | Department of Pathology, University of Cambridge | COVID-19 Genomics UK (COG-UK) Consortium | Luke W Meredith, M. Estee Torok , Myra Hosmillo, William L. Hamilton, Martin D. Curran, Theresa Feltwell, Grant Hall, Anna Yakovleva, Fahad A Khokhar, Charlotte J. Houldcroft, Laura G Caller, Aminu S. Jahun, Sarah L. Caddy, Ian Goodfellow |
| EPI_ISL_433713 | hCoV-19/England/CAMB-782D0/2020 | Europe / United Kingdom / England | 2020-04-04 | Department of Pathology, University of Cambridge | COVID-19 Genomics UK (COG-UK) Consortium | Luke W Meredith, M. Estee Torok , Myra Hosmillo, William L. Hamilton, Martin D. Curran, Theresa Feltwell, Grant Hall, Anna Yakovleva, Fahad A Khokhar, Charlotte J. Houldcroft, Laura G Caller, Aminu S. Jahun, Sarah L. Caddy, Ian Goodfellow |
| EPI_ISL_433714 | hCoV-19/England/CAMB-782EF/2020 | Europe / United Kingdom / England | 2020-04-05 | Department of Pathology, University of Cambridge | COVID-19 Genomics UK (COG-UK) Consortium | Luke W Meredith, M. Estee Torok , Myra Hosmillo, William L. Hamilton, Martin D. Curran, Theresa Feltwell, Grant Hall, Anna Yakovleva, Fahad A Khokhar, Charlotte J. Houldcroft, Laura G Caller, Aminu S. Jahun, Sarah L. Caddy, Ian Goodfellow |
| EPI_ISL_433716 | hCoV-19/England/CAMB-7830A/2020 | Europe / United Kingdom / England | 2020 | Department of Pathology, University of Cambridge | COVID-19 Genomics UK (COG-UK) Consortium | Luke W Meredith, M. Estee Torok , Myra Hosmillo, William L. Hamilton, Martin D. Curran, Theresa Feltwell, Grant Hall, Anna Yakovleva, Fahad A Khokhar, Charlotte J. Houldcroft, Laura G Caller, Aminu S. Jahun, Sarah L. Caddy, Ian Goodfellow |
| EPI_ISL_433717 | hCoV-19/England/CAMB-78319/2020 | Europe / United Kingdom / England | 2020-04-04 | Department of Pathology, University of Cambridge | COVID-19 Genomics UK (COG-UK) Consortium | Luke W Meredith, M. Estee Torok , Myra Hosmillo, William L. Hamilton, Martin D. Curran, Theresa Feltwell, Grant Hall, Anna Yakovleva, Fahad A Khokhar, Charlotte J. Houldcroft, Laura G Caller, Aminu S. Jahun, Sarah L. Caddy, Ian Goodfellow |
| EPI_ISL_433720 | hCoV-19/England/CAMB-78346/2020 | Europe / United Kingdom / England | 2020-04-03 | Department of Pathology, University of Cambridge | COVID-19 Genomics UK (COG-UK) Consortium | Luke W Meredith, M. Estee Torok , Myra Hosmillo, William L. Hamilton, Martin D. Curran, Theresa Feltwell, Grant Hall, Anna Yakovleva, Fahad A Khokhar, Charlotte J. Houldcroft, Laura G Caller, Aminu S. Jahun, Sarah L. Caddy, Ian Goodfellow |
| EPI_ISL_433732 | hCoV-19/England/CAMB-78416/2020 | Europe / United Kingdom / England | 2020-04-05 | Department of Pathology, University of Cambridge | COVID-19 Genomics UK (COG-UK) Consortium | Luke W Meredith, M. Estee Torok , Myra Hosmillo, William L. Hamilton, Martin D. Curran, Theresa Feltwell, Grant Hall, Anna Yakovleva, Fahad A Khokhar, Charlotte J. Houldcroft, Laura G Caller, Aminu S. Jahun, Sarah L. Caddy, Ian Goodfellow |
| EPI_ISL_433734 | hCoV-19/England/CAMB-78452/2020 | Europe / United Kingdom / England | 2020-04-05 | Department of Pathology, University of Cambridge | COVID-19 Genomics UK (COG-UK) Consortium | Luke W Meredith, M. Estee Torok , Myra Hosmillo, William L. Hamilton, Martin D. Curran, Theresa Feltwell, Grant Hall, Anna Yakovleva, Fahad A Khokhar, Charlotte J. Houldcroft, Laura G Caller, Aminu S. Jahun, Sarah L. Caddy, Ian Goodfellow |
| EPI_ISL_433735 | hCoV-19/England/CAMB-78461/2020 | Europe / United Kingdom / England | 2020-04-04 | Department of Pathology, University of Cambridge | COVID-19 Genomics UK (COG-UK) Consortium | Luke W Meredith, M. Estee Torok , Myra Hosmillo, William L. Hamilton, Martin D. Curran, Theresa Feltwell, Grant Hall, Anna Yakovleva, Fahad A Khokhar, Charlotte J. Houldcroft, Laura G Caller, Aminu S. Jahun, Sarah L. Caddy, Ian Goodfellow |
| EPI_ISL_433736 | hCoV-19/England/CAMB-78470/2020 | Europe / United Kingdom / England | 2020-04-05 | Department of Pathology, University of Cambridge | COVID-19 Genomics UK (COG-UK) Consortium | Luke W Meredith, M. Estee Torok , Myra Hosmillo, William L. Hamilton, Martin D. Curran, Theresa Feltwell, Grant Hall, Anna Yakovleva, Fahad A Khokhar, Charlotte J. Houldcroft, Laura G Caller, Aminu S. Jahun, Sarah L. Caddy, Ian Goodfellow |
| EPI_ISL_433737 | hCoV-19/England/CAMB-7848F/2020 | Europe / United Kingdom / England | 2020-04-04 | Department of Pathology, University of Cambridge | COVID-19 Genomics UK (COG-UK) Consortium | Luke W Meredith, M. Estee Torok , Myra Hosmillo, William L. Hamilton, Martin D. Curran, Theresa Feltwell, Grant Hall, Anna Yakovleva, Fahad A Khokhar, Charlotte J. Houldcroft, Laura G Caller, Aminu S. Jahun, Sarah L. Caddy, Ian Goodfellow |
| EPI_ISL_433738 | hCoV-19/England/CAMB-7849E/2020 | Europe / United Kingdom / England | 2020-04-06 | Department of Pathology, University of Cambridge | COVID-19 Genomics UK (COG-UK) Consortium | Luke W Meredith, M. Estee Torok , Myra Hosmillo, William L. Hamilton, Martin D. Curran, Theresa Feltwell, Grant Hall, Anna Yakovleva, Fahad A Khokhar, Charlotte J. Houldcroft, Laura G Caller, Aminu S. Jahun, Sarah L. Caddy, Ian Goodfellow |
| EPI_ISL_433739 | hCoV-19/England/CAMB-784AD/2020 | Europe / United Kingdom / England | 2020-03-26 | Department of Pathology, University of Cambridge | COVID-19 Genomics UK (COG-UK) Consortium | Luke W Meredith, M. Estee Torok , Myra Hosmillo, William L. Hamilton, Martin D. Curran, Theresa Feltwell, Grant Hall, Anna Yakovleva, Fahad A Khokhar, Charlotte J. Houldcroft, Laura G Caller, Aminu S. Jahun, Sarah L. Caddy, Ian Goodfellow |
| EPI_ISL_433740 | hCoV-19/England/CAMB-784BC/2020 | Europe / United Kingdom / England | 2020 | Department of Pathology, University of Cambridge | COVID-19 Genomics UK (COG-UK) Consortium | Luke W Meredith, M. Estee Torok , Myra Hosmillo, William L. Hamilton, Martin D. Curran, Theresa Feltwell, Grant Hall, Anna Yakovleva, Fahad A Khokhar, Charlotte J. Houldcroft, Laura G Caller, Aminu S. Jahun, Sarah L. Caddy, Ian Goodfellow |
| EPI_ISL_433742 | hCoV-19/England/CAMB-784DA/2020 | Europe / United Kingdom / England | 2020-04-06 | Department of Pathology, University of Cambridge | COVID-19 Genomics UK (COG-UK) Consortium | Luke W Meredith, M. Estee Torok , Myra Hosmillo, William L. Hamilton, Martin D. Curran, Theresa Feltwell, Grant Hall, Anna Yakovleva, Fahad A Khokhar, Charlotte J. Houldcroft, Laura G Caller, Aminu S. Jahun, Sarah L. Caddy, Ian Goodfellow |
| EPI_ISL_433765 | hCoV-19/England/CAMB-7870E/2020 | Europe / United Kingdom / England | 2020-04-05 | Department of Pathology, University of Cambridge | COVID-19 Genomics UK (COG-UK) Consortium | Luke W Meredith, M. Estee Torok , Myra Hosmillo, William L. Hamilton, Martin D. Curran, Theresa Feltwell, Grant Hall, Anna Yakovleva, Fahad A Khokhar, Charlotte J. Houldcroft, Laura G Caller, Aminu S. Jahun, Sarah L. Caddy, Ian Goodfellow |
| EPI_ISL_433766 | hCoV-19/England/CAMB-7872C/2020 | Europe / United Kingdom / England | 2020 | Department of Pathology, University of Cambridge | COVID-19 Genomics UK (COG-UK) Consortium | Luke W Meredith, M. Estee Torok , Myra Hosmillo, William L. Hamilton, Martin D. Curran, Theresa Feltwell, Grant Hall, Anna Yakovleva, Fahad A Khokhar, Charlotte J. Houldcroft, Laura G Caller, Aminu S. Jahun, Sarah L. Caddy, Ian Goodfellow |
| EPI_ISL_433767 | hCoV-19/England/CAMB-7873B/2020 | Europe / United Kingdom / England | 2020-04-06 | Department of Pathology, University of Cambridge | COVID-19 Genomics UK (COG-UK) Consortium | Luke W Meredith, M. Estee Torok , Myra Hosmillo, William L. Hamilton, Martin D. Curran, Theresa Feltwell, Grant Hall, Anna Yakovleva, Fahad A Khokhar, Charlotte J. Houldcroft, Laura G Caller, Aminu S. Jahun, Sarah L. Caddy, Ian Goodfellow |
| EPI_ISL_433768 | hCoV-19/England/CAMB-7874A/2020 | Europe / United Kingdom / England | 2020-04-06 | Department of Pathology, University of Cambridge | COVID-19 Genomics UK (COG-UK) Consortium | Luke W Meredith, M. Estee Torok , Myra Hosmillo, William L. Hamilton, Martin D. Curran, Theresa Feltwell, Grant Hall, Anna Yakovleva, Fahad A Khokhar, Charlotte J. Houldcroft, Laura G Caller, Aminu S. Jahun, Sarah L. Caddy, Ian Goodfellow |
| EPI_ISL_433770 | hCoV-19/England/CAMB-78786/2020 | Europe / United Kingdom / England | 2020-04-05 | Department of Pathology, University of Cambridge | COVID-19 Genomics UK (COG-UK) Consortium | Luke W Meredith, M. Estee Torok , Myra Hosmillo, William L. Hamilton, Martin D. Curran, Theresa Feltwell, Grant Hall, Anna Yakovleva, Fahad A Khokhar, Charlotte J. Houldcroft, Laura G Caller, Aminu S. Jahun, Sarah L. Caddy, Ian Goodfellow |
| EPI_ISL_433771 | hCoV-19/England/CAMB-787A4/2020 | Europe / United Kingdom / England | 2020-04-05 | Department of Pathology, University of Cambridge | COVID-19 Genomics UK (COG-UK) Consortium | Luke W Meredith, M. Estee Torok , Myra Hosmillo, William L. Hamilton, Martin D. Curran, Theresa Feltwell, Grant Hall, Anna Yakovleva, Fahad A Khokhar, Charlotte J. Houldcroft, Laura G Caller, Aminu S. Jahun, Sarah L. Caddy, Ian Goodfellow |
| EPI_ISL_433772 | hCoV-19/England/CAMB-787B3/2020 | Europe / United Kingdom / England | 2020-04-08 | Department of Pathology, University of Cambridge | COVID-19 Genomics UK (COG-UK) Consortium | Luke W Meredith, M. Estee Torok , Myra Hosmillo, William L. Hamilton, Martin D. Curran, Theresa Feltwell, Grant Hall, Anna Yakovleva, Fahad A Khokhar, Charlotte J. Houldcroft, Laura G Caller, Aminu S. Jahun, Sarah L. Caddy, Ian Goodfellow |
| EPI_ISL_433775 | hCoV-19/England/CAMB-787E0/2020 | Europe / United Kingdom / England | 2020-04-07 | Department of Pathology, University of Cambridge | COVID-19 Genomics UK (COG-UK) Consortium | Luke W Meredith, M. Estee Torok , Myra Hosmillo, William L. Hamilton, Martin D. Curran, Theresa Feltwell, Grant Hall, Anna Yakovleva, Fahad A Khokhar, Charlotte J. Houldcroft, Laura G Caller, Aminu S. Jahun, Sarah L. Caddy, Ian Goodfellow |
| EPI_ISL_433776 | hCoV-19/England/CAMB-7880B/2020 | Europe / United Kingdom / England | 2020-04-07 | Department of Pathology, University of Cambridge | COVID-19 Genomics UK (COG-UK) Consortium | Luke W Meredith, M. Estee Torok , Myra Hosmillo, William L. Hamilton, Martin D. Curran, Theresa Feltwell, Grant Hall, Anna Yakovleva, Fahad A Khokhar, Charlotte J. Houldcroft, Laura G Caller, Aminu S. Jahun, Sarah L. Caddy, Ian Goodfellow |
| EPI_ISL_433778 | hCoV-19/England/CAMB-78829/2020 | Europe / United Kingdom / England | 2020-04-07 | Department of Pathology, University of Cambridge | COVID-19 Genomics UK (COG-UK) Consortium | Luke W Meredith, M. Estee Torok , Myra Hosmillo, William L. Hamilton, Martin D. Curran, Theresa Feltwell, Grant Hall, Anna Yakovleva, Fahad A Khokhar, Charlotte J. Houldcroft, Laura G Caller, Aminu S. Jahun, Sarah L. Caddy, Ian Goodfellow |
| EPI_ISL_433779 | hCoV-19/England/CAMB-78838/2020 | Europe / United Kingdom / England | 2020-04-08 | Department of Pathology, University of Cambridge | COVID-19 Genomics UK (COG-UK) Consortium | Luke W Meredith, M. Estee Torok , Myra Hosmillo, William L. Hamilton, Martin D. Curran, Theresa Feltwell, Grant Hall, Anna Yakovleva, Fahad A Khokhar, Charlotte J. Houldcroft, Laura G Caller, Aminu S. Jahun, Sarah L. Caddy, Ian Goodfellow |
| EPI_ISL_433780 | hCoV-19/England/CAMB-78847/2020 | Europe / United Kingdom / England | 2020-04-06 | Department of Pathology, University of Cambridge | COVID-19 Genomics UK (COG-UK) Consortium | Luke W Meredith, M. Estee Torok , Myra Hosmillo, William L. Hamilton, Martin D. Curran, Theresa Feltwell, Grant Hall, Anna Yakovleva, Fahad A Khokhar, Charlotte J. Houldcroft, Laura G Caller, Aminu S. Jahun, Sarah L. Caddy, Ian Goodfellow |
| EPI_ISL_433781 | hCoV-19/England/CAMB-78865/2020 | Europe / United Kingdom / England | 2020-04-06 | Department of Pathology, University of Cambridge | COVID-19 Genomics UK (COG-UK) Consortium | Luke W Meredith, M. Estee Torok , Myra Hosmillo, William L. Hamilton, Martin D. Curran, Theresa Feltwell, Grant Hall, Anna Yakovleva, Fahad A Khokhar, Charlotte J. Houldcroft, Laura G Caller, Aminu S. Jahun, Sarah L. Caddy, Ian Goodfellow |
| EPI_ISL_433782 | hCoV-19/England/CAMB-78874/2020 | Europe / United Kingdom / England | 2020-04-06 | Department of Pathology, University of Cambridge | COVID-19 Genomics UK (COG-UK) Consortium | Luke W Meredith, M. Estee Torok , Myra Hosmillo, William L. Hamilton, Martin D. Curran, Theresa Feltwell, Grant Hall, Anna Yakovleva, Fahad A Khokhar, Charlotte J. Houldcroft, Laura G Caller, Aminu S. Jahun, Sarah L. Caddy, Ian Goodfellow |
| EPI_ISL_433783 | hCoV-19/England/CAMB-78883/2020 | Europe / United Kingdom / England | 2020-04-06 | Department of Pathology, University of Cambridge | COVID-19 Genomics UK (COG-UK) Consortium | Luke W Meredith, M. Estee Torok , Myra Hosmillo, William L. Hamilton, Martin D. Curran, Theresa Feltwell, Grant Hall, Anna Yakovleva, Fahad A Khokhar, Charlotte J. Houldcroft, Laura G Caller, Aminu S. Jahun, Sarah L. Caddy, Ian Goodfellow |
| EPI_ISL_433784 | hCoV-19/England/CAMB-78892/2020 | Europe / United Kingdom / England | 2020-04-07 | Department of Pathology, University of Cambridge | COVID-19 Genomics UK (COG-UK) Consortium | Luke W Meredith, M. Estee Torok , Myra Hosmillo, William L. Hamilton, Martin D. Curran, Theresa Feltwell, Grant Hall, Anna Yakovleva, Fahad A Khokhar, Charlotte J. Houldcroft, Laura G Caller, Aminu S. Jahun, Sarah L. Caddy, Ian Goodfellow |
| EPI_ISL_433785 | hCoV-19/England/CAMB-788A1/2020 | Europe / United Kingdom / England | 2020-04-06 | Department of Pathology, University of Cambridge | COVID-19 Genomics UK (COG-UK) Consortium | Luke W Meredith, M. Estee Torok , Myra Hosmillo, William L. Hamilton, Martin D. Curran, Theresa Feltwell, Grant Hall, Anna Yakovleva, Fahad A Khokhar, Charlotte J. Houldcroft, Laura G Caller, Aminu S. Jahun, Sarah L. Caddy, Ian Goodfellow |
| EPI_ISL_433786 | hCoV-19/England/CAMB-788B0/2020 | Europe / United Kingdom / England | 2020-04-07 | Department of Pathology, University of Cambridge | COVID-19 Genomics UK (COG-UK) Consortium | Luke W Meredith, M. Estee Torok , Myra Hosmillo, William L. Hamilton, Martin D. Curran, Theresa Feltwell, Grant Hall, Anna Yakovleva, Fahad A Khokhar, Charlotte J. Houldcroft, Laura G Caller, Aminu S. Jahun, Sarah L. Caddy, Ian Goodfellow |
| EPI_ISL_433787 | hCoV-19/England/CAMB-788CF/2020 | Europe / United Kingdom / England | 2020-04-06 | Department of Pathology, University of Cambridge | COVID-19 Genomics UK (COG-UK) Consortium | Luke W Meredith, M. Estee Torok , Myra Hosmillo, William L. Hamilton, Martin D. Curran, Theresa Feltwell, Grant Hall, Anna Yakovleva, Fahad A Khokhar, Charlotte J. Houldcroft, Laura G Caller, Aminu S. Jahun, Sarah L. Caddy, Ian Goodfellow |
| EPI_ISL_433789 | hCoV-19/England/CAMB-788ED/2020 | Europe / United Kingdom / England | 2020-04-07 | Department of Pathology, University of Cambridge | COVID-19 Genomics UK (COG-UK) Consortium | Luke W Meredith, M. Estee Torok , Myra Hosmillo, William L. Hamilton, Martin D. Curran, Theresa Feltwell, Grant Hall, Anna Yakovleva, Fahad A Khokhar, Charlotte J. Houldcroft, Laura G Caller, Aminu S. Jahun, Sarah L. Caddy, Ian Goodfellow |
| EPI_ISL_433790 | hCoV-19/England/CAMB-788FC/2020 | Europe / United Kingdom / England | 2020-04-07 | Department of Pathology, University of Cambridge | COVID-19 Genomics UK (COG-UK) Consortium | Luke W Meredith, M. Estee Torok , Myra Hosmillo, William L. Hamilton, Martin D. Curran, Theresa Feltwell, Grant Hall, Anna Yakovleva, Fahad A Khokhar, Charlotte J. Houldcroft, Laura G Caller, Aminu S. Jahun, Sarah L. Caddy, Ian Goodfellow |
| EPI_ISL_433791 | hCoV-19/England/CAMB-78917/2020 | Europe / United Kingdom / England | 2020-04-08 | Department of Pathology, University of Cambridge | COVID-19 Genomics UK (COG-UK) Consortium | Luke W Meredith, M. Estee Torok , Myra Hosmillo, William L. Hamilton, Martin D. Curran, Theresa Feltwell, Grant Hall, Anna Yakovleva, Fahad A Khokhar, Charlotte J. Houldcroft, Laura G Caller, Aminu S. Jahun, Sarah L. Caddy, Ian Goodfellow |
| EPI_ISL_433792 | hCoV-19/England/CAMB-78926/2020 | Europe / United Kingdom / England | 2020-04-07 | Department of Pathology, University of Cambridge | COVID-19 Genomics UK (COG-UK) Consortium | Luke W Meredith, M. Estee Torok , Myra Hosmillo, William L. Hamilton, Martin D. Curran, Theresa Feltwell, Grant Hall, Anna Yakovleva, Fahad A Khokhar, Charlotte J. Houldcroft, Laura G Caller, Aminu S. Jahun, Sarah L. Caddy, Ian Goodfellow |
| EPI_ISL_433793 | hCoV-19/England/CAMB-78935/2020 | Europe / United Kingdom / England | 2020-04-06 | Department of Pathology, University of Cambridge | COVID-19 Genomics UK (COG-UK) Consortium | Luke W Meredith, M. Estee Torok , Myra Hosmillo, William L. Hamilton, Martin D. Curran, Theresa Feltwell, Grant Hall, Anna Yakovleva, Fahad A Khokhar, Charlotte J. Houldcroft, Laura G Caller, Aminu S. Jahun, Sarah L. Caddy, Ian Goodfellow |
| EPI_ISL_433794 | hCoV-19/England/CAMB-78944/2020 | Europe / United Kingdom / England | 2020-04-06 | Department of Pathology, University of Cambridge | COVID-19 Genomics UK (COG-UK) Consortium | Luke W Meredith, M. Estee Torok , Myra Hosmillo, William L. Hamilton, Martin D. Curran, Theresa Feltwell, Grant Hall, Anna Yakovleva, Fahad A Khokhar, Charlotte J. Houldcroft, Laura G Caller, Aminu S. Jahun, Sarah L. Caddy, Ian Goodfellow |
| EPI_ISL_433795 | hCoV-19/England/CAMB-78953/2020 | Europe / United Kingdom / England | 2020-04-07 | Department of Pathology, University of Cambridge | COVID-19 Genomics UK (COG-UK) Consortium | Luke W Meredith, M. Estee Torok , Myra Hosmillo, William L. Hamilton, Martin D. Curran, Theresa Feltwell, Grant Hall, Anna Yakovleva, Fahad A Khokhar, Charlotte J. Houldcroft, Laura G Caller, Aminu S. Jahun, Sarah L. Caddy, Ian Goodfellow |
| EPI_ISL_433796 | hCoV-19/England/CAMB-78962/2020 | Europe / United Kingdom / England | 2020-04-07 | Department of Pathology, University of Cambridge | COVID-19 Genomics UK (COG-UK) Consortium | Luke W Meredith, M. Estee Torok , Myra Hosmillo, William L. Hamilton, Martin D. Curran, Theresa Feltwell, Grant Hall, Anna Yakovleva, Fahad A Khokhar, Charlotte J. Houldcroft, Laura G Caller, Aminu S. Jahun, Sarah L. Caddy, Ian Goodfellow |
| EPI_ISL_433797 | hCoV-19/England/CAMB-78971/2020 | Europe / United Kingdom / England | 2020-04-07 | Department of Pathology, University of Cambridge | COVID-19 Genomics UK (COG-UK) Consortium | Luke W Meredith, M. Estee Torok , Myra Hosmillo, William L. Hamilton, Martin D. Curran, Theresa Feltwell, Grant Hall, Anna Yakovleva, Fahad A Khokhar, Charlotte J. Houldcroft, Laura G Caller, Aminu S. Jahun, Sarah L. Caddy, Ian Goodfellow |
| EPI_ISL_433798 | hCoV-19/England/CAMB-78980/2020 | Europe / United Kingdom / England | 2020-04-07 | Department of Pathology, University of Cambridge | COVID-19 Genomics UK (COG-UK) Consortium | Luke W Meredith, M. Estee Torok , Myra Hosmillo, William L. Hamilton, Martin D. Curran, Theresa Feltwell, Grant Hall, Anna Yakovleva, Fahad A Khokhar, Charlotte J. Houldcroft, Laura G Caller, Aminu S. Jahun, Sarah L. Caddy, Ian Goodfellow |
| EPI_ISL_434100 | hCoV-19/USA/WA-S340/2020 | North America / USA / Washington / Yaki | 2020-04-04 | Washington State Department of Health | Seattle Flu Study | Chu et al |
| EPI_ISL_434101 | hCoV-19/USA/WA-S341/2020 | North America / USA / Washington / Yaki | 2020-04-02 | Washington State Department of Health | Seattle Flu Study | Chu et al |
| EPI_ISL_434102 | hCoV-19/USA/WA-S342/2020 | North America / USA / Washington / Yaki | 2020-04-02 | Washington State Department of Health | Seattle Flu Study | Chu et al |
| EPI_ISL_434103 | hCoV-19/USA/WA-S343/2020 | North America / USA / Washington / Yaki | 2020-04-02 | Washington State Department of Health | Seattle Flu Study | Chu et al |
| EPI_ISL_434104 | hCoV-19/USA/WA-S344/2020 | North America / USA / Washington / Yaki | 2020-04-02 | Washington State Department of Health | Seattle Flu Study | Chu et al |
| EPI_ISL_434105 | hCoV-19/USA/WA-S345/2020 | North America / USA / Washington / Yaki | 2020-04-05 | Washington State Department of Health | Seattle Flu Study | Chu et al |
| EPI_ISL_434106 | hCoV-19/USA/WA-S346/2020 | North America / USA / Washington / Yaki | 2020-04-04 | Washington State Department of Health | Seattle Flu Study | Chu et al |
| EPI_ISL_434107 | hCoV-19/USA/WA-S347/2020 | North America / USA / Washington / Yaki | 2020-04-01 | Washington State Department of Health | Seattle Flu Study | Chu et al |
| EPI_ISL_434108 | hCoV-19/USA/WA-S348/2020 | North America / USA / Washington / Asot | 2020-04-01 | Washington State Department of Health | Seattle Flu Study | Chu et al |
| EPI_ISL_434109 | hCoV-19/USA/WA-S349/2020 | North America / USA / Washington / Mas | 2020-04-02 | Washington State Department of Health | Seattle Flu Study | Chu et al |
| EPI_ISL_434110 | hCoV-19/USA/WA-S350/2020 | North America / USA / Washington / Yaki | 2020-04-02 | Washington State Department of Health | Seattle Flu Study | Chu et al |
| EPI_ISL_434111 | hCoV-19/USA/WA-S351/2020 | North America / USA / Washington / Yaki | 2020-03-25 | Washington State Department of Health | Seattle Flu Study | Chu et al |
| EPI_ISL_434112 | hCoV-19/USA/WA-S352/2020 | North America / USA / Washington / Yaki | 2020-03-25 | Washington State Department of Health | Seattle Flu Study | Chu et al |
| EPI_ISL_434113 | hCoV-19/USA/WA-S353/2020 | North America / USA / Washington / Yaki | 2020-03-25 | Washington State Department of Health | Seattle Flu Study | Chu et al |
| EPI_ISL_434114 | hCoV-19/USA/WA-S354/2020 | North America / USA / Washington / Yaki | 2020-03-25 | Washington State Department of Health | Seattle Flu Study | Chu et al |
| EPI_ISL_434115 | hCoV-19/USA/WA-S355/2020 | North America / USA / Washington / Yaki | 2020-03-25 | Washington State Department of Health | Seattle Flu Study | Chu et al |
| EPI_ISL_434116 | hCoV-19/USA/WA-S356/2020 | North America / USA / Washington / Yaki | 2020-03-25 | Washington State Department of Health | Seattle Flu Study | Chu et al |
| EPI_ISL_434117 | hCoV-19/USA/WA-S357/2020 | North America / USA / Washington / Yaki | 2020-03-25 | Washington State Department of Health | Seattle Flu Study | Chu et al |
| EPI_ISL_434118 | hCoV-19/USA/WA-S358/2020 | North America / USA / Washington / Yaki | 2020-03-25 | Washington State Department of Health | Seattle Flu Study | Chu et al |
| EPI_ISL_434119 | hCoV-19/USA/WA-S359/2020 | North America / USA / Washington / Yaki | 2020-03-25 | Washington State Department of Health | Seattle Flu Study | Chu et al |
| EPI_ISL_434120 | hCoV-19/USA/WA-S360/2020 | North America / USA / Washington / Yaki | 2020-03-25 | Washington State Department of Health | Seattle Flu Study | Chu et al |
| EPI_ISL_434121 | hCoV-19/USA/WA-S361/2020 | North America / USA / Washington / Wha | 2020-03-26 | Washington State Department of Health | Seattle Flu Study | Chu et al |
| EPI_ISL_434122 | hCoV-19/USA/WA-S362/2020 | North America / USA / Washington / Wha | 2020-03-26 | Washington State Department of Health | Seattle Flu Study | Chu et al |
| EPI_ISL_434123 | hCoV-19/USA/WA-S363/2020 | North America / USA / Washington / Wha | 2020-03-26 | Washington State Department of Health | Seattle Flu Study | Chu et al |
| EPI_ISL_434124 | hCoV-19/USA/WA-S364/2020 | North America / USA / Washington / Wha | 2020-03-26 | Washington State Department of Health | Seattle Flu Study | Chu et al |
| EPI_ISL_434125 | hCoV-19/USA/WA-S365/2020 | North America / USA / Washington | 2020-03-10 | Washington State Department of Health | Seattle Flu Study | Chu et al |
| EPI_ISL_434126 | hCoV-19/USA/WA-S366/2020 | North America / USA / Washington | 2020-03-09 | Washington State Department of Health | Seattle Flu Study | Chu et al |
| EPI_ISL_434127 | hCoV-19/USA/WA-S367/2020 | North America / USA / Washington | 2020-03-08 | Washington State Department of Health | Seattle Flu Study | Chu et al |
| EPI_ISL_434128 | hCoV-19/USA/WA-S368/2020 | North America / USA / Washington / Yaki | 2020-03-26 | Washington State Department of Health | Seattle Flu Study | Chu et al |
| EPI_ISL_434129 | hCoV-19/USA/WA-S369/2020 | North America / USA / Washington / Yaki | 2020-03-26 | Washington State Department of Health | Seattle Flu Study | Chu et al |
| EPI_ISL_434130 | hCoV-19/USA/WA-S370/2020 | North America / USA / Washington / Yaki | 2020-03-28 | Washington State Department of Health | Seattle Flu Study | Chu et al |
| EPI_ISL_434131 | hCoV-19/USA/WA-S371/2020 | North America / USA / Washington / Skag | 2020-03-27 | Washington State Department of Health | Seattle Flu Study | Chu et al |
| EPI_ISL_434132 | hCoV-19/USA/WA-S372/2020 | North America / USA / Washington / Yaki | 2020-03-26 | Washington State Department of Health | Seattle Flu Study | Chu et al |
| EPI_ISL_434133 | hCoV-19/USA/WA-S373/2020 | North America / USA / Washington / Yaki | 2020-03-28 | Washington State Department of Health | Seattle Flu Study | Chu et al |
| EPI_ISL_434134 | hCoV-19/USA/WA-S374/2020 | North America / USA / Washington / Yaki | 2020-03-26 | Washington State Department of Health | Seattle Flu Study | Chu et al |
| EPI_ISL_434135 | hCoV-19/USA/WA-S375/2020 | North America / USA / Washington / Yaki | 2020-03-26 | Washington State Department of Health | Seattle Flu Study | Chu et al |
| EPI_ISL_434136 | hCoV-19/USA/WA-S376/2020 | North America / USA / Washington / Yaki | 2020-03-27 | Washington State Department of Health | Seattle Flu Study | Chu et al |
| EPI_ISL_434137 | hCoV-19/USA/WA-S377/2020 | North America / USA / Washington / Yaki | 2020-03-26 | Washington State Department of Health | Seattle Flu Study | Chu et al |
| EPI_ISL_434138 | hCoV-19/USA/WA-S378/2020 | North America / USA / Washington / Skag | 2020-03-27 | Washington State Department of Health | Seattle Flu Study | Chu et al |
| EPI_ISL_434139 | hCoV-19/USA/WA-S379/2020 | North America / USA / Washington / Skag | 2020-03-27 | Washington State Department of Health | Seattle Flu Study | Chu et al |
| EPI_ISL_434140 | hCoV-19/USA/WA-S380/2020 | North America / USA / Washington / Fran | 2020-03-27 | Washington State Department of Health | Seattle Flu Study | Chu et al |
| EPI_ISL_434141 | hCoV-19/USA/WA-S381/2020 | North America / USA / Washington / Skag | 2020-03-29 | Washington State Department of Health | Seattle Flu Study | Chu et al |
| EPI_ISL_434142 | hCoV-19/USA/WA-S382/2020 | North America / USA / Washington / Skag | 2020-03-27 | Washington State Department of Health | Seattle Flu Study | Chu et al |
| EPI_ISL_434143 | hCoV-19/USA/WA-S383/2020 | North America / USA / Washington / Adar | 2020-03-27 | Washington State Department of Health | Seattle Flu Study | Chu et al |
| EPI_ISL_434144 | hCoV-19/USA/WA-S384/2020 | North America / USA / Washington / Skag | 2020-03-27 | Washington State Department of Health | Seattle Flu Study | Chu et al |
| EPI_ISL_434145 | hCoV-19/USA/WA-S385/2020 | North America / USA / Washington / Pien | 2020-03-28 | Washington State Department of Health | Seattle Flu Study | Chu et al |
| EPI_ISL_434146 | hCoV-19/USA/WA-S386/2020 | North America / USA / Washington / Yaki | 2020-03-26 | Washington State Department of Health | Seattle Flu Study | Chu et al |
| EPI_ISL_434147 | hCoV-19/USA/WA-S387/2020 | North America / USA / Washington / Skag | 2020-03-27 | Washington State Department of Health | Seattle Flu Study | Chu et al |
| EPI_ISL_434148 | hCoV-19/USA/WA-S388/2020 | North America / USA / Washington / Skag | 2020-03-27 | Washington State Department of Health | Seattle Flu Study | Chu et al |
| EPI_ISL_434149 | hCoV-19/USA/WA-S389/2020 | North America / USA / Washington / Skag | 2020-03-28 | Washington State Department of Health | Seattle Flu Study | Chu et al |
| EPI_ISL_434150 | hCoV-19/USA/WA-S390/2020 | North America / USA / Washington / Skag | 2020-03-27 | Washington State Department of Health | Seattle Flu Study | Chu et al |
| EPI_ISL_434151 | hCoV-19/USA/WA-S391/2020 | North America / USA / Washington / Yaki | 2020-03-26 | Washington State Department of Health | Seattle Flu Study | Chu et al |
| EPI_ISL_434152 | hCoV-19/USA/WA-S392/2020 | North America / USA / Washington / King | 2020-03-30 | Washington State Department of Health | Seattle Flu Study | Chu et al |
| EPI_ISL_434153 | hCoV-19/USA/WA-S393/2020 | North America / USA / Washington / Yaki | 2020-03-28 | Washington State Department of Health | Seattle Flu Study | Chu et al |
| EPI_ISL_434154 | hCoV-19/USA/WA-S394/2020 | North America / USA / Washington / Yaki | 2020-03-30 | Washington State Department of Health | Seattle Flu Study | Chu et al |
| EPI_ISL_434155 | hCoV-19/USA/WA-S395/2020 | North America / USA / Washington / Snol | 2020-03-30 | Washington State Department of Health | Seattle Flu Study | Chu et al |
| EPI_ISL_434156 | hCoV-19/USA/WA-S396/2020 | North America / USA / Washington / Yaki | 2020-03-25 | Washington State Department of Health | Seattle Flu Study | Chu et al |
| EPI_ISL_434157 | hCoV-19/USA/WA-S397/2020 | North America / USA / Washington / Yaki | 2020-03-25 | Washington State Department of Health | Seattle Flu Study | Chu et al |
| EPI_ISL_434158 | hCoV-19/USA/WA-S398/2020 | North America / USA / Washington / King | 2020-03-30 | Washington State Department of Health | Seattle Flu Study | Chu et al |
| EPI_ISL_434159 | hCoV-19/USA/WA-S399/2020 | North America / USA / Washington / Yaki | 2020-03-28 | Washington State Department of Health | Seattle Flu Study | Chu et al |
| EPI_ISL_434160 | hCoV-19/USA/WA-S400/2020 | North America / USA / Washington / Yaki | 2020-03-28 | Washington State Department of Health | Seattle Flu Study | Chu et al |
| EPI_ISL_434161 | hCoV-19/USA/WA-S401/2020 | North America / USA / Washington / Yaki | 2020-03-28 | Washington State Department of Health | Seattle Flu Study | Chu et al |
| EPI_ISL_434162 | hCoV-19/USA/WA-S402/2020 | North America / USA / Washington / Yaki | 2020-03-28 | Washington State Department of Health | Seattle Flu Study | Chu et al |
| EPI_ISL_434163 | hCoV-19/USA/WA-S403/2020 | North America / USA / Washington / King | 2020-03-26 | Washington State Department of Health | Seattle Flu Study | Chu et al |
| EPI_ISL_434164 | hCoV-19/USA/WA-S404/2020 | North America / USA / Washington / Yaki | 2020-03-26 | Washington State Department of Health | Seattle Flu Study | Chu et al |
| EPI_ISL_434165 | hCoV-19/USA/WA-S405/2020 | North America / USA / Washington / Yaki | 2020-03-25 | Washington State Department of Health | Seattle Flu Study | Chu et al |
| EPI_ISL_434166 | hCoV-19/USA/WA-S406/2020 | North America / USA / Washington / Yaki | 2020-03-26 | Washington State Department of Health | Seattle Flu Study | Chu et al |
| EPI_ISL_434167 | hCoV-19/USA/WA-S407/2020 | North America / USA / Washington / Yaki | 2020-03-28 | Washington State Department of Health | Seattle Flu Study | Chu et al |
| EPI_ISL_434169 | hCoV-19/USA/WA-S409/2020 | North America / USA / Washington / Yaki | 2020-03-28 | Washington State Department of Health | Seattle Flu Study | Chu et al |
| EPI_ISL_434170 | hCoV-19/USA/WA-S410/2020 | North America / USA / Washington / Yaki | 2020-03-29 | Washington State Department of Health | Seattle Flu Study | Chu et al |
| EPI_ISL_434171 | hCoV-19/USA/WA-S411/2020 | North America / USA / Washington / Yaki | 2020-03-28 | Washington State Department of Health | Seattle Flu Study | Chu et al |
| EPI_ISL_434172 | hCoV-19/USA/WA-S412/2020 | North America / USA / Washington / King | 2020-03-30 | Washington State Department of Health | Seattle Flu Study | Chu et al |
| EPI_ISL_434173 | hCoV-19/USA/WA-S413/2020 | North America / USA / Washington / Fran | 2020-03-26 | Washington State Department of Health | Seattle Flu Study | Chu et al |
| EPI_ISL_434175 | hCoV-19/USA/WA-S415/2020 | North America / USA / Washington / Yaki | 2020-03-28 | Washington State Department of Health | Seattle Flu Study | Chu et al |
| EPI_ISL_434176 | hCoV-19/USA/WA-S416/2020 | North America / USA / Washington / Yaki | 2020-03-29 | Washington State Department of Health | Seattle Flu Study | Chu et al |
| EPI_ISL_434177 | hCoV-19/USA/WA-S417/2020 | North America / USA / Washington / Yaki | 2020-03-28 | Washington State Department of Health | Seattle Flu Study | Chu et al |
| EPI_ISL_434178 | hCoV-19/USA/WA-S418/2020 | North America / USA / Washington / Yaki | 2020-03-25 | Washington State Department of Health | Seattle Flu Study | Chu et al |
| EPI_ISL_434179 | hCoV-19/USA/WA-S419/2020 | North America / USA / Washington / Yaki | 2020-03-28 | Washington State Department of Health | Seattle Flu Study | Chu et al |
| EPI_ISL_434180 | hCoV-19/USA/WA-S420/2020 | North America / USA / Washington / Yaki | 2020-03-28 | Washington State Department of Health | Seattle Flu Study | Chu et al |
| EPI_ISL_434182 | hCoV-19/USA/WA-S422/2020 | North America / USA / Washington / Yaki | 2020-03-25 | Washington State Department of Health | Seattle Flu Study | Chu et al |
| EPI_ISL_434183 | hCoV-19/USA/WA-S423/2020 | North America / USA / Washington / Yaki | 2020-03-28 | Washington State Department of Health | Seattle Flu Study | Chu et al |

| | | | | | | |
|---|---|---|---|---|---|---|
| EPI_ISL_434184 | hCoV-19/USA/WA-S424/2020 | North America / USA / Washington / Yaki | 2020-03-28 | Washington State Department of Health | Seattle Flu Study | Chu et al |
| EPI_ISL_434185 | hCoV-19/USA/WA-S425/2020 | North America / USA / Washington / Yaki | 2020-03-28 | Washington State Department of Health | Seattle Flu Study | Chu et al |
| EPI_ISL_434186 | hCoV-19/USA/WA-S426/2020 | North America / USA / Washington / Yaki | 2020-03-28 | Washington State Department of Health | Seattle Flu Study | Chu et al |
| EPI_ISL_434187 | hCoV-19/USA/WA-S427/2020 | North America / USA / Washington / Yaki | 2020-03-28 | Washington State Department of Health | Seattle Flu Study | Chu et al |
| EPI_ISL_434188 | hCoV-19/USA/WA-S428/2020 | North America / USA / Washington / Yaki | 2020-03-28 | Washington State Department of Health | Seattle Flu Study | Chu et al |
| EPI_ISL_434189 | hCoV-19/USA/WA-S429/2020 | North America / USA / Washington / Yaki | 2020-03-25 | Washington State Department of Health | Seattle Flu Study | Chu et al |
| EPI_ISL_434190 | hCoV-19/USA/WA-S430/2020 | North America / USA / Washington / Yaki | 2020-03-25 | Washington State Department of Health | Seattle Flu Study | Chu et al |
| EPI_ISL_434191 | hCoV-19/USA/WA-S431/2020 | North America / USA / Washington / Snol | 2020-03-28 | Washington State Department of Health | Seattle Flu Study | Chu et al |
| EPI_ISL_434192 | hCoV-19/USA/WA-S432/2020 | North America / USA / Washington / Yaki | 2020-03-28 | Washington State Department of Health | Seattle Flu Study | Chu et al |
| EPI_ISL_434193 | hCoV-19/USA/WA-S433/2020 | North America / USA / Washington / Yaki | 2020-03-28 | Washington State Department of Health | Seattle Flu Study | Chu et al |
| EPI_ISL_434194 | hCoV-19/USA/WA-S434/2020 | North America / USA / Washington / Snol | 2020-03-28 | Washington State Department of Health | Seattle Flu Study | Chu et al |
| EPI_ISL_434195 | hCoV-19/USA/WA-S435/2020 | North America / USA / Washington | 2020-03-30 | Washington State Department of Health | Seattle Flu Study | Chu et al |
| EPI_ISL_434196 | hCoV-19/USA/WA-S436/2020 | North America / USA / Washington / Yaki | 2020-03-31 | Washington State Department of Health | Seattle Flu Study | Chu et al |
| EPI_ISL_434197 | hCoV-19/USA/WA-S437/2020 | North America / USA / Washington / Yaki | 2020-03-31 | Washington State Department of Health | Seattle Flu Study | Chu et al |
| EPI_ISL_434199 | hCoV-19/USA/WA-S439/2020 | North America / USA / Washington / Pien | 2020-04-02 | Washington State Department of Health | Seattle Flu Study | Chu et al |
| EPI_ISL_434656 | hCoV-19/Sweden/20-06503/2020 | Europe / Sweden / Stockholm | 2020-03-30 | Trollbackens VC | The Public Health Agency of Sweden | Amelie Holmqvist, Oskar Karlsson Lindsjo, Maria Lind Karlberg, Anna-Malin Linde, Olov Svartstrom, Anna Risberg, Theresa Enkirch, Mia Brytting, Karin Tegmark-Wisell |
| EPI_ISL_434657 | hCoV-19/Sweden/20-06813/2020 | Europe / Sweden / Skane | 2020-04-01 | Kristianstadkliniken | The Public Health Agency of Sweden | Mia Settergren Hammer, Oskar Karlsson Lindsjo, Maria Lind Karlberg, Anna-Malin Linde, Olov Svartstrom, Anna Risberg, Theresa Enkirch, Mia Brytting, Karin Tegmark-Wisell |
| EPI_ISL_434658 | hCoV-19/Sweden/20-06883/2020 | Europe / Sweden / Skane | 2020-04-03 | Lundens VC | The Public Health Agency of Sweden | Marita Dagner, Oskar Karlsson Lindsjo, Maria Lind Karlberg, Anna-Malin Linde, Olov Svartstrom, Anna Risberg, Theresa Enkirch, Mia Brytting, Karin Tegmark-Wisell |
| EPI_ISL_434659 | hCoV-19/Sweden/20-06908/2020 | Europe / Sweden / Uppsala | 2020-04-06 | Knivsta VC | The Public Health Agency of Sweden | Johanna Carlson, Oskar Karlsson Lindsjo, Maria Lind Karlberg, Anna-Malin Linde, Olov Svartstrom, Anna Risberg, Theresa Enkirch, Mia Brytting, Karin Tegmark-Wisell |
| EPI_ISL_434660 | hCoV-19/Sweden/20-06928/2020 | Europe / Sweden / Vastra Gotaland | 2020-04-06 | Narhalsan Backa vardcentral | The Public Health Agency of Sweden | Mats Olsson, Oskar Karlsson Lindsjo, Maria Lind Karlberg, Anna-Malin Linde, Olov Svartstrom, Anna Risberg, Theresa Enkirch, Mia Brytting, Karin Tegmark-Wisell |
| EPI_ISL_434661 | hCoV-19/Sweden/20-07010/2020 | Europe / Sweden / Vastra Gotaland | 2020-04-07 | Omtanken Grimmered | The Public Health Agency of Sweden | Bernd Sengpiel, Oskar Karlsson Lindsjo, Maria Lind Karlberg, Anna-Malin Linde, Olov Svartstrom, Anna Risberg, Theresa Enkirch, Mia Brytting, Karin Tegmark-Wisell |
| EPI_ISL_434662 | hCoV-19/Sweden/20-07013/2020 | Europe / Sweden / Vastra Gotaland | 2020-04-07 | Omtanken Grimmered | The Public Health Agency of Sweden | Bernd Sengpiel, Oskar Karlsson Lindsjo, Maria Lind Karlberg, Anna-Malin Linde, Olov Svartstrom, Anna Risberg, Theresa Enkirch, Mia Brytting, Karin Tegmark-Wisell |
| EPI_ISL_434663 | hCoV-19/Sweden/20-07025/2020 | Europe / Sweden / Stockholm | 2020-04-06 | Surbrunns VC | The Public Health Agency of Sweden | Erik Embring, Oskar Karlsson Lindsjo, Maria Lind Karlberg, Anna-Malin Linde, Olov Svartstrom, Anna Risberg, Theresa Enkirch, Mia Brytting, Karin Tegmark-Wisell |
| EPI_ISL_434664 | hCoV-19/Sweden/20-07034/2020 | Europe / Sweden / Vastra Gotaland | 2020-04-06 | Omtanken Grimmered | The Public Health Agency of Sweden | Bernd Sengpiel, Oskar Karlsson Lindsjo, Maria Lind Karlberg, Anna-Malin Linde, Olov Svartstrom, Anna Risberg, Theresa Enkirch, Mia Brytting, Karin Tegmark-Wisell |
| EPI_ISL_434665 | hCoV-19/Sweden/20-07042/2020 | Europe / Sweden / Uppsala | 2020-04-04 | Uppsala Narakut Aleris | The Public Health Agency of Sweden | Annika Nilsson, Oskar Karlsson Lindsjo, Maria Lind Karlberg, Anna-Malin Linde, Olov Svartstrom, Anna Risberg, Theresa Enkirch, Mia Brytting, Karin Tegmark-Wisell |
| EPI_ISL_434666 | hCoV-19/Sweden/20-07171/2020 | Europe / Sweden / Vasterbotten | 2020-04-08 | Hornefors Halsocentral | The Public Health Agency of Sweden | Camilla Elback, Oskar Karlsson Lindsjo, Maria Lind Karlberg, Anna-Malin Linde, Olov Svartstrom, Anna Risberg, Theresa Enkirch, Mia Brytting, Karin Tegmark-Wisell |
| EPI_ISL_434667 | hCoV-19/Sweden/20-07298/2020 | Europe / Sweden / Uppsala | 2020-04-14 | Ulltuna Vardcentral | The Public Health Agency of Sweden | Heidi Lindback, Oskar Karlsson Lindsjo, Maria Lind Karlberg, Anna-Malin Linde, Olov Svartstrom, Anna Risberg, Theresa Enkirch, Mia Brytting, Karin Tegmark-Wisell |
| EPI_ISL_434668 | hCoV-19/Sweden/20-07389/2020 | Europe / Sweden / Vastmanland | 2020-04-15 | Kungsors VC | The Public Health Agency of Sweden | Jessica Karlsson, Oskar Karlsson Lindsjo, Maria Lind Karlberg, Anna-Malin Linde, Olov Svartstrom, Anna Risberg, Theresa Enkirch, Mia Brytting, Karin Tegmark-Wisell |
| EPI_ISL_434669 | hCoV-19/Sweden/20-07390/2020 | Europe / Sweden / Vastmanland | 2020-04-15 | Lakargruppen | The Public Health Agency of Sweden | Boris Klanger, Oskar Karlsson Lindsjo, Maria Lind Karlberg, Anna-Malin Linde, Olov Svartstrom, Anna Risberg, Theresa Enkirch, Mia Brytting, Karin Tegmark-Wisell |
| EPI_ISL_434670 | hCoV-19/Sweden/20-07439/2020 | Europe / Sweden / Vastra Gotaland | 2020-04-17 | Narhalsan Molnlycke, Barn och ungdomsmedicin | The Public Health Agency of Sweden | Mats Reimer, Oskar Karlsson Lindsjo, Maria Lind Karlberg, Anna-Malin Linde, Olov Svartstrom, Anna Risberg, Theresa Enkirch, Mia Brytting, Karin Tegmark-Wisell |
| EPI_ISL_434671 | hCoV-19/Sweden/20-07440/2020 | Europe / Sweden / Vastra Gotaland | 2020-04-16 | Narhalsan Sjobo vardcentral | The Public Health Agency of Sweden | Lovisa Hjerten, Oskar Karlsson Lindsjo, Maria Lind Karlberg, Anna-Malin Linde, Olov Svartstrom, Anna Risberg, Theresa Enkirch, Mia Brytting, Karin Tegmark-Wisell |
| EPI_ISL_434672 | hCoV-19/Sweden/20-07459/2020 | Europe / Sweden / Stockholm | 2020-04-16 | Surbrunns VC | The Public Health Agency of Sweden | Erik Embring, Oskar Karlsson Lindsjo, Maria Lind Karlberg, Anna-Malin Linde, Olov Svartstrom, Anna Risberg, Theresa Enkirch, Mia Brytting, Karin Tegmark-Wisell |
| EPI_ISL_434673 | hCoV-19/Sweden/20-07480/2020 | Europe / Sweden / Vastra Gotaland | 2020-04-16 | Omtanken Grimmered | The Public Health Agency of Sweden | Bernd Sengpiel, Oskar Karlsson Lindsjo, Maria Lind Karlberg, Anna-Malin Linde, Olov Svartstrom, Anna Risberg, Theresa Enkirch, Mia Brytting, Karin Tegmark-Wisell |
| EPI_ISL_434674 | hCoV-19/Sweden/20-07572/2020 | Europe / Sweden / Vastra Gotaland | 2020-04-21 | Narhalsan Backa vardcentral | The Public Health Agency of Sweden | Mats Olsson, Oskar Karlsson Lindsjo, Maria Lind Karlberg, Anna-Malin Linde, Olov Svartstrom, Anna Risberg, Theresa Enkirch, Mia Brytting, Karin Tegmark-Wisell |
| EPI_ISL_434675 | hCoV-19/Sweden/20-07607/2020 | Europe / Sweden / Stockholm | 2020-04-20 | Surbrunns VC | The Public Health Agency of Sweden | Erik Embring, Oskar Karlsson Lindsjo, Maria Lind Karlberg, Anna-Malin Linde, Olov Svartstrom, Anna Risberg, Theresa Enkirch, Mia Brytting, Karin Tegmark-Wisell |
| EPI_ISL_434676 | hCoV-19/Sweden/20-07616/2020 | Europe / Sweden / Stockholm | 2020-04-20 | Surbrunns VC | The Public Health Agency of Sweden | Erik Embring, Oskar Karlsson Lindsjo, Maria Lind Karlberg, Anna-Malin Linde, Olov Svartstrom, Anna Risberg, Theresa Enkirch, Mia Brytting, Karin Tegmark-Wisell |
| EPI_ISL_434677 | hCoV-19/env/USA/FL-UF3/2020 | North America / USA / Florida | 2020-03-25 | Lednicky Laboratory at Emerging Pathogens Institute | Lednicky Laboratory at Emerging Pathogens Institute | Shankar,S.N., Wu,C.-Y., Clugston,J.R., Elbadry,M.A., Morris,J.G. Jr. and Lednicky,J.A. |
| EPI_ISL_426300 | hCoV-19/USA/NY-Wadsworth-26088-01/2020 | North America / USA / New York / Sarato | 2020-03-21 | Wadsworth Center, New York State Department of Health | Wadsworth Center, New York State Department of Health | Kirsten St. George, Daryl M. Lamson, Sara Griesemer, Jonathan Plitnick, Navjot Singh, Matthew D. Shudt, Erica Lasek-Nesselquist |
| EPI_ISL_426301 | hCoV-19/USA/NY-Wadsworth-31302-01/2020 | North America / USA / New York / Ulster | 2020-03-27 | Wadsworth Center, New York State Department of Health | Wadsworth Center, New York State Department of Health | Kirsten St. George, Daryl M. Lamson, Sara Griesemer, Jonathan Plitnick, Navjot Singh, Matthew D. Shudt, Erica Lasek-Nesselquist |
| EPI_ISL_426302 | hCoV-19/USA/NY-Wadsworth-26856-01/2020 | North America / USA / New York / Putnar | 2020-03-21 | Wadsworth Center, New York State Department of Health | Wadsworth Center, New York State Department of Health | Kirsten St. George, Daryl M. Lamson, Sara Griesemer, Jonathan Plitnick, Navjot Singh, Matthew D. Shudt, Erica Lasek-Nesselquist |
| EPI_ISL_426303 | hCoV-19/USA/NY-Wadsworth-31364-01/2020 | North America / USA / New York / Cayug | 2020-03-27 | Wadsworth Center, New York State Department of Health | Wadsworth Center, New York State Department of Health | Kirsten St. George, Daryl M. Lamson, Sara Griesemer, Jonathan Plitnick, Navjot Singh, Matthew D. Shudt, Erica Lasek-Nesselquist |
| EPI_ISL_426304 | hCoV-19/USA/NY-Wadsworth-27162-01/2020 | North America / USA / New York / Jeffers | 2020-03-23 | Wadsworth Center, New York State Department of Health | Wadsworth Center, New York State Department of Health | Kirsten St. George, Daryl M. Lamson, Sara Griesemer, Jonathan Plitnick, Navjot Singh, Matthew D. Shudt, Erica Lasek-Nesselquist |
| EPI_ISL_426305 | hCoV-19/USA/NY-Wadsworth-31930-01/2020 | North America / USA / New York / Washi | 2020-03-28 | Wadsworth Center, New York State Department of Health | Wadsworth Center, New York State Department of Health | Kirsten St. George, Daryl M. Lamson, Sara Griesemer, Jonathan Plitnick, Navjot Singh, Matthew D. Shudt, Erica Lasek-Nesselquist |
| EPI_ISL_426306 | hCoV-19/USA/NY-Wadsworth-16004-01/2020 | North America / USA / New York / Richm | 2020-03-15 | Wadsworth Center, New York State Department of Health | Wadsworth Center, New York State Department of Health | Kirsten St. George, Daryl M. Lamson, Sara Griesemer, Jonathan Plitnick, Navjot Singh, Matthew D. Shudt, Erica Lasek-Nesselquist |
| EPI_ISL_426307 | hCoV-19/USA/NY-Wadsworth-30164-01/2020 | North America / USA / New York / Onond | 2020-03-26 | Wadsworth Center, New York State Department of Health | Wadsworth Center, New York State Department of Health | Kirsten St. George, Daryl M. Lamson, Sara Griesemer, Jonathan Plitnick, Navjot Singh, Matthew D. Shudt, Erica Lasek-Nesselquist |
| EPI_ISL_426308 | hCoV-19/USA/NY-Wadsworth-30246-01/2020 | North America / USA / New York / Montg | 2020-03-26 | Wadsworth Center, New York State Department of Health | Wadsworth Center, New York State Department of Health | Kirsten St. George, Daryl M. Lamson, Sara Griesemer, Jonathan Plitnick, Navjot Singh, Matthew D. Shudt, Erica Lasek-Nesselquist |
| EPI_ISL_426309 | hCoV-19/USA/NY-Wadsworth-32104-01/2020 | North America / USA / New York / Orang | 2020-03-28 | Wadsworth Center, New York State Department of Health | Wadsworth Center, New York State Department of Health | Kirsten St. George, Daryl M. Lamson, Sara Griesemer, Jonathan Plitnick, Navjot Singh, Matthew D. Shudt, Erica Lasek-Nesselquist |
| EPI_ISL_426310 | hCoV-19/USA/NY-Wadsworth-15616-01/2020 | North America / USA / New York / Schen | 2020-03-14 | Wadsworth Center, New York State Department of Health | Wadsworth Center, New York State Department of Health | Kirsten St. George, Daryl M. Lamson, Sara Griesemer, Jonathan Plitnick, Navjot Singh, Matthew D. Shudt, Erica Lasek-Nesselquist |
| EPI_ISL_426311 | hCoV-19/USA/NY-Wadsworth-11736-02/2020 | North America / USA / New York / Bronx | 2020-03-08 | Wadsworth Center, New York State Department of Health | Wadsworth Center, New York State Department of Health | Kirsten St. George, Daryl M. Lamson, Sara Griesemer, Jonathan Plitnick, Navjot Singh, Matthew D. Shudt, Erica Lasek-Nesselquist |
| EPI_ISL_426312 | hCoV-19/USA/NY-Wadsworth-30878-01/2020 | North America / USA / New York / Suffolk | 2020-03-27 | Wadsworth Center, New York State Department of Health | Wadsworth Center, New York State Department of Health | Kirsten St. George, Daryl M. Lamson, Sara Griesemer, Jonathan Plitnick, Navjot Singh, Matthew D. Shudt, Erica Lasek-Nesselquist |
| EPI_ISL_426313 | hCoV-19/USA/NY-Wadsworth-22503-01/2020 | North America / USA / New York / Clintor | 2020-03-19 | Wadsworth Center, New York State Department of Health | Wadsworth Center, New York State Department of Health | Kirsten St. George, Daryl M. Lamson, Sara Griesemer, Jonathan Plitnick, Navjot Singh, Matthew D. Shudt, Erica Lasek-Nesselquist |
| EPI_ISL_426314 | hCoV-19/USA/NY-Wadsworth-29275-01/2020 | North America / USA / New York / Albany | 2020-03-25 | Wadsworth Center, New York State Department of Health | Wadsworth Center, New York State Department of Health | Kirsten St. George, Daryl M. Lamson, Sara Griesemer, Jonathan Plitnick, Navjot Singh, Matthew D. Shudt, Erica Lasek-Nesselquist |
| EPI_ISL_426315 | hCoV-19/USA/NY-Wadsworth-22874-01/2020 | North America / USA / New York / Onond | 2020-03-19 | Wadsworth Center, New York State Department of Health | Wadsworth Center, New York State Department of Health | Kirsten St. George, Daryl M. Lamson, Sara Griesemer, Jonathan Plitnick, Navjot Singh, Matthew D. Shudt, Erica Lasek-Nesselquist |
| EPI_ISL_426316 | hCoV-19/USA/NY-Wadsworth-16833-01/2020 | North America / USA / New York / Rockla | 2020-03-15 | Wadsworth Center, New York State Department of Health | Wadsworth Center, New York State Department of Health | Kirsten St. George, Daryl M. Lamson, Sara Griesemer, Jonathan Plitnick, Navjot Singh, Matthew D. Shudt, Erica Lasek-Nesselquist |
| EPI_ISL_426317 | hCoV-19/USA/NY-Wadsworth-16480-01/2020 | North America / USA / New York / New Y | 2020-03-15 | Wadsworth Center, New York State Department of Health | Wadsworth Center, New York State Department of Health | Kirsten St. George, Daryl M. Lamson, Sara Griesemer, Jonathan Plitnick, Navjot Singh, Matthew D. Shudt, Erica Lasek-Nesselquist |
| EPI_ISL_426318 | hCoV-19/USA/NY-Wadsworth-26551-01/2020 | North America / USA / New York / Ulster | 2020-03-22 | Wadsworth Center, New York State Department of Health | Wadsworth Center, New York State Department of Health | Kirsten St. George, Daryl M. Lamson, Sara Griesemer, Jonathan Plitnick, Navjot Singh, Matthew D. Shudt, Erica Lasek-Nesselquist |
| EPI_ISL_426319 | hCoV-19/USA/NY-Wadsworth-16166-01/2020 | North America / USA / New York / Bronx | 2020-03-15 | Wadsworth Center, New York State Department of Health | Wadsworth Center, New York State Department of Health | Kirsten St. George, Daryl M. Lamson, Sara Griesemer, Jonathan Plitnick, Navjot Singh, Matthew D. Shudt, Erica Lasek-Nesselquist |
| EPI_ISL_426320 | hCoV-19/USA/NY-Wadsworth-25860-01/2020 | North America / USA / New York / Broom | 2020-03-20 | Wadsworth Center, New York State Department of Health | Wadsworth Center, New York State Department of Health | Kirsten St. George, Daryl M. Lamson, Sara Griesemer, Jonathan Plitnick, Navjot Singh, Matthew D. Shudt, Erica Lasek-Nesselquist |
| EPI_ISL_426321 | hCoV-19/USA/NY-Wadsworth-31617-01/2020 | North America / USA / New York / Chena | 2020-03-27 | Wadsworth Center, New York State Department of Health | Wadsworth Center, New York State Department of Health | Kirsten St. George, Daryl M. Lamson, Sara Griesemer, Jonathan Plitnick, Navjot Singh, Matthew D. Shudt, Erica Lasek-Nesselquist |
| EPI_ISL_426322 | hCoV-19/USA/NY-Wadsworth-14726-02/2020 | North America / USA / New York / Nassa | 2020-03-12 | Wadsworth Center, New York State Department of Health | Wadsworth Center, New York State Department of Health | Kirsten St. George, Daryl M. Lamson, Sara Griesemer, Jonathan Plitnick, Navjot Singh, Matthew D. Shudt, Erica Lasek-Nesselquist |
| EPI_ISL_426323 | hCoV-19/USA/NY-Wadsworth-16766-01/2020 | North America / USA / New York / Dutche | 2020-03-16 | Wadsworth Center, New York State Department of Health | Wadsworth Center, New York State Department of Health | Kirsten St. George, Daryl M. Lamson, Sara Griesemer, Jonathan Plitnick, Navjot Singh, Matthew D. Shudt, Erica Lasek-Nesselquist |
| EPI_ISL_426324 | hCoV-19/USA/NY-Wadsworth-11190-01/2020 | North America / USA / New York / Westcl | 2020-03-06 | Wadsworth Center, New York State Department of Health | Wadsworth Center, New York State Department of Health | Kirsten St. George, Daryl M. Lamson, Sara Griesemer, Jonathan Plitnick, Navjot Singh, Matthew D. Shudt, Erica Lasek-Nesselquist |
| EPI_ISL_426325 | hCoV-19/USA/NY-Wadsworth-11191-01/2020 | North America / USA / New York / Westcl | 2020-03-06 | Wadsworth Center, New York State Department of Health | Wadsworth Center, New York State Department of Health | Kirsten St. George, Daryl M. Lamson, Sara Griesemer, Jonathan Plitnick, Navjot Singh, Matthew D. Shudt, Erica Lasek-Nesselquist |
| EPI_ISL_426326 | hCoV-19/USA/NY-Wadsworth-11245-01/2020 | North America / USA / New York / Westcl | 2020-03-06 | Wadsworth Center, New York State Department of Health | Wadsworth Center, New York State Department of Health | Kirsten St. George, Daryl M. Lamson, Sara Griesemer, Jonathan Plitnick, Navjot Singh, Matthew D. Shudt, Erica Lasek-Nesselquist |
| EPI_ISL_426327 | hCoV-19/USA/NY-Wadsworth-11249-01/2020 | North America / USA / New York / Westcl | 2020-03-06 | Wadsworth Center, New York State Department of Health | Wadsworth Center, New York State Department of Health | Kirsten St. George, Daryl M. Lamson, Sara Griesemer, Jonathan Plitnick, Navjot Singh, Matthew D. Shudt, Erica Lasek-Nesselquist |
| EPI_ISL_426328 | hCoV-19/USA/NY-Wadsworth-11330-01/2020 | North America / USA / New York / Westcl | 2020-03-07 | Wadsworth Center, New York State Department of Health | Wadsworth Center, New York State Department of Health | Kirsten St. George, Daryl M. Lamson, Sara Griesemer, Jonathan Plitnick, Navjot Singh, Matthew D. Shudt, Erica Lasek-Nesselquist |
| EPI_ISL_426356 | hCoV-19/Czech Republic/Seq3/2020 | Europe / Czech Republic / Central Czech | 2020-03-21 | Laboratory of Molecular Genetics, 2nd Faculty of Medicine, Charles University in Prague, Prague, Czech Republic | Laboratory of Molecular Genetics, 2nd Faculty of Medicine, Charles University in Prague, Prague, Czech Republic | Lenka Kramna, Katerina Polackova, Ondrej Cinek |
| EPI_ISL_426357 | hCoV-19/Czech Republic/Seq4/2020 | Europe / Czech Republic / Central Czech | 2020-03-18 | Laboratory of Molecular Genetics, 2nd Faculty of Medicine, Charles University in Prague, Prague, Czech Republic | Laboratory of Molecular Genetics, 2nd Faculty of Medicine, Charles University in Prague, Prague, Czech Republic | Lenka Kramna, Katerina Polackova, Ondrej Cinek |
| EPI_ISL_426358 | hCoV-19/Czech Republic/Seq5/2020 | Europe / Czech Republic / Central Czech | 2020-03-19 | Laboratory of Molecular Genetics, 2nd Faculty of Medicine, Charles University in Prague, Prague, Czech Republic | Laboratory of Molecular Genetics, 2nd Faculty of Medicine, Charles University in Prague, Prague, Czech Republic | Lenka Kramna, Katerina Polackova, Ondrej Cinek |
| EPI_ISL_426359 | hCoV-19/Czech Republic/Seq6/2020 | Europe / Czech Republic / Central Czech | 2020-03-17 | Laboratory of Molecular Genetics, 2nd Faculty of Medicine, Charles University in Prague, Prague, Czech Republic | Laboratory of Molecular Genetics, 2nd Faculty of Medicine, Charles University in Prague, Prague, Czech Republic | Lenka Kramna, Katerina Polackova, Ondrej Cinek |
| EPI_ISL_426360 | hCoV-19/Czech Republic/Seq7/2020 | Europe / Czech Republic / Central Czech | 2020-03-18 | Laboratory of Molecular Genetics, 2nd Faculty of Medicine, Charles University in Prague, Prague, Czech Republic | Laboratory of Molecular Genetics, 2nd Faculty of Medicine, Charles University in Prague, Prague, Czech Republic | Lenka Kramna, Katerina Polackova, Ondrej Cinek |
| EPI_ISL_426361 | hCoV-19/Mexico/CDMX-INCMNSZ_01/2020 | North America / Mexico / Mexico City | 2020-03-12 | Instituto Nacional de Ciencias Medicas y Nutricion Salvador Zubiran | Instituto Nacional de Ciencias Medicas y Nutricion Salvador Zubiran | Guillermo M. Ruiz-Palacios, Pilar Ramos Cervantes, Violeta Ibarra Gonzalez, Fernando Ledesma Barrientos, Luis Alberto García Andrade, Alfredo Ponce de León Garduño, Irma López Martínez, Lucia Hernández Rivas, Gisela Barrera Badillo, Edgar Mendieta Condado, Fabiola Garcés Ayala, Adnan Araiza Rodríguez, José Ernesto Ramírez González, Celia Boukadida, Santiago Avila Ríos, Mario Mújica Sánchez, José Arturo Martínez Orozco, Eduardo Becerril Vargas, Joel Armando Vázquez Pérez, Víctor Hugo Borja Aburto, Concepción Grajales Muñiz, Cesar Raúl González Bonilla, Carolina González Torres, Francisco Javier Gaytán Cervantes, José Esteban Muñoz Medina, Blanca Taboada, Alejandro Sánchez, Pavel Isa, Ricardo Grande, Gloria Vázquez, Francisco Pulido, Carlos F. Arias |
| EPI_ISL_426362 | hCoV-19/Mexico/CDMX-INCMNSZ_02/2020 | North America / Mexico / Mexico City | 2020-03-10 | Instituto Nacional de Ciencias Medicas y Nutricion Salvador Zubiran | Instituto Nacional de Ciencias Medicas y Nutricion Salvador Zubiran | Guillermo M. Ruiz-Palacios, Pilar Ramos Cervantes, Violeta Ibarra Gonzalez, Fernando Ledesma Barrientos, Luis Alberto García Andrade, Alfredo Ponce de León Garduño, Irma López Martínez, Lucia Hernández Rivas, Gisela Barrera Badillo, Edgar Mendieta Condado, Fabiola Garcés Ayala, Adnan Araiza Rodríguez, José Ernesto Ramírez González, Celia Boukadida, Santiago Avila Ríos, Mario Mújica Sánchez, José Arturo Martínez Orozco, Eduardo Becerril Vargas, Joel Armando Vázquez Pérez, Víctor Hugo Borja Aburto, Concepción Grajales Muñiz, Cesar Raúl González Bonilla, Carolina González Torres, Francisco Javier Gaytán Cervantes, José Esteban Muñoz Medina, Blanca Taboada, Alejandro Sánchez, Pavel Isa, Ricardo Grande, Gloria Vázquez, Francisco Pulido, Carlos F. Arias |
| EPI_ISL_426363 | hCoV-19/Mexico/CDMX-INCMNSZ_03/2020 | North America / Mexico / Mexico City | 2020-03-12 | Instituto Nacional de Ciencias Medicas y Nutricion Salvador Zubiran | Instituto Nacional de Ciencias Medicas y Nutricion Salvador Zubiran | Guillermo M. Ruiz-Palacios, Pilar Ramos Cervantes, Violeta Ibarra Gonzalez, Fernando Ledesma Barrientos, Luis Alberto García Andrade, Alfredo Ponce de León Garduño, Irma López Martínez, Lucia Hernández Rivas, Gisela Barrera Badillo, Edgar Mendieta Condado, Fabiola Garcés Ayala, Adnan Araiza Rodríguez, José Ernesto Ramírez González, Celia Boukadida, Santiago Avila Ríos, Mario Mújica Sánchez, José Arturo Martínez Orozco, Eduardo Becerril Vargas, Joel Armando Vázquez Pérez, Víctor Hugo Borja Aburto, Concepción Grajales Muñiz, Cesar Raúl González Bonilla, Carolina González Torres, Francisco Javier Gaytán Cervantes, José Esteban Muñoz Medina, Blanca Taboada, Alejandro Sánchez, Pavel Isa, Ricardo Grande, Gloria Vázquez, Francisco Pulido, Carlos F. Arias |
| EPI_ISL_426364 | hCoV-19/Mexico/CDMX-INCMNSZ_04/2020 | North America / Mexico / Mexico City | 2020-03-12 | Instituto Nacional de Ciencias Medicas y Nutricion Salvador Zubiran | Instituto Nacional de Ciencias Medicas y Nutricion | Guillermo M. Ruiz-Palacios, Pilar Ramos Cervantes, Violeta Ibarra Gonzalez, Fernando Ledesma Barrientos, Luis Alberto García Andrade, Alfredo Ponce de León Garduño, Irma López Martínez, Lucia Hernández Rivas, Gisela Barrera Badillo, Edgar Mendieta Condado, Fabiola Garcés Ayala, Adnan Araiza Rodríguez, José Ernesto Ramírez González, Celia Boukadida, Santiago Avila Ríos, Mario Mújica Sánchez, José Arturo Martínez Orozco, Eduardo Becerril Vargas, Joel Armando Vázquez Pérez, Víctor Hugo Borja Aburto, Concepción Grajales Muñiz, Cesar Raúl González Bonilla, Carolina González Torres, Francisco Javier Gaytán Cervantes, José Esteban Muñoz Medina, Blanca Taboada, Alejandro Sánchez, Pavel Isa, Ricardo Grande, Gloria Vázquez, Francisco Pulido, Carlos F. Arias |
| EPI_ISL_426365 | hCoV-19/Mexico/CDMX-INCMNSZ_05/2020 | North America / Mexico / Mexico City | 2020-03-12 | Instituto Nacional de Ciencias Medicas y Nutricion Salvador Zubiran | Instituto Nacional de Ciencias Medicas y Nutricion Salvador Zubiran | Guillermo M. Ruiz-Palacios, Pilar Ramos Cervantes, Violeta Ibarra Gonzalez, Fernando Ledesma Barrientos, Luis Alberto García Andrade, Alfredo Ponce de León Garduño, Irma López Martínez, Lucia Hernández Rivas, Gisela Barrera Badillo, Edgar Mendieta Condado, Fabiola Garcés Ayala, Adnan Araiza Rodríguez, José Ernesto Ramírez González, Celia Boukadida, Santiago Avila Ríos, Mario Mújica Sánchez, José Arturo Martínez Orozco, Eduardo Becerril Vargas, Joel Armando Vázquez Pérez, Víctor Hugo Borja Aburto, Concepción Grajales Muñiz, Cesar Raúl González Bonilla, Carolina González Torres, Francisco Javier Gaytán Cervantes, José Esteban Muñoz Medina, Blanca Taboada, Alejandro Sánchez, Pavel Isa, Ricardo Grande, Gloria Vázquez, Francisco Pulido, Carlos F. Arias |
| EPI_ISL_426379 | hCoV-19/Slovenia/MB3329/2020 | Europe / Slovenia | 2020-03-29 | The National Laboratory of Health, Environment and Food, Maribor, Slovenia | The National Laboratory of Health, Environment and Food, Maribor, Slovenia | Mahnic A., Hedzet S., Janezic S., Duh D., Zavrsnik J., Blazun Vosner H., Rupnik M. |
| EPI_ISL_427600 | hCoV-19/USA/NY-WCMP12C01P/2020 | North America / USA / New York / Manha | 2020-03-16 | NewYork-Presbyterian & Mason Lab | Mason Lab | Daniel J. Butler, Christopher Mozsary, Cem Meydan, David Danko, Jonathan Foox, Joel Rosiene, Alon Shaiber, Matthew MacKay, Ebrahim Afshinnekoo, Fritz J. Sedlazeck, Nikolay A. Ivanov, Maria Sierra, Craig D. Westover, Krista Ryon, Benjamin Young, Chandrima Bhattacharya, Phyllis Ruggiero, Justyna Gawrys, Iman Hajirasouliha, Dmitry Meleshko, Mirella Salvatore, Dong Xu, Jenny Xiang, John Sipley, Lin Cong, Arryn Craney, Priya Velu, Lars F. Westblade, Massimo Loda, Shawn Levy, Melissa Cushing, Marcin Imielinski, Hanna Rennert, Christopher E. Mason |
| EPI_ISL_427601 | hCoV-19/USA/NY-WCMP12C04P/2020 | North America / USA / New York / Manha | 2020-03-16 | NewYork-Presbyterian & Mason Lab | Mason Lab | Daniel J. Butler, Christopher Mozsary, Cem Meydan, David Danko, Jonathan Foox, Joel Rosiene, Alon Shaiber, Matthew MacKay, Ebrahim Afshinnekoo, Fritz J. Sedlazeck, Nikolay A. Ivanov, Maria Sierra, Craig D. Westover, Krista Ryon, Benjamin Young, Chandrima Bhattacharya, Phyllis Ruggiero, Justyna Gawrys, Iman Hajirasouliha, Dmitry Meleshko, Mirella Salvatore, Dong Xu, Jenny Xiang, John Sipley, Lin Cong, Arryn Craney, Priya Velu, Lars F. Westblade, Massimo Loda, Shawn Levy, Melissa Cushing, Marcin Imielinski, Hanna Rennert, Christopher E. Mason |
| EPI_ISL_427602 | hCoV-19/USA/NY-WCMP12C09P/2020 | North America / USA / New York / Manha | 2020-03-16 | NewYork-Presbyterian & Mason Lab | Mason Lab | Daniel J. Butler, Christopher Mozsary, Cem Meydan, David Danko, Jonathan Foox, Joel Rosiene, Alon Shaiber, Matthew MacKay, Ebrahim Afshinnekoo, Fritz J. Sedlazeck, Nikolay A. Ivanov, Maria Sierra, Craig D. Westover, Krista Ryon, Benjamin Young, Chandrima Bhattacharya, Phyllis Ruggiero, Justyna Gawrys, Iman Hajirasouliha, Dmitry Meleshko, Mirella Salvatore, Dong Xu, Jenny Xiang, John Sipley, Lin Cong, Arryn Craney, Priya Velu, Lars F. Westblade, Massimo Loda, Shawn Levy, Melissa Cushing, Marcin Imielinski, Hanna Rennert, Christopher E. Mason |
| EPI_ISL_427603 | hCoV-19/USA/NY-WCMP12D02P/2020 | North America / USA / New York / Manha | 2020-03-16 | NewYork-Presbyterian & Mason Lab | Mason Lab | Daniel J. Butler, Christopher Mozsary, Cem Meydan, David Danko, Jonathan Foox, Joel Rosiene, Alon Shaiber, Matthew MacKay, Ebrahim Afshinnekoo, Fritz J. Sedlazeck, Nikolay A. Ivanov, Maria Sierra, Craig D. Westover, Krista Ryon, Benjamin Young, Chandrima Bhattacharya, Phyllis Ruggiero, Justyna Gawrys, Iman Hajirasouliha, Dmitry Meleshko, Mirella Salvatore, Dong Xu, Jenny Xiang, John Sipley, Lin Cong, Arryn Craney, Priya Velu, Lars F. Westblade, Massimo Loda, Shawn Levy, Melissa Cushing, Marcin Imielinski, Hanna Rennert, Christopher E. Mason |
| EPI_ISL_427604 | hCoV-19/USA/NY-WCMP12D08P/2020 | North America / USA / New York / Manha | 2020-03-16 | NewYork-Presbyterian & Mason Lab | Mason Lab | Daniel J. Butler, Christopher Mozsary, Cem Meydan, David Danko, Jonathan Foox, Joel Rosiene, Alon Shaiber, Matthew MacKay, Ebrahim Afshinnekoo, Fritz J. Sedlazeck, Nikolay A. Ivanov, Maria Sierra, Craig D. Westover, Krista Ryon, Benjamin Young, Chandrima Bhattacharya, Phyllis Ruggiero, Justyna Gawrys, Iman Hajirasouliha, Dmitry Meleshko, Mirella Salvatore, Dong Xu, Jenny Xiang, John Sipley, Lin Cong, Arryn Craney, Priya Velu, Lars F. Westblade, Massimo Loda, Shawn Levy, Melissa Cushing, Marcin Imielinski, Hanna Rennert, Christopher E. Mason |
| EPI_ISL_427605 | hCoV-19/USA/NY-WCMP12D09P/2020 | North America / USA / New York / Manha | 2020-03-16 | NewYork-Presbyterian & Mason Lab | Mason Lab | Daniel J. Butler, Christopher Mozsary, Cem Meydan, David Danko, Jonathan Foox, Joel Rosiene, Alon Shaiber, Matthew MacKay, Ebrahim Afshinnekoo, Fritz J. Sedlazeck, Nikolay A. Ivanov, Maria Sierra, Craig D. Westover, Krista Ryon, Benjamin Young, Chandrima Bhattacharya, Phyllis Ruggiero, Justyna Gawrys, Iman Hajirasouliha, Dmitry Meleshko, Mirella Salvatore, Dong Xu, Jenny Xiang, John Sipley, Lin Cong, Arryn Craney, Priya Velu, Lars F. Westblade, Massimo Loda, Shawn Levy, Melissa Cushing, Marcin Imielinski, Hanna Rennert, Christopher E. Mason |
| EPI_ISL_427606 | hCoV-19/USA/NY-WCMP12E01P/2020 | North America / USA / New York / Manha | 2020-03-16 | NewYork-Presbyterian & Mason Lab | Mason Lab | Daniel J. Butler, Christopher Mozsary, Cem Meydan, David Danko, Jonathan Foox, Joel Rosiene, Alon Shaiber, Matthew MacKay, Ebrahim Afshinnekoo, Fritz J. Sedlazeck, Nikolay A. Ivanov, Maria Sierra, Craig D. Westover, Krista Ryon, Benjamin Young, Chandrima Bhattacharya, Phyllis Ruggiero, Justyna Gawrys, Iman Hajirasouliha, Dmitry Meleshko, Mirella Salvatore, Dong Xu, Jenny Xiang, John Sipley, Lin Cong, Arryn Craney, Priya Velu, Lars F. Westblade, Massimo Loda, Shawn Levy, Melissa Cushing, Marcin Imielinski, Hanna Rennert, Christopher E. Mason |
| EPI_ISL_427607 | hCoV-19/USA/NY-WCMP12E06P/2020 | North America / USA / New York / Manha | 2020-03-16 | NewYork-Presbyterian & Mason Lab | Mason Lab | Daniel J. Butler, Christopher Mozsary, Cem Meydan, David Danko, Jonathan Foox, Joel Rosiene, Alon Shaiber, Matthew MacKay, Ebrahim Afshinnekoo, Fritz J. Sedlazeck, Nikolay A. Ivanov, Maria Sierra, Craig D. Westover, Krista Ryon, Benjamin Young, Chandrima Bhattacharya, Phyllis Ruggiero, Justyna Gawrys, Iman Hajirasouliha, Dmitry Meleshko, Mirella Salvatore, Dong Xu, Jenny Xiang, John Sipley, Lin Cong, Arryn Craney, Priya Velu, Lars F. Westblade, Massimo Loda, Shawn Levy, Melissa Cushing, Marcin Imielinski, Hanna Rennert, Christopher E. Mason |
| EPI_ISL_427608 | hCoV-19/USA/NY-WCMP12E09P/2020 | North America / USA / New York / Manha | 2020-03-16 | NewYork-Presbyterian & Mason Lab | Mason Lab | Daniel J. Butler, Christopher Mozsary, Cem Meydan, David Danko, Jonathan Foox, Joel Rosiene, Alon Shaiber, Matthew MacKay, Ebrahim Afshinnekoo, Fritz J. Sedlazeck, Nikolay A. Ivanov, Maria Sierra, Craig D. Westover, Krista Ryon, Benjamin Young, Chandrima Bhattacharya, Phyllis Ruggiero, Justyna Gawrys, Iman Hajirasouliha, Dmitry Meleshko, Mirella Salvatore, Dong Xu, Jenny Xiang, John Sipley, Lin Cong, Arryn Craney, Priya Velu, Lars F. Westblade, Massimo Loda, Shawn Levy, Melissa Cushing, Marcin Imielinski, Hanna Rennert, Christopher E. Mason |
| EPI_ISL_427609 | hCoV-19/USA/NY-WCMP12F01P/2020 | North America / USA / New York / Manha | 2020-03-16 | NewYork-Presbyterian & Mason Lab | Mason Lab | Daniel J. Butler, Christopher Mozsary, Cem Meydan, David Danko, Jonathan Foox, Joel Rosiene, Alon Shaiber, Matthew MacKay, Ebrahim Afshinnekoo, Fritz J. Sedlazeck, Nikolay A. Ivanov, Maria Sierra, Craig D. Westover, Krista Ryon, Benjamin Young, Chandrima Bhattacharya, Phyllis Ruggiero, Justyna Gawrys, Iman Hajirasouliha, Dmitry Meleshko, Mirella Salvatore, Dong Xu, Jenny Xiang, John Sipley, Lin Cong, Arryn Craney, Priya Velu, Lars F. Westblade, Massimo Loda, Shawn Levy, Melissa Cushing, Marcin Imielinski, Hanna Rennert, Christopher E. Mason |
| EPI_ISL_427610 | hCoV-19/USA/NY-WCMP12F02P/2020 | North America / USA / New York / Manha | 2020-03-16 | NewYork-Presbyterian & Mason Lab | Mason Lab | Daniel J. Butler, Christopher Mozsary, Cem Meydan, David Danko, Jonathan Foox, Joel Rosiene, Alon Shaiber, Matthew MacKay, Ebrahim Afshinnekoo, Fritz J. Sedlazeck, Nikolay A. Ivanov, Maria Sierra, Craig D. Westover, Krista Ryon, Benjamin Young, Chandrima Bhattacharya, Phyllis Ruggiero, Justyna Gawrys, Iman Hajirasouliha, Dmitry Meleshko, Mirella Salvatore, Dong Xu, Jenny Xiang, John Sipley, Lin Cong, Arryn Craney, Priya Velu, Lars F. Westblade, Massimo Loda, Shawn Levy, Melissa Cushing, Marcin Imielinski, Hanna Rennert, Christopher E. Mason |
| EPI_ISL_427611 | hCoV-19/USA/NY-WCMP12F03P/2020 | North America / USA / New York / Manha | 2020-03-16 | NewYork-Presbyterian & Mason Lab | Mason Lab | Daniel J. Butler, Christopher Mozsary, Cem Meydan, David Danko, Jonathan Foox, Joel Rosiene, Alon Shaiber, Matthew MacKay, Ebrahim Afshinnekoo, Fritz J. Sedlazeck, Nikolay A. Ivanov, Maria Sierra, Craig D. Westover, Krista Ryon, Benjamin Young, Chandrima Bhattacharya, Phyllis Ruggiero, Justyna Gawrys, Iman Hajirasouliha, Dmitry Meleshko, Mirella Salvatore, Dong Xu, Jenny Xiang, John Sipley, Lin Cong, Arryn Craney, Priya Velu, Lars F. Westblade, Massimo Loda, Shawn Levy, Melissa Cushing, Marcin Imielinski, Hanna Rennert, Christopher E. Mason |
| EPI_ISL_427612 | hCoV-19/USA/NY-WCMP12F07P/2020 | North America / USA / New York / Manha | 2020-03-16 | NewYork-Presbyterian & Mason Lab | Mason Lab | Daniel J. Butler, Christopher Mozsary, Cem Meydan, David Danko, Jonathan Foox, Joel Rosiene, Alon Shaiber, Matthew MacKay, Ebrahim Afshinnekoo, Fritz J. Sedlazeck, Nikolay A. Ivanov, Maria Sierra, Craig D. Westover, Krista Ryon, Benjamin Young, Chandrima Bhattacharya, Phyllis Ruggiero, Justyna Gawrys, Iman Hajirasouliha, Dmitry Meleshko, Mirella Salvatore, Dong Xu, Jenny Xiang, John Sipley, Lin Cong, Arryn Craney, Priya Velu, Lars F. Westblade, Massimo Loda, Shawn Levy, Melissa Cushing, Marcin Imielinski, Hanna Rennert, Christopher E. Mason |
| EPI_ISL_427613 | hCoV-19/USA/NY-WCMP12G01P/2020 | North America / USA / New York / Manha | 2020-03-16 | NewYork-Presbyterian & Mason Lab | Mason Lab | Daniel J. Butler, Christopher Mozsary, Cem Meydan, David Danko, Jonathan Foox, Joel Rosiene, Alon Shaiber, Matthew MacKay, Ebrahim Afshinnekoo, Fritz J. Sedlazeck, Nikolay A. Ivanov, Maria Sierra, Craig D. Westover, Krista Ryon, Benjamin Young, Chandrima Bhattacharya, Phyllis Ruggiero, Justyna Gawrys, Iman Hajirasouliha, Dmitry Meleshko, Mirella Salvatore, Dong Xu, Jenny Xiang, John Sipley, Lin Cong, Arryn Craney, Priya Velu, Lars F. Westblade, Massimo Loda, Shawn Levy, Melissa Cushing, Marcin Imielinski, Hanna Rennert, Christopher E. Mason |
| EPI_ISL_427614 | hCoV-19/USA/NY-WCMP12G02P/2020 | North America / USA / New York / Manha | 2020-03-16 | NewYork-Presbyterian & Mason Lab | Mason Lab | Daniel J. Butler, Christopher Mozsary, Cem Meydan, David Danko, Jonathan Foox, Joel Rosiene, Alon Shaiber, Matthew MacKay, Ebrahim Afshinnekoo, Fritz J. Sedlazeck, Nikolay A. Ivanov, Maria Sierra, Craig D. Westover, Krista Ryon, Benjamin Young, Chandrima Bhattacharya, Phyllis Ruggiero, Justyna Gawrys, Iman Hajirasouliha, Dmitry Meleshko, Mirella Salvatore, Dong Xu, Jenny Xiang, John Sipley, Lin Cong, Arryn Craney, Priya Velu, Lars F. Westblade, Massimo Loda, Shawn Levy, Melissa Cushing, Marcin Imielinski, Hanna Rennert, Christopher E. Mason |
| EPI_ISL_427615 | hCoV-19/USA/NY-WCMP12G03P/2020 | North America / USA / New York / Manha | 2020-03-16 | NewYork-Presbyterian & Mason Lab | Mason Lab | Daniel J. Butler, Christopher Mozsary, Cem Meydan, David Danko, Jonathan Foox, Joel Rosiene, Alon Shaiber, Matthew MacKay, Ebrahim Afshinnekoo, Fritz J. Sedlazeck, Nikolay A. Ivanov, Maria Sierra, Craig D. Westover, Krista Ryon, Benjamin Young, Chandrima Bhattacharya, Phyllis Ruggiero, Justyna Gawrys, Iman Hajirasouliha, Dmitry Meleshko, Mirella Salvatore, Dong Xu, Jenny Xiang, John Sipley, Lin Cong, Arryn Craney, Priya Velu, Lars F. Westblade, Massimo Loda, Shawn Levy, Melissa Cushing, Marcin Imielinski, Hanna Rennert, Christopher E. Mason |
| EPI_ISL_427616 | hCoV-19/USA/NY-WCMP12G06P/2020 | North America / USA / New York / Manha | 2020-03-16 | NewYork-Presbyterian & Mason Lab | Mason Lab | Daniel J. Butler, Christopher Mozsary, Cem Meydan, David Danko, Jonathan Foox, Joel Rosiene, Alon Shaiber, Matthew MacKay, Ebrahim Afshinnekoo, Fritz J. Sedlazeck, Nikolay A. Ivanov, Maria Sierra, Craig D. Westover, Krista Ryon, Benjamin Young, Chandrima Bhattacharya, Phyllis Ruggiero, Justyna Gawrys, Iman Hajirasouliha, Dmitry Meleshko, Mirella Salvatore, Dong Xu, Jenny Xiang, John Sipley, Lin Cong, Arryn Craney, Priya Velu, Lars F. Westblade, Massimo Loda, Shawn Levy, Melissa Cushing, Marcin Imielinski, Hanna Rennert, Christopher E. Mason |
| EPI_ISL_427617 | hCoV-19/USA/NY-WCMP12G08P/2020 | North America / USA / New York / Manha | 2020-03-16 | NewYork-Presbyterian & Mason Lab | Mason Lab | Daniel J. Butler, Christopher Mozsary, Cem Meydan, David Danko, Jonathan Foox, Joel Rosiene, Alon Shaiber, Matthew MacKay, Ebrahim Afshinnekoo, Fritz J. Sedlazeck, Nikolay A. Ivanov, Maria Sierra, Craig D. Westover, Krista Ryon, Benjamin Young, Chandrima Bhattacharya, Phyllis Ruggiero, Justyna Gawrys, Iman Hajirasouliha, Dmitry Meleshko, Mirella Salvatore, Dong Xu, Jenny Xiang, John Sipley, Lin Cong, Arryn Craney, Priya Velu, Lars F. Westblade, Massimo Loda, Shawn Levy, Melissa Cushing, Marcin Imielinski, Hanna Rennert, Christopher E. Mason |
| EPI_ISL_427618 | hCoV-19/USA/NY-WCMP12H01P/2020 | North America / USA / New York / Manha | 2020-03-16 | NewYork-Presbyterian & Mason Lab | Mason Lab | Daniel J. Butler, Christopher Mozsary, Cem Meydan, David Danko, Jonathan Foox, Joel Rosiene, Alon Shaiber, Matthew MacKay, Ebrahim Afshinnekoo, Fritz J. Sedlazeck, Nikolay A. Ivanov, Maria Sierra, Craig D. Westover, Krista Ryon, Benjamin Young, Chandrima Bhattacharya, Phyllis Ruggiero, Justyna Gawrys, Iman Hajirasouliha, Dmitry Meleshko, Mirella Salvatore, Dong Xu, Jenny Xiang, John Sipley, Lin Cong, Arryn Craney, Priya Velu, Lars F. Westblade, Massimo Loda, Shawn Levy, Melissa Cushing, Marcin Imielinski, Hanna Rennert, Christopher E. Mason |
| EPI_ISL_427619 | hCoV-19/USA/AK-PHL092/2020 | North America / USA / Alaska / Interior | 2020-04-02 | Alaska State Virology Laboratory | Alaska State Virology Laboratory | Chen, J. |
| EPI_ISL_427620 | hCoV-19/USA/AK-PHL101/2020 | North America / USA / Alaska / Interior | 2020-04-04 | Alaska State Virology Laboratory | Alaska State Virology Laboratory | Chen, J. |
| EPI_ISL_427621 | hCoV-19/USA/AK-PHL115/2020 | North America / USA / Alaska / Interior | 2020-03-25 | Alaska State Virology Laboratory | Alaska State Virology Laboratory | Chen, J. |
| EPI_ISL_427622 | hCoV-19/USA/AK-PHL118/2020 | North America / USA / Alaska / Interior | 2020-03-24 | Alaska State Virology Laboratory | Alaska State Virology Laboratory | Chen, J. |
| EPI_ISL_427627 | hCoV-19/USA/NY-NYUMC227/2020 | North America / USA / New York / Brookl | 2020-04-01 | NYU Langone Health | Departments of Pathology and Medicine, New York University School of Medicine | Maria Aguero-Rosenfeld, Brendan Belovarac, Margaret Black, Ludovic Boytard, John Cadley, Paolo Cotzia, John Chen, Dacia Dimartino, Xiaojun Feng, Tatyana Gindin, Emily Guzman, Adriana Heguy, Megan Hogan, Emily Huang, George Jour, Andrew Lytle, Christian Marier, Matthew T. Maurano, Mark J. Mulligan, Peter Meyn, Iman Osman, Jared Pinnell, Vanessa Raabe, Sitharam Ramaswami, Amy Rapkiewicz, Marie Samanovic-Golden, Antonio Serrano, Guomiao Shen, Matija Snuderl, Theodore Vougiouklakis, Nick Vulpescu, Gael Westby, Paul Zappile, Yutong Zhang |
| EPI_ISL_427628 | hCoV-19/USA/NY-NYUMC228/2020 | North America / USA / New York / Brookl | 2020-04-03 | NYU Langone Health | Departments of Pathology and Medicine, New York University School of Medicine | Maria Aguero-Rosenfeld, Brendan Belovarac, Margaret Black, Ludovic Boytard, John Cadley, Paolo Cotzia, John Chen, Dacia Dimartino, Xiaojun Feng, Tatyana Gindin, Emily Guzman, Adriana Heguy, Megan Hogan, Emily Huang, George Jour, Andrew Lytle, Christian Marier, Matthew T. Maurano, Mark J. Mulligan, Peter Meyn, Iman Osman, Jared Pinnell, Vanessa Raabe, Sitharam Ramaswami, Amy Rapkiewicz, Marie Samanovic-Golden, Antonio Serrano, Guomiao Shen, Matija Snuderl, Theodore Vougiouklakis, Nick Vulpescu, Gael Westby, Paul Zappile, Yutong Zhang |
| EPI_ISL_427629 | hCoV-19/USA/NY-NYUMC229/2020 | North America / USA / New York / Queen | 2020-03-30 | NYU Langone Health | Departments of Pathology and Medicine, New York University School of Medicine | Maria Aguero-Rosenfeld, Brendan Belovarac, Margaret Black, Ludovic Boytard, John Cadley, Paolo Cotzia, John Chen, Dacia Dimartino, Xiaojun Feng, Tatyana Gindin, Emily Guzman, Adriana Heguy, Megan Hogan, Emily Huang, George Jour, Andrew Lytle, Christian Marier, Matthew T. Maurano, Mark J. Mulligan, Peter Meyn, Iman Osman, Jared Pinnell, Vanessa Raabe, Sitharam Ramaswami, Amy Rapkiewicz, Marie Samanovic-Golden, Antonio Serrano, Guomiao Shen, Matija Snuderl, Theodore Vougiouklakis, Nick Vulpescu, Gael Westby, Paul Zappile, Yutong Zhang |
| EPI_ISL_427630 | hCoV-19/USA/NY-NYUMC230/2020 | North America / USA / New York / Staten | 2020-04-06 | NYU Langone Health | Departments of Pathology and Medicine, New York University School of Medicine | Maria Aguero-Rosenfeld, Brendan Belovarac, Margaret Black, Ludovic Boytard, John Cadley, Paolo Cotzia, John Chen, Dacia Dimartino, Xiaojun Feng, Tatyana Gindin, Emily Guzman, Adriana Heguy, Megan Hogan, Emily Huang, George Jour, Andrew Lytle, Christian Marier, Matthew T. Maurano, Mark J. Mulligan, Peter Meyn, Iman Osman, Jared Pinnell, Vanessa Raabe, Sitharam Ramaswami, Amy Rapkiewicz, Marie Samanovic-Golden, Antonio Serrano, Guomiao Shen, Matija Snuderl, Theodore Vougiouklakis, Nick Vulpescu, Gael Westby, Paul Zappile, Yutong Zhang |
| EPI_ISL_427631 | hCoV-19/USA/NY-NYUMC231/2020 | North America / USA / New York / Staten | 2020-04-06 | NYU Langone Health | Departments of Pathology and Medicine, New York University School of Medicine | Maria Aguero-Rosenfeld, Brendan Belovarac, Margaret Black, Ludovic Boytard, John Cadley, Paolo Cotzia, John Chen, Dacia Dimartino, Xiaojun Feng, Tatyana Gindin, Emily Guzman, Adriana Heguy, Megan Hogan, Emily Huang, George Jour, Andrew Lytle, Christian Marier, Matthew T. Maurano, Mark J. Mulligan, Peter Meyn, Iman Osman, Jared Pinnell, Vanessa Raabe, Sitharam Ramaswami, Amy Rapkiewicz, Marie Samanovic-Golden, Antonio Serrano, Guomiao Shen, Matija Snuderl, Theodore Vougiouklakis, Nick Vulpescu, Gael Westby, Paul Zappile, Yutong Zhang |
| EPI_ISL_427632 | hCoV-19/USA/NY-NYUMC232/2020 | North America / USA / New York / Queen | 2020-04-06 | NYU Langone Health | Departments of Pathology and Medicine, New York University School of Medicine | Maria Aguero-Rosenfeld, Brendan Belovarac, Margaret Black, Ludovic Boytard, John Cadley, Paolo Cotzia, John Chen, Dacia Dimartino, Xiaojun Feng, Tatyana Gindin, Emily Guzman, Adriana Heguy, Megan Hogan, Emily Huang, George Jour, Andrew Lytle, Christian Marier, Matthew T. Maurano, Mark J. Mulligan, Peter Meyn, Iman Osman, Jared Pinnell, Vanessa Raabe, Sitharam Ramaswami, Amy Rapkiewicz, Marie Samanovic-Golden, Antonio Serrano, Guomiao Shen, Matija Snuderl, Theodore Vougiouklakis, Nick Vulpescu, Gael Westby, Paul Zappile, Yutong Zhang |
| EPI_ISL_427633 | hCoV-19/USA/NY-NYUMC233/2020 | North America / USA / New York / Brookl | 2020-04-06 | NYU Langone Health | Departments of Pathology and Medicine, New York University School of Medicine | Maria Aguero-Rosenfeld, Brendan Belovarac, Margaret Black, Ludovic Boytard, John Cadley, Paolo Cotzia, John Chen, Dacia Dimartino, Xiaojun Feng, Tatyana Gindin, Emily Guzman, Adriana Heguy, Megan Hogan, Emily Huang, George Jour, Andrew Lytle, Christian Marier, Matthew T. Maurano, Mark J. Mulligan, Peter Meyn, Iman Osman, Jared Pinnell, Vanessa Raabe, Sitharam Ramaswami, Amy Rapkiewicz, Marie Samanovic-Golden, Antonio Serrano, Guomiao Shen, Matija Snuderl, Theodore Vougiouklakis, Nick Vulpescu, Gael Westby, Paul Zappile, Yutong Zhang |
| EPI_ISL_427634 | hCoV-19/USA/NY-NYUMC235/2020 | North America / USA / New York / Brookl | 2020-04-06 | NYU Langone Health | Departments of Pathology and Medicine, New York University School of Medicine | Maria Aguero-Rosenfeld, Brendan Belovarac, Margaret Black, Ludovic Boytard, John Cadley, Paolo Cotzia, John Chen, Dacia Dimartino, Xiaojun Feng, Tatyana Gindin, Emily Guzman, Adriana Heguy, Megan Hogan, Emily Huang, George Jour, Andrew Lytle, Christian Marier, Matthew T. Maurano, Mark J. Mulligan, Peter Meyn, Iman Osman, Jared Pinnell, Vanessa Raabe, Sitharam Ramaswami, Amy Rapkiewicz, Marie Samanovic-Golden, Antonio Serrano, Guomiao Shen, Matija Snuderl, Theodore Vougiouklakis, Nick Vulpescu, Gael Westby, Paul Zappile, Yutong Zhang |
| EPI_ISL_427635 | hCoV-19/USA/NY-NYUMC236/2020 | North America / USA / New York / Brookl | 2020-04-05 | NYU Langone Health | Departments of Pathology and Medicine, New York University School of Medicine | Maria Aguero-Rosenfeld, Brendan Belovarac, Margaret Black, Ludovic Boytard, John Cadley, Paolo Cotzia, John Chen, Dacia Dimartino, Xiaojun Feng, Tatyana Gindin, Emily Guzman, Adriana Heguy, Megan Hogan, Emily Huang, George Jour, Andrew Lytle, Christian Marier, Matthew T. Maurano, Mark J. Mulligan, Peter Meyn, Iman Osman, Jared Pinnell, Vanessa Raabe, Sitharam Ramaswami, Amy Rapkiewicz, Marie Samanovic-Golden, Antonio Serrano, Guomiao Shen, Matija Snuderl, Theodore Vougiouklakis, Nick Vulpescu, Gael Westby, Paul Zappile, Yutong Zhang |
| EPI_ISL_427636 | hCoV-19/USA/NY-NYUMC237/2020 | North America / USA / New York / Brookl | 2020-04-05 | NYU Langone Health | Departments of Pathology and Medicine, New York University School of Medicine | Maria Aguero-Rosenfeld, Brendan Belovarac, Margaret Black, Ludovic Boytard, John Cadley, Paolo Cotzia, John Chen, Dacia Dimartino, Xiaojun Feng, Tatyana Gindin, Emily Guzman, Adriana Heguy, Megan Hogan, Emily Huang, George Jour, Andrew Lytle, Christian Marier, Matthew T. Maurano, Mark J. Mulligan, Peter Meyn, Iman Osman, Jared Pinnell, Vanessa Raabe, Sitharam Ramaswami, Amy Rapkiewicz, Marie Samanovic-Golden, Antonio Serrano, Guomiao Shen, Matija Snuderl, Theodore Vougiouklakis, Nick Vulpescu, Gael Westby, Paul Zappile, Yutong Zhang |
| EPI_ISL_427637 | hCoV-19/USA/NY-NYUMC238/2020 | North America / USA / New York / Queen | 2020-04-06 | NYU Langone Health | Departments of Pathology and Medicine, New York University School of Medicine | Maria Aguero-Rosenfeld, Brendan Belovarac, Margaret Black, Ludovic Boytard, John Cadley, Paolo Cotzia, John Chen, Dacia Dimartino, Xiaojun Feng, Tatyana Gindin, Emily Guzman, Adriana Heguy, Megan Hogan, Emily Huang, George Jour, Andrew Lytle, Christian Marier, Matthew T. Maurano, Mark J. Mulligan, Peter Meyn, Iman Osman, Jared Pinnell, Vanessa Raabe, Sitharam Ramaswami, Amy Rapkiewicz, Marie Samanovic-Golden, Antonio Serrano, Guomiao Shen, Matija Snuderl, Theodore Vougiouklakis, Nick Vulpescu, Gael Westby, Paul Zappile, Yutong Zhang |
| EPI_ISL_427638 | hCoV-19/USA/NY-NYUMC239/2020 | North America / USA / New York / Brookl | 2020-04-06 | NYU Langone Health | Departments of Pathology and Medicine, New York University School of Medicine | Maria Aguero-Rosenfeld, Brendan Belovarac, Margaret Black, Ludovic Boytard, John Cadley, Paolo Cotzia, John Chen, Dacia Dimartino, Xiaojun Feng, Tatyana Gindin, Emily Guzman, Adriana Heguy, Megan Hogan, Emily Huang, George Jour, Andrew Lytle, Christian Marier, Matthew T. Maurano, Mark J. Mulligan, Peter Meyn, Iman Osman, Jared Pinnell, Vanessa Raabe, Sitharam Ramaswami, Amy Rapkiewicz, Marie Samanovic-Golden, Antonio Serrano, Guomiao Shen, Matija Snuderl, Theodore Vougiouklakis, Nick Vulpescu, Gael Westby, Paul Zappile, Yutong Zhang |
| EPI_ISL_427639 | hCoV-19/USA/NY-NYUMC240/2020 | North America / USA / New York / Queen | 2020-04-06 | NYU Langone Health | Departments of Pathology and Medicine, New York University School of Medicine | Maria Aguero-Rosenfeld, Brendan Belovarac, Margaret Black, Ludovic Boytard, John Cadley, Paolo Cotzia, John Chen, Dacia Dimartino, Xiaojun Feng, Tatyana Gindin, Emily Guzman, Adriana Heguy, Megan Hogan, Emily Huang, George Jour, Andrew Lytle, Christian Marier, Matthew T. Maurano, Mark J. Mulligan, Peter Meyn, Iman Osman, Jared Pinnell, Vanessa Raabe, Sitharam Ramaswami, Amy Rapkiewicz, Marie Samanovic-Golden, Antonio Serrano, Guomiao Shen, Matija Snuderl, Theodore Vougiouklakis, Nick Vulpescu, Gael Westby, Paul Zappile, Yutong Zhang |
| EPI_ISL_427640 | hCoV-19/USA/NY-NYUMC241/2020 | North America / USA / New York / Brookl | 2020-04-05 | NYU Langone Health | Departments of Pathology and Medicine, New York University School of Medicine | Maria Aguero-Rosenfeld, Brendan Belovarac, Margaret Black, Ludovic Boytard, John Cadley, Paolo Cotzia, John Chen, Dacia Dimartino, Xiaojun Feng, Tatyana Gindin, Emily Guzman, Adriana Heguy, Megan Hogan, Emily Huang, George Jour, Andrew Lytle, Christian Marier, Matthew T. Maurano, Mark J. Mulligan, Peter Meyn, Iman Osman, Jared Pinnell, Vanessa Raabe, Sitharam Ramaswami, Amy Rapkiewicz, Marie Samanovic-Golden, Antonio Serrano, Guomiao Shen, Matija Snuderl, Theodore Vougiouklakis, Nick Vulpescu, Gael Westby, Paul Zappile, Yutong Zhang |
| EPI_ISL_427641 | hCoV-19/USA/NY-NYUMC242/2020 | North America / USA / New York / Brookl | 2020-04-06 | NYU Langone Health | Departments of Pathology and Medicine, New York University School of Medicine | Maria Aguero-Rosenfeld, Brendan Belovarac, Margaret Black, Ludovic Boytard, John Cadley, Paolo Cotzia, John Chen, Dacia Dimartino, Xiaojun Feng, Tatyana Gindin, Emily Guzman, Adriana Heguy, Megan Hogan, Emily Huang, George Jour, Andrew Lytle, Christian Marier, Matthew T. Maurano, Mark J. Mulligan, Peter Meyn, Iman Osman, Jared Pinnell, Vanessa Raabe, Sitharam Ramaswami, Amy Rapkiewicz, Marie Samanovic-Golden, Antonio Serrano, Guomiao Shen, Matija Snuderl, Theodore Vougiouklakis, Nick Vulpescu, Gael Westby, Paul Zappile, Yutong Zhang |
| EPI_ISL_427643 | hCoV-19/Australia/NSW219/2020 | Oceania / Australia / New South Wales / | 2020-02-28 | Centre for Infectious Diseases and Microbiology Public Health | NSW Health Pathology - Institute of Clinical Pathology and Medical Research; Westmead Hospital; University of Sydney | Timms V, Gall M, Arnott A, Sadsad R, Draper J, Sim E, Bachmann N, Rockett R, Lam C, Gray K, Carter I, Holmes EC, O'Sullivan MV, Byun R, Sintchenko V, Chen SC, Eden JS, Maddocks S, Kok J, Propenko M, Sorrell T, Chang S, Basile K, Dwyer DE for the 2019-nCoV Study Group |
| EPI_ISL_427644 | hCoV-19/Australia/NSW217/2020 | Oceania / Australia / New South Wales / | 2020-03-02 | Centre for Infectious Diseases and Microbiology Public Health | NSW Health Pathology - Institute of Clinical Pathology and Medical Research; Westmead Hospital; University of Sydney | Rockett R, Lam C, Gray K, Timms V, Gall M, Arnott A, Sadsad R, Draper J, Sim E, Bachmann N, Carter I, Holmes EC, O'Sullivan MV, Byun R, Sintchenko V, Chen SC, Eden JS, Maddocks S, Kok J, Propenko M, Sorrell T, Chang S, Basile K, Dwyer DE for the 2019-nCoV Study Group |
| EPI_ISL_427645 | hCoV-19/Australia/NSW65/2020 | Oceania / Australia / New South Wales / | 2020-03-05 | Centre for Infectious Diseases and Microbiology Public Health | NSW Health Pathology - Institute of Clinical Pathology and Medical Research; Westmead Hospital; University of Sydney | Gray K, Timms V, Gall M, Arnott A, Sadsad R, Draper J, Sim E, Bachmann N, Rockett R, Lam C, Carter I, Holmes EC, O'Sullivan MV, Byun R, Sintchenko V, Chen SC, Eden JS, Maddocks S, Kok J, Propenko M, Sorrell T, Chang S, Basile K, Dwyer DE for the 2019-nCoV Study Group |
| EPI_ISL_427646 | hCoV-19/Australia/NSW66/2020 | Oceania / Australia / New South Wales / | 2020-03-04 | Centre for Infectious Diseases and Microbiology Public Health | NSW Health Pathology - Institute of Clinical Pathology and Medical Research; Westmead Hospital; University of Sydney | Timms V, Gall M, Arnott A, Sadsad R, Draper J, Sim E, Bachmann N, Rockett R, Lam C, Gray K, Carter I, Holmes EC, O'Sullivan MV, Byun R, Sintchenko V, Chen SC, Eden JS, Maddocks S, Kok J, Propenko M, Sorrell T, Chang S, Basile K, Dwyer DE for the 2019-nCoV Study Group |
| EPI_ISL_427647 | hCoV-19/Australia/NSW155/2020 | Oceania / Australia / New South Wales / | 2020-03-06 | Centre for Infectious Diseases and Microbiology Public Health | NSW Health Pathology - Institute of Clinical Pathology and Medical Research; Westmead Hospital; University of Sydney | Gray K, Timms V, Gall M, Arnott A, Sadsad R, Draper J, Sim E, Bachmann N, Rockett R, Lam C, Carter I, Holmes EC, O'Sullivan MV, Byun R, Sintchenko V, Chen SC, Eden JS, Maddocks S, Kok J, Propenko M, Sorrell T, Chang S, Basile K, Dwyer DE for the 2019-nCoV Study Group |
| EPI_ISL_427648 | hCoV-19/Australia/NSW152/2020 | Oceania / Australia / New South Wales / | 2020-03-06 | Centre for Infectious Diseases and Microbiology Public Health | NSW Health Pathology - Institute of Clinical Pathology and Medical Research; Westmead Hospital; University of Sydney | Bachmann N, Rockett R, Lam C, Gray K, Timms V, Gall M, Arnott A, Sadsad R, Draper J, Sim E, Carter I, Holmes EC, O'Sullivan MV, Byun R, Sintchenko V, Chen SC, Eden JS, Maddocks S, Kok J, Propenko M, Sorrell T, Chang S, Basile K, Dwyer DE for the 2019-nCoV Study Group |
| EPI_ISL_427649 | hCoV-19/Australia/NSW22/2020 | Oceania / Australia / New South Wales / | 2020-03-03 | Centre for Infectious Diseases and Microbiology Public Health | NSW Health Pathology - Institute of Clinical Pathology and Medical Research; Westmead Hospital; University of Sydney | Timms V, Gall M, Arnott A, Sadsad R, Draper J, Sim E, Bachmann N, Rockett R, Lam C, Gray K, Carter I, Holmes EC, O'Sullivan MV, Byun R, Sintchenko V, Chen SC, Eden JS, Maddocks S, Kok J, Propenko M, Sorrell T, Chang S, Basile K, Dwyer DE for the 2019-nCoV Study Group |
| EPI_ISL_427650 | hCoV-19/Australia/NSW23/2020 | Oceania / Australia / New South Wales / | 2020-03-04 | Centre for Infectious Diseases and Microbiology Public Health | NSW Health Pathology - Institute of Clinical Pathology and Medical Research; Westmead Hospital; University of Sydney | Gall M, Arnott A, Sadsad R, Draper J, Sim E, Bachmann N, Rockett R, Lam C, Gray K, Timms V, Carter I, Holmes EC, O'Sullivan MV, Byun R, Sintchenko V, Chen SC, Eden JS, Maddocks S, Kok J, Propenko M, Sorrell T, Chang S, Basile K, Dwyer DE for the 2019-nCoV Study Group |
| EPI_ISL_427651 | hCoV-19/Australia/NSW29/2020 | Oceania / Australia / New South Wales / | 2020-03-08 | Centre for Infectious Diseases and Microbiology Public Health | NSW Health Pathology - Institute of Clinical Pathology and Medical Research; Westmead Hospital; University of Sydney | Arnott A, Sadsad R, Draper J, Sim E, Bachmann N, Rockett R, Lam C, Gray K, Timms V, Gall M, Carter I, Holmes EC, O'Sullivan MV, Byun R, Sintchenko V, Chen SC, Eden JS, Maddocks S, Kok J, Propenko M, Sorrell T, Chang S, Basile K, Dwyer DE for the 2019-nCoV Study Group |
| EPI_ISL_427652 | hCoV-19/Australia/NSW154/2020 | Oceania / Australia / New South Wales / | 2020-03-09 | Centre for Infectious Diseases and Microbiology Public Health | NSW Health Pathology - Institute of Clinical Pathology and Medical Research; Westmead Hospital; University of Sydney | Lam C, Gray K, Timms, V, Gall M, Arnott A, Sadsad R, Draper J, Sim E, Bachmann N, Rockett R, Carter I, Holmes EC, O'Sullivan MV, Byun R, Sintchenko V, Chen SC, Eden JS, Maddocks S, Kok J, Propenko M, Sorrell T, Chang S, Basile K, Dwyer DE for the 2019-nCoV Study Group |
| EPI_ISL_427653 | hCoV-19/Australia/NSW153/2020 | Oceania / Australia / New South Wales / | 2020-03-08 | Centre for Infectious Diseases and Microbiology Public Health | NSW Health Pathology - Institute of Clinical Pathology and Medical Research; Westmead Hospital; University of Sydney | Rockett R, Lam C, Gray K, Timms V, Gall M, Arnott A, Sadsad R, Draper J, Sim E, Bachmann N, Carter I, Holmes EC, O'Sullivan MV, Byun R, Sintchenko V, Chen SC, Eden JS, Maddocks S, Kok J, Propenko M, Sorrell T, Chang S, Basile K, Dwyer DE for the 2019-nCoV Study Group |
| EPI_ISL_427654 | hCoV-19/Australia/NSW33/2020 | Oceania / Australia / New South Wales / | 2020-03-06 | Centre for Infectious Diseases and Microbiology Public Health | NSW Health Pathology - Institute of Clinical Pathology and Medical Research; Westmead Hospital; University of Sydney | Sadsad R, Draper J, Sim E, Bachmann N, Rockett R, Lam C, Gray K, Timms V, Gall M, Arnott A, Carter I, Holmes EC, O'Sullivan MV, Byun R, Sintchenko V, Chen SC, Eden JS, Maddocks S, Kok J, Propenko M, Sorrell T, Chang S, Basile K, Dwyer DE for the 2019-nCoV Study Group |
| EPI_ISL_427655 | hCoV-19/Australia/NSW34/2020 | Oceania / Australia / New South Wales / | 2020-03-11 | Centre for Infectious Diseases and Microbiology Public Health | NSW Health Pathology - Institute of Clinical Pathology and Medical Research; Westmead Hospital; University of Sydney | Draper J, Sim E, Bachmann N, Rockett R, Lam C, Gray K, Timms V, Gall M, Arnott A, Sadsad R, Carter I, Holmes EC, O'Sullivan MV, Byun R, Sintchenko V, Chen SC, Eden JS, Maddocks S, Kok J, Propenko M, Sorrell T, Chang S, Basile K, Dwyer DE for the 2019-nCoV Study Group |
| EPI_ISL_427656 | hCoV-19/Australia/NSW102/2020 | Oceania / Australia / New South Wales / | 2020-03-11 | Centre for Infectious Diseases and Microbiology Public Health | NSW Health Pathology - Institute of Clinical Pathology and Medical Research; Westmead Hospital; University of Sydney | Bachmann N, Rockett R, Lam C, Gray K, Timms V, Gall M, Arnott A, Sadsad R, Draper J, Sim E, Carter I, Holmes EC, O'Sullivan MV, Byun R, Sintchenko V, Chen SC, Eden JS, Maddocks S, Kok J, Propenko M, Sorrell T, Chang S, Basile K, Dwyer DE for the 2019-nCoV Study Group |
| EPI_ISL_427657 | hCoV-19/Australia/NSW103/2020 | Oceania / Australia / New South Wales / | 2020-03-10 | Centre for Infectious Diseases and Microbiology Public Health | NSW Health Pathology - Institute of Clinical Pathology and Medical Research; Westmead Hospital; University of Sydney | Rockett R, Lam C, Gray K, Timms V, Gall M, Arnott A, Sadsad R, Draper J, Sim E, Bachmann N, Carter I, Holmes EC, O'Sullivan MV, Byun R, Sintchenko V, Chen SC, Eden JS, Maddocks S, Kok J, Propenko M, Sorrell T, Chang S, Basile K, Dwyer DE for the 2019-nCoV Study Group |
| EPI_ISL_427658 | hCoV-19/Australia/NSW46/2020 | Oceania / Australia / New South Wales / | 2020-03-10 | Centre for Infectious Diseases and Microbiology Public Health | NSW Health Pathology - Institute of Clinical Pathology and Medical Research; Westmead Hospital; University of Sydney | Sim E, Bachmann N, Rockett R, Lam C, Gray K, Timms V, Gall M, Arnott A, Sadsad R, Draper J, Carter I, Holmes EC, O'Sullivan MV, Byun R, Sintchenko V, Chen SC, Eden JS, Maddocks S, Kok J, Propenko M, Sorrell T, Chang S, Basile K, Dwyer DE for the 2019-nCoV Study Group |
| EPI_ISL_427659 | hCoV-19/Australia/NSW49/2020 | Oceania / Australia / New South Wales / | 2020-03-10 | Centre for Infectious Diseases and Microbiology Public Health | NSW Health Pathology - Institute of Clinical Pathology and Medical Research; Westmead Hospital; University of Sydney | Bachmann N, Rockett R, Lam C, Gray K, Timms V, Gall M, Arnott A, Sadsad R, Draper J, Sim E, Carter I, Holmes EC, O'Sullivan MV, Byun R, Sintchenko V, Chen SC, Eden JS, Maddocks S, Kok J, Propenko M, Sorrell T, Chang S, Basile K, Dwyer DE for the 2019-nCoV Study Group |
| EPI_ISL_427660 | hCoV-19/Australia/NSW51/2020 | Oceania / Australia / New South Wales / | 2020-03-10 | Centre for Infectious Diseases and Microbiology Public Health | NSW Health Pathology - Institute of Clinical Pathology and Medical Research; Westmead Hospital; University of Sydney | Rockett R, Lam C, Gray K, Timms V, Gall M, Arnott A, Sadsad R, Draper J, Sim E, Bachmann N, Carter I, Holmes EC, O'Sullivan MV, Byun R, Sintchenko V, Chen SC, Eden JS, Maddocks S, Kok J, Propenko M, Sorrell T, Chang S, Basile K, Dwyer DE for the 2019-nCoV Study Group |
| EPI_ISL_427661 | hCoV-19/Australia/NSW111/2020 | Oceania / Australia / New South Wales / | 2020-03-16 | Centre for Infectious Diseases and Microbiology Public Health | NSW Health Pathology - Institute of Clinical Pathology and Medical Research; Westmead Hospital; University of Sydney | Sim E, Bachmann N, Rockett R, Lam C, Gray K, Timms V, Gall M, Arnott A, Sadsad R, Draper J, Carter I, Holmes EC, O'Sullivan MV, Byun R, Sintchenko V, Chen SC, Eden JS, Maddocks S, Kok J, Propenko M, Sorrell T, Chang S, Basile K, Dwyer DE for the 2019-nCoV Study Group |
| EPI_ISL_427662 | hCoV-19/Australia/NSW59/2020 | Oceania / Australia / New South Wales / | 2020-03-17 | Centre for Infectious Diseases and Microbiology Public Health | NSW Health Pathology - Institute of Clinical Pathology and Medical Research; Westmead Hospital; University of Sydney | Lam C, Gray K, Timms, V, Gall M, Arnott A, Sadsad R, Draper J, Sim E, Bachmann N, Rockett R, Carter I, Holmes EC, O'Sullivan MV, Byun R, Sintchenko V, Chen SC, Eden JS, Maddocks S, Kok J, Propenko M, Sorrell T, Chang S, Basile K, Dwyer DE for the 2019-nCoV Study Group |
| EPI_ISL_427663 | hCoV-19/Australia/NSW60/2020 | Oceania / Australia / New South Wales / | 2020-03-17 | Centre for Infectious Diseases and Microbiology Public Health | NSW Health Pathology - Institute of Clinical Pathology and Medical Research; Westmead Hospital; University of Sydney | Timms V, Gall M, Arnott A, Sadsad R, Draper J, Sim E, Bachmann N, Rockett R, Lam C, Gray K, Carter I, Holmes EC, O'Sullivan MV, Byun R, Sintchenko V, Chen SC, Eden JS, Maddocks S, Kok J, Propenko M, Sorrell T, Chang S, Basile K, Dwyer DE for the 2019-nCoV Study Group |
| EPI_ISL_427664 | hCoV-19/Australia/NSW63/2020 | Oceania / Australia / New South Wales / | 2020-03-17 | Centre for Infectious Diseases and Microbiology Public Health | NSW Health Pathology - Institute of Clinical Pathology and Medical Research; Westmead Hospital; University of Sydney | Timms V, Gall M, Arnott A, Sadsad R, Draper J, Sim E, Bachmann N, Rockett R, Lam C, Gray K, Carter I, Holmes EC, O'Sullivan MV, Byun R, Sintchenko V, Chen SC, Eden JS, Maddocks S, Kok J, Propenko M, Sorrell T, Chang S, Basile K, Dwyer DE for the 2019-nCoV Study Group |
| EPI_ISL_427665 | hCoV-19/Australia/NSW218/2020 | Oceania / Australia / New South Wales / | 2020-03-11 | Centre for Infectious Diseases and Microbiology Public Health | NSW Health Pathology - Institute of Clinical Pathology and Medical Research; Westmead Hospital; University of Sydney | Timms V, Gall M, Arnott A, Sadsad R, Draper J, Sim E, Bachmann N, Rockett R, Lam C, Gray K, Carter I, Holmes EC, O'Sullivan MV, Byun R, Sintchenko V, Chen SC, Eden JS, Maddocks S, Kok J, Propenko M, Sorrell T, Chang S, Basile K, Dwyer DE for the 2019-nCoV Study Group |
| EPI_ISL_427666 | hCoV-19/Australia/NSW112/2020 | Oceania / Australia / New South Wales / | 2020-03-15 | Centre for Infectious Diseases and Microbiology Public Health | NSW Health Pathology - Institute of Clinical Pathology and Medical Research; Westmead Hospital; University of Sydney | Bachmann N, Rockett R, Lam C, Gray K, Timms V, Gall M, Arnott A, Sadsad R, Draper J, Sim E, Carter I, Holmes EC, O'Sullivan MV, Byun R, Sintchenko V, Chen SC, Eden JS, Maddocks S, Kok J, Propenko M, Sorrell T, Chang S, Basile K, Dwyer DE for the 2019-nCoV Study Group |
| EPI_ISL_427667 | hCoV-19/Australia/NSW177/2020 | Oceania / Australia / New South Wales / | 2020-03-23 | Centre for Infectious Diseases and Microbiology Public Health | NSW Health Pathology - Institute of Clinical Pathology and Medical Research; Westmead Hospital; University of Sydney | Gall M, Arnott A, Sadsad R, Draper J, Sim E, Bachmann N, Rockett R, Lam C, Gray K, Timms V, Carter I, Holmes EC, O'Sullivan MV, Byun R, Sintchenko V, Chen SC, Eden JS, Maddocks S, Kok J, Propenko M, Sorrell T, Chang S, Basile K, Dwyer DE for the 2019-nCoV Study Group |
| EPI_ISL_427668 | hCoV-19/Australia/NSW174/2020 | Oceania / Australia / New South Wales / | 2020-03-21 | Centre for Infectious Diseases and Microbiology Public Health | NSW Health Pathology - Institute of Clinical Pathology and Medical Research; Westmead Hospital; University of Sydney | Lam C, Gray K, Timms, V, Gall M, Arnott A, Sadsad R, Draper J, Sim E, Bachmann N, Rockett R, Carter I, Holmes EC, O'Sullivan MV, Byun R, Sintchenko V, Chen SC, Eden JS, Maddocks S, Kok J, Propenko M, Sorrell T, Chang S, Basile K, Dwyer DE for the 2019-nCoV Study Group |
| EPI_ISL_427669 | hCoV-19/Australia/NSW119/2020 | Oceania / Australia / New South Wales / | 2020-03-22 | Centre for Infectious Diseases and Microbiology Public Health | NSW Health Pathology - Institute of Clinical Pathology and Medical Research; Westmead Hospital; University of Sydney | Sadsad R, Draper J, Sim E, Bachmann N, Rockett R, Lam C, Gray K, Timms V, Gall M, Arnott A, Carter I, Holmes EC, O'Sullivan MV, Byun R, Sintchenko V, Chen SC, Eden JS, Maddocks S, Kok J, Propenko M, Sorrell T, Chang S, Basile K, Dwyer DE for the 2019-nCoV Study Group |
| EPI_ISL_427670 | hCoV-19/Australia/NSW178/2020 | Oceania / Australia / New South Wales / | 2020-03-21 | Centre for Infectious Diseases and Microbiology Public Health | NSW Health Pathology - Institute of Clinical Pathology and Medical Research; Westmead Hospital; University of Sydney | Arnott A, Sadsad R, Draper J, Sim E, Bachmann N, Rockett R, Lam C, Gray K, Timms V, Gall M, Carter I, Holmes EC, O'Sullivan MV, Byun R, Sintchenko V, Chen SC, Eden JS, Maddocks S, Kok J, Propenko M, Sorrell T, Chang S, Basile K, Dwyer DE for the 2019-nCoV Study Group |
| EPI_ISL_427671 | hCoV-19/Australia/NSW212/2020 | Oceania / Australia / New South Wales / | 2020-03-21 | Centre for Infectious Diseases and Microbiology Public Health | NSW Health Pathology - Institute of Clinical Pathology and Medical Research; Westmead Hospital; University of Sydney | Bachmann N, Rockett R, Lam C, Gray K, Timms V, Gall M, Arnott A, Sadsad R, Draper J, Sim E, Carter I, Holmes EC, O'Sullivan MV, Byun R, Sintchenko V, Chen SC, Eden JS, Maddocks S, Kok J, Propenko M, Sorrell T, Chang S, Basile K, Dwyer DE for the 2019-nCoV Study Group |
| EPI_ISL_427672 | hCoV-19/Australia/NSW95/2020 | Oceania / Australia / New South Wales / | 2020-03-23 | Centre for Infectious Diseases and Microbiology Public Health | NSW Health Pathology - Institute of Clinical Pathology and Medical Research; Westmead Hospital; University of Sydney | Gray K, Timms V, Gall M, Arnott A, Sadsad R, Draper J, Sim E, Bachmann N, Rockett R, Lam C, Carter I, Holmes EC, O'Sullivan MV, Byun R, Sintchenko V, Chen SC, Eden JS, Maddocks S, Kok J, Propenko M, Sorrell T, Chang S, Basile K, Dwyer DE for the 2019-nCoV Study Group |
| EPI_ISL_427673 | hCoV-19/Australia/NSW135/2020 | Oceania / Australia / New South Wales / | 2020-03-22 | Centre for Infectious Diseases and Microbiology Public Health | NSW Health Pathology - Institute of Clinical Pathology and Medical Research; Westmead Hospital; University of Sydney | Gray K, Timms V, Gall M, Arnott A, Sadsad R, Draper J, Sim E, Bachmann N, Rockett R, Lam C, Carter I, Holmes EC, O'Sullivan MV, Byun R, Sintchenko V, Chen SC, Eden JS, Maddocks S, Kok J, Propenko M, Sorrell T, Chang S, Basile K, Dwyer DE for the 2019-nCoV Study Group |
| EPI_ISL_427674 | hCoV-19/Australia/NSW157/2020 | Oceania / Australia / New South Wales / | 2020-03-22 | Centre for Infectious Diseases and Microbiology Public Health | NSW Health Pathology - Institute of Clinical Pathology and Medical Research; Westmead Hospital; University of Sydney | Gall M, Arnott A, Sadsad R, Draper J, Sim E, Bachmann N, Rockett R, Lam C, Gray K, Timms V, Carter I, Holmes EC, O'Sullivan MV, Byun R, Sintchenko V, Chen SC, Eden JS, Maddocks S, Kok J, Propenko M, Sorrell T, Chang S, Basile K, Dwyer DE for the 2019-nCoV Study Group |
| EPI_ISL_427675 | hCoV-19/Australia/NSW92/2020 | Oceania / Australia / New South Wales / | 2020-03-17 | Centre for Infectious Diseases and Microbiology Public Health | NSW Health Pathology - Institute of Clinical Pathology and Medical Research; Westmead Hospital; University of Sydney | Bachmann N, Rockett R, Lam C, Gray K, Timms V, Gall M, Arnott A, Sadsad R, Draper J, Sim E, Carter I, Holmes EC, O'Sullivan MV, Byun R, Sintchenko V, Chen SC, Eden JS, Maddocks S, Kok J, Propenko M, Sorrell T, Chang S, Basile K, Dwyer DE for the 2019-nCoV Study Group |
| EPI_ISL_427676 | hCoV-19/Australia/NSW170/2020 | Oceania / Australia / New South Wales / | 2020-03-22 | Centre for Infectious Diseases and Microbiology Public Health | NSW Health Pathology - Institute of Clinical Pathology and Medical Research; Westmead Hospital; University of Sydney | Draper J, Sim E, Bachmann N, Rockett R, Lam C, Gray K, Timms V, Gall M, Arnott A, Sadsad R, Carter I, Holmes EC, O'Sullivan MV, Byun R, Sintchenko V, Chen SC, Eden JS, Maddocks S, Kok J, Propenko M, Sorrell T, Chang S, Basile K, Dwyer DE for the 2019-nCoV Study Group |
| EPI_ISL_427677 | hCoV-19/Australia/NSW104/2020 | Oceania / Australia / New South Wales / | 2020-03-16 | Centre for Infectious Diseases and Microbiology Public Health | NSW Health Pathology - Institute of Clinical Pathology and Medical Research; Westmead Hospital; University of Sydney | Lam C, Gray K, Timms, V, Gall M, Arnott A, Sadsad R, Draper J, Sim E, Bachmann N, Rockett R, Carter I, Holmes EC, O'Sullivan MV, Byun R, Sintchenko V, Chen SC, Eden JS, Maddocks S, Kok J, Propenko M, Sorrell T, Chang S, Basile K, Dwyer DE for the 2019-nCoV Study Group |

| | | | | | | |
|---|---|---|---|---|---|---|
| EPI_ISL_427678 | hCoV-19/Australia/NSW96/2020 | Oceania / Australia / New South Wales / ! | 2020-03-23 | Centre for Infectious Diseases and Microbiology Public Health | NSW Health Pathology - Institute of Clinical Pathology and Medical Research; Westmead Hospital; University of Sydney | Timms V, Gall M, Arnott A, Sadsad R, Draper J, Sim E, Bachmann N, Rockett R, Lam C, Gray K, Carter I, Holmes EC, O'Sullivan MV, Byun R, Sintchenko V, Chen SC, Eden JS, Maddocks S, Kok J, Propenko M, Sorrell T, Chang S, Basile K, Dwyer DE for the 2019-nCoV Study Group |
| EPI_ISL_427679 | hCoV-19/Australia/NSW120/2020 | Oceania / Australia / New South Wales / ! | 2020-03-22 | Centre for Infectious Diseases and Microbiology Public Health | NSW Health Pathology - Institute of Clinical Pathology and Medical Research; Westmead Hospital; University of Sydney | Draper J, Sim E, Bachmann N, Rockett R, Lam C, Gray K, Timms V, Gall M, Arnott A, Sadsad R, Carter I, Holmes EC, O'Sullivan MV, Byun R, Sintchenko V, Chen SC, Eden JS, Maddocks S, Kok J, Propenko M, Sorrell T, Chang S, Basile K, Dwyer DE for the 2019-nCoV Study Group |
| EPI_ISL_427680 | hCoV-19/Australia/NSW90/2020 | Oceania / Australia / New South Wales / ! | 2020-03-22 | Centre for Infectious Diseases and Microbiology Public Health | NSW Health Pathology - Institute of Clinical Pathology and Medical Research; Westmead Hospital; University of Sydney | Draper J, Sim E, Bachmann N, Rockett R, Lam C, Gray K, Timms V, Gall M, Arnott A, Sadsad R, Carter I, Holmes EC, O'Sullivan MV, Byun R, Sintchenko V, Chen SC, Eden JS, Maddocks S, Kok J, Propenko M, Sorrell T, Chang S, Basile K, Dwyer DE for the 2019-nCoV Study Group |
| EPI_ISL_427681 | hCoV-19/Australia/NSW93/2020 | Oceania / Australia / New South Wales / ! | 2020-03-19 | Centre for Infectious Diseases and Microbiology Public Health | NSW Health Pathology - Institute of Clinical Pathology and Medical Research; Westmead Hospital; University of Sydney | Rockett R, Lam C, Gray K, Timms V, Gall M, Arnott A, Sadsad R, Draper J, Sim E, Bachmann N, Carter I, Holmes EC, O'Sullivan MV, Byun R, Sintchenko V, Chen SC, Eden JS, Maddocks S, Kok J, Propenko M, Sorrell T, Chang S, Basile K, Dwyer DE for the 2019-nCoV Study Group |
| EPI_ISL_427682 | hCoV-19/Australia/NSW146/2020 | Oceania / Australia / New South Wales / ! | 2020-03-23 | Centre for Infectious Diseases and Microbiology Public Health | NSW Health Pathology - Institute of Clinical Pathology and Medical Research; Westmead Hospital; University of Sydney | Timms V, Gall M, Arnott A, Sadsad R, Draper J, Sim E, Bachmann N, Rockett R, Lam C, Gray K, Carter I, Holmes EC, O'Sullivan MV, Byun R, Sintchenko V, Chen SC, Eden JS, Maddocks S, Kok J, Propenko M, Sorrell T, Chang S, Basile K, Dwyer DE for the 2019-nCoV Study Group |
| EPI_ISL_427683 | hCoV-19/Australia/NSW148/2020 | Oceania / Australia / New South Wales / ! | 2020-03-23 | Centre for Infectious Diseases and Microbiology Public Health | NSW Health Pathology - Institute of Clinical Pathology and Medical Research; Westmead Hospital; University of Sydney | Arnott A, Sadsad R, Draper J, Sim E, Bachmann N, Rockett R, Lam C, Gray K, Timms V, Gall M, Carter I, Holmes EC, O'Sullivan MV, Byun R, Sintchenko V, Chen SC, Eden JS, Maddocks S, Kok J, Propenko M, Sorrell T, Chang S, Basile K, Dwyer DE for the 2019-nCoV Study Group |
| EPI_ISL_427684 | hCoV-19/Australia/NSW200/2020 | Oceania / Australia / New South Wales / ! | 2020-03-23 | Centre for Infectious Diseases and Microbiology Public Health | NSW Health Pathology - Institute of Clinical Pathology and Medical Research; Westmead Hospital; University of Sydney | Draper J, Sim E, Bachmann N, Rockett R, Lam C, Gray K, Timms V, Gall M, Arnott A, Sadsad R, Carter I, Holmes EC, O'Sullivan MV, Byun R, Sintchenko V, Chen SC, Eden JS, Maddocks S, Kok J, Propenko M, Sorrell T, Chang S, Basile K, Dwyer DE for the 2019-nCoV Study Group |
| EPI_ISL_427685 | hCoV-19/Australia/NSW179/2020 | Oceania / Australia / New South Wales / ! | 2020-03-23 | Centre for Infectious Diseases and Microbiology Public Health | NSW Health Pathology - Institute of Clinical Pathology and Medical Research; Westmead Hospital; University of Sydney | Sadsad R, Draper J, Sim E, Bachmann N, Rockett R, Lam C, Gray K, Timms V, Gall M, Arnott A, Carter I, Holmes EC, O'Sullivan MV, Byun R, Sintchenko V, Chen SC, Eden JS, Maddocks S, Kok J, Propenko M, Sorrell T, Chang S, Basile K, Dwyer DE for the 2019-nCoV Study Group |
| EPI_ISL_427686 | hCoV-19/Australia/NSW105/2020 | Oceania / Australia / New South Wales / ! | 2020-03-23 | Centre for Infectious Diseases and Microbiology Public Health | NSW Health Pathology - Institute of Clinical Pathology and Medical Research; Westmead Hospital; University of Sydney | Gray K, Timms V, Gall M, Arnott A, Sadsad R, Draper J, Sim E, Bachmann N, Rockett R, Lam C, Carter I, Holmes EC, O'Sullivan MV, Byun R, Sintchenko V, Chen SC, Eden JS, Maddocks S, Kok J, Propenko M, Sorrell T, Chang S, Basile K, Dwyer DE for the 2019-nCoV Study Group |
| EPI_ISL_427687 | hCoV-19/Australia/NSW175/2020 | Oceania / Australia / New South Wales / ! | 2020-03-18 | Centre for Infectious Diseases and Microbiology Public Health | NSW Health Pathology - Institute of Clinical Pathology and Medical Research; Westmead Hospital; University of Sydney | Gray K, Timms V, Gall M, Arnott A, Sadsad R, Draper J, Sim E, Bachmann N, Rockett R, Lam C, Carter I, Holmes EC, O'Sullivan MV, Byun R, Sintchenko V, Chen SC, Eden JS, Maddocks S, Kok J, Propenko M, Sorrell T, Chang S, Basile K, Dwyer DE for the 2019-nCoV Study Group |
| EPI_ISL_427688 | hCoV-19/Australia/NSW121/2020 | Oceania / Australia / New South Wales / ! | 2020-03-22 | Centre for Infectious Diseases and Microbiology Public Health | NSW Health Pathology - Institute of Clinical Pathology and Medical Research; Westmead Hospital; University of Sydney | Sim E, Bachmann N, Rockett R, Lam C, Gray K, Timms V, Gall M, Arnott A, Sadsad R, Draper J, Carter I, Holmes EC, O'Sullivan MV, Byun R, Sintchenko V, Chen SC, Eden JS, Maddocks S, Kok J, Propenko M, Sorrell T, Chang S, Basile K, Dwyer DE for the 2019-nCoV Study Group |
| EPI_ISL_427689 | hCoV-19/Australia/NSW128/2020 | Oceania / Australia / New South Wales / ! | 2020-03-22 | Centre for Infectious Diseases and Microbiology Public Health | NSW Health Pathology - Institute of Clinical Pathology and Medical Research; Westmead Hospital; University of Sydney | Arnott A, Sadsad R, Draper J, Sim E, Bachmann N, Rockett R, Lam C, Gray K, Timms V, Gall M, Carter I, Holmes EC, O'Sullivan MV, Byun R, Sintchenko V, Chen SC, Eden JS, Maddocks S, Kok J, Propenko M, Sorrell T, Chang S, Basile K, Dwyer DE for the 2019-nCoV Study Group |
| EPI_ISL_427690 | hCoV-19/Australia/NSW80/2020 | Oceania / Australia / New South Wales / ! | 2020-03-21 | Centre for Infectious Diseases and Microbiology Public Health | NSW Health Pathology - Institute of Clinical Pathology and Medical Research; Westmead Hospital; University of Sydney | Draper J, Sim E, Bachmann N, Rockett R, Lam C, Gray K, Timms V, Gall M, Arnott A, Sadsad R, Carter I, Holmes EC, O'Sullivan MV, Byun R, Sintchenko V, Chen SC, Eden JS, Maddocks S, Kok J, Propenko M, Sorrell T, Chang S, Basile K, Dwyer DE for the 2019-nCoV Study Group |
| EPI_ISL_427691 | hCoV-19/Australia/NSW187/2020 | Oceania / Australia / New South Wales / ! | 2020-03-22 | Centre for Infectious Diseases and Microbiology Public Health | NSW Health Pathology - Institute of Clinical Pathology and Medical Research; Westmead Hospital; University of Sydney | Gall M, Arnott A, Sadsad R, Draper J, Sim E, Bachmann N, Rockett R, Lam C, Gray K, Timms V, Carter I, Holmes EC, O'Sullivan MV, Byun R, Sintchenko V, Chen SC, Eden JS, Maddocks S, Kok J, Propenko M, Sorrell T, Chang S, Basile K, Dwyer DE for the 2019-nCoV Study Group |
| EPI_ISL_427692 | hCoV-19/Australia/NSW136/2020 | Oceania / Australia / New South Wales / ! | 2020-03-23 | Centre for Infectious Diseases and Microbiology Public Health | NSW Health Pathology - Institute of Clinical Pathology and Medical Research; Westmead Hospital; University of Sydney | Timms V, Gall M, Arnott A, Sadsad R, Draper J, Sim E, Bachmann N, Rockett R, Lam C, Gray K, Carter I, Holmes EC, O'Sullivan MV, Byun R, Sintchenko V, Chen SC, Eden JS, Maddocks S, Kok J, Propenko M, Sorrell T, Chang S, Basile K, Dwyer DE for the 2019-nCoV Study Group |
| EPI_ISL_427693 | hCoV-19/Australia/NSW118/2020 | Oceania / Australia / New South Wales / ! | 2020-03-22 | Centre for Infectious Diseases and Microbiology Public Health | NSW Health Pathology - Institute of Clinical Pathology and Medical Research; Westmead Hospital; University of Sydney | Arnott A, Sadsad R, Draper J, Sim E, Bachmann N, Rockett R, Lam C, Gray K, Timms V, Gall M, Carter I, Holmes EC, O'Sullivan MV, Byun R, Sintchenko V, Chen SC, Eden JS, Maddocks S, Kok J, Propenko M, Sorrell T, Chang S, Basile K, Dwyer DE for the 2019-nCoV Study Group |
| EPI_ISL_427694 | hCoV-19/Australia/NSW188/2020 | Oceania / Australia / New South Wales / ! | 2020-03-22 | Centre for Infectious Diseases and Microbiology Public Health | NSW Health Pathology - Institute of Clinical Pathology and Medical Research; Westmead Hospital; University of Sydney | Arnott A, Sadsad R, Draper J, Sim E, Bachmann N, Rockett R, Lam C, Gray K, Timms V, Gall M, Carter I, Holmes EC, O'Sullivan MV, Byun R, Sintchenko V, Chen SC, Eden JS, Maddocks S, Kok J, Propenko M, Sorrell T, Chang S, Basile K, Dwyer DE for the 2019-nCoV Study Group |
| EPI_ISL_427695 | hCoV-19/Australia/NSW206/2020 | Oceania / Australia / New South Wales / ! | 2020-03-22 | Centre for Infectious Diseases and Microbiology Public Health | NSW Health Pathology - Institute of Clinical Pathology and Medical Research; Westmead Hospital; University of Sydney | Timms V, Gall M, Arnott A, Sadsad R, Draper J, Sim E, Bachmann N, Rockett R, Lam C, Gray K, Carter I, Holmes EC, O'Sullivan MV, Byun R, Sintchenko V, Chen SC, Eden JS, Maddocks S, Kok J, Propenko M, Sorrell T, Chang S, Basile K, Dwyer DE for the 2019-nCoV Study Group |
| EPI_ISL_427696 | hCoV-19/Australia/NSW115/2020 | Oceania / Australia / New South Wales / ! | 2020-03-21 | Centre for Infectious Diseases and Microbiology Public Health | NSW Health Pathology - Institute of Clinical Pathology and Medical Research; Westmead Hospital; University of Sydney | Gray K, Timms V, Gall M, Arnott A, Sadsad R, Draper J, Sim E, Bachmann N, Rockett R, Lam C, Carter I, Holmes EC, O'Sullivan MV, Byun R, Sintchenko V, Chen SC, Eden JS, Maddocks S, Kok J, Propenko M, Sorrell T, Chang S, Basile K, Dwyer DE for the 2019-nCoV Study Group |
| EPI_ISL_427697 | hCoV-19/Australia/NSW167/2020 | Oceania / Australia / New South Wales / ! | 2020-03-22 | Centre for Infectious Diseases and Microbiology Public Health | NSW Health Pathology - Institute of Clinical Pathology and Medical Research; Westmead Hospital; University of Sydney | Gall M, Arnott A, Sadsad R, Draper J, Sim E, Bachmann N, Rockett R, Lam C, Gray K, Timms V, Carter I, Holmes EC, O'Sullivan MV, Byun R, Sintchenko V, Chen SC, Eden JS, Maddocks S, Kok J, Propenko M, Sorrell T, Chang S, Basile K, Dwyer DE for the 2019-nCoV Study Group |
| EPI_ISL_427698 | hCoV-19/Australia/NSW116/2020 | Oceania / Australia / New South Wales / ! | 2020-03-21 | Centre for Infectious Diseases and Microbiology Public Health | NSW Health Pathology - Institute of Clinical Pathology and Medical Research; Westmead Hospital; University of Sydney | Timms V, Gall M, Arnott A, Sadsad R, Draper J, Sim E, Bachmann N, Rockett R, Lam C, Gray K, Carter I, Holmes EC, O'Sullivan MV, Byun R, Sintchenko V, Chen SC, Eden JS, Maddocks S, Kok J, Propenko M, Sorrell T, Chang S, Basile K, Dwyer DE for the 2019-nCoV Study Group |
| EPI_ISL_427699 | hCoV-19/Australia/NSW211/2020 | Oceania / Australia / New South Wales / ! | 2020-03-21 | Centre for Infectious Diseases and Microbiology Public Health | NSW Health Pathology - Institute of Clinical Pathology and Medical Research; Westmead Hospital; University of Sydney | Sim E, Bachmann N, Rockett R, Lam C, Gray K, Timms V, Gall M, Arnott A, Sadsad R, Draper J, Carter I, Holmes EC, O'Sullivan MV, Byun R, Sintchenko V, Chen SC, Eden JS, Maddocks S, Kok J, Propenko M, Sorrell T, Chang S, Basile K, Dwyer DE for the 2019-nCoV Study Group |
| EPI_ISL_428148 | hCoV-19/Sweden/RV-FOI-4/2020 | Europe / Sweden / Västerbotten | 2020-03-18 | Klinisk mikrobiologi, Region Västerbotten | Unit for Biological Agents, Department for CBRN Defence and Security, Swedish Defence Research Agency | FOI bioinformatics team |
| EPI_ISL_428800 | hCoV-19/USA/NY-NYUMC286/2020 | North America / USA / New York / Brookl | 2020-04-05 | NYU Langone Health | Departments of Pathology and Medicine, New York University School of Medicine | Maria Aguero-Rosenfeld, Brendan Belovarac, Margaret Black, Ludovic Boytard, John Cadley, Paolo Cotzia, John Chen, Dacia Dimartino, Xiaojun Feng, Tatyana Gindin, Emily Guzman, Adriana Heguy, Megan Hogan, Emily Huang, George Jour, Andrew Lytle, Christian Marier, Matthew T. Maurano, Mark J. Mulligan, Peter Meyn, Iman Osman, Jared Pinnell, Vanessa Raabe, Sitharam Ramaswami, Amy Rapkiewicz, Marie Samanovic-Golden, Antonio Serrano, Guomiao Shen, Matija Snuderl, Theodore Vougiouklakis, Nick Vulpescu, Gael Westby, Paul Zappile, Yutong Zhang |
| EPI_ISL_428801 | hCoV-19/USA/NY-NYUMC287/2020 | North America / USA / New York / Brookl | 2020-04-05 | NYU Langone Health | Departments of Pathology and Medicine, New York University School of Medicine | Maria Aguero-Rosenfeld, Brendan Belovarac, Margaret Black, Ludovic Boytard, John Cadley, Paolo Cotzia, John Chen, Dacia Dimartino, Xiaojun Feng, Tatyana Gindin, Emily Guzman, Adriana Heguy, Megan Hogan, Emily Huang, George Jour, Andrew Lytle, Christian Marier, Matthew T. Maurano, Mark J. Mulligan, Peter Meyn, Iman Osman, Jared Pinnell, Vanessa Raabe, Sitharam Ramaswami, Amy Rapkiewicz, Marie Samanovic-Golden, Antonio Serrano, Guomiao Shen, Matija Snuderl, Theodore Vougiouklakis, Nick Vulpescu, Gael Westby, Paul Zappile, Yutong Zhang |
| EPI_ISL_428804 | hCoV-19/USA/NY-NYUMC290/2020 | North America / USA / New York | 2020-04-06 | NYU Langone Health | Departments of Pathology and Medicine, New York University School of Medicine | Maria Aguero-Rosenfeld, Brendan Belovarac, Margaret Black, Ludovic Boytard, John Cadley, Paolo Cotzia, John Chen, Dacia Dimartino, Xiaojun Feng, Tatyana Gindin, Emily Guzman, Adriana Heguy, Megan Hogan, Emily Huang, George Jour, Andrew Lytle, Christian Marier, Matthew T. Maurano, Mark J. Mulligan, Peter Meyn, Iman Osman, Jared Pinnell, Vanessa Raabe, Sitharam Ramaswami, Amy Rapkiewicz, Marie Samanovic-Golden, Antonio Serrano, Guomiao Shen, Matija Snuderl, Theodore Vougiouklakis, Nick Vulpescu, Gael Westby, Paul Zappile, Yutong Zhang |
| EPI_ISL_428805 | hCoV-19/USA/NY-NYUMC291/2020 | North America / USA / New York / Queen | 2020-04-06 | NYU Langone Health | Departments of Pathology and Medicine, New York University School of Medicine | Maria Aguero-Rosenfeld, Brendan Belovarac, Margaret Black, Ludovic Boytard, John Cadley, Paolo Cotzia, John Chen, Dacia Dimartino, Xiaojun Feng, Tatyana Gindin, Emily Guzman, Adriana Heguy, Megan Hogan, Emily Huang, George Jour, Andrew Lytle, Christian Marier, Matthew T. Maurano, Mark J. Mulligan, Peter Meyn, Iman Osman, Jared Pinnell, Vanessa Raabe, Sitharam Ramaswami, Amy Rapkiewicz, Marie Samanovic-Golden, Antonio Serrano, Guomiao Shen, Matija Snuderl, Theodore Vougiouklakis, Nick Vulpescu, Gael Westby, Paul Zappile, Yutong Zhang |
| EPI_ISL_428900 | hCoV-19/Russia/Moscow-80112/2020 | Europe / Russia / Moscow Region | 2020-03-23 | State Research Center of Virology and Biotechnology VECTOR, Department of Collection of Microorganisms | State Research Center of Virology and Biotechnology VECTOR, Department of Collection of Microorganisms | Sergey A. Bodnev, Oleg V. Pyankov, Tatyana V. Tregubchak, Alexander N. Shvalov, Elena V. Gavrilova, Rinat A. Maksyutov |
| EPI_ISL_428901 | hCoV-19/Russia/Crymea-80603/2020 | Europe / Russia / Republic of Crymea | 2020-03-23 | State Research Center of Virology and Biotechnology VECTOR, Department of Collection of Microorganisms | State Research Center of Virology and Biotechnology VECTOR, Department of Collection of Microorganisms | Sergey A. Bodnev, Oleg V. Pyankov, Tatyana V. Tregubchak, Alexander N. Shvalov, Elena V. Gavrilova, Rinat A. Maksyutov |
| EPI_ISL_428902 | hCoV-19/Russia/Orenburg-80305/2020 | Europe / Russia / Orenburg Region | 2020-03-23 | State Research Center of Virology and Biotechnology VECTOR, Department of Collection of Microorganisms | State Research Center of Virology and Biotechnology VECTOR, Department of Collection of Microorganisms | Sergey A. Bodnev, Oleg V. Pyankov, Tatyana V. Tregubchak, Alexander N. Shvalov, Elena V. Gavrilova, Rinat A. Maksyutov |
| EPI_ISL_428903 | hCoV-19/Russia/Primorskiy-79001/2020 | Europe / Russia / Primorskiy Region | 2020-03-22 | State Research Center of Virology and Biotechnology VECTOR, Department of Collection of Microorganisms | State Research Center of Virology and Biotechnology VECTOR, Department of Collection of Microorganisms | Sergey A. Bodnev, Oleg V. Pyankov, Tatyana V. Tregubchak, Alexander N. Shvalov, Elena V. Gavrilova, Rinat A. Maksyutov |
| EPI_ISL_428904 | hCoV-19/Russia/Vector_77628/2020 | Europe / Russia / Moscow | 2020-03-22 | State Research Center of Virology and Biotechnology VECTOR, Department of Collection of Microorganisms | State Research Center of Virology and Biotechnology VECTOR, Department of Collection of Microorganisms | Sergey A. Bodnev, Oleg V. Pyankov, Tatyana V. Tregubchak, Alexander N. Shvalov, Elena V. Gavrilova, Rinat A. Maksyutov |
| EPI_ISL_428905 | hCoV-19/Russia/Moscow-80402/2020 | Europe / Russia / Moscow | 2020-03-23 | State Research Center of Virology and Biotechnology VECTOR, Department of Collection of Microorganisms | State Research Center of Virology and Biotechnology VECTOR, Department of Collection of Microorganisms | Sergey A. Bodnev, Oleg V. Pyankov, Tatyana V. Tregubchak, Alexander N. Shvalov, Elena V. Gavrilova, Rinat A. Maksyutov |
| EPI_ISL_428906 | hCoV-19/Russia/Buryatia-84504/2020 | Europe / Russia / Republic of Buryatia | 2020-03-27 | State Research Center of Virology and Biotechnology VECTOR, Department of Collection of Microorganisms | State Research Center of Virology and Biotechnology VECTOR, Department of Collection of Microorganisms | Oleg V. Pyankov, Sergey A. Bodnev, Tatyana V. Tregubchak, Alexander N. Shvalov, Elena V. Gavrilova, Rinat A. Maksyutov |
| EPI_ISL_428907 | hCoV-19/Russia/Sakha-89702/2020 | Europe / Russia / Republic of Sakha (Yak | 2020-04-01 | State Research Center of Virology and Biotechnology VECTOR, Department of Collection of Microorganisms | State Research Center of Virology and Biotechnology VECTOR, Department of Collection of Microorganisms | Oleg V. Pyankov, Sergey A. Bodnev, Tatyana V. Tregubchak, Alexander N. Shvalov, Elena V. Gavrilova, Rinat A. Maksyutov |
| EPI_ISL_428908 | hCoV-19/Russia/Pskov-83602/2020 | Europe / Russia / Pskov Region | 2020-03-25 | State Research Center of Virology and Biotechnology VECTOR, Department of Collection of Microorganisms | State Research Center of Virology and Biotechnology VECTOR, Department of Collection of Microorganisms | Oleg V. Pyankov, Sergey A. Bodnev, Tatyana V. Tregubchak, Alexander N. Shvalov, Elena V. Gavrilova, Rinat A. Maksyutov |
| EPI_ISL_428909 | hCoV-19/Russia/Chechenya-83801/2020 | Europe / Russia / Chechen Republic | 2020-03-26 | State Research Center of Virology and Biotechnology VECTOR, Department of Collection of Microorganisms | State Research Center of Virology and Biotechnology VECTOR, Department of Collection of Microorganisms | Oleg V. Pyankov, Sergey A. Bodnev, Tatyana V. Tregubchak, Alexander N. Shvalov, Elena V. Gavrilova, Rinat A. Maksyutov |
| EPI_ISL_428910 | hCoV-19/Russia/Khabarovsk-88201/2020 | Europe / Russia / Khabarovsk Region | 2020-03-31 | State Research Center of Virology and Biotechnology VECTOR, Department of Collection of Microorganisms | State Research Center of Virology and Biotechnology VECTOR, Department of Collection of Microorganisms | Oleg V. Pyankov, Sergey A. Bodnev, Tatyana V. Tregubchak, Alexander N. Shvalov, Elena V. Gavrilova, Rinat A. Maksyutov |
| EPI_ISL_428911 | hCoV-19/Russia/Buryatia-88903/2020 | Europe / Russia / Republic of Buryatia | 2020-04-01 | State Research Center of Virology and Biotechnology VECTOR, Department of Collection of Microorganisms | State Research Center of Virology and Biotechnology VECTOR, Department of Collection of Microorganisms | Oleg V. Pyankov, Sergey A. Bodnev, Tatyana V. Tregubchak, Alexander N. Shvalov, Elena V. Gavrilova, Rinat A. Maksyutov |
| EPI_ISL_428912 | hCoV-19/Russia/Sakhalin-84401/2020 | Europe / Russia / Sakhalin Region | 2020-03-27 | State Research Center of Virology and Biotechnology VECTOR, Department of Collection of Microorganisms | State Research Center of Virology and Biotechnology VECTOR, Department of Collection of Microorganisms | Oleg V. Pyankov, Sergey A. Bodnev, Tatyana V. Tregubchak, Alexander N. Shvalov, Elena V. Gavrilova, Rinat A. Maksyutov |
| EPI_ISL_428913 | hCoV-19/Russia/Buryatia-84508/2020 | Europe / Russia / Republic of Buryatia | 2020-03-27 | State Research Center of Virology and Biotechnology VECTOR, Department of Collection of Microorganisms | State Research Center of Virology and Biotechnology VECTOR, Department of Collection of Microorganisms | Oleg V. Pyankov, Sergey A. Bodnev, Tatyana V. Tregubchak, Alexander N. Shvalov, Elena V. Gavrilova, Rinat A. Maksyutov |
| EPI_ISL_428914 | hCoV-19/Russia/Buryatia-87104/2020 | Europe / Russia / Republic of Buryatia | 2020-03-30 | State Research Center of Virology and Biotechnology VECTOR, Department of Collection of Microorganisms | State Research Center of Virology and Biotechnology VECTOR, Department of Collection of Microorganisms | Oleg V. Pyankov, Sergey A. Bodnev, Tatyana V. Tregubchak, Alexander N. Shvalov, Elena V. Gavrilova, Rinat A. Maksyutov |
| EPI_ISL_428915 | hCoV-19/Russia/Sverdlovsk-82904/2020 | Europe / Russia / Sverdlovsk Region | 2020-03-25 | State Research Center of Virology and Biotechnology VECTOR, Department of Collection of Microorganisms | State Research Center of Virology and Biotechnology VECTOR, Department of Collection of Microorganisms | Oleg V. Pyankov, Sergey A. Bodnev, Tatyana V. Tregubchak, Alexander N. Shvalov, Elena V. Gavrilova, Rinat A. Maksyutov |
| EPI_ISL_428916 | hCoV-19/Russia/Chechenya-83803/2020 | Europe / Russia / Chechen Republic | 2020-03-26 | State Research Center of Virology and Biotechnology VECTOR, Department of Collection of Microorganisms | State Research Center of Virology and Biotechnology VECTOR, Department of Collection of Microorganisms | Oleg V. Pyankov, Sergey A. Bodnev, Tatyana V. Tregubchak, Alexander N. Shvalov, Elena V. Gavrilova, Rinat A. Maksyutov |
| EPI_ISL_428917 | hCoV-19/Russia/Tatarstan-82208/2020 | Europe / Russia / Tatarstan | 2020-03-24 | State Research Center of Virology and Biotechnology VECTOR, Department of Collection of Microorganisms | State Research Center of Virology and Biotechnology VECTOR, Department of Collection of Microorganisms | Sergey A. Bodnev, Oleg V. Pyankov, Tatyana V. Tregubchak, Alexander N. Shvalov, Elena V. Gavrilova, Rinat A. Maksyutov |
| EPI_ISL_428918 | hCoV-19/Russia/Novosibirsk-84101/2020 | Europe / Russia / Novosibirsk Region | 2020-03-27 | State Research Center of Virology and Biotechnology VECTOR, Department of Collection of Microorganisms | State Research Center of Virology and Biotechnology VECTOR, Department of Collection of Microorganisms | Sergey A. Bodnev, Oleg V. Pyankov, Tatyana V. Tregubchak, Alexander N. Shvalov, Elena V. Gavrilova, Rinat A. Maksyutov |
| EPI_ISL_428919 | hCoV-19/Russia/Buryatia-87106/2020 | Europe / Russia / Republic of Buryatia | 2020-03-30 | State Research Center of Virology and Biotechnology VECTOR, Department of Collection of Microorganisms | State Research Center of Virology and Biotechnology VECTOR, Department of Collection of Microorganisms | Sergey A. Bodnev, Oleg V. Pyankov, Tatyana V. Tregubchak, Alexander N. Shvalov, Elena V. Gavrilova, Rinat A. Maksyutov |
| EPI_ISL_428920 | hCoV-19/Russia/Buryatia-87105/2020 | Europe / Russia / Republic of Buryatia | 2020-03-30 | State Research Center of Virology and Biotechnology VECTOR, Department of Collection of Microorganisms | State Research Center of Virology and Biotechnology VECTOR, Department of Collection of Microorganisms | Sergey A. Bodnev, Oleg V. Pyankov, Tatyana V. Tregubchak, Alexander N. Shvalov, Elena V. Gavrilova, Rinat A. Maksyutov |
| EPI_ISL_428921 | hCoV-19/Russia/Tver-72601/2020 | Europe / Russia / Tver Region | 2020-03-19 | State Research Center of Virology and Biotechnology VECTOR, Department of Collection of Microorganisms | State Research Center of Virology and Biotechnology VECTOR, Department of Collection of Microorganisms | Sergey A. Bodnev, Oleg V. Pyankov, Tatyana V. Tregubchak, Alexander N. Shvalov, Elena V. Gavrilova, Rinat A. Maksyutov |
| EPI_ISL_428922 | hCoV-19/Russia/Sverdlovsk-86704/2020 | Europe / Russia / Sverdlovsk Region | 2020-03-30 | State Research Center of Virology and Biotechnology VECTOR, Department of Collection of Microorganisms | State Research Center of Virology and Biotechnology VECTOR, Department of Collection of Microorganisms | Sergey A. Bodnev, Oleg V. Pyankov, Tatyana V. Tregubchak, Alexander N. Shvalov, Elena V. Gavrilova, Rinat A. Maksyutov |
| EPI_ISL_428923 | hCoV-19/Russia/Sverdlovsk-84302/2020 | Europe / Russia / Sverdlovsk Region | 2020-03-27 | State Research Center of Virology and Biotechnology VECTOR, Department of Collection of Microorganisms | State Research Center of Virology and Biotechnology VECTOR, Department of Collection of Microorganisms | Sergey A. Bodnev, Oleg V. Pyankov, Tatyana V. Tregubchak, Alexander N. Shvalov, Elena V. Gavrilova, Rinat A. Maksyutov |
| EPI_ISL_428924 | hCoV-19/Russia/Chelyabinsk-85311/2020 | Europe / Russia / Chelyabinsk Region | 2020-03-28 | State Research Center of Virology and Biotechnology VECTOR, Department of Collection of Microorganisms | State Research Center of Virology and Biotechnology VECTOR, Department of Collection of Microorganisms | Sergey A. Bodnev, Oleg V. Pyankov, Tatyana V. Tregubchak, Alexander N. Shvalov, Elena V. Gavrilova, Rinat A. Maksyutov |
| EPI_ISL_428925 | hCoV-19/Poland/PL_P2/2020 | Europe / Poland / Malopolskie | 2020-03-28 | ViroGenetics - BSL3 Laboratory of Virology; Human Genome Variation Research Group & Genomics Centre MCB; Bioinformatics Research Group; Wojewódzka Stacja Sanitarno-Epidemiologiczna w Krakowie | ViroGenetics - BSL3 Laboratory of Virology; Human Genome Variation Research Group & Genomics Centre MCB; Bioinformatics Research Group | Wojciech Branicki, Ewelina Pośpiech, Michał Kowalski, Agnieszka Starowicz, Adrianna Klajmon, Aleksandra Pisarek, Danuta Piniewska-Róg, Kamila Marszałek, Tomasz Gromowski, Katarzyna Kopera, Katarzyna Dudek, Inga Drebot, Katarzyna Guła, Magda Pachota, Aleksandra Synowiec, Marek Sanak, Jarosław Foremny, Paweł P Łabaj, Krzysztof Pyrć |
| EPI_ISL_428926 | hCoV-19/Poland/PL_P3/2020 | Europe / Poland / Malopolskie | 2020-03-28 | ViroGenetics - BSL3 Laboratory of Virology; Human Genome Variation Research Group & Genomics Centre MCB; Bioinformatics Research Group; Wojewódzka Stacja Sanitarno-Epidemiologiczna w Krakowie | ViroGenetics - BSL3 Laboratory of Virology; Human Genome Variation Research Group & Genomics Centre MCB; Bioinformatics Research Group | Wojciech Branicki, Ewelina Pośpiech, Michał Kowalski, Agnieszka Starowicz, Adrianna Klajmon, Aleksandra Pisarek, Danuta Piniewska-Róg, Kamila Marszałek, Tomasz Gromowski, Katarzyna Kopera, Katarzyna Dudek, Inga Drebot, Katarzyna Guła, Magda Pachota, Aleksandra Synowiec, Marek Sanak, Jarosław Foremny, Paweł P Łabaj, Krzysztof Pyrć |
| EPI_ISL_428927 | hCoV-19/Poland/PL_P4/2020 | Europe / Poland / Malopolskie | 2020-03-28 | ViroGenetics - BSL3 Laboratory of Virology; Human Genome Variation Research Group & Genomics Centre MCB; Bioinformatics Research Group; Wojewódzka Stacja Sanitarno-Epidemiologiczna w Krakowie | ViroGenetics - BSL3 Laboratory of Virology; Human Genome Variation Research Group & Genomics Centre MCB; Bioinformatics Research Group | Wojciech Branicki, Ewelina Pośpiech, Michał Kowalski, Agnieszka Starowicz, Adrianna Klajmon, Aleksandra Pisarek, Danuta Piniewska-Róg, Kamila Marszałek, Tomasz Gromowski, Katarzyna Kopera, Katarzyna Dudek, Inga Drebot, Katarzyna Guła, Magda Pachota, Aleksandra Synowiec, Marek Sanak, Jarosław Foremny, Paweł P Łabaj, Krzysztof Pyrć |
| EPI_ISL_428928 | hCoV-19/Poland/PL_P5/2020 | Europe / Poland / Malopolskie | 2020-03-30 | ViroGenetics - BSL3 Laboratory of Virology; Human Genome Variation Research Group & Genomics Centre MCB; Bioinformatics Research Group; Wojewódzka Stacja Sanitarno-Epidemiologiczna w Krakowie | ViroGenetics - BSL3 Laboratory of Virology; Human Genome Variation Research Group & Genomics Centre MCB; Bioinformatics Research Group | Wojciech Branicki, Ewelina Pośpiech, Michał Kowalski, Agnieszka Starowicz, Adrianna Klajmon, Aleksandra Pisarek, Danuta Piniewska-Róg, Kamila Marszałek, Tomasz Gromowski, Katarzyna Kopera, Katarzyna Dudek, Inga Drebot, Katarzyna Guła, Magda Pachota, Aleksandra Synowiec, Marek Sanak, Jarosław Foremny, Paweł P Łabaj, Krzysztof Pyrć |
| EPI_ISL_428929 | hCoV-19/Poland/PL_P6/2020 | Europe / Poland / Malopolskie | 2020-03-29 | ViroGenetics - BSL3 Laboratory of Virology; Human Genome Variation Research Group & Genomics Centre MCB; Bioinformatics Research Group; Wojewódzka Stacja Sanitarno-Epidemiologiczna w Krakowie | ViroGenetics - BSL3 Laboratory of Virology; Human Genome Variation Research Group & Genomics Centre MCB; Bioinformatics Research Group | Wojciech Branicki, Ewelina Pośpiech, Michał Kowalski, Agnieszka Starowicz, Adrianna Klajmon, Aleksandra Pisarek, Danuta Piniewska-Róg, Kamila Marszałek, Tomasz Gromowski, Katarzyna Kopera, Katarzyna Dudek, Inga Drebot, Katarzyna Guła, Magda Pachota, Aleksandra Synowiec, Marek Sanak, Jarosław Foremny, Paweł P Łabaj, Krzysztof Pyrć |
| EPI_ISL_428930 | hCoV-19/Poland/PL_P7/2020 | Europe / Poland / Malopolskie | 2020-03-29 | ViroGenetics - BSL3 Laboratory of Virology; Human Genome Variation Research Group & Genomics Centre MCB; Bioinformatics Research Group; Wojewódzka Stacja Sanitarno-Epidemiologiczna w Krakowie | ViroGenetics - BSL3 Laboratory of Virology; Human Genome Variation Research Group & Genomics Centre MCB; Bioinformatics Research Group | Wojciech Branicki, Ewelina Pośpiech, Michał Kowalski, Agnieszka Starowicz, Adrianna Klajmon, Aleksandra Pisarek, Danuta Piniewska-Róg, Kamila Marszałek, Tomasz Gromowski, Katarzyna Kopera, Katarzyna Dudek, Inga Drebot, Katarzyna Guła, Magda Pachota, Aleksandra Synowiec, Marek Sanak, Jarosław Foremny, Paweł P Łabaj, Krzysztof Pyrć |
| EPI_ISL_428931 | hCoV-19/Poland/PL_P8/2020 | Europe / Poland / Malopolskie | 2020-03-28 | ViroGenetics - BSL3 Laboratory of Virology; Human Genome Variation Research Group & Genomics Centre MCB; Bioinformatics Research Group; Wojewódzka Stacja Sanitarno-Epidemiologiczna w Krakowie | ViroGenetics - BSL3 Laboratory of Virology; Human Genome Variation Research Group & Genomics Centre MCB; Bioinformatics Research Group | Wojciech Branicki, Ewelina Pośpiech, Michał Kowalski, Agnieszka Starowicz, Adrianna Klajmon, Aleksandra Pisarek, Danuta Piniewska-Róg, Kamila Marszałek, Tomasz Gromowski, Katarzyna Kopera, Katarzyna Dudek, Inga Drebot, Katarzyna Guła, Magda Pachota, Aleksandra Synowiec, Marek Sanak, Jarosław Foremny, Paweł P Łabaj, Krzysztof Pyrć |
| EPI_ISL_428932 | hCoV-19/Poland/PL_P9/2020 | Europe / Poland / Malopolskie | 2020-03-28 | ViroGenetics - BSL3 Laboratory of Virology; Human Genome Variation Research Group & Genomics Centre MCB; Bioinformatics Research Group; Wojewódzka Stacja Sanitarno-Epidemiologiczna w Krakowie | ViroGenetics - BSL3 Laboratory of Virology; Human Genome Variation Research Group & Genomics Centre MCB; Bioinformatics Research Group | Wojciech Branicki, Ewelina Pośpiech, Michał Kowalski, Agnieszka Starowicz, Adrianna Klajmon, Aleksandra Pisarek, Danuta Piniewska-Róg, Kamila Marszałek, Tomasz Gromowski, Katarzyna Kopera, Katarzyna Dudek, Inga Drebot, Katarzyna Guła, Magda Pachota, Aleksandra Synowiec, Marek Sanak, Jarosław Foremny, Paweł P Łabaj, Krzysztof Pyrć |
| EPI_ISL_428935 | hCoV-19/USA/WI-UW-259/2020 | North America / USA / Wisconsin / Madis | 2020-04-17 | University of Wisconsin-Madison AIDS Vaccine Research Laboratories | University of Wisconsin-Madison AIDS Vaccine Research Laboratories | Gage Moreno, Katarina Braun, et al. AIDS Vaccine Research Laboratories |
| EPI_ISL_428936 | hCoV-19/USA/WI-UW-260/2020 | North America / USA / Wisconsin / Madis | 2020-04-18 | University of Wisconsin-Madison AIDS Vaccine Research Laboratories | University of Wisconsin-Madison AIDS Vaccine Research Laboratories | Gage Moreno, Katarina Braun, et al. AIDS Vaccine Research Laboratories |
| EPI_ISL_428939 | hCoV-19/Luxembourg/LNS3259987/2020 | Europe / Luxembourg | 2020-04-01 | Laboratoire National de Sante, Microbiology, Virology | Laboratoire National de Sante, Microbiology, Epidemiology and Microbial Genomics | Anke Wienecke-Baldacchino, Ardashel Latsuzbaia, Jessica Tapp, Catherine Ragimbeau, Guillaume Fournier, Tamir Abdelrahman, Trung Nguyen Nguyen, Joel Mossong |
| EPI_ISL_428940 | hCoV-19/Luxembourg/LNS7841004/2020 | Europe / Luxembourg | 2020-04-02 | Laboratoire National de Sante, Microbiology, Virology | Laboratoire National de Sante, Microbiology, Epidemiology and Microbial Genomics | Anke Wienecke-Baldacchino, Ardashel Latsuzbaia, Jessica Tapp, Catherine Ragimbeau, Guillaume Fournier, Tamir Abdelrahman, Trung Nguyen Nguyen, Joel Mossong |
| EPI_ISL_428942 | hCoV-19/Luxembourg/LNS0113568/2020 | Europe / Luxembourg | 2020-04-04 | Laboratoire National de Sante, Microbiology, Virology | Laboratoire National de Sante, Microbiology, Epidemiology and Microbial Genomics | Anke Wienecke-Baldacchino, Ardashel Latsuzbaia, Jessica Tapp, Catherine Ragimbeau, Guillaume Fournier, Tamir Abdelrahman, Trung Nguyen Nguyen, Joel Mossong |
| EPI_ISL_428943 | hCoV-19/Luxembourg/LNS6969569/2020 | Europe / Luxembourg | 2020-04-01 | Laboratoire National de Sante, Microbiology, Virology | Laboratoire National de Sante, Microbiology, Epidemiology and Microbial Genomics | Anke Wienecke-Baldacchino, Ardashel Latsuzbaia, Jessica Tapp, Catherine Ragimbeau, Guillaume Fournier, Tamir Abdelrahman, Trung Nguyen Nguyen, Joel Mossong |
| EPI_ISL_428944 | hCoV-19/Luxembourg/LNS1387711/2020 | Europe / Luxembourg | 2020-04-01 | Laboratoire National de Sante, Microbiology, Virology | Laboratoire National de Sante, Microbiology, Epidemiology and Microbial Genomics | Anke Wienecke-Baldacchino, Ardashel Latsuzbaia, Jessica Tapp, Catherine Ragimbeau, Guillaume Fournier, Tamir Abdelrahman, Trung Nguyen Nguyen, Joel Mossong |
| EPI_ISL_428945 | hCoV-19/Luxembourg/LNS3702558/2020 | Europe / Luxembourg | 2020-04-03 | Laboratoire National de Sante, Microbiology, Virology | Laboratoire National de Sante, Microbiology, Epidemiology and Microbial Genomics | Anke Wienecke-Baldacchino, Ardashel Latsuzbaia, Jessica Tapp, Catherine Ragimbeau, Guillaume Fournier, Tamir Abdelrahman, Trung Nguyen Nguyen, Joel Mossong |
| EPI_ISL_428946 | hCoV-19/Luxembourg/LNS1222966/2020 | Europe / Luxembourg | 2020-04-02 | Laboratoire National de Sante, Microbiology, Virology | Laboratoire National de Sante, Microbiology, Epidemiology and Microbial Genomics | Anke Wienecke-Baldacchino, Ardashel Latsuzbaia, Jessica Tapp, Catherine Ragimbeau, Guillaume Fournier, Tamir Abdelrahman, Trung Nguyen Nguyen, Joel Mossong |
| EPI_ISL_428948 | hCoV-19/Luxembourg/LNS0522318/2020 | Europe / Luxembourg | 2020-04-04 | Laboratoire National de Sante, Microbiology, Virology | Laboratoire National de Sante, Microbiology, Epidemiology and Microbial Genomics | Anke Wienecke-Baldacchino, Ardashel Latsuzbaia, Jessica Tapp, Catherine Ragimbeau, Guillaume Fournier, Tamir Abdelrahman, Trung Nguyen Nguyen, Joel Mossong |
| EPI_ISL_428949 | hCoV-19/Luxembourg/LNS0078676/2020 | Europe / Luxembourg | 2020-04-03 | Laboratoire National de Sante, Microbiology, Virology | Laboratoire National de Sante, Microbiology, Epidemiology and Microbial Genomics | Anke Wienecke-Baldacchino, Ardashel Latsuzbaia, Jessica Tapp, Catherine Ragimbeau, Guillaume Fournier, Tamir Abdelrahman, Trung Nguyen Nguyen, Joel Mossong |
| EPI_ISL_428954 | hCoV-19/Luxembourg/LNS5971805/2020 | Europe / Luxembourg | 2020-04-05 | Laboratoire National de Sante, Microbiology, Virology | Laboratoire National de Sante, Microbiology, Epidemiology and Microbial Genomics | Anke Wienecke-Baldacchino, Ardashel Latsuzbaia, Jessica Tapp, Catherine Ragimbeau, Guillaume Fournier, Tamir Abdelrahman, Trung Nguyen Nguyen, Joel Mossong |
| EPI_ISL_428956 | hCoV-19/Luxembourg/LNS6854244/2020 | Europe / Luxembourg | 2020-04-01 | Laboratoire National de Sante, Microbiology, Virology | Laboratoire National de Sante, Microbiology, Epidemiology and Microbial Genomics | Anke Wienecke-Baldacchino, Ardashel Latsuzbaia, Jessica Tapp, Catherine Ragimbeau, Guillaume Fournier, Tamir Abdelrahman, Trung Nguyen Nguyen, Joel Mossong |
| EPI_ISL_428959 | hCoV-19/Luxembourg/LNS4295690/2020 | Europe / Luxembourg | 2020-04-01 | Laboratoire National de Sante, Microbiology, Virology | Laboratoire National de Sante, Microbiology, Epidemiology and Microbial Genomics | Anke Wienecke-Baldacchino, Ardashel Latsuzbaia, Jessica Tapp, Catherine Ragimbeau, Guillaume Fournier, Tamir Abdelrahman, Trung Nguyen Nguyen, Joel Mossong |
| EPI_ISL_428960 | hCoV-19/Luxembourg/LNS6326231/2020 | Europe / Luxembourg | 2020-04-02 | Laboratoire National de Sante, Microbiology, Virology | Laboratoire National de Sante, Microbiology, Epidemiology and Microbial Genomics | Anke Wienecke-Baldacchino, Ardashel Latsuzbaia, Jessica Tapp, Catherine Ragimbeau, Guillaume Fournier, Tamir Abdelrahman, Trung Nguyen Nguyen, Joel Mossong |
| EPI_ISL_428961 | hCoV-19/Luxembourg/LNS9322962/2020 | Europe / Luxembourg | 2020-04-03 | Laboratoire National de Sante, Microbiology, Virology | Laboratoire National de Sante, Microbiology, Epidemiology and Microbial Genomics | Anke Wienecke-Baldacchino, Ardashel Latsuzbaia, Jessica Tapp, Catherine Ragimbeau, Guillaume Fournier, Tamir Abdelrahman, Trung Nguyen Nguyen, Joel Mossong |
| EPI_ISL_428962 | hCoV-19/Luxembourg/LNS6274573/2020 | Europe / Luxembourg | 2020-04-06 | Laboratoire National de Sante, Microbiology, Virology | Laboratoire National de Sante, Microbiology, Epidemiology and Microbial Genomics | Anke Wienecke-Baldacchino, Ardashel Latsuzbaia, Jessica Tapp, Catherine Ragimbeau, Guillaume Fournier, Tamir Abdelrahman, Trung Nguyen Nguyen, Joel Mossong |
| EPI_ISL_428990 | hCoV-19/USA/CA-CZB0104/2020 | North America / USA / California / San Fr | 2020-03-28 | UCSF Clinical Microbiology Laboratory | Chan-Zuckerberg Biohub | CZB Cliahub Consortium |
| EPI_ISL_428991 | hCoV-19/USA/CA-CZB071/2020 | North America / USA / California / San Fr | 2020-03-28 | UCSF Clinical Microbiology Laboratory | Chan-Zuckerberg Biohub | CZB Cliahub Consortium |
| EPI_ISL_428992 | hCoV-19/USA/CA-CZB0138/2020 | North America / USA / California / San Fr | 2020-04-08 | UCSF Clinical Microbiology Laboratory | Chan-Zuckerberg Biohub | CZB Cliahub Consortium |
| EPI_ISL_428993 | hCoV-19/USA/CA-CZB0157/2020 | North America / USA / California / San Fr | 2020-04-08 | UCSF Clinical Microbiology Laboratory | Chan-Zuckerberg Biohub | CZB Cliahub Consortium |
| EPI_ISL_428994 | hCoV-19/USA/CA-CZB0703/2020 | North America / USA / California / San Fr | 2020-03-23 | UCSF Clinical Microbiology Laboratory | Chan-Zuckerberg Biohub | CZB Cliahub Consortium |
| EPI_ISL_428995 | hCoV-19/USA/CA-CZB0133/2020 | North America / USA / California / San Fr | 2020-04-08 | UCSF Clinical Microbiology Laboratory | Chan-Zuckerberg Biohub | CZB Cliahub Consortium |
| EPI_ISL_428996 | hCoV-19/USA/CA-CZB077/2020 | North America / USA / California / San Fr | 2020-03-28 | UCSF Clinical Microbiology Laboratory | Chan-Zuckerberg Biohub | CZB Cliahub Consortium |
| EPI_ISL_428997 | hCoV-19/USA/CA-CZB015b/2020 | North America / USA / California / San Fr | 2020-03-25 | UCSF Clinical Microbiology Laboratory | Chan-Zuckerberg Biohub | CZB Cliahub Consortium |
| EPI_ISL_428998 | hCoV-19/USA/CA-CZB084/2020 | North America / USA / California / San Fr | 2020-03-28 | UCSF Clinical Microbiology Laboratory | Chan-Zuckerberg Biohub | CZB Cliahub Consortium |
| EPI_ISL_428999 | hCoV-19/USA/CA-CZB069/2020 | North America / USA / California / San Fr | 2020-03-28 | UCSF Clinical Microbiology Laboratory | Chan-Zuckerberg Biohub | CZB Cliahub Consortium |
| EPI_ISL_429402 | hCoV-19/Denmark/ALAB-SSI204/2020 | Europe / Denmark / Unknown | 2020-03-09 | Department of Virus and Microbiological Special Diagnostics, Statens Serum Institut, Copenhagen, Denmark, Artillerivej 5, 2300 Copenahgen S | Albertsen lab, Department of Chemistry and Bioscience, Aalborg University, Denmark | Rasmus Kirkegaard |
| EPI_ISL_429404 | hCoV-19/Denmark/ALAB-SSI207/2020 | Europe / Denmark / Unknown | 2020-03-10 | Department of Virus and Microbiological Special Diagnostics, Statens Serum Institut, Copenhagen, Denmark, Artillerivej 5, 2300 Copenahgen S | Albertsen lab, Department of Chemistry and Bioscience, Aalborg University, Denmark | Rasmus Kirkegaard |
| EPI_ISL_429405 | hCoV-19/Denmark/ALAB-SSI209/2020 | Europe / Denmark / Unknown | 2020-03-10 | Department of Virus and Microbiological Special Diagnostics, Statens Serum Institut, Copenhagen, Denmark, Artillerivej 5, 2300 Copenahgen S | Albertsen lab, Department of Chemistry and Bioscience, Aalborg University, Denmark | Rasmus Kirkegaard |
| EPI_ISL_429406 | hCoV-19/Denmark/ALAB-SSI213/2020 | Europe / Denmark / Unknown | 2020-03-11 | Department of Virus and Microbiological Special Diagnostics, Statens Serum Institut, Copenhagen, Denmark, Artillerivej 5, 2300 Copenahgen S | Albertsen lab, Department of Chemistry and Bioscience, Aalborg University, Denmark | Rasmus Kirkegaard |
| EPI_ISL_429407 | hCoV-19/Denmark/ALAB-SSI215/2020 | Europe / Denmark / Unknown | 2020-03-11 | Department of Virus and Microbiological Special Diagnostics, Statens Serum Institut, Copenhagen, Denmark, Artillerivej 5, 2300 Copenahgen S | Albertsen lab, Department of Chemistry and Bioscience, Aalborg University, Denmark | Rasmus Kirkegaard |
| EPI_ISL_429408 | hCoV-19/Denmark/ALAB-SSI216/2020 | Europe / Denmark / Unknown | 2020-03-10 | Department of Virus and Microbiological Special Diagnostics, Statens Serum Institut, Copenhagen, Denmark, Artillerivej 5, 2300 Copenahgen S | Albertsen lab, Department of Chemistry and Bioscience, Aalborg University, Denmark | Rasmus Kirkegaard |
| EPI_ISL_429409 | hCoV-19/Denmark/ALAB-SSI219/2020 | Europe / Denmark / Unknown | 2020-03-10 | Department of Virus and Microbiological Special Diagnostics, Statens Serum Institut, Copenhagen, Denmark, Artillerivej 5, 2300 Copenahgen S | Albertsen lab, Department of Chemistry and Bioscience, Aalborg University, Denmark | Rasmus Kirkegaard |
| EPI_ISL_429410 | hCoV-19/Denmark/ALAB-SSI221/2020 | Europe / Denmark / Unknown | 2020-03-10 | Department of Virus and Microbiological Special Diagnostics, Statens Serum Institut, Copenhagen, Denmark, Artillerivej 5, 2300 Copenahgen S | Albertsen lab, Department of Chemistry and Bioscience, Aalborg University, Denmark | Rasmus Kirkegaard |
| EPI_ISL_429413 | hCoV-19/Denmark/ALAB-SSI234/2020 | Europe / Denmark / Unknown | 2020-03-10 | Department of Virus and Microbiological Special Diagnostics, Statens Serum Institut, Copenhagen, Denmark, Artillerivej 5, 2300 Copenahgen S | Albertsen lab, Department of Chemistry and Bioscience, Aalborg University, Denmark | Rasmus Kirkegaard |
| EPI_ISL_429414 | hCoV-19/Denmark/ALAB-SSI235/2020 | Europe / Denmark / Unknown | 2020-03-09 | Department of Virus and Microbiological Special Diagnostics, Statens Serum Institut, Copenhagen, Denmark, Artillerivej 5, 2300 Copenahgen S | Albertsen lab, Department of Chemistry and Bioscience, Aalborg University, Denmark | Rasmus Kirkegaard |
| EPI_ISL_429415 | hCoV-19/Denmark/ALAB-SSI237/2020 | Europe / Denmark / Unknown | 2020-03-10 | Department of Virus and Microbiological Special Diagnostics, Statens Serum Institut, Copenhagen, Denmark, Artillerivej 5, 2300 Copenahgen S | Albertsen lab, Department of Chemistry and Bioscience, Aalborg University, Denmark | Rasmus Kirkegaard |
| EPI_ISL_429420 | hCoV-19/Denmark/ALAB-SSI258/2020 | Europe / Denmark / Unknown | 2020-03-10 | Department of Virus and Microbiological Special Diagnostics, Statens Serum Institut, Copenhagen, Denmark, Artillerivej 5, 2300 Copenahgen S | Albertsen lab, Department of Chemistry and Bioscience, Aalborg University, Denmark | Rasmus Kirkegaard |
| EPI_ISL_429421 | hCoV-19/Denmark/ALAB-SSI259/2020 | Europe / Denmark / Unknown | 2020-03-10 | Department of Virus and Microbiological Special Diagnostics, Statens Serum Institut, Copenhagen, Denmark, Artillerivej 5, 2300 Copenahgen S | Albertsen lab, Department of Chemistry and Bioscience, Aalborg University, Denmark | Rasmus Kirkegaard |
| EPI_ISL_429424 | hCoV-19/Denmark/ALAB-SSI262/2020 | Europe / Denmark / Unknown | 2020-03-10 | Department of Virus and Microbiological Special Diagnostics, Statens Serum Institut, Copenhagen, Denmark, Artillerivej 5, 2300 Copenahgen S | Albertsen lab, Department of Chemistry and Bioscience, Aalborg University, Denmark | Rasmus Kirkegaard |
| EPI_ISL_429425 | hCoV-19/Denmark/ALAB-SSI263/2020 | Europe / Denmark / Unknown | 2020-03-10 | Department of Virus and Microbiological Special Diagnostics, Statens Serum Institut, Copenhagen, Denmark, Artillerivej 5, 2300 Copenahgen S | Albertsen lab, Department of Chemistry and Bioscience, Aalborg University, Denmark | Rasmus Kirkegaard |
| EPI_ISL_429426 | hCoV-19/Denmark/ALAB-SSI264/2020 | Europe / Denmark / Unknown | 2020-03-10 | Department of Virus and Microbiological Special Diagnostics, Statens Serum Institut, Copenhagen, Denmark, Artillerivej 5, 2300 Copenahgen S | Albertsen lab, Department of Chemistry and Bioscience, Aalborg University, Denmark | Rasmus Kirkegaard |
| EPI_ISL_429427 | hCoV-19/Denmark/ALAB-SSI265/2020 | Europe / Denmark / Unknown | 2020-03-10 | Department of Virus and Microbiological Special Diagnostics, Statens Serum Institut, Copenhagen, Denmark, Artillerivej 5, 2300 Copenahgen S | Albertsen lab, Department of Chemistry and Bioscience, Aalborg University, Denmark | Rasmus Kirkegaard |
| EPI_ISL_429428 | hCoV-19/Denmark/ALAB-SSI268/2020 | Europe / Denmark / Unknown | 2020-03-10 | Department of Virus and Microbiological Special Diagnostics, Statens Serum Institut, Copenhagen, Denmark, Artillerivej 5, 2300 Copenahgen S | Albertsen lab, Department of Chemistry and Bioscience, Aalborg University, Denmark | Rasmus Kirkegaard |
| EPI_ISL_429431 | hCoV-19/Denmark/ALAB-SSI271/2020 | Europe / Denmark / Unknown | 2020-03-10 | Department of Virus and Microbiological Special Diagnostics, Statens Serum Institut, Copenhagen, Denmark, Artillerivej 5, 2300 Copenahgen S | Albertsen lab, Department of Chemistry and Bioscience, Aalborg University, Denmark | Rasmus Kirkegaard |
| EPI_ISL_429433 | hCoV-19/Denmark/ALAB-SSI272/2020 | Europe / Denmark / Unknown | 2020-03-10 | Department of Virus and Microbiological Special Diagnostics, Statens Serum Institut, Copenhagen, Denmark, Artillerivej 5, 2300 Copenahgen S | Albertsen lab, Department of Chemistry and Bioscience, Aalborg University, Denmark | Rasmus Kirkegaard |
| EPI_ISL_429434 | hCoV-19/Denmark/ALAB-SSI273/2020 | Europe / Denmark / Unknown | 2020-03-10 | Department of Virus and Microbiological Special Diagnostics, Statens Serum Institut, Copenhagen, Denmark, Artillerivej 5, 2300 Copenahgen S | Albertsen lab, Department of Chemistry and Bioscience, Aalborg University, Denmark | Rasmus Kirkegaard |
| EPI_ISL_429439 | hCoV-19/Denmark/ALAB-SSI279/2020 | Europe / Denmark / Unknown | 2020-03-10 | Department of Virus and Microbiological Special Diagnostics, Statens Serum Institut, Copenhagen, Denmark, Artillerivej 5, 2300 Copenahgen S | Albertsen lab, Department of Chemistry and Bioscience, Aalborg University, Denmark | Rasmus Kirkegaard |
| EPI_ISL_429440 | hCoV-19/Denmark/ALAB-SSI280/2020 | Europe / Denmark / Unknown | 2020-03-10 | Department of Virus and Microbiological Special Diagnostics, Statens Serum Institut, Copenhagen, Denmark, Artillerivej 5, 2300 Copenahgen S | Albertsen lab, Department of Chemistry and Bioscience, Aalborg University, Denmark | Rasmus Kirkegaard |
| EPI_ISL_429445 | hCoV-19/Denmark/ALAB-SSI287/2020 | Europe / Denmark / Unknown | 2020-03-10 | Department of Virus and Microbiological Special Diagnostics, Statens Serum Institut, Copenhagen, Denmark, Artillerivej 5, 2300 Copenahgen S | Albertsen lab, Department of Chemistry and Bioscience, Aalborg University, Denmark | Rasmus Kirkegaard |
| EPI_ISL_429447 | hCoV-19/Denmark/ALAB-SSI29/2020 | Europe / Denmark / Unknown | 2020-03-03 | Department of Virus and Microbiological Special Diagnostics, Statens Serum Institut, Copenhagen, Denmark, Artillerivej 5, 2300 Copenahgen S | Albertsen lab, Department of Chemistry and Bioscience, Aalborg University, Denmark | Rasmus Kirkegaard |
| EPI_ISL_429448 | hCoV-19/Denmark/ALAB-SSI293/2020 | Europe / Denmark / Unknown | 2020-03-10 | Department of Virus and Microbiological Special Diagnostics, Statens Serum Institut, Copenhagen, Denmark, Artillerivej 5, 2300 Copenahgen S | Albertsen lab, Department of Chemistry and Bioscience, Aalborg University, Denmark | Rasmus Kirkegaard |
| EPI_ISL_429449 | hCoV-19/Denmark/ALAB-SSI298/2020 | Europe / Denmark / Unknown | 2020-03-10 | Department of Virus and Microbiological Special Diagnostics, Statens Serum Institut, Copenhagen, Denmark, Artillerivej 5, 2300 Copenahgen S | Albertsen lab, Department of Chemistry and Bioscience, Aalborg University, Denmark | Rasmus Kirkegaard |
| EPI_ISL_429450 | hCoV-19/Denmark/ALAB-SSI299/2020 | Europe / Denmark / Unknown | 2020-03-10 | Department of Virus and Microbiological Special Diagnostics, Statens Serum Institut, Copenhagen, Denmark, Artillerivej 5, 2300 Copenahgen S | Albertsen lab, Department of Chemistry and Bioscience, Aalborg University, Denmark | Rasmus Kirkegaard |
| EPI_ISL_429451 | hCoV-19/Denmark/ALAB-SSI300/2020 | Europe / Denmark / Unknown | 2020-03-10 | Department of Virus and Microbiological Special Diagnostics, Statens Serum Institut, Copenhagen, Denmark, Artillerivej 5, 2300 Copenahgen S | Albertsen lab, Department of Chemistry and Bioscience, Aalborg University, Denmark | Rasmus Kirkegaard |
| EPI_ISL_429452 | hCoV-19/Denmark/ALAB-SSI301/2020 | Europe / Denmark / Unknown | 2020-03-10 | Department of Virus and Microbiological Special Diagnostics, Statens Serum Institut, Copenhagen, Denmark, Artillerivej 5, 2300 Copenahgen S | Albertsen lab, Department of Chemistry and Bioscience, Aalborg University, Denmark | Rasmus Kirkegaard |
| EPI_ISL_429453 | hCoV-19/Denmark/ALAB-SSI302/2020 | Europe / Denmark / Unknown | 2020-03-10 | Department of Virus and Microbiological Special Diagnostics, Statens Serum Institut, Copenhagen, Denmark, Artillerivej 5, 2300 Copenahgen S | Albertsen lab, Department of Chemistry and Bioscience, Aalborg University, Denmark | Rasmus Kirkegaard |
| EPI_ISL_429457 | hCoV-19/Denmark/ALAB-SSI330/2020 | Europe / Denmark / Unknown | 2020-03-09 | Department of Virus and Microbiological Special Diagnostics, Statens Serum Institut, Copenhagen, Denmark, Artillerivej 5, 2300 Copenahgen S | Albertsen lab, Department of Chemistry and Bioscience, Aalborg University, Denmark | Rasmus Kirkegaard |
| EPI_ISL_429459 | hCoV-19/Denmark/ALAB-SSI337/2020 | Europe / Denmark / Unknown | 2020-03-09 | Department of Virus and Microbiological Special Diagnostics, Statens Serum Institut, Copenhagen, Denmark, Artillerivej 5, 2300 Copenahgen S | Albertsen lab, Department of Chemistry and Bioscience, Aalborg University, Denmark | Rasmus Kirkegaard |
| EPI_ISL_429460 | hCoV-19/Denmark/ALAB-SSI346/2020 | Europe / Denmark / Unknown | 2020-03-23 | Department of Virus and Microbiological Special Diagnostics, Statens Serum Institut, Copenhagen, Denmark, Artillerivej 5, 2300 Copenahgen S | Albertsen lab, Department of Chemistry and Bioscience, Aalborg University, Denmark | Rasmus Kirkegaard |
| EPI_ISL_429461 | hCoV-19/Denmark/ALAB-SSI347/2020 | Europe / Denmark / Unknown | 2020-03-23 | Department of Virus and Microbiological Special Diagnostics, Statens Serum Institut, Copenhagen, Denmark, Artillerivej 5, 2300 Copenahgen S | Albertsen lab, Department of Chemistry and Bioscience, Aalborg University, Denmark | Rasmus Kirkegaard |
| EPI_ISL_429462 | hCoV-19/Denmark/ALAB-SSI348/2020 | Europe / Denmark / Unknown | 2020-03-23 | Department of Virus and Microbiological Special Diagnostics, Statens Serum Institut, Copenhagen, Denmark, Artillerivej 5, 2300 Copenahgen S | Albertsen lab, Department of Chemistry and Bioscience, Aalborg University, Denmark | Rasmus Kirkegaard |
| EPI_ISL_429463 | hCoV-19/Denmark/ALAB-SSI349/2020 | Europe / Denmark / Unknown | 2020-03-23 | Department of Virus and Microbiological Special Diagnostics, Statens Serum Institut, Copenhagen, Denmark, Artillerivej 5, 2300 Copenahgen S | Albertsen lab, Department of Chemistry and Bioscience, Aalborg University, Denmark | Rasmus Kirkegaard |
| EPI_ISL_429464 | hCoV-19/Denmark/ALAB-SSI351/2020 | Europe / Denmark / Unknown | 2020-03-23 | Department of Virus and Microbiological Special Diagnostics, Statens Serum Institut, Copenhagen, Denmark, Artillerivej 5, 2300 Copenahgen S | Albertsen lab, Department of Chemistry and Bioscience, Aalborg University, Denmark | Rasmus Kirkegaard |
| EPI_ISL_429465 | hCoV-19/Denmark/ALAB-SSI352/2020 | Europe / Denmark / Unknown | 2020-03-23 | Department of Virus and Microbiological Special Diagnostics, Statens Serum Institut, Copenhagen, Denmark, Artillerivej 5, 2300 Copenahgen S | Albertsen lab, Department of Chemistry and Bioscience, Aalborg University, Denmark | Rasmus Kirkegaard |
| EPI_ISL_429466 | hCoV-19/Denmark/ALAB-SSI354/2020 | Europe / Denmark / Unknown | 2020-03-23 | Department of Virus and Microbiological Special Diagnostics, Statens Serum Institut, Copenhagen, Denmark, Artillerivej 5, 2300 Copenahgen S | Albertsen lab, Department of Chemistry and Bioscience, Aalborg University, Denmark | Rasmus Kirkegaard |
| EPI_ISL_429468 | hCoV-19/Denmark/ALAB-SSI357/2020 | Europe / Denmark / Unknown | 2020-03-23 | Department of Virus and Microbiological Special Diagnostics, Statens Serum Institut, Copenhagen, Denmark, Artillerivej 5, 2300 Copenahgen S | Albertsen lab, Department of Chemistry and Bioscience, Aalborg University, Denmark | Rasmus Kirkegaard |
| EPI_ISL_429469 | hCoV-19/Denmark/ALAB-SSI358/2020 | Europe / Denmark / Unknown | 2020-03-24 | Department of Virus and Microbiological Special Diagnostics, Statens Serum Institut, Copenhagen, Denmark, Artillerivej 5, 2300 Copenahgen S | Albertsen lab, Department of Chemistry and Bioscience, Aalborg University, Denmark | Rasmus Kirkegaard |
| EPI_ISL_429470 | hCoV-19/Denmark/ALAB-SSI359/2020 | Europe / Denmark / Unknown | 2020-03-23 | Department of Virus and Microbiological Special Diagnostics, Statens Serum Institut, Copenhagen, Denmark, Artillerivej 5, 2300 Copenahgen S | Albertsen lab, Department of Chemistry and Bioscience, Aalborg University, Denmark | Rasmus Kirkegaard |
| EPI_ISL_429471 | hCoV-19/Denmark/ALAB-SSI361/2020 | Europe / Denmark / Unknown | 2020-03-23 | Department of Virus and Microbiological Special Diagnostics, Statens Serum Institut, Copenhagen, Denmark, Artillerivej 5, 2300 Copenahgen S | Albertsen lab, Department of Chemistry and Bioscience, Aalborg University, Denmark | Rasmus Kirkegaard |
| EPI_ISL_429472 | hCoV-19/Denmark/ALAB-SSI364/2020 | Europe / Denmark / Unknown | 2020-03-23 | Department of Virus and Microbiological Special Diagnostics, Statens Serum Institut, Copenhagen, Denmark, Artillerivej 5, 2300 Copenahgen S | Albertsen lab, Department of Chemistry and Bioscience, Aalborg University, Denmark | Rasmus Kirkegaard |
| EPI_ISL_429473 | hCoV-19/Denmark/ALAB-SSI365/2020 | Europe / Denmark / Unknown | 2020-03-23 | Department of Virus and Microbiological Special Diagnostics, Statens Serum Institut, Copenhagen, Denmark, Artillerivej 5, 2300 Copenahgen S | Albertsen lab, Department of Chemistry and Bioscience, Aalborg University, Denmark | Rasmus Kirkegaard |
| EPI_ISL_429474 | hCoV-19/Denmark/ALAB-SSI366/2020 | Europe / Denmark / Unknown | 2020-03-23 | Department of Virus and Microbiological Special Diagnostics, Statens Serum Institut, Copenhagen, Denmark, Artillerivej 5, 2300 Copenahgen S | Albertsen lab, Department of Chemistry and Bioscience, Aalborg University, Denmark | Rasmus Kirkegaard |
| EPI_ISL_429476 | hCoV-19/Denmark/ALAB-SSI370/2020 | Europe / Denmark / Unknown | 2020-03-23 | Department of Virus and Microbiological Special Diagnostics, Statens Serum Institut, Copenhagen, Denmark, Artillerivej 5, 2300 Copenahgen S | Albertsen lab, Department of Chemistry and Bioscience, Aalborg University, Denmark | Rasmus Kirkegaard |
| EPI_ISL_429477 | hCoV-19/Denmark/ALAB-SSI371/2020 | Europe / Denmark / Unknown | 2020-03-23 | Department of Virus and Microbiological Special Diagnostics, Statens Serum Institut, Copenhagen, Denmark, Artillerivej 5, 2300 Copenahgen S | Albertsen lab, Department of Chemistry and Bioscience, Aalborg University, Denmark | Rasmus Kirkegaard |
| EPI_ISL_429478 | hCoV-19/Denmark/ALAB-SSI372/2020 | Europe / Denmark / Unknown | 2020-03-23 | Department of Virus and Microbiological Special Diagnostics, Statens Serum Institut, Copenhagen, Denmark, Artillerivej 5, 2300 Copenahgen S | Albertsen lab, Department of Chemistry and Bioscience, Aalborg University, Denmark | Rasmus Kirkegaard |
| EPI_ISL_429479 | hCoV-19/Denmark/ALAB-SSI374/2020 | Europe / Denmark / Unknown | 2020-03-22 | Department of Virus and Microbiological Special Diagnostics, Statens Serum Institut, Copenhagen, Denmark, Artillerivej 5, 2300 Copenahgen S | Albertsen lab, Department of Chemistry and Bioscience, Aalborg University, Denmark | Rasmus Kirkegaard |
| EPI_ISL_429480 | hCoV-19/Denmark/ALAB-SSI375/2020 | Europe / Denmark / Unknown | 2020-03-22 | Department of Virus and Microbiological Special Diagnostics, Statens Serum Institut, Copenhagen, Denmark, Artillerivej 5, 2300 Copenahgen S | Albertsen lab, Department of Chemistry and Bioscience, Aalborg University, Denmark | Rasmus Kirkegaard |
| EPI_ISL_429481 | hCoV-19/Denmark/ALAB-SSI376/2020 | Europe / Denmark / Unknown | 2020-03-22 | Department of Virus and Microbiological Special Diagnostics, Statens Serum Institut, Copenhagen, Denmark, Artillerivej 5, 2300 Copenahgen S | Albertsen lab, Department of Chemistry and Bioscience, Aalborg University, Denmark | Rasmus Kirkegaard |
| EPI_ISL_429482 | hCoV-19/Denmark/ALAB-SSI377/2020 | Europe / Denmark / Unknown | 2020-03-22 | Department of Virus and Microbiological Special Diagnostics, Statens Serum Institut, Copenhagen, Denmark, Artillerivej 5, 2300 Copenahgen S | Albertsen lab, Department of Chemistry and Bioscience, Aalborg University, Denmark | Rasmus Kirkegaard |
| EPI_ISL_429483 | hCoV-19/Denmark/ALAB-SSI383/2020 | Europe / Denmark / Unknown | 2020-03-24 | Department of Virus and Microbiological Special Diagnostics, Statens Serum Institut, Copenhagen, Denmark, Artillerivej 5, 2300 Copenahgen S | Albertsen lab, Department of Chemistry and Bioscience, Aalborg University, Denmark | Rasmus Kirkegaard |
| EPI_ISL_429484 | hCoV-19/Denmark/ALAB-SSI384/2020 | Europe / Denmark / Unknown | 2020-03-24 | Department of Virus and Microbiological Special Diagnostics, Statens Serum Institut, Copenhagen, Denmark, Artillerivej 5, 2300 Copenahgen S | Albertsen lab, Department of Chemistry and Bioscience, Aalborg University, Denmark | Rasmus Kirkegaard |
| EPI_ISL_429485 | hCoV-19/Denmark/ALAB-SSI385/2020 | Europe / Denmark / Unknown | 2020-03-24 | Department of Virus and Microbiological Special Diagnostics, Statens Serum Institut, Copenhagen, Denmark, Artillerivej 5, 2300 Copenahgen S | Albertsen lab, Department of Chemistry and Bioscience, Aalborg University, Denmark | Rasmus Kirkegaard |
| EPI_ISL_429486 | hCoV-19/Denmark/ALAB-SSI390/2020 | Europe / Denmark / Unknown | 2020-03-24 | Department of Virus and Microbiological Special Diagnostics, Statens Serum Institut, Copenhagen, Denmark, Artillerivej 5, 2300 Copenahgen S | Albertsen lab, Department of Chemistry and Bioscience, Aalborg University, Denmark | Rasmus Kirkegaard |
| EPI_ISL_429488 | hCoV-19/Denmark/ALAB-SSI393/2020 | Europe / Denmark / Unknown | 2020-03-24 | Department of Virus and Microbiological Special Diagnostics, Statens Serum Institut, Copenhagen, Denmark, Artillerivej 5, 2300 Copenahgen S | Albertsen lab, Department of Chemistry and Bioscience, Aalborg University, Denmark | Rasmus Kirkegaard |
| EPI_ISL_429489 | hCoV-19/Denmark/ALAB-SSI394/2020 | Europe / Denmark / Unknown | 2020-03-24 | Department of Virus and Microbiological Special Diagnostics, Statens Serum Institut, Copenhagen, Denmark, Artillerivej 5, 2300 Copenahgen S | Albertsen lab, Department of Chemistry and Bioscience, Aalborg University, Denmark | Rasmus Kirkegaard |
| EPI_ISL_429490 | hCoV-19/Denmark/ALAB-SSI395/2020 | Europe / Denmark / Unknown | 2020-03-23 | Department of Virus and Microbiological Special Diagnostics, Statens Serum Institut, Copenhagen, Denmark, Artillerivej 5, 2300 Copenahgen S | Albertsen lab, Department of Chemistry and Bioscience, Aalborg University, Denmark | Rasmus Kirkegaard |
| EPI_ISL_429492 | hCoV-19/Denmark/ALAB-SSI399/2020 | Europe / Denmark / Unknown | 2020-03-23 | Department of Virus and Microbiological Special Diagnostics, Statens Serum Institut, Copenhagen, Denmark, Artillerivej 5, 2300 Copenahgen S | Albertsen lab, Department of Chemistry and Bioscience, Aalborg University, Denmark | Rasmus Kirkegaard |
| EPI_ISL_429493 | hCoV-19/Denmark/ALAB-SSI400/2020 | Europe / Denmark / Unknown | 2020-03-23 | Department of Virus and Microbiological Special Diagnostics, Statens Serum Institut, Copenhagen, Denmark, Artillerivej 5, 2300 Copenahgen S | Albertsen lab, Department of Chemistry and Bioscience, Aalborg University, Denmark | Rasmus Kirkegaard |
| EPI_ISL_429495 | hCoV-19/Denmark/ALAB-SSI402/2020 | Europe / Denmark / Unknown | 2020-03-09 | Department of Virus and Microbiological Special Diagnostics, Statens Serum Institut, Copenhagen, Denmark, Artillerivej 5, 2300 Copenahgen S | Albertsen lab, Department of Chemistry and Bioscience, Aalborg University, Denmark | Rasmus Kirkegaard |
| EPI_ISL_429496 | hCoV-19/Denmark/ALAB-SSI403A/2020 | Europe / Denmark / Unknown | 2020-03-23 | Department of Virus and Microbiological Special Diagnostics, Statens Serum Institut, Copenhagen, Denmark, Artillerivej 5, 2300 Copenahgen S | Albertsen lab, Department of Chemistry and Bioscience, Aalborg University, Denmark | Rasmus Kirkegaard |
| EPI_ISL_429497 | hCoV-19/Denmark/ALAB-SSI404/2020 | Europe / Denmark / Unknown | 2020-03-08 | Department of Virus and Microbiological Special Diagnostics, Statens Serum Institut, Copenhagen, Denmark, Artillerivej 5, 2300 Copenahgen S | Albertsen lab, Department of Chemistry and Bioscience, Aalborg University, Denmark | Rasmus Kirkegaard |
| EPI_ISL_429498 | hCoV-19/Denmark/ALAB-SSI404A/2020 | Europe / Denmark / Unknown | 2020-03-23 | Department of Virus and Microbiological Special Diagnostics, Statens Serum Institut, Copenhagen, Denmark, Artillerivej 5, 2300 Copenahgen S | Albertsen lab, Department of Chemistry and Bioscience, Aalborg University, Denmark | Rasmus Kirkegaard |
| EPI_ISL_429499 | hCoV-19/Denmark/ALAB-SSI405/2020 | Europe / Denmark / Unknown | 2020-03-08 | Department of Virus and Microbiological Special Diagnostics, Statens Serum Institut, Copenhagen, Denmark, Artillerivej 5, 2300 Copenahgen S | Albertsen lab, Department of Chemistry and Bioscience, Aalborg University, Denmark | Rasmus Kirkegaard |
| EPI_ISL_430400 | hCoV-19/USA/NY-NYUMC373/2020 | North America / USA / New York / Brookl | 2020-04-10 | NYU Langone Health | Departments of Pathology and Medicine, New York University School of Medicine | Maria Aguero-Rosenfeld, Brendan Belovarac, Margaret Black, Ludovic Boytard, John Cadley, Paolo Cotzia, John Chen, Dacia Dimartino, Xiaojun Feng, Tatyana Gindin, Emily Guzman, Adriana Heguy, Megan Hogan, Emily Huang, George Jour, Lawrence H. Lin, Raven Luther, Andrew Lytle, Christian Marier, Matthew T. Maurano, Mark J. Mulligan, Peter Meyn, Raquel Ordonez Ciriza, Iman Osman, Jared Pinnell, Vanessa Raabe, Sitharam Ramaswami, Amy Rapkiewicz, Andre M. Ribeiro-dos-Santos, Marie Samanovic-Golden, Antonio Serrano, Guomiao Shen, Matija Snuderl, Theodore Vougiouklakis, Nick Vulpescu, Gael Westby, Paul Zappile, Yutong Zhang |
| EPI_ISL_430401 | hCoV-19/USA/CT-NYUMC374/2020 | North America / USA / Connecticut / Fairf | 2020-04-10 | NYU Langone Health | Departments of Pathology and Medicine, New York University School of Medicine | Maria Aguero-Rosenfeld, Brendan Belovarac, Margaret Black, Ludovic Boytard, John Cadley, Paolo Cotzia, John Chen, Dacia Dimartino, Xiaojun Feng, Tatyana Gindin, Emily Guzman, Adriana Heguy, Megan Hogan, Emily Huang, George Jour, Lawrence H. Lin, Raven Luther, Andrew Lytle, Christian Marier, Matthew T. Maurano, Mark J. Mulligan, Peter Meyn, Raquel Ordonez Ciriza, Iman Osman, Jared Pinnell, Vanessa Raabe, Sitharam Ramaswami, Amy Rapkiewicz, Andre M. Ribeiro-dos-Santos, Marie Samanovic-Golden, Antonio Serrano, Guomiao Shen, Matija Snuderl, Theodore Vougiouklakis, Nick Vulpescu, Gael Westby, Paul Zappile, Yutong Zhang |
| EPI_ISL_430402 | hCoV-19/USA/NY-NYUMC375/2020 | North America / USA / New York / Brookl | 2020-04-10 | NYU Langone Health | Departments of Pathology and Medicine, New York University School of Medicine | Maria Aguero-Rosenfeld, Brendan Belovarac, Margaret Black, Ludovic Boytard, John Cadley, Paolo Cotzia, John Chen, Dacia Dimartino, Xiaojun Feng, Tatyana Gindin, Emily Guzman, Adriana Heguy, Megan Hogan, Emily Huang, George Jour, Lawrence H. Lin, Raven Luther, Andrew Lytle, Christian Marier, Matthew T. Maurano, Mark J. Mulligan, Peter Meyn, Raquel Ordonez Ciriza, Iman Osman, Jared Pinnell, Vanessa Raabe, Sitharam Ramaswami, Amy Rapkiewicz, Andre M. Ribeiro-dos-Santos, Marie Samanovic-Golden, Antonio Serrano, Guomiao Shen, Matija Snuderl, Theodore Vougiouklakis, Nick Vulpescu, Gael Westby, Paul Zappile, Yutong Zhang |
| EPI_ISL_430403 | hCoV-19/USA/NJ-NYUMC376/2020 | North America / USA / New York | 2020-04-10 | NYU Langone Health | Departments of Pathology and Medicine, New York University School of Medicine | Maria Aguero-Rosenfeld, Brendan Belovarac, Margaret Black, Ludovic Boytard, John Cadley, Paolo Cotzia, John Chen, Dacia Dimartino, Xiaojun Feng, Tatyana Gindin, Emily Guzman, Adriana Heguy, Megan Hogan, Emily Huang, George Jour, Lawrence H. Lin, Raven Luther, Andrew Lytle, Christian Marier, Matthew T. Maurano, Mark J. Mulligan, Peter Meyn, Raquel Ordonez Ciriza, Iman Osman, Jared Pinnell, Vanessa Raabe, Sitharam Ramaswami, Amy Rapkiewicz, Andre M. Ribeiro-dos-Santos, Marie Samanovic-Golden, Antonio Serrano, Guomiao Shen, Matija Snuderl, Theodore Vougiouklakis, Nick Vulpescu, Gael Westby, Paul Zappile, Yutong Zhang |
| EPI_ISL_430404 | hCoV-19/USA/NY-NYUMC377/2020 | North America / USA / New York / Brookl | 2020-04-10 | NYU Langone Health | Departments of Pathology and Medicine, New York University School of Medicine | Maria Aguero-Rosenfeld, Brendan Belovarac, Margaret Black, Ludovic Boytard, John Cadley, Paolo Cotzia, John Chen, Dacia Dimartino, Xiaojun Feng, Tatyana Gindin, Emily Guzman, Adriana Heguy, Megan Hogan, Emily Huang, George Jour, Lawrence H. Lin, Raven Luther, Andrew Lytle, Christian Marier, Matthew T. Maurano, Mark J. Mulligan, Peter Meyn, Raquel Ordonez Ciriza, Iman Osman, Jared Pinnell, Vanessa Raabe, Sitharam Ramaswami, Amy Rapkiewicz, Andre M. Ribeiro-dos-Santos, Marie Samanovic-Golden, Antonio Serrano, Guomiao Shen, Matija Snuderl, Theodore Vougiouklakis, Nick Vulpescu, Gael Westby, Paul Zappile, Yutong Zhang |
| EPI_ISL_430405 | hCoV-19/USA/NJ-NYUMC378/2020 | North America / USA / New Jersey / Huds | 2020-04-10 | NYU Langone Health | Departments of Pathology and Medicine, New York University School of Medicine | Maria Aguero-Rosenfeld, Brendan Belovarac, Margaret Black, Ludovic Boytard, John Cadley, Paolo Cotzia, John Chen, Dacia Dimartino, Xiaojun Feng, Tatyana Gindin, Emily Guzman, Adriana Heguy, Megan Hogan, Emily Huang, George Jour, Lawrence H. Lin, Raven Luther, Andrew Lytle, Christian Marier, Matthew T. Maurano, Mark J. Mulligan, Peter Meyn, Raquel Ordonez Ciriza, Iman Osman, Jared Pinnell, Vanessa Raabe, Sitharam Ramaswami, Amy Rapkiewicz, Andre M. Ribeiro-dos-Santos, Marie Samanovic-Golden, Antonio Serrano, Guomiao Shen, Matija Snuderl, Theodore Vougiouklakis, Nick Vulpescu, Gael Westby, Paul Zappile, Yutong Zhang |
| EPI_ISL_430406 | hCoV-19/USA/NY-NYUMC379/2020 | North America / USA / New York / Manha | 2020-04-10 | NYU Langone Health | Departments of Pathology and Medicine, New York University School of Medicine | Maria Aguero-Rosenfeld, Brendan Belovarac, Margaret Black, Ludovic Boytard, John Cadley, Paolo Cotzia, John Chen, Dacia Dimartino, Xiaojun Feng, Tatyana Gindin, Emily Guzman, Adriana Heguy, Megan Hogan, Emily Huang, George Jour, Lawrence H. Lin, Raven Luther, Andrew Lytle, Christian Marier, Matthew T. Maurano, Mark J. Mulligan, Peter Meyn, Raquel Ordonez Ciriza, Iman Osman, Jared Pinnell, Vanessa Raabe, Sitharam Ramaswami, Amy Rapkiewicz, Andre M. Ribeiro-dos-Santos, Marie Samanovic-Golden, Antonio Serrano, Guomiao Shen, Matija Snuderl, Theodore Vougiouklakis, Nick Vulpescu, Gael Westby, Paul Zappile, Yutong Zhang |
| EPI_ISL_430407 | hCoV-19/USA/NY-NYUMC380/2020 | North America / USA / New York / Brookl | 2020-04-10 | NYU Langone Health | Departments of Pathology and Medicine, New York University School of Medicine | Maria Aguero-Rosenfeld, Brendan Belovarac, Margaret Black, Ludovic Boytard, John Cadley, Paolo Cotzia, John Chen, Dacia Dimartino, Xiaojun Feng, Tatyana Gindin, Emily Guzman, Adriana Heguy, Megan Hogan, Emily Huang, George Jour, Lawrence H. Lin, Raven Luther, Andrew Lytle, Christian Marier, Matthew T. Maurano, Mark J. Mulligan, Peter Meyn, Raquel Ordonez Ciriza, Iman Osman, Jared Pinnell, Vanessa Raabe, Sitharam Ramaswami, Amy Rapkiewicz, Andre M. Ribeiro-dos-Santos, Marie Samanovic-Golden, Antonio Serrano, Guomiao Shen, Matija Snuderl, Theodore Vougiouklakis, Nick Vulpescu, Gael Westby, Paul Zappile, Yutong Zhang |
| EPI_ISL_430408 | hCoV-19/USA/NY-NYUMC381/2020 | North America / USA / New York / Nassa | 2020-04-09 | NYU Langone Health | Departments of Pathology and Medicine, New York University School of Medicine | Maria Aguero-Rosenfeld, Brendan Belovarac, Margaret Black, Ludovic Boytard, John Cadley, Paolo Cotzia, John Chen, Dacia Dimartino, Xiaojun Feng, Tatyana Gindin, Emily Guzman, Adriana Heguy, Megan Hogan, Emily Huang, George Jour, Lawrence H. Lin, Raven Luther, Andrew Lytle, Christian Marier, Matthew T. Maurano, Mark J. Mulligan, Peter Meyn, Raquel Ordonez Ciriza, Iman Osman, Jared Pinnell, Vanessa Raabe, Sitharam Ramaswami, Amy Rapkiewicz, Andre M. Ribeiro-dos-Santos, Marie Samanovic-Golden, Antonio Serrano, Guomiao Shen, Matija Snuderl, Theodore Vougiouklakis, Nick Vulpescu, Gael Westby, Paul Zappile, Yutong Zhang |
| EPI_ISL_430409 | hCoV-19/USA/NY-NYUMC382/2020 | North America / USA / New York / Brookl | 2020-04-10 | NYU Langone Health | Departments of Pathology and Medicine, New York University School of Medicine | Maria Aguero-Rosenfeld, Brendan Belovarac, Margaret Black, Ludovic Boytard, John Cadley, Paolo Cotzia, John Chen, Dacia Dimartino, Xiaojun Feng, Tatyana Gindin, Emily Guzman, Adriana Heguy, Megan Hogan, Emily Huang, George Jour, Lawrence H. Lin, Raven Luther, Andrew Lytle, Christian Marier, Matthew T. Maurano, Mark J. Mulligan, Peter Meyn, Raquel Ordonez Ciriza, Iman Osman, Jared Pinnell, Vanessa Raabe, Sitharam Ramaswami, Amy Rapkiewicz, Andre M. Ribeiro-dos-Santos, Marie Samanovic-Golden, Antonio Serrano, Guomiao Shen, Matija Snuderl, Theodore Vougiouklakis, Nick Vulpescu, Gael Westby, Paul Zappile, Yutong Zhang |
| EPI_ISL_430410 | hCoV-19/USA/NY-NYUMC383/2020 | North America / USA / New York / Brookl | 2020-04-10 | NYU Langone Health | Departments of Pathology and Medicine, New York University School of Medicine | Maria Aguero-Rosenfeld, Brendan Belovarac, Margaret Black, Ludovic Boytard, John Cadley, Paolo Cotzia, John Chen, Dacia Dimartino, Xiaojun Feng, Tatyana Gindin, Emily Guzman, Adriana Heguy, Megan Hogan, Emily Huang, George Jour, Lawrence H. Lin, Raven Luther, Andrew Lytle, Christian Marier, Matthew T. Maurano, Mark J. Mulligan, Peter Meyn, Raquel Ordonez Ciriza, Iman Osman, Jared Pinnell, Vanessa Raabe, Sitharam Ramaswami, Amy Rapkiewicz, Andre M. Ribeiro-dos-Santos, Marie Samanovic-Golden, Antonio Serrano, Guomiao Shen, Matija Snuderl, Theodore Vougiouklakis, Nick Vulpescu, Gael Westby, Paul Zappile, Yutong Zhang |

| | | | | | | |
|---|---|---|---|---|---|---|
| EPI_ISL_430411 | hCoV-19/USA/NY-NYUMC384/2020 | North America / USA / New York / Brookl | 2020-04-09 | NYU Langone Health | Departments of Pathology and Medicine, New York University School of Medicine | Maria Aguero-Rosenfeld, Brendan Belovarac, Margaret Black, Ludovic Boytard, John Cadley, Paolo Cotzia, John Chen, Dacia Dimartino, Xiaojun Feng, Tatyana Gindin, Emily Guzman, Adriana Heguy, Megan Hogan, Emily Huang, George Jour, Lawrence H. Lin, Raven Luther, Andrew Lytle, Christian Marier, Matthew T. Maurano, Mark J. Mulligan, Peter Meyn, Raquel Ordonez Ciriza, Iman Osman, Jared Pinnell, Vanessa Raabe, Sitharam Ramaswami, Amy Rapkiewicz, Andre M. Ribeiro-dos-Santos, Marie Samanovic-Golden, Antonio Serrano, Guomiao Shen, Matija Snuderl, Theodore Vougiouklakis, Nick Vulpescu, Gael Westby, Paul Zappile, Yutong Zhang |
| EPI_ISL_430412 | hCoV-19/USA/NY-NYUMC385/2020 | North America / USA / New York / Staten | 2020-04-10 | NYU Langone Health | Departments of Pathology and Medicine, New York University School of Medicine | Maria Aguero-Rosenfeld, Brendan Belovarac, Margaret Black, Ludovic Boytard, John Cadley, Paolo Cotzia, John Chen, Dacia Dimartino, Xiaojun Feng, Tatyana Gindin, Emily Guzman, Adriana Heguy, Megan Hogan, Emily Huang, George Jour, Lawrence H. Lin, Raven Luther, Andrew Lytle, Christian Marier, Matthew T. Maurano, Mark J. Mulligan, Peter Meyn, Raquel Ordonez Ciriza, Iman Osman, Jared Pinnell, Vanessa Raabe, Sitharam Ramaswami, Amy Rapkiewicz, Andre M. Ribeiro-dos-Santos, Marie Samanovic-Golden, Antonio Serrano, Guomiao Shen, Matija Snuderl, Theodore Vougiouklakis, Nick Vulpescu, Gael Westby, Paul Zappile, Yutong Zhang |
| EPI_ISL_430413 | hCoV-19/USA/NY-NYUMC386/2020 | North America / USA / New York / Nassa | 2020-04-09 | NYU Langone Health | Departments of Pathology and Medicine, New York University School of Medicine | Maria Aguero-Rosenfeld, Brendan Belovarac, Margaret Black, Ludovic Boytard, John Cadley, Paolo Cotzia, John Chen, Dacia Dimartino, Xiaojun Feng, Tatyana Gindin, Emily Guzman, Adriana Heguy, Megan Hogan, Emily Huang, George Jour, Lawrence H. Lin, Raven Luther, Andrew Lytle, Christian Marier, Matthew T. Maurano, Mark J. Mulligan, Peter Meyn, Raquel Ordonez Ciriza, Iman Osman, Jared Pinnell, Vanessa Raabe, Sitharam Ramaswami, Amy Rapkiewicz, Andre M. Ribeiro-dos-Santos, Marie Samanovic-Golden, Antonio Serrano, Guomiao Shen, Matija Snuderl, Theodore Vougiouklakis, Nick Vulpescu, Gael Westby, Paul Zappile, Yutong Zhang |
| EPI_ISL_430414 | hCoV-19/USA/NY-NYUMC387/2020 | North America / USA / New York / Rockla | 2020-04-10 | NYU Langone Health | Departments of Pathology and Medicine, New York University School of Medicine | Maria Aguero-Rosenfeld, Brendan Belovarac, Margaret Black, Ludovic Boytard, John Cadley, Paolo Cotzia, John Chen, Dacia Dimartino, Xiaojun Feng, Tatyana Gindin, Emily Guzman, Adriana Heguy, Megan Hogan, Emily Huang, George Jour, Lawrence H. Lin, Raven Luther, Andrew Lytle, Christian Marier, Matthew T. Maurano, Mark J. Mulligan, Peter Meyn, Raquel Ordonez Ciriza, Iman Osman, Jared Pinnell, Vanessa Raabe, Sitharam Ramaswami, Amy Rapkiewicz, Andre M. Ribeiro-dos-Santos, Marie Samanovic-Golden, Antonio Serrano, Guomiao Shen, Matija Snuderl, Theodore Vougiouklakis, Nick Vulpescu, Gael Westby, Paul Zappile, Yutong Zhang |
| EPI_ISL_430415 | hCoV-19/USA/NY-NYUMC388/2020 | North America / USA / New York / Manha | 2020-04-10 | NYU Langone Health | Departments of Pathology and Medicine, New York University School of Medicine | Maria Aguero-Rosenfeld, Brendan Belovarac, Margaret Black, Ludovic Boytard, John Cadley, Paolo Cotzia, John Chen, Dacia Dimartino, Xiaojun Feng, Tatyana Gindin, Emily Guzman, Adriana Heguy, Megan Hogan, Emily Huang, George Jour, Lawrence H. Lin, Raven Luther, Andrew Lytle, Christian Marier, Matthew T. Maurano, Mark J. Mulligan, Peter Meyn, Raquel Ordonez Ciriza, Iman Osman, Jared Pinnell, Vanessa Raabe, Sitharam Ramaswami, Amy Rapkiewicz, Andre M. Ribeiro-dos-Santos, Marie Samanovic-Golden, Antonio Serrano, Guomiao Shen, Matija Snuderl, Theodore Vougiouklakis, Nick Vulpescu, Gael Westby, Paul Zappile, Yutong Zhang |
| EPI_ISL_430416 | hCoV-19/USA/NY-NYUMC389/2020 | North America / USA / New York / Manha | 2020-04-13 | NYU Langone Health | Departments of Pathology and Medicine, New York University School of Medicine | Maria Aguero-Rosenfeld, Brendan Belovarac, Margaret Black, Ludovic Boytard, John Cadley, Paolo Cotzia, John Chen, Dacia Dimartino, Xiaojun Feng, Tatyana Gindin, Emily Guzman, Adriana Heguy, Megan Hogan, Emily Huang, George Jour, Lawrence H. Lin, Raven Luther, Andrew Lytle, Christian Marier, Matthew T. Maurano, Mark J. Mulligan, Peter Meyn, Raquel Ordonez Ciriza, Iman Osman, Jared Pinnell, Vanessa Raabe, Sitharam Ramaswami, Amy Rapkiewicz, Andre M. Ribeiro-dos-Santos, Marie Samanovic-Golden, Antonio Serrano, Guomiao Shen, Matija Snuderl, Theodore Vougiouklakis, Nick Vulpescu, Gael Westby, Paul Zappile, Yutong Zhang |
| EPI_ISL_430417 | hCoV-19/USA/NY-NYUMC390/2020 | North America / USA / New York / Nassa | 2020-04-13 | NYU Langone Health | Departments of Pathology and Medicine, New York University School of Medicine | Maria Aguero-Rosenfeld, Brendan Belovarac, Margaret Black, Ludovic Boytard, John Cadley, Paolo Cotzia, John Chen, Dacia Dimartino, Xiaojun Feng, Tatyana Gindin, Emily Guzman, Adriana Heguy, Megan Hogan, Emily Huang, George Jour, Lawrence H. Lin, Raven Luther, Andrew Lytle, Christian Marier, Matthew T. Maurano, Mark J. Mulligan, Peter Meyn, Raquel Ordonez Ciriza, Iman Osman, Jared Pinnell, Vanessa Raabe, Sitharam Ramaswami, Amy Rapkiewicz, Andre M. Ribeiro-dos-Santos, Marie Samanovic-Golden, Antonio Serrano, Guomiao Shen, Matija Snuderl, Theodore Vougiouklakis, Nick Vulpescu, Gael Westby, Paul Zappile, Yutong Zhang |
| EPI_ISL_430418 | hCoV-19/USA/NY-NYUMC391/2020 | North America / USA / New York / Nassa | 2020-04-13 | NYU Langone Health | Departments of Pathology and Medicine, New York University School of Medicine | Maria Aguero-Rosenfeld, Brendan Belovarac, Margaret Black, Ludovic Boytard, John Cadley, Paolo Cotzia, John Chen, Dacia Dimartino, Xiaojun Feng, Tatyana Gindin, Emily Guzman, Adriana Heguy, Megan Hogan, Emily Huang, George Jour, Lawrence H. Lin, Raven Luther, Andrew Lytle, Christian Marier, Matthew T. Maurano, Mark J. Mulligan, Peter Meyn, Raquel Ordonez Ciriza, Iman Osman, Jared Pinnell, Vanessa Raabe, Sitharam Ramaswami, Amy Rapkiewicz, Andre M. Ribeiro-dos-Santos, Marie Samanovic-Golden, Antonio Serrano, Guomiao Shen, Matija Snuderl, Theodore Vougiouklakis, Nick Vulpescu, Gael Westby, Paul Zappile, Yutong Zhang |
| EPI_ISL_430419 | hCoV-19/USA/NY-NYUMC392/2020 | North America / USA / New York / Nassa | 2020-04-13 | NYU Langone Health | Departments of Pathology and Medicine, New York University School of Medicine | Maria Aguero-Rosenfeld, Brendan Belovarac, Margaret Black, Ludovic Boytard, John Cadley, Paolo Cotzia, John Chen, Dacia Dimartino, Xiaojun Feng, Tatyana Gindin, Emily Guzman, Adriana Heguy, Megan Hogan, Emily Huang, George Jour, Lawrence H. Lin, Raven Luther, Andrew Lytle, Christian Marier, Matthew T. Maurano, Mark J. Mulligan, Peter Meyn, Raquel Ordonez Ciriza, Iman Osman, Jared Pinnell, Vanessa Raabe, Sitharam Ramaswami, Amy Rapkiewicz, Andre M. Ribeiro-dos-Santos, Marie Samanovic-Golden, Antonio Serrano, Guomiao Shen, Matija Snuderl, Theodore Vougiouklakis, Nick Vulpescu, Gael Westby, Paul Zappile, Yutong Zhang |
| EPI_ISL_430420 | hCoV-19/USA/NY-NYUMC393/2020 | North America / USA / New York / Staten | 2020-04-13 | NYU Langone Health | Departments of Pathology and Medicine, New York University School of Medicine | Maria Aguero-Rosenfeld, Brendan Belovarac, Margaret Black, Ludovic Boytard, John Cadley, Paolo Cotzia, John Chen, Dacia Dimartino, Xiaojun Feng, Tatyana Gindin, Emily Guzman, Adriana Heguy, Megan Hogan, Emily Huang, George Jour, Lawrence H. Lin, Raven Luther, Andrew Lytle, Christian Marier, Matthew T. Maurano, Mark J. Mulligan, Peter Meyn, Raquel Ordonez Ciriza, Iman Osman, Jared Pinnell, Vanessa Raabe, Sitharam Ramaswami, Amy Rapkiewicz, Andre M. Ribeiro-dos-Santos, Marie Samanovic-Golden, Antonio Serrano, Guomiao Shen, Matija Snuderl, Theodore Vougiouklakis, Nick Vulpescu, Gael Westby, Paul Zappile, Yutong Zhang |
| EPI_ISL_430421 | hCoV-19/USA/NY-NYUMC394/2020 | North America / USA / New York / Brookl | 2020-04-13 | NYU Langone Health | Departments of Pathology and Medicine, New York University School of Medicine | Maria Aguero-Rosenfeld, Brendan Belovarac, Margaret Black, Ludovic Boytard, John Cadley, Paolo Cotzia, John Chen, Dacia Dimartino, Xiaojun Feng, Tatyana Gindin, Emily Guzman, Adriana Heguy, Megan Hogan, Emily Huang, George Jour, Lawrence H. Lin, Raven Luther, Andrew Lytle, Christian Marier, Matthew T. Maurano, Mark J. Mulligan, Peter Meyn, Raquel Ordonez Ciriza, Iman Osman, Jared Pinnell, Vanessa Raabe, Sitharam Ramaswami, Amy Rapkiewicz, Andre M. Ribeiro-dos-Santos, Marie Samanovic-Golden, Antonio Serrano, Guomiao Shen, Matija Snuderl, Theodore Vougiouklakis, Nick Vulpescu, Gael Westby, Paul Zappile, Yutong Zhang |
| EPI_ISL_430422 | hCoV-19/USA/NY-NYUMC395/2020 | North America / USA / New York / Queen | 2020-04-13 | NYU Langone Health | Departments of Pathology and Medicine, New York University School of Medicine | Maria Aguero-Rosenfeld, Brendan Belovarac, Margaret Black, Ludovic Boytard, John Cadley, Paolo Cotzia, John Chen, Dacia Dimartino, Xiaojun Feng, Tatyana Gindin, Emily Guzman, Adriana Heguy, Megan Hogan, Emily Huang, George Jour, Lawrence H. Lin, Raven Luther, Andrew Lytle, Christian Marier, Matthew T. Maurano, Mark J. Mulligan, Peter Meyn, Raquel Ordonez Ciriza, Iman Osman, Jared Pinnell, Vanessa Raabe, Sitharam Ramaswami, Amy Rapkiewicz, Andre M. Ribeiro-dos-Santos, Marie Samanovic-Golden, Antonio Serrano, Guomiao Shen, Matija Snuderl, Theodore Vougiouklakis, Nick Vulpescu, Gael Westby, Paul Zappile, Yutong Zhang |
| EPI_ISL_430423 | hCoV-19/USA/NY-NYUMC396/2020 | North America / USA / New York / Queen | 2020-04-11 | NYU Langone Health | Departments of Pathology and Medicine, New York University School of Medicine | Maria Aguero-Rosenfeld, Brendan Belovarac, Margaret Black, Ludovic Boytard, John Cadley, Paolo Cotzia, John Chen, Dacia Dimartino, Xiaojun Feng, Tatyana Gindin, Emily Guzman, Adriana Heguy, Megan Hogan, Emily Huang, George Jour, Lawrence H. Lin, Raven Luther, Andrew Lytle, Christian Marier, Matthew T. Maurano, Mark J. Mulligan, Peter Meyn, Raquel Ordonez Ciriza, Iman Osman, Jared Pinnell, Vanessa Raabe, Sitharam Ramaswami, Amy Rapkiewicz, Andre M. Ribeiro-dos-Santos, Marie Samanovic-Golden, Antonio Serrano, Guomiao Shen, Matija Snuderl, Theodore Vougiouklakis, Nick Vulpescu, Gael Westby, Paul Zappile, Yutong Zhang |
| EPI_ISL_430424 | hCoV-19/USA/NY-NYUMC397/2020 | North America / USA / New York / Brookl | 2020-04-13 | NYU Langone Health | Departments of Pathology and Medicine, New York University School of Medicine | Maria Aguero-Rosenfeld, Brendan Belovarac, Margaret Black, Ludovic Boytard, John Cadley, Paolo Cotzia, John Chen, Dacia Dimartino, Xiaojun Feng, Tatyana Gindin, Emily Guzman, Adriana Heguy, Megan Hogan, Emily Huang, George Jour, Lawrence H. Lin, Raven Luther, Andrew Lytle, Christian Marier, Matthew T. Maurano, Mark J. Mulligan, Peter Meyn, Raquel Ordonez Ciriza, Iman Osman, Jared Pinnell, Vanessa Raabe, Sitharam Ramaswami, Amy Rapkiewicz, Andre M. Ribeiro-dos-Santos, Marie Samanovic-Golden, Antonio Serrano, Guomiao Shen, Matija Snuderl, Theodore Vougiouklakis, Nick Vulpescu, Gael Westby, Paul Zappile, Yutong Zhang |
| EPI_ISL_430425 | hCoV-19/USA/NY-NYUMC398/2020 | North America / USA / New York / Manha | 2020-04-13 | NYU Langone Health | Departments of Pathology and Medicine, New York University School of Medicine | Maria Aguero-Rosenfeld, Brendan Belovarac, Margaret Black, Ludovic Boytard, John Cadley, Paolo Cotzia, John Chen, Dacia Dimartino, Xiaojun Feng, Tatyana Gindin, Emily Guzman, Adriana Heguy, Megan Hogan, Emily Huang, George Jour, Lawrence H. Lin, Raven Luther, Andrew Lytle, Christian Marier, Matthew T. Maurano, Mark J. Mulligan, Peter Meyn, Raquel Ordonez Ciriza, Iman Osman, Jared Pinnell, Vanessa Raabe, Sitharam Ramaswami, Amy Rapkiewicz, Andre M. Ribeiro-dos-Santos, Marie Samanovic-Golden, Antonio Serrano, Guomiao Shen, Matija Snuderl, Theodore Vougiouklakis, Nick Vulpescu, Gael Westby, Paul Zappile, Yutong Zhang |
| EPI_ISL_430426 | hCoV-19/USA/NY-NYUMC399/2020 | North America / USA / New York / Manha | 2020-04-13 | NYU Langone Health | Departments of Pathology and Medicine, New York University School of Medicine | Maria Aguero-Rosenfeld, Brendan Belovarac, Margaret Black, Ludovic Boytard, John Cadley, Paolo Cotzia, John Chen, Dacia Dimartino, Xiaojun Feng, Tatyana Gindin, Emily Guzman, Adriana Heguy, Megan Hogan, Emily Huang, George Jour, Lawrence H. Lin, Raven Luther, Andrew Lytle, Christian Marier, Matthew T. Maurano, Mark J. Mulligan, Peter Meyn, Raquel Ordonez Ciriza, Iman Osman, Jared Pinnell, Vanessa Raabe, Sitharam Ramaswami, Amy Rapkiewicz, Andre M. Ribeiro-dos-Santos, Marie Samanovic-Golden, Antonio Serrano, Guomiao Shen, Matija Snuderl, Theodore Vougiouklakis, Nick Vulpescu, Gael Westby, Paul Zappile, Yutong Zhang |
| EPI_ISL_430427 | hCoV-19/USA/NY-NYUMC400/2020 | North America / USA / New York / Nassa | 2020-04-13 | NYU Langone Health | Departments of Pathology and Medicine, New York University School of Medicine | Maria Aguero-Rosenfeld, Brendan Belovarac, Margaret Black, Ludovic Boytard, John Cadley, Paolo Cotzia, John Chen, Dacia Dimartino, Xiaojun Feng, Tatyana Gindin, Emily Guzman, Adriana Heguy, Megan Hogan, Emily Huang, George Jour, Lawrence H. Lin, Raven Luther, Andrew Lytle, Christian Marier, Matthew T. Maurano, Mark J. Mulligan, Peter Meyn, Raquel Ordonez Ciriza, Iman Osman, Jared Pinnell, Vanessa Raabe, Sitharam Ramaswami, Amy Rapkiewicz, Andre M. Ribeiro-dos-Santos, Marie Samanovic-Golden, Antonio Serrano, Guomiao Shen, Matija Snuderl, Theodore Vougiouklakis, Nick Vulpescu, Gael Westby, Paul Zappile, Yutong Zhang |
| EPI_ISL_430428 | hCoV-19/USA/NJ-NYUMC401/2020 | North America / USA / New Jersey / Hud | 2020-04-13 | NYU Langone Health | Departments of Pathology and Medicine, New York University School of Medicine | Maria Aguero-Rosenfeld, Brendan Belovarac, Margaret Black, Ludovic Boytard, John Cadley, Paolo Cotzia, John Chen, Dacia Dimartino, Xiaojun Feng, Tatyana Gindin, Emily Guzman, Adriana Heguy, Megan Hogan, Emily Huang, George Jour, Lawrence H. Lin, Raven Luther, Andrew Lytle, Christian Marier, Matthew T. Maurano, Mark J. Mulligan, Peter Meyn, Raquel Ordonez Ciriza, Iman Osman, Jared Pinnell, Vanessa Raabe, Sitharam Ramaswami, Amy Rapkiewicz, Andre M. Ribeiro-dos-Santos, Marie Samanovic-Golden, Antonio Serrano, Guomiao Shen, Matija Snuderl, Theodore Vougiouklakis, Nick Vulpescu, Gael Westby, Paul Zappile, Yutong Zhang |
| EPI_ISL_430429 | hCoV-19/USA/NY-NYUMC402/2020 | North America / USA / New York / Brookl | 2020-04-13 | NYU Langone Health | Departments of Pathology and Medicine, New York University School of Medicine | Maria Aguero-Rosenfeld, Brendan Belovarac, Margaret Black, Ludovic Boytard, John Cadley, Paolo Cotzia, John Chen, Dacia Dimartino, Xiaojun Feng, Tatyana Gindin, Emily Guzman, Adriana Heguy, Megan Hogan, Emily Huang, George Jour, Lawrence H. Lin, Raven Luther, Andrew Lytle, Christian Marier, Matthew T. Maurano, Mark J. Mulligan, Peter Meyn, Raquel Ordonez Ciriza, Iman Osman, Jared Pinnell, Vanessa Raabe, Sitharam Ramaswami, Amy Rapkiewicz, Andre M. Ribeiro-dos-Santos, Marie Samanovic-Golden, Antonio Serrano, Guomiao Shen, Matija Snuderl, Theodore Vougiouklakis, Nick Vulpescu, Gael Westby, Paul Zappile, Yutong Zhang |
| EPI_ISL_430430 | hCoV-19/USA/NY-NYUMC403/2020 | North America / USA / New York / Bronx | 2020-04-13 | NYU Langone Health | Departments of Pathology and Medicine, New York University School of Medicine | Maria Aguero-Rosenfeld, Brendan Belovarac, Margaret Black, Ludovic Boytard, John Cadley, Paolo Cotzia, John Chen, Dacia Dimartino, Xiaojun Feng, Tatyana Gindin, Emily Guzman, Adriana Heguy, Megan Hogan, Emily Huang, George Jour, Lawrence H. Lin, Raven Luther, Andrew Lytle, Christian Marier, Matthew T. Maurano, Mark J. Mulligan, Peter Meyn, Raquel Ordonez Ciriza, Iman Osman, Jared Pinnell, Vanessa Raabe, Sitharam Ramaswami, Amy Rapkiewicz, Andre M. Ribeiro-dos-Santos, Marie Samanovic-Golden, Antonio Serrano, Guomiao Shen, Matija Snuderl, Theodore Vougiouklakis, Nick Vulpescu, Gael Westby, Paul Zappile, Yutong Zhang |
| EPI_ISL_430431 | hCoV-19/USA/NY-NYUMC404/2020 | North America / USA / New York / Nassa | 2020-04-13 | NYU Langone Health | Departments of Pathology and Medicine, New York University School of Medicine | Maria Aguero-Rosenfeld, Brendan Belovarac, Margaret Black, Ludovic Boytard, John Cadley, Paolo Cotzia, John Chen, Dacia Dimartino, Xiaojun Feng, Tatyana Gindin, Emily Guzman, Adriana Heguy, Megan Hogan, Emily Huang, George Jour, Lawrence H. Lin, Raven Luther, Andrew Lytle, Christian Marier, Matthew T. Maurano, Mark J. Mulligan, Peter Meyn, Raquel Ordonez Ciriza, Iman Osman, Jared Pinnell, Vanessa Raabe, Sitharam Ramaswami, Amy Rapkiewicz, Andre M. Ribeiro-dos-Santos, Marie Samanovic-Golden, Antonio Serrano, Guomiao Shen, Matija Snuderl, Theodore Vougiouklakis, Nick Vulpescu, Gael Westby, Paul Zappile, Yutong Zhang |
| EPI_ISL_430432 | hCoV-19/USA/NY-NYUMC405/2020 | North America / USA / New York / Staten | 2020-04-13 | NYU Langone Health | Departments of Pathology and Medicine, New York University School of Medicine | Maria Aguero-Rosenfeld, Brendan Belovarac, Margaret Black, Ludovic Boytard, John Cadley, Paolo Cotzia, John Chen, Dacia Dimartino, Xiaojun Feng, Tatyana Gindin, Emily Guzman, Adriana Heguy, Megan Hogan, Emily Huang, George Jour, Lawrence H. Lin, Raven Luther, Andrew Lytle, Christian Marier, Matthew T. Maurano, Mark J. Mulligan, Peter Meyn, Raquel Ordonez Ciriza, Iman Osman, Jared Pinnell, Vanessa Raabe, Sitharam Ramaswami, Amy Rapkiewicz, Andre M. Ribeiro-dos-Santos, Marie Samanovic-Golden, Antonio Serrano, Guomiao Shen, Matija Snuderl, Theodore Vougiouklakis, Nick Vulpescu, Gael Westby, Paul Zappile, Yutong Zhang |
| EPI_ISL_430433 | hCoV-19/USA/NY-NYUMC406/2020 | North America / USA / New York / Queen | 2020-04-13 | NYU Langone Health | Departments of Pathology and Medicine, New York University School of Medicine | Maria Aguero-Rosenfeld, Brendan Belovarac, Margaret Black, Ludovic Boytard, John Cadley, Paolo Cotzia, John Chen, Dacia Dimartino, Xiaojun Feng, Tatyana Gindin, Emily Guzman, Adriana Heguy, Megan Hogan, Emily Huang, George Jour, Lawrence H. Lin, Raven Luther, Andrew Lytle, Christian Marier, Matthew T. Maurano, Mark J. Mulligan, Peter Meyn, Raquel Ordonez Ciriza, Iman Osman, Jared Pinnell, Vanessa Raabe, Sitharam Ramaswami, Amy Rapkiewicz, Andre M. Ribeiro-dos-Santos, Marie Samanovic-Golden, Antonio Serrano, Guomiao Shen, Matija Snuderl, Theodore Vougiouklakis, Nick Vulpescu, Gael Westby, Paul Zappile, Yutong Zhang |
| EPI_ISL_430434 | hCoV-19/USA/NY-NYUMC407/2020 | North America / USA / New York / Queen | 2020-04-12 | NYU Langone Health | Departments of Pathology and Medicine, New York University School of Medicine | Maria Aguero-Rosenfeld, Brendan Belovarac, Margaret Black, Ludovic Boytard, John Cadley, Paolo Cotzia, John Chen, Dacia Dimartino, Xiaojun Feng, Tatyana Gindin, Emily Guzman, Adriana Heguy, Megan Hogan, Emily Huang, George Jour, Lawrence H. Lin, Raven Luther, Andrew Lytle, Christian Marier, Matthew T. Maurano, Mark J. Mulligan, Peter Meyn, Raquel Ordonez Ciriza, Iman Osman, Jared Pinnell, Vanessa Raabe, Sitharam Ramaswami, Amy Rapkiewicz, Andre M. Ribeiro-dos-Santos, Marie Samanovic-Golden, Antonio Serrano, Guomiao Shen, Matija Snuderl, Theodore Vougiouklakis, Nick Vulpescu, Gael Westby, Paul Zappile, Yutong Zhang |
| EPI_ISL_430439 | hCoV-19/Malaysia/IMR_WC1177/2020 | Asia / Malaysia | 2020-03-05 | Institute for Medical Research, Infectious Disease Research Centre, National Institutes of Health, Ministry of Health Malaysia | Institute for Medical Research Infectious Disease Research Centre, National Institutes of Health, Ministry of Health Malaysia | Suppiah.J, Mohd-Zawawi.Z, Kalyanasundram.J, Azizan.M-A, Mat-Sharani.S, Hisham.H-A, Tan.L-P, Abdul-Wahid.M-Z, Mohd-Zain.R, Ahmad.N, Thayan.R |
| EPI_ISL_430440 | hCoV-19/Malaysia/IMR_WC1170/2020 | Asia / Malaysia | 2020-03-05 | Institute for Medical Research, Infectious Disease Research Centre, National Institutes of Health, Ministry of Health Malaysia | Institute for Medical Research, Infectious Disease Research Centre, National Institutes of Health, Ministry of Health Malaysia | Suppiah.J, Mohd-Zawawi.Z, Kalyanasundram.J, Azizan.M-A, Mat-Sharani.S, Hisham.H-A, Tan.L-P, Abdul-Wahid.M-Z, Tengku-Abd-Rashid.T-R, Mohd-Zain.R, Ahmad.N, Thayan.R |
| EPI_ISL_430441 | hCoV-19/Malaysia/IMR_WC1097/2020 | Asia / Malaysia | 2020-02-29 | Institute for Medical Research, Infectious Disease Research Centre, National Institutes of Health, Ministry of Health Malaysia | Institute for Medical Research, Infectious Disease Research Centre, National Institutes of Health, Ministry of Health Malaysia | Suppiah.J, Mohd-Zawawi.Z, Kalyanasundram.J, Azizan.M-A, Mat-Sharani.S, Hisham.H-A, Tan.L-P, Abdul-Wahid.M-Z, Tengku-Rogayah.TAR, Mohd-Zain.R, Ahmad.N, Thayan.R |
| EPI_ISL_430442 | hCoV-19/Malaysia/IMR_WC1098/2020 | Asia / Malaysia | 2020-02-29 | Institute for Medical Research, Infectious Disease Research Centre, National Institutes of Health, Ministry of Health Malaysia | Institute for Medical Research, Infectious Disease Research Centre, National Institutes of Health, Ministry of Health Malaysia | Suppiah.J, Mohd-Zawawi.Z, Kalyanasundram.J, Azizan.M-A, Mat-Sharani.S, Hisham.H-A, Tan.L-P, Abdul-Wahid.M-Z, Tengku-Rogayah.TAR, Mohd-Zain.R, Ahmad.N, Thayan.R |
| EPI_ISL_430443 | hCoV-19/Malaysia/IMR_WC085/2020 | Asia / Malaysia | 2020-01-28 | Institute for Medical Research, Infectious Disease Research Centre, National Institutes of Health, Ministry of Health Malaysia | Institute for Medical Research, Infectious Disease Research Centre, National Institutes of Health, Ministry of Health Malaysia | Suppiah.J, Mohd-Zawawi.Z, Kalyanasundram.J, Azizan.M-A, Mat-Sharani.S, Hisham.H-A, Tan.L-P, Abdul-Wahid.M-Z, Tengku-Rogayah.TAR, Mohd-Zain.R, Ahmad.N, Thayan.R |
| EPI_ISL_430444 | hCoV-19/Malaysia/IMR_WC627/2020 | Asia / Malaysia | 2020-02-12 | Institute for Medical Research, Infectious Disease Research Centre, National Institutes of Health, Ministry of Health Malaysia | Institute for Medical Research, Infectious Disease Research Centre, National Institutes of Health, Ministry of Health Malaysia | Suppiah.J, Mohd-Zawawi.Z, Kalyanasundram.J, Azizan.M-A, Mat-Sharani.S, Hisham.H-A, Tan.L-P, Abdul-Wahid.M-Z, Tengku-Rogayah.TAR, Mohd-Zain.R, Ahmad.N, Thayan.R |
| EPI_ISL_430464 | hCoV-19/India/S3/2020 | Asia / India / West Bengal / Kolkata | 2020-03-21 | ICMR-National Institute of Cholera and Enteric Diseases | National Institute of Biomedical Genomics | Arindam Maitra, Mamta Chawla Sarkar, Sreedhar Chinnaswamy, Hasina Banu, Ananya Chatterjee, Shanta Dutta, Saumitra Das |
| EPI_ISL_430465 | hCoV-19/India/S5/2020 | Asia / India / West Bengal / Darjeeling | 2020-03-28 | ICMR-National Institute of Cholera and Enteric Diseases | National Institute of Biomedical Genomics | Arindam Maitra, Mamta Chawla Sarkar, Sreedhar Chinnaswamy, Hasina Banu, Ananya Chatterjee, Shanta Dutta, Saumitra Das |
| EPI_ISL_430466 | hCoV-19/India/S6/2020 | Asia / India / West Bengal / Tehatta | 2020-03-26 | ICMR-National Institute of Cholera and Enteric Diseases | National Institute of Biomedical Genomics | Arindam Maitra, Mamta Chawla Sarkar, Sreedhar Chinnaswamy, Hasina Banu, Ananya Chatterjee, Shanta Dutta, Saumitra Das |
| EPI_ISL_430467 | hCoV-19/India/S11/2020 | Asia / India / West Bengal / East Medinip | 2020-04-03 | ICMR-National Institute of Cholera and Enteric Diseases | National Institute of Biomedical Genomics | Arindam Maitra, Mamta Chawla Sarkar, Sreedhar Chinnaswamy, Hasina Banu, Ananya Chatterjee, Shanta Dutta, Saumitra Das |
| EPI_ISL_430468 | hCoV-19/India/S2/2020 | Asia / India / West_Bengal / Kolkata | 2020-03-21 | ICMR-National Institute of Cholera and Enteric Diseases | National Institute of Biomedical Genomics | Arindam Maitra, Mamta Chawla Sarkar, Sreedhar Chinnaswamy, Hasina Banu, Ananya Chatterjee, Shanta Dutta, Saumitra Das |
| EPI_ISL_430469 | hCoV-19/Greece/127_HPI/2020 | Europe / Greece / Athens | 2020-02-29 | Hellenic Pasteur Institute, Public Health Laboratories | Hellenic Pasteur Institute, Public Health Laboratories, Unit of Bioinformatics and Applied Genomics | Vassiliki Pogka, Timokratis Karamitros, Athanasios Kossyvakis, Antonios Kalliaropoulos, Horefti Elina, Evangelidou Maria, Androniki Voulgari-Kokota, Aspasia Kontou, Andreas Mentis |
| EPI_ISL_430471 | hCoV-19/Australia/VIC935/2020 | Oceania / Australia / Victoria | 2020-03-25 | Microbiological Diagnostic Unit Public Health Laboratory | Microbiological Diagnostic Unit Public Health Laboratory | Seemann T., Schultz M., Sait, M., Sherry, N. |
| EPI_ISL_430472 | hCoV-19/Australia/VIC938/2020 | Oceania / Australia / Victoria | 2020-03-25 | Microbiological Diagnostic Unit Public Health Laboratory | Microbiological Diagnostic Unit Public Health Laboratory | Seemann T., Schultz M., Sait, M., Sherry, N. |
| EPI_ISL_430473 | hCoV-19/Australia/VIC1008/2020 | Oceania / Australia / Victoria | 2020-03-31 | Victorian Infectious Diseases Reference Laboratory (VIDRL) | Microbiological Diagnostic Unit Public Health Laboratory and Victorian Infectious Diseases Reference Laboratory, The Peter Doherty Institute for Infection and Immunity | Caly L., Seemann T., Sait, M., Schultz M., Druce J., Sherry, N. |
| EPI_ISL_430476 | hCoV-19/Australia/VIC1016/2020 | Oceania / Australia / Victoria | 2020-04-01 | Victorian Infectious Diseases Reference Laboratory (VIDRL) | Microbiological Diagnostic Unit Public Health Laboratory and Victorian Infectious Diseases Reference Laboratory, The Peter Doherty Institute for Infection and Immunity | Caly L., Seemann T., Sait, M., Schultz M., Druce J., Sherry, N. |
| EPI_ISL_430478 | hCoV-19/Australia/VIC994/2020 | Oceania / Australia / Victoria | 2020-03-31 | Victorian Infectious Diseases Reference Laboratory (VIDRL) | Microbiological Diagnostic Unit Public Health Laboratory and Victorian Infectious Diseases Reference Laboratory, The Peter Doherty Institute for Infection and Immunity | Caly L., Seemann T., Sait, M., Schultz M., Druce J., Sherry, N. |
| EPI_ISL_430479 | hCoV-19/Australia/VIC995/2020 | Oceania / Australia / Victoria | 2020-03-31 | Victorian Infectious Diseases Reference Laboratory (VIDRL) | Microbiological Diagnostic Unit Public Health Laboratory and Victorian Infectious Diseases Reference Laboratory, The Peter Doherty Institute for Infection and Immunity | Caly L., Seemann T., Sait, M., Schultz M., Druce J., Sherry, N. |
| EPI_ISL_430481 | hCoV-19/Australia/VIC997/2020 | Oceania / Australia / Victoria | 2020-03-31 | Victorian Infectious Diseases Reference Laboratory (VIDRL) | Microbiological Diagnostic Unit Public Health Laboratory and Victorian Infectious Diseases Reference Laboratory, The Peter Doherty Institute for Infection and Immunity | Caly L., Seemann T., Sait, M., Schultz M., Druce J., Sherry, N. |
| EPI_ISL_430482 | hCoV-19/Australia/VIC998/2020 | Oceania / Australia / Victoria | 2020-03-31 | Victorian Infectious Diseases Reference Laboratory (VIDRL) | Microbiological Diagnostic Unit Public Health Laboratory and Victorian Infectious Diseases Reference Laboratory, The Peter Doherty Institute for Infection and Immunity | Caly L., Seemann T., Sait, M., Schultz M., Druce J., Sherry, N. |
| EPI_ISL_430484 | hCoV-19/Australia/VIC1000/2020 | Oceania / Australia / Victoria | 2020-03-31 | Victorian Infectious Diseases Reference Laboratory (VIDRL) | Microbiological Diagnostic Unit Public Health Laboratory and Victorian Infectious Diseases Reference Laboratory, The Peter Doherty Institute for Infection and Immunity | Caly L., Seemann T., Sait, M., Schultz M., Druce J., Sherry, N. |
| EPI_ISL_430486 | hCoV-19/Australia/VIC1002/2020 | Oceania / Australia / Victoria | 2020-03-31 | Victorian Infectious Diseases Reference Laboratory (VIDRL) | Microbiological Diagnostic Unit Public Health Laboratory and Victorian Infectious Diseases Reference Laboratory, The Peter Doherty Institute for Infection and Immunity | Caly L., Seemann T., Sait, M., Schultz M., Druce J., Sherry, N. |
| EPI_ISL_430490 | hCoV-19/Australia/VIC1018/2020 | Oceania / Australia / Victoria | 2020-04-01 | Victorian Infectious Diseases Reference Laboratory (VIDRL) | Microbiological Diagnostic Unit Public Health Laboratory and Victorian Infectious Diseases Reference Laboratory, The Peter Doherty Institute for Infection and Immunity | Caly L., Seemann T., Sait, M., Schultz M., Druce J., Sherry, N. |
| EPI_ISL_430498 | hCoV-19/Australia/VIC1208/2020 | Oceania / Australia / Victoria | 2020-04-01 | Victorian Infectious Diseases Reference Laboratory (VIDRL) | Microbiological Diagnostic Unit Public Health Laboratory and Victorian Infectious Diseases Reference Laboratory, The Peter Doherty Institute for Infection and Immunity | Caly L., Seemann T., Sait, M., Schultz M., Druce J., Sherry, N. |
| EPI_ISL_431778 | hCoV-19/mink/Netherlands/1/2020 | Europe / Netherlands / Milheeze | 2020-04-24 | unknown | Virology, Wageningen Bioveterinary Research | Oreshkova,N., Vreman,S., Molenaar,R.J., Harders,F., Hakze van der Honing,R.W., Gerhards,N., Bouwstra,R., Hissink,H., Smit,L., Tacken,M., Weesendorp,E., Stegeman,A. and van der Poel,W.H.M. |
| EPI_ISL_431782 | hCoV-19/Fujian/IM3520014T/2020 | Asia / China / Fujian | 2020-03-22 | Fujian Center for Disease Control and Prevention | Fujian Center for Disease Control and Prevention | Lin Qi, Huang Zhimiao, Zhang Yanhua, Weng Yuwei |
| EPI_ISL_431783 | hCoV-19/Fujian/IM3520016T/2020 | Asia / China / Fujian | 2020-03-19 | Fujian Center for Disease Control and Prevention | Fujian Center for Disease Control and Prevention | Lin Qi, Huang Zhimiao, Zhang Yanhua, Weng Yuwei |
| EPI_ISL_431784 | hCoV-19/Fujian/IM3520017T/2020 | Asia / China / Fujian | 2020-03-18 | Fujian Center for Disease Control and Prevention | Fujian Center for Disease Control and Prevention | Lin Qi, Huang Zhimiao, Zhang Yanhua, Weng Yuwei |
| EPI_ISL_431785 | hCoV-19/Fujian/3520022Y/2020 | Asia / China / Fujian | 2020-01-23 | Fujian Center for Disease Control and Prevention | Fujian Center for Disease Control and Prevention | Lin Qi, Huang Zhimiao, Zhang Yanhua, Weng Yuwei |
| EPI_ISL_432176 | hCoV-19/Wales/PHWC-2771D/2020 | Europe / United Kingdom / Wales | 2020-04-01 | Wales Specialist Virology Centre | Public Health Wales Microbiology Cardiff | Catherine Moore, Johnathan Evans, Malorie Perry, Simon Cottrell, Alec Birchley, Alexander Adams, Amy Gaskin, Bree Gatica-Wilcox, Jason Coombes, Lauren Gilbert, Lee Graham, Nicole Pacchiarini, Sara Kumziene-Summerhayes, Sarah Taylor, Sophie Jones, Sara Rey, Matthew Bull, Joanne Watkins, Sally Corden, Tom Connor |
| EPI_ISL_432177 | hCoV-19/Wales/PHWC-252F0/2020 | Europe / United Kingdom / Wales | 2020-03-22 | Wales Specialist Virology Centre | Public Health Wales Microbiology Cardiff | Catherine Moore, Johnathan Evans, Malorie Perry, Simon Cottrell, Alec Birchley, Alexander Adams, Amy Gaskin, Bree Gatica-Wilcox, Jason Coombes, Lauren Gilbert, Lee Graham, Nicole Pacchiarini, Sara Kumziene-Summerhayes, Sarah Taylor, Sophie Jones, Sara Rey, Matthew Bull, Joanne Watkins, Sally Corden, Tom Connor |
| EPI_ISL_432178 | hCoV-19/Wales/PHWC-2A12B/2020 | Europe / United Kingdom / Wales | 2020-04-04 | Wales Specialist Virology Centre | Public Health Wales Microbiology Cardiff | Catherine Moore, Johnathan Evans, Malorie Perry, Simon Cottrell, Alec Birchley, Alexander Adams, Amy Gaskin, Bree Gatica-Wilcox, Jason Coombes, Lauren Gilbert, Lee Graham, Nicole Pacchiarini, Sara Kumziene-Summerhayes, Sarah Taylor, Sophie Jones, Sara Rey, Matthew Bull, Joanne Watkins, Sally Corden, Tom Connor |
| EPI_ISL_432179 | hCoV-19/Wales/PHWC-25384/2020 | Europe / United Kingdom / Wales | 2020-03-23 | Wales Specialist Virology Centre | Public Health Wales Microbiology Cardiff | Catherine Moore, Johnathan Evans, Malorie Perry, Simon Cottrell, Alec Birchley, Alexander Adams, Amy Gaskin, Bree Gatica-Wilcox, Jason Coombes, Lauren Gilbert, Lee Graham, Nicole Pacchiarini, Sara Kumziene-Summerhayes, Sarah Taylor, Sophie Jones, Sara Rey, Matthew Bull, Joanne Watkins, Sally Corden, Tom Connor |
| EPI_ISL_432180 | hCoV-19/Wales/PHWC-27A9C/2020 | Europe / United Kingdom / Wales | 2020-03-30 | Wales Specialist Virology Centre | Public Health Wales Microbiology Cardiff | Catherine Moore, Johnathan Evans, Malorie Perry, Simon Cottrell, Alec Birchley, Alexander Adams, Amy Gaskin, Bree Gatica-Wilcox, Jason Coombes, Lauren Gilbert, Lee Graham, Nicole Pacchiarini, Sara Kumziene-Summerhayes, Sarah Taylor, Sophie Jones, Sara Rey, Matthew Bull, Joanne Watkins, Sally Corden, Tom Connor |
| EPI_ISL_432181 | hCoV-19/Wales/PHWC-29E16/2020 | Europe / United Kingdom / Wales | 2020-04-04 | Wales Specialist Virology Centre | Public Health Wales Microbiology Cardiff | Catherine Moore, Johnathan Evans, Malorie Perry, Simon Cottrell, Alec Birchley, Alexander Adams, Amy Gaskin, Bree Gatica-Wilcox, Jason Coombes, Lauren Gilbert, Lee Graham, Nicole Pacchiarini, Sara Kumziene-Summerhayes, Sarah Taylor, Sophie Jones, Sara Rey, Matthew Bull, Joanne Watkins, Sally Corden, Tom Connor |
| EPI_ISL_432182 | hCoV-19/Wales/PHWC-2531B/2020 | Europe / United Kingdom / Wales | 2020-03-22 | Wales Specialist Virology Centre | Public Health Wales Microbiology Cardiff | Catherine Moore, Johnathan Evans, Malorie Perry, Simon Cottrell, Alec Birchley, Alexander Adams, Amy Gaskin, Bree Gatica-Wilcox, Jason Coombes, Lauren Gilbert, Lee Graham, Nicole Pacchiarini, Sara Kumziene-Summerhayes, Sarah Taylor, Sophie Jones, Sara Rey, Matthew Bull, Joanne Watkins, Sally Corden, Tom Connor |
| EPI_ISL_432183 | hCoV-19/Wales/PHWC-25366/2020 | Europe / United Kingdom / Wales | 2020-03-24 | Wales Specialist Virology Centre | Public Health Wales Microbiology Cardiff | Catherine Moore, Johnathan Evans, Malorie Perry, Simon Cottrell, Alec Birchley, Alexander Adams, Amy Gaskin, Bree Gatica-Wilcox, Jason Coombes, Lauren Gilbert, Lee Graham, Nicole Pacchiarini, Sara Kumziene-Summerhayes, Sarah Taylor, Sophie Jones, Sara Rey, Matthew Bull, Joanne Watkins, Sally Corden, Tom Connor |
| EPI_ISL_432184 | hCoV-19/Wales/PHWC-2A167/2020 | Europe / United Kingdom / Wales | 2020-04-05 | Wales Specialist Virology Centre | Public Health Wales Microbiology Cardiff | Catherine Moore, Johnathan Evans, Malorie Perry, Simon Cottrell, Alec Birchley, Alexander Adams, Amy Gaskin, Bree Gatica-Wilcox, Jason Coombes, Lauren Gilbert, Lee Graham, Nicole Pacchiarini, Sara Kumziene-Summerhayes, Sarah Taylor, Sophie Jones, Sara Rey, Matthew Bull, Joanne Watkins, Sally Corden, Tom Connor |
| EPI_ISL_432185 | hCoV-19/Wales/PHWC-27CA5/2020 | Europe / United Kingdom / Wales | 2020-04-01 | Wales Specialist Virology Centre | Public Health Wales Microbiology Cardiff | Catherine Moore, Johnathan Evans, Malorie Perry, Simon Cottrell, Alec Birchley, Alexander Adams, Amy Gaskin, Bree Gatica-Wilcox, Jason Coombes, Lauren Gilbert, Lee Graham, Nicole Pacchiarini, Sara Kumziene-Summerhayes, Sarah Taylor, Sophie Jones, Sara Rey, Matthew Bull, Joanne Watkins, Sally Corden, Tom Connor |
| EPI_ISL_432186 | hCoV-19/Wales/PHWC-29E61/2020 | Europe / United Kingdom / Wales | 2020-04-04 | Wales Specialist Virology Centre | Public Health Wales Microbiology Cardiff | Catherine Moore, Johnathan Evans, Malorie Perry, Simon Cottrell, Alec Birchley, Alexander Adams, Amy Gaskin, Bree Gatica-Wilcox, Jason Coombes, Lauren Gilbert, Lee Graham, Nicole Pacchiarini, Sara Kumziene-Summerhayes, Sarah Taylor, Sophie Jones, Sara Rey, Matthew Bull, Joanne Watkins, Sally Corden, Tom Connor |
| EPI_ISL_432187 | hCoV-19/Wales/PHWC-2A06A/2020 | Europe / United Kingdom / Wales | 2020-04-04 | Wales Specialist Virology Centre | Public Health Wales Microbiology Cardiff | Catherine Moore, Johnathan Evans, Malorie Perry, Simon Cottrell, Alec Birchley, Alexander Adams, Amy Gaskin, Bree Gatica-Wilcox, Jason Coombes, Lauren Gilbert, Lee Graham, Nicole Pacchiarini, Sara Kumziene-Summerhayes, Sarah Taylor, Sophie Jones, Sara Rey, Matthew Bull, Joanne Watkins, Sally Corden, Tom Connor |
| EPI_ISL_432188 | hCoV-19/Wales/PHWC-25357/2020 | Europe / United Kingdom / Wales | 2020-03-23 | Wales Specialist Virology Centre | Public Health Wales Microbiology Cardiff | Catherine Moore, Johnathan Evans, Malorie Perry, Simon Cottrell, Alec Birchley, Alexander Adams, Amy Gaskin, Bree Gatica-Wilcox, Jason Coombes, Lauren Gilbert, Lee Graham, Nicole Pacchiarini, Sara Kumziene-Summerhayes, Sarah Taylor, Sophie Jones, Sara Rey, Matthew Bull, Joanne Watkins, Sally Corden, Tom Connor |
| EPI_ISL_432189 | hCoV-19/Wales/PHWC-29C76/2020 | Europe / United Kingdom / Wales | 2020-04-05 | Wales Specialist Virology Centre | Public Health Wales Microbiology Cardiff | Catherine Moore, Johnathan Evans, Malorie Perry, Simon Cottrell, Alec Birchley, Alexander Adams, Amy Gaskin, Bree Gatica-Wilcox, Jason Coombes, Lauren Gilbert, Lee Graham, Nicole Pacchiarini, Sara Kumziene-Summerhayes, Sarah Taylor, Sophie Jones, Sara Rey, Matthew Bull, Joanne Watkins, Sally Corden, Tom Connor |
| EPI_ISL_432190 | hCoV-19/Wales/PHWC-25393/2020 | Europe / United Kingdom / Wales | 2020-03-23 | Wales Specialist Virology Centre | Public Health Wales Microbiology Cardiff | Catherine Moore, Johnathan Evans, Malorie Perry, Simon Cottrell, Alec Birchley, Alexander Adams, Amy Gaskin, Bree Gatica-Wilcox, Jason Coombes, Lauren Gilbert, Lee Graham, Nicole Pacchiarini, Sara Kumziene-Summerhayes, Sarah Taylor, Sophie Jones, Sara Rey, Matthew Bull, Joanne Watkins, Sally Corden, Tom Connor |
| EPI_ISL_432191 | hCoV-19/Wales/PHWC-2B2BE/2020 | Europe / United Kingdom / Wales | 2020-04-08 | Wales Specialist Virology Centre | Public Health Wales Microbiology Cardiff | Catherine Moore, Johnathan Evans, Malorie Perry, Simon Cottrell, Alec Birchley, Alexander Adams, Amy Gaskin, Bree Gatica-Wilcox, Jason Coombes, Lauren Gilbert, Lee Graham, Nicole Pacchiarini, Sara Kumziene-Summerhayes, Sarah Taylor, Sophie Jones, Sara Rey, Matthew Bull, Joanne Watkins, Sally Corden, Tom Connor |
| EPI_ISL_432192 | hCoV-19/Wales/PHWC-27CD2/2020 | Europe / United Kingdom / Wales | 2020-04-01 | Wales Specialist Virology Centre | Public Health Wales Microbiology Cardiff | Catherine Moore, Johnathan Evans, Malorie Perry, Simon Cottrell, Alec Birchley, Alexander Adams, Amy Gaskin, Bree Gatica-Wilcox, Jason Coombes, Lauren Gilbert, Lee Graham, Nicole Pacchiarini, Sara Kumziene-Summerhayes, Sarah Taylor, Sophie Jones, Sara Rey, Matthew Bull, Joanne Watkins, Sally Corden, Tom Connor |
| EPI_ISL_432193 | hCoV-19/Wales/PHWC-29BC4/2020 | Europe / United Kingdom / Wales | 2020-04-04 | Wales Specialist Virology Centre | Public Health Wales Microbiology Cardiff | Catherine Moore, Johnathan Evans, Malorie Perry, Simon Cottrell, Alec Birchley, Alexander Adams, Amy Gaskin, Bree Gatica-Wilcox, Jason Coombes, Lauren Gilbert, Lee Graham, Nicole Pacchiarini, Sara Kumziene-Summerhayes, Sarah Taylor, Sophie Jones, Sara Rey, Matthew Bull, Joanne Watkins, Sally Corden, Tom Connor |
| EPI_ISL_432194 | hCoV-19/Wales/PHWC-29E43/2020 | Europe / United Kingdom / Wales | 2020-04-04 | Wales Specialist Virology Centre | Public Health Wales Microbiology Cardiff | Catherine Moore, Johnathan Evans, Malorie Perry, Simon Cottrell, Alec Birchley, Alexander Adams, Amy Gaskin, Bree Gatica-Wilcox, Jason Coombes, Lauren Gilbert, Lee Graham, Nicole Pacchiarini, Sara Kumziene-Summerhayes, Sarah Taylor, Sophie Jones, Sara Rey, Matthew Bull, Joanne Watkins, Sally Corden, Tom Connor |
| EPI_ISL_432195 | hCoV-19/Wales/PHWC-29C3A/2020 | Europe / United Kingdom / Wales | 2020-04-04 | Wales Specialist Virology Centre | Public Health Wales Microbiology Cardiff | Catherine Moore, Johnathan Evans, Malorie Perry, Simon Cottrell, Alec Birchley, Alexander Adams, Amy Gaskin, Bree Gatica-Wilcox, Jason Coombes, Lauren Gilbert, Lee Graham, Nicole Pacchiarini, Sara Kumziene-Summerhayes, Sarah Taylor, Sophie Jones, Sara Rey, Matthew Bull, Joanne Watkins, Sally Corden, Tom Connor |
| EPI_ISL_432196 | hCoV-19/Wales/PHWC-29C49/2020 | Europe / United Kingdom / Wales | 2020-04-05 | Wales Specialist Virology Centre | Public Health Wales Microbiology Cardiff | Catherine Moore, Johnathan Evans, Malorie Perry, Simon Cottrell, Alec Birchley, Alexander Adams, Amy Gaskin, Bree Gatica-Wilcox, Jason Coombes, Lauren Gilbert, Lee Graham, Nicole Pacchiarini, Sara Kumziene-Summerhayes, Sarah Taylor, Sophie Jones, Sara Rey, Matthew Bull, Joanne Watkins, Sally Corden, Tom Connor |
| EPI_ISL_432197 | hCoV-19/Wales/PHWC-29DDD/2020 | Europe / United Kingdom / Wales | 2020-04-04 | Wales Specialist Virology Centre | Public Health Wales Microbiology Cardiff | Catherine Moore, Johnathan Evans, Malorie Perry, Simon Cottrell, Alec Birchley, Alexander Adams, Amy Gaskin, Bree Gatica-Wilcox, Jason Coombes, Lauren Gilbert, Lee Graham, Nicole Pacchiarini, Sara Kumziene-Summerhayes, Sarah Taylor, Sophie Jones, Sara Rey, Matthew Bull, Joanne Watkins, Sally Corden, Tom Connor |
| EPI_ISL_432198 | hCoV-19/Wales/PHWC-27ABA/2020 | Europe / United Kingdom / Wales | 2020-03-30 | Wales Specialist Virology Centre | Public Health Wales Microbiology Cardiff | Catherine Moore, Johnathan Evans, Malorie Perry, Simon Cottrell, Alec Birchley, Alexander Adams, Amy Gaskin, Bree Gatica-Wilcox, Jason Coombes, Lauren Gilbert, Lee Graham, Nicole Pacchiarini, Sara Kumziene-Summerhayes, Sarah Taylor, Sophie Jones, Sara Rey, Matthew Bull, Joanne Watkins, Sally Corden, Tom Connor |
| EPI_ISL_432199 | hCoV-19/Wales/PHWC-29F13/2020 | Europe / United Kingdom / Wales | 2020-04-01 | Wales Specialist Virology Centre | Public Health Wales Microbiology Cardiff | Catherine Moore, Johnathan Evans, Malorie Perry, Simon Cottrell, Alec Birchley, Alexander Adams, Amy Gaskin, Bree Gatica-Wilcox, Jason Coombes, Lauren Gilbert, Lee Graham, Nicole Pacchiarini, Sara Kumziene-Summerhayes, Sarah Taylor, Sophie Jones, Sara Rey, Matthew Bull, Joanne Watkins, Sally Corden, Tom Connor |
| EPI_ISL_432200 | hCoV-19/Wales/PHWC-276C5/2020 | Europe / United Kingdom / Wales | 2020-03-31 | Wales Specialist Virology Centre | Public Health Wales Microbiology Cardiff | Catherine Moore, Johnathan Evans, Malorie Perry, Simon Cottrell, Alec Birchley, Alexander Adams, Amy Gaskin, Bree Gatica-Wilcox, Jason Coombes, Lauren Gilbert, Lee Graham, Nicole Pacchiarini, Sara Kumziene-Summerhayes, Sarah Taylor, Sophie Jones, Sara Rey, Matthew Bull, Joanne Watkins, Sally Corden, Tom Connor |
| EPI_ISL_432201 | hCoV-19/Wales/PHWC-2A0E2/2020 | Europe / United Kingdom / Wales | 2020-04-04 | Wales Specialist Virology Centre | Public Health Wales Microbiology Cardiff | Catherine Moore, Johnathan Evans, Malorie Perry, Simon Cottrell, Alec Birchley, Alexander Adams, Amy Gaskin, Bree Gatica-Wilcox, Jason Coombes, Lauren Gilbert, Lee Graham, Nicole Pacchiarini, Sara Kumziene-Summerhayes, Sarah Taylor, Sophie Jones, Sara Rey, Matthew Bull, Joanne Watkins, Sally Corden, Tom Connor |
| EPI_ISL_432202 | hCoV-19/Wales/PHWC-29E8F/2020 | Europe / United Kingdom / Wales | 2020-04-05 | Wales Specialist Virology Centre | Public Health Wales Microbiology Cardiff | Catherine Moore, Johnathan Evans, Malorie Perry, Simon Cottrell, Alec Birchley, Alexander Adams, Amy Gaskin, Bree Gatica-Wilcox, Jason Coombes, Lauren Gilbert, Lee Graham, Nicole Pacchiarini, Sara Kumziene-Summerhayes, Sarah Taylor, Sophie Jones, Sara Rey, Matthew Bull, Joanne Watkins, Sally Corden, Tom Connor |
| EPI_ISL_432203 | hCoV-19/Wales/PHWC-29E52/2020 | Europe / United Kingdom / Wales | 2020-04-04 | Wales Specialist Virology Centre | Public Health Wales Microbiology Cardiff | Catherine Moore, Johnathan Evans, Malorie Perry, Simon Cottrell, Alec Birchley, Alexander Adams, Amy Gaskin, Bree Gatica-Wilcox, Jason Coombes, Lauren Gilbert, Lee Graham, Nicole Pacchiarini, Sara Kumziene-Summerhayes, Sarah Taylor, Sophie Jones, Sara Rey, Matthew Bull, Joanne Watkins, Sally Corden, Tom Connor |
| EPI_ISL_432204 | hCoV-19/Wales/PHWC-2A10D/2020 | Europe / United Kingdom / Wales | 2020-04-04 | Wales Specialist Virology Centre | Public Health Wales Microbiology Cardiff | Catherine Moore, Johnathan Evans, Malorie Perry, Simon Cottrell, Alec Birchley, Alexander Adams, Amy Gaskin, Bree Gatica-Wilcox, Jason Coombes, Lauren Gilbert, Lee Graham, Nicole Pacchiarini, Sara Kumziene-Summerhayes, Sarah Taylor, Sophie Jones, Sara Rey, Matthew Bull, Joanne Watkins, Sally Corden, Tom Connor |
| EPI_ISL_432205 | hCoV-19/Wales/PHWC-29E34/2020 | Europe / United Kingdom / Wales | 2020-04-04 | Wales Specialist Virology Centre | Public Health Wales Microbiology Cardiff | Catherine Moore, Johnathan Evans, Malorie Perry, Simon Cottrell, Alec Birchley, Alexander Adams, Amy Gaskin, Bree Gatica-Wilcox, Jason Coombes, Lauren Gilbert, Lee Graham, Nicole Pacchiarini, Sara Kumziene-Summerhayes, Sarah Taylor, Sophie Jones, Sara Rey, Matthew Bull, Joanne Watkins, Sally Corden, Tom Connor |
| EPI_ISL_432206 | hCoV-19/Wales/PHWC-27698/2020 | Europe / United Kingdom / Wales | 2020-03-31 | Wales Specialist Virology Centre | Public Health Wales Microbiology Cardiff | Catherine Moore, Johnathan Evans, Malorie Perry, Simon Cottrell, Alec Birchley, Alexander Adams, Amy Gaskin, Bree Gatica-Wilcox, Jason Coombes, Lauren Gilbert, Lee Graham, Nicole Pacchiarini, Sara Kumziene-Summerhayes, Sarah Taylor, Sophie Jones, Sara Rey, Matthew Bull, Joanne Watkins, Sally Corden, Tom Connor |
| EPI_ISL_432207 | hCoV-19/Wales/PHWC-2530C/2020 | Europe / United Kingdom / Wales | 2020-03-23 | Wales Specialist Virology Centre | Public Health Wales Microbiology Cardiff | Catherine Moore, Johnathan Evans, Malorie Perry, Simon Cottrell, Alec Birchley, Alexander Adams, Amy Gaskin, Bree Gatica-Wilcox, Jason Coombes, Lauren Gilbert, Lee Graham, Nicole Pacchiarini, Sara Kumziene-Summerhayes, Sarah Taylor, Sophie Jones, Sara Rey, Matthew Bull, Joanne Watkins, Sally Corden, Tom Connor |
| EPI_ISL_432208 | hCoV-19/Wales/PHWC-278A1/2020 | Europe / United Kingdom / Wales | 2020-03-31 | Wales Specialist Virology Centre | Public Health Wales Microbiology Cardiff | Catherine Moore, Johnathan Evans, Malorie Perry, Simon Cottrell, Alec Birchley, Alexander Adams, Amy Gaskin, Bree Gatica-Wilcox, Jason Coombes, Lauren Gilbert, Lee Graham, Nicole Pacchiarini, Sara Kumziene-Summerhayes, Sarah Taylor, Sophie Jones, Sara Rey, Matthew Bull, Joanne Watkins, Sally Corden, Tom Connor |
| EPI_ISL_432209 | hCoV-19/Wales/PHWC-27A41/2020 | Europe / United Kingdom / Wales | 2020-04-02 | Wales Specialist Virology Centre | Public Health Wales Microbiology Cardiff | Catherine Moore, Johnathan Evans, Malorie Perry, Simon Cottrell, Alec Birchley, Alexander Adams, Amy Gaskin, Bree Gatica-Wilcox, Jason Coombes, Lauren Gilbert, Lee Graham, Nicole Pacchiarini, Sara Kumziene-Summerhayes, Sarah Taylor, Sophie Jones, Sara Rey, Matthew Bull, Joanne Watkins, Sally Corden, Tom Connor |
| EPI_ISL_432210 | hCoV-19/Wales/PHWC-277B3/2020 | Europe / United Kingdom / Wales | 2020-04-01 | Wales Specialist Virology Centre | Public Health Wales Microbiology Cardiff | Catherine Moore, Johnathan Evans, Malorie Perry, Simon Cottrell, Alec Birchley, Alexander Adams, Amy Gaskin, Bree Gatica-Wilcox, Jason Coombes, Lauren Gilbert, Lee Graham, Nicole Pacchiarini, Sara Kumziene-Summerhayes, Sarah Taylor, Sophie Jones, Sara Rey, Matthew Bull, Joanne Watkins, Sally Corden, Tom Connor |
| EPI_ISL_432211 | hCoV-19/Wales/PHWC-2764D/2020 | Europe / United Kingdom / Wales | 2020-03-30 | Wales Specialist Virology Centre | Public Health Wales Microbiology Cardiff | Catherine Moore, Johnathan Evans, Malorie Perry, Simon Cottrell, Alec Birchley, Alexander Adams, Amy Gaskin, Bree Gatica-Wilcox, Jason Coombes, Lauren Gilbert, Lee Graham, Nicole Pacchiarini, Sara Kumziene-Summerhayes, Sarah Taylor, Sophie Jones, Sara Rey, Matthew Bull, Joanne Watkins, Sally Corden, Tom Connor |
| EPI_ISL_432212 | hCoV-19/Wales/PHWC-27944/2020 | Europe / United Kingdom / Wales | 2020-04-01 | Wales Specialist Virology Centre | Public Health Wales Microbiology Cardiff | Catherine Moore, Johnathan Evans, Malorie Perry, Simon Cottrell, Alec Birchley, Alexander Adams, Amy Gaskin, Bree Gatica-Wilcox, Jason Coombes, Lauren Gilbert, Lee Graham, Nicole Pacchiarini, Sara Kumziene-Summerhayes, Sarah Taylor, Sophie Jones, Sara Rey, Matthew Bull, Joanne Watkins, Sally Corden, Tom Connor |
| EPI_ISL_432213 | hCoV-19/Wales/PHWC-2AF5E/2020 | Europe / United Kingdom / Wales | 2020-04-07 | Wales Specialist Virology Centre | Public Health Wales Microbiology Cardiff | Catherine Moore, Johnathan Evans, Malorie Perry, Simon Cottrell, Alec Birchley, Alexander Adams, Amy Gaskin, Bree Gatica-Wilcox, Jason Coombes, Lauren Gilbert, Lee Graham, Nicole Pacchiarini, Sara Kumziene-Summerhayes, Sarah Taylor, Sophie Jones, Sara Rey, Matthew Bull, Joanne Watkins, Sally Corden, Tom Connor |
| EPI_ISL_432214 | hCoV-19/Wales/PHWC-273CE/2020 | Europe / United Kingdom / Wales | 2020-03-31 | Wales Specialist Virology Centre | Public Health Wales Microbiology Cardiff | Catherine Moore, Johnathan Evans, Malorie Perry, Simon Cottrell, Alec Birchley, Alexander Adams, Amy Gaskin, Bree Gatica-Wilcox, Jason Coombes, Lauren Gilbert, Lee Graham, Nicole Pacchiarini, Sara Kumziene-Summerhayes, Sarah Taylor, Sophie Jones, Sara Rey, Matthew Bull, Joanne Watkins, Sally Corden, Tom Connor |
| EPI_ISL_432215 | hCoV-19/Wales/PHWC-29DCE/2020 | Europe / United Kingdom / Wales | 2020-04-04 | Wales Specialist Virology Centre | Public Health Wales Microbiology Cardiff | Catherine Moore, Johnathan Evans, Malorie Perry, Simon Cottrell, Alec Birchley, Alexander Adams, Amy Gaskin, Bree Gatica-Wilcox, Jason Coombes, Lauren Gilbert, Lee Graham, Nicole Pacchiarini, Sara Kumziene-Summerhayes, Sarah Taylor, Sophie Jones, Sara Rey, Matthew Bull, Joanne Watkins, Sally Corden, Tom Connor |
| EPI_ISL_432216 | hCoV-19/Wales/PHWC-2AF03/2020 | Europe / United Kingdom / Wales | 2020-04-06 | Wales Specialist Virology Centre | Public Health Wales Microbiology Cardiff | Catherine Moore, Johnathan Evans, Malorie Perry, Simon Cottrell, Alec Birchley, Alexander Adams, Amy Gaskin, Bree Gatica-Wilcox, Jason Coombes, Lauren Gilbert, Lee Graham, Nicole Pacchiarini, Sara Kumziene-Summerhayes, Sarah Taylor, Sophie Jones, Sara Rey, Matthew Bull, Joanne Watkins, Sally Corden, Tom Connor |
| EPI_ISL_432217 | hCoV-19/Wales/PHWC-27A32/2020 | Europe / United Kingdom / Wales | 2020-04-01 | Wales Specialist Virology Centre | Public Health Wales Microbiology Cardiff | Catherine Moore, Johnathan Evans, Malorie Perry, Simon Cottrell, Alec Birchley, Alexander Adams, Amy Gaskin, Bree Gatica-Wilcox, Jason Coombes, Lauren Gilbert, Lee Graham, Nicole Pacchiarini, Sara Kumziene-Summerhayes, Sarah Taylor, Sophie Jones, Sara Rey, Matthew Bull, Joanne Watkins, Sally Corden, Tom Connor |
| EPI_ISL_432218 | hCoV-19/Wales/PHWC-29BB5/2020 | Europe / United Kingdom / Wales | 2020-03-31 | Wales Specialist Virology Centre | Public Health Wales Microbiology Cardiff | Catherine Moore, Johnathan Evans, Malorie Perry, Simon Cottrell, Alec Birchley, Alexander Adams, Amy Gaskin, Bree Gatica-Wilcox, Jason Coombes, Lauren Gilbert, Lee Graham, Nicole Pacchiarini, Sara Kumziene-Summerhayes, Sarah Taylor, Sophie Jones, Sara Rey, Matthew Bull, Joanne Watkins, Sally Corden, Tom Connor |
| EPI_ISL_432219 | hCoV-19/Wales/PHWC-29C0D/2020 | Europe / United Kingdom / Wales | 2020-04-04 | Wales Specialist Virology Centre | Public Health Wales Microbiology Cardiff | Catherine Moore, Johnathan Evans, Malorie Perry, Simon Cottrell, Alec Birchley, Alexander Adams, Amy Gaskin, Bree Gatica-Wilcox, Jason Coombes, Lauren Gilbert, Lee Graham, Nicole Pacchiarini, Sara Kumziene-Summerhayes, Sarah Taylor, Sophie Jones, Sara Rey, Matthew Bull, Joanne Watkins, Sally Corden, Tom Connor |
| EPI_ISL_432220 | hCoV-19/Wales/PHWC-25375/2020 | Europe / United Kingdom / Wales | 2020-03-23 | Wales Specialist Virology Centre | Public Health Wales Microbiology Cardiff | Catherine Moore, Johnathan Evans, Malorie Perry, Simon Cottrell, Alec Birchley, Alexander Adams, Amy Gaskin, Bree Gatica-Wilcox, Jason Coombes, Lauren Gilbert, Lee Graham, Nicole Pacchiarini, Sara Kumziene-Summerhayes, Sarah Taylor, Sophie Jones, Sara Rey, Matthew Bull, Joanne Watkins, Sally Corden, Tom Connor |
| EPI_ISL_432221 | hCoV-19/Wales/PHWC-27917/2020 | Europe / United Kingdom / Wales | 2020-04-01 | Wales Specialist Virology Centre | Public Health Wales Microbiology Cardiff | Catherine Moore, Johnathan Evans, Malorie Perry, Simon Cottrell, Alec Birchley, Alexander Adams, Amy Gaskin, Bree Gatica-Wilcox, Jason Coombes, Lauren Gilbert, Lee Graham, Nicole Pacchiarini, Sara Kumziene-Summerhayes, Sarah Taylor, Sophie Jones, Sara Rey, Matthew Bull, Joanne Watkins, Sally Corden, Tom Connor |
| EPI_ISL_432222 | hCoV-19/Wales/PHWC-27B3F/2020 | Europe / United Kingdom / Wales | 2020-03-30 | Wales Specialist Virology Centre | Public Health Wales Microbiology Cardiff | Catherine Moore, Johnathan Evans, Malorie Perry, Simon Cottrell, Alec Birchley, Alexander Adams, Amy Gaskin, Bree Gatica-Wilcox, Jason Coombes, Lauren Gilbert, Lee Graham, Nicole Pacchiarini, Sara Kumziene-Summerhayes, Sarah Taylor, Sophie Jones, Sara Rey, Matthew Bull, Joanne Watkins, Sally Corden, Tom Connor |
| EPI_ISL_432223 | hCoV-19/Wales/PHWC-27443/2020 | Europe / United Kingdom / Wales | 2020-03-31 | Wales Specialist Virology Centre | Public Health Wales Microbiology Cardiff | Catherine Moore, Johnathan Evans, Malorie Perry, Simon Cottrell, Alec Birchley, Alexander Adams, Amy Gaskin, Bree Gatica-Wilcox, Jason Coombes, Lauren Gilbert, Lee Graham, Nicole Pacchiarini, Sara Kumziene-Summerhayes, Sarah Taylor, Sophie Jones, Sara Rey, Matthew Bull, Joanne Watkins, Sally Corden, Tom Connor |
| EPI_ISL_432224 | hCoV-19/Wales/PHWC-27786/2020 | Europe / United Kingdom / Wales | 2020-03-31 | Wales Specialist Virology Centre | Public Health Wales Microbiology Cardiff | Catherine Moore, Johnathan Evans, Malorie Perry, Simon Cottrell, Alec Birchley, Alexander Adams, Amy Gaskin, Bree Gatica-Wilcox, Jason Coombes, Lauren Gilbert, Lee Graham, Nicole Pacchiarini, Sara Kumziene-Summerhayes, Sarah Taylor, Sophie Jones, Sara Rey, Matthew Bull, Joanne Watkins, Sally Corden, Tom Connor |
| EPI_ISL_432225 | hCoV-19/Wales/PHWC-2B254/2020 | Europe / United Kingdom / Wales | 2020-04-08 | Wales Specialist Virology Centre | Public Health Wales Microbiology Cardiff | Catherine Moore, Johnathan Evans, Malorie Perry, Simon Cottrell, Alec Birchley, Alexander Adams, Amy Gaskin, Bree Gatica-Wilcox, Jason Coombes, Lauren Gilbert, Lee Graham, Nicole Pacchiarini, Sara Kumziene-Summerhayes, Sarah Taylor, Sophie Jones, Sara Rey, Matthew Bull, Joanne Watkins, Sally Corden, Tom Connor |
| EPI_ISL_432226 | hCoV-19/Wales/PHWC-29DEC/2020 | Europe / United Kingdom / Wales | 2020-04-04 | Wales Specialist Virology Centre | Public Health Wales Microbiology Cardiff | Catherine Moore, Johnathan Evans, Malorie Perry, Simon Cottrell, Alec Birchley, Alexander Adams, Amy Gaskin, Bree Gatica-Wilcox, Jason Coombes, Lauren Gilbert, Lee Graham, Nicole Pacchiarini, Sara Kumziene-Summerhayes, Sarah Taylor, Sophie Jones, Sara Rey, Matthew Bull, Joanne Watkins, Sally Corden, Tom Connor |
| EPI_ISL_432227 | hCoV-19/Wales/PHWC-27AAB/2020 | Europe / United Kingdom / Wales | 2020-03-30 | Wales Specialist Virology Centre | Public Health Wales Microbiology Cardiff | Catherine Moore, Johnathan Evans, Malorie Perry, Simon Cottrell, Alec Birchley, Alexander Adams, Amy Gaskin, Bree Gatica-Wilcox, Jason Coombes, Lauren Gilbert, Lee Graham, Nicole Pacchiarini, Sara Kumziene-Summerhayes, Sarah Taylor, Sophie Jones, Sara Rey, Matthew Bull, Joanne Watkins, Sally Corden, Tom Connor |
| EPI_ISL_432228 | hCoV-19/Wales/PHWC-2B281/2020 | Europe / United Kingdom / Wales | 2020-04-08 | Wales Specialist Virology Centre | Public Health Wales Microbiology Cardiff | Catherine Moore, Johnathan Evans, Malorie Perry, Simon Cottrell, Alec Birchley, Alexander Adams, Amy Gaskin, Bree Gatica-Wilcox, Jason Coombes, Lauren Gilbert, Lee Graham, Nicole Pacchiarini, Sara Kumziene-Summerhayes, Sarah Taylor, Sophie Jones, Sara Rey, Matthew Bull, Joanne Watkins, Sally Corden, Tom Connor |
| EPI_ISL_432229 | hCoV-19/Wales/PHWC-29D91/2020 | Europe / United Kingdom / Wales | 2020-04-04 | Wales Specialist Virology Centre | Public Health Wales Microbiology Cardiff | Catherine Moore, Johnathan Evans, Malorie Perry, Simon Cottrell, Alec Birchley, Alexander Adams, Amy Gaskin, Bree Gatica-Wilcox, Jason Coombes, Lauren Gilbert, Lee Graham, Nicole Pacchiarini, Sara Kumziene-Summerhayes, Sarah Taylor, Sophie Jones, Sara Rey, Matthew Bull, Joanne Watkins, Sally Corden, Tom Connor |
| EPI_ISL_432230 | hCoV-19/Wales/PHWC-2A0B5/2020 | Europe / United Kingdom / Wales | 2020-04-04 | Wales Specialist Virology Centre | Public Health Wales Microbiology Cardiff | Catherine Moore, Johnathan Evans, Malorie Perry, Simon Cottrell, Alec Birchley, Alexander Adams, Amy Gaskin, Bree Gatica-Wilcox, Jason Coombes, Lauren Gilbert, Lee Graham, Nicole Pacchiarini, Sara Kumziene-Summerhayes, Sarah Taylor, Sophie Jones, Sara Rey, Matthew Bull, Joanne Watkins, Sally Corden, Tom Connor |
| EPI_ISL_432231 | hCoV-19/Wales/PHWC-2532A/2020 | Europe / United Kingdom / Wales | 2020-03-22 | Wales Specialist Virology Centre | Public Health Wales Microbiology Cardiff | Catherine Moore, Johnathan Evans, Malorie Perry, Simon Cottrell, Alec Birchley, Alexander Adams, Amy Gaskin, Bree Gatica-Wilcox, Jason Coombes, Lauren Gilbert, Lee Graham, Nicole Pacchiarini, Sara Kumziene-Summerhayes, Sarah Taylor, Sophie Jones, Sara Rey, Matthew Bull, Joanne Watkins, Sally Corden, Tom Connor |
| EPI_ISL_432232 | hCoV-19/Wales/PHWC-27391/2020 | Europe / United Kingdom / Wales | 2020-03-31 | Wales Specialist Virology Centre | Public Health Wales Microbiology Cardiff | Catherine Moore, Johnathan Evans, Malorie Perry, Simon Cottrell, Alec Birchley, Alexander Adams, Amy Gaskin, Bree Gatica-Wilcox, Jason Coombes, Lauren Gilbert, Lee Graham, Nicole Pacchiarini, Sara Kumziene-Summerhayes, Sarah Taylor, Sophie Jones, Sara Rey, Matthew Bull, Joanne Watkins, Sally Corden, Tom Connor |
| EPI_ISL_432233 | hCoV-19/Wales/PHWC-275D7/2020 | Europe / United Kingdom / Wales | 2020-04-01 | Wales Specialist Virology Centre | Public Health Wales Microbiology Cardiff | Catherine Moore, Johnathan Evans, Malorie Perry, Simon Cottrell, Alec Birchley, Alexander Adams, Amy Gaskin, Bree Gatica-Wilcox, Jason Coombes, Lauren Gilbert, Lee Graham, Nicole Pacchiarini, Sara Kumziene-Summerhayes, Sarah Taylor, Sophie Jones, Sara Rey, Matthew Bull, Joanne Watkins, Sally Corden, Tom Connor |
| EPI_ISL_432234 | hCoV-19/Wales/PHWC-2A097/2020 | Europe / United Kingdom / Wales | 2020-04-04 | Wales Specialist Virology Centre | Public Health Wales Microbiology Cardiff | Catherine Moore, Johnathan Evans, Malorie Perry, Simon Cottrell, Alec Birchley, Alexander Adams, Amy Gaskin, Bree Gatica-Wilcox, Jason Coombes, Lauren Gilbert, Lee Graham, Nicole Pacchiarini, Sara Kumziene-Summerhayes, Sarah Taylor, Sophie Jones, Sara Rey, Matthew Bull, Joanne Watkins, Sally Corden, Tom Connor |
| EPI_ISL_432235 | hCoV-19/Wales/PHWC-272FE/2020 | Europe / United Kingdom / Wales | 2020-03-30 | Wales Specialist Virology Centre | Public Health Wales Microbiology Cardiff | Catherine Moore, Johnathan Evans, Malorie Perry, Simon Cottrell, Alec Birchley, Alexander Adams, Amy Gaskin, Bree Gatica-Wilcox, Jason Coombes, Lauren Gilbert, Lee Graham, Nicole Pacchiarini, Sara Kumziene-Summerhayes, Sarah Taylor, Sophie Jones, Sara Rey, Matthew Bull, Joanne Watkins, Sally Corden, Tom Connor |
| EPI_ISL_433370 | hCoV-19/Scotland/CVR977/2020 | Europe / United Kingdom / Scotland | 2020-03-27 | West of Scotland Specialist Virology Centre, NHSGGC / MRC-University of Glasgow Centre for Virus Research | COVID-19 Genomics UK (COG-UK) Consortium | Ana da Silva Filipe, Natasha Johnson, Kathy Smollett, Daniel Mair, Stephen Carmichael, Lily Tong, Jenna Nichols, Elihu Aranday-Cortes, Kirstyn Brunker, Yasmin Parr, Kyriaki Nomikou; Sarah McDonald, Marc Niebel, Patawee Asamaphan; Richard Orton, Joseph Hughes, Sreenu Vattipally, David L Robertson; Alasdair MacLean, Rory Gunson; Kathy Li, Natasha Jesudason, Rajiv Shah, James Shepherd, Antonia Ho, Emma Thomson |
| EPI_ISL_433371 | hCoV-19/Scotland/CVR987/2020 | Europe / United Kingdom / Scotland | 2020-03-29 | West of Scotland Specialist Virology Centre, NHSGGC / MRC-University of Glasgow Centre for Virus Research | COVID-19 Genomics UK (COG-UK) Consortium | Ana da Silva Filipe, Natasha Johnson, Kathy Smollett, Daniel Mair, Stephen Carmichael, Lily Tong, Jenna Nichols, Elihu Aranday-Cortes, Kirstyn Brunker, Yasmin Parr, Kyriaki Nomikou; Sarah McDonald, Marc Niebel, Patawee Asamaphan; Richard Orton, Joseph Hughes, Sreenu Vattipally, David L Robertson; Alasdair MacLean, Rory Gunson; Kathy Li, Natasha Jesudason, Rajiv Shah, James Shepherd, Antonia Ho, Emma Thomson |
| EPI_ISL_433372 | hCoV-19/Scotland/CVR997/2020 | Europe / United Kingdom / Scotland | 2020-03-28 | West of Scotland Specialist Virology Centre, NHSGGC / MRC-University of Glasgow Centre for Virus Research | COVID-19 Genomics UK (COG-UK) Consortium | Ana da Silva Filipe, Natasha Johnson, Kathy Smollett, Daniel Mair, Stephen Carmichael, Lily Tong, Jenna Nichols, Elihu Aranday-Cortes, Kirstyn Brunker, Yasmin Parr, Kyriaki Nomikou; Sarah McDonald, Marc Niebel, Patawee Asamaphan; Richard Orton, Joseph Hughes, Sreenu Vattipally, David L Robertson; Alasdair MacLean, Rory Gunson; Kathy Li, Natasha Jesudason, Rajiv Shah, James Shepherd, Antonia Ho, Emma Thomson |
| EPI_ISL_433374 | hCoV-19/Scotland/EDB1001/2020 | Europe / United Kingdom / Scotland | 2020-04-06 | Virology Department, Royal Infirmary of Edinburgh, NHS Lothian / School of Biological Sciences, University of Edinburgh / Institute of Genetics and Molecular Medicine, University of Edinburgh | COVID-19 Genomics UK (COG-UK) Consortium | McHugh M, Dewar R, Rooke S, Gallagher M, Balcaza C, O'Toole A, Hill V, McCrone JT, Colquhoun R, Yu X, Jackson B, Rambaut A, Williams TC, Templeton K |
| EPI_ISL_433375 | hCoV-19/Scotland/EDB1005/2020 | Europe / United Kingdom / Scotland | 2020-04-06 | Virology Department, Royal Infirmary of Edinburgh, NHS Lothian / School of Biological Sciences, University of Edinburgh / Institute of Genetics and Molecular Medicine, University of Edinburgh | COVID-19 Genomics UK (COG-UK) Consortium | McHugh M, Dewar R, Rooke S, Gallagher M, Balcaza C, O'Toole A, Hill V, McCrone JT, Colquhoun R, Yu X, Jackson B, Rambaut A, Williams TC, Templeton K |
| EPI_ISL_433376 | hCoV-19/Scotland/EDB1037/2020 | Europe / United Kingdom / Scotland | 2020-04-06 | Virology Department, Royal Infirmary of Edinburgh, NHS Lothian / School of Biological Sciences, University of Edinburgh / Institute of Genetics and Molecular Medicine, University of Edinburgh | COVID-19 Genomics UK (COG-UK) Consortium | McHugh M, Dewar R, Rooke S, Gallagher M, Balcaza C, O'Toole A, Hill V, McCrone JT, Colquhoun R, Yu X, Jackson B, Rambaut A, Williams TC, Templeton K |
| EPI_ISL_433377 | hCoV-19/Scotland/EDB1040/2020 | Europe / United Kingdom / Scotland | 2020-04-06 | Virology Department, Royal Infirmary of Edinburgh, NHS Lothian / School of Biological Sciences, University of Edinburgh / Institute of Genetics and Molecular Medicine, University of Edinburgh | COVID-19 Genomics UK (COG-UK) Consortium | McHugh M, Dewar R, Rooke S, Gallagher M, Balcaza C, O'Toole A, Hill V, McCrone JT, Colquhoun R, Yu X, Jackson B, Rambaut A, Williams TC, Templeton K |
| EPI_ISL_433378 | hCoV-19/Scotland/EDB1046/2020 | Europe / United Kingdom / Scotland | 2020-04-06 | Virology Department, Royal Infirmary of Edinburgh, NHS Lothian / School of Biological Sciences, University of Edinburgh / Institute of Genetics and Molecular Medicine, University of Edinburgh | COVID-19 Genomics UK (COG-UK) Consortium | McHugh M, Dewar R, Rooke S, Gallagher M, Balcaza C, O'Toole A, Hill V, McCrone JT, Colquhoun R, Yu X, Jackson B, Rambaut A, Williams TC, Templeton K |
| EPI_ISL_433379 | hCoV-19/Scotland/EDB1062/2020 | Europe / United Kingdom / Scotland | 2020-04-06 | Virology Department, Royal Infirmary of Edinburgh, NHS Lothian / School of Biological Sciences, University of Edinburgh / Institute of Genetics and Molecular Medicine, University of Edinburgh | COVID-19 Genomics UK (COG-UK) Consortium | McHugh M, Dewar R, Rooke S, Gallagher M, Balcaza C, O'Toole A, Hill V, McCrone JT, Colquhoun R, Yu X, Jackson B, Rambaut A, Williams TC, Templeton K |
| EPI_ISL_433380 | hCoV-19/Scotland/EDB1070/2020 | Europe / United Kingdom / Scotland | 2020-04-06 | Virology Department, Royal Infirmary of Edinburgh, NHS Lothian / School of Biological Sciences, University of Edinburgh / Institute of Genetics and Molecular Medicine, University of Edinburgh | COVID-19 Genomics UK (COG-UK) Consortium | McHugh M, Dewar R, Rooke S, Gallagher M, Balcaza C, O'Toole A, Hill V, McCrone JT, Colquhoun R, Yu X, Jackson B, Rambaut A, Williams TC, Templeton K |
| EPI_ISL_433381 | hCoV-19/Scotland/EDB1071/2020 | Europe / United Kingdom / Scotland | 2020-04-06 | Virology Department, Royal Infirmary of Edinburgh, NHS Lothian / School of Biological Sciences, University of Edinburgh / Institute of Genetics and Molecular Medicine, University of Edinburgh | COVID-19 Genomics UK (COG-UK) Consortium | McHugh M, Dewar R, Rooke S, Gallagher M, Balcaza C, O'Toole A, Hill V, McCrone JT, Colquhoun R, Yu X, Jackson B, Rambaut A, Williams TC, Templeton K |
| EPI_ISL_433382 | hCoV-19/Scotland/EDB1072/2020 | Europe / United Kingdom / Scotland | 2020-04-06 | Virology Department, Royal Infirmary of Edinburgh, NHS Lothian / School of Biological Sciences, University of Edinburgh / Institute of Genetics and Molecular Medicine, University of Edinburgh | COVID-19 Genomics UK (COG-UK) Consortium | McHugh M, Dewar R, Rooke S, Gallagher M, Balcaza C, O'Toole A, Hill V, McCrone JT, Colquhoun R, Yu X, Jackson B, Rambaut A, Williams TC, Templeton K |
| EPI_ISL_433383 | hCoV-19/Scotland/EDB1073/2020 | Europe / United Kingdom / Scotland | 2020-04-06 | Virology Department, Royal Infirmary of Edinburgh, NHS Lothian / School of Biological Sciences, University of Edinburgh / Institute of Genetics and Molecular Medicine, University of Edinburgh | COVID-19 Genomics UK (COG-UK) Consortium | McHugh M, Dewar R, Rooke S, Gallagher M, Balcaza C, O'Toole A, Hill V, McCrone JT, Colquhoun R, Yu X, Jackson B, Rambaut A, Williams TC, Templeton K |
| EPI_ISL_433384 | hCoV-19/Scotland/EDB1076/2020 | Europe / United Kingdom / Scotland | 2020-04-07 | Virology Department, Royal Infirmary of Edinburgh, NHS Lothian / School of Biological Sciences, University of Edinburgh / Institute of Genetics and Molecular Medicine, University of Edinburgh | COVID-19 Genomics UK (COG-UK) Consortium | McHugh M, Dewar R, Rooke S, Gallagher M, Balcaza C, O'Toole A, Hill V, McCrone JT, Colquhoun R, Yu X, Jackson B, Rambaut A, Williams TC, Templeton K |
| EPI_ISL_433385 | hCoV-19/Scotland/EDB1077/2020 | Europe / United Kingdom / Scotland | 2020-04-07 | Virology Department, Royal Infirmary of Edinburgh, NHS Lothian / School of Biological Sciences, University of Edinburgh / Institute of Genetics and Molecular Medicine, University of Edinburgh | COVID-19 Genomics UK (COG-UK) Consortium | McHugh M, Dewar R, Rooke S, Gallagher M, Balcaza C, O'Toole A, Hill V, McCrone JT, Colquhoun R, Yu X, Jackson B, Rambaut A, Williams TC, Templeton K |
| EPI_ISL_433386 | hCoV-19/Scotland/EDB1078/2020 | Europe / United Kingdom / Scotland | 2020-04-07 | Virology Department, Royal Infirmary of Edinburgh, NHS Lothian / School of Biological Sciences, University of Edinburgh / Institute of Genetics and Molecular Medicine, University of Edinburgh | COVID-19 Genomics UK (COG-UK) Consortium | McHugh M, Dewar R, Rooke S, Gallagher M, Balcaza C, O'Toole A, Hill V, McCrone JT, Colquhoun R, Yu X, Jackson B, Rambaut A, Williams TC, Templeton K |
| EPI_ISL_433387 | hCoV-19/Scotland/EDB1079/2020 | Europe / United Kingdom / Scotland | 2020-04-07 | Virology Department, Royal Infirmary of Edinburgh, NHS Lothian / School of Biological Sciences, University of Edinburgh / Institute of Genetics and Molecular Medicine, University of Edinburgh | COVID-19 Genomics UK (COG-UK) Consortium | McHugh M, Dewar R, Rooke S, Gallagher M, Balcaza C, O'Toole A, Hill V, McCrone JT, Colquhoun R, Yu X, Jackson B, Rambaut A, Williams TC, Templeton K |

| | | | | | | |
|---|---|---|---|---|---|---|
| EPI_ISL_433388 | hCoV-19/Scotland/EDB1081/2020 | Europe / United Kingdom / Scotland | 2020-04-07 | Virology Department, Royal Infirmary of Edinburgh, NHS Lothian / School of Biological Sciences, University of Edinburgh / Institute of Genetics and Molecular Medicine, University of Edinburgh | COVID-19 Genomics UK (COG-UK) Consortium | McHugh M, Dewar R, Rooke S, Gallagher M, Balcaza C, O'Toole A, Hill V, McCrone JT, Colquhoun R, Yu X, Jackson B, Rambaut A, Williams TC, Templeton K |
| EPI_ISL_433389 | hCoV-19/Scotland/EDB1093/2020 | Europe / United Kingdom / Scotland | 2020-04-07 | Virology Department, Royal Infirmary of Edinburgh, NHS Lothian / School of Biological Sciences, University of Edinburgh / Institute of Genetics and Molecular Medicine, University of Edinburgh | COVID-19 Genomics UK (COG-UK) Consortium | McHugh M, Dewar R, Rooke S, Gallagher M, Balcaza C, O'Toole A, Hill V, McCrone JT, Colquhoun R, Yu X, Jackson B, Rambaut A, Williams TC, Templeton K |
| EPI_ISL_433390 | hCoV-19/Scotland/EDB1094/2020 | Europe / United Kingdom / Scotland | 2020-04-07 | Virology Department, Royal Infirmary of Edinburgh, NHS Lothian / School of Biological Sciences, University of Edinburgh / Institute of Genetics and Molecular Medicine, University of Edinburgh | COVID-19 Genomics UK (COG-UK) Consortium | McHugh M, Dewar R, Rooke S, Gallagher M, Balcaza C, O'Toole A, Hill V, McCrone JT, Colquhoun R, Yu X, Jackson B, Rambaut A, Williams TC, Templeton K |
| EPI_ISL_433391 | hCoV-19/Scotland/EDB1095/2020 | Europe / United Kingdom / Scotland | 2020-04-07 | Virology Department, Royal Infirmary of Edinburgh, NHS Lothian / School of Biological Sciences, University of Edinburgh / Institute of Genetics and Molecular Medicine, University of Edinburgh | COVID-19 Genomics UK (COG-UK) Consortium | McHugh M, Dewar R, Rooke S, Gallagher M, Balcaza C, O'Toole A, Hill V, McCrone JT, Colquhoun R, Yu X, Jackson B, Rambaut A, Williams TC, Templeton K |
| EPI_ISL_433392 | hCoV-19/Scotland/EDB1096/2020 | Europe / United Kingdom / Scotland | 2020-04-07 | Virology Department, Royal Infirmary of Edinburgh, NHS Lothian / School of Biological Sciences, University of Edinburgh / Institute of Genetics and Molecular Medicine, University of Edinburgh | COVID-19 Genomics UK (COG-UK) Consortium | McHugh M, Dewar R, Rooke S, Gallagher M, Balcaza C, O'Toole A, Hill V, McCrone JT, Colquhoun R, Yu X, Jackson B, Rambaut A, Williams TC, Templeton K |
| EPI_ISL_433395 | hCoV-19/Scotland/EDB1102/2020 | Europe / United Kingdom / Scotland | 2020-04-08 | Virology Department, Royal Infirmary of Edinburgh, NHS Lothian / School of Biological Sciences, University of Edinburgh / Institute of Genetics and Molecular Medicine, University of Edinburgh | COVID-19 Genomics UK (COG-UK) Consortium | McHugh M, Dewar R, Rooke S, Gallagher M, Balcaza C, O'Toole A, Hill V, McCrone JT, Colquhoun R, Yu X, Jackson B, Rambaut A, Williams TC, Templeton K |
| EPI_ISL_433396 | hCoV-19/Scotland/EDB1105/2020 | Europe / United Kingdom / Scotland | 2020-04-08 | Virology Department, Royal Infirmary of Edinburgh, NHS Lothian / School of Biological Sciences, University of Edinburgh / Institute of Genetics and Molecular Medicine, University of Edinburgh | COVID-19 Genomics UK (COG-UK) Consortium | McHugh M, Dewar R, Rooke S, Gallagher M, Balcaza C, O'Toole A, Hill V, McCrone JT, Colquhoun R, Yu X, Jackson B, Rambaut A, Williams TC, Templeton K |
| EPI_ISL_433397 | hCoV-19/Scotland/EDB1106/2020 | Europe / United Kingdom / Scotland | 2020-04-08 | Virology Department, Royal Infirmary of Edinburgh, NHS Lothian / School of Biological Sciences, University of Edinburgh / Institute of Genetics and Molecular Medicine, University of Edinburgh | COVID-19 Genomics UK (COG-UK) Consortium | McHugh M, Dewar R, Rooke S, Gallagher M, Balcaza C, O'Toole A, Hill V, McCrone JT, Colquhoun R, Yu X, Jackson B, Rambaut A, Williams TC, Templeton K |
| EPI_ISL_433398 | hCoV-19/Scotland/EDB1107/2020 | Europe / United Kingdom / Scotland | 2020-04-08 | Virology Department, Royal Infirmary of Edinburgh, NHS Lothian / School of Biological Sciences, University of Edinburgh / Institute of Genetics and Molecular Medicine, University of Edinburgh | COVID-19 Genomics UK (COG-UK) Consortium | McHugh M, Dewar R, Rooke S, Gallagher M, Balcaza C, O'Toole A, Hill V, McCrone JT, Colquhoun R, Yu X, Jackson B, Rambaut A, Williams TC, Templeton K |
| EPI_ISL_433446 | hCoV-19/Scotland/EDB1433/2020 | Europe / United Kingdom / Scotland | 2020-04-12 | Virology Department, Royal Infirmary of Edinburgh, NHS Lothian / School of Biological Sciences, University of Edinburgh / Institute of Genetics and Molecular Medicine, University of Edinburgh | COVID-19 Genomics UK (COG-UK) Consortium | McHugh M, Dewar R, Rooke S, Gallagher M, Balcaza C, O'Toole A, Hill V, McCrone JT, Colquhoun R, Yu X, Jackson B, Rambaut A, Williams TC, Templeton K |
| EPI_ISL_433447 | hCoV-19/Scotland/EDB1436/2020 | Europe / United Kingdom / Scotland | 2020-04-12 | Virology Department, Royal Infirmary of Edinburgh, NHS Lothian / School of Biological Sciences, University of Edinburgh / Institute of Genetics and Molecular Medicine, University of Edinburgh | COVID-19 Genomics UK (COG-UK) Consortium | McHugh M, Dewar R, Rooke S, Gallagher M, Balcaza C, O'Toole A, Hill V, McCrone JT, Colquhoun R, Yu X, Jackson B, Rambaut A, Williams TC, Templeton K |
| EPI_ISL_433448 | hCoV-19/Scotland/EDB1437/2020 | Europe / United Kingdom / Scotland | 2020-04-12 | Virology Department, Royal Infirmary of Edinburgh, NHS Lothian / School of Biological Sciences, University of Edinburgh / Institute of Genetics and Molecular Medicine, University of Edinburgh | COVID-19 Genomics UK (COG-UK) Consortium | McHugh M, Dewar R, Rooke S, Gallagher M, Balcaza C, O'Toole A, Hill V, McCrone JT, Colquhoun R, Yu X, Jackson B, Rambaut A, Williams TC, Templeton K |
| EPI_ISL_433449 | hCoV-19/Scotland/EDB1444/2020 | Europe / United Kingdom / Scotland | 2020-04-13 | Virology Department, Royal Infirmary of Edinburgh, NHS Lothian / School of Biological Sciences, University of Edinburgh / Institute of Genetics and Molecular Medicine, University of Edinburgh | COVID-19 Genomics UK (COG-UK) Consortium | McHugh M, Dewar R, Rooke S, Gallagher M, Balcaza C, O'Toole A, Hill V, McCrone JT, Colquhoun R, Yu X, Jackson B, Rambaut A, Williams TC, Templeton K |
| EPI_ISL_433451 | hCoV-19/Scotland/EDB1448/2020 | Europe / United Kingdom / Scotland | 2020-04-12 | Virology Department, Royal Infirmary of Edinburgh, NHS Lothian / School of Biological Sciences, University of Edinburgh / Institute of Genetics and Molecular Medicine, University of Edinburgh | COVID-19 Genomics UK (COG-UK) Consortium | McHugh M, Dewar R, Rooke S, Gallagher M, Balcaza C, O'Toole A, Hill V, McCrone JT, Colquhoun R, Yu X, Jackson B, Rambaut A, Williams TC, Templeton K |
| EPI_ISL_433452 | hCoV-19/Scotland/EDB1450/2020 | Europe / United Kingdom / Scotland | 2020-04-13 | Virology Department, Royal Infirmary of Edinburgh, NHS Lothian / School of Biological Sciences, University of Edinburgh / Institute of Genetics and Molecular Medicine, University of Edinburgh | COVID-19 Genomics UK (COG-UK) Consortium | McHugh M, Dewar R, Rooke S, Gallagher M, Balcaza C, O'Toole A, Hill V, McCrone JT, Colquhoun R, Yu X, Jackson B, Rambaut A, Williams TC, Templeton K |
| EPI_ISL_433453 | hCoV-19/Scotland/EDB1452/2020 | Europe / United Kingdom / Scotland | 2020-04-13 | Virology Department, Royal Infirmary of Edinburgh, NHS Lothian / School of Biological Sciences, University of Edinburgh / Institute of Genetics and Molecular Medicine, University of Edinburgh | COVID-19 Genomics UK (COG-UK) Consortium | McHugh M, Dewar R, Rooke S, Gallagher M, Balcaza C, O'Toole A, Hill V, McCrone JT, Colquhoun R, Yu X, Jackson B, Rambaut A, Williams TC, Templeton K |
| EPI_ISL_433454 | hCoV-19/Scotland/EDB1453/2020 | Europe / United Kingdom / Scotland | 2020-04-13 | Virology Department, Royal Infirmary of Edinburgh, NHS Lothian / School of Biological Sciences, University of Edinburgh / Institute of Genetics and Molecular Medicine, University of Edinburgh | COVID-19 Genomics UK (COG-UK) Consortium | McHugh M, Dewar R, Rooke S, Gallagher M, Balcaza C, O'Toole A, Hill V, McCrone JT, Colquhoun R, Yu X, Jackson B, Rambaut A, Williams TC, Templeton K |
| EPI_ISL_433455 | hCoV-19/Scotland/EDB1457/2020 | Europe / United Kingdom / Scotland | 2020-04-11 | Virology Department, Royal Infirmary of Edinburgh, NHS Lothian / School of Biological Sciences, University of Edinburgh / Institute of Genetics and Molecular Medicine, University of Edinburgh | COVID-19 Genomics UK (COG-UK) Consortium | McHugh M, Dewar R, Rooke S, Gallagher M, Balcaza C, O'Toole A, Hill V, McCrone JT, Colquhoun R, Yu X, Jackson B, Rambaut A, Williams TC, Templeton K |
| EPI_ISL_433456 | hCoV-19/Scotland/EDB1459/2020 | Europe / United Kingdom / Scotland | 2020-04-11 | Virology Department, Royal Infirmary of Edinburgh, NHS Lothian / School of Biological Sciences, University of Edinburgh / Institute of Genetics and Molecular Medicine, University of Edinburgh | COVID-19 Genomics UK (COG-UK) Consortium | McHugh M, Dewar R, Rooke S, Gallagher M, Balcaza C, O'Toole A, Hill V, McCrone JT, Colquhoun R, Yu X, Jackson B, Rambaut A, Williams TC, Templeton K |
| EPI_ISL_433459 | hCoV-19/Scotland/EDB1469/2020 | Europe / United Kingdom / Scotland | 2020-04-11 | Virology Department, Royal Infirmary of Edinburgh, NHS Lothian / School of Biological Sciences, University of Edinburgh / Institute of Genetics and Molecular Medicine, University of Edinburgh | COVID-19 Genomics UK (COG-UK) Consortium | McHugh M, Dewar R, Rooke S, Gallagher M, Balcaza C, O'Toole A, Hill V, McCrone JT, Colquhoun R, Yu X, Jackson B, Rambaut A, Williams TC, Templeton K |
| EPI_ISL_433460 | hCoV-19/Scotland/EDB1470/2020 | Europe / United Kingdom / Scotland | 2020-04-11 | Virology Department, Royal Infirmary of Edinburgh, NHS Lothian / School of Biological Sciences, University of Edinburgh / Institute of Genetics and Molecular Medicine, University of Edinburgh | COVID-19 Genomics UK (COG-UK) Consortium | McHugh M, Dewar R, Rooke S, Gallagher M, Balcaza C, O'Toole A, Hill V, McCrone JT, Colquhoun R, Yu X, Jackson B, Rambaut A, Williams TC, Templeton K |
| EPI_ISL_433461 | hCoV-19/Scotland/EDB1471/2020 | Europe / United Kingdom / Scotland | 2020-04-11 | Virology Department, Royal Infirmary of Edinburgh, NHS Lothian / School of Biological Sciences, University of Edinburgh / Institute of Genetics and Molecular Medicine, University of Edinburgh | COVID-19 Genomics UK (COG-UK) Consortium | McHugh M, Dewar R, Rooke S, Gallagher M, Balcaza C, O'Toole A, Hill V, McCrone JT, Colquhoun R, Yu X, Jackson B, Rambaut A, Williams TC, Templeton K |
| EPI_ISL_433462 | hCoV-19/Scotland/EDB1507/2020 | Europe / United Kingdom / Scotland | 2020-04-13 | Virology Department, Royal Infirmary of Edinburgh, NHS Lothian / School of Biological Sciences, University of Edinburgh / Institute of Genetics and Molecular Medicine, University of Edinburgh | COVID-19 Genomics UK (COG-UK) Consortium | McHugh M, Dewar R, Rooke S, Gallagher M, Balcaza C, O'Toole A, Hill V, McCrone JT, Colquhoun R, Yu X, Jackson B, Rambaut A, Williams TC, Templeton K |
| EPI_ISL_433463 | hCoV-19/Scotland/EDB1510/2020 | Europe / United Kingdom / Scotland | 2020-04-12 | Virology Department, Royal Infirmary of Edinburgh, NHS Lothian / School of Biological Sciences, University of Edinburgh / Institute of Genetics and Molecular Medicine, University of Edinburgh | COVID-19 Genomics UK (COG-UK) Consortium | McHugh M, Dewar R, Rooke S, Gallagher M, Balcaza C, O'Toole A, Hill V, McCrone JT, Colquhoun R, Yu X, Jackson B, Rambaut A, Williams TC, Templeton K |
| EPI_ISL_433464 | hCoV-19/Scotland/EDB1511/2020 | Europe / United Kingdom / Scotland | 2020-04-13 | Virology Department, Royal Infirmary of Edinburgh, NHS Lothian / School of Biological Sciences, University of Edinburgh / Institute of Genetics and Molecular Medicine, University of Edinburgh | COVID-19 Genomics UK (COG-UK) Consortium | McHugh M, Dewar R, Rooke S, Gallagher M, Balcaza C, O'Toole A, Hill V, McCrone JT, Colquhoun R, Yu X, Jackson B, Rambaut A, Williams TC, Templeton K |
| EPI_ISL_433467 | hCoV-19/England/CAMB-7E2E9/2020 | Europe / United Kingdom / England | 2020-04-17 | Department of Pathology, University of Cambridge | COVID-19 Genomics UK (COG-UK) Consortium | Luke W Meredith, M. Estee Torok , Myra Hosmillo, William L. Hamilton, Martin D. Curran, Theresa Feltwell, Grant Hall, Anna Yakovleva, Fahad A Khokhar, Charlotte J. Houldcroft, Laura G Caller, Aminu S. Jahun, Sarah L. Caddy, Ian Goodfellow |
| EPI_ISL_433469 | hCoV-19/England/CAMB-7E304/2020 | Europe / United Kingdom / England | 2020-04-17 | Department of Pathology, University of Cambridge | COVID-19 Genomics UK (COG-UK) Consortium | Luke W Meredith, M. Estee Torok , Myra Hosmillo, William L. Hamilton, Martin D. Curran, Theresa Feltwell, Grant Hall, Anna Yakovleva, Fahad A Khokhar, Charlotte J. Houldcroft, Laura G Caller, Aminu S. Jahun, Sarah L. Caddy, Ian Goodfellow |
| EPI_ISL_433470 | hCoV-19/England/CAMB-7E313/2020 | Europe / United Kingdom / England | 2020-04-17 | Department of Pathology, University of Cambridge | COVID-19 Genomics UK (COG-UK) Consortium | Luke W Meredith, M. Estee Torok , Myra Hosmillo, William L. Hamilton, Martin D. Curran, Theresa Feltwell, Grant Hall, Anna Yakovleva, Fahad A Khokhar, Charlotte J. Houldcroft, Laura G Caller, Aminu S. Jahun, Sarah L. Caddy, Ian Goodfellow |
| EPI_ISL_433471 | hCoV-19/England/CAMB-7E322/2020 | Europe / United Kingdom / England | 2020-04-17 | Department of Pathology, University of Cambridge | COVID-19 Genomics UK (COG-UK) Consortium | Luke W Meredith, M. Estee Torok , Myra Hosmillo, William L. Hamilton, Martin D. Curran, Theresa Feltwell, Grant Hall, Anna Yakovleva, Fahad A Khokhar, Charlotte J. Houldcroft, Laura G Caller, Aminu S. Jahun, Sarah L. Caddy, Ian Goodfellow |
| EPI_ISL_433472 | hCoV-19/England/CAMB-7E331/2020 | Europe / United Kingdom / England | 2020-04-16 | Department of Pathology, University of Cambridge | COVID-19 Genomics UK (COG-UK) Consortium | Luke W Meredith, M. Estee Torok , Myra Hosmillo, William L. Hamilton, Martin D. Curran, Theresa Feltwell, Grant Hall, Anna Yakovleva, Fahad A Khokhar, Charlotte J. Houldcroft, Laura G Caller, Aminu S. Jahun, Sarah L. Caddy, Ian Goodfellow |
| EPI_ISL_433478 | hCoV-19/England/CAMB-7F8E6/2020 | Europe / United Kingdom / England | 2020-04-18 | Department of Pathology, University of Cambridge | COVID-19 Genomics UK (COG-UK) Consortium | Luke W Meredith, M. Estee Torok , Myra Hosmillo, William L. Hamilton, Martin D. Curran, Theresa Feltwell, Grant Hall, Anna Yakovleva, Fahad A Khokhar, Charlotte J. Houldcroft, Laura G Caller, Aminu S. Jahun, Sarah L. Caddy, Ian Goodfellow |
| EPI_ISL_433480 | hCoV-19/England/CAMB-7FB47/2020 | Europe / United Kingdom / England | 2020-04-17 | Department of Pathology, University of Cambridge | COVID-19 Genomics UK (COG-UK) Consortium | Luke W Meredith, M. Estee Torok , Myra Hosmillo, William L. Hamilton, Martin D. Curran, Theresa Feltwell, Grant Hall, Anna Yakovleva, Fahad A Khokhar, Charlotte J. Houldcroft, Laura G Caller, Aminu S. Jahun, Sarah L. Caddy, Ian Goodfellow |
| EPI_ISL_433481 | hCoV-19/England/CAMB-7FB56/2020 | Europe / United Kingdom / England | 2020-04-17 | Department of Pathology, University of Cambridge | COVID-19 Genomics UK (COG-UK) Consortium | Luke W Meredith, M. Estee Torok , Myra Hosmillo, William L. Hamilton, Martin D. Curran, Theresa Feltwell, Grant Hall, Anna Yakovleva, Fahad A Khokhar, Charlotte J. Houldcroft, Laura G Caller, Aminu S. Jahun, Sarah L. Caddy, Ian Goodfellow |
| EPI_ISL_433482 | hCoV-19/England/CAMB-7FB65/2020 | Europe / United Kingdom / England | 2020-04-17 | Department of Pathology, University of Cambridge | COVID-19 Genomics UK (COG-UK) Consortium | Luke W Meredith, M. Estee Torok , Myra Hosmillo, William L. Hamilton, Martin D. Curran, Theresa Feltwell, Grant Hall, Anna Yakovleva, Fahad A Khokhar, Charlotte J. Houldcroft, Laura G Caller, Aminu S. Jahun, Sarah L. Caddy, Ian Goodfellow |
| EPI_ISL_433483 | hCoV-19/England/CAMB-7FB74/2020 | Europe / United Kingdom / England | 2020-04-17 | Department of Pathology, University of Cambridge | COVID-19 Genomics UK (COG-UK) Consortium | Luke W Meredith, M. Estee Torok , Myra Hosmillo, William L. Hamilton, Martin D. Curran, Theresa Feltwell, Grant Hall, Anna Yakovleva, Fahad A Khokhar, Charlotte J. Houldcroft, Laura G Caller, Aminu S. Jahun, Sarah L. Caddy, Ian Goodfellow |
| EPI_ISL_433484 | hCoV-19/England/CAMB-7FB83/2020 | Europe / United Kingdom / England | 2020-04-17 | Department of Pathology, University of Cambridge | COVID-19 Genomics UK (COG-UK) Consortium | Luke W Meredith, M. Estee Torok , Myra Hosmillo, William L. Hamilton, Martin D. Curran, Theresa Feltwell, Grant Hall, Anna Yakovleva, Fahad A Khokhar, Charlotte J. Houldcroft, Laura G Caller, Aminu S. Jahun, Sarah L. Caddy, Ian Goodfellow |
| EPI_ISL_433485 | hCoV-19/England/CAMB-7FB92/2020 | Europe / United Kingdom / England | 2020-04-17 | Department of Pathology, University of Cambridge | COVID-19 Genomics UK (COG-UK) Consortium | Luke W Meredith, M. Estee Torok , Myra Hosmillo, William L. Hamilton, Martin D. Curran, Theresa Feltwell, Grant Hall, Anna Yakovleva, Fahad A Khokhar, Charlotte J. Houldcroft, Laura G Caller, Aminu S. Jahun, Sarah L. Caddy, Ian Goodfellow |
| EPI_ISL_433486 | hCoV-19/England/CAMB-7FBA1/2020 | Europe / United Kingdom / England | 2020-04-17 | Department of Pathology, University of Cambridge | COVID-19 Genomics UK (COG-UK) Consortium | Luke W Meredith, M. Estee Torok , Myra Hosmillo, William L. Hamilton, Martin D. Curran, Theresa Feltwell, Grant Hall, Anna Yakovleva, Fahad A Khokhar, Charlotte J. Houldcroft, Laura G Caller, Aminu S. Jahun, Sarah L. Caddy, Ian Goodfellow |
| EPI_ISL_433487 | hCoV-19/England/CAMB-7FBB0/2020 | Europe / United Kingdom / England | 2020-04-16 | Department of Pathology, University of Cambridge | COVID-19 Genomics UK (COG-UK) Consortium | Luke W Meredith, M. Estee Torok , Myra Hosmillo, William L. Hamilton, Martin D. Curran, Theresa Feltwell, Grant Hall, Anna Yakovleva, Fahad A Khokhar, Charlotte J. Houldcroft, Laura G Caller, Aminu S. Jahun, Sarah L. Caddy, Ian Goodfellow |
| EPI_ISL_433488 | hCoV-19/England/CAMB-7FBCF/2020 | Europe / United Kingdom / England | 2020 | Department of Pathology, University of Cambridge | COVID-19 Genomics UK (COG-UK) Consortium | Luke W Meredith, M. Estee Torok , Myra Hosmillo, William L. Hamilton, Martin D. Curran, Theresa Feltwell, Grant Hall, Anna Yakovleva, Fahad A Khokhar, Charlotte J. Houldcroft, Laura G Caller, Aminu S. Jahun, Sarah L. Caddy, Ian Goodfellow |
| EPI_ISL_433489 | hCoV-19/England/CAMB-7FBDE/2020 | Europe / United Kingdom / England | 2020-04-16 | Department of Pathology, University of Cambridge | COVID-19 Genomics UK (COG-UK) Consortium | Luke W Meredith, M. Estee Torok , Myra Hosmillo, William L. Hamilton, Martin D. Curran, Theresa Feltwell, Grant Hall, Anna Yakovleva, Fahad A Khokhar, Charlotte J. Houldcroft, Laura G Caller, Aminu S. Jahun, Sarah L. Caddy, Ian Goodfellow |
| EPI_ISL_433490 | hCoV-19/England/CAMB-7FBFC/2020 | Europe / United Kingdom / England | 2020-04-16 | Department of Pathology, University of Cambridge | COVID-19 Genomics UK (COG-UK) Consortium | Luke W Meredith, M. Estee Torok , Myra Hosmillo, William L. Hamilton, Martin D. Curran, Theresa Feltwell, Grant Hall, Anna Yakovleva, Fahad A Khokhar, Charlotte J. Houldcroft, Laura G Caller, Aminu S. Jahun, Sarah L. Caddy, Ian Goodfellow |
| EPI_ISL_433491 | hCoV-19/England/CAMB-7FC08/2020 | Europe / United Kingdom / England | 2020-04-16 | Department of Pathology, University of Cambridge | COVID-19 Genomics UK (COG-UK) Consortium | Luke W Meredith, M. Estee Torok , Myra Hosmillo, William L. Hamilton, Martin D. Curran, Theresa Feltwell, Grant Hall, Anna Yakovleva, Fahad A Khokhar, Charlotte J. Houldcroft, Laura G Caller, Aminu S. Jahun, Sarah L. Caddy, Ian Goodfellow |
| EPI_ISL_433492 | hCoV-19/England/CAMB-7FC17/2020 | Europe / United Kingdom / England | 2020-04-16 | Department of Pathology, University of Cambridge | COVID-19 Genomics UK (COG-UK) Consortium | Luke W Meredith, M. Estee Torok , Myra Hosmillo, William L. Hamilton, Martin D. Curran, Theresa Feltwell, Grant Hall, Anna Yakovleva, Fahad A Khokhar, Charlotte J. Houldcroft, Laura G Caller, Aminu S. Jahun, Sarah L. Caddy, Ian Goodfellow |
| EPI_ISL_433493 | hCoV-19/England/CAMB-7FC26/2020 | Europe / United Kingdom / England | 2020-04-16 | Department of Pathology, University of Cambridge | COVID-19 Genomics UK (COG-UK) Consortium | Luke W Meredith, M. Estee Torok , Myra Hosmillo, William L. Hamilton, Martin D. Curran, Theresa Feltwell, Grant Hall, Anna Yakovleva, Fahad A Khokhar, Charlotte J. Houldcroft, Laura G Caller, Aminu S. Jahun, Sarah L. Caddy, Ian Goodfellow |
| EPI_ISL_433494 | hCoV-19/England/CAMB-7FC35/2020 | Europe / United Kingdom / England | 2020-04-16 | Department of Pathology, University of Cambridge | COVID-19 Genomics UK (COG-UK) Consortium | Luke W Meredith, M. Estee Torok , Myra Hosmillo, William L. Hamilton, Martin D. Curran, Theresa Feltwell, Grant Hall, Anna Yakovleva, Fahad A Khokhar, Charlotte J. Houldcroft, Laura G Caller, Aminu S. Jahun, Sarah L. Caddy, Ian Goodfellow |
| EPI_ISL_433495 | hCoV-19/England/CAMB-7FC44/2020 | Europe / United Kingdom / England | 2020-04-16 | Department of Pathology, University of Cambridge | COVID-19 Genomics UK (COG-UK) Consortium | Luke W Meredith, M. Estee Torok , Myra Hosmillo, William L. Hamilton, Martin D. Curran, Theresa Feltwell, Grant Hall, Anna Yakovleva, Fahad A Khokhar, Charlotte J. Houldcroft, Laura G Caller, Aminu S. Jahun, Sarah L. Caddy, Ian Goodfellow |
| EPI_ISL_433496 | hCoV-19/England/CAMB-7FC53/2020 | Europe / United Kingdom / England | 2020-04-16 | Department of Pathology, University of Cambridge | COVID-19 Genomics UK (COG-UK) Consortium | Luke W Meredith, M. Estee Torok , Myra Hosmillo, William L. Hamilton, Martin D. Curran, Theresa Feltwell, Grant Hall, Anna Yakovleva, Fahad A Khokhar, Charlotte J. Houldcroft, Laura G Caller, Aminu S. Jahun, Sarah L. Caddy, Ian Goodfellow |
| EPI_ISL_433497 | hCoV-19/England/CAMB-7FC62/2020 | Europe / United Kingdom / England | 2020-04-16 | Department of Pathology, University of Cambridge | COVID-19 Genomics UK (COG-UK) Consortium | Luke W Meredith, M. Estee Torok , Myra Hosmillo, William L. Hamilton, Martin D. Curran, Theresa Feltwell, Grant Hall, Anna Yakovleva, Fahad A Khokhar, Charlotte J. Houldcroft, Laura G Caller, Aminu S. Jahun, Sarah L. Caddy, Ian Goodfellow |
| EPI_ISL_433498 | hCoV-19/Scotland/CVR1000/2020 | Europe / United Kingdom / Scotland | 2020-03-29 | West of Scotland Specialist Virology Centre, NHSGGC / MRC-University of Glasgow Centre for Virus Research | COVID-19 Genomics UK (COG-UK) Consortium | Ana da Silva Filipe, Natasha Johnson, Kathy Smollett, Daniel Mair, Stephen Carmichael, Lily Tong, Jenna Nichols, Elihu Aranday-Cortes, Kirstyn Brunker, Yasmin Parr, Kyriaki Nomikou; Sarah McDonald, Marc Niebel, Patawee Asamaphan; Richard Orton, Joseph Hughes, Sreenu Vattipally, David L Robertson; Alasdair MacLean, Rory Gunson; Kathy Li, Natasha Jesudason, Rajiv Shah, James Shepherd, Antonia Ho, Emma Thomson |
| EPI_ISL_433499 | hCoV-19/Scotland/CVR1022/2020 | Europe / United Kingdom / Scotland | 2020-03-30 | West of Scotland Specialist Virology Centre, NHSGGC / MRC-University of Glasgow Centre for Virus Research | COVID-19 Genomics UK (COG-UK) Consortium | Ana da Silva Filipe, Natasha Johnson, Kathy Smollett, Daniel Mair, Stephen Carmichael, Lily Tong, Jenna Nichols, Elihu Aranday-Cortes, Kirstyn Brunker, Yasmin Parr, Kyriaki Nomikou; Sarah McDonald, Marc Niebel, Patawee Asamaphan; Richard Orton, Joseph Hughes, Sreenu Vattipally, David L Robertson; Alasdair MacLean, Rory Gunson; Kathy Li, Natasha Jesudason, Rajiv Shah, James Shepherd, Antonia Ho, Emma Thomson |
| EPI_ISL_433501 | hCoV-19/Scotland/CVR1053/2020 | Europe / United Kingdom / Scotland | 2020-03-28 | West of Scotland Specialist Virology Centre, NHSGGC / MRC-University of Glasgow Centre for Virus Research | COVID-19 Genomics UK (COG-UK) Consortium | Ana da Silva Filipe, Natasha Johnson, Kathy Smollett, Daniel Mair, Stephen Carmichael, Lily Tong, Jenna Nichols, Elihu Aranday-Cortes, Kirstyn Brunker, Yasmin Parr, Kyriaki Nomikou; Sarah McDonald, Marc Niebel, Patawee Asamaphan; Richard Orton, Joseph Hughes, Sreenu Vattipally, David L Robertson; Alasdair MacLean, Rory Gunson; Kathy Li, Natasha Jesudason, Rajiv Shah, James Shepherd, Antonia Ho, Emma Thomson |
| EPI_ISL_433502 | hCoV-19/Scotland/CVR1056/2020 | Europe / United Kingdom / Scotland | 2020-03-28 | West of Scotland Specialist Virology Centre, NHSGGC / MRC-University of Glasgow Centre for Virus Research | COVID-19 Genomics UK (COG-UK) Consortium | Ana da Silva Filipe, Natasha Johnson, Kathy Smollett, Daniel Mair, Stephen Carmichael, Lily Tong, Jenna Nichols, Elihu Aranday-Cortes, Kirstyn Brunker, Yasmin Parr, Kyriaki Nomikou; Sarah McDonald, Marc Niebel, Patawee Asamaphan; Richard Orton, Joseph Hughes, Sreenu Vattipally, David L Robertson; Alasdair MacLean, Rory Gunson; Kathy Li, Natasha Jesudason, Rajiv Shah, James Shepherd, Antonia Ho, Emma Thomson |
| EPI_ISL_433505 | hCoV-19/Scotland/CVR1068/2020 | Europe / United Kingdom / Scotland | 2020-03-29 | West of Scotland Specialist Virology Centre, NHSGGC / MRC-University of Glasgow Centre for Virus Research | COVID-19 Genomics UK (COG-UK) Consortium | Ana da Silva Filipe, Natasha Johnson, Kathy Smollett, Daniel Mair, Stephen Carmichael, Lily Tong, Jenna Nichols, Elihu Aranday-Cortes, Kirstyn Brunker, Yasmin Parr, Kyriaki Nomikou; Sarah McDonald, Marc Niebel, Patawee Asamaphan; Richard Orton, Joseph Hughes, Sreenu Vattipally, David L Robertson; Alasdair MacLean, Rory Gunson; Kathy Li, Natasha Jesudason, Rajiv Shah, James Shepherd, Antonia Ho, Emma Thomson |
| EPI_ISL_433507 | hCoV-19/Scotland/CVR1081/2020 | Europe / United Kingdom / Scotland | 2020-03-29 | West of Scotland Specialist Virology Centre, NHSGGC / MRC-University of Glasgow Centre for Virus Research | COVID-19 Genomics UK (COG-UK) Consortium | Ana da Silva Filipe, Natasha Johnson, Kathy Smollett, Daniel Mair, Stephen Carmichael, Lily Tong, Jenna Nichols, Elihu Aranday-Cortes, Kirstyn Brunker, Yasmin Parr, Kyriaki Nomikou; Sarah McDonald, Marc Niebel, Patawee Asamaphan; Richard Orton, Joseph Hughes, Sreenu Vattipally, David L Robertson; Alasdair MacLean, Rory Gunson; Kathy Li, Natasha Jesudason, Rajiv Shah, James Shepherd, Antonia Ho, Emma Thomson |
| EPI_ISL_433508 | hCoV-19/Scotland/CVR1085/2020 | Europe / United Kingdom / Scotland | 2020-03-29 | West of Scotland Specialist Virology Centre, NHSGGC / MRC-University of Glasgow Centre for Virus Research | COVID-19 Genomics UK (COG-UK) Consortium | Ana da Silva Filipe, Natasha Johnson, Kathy Smollett, Daniel Mair, Stephen Carmichael, Lily Tong, Jenna Nichols, Elihu Aranday-Cortes, Kirstyn Brunker, Yasmin Parr, Kyriaki Nomikou; Sarah McDonald, Marc Niebel, Patawee Asamaphan; Richard Orton, Joseph Hughes, Sreenu Vattipally, David L Robertson; Alasdair MacLean, Rory Gunson; Kathy Li, Natasha Jesudason, Rajiv Shah, James Shepherd, Antonia Ho, Emma Thomson |
| EPI_ISL_433509 | hCoV-19/Scotland/CVR1093/2020 | Europe / United Kingdom / Scotland | 2020-03-29 | West of Scotland Specialist Virology Centre, NHSGGC / MRC-University of Glasgow Centre for Virus Research | COVID-19 Genomics UK (COG-UK) Consortium | Ana da Silva Filipe, Natasha Johnson, Kathy Smollett, Daniel Mair, Stephen Carmichael, Lily Tong, Jenna Nichols, Elihu Aranday-Cortes, Kirstyn Brunker, Yasmin Parr, Kyriaki Nomikou; Sarah McDonald, Marc Niebel, Patawee Asamaphan; Richard Orton, Joseph Hughes, Sreenu Vattipally, David L Robertson; Alasdair MacLean, Rory Gunson; Kathy Li, Natasha Jesudason, Rajiv Shah, James Shepherd, Antonia Ho, Emma Thomson |
| EPI_ISL_433510 | hCoV-19/Scotland/CVR1101/2020 | Europe / United Kingdom / Scotland | 2020-03-29 | West of Scotland Specialist Virology Centre, NHSGGC / MRC-University of Glasgow Centre for Virus Research | COVID-19 Genomics UK (COG-UK) Consortium | Ana da Silva Filipe, Natasha Johnson, Kathy Smollett, Daniel Mair, Stephen Carmichael, Lily Tong, Jenna Nichols, Elihu Aranday-Cortes, Kirstyn Brunker, Yasmin Parr, Kyriaki Nomikou; Sarah McDonald, Marc Niebel, Patawee Asamaphan; Richard Orton, Joseph Hughes, Sreenu Vattipally, David L Robertson; Alasdair MacLean, Rory Gunson; Kathy Li, Natasha Jesudason, Rajiv Shah, James Shepherd, Antonia Ho, Emma Thomson |
| EPI_ISL_433511 | hCoV-19/Scotland/CVR1115/2020 | Europe / United Kingdom / Scotland | 2020-03-29 | West of Scotland Specialist Virology Centre, NHSGGC / MRC-University of Glasgow Centre for Virus Research | COVID-19 Genomics UK (COG-UK) Consortium | Ana da Silva Filipe, Natasha Johnson, Kathy Smollett, Daniel Mair, Stephen Carmichael, Lily Tong, Jenna Nichols, Elihu Aranday-Cortes, Kirstyn Brunker, Yasmin Parr, Kyriaki Nomikou; Sarah McDonald, Marc Niebel, Patawee Asamaphan; Richard Orton, Joseph Hughes, Sreenu Vattipally, David L Robertson; Alasdair MacLean, Rory Gunson; Kathy Li, Natasha Jesudason, Rajiv Shah, James Shepherd, Antonia Ho, Emma Thomson |
| EPI_ISL_433515 | hCoV-19/Scotland/CVR1133b/2020 | Europe / United Kingdom / Scotland | 2020-03-29 | West of Scotland Specialist Virology Centre, NHSGGC / MRC-University of Glasgow Centre for Virus Research | COVID-19 Genomics UK (COG-UK) Consortium | Ana da Silva Filipe, Natasha Johnson, Kathy Smollett, Daniel Mair, Stephen Carmichael, Lily Tong, Jenna Nichols, Elihu Aranday-Cortes, Kirstyn Brunker, Yasmin Parr, Kyriaki Nomikou; Sarah McDonald, Marc Niebel, Patawee Asamaphan; Richard Orton, Joseph Hughes, Sreenu Vattipally, David L Robertson; Alasdair MacLean, Rory Gunson; Kathy Li, Natasha Jesudason, Rajiv Shah, James Shepherd, Antonia Ho, Emma Thomson |
| EPI_ISL_433517 | hCoV-19/Scotland/CVR1140/2020 | Europe / United Kingdom / Scotland | 2020-03-29 | West of Scotland Specialist Virology Centre, NHSGGC / MRC-University of Glasgow Centre for Virus Research | COVID-19 Genomics UK (COG-UK) Consortium | Ana da Silva Filipe, Natasha Johnson, Kathy Smollett, Daniel Mair, Stephen Carmichael, Lily Tong, Jenna Nichols, Elihu Aranday-Cortes, Kirstyn Brunker, Yasmin Parr, Kyriaki Nomikou; Sarah McDonald, Marc Niebel, Patawee Asamaphan; Richard Orton, Joseph Hughes, Sreenu Vattipally, David L Robertson; Alasdair MacLean, Rory Gunson; Kathy Li, Natasha Jesudason, Rajiv Shah, James Shepherd, Antonia Ho, Emma Thomson |
| EPI_ISL_433518 | hCoV-19/Scotland/CVR1142/2020 | Europe / United Kingdom / Scotland | 2020-03-29 | West of Scotland Specialist Virology Centre, NHSGGC / MRC-University of Glasgow Centre for Virus Research | COVID-19 Genomics UK (COG-UK) Consortium | Ana da Silva Filipe, Natasha Johnson, Kathy Smollett, Daniel Mair, Stephen Carmichael, Lily Tong, Jenna Nichols, Elihu Aranday-Cortes, Kirstyn Brunker, Yasmin Parr, Kyriaki Nomikou; Sarah McDonald, Marc Niebel, Patawee Asamaphan; Richard Orton, Joseph Hughes, Sreenu Vattipally, David L Robertson; Alasdair MacLean, Rory Gunson; Kathy Li, Natasha Jesudason, Rajiv Shah, James Shepherd, Antonia Ho, Emma Thomson |
| EPI_ISL_433521 | hCoV-19/Scotland/CVR1159/2020 | Europe / United Kingdom / Scotland | 2020-03-29 | West of Scotland Specialist Virology Centre, NHSGGC / MRC-University of Glasgow Centre for Virus Research | COVID-19 Genomics UK (COG-UK) Consortium | Ana da Silva Filipe, Natasha Johnson, Kathy Smollett, Daniel Mair, Stephen Carmichael, Lily Tong, Jenna Nichols, Elihu Aranday-Cortes, Kirstyn Brunker, Yasmin Parr, Kyriaki Nomikou; Sarah McDonald, Marc Niebel, Patawee Asamaphan; Richard Orton, Joseph Hughes, Sreenu Vattipally, David L Robertson; Alasdair MacLean, Rory Gunson; Kathy Li, Natasha Jesudason, Rajiv Shah, James Shepherd, Antonia Ho, Emma Thomson |
| EPI_ISL_433522 | hCoV-19/Scotland/CVR1204/2020 | Europe / United Kingdom / Scotland | 2020-03-30 | West of Scotland Specialist Virology Centre, NHSGGC / MRC-University of Glasgow Centre for Virus Research | COVID-19 Genomics UK (COG-UK) Consortium | Ana da Silva Filipe, Natasha Johnson, Kathy Smollett, Daniel Mair, Stephen Carmichael, Lily Tong, Jenna Nichols, Elihu Aranday-Cortes, Kirstyn Brunker, Yasmin Parr, Kyriaki Nomikou; Sarah McDonald, Marc Niebel, Patawee Asamaphan; Richard Orton, Joseph Hughes, Sreenu Vattipally, David L Robertson; Alasdair MacLean, Rory Gunson; Kathy Li, Natasha Jesudason, Rajiv Shah, James Shepherd, Antonia Ho, Emma Thomson |
| EPI_ISL_433525 | hCoV-19/Scotland/CVR1261/2020 | Europe / United Kingdom / Scotland | 2020-03-31 | West of Scotland Specialist Virology Centre, NHSGGC / MRC-University of Glasgow Centre for Virus Research | COVID-19 Genomics UK (COG-UK) Consortium | Ana da Silva Filipe, Natasha Johnson, Kathy Smollett, Daniel Mair, Stephen Carmichael, Lily Tong, Jenna Nichols, Elihu Aranday-Cortes, Kirstyn Brunker, Yasmin Parr, Kyriaki Nomikou; Sarah McDonald, Marc Niebel, Patawee Asamaphan; Richard Orton, Joseph Hughes, Sreenu Vattipally, David L Robertson; Alasdair MacLean, Rory Gunson; Kathy Li, Natasha Jesudason, Rajiv Shah, James Shepherd, Antonia Ho, Emma Thomson |
| EPI_ISL_433527 | hCoV-19/Scotland/CVR1267/2020 | Europe / United Kingdom / Scotland | 2020-03-31 | West of Scotland Specialist Virology Centre, NHSGGC / MRC-University of Glasgow Centre for Virus Research | COVID-19 Genomics UK (COG-UK) Consortium | Ana da Silva Filipe, Natasha Johnson, Kathy Smollett, Daniel Mair, Stephen Carmichael, Lily Tong, Jenna Nichols, Elihu Aranday-Cortes, Kirstyn Brunker, Yasmin Parr, Kyriaki Nomikou; Sarah McDonald, Marc Niebel, Patawee Asamaphan; Richard Orton, Joseph Hughes, Sreenu Vattipally, David L Robertson; Alasdair MacLean, Rory Gunson; Kathy Li, Natasha Jesudason, Rajiv Shah, James Shepherd, Antonia Ho, Emma Thomson |
| EPI_ISL_433529 | hCoV-19/Scotland/CVR1271/2020 | Europe / United Kingdom / Scotland | 2020-03-31 | West of Scotland Specialist Virology Centre, NHSGGC / MRC-University of Glasgow Centre for Virus Research | COVID-19 Genomics UK (COG-UK) Consortium | Ana da Silva Filipe, Natasha Johnson, Kathy Smollett, Daniel Mair, Stephen Carmichael, Lily Tong, Jenna Nichols, Elihu Aranday-Cortes, Kirstyn Brunker, Yasmin Parr, Kyriaki Nomikou; Sarah McDonald, Marc Niebel, Patawee Asamaphan; Richard Orton, Joseph Hughes, Sreenu Vattipally, David L Robertson; Alasdair MacLean, Rory Gunson; Kathy Li, Natasha Jesudason, Rajiv Shah, James Shepherd, Antonia Ho, Emma Thomson |
| EPI_ISL_433534 | hCoV-19/Scotland/CVR1305/2020 | Europe / United Kingdom / Scotland | 2020-03-30 | West of Scotland Specialist Virology Centre, NHSGGC / MRC-University of Glasgow Centre for Virus Research | COVID-19 Genomics UK (COG-UK) Consortium | Ana da Silva Filipe, Natasha Johnson, Kathy Smollett, Daniel Mair, Stephen Carmichael, Lily Tong, Jenna Nichols, Elihu Aranday-Cortes, Kirstyn Brunker, Yasmin Parr, Kyriaki Nomikou; Sarah McDonald, Marc Niebel, Patawee Asamaphan; Richard Orton, Joseph Hughes, Sreenu Vattipally, David L Robertson; Alasdair MacLean, Rory Gunson; Kathy Li, Natasha Jesudason, Rajiv Shah, James Shepherd, Antonia Ho, Emma Thomson |
| EPI_ISL_433535 | hCoV-19/Scotland/CVR1308/2020 | Europe / United Kingdom / Scotland | 2020-03-26 | West of Scotland Specialist Virology Centre, NHSGGC / MRC-University of Glasgow Centre for Virus Research | COVID-19 Genomics UK (COG-UK) Consortium | Ana da Silva Filipe, Natasha Johnson, Kathy Smollett, Daniel Mair, Stephen Carmichael, Lily Tong, Jenna Nichols, Elihu Aranday-Cortes, Kirstyn Brunker, Yasmin Parr, Kyriaki Nomikou; Sarah McDonald, Marc Niebel, Patawee Asamaphan; Richard Orton, Joseph Hughes, Sreenu Vattipally, David L Robertson; Alasdair MacLean, Rory Gunson; Kathy Li, Natasha Jesudason, Rajiv Shah, James Shepherd, Antonia Ho, Emma Thomson |
| EPI_ISL_433536 | hCoV-19/Scotland/CVR1309/2020 | Europe / United Kingdom / Scotland | 2020-03-25 | West of Scotland Specialist Virology Centre, NHSGGC / MRC-University of Glasgow Centre for Virus Research | COVID-19 Genomics UK (COG-UK) Consortium | Ana da Silva Filipe, Natasha Johnson, Kathy Smollett, Daniel Mair, Stephen Carmichael, Lily Tong, Jenna Nichols, Elihu Aranday-Cortes, Kirstyn Brunker, Yasmin Parr, Kyriaki Nomikou; Sarah McDonald, Marc Niebel, Patawee Asamaphan; Richard Orton, Joseph Hughes, Sreenu Vattipally, David L Robertson; Alasdair MacLean, Rory Gunson; Kathy Li, Natasha Jesudason, Rajiv Shah, James Shepherd, Antonia Ho, Emma Thomson |
| EPI_ISL_433537 | hCoV-19/Scotland/CVR1311/2020 | Europe / United Kingdom / Scotland | 2020-03-27 | West of Scotland Specialist Virology Centre, NHSGGC / MRC-University of Glasgow Centre for Virus Research | COVID-19 Genomics UK (COG-UK) Consortium | Ana da Silva Filipe, Natasha Johnson, Kathy Smollett, Daniel Mair, Stephen Carmichael, Lily Tong, Jenna Nichols, Elihu Aranday-Cortes, Kirstyn Brunker, Yasmin Parr, Kyriaki Nomikou; Sarah McDonald, Marc Niebel, Patawee Asamaphan; Richard Orton, Joseph Hughes, Sreenu Vattipally, David L Robertson; Alasdair MacLean, Rory Gunson; Kathy Li, Natasha Jesudason, Rajiv Shah, James Shepherd, Antonia Ho, Emma Thomson |
| EPI_ISL_433538 | hCoV-19/Scotland/CVR1312/2020 | Europe / United Kingdom / Scotland | 2020-03-30 | West of Scotland Specialist Virology Centre, NHSGGC / MRC-University of Glasgow Centre for Virus Research | COVID-19 Genomics UK (COG-UK) Consortium | Ana da Silva Filipe, Natasha Johnson, Kathy Smollett, Daniel Mair, Stephen Carmichael, Lily Tong, Jenna Nichols, Elihu Aranday-Cortes, Kirstyn Brunker, Yasmin Parr, Kyriaki Nomikou; Sarah McDonald, Marc Niebel, Patawee Asamaphan; Richard Orton, Joseph Hughes, Sreenu Vattipally, David L Robertson; Alasdair MacLean, Rory Gunson; Kathy Li, Natasha Jesudason, Rajiv Shah, James Shepherd, Antonia Ho, Emma Thomson |
| EPI_ISL_433539 | hCoV-19/Scotland/CVR1314/2020 | Europe / United Kingdom / Scotland | 2020-03-31 | West of Scotland Specialist Virology Centre, NHSGGC / MRC-University of Glasgow Centre for Virus Research | COVID-19 Genomics UK (COG-UK) Consortium | Ana da Silva Filipe, Natasha Johnson, Kathy Smollett, Daniel Mair, Stephen Carmichael, Lily Tong, Jenna Nichols, Elihu Aranday-Cortes, Kirstyn Brunker, Yasmin Parr, Kyriaki Nomikou; Sarah McDonald, Marc Niebel, Patawee Asamaphan; Richard Orton, Joseph Hughes, Sreenu Vattipally, David L Robertson; Alasdair MacLean, Rory Gunson; Kathy Li, Natasha Jesudason, Rajiv Shah, James Shepherd, Antonia Ho, Emma Thomson |
| EPI_ISL_433541 | hCoV-19/Scotland/CVR1325/2020 | Europe / United Kingdom / Scotland | 2020-03-31 | West of Scotland Specialist Virology Centre, NHSGGC / MRC-University of Glasgow Centre for Virus Research | COVID-19 Genomics UK (COG-UK) Consortium | Ana da Silva Filipe, Natasha Johnson, Kathy Smollett, Daniel Mair, Stephen Carmichael, Lily Tong, Jenna Nichols, Elihu Aranday-Cortes, Kirstyn Brunker, Yasmin Parr, Kyriaki Nomikou; Sarah McDonald, Marc Niebel, Patawee Asamaphan; Richard Orton, Joseph Hughes, Sreenu Vattipally, David L Robertson; Alasdair MacLean, Rory Gunson; Kathy Li, Natasha Jesudason, Rajiv Shah, James Shepherd, Antonia Ho, Emma Thomson |
| EPI_ISL_433542 | hCoV-19/Scotland/CVR1329/2020 | Europe / United Kingdom / Scotland | 2020-03-31 | West of Scotland Specialist Virology Centre, NHSGGC / MRC-University of Glasgow Centre for Virus Research | COVID-19 Genomics UK (COG-UK) Consortium | Ana da Silva Filipe, Natasha Johnson, Kathy Smollett, Daniel Mair, Stephen Carmichael, Lily Tong, Jenna Nichols, Elihu Aranday-Cortes, Kirstyn Brunker, Yasmin Parr, Kyriaki Nomikou; Sarah McDonald, Marc Niebel, Patawee Asamaphan; Richard Orton, Joseph Hughes, Sreenu Vattipally, David L Robertson; Alasdair MacLean, Rory Gunson; Kathy Li, Natasha Jesudason, Rajiv Shah, James Shepherd, Antonia Ho, Emma Thomson |
| EPI_ISL_433543 | hCoV-19/Scotland/CVR1331/2020 | Europe / United Kingdom / Scotland | 2020-03-31 | West of Scotland Specialist Virology Centre, NHSGGC / MRC-University of Glasgow Centre for Virus Research | COVID-19 Genomics UK (COG-UK) Consortium | Ana da Silva Filipe, Natasha Johnson, Kathy Smollett, Daniel Mair, Stephen Carmichael, Lily Tong, Jenna Nichols, Elihu Aranday-Cortes, Kirstyn Brunker, Yasmin Parr, Kyriaki Nomikou; Sarah McDonald, Marc Niebel, Patawee Asamaphan; Richard Orton, Joseph Hughes, Sreenu Vattipally, David L Robertson; Alasdair MacLean, Rory Gunson; Kathy Li, Natasha Jesudason, Rajiv Shah, James Shepherd, Antonia Ho, Emma Thomson |
| EPI_ISL_433544 | hCoV-19/Scotland/CVR1334/2020 | Europe / United Kingdom / Scotland | 2020-03-31 | West of Scotland Specialist Virology Centre, NHSGGC / MRC-University of Glasgow Centre for Virus Research | COVID-19 Genomics UK (COG-UK) Consortium | Ana da Silva Filipe, Natasha Johnson, Kathy Smollett, Daniel Mair, Stephen Carmichael, Lily Tong, Jenna Nichols, Elihu Aranday-Cortes, Kirstyn Brunker, Yasmin Parr, Kyriaki Nomikou; Sarah McDonald, Marc Niebel, Patawee Asamaphan; Richard Orton, Joseph Hughes, Sreenu Vattipally, David L Robertson; Alasdair MacLean, Rory Gunson; Kathy Li, Natasha Jesudason, Rajiv Shah, James Shepherd, Antonia Ho, Emma Thomson |
| EPI_ISL_433545 | hCoV-19/Scotland/CVR1336/2020 | Europe / United Kingdom / Scotland | 2020-03-31 | West of Scotland Specialist Virology Centre, NHSGGC / MRC-University of Glasgow Centre for Virus Research | COVID-19 Genomics UK (COG-UK) Consortium | Ana da Silva Filipe, Natasha Johnson, Kathy Smollett, Daniel Mair, Stephen Carmichael, Lily Tong, Jenna Nichols, Elihu Aranday-Cortes, Kirstyn Brunker, Yasmin Parr, Kyriaki Nomikou; Sarah McDonald, Marc Niebel, Patawee Asamaphan; Richard Orton, Joseph Hughes, Sreenu Vattipally, David L Robertson; Alasdair MacLean, Rory Gunson; Kathy Li, Natasha Jesudason, Rajiv Shah, James Shepherd, Antonia Ho, Emma Thomson |
| EPI_ISL_433546 | hCoV-19/Scotland/CVR1337/2020 | Europe / United Kingdom / Scotland | 2020-03-31 | West of Scotland Specialist Virology Centre, NHSGGC / MRC-University of Glasgow Centre for Virus Research | COVID-19 Genomics UK (COG-UK) Consortium | Ana da Silva Filipe, Natasha Johnson, Kathy Smollett, Daniel Mair, Stephen Carmichael, Lily Tong, Jenna Nichols, Elihu Aranday-Cortes, Kirstyn Brunker, Yasmin Parr, Kyriaki Nomikou; Sarah McDonald, Marc Niebel, Patawee Asamaphan; Richard Orton, Joseph Hughes, Sreenu Vattipally, David L Robertson; Alasdair MacLean, Rory Gunson; Kathy Li, Natasha Jesudason, Rajiv Shah, James Shepherd, Antonia Ho, Emma Thomson |
| EPI_ISL_433547 | hCoV-19/Scotland/CVR1343/2020 | Europe / United Kingdom / Scotland | 2020-03-31 | West of Scotland Specialist Virology Centre, NHSGGC / MRC-University of Glasgow Centre for Virus Research | COVID-19 Genomics UK (COG-UK) Consortium | Ana da Silva Filipe, Natasha Johnson, Kathy Smollett, Daniel Mair, Stephen Carmichael, Lily Tong, Jenna Nichols, Elihu Aranday-Cortes, Kirstyn Brunker, Yasmin Parr, Kyriaki Nomikou; Sarah McDonald, Marc Niebel, Patawee Asamaphan; Richard Orton, Joseph Hughes, Sreenu Vattipally, David L Robertson; Alasdair MacLean, Rory Gunson; Kathy Li, Natasha Jesudason, Rajiv Shah, James Shepherd, Antonia Ho, Emma Thomson |
| EPI_ISL_433548 | hCoV-19/Scotland/CVR1345/2020 | Europe / United Kingdom / Scotland | 2020-03-31 | West of Scotland Specialist Virology Centre, NHSGGC / MRC-University of Glasgow Centre for Virus Research | COVID-19 Genomics UK (COG-UK) Consortium | Ana da Silva Filipe, Natasha Johnson, Kathy Smollett, Daniel Mair, Stephen Carmichael, Lily Tong, Jenna Nichols, Elihu Aranday-Cortes, Kirstyn Brunker, Yasmin Parr, Kyriaki Nomikou; Sarah McDonald, Marc Niebel, Patawee Asamaphan; Richard Orton, Joseph Hughes, Sreenu Vattipally, David L Robertson; Alasdair MacLean, Rory Gunson; Kathy Li, Natasha Jesudason, Rajiv Shah, James Shepherd, Antonia Ho, Emma Thomson |
| EPI_ISL_433549 | hCoV-19/Scotland/CVR1351/2020 | Europe / United Kingdom / Scotland | 2020-04-01 | West of Scotland Specialist Virology Centre, NHSGGC / MRC-University of Glasgow Centre for Virus Research | COVID-19 Genomics UK (COG-UK) Consortium | Ana da Silva Filipe, Natasha Johnson, Kathy Smollett, Daniel Mair, Stephen Carmichael, Lily Tong, Jenna Nichols, Elihu Aranday-Cortes, Kirstyn Brunker, Yasmin Parr, Kyriaki Nomikou; Sarah McDonald, Marc Niebel, Patawee Asamaphan; Richard Orton, Joseph Hughes, Sreenu Vattipally, David L Robertson; Alasdair MacLean, Rory Gunson; Kathy Li, Natasha Jesudason, Rajiv Shah, James Shepherd, Antonia Ho, Emma Thomson |
| EPI_ISL_433550 | hCoV-19/Scotland/CVR1352/2020 | Europe / United Kingdom / Scotland | 2020-04-01 | West of Scotland Specialist Virology Centre, NHSGGC / MRC-University of Glasgow Centre for Virus Research | COVID-19 Genomics UK (COG-UK) Consortium | Ana da Silva Filipe, Natasha Johnson, Kathy Smollett, Daniel Mair, Stephen Carmichael, Lily Tong, Jenna Nichols, Elihu Aranday-Cortes, Kirstyn Brunker, Yasmin Parr, Kyriaki Nomikou; Sarah McDonald, Marc Niebel, Patawee Asamaphan; Richard Orton, Joseph Hughes, Sreenu Vattipally, David L Robertson; Alasdair MacLean, Rory Gunson; Kathy Li, Natasha Jesudason, Rajiv Shah, James Shepherd, Antonia Ho, Emma Thomson |
| EPI_ISL_433551 | hCoV-19/Scotland/CVR1356/2020 | Europe / United Kingdom / Scotland | 2020-03-31 | West of Scotland Specialist Virology Centre, NHSGGC / MRC-University of Glasgow Centre for Virus Research | COVID-19 Genomics UK (COG-UK) Consortium | Ana da Silva Filipe, Natasha Johnson, Kathy Smollett, Daniel Mair, Stephen Carmichael, Lily Tong, Jenna Nichols, Elihu Aranday-Cortes, Kirstyn Brunker, Yasmin Parr, Kyriaki Nomikou; Sarah McDonald, Marc Niebel, Patawee Asamaphan; Richard Orton, Joseph Hughes, Sreenu Vattipally, David L Robertson; Alasdair MacLean, Rory Gunson; Kathy Li, Natasha Jesudason, Rajiv Shah, James Shepherd, Antonia Ho, Emma Thomson |
| EPI_ISL_433552 | hCoV-19/Scotland/CVR1358/2020 | Europe / United Kingdom / Scotland | 2020-03-31 | West of Scotland Specialist Virology Centre, NHSGGC / MRC-University of Glasgow Centre for Virus Research | COVID-19 Genomics UK (COG-UK) Consortium | Ana da Silva Filipe, Natasha Johnson, Kathy Smollett, Daniel Mair, Stephen Carmichael, Lily Tong, Jenna Nichols, Elihu Aranday-Cortes, Kirstyn Brunker, Yasmin Parr, Kyriaki Nomikou; Sarah McDonald, Marc Niebel, Patawee Asamaphan; Richard Orton, Joseph Hughes, Sreenu Vattipally, David L Robertson; Alasdair MacLean, Rory Gunson; Kathy Li, Natasha Jesudason, Rajiv Shah, James Shepherd, Antonia Ho, Emma Thomson |
| EPI_ISL_433554 | hCoV-19/Scotland/CVR1361/2020 | Europe / United Kingdom / Scotland | 2020-04-01 | West of Scotland Specialist Virology Centre, NHSGGC / MRC-University of Glasgow Centre for Virus Research | COVID-19 Genomics UK (COG-UK) Consortium | Ana da Silva Filipe, Natasha Johnson, Kathy Smollett, Daniel Mair, Stephen Carmichael, Lily Tong, Jenna Nichols, Elihu Aranday-Cortes, Kirstyn Brunker, Yasmin Parr, Kyriaki Nomikou; Sarah McDonald, Marc Niebel, Patawee Asamaphan; Richard Orton, Joseph Hughes, Sreenu Vattipally, David L Robertson; Alasdair MacLean, Rory Gunson; Kathy Li, Natasha Jesudason, Rajiv Shah, James Shepherd, Antonia Ho, Emma Thomson |
| EPI_ISL_433557 | hCoV-19/Scotland/CVR1366/2020 | Europe / United Kingdom / Scotland | 2020-04-01 | West of Scotland Specialist Virology Centre, NHSGGC / MRC-University of Glasgow Centre for Virus Research | COVID-19 Genomics UK (COG-UK) Consortium | Ana da Silva Filipe, Natasha Johnson, Kathy Smollett, Daniel Mair, Stephen Carmichael, Lily Tong, Jenna Nichols, Elihu Aranday-Cortes, Kirstyn Brunker, Yasmin Parr, Kyriaki Nomikou; Sarah McDonald, Marc Niebel, Patawee Asamaphan; Richard Orton, Joseph Hughes, Sreenu Vattipally, David L Robertson; Alasdair MacLean, Rory Gunson; Kathy Li, Natasha Jesudason, Rajiv Shah, James Shepherd, Antonia Ho, Emma Thomson |
| EPI_ISL_433559 | hCoV-19/Scotland/CVR1371/2020 | Europe / United Kingdom / Scotland | 2020-03-31 | West of Scotland Specialist Virology Centre, NHSGGC / MRC-University of Glasgow Centre for Virus Research | COVID-19 Genomics UK (COG-UK) Consortium | Ana da Silva Filipe, Natasha Johnson, Kathy Smollett, Daniel Mair, Stephen Carmichael, Lily Tong, Jenna Nichols, Elihu Aranday-Cortes, Kirstyn Brunker, Yasmin Parr, Kyriaki Nomikou; Sarah McDonald, Marc Niebel, Patawee Asamaphan; Richard Orton, Joseph Hughes, Sreenu Vattipally, David L Robertson; Alasdair MacLean, Rory Gunson; Kathy Li, Natasha Jesudason, Rajiv Shah, James Shepherd, Antonia Ho, Emma Thomson |
| EPI_ISL_433560 | hCoV-19/Scotland/CVR1373/2020 | Europe / United Kingdom / Scotland | 2020-04-01 | West of Scotland Specialist Virology Centre, NHSGGC / MRC-University of Glasgow Centre for Virus Research | COVID-19 Genomics UK (COG-UK) Consortium | Ana da Silva Filipe, Natasha Johnson, Kathy Smollett, Daniel Mair, Stephen Carmichael, Lily Tong, Jenna Nichols, Elihu Aranday-Cortes, Kirstyn Brunker, Yasmin Parr, Kyriaki Nomikou; Sarah McDonald, Marc Niebel, Patawee Asamaphan; Richard Orton, Joseph Hughes, Sreenu Vattipally, David L Robertson; Alasdair MacLean, Rory Gunson; Kathy Li, Natasha Jesudason, Rajiv Shah, James Shepherd, Antonia Ho, Emma Thomson |
| EPI_ISL_433561 | hCoV-19/Scotland/CVR1377/2020 | Europe / United Kingdom / Scotland | 2020-04-01 | West of Scotland Specialist Virology Centre, NHSGGC / MRC-University of Glasgow Centre for Virus Research | COVID-19 Genomics UK (COG-UK) Consortium | Ana da Silva Filipe, Natasha Johnson, Kathy Smollett, Daniel Mair, Stephen Carmichael, Lily Tong, Jenna Nichols, Elihu Aranday-Cortes, Kirstyn Brunker, Yasmin Parr, Kyriaki Nomikou; Sarah McDonald, Marc Niebel, Patawee Asamaphan; Richard Orton, Joseph Hughes, Sreenu Vattipally, David L Robertson; Alasdair MacLean, Rory Gunson; Kathy Li, Natasha Jesudason, Rajiv Shah, James Shepherd, Antonia Ho, Emma Thomson |
| EPI_ISL_433562 | hCoV-19/Scotland/CVR1379/2020 | Europe / United Kingdom / Scotland | 2020-04-01 | West of Scotland Specialist Virology Centre, NHSGGC / MRC-University of Glasgow Centre for Virus Research | COVID-19 Genomics UK (COG-UK) Consortium | Ana da Silva Filipe, Natasha Johnson, Kathy Smollett, Daniel Mair, Stephen Carmichael, Lily Tong, Jenna Nichols, Elihu Aranday-Cortes, Kirstyn Brunker, Yasmin Parr, Kyriaki Nomikou; Sarah McDonald, Marc Niebel, Patawee Asamaphan; Richard Orton, Joseph Hughes, Sreenu Vattipally, David L Robertson; Alasdair MacLean, Rory Gunson; Kathy Li, Natasha Jesudason, Rajiv Shah, James Shepherd, Antonia Ho, Emma Thomson |
| EPI_ISL_433563 | hCoV-19/Scotland/CVR1384/2020 | Europe / United Kingdom / Scotland | 2020-04-01 | West of Scotland Specialist Virology Centre, NHSGGC / MRC-University of Glasgow Centre for Virus Research | COVID-19 Genomics UK (COG-UK) Consortium | Ana da Silva Filipe, Natasha Johnson, Kathy Smollett, Daniel Mair, Stephen Carmichael, Lily Tong, Jenna Nichols, Elihu Aranday-Cortes, Kirstyn Brunker, Yasmin Parr, Kyriaki Nomikou; Sarah McDonald, Marc Niebel, Patawee Asamaphan; Richard Orton, Joseph Hughes, Sreenu Vattipally, David L Robertson; Alasdair MacLean, Rory Gunson; Kathy Li, Natasha Jesudason, Rajiv Shah, James Shepherd, Antonia Ho, Emma Thomson |
| EPI_ISL_433565 | hCoV-19/Scotland/CVR1387/2020 | Europe / United Kingdom / Scotland | 2020-04-01 | West of Scotland Specialist Virology Centre, NHSGGC / MRC-University of Glasgow Centre for Virus Research | COVID-19 Genomics UK (COG-UK) Consortium | Ana da Silva Filipe, Natasha Johnson, Kathy Smollett, Daniel Mair, Stephen Carmichael, Lily Tong, Jenna Nichols, Elihu Aranday-Cortes, Kirstyn Brunker, Yasmin Parr, Kyriaki Nomikou; Sarah McDonald, Marc Niebel, Patawee Asamaphan; Richard Orton, Joseph Hughes, Sreenu Vattipally, David L Robertson; Alasdair MacLean, Rory Gunson; Kathy Li, Natasha Jesudason, Rajiv Shah, James Shepherd, Antonia Ho, Emma Thomson |
| EPI_ISL_433566 | hCoV-19/Scotland/CVR1393/2020 | Europe / United Kingdom / Scotland | 2020-03-31 | West of Scotland Specialist Virology Centre, NHSGGC / MRC-University of Glasgow Centre for Virus Research | COVID-19 Genomics UK (COG-UK) Consortium | Ana da Silva Filipe, Natasha Johnson, Kathy Smollett, Daniel Mair, Stephen Carmichael, Lily Tong, Jenna Nichols, Elihu Aranday-Cortes, Kirstyn Brunker, Yasmin Parr, Kyriaki Nomikou; Sarah McDonald, Marc Niebel, Patawee Asamaphan; Richard Orton, Joseph Hughes, Sreenu Vattipally, David L Robertson; Alasdair MacLean, Rory Gunson; Kathy Li, Natasha Jesudason, Rajiv Shah, James Shepherd, Antonia Ho, Emma Thomson |
